# Huntingtin loss-of-function contributes to transcriptional deregulation in Huntington’s disease

**DOI:** 10.1101/2024.05.20.594947

**Authors:** Kozłowska Emilia, Ciołak Agata, Adamek Grażyna, Szcześniak Julia, Fiszer Agnieszka

**Author notes:** Corresponding author, Institute of Bioorganic Chemistry, Polish Academy of Sciences, Noskowskiego 12/14 Str., 61-704 Poznań, Poland; (ext. 1450).

## Abstract

Huntington’s disease (HD) is a fatal neurodegenerative disorder that is caused by the expansion of CAG repeats in the *HTT* gene, which results in a long polyglutamine (polyQ) tract in the huntingtin protein (HTT). In this study, we searched for networks of deregulated RNAs that contribute to initial transcriptional changes in HD neuronal cells and HTT-deficient cells.

We used RNA-seq (including small RNA sequencing) to analyze a set of isogenic, human induced pluripotent stem cell (iPSC)-derived neural stem cells (NSCs); and we observed numerous changes in gene expression and substantial dysregulation of miRNA expression in HD and *HTT*-knockout (*HTT*-KO) cell lines. The gene set that was upregulated in both HD and *HTT*-KO cells was enriched in genes that are associated with DNA binding and regulation of transcription. For both of these models, we confirmed the substantial upregulation of the transcription factors (TFs) *TWIST1, SIX1, TBX1, TBX15, MSX2, MEOX2* and *FOXD1* in NSCs and medium spiny neuron (MSN)-like cells. Moreover, we identified miRNAs that were consistently deregulated in HD and *HTT*-KO NSCs and MSN-like cells, including miR-214, miR-199, and miR-9. We suggest that these miRNAs function in the network that regulates *TWIST1* and *HTT* expression *via* regulatory feed-forward loop (FFL) in HD. Additionally, we reported that the expression of selected TFs and miRNAs tended to progressively change during the neural differentiation of HD cells, what was not observed in *HTT*-KO model. Based on comparing the HD and *HTT*-KO cell lines, we propose that early transcriptional deregulation in HD is largely caused by loss of HTT function.

## Introduction

Huntington’s disease (HD) is an autosomal dominant neurodegenerative disorder that is caused by expansion of the CAG repeat in the huntingtin *(HTT)* gene [1, 2]. This mutation results in an abnormally long polyglutamine (polyQ) sequence in a large, multifunctional protein called huntingtin (HTT). Mutant HTT (mutHTT) exhibits toxic properties and causes the dysfunction and death of neurons, especially medium spiny neurons (MSNs) in the striatum and, in the later stages of HD, neurons in the cerebral cortex. Due to this particular pattern of neurodegeneration, HD is characterized primarily by cognitive and behavioral disturbances and motor dysfunctions like chorea. The disease manifests at a mean age of 40 years and causes death 15–20 years after the appearance of the first motor symptoms [3].

The CAG tract, which is located in the first exon of *HTT,* exhibits polymorphisms in length in the population [4]. Normal alleles contain <35 CAG repeats, whereas full penetrance alleles contain more than 39 CAG repeats and are associated with the development of clinical signs and symptoms of HD. Patients with more than 56 CAG repeats in *HTT* usually develop juvenile HD characterized by the onset before 20 years of age [5].

HTT is a protein that is expressed in various cell types and localizes to various subcellular compartments. However, its expression levels vary depending on the cell compartment or cell type. Enrichment of HTT in the nucleus has been observed during embryogenesis and in brain cells [6]. The normal functions of HTT are still being defined. Wild-type huntingtin (wtHTT) is a multifunctional protein that plays roles in apoptosis, endocytosis, vesicle recycling, endosomal trafficking, tissue maintenance, cell morphology, neurogenesis and postsynaptic signaling [7]. Moreover, wtHTT interacts with numerous protein partners that are involved in processes such as signal transduction, cellular homeostasis, and transcription [8, 9].

HD patients usually have one normal and one mutant *HTT* allele but there are rare cases of biallelic mutation [10, 11]. Although the mutHTT gain-of-function (GoF) mechanism is well recognized [12–14], the loss of the WT protein may significantly contribute to the pathogenesis of HD [15] due to large amount of data indicating that wtHTT plays a role in neuronal survival [7, 16]. HTT loss-of-function (LoF) mechanisms have been widely studied in mice, and the crucial role of Htt in the embryonic stage and its important functions in adult animals have been proven. The complete knockout of the mouse *Htt* gene caused embryonic death before gastrulation and nervous system formation [17, 18], and mice with a 50% reduction in the level of endogenous *Htt* exhibited strong malformations of the cortex and striatum [17, 19]. Additionally, mice with substantially decreased levels of wtHtt from the embryonic stage showed motor abnormalities and neurodegenerative changes [20, 21] or developed seizure disorder [22]. Elimination of Htt in the developing or early postnatal mouse brain caused apoptosis and axon degeneration in neurons, resulting in striatum and cortex degradation and death within 13 months [23]. In adult mice, inactivation of wt*Htt* causes severe motor function and behavioral impairment, progressive brain atrophy, bilateral thalamic calcification, and disrupted brain iron homeostasis [24].

Transcriptional dysregulation has been proposed to be one of the earliest and central molecular mechanisms underlying HD pathogenesis [25–27]. Experimental evidence from human tissues and *in vivo* and *in vitro* HD models has demonstrated massive changes in the levels of protein-coding and noncoding RNAs, such as miRNAs [28–31]. A variety of mechanisms have been suggested to explain how mutHTT or loss of wtHTT causes transcriptional dysregulation, including altered interactions with positive or negative regulators of transcription [27]. In this study, we searched for networks of deregulated RNAs, which include transcription factors (TFs) and miRNAs, that contribute to the initial molecular disruptions in HD neuronal cells. Based on a comparison of neural HD and *HTT*-KO cell lines, with reference to isogenic control lines, we propose that a deficiency of properly functioning HTT substantially contributes to the transcriptional deregulation that is observed in HD.

## Materials and methods

### iPSC lines

The human HD iPSC line ND42222 (with 19/109 CAG repeats in *HTT)* was obtained from the NINDS Repository. The generation of two isogenic control lines (C105 and C39) and an *HTT*-knockout line (C37) *via* CRISPR-Cas9 editing of the ND42222 cell line was previously described [32]. For clarity, in this study, the ND42222 line is called HD, C105 is called IC1, C39 is called IC2 and C37 is called KO (Fig. 1 a). IC1 was initially intended to be used as an isogenic control for all the analyses in this study but, in the meantime, we observed that the expression of the corrected allele was inhibited in this clone, resulting in the monoallelic expression of *HTT* [33]. The experiment descriptions and figure legends indicate whether IC1 or IC2 was used.

**Figure 1.**
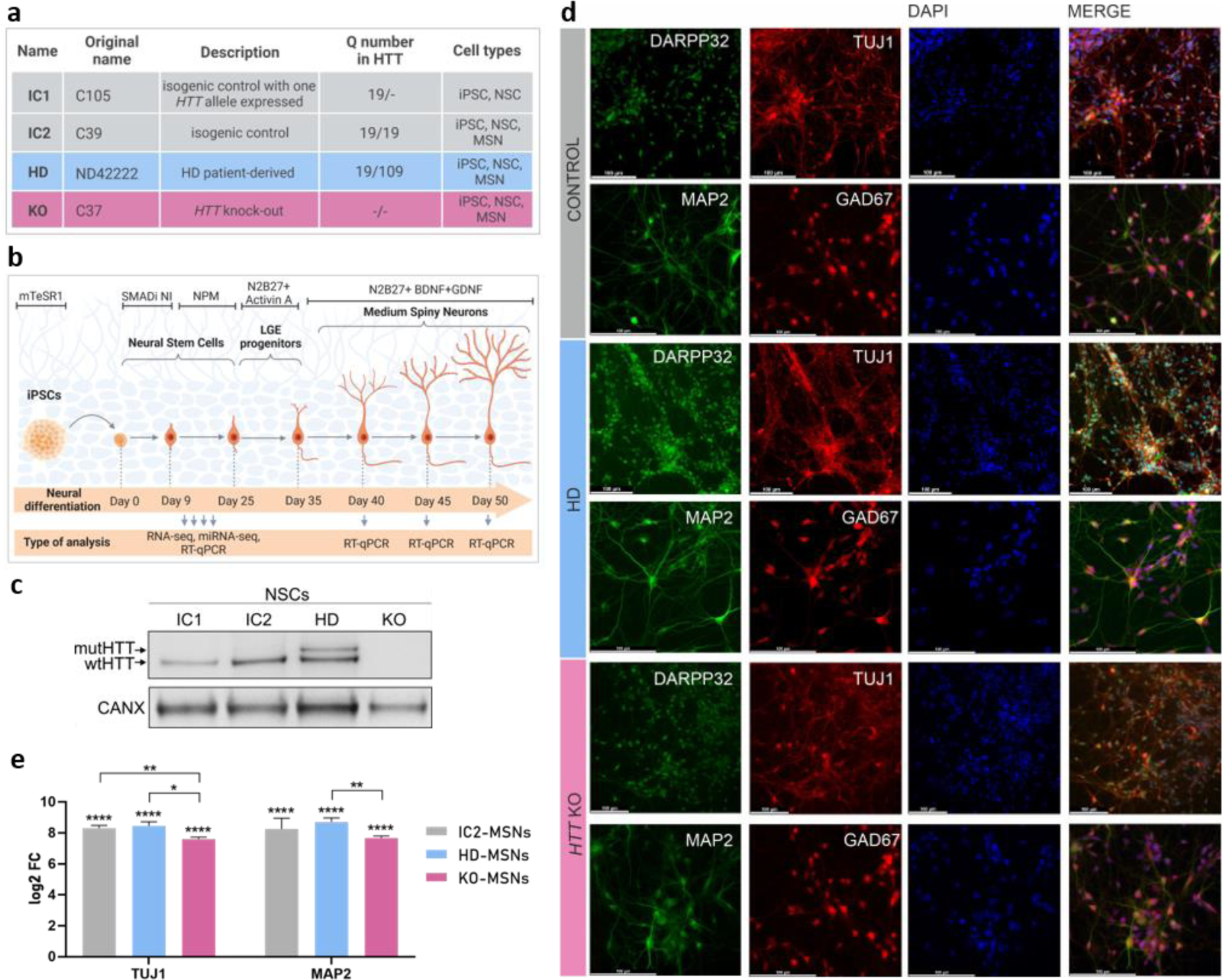
Neuronal differentiation of a set of isogenic iPSC lines. (a) Table with information about the investigated isogenic cell lines, including the nomenclature used, the number of CAG repeats in the expressed *HTT* alleles, and the cell types used in the experiments. (b) Schedule of neuronal differentiation. The arrows at the bottom indicate the points at which the cells were harvested for RNA isolation. (**c**) Western blotting analysis of huntingtin levels in iPSC-derived IC1-NSCs, IC2-NSCs, HD-NSCs and KO-NSCs. Calnexin was used as a reference protein. (**d**) Representative images of ICC staining for the markers MAP2, TUJ1, GAD67 and DARPP32 in the obtained neurons. Scale bar = 100 µm (**e**) Relative expression of the neuronal markers *TUJ1* and *MAP2* in iPSC-derived IC2-MSNs, HD-MSNs and KO-MSNs was assessed by RT‒qPCR. FC was calculated relative to that of iPSCs using the *delta*-*delta Ct* method and is shown as log2. Reference gene: *RPLP0*. Statistical analysis was performed using multiple *t* tests. *0.01<p<0,05; **p<0.01; ***p<0.001; ****p <0.0001.

### Differentiation of iPSCs into NSCs

A set of iPSC lines, namely, HD, IC1, IC2 and KO cell lines, was differentiated into NSCs using a STEMdiff SMADi Neural Induction Kit (STEMCELL Technologies) and a monolayer protocol following the manufacturer’s instructions. Briefly, iPSCs were grown in StemFlex (Gibco) medium on a Geltrex (Gibco)-coated 6-well plate until they reached 70–80% confluence. Then, the iPSCs were dissociated into single-cell suspensions by incubation with Accutase (STEMCELL Technologies) for 5-7 min (at 37°C). The cells were counted using a TC20 Automated Cell Counter (Bio-Rad), resuspended at a concentration of 1×10^6^ cells/mL, and seeded on Geltrex-coated plates in STEMdiff Neural Induction Medium supplemented with SMADi and 10 nM Y-27632 (a ROCK inhibitor) (all reagents from STEMCELL Technologies). For further cultivation, the cells were detached using Accutase, and after the third passage, they were grown in STEMdiff Neural Progenitor Medium (STEMCELL Technologies). After the fourth passage, the expression of the markers *SOX1, SOX2, PAX6*, and *NES* was confirmed by immunocytochemistry ICC (data not shown).

### Differentiation of NSCs into neuronal cells

The neural differentiation of NSCs was performed according to the protocol published by M. Fjodorova and M. Li [34] with slight modifications. Briefly, NSCs were grown in STEMdiff Neural Progenitor Medium (NPM) until they reached full confluence and then dissociated into single-cell suspensions by incubation with Accutase. The collected cells were passaged at a 1:5 ratio onto a Geltrex-coated 6-well plate in NPM supplemented with 10 nM Y-27632. The next day, the NPM was exchanged for lateral ganglionic eminence (LGE) pattering medium (N2B27 supplemented with 25 ng/mL recombinant human activin A) (STEMCELL Technologies). The medium was changed every day until day 10 when the LGE progenitors were passaged for terminal differentiation into MSNs. Then, the cells were passaged with Accutase at a ratio of 1:5 onto PDL/laminin-coated 6-well and 12-well plates (for ICC) in LGE pattering medium supplemented with 10 nM Y-27632. The next day, the LGE pattering medium was exchanged for terminal differentiation medium (N2B27 supplemented with 10 ng/mL recombinant human BDNF and 10 ng/mL recombinant human GDNF) (both from STEMCELL Technologies). The neuronal medium was half-replaced every 2 days until day 50 of differentiation.

### ICC

Neuronal cells were cultured on PDL/laminin-coated glass coverslips. First, the cells were fixed by adding 2% PFA directly to the cell culture medium and culturing the cells for 5 min, and then, the cells were gently washed with PBS and fixed with 4% PFA in PBS for 10 min. The cells were permeabilized with 0.5% Tween, blocked with 1% BSA, and then incubated with primary antibodies and fluorescent dye-conjugated secondary antibodies (all listed in Table S1). DAPI was used for nuclear staining. Images were captured with a Leica DMI6000 microscope.

### Protein isolation and western blotting

For protein isolation, cells were detached with Accutase and centrifuged at 300 rpm for 4 min. The pellets were washed with PBS, lysed in PB buffer (60 mM Tris-base, 2% SDS, 10% sucrose, 2 mM PMSF) and incubated at 95°C for 5 min. The protein concentrations were determined by measuring the absorbance at 280 nm using a DeNovix spectrophotometer. Next, 30 μg of total protein was diluted in 4x SDS loading buffer, denatured at 95°C for 5 min and run on 3–8% NuPAGE Tris-acetate gels in NuPAGE Tris-acetate SDS Running Buffer (20x) (Thermo Fisher Scientific) at 4°C. The proteins were wet transferred overnight onto a 0.45-μm nitrocellulose membrane (GE Healthcare) in ice-cold Towbin buffer at 4°C. Then, specific primary and secondary antibodies, which are listed in Table S1, were added and incubated with the membranes. Immunodetection was performed using Westar Supernova XLS3 (Cyanagen). The chemiluminescent signals on the membranes were scanned using a G:BOX documentation system (Syngene).

### RNA isolation

Total RNA was isolated from cultured cells with Direct-zol RNA Microprep (ZYMO RESEARCH) according to the manufacturer’s protocol. Due to the lower number of cells, total RNA was isolated from MSNs and NSCs after electroporation and transfection with an Arcturus PicoPure RNA Isolation Kit (Thermo Fisher Scientific). The concentration of isolated total RNA was calculated by measuring the absorbance at 260 nm using a DeNovix spectrophotometer. For RNA-seq, RNA quality was assessed using an RNA Pico 6000 kit (Agilent) and a Bioanalyzer 2100 (Agilent).

### RNA-seq

Total RNA was isolated from 3 biological replicates of control NSCs (IC1; 4th to 6th passage) and 4 biological replicates of HD and *HTT*-KO NSC lines (4th to 7th passage). RNA-seq was performed by The Genomics Core Facility in The Centre of New Technologies, University of Warsaw, Poland. Sequencing libraries were generated using the KAPA RNA HyperPrep Kit with RiboErase (HMR) (Kapa Biosciences) and IDT adapters for Illumina TruSeq DNA UD Indexes (96 Indexes, 96 Samples) for total RNA or using the TruSeq Small RNA Library Prep (Illumina) for miRNAs. The quality and quantity of the sequencing libraries were analyzed using a Bioanalyzer 2100 (Agilent) with a High Sensitivity DNA Kit (Agilent) and a Kapa Library Quantification Kit (Kapa Biosciences).

The samples were sequenced with the NovaSeq 6000 system (Illumina) with a paired-end 2x100 cycle procedure, 50 MR/sample for total RNA and 10 MR/sample for miRNAs. Bioinformatics analysis was performed by the IDEAS4BIOLOGY company. Quality reports were generated with FastQC v0.11.9. The reads were subjected to quality filtering and adapter trimming using BBDUK 2 v. 37.02. rRNAs mapped reads were removed following alignment with bowtie2 v. 2.3.5.1, with the -X 1000 parameter (for mapped read distance) and the *--un-conc* parameter. The expression values were obtained with RSEM v. 1.3.1, using default settings and bowtie2 as an alignment method. The ENSEMBL v. 102 and GRCh38 genome assembly were used as references. Differential expression analysis was performed using DESeq2 v. 1.30.0.

PANTHER v.17 [35] was used for Gene Ontology (GO) analysis of the DEGs (|log2FC| >1.5; padj < 0.05) in the HD and *HTT*-KO cell lines. Significant GO terms were determined by Fisher’s exact test after FDR correction at p < 0.05 and sorted by fold enrichment.

The expression correlation of selected genes and miRNAs was calculated using the Spearman method. The correlation matrix generated with *cor* function (*stats* R package) was visualized with corrplot v. 0.92, employing hierarchical clustering ordering.

Full RNA-seq data will be available on GEO with a peer-reviewed publication.

### Bioinformatics analysis of genes whose expression increased or decreased over time

To select genes whose expression increased or decreased with time in NSCs according to RNA-seq data, we used the following criteria: (I) genes with TPM values below 1 at each time point were excluded; (II) genes that were classified as “increasing” or “decreasing” showed a consequent increase or decrease in TPM values, respectively, at subsequent passages; and (III) genes whose expression exhibited at least a 25% change in expression between the first and last time points were included. The percentage change in expression was calculated as the difference in the expression values at the first and last time points divided by the initial expression value and multiplied by 100%. The genes were sorted starting from the genes with the greatest changes in expression. GO enrichment analysis was conducted on selected sets of “increasing” and “decreasing” genes using the *enrichGO* function from the clusterProfiler v. 4.10.0 R package. Significantly enriched GO terms were identified through Benjamini–Hochberg correction (cutoff padj < 0.05, q value < 0.05). The results were sorted by GeneRatio and visualized using a *dotplot* (enrichplot v. 1.22.0 R package).

### RT–qPCR

To analyze mRNA levels, reverse transcription (RT) was performed using a High-Capacity cDNA Reverse Transcription Kit (Applied Biosystems) with random primers according to the manufacturer’s protocols. RT‒qPCR was performed using SsoAdvanced Universal SYBR Green Supermix (Bio-Rad) and a CFX Connect Real-Time System (Bio-Rad) according to the manufacturer’s protocols and established RT‒qPCR guidelines. Based on our RNA-seq data, we selected the genes with the most stable expression, namely, *RPLP0* and *EEF2,* as endogenous controls for data normalization. A list of primers is provided in Table S2.

RT for miRNA analysis was performed using the TaqMan Advanced miRNA cDNA Synthesis Kit (Applied Biosystems). RT‒qPCR was performed using TaqMan Advanced miRNA Assays (Applied Biosystems) according to the manufacturer’s instructions. The levels of miR-92a and miR-16 were utilized as endogenous controls to normalize the data.

All the RT‒qPCRs were performed in triplicate. The comparative cycle threshold (CT) 2^−ΔΔCT^ method was used to calculate the relative expression of mRNAs and miRNAs.

### Electroporation of NSCs with plasmids

To prepare plasmids expressing HTT-19Q, HTT-109Q and GFP (pCD, cat #CD550A-1), bacterial cultures (*E.coli DH5ⲁ*) from stabs were spread on LB agar plates supplemented with 100 µg/mL ampicillin and incubated overnight at 37°C. Then, single colonies were picked, inoculated in 5 ml of LB supplemented with 100 µg/mL ampicillin, and incubated for 8 h at 37°C with shaking at 250 rpm. Next, starter cultures were diluted 1:1000 in LB medium supplemented with 100 µg/mL ampicillin and incubated at 37°C with shaking at 250 rpm. Then, plasmids were isolated using the GeneJet Plasmid Maxiprep Kit (Thermo Scientific). KO-NSCs between the 7th and 10th passages were dissociated into single-cell suspensions using Accutase and counted using a TC20 Automated Cell Counter (Bio-Rad). KO-NSCs were electroporated with the Neon Transfection System (Invitrogen). A total of 6 × 10^5^ cells were resuspended in buffer R and combined with 50 μg of the HTT-19Q plasmid, 50 μg of the HTT-109Q plasmid or 35 μg of the GFP plasmid. Cells were electroporated in 10 μL tips using the following conditions: 1150 V, 10 ms, 3 pulses. After electroporation, the cells were seeded on Geltrex-coated 12-well plates in STEMdiff Neural Progenitor Medium (STEMCELL Technologies) supplemented with 10 nM Y-27632. 48 h after electroporation, the cells were harvested for total RNA and protein isolation. HTT-19Q and HTT-109Q plasmids were precisely pBacMam2-DiEx-LIC-C-flag_huntingtin_full-length_Q19 and pBacMam2-DiEx-LIC-C-flag_huntingtin_full-length_Q109, respectively, and were a gift from Cheryl Arrowsmith (Addgene plasmid #111741; http://n2t.net/addgene:111741; RRID:Addgene_111741 and Addgene plasmid #111730; http://n2t.net/addgene:111730; RRID:Addgene_111730) [35].

### Transfection of NSCs with the miR mimic

For miR-9 overexpression, NSCs (after 4th passage) were cultured on Geltrex-treated 12-well plates in NPM for 24 h until they reached 30 to 50% confluence. Lipofectamine 2000 (Thermo Fisher) transfection reagent was mixed with 1 nM, 10 nM, or 30 nM miR-9 mimic (miRVana; cat# 4464066) or 20 nM control fluorescent BlockIT siRNA (Thermo Fisher) to monitor the transfection efficiency. After 20 min, the mixture was applied to the NSC cultures. After 4 h, the medium was replaced with fresh medium supplemented with 10 nM Y-27632, and after another 24 h, the medium was replaced with medium without Y-27632. The cells were harvested after 48 h for total RNA isolation.

### Statistical analysis

The experiments were repeated at least three times. The graphs presenting the values and error bars (mean ±SEM) were generated using GraphPad Prism 8 software. The statistical significance was calculated using multiple *t* tests after checking that the data followed a normal distribution. p values < 0.05 were considered to indicate a significant difference.

The representation factor (RF) was calculated using the tool provided by http://nemates.org/MA/progs/overlap_stats.html and is defined as the number of overlapping genes divided by the expected number of overlapping genes drawn from two independent groups. An RF > 1 indicates more overlap than random (expected from two independent groups), and an RF < 1 indicates less overlap than random. The probability of each overlap was determined using the hypergeometric probability formula.

## Results

### Neuronal differentiation of a set of isogenic cell lines with different *HTT* variants

In this study, we aimed to reveal and compare networks of deregulated RNAs during the neural differentiation of a set of isogenic human cell lines in the context of HTT function and dysfunction. To investigate both mechanisms underlying HD, namely, mutHTT GoF and wtHTT LoF, we used an HD cell line endogenously expressing *mutHTT* (together with the *wtHTT* allele), control lines expressing normal *HTT* (named IC1 and IC2, which express one and two *wtHTT* alleles, respectively), and an *HTT-* knockout line (named KO) (Fig. 1 a). The generation of a set of isogenic iPSC lines *via* CRISPR-Cas9 editing of the HD line was previously described [32]. To model neural cells from the region that is most affected in HD brain, the striatum, we differentiated the control, HD and KO iPSCs into NSCs, then to striatal progenitors, and subsequently to MSN-like cells (Fig. 1 b). We observed lower protein levels of wtHTT in IC1-NSCs than in IC2-NSCs (Fig. 1 c), which is consistent with the RT-ddPCR results of transcription in these cell lines [33]; this result is the consequence of one *HTT* allele being inactive in IC1 cells. As expected, HTT was not detected in KO-NSCs (Fig. 1 c). We confirmed the neuronal state of the obtained MSNs by ICC staining for striatal markers, DARPP-32 and GAD67, and other neuronal markers, TUJ1 and MAP2 (Fig. 1 d). Moreover, *TUJ1* and *MAP2* expression was quantified using RT‒ qPCR, and a more than 150-fold change (FC, presented as log2 FC) in their expression was observed in all the MSN lines compared to iPSCs (Fig. 1 e). However, *TUJ1* expression was significantly lower in the obtained KO-MSNs than in other MSN lines, and *MAP2* expression was lower in the KO-MSNs than in the HD-MSNs. Generally, no clear differences in the cell morphology of control, HD and KO cells were observed; however, the KO line was unstable (KO-iPSCs tended to spontaneously differentiate into nondefined cell types) and was relatively difficult to differentiate (some neural differentiation experiments with this cell line were unsuccessful).

### Overlap of transcriptional changes in mut*HTT*-expressing and *HTT*-knockout neuronal cells

RNA was isolated from IC1, HD, and KO NSC lines at subsequent passages, and then RNA sequencing was performed to assess differences in the transcriptome between the cell lines. Hierarchical clustering showed a clear separation of the HD and KO datasets from the control NSC dataset (Fig. S1). RNA-seq reads from the HD and KO lines were compared with those from the IC1 line, and the analysis revealed a substantially higher number of differentially expressed genes (DEGs) in the KO line than in the HD line (Fig. 2 a-c). Specifically, we identified 1401 DEGs in KO-NSCs and 822 DEGs in HD-NSCs (|log2FC| > 1.5; padj < 0.05) (Fig. 2 c; a full list of identified genes is available in Table S3). A highly significant overlap of 331 DEGs was confirmed by calculating the RF, which equaled 9; this suggested a much larger group of common DEGs between KO-NSCs and HD-NSCs than random (Fig. 2 c). In the HD-NSC model, a similar number of genes, 387 and 435, were upregulated and downregulated, respectively (Fig. 2 d, e). In the case of KO-NSCs, we observed a large disproportion, which included 1231 upregulated genes and only 170 downregulated genes (Fig. 2 d, e). The HD and KO lines shared 280 upregulated genes (Fig. 2 d) and 40 downregulated genes (Fig. 2 e), and these overlaps were characterized by high statistically significant RFs (∼18 and ∼17, respectively). The overlapping results were especially striking in the HD line, as the majority (∼70%) of upregulated genes in this model were also upregulated in the KO model (Fig. 2 d).

**Figure 2.**
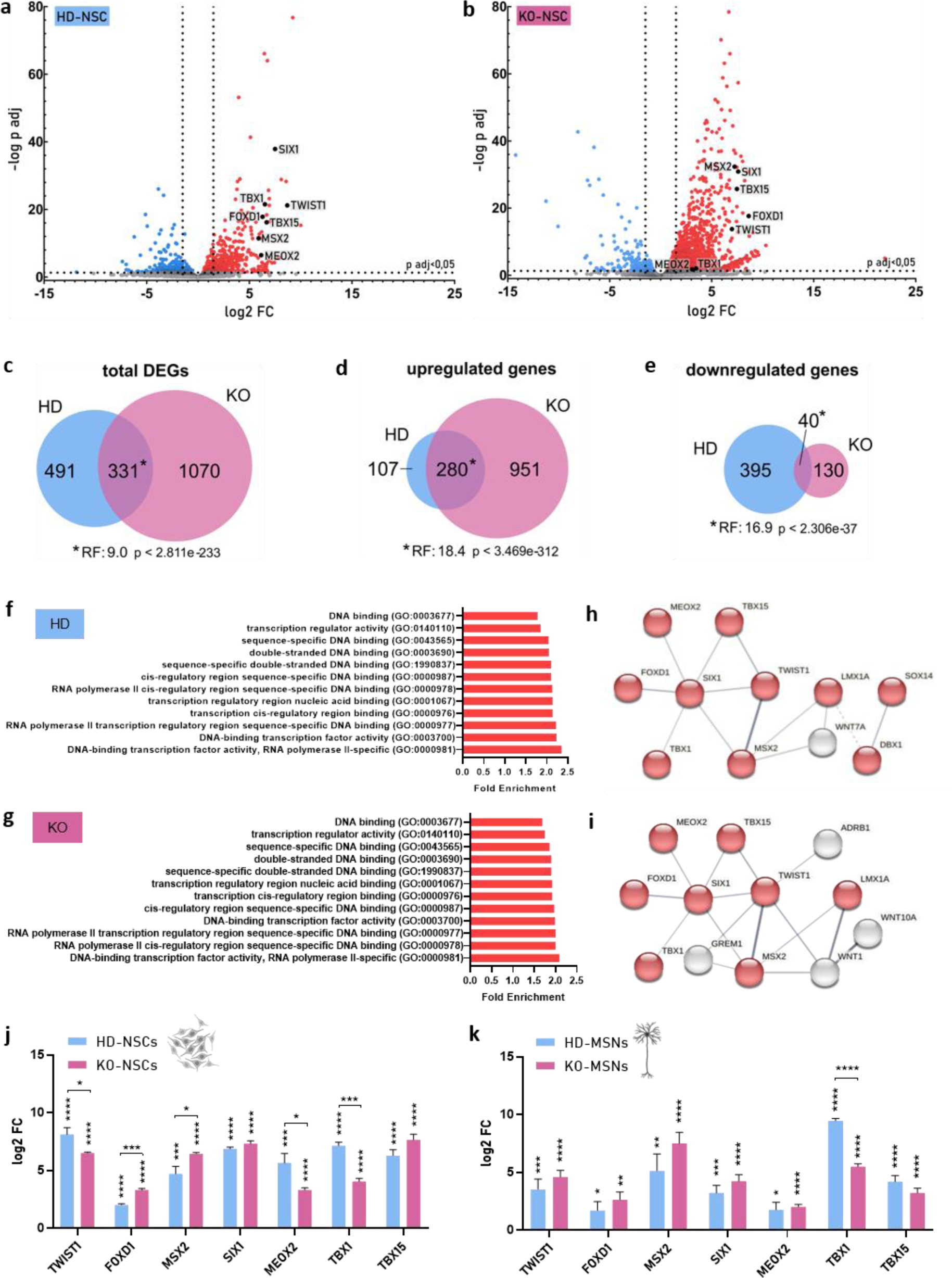
Summary of the RNA-seq results for the set of isogenic NSCs and deregulation of TFs in HD and KO neural cells. (**a**, **b**) Volcano plot points indicate genes with significantly increased (red dots) or decreased (blue dots) expression in HD-NSCs (a) and KO-NSCs (b) compared with IC1-NSCs. The x-axis shows log2 of FCs in expression (vertical lines indicate a cutoff of |1.5| in this value), and the y-axis presents the –log10 of p values adjusted TFs that were selected for validation in the later part of this study are indicated as black dots. (**c, d, e**) Venn diagrams showing the numbers of genes that were differentially expressed in HD-NSCs and KO-NSCs with total DEGs (c) and the separation of upregulated (d) and downregulated genes (e). (**f**, **g**) GO Panther enrichment analysis of DEGs in HD (f) and KO (g) NSCs. Selected significantly enriched GO terms are shown with false discovery rate (FDR)-corrected fold enrichment (p values <0.05). (**h, i**) Prediction of the PPI network of deregulated TFs in HD (h) and KO (i) NSCs. Lines represent interactions of proteins in networks, and nodes represent proteins. Red indicates TFs related to RNA polymerase II (GO: 0000981). The FDR-corrected p values were 1.90E-04 for (g) and 5.22e-05 for (i). The figure of interaction networks was adapted from STRING v11.5. (**j, k**) Relative expression of TFs in HD-NSCs and KO-NSCs (vs. IC1-NSCs) (j) and HD-MSNs and KO-MSNs (vs. IC2-MSNs) (k) was analyzed using RT‒qPCR. FC was calculated relative to that of IC lines using the *delta*-*delta Ct* method and is shown as log2. Reference genes: *EEF2* and *RPLP0*. Statistical analysis for (j) and (k) was performed using multiple *t* tests. *0.01<p<0,05; **p<0.01; ***p<0.001; ****p <0.0001.

To classify and cluster DEGs in HD-NSCs and KO-NSCs, we performed GO enrichment analysis using the PANTHER GO-Slim algorithm [36]. Analyses in the molecular function (MF) category revealed many disrupted pathways in both cellular models (Table S4). Among the DEGs in both cellular models, we observed significant enrichment of DNA binding factors, mainly TFs that are associated with RNA polymerase II (Fig. 2 f, g). We also conducted an analysis using the STRING v11.5 database to study the protein‒protein interaction (PPI) networks of the DEGs, and we also observed enrichment in the TF network (Fig. 2 h, i). Therefore, we selected seven TFs, *TWIST1, FOXD1, MSX2, SIX1, MEOX2, TBX1,* and *TBX15*, which were all highly upregulated in both the HD and KO models (FC ranged from ∼6 to ∼500; most TFs had FC>50) for validation (Fig. 2 h, i and Fig. 2 a, b; precise values are given in Table S3). *LMX1A, SOX14* and *DBX1,* which were also present in the identified PPI networks (Fig. 2 h, i), were excluded from validation due to low expression levels (TPM<1 in HD-NSCs). All the genes, except *TBX1*, that were selected for further analysis were also reported to be upregulated in brain tissues from HD patients [37], and one of them, namely, *TWIST1*, has already been described in the context of HD [38, 39]. RT‒qPCR confirmed strong upregulation, similar to the RNA-seq results, of all 7 genes (*TWIST1, FOXD1, MSX2, SIX1, MEOX2, TBX1,* and *TBX15)* in both HD-NSCs and KO-NSCs (Fig. 2 j). Next, we analyzed the expression levels of the selected TFs in MSN-like cells that were differentiated from NSCs. We have shown strong upregulation of all the studied TFs in HD and KO neurons, as compared to IC2-MSNs, with a FC of at least 8 but usually much higher (data shown as log2FC, Fig. 2 k). Overall, no clear trend towards stronger TFs deregulation was observed in the HD or KO models.

### Rescue of *HTT* expression resulted in TFs downregulation in KO-NSCs

To confirm the impact of wtHTT or mutHTT on the expression of the selected TFs, we rescued *HTT* expression by the electroporation of KO-NSCs with plasmids expressing the full-length cDNA of human *HTT* with 19 or 109 CAG repeats [35]. After the optimization of the electroporation conditions for plasmid delivery (Fig. 3 a), we confirmed the expression of *HTT* (Fig. 3 b). The delivery of plasmid encoding normal HTT (HTT-19Q) into NSCs without endogenous *HTT* expression (KO-NSCs) resulted in significant downregulation of all the investigated TFs, specifically *TWIST1, FOXD1, MSX2, SIX1, MEOX2, TBX1* and *TBX15* (Fig. 3 c). The mRNA levels were decreased ∼40-60% of the initial levels in the KO-NSCs. Moreover, after overexpression of mutHTT from a plasmid (HTT-109Q), we observed significant downregulation of *TWIST1, MEOX2* and *TBX15* to ∼60-80% of the initial level in KO-NSCs. The less substantial decrease in the mRNA levels of the investigated factors caused by HTT-109Q, compared to that caused by HTT-19Q, may indicate that mutHTT exerts weaker effects than wtHTT but the impact of lower expression of mutHTT from the plasmid cannot be excluded (Fig. 3 b).

**Figure 3.**
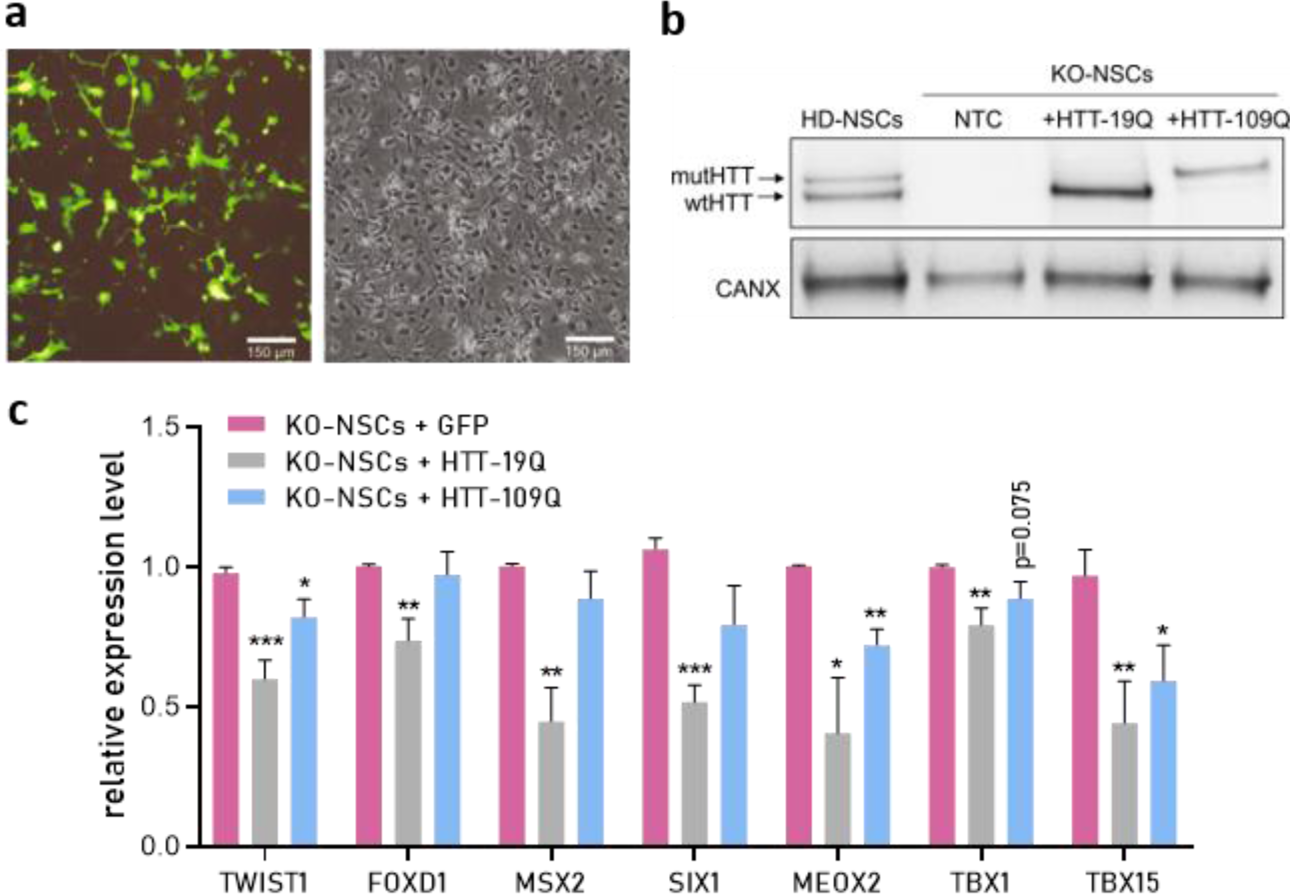
Effect of HTT rescue on the expression of selected TFs in KO-NSCs. (**a**) Representative images of KO-NSCs 48 h after electroporation with a plasmid encoding GFP. Scale bar = 150 µm. (**b**) Western blotting analysis of HTT levels in KO-NSCs 48 h after electroporation with the HTT-109Q or HTT-19Q plasmid. A sample from HD-NSCs was also loaded, and calnexin was used as a reference protein. NTC – nontreated cells. (**c**) Relative expression of TFs in KO-NSCs 48 h after electroporation with the HTT-109Q or HTT-19Q plasmid (for HTT-109Q, n=5; for HTT-19Q, n=4). FC was calculated relative to GFP-electroporated cells using the *delta*-*delta Ct* method and is shown as log2. Reference genes: *EEF2 or RPLP0*. Statistical analysis was performed using multiple *t* tests. *0.01<p<0,05; **p<0.01; ***p<0.001; ****p <0.0001.

### Overlapping changes in the miRNome of HD and *HTT*-KO neural cells

Strong evidence indicates that miRNA deregulation plays an important role in the pathogenesis of HD [30, 40–44]. Using small RNA sequencing, we profiled the miRNome of HD and KO-NSCs and compared these miRNomes to that of IC1-NSCs (Fig. S2). We found 16 and 68 significantly deregulated miRNAs (|log2FC|>1.5; padj<0.05; the full list of identified miRNAs is available in Table S5) in HD- and KO-NSCs, respectively (Fig. 4 a-c). As many as 12 deregulated miRNAs were common to HD and KO cells, with a significant RF of 7.4 (Fig. 4 c). In HD-NSCs, we identified 15 upregulated and only 1 downregulated miRNA (Fig. 4 d, e). Similar to the mRNA analysis, we identified more changes in KO-NSCs compared with HD-NSCs; in KO-NSCs, we identified 53 upregulated and 15 downregulated miRNAs (Fig. 4 d, e). Subsequently, we confirmed that the deregulation of miRNAs in the HD and KO cell lines was unrelated to the deregulation of genes associated with the biogenesis and functioning of miRNAs (Table S6). To validate the RNA-seq results with RT-qPCR, we selected five of the most strongly deregulated miRNAs, four of which were upregulated—miR-214-3p, miR-199a/b-3p, miR-199a-5p, and miR-143-3p— both in HD- and KO-NSCs, while miR-9-5p was downregulated in the HD cell line. It has already been shown that miR-143 and miR-199 are upregulated in HD, in the striatal tissues of patients [30] and in the striatal cells of mice [45], respectively. miR-214 is also known to be a posttranscriptional regulator of *HTT* expression [46, 47]. Moreover, the miR-214/199a cluster is regulated by TWIST1 [48]. Additionally, miR-9 is known to be a neuronal-specific miRNA that is downregulated in the brains of HD patients [49]. Other miRNAs that we selected for validation were reported to be broadly expressed in human tissues. RT‒qPCR confirmed the strong upregulation of miR-214-3p, miR-199a/b-3p, miR-199a-5p and miR-143-3p in HD-NSCs and KO-NSCs and the downregulation of miR-9-5p in HD-NSCs. Moreover, based on RT-qPCR results, miR-9-5p was also downregulated in KO-NSCs (Fig. 4 f).

**Figure 4.**
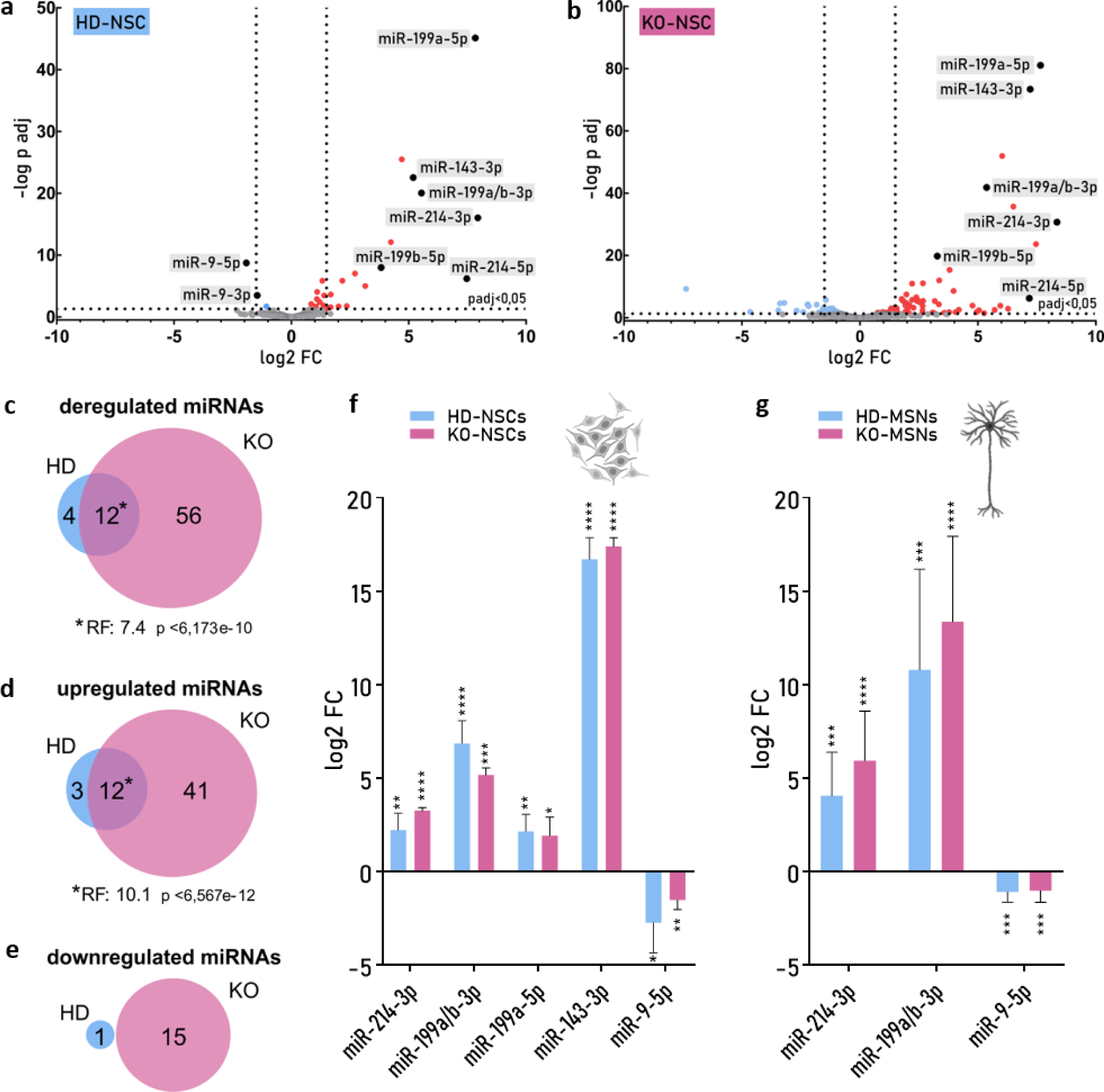
Deregulation of miRNA expression in HD and KO neural cells. (**a**, **b**) Volcano plot points indicate the miRNAs with significantly increased (red dots) or decreased (blue dots) expression in HD-NSCs (a) and KO-NSCs (b) versus IC1-NSCs. The x-axis shows log2 of FCs in expression (vertical lines indicate a cutoff of |1.5| in this value), and the y-axis presents the – log10 of p values adjusted. miRNAs that were selected for validation are indicated as black dots. (**c, d, e**) Venn diagrams showing the numbers of miRNAs that were differentially expressed in HD-NSCs and KO-NSCs and the total number of differentially expressed miRNAs (c) and the number of upregulated (d) and downregulated (e) miRNAs. (**f**) Validation of RNA-seq data using RT‒qPCR to analyze the expression of selected miRNAs in HD-NSCs and KO-NSCs relative to IC1-NSCs. (**g**) Relative expression of selected miRNAs in HD-MSNs and KO-MSNs was assessed using RT‒qPCR (vs. IC2-MSNs). FC (f, g) was calculated relative to that of IC lines using the *delta*-*delta Ct* method and is shown as log2. Reference miRNAs: miR-16 and miR-92a. Statistical analysis for (f) and (g) was performed using multiple *t* tests. *0.01<p<0,05; **p<0.01; ***p<0.001; ****p <0.0001.

Next, we analyzed the levels of selected miRNAs in MSNs. Similar to NSCs, we observed strong upregulation of miR-214-3p and miR-199a/b-3p and downregulation of miR-9-5p in HD-MSNs and KO-MSNs compared to IC2-MSNs (Fig. 4 g). Also, we observed a tendency toward stronger upregulation of the miR-214/199a cluster in KO neurons than in HD neurons. The expression levels of miR-143-3p and miR-199a-5p were too low to accurately detect those miRNAs in all MSN lines.

### Overexpression of miR-9-5p led to downregulation of selected TFs in HD-NSCs

According to correlation analysis of our RNA-seq data from NSCs, *TWIST1*, *TBX1* and *MEOX2* expression levels were strongly positively correlated with each other, and the levels of these TFs were negatively correlated with those of miR-9 (Fig. 5 a, Fig. S3). The other TFs, i.e., *MSX2*, *SIX1*, *FOXD1*, and *TBX15*, also tended to form a positively correlated network in terms of their expression levels, and the expression of these TFs was significantly correlated with that of miR-143-3p. As expected, the expression levels of miRNAs from cluster miR199a/214 were also positively correlated.

**Figure 5.**
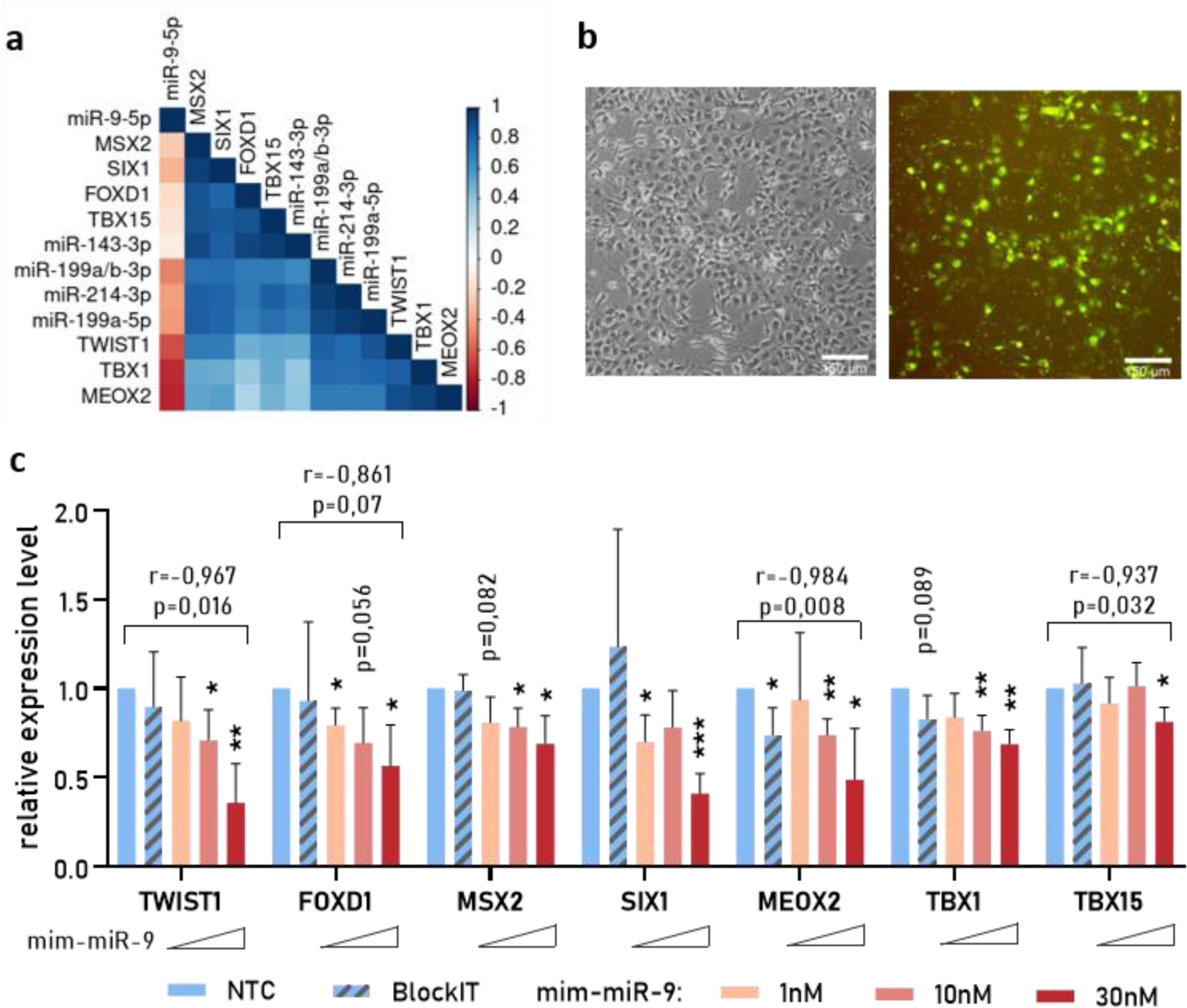
Effect of miR-9 on the expression of selected TFs in HD-NSCs. (a) Analysis of expression correlation of selected TFs and miRNAs. A heatmap displaying Spearman correlation coefficients calculated based on TPM expression values from IC1, KO and HD NSC models. (b) Representative images of HD-NSCs 24 h after transfection with 20 nM BlockIT control oligonucleotide. Scale bar = 150 µm. (**c**) Relative expression of selected TFs in HD-NSCs after transfection with 1 nM, 10 nM or 30 nM miR-9 mimic. FC was calculated relative to that of nontreated cells (NTC) using the *delta*-*delta Ct* method. Reference genes: *EEF2* and *RPLP0*. Statistical analysis was performed using multiple *t* tests. *0.01<p<0,05; **p<0.01; ***p<0.001; ****p <0.0001. r - Pearson correlation coefficient.

We decided to validate regulation of TFs expression by miR-9 in our models as downregulation of this brain-enriched miRNA was found important in HD brain [49]. miR-9-5p is also known to directly regulate the expression of TFs, including *TWIST1* [50]. We investigated whether the deregulation of TFs in HD-NSCs may be related to the miR-9-5p level by delivering a miR-9 mimic. We monitored the transfection efficiency using the FITC-labeled oligonucleotide BlockIT (Fig. 5 b). HD-NSCs were transfected with three concentrations of the miR-9-5p mimic. We observed significant downregulation of all 7 investigated TFs after mim-miR-9 delivery compared to their expression levels in nontreated cells (NTCs) (Fig. 5 c). Transfection with 30 nM mim-miR-9 resulted in a decrease in the expression level of *TWIST1, FOXD1, MSX2, SIX1, MEOX2, TBX1,* and *TBX15* to 35-80%, compared to their expression level in NTCs. The level of downregulation of *TWIST1, MEOX2* and *TBX15* and the concentration of delivered mim-miR-9 tended to be negatively correlated (the Pearson correlation coefficients (r) were *-0,967, -0,984* and *–0,937,* respectively) (Fig. 5 c).

### A trend for mRNA and miRNA levels to increase over time in HD neuronal cells

There are suggestions of increasing dysregulation of *TWIST1* and miR-9 as HD progresses [39, 49]. Our RNA-seq results revealed interesting differences between cells with *HTT* mutation and those without *HTT* expression, i.e., in HD-NSCs but not in KO-NSCs there was a tendency for the expression of several TFs (*TWIST1, MSX2, MEOX2,* and *TBX1*) to increase over time (with subsequent passages of NSCs) (Table S7 a). This observation was confirmed by RT‒qPCR for these TFs (Table S7 b). To further explore this phenomenon, we performed an additional bioinformatics analysis of our RNA-seq data, which revealed genes whose expression increased or decreased over time in HD, KO and control NSCs (lists of genes are provided in Table S8 and S9). For each cell line, differences in the TPM values were calculated between three (for IC1) or four (for HD and KO-NSC) time points, and specific criteria (described in Materials and Methods) were applied to categorize genes as “increasing” or “decreasing”.

The HD model exhibited the highest number of “increasing” genes (2835, representing 67% of ”increasing” genes in all NSC lines), including 2508 genes that were unique to HD-NSCs (which constituted 88.5% of “increasing” genes in this cell line) (Fig. 6 a). The number of genes whose expression increased in the IC1-NSCs and KO-NSCs was considerably lower than that in the cells with mutation (722 and 1036, respectively). As expected, GO analysis revealed that in control cells (IC1-NSCs), “increasing” genes were associated mainly with neural differentiation (Fig. S4). The overlap between KO or HD and IC1 was relatively small (5.2% of HD and 5.3% of KO “increasing” genes), which might indicate disruptions of neural differentiation in cells with mutation and in those without *HTT* expression (Fig. 6 a). To classify genes whose expression increased specifically in HD cells, we performed GO enrichment analysis on a set of 2508 genes specific to HD-NSCs. Among those genes, we found enrichment in 274 different biological processes (the top 15 are plotted in Fig. 6 b; all the GO terms are presented in Table S10), i.e., processing different types of RNAs, posttranslational modifications of proteins, and Wnt pathway regulation. Moreover, PPI analysis (STRING) revealed enrichment of TFs that are associated with polymerase II, which confirmed our previous observation (Fig. S5).

**Figure 6.**
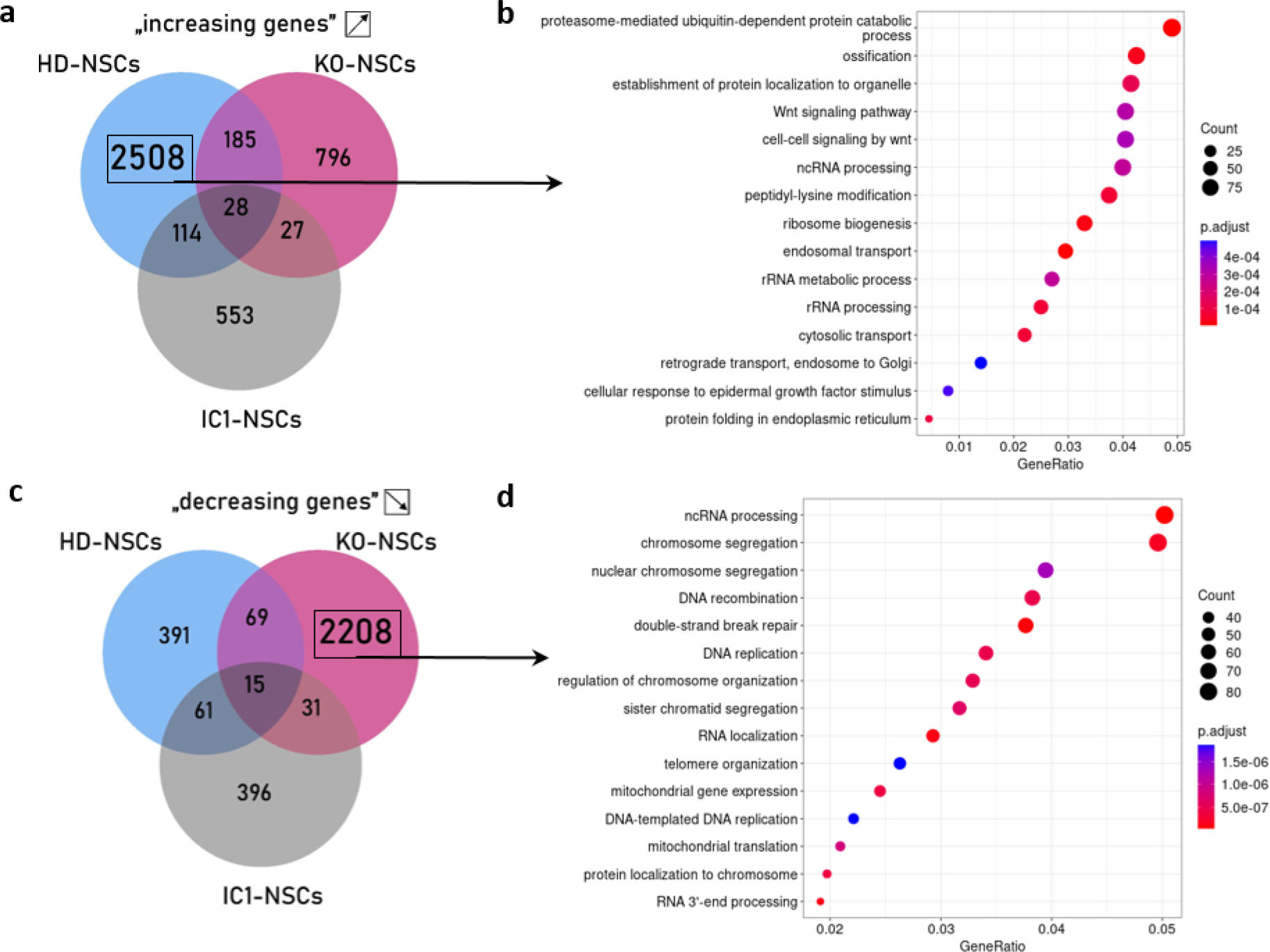
Increasing and decreasing gene expression over time in NSC culture. (**a**) Venn diagram showing the number of genes with increasing expression in IC1, HD and KO cells based on RNA-seq data from 3 (for IC) or 4 (for HD and KO) time points (subsequent passages) of NSC culture. (**b**) GO biological process (GO-BP) enrichment analysis for genes classified as “increasing” only in HD-NSCs. (**c**) Venn diagram analogous as in (a) for genes with decreasing expression. (**d**) GO-BP enrichment analysis for genes classified as classified as “decreasing” only in KO-NSCs.

Regarding “decreasing” genes, we identified the highest number of these genes in the KO-NSC model. Precisely, we found 2323 genes with decreasing expression in KO-NSCs (which constituted 73% of “decreasing” genes in all cell lines) (Fig. 6 c). Most of these genes were unique to the KO model (2208 genes which accounted for 70% of “decreasing” genes in this cell line). In the IC1 and HD models, a substantially lower number of genes with decreasing expression was identified (503 and 536, respectively) (Fig. 6 c). In the KO model, “decreasing” genes were classified as being associated with ncRNA processing, chromosome segregation and DNA recombination (Fig. 6 d, all the GO terms are presented in Table S10). There was no substantial overlap between the two most numerous groups, “increasing” genes in HD-NSCs and “decreasing” genes in KO-NSCs (RF = 1.1, p <0,091), indicating significant differences in progressive changes in gene expression in these models.

To further study the time-dependent increase of the selected TFs expression identified in HD-NSCs, we analyzed their levels in MSN-like cells at three time points of differentiation (Fig. 1 b). In three independent differentiation experiments, we observed a trend toward increasing expression of *FOXD1, MEOX2* and *TBX1* (as well as a result that was close to statistically significance for *TWIST1*) in HD-MSNs but not in KO-MSNs (Fig. 7 a, b). Next, we evaluated miRNA expression in the same neuronal samples and observed a trend toward increasing upregulation of miR-214 and miR-199 (high R but nonsignificant for both miRNAs) in HD-MSNs (Fig. 7 c). No such trend was observed in the KO-MSNs (Fig. 7 d).

**Figure 7.**
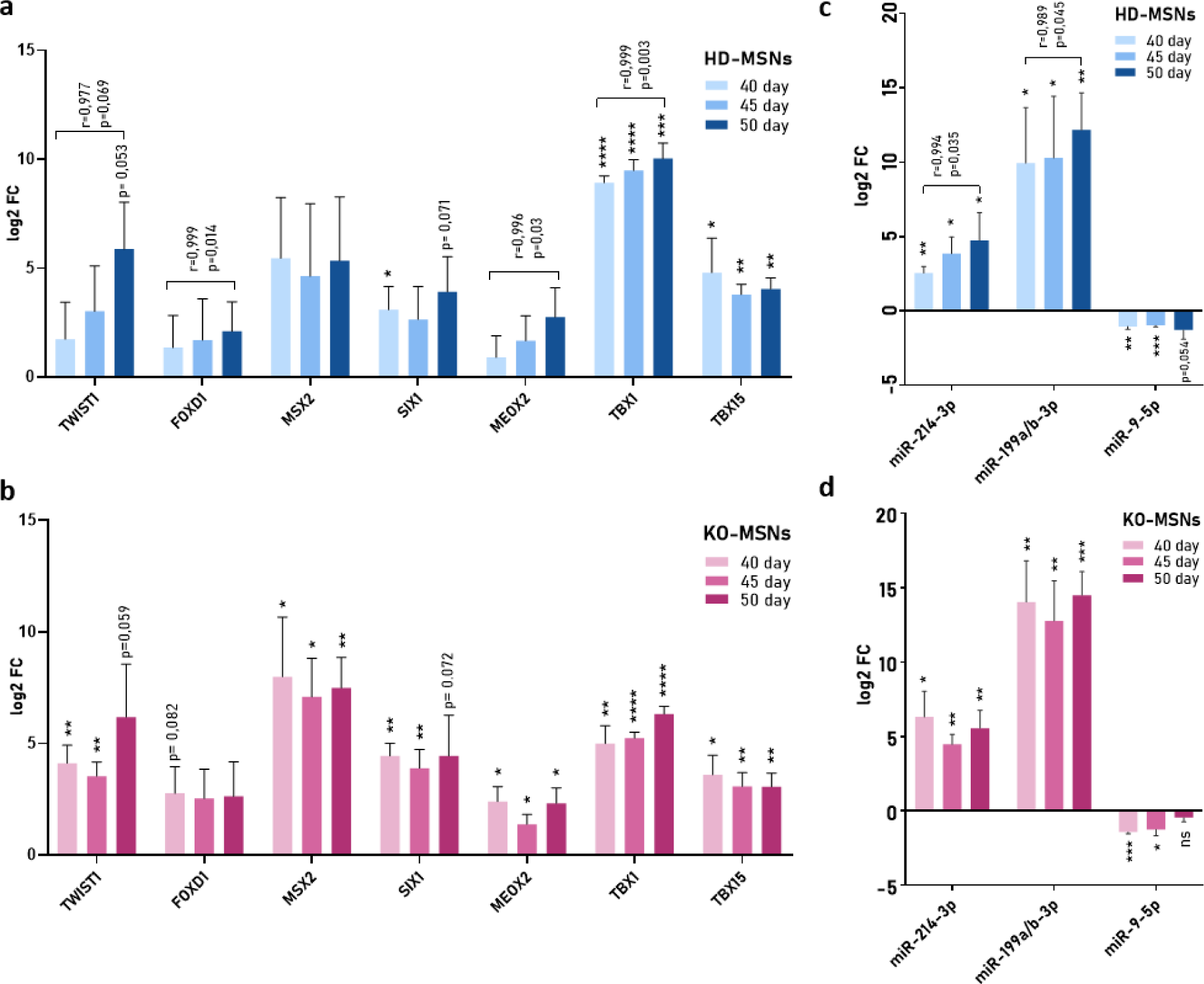
Deregulation of selected TFs and miRNAs during the differentiation of HD-MSNs and KO-MSNs. (**a**, **b**) Relative expression of the selected TFs *TWIST1, FOXD1, MSX2, SIX1, MEOX2, TBX1* and *TBX15* in 40-, 45- and 50-day-old HD-MSNs (a) and KO-MSNs (b) was analyzed using RT‒qPCR. Reference genes: *EEF2* and *RPLP0*. (**c**, **d**) Relative expression of miR-214-3p, miR-199a/b-3p and miR-9-5p in 40-, 45- and 50-day-old HD-MSNs (c) and KO-MSNs (d) was assessed using RT‒qPCR. Reference miRNAs: miR-16 and miR-92a. FC was calculated using the *delta-delta Ct* method and is shown as the log2 relative to that of IC2-MSNs at each of the time points; for example, 40-day HD-MSNs/KO-MSNs were referred to as 40-day IC2-MSNs. Statistical analysis was performed using multiple *t* tests. *0.01<p<0,05; **p<0.01; ***p<0.001; ****p <0.0001. r-Pearson correlation coefficient.

## Discussion

In the last several years, HD has begun to be recognized as a neurodevelopmental disorder [51, 52]. Early neurodevelopmental defects occur in HD and may contribute to the neurodegeneration phenotype in adults [31, 53–55]. Mutation of *HTT* leads to many changes in molecular processes that, although initially subtle, have already been observed in HD patient-derived iPSCs or embryonic stem cells (ESCs) [56–59]. At later steps of cell differentiation, these changes lead to disruptions that have significant implications for the functions of neural cells [31, 56, 60–65]. Stem cell-derived neural lines are models that are widely used in HD research to investigate aspects of transcriptional dysregulation [63, 66, 67], but the underlying mechanisms appear to be diverse and complex due to the interplay of the factors involved [26].

In this study, we used a unique set of human neural cell lines with the same genetic background in the context of HTT function (KO model) and dysfunction (HD model). A straightforward comparison of three cell lines (control, *HTT*-KO and HD) provided insights into the effects of HTT LoF and GoF. We focused on global transcriptional deregulation in the iPSC-derived neuronal HD and *HTT*-KO lines and investigated selected deregulated RNAs in more detail. We studied particular mRNAs (TFs) and miRNAs, both of which play key roles in the regulation of gene expression at the transcription and posttranscription levels, respectively. Moreover, TFs and miRNAs expression is directly interlinked, as TFs regulate pri-miRNA transcription, and mature miRNAs modulate the expression levels of TFs [68, 69]. Additionally, there are indirect links between TFs and miRNAs levels that involve other cellular factors. The network of these interactions may easily become deregulated in the presence of an abnormal molecule, such as mutHTT, in the case of HD, or in the absence of a specific important protein, as in the case of the *HTT*-KO model. Importantly, crucial effects of mutHTT GoF are linked to the nuclear localization of HTT or its fragments [12, 70]. wtHTT is involved in transcriptional regulation, acts as a scaffold for regulatory protein complexes in the nucleus, and interacts directly with various TFs and DNA in promoter regions [71, 72]. All these interactions could be altered as a result of polyQ tract expansion [12, 72–74]. It was observed that wtHTT may be transported to the nucleus to act as both a transcriptional activator [75] and a transcriptional repressor [71]. Interestingly, in some interactions, mutHTT can function as a repressor until it is truncated by proteases [71]. Generally, changes in transcription in HD that are directly related to HTT may result from the improper function of mutHTT in particular interactions (with TF or chromatin), the occurrence of novel interactions, or a deficiency in wtHTT, which normally regulates transcription [12, 76]. These individual mechanisms may be specific to a particular HTT-TF interaction, to a cell type, or developmental stage.

In our study, we demonstrated massive transcriptional changes in the HD and *HTT*-KO NSC lines, which surprisingly substantially overlapped (Fig. 2 d). Among the genes whose expression was upregulated in HD and KO cells, we found enrichment in TFs (Fig. 2 f-i). The substantial increase in the expression levels of the TFs that were validated in our study, namely, *TWIST1, FOXD1, MSX2, SIX1, MEOX2,* and *TBX15,* is consistent with reported changes observed in the prefrontal cortex of brain tissue of HD patients [37]. Moreover, *TWIST1, SIX1, TBX1* and *TBX15* were also upregulated in a set of nonisogenic HD neural differentiated iPSCs [31]. Upregulation of numerous TFs, including *TWIST1, FOXD1* and *SIX1,* was shown in mice in which polycomb repressive complex 2 (PRC2) was depleted in striatal neurons [77]. PRC2 is known to support neuronal differentiation through negative gene regulation, and its deficiency caused upregulation of genes that are usually suppressed in MSNs [77]. Moreover, *MEOX2* was suggested to be a suppressor gene of mutHTT toxicity in mouse ESCs that express the N-terminal fragment of *HTT* with 128 CAG repeats [59]. However, among the genes that were validated in our study, only *TWIST1* has been described in detail in the context of HD [38, 39]. TWIST1, which is a basic helix-loop-helix TF, is a highly conserved, antiapoptotic protein that participates in neuronal development, neural crest formation and many other processes during embryogenesis [78]. In somatic cells, the expression of *TWIST1* is very low, which we also observed in our control NSC and MSN lines. There are two different proposed mechanisms underlying *TWIST1* upregulation in HD. The results of two studies led to opposite conclusions, and neurotoxic [38] or neuroprotective [39] functions of TWIST1 were indicated. Pan et al. suggested that in cortical mouse neurons, increased levels of H3K4me3 in the *Twist1* promoter were directly mediated by mutHtt [38]. On the other hand, Jen et al. proposed that in HD striatal progenitor cells, *Twist1* expression is upregulated by mutHtt through a signal transducer and activator of transcription 3 (STAT3)-mediated pathway [39]. RNA-seq analysis demonstrated that *Twist1* knockdown partially rescued the expression of a set of DEGs in HD cortical neurons, suggesting that upregulation of *Twist1* was a deleterious effect of *HTT* mutation in these types of neurons [38]. On the other hand, in striatal cells, Twist1 might act as a neuroprotective, antiapoptotic factor, as decreased Twist1 levels in HD cells resulted in significantly increased neuronal death [39]. These data suggest that in different types of cells, we might observe different mechanisms of neurodegeneration and that the role of TWIST1 in HD should be further evaluated.

In our HD and KO cellular models, we observed strong upregulation of *TWIST1* in both the early (NSCs) and late (MSNs) stages of neural differentiation. In HD cells, we observed a gradual increase in the expression level of *TWIST1* in the HD model during neuronal differentiation, but no such progressive upregulation was observed in the *HTT*-KO cells, in which the *TWIST1* mRNA level was high beginning at the first investigated time point of differentiation (Fig. 7 a, b). These findings indicate that a progressive loss of the repressor function of HTT could be responsible for *TWIST1* upregulation in our HD cellular models. Moreover, in *HTT*-KO cells, the overexpression of wt*HTT* resulted in the downregulation of the expression of all the investigated TFs (Fig. 3 c). Surprisingly, mut*HTT* overexpression also exerted this effect on three TFs, *TWIST1, MEOX2,* and *TBX15* (with a trend in *TBX1;* p value of 0.075), suggesting that, at least for some of the investigated genes, including *TWIST1*, mutHTT is able, to some extent, to function in the regulation of their expression levels. Similar findings were obtained in a human ESC model in which exogenous expression of mut*HTT* in HTT-KO ESCs led to partial rescue of the HD-like phenotype [79].

Many brain-specific miRNAs are downregulated in HD [49, 80, 81]. One of these miRNAs is miR-9, which was downregulated in our both cellular models, HD and *HTT*-KO. miR-9 is also known to directly regulate *TWIST1* expression [50], which is consistent with our RNA-seq data that revealed a negative correlation between the expression of miR-9 and *TWIST1* in NSCs (Fig. 5 a). Moreover, when we overexpressed this miRNA in HD-NSCs, we observed dose-dependent downregulation of TFs, especially *TWIST1* (Fig. 5 c). Furthermore, TWIST1 and the miR-199a/214 cluster may regulate the expression of one another [48], and we demonstrated a significant positive correlation between these expression levels in NSCs (Fig. 5 a). On the other hand, miR-214 regulates *HTT* expression [46, 47]. Therefore, we proposed a network of gene regulation by TFs and miRNAs involving a regulatory feed-forward loop (FFL) that occurs in HD (Fig. 8) due to wtHTT deficiency. In healthy neurons, miR-9 contributes to the inhibition of *TWIST1* expression, and a low level of this TF leads to low expression of miR-214 and miR-199a. However, there are likely other associations between the expression of these TFs and miRNAs. Additionally, we observed a significant positive correlation between the expression of the miRNAs miR-199a/214 cluster and miR-143 and also selected TFs: *SIX1*, *MSX2*, *TBX15* and *FOXD1*, which could be further investigated as not reported previously.

**Figure 8.**
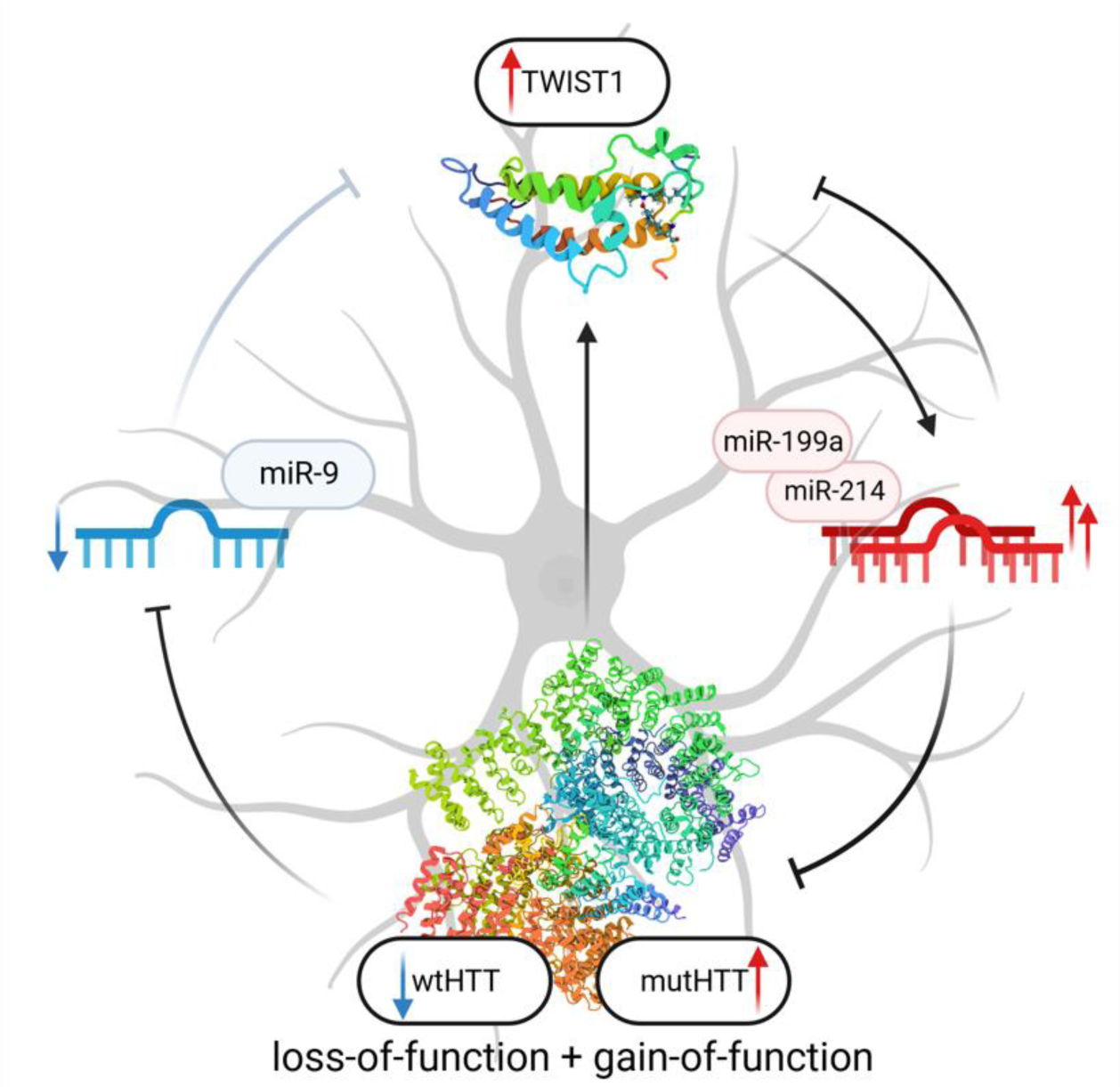
Model of the molecular network implicated in HD pathogenesis. Proposed functional network in HD neuronal cells: crosstalk mediated by a regulatory feed-forward loop (FFL) involving HTT (due to its deficiency or dysfunction), TWIST1 TF, miR-214/199a and miR-9. For more details, see the text.

It is challenging to investigate aspects of mutHTT GoF and wtHTT partial LoF, as these mechanisms occur simultaneously in HD, and they influence and enhance each other. Therefore, it can be hypothesized that separated HTT GoF and partial LoF mechanisms would be less pronounced. The LoF of HTT may be easily separated and analyzed in *HTT*-KO models. In our study, we found a more severe phenotype in the *HTT*-KO cell line than in the HD cell line, as indicated by the markedly higher number of upregulated genes in the KO-NSCs compared with the HD-NSCs (Fig. 2 b, d). This finding suggested that complete loss of the repressor function of HTT had prominent effects in the KO model. Moreover, we observed difficulties in neural differentiation in the KO line, which was also indicated by the lower expression of *TUJ1* in the KO-MSNs than in the other lines (Fig. 1 e) and supported by the lower *PAX6* expression in the KO-NSCs than in the IC1 or HD lines according to the RNA-seq data (log2FC = -2.38; KO vs. IC1, Table S3). Taken together, these results confirm the important role of wtHTT in neuronal differentiation and highlight a spectrum of deregulation events that are induced by a lack of HTT in neural cells. Our findings regarding the *HTT*-KO model, although reported at the cell culture level, are consistent with the more severe phenotype observed in ultra-rare patients with very low wtHTT levels. Mutations in both *HTT* alleles (other than CAG expansion) cause a severe neurodevelopmental disorder called Lopes–Maciel–Rodan syndrome (LOMARS) [82–84]. LOMARS has been reported thus far only for several patients who suffered from severe neurological symptoms already in early childhood. Similarly, a lower *wtHTT* expression as a result of a SNP (rs13102260: G>A) in the *HTT* promoter region was correlated with an earlier age of HD onset in patients [85]. Investigation of mutation homozygosity in HD patients is difficult, as such cases are extremely rare, and therefore data are very limited and not fully conclusive. Two studies suggested rather similar age of onset for homozygous and heterozygous HD patients [10, 86]. Interestingly, in case of biallelic *HTT* mutation in patients, more rapid disease progression in homozygous patients was suggested in one of these studies [10], and in the second study, some worse parameters in BMI and gait were reported [11]. Similarly, in different knock-in mouse models of HD, more aggressive and earlier phenotypes are observed in homozygous than in heterozygous animals [87–90]. Nevertheless, it cannot be excluded that a worse course of the disease is caused by a greater mutation effect due to the expression of two mutant alleles.

The mechanism underlying HTT LoF in HD may be nonclassical, as a reduced level of wtHTT (due to the expression of one wt allele) is not the only cause. An interesting general hypothesis is that the dominant negative effect of the mutated protein on the wt protein is mediated by interference with the normal functions [91, 92]. The dominant role of mutHTT in HD has been proven, e.g., studies showing that increasing wtHTT levels have limited effectiveness in correcting the HD phenotype [79, 93]. Regarding the effect of HTT LoF on HD, it can be hypothesized that the already limited function of wtHTT becomes even more limited in the presence of mutHTT. Our results support this suggestion, as more than 70% of upregulated genes in HD-NSCs were common to those upregulated in KO-NSCs (Fig. 2 d). This indicates that increased expression of genes in HD-NSCs was mainly a direct or indirect result of a deficiency of properly functioning HTT in the cellular regulatory network. In a recent study, a set of isogenic hESC lines, similar to those used in our study, including HD, control and HTT-KO hESC lines, was assessed in activin A-stimulation tests and neuroloid formation assays [79]. Although different features were analyzed, the authors also found that *HTT*-KO resulted in an HD signature phenotype, suggesting a dominant negative mechanism of mutHTT in HD.

The investigation of the LoF of wtHTT in HD has additional important implications for the design of therapeutic strategies. Approaches that aim to diminish mut*HTT* expression are currently considered the most promising strategies for eliminating the dominant effects of mutHTT GoF [94, 95]. These strategies are mainly based on the use of short-interfering siRNAs (siRNAs) or antisense oligonucleotides (ASOs) that promote mut*HTT* RNA degradation [96]. Despite advances in clinical trials, e.g., with Tominersen [97], there are still important considerations regarding many issues that are related to the LoF of HTT in HD [98, 99]. First, there are concerns about brain regions that should be specifically targeted in HTT-lowering strategies. It might be crucial to downregulate HTT in specific brain cells in HD, but in other cell types, it might be better to retain both wtHTT and mutHTT, as they function relatively well. Second, there are deliberations about the patient age at which treatment should be administered. HTT plays an important role in early life, which indicates that HTT should not be downregulated before adulthood; however, turning off the expression of mut*Htt* at an early age in mice (at 21 postnatal day) did not reduce the later development of the HD phenotype [100]. Third, the requirement for allele selectivity in the downregulation of *HTT* is under debate. Although generally considered safer, allele-selective strategies are more complex in terms of design and universality for all patients. The approaches for preferential silencing mut*HTT* expression are based mainly on targeting SNP regions or direct mutation sites (CAG repeat expansion) [96]. Currently, the most advanced clinical trial involving allele-selective *HTT* targeting is SELECT HD (clinicaltrials.gov NCT05032196), which uses ASOs to target regions of specific SNPs [101]. In contrast to the development of allele-selective strategies, several studies in rodents and primates concluded that wtHTT levels may decrease to ∼50% in adulthood without any phenotypic alterations [98]. The same conclusion may be drawn based on the lack of dysfunction in adult people whose one *HTT* allele is inactive [82]. Nevertheless, it should be assumed that any downregulation of wtHTT may have more prominent adverse effects when it is accompanied by the presence of mutHTT in HD patients. Further research should re-examine the potential risk of non-allele-selective therapies, and we believe that a focus on allele-selective therapies will be safer for HD patients.

## Author Contributions

Conceptualization: EK, AF. Experiments design: EK, AC, AF. Cell culture: EK, AC, JS. Neural differentiation: EK, AC. RNA-seq results analysis: EK. RT-qPCR for mRNAs: AC. RT-qPCR for miRNAs: EK, JS. Immunocytochemistry: EK, AC. Bioinformatics analyses: GA. NSC transfection: EK. NSC electroporation: AC. Figures: EK, AC, AF. Manuscript writing: EK, AF with the input and revision from AC, GA. Supervision: AF. All authors read and approved the final manuscript.

## Acknowledgments

Microscopic images were obtained in the Laboratory of Subcellular Structures Analyses, IBCH PAS. The authors would like to thank Julia Misiorek and Eliza Walczak for contributions to some experiments in frame of this study. Figures were created with BioRender.com and GraphPad Prism.

## Availability of data and materials

Raw data (RCC files) from total and small RNA-seq analysis will be available on Gene Expression Omnibus (GEO) upon publication.

## Abbreviations

DEG: differentially expressed genes
FC: fold change
FFL: feed-forward loop
GO: gene ontology
GoF: gain-of-function
HD: Huntington’s disease
HTT: huntingtin
IC: isogenic control
ICC: immunocytochemistry
iPSC: induced pluripotent stem cell
KO: knock out
LoF: loss-of-function
LOMARS: Lopes-Maciel-Rodan syndrome
MSN: medium spiny neuron
NSC: neural stem cell
NTC: nontreated control
mutHTT: mutant huntingtin
polyQ: polyglutamine
RF: representation factor
PPI: protein-protein interaction
PRC2: polycomb repressive complex 2
Q: glutamine
TF: transcription factor
TPM: transcript per million
wtHTT: wild-type huntingtin

## Supplementary files

Supplementary Figure 1. Heatmaps of deregulated genes in NSCs based on RNA-seq results

Supplementary Figure 2. Heatmaps of deregulated miRNAs in NSCs based on miRNA-seq results

Supplementary Figure 3. Correlation of expression of selected TFs and miRNAs in IC1, HD and KO NSCs

Supplementary Figure 4. GO enrichment analysis for genes classified as “increasing” in control NSCs

Supplementary Figure 5. Enrichment of TFs that are associated with polymerase II in HD among „increasing genes” unique for HD-NSCs

Supplementary Table 1. A list of primary and secondary antibodies with their dilutions used in immunocytochemistry and western blot

Supplementary Table 2. A list of primers used for RT-qPCR

Supplementary Table 3. Lists of DEGs in HD and KO-NSCs

Supplementary Table 4. GO enrichment analysis of DEGs in HD and KO-NSCs

Supplementary Table 5. Lists of deregulated miRNAs in HD and KO-NSCs

Supplementary Table 6. Analysis of miRNA biogenesis and function genes in HD-NSCs

Supplementary Table 7. Correlation analysis of expression level of selected TFs with subsequent passages of HD and KO-NSCs

Supplementary Table 8. List of genes with expression decreasing with time in IC1, HD and KO-NSCs

Supplementary Table 9. List of genes with expression increasing with time in IC1, HD and KO-NSCs

Supplementary Table 10. GO analysis of genes increasing and decreasing in time in HD and KO- NSCs

