## Supplementary Figures for "Huntingtin loss-of-function contributes to transcriptional deregulation in Huntington’s disease"

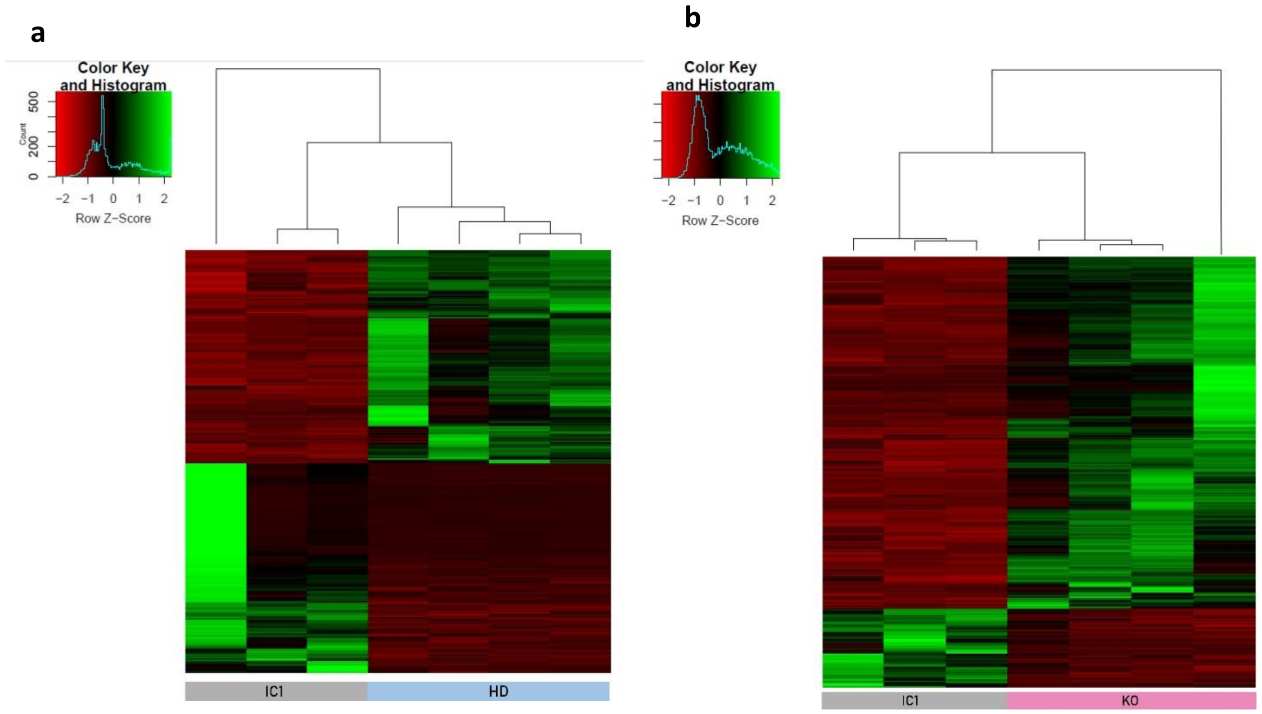


**Supplementary Figure 1. Heatmaps of deregulated genes in NSCs based on RNA-seq results**
Heatmap representing relative expression levels of the DEGs common to control (IC1) versus HD (a) and KO (b), with corrected padj < 0.05. Samples (in columns) and genes (in rows) are clustered by similarity. Shades of green represent upregulation, shades of red represent downregulation.

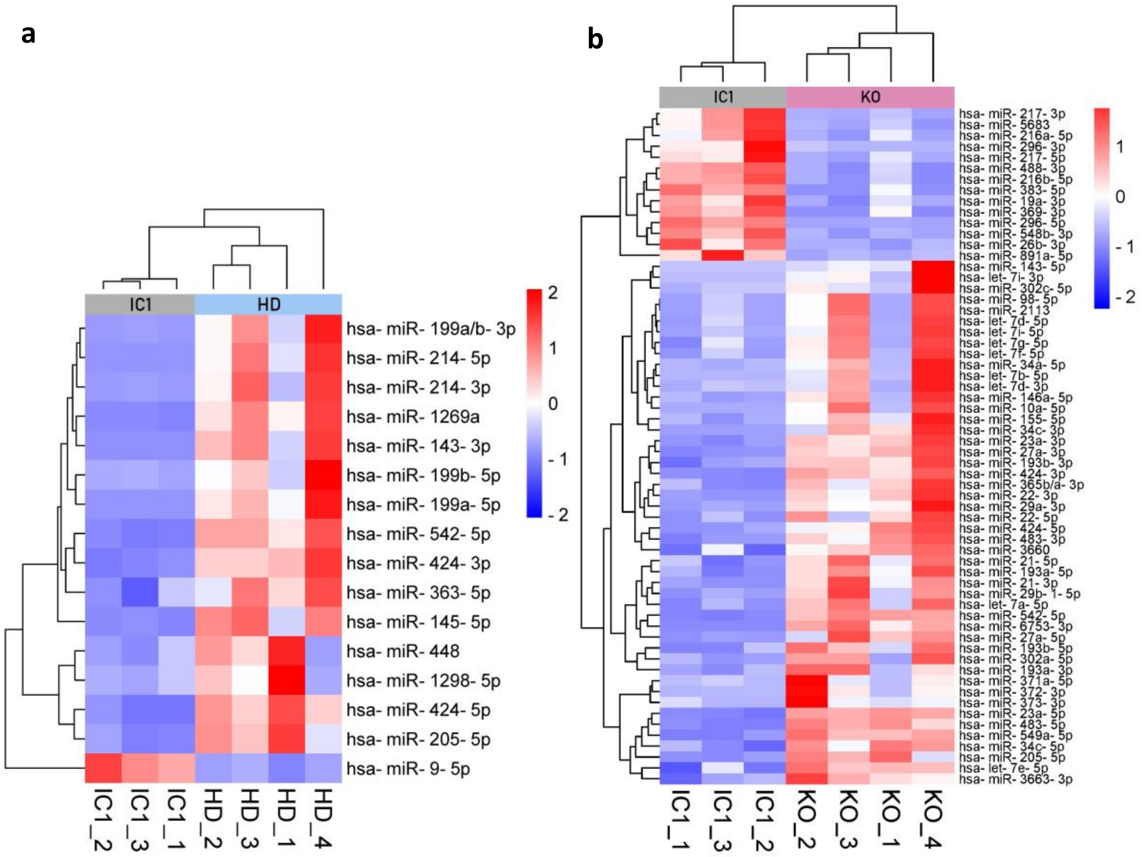


**Supplementary Figure 2. Heatmaps of deregulated miRNAs in NSCs based on miRNA-seq results**

Heatmap representing relative expression levels of the DE miRNAs common to control (IC1) versus HD (a) and KO (b), with log2FC| >1.5 and corrected padj < 0.05. Samples (in columns) and genes (in rows) are clustered by similarity. Shades of red represent upregulation, shades of blue represent downregulation.

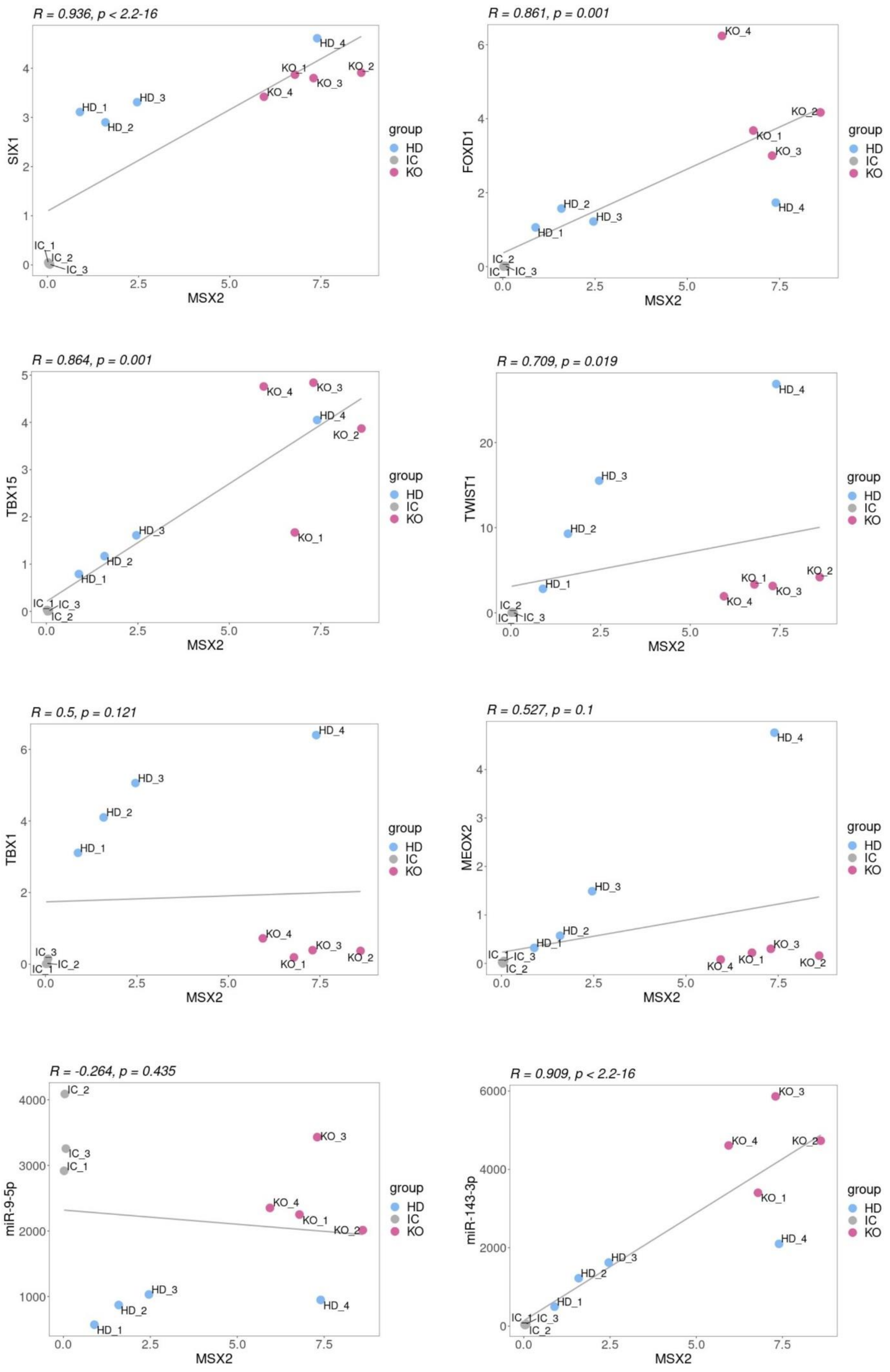
R = 0.936, p < 2.2-16
SIX1
MSX2
R = 0.861, p = 0.001
FOXD1
MSX2
R = 0.864, p = 0.001
TBX15
MSX2
R = 0.709, p = 0.019
TWIST1
MSX2
R = 0.5, p = 0.121
TBX1
MSX2
R = 0.527, p = 0.1
MEOX2
MSX2
R = -0.264, p = 0.435
miR-9-5p
MSX2
R = 0.909, p < 2.2-16
miR-143-3p
MSX2
group
HD
IC
KO

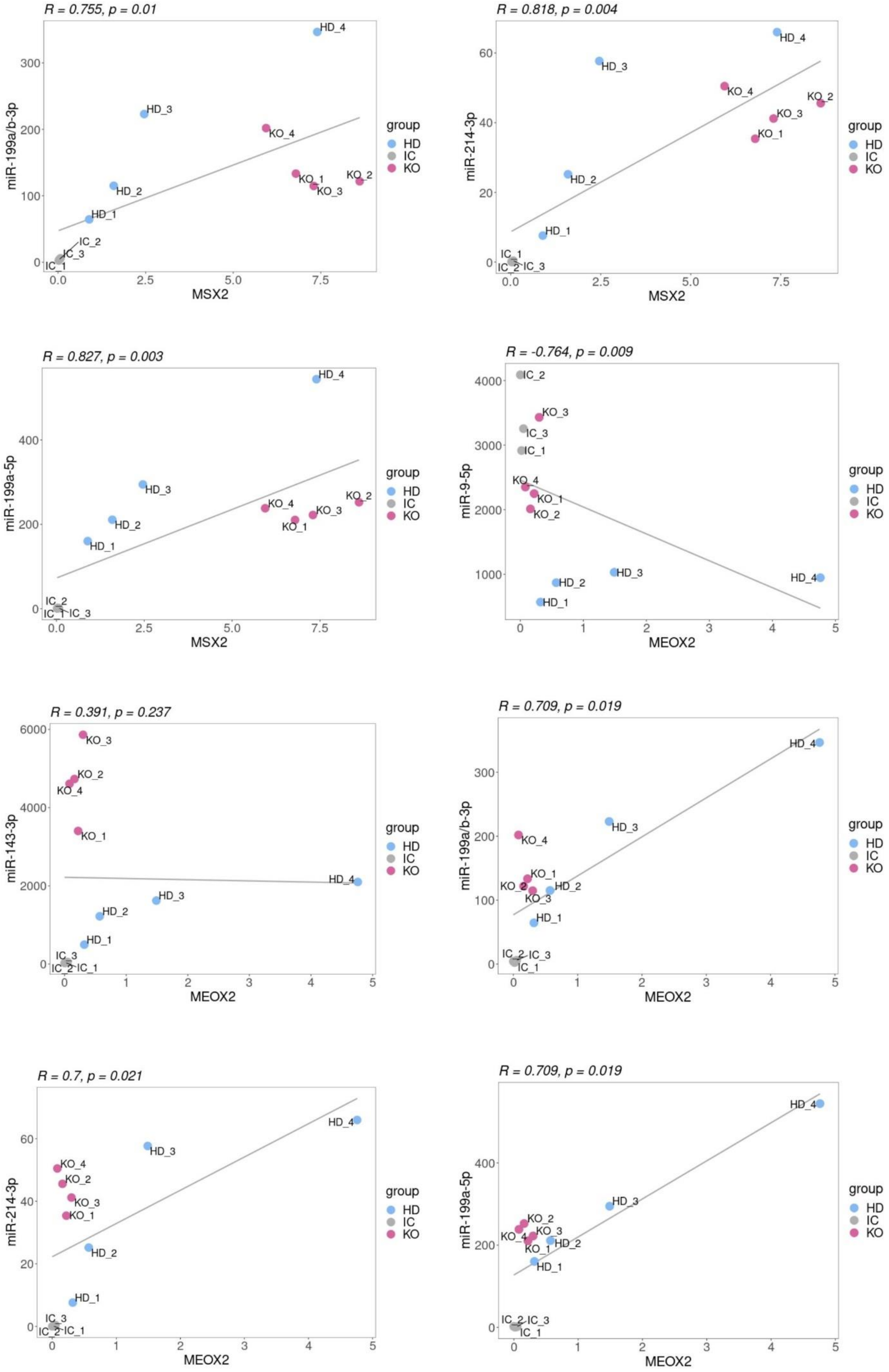
R = 0.755, p = 0.01
miR-199a/b-3p
MSX2
R = 0.818, p = 0.004
miR-214-3p
MSX2
R = 0.827, p = 0.003
miR-199a-5p
MSX2
R = -0.764, p = 0.009
miR-9-5p
MEOX2
R = 0.391, p = 0.237
miR-143-3p
MEOX2
R = 0.709, p = 0.019
miR-199a/b-3p
MEOX2
R = 0.7, p = 0.021
miR-214-3p
MEOX2
R = 0.709, p = 0.019
miR-199a-5p
MEOX2
group
HD
IC
KO
HD_1
HD_2
HD_3
HD_4
IC_1
IC_2
IC_3
KO_1
KO_2
KO_3
KO_4

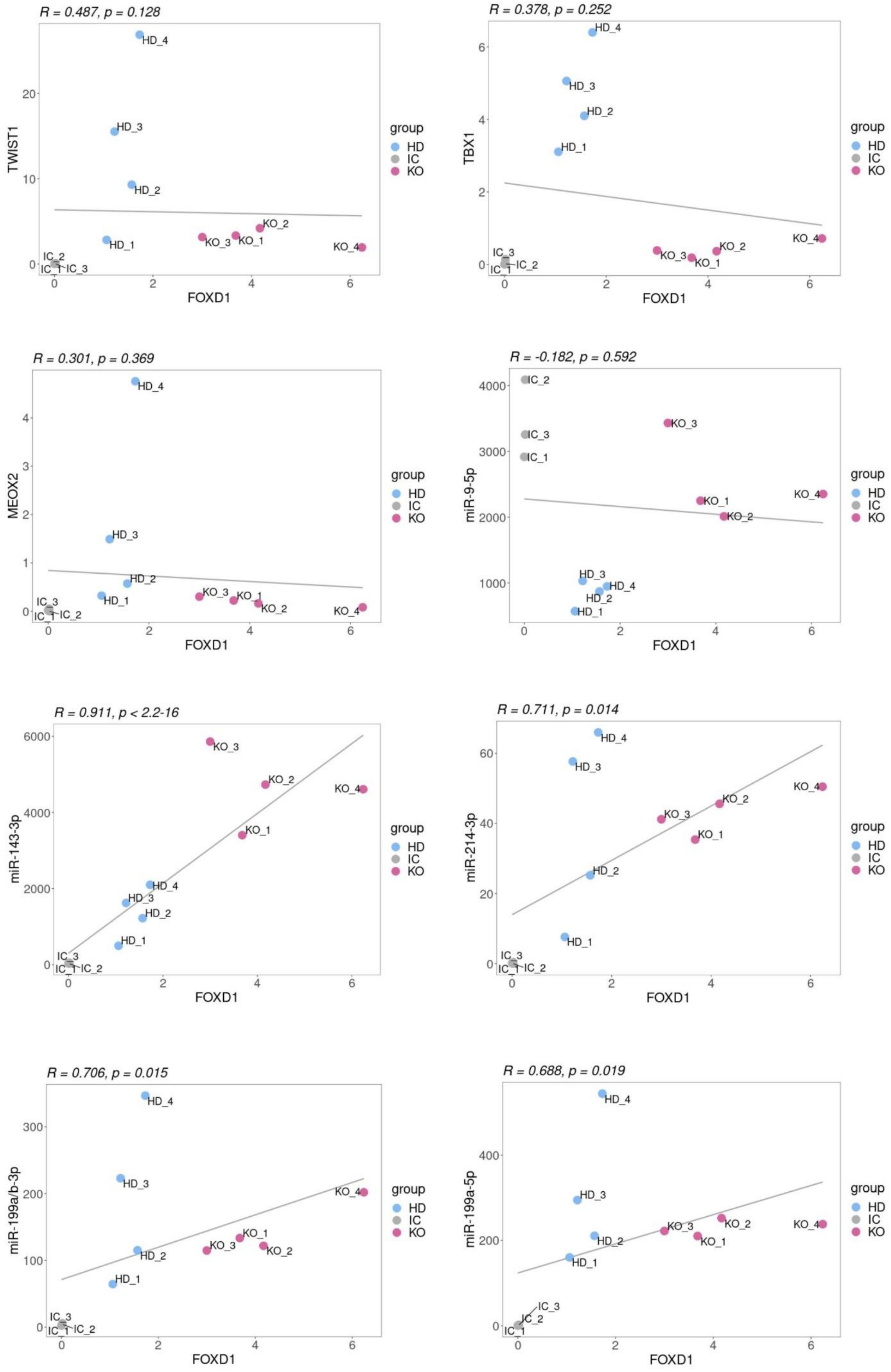
R = 0.487, p = 0.128
TWIST1
FOXD1
R = 0.378, p = 0.252
TBX1
FOXD1
R = 0.301, p = 0.369
MEOX2
FOXD1
R = -0.182, p = 0.592
miR-9-5p
FOXD1
R = 0.911, p < 2.2-16
miR-143-3p
FOXD1
R = 0.711, p = 0.014
miR-214-3p
FOXD1
R = 0.706, p = 0.015
miR-199a/b-3p
FOXD1
R = 0.688, p = 0.019
miR-199a-5p
FOXD1
group
HD
IC
KO
HD_1
HD_2
HD_3
HD_4
IC_1
IC_2
IC_3
KO_1
KO_2
KO_3
KO_4

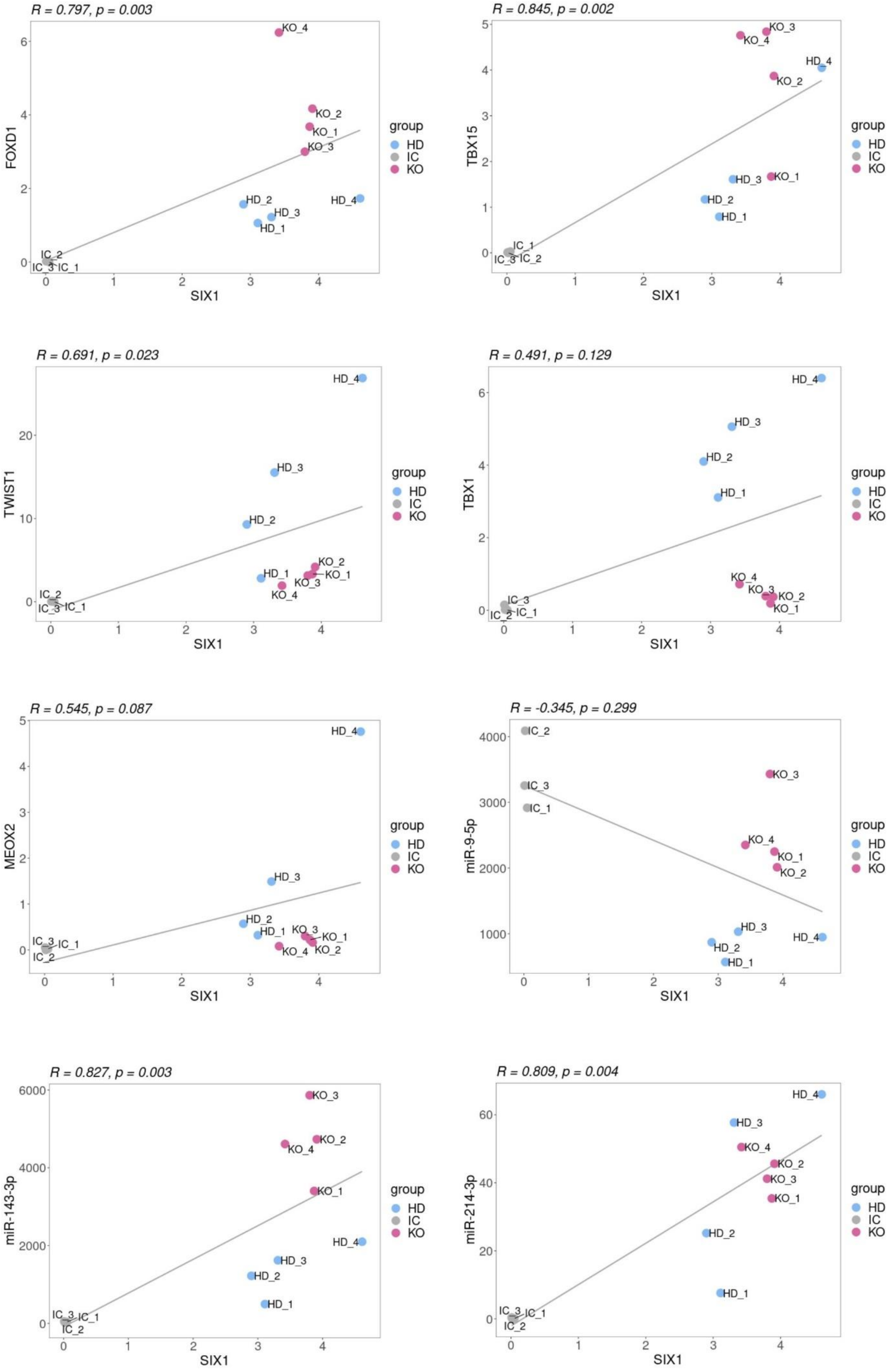

R = 0.797, p = 0.003
FOXD1
SIX1
KO_4
KO_2
KO_1
KO_3
HD_2
HD_3
HD_1
HD_4
IC_2
IC_3
IC_1
group
HD
IC
KO
R = 0.845, p = 0.002
TBX15
SIX1
KO_3
KO_4
HD_4
KO_2
HD_3
KO_1
HD_2
HD_1
IC_1
IC_3
IC_2
group
HD
IC
KO
R = 0.691, p = 0.023
TWIST1
SIX1
HD_4
HD_3
HD_2
KO_2
HD_1
KO_1
KO_3
KO_4
IC_2
IC_3
IC_1
group
HD
IC
KO
R = 0.491, p = 0.129
TBX1
SIX1
HD_4
HD_3
HD_2
HD_1
KO_4
KO_3
KO_2
KO_1
IC_3
IC_2
IC_1
group
HD
IC
KO
R = 0.545, p = 0.087
MEOX2
SIX1
HD_4
HD_3
HD_2
HD_1
KO_3
KO_1
KO_4
KO_2
IC_3
IC_1
IC_2
group
HD
IC
KO
R = -0.345, p = 0.299
miR-9-5p
SIX1
IC_2
IC_3
IC_1
KO_3
KO_4
KO_1
KO_2
HD_3
HD_2
HD_1
HD_4
group
HD
IC
KO
R = 0.827, p = 0.003
miR-143-3p
SIX1
KO_3
KO_2
KO_4
KO_1
HD_4
HD_3
HD_2
HD_1
IC_3
IC_1
IC_2
group
HD
IC
KO
R = 0.809, p = 0.004
miR-214-3p
SIX1
HD_4
HD_3
KO_4
KO_2
KO_3
KO_1
HD_2
HD_1
IC_3
IC_1
IC_2
group
HD
IC
KO

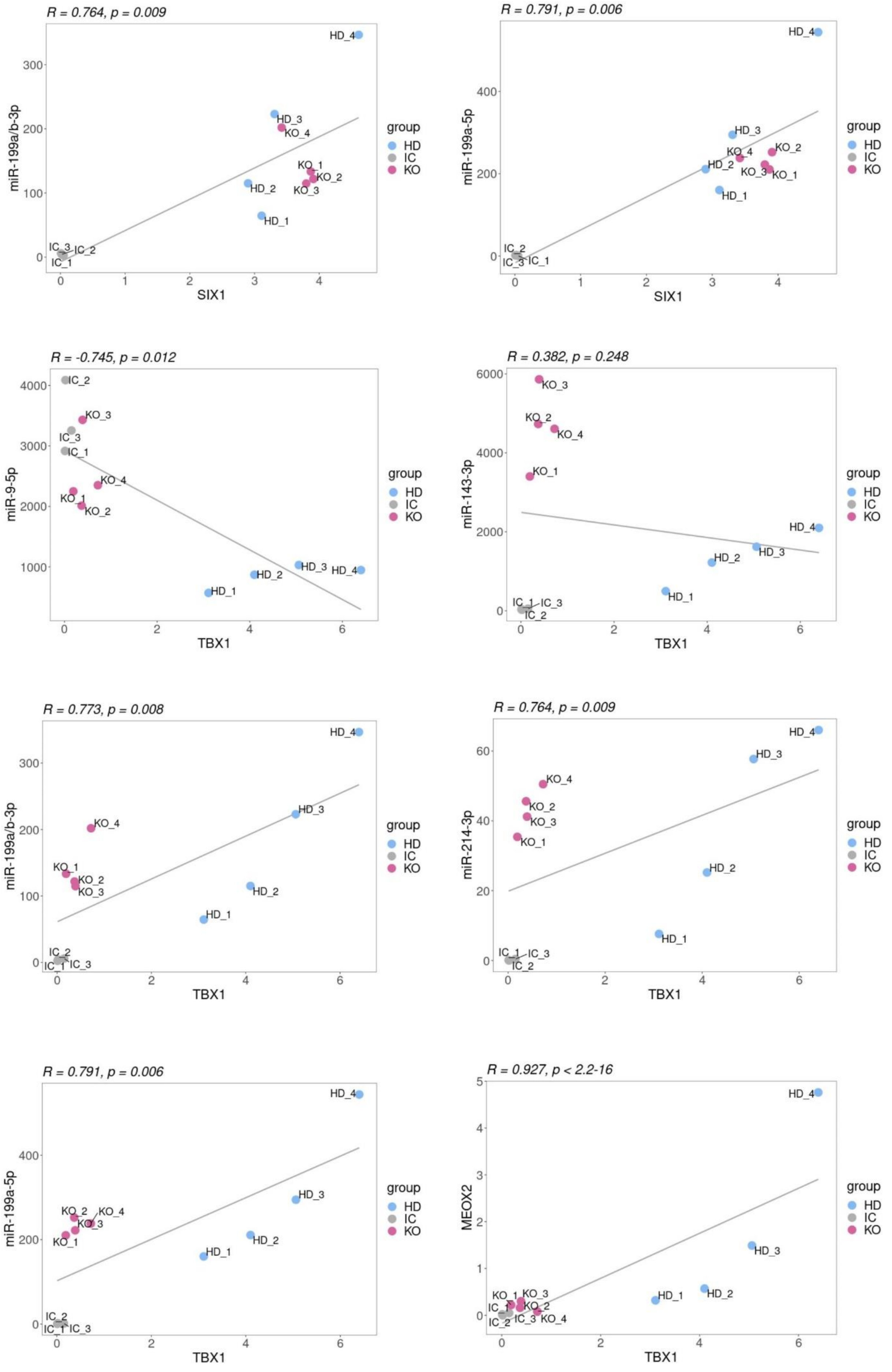
R = 0.764, p = 0.009
miR-199a/b-3p
SIX1
HD_4
HD_3
KO_4
KO_1
KO_2
KO_3
HD_2
HD_1
IC_3
IC_2
IC_1
group
HD
IC
KO
R = 0.791, p = 0.006
miR-199a-5p
SIX1
HD_4
HD_3
KO_4
KO_2
HD_2
KO_3
KO_1
HD_1
IC_2
IC_3
IC_1
group
HD
IC
KO
R = -0.745, p = 0.012
miR-9-5p
TBX1
IC_2
KO_3
IC_3
IC_1
KO_4
KO_1
KO_2
HD_3
HD_4
HD_2
HD_1
group
HD
IC
KO
R = 0.382, p = 0.248
miR-143-3p
TBX1
KO_3
KO_2
KO_4
KO_1
HD_4
HD_3
HD_2
HD_1
IC_1
IC_3
IC_2
group
HD
IC
KO
R = 0.773, p = 0.008
miR-199a/b-3p
TBX1
HD_4
HD_3
KO_4
KO_1
KO_2
KO_3
HD_2
HD_1
IC_2
IC_1
IC_3
group
HD
IC
KO
R = 0.764, p = 0.009
miR-214-3p
TBX1
HD_4
HD_3
KO_4
KO_2
KO_3
KO_1
HD_2
HD_1
IC_1
IC_3
IC_2
group
HD
IC
KO
R = 0.791, p = 0.006
miR-199a-5p
TBX1
HD_4
HD_3
KO_2
KO_4
KO_3
KO_1
HD_2
HD_1
IC_2
IC_1
IC_3
group
HD
IC
KO
R = 0.927, p < 2.2-16
MEOX2
TBX1
HD_4
HD_3
HD_2
HD_1
KO_1
KO_3
IC_1
KO_2
IC_2
IC_3
KO_4
group
HD
IC
KO

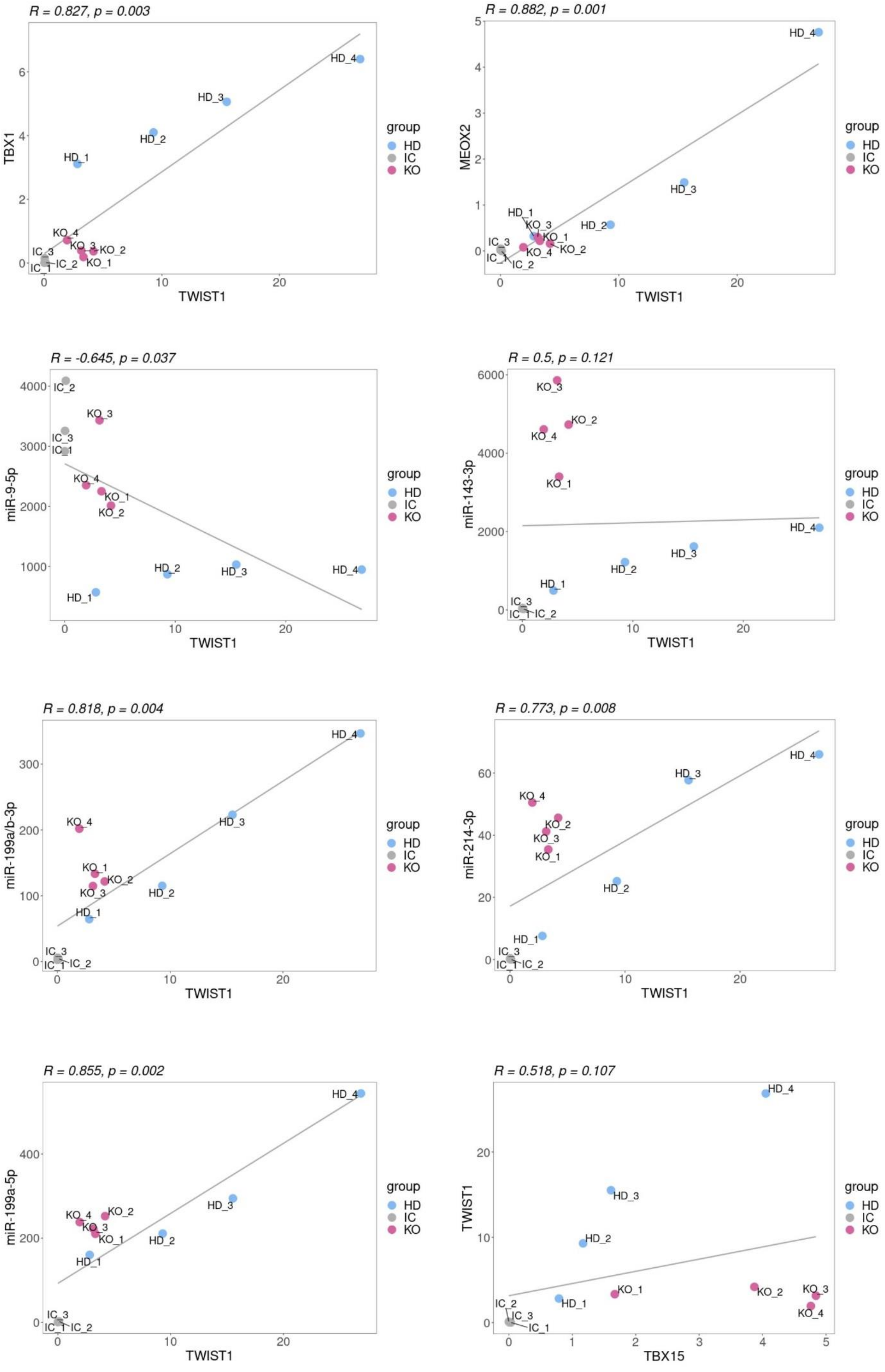
R = 0.827, p = 0.003
TBX1
TWIST1
R = 0.882, p = 0.001
MEOX2
TWIST1
R = -0.645, p = 0.037
miR-9-5p
TWIST1
R = 0.5, p = 0.121
miR-143-3p
TWIST1
R = 0.818, p = 0.004
miR-199a/b-3p
TWIST1
R = 0.773, p = 0.008
miR-214-3p
TWIST1
R = 0.855, p = 0.002
miR-199a-5p
TWIST1
R = 0.518, p = 0.107
TWIST1
TBX15
group
HD
IC
KO

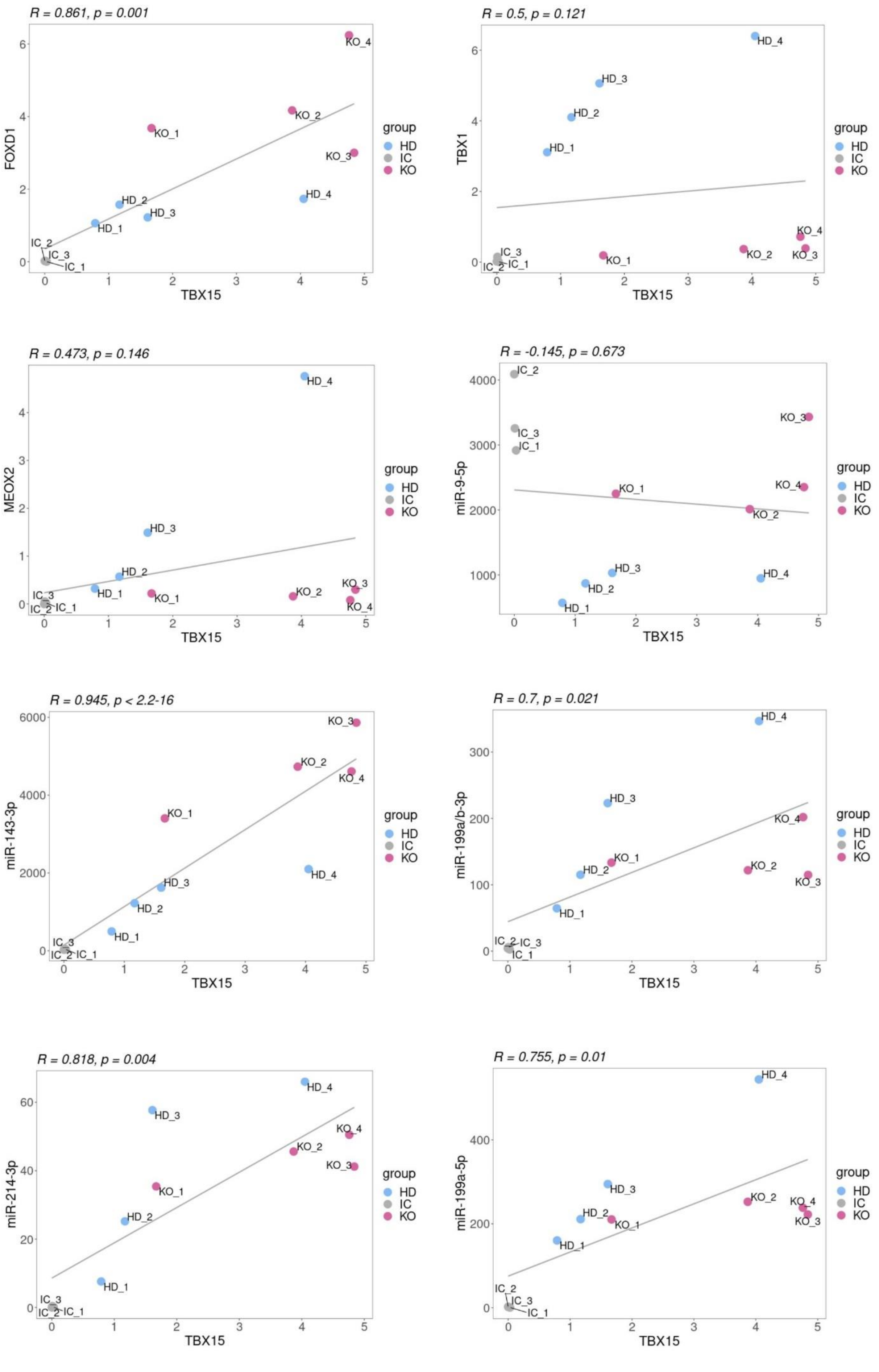
R = 0.861, p = 0.001
FOXD1
TBX15
R = 0.5, p = 0.121
TBX1
TBX15
R = 0.473, p = 0.146
MEOX2
TBX15
R = -0.145, p = 0.673
miR-9-5p
TBX15
R = 0.945, p < 2.2-16
miR-143-3p
TBX15
R = 0.7, p = 0.021
miR-199a/b-3p
TBX15
R = 0.818, p = 0.004
miR-214-3p
TBX15
R = 0.755, p = 0.01
miR-199a-5p
TBX15
group
HD
IC
KO

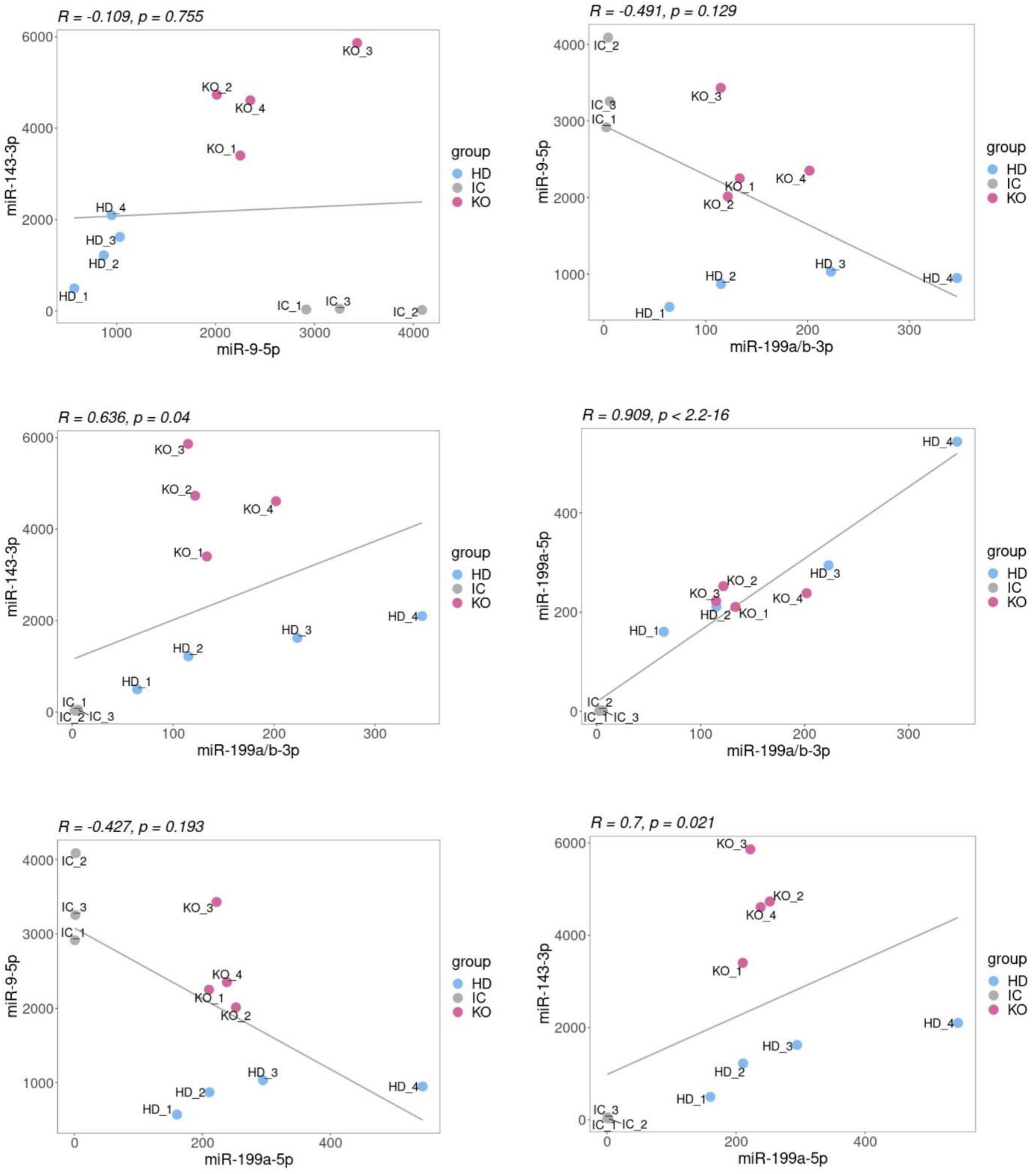
R = -0.109, p = 0.755
miR-143-3p
miR-9-5p
KO_3
KO_2
KO_4
KO_1
HD_4
HD_3
HD_2
HD_1
IC_1
IC_3
IC_2
group
HD
IC
KO
R = -0.491, p = 0.129
miR-9-5p
miR-199a/b-3p
R = 0.636, p = 0.04
miR-143-3p
miR-199a/b-3p
R = 0.909, p < 2.2-16
miR-199a-5p
miR-199a/b-3p
R = -0.427, p = 0.193
miR-9-5p
miR-199a-5p
R = 0.7, p = 0.021
miR-143-3p
miR-199a-5p

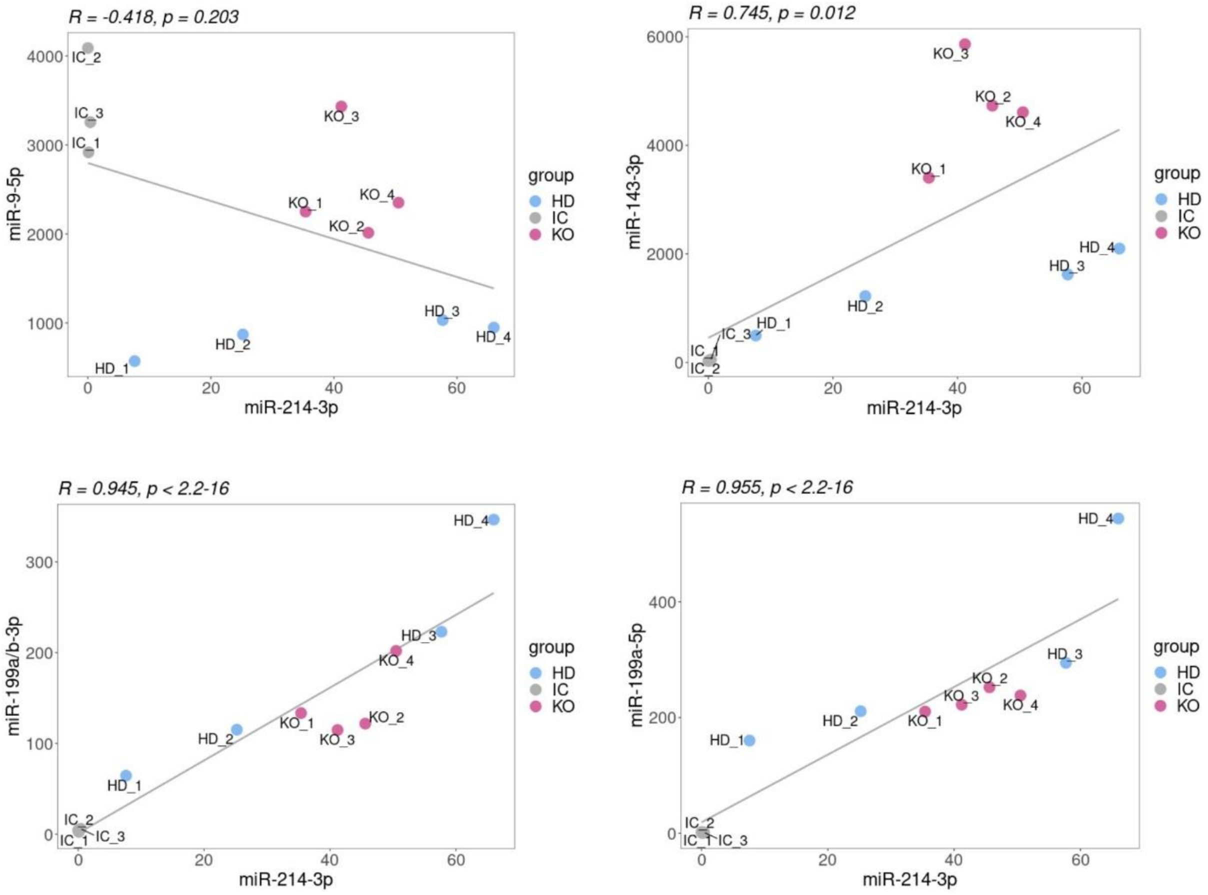


**Supplementary Figure 3. Correlation of expression of selected TFs and miRNAs in IC1, HD and KO NSCs**

Correlation plots for selected gene pairs were generated using ggplot2 v. 3.4.4 R package. Scatter plots present TPM expression values across all samples for selected gene pairs. Presented regression lines were added using the *geom_smooth* function (ggplot2 package) and Spearman correlation coefficients were calculated using the *cor.test* function (*stats* R package).

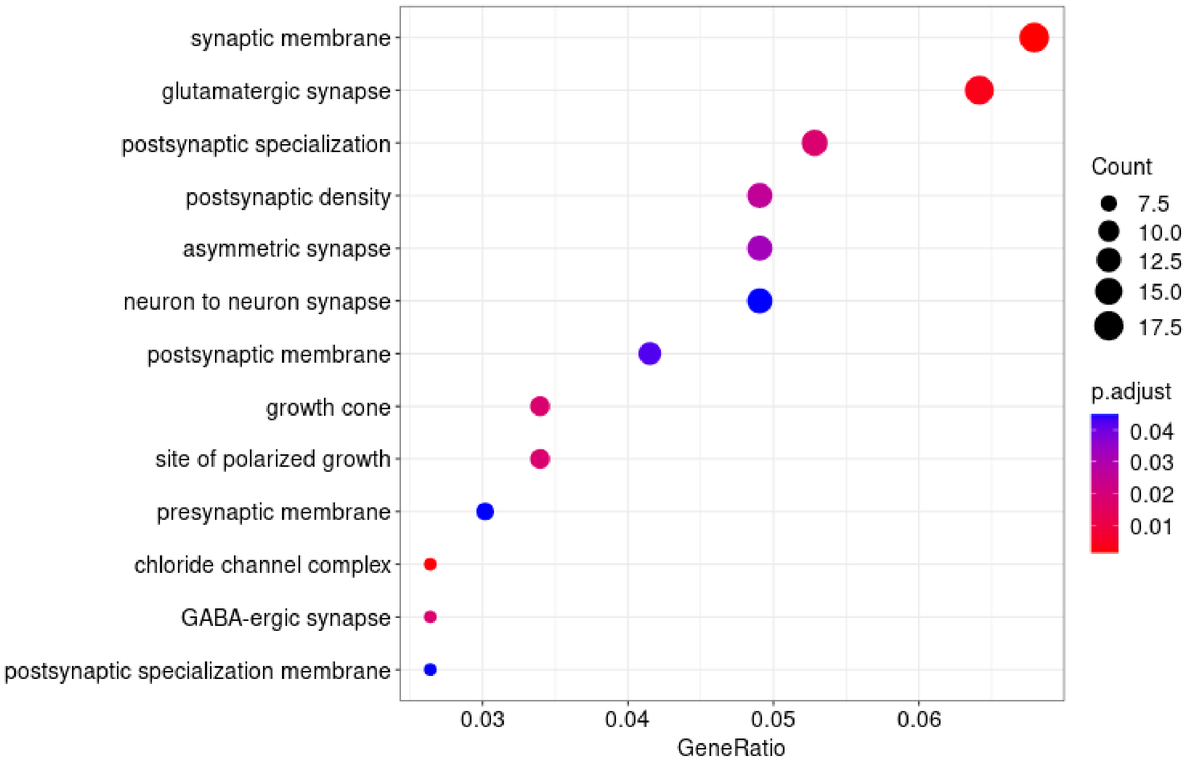


**Supplementary Figure 4. GO enrichment analysis for genes classified as "increasing" in control NSCs**

GO biological processes (BP) enrichment analysis was performed using a clusterProfiler R package. The significantly enriched GO-BP categories are listed and sorted by significance.

The y-axis is GO-BP term, and the x-axis shows the gene ratio representing the proportion of enriched genes in a GO term over the number of genes in the inputted gene list.

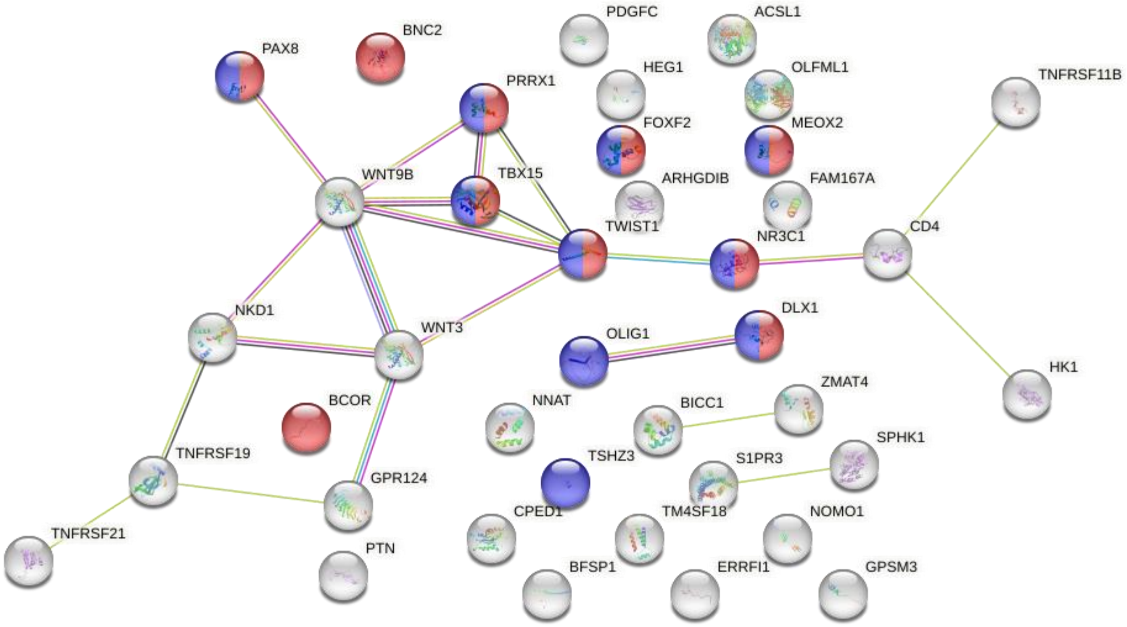


| GO-term | description | FDR |
|---|---|---|
| GO:0000977 (red) | RNA polymerase II transcription regulatory region sequence-specific DNA binding | 0,0215 |
| GO:0000981 (blue) | DNA-binding transcription factor activity, RNA polymerase II-specific | 0,0265 |

**Supplementary Figure 5. Enrichment of TFs that are associated with polymerase II in HD among „increasing genes" unique for HD-NSCs**

Prediction of the PPI network was conducted using the STRING v11.5 database
