## Supplementary Table 1 for "Huntingtin loss-of-function contributes to transcriptional deregulation in Huntington’s disease"

**Supplementary Table 1.** A list of primary and secondary antibodies with their dilutions used in immunocytochemistry and western blot

| **Primary antibodies** | **Dilution** | **Manufacturer** |
|---|---|---|
| anti-DARPP-32 (19A3) | 1:400 | Cell Signaling Technology (Cat. No. 2306) |
| anti-GAD67 | 1:50 | Santa Cruz Biotechnology (Cat. No. SC-28376) |
| anti-MAP2 | 1:200 | Cell Signaling Technology (Cat. No. 4542) |
| anti-TUJ1 | 1:500 | BioLegend (Cat. No. MMS-435P) |
| anti-HTT | 1:1000 | MAB2166 (Sigma-Aldrich/Merck) |
| anti-CANX | 1:1000 | C4731 (Sigma-Aldrich/Merck) |
| **Secondary antibodies** | **Dilution** | **Manufacturer** |
| anti-rabbit Alexa Fluor 488 | 1:1000 | Jackson ImmunoResearch (711-546-152) |
| anti-mouse Alexa Fluor 594 | 1:1000 | Jackson ImmunoResearch (715-586-151) |
| anti-rabbit | 1:1000 | Jackson ImmunoResearch (711-035-152) |
| anti-mouse | 1:1000 | A9917 (Sigma-Aldrich/Merck) |
