## Supplementary Table 2 for "Huntingtin loss-of-function contributes to transcriptional deregulation in Huntington’s disease"

**Supplementary Table 2.** A list of primers used for RT-qPCR

| Name | Forward (5'-3') | Reverse (5'-3') |
|---|---|---|
| *PAX6* | TGCTCCGGCATGAAATATACTA | GTCTCCAAATGTGCAGCAAC |
| *SOX1* | ACCAGGCCATGGATGAAG | CTTAATTGCTGGGGAATTGG |
| *SOX2* | CAAAAATGGCCATGCAGGTT | AGTTGGGATCGAACAAAAGCTATT |
| *RPLP0* | CATATCCGGGGGAATGTGGG | CAGCAGCTGGCACCTTATTG |
| *EEF2* | TCATCGAGGAGTCGGGAGAG | ACGACCGGGTCAGATTTCTTG |
| *TWIST1* | TACGCCTTCTCGGTCTGGAG | TTCTCTGGAAACAATGACATCTAGG |
| *FOXD1* | CGCTCGAGGAAGAAGGTAGG | GAGGAGCGAACAAAACACCG |
| *SIX1* | AGGTCAGCAACTGGTTTAAGAACC | GAGGAGAGAGTTGGTTCTGCTTG |
| *MSX2* | CGGAAAATTCAGAAGATGGAGCG | CGGCTTCCGATTGGTCTTGTGT |
| *MEOX2* | TCTCACCAGACTGAGGCGATAC | TCCACTTCATCCGCCTGTTTTGG |
| *TBX1* | TGGACCCACGCAAAGATAGC | TGAGCTGCGTGATCCGATG |
| *TBX15* | CTGGATGAGACAGGTGGTCAGT | GGTGAAAGGTCACTGCTGAAGTC |
