## Supplementary Table 3 for "Huntingtin loss-of-function contributes to transcriptional deregulation in Huntington’s disease"

**Supplementary Table 3. Lists of DEGs in HD and KO-NSCs**

**HD-NSC vs IC1-NSC**

| gene_ID | Gene_name | log2FoldChange | Fold_change | P-value | P_adjusted | Description |
|---|---|---|---|---|---|---|
| ENSG00000069122 | ADGRF5 | 10,01748938 | 1036,489209 | 7,48E-19 | 5,25E-16 | adhesion G protein-coupled receptor F5 |
| ENSG00000277649 | TIMM22 | 9,553103155 | 751,225984 | 0,00941 | 0,12417731 | translocase of inner mitochondrial membrane 22 |
| ENSG00000278501 | TIMM22 | 9,553103155 | 751,225984 | 0,00941 | 0,12417731 | translocase of inner mitochondrial membrane 22 |
| ENSG00000287080 | H3C3 | 9,222647254 | 597,4388537 | 6,48E-82 | 1,82E-77 | H3 clustered histone 3 |
| ENSG00000122691 | TWIST1 | 8,694692813 | 414,3461804 | 4,25E-25 | 6,30E-22 | twist family bHLH transcription factor 1 |
| ENSG00000229807 | XIST | 8,58302298 | 383,4841074 | 1,96E-32 | 5,49E-29 | X inactive specific transcript |
| ENSG00000278444 | CDC42EP5 | 8,362411405 | 329,1066584 | 0,00311 | 0,06649378 | CDC42 effector protein 5 |
| ENSG00000276410 | H2BC3 | 8,113259472 | 276,9073375 | 4,08E-33 | 1,43E-29 | H2B clustered histone 3 |
| ENSG00000163599 | CTLA4 | 7,997303853 | 255,5220272 | 0,00752 | 0,11001711 | cytotoxic T-lymphocyte associated protein 4 |
| ENSG00000232821 | AC003986,2 | 7,98026835 | 252,5225387 | 9,99E-09 | 1,82E-06 | novel transcript |
| ENSG00000287996 | AL359208,1 | 7,829342642 | 227,4400707 | 8,71E-09 | 1,63E-06 | novel transcript |
| ENSG00000205810 | KLRC3 | 7,797664528 | 222,500463 | 1,40E-08 | 2,44E-06 | killer cell lectin like receptor C3 |
| ENSG00000126778 | SIX1 | 7,506493967 | 181,8359904 | 2,81E-42 | 1,32E-38 | SIX homeobox 1 |
| ENSG00000143196 | DPT | 7,443323433 | 174,0458293 | 4,17E-07 | 5,27E-05 | dermatopontin |
| ENSG00000274847 | MAFIP | 7,201983644 | 147,2356931 | 5,54E-07 | 6,61E-05 | MAFF interacting protein (pseudogene) |
| ENSG00000019186 | CYP24A1 | 7,04078668 | 131,6703481 | 0,0028 | 0,06237385 | cytochrome P450 family 24 subfamily A member 1 |
| ENSG00000011465 | DCN | 7,010945526 | 128,9748113 | 4,97E-20 | 3,98E-17 | decorin |
| ENSG00000278299 | TBC1D3C | 7,005489679 | 128,4879878 | 0,05701 | 0,31850415 | TBC1 domain family member 3C |
| ENSG00000285032 | MTMR9 | 6,947609608 | 123,4351617 | 0,0591 | 0,32424653 | myotubularin related protein 9 |
| ENSG00000106034 | CPED1 | 6,946196079 | 123,3142811 | 3,27E-27 | 6,12E-24 | cadherin like and PC-esterase domain containing 1 |
| ENSG00000205809 | KLRC2 | 6,945259778 | 123,2342769 | 1,08E-07 | 1,52E-05 | killer cell lectin like receptor C2 |
| ENSG00000139329 | LUM | 6,941670817 | 122,9280906 | 4,26E-25 | 6,30E-22 | lumican |
| ENSG00000000005 | TNMD | 6,878134234 | 117,6317935 | 1,00E-07 | 1,42E-05 | tenomodulin |
| ENSG00000111799 | COL12A1 | 6,872037667 | 117,1357518 | 1,05E-29 | 2,27E-26 | collagen type XII alpha 1 chain |
| ENSG00000212643 | ZRSR2P1 | 6,788874927 | 110,5745015 | 0,00151 | 0,04166341 | ZRSR2 pseudogene 1 |
| ENSG00000182492 | BGN | 6,757104889 | 108,166121 | 1,14E-68 | 1,07E-64 | biglycan |
| ENSG00000255641 | AC068775,1 | 6,749586679 | 107,6039091 | 0,00028 | 0,01368428 | novel protein |
| ENSG00000285330 | AC126283,2 | 6,748550725 | 107,5266698 | 0,00025 | 0,01271239 | novel protein |
| ENSG00000071991 | CDH19 | 6,722576575 | 105,6080922 | 9,73E-07 | 0,00011184 | cadherin 19 |
| ENSG00000255819 | KLRC4-KLRK1 | 6,714334641 | 105,0064866 | 7,88E-07 | 9,25E-05 | KLRC4-KLRK1 readthrough |

| | | | | | | |
|---|---|---|---|---|---|---|
| ENSG00000130300 | PLVAP | 6,690902047 | 103,3147227 | 3,97E-22 | 4,13E-19 | plasmalemma vesicle associated protein |
| ENSG00000092607 | TBX15 | 6,684193389 | 102,8354154 | 8,55E-20 | 6,49E-17 | T-box transcription factor 15 |
| ENSG00000277726 | AL109811,3 | 6,6453891 | 100,1063097 | 9,53E-06 | 0,00080313 | novel protein |
| ENSG00000258952 | SALRNA1 | 6,636308709 | 99,47821456 | 7,70E-06 | 0,00067131 | senescence associated long non-coding RNA 1 |
| ENSG00000251254 | GTF2F2P1 | 6,614496218 | 97,98548978 | 3,69E-05 | 0,00259353 | general transcription factor IIF subunit 2 pseudogene 1 |
| ENSG00000159184 | HOXB13 | 6,540431199 | 93,0820577 | 0,0003 | 0,01422985 | homeobox B13 |
| ENSG00000288297 | ECSCR | 6,522657847 | 91,94236344 | 5,46E-06 | 0,00049451 | endothelial cell surface expressed chemotaxis and apoptosis regulator |
| ENSG00000249751 | ECSCR | 6,522657847 | 91,94236344 | 5,46E-06 | 0,00049451 | endothelial cell surface expressed chemotaxis and apoptosis regulator |
| ENSG00000184058 | TBX1 | 6,514183688 | 91,40389176 | 1,74E-25 | 2,87E-22 | T-box transcription factor 1 |
| ENSG00000145681 | HAPLN1 | 6,455927456 | 87,78651615 | 6,14E-71 | 8,61E-67 | hyaluronan and proteoglycan link protein 1 |
| ENSG00000287496 | AC093227,3 | 6,452972044 | 87,60686644 | 0,00027 | 0,01321807 | novel transcript |
| ENSG00000162761 | LMX1A | 6,434326367 | 86,48190316 | 1,76E-06 | 0,00018061 | LIM homeobox transcription factor 1 alpha |
| ENSG00000123610 | TNFAIP6 | 6,421969201 | 85,74432006 | 4,29E-05 | 0,00295839 | TNF alpha induced protein 6 |
| ENSG00000251493 | FOXD1 | 6,288264831 | 78,15492215 | 1,62E-21 | 1,56E-18 | forkhead box D1 |
| ENSG00000215912 | TTC34 | 6,230244434 | 75,07415539 | 3,78E-07 | 4,84E-05 | tetratricopeptide repeat domain 34 |
| ENSG00000008196 | TFAP2B | 6,207603045 | 73,90515253 | 8,36E-17 | 4,60E-14 | transcription factor AP-2 beta |
| ENSG00000268357 | VN1R81P | 6,194960235 | 73,26032698 | 0,0002 | 0,01057544 | vomeronasal 1 receptor 81 pseudogene |
| ENSG00000234475 | HSPA1A | 6,183176206 | 72,664369 | 0,01415 | 0,15611849 | heat shock protein family A (Hsp70) member 1A |
| ENSG00000106511 | MEOX2 | 6,154190052 | 71,21898907 | 1,60E-09 | 3,43E-07 | mesenchyme homeobox 2 |
| ENSG00000284811 | AC270285,1 | 6,128224064 | 69,94863812 | 8,51E-06 | 0,00072613 | |
| ENSG00000168497 | CAVIN2 | 6,101926376 | 68,68515307 | 7,87E-15 | 3,45E-12 | caveolae associated protein 2 |
| ENSG00000226268 | AC135977,1 | 6,069223706 | 67,14572784 | 0,07338 | 0,35662549 | solute carrier family 25, member 33 (SLC25A33) pseudogene |
| ENSG00000223618 | VPS52 | 6,058472045 | 66,64718544 | 0,01609 | 0,16620294 | VPS52 subunit of GARP complex |
| ENSG00000274741 | AC138827,3 | 6,056119121 | 66,53857763 | 0,00049 | 0,02017094 | occludin (OCLN) pseudogene |
| ENSG00000262456 | AC006435,1 | 6,026378912 | 65,18096977 | 0,01683 | 0,1704098 | novel transcript, antisense to MNT |
| ENSG00000198851 | CD3E | 5,979392621 | 63,09232544 | 0,00433 | 0,08098807 | CD3e molecule |
| ENSG00000157554 | ERG | 5,972259523 | 62,7811488 | 3,72E-16 | 1,91E-13 | ETS transcription factor ERG |
| ENSG00000163081 | CCDC140 | 5,897899149 | 59,6272191 | 9,19E-06 | 0,00077656 | CCDC140 long non-coding RNA |
| ENSG00000120149 | MSX2 | 5,896931316 | 59,58723156 | 8,19E-15 | 3,54E-12 | msh homeobox 2 |
| ENSG00000227191 | TRGC2 | 5,881623489 | 58,95831913 | 7,17E-05 | 0,00458395 | T cell receptor gamma constant 2 |

| | | | | | | |
|---|---|---|---|---|---|---|
| ENSG00000183542 | KLRC4 | 5,880895654 | 58,92858233 | 0,00092 | 0,03058422 | killer cell lectin like receptor C4 |
| ENSG00000240821 | RPL9P25 | 5,872568081 | 58,5894126 | 0,00022 | 0,01145716 | ribosomal protein L9 pseudogene 25 |
| ENSG00000287554 | AC006364,1 | 5,862510351 | 58,18237774 | 0,0004 | 0,0176355 | novel transcript |
| ENSG00000276886 | GREM1 | 5,859616771 | 58,06579964 | 1,33E-08 | 2,35E-06 | gremlin 1, DAN family BMP antagonist |
| ENSG00000288370 | CTAGE4 | 5,744669631 | 53,6188968 | 0,00043 | 0,018296 | CTAGE family member 4 |
| ENSG00000180938 | ZNF572 | 5,648608692 | 50,16498077 | 1,96E-09 | 4,11E-07 | zinc finger protein 572 |
| ENSG00000133110 | POSTN | 5,642817564 | 49,96401692 | 7,01E-20 | 5,47E-17 | periostin |
| ENSG00000237070 | AC005550,2 | 5,580918828 | 47,86565107 | 0,00038 | 0,01693013 | novel transcript, antisense to MEOX2 |
| ENSG00000234816 | H2AC5P | 5,447800608 | 43,64669798 | 6,69E-05 | 0,00431491 | H2A clustered histone 5, pseudogene |
| ENSG00000158055 | GRHL3 | 5,428221009 | 43,05834636 | 9,19E-10 | 2,11E-07 | grainyhead like transcription factor 3 |
| ENSG00000138207 | RBP4 | 5,371257961 | 41,39136614 | 0,00078 | 0,02733095 | retinol binding protein 4 |
| ENSG00000287352 | AC011029,1 | 5,354506148 | 40,91353101 | 0,00028 | 0,01358105 | novel transcript |
| ENSG00000266524 | GDF10 | 5,353545915 | 40,88630873 | 2,21E-24 | 3,10E-21 | growth differentiation factor 10 |
| ENSG00000134853 | PDGFRA | 5,350096481 | 40,78866774 | 2,68E-24 | 3,59E-21 | platelet derived growth factor receptor alpha |
| ENSG00000160180 | TFF3 | 5,344365702 | 40,62696539 | 0,00041 | 0,01789203 | trefoil factor 3 |
| ENSG00000172987 | HPSE2 | 5,321870148 | 39,99839341 | 9,62E-15 | 4,09E-12 | heparanase 2 (inactive) |
| ENSG00000186564 | FOXD2 | 5,257326539 | 38,24837469 | 0,0001 | 0,00619321 | forkhead box D2 |
| ENSG00000230630 | DNM3OS | 5,235421452 | 37,67201905 | 5,03E-16 | 2,48E-13 | DNM3 opposite strand/antisense RNA |
| ENSG00000106991 | ENG | 5,182997341 | 36,32768031 | 3,23E-17 | 1,81E-14 | endoglin |
| ENSG00000141052 | MYOCD | 5,168582567 | 35,96651742 | 1,29E-09 | 2,85E-07 | myocardin |
| ENSG00000137273 | FOXF2 | 5,150140839 | 35,50968946 | 3,18E-18 | 2,08E-15 | forkhead box F2 |
| ENSG00000178726 | THBD | 5,149539069 | 35,49488094 | 3,59E-10 | 8,82E-08 | thrombomodulin |
| ENSG00000164741 | DLC1 | 5,112722495 | 34,60053632 | 7,87E-46 | 4,42E-42 | DLC1 Rho GTPase activating protein |
| ENSG00000174600 | CMKLR1 | 4,94566602 | 30,81724566 | 3,19E-11 | 9,03E-09 | chemerin chemokine-like receptor 1 |
| ENSG00000125618 | PAX8 | 4,87647768 | 29,3742005 | 6,68E-23 | 8,15E-20 | paired box 8 |
| ENSG00000150048 | CLEC1A | 4,853835332 | 28,91678652 | 1,27E-10 | 3,30E-08 | C-type lectin domain family 1 member A |
| ENSG00000151617 | EDNRA | 4,822188724 | 28,28938139 | 6,99E-21 | 6,54E-18 | endothelin receptor type A |
| ENSG00000107562 | CXCL12 | 4,812006464 | 28,09042339 | 2,77E-20 | 2,35E-17 | C-X-C motif chemokine ligand 12 |
| ENSG00000248572 | EGFLAM-AS2 | 4,778038083 | 27,43675741 | 0,00076 | 0,02699099 | EGFLAM antisense RNA 2 |
| ENSG00000288673 | AC270289,4 | 4,727182592 | 26,4864501 | 9,65E-09 | 1,77E-06 | DLC1 Rho GTPase activating protein |
| ENSG00000120708 | TGFBI | 4,709620637 | 26,16598451 | 4,20E-08 | 6,59E-06 | transforming growth factor beta induced |
| ENSG00000187140 | FOXD3 | 4,679090384 | 25,61807884 | 3,35E-06 | 0,00032172 | forkhead box D3 |
| ENSG00000163453 | IGFBP7 | 4,58706234 | 24,03495738 | 8,61E-22 | 8,63E-19 | insulin like growth factor binding protein 7 |

| | | | | | | |
|---|---|---|---|---|---|---|
| ENSG00000104415 | CCN4 | 4,585758638 | 24,01324782 | 1,41E-06 | 0,00015278 | cellular communication network factor 4 |
| ENSG00000155011 | DKK2 | 4,576285684 | 23,85608973 | 1,85E-18 | 1,23E-15 | dickkopf WNT signaling pathway inhibitor 2 |
| ENSG00000172819 | RARG | 4,420129655 | 21,40876479 | 9,24E-12 | 2,82E-09 | retinoic acid receptor gamma |
| ENSG00000109132 | PHOX2B | 4,400746958 | 21,12306024 | 0,00123 | 0,03702164 | paired like homeobox 2B |
| ENSG00000112837 | TBX18 | 4,390342692 | 20,97127531 | 1,40E-19 | 1,03E-16 | T-box transcription factor 18 |
| ENSG00000248498 | ASNSP1 | 4,38911132 | 20,9533835 | 0,00011 | 0,00631075 | asparagine synthetase pseudogene 1 |
| ENSG00000005102 | MEOX1 | 4,376944748 | 20,7774219 | 1,05E-13 | 3,93E-11 | mesenchyme homeobox 1 |
| ENSG00000258815 | LINC02820 | 4,356421972 | 20,48394907 | 1,29E-09 | 2,85E-07 | long intergenic non-protein coding RNA 2820 |
| ENSG00000165694 | FRMD7 | 4,3488593 | 20,37685223 | 1,55E-17 | 9,25E-15 | FERM domain containing 7 |
| ENSG00000184254 | ALDH1A3 | 4,341881902 | 20,27854029 | 5,66E-08 | 8,67E-06 | aldehyde dehydrogenase 1 family member A3 |
| ENSG00000111432 | FZD10 | 4,335444284 | 20,18825463 | 1,82E-11 | 5,42E-09 | frizzled class receptor 10 |
| ENSG00000175899 | A2M | 4,285752594 | 19,50473638 | 1,54E-22 | 1,80E-19 | alpha-2-macroglobulin |
| ENSG00000104313 | EYA1 | 4,258177594 | 19,13547215 | 2,01E-20 | 1,77E-17 | EYA transcriptional coactivator and phosphatase 1 |
| ENSG00000054598 | FOXC1 | 4,227847409 | 18,73738098 | 1,53E-23 | 1,96E-20 | forkhead box C1 |
| ENSG00000112175 | BMP5 | 4,214158316 | 18,56043111 | 7,22E-16 | 3,43E-13 | bone morphogenetic protein 5 |
| ENSG00000248215 | LINC02505 | 4,173296873 | 18,04211886 | 0,00107 | 0,03382055 | long intergenic non-protein coding RNA 2505 |
| ENSG00000188783 | PRELP | 4,167965281 | 17,97556585 | 7,65E-09 | 1,46E-06 | proline and arginine rich end leucine rich repeat protein |
| ENSG00000189058 | APOD | 4,153892205 | 17,80107182 | 0,00256 | 0,05892196 | apolipoprotein D |
| ENSG00000145908 | ZNF300 | 4,094517523 | 17,08333244 | 6,92E-14 | 2,69E-11 | zinc finger protein 300 |
| ENSG00000124491 | F13A1 | 4,081286804 | 16,9273802 | 2,46E-33 | 9,84E-30 | coagulation factor XIII A chain |
| ENSG00000064300 | NGFR | 4,066327753 | 16,75277 | 6,70E-08 | 1,01E-05 | nerve growth factor receptor |
| ENSG00000164318 | EGFLAM | 4,063479221 | 16,71972509 | 3,83E-16 | 1,92E-13 | EGF like, fibronectin type III and laminin G domains |
| ENSG00000041982 | TNC | 4,054431091 | 16,61519237 | 1,39E-09 | 3,03E-07 | tenascin C |
| ENSG00000166825 | ANPEP | 4,041548377 | 16,46748552 | 9,02E-09 | 1,68E-06 | alanyl aminopeptidase, membrane |
| ENSG00000282416 | AC092299,1 | 4,0197037 | 16,22002007 | 0,25451 | 0,55944852 | novel pseudogene |
| ENSG00000234451 | BX005266,3 | 4,018090596 | 16,2018943 | 0,00233 | 0,05562927 | G protein-coupled receptor, family C, group 5,member B (GPRC5B) pseudogene |
| ENSG00000280916 | FOXCUT | 4,018051094 | 16,20145069 | 0,0003 | 0,01421667 | FOXC1 upstream transcript |
| ENSG00000184584 | STING1 | 3,987520843 | 15,86219856 | 4,02E-06 | 0,000377 | stimulator of interferon response cGAMP interactor 1 |
| ENSG00000163017 | ACTG2 | 3,968245452 | 15,65167824 | 7,74E-14 | 2,98E-11 | actin gamma 2, smooth muscle |
| ENSG00000135903 | PAX3 | 3,957702719 | 15,53771791 | 1,10E-57 | 7,68E-54 | paired box 3 |
| ENSG00000169071 | ROR2 | 3,949846977 | 15,45334208 | 4,70E-18 | 3,00E-15 | receptor tyrosine kinase like orphan receptor 2 |
| ENSG00000170961 | HAS2 | 3,941498514 | 15,36417625 | 2,57E-06 | 0,00025343 | hyaluronan synthase 2 |

| | | | | | | |
|---|---|---|---|---|---|---|
| ENSG00000135318 | NT5E | 3,938823045 | 15,33570989 | 3,96E-20 | 3,27E-17 | 5'-nucleotidase ecto |
| ENSG00000285579 | AC015687,1 | 3,911487901 | 15,04787538 | 9,66E-06 | 0,00080972 | novel transcript |
| ENSG00000111348 | ARHGDIB | 3,910640141 | 15,03903551 | 9,56E-08 | 1,38E-05 | Rho GDP dissociation inhibitor beta |
| ENSG00000184374 | COLEC10 | 3,908483204 | 15,01656783 | 3,39E-06 | 0,00032451 | collectin subfamily member 10 |
| ENSG00000047457 | CP | 3,904684266 | 14,97707787 | 8,79E-12 | 2,71E-09 | ceruloplasmin |
| ENSG00000196867 | ZFP28 | 3,890797933 | 14,83361096 | 7,73E-18 | 4,72E-15 | ZFP28 zinc finger protein |
| ENSG00000164035 | EMCN | 3,875878179 | 14,6809984 | 9,62E-05 | 0,00581492 | endomucin |
| ENSG00000277400 | AC145212,1 | 3,871070963 | 14,63216113 | 1,15E-08 | 2,06E-06 | MaFF-interacting protein |
| ENSG00000087303 | NID2 | 3,853502019 | 14,45505326 | 1,75E-32 | 5,46E-29 | nidogen 2 |
| ENSG00000122861 | PLAU | 3,84978405 | 14,4178491 | 1,12E-09 | 2,51E-07 | plasminogen activator, urokinase |
| ENSG00000170965 | PLAC1 | 3,825227162 | 14,174512 | 0,00788 | 0,11292581 | placenta enriched 1 |
| ENSG00000197467 | COL13A1 | 3,803248134 | 13,96020414 | 4,99E-30 | 1,17E-26 | collagen type XIII alpha 1 chain |
| ENSG00000275571 | TTC34 | 3,77895469 | 13,72709736 | 6,06E-08 | 9,24E-06 | tetratricopeptide repeat domain 34 |
| ENSG00000145824 | CXCL14 | 3,724097049 | 13,21493157 | 3,73E-10 | 9,10E-08 | C-X-C motif chemokine ligand 14 |
| ENSG00000108691 | CCL2 | 3,722186819 | 13,19744565 | 7,98E-06 | 0,00068866 | C-C motif chemokine ligand 2 |
| ENSG00000189223 | PAX8-AS1 | 3,721939756 | 13,19518576 | 1,03E-10 | 2,73E-08 | PAX8 antisense RNA 1 |
| ENSG00000168542 | COL3A1 | 3,693689459 | 12,93931608 | 2,70E-11 | 7,82E-09 | collagen type III alpha 1 chain |
| ENSG00000211689 | TRGC1 | 3,68771435 | 12,88583704 | 6,30E-07 | 7,46E-05 | T cell receptor gamma constant 1 |
| ENSG00000138061 | CYP1B1 | 3,660227488 | 12,64265436 | 5,88E-27 | 1,03E-23 | cytochrome P450 family 1 subfamily B member 1 |
| ENSG00000151468 | CCDC3 | 3,640778379 | 12,47336123 | 6,36E-14 | 2,51E-11 | coiled-coil domain containing 3 |
| ENSG00000176692 | FOXC2 | 3,637883836 | 12,44836045 | 2,51E-10 | 6,24E-08 | forkhead box C2 |
| ENSG00000079689 | SCGN | 3,637102676 | 12,44162201 | 9,26E-07 | 0,00010783 | secretagogin, EF-hand calcium binding protein |
| ENSG00000228592 | D21S2088E | 3,616767187 | 12,26748147 | 1,18E-06 | 0,00013278 | D21S2088E |
| ENSG00000288335 | TWIST2 | 3,591214303 | 12,05211387 | 1,75E-06 | 0,00017951 | twist family bHLH transcription factor 2 |
| ENSG00000233608 | TWIST2 | 3,591214303 | 12,05211387 | 1,75E-06 | 0,00017951 | twist family bHLH transcription factor 2 |
| ENSG00000135636 | DYSF | 3,586558443 | 12,01328202 | 1,34E-16 | 7,23E-14 | dysferlin |
| ENSG00000017483 | SLC38A5 | 3,585825581 | 12,00718105 | 2,75E-09 | 5,64E-07 | solute carrier family 38 member 5 |
| ENSG00000223345 | H2BP1 | 3,573599025 | 11,90585253 | 0,00049 | 0,01999623 | H2B histone pseudogene 1 |
| ENSG00000213654 | GPSM3 | 3,557890442 | 11,77692053 | 0,00046 | 0,01934795 | G protein signaling modulator 3 |
| ENSG00000247993 | FOXD1-AS1 | 3,546096641 | 11,68103853 | 6,29E-07 | 7,46E-05 | FOXD1 antisense RNA 1 |
| ENSG00000277297 | ATP5F1AP10 | 3,539312102 | 11,62623526 | 0,00055 | 0,02167479 | ATP synthase F1 subunit alpha pseudogene 10 |
| ENSG00000100234 | TIMP3 | 3,537904698 | 11,61489895 | 1,42E-08 | 2,45E-06 | TIMP metallopeptidase inhibitor 3 |
| ENSG00000013297 | CLDN11 | 3,47672771 | 11,13266979 | 1,05E-13 | 3,93E-11 | claudin 11 |

| | | | | | | |
|---|---|---|---|---|---|---|
| ENSG00000151702 | FLI1 | 3,450211357 | 10,92992319 | 3,76E-07 | 4,84E-05 | Fli-1 proto-oncogene, ETS transcription factor |
| ENSG00000186895 | FGF3 | 3,444372181 | 10,88577473 | 0,01189 | 0,14209957 | fibroblast growth factor 3 |
| ENSG00000275018 | AL121581,1 | 3,442295042 | 10,87011307 | 0,06061 | 0,32697886 | novel transcript |
| ENSG00000267120 | AD000671,2 | 3,425458904 | 10,74399702 | 0,0021 | 0,05165212 | novel transcript |
| ENSG00000177103 | DSCAML1 | 3,42395198 | 10,73278055 | 0,00012 | 0,00687924 | DS cell adhesion molecule like 1 |
| ENSG00000267601 | AC022966,1 | 3,418233985 | 10,69032631 | 0,06646 | 0,34124715 | novel transcript, antisense to TIMP2 |
| ENSG00000274801 | AC091304,4 | 3,40610783 | 10,60084844 | 0,05542 | 0,31484345 | cholinergic receptor, nicotinic, beta 4 (CHRNB4) pseudogene |
| ENSG00000261189 | AL031058,1 | 3,391363143 | 10,49305701 | 2,79E-05 | 0,00204716 | novel transcript, antisense to DSP |
| ENSG00000147256 | ARHGAP36 | 3,387920717 | 10,46804929 | 3,22E-05 | 0,00229769 | Rho GTPase activating protein 36 |
| ENSG00000270249 | AC093668,1 | 3,380151969 | 10,41183153 | 0,00673 | 0,10439701 | novel protein |
| ENSG00000124205 | EDN3 | 3,377511445 | 10,39279249 | 4,27E-07 | 5,38E-05 | endothelin 3 |
| ENSG00000020181 | ADGRA2 | 3,376202918 | 10,38337048 | 3,53E-12 | 1,13E-09 | adhesion G protein-coupled receptor A2 |
| ENSG00000204849 | SPATA31A1 | 3,346995578 | 10,17527287 | 0,00073 | 0,02639057 | SPATA31 subfamily A member 1 |
| ENSG00000143867 | OSR1 | 3,339287904 | 10,12105589 | 3,07E-08 | 5,13E-06 | odd-skipped related transcription factor 1 |
| ENSG00000102760 | RGCC | 3,328007899 | 10,04223091 | 1,30E-06 | 0,00014394 | regulator of cell cycle |
| ENSG00000133115 | STOML3 | 3,326821948 | 10,03397921 | 0,00873 | 0,11841612 | stomatin like 3 |
| ENSG00000259807 | AC009093,1 | 3,326778276 | 10,03367547 | 3,88E-06 | 0,00036652 | novel transcript |
| ENSG00000229970 | AC007128,1 | 3,319061585 | 9,980150592 | 9,09E-05 | 0,00555669 | novel transcript, antisense to ICA1 |
| ENSG00000019549 | SNAI2 | 3,299075546 | 9,842846163 | 5,21E-09 | 1,01E-06 | snail family transcriptional repressor 2 |
| ENSG00000096696 | DSP | 3,287022803 | 9,760958359 | 3,94E-10 | 9,53E-08 | desmoplakin |
| ENSG00000102359 | SRPX2 | 3,277853851 | 9,699119952 | 1,82E-22 | 2,04E-19 | sushi repeat containing protein X-linked 2 |
| ENSG00000164761 | TNFRSF11B | 3,270590843 | 9,650414052 | 1,40E-08 | 2,44E-06 | TNF receptor superfamily member 11b |
| ENSG00000167157 | PRRX2 | 3,259779459 | 9,578365308 | 3,90E-09 | 7,88E-07 | paired related homeobox 2 |
| ENSG00000105880 | DLX5 | 3,248956535 | 9,506778418 | 5,38E-10 | 1,26E-07 | distal-less homeobox 5 |
| ENSG00000231245 | C1DP1 | 3,235644991 | 9,419464152 | 0,01532 | 0,16232837 | C1D nuclear receptor corepressor pseudogene 1 |
| ENSG00000280944 | AC241851,2 | 3,220345126 | 9,320098006 | 0,00012 | 0,00671805 | novel transcript |
| ENSG00000288508 | ISLR | 3,217820052 | 9,303799794 | 9,69E-08 | 1,38E-05 | immunoglobulin superfamily containing leucine rich repeat |
| ENSG00000129009 | ISLR | 3,217820052 | 9,303799794 | 9,69E-08 | 1,38E-05 | immunoglobulin superfamily containing leucine rich repeat |
| ENSG00000261757 | AC005592,1 | 3,21240858 | 9,268967109 | 0,03972 | 0,26947142 | novel transcript |
| ENSG00000132854 | KANK4 | 3,197989648 | 9,176790344 | 7,95E-09 | 1,51E-06 | KN motif and ankyrin repeat domains 4 |

| | | | | | | |
|---|---|---|---|---|---|---|
| ENSG00000288550 | AC018809,3 | 3,164510882 | 8,96628826 | 0,16169 | 0,48914394 | novel protein |
| ENSG00000183801 | OLFML1 | 3,163500529 | 8,960011158 | 3,68E-08 | 5,89E-06 | olfactomedin like 1 |
| ENSG00000250208 | FZD10-AS1 | 3,159517043 | 8,935305417 | 2,20E-07 | 2,95E-05 | FZD10 antisense divergent transcript |
| ENSG00000275300 | MCCC2 | 3,149524541 | 8,873630877 | 0,00802 | 0,11389729 | methylcrotonoyl-CoA carboxylase 2 |
| ENSG00000288539 | AC022133,2 | 3,139801033 | 8,81402527 | 0,0003 | 0,014483 | novel transcript |
| ENSG00000170624 | SGCD | 3,139460775 | 8,811946739 | 5,12E-06 | 0,00046678 | sarcoglycan delta |
| ENSG00000123572 | NRK | 3,138997271 | 8,809116121 | 4,41E-12 | 1,38E-09 | Nik related kinase |
| ENSG00000232392 | GOT2P7 | 3,13656725 | 8,794290868 | 0,01774 | 0,17544816 | GOT2 pseudogene 7 |
| ENSG00000163132 | MSX1 | 3,1272969 | 8,737962411 | 3,11E-12 | 1,00E-09 | msh homeobox 1 |
| ENSG00000147869 | CER1 | 3,122217073 | 8,707249558 | 0,00043 | 0,01841098 | cerberus 1, DAN family BMP antagonist |
| ENSG00000179772 | FOXS1 | 3,121577499 | 8,703390324 | 0,01906 | 0,18296265 | forkhead box S1 |
| ENSG00000157502 | PWWP3B | 3,121415909 | 8,702415549 | 6,21E-09 | 1,20E-06 | PWWP domain containing 3B |
| ENSG00000179256 | SMCO3 | 3,119039238 | 8,688091146 | 0,00409 | 0,07837165 | single-pass membrane protein with coiled-coil domains 3 |
| ENSG00000116774 | OLFML3 | 3,10519873 | 8,605140393 | 1,64E-14 | 6,75E-12 | olfactomedin like 3 |
| ENSG00000285314 | AC002470,2 | 3,09570295 | 8,548687579 | 0,01146 | 0,1389947 | novel transcript |
| ENSG00000179855 | GIPC3 | 3,091534208 | 8,524021358 | 0,00029 | 0,01402048 | GIPC PDZ domain containing family member 3 |
| ENSG00000277646 | AL353684,1 | 3,091126251 | 8,521611328 | 0,00085 | 0,02895387 | pyruvate dehydrogenase kinase, isozyme 1 (PDK1) pseudogene |
| ENSG00000234109 | RPL7P36 | 3,088993161 | 8,509021046 | 0,01007 | 0,12883758 | ribosomal protein L7 pseudogene 36 |
| ENSG00000050555 | LAMC3 | 3,076660615 | 8,436593649 | 2,64E-06 | 0,00025849 | laminin subunit gamma 3 |
| ENSG00000106852 | LHX6 | 3,058828191 | 8,33295501 | 0,03266 | 0,24385689 | LIM homeobox 6 |
| ENSG00000261040 | WFDC21P | 3,054341944 | 8,307082856 | 0,02202 | 0,1972903 | WAP four-disulfide core domain 21, pseudogene |
| ENSG00000092068 | SLC7A8 | 3,044739352 | 8,251974437 | 5,50E-11 | 1,54E-08 | solute carrier family 7 member 8 |
| ENSG00000185885 | IFITM1 | 3,041106476 | 8,231221136 | 1,55E-08 | 2,66E-06 | interferon induced transmembrane protein 1 |
| ENSG00000164283 | ESM1 | 3,022090841 | 8,123440294 | 0,00608 | 0,09842873 | endothelial cell specific molecule 1 |
| ENSG00000111262 | KCNA1 | 3,014310993 | 8,079751896 | 0,00092 | 0,03058422 | potassium voltage-gated channel subfamily A member 1 |
| ENSG00000284115 | AC113189,9 | 3,006776068 | 8,03766288 | 0,01432 | 0,15653473 | novel transcript |
| ENSG00000272884 | AC113189,4 | 3,006776068 | 8,03766288 | 0,01432 | 0,15653473 | novel transcript |
| ENSG00000263244 | AC087190,3 | 2,973824069 | 7,856158674 | 0,30002 | 0,59160746 | novel transcript, 3' overlapping chromosome 16 open reading frame 72 |
| ENSG00000178394 | HTR1A | 2,9693849 | 7,832022448 | 0,00048 | 0,01988165 | 5-hydroxytryptamine receptor 1A |
| ENSG00000000971 | CFH | 2,968658901 | 7,828082174 | 0,00019 | 0,01030113 | complement factor H |
| ENSG00000249899 | AC025180,1 | 2,964481724 | 7,805449543 | 2,96E-06 | 0,00028624 | novel transcript |

| | | | | | | |
|---|---|---|---|---|---|---|
| ENSG00000237718 | AC009095,1 | 2,957617509 | 7,768400113 | 0,07872 | 0,36645437 | novel transcript |
| ENSG00000177468 | OLIG3 | 2,940820904 | 7,678480824 | 0,00034 | 0,01565603 | oligodendrocyte transcription factor 3 |
| ENSG00000173918 | C1QTNF1 | 2,937304825 | 7,659789918 | 3,55E-08 | 5,73E-06 | C1q and TNF related 1 |
| ENSG00000124216 | SNAI1 | 2,931686147 | 7,63001634 | 1,16E-14 | 4,86E-12 | snail family transcriptional repressor 1 |
| ENSG00000240032 | LNCSRLR | 2,925031032 | 7,59490033 | 0,00754 | 0,11001711 | lncRNA sorafenib resistance in renal cell carcinoma associated |
| ENSG00000214286 | PDCL3P3 | 2,920522356 | 7,571201977 | 0,00024 | 0,01236909 | PDCL3 pseudogene 3 |
| ENSG00000124253 | PCK1 | 2,91604427 | 7,547737613 | 0,00085 | 0,02896138 | phosphoenolpyruvate carboxykinase 1 |
| ENSG00000259856 | RAB43P1 | 2,91500045 | 7,542278642 | 0,01293 | 0,14886653 | RAB43 pseudogene 1 |
| ENSG00000168122 | ZNF355P | 2,903255364 | 7,481125659 | 0,00524 | 0,09023148 | zinc finger protein 355, pseudogene |
| ENSG00000113389 | NPR3 | 2,900447019 | 7,466577093 | 4,84E-09 | 9,49E-07 | natriuretic peptide receptor 3 |
| ENSG00000177409 | SAMD9L | 2,891672103 | 7,421300898 | 3,37E-07 | 4,35E-05 | sterile alpha motif domain containing 9 like |
| ENSG00000261127 | AC133548,2 | 2,891273171 | 7,419249058 | 0,00126 | 0,03785412 | protein phosphatase 2, regulatory subunit B'', beta (PPP2R3B) pseudogene |
| ENSG00000180287 | PLD5 | 2,884127058 | 7,382590095 | 2,87E-07 | 3,76E-05 | phospholipase D family member 5 |
| ENSG00000078098 | FAP | 2,876429834 | 7,343306584 | 0,01773 | 0,1753902 | fibroblast activation protein alpha |
| ENSG00000254035 | AC104117,3 | 2,864683208 | 7,283759098 | 0,00611 | 0,098625 | novel transcript |
| ENSG00000205927 | OLIG2 | 2,863105651 | 7,275798813 | 0,00101 | 0,03265524 | oligodendrocyte transcription factor 2 |
| ENSG00000226496 | LINC00323 | 2,862915666 | 7,274840743 | 0,00764 | 0,11109228 | long intergenic non-protein coding RNA 323 |
| ENSG00000105851 | PIK3CG | 2,861262384 | 7,266508787 | 0,00574 | 0,09484162 | phosphatidylinositol-4,5-bisphosphate 3-kinase catalytic subunit gamma |
| ENSG00000176678 | FOXL1 | 2,856606991 | 7,243098481 | 0,00011 | 0,00665113 | forkhead box L1 |
| ENSG00000163359 | COL6A3 | 2,841291478 | 7,166613139 | 0,00413 | 0,07872305 | collagen type VI alpha 3 chain |
| ENSG00000197057 | DTHD1 | 2,827783272 | 7,099824054 | 0,04262 | 0,2782038 | death domain containing 1 |
| ENSG00000182168 | UNC5C | 2,819474927 | 7,05905434 | 1,78E-08 | 3,02E-06 | unc-5 netrin receptor C |
| ENSG00000185614 | INKA1 | 2,816428657 | 7,044164777 | 5,02E-06 | 0,00046033 | inka box actin regulator 1 |
| ENSG00000107831 | FGF8 | 2,808504201 | 7,005578559 | 2,58E-05 | 0,00190142 | fibroblast growth factor 8 |
| ENSG00000258548 | LINC00645 | 2,800542968 | 6,967026099 | 8,06E-06 | 0,00069151 | long intergenic non-protein coding RNA 645 |
| ENSG00000267340 | AC060780,2 | 2,799706096 | 6,962985874 | 0,029 | 0,22977124 | ribosomal protein, large, P1 (RPLP1) pseudogene |
| ENSG00000064218 | DMRT3 | 2,797520849 | 6,952447034 | 9,55E-05 | 0,00580109 | doublesex and mab-3 related transcription factor 3 |
| ENSG00000184221 | OLIG1 | 2,790195877 | 6,917236952 | 0,00159 | 0,04261325 | oligodendrocyte transcription factor 1 |
| ENSG00000010610 | CD4 | 2,78658925 | 6,899965987 | 1,32E-09 | 2,90E-07 | CD4 molecule |
| ENSG00000243089 | AC107029,1 | 2,773640997 | 6,838315551 | 0,00183 | 0,04744039 | novel transcript |

| | | | | | | |
|---|---|---|---|---|---|---|
| ENSG00000282278 | AC058822,1 | 2,768262067 | 6,812867113 | 0,07308 | 0,35640071 | novel FIP1L1-PDGFRA fusion protein |
| ENSG00000154342 | WNT3A | 2,760282097 | 6,775287174 | 0,0002 | 0,01056153 | Wnt family member 3A |
| ENSG00000184232 | OAF | 2,758771315 | 6,768195844 | 5,64E-15 | 2,51E-12 | out at first homolog |
| ENSG00000113356 | POLR3G | 2,75168528 | 6,735034241 | 5,97E-05 | 0,0039327 | RNA polymerase III subunit G |
| ENSG00000261857 | MIA | 2,750979608 | 6,731740708 | 0,00106 | 0,03368549 | MIA SH3 domain containing |
| ENSG00000282435 | AC246817,4 | 2,750065017 | 6,7274745 | 0,00956 | 0,12500859 | novel transcript |
| ENSG00000196954 | CASP4 | 2,749690456 | 6,725728097 | 0,03163 | 0,24001172 | caspase 4 |
| ENSG00000271454 | AL353729,2 | 2,746368816 | 6,710260694 | 1,47E-06 | 0,00015662 | CAS1 domain containing 1 (CASD1) pseudogene |
| ENSG00000277013 | AC008556,1 | 2,730278281 | 6,635836226 | 5,29E-07 | 6,37E-05 | novel transcript |
| ENSG00000287460 | AL442636,1 | 2,72862056 | 6,628215733 | 3,90E-07 | 4,96E-05 | novel transcript |
| ENSG00000236013 | AL357146,1 | 2,727184377 | 6,621620721 | 0,00117 | 0,03594634 | novel transcript |
| ENSG00000162849 | KIF26B | 2,718522115 | 6,581982154 | 2,08E-12 | 6,79E-10 | kinesin family member 26B |
| ENSG00000118777 | ABCG2 | 2,713570279 | 6,559429207 | 5,50E-06 | 0,00049614 | ATP binding cassette subfamily G member 2 (Junior blood group) |
| ENSG00000073756 | PTGS2 | 2,707230616 | 6,53066822 | 0,00568 | 0,09412963 | prostaglandin-endoperoxide synthase 2 |
| ENSG00000180347 | ITPRID1 | 2,705924243 | 6,524757316 | 0,01092 | 0,13517515 | ITPR interacting domain containing 1 |
| ENSG00000158525 | CPA5 | 2,680278942 | 6,409798223 | 0,01417 | 0,15631436 | carboxypeptidase A5 |
| ENSG00000166073 | GPR176 | 2,679226339 | 6,405123284 | 4,99E-07 | 6,11E-05 | G protein-coupled receptor 176 |
| ENSG00000288520 | AL357075,4 | 2,652124712 | 6,285923488 | 0,17184 | 0,49710979 | CNK3/IPCEF1 fusion protein |
| ENSG00000282046 | GREM1 | 2,651526673 | 6,28331833 | 0,00058 | 0,02257221 | gremlin 1, DAN family BMP antagonist |
| ENSG00000223867 | AC098592,1 | 2,646116474 | 6,259799609 | 0,10473 | 0,41326267 | pseudogene similar to part of serine/threonine kinase |
| ENSG00000126785 | RHOJ | 2,644568634 | 6,253087191 | 1,71E-06 | 0,0001789 | ras homolog family member J |
| ENSG00000131196 | NFATC1 | 2,64230351 | 6,243277146 | 4,91E-10 | 1,16E-07 | nuclear factor of activated T cells 1 |
| ENSG00000113721 | PDGFRB | 2,639465206 | 6,23100643 | 1,95E-20 | 1,76E-17 | platelet derived growth factor receptor beta |
| ENSG00000259656 | AC013452,2 | 2,634331517 | 6,208873408 | 0,02896 | 0,22977124 | novel transcript |
| ENSG00000127863 | TNFRSF19 | 2,632726447 | 6,20196957 | 2,91E-06 | 0,00028253 | TNF receptor superfamily member 19 |
| ENSG00000177301 | KCNA2 | 2,629957938 | 6,190079499 | 0,00055 | 0,02167479 | potassium voltage-gated channel subfamily A member 2 |
| ENSG00000258856 | AL136520,1 | 2,625291353 | 6,170089228 | 0,01179 | 0,14150455 | serine/arginine-rich splicing factor 10 (SRSF10) pseudogene |
| ENSG00000197550 | AL359955,1 | 2,622608374 | 6,158625381 | 9,86E-06 | 0,00082058 | zinc finger pseudogene |
| ENSG00000250420 | AACSP1 | 2,616156948 | 6,131146801 | 1,05E-05 | 0,00087065 | acetoacetyl-CoA synthetase pseudogene 1 |
| ENSG00000253819 | LINC01151 | 2,612027673 | 6,113623352 | 0,05066 | 0,30278705 | long intergenic non-protein coding RNA 1151 |
| ENSG00000162878 | PKDCC | 2,611996686 | 6,113492039 | 2,52E-17 | 1,44E-14 | protein kinase domain containing, cytoplasmic |

| | | | | | | |
|---|---|---|---|---|---|---|
| ENSG00000171992 | SYNPO | 2,606989618 | 6,092311116 | 6,00E-11 | 1,67E-08 | synaptopodin |
| ENSG00000285585 | AC069444,2 | 2,600919455 | 6,066731468 | 0,13696 | 0,46423633 | novel protein |
| ENSG00000189108 | IL1RAPL2 | 2,596518597 | 6,04825345 | 0,00094 | 0,03101374 | interleukin 1 receptor accessory protein like 2 |
| ENSG00000223749 | MIR503HG | 2,590549542 | 6,023280903 | 5,02E-13 | 1,73E-10 | MIR503 host gene |
| ENSG00000169129 | AFAP1L2 | 2,590079472 | 6,021318668 | 1,75E-08 | 2,99E-06 | actin filament associated protein 1 like 2 |
| ENSG00000215612 | HMX1 | 2,577320969 | 5,96830378 | 0,00797 | 0,11367829 | H6 family homeobox 1 |
| ENSG00000288204 | HMX1 | 2,577320969 | 5,96830378 | 0,00797 | 0,11367829 | H6 family homeobox 1 |
| ENSG00000139926 | FRMD6 | 2,576064662 | 5,963108811 | 1,81E-08 | 3,06E-06 | FERM domain containing 6 |
| ENSG00000019991 | HGF | 2,570364829 | 5,939596099 | 0,00135 | 0,03912393 | hepatocyte growth factor |
| ENSG00000187800 | PEAR1 | 2,568471904 | 5,931808011 | 4,87E-10 | 1,16E-07 | platelet endothelial aggregation receptor 1 |
| ENSG00000285304 | Z83844,2 | 2,557598088 | 5,887267121 | 0,22985 | 0,54324957 | novel protein |
| ENSG00000166743 | ACSM1 | 2,556018288 | 5,880823893 | 0,00342 | 0,07051554 | acyl-CoA synthetase medium chain family member 1 |
| ENSG00000167600 | CYP2S1 | 2,54914384 | 5,8528684 | 4,53E-07 | 5,62E-05 | cytochrome P450 family 2 subfamily S member 1 |
| ENSG00000125378 | BMP4 | 2,547865717 | 5,847685479 | 4,79E-07 | 5,92E-05 | bone morphogenetic protein 4 |
| ENSG00000217236 | SP9 | 2,546588371 | 5,842510293 | 5,05E-07 | 6,11E-05 | Sp9 transcription factor |
| ENSG00000050767 | COL23A1 | 2,54302968 | 5,828116352 | 0,00384 | 0,07538386 | collagen type XXIII alpha 1 chain |
| ENSG00000091136 | LAMB1 | 2,529130139 | 5,772235409 | 9,80E-11 | 2,62E-08 | laminin subunit beta 1 |
| ENSG00000145536 | ADAMTS16 | 2,526813613 | 5,762974408 | 1,95E-07 | 2,66E-05 | ADAM metallopeptidase with thrombospondin type 1 motif 16 |
| ENSG00000150556 | LYPD6B | 2,524395724 | 5,753324023 | 0,02941 | 0,23078264 | LY6/PLAUR domain containing 6B |
| ENSG00000106333 | PCOLCE | 2,522982175 | 5,747689691 | 1,09E-13 | 4,03E-11 | procollagen C-endopeptidase enhancer |
| ENSG00000220256 | AC093802,1 | 2,522501183 | 5,745773741 | 0,00595 | 0,09722621 | novel transcript |
| ENSG00000144355 | DLX1 | 2,516356909 | 5,721355178 | 2,26E-09 | 4,65E-07 | distal-less homeobox 1 |
| ENSG00000183246 | RIMBP3C | 2,512660591 | 5,706715302 | 0,08896 | 0,3857873 | RIMS binding protein 3C |
| ENSG00000157613 | CREB3L1 | 2,510757214 | 5,69919128 | 1,47E-06 | 0,00015662 | cAMP responsive element binding protein 3 like 1 |
| ENSG00000205413 | SAMD9 | 2,507582454 | 5,686663566 | 0,00103 | 0,03289327 | sterile alpha motif domain containing 9 |
| ENSG00000181234 | TMEM132C | 2,506870381 | 5,683857484 | 9,58E-08 | 1,38E-05 | transmembrane protein 132C |
| ENSG00000261600 | AC233266,2 | 2,504652771 | 5,675127365 | 0,02797 | 0,22566447 | novel transcript |
| ENSG00000153071 | DAB2 | 2,486046519 | 5,602405929 | 1,45E-06 | 0,00015595 | DAB adaptor protein 2 |
| ENSG00000228582 | TAP2 | 2,482006028 | 5,586737472 | 0,00306 | 0,06600584 | transporter 2, ATP binding cassette subfamily B member |
| ENSG00000259727 | AC103740,2 | 2,475938631 | 5,563291232 | 0,00015 | 0,00831137 | novel transcript |
| ENSG00000173068 | BNC2 | 2,472860585 | 5,551434388 | 9,48E-07 | 0,00010966 | basonuclin 2 |
| ENSG00000183770 | FOXL2 | 2,471996603 | 5,548110812 | 0,00233 | 0,05562927 | forkhead box L2 |

| | | | | | | |
|---|---|---|---|---|---|---|
| ENSG00000159167 | STC1 | 2,466167637 | 5,525739832 | 4,03E-09 | 8,07E-07 | stanniocalcin 1 |
| ENSG00000134817 | APLNR | 2,462464626 | 5,511574926 | 0,0004 | 0,01762658 | apelin receptor |
| ENSG00000243695 | AL353729,1 | 2,462316663 | 5,511009687 | 1,06E-06 | 0,00012011 | TGF beta-inducible nuclear protein 1 (TINP1) pseudogene |
| ENSG00000110799 | VWF | 2,46068732 | 5,504789207 | 1,27E-06 | 0,00014151 | von Willebrand factor |
| ENSG00000163762 | TM4SF18 | 2,449881331 | 5,463711589 | 6,19E-06 | 0,00055276 | transmembrane 4 L six family member 18 |
| ENSG00000285283 | AL035078,4 | 2,44820757 | 5,45737647 | 0,35404 | 0,62877376 | novel protein |
| ENSG00000285479 | CACNA1C | 2,44742463 | 5,454415597 | 2,82E-05 | 0,00204966 | calcium voltage-gated channel subunit alpha1 C |
| ENSG00000151067 | CACNA1C | 2,44742463 | 5,454415597 | 2,82E-05 | 0,00204966 | calcium voltage-gated channel subunit alpha1 C |
| ENSG00000213949 | ITGA1 | 2,444537544 | 5,443511268 | 9,50E-07 | 0,00010966 | integrin subunit alpha 1 |
| ENSG00000115594 | IL1R1 | 2,44265171 | 5,436400372 | 2,12E-06 | 0,00021347 | interleukin 1 receptor type 1 |
| ENSG00000123104 | ITPR2 | 2,441483478 | 5,431999992 | 1,16E-06 | 0,00013103 | inositol 1,4,5-trisphosphate receptor type 2 |
| ENSG00000108821 | COL1A1 | 2,439899845 | 5,426040609 | 2,51E-05 | 0,00186998 | collagen type I alpha 1 chain |
| ENSG00000112769 | LAMA4 | 2,436411301 | 5,412935889 | 3,26E-05 | 0,00231818 | laminin subunit alpha 4 |
| ENSG00000285633 | AL132633,1 | 2,435357702 | 5,408984269 | 0,00181 | 0,04716007 | novel transcript |
| ENSG00000133710 | SPINK5 | 2,417036039 | 5,340726623 | 1,51E-11 | 4,54E-09 | serine peptidase inhibitor Kazal type 5 |
| ENSG00000256630 | AC090424,1 | 2,416987966 | 5,340548666 | 0,00951 | 0,12488553 | ribosomal protein S6 (RPS6) pseudogene |
| ENSG00000244578 | LINC01391 | 2,416969919 | 5,340481859 | 0,03785 | 0,26339938 | long intergenic non-protein coding RNA 1391 |
| ENSG00000236714 | LINC01844 | 2,405996273 | 5,300014376 | 0,06716 | 0,34321767 | long intergenic non-protein coding RNA 1844 |
| ENSG00000103316 | CRYM | 2,404850595 | 5,295807181 | 7,35E-05 | 0,00463465 | crystallin mu |
| ENSG00000112936 | C7 | 2,403908453 | 5,292349921 | 0,00181 | 0,04700241 | complement C7 |
| ENSG00000166923 | GREM1 | 2,402912376 | 5,288697196 | 0,00114 | 0,03540527 | gremlin 1, DAN family BMP antagonist |
| ENSG00000163191 | S100A11 | 2,397134186 | 5,267557604 | 5,30E-14 | 2,12E-11 | S100 calcium binding protein A11 |
| ENSG00000171234 | UGT2B7 | 2,3971115 | 5,267474774 | 0,00417 | 0,07917545 | UDP glucuronosyltransferase family 2 member B7 |
| ENSG00000286235 | AL035461,3 | 2,39382855 | 5,255501906 | 0,23626 | 0,54725688 | novel protein |
| ENSG00000101825 | MXRA5 | 2,392919387 | 5,252191017 | 3,37E-08 | 5,53E-06 | matrix remodeling associated 5 |
| ENSG00000159247 | TUBBP5 | 2,392069614 | 5,249098296 | 3,80E-08 | 6,06E-06 | tubulin beta pseudogene 5 |
| ENSG00000166250 | CLMP | 2,386541449 | 5,22902312 | 2,28E-05 | 0,00173528 | CXADR like membrane protein |
| ENSG00000180787 | ZFP3 | 2,385367419 | 5,224769599 | 7,14E-08 | 1,07E-05 | ZFP3 zinc finger protein |
| ENSG00000169908 | TM4SF1 | 2,384983108 | 5,223377989 | 0,01986 | 0,18711351 | transmembrane 4 L six family member 1 |
| ENSG00000239865 | RPP21 | 2,384466169 | 5,221506712 | 0,1744 | 0,49877327 | ribonuclease P/MRP subunit p21 |
| ENSG00000250155 | AC008957,1 | 2,383925711 | 5,219551011 | 0,00022 | 0,01144572 | novel transcript |
| ENSG00000136541 | ERMN | 2,371772396 | 5,175766008 | 0,04088 | 0,2735803 | ermin |
| ENSG00000211448 | DIO2 | 2,367629862 | 5,160925703 | 0,0041 | 0,07854299 | iodothyronine deiodinase 2 |

| | | | | | | |
|---|---|---|---|---|---|---|
| ENSG00000113580 | NR3C1 | 2,352445617 | 5,106892241 | 8,86E-05 | 0,0054472 | nuclear receptor subfamily 3 group C member 1 |
| ENSG00000255690 | TRIL | 2,348245987 | 5,092047886 | 2,08E-05 | 0,0016108 | TLR4 interactor with leucine rich repeats |
| ENSG00000259757 | AC091891,1 | 2,344613896 | 5,079244406 | 0,01803 | 0,17729131 | novel transcript |
| ENSG00000183239 | AL109615,1 | 2,327193719 | 5,018282607 | 0,0249 | 0,21153891 | ribosomal protein L29 (RPL29) pseudogene |
| ENSG00000225028 | AC096541,1 | 2,321737338 | 4,999338929 | 0,01636 | 0,16750744 | novel transcript |
| ENSG00000253706 | AC011632,1 | 2,30655471 | 4,947002778 | 0,001 | 0,03235839 | novel transcript, antisense to PI15 |
| ENSG00000274441 | AC008957,2 | 2,304553929 | 4,940146854 | 0,00046 | 0,01934795 | novel transcript, antisense SLC1A3 |
| ENSG00000233854 | POU6F2-AS2 | 2,299006559 | 4,921187751 | 0,00534 | 0,09086835 | POU6F2 antisense RNA 2 |
| ENSG00000286783 | AC093151,6 | 2,298733882 | 4,92025771 | 0,05253 | 0,30766476 | SET domain and mariner transposase fusion gene (SETMAR) pseudogene |
| ENSG00000286640 | AC093151,5 | 2,298733882 | 4,92025771 | 0,05253 | 0,30766476 | SET domain and mariner transposase fusion gene (SETMAR) pseudogene |
| ENSG00000227365 | AC008060,2 | 2,297138919 | 4,914821153 | 0,13736 | 0,46438162 | novel transcript, antisense to CNPY1 |
| ENSG00000163520 | FBLN2 | 2,291935717 | 4,897127362 | 8,11E-09 | 1,53E-06 | fibulin 2 |
| ENSG00000160307 | S100B | 2,284250455 | 4,871109637 | 0,00441 | 0,08186623 | S100 calcium binding protein B |
| ENSG00000137809 | ITGA11 | 2,281945543 | 4,863333556 | 7,70E-11 | 2,10E-08 | integrin subunit alpha 11 |
| ENSG00000258930 | AC005230,1 | 2,279693878 | 4,855749099 | 0,03818 | 0,26456066 | tetratricopeptide repeat domain 7B (TTC7B) pseudogene |
| ENSG00000170017 | ALCAM | 2,274866526 | 4,839528598 | 1,02E-12 | 3,42E-10 | activated leukocyte cell adhesion molecule |
| ENSG00000287818 | AL512283,3 | 2,270765855 | 4,825792405 | 0,01267 | 0,14729065 | novel transcript, antisense to PKIB |
| ENSG00000275013 | PRAME | 2,264618817 | 4,805274401 | 0,00351 | 0,07177624 | PRAME nuclear receptor transcriptional regulator |
| ENSG00000007350 | TKTL1 | 2,263416512 | 4,801271476 | 1,60E-06 | 0,00016973 | transketolase like 1 |
| ENSG00000121297 | TSHZ3 | 2,261865935 | 4,796113948 | 1,53E-07 | 2,14E-05 | teashirt zinc finger homeobox 3 |
| ENSG00000171227 | TMEM37 | 2,26170606 | 4,795582486 | 2,57E-06 | 0,00025343 | transmembrane protein 37 |
| ENSG00000113430 | IRX4 | 2,26072979 | 4,792338418 | 0,08358 | 0,37573324 | iroquois homeobox 4 |
| ENSG00000226816 | AC005082,1 | 2,259312039 | 4,787631253 | 0,00082 | 0,02838869 | novel transcript |
| ENSG00000134548 | SPX | 2,257223633 | 4,780705821 | 0,00032 | 0,01491513 | spexin hormone |
| ENSG00000158104 | HPD | 2,256680095 | 4,77890502 | 0,01375 | 0,15390159 | 4-hydroxyphenylpyruvate dioxygenase |
| ENSG00000144668 | ITGA9 | 2,254307859 | 4,771053483 | 7,88E-11 | 2,13E-08 | integrin subunit alpha 9 |
| ENSG00000278527 | SNORD22 | 2,25429259 | 4,77100299 | 0,02485 | 0,21147695 | Small nucleolar RNA SNORD22 |
| ENSG00000287059 | AC090004,2 | 2,252795762 | 4,766055533 | 0,00542 | 0,09156474 | novel transcript |
| ENSG00000258610 | AF111169,1 | 2,252606913 | 4,765431699 | 0,00771 | 0,11160063 | novel transcript |
| ENSG00000249906 | AC006487,1 | 2,250402939 | 4,75815721 | 0,07181 | 0,35363591 | novel transcript |
| ENSG00000104371 | DKK4 | 2,248253382 | 4,751073028 | 0,04078 | 0,27303021 | dickkopf WNT signaling pathway inhibitor 4 |

| | | | | | | |
|---|---|---|---|---|---|---|
| ENSG00000213383 | AC104297,1 | 2,241210632 | 4,727936411 | 0,00028 | 0,01358105 | pyrophosphatase (inorganic) 1 (PPA1) pseudogene |
| ENSG00000234779 | BNC2-AS1 | 2,239069314 | 4,720924185 | 0,01 | 0,12811599 | BNC2 antisense RNA 1 |
| ENSG00000185112 | FAM43A | 2,238303043 | 4,718417383 | 4,33E-06 | 0,00040072 | family with sequence similarity 43 member A |
| ENSG00000140937 | CDH11 | 2,235394404 | 4,708914096 | 1,77E-09 | 3,73E-07 | cadherin 11 |
| ENSG00000075035 | WSCD2 | 2,233451126 | 4,702575562 | 0,01408 | 0,15573084 | WSC domain containing 2 |
| ENSG00000231179 | MICB | 2,230682149 | 4,693558525 | 0,00816 | 0,11479236 | MHC class I polypeptide-related sequence B |
| ENSG00000204516 | MICB | 2,230682149 | 4,693558525 | 0,00816 | 0,11479236 | MHC class I polypeptide-related sequence B |
| ENSG00000271474 | AC106881,1 | 2,229659604 | 4,690233032 | 1,77E-05 | 0,00138668 | novel transcript, antisense to UNC5C |
| ENSG00000050628 | PTGER3 | 2,226463405 | 4,679853621 | 0,0013 | 0,03846063 | prostaglandin E receptor 3 |
| ENSG00000277829 | AL031651,2 | 2,224530364 | 4,673587368 | 0,00031 | 0,01478522 | novel transcript, sense intronic to KIAA1755 |
| ENSG00000225826 | LINC00626 | 2,222107826 | 4,66574618 | 0,00443 | 0,08207522 | long intergenic non-protein coding RNA 626 |
| ENSG00000122121 | XPNPEP2 | 2,220303869 | 4,65991574 | 3,01E-05 | 0,00217789 | X-prolyl aminopeptidase 2 |
| ENSG00000224057 | EGFR-AS1 | 2,215690822 | 4,645039363 | 0,00535 | 0,09094215 | EGFR antisense RNA 1 |
| ENSG00000285897 | AC004687,2 | 2,213401864 | 4,637675456 | 0,02104 | 0,19259837 | novel transcript |
| ENSG00000171522 | PTGER4 | 2,211598766 | 4,631882853 | 3,36E-07 | 4,35E-05 | prostaglandin E receptor 4 |
| ENSG00000196569 | LAMA2 | 2,207547605 | 4,618894535 | 0,00155 | 0,04237695 | laminin subunit alpha 2 |
| ENSG00000213860 | RPL21P75 | 2,205714366 | 4,613029011 | 0,40667 | 0,66608043 | ribosomal protein L21 pseudogene 75 |
| ENSG00000081189 | MEF2C | 2,205414587 | 4,612070568 | 7,34E-08 | 1,09E-05 | myocyte enhancer factor 2C |
| ENSG00000133636 | NTS | 2,203029796 | 4,604453059 | 3,91E-05 | 0,0027316 | neurotensin |
| ENSG00000105889 | STEAP1B | 2,202362849 | 4,602324947 | 0,00021 | 0,01078094 | STEAP family member 1B |
| ENSG00000261312 | AC002550,1 | 2,200508001 | 4,596411621 | 2,33E-05 | 0,00175493 | novel transcript, antisense to C16orf88 |
| ENSG00000244721 | HLA-L | 2,197261561 | 4,586080125 | 0,4486 | 0,6944286 | major histocompatibility complex, class I, L (pseudogene) |
| ENSG00000082497 | SERTAD4 | 2,189672421 | 4,562018891 | 4,32E-07 | 5,38E-05 | SERTA domain containing 4 |
| ENSG00000281230 | SERTAD4 | 2,189672421 | 4,562018891 | 4,32E-07 | 5,38E-05 | SERTA domain containing 4 |
| ENSG00000069702 | TGFBR3 | 2,187065689 | 4,553783454 | 2,79E-06 | 0,0002725 | transforming growth factor beta receptor 3 |
| ENSG00000143369 | ECM1 | 2,185293462 | 4,548192958 | 3,26E-15 | 1,47E-12 | extracellular matrix protein 1 |
| ENSG00000164438 | TLX3 | 2,177752397 | 4,524481268 | 0,08112 | 0,37180968 | T cell leukemia homeobox 3 |
| ENSG00000237181 | PRKAR1B-AS1 | 2,17572035 | 4,518112987 | 0,07497 | 0,3592137 | PRKAR1B antisense RNA 1 |
| ENSG00000167244 | IGF2 | 2,17355343 | 4,511331901 | 2,06E-06 | 0,00020928 | insulin like growth factor 2 |
| ENSG00000154229 | PRKCA | 2,166623837 | 4,489714904 | 1,08E-05 | 0,00088915 | protein kinase C alpha |
| ENSG00000260461 | AL133355,1 | 2,166330321 | 4,488801565 | 0,01078 | 0,13435594 | novel transcript, overlapping to OBFC1 |
| ENSG00000226625 | RBM17P1 | 2,161765651 | 4,474621495 | 1,42E-05 | 0,00114173 | RNA binding motif protein 17 pseudogene 1 |
| ENSG00000078401 | EDN1 | 2,160007926 | 4,469173104 | 0,00649 | 0,1020586 | endothelin 1 |

| | | | | | | |
|---|---|---|---|---|---|---|
| ENSG00000122176 | FMOD | 2,15801376 | 4,462999855 | 0,00285 | 0,06337125 | fibromodulin |
| ENSG00000106714 | CNTNAP3 | 2,157950306 | 4,462803564 | 3,21E-14 | 1,30E-11 | contactin associated protein family member 3 |
| ENSG00000145626 | UGT3A1 | 2,156379017 | 4,457945616 | 0,10484 | 0,41357626 | UDP glycosyltransferase family 3 member A1 |
| ENSG00000182508 | LHFPL1 | 2,153817573 | 4,450037744 | 5,62E-05 | 0,00374527 | LHFPL tetraspan subfamily member 1 |
| ENSG00000180353 | HCLS1 | 2,147209703 | 4,429702171 | 0,00898 | 0,12020467 | hematopoietic cell-specific Lyn substrate 1 |
| ENSG00000181649 | PHLDA2 | 2,137631923 | 4,400391616 | 0,0022 | 0,05331646 | pleckstrin homology like domain family A member 2 |
| ENSG00000243914 | RPL5P14 | 2,136237466 | 4,39614041 | 0,03994 | 0,27022885 | ribosomal protein L5 pseudogene 14 |
| ENSG00000079308 | TNS1 | 2,135863193 | 4,395000084 | 8,33E-05 | 0,00520428 | tensin 1 |
| ENSG00000077238 | IL4R | 2,134519802 | 4,390909507 | 3,17E-08 | 5,23E-06 | interleukin 4 receptor |
| ENSG00000113924 | HGD | 2,133898446 | 4,389018787 | 0,01593 | 0,16534892 | homogentisate 1,2-dioxygenase |
| ENSG00000096433 | ITPR3 | 2,133226567 | 4,386975248 | 8,62E-05 | 0,00533908 | inositol 1,4,5-trisphosphate receptor type 3 |
| ENSG00000065371 | ROPN1 | 2,130407695 | 4,378411937 | 0,0436 | 0,28186685 | rhophilin associated tail protein 1 |
| ENSG00000248187 | AC078850,1 | 2,128485572 | 4,3725824 | 0,02751 | 0,22359982 | novel transcript |
| ENSG00000115085 | ZAP70 | 2,127880838 | 4,370749931 | 0,02139 | 0,19418806 | zeta chain of T cell receptor associated protein kinase 70 |
| ENSG00000152583 | SPARCL1 | 2,122856151 | 4,355553754 | 0,02253 | 0,19993459 | SPARC like 1 |
| ENSG00000286156 | AC026273,1 | 2,120051177 | 4,347093652 | 0,00084 | 0,02886269 | BMS1, ribosome biogenesis factor (BMS1) pseudogene |
| ENSG00000009709 | PAX7 | 2,118007698 | 4,340940651 | 4,11E-05 | 0,00284169 | paired box 7 |
| ENSG00000274049 | INO80B-WBP1 | 2,117800317 | 4,340316705 | 0,04342 | 0,28108864 | INO80B-WBP1 readthrough (NMD candidate) |
| ENSG00000159231 | CBR3 | 2,117412616 | 4,33915047 | 0,00594 | 0,09720154 | carbonyl reductase 3 |
| ENSG00000226440 | LAMA4-AS1 | 2,11526026 | 4,33268172 | 0,02116 | 0,19303449 | LAMA4 antisense RNA 1 |
| ENSG00000231764 | DLX6-AS1 | 2,115179649 | 4,332439638 | 0,00056 | 0,02194222 | DLX6 antisense RNA 1 |
| ENSG00000162998 | FRZB | 2,112594842 | 4,324684365 | 7,22E-11 | 1,98E-08 | frizzled related protein |
| ENSG00000256566 | AL049650,1 | 2,107805295 | 4,310350819 | 0,05431 | 0,3118853 | novel protein |
| ENSG00000169594 | BNC1 | 2,101530466 | 4,291644176 | 0,00022 | 0,01111111 | basonuclin 1 |
| ENSG00000163710 | PCOLCE2 | 2,10038391 | 4,288234825 | 3,98E-05 | 0,00276392 | procollagen C-endopeptidase enhancer 2 |
| ENSG00000185652 | NTF3 | 2,098786182 | 4,2834884 | 0,00014 | 0,00790631 | neurotrophin 3 |
| ENSG00000164756 | SLC30A8 | 2,098143929 | 4,281581921 | 0,00905 | 0,12067214 | solute carrier family 30 member 8 |
| ENSG00000234288 | OR1AA1P | 2,097483024 | 4,279620958 | 0,02154 | 0,19517279 | olfactory receptor family 1 subfamily AA member 1 pseudogene |
| ENSG00000152952 | PLOD2 | 2,095043293 | 4,272389839 | 1,10E-05 | 0,0008999 | procollagen-lysine,2-oxoglutarate 5-dioxygenase 2 |
| ENSG00000264630 | PRKCA-AS1 | 2,085881242 | 4,245343359 | 0,02036 | 0,18949014 | PRKCA antisense RNA 1 |
| ENSG00000234702 | VDAC1P3 | 2,083472955 | 4,23826253 | 0,008 | 0,11389729 | voltage dependent anion channel 1 pseudogene 3 |
| ENSG00000164125 | GASK1B | 2,083114634 | 4,237210005 | 4,43E-08 | 6,91E-06 | golgi associated kinase 1B |

| | | | | | | |
|---|---|---|---|---|---|---|
| ENSG00000286546 | AL079338,1 | 2,081621911 | 4,232828131 | 0,02164 | 0,19544769 | novel transcript, antisense to MACROD2 |
| ENSG00000239263 | RBM43P1 | 2,078321348 | 4,223155436 | 0,01169 | 0,14072598 | RNA binding motif protein 43 pseudogene 1 |
| ENSG00000153064 | BANK1 | 2,07474101 | 4,212687821 | 0,00308 | 0,06617279 | B cell scaffold protein with ankyrin repeats 1 |
| ENSG00000278212 | AC134878,2 | 2,072604453 | 4,206453666 | 0,01339 | 0,15158202 | MAFF interacting protein (pseudogene) |
| ENSG00000197461 | PDGFA | 2,072391333 | 4,20583232 | 0,00019 | 0,00993952 | platelet derived growth factor subunit A |
| ENSG00000276539 | PLPPR1 | 2,06940275 | 4,197128837 | 0,0702 | 0,34975886 | phospholipid phosphatase related 1 |
| ENSG00000251634 | NAIPP4 | 2,067601549 | 4,191892003 | 0,12013 | 0,43868155 | NAIP pseudogene 4 |
| ENSG00000250722 | SELENOP | 2,066724821 | 4,18934536 | 4,26E-06 | 0,0003954 | selenoprotein P |
| ENSG00000180801 | ARSJ | 2,064360697 | 4,182485958 | 6,44E-05 | 0,00418528 | arylsulfatase family member J |
| ENSG00000164093 | PITX2 | 2,062452421 | 4,176957373 | 0,02682 | 0,22097601 | paired like homeodomain 2 |
| ENSG00000116745 | RPE65 | 2,061063778 | 4,172938844 | 0,07382 | 0,35713141 | retinoid isomerohydrolase RPE65 |
| ENSG00000187823 | RTL4 | 2,060492337 | 4,171286302 | 0,00428 | 0,08037599 | retrotransposon Gag like 4 |
| ENSG00000206340 | C4A | 2,056503724 | 4,15976989 | 0,07559 | 0,36093649 | complement C4A (Rodgers blood group) |
| ENSG00000134954 | ETS1 | 2,055960181 | 4,158202972 | 1,57E-06 | 0,00016635 | ETS proto-oncogene 1, transcription factor |
| ENSG00000274630 | AC125257,2 | 2,045566644 | 4,128353884 | 0,06792 | 0,34439007 | novel transcript, antisense to ACLY |
| ENSG00000109099 | PMP22 | 2,045233078 | 4,127399476 | 1,71E-05 | 0,00135257 | peripheral myelin protein 22 |
| ENSG00000261051 | AC107021,2 | 2,039727918 | 4,1116798 | 0,01575 | 0,16461429 | novel transcript, overlapping to PLOD2 |
| ENSG00000104321 | TRPA1 | 2,031987955 | 4,089679977 | 0,0044 | 0,08180866 | transient receptor potential cation channel subfamily A member 1 |
| ENSG00000137573 | SULF1 | 2,029748298 | 4,083336035 | 6,68E-06 | 0,00059512 | sulfatase 1 |
| ENSG00000188580 | NKAIN2 | 2,029225324 | 4,081856102 | 0,00163 | 0,04346965 | sodium/potassium transporting ATPase interacting 2 |
| ENSG00000278139 | P3R3URF-PIK3 | 2,024103845 | 4,067391453 | 0,36405 | 0,63490528 | P3R3URF-PIK3R3 readthrough |
| ENSG00000152207 | CYSLTR2 | 2,020471617 | 4,057163989 | 0,02634 | 0,21861252 | cysteinyl leukotriene receptor 2 |
| ENSG00000228271 | LINC02048 | 2,017396217 | 4,048524525 | 0,05591 | 0,31600461 | long intergenic non-protein coding RNA 2048 |
| ENSG00000230457 | PA2G4P4 | 2,017027269 | 4,047489306 | 0,00253 | 0,05846996 | proliferation-associated 2G4 pseudogene 4 |
| ENSG00000260244 | AC104083,1 | 2,014819882 | 4,041301206 | 0,39187 | 0,65534953 | novel transcript, overlapping GUCY1A3 |
| ENSG00000282304 | AC140479,4 | 2,013628031 | 4,03796395 | 0,03827 | 0,26488494 | novel transcript |
| ENSG00000253831 | ETV3L | 2,013284221 | 4,037001773 | 0,1037 | 0,41186474 | ETS variant transcription factor 3 like |
| ENSG00000283213 | AC130466,1 | 2,01271372 | 4,035405694 | 0,00972 | 0,12625914 | novel transcript |
| ENSG00000282053 | AC019257,1 | 2,011880225 | 4,033074973 | 0,05114 | 0,30402651 | novel transcript |
| ENSG00000166033 | HTRA1 | 2,00182428 | 4,005061176 | 5,65E-05 | 0,00375785 | HtrA serine peptidase 1 |
| ENSG00000188770 | OPTC | 2,001602229 | 4,00444479 | 0,0311 | 0,23762196 | opticin |
| ENSG00000161249 | DMKN | 2,001411994 | 4,003916796 | 0,0125 | 0,14623734 | dermokine |

| | | | | | | |
|---|---|---|---|---|---|---|
| ENSG00000256824 | AP000721,2 | 2,001375579 | 4,003815732 | 0,10308 | 0,41078404 | novel transcript, antisense to MACROD1 |
| ENSG00000145642 | SHISAL2B | 2,000335462 | 4,000930207 | 0,0152 | 0,16197635 | shisa like 2B |
| ENSG00000255337 | AP001830,1 | 1,999764885 | 3,999348177 | 0,05971 | 0,32521812 | novel transcript |
| ENSG00000225953 | SATB2-AS1 | 1,999246517 | 3,997911448 | 0,01201 | 0,14273718 | SATB2 antisense RNA 1 |
| ENSG00000258373 | SLC25A3P2 | 1,994877329 | 3,985822126 | 0,05229 | 0,30699657 | solute carrier family 25 member 3 pseudogene 2 |
| ENSG00000224864 | AC011447,1 | 1,994177506 | 3,98388915 | 0,19608 | 0,51693058 | ribosomal protein S16 (RPS16) pseudogene |
| ENSG00000162733 | DDR2 | 1,993198681 | 3,981187119 | 2,45E-05 | 0,00183951 | discoidin domain receptor tyrosine kinase 2 |
| ENSG00000162618 | ADGRL4 | 1,99308301 | 3,980867933 | 0,00076 | 0,02699099 | adhesion G protein-coupled receptor L4 |
| ENSG00000174807 | CD248 | 1,992683792 | 3,979766512 | 5,07E-13 | 1,73E-10 | CD248 molecule |
| ENSG00000286911 | AC009533,3 | 1,990755649 | 3,974451158 | 0,05917 | 0,32424653 | novel transcript |
| ENSG00000165757 | JCAD | 1,987006809 | 3,964136962 | 0,00018 | 0,00974504 | junctional cadherin 5 associated |
| ENSG00000238072 | CDC26P1 | 1,985644867 | 3,96039648 | 0,14394 | 0,47144689 | cell division cycle 26 pseudogene 1 |
| ENSG00000170577 | SIX2 | 1,981647525 | 3,949438414 | 0,00532 | 0,09086355 | SIX homeobox 2 |
| ENSG00000276600 | RAB7B | 1,977709658 | 3,938673035 | 0,00077 | 0,02718618 | RAB7B, member RAS oncogene family |
| ENSG00000237424 | FOXD2-AS1 | 1,976460644 | 3,935264603 | 0,0037 | 0,07414958 | FOXD2 adjacent opposite strand RNA 1 |
| ENSG00000121068 | TBX2 | 1,974478968 | 3,92986286 | 0,05643 | 0,31712073 | T-box transcription factor 2 |
| ENSG00000137507 | LRRC32 | 1,970926084 | 3,920196806 | 0,00018 | 0,00957024 | leucine rich repeat containing 32 |
| ENSG00000257986 | LINC02306 | 1,963476253 | 3,900005755 | 0,0085 | 0,11637365 | long intergenic non-protein coding RNA 2306 |
| ENSG00000165424 | ZCCHC24 | 1,947844741 | 3,857977531 | 2,70E-13 | 9,60E-11 | zinc finger CCHC-type containing 24 |
| ENSG00000108771 | DHX58 | 1,943179475 | 3,845522076 | 0,04225 | 0,27788591 | DExH-box helicase 58 |
| ENSG00000024422 | EHD2 | 1,940734929 | 3,839011625 | 0,00174 | 0,0457836 | EH domain containing 2 |
| ENSG00000107796 | ACTA2 | 1,939300471 | 3,83519643 | 4,36E-05 | 0,00299152 | actin alpha 2, smooth muscle |
| ENSG00000287988 | FO082814,1 | 1,937321856 | 3,829940173 | 0,18728 | 0,5105374 | methylenetetrahydrofolate dehydrogenase (NADP+ dependent) 1 like (MTHFD1L) pseudogene |
| ENSG00000145808 | ADAMTS19 | 1,933032227 | 3,818569355 | 0,00116 | 0,03561475 | ADAM metallopeptidase with thrombospondin type 1 motif 19 |
| ENSG00000229415 | SFTA3 | 1,931951498 | 3,815709919 | 0,01097 | 0,1353078 | surfactant associated 3 |
| ENSG00000154127 | UBASH3B | 1,931276112 | 3,813924044 | 1,31E-06 | 0,00014394 | ubiquitin associated and SH3 domain containing B |
| ENSG00000234571 | H2BP2 | 1,93058523 | 3,812098058 | 0,00482 | 0,08564971 | H2B histone pseudogene 2 |
| ENSG00000256116 | AP001453,1 | 1,929431557 | 3,809050874 | 0,06523 | 0,33808525 | novel transcript |
| ENSG00000234218 | MICB | 1,927613593 | 3,804254048 | 0,02589 | 0,21707964 | MHC class I polypeptide-related sequence B |
| ENSG00000115232 | ITGA4 | 1,927009407 | 3,802661201 | 8,14E-08 | 1,19E-05 | integrin subunit alpha 4 |
| ENSG00000260272 | AC093525,2 | 1,926398042 | 3,801050102 | 0,11737 | 0,4347145 | novel protein |

| | | | | | | |
|---|---|---|---|---|---|---|
| ENSG00000153233 | PTPRR | 1,924823892 | 3,796904972 | 0,00213 | 0,05214547 | protein tyrosine phosphatase receptor type R |
| ENSG00000147257 | GPC3 | 1,924695008 | 3,796565788 | 8,91E-06 | 0,00075488 | glypican 3 |
| ENSG00000235308 | AL445991,1 | 1,923668046 | 3,79386422 | 0,03604 | 0,25711935 | novel pseudogene |
| ENSG00000276592 | PHBP6 | 1,922302231 | 3,790274227 | 0,16271 | 0,48945635 | prohibitin pseudogene 6 |
| ENSG00000116132 | PRRX1 | 1,919441579 | 3,782766117 | 1,17E-05 | 0,00094692 | paired related homeobox 1 |
| ENSG00000173213 | TUBB8B | 1,918405183 | 3,780049647 | 0,00204 | 0,05084299 | tubulin beta 8B |
| ENSG00000170374 | SP7 | 1,917199541 | 3,776892027 | 0,00775 | 0,11195452 | Sp7 transcription factor |
| ENSG00000272459 | AC139795,3 | 1,907554628 | 3,751726417 | 0,00837 | 0,11551322 | novel transcript |
| ENSG00000240801 | AC132217,1 | 1,905876213 | 3,747364239 | 0,0803 | 0,37007763 | novel transcript, 3' overlapping ncRNA IGF2 |
| ENSG00000153165 | RGPD3 | 1,904686953 | 3,744276441 | 0,0125 | 0,14623734 | RANBP2 like and GRIP domain containing 3 |
| ENSG00000277626 | WNT3 | 1,902423965 | 3,738407832 | 0,00533 | 0,09086355 | Wnt family member 3 |
| ENSG00000249413 | AC116049,1 | 1,898937546 | 3,729384497 | 0,18905 | 0,51243448 | novel transcript |
| ENSG00000269987 | AC004542,2 | 1,897370851 | 3,725336771 | 0,41004 | 0,66821634 | novel transcript |
| ENSG00000102445 | RUBCNL | 1,89712455 | 3,724700824 | 1,46E-09 | 3,15E-07 | rubicon like autophagy enhancer |
| ENSG00000165061 | ZMAT4 | 1,895698617 | 3,721021218 | 0,00442 | 0,081876 | zinc finger matrin-type 4 |
| ENSG00000065717 | TLE2 | 1,895658016 | 3,7209165 | 0,02031 | 0,18924656 | TLE family member 2, transcriptional corepressor |
| ENSG00000259848 | AC097374,1 | 1,893337179 | 3,714935544 | 0,00595 | 0,09722621 | pseudogene similar to part of asparagine synthetase (ASNS) |
| ENSG00000226491 | FTOP1 | 1,886523585 | 3,697431924 | 0,01087 | 0,1349201 | FTO pseudogene 1 |
| ENSG00000176170 | SPHK1 | 1,886442156 | 3,697223238 | 0,0029 | 0,06396346 | sphingosine kinase 1 |
| ENSG00000006377 | DLX6 | 1,881741782 | 3,685197099 | 0,00043 | 0,01859241 | distal-less homeobox 6 |
| ENSG00000274840 | AC132807,2 | 1,880204273 | 3,6812718 | 0,02471 | 0,21047862 | novel transcript |
| ENSG00000240755 | ERLEC1P1 | 1,87745538 | 3,67426423 | 0,59306 | 0,79148977 | endoplasmic reticulum lectin 1 pseudogene 1 |
| ENSG00000166763 | STRCP1 | 1,876634263 | 3,672173599 | 0,08279 | 0,37466043 | stereocilin pseudogene 1 |
| ENSG00000254894 | NAV2-AS1 | 1,876239011 | 3,67116768 | 0,2915 | 0,58655161 | NAV2 antisense RNA 1 |
| ENSG00000084636 | COL16A1 | 1,875277962 | 3,668722952 | 0,00219 | 0,05325085 | collagen type XVI alpha 1 chain |
| ENSG00000152779 | SLC16A12 | 1,874660077 | 3,667152029 | 0,00164 | 0,04367867 | solute carrier family 16 member 12 |
| ENSG00000138316 | ADAMTS14 | 1,871896962 | 3,660135254 | 0,06347 | 0,33385402 | ADAM metallopeptidase with thrombospondin type 1 motif 14 |
| ENSG00000135905 | DOCK10 | 1,8675034 | 3,649005687 | 5,51E-05 | 0,00367877 | dedicator of cytokinesis 10 |
| ENSG00000078596 | ITM2A | 1,865481803 | 3,643896047 | 0,00552 | 0,09228058 | integral membrane protein 2A |
| ENSG00000269113 | TRABD2B | 1,865384264 | 3,643649696 | 1,75E-07 | 2,42E-05 | TraB domain containing 2B |
| ENSG00000258385 | AL391516,1 | 1,859365682 | 3,628480917 | 0,03103 | 0,23730035 | WD repeat-containing protein 75 (WDR75) pseudogene |

| | | | | | | |
|---|---|---|---|---|---|---|
| ENSG00000196196 | HRCT1 | 1,859176902 | 3,628006151 | 0,06403 | 0,33545921 | histidine rich carboxyl terminus 1 |
| ENSG00000287804 | Z98745,2 | 1,854965219 | 3,6174303 | 0,2093 | 0,52826608 | novel transcript |
| ENSG00000171956 | FOXB1 | 1,853108389 | 3,612777458 | 0,00417 | 0,07917545 | forkhead box B1 |
| ENSG00000288029 | AC115282,2 | 1,851924362 | 3,609813649 | 0,12905 | 0,45247986 | novel transcript |
| ENSG00000112319 | EYA4 | 1,850576662 | 3,606443099 | 1,39E-05 | 0,00112108 | EYA transcriptional coactivator and phosphatase 4 |
| ENSG00000137252 | HCRTR2 | 1,847303386 | 3,598269853 | 0,04106 | 0,27450533 | hypocretin receptor 2 |
| ENSG00000239801 | DENND6A-AS: | 1,84678539 | 3,596978136 | 0,2204 | 0,53750545 | DENND6A antisense RNA 1 |
| ENSG00000218109 | KIRREL3-AS3 | 1,846635022 | 3,596603251 | 0,05216 | 0,30680362 | KIRREL3 antisense RNA 3 |
| ENSG00000007312 | CD79B | 1,8428479 | 3,587174431 | 0,03343 | 0,24712306 | CD79b molecule |
| ENSG00000250072 | SH3TC2-DT | 1,842836342 | 3,587145692 | 0,02061 | 0,19045384 | SH3TC2 divergent transcript |
| ENSG00000008516 | MMP25 | 1,839023985 | 3,577679087 | 3,97E-05 | 0,00276043 | matrix metallopeptidase 25 |
| ENSG00000272479 | AC010857,1 | 1,837826537 | 3,574710818 | 0,07777 | 0,36490452 | novel transcript |
| ENSG00000167994 | RAB3IL1 | 1,832777811 | 3,562222943 | 0,00246 | 0,05751537 | RAB3A interacting protein like 1 |
| ENSG00000224292 | AF196972,1 | 1,832286772 | 3,561010704 | 0,04947 | 0,29935562 | novel transcript, antisense to PORCN, EBP |
| ENSG00000064195 | DLX3 | 1,831026156 | 3,557900479 | 0,0136 | 0,1529196 | distal-less homeobox 3 |
| ENSG00000149090 | PAMR1 | 1,830615701 | 3,556888379 | 0,00873 | 0,11840155 | peptidase domain containing associated with muscle regeneration 1 |
| ENSG00000186867 | QRFPR | 1,829470432 | 3,554065899 | 0,01424 | 0,15639135 | pyroglutamylated RFamide peptide receptor |
| ENSG00000232691 | ZNRD1ASP | 1,827501983 | 3,54921995 | 0,094 | 0,39609058 | zinc ribbon domain containing 1 antisense, pseudogene |
| ENSG00000250765 | AC138035,2 | 1,826559411 | 3,546901857 | 0,19024 | 0,51331106 | novel transcript |
| ENSG00000120820 | GLT8D2 | 1,823891554 | 3,540348923 | 2,15E-06 | 0,00021574 | glycosyltransferase 8 domain containing 2 |
| ENSG00000146648 | EGFR | 1,821785708 | 3,535184983 | 7,75E-06 | 0,00067314 | epidermal growth factor receptor |
| ENSG00000273796 | BX322562,1 | 1,819473196 | 3,529522936 | 0,0424 | 0,2781925 | novel transcript |
| ENSG00000131914 | LIN28A | 1,812101534 | 3,511534316 | 0,00141 | 0,04015868 | lin-28 homolog A |
| ENSG00000146910 | CNPY1 | 1,81109509 | 3,509085477 | 0,0645 | 0,33665657 | canopy FGF signaling regulator 1 |
| ENSG00000216636 | RPL7P25 | 1,808977914 | 3,503939618 | 0,0243 | 0,20836485 | ribosomal protein L7 pseudogene 25 |
| ENSG00000254066 | LINC01938 | 1,808922909 | 3,503806027 | 0,04248 | 0,2782038 | long intergenic non-protein coding RNA 1938 |
| ENSG00000160963 | COL26A1 | 1,808503394 | 3,502787319 | 3,29E-07 | 4,29E-05 | collagen type XXVI alpha 1 chain |
| ENSG00000113578 | FGF1 | 1,808125114 | 3,501868996 | 0,01977 | 0,18665735 | fibroblast growth factor 1 |
| ENSG00000276298 | AC234782,1 | 1,80685369 | 3,498784216 | 0,0014 | 0,04004246 | HYDIN, axonemal central pair apparatus protein (HYDIN) pseudogene |
| ENSG00000278565 | CEACAM21 | 1,806191039 | 3,497177541 | 0,00055 | 0,02167479 | CEA cell adhesion molecule 21 |
| ENSG00000189221 | MAOA | 1,804546876 | 3,493194263 | 0,00019 | 0,0101066 | monoamine oxidase A |

| | | | | | | |
|---|---|---|---|---|---|---|
| ENSG00000282540 | WHAMMP2 | 1,803708058 | 3,491163824 | 0,02087 | 0,19166705 | WAS protein homolog associated with actin, golgi membranes and microtubules pseudogene 2 |
| ENSG00000161896 | IP6K3 | 1,801589649 | 3,48604127 | 0,1472 | 0,47471707 | inositol hexakisphosphate kinase 3 |
| ENSG00000204381 | LAYN | 1,797301455 | 3,475694921 | 0,00268 | 0,06058562 | layilin |
| ENSG00000268643 | AC006486,1 | 1,796397292 | 3,473517322 | 0,0922 | 0,39260493 | novel protein |
| ENSG00000287248 | AC009271,2 | 1,794676604 | 3,469376963 | 0,02982 | 0,23250598 | novel transcript |
| ENSG00000146070 | PLA2G7 | 1,791574298 | 3,461924586 | 3,96E-06 | 0,00037236 | phospholipase A2 group VII |
| ENSG00000228158 | TLE1P1 | 1,790675696 | 3,459768952 | 0,04249 | 0,2782038 | TLE family member 1 pseudogene 1 |
| ENSG00000216867 | RPL22P12 | 1,789401237 | 3,456713984 | 0,06395 | 0,33512602 | ribosomal protein L22 pseudogene 12 |
| ENSG00000272855 | AC104458,1 | 1,786410846 | 3,449556393 | 0,07469 | 0,35882064 | novel transcript |
| ENSG00000287542 | AC098582,1 | 1,786152268 | 3,448938176 | 0,07333 | 0,35655334 | HECT and RLD domain containing E3 ubiquitin protein ligase 3 |
| ENSG00000076662 | ICAM3 | 1,783995164 | 3,443785209 | 0,00444 | 0,08217356 | intercellular adhesion molecule 3 |
| ENSG00000235649 | MXRA5Y | 1,782751798 | 3,440818511 | 0,00146 | 0,04079507 | matrix remodeling associated 5 Y-linked (pseudogene) |
| ENSG00000284116 | AL772307,1 | 1,780487095 | 3,435421447 | 7,34E-05 | 0,00463465 | novel transcript |
| ENSG00000166289 | PLEKHF1 | 1,779937391 | 3,434112712 | 0,0007 | 0,02564289 | pleckstrin homology and FYVE domain containing 1 |
| ENSG00000288147 | MG910336,1 | 1,777304421 | 3,427851049 | 0,05014 | 0,3009578 | uncharacterized LOC389831 |
| ENSG00000154269 | ENPP3 | 1,775790928 | 3,424256866 | 0,04735 | 0,29242051 | ectonucleotide pyrophosphatase/phosphodiesterase 3 |
| ENSG00000269145 | AC007192,2 | 1,774836779 | 3,421992931 | 0,17416 | 0,49867554 | novel transcript, antisense to MAST3 |
| ENSG00000166501 | PRKCB | 1,774833696 | 3,421985617 | 0,0003 | 0,014483 | protein kinase C beta |
| ENSG00000027075 | PRKCH | 1,774369373 | 3,420884447 | 0,00399 | 0,07731709 | protein kinase C eta |
| ENSG00000268027 | AC243960,1 | 1,772271696 | 3,4159141 | 0,04546 | 0,2880569 | novel transcript, sense intronic to CEACAM21 |
| ENSG00000227954 | TARID | 1,772257165 | 3,415879695 | 0,00517 | 0,08972908 | TCF21 antisense RNA inducing promoter demethylation |
| ENSG00000247809 | NR2F2-AS1 | 1,770350642 | 3,411368588 | 0,00083 | 0,02863467 | NR2F2 antisense RNA 1 |
| ENSG00000269823 | AL513523,9 | 1,7687184 | 3,407511202 | 0,07398 | 0,35740873 | |
| ENSG00000237260 | PPIAP67 | 1,76803125 | 3,405888605 | 0,089 | 0,38586424 | peptidylprolyl isomerase A pseudogene 67 |
| ENSG00000251076 | AC104126,1 | 1,766074594 | 3,401272497 | 0,03311 | 0,24583794 | novel transcript |
| ENSG00000115844 | DLX2 | 1,761547427 | 3,390616056 | 9,90E-06 | 0,00082202 | distal-less homeobox 2 |
| ENSG00000283563 | AC098650,1 | 1,75714383 | 3,380282518 | 1,74E-06 | 0,00017951 | novel protein |
| ENSG00000140563 | MCTP2 | 1,756887692 | 3,379682434 | 0,04632 | 0,28930497 | multiple C2 and transmembrane domain containing 2 |
| ENSG00000154856 | APCDD1 | 1,756362063 | 3,37845131 | 6,47E-05 | 0,00419052 | APC down-regulated 1 |
| ENSG00000169418 | NPR1 | 1,753490374 | 3,37173318 | 0,00465 | 0,08400694 | natriuretic peptide receptor 1 |
| ENSG00000277585 | MUC4 | 1,753128092 | 3,370886594 | 0,24472 | 0,55280101 | mucin 4, cell surface associated |

| | | | | | | |
|---|---|---|---|---|---|---|
| ENSG00000135046 | ANXA1 | 1,748670679 | 3,360487829 | 9,17E-07 | 0,00010719 | annexin A1 |
| ENSG00000108379 | WNT3 | 1,747483241 | 3,357723053 | 0,0855 | 0,37899774 | Wnt family member 3 |
| ENSG00000158887 | MPZ | 1,746378743 | 3,355153432 | 7,44E-05 | 0,00468038 | myelin protein zero |
| ENSG00000256007 | ARAP1-AS1 | 1,744062381 | 3,349770789 | 0,23058 | 0,54366934 | ARAP1 antisense RNA 1 |
| ENSG00000214415 | GNAT3 | 1,743866007 | 3,349314861 | 0,14347 | 0,47083258 | G protein subunit alpha transducin 3 |
| ENSG00000287134 | AC106845,1 | 1,742616334 | 3,346414916 | 0,11625 | 0,43279558 | novel transcript, antisense to DLC1 |
| ENSG00000249421 | ADAMTS19-A: | 1,741333133 | 3,343439782 | 0,03538 | 0,25487994 | ADAMTS19 antisense RNA 1 |
| ENSG00000237321 | AL354936,1 | 1,73839703 | 3,336642293 | 0,18197 | 0,50521773 | novel transcript |
| ENSG00000165929 | TC2N | 1,730010858 | 3,31730315 | 0,01322 | 0,15023876 | tandem C2 domains, nuclear |
| ENSG00000226747 | FSIP2-AS2 | 1,7265655 | 3,309390418 | 0,02193 | 0,19698513 | FSIP2 antisense RNA 2 |
| ENSG00000156510 | HKDC1 | 1,725121473 | 3,306079629 | 0,05403 | 0,31168784 | hexokinase domain containing 1 |
| ENSG00000277527 | LINC01796 | 1,724404986 | 3,304438134 | 0,11591 | 0,43214457 | long intergenic non-protein coding RNA 1796 |
| ENSG00000233123 | LINC01007 | 1,723483364 | 3,302327869 | 0,08815 | 0,38442787 | long intergenic non-protein coding RNA 1007 |
| ENSG00000185811 | IKZF1 | 1,718446832 | 3,29081935 | 0,00015 | 0,00860562 | IKAROS family zinc finger 1 |
| ENSG00000280114 | AL021920,2 | 1,716673989 | 3,286777941 | 0,00436 | 0,08120816 | phosphodiesterase 4D interacting protein (myomegalin) (PDE4DIP) pseudogene |
| ENSG00000163092 | XIRP2 | 1,716611307 | 3,28663514 | 0,10929 | 0,42113687 | xin actin binding repeat containing 2 |
| ENSG00000065361 | ERBB3 | 1,715890296 | 3,284992998 | 0,00127 | 0,03791245 | erb-b2 receptor tyrosine kinase 3 |
| ENSG00000203952 | CCDC160 | 1,714482583 | 3,281789223 | 8,04E-08 | 1,19E-05 | coiled-coil domain containing 160 |
| ENSG00000109158 | GABRA4 | 1,714020667 | 3,280738643 | 0,09878 | 0,40461924 | gamma-aminobutyric acid type A receptor subunit alpha4 |
| ENSG00000285816 | AP000944,5 | 1,712997941 | 3,278413753 | 0,01303 | 0,14938099 | novel protein, POLA2-CDC42EP2 readthrough |
| ENSG00000273403 | AC107294,3 | 1,712901619 | 3,278194875 | 0,10383 | 0,4122061 | novel transcript |
| ENSG00000072657 | TRHDE | 1,711709174 | 3,275486435 | 1,17E-08 | 2,09E-06 | thyrotropin releasing hormone degrading enzyme |
| ENSG00000186377 | CYP4X1 | 1,710851349 | 3,273539413 | 0,02925 | 0,23044387 | cytochrome P450 family 4 subfamily X member 1 |
| ENSG00000238213 | TARDBPP2 | 1,709729396 | 3,270994641 | 0,08272 | 0,37456114 | TAR DNA binding protein pseudogene 2 |
| ENSG00000172985 | SH3RF3 | 1,703386028 | 3,256644017 | 0,00016 | 0,0087027 | SH3 domain containing ring finger 3 |
| ENSG00000254405 | MSH5-SAPCD: | 1,703065395 | 3,255920323 | 0,03568 | 0,25626339 | MSH5-SAPCD1 readthrough (NMD candidate) |
| ENSG00000143341 | HMCN1 | 1,701334686 | 3,252016744 | 1,74E-06 | 0,00017951 | hemicentin 1 |
| ENSG00000185758 | CLDN24 | 1,701223373 | 3,251765842 | 0,05762 | 0,32039326 | claudin 24 |
| ENSG00000107821 | KAZALD1 | 1,698640734 | 3,245949904 | 2,58E-07 | 3,42E-05 | Kazal type serine peptidase inhibitor domain 1 |
| ENSG00000213303 | AC008481,1 | 1,698637065 | 3,245941648 | 0,08827 | 0,38452862 | ribosomal protein S12 (RPS12) pseudogene |
| ENSG00000145335 | SNCA | 1,697929573 | 3,24435024 | 1,03E-08 | 1,86E-06 | synuclein alpha |
| ENSG00000005187 | ACSM3 | 1,69639461 | 3,240900233 | 0,0049 | 0,0866911 | acyl-CoA synthetase medium chain family member 3 |

| | | | | | | |
|---|---|---|---|---|---|---|
| ENSG00000158815 | FGF17 | 1,696145119 | 3,24033982 | 0,00013 | 0,00711881 | fibroblast growth factor 17 |
| ENSG00000284933 | ARHGEF5 | 1,695229131 | 3,238283134 | 0,00166 | 0,04403455 | Rho guanine nucleotide exchange factor 5 |
| ENSG00000261235 | AC092142,1 | 1,693772193 | 3,235014533 | 0,13048 | 0,45477153 | novel transcript, antisense to HSD17B2 |
| ENSG00000288013 | AP003486,2 | 1,693539227 | 3,234492184 | 0,01547 | 0,16309786 | novel transcript |
| ENSG00000285602 | AC097625,1 | 1,692832701 | 3,232908555 | 0,07914 | 0,36738337 | novel protein |
| ENSG00000170356 | OR2A20P | 1,690473802 | 3,227626862 | 0,00396 | 0,07705722 | olfactory receptor family 2 subfamily A member 20 pseudogene |
| ENSG00000257225 | AC079601,1 | 1,685408178 | 3,216313819 | 0,00788 | 0,11292581 | novel transcript, antisense to PRICKLE1 |
| ENSG00000179930 | ZNF648 | 1,682524762 | 3,209892011 | 0,00303 | 0,06571644 | zinc finger protein 648 |
| ENSG00000088756 | ARHGAP28 | 1,680136013 | 3,204581615 | 1,18E-06 | 0,00013244 | Rho GTPase activating protein 28 |
| ENSG00000154654 | NCAM2 | 1,679627945 | 3,203453268 | 0,00078 | 0,02733095 | neural cell adhesion molecule 2 |
| ENSG00000113083 | LOX | 1,679459437 | 3,203079125 | 0,00045 | 0,01917619 | lysyl oxidase |
| ENSG00000232650 | LINC01780 | 1,67830718 | 3,200521898 | 0,07991 | 0,36897172 | long intergenic non-protein coding RNA 1780 |
| ENSG00000287086 | AC034193,1 | 1,67470094 | 3,192531688 | 0,10836 | 0,41899991 | novel transcript, antisense to VHL |
| ENSG00000134107 | BHLHE40 | 1,673770396 | 3,190473156 | 0,00145 | 0,04070054 | basic helix-loop-helix family member e40 |
| ENSG00000259442 | AC105339,3 | 1,673160496 | 3,189124669 | 0,23482 | 0,5463188 | novel transcript, antisense FSD2 |
| ENSG00000130052 | STARD8 | 1,670316794 | 3,182844762 | 0,00061 | 0,02348863 | StAR related lipid transfer domain containing 8 |
| ENSG00000231389 | HLA-DPA1 | 1,667177775 | 3,17592705 | 0,05897 | 0,3241112 | major histocompatibility complex, class II, DP alpha 1 |
| ENSG00000142178 | SIK1 | 1,666509313 | 3,17445585 | 2,80E-06 | 0,00027319 | salt inducible kinase 1 |
| ENSG00000006283 | CACNA1G | 1,664491567 | 3,170019175 | 0,00621 | 0,09929843 | calcium voltage-gated channel subunit alpha1 G |
| ENSG00000270106 | TSNAX-DISC1 | 1,662964255 | 3,166665003 | 0,02628 | 0,21852536 | TSNAX-DISC1 readthrough (NMD candidate) |
| ENSG00000271811 | Z97200,1 | 1,662722112 | 3,166133553 | 0,00099 | 0,0320617 | novel transcript |
| ENSG00000250669 | AC112206,4 | 1,662710588 | 3,166108262 | 0,05552 | 0,31503413 | mitochondrial ribosomal protein L39 (MRPL39) pseudogene |
| ENSG00000215146 | BX322639,1 | 1,660689949 | 3,161676914 | 0,02382 | 0,20612209 | zinc finger protein pseudogene |
| ENSG00000273483 | AL354760,1 | 1,660232847 | 3,160675331 | 0,00086 | 0,02908139 | novel transcript, antisense to WNT2B |
| ENSG00000225408 | AL136980,1 | 1,658717021 | 3,157356183 | 0,09166 | 0,39158946 | novel transcript |
| ENSG00000171517 | LPAR3 | 1,657155655 | 3,153940962 | 0,04328 | 0,28034368 | lysophosphatidic acid receptor 3 |
| ENSG00000270661 | Z99289,2 | 1,655325357 | 3,149942201 | 0,08877 | 0,38517778 | novel transcript, sense intronic LAMA4 |
| ENSG00000277782 | AC068870,2 | 1,655188165 | 3,149642674 | 0,00339 | 0,07021313 | novel transcript |
| ENSG00000205562 | AL049775,1 | 1,652270511 | 3,143279388 | 0,00046 | 0,01934795 | novel transcript |
| ENSG00000267280 | TBX2-AS1 | 1,650716759 | 3,139895966 | 0,13888 | 0,46622186 | TBX2 antisense RNA 1 |
| ENSG00000136352 | NKX2-1 | 1,646549971 | 3,130840411 | 0,00411 | 0,07854299 | NK2 homeobox 1 |

| | | | | | | |
|---|---|---|---|---|---|---|
| ENSG00000269952 | AL117336,1 | 1,645876705 | 3,129379673 | 0,09058 | 0,38933013 | novel transcript, sense intronic to CREM |
| ENSG00000165092 | ALDH1A1 | 1,644037311 | 3,125392348 | 0,18533 | 0,50876814 | aldehyde dehydrogenase 1 family member A1 |
| ENSG00000251467 | AC020893,1 | 1,643385778 | 3,123981214 | 0,08179 | 0,37300773 | dullard homolog (Xenopus laevis) (DULLARD) pseudogene |
| ENSG00000260492 | CISTR | 1,642683403 | 3,122460677 | 0,02599 | 0,21717729 | chondrogenesis-associated transcript |
| ENSG00000125148 | MT2A | 1,640768031 | 3,11831794 | 0,00039 | 0,01730459 | metallothionein 2A |
| ENSG00000227403 | LINC01806 | 1,639097704 | 3,114709695 | 0,11488 | 0,43031399 | long intergenic non-protein coding RNA 1806 |
| ENSG00000237819 | CDK6-AS1 | 1,637528953 | 3,111324677 | 0,00431 | 0,0807188 | CDK6 antisense RNA 1 |
| ENSG00000145075 | CCDC39 | 1,634130988 | 3,10400523 | 0,24979 | 0,55657567 | coiled-coil domain containing 39 |
| ENSG00000233355 | CHRM3-AS2 | 1,632804328 | 3,10115219 | 0,00476 | 0,08508173 | CHRM3 antisense RNA 2 |
| ENSG00000270137 | AF230666,2 | 1,630015764 | 3,095163807 | 0,02787 | 0,22522567 | novel transcript, intronic to PHF20L1 |
| ENSG00000224509 | MRPS9-AS2 | 1,628255835 | 3,09139035 | 0,10394 | 0,41230119 | MRPS9 antisense RNA 2 |
| ENSG00000264164 | AC090616,2 | 1,624818254 | 3,08403311 | 0,085 | 0,37843844 | bromodomain PHD finger transcription factor (BPTF) pseudogene |
| ENSG00000261456 | TUBB8 | 1,622223241 | 3,078490766 | 0,00032 | 0,014904 | tubulin beta 8 class VIII |
| ENSG00000171291 | ZNF439 | 1,620104663 | 3,073973362 | 0,00045 | 0,01917619 | zinc finger protein 439 |
| ENSG00000248429 | FAM198B-AS1 | 1,620051844 | 3,073860821 | 0,14017 | 0,46710628 | FAM198B antisense RNA 1 |
| ENSG00000249669 | CARMN | 1,619334114 | 3,072331979 | 0,07412 | 0,35756831 | cardiac mesoderm enhancer-associated non-coding RNA |
| ENSG00000148908 | RGS10 | 1,617735546 | 3,068929588 | 0,01903 | 0,18296265 | regulator of G protein signaling 10 |
| ENSG00000170775 | GPR37 | 1,617575713 | 3,068589608 | 5,29E-08 | 8,20E-06 | G protein-coupled receptor 37 |
| ENSG00000278202 | AC243919,2 | 1,613864968 | 3,060707053 | 0,05351 | 0,31052177 | novel transcript |
| ENSG00000123095 | BHLHE41 | 1,613622862 | 3,060193463 | 0,01069 | 0,13371001 | basic helix-loop-helix family member e41 |
| ENSG00000177098 | SCN4B | 1,612560166 | 3,05794014 | 0,01992 | 0,18715502 | sodium voltage-gated channel beta subunit 4 |
| ENSG00000285864 | AP000593,4 | 1,612155593 | 3,057082726 | 0,13038 | 0,45477153 | novel transcript |
| ENSG00000228605 | GPANK1 | 1,610523753 | 3,053626799 | 0,023 | 0,20221357 | G-patch domain and ankyrin repeats 1 |
| ENSG00000206408 | GPANK1 | 1,610523753 | 3,053626799 | 0,023 | 0,20221357 | G-patch domain and ankyrin repeats 1 |
| ENSG00000223772 | GEMIN8P3 | 1,609284451 | 3,051004804 | 0,25887 | 0,56260889 | gem nuclear organelle associated protein 8 pseudogene 3 |
| ENSG00000137491 | SLCO2B1 | 1,605390869 | 3,042781775 | 0,03865 | 0,26598457 | solute carrier organic anion transporter family member 2B1 |
| ENSG00000226107 | AC004383,1 | 1,602885878 | 3,037503093 | 0,04188 | 0,27720117 | ribosomal protein L36a (RPL36A) pseudogene |
| ENSG00000275993 | SIK1B | 1,601412095 | 3,034401722 | 4,14E-08 | 6,53E-06 | salt inducible kinase 1B (putative) |
| ENSG00000196549 | MME | 1,597002642 | 3,025141535 | 8,99E-05 | 0,00550571 | membrane metalloendopeptidase |
| ENSG00000183036 | PCP4 | 1,596465907 | 3,024016282 | 0,00075 | 0,02677782 | Purkinje cell protein 4 |
| ENSG00000141738 | GRB7 | 1,594802892 | 3,020532463 | 0,04873 | 0,29677012 | growth factor receptor bound protein 7 |
| ENSG00000144331 | ZNF385B | 1,590414081 | 3,011357687 | 0,00813 | 0,11477829 | zinc finger protein 385B |

| | | | | | | |
|---|---|---|---|---|---|---|
| ENSG00000181552 | EDDM3B | 1,587651752 | 3,005597356 | 0,05523 | 0,31443527 | epididymal protein 3B |
| ENSG00000079215 | SLC1A3 | 1,587017978 | 3,00427729 | 0,00815 | 0,11479236 | solute carrier family 1 member 3 |
| ENSG00000132031 | MATN3 | 1,586679885 | 3,003573326 | 6,87E-06 | 0,00060953 | matrilin 3 |
| ENSG00000101888 | NXT2 | 1,586548155 | 3,003299087 | 0,006 | 0,09769652 | nuclear transport factor 2 like export factor 2 |
| ENSG00000204767 | INSYN2B | 1,586458253 | 3,003111942 | 0,01419 | 0,15631436 | inhibitory synaptic factor family member 2B |
| ENSG00000265939 | UBE2CP2 | 1,583265034 | 2,996472294 | 0,06241 | 0,33099652 | ubiquitin conjugating enzyme E2 C pseudogene 2 |
| ENSG00000144339 | TMEFF2 | 1,58210965 | 2,994073526 | 2,13E-05 | 0,00164156 | transmembrane protein with EGF like and two follistatin like domains 2 |
| ENSG00000169174 | PCSK9 | 1,581937668 | 2,993716626 | 0,00437 | 0,08120816 | proprotein convertase subtilisin/kexin type 9 |
| ENSG00000285446 | Z84488,1 | 1,581551757 | 2,992915935 | 0,29187 | 0,58674733 | novel protein |
| ENSG00000254835 | RNF185-AS1 | 1,579570031 | 2,988807604 | 0,20577 | 0,52520068 | RNF185 antisense RNA 1 |
| ENSG00000273622 | CDC42EP5 | 1,578338442 | 2,986257231 | 0,05668 | 0,31746133 | CDC42 effector protein 5 |
| ENSG00000138080 | EMILIN1 | 1,577331045 | 2,984172732 | 0,00121 | 0,03678152 | elastin microfibril interfacer 1 |
| ENSG00000189120 | SP6 | 1,575830982 | 2,981071508 | 0,04094 | 0,2738393 | Sp6 transcription factor |
| ENSG00000246283 | AC090510,1 | 1,575740574 | 2,980884702 | 0,42673 | 0,68012899 | novel transcript |
| ENSG00000254122 | PCDHGB7 | 1,572255622 | 2,973692812 | 6,41E-16 | 3,10E-13 | protocadherin gamma subfamily B, 7 |
| ENSG00000226247 | SUPT4H1P1 | 1,569621194 | 2,968267666 | 0,1427 | 0,47017263 | SPT4 homolog, DSIF elongation factor subunit pseudogene |
| ENSG00000258964 | AL137129,1 | 1,567910774 | 2,964750653 | 0,24683 | 0,55453071 | novel transcript |
| ENSG00000111110 | PPM1H | 1,567708362 | 2,964334725 | 4,09E-06 | 0,00038088 | protein phosphatase, Mg2+/Mn2+ dependent 1H |
| ENSG00000267026 | AL136084,1 | 1,567205096 | 2,963300833 | 0,14078 | 0,46780719 | novel transcript, antisense to GALNT12 |
| ENSG00000170891 | CYTL1 | 1,564508501 | 2,95776719 | 0,05679 | 0,3177005 | cytokine like 1 |
| ENSG00000139567 | ACVRL1 | 1,56204657 | 2,952724123 | 0,00402 | 0,07768614 | activin A receptor like type 1 |
| ENSG00000071205 | ARHGAP10 | 1,561783939 | 2,952186653 | 5,91E-05 | 0,00390694 | Rho GTPase activating protein 10 |
| ENSG00000256500 | AL139300,1 | 1,559824561 | 2,948179899 | 0,31976 | 0,60470232 | novel protein |
| ENSG00000125398 | SOX9 | 1,55957489 | 2,947669734 | 9,67E-06 | 0,00080972 | SRY-box transcription factor 9 |
| ENSG00000139780 | METTL21C | 1,559120293 | 2,946741064 | 0,11899 | 0,43696075 | methyltransferase like 21C |
| ENSG00000261586 | AC068987,3 | 1,5584384 | 2,945348607 | 0,13017 | 0,45457323 | novel transcript |
| ENSG00000276386 | CNTNAP3P2 | 1,555114927 | 2,938571347 | 1,53E-10 | 3,93E-08 | CNTNAP3 pseudogene 2 |
| ENSG00000277575 | AL162233,1 | 1,554719911 | 2,937766865 | 0,02715 | 0,22224238 | CAS1 domain containing 1 (CASD1) pseudogene |
| ENSG00000162267 | ITIH3 | 1,553704637 | 2,935700185 | 0,02074 | 0,19107121 | inter-alpha-trypsin inhibitor heavy chain 3 |
| ENSG00000120051 | CFAP58 | 1,549264464 | 2,92667889 | 0,02995 | 0,23316056 | cilia and flagella associated protein 58 |
| ENSG00000228960 | OR2A9P | 1,547556306 | 2,923215738 | 7,16E-06 | 0,00062946 | olfactory receptor family 2 subfamily A member 9 pseudogene |

| | | | | | | |
|---|---|---|---|---|---|---|
| ENSG00000110195 | FOLR1 | 1,546426298 | 2,92092699 | 0,0599 | 0,32541248 | folate receptor alpha |
| ENSG00000254783 | AP003084,1 | 1,543033699 | 2,914066295 | 0,04002 | 0,27049438 | immunoglobin superfamily, member 21 (IGSF21) pseudogene |
| ENSG00000258325 | ITFG2-AS1 | 1,53904617 | 2,906023098 | 0,0956 | 0,39857837 | ITFG2 antisense RNA 1 |
| ENSG00000116147 | TNR | 1,53856863 | 2,905061347 | 0,14104 | 0,4679289 | tenascin R |
| ENSG00000135333 | EPHA7 | 1,537856613 | 2,90362796 | 0,02424 | 0,20804286 | EPH receptor A7 |
| ENSG00000174145 | NWD2 | 1,53611876 | 2,90013239 | 0,0206 | 0,19042717 | NACHT and WD repeat domain containing 2 |
| ENSG00000215720 | MFFP1 | 1,534798996 | 2,897480589 | 0,08147 | 0,37244139 | MFF pseudogene 1 |
| ENSG00000166426 | CRABP1 | 1,533079667 | 2,894029579 | 4,91E-07 | 6,04E-05 | cellular retinoic acid binding protein 1 |
| ENSG00000274829 | PARN | 1,531199193 | 2,890259826 | 0,04912 | 0,29843204 | poly(A)-specific ribonuclease |
| ENSG00000187474 | FPR3 | 1,529994421 | 2,887847224 | 0,01559 | 0,16367661 | formyl peptide receptor 3 |
| ENSG00000124749 | COL21A1 | 1,529404867 | 2,886667353 | 0,32876 | 0,60992716 | collagen type XXI alpha 1 chain |
| ENSG00000184838 | PRR16 | 1,528770153 | 2,885397641 | 0,00382 | 0,0752235 | proline rich 16 |
| ENSG00000150361 | KLHL1 | 1,528370124 | 2,884597693 | 0,02062 | 0,19045384 | kelch like family member 1 |
| ENSG00000118733 | OLFM3 | 1,526197222 | 2,880256352 | 0,0143 | 0,15653473 | olfactomedin 3 |
| ENSG00000064489 | BORCS8-MEF2 | 1,52385638 | 2,875586787 | 0,11064 | 0,42326288 | BORCS8-MEF2B readthrough |
| ENSG00000229685 | HLA-DPA1 | 1,522361298 | 2,872608326 | 0,05105 | 0,30386013 | major histocompatibility complex, class II, DP alpha 1 |
| ENSG00000234073 | AC011816,1 | 1,517475181 | 2,862895838 | 0,17943 | 0,5038121 | chromobox homolog 3 (CBX3) pseudogene |
| ENSG00000282397 | AC104021,1 | 1,515335938 | 2,858653853 | 0,10791 | 0,41847368 | novel transcript |
| ENSG00000282033 | AC074387,1 | 1,513764661 | 2,855542111 | 0,01706 | 0,17158055 | uncharacterized LOC105373553 |
| ENSG00000285665 | TOP1P1 | 1,51253908 | 2,853117336 | 0,18439 | 0,50822347 | DNA topoisomerase I pseudogene 1 |
| ENSG00000269397 | AC011503,2 | 1,512165052 | 2,852377743 | 2,53E-05 | 0,00187986 | novel transcript, sense intronic to ZNF726 |
| ENSG00000162511 | LAPTM5 | 1,510083692 | 2,848265617 | 0,06493 | 0,33778346 | lysosomal protein transmembrane 5 |
| ENSG00000269983 | AC146944,2 | 1,508506504 | 2,845153528 | 0,1143 | 0,42980219 | novel transcript |
| ENSG00000176134 | AL445665,1 | 1,507433336 | 2,843037909 | 5,08E-06 | 0,00046411 | methylenetetrahydrofolate dehydrogenase (NADP+ dependent) 1-like (MTHFD1L) pseudogene |
| ENSG00000128918 | ALDH1A2 | 1,506849887 | 2,841888372 | 0,05266 | 0,30807312 | aldehyde dehydrogenase 1 family member A2 |
| ENSG00000277734 | TRAC | 1,506538682 | 2,841275412 | 0,15413 | 0,48239691 | T cell receptor alpha constant |
| ENSG00000163251 | FZD5 | 1,505262854 | 2,838763878 | 0,0944 | 0,39663526 | frizzled class receptor 5 |
| ENSG00000234303 | CLYBL-AS1 | 1,504583158 | 2,837426768 | 0,15186 | 0,48031011 | CLYBL antisense RNA 1 |
| ENSG00000205359 | SLCO6A1 | 1,504494636 | 2,837252672 | 0,01101 | 0,13537791 | solute carrier organic anion transporter family member 6A1 |
| ENSG00000226245 | ZNF32-AS1 | 1,504035931 | 2,836350711 | 0,1574 | 0,48457045 | ZNF32 antisense RNA 1 |

| | | | | | | |
|---|---|---|---|---|---|---|
| ENSG00000230916 | MTCO1P53 | 1,503883273 | 2,836050601 | 0,09558 | 0,3985545 | MT-CO1 pseudogene 53 |
| ENSG00000163492 | CCDC141 | 1,502732181 | 2,833788686 | 0,00379 | 0,0751395 | coiled-coil domain containing 141 |
| ENSG00000145708 | CRHBP | 1,502662629 | 2,833652073 | 0,08258 | 0,37428581 | corticotropin releasing hormone binding protein |
| ENSG00000263677 | AC012588,1 | 1,50236204 | 2,833061735 | 0,0161 | 0,16621463 | novel transcript |
| ENSG00000223872 | AC006372,1 | 1,501934344 | 2,832221981 | 0,19637 | 0,51741432 | novel transcript |
| ENSG00000172238 | ATOH1 | 1,501370794 | 2,831115867 | 0,28101 | 0,57839016 | atonal bHLH transcription factor 1 |
| ENSG00000277481 | PKD1L3 | 1,501321169 | 2,831018484 | 0,07291 | 0,35631792 | polycystin 1 like 3, transient receptor potential channel interacting |
| ENSG00000274654 | AC022467,1 | 1,500319928 | 2,829054419 | 0,08307 | 0,37504927 | novel transcript, sense intronic to FBN1 |
| ENSG00000007129 | CEACAM21 | 1,499768123 | 2,827972563 | 0,00046 | 0,01932628 | CEA cell adhesion molecule 21 |
| ENSG00000197085 | NPSR1-AS1 | 1,495452164 | 2,819525056 | 0,10632 | 0,41594715 | NPSR1 antisense RNA 1 |
| ENSG00000197442 | MAP3K5 | 1,494983692 | 2,818609648 | 0,0556 | 0,31505767 | mitogen-activated protein kinase kinase kinase 5 |
| ENSG00000151655 | ITIH2 | 1,494384533 | 2,817439309 | 0,12289 | 0,44269063 | inter-alpha-trypsin inhibitor heavy chain 2 |
| ENSG00000185686 | PRAME | 1,493635475 | 2,815976853 | 0,06335 | 0,33340778 | PRAME nuclear receptor transcriptional regulator |
| ENSG00000249302 | FTH1P24 | 1,49260493 | 2,813966064 | 0,09282 | 0,39343521 | ferritin heavy chain 1 pseudogene 24 |
| ENSG00000242288 | BMS1P4-AGA | 1,491085813 | 2,811004596 | 0,05867 | 0,32370576 | BMS1P4-AGAP5 readthough |
| ENSG00000239926 | PRDX3P4 | 1,490821346 | 2,810489346 | 0,1235 | 0,44415944 | peroxiredoxin 3 pseudogene 4 |
| ENSG00000166387 | PPFIBP2 | 1,490763006 | 2,810375697 | 0,0051 | 0,08908575 | PPFIA binding protein 2 |
| ENSG00000148848 | ADAM12 | 1,490577979 | 2,810015287 | 7,22E-08 | 1,08E-05 | ADAM metallopeptidase domain 12 |
| ENSG00000266869 | AC005993,1 | 1,487738852 | 2,804490804 | 0,07037 | 0,35012629 | novel transcript |
| ENSG00000181019 | NQO1 | 1,486588284 | 2,802255077 | 3,15E-08 | 5,22E-06 | NAD(P)H quinone dehydrogenase 1 |
| ENSG00000117228 | GBP1 | 1,486447578 | 2,801981788 | 0,0852 | 0,37857645 | guanylate binding protein 1 |
| ENSG00000251363 | LINC02315 | 1,486137707 | 2,801380024 | 4,36E-05 | 0,00299152 | long intergenic non-protein coding RNA 2315 |
| ENSG00000237928 | NFIA-AS2 | 1,485472338 | 2,80008833 | 0,00531 | 0,09082808 | NFIA antisense RNA 2 |
| ENSG00000266441 | AP005205,2 | 1,484420846 | 2,798048261 | 0,00976 | 0,12641099 | novel transcript, antisense to ARHGAP28, |
| ENSG00000137747 | TMPRSS13 | 1,483068774 | 2,795427201 | 0,22372 | 0,53967355 | transmembrane serine protease 13 |
| ENSG00000213014 | VN2R17P | 1,48179384 | 2,792957926 | 0,09059 | 0,38933013 | vomeronasal 2 receptor 17, pseudogene |
| ENSG00000264272 | AC100786,1 | 1,481608614 | 2,792599362 | 0,30708 | 0,59617071 | novel transcript, antisense to TTYH2 |
| ENSG00000215089 | KRT18P11 | 1,478734004 | 2,787040572 | 0,12275 | 0,44256265 | keratin 18 pseudogene 11 |
| ENSG00000115414 | FN1 | 1,478401574 | 2,786398447 | 3,15E-05 | 0,00225882 | fibronectin 1 |
| ENSG00000279413 | AC112497,2 | 1,477750387 | 2,785141038 | 0,0015 | 0,04149555 | |
| ENSG00000077009 | NMRK2 | 1,477732957 | 2,78510739 | 0,08776 | 0,38337966 | nicotinamide riboside kinase 2 |
| ENSG00000283128 | AC009403,2 | 1,476700423 | 2,783114808 | 0,11174 | 0,42501069 | novel transcript |

| | | | | | | |
|---|---|---|---|---|---|---|
| ENSG00000279263 | OR2L8 | 1,474705003 | 2,779268086 | 0,21726 | 0,53520949 | olfactory receptor family 2 subfamily L member 8 |
| ENSG00000204136 | GGTA1 | 1,474098465 | 2,778099872 | 0,07305 | 0,35640071 | glycoprotein alpha-galactosyltransferase 1 (inactive) |
| ENSG00000124942 | AHNAK | 1,47336643 | 2,7766906 | 0,00264 | 0,059998 | AHNAK nucleoprotein |
| ENSG00000213694 | S1PR3 | 1,472995217 | 2,775976235 | 0,00023 | 0,01156464 | sphingosine-1-phosphate receptor 3 |
| ENSG00000287672 | AC003086,1 | 1,472862349 | 2,775720587 | 0,15153 | 0,47966555 | novel transcript |
| ENSG00000233817 | AL162727,1 | 1,471304037 | 2,77272404 | 0,15256 | 0,48061452 | novel transcript |
| ENSG00000204161 | TMEM273 | 1,471192218 | 2,772509143 | 0,03216 | 0,24193883 | transmembrane protein 273 |
| ENSG00000286606 | AC097468,3 | 1,469978425 | 2,770177508 | 0,03293 | 0,24506146 | novel transcript |
| ENSG00000243978 | RTL9 | 1,469677978 | 2,769600668 | 8,70E-05 | 0,00536441 | retrotransposon Gag like 9 |
| ENSG00000277170 | AC012676,3 | 1,469320069 | 2,768913662 | 0,27623 | 0,57538841 | novel transcript, sense intronic with DNAJA3 |
| ENSG00000242610 | OR5BH1P | 1,466844679 | 2,764166807 | 0,07486 | 0,35906132 | olfactory receptor family 5 subfamily BH member 1 pseudogene |
| ENSG00000162624 | LHX8 | 1,466721311 | 2,763930445 | 0,13918 | 0,46648276 | LIM homeobox 8 |
| ENSG00000205578 | POM121B | 1,466145929 | 2,762828343 | 0,10716 | 0,41721821 | POM121 transmembrane nucleoporin B (pseudogene) |
| ENSG00000111145 | ELK3 | 1,46545198 | 2,761499717 | 0,00186 | 0,04794759 | ETS transcription factor ELK3 |
| ENSG00000166147 | FBN1 | 1,463977942 | 2,758679664 | 5,85E-10 | 1,36E-07 | fibrillin 1 |
| ENSG00000141753 | IGFBP4 | 1,461784073 | 2,754487801 | 6,56E-13 | 2,22E-10 | insulin like growth factor binding protein 4 |
| ENSG00000148841 | ITPRIP | 1,461096044 | 2,753174484 | 0,00133 | 0,03905237 | inositol 1,4,5-trisphosphate receptor interacting protein |
| ENSG00000229502 | AL391863,1 | 1,461077229 | 2,753138577 | 0,00802 | 0,11389729 | novel transcript |
| ENSG00000166106 | ADAMTS15 | 1,460752256 | 2,752518492 | 1,58E-05 | 0,00125586 | ADAM metallopeptidase with thrombospondin type 1 motif 15 |
| ENSG00000172752 | COL6A5 | 1,458988668 | 2,749155797 | 0,11258 | 0,42638197 | collagen type VI alpha 5 chain |
| ENSG00000150893 | FREM2 | 1,458839189 | 2,748870969 | 0,00029 | 0,01407223 | FRAS1 related extracellular matrix 2 |
| ENSG00000235621 | LINC00494 | 1,456106628 | 2,743669351 | 0,17433 | 0,49875953 | long intergenic non-protein coding RNA 494 |
| ENSG00000259203 | AC016044,1 | 1,45552459 | 2,742562675 | 0,02945 | 0,23083495 | novel transcript |
| ENSG00000100599 | RIN3 | 1,453557483 | 2,738825752 | 1,36E-06 | 0,00014891 | Ras and Rab interactor 3 |
| ENSG00000253223 | AC110998,1 | 1,452461208 | 2,736745364 | 0,02498 | 0,21208552 | PRP3 pre-mRNA processing factor 3 homolog (S, cerevisiae) (PRPF3) pseudogene |
| ENSG00000238289 | MICB | 1,452394283 | 2,736618412 | 0,04607 | 0,28930497 | MHC class I polypeptide-related sequence B |
| ENSG00000187678 | SPRY4 | 1,451824523 | 2,73553786 | 0,0033 | 0,06920094 | sprouty RTK signaling antagonist 4 |
| ENSG00000267303 | AC011511,4 | 1,450624025 | 2,733262507 | 0,17985 | 0,50407792 | novel transcript |
| ENSG00000271871 | AC005740,4 | 1,44904585 | 2,730274205 | 0,09575 | 0,39858189 | novel transcript, antisense to PCDH12 |
| ENSG00000255513 | AC005363,1 | 1,448894051 | 2,729986943 | 0,2654 | 0,56822387 | ribosomal protein L26 (RPL26) pseudogene |

| | | | | | | |
|---|---|---|---|---|---|---|
| ENSG00000254718 | AL157756,1 | 1,448683907 | 2,72958932 | 0,06765 | 0,34407703 | novel transcript, antisense to PPM1A |
| ENSG00000122870 | BICC1 | 1,448407147 | 2,729065739 | 0,01023 | 0,1300932 | BicC family RNA binding protein 1 |
| ENSG00000151812 | SLC35F4 | 1,447917898 | 2,728140411 | 0,00247 | 0,05755506 | solute carrier family 35 member F4 |
| ENSG00000136630 | HLX | 1,446662898 | 2,725768233 | 0,06534 | 0,3384921 | H2,0 like homeobox |
| ENSG00000171488 | LRRC8C | 1,4460102 | 2,724535333 | 0,00058 | 0,02257221 | leucine rich repeat containing 8 VRAC subunit C |
| ENSG00000185070 | FLRT2 | 1,444844271 | 2,72233436 | 2,16E-05 | 0,00166063 | fibronectin leucine rich transmembrane protein 2 |
| ENSG00000233077 | LINC01271 | 1,442495393 | 2,717905683 | 0,10553 | 0,41477267 | long intergenic non-protein coding RNA 1271 |
| ENSG00000154864 | PIEZO2 | 1,439794877 | 2,712822917 | 2,55E-05 | 0,00188567 | piezo type mechanosensitive ion channel component 2 |
| ENSG00000283809 | AC007326,4 | 1,439460632 | 2,712194481 | 0,44198 | 0,68994635 | novel protein |
| ENSG00000213809 | KLRK1 | 1,438443316 | 2,710282651 | 0,11799 | 0,43562845 | killer cell lectin like receptor K1 |
| ENSG00000231617 | DAXX | 1,438292729 | 2,709999771 | 0,27019 | 0,57192517 | death domain associated protein |
| ENSG00000154529 | CNTNAP3B | 1,437669832 | 2,708829954 | 5,64E-08 | 8,67E-06 | contactin associated protein family member 3B |
| ENSG00000075651 | PLD1 | 1,436483459 | 2,706603315 | 0,00691 | 0,10601066 | phospholipase D1 |
| ENSG00000100146 | SOX10 | 1,435211373 | 2,704217839 | 0,01578 | 0,16470677 | SRY-box transcription factor 10 |
| ENSG00000111817 | DSE | 1,434853876 | 2,703547822 | 0,00034 | 0,01573727 | dermatan sulfate epimerase |
| ENSG00000094963 | FMO2 | 1,434555192 | 2,702988159 | 0,15624 | 0,4840074 | flavin containing dimethylaniline monoxygenase 2 |
| ENSG00000152580 | IGSF10 | 1,434093294 | 2,7021229 | 0,01253 | 0,1464509 | immunoglobulin superfamily member 10 |
| ENSG00000176928 | GCNT4 | 1,432376916 | 2,69891009 | 0,00376 | 0,07477699 | glucosaminyl (N-acetyl) transferase 4 |
| ENSG00000285508 | AL034430,1 | 1,431705029 | 2,697653454 | 0,14703 | 0,47471707 | novel protein |
| ENSG00000178033 | CALHM5 | 1,431643871 | 2,6975391 | 0,003 | 0,0654025 | calcium homeostasis modulator family member 5 |
| ENSG00000084453 | SLCO1A2 | 1,429823818 | 2,694138124 | 1,42E-06 | 0,00015339 | solute carrier organic anion transporter family member 1A2 |
| ENSG00000092969 | TGFB2 | 1,429743955 | 2,693988991 | 0,02219 | 0,19806119 | transforming growth factor beta 2 |
| ENSG00000182022 | CHST15 | 1,429730269 | 2,693963435 | 0,0077 | 0,11157538 | carbohydrate sulfotransferase 15 |
| ENSG00000139211 | AMIGO2 | 1,427980192 | 2,690697473 | 0,00297 | 0,06492836 | adhesion molecule with Ig like domain 2 |
| ENSG00000264982 | AC015563,2 | 1,42711272 | 2,689080082 | 0,14863 | 0,47609532 | novel transcript, antisense to FAM59A |
| ENSG00000278494 | RRN3 | 1,426967276 | 2,688808999 | 0,10887 | 0,42013883 | RRN3 homolog, RNA polymerase I transcription factor |
| ENSG00000254432 | AC113143,1 | 1,426914827 | 2,688711249 | 0,06163 | 0,32867767 | novel transcript |
| ENSG00000283228 | AC068547,1 | 1,426169159 | 2,687321928 | 0,16481 | 0,49092529 | novel transcript |
| ENSG00000099260 | PALMD | 1,426048429 | 2,687097052 | 0,00202 | 0,05060501 | palmdelphin |
| ENSG00000236936 | AL031005,1 | 1,425420659 | 2,68592805 | 0,17222 | 0,49710979 | novel transcript |
| ENSG00000285160 | THSD1P1 | 1,424003534 | 2,683291023 | 0,1997 | 0,52077515 | thrombospondin type 1 domain containing 1 pseudogene 1 |
| ENSG00000178718 | RPP25 | 1,423436636 | 2,682236848 | 1,41E-05 | 0,00113396 | ribonuclease P and MRP subunit p25 |

| | | | | | | |
|---|---|---|---|---|---|---|
| ENSG00000269994 | AL513318,1 | 1,420823625 | 2,677383174 | 1,87E-07 | 2,57E-05 | novel transcript |
| ENSG00000228639 | ROCR | 1,420312505 | 2,676434795 | 0,08173 | 0,37295776 | regulator of chondrogenesis RNA |
| ENSG00000244021 | AC093591,1 | 1,419916856 | 2,675700902 | 0,05525 | 0,31443527 | ribosomal protein L21 (RPL21) pseudogene |
| ENSG00000213145 | CRIP1 | 1,418182382 | 2,672485985 | 0,0028 | 0,06237064 | cysteine rich protein 1 |
| ENSG00000283378 | CNTNAP3C | 1,417866292 | 2,671900516 | 0,0019 | 0,0485619 | contactin associated protein family member 3C |
| ENSG00000136237 | RAPGEF5 | 1,417374631 | 2,670990103 | 0,00625 | 0,09968224 | Rap guanine nucleotide exchange factor 5 |
| ENSG00000095637 | SORBS1 | 1,415262935 | 2,667083393 | 7,35E-05 | 0,00463465 | sorbin and SH3 domain containing 1 |
| ENSG00000263535 | AK4P1 | 1,414645323 | 2,665941869 | 0,05 | 0,30089233 | adenylate kinase 4 pseudogene 1 |
| ENSG00000012211 | PRICKLE3 | 1,414571805 | 2,665806019 | 0,00166 | 0,04403004 | prickle planar cell polarity protein 3 |
| ENSG00000235174 | RPL39P3 | 1,41417133 | 2,665066127 | 0,34082 | 0,61918628 | ribosomal protein L39 pseudogene 3 |
| ENSG00000253463 | HMGB1P19 | 1,408767565 | 2,655102515 | 0,15054 | 0,47863728 | high mobility group box 1 pseudogene 19 |
| ENSG00000105894 | PTN | 1,408330426 | 2,654298135 | 0,00156 | 0,04237714 | pleiotrophin |
| ENSG00000283239 | AC019257,8 | 1,406016387 | 2,650044135 | 0,11767 | 0,43523522 | novel transcript |
| ENSG00000038427 | VCAN | 1,405279904 | 2,648691657 | 0,00329 | 0,06904163 | versican |
| ENSG00000226476 | LINC01748 | 1,405077636 | 2,648320331 | 8,16E-08 | 1,19E-05 | long intergenic non-protein coding RNA 1748 |
| ENSG00000225080 | PFN1P4 | 1,403293806 | 2,645047822 | 0,10261 | 0,41040107 | profilin 1 pseudogene 4 |
| ENSG00000163513 | TGFBR2 | 1,401334553 | 2,64145815 | 0,00078 | 0,02723246 | transforming growth factor beta receptor 2 |
| ENSG00000173867 | AC013489,1 | 1,400255304 | 2,639482871 | 0,0007 | 0,02564289 | novel transcript |
| ENSG00000236489 | AC133473,1 | 1,399650234 | 2,638376098 | 0,10401 | 0,41241152 | ribosomal protein L34 (RPL34) pseudogene |
| ENSG00000165704 | HPRT1 | 1,398046216 | 2,635444327 | 4,43E-05 | 0,00301829 | hypoxanthine phosphoribosyltransferase 1 |
| ENSG00000259658 | AC027559,1 | 1,396259863 | 2,632183125 | 0,042 | 0,27753946 | ubiquitin-conjugating enzyme E2Q family member 2 (UBE2Q2) pseudogene |
| ENSG00000288605 | AC007461,1 | 1,395590106 | 2,630961444 | 0,1728 | 0,49764744 | novel transcript |
| ENSG00000273398 | AC017083,3 | 1,394365614 | 2,628729354 | 0,04383 | 0,28236923 | novel protein |
| ENSG00000198053 | SIRPA | 1,392634992 | 2,625577884 | 0,00033 | 0,01519823 | signal regulatory protein alpha |
| ENSG00000275719 | AC008622,2 | 1,39215413 | 2,624702903 | 0,18518 | 0,50876814 | novel transcript, antisense to SLC1A5 |
| ENSG00000140470 | ADAMTS17 | 1,392123601 | 2,624647363 | 0,00224 | 0,05388971 | ADAM metallopeptidase with thrombospondin type 1 motif 17 |
| ENSG00000242134 | RPL5P13 | 1,390954877 | 2,622522002 | 0,20883 | 0,52791772 | ribosomal protein L5 pseudogene 13 |
| ENSG00000253953 | PCDHGB4 | 1,390726661 | 2,622107186 | 3,70E-09 | 7,52E-07 | protocadherin gamma subfamily B, 4 |
| ENSG00000184845 | DRD1 | 1,390347251 | 2,621417696 | 0,01334 | 0,15119996 | dopamine receptor D1 |
| ENSG00000153820 | SPHKAP | 1,388817034 | 2,618638723 | 0,32785 | 0,60960585 | SPHK1 interactor, AKAP domain containing |
| ENSG00000150907 | FOXO1 | 1,388315655 | 2,617728827 | 0,00337 | 0,07000131 | forkhead box O1 |

| | | | | | | |
|---|---|---|---|---|---|---|
| ENSG00000174640 | SLCO2A1 | 1,38556991 | 2,612751489 | 0,13814 | 0,46543358 | solute carrier organic anion transporter family member 2A1 |
| ENSG00000169242 | EFNA1 | 1,384817565 | 2,611389332 | 0,0002 | 0,01061649 | ephrin A1 |
| ENSG00000259436 | AC010247,2 | 1,38286223 | 2,607852423 | 0,00771 | 0,11160063 | novel transcript, antisense to POU2F2 |
| ENSG00000277639 | AC007906,2 | 1,382668715 | 2,607502642 | 0,00277 | 0,06186307 | novel protein |
| ENSG00000159200 | RCAN1 | 1,38208556 | 2,606448871 | 3,98E-10 | 9,53E-08 | regulator of calcineurin 1 |
| ENSG00000172986 | GXYLT2 | 1,380096033 | 2,602856964 | 0,00152 | 0,04177696 | glucoside xylosyltransferase 2 |
| ENSG00000247877 | AC021086,1 | 1,379987216 | 2,602660647 | 0,00175 | 0,0459021 | novel transcript |
| ENSG00000187773 | DIPK1C | 1,379893289 | 2,602491206 | 0,21702 | 0,53505085 | divergent protein kinase domain 1C |
| ENSG00000147246 | HTR2C | 1,379733297 | 2,602202611 | 0,08368 | 0,37585132 | 5-hydroxytryptamine receptor 2C |
| ENSG00000114200 | BCHE | 1,379650935 | 2,602054059 | 0,01656 | 0,16855868 | butyrylcholinesterase |
| ENSG00000180071 | ANKRD18A | 1,379447923 | 2,601687931 | 0,00918 | 0,12211443 | ankyrin repeat domain 18A |
| ENSG00000285245 | AL162417,1 | 1,378734427 | 2,600401564 | 0,36194 | 0,63352908 | novel protein |
| ENSG00000233766 | CAVIN2-AS1 | 1,378487073 | 2,599955757 | 0,00674 | 0,10442057 | CAVIN2 and TMEFF2 antisense RNA 1 |
| ENSG00000162882 | HAAO | 1,377796987 | 2,598712414 | 4,87E-06 | 0,00044851 | 3-hydroxyanthranilate 3,4-dioxygenase |
| ENSG00000077943 | ITGA8 | 1,376349124 | 2,596105701 | 0,00118 | 0,03607441 | integrin subunit alpha 8 |
| ENSG00000272293 | AC083964,1 | 1,375004689 | 2,593687538 | 0,05393 | 0,31146902 | novel transcript, antisense to FBXO25 |
| ENSG00000257921 | AC025165,3 | 1,373266167 | 2,590563894 | 0,20454 | 0,52439888 | novel protein |
| ENSG00000249069 | LINC01033 | 1,372867005 | 2,589847242 | 0,1659 | 0,49195088 | long intergenic non-protein coding RNA 1033 |
| ENSG00000249193 | HSPD1P5 | 1,371926083 | 2,588158701 | 0,12187 | 0,44093327 | heat shock protein family D (Hsp60) member 1 pseudogene 5 |
| ENSG00000230798 | FOXD3-AS1 | 1,371510469 | 2,587413207 | 0,10454 | 0,41308286 | FOXD3 antisense RNA 1 |
| ENSG00000168671 | UGT3A2 | 1,371089783 | 2,586658834 | 0,0589 | 0,3241112 | UDP glycosyltransferase family 3 member A2 |
| ENSG00000254553 | AL033529,1 | 1,370566589 | 2,58572095 | 0,55259 | 0,76595895 | novel transcript |
| ENSG00000267325 | LINC01415 | 1,37032545 | 2,585288797 | 0,08994 | 0,38780297 | long intergenic non-protein coding RNA 1415 |
| ENSG00000273073 | AC073869,1 | 1,369732961 | 2,584227283 | 0,15908 | 0,48655958 | novel transcript |
| ENSG00000166450 | PRTG | 1,369338948 | 2,583521603 | 0,00666 | 0,10355399 | protogenin |
| ENSG00000272255 | AC113361,1 | 1,368241919 | 2,581557834 | 0,07779 | 0,36490452 | novel transcript |
| ENSG00000205129 | C4orf47 | 1,367707587 | 2,580601878 | 0,09193 | 0,39202624 | chromosome 4 open reading frame 47 |
| ENSG00000200488 | RN7SKP203 | 1,366141056 | 2,577801288 | 0,06271 | 0,3313139 | RN7SK pseudogene 203 |
| ENSG00000272760 | AC093726,1 | 1,365564476 | 2,576771262 | 0,34927 | 0,62541638 | novel transcript, antisense to PAXIP1 |
| ENSG00000154258 | ABCA9 | 1,365558635 | 2,57676083 | 9,94E-05 | 0,00595921 | ATP binding cassette subfamily A member 9 |
| ENSG00000172640 | OR10AD1 | 1,364355123 | 2,574612164 | 0,01712 | 0,17198199 | olfactory receptor family 10 subfamily AD member 1 |

| | | | | | | |
|---|---|---|---|---|---|---|
| ENSG00000166292 | TMEM100 | 1,363214494 | 2,572577419 | 0,04191 | 0,2772815 | transmembrane protein 100 |
| ENSG00000259308 | FABP5P8 | 1,362931644 | 2,572073098 | 0,09051 | 0,38920435 | fatty acid binding protein 5 pseudogene 8 |
| ENSG00000266442 | MSH5-SAPCD: | 1,361852543 | 2,570149968 | 0,2359 | 0,5471053 | MSH5-SAPCD1 readthrough (NMD candidate) |
| ENSG00000230872 | MFSD13B | 1,360887122 | 2,568430653 | 0,19687 | 0,51812824 | major facilitator superfamily domain containing 13B (pseudogene) |
| ENSG00000233405 | LINC01046 | 1,360446401 | 2,567646157 | 0,17022 | 0,49606609 | long intergenic non-protein coding RNA 1046 |
| ENSG00000149150 | SLC43A1 | 1,360152278 | 2,567122744 | 0,00026 | 0,01310355 | solute carrier family 43 member 1 |
| ENSG00000273853 | AC023158,2 | 1,359536679 | 2,566027583 | 0,20872 | 0,52791772 | novel transcript, sense intronic to SYT10 |
| ENSG00000240463 | RPS19P3 | 1,359164959 | 2,565366516 | 0,12589 | 0,448127 | ribosomal protein S19 pseudogene 3 |
| ENSG00000101230 | ISM1 | 1,358413416 | 2,564030487 | 0,0018 | 0,04684971 | isthmin 1 |
| ENSG00000138722 | MMRN1 | 1,356304461 | 2,560285083 | 0,01881 | 0,18170309 | multimerin 1 |
| ENSG00000267809 | NDUFV2P1 | 1,356188839 | 2,560079902 | 0,05991 | 0,32541248 | NADH:ubiquinone oxidoreductase core subunit V2 pseudogene 1 |
| ENSG00000135862 | LAMC1 | 1,355034804 | 2,558032873 | 0,00028 | 0,01368428 | laminin subunit gamma 1 |
| ENSG00000111405 | ENDOU | 1,35191659 | 2,55250996 | 0,14129 | 0,46803321 | endonuclease, poly(U) specific |
| ENSG00000287561 | AL121899,4 | 1,351779367 | 2,552267186 | 0,09166 | 0,39158946 | novel transcript |
| ENSG00000134058 | CDK7 | 1,350570171 | 2,550128898 | 0,05085 | 0,30326674 | cyclin dependent kinase 7 |
| ENSG00000085741 | WNT11 | 1,348754963 | 2,546922328 | 0,08242 | 0,37394502 | Wnt family member 11 |
| ENSG00000276573 | AL442067,1 | 1,3475274 | 2,544756119 | 0,23929 | 0,5491566 | novel transcript, sense intronic to RAP2A |
| ENSG00000223631 | LINC01120 | 1,346259265 | 2,542520251 | 0,16973 | 0,49597212 | long intergenic non-protein coding RNA 1120 |
| ENSG00000166863 | TAC3 | 1,345627349 | 2,541406843 | 0,07782 | 0,36497315 | tachykinin precursor 3 |
| ENSG00000181201 | H2BU2P | 1,345509332 | 2,541198956 | 0,07308 | 0,35640071 | H2B,U histone 2, pseudogene |
| ENSG00000285234 | H2BU2P | 1,345509332 | 2,541198956 | 0,07308 | 0,35640071 | H2B,U histone 2, pseudogene |
| ENSG00000270935 | AL021808,1 | 1,344381364 | 2,539212902 | 0,22576 | 0,5409527 | cell division cycle associated 7 (CDCA7) pseudogene |
| ENSG00000282142 | AC245187,2 | 1,344334197 | 2,539129888 | 0,06 | 0,32549025 | novel transcript |
| ENSG00000255282 | WTAPP1 | 1,342725555 | 2,53630027 | 0,1013 | 0,40805419 | Wilms tumor 1 associated protein pseudogene 1 |
| ENSG00000256463 | SALL3 | 1,339840669 | 2,531233624 | 0,00803 | 0,11389729 | spalt like transcription factor 3 |
| ENSG00000277015 | SALL3 | 1,339840669 | 2,531233624 | 0,00803 | 0,11389729 | spalt like transcription factor 3 |
| ENSG00000134508 | CABLES1 | 1,338964022 | 2,529695997 | 0,00073 | 0,0264624 | Cdk5 and Abl enzyme substrate 1 |
| ENSG00000156427 | FGF18 | 1,337045801 | 2,526334725 | 0,00972 | 0,12625914 | fibroblast growth factor 18 |
| ENSG00000102271 | KLHL4 | 1,335092841 | 2,522917168 | 0,00942 | 0,1242251 | kelch like family member 4 |
| ENSG00000269421 | AC011503,3 | 1,333085464 | 2,519409204 | 0,06166 | 0,32868677 | zinc finger protein 92 pseudogene 3 |
| ENSG00000156687 | UNC5D | 1,331506272 | 2,516652936 | 0,01482 | 0,15970341 | unc-5 netrin receptor D |

| | | | | | | |
|---|---|---|---|---|---|---|
| ENSG00000064309 | CDON | 1,331494985 | 2,516633247 | 2,40E-05 | 0,00180654 | cell adhesion associated, oncogene regulated |
| ENSG00000243896 | OR2A7 | 1,330625136 | 2,515116341 | 0,0678 | 0,34426294 | olfactory receptor family 2 subfamily A member 7 |
| ENSG00000267561 | AC093155,3 | 1,32948258 | 2,513125259 | 0,19286 | 0,51493916 | novel protein |
| ENSG00000167779 | IGFBP6 | 1,329411972 | 2,513002266 | 0,1526 | 0,48061452 | insulin like growth factor binding protein 6 |
| ENSG00000031081 | ARHGAP31 | 1,329225181 | 2,512676919 | 0,01204 | 0,14273718 | Rho GTPase activating protein 31 |
| ENSG00000138829 | FBN2 | 1,328924206 | 2,51215278 | 4,88E-05 | 0,00329863 | fibrillin 2 |
| ENSG00000122025 | FLT3 | 1,328281054 | 2,511033114 | 0,03364 | 0,24775268 | fms related receptor tyrosine kinase 3 |
| ENSG00000255104 | AC005324,3 | 1,327986475 | 2,510520448 | 0,00359 | 0,07292667 | novel transcript |
| ENSG00000139174 | PRICKLE1 | 1,327963322 | 2,510480158 | 1,79E-10 | 4,52E-08 | prickle planar cell polarity protein 1 |
| ENSG00000112559 | MDFI | 1,326041439 | 2,507138052 | 0,00028 | 0,01366998 | MyoD family inhibitor |
| ENSG00000225932 | CTAGE4 | 1,325809198 | 2,506734494 | 0,04003 | 0,27049438 | CTAGE family member 4 |
| ENSG00000183160 | TMEM119 | 1,325210362 | 2,505694211 | 0,1198 | 0,43834197 | transmembrane protein 119 |
| ENSG00000187772 | LIN28B | 1,324502527 | 2,504465134 | 0,00449 | 0,08275517 | lin-28 homolog B |
| ENSG00000229190 | AC069542,1 | 1,324473602 | 2,504414922 | 0,07682 | 0,36335899 | novel transcript |
| ENSG00000248192 | ATG10-AS1 | 1,324315829 | 2,504141054 | 0,16703 | 0,49293221 | ATG10 antisense RNA 1 |
| ENSG00000116962 | NID1 | 1,324283436 | 2,504084829 | 0,00033 | 0,01523174 | nidogen 1 |
| ENSG00000228643 | AC079779,2 | 1,323720959 | 2,503108727 | 0,07765 | 0,36468824 | novel transcript |
| ENSG00000271659 | AL161729,4 | 1,323719453 | 2,503106115 | 0,15119 | 0,47956749 | novel transcript, antisense to PTCH1 |
| ENSG00000003436 | TFPI | 1,318922016 | 2,494796286 | 1,65E-06 | 0,00017332 | tissue factor pathway inhibitor |
| ENSG00000273291 | AC092042,3 | 1,318892938 | 2,494746002 | 0,29315 | 0,58745839 | novel protein |
| ENSG00000226386 | PARD3-AS1 | 1,318543843 | 2,494142411 | 0,20822 | 0,52760275 | PARD3 antisense RNA 1 |
| ENSG00000167173 | C15orf39 | 1,317174501 | 2,491776205 | 4,88E-06 | 0,00044851 | chromosome 15 open reading frame 39 |
| ENSG00000124835 | AC105760,1 | 1,316793319 | 2,491117926 | 0,0616 | 0,32867767 | novel transcript |
| ENSG00000260725 | AC005307,1 | 1,31584435 | 2,48947987 | 0,0666 | 0,34143033 | novel transcript |
| ENSG00000151364 | KCTD14 | 1,315834757 | 2,489463318 | 0,01199 | 0,14264319 | potassium channel tetramerization domain containing 14 |
| ENSG00000242156 | AC000041,1 | 1,314989188 | 2,48800466 | 0,14986 | 0,47782859 | adaptor-related protein complex 1, beta 1 subunit (AP1B1) pseudogene |
| ENSG00000247516 | MIR4458HG | 1,313432959 | 2,485322308 | 0,28325 | 0,58053323 | MIR4458 host gene |
| ENSG00000179546 | HTR1D | 1,313053533 | 2,484668758 | 0,00123 | 0,03698156 | 5-hydroxytryptamine receptor 1D |
| ENSG00000197943 | PLCG2 | 1,312843684 | 2,484307374 | 0,00563 | 0,09352688 | phospholipase C gamma 2 |
| ENSG00000146072 | TNFRSF21 | 1,31232951 | 2,48342213 | 3,91E-07 | 4,96E-05 | TNF receptor superfamily member 21 |
| ENSG00000185924 | RTN4RL1 | 1,311084008 | 2,481279076 | 0,00079 | 0,02756138 | reticulon 4 receptor like 1 |
| ENSG00000280689 | BORCS5 | 1,31055824 | 2,480374977 | 0,11639 | 0,43288152 | BLOC-1 related complex subunit 5 |

| | | | | | | |
|---|---|---|---|---|---|---|
| ENSG00000257718 | CPNE8-AS1 | 1,309345869 | 2,478291465 | 0,10908 | 0,42061552 | CPNE8 antisense RNA 1 |
| ENSG00000125864 | BFSP1 | 1,308488611 | 2,476819288 | 0,01752 | 0,17446742 | beaded filament structural protein 1 |
| ENSG00000183055 | FAM133CP | 1,308293861 | 2,476484962 | 0,09869 | 0,40456111 | family with sequence similarity 133 member C, pseudogene |
| ENSG00000229653 | ZNRD1ASP | 1,308069529 | 2,476099911 | 0,0844 | 0,37731853 | zinc ribbon domain containing 1 antisense, pseudogene |
| ENSG00000137871 | ZNF280D | 1,306714843 | 2,473775953 | 0,00917 | 0,12205788 | zinc finger protein 280D |
| ENSG00000088387 | DOCK9 | 1,306629002 | 2,473628767 | 1,33E-06 | 0,00014608 | dedicator of cytokinesis 9 |
| ENSG00000248231 | KRT8P4 | 1,305612558 | 2,471886597 | 0,04796 | 0,29426648 | keratin 8 pseudogene 4 |
| ENSG00000130303 | BST2 | 1,304608846 | 2,470167454 | 0,05523 | 0,31443527 | bone marrow stromal cell antigen 2 |
| ENSG00000058335 | RASGRF1 | 1,304585041 | 2,470126695 | 0,117 | 0,43438325 | Ras protein specific guanine nucleotide releasing factor 1 |
| ENSG00000223501 | VPS52 | 1,303191415 | 2,467741735 | 0,08286 | 0,37473201 | VPS52 subunit of GARP complex |
| ENSG00000187546 | AGMO | 1,30258138 | 2,466698486 | 0,24831 | 0,55546558 | alkylglycerol monooxygenase |
| ENSG00000254245 | PCDHGA3 | 1,30251475 | 2,466584566 | 0,01526 | 0,16213925 | protocadherin gamma subfamily A, 3 |
| ENSG00000090989 | EXOC1 | 1,300814097 | 2,463678661 | 0,00102 | 0,03289039 | exocyst complex component 1 |
| ENSG00000231252 | AC099792,1 | 1,300097182 | 2,462454695 | 0,00081 | 0,02825371 | novel transcript |
| ENSG00000165646 | SLC18A2 | 1,299858607 | 2,462047519 | 0,03265 | 0,24385689 | solute carrier family 18 member A2 |
| ENSG00000163659 | TIPARP | 1,298154853 | 2,459141674 | 0,0042 | 0,07947406 | TCDD inducible poly(ADP-ribose) polymerase |
| ENSG00000186815 | TPCN1 | 1,296400214 | 2,456152628 | 0,00016 | 0,0090156 | two pore segment channel 1 |
| ENSG00000259144 | RANBP20P | 1,295541546 | 2,454691201 | 0,13777 | 0,46481656 | RAN binding protein 20 pseudogene |
| ENSG00000259223 | AC009654,1 | 1,295451298 | 2,454537653 | 0,01248 | 0,14623734 | novel transcript |
| ENSG00000148926 | ADM | 1,294416037 | 2,452776938 | 0,00026 | 0,01313468 | adrenomedullin |
| ENSG00000250111 | AC107982,1 | 1,292178848 | 2,44897636 | 0,19055 | 0,51358916 | coiled-coil domain containing 144B (pseudogene) (CCDC144B) pseudogene |
| ENSG00000100558 | PLEK2 | 1,291961919 | 2,448608151 | 0,11902 | 0,43696075 | pleckstrin 2 |
| ENSG00000258466 | AL049779,1 | 1,291692478 | 2,448150886 | 0,15216 | 0,48061452 | novel protein |
| ENSG00000148200 | NR6A1 | 1,291306677 | 2,447496297 | 0,00025 | 0,01250193 | nuclear receptor subfamily 6 group A member 1 |
| ENSG00000135547 | HEY2 | 1,290622603 | 2,446336058 | 0,03037 | 0,23517553 | hes related family bHLH transcription factor with YRPW motif 2 |
| ENSG00000174652 | ZNF266 | 1,289864631 | 2,445051125 | 2,98E-11 | 8,52E-09 | zinc finger protein 266 |
| ENSG00000140092 | FBLN5 | 1,288793635 | 2,443236694 | 3,12E-06 | 0,00030074 | fibulin 5 |
| ENSG00000231804 | AL353748,1 | 1,288769542 | 2,443195893 | 0,1731 | 0,49783873 | ribosomal protein L21 (RPL21) pseudogene |
| ENSG00000272936 | AC096586,1 | 1,287809252 | 2,441570188 | 0,58209 | 0,78454692 | novel transcript, antisense to GNPDA2 |
| ENSG00000198019 | FCGR1B | 1,286477972 | 2,439318212 | 0,05552 | 0,31503413 | Fc fragment of IgG receptor Ib |

| | | | | | | |
|---|---|---|---|---|---|---|
| ENSG00000226876 | SMYD3-AS1 | 1,286325911 | 2,43906112 | 0,07952 | 0,36821728 | SMYD3 antisense RNA 1 |
| ENSG00000149633 | KIAA1755 | 1,285359751 | 2,437428249 | 0,00223 | 0,05379687 | KIAA1755 |
| ENSG00000231945 | VARS1 | 1,284343566 | 2,435712013 | 0,10576 | 0,41527838 | valyl-tRNA synthetase 1 |
| ENSG00000260118 | AL157700,1 | 1,281820391 | 2,431455842 | 0,06497 | 0,33787601 | novel transcript |
| ENSG00000284858 | AC091185,2 | 1,281730847 | 2,431304933 | 0,18277 | 0,50616031 | novel oocyte-secreted protein family member pseudogene |
| ENSG00000151090 | THRB | 1,281316275 | 2,430606375 | 0,07059 | 0,3507077 | thyroid hormone receptor beta |
| ENSG00000186973 | FAM183A | 1,280203866 | 2,428732947 | 0,0018 | 0,0469168 | family with sequence similarity 183 member A |
| ENSG00000136826 | KLF4 | 1,279593777 | 2,427706097 | 0,01541 | 0,16264924 | Kruppel like factor 4 |
| ENSG00000265490 | AP001178,2 | 1,278694839 | 2,426193874 | 0,35806 | 0,63185282 | novel transcript, antisense to ENOSF1 |
| ENSG00000243663 | RPS4XP14 | 1,277777197 | 2,424651157 | 0,09285 | 0,39343521 | ribosomal protein S4X pseudogene 14 |
| ENSG00000284817 | RPS4XP14 | 1,277777197 | 2,424651157 | 0,09285 | 0,39343521 | ribosomal protein S4X pseudogene 14 |
| ENSG00000235844 | HLA-DPA1 | 1,276607369 | 2,422685895 | 0,13501 | 0,46159318 | major histocompatibility complex, class II, DP alpha 1 |
| ENSG00000168384 | HLA-DPA1 | 1,276607369 | 2,422685895 | 0,13501 | 0,46159318 | major histocompatibility complex, class II, DP alpha 1 |
| ENSG00000235394 | ZNRD1ASP | 1,27651949 | 2,422538326 | 0,21904 | 0,5365253 | zinc ribbon domain containing 1 antisense, pseudogene |
| ENSG00000277945 | AC107308,1 | 1,273392774 | 2,417293706 | 0,1619 | 0,48914394 | novel transcript, sense intronic to HMGA2 |
| ENSG00000228638 | FCF1P2 | 1,273383925 | 2,417278879 | 0,22985 | 0,54324957 | FCF1 pseudogene 2 |
| ENSG00000008441 | NFIX | 1,27288111 | 2,416436544 | 0,17759 | 0,50200419 | nuclear factor I X |
| ENSG00000260382 | AC087481,1 | 1,271196484 | 2,413616532 | 0,13847 | 0,46575418 | transmembrane protein 183A (TMEM183A) pseudogene |
| ENSG00000183876 | ARSI | 1,270276762 | 2,412078337 | 0,03286 | 0,24470884 | arylsulfatase family member I |
| ENSG00000022355 | GABRA1 | 1,269454986 | 2,410704779 | 0,25488 | 0,55944852 | gamma-aminobutyric acid type A receptor subunit alpha1 |
| ENSG00000241506 | PSMC1P1 | 1,268326827 | 2,408820392 | 0,2758 | 0,57533851 | proteasome 26S subunit, ATPase 1 pseudogene 1 |
| ENSG00000162105 | SHANK2 | 1,267289261 | 2,407088625 | 0,0004 | 0,0176948 | SH3 and multiple ankyrin repeat domains 2 |
| ENSG00000287091 | AC009961,3 | 1,267120195 | 2,406806561 | 0,13288 | 0,45832984 | novel transcript, sense intronic to CD302and LY75-CD302 |
| ENSG00000185903 | OR11N1P | 1,26693181 | 2,406492305 | 0,14395 | 0,47144689 | olfactory receptor family 11 subfamily N member 1 pseudogene |
| ENSG00000263155 | MYZAP | 1,266369593 | 2,405554679 | 0,34844 | 0,62472936 | myocardial zonula adherens protein |
| ENSG00000287844 | AC018953,2 | 1,266074461 | 2,405062625 | 0,237 | 0,54798297 | novel transcript |
| ENSG00000178882 | RFLNA | 1,265670867 | 2,404389904 | 0,01851 | 0,17984912 | refilin A |
| ENSG00000162039 | MEIOB | 1,2651773 | 2,403567467 | 0,20104 | 0,52181085 | meiosis specific with OB-fold |
| ENSG00000274303 | 7SK | 1,26465015 | 2,402689382 | 0,25565 | 0,56008719 | 7SK RNA |
| ENSG00000215808 | LINC01139 | 1,263890371 | 2,401424366 | 0,01216 | 0,14324383 | long intergenic non-protein coding RNA 1139 |
| ENSG00000236252 | AL512605,1 | 1,263755267 | 2,401199491 | 0,05082 | 0,30326674 | uncharacterized LOC101928381 |

| | | | | | | |
|---|---|---|---|---|---|---|
| ENSG00000113739 | STC2 | 1,263559431 | 2,400873566 | 0,09091 | 0,390226 | stanniocalcin 2 |
| ENSG00000104332 | SFRP1 | 1,263228326 | 2,400322619 | 0,00078 | 0,02733095 | secreted frizzled related protein 1 |
| ENSG00000118322 | ATP10B | 1,262602082 | 2,399280915 | 0,01808 | 0,17756463 | ATPase phospholipid transporting 10B (putative) |
| ENSG00000095303 | PTGS1 | 1,261622314 | 2,39765206 | 0,00537 | 0,09125239 | prostaglandin-endoperoxide synthase 1 |
| ENSG00000178695 | KCTD12 | 1,261075418 | 2,396743331 | 0,00224 | 0,05388971 | potassium channel tetramerization domain containing 12 |
| ENSG00000187536 | TPM3P7 | 1,259127835 | 2,393510003 | 0,07196 | 0,3539609 | tropomyosin 3 pseudogene 7 |
| ENSG00000285107 | CACNA1C-AS4 | 1,258400748 | 2,392304029 | 0,19173 | 0,51436129 | CACNA1C antisense RNA 4 |
| ENSG00000256025 | CACNA1C-AS4 | 1,258400748 | 2,392304029 | 0,19173 | 0,51436129 | CACNA1C antisense RNA 4 |
| ENSG00000256232 | LINC02387 | 1,257010276 | 2,389999432 | 0,13326 | 0,4592636 | long intergenic non-protein coding RNA 2387 |
| ENSG00000231453 | LINC01305 | 1,254917731 | 2,386535391 | 0,32249 | 0,60577115 | long intergenic non-protein coding RNA 1305 |
| ENSG00000165197 | VEGFD | 1,254613097 | 2,386031513 | 0,11364 | 0,42841156 | vascular endothelial growth factor D |
| ENSG00000124134 | KCNS1 | 1,254512093 | 2,385864471 | 0,00068 | 0,02516175 | potassium voltage-gated channel modifier subfamily S member 1 |
| ENSG00000157557 | ETS2 | 1,25444942 | 2,385760828 | 3,73E-05 | 0,00261377 | ETS proto-oncogene 2, transcription factor |
| ENSG00000205946 | USP17L6P | 1,253857139 | 2,384781583 | 0,21626 | 0,53480193 | ubiquitin specific peptidase 17 like family member 6, pseudogene |
| ENSG00000273541 | AL137060,1 | 1,253635105 | 2,384414588 | 0,23063 | 0,54367096 | |
| ENSG00000068366 | ACSL4 | 1,253012934 | 2,383386517 | 0,02094 | 0,19206619 | acyl-CoA synthetase long chain family member 4 |
| ENSG00000253722 | AC010834,1 | 1,252944095 | 2,383272794 | 0,49065 | 0,72477304 | novel transcript |
| ENSG00000101938 | CHRDL1 | 1,250214767 | 2,378768319 | 0,02313 | 0,20293923 | chordin like 1 |
| ENSG00000232176 | AL161909,1 | 1,24799304 | 2,375107875 | 0,08369 | 0,37585132 | ribosomal protein S6 (RPS6) pseudogene |
| ENSG00000227060 | LINC00629 | 1,246930718 | 2,373359617 | 0,07064 | 0,3507077 | long intergenic non-protein coding RNA 629 |
| ENSG00000162595 | DIRAS3 | 1,246760078 | 2,373078916 | 0,00825 | 0,11480072 | DIRAS family GTPase 3 |
| ENSG00000269696 | AC005498,3 | 1,246026983 | 2,371873359 | 0,00158 | 0,04261325 | novel transcript, antisense to ZFP28 |
| ENSG00000134343 | ANO3 | 1,245876193 | 2,371625465 | 0,02154 | 0,19517279 | anoctamin 3 |
| ENSG00000271889 | AC016747,2 | 1,244642548 | 2,369598362 | 0,17363 | 0,49838936 | novel transcript |
| ENSG00000277277 | FAM243B | 1,244538441 | 2,369427375 | 0,16522 | 0,49131595 | family with sequence similarity 243 member B |
| ENSG00000213659 | RSU1P3 | 1,244407547 | 2,369212408 | 0,11537 | 0,43106723 | Ras suppressor protein 1 pseudogene 3 |
| ENSG00000197454 | OR2L5 | 1,243148669 | 2,367145964 | 0,1025 | 0,41028143 | olfactory receptor family 2 subfamily L member 5 |
| ENSG00000259627 | AC079328,2 | 1,242887655 | 2,366717736 | 0,09819 | 0,40370642 | novel transcript, antisense to TPM1 |
| ENSG00000213468 | FIRRE | 1,242860214 | 2,36667272 | 0,03965 | 0,26944441 | firre intergenic repeating RNA element |
| ENSG00000281358 | RASSF1-AS1 | 1,242457756 | 2,366012598 | 0,13242 | 0,45752148 | RASSF1 antisense RNA 1 |
| ENSG00000130720 | FIBCD1 | 1,24229304 | 2,365742482 | 0,08502 | 0,37844947 | fibrinogen C domain containing 1 |

| | | | | | | |
|---|---|---|---|---|---|---|
| ENSG00000145741 | BTF3 | 1,238403002 | 2,359372158 | 2,22E-09 | 4,61E-07 | basic transcription factor 3 |
| ENSG00000165807 | PPP1R36 | 1,237834917 | 2,358443299 | 0,11781 | 0,43532264 | protein phosphatase 1 regulatory subunit 36 |
| ENSG00000287219 | AL021707,9 | 1,237262401 | 2,357507566 | 0,16641 | 0,49238711 | novel transcript |
| ENSG00000276077 | CU633904,1 | 1,23686735 | 2,356862101 | 0,16958 | 0,49580293 | uncharacterized LOC102724951 |
| ENSG00000182795 | C1orf116 | 1,236294074 | 2,355925754 | 0,09266 | 0,39335125 | chromosome 1 open reading frame 116 |
| ENSG00000258819 | LINC02289 | 1,235887276 | 2,355261545 | 0,00717 | 0,10764638 | long intergenic non-protein coding RNA 2289 |
| ENSG00000227536 | SOCS5P4 | 1,234952715 | 2,353736328 | 0,05654 | 0,31727683 | suppressor of cytokine signaling 5 pseudogene 4 |
| ENSG00000004948 | CALCR | 1,234559238 | 2,353094463 | 0,25782 | 0,56199574 | calcitonin receptor |
| ENSG00000240563 | L1TD1 | 1,233708394 | 2,351707111 | 0,02006 | 0,18786958 | LINE1 type transposase domain containing 1 |
| ENSG00000152661 | GJA1 | 1,233636452 | 2,351589843 | 0,01975 | 0,1865312 | gap junction protein alpha 1 |
| ENSG00000179071 | CCDC89 | 1,233365732 | 2,351148612 | 0,14196 | 0,46851861 | coiled-coil domain containing 89 |
| ENSG00000276931 | AC009041,3 | 1,233131379 | 2,350766719 | 0,05675 | 0,31760957 | novel transcript, sense intronic to LMF1 |
| ENSG00000186439 | TRDN | 1,232547254 | 2,349815123 | 0,15965 | 0,48693054 | triadin |
| ENSG00000104611 | SH2D4A | 1,232479341 | 2,349704511 | 0,02562 | 0,21563704 | SH2 domain containing 4A |
| ENSG00000234243 | GPSM3 | 1,232308951 | 2,349427014 | 0,18467 | 0,50835796 | G protein signaling modulator 3 |
| ENSG00000236697 | GPSM3 | 1,232308951 | 2,349427014 | 0,18467 | 0,50835796 | G protein signaling modulator 3 |
| ENSG00000237052 | GPSM3 | 1,232308951 | 2,349427014 | 0,18467 | 0,50835796 | G protein signaling modulator 3 |
| ENSG00000206314 | GPSM3 | 1,232308951 | 2,349427014 | 0,18467 | 0,50835796 | G protein signaling modulator 3 |
| ENSG00000234508 | GPSM3 | 1,232308951 | 2,349427014 | 0,18467 | 0,50835796 | G protein signaling modulator 3 |
| ENSG00000177144 | NUDT4B | 1,229918708 | 2,345537731 | 0,22071 | 0,53767462 | nudix hydrolase 4B |
| ENSG00000100985 | MMP9 | 1,229881369 | 2,345477025 | 0,02312 | 0,20293306 | matrix metallopeptidase 9 |
| ENSG00000286549 | AP001631,2 | 1,229302727 | 2,34453648 | 0,16179 | 0,48914394 | novel transcript, sense intronic to MX2 |
| ENSG00000234832 | NXT1-AS1 | 1,228847437 | 2,343796701 | 0,14051 | 0,46742953 | NXT1 antisense RNA 1 |
| ENSG00000235248 | OR13C1P | 1,228426227 | 2,343112504 | 0,24269 | 0,55158827 | olfactory receptor family 13 subfamily C member 1 pseudogene |
| ENSG00000273396 | LINC01396 | 1,228376259 | 2,343031352 | 0,15276 | 0,48079669 | long intergenic non-protein coding RNA 1396 |
| ENSG00000283149 | AC068631,2 | 1,227564554 | 2,341713461 | 0,2388 | 0,54892264 | novel protein |
| ENSG00000275263 | AC135048,3 | 1,227317199 | 2,341312 | 0,08991 | 0,38777461 | novel transcript |
| ENSG00000277196 | AC007325,2 | 1,226485742 | 2,339963039 | 0,0799 | 0,36897172 | proline dehydrogenase 1, mitochondrial |
| ENSG00000254224 | AP003465,1 | 1,226483775 | 2,339959849 | 0,08101 | 0,37153397 | novel transcript |
| ENSG00000253307 | AC011676,1 | 1,2259168 | 2,339040433 | 0,23684 | 0,54768226 | novel transcript |
| ENSG00000106355 | LSM5 | 1,223976199 | 2,335896253 | 0,00084 | 0,02886269 | LSM5 homolog, U6 small nuclear RNA and mRNA degradation associated |

| | | | | | | |
|---|---|---|---|---|---|---|
| ENSG00000102048 | ASB9 | 1,222263605 | 2,333125002 | 0,06111 | 0,32798786 | ankyrin repeat and SOCS box containing 9 |
| ENSG00000236739 | CLIC4P1 | 1,221874023 | 2,332495056 | 0,17491 | 0,49911246 | chloride intracellular channel 4 pseudogene 1 |
| ENSG00000131668 | BARX1 | 1,221735078 | 2,332270425 | 0,02707 | 0,22193038 | BARX homeobox 1 |
| ENSG00000128606 | LRRC17 | 1,221575218 | 2,332012009 | 0,01799 | 0,17701257 | leucine rich repeat containing 17 |
| ENSG00000273999 | RBM17P2 | 1,221282522 | 2,331538936 | 5,69E-06 | 0,00051205 | RNA binding motif protein 17 pseudogene 2 |
| ENSG00000284644 | AC074386,1 | 1,220995557 | 2,331075218 | 0,06073 | 0,32716307 | novel transcript, antisense to OR2A1 and ARHGEF5 |
| ENSG00000103489 | XYLT1 | 1,220053633 | 2,329553774 | 0,00019 | 0,01011637 | xylosyltransferase 1 |
| ENSG00000182621 | PLCB1 | 1,220000887 | 2,329468605 | 0,00044 | 0,01870943 | phospholipase C beta 1 |
| ENSG00000281742 | MCCC2 | 1,219766632 | 2,329090393 | 0,2331 | 0,54557342 | methylcrotonoyl-CoA carboxylase 2 |
| ENSG00000189184 | PCDH18 | 1,219687429 | 2,32896253 | 0,06711 | 0,34304078 | protocadherin 18 |
| ENSG00000182240 | BACE2 | 1,21936707 | 2,328445428 | 0,00208 | 0,05143262 | beta-secretase 2 |
| ENSG00000256045 | MTRNR2L10 | 1,219097008 | 2,328009601 | 0,33118 | 0,61131843 | MT-RNR2 like 10 |
| ENSG00000153179 | RASSF3 | 1,217962822 | 2,326180136 | 2,29E-07 | 3,06E-05 | Ras association domain family member 3 |
| ENSG00000101935 | AMMECR1 | 1,217877136 | 2,326041982 | 0,00012 | 0,00685911 | AMMECR nuclear protein 1 |
| ENSG00000198759 | EGFL6 | 1,215283444 | 2,321863957 | 0,21434 | 0,53334744 | EGF like domain multiple 6 |
| ENSG00000159289 | GOLGA6A | 1,214603661 | 2,320770176 | 0,14177 | 0,46830089 | golgin A6 family member A |
| ENSG00000130707 | ASS1 | 1,2141393 | 2,32002331 | 0,02359 | 0,20499484 | argininosuccinate synthase 1 |
| ENSG00000108244 | KRT23 | 1,212701383 | 2,317712121 | 0,23602 | 0,5471053 | keratin 23 |
| ENSG00000147255 | IGSF1 | 1,210783985 | 2,314633837 | 0,01553 | 0,16340375 | immunoglobulin superfamily member 1 |
| ENSG00000231233 | CFAP58-DT | 1,210109953 | 2,313552686 | 0,19413 | 0,51562158 | CFAP58 divergent transcript |
| ENSG00000239483 | RPS15AP16 | 1,209675508 | 2,312856101 | 0,10461 | 0,41313952 | ribosomal protein S15a pseudogene 16 |
| ENSG00000223486 | AC092198,1 | 1,20963007 | 2,312783257 | 0,01878 | 0,18160589 | novel transcript |
| ENSG00000242198 | AC008897,1 | 1,208745505 | 2,311365647 | 0,1708 | 0,49641416 | ribosomal protein S26 (RPS26) pseudogene |
| ENSG00000214967 | NPIPA7 | 1,20868518 | 2,311269001 | 0,02159 | 0,19542707 | nuclear pore complex interacting protein family member A7 |
| ENSG00000259250 | AC018904,1 | 1,208516574 | 2,310998902 | 0,09964 | 0,40615251 | novel transcript, antisense to ADAM10 |
| ENSG00000254656 | RTL1 | 1,208486012 | 2,310949946 | 0,06278 | 0,33160043 | retrotransposon Gag like 1 |
| ENSG00000123096 | SSPN | 1,207803421 | 2,309856811 | 0,01239 | 0,14545533 | sarcospan |
| ENSG00000230836 | LINC01293 | 1,205804518 | 2,306658642 | 0,09275 | 0,39337162 | long intergenic non-protein coding RNA 1293 |
| ENSG00000251224 | CNOT10-AS1 | 1,205334763 | 2,305907695 | 0,13936 | 0,46666406 | CNOT10 antisense RNA 1 |
| ENSG00000137558 | PI15 | 1,205172589 | 2,3056485 | 0,15649 | 0,48425071 | peptidase inhibitor 15 |
| ENSG00000182107 | TMEM30B | 1,204390464 | 2,304398883 | 0,1163 | 0,43279558 | transmembrane protein 30B |
| ENSG00000249532 | MIR302CHG | 1,20292788 | 2,3020639 | 0,12381 | 0,4446254 | miR-302/367 cluster host gene |

| | | | | | | |
|---|---|---|---|---|---|---|
| ENSG00000231646 | FSIP2-AS1 | 1,202327331 | 2,301105822 | 0,00602 | 0,09782483 | FSIP2 antisense RNA 1 |
| ENSG00000227094 | AL512785,1 | 1,201853325 | 2,300349904 | 0,36205 | 0,63354671 | novel transcript |
| ENSG00000200997 | RNVU1-34 | 1,201594135 | 2,299936668 | 0,19833 | 0,51980353 | RNA, variant U1 small nuclear 34 |
| ENSG00000100292 | HMOX1 | 1,200245883 | 2,297788297 | 2,25E-06 | 0,00022506 | heme oxygenase 1 |
| ENSG00000237225 | OR13K1P | 1,199376714 | 2,296404383 | 0,01385 | 0,15453657 | olfactory receptor family 13 subfamily K member 1 pseudogene |
| ENSG00000105974 | CAV1 | 1,196804531 | 2,292313768 | 0,15883 | 0,48624756 | caveolin 1 |
| ENSG00000248690 | HAS2-AS1 | 1,195049441 | 2,289526781 | 0,01454 | 0,15764244 | HAS2 antisense RNA 1 |
| ENSG00000250882 | AC010261,1 | 1,193854718 | 2,287631567 | 0,32814 | 0,60971379 | novel transcript |
| ENSG00000286338 | AC099487,2 | 1,193584091 | 2,287202482 | 0,18966 | 0,51284737 | novel transcript |
| ENSG00000273047 | AL121845,2 | 1,193532646 | 2,287120924 | 0,40154 | 0,66303726 | novel transcript, LIME1-SLC2A4RG readthrough |
| ENSG00000196656 | AC004057,1 | 1,192812365 | 2,285979339 | 0,43965 | 0,68844641 | ribosomal protein S26 (RPS26) pseudogene |
| ENSG00000270909 | CR788268,1 | 1,192327573 | 2,285211306 | 8,64E-05 | 0,00533996 | CAS1 domain containing 1 (CASD1) pseudogene |
| ENSG00000226683 | PWWP2AP1 | 1,191947524 | 2,284609392 | 0,27523 | 0,57491355 | PWWP domain containing 2A pseudogene 1 |
| ENSG00000250892 | AC108475,1 | 1,191423397 | 2,283779551 | 0,28931 | 0,58494545 | pseudogene similar to part of WEE1 homolog (WEE1) |
| ENSG00000143344 | RGL1 | 1,190999778 | 2,283109064 | 0,00424 | 0,07987039 | ral guanine nucleotide dissociation stimulator like 1 |
| ENSG00000272167 | AL606537,1 | 1,190481033 | 2,282288281 | 0,22659 | 0,54144 | PROX1 antisense RNA 1 |
| ENSG00000114854 | TNNC1 | 1,188140706 | 2,278588974 | 0,05607 | 0,31627719 | troponin C1, slow skeletal and cardiac type |
| ENSG00000286172 | RNVU1-8 | 1,188012839 | 2,278387031 | 0,09531 | 0,39794903 | RNA, variant U1 small nuclear 8 |
| ENSG00000198807 | PAX9 | 1,187503251 | 2,277582402 | 0,20342 | 0,52321822 | paired box 9 |
| ENSG00000251680 | AC008591,1 | 1,186811846 | 2,276491143 | 0,02403 | 0,20719534 | novel transcript |
| ENSG00000149218 | ENDOD1 | 1,185715314 | 2,274761534 | 0,00248 | 0,05771514 | endonuclease domain containing 1 |
| ENSG00000214190 | RNF152P1 | 1,185455099 | 2,274351279 | 0,21732 | 0,53520949 | ring finger protein 152 pseudogene 1 |
| ENSG00000225401 | TGIF2P1 | 1,185436566 | 2,274322063 | 0,17308 | 0,49783873 | TGFB induced factor homeobox 2 pseudogene 1 |
| ENSG00000244733 | AL132656,2 | 1,184858974 | 2,273411705 | 0,05905 | 0,32417102 | novel transcript |
| ENSG00000176076 | KCNE5 | 1,184017495 | 2,272086082 | 0,00988 | 0,12735196 | potassium voltage-gated channel subfamily E regulatory subunit 5 |
| ENSG00000226437 | TRIM39 | 1,182682344 | 2,269984339 | 0,20167 | 0,52214043 | tripartite motif containing 39 |
| ENSG00000264186 | SNRPCP4 | 1,181767291 | 2,268545021 | 0,20233 | 0,52249745 | small nuclear ribonucleoprotein polypeptide C pseudogene 4 |
| ENSG00000261349 | AL031432,2 | 1,181745401 | 2,268510601 | 0,27428 | 0,57420712 | novel pseudogene |
| ENSG00000225558 | UBE2D3P2 | 1,181695807 | 2,26843262 | 0,28022 | 0,57784691 | ubiquitin conjugating enzyme E2 D3 pseudogene 2 |
| ENSG00000288299 | LDHA | 1,180812545 | 2,26704424 | 0,00035 | 0,01612354 | lactate dehydrogenase A |

| | | | | | | |
|---|---|---|---|---|---|---|
| ENSG00000230162 | CT45A11P | 1,18035042 | 2,266318175 | 0,29981 | 0,59160746 | cancer/testis antigen family 45 member A11, pseudogene |
| ENSG00000181577 | C6orf223 | 1,180313256 | 2,266259795 | 0,20528 | 0,52495117 | chromosome 6 open reading frame 223 |
| ENSG00000077274 | CAPN6 | 1,178599679 | 2,263569619 | 0,03615 | 0,25739586 | calpain 6 |
| ENSG00000198947 | DMD | 1,178588041 | 2,263551359 | 0,00461 | 0,08360185 | dystrophin |
| ENSG00000111077 | TNS2 | 1,177350147 | 2,261609968 | 0,02125 | 0,19328293 | tensin 2 |
| ENSG00000233786 | CDC27P1 | 1,177220615 | 2,26140692 | 0,34107 | 0,6192144 | cell division cycle 27 pseudogene 1 |
| ENSG00000261554 | AC037471,2 | 1,171670834 | 2,252724416 | 0,2224 | 0,53859118 | vomeronasal 1 receptor (VN1R) pseudogene |
| ENSG00000168938 | PPIC | 1,170774104 | 2,251324634 | 0,00507 | 0,08898421 | peptidylprolyl isomerase C |
| ENSG00000259182 | AC019254,1 | 1,170691936 | 2,251196414 | 0,11359 | 0,42836024 | novel transcript |
| ENSG00000130203 | APOE | 1,169952657 | 2,250043131 | 0,0162 | 0,16670888 | apolipoprotein E |
| ENSG00000143320 | CRABP2 | 1,16949533 | 2,249329993 | 0,01762 | 0,17483597 | cellular retinoic acid binding protein 2 |
| ENSG00000227210 | AC079145,1 | 1,16903627 | 2,248614379 | 0,00319 | 0,06768121 | novel transcript |
| ENSG00000233013 | FAM157B | 1,168699989 | 2,248090305 | 0,20271 | 0,52269757 | family with sequence similarity 157 member B |
| ENSG00000221970 | OR2A1 | 1,168685804 | 2,2480682 | 0,04609 | 0,28930497 | olfactory receptor family 2 subfamily A member 1 |
| ENSG00000256950 | AC069503,2 | 1,167469508 | 2,246173715 | 0,29558 | 0,58930301 | novel transcript |
| ENSG00000176971 | FIBIN | 1,167429393 | 2,24611126 | 0,05661 | 0,31740666 | fin bud initiation factor homolog |
| ENSG00000184005 | ST6GALNAC3 | 1,166490113 | 2,244649385 | 0,01295 | 0,14897258 | ST6 N-acetylgalactosaminide alpha-2,6-sialyltransferase 3 |
| ENSG00000240793 | UBA52P8 | 1,166270593 | 2,244307866 | 0,14963 | 0,47763598 | ubiquitin A-52 residue ribosomal protein fusion product 1 pseudogene 8 |
| ENSG00000188707 | ZBED6CL | 1,165621145 | 2,243297788 | 0,06923 | 0,34753175 | ZBED6 C-terminal like |
| ENSG00000265752 | AC010754,1 | 1,165423126 | 2,242989902 | 0,25425 | 0,55925777 | novel transcript |
| ENSG00000223838 | AC007091,1 | 1,165251048 | 2,242722386 | 0,17854 | 0,50321732 | novel transcript |
| ENSG00000168454 | TXNDC2 | 1,164035606 | 2,240833732 | 0,12438 | 0,44539699 | thioredoxin domain containing 2 |
| ENSG00000142173 | COL6A2 | 1,16354701 | 2,24007496 | 0,04906 | 0,29818993 | collagen type VI alpha 2 chain |
| ENSG00000274351 | AL137000,1 | 1,162140395 | 2,23789197 | 0,24345 | 0,55221671 | protein-kinase, interferon-inducible double stranded RNA dependent inhibitor, repressor of (P58 repressor) PRKRIR |
| ENSG00000285920 | AC087721,2 | 1,16213281 | 2,237880206 | 0,51821 | 0,74307441 | novel protein |
| ENSG00000248746 | ACTN3 | 1,161616971 | 2,237080188 | 0,16284 | 0,48945635 | actinin alpha 3 |
| ENSG00000149573 | MPZL2 | 1,160958339 | 2,23605913 | 0,15885 | 0,48624756 | myelin protein zero like 2 |
| ENSG00000258215 | AC078886,1 | 1,160288104 | 2,235020562 | 0,15441 | 0,48266733 | tripartite motif containing 61 (TRIM61) pseudogene |
| ENSG00000161544 | CYGB | 1,159991721 | 2,234561453 | 0,02912 | 0,23027064 | cytoglobin |
| ENSG00000287572 | AC270284,1 | 1,15939436 | 2,233636404 | 0,25982 | 0,56377449 | novel transcript |
| ENSG00000101928 | MOSPD1 | 1,158063745 | 2,231577245 | 0,01346 | 0,15210212 | motile sperm domain containing 1 |

| | | | | | | |
|---|---|---|---|---|---|---|
| ENSG00000225991 | RPL23AP34 | 1,157913034 | 2,231344134 | 0,52606 | 0,7497214 | ribosomal protein L23a pseudogene 34 |
| ENSG00000132481 | TRIM47 | 1,157717352 | 2,231041503 | 0,16571 | 0,49182669 | tripartite motif containing 47 |
| ENSG00000285154 | AC243837,4 | 1,157362823 | 2,230493313 | 0,34008 | 0,61888023 | olfactory receptor 2A1/2A42-like |
| ENSG00000180921 | FAM83H | 1,156313594 | 2,228871732 | 0,04832 | 0,29520031 | family with sequence similarity 83 member H |
| ENSG00000165072 | MAMDC2 | 1,155949924 | 2,228309955 | 0,07607 | 0,36217039 | MAM domain containing 2 |
| ENSG00000189127 | ANKRD34B | 1,155765277 | 2,228024778 | 0,15822 | 0,48525382 | ankyrin repeat domain 34B |
| ENSG00000236333 | TRHDE-AS1 | 1,155610588 | 2,227785897 | 2,21E-05 | 0,00169472 | TRHDE antisense RNA 1 |
| ENSG00000236166 | AL021408,1 | 1,155370964 | 2,227415905 | 0,09515 | 0,39774693 | novel transcript |
| ENSG00000167588 | GPD1 | 1,155199988 | 2,227151945 | 0,18656 | 0,509961 | glycerol-3-phosphate dehydrogenase 1 |
| ENSG00000272805 | TAS2R10 | 1,153678281 | 2,224804058 | 0,13769 | 0,46480929 | taste 2 receptor member 10 |
| ENSG00000121318 | TAS2R10 | 1,153678281 | 2,224804058 | 0,13769 | 0,46480929 | taste 2 receptor member 10 |
| ENSG00000277238 | TAS2R10 | 1,153678281 | 2,224804058 | 0,13769 | 0,46480929 | taste 2 receptor member 10 |
| ENSG00000225066 | MDM4P1 | 1,153550065 | 2,224606342 | 0,15939 | 0,48684894 | MDM4 pseudogene 1 |
| ENSG00000232024 | LSM12P1 | 1,15287812 | 2,223570458 | 0,43765 | 0,68737508 | LSM12 pseudogene 1 |
| ENSG00000173546 | CSPG4 | 1,152791321 | 2,223436683 | 0,01647 | 0,16805575 | chondroitin sulfate proteoglycan 4 |
| ENSG00000184672 | RALYL | 1,152721219 | 2,223328646 | 0,06762 | 0,34407703 | RALY RNA binding protein like |
| ENSG00000127399 | LRRC61 | 1,151971279 | 2,222173218 | 0,22774 | 0,54184325 | leucine rich repeat containing 61 |
| ENSG00000168491 | CCDC110 | 1,15167234 | 2,221712812 | 0,11967 | 0,43818503 | coiled-coil domain containing 110 |
| ENSG00000152518 | ZFP36L2 | 1,151334049 | 2,221191913 | 0,00065 | 0,02447582 | ZFP36 ring finger protein like 2 |
| ENSG00000258677 | AC022826,2 | 1,151322925 | 2,221174787 | 0,24817 | 0,55542983 | novel protein, UBE2W-STAU2 readthrough |
| ENSG00000274667 | AC090517,2 | 1,150998844 | 2,220675889 | 0,19375 | 0,51547251 | novel transcript, sense intronic to |
| ENSG00000172380 | GNG12 | 1,150884271 | 2,220499538 | 0,00208 | 0,05143262 | G protein subunit gamma 12 |
| ENSG00000256040 | PAPPA-AS1 | 1,149560148 | 2,218462471 | 0,59242 | 0,79117578 | PAPPA antisense RNA 1 |
| ENSG00000258388 | PPT2-EGFL8 | 1,148531989 | 2,216882012 | 0,25503 | 0,55951106 | PPT2-EGFL8 readthrough (NMD candidate) |
| ENSG00000284838 | C1QTNF4 | 1,14790143 | 2,215913292 | 0,00507 | 0,08898421 | C1q and TNF related 4 |
| ENSG00000172247 | C1QTNF4 | 1,14790143 | 2,215913292 | 0,00507 | 0,08898421 | C1q and TNF related 4 |
| ENSG00000237105 | HCP5 | 1,145667961 | 2,212485439 | 0,54484 | 0,76084582 | HLA complex P5 |
| ENSG00000177459 | ERICH5 | 1,14558673 | 2,212360869 | 0,31824 | 0,60349082 | glutamate rich 5 |
| ENSG00000109610 | SOD3 | 1,144971951 | 2,211418311 | 0,10589 | 0,4154675 | superoxide dismutase 3 |
| ENSG00000162643 | DNAI3 | 1,144731771 | 2,211050185 | 0,02638 | 0,2188126 | dynein axonemal intermediate chain 3 |
| ENSG00000175556 | LONRF3 | 1,144708723 | 2,211014862 | 0,06101 | 0,32783036 | LON peptidase N-terminal domain and ring finger 3 |
| ENSG00000279463 | AL159972,1 | 1,14456726 | 2,210798073 | 0,19501 | 0,51588752 | novel transcript, sense intronic to KLF12 |
| ENSG00000206457 | CCHCR1 | 1,142821827 | 2,208124974 | 0,32778 | 0,60960585 | coiled-coil alpha-helical rod protein 1 |

| | | | | | | |
|---|---|---|---|---|---|---|
| ENSG00000188959 | C9orf152 | 1,141785279 | 2,20653905 | 0,11908 | 0,43696075 | chromosome 9 open reading frame 152 |
| ENSG00000264743 | DPRXP4 | 1,140583857 | 2,204702292 | 0,16721 | 0,49323767 | divergent-paired related homeobox pseudogene 4 |
| ENSG00000134438 | RAX | 1,138459999 | 2,201459036 | 0,09996 | 0,40649331 | retina and anterior neural fold homeobox |
| ENSG00000162520 | SYNC | 1,137352329 | 2,199769451 | 0,10386 | 0,41222634 | syncoilin, intermediate filament protein |
| ENSG00000235387 | SPAAR | 1,136905352 | 2,199088023 | 0,32243 | 0,60577115 | small regulatory polypeptide of amino acid response |
| ENSG00000172889 | EGFL7 | 1,136895479 | 2,199072973 | 0,00337 | 0,07000131 | EGF like domain multiple 7 |
| ENSG00000171345 | KRT19 | 1,136819573 | 2,198957274 | 0,01896 | 0,18263335 | keratin 19 |
| ENSG00000281766 | RYBP | 1,136316058 | 2,19818995 | 0,01211 | 0,14307474 | RING1 and YY1 binding protein |
| ENSG00000072201 | LNX1 | 1,134745015 | 2,195797503 | 0,06407 | 0,33563563 | ligand of numb-protein X 1 |
| ENSG00000158270 | COLEC12 | 1,1343266 | 2,195160764 | 0,17057 | 0,49641416 | collectin subfamily member 12 |
| ENSG00000134333 | LDHA | 1,133420972 | 2,193783221 | 0,00011 | 0,00668974 | lactate dehydrogenase A |
| ENSG00000008394 | MGST1 | 1,131689304 | 2,1911516 | 0,01922 | 0,18350688 | microsomal glutathione S-transferase 1 |
| ENSG00000276920 | NKD2 | 1,131265988 | 2,190508765 | 0,06745 | 0,34371006 | NKD inhibitor of WNT signaling pathway 2 |
| ENSG00000116704 | SLC35D1 | 1,129436563 | 2,187732828 | 0,0019 | 0,04858818 | solute carrier family 35 member D1 |
| ENSG00000147883 | CDKN2B | 1,129373383 | 2,187637023 | 0,01608 | 0,16620111 | cyclin dependent kinase inhibitor 2B |
| ENSG00000226493 | RPL26P27 | 1,128656043 | 2,186549551 | 0,16848 | 0,49474143 | ribosomal protein L26 pseudogene 27 |
| ENSG00000145675 | PIK3R1 | 1,127320707 | 2,184526651 | 0,0034 | 0,07030692 | phosphoinositide-3-kinase regulatory subunit 1 |
| ENSG00000173261 | PLAC8L1 | 1,126781552 | 2,183710417 | 0,06255 | 0,33112961 | PLAC8 like 1 |
| ENSG00000256162 | SMLR1 | 1,12612315 | 2,182714065 | 0,09652 | 0,40027716 | small leucine rich protein 1 |
| ENSG00000052850 | ALX4 | 1,125593973 | 2,181913598 | 0,10403 | 0,41241152 | ALX homeobox 4 |
| ENSG00000131471 | AOC3 | 1,124839694 | 2,180773134 | 0,03621 | 0,25754934 | amine oxidase copper containing 3 |
| ENSG00000231965 | AF131215,1 | 1,124836253 | 2,180767932 | 0,12391 | 0,44467458 | ribosomal protein L17 (RPL17) pseudogene |
| ENSG00000285026 | AC270301,5 | 1,124836253 | 2,180767932 | 0,12391 | 0,44467458 | ribosomal protein L17 (RPL17) pseudogene |
| ENSG00000257185 | LINC02293 | 1,124623722 | 2,180446696 | 0,07882 | 0,36667857 | long intergenic non-protein coding RNA 2293 |
| ENSG00000264714 | KIAA0895LP1 | 1,124434221 | 2,180160308 | 0,17745 | 0,50191142 | KIAA00895L pseudogene 1 |
| ENSG00000213856 | VDAC1P2 | 1,124351801 | 2,180035761 | 0,16035 | 0,48762464 | voltage dependent anion channel 1 pseudogene 2 |
| ENSG00000219547 | RPL17P25 | 1,123156027 | 2,178229594 | 0,37353 | 0,6422142 | ribosomal protein L17 pseudogene 25 |
| ENSG00000204814 | AL935212,1 | 1,122554451 | 2,177321503 | 0,20403 | 0,52370308 | novel transcript |
| ENSG00000215695 | RSC1A1 | 1,122462309 | 2,177182446 | 0,44491 | 0,69205935 | regulator of solute carriers 1 |
| ENSG00000270008 | AL137247,2 | 1,122397341 | 2,177084404 | 0,55219 | 0,76565817 | novel transcript |
| ENSG00000221874 | ZNF816-ZNF3: | 1,122019111 | 2,176513715 | 0,22263 | 0,53875487 | ZNF816-ZNF321P readthrough |
| ENSG00000115183 | TANC1 | 1,121558349 | 2,1758187 | 0,00191 | 0,04876303 | tetratricopeptide repeat, ankyrin repeat and coiled-coil containing 1 |

| | | | | | | |
|---|---|---|---|---|---|---|
| ENSG00000073060 | SCARB1 | 1,121520603 | 2,175761774 | 0,00026 | 0,0130421 | scavenger receptor class B member 1 |
| ENSG00000198948 | MFAP3L | 1,121505871 | 2,175739556 | 0,0147 | 0,15872563 | microfibril associated protein 3 like |
| ENSG00000184185 | KCNJ12 | 1,120497409 | 2,174219218 | 0,0128 | 0,14802902 | potassium inwardly rectifying channel subfamily J member 12 |
| ENSG00000232273 | FTH1P1 | 1,120027637 | 2,173511361 | 0,11023 | 0,42280938 | ferritin heavy chain 1 pseudogene 1 |
| ENSG00000271947 | AC017076,1 | 1,119896149 | 2,173313275 | 0,13968 | 0,4668467 | novel transcript |
| ENSG00000220842 | RPL21P16 | 1,118972974 | 2,171923026 | 0,51372 | 0,73999943 | ribosomal protein L21 pseudogene 16 |
| ENSG00000184697 | CLDN6 | 1,118630346 | 2,171407274 | 0,08461 | 0,37769282 | claudin 6 |
| ENSG00000238090 | AC006195,1 | 1,118414447 | 2,171082348 | 0,22316 | 0,53927144 | RAB3A interacting protein (rabin3)-like 1 (RAB3IL1) pseudogene |
| ENSG00000154133 | ROBO4 | 1,116194378 | 2,167743979 | 0,04155 | 0,27616337 | roundabout guidance receptor 4 |
| ENSG00000188483 | IER5L | 1,114310652 | 2,164915405 | 1,12E-05 | 0,00091107 | immediate early response 5 like |
| ENSG00000164692 | COL1A2 | 1,113974131 | 2,164410479 | 0,02571 | 0,21626994 | collagen type I alpha 2 chain |
| ENSG00000162885 | B3GALNT2 | 1,112204509 | 2,161757223 | 0,00302 | 0,06570434 | beta-1,3-N-acetylgalactosaminyltransferase 2 |
| ENSG00000123405 | NFE2 | 1,110878856 | 2,159771755 | 0,03849 | 0,26548082 | nuclear factor, erythroid 2 |
| ENSG00000238186 | AL603839,2 | 1,10872769 | 2,156553774 | 0,31624 | 0,60226233 | novel transcript |
| ENSG00000157399 | ARSL | 1,108534572 | 2,156265119 | 0,00109 | 0,03441486 | arylsulfatase L |
| ENSG00000278083 | AC013476,1 | 1,107444073 | 2,154635864 | 0,08484 | 0,37817642 | zinc finger protein 354B (ZNF354B) pseudogene |
| ENSG00000284686 | AC119674,2 | 1,107052256 | 2,154050773 | 0,1969 | 0,51817259 | novel transcript |
| ENSG00000263563 | UBBP4 | 1,106613362 | 2,153395572 | 0,05383 | 0,31110058 | ubiquitin B pseudogene 4 |
| ENSG00000143248 | RGS5 | 1,106587957 | 2,153357652 | 0,00533 | 0,09086355 | regulator of G protein signaling 5 |
| ENSG00000284320 | SHMT1 | 1,105855217 | 2,152264246 | 0,48939 | 0,72387044 | serine hydroxymethyltransferase 1 |
| ENSG00000234953 | AL138799,3 | 1,10514534 | 2,151205487 | 0,3772 | 0,6456352 | novel transcript |
| ENSG00000106018 | VIPR2 | 1,105044986 | 2,151055854 | 0,08132 | 0,37209365 | vasoactive intestinal peptide receptor 2 |
| ENSG00000257681 | AC025265,1 | 1,104125554 | 2,14968542 | 0,23519 | 0,54667863 | novel transcript, antisense to STAB2 |
| ENSG00000260880 | HCCAT5 | 1,103718708 | 2,149079285 | 0,17405 | 0,49860418 | hepatocellular carcinoma associated transcript 5 |
| ENSG00000276408 | AC025287,2 | 1,103696145 | 2,149045674 | 0,1064 | 0,41609301 | novel transcript |
| ENSG00000270553 | AC011921,3 | 1,102572099 | 2,147371942 | 0,29231 | 0,58692263 | splicing factor 3B, 14 kDa subunit (SF3B14) pseudogene |
| ENSG00000232994 | RPL7P14 | 1,102122864 | 2,146703384 | 0,36523 | 0,63543797 | ribosomal protein L7 pseudogene 14 |
| ENSG00000225406 | ELF2P4 | 1,101969024 | 2,146474486 | 0,06451 | 0,33665657 | ELF2 pseudogene 4 |
| ENSG00000134996 | OSTF1 | 1,100712898 | 2,144606406 | 0,03824 | 0,26482463 | osteoclast stimulating factor 1 |
| ENSG00000177452 | AL161742,1 | 1,100119826 | 2,143724969 | 0,06032 | 0,32601555 | ribosomal protein L19 (RPL19) pseudogene |
| ENSG00000144369 | FAM171B | 1,098255349 | 2,1409563 | 0,03544 | 0,25505326 | family with sequence similarity 171 member B |

| | | | | | | |
|---|---|---|---|---|---|---|
| ENSG00000164330 | EBF1 | 1,097922085 | 2,140461794 | 0,00412 | 0,07872305 | EBF transcription factor 1 |
| ENSG00000165495 | PKNOX2 | 1,097921438 | 2,140460835 | 1,02E-06 | 0,0001164 | PBX/knotted 1 homeobox 2 |
| ENSG00000148700 | ADD3 | 1,097655071 | 2,140065675 | 0,02049 | 0,18981557 | adducin 3 |
| ENSG00000134716 | CYP2J2 | 1,097468009 | 2,139788209 | 0,25882 | 0,56260423 | cytochrome P450 family 2 subfamily J member 2 |
| ENSG00000161638 | ITGA5 | 1,097464292 | 2,139782696 | 0,00015 | 0,00833587 | integrin subunit alpha 5 |
| ENSG00000118971 | CCND2 | 1,097254183 | 2,139471087 | 0,00598 | 0,09746931 | cyclin D2 |
| ENSG00000224103 | HLA-DPA1 | 1,097047587 | 2,139164735 | 0,35627 | 0,63039976 | major histocompatibility complex, class II, DP alpha 1 |
| ENSG00000174715 | PPIAP72 | 1,09588567 | 2,137442588 | 0,19673 | 0,51793033 | peptidylprolyl isomerase A pseudogene 72 |
| ENSG00000174498 | IGDCC3 | 1,095776514 | 2,137280874 | 0,00012 | 0,00687924 | immunoglobulin superfamily DCC subclass member 3 |
| ENSG00000176054 | RPL23P2 | 1,095459334 | 2,136811039 | 0,1543 | 0,48266239 | ribosomal protein L23 pseudogene 2 |
| ENSG00000123575 | FAM199X | 1,095447677 | 2,136793773 | 0,01832 | 0,1790509 | family with sequence similarity 199, X-linked |
| ENSG00000278787 | UBR7 | 1,095011179 | 2,136147369 | 0,02606 | 0,21736588 | ubiquitin protein ligase E3 component n-recognin 7 |
| ENSG00000272825 | AL844908,1 | 1,094148544 | 2,134870477 | 0,03575 | 0,25654732 | novel transcript |
| ENSG00000161021 | MAML1 | 1,093405383 | 2,133771046 | 0,16381 | 0,49006042 | mastermind like transcriptional coactivator 1 |
| ENSG00000288100 | AL161663,2 | 1,092937771 | 2,133079552 | 0,23375 | 0,54563244 | novel transcript, antisense to STXBP6 |
| ENSG00000227117 | HORMAD2-AS | 1,092935875 | 2,133076749 | 0,24394 | 0,55250543 | HORMAD2 and MTMR3 antisense RNA 1 |
| ENSG00000228956 | SATB1-AS1 | 1,09232266 | 2,13217028 | 0,05948 | 0,32472914 | SATB1 antisense RNA 1 |
| ENSG00000277368 | AL138966,2 | 1,09095333 | 2,130147497 | 0,26449 | 0,56726601 | novel transcript, antisense to ring finger protein (C3H2C3 type) 6 |
| ENSG00000240861 | AC009153,1 | 1,090291335 | 2,129170282 | 0,22157 | 0,53805234 | ribosomal protein S10 (RPS10) pseudogene |
| ENSG00000202406 | RN7SKP187 | 1,089558277 | 2,128088689 | 0,26901 | 0,57085484 | RN7SK pseudogene 187 |
| ENSG00000160867 | FGFR4 | 1,088913052 | 2,127137144 | 0,06237 | 0,33099652 | fibroblast growth factor receptor 4 |
| ENSG00000168899 | VAMP5 | 1,088675788 | 2,126787346 | 0,03326 | 0,24649377 | vesicle associated membrane protein 5 |
| ENSG00000171621 | SPSB1 | 1,088396377 | 2,126375485 | 0,00564 | 0,09352688 | splA/ryanodine receptor domain and SOCS box containing 1 |
| ENSG00000099219 | ERMP1 | 1,088336038 | 2,126286554 | 0,00021 | 0,01087967 | endoplasmic reticulum metallopeptidase 1 |
| ENSG00000139239 | RPL14P1 | 1,085790627 | 2,12253836 | 0,23572 | 0,5471053 | ribosomal protein L14 pseudogene 1 |
| ENSG00000183153 | GJD3 | 1,084236489 | 2,120253094 | 0,32882 | 0,60992716 | gap junction protein delta 3 |
| ENSG00000188883 | KLRG2 | 1,083950089 | 2,119832229 | 0,37435 | 0,64275601 | killer cell lectin like receptor G2 |
| ENSG00000239215 | RPL27P12 | 1,082665052 | 2,117944892 | 0,30893 | 0,59771457 | ribosomal protein L27 pseudogene 12 |
| ENSG00000283924 | TNPO1P2 | 1,082635937 | 2,117902152 | 0,22049 | 0,53750545 | transportin 1 pseudogene 2 |
| ENSG00000230662 | TNPO1P2 | 1,082635937 | 2,117902152 | 0,22049 | 0,53750545 | transportin 1 pseudogene 2 |
| ENSG00000287038 | AL162388,2 | 1,082525736 | 2,11774038 | 0,01275 | 0,14784043 | novel transcript |

| | | | | | | |
|---|---|---|---|---|---|---|
| ENSG00000156504 | FAM122B | 1,082033541 | 2,117018007 | 0,00119 | 0,03622756 | family with sequence similarity 122B |
| ENSG00000152953 | STK32B | 1,08116518 | 2,115744153 | 0,10086 | 0,40770184 | serine/threonine kinase 32B |
| ENSG00000259999 | AC009054,1 | 1,079766688 | 2,113694228 | 0,3662 | 0,63617598 | novel transcript, sense intronic to CFDP1 |
| ENSG00000203706 | SERTAD4-AS1 | 1,078647925 | 2,112055762 | 0,01522 | 0,16205591 | SERTAD4 antisense RNA 1 |
| ENSG00000280963 | SERTAD4-AS1 | 1,078647925 | 2,112055762 | 0,01522 | 0,16205591 | SERTAD4 antisense RNA 1 |
| ENSG00000225178 | RPSAP58 | 1,077715788 | 2,110691586 | 0,01727 | 0,17300352 | ribosomal protein SA pseudogene 58 |
| ENSG00000228760 | BAG6 | 1,077210138 | 2,109951939 | 0,01652 | 0,16838361 | BAG cochaperone 6 |
| ENSG00000091106 | NLRC4 | 1,076706949 | 2,109216151 | 0,15754 | 0,48464679 | NLR family CARD domain containing 4 |
| ENSG00000188488 | SERPINA5 | 1,07591217 | 2,108054507 | 0,03399 | 0,248958 | serpin family A member 5 |
| ENSG00000287642 | AC093599,2 | 1,075787448 | 2,107872271 | 0,13491 | 0,46159318 | novel transcript, antisense to FHDC1 |
| ENSG00000250993 | AC020551,1 | 1,075691891 | 2,107732661 | 0,35279 | 0,62759135 | novel transcript |
| ENSG00000136859 | ANGPTL2 | 1,075427565 | 2,107346524 | 0,00209 | 0,05143262 | angiopoietin like 2 |
| ENSG00000165168 | CYBB | 1,075354499 | 2,107239799 | 0,08449 | 0,37732439 | cytochrome b-245 beta chain |
| ENSG00000250786 | SNHG18 | 1,075072858 | 2,106828467 | 0,01188 | 0,14209957 | small nucleolar RNA host gene 18 |
| ENSG00000249364 | AC112206,2 | 1,072971129 | 2,103761459 | 0,03396 | 0,24883097 | novel transcript |
| ENSG00000125968 | ID1 | 1,072124174 | 2,102526777 | 0,05158 | 0,3052902 | inhibitor of DNA binding 1, HLH protein |
| ENSG00000231114 | AC078842,2 | 1,072122899 | 2,102524918 | 0,00955 | 0,12499921 | novel transcript |
| ENSG00000280571 | AC006059,2 | 1,071744979 | 2,101974226 | 0,07623 | 0,36242671 | novel protein |
| ENSG00000223932 | GPANK1 | 1,071657176 | 2,101846302 | 0,08753 | 0,38311781 | G-patch domain and ankyrin repeats 1 |
| ENSG00000179388 | EGR3 | 1,070321125 | 2,099900726 | 0,32299 | 0,60584543 | early growth response 3 |
| ENSG00000010932 | FMO1 | 1,069713246 | 2,09901612 | 0,10966 | 0,42180924 | flavin containing dimethylaniline monoxygenase 1 |
| ENSG00000274092 | AC106739,1 | 1,069285001 | 2,098393146 | 0,27471 | 0,57463533 | novel transcript, antisense to IL4R |
| ENSG00000254198 | AC113191,1 | 1,069153128 | 2,098201347 | 0,01361 | 0,1529196 | Family with sequence similarity 9, member C (FAM9C) pseudogene |
| ENSG00000137310 | TCF19 | 1,066998876 | 2,095070621 | 0,45445 | 0,69856079 | transcription factor 19 |
| ENSG00000122707 | RECK | 1,066621401 | 2,094522528 | 0,01486 | 0,16001992 | reversion inducing cysteine rich protein with kazal motifs |
| ENSG00000258646 | AL049780,1 | 1,066283715 | 2,094032328 | 0,29456 | 0,58857672 | novel transcript, antisense to EIF2B2 |
| ENSG00000172572 | PDE3A | 1,065548882 | 2,092966009 | 0,01518 | 0,1619728 | phosphodiesterase 3A |
| ENSG00000286334 | AL161430,1 | 1,065276046 | 2,092570235 | 0,29633 | 0,58968627 | novel transcript, antisense to MBNL2 |
| ENSG00000241155 | ARHGAP31-AS | 1,064597628 | 2,091586449 | 0,14661 | 0,47448174 | ARHGAP31 antisense RNA 1 |
| ENSG00000163430 | FSTL1 | 1,064408828 | 2,091312748 | 0,00014 | 0,00778966 | follistatin like 1 |
| ENSG00000164308 | ERAP2 | 1,063796596 | 2,090425452 | 0,01815 | 0,17819547 | endoplasmic reticulum aminopeptidase 2 |
| ENSG00000281641 | SAMD12-AS1 | 1,063542946 | 2,090057953 | 0,34062 | 0,61906935 | SAMD12 antisense RNA 1 |

| | | | | | | |
|---|---|---|---|---|---|---|
| ENSG00000288235 | FAM106C | 1,063340443 | 2,089764604 | 0,18781 | 0,51107563 | family with sequence similarity 106 member C |
| ENSG00000135697 | BCO1 | 1,062533132 | 2,088595529 | 0,2116 | 0,53038627 | beta-carotene oxygenase 1 |
| ENSG00000224401 | RPL7P57 | 1,062332187 | 2,088304642 | 0,17345 | 0,49804988 | ribosomal protein L7 pseudogene 57 |
| ENSG00000224216 | AC234781,1 | 1,061850032 | 2,087606837 | 0,22738 | 0,54168591 | novel transcript |
| ENSG00000236212 | AC009262,1 | 1,061416503 | 2,086979607 | 0,18277 | 0,50616031 | novel transcript |
| ENSG00000138944 | SHISAL1 | 1,061269319 | 2,086766704 | 0,00403 | 0,07780095 | shisa like 1 |
| ENSG00000116525 | TRIM62 | 1,060802523 | 2,086091622 | 7,96E-06 | 0,00068866 | tripartite motif containing 62 |
| ENSG00000229896 | AL157373,2 | 1,060776784 | 2,086054404 | 0,25872 | 0,56260205 | novel transcript |
| ENSG00000106688 | SLC1A1 | 1,059926923 | 2,084825917 | 0,0412 | 0,27486753 | solute carrier family 1 member 1 |
| ENSG00000225210 | DUXAP9 | 1,059577839 | 2,08432152 | 0,14178 | 0,46830089 | double homeobox A pseudogene 9 |
| ENSG00000256073 | URB1-AS1 | 1,059308811 | 2,08393288 | 0,02936 | 0,23068546 | URB1 antisense RNA 1 (head to head) |
| ENSG00000235631 | RNF148 | 1,059218181 | 2,083801971 | 0,10148 | 0,40833508 | ring finger protein 148 |
| ENSG00000088882 | CPXM1 | 1,058852989 | 2,083274562 | 0,00169 | 0,04470397 | carboxypeptidase X, M14 family member 1 |
| ENSG00000233994 | GDI2P2 | 1,058561455 | 2,082853624 | 0,23388 | 0,5456699 | GDP dissociation inhibitor 2 pseudogene 2 |
| ENSG00000262533 | AC090617,3 | 1,057825134 | 2,08179085 | 0,49087 | 0,72482729 | novel transcript, antisense to DPH1 and OVCA2 |
| ENSG00000288630 | AC009093,11 | 1,05778313 | 2,081730241 | 0,4664 | 0,70665143 | novel transcript |
| ENSG00000139832 | RAB20 | 1,057510903 | 2,081337469 | 0,09358 | 0,39524231 | RAB20, member RAS oncogene family |
| ENSG00000238246 | AC069549,1 | 1,057373502 | 2,081139253 | 0,30305 | 0,59337193 | novel transcript |
| ENSG00000226101 | LINC02097 | 1,057189273 | 2,080873513 | 0,01549 | 0,16309786 | long intergenic non-protein coding RNA 2097 |
| ENSG00000240775 | AC021205,1 | 1,056784673 | 2,08029002 | 0,17792 | 0,50233162 | ribosomal protein S26 (RPS26) pseudogene |
| ENSG00000228124 | AL096678,1 | 1,055028998 | 2,077759969 | 0,25247 | 0,55796133 | novel transcript |
| ENSG00000119147 | ECRG4 | 1,05480384 | 2,077435724 | 0,02077 | 0,19121359 | ECRG4 augurin precursor |
| ENSG00000231794 | AC009542,1 | 1,054655948 | 2,077222775 | 0,18779 | 0,51107563 | novel transcript, antisense to WDR91 |
| ENSG00000164659 | ELAPOR2 | 1,053879459 | 2,07610507 | 0,0003 | 0,014483 | endosome-lysosome associated apoptosis and autophagy regulator family member 2 |
| ENSG00000148175 | STOM | 1,053550466 | 2,075631687 | 0,00552 | 0,09228058 | stomatin |
| ENSG00000134569 | LRP4 | 1,052828927 | 2,074593855 | 0,0274 | 0,22324969 | LDL receptor related protein 4 |
| ENSG00000263505 | SLC25A3P3 | 1,051321641 | 2,072427512 | 0,05786 | 0,32112994 | solute carrier family 25 member 3 pseudogene 3 |
| ENSG00000268995 | VN1R82P | 1,051109353 | 2,072122583 | 0,47107 | 0,71051384 | vomeronasal 1 receptor 82 pseudogene |
| ENSG00000233795 | POLR1H | 1,049989723 | 2,070515098 | 0,49073 | 0,72478928 | RNA polymerase I subunit H |
| ENSG00000106536 | POU6F2 | 1,049775559 | 2,070207759 | 1,56E-05 | 0,0012439 | POU class 6 homeobox 2 |
| ENSG00000261026 | AC105046,1 | 1,048824279 | 2,068843161 | 0,37304 | 0,64180957 | novel transcript, overlapping to EGR3 |
| ENSG00000151491 | EPS8 | 1,048198506 | 2,06794599 | 0,00262 | 0,05977985 | epidermal growth factor receptor pathway substrate 8 |

| | | | | | | |
|---|---|---|---|---|---|---|
| ENSG00000266601 | AC018521,6 | 1,046923514 | 2,066119235 | 0,23487 | 0,54635524 | novel transcript, antisense to SP2 |
| ENSG00000234343 | NEU1 | 1,046293552 | 2,065217248 | 0,30162 | 0,59280146 | neuraminidase 1 |
| ENSG00000227440 | ATP5MC1P4 | 1,0459654 | 2,064747552 | 0,26737 | 0,56973003 | ATP synthase membrane subunit c locus 1 pseudogene 4 |
| ENSG00000272410 | AC022384,1 | 1,045930723 | 2,064697923 | 0,48485 | 0,72052946 | novel protein |
| ENSG00000219294 | PIP5K1P1 | 1,044654946 | 2,062872915 | 0,10008 | 0,40666505 | phosphatidylinositol-4-phosphate 5-kinase type 1 pseudogene 1 |
| ENSG00000273061 | CDC37L1-DT | 1,044545603 | 2,062716574 | 0,13051 | 0,45477153 | CDC37L1 divergent transcript |
| ENSG00000240668 | KRT8P36 | 1,042547312 | 2,059861464 | 0,22913 | 0,54270085 | keratin 8 pseudogene 36 |
| ENSG00000157514 | TSC22D3 | 1,042206703 | 2,059375204 | 1,31E-06 | 0,00014394 | TSC22 domain family member 3 |
| ENSG00000272670 | DOC2B | 1,041980423 | 2,059052226 | 0,63411 | 0,81555826 | double C2 domain beta |
| ENSG00000218358 | RAET1K | 1,041787045 | 2,05877625 | 0,04898 | 0,29775794 | retinoic acid early transcript 1K pseudogene |
| ENSG00000198865 | CCDC152 | 1,039346782 | 2,055296853 | 0,07962 | 0,36831832 | coiled-coil domain containing 152 |
| ENSG00000223503 | AL353807,1 | 1,038880511 | 2,0546327 | 0,29335 | 0,58775703 | claudin 6 (CLDN6) pseudogene |
| ENSG00000265943 | AC090912,1 | 1,038079388 | 2,053492087 | 0,14903 | 0,47664156 | novel transcript, antisense to RBBP8 |
| ENSG00000070759 | TESK2 | 1,037238484 | 2,052295516 | 0,00158 | 0,04261325 | testis associated actin remodelling kinase 2 |
| ENSG00000086717 | PPEF1 | 1,037107572 | 2,052109297 | 0,06728 | 0,34325886 | protein phosphatase with EF-hand domain 1 |
| ENSG00000165182 | CXorf58 | 1,037063763 | 2,052046983 | 0,40216 | 0,66350593 | chromosome X open reading frame 58 |
| ENSG00000100504 | PYGL | 1,036716991 | 2,051553804 | 0,00479 | 0,08544369 | glycogen phosphorylase L |
| ENSG00000198682 | PAPSS2 | 1,036496164 | 2,051239806 | 0,01562 | 0,16386137 | 3'-phosphoadenosine 5'-phosphosulfate synthase 2 |
| ENSG00000137766 | UNC13C | 1,035700105 | 2,050108272 | 0,02335 | 0,20398797 | unc-13 homolog C |
| ENSG00000216657 | GLRX3P2 | 1,035378748 | 2,049651667 | 0,19582 | 0,51663034 | glutaredoxin 3 pseudogene 2 |
| ENSG00000151790 | TDO2 | 1,034614634 | 2,048566369 | 0,27789 | 0,5762948 | tryptophan 2,3-dioxygenase |
| ENSG00000204209 | DAXX | 1,032982772 | 2,046250503 | 0,10657 | 0,41635243 | death domain associated protein |
| ENSG00000233401 | PRKAR1AP1 | 1,032860762 | 2,046077458 | 0,25874 | 0,56260205 | protein kinase cAMP-dependent type I regulatory subunit alpha pseudogene 1 |
| ENSG00000145757 | SPATA9 | 1,032144951 | 2,045062524 | 0,27984 | 0,57751109 | spermatogenesis associated 9 |
| ENSG00000283453 | AC244258,1 | 1,03202421 | 2,044891376 | 0,05927 | 0,32445511 | primase, DNA, polypeptide 2 (58kDa) (PRIM2) pseudogene |
| ENSG00000225230 | AC008937,1 | 1,032008617 | 2,044869274 | 0,06205 | 0,32975861 | novel transcript |
| ENSG00000231588 | SHQ1P1 | 1,031414378 | 2,044027177 | 0,25475 | 0,55944852 | SHQ1, H/ACA ribonucleoprotein assembly factor pseudogene 1 |
| ENSG00000144677 | CTDSPL | 1,030720502 | 2,043044322 | 0,00156 | 0,04237714 | CTD small phosphatase like |
| ENSG00000229243 | LINC01981 | 1,029895649 | 2,041876555 | 0,03087 | 0,2370319 | long intergenic non-protein coding RNA 1981 |
| ENSG00000115738 | ID2 | 1,029200853 | 2,040893433 | 0,15119 | 0,47956749 | inhibitor of DNA binding 2 |

| | | | | | | |
|---|---|---|---|---|---|---|
| ENSG00000138271 | GPR87 | 1,029180561 | 2,040864728 | 0,23781 | 0,54864017 | G protein-coupled receptor 87 |
| ENSG00000163909 | HEYL | 1,028209454 | 2,039491442 | 0,01634 | 0,16750744 | hes related family bHLH transcription factor with YRPW motif like |
| ENSG00000232467 | TMA16P2 | 1,028206993 | 2,039487963 | 0,18058 | 0,50407792 | translation machinery associated 16 homolog pseudogene 2 |
| ENSG00000257379 | AC023509,1 | 1,028175577 | 2,039443553 | 0,33537 | 0,61502114 | novel transcript |
| ENSG00000277299 | AC084876,1 | 1,027447637 | 2,038414771 | 0,24132 | 0,55030444 | novel transcript, antisense to GIT2 |
| ENSG00000087245 | MMP2 | 1,026765231 | 2,037450813 | 2,05E-07 | 2,77E-05 | matrix metallopeptidase 2 |
| ENSG00000163739 | CXCL1 | 1,026292349 | 2,036783093 | 0,25008 | 0,55657567 | C-X-C motif chemokine ligand 1 |
| ENSG00000263020 | AL662899,2 | 1,025490265 | 2,035651033 | 0,27357 | 0,57362818 | novel protein |
| ENSG00000196562 | SULF2 | 1,025333468 | 2,035429804 | 0,05353 | 0,31055447 | sulfatase 2 |
| ENSG00000232305 | CLCP1 | 1,024087043 | 2,033672042 | 0,26048 | 0,56388142 | Charcot-Leyden crystal protein pseudogene 1 |
| ENSG00000226877 | AL354733,1 | 1,020854235 | 2,029120067 | 0,40501 | 0,66481282 | novel transcript, antisense to KIF27 |
| ENSG00000257880 | AC078789,1 | 1,020803909 | 2,029049286 | 0,40919 | 0,66785537 | novel transcript, sense intronic to FAM19A2 |
| ENSG00000185551 | NR2F2 | 1,017256552 | 2,024066306 | 0,01597 | 0,16547046 | nuclear receptor subfamily 2 group F member 2 |
| ENSG00000276725 | CEP170 | 1,017157456 | 2,023927282 | 0,2732 | 0,57344244 | centrosomal protein 170 |
| ENSG00000267243 | AC005381,1 | 1,016892176 | 2,02355516 | 0,30394 | 0,59407983 | novel transcript |
| ENSG00000135312 | HTR1B | 1,016663937 | 2,023235053 | 0,0857 | 0,37966806 | 5-hydroxytryptamine receptor 1B |
| ENSG00000128610 | FEZF1 | 1,016339492 | 2,022780102 | 0,05999 | 0,32549025 | FEZ family zinc finger 1 |
| ENSG00000256975 | AC092821,2 | 1,016335704 | 2,022774791 | 0,27061 | 0,5723494 | RAN binding protein (RANBP) pseudogene |
| ENSG00000278496 | AL158062,1 | 1,015627772 | 2,021782456 | 0,34075 | 0,61914946 | retinoblastoma binding protein 8 (RBBP8) pseudogene |
| ENSG00000145649 | GZMA | 1,0153875 | 2,021445769 | 0,14341 | 0,47083258 | granzyme A |
| ENSG00000253853 | AC246817,1 | 1,014649193 | 2,020411548 | 0,05698 | 0,31838529 | novel transcript |
| ENSG00000287652 | AC000082,1 | 1,013817297 | 2,019246861 | 0,48961 | 0,72395897 | novel transcript, antisense to DGCR8and CDC45 |
| ENSG00000114279 | FGF12 | 1,013077985 | 2,018212359 | 0,00234 | 0,05563463 | fibroblast growth factor 12 |
| ENSG00000275857 | AC009133,4 | 1,012961454 | 2,018049349 | 0,77273 | 0,89496764 | novel transcript, antisense to KIF22 |
| ENSG00000277454 | AC245078,1 | 1,012123589 | 2,016877679 | 0,29072 | 0,58579089 | PRH1-PRR4 readthrough |
| ENSG00000241991 | AC114982,1 | 1,011971041 | 2,016664429 | 0,27109 | 0,57254873 | ribosomal protein L31 (RPL31) pseudogene |
| ENSG00000057704 | TMCC3 | 1,011455617 | 2,015944075 | 0,00135 | 0,03911553 | transmembrane and coiled-coil domain family 3 |
| ENSG00000178093 | TSSK6 | 1,010585366 | 2,014728399 | 0,0161 | 0,16621463 | testis specific serine kinase 6 |
| ENSG00000171729 | TMEM51 | 1,009639554 | 2,013408003 | 0,00153 | 0,04189512 | transmembrane protein 51 |
| ENSG00000266289 | AC012213,3 | 1,009336802 | 2,012985531 | 0,11826 | 0,43606269 | novel transcript |
| ENSG00000267289 | AC008752,2 | 1,009311473 | 2,012950189 | 0,34913 | 0,62524308 | novel transcript |

| | | | | | | |
|---|---|---|---|---|---|---|
| ENSG00000270955 | AC135731,2 | 1,009198046 | 2,012791934 | 0,23219 | 0,54498326 | NEDD4 binding protein 2 (N4BP2) pseudogene |
| ENSG00000244060 | RPS2P41 | 1,008800052 | 2,012236745 | 0,21601 | 0,53473714 | ribosomal protein S2 pseudogene 41 |
| ENSG00000249664 | AC027338,1 | 1,007368022 | 2,010240375 | 0,30252 | 0,59306846 | novel transcript |
| ENSG00000152137 | HSPB8 | 1,007086145 | 2,009847648 | 0,0122 | 0,14356185 | heat shock protein family B (small) member 8 |
| ENSG00000005243 | COPZ2 | 1,006869564 | 2,009545947 | 0,02487 | 0,21153603 | COPI coat complex subunit zeta 2 |
| ENSG00000283567 | C19orf85 | 1,006792151 | 2,00943812 | 0,2964 | 0,58976635 | chromosome 19 open reading frame 85 |
| ENSG00000261771 | DNAAF4-CCPC | 1,005589991 | 2,007764406 | 0,39595 | 0,65863599 | DNAAF4-CCPG1 readthrough (NMD candidate) |
| ENSG00000010030 | ETV7 | 1,004800134 | 2,006665482 | 0,33726 | 0,61706507 | ETS variant transcription factor 7 |
| ENSG00000090339 | ICAM1 | 1,004554997 | 2,006324546 | 0,01003 | 0,12839387 | intercellular adhesion molecule 1 |
| ENSG00000187527 | ATP13A5 | 1,004543312 | 2,006308295 | 0,25651 | 0,56092974 | ATPase 13A5 |
| ENSG00000101017 | CD40 | 1,003320246 | 2,004608139 | 0,13301 | 0,45866131 | CD40 molecule |
| ENSG00000250295 | RDH10-AS1 | 1,003271345 | 2,004540193 | 0,09485 | 0,39735025 | RDH10 antisense RNA 1 |
| ENSG00000198881 | ASB12 | 1,002520674 | 2,00349745 | 0,28635 | 0,58267001 | ankyrin repeat and SOCS box containing 12 |
| ENSG00000284297 | AC208162,3 | 1,00249168 | 2,003457186 | 0,36043 | 0,63269368 | novel transcript |
| ENSG00000104953 | TLE6 | 1,001960551 | 2,002719748 | 0,17063 | 0,49641416 | TLE family member 6, subcortical maternal complex member |
| ENSG00000007171 | NOS2 | 1,001582187 | 2,002194581 | 0,18615 | 0,50956086 | nitric oxide synthase 2 |
| ENSG00000159197 | KCNE2 | 1,001407697 | 2,001952435 | 0,38746 | 0,65234549 | potassium voltage-gated channel subfamily E regulatory subunit 2 |
| ENSG00000272010 | AC100814,1 | 1,001359954 | 2,001886186 | 0,26066 | 0,56400103 | novel transcript |
| ENSG00000115884 | SDC1 | 1,001339501 | 2,001857805 | 0,00049 | 0,01999623 | syndecan 1 |
| ENSG00000233063 | SAR1AP4 | 1,001256282 | 2,001742336 | 0,39412 | 0,65707455 | secretion associated Ras related GTPase 1A pseudogene 4 |
| ENSG00000226710 | SNHG32 | 1,001033999 | 2,001433941 | 0,01451 | 0,15751168 | small nucleolar RNA host gene 32 |
| ENSG00000268903 | AL627309,6 | 1,000684892 | 2,000949688 | 0,16336 | 0,48977654 | novel pseudogene |
| ENSG00000284369 | COG5 | 1,000482455 | 2,000668936 | 0,08238 | 0,37394502 | component of oligomeric golgi complex 5 |
| ENSG00000225265 | TAF1A-AS1 | 1,000116168 | 2,00016105 | 0,03227 | 0,2425006 | TAF1A antisense RNA 1 |
| ENSG00000182158 | CREB3L2 | 0,999376013 | 1,999135157 | 4,71E-09 | 9,32E-07 | cAMP responsive element binding protein 3 like 2 |
| ENSG00000276040 | SPATA31A7 | 0,998739469 | 1,998253296 | 0,4711 | 0,71051384 | SPATA31 subfamily A member 7 |
| ENSG00000239828 | AC063944,1 | 0,998422372 | 1,997814139 | 0,08868 | 0,38505941 | novel transcript |
| ENSG00000271833 | AL445222,1 | 0,997316545 | 1,996283399 | 0,271 | 0,57248369 | novel transcript |
| ENSG00000158516 | CPA2 | 0,997114794 | 1,996004252 | 0,18725 | 0,5105374 | carboxypeptidase A2 |
| ENSG00000275426 | AC253576,2 | 0,997109647 | 1,995997131 | 0,05903 | 0,32417102 | novel transcript |
| ENSG00000227245 | AC092042,1 | 0,996903199 | 1,995711526 | 0,38061 | 0,64818855 | novel transcript |

| | | | | | | |
|---|---|---|---|---|---|---|
| ENSG00000225778 | PROSER2-AS1 | 0,996568978 | 1,995249245 | 0,1469 | 0,47471707 | PROSER2 antisense RNA 1 |
| ENSG00000226468 | AC018641,1 | 0,995711044 | 1,994063073 | 0,41073 | 0,66856387 | phosphoribosyl pyrophosphate synthetase 2 (PRPS2) pseudogene |
| ENSG00000248309 | MEF2C-AS1 | 0,995094273 | 1,993210769 | 0,15546 | 0,48334025 | MEF2C antisense RNA 1 |
| ENSG00000273474 | AL157392,4 | 0,994796415 | 1,992799294 | 0,20231 | 0,52249745 | novel transcript, antisense to FRMD4A |
| ENSG00000157227 | MMP14 | 0,994742724 | 1,992725131 | 0,00552 | 0,09228058 | matrix metallopeptidase 14 |
| ENSG00000235426 | AL133481,1 | 0,994519527 | 1,992416863 | 0,43807 | 0,68754225 | novel transcript |
| ENSG00000273294 | C1QTNF3-AM | 0,994492804 | 1,992379958 | 0,38051 | 0,64814735 | C1QTNF3-AMACR readthrough (NMD candidate) |
| ENSG00000180964 | TCEAL8 | 0,99341815 | 1,990896401 | 0,02629 | 0,21852536 | transcription elongation factor A like 8 |
| ENSG00000214575 | CPEB1 | 0,992787091 | 1,990025739 | 0,01171 | 0,14084308 | cytoplasmic polyadenylation element binding protein 1 |
| ENSG00000149571 | KIRREL3 | 0,992466912 | 1,98958414 | 0,00888 | 0,11966089 | kirre like nephrin family adhesion molecule 3 |
| ENSG00000288564 | AL355385,2 | 0,99243302 | 1,989537401 | 0,50006 | 0,73078231 | novel protein |
| ENSG00000255624 | C10orf88B | 0,991975415 | 1,988906444 | 0,172 | 0,49710979 | C10orf88B (pseudogene) |
| ENSG00000274769 | AC016747,4 | 0,991485183 | 1,988230723 | 0,19944 | 0,52039716 | novel transcript |
| ENSG00000100979 | PLTP | 0,991105384 | 1,987707377 | 0,00216 | 0,052624 | phospholipid transfer protein |
| ENSG00000102098 | SCML2 | 0,990671247 | 1,987109325 | 0,00039 | 0,01742594 | Scm polycomb group protein like 2 |
| ENSG00000176399 | DMRTA1 | 0,990405178 | 1,986742885 | 0,07374 | 0,35703856 | DMRT like family A1 |
| ENSG00000127824 | TUBA4A | 0,990302673 | 1,98660173 | 0,02082 | 0,19144697 | tubulin alpha 4a |
| ENSG00000278041 | AL133325,3 | 0,990171685 | 1,986421367 | 0,16112 | 0,48841163 | novel transcript |
| ENSG00000215875 | ST13P20 | 0,990152845 | 1,986395426 | 0,32149 | 0,60577115 | ST13, Hsp70 interacting protein pseudogene 20 |
| ENSG00000274340 | AC032011,1 | 0,98883855 | 1,984586645 | 0,08728 | 0,38244677 | novel transcript |
| ENSG00000266127 | ZNF415P1 | 0,988749425 | 1,984464047 | 0,27762 | 0,57600845 | zinc finger protein 415 pseudogene 1 |
| ENSG00000234210 | AC006372,3 | 0,987733442 | 1,983067028 | 0,28423 | 0,58107964 | novel transcript |
| ENSG00000257550 | AC023509,2 | 0,986551133 | 1,981442542 | 0,11591 | 0,43214457 | novel transcript, antisense to ATF7 |
| ENSG00000099860 | GADD45B | 0,98642694 | 1,98127198 | 0,03381 | 0,24846467 | growth arrest and DNA damage inducible beta |
| ENSG00000168461 | RAB31 | 0,986346175 | 1,981161067 | 0,023 | 0,20221357 | RAB31, member RAS oncogene family |
| ENSG00000231181 | AL954705,1 | 0,986248112 | 1,981026408 | 0,31853 | 0,6036094 | ribosomal protein L9 (RPL9) pseudogene |
| ENSG00000253626 | EIF5AL1 | 0,986234585 | 1,981007833 | 0,21106 | 0,52983259 | eukaryotic translation initiation factor 5A like 1 |
| ENSG00000156531 | PHF6 | 0,985945404 | 1,98061079 | 0,00297 | 0,06492836 | PHD finger protein 6 |
| ENSG00000129521 | EGLN3 | 0,985743962 | 1,980334258 | 0,04686 | 0,2912788 | egl-9 family hypoxia inducible factor 3 |
| ENSG00000072315 | TRPC5 | 0,985452852 | 1,979934703 | 0,03014 | 0,23425595 | transient receptor potential cation channel subfamily C member 5 |
| ENSG00000273363 | AL353801,3 | 0,985113031 | 1,979468392 | 0,20443 | 0,52437248 | novel transcript |

| | | | | | | |
|---|---|---|---|---|---|---|
| ENSG00000285909 | AP002762,2 | 0,985099274 | 1,979449518 | 0,20206 | 0,52230471 | novel transcript, antisense to CRTAM |
| ENSG00000272137 | AL451064,1 | 0,984888931 | 1,979160937 | 0,09078 | 0,38999312 | novel transcript |
| ENSG00000242595 | RPL26P26 | 0,984424601 | 1,978524049 | 0,30395 | 0,59407983 | ribosomal protein L26 pseudogene 26 |
| ENSG00000166025 | AMOTL1 | 0,984256506 | 1,978293535 | 0,00207 | 0,05132911 | angiomotin like 1 |
| ENSG00000286231 | AL445423,3 | 0,983553183 | 1,977329339 | 0,1216 | 0,44093327 | novel protein |
| ENSG00000236813 | BTF3P8 | 0,983131928 | 1,976752059 | 0,26475 | 0,56748133 | basic transcription factor 3 pseudogene 8 |
| ENSG00000010379 | SLC6A13 | 0,982793612 | 1,97628856 | 0,15837 | 0,48549236 | solute carrier family 6 member 13 |
| ENSG00000224729 | PCOLCE-AS1 | 0,982787724 | 1,976280495 | 0,24086 | 0,55001001 | PCOLCE antisense RNA 1 |
| ENSG00000188868 | ZNF563 | 0,981377581 | 1,97434975 | 0,00197 | 0,04968365 | zinc finger protein 563 |
| ENSG00000250742 | LINC02381 | 0,981296443 | 1,974238713 | 0,04178 | 0,27685414 | long intergenic non-protein coding RNA 2381 |
| ENSG00000115170 | ACVR1 | 0,979947899 | 1,972394178 | 0,00113 | 0,03504073 | activin A receptor type 1 |
| ENSG00000239557 | PPP2R5CP | 0,979490628 | 1,971769114 | 0,31401 | 0,60070179 | protein phosphatase 2 regulatory subunit B'gamma pseudogene |
| ENSG00000251689 | AC079140,5 | 0,979460794 | 1,971728339 | 0,10543 | 0,4147036 | novel zinc finger protein pseudogene |
| ENSG00000135549 | PKIB | 0,978783308 | 1,970802639 | 0,03177 | 0,24047554 | cAMP-dependent protein kinase inhibitor beta |
| ENSG00000283108 | AC011451,3 | 0,978616296 | 1,970574503 | 0,30055 | 0,5918686 | novel transcript |
| ENSG00000275374 | AC113385,3 | 0,978136481 | 1,969919234 | 0,11537 | 0,43106723 | glucuronidase, beta (GUSB) pseudogene |
| ENSG00000173930 | SLCO4C1 | 0,977931317 | 1,969639115 | 0,11843 | 0,43614969 | solute carrier organic anion transporter family member 4C1 |
| ENSG00000235330 | RPL12P25 | 0,977549259 | 1,969117579 | 0,22138 | 0,53805234 | ribosomal protein L12 pseudogene 25 |
| ENSG00000053108 | FSTL4 | 0,977091562 | 1,968492973 | 0,02021 | 0,18878814 | follistatin like 4 |
| ENSG00000225078 | AL365338,1 | 0,976594356 | 1,967814673 | 0,29963 | 0,59157893 | COP9 constitutive photomorphogenic homolog subunit 2 (Arabidopsis) (COPS2) pseudogene |
| ENSG00000162552 | WNT4 | 0,975496377 | 1,966317616 | 0,02929 | 0,23059905 | Wnt family member 4 |
| ENSG00000123612 | ACVR1C | 0,974811924 | 1,965384964 | 0,32299 | 0,60584543 | activin A receptor type 1C |
| ENSG00000236808 | POLR1H | 0,974512882 | 1,964977621 | 0,52289 | 0,7470043 | RNA polymerase I subunit H |
| ENSG00000233490 | GPSM3 | 0,973943331 | 1,964202036 | 0,01322 | 0,15023876 | G protein signaling modulator 3 |
| ENSG00000272991 | AF129408,1 | 0,973889001 | 1,964128068 | 0,40948 | 0,66797746 | novel transcript, antisense to BRWD1 |
| ENSG00000260537 | AC012184,2 | 0,971063492 | 1,960285099 | 0,31678 | 0,60267293 | novel protein, DDX19B and DDX19A readthrough |
| ENSG00000119042 | SATB2 | 0,970932436 | 1,960107033 | 8,87E-05 | 0,0054472 | SATB homeobox 2 |
| ENSG00000285684 | AL137781,1 | 0,970745848 | 1,959853543 | 0,15805 | 0,48506908 | novel transcript |
| ENSG00000236177 | HLA-DPA1 | 0,97011853 | 1,959001538 | 0,26371 | 0,56629809 | major histocompatibility complex, class II, DP alpha 1 |
| ENSG00000172159 | FRMD3 | 0,969703296 | 1,958437782 | 0,02122 | 0,19328293 | FERM domain containing 3 |
| ENSG00000117298 | ECE1 | 0,969394262 | 1,958018318 | 9,34E-08 | 1,36E-05 | endothelin converting enzyme 1 |

| | | | | | | |
|---|---|---|---|---|---|---|
| ENSG00000143167 | GPA33 | 0,969002031 | 1,957486057 | 0,0367 | 0,2596357 | glycoprotein A33 |
| ENSG00000105967 | TFEC | 0,968773517 | 1,957176028 | 0,11102 | 0,42382951 | transcription factor EC |
| ENSG00000115290 | GRB14 | 0,968604652 | 1,956946957 | 0,02694 | 0,22168572 | growth factor receptor bound protein 14 |
| ENSG00000135535 | CD164 | 0,968453672 | 1,95674217 | 0,01099 | 0,1353263 | CD164 molecule |
| ENSG00000141338 | ABCA8 | 0,968047271 | 1,956191042 | 0,10472 | 0,41326267 | ATP binding cassette subfamily A member 8 |
| ENSG00000116729 | WLS | 0,967652024 | 1,955655189 | 0,00242 | 0,05669968 | Wnt ligand secretion mediator |
| ENSG00000237207 | RBM17P3 | 0,967044317 | 1,954831581 | 0,15567 | 0,48345635 | RNA binding motif protein 17 pseudogene 3 |
| ENSG00000287834 | AP005210,2 | 0,967005242 | 1,954778635 | 0,02518 | 0,21337619 | novel transcript, antisense to ARHGAP28 |
| ENSG00000260661 | AC116903,2 | 0,966997834 | 1,954768598 | 0,24815 | 0,55542983 | novel transcript, antisense to SLCO3A1 |
| ENSG00000157404 | KIT | 0,966969618 | 1,954730367 | 0,03325 | 0,24649377 | KIT proto-oncogene, receptor tyrosine kinase |
| ENSG00000183421 | RIPK4 | 0,966305438 | 1,953830667 | 0,37887 | 0,64679625 | receptor interacting serine/threonine kinase 4 |
| ENSG00000269506 | AC110792,2 | 0,966060373 | 1,953498806 | 0,01667 | 0,16947852 | novel transcript, antisense to CHIC2 |
| ENSG00000270659 | AC079610,1 | 0,965674261 | 1,952976055 | 0,1905 | 0,51357771 | novel transcript |
| ENSG00000240499 | AC004594,1 | 0,965628256 | 1,952913779 | 0,07791 | 0,36517053 | novel transcript, antisense to CADPS2 |
| ENSG00000260914 | AC026464,4 | 0,965473664 | 1,952704526 | 0,20125 | 0,52181085 | novel protein |
| ENSG00000065308 | TRAM2 | 0,965051646 | 1,952133403 | 1,59E-05 | 0,00126474 | translocation associated membrane protein 2 |
| ENSG00000270181 | BIVM-ERCC5 | 0,964825555 | 1,9518275 | 0,1474 | 0,47490365 | BIVM-ERCC5 readthrough |
| ENSG00000268015 | AC010320,2 | 0,964588395 | 1,951506672 | 0,22838 | 0,54241083 | novel transcript, antisense to PPP2R1A |
| ENSG00000254233 | LINC02365 | 0,96406851 | 1,95080356 | 0,40904 | 0,66785537 | long intergenic non-protein coding RNA 2365 |
| ENSG00000283060 | ID3 | 0,963765174 | 1,950393434 | 0,09891 | 0,40481169 | inhibitor of DNA binding 3, HLH protein |
| ENSG00000117318 | ID3 | 0,963765174 | 1,950393434 | 0,09891 | 0,40481169 | inhibitor of DNA binding 3, HLH protein |
| ENSG00000235779 | AC079779,3 | 0,9637533 | 1,950377381 | 0,19387 | 0,51547251 | novel transcript |
| ENSG00000165568 | AKR1E2 | 0,963297696 | 1,949761548 | 0,00942 | 0,1242017 | aldo-keto reductase family 1 member E2 |
| ENSG00000047365 | ARAP2 | 0,963130764 | 1,949535957 | 0,05473 | 0,31318759 | ArfGAP with RhoGAP domain, ankyrin repeat and PH domain 2 |
| ENSG00000249379 | AL033397,2 | 0,962696341 | 1,948949004 | 0,16257 | 0,48945635 | novel transcript |
| ENSG00000149575 | SCN2B | 0,962565238 | 1,948771903 | 0,09993 | 0,40649331 | sodium voltage-gated channel beta subunit 2 |
| ENSG00000268266 | AC003005,2 | 0,961931942 | 1,947916643 | 0,28907 | 0,58469988 | novel transcript |
| ENSG00000234476 | LINC02765 | 0,961843505 | 1,94779724 | 0,24867 | 0,55591283 | long intergenic non-protein coding RNA 2765 |
| ENSG00000277067 | CU634019,1 | 0,961470988 | 1,947294367 | 0,14827 | 0,47551028 | uncharacterized LOC102724843 |
| ENSG00000164124 | TMEM144 | 0,96126532 | 1,947016783 | 0,07229 | 0,35498117 | transmembrane protein 144 |
| ENSG00000224985 | AL590714,1 | 0,960241082 | 1,945634993 | 0,02554 | 0,21502349 | novel transcript |
| ENSG00000047648 | ARHGAP6 | 0,95921474 | 1,944251349 | 0,00073 | 0,02636261 | Rho GTPase activating protein 6 |

| | | | | | | |
|---|---|---|---|---|---|---|
| ENSG00000261468 | AC096921,2 | 0,959181176 | 1,944206117 | 0,12057 | 0,43944057 | novel transcript |
| ENSG00000272416 | AC025175,1 | 0,958918749 | 1,943852497 | 0,36154 | 0,63347405 | novel transcript |
| ENSG00000251199 | AC015631,1 | 0,958840193 | 1,943746656 | 0,25642 | 0,56092974 | novel transcript |
| ENSG00000283109 | AC116049,2 | 0,958380937 | 1,943127997 | 0,62278 | 0,80873034 | novel transcript |
| ENSG00000185559 | DLK1 | 0,956979619 | 1,941241515 | 0,22582 | 0,54105066 | delta like non-canonical Notch ligand 1 |
| ENSG00000240583 | AQP1 | 0,955657745 | 1,939463661 | 0,13968 | 0,4668467 | aquaporin 1 (Colton blood group) |
| ENSG00000185222 | TCEAL9 | 0,955098048 | 1,938711387 | 0,00011 | 0,00628279 | transcription elongation factor A like 9 |
| ENSG00000227073 | SDHDP2 | 0,954966353 | 1,938534423 | 0,45339 | 0,69750019 | succinate dehydrogenase complex subunit D pseudogene 2 |
| ENSG00000183337 | BCOR | 0,954098651 | 1,937368851 | 0,00324 | 0,06836331 | BCL6 corepressor |
| ENSG00000185201 | IFITM2 | 0,95399146 | 1,937224911 | 9,69E-05 | 0,00584194 | interferon induced transmembrane protein 2 |
| ENSG00000231528 | FAM225A | 0,953880157 | 1,937075461 | 0,41344 | 0,67078587 | family with sequence similarity 225 member A |
| ENSG00000088970 | KIZ | 0,95369623 | 1,936828523 | 0,03252 | 0,2434582 | kizuna centrosomal protein |
| ENSG00000232312 | GPANK1 | 0,953153987 | 1,936100694 | 0,10257 | 0,41028143 | G-patch domain and ankyrin repeats 1 |
| ENSG00000255693 | LINC02389 | 0,953113332 | 1,936046136 | 0,02921 | 0,2303054 | long intergenic non-protein coding RNA 2389 |
| ENSG00000135525 | MAP7 | 0,953038153 | 1,93594525 | 0,26012 | 0,56382871 | microtubule associated protein 7 |
| ENSG00000278493 | AC039056,2 | 0,952704281 | 1,935497281 | 0,30187 | 0,59294633 | novel transcript, sense intronic to EHD4 |
| ENSG00000210127 | MT-TA | 0,952684035 | 1,935470119 | 0,08211 | 0,37331809 | mitochondrially encoded tRNA-Ala (GCN) |
| ENSG00000108604 | SMARCD2 | 0,950852677 | 1,933014791 | 1,97E-05 | 0,00152467 | SWI/SNF related, matrix associated, actin dependent regulator of chromatin, subfamily d, member 2 |
| ENSG00000273703 | H2BC14 | 0,950693591 | 1,932801649 | 0,19044 | 0,51350457 | H2B clustered histone 14 |
| ENSG00000110719 | TCIRG1 | 0,950140881 | 1,932061317 | 0,04191 | 0,2772815 | T cell immune regulator 1, ATPase H+ transporting V0 subunit a3 |
| ENSG00000121904 | CSMD2 | 0,949769506 | 1,931564035 | 0,01337 | 0,15144828 | CUB and Sushi multiple domains 2 |
| ENSG00000165874 | SHLD2P1 | 0,949718014 | 1,931495096 | 0,0563 | 0,31687263 | shieldin complex subunit 2 pseudogene 1 |
| ENSG00000131724 | IL13RA1 | 0,949609914 | 1,931350376 | 0,00501 | 0,08821489 | interleukin 13 receptor subunit alpha 1 |
| ENSG00000281566 | AL157778,1 | 0,949609158 | 1,931349364 | 0,00368 | 0,07408435 | novel transcript |
| ENSG00000275146 | AP003499,3 | 0,948650667 | 1,930066648 | 0,30158 | 0,59277108 | |
| ENSG00000240395 | RPL5P23 | 0,948476041 | 1,929833044 | 0,1566 | 0,48425071 | ribosomal protein L5 pseudogene 23 |
| ENSG00000286154 | AC009974,2 | 0,947851994 | 1,928998464 | 0,14038 | 0,46721845 | novel transcript, antisense to CYP27A1 |
| ENSG00000164220 | F2RL2 | 0,947595183 | 1,928655117 | 0,00794 | 0,11357366 | coagulation factor II thrombin receptor like 2 |
| ENSG00000223957 | NEU1 | 0,947571587 | 1,928623573 | 0,10946 | 0,42142456 | neuraminidase 1 |
| ENSG00000282733 | NSFP1 | 0,946984748 | 1,927839234 | 0,37566 | 0,64399236 | N-ethylmaleimide-sensitive factor pseudogene 1 |

| | | | | | | |
|---|---|---|---|---|---|---|
| ENSG00000182326 | C1S | 0,94674689 | 1,927521416 | 0,00167 | 0,04426888 | complement C1s |
| ENSG00000265204 | AC090772,2 | 0,946479892 | 1,927164726 | 0,2192 | 0,53667844 | novel transcript, antisense to TTC39C |
| ENSG00000198108 | CHSY3 | 0,945654125 | 1,926061974 | 0,00055 | 0,02176177 | chondroitin sulfate synthase 3 |
| ENSG00000196932 | TMEM26 | 0,944833046 | 1,924966109 | 0,27798 | 0,57629868 | transmembrane protein 26 |
| ENSG00000287381 | AC012433,2 | 0,944705354 | 1,924795739 | 0,13108 | 0,45580833 | novel transcript, antisense to MAPK4 |
| ENSG00000117791 | MTARC2 | 0,943720376 | 1,923482063 | 0,04385 | 0,28238629 | mitochondrial amidoxime reducing component 2 |
| ENSG00000237234 | Z99289,1 | 0,94364624 | 1,923383224 | 0,27735 | 0,57586315 | novel transcript |
| ENSG00000254715 | OR7E154P | 0,943197761 | 1,92278541 | 0,61971 | 0,80726991 | olfactory receptor family 7 subfamily E member 154 pseudogene |
| ENSG00000143387 | CTSK | 0,942862188 | 1,922338219 | 0,08448 | 0,37732439 | cathepsin K |
| ENSG00000282978 | AC110994,2 | 0,942479771 | 1,92182873 | 0,21939 | 0,53676938 | basic transcription factor 3 (BTF3) pseudogene |
| ENSG00000213189 | BTF3L4P2 | 0,94196069 | 1,92113738 | 0,38013 | 0,64784628 | basic transcription factor 3 like 4 pseudogene 2 |
| ENSG00000276194 | HNF1B | 0,941772824 | 1,920887229 | 0,43052 | 0,68295004 | HNF1 homeobox B |
| ENSG00000259414 | AC242376,1 | 0,941324181 | 1,920289973 | 0,42034 | 0,67570208 | hect domain and RLD 2 pseudogene 7 |
| ENSG00000106153 | CHCHD2 | 0,941310041 | 1,920271152 | 2,41E-06 | 0,00023976 | coiled-coil-helix-coiled-coil-helix domain containing 2 |
| ENSG00000099139 | PCSK5 | 0,941274005 | 1,920223188 | 0,01568 | 0,16409525 | proprotein convertase subtilisin/kexin type 5 |
| ENSG00000232754 | AL035658,1 | 0,941071374 | 1,919953505 | 0,41496 | 0,67178888 | novel transcript, antisense to EP300 |
| ENSG00000231995 | AL590399,4 | 0,940626348 | 1,919361351 | 0,23367 | 0,54563244 | pseudogene similar to part of ribosomal protein L7a (RPL7A) |
| ENSG00000113494 | PRLR | 0,940278112 | 1,918898114 | 0,03633 | 0,25798936 | prolactin receptor |
| ENSG00000254481 | PTP4A2P2 | 0,940017798 | 1,918551906 | 0,05816 | 0,32183657 | PTP4A2 pseudogene 2 |
| ENSG00000204267 | TAP2 | 0,937823944 | 1,915636652 | 0,28538 | 0,58187605 | transporter 2, ATP binding cassette subfamily B member |
| ENSG00000145794 | MEGF10 | 0,937784801 | 1,915584677 | 0,09587 | 0,39884178 | multiple EGF like domains 10 |
| ENSG00000197386 | HTT | 0,93737425 | 1,915039633 | 2,47E-05 | 0,00184973 | huntingtin |
| ENSG00000258337 | AC130895,1 | 0,937124678 | 1,914708379 | 0,24883 | 0,5559185 | novel transcript, sense intronic to MED13L |
| ENSG00000215512 | AP005901,1 | 0,936528095 | 1,913916772 | 0,32662 | 0,60877026 | ankyrin repeat domain 30A (ANKRD30A) pseudogene |
| ENSG00000187475 | H1-6 | 0,936332435 | 1,913657222 | 0,10609 | 0,41557034 | H1,6 linker histone, cluster member |
| ENSG00000285327 | XKR5 | 0,935588843 | 1,912671141 | 0,18658 | 0,509961 | XK related 5 |
| ENSG00000108176 | DNAJC12 | 0,93464598 | 1,911421537 | 0,00077 | 0,02721863 | DnaJ heat shock protein family (Hsp40) member C12 |
| ENSG00000253385 | AP003696,1 | 0,934351184 | 1,911031004 | 0,18032 | 0,50407792 | novel transcript |
| ENSG00000288542 | AL133318,1 | 0,933418175 | 1,909795516 | 0,79017 | 0,9027893 | novel transcript |
| ENSG00000238064 | HCG25 | 0,933300204 | 1,909639356 | 0,34778 | 0,62427624 | HLA complex group 25 |
| ENSG00000225447 | RPS15AP10 | 0,932687835 | 1,908828959 | 0,12591 | 0,448127 | ribosomal protein S15a pseudogene 10 |

| | | | | | | |
|---|---|---|---|---|---|---|
| ENSG00000108039 | XPNPEP1 | 0,932420913 | 1,908475827 | 0,00122 | 0,03692127 | X-prolyl aminopeptidase 1 |
| ENSG00000122035 | RASL11A | 0,93207074 | 1,908012654 | 0,22931 | 0,54277792 | RAS like family 11 member A |
| ENSG00000224481 | AC245100,1 | 0,931998243 | 1,907916778 | 0,27864 | 0,57683444 | novel transcript |
| ENSG00000272361 | AC005014,2 | 0,931159907 | 1,906808428 | 0,27179 | 0,57305058 | novel transcript |
| ENSG00000146373 | RNF217 | 0,931058695 | 1,90667466 | 0,00619 | 0,09912441 | ring finger protein 217 |
| ENSG00000171243 | SOSTDC1 | 0,930277035 | 1,905641893 | 0,1319 | 0,45708171 | sclerostin domain containing 1 |
| ENSG00000219747 | AL133260,1 | 0,929847833 | 1,905075049 | 0,37995 | 0,64769466 | 60S ribosomal protein L32 (RPL32) pseudogene |
| ENSG00000121931 | LRIF1 | 0,929817962 | 1,905035605 | 0,03625 | 0,25760275 | ligand dependent nuclear receptor interacting factor 1 |
| ENSG00000275759 | AC026367,3 | 0,929783634 | 1,904990276 | 0,23385 | 0,54565913 | novel transcript |
| ENSG00000157600 | TMEM164 | 0,929608251 | 1,904758709 | 6,24E-09 | 1,20E-06 | transmembrane protein 164 |
| ENSG00000100101 | Z83844,1 | 0,928991928 | 1,903945164 | 0,69997 | 0,85510522 | novel transcript |
| ENSG00000283753 | RPL21P121 | 0,928807194 | 1,903701384 | 0,29495 | 0,58868965 | ribosomal protein L21 pseudogene 121 |
| ENSG00000237382 | RPL21P121 | 0,928807194 | 1,903701384 | 0,29495 | 0,58868965 | ribosomal protein L21 pseudogene 121 |
| ENSG00000228366 | AL592437,1 | 0,928534711 | 1,903341864 | 0,22602 | 0,54121239 | FUS interacting protein (serine-arginine rich) 1 (FUSIP1) pseudogene |
| ENSG00000131459 | GFPT2 | 0,928139993 | 1,902821185 | 0,0002 | 0,01057544 | glutamine-fructose-6-phosphate transaminase 2 |
| ENSG00000277526 | AC245123,1 | 0,9276707 | 1,902202319 | 0,24367 | 0,55235189 | novel transcript |
| ENSG00000228988 | AL353771,1 | 0,927442482 | 1,901901435 | 0,41921 | 0,67514348 | novel transcript |
| ENSG00000163803 | PLB1 | 0,927380991 | 1,901820374 | 0,05802 | 0,32139417 | phospholipase B1 |
| ENSG00000255062 | AP001318,2 | 0,926973417 | 1,901283169 | 0,09212 | 0,39242415 | novel transcript, antisense to DCPS and TIRAP |
| ENSG00000165675 | ENOX2 | 0,92667819 | 1,900894139 | 0,00143 | 0,04046504 | ecto-NOX disulfide-thiol exchanger 2 |
| ENSG00000165895 | ARHGAP42 | 0,926464445 | 1,900612529 | 0,02122 | 0,19328293 | Rho GTPase activating protein 42 |
| ENSG00000284652 | BX323043,1 | 0,926386093 | 1,90050931 | 0,21318 | 0,53226622 | novel transcript |
| ENSG00000010438 | PRSS3 | 0,926057955 | 1,900077092 | 0,23285 | 0,54547822 | serine protease 3 |
| ENSG00000224599 | BMS1P12 | 0,925882228 | 1,899845668 | 0,02037 | 0,18949014 | BMS1 pseudogene 12 |
| ENSG00000185483 | ROR1 | 0,92575779 | 1,899681807 | 9,70E-05 | 0,00584194 | receptor tyrosine kinase like orphan receptor 1 |
| ENSG00000197665 | AC007952,1 | 0,925351553 | 1,899146966 | 0,2864 | 0,58267699 | novel transcript, antisense to GRAPL |
| ENSG00000164176 | EDIL3 | 0,925289916 | 1,89906583 | 0,00658 | 0,10287723 | EGF like repeats and discoidin domains 3 |
| ENSG00000171054 | OR13H1 | 0,924718403 | 1,898313678 | 0,06593 | 0,33989895 | olfactory receptor family 13 subfamily H member 1 |
| ENSG00000007314 | SCN4A | 0,924324301 | 1,897795185 | 0,11151 | 0,42449053 | sodium voltage-gated channel alpha subunit 4 |
| ENSG00000260645 | AL359715,2 | 0,92406078 | 1,897448567 | 0,04637 | 0,2893324 | novel transcript |
| ENSG00000162551 | ALPL | 0,923770782 | 1,897067196 | 0,04563 | 0,2883604 | alkaline phosphatase, biomineralization associated |
| ENSG00000242612 | DECR2 | 0,923055811 | 1,896127281 | 0,00157 | 0,04253303 | 2,4-dienoyl-CoA reductase 2 |

| | | | | | | |
|---|---|---|---|---|---|---|
| ENSG00000257761 | AC078860,2 | 0,923016289 | 1,896075339 | 0,35716 | 0,63118399 | novel transcript, antisense to LGR5 |
| ENSG00000184494 | NEU1 | 0,922447437 | 1,895327866 | 0,09355 | 0,39523994 | neuraminidase 1 |
| ENSG00000240751 | AC026348,1 | 0,921661699 | 1,894295891 | 0,17303 | 0,4978037 | Rab geranylgeranyltransferase, beta subunit (RABGGTB) pseudogene |
| ENSG00000174482 | LINGO2 | 0,921302296 | 1,893824045 | 0,09272 | 0,39335125 | leucine rich repeat and Ig domain containing 2 |
| ENSG00000139117 | CPNE8 | 0,92088847 | 1,893280894 | 0,0144 | 0,1569305 | copine 8 |
| ENSG00000283352 | AL021368,4 | 0,920250881 | 1,892444356 | 0,33011 | 0,61103391 | novel transcript |
| ENSG00000014257 | ACP3 | 0,920226138 | 1,892411899 | 0,46255 | 0,70358709 | acid phosphatase 3 |
| ENSG00000235030 | IER3 | 0,920053258 | 1,892185143 | 0,00879 | 0,11877365 | immediate early response 3 |
| ENSG00000230128 | IER3 | 0,920053258 | 1,892185143 | 0,00879 | 0,11877365 | immediate early response 3 |
| ENSG00000237155 | IER3 | 0,920053258 | 1,892185143 | 0,00879 | 0,11877365 | immediate early response 3 |
| ENSG00000227231 | IER3 | 0,920053258 | 1,892185143 | 0,00879 | 0,11877365 | immediate early response 3 |
| ENSG00000115705 | TPO | 0,91944345 | 1,891385512 | 0,44696 | 0,69334376 | thyroid peroxidase |
| ENSG00000142733 | MAP3K6 | 0,919383254 | 1,891306595 | 0,00014 | 0,00801207 | mitogen-activated protein kinase kinase kinase 6 |
| ENSG00000287141 | AC093675,2 | 0,919320393 | 1,891224189 | 0,24156 | 0,55039378 | novel transcript, antisense to TMEM87B |
| ENSG00000081803 | CADPS2 | 0,919121787 | 1,890963855 | 0,00182 | 0,0471652 | calcium dependent secretion activator 2 |
| ENSG00000286060 | AL365256,1 | 0,918962672 | 1,890755312 | 0,21581 | 0,53440695 | novel transcript, antisense to IGSF1 |
| ENSG00000270248 | AC011468,4 | 0,917865601 | 1,889318068 | 0,30469 | 0,59445929 | transmembrane protein 50B (TMEM50B) pseudogene |
| ENSG00000249068 | AC008417,1 | 0,917832165 | 1,889274282 | 0,21849 | 0,53578053 | cancer/testis antigen family 45, member A5 pseudogene |
| ENSG00000273261 | AC092953,2 | 0,917382552 | 1,888685585 | 0,37469 | 0,6431311 | novel transcript, antisense to ATP11B |
| ENSG00000224014 | AL390728,3 | 0,917242479 | 1,88850222 | 0,25629 | 0,56078582 | SMT3 suppressor of mif two 3 homolog 3 (yeast) (SUMO3) pseudogene |
| ENSG00000200966 | RN7SKP87 | 0,917142486 | 1,888371331 | 0,34137 | 0,61943649 | RN7SK pseudogene 87 |
| ENSG00000276461 | TCF20 | 0,917087302 | 1,888299102 | 0,1442 | 0,47175088 | transcription factor 20 |
| ENSG00000262024 | TCF20 | 0,917087302 | 1,888299102 | 0,1442 | 0,47175088 | transcription factor 20 |
| ENSG00000118520 | ARG1 | 0,917024042 | 1,888216304 | 0,2414 | 0,55033747 | arginase 1 |
| ENSG00000204711 | C9orf135 | 0,916284246 | 1,887248299 | 0,11184 | 0,42531464 | chromosome 9 open reading frame 135 |
| ENSG00000219902 | RPL35P3 | 0,915623551 | 1,886384215 | 0,35187 | 0,62699036 | ribosomal protein L35 pseudogene 3 |
| ENSG00000263797 | AP005210,1 | 0,915454313 | 1,886162942 | 0,13765 | 0,46480929 | novel transcript, antisense to ARHGAP28 |
| ENSG00000169504 | CLIC4 | 0,915351177 | 1,886028108 | 0,06623 | 0,3406271 | chloride intracellular channel 4 |
| ENSG00000271874 | AC025754,2 | 0,914909488 | 1,885450778 | 0,42281 | 0,67683766 | novel transcript |
| ENSG00000154319 | FAM167A | 0,914892232 | 1,885428227 | 0,02991 | 0,23288546 | family with sequence similarity 167 member A |
| ENSG00000206256 | PPT2 | 0,914724206 | 1,885208649 | 0,00639 | 0,10091712 | palmitoyl-protein thioesterase 2 |

| | | | | | | |
|---|---|---|---|---|---|---|
| ENSG00000042781 | USH2A | 0,914374271 | 1,884751436 | 0,119 | 0,43696075 | usherin |
| ENSG00000249787 | AC113385,1 | 0,914345132 | 1,884713369 | 0,2171 | 0,53509398 | novel transcript |
| ENSG00000266648 | SETP3 | 0,914121158 | 1,884420795 | 0,21261 | 0,53153879 | SET pseudogene 3 |
| ENSG00000288438 | MG910338,3 | 0,913995354 | 1,88425648 | 0,29412 | 0,58841393 | uncharacterized LOC102723360 |
| ENSG00000170561 | IRX2 | 0,913613873 | 1,883758305 | 0,07891 | 0,36683128 | iroquois homeobox 2 |
| ENSG00000165338 | HECTD2 | 0,913573962 | 1,883706194 | 0,03647 | 0,25869664 | HECT domain E3 ubiquitin protein ligase 2 |
| ENSG00000125845 | BMP2 | 0,913157205 | 1,883162119 | 0,00396 | 0,07705722 | bone morphogenetic protein 2 |
| ENSG00000124813 | RUNX2 | 0,912948083 | 1,88288917 | 0,05987 | 0,32540929 | RUNX family transcription factor 2 |
| ENSG00000275935 | LENG1 | 0,91271925 | 1,88259054 | 0,5565 | 0,76843732 | leukocyte receptor cluster member 1 |
| ENSG00000277457 | LENG1 | 0,91271925 | 1,88259054 | 0,5565 | 0,76843732 | leukocyte receptor cluster member 1 |
| ENSG00000276628 | LENG1 | 0,91271925 | 1,88259054 | 0,5565 | 0,76843732 | leukocyte receptor cluster member 1 |
| ENSG00000278257 | LENG1 | 0,91271925 | 1,88259054 | 0,5565 | 0,76843732 | leukocyte receptor cluster member 1 |
| ENSG00000278426 | LENG1 | 0,91271925 | 1,88259054 | 0,5565 | 0,76843732 | leukocyte receptor cluster member 1 |
| ENSG00000276774 | LENG1 | 0,91271925 | 1,88259054 | 0,5565 | 0,76843732 | leukocyte receptor cluster member 1 |
| ENSG00000214185 | XPOTP1 | 0,912364933 | 1,882128243 | 0,39414 | 0,65707455 | exportin for tRNA pseudogene 1 |
| ENSG00000104213 | PDGFRL | 0,911668527 | 1,881219937 | 0,11205 | 0,42549415 | platelet derived growth factor receptor like |
| ENSG00000243422 | RPL23AP49 | 0,911574647 | 1,881097525 | 0,06055 | 0,3267765 | ribosomal protein L23a pseudogene 49 |
| ENSG00000248896 | AC105001,1 | 0,910983954 | 1,880327491 | 0,15802 | 0,48506908 | novel transcript, antisense to SOX7 and PINX1 |
| ENSG00000189152 | GRAPL | 0,910981736 | 1,8803246 | 0,25899 | 0,56278415 | GRB2 related adaptor protein like |
| ENSG00000270933 | AC010719,1 | 0,910847332 | 1,880149435 | 0,21143 | 0,53032236 | novel transcript |
| ENSG00000242361 | HLA-DMA | 0,910844901 | 1,880146266 | 0,03033 | 0,23505966 | major histocompatibility complex, class II, DM alpha |
| ENSG00000075223 | SEMA3C | 0,910680631 | 1,879932199 | 0,01713 | 0,17198199 | semaphorin 3C |
| ENSG00000253523 | AC104350,1 | 0,9106558 | 1,879899843 | 0,25175 | 0,55773568 | novel transcript |
| ENSG00000157782 | CABP1 | 0,910285149 | 1,879416929 | 0,12932 | 0,45283544 | calcium binding protein 1 |
| ENSG00000206557 | TRIM71 | 0,91022054 | 1,879332764 | 0,01204 | 0,14273718 | tripartite motif containing 71 |
| ENSG00000120519 | SLC10A7 | 0,910156727 | 1,879249639 | 0,07327 | 0,35655334 | solute carrier family 10 member 7 |
| ENSG00000238133 | MAP3K20-AS1 | 0,909561668 | 1,878474677 | 0,1295 | 0,45306297 | MAP3K20 antisense RNA 1 |
| ENSG00000235619 | RPL36AP33 | 0,908515313 | 1,877112755 | 0,36839 | 0,63803074 | ribosomal protein L36a pseudogene 33 |
| ENSG00000138119 | MYOF | 0,908417952 | 1,876986082 | 0,07708 | 0,36384125 | myoferlin |
| ENSG00000269903 | AC025165,4 | 0,908255057 | 1,876774162 | 0,17772 | 0,50205238 | novel transcript |
| ENSG00000111199 | TRPV4 | 0,908090539 | 1,876560157 | 0,22116 | 0,53805234 | transient receptor potential cation channel subfamily V member 4 |
| ENSG00000197329 | PELI1 | 0,908090341 | 1,876559898 | 0,02641 | 0,21889802 | pellino E3 ubiquitin protein ligase 1 |

| | | | | | | |
|---|---|---|---|---|---|---|
| ENSG00000104081 | BMF | 0,908069932 | 1,876533353 | 0,03233 | 0,24253636 | Bcl2 modifying factor |
| ENSG00000283681 | TCF20 | 0,908039744 | 1,876494087 | 0,14524 | 0,47289051 | transcription factor 20 |
| ENSG00000283026 | TCF20 | 0,908039744 | 1,876494087 | 0,14524 | 0,47289051 | transcription factor 20 |
| ENSG00000281897 | TCF20 | 0,908039744 | 1,876494087 | 0,14524 | 0,47289051 | transcription factor 20 |
| ENSG00000253521 | HPYR1 | 0,90792062 | 1,87633915 | 0,20443 | 0,52437248 | Helicobacter pylori responsive 1 |
| ENSG00000204446 | LINC02872 | 0,907522308 | 1,875821185 | 0,25648 | 0,56092974 | long intergenic non-protein coding RNA 2872 |
| ENSG00000123268 | ATF1 | 0,907517738 | 1,875815243 | 0,02616 | 0,21795549 | activating transcription factor 1 |
| ENSG00000198153 | ZNF849P | 0,907085666 | 1,87525354 | 0,31809 | 0,60349082 | zinc finger protein 849, pseudogene |
| ENSG00000125510 | OPRL1 | 0,906570131 | 1,874583554 | 0,00341 | 0,07036709 | opioid related nociceptin receptor 1 |
| ENSG00000270115 | AL513327,2 | 0,906106606 | 1,873981363 | 0,06651 | 0,34124715 | novel transcript |
| ENSG00000138639 | ARHGAP24 | 0,905934491 | 1,873757809 | 0,0666 | 0,34143033 | Rho GTPase activating protein 24 |
| ENSG00000182752 | PAPPA | 0,905290021 | 1,872920965 | 0,06933 | 0,34785467 | pappalysin 1 |
| ENSG00000140968 | IRF8 | 0,904813808 | 1,872302843 | 0,40279 | 0,66396567 | interferon regulatory factor 8 |
| ENSG00000179715 | PCED1B | 0,903988039 | 1,871231481 | 0,09949 | 0,40608287 | PC-esterase domain containing 1B |
| ENSG00000213185 | FAM24B | 0,903355367 | 1,870411061 | 0,29749 | 0,59034513 | family with sequence similarity 24 member B |
| ENSG00000282892 | TCF20 | 0,902661047 | 1,869511113 | 0,14118 | 0,4679289 | transcription factor 20 |
| ENSG00000280467 | TCF20 | 0,902661047 | 1,869511113 | 0,14118 | 0,4679289 | transcription factor 20 |
| ENSG00000141540 | TTYH2 | 0,902514594 | 1,869321342 | 0,00069 | 0,02538363 | tweety family member 2 |
| ENSG00000122477 | LRRC39 | 0,902199715 | 1,868913392 | 0,07661 | 0,36266919 | leucine rich repeat containing 39 |
| ENSG00000235947 | EGOT | 0,901586442 | 1,868119109 | 0,37938 | 0,64719976 | eosinophil granule ontogeny transcript |
| ENSG00000276542 | AC097478,2 | 0,901583374 | 1,868115135 | 0,19341 | 0,51546367 | novel transcript, antisense to SNCA |
| ENSG00000205832 | C16orf96 | 0,901436247 | 1,867924633 | 0,01039 | 0,13135977 | chromosome 16 open reading frame 96 |
| ENSG00000127528 | KLF2 | 0,900696411 | 1,866966979 | 0,33391 | 0,61382024 | Kruppel like factor 2 |
| ENSG00000136960 | ENPP2 | 0,900680125 | 1,866945904 | 0,07881 | 0,36667857 | ectonucleotide pyrophosphatase/phosphodiesterase 2 |
| ENSG00000288243 | STING1 | 0,900542345 | 1,866767616 | 0,35894 | 0,63196414 | stimulator of interferon response cGAMP interactor 1 |
| ENSG00000154608 | CEP170P1 | 0,900170611 | 1,866286675 | 0,11635 | 0,43288152 | centrosomal protein 170 pseudogene 1 |
| ENSG00000163666 | HESX1 | 0,899848617 | 1,865870185 | 0,08047 | 0,37018111 | HESX homeobox 1 |
| ENSG00000156103 | MMP16 | 0,898611364 | 1,864270705 | 0,04575 | 0,28848581 | matrix metallopeptidase 16 |
| ENSG00000178522 | AMBN | 0,898554193 | 1,864196829 | 0,3205 | 0,6053879 | ameloblastin |
| ENSG00000103710 | RASL12 | 0,898521375 | 1,864154423 | 0,15894 | 0,48628471 | RAS like family 12 |
| ENSG00000227001 | NBPF2P | 0,897348109 | 1,862639022 | 0,25125 | 0,55719434 | NBPF member 2, pseudogene |
| ENSG00000233904 | HLA-E | 0,89719114 | 1,862436374 | 0,16625 | 0,49212363 | major histocompatibility complex, class I, E |
| ENSG00000229252 | HLA-E | 0,89719114 | 1,862436374 | 0,16625 | 0,49212363 | major histocompatibility complex, class I, E |

| | | | | | | |
|---|---|---|---|---|---|---|
| ENSG00000259763 | LINC02157 | 0,897097954 | 1,86231608 | 0,30923 | 0,59774312 | long intergenic non-protein coding RNA 2157 |
| ENSG00000288198 | PPFIA1 | 0,896788044 | 1,861916073 | 0,57742 | 0,78140182 | PTPRF interacting protein alpha 1 |
| ENSG00000258631 | AC110023,1 | 0,896456373 | 1,861488073 | 0,13943 | 0,46666406 | novel transcript |
| ENSG00000271156 | AC090578,2 | 0,895778023 | 1,860613014 | 0,42352 | 0,67732256 | novel transcript |
| ENSG00000277763 | AL138995,1 | 0,895631639 | 1,860424235 | 0,35584 | 0,63013888 | novel transcript |
| ENSG00000266921 | AC006213,1 | 0,89505626 | 1,859682405 | 0,17962 | 0,5039322 | novel transcript, antisense to ZNF155 |
| ENSG00000183873 | SCN5A | 0,89435062 | 1,858773033 | 0,17414 | 0,49867554 | sodium voltage-gated channel alpha subunit 5 |
| ENSG00000137767 | SQOR | 0,894231814 | 1,85861997 | 0,23335 | 0,54563244 | sulfide quinone oxidoreductase |
| ENSG00000240875 | LINC00886 | 0,894210119 | 1,858592021 | 0,08958 | 0,38705587 | long intergenic non-protein coding RNA 886 |
| ENSG00000232192 | KIF26B-AS1 | 0,894193819 | 1,858571022 | 0,27107 | 0,57254873 | KIF26B antisense RNA 1 |
| ENSG00000107159 | CA9 | 0,893782623 | 1,858041368 | 0,10407 | 0,41243906 | carbonic anhydrase 9 |
| ENSG00000225505 | AC104332,1 | 0,893518176 | 1,857700819 | 0,10404 | 0,41241152 | novel pseudogene |
| ENSG00000105810 | CDK6 | 0,893373132 | 1,857514062 | 0,02711 | 0,22199012 | cyclin dependent kinase 6 |
| ENSG00000234450 | AC005534,1 | 0,893039368 | 1,85708438 | 0,33436 | 0,6142045 | ribosomal protein L26-like 1 (RPL26L1) pseudogene |
| ENSG00000248936 | AC027607,1 | 0,892658095 | 1,856593658 | 0,2774 | 0,57586315 | putatitive novel transcript |
| ENSG00000175746 | C15orf54 | 0,892570709 | 1,856481204 | 0,30449 | 0,59430086 | chromosome 15 putative open reading frame 54 |
| ENSG00000108798 | ABI3 | 0,892196254 | 1,855999413 | 0,13703 | 0,4642578 | ABI family member 3 |
| ENSG00000145703 | IQGAP2 | 0,892076525 | 1,85584539 | 0,01249 | 0,14623734 | IQ motif containing GTPase activating protein 2 |
| ENSG00000272525 | AC099522,2 | 0,891613549 | 1,855249925 | 0,05657 | 0,31735962 | novel transcript |
| ENSG00000230254 | HLA-E | 0,891370853 | 1,854937854 | 0,16201 | 0,48916554 | major histocompatibility complex, class I, E |
| ENSG00000256030 | CBX3P4 | 0,891347185 | 1,854907423 | 0,25993 | 0,56377449 | chromobox 3 pseudogene 4 |
| ENSG00000121690 | DEPDC7 | 0,890756108 | 1,854147617 | 0,03143 | 0,23928335 | DEP domain containing 7 |
| ENSG00000166391 | MOGAT2 | 0,890550913 | 1,85388392 | 0,44978 | 0,69538334 | monoacylglycerol O-acyltransferase 2 |
| ENSG00000162946 | DISC1 | 0,890515369 | 1,853838246 | 0,00022 | 0,01111111 | DISC1 scaffold protein |
| ENSG00000282840 | AC004554,1 | 0,890002688 | 1,853179577 | 0,25547 | 0,55995963 | novel transcript |
| ENSG00000166265 | CYYR1 | 0,889538738 | 1,852583716 | 0,05796 | 0,32124805 | cysteine and tyrosine rich 1 |
| ENSG00000240350 | SOCAR | 0,889527122 | 1,852568801 | 0,10925 | 0,42109719 | serous ovarian cancer associated RNA |
| ENSG00000266469 | AC005288,1 | 0,889304881 | 1,852283443 | 0,25996 | 0,56377449 | novel transcript, antisense to MED1 & FBXL20 |
| ENSG00000271980 | AC012640,4 | 0,889243726 | 1,852204927 | 0,19492 | 0,51588752 | novel transcript, antisense to CCT5 |
| ENSG00000188649 | CC2D2B | 0,889182033 | 1,852125724 | 0,28366 | 0,58072192 | coiled-coil and C2 domain containing 2B |
| ENSG00000139289 | PHLDA1 | 0,888933525 | 1,851806717 | 0,0021 | 0,05163115 | pleckstrin homology like domain family A member 1 |
| ENSG00000149948 | HMGA2 | 0,888604799 | 1,851384821 | 0,01286 | 0,14834544 | high mobility group AT-hook 2 |
| ENSG00000258565 | BLZF2P | 0,888485859 | 1,851232194 | 0,34036 | 0,61897407 | basic leucine zipper nuclear factor 2, pseudogene |

| | | | | | | |
|---|---|---|---|---|---|---|
| ENSG00000270426 | AC099343,2 | 0,888209282 | 1,850877331 | 0,13216 | 0,45719971 | novel transcript, sense intronic to IRF2 |
| ENSG00000288516 | PRKACA | 0,887474738 | 1,849935202 | 0,04466 | 0,28548113 | protein kinase cAMP-activated catalytic subunit alpha |
| ENSG00000241014 | AC114490,1 | 0,88723629 | 1,84962947 | 0,14235 | 0,46945664 | novel platelet-activating factor receptor family pseudogene |
| ENSG00000274001 | AL512506,1 | 0,887165453 | 1,849538655 | 0,381 | 0,64851669 | novel transcript, sense intronic to CCDC122 |
| ENSG00000115935 | WIPF1 | 0,886959203 | 1,849274261 | 0,00374 | 0,07458671 | WAS/WASL interacting protein family member 1 |
| ENSG00000274598 | AC087893,2 | 0,886252727 | 1,848368908 | 0,17482 | 0,49908125 | novel transcript, sense intronic to SSH1 |
| ENSG00000286882 | AC116428,1 | 0,886235121 | 1,848346352 | 0,32587 | 0,60842402 | novel transcript, antisense to FER |
| ENSG00000106546 | AHR | 0,886168183 | 1,848260594 | 0,07671 | 0,362924 | aryl hydrocarbon receptor |
| ENSG00000251095 | AC093866,1 | 0,8861211 | 1,848200277 | 0,14742 | 0,47491949 | novel transcript |
| ENSG00000103335 | PIEZO1 | 0,885329179 | 1,847186045 | 0,00818 | 0,11480072 | piezo type mechanosensitive ion channel component 1 |
| ENSG00000183785 | TUBA8 | 0,885061721 | 1,846843631 | 0,16174 | 0,48914394 | tubulin alpha 8 |
| ENSG00000105289 | TJP3 | 0,884737688 | 1,846428872 | 0,15165 | 0,47995077 | tight junction protein 3 |
| ENSG00000286506 | FP325317,2 | 0,884335551 | 1,845914269 | 0,45762 | 0,70082333 | methylenetetrahydrofolate dehydrogenase (NADP+ dependent) 1 like (MTHFD1L) pseudogene |
| ENSG00000225507 | AC069282,1 | 0,882690015 | 1,843810023 | 0,22004 | 0,53747564 | pseudogene similar to part of TTK protein kinase TTK |
| ENSG00000090104 | RGS1 | 0,88244515 | 1,843497104 | 0,424 | 0,67778738 | regulator of G protein signaling 1 |
| ENSG00000226435 | ANKRD18DP | 0,882004862 | 1,842934583 | 0,4274 | 0,68046596 | ankyrin repeat domain 18D, pseudogene |
| ENSG00000234683 | AL138799,2 | 0,88188155 | 1,842777069 | 0,52633 | 0,74973856 | ribosomal protein S20 (RPS20) pseudogene |
| ENSG00000149177 | PTPRJ | 0,881778966 | 1,842646041 | 0,0045 | 0,08275926 | protein tyrosine phosphatase receptor type J |
| ENSG00000155090 | KLF10 | 0,881297628 | 1,842031366 | 0,10006 | 0,40666505 | Kruppel like factor 10 |
| ENSG00000196628 | TCF4 | 0,881216604 | 1,841927917 | 0,00256 | 0,05892196 | transcription factor 4 |
| ENSG00000113248 | PCDHB15 | 0,881082734 | 1,84175701 | 0,00079 | 0,0274628 | protocadherin beta 15 |
| ENSG00000177359 | AC024940,1 | 0,880568253 | 1,841100336 | 0,21692 | 0,53505085 | ovostatin 2 |
| ENSG00000204568 | MRPS18B | 0,880274503 | 1,840725504 | 0,23584 | 0,5471053 | mitochondrial ribosomal protein S18B |
| ENSG00000285162 | AC004593,2 | 0,879975422 | 1,840343948 | 0,32234 | 0,60577115 | chimerin 2 |
| ENSG00000108384 | RAD51C | 0,879608492 | 1,839875941 | 5,53E-07 | 6,61E-05 | RAD51 paralog C |
| ENSG00000049246 | PER3 | 0,878519438 | 1,838487589 | 0,00162 | 0,0432168 | period circadian regulator 3 |
| ENSG00000258790 | AL121594,1 | 0,877433968 | 1,83710485 | 0,14533 | 0,47300165 | novel transcript |
| ENSG00000101438 | SLC32A1 | 0,877338384 | 1,836983139 | 0,20175 | 0,52216179 | solute carrier family 32 member 1 |
| ENSG00000255970 | LINC02421 | 0,877304622 | 1,83694015 | 0,17772 | 0,50205238 | long intergenic non-protein coding RNA 2421 |
| ENSG00000233330 | AL078581,2 | 0,87716826 | 1,836766533 | 0,25938 | 0,56317998 | novel transcript |
| ENSG00000148482 | SLC39A12 | 0,877141982 | 1,836733077 | 0,32995 | 0,6108747 | solute carrier family 39 member 12 |

| | | | | | | |
|---|---|---|---|---|---|---|
| ENSG00000142089 | IFITM3 | 0,876359268 | 1,835736853 | 0,00321 | 0,06789583 | interferon induced transmembrane protein 3 |
| ENSG00000227267 | AC072039,1 | 0,875394417 | 1,834509552 | 0,31223 | 0,59925736 | downstream neighbor of SON (DONSON) pseudogene |
| ENSG00000174080 | CTSF | 0,875214773 | 1,834281134 | 0,00058 | 0,02257221 | cathepsin F |
| ENSG00000261737 | CLCA4-AS1 | 0,875001814 | 1,834010393 | 0,24995 | 0,55657567 | CLCA4 antisense RNA 1 |
| ENSG00000144218 | AFF3 | 0,874945428 | 1,833938714 | 0,00778 | 0,11221258 | AF4/FMR2 family member 3 |
| ENSG00000185339 | TCN2 | 0,874167281 | 1,832949808 | 0,00046 | 0,01927018 | transcobalamin 2 |
| ENSG00000188985 | DHFRP1 | 0,873809602 | 1,832495432 | 0,56339 | 0,7724391 | dihydrofolate reductase pseudogene 1 |
| ENSG00000255760 | LINC02422 | 0,873665552 | 1,83231247 | 0,24824 | 0,55542983 | long intergenic non-protein coding RNA 2422 |
| ENSG00000148935 | GAS2 | 0,873553554 | 1,832170232 | 0,04823 | 0,29503004 | growth arrest specific 2 |
| ENSG00000184564 | SLITRK6 | 0,873267123 | 1,83180651 | 0,12525 | 0,44701864 | SLIT and NTRK like family member 6 |
| ENSG00000231007 | CDC20P1 | 0,87312792 | 1,831629772 | 0,04792 | 0,29413956 | cell division cycle 20 pseudogene 1 |
| ENSG00000236430 | KRT8P29 | 0,872980391 | 1,83144248 | 0,24347 | 0,55221671 | keratin 8 pseudogene 29 |
| ENSG00000251111 | FCF1P8 | 0,872798469 | 1,831211552 | 0,18913 | 0,51249536 | FCF1 pseudogene 8 |
| ENSG00000183386 | FHL3 | 0,872550435 | 1,83089675 | 0,00116 | 0,03569587 | four and a half LIM domains 3 |
| ENSG00000179460 | EEF1A1P27 | 0,872108814 | 1,830336382 | 0,48082 | 0,71789648 | eukaryotic translation elongation factor 1 alpha 1 pseudogene 27 |
| ENSG00000231573 | HCG25 | 0,872100022 | 1,830325228 | 0,37782 | 0,64626684 | HLA complex group 25 |
| ENSG00000262171 | AC130651,1 | 0,871894485 | 1,830064485 | 0,25499 | 0,55947845 | novel transcript, antisense to C16orf63 |
| ENSG00000282412 | AC243629,3 | 0,871894485 | 1,830064485 | 0,25499 | 0,55947845 | novel transcript, antisense to C16orf63 |
| ENSG00000182263 | FIGN | 0,870946564 | 1,828862439 | 0,04991 | 0,30075789 | fidgetin, microtubule severing factor |
| ENSG00000262292 | AC007952,5 | 0,870590773 | 1,828411468 | 0,23795 | 0,54864017 | novel transcript |
| ENSG00000272248 | AL138831,2 | 0,870313977 | 1,828060702 | 0,17636 | 0,50050439 | novel transcript |
| ENSG00000140287 | HDC | 0,869984931 | 1,827643811 | 0,16444 | 0,49065086 | histidine decarboxylase |
| ENSG00000223864 | NPM1P19 | 0,86990676 | 1,827544785 | 0,21878 | 0,53630871 | nucleophosmin 1 pseudogene 19 |
| ENSG00000105849 | POLR1F | 0,869758251 | 1,827356669 | 0,07272 | 0,35572609 | RNA polymerase I subunit F |
| ENSG00000285129 | RPS3AP30 | 0,869749548 | 1,827345645 | 0,30754 | 0,59664516 | RPS3A pseudogene 30 |
| ENSG00000274943 | AC079684,1 | 0,868525123 | 1,825795423 | 0,26072 | 0,56400103 | novel transcript, sense intronic to PPHLN1 |
| ENSG00000266306 | AC061975,5 | 0,868397003 | 1,825633288 | 0,39705 | 0,65962661 | leucine rich repeat containing 37, member A3 (LRRC37A3) pseudogene |
| ENSG00000272472 | AL512283,1 | 0,868179914 | 1,825358597 | 0,26269 | 0,56575908 | novel transcript |
| ENSG00000140416 | TPM1 | 0,867111909 | 1,824007813 | 0,0041 | 0,07854299 | tropomyosin 1 |
| ENSG00000286370 | AC064856,1 | 0,867072913 | 1,823958511 | 0,01572 | 0,16443194 | novel transcript |
| ENSG00000197837 | H4-16 | 0,866933593 | 1,823782381 | 0,05107 | 0,30386013 | H4 histone 16 |

| | | | | | | |
|---|---|---|---|---|---|---|
| ENSG00000237719 | Z95152,1 | 0,866565055 | 1,823316553 | 0,18887 | 0,51222043 | nudix (nucleoside diphosphate linked moiety X) type 5 motif (NUDT5) pseudogene |
| ENSG00000147010 | SH3KBP1 | 0,865889534 | 1,822463012 | 0,01605 | 0,16606232 | SH3 domain containing kinase binding protein 1 |
| ENSG00000170854 | RIOX2 | 0,865777702 | 1,822321747 | 0,07498 | 0,3592137 | ribosomal oxygenase 2 |
| ENSG00000072952 | IRAG1 | 0,865256402 | 1,821663392 | 0,01565 | 0,164013 | inositol 1,4,5-triphosphate receptor associated 1 |
| ENSG00000125810 | CD93 | 0,864736501 | 1,821007042 | 0,20716 | 0,52670822 | CD93 molecule |
| ENSG00000250102 | LINC02377 | 0,864641012 | 1,820886517 | 0,28056 | 0,57800643 | long intergenic non-protein coding RNA 2377 |
| ENSG00000151726 | ACSL1 | 0,864426602 | 1,820615921 | 0,15994 | 0,48710442 | acyl-CoA synthetase long chain family member 1 |
| ENSG00000226797 | AC005828,1 | 0,86407753 | 1,820175462 | 0,33184 | 0,61186662 | novel transcript, antisense to TANC2 |
| ENSG00000266680 | AL135905,1 | 0,864024703 | 1,820108814 | 0,10966 | 0,42180924 | novel transcript, antisense to PTP4A1 |
| ENSG00000185674 | LYG2 | 0,863931721 | 1,81999151 | 0,30218 | 0,59294633 | lysozyme g2 |
| ENSG00000234546 | LNCTAM34A | 0,86380382 | 1,819830168 | 0,37952 | 0,64731477 | long non coding transcriptional activator of miR34a |
| ENSG00000106078 | COBL | 0,86359285 | 1,819564068 | 0,0521 | 0,30670474 | cordon-bleu WH2 repeat protein |
| ENSG00000251188 | AC079140,3 | 0,863423795 | 1,819350864 | 0,18135 | 0,50489668 | novel zinc finger protein pseudogene |
| ENSG00000188820 | CALHM6 | 0,863140417 | 1,818993537 | 0,24617 | 0,55394307 | calcium homeostasis modulator family member 6 |
| ENSG00000258471 | AL161668,3 | 0,862652968 | 1,81837905 | 0,196 | 0,51680811 | novel transcript, antisense to SLC39A2 |
| ENSG00000101955 | SRPX | 0,861050978 | 1,816361016 | 0,04963 | 0,29985549 | sushi repeat containing protein X-linked |
| ENSG00000260808 | AP003096,1 | 0,861031661 | 1,816336695 | 0,19361 | 0,51547251 | novel transcript |
| ENSG00000218996 | ARL4AP5 | 0,860524994 | 1,81569892 | 0,13597 | 0,46294575 | ADP ribosylation factor like GTPase 4A pseudogene 5 |
| ENSG00000107736 | CDH23 | 0,860088942 | 1,815150211 | 0,01438 | 0,15682082 | cadherin related 23 |
| ENSG00000106366 | SERPINE1 | 0,85955058 | 1,814472988 | 0,29299 | 0,58745499 | serpin family E member 1 |
| ENSG00000126458 | RRAS | 0,859547932 | 1,814469658 | 0,063 | 0,33231464 | RAS related |
| ENSG00000155657 | TTN | 0,858977372 | 1,81375221 | 0,03604 | 0,25711935 | titin |
| ENSG00000119715 | ESRRB | 0,858951511 | 1,813719698 | 0,15675 | 0,48425071 | estrogen related receptor beta |
| ENSG00000283852 | BCAP29 | 0,858576342 | 1,813248105 | 0,42636 | 0,67986793 | B cell receptor associated protein 29 |
| ENSG00000185681 | MORN5 | 0,858125423 | 1,812681459 | 0,4649 | 0,70550186 | MORN repeat containing 5 |
| ENSG00000214688 | C10orf105 | 0,857897066 | 1,812394561 | 0,08762 | 0,38323183 | chromosome 10 open reading frame 105 |
| ENSG00000254221 | PCDHGB1 | 0,857422709 | 1,811798744 | 0,00147 | 0,04107269 | protocadherin gamma subfamily B, 1 |
| ENSG00000103472 | RRN3P2 | 0,857411243 | 1,811784344 | 0,15448 | 0,48266733 | RRN3 pseudogene 2 |
| ENSG00000185933 | CALHM1 | 0,857292824 | 1,811635636 | 0,22569 | 0,54090885 | calcium homeostasis modulator 1 |
| ENSG00000219529 | AP000580,1 | 0,856844402 | 1,811072627 | 0,23457 | 0,54623956 | ribosomal protein S20 (RPS20) pseudogene |
| ENSG00000273841 | TAF9 | 0,856373824 | 1,810481989 | 0,13426 | 0,46055924 | TATA-box binding protein associated factor 9 |
| ENSG00000016082 | ISL1 | 0,85621255 | 1,810279611 | 0,18045 | 0,50407792 | ISL LIM homeobox 1 |

| | | | | | | |
|---|---|---|---|---|---|---|
| ENSG00000139800 | ZIC5 | 0,856144636 | 1,810194396 | 0,02006 | 0,18786958 | Zic family member 5 |
| ENSG00000260192 | LINC02240 | 0,855959339 | 1,809961914 | 0,27249 | 0,57315552 | long intergenic non-protein coding RNA 2240 |
| ENSG00000275038 | AC091980,2 | 0,85452529 | 1,808163692 | 0,41891 | 0,67491438 | novel transcript, antisense to TLX3 |
| ENSG00000286634 | AC146507,3 | 0,854303903 | 1,807886244 | 0,11436 | 0,42986344 | novel transcript, sense intronic to NSD1 |
| ENSG00000272755 | AC245297,2 | 0,854176423 | 1,807726502 | 0,10532 | 0,41450376 | novel transcript |
| ENSG00000261019 | AC010132,4 | 0,853664323 | 1,807084944 | 0,11708 | 0,43438325 | novel transcript |
| ENSG00000271597 | AC112230,1 | 0,853048702 | 1,806313996 | 0,35614 | 0,63035942 | golgi reassembly stacking protein 2, 55kDa (GORASP2) pseudogene |
| ENSG00000116701 | NCF2 | 0,85282146 | 1,806029502 | 0,08976 | 0,38749519 | neutrophil cytosolic factor 2 |
| ENSG00000267264 | AC006504,3 | 0,851929947 | 1,804913812 | 0,30427 | 0,59424503 | vomeronasal receptor pseudogene |
| ENSG00000145864 | GABRB2 | 0,850865931 | 1,803583143 | 0,04119 | 0,27486753 | gamma-aminobutyric acid type A receptor subunit beta2 |
| ENSG00000265533 | AC114689,3 | 0,850844626 | 1,803556508 | 0,35494 | 0,62943538 | novel transcript |
| ENSG00000100097 | LGALS1 | 0,850440124 | 1,803050899 | 0,00444 | 0,08217356 | galectin 1 |
| ENSG00000171132 | PRKCE | 0,849973052 | 1,802467257 | 0,00261 | 0,05961957 | protein kinase C epsilon |
| ENSG00000235105 | AL356968,2 | 0,849968637 | 1,802461741 | 0,12644 | 0,44907902 | amyotrophic lateral sclerosis 2 (juvenile) chromosome region, candidate 2 (ALS2CR2) pseudogene |
| ENSG00000259937 | AC023158,1 | 0,849474867 | 1,801844945 | 0,41596 | 0,67284446 | novel transcript, sense intronic to SYT10 |
| ENSG00000226964 | RHEBP2 | 0,849412617 | 1,8017672 | 0,08296 | 0,37497922 | RHEB pseudogene 2 |
| ENSG00000049245 | VAMP3 | 0,849214501 | 1,801519792 | 0,03365 | 0,24782119 | vesicle associated membrane protein 3 |
| ENSG00000238120 | LINC01589 | 0,84916013 | 1,801451898 | 0,18373 | 0,50744379 | long intergenic non-protein coding RNA 1589 |
| ENSG00000274750 | H3C6 | 0,8489534 | 1,801193779 | 0,10052 | 0,40713342 | H3 clustered histone 6 |
| ENSG00000231875 | AL359885,1 | 0,848855304 | 1,801071311 | 0,3675 | 0,63748617 | novel pseudogene |
| ENSG00000173890 | GPR160 | 0,848133537 | 1,800170477 | 0,1486 | 0,47609532 | G protein-coupled receptor 160 |
| ENSG00000183908 | LRRC55 | 0,84774086 | 1,799680569 | 0,16067 | 0,48807056 | leucine rich repeat containing 55 |
| ENSG00000118412 | CASP8AP2 | 0,846405636 | 1,798015722 | 0,209 | 0,5280176 | caspase 8 associated protein 2 |
| ENSG00000258653 | AC005520,1 | 0,845923539 | 1,797414991 | 0,11465 | 0,43014087 | novel protein |
| ENSG00000118985 | ELL2 | 0,845808715 | 1,797271939 | 0,04925 | 0,29853148 | elongation factor for RNA polymerase II 2 |
| ENSG00000237441 | RGL2 | 0,845291303 | 1,796627477 | 0,09985 | 0,40629454 | ral guanine nucleotide dissociation stimulator like 2 |
| ENSG00000233836 | AC139769,1 | 0,845082154 | 1,796367037 | 0,04534 | 0,28773055 | zinc finger protein 726 pseudogene 1 |
| ENSG00000136111 | TBC1D4 | 0,844905715 | 1,796147358 | 0,00228 | 0,05474871 | TBC1 domain family member 4 |
| ENSG00000274849 | AC023043,4 | 0,844645348 | 1,795823231 | 0,285 | 0,58171017 | novel transcript, sense intronic to MOCOS |
| ENSG00000164142 | FAM160A1 | 0,844241221 | 1,795320257 | 0,03401 | 0,24897185 | family with sequence similarity 160 member A1 |
| ENSG00000246898 | LINC00920 | 0,844232322 | 1,795309183 | 0,23336 | 0,54563244 | long intergenic non-protein coding RNA 920 |

| | | | | | | |
|---|---|---|---|---|---|---|
| ENSG00000187017 | ESPN | 0,84414624 | 1,795202065 | 0,07583 | 0,36167805 | espin |
| ENSG00000168306 | ACOX2 | 0,843809849 | 1,794783529 | 0,13985 | 0,46689302 | acyl-CoA oxidase 2 |
| ENSG00000229132 | EIF4A1P10 | 0,843783881 | 1,794751223 | 0,47515 | 0,71372446 | eukaryotic translation initiation factor 4A1 pseudogene 10 |
| ENSG00000224333 | GAPDHP20 | 0,843761772 | 1,79472372 | 0,17925 | 0,50368273 | glyceraldehyde 3 phosphate dehydrogenase pseudogene 20 |
| ENSG00000248909 | HMGB1P21 | 0,843141067 | 1,793951724 | 0,12886 | 0,45214857 | high mobility group box 1 pseudogene 21 |
| ENSG00000157654 | PALM2AKAP2 | 0,842991571 | 1,793765839 | 0,00511 | 0,08910452 | PALM2 and AKAP2 fusion |
| ENSG00000227383 | RPL35AP2 | 0,842640033 | 1,79332881 | 0,27775 | 0,57618023 | ribosomal protein L35a pseudogene 2 |
| ENSG00000150961 | SEC24D | 0,842633804 | 1,793321067 | 0,0614 | 0,32865897 | SEC24 homolog D, COPII coat complex component |
| ENSG00000262795 | IFNGR2 | 0,842588666 | 1,79326496 | 0,38313 | 0,64987805 | interferon gamma receptor 2 |
| ENSG00000255710 | AP003170,1 | 0,842507032 | 1,793163491 | 0,09552 | 0,39847312 | interferon stimulated exonuclease gene 20kDa-like 2 (ISG20L2) pseudogene |
| ENSG00000141524 | TMC6 | 0,842066388 | 1,792615887 | 0,06983 | 0,34878306 | transmembrane channel like 6 |
| ENSG00000274419 | TBC1D3D | 0,841877799 | 1,792381573 | 0,2724 | 0,57315552 | TBC1 domain family member 3D |
| ENSG00000229060 | AL451124,1 | 0,841085922 | 1,791398026 | 0,40072 | 0,66269448 | novel pseudogene |
| ENSG00000259719 | LINC02284 | 0,84080843 | 1,791053498 | 0,31895 | 0,60399503 | long intergenic non-protein coding RNA 2284 |
| ENSG00000178425 | NT5DC1 | 0,840796823 | 1,791039087 | 0,07916 | 0,36739479 | 5'-nucleotidase domain containing 1 |
| ENSG00000163624 | CDS1 | 0,840580166 | 1,790770138 | 0,03962 | 0,26931682 | CDP-diacylglycerol synthase 1 |
| ENSG00000084093 | REST | 0,840120998 | 1,790200279 | 0,02891 | 0,22977124 | RE1 silencing transcription factor |
| ENSG00000278308 | NPIPA3 | 0,840119156 | 1,790197993 | 0,19735 | 0,51866271 | nuclear pore complex interacting protein family member A3 |
| ENSG00000274618 | H4C6 | 0,839849854 | 1,789863856 | 0,18027 | 0,50407792 | H4 clustered histone 6 |
| ENSG00000287787 | AC092275,1 | 0,839670724 | 1,789641634 | 0,30452 | 0,59430086 | novel transcript, sense intronic to GSE1 |
| ENSG00000124743 | KLHL31 | 0,839469559 | 1,789392109 | 0,16723 | 0,49323767 | kelch like family member 31 |
| ENSG00000112902 | SEMA5A | 0,838433963 | 1,788108107 | 0,01169 | 0,14074952 | semaphorin 5A |
| ENSG00000281980 | AC246817,3 | 0,838230512 | 1,787855963 | 0,09587 | 0,39884178 | novel transcript |
| ENSG00000165379 | LRFN5 | 0,837975584 | 1,787540073 | 0,02787 | 0,22522567 | leucine rich repeat and fibronectin type III domain containing 5 |
| ENSG00000162888 | C1orf147 | 0,8377082 | 1,787208807 | 0,16087 | 0,48815237 | chromosome 1 open reading frame 147 |
| ENSG00000180440 | SERTM1 | 0,837663608 | 1,787153567 | 0,27854 | 0,57673408 | serine rich and transmembrane domain containing 1 |
| ENSG00000123562 | MORF4L2 | 0,837447852 | 1,786886317 | 0,01423 | 0,15639135 | mortality factor 4 like 2 |
| ENSG00000261934 | PCDHGA9 | 0,836714901 | 1,785978733 | 0,05306 | 0,30931591 | protocadherin gamma subfamily A, 9 |
| ENSG00000272477 | AC144521,1 | 0,836588813 | 1,78582265 | 0,27467 | 0,57463533 | novel transcript, sense intronic to SATB1 |

| | | | | | | |
|---|---|---|---|---|---|---|
| ENSG00000122254 | HS3ST2 | 0,836454753 | 1,785656712 | 0,16981 | 0,49606609 | heparan sulfate-glucosamine 3-sulfotransferase 2 |
| ENSG00000125409 | TEKT3 | 0,836364753 | 1,78554532 | 0,12983 | 0,45396158 | tektin 3 |
| ENSG00000254535 | PABPC4L | 0,836165057 | 1,785298185 | 0,02633 | 0,21861252 | poly(A) binding protein cytoplasmic 4 like |
| ENSG00000214814 | FER1L6 | 0,836118059 | 1,785240027 | 0,03377 | 0,24838006 | fer-1 like family member 6 |
| ENSG00000283886 | BX664615,2 | 0,835533222 | 1,784516476 | 0,24868 | 0,55591283 | Uncharacterized protein FLJ76381 |
| ENSG00000200673 | RN7SKP174 | 0,834441507 | 1,78316661 | 0,29062 | 0,58566807 | RN7SK pseudogene 174 |
| ENSG00000219608 | HIGD1AP16 | 0,83394429 | 1,782552157 | 0,38742 | 0,6523395 | HIG1 hypoxia inducible domain family member 1A pseudogene 16 |
| ENSG00000267022 | AC067968,1 | 0,832712902 | 1,781031338 | 0,12441 | 0,44539699 | novel protein |
| ENSG00000277401 | TJP1 | 0,832398396 | 1,780643118 | 0,15553 | 0,48345606 | tight junction protein 1 |
| ENSG00000149591 | TAGLN | 0,832306205 | 1,780529335 | 0,07712 | 0,36384125 | transgelin |
| ENSG00000274031 | AC092140,2 | 0,8321666 | 1,780357047 | 0,32169 | 0,60577115 | novel transcript, antisense to OGFOD1 |
| ENSG00000148680 | HTR7 | 0,832141398 | 1,780325946 | 0,39864 | 0,66118205 | 5-hydroxytryptamine receptor 7 |
| ENSG00000198691 | ABCA4 | 0,831840387 | 1,779954529 | 0,12843 | 0,45191041 | ATP binding cassette subfamily A member 4 |
| ENSG00000197147 | LRRC8B | 0,831706078 | 1,779788831 | 0,00837 | 0,11551322 | leucine rich repeat containing 8 VRAC subunit B |
| ENSG00000224881 | AC068279,1 | 0,831624261 | 1,7796879 | 0,17783 | 0,50222458 | novel pseudogene |
| ENSG00000111885 | MAN1A1 | 0,831606995 | 1,779666601 | 0,00052 | 0,02082753 | mannosidase alpha class 1A member 1 |
| ENSG00000257767 | AC002996,1 | 0,830958106 | 1,77886633 | 0,41713 | 0,6738387 | novel protein |
| ENSG00000197705 | KLHL14 | 0,830489124 | 1,778288162 | 0,09106 | 0,39033853 | kelch like family member 14 |
| ENSG00000111907 | TPD52L1 | 0,830080087 | 1,777784049 | 0,02422 | 0,20804286 | TPD52 like 1 |
| ENSG00000237027 | AL159174,1 | 0,830037533 | 1,777731611 | 0,51182 | 0,7388352 | novel transcript |
| ENSG00000271741 | AC114490,2 | 0,82990924 | 1,777573532 | 0,25339 | 0,55846055 | novel transcript, ZMYM6-ZMYM6NB readthrough |
| ENSG00000272617 | AC026464,6 | 0,829810348 | 1,777451688 | 0,3612 | 0,63317043 | novel protein, COG8-PDF readthrough |
| ENSG00000282105 | RPL41P2 | 0,829528049 | 1,77710392 | 0,26211 | 0,56501119 | ribosomal protein L41 pseudogene 2 |
| ENSG00000204860 | FAM201A | 0,829460054 | 1,777020166 | 0,18941 | 0,51258824 | family with sequence similarity 201 member A |
| ENSG00000184675 | AMER1 | 0,829022047 | 1,776480739 | 0,00804 | 0,11395828 | APC membrane recruitment protein 1 |
| ENSG00000131773 | KHDRBS3 | 0,828928795 | 1,776365916 | 0,01859 | 0,18025307 | KH RNA binding domain containing, signal transduction associated 3 |
| ENSG00000288257 | STRA6 | 0,828658475 | 1,776033107 | 0,00408 | 0,07836207 | signaling receptor and transporter of retinol STRA6 |
| ENSG00000136997 | MYC | 0,828246025 | 1,775525431 | 0,0339 | 0,24870776 | MYC proto-oncogene, bHLH transcription factor |
| ENSG00000272209 | AL023583,1 | 0,82803675 | 1,775267896 | 0,44936 | 0,69503754 | novel transcript |
| ENSG00000248557 | AC024558,1 | 0,827856522 | 1,775046135 | 0,29094 | 0,58593792 | methyltransferase like 5 (METTL5) pseudogene |
| ENSG00000223945 | AC244033,1 | 0,827575097 | 1,774699913 | 0,18121 | 0,50471399 | ribosomal protein L6 (RPL6) pseudogene |

| | | | | | | |
|---|---|---|---|---|---|---|
| ENSG00000286707 | AC010896,1 | 0,827565473 | 1,774688074 | 0,10774 | 0,41834811 | novel transcript |
| ENSG00000285397 | AC270273,5 | 0,827477856 | 1,774580298 | 0,22498 | 0,54037642 | family with sequence similarity 86 member B1 |
| ENSG00000267576 | AC011472,3 | 0,827218015 | 1,774260711 | 0,4229 | 0,67683766 | novel transcript, sense intronic to TSPAN16 |
| ENSG00000226874 | AC005154,1 | 0,826793168 | 1,7737383 | 0,32724 | 0,6090604 | pseudogene similar to part of transient receptor potential cation channel, subfamily C, member 6 TRPC6 |
| ENSG00000266934 | AC005746,1 | 0,826788739 | 1,773732855 | 0,23584 | 0,5471053 | novel transcript, sense intronic to BCAS3 |
| ENSG00000079150 | FKBP7 | 0,826730082 | 1,77366074 | 0,13358 | 0,45946836 | FKBP prolyl isomerase 7 |
| ENSG00000227692 | MED28P3 | 0,826199677 | 1,773008776 | 0,36024 | 0,63261878 | mediator complex subunit 28 pseudogene 3 |
| ENSG00000130829 | DUSP9 | 0,825893217 | 1,772632189 | 0,2284 | 0,54241083 | dual specificity phosphatase 9 |
| ENSG00000230417 | LINC00595 | 0,825852595 | 1,772582279 | 0,34817 | 0,62452278 | long intergenic non-protein coding RNA 595 |
| ENSG00000270533 | CR382285,1 | 0,825651761 | 1,772335539 | 0,45338 | 0,69750019 | ribosomal protein L7a (RPL7A) pseudogene |
| ENSG00000169193 | CCDC126 | 0,825477857 | 1,772121913 | 0,08516 | 0,37857645 | coiled-coil domain containing 126 |
| ENSG00000236264 | RPL26P30 | 0,824907593 | 1,771421572 | 0,35484 | 0,62936991 | ribosomal protein L26 pseudogene 30 |
| ENSG00000116815 | CD58 | 0,824726782 | 1,771199577 | 0,12633 | 0,44903997 | CD58 molecule |
| ENSG00000285752 | CDC42-AS1 | 0,824594747 | 1,771037483 | 0,00772 | 0,11169557 | CDC42 antisense RNA 1 |
| ENSG00000161791 | FMNL3 | 0,824554054 | 1,77098753 | 3,75E-06 | 0,00035686 | formin like 3 |
| ENSG00000116194 | ANGPTL1 | 0,824364916 | 1,770755368 | 0,09083 | 0,39006208 | angiopoietin like 1 |
| ENSG00000137965 | IFI44 | 0,824296334 | 1,770671193 | 0,3957 | 0,65846795 | interferon induced protein 44 |
| ENSG00000286645 | AC013267,1 | 0,824057876 | 1,770378549 | 0,46043 | 0,70227822 | novel transcript |
| ENSG00000174370 | C11orf45 | 0,823987265 | 1,770291902 | 0,22297 | 0,53908993 | chromosome 11 open reading frame 45 |
| ENSG00000178878 | APOLD1 | 0,823668479 | 1,769900772 | 0,01762 | 0,17483597 | apolipoprotein L domain containing 1 |
| ENSG00000268521 | VN1R83P | 0,823452909 | 1,769636329 | 0,26606 | 0,56902849 | vomeronasal 1 receptor 83 pseudogene |
| ENSG00000143153 | ATP1B1 | 0,823352228 | 1,769512838 | 0,01152 | 0,13944196 | ATPase Na+/K+ transporting subunit beta 1 |
| ENSG00000185338 | SOCS1 | 0,823271266 | 1,769413538 | 0,18189 | 0,50521394 | suppressor of cytokine signaling 1 |
| ENSG00000147231 | RADX | 0,823150124 | 1,769264968 | 0,00718 | 0,10764638 | RPA1 related single stranded DNA binding protein, X-linked |
| ENSG00000245067 | IGFBP7-AS1 | 0,822531272 | 1,768506195 | 0,39958 | 0,66181917 | IGFBP7 antisense RNA 1 |
| ENSG00000022567 | SLC45A4 | 0,821015156 | 1,766648663 | 0,00025 | 0,01256697 | solute carrier family 45 member 4 |
| ENSG00000227253 | AL158834,2 | 0,82038762 | 1,765880381 | 0,21623 | 0,53480193 | novel transcript |
| ENSG00000129595 | EPB41L4A | 0,820192989 | 1,765642166 | 0,05294 | 0,30886444 | erythrocyte membrane protein band 4,1 like 4A |
| ENSG00000284989 | AL451062,2 | 0,819679469 | 1,765013807 | 0,40338 | 0,66399303 | novel transcript |
| ENSG00000128045 | RASL11B | 0,819491502 | 1,76478386 | 0,00367 | 0,07405288 | RAS like family 11 member B |
| ENSG00000232143 | GNL1 | 0,818943972 | 1,764114219 | 0,41864 | 0,67477592 | G protein nucleolar 1 (putative) |

| | | | | | | |
|---|---|---|---|---|---|---|
| ENSG00000228581 | GNL1 | 0,818943972 | 1,764114219 | 0,41864 | 0,67477592 | G protein nucleolar 1 (putative) |
| ENSG00000204590 | GNL1 | 0,818943972 | 1,764114219 | 0,41864 | 0,67477592 | G protein nucleolar 1 (putative) |
| ENSG00000273123 | AC020634,2 | 0,818784244 | 1,763918915 | 0,38054 | 0,6481589 | novel transcript, antisense to MYLK |
| ENSG00000018625 | ATP1A2 | 0,81857985 | 1,76366903 | 0,00097 | 0,03167816 | ATPase Na+/K+ transporting subunit alpha 2 |
| ENSG00000183783 | KCTD8 | 0,818456695 | 1,763518481 | 0,14932 | 0,47695763 | potassium channel tetramerization domain containing 8 |
| ENSG00000267890 | AC010624,2 | 0,818165488 | 1,763162552 | 0,22787 | 0,54197226 | novel transcript, similar to SYT3 |
| ENSG00000236908 | LINC02827 | 0,818111271 | 1,763096294 | 0,35542 | 0,62972424 | long intergenic non-protein coding RNA 2827 |
| ENSG00000147082 | CCNB3 | 0,818022929 | 1,762988335 | 0,09014 | 0,38834203 | cyclin B3 |
| ENSG00000271075 | AL139317,4 | 0,817813328 | 1,762732219 | 0,13542 | 0,46210588 | golgi membrane protein 1 (GOLM1) pseudogene |
| ENSG00000273035 | AC007684,2 | 0,817785221 | 1,762697877 | 0,4006 | 0,66264525 | novel transcript, sense intronic to MAP4K3 |
| ENSG00000095203 | EPB41L4B | 0,817618531 | 1,762494225 | 0,03509 | 0,2533325 | erythrocyte membrane protein band 4,1 like 4B |
| ENSG00000258066 | AC138331,1 | 0,817588579 | 1,762457635 | 0,38243 | 0,64928247 | novel transcript |
| ENSG00000276232 | AC006064,5 | 0,817465791 | 1,762307638 | 0,25002 | 0,55657567 | small Cajal body-specific RNA 10 |
| ENSG00000272568 | AC005162,2 | 0,816659565 | 1,761323077 | 0,13935 | 0,46666406 | novel transcript, antisense to CPVL |
| ENSG00000136810 | TXN | 0,816504512 | 1,76113379 | 0,19118 | 0,51406851 | thioredoxin |
| ENSG00000250474 | WBP1LP2 | 0,816107512 | 1,760649229 | 0,34531 | 0,62249219 | WBP1L pseudogene 2 |
| ENSG00000189190 | ZNF600 | 0,815842592 | 1,760325953 | 0,03744 | 0,26177644 | zinc finger protein 600 |
| ENSG00000226937 | CEP164P1 | 0,815623355 | 1,760058468 | 0,01486 | 0,16001992 | centrosomal protein 164 pseudogene 1 |
| ENSG00000236340 | GRM8-AS1 | 0,815311998 | 1,75967866 | 0,41773 | 0,67424452 | GRM8 antisense RNA 1 |
| ENSG00000095370 | SH2D3C | 0,814827456 | 1,759087755 | 0,02708 | 0,22194635 | SH2 domain containing 3C |
| ENSG00000258655 | ARHGAP5-AS1 | 0,814166382 | 1,758281888 | 0,07962 | 0,36831832 | ARHGAP5 antisense RNA 1 (head to head) |
| ENSG00000131982 | UBE2L2 | 0,813737328 | 1,757759057 | 0,32976 | 0,61069811 | ubiquitin conjugating enzyme E2 L2 (pseudogene) |
| ENSG00000226766 | FABP7P1 | 0,813665633 | 1,757671706 | 0,19052 | 0,51357771 | fatty acid binding protein 7 pseudogene 1 |
| ENSG00000227560 | RPS15AP30 | 0,813549891 | 1,757530701 | 0,16087 | 0,48815237 | ribosomal protein S15a pseudogene 30 |
| ENSG00000286285 | AL138778,1 | 0,813541544 | 1,757520532 | 0,45817 | 0,70115011 | novel transcript |
| ENSG00000111846 | GCNT2 | 0,813483232 | 1,757449497 | 0,1942 | 0,51562158 | glucosaminyl (N-acetyl) transferase 2 (I blood group) |
| ENSG00000212807 | OR2A42 | 0,813340562 | 1,757275709 | 0,04573 | 0,28843251 | olfactory receptor family 2 subfamily A member 42 |
| ENSG00000101443 | WFDC2 | 0,813227179 | 1,757137608 | 0,06175 | 0,32900116 | WAP four-disulfide core domain 2 |
| ENSG00000226288 | OR52I2 | 0,812608194 | 1,756383874 | 0,32098 | 0,60568214 | olfactory receptor family 52 subfamily I member 2 |
| ENSG00000234882 | EIF3EP1 | 0,811882418 | 1,755500513 | 0,56158 | 0,77136826 | eukaryotic translation initiation factor 3 subunit E pseudogene 1 |
| ENSG00000251370 | SEMA5A-AS1 | 0,811689757 | 1,755266096 | 0,04044 | 0,27194236 | SEMA5A antisense RNA 1 |
| ENSG00000111796 | KLRB1 | 0,811620569 | 1,755181919 | 0,19366 | 0,51547251 | killer cell lectin like receptor B1 |

| | | | | | | |
|---|---|---|---|---|---|---|
| ENSG00000269289 | AC011503,1 | 0,811264356 | 1,754748603 | 0,06286 | 0,33177754 | novel transcript, antisense to ZNF726 |
| ENSG00000255663 | AP002373,1 | 0,811166136 | 1,754629143 | 0,70152 | 0,85609945 | novel transcript, RBM7-REXO2 readthrough |
| ENSG00000188517 | COL25A1 | 0,811097922 | 1,754546182 | 0,09255 | 0,39322559 | collagen type XXV alpha 1 chain |
| ENSG00000229246 | LINC00377 | 0,810900396 | 1,754305976 | 0,43693 | 0,68701656 | long intergenic non-protein coding RNA 377 |
| ENSG00000283900 | TPTEP2-CSNK | 0,810624275 | 1,753970247 | 0,12007 | 0,43868155 | TPTEP2-CSNK1E readthrough |
| ENSG00000267924 | AC139769,2 | 0,809902353 | 1,753092782 | 0,03096 | 0,23719337 | novel transcript |
| ENSG00000259663 | AC010478,1 | 0,80975976 | 1,752919519 | 0,22668 | 0,54144 | novel transcript, antisense to ISL1 |
| ENSG00000169126 | ARMC4 | 0,809580293 | 1,752701475 | 0,02709 | 0,22195182 | armadillo repeat containing 4 |
| ENSG00000256238 | SUPT16HP1 | 0,80943613 | 1,752526343 | 0,17585 | 0,49979999 | SPT16 homolog, facilitates chromatin remodeling subunit pseudogene 1 |
| ENSG00000163932 | PRKCD | 0,809247806 | 1,75229759 | 0,00697 | 0,10629039 | protein kinase C delta |
| ENSG00000182247 | UBE2E2 | 0,809246003 | 1,7522954 | 0,0341 | 0,24932302 | ubiquitin conjugating enzyme E2 E2 |
| ENSG00000186312 | CA5BP1 | 0,808480393 | 1,751365738 | 0,00065 | 0,02442894 | carbonic anhydrase 5B pseudogene 1 |
| ENSG00000272509 | AC087752,4 | 0,808278077 | 1,751120152 | 0,30428 | 0,59424503 | novel transcript, antisense to CCNE2 |
| ENSG00000266885 | AC132825,4 | 0,808231927 | 1,751064137 | 0,0397 | 0,26947142 | chromosome 17 open reading frame 51 (C17orf51) pseudogene |
| ENSG00000225871 | AC245100,2 | 0,808231656 | 1,751063808 | 0,13491 | 0,46159318 | family with sequence similarity 91, member A1 (FAM91A1) pseudogene |
| ENSG00000163631 | ALB | 0,807447908 | 1,750112796 | 0,29234 | 0,58692263 | albumin |
| ENSG00000122873 | CISD1 | 0,806595501 | 1,749079058 | 0,06268 | 0,33130676 | CDGSH iron sulfur domain 1 |
| ENSG00000171246 | NPTX1 | 0,806568948 | 1,749046868 | 0,05107 | 0,30386013 | neuronal pentraxin 1 |
| ENSG00000280279 | LINC02887 | 0,806450037 | 1,748902712 | 0,10424 | 0,41262354 | long intergenic non-protein coding RNA 2887 |
| ENSG00000282235 | LINC02887 | 0,806450037 | 1,748902712 | 0,10424 | 0,41262354 | long intergenic non-protein coding RNA 2887 |
| ENSG00000164778 | EN2 | 0,806269957 | 1,748684424 | 0,41709 | 0,6738387 | engrailed homeobox 2 |
| ENSG00000253811 | AC136424,2 | 0,805623386 | 1,747900893 | 0,3735 | 0,64220148 | novel transcript |
| ENSG00000278588 | H2BC10 | 0,805580262 | 1,747848648 | 0,18247 | 0,50597301 | H2B clustered histone 10 |
| ENSG00000269604 | AC005523,2 | 0,805155747 | 1,747334417 | 0,62723 | 0,81132445 | novel transcript, antisense to FEM1A |
| ENSG00000211683 | AP000346,1 | 0,804903352 | 1,747028752 | 0,2356 | 0,54704325 | novel transcript |
| ENSG00000155254 | MARVELD1 | 0,804863913 | 1,746980994 | 3,85E-05 | 0,00269251 | MARVEL domain containing 1 |
| ENSG00000261994 | WEE2-AS1 | 0,804797739 | 1,746900865 | 0,26747 | 0,56973003 | WEE2 antisense RNA 1 |
| ENSG00000112715 | VEGFA | 0,803618246 | 1,745473249 | 0,00024 | 0,01233984 | vascular endothelial growth factor A |
| ENSG00000174132 | FAM174A | 0,802855948 | 1,744551211 | 0,01158 | 0,13993374 | family with sequence similarity 174 member A |
| ENSG00000278727 | AC000403,1 | 0,802456591 | 1,744068363 | 0,19254 | 0,51490739 | novel transcript |

| | | | | | | |
|---|---|---|---|---|---|---|
| ENSG00000081248 | CACNA1S | 0,802058584 | 1,743587281 | 0,37865 | 0,64679625 | calcium voltage-gated channel subunit alpha1 S |
| ENSG00000261578 | AP003119,3 | 0,801713796 | 1,743170632 | 0,00802 | 0,11389729 | novel transcript, overlapping to TSKU |
| ENSG00000276809 | AL138955,1 | 0,801696888 | 1,743150203 | 0,35427 | 0,62895806 | novel transcript, sense intronic to DNAJC3 |
| ENSG00000274286 | ADRA2B | 0,80168053 | 1,743130438 | 0,07367 | 0,35701531 | adrenoceptor alpha 2B |
| ENSG00000172554 | SNTG2 | 0,801651278 | 1,743095095 | 0,16042 | 0,48768584 | syntrophin gamma 2 |
| ENSG00000232457 | SLC16A6P1 | 0,801401343 | 1,742793144 | 0,09473 | 0,39724142 | SLC16A6 pseudogene 1 |
| ENSG00000144655 | CSRNP1 | 0,800393185 | 1,741575703 | 0,00773 | 0,11178616 | cysteine and serine rich nuclear protein 1 |
| ENSG00000122786 | CALD1 | 0,80012023 | 1,741246231 | 0,00068 | 0,02516175 | caldesmon 1 |
| ENSG00000284468 | SLC25A24 | 0,79998282 | 1,741080393 | 0,03586 | 0,25662951 | solute carrier family 25 member 24 |
| ENSG00000085491 | SLC25A24 | 0,79998282 | 1,741080393 | 0,03586 | 0,25662951 | solute carrier family 25 member 24 |
| ENSG00000250473 | DUTP7 | 0,79907665 | 1,739987148 | 0,16397 | 0,49027508 | deoxyuridine triphosphatase pseudogene 7 |
| ENSG00000232486 | DEPDC1P2 | 0,799072134 | 1,739981701 | 0,04521 | 0,2876168 | DEP domain containing 1 pseudogene 2 |
| ENSG00000272842 | AL391834,1 | 0,799068301 | 1,739977078 | 0,22545 | 0,5407493 | novel transcript, antisense to DENND4C |
| ENSG00000249713 | AC026725,1 | 0,798863702 | 1,739730338 | 0,28908 | 0,58469988 | novel transcript |
| ENSG00000267343 | ZNF833P | 0,798605204 | 1,739418645 | 0,40515 | 0,6648439 | zinc finger protein 833, pseudogene |
| ENSG00000171914 | TLN2 | 0,798573019 | 1,739379842 | 0,00157 | 0,04253303 | talin 2 |
| ENSG00000261436 | AC009081,2 | 0,798565817 | 1,739371158 | 0,25272 | 0,55796133 | novel transcript |
| ENSG00000166710 | B2M | 0,798395997 | 1,739166428 | 0,11138 | 0,42416781 | beta-2-microglobulin |
| ENSG00000230678 | BRD2 | 0,798364854 | 1,739128886 | 0,13284 | 0,45832113 | bromodomain containing 2 |
| ENSG00000278961 | SMIM34B | 0,798311827 | 1,739064964 | 0,33405 | 0,61387439 | small integral membrane protein 34B |
| ENSG00000136261 | BZW2 | 0,798259656 | 1,739002077 | 0,03233 | 0,24253636 | basic leucine zipper and W2 domains 2 |
| ENSG00000079819 | EPB41L2 | 0,79736377 | 1,737922525 | 0,03168 | 0,24008374 | erythrocyte membrane protein band 4,1 like 2 |
| ENSG00000076716 | GPC4 | 0,797177918 | 1,737698657 | 0,01526 | 0,16213925 | glypican 4 |
| ENSG00000106819 | ASPN | 0,797132691 | 1,737644182 | 0,08269 | 0,37450725 | asporin |
| ENSG00000250227 | TRIM60P14 | 0,797086495 | 1,737588542 | 0,21768 | 0,53547632 | tripartite motif containing 60 pseudogene 14 |
| ENSG00000248931 | MTCYBP40 | 0,796496651 | 1,736878276 | 0,48192 | 0,71869839 | MT-CYB pseudogene 40 |
| ENSG00000241217 | RN7SL809P | 0,796386918 | 1,736746172 | 0,32785 | 0,60960585 | RNA, 7SL, cytoplasmic 809, pseudogene |
| ENSG00000105329 | TGFB1 | 0,796271741 | 1,736607525 | 0,21364 | 0,53275614 | transforming growth factor beta 1 |
| ENSG00000154642 | C21orf91 | 0,795781262 | 1,736017224 | 0,10552 | 0,41477267 | chromosome 21 open reading frame 91 |
| ENSG00000085276 | MECOM | 0,795714613 | 1,735937027 | 0,13153 | 0,45658336 | MDS1 and EVI1 complex locus |
| ENSG00000149054 | ZNF215 | 0,79512439 | 1,735226981 | 0,01831 | 0,17903469 | zinc finger protein 215 |
| ENSG00000254087 | LYN | 0,795080087 | 1,735173694 | 0,03678 | 0,25987876 | LYN proto-oncogene, Src family tyrosine kinase |
| ENSG00000174827 | PDZK1 | 0,794940635 | 1,73500598 | 0,05905 | 0,32417102 | PDZ domain containing 1 |

| | | | | | | |
|---|---|---|---|---|---|---|
| ENSG00000180530 | NRIP1 | 0,794813765 | 1,734853411 | 0,03096 | 0,23719337 | nuclear receptor interacting protein 1 |
| ENSG00000274346 | AC016999,2 | 0,793973433 | 1,733843198 | 0,40135 | 0,6630277 | novel zinc finger protein pseudogene |
| ENSG00000254560 | BBOX1-AS1 | 0,793848953 | 1,733693604 | 0,08866 | 0,38505257 | BBOX1 antisense RNA 1 |
| ENSG00000287886 | AC005914,1 | 0,793586608 | 1,733378371 | 0,25294 | 0,55808134 | novel transcript |
| ENSG00000162944 | RFTN2 | 0,793344413 | 1,733087402 | 0,00033 | 0,01523174 | raftlin family member 2 |
| ENSG00000248926 | AC114801,2 | 0,793053043 | 1,732737418 | 0,43771 | 0,68744243 | FUS interacting protein (serine/arginine-rich) 1 (FUSIP1) pseudogene |
| ENSG00000166888 | STAT6 | 0,793006673 | 1,732681727 | 6,26E-05 | 0,00410381 | signal transducer and activator of transcription 6 |
| ENSG00000284842 | FAM91A2P | 0,792998773 | 1,732672239 | 0,49641 | 0,72831792 | family with sequence similarity 91 member A2, pseudogene |
| ENSG00000236297 | AC048351,1 | 0,792832848 | 1,732472975 | 0,46253 | 0,70358709 | eukaryotic translation elongation factor 1 alpha 1 (EEF1A1) pseudogene |
| ENSG00000283782 | AC116366,2 | 0,792679605 | 1,732288962 | 0,62396 | 0,80955861 | novel protein |
| ENSG00000260625 | AC026471,2 | 0,792449659 | 1,732012881 | 0,08692 | 0,38197222 | novel transcript, antisense to C16orf58 |
| ENSG00000260772 | AC012321,1 | 0,792234632 | 1,731754752 | 0,40852 | 0,66729874 | novel transcript |
| ENSG00000114698 | PLSCR4 | 0,792199986 | 1,731713165 | 0,04392 | 0,28268806 | phospholipid scramblase 4 |
| ENSG00000171451 | DSEL | 0,792027984 | 1,731506717 | 0,0598 | 0,32535617 | dermatan sulfate epimerase like |
| ENSG00000244676 | AL109761,1 | 0,791927414 | 1,731386018 | 0,14722 | 0,47471707 | novel transcript, antisense to C21orf91 |
| ENSG00000119686 | FLVCR2 | 0,791176541 | 1,730485126 | 0,05155 | 0,30525036 | FLVCR heme transporter 2 |
| ENSG00000233623 | PGAM1P11 | 0,791166224 | 1,730472751 | 0,37302 | 0,64180957 | phosphoglycerate mutase 1 pseudogene 11 |
| ENSG00000007944 | MYLIP | 0,791072313 | 1,730360111 | 0,03217 | 0,24200947 | myosin regulatory light chain interacting protein |
| ENSG00000118972 | FGF23 | 0,790515997 | 1,729692998 | 0,1997 | 0,52077515 | fibroblast growth factor 23 |
| ENSG00000158195 | WASF2 | 0,790270331 | 1,729398485 | 6,99E-05 | 0,004476 | WASP family member 2 |
| ENSG00000074695 | LMAN1 | 0,79014435 | 1,729247476 | 0,06092 | 0,32753264 | lectin, mannose binding 1 |
| ENSG00000206052 | DOK6 | 0,789843484 | 1,728886888 | 0,03412 | 0,24936671 | docking protein 6 |
| ENSG00000118515 | SGK1 | 0,789527304 | 1,728508028 | 0,00973 | 0,12625914 | serum/glucocorticoid regulated kinase 1 |
| ENSG00000204103 | MAFB | 0,789427817 | 1,728388836 | 0,05981 | 0,32535617 | MAF bZIP transcription factor B |
| ENSG00000279375 | AC244517,7 | 0,789065176 | 1,727954436 | 0,39279 | 0,6559444 | novel transcript, antisense to PCDHB10 |
| ENSG00000101187 | SLCO4A1 | 0,788710046 | 1,72752914 | 0,02029 | 0,18921073 | solute carrier organic anion transporter family member 4A1 |
| ENSG00000227694 | RPL23AP74 | 0,788281511 | 1,727016075 | 0,33149 | 0,61161075 | ribosomal protein L23a pseudogene 74 |
| ENSG00000042980 | ADAM28 | 0,788185766 | 1,726901465 | 0,25359 | 0,55857654 | ADAM metallopeptidase domain 28 |
| ENSG00000204435 | CSNK2B | 0,788146366 | 1,726854303 | 0,22113 | 0,53805234 | casein kinase 2 beta |

| | | | | | | |
|---|---|---|---|---|---|---|
| ENSG00000248115 | AC023154,1 | 0,787753974 | 1,726384687 | 0,18708 | 0,5105374 | novel transcript |
| ENSG00000239264 | TXNDC5 | 0,787416641 | 1,72598107 | 0,00414 | 0,0788858 | thioredoxin domain containing 5 |
| ENSG00000275481 | AC025031,4 | 0,787360805 | 1,725914271 | 0,11626 | 0,43279558 | novel transcript |
| ENSG00000264345 | LINC01894 | 0,786941992 | 1,725413312 | 0,11413 | 0,42945959 | long intergenic non-protein coding RNA 1894 |
| ENSG00000231154 | MORF4L2-AS1 | 0,786883962 | 1,725343911 | 0,0131 | 0,14972794 | MORF4L2 antisense RNA 1 |
| ENSG00000177932 | ZNF354C | 0,786790821 | 1,725232527 | 0,01588 | 0,16502318 | zinc finger protein 354C |
| ENSG00000277610 | RNVU1-4 | 0,786646311 | 1,725059724 | 0,33972 | 0,61872249 | RNA, variant U1 small nuclear 4 |
| ENSG00000251356 | RAB5CP2 | 0,786207548 | 1,724535166 | 0,3823 | 0,64928247 | RAB5C, member RAS oncogene family pseudogene 2 |
| ENSG00000259125 | LRP1-AS | 0,786190784 | 1,724515127 | 0,46293 | 0,70406535 | LRP1 antisense RNA |
| ENSG00000286863 | AL513166,1 | 0,786041584 | 1,724336791 | 0,05007 | 0,30089233 | novel transcript |
| ENSG00000196653 | ZNF502 | 0,78579586 | 1,724043122 | 0,03982 | 0,26986107 | zinc finger protein 502 |
| ENSG00000281448 | ZNF502 | 0,78579586 | 1,724043122 | 0,03982 | 0,26986107 | zinc finger protein 502 |
| ENSG00000198406 | BZW1P2 | 0,785007252 | 1,723100981 | 0,63898 | 0,81893471 | basic leucine zipper and W2 domains 1 pseudogene 2 |
| ENSG00000250565 | ATP6V1E2 | 0,784678836 | 1,722708778 | 0,00693 | 0,10609569 | ATPase H+ transporting V1 subunit E2 |
| ENSG00000278058 | AC009159,3 | 0,784225409 | 1,72216743 | 0,21413 | 0,53334744 | novel transcript |
| ENSG00000138735 | PDE5A | 0,783904063 | 1,721783877 | 0,14451 | 0,47209141 | phosphodiesterase 5A |
| ENSG00000231201 | AF127577,2 | 0,78381762 | 1,721680715 | 0,10431 | 0,41269448 | novel transcript |
| ENSG00000130208 | APOC1 | 0,783733846 | 1,721580744 | 0,0309 | 0,23705504 | apolipoprotein C1 |
| ENSG00000126368 | NR1D1 | 0,783573078 | 1,721388908 | 0,12458 | 0,44559634 | nuclear receptor subfamily 1 group D member 1 |
| ENSG00000154309 | DISP1 | 0,783524806 | 1,721331312 | 0,00299 | 0,06528619 | dispatched RND transporter family member 1 |
| ENSG00000125868 | DSTN | 0,783431117 | 1,721219532 | 0,04263 | 0,27820551 | destrin, actin depolymerizing factor |
| ENSG00000117151 | CTBS | 0,783400526 | 1,721183035 | 0,11239 | 0,42609972 | chitobiase |
| ENSG00000233170 | AC138356,2 | 0,783319892 | 1,721086839 | 0,16844 | 0,49467529 | chromosome X open reading frame 56 (CXorf56) pseudogene |
| ENSG00000162576 | MXRA8 | 0,782803121 | 1,720470459 | 0,03233 | 0,24253636 | matrix remodeling associated 8 |
| ENSG00000137868 | STRA6 | 0,782775675 | 1,72043773 | 0,00169 | 0,04470397 | signaling receptor and transporter of retinol STRA6 |
| ENSG00000165816 | VWA2 | 0,782743962 | 1,720399911 | 0,20836 | 0,5276952 | von Willebrand factor A domain containing 2 |
| ENSG00000274290 | H2BC6 | 0,782727833 | 1,720380678 | 0,25358 | 0,55857654 | H2B clustered histone 6 |
| ENSG00000265222 | AC079336,2 | 0,78265718 | 1,720296428 | 0,45078 | 0,69592991 | novel transcript, sense intronic MYO1D |
| ENSG00000158125 | XDH | 0,781672823 | 1,719123062 | 0,03081 | 0,23691425 | xanthine dehydrogenase |
| ENSG00000121039 | RDH10 | 0,781603629 | 1,719040613 | 0,13113 | 0,45586859 | retinol dehydrogenase 10 |
| ENSG00000244018 | RPL35P6 | 0,781455289 | 1,718863868 | 0,20011 | 0,5209612 | ribosomal protein L35 pseudogene 6 |

| | | | | | | |
|---|---|---|---|---|---|---|
| ENSG00000254593 | OR7E126P | 0,781412883 | 1,718813344 | 0,14771 | 0,47507961 | olfactory receptor family 7 subfamily E member 126 pseudogene |
| ENSG00000204778 | CBWD4P | 0,781267246 | 1,718639843 | 0,3947 | 0,65761928 | COBW domain containing 4, pseudogene |
| ENSG00000280693 | SH3PXD2A-AS | 0,780855351 | 1,718149234 | 0,13593 | 0,46291909 | SH3PXD2A antisense RNA 1 |
| ENSG00000198478 | SH3BGRL2 | 0,780828016 | 1,71811668 | 0,07147 | 0,35325412 | SH3 domain binding glutamate rich protein like 2 |
| ENSG00000181472 | ZBTB2 | 0,78081063 | 1,718095976 | 0,1098 | 0,42203441 | zinc finger and BTB domain containing 2 |
| ENSG00000167550 | RHEBL1 | 0,780287412 | 1,717472992 | 0,03936 | 0,26861041 | RHEB like 1 |
| ENSG00000278693 | ASB2 | 0,780223942 | 1,717397435 | 0,50287 | 0,73217708 | ankyrin repeat and SOCS box containing 2 |
| ENSG00000272009 | AL121944,1 | 0,779748094 | 1,716831074 | 0,28831 | 0,58436774 | novel transcript, antisense to ZNF165 |
| ENSG00000121769 | FABP3 | 0,779672094 | 1,716740636 | 0,02917 | 0,2303054 | fatty acid binding protein 3 |
| ENSG00000134049 | IER3IP1 | 0,779622678 | 1,716681834 | 0,16812 | 0,49421665 | immediate early response 3 interacting protein 1 |
| ENSG00000116874 | WARS2 | 0,779438475 | 1,716462662 | 0,10111 | 0,40780603 | tryptophanyl tRNA synthetase 2, mitochondrial |
| ENSG00000185585 | OLFML2A | 0,779160089 | 1,716131482 | 0,25301 | 0,55809014 | olfactomedin like 2A |
| ENSG00000132294 | EFR3A | 0,778964131 | 1,715898398 | 0,05423 | 0,31168784 | EFR3 homolog A |
| ENSG00000200959 | SNORA74A | 0,778792112 | 1,715693817 | 0,05289 | 0,30865611 | small nucleolar RNA, H/ACA box 74A |
| ENSG00000137962 | ARHGAP29 | 0,778674589 | 1,715554061 | 0,08381 | 0,37610661 | Rho GTPase activating protein 29 |
| ENSG00000286952 | AC093928,1 | 0,77858275 | 1,715444855 | 0,48497 | 0,72063669 | novel transcript, antisense to DNAH12 |
| ENSG00000150722 | PPP1R1C | 0,778556693 | 1,715413873 | 0,17865 | 0,50323303 | protein phosphatase 1 regulatory inhibitor subunit 1C |
| ENSG00000285850 | AC009152,3 | 0,778257695 | 1,71505839 | 0,26117 | 0,56416552 | novel transcript |
| ENSG00000285903 | AC009152,4 | 0,778257695 | 1,71505839 | 0,26117 | 0,56416552 | novel transcript |
| ENSG00000277773 | SMN2 | 0,778209989 | 1,715001679 | 0,15506 | 0,48296753 | survival of motor neuron 2, centromeric |
| ENSG00000169064 | ZBBX | 0,777999132 | 1,714751041 | 0,14547 | 0,47300165 | zinc finger B-box domain containing |
| ENSG00000239280 | AC108693,1 | 0,777618381 | 1,71429855 | 0,05388 | 0,31122212 | pseudogene similar to part of dihydrolipoamide branched chain transacylase E2 DBT |
| ENSG00000137880 | GCHFR | 0,777298167 | 1,713918094 | 0,19248 | 0,51490739 | GTP cyclohydrolase I feedback regulator |
| ENSG00000138604 | GLCE | 0,777232993 | 1,713840669 | 0,101 | 0,40770184 | glucuronic acid epimerase |
| ENSG00000144642 | RBMS3 | 0,777090673 | 1,71367161 | 0,24494 | 0,55288356 | RNA binding motif single stranded interacting protein 3 |
| ENSG00000100276 | RASL10A | 0,776897367 | 1,713442011 | 0,1463 | 0,47433542 | RAS like family 10 member A |
| ENSG00000250568 | AC098591,3 | 0,776826661 | 1,713358038 | 0,28893 | 0,58464073 | poly(A) binding protein, cytoplasmic 1 (PABPC1) pseudogene |
| ENSG00000224393 | MPIG6B | 0,775359452 | 1,711616452 | 0,49329 | 0,72631 | megakaryocyte and platelet inhibitory receptor G6b |
| ENSG00000163638 | ADAMTS9 | 0,774999544 | 1,71118951 | 0,04146 | 0,27595476 | ADAM metallopeptidase with thrombospondin type 1 motif 9 |

| | | | | | | |
|---|---|---|---|---|---|---|
| ENSG00000230325 | AL359921,1 | 0,774648798 | 1,710773539 | 0,25254 | 0,55796133 | novel transcript |
| ENSG00000179776 | CDH5 | 0,774481211 | 1,710574823 | 0,20814 | 0,52756478 | cadherin 5 |
| ENSG00000273184 | AC010655,4 | 0,77380732 | 1,70977599 | 0,14255 | 0,46989963 | novel transcript, readthrough between HILPDA-METTL2B |
| ENSG00000171428 | NAT1 | 0,773511321 | 1,70942523 | 0,17996 | 0,50407792 | N-acetyltransferase 1 |
| ENSG00000126500 | FLRT1 | 0,77336465 | 1,70925145 | 0,04187 | 0,27720117 | fibronectin leucine rich transmembrane protein 1 |
| ENSG00000243312 | AC093827,1 | 0,773081543 | 1,708916069 | 0,15619 | 0,4840074 | ribosomal protein L6 (RPL6) pseudogene |
| ENSG00000105173 | CCNE1 | 0,773036501 | 1,708862717 | 0,06799 | 0,34452206 | cyclin E1 |
| ENSG00000232586 | KIAA1614-AS1 | 0,772811951 | 1,708596759 | 0,21976 | 0,5371755 | KIAA1614 antisense RNA 1 |
| ENSG00000237263 | MAPK6P3 | 0,772357496 | 1,708058629 | 0,05786 | 0,32112994 | mitogen-activated protein kinase 6 pseudogene 3 |
| ENSG00000236729 | HCG17 | 0,772225198 | 1,707902003 | 0,60302 | 0,7971742 | HLA complex group 17 |
| ENSG00000166368 | OR2D2 | 0,771920229 | 1,707541011 | 0,28542 | 0,58187605 | olfactory receptor family 2 subfamily D member 2 |
| ENSG00000287686 | AC062029,1 | 0,771827423 | 1,707431171 | 0,11267 | 0,42643639 | novel transcript |
| ENSG00000287932 | AC000065,2 | 0,771589892 | 1,707150077 | 0,1999 | 0,52088599 | novel transcript, sense intronic to CDK6 |
| ENSG00000133124 | IRS4 | 0,771090137 | 1,706558815 | 0,16672 | 0,49257009 | insulin receptor substrate 4 |
| ENSG00000236529 | AC093582,1 | 0,770821884 | 1,70624153 | 0,27636 | 0,57542877 | novel transcript |
| ENSG00000285911 | AL391834,3 | 0,770453335 | 1,705805711 | 0,60275 | 0,7971742 | Novel transcript |
| ENSG00000188906 | LRRK2 | 0,770357686 | 1,705692622 | 0,03968 | 0,26947142 | leucine rich repeat kinase 2 |
| ENSG00000272345 | AL031775,1 | 0,770353344 | 1,705687488 | 0,22814 | 0,54221266 | novel transcript, antisense to ACOT13 |
| ENSG00000257524 | AL157935,2 | 0,769875664 | 1,705122824 | 0,23016 | 0,54335232 | novel protein |
| ENSG00000163584 | RPL22L1 | 0,769773381 | 1,70500194 | 0,18884 | 0,5121977 | ribosomal protein L22 like 1 |
| ENSG00000009790 | TRAF3IP3 | 0,769749124 | 1,704973273 | 0,32145 | 0,60577115 | TRAF3 interacting protein 3 |
| ENSG00000244398 | AC116533,1 | 0,769688174 | 1,704901244 | 0,14144 | 0,46805553 | ribosomal protein L36a (RPL36A) pseudogene |
| ENSG00000112984 | KIF20A | 0,769546092 | 1,704733348 | 0,10142 | 0,40833508 | kinesin family member 20A |
| ENSG00000198743 | SLC5A3 | 0,769199063 | 1,704323336 | 0,02339 | 0,2040793 | solute carrier family 5 member 3 |
| ENSG00000271369 | AC087783,1 | 0,768279032 | 1,703236807 | 0,08663 | 0,38130911 | NADH dehydrogenase (ubiquinone) 1 beta subcomplex, 8, 19kDa, pseudogene |
| ENSG00000258932 | AL390334,1 | 0,768171717 | 1,703110116 | 0,21578 | 0,53440695 | ribosomal protein S27a (RPS27A) pseudogene |
| ENSG00000234345 | ELF2P1 | 0,767834181 | 1,7027117 | 0,03663 | 0,25948322 | ELF2 pseudogene 1 |
| ENSG00000232093 | DCST1-AS1 | 0,767499733 | 1,70231702 | 0,18611 | 0,50956086 | DCST1 antisense RNA 1 |
| ENSG00000237732 | CT75 | 0,767010449 | 1,701739784 | 0,1744 | 0,49877327 | cancer/testis associated transcript 75 |
| ENSG00000127334 | DYRK2 | 0,766813882 | 1,701507938 | 0,01567 | 0,16407493 | dual specificity tyrosine phosphorylation regulated kinase 2 |
| ENSG00000170190 | SLC16A5 | 0,766329038 | 1,700936211 | 0,01911 | 0,18296265 | solute carrier family 16 member 5 |

| | | | | | | |
|---|---|---|---|---|---|---|
| ENSG00000284330 | AC256236,1 | 0,765954032 | 1,700494137 | 0,8266 | 0,91973568 | |
| ENSG00000151224 | MAT1A | 0,765952147 | 1,700491915 | 0,50622 | 0,73467375 | methionine adenosyltransferase 1A |
| ENSG00000236723 | AL606760,2 | 0,765523392 | 1,699986621 | 0,21507 | 0,53378006 | novel transcript, antisense to CPT2 |
| ENSG00000152684 | PELO | 0,765476105 | 1,699930902 | 0,0169 | 0,17086866 | pelota mRNA surveillance and ribosome rescue factor |
| ENSG00000050327 | ARHGEF5 | 0,764860034 | 1,699205138 | 0,17553 | 0,49954066 | Rho guanine nucleotide exchange factor 5 |
| ENSG00000140285 | FGF7 | 0,764409426 | 1,698674496 | 0,16731 | 0,49337433 | fibroblast growth factor 7 |
| ENSG00000164300 | SERINC5 | 0,764140218 | 1,698357551 | 0,02158 | 0,19535533 | serine incorporator 5 |
| ENSG00000238249 | HMGN2P17 | 0,764098659 | 1,698308629 | 0,45627 | 0,70003941 | high mobility group nucleosomal binding domain 2 pseudogene 17 |
| ENSG00000213443 | NPM1P7 | 0,763878849 | 1,698049892 | 0,1206 | 0,43944057 | nucleophosmin 1 pseudogene 7 |
| ENSG00000204816 | FGF7P5 | 0,763426603 | 1,697517683 | 0,49222 | 0,72545279 | fibroblast growth factor 7 pseudogene 5 |
| ENSG00000163235 | TGFA | 0,763386172 | 1,697470111 | 0,28892 | 0,58464073 | transforming growth factor alpha |
| ENSG00000170270 | GON7 | 0,763331675 | 1,697405991 | 0,40941 | 0,66795766 | GON7 subunit of KEOPS complex |
| ENSG00000237227 | AL162739,1 | 0,76318411 | 1,697232383 | 0,60024 | 0,79579686 | ribosomal protein L36 (RPL36) pseudogene |
| ENSG00000147224 | PRPS1 | 0,762624248 | 1,696573871 | 0,00016 | 0,00880308 | phosphoribosyl pyrophosphate synthetase 1 |
| ENSG00000262145 | ATP23 | 0,762514719 | 1,696445073 | 0,3286 | 0,60992716 | ATP23 metallopeptidase and ATP synthase assembly factor homolog |
| ENSG00000087842 | PIR | 0,762289558 | 1,696180329 | 0,0204 | 0,18959706 | pirin |
| ENSG00000234495 | TRIM27 | 0,761229134 | 1,694934044 | 0,07342 | 0,35662549 | tripartite motif containing 27 |
| ENSG00000228830 | AL160408,2 | 0,761148879 | 1,69483976 | 0,48461 | 0,72037164 | novel transcript |
| ENSG00000144354 | CDCA7 | 0,76114045 | 1,694829858 | 0,03394 | 0,24882853 | cell division cycle associated 7 |
| ENSG00000282413 | AL133461,1 | 0,760993345 | 1,694657052 | 0,0782 | 0,36540845 | VMA21 vacuolar H+-ATPase homolog (S, cerevisiae) (VMA21) pseudogene |
| ENSG00000179833 | SERTAD2 | 0,76025315 | 1,693787808 | 0,01264 | 0,14722977 | SERTA domain containing 2 |
| ENSG00000138315 | OIT3 | 0,760210483 | 1,693737716 | 0,40972 | 0,6680455 | oncoprotein induced transcript 3 |
| ENSG00000275004 | ZNF280B | 0,760191695 | 1,693715659 | 0,26164 | 0,56452671 | zinc finger protein 280B |
| ENSG00000214944 | ARHGEF28 | 0,759901629 | 1,693375157 | 0,02576 | 0,21654781 | Rho guanine nucleotide exchange factor 28 |
| ENSG00000024526 | DEPDC1 | 0,759540777 | 1,692951657 | 0,15135 | 0,47956749 | DEP domain containing 1 |
| ENSG00000270802 | AC005776,1 | 0,758987464 | 1,692302488 | 0,49405 | 0,72659022 | ribosomal protein L21 (RPL21) pseudogene |
| ENSG00000163602 | RYBP | 0,758509569 | 1,691742003 | 0,1177 | 0,43523522 | RING1 and YY1 binding protein |
| ENSG00000286347 | AC114802,1 | 0,758353776 | 1,691559327 | 0,20604 | 0,52555013 | novel transcript, antisense to ANTXR1 |
| ENSG00000137501 | SYTL2 | 0,758353159 | 1,691558603 | 0,00336 | 0,07000131 | synaptotagmin like 2 |
| ENSG00000119547 | ONECUT2 | 0,758344662 | 1,69154864 | 0,11404 | 0,42916778 | one cut homeobox 2 |

| | | | | | | |
|---|---|---|---|---|---|---|
| ENSG00000273897 | PMS2P8 | 0,758220115 | 1,691402615 | 0,42506 | 0,67871429 | PMS1 homolog 2, mismatch repair system component pseudogene 8 |
| ENSG00000108932 | SLC16A6 | 0,757737098 | 1,690836425 | 0,15021 | 0,47830235 | solute carrier family 16 member 6 |
| ENSG00000260265 | LINC02562 | 0,75767082 | 1,69075875 | 0,11467 | 0,43014087 | long intergenic non-protein coding RNA 2562 |
| ENSG00000126561 | STAT5A | 0,757635449 | 1,690717298 | 0,1431 | 0,47072472 | signal transducer and activator of transcription 5A |
| ENSG00000233673 | ANAPC1P1 | 0,757558094 | 1,690626647 | 0,45318 | 0,69750019 | anaphase promoting complex subunit 1 pseudogene 1 |
| ENSG00000281468 | AC006504,7 | 0,757412384 | 1,690455904 | 0,29006 | 0,58531219 | novel transcript, sense intronic to novel transcript |
| ENSG00000151287 | TEX30 | 0,757231231 | 1,690243655 | 0,15631 | 0,48409235 | testis expressed 30 |
| ENSG00000188338 | SLC38A3 | 0,756936691 | 1,689898611 | 0,06138 | 0,32865897 | solute carrier family 38 member 3 |
| ENSG00000242338 | BMS1P4 | 0,756439911 | 1,689316808 | 0,0565 | 0,31717392 | BMS1 pseudogene 4 |
| ENSG00000275079 | LPCAT1 | 0,756364326 | 1,689228304 | 0,07811 | 0,36540632 | lysophosphatidylcholine acyltransferase 1 |
| ENSG00000163071 | SPATA18 | 0,756301461 | 1,689154698 | 0,1445 | 0,47209141 | spermatogenesis associated 18 |
| ENSG00000228450 | AL049734,1 | 0,756222495 | 1,689062245 | 0,41426 | 0,67104294 | pseudogene similar to part of PYRIN-containing Apaf1-like protein 3 (PYPAF3) |
| ENSG00000287548 | AC021269,3 | 0,756157045 | 1,68898562 | 0,14503 | 0,47264896 | novel transcript |
| ENSG00000260799 | KRT8P50 | 0,755810651 | 1,68858014 | 0,40656 | 0,66597268 | keratin 8 pseudogene 50 |
| ENSG00000060718 | COL11A1 | 0,755712985 | 1,688465832 | 0,05372 | 0,31090219 | collagen type XI alpha 1 chain |
| ENSG00000286996 | AC099332,2 | 0,755384235 | 1,688081122 | 0,44898 | 0,69460209 | novel transcript |
| ENSG00000230397 | SPTLC1P1 | 0,754661693 | 1,687235894 | 0,25063 | 0,55690541 | serine palmitoyltransferase long chain base subunit 1 pseudogene 1 |
| ENSG00000256682 | AC006518,2 | 0,75454577 | 1,687100328 | 0,33167 | 0,61166253 | taste receptor, type 2, member 12 pseudogene |
| ENSG00000282363 | AC006518,8 | 0,75454577 | 1,687100328 | 0,33167 | 0,61166253 | taste receptor, type 2, member 12 pseudogene |
| ENSG00000272751 | AC006518,4 | 0,75454577 | 1,687100328 | 0,33167 | 0,61166253 | taste receptor, type 2, member 12 pseudogene |
| ENSG00000171700 | RGS19 | 0,754339093 | 1,686858656 | 0,00027 | 0,0132952 | regulator of G protein signaling 19 |
| ENSG00000172349 | IL16 | 0,754031526 | 1,686499073 | 0,1283 | 0,45171208 | interleukin 16 |
| ENSG00000205835 | GMNC | 0,753980259 | 1,686439144 | 0,23968 | 0,5491566 | geminin coiled-coil domain containing |
| ENSG00000278766 | AC092683,2 | 0,753735744 | 1,686153343 | 0,16789 | 0,49407144 | novel transcript |
| ENSG00000231500 | RPS18 | 0,753619487 | 1,686017472 | 0,19463 | 0,51562158 | ribosomal protein S18 |
| ENSG00000235033 | DAAM2-AS1 | 0,753278219 | 1,685618694 | 0,01635 | 0,16750744 | DAAM2 antisense RNA 1 |
| ENSG00000247595 | SPTY2D1OS | 0,75322764 | 1,685559599 | 0,04366 | 0,28188832 | SPTY2D1 opposite strand |
| ENSG00000088836 | SLC4A11 | 0,752426302 | 1,684623623 | 0,15499 | 0,48292767 | solute carrier family 4 member 11 |
| ENSG00000263567 | AC007923,1 | 0,752312257 | 1,684490459 | 0,41993 | 0,67551392 | novel transcript |
| ENSG00000256603 | AP003170,4 | 0,75181316 | 1,683907814 | 0,29127 | 0,5862741 | novel transcript |

| | | | | | | |
|---|---|---|---|---|---|---|
| ENSG00000181773 | GPR3 | 0,751644627 | 1,683711115 | 0,00254 | 0,0585725 | G protein-coupled receptor 3 |
| ENSG00000256947 | AP002518,2 | 0,751214565 | 1,683209281 | 0,27991 | 0,57755546 | novel transcript |
| ENSG00000135723 | FHOD1 | 0,751035199 | 1,683000026 | 0,00522 | 0,09023148 | formin homology 2 domain containing 1 |
| ENSG00000241409 | AC064852,1 | 0,750974804 | 1,682929573 | 0,34411 | 0,62149652 | novel transcript |
| ENSG00000256966 | AL513165,2 | 0,750887108 | 1,682827276 | 0,14988 | 0,47785837 | novel transcript |
| ENSG00000107566 | ERLIN1 | 0,750429042 | 1,682293052 | 0,04609 | 0,28930497 | ER lipid raft associated 1 |
| ENSG00000277806 | AC006213,4 | 0,749822149 | 1,681585517 | 0,28634 | 0,58267001 | novel transcript, sense intronic to ZNF155 |
| ENSG00000185304 | RGPD2 | 0,749791327 | 1,681549591 | 0,67693 | 0,84109542 | RANBP2 like and GRIP domain containing 2 |
| ENSG00000102287 | GABRE | 0,749291071 | 1,680966613 | 0,00722 | 0,10783655 | gamma-aminobutyric acid type A receptor subunit epsilon |
| ENSG00000165588 | OTX2 | 0,748568189 | 1,680124554 | 0,26594 | 0,56887049 | orthodenticle homeobox 2 |
| ENSG00000272599 | AC016394,1 | 0,748197888 | 1,679693367 | 0,10091 | 0,40770184 | novel transcript to NUDT13 |
| ENSG00000131634 | TMEM204 | 0,748124234 | 1,679607614 | 0,08427 | 0,37695424 | transmembrane protein 204 |
| ENSG00000100191 | SLC5A4 | 0,748039717 | 1,679509222 | 0,44609 | 0,69284796 | solute carrier family 5 member 4 |
| ENSG00000136110 | CNMD | 0,747293134 | 1,678640314 | 0,19565 | 0,51643293 | chondromodulin |
| ENSG00000112893 | MAN2A1 | 0,747260863 | 1,678602766 | 0,00808 | 0,11424852 | mannosidase alpha class 2A member 1 |
| ENSG00000241627 | UBQLN4P1 | 0,747113542 | 1,678431364 | 0,49402 | 0,72659022 | ubiquilin 4 pseudogene 1 |
| ENSG00000164442 | CITED2 | 0,746754062 | 1,678013196 | 0,00892 | 0,11991469 | Cbp/p300 interacting transactivator with Glu/Asp rich carboxy-terminal domain 2 |
| ENSG00000182379 | NXPH4 | 0,746651539 | 1,677893956 | 0,10451 | 0,41305105 | neurexophilin 4 |
| ENSG00000286181 | AC232271,3 | 0,746520765 | 1,677741868 | 0,14396 | 0,47144689 | FOXP3 regulating long intergenic non-coding RNA |
| ENSG00000232149 | FERP1 | 0,746431327 | 1,677637863 | 0,39295 | 0,65606871 | FER tyrosine kinase pseudogene 1 |
| ENSG00000280036 | AC020661,4 | 0,746419056 | 1,677623593 | 0,07112 | 0,35204529 | novel transcript, sense intronic to INO80 |
| ENSG00000216639 | AL133406,1 | 0,745821625 | 1,67692902 | 0,26649 | 0,56917012 | ribosomal protein L7 (RPL7) pseudogene |
| ENSG00000251667 | BRCC3P1 | 0,745669197 | 1,676751853 | 0,13269 | 0,45817003 | BRCA1/BRCA2-containing complex subunit 3 pseudogene 1 |
| ENSG00000225630 | MTND2P28 | 0,745659911 | 1,676741061 | 0,23545 | 0,54683669 | MT-ND2 pseudogene 28 |
| ENSG00000011304 | PTBP1 | 0,745573044 | 1,676640105 | 0,00073 | 0,02639057 | polypyrimidine tract binding protein 1 |
| ENSG00000166831 | RBPMS2 | 0,74478418 | 1,67572357 | 0,00074 | 0,02670595 | RNA binding protein, mRNA processing factor 2 |
| ENSG00000126010 | GRPR | 0,744704597 | 1,675631135 | 0,10573 | 0,41526993 | gastrin releasing peptide receptor |
| ENSG00000157107 | FCHO2 | 0,744591098 | 1,675499317 | 0,06567 | 0,33945082 | FCH and mu domain containing endocytic adaptor 2 |
| ENSG00000112343 | TRIM38 | 0,744544616 | 1,675445334 | 0,01915 | 0,18320213 | tripartite motif containing 38 |
| ENSG00000211829 | TRDC | 0,744375806 | 1,675249302 | 0,57109 | 0,77786458 | T cell receptor delta constant |
| ENSG00000278719 | MCM8-AS1 | 0,744102539 | 1,674932016 | 0,31479 | 0,601382 | MCM8 antisense RNA 1 |

| | | | | | | |
|---|---|---|---|---|---|---|
| | | | | | | mitochondrially encoded cytochrome c oxidase (MT-CO1) |
| ENSG00000242152 | AC108729,1 | 0,744101213 | 1,674930477 | 0,51529 | 0,74138689 | pseudogene |
| ENSG00000223791 | UBE2E1-AS1 | 0,743994225 | 1,674806271 | 0,33499 | 0,61467789 | UBE2E1 antisense RNA 1 |
| ENSG00000114529 | C3orf52 | 0,74386615 | 1,674657597 | 0,07321 | 0,35655334 | chromosome 3 open reading frame 52 |
| ENSG00000278091 | ZNF85 | 0,743409159 | 1,674127213 | 0,24289 | 0,55182091 | zinc finger protein 85 |
| ENSG00000165731 | RET | 0,743398659 | 1,674115029 | 0,07341 | 0,35662549 | ret proto-oncogene |
| ENSG00000274636 | GLIS2 | 0,743199132 | 1,673883512 | 0,05038 | 0,30188271 | GLIS family zinc finger 2 |
| ENSG00000228824 | MIR4500HG | 0,742973436 | 1,673621669 | 0,16029 | 0,48753847 | MIR4500 host gene |
| ENSG00000092531 | SNAP23 | 0,742810154 | 1,673432262 | 0,00377 | 0,07484606 | synaptosome associated protein 23 |
| ENSG00000253915 | MAPRE1P1 | 0,742479563 | 1,673048842 | 0,10294 | 0,41078404 | MAPRE1 pseudogene 1 |
| ENSG00000288077 | AC027129,1 | 0,742266731 | 1,672802045 | 0,35839 | 0,63185282 | novel transcript, antisense to COLQ |
| ENSG00000100385 | IL2RB | 0,742144838 | 1,672660717 | 0,19704 | 0,51824743 | interleukin 2 receptor subunit beta |
| ENSG00000267781 | SLC25A36P1 | 0,742100479 | 1,672609287 | 0,5345 | 0,75405583 | SLC25A36 pseudogene 1 |
| ENSG00000271870 | AC024060,2 | 0,742098921 | 1,672607481 | 0,40107 | 0,66295784 | novel transcript, antisense to CRBN |
| ENSG00000275591 | XKR5 | 0,741688363 | 1,672131563 | 0,0302 | 0,23452348 | XK related 5 |
| ENSG00000271303 | SRXN1 | 0,741631472 | 1,672065626 | 0,01688 | 0,17073473 | sulfiredoxin 1 |
| ENSG00000181856 | SLC2A4 | 0,74103637 | 1,671376053 | 0,30743 | 0,59658791 | solute carrier family 2 member 4 |
| ENSG00000137473 | TTC29 | 0,740995884 | 1,67132915 | 0,16291 | 0,48945635 | tetratricopeptide repeat domain 29 |
| ENSG00000132640 | BTBD3 | 0,740474496 | 1,670725243 | 0,04517 | 0,2876168 | BTB domain containing 3 |
| ENSG00000254154 | CRYZL2P-SEC1 | 0,740028834 | 1,67020922 | 0,1642 | 0,49065086 | CRYZL2P-SEC16B readthrough |
| ENSG00000288658 | AC010980,1 | 0,739878992 | 1,670035757 | 0,07806 | 0,36538634 | novel protein, ortholog of Gm2102 (M, musculus) |
| ENSG00000261528 | CCNYL7 | 0,739425271 | 1,669510621 | 0,15457 | 0,48266733 | cyclin Y like 7 (pseudogene) |
| ENSG00000101333 | PLCB4 | 0,739391224 | 1,669471222 | 0,11481 | 0,43027964 | phospholipase C beta 4 |
| ENSG00000095015 | MAP3K1 | 0,739076788 | 1,669107399 | 0,05336 | 0,31018511 | mitogen-activated protein kinase kinase kinase 1 |
| ENSG00000285532 | LINC02788 | 0,738352496 | 1,668269648 | 0,30628 | 0,59519791 | long intergenic non-protein coding RNA 2788 |
| ENSG00000234997 | AC016745,1 | 0,738095784 | 1,667972825 | 0,26865 | 0,57070906 | novel transcript |
| ENSG00000266744 | AC005304,3 | 0,737912385 | 1,667760801 | 0,34746 | 0,62411645 | novel transcript |
| ENSG00000111328 | CDK2AP1 | 0,73788247 | 1,66772622 | 0,10101 | 0,40770184 | cyclin dependent kinase 2 associated protein 1 |
| ENSG00000173535 | TNFRSF10C | 0,737517099 | 1,667303912 | 0,2906 | 0,58565836 | TNF receptor superfamily member 10c |
| ENSG00000212122 | TSSK1B | 0,737167478 | 1,666899908 | 0,60462 | 0,79801151 | testis specific serine kinase 1B |
| ENSG00000256417 | AC006206,2 | 0,737013089 | 1,666721535 | 0,14242 | 0,46962982 | novel transcript |
| ENSG00000282844 | ACTN4 | 0,736851741 | 1,666535144 | 0,03027 | 0,23475421 | actinin alpha 4 |
| ENSG00000113328 | CCNG1 | 0,736274576 | 1,665868563 | 0,16148 | 0,48899841 | cyclin G1 |

| | | | | | | |
|---|---|---|---|---|---|---|
| ENSG00000131759 | RARA | 0,736125059 | 1,665695925 | 0,00721 | 0,10778628 | retinoic acid receptor alpha |
| ENSG00000120549 | KIAA1217 | 0,736015337 | 1,665569248 | 0,06411 | 0,33569301 | KIAA1217 |
| ENSG00000285900 | AC079177,1 | 0,735484257 | 1,664956238 | 0,14542 | 0,47300165 | novel transcript, antisense to DMD |
| ENSG00000270571 | AC007681,1 | 0,735066088 | 1,664473715 | 0,21279 | 0,53180216 | novel transcript, antisense to TACR1 |
| ENSG00000242242 | NECTIN3-AS1 | 0,734982036 | 1,664376745 | 0,0744 | 0,35811443 | NECTIN3 antisense RNA 1 |
| ENSG00000166268 | MYRFL | 0,733971411 | 1,663211238 | 0,22294 | 0,53908993 | myelin regulatory factor like |
| ENSG00000258776 | AL161757,4 | 0,73395458 | 1,663191834 | 0,07567 | 0,36117248 | novel transcript |
| ENSG00000182771 | GRID1 | 0,733105329 | 1,662213075 | 0,13362 | 0,45953428 | glutamate ionotropic receptor delta type subunit 1 |
| ENSG00000221914 | PPP2R2A | 0,733073146 | 1,662175995 | 0,06871 | 0,34648746 | protein phosphatase 2 regulatory subunit Balpha |
| ENSG00000260807 | CEROX1 | 0,732790443 | 1,661850316 | 0,06967 | 0,34826564 | cytoplasmic endogenous regulator of oxidative phosphorylation 1 |
| ENSG00000229082 | OR5AW1P | 0,732075202 | 1,661026629 | 0,31941 | 0,60429866 | olfactory receptor family 5 subfamily AW member 1 pseudogene |
| ENSG00000277959 | AL162274,2 | 0,731024013 | 1,659816798 | 0,37472 | 0,6431311 | novel transcript, antisense to BNIP3 |
| ENSG00000250820 | AC138965,1 | 0,730570164 | 1,659294728 | 0,48372 | 0,71988524 | novel transcript, sense overlapping to THOC3 |
| ENSG00000102313 | ITIH6 | 0,730507338 | 1,659222471 | 0,51305 | 0,7394947 | inter-alpha-trypsin inhibitor heavy chain family member 6 |
| ENSG00000184347 | SLIT3 | 0,730392527 | 1,659090434 | 0,02328 | 0,20394642 | slit guidance ligand 3 |
| ENSG00000261594 | TPBGL | 0,730340299 | 1,659030373 | 0,24929 | 0,55642837 | trophoblast glycoprotein like |
| ENSG00000271379 | AC023513,1 | 0,730220897 | 1,658893073 | 0,41043 | 0,66846487 | family with sequence similarity 173, member B (FAM173B) pseudogene |
| ENSG00000184160 | ADRA2C | 0,730140883 | 1,65880107 | 0,34733 | 0,6240601 | adrenoceptor alpha 2C |
| ENSG00000170340 | B3GNT2 | 0,730034595 | 1,658678866 | 0,01023 | 0,1300932 | UDP-GlcNAc:betaGal beta-1,3-N-acetylglucosaminyltransferase 2 |
| ENSG00000101608 | MYL12A | 0,729862427 | 1,658480934 | 0,07684 | 0,3633846 | myosin light chain 12A |
| ENSG00000137801 | THBS1 | 0,729817711 | 1,658429531 | 0,00832 | 0,11538752 | thrombospondin 1 |
| ENSG00000183010 | PYCR1 | 0,729595989 | 1,658174673 | 0,01255 | 0,14662712 | pyrroline-5-carboxylate reductase 1 |
| ENSG00000079337 | RAPGEF3 | 0,729535607 | 1,658105274 | 0,08859 | 0,38491339 | Rap guanine nucleotide exchange factor 3 |
| ENSG00000254090 | MTND2P32 | 0,729499884 | 1,658064217 | 0,21736 | 0,53526451 | MT-ND2 pseudogene 32 |
| ENSG00000130429 | ARPC1B | 0,729359745 | 1,657903166 | 0,00539 | 0,09144669 | actin related protein 2/3 complex subunit 1B |
| ENSG00000217527 | RPS16P5 | 0,728804477 | 1,657265191 | 0,31121 | 0,59885232 | ribosomal protein S16 pseudogene 5 |
| ENSG00000155542 | SETD9 | 0,728409314 | 1,656811318 | 0,11469 | 0,43014087 | SET domain containing 9 |
| ENSG00000254319 | AC246817,2 | 0,728405958 | 1,656807464 | 0,11389 | 0,42900168 | novel transcript |
| ENSG00000109107 | ALDOC | 0,728127566 | 1,656487787 | 0,06107 | 0,32783036 | aldolase, fructose-bisphosphate C |

| | | | | | | |
|---|---|---|---|---|---|---|
| ENSG00000124575 | H1-3 | 0,727953885 | 1,656288379 | 0,00803 | 0,11389729 | H1,3 linker histone, cluster member |
| ENSG00000011566 | MAP4K3 | 0,727901851 | 1,656228643 | 0,03389 | 0,24867791 | mitogen-activated protein kinase kinase kinase kinase 3 |
| ENSG00000275322 | AC103746,1 | 0,72759414 | 1,655875426 | 0,26581 | 0,56871984 | novel transcript |
| ENSG00000182722 | SEPHS1P1 | 0,727133369 | 1,655346652 | 0,3727 | 0,6416226 | selenophosphate synthetase 1 pseudogene 1 |
| ENSG00000269897 | COMMD3-BM | 0,727087819 | 1,65529439 | 0,5058 | 0,73418581 | COMMD3-BMI1 readthrough |
| ENSG00000267699 | AC091551,1 | 0,726849771 | 1,655021285 | 0,22737 | 0,54168591 | novel protein |
| ENSG00000244219 | TMEM225B | 0,726592989 | 1,654726737 | 0,25398 | 0,55914254 | transmembrane protein 225B |
| ENSG00000115850 | LCT | 0,726137343 | 1,654204207 | 0,39988 | 0,66197344 | lactase |
| ENSG00000198771 | RCSD1 | 0,725616405 | 1,653607004 | 0,44327 | 0,69084105 | RCSD domain containing 1 |
| ENSG00000177105 | RHOG | 0,72539173 | 1,653349504 | 0,06506 | 0,33804854 | ras homolog family member G |
| ENSG00000255222 | SETP17 | 0,724530574 | 1,652362901 | 0,49984 | 0,73058737 | SET pseudogene 17 |
| ENSG00000121316 | PLBD1 | 0,724452973 | 1,652274024 | 0,27712 | 0,57572679 | phospholipase B domain containing 1 |
| ENSG00000242114 | MTFP1 | 0,724434646 | 1,652253035 | 0,01319 | 0,15012165 | mitochondrial fission process 1 |
| ENSG00000232023 | LINC01807 | 0,724311451 | 1,652111952 | 0,35801 | 0,63185282 | long intergenic non-protein coding RNA 1807 |
| ENSG00000225674 | IPO7P2 | 0,723352347 | 1,651013992 | 0,10302 | 0,41078404 | importin 7 pseudogene 2 |
| ENSG00000261728 | AL138690,1 | 0,722866482 | 1,650458064 | 0,44582 | 0,69284796 | novel transcript |
| ENSG00000267402 | TCF4-AS2 | 0,722818698 | 1,650403399 | 0,16475 | 0,49092529 | TCF4 antisense RNA 2 |
| ENSG00000196796 | NPIPB10P | 0,722408264 | 1,649933941 | 0,45334 | 0,69750019 | nuclear pore complex interacting protein family, member B10, pseudogene |
| ENSG00000286132 | AC022415,2 | 0,722145045 | 1,649632939 | 0,29023 | 0,58540802 | novel protein |
| ENSG00000196531 | NACA | 0,721977088 | 1,649440902 | 0,15347 | 0,48185226 | nascent polypeptide associated complex subunit alpha |
| ENSG00000044524 | EPHA3 | 0,72196984 | 1,649432615 | 0,27686 | 0,57549599 | EPH receptor A3 |
| ENSG00000275947 | TMEM251 | 0,721825729 | 1,649267861 | 0,29647 | 0,58978642 | transmembrane protein 251 |
| ENSG00000142676 | RPL11 | 0,721689777 | 1,649112451 | 0,14338 | 0,47083258 | ribosomal protein L11 |
| ENSG00000276787 | FAN1 | 0,721583641 | 1,648991133 | 0,26719 | 0,56958284 | FANCD2 and FANCI associated nuclease 1 |
| ENSG00000124610 | H1-1 | 0,72143082 | 1,648816469 | 0,52235 | 0,74653683 | H1,1 linker histone, cluster member |
| ENSG00000101445 | PPP1R16B | 0,721253283 | 1,648613579 | 0,21661 | 0,53490925 | protein phosphatase 1 regulatory subunit 16B |
| ENSG00000233651 | AL590399,5 | 0,721219919 | 1,648575453 | 0,65001 | 0,8246264 | novel transcript |
| ENSG00000158806 | NPM2 | 0,721131404 | 1,64847431 | 0,12231 | 0,44185689 | nucleophosmin/nucleoplasmin 2 |
| ENSG00000100678 | SLC8A3 | 0,720924161 | 1,648237524 | 0,02488 | 0,21153603 | solute carrier family 8 member A3 |
| ENSG00000121644 | DESI2 | 0,7208499 | 1,648152685 | 0,07268 | 0,35563034 | desumoylating isopeptidase 2 |
| ENSG00000176018 | LYSMD3 | 0,720836628 | 1,648137523 | 0,22844 | 0,54241083 | LysM domain containing 3 |
| ENSG00000285278 | TFAP2A-AS2 | 0,720717072 | 1,648000947 | 0,05321 | 0,30971589 | TFAP2A antisense RNA 2 |

| | | | | | | |
|---|---|---|---|---|---|---|
| ENSG00000169641 | LUZP1 | 0,720455775 | 1,647702493 | 0,00229 | 0,05480145 | leucine zipper protein 1 |
| ENSG00000076770 | MBNL3 | 0,720426076 | 1,647668575 | 0,10308 | 0,41078404 | muscleblind like splicing regulator 3 |
| ENSG00000214041 | PGAM1P4 | 0,720374898 | 1,647610126 | 0,40846 | 0,66723037 | phosphoglycerate mutase 1 pseudogene 4 |
| ENSG00000203814 | H2BC18 | 0,72028356 | 1,647505818 | 0,2388 | 0,54892264 | H2B clustered histone 18 |
| ENSG00000280994 | LINC01347 | 0,720277995 | 1,647499463 | 0,45791 | 0,70103159 | long intergenic non-protein coding RNA 1347 |
| ENSG00000214110 | LDHAP4 | 0,720124273 | 1,647323928 | 0,11912 | 0,43696075 | lactate dehydrogenase A pseudogene 4 |
| ENSG00000273523 | AL139082,1 | 0,720025919 | 1,647211628 | 0,18585 | 0,50915451 | novel transcript, antisense to NEK3 & NEK5 |
| ENSG00000186280 | KDM4D | 0,719981598 | 1,647161025 | 0,04917 | 0,29853148 | lysine demethylase 4D |
| ENSG00000233029 | AC244453,2 | 0,719976727 | 1,647155463 | 0,69126 | 0,84945675 | novel transcript |
| ENSG00000144035 | NAT8 | 0,719866142 | 1,647029211 | 0,21687 | 0,53505085 | N-acetyltransferase 8 (putative) |
| ENSG00000272688 | AP005329,3 | 0,719857607 | 1,647019467 | 0,1661 | 0,492091 | novel transcript |
| ENSG00000148154 | UGCG | 0,719840297 | 1,646999705 | 0,06333 | 0,33335893 | UDP-glucose ceramide glucosyltransferase |
| ENSG00000145192 | AHSG | 0,719631784 | 1,646761681 | 0,35336 | 0,62813516 | alpha 2-HS glycoprotein |
| ENSG00000255647 | AC093510,1 | 0,719565651 | 1,646686195 | 0,228 | 0,54213739 | novel transcript |
| ENSG00000285600 | AC023593,1 | 0,719032637 | 1,646077928 | 0,07274 | 0,35572609 | Novel transcript, antisense to MCM2 |
| ENSG00000150867 | PIP4K2A | 0,719011815 | 1,646054171 | 0,08941 | 0,38670267 | phosphatidylinositol-5-phosphate 4-kinase type 2 alpha |
| ENSG00000224892 | RPS4XP16 | 0,718957895 | 1,645992652 | 0,41377 | 0,67085666 | ribosomal protein S4X pseudogene 16 |
| ENSG00000236494 | AC008080,4 | 0,718890937 | 1,64591626 | 0,41722 | 0,6738387 | novel transcript |
| ENSG00000152104 | PTPN14 | 0,718864283 | 1,645885852 | 0,02734 | 0,22320635 | protein tyrosine phosphatase non-receptor type 14 |
| ENSG00000110944 | IL23A | 0,718808061 | 1,645821712 | 0,34903 | 0,62519057 | interleukin 23 subunit alpha |
| ENSG00000091428 | RAPGEF4 | 0,718566762 | 1,645546462 | 0,04745 | 0,29254611 | Rap guanine nucleotide exchange factor 4 |
| ENSG00000156466 | GDF6 | 0,718565319 | 1,645544817 | 0,08133 | 0,37209365 | growth differentiation factor 6 |
| ENSG00000213881 | NPM1P6 | 0,718398427 | 1,64535447 | 0,27333 | 0,57344244 | nucleophosmin 1 pseudogene 6 |
| ENSG00000183508 | TENT5C | 0,718319505 | 1,645264464 | 0,02232 | 0,19848029 | terminal nucleotidyltransferase 5C |
| ENSG00000198756 | COLGALT2 | 0,718263286 | 1,645200352 | 0,06267 | 0,33130676 | collagen beta(1-O)galactosyltransferase 2 |
| ENSG00000106565 | TMEM176B | 0,717940997 | 1,644832866 | 0,36111 | 0,63316311 | transmembrane protein 176B |
| ENSG00000277152 | AC110048,2 | 0,717261411 | 1,644058244 | 0,20799 | 0,52756478 | novel transcript |
| ENSG00000121577 | POPDC2 | 0,717242428 | 1,644036612 | 0,21815 | 0,53569444 | popeye domain containing 2 |
| ENSG00000095951 | HIVEP1 | 0,717136199 | 1,643915562 | 0,05036 | 0,30181338 | HIVEP zinc finger 1 |
| ENSG00000153094 | BCL2L11 | 0,717119569 | 1,643896613 | 0,03892 | 0,26698555 | BCL2 like 11 |
| ENSG00000220867 | HSPE1P26 | 0,716803988 | 1,64353706 | 0,29408 | 0,58837921 | heat shock protein family E (Hsp10) member 1 pseudogene 26 |
| ENSG00000129084 | PSMA1 | 0,716627162 | 1,643335629 | 0,18481 | 0,50844776 | proteasome 20S subunit alpha 1 |

| | | | | | | |
|---|---|---|---|---|---|---|
| ENSG00000137960 | GIPC2 | 0,716554342 | 1,643252684 | 0,25492 | 0,5594539 | GIPC PDZ domain containing family member 2 |
| ENSG00000230427 | ALG14-AS1 | 0,716420711 | 1,643100484 | 0,35277 | 0,62759135 | ALG14 antisense RNA 1 |
| ENSG00000105281 | SLC1A5 | 0,716327674 | 1,642994526 | 0,00653 | 0,10244112 | solute carrier family 1 member 5 |
| ENSG00000231822 | SMC3P1 | 0,716279191 | 1,642939313 | 0,1666 | 0,49255702 | SMC3 pseudogene 1 |
| ENSG00000015413 | DPEP1 | 0,715744709 | 1,642330758 | 0,48254 | 0,71901075 | dipeptidase 1 |
| ENSG00000163378 | EOGT | 0,715654749 | 1,642228353 | 0,03071 | 0,23637289 | EGF domain specific O-linked N-acetylglucosamine transferase |
| ENSG00000070882 | OSBPL3 | 0,715058945 | 1,641550287 | 0,14177 | 0,46830089 | oxysterol binding protein like 3 |
| ENSG00000172795 | DCP2 | 0,714962805 | 1,641440898 | 0,04324 | 0,28025067 | decapping mRNA 2 |
| ENSG00000288632 | AC133555,6 | 0,714885335 | 1,641352758 | 0,11493 | 0,43039322 | novel transcript |
| ENSG00000259547 | CYCSP2 | 0,714803479 | 1,641259633 | 0,57653 | 0,78091994 | CYCS pseudogene 2 |
| ENSG00000176055 | MBLAC2 | 0,714796855 | 1,641252097 | 0,06631 | 0,34094389 | metallo-beta-lactamase domain containing 2 |
| ENSG00000241438 | TDGF1P6 | 0,714744621 | 1,641192676 | 0,43314 | 0,68455289 | teratocarcinoma-derived growth factor 1 pseudogene 6 |
| ENSG00000270720 | AC104785,1 | 0,714306595 | 1,640694459 | 0,29393 | 0,58836931 | novel transcript |
| ENSG00000254838 | GVINP1 | 0,714303878 | 1,640691369 | 0,03219 | 0,24204878 | GTPase, very large interferon inducible pseudogene 1 |
| ENSG00000205795 | CYS1 | 0,71397504 | 1,640317443 | 0,40151 | 0,6630277 | cystin 1 |
| ENSG00000286939 | U82671,1 | 0,713800688 | 1,640119221 | 0,46187 | 0,70317543 | novel transcript, antisense to ZNF185 |
| ENSG00000259803 | SLC22A31 | 0,713657127 | 1,639956022 | 0,35758 | 0,63140243 | solute carrier family 22 member 31 |
| ENSG00000104808 | DHDH | 0,713525274 | 1,639806147 | 0,35158 | 0,62675995 | dihydrodiol dehydrogenase |
| ENSG00000260608 | AC024270,3 | 0,71346706 | 1,639739981 | 0,41333 | 0,67078587 | novel transcript, antisense to BTBD1 |
| ENSG00000214013 | GANC | 0,713379621 | 1,639640603 | 0,03494 | 0,25281694 | glucosidase alpha, neutral C |
| ENSG00000286070 | AP000356,5 | 0,71336033 | 1,639618678 | 0,66165 | 0,831498 | novel protein |
| ENSG00000235092 | ID2-AS1 | 0,713257426 | 1,639501732 | 0,06913 | 0,34731919 | ID2 antisense RNA 1 |
| ENSG00000141179 | PCTP | 0,713232541 | 1,639473452 | 0,00429 | 0,0804932 | phosphatidylcholine transfer protein |
| ENSG00000180152 | XIAPP3 | 0,71292227 | 1,6391209 | 0,38241 | 0,64928247 | X-linked inhibitor of apoptosis pseudogene 3 |
| ENSG00000036448 | MYOM2 | 0,712854144 | 1,6390435 | 0,44235 | 0,69006979 | myomesin 2 |
| ENSG00000234851 | RPL23AP42 | 0,712659385 | 1,638822249 | 0,14619 | 0,47428193 | ribosomal protein L23a pseudogene 42 |
| ENSG00000254615 | AC027031,2 | 0,712657172 | 1,638819736 | 0,02152 | 0,19510325 | novel transcript |
| ENSG00000234769 | WASH4P | 0,712290318 | 1,638403063 | 0,292 | 0,5868753 | WASP family homolog 4, pseudogene |
| ENSG00000259080 | AC074091,2 | 0,712250756 | 1,638358135 | 0,46635 | 0,70665143 | novel transcript |
| ENSG00000274588 | DGKK | 0,712034048 | 1,638112055 | 0,14911 | 0,47668027 | diacylglycerol kinase kappa |
| ENSG00000095777 | MYO3A | 0,711433816 | 1,637430661 | 0,11845 | 0,43614969 | myosin IIIA |
| ENSG00000066382 | MPPED2 | 0,711182958 | 1,637145968 | 0,01452 | 0,15751905 | metallophosphoesterase domain containing 2 |

| | | | | | | |
|---|---|---|---|---|---|---|
| ENSG00000243697 | AC009108,1 | 0,711069759 | 1,637017517 | 0,29949 | 0,59139511 | ribosomal protein L7a (RPL7A) pseudogene |
| ENSG00000186395 | KRT10 | 0,710889434 | 1,636812916 | 0,22109 | 0,53805234 | keratin 10 |
| ENSG00000288185 | IL27RA | 0,710608088 | 1,636493745 | 0,05511 | 0,31415946 | interleukin 27 receptor subunit alpha |
| ENSG00000104998 | IL27RA | 0,710608088 | 1,636493745 | 0,05511 | 0,31415946 | interleukin 27 receptor subunit alpha |
| ENSG00000180257 | ZNF816 | 0,710606174 | 1,636491574 | 0,13729 | 0,4642578 | zinc finger protein 816 |
| ENSG00000128059 | PPAT | 0,710303956 | 1,636148795 | 0,05063 | 0,30274123 | phosphoribosyl pyrophosphate amidotransferase |
| ENSG00000267436 | AC005786,3 | 0,710235626 | 1,636071304 | 0,19431 | 0,51562158 | novel transcript, antisense to MFSD12 |
| ENSG00000225386 | AC099754,1 | 0,709995746 | 1,635799294 | 0,20226 | 0,5224664 | novel transcript |
| ENSG00000139946 | PELI2 | 0,709885279 | 1,635674045 | 0,08082 | 0,37094893 | pellino E3 ubiquitin protein ligase family member 2 |
| ENSG00000221974 | GTF2H4 | 0,709828708 | 1,635609908 | 0,21571 | 0,53440695 | general transcription factor IIH subunit 4 |
| ENSG00000233149 | GTF2H4 | 0,709828708 | 1,635609908 | 0,21571 | 0,53440695 | general transcription factor IIH subunit 4 |
| ENSG00000100344 | PNPLA3 | 0,709725228 | 1,635492595 | 0,00118 | 0,03613087 | patatin like phospholipase domain containing 3 |
| ENSG00000241399 | CD302 | 0,709475378 | 1,635209381 | 0,013 | 0,14927519 | CD302 molecule |
| ENSG00000266573 | AC018697,1 | 0,709397234 | 1,635120812 | 0,1 | 0,40656692 | novel transcript |
| ENSG00000151414 | NEK7 | 0,709219896 | 1,634919833 | 0,0962 | 0,39965673 | NIMA related kinase 7 |
| ENSG00000224549 | AC017067,1 | 0,709144673 | 1,63483459 | 0,34829 | 0,62466592 | novel transcript |
| ENSG00000228502 | EEF1A1P11 | 0,709096164 | 1,634779621 | 0,44441 | 0,69184225 | eukaryotic translation elongation factor 1 alpha 1 pseudogene 11 |
| ENSG00000288246 | AC140725,6 | 0,708984594 | 1,634653201 | 0,27081 | 0,57239492 | novel transcript |
| ENSG00000259553 | AC140725,1 | 0,708984594 | 1,634653201 | 0,27081 | 0,57239492 | novel transcript |
| ENSG00000162702 | ZNF281 | 0,708917937 | 1,634577677 | 0,13245 | 0,45752148 | zinc finger protein 281 |
| ENSG00000142856 | ITGB3BP | 0,708851492 | 1,634502396 | 0,17021 | 0,49606609 | integrin subunit beta 3 binding protein |
| ENSG00000101096 | NFATC2 | 0,708729695 | 1,634364411 | 0,06146 | 0,32865897 | nuclear factor of activated T cells 2 |
| ENSG00000213735 | ANAPC10P1 | 0,708544829 | 1,634154999 | 0,38936 | 0,65372387 | anaphase promoting complex subunit 10 pseudogene 1 |
| ENSG00000260238 | PMF1-BGLAP | 0,708412745 | 1,634005393 | 0,25173 | 0,55773568 | PMF1-BGLAP readthrough |
| ENSG00000066813 | ACSM2B | 0,708185017 | 1,633747488 | 0,45241 | 0,69702004 | acyl-CoA synthetase medium chain family member 2B |
| ENSG00000134709 | HOOK1 | 0,707767686 | 1,633274958 | 0,21353 | 0,53275614 | hook microtubule tethering protein 1 |
| ENSG00000229855 | AC008568,1 | 0,707701082 | 1,633199558 | 0,2235 | 0,53953208 | novel transcript |
| ENSG00000180596 | H2BC4 | 0,707673757 | 1,633168625 | 0,25059 | 0,55690541 | H2B clustered histone 4 |
| ENSG00000229647 | MYOSLID | 0,707206607 | 1,632639884 | 0,39144 | 0,65510995 | myocardin-induced smooth muscle lncRNA, inducer of differentiation |
| ENSG00000237481 | AL117350,1 | 0,707060653 | 1,632474722 | 0,30002 | 0,59160746 | novel transcript |
| ENSG00000187554 | TLR5 | 0,707001032 | 1,63240726 | 0,13671 | 0,46406381 | toll like receptor 5 |

| | | | | | | |
|---|---|---|---|---|---|---|
| ENSG00000126247 | CAPNS1 | 0,706791473 | 1,632170162 | 0,11774 | 0,43532264 | calpain small subunit 1 |
| ENSG00000105835 | NAMPT | 0,706537223 | 1,631882546 | 0,12858 | 0,45191041 | nicotinamide phosphoribosyltransferase |
| ENSG00000198889 | DCAF12L1 | 0,706004989 | 1,631280628 | 0,21763 | 0,53547632 | DDB1 and CUL4 associated factor 12 like 1 |
| ENSG00000169548 | ZNF280A | 0,705937909 | 1,631204782 | 0,50036 | 0,73081513 | zinc finger protein 280A |
| ENSG00000275499 | ZNF280A | 0,705937909 | 1,631204782 | 0,50036 | 0,73081513 | zinc finger protein 280A |
| ENSG00000253773 | C8orf37-AS1 | 0,705757903 | 1,631001267 | 0,28775 | 0,58398675 | C8orf37 antisense RNA 1 |
| ENSG00000197465 | GYPE | 0,70529683 | 1,630480096 | 0,3665 | 0,63640899 | glycophorin E (MNS blood group) |
| ENSG00000080608 | PUM3 | 0,705113629 | 1,630273063 | 0,09132 | 0,3908781 | pumilio RNA binding family member 3 |
| ENSG00000108958 | AC130689,1 | 0,704850657 | 1,629975926 | 0,36823 | 0,63803074 | succinate dehydrogenase complex, subunit C, integral membrane protein, 15kDa (SDHC) pseudogene |
| ENSG00000253762 | AC069133,1 | 0,704737806 | 1,629848431 | 0,12835 | 0,45177856 | novel transcript |
| ENSG00000151012 | SLC7A11 | 0,704628884 | 1,629725383 | 0,04613 | 0,28930497 | solute carrier family 7 member 11 |
| ENSG00000263322 | AC110597,2 | 0,704383156 | 1,629447824 | 0,31284 | 0,59981841 | novel transcript |
| ENSG00000161970 | RPL26 | 0,704350719 | 1,629411187 | 0,10257 | 0,41028143 | ribosomal protein L26 |
| ENSG00000111057 | KRT18 | 0,704127479 | 1,629159075 | 0,23189 | 0,5446505 | keratin 18 |
| ENSG00000086827 | ZW10 | 0,70403796 | 1,629057989 | 0,12346 | 0,44412763 | zw10 kinetochore protein |
| ENSG00000237440 | ZNF737 | 0,703993939 | 1,629008283 | 0,00131 | 0,03846063 | zinc finger protein 737 |
| ENSG00000239830 | RPS4XP22 | 0,703761206 | 1,628745515 | 0,39668 | 0,65929197 | ribosomal protein S4X pseudogene 22 |
| ENSG00000178358 | OR2D3 | 0,703695031 | 1,628670808 | 0,19793 | 0,51960355 | olfactory receptor family 2 subfamily D member 3 |
| ENSG00000237238 | BMS1P10 | 0,703539165 | 1,628494859 | 0,07722 | 0,36386039 | BMS1 pseudogene 10 |
| ENSG00000166750 | SLFN5 | 0,703380239 | 1,628315475 | 0,04727 | 0,29217708 | schlafen family member 5 |
| ENSG00000188487 | INSC | 0,703175991 | 1,628084964 | 0,54938 | 0,76407481 | INSC spindle orientation adaptor protein |
| ENSG00000179344 | HLA-DQB1 | 0,703071309 | 1,627966835 | 0,33805 | 0,61769428 | major histocompatibility complex, class II, DQ beta 1 |
| ENSG00000275713 | H2BC9 | 0,703031728 | 1,627922171 | 0,2324 | 0,54513458 | H2B clustered histone 9 |
| ENSG00000274349 | ZNF658 | 0,70300414 | 1,627891041 | 0,19464 | 0,51562158 | zinc finger protein 658 |
| ENSG00000287985 | AC073270,2 | 0,702599171 | 1,627434151 | 0,38825 | 0,6529693 | novel transcript |
| ENSG00000173706 | HEG1 | 0,702472621 | 1,627291403 | 0,00683 | 0,10540108 | heart development protein with EGF like domains 1 |
| ENSG00000235946 | H2BW3P | 0,70212959 | 1,626904526 | 0,07721 | 0,36386039 | H2B,W histone 3, pseudogene |
| ENSG00000174028 | FAM3C2P | 0,701946043 | 1,626697557 | 0,28867 | 0,58455903 | family with sequence similarity 3 member C2, pseudogene |
| ENSG00000213033 | AURKAP1 | 0,701703034 | 1,626423577 | 0,10413 | 0,41251856 | aurora kinase A pseudogene 1 |
| ENSG00000185130 | H2BC13 | 0,701619502 | 1,626329409 | 0,14444 | 0,47208992 | H2B clustered histone 13 |
| ENSG00000234324 | RPL9P2 | 0,701429538 | 1,62611528 | 0,36816 | 0,6380264 | ribosomal protein L9 pseudogene 2 |

| | | | | | | |
|---|---|---|---|---|---|---|
| ENSG00000155438 | NIFK | 0,701395541 | 1,626076961 | 0,10097 | 0,40770184 | nucleolar protein interacting with the FHA domain of MKI67 |
| ENSG00000198331 | HYLS1 | 0,701375589 | 1,626054473 | 0,04619 | 0,28930497 | HYLS1 centriolar and ciliogenesis associated |
| ENSG00000112245 | PTP4A1 | 0,701256605 | 1,625920372 | 0,24878 | 0,55591283 | protein tyrosine phosphatase 4A1 |
| ENSG00000030304 | MUSK | 0,701172709 | 1,625825824 | 0,25999 | 0,56379428 | muscle associated receptor tyrosine kinase |
| ENSG00000233164 | C6orf136 | 0,701114737 | 1,625760495 | 0,15332 | 0,48157724 | chromosome 6 open reading frame 136 |
| ENSG00000049130 | KITLG | 0,700939095 | 1,625562578 | 0,09445 | 0,39676397 | KIT ligand |
| ENSG00000272672 | AL391497,1 | 0,700924295 | 1,625545901 | 0,22179 | 0,53818364 | novel transcript |
| ENSG00000235251 | AC093155,2 | 0,700400782 | 1,624956144 | 0,33913 | 0,61812227 | pseudogene similar to part of SUMO-1 activating enzyme subunit 2 (UBA2) |
| ENSG00000175820 | CCDC168 | 0,700387611 | 1,624941309 | 0,16798 | 0,49417064 | coiled-coil domain containing 168 |
| ENSG00000223518 | CSNK1A1P1 | 0,700294005 | 1,624835882 | 0,21416 | 0,53334744 | casein kinase 1 alpha 1 pseudogene 1 |
| ENSG00000163347 | CLDN1 | 0,699910723 | 1,624404268 | 0,31697 | 0,60268749 | claudin 1 |
| ENSG00000274746 | ZNF100 | 0,699716136 | 1,624185187 | 0,04498 | 0,28673676 | zinc finger protein 100 |
| ENSG00000224885 | EIPR1-IT1 | 0,699481475 | 1,623921028 | 0,30609 | 0,59513628 | EIPR1 intronic transcript 1 |
| ENSG00000273786 | AC020658,4 | 0,699468348 | 1,623906252 | 0,20689 | 0,52654093 | novel transcript, sense intronic to novel protein |
| ENSG00000259609 | AC027807,1 | 0,699164749 | 1,623564555 | 0,5666 | 0,77458849 | family with sequence similarity 154, member B (Fam154b) pseudogene |
| ENSG00000260033 | AC023813,2 | 0,698970075 | 1,62334549 | 0,31059 | 0,59852818 | ribosomal L24 domain containing 1 (RSL24D1) pseudogene |
| ENSG00000168589 | DYNLRB2 | 0,698511653 | 1,622829747 | 0,46335 | 0,70419128 | dynein light chain roadblock-type 2 |
| ENSG00000137478 | FCHSD2 | 0,698212639 | 1,622493433 | 0,02755 | 0,22376406 | FCH and double SH3 domains 2 |
| ENSG00000277954 | AC092376,2 | 0,697718255 | 1,621937531 | 0,15107 | 0,47946774 | novel transcript, antisense to WWOX |
| ENSG00000284902 | AC074008,1 | 0,697405753 | 1,621586242 | 0,17916 | 0,50364038 | novel transcript, sense intronic to ALMS1 |
| ENSG00000004399 | PLXND1 | 0,697311689 | 1,621480516 | 0,00753 | 0,11001711 | plexin D1 |
| ENSG00000286394 | AC117373,2 | 0,697311128 | 1,621479886 | 0,53192 | 0,75280487 | novel transcript, antisense to TMEM132D |
| ENSG00000171174 | RBKS | 0,697089726 | 1,621231067 | 0,12718 | 0,45032932 | ribokinase |
| ENSG00000082213 | C5orf22 | 0,696752756 | 1,62085244 | 0,05967 | 0,32521754 | chromosome 5 open reading frame 22 |
| ENSG00000112210 | RAB23 | 0,696726955 | 1,620823453 | 0,03566 | 0,2562416 | RAB23, member RAS oncogene family |
| ENSG00000137628 | DDX60 | 0,696406707 | 1,620463704 | 0,16694 | 0,49293221 | DExD/H-box helicase 60 |
| ENSG00000276649 | AL117335,1 | 0,696104746 | 1,620124571 | 0,20995 | 0,52885604 | novel transcript, antisense to SIRPA |
| ENSG00000235531 | MSC-AS1 | 0,695845686 | 1,619833677 | 0,17457 | 0,49884018 | MSC antisense RNA 1 |
| ENSG00000233892 | PAIP1P1 | 0,695546262 | 1,619497524 | 0,40362 | 0,66399439 | poly(A) binding protein interacting protein 1 pseudogene 1 |
| ENSG00000116285 | ERRFI1 | 0,695528713 | 1,619477824 | 0,01615 | 0,16645939 | ERBB receptor feedback inhibitor 1 |

| | | | | | | |
|---|---|---|---|---|---|---|
| ENSG00000231401 | AC023481,1 | 0,695104979 | 1,619002237 | 0,17066 | 0,49641416 | novel transcript |
| ENSG00000095464 | PDE6C | 0,694513708 | 1,618338844 | 0,33877 | 0,61792351 | phosphodiesterase 6C |
| ENSG00000163528 | CHCHD4 | 0,694175988 | 1,617960053 | 0,14135 | 0,46803321 | coiled-coil-helix-coiled-coil-helix domain containing 4 |
| ENSG00000231310 | TBL1XR1-AS1 | 0,693933251 | 1,617687849 | 0,33339 | 0,61335912 | TBL1XR1 antisense RNA 1 |
| ENSG00000101680 | LAMA1 | 0,693906902 | 1,617658305 | 0,00298 | 0,06507521 | laminin subunit alpha 1 |
| ENSG00000107551 | RASSF4 | 0,693840821 | 1,617584212 | 0,01386 | 0,15454988 | Ras association domain family member 4 |
| ENSG00000109189 | USP46 | 0,693701885 | 1,617428441 | 0,08677 | 0,38157196 | ubiquitin specific peptidase 46 |
| ENSG00000177954 | RPS27 | 0,693639091 | 1,617358043 | 0,27318 | 0,57344244 | ribosomal protein S27 |
| ENSG00000255220 | DDX18P5 | 0,693180707 | 1,616844245 | 0,24249 | 0,55137204 | DEAD-box helicase 18 pseudogene 5 |
| ENSG00000118276 | B4GALT6 | 0,69312208 | 1,616778543 | 0,16451 | 0,49065086 | beta-1,4-galactosyltransferase 6 |
| ENSG00000181227 | DLSTP1 | 0,693031854 | 1,616677432 | 0,13955 | 0,46677981 | dihydrolipoamide S-succinyltransferase pseudogene 1 |
| ENSG00000101986 | ABCD1 | 0,692644898 | 1,61624387 | 0,00087 | 0,02936196 | ATP binding cassette subfamily D member 1 |
| ENSG00000107798 | LIPA | 0,692430199 | 1,616003361 | 0,15695 | 0,48427418 | lipase A, lysosomal acid type |
| ENSG00000147003 | CLTRN | 0,69221528 | 1,615762642 | 0,14765 | 0,47507961 | collectrin, amino acid transport regulator |
| ENSG00000229751 | AC044781,1 | 0,692201195 | 1,615746868 | 0,54021 | 0,75814808 | novel transcript |
| ENSG00000176142 | TMEM39A | 0,692064982 | 1,615594323 | 0,07338 | 0,35662549 | transmembrane protein 39A |
| ENSG00000136897 | MRPL50 | 0,692038602 | 1,615564782 | 0,18347 | 0,50703725 | mitochondrial ribosomal protein L50 |
| ENSG00000245857 | GS1-24F4,2 | 0,691709653 | 1,615196459 | 0,309 | 0,59774312 | uncharacterized LOC100652791 |
| ENSG00000118640 | VAMP8 | 0,691640135 | 1,615118631 | 0,42594 | 0,67940091 | vesicle associated membrane protein 8 |
| ENSG00000186522 | SEPTIN10 | 0,691242965 | 1,614674054 | 0,11512 | 0,43076625 | septin 10 |
| ENSG00000150760 | DOCK1 | 0,691123487 | 1,614540339 | 0,04246 | 0,2782038 | dedicator of cytokinesis 1 |
| ENSG00000117586 | TNFSF4 | 0,690896714 | 1,614286574 | 0,23076 | 0,54378056 | TNF superfamily member 4 |
| ENSG00000133119 | RFC3 | 0,690563169 | 1,613913401 | 0,23971 | 0,5491566 | replication factor C subunit 3 |
| ENSG00000285608 | AL161665,1 | 0,690040417 | 1,613328715 | 0,37549 | 0,6438135 | novel transcript, antisense to ARHGAP5 |
| ENSG00000270105 | AC136475,8 | 0,689799121 | 1,613058902 | 0,58264 | 0,7847455 | novel transcript |
| ENSG00000151332 | MBIP | 0,689767924 | 1,613024022 | 0,23718 | 0,54813266 | MAP3K12 binding inhibitory protein 1 |
| ENSG00000259692 | AC104041,1 | 0,689706659 | 1,612955525 | 0,01405 | 0,15563367 | novel transcript |
| ENSG00000273375 | AC055764,2 | 0,689692824 | 1,612940058 | 0,31949 | 0,60429866 | novel transcript |
| ENSG00000198093 | ZNF649 | 0,689662623 | 1,612906294 | 0,11821 | 0,43597399 | zinc finger protein 649 |
| ENSG00000141905 | NFIC | 0,689595746 | 1,612831528 | 0,02511 | 0,21287111 | nuclear factor I C |
| ENSG00000107738 | VSIR | 0,689191775 | 1,61237998 | 0,29985 | 0,59160746 | V-set immunoregulatory receptor |
| ENSG00000255107 | AC079089,2 | 0,689026661 | 1,612195456 | 0,47885 | 0,7165637 | BAG family molecular chaperone regulator 4 (BAG4) pseudogene |

| | | | | | | |
|---|---|---|---|---|---|---|
| ENSG00000219102 | HNRNPA3P12 | 0,689024404 | 1,612192933 | 0,22202 | 0,53851366 | heterogeneous nuclear ribonucleoprotein A3 pseudogene 12 |
| ENSG00000058668 | ATP2B4 | 0,688989644 | 1,61215409 | 0,05581 | 0,31585374 | ATPase plasma membrane Ca2+ transporting 4 |
| ENSG00000230928 | AL139241,1 | 0,68897514 | 1,612137883 | 0,15841 | 0,48554971 | novel transcript |
| ENSG00000235934 | AC007405,2 | 0,68895104 | 1,612110953 | 0,3862 | 0,65163285 | novel transcript |
| ENSG00000136068 | FLNB | 0,688715672 | 1,611847966 | 0,01373 | 0,15370548 | filamin B |
| ENSG00000225890 | HLA-DQA1 | 0,688592391 | 1,611710237 | 0,23123 | 0,54417127 | major histocompatibility complex, class II, DQ alpha 1 |
| ENSG00000068438 | FTSJ1 | 0,688512102 | 1,611620545 | 0,02679 | 0,2209591 | FtsJ RNA 2'-O-methyltransferase 1 |
| ENSG00000134873 | CLDN10 | 0,688343294 | 1,611431982 | 0,2956 | 0,58930301 | claudin 10 |
| ENSG00000131473 | ACLY | 0,687893319 | 1,610929457 | 0,04976 | 0,30012887 | ATP citrate lyase |
| ENSG00000197056 | ZMYM1 | 0,687233259 | 1,610192595 | 0,21064 | 0,52920077 | zinc finger MYM-type containing 1 |
| ENSG00000138801 | PAPSS1 | 0,687212504 | 1,61016943 | 0,06794 | 0,34439007 | 3'-phosphoadenosine 5'-phosphosulfate synthase 1 |
| ENSG00000197043 | ANXA6 | 0,687212315 | 1,61016922 | 0,00513 | 0,08937569 | annexin A6 |
| ENSG00000272002 | AC010904,2 | 0,687009223 | 1,609942567 | 0,32368 | 0,60638344 | novel transcript |
| ENSG00000153902 | LGI4 | 0,686677783 | 1,609572747 | 0,08592 | 0,38009386 | leucine rich repeat LGI family member 4 |
| ENSG00000283515 | AC020915,4 | 0,686522351 | 1,609399346 | 0,38139 | 0,64874863 | novel protein |
| ENSG00000067955 | CBFB | 0,686424555 | 1,609290253 | 0,01651 | 0,16833274 | core-binding factor subunit beta |
| ENSG00000286767 | AL023877,1 | 0,686205974 | 1,609046449 | 0,44408 | 0,6916877 | novel transcript |
| ENSG00000287261 | AL139243,1 | 0,68613437 | 1,608966591 | 0,35616 | 0,63035942 | novel transcript, antisense to HPS1 |
| ENSG00000132842 | AP3B1 | 0,686048555 | 1,608870888 | 0,11007 | 0,4224508 | adaptor related protein complex 3 subunit beta 1 |
| ENSG00000163510 | CWC22 | 0,6859133 | 1,608720062 | 0,16422 | 0,49065086 | CWC22 spliceosome associated protein homolog |
| ENSG00000270765 | GAS2L2 | 0,685708855 | 1,608492105 | 0,31548 | 0,60172912 | growth arrest specific 2 like 2 |
| ENSG00000128052 | KDR | 0,685593711 | 1,608363734 | 0,21433 | 0,53334744 | kinase insert domain receptor |
| ENSG00000233010 | RPEP4 | 0,685376818 | 1,608121952 | 0,31699 | 0,60268749 | ribulose-5-phosphate-3-epimerase pseudogene 4 |
| ENSG00000083444 | PLOD1 | 0,685323005 | 1,60806197 | 0,00547 | 0,0920741 | procollagen-lysine,2-oxoglutarate 5-dioxygenase 1 |
| ENSG00000273175 | BX323046,1 | 0,685058655 | 1,607767346 | 0,27783 | 0,57629036 | novel transcript |
| ENSG00000288172 | AL356419,1 | 0,684905202 | 1,607596344 | 0,17811 | 0,50236177 | novel transcript |
| ENSG00000262525 | TAS2R46 | 0,684804946 | 1,607484634 | 0,27178 | 0,57305058 | taste 2 receptor member 46 |
| ENSG00000226761 | TAS2R46 | 0,684804946 | 1,607484634 | 0,27178 | 0,57305058 | taste 2 receptor member 46 |
| ENSG00000224280 | AC005014,1 | 0,684736034 | 1,607407851 | 0,3866 | 0,65194712 | ataxin 2-like (ATXN2L) pseudogene |
| ENSG00000198938 | MT-CO3 | 0,684674062 | 1,607338806 | 0,17743 | 0,50191142 | mitochondrially encoded cytochrome c oxidase III |
| ENSG00000184867 | ARMCX2 | 0,684046361 | 1,606639623 | 0,03469 | 0,25183223 | armadillo repeat containing X-linked 2 |

| | | | | | | |
|---|---|---|---|---|---|---|
| ENSG00000170222 | ADPRM | 0,683692962 | 1,606246112 | 0,28601 | 0,58240438 | ADP-ribose/CDP-alcohol diphosphatase, manganese dependent |
| ENSG00000232814 | COL4A2-AS1 | 0,683520585 | 1,606054205 | 0,31197 | 0,59925736 | COL4A2 antisense RNA 1 |
| ENSG00000164163 | ABCE1 | 0,683451429 | 1,60597722 | 0,1044 | 0,41285329 | ATP binding cassette subfamily E member 1 |
| ENSG00000272156 | AC008280,2 | 0,683333048 | 1,605845446 | 0,34129 | 0,61939402 | novel transcript |
| ENSG00000085871 | MGST2 | 0,683224679 | 1,605724827 | 0,18002 | 0,50407792 | microsomal glutathione S-transferase 2 |
| ENSG00000224121 | ATG12P2 | 0,682985337 | 1,60545846 | 0,26217 | 0,56507214 | autophagy related 12 pseudogene 2 |
| ENSG00000263606 | CHORDC1P4 | 0,682798898 | 1,605251001 | 0,1443 | 0,47189017 | CHORDC1 pseudogene 4 |
| ENSG00000091127 | PUS7 | 0,682700748 | 1,605141795 | 0,09181 | 0,39174496 | pseudouridine synthase 7 |
| ENSG00000188313 | PLSCR1 | 0,682276642 | 1,604670005 | 0,08143 | 0,37234025 | phospholipid scramblase 1 |
| ENSG00000228056 | CFL1P3 | 0,68219602 | 1,604580333 | 0,30128 | 0,59258306 | cofilin 1 pseudogene 3 |
| ENSG00000259399 | TGIF2-RAB5IF | 0,682059286 | 1,604428264 | 0,2574 | 0,56160845 | TGIF2-RAB5IF readthrough |
| ENSG00000274569 | AC024884,1 | 0,681983849 | 1,604344372 | 0,47385 | 0,71294959 | apolipoprotein F (APOF) pseudogene |
| ENSG00000248866 | USP46-DT | 0,681871416 | 1,604219346 | 0,01042 | 0,13153018 | USP46 divergent transcript |
| ENSG00000276704 | AL442067,2 | 0,681312261 | 1,603597709 | 0,31017 | 0,5983747 | novel transcript, sense intronic to RAP2A |
| ENSG00000230581 | ACTG1P14 | 0,681303945 | 1,603588465 | 0,58263 | 0,7847455 | actin gamma 1 pseudogene 14 |
| ENSG00000249853 | HS3ST5 | 0,681227879 | 1,603503919 | 0,00894 | 0,11991469 | heparan sulfate-glucosamine 3-sulfotransferase 5 |
| ENSG00000068697 | LAPTM4A | 0,681155661 | 1,603423653 | 0,01738 | 0,17367715 | lysosomal protein transmembrane 4 alpha |
| ENSG00000236426 | AL583824,1 | 0,68094197 | 1,603186172 | 0,38893 | 0,65351197 | novel transcript |
| ENSG00000213453 | FTH1P3 | 0,680914375 | 1,603155508 | 0,43365 | 0,68473723 | ferritin heavy chain 1 pseudogene 3 |
| ENSG00000159712 | ANKRD18CP | 0,680270499 | 1,602440178 | 0,10376 | 0,4119916 | ankyrin repeat domain 18C, pseudogene |
| ENSG00000156017 | CARNMT1 | 0,680242611 | 1,602409201 | 0,06164 | 0,32867767 | carnosine N-methyltransferase 1 |
| ENSG00000162757 | C1orf74 | 0,680052007 | 1,602197511 | 0,16689 | 0,49287023 | chromosome 1 open reading frame 74 |
| ENSG00000165621 | OXGR1 | 0,679762162 | 1,601875654 | 0,32248 | 0,60577115 | oxoglutarate receptor 1 |
| ENSG00000236156 | CHCHD4P3 | 0,679508875 | 1,601594444 | 0,3571 | 0,63118328 | coiled-coil-helix-coiled-coil-helix domain containing 4 pseudogene 3 |
| ENSG00000069956 | MAPK6 | 0,67928427 | 1,601345121 | 0,13507 | 0,46159318 | mitogen-activated protein kinase 6 |
| ENSG00000204899 | MZT1 | 0,679182713 | 1,601232399 | 0,15147 | 0,47963227 | mitotic spindle organizing protein 1 |
| ENSG00000164294 | GPX8 | 0,678944997 | 1,600968583 | 0,03103 | 0,23730035 | glutathione peroxidase 8 (putative) |
| ENSG00000176994 | SMCR8 | 0,678890616 | 1,600908237 | 0,46047 | 0,70229998 | SMCR8-C9orf72 complex subunit |
| ENSG00000160973 | FOXH1 | 0,678702857 | 1,600699901 | 0,49183 | 0,72530727 | forkhead box H1 |
| ENSG00000109113 | RAB34 | 0,678666038 | 1,600659051 | 0,00375 | 0,07474822 | RAB34, member RAS oncogene family |
| ENSG00000169122 | FAM110B | 0,678645376 | 1,600636126 | 0,00432 | 0,08088101 | family with sequence similarity 110 member B |

| | | | | | | |
|---|---|---|---|---|---|---|
| ENSG00000005022 | SLC25A5 | 0,67846039 | 1,600430901 | 0,11806 | 0,43562845 | solute carrier family 25 member 5 |
| ENSG00000186288 | PABPC1L2A | 0,678097616 | 1,600028515 | 0,53875 | 0,75715394 | poly(A) binding protein cytoplasmic 1 like 2A |
| ENSG00000189171 | S100A13 | 0,677960409 | 1,599876351 | 0,03949 | 0,2689738 | S100 calcium binding protein A13 |
| ENSG00000264217 | RPL35AP23 | 0,677421723 | 1,599279087 | 0,43052 | 0,68295004 | ribosomal protein L35a pseudogene 23 |
| ENSG00000273645 | KBTBD11 | 0,677320705 | 1,59916711 | 0,69514 | 0,85206566 | kelch repeat and BTB domain containing 11 |
| ENSG00000248740 | LINC02428 | 0,677173735 | 1,599004207 | 0,02383 | 0,20614699 | long intergenic non-protein coding RNA 2428 |
| ENSG00000196636 | SDHAF3 | 0,67664708 | 1,598420598 | 0,25279 | 0,55796133 | succinate dehydrogenase complex assembly factor 3 |
| ENSG00000276663 | AC009090,3 | 0,676593019 | 1,598360703 | 0,29205 | 0,5868753 | novel transcript, antisense to ARL2BP |
| ENSG00000182347 | PDSS1P1 | 0,676367492 | 1,598110862 | 0,20747 | 0,52696827 | decaprenyl diphosphate synthase subunit 1 pseudogene 1 |
| ENSG00000142556 | ZNF614 | 0,676036086 | 1,597743797 | 0,2073 | 0,52677042 | zinc finger protein 614 |
| ENSG00000148942 | SLC5A12 | 0,6756559 | 1,597322807 | 0,24705 | 0,55476618 | solute carrier family 5 member 12 |
| ENSG00000107036 | RIC1 | 0,675338814 | 1,596971774 | 0,04153 | 0,27616337 | RIC1 homolog, RAB6A GEF complex partner 1 |
| ENSG00000060749 | QSER1 | 0,675087398 | 1,596693497 | 0,19017 | 0,51331106 | glutamine and serine rich 1 |
| ENSG00000273387 | AC005005,3 | 0,67505559 | 1,596658294 | 0,09556 | 0,3985545 | novel transcript, antisense to SMTN |
| ENSG00000257949 | TEN1 | 0,675027006 | 1,59662666 | 0,31204 | 0,59925736 | TEN1 subunit of CST complex |
| ENSG00000198416 | ZNF658B | 0,675007794 | 1,596605399 | 0,16276 | 0,48945635 | zinc finger protein 658B (pseudogene) |
| ENSG00000257964 | AC020612,3 | 0,674933199 | 1,596522847 | 0,2432 | 0,55199251 | novel transcript, antisense to PRPF40B |
| ENSG00000112246 | SIM1 | 0,674336527 | 1,595862692 | 0,18615 | 0,50956086 | SIM bHLH transcription factor 1 |
| ENSG00000237754 | AC093655,2 | 0,67415607 | 1,595663088 | 0,44632 | 0,69300607 | septin 13 (SEPT13) pseudogene |
| ENSG00000166002 | SMCO4 | 0,673962398 | 1,595448896 | 0,00668 | 0,10370681 | single-pass membrane protein with coiled-coil domains 4 |
| ENSG00000277050 | AL122125,1 | 0,673775314 | 1,595242017 | 0,38545 | 0,65112398 | novel transcript, antisense to FRMD6 |
| ENSG00000228915 | OR7E128P | 0,673716453 | 1,595176933 | 0,1372 | 0,4642578 | olfactory receptor family 7 subfamily E member 128 pseudogene |
| ENSG00000138750 | NUP54 | 0,673702221 | 1,595161197 | 0,20574 | 0,52518691 | nucleoporin 54 |
| ENSG00000138772 | ANXA3 | 0,672989597 | 1,594373456 | 0,13745 | 0,464516 | annexin A3 |
| ENSG00000119457 | SLC46A2 | 0,672875363 | 1,594247218 | 0,56348 | 0,7724837 | solute carrier family 46 member 2 |
| ENSG00000136603 | SKIL | 0,672747028 | 1,594105407 | 0,14678 | 0,47470036 | SKI like proto-oncogene |
| ENSG00000115128 | SF3B6 | 0,672356382 | 1,593673822 | 0,24864 | 0,55591283 | splicing factor 3b subunit 6 |
| ENSG00000234444 | ZNF736 | 0,672311571 | 1,593624322 | 0,01934 | 0,18420145 | zinc finger protein 736 |
| ENSG00000251332 | PSME2P4 | 0,672182173 | 1,593481393 | 0,31016 | 0,5983747 | proteasome activator subunit 2 pseudogene 4 |
| ENSG00000175868 | CALCB | 0,67215289 | 1,593449049 | 0,34435 | 0,62180651 | calcitonin related polypeptide beta |
| ENSG00000184908 | CLCNKB | 0,671840118 | 1,593103632 | 0,20855 | 0,52778377 | chloride voltage-gated channel Kb |
| ENSG00000182132 | KCNIP1 | 0,671362262 | 1,592576044 | 0,09548 | 0,39838572 | potassium voltage-gated channel interacting protein 1 |

| | | | | | | |
|---|---|---|---|---|---|---|
| ENSG00000160838 | LRRC71 | 0,671121576 | 1,592310376 | 0,33159 | 0,61165554 | leucine rich repeat containing 71 |
| ENSG00000232528 | AL109809,1 | 0,670980067 | 1,592154199 | 0,41362 | 0,67085666 | pseudogene similar to F830045P16Rik |
| ENSG00000229644 | NAMPTP1 | 0,670792046 | 1,591946713 | 0,32529 | 0,60778059 | nicotinamide phosphoribosyltransferase pseudogene 1 |
| ENSG00000070785 | EIF2B3 | 0,670724412 | 1,591872085 | 0,12093 | 0,43998535 | eukaryotic translation initiation factor 2B subunit gamma |
| ENSG00000170266 | GLB1 | 0,670508079 | 1,5916334 | 0,08795 | 0,3838874 | galactosidase beta 1 |
| ENSG00000241127 | YAE1 | 0,670354684 | 1,591464178 | 0,24815 | 0,55542983 | YAE1 maturation factor of ABCE1 |
| ENSG00000268894 | PLCE1-AS1 | 0,670095397 | 1,591178179 | 0,24649 | 0,5541223 | PLCE1 antisense RNA 1 |
| ENSG00000259865 | AL390728,6 | 0,669876547 | 1,590936823 | 0,24322 | 0,55199251 | novel transcript |
| ENSG00000233297 | RASA4DP | 0,669722088 | 1,590766502 | 0,52452 | 0,74828894 | RAS p21 protein activator 4CD, pseudogene |
| ENSG00000169413 | RNASE6 | 0,669673496 | 1,590712924 | 0,41615 | 0,67303919 | ribonuclease A family member k6 |
| ENSG00000220695 | Z86062,1 | 0,669583609 | 1,590613818 | 0,54215 | 0,75894841 | RP42 homolog (RP42) pseudogene |
| ENSG00000247746 | USP51 | 0,669533184 | 1,590558224 | 0,05866 | 0,3237033 | ubiquitin specific peptidase 51 |
| ENSG00000114423 | CBLB | 0,669511642 | 1,590534474 | 0,0429 | 0,27914798 | Cbl proto-oncogene B |
| ENSG00000235677 | NPM1P26 | 0,669183615 | 1,590172874 | 0,27085 | 0,57239492 | nucleophosmin 1 pseudogene 26 |
| ENSG00000176222 | ZNF404 | 0,669056203 | 1,590032444 | 0,19012 | 0,51331106 | zinc finger protein 404 |
| ENSG00000172901 | LVRN | 0,668954117 | 1,589919936 | 0,44449 | 0,69185302 | laeverin |
| ENSG00000088179 | PTPN4 | 0,668832805 | 1,58978625 | 0,1059 | 0,4154675 | protein tyrosine phosphatase non-receptor type 4 |
| ENSG00000234193 | ACSL3-AS1 | 0,668201556 | 1,589090794 | 0,5029 | 0,73217708 | ACSL3 antisense RNA 1 |
| ENSG00000285292 | ABCF2-H2BE1 | 0,668126081 | 1,589007662 | 0,11641 | 0,43288152 | ABCF2-H2BE1 readthrough |
| ENSG00000260920 | AL031985,3 | 0,668037828 | 1,588910462 | 0,08237 | 0,37394502 | novel transcript, sense overlapping with ZNF643 |
| ENSG00000070669 | ASNS | 0,667948174 | 1,588811724 | 0,2248 | 0,54036191 | asparagine synthetase (glutamine-hydrolyzing) |
| ENSG00000122877 | EGR2 | 0,667927912 | 1,58878941 | 0,33806 | 0,61769428 | early growth response 2 |
| ENSG00000268869 | ESPNP | 0,667702546 | 1,588541242 | 0,11293 | 0,42702832 | espin pseudogene |
| ENSG00000255139 | AP000442,1 | 0,667612996 | 1,588442643 | 0,31257 | 0,59946361 | novel transcript |
| ENSG00000273369 | AC096586,2 | 0,667409102 | 1,588218166 | 0,24818 | 0,55542983 | novel transcript, antisense to GUF1 |
| ENSG00000041357 | PSMA4 | 0,667400816 | 1,588209044 | 0,24028 | 0,54948131 | proteasome 20S subunit alpha 4 |
| ENSG00000126733 | DACH2 | 0,667257511 | 1,588051293 | 0,30106 | 0,59248675 | dachshund family transcription factor 2 |
| ENSG00000277544 | AC104073,3 | 0,667172312 | 1,587957513 | 0,24906 | 0,55619875 | novel zinc finger protein pseudogene |
| ENSG00000267751 | AC009005,1 | 0,667161128 | 1,587945203 | 0,10463 | 0,41313952 | novel transcript, antisense to BSG |
| ENSG00000104419 | NDRG1 | 0,66674044 | 1,587482227 | 0,04029 | 0,27141252 | N-myc downstream regulated 1 |
| ENSG00000241369 | LINC01192 | 0,666726864 | 1,587467289 | 0,41318 | 0,67067821 | long intergenic non-protein coding RNA 1192 |
| ENSG00000107833 | NPM3 | 0,666525043 | 1,587245231 | 0,01911 | 0,18296265 | nucleophosmin/nucleoplasmin 3 |
| ENSG00000282486 | LRRC37A9P | 0,666495711 | 1,58721296 | 0,51293 | 0,73946844 | leucine rich repeat containing 37 member A9, pseudogene |

| | | | | | | |
|---|---|---|---|---|---|---|
| ENSG00000197977 | ELOVL2 | 0,66611839 | 1,586797897 | 0,22459 | 0,54036191 | ELOVL fatty acid elongase 2 |
| ENSG00000128595 | CALU | 0,666038647 | 1,586710191 | 0,09711 | 0,40152057 | calumenin |
| ENSG00000233178 | AL161457,2 | 0,665992529 | 1,58665947 | 0,0241 | 0,20752957 | novel transcript, antisense to PGM5 |
| ENSG00000220392 | FCF1P5 | 0,665794318 | 1,586441495 | 0,29621 | 0,58961035 | FCF1 pseudogene 5 |
| ENSG00000256167 | ATF4P4 | 0,665643044 | 1,586275157 | 0,45872 | 0,70133296 | activating transcription factor 4 pseudogene 4 |
| ENSG00000273245 | AC092653,1 | 0,665609827 | 1,586238635 | 0,46507 | 0,70560072 | novel transcript |
| ENSG00000125686 | MED1 | 0,665384513 | 1,585990921 | 0,10921 | 0,42097623 | mediator complex subunit 1 |
| ENSG00000131462 | TUBG1 | 0,665216261 | 1,585805969 | 0,08143 | 0,37234025 | tubulin gamma 1 |
| ENSG00000100129 | EIF3L | 0,664999781 | 1,585568032 | 0,24159 | 0,55039378 | eukaryotic translation initiation factor 3 subunit L |
| ENSG00000255396 | AP000857,1 | 0,664872126 | 1,585427742 | 0,4699 | 0,70928807 | ras homolog gene family, member B (RHOB) pseudogene |
| ENSG00000180346 | TIGD2 | 0,664473842 | 1,584990114 | 0,20204 | 0,52230471 | tigger transposable element derived 2 |
| ENSG00000230409 | TCEA1P2 | 0,664464714 | 1,584980086 | 0,32084 | 0,60558211 | transcription elongation factor A1 pseudogene 2 |
| ENSG00000186063 | AIDA | 0,66445902 | 1,58497383 | 0,05259 | 0,30799404 | axin interactor, dorsalization associated |
| ENSG00000273353 | AL008718,3 | 0,664372297 | 1,584878557 | 0,54153 | 0,75882662 | novel transcript, sense intronic to NUP50 |
| ENSG00000273275 | AC017083,2 | 0,664261789 | 1,584757163 | 0,8491 | 0,93130502 | novel transcript |
| ENSG00000224903 | RNF32-AS1 | 0,66421071 | 1,584701055 | 0,13077 | 0,45532399 | RNF32 antisense RNA 1 |
| ENSG00000231185 | SPRY4-AS1 | 0,664206539 | 1,584696474 | 0,38921 | 0,65363949 | SPRY4 antisense RNA 1 |
| ENSG00000083857 | FAT1 | 0,664160043 | 1,584645402 | 0,06749 | 0,34374257 | FAT atypical cadherin 1 |
| ENSG00000068650 | ATP11A | 0,664084855 | 1,584562818 | 0,02884 | 0,22968658 | ATPase phospholipid transporting 11A |
| ENSG00000206190 | ATP10A | 0,663714248 | 1,58415582 | 0,00543 | 0,0916301 | ATPase phospholipid transporting 10A (putative) |
| ENSG00000156006 | NAT2 | 0,663602313 | 1,584032914 | 0,38803 | 0,65289784 | N-acetyltransferase 2 |
| ENSG00000231799 | PA2G4P6 | 0,662403664 | 1,582717382 | 0,30934 | 0,59780895 | proliferation-associated 2G4 pseudogene 6 |
| ENSG00000144713 | RPL32 | 0,662294769 | 1,582597923 | 0,16703 | 0,49293221 | ribosomal protein L32 |
| ENSG00000286615 | AC011416,4 | 0,662040408 | 1,582318921 | 0,31344 | 0,6002104 | novel transcript |
| ENSG00000116044 | NFE2L2 | 0,661940432 | 1,582209272 | 0,11881 | 0,43692176 | nuclear factor, erythroid 2 like 2 |
| ENSG00000138326 | RPS24 | 0,661928832 | 1,582196551 | 0,19067 | 0,51369186 | ribosomal protein S24 |
| ENSG00000235655 | H3P6 | 0,661850476 | 1,58211062 | 0,70161 | 0,85613754 | H3 histone pseudogene 6 |
| ENSG00000120798 | NR2C1 | 0,661718555 | 1,581965957 | 0,08445 | 0,37732439 | nuclear receptor subfamily 2 group C member 1 |
| ENSG00000143819 | EPHX1 | 0,661418552 | 1,581637029 | 0,00704 | 0,10676605 | epoxide hydrolase 1 |
| ENSG00000275454 | AC105020,6 | 0,661358792 | 1,581571515 | 0,19891 | 0,52005179 | novel transcript, antisense to IMP3 |
| ENSG00000169019 | COMMD8 | 0,661346314 | 1,581557835 | 0,22699 | 0,54168591 | COMM domain containing 8 |
| ENSG00000204392 | LSM2 | 0,661211319 | 1,581409854 | 0,01685 | 0,17053963 | LSM2 homolog, U6 small nuclear RNA and mRNA degradation associated |

| | | | | | | |
|---|---|---|---|---|---|---|
| ENSG00000120800 | UTP20 | 0,66098657 | 1,581163514 | 0,062 | 0,32961785 | UTP20 small subunit processome component |
| ENSG00000185267 | CDNF | 0,660873579 | 1,581039683 | 0,1502 | 0,47830235 | cerebral dopamine neurotrophic factor |
| ENSG00000231296 | AL050341,1 | 0,660855158 | 1,581019496 | 0,42689 | 0,6801859 | pseudogene similar to part of RIO kinase 2 (yeast) (RIOK2) |
| ENSG00000176907 | TCIM | 0,660585545 | 1,58072406 | 0,17887 | 0,50335172 | transcriptional and immune response regulator |
| ENSG00000223379 | AL391987,3 | 0,660427822 | 1,580551256 | 0,332 | 0,61202931 | novel transcript |
| ENSG00000254473 | AL354920,1 | 0,660359817 | 1,580476756 | 0,10196 | 0,40933045 | novel transcript, antisense to UBQLN1 |
| ENSG00000170802 | FOXN2 | 0,659816573 | 1,579881741 | 0,22338 | 0,53953208 | forkhead box N2 |
| ENSG00000105640 | RPL18A | 0,659639815 | 1,579688188 | 0,2397 | 0,5491566 | ribosomal protein L18a |
| ENSG00000145781 | COMMD10 | 0,659337873 | 1,579357609 | 0,29965 | 0,59157893 | COMM domain containing 10 |
| ENSG00000215093 | EEF1A1P29 | 0,65928014 | 1,579294408 | 0,49841 | 0,72960863 | eukaryotic translation elongation factor 1 alpha 1 pseudogene 29 |
| ENSG00000130227 | XPO7 | 0,659259429 | 1,579271736 | 0,12024 | 0,43879231 | exportin 7 |
| ENSG00000234473 | ZNF101P2 | 0,659197943 | 1,579204431 | 0,27232 | 0,57315552 | zinc finger protein 101 pseudogene 2 |
| ENSG00000258285 | TESC-AS1 | 0,658906663 | 1,578885622 | 0,49601 | 0,72828309 | TESC antisense RNA 1 (head to head) |
| ENSG00000285023 | CHORDC1 | 0,658900658 | 1,57887905 | 0,0984 | 0,40390309 | cysteine and histidine rich domain containing 1 |
| ENSG00000189398 | OR7E12P | 0,658827785 | 1,5787993 | 0,16169 | 0,48914394 | olfactory receptor family 7 subfamily E member 12 pseudogene |
| ENSG00000106066 | CPVL | 0,658752161 | 1,578716544 | 0,14599 | 0,47387591 | carboxypeptidase vitellogenic like |
| ENSG00000188312 | CENPP | 0,658707476 | 1,578667646 | 0,07021 | 0,34975886 | centromere protein P |
| ENSG00000058804 | NDC1 | 0,658702406 | 1,578662099 | 0,20226 | 0,5224664 | NDC1 transmembrane nucleoporin |
| ENSG00000048392 | RRM2B | 0,658522909 | 1,578465698 | 0,06544 | 0,33888576 | ribonucleotide reductase regulatory TP53 inducible subunit M2B |
| ENSG00000172575 | RASGRP1 | 0,658406812 | 1,57833868 | 0,17582 | 0,49976397 | RAS guanyl releasing protein 1 |
| ENSG00000286113 | AC022868,1 | 0,657640029 | 1,577500026 | 0,37379 | 0,6423379 | long intergenic non-protein coding RNA 1617 |
| ENSG00000258704 | SRP54-AS1 | 0,657556836 | 1,577409063 | 0,24312 | 0,55199251 | SRP54 antisense RNA 1 (head to head) |
| ENSG00000091073 | DTX2 | 0,657487373 | 1,577333115 | 0,00287 | 0,06351531 | deltex E3 ubiquitin ligase 2 |
| ENSG00000123836 | PFKFB2 | 0,657265813 | 1,577090897 | 0,02123 | 0,19328293 | 6-phosphofructo-2-kinase/fructose-2,6-biphosphatase 2 |
| ENSG00000276262 | NSF | 0,657163157 | 1,576978681 | 0,10058 | 0,40713342 | N-ethylmaleimide sensitive factor, vesicle fusing ATPase |
| ENSG00000073969 | NSF | 0,657163157 | 1,576978681 | 0,10058 | 0,40713342 | N-ethylmaleimide sensitive factor, vesicle fusing ATPase |
| ENSG00000137947 | GTF2B | 0,657070975 | 1,576877923 | 0,17956 | 0,50389568 | general transcription factor IIB |
| ENSG00000139514 | SLC7A1 | 0,656776003 | 1,576555549 | 0,01583 | 0,16482188 | solute carrier family 7 member 1 |
| ENSG00000165164 | CFAP47 | 0,656630639 | 1,576396705 | 0,17122 | 0,49650902 | cilia and flagella associated protein 47 |
| ENSG00000249098 | AC055733,1 | 0,656311039 | 1,576047524 | 0,47978 | 0,71717676 | cysteine and glycine-rich protein 2 pseudogene |

| | | | | | | |
|---|---|---|---|---|---|---|
| ENSG00000198842 | STYXL2 | 0,656203872 | 1,575930456 | 0,2623 | 0,56516131 | serine/threonine/tyrosine interacting like 2 |
| ENSG00000035403 | VCL | 0,655355905 | 1,57500445 | 0,00098 | 0,0320617 | vinculin |
| ENSG00000122585 | NPY | 0,655225094 | 1,574861649 | 0,52208 | 0,7463592 | neuropeptide Y |
| ENSG00000287110 | AC112484,5 | 0,655171263 | 1,574802888 | 0,13346 | 0,4594073 | novel transcript |
| ENSG00000197312 | DDI2 | 0,655126313 | 1,574753823 | 0,00013 | 0,00758921 | DNA damage inducible 1 homolog 2 |
| ENSG00000261335 | AC005837,1 | 0,655123932 | 1,574751223 | 0,11078 | 0,42340354 | novel transcript, antisense to MXRA7 |
| ENSG00000135248 | FAM71F1 | 0,654986058 | 1,574600737 | 0,41515 | 0,67201735 | family with sequence similarity 71 member F1 |
| ENSG00000130340 | SNX9 | 0,654976905 | 1,574590747 | 0,01777 | 0,17559058 | sorting nexin 9 |
| ENSG00000196358 | NTNG2 | 0,654955183 | 1,574567039 | 0,20313 | 0,52299224 | netrin G2 |
| ENSG00000274372 | LINC02804 | 0,654896686 | 1,574503196 | 0,36465 | 0,63504684 | long intergenic non-protein coding RNA 2804 |
| ENSG00000283638 | AC002407,1 | 0,654841286 | 1,574442736 | 0,09836 | 0,40379437 | novel transcript |
| ENSG00000121481 | RNF2 | 0,654723059 | 1,574313718 | 0,1661 | 0,492091 | ring finger protein 2 |
| ENSG00000196305 | IARS1 | 0,654709948 | 1,57429941 | 0,08719 | 0,38232685 | isoleucyl-tRNA synthetase 1 |
| ENSG00000138821 | SLC39A8 | 0,654524314 | 1,574096856 | 0,09449 | 0,39680104 | solute carrier family 39 member 8 |
| ENSG00000183291 | SELENOF | 0,654238617 | 1,573785168 | 0,13706 | 0,4642578 | selenoprotein F |
| ENSG00000250616 | AC012645,1 | 0,654219219 | 1,573764007 | 0,17754 | 0,50196932 | novel transcript, antisense to YPEL3 |
| ENSG00000241158 | ADAMTS9-AS: | 0,654008042 | 1,573533662 | 0,07462 | 0,35872988 | ADAMTS9 antisense RNA 1 |
| ENSG00000228522 | AL845321,1 | 0,653768761 | 1,573272702 | 0,52201 | 0,7463592 | methylenetetrahydrofolate dehydrogenase (NADP+ dependent) 1-like (MTHFD1L) pseudogene |
| ENSG00000109929 | SC5D | 0,65372017 | 1,573219715 | 0,11281 | 0,42686158 | sterol-C5-desaturase |
| ENSG00000271551 | AL355297,3 | 0,653585748 | 1,573073138 | 0,20111 | 0,52181085 | novel transcript, antisense to ARID1B |
| ENSG00000173250 | GPR151 | 0,653565175 | 1,573050706 | 0,26956 | 0,57134267 | G protein-coupled receptor 151 |
| ENSG00000119285 | HEATR1 | 0,653216673 | 1,57267076 | 0,12376 | 0,4446254 | HEAT repeat containing 1 |
| ENSG00000164961 | WASHC5 | 0,653124551 | 1,572570342 | 0,14199 | 0,46856898 | WASH complex subunit 5 |
| ENSG00000223367 | RPS18 | 0,653032923 | 1,572470469 | 0,16809 | 0,49417662 | ribosomal protein S18 |
| ENSG00000235650 | RPS18 | 0,653032923 | 1,572470469 | 0,16809 | 0,49417662 | ribosomal protein S18 |
| ENSG00000170425 | ADORA2B | 0,652910102 | 1,572336605 | 0,13125 | 0,4561992 | adenosine A2b receptor |
| ENSG00000004961 | HCCS | 0,652677445 | 1,572083061 | 0,03194 | 0,24116556 | holocytochrome c synthase |
| ENSG00000254946 | LINC02751 | 0,652319553 | 1,571693121 | 0,5739 | 0,77926229 | long intergenic non-protein coding RNA 2751 |
| ENSG00000145860 | RNF145 | 0,651846845 | 1,57117823 | 0,01768 | 0,17511677 | ring finger protein 145 |
| ENSG00000227176 | NSA2P5 | 0,651353705 | 1,570641264 | 0,33336 | 0,61333573 | NSA2 pseudogene 5 |
| ENSG00000133935 | ERG28 | 0,650872434 | 1,570117398 | 0,08131 | 0,37209365 | ergosterol biosynthesis 28 homolog |
| ENSG00000182459 | TEX19 | 0,650860577 | 1,570104494 | 0,5416 | 0,75882662 | testis expressed 19 |

| | | | | | | |
|---|---|---|---|---|---|---|
| ENSG00000105048 | TNNT1 | 0,650859014 | 1,570102793 | 0,17902 | 0,50343511 | troponin T1, slow skeletal type |
| ENSG00000138161 | CUZD1 | 0,650529511 | 1,569744232 | 0,42181 | 0,67643128 | CUB and zona pellucida like domains 1 |
| ENSG00000089472 | HEPH | 0,650511101 | 1,569724201 | 0,04251 | 0,2782038 | hephaestin |
| ENSG00000155380 | SLC16A1 | 0,650415787 | 1,569620498 | 0,0237 | 0,20555253 | solute carrier family 16 member 1 |
| ENSG00000270872 | SRGAP2D | 0,650093823 | 1,569270247 | 0,22268 | 0,53876885 | SLIT-ROBO Rho GTPase activating protein 2D (pseudogene) |
| ENSG00000134245 | WNT2B | 0,649822032 | 1,568974639 | 0,06105 | 0,32783036 | Wnt family member 2B |
| ENSG00000228251 | AC012442,1 | 0,649699048 | 1,568840895 | 0,54542 | 0,76113525 | novel transcript |
| ENSG00000144847 | IGSF11 | 0,649578136 | 1,568709417 | 0,07189 | 0,3538103 | immunoglobulin superfamily member 11 |
| ENSG00000204745 | AC083899,1 | 0,649469211 | 1,568590981 | 0,08445 | 0,37732439 | anaphase promoting complex subunit 1 (ANAPC1) pseudogene |
| ENSG00000026103 | FAS | 0,649313377 | 1,568421559 | 0,10582 | 0,41542189 | Fas cell surface death receptor |
| ENSG00000198160 | MIER1 | 0,64914712 | 1,568240822 | 0,12913 | 0,45251045 | MIER1 transcriptional regulator |
| ENSG00000237971 | AL445523,1 | 0,648935592 | 1,568010904 | 0,27594 | 0,57533851 | karyopherin alpha 2 (RAG cohort 1, importin alpha 1) (KPNA2) pseudogene |
| ENSG00000132313 | MRPL35 | 0,648662555 | 1,567714178 | 0,18045 | 0,50407792 | mitochondrial ribosomal protein L35 |
| ENSG00000234840 | LINC01239 | 0,648627543 | 1,567676133 | 0,03677 | 0,25984282 | long intergenic non-protein coding RNA 1239 |
| ENSG00000116489 | CAPZA1 | 0,648200931 | 1,567212632 | 0,12746 | 0,45071101 | capping actin protein of muscle Z-line subunit alpha 1 |
| ENSG00000184575 | XPOT | 0,648197118 | 1,56720849 | 0,06482 | 0,33757437 | exportin for tRNA |
| ENSG00000238018 | AC093110,1 | 0,648140964 | 1,56714749 | 0,28589 | 0,58229607 | novel transcript, antisense to SPTBN1 |
| ENSG00000187840 | EIF4EBP1 | 0,648095721 | 1,567098345 | 0,05879 | 0,32384911 | eukaryotic translation initiation factor 4E binding protein 1 |
| ENSG00000268658 | LINC00664 | 0,647925936 | 1,566913931 | 0,05377 | 0,31096353 | long intergenic non-protein coding RNA 664 |
| ENSG00000260526 | AC109347,1 | 0,647811983 | 1,566790172 | 0,24432 | 0,55253387 | novel transcript, antisense to AP1AR |
| ENSG00000034693 | PEX3 | 0,647685504 | 1,566652818 | 0,1648 | 0,49092529 | peroxisomal biogenesis factor 3 |
| ENSG00000181827 | RFX7 | 0,64757885 | 1,566537005 | 0,15192 | 0,48035862 | regulatory factor X7 |
| ENSG00000089009 | RPL6 | 0,647453702 | 1,566401121 | 0,18029 | 0,50407792 | ribosomal protein L6 |
| ENSG00000213041 | AC093422,1 | 0,6473653 | 1,566305141 | 0,40949 | 0,66797746 | ribosomal protein L17 (RPL17) pseudogene |
| ENSG00000180667 | YOD1 | 0,647065606 | 1,565979803 | 0,18719 | 0,5105374 | YOD1 deubiquitinase |
| ENSG00000181826 | RELL1 | 0,646793314 | 1,56568427 | 0,05722 | 0,31908628 | RELT like 1 |
| ENSG00000274779 | NOMO1 | 0,64675832 | 1,565646294 | 0,07542 | 0,36063909 | NODAL modulator 1 |
| ENSG00000153317 | ASAP1 | 0,646368179 | 1,565222961 | 0,13238 | 0,45750519 | ArfGAP with SH3 domain, ankyrin repeat and PH domain 1 |
| ENSG00000232139 | LINC00867 | 0,64620349 | 1,565044295 | 0,32452 | 0,60706463 | long intergenic non-protein coding RNA 867 |
| ENSG00000145391 | SETD7 | 0,645977044 | 1,564798664 | 1,87E-05 | 0,0014596 | SET domain containing 7, histone lysine methyltransferase |
| ENSG00000160691 | SHC1 | 0,645969337 | 1,564790305 | 0,00403 | 0,07780095 | SHC adaptor protein 1 |

| | | | | | | |
|---|---|---|---|---|---|---|
| ENSG00000179304 | FAM156B | 0,645701764 | 1,564500114 | 0,48765 | 0,72265932 | family with sequence similarity 156 member B |
| ENSG00000259751 | TUBAP11 | 0,645448241 | 1,564225211 | 0,25486 | 0,55944852 | tubulin alpha pseudogene 11 |
| ENSG00000165695 | AK8 | 0,645431657 | 1,564207229 | 0,09379 | 0,39568176 | adenylate kinase 8 |
| ENSG00000073849 | ST6GAL1 | 0,645387523 | 1,564159379 | 0,03739 | 0,26168679 | ST6 beta-galactoside alpha-2,6-sialyltransferase 1 |
| ENSG00000267283 | AC005306,1 | 0,645105296 | 1,563853421 | 0,67144 | 0,83732104 | novel transcript, antisense to BTBD2 |
| ENSG00000228886 | AL138963,1 | 0,644956917 | 1,56369259 | 0,26756 | 0,56973105 | novel transcript |
| ENSG00000259775 | AL138976,2 | 0,644790142 | 1,563511837 | 0,10845 | 0,41905863 | novel transcript, antisense to EIF5 |
| ENSG00000284153 | RPS4XP17 | 0,644751206 | 1,563469642 | 0,42287 | 0,67683766 | ribosomal protein S4X pseudogene 17 |
| ENSG00000244097 | RPS4XP17 | 0,644751206 | 1,563469642 | 0,42287 | 0,67683766 | ribosomal protein S4X pseudogene 17 |
| ENSG00000236251 | HSPA1L | 0,644745312 | 1,563463254 | 0,61642 | 0,80495273 | heat shock protein family A (Hsp70) member 1 like |
| ENSG00000206383 | HSPA1L | 0,644745312 | 1,563463254 | 0,61642 | 0,80495273 | heat shock protein family A (Hsp70) member 1 like |
| ENSG00000157978 | LDLRAP1 | 0,644743234 | 1,563461002 | 0,00066 | 0,0248432 | low density lipoprotein receptor adaptor protein 1 |
| ENSG00000198744 | MTCO3P12 | 0,644649016 | 1,5633589 | 0,21821 | 0,53571161 | MT-CO3 pseudogene 12 |
| ENSG00000145246 | ATP10D | 0,644172623 | 1,562842748 | 0,06283 | 0,33170499 | ATPase phospholipid transporting 10D (putative) |
| ENSG00000111058 | ACSS3 | 0,644083664 | 1,562746384 | 0,01467 | 0,15857384 | acyl-CoA synthetase short chain family member 3 |
| ENSG00000078328 | RBFOX1 | 0,644027978 | 1,562686065 | 0,19041 | 0,51350457 | RNA binding fox-1 homolog 1 |
| ENSG00000123560 | PLP1 | 0,643856061 | 1,562499861 | 0,09778 | 0,4029082 | proteolipid protein 1 |
| ENSG00000072274 | TFRC | 0,643833971 | 1,562475936 | 0,03992 | 0,27022885 | transferrin receptor |
| ENSG00000119681 | LTBP2 | 0,64309057 | 1,561671021 | 0,07384 | 0,35713141 | latent transforming growth factor beta binding protein 2 |
| ENSG00000269955 | FMC1-LUC7L2 | 0,642936031 | 1,561503746 | 0,69642 | 0,85299392 | FMC1-LUC7L2 readthrough |
| ENSG00000155714 | PDZD9 | 0,642836944 | 1,561396503 | 0,35141 | 0,62674766 | PDZ domain containing 9 |
| ENSG00000224587 | MDC1 | 0,642512951 | 1,561045893 | 0,08768 | 0,38329529 | mediator of DNA damage checkpoint 1 |
| ENSG00000237095 | MDC1 | 0,642512951 | 1,561045893 | 0,08768 | 0,38329529 | mediator of DNA damage checkpoint 1 |
| ENSG00000135069 | PSAT1 | 0,642279872 | 1,560793713 | 0,02332 | 0,20398797 | phosphoserine aminotransferase 1 |
| ENSG00000138615 | CILP | 0,642273371 | 1,56078668 | 0,31116 | 0,59885232 | cartilage intermediate layer protein |
| ENSG00000265681 | RPL17 | 0,642250725 | 1,560762181 | 0,19104 | 0,51406315 | ribosomal protein L17 |
| ENSG00000064102 | INTS13 | 0,641946346 | 1,560432927 | 0,13638 | 0,46359081 | integrator complex subunit 13 |
| ENSG00000134909 | ARHGAP32 | 0,641889208 | 1,560371126 | 0,0835 | 0,37567721 | Rho GTPase activating protein 32 |
| ENSG00000164976 | MYORG | 0,641655053 | 1,560117893 | 0,05361 | 0,31071932 | myogenesis regulating glycosidase (putative) |
| ENSG00000197223 | C1D | 0,641513395 | 1,559964712 | 0,13444 | 0,46059969 | C1D nuclear receptor corepressor |
| ENSG00000278666 | AC106795,4 | 0,641158736 | 1,559581273 | 0,13714 | 0,4642578 | THO complex 3 (THOC3) pseudogene |
| ENSG00000170915 | PAQR8 | 0,641150557 | 1,559572431 | 0,02735 | 0,22320635 | progestin and adipoQ receptor family member 8 |
| ENSG00000108091 | CCDC6 | 0,64109006 | 1,559507034 | 0,12311 | 0,44322266 | coiled-coil domain containing 6 |

| | | | | | | |
|---|---|---|---|---|---|---|
| ENSG00000008513 | ST3GAL1 | 0,641059326 | 1,559473813 | 0,00683 | 0,10536721 | ST3 beta-galactoside alpha-2,3-sialyltransferase 1 |
| ENSG00000197958 | RPL12 | 0,640691979 | 1,55907678 | 0,17623 | 0,50038183 | ribosomal protein L12 |
| ENSG00000127870 | RNF6 | 0,640671929 | 1,559055114 | 0,18577 | 0,50915451 | ring finger protein 6 |
| ENSG00000103512 | NOMO1 | 0,640375938 | 1,558735282 | 0,52222 | 0,74649491 | NODAL modulator 1 |
| ENSG00000107560 | RAB11FIP2 | 0,640211973 | 1,558558139 | 0,04519 | 0,2876168 | RAB11 family interacting protein 2 |
| ENSG00000277785 | SNORD116-21 | 0,639892402 | 1,558212942 | 0,55742 | 0,76914186 | small nucleolar RNA, C/D box 116-21 |
| ENSG00000227082 | LINC02798 | 0,639726349 | 1,558033602 | 0,08159 | 0,37274873 | long intergenic non-protein coding RNA 2798 |
| ENSG00000254876 | SUGT1P4-STR | 0,639660542 | 1,557962536 | 0,08836 | 0,38462876 | SUGT1P4-STRA6LP readthrough |
| ENSG00000184933 | OR6A2 | 0,639631709 | 1,5579314 | 0,29377 | 0,58826777 | olfactory receptor family 6 subfamily A member 2 |
| ENSG00000280109 | PLAC4 | 0,639507526 | 1,557797303 | 0,11733 | 0,43468366 | placenta enriched 4 |
| ENSG00000283486 | FAM95C | 0,639332765 | 1,557608611 | 0,41191 | 0,66967031 | family with sequence similarity 95 member C |
| ENSG00000081237 | PTPRC | 0,639307059 | 1,557580858 | 0,50371 | 0,73293685 | protein tyrosine phosphatase receptor type C |
| ENSG00000137168 | PPIL1 | 0,639240328 | 1,557508815 | 0,22664 | 0,54144 | peptidylprolyl isomerase like 1 |
| ENSG00000137693 | YAP1 | 0,639045325 | 1,557298306 | 0,07714 | 0,36384125 | Yes1 associated transcriptional regulator |
| ENSG00000242154 | AC004884,2 | 0,638820749 | 1,55705591 | 0,32639 | 0,60866572 | novel transcript |
| ENSG00000273675 | AL118556,2 | 0,638259262 | 1,556450033 | 0,28272 | 0,58009943 | zinc finger protein 124 (ZNF124) pseudogene |
| ENSG00000073712 | FERMT2 | 0,638254478 | 1,556444871 | 0,00733 | 0,10869257 | fermitin family member 2 |
| ENSG00000148737 | TCF7L2 | 0,638209019 | 1,556395828 | 0,05414 | 0,31168784 | transcription factor 7 like 2 |
| ENSG00000249174 | AC106744,1 | 0,63809987 | 1,556278082 | 0,29734 | 0,59031059 | novel transcript |
| ENSG00000198088 | NUP62CL | 0,637658031 | 1,555801531 | 0,18658 | 0,509961 | nucleoporin 62 C-terminal like |
| ENSG00000231249 | ITPR1-DT | 0,637326356 | 1,555443893 | 0,37161 | 0,64073742 | ITPR1 divergent transcript |
| ENSG00000131469 | RPL27 | 0,637286758 | 1,555401201 | 0,21196 | 0,53079538 | ribosomal protein L27 |
| ENSG00000125354 | SEPTIN6 | 0,637249556 | 1,555361094 | 0,0522 | 0,30681232 | septin 6 |
| ENSG00000214062 | RPL7P17 | 0,636718836 | 1,554789033 | 0,27306 | 0,57344244 | ribosomal protein L7 pseudogene 17 |
| ENSG00000282269 | PRR4 | 0,636139795 | 1,554165126 | 0,32115 | 0,60573903 | proline rich 4 |
| ENSG00000174442 | ZWILCH | 0,636049814 | 1,554068196 | 0,18584 | 0,50915451 | zwilch kinetochore protein |
| ENSG00000278087 | NOMO3 | 0,636001219 | 1,55401585 | 0,18872 | 0,5120698 | NODAL modulator 3 |
| ENSG00000138386 | NAB1 | 0,636000834 | 1,554015436 | 0,06988 | 0,34890292 | NGFI-A binding protein 1 |
| ENSG00000254996 | ANKHD1-EIF4 | 0,635950076 | 1,553960763 | 0,15304 | 0,48120112 | ANKHD1-EIF4EBP3 readthrough |
| ENSG00000272129 | AL359715,3 | 0,635918634 | 1,553926896 | 0,39711 | 0,65966023 | novel transcript |
| ENSG00000066455 | GOLGA5 | 0,635852303 | 1,553855453 | 0,17008 | 0,49606609 | golgin A5 |
| ENSG00000150753 | CCT5 | 0,63583602 | 1,553837915 | 0,10149 | 0,40833508 | chaperonin containing TCP1 subunit 5 |
| ENSG00000181751 | MACIR | 0,6357117 | 1,553704023 | 0,16064 | 0,48804 | macrophage immunometabolism regulator |

| | | | | | | |
|---|---|---|---|---|---|---|
| ENSG00000111906 | HDDC2 | 0,635380892 | 1,553347802 | 0,1002 | 0,4067769 | HD domain containing 2 |
| ENSG00000123870 | ZNF137P | 0,635344906 | 1,553309056 | 0,28656 | 0,58289163 | zinc finger protein 137, pseudogene |
| ENSG00000185127 | C6orf120 | 0,635287122 | 1,553246843 | 0,06614 | 0,3404567 | chromosome 6 open reading frame 120 |
| ENSG00000235701 | PCBP2P1 | 0,635080041 | 1,55302391 | 0,24935 | 0,55642837 | poly(rC) binding protein 2 pseudogene 1 |
| ENSG00000196092 | PAX5 | 0,635030453 | 1,552970531 | 0,26228 | 0,56516131 | paired box 5 |
| ENSG00000130962 | PRRG1 | 0,634790839 | 1,552712622 | 0,13859 | 0,46578766 | proline rich and Gla domain 1 |
| ENSG00000227540 | DNAJC9-AS1 | 0,634781513 | 1,552702585 | 0,29571 | 0,5893128 | DNAJC9 antisense RNA 1 |
| ENSG00000167306 | MYO5B | 0,634759315 | 1,552678694 | 0,13153 | 0,45658336 | myosin VB |
| ENSG00000261655 | AC100803,3 | 0,634578608 | 1,552484224 | 0,42721 | 0,680351 | novel transcript, overlapping to GPR20 |
| ENSG00000282573 | AC100803,3 | 0,634578608 | 1,552484224 | 0,42721 | 0,680351 | novel transcript, overlapping to GPR20 |
| ENSG00000186197 | EDARADD | 0,63428044 | 1,552163399 | 0,16742 | 0,49362211 | EDAR associated death domain |
| ENSG00000106591 | MRPL32 | 0,634257167 | 1,552138359 | 0,21051 | 0,5290538 | mitochondrial ribosomal protein L32 |
| ENSG00000253251 | SHLD3 | 0,633903848 | 1,551758284 | 0,35586 | 0,63013888 | shieldin complex subunit 3 |
| ENSG00000260830 | AL135744,1 | 0,633818435 | 1,551666417 | 0,31048 | 0,59844173 | novel transcript, antisense to CHD8 and SUPT16H |
| ENSG00000204304 | PBX2 | 0,633764447 | 1,551608352 | 0,43095 | 0,68325486 | PBX homeobox 2 |
| ENSG00000277595 | AC007546,1 | 0,633623566 | 1,551456843 | 0,37885 | 0,64679625 | novel transcript, sense intronic to ANKRD13A |
| ENSG00000286185 | AC242842,3 | 0,633523847 | 1,551349611 | 0,22625 | 0,54121239 | novel protein, identical to neuroblastoma breakpoint family, member 19 NBPF19 |
| ENSG00000249550 | LINC01234 | 0,633409992 | 1,551227185 | 0,17858 | 0,50323303 | long intergenic non-protein coding RNA 1234 |
| ENSG00000101132 | PFDN4 | 0,633216964 | 1,55101965 | 0,17933 | 0,50368273 | prefoldin subunit 4 |
| ENSG00000251201 | TMED7-TICAI | 0,63320075 | 1,551002218 | 0,03211 | 0,24164117 | TMED7-TICAM2 readthrough |
| ENSG00000288074 | AC026341,3 | 0,633058363 | 1,550849149 | 0,40478 | 0,66474345 | novel transcript |
| ENSG00000001167 | NFYA | 0,632913673 | 1,55069362 | 0,16832 | 0,49453836 | nuclear transcription factor Y subunit alpha |
| ENSG00000285708 | AC097634,4 | 0,632832551 | 1,550606428 | 0,32177 | 0,60577115 | novel protein |
| ENSG00000146374 | RSPO3 | 0,632754316 | 1,550522344 | 0,22975 | 0,54324957 | R-spondin 3 |
| ENSG00000250988 | SNHG21 | 0,632627905 | 1,550386491 | 0,29769 | 0,59036937 | small nucleolar RNA host gene 21 |
| ENSG00000273673 | AC011295,1 | 0,632547861 | 1,550300474 | 0,20099 | 0,52181085 | ribosomal protein L9 pseudogene 9 |
| ENSG00000171408 | PDE7B | 0,63253795 | 1,550289824 | 0,31306 | 0,59987747 | phosphodiesterase 7B |
| ENSG00000173207 | CKS1B | 0,632517049 | 1,550267364 | 0,07415 | 0,35759867 | CDC28 protein kinase regulatory subunit 1B |
| ENSG00000104635 | SLC39A14 | 0,632396193 | 1,550137503 | 0,05962 | 0,3251502 | solute carrier family 39 member 14 |
| ENSG00000125485 | DDX31 | 0,632384392 | 1,550124823 | 0,08534 | 0,37867628 | DEAD-box helicase 31 |
| ENSG00000271367 | AL034374,1 | 0,632255579 | 1,549986424 | 0,58442 | 0,78572744 | novel transcript |
| ENSG00000226284 | ARPC3P1 | 0,632228007 | 1,549956801 | 0,41642 | 0,67315741 | actin related protein 2/3 complex subunit 3 pseudogene 1 |

| | | | | | | |
|---|---|---|---|---|---|---|
| ENSG00000273173 | SNURF | 0,632088431 | 1,549806856 | 0,18159 | 0,50490902 | SNRPN upstream reading frame |
| ENSG00000100078 | PLA2G3 | 0,632010752 | 1,549723412 | 0,07132 | 0,3527929 | phospholipase A2 group III |
| ENSG00000136371 | MTHFS | 0,631979353 | 1,549689684 | 0,02079 | 0,19127841 | methenyltetrahydrofolate synthetase |
| ENSG00000168803 | ADAL | 0,631977885 | 1,549688106 | 0,09096 | 0,39027377 | adenosine deaminase like |
| ENSG00000142937 | RPS8 | 0,631921027 | 1,549627033 | 0,14943 | 0,4771595 | ribosomal protein S8 |
| ENSG00000242428 | CFAP20DC-AS | 0,63188826 | 1,549591837 | 0,35571 | 0,63008438 | CFAP20DC antisense RNA 1 |
| ENSG00000273011 | AC092681,3 | 0,631856161 | 1,54955736 | 0,41102 | 0,6688445 | novel transcript |
| ENSG00000225133 | MORF4L1P4 | 0,631846089 | 1,549546543 | 0,43383 | 0,68475933 | mortality factor 4 like 1 pseudogene 4 |
| ENSG00000287960 | AC008567,3 | 0,631701075 | 1,549390797 | 0,09177 | 0,39169294 | novel transcript |
| ENSG00000258643 | BCL2L2-PABPI | 0,631496348 | 1,549170945 | 0,24819 | 0,55542983 | BCL2L2-PABPN1 readthrough |
| ENSG00000141429 | GALNT1 | 0,631354694 | 1,549018844 | 0,21422 | 0,53334744 | polypeptide N-acetylgalactosaminyltransferase 1 |
| ENSG00000143632 | ACTA1 | 0,631290647 | 1,548950078 | 0,40081 | 0,66280184 | actin alpha 1, skeletal muscle |
| ENSG00000269743 | SLC25A53 | 0,631237733 | 1,548893268 | 0,00054 | 0,02167371 | solute carrier family 25 member 53 |
| ENSG00000172086 | KRCC1 | 0,631085108 | 1,548729416 | 0,20084 | 0,52177529 | lysine rich coiled-coil 1 |
| ENSG00000087302 | RTRAF | 0,630995993 | 1,548633754 | 0,21212 | 0,5308403 | RNA transcription, translation and transport factor |
| ENSG00000184983 | NDUFA6 | 0,630802559 | 1,548426131 | 0,11042 | 0,42308928 | NADH:ubiquinone oxidoreductase subunit A6 |
| ENSG00000151572 | ANO4 | 0,630772372 | 1,548393731 | 0,17802 | 0,50235771 | anoctamin 4 |
| ENSG00000111596 | CNOT2 | 0,630753867 | 1,548373871 | 0,16321 | 0,48972962 | CCR4-NOT transcription complex subunit 2 |
| ENSG00000228575 | MDC1 | 0,630520221 | 1,548123131 | 0,0953 | 0,39794903 | mediator of DNA damage checkpoint 1 |
| ENSG00000184349 | EFNA5 | 0,630448999 | 1,548046705 | 0,01495 | 0,16037017 | ephrin A5 |
| ENSG00000181690 | PLAG1 | 0,630406997 | 1,548001636 | 0,18257 | 0,50600317 | PLAG1 zinc finger |
| ENSG00000166562 | SEC11C | 0,63031985 | 1,547908131 | 0,08072 | 0,37077071 | SEC11 homolog C, signal peptidase complex subunit |
| ENSG00000237766 | GGTA2P | 0,630167386 | 1,547744557 | 0,39619 | 0,65876705 | glycoprotein alpha-galactosyltransferase 2, pseudogene |
| ENSG00000110713 | NUP98 | 0,630147628 | 1,547723361 | 0,13992 | 0,46689302 | nucleoporin 98 and 96 precursor |
| ENSG00000277060 | NLRP2 | 0,630083541 | 1,54765461 | 0,26299 | 0,56587311 | NLR family pyrin domain containing 2 |
| ENSG00000260941 | LINC00622 | 0,62978499 | 1,547334371 | 0,17443 | 0,498816 | long intergenic non-protein coding RNA 622 |
| ENSG00000226383 | LINC01876 | 0,629754249 | 1,547301401 | 0,0331 | 0,24582875 | long intergenic non-protein coding RNA 1876 |
| ENSG00000067167 | TRAM1 | 0,629666481 | 1,547207272 | 0,20961 | 0,52860172 | translocation associated membrane protein 1 |
| ENSG00000100181 | TPTEP1 | 0,629658199 | 1,54719839 | 0,06328 | 0,33320544 | TPTE pseudogene 1 |
| ENSG00000248610 | HSPA8P4 | 0,629479716 | 1,547006991 | 0,49394 | 0,72657717 | heat shock protein family A (Hsp70) member 8 pseudogene 4 |
| ENSG00000234937 | AL139128,1 | 0,629399097 | 1,546920544 | 0,21814 | 0,53569444 | proteasome (prosome, macropain) 26S subunit, non-ATPase, 7 (Mov34 homolog) (PSMD7) pseudogene |

| | | | | | | |
|---|---|---|---|---|---|---|
| ENSG00000125691 | RPL23 | 0,629274904 | 1,546787385 | 0,13984 | 0,46689302 | ribosomal protein L23 |
| ENSG00000171862 | PTEN | 0,629256756 | 1,546767929 | 0,26339 | 0,56608528 | phosphatase and tensin homolog |
| ENSG00000132963 | POMP | 0,629160749 | 1,546664999 | 0,15014 | 0,47828953 | proteasome maturation protein |
| ENSG00000171492 | LRRC8D | 0,629083034 | 1,546581686 | 0,11456 | 0,42993937 | leucine rich repeat containing 8 VRAC subunit D |
| ENSG00000241634 | AC069499,1 | 0,628719104 | 1,546191598 | 0,17224 | 0,49710979 | ribosomal protein L13 (RPL13) pseudogene |
| ENSG00000135074 | ADAM19 | 0,628575633 | 1,546037843 | 0,00026 | 0,01300234 | ADAM metallopeptidase domain 19 |
| ENSG00000078140 | UBE2K | 0,628478768 | 1,545934043 | 0,10596 | 0,4154675 | ubiquitin conjugating enzyme E2 K |
| ENSG00000188419 | CHM | 0,628416235 | 1,545867037 | 0,08197 | 0,37306719 | CHM Rab escort protein |
| ENSG00000234841 | AC017007,2 | 0,628099925 | 1,545528143 | 0,4317 | 0,68379763 | TAF4b RNA polymerase II, TATA box binding protein (TBP)-associated factor, 105kDa (TAF4B) pseudogene |
| ENSG00000287811 | AL391840,3 | 0,628046646 | 1,545471068 | 0,18128 | 0,50481327 | novel transcript |
| ENSG00000248780 | ARL4AP2 | 0,627998906 | 1,545419928 | 0,40796 | 0,66699097 | ADP ribosylation factor like GTPase 4A pseudogene 2 |
| ENSG00000223865 | HLA-DPB1 | 0,627884415 | 1,545297289 | 0,04693 | 0,29141554 | major histocompatibility complex, class II, DP beta 1 |
| ENSG00000242931 | RPL7P49 | 0,62749879 | 1,544884295 | 0,34576 | 0,62283284 | ribosomal protein L7 pseudogene 49 |
| ENSG00000164211 | STARD4 | 0,627291499 | 1,544662336 | 0,11881 | 0,43692176 | StAR related lipid transfer domain containing 4 |
| ENSG00000106524 | ANKMY2 | 0,627249397 | 1,544617259 | 0,17068 | 0,49641416 | ankyrin repeat and MYND domain containing 2 |
| ENSG00000135916 | ITM2C | 0,62720476 | 1,544569469 | 0,01792 | 0,17654822 | integral membrane protein 2C |
| ENSG00000108094 | CUL2 | 0,627185715 | 1,54454908 | 0,19449 | 0,51562158 | cullin 2 |
| ENSG00000275560 | AC008115,3 | 0,627008487 | 1,544359352 | 0,25069 | 0,55690541 | novel transcript |
| ENSG00000146109 | ABT1 | 0,626857553 | 1,54419779 | 0,05526 | 0,31443768 | activator of basal transcription 1 |
| ENSG00000267724 | AC012254,3 | 0,626758018 | 1,544091256 | 0,14282 | 0,47034016 | novel transcript, antisense to HDHD2 |
| ENSG00000231789 | PIK3CD-AS2 | 0,626481375 | 1,543795198 | 0,24226 | 0,55112431 | PIK3CD antisense RNA 2 |
| ENSG00000136536 | MARCHF7 | 0,626481019 | 1,543794817 | 0,21633 | 0,53480193 | membrane associated ring-CH-type finger 7 |
| ENSG00000172186 | HMGN1P35 | 0,626442592 | 1,543753697 | 0,53444 | 0,75405583 | high mobility group nucleosome binding domain 1 pseudogene 35 |
| ENSG00000196126 | HLA-DRB1 | 0,62593101 | 1,543206377 | 0,39612 | 0,65876705 | major histocompatibility complex, class II, DR beta 1 |
| ENSG00000278384 | AL354822,1 | 0,625716166 | 1,542976582 | 0,16053 | 0,48780354 | |
| ENSG00000178996 | SNX18 | 0,625557355 | 1,542806742 | 0,02078 | 0,19127841 | sorting nexin 18 |
| ENSG00000137760 | ALKBH8 | 0,625518112 | 1,542764776 | 0,10576 | 0,41527838 | alkB homolog 8, tRNA methyltransferase |
| ENSG00000100479 | POLE2 | 0,625497021 | 1,542742222 | 0,18943 | 0,51258824 | DNA polymerase epsilon 2, accessory subunit |
| ENSG00000089682 | RBM41 | 0,625474337 | 1,542717965 | 0,03602 | 0,25706681 | RNA binding motif protein 41 |
| ENSG00000286313 | AL360178,1 | 0,625208591 | 1,542433821 | 0,60058 | 0,79589975 | novel transcript, antisense to PDE7B |
| ENSG00000134602 | STK26 | 0,625171219 | 1,542393866 | 0,26112 | 0,56416552 | serine/threonine kinase 26 |

| | | | | | | |
|---|---|---|---|---|---|---|
| ENSG00000176273 | SLC35G1 | 0,624996808 | 1,542207413 | 0,12185 | 0,44093327 | solute carrier family 35 member G1 |
| ENSG00000118922 | KLF12 | 0,624773073 | 1,541968264 | 0,21015 | 0,52904355 | Kruppel like factor 12 |
| ENSG00000108561 | C1QBP | 0,624691181 | 1,54188074 | 0,04222 | 0,27788591 | complement C1q binding protein |
| ENSG00000235920 | THRAP3P3 | 0,624598631 | 1,54178183 | 0,26186 | 0,56477027 | THRAP3 pseudogene 3 |
| ENSG00000198034 | RPS4X | 0,6244631 | 1,541636997 | 0,20974 | 0,52867318 | ribosomal protein S4 X-linked |
| ENSG00000123338 | NCKAP1L | 0,624434 | 1,541605902 | 0,0595 | 0,32479124 | NCK associated protein 1 like |
| ENSG00000115761 | NOL10 | 0,624372146 | 1,541539808 | 0,1283 | 0,45171208 | nucleolar protein 10 |
| ENSG00000131931 | THAP1 | 0,62423034 | 1,541388295 | 0,15548 | 0,48334025 | THAP domain containing 1 |
| ENSG00000131069 | ACSS2 | 0,624109629 | 1,541259331 | 0,02388 | 0,20636344 | acyl-CoA synthetase short chain family member 2 |
| ENSG00000262446 | GBA | 0,623893612 | 1,541028573 | 0,19154 | 0,51435195 | glucosylceramidase beta |
| ENSG00000122884 | P4HA1 | 0,623855125 | 1,540987464 | 0,03906 | 0,26753929 | prolyl 4-hydroxylase subunit alpha 1 |
| ENSG00000285943 | AC112128,1 | 0,623568141 | 1,540680958 | 0,02046 | 0,18969631 | novel protein |
| ENSG00000276368 | H2AC14 | 0,623495278 | 1,540603148 | 0,24295 | 0,55192089 | H2A clustered histone 14 |
| ENSG00000255382 | LINC02734 | 0,623410909 | 1,540513055 | 0,56171 | 0,77136826 | long intergenic non-protein coding RNA 2734 |
| ENSG00000115159 | GPD2 | 0,623297268 | 1,540391714 | 0,14036 | 0,46721845 | glycerol-3-phosphate dehydrogenase 2 |
| ENSG00000286650 | AL139234,1 | 0,623181608 | 1,540268227 | 0,28029 | 0,57784691 | novel transcript |
| ENSG00000163634 | THOC7 | 0,623171605 | 1,540257547 | 0,1236 | 0,4443454 | THO complex 7 |
| ENSG00000144893 | MED12L | 0,623164749 | 1,540250228 | 0,08165 | 0,37277532 | mediator complex subunit 12L |
| ENSG00000237992 | LINC01808 | 0,623147967 | 1,540232311 | 0,54604 | 0,76166626 | long intergenic non-protein coding RNA 1808 |
| ENSG00000090054 | SPTLC1 | 0,623147298 | 1,540231596 | 0,08284 | 0,37470115 | serine palmitoyltransferase long chain base subunit 1 |
| ENSG00000131951 | LRRC9 | 0,623044981 | 1,540122367 | 0,13947 | 0,46666406 | leucine rich repeat containing 9 |
| ENSG00000177602 | HASPIN | 0,623012792 | 1,540088004 | 0,09948 | 0,40608287 | histone H3 associated protein kinase |
| ENSG00000227097 | RPS28P7 | 0,622878323 | 1,539944464 | 0,4645 | 0,70506031 | ribosomal protein S28 pseudogene 7 |
| ENSG00000120370 | GORAB | 0,622835759 | 1,539899031 | 0,12571 | 0,44776504 | golgin, RAB6 interacting |
| ENSG00000185684 | EP400P1 | 0,622737783 | 1,539794457 | 0,27748 | 0,57588377 | EP400 pseudogene 1 |
| ENSG00000151233 | GXYLT1 | 0,622619585 | 1,53966831 | 0,0109 | 0,13508104 | glucoside xylosyltransferase 1 |
| ENSG00000178074 | C2orf69 | 0,622470456 | 1,539509165 | 0,14814 | 0,47527573 | chromosome 2 open reading frame 69 |
| ENSG00000165462 | PHOX2A | 0,622155608 | 1,539173226 | 0,50634 | 0,73476122 | paired like homeobox 2A |
| ENSG00000284669 | AC092053,3 | 0,621940033 | 1,538943251 | 0,17337 | 0,49804988 | novel transcript, antisense CSRNP1 |
| ENSG00000180581 | SRP9P1 | 0,621931016 | 1,538933632 | 0,11455 | 0,42992744 | signal recognition particle 9 pseudogene 1 |
| ENSG00000240175 | MAGI1-AS1 | 0,621678023 | 1,538663787 | 0,45878 | 0,70137313 | MAGI1 antisense RNA 1 |
| ENSG00000134250 | NOTCH2 | 0,621623832 | 1,538605992 | 0,02242 | 0,19922746 | notch receptor 2 |
| ENSG00000172216 | CEBPB | 0,621474314 | 1,538446543 | 0,03969 | 0,26947142 | CCAAT enhancer binding protein beta |

| | | | | | | |
|---|---|---|---|---|---|---|
| ENSG00000158560 | DYNC1I1 | 0,621312277 | 1,538273761 | 0,12066 | 0,4394821 | dynein cytoplasmic 1 intermediate chain 1 |
| ENSG00000123570 | RAB9B | 0,621160985 | 1,538112454 | 0,12566 | 0,44769898 | RAB9B, member RAS oncogene family |
| ENSG00000270890 | AL049844,2 | 0,621106219 | 1,538054067 | 0,26974 | 0,57140742 | synaptophysin-like 1 (SYPL1) pseudogene |
| ENSG00000181481 | RNF135 | 0,621038447 | 1,537981817 | 0,03386 | 0,24867791 | ring finger protein 135 |
| ENSG00000272791 | AC073389,3 | 0,620991429 | 1,537931694 | 0,21715 | 0,53517619 | novel transcript |
| ENSG00000114738 | MAPKAPK3 | 0,620714777 | 1,537636808 | 0,04744 | 0,29251695 | MAPK activated protein kinase 3 |
| ENSG00000164039 | BDH2 | 0,620578629 | 1,537491707 | 0,22029 | 0,53750545 | 3-hydroxybutyrate dehydrogenase 2 |
| ENSG00000237863 | AL035425,1 | 0,620501057 | 1,53740904 | 0,22567 | 0,54090479 | novel transcript, antisense to IRS4 |
| ENSG00000276068 | NAIP | 0,620413296 | 1,53731552 | 0,10189 | 0,40926385 | NLR family apoptosis inhibitory protein |
| ENSG00000168505 | GBX2 | 0,620412763 | 1,537314953 | 0,32561 | 0,60804921 | gastrulation brain homeobox 2 |
| ENSG00000286682 | AC025262,3 | 0,620181167 | 1,537068188 | 0,42582 | 0,67925635 | novel transcript, antisense to TBC1D30 |
| ENSG00000138650 | PCDH10 | 0,620135685 | 1,537019731 | 0,24313 | 0,55199251 | protocadherin 10 |
| ENSG00000269378 | ITGB1P1 | 0,620119038 | 1,537001995 | 0,70071 | 0,85544787 | integrin subunit beta 1 pseudogene 1 |
| ENSG00000081853 | PCDHGA2 | 0,619555427 | 1,536401659 | 0,12005 | 0,43868155 | protocadherin gamma subfamily A, 2 |
| ENSG00000138448 | ITGAV | 0,61953979 | 1,536385007 | 0,10465 | 0,41313952 | integrin subunit alpha V |
| ENSG00000276411 | AC106875,2 | 0,619300169 | 1,536129846 | 0,36474 | 0,6350493 | novel transcript |
| ENSG00000154723 | ATP5PF | 0,61926043 | 1,536087534 | 0,25622 | 0,56072859 | ATP synthase peripheral stalk subunit F6 |
| ENSG00000135045 | C9orf40 | 0,619141017 | 1,535960396 | 0,01638 | 0,16750744 | chromosome 9 open reading frame 40 |
| ENSG00000240828 | RPL21P4 | 0,618932522 | 1,535738439 | 0,36757 | 0,63756038 | ribosomal protein L21 pseudogene 4 |
| ENSG00000173517 | PEAK1 | 0,618597388 | 1,535381732 | 0,00852 | 0,11640065 | pseudopodium enriched atypical kinase 1 |
| ENSG00000188375 | H3-5 | 0,618333267 | 1,535100668 | 0,21151 | 0,53032236 | H3,5 histone |
| ENSG00000226828 | LINC02774 | 0,617967402 | 1,534711418 | 0,47842 | 0,71617224 | long intergenic non-protein coding RNA 2774 |
| ENSG00000255423 | EBLN2 | 0,617729848 | 1,534458734 | 0,08177 | 0,3729912 | endogenous Bornavirus like nucleoprotein 2 |
| ENSG00000124207 | CSE1L | 0,617576573 | 1,534295718 | 0,20498 | 0,52453221 | chromosome segregation 1 like |
| ENSG00000257763 | OR5BK1P | 0,617564937 | 1,534283344 | 0,58403 | 0,78572744 | olfactory receptor family 5 subfamily BK member 1 pseudogene |
| ENSG00000176014 | TUBB6 | 0,617563827 | 1,534282164 | 0,04039 | 0,27186146 | tubulin beta 6 class V |
| ENSG00000136560 | TANK | 0,617363613 | 1,534069253 | 0,18509 | 0,50871188 | TRAF family member associated NFKB activator |
| ENSG00000165124 | SVEP1 | 0,617267085 | 1,533966616 | 0,23806 | 0,54864017 | sushi, von Willebrand factor type A, EGF and pentraxin domain containing 1 |
| ENSG00000139436 | GIT2 | 0,617139631 | 1,533831105 | 0,1943 | 0,51562158 | GIT ArfGAP 2 |
| ENSG00000281556 | AC243960,18 | 0,61710977 | 1,533799357 | 0,32441 | 0,60694578 | novel transcript, sense intronic to CEACAM21 |
| ENSG00000276853 | AC026124,2 | 0,617098343 | 1,533787208 | 0,461 | 0,70277253 | novel transcript, sense intronic to LEMD3 |

| | | | | | | |
|---|---|---|---|---|---|---|
| ENSG00000277692 | AL121583,1 | 0,617098334 | 1,533787199 | 0,4731 | 0,71242337 | novel transcript |
| ENSG00000129173 | E2F8 | 0,616883946 | 1,533559292 | 0,1601 | 0,48727733 | E2F transcription factor 8 |
| ENSG00000274166 | GOLGA6L10 | 0,616855474 | 1,533529026 | 0,19811 | 0,51977271 | golgin A6 family like 10 |
| ENSG00000089195 | TRMT6 | 0,616740575 | 1,533406899 | 0,14935 | 0,47696569 | tRNA methyltransferase 6 |
| ENSG00000147044 | CASK | 0,616670084 | 1,533331977 | 0,00975 | 0,12637434 | calcium/calmodulin dependent serine protein kinase |
| ENSG00000124279 | FASTKD3 | 0,616566971 | 1,53322239 | 0,31125 | 0,59885232 | FAST kinase domains 3 |
| ENSG00000146776 | ATXN7L1 | 0,616210331 | 1,532843417 | 0,05665 | 0,31746133 | ataxin 7 like 1 |
| ENSG00000279058 | AGAP14P | 0,616187648 | 1,532819318 | 0,30741 | 0,59658791 | ArfGAP with GTPase domain, ankyrin repeat and PH domain 14, pseudogene |
| ENSG00000285851 | AL359762,3 | 0,616074934 | 1,532699567 | 0,16661 | 0,49255702 | novel transcript |
| ENSG00000243199 | AC115223,1 | 0,615950511 | 1,532567388 | 0,59917 | 0,79538547 | ribosomal protein L6 (RPL6) pseudogene |
| ENSG00000137225 | CAPN11 | 0,615875863 | 1,532488092 | 0,49664 | 0,72841371 | calpain 11 |
| ENSG00000177565 | TBL1XR1 | 0,615484003 | 1,532071899 | 0,14075 | 0,46776292 | TBL1X receptor 1 |
| ENSG00000162772 | ATF3 | 0,615249958 | 1,531823375 | 0,04786 | 0,29380293 | activating transcription factor 3 |
| ENSG00000253119 | HNRNPA3P7 | 0,615239479 | 1,531812248 | 0,2466 | 0,55423575 | heterogeneous nuclear ribonucleoprotein A3 pseudogene 7 |
| ENSG00000167414 | GNG8 | 0,615238425 | 1,531811128 | 0,58899 | 0,7885434 | G protein subunit gamma 8 |
| ENSG00000136205 | TNS3 | 0,614971348 | 1,53152758 | 0,00831 | 0,11527851 | tensin 3 |
| ENSG00000153956 | CACNA2D1 | 0,614951147 | 1,531506135 | 0,0675 | 0,34374257 | calcium voltage-gated channel auxiliary subunit alpha2delta 1 |
| ENSG00000116157 | GPX7 | 0,614901242 | 1,531453159 | 0,20695 | 0,52659911 | glutathione peroxidase 7 |
| ENSG00000111711 | GOLT1B | 0,614834353 | 1,531382157 | 0,19606 | 0,51693058 | golgi transport 1B |
| ENSG00000129636 | ITFG1 | 0,614832701 | 1,531380404 | 0,2063 | 0,5259882 | integrin alpha FG-GAP repeat containing 1 |
| ENSG00000091483 | FH | 0,61463809 | 1,531173844 | 0,11244 | 0,42619744 | fumarate hydratase |
| ENSG00000198301 | SDAD1 | 0,614614666 | 1,531148983 | 0,08425 | 0,37695424 | SDA1 domain containing 1 |
| ENSG00000261135 | AL137802,2 | 0,614613837 | 1,531148103 | 0,27838 | 0,57651947 | novel transcript |
| ENSG00000153214 | TMEM87B | 0,614403469 | 1,530924854 | 0,08507 | 0,37850509 | transmembrane protein 87B |
| ENSG00000287502 | AC092418,2 | 0,614286643 | 1,530800888 | 0,55805 | 0,76971239 | novel transcript |
| ENSG00000237889 | DDX39B | 0,614201156 | 1,530710183 | 0,20879 | 0,52791772 | DExD-box helicase 39B |
| ENSG00000166197 | NOLC1 | 0,613936902 | 1,530429833 | 0,10554 | 0,41478571 | nucleolar and coiled-body phosphoprotein 1 |
| ENSG00000182631 | RXFP3 | 0,613797421 | 1,530281877 | 0,47432 | 0,71307604 | relaxin family peptide receptor 3 |
| ENSG00000102001 | CACNA1F | 0,613781184 | 1,530264655 | 0,30656 | 0,59558906 | calcium voltage-gated channel subunit alpha1 F |
| ENSG00000154727 | GABPA | 0,613515868 | 1,52998326 | 0,15187 | 0,48031011 | GA binding protein transcription factor subunit alpha |

| | | | | | | |
|---|---|---|---|---|---|---|
| ENSG00000171204 | TMEM126B | 0,613507706 | 1,529974604 | 0,26915 | 0,57086439 | transmembrane protein 126B |
| ENSG00000185053 | SGCZ | 0,613456011 | 1,529919783 | 0,29736 | 0,59031059 | sarcoglycan zeta |
| ENSG00000287387 | AC010883,3 | 0,613359376 | 1,529817308 | 0,18022 | 0,50407792 | novel transcript, antisense to ZFP36L2 |
| ENSG00000280618 | CYFIP1 | 0,613330377 | 1,529786558 | 0,15453 | 0,48266733 | cytoplasmic FMR1 interacting protein 1 |
| ENSG00000277517 | AC212493,1 | 0,613065796 | 1,529506031 | 0,32069 | 0,60555138 | novel transcript, antisense to MAMDC2 |
| ENSG00000272906 | AL353708,3 | 0,613006867 | 1,529443557 | 0,27077 | 0,57239492 | novel transcript, antisense to TOR1AIP1 |
| ENSG00000154310 | TNIK | 0,612990462 | 1,529426167 | 0,13426 | 0,46055924 | TRAF2 and NCK interacting kinase |
| ENSG00000041353 | RAB27B | 0,612976587 | 1,529411457 | 0,26451 | 0,5672714 | RAB27B, member RAS oncogene family |
| ENSG00000136146 | MED4 | 0,612955824 | 1,529389446 | 0,21929 | 0,53676938 | mediator complex subunit 4 |
| ENSG00000276282 | AC022960,2 | 0,612816744 | 1,529242016 | 0,22652 | 0,54144 | novel transcript, sense intronic to TRAPPC8 |
| ENSG00000120690 | ELF1 | 0,612792666 | 1,529216494 | 0,00654 | 0,1024996 | E74 like ETS transcription factor 1 |
| ENSG00000131236 | CAP1 | 0,61221364 | 1,528602865 | 0,14641 | 0,47433542 | cyclase associated actin cytoskeleton regulatory protein 1 |
| ENSG00000134910 | STT3A | 0,612195312 | 1,528583447 | 0,10681 | 0,41660052 | STT3 oligosaccharyltransferase complex catalytic subunit A |
| ENSG00000250682 | LINC00491 | 0,612140337 | 1,528525199 | 0,03183 | 0,24060936 | long intergenic non-protein coding RNA 491 |
| ENSG00000263235 | AC006111,2 | 0,612025191 | 1,528403208 | 0,2605 | 0,56389648 | novel transcript, sense overlapping TRAP1 |
| ENSG00000233450 | KIFC1 | 0,611789275 | 1,528153297 | 0,43041 | 0,68295004 | kinesin family member C1 |
| ENSG00000182400 | TRAPPC6B | 0,611679571 | 1,528037099 | 0,14857 | 0,47609532 | trafficking protein particle complex 6B |
| ENSG00000135638 | EMX1 | 0,611617104 | 1,527970938 | 0,19995 | 0,52092803 | empty spiracles homeobox 1 |
| ENSG00000111196 | MAGOHB | 0,611538337 | 1,527887518 | 0,20555 | 0,52513499 | mago homolog B, exon junction complex subunit |
| ENSG00000277564 | RBFOX2 | 0,611190747 | 1,527519447 | 0,06585 | 0,33965348 | RNA binding fox-1 homolog 2 |
| ENSG00000287723 | AP005901,6 | 0,611167869 | 1,527495224 | 0,61791 | 0,80610497 | novel transcript |
| ENSG00000123892 | RAB38 | 0,611112861 | 1,527436983 | 0,34679 | 0,62352276 | RAB38, member RAS oncogene family |
| ENSG00000132953 | XPO4 | 0,611102656 | 1,527426179 | 0,1444 | 0,47200462 | exportin 4 |
| ENSG00000055208 | TAB2 | 0,611001644 | 1,527319239 | 0,10973 | 0,42183783 | TGF-beta activated kinase 1 (MAP3K7) binding protein 2 |
| ENSG00000172845 | SP3 | 0,610575183 | 1,526867829 | 0,11253 | 0,42637009 | Sp3 transcription factor |
| ENSG00000233858 | LINC02599 | 0,610503287 | 1,52679174 | 0,45564 | 0,69957443 | long intergenic non-protein coding RNA 2599 |
| ENSG00000115514 | TXNDC9 | 0,610466184 | 1,526752475 | 0,31825 | 0,60349082 | thioredoxin domain containing 9 |
| ENSG00000115841 | RMDN2 | 0,610427461 | 1,526711497 | 0,12645 | 0,44907902 | regulator of microtubule dynamics 2 |
| ENSG00000170522 | ELOVL6 | 0,610372535 | 1,526653373 | 0,17588 | 0,49983618 | ELOVL fatty acid elongase 6 |
| ENSG00000243250 | RPS6P16 | 0,610323375 | 1,526601353 | 0,39824 | 0,66084743 | ribosomal protein S6 pseudogene 16 |
| ENSG00000101812 | H2BW2 | 0,610279368 | 1,526554788 | 0,1134 | 0,42803427 | H2B,W histone 2 |
| ENSG00000229061 | ATAT1 | 0,610254296 | 1,526528258 | 0,15929 | 0,4867709 | alpha tubulin acetyltransferase 1 |
| ENSG00000206488 | ATAT1 | 0,610254296 | 1,526528258 | 0,15929 | 0,4867709 | alpha tubulin acetyltransferase 1 |

| | | | | | | |
|---|---|---|---|---|---|---|
| ENSG00000234549 | ATAT1 | 0,610254296 | 1,526528258 | 0,15929 | 0,4867709 | alpha tubulin acetyltransferase 1 |
| ENSG00000172215 | CXCR6 | 0,610170955 | 1,526440077 | 0,25284 | 0,55796133 | C-X-C motif chemokine receptor 6 |
| ENSG00000139263 | LRIG3 | 0,610147059 | 1,526414793 | 0,1904 | 0,51350457 | leucine rich repeats and immunoglobulin like domains 3 |
| ENSG00000136698 | CFC1 | 0,609957015 | 1,526213735 | 0,64661 | 0,82303578 | cripto, FRL-1, cryptic family 1 |
| ENSG00000116017 | ARID3A | 0,609910335 | 1,526164353 | 0,13093 | 0,45558936 | AT-rich interaction domain 3A |
| ENSG00000224020 | MIR181A2HG | 0,609764573 | 1,526010165 | 0,11604 | 0,4324276 | MIR181A2 host gene |
| ENSG00000182199 | SHMT2 | 0,60970131 | 1,525943251 | 0,01805 | 0,17733967 | serine hydroxymethyltransferase 2 |
| ENSG00000167862 | MRPL58 | 0,609630933 | 1,525868815 | 0,26145 | 0,56450372 | mitochondrial ribosomal protein L58 |
| ENSG00000281333 | AC024941,2 | 0,609604503 | 1,525840861 | 0,36337 | 0,6341603 | novel transcript, antisense to KITLG |
| ENSG00000266964 | FXYD1 | 0,609569146 | 1,525803467 | 0,21804 | 0,5356729 | FXYD domain containing ion transport regulator 1 |
| ENSG00000217495 | AL136116,2 | 0,609430073 | 1,52565639 | 0,50142 | 0,73136997 | actin protein pseudogene |
| ENSG00000249395 | CASC9 | 0,609177533 | 1,525389351 | 0,04396 | 0,28274932 | cancer susceptibility 9 |
| ENSG00000285053 | AL357556,3 | 0,609024009 | 1,525227036 | 0,37389 | 0,64236763 | Tubulin-specific chaperone E |
| ENSG00000027869 | SH2D2A | 0,608996122 | 1,525197553 | 0,44418 | 0,691712 | SH2 domain containing 2A |
| ENSG00000273284 | AP001033,2 | 0,608955215 | 1,525154308 | 0,35523 | 0,62970025 | novel transcript, sense intronic to ANKRD12 |
| ENSG00000222112 | RN7SKP16 | 0,60854695 | 1,524722769 | 0,34441 | 0,62187731 | RN7SK pseudogene 16 |
| ENSG00000286811 | AL353138,1 | 0,608417261 | 1,524585713 | 0,41588 | 0,67283255 | novel transcript |
| ENSG00000285395 | XYLT1 | 0,608344618 | 1,524508948 | 0,08127 | 0,37209365 | xylosyltransferase 1 |
| ENSG00000168818 | STX18 | 0,608294773 | 1,524456277 | 0,06427 | 0,33593883 | syntaxin 18 |
| ENSG00000145431 | PDGFC | 0,608244283 | 1,524402927 | 0,11075 | 0,42337179 | platelet derived growth factor C |
| ENSG00000072422 | RHOBTB1 | 0,607872666 | 1,524010313 | 0,12224 | 0,44171717 | Rho related BTB domain containing 1 |
| ENSG00000257207 | AC112229,3 | 0,607849399 | 1,523985736 | 0,7094 | 0,85983746 | novel transcript |
| ENSG00000253368 | TRNP1 | 0,607840606 | 1,523976447 | 0,12257 | 0,44233393 | TMF1 regulated nuclear protein 1 |
| ENSG00000183087 | GAS6 | 0,607767957 | 1,523899707 | 0,05515 | 0,31428901 | growth arrest specific 6 |
| ENSG00000123983 | ACSL3 | 0,607737157 | 1,523867173 | 0,09671 | 0,40087593 | acyl-CoA synthetase long chain family member 3 |
| ENSG00000120555 | SEPTIN7P9 | 0,607727006 | 1,523856451 | 0,46737 | 0,70725568 | septin 7 pseudogene 9 |
| ENSG00000288167 | SEPTIN7P9 | 0,607727006 | 1,523856451 | 0,46737 | 0,70725568 | septin 7 pseudogene 9 |
| ENSG00000134970 | TMED7 | 0,607638016 | 1,523762458 | 0,21841 | 0,53578053 | transmembrane p24 trafficking protein 7 |
| ENSG00000166960 | CCDC178 | 0,607550095 | 1,523669599 | 0,32362 | 0,60635905 | coiled-coil domain containing 178 |
| ENSG00000255203 | OR7E2P | 0,607298939 | 1,52340437 | 0,29936 | 0,59128796 | olfactory receptor family 7 subfamily E member 2 pseudogene |
| ENSG00000163655 | GMPS | 0,607259816 | 1,523363059 | 0,08545 | 0,37891248 | guanine monophosphate synthase |
| ENSG00000150457 | LATS2 | 0,607040403 | 1,523131394 | 3,38E-05 | 0,00239114 | large tumor suppressor kinase 2 |

| | | | | | | |
|---|---|---|---|---|---|---|
| ENSG00000185164 | NOMO2 | 0,606930144 | 1,523014993 | 0,17947 | 0,50387897 | NODAL modulator 2 |
| ENSG00000255189 | GLYATL1P1 | 0,606818442 | 1,522897076 | 0,59712 | 0,79434997 | glycine-N-acyltransferase like 1 pseudogene 1 |
| ENSG00000073417 | PDE8A | 0,606769825 | 1,522845758 | 0,03156 | 0,23991209 | phosphodiesterase 8A |
| ENSG00000275180 | AC048341,2 | 0,606577174 | 1,522642417 | 0,28358 | 0,58072192 | novel transcript |
| ENSG00000138802 | SEC24B | 0,606516612 | 1,5225785 | 0,14862 | 0,47609532 | SEC24 homolog B, COPII coat complex component |
| ENSG00000101361 | NOP56 | 0,606098647 | 1,522137457 | 0,10031 | 0,4067769 | NOP56 ribonucleoprotein |
| ENSG00000154719 | MRPL39 | 0,605970505 | 1,522002264 | 0,1993 | 0,52034238 | mitochondrial ribosomal protein L39 |
| ENSG00000285361 | AL844170,1 | 0,605436536 | 1,521439047 | 0,46306 | 0,70415312 | novel transcript |
| ENSG00000214067 | AL360182,1 | 0,60540284 | 1,521403511 | 0,42243 | 0,67663831 | ribosomal protein L21 (RPL21) pseudogene |
| ENSG00000206530 | CFAP44 | 0,605185842 | 1,521174692 | 0,04926 | 0,29853148 | cilia and flagella associated protein 44 |
| ENSG00000127366 | TAS2R5 | 0,605084585 | 1,521067931 | 0,24434 | 0,55253387 | taste 2 receptor member 5 |
| ENSG00000141101 | NOB1 | 0,604870662 | 1,520842404 | 0,13681 | 0,4641716 | NIN1 (RPN12) binding protein 1 homolog |
| ENSG00000196323 | ZBTB44 | 0,60470704 | 1,520669928 | 0,10606 | 0,41557034 | zinc finger and BTB domain containing 44 |
| ENSG00000143149 | ALDH9A1 | 0,604675202 | 1,520636369 | 0,17224 | 0,49710979 | aldehyde dehydrogenase 9 family member A1 |
| ENSG00000211460 | TSN | 0,604647628 | 1,520607307 | 0,04552 | 0,28824362 | translin |
| ENSG00000166004 | CEP295 | 0,604626576 | 1,520585117 | 0,23612 | 0,54721765 | centrosomal protein 295 |
| ENSG00000143036 | SLC44A3 | 0,604370405 | 1,52031514 | 0,17976 | 0,50407792 | solute carrier family 44 member 3 |
| ENSG00000181562 | EDDM3A | 0,604281595 | 1,520221555 | 0,52875 | 0,75099628 | epididymal protein 3A |
| ENSG00000171812 | COL8A2 | 0,604000109 | 1,519924971 | 0,23568 | 0,5471053 | collagen type VIII alpha 2 chain |
| ENSG00000287977 | AC068630,1 | 0,603889232 | 1,519808163 | 0,32904 | 0,61007355 | tigger transposable element derived 1 (TIGD1) pseudogene |
| ENSG00000288424 | AGAP12P | 0,60378621 | 1,519699638 | 0,56776 | 0,77563329 | ArfGAP with GTPase domain, ankyrin repeat and PH domain 12, pseudogene |
| ENSG00000265018 | AGAP12P | 0,60378621 | 1,519699638 | 0,56776 | 0,77563329 | ArfGAP with GTPase domain, ankyrin repeat and PH domain 12, pseudogene |
| ENSG00000006282 | SPATA20 | 0,603435935 | 1,519330712 | 0,01319 | 0,15012165 | spermatogenesis associated 20 |
| ENSG00000240758 | AC010655,2 | 0,603432328 | 1,519326913 | 0,22059 | 0,53759541 | novel transcript |
| ENSG00000115415 | STAT1 | 0,603428359 | 1,519322734 | 0,16544 | 0,49168875 | signal transducer and activator of transcription 1 |
| ENSG00000267508 | ZNF285 | 0,603117117 | 1,518994996 | 0,00967 | 0,12594734 | zinc finger protein 285 |
| ENSG00000273783 | AL136040,1 | 0,603036882 | 1,51891052 | 0,10557 | 0,4148224 | novel transcript, antisense to GTF2A1 |
| ENSG00000138382 | METTL5 | 0,602908857 | 1,518775738 | 0,12247 | 0,4421901 | methyltransferase like 5 |
| ENSG00000126790 | L3HYPDH | 0,602871717 | 1,518736639 | 0,16303 | 0,48967932 | trans-L-3-hydroxyproline dehydratase |
| ENSG00000269921 | AC068620,1 | 0,602499497 | 1,51834485 | 0,25455 | 0,55944852 | novel transcript |

| | | | | | | |
|---|---|---|---|---|---|---|
| ENSG00000176890 | TYMS | 0,602317818 | 1,518153657 | 0,1125 | 0,4263539 | thymidylate synthetase |
| ENSG00000114023 | FAM162A | 0,602288188 | 1,518122478 | 0,06804 | 0,34462459 | family with sequence similarity 162 member A |
| ENSG00000189014 | SHLD2P3 | 0,60222382 | 1,518054746 | 0,5264 | 0,74978877 | shieldin complex subunit 2 pseudogene 3 |
| ENSG00000260922 | AC009139,1 | 0,602169548 | 1,51799764 | 0,5627 | 0,77194048 | novel transcript, antisense to ADAMTS18 |
| ENSG00000287054 | AC113410,5 | 0,602134906 | 1,517961191 | 0,50514 | 0,73378796 | novel transcript, antisense to ZCCHC10 |
| ENSG00000148180 | GSN | 0,6021253 | 1,517951083 | 0,02053 | 0,19002626 | gelsolin |
| ENSG00000229117 | RPL41 | 0,602099535 | 1,517923974 | 0,10604 | 0,41557034 | ribosomal protein L41 |
| ENSG00000163376 | KBTBD8 | 0,601943899 | 1,517760232 | 0,31962 | 0,60451366 | kelch repeat and BTB domain containing 8 |
| ENSG00000145147 | SLIT2 | 0,601820263 | 1,517630168 | 0,08643 | 0,38089807 | slit guidance ligand 2 |
| ENSG00000250483 | PPM1AP1 | 0,601718679 | 1,517523312 | 0,40654 | 0,66597268 | protein phosphatase, Mg2+/Mn2+ dependent 1A pseudogene 1 |
| ENSG00000010219 | DYRK4 | 0,601609493 | 1,517408467 | 0,26754 | 0,56973003 | dual specificity tyrosine phosphorylation regulated kinase 4 |
| ENSG00000198589 | LRBA | 0,601541157 | 1,517336594 | 0,12656 | 0,44922932 | LPS responsive beige-like anchor protein |
| ENSG00000217783 | LDHAL6FP | 0,60147602 | 1,517268088 | 0,46874 | 0,70833955 | lactate dehydrogenase A like 6F, pseudogene |
| ENSG00000134987 | WDR36 | 0,60144038 | 1,517230606 | 0,19289 | 0,51495054 | WD repeat domain 36 |
| ENSG00000286757 | AL137139,2 | 0,60135665 | 1,517142553 | 0,12824 | 0,45171208 | novel transcript, antisense to CLYBL |
| ENSG00000265399 | AP005329,2 | 0,601226159 | 1,517005335 | 0,2075 | 0,5269766 | novel transcript, antisense to MYOM1 |
| ENSG00000114779 | ABHD14B | 0,601048942 | 1,516819001 | 0,0134 | 0,15158202 | abhydrolase domain containing 14B |
| ENSG00000141384 | TAF4B | 0,600953675 | 1,516718843 | 0,16009 | 0,48727733 | TATA-box binding protein associated factor 4b |
| ENSG00000124508 | BTN2A2 | 0,600651767 | 1,516401477 | 0,02703 | 0,22193038 | butyrophilin subfamily 2 member A2 |
| ENSG00000184500 | PROS1 | 0,600505431 | 1,516247673 | 0,19363 | 0,51547251 | protein S |
| ENSG00000156076 | WIF1 | 0,600415091 | 1,51615273 | 0,55937 | 0,77045525 | WNT inhibitory factor 1 |
| ENSG00000233912 | AC026202,2 | 0,600383507 | 1,516119538 | 0,5121 | 0,73906991 | novel transcript, antisense to ARL8B |
| ENSG00000276618 | RAD17 | 0,600359506 | 1,516094316 | 0,23891 | 0,54892264 | RAD17 checkpoint clamp loader component |
| ENSG00000149308 | NPAT | 0,600233835 | 1,515962256 | 0,22649 | 0,54144 | nuclear protein, coactivator of histone transcription |
| ENSG00000280828 | AC090114,3 | 0,600161558 | 1,515886311 | 0,1727 | 0,49752358 | pseudogene similar to part of tousled-like kinase 2 (TLK2) |
| ENSG00000231728 | TMSB15B-AS1 | 0,599995908 | 1,515712267 | 0,00264 | 0,059998 | TMSB15B antisense RNA 1 |
| ENSG00000242479 | AC109992,1 | 0,599936969 | 1,515650347 | 0,4256 | 0,67918049 | eukaryotic translation initiation factor 2-alpha kinase 1 (EIF2AK1) pseudogene |
| ENSG00000166225 | FRS2 | 0,599674445 | 1,515374572 | 0,22585 | 0,54106597 | fibroblast growth factor receptor substrate 2 |
| ENSG00000244405 | ETV5 | 0,599507734 | 1,515199473 | 0,1723 | 0,49714741 | ETS variant transcription factor 5 |

| | | | | | | |
|---|---|---|---|---|---|---|
| ENSG00000173166 | RAPH1 | 0,599445044 | 1,515133634 | 0,15189 | 0,48031011 | Ras association (RalGDS/AF-6) and pleckstrin homology domains 1 |
| ENSG00000188467 | SLC24A5 | 0,59939205 | 1,51507798 | 0,39182 | 0,65534953 | solute carrier family 24 member 5 |
| ENSG00000187999 | HNRNPA1P61 | 0,59930831 | 1,514990041 | 0,49062 | 0,72476953 | heterogeneous nuclear ribonucleoprotein A1 pseudogene 61 |
| ENSG00000112378 | PERP | 0,599209797 | 1,514886595 | 0,21055 | 0,52907125 | p53 apoptosis effector related to PMP22 |
| ENSG00000259820 | AC083843,2 | 0,598926846 | 1,514589514 | 0,22765 | 0,54168591 | novel transcript |
| ENSG00000134901 | POGLUT2 | 0,598829105 | 1,514486906 | 0,10289 | 0,41078404 | protein O-glucosyltransferase 2 |
| ENSG00000122194 | PLG | 0,598555994 | 1,514200231 | 0,52537 | 0,74902159 | plasminogen |
| ENSG00000285187 | AC270290,1 | 0,598339603 | 1,513973133 | 0,63141 | 0,81403245 | novel transcript, antisense to SOX7 and PINX1 |
| ENSG00000241563 | CORT | 0,598194174 | 1,513820526 | 0,25662 | 0,56097949 | cortistatin |
| ENSG00000177879 | AP3S1 | 0,598074659 | 1,513695124 | 0,19587 | 0,51672115 | adaptor related protein complex 3 subunit sigma 1 |
| ENSG00000158470 | B4GALT5 | 0,59798573 | 1,513601822 | 0,027 | 0,22188397 | beta-1,4-galactosyltransferase 5 |
| ENSG00000159111 | MRPL10 | 0,59798116 | 1,513597027 | 0,11641 | 0,43288152 | mitochondrial ribosomal protein L10 |
| ENSG00000250421 | AC106798,1 | 0,597954739 | 1,513569308 | 0,43429 | 0,68505296 | novel transcript |
| ENSG00000165672 | PRDX3 | 0,597758827 | 1,513363785 | 0,15566 | 0,48345635 | peroxiredoxin 3 |
| ENSG00000187753 | C9orf153 | 0,597552886 | 1,513147772 | 0,26679 | 0,56923396 | chromosome 9 open reading frame 153 |
| ENSG00000234284 | ZNF879 | 0,597544653 | 1,513139137 | 0,15234 | 0,48061452 | zinc finger protein 879 |
| ENSG00000148516 | ZEB1 | 0,597377114 | 1,512963428 | 0,20041 | 0,52108722 | zinc finger E-box binding homeobox 1 |
| ENSG00000137393 | RNF144B | 0,597366944 | 1,512952762 | 0,30781 | 0,59671558 | ring finger protein 144B |
| ENSG00000183423 | LRIT3 | 0,597227605 | 1,512806644 | 0,48903 | 0,7236969 | leucine rich repeat, Ig-like and transmembrane domains 3 |
| ENSG00000060982 | BCAT1 | 0,596912363 | 1,512476119 | 0,12922 | 0,45265016 | branched chain amino acid transaminase 1 |
| ENSG00000285713 | AC098588,2 | 0,596833897 | 1,512393859 | 0,42459 | 0,67841584 | novel transcript |
| ENSG00000136098 | NEK3 | 0,596695053 | 1,512248315 | 0,03083 | 0,23697329 | NIMA related kinase 3 |
| ENSG00000166444 | DENND2B | 0,596648436 | 1,512199451 | 0,00142 | 0,040295 | DENN domain containing 2B |
| ENSG00000260619 | AC068870,1 | 0,596515359 | 1,512059969 | 0,35091 | 0,62636083 | novel transcript |
| ENSG00000101542 | CDH20 | 0,596275141 | 1,511808223 | 0,02696 | 0,22175312 | cadherin 20 |
| ENSG00000164932 | CTHRC1 | 0,596186129 | 1,511714949 | 0,13158 | 0,45658562 | collagen triple helix repeat containing 1 |
| ENSG00000169760 | NLGN1 | 0,595880094 | 1,511394308 | 0,143 | 0,47061399 | neuroligin 1 |
| ENSG00000170370 | EMX2 | 0,595845918 | 1,511358504 | 0,19714 | 0,51841425 | empty spiracles homeobox 2 |
| ENSG00000133112 | TPT1 | 0,595582743 | 1,511082829 | 0,16376 | 0,49000347 | tumor protein, translationally-controlled 1 |
| ENSG00000187325 | TAF9B | 0,595406444 | 1,510898184 | 0,28285 | 0,58009943 | TATA-box binding protein associated factor 9b |
| ENSG00000055955 | ITIH4 | 0,595241321 | 1,510725265 | 0,15467 | 0,48266733 | inter-alpha-trypsin inhibitor heavy chain 4 |

| | | | | | | |
|---|---|---|---|---|---|---|
| ENSG00000107968 | MAP3K8 | 0,595116491 | 1,510594554 | 0,05929 | 0,32445511 | mitogen-activated protein kinase kinase kinase 8 |
| ENSG00000153561 | RMND5A | 0,59508307 | 1,510559561 | 0,04717 | 0,29216937 | required for meiotic nuclear division 5 homolog A |
| ENSG00000172340 | SUCLG2 | 0,59489374 | 1,510361338 | 0,18033 | 0,50407792 | succinate-CoA ligase GDP-forming subunit beta |
| ENSG00000228816 | AK3P5 | 0,594803714 | 1,510267092 | 0,22167 | 0,53805234 | adenylate kinase 3 pseudogene 5 |
| ENSG00000273607 | CHCHD10 | 0,594704631 | 1,510163372 | 0,08591 | 0,38009386 | coiled-coil-helix-coiled-coil-helix domain containing 10 |
| ENSG00000164323 | CFAP97 | 0,594677832 | 1,51013532 | 0,25016 | 0,55657567 | cilia and flagella associated protein 97 |
| ENSG00000177125 | ZBTB34 | 0,594579922 | 1,510032837 | 0,13011 | 0,45452604 | zinc finger and BTB domain containing 34 |
| ENSG00000143368 | SF3B4 | 0,594551709 | 1,510003307 | 0,15954 | 0,48693054 | splicing factor 3b subunit 4 |
| ENSG00000196793 | ZNF239 | 0,594484607 | 1,509933076 | 0,16879 | 0,49492999 | zinc finger protein 239 |
| ENSG00000235443 | POLR1H | 0,59431375 | 1,509754266 | 0,69832 | 0,85390309 | RNA polymerase I subunit H |
| ENSG00000236949 | POLR1H | 0,59431375 | 1,509754266 | 0,69832 | 0,85390309 | RNA polymerase I subunit H |
| ENSG00000136161 | RCBTB2 | 0,594207954 | 1,509643557 | 0,25531 | 0,55990881 | RCC1 and BTB domain containing protein 2 |
| ENSG00000170275 | CRTAP | 0,594087036 | 1,509517033 | 0,00473 | 0,08480131 | cartilage associated protein |
| ENSG00000153347 | FAM81B | 0,594048382 | 1,509476589 | 0,52274 | 0,74689824 | family with sequence similarity 81 member B |
| ENSG00000124215 | CDH26 | 0,594005768 | 1,509432004 | 0,25709 | 0,56155289 | cadherin 26 |
| ENSG00000100138 | SNU13 | 0,593972544 | 1,509397243 | 0,12909 | 0,45248057 | small nuclear ribonucleoprotein 13 |
| ENSG00000072042 | RDH11 | 0,593929852 | 1,509352578 | 0,09535 | 0,39794903 | retinol dehydrogenase 11 |
| ENSG00000100567 | PSMA3 | 0,59388011 | 1,509300538 | 0,18255 | 0,5059851 | proteasome 20S subunit alpha 3 |
| ENSG00000231321 | RXRB | 0,593806541 | 1,509223575 | 0,24128 | 0,55027101 | retinoid X receptor beta |
| ENSG00000277103 | AL445670,1 | 0,593803074 | 1,509219948 | 0,27798 | 0,57629868 | sorting nexin 18 (SNX18) pseudogene |
| ENSG00000174842 | GLMN | 0,593761907 | 1,509176884 | 0,19179 | 0,51436129 | glomulin, FKBP associated protein |
| ENSG00000264058 | AC073508,2 | 0,593575884 | 1,508982301 | 0,54098 | 0,75863636 | novel transcript |
| ENSG00000068305 | MEF2A | 0,593443171 | 1,508843496 | 0,13195 | 0,45708171 | myocyte enhancer factor 2A |
| ENSG00000214273 | AGGF1P1 | 0,593410578 | 1,508809409 | 0,34937 | 0,62549486 | angiogenic factor with G-patch and FHA domains 1 pseudogene 1 |
| ENSG00000112419 | PHACTR2 | 0,592893157 | 1,508268373 | 0,10305 | 0,41078404 | phosphatase and actin regulator 2 |
| ENSG00000182667 | NTM | 0,59284066 | 1,508213491 | 0,09921 | 0,40549509 | neurotrimin |
| ENSG00000172476 | RAB40A | 0,592800708 | 1,508171725 | 0,04294 | 0,27922774 | RAB40A, member RAS oncogene family |
| ENSG00000144895 | EIF2A | 0,592710419 | 1,508077342 | 0,12736 | 0,45051698 | eukaryotic translation initiation factor 2A |
| ENSG00000000419 | DPM1 | 0,592496634 | 1,507853885 | 0,22453 | 0,54035379 | dolichyl-phosphate mannosyltransferase subunit 1, catalytic |
| ENSG00000166145 | SPINT1 | 0,592389635 | 1,507742057 | 0,19837 | 0,51980353 | serine peptidase inhibitor, Kunitz type 1 |
| ENSG00000136122 | BORA | 0,592374596 | 1,50772634 | 0,31524 | 0,60158701 | BORA aurora kinase A activator |

| | | | | | | |
|---|---|---|---|---|---|---|
| ENSG00000136156 | ITM2B | 0,592260476 | 1,507607081 | 0,19889 | 0,52005179 | integral membrane protein 2B |
| ENSG00000162244 | RPL29 | 0,591919419 | 1,507250721 | 0,24186 | 0,55068342 | ribosomal protein L29 |
| ENSG00000113396 | SLC27A6 | 0,591589794 | 1,506906386 | 0,36209 | 0,63354671 | solute carrier family 27 member 6 |
| ENSG00000185591 | SP1 | 0,59148354 | 1,506795407 | 0,09512 | 0,39772599 | Sp1 transcription factor |
| ENSG00000007237 | GAS7 | 0,591480999 | 1,506792752 | 0,02367 | 0,20544458 | growth arrest specific 7 |
| ENSG00000138756 | BMP2K | 0,59134303 | 1,50664866 | 0,15322 | 0,48147318 | BMP2 inducible kinase |
| ENSG00000281221 | PSMC4 | 0,591316831 | 1,506621301 | 0,14081 | 0,46780719 | proteasome 26S subunit, ATPase 4 |
| ENSG00000013275 | PSMC4 | 0,591316831 | 1,506621301 | 0,14081 | 0,46780719 | proteasome 26S subunit, ATPase 4 |
| ENSG00000272076 | AC090186,1 | 0,59131496 | 1,506619346 | 0,56224 | 0,77176502 | novel transcript |
| ENSG00000171863 | RPS7 | 0,591309924 | 1,506614088 | 0,15803 | 0,48506908 | ribosomal protein S7 |
| ENSG00000247317 | LY6E-DT | 0,591158312 | 1,506455766 | 0,6173 | 0,8057177 | LY6E divergent transcript |
| ENSG00000088930 | XRN2 | 0,59105449 | 1,50634736 | 0,2096 | 0,52860172 | 5'-3' exoribonuclease 2 |
| ENSG00000097046 | CDC7 | 0,591030493 | 1,506322304 | 0,20257 | 0,52264621 | cell division cycle 7 |
| ENSG00000133773 | CCDC59 | 0,59101521 | 1,506306347 | 0,22881 | 0,54259385 | coiled-coil domain containing 59 |
| ENSG00000274819 | AC018659,3 | 0,590977747 | 1,506267233 | 0,57547 | 0,78019408 | |
| ENSG00000223409 | RPL17P17 | 0,590969904 | 1,506259044 | 0,3927 | 0,65593262 | ribosomal protein L17 pseudogene 17 |
| ENSG00000067248 | DHX29 | 0,590942256 | 1,506230178 | 0,17498 | 0,49911246 | DExH-box helicase 29 |
| ENSG00000133863 | TEX15 | 0,590724159 | 1,506002494 | 0,34302 | 0,62060539 | testis expressed 15, meiosis and synapsis associated |
| ENSG00000163468 | CCT3 | 0,590583706 | 1,505855885 | 0,18835 | 0,51161341 | chaperonin containing TCP1 subunit 3 |
| ENSG00000125249 | RAP2A | 0,59047737 | 1,505744898 | 0,1827 | 0,50615118 | RAP2A, member of RAS oncogene family |
| ENSG00000226965 | AC073114,1 | 0,590453728 | 1,505720223 | 0,33917 | 0,61815389 | novel transcript |
| ENSG00000283802 | ADAMTS2 | 0,590323793 | 1,505584617 | 0,03249 | 0,24332854 | ADAM metallopeptidase with thrombospondin type 1 motif 2 |
| ENSG00000146233 | CYP39A1 | 0,590320347 | 1,505581021 | 0,1507 | 0,47887343 | cytochrome P450 family 39 subfamily A member 1 |
| ENSG00000079387 | SENP1 | 0,590215485 | 1,505471592 | 0,16377 | 0,49000347 | SUMO specific peptidase 1 |
| ENSG00000239763 | AC009120,1 | 0,590137188 | 1,50538989 | 0,33736 | 0,61706766 | ribosomal protein L21 (RPL21) pseudogene |
| ENSG00000151135 | TMEM263 | 0,590072316 | 1,505322201 | 0,13156 | 0,45658491 | transmembrane protein 263 |
| ENSG00000231665 | OGFOD1P1 | 0,589657847 | 1,504889801 | 0,21173 | 0,53061996 | 2-oxoglutarate and iron dependent oxygenase domain containing 1 pseudogene 1 |
| ENSG00000196950 | SLC39A10 | 0,589313216 | 1,504530356 | 0,22751 | 0,54168591 | solute carrier family 39 member 10 |
| ENSG00000146530 | VWDE | 0,588944143 | 1,504145514 | 0,39843 | 0,66102743 | von Willebrand factor D and EGF domains |
| ENSG00000223393 | AL118511,1 | 0,588798837 | 1,503994027 | 0,06153 | 0,32865897 | novel transcript |
| ENSG00000250329 | KDELC1P1 | 0,588668197 | 1,503857842 | 0,36433 | 0,6349692 | KDEL motif containing 1 pseudogene 1 |

| | | | | | | |
|---|---|---|---|---|---|---|
| ENSG00000231787 | H2BP9 | 0,588585831 | 1,503771987 | 0,37547 | 0,6438135 | H2B histone pseudogene 9 |
| ENSG00000139209 | SLC38A4 | 0,588562662 | 1,503747837 | 0,40679 | 0,66618707 | solute carrier family 38 member 4 |
| ENSG00000127252 | PLAAT1 | 0,58824101 | 1,503412611 | 0,12849 | 0,45191041 | phospholipase A and acyltransferase 1 |
| ENSG00000150456 | EEF1AKMT1 | 0,588005825 | 1,503167548 | 0,23889 | 0,54892264 | EEF1A lysine methyltransferase 1 |
| ENSG00000133135 | RNF128 | 0,587779375 | 1,502931624 | 0,58341 | 0,78522606 | ring finger protein 128 |
| ENSG00000197157 | SND1 | 0,587723735 | 1,502873662 | 0,06322 | 0,33302692 | staphylococcal nuclease and tudor domain containing 1 |
| ENSG00000285253 | AC090517,4 | 0,587406699 | 1,502543437 | 0,56559 | 0,773864 | zinc finger protein 280D |
| ENSG00000236977 | ANKRD44-IT1 | 0,587354965 | 1,502489558 | 0,50088 | 0,73097006 | ANKRD44 intronic transcript 1 |
| ENSG00000117519 | CNN3 | 0,587223908 | 1,502353076 | 0,0661 | 0,34035709 | calponin 3 |
| ENSG00000155329 | ZCCHC10 | 0,587120123 | 1,502245003 | 0,19339 | 0,51546367 | zinc finger CCHC-type containing 10 |
| ENSG00000112699 | GMDS | 0,587063244 | 1,502185778 | 0,01297 | 0,14899372 | GDP-mannose 4,6-dehydratase |
| ENSG00000183527 | PSMG1 | 0,586782461 | 1,501893445 | 0,01987 | 0,18711351 | proteasome assembly chaperone 1 |
| ENSG00000186352 | ANKRD37 | 0,586766625 | 1,501876959 | 0,18277 | 0,50616031 | ankyrin repeat domain 37 |
| ENSG00000242474 | AC093627,5 | 0,586516107 | 1,501616186 | 0,13434 | 0,46058584 | novel transcript |
| ENSG00000288472 | AC093627,29 | 0,586516107 | 1,501616186 | 0,13434 | 0,46058584 | novel transcript |
| ENSG00000064989 | CALCRL | 0,586513758 | 1,501613741 | 0,06161 | 0,32867767 | calcitonin receptor like receptor |
| ENSG00000107957 | SH3PXD2A | 0,586471736 | 1,501570004 | 0,08965 | 0,38717603 | SH3 and PX domains 2A |
| ENSG00000011426 | ANLN | 0,586399415 | 1,501494734 | 0,17434 | 0,49875953 | anillin actin binding protein |
| ENSG00000099958 | DERL3 | 0,586399234 | 1,501494545 | 0,1536 | 0,48191866 | derlin 3 |
| ENSG00000259845 | HERC2P10 | 0,586328603 | 1,501421038 | 0,11847 | 0,43614969 | hect domain and RLD 2 pseudogene 10 |
| ENSG00000023697 | DERA | 0,58625784 | 1,501347396 | 0,23885 | 0,54892264 | deoxyribose-phosphate aldolase |
| ENSG00000159593 | NAE1 | 0,586124739 | 1,50120889 | 0,1929 | 0,51495054 | NEDD8 activating enzyme E1 subunit 1 |
| ENSG00000232006 | AC005537,1 | 0,586123971 | 1,501208091 | 0,17814 | 0,50236177 | novel transcript |
| ENSG00000151883 | PARP8 | 0,586113597 | 1,501197296 | 0,18037 | 0,50407792 | poly(ADP-ribose) polymerase family member 8 |
| ENSG00000024862 | CCDC28A | 0,586097858 | 1,501180919 | 0,20274 | 0,52269757 | coiled-coil domain containing 28A |
| ENSG00000077063 | CTTNBP2 | 0,585874669 | 1,500948701 | 0,05033 | 0,30174609 | cortactin binding protein 2 |
| ENSG00000170515 | PA2G4 | 0,585704229 | 1,500771389 | 0,09339 | 0,39491488 | proliferation-associated 2G4 |
| ENSG00000129518 | EAPP | 0,585703084 | 1,500770197 | 0,22185 | 0,53821014 | E2F associated phosphoprotein |
| ENSG00000248278 | SUMO2P17 | 0,585599976 | 1,500662943 | 0,41904 | 0,67509192 | SUMO2 pseudogene 17 |
| ENSG00000109762 | SNX25 | 0,585585641 | 1,500648032 | 0,01992 | 0,18715502 | sorting nexin 25 |
| ENSG00000261603 | PRSS46P | 0,585265245 | 1,500314802 | 0,48829 | 0,72323505 | serine protease 46, pseudogene |
| ENSG00000241318 | WDR82P2 | 0,584907847 | 1,499943176 | 0,44063 | 0,68910892 | WD repeat domain 82 pseudogene 2 |
| ENSG00000198492 | YTHDF2 | 0,584840625 | 1,499873289 | 0,15651 | 0,48425071 | YTH N6-methyladenosine RNA binding protein 2 |

| | | | | | | |
|---|---|---|---|---|---|---|
| ENSG00000162385 | MAGOH | 0,584833836 | 1,49986623 | 0,17892 | 0,5033932 | mago homolog, exon junction complex subunit |
| ENSG00000256646 | AC010132,3 | 0,584795859 | 1,499826749 | 0,31552 | 0,60172912 | UPF0415 protein C7orf25 |
| ENSG00000103064 | SLC7A6 | 0,584755538 | 1,499784832 | 0,08725 | 0,38241534 | solute carrier family 7 member 6 |
| ENSG00000198825 | INPP5F | 0,584747116 | 1,499776077 | 0,11668 | 0,43365709 | inositol polyphosphate-5-phosphatase F |
| ENSG00000111961 | SASH1 | 0,58470612 | 1,499733459 | 0,09426 | 0,39648446 | SAM and SH3 domain containing 1 |
| ENSG00000170231 | FABP6 | 0,584625951 | 1,499650124 | 0,53247 | 0,75300764 | fatty acid binding protein 6 |
| ENSG00000182118 | FAM89A | 0,584603411 | 1,499626693 | 0,10973 | 0,42183783 | family with sequence similarity 89 member A |
| ENSG00000198298 | ZNF485 | 0,584388196 | 1,499403002 | 0,22676 | 0,54154849 | zinc finger protein 485 |
| ENSG00000096968 | JAK2 | 0,584317285 | 1,499329305 | 0,13922 | 0,46655993 | Janus kinase 2 |
| ENSG00000023445 | BIRC3 | 0,584057083 | 1,499058914 | 0,38708 | 0,65213944 | baculoviral IAP repeat containing 3 |
| ENSG00000130935 | NOL11 | 0,583923331 | 1,498919943 | 0,21501 | 0,53373528 | nucleolar protein 11 |
| ENSG00000278845 | MRPL45 | 0,583897842 | 1,49889346 | 0,03995 | 0,27022885 | mitochondrial ribosomal protein L45 |
| ENSG00000124802 | EEF1E1 | 0,583871742 | 1,498866344 | 0,16998 | 0,49606609 | eukaryotic translation elongation factor 1 epsilon 1 |
| ENSG00000226453 | LINC02542 | 0,583733532 | 1,49872276 | 0,42132 | 0,67634721 | long intergenic non-protein coding RNA 2542 |
| ENSG00000287606 | LINC02542 | 0,583733532 | 1,49872276 | 0,42132 | 0,67634721 | long intergenic non-protein coding RNA 2542 |
| ENSG00000155846 | PPARGC1B | 0,583701077 | 1,498689045 | 0,04722 | 0,29217708 | PPARG coactivator 1 beta |
| ENSG00000288079 | AC026329,1 | 0,583432694 | 1,498410271 | 0,08396 | 0,37651068 | novel transcript, antisense to BOC |
| ENSG00000281617 | AC073135,6 | 0,583315389 | 1,498288441 | 0,35738 | 0,63130076 | Ankyrin repeat domain 18A (ANKRD18A) pseudogene |
| ENSG00000183853 | KIRREL1 | 0,583095929 | 1,498060542 | 0,02764 | 0,22402593 | kirre like nephrin family adhesion molecule 1 |
| ENSG00000152359 | POC5 | 0,582993248 | 1,497953924 | 0,23804 | 0,54864017 | POC5 centriolar protein |
| ENSG00000077942 | FBLN1 | 0,582881677 | 1,497838083 | 0,02743 | 0,22348662 | fibulin 1 |
| ENSG00000223899 | SEC13P1 | 0,582873352 | 1,49782944 | 0,29913 | 0,59122003 | SEC13 homolog, nuclear pore and COPII coat complex component pseudogene 1 |
| ENSG00000005175 | RPAP3 | 0,582804506 | 1,497757966 | 0,19243 | 0,514872 | RNA polymerase II associated protein 3 |
| ENSG00000203737 | GPR52 | 0,582717127 | 1,497667254 | 0,21454 | 0,53351409 | G protein-coupled receptor 52 |
| ENSG00000187950 | OVCH1 | 0,582703592 | 1,497653204 | 0,43592 | 0,68646498 | ovochymase 1 |
| ENSG00000262826 | INTS3 | 0,582494916 | 1,497436594 | 0,04962 | 0,29985549 | integrator complex subunit 3 |
| ENSG00000189266 | PNRC2 | 0,582374436 | 1,497311547 | 0,1687 | 0,49492999 | proline rich nuclear receptor coactivator 2 |
| ENSG00000224973 | LARGE-AS1 | 0,582175296 | 1,497104883 | 0,20874 | 0,52791772 | LARGE antisense RNA 1 |
| ENSG00000151689 | INPP1 | 0,582160652 | 1,497089687 | 0,14654 | 0,47441255 | inositol polyphosphate-1-phosphatase |
| ENSG00000286672 | AL096712,2 | 0,582138239 | 1,497066428 | 0,21615 | 0,53480193 | novel transcript, antisense to TBC1D22B |
| ENSG00000258578 | AC008015,1 | 0,582098802 | 1,497025506 | 0,58506 | 0,7860906 | ubiquitin-conjugating enzyme E2L 3 (UBE2L3) pseudogene |
| ENSG00000130695 | CEP85 | 0,581999521 | 1,49692249 | 0,0801 | 0,36938412 | centrosomal protein 85 |

| | | | | | | |
|---|---|---|---|---|---|---|
| ENSG00000120805 | ARL1 | 0,581902895 | 1,496822236 | 0,1805 | 0,50407792 | ADP ribosylation factor like GTPase 1 |
| ENSG00000100320 | RBFOX2 | 0,581901771 | 1,49682107 | 0,05183 | 0,30603373 | RNA binding fox-1 homolog 2 |
| ENSG00000161551 | ZNF577 | 0,581812663 | 1,496728621 | 0,12842 | 0,45191041 | zinc finger protein 577 |
| ENSG00000183336 | BOLA2 | 0,581792866 | 1,496708083 | 0,3271 | 0,60902638 | bolA family member 2 |
| ENSG00000125703 | ATG4C | 0,581567118 | 1,496473901 | 0,23825 | 0,54870569 | autophagy related 4C cysteine peptidase |
| ENSG00000135912 | TTLL4 | 0,581566363 | 1,496473117 | 0,01191 | 0,14219678 | tubulin tyrosine ligase like 4 |
| ENSG00000168118 | RAB4A | 0,581530208 | 1,496435615 | 0,16829 | 0,49453836 | RAB4A, member RAS oncogene family |
| ENSG00000263050 | AC090617,5 | 0,58140226 | 1,496302907 | 0,46142 | 0,7028392 | novel transcript |
| ENSG00000146411 | SLC2A12 | 0,581314099 | 1,496211473 | 0,26401 | 0,56658292 | solute carrier family 2 member 12 |
| ENSG00000277161 | PIGW | 0,581291805 | 1,496188352 | 0,08257 | 0,37428581 | phosphatidylinositol glycan anchor biosynthesis class W |
| ENSG00000111785 | RIC8B | 0,581226718 | 1,496120853 | 0,19013 | 0,51331106 | RIC8 guanine nucleotide exchange factor B |
| ENSG00000251682 | AC122718,2 | 0,581218401 | 1,496112229 | 0,31836 | 0,60349835 | DEAD (Asp-Glu-Ala-Asp) box polypeptide 25 (DDX25) pseudogene |
| ENSG00000286427 | AC009486,2 | 0,581111544 | 1,49600142 | 0,52919 | 0,75112358 | novel transcript |
| ENSG00000182287 | AP1S2 | 0,581084276 | 1,495973144 | 0,26478 | 0,56748133 | adaptor related protein complex 1 subunit sigma 2 |
| ENSG00000182718 | ANXA2 | 0,58072577 | 1,495601445 | 0,09334 | 0,39489949 | annexin A2 |
| ENSG00000133193 | FAM104A | 0,580425467 | 1,495290161 | 0,17037 | 0,49641416 | family with sequence similarity 104 member A |
| ENSG00000066583 | ISOC1 | 0,580328519 | 1,495189682 | 0,17401 | 0,49860418 | isochorismatase domain containing 1 |
| ENSG00000064703 | DDX20 | 0,580213644 | 1,495070633 | 0,15571 | 0,48346026 | DEAD-box helicase 20 |
| ENSG00000197713 | RPE | 0,580158027 | 1,495012997 | 0,20331 | 0,52321009 | ribulose-5-phosphate-3-epimerase |
| ENSG00000160710 | ADAR | 0,580124111 | 1,494977852 | 0,03765 | 0,26251324 | adenosine deaminase RNA specific |
| ENSG00000108590 | MED31 | 0,580017677 | 1,494867565 | 0,21006 | 0,52899327 | mediator complex subunit 31 |
| ENSG00000242852 | ZNF709 | 0,579951696 | 1,494799199 | 0,32478 | 0,60732633 | zinc finger protein 709 |
| ENSG00000138039 | LHCGR | 0,579904071 | 1,494749855 | 0,39083 | 0,65464997 | luteinizing hormone/choriogonadotropin receptor |
| ENSG00000165997 | ARL5B | 0,579830392 | 1,49467352 | 0,15559 | 0,48345635 | ADP ribosylation factor like GTPase 5B |
| ENSG00000058673 | ZC3H11A | 0,579813555 | 1,494656077 | 0,15169 | 0,48001435 | zinc finger CCCH-type containing 11A |
| ENSG00000120675 | DNAJC15 | 0,579657496 | 1,494494405 | 0,17154 | 0,49702944 | DnaJ heat shock protein family (Hsp40) member C15 |
| ENSG00000120063 | GNA13 | 0,579628488 | 1,494464355 | 0,19213 | 0,51468713 | G protein subunit alpha 13 |
| ENSG00000078114 | NEBL | 0,579622232 | 1,494457876 | 0,25331 | 0,55839162 | nebulette |
| ENSG00000131849 | ZNF132 | 0,579392772 | 1,494220202 | 0,1103 | 0,42293017 | zinc finger protein 132 |
| ENSG00000117751 | PPP1R8 | 0,579312954 | 1,494137536 | 0,13723 | 0,4642578 | protein phosphatase 1 regulatory subunit 8 |
| ENSG00000127362 | TAS2R3 | 0,579252435 | 1,494074859 | 0,19814 | 0,51977271 | taste 2 receptor member 3 |
| ENSG00000155629 | PIK3AP1 | 0,579193021 | 1,494013331 | 0,34614 | 0,62331777 | phosphoinositide-3-kinase adaptor protein 1 |

| | | | | | | |
|---|---|---|---|---|---|---|
| ENSG00000170855 | TRIAP1 | 0,579109646 | 1,493926992 | 0,19816 | 0,51977271 | TP53 regulated inhibitor of apoptosis 1 |
| ENSG00000151151 | IPMK | 0,579022814 | 1,493837079 | 0,08842 | 0,38471382 | inositol polyphosphate multikinase |
| ENSG00000175093 | SPSB4 | 0,578672103 | 1,49347398 | 0,02136 | 0,19404207 | splA/ryanodine receptor domain and SOCS box containing 4 |
| ENSG00000154330 | PGM5 | 0,578659065 | 1,493460484 | 0,36149 | 0,63347405 | phosphoglucomutase 5 |
| ENSG00000123094 | RASSF8 | 0,57840086 | 1,493193217 | 0,25012 | 0,55657567 | Ras association domain family member 8 |
| ENSG00000273257 | AC069200,1 | 0,578322095 | 1,493111697 | 0,45868 | 0,70133296 | novel transcript |
| ENSG00000198189 | HSD17B11 | 0,578300076 | 1,493088909 | 0,13979 | 0,46689302 | hydroxysteroid 17-beta dehydrogenase 11 |
| ENSG00000274297 | AC009269,5 | 0,578248725 | 1,493035765 | 0,51771 | 0,74272749 | novel transcript, sense intronic to UACA |
| ENSG00000226604 | PAPPA-AS2 | 0,578199493 | 1,492984816 | 0,23064 | 0,54367096 | PAPPA antisense RNA 2 |
| ENSG00000223583 | AL513365,1 | 0,5781956 | 1,492980787 | 0,44189 | 0,68994635 | ribosomal protein L17 (RPL17) pseudogene |
| ENSG00000186871 | ERCC6L | 0,578135476 | 1,492918569 | 0,19756 | 0,51902178 | ERCC excision repair 6 like, spindle assembly checkpoint helicase |
| ENSG00000169612 | RAMAC | 0,578083846 | 1,492865142 | 0,25073 | 0,55690541 | RNA guanine-7 methyltransferase activating subunit |
| ENSG00000108582 | CPD | 0,577866627 | 1,492640387 | 0,12297 | 0,44289305 | carboxypeptidase D |
| ENSG00000235784 | HNRNPA1P29 | 0,577824475 | 1,492596776 | 0,26575 | 0,56866772 | heterogeneous nuclear ribonucleoprotein A1 pseudogene 29 |
| ENSG00000224859 | POLR1H | 0,577779331 | 1,492550071 | 0,06886 | 0,34681313 | RNA polymerase I subunit H |
| ENSG00000276867 | AC074050,4 | 0,577662742 | 1,492429458 | 0,48355 | 0,71980944 | novel transcript |
| ENSG00000109458 | GAB1 | 0,577632116 | 1,492397777 | 0,09121 | 0,39063588 | GRB2 associated binding protein 1 |
| ENSG00000156876 | SASS6 | 0,57743959 | 1,492198631 | 0,22991 | 0,54327768 | SAS-6 centriolar assembly protein |
| ENSG00000258448 | AL442663,2 | 0,577400644 | 1,492158349 | 0,36404 | 0,63490528 | required for meiotic nuclear division 1 homolog (S, cerevisiae) (RMND1) pseudogene |
| ENSG00000198612 | COPS8 | 0,577155574 | 1,491904898 | 0,31213 | 0,59925736 | COP9 signalosome subunit 8 |
| ENSG00000225215 | SMARCE1P1 | 0,577124168 | 1,491872422 | 0,28969 | 0,58522301 | SWI/SNF related, matrix associated, actin dependent regulator of chromatin, subfamily e, member 1 pseudogene 1 |
| ENSG00000073921 | PICALM | 0,576904916 | 1,491645713 | 0,22562 | 0,54087803 | phosphatidylinositol binding clathrin assembly protein |
| ENSG00000173276 | ZBTB21 | 0,576859525 | 1,491598783 | 0,17043 | 0,49641416 | zinc finger and BTB domain containing 21 |
| ENSG00000274929 | AL157813,1 | 0,576848466 | 1,491587349 | 0,23848 | 0,5488078 | novel transcript, sense intronic to RCBTB2 |
| ENSG00000265354 | TIMM23 | 0,576799793 | 1,491537027 | 0,16284 | 0,48945635 | translocase of inner mitochondrial membrane 23 |
| ENSG00000172728 | FUT10 | 0,576771184 | 1,49150745 | 0,11486 | 0,43031399 | fucosyltransferase 10 |
| ENSG00000271092 | TLCD4-RWDD | 0,57674184 | 1,491477113 | 0,28705 | 0,58349114 | TLCD4-RWDD3 readthrough |

| | | | | | | |
|---|---|---|---|---|---|---|
| ENSG00000144935 | TRPC1 | 0,576719179 | 1,491453687 | 0,15771 | 0,4847949 | transient receptor potential cation channel subfamily C member 1 |
| ENSG00000119777 | TMEM214 | 0,576551846 | 1,491280708 | 0,04385 | 0,28238629 | transmembrane protein 214 |
| ENSG00000096080 | MRPS18A | 0,576453606 | 1,491179163 | 0,13081 | 0,45538979 | mitochondrial ribosomal protein S18A |
| ENSG00000227129 | NEU1 | 0,57635392 | 1,491076131 | 0,04138 | 0,27554991 | neuraminidase 1 |
| ENSG00000168439 | STIP1 | 0,576296774 | 1,49101707 | 0,12441 | 0,44539699 | stress induced phosphoprotein 1 |
| ENSG00000206560 | ANKRD28 | 0,576021441 | 1,490732542 | 0,12431 | 0,44533764 | ankyrin repeat domain 28 |
| ENSG00000188803 | SHISA6 | 0,575995758 | 1,490706004 | 0,01635 | 0,16750744 | shisa family member 6 |
| ENSG00000160606 | TLCD1 | 0,575966378 | 1,490675646 | 0,08617 | 0,38054156 | TLC domain containing 1 |
| ENSG00000162368 | CMPK1 | 0,575652615 | 1,490351483 | 0,19896 | 0,52006724 | cytidine/uridine monophosphate kinase 1 |
| ENSG00000052723 | SIKE1 | 0,57560634 | 1,490303681 | 0,14748 | 0,47493724 | suppressor of IKBKE 1 |
| ENSG00000106392 | C1GALT1 | 0,57549453 | 1,490188185 | 0,17209 | 0,49710979 | core 1 synthase, glycoprotein-N-acetylgalactosamine 3-beta-galactosyltransferase 1 |
| ENSG00000145248 | SLC10A4 | 0,575437574 | 1,490129355 | 0,10738 | 0,41771747 | solute carrier family 10 member 4 |
| ENSG00000230569 | AC114763,2 | 0,57542983 | 1,490121357 | 0,57727 | 0,78134056 | novel transcript |
| ENSG00000117450 | PRDX1 | 0,575391721 | 1,490081996 | 0,04953 | 0,29942963 | peroxiredoxin 1 |
| ENSG00000137098 | SPAG8 | 0,575250487 | 1,489936129 | 0,04282 | 0,27885381 | sperm associated antigen 8 |
| ENSG00000140105 | WARS1 | 0,57496785 | 1,489644267 | 0,18298 | 0,50649341 | tryptophanyl-tRNA synthetase 1 |
| ENSG00000277222 | BTBD7 | 0,574859148 | 1,489532032 | 0,07942 | 0,3679672 | BTB domain containing 7 |
| ENSG00000109846 | CRYAB | 0,574850082 | 1,489522671 | 0,5417 | 0,75882662 | crystallin alpha B |
| ENSG00000120437 | ACAT2 | 0,574754246 | 1,489423727 | 0,09852 | 0,40417627 | acetyl-CoA acetyltransferase 2 |
| ENSG00000108298 | RPL19 | 0,574692808 | 1,489360301 | 0,16909 | 0,49515165 | ribosomal protein L19 |
| ENSG00000285226 | PAK1IP1 | 0,574691001 | 1,489358435 | 0,16886 | 0,49495623 | PAK1 interacting protein 1 |
| ENSG00000111845 | PAK1IP1 | 0,574691001 | 1,489358435 | 0,16886 | 0,49495623 | PAK1 interacting protein 1 |
| ENSG00000139679 | LPAR6 | 0,574615458 | 1,489280451 | 0,31487 | 0,60143663 | lysophosphatidic acid receptor 6 |
| ENSG00000134419 | RPS15A | 0,574563192 | 1,489226499 | 0,18317 | 0,50661194 | ribosomal protein S15a |
| ENSG00000145331 | TRMT10A | 0,574545239 | 1,489207967 | 0,19721 | 0,5184837 | tRNA methyltransferase 10A |
| ENSG00000248491 | AC093772,1 | 0,574537047 | 1,489199511 | 0,58214 | 0,78454692 | novel transcript |
| ENSG00000279389 | GARRE1 | 0,574442322 | 1,489101736 | 0,11999 | 0,43868155 | granule associated Rac and RHOG effector 1 |
| ENSG00000112149 | CD83 | 0,574397427 | 1,489055397 | 0,18063 | 0,50411652 | CD83 molecule |
| ENSG00000178826 | TMEM139 | 0,574349786 | 1,489006226 | 0,22142 | 0,53805234 | transmembrane protein 139 |
| ENSG00000185480 | PARPBP | 0,574075585 | 1,488723249 | 0,21635 | 0,53480193 | PARP1 binding protein |
| ENSG00000116106 | EPHA4 | 0,573598315 | 1,488230835 | 0,24447 | 0,55253387 | EPH receptor A4 |

| | | | | | | |
|---|---|---|---|---|---|---|
| ENSG00000186787 | SPIN2B | 0,573544098 | 1,488174907 | 0,25823 | 0,5622185 | spindlin family member 2B |
| ENSG00000254206 | NPIPB11 | 0,573477651 | 1,488106367 | 0,15991 | 0,48710442 | nuclear pore complex interacting protein family member B11 |
| ENSG00000230894 | AL133284,1 | 0,573383249 | 1,488008996 | 0,31107 | 0,59885232 | novel transcript |
| ENSG00000260296 | AC095057,3 | 0,573054663 | 1,487670129 | 0,27906 | 0,5771769 | novel transcript, overlapping N4BP2 |
| ENSG00000231206 | HNRNPA1P25 | 0,573029434 | 1,487644113 | 0,40015 | 0,66233089 | heterogeneous nuclear ribonucleoprotein A1 pseudogene 25 |
| ENSG00000095209 | TMEM38B | 0,572693508 | 1,487297761 | 0,16946 | 0,49561922 | transmembrane protein 38B |
| ENSG00000189129 | PLAC9 | 0,572659741 | 1,48726295 | 0,16205 | 0,48917419 | placenta associated 9 |
| ENSG00000286830 | AC105052,5 | 0,572632155 | 1,487234512 | 0,14776 | 0,47507961 | novel transcript |
| ENSG00000229591 | AC006017,1 | 0,572569864 | 1,4871703 | 0,29634 | 0,58968627 | novel transcript |
| ENSG00000135336 | ORC3 | 0,572549535 | 1,487149344 | 0,29378 | 0,58826777 | origin recognition complex subunit 3 |
| ENSG00000234197 | ETV5-AS1 | 0,572518841 | 1,487117705 | 0,34168 | 0,6197312 | ETV5 antisense RNA 1 |
| ENSG00000139324 | TMTC3 | 0,572365897 | 1,48696006 | 0,18476 | 0,50844008 | transmembrane O-mannosyltransferase targeting cadherins 3 |
| ENSG00000204956 | PCDHGA1 | 0,572175996 | 1,486764346 | 0,15827 | 0,48529549 | protocadherin gamma subfamily A, 1 |
| ENSG00000146476 | ARMT1 | 0,572127511 | 1,486714381 | 0,19978 | 0,52077977 | acidic residue methyltransferase 1 |
| ENSG00000173848 | NET1 | 0,571967332 | 1,486549323 | 0,04036 | 0,27171411 | neuroepithelial cell transforming 1 |
| ENSG00000275176 | ACACA | 0,571905548 | 1,486485663 | 0,19203 | 0,5145257 | acetyl-CoA carboxylase alpha |
| ENSG00000275835 | TUBGCP5 | 0,571900147 | 1,486480098 | 0,16365 | 0,49000031 | tubulin gamma complex associated protein 5 |
| ENSG00000182197 | EXT1 | 0,571853763 | 1,486432307 | 0,10645 | 0,41619134 | exostosin glycosyltransferase 1 |
| ENSG00000240549 | THOC7-AS1 | 0,571829521 | 1,48640733 | 0,3077 | 0,59671558 | THOC7 antisense RNA 1 |
| ENSG00000140006 | WDR89 | 0,571777743 | 1,486353984 | 0,22132 | 0,53805234 | WD repeat domain 89 |
| ENSG00000206448 | PPIAP30 | 0,571688767 | 1,486262318 | 0,44496 | 0,69205935 | peptidylprolyl isomerase A pseudogene 30 |
| ENSG00000117009 | KMO | 0,571630349 | 1,486202137 | 0,34203 | 0,61989127 | kynurenine 3-monooxygenase |
| ENSG00000094804 | CDC6 | 0,57153828 | 1,486107295 | 0,02842 | 0,22802307 | cell division cycle 6 |
| ENSG00000188460 | ACTBP11 | 0,571502846 | 1,486070795 | 0,19048 | 0,51355616 | ACTB pseudogene 11 |
| ENSG00000234737 | KRT18P15 | 0,57101585 | 1,485569242 | 0,55178 | 0,76553533 | keratin 18 pseudogene 15 |
| ENSG00000181333 | HEPHL1 | 0,570998198 | 1,485551065 | 0,16084 | 0,48815237 | hephaestin like 1 |
| ENSG00000139131 | YARS2 | 0,570951964 | 1,485503458 | 0,1031 | 0,41078404 | tyrosyl-tRNA synthetase 2 |
| ENSG00000143575 | HAX1 | 0,57088678 | 1,485436342 | 0,10534 | 0,41451879 | HCLS1 associated protein X-1 |
| ENSG00000174371 | EXO1 | 0,570826102 | 1,485373867 | 0,1985 | 0,51980353 | exonuclease 1 |
| ENSG00000078246 | TULP3 | 0,570824846 | 1,485372574 | 0,14807 | 0,47519813 | TUB like protein 3 |

| | | | | | | |
|---|---|---|---|---|---|---|
| ENSG00000197893 | NRAP | 0,570778457 | 1,485324814 | 0,30622 | 0,59517372 | nebulin related anchoring protein |
| ENSG00000288065 | AC007881,4 | 0,570763321 | 1,485309231 | 0,32423 | 0,60681275 | novel transcript, antisense to MCEE |
| ENSG00000258227 | CLEC5A | 0,570679508 | 1,485222944 | 0,49425 | 0,72675894 | C-type lectin domain containing 5A |
| ENSG00000250486 | FAM218A | 0,570650447 | 1,485193027 | 0,24089 | 0,55001001 | family with sequence similarity 218 member A |
| ENSG00000070371 | CLTCL1 | 0,570464401 | 1,485001513 | 0,02632 | 0,21861252 | clathrin heavy chain like 1 |
| ENSG00000065559 | MAP2K4 | 0,570344156 | 1,484877748 | 0,23631 | 0,54725688 | mitogen-activated protein kinase kinase 4 |
| ENSG00000205693 | MANSC4 | 0,570243306 | 1,484773952 | 0,48133 | 0,71822668 | MANSC domain containing 4 |
| ENSG00000197557 | TTC30A | 0,570207605 | 1,484737211 | 0,15965 | 0,48693054 | tetratricopeptide repeat domain 30A |
| ENSG00000178035 | IMPDH2 | 0,569982076 | 1,484505127 | 0,1344 | 0,46058584 | inosine monophosphate dehydrogenase 2 |
| ENSG00000169282 | KCNAB1 | 0,569980625 | 1,484503634 | 0,27599 | 0,57533851 | potassium voltage-gated channel subfamily A member regulatory beta subunit 1 |
| ENSG00000240972 | MIF | 0,569956649 | 1,484478963 | 0,24931 | 0,55642837 | macrophage migration inhibitory factor |
| ENSG00000278842 | AC008147,4 | 0,569924736 | 1,484446127 | 0,49147 | 0,72510994 | novel pseudogene |
| ENSG00000117036 | ETV3 | 0,569873741 | 1,484393656 | 0,05918 | 0,32424653 | ETS variant transcription factor 3 |
| ENSG00000206487 | C6orf136 | 0,569807281 | 1,484325278 | 0,13069 | 0,45508994 | chromosome 6 open reading frame 136 |
| ENSG00000147592 | LACTB2 | 0,569653113 | 1,484166669 | 0,27216 | 0,57306433 | lactamase beta 2 |
| ENSG00000286810 | AL513128,3 | 0,569620797 | 1,484133425 | 0,13281 | 0,45830601 | novel transcript |
| ENSG00000121578 | B4GALT4 | 0,569454389 | 1,483962247 | 0,04541 | 0,2878744 | beta-1,4-galactosyltransferase 4 |
| ENSG00000213049 | HNRNPA1P34 | 0,569339527 | 1,483844104 | 0,48838 | 0,72325321 | heterogeneous nuclear ribonucleoprotein A1 pseudogene 34 |
| ENSG00000261296 | AC126323,6 | 0,5692839 | 1,483786891 | 0,19977 | 0,52077977 | novel transcript |
| ENSG00000213976 | AC010615,1 | 0,569135156 | 1,483633919 | 0,21139 | 0,53032236 | zinc finger protein pseudogene |
| ENSG00000235754 | APOM | 0,569076027 | 1,483573113 | 0,08038 | 0,37007763 | apolipoprotein M |
| ENSG00000100439 | ABHD4 | 0,568820477 | 1,483310346 | 0,13176 | 0,4570124 | abhydrolase domain containing 4, N-acyl phospholipase B |
| ENSG00000182774 | RPS17 | 0,568809358 | 1,483298914 | 0,18379 | 0,50744379 | ribosomal protein S17 |
| ENSG00000256146 | AC006206,1 | 0,568778382 | 1,483267066 | 0,46133 | 0,70278869 | novel transcript |
| ENSG00000111144 | LTA4H | 0,568759119 | 1,483247261 | 0,0603 | 0,32601555 | leukotriene A4 hydrolase |
| ENSG00000117859 | OSBPL9 | 0,568619927 | 1,483104164 | 0,14735 | 0,47490238 | oxysterol binding protein like 9 |
| ENSG00000230092 | AL669831,4 | 0,568561048 | 1,483043637 | 0,27128 | 0,57271346 | F-box only protein 25 (FBXO25) pseudogene |
| ENSG00000162398 | LEXM | 0,568517522 | 1,482998894 | 0,65626 | 0,82829288 | lymphocyte expansion molecule |
| ENSG00000110700 | RPS13 | 0,568496061 | 1,482976834 | 0,19368 | 0,51547251 | ribosomal protein S13 |
| ENSG00000102572 | STK24 | 0,56843552 | 1,482914604 | 0,10206 | 0,40934973 | serine/threonine kinase 24 |
| ENSG00000164128 | NPY1R | 0,56841156 | 1,482889976 | 0,38895 | 0,65351197 | neuropeptide Y receptor Y1 |

| | | | | | | |
|---|---|---|---|---|---|---|
| ENSG00000004864 | SLC25A13 | 0,568321751 | 1,482797668 | 0,08737 | 0,38267915 | solute carrier family 25 member 13 |
| ENSG00000124198 | ARFGEF2 | 0,568321378 | 1,482797284 | 0,1383 | 0,46546655 | ADP ribosylation factor guanine nucleotide exchange factor 2 |
| ENSG00000153107 | ANAPC1 | 0,568105907 | 1,48257584 | 0,10509 | 0,41394479 | anaphase promoting complex subunit 1 |
| ENSG00000136891 | TEX10 | 0,568065767 | 1,482534592 | 0,10825 | 0,41895524 | testis expressed 10 |
| ENSG00000126814 | TRMT5 | 0,568003388 | 1,482470491 | 0,18032 | 0,50407792 | tRNA methyltransferase 5 |
| ENSG00000118508 | RAB32 | 0,567953096 | 1,482418813 | 0,15386 | 0,48214138 | RAB32, member RAS oncogene family |
| ENSG00000224186 | C5orf66 | 0,567895255 | 1,482359381 | 0,32831 | 0,60972026 | chromosome 5 open reading frame 66 |
| ENSG00000165304 | MELK | 0,567796276 | 1,482257684 | 0,2449 | 0,55288356 | maternal embryonic leucine zipper kinase |
| ENSG00000242553 | AP001432,1 | 0,567647309 | 1,48210464 | 0,30191 | 0,59294633 | novel transcript, antisense to DSCR3 |
| ENSG00000099341 | PSMD8 | 0,567574262 | 1,4820296 | 0,16591 | 0,49195088 | proteasome 26S subunit, non-ATPase 8 |
| ENSG00000165617 | DACT1 | 0,567516926 | 1,481970702 | 0,07273 | 0,35572609 | dishevelled binding antagonist of beta catenin 1 |
| ENSG00000138434 | ITPRID2 | 0,567513493 | 1,481967175 | 0,13295 | 0,45853196 | ITPR interacting domain containing 2 |
| ENSG00000167705 | RILP | 0,567415061 | 1,481866067 | 0,31692 | 0,60268749 | Rab interacting lysosomal protein |
| ENSG00000197763 | TXNRD3 | 0,567393888 | 1,481844319 | 0,03524 | 0,25425072 | thioredoxin reductase 3 |
| ENSG00000100528 | CNIH1 | 0,567375172 | 1,481825095 | 0,25482 | 0,55944852 | cornichon family AMPA receptor auxiliary protein 1 |
| ENSG00000113441 | LNPEP | 0,567336032 | 1,481784894 | 0,09844 | 0,40393831 | leucyl and cystinyl aminopeptidase |
| ENSG00000164162 | ANAPC10 | 0,567293757 | 1,481741475 | 0,16703 | 0,49293221 | anaphase promoting complex subunit 10 |
| ENSG00000077152 | UBE2T | 0,567280918 | 1,481728288 | 0,30144 | 0,59262328 | ubiquitin conjugating enzyme E2 T |
| ENSG00000277823 | PRH1 | 0,567231285 | 1,481677314 | 0,5382 | 0,75658794 | proline rich protein HaeIII subfamily 1 |
| ENSG00000176731 | RBIS | 0,567225007 | 1,481670866 | 0,33857 | 0,61774316 | ribosomal biogenesis factor |
| ENSG00000164647 | STEAP1 | 0,567113237 | 1,48155608 | 0,27282 | 0,57322782 | STEAP family member 1 |
| ENSG00000156642 | NPTN | 0,56708756 | 1,481529712 | 0,07748 | 0,36443202 | neuroplastin |
| ENSG00000143476 | DTL | 0,566963589 | 1,48140241 | 0,15995 | 0,48710442 | denticleless E3 ubiquitin protein ligase homolog |
| ENSG00000166333 | ILK | 0,566933381 | 1,481371391 | 0,15229 | 0,48061452 | integrin linked kinase |
| ENSG00000250284 | AC109439,1 | 0,566814279 | 1,481249102 | 0,12825 | 0,45171208 | novel transcript |
| ENSG00000170191 | NANP | 0,566769411 | 1,481203035 | 0,04439 | 0,28421599 | N-acetylneuraminic acid phosphatase |
| ENSG00000117748 | RPA2 | 0,566752329 | 1,481185497 | 0,23508 | 0,54665183 | replication protein A2 |
| ENSG00000136867 | SLC31A2 | 0,566737961 | 1,481170746 | 0,55032 | 0,76457697 | solute carrier family 31 member 2 |
| ENSG00000272414 | FAM47E-STBD | 0,566729287 | 1,481161841 | 0,45632 | 0,70003941 | FAM47E-STBD1 readthrough |
| ENSG00000234257 | SOD2P1 | 0,566662337 | 1,481093108 | 0,46783 | 0,70757147 | superoxide dismutase 2 pseudogene 1 |
| ENSG00000197111 | PCBP2 | 0,566618768 | 1,481048379 | 0,12902 | 0,45241788 | poly(rC) binding protein 2 |
| ENSG00000189362 | NEMP2 | 0,566526254 | 1,480953409 | 0,18511 | 0,50871969 | nuclear envelope integral membrane protein 2 |

| | | | | | | |
|---|---|---|---|---|---|---|
| ENSG00000116191 | RALGPS2 | 0,566113215 | 1,480529477 | 0,22342 | 0,53953208 | Ral GEF with PH domain and SH3 binding motif 2 |
| ENSG00000282977 | PCBP2-OT1 | 0,565863368 | 1,4802731 | 0,07402 | 0,35746749 | PCBP2 overlapping transcript 1 |
| ENSG00000261997 | AC007336,1 | 0,565746839 | 1,480153541 | 0,31102 | 0,59883009 | novel transcript |
| ENSG00000249590 | AC004832,3 | 0,565655126 | 1,48005945 | 0,70452 | 0,85756851 | novel SEC14-like 2 (S, cerevisiae) (SEC14L2) and mitochondrial protein 18 kDa (MTP18) protein |
| ENSG00000157224 | CLDN12 | 0,565601805 | 1,480004749 | 0,27935 | 0,57742338 | claudin 12 |
| ENSG00000181004 | BBS12 | 0,565552615 | 1,479954288 | 0,32296 | 0,60584543 | Bardet-Biedl syndrome 12 |
| ENSG00000171004 | HS6ST2 | 0,56539805 | 1,47979574 | 0,19145 | 0,51435195 | heparan sulfate 6-O-sulfotransferase 2 |
| ENSG00000225156 | AC012354,1 | 0,565359424 | 1,479756121 | 0,17025 | 0,49610888 | novel transcript |
| ENSG00000132773 | TOE1 | 0,565266814 | 1,479661135 | 0,16417 | 0,49061845 | target of EGR1, exonuclease |
| ENSG00000173218 | VANGL1 | 0,565135617 | 1,479526582 | 0,07341 | 0,35662549 | VANGL planar cell polarity protein 1 |
| ENSG00000224858 | RPL29P11 | 0,565093097 | 1,479482977 | 0,57923 | 0,78270914 | ribosomal protein L29 pseudogene 11 |
| ENSG00000139155 | SLCO1C1 | 0,565078819 | 1,479468335 | 0,54276 | 0,75956237 | solute carrier organic anion transporter family member 1C1 |
| ENSG00000002549 | LAP3 | 0,564898667 | 1,479283603 | 0,01356 | 0,15268762 | leucine aminopeptidase 3 |
| ENSG00000238063 | LINC01685 | 0,564774493 | 1,479156285 | 0,51866 | 0,74352557 | long intergenic non-protein coding RNA 1685 |
| ENSG00000163116 | STPG2 | 0,564712786 | 1,47909302 | 0,30131 | 0,59258306 | sperm tail PG-rich repeat containing 2 |
| ENSG00000131100 | ATP6V1E1 | 0,564647054 | 1,479025631 | 0,1178 | 0,43532264 | ATPase H+ transporting V1 subunit E1 |
| ENSG00000286136 | AC107071,1 | 0,564577132 | 1,47895395 | 0,2908 | 0,58580853 | novel transcript |
| ENSG00000146376 | ARHGAP18 | 0,564539161 | 1,478915025 | 0,23511 | 0,54666157 | Rho GTPase activating protein 18 |
| ENSG00000147604 | RPL7 | 0,564451593 | 1,478825261 | 0,16237 | 0,48945635 | ribosomal protein L7 |
| ENSG00000274265 | AC245297,3 | 0,563934818 | 1,478295639 | 0,06834 | 0,34555414 | novel transcript |
| ENSG00000230314 | ELOVL2-AS1 | 0,563860814 | 1,478219811 | 0,215 | 0,53373528 | ELOVL2 antisense RNA 1 |
| ENSG00000186777 | ZNF732 | 0,563834762 | 1,478193118 | 0,08009 | 0,3693792 | zinc finger protein 732 |
| ENSG00000280165 | PCDH20 | 0,563677119 | 1,478031605 | 0,28636 | 0,58267001 | protocadherin 20 |
| ENSG00000028839 | TBPL1 | 0,563557806 | 1,477909375 | 0,25007 | 0,55657567 | TATA-box binding protein like 1 |
| ENSG00000109667 | SLC2A9 | 0,563491774 | 1,477841732 | 0,48397 | 0,71999322 | solute carrier family 2 member 9 |
| ENSG00000165480 | SKA3 | 0,563012074 | 1,477350427 | 0,17159 | 0,49704622 | spindle and kinetochore associated complex subunit 3 |
| ENSG00000197279 | ZNF165 | 0,562934952 | 1,477271455 | 0,19178 | 0,51436129 | zinc finger protein 165 |
| ENSG00000105185 | PDCD5 | 0,562869496 | 1,477204432 | 0,23376 | 0,54563244 | programmed cell death 5 |
| ENSG00000259005 | AC005479,1 | 0,562857122 | 1,477191762 | 0,43821 | 0,68763409 | novel transcript |
| ENSG00000213713 | PIGCP1 | 0,562839243 | 1,477173456 | 0,24188 | 0,5506887 | phosphatidylinositol glycan anchor biosynthesis class C pseudogene 1 |
| ENSG00000179021 | C3orf38 | 0,562669417 | 1,476999581 | 0,29429 | 0,58857672 | chromosome 3 open reading frame 38 |

| | | | | | | |
|---|---|---|---|---|---|---|
| ENSG00000088038 | CNOT3 | 0,562604889 | 1,47693352 | 0,10885 | 0,42013578 | CCR4-NOT transcription complex subunit 3 |
| ENSG00000227739 | TUBB | 0,562545 | 1,476872211 | 0,11473 | 0,43020176 | tubulin beta class I |
| ENSG00000239672 | NME1 | 0,562529564 | 1,47685641 | 0,17553 | 0,49954066 | NME/NM23 nucleoside diphosphate kinase 1 |
| ENSG00000180855 | ZNF443 | 0,562460713 | 1,47678593 | 0,28088 | 0,57822547 | zinc finger protein 443 |
| ENSG00000144118 | RALB | 0,56239794 | 1,476721675 | 0,11736 | 0,4347145 | RAS like proto-oncogene B |
| ENSG00000027697 | IFNGR1 | 0,562059719 | 1,476375517 | 0,21765 | 0,53547632 | interferon gamma receptor 1 |
| ENSG00000150316 | CWC15 | 0,561956953 | 1,476270356 | 0,13347 | 0,4594073 | CWC15 spliceosome associated protein homolog |
| ENSG00000136141 | LRCH1 | 0,561920475 | 1,47623303 | 0,08485 | 0,37817642 | leucine rich repeats and calponin homology domain containing 1 |
| ENSG00000196850 | PPTC7 | 0,561911408 | 1,476223752 | 0,15407 | 0,48233496 | protein phosphatase targeting COQ7 |
| ENSG00000120696 | KBTBD7 | 0,561789382 | 1,476098896 | 0,17223 | 0,49710979 | kelch repeat and BTB domain containing 7 |
| ENSG00000173926 | MARCHF3 | 0,561740037 | 1,476048409 | 0,18267 | 0,50615118 | membrane associated ring-CH-type finger 3 |
| ENSG00000287645 | AC092939,1 | 0,561705008 | 1,47601257 | 0,57647 | 0,78087449 | novel transcript, antisense to PEX5L |
| ENSG00000027001 | MIPEP | 0,561363756 | 1,475663479 | 0,08076 | 0,37083982 | mitochondrial intermediate peptidase |
| ENSG00000112742 | TTK | 0,561245874 | 1,475542908 | 0,30824 | 0,59696396 | TTK protein kinase |
| ENSG00000261105 | LMO7-AS1 | 0,561120434 | 1,475414618 | 0,22507 | 0,54049887 | LMO7 antisense RNA 1 |
| ENSG00000198795 | ZNF521 | 0,560918541 | 1,47520816 | 0,20036 | 0,52107152 | zinc finger protein 521 |
| ENSG00000133302 | SLF1 | 0,56083267 | 1,475120356 | 0,2363 | 0,54725688 | SMC5-SMC6 complex localization factor 1 |
| ENSG00000142534 | RPS11 | 0,560742806 | 1,475028475 | 0,21526 | 0,53398403 | ribosomal protein S11 |
| ENSG00000176463 | SLCO3A1 | 0,560637615 | 1,474920931 | 0,0035 | 0,07173085 | solute carrier organic anion transporter family member 3A1 |
| ENSG00000173275 | ZNF449 | 0,560629187 | 1,474912315 | 0,25017 | 0,55657567 | zinc finger protein 449 |
| ENSG00000235410 | AL157896,1 | 0,560588227 | 1,47487044 | 0,4324 | 0,68411566 | novel transcript |
| ENSG00000101190 | TCFL5 | 0,560536697 | 1,474817762 | 0,02945 | 0,23083495 | transcription factor like 5 |
| ENSG00000129562 | DAD1 | 0,560213457 | 1,474487362 | 0,20717 | 0,52670822 | defender against cell death 1 |
| ENSG00000123728 | RAP2C | 0,56012988 | 1,474401946 | 0,09796 | 0,40323803 | RAP2C, member of RAS oncogene family |
| ENSG00000117133 | RPF1 | 0,56011202 | 1,474383693 | 0,14362 | 0,47123212 | ribosome production factor 1 homolog |
| ENSG00000197798 | FAM118B | 0,559673131 | 1,473935233 | 0,14914 | 0,47671235 | family with sequence similarity 118 member B |
| ENSG00000152558 | TMEM123 | 0,559536919 | 1,473796078 | 0,18025 | 0,50407792 | transmembrane protein 123 |
| ENSG00000135972 | MRPS9 | 0,559308468 | 1,47356272 | 0,19509 | 0,51588752 | mitochondrial ribosomal protein S9 |
| ENSG00000071539 | TRIP13 | 0,559150207 | 1,473401082 | 0,18526 | 0,50876814 | thyroid hormone receptor interactor 13 |
| ENSG00000118705 | RPN2 | 0,559091652 | 1,473341281 | 0,15234 | 0,48061452 | ribophorin II |
| ENSG00000262664 | OVCA2 | 0,559084731 | 1,473334214 | 0,23435 | 0,54616925 | OVCA2 serine hydrolase domain containing |

| | | | | | | |
|---|---|---|---|---|---|---|
| ENSG00000171469 | ZNF561 | 0,559011709 | 1,473259643 | 0,20131 | 0,52181085 | zinc finger protein 561 |
| ENSG00000164185 | ZNF474 | 0,558965916 | 1,47321288 | 0,3001 | 0,59160746 | zinc finger protein 474 |
| ENSG00000158636 | EMSY | 0,558861685 | 1,473106448 | 0,22374 | 0,53967355 | EMSY transcriptional repressor, BRCA2 interacting |
| ENSG00000125954 | CHURC1-FNTE | 0,558815182 | 1,473058966 | 0,62723 | 0,81132445 | CHURC1-FNTB readthrough |
| ENSG00000102580 | DNAJC3 | 0,558777665 | 1,47302066 | 0,03588 | 0,25662951 | DnaJ heat shock protein family (Hsp40) member C3 |
| ENSG00000253456 | AC136424,1 | 0,558767364 | 1,473010143 | 0,5223 | 0,74653075 | novel transcript |
| ENSG00000119421 | NDUFA8 | 0,558614657 | 1,472854235 | 0,12097 | 0,44006069 | NADH:ubiquinone oxidoreductase subunit A8 |
| ENSG00000116212 | LRRC42 | 0,558563157 | 1,47280166 | 0,06428 | 0,33593883 | leucine rich repeat containing 42 |
| ENSG00000163041 | H3-3A | 0,558530781 | 1,472768608 | 0,17199 | 0,49710979 | H3,3 histone A |
| ENSG00000117013 | KCNQ4 | 0,558516141 | 1,472753662 | 0,22619 | 0,54121239 | potassium voltage-gated channel subfamily Q member 4 |
| ENSG00000204789 | ZNF204P | 0,55841323 | 1,472648611 | 0,31307 | 0,59987747 | zinc finger protein 204, pseudogene |
| ENSG00000280810 | NSFP1 | 0,558374276 | 1,472608849 | 0,22078 | 0,53776578 | N-ethylmaleimide-sensitive factor pseudogene 1 |
| ENSG00000004700 | RECQL | 0,558333317 | 1,472567042 | 0,24903 | 0,55617627 | RecQ like helicase |
| ENSG00000173039 | RELA | 0,558287623 | 1,472520402 | 0,12692 | 0,44966573 | RELA proto-oncogene, NF-kB subunit |
| ENSG00000228199 | PAIP1P1 | 0,558114341 | 1,472343549 | 0,68988 | 0,84877222 | poly(A) binding protein interacting protein 1 pseudogene 1 |
| ENSG00000140463 | BBS4 | 0,558067692 | 1,472295941 | 0,17231 | 0,49714741 | Bardet-Biedl syndrome 4 |
| ENSG00000145685 | LHFPL2 | 0,558053534 | 1,472281493 | 0,01295 | 0,14897258 | LHFPL tetraspan subfamily member 2 |
| ENSG00000171016 | PYGO1 | 0,557926004 | 1,472151354 | 0,08222 | 0,37359506 | pygopus family PHD finger 1 |
| ENSG00000121211 | MND1 | 0,557801967 | 1,472024789 | 0,30774 | 0,59671558 | meiotic nuclear divisions 1 |
| ENSG00000254837 | AP001372,2 | 0,557783197 | 1,472005637 | 0,22381 | 0,53967355 | uncharacterized LOC100287896 |
| ENSG00000136930 | PSMB7 | 0,557757406 | 1,471979323 | 0,19775 | 0,51937526 | proteasome 20S subunit beta 7 |
| ENSG00000159479 | MED8 | 0,557639952 | 1,47185949 | 0,14646 | 0,47433542 | mediator complex subunit 8 |
| ENSG00000163104 | SMARCAD1 | 0,557411737 | 1,47162668 | 0,21359 | 0,53275614 | SWI/SNF-related, matrix-associated actin-dependent regulator of chromatin, subfamily a, containing DEAD/H box 1 |
| ENSG00000014914 | MTMR11 | 0,557365399 | 1,471579414 | 0,04262 | 0,2782038 | myotubularin related protein 11 |
| ENSG00000055147 | FAM114A2 | 0,557185739 | 1,471396168 | 0,18771 | 0,51107563 | family with sequence similarity 114 member A2 |
| ENSG00000266490 | AC127024,5 | 0,556992046 | 1,471198634 | 0,32945 | 0,61046296 | novel transcript |
| ENSG00000286330 | AL353660,1 | 0,55696013 | 1,471166088 | 0,20413 | 0,52388504 | novel transcript |
| ENSG00000100938 | GMPR2 | 0,556850476 | 1,471054275 | 0,11565 | 0,43162108 | guanosine monophosphate reductase 2 |
| ENSG00000167881 | SRP68 | 0,556740224 | 1,47094186 | 0,10419 | 0,4125943 | signal recognition particle 68 |
| ENSG00000259661 | AC068831,3 | 0,556739487 | 1,470941108 | 0,58619 | 0,78664449 | novel transcript, antisense to MAN2A2 |
| ENSG00000102290 | PCDH11X | 0,556721011 | 1,47092227 | 0,23159 | 0,54440987 | protocadherin 11 X-linked |

| | | | | | | |
|---|---|---|---|---|---|---|
| ENSG00000262179 | MYMX | 0,556709239 | 1,470910269 | 0,41267 | 0,67028426 | myomixer, myoblast fusion factor |
| ENSG00000034533 | ASTE1 | 0,556613546 | 1,470812708 | 0,26491 | 0,56757563 | asteroid homolog 1 |
| ENSG00000276702 | AC010809,3 | 0,556560962 | 1,470759099 | 0,4429 | 0,69042854 | novel transcript, antisense to RYR3 |
| ENSG00000236693 | HLA-DPB1 | 0,556555637 | 1,470753671 | 0,20863 | 0,52785962 | major histocompatibility complex, class II, DP beta 1 |
| ENSG00000230763 | HLA-DPB1 | 0,556555637 | 1,470753671 | 0,20863 | 0,52785962 | major histocompatibility complex, class II, DP beta 1 |
| ENSG00000138640 | FAM13A | 0,556339687 | 1,470533537 | 0,09441 | 0,39663526 | family with sequence similarity 13 member A |
| ENSG00000112306 | RPS12 | 0,55633052 | 1,470524194 | 0,21256 | 0,53151956 | ribosomal protein S12 |
| ENSG00000109787 | KLF3 | 0,556190053 | 1,470381024 | 0,16289 | 0,48945635 | Kruppel like factor 3 |
| ENSG00000134852 | CLOCK | 0,556163921 | 1,47035439 | 0,23995 | 0,54936679 | clock circadian regulator |
| ENSG00000184752 | NDUFA12 | 0,555998688 | 1,470185999 | 0,28988 | 0,58531219 | NADH:ubiquinone oxidoreductase subunit A12 |
| ENSG00000116750 | UCHL5 | 0,555817701 | 1,470001575 | 0,17608 | 0,50015171 | ubiquitin C-terminal hydrolase L5 |
| ENSG00000250472 | TRIM36-IT1 | 0,555704095 | 1,469885824 | 0,59791 | 0,79484486 | TRIM36 intronic transcript 1 |
| ENSG00000113583 | C5orf15 | 0,555659105 | 1,469839986 | 0,16449 | 0,49065086 | chromosome 5 open reading frame 15 |
| ENSG00000272263 | AC034198,2 | 0,555601914 | 1,469781721 | 0,3709 | 0,64013451 | novel transcript, antisense to CAND2 |
| ENSG00000120705 | ETF1 | 0,555585329 | 1,469764824 | 0,14326 | 0,47082074 | eukaryotic translation termination factor 1 |
| ENSG00000197061 | H4C3 | 0,555420526 | 1,469596938 | 0,04672 | 0,29062603 | H4 clustered histone 3 |
| ENSG00000276776 | TC2N | 0,555319053 | 1,469493577 | 0,40151 | 0,6630277 | tandem C2 domains, nuclear |
| ENSG00000262860 | LSM14A | 0,555314002 | 1,469488432 | 0,14766 | 0,47507961 | LSM14A mRNA processing body assembly factor |
| ENSG00000169299 | PGM2 | 0,55529567 | 1,46946976 | 0,19545 | 0,51643293 | phosphoglucomutase 2 |
| ENSG00000058056 | USP13 | 0,555193137 | 1,469365328 | 0,0514 | 0,30462001 | ubiquitin specific peptidase 13 |
| ENSG00000124429 | POF1B | 0,555179535 | 1,469351474 | 0,37861 | 0,64679625 | POF1B actin binding protein |
| ENSG00000167461 | RAB8A | 0,555166672 | 1,469338374 | 0,20714 | 0,52670822 | RAB8A, member RAS oncogene family |
| ENSG00000287414 | AC093787,2 | 0,55515743 | 1,469328961 | 0,30552 | 0,59501105 | novel transcript |
| ENSG00000224830 | OR2X1P | 0,555149018 | 1,469320393 | 0,57284 | 0,77868876 | olfactory receptor family 2 subfamily X member 1 pseudogene |
| ENSG00000106803 | SEC61B | 0,555097361 | 1,469267784 | 0,29544 | 0,58916761 | SEC61 translocon subunit beta |
| ENSG00000119048 | UBE2B | 0,555016466 | 1,469185402 | 0,2448 | 0,55282045 | ubiquitin conjugating enzyme E2 B |
| ENSG00000285005 | PLEKHM3 | 0,554937067 | 1,469104547 | 0,43777 | 0,68744895 | pleckstrin homology domain containing M3 |
| ENSG00000134874 | DZIP1 | 0,554904693 | 1,469071581 | 0,04374 | 0,28225644 | DAZ interacting zinc finger protein 1 |
| ENSG00000126883 | NUP214 | 0,554794294 | 1,468959168 | 0,11121 | 0,42416781 | nucleoporin 214 |
| ENSG00000109805 | NCAPG | 0,554513652 | 1,468673445 | 0,19515 | 0,51597848 | non-SMC condensin I complex subunit G |
| ENSG00000153975 | ZUP1 | 0,554355404 | 1,468512356 | 0,25453 | 0,55944852 | zinc finger containing ubiquitin peptidase 1 |
| ENSG00000187959 | CPSF4L | 0,554298459 | 1,468454392 | 0,51274 | 0,73941533 | cleavage and polyadenylation specific factor 4 like |

| | | | | | | |
|---|---|---|---|---|---|---|
| ENSG00000224243 | SOX1-OT | 0,554258835 | 1,468414062 | 0,11613 | 0,43264096 | SOX1 overlapping transcript |
| ENSG00000137804 | NUSAP1 | 0,554221236 | 1,468375793 | 0,30293 | 0,59333137 | nucleolar and spindle associated protein 1 |
| ENSG00000234883 | MIR155HG | 0,55418415 | 1,468338048 | 0,44714 | 0,69341308 | MIR155 host gene |
| ENSG00000066136 | NFYC | 0,554043168 | 1,468194567 | 0,07974 | 0,36858935 | nuclear transcription factor Y subunit gamma |
| ENSG00000270585 | SNX2P1 | 0,553979411 | 1,468129684 | 0,34015 | 0,61889407 | sorting nexin 2 pseudogene 1 |
| ENSG00000232623 | MRAP-AS1 | 0,553978123 | 1,468128373 | 0,64459 | 0,82154386 | MRAP antisense RNA 1 |
| ENSG00000147654 | EBAG9 | 0,553879312 | 1,468027824 | 0,23971 | 0,5491566 | estrogen receptor binding site associated antigen 9 |
| ENSG00000272744 | AC107214,2 | 0,553843275 | 1,467991154 | 0,25737 | 0,56160845 | novel transcript |
| ENSG00000145425 | RPS3A | 0,553763671 | 1,467910157 | 0,17389 | 0,49854855 | ribosomal protein S3A |
| ENSG00000167004 | PDIA3 | 0,553569311 | 1,467712413 | 0,03623 | 0,25760275 | protein disulfide isomerase family A member 3 |
| ENSG00000093100 | AC016026,1 | 0,55335258 | 1,46749194 | 0,17102 | 0,49641416 | novel transcript |
| ENSG00000122958 | VPS26A | 0,553325018 | 1,467463905 | 0,2372 | 0,54814212 | VPS26, retromer complex component A |
| ENSG00000204149 | AGAP6 | 0,553217869 | 1,46735492 | 0,03251 | 0,24342027 | ArfGAP with GTPase domain, ankyrin repeat and PH domain 6 |
| ENSG00000175054 | ATR | 0,553156029 | 1,467292025 | 0,20199 | 0,52230208 | ATR serine/threonine kinase |
| ENSG00000095319 | NUP188 | 0,553135373 | 1,467271017 | 0,07691 | 0,36356213 | nucleoporin 188 |
| ENSG00000269427 | AC024075,3 | 0,553135058 | 1,467270697 | 0,57377 | 0,77921182 | novel transcript, antisense C19orf42 |
| ENSG00000040731 | CDH10 | 0,553128086 | 1,467263606 | 0,24732 | 0,5549105 | cadherin 10 |
| ENSG00000103599 | IQCH | 0,553082474 | 1,467217218 | 0,23054 | 0,54361941 | IQ motif containing H |
| ENSG00000133818 | RRAS2 | 0,552991987 | 1,467125195 | 0,19108 | 0,51406315 | RAS related 2 |
| ENSG00000258484 | SPESP1 | 0,552862008 | 1,466993022 | 0,36112 | 0,63316311 | sperm equatorial segment protein 1 |
| ENSG00000182552 | RWDD4 | 0,552842663 | 1,466973351 | 0,08992 | 0,38777461 | RWD domain containing 4 |
| ENSG00000101407 | TTI1 | 0,552513217 | 1,466638399 | 0,17957 | 0,50389568 | TELO2 interacting protein 1 |
| ENSG00000165389 | SPTSSA | 0,552408232 | 1,466531676 | 0,16625 | 0,49212363 | serine palmitoyltransferase small subunit A |
| ENSG00000257261 | AC008014,1 | 0,552299543 | 1,466421194 | 0,17115 | 0,49641416 | novel transcript |
| ENSG00000287279 | AL121932,2 | 0,552249998 | 1,466370836 | 0,29967 | 0,59157893 | novel transcript |
| ENSG00000287858 | AC020550,2 | 0,552118322 | 1,466237005 | 0,40317 | 0,66399303 | novel transcript, antisense to RFTN2 |
| ENSG00000259589 | AC073167,1 | 0,552117359 | 1,466236027 | 0,50453 | 0,73328552 | novel transcript, antisense to HERC1 |
| ENSG00000124333 | VAMP7 | 0,552098879 | 1,466217245 | 0,2511 | 0,55719428 | vesicle associated membrane protein 7 |
| ENSG00000003056 | M6PR | 0,551982568 | 1,466099042 | 0,11898 | 0,43696075 | mannose-6-phosphate receptor, cation dependent |
| ENSG00000119041 | GTF3C3 | 0,551974473 | 1,466090816 | 0,13437 | 0,46058584 | general transcription factor IIIC subunit 3 |
| ENSG00000162396 | PARS2 | 0,551928549 | 1,466044148 | 0,2101 | 0,52904355 | prolyl-tRNA synthetase 2, mitochondrial |
| ENSG00000215057 | PKMP5 | 0,551807661 | 1,465921309 | 0,35239 | 0,62742225 | pyruvate kinase M1/2 pseudogene 5 |

| | | | | | | |
|---|---|---|---|---|---|---|
| ENSG00000119326 | CTNNAL1 | 0,551739778 | 1,465852334 | 0,12064 | 0,43944057 | catenin alpha like 1 |
| ENSG00000165609 | NUDT5 | 0,551702901 | 1,465814866 | 0,20354 | 0,52337667 | nudix hydrolase 5 |
| ENSG00000103194 | USP10 | 0,551547666 | 1,465657151 | 0,10694 | 0,41675015 | ubiquitin specific peptidase 10 |
| ENSG00000174173 | TRMT10C | 0,551530185 | 1,465639392 | 0,21485 | 0,53373102 | tRNA methyltransferase 10C, mitochondrial RNase P subunit |
| ENSG00000143110 | C1orf162 | 0,551416584 | 1,46552399 | 0,56326 | 0,77233365 | chromosome 1 open reading frame 162 |
| ENSG00000230530 | LIMD1-AS1 | 0,551405802 | 1,465513037 | 0,11977 | 0,4383162 | LIMD1 antisense RNA 1 |
| ENSG00000119938 | PPP1R3C | 0,551387105 | 1,465494044 | 0,23557 | 0,54701683 | protein phosphatase 1 regulatory subunit 3C |
| ENSG00000282318 | DLGAP2 | 0,551367102 | 1,465473726 | 0,68762 | 0,8478032 | DLG associated protein 2 |
| ENSG00000162694 | EXTL2 | 0,551020469 | 1,465121662 | 0,2153 | 0,53398925 | exostosin like glycosyltransferase 2 |
| ENSG00000206289 | RXRB | 0,550995568 | 1,465096374 | 0,2001 | 0,5209612 | retinoid X receptor beta |
| ENSG00000083642 | PDS5B | 0,550927389 | 1,465027138 | 0,22786 | 0,54197226 | PDS5 cohesin associated factor B |
| ENSG00000261868 | MFSD1P1 | 0,550669254 | 1,46476503 | 0,42461 | 0,67841584 | major facilitator superfamily domain containing 1 pseudogene 1 |
| ENSG00000155304 | HSPA13 | 0,550586758 | 1,464681275 | 0,28399 | 0,58090552 | heat shock protein family A (Hsp70) member 13 |
| ENSG00000077312 | SNRPA | 0,550507165 | 1,464600471 | 0,17749 | 0,50192264 | small nuclear ribonucleoprotein polypeptide A |
| ENSG00000167085 | PHB | 0,55043131 | 1,464523466 | 0,02658 | 0,21992823 | prohibitin |
| ENSG00000033800 | PIAS1 | 0,550417104 | 1,464509046 | 0,15667 | 0,48425071 | protein inhibitor of activated STAT 1 |
| ENSG00000136732 | GYPC | 0,550398057 | 1,464489711 | 0,03885 | 0,26677873 | glycophorin C (Gerbich blood group) |
| ENSG00000104951 | IL4I1 | 0,550366348 | 1,464457523 | 0,46625 | 0,70659219 | interleukin 4 induced 1 |
| ENSG00000168556 | ING2 | 0,550360169 | 1,464451251 | 0,23362 | 0,54563244 | inhibitor of growth family member 2 |
| ENSG00000079257 | LXN | 0,549747371 | 1,463829343 | 0,26483 | 0,56748133 | latexin |
| ENSG00000163629 | PTPN13 | 0,549706766 | 1,463788145 | 0,11295 | 0,42702832 | protein tyrosine phosphatase non-receptor type 13 |
| ENSG00000278243 | GFUS | 0,549668089 | 1,463748903 | 0,30405 | 0,59411013 | GDP-L-fucose synthase |
| ENSG00000151929 | BAG3 | 0,549567252 | 1,463646598 | 0,02808 | 0,22614034 | BAG cochaperone 3 |
| ENSG00000259632 | AC090971,4 | 0,549565056 | 1,46364437 | 0,51481 | 0,74111557 | UTP6, small subunit (SSU) processome component, homolog (yeast) (UTP6) pseudogene |
| ENSG00000119699 | TGFB3 | 0,549349784 | 1,463425988 | 0,14327 | 0,47082074 | transforming growth factor beta 3 |
| ENSG00000221818 | EBF2 | 0,549338423 | 1,463414464 | 0,05064 | 0,30274123 | EBF transcription factor 2 |
| ENSG00000251485 | NRBF2P6 | 0,549332578 | 1,463408535 | 0,18401 | 0,50777976 | NRBF2 pseudogene 6 |
| ENSG00000089335 | ZNF302 | 0,549327333 | 1,463403215 | 0,09625 | 0,39974205 | zinc finger protein 302 |
| ENSG00000144034 | TPRKB | 0,549316364 | 1,463392088 | 0,32517 | 0,60763418 | TP53RK binding protein |
| ENSG00000086200 | IPO11 | 0,549293514 | 1,463368911 | 0,23683 | 0,54768226 | importin 11 |

| | | | | | | |
|---|---|---|---|---|---|---|
| ENSG00000123473 | STIL | 0,549285701 | 1,463360986 | 0,17538 | 0,49944679 | STIL centriolar assembly protein |
| ENSG00000166681 | BEX3 | 0,549266652 | 1,463341664 | 0,01084 | 0,13471553 | brain expressed X-linked 3 |
| ENSG00000232303 | DFFBP1 | 0,549219445 | 1,463293782 | 0,4467 | 0,69321646 | DNA fragmentation factor subunit beta pseudogene 1 |
| ENSG00000169564 | PCBP1 | 0,549219047 | 1,463293379 | 0,12173 | 0,44093327 | poly(rC) binding protein 1 |
| ENSG00000116731 | PRDM2 | 0,549211727 | 1,463285954 | 0,09883 | 0,40464633 | PR/SET domain 2 |
| ENSG00000273855 | AC020658,5 | 0,549173467 | 1,463247148 | 0,56305 | 0,77224059 | novel transcript, sense intronic to PLCB2 |
| ENSG00000198909 | MAP3K3 | 0,54917245 | 1,463246117 | 0,00783 | 0,11257373 | mitogen-activated protein kinase kinase kinase 3 |
| ENSG00000018699 | TTC27 | 0,549169728 | 1,463243356 | 0,20212 | 0,52239826 | tetratricopeptide repeat domain 27 |
| ENSG00000163755 | HPS3 | 0,549036287 | 1,463108021 | 0,16816 | 0,49426916 | HPS3 biogenesis of lysosomal organelles complex 2 subunit 1 |
| ENSG00000178021 | TSPYL6 | 0,548940554 | 1,463010936 | 0,48394 | 0,71999322 | TSPY like 6 |
| ENSG00000120686 | UFM1 | 0,548918906 | 1,462988983 | 0,26131 | 0,56432558 | ubiquitin fold modifier 1 |
| ENSG00000260633 | AC010207,1 | 0,548907964 | 1,462977887 | 0,1367 | 0,46406381 | novel transcript |
| ENSG00000124789 | NUP153 | 0,548760462 | 1,46282832 | 0,08461 | 0,37769282 | nucleoporin 153 |
| ENSG00000179083 | FAM133A | 0,548627258 | 1,462693263 | 0,16991 | 0,49606609 | family with sequence similarity 133 member A |
| ENSG00000185634 | SHC4 | 0,548489017 | 1,462553113 | 0,05227 | 0,30698377 | SHC adaptor protein 4 |
| ENSG00000108175 | ZMIZ1 | 0,548478599 | 1,462542551 | 0,04471 | 0,28565589 | zinc finger MIZ-type containing 1 |
| ENSG00000171067 | C11orf24 | 0,548464433 | 1,462528191 | 0,13383 | 0,45985981 | chromosome 11 open reading frame 24 |
| ENSG00000155115 | GTF3C6 | 0,548452792 | 1,462516389 | 0,27939 | 0,57742338 | general transcription factor IIIC subunit 6 |
| ENSG00000118680 | MYL12B | 0,548356663 | 1,462418943 | 0,28231 | 0,5797164 | myosin light chain 12B |
| ENSG00000166200 | COPS2 | 0,548314741 | 1,462376448 | 0,22784 | 0,54197226 | COP9 signalosome subunit 2 |
| ENSG00000236478 | AC012513,2 | 0,548281104 | 1,462342353 | 0,24018 | 0,54948131 | heterogeneous nuclear ribonucleoprotein A3 (HNRNPA3) pseudogene |
| ENSG00000096654 | ZNF184 | 0,548111214 | 1,46217016 | 0,2541 | 0,55925777 | zinc finger protein 184 |
| ENSG00000106299 | WASL | 0,54806583 | 1,462124163 | 0,18559 | 0,50901954 | WASP like actin nucleation promoting factor |
| ENSG00000112309 | B3GAT2 | 0,548040626 | 1,462098621 | 0,26372 | 0,56629809 | beta-1,3-glucuronyltransferase 2 |
| ENSG00000009307 | CSDE1 | 0,547966162 | 1,462023157 | 0,19913 | 0,52017864 | cold shock domain containing E1 |
| ENSG00000237289 | CKMT1B | 0,547885309 | 1,461941223 | 0,14836 | 0,47569562 | creatine kinase, mitochondrial 1B |
| ENSG00000137073 | UBAP2 | 0,547752441 | 1,461806589 | 0,12217 | 0,44167604 | ubiquitin associated protein 2 |
| ENSG00000051382 | PIK3CB | 0,547662424 | 1,461715382 | 0,20382 | 0,52356486 | phosphatidylinositol-4,5-bisphosphate 3-kinase catalytic subunit beta |
| ENSG00000213071 | LPAL2 | 0,547537644 | 1,461588963 | 0,04411 | 0,28306891 | lipoprotein(a) like 2, pseudogene |
| ENSG00000237273 | RSL24D1P8 | 0,54751214 | 1,461563125 | 0,23598 | 0,5471053 | ribosomal L24 domain containing 1 pseudogene 8 |

| | | | | | | |
|---|---|---|---|---|---|---|
| ENSG00000149679 | CABLES2 | 0,547415283 | 1,461465004 | 0,08738 | 0,38267915 | Cdk5 and Abl enzyme substrate 2 |
| ENSG00000188229 | TUBB4B | 0,547012097 | 1,461056629 | 0,1115 | 0,42449053 | tubulin beta 4B class IVb |
| ENSG00000083312 | TNPO1 | 0,546905847 | 1,460949031 | 0,16076 | 0,48807078 | transportin 1 |
| ENSG00000061918 | GUCY1B1 | 0,546878023 | 1,460920855 | 0,20253 | 0,52259159 | guanylate cyclase 1 soluble subunit beta 1 |
| ENSG00000278936 | AC244517,4 | 0,546691466 | 1,460731954 | 0,59913 | 0,79538547 | novel transcript, antisense to PCDHB16, PCDHB9, PCDHB10 and PCDHB11 |
| ENSG00000228623 | ZNF883 | 0,546679717 | 1,460720058 | 0,20326 | 0,5231305 | zinc finger protein 883 |
| ENSG00000112685 | EXOC2 | 0,546604748 | 1,460644154 | 0,1668 | 0,49266358 | exocyst complex component 2 |
| ENSG00000168661 | ZNF30 | 0,546584745 | 1,460623902 | 0,29467 | 0,58857672 | zinc finger protein 30 |
| ENSG00000126821 | SGPP1 | 0,546580643 | 1,460619749 | 0,08224 | 0,37359506 | sphingosine-1-phosphate phosphatase 1 |
| ENSG00000285281 | SGPP1 | 0,546580643 | 1,460619749 | 0,08224 | 0,37359506 | sphingosine-1-phosphate phosphatase 1 |
| ENSG00000228653 | HNRNPCP7 | 0,546418527 | 1,460455629 | 0,25015 | 0,55657567 | heterogeneous nuclear ribonucleoprotein C pseudogene 7 |
| ENSG00000120992 | LYPLA1 | 0,546404752 | 1,460441684 | 0,18471 | 0,50835796 | lysophospholipase 1 |
| ENSG00000230090 | AC108025,1 | 0,545945163 | 1,459976515 | 0,43202 | 0,68400977 | novel transcript |
| ENSG00000082515 | MRPL22 | 0,545940469 | 1,459971765 | 0,25372 | 0,55878634 | mitochondrial ribosomal protein L22 |
| ENSG00000205208 | C4orf46 | 0,545936229 | 1,459967475 | 0,13763 | 0,46480929 | chromosome 4 open reading frame 46 |
| ENSG00000258026 | AC078955,1 | 0,545916577 | 1,459947588 | 0,66346 | 0,83242946 | novel transcript, antisense to ACSS3 |
| ENSG00000236896 | AL354726,1 | 0,545905573 | 1,459936452 | 0,56493 | 0,77334048 | novel transcript |
| ENSG00000146587 | RBAK | 0,545683943 | 1,45971219 | 0,26364 | 0,56629809 | RB associated KRAB zinc finger |
| ENSG00000100601 | ALKBH1 | 0,545612406 | 1,459639812 | 0,09342 | 0,39496269 | alkB homolog 1, histone H2A dioxygenase |
| ENSG00000212719 | LINC02693 | 0,545269419 | 1,459292838 | 0,02183 | 0,19644572 | long intergenic non-protein coding RNA 2693 |
| ENSG00000231381 | RNF2P1 | 0,545231266 | 1,459254246 | 0,4934 | 0,72634709 | ring finger protein 2 pseudogene 1 |
| ENSG00000065809 | FAM107B | 0,545224503 | 1,459247405 | 0,13719 | 0,4642578 | family with sequence similarity 107 member B |
| ENSG00000113013 | HSPA9 | 0,544998397 | 1,459018723 | 0,13531 | 0,46182547 | heat shock protein family A (Hsp70) member 9 |
| ENSG00000250853 | RNF138P1 | 0,544961215 | 1,458981121 | 0,45733 | 0,70074793 | ring finger protein 138 pseudogene 1 |
| ENSG00000132507 | EIF5A | 0,54490392 | 1,45892318 | 0,03587 | 0,25662951 | eukaryotic translation initiation factor 5A |
| ENSG00000288145 | EIF5A | 0,54490392 | 1,45892318 | 0,03587 | 0,25662951 | eukaryotic translation initiation factor 5A |
| ENSG00000251349 | MSANTD3-TM | 0,544570484 | 1,458586033 | 0,55434 | 0,76737943 | MSANTD3-TMEFF1 readthrough |
| ENSG00000236308 | AL138921,2 | 0,544545104 | 1,458560373 | 0,44839 | 0,69432421 | novel transcript, antisense to CHUK |
| ENSG00000143079 | CTTNBP2NL | 0,544425764 | 1,458439726 | 0,12689 | 0,44963011 | CTTNBP2 N-terminal like |
| ENSG00000170954 | ZNF415 | 0,544217748 | 1,458229455 | 0,22963 | 0,54315596 | zinc finger protein 415 |
| ENSG00000284052 | AC006460,2 | 0,54415133 | 1,458162324 | 0,01735 | 0,17351217 | novel transcript |
| ENSG00000158079 | PTPDC1 | 0,544143963 | 1,458154877 | 0,12414 | 0,44504227 | protein tyrosine phosphatase domain containing 1 |

| | | | | | | |
|---|---|---|---|---|---|---|
| ENSG00000286223 | AC064836,3 | 0,544107004 | 1,458117523 | 0,16658 | 0,49255702 | novel transcript, antisense to NOP58 |
| ENSG00000126261 | UBA2 | 0,544103884 | 1,45811437 | 0,16446 | 0,49065086 | ubiquitin like modifier activating enzyme 2 |
| ENSG00000163931 | TKT | 0,544099239 | 1,458109675 | 0,07943 | 0,3679672 | transketolase |
| ENSG00000132432 | SEC61G | 0,543960645 | 1,457969607 | 0,2397 | 0,5491566 | SEC61 translocon subunit gamma |
| ENSG00000277118 | Metazoa_SRP | 0,543951649 | 1,457960515 | 0,3557 | 0,63008438 | Metazoan signal recognition particle RNA |
| ENSG00000285953 | AC000120,3 | 0,543948029 | 1,457956857 | 0,50879 | 0,73694212 | novel protein |
| ENSG00000143183 | TMCO1 | 0,543798864 | 1,457806122 | 0,20951 | 0,52856218 | transmembrane and coiled-coil domains 1 |
| ENSG00000226715 | LINC01709 | 0,543788128 | 1,457795273 | 0,10027 | 0,4067769 | long intergenic non-protein coding RNA 1709 |
| ENSG00000117533 | VAMP4 | 0,543309791 | 1,45731201 | 0,19669 | 0,51792163 | vesicle associated membrane protein 4 |
| ENSG00000236498 | AC107081,1 | 0,543266787 | 1,457268571 | 0,38833 | 0,6529693 | novel transcript |
| ENSG00000144566 | RAB5A | 0,543154316 | 1,457154968 | 0,24153 | 0,55039174 | RAB5A, member RAS oncogene family |
| ENSG00000235072 | ARNILA | 0,543116669 | 1,457116945 | 0,28868 | 0,58455903 | androgen receptor negatively regulated lncRNA |
| ENSG00000225695 | HNRNPA1P35 | 0,543074047 | 1,457073897 | 0,39906 | 0,66150777 | heterogeneous nuclear ribonucleoprotein A1 pseudogene 35 |
| ENSG00000282853 | ECH1 | 0,543068834 | 1,457068632 | 0,10038 | 0,4067769 | enoyl-CoA hydratase 1 |
| ENSG00000104823 | ECH1 | 0,543068834 | 1,457068632 | 0,10038 | 0,4067769 | enoyl-CoA hydratase 1 |
| ENSG00000101557 | USP14 | 0,542931197 | 1,456929631 | 0,09391 | 0,39593973 | ubiquitin specific peptidase 14 |
| ENSG00000227409 | ZMYM4-AS1 | 0,542861397 | 1,456859144 | 0,57757 | 0,78149551 | ZMYM4 antisense RNA 1 |
| ENSG00000226067 | LINC00623 | 0,542830953 | 1,4568284 | 0,05188 | 0,30606616 | long intergenic non-protein coding RNA 623 |
| ENSG00000079931 | MOXD1 | 0,542808918 | 1,45680615 | 0,15949 | 0,48693054 | monooxygenase DBH like 1 |
| ENSG00000253607 | AC104316,1 | 0,542706017 | 1,456702247 | 0,41616 | 0,67303919 | novel transcript, antisense to DERL1 |
| ENSG00000166349 | RAG1 | 0,542703511 | 1,456699716 | 0,32066 | 0,60553919 | recombination activating 1 |
| ENSG00000035681 | NSMAF | 0,542625892 | 1,456621346 | 0,14784 | 0,4751727 | neutral sphingomyelinase activation associated factor |
| ENSG00000101574 | METTL4 | 0,542580111 | 1,456575124 | 0,21556 | 0,53437517 | methyltransferase like 4 |
| ENSG00000168564 | CDKN2AIP | 0,542465061 | 1,456458971 | 0,24141 | 0,55033747 | CDKN2A interacting protein |
| ENSG00000126878 | AIF1L | 0,54227014 | 1,456262203 | 0,02772 | 0,22434063 | allograft inflammatory factor 1 like |
| ENSG00000274995 | AC013564,1 | 0,542112704 | 1,456103296 | 0,54289 | 0,75963423 | novel transcript, sense intronic to SMAD6 |
| ENSG00000254285 | KRT8P3 | 0,542093578 | 1,456083992 | 0,59794 | 0,79484486 | keratin 8 pseudogene 3 |
| ENSG00000130449 | ZSWIM6 | 0,541970187 | 1,455959461 | 0,26073 | 0,56400103 | zinc finger SWIM-type containing 6 |
| ENSG00000163421 | PROK2 | 0,541896461 | 1,45588506 | 0,22041 | 0,53750545 | prokineticin 2 |
| ENSG00000259993 | AC135731,1 | 0,5418241 | 1,455812039 | 0,41289 | 0,67053125 | hyperpolarization activated cyclic nucleotide-gated potassium channel 2 (HCN2) pseudogene |
| ENSG00000238276 | AL354863,1 | 0,541503896 | 1,45548896 | 0,21438 | 0,53334744 | uncharacterized LOC101927760 |

| | | | | | | |
|---|---|---|---|---|---|---|
| ENSG00000179933 | C14orf119 | 0,541487944 | 1,455472866 | 0,18707 | 0,5105374 | chromosome 14 open reading frame 119 |
| ENSG00000125107 | CNOT1 | 0,541463566 | 1,455448272 | 0,21186 | 0,53070055 | CCR4-NOT transcription complex subunit 1 |
| ENSG00000174547 | MRPL11 | 0,541414915 | 1,455399192 | 0,15995 | 0,48710442 | mitochondrial ribosomal protein L11 |
| ENSG00000116752 | BCAS2 | 0,541225191 | 1,45520781 | 0,16454 | 0,49065086 | BCAS2 pre-mRNA processing factor |
| ENSG00000100749 | VRK1 | 0,541101326 | 1,455082876 | 0,26473 | 0,56748133 | VRK serine/threonine kinase 1 |
| ENSG00000185728 | YTHDF3 | 0,541044293 | 1,455025355 | 0,24834 | 0,55546558 | YTH N6-methyladenosine RNA binding protein 3 |
| ENSG00000241020 | AP000873,1 | 0,541007727 | 1,454988477 | 0,56415 | 0,77273028 | ribosomal protein L32 (RPL32) pseudogene |
| ENSG00000008282 | SYPL1 | 0,540968955 | 1,454949375 | 0,25539 | 0,55992862 | synaptophysin like 1 |
| ENSG00000135801 | TAF5L | 0,540952337 | 1,454932616 | 0,17202 | 0,49710979 | TATA-box binding protein associated factor 5 like |
| ENSG00000108021 | TASOR2 | 0,540870451 | 1,454850037 | 0,11489 | 0,43031399 | transcription activation suppressor family member 2 |
| ENSG00000116690 | PRG4 | 0,540783243 | 1,454762097 | 0,48229 | 0,71892541 | proteoglycan 4 |
| ENSG00000138768 | USO1 | 0,540698582 | 1,454676731 | 0,29013 | 0,58531219 | USO1 vesicle transport factor |
| ENSG00000067082 | KLF6 | 0,540658052 | 1,454635865 | 0,1678 | 0,49395055 | Kruppel like factor 6 |
| ENSG00000251288 | AC018797,2 | 0,54063442 | 1,454612037 | 0,30498 | 0,59446033 | WD repeat domain 12 (WDR12) pseudogene |
| ENSG00000116459 | ATP5PB | 0,540624219 | 1,454601752 | 0,18727 | 0,5105374 | ATP synthase peripheral stalk-membrane subunit b |
| ENSG00000181804 | SLC9A9 | 0,540405232 | 1,454380975 | 0,31752 | 0,60289482 | solute carrier family 9 member A9 |
| ENSG00000150594 | ADRA2A | 0,540302439 | 1,454277352 | 0,65935 | 0,82984259 | adrenoceptor alpha 2A |
| ENSG00000162613 | FUBP1 | 0,540244702 | 1,454219153 | 0,27288 | 0,57325456 | far upstream element binding protein 1 |
| ENSG00000249709 | ZNF564 | 0,540210787 | 1,454184967 | 0,21493 | 0,53373528 | zinc finger protein 564 |
| ENSG00000113161 | HMGCR | 0,540183853 | 1,45415782 | 0,1922 | 0,51474669 | 3-hydroxy-3-methylglutaryl-CoA reductase |
| ENSG00000170421 | KRT8 | 0,53990527 | 1,45387705 | 0,37651 | 0,6449012 | keratin 8 |
| ENSG00000114942 | EEF1B2 | 0,539864615 | 1,45383608 | 0,14468 | 0,47224655 | eukaryotic translation elongation factor 1 beta 2 |
| ENSG00000283391 | EEF1B2 | 0,539864615 | 1,45383608 | 0,14468 | 0,47224655 | eukaryotic translation elongation factor 1 beta 2 |
| ENSG00000150768 | DLAT | 0,539857176 | 1,453828584 | 0,10691 | 0,41675015 | dihydrolipoamide S-acetyltransferase |
| ENSG00000270048 | AC068790,4 | 0,539802314 | 1,4537733 | 0,39947 | 0,66176242 | novel transcript, sense intronic to CCDC92 |
| ENSG00000138663 | COPS4 | 0,539800406 | 1,453771377 | 0,22888 | 0,54262488 | COP9 signalosome subunit 4 |
| ENSG00000180611 | MB21D2 | 0,539798729 | 1,453769687 | 0,2173 | 0,53520949 | Mab-21 domain containing 2 |
| ENSG00000287275 | AC011451,4 | 0,539766333 | 1,453737043 | 0,05121 | 0,30437785 | novel transcript |
| ENSG00000183621 | ZNF438 | 0,539696405 | 1,453666581 | 0,31111 | 0,59885232 | zinc finger protein 438 |
| ENSG00000185947 | ZNF267 | 0,539692044 | 1,453662187 | 0,22906 | 0,54263903 | zinc finger protein 267 |
| ENSG00000213588 | ZBTB9 | 0,539659134 | 1,453629028 | 0,2625 | 0,56541759 | zinc finger and BTB domain containing 9 |
| ENSG00000066032 | CTNNA2 | 0,539651176 | 1,453621009 | 0,16581 | 0,49189706 | catenin alpha 2 |
| ENSG00000163636 | PSMD6 | 0,539500006 | 1,453468702 | 0,18701 | 0,5105217 | proteasome 26S subunit, non-ATPase 6 |

| | | | | | | |
|---|---|---|---|---|---|---|
| ENSG00000275495 | AC114936,2 | 0,539481696 | 1,453450256 | 0,10242 | 0,4100442 | THO complex 3 (THOC3) pseudogene |
| ENSG00000166024 | R3HCC1L | 0,539423244 | 1,453391369 | 0,1616 | 0,48914394 | R3H domain and coiled-coil containing 1 like |
| ENSG00000034152 | MAP2K3 | 0,539420742 | 1,453388848 | 0,02938 | 0,23068546 | mitogen-activated protein kinase kinase 3 |
| ENSG00000281930 | SMIM10L1 | 0,539345256 | 1,453312805 | 0,11588 | 0,43214457 | small integral membrane protein 10 like 1 |
| ENSG00000233337 | UBE2FP3 | 0,539304436 | 1,453271685 | 0,52344 | 0,74736255 | UBE2F pseudogene 3 |
| ENSG00000166012 | TAF1D | 0,539207052 | 1,45317359 | 0,11569 | 0,43162108 | TATA-box binding protein associated factor, RNA polymerase I subunit D |
| ENSG00000100852 | ARHGAP5 | 0,539181195 | 1,453147546 | 0,16303 | 0,48967932 | Rho GTPase activating protein 5 |
| ENSG00000253167 | WASHC5-AS1 | 0,539161841 | 1,453128052 | 0,44123 | 0,68956601 | WASHC5 antisense RNA 1 |
| ENSG00000174808 | BTC | 0,539141743 | 1,453107809 | 0,40555 | 0,66526837 | betacellulin |
| ENSG00000182890 | GLUD2 | 0,538940463 | 1,45290509 | 0,14623 | 0,47428193 | glutamate dehydrogenase 2 |
| ENSG00000288118 | GLUD2 | 0,538940463 | 1,45290509 | 0,14623 | 0,47428193 | glutamate dehydrogenase 2 |
| ENSG00000109971 | HSPA8 | 0,538868441 | 1,45283256 | 0,1733 | 0,4980379 | heat shock protein family A (Hsp70) member 8 |
| ENSG00000273749 | CYFIP1 | 0,538686337 | 1,452649188 | 0,26742 | 0,56973003 | cytoplasmic FMR1 interacting protein 1 |
| ENSG00000205885 | C1RL-AS1 | 0,53865842 | 1,452621079 | 0,22845 | 0,54241083 | C1RL antisense RNA 1 |
| ENSG00000144285 | SCN1A | 0,538486487 | 1,452447973 | 0,34025 | 0,61897407 | sodium voltage-gated channel alpha subunit 1 |
| ENSG00000144744 | UBA3 | 0,538484391 | 1,452445862 | 0,17015 | 0,49606609 | ubiquitin like modifier activating enzyme 3 |
| ENSG00000288170 | DDX11L9 | 0,538335178 | 1,452295649 | 0,28869 | 0,58455903 | DEAD/H-box helicase 11 like 9 (pseudogene) |
| ENSG00000248472 | DDX11L9 | 0,538335178 | 1,452295649 | 0,28869 | 0,58455903 | DEAD/H-box helicase 11 like 9 (pseudogene) |
| ENSG00000085274 | MYNN | 0,538315673 | 1,452276014 | 0,29859 | 0,59089801 | myoneurin |
| ENSG00000124493 | GRM4 | 0,538311926 | 1,452272243 | 0,36712 | 0,63709691 | glutamate metabotropic receptor 4 |
| ENSG00000115966 | ATF2 | 0,538266303 | 1,452226317 | 0,25265 | 0,55796133 | activating transcription factor 2 |
| ENSG00000251600 | AC107223,1 | 0,538237642 | 1,452197468 | 0,25619 | 0,56069543 | novel transcript, antisense to GYPA |
| ENSG00000286693 | AC105230,1 | 0,538186542 | 1,452146032 | 0,44475 | 0,69205935 | novel transcript, antisense to UNC5D |
| ENSG00000198056 | PRIM1 | 0,538155469 | 1,452114756 | 0,17578 | 0,49972993 | DNA primase subunit 1 |
| ENSG00000241735 | FABP5P3 | 0,538043233 | 1,452001791 | 0,41238 | 0,67001327 | fatty acid binding protein 5 pseudogene 3 |
| ENSG00000141655 | TNFRSF11A | 0,537902131 | 1,451859786 | 0,28792 | 0,5840701 | TNF receptor superfamily member 11a |
| ENSG00000241674 | HLA-DMB | 0,537824684 | 1,451781849 | 0,4837 | 0,71988524 | major histocompatibility complex, class II, DM beta |
| ENSG00000106211 | HSPB1 | 0,537820143 | 1,45177728 | 0,08608 | 0,3803347 | heat shock protein family B (small) member 1 |
| ENSG00000180957 | PITPNB | 0,537795821 | 1,451752805 | 0,14103 | 0,4679289 | phosphatidylinositol transfer protein beta |
| ENSG00000113194 | FAF2 | 0,537714928 | 1,451671406 | 0,14709 | 0,47471707 | Fas associated factor family member 2 |
| ENSG00000287708 | AL137017,1 | 0,537562534 | 1,451518072 | 0,57315 | 0,77878548 | novel transcript, antisense to NFIB |
| ENSG00000022267 | FHL1 | 0,537407657 | 1,451362256 | 0,14298 | 0,47060958 | four and a half LIM domains 1 |

| | | | | | | |
|---|---|---|---|---|---|---|
| ENSG00000170782 | OR10A4 | 0,537347748 | 1,451301988 | 0,48243 | 0,71895682 | olfactory receptor family 10 subfamily A member 4 |
| ENSG00000090857 | PDPR | 0,537307399 | 1,451261399 | 0,03143 | 0,23928335 | pyruvate dehydrogenase phosphatase regulatory subunit |
| ENSG00000242779 | ZNF702P | 0,537284174 | 1,451238037 | 0,19548 | 0,51643293 | zinc finger protein 702, pseudogene |
| ENSG00000047315 | POLR2B | 0,537124259 | 1,451077183 | 0,1626 | 0,48945635 | RNA polymerase II subunit B |
| ENSG00000152240 | HAUS1 | 0,537025405 | 1,450977758 | 0,22564 | 0,54087803 | HAUS augmin like complex subunit 1 |
| ENSG00000081870 | HSPB11 | 0,53701213 | 1,450964407 | 0,03788 | 0,26341726 | heat shock protein family B (small) member 11 |
| ENSG00000166181 | API5 | 0,537008018 | 1,450960272 | 0,10447 | 0,41292721 | apoptosis inhibitor 5 |
| ENSG00000138095 | LRPPRC | 0,537002745 | 1,450954969 | 0,15479 | 0,48266733 | leucine rich pentatricopeptide repeat containing |
| ENSG00000129515 | SNX6 | 0,536822712 | 1,450773917 | 0,21653 | 0,53490925 | sorting nexin 6 |
| ENSG00000116678 | LEPR | 0,536630436 | 1,450580576 | 0,1841 | 0,50785017 | leptin receptor |
| ENSG00000288362 | CYP11A1 | 0,536510361 | 1,45045985 | 0,34484 | 0,62213186 | cytochrome P450 family 11 subfamily A member 1 |
| ENSG00000140459 | CYP11A1 | 0,536510361 | 1,45045985 | 0,34484 | 0,62213186 | cytochrome P450 family 11 subfamily A member 1 |
| ENSG00000141367 | CLTC | 0,536509306 | 1,450458789 | 0,18048 | 0,50407792 | clathrin heavy chain |
| ENSG00000284967 | FDFT1 | 0,536452385 | 1,450401564 | 0,09234 | 0,39278463 | farnesyl-diphosphate farnesyltransferase 1 |
| ENSG00000070761 | CFAP20 | 0,536417279 | 1,45036627 | 0,20563 | 0,52513499 | cilia and flagella associated protein 20 |
| ENSG00000107854 | TNKS2 | 0,53621231 | 1,450160225 | 0,13857 | 0,46578766 | tankyrase 2 |
| ENSG00000270147 | AC068620,2 | 0,536197086 | 1,450144923 | 0,45631 | 0,70003941 | novel transcript |
| ENSG00000019485 | PRDM11 | 0,536132453 | 1,450079958 | 0,00688 | 0,10575605 | PR/SET domain 11 |
| ENSG00000270362 | HMGN3-AS1 | 0,536109562 | 1,45005695 | 0,14118 | 0,4679289 | HMGN3 antisense RNA 1 |
| ENSG00000134013 | LOXL2 | 0,535897747 | 1,44984407 | 0,01197 | 0,14255836 | lysyl oxidase like 2 |
| ENSG00000251126 | AC017007,5 | 0,535782726 | 1,449728483 | 0,57999 | 0,78332989 | novel transcript, antisense to ANK2 |
| ENSG00000257315 | ZBED6 | 0,535649771 | 1,449594886 | 0,07959 | 0,36831832 | zinc finger BED-type containing 6 |
| ENSG00000277075 | H2AC8 | 0,535446764 | 1,449390923 | 0,32379 | 0,60650764 | H2A clustered histone 8 |
| ENSG00000101346 | POFUT1 | 0,535341218 | 1,449284891 | 0,01273 | 0,14778337 | protein O-fucosyltransferase 1 |
| ENSG00000185010 | F8 | 0,535231954 | 1,449175132 | 0,19631 | 0,51733996 | coagulation factor VIII |
| ENSG00000100387 | RBX1 | 0,534827949 | 1,448769368 | 0,25061 | 0,55690541 | ring-box 1 |
| ENSG00000152894 | PTPRK | 0,5347857 | 1,448726942 | 0,14642 | 0,47433542 | protein tyrosine phosphatase receptor type K |
| ENSG00000133641 | C12orf29 | 0,534742033 | 1,448683094 | 0,32688 | 0,60888005 | chromosome 12 open reading frame 29 |
| ENSG00000086475 | SEPHS1 | 0,534700537 | 1,448641427 | 0,24663 | 0,55425129 | selenophosphate synthetase 1 |
| ENSG00000162736 | NCSTN | 0,534661637 | 1,448602366 | 0,11549 | 0,43134353 | nicastrin |
| ENSG00000075188 | NUP37 | 0,534609274 | 1,44854979 | 0,23952 | 0,5491566 | nucleoporin 37 |
| ENSG00000162669 | HFM1 | 0,534506775 | 1,448446878 | 0,26886 | 0,57085484 | helicase for meiosis 1 |
| ENSG00000167105 | TMEM92 | 0,534476559 | 1,448416542 | 0,51943 | 0,74396498 | transmembrane protein 92 |

| | | | | | | |
|---|---|---|---|---|---|---|
| ENSG00000231628 | AL133406,2 | 0,534409144 | 1,448348862 | 0,19369 | 0,51547251 | novel transcript |
| ENSG00000114251 | WNT5A | 0,534325774 | 1,448265167 | 0,3642 | 0,634951 | Wnt family member 5A |
| ENSG00000152601 | MBNL1 | 0,534270415 | 1,448209595 | 0,2329 | 0,54547822 | muscleblind like splicing regulator 1 |
| ENSG00000106397 | PLOD3 | 0,534168614 | 1,448107409 | 0,0072 | 0,10776454 | procollagen-lysine,2-oxoglutarate 5-dioxygenase 3 |
| ENSG00000230280 | HNRNPA1P59 | 0,534055801 | 1,447994177 | 0,17345 | 0,49804988 | heterogeneous nuclear ribonucleoprotein A1 pseudogene 59 |
| ENSG00000162636 | FAM102B | 0,534047305 | 1,44798565 | 0,2012 | 0,52181085 | family with sequence similarity 102 member B |
| ENSG00000143870 | PDIA6 | 0,534030832 | 1,447969116 | 0,11396 | 0,42904457 | protein disulfide isomerase family A member 6 |
| ENSG00000140740 | UQCRC2 | 0,53396345 | 1,44790149 | 0,27884 | 0,57700804 | ubiquinol-cytochrome c reductase core protein 2 |
| ENSG00000176024 | ZNF613 | 0,533915315 | 1,447853182 | 0,21865 | 0,53608242 | zinc finger protein 613 |
| ENSG00000159579 | RSPRY1 | 0,53388018 | 1,447817922 | 0,14696 | 0,47471707 | ring finger and SPRY domain containing 1 |
| ENSG00000109794 | FAM149A | 0,533863706 | 1,447801389 | 0,02173 | 0,19584506 | family with sequence similarity 149 member A |
| ENSG00000198888 | MT-ND1 | 0,533710226 | 1,447647375 | 0,15023 | 0,47830235 | mitochondrially encoded NADH:ubiquinone oxidoreductase core subunit 1 |
| ENSG00000225282 | AP000350,2 | 0,533706977 | 1,447644115 | 0,33395 | 0,61382024 | kelch-like 5 (Drosophila) (KLHL5) pseudogene |
| ENSG00000281900 | AP000350,17 | 0,533706977 | 1,447644115 | 0,33395 | 0,61382024 | kelch-like 5 (Drosophila) (KLHL5) pseudogene |
| ENSG00000117153 | KLHL12 | 0,533689489 | 1,447626567 | 0,18265 | 0,50615118 | kelch like family member 12 |
| ENSG00000273633 | CDC42EP5 | 0,533669661 | 1,447606671 | 0,44488 | 0,69205935 | CDC42 effector protein 5 |
| ENSG00000111229 | ARPC3 | 0,533605021 | 1,447541813 | 0,28517 | 0,58171017 | actin related protein 2/3 complex subunit 3 |
| ENSG00000183054 | RGPD6 | 0,533554447 | 1,447491069 | 0,34835 | 0,62469429 | RANBP2 like and GRIP domain containing 6 |
| ENSG00000136108 | CKAP2 | 0,533524347 | 1,447460869 | 0,31173 | 0,59912688 | cytoskeleton associated protein 2 |
| ENSG00000197165 | SULT1A2 | 0,533505044 | 1,447441503 | 0,37381 | 0,6423379 | sulfotransferase family 1A member 2 |
| ENSG00000136518 | ACTL6A | 0,533415771 | 1,447351938 | 0,15219 | 0,48061452 | actin like 6A |
| ENSG00000148943 | LIN7C | 0,533284559 | 1,447220309 | 0,15251 | 0,48061452 | lin-7 homolog C, crumbs cell polarity complex component |
| ENSG00000275895 | U2AF1L5 | 0,533198813 | 1,447134297 | 0,10841 | 0,41904372 | U2 small nuclear RNA auxiliary factor 1 like 5 |
| ENSG00000258210 | AC144548,1 | 0,533172153 | 1,447107556 | 0,09946 | 0,40605358 | novel transcript |
| ENSG00000266598 | AC037487,2 | 0,533168706 | 1,447104098 | 0,57922 | 0,78270914 | novel transcript, antisense to GNA13 |
| ENSG00000104369 | JPH1 | 0,533166275 | 1,447101659 | 0,09399 | 0,39609058 | junctophilin 1 |
| ENSG00000082258 | CCNT2 | 0,53285543 | 1,446789898 | 0,2601 | 0,56382871 | cyclin T2 |
| ENSG00000266126 | AC005730,3 | 0,532807563 | 1,446741896 | 0,4589 | 0,70147844 | novel transcript |
| ENSG00000135338 | LCA5 | 0,532700028 | 1,446634064 | 0,20182 | 0,52216179 | lebercilin LCA5 |
| ENSG00000270757 | HSPE1-MOB4 | 0,532658037 | 1,446591958 | 0,62714 | 0,81127419 | HSPE1-MOB4 readthrough |
| ENSG00000148110 | MFSD14B | 0,532507609 | 1,446441132 | 0,10362 | 0,41173831 | major facilitator superfamily domain containing 14B |

| | | | | | | |
|---|---|---|---|---|---|---|
| ENSG00000282572 | AC215522,2 | 0,5324612 | 1,446394603 | 0,54099 | 0,75863636 | novel transcript similar to family member with sequence similarity 157 |
| ENSG00000158164 | TMSB15A | 0,53231389 | 1,446246923 | 0,35956 | 0,6323445 | thymosin beta 15a |
| ENSG00000169679 | BUB1 | 0,532267512 | 1,446200432 | 0,31319 | 0,59995613 | BUB1 mitotic checkpoint serine/threonine kinase |
| ENSG00000287149 | AC092640,1 | 0,532245077 | 1,446177942 | 0,53253 | 0,75301809 | novel transcript, antisense to CCDC141and TTN |
| ENSG00000114742 | WDR48 | 0,532124535 | 1,446057115 | 0,2315 | 0,54440987 | WD repeat domain 48 |
| ENSG00000262681 | AC005722,3 | 0,532056861 | 1,445989284 | 0,57381 | 0,77921182 | novel transcript |
| ENSG00000089597 | GANAB | 0,531765428 | 1,445697215 | 0,05298 | 0,30901632 | glucosidase II alpha subunit |
| ENSG00000160124 | CCDC58 | 0,531704333 | 1,445635995 | 0,20131 | 0,52181085 | coiled-coil domain containing 58 |
| ENSG00000165501 | LRR1 | 0,531679836 | 1,445611448 | 0,19382 | 0,51547251 | leucine rich repeat protein 1 |
| ENSG00000197965 | MPZL1 | 0,531661505 | 1,44559308 | 0,11337 | 0,42803427 | myelin protein zero like 1 |
| ENSG00000179133 | C10orf67 | 0,531637034 | 1,44556856 | 0,18726 | 0,5105374 | chromosome 10 open reading frame 67 |
| ENSG00000112320 | SOBP | 0,531501052 | 1,445432313 | 0,11103 | 0,42382951 | sine oculis binding protein homolog |
| ENSG00000278715 | SNORD116-20 | 0,531452083 | 1,445383253 | 0,63205 | 0,81450663 | small nucleolar RNA, C/D box 116-20 |
| ENSG00000035141 | FAM136A | 0,531426493 | 1,445357615 | 0,20761 | 0,52713166 | family with sequence similarity 136 member A |
| ENSG00000136888 | ATP6V1G1 | 0,531393577 | 1,445324638 | 0,18488 | 0,50844776 | ATPase H+ transporting V1 subunit G1 |
| ENSG00000198856 | OSTC | 0,531293499 | 1,445224381 | 0,15924 | 0,4867709 | oligosaccharyltransferase complex non-catalytic subunit |
| ENSG00000156675 | RAB11FIP1 | 0,531143234 | 1,445073861 | 0,29708 | 0,59016006 | RAB11 family interacting protein 1 |
| ENSG00000038274 | MAT2B | 0,531056571 | 1,444987058 | 0,25659 | 0,56097949 | methionine adenosyltransferase 2B |
| ENSG00000100532 | CGRRF1 | 0,53101165 | 1,444942066 | 0,24374 | 0,55240702 | cell growth regulator with ring finger domain 1 |
| ENSG00000105879 | CBLL1 | 0,530966186 | 1,444896532 | 0,19305 | 0,51505487 | Cbl proto-oncogene like 1 |
| ENSG00000196970 | NXF4 | 0,530867262 | 1,44479746 | 0,57347 | 0,77895239 | nuclear RNA export factor 4 pseudogene |
| ENSG00000261241 | LINC02128 | 0,530842803 | 1,444772966 | 0,66126 | 0,83123598 | long intergenic non-protein coding RNA 2128 |
| ENSG00000174718 | RESF1 | 0,530830596 | 1,444760741 | 0,31228 | 0,59926552 | retroelement silencing factor 1 |
| ENSG00000189089 | RIMKLBP1 | 0,530739544 | 1,444669562 | 0,30887 | 0,59767335 | ribosomal modification protein rimK like family member B pseudogene 1 |
| ENSG00000143761 | ARF1 | 0,530724923 | 1,444654921 | 0,07997 | 0,36911886 | ADP ribosylation factor 1 |
| ENSG00000167553 | TUBA1C | 0,530635372 | 1,444565251 | 0,28442 | 0,58127794 | tubulin alpha 1c |
| ENSG00000176340 | COX8A | 0,530578155 | 1,444507961 | 0,14548 | 0,47300165 | cytochrome c oxidase subunit 8A |
| ENSG00000163219 | ARHGAP25 | 0,530464261 | 1,444393929 | 0,10107 | 0,40780603 | Rho GTPase activating protein 25 |
| ENSG00000256473 | AC087260,1 | 0,53046197 | 1,444391635 | 0,40732 | 0,66661761 | novel transcript, antisense to SOX5 |
| ENSG00000143190 | POU2F1 | 0,530389901 | 1,444319483 | 0,04996 | 0,30088335 | POU class 2 homeobox 1 |
| ENSG00000158710 | TAGLN2 | 0,530253433 | 1,444182868 | 0,18561 | 0,50901954 | transgelin 2 |

| | | | | | | |
|---|---|---|---|---|---|---|
| ENSG00000279342 | AP000866,6 | 0,530153838 | 1,444083174 | 0,27706 | 0,57569532 | novel transcript, sense intronic to MSANTD2 |
| ENSG00000185624 | P4HB | 0,529885834 | 1,443814936 | 0,04897 | 0,29775453 | prolyl 4-hydroxylase subunit beta |
| ENSG00000143418 | CERS2 | 0,529855008 | 1,443784086 | 0,10355 | 0,41160135 | ceramide synthase 2 |
| ENSG00000285547 | AL133500,1 | 0,529848561 | 1,443777635 | 0,72296 | 0,86768626 | novel protein |
| ENSG00000065833 | ME1 | 0,529716572 | 1,443645553 | 0,23275 | 0,54542294 | malic enzyme 1 |
| ENSG00000118197 | DDX59 | 0,529671063 | 1,443600015 | 0,08609 | 0,3803347 | DEAD-box helicase 59 |
| ENSG00000275536 | AC118459,1 | 0,529501246 | 1,443430101 | 0,59703 | 0,79434997 | novel transcript |
| ENSG00000167202 | TBC1D2B | 0,529369952 | 1,443298746 | 0,06516 | 0,33804854 | TBC1 domain family member 2B |
| ENSG00000089063 | TMEM230 | 0,529353003 | 1,443281791 | 0,21832 | 0,535736 | transmembrane protein 230 |
| ENSG00000286599 | AC110611,1 | 0,529304527 | 1,443233295 | 0,14766 | 0,47507961 | novel transcript, antisense to EXOC1 |
| ENSG00000253908 | AC104115,2 | 0,529116744 | 1,443045454 | 0,36477 | 0,6350493 | zinc finger protein pseudogene |
| ENSG00000152944 | MED21 | 0,528979001 | 1,442907685 | 0,27098 | 0,57248186 | mediator complex subunit 21 |
| ENSG00000116128 | BCL9 | 0,528935449 | 1,442864127 | 0,10289 | 0,41078404 | BCL9 transcription coactivator |
| ENSG00000131238 | PPT1 | 0,528924081 | 1,442852757 | 0,03583 | 0,25662951 | palmitoyl-protein thioesterase 1 |
| ENSG00000139684 | ESD | 0,528887598 | 1,442816271 | 0,33932 | 0,61833676 | esterase D |
| ENSG00000164815 | ORC5 | 0,528791491 | 1,442720159 | 0,24529 | 0,55295131 | origin recognition complex subunit 5 |
| ENSG00000123360 | PDE1B | 0,528752202 | 1,44268087 | 0,03744 | 0,26177644 | phosphodiesterase 1B |
| ENSG00000117592 | PRDX6 | 0,528713262 | 1,442641931 | 0,17627 | 0,50045059 | peroxiredoxin 6 |
| ENSG00000169375 | SIN3A | 0,528603515 | 1,442532192 | 0,04663 | 0,29026543 | SIN3 transcription regulator family member A |
| ENSG00000174607 | UGT8 | 0,528495905 | 1,442424598 | 0,11446 | 0,42986344 | UDP glycosyltransferase 8 |
| ENSG00000100575 | TIMM9 | 0,528476491 | 1,442405188 | 0,32569 | 0,60814905 | translocase of inner mitochondrial membrane 9 |
| ENSG00000114790 | ARHGEF26 | 0,528439051 | 1,442367756 | 0,07598 | 0,36203962 | Rho guanine nucleotide exchange factor 26 |
| ENSG00000243402 | AC022973,2 | 0,528392962 | 1,442321679 | 0,61308 | 0,8035281 | ribosomal protein L15 (RPL15) pseudogene |
| ENSG00000236398 | TAS2R39 | 0,528323569 | 1,442252305 | 0,58821 | 0,78810304 | taste 2 receptor member 39 |
| ENSG00000135763 | URB2 | 0,528309519 | 1,442238259 | 0,10741 | 0,41771747 | URB2 ribosome biogenesis homolog |
| ENSG00000198182 | ZNF607 | 0,528279777 | 1,442208527 | 0,15955 | 0,48693054 | zinc finger protein 607 |
| ENSG00000286271 | AC008945,2 | 0,52824476 | 1,442173523 | 0,2453 | 0,55295131 | novel transcript, antisense to SEPP1 |
| ENSG00000144591 | GMPPA | 0,528133831 | 1,442062637 | 0,00589 | 0,09677469 | GDP-mannose pyrophosphorylase A |
| ENSG00000165355 | FBXO33 | 0,528015185 | 1,441944048 | 0,17508 | 0,49919753 | F-box protein 33 |
| ENSG00000134255 | CEPT1 | 0,527750569 | 1,441679595 | 0,14329 | 0,47082074 | choline/ethanolamine phosphotransferase 1 |
| ENSG00000131778 | CHD1L | 0,527747135 | 1,441676163 | 0,04455 | 0,28482141 | chromodomain helicase DNA binding protein 1 like |
| ENSG00000149212 | SESN3 | 0,527743949 | 1,441672979 | 0,13035 | 0,45477153 | sestrin 3 |
| ENSG00000100522 | GNPNAT1 | 0,527592212 | 1,441521357 | 0,00308 | 0,06617279 | glucosamine-phosphate N-acetyltransferase 1 |

| | | | | | | |
|---|---|---|---|---|---|---|
| ENSG00000140987 | ZSCAN32 | 0,527538952 | 1,441468142 | 0,16191 | 0,48914394 | zinc finger and SCAN domain containing 32 |
| ENSG00000137312 | FLOT1 | 0,527523547 | 1,44145275 | 0,39037 | 0,65441492 | flotillin 1 |
| ENSG00000069275 | NUCKS1 | 0,527476736 | 1,44140598 | 0,04485 | 0,28619498 | nuclear casein kinase and cyclin dependent kinase substrate 1 |
| ENSG00000166974 | MAPRE2 | 0,527451269 | 1,441380537 | 0,15359 | 0,48191866 | microtubule associated protein RP/EB family member 2 |
| ENSG00000247134 | AC090204,1 | 0,527450545 | 1,441379812 | 0,33027 | 0,61114402 | novel transcript |
| ENSG00000128513 | POT1 | 0,527402113 | 1,441331426 | 0,24144 | 0,55035777 | protection of telomeres 1 |
| ENSG00000134153 | EMC7 | 0,527387744 | 1,441317071 | 0,14561 | 0,47304491 | ER membrane protein complex subunit 7 |
| ENSG00000108953 | YWHAE | 0,527353679 | 1,441283038 | 0,19166 | 0,51436129 | tyrosine 3-monooxygenase/tryptophan 5-monooxygenase activation protein epsilon |
| ENSG00000229021 | AL450992,1 | 0,527276142 | 1,441205579 | 0,11339 | 0,42803427 | novel transcript |
| ENSG00000204356 | NELFE | 0,52711124 | 1,441040857 | 0,17058 | 0,49641416 | negative elongation factor complex member E |
| ENSG00000285335 | CACNA1C-AS1 | 0,526951436 | 1,440881245 | 0,50269 | 0,73209345 | CACNA1C antisense RNA 1 |
| ENSG00000246627 | CACNA1C-AS1 | 0,526951436 | 1,440881245 | 0,50269 | 0,73209345 | CACNA1C antisense RNA 1 |
| ENSG00000163811 | WDR43 | 0,526843791 | 1,44077374 | 0,047 | 0,29158646 | WD repeat domain 43 |
| ENSG00000131844 | MCCC2 | 0,526772432 | 1,440702478 | 0,02043 | 0,18959706 | methylcrotonoyl-CoA carboxylase 2 |
| ENSG00000254595 | AC084337,1 | 0,526624076 | 1,440554334 | 0,3928 | 0,6559444 | voltage-dependent anion channel 1(VDAC1) pseudogene |
| ENSG00000069849 | ATP1B3 | 0,526604646 | 1,440534933 | 0,18391 | 0,50761591 | ATPase Na+/K+ transporting subunit beta 3 |
| ENSG00000136167 | LCP1 | 0,526433432 | 1,440363986 | 0,40638 | 0,66593291 | lymphocyte cytosolic protein 1 |
| ENSG00000198712 | MT-CO2 | 0,526347301 | 1,440277996 | 0,19509 | 0,51588752 | mitochondrially encoded cytochrome c oxidase II |
| ENSG00000184731 | FAM110C | 0,526292437 | 1,440223225 | 0,20777 | 0,52736343 | family with sequence similarity 110 member C |
| ENSG00000198804 | MT-CO1 | 0,52620792 | 1,440138855 | 0,20648 | 0,52609885 | mitochondrially encoded cytochrome c oxidase I |
| ENSG00000277879 | AL391988,1 | 0,526140266 | 1,440071323 | 0,2969 | 0,58997679 | novel transcript, antisense to SLC18A2 |
| ENSG00000101605 | MYOM1 | 0,52587429 | 1,439805855 | 0,20812 | 0,52756478 | myomesin 1 |
| ENSG00000253537 | PCDHGA7 | 0,525761349 | 1,439693144 | 0,13709 | 0,4642578 | protocadherin gamma subfamily A, 7 |
| ENSG00000170085 | SIMC1 | 0,525640087 | 1,439572139 | 0,15763 | 0,48475003 | SUMO interacting motifs containing 1 |
| ENSG00000196922 | ZNF252P | 0,525576573 | 1,439508764 | 0,15744 | 0,48457045 | zinc finger protein 252, pseudogene |
| ENSG00000243836 | WDR86-AS1 | 0,525516837 | 1,439449162 | 0,56949 | 0,77669901 | WDR86 antisense RNA 1 |
| ENSG00000256262 | USP30-AS1 | 0,52519806 | 1,439131137 | 0,68245 | 0,84436596 | USP30 antisense RNA 1 |
| ENSG00000198369 | SPRED2 | 0,525190382 | 1,439123478 | 0,03439 | 0,25062659 | sprouty related EVH1 domain containing 2 |
| ENSG00000255856 | AC069503,1 | 0,525172413 | 1,439105554 | 0,49982 | 0,73058737 | novel transcript |
| ENSG00000121350 | PYROXD1 | 0,52494064 | 1,438874376 | 0,25422 | 0,55925777 | pyridine nucleotide-disulphide oxidoreductase domain 1 |
| ENSG00000186130 | ZBTB6 | 0,524705415 | 1,438639793 | 0,22466 | 0,54036191 | zinc finger and BTB domain containing 6 |

| | | | | | | |
|---|---|---|---|---|---|---|
| ENSG00000213782 | DDX47 | 0,524672886 | 1,438607356 | 0,1273 | 0,45051698 | DEAD-box helicase 47 |
| ENSG00000113719 | ERGIC1 | 0,524584262 | 1,438518985 | 0,12313 | 0,44322266 | endoplasmic reticulum-golgi intermediate compartment 1 |
| ENSG00000175279 | CENPS | 0,524550413 | 1,438485235 | 0,13918 | 0,46648276 | centromere protein S |
| ENSG00000226916 | WDR46 | 0,524542157 | 1,438477003 | 0,13874 | 0,46603966 | WD repeat domain 46 |
| ENSG00000204221 | WDR46 | 0,524542157 | 1,438477003 | 0,13874 | 0,46603966 | WD repeat domain 46 |
| ENSG00000204904 | LINC01545 | 0,524463702 | 1,43839878 | 0,63196 | 0,81447173 | long intergenic non-protein coding RNA 1545 |
| ENSG00000175449 | RFESD | 0,524439414 | 1,438374564 | 0,31504 | 0,60145932 | Rieske Fe-S domain containing |
| ENSG00000140575 | IQGAP1 | 0,524422012 | 1,438357215 | 0,17613 | 0,50021044 | IQ motif containing GTPase activating protein 1 |
| ENSG00000271843 | AC012557,1 | 0,524387044 | 1,438322352 | 0,32087 | 0,60558211 | novel transcript |
| ENSG00000170549 | IRX1 | 0,524383035 | 1,438318356 | 0,25284 | 0,55796133 | iroquois homeobox 1 |
| ENSG00000122376 | SHLD2 | 0,524374432 | 1,438309779 | 0,10226 | 0,40974995 | shieldin complex subunit 2 |
| ENSG00000248496 | ZBED9 | 0,524370062 | 1,438305422 | 0,20036 | 0,52107152 | zinc finger BED-type containing 9 |
| ENSG00000157456 | CCNB2 | 0,524363002 | 1,438298383 | 0,31761 | 0,60294242 | cyclin B2 |
| ENSG00000145780 | FEM1C | 0,524332227 | 1,438267702 | 0,32807 | 0,60971379 | fem-1 homolog C |
| ENSG00000270140 | AC005520,3 | 0,524272456 | 1,438208116 | 0,48906 | 0,7236969 | novel transcript |
| ENSG00000159592 | GPBP1L1 | 0,524157724 | 1,438093746 | 0,15897 | 0,48633228 | GC-rich promoter binding protein 1 like 1 |
| ENSG00000185305 | ARL15 | 0,524157244 | 1,438093267 | 0,32065 | 0,60553919 | ADP ribosylation factor like GTPase 15 |
| ENSG00000230782 | AL807761,4 | 0,524145007 | 1,438081069 | 0,17609 | 0,50015171 | novel transcript |
| ENSG00000198155 | ZNF876P | 0,524134564 | 1,438070659 | 0,07249 | 0,35549468 | zinc finger protein 876, pseudogene |
| ENSG00000088986 | DYNLL1 | 0,52391796 | 1,437854766 | 0,2566 | 0,56097949 | dynein light chain LC8-type 1 |
| ENSG00000114120 | SLC25A36 | 0,523901803 | 1,437838663 | 0,23744 | 0,54846286 | solute carrier family 25 member 36 |
| ENSG00000169139 | UBE2V2 | 0,523766616 | 1,437703938 | 0,16427 | 0,49065086 | ubiquitin conjugating enzyme E2 V2 |
| ENSG00000180263 | FGD6 | 0,523635694 | 1,437573475 | 0,1071 | 0,41711412 | FYVE, RhoGEF and PH domain containing 6 |
| ENSG00000155368 | DBI | 0,523599167 | 1,437537077 | 0,14909 | 0,47668027 | diazepam binding inhibitor, acyl-CoA binding protein |
| ENSG00000223685 | LINC00571 | 0,523454141 | 1,437392577 | 0,35821 | 0,63185282 | long intergenic non-protein coding RNA 571 |
| ENSG00000122406 | RPL5 | 0,523318444 | 1,437257385 | 0,12846 | 0,45191041 | ribosomal protein L5 |
| ENSG00000174628 | IQCK | 0,523226035 | 1,437165327 | 0,22755 | 0,54168591 | IQ motif containing K |
| ENSG00000147687 | TATDN1 | 0,523223786 | 1,437163087 | 0,1171 | 0,43439803 | TatD DNase domain containing 1 |
| ENSG00000101849 | TBL1X | 0,523181819 | 1,437121282 | 0,01584 | 0,16482188 | transducin beta like 1 X-linked |
| ENSG00000069667 | RORA | 0,522893684 | 1,436834288 | 0,21933 | 0,53676938 | RAR related orphan receptor A |
| ENSG00000155893 | PXYLP1 | 0,522814467 | 1,436755395 | 0,19097 | 0,51398916 | 2-phosphoxylose phosphatase 1 |
| ENSG00000259192 | AC109631,1 | 0,522744233 | 1,436685453 | 0,22533 | 0,54070011 | CTD (carboxy-terminal domain, RNA polymerase II, polypeptide A) small phosphatase 2 (CTDSP2) pseudogene |

| | | | | | | |
|---|---|---|---|---|---|---|
| ENSG00000286281 | AL353152,2 | 0,522627474 | 1,436569185 | 0,48384 | 0,719985 | novel transcript |
| ENSG00000272402 | AL031775,2 | 0,52254438 | 1,436486446 | 0,54078 | 0,75852611 | novel transcript, antisense to C6orf62 |
| ENSG00000261402 | AL591222,1 | 0,522428 | 1,436370571 | 0,56119 | 0,7713106 | novel transcript |
| ENSG00000145623 | OSMR | 0,52242257 | 1,436365165 | 0,23684 | 0,54768226 | oncostatin M receptor |
| ENSG00000287731 | AL035603,1 | 0,522391666 | 1,436334396 | 0,4489 | 0,69460209 | novel transcript |
| ENSG00000183479 | TREX2 | 0,522375701 | 1,436318502 | 0,56204 | 0,77155921 | three prime repair exonuclease 2 |
| ENSG00000054277 | OPN3 | 0,522197075 | 1,436140677 | 0,17579 | 0,49972993 | opsin 3 |
| ENSG00000110080 | ST3GAL4 | 0,521928332 | 1,435873179 | 0,06314 | 0,3327742 | ST3 beta-galactoside alpha-2,3-sialyltransferase 4 |
| ENSG00000183281 | PLGLB1 | 0,521835755 | 1,435781043 | 0,23574 | 0,5471053 | plasminogen like B1 |
| ENSG00000125775 | SDCBP2 | 0,52179441 | 1,435739898 | 0,37882 | 0,64679625 | syndecan binding protein 2 |
| ENSG00000132623 | ANKEF1 | 0,521705655 | 1,435651572 | 0,25775 | 0,56191608 | ankyrin repeat and EF-hand domain containing 1 |
| ENSG00000240180 | RPL12P28 | 0,521655295 | 1,43560146 | 0,62264 | 0,80866078 | ribosomal protein L12 pseudogene 28 |
| ENSG00000213090 | SCYL2P1 | 0,521651211 | 1,435597396 | 0,35892 | 0,63196414 | SCYL2 pseudogene 1 |
| ENSG00000109814 | UGDH | 0,52155992 | 1,435506557 | 0,02393 | 0,20662493 | UDP-glucose 6-dehydrogenase |
| ENSG00000219545 | UMAD1 | 0,521463318 | 1,435410439 | 0,20701 | 0,52670763 | UBAP1-MVB12-associated (UMA) domain containing 1 |
| ENSG00000124767 | GLO1 | 0,521434128 | 1,435381397 | 0,24819 | 0,55542983 | glyoxalase I |
| ENSG00000135929 | CYP27A1 | 0,521349792 | 1,435297491 | 0,06254 | 0,33112961 | cytochrome P450 family 27 subfamily A member 1 |
| ENSG00000179119 | SPTY2D1 | 0,521334476 | 1,435282253 | 0,06738 | 0,34363554 | SPT2 chromatin protein domain containing 1 |
| ENSG00000122126 | OCRL | 0,521314868 | 1,435262746 | 0,19063 | 0,51369186 | OCRL inositol polyphosphate-5-phosphatase |
| ENSG00000134986 | NREP | 0,521121857 | 1,435070742 | 0,15404 | 0,48233496 | neuronal regeneration related protein |
| ENSG00000099204 | ABLIM1 | 0,521098935 | 1,435047942 | 0,17007 | 0,49606609 | actin binding LIM protein 1 |
| ENSG00000155463 | OXA1L | 0,520991557 | 1,434941137 | 0,09421 | 0,39648446 | OXA1L mitochondrial inner membrane protein |
| ENSG00000272343 | AC107952,2 | 0,52087187 | 1,434822098 | 0,3663 | 0,63627876 | novel transcript |
| ENSG00000182899 | RPL35A | 0,520771138 | 1,43472192 | 0,24305 | 0,55199251 | ribosomal protein L35a |
| ENSG00000214265 | AC124312,1 | 0,52066896 | 1,434620309 | 0,58827 | 0,78810304 | novel protein |
| ENSG00000143353 | LYPLAL1 | 0,520642954 | 1,434594449 | 0,35947 | 0,63230932 | lysophospholipase like 1 |
| ENSG00000123975 | CKS2 | 0,520528237 | 1,434480381 | 0,44949 | 0,69516404 | CDC28 protein kinase regulatory subunit 2 |
| ENSG00000155561 | NUP205 | 0,520482471 | 1,434434877 | 0,1433 | 0,47082074 | nucleoporin 205 |
| ENSG00000145220 | LYAR | 0,520382149 | 1,434335132 | 0,08523 | 0,37859143 | Ly1 antibody reactive |
| ENSG00000256667 | KLRA1P | 0,520373045 | 1,434326081 | 0,24146 | 0,55035937 | killer cell lectin like receptor A1, pseudogene |
| ENSG00000173226 | IQCB1 | 0,520356385 | 1,434309518 | 0,2496 | 0,55657567 | IQ motif containing B1 |
| ENSG00000117000 | RLF | 0,52030443 | 1,434257866 | 0,15323 | 0,48147318 | RLF zinc finger |
| ENSG00000135341 | MAP3K7 | 0,520255046 | 1,434208772 | 0,12737 | 0,45051698 | mitogen-activated protein kinase kinase kinase 7 |

| | | | | | | |
|---|---|---|---|---|---|---|
| ENSG00000156171 | DRAM2 | 0,519970963 | 1,433926387 | 0,23562 | 0,54704432 | DNA damage regulated autophagy modulator 2 |
| ENSG00000133657 | ATP13A3 | 0,519967101 | 1,433922549 | 0,13999 | 0,46694389 | ATPase 13A3 |
| ENSG00000232888 | RPS11P5 | 0,519879721 | 1,433835703 | 0,34187 | 0,6197312 | ribosomal protein S11 pseudogene 5 |
| ENSG00000151388 | ADAMTS12 | 0,519799589 | 1,433756065 | 0,17982 | 0,50407792 | ADAM metallopeptidase with thrombospondin type 1 motif 12 |
| ENSG00000037241 | RPL26L1 | 0,519644883 | 1,433602326 | 0,3586 | 0,63185282 | ribosomal protein L26 like 1 |
| ENSG00000286742 | AC000065,1 | 0,519617253 | 1,43357487 | 0,16435 | 0,49065086 | novel transcript, antisense to CDK6 |
| ENSG00000071082 | RPL31 | 0,519501865 | 1,433460216 | 0,20275 | 0,52269757 | ribosomal protein L31 |
| ENSG00000133111 | RFXAP | 0,519306035 | 1,433265653 | 0,17496 | 0,49911246 | regulatory factor X associated protein |
| ENSG00000077458 | FAM76B | 0,519203915 | 1,433164204 | 0,26728 | 0,56973003 | family with sequence similarity 76 member B |
| ENSG00000117154 | IGSF21 | 0,51910864 | 1,433069561 | 0,19256 | 0,51490739 | immunoglobin superfamily member 21 |
| ENSG00000227811 | INKA2-AS1 | 0,519086139 | 1,433047211 | 0,3759 | 0,64417233 | INKA2 antisense RNA 1 |
| ENSG00000090061 | CCNK | 0,519069127 | 1,433030313 | 0,14038 | 0,46721845 | cyclin K |
| ENSG00000224083 | MTCO1P11 | 0,518905518 | 1,432867809 | 0,5305 | 0,75210231 | MT-CO1 pseudogene 11 |
| ENSG00000184432 | COPB2 | 0,518869143 | 1,432831682 | 0,17177 | 0,49710979 | COPI coat complex subunit beta 2 |
| ENSG00000141380 | SS18 | 0,518780721 | 1,432743868 | 0,07724 | 0,36386039 | SS18 subunit of BAF chromatin remodeling complex |
| ENSG00000271711 | AC069236,1 | 0,518761373 | 1,432724653 | 0,31817 | 0,60349082 | Sin3A-associated protein, 30kDa, pseudogene, |
| ENSG00000147677 | EIF3H | 0,518759736 | 1,432723027 | 0,22901 | 0,54263903 | eukaryotic translation initiation factor 3 subunit H |
| ENSG00000138018 | SELENOI | 0,518575294 | 1,432539872 | 0,08333 | 0,37550713 | selenoprotein I |
| ENSG00000134900 | TPP2 | 0,518561158 | 1,432525836 | 0,20011 | 0,5209612 | tripeptidyl peptidase 2 |
| ENSG00000101898 | MCTS2P | 0,518504934 | 1,432470009 | 0,61024 | 0,80221019 | MCTS family member 2, pseudogene |
| ENSG00000156261 | CCT8 | 0,518493842 | 1,432458996 | 0,097 | 0,40141222 | chaperonin containing TCP1 subunit 8 |
| ENSG00000182957 | SPATA13 | 0,518319533 | 1,432285933 | 0,18332 | 0,50679418 | spermatogenesis associated 13 |
| ENSG00000058262 | SEC61A1 | 0,518312143 | 1,432278597 | 0,12545 | 0,44729629 | SEC61 translocon subunit alpha 1 |
| ENSG00000182934 | SRPRA | 0,518266511 | 1,432233296 | 0,10864 | 0,41972109 | SRP receptor subunit alpha |
| ENSG00000100290 | BIK | 0,518260009 | 1,432226841 | 0,46109 | 0,70278639 | BCL2 interacting killer |
| ENSG00000126088 | UROD | 0,518185383 | 1,432152758 | 0,03168 | 0,24008374 | uroporphyrinogen decarboxylase |
| ENSG00000118689 | FOXO3 | 0,517993386 | 1,431962177 | 0,00075 | 0,02678818 | forkhead box O3 |
| ENSG00000241261 | RPL17P19 | 0,517984985 | 1,431953838 | 0,43382 | 0,68475933 | ribosomal protein L17 pseudogene 19 |
| ENSG00000246422 | DIAPH1-AS1 | 0,517878862 | 1,431848509 | 0,18782 | 0,51107563 | DIAPH1 antisense RNA 1 |
| ENSG00000131467 | PSME3 | 0,517848586 | 1,431818461 | 0,04656 | 0,29026341 | proteasome activator subunit 3 |
| ENSG00000140807 | NKD1 | 0,517796532 | 1,4317668 | 0,16524 | 0,49131595 | NKD inhibitor of WNT signaling pathway 1 |
| ENSG00000244026 | FAM86DP | 0,51779048 | 1,431760794 | 0,0216 | 0,19543346 | family with sequence similarity 86 member D, pseudogene |

| | | | | | | |
|---|---|---|---|---|---|---|
| ENSG00000259129 | LINC00648 | 0,517745444 | 1,4317161 | 0,09358 | 0,39524231 | long intergenic non-protein coding RNA 648 |
| ENSG00000143061 | IGSF3 | 0,517727589 | 1,431698381 | 0,02536 | 0,21419157 | immunoglobulin superfamily member 3 |
| ENSG00000124164 | VAPB | 0,517581817 | 1,431553727 | 0,08387 | 0,3763216 | VAMP associated protein B and C |
| ENSG00000154845 | PPP4R1 | 0,517513585 | 1,431486024 | 0,20317 | 0,52304241 | protein phosphatase 4 regulatory subunit 1 |
| ENSG00000012660 | ELOVL5 | 0,517475293 | 1,43144803 | 0,27146 | 0,57285347 | ELOVL fatty acid elongase 5 |
| ENSG00000095139 | ARCN1 | 0,517415732 | 1,431388935 | 0,04363 | 0,28187766 | archain 1 |
| ENSG00000237172 | B3GNT9 | 0,517311358 | 1,431285383 | 0,32441 | 0,60694578 | UDP-GlcNAc:betaGal beta-1,3-N-acetylglucosaminyltransferase 9 |
| ENSG00000006625 | GGCT | 0,517307941 | 1,431281993 | 0,12942 | 0,45292332 | gamma-glutamylcyclotransferase |
| ENSG00000107164 | FUBP3 | 0,51724192 | 1,431216495 | 0,21302 | 0,53205064 | far upstream element binding protein 3 |
| ENSG00000141030 | COPS3 | 0,51721342 | 1,431188222 | 0,25445 | 0,55944852 | COP9 signalosome subunit 3 |
| ENSG00000234369 | TATDN1P1 | 0,517179178 | 1,431154254 | 0,44475 | 0,69205935 | TatD DNase domain containing 1 pseudogene 1 |
| ENSG00000267532 | MIR497HG | 0,516979676 | 1,430956361 | 0,36558 | 0,6357945 | mir-497-195 cluster host gene |
| ENSG00000254929 | AL591684,2 | 0,516965159 | 1,430941963 | 0,3582 | 0,63185282 | novel transcript |
| ENSG00000116161 | CACYBP | 0,516930748 | 1,430907832 | 0,05093 | 0,30366564 | calcyclin binding protein |
| ENSG00000231063 | RPL6P6 | 0,516804088 | 1,430782213 | 0,30582 | 0,59512079 | ribosomal protein L6 pseudogene 6 |
| ENSG00000095002 | MSH2 | 0,516753186 | 1,430731731 | 0,16857 | 0,49492866 | mutS homolog 2 |
| ENSG00000115380 | EFEMP1 | 0,516648665 | 1,430628082 | 0,47643 | 0,71465252 | EGF containing fibulin extracellular matrix protein 1 |
| ENSG00000196890 | H2BU1 | 0,516565006 | 1,430545124 | 0,37118 | 0,64038503 | H2B,U histone 1 |
| ENSG00000168214 | RBPJ | 0,516521969 | 1,43050245 | 0,21469 | 0,53362255 | recombination signal binding protein for immunoglobulin kappa J region |
| ENSG00000138246 | DNAJC13 | 0,516428349 | 1,430409624 | 0,18147 | 0,50490902 | DnaJ heat shock protein family (Hsp40) member C13 |
| ENSG00000280837 | CPS1-IT1 | 0,516427967 | 1,430409246 | 0,5528 | 0,7660829 | CPS1 intronic transcript 1 |
| ENSG00000000460 | C1orf112 | 0,516301506 | 1,430283867 | 0,21606 | 0,53474184 | chromosome 1 open reading frame 112 |
| ENSG00000277462 | ZNF670 | 0,516198499 | 1,43018175 | 0,13975 | 0,4668467 | zinc finger protein 670 |
| ENSG00000236895 | GTF2H4 | 0,516195031 | 1,430178312 | 0,37382 | 0,6423379 | general transcription factor IIH subunit 4 |
| ENSG00000137936 | BCAR3 | 0,516169541 | 1,430153044 | 0,07638 | 0,36251343 | BCAR3 adaptor protein, NSP family member |
| ENSG00000163297 | ANTXR2 | 0,516161495 | 1,430145067 | 0,05489 | 0,31349896 | ANTXR cell adhesion molecule 2 |
| ENSG00000184678 | H2BC21 | 0,516150477 | 1,430134146 | 0,3589 | 0,63196414 | H2B clustered histone 21 |
| ENSG00000115652 | UXS1 | 0,516098556 | 1,430082678 | 0,11138 | 0,42416781 | UDP-glucuronate decarboxylase 1 |
| ENSG00000131373 | HACL1 | 0,516063642 | 1,430048069 | 0,13215 | 0,45719971 | 2-hydroxyacyl-CoA lyase 1 |
| ENSG00000057663 | ATG5 | 0,516040217 | 1,430024849 | 0,09573 | 0,39858189 | autophagy related 5 |
| ENSG00000125962 | ARMCX5 | 0,515853561 | 1,429839845 | 0,02897 | 0,22977124 | armadillo repeat containing X-linked 5 |

| | | | | | | |
|---|---|---|---|---|---|---|
| ENSG00000139793 | MBNL2 | 0,515797874 | 1,429784655 | 0,20296 | 0,52282418 | muscleblind like splicing regulator 2 |
| ENSG00000136738 | STAM | 0,515775924 | 1,429762902 | 0,19673 | 0,51793033 | signal transducing adaptor molecule |
| ENSG00000173915 | ATP5MD | 0,515766118 | 1,429753183 | 0,38983 | 0,65396251 | ATP synthase membrane subunit DAPIT |
| ENSG00000229598 | PRDX3P1 | 0,515740062 | 1,429727361 | 0,40086 | 0,66282495 | peroxiredoxin 3 pseudogene 1 |
| ENSG00000270882 | H4C14 | 0,515645557 | 1,429633709 | 0,30226 | 0,59294633 | H4 clustered histone 14 |
| ENSG00000143603 | KCNN3 | 0,515618362 | 1,429606761 | 0,1892 | 0,51249536 | potassium calcium-activated channel subfamily N member 3 |
| ENSG00000232280 | FLOT1 | 0,515576634 | 1,429565411 | 0,07202 | 0,35416676 | flotillin 1 |
| ENSG00000128699 | ORMDL1 | 0,515493108 | 1,429482649 | 0,3515 | 0,62674766 | ORMDL sphingolipid biosynthesis regulator 1 |
| ENSG00000141682 | PMAIP1 | 0,515488232 | 1,429477817 | 0,15261 | 0,48061452 | phorbol-12-myristate-13-acetate-induced protein 1 |
| ENSG00000263647 | BPTFP1 | 0,515396762 | 1,429387188 | 0,46451 | 0,70506031 | bromodomain PHD finger transcription factor pseudogene 1 |
| ENSG00000162630 | B3GALT2 | 0,515389817 | 1,429380307 | 0,26653 | 0,56917012 | beta-1,3-galactosyltransferase 2 |
| ENSG00000061455 | PRDM6 | 0,515384063 | 1,429374606 | 0,3866 | 0,65194712 | PR/SET domain 6 |
| ENSG00000141428 | C18orf21 | 0,515333025 | 1,42932404 | 0,29683 | 0,58995143 | chromosome 18 open reading frame 21 |
| ENSG00000103187 | COTL1 | 0,515316258 | 1,429307429 | 0,16942 | 0,49554671 | coactosin like F-actin binding protein 1 |
| ENSG00000276148 | AC084824,5 | 0,515309915 | 1,429301145 | 0,50921 | 0,73706413 | novel transcript, antisense to DNM1L |
| ENSG00000275779 | NTAN1 | 0,515216085 | 1,429208189 | 0,0716 | 0,35351167 | N-terminal asparagine amidase |
| ENSG00000186480 | INSIG1 | 0,515199847 | 1,429192102 | 0,00364 | 0,07353585 | insulin induced gene 1 |
| ENSG00000273055 | AC005046,1 | 0,515183579 | 1,429175987 | 0,46121 | 0,70278639 | novel transcript, antisense to LAMB1 |
| ENSG00000005100 | DHX33 | 0,515142286 | 1,429135081 | 0,02535 | 0,21419157 | DEAH-box helicase 33 |
| ENSG00000112695 | COX7A2 | 0,514974844 | 1,428969223 | 0,24881 | 0,55591283 | cytochrome c oxidase subunit 7A2 |
| ENSG00000108107 | RPL28 | 0,514944494 | 1,428939162 | 0,12218 | 0,44167604 | ribosomal protein L28 |
| ENSG00000166266 | CUL5 | 0,514819197 | 1,428815065 | 0,22488 | 0,5403718 | cullin 5 |
| ENSG00000204370 | SDHD | 0,514744847 | 1,428741432 | 0,08904 | 0,38596521 | succinate dehydrogenase complex subunit D |
| ENSG00000119446 | RBM18 | 0,514723855 | 1,428720643 | 0,24427 | 0,55253387 | RNA binding motif protein 18 |
| ENSG00000141376 | BCAS3 | 0,514400547 | 1,428400503 | 0,09373 | 0,39568176 | BCAS3 microtubule associated cell migration factor |
| ENSG00000122420 | PTGFR | 0,514140489 | 1,428143045 | 0,30368 | 0,59385655 | prostaglandin F receptor |
| ENSG00000101665 | SMAD7 | 0,514130926 | 1,428133579 | 0,13237 | 0,45750519 | SMAD family member 7 |
| ENSG00000275903 | PSMB3 | 0,51412624 | 1,428128939 | 0,16461 | 0,49065086 | proteasome 20S subunit beta 3 |
| ENSG00000132823 | OSER1 | 0,514097787 | 1,428100774 | 0,30257 | 0,59306846 | oxidative stress responsive serine rich 1 |
| ENSG00000013375 | PGM3 | 0,51403835 | 1,42804194 | 0,27831 | 0,57645922 | phosphoglucomutase 3 |
| ENSG00000163291 | PAQR3 | 0,513931216 | 1,427935898 | 0,30975 | 0,59807209 | progestin and adipoQ receptor family member 3 |

| | | | | | | |
|---|---|---|---|---|---|---|
| ENSG00000115368 | WDR75 | 0,51380581 | 1,42781178 | 0,12262 | 0,44239528 | WD repeat domain 75 |
| ENSG00000143569 | UBAP2L | 0,513773762 | 1,427780063 | 0,09644 | 0,40027371 | ubiquitin associated protein 2 like |
| ENSG00000162980 | ARL5A | 0,51371569 | 1,427722592 | 0,14183 | 0,46831266 | ADP ribosylation factor like GTPase 5A |
| ENSG00000113805 | CNTN3 | 0,513704139 | 1,427711162 | 0,14039 | 0,46721845 | contactin 3 |
| ENSG00000184743 | ATL3 | 0,513632265 | 1,427640036 | 0,01283 | 0,14814513 | atlastin GTPase 3 |
| ENSG00000164151 | ICE1 | 0,513581949 | 1,427590246 | 0,14233 | 0,46945051 | interactor of little elongation complex ELL subunit 1 |
| ENSG00000158296 | SLC13A3 | 0,513546137 | 1,427554809 | 0,25009 | 0,55657567 | solute carrier family 13 member 3 |
| ENSG00000129315 | CCNT1 | 0,513259819 | 1,427271524 | 0,19975 | 0,52077515 | cyclin T1 |
| ENSG00000149257 | SERPINH1 | 0,513203835 | 1,42721614 | 0,08334 | 0,37550713 | serpin family H member 1 |
| ENSG00000171448 | ZBTB26 | 0,513184167 | 1,427196682 | 0,21752 | 0,53541589 | zinc finger and BTB domain containing 26 |
| ENSG00000270587 | AC046185,2 | 0,513181409 | 1,427193954 | 0,42139 | 0,67641736 | family with sequence similarity 136, member A (FAM136A) pseudogene |
| ENSG00000093009 | CDC45 | 0,51312775 | 1,427140873 | 0,08584 | 0,38002516 | cell division cycle 45 |
| ENSG00000164024 | METAP1 | 0,513032731 | 1,427046882 | 0,19298 | 0,51505487 | methionyl aminopeptidase 1 |
| ENSG00000146463 | ZMYM4 | 0,512998574 | 1,427013095 | 0,22244 | 0,53859118 | zinc finger MYM-type containing 4 |
| ENSG00000124275 | MTRR | 0,512968773 | 1,426983619 | 0,1429 | 0,47049576 | 5-methyltetrahydrofolate-homocysteine methyltransferase reductase |
| ENSG00000121152 | NCAPH | 0,512892571 | 1,426908248 | 0,13049 | 0,45477153 | non-SMC condensin I complex subunit H |
| ENSG00000101544 | ADNP2 | 0,512747474 | 1,426764746 | 0,12658 | 0,44926656 | ADNP homeobox 2 |
| ENSG00000126583 | PRKCG | 0,512728283 | 1,426745767 | 0,17747 | 0,50191142 | protein kinase C gamma |
| ENSG00000233578 | EIF4EP1 | 0,512696626 | 1,426714461 | 0,26076 | 0,56400103 | eukaryotic translation initiation factor 4E pseudogene 1 |
| ENSG00000120254 | MTHFD1L | 0,512680762 | 1,426698773 | 0,1236 | 0,4443454 | methylenetetrahydrofolate dehydrogenase (NADP+ dependent) 1 like |
| ENSG00000168813 | ZNF507 | 0,512605963 | 1,426624805 | 0,14748 | 0,47493724 | zinc finger protein 507 |
| ENSG00000166394 | CYB5R2 | 0,512525227 | 1,42654497 | 0,08072 | 0,37077071 | cytochrome b5 reductase 2 |
| ENSG00000080503 | SMARCA2 | 0,51244852 | 1,426469124 | 0,32106 | 0,6057178 | SWI/SNF related, matrix associated, actin dependent regulator of chromatin, subfamily a, member 2 |
| ENSG00000136715 | SAP130 | 0,5123943 | 1,426415514 | 0,18294 | 0,50641825 | Sin3A associated protein 130 |
| ENSG00000215784 | FAM72D | 0,512323 | 1,426345021 | 0,17464 | 0,49886732 | family with sequence similarity 72 member D |
| ENSG00000163491 | NEK10 | 0,512181732 | 1,426205361 | 0,12248 | 0,4421901 | NIMA related kinase 10 |
| ENSG00000176444 | CLK2 | 0,512138436 | 1,42616256 | 0,10137 | 0,40826418 | CDC like kinase 2 |
| ENSG00000215190 | LINC00680 | 0,512123133 | 1,426147433 | 0,15312 | 0,48122632 | long intergenic non-protein coding RNA 680 |
| ENSG00000091527 | CDV3 | 0,512047624 | 1,426072792 | 0,11628 | 0,43279558 | CDV3 homolog |

| | | | | | | |
|---|---|---|---|---|---|---|
| ENSG00000060688 | SNRNP40 | 0,511998675 | 1,426024408 | 0,15037 | 0,47854874 | small nuclear ribonucleoprotein U5 subunit 40 |
| ENSG00000138614 | INTS14 | 0,511957783 | 1,425983989 | 0,17318 | 0,49786688 | integrator complex subunit 14 |
| ENSG00000198216 | CACNA1E | 0,511894783 | 1,42592172 | 0,28402 | 0,58092186 | calcium voltage-gated channel subunit alpha1 E |
| ENSG00000118961 | LDAH | 0,511706571 | 1,425735709 | 0,17345 | 0,49804988 | lipid droplet associated hydrolase |
| ENSG00000277496 | AL357033,4 | 0,511688848 | 1,425718193 | 0,52122 | 0,74558236 | novel transcript, antisense to SLCO4A1 |
| ENSG00000164116 | GUCY1A1 | 0,511620637 | 1,425650787 | 0,35607 | 0,6303154 | guanylate cyclase 1 soluble subunit alpha 1 |
| ENSG00000066654 | THUMPD1 | 0,511387994 | 1,425420911 | 0,15679 | 0,48425071 | THUMP domain containing 1 |
| ENSG00000231419 | LINC00689 | 0,511375841 | 1,425408903 | 0,4013 | 0,6630277 | long intergenic non-protein coding RNA 689 |
| ENSG00000170348 | TMED10 | 0,51133145 | 1,425365045 | 0,09198 | 0,39215144 | transmembrane p24 trafficking protein 10 |
| ENSG00000101327 | PDYN | 0,511296501 | 1,425330517 | 0,43872 | 0,6878469 | prodynorphin |
| ENSG00000185875 | THNSL1 | 0,511095303 | 1,425131754 | 0,10334 | 0,41114168 | threonine synthase like 1 |
| ENSG00000187097 | ENTPD5 | 0,511078603 | 1,425115258 | 0,10773 | 0,41834811 | ectonucleoside triphosphate diphosphohydrolase 5 (inactive) |
| ENSG00000250021 | ARPIN-AP3S2 | 0,510985327 | 1,42502312 | 0,36421 | 0,634951 | ARPIN-AP3S2 readthrough |
| ENSG00000162972 | MAIP1 | 0,51085157 | 1,424891009 | 0,26245 | 0,56537068 | matrix AAA peptidase interacting protein 1 |
| ENSG00000057608 | GDI2 | 0,510790225 | 1,424830422 | 0,23514 | 0,54666157 | GDP dissociation inhibitor 2 |
| ENSG00000104320 | NBN | 0,510712526 | 1,424753687 | 0,28544 | 0,58187705 | nibrin |
| ENSG00000124356 | STAMBP | 0,510344359 | 1,424390145 | 0,0428 | 0,27879697 | STAM binding protein |
| ENSG00000074935 | TUBE1 | 0,510134309 | 1,424182775 | 0,3055 | 0,59500307 | tubulin epsilon 1 |
| ENSG00000273761 | DDX24 | 0,510111017 | 1,424159782 | 0,21264 | 0,53156238 | DEAD-box helicase 24 |
| ENSG00000143179 | UCK2 | 0,510091671 | 1,424140685 | 0,02298 | 0,20221357 | uridine-cytidine kinase 2 |
| ENSG00000094841 | UPRT | 0,510071152 | 1,42412043 | 0,23324 | 0,54562736 | uracil phosphoribosyltransferase homolog |
| ENSG00000270084 | GAS5-AS1 | 0,509820872 | 1,423873394 | 0,07976 | 0,36858935 | GAS5 antisense RNA 1 |
| ENSG00000170312 | CDK1 | 0,509725205 | 1,423778978 | 0,28339 | 0,58063542 | cyclin dependent kinase 1 |
| ENSG00000243989 | ACY1 | 0,509482253 | 1,423539231 | 0,01306 | 0,14957757 | aminoacylase 1 |
| ENSG00000172270 | BSG | 0,509430225 | 1,423487895 | 0,06242 | 0,33099652 | basigin (Ok blood group) |
| ENSG00000163507 | CIP2A | 0,509230612 | 1,423290954 | 0,29278 | 0,58724158 | cellular inhibitor of PP2A |
| ENSG00000150093 | ITGB1 | 0,509084287 | 1,423146604 | 0,20794 | 0,52755008 | integrin subunit beta 1 |
| ENSG00000158092 | NCK1 | 0,508906163 | 1,422970904 | 0,21354 | 0,53275614 | NCK adaptor protein 1 |
| ENSG00000220685 | AL139094,1 | 0,5086403 | 1,4227087 | 0,33123 | 0,61131843 | transcription factor B2, mitochondrial (TFB2M) pseudogene |
| ENSG00000174231 | PRPF8 | 0,508606874 | 1,422675738 | 0,18676 | 0,51023628 | pre-mRNA processing factor 8 |
| ENSG00000134690 | CDCA8 | 0,508576204 | 1,422645493 | 0,23853 | 0,5488078 | cell division cycle associated 8 |
| ENSG00000163466 | ARPC2 | 0,508537225 | 1,422607056 | 0,17183 | 0,49710979 | actin related protein 2/3 complex subunit 2 |

| | | | | | | |
|---|---|---|---|---|---|---|
| ENSG00000115760 | BIRC6 | 0,508407005 | 1,422478656 | 0,23648 | 0,54754423 | baculoviral IAP repeat containing 6 |
| ENSG00000230224 | PHBP9 | 0,508223505 | 1,422297738 | 0,30482 | 0,59446033 | prohibitin pseudogene 9 |
| ENSG00000109689 | STIM2 | 0,508222578 | 1,422296825 | 0,19911 | 0,52017864 | stromal interaction molecule 2 |
| ENSG00000142657 | PGD | 0,508144487 | 1,42221984 | 0,07409 | 0,35756831 | phosphogluconate dehydrogenase |
| ENSG00000143514 | TP53BP2 | 0,508076448 | 1,422152768 | 0,13245 | 0,45752148 | tumor protein p53 binding protein 2 |
| ENSG00000121274 | TENT4B | 0,508064774 | 1,42214126 | 0,21485 | 0,53373102 | terminal nucleotidyltransferase 4B |
| ENSG00000100523 | DDHD1 | 0,508063642 | 1,422140145 | 0,11785 | 0,43532267 | DDHD domain containing 1 |
| ENSG00000089486 | CDIP1 | 0,507935545 | 1,422013878 | 0,06917 | 0,34742129 | cell death inducing p53 target 1 |
| ENSG00000169851 | PCDH7 | 0,507820048 | 1,421900041 | 0,10301 | 0,41078404 | protocadherin 7 |
| ENSG00000038382 | TRIO | 0,507770006 | 1,421850721 | 0,02686 | 0,22118271 | trio Rho guanine nucleotide exchange factor |
| ENSG00000107789 | MINPP1 | 0,507672568 | 1,421754694 | 0,17867 | 0,50323303 | multiple inositol-polyphosphate phosphatase 1 |
| ENSG00000138698 | RAP1GDS1 | 0,507650522 | 1,421732968 | 0,24637 | 0,55403311 | Rap1 GTPase-GDP dissociation stimulator 1 |
| ENSG00000110955 | ATP5F1B | 0,507588066 | 1,421671422 | 0,15719 | 0,48443207 | ATP synthase F1 subunit beta |
| ENSG00000114054 | PCCB | 0,507581229 | 1,421664684 | 0,05777 | 0,32104013 | propionyl-CoA carboxylase subunit beta |
| ENSG00000145725 | PPIP5K2 | 0,507186965 | 1,421276221 | 0,12166 | 0,44093327 | diphosphoinositol pentakisphosphate kinase 2 |
| ENSG00000169813 | HNRNPF | 0,507176855 | 1,42126626 | 0,20584 | 0,5253024 | heterogeneous nuclear ribonucleoprotein F |
| ENSG00000113593 | PPWD1 | 0,506995145 | 1,42108726 | 0,1737 | 0,49846579 | peptidylprolyl isomerase domain and WD repeat containing 1 |
| ENSG00000052841 | TTC17 | 0,506992177 | 1,421084337 | 0,19457 | 0,51562158 | tetratricopeptide repeat domain 17 |
| ENSG00000163006 | CCDC138 | 0,506969473 | 1,421061973 | 0,20417 | 0,52394055 | coiled-coil domain containing 138 |
| ENSG00000104231 | ZFAND1 | 0,506850448 | 1,420944738 | 0,3034 | 0,59367791 | zinc finger AN1-type containing 1 |
| ENSG00000060709 | RIMBP2 | 0,506689428 | 1,420786155 | 0,04174 | 0,27670157 | RIMS binding protein 2 |
| ENSG00000166670 | MMP10 | 0,506662595 | 1,420759729 | 0,6688 | 0,83569335 | matrix metallopeptidase 10 |
| ENSG00000119321 | FKBP15 | 0,50656287 | 1,420661524 | 0,24878 | 0,55591283 | FKBP prolyl isomerase 15 |
| ENSG00000119314 | PTBP3 | 0,50651596 | 1,420615331 | 0,16853 | 0,49484328 | polypyrimidine tract binding protein 3 |
| ENSG00000163002 | NUP35 | 0,506483633 | 1,420583499 | 0,28245 | 0,57981197 | nucleoporin 35 |
| ENSG00000134825 | TMEM258 | 0,506451165 | 1,420551529 | 0,18937 | 0,5125846 | transmembrane protein 258 |
| ENSG00000284752 | GMPR2 | 0,506421602 | 1,420522421 | 0,15451 | 0,48266733 | guanosine monophosphate reductase 2 |
| ENSG00000237423 | LINC01522 | 0,506390276 | 1,420491577 | 0,48975 | 0,72409833 | long intergenic non-protein coding RNA 1522 |
| ENSG00000081181 | ARG2 | 0,506305729 | 1,420408333 | 0,40681 | 0,66618707 | arginase 2 |
| ENSG00000168288 | MMADHC | 0,506038833 | 1,420145584 | 0,18789 | 0,51107563 | metabolism of cobalamin associated D |
| ENSG00000244270 | RPL32P29 | 0,505951596 | 1,420059713 | 0,6219 | 0,80819595 | ribosomal protein L32 pseudogene 29 |
| ENSG00000262280 | LINC01896 | 0,505754822 | 1,41986604 | 0,45947 | 0,7017037 | long intergenic non-protein coding RNA 1896 |

| | | | | | | |
|---|---|---|---|---|---|---|
| ENSG00000164393 | ADGRF2 | 0,505715429 | 1,419827271 | 0,62387 | 0,80952593 | adhesion G protein-coupled receptor F2 |
| ENSG00000058091 | CDK14 | 0,505689749 | 1,419801998 | 0,01885 | 0,1820155 | cyclin dependent kinase 14 |
| ENSG00000186416 | NKRF | 0,505620623 | 1,41973397 | 0,09976 | 0,40626368 | NFKB repressing factor |
| ENSG00000104064 | GABPB1 | 0,505543212 | 1,419657793 | 0,24978 | 0,55657567 | GA binding protein transcription factor subunit beta 1 |
| ENSG00000096063 | SRPK1 | 0,505443678 | 1,419559852 | 0,2004 | 0,52108722 | SRSF protein kinase 1 |
| ENSG00000263528 | IKBKE | 0,505396613 | 1,419513543 | 0,0609 | 0,32750733 | inhibitor of nuclear factor kappa B kinase subunit epsilon |
| ENSG00000268870 | AC008758,5 | 0,505309421 | 1,419427755 | 0,44752 | 0,69342177 | novel zinc finger protein |
| ENSG00000196976 | LAGE3 | 0,505304335 | 1,419422751 | 0,07784 | 0,36497628 | L antigen family member 3 |
| ENSG00000115806 | GORASP2 | 0,505299695 | 1,419418185 | 0,15682 | 0,48425071 | golgi reassembly stacking protein 2 |
| ENSG00000196151 | WDSUB1 | 0,505295617 | 1,419414173 | 0,09134 | 0,3908781 | WD repeat, sterile alpha motif and U-box domain containing 1 |
| ENSG00000125631 | HTR5BP | 0,505238762 | 1,419358237 | 0,57712 | 0,78134056 | 5-hydroxytryptamine receptor 5B, pseudogene |
| ENSG00000172935 | MRGPRF | 0,505170467 | 1,419291048 | 0,45073 | 0,69592991 | MAS related GPR family member F |
| ENSG00000151304 | SRFBP1 | 0,505150368 | 1,419271275 | 0,09966 | 0,40615251 | serum response factor binding protein 1 |
| ENSG00000156206 | CFAP161 | 0,505141458 | 1,419262509 | 0,58193 | 0,78454692 | cilia and flagella associated protein 161 |
| ENSG00000100632 | ERH | 0,505103732 | 1,419225397 | 0,27807 | 0,57635142 | ERH mRNA splicing and mitosis factor |
| ENSG00000288558 | DUS4L-BCAP2 | 0,504969976 | 1,419093823 | 0,2659 | 0,56882152 | DUS4L-BCAP29 readthrough |
| ENSG00000129048 | ACKR4 | 0,504582605 | 1,41871284 | 0,49164 | 0,72520694 | atypical chemokine receptor 4 |
| ENSG00000165490 | DDIAS | 0,504551972 | 1,418682716 | 0,36687 | 0,6368162 | DNA damage induced apoptosis suppressor |
| ENSG00000270387 | AC115621,1 | 0,504537287 | 1,418668277 | 0,62446 | 0,80979211 | ret finger protein-like 4A (RFPL4A) pseudogene |
| ENSG00000232909 | AL157823,2 | 0,504506629 | 1,418638129 | 0,43092 | 0,68325486 | novel transcript |
| ENSG00000174021 | GNG5 | 0,50450304 | 1,4186346 | 0,31849 | 0,6036094 | G protein subunit gamma 5 |
| ENSG00000163689 | CFAP20DC | 0,50447649 | 1,418608493 | 0,01612 | 0,1663376 | CFAP20 domain containing |
| ENSG00000113282 | CLINT1 | 0,50441048 | 1,418543587 | 0,10942 | 0,42140241 | clathrin interactor 1 |
| ENSG00000181450 | ZNF678 | 0,504357951 | 1,418491938 | 0,29067 | 0,58571697 | zinc finger protein 678 |
| ENSG00000036549 | ZZZ3 | 0,504337491 | 1,418471821 | 0,17514 | 0,49924834 | zinc finger ZZ-type containing 3 |
| ENSG00000182117 | NOP10 | 0,504123996 | 1,418261926 | 0,18362 | 0,50736104 | NOP10 ribonucleoprotein |
| ENSG00000084073 | ZMPSTE24 | 0,504097586 | 1,418235964 | 0,08445 | 0,37732439 | zinc metallopeptidase STE24 |
| ENSG00000021645 | NRXN3 | 0,50409626 | 1,41823466 | 0,09572 | 0,39858189 | neurexin 3 |
| ENSG00000153563 | CD8A | 0,503963452 | 1,41810411 | 0,43879 | 0,6878469 | CD8a molecule |
| ENSG00000183814 | LIN9 | 0,503920456 | 1,418061847 | 0,2333 | 0,54562736 | lin-9 DREAM MuvB core complex component |
| ENSG00000243725 | TTC4 | 0,50386828 | 1,418010563 | 0,03692 | 0,26058168 | tetratricopeptide repeat domain 4 |
| ENSG00000255529 | POLR2M | 0,503724764 | 1,41786951 | 0,02277 | 0,20142881 | RNA polymerase II subunit M |

| | | | | | | |
|---|---|---|---|---|---|---|
| ENSG00000095574 | IKZF5 | 0,503665405 | 1,417811174 | 0,23087 | 0,54390394 | IKAROS family zinc finger 5 |
| ENSG00000125870 | SNRPB2 | 0,503632023 | 1,417778368 | 0,16721 | 0,49323767 | small nuclear ribonucleoprotein polypeptide B2 |
| ENSG00000106100 | NOD1 | 0,503556252 | 1,417703908 | 0,05266 | 0,30807312 | nucleotide binding oligomerization domain containing 1 |
| ENSG00000273909 | AC138827,2 | 0,50340582 | 1,417556089 | 0,5617 | 0,77136826 | novel transcript |
| ENSG00000101911 | PRPS2 | 0,503391119 | 1,417541645 | 0,3091 | 0,59774312 | phosphoribosyl pyrophosphate synthetase 2 |
| ENSG00000215641 | TRIM27 | 0,503180744 | 1,417334952 | 0,1456 | 0,47304491 | tripartite motif containing 27 |
| ENSG00000286458 | AC083870,1 | 0,503108385 | 1,417263867 | 0,24949 | 0,55657567 | novel transcript, antisense to AGBL3and CALD1 |
| ENSG00000139133 | ALG10 | 0,502809578 | 1,416970358 | 0,20636 | 0,52602996 | ALG10 alpha-1,2-glucosyltransferase |
| ENSG00000156735 | BAG4 | 0,50272456 | 1,416886858 | 0,06813 | 0,34500646 | BAG cochaperone 4 |
| ENSG00000240303 | ACAD11 | 0,502708673 | 1,416871255 | 0,21566 | 0,53440695 | acyl-CoA dehydrogenase family member 11 |
| ENSG00000076706 | MCAM | 0,502613143 | 1,416777438 | 0,03726 | 0,26113308 | melanoma cell adhesion molecule |
| ENSG00000112697 | TMEM30A | 0,502607129 | 1,416771533 | 0,19524 | 0,51608421 | transmembrane protein 30A |
| ENSG00000282789 | CRYZP1 | 0,50257412 | 1,416739117 | 0,3898 | 0,65396251 | crystallin zeta pseudogene 1 |
| ENSG00000233025 | CRYZP1 | 0,50257412 | 1,416739117 | 0,3898 | 0,65396251 | crystallin zeta pseudogene 1 |
| ENSG00000272508 | AL136982,6 | 0,502494366 | 1,4166608 | 0,46225 | 0,70351872 | novel transcript |
| ENSG00000137055 | PLAA | 0,502448916 | 1,416616171 | 0,2091 | 0,52816443 | phospholipase A2 activating protein |
| ENSG00000110880 | CORO1C | 0,502364741 | 1,416533519 | 0,12642 | 0,44907109 | coronin 1C |
| ENSG00000134294 | SLC38A2 | 0,502339252 | 1,416508494 | 0,02908 | 0,2300184 | solute carrier family 38 member 2 |
| ENSG00000253729 | PRKDC | 0,502117813 | 1,416291091 | 0,17651 | 0,50072935 | protein kinase, DNA-activated, catalytic subunit |
| ENSG00000185246 | PRPF39 | 0,502113162 | 1,416286525 | 0,33907 | 0,61804482 | pre-mRNA processing factor 39 |
| ENSG00000104299 | INTS9 | 0,502047121 | 1,416221694 | 0,16377 | 0,49000347 | integrator complex subunit 9 |
| ENSG00000163281 | GNPDA2 | 0,501913268 | 1,416090303 | 0,30666 | 0,59571748 | glucosamine-6-phosphate deaminase 2 |
| ENSG00000286219 | NOTCH2NLC | 0,501876033 | 1,416053755 | 0,07547 | 0,36080119 | notch 2 N-terminal like C |
| ENSG00000212952 | BX088651,1 | 0,50172372 | 1,415904263 | 0,41592 | 0,67283439 | novel pseudogene |
| ENSG00000171150 | SOCS5 | 0,501578968 | 1,415762206 | 0,14629 | 0,47433542 | suppressor of cytokine signaling 5 |
| ENSG00000115339 | GALNT3 | 0,50152006 | 1,415704398 | 0,34489 | 0,62213186 | polypeptide N-acetylgalactosaminyltransferase 3 |
| ENSG00000070950 | RAD18 | 0,501471501 | 1,415656749 | 0,25843 | 0,56243993 | RAD18 E3 ubiquitin protein ligase |
| ENSG00000122033 | MTIF3 | 0,501457758 | 1,415643264 | 0,22282 | 0,53904413 | mitochondrial translational initiation factor 3 |
| ENSG00000129116 | PALLD | 0,501398289 | 1,415584911 | 0,10281 | 0,41062092 | palladin, cytoskeletal associated protein |
| ENSG00000136143 | SUCLA2 | 0,501344868 | 1,415532495 | 0,15787 | 0,48490111 | succinate-CoA ligase ADP-forming subunit beta |
| ENSG00000165912 | PACSIN3 | 0,501339792 | 1,415527515 | 0,42178 | 0,67643128 | protein kinase C and casein kinase substrate in neurons 3 |
| ENSG00000014123 | UFL1 | 0,501222059 | 1,415412003 | 0,28421 | 0,58107964 | UFM1 specific ligase 1 |

| | | | | | | |
|---|---|---|---|---|---|---|
| ENSG00000203880 | PCMTD2 | 0,500994715 | 1,415188977 | 0,22249 | 0,53862734 | protein-L-isoaspartate (D-aspartate) O-methyltransferase domain containing 2 |
| ENSG00000156574 | NODAL | 0,500985379 | 1,415179818 | 0,21201 | 0,53079538 | nodal growth differentiation factor |
| ENSG00000176087 | SLC35A4 | 0,500960758 | 1,415155667 | 0,01498 | 0,16040868 | solute carrier family 35 member A4 |
| ENSG00000166788 | SAAL1 | 0,500928307 | 1,415123836 | 0,151 | 0,47929859 | serum amyloid A like 1 |
| ENSG00000107438 | PDLIM1 | 0,500747947 | 1,414946933 | 0,21362 | 0,53275614 | PDZ and LIM domain 1 |
| ENSG00000241468 | ATP5MF | 0,5007101 | 1,414909815 | 0,24724 | 0,55483158 | ATP synthase membrane subunit f |
| ENSG00000274333 | CU633967,1 | 0,500702784 | 1,41490264 | 0,17096 | 0,49641416 | uncharacterized LOC102724219 |
| ENSG00000151239 | TWF1 | 0,500677811 | 1,414878149 | 0,42322 | 0,67699936 | twinfilin actin binding protein 1 |
| ENSG00000136754 | ABI1 | 0,50060088 | 1,414802702 | 0,23236 | 0,54512385 | abl interactor 1 |
| ENSG00000129484 | PARP2 | 0,500510632 | 1,414714202 | 0,25604 | 0,56063827 | poly(ADP-ribose) polymerase 2 |
| ENSG00000196678 | ERI2 | 0,500482298 | 1,414686418 | 0,16454 | 0,49065086 | ERI1 exoribonuclease family member 2 |
| ENSG00000248919 | ATP5MF-PTCD | 0,500467976 | 1,414672374 | 0,28254 | 0,57996606 | ATP5MF-PTCD1 readthrough |
| ENSG00000242110 | AMACR | 0,500403868 | 1,414609513 | 0,18432 | 0,50808994 | alpha-methylacyl-CoA racemase |
| ENSG00000110107 | PRPF19 | 0,50040048 | 1,414606191 | 0,0578 | 0,32108765 | pre-mRNA processing factor 19 |
| ENSG00000124786 | SLC35B3 | 0,500349587 | 1,41455629 | 0,19202 | 0,5145257 | solute carrier family 35 member B3 |
| ENSG00000122482 | ZNF644 | 0,500319652 | 1,414526939 | 0,16571 | 0,49182669 | zinc finger protein 644 |
| ENSG00000132286 | TIMM10B | 0,500309236 | 1,414516726 | 0,12738 | 0,45051698 | translocase of inner mitochondrial membrane 10B |
| ENSG00000197301 | HMGA2-AS1 | 0,500256895 | 1,414465408 | 0,07215 | 0,35442193 | HMGA2 antisense RNA 1 |
| ENSG00000125245 | GPR18 | 0,500195453 | 1,414405169 | 0,30149 | 0,59266609 | G protein-coupled receptor 18 |
| ENSG00000185896 | LAMP1 | 0,500183311 | 1,414393266 | 0,05993 | 0,32541248 | lysosomal associated membrane protein 1 |
| ENSG00000131435 | PDLIM4 | 0,500132353 | 1,414343309 | 0,09605 | 0,39928497 | PDZ and LIM domain 4 |
| ENSG00000177889 | UBE2N | 0,500075681 | 1,414287751 | 0,25733 | 0,56160845 | ubiquitin conjugating enzyme E2 N |
| ENSG00000250071 | AC093214,1 | 0,500014859 | 1,414228128 | 0,543 | 0,75965603 | actin related protein 2/3 complex, subunit 1A, 41kDa (ARPC1A) pseudogene |
| ENSG00000148572 | NRBF2 | 0,499944196 | 1,414158861 | 0,20954 | 0,52856218 | nuclear receptor binding factor 2 |
| ENSG00000067798 | NAV3 | 0,499846801 | 1,414063396 | 0,23051 | 0,54361941 | neuron navigator 3 |
| ENSG00000273306 | AC018690,1 | 0,499835474 | 1,414052293 | 0,18333 | 0,50679418 | novel transcript |
| ENSG00000205302 | SNX2 | 0,49983359 | 1,414050447 | 0,11036 | 0,42294747 | sorting nexin 2 |
| ENSG00000117862 | TXNDC12 | 0,499793425 | 1,41401108 | 0,16078 | 0,48808991 | thioredoxin domain containing 12 |
| ENSG00000141458 | NPC1 | 0,499741589 | 1,413960276 | 0,11186 | 0,42531464 | NPC intracellular cholesterol transporter 1 |
| ENSG00000115363 | EVA1A | 0,499726006 | 1,413945003 | 0,28964 | 0,58522301 | eva-1 homolog A, regulator of programmed cell death |
| ENSG00000254959 | INMT-MINDY | 0,499702061 | 1,413921535 | 0,88657 | 0,94987292 | INMT-MINDY4 readthrough (NMD candidate) |

| | | | | | | |
|---|---|---|---|---|---|---|
| ENSG00000162434 | JAK1 | 0,499563547 | 1,41378579 | 0,17507 | 0,49919753 | Janus kinase 1 |
| ENSG00000180764 | PIPSL | 0,499467361 | 1,413691535 | 0,24693 | 0,5545754 | PIP5K1A and PSMD4 like (pseudogene) |
| ENSG00000124784 | RIOK1 | 0,49946327 | 1,413687526 | 0,18875 | 0,5120892 | RIO kinase 1 |
| ENSG00000256223 | ZNF10 | 0,499447137 | 1,413671718 | 0,17001 | 0,49606609 | zinc finger protein 10 |
| ENSG00000197771 | MCMBP | 0,499441733 | 1,413666423 | 0,10866 | 0,41972109 | minichromosome maintenance complex binding protein |
| ENSG00000145817 | YIPF5 | 0,499346286 | 1,413572899 | 0,27052 | 0,5723494 | Yip1 domain family member 5 |
| ENSG00000175105 | ZNF654 | 0,499328404 | 1,413555378 | 0,26519 | 0,56799073 | zinc finger protein 654 |
| ENSG00000215388 | ACTG1P3 | 0,49926663 | 1,413494853 | 0,40429 | 0,6642832 | actin gamma 1 pseudogene 3 |
| ENSG00000175582 | RAB6A | 0,49916638 | 1,413396636 | 0,23658 | 0,54760364 | RAB6A, member RAS oncogene family |
| ENSG00000249319 | AC068533,4 | 0,49909859 | 1,413330224 | 0,62197 | 0,80819595 | novel protein |
| ENSG00000234187 | AIMP1P1 | 0,498913048 | 1,41314847 | 0,50167 | 0,73146914 | aminoacyl tRNA synthetase complex interacting multifunctional protein 1 pseudogene 1 |
| ENSG00000090020 | SLC9A1 | 0,498909473 | 1,413144968 | 0,00733 | 0,10873533 | solute carrier family 9 member A1 |
| ENSG00000164031 | DNAJB14 | 0,498901992 | 1,413137641 | 0,174 | 0,49860418 | DnaJ heat shock protein family (Hsp40) member B14 |
| ENSG00000085449 | WDFY1 | 0,498723639 | 1,412962952 | 0,22027 | 0,53750545 | WD repeat and FYVE domain containing 1 |
| ENSG00000145975 | FAM217A | 0,498692039 | 1,412932004 | 0,49544 | 0,72768069 | family with sequence similarity 217 member A |
| ENSG00000280651 | AC156455,2 | 0,498686067 | 1,412926155 | 0,20234 | 0,52249745 | novel transcript |
| ENSG00000067064 | IDI1 | 0,498563829 | 1,412806445 | 0,29458 | 0,58857672 | isopentenyl-diphosphate delta isomerase 1 |
| ENSG00000284548 | METTL9 | 0,498454834 | 1,412699712 | 0,76127 | 0,88879633 | methyltransferase like 9 |
| ENSG00000243849 | CFAP44-AS1 | 0,498445008 | 1,41269009 | 0,49735 | 0,72869621 | CFAP44 antisense RNA 1 |
| ENSG00000117479 | SLC19A2 | 0,498209793 | 1,412459786 | 0,14005 | 0,46694389 | solute carrier family 19 member 2 |
| ENSG00000168090 | COPS6 | 0,498178938 | 1,412429578 | 0,16951 | 0,49565021 | COP9 signalosome subunit 6 |
| ENSG00000243101 | RPS3P7 | 0,498126273 | 1,412378018 | 0,41801 | 0,67432365 | ribosomal protein S3 pseudogene 7 |
| ENSG00000135913 | USP37 | 0,49805284 | 1,412306131 | 0,22144 | 0,53805234 | ubiquitin specific peptidase 37 |
| ENSG00000214999 | AC129492,1 | 0,497986791 | 1,412241474 | 0,56616 | 0,77421829 | novel transcript, antisense to ALOX12B |
| ENSG00000274320 | GOLGA6L9 | 0,497863249 | 1,412120545 | 0,39403 | 0,65702282 | golgin A6 family like 9 |
| ENSG00000029363 | BCLAF1 | 0,497608999 | 1,411871705 | 0,21783 | 0,53550245 | BCL2 associated transcription factor 1 |
| ENSG00000156875 | MFSD14A | 0,497454266 | 1,411720287 | 0,1887 | 0,5120698 | major facilitator superfamily domain containing 14A |
| ENSG00000278811 | LINC00624 | 0,497441381 | 1,411707678 | 0,31078 | 0,5987267 | long intergenic non-protein coding RNA 624 |
| ENSG00000077092 | RARB | 0,497438849 | 1,411705201 | 0,30316 | 0,59350995 | retinoic acid receptor beta |
| ENSG00000118939 | UCHL3 | 0,49741757 | 1,411684379 | 0,2857 | 0,58214927 | ubiquitin C-terminal hydrolase L3 |
| ENSG00000179241 | LDLRAD3 | 0,497342726 | 1,411611146 | 0,06846 | 0,3457466 | low density lipoprotein receptor class A domain containing 3 |

| | | | | | | |
|---|---|---|---|---|---|---|
| ENSG00000131876 | SNRPA1 | 0,497309148 | 1,411578291 | 0,17733 | 0,50187931 | small nuclear ribonucleoprotein polypeptide A' |
| ENSG00000238161 | OR7E117P | 0,497297802 | 1,41156719 | 0,53286 | 0,75333268 | olfactory receptor family 7 subfamily E member 117 pseudogene |
| ENSG00000248243 | LINC02014 | 0,497210872 | 1,411482139 | 0,35385 | 0,62865377 | long intergenic non-protein coding RNA 2014 |
| ENSG00000119771 | KLHL29 | 0,497095615 | 1,41136938 | 0,11102 | 0,42382951 | kelch like family member 29 |
| ENSG00000102753 | KPNA3 | 0,497049218 | 1,41132399 | 0,36306 | 0,63401731 | karyopherin subunit alpha 3 |
| ENSG00000257657 | AC079950,1 | 0,497016539 | 1,411292022 | 0,50583 | 0,73418581 | novel transcript |
| ENSG00000153339 | TRAPPC8 | 0,497002589 | 1,411278376 | 0,30077 | 0,59208755 | trafficking protein particle complex 8 |
| ENSG00000275092 | AL031710,2 | 0,496957327 | 1,411234101 | 0,32211 | 0,60577115 | novel transcript, sense intronic to MAPK8IP3 |
| ENSG00000159596 | TMEM69 | 0,496939702 | 1,41121686 | 0,19496 | 0,51588752 | transmembrane protein 69 |
| ENSG00000257181 | AC025423,4 | 0,496914646 | 1,411192351 | 0,34743 | 0,62411645 | novel transcript |
| ENSG00000158402 | CDC25C | 0,496808857 | 1,411088875 | 0,23276 | 0,54542294 | cell division cycle 25C |
| ENSG00000175029 | CTBP2 | 0,496808486 | 1,411088513 | 0,154 | 0,48230757 | C-terminal binding protein 2 |
| ENSG00000125871 | MGME1 | 0,496678894 | 1,410961765 | 0,26285 | 0,56582568 | mitochondrial genome maintenance exonuclease 1 |
| ENSG00000173726 | TOMM20 | 0,496665407 | 1,410948575 | 0,13148 | 0,45657505 | translocase of outer mitochondrial membrane 20 |
| ENSG00000230870 | FBXW11P1 | 0,49666257 | 1,4109458 | 0,4472 | 0,69342177 | F-box and WD repeat domain containing 11 pseudogene 1 |
| ENSG00000256056 | AC087257,2 | 0,496630727 | 1,410914658 | 0,45742 | 0,70074793 | chromosome 12 open reading frame 71 (C12orf71) pseudogene |
| ENSG00000270469 | AC006557,5 | 0,496627057 | 1,41091107 | 0,60245 | 0,79709598 | non-SMC element 4 homolog A (S, cerevisiae), pseudogene |
| ENSG00000255491 | AC100858,2 | 0,496625653 | 1,410909696 | 0,41218 | 0,66995975 | uncharacterized LOC105375743 |
| ENSG00000102409 | BEX4 | 0,496559935 | 1,410845428 | 0,22667 | 0,54144 | brain expressed X-linked 4 |
| ENSG00000189043 | NDUFA4 | 0,496482072 | 1,410769286 | 0,38377 | 0,65041894 | NDUFA4 mitochondrial complex associated |
| ENSG00000258469 | CHMP4BP1 | 0,496472785 | 1,410760204 | 0,40655 | 0,66597268 | charged multivesicular body protein 4B pseudogene 1 |
| ENSG00000168398 | BDKRB2 | 0,496455889 | 1,410743682 | 0,46016 | 0,70210714 | bradykinin receptor B2 |
| ENSG00000120656 | TAF12 | 0,496443178 | 1,410731253 | 0,1363 | 0,46342304 | TATA-box binding protein associated factor 12 |
| ENSG00000228267 | C4B | 0,496374727 | 1,410664321 | 0,54478 | 0,76084582 | complement C4B (Chido blood group) |
| ENSG00000270091 | AC015726,1 | 0,496260932 | 1,410553056 | 0,31562 | 0,60172912 | novel transcript |
| ENSG00000170571 | EMB | 0,496220649 | 1,410513671 | 0,39615 | 0,65876705 | embigin |
| ENSG00000157181 | ODR4 | 0,496212361 | 1,410505568 | 0,22938 | 0,54286041 | odr-4 GPCR localization factor homolog |
| ENSG00000179941 | BBS10 | 0,496122395 | 1,410417612 | 0,35836 | 0,63185282 | Bardet-Biedl syndrome 10 |
| ENSG00000206286 | VPS52 | 0,496082238 | 1,410378354 | 0,22404 | 0,53995756 | VPS52 subunit of GARP complex |
| ENSG00000277224 | H2BC7 | 0,495959865 | 1,410258727 | 0,22164 | 0,53805234 | H2B clustered histone 7 |

| | | | | | | |
|---|---|---|---|---|---|---|
| ENSG00000243813 | AC078857,1 | 0,495668113 | 1,409973563 | 0,532 | 0,75285051 | zinc finger protein pseudogene |
| ENSG00000228547 | OR7E26P | 0,495585598 | 1,409892923 | 0,60366 | 0,79770754 | olfactory receptor family 7 subfamily E member 26 pseudogene |
| ENSG00000083937 | CHMP2B | 0,495522236 | 1,409831003 | 0,09798 | 0,40323803 | charged multivesicular body protein 2B |
| ENSG00000204228 | HSD17B8 | 0,495518547 | 1,409827398 | 0,31537 | 0,60163243 | hydroxysteroid 17-beta dehydrogenase 8 |
| ENSG00000198553 | KCNRG | 0,495465517 | 1,409775577 | 0,28322 | 0,58053323 | potassium channel regulator |
| ENSG00000229334 | AC046143,1 | 0,495167497 | 1,409484387 | 0,25594 | 0,56050326 | novel transcript |
| ENSG00000183098 | GPC6 | 0,495150344 | 1,409467629 | 0,32177 | 0,60577115 | glypican 6 |
| ENSG00000084754 | HADHA | 0,495136773 | 1,40945437 | 0,16623 | 0,49212363 | hydroxyacyl-CoA dehydrogenase trifunctional multienzyme complex subunit alpha |
| ENSG00000102908 | NFAT5 | 0,495060614 | 1,409379968 | 0,06619 | 0,34056415 | nuclear factor of activated T cells 5 |
| ENSG00000113048 | MRPS27 | 0,494991485 | 1,409312437 | 0,11312 | 0,42753722 | mitochondrial ribosomal protein S27 |
| ENSG00000124370 | MCEE | 0,494957815 | 1,409279547 | 0,05596 | 0,31610544 | methylmalonyl-CoA epimerase |
| ENSG00000067704 | IARS2 | 0,494916191 | 1,409238887 | 0,10932 | 0,42119293 | isoleucyl-tRNA synthetase 2, mitochondrial |
| ENSG00000243927 | MRPS6 | 0,494893862 | 1,409217077 | 0,10756 | 0,41807472 | mitochondrial ribosomal protein S6 |
| ENSG00000170485 | NPAS2 | 0,494857685 | 1,40918174 | 0,12885 | 0,45214857 | neuronal PAS domain protein 2 |
| ENSG00000271784 | AL031055,1 | 0,494837036 | 1,40916157 | 0,54302 | 0,75965603 | novel transcript, antisense to TP53RK |
| ENSG00000273784 | AL137058,2 | 0,494699199 | 1,409026943 | 0,22702 | 0,54168591 | novel transcript |
| ENSG00000135837 | CEP350 | 0,494580527 | 1,408911045 | 0,19651 | 0,51769123 | centrosomal protein 350 |
| ENSG00000151725 | CENPU | 0,494504503 | 1,408836804 | 0,31716 | 0,6027392 | centromere protein U |
| ENSG00000269867 | AC010326,3 | 0,494361623 | 1,408697284 | 0,4841 | 0,72005601 | novel transcript, sense intronic to ZNF814 |
| ENSG00000198464 | ZNF480 | 0,494282129 | 1,408619665 | 0,37372 | 0,6423379 | zinc finger protein 480 |
| ENSG00000109475 | RPL34 | 0,494264215 | 1,408602174 | 0,14798 | 0,4751727 | ribosomal protein L34 |
| ENSG00000264007 | AC104564,2 | 0,494243822 | 1,408582263 | 0,62146 | 0,80788604 | novel transcript |
| ENSG00000107960 | STN1 | 0,494227826 | 1,408566646 | 0,24885 | 0,55592231 | STN1 subunit of CST complex |
| ENSG00000277245 | AC084782,3 | 0,49422608 | 1,408564941 | 0,55716 | 0,76897426 | novel transcript, antisense to MNS1 |
| ENSG00000173947 | PIFO | 0,494219328 | 1,408558349 | 0,22034 | 0,53750545 | primary cilia formation |
| ENSG00000244462 | RBM12 | 0,494219193 | 1,408558218 | 0,20178 | 0,52216179 | RNA binding motif protein 12 |
| ENSG00000100934 | SEC23A | 0,494195831 | 1,408535408 | 0,0348 | 0,25226698 | Sec23 homolog A, COPII coat complex component |
| ENSG00000275949 | AC139677,1 | 0,494149967 | 1,408490631 | 0,27617 | 0,57535709 | ADP-ribosylation factor-like protein 17 |
| ENSG00000105193 | RPS16 | 0,49410978 | 1,408451397 | 0,14553 | 0,47304491 | ribosomal protein S16 |
| ENSG00000058729 | RIOK2 | 0,493975305 | 1,40832012 | 0,23999 | 0,54936679 | RIO kinase 2 |

| | | | | | | |
|---|---|---|---|---|---|---|
| ENSG00000255566 | AC135279,2 | 0,493947573 | 1,408293049 | 0,59343 | 0,7917651 | mitochondrial ribosomal protein S25 (MRPS25) pseudogene |
| ENSG00000107443 | CCNJ | 0,493829488 | 1,408177785 | 0,23143 | 0,54440536 | cyclin J |
| ENSG00000272518 | AC036214,2 | 0,493827823 | 1,40817616 | 0,18413 | 0,50785017 | novel transcript, sense intronic to MRPS28 |
| ENSG00000242290 | ZBTB20-AS5 | 0,493788245 | 1,408137529 | 0,4826 | 0,71901075 | ZBTB20 antisense RNA 5 |
| ENSG00000256742 | KDM2B-DT | 0,493763619 | 1,408113493 | 0,29766 | 0,59036937 | KDM2B divergent transcript |
| ENSG00000173409 | ARV1 | 0,493650813 | 1,408003396 | 0,24495 | 0,55288356 | ARV1 homolog, fatty acid homeostasis modulator |
| ENSG00000261202 | Z83847,1 | 0,49362116 | 1,407974455 | 0,3779 | 0,64628549 | novel transcript, antisense to FAM83F |
| ENSG00000274332 | LRRC37A2 | 0,493447534 | 1,407805019 | 0,31464 | 0,60128422 | leucine rich repeat containing 37 member A2 |
| ENSG00000147324 | MFHAS1 | 0,49339887 | 1,407757532 | 0,02037 | 0,18949014 | malignant fibrous histiocytoma amplified sequence 1 |
| ENSG00000213842 | SUGT1P2 | 0,493366962 | 1,407726397 | 0,24996 | 0,55657567 | SUGT1 pseudogene 2 |
| ENSG00000143578 | CREB3L4 | 0,493347529 | 1,407707436 | 0,14778 | 0,47507961 | cAMP responsive element binding protein 3 like 4 |
| ENSG00000111843 | TMEM14C | 0,493280172 | 1,407641714 | 0,18312 | 0,50652733 | transmembrane protein 14C |
| ENSG00000284936 | TMEM14C | 0,493280172 | 1,407641714 | 0,18312 | 0,50652733 | transmembrane protein 14C |
| ENSG00000117597 | UTP25 | 0,4932476 | 1,407609934 | 0,00651 | 0,10236887 | UTP25 small subunit processor component |
| ENSG00000079246 | XRCC5 | 0,493159546 | 1,407524023 | 0,22811 | 0,54221266 | X-ray repair cross complementing 5 |
| ENSG00000165512 | ZNF22 | 0,493153235 | 1,407517866 | 0,26846 | 0,57056686 | zinc finger protein 22 |
| ENSG00000148481 | MINDY3 | 0,493148124 | 1,40751288 | 0,23814 | 0,54870569 | MINDY lysine 48 deubiquitinase 3 |
| ENSG00000104388 | RAB2A | 0,493070251 | 1,407436908 | 0,30452 | 0,59430086 | RAB2A, member RAS oncogene family |
| ENSG00000168685 | IL7R | 0,49305286 | 1,407419942 | 0,52923 | 0,75112358 | interleukin 7 receptor |
| ENSG00000152464 | RPP38 | 0,493013432 | 1,407381478 | 0,15249 | 0,48061452 | ribonuclease P/MRP subunit p38 |
| ENSG00000236184 | TCEA1P4 | 0,49298878 | 1,40735743 | 0,32483 | 0,60735626 | transcription elongation factor A1 pseudogene 4 |
| ENSG00000119608 | PROX2 | 0,492926743 | 1,407296914 | 0,32161 | 0,60577115 | prospero homeobox 2 |
| ENSG00000155313 | USP25 | 0,492884792 | 1,407255993 | 0,17977 | 0,50407792 | ubiquitin specific peptidase 25 |
| ENSG00000272192 | AC100812,1 | 0,492873479 | 1,407244958 | 0,50495 | 0,73369183 | novel transcript |
| ENSG00000183726 | TMEM50A | 0,492859884 | 1,407231697 | 0,24161 | 0,55039378 | transmembrane protein 50A |
| ENSG00000171365 | CLCN5 | 0,492742957 | 1,407117649 | 0,01699 | 0,17104675 | chloride voltage-gated channel 5 |
| ENSG00000135914 | HTR2B | 0,492647333 | 1,407024386 | 0,32597 | 0,6085098 | 5-hydroxytryptamine receptor 2B |
| ENSG00000240661 | AC063952,1 | 0,492620022 | 1,406997751 | 0,13992 | 0,46689302 | novel V-set domain-containing pseudogene |
| ENSG00000117399 | CDC20 | 0,492615175 | 1,406993023 | 0,12058 | 0,43944057 | cell division cycle 20 |
| ENSG00000126524 | SBDS | 0,492434437 | 1,40681677 | 0,21181 | 0,53070055 | SBDS ribosome maturation factor |
| ENSG00000158290 | CUL4B | 0,492370374 | 1,406754301 | 0,29917 | 0,59122555 | cullin 4B |
| ENSG00000278540 | ACACA | 0,492269063 | 1,406655517 | 0,1352 | 0,46165567 | acetyl-CoA carboxylase alpha |

| | | | | | | |
|---|---|---|---|---|---|---|
| ENSG00000198128 | OR2L3 | 0,492143709 | 1,406533299 | 0,62296 | 0,80886176 | olfactory receptor family 2 subfamily L member 3 |
| ENSG00000132541 | RIDA | 0,492136496 | 1,406526268 | 0,43096 | 0,68325486 | reactive intermediate imine deaminase A homolog |
| ENSG00000075420 | FNDC3B | 0,492014786 | 1,406407614 | 0,06044 | 0,32647604 | fibronectin type III domain containing 3B |
| ENSG00000230793 | SMARCE1P5 | 0,491950927 | 1,406345362 | 0,40531 | 0,66502482 | SWI/SNF related, matrix associated, actin dependent regulator of chromatin, subfamily e, member 1 pseudogene 5 |
| ENSG00000278590 | RN7SL113P | 0,491947085 | 1,406341617 | 0,53072 | 0,75212511 | RNA, 7SL, cytoplasmic 113, pseudogene |
| ENSG00000272182 | AC135507,1 | 0,491906936 | 1,40630248 | 0,35223 | 0,62740642 | novel transcript, antisense to PDHB |
| ENSG00000181061 | HIGD1A | 0,491843948 | 1,406241083 | 0,26791 | 0,56999323 | HIG1 hypoxia inducible domain family member 1A |
| ENSG00000184226 | PCDH9 | 0,491730466 | 1,406130472 | 0,23775 | 0,54863436 | protocadherin 9 |
| ENSG00000287456 | AL157369,1 | 0,491683958 | 1,406085144 | 0,35535 | 0,62970025 | novel transcript, antisense to SUCLA2 |
| ENSG00000230042 | AK3P3 | 0,491627648 | 1,406030264 | 0,43812 | 0,68755048 | adenylate kinase 3 pseudogene 3 |
| ENSG00000183431 | SF3A3 | 0,491544074 | 1,405948817 | 0,04491 | 0,28641225 | splicing factor 3a subunit 3 |
| ENSG00000112561 | TFEB | 0,491512969 | 1,405918504 | 0,30955 | 0,59799482 | transcription factor EB |
| ENSG00000167447 | SMG8 | 0,491337527 | 1,405747544 | 0,06041 | 0,32637172 | SMG8 nonsense mediated mRNA decay factor |
| ENSG00000014824 | SLC30A9 | 0,491168553 | 1,405582908 | 0,23865 | 0,54890467 | solute carrier family 30 member 9 |
| ENSG00000077514 | POLD3 | 0,491155935 | 1,405570614 | 0,12834 | 0,45177856 | DNA polymerase delta 3, accessory subunit |
| ENSG00000262576 | PCDHGA4 | 0,491152726 | 1,405567488 | 0,18565 | 0,5090363 | protocadherin gamma subfamily A, 4 |
| ENSG00000114209 | PDCD10 | 0,491151696 | 1,405566485 | 0,23109 | 0,54397066 | programmed cell death 10 |
| ENSG00000155966 | AFF2 | 0,491130034 | 1,40554538 | 0,14002 | 0,46694389 | AF4/FMR2 family member 2 |
| ENSG00000164332 | UBLCP1 | 0,490965732 | 1,405385318 | 0,14178 | 0,46830089 | ubiquitin like domain containing CTD phosphatase 1 |
| ENSG00000204920 | ZNF155 | 0,49095907 | 1,405378828 | 0,24117 | 0,55024202 | zinc finger protein 155 |
| ENSG00000273148 | LINC00653 | 0,490872079 | 1,405294089 | 0,31196 | 0,59925736 | long intergenic non-protein coding RNA 653 |
| ENSG00000278603 | AC245033,3 | 0,490696392 | 1,405122968 | 0,61888 | 0,80669814 | dynamin 1 (DNM1) pseudogene |
| ENSG00000260475 | AL353719,1 | 0,490680216 | 1,405107213 | 0,45781 | 0,70096886 | novel transcript |
| ENSG00000171988 | JMJD1C | 0,490360263 | 1,404795631 | 0,24769 | 0,55509241 | jumonji domain containing 1C |
| ENSG00000094880 | CDC23 | 0,490323059 | 1,404759405 | 0,11517 | 0,43077792 | cell division cycle 23 |
| ENSG00000244730 | AC139451,1 | 0,490287245 | 1,404724533 | 0,42549 | 0,67907903 | PHD finger protein 5A (PHF5A) pseudogene |
| ENSG00000249741 | AC093890,1 | 0,490084821 | 1,40452745 | 0,54662 | 0,76230405 | salvador homolog 1 (Drosophila) (SAV1) pseudogene |
| ENSG00000184182 | UBE2F | 0,489998113 | 1,404443039 | 0,19597 | 0,51680811 | ubiquitin conjugating enzyme E2 F (putative) |
| ENSG00000179055 | OR13D1 | 0,489946866 | 1,404393152 | 0,68156 | 0,84375324 | olfactory receptor family 13 subfamily D member 1 |
| ENSG00000143614 | GATAD2B | 0,489936333 | 1,404382898 | 0,06518 | 0,33804854 | GATA zinc finger domain containing 2B |
| ENSG00000102316 | MAGED2 | 0,489812232 | 1,404262098 | 0,16659 | 0,49255702 | MAGE family member D2 |

| | | | | | | |
|---|---|---|---|---|---|---|
| ENSG00000170989 | S1PR1 | 0,489786232 | 1,404236791 | 0,32719 | 0,60902638 | sphingosine-1-phosphate receptor 1 |
| ENSG00000213585 | VDAC1 | 0,489703213 | 1,404155987 | 0,12204 | 0,44133251 | voltage dependent anion channel 1 |
| ENSG00000170677 | SOCS6 | 0,489635922 | 1,404090496 | 0,32221 | 0,60577115 | suppressor of cytokine signaling 6 |
| ENSG00000143756 | FBXO28 | 0,489605374 | 1,404060765 | 0,22743 | 0,54168591 | F-box protein 28 |
| ENSG00000006634 | DBF4 | 0,489392371 | 1,403853481 | 0,29775 | 0,59036937 | DBF4 zinc finger |
| ENSG00000273890 | AC034102,8 | 0,489345147 | 1,403807529 | 0,57138 | 0,77809338 | novel transcript, antisence to IKZF4 |
| ENSG00000233822 | H2BC15 | 0,489343586 | 1,403806011 | 0,09402 | 0,39609058 | H2B clustered histone 15 |
| ENSG00000224488 | SAMM50P1 | 0,489308134 | 1,403771515 | 0,48949 | 0,72394433 | SAMM50 sorting and assembly machinery component pseudogene 1 |
| ENSG00000198805 | PNP | 0,489213882 | 1,403679809 | 0,10908 | 0,42061552 | purine nucleoside phosphorylase |
| ENSG00000227056 | RPL6P2 | 0,489176901 | 1,403643828 | 0,52445 | 0,74828894 | ribosomal protein L6 pseudogene 2 |
| ENSG00000174429 | ABRA | 0,48914137 | 1,403609259 | 0,54616 | 0,76176608 | actin binding Rho activating protein |
| ENSG00000271161 | BOLA2P2 | 0,489132422 | 1,403600554 | 0,36605 | 0,63617598 | bolA family member 2 pseudogene 2 |
| ENSG00000172878 | METAP1D | 0,489090117 | 1,403559395 | 0,14752 | 0,47501835 | methionyl aminopeptidase type 1D, mitochondrial |
| ENSG00000233184 | AC093157,1 | 0,48906632 | 1,403536245 | 0,07051 | 0,35057542 | novel transcript |
| ENSG00000112078 | KCTD20 | 0,489063578 | 1,403533577 | 0,08359 | 0,37573324 | potassium channel tetramerization domain containing 20 |
| ENSG00000087301 | TXNDC16 | 0,489057123 | 1,403527298 | 0,20005 | 0,5209612 | thioredoxin domain containing 16 |
| ENSG00000246323 | AC113382,1 | 0,489056934 | 1,403527113 | 0,57933 | 0,78270914 | novel transcript, antisense to FAM13B |
| ENSG00000055130 | CUL1 | 0,488996181 | 1,403468011 | 0,22518 | 0,54058261 | cullin 1 |
| ENSG00000148248 | SURF4 | 0,488878606 | 1,403353637 | 0,0652 | 0,33804854 | surfeit 4 |
| ENSG00000280951 | SURF4 | 0,488878606 | 1,403353637 | 0,0652 | 0,33804854 | surfeit 4 |
| ENSG00000163818 | LZTFL1 | 0,488782129 | 1,403259794 | 0,26644 | 0,56917012 | leucine zipper transcription factor like 1 |
| ENSG00000162437 | RAVER2 | 0,488766811 | 1,403244895 | 0,25263 | 0,55796133 | ribonucleoprotein, PTB binding 2 |
| ENSG00000166260 | COX11 | 0,488755819 | 1,403234204 | 0,19856 | 0,51980353 | cytochrome c oxidase copper chaperone COX11 |
| ENSG00000137460 | FHDC1 | 0,48871981 | 1,40319918 | 0,20205 | 0,52230471 | FH2 domain containing 1 |
| ENSG00000185220 | PGBD2 | 0,488709925 | 1,403189566 | 0,33819 | 0,6177255 | piggyBac transposable element derived 2 |
| ENSG00000103274 | NUBP1 | 0,488575316 | 1,403058649 | 0,17885 | 0,50335172 | nucleotide binding protein 1 |
| ENSG00000025772 | TOMM34 | 0,488505887 | 1,402991129 | 0,13387 | 0,45992327 | translocase of outer mitochondrial membrane 34 |
| ENSG00000186468 | RPS23 | 0,488435654 | 1,40292283 | 0,20728 | 0,52674983 | ribosomal protein S23 |
| ENSG00000149925 | ALDOA | 0,488398616 | 1,402886815 | 0,04535 | 0,28773055 | aldolase, fructose-bisphosphate A |
| ENSG00000004866 | ST7 | 0,488322142 | 1,402812452 | 0,18044 | 0,50407792 | suppression of tumorigenicity 7 |
| ENSG00000264943 | SH3GL1P2 | 0,488301485 | 1,402792366 | 0,26641 | 0,56917012 | SH3 domain containing GRB2 like 1, endophilin A2 pseudogene 2 |

| | | | | | | |
|---|---|---|---|---|---|---|
| ENSG00000244687 | UBE2V1 | 0,4882914 | 1,40278256 | 0,13626 | 0,46338195 | ubiquitin conjugating enzyme E2 V1 |
| ENSG00000102678 | FGF9 | 0,488287411 | 1,402778682 | 0,1411 | 0,4679289 | fibroblast growth factor 9 |
| ENSG00000075303 | SLC25A40 | 0,48824901 | 1,402741344 | 0,18672 | 0,51019454 | solute carrier family 25 member 40 |
| ENSG00000249773 | AC092647,5 | 0,488219148 | 1,402712309 | 0,35315 | 0,6278392 | novel zinc finger protein 713 (ZNF713) and mitochondrial ribosomal protein S17 (MRPS17) protein |
| ENSG00000185798 | WDR53 | 0,488180691 | 1,402674919 | 0,10023 | 0,4067769 | WD repeat domain 53 |
| ENSG00000196812 | ZSCAN16 | 0,488168892 | 1,402663447 | 0,25607 | 0,56065266 | zinc finger and SCAN domain containing 16 |
| ENSG00000204713 | TRIM27 | 0,488161958 | 1,402656705 | 0,47504 | 0,71364583 | tripartite motif containing 27 |
| ENSG00000227460 | SYNGAP1 | 0,488024723 | 1,402523285 | 0,15234 | 0,48061452 | synaptic Ras GTPase activating protein 1 |
| ENSG00000133104 | SPART | 0,488007043 | 1,402506098 | 0,0764 | 0,36251343 | spartin |
| ENSG00000000003 | TSPAN6 | 0,487926089 | 1,402427401 | 0,18067 | 0,50417319 | tetraspanin 6 |
| ENSG00000161813 | LARP4 | 0,487904354 | 1,402406273 | 0,30055 | 0,5918686 | La ribonucleoprotein 4 |
| ENSG00000137265 | IRF4 | 0,487740613 | 1,402247113 | 0,51154 | 0,73860667 | interferon regulatory factor 4 |
| ENSG00000196437 | ZNF569 | 0,487724183 | 1,402231144 | 0,32164 | 0,60577115 | zinc finger protein 569 |
| ENSG00000085377 | PREP | 0,487672899 | 1,402181299 | 0,02257 | 0,20017681 | prolyl endopeptidase |
| ENSG00000273213 | H3-2 | 0,487624854 | 1,402134604 | 0,40743 | 0,66661764 | H3,2 histone (putative) |
| ENSG00000139637 | MYG1 | 0,487550996 | 1,402062825 | 0,0292 | 0,2303054 | MYG1 exonuclease |
| ENSG00000136628 | EPRS1 | 0,487490155 | 1,402003698 | 0,21961 | 0,53699318 | glutamyl-prolyl-tRNA synthetase 1 |
| ENSG00000163125 | RPRD2 | 0,487473509 | 1,401987522 | 0,15536 | 0,48334025 | regulation of nuclear pre-mRNA domain containing 2 |
| ENSG00000083544 | TDRD3 | 0,487456454 | 1,401970948 | 0,22311 | 0,53927144 | tudor domain containing 3 |
| ENSG00000262771 | SSBP1 | 0,48744475 | 1,401959574 | 0,09355 | 0,39523994 | single stranded DNA binding protein 1 |
| ENSG00000136152 | COG3 | 0,487442164 | 1,401957061 | 0,20397 | 0,52366556 | component of oligomeric golgi complex 3 |
| ENSG00000220517 | ASS1P1 | 0,487422987 | 1,401938427 | 0,68853 | 0,84821636 | argininosuccinate synthetase 1 pseudogene 1 |
| ENSG00000124613 | ZNF391 | 0,487416909 | 1,40193252 | 0,10195 | 0,40933045 | zinc finger protein 391 |
| ENSG00000261167 | AC107027,3 | 0,487395054 | 1,401911283 | 0,23163 | 0,54440987 | novel transcript, overlapping to MRPL3 |
| ENSG00000103035 | PSMD7 | 0,487356587 | 1,401873904 | 0,26964 | 0,57135887 | proteasome 26S subunit, non-ATPase 7 |
| ENSG00000135070 | ISCA1 | 0,487312817 | 1,401831373 | 0,21893 | 0,53639845 | iron-sulfur cluster assembly 1 |
| ENSG00000214114 | MYCBP | 0,487142289 | 1,401665685 | 0,20073 | 0,52160583 | MYC binding protein |
| ENSG00000174738 | NR1D2 | 0,486979486 | 1,401507521 | 0,16182 | 0,48914394 | nuclear receptor subfamily 1 group D member 2 |
| ENSG00000174780 | SRP72 | 0,4868182 | 1,401350848 | 0,25044 | 0,55681669 | signal recognition particle 72 |
| ENSG00000197149 | AC107956,1 | 0,486764122 | 1,40129832 | 0,61286 | 0,8035281 | ribosomal protein S2 (RPS2) pseudogene |
| ENSG00000250317 | SMIM20 | 0,486655575 | 1,401192892 | 0,36436 | 0,6349692 | small integral membrane protein 20 |
| ENSG00000162607 | USP1 | 0,486554725 | 1,401094947 | 0,18159 | 0,50490902 | ubiquitin specific peptidase 1 |

| | | | | | | |
|---|---|---|---|---|---|---|
| ENSG00000123349 | PFDN5 | 0,486511786 | 1,401053246 | 0,10814 | 0,41884147 | prefoldin subunit 5 |
| ENSG00000259216 | AC084757,1 | 0,486440625 | 1,400984141 | 0,55584 | 0,7681354 | oxidation resistance 1 (OXR1) pseudogene |
| ENSG00000116954 | RRAGC | 0,486423277 | 1,400967295 | 0,28888 | 0,58464073 | Ras related GTP binding C |
| ENSG00000154678 | PDE1C | 0,486308554 | 1,400855895 | 0,32647 | 0,60866572 | phosphodiesterase 1C |
| ENSG00000166478 | ZNF143 | 0,486262434 | 1,400811113 | 0,23314 | 0,54557342 | zinc finger protein 143 |
| ENSG00000137076 | TLN1 | 0,48624739 | 1,400796506 | 0,02229 | 0,19837076 | talin 1 |
| ENSG00000136457 | CHAD | 0,486240179 | 1,400789504 | 0,58469 | 0,78582649 | chondroadherin |
| ENSG00000215032 | GNL3LP1 | 0,486159543 | 1,400711213 | 0,20277 | 0,52269757 | G protein nucleolar 3 like pseudogene 1 |
| ENSG00000224430 | MKRN5P | 0,485970105 | 1,400527299 | 0,33476 | 0,61445802 | makorin ring finger protein 5, pseudogene |
| ENSG00000125863 | MKKS | 0,485958059 | 1,400515605 | 0,39269 | 0,65593262 | McKusick-Kaufman syndrome |
| ENSG00000106086 | PLEKHA8 | 0,485860906 | 1,400421296 | 0,17102 | 0,49641416 | pleckstrin homology domain containing A8 |
| ENSG00000175536 | LIPT2 | 0,48570035 | 1,400265453 | 0,1707 | 0,49641416 | lipoyl(octanoyl) transferase 2 |
| ENSG00000196455 | PIK3R4 | 0,485608932 | 1,400176727 | 0,20521 | 0,52486901 | phosphoinositide-3-kinase regulatory subunit 4 |
| ENSG00000188828 | GLRA4 | 0,485578725 | 1,40014741 | 0,20162 | 0,52214043 | glycine receptor alpha 4 (pseudogene) |
| ENSG00000086065 | CHMP5 | 0,48552468 | 1,400094961 | 0,15941 | 0,48684894 | charged multivesicular body protein 5 |
| ENSG00000178764 | ZHX2 | 0,485507531 | 1,400078317 | 0,13727 | 0,4642578 | zinc fingers and homeoboxes 2 |
| ENSG00000198522 | GPN1 | 0,485485908 | 1,400057333 | 0,21299 | 0,53201658 | GPN-loop GTPase 1 |
| ENSG00000251555 | AC096711,2 | 0,48543589 | 1,400008794 | 0,5311 | 0,75245598 | novel transcript |
| ENSG00000285646 | AL021155,4 | 0,485390657 | 1,3999649 | 0,03668 | 0,25960341 | novel transcript |
| ENSG00000188869 | TMC3 | 0,485278682 | 1,399856247 | 0,27601 | 0,57533851 | transmembrane channel like 3 |
| ENSG00000125743 | SNRPD2 | 0,485260064 | 1,399838181 | 0,26747 | 0,56973003 | small nuclear ribonucleoprotein D2 polypeptide |
| ENSG00000213068 | AL451050,1 | 0,48525654 | 1,399834762 | 0,67825 | 0,84162176 | ribosomal protein S17 (RPS17) pseudogene |
| ENSG00000276390 | AC004241,3 | 0,485229889 | 1,399808903 | 0,33812 | 0,6177255 | novel transcript, sense intronic to RPAP3 |
| ENSG00000141232 | TOB1 | 0,485171819 | 1,39975256 | 0,0825 | 0,37413032 | transducer of ERBB2, 1 |
| ENSG00000115419 | GLS | 0,484923966 | 1,399512106 | 0,11666 | 0,43361843 | glutaminase |
| ENSG00000165732 | DDX21 | 0,484885322 | 1,399474619 | 0,11372 | 0,42855874 | DExD-box helicase 21 |
| ENSG00000156273 | BACH1 | 0,484647531 | 1,39924397 | 0,22388 | 0,53971809 | BTB domain and CNC homolog 1 |
| ENSG00000234224 | TMEM229A | 0,484529221 | 1,399129228 | 0,26679 | 0,56923396 | transmembrane protein 229A |
| ENSG00000156299 | TIAM1 | 0,484504414 | 1,39910517 | 0,06887 | 0,34681313 | TIAM Rac1 associated GEF 1 |
| ENSG00000134532 | SOX5 | 0,484457057 | 1,399059245 | 0,15053 | 0,47863728 | SRY-box transcription factor 5 |
| ENSG00000108671 | PSMD11 | 0,484432933 | 1,399035851 | 0,07011 | 0,34975886 | proteasome 26S subunit, non-ATPase 11 |
| ENSG00000121486 | TRMT1L | 0,484417973 | 1,399021344 | 0,05071 | 0,30287257 | tRNA methyltransferase 1 like |
| ENSG00000248121 | SMURF2P1 | 0,484325921 | 1,398932082 | 0,40979 | 0,6680777 | SMAD specific E3 ubiquitin protein ligase 2 pseudogene 1 |

| | | | | | | |
|---|---|---|---|---|---|---|
| ENSG00000287806 | AL135784,1 | 0,484308262 | 1,398914959 | 0,57422 | 0,77939939 | novel transcript, antisense to IGSF1 |
| ENSG00000157500 | APPL1 | 0,484219514 | 1,398828906 | 0,2488 | 0,55591283 | adaptor protein, phosphotyrosine interacting with PH domain and leucine zipper 1 |
| ENSG00000111445 | RFC5 | 0,484210351 | 1,398820022 | 0,24193 | 0,55071239 | replication factor C subunit 5 |
| ENSG00000124214 | STAU1 | 0,484194411 | 1,398804566 | 0,09703 | 0,40143759 | staufen double-stranded RNA binding protein 1 |
| ENSG00000275805 | AC090206,1 | 0,484131489 | 1,39874356 | 0,33951 | 0,61852579 | novel transcript, sense intronic to KCTD1 |
| ENSG00000166716 | ZNF592 | 0,484118231 | 1,398730706 | 0,0618 | 0,32905359 | zinc finger protein 592 |
| ENSG00000101126 | ADNP | 0,484111922 | 1,398724589 | 0,23066 | 0,54367096 | activity dependent neuroprotector homeobox |
| ENSG00000156026 | MCU | 0,4841119 | 1,398724568 | 0,20558 | 0,52513499 | mitochondrial calcium uniporter |
| ENSG00000254999 | BRK1 | 0,484081447 | 1,398695043 | 0,24416 | 0,55253387 | BRICK1 subunit of SCAR/WAVE actin nucleating complex |
| ENSG00000196104 | SPOCK3 | 0,484022687 | 1,398638077 | 0,37131 | 0,64047773 | SPARC (osteonectin), cwcv and kazal like domains proteoglycan 3 |
| ENSG00000005700 | IBTK | 0,483958026 | 1,398575392 | 0,22488 | 0,5403718 | inhibitor of Bruton tyrosine kinase |
| ENSG00000116819 | TFAP2E | 0,483716795 | 1,398341557 | 0,29155 | 0,58655792 | transcription factor AP-2 epsilon |
| ENSG00000259326 | AC116158,1 | 0,483685638 | 1,398311359 | 0,22348 | 0,53953208 | novel transcript, antisense to RASGRP1 |
| ENSG00000282619 | AF186192,6 | 0,483676536 | 1,398302536 | 0,37117 | 0,64038503 | zinc finger protein 154 (ZNF154) pseudogene |
| ENSG00000255085 | AF186192,2 | 0,483676536 | 1,398302536 | 0,37117 | 0,64038503 | zinc finger protein 154 (ZNF154) pseudogene |
| ENSG00000124782 | RREB1 | 0,483637013 | 1,39826423 | 0,04802 | 0,29426648 | ras responsive element binding protein 1 |
| ENSG00000156471 | PTDSS1 | 0,483591939 | 1,398220546 | 0,18501 | 0,50863955 | phosphatidylserine synthase 1 |
| ENSG00000130177 | CDC16 | 0,483538967 | 1,398169207 | 0,22239 | 0,53859118 | cell division cycle 16 |
| ENSG00000075407 | ZNF37A | 0,483514541 | 1,398145535 | 0,11253 | 0,42637009 | zinc finger protein 37A |
| ENSG00000187240 | DYNC2H1 | 0,4835038 | 1,398135126 | 0,15067 | 0,47887343 | dynein cytoplasmic 2 heavy chain 1 |
| ENSG00000063587 | ZNF275 | 0,483498063 | 1,398129567 | 0,13577 | 0,462686 | zinc finger protein 275 |
| ENSG00000280475 | AC007537,2 | 0,483461938 | 1,398094557 | 0,5404 | 0,75829455 | ribosomal protein L21 (RPL21) pseudogene |
| ENSG00000242405 | AC007537,1 | 0,483461938 | 1,398094557 | 0,5404 | 0,75829455 | ribosomal protein L21 (RPL21) pseudogene |
| ENSG00000181652 | ATG9B | 0,483406044 | 1,398040392 | 0,12815 | 0,45171208 | autophagy related 9B |
| ENSG00000233460 | RPL35AP31 | 0,483381873 | 1,39801697 | 0,47775 | 0,71547332 | ribosomal protein L35a pseudogene 31 |
| ENSG00000108443 | RPS6KB1 | 0,483365886 | 1,398001478 | 0,23715 | 0,54813266 | ribosomal protein S6 kinase B1 |
| ENSG00000277566 | AC089999,2 | 0,483355567 | 1,397991478 | 0,55987 | 0,77074768 | novel transcript, sense intronic to TPCN1 |
| ENSG00000181544 | FANCB | 0,483334243 | 1,397970816 | 0,19801 | 0,51977271 | FA complementation group B |
| ENSG00000238207 | AC009312,1 | 0,483230837 | 1,397870619 | 0,45158 | 0,69662476 | novel transcript |
| ENSG00000196757 | ZNF700 | 0,483155444 | 1,39779757 | 0,16877 | 0,49492999 | zinc finger protein 700 |
| ENSG00000008086 | CDKL5 | 0,483127741 | 1,39777073 | 0,09335 | 0,39489949 | cyclin dependent kinase like 5 |

| | | | | | | |
|---|---|---|---|---|---|---|
| ENSG00000134108 | ARL8B | 0,483020949 | 1,397667266 | 0,18464 | 0,50835796 | ADP ribosylation factor like GTPase 8B |
| ENSG00000168301 | KCTD6 | 0,48279517 | 1,397448552 | 0,14155 | 0,46809977 | potassium channel tetramerization domain containing 6 |
| ENSG00000159217 | IGF2BP1 | 0,482741325 | 1,397396396 | 0,04007 | 0,27058857 | insulin like growth factor 2 mRNA binding protein 1 |
| ENSG00000235272 | RAMACL | 0,482726266 | 1,397381811 | 0,23737 | 0,54833864 | RNA guanine-7 methyltransferase activating subunit like (pseudogene) |
| ENSG00000271730 | AL390208,1 | 0,482711924 | 1,397367919 | 0,64256 | 0,82082347 | novel transcript, antisense to SESN1 |
| ENSG00000160801 | PTH1R | 0,482633766 | 1,397292219 | 0,29686 | 0,58996365 | parathyroid hormone 1 receptor |
| ENSG00000087076 | HSD17B14 | 0,482609819 | 1,397269025 | 0,18426 | 0,50802406 | hydroxysteroid 17-beta dehydrogenase 14 |
| ENSG00000273338 | AC103591,3 | 0,482479431 | 1,397142748 | 0,49152 | 0,72510994 | novel transcript, antisense to DNAJB4 |
| ENSG00000272181 | AC012557,2 | 0,482397884 | 1,397063779 | 0,66175 | 0,83154613 | novel transcript, antisense to PSMD6 |
| ENSG00000285317 | DNAJC8 | 0,482313451 | 1,396982018 | 0,18921 | 0,51249536 | DnaJ heat shock protein family (Hsp40) member C8 |
| ENSG00000126698 | DNAJC8 | 0,482313451 | 1,396982018 | 0,18921 | 0,51249536 | DnaJ heat shock protein family (Hsp40) member C8 |
| ENSG00000091140 | DLD | 0,482296419 | 1,396965526 | 0,22626 | 0,54121239 | dihydrolipoamide dehydrogenase |
| ENSG00000146386 | ABRACL | 0,482246327 | 1,396917023 | 0,2387 | 0,54892264 | ABRA C-terminal like |
| ENSG00000162852 | CNST | 0,482170514 | 1,396843617 | 0,22018 | 0,53750545 | consortin, connexin sorting protein |
| ENSG00000178741 | COX5A | 0,482102438 | 1,396777707 | 0,28771 | 0,58395529 | cytochrome c oxidase subunit 5A |
| ENSG00000137944 | KYAT3 | 0,482067785 | 1,396744157 | 0,14002 | 0,46694389 | kynurenine aminotransferase 3 |
| ENSG00000149474 | KAT14 | 0,482012069 | 1,396690216 | 0,07577 | 0,36145964 | lysine acetyltransferase 14 |
| ENSG00000161960 | EIF4A1 | 0,481938248 | 1,396618752 | 0,17433 | 0,49875953 | eukaryotic translation initiation factor 4A1 |
| ENSG00000147403 | RPL10 | 0,481933168 | 1,396613833 | 0,29013 | 0,58531219 | ribosomal protein L10 |
| ENSG00000244332 | AL138759,1 | 0,481916302 | 1,396597506 | 0,49071 | 0,72478928 | novel transcript |
| ENSG00000115484 | CCT4 | 0,48184435 | 1,396527855 | 0,18621 | 0,50957623 | chaperonin containing TCP1 subunit 4 |
| ENSG00000112031 | MTRF1L | 0,481822821 | 1,396507016 | 0,23547 | 0,54683669 | mitochondrial translational release factor 1 like |
| ENSG00000198042 | MAK16 | 0,481661402 | 1,396350773 | 0,09939 | 0,40593997 | MAK16 homolog |
| ENSG00000136197 | C7orf25 | 0,481643423 | 1,396333371 | 0,11487 | 0,43031399 | chromosome 7 open reading frame 25 |
| ENSG00000047597 | XK | 0,481635257 | 1,396325468 | 0,38431 | 0,65052153 | X-linked Kx blood group |
| ENSG00000167005 | NUDT21 | 0,481540601 | 1,396233857 | 0,21664 | 0,53492477 | nudix hydrolase 21 |
| ENSG00000215417 | MIR17HG | 0,48152325 | 1,396217065 | 0,07262 | 0,35556123 | miR-17-92a-1 cluster host gene |
| ENSG00000197782 | ZNF780A | 0,481483592 | 1,396178685 | 0,27192 | 0,57305058 | zinc finger protein 780A |
| ENSG00000280568 | ZNF780A | 0,481483592 | 1,396178685 | 0,27192 | 0,57305058 | zinc finger protein 780A |
| ENSG00000281380 | AC244635,2 | 0,481479952 | 1,396175163 | 0,65135 | 0,82534198 | NLR family, apoptosis inhibitory protein pseudogene |
| ENSG00000179978 | NAIPP2 | 0,481479952 | 1,396175163 | 0,65135 | 0,82534198 | NAIP pseudogene 2 |
| ENSG00000075336 | TIMM21 | 0,481413089 | 1,396110458 | 0,17261 | 0,49744497 | translocase of inner mitochondrial membrane 21 |

| | | | | | | |
|---|---|---|---|---|---|---|
| ENSG00000100281 | HMGXB4 | 0,481406701 | 1,396104276 | 0,16713 | 0,4931666 | HMG-box containing 4 |
| ENSG00000198680 | TUSC1 | 0,481357308 | 1,396056479 | 0,20725 | 0,52674983 | tumor suppressor candidate 1 |
| ENSG00000267655 | AC125437,1 | 0,481208751 | 1,395912732 | 0,30357 | 0,59380161 | novel transcript, sense intronic to ATP9B |
| ENSG00000001617 | SEMA3F | 0,481152773 | 1,39585857 | 0,13111 | 0,45585953 | semaphorin 3F |
| ENSG00000276497 | APBA1 | 0,481128853 | 1,395835426 | 0,13418 | 0,46036768 | amyloid beta precursor protein binding family A member 1 |
| ENSG00000101146 | RAE1 | 0,481102745 | 1,395810167 | 0,16761 | 0,49386723 | ribonucleic acid export 1 |
| ENSG00000153250 | RBMS1 | 0,481056008 | 1,395764949 | 0,12416 | 0,44504227 | RNA binding motif single stranded interacting protein 1 |
| ENSG00000275333 | FAM120B | 0,480946899 | 1,395659393 | 0,22927 | 0,54273374 | family with sequence similarity 120B |
| ENSG00000112584 | FAM120B | 0,480946899 | 1,395659393 | 0,22927 | 0,54273374 | family with sequence similarity 120B |
| ENSG00000184178 | SCFD2 | 0,480625322 | 1,395348336 | 0,09102 | 0,39033853 | sec1 family domain containing 2 |
| ENSG00000256538 | AC046130,2 | 0,480565716 | 1,395290686 | 0,49702 | 0,72862056 | novel transcript |
| ENSG00000165376 | CLDN2 | 0,480454258 | 1,395182895 | 0,30112 | 0,59252799 | claudin 2 |
| ENSG00000151503 | NCAPD3 | 0,480441032 | 1,395170105 | 0,12852 | 0,45191041 | non-SMC condensin II complex subunit D3 |
| ENSG00000236492 | DDR1-DT | 0,480338138 | 1,395070604 | 0,84076 | 0,92746594 | DDR1 divergent transcript |
| ENSG00000135387 | CAPRIN1 | 0,480298859 | 1,395032622 | 0,21744 | 0,53536715 | cell cycle associated protein 1 |
| ENSG00000178163 | ZNF518B | 0,480289706 | 1,395023771 | 0,2084 | 0,52769715 | zinc finger protein 518B |
| ENSG00000156931 | VPS8 | 0,480262124 | 1,394997101 | 0,22239 | 0,53859118 | VPS8 subunit of CORVET complex |
| ENSG00000181789 | COPG1 | 0,480201106 | 1,394938101 | 0,03826 | 0,26488494 | COPI coat complex subunit gamma 1 |
| ENSG00000136144 | RCBTB1 | 0,480151923 | 1,394890548 | 0,08861 | 0,38491339 | RCC1 and BTB domain containing protein 1 |
| ENSG00000128886 | ELL3 | 0,480084203 | 1,394825073 | 0,36065 | 0,63299251 | elongation factor for RNA polymerase II 3 |
| ENSG00000115041 | KCNIP3 | 0,480066387 | 1,394807849 | 0,20648 | 0,52609885 | potassium voltage-gated channel interacting protein 3 |
| ENSG00000240210 | AL122013,1 | 0,480040573 | 1,394782891 | 0,57877 | 0,78253224 | ribosomal protein L7a (RPL7A) pseudogene |
| ENSG00000112335 | SNX3 | 0,480039746 | 1,394782092 | 0,23931 | 0,5491566 | sorting nexin 3 |
| ENSG00000153046 | CDYL | 0,480021264 | 1,394764223 | 0,1122 | 0,42578355 | chromodomain Y like |
| ENSG00000165660 | ABRAXAS2 | 0,479950412 | 1,394695727 | 0,26672 | 0,56923396 | abraxas 2, BRISC complex subunit |
| ENSG00000288617 | AL049777,1 | 0,479947323 | 1,394692741 | 0,43852 | 0,6877408 | FOXG1-C14orf23 readthrough |
| ENSG00000004897 | CDC27 | 0,479750044 | 1,394502039 | 0,34458 | 0,62200601 | cell division cycle 27 |
| ENSG00000158955 | WNT9B | 0,479530051 | 1,394289411 | 0,58249 | 0,7847455 | Wnt family member 9B |
| ENSG00000005075 | POLR2J | 0,479481236 | 1,394242235 | 0,35153 | 0,62674766 | RNA polymerase II subunit J |
| ENSG00000260780 | AC087463,1 | 0,479447103 | 1,394209248 | 0,51081 | 0,73828855 | novel transcript |
| ENSG00000256546 | AC156455,1 | 0,479361848 | 1,394126861 | 0,21845 | 0,53578053 | novel transcript |
| ENSG00000257489 | AC010203,1 | 0,479354789 | 1,39412004 | 0,27589 | 0,57533851 | TBC1 domain family, member 2B (TBC1D2B) pseudogene |
| ENSG00000265474 | AC010761,4 | 0,479226876 | 1,39399644 | 0,41913 | 0,67514348 | novel transcript, antisense to TRAF4 |

| | | | | | | |
|---|---|---|---|---|---|---|
| ENSG00000196449 | YRDC | 0,479176215 | 1,393947489 | 0,06084 | 0,32750733 | yrdC N6-threonylcarbamoyltransferase domain containing |
| ENSG00000282899 | E2F2 | 0,478705349 | 1,393492607 | 0,16328 | 0,48972962 | E2F transcription factor 2 |
| ENSG00000156239 | N6AMT1 | 0,47864185 | 1,393431275 | 0,27075 | 0,57239492 | N-6 adenine-specific DNA methyltransferase 1 |
| ENSG00000182712 | CMC4 | 0,478632939 | 1,393422669 | 0,18627 | 0,50961491 | C-X9-C motif containing 4 |
| ENSG00000118482 | PHF3 | 0,478577019 | 1,39336866 | 0,14862 | 0,47609532 | PHD finger protein 3 |
| ENSG00000260930 | LINC01416 | 0,47851191 | 1,393305778 | 0,56103 | 0,77125979 | long intergenic non-protein coding RNA 1416 |
| ENSG00000196659 | TTC30B | 0,47844113 | 1,393237423 | 0,28176 | 0,57909527 | tetratricopeptide repeat domain 30B |
| ENSG00000283156 | AC068620,3 | 0,478373631 | 1,39317224 | 0,12619 | 0,44887364 | novel Myb/SANT-like DNA-binding domain containing pseudogene |
| ENSG00000168566 | SNRNP48 | 0,478368445 | 1,393167232 | 0,14312 | 0,47072472 | small nuclear ribonucleoprotein U11/U12 subunit 48 |
| ENSG00000196074 | SYCP2 | 0,478323022 | 1,393123369 | 0,38407 | 0,65052153 | synaptonemal complex protein 2 |
| ENSG00000234817 | ECI2-DT | 0,478297673 | 1,393098891 | 0,67974 | 0,84268708 | ECI2 divergent transcript |
| ENSG00000165410 | CFL2 | 0,478279074 | 1,393080931 | 0,36148 | 0,63347405 | cofilin 2 |
| ENSG00000129128 | SPCS3 | 0,478254515 | 1,393057217 | 0,10992 | 0,42220696 | signal peptidase complex subunit 3 |
| ENSG00000178127 | NDUFV2 | 0,478183945 | 1,392989077 | 0,26041 | 0,56383537 | NADH:ubiquinone oxidoreductase core subunit V2 |
| ENSG00000157212 | PAXIP1 | 0,478167648 | 1,392973342 | 0,05832 | 0,32231869 | PAX interacting protein 1 |
| ENSG00000232613 | LINC02576 | 0,478162354 | 1,39296823 | 0,5842 | 0,78572744 | long intergenic non-protein coding RNA 2576 |
| ENSG00000065150 | IPO5 | 0,478075525 | 1,392884396 | 0,24996 | 0,55657567 | importin 5 |
| ENSG00000275413 | AC002553,2 | 0,478020003 | 1,392830792 | 0,49712 | 0,72865531 | novel transcript, antisense to TTC19 |
| ENSG00000255893 | AP000786,1 | 0,477857488 | 1,392673903 | 0,30881 | 0,59767335 | novel transcript |
| ENSG00000165119 | HNRNPK | 0,477784665 | 1,392603607 | 0,21366 | 0,53275614 | heterogeneous nuclear ribonucleoprotein K |
| ENSG00000113387 | SUB1 | 0,477773676 | 1,392593 | 0,28264 | 0,58004357 | SUB1 regulator of transcription |
| ENSG00000147669 | POLR2K | 0,477767242 | 1,392586789 | 0,30066 | 0,59200188 | RNA polymerase II, I and III subunit K |
| ENSG00000282207 | AC254944,1 | 0,47775758 | 1,392577462 | 0,50172 | 0,73146914 | novel transcript, antisense to AC026150,11 |
| ENSG00000270016 | AC026150,3 | 0,47775758 | 1,392577462 | 0,50172 | 0,73146914 | novel transcript, antisense to AC026150,11 |
| ENSG00000122643 | NT5C3A | 0,477750531 | 1,392570658 | 0,28908 | 0,58469988 | 5'-nucleotidase, cytosolic IIIA |
| ENSG00000026950 | BTN3A1 | 0,477746426 | 1,392566696 | 0,18255 | 0,5059851 | butyrophilin subfamily 3 member A1 |
| ENSG00000207501 | RNVU1-14 | 0,477597745 | 1,392423189 | 0,48737 | 0,72237072 | RNA, variant U1 small nuclear 14 |
| ENSG00000235036 | AL035456,1 | 0,477582267 | 1,392408249 | 0,22467 | 0,54036191 | pseudogene similar to DNAJ protein |
| ENSG00000198677 | TTC37 | 0,477572385 | 1,392398712 | 0,27061 | 0,5723494 | tetratricopeptide repeat domain 37 |
| ENSG00000275854 | AC084824,4 | 0,47734532 | 1,392179581 | 0,41353 | 0,67078864 | novel transcript, sense intronic to YARS2 |
| ENSG00000204291 | COL15A1 | 0,477316531 | 1,3921518 | 0,14688 | 0,47471707 | collagen type XV alpha 1 chain |
| ENSG00000184260 | H2AC20 | 0,47730246 | 1,392138222 | 0,30849 | 0,5973408 | H2A clustered histone 20 |

| | | | | | | |
|---|---|---|---|---|---|---|
| ENSG00000004975 | DVL2 | 0,477211327 | 1,392050286 | 0,11748 | 0,43477245 | dishevelled segment polarity protein 2 |
| ENSG00000267711 | AC060766,6 | 0,477200335 | 1,392039679 | 0,54482 | 0,76084582 | novel transcript |
| ENSG00000138303 | ASCC1 | 0,477182047 | 1,392022033 | 0,06458 | 0,33679627 | activating signal cointegrator 1 complex subunit 1 |
| ENSG00000196139 | AKR1C3 | 0,47709225 | 1,391935393 | 0,51833 | 0,74319811 | aldo-keto reductase family 1 member C3 |
| ENSG00000256235 | SMIM3 | 0,477083173 | 1,391926636 | 0,135 | 0,46159318 | small integral membrane protein 3 |
| ENSG00000151623 | NR3C2 | 0,477031919 | 1,391877186 | 0,29314 | 0,58745839 | nuclear receptor subfamily 3 group C member 2 |
| ENSG00000142168 | SOD1 | 0,477028606 | 1,39187399 | 0,24878 | 0,55591283 | superoxide dismutase 1 |
| ENSG00000085511 | MAP3K4 | 0,476846939 | 1,391698733 | 0,12268 | 0,44246664 | mitogen-activated protein kinase kinase kinase 4 |
| ENSG00000213917 | RPL5P8 | 0,476686678 | 1,391544146 | 0,34617 | 0,62332894 | ribosomal protein L5 pseudogene 8 |
| ENSG00000138115 | CYP2C8 | 0,476649875 | 1,391508648 | 0,18053 | 0,50407792 | cytochrome P450 family 2 subfamily C member 8 |
| ENSG00000181817 | LSM10 | 0,476636746 | 1,391495985 | 0,26612 | 0,56908449 | LSM10, U7 small nuclear RNA associated |
| ENSG00000155099 | PIP4P2 | 0,476630692 | 1,391490146 | 0,4621 | 0,70341975 | phosphatidylinositol-4,5-bisphosphate 4-phosphatase 2 |
| ENSG00000151465 | CDC123 | 0,476612468 | 1,391472569 | 0,1246 | 0,4456051 | cell division cycle 123 |
| ENSG00000235859 | AC006978,1 | 0,476493651 | 1,391357975 | 0,23529 | 0,54677606 | proteasome (prosome, macropain) 26S subunit, ATPase, 5 (PSMC5) pseudogene |
| ENSG00000165288 | BRWD3 | 0,476432767 | 1,391299259 | 0,15742 | 0,48457045 | bromodomain and WD repeat domain containing 3 |
| ENSG00000268584 | AC073389,1 | 0,476424024 | 1,391290827 | 0,20579 | 0,52520068 | novel transcript, antisense to USP54 |
| ENSG00000149262 | INTS4 | 0,476351703 | 1,391221085 | 0,23781 | 0,54864017 | integrator complex subunit 4 |
| ENSG00000066027 | PPP2R5A | 0,476312466 | 1,391183248 | 0,02899 | 0,22977124 | protein phosphatase 2 regulatory subunit B'alpha |
| ENSG00000025156 | HSF2 | 0,476298719 | 1,391169992 | 0,29991 | 0,59160746 | heat shock transcription factor 2 |
| ENSG00000164111 | ANXA5 | 0,476292952 | 1,391164431 | 0,0861 | 0,3803347 | annexin A5 |
| ENSG00000164841 | TMEM74 | 0,476148857 | 1,39102549 | 0,44222 | 0,69006952 | transmembrane protein 74 |
| ENSG00000213109 | AL365223,1 | 0,476126224 | 1,391003667 | 0,3619 | 0,63352908 | ribosomal protein L7a (RPL7A) pseudogene |
| ENSG00000163611 | SPICE1 | 0,476123059 | 1,391000617 | 0,29888 | 0,59104323 | spindle and centriole associated protein 1 |
| ENSG00000129219 | PLD2 | 0,476105985 | 1,390984154 | 0,01748 | 0,17422253 | phospholipase D2 |
| ENSG00000256043 | CTSO | 0,476093266 | 1,390971891 | 0,66519 | 0,83337994 | cathepsin O |
| ENSG00000198312 | BMS1P9 | 0,476067591 | 1,390947137 | 0,30582 | 0,59512079 | BMS1 pseudogene 9 |
| ENSG00000275145 | FRG1 | 0,475998665 | 1,390880685 | 0,23653 | 0,54760364 | FSHD region gene 1 |
| ENSG00000216802 | AL590617,1 | 0,475780343 | 1,390670219 | 0,44813 | 0,69411284 | mitochondrial carrier homolog 1 (MTCH1) pseudogene |
| ENSG00000169083 | AR | 0,475745578 | 1,390636709 | 0,08642 | 0,38089807 | androgen receptor |
| ENSG00000083635 | NUFIP1 | 0,475737474 | 1,390628897 | 0,19808 | 0,51977271 | nuclear FMR1 interacting protein 1 |
| ENSG00000203740 | METTL11B | 0,475704749 | 1,390597354 | 0,53751 | 0,75614307 | methyltransferase like 11B |
| ENSG00000280831 | RPS25 | 0,475678753 | 1,390572296 | 0,27197 | 0,573058 | ribosomal protein S25 |

| | | | | | | |
|---|---|---|---|---|---|---|
| ENSG00000118181 | RPS25 | 0,475678753 | 1,390572296 | 0,27197 | 0,573058 | ribosomal protein S25 |
| ENSG00000091164 | TXNL1 | 0,475645389 | 1,390540138 | 0,19911 | 0,52017864 | thioredoxin like 1 |
| ENSG00000245598 | DACT3-AS1 | 0,475595687 | 1,390492234 | 0,50791 | 0,73602279 | DACT3 antisense RNA 1 |
| ENSG00000225774 | SIRPAP1 | 0,47550735 | 1,390407096 | 0,41964 | 0,67535925 | signal regulatory protein alpha pseudogene 1 |
| ENSG00000056050 | HPF1 | 0,475487303 | 1,390387775 | 0,39543 | 0,6582534 | histone PARylation factor 1 |
| ENSG00000165487 | MICU2 | 0,475468234 | 1,390369398 | 0,22898 | 0,54262488 | mitochondrial calcium uptake 2 |
| ENSG00000217165 | ANKRD18EP | 0,475343931 | 1,390249609 | 0,50047 | 0,73086101 | ankyrin repeat domain 18E, pseudogene |
| ENSG00000144959 | NCEH1 | 0,475237612 | 1,390147158 | 0,29775 | 0,59036937 | neutral cholesterol ester hydrolase 1 |
| ENSG00000112365 | ZBTB24 | 0,474947677 | 1,389867812 | 0,05409 | 0,31168784 | zinc finger and BTB domain containing 24 |
| ENSG00000253516 | HMGB1P41 | 0,474816336 | 1,389741286 | 0,41839 | 0,67471324 | high mobility group box 1 pseudogene 41 |
| ENSG00000188092 | GPR89B | 0,474806488 | 1,389731799 | 0,18774 | 0,51107563 | G protein-coupled receptor 89B |
| ENSG00000116984 | MTR | 0,474805265 | 1,389730621 | 0,0532 | 0,30971589 | 5-methyltetrahydrofolate-homocysteine methyltransferase |
| ENSG00000108518 | PFN1 | 0,474645785 | 1,389577005 | 0,09249 | 0,39312083 | profilin 1 |
| ENSG00000248098 | BCKDHA | 0,474607346 | 1,389539982 | 0,26828 | 0,5704222 | branched chain keto acid dehydrogenase E1 subunit alpha |
| ENSG00000120647 | CCDC77 | 0,474568036 | 1,38950212 | 0,10309 | 0,41078404 | coiled-coil domain containing 77 |
| ENSG00000115904 | SOS1 | 0,474516712 | 1,389452689 | 0,23886 | 0,54892264 | SOS Ras/Rac guanine nucleotide exchange factor 1 |
| ENSG00000237361 | TUSC8 | 0,474448364 | 1,389386866 | 0,45601 | 0,69996535 | tumor suppressor candidate 8 |
| ENSG00000123485 | HJURP | 0,47441498 | 1,389354715 | 0,16072 | 0,48807078 | Holliday junction recognition protein |
| ENSG00000287921 | AL713868,1 | 0,474411455 | 1,389351321 | 0,80238 | 0,90900521 | novel transcript, antisense to SLC35F3 |
| ENSG00000101282 | RSPO4 | 0,474394638 | 1,389335126 | 0,48882 | 0,72350189 | R-spondin 4 |
| ENSG00000171103 | TRMT61B | 0,474308981 | 1,38925264 | 0,20569 | 0,52518691 | tRNA methyltransferase 61B |
| ENSG00000180332 | KCTD4 | 0,47430572 | 1,389249499 | 0,3451 | 0,62231657 | potassium channel tetramerization domain containing 4 |
| ENSG00000164167 | LSM6 | 0,474295329 | 1,389239493 | 0,15703 | 0,48438698 | LSM6 homolog, U6 small nuclear RNA and mRNA degradation associated |
| ENSG00000164654 | MIOS | 0,474160799 | 1,389109954 | 0,15982 | 0,48710442 | meiosis regulator for oocyte development |
| ENSG00000144843 | ADPRH | 0,474155091 | 1,389104458 | 0,21096 | 0,52978958 | ADP-ribosylarginine hydrolase |
| ENSG00000242419 | PCDHGC4 | 0,474129792 | 1,389080099 | 0,09127 | 0,39075461 | protocadherin gamma subfamily C, 4 |
| ENSG00000158773 | USF1 | 0,474112288 | 1,389063245 | 0,23539 | 0,54683669 | upstream transcription factor 1 |
| ENSG00000184634 | MED12 | 0,474069278 | 1,389021835 | 0,15302 | 0,48120112 | mediator complex subunit 12 |
| ENSG00000234028 | EIF2AK3-DT | 0,474065545 | 1,38901824 | 0,21178 | 0,53069991 | EIF2AK3 divergent transcript |
| ENSG00000140941 | MAP1LC3B | 0,474012589 | 1,388967256 | 0,22426 | 0,54011527 | microtubule associated protein 1 light chain 3 beta |
| ENSG00000088451 | TGDS | 0,473812337 | 1,388774475 | 0,35487 | 0,62936991 | TDP-glucose 4,6-dehydratase |

| | | | | | | |
|---|---|---|---|---|---|---|
| ENSG00000165629 | ATP5F1C | 0,473759124 | 1,388723252 | 0,24797 | 0,55537155 | ATP synthase F1 subunit gamma |
| ENSG00000182551 | ADI1 | 0,473645545 | 1,388613926 | 0,04013 | 0,27065903 | acireductone dioxygenase 1 |
| ENSG00000072041 | SLC6A15 | 0,473615134 | 1,388584655 | 0,3193 | 0,60429866 | solute carrier family 6 member 15 |
| ENSG00000253910 | PCDHGB2 | 0,473505584 | 1,388479218 | 0,38841 | 0,6529693 | protocadherin gamma subfamily B, 2 |
| ENSG00000229539 | PANK2-AS1 | 0,473499074 | 1,388472952 | 0,42221 | 0,67663014 | PANK2 antisense RNA 1 |
| ENSG00000168411 | RFWD3 | 0,473465519 | 1,388440659 | 0,05214 | 0,30680362 | ring finger and WD repeat domain 3 |
| ENSG00000173093 | CCDC63 | 0,473427941 | 1,388404495 | 0,66838 | 0,83536064 | coiled-coil domain containing 63 |
| ENSG00000123066 | MED13L | 0,473401531 | 1,388379079 | 0,00758 | 0,11039804 | mediator complex subunit 13L |
| ENSG00000136240 | KDELR2 | 0,473315402 | 1,388296195 | 0,19141 | 0,51435195 | KDEL endoplasmic reticulum protein retention receptor 2 |
| ENSG00000088205 | DDX18 | 0,47326912 | 1,388251659 | 0,21574 | 0,53440695 | DEAD-box helicase 18 |
| ENSG00000287978 | AC245407,2 | 0,473179726 | 1,388165641 | 0,16995 | 0,49606609 | novel transcript |
| ENSG00000070756 | PABPC1 | 0,473120906 | 1,388109046 | 0,06584 | 0,33965348 | poly(A) binding protein cytoplasmic 1 |
| ENSG00000152382 | TADA1 | 0,473113879 | 1,388102285 | 0,29787 | 0,59037822 | transcriptional adaptor 1 |
| ENSG00000259863 | SH3RF3-AS1 | 0,472994363 | 1,387987296 | 0,52704 | 0,75012908 | SH3RF3 antisense RNA 1 |
| ENSG00000243789 | JMJD7 | 0,472992553 | 1,387985554 | 0,35443 | 0,62908147 | jumonji domain containing 7 |
| ENSG00000166226 | CCT2 | 0,472909197 | 1,387905362 | 0,12982 | 0,4539604 | chaperonin containing TCP1 subunit 2 |
| ENSG00000087053 | MTMR2 | 0,472868359 | 1,387866075 | 0,17678 | 0,50110131 | myotubularin related protein 2 |
| ENSG00000213780 | GTF2H4 | 0,472864743 | 1,387862596 | 0,63929 | 0,81902626 | general transcription factor IIH subunit 4 |
| ENSG00000110975 | SYT10 | 0,472823742 | 1,387823154 | 0,26098 | 0,56410815 | synaptotagmin 10 |
| ENSG00000163918 | RFC4 | 0,472738457 | 1,387741115 | 0,21381 | 0,53287453 | replication factor C subunit 4 |
| ENSG00000189079 | ARID2 | 0,472699702 | 1,387703837 | 0,23212 | 0,54496653 | AT-rich interaction domain 2 |
| ENSG00000162526 | TSSK3 | 0,472666941 | 1,387672325 | 0,26465 | 0,56739986 | testis specific serine kinase 3 |
| ENSG00000188343 | CIBAR1 | 0,472658255 | 1,387663971 | 0,33438 | 0,6142045 | CBY1 interacting BAR domain containing 1 |
| ENSG00000168374 | ARF4 | 0,472655374 | 1,3876612 | 0,12432 | 0,44533764 | ADP ribosylation factor 4 |
| ENSG00000215251 | FASTKD5 | 0,472640679 | 1,387647065 | 0,29889 | 0,59104323 | FAST kinase domains 5 |
| ENSG00000105819 | PMPCB | 0,472527239 | 1,387537958 | 0,2615 | 0,56452671 | peptidase, mitochondrial processing subunit beta |
| ENSG00000198246 | SLC29A3 | 0,472523199 | 1,387534072 | 0,12684 | 0,44957921 | solute carrier family 29 member 3 |
| ENSG00000116133 | DHCR24 | 0,472404967 | 1,387420366 | 0,05018 | 0,30107555 | 24-dehydrocholesterol reductase |
| ENSG00000166845 | C18orf54 | 0,472376361 | 1,387392856 | 0,34087 | 0,61918628 | chromosome 18 open reading frame 54 |
| ENSG00000198915 | RASGEF1A | 0,472369177 | 1,387385948 | 0,38744 | 0,6523395 | RasGEF domain family member 1A |
| ENSG00000224660 | SH3BP5-AS1 | 0,472310444 | 1,387329467 | 0,26644 | 0,56917012 | SH3BP5 antisense RNA 1 |
| ENSG00000117266 | CDK18 | 0,472291098 | 1,387310863 | 0,34007 | 0,61888023 | cyclin dependent kinase 18 |
| ENSG00000188163 | FAM166A | 0,472285815 | 1,387305784 | 0,43832 | 0,68766712 | family with sequence similarity 166 member A |

| | | | | | | |
|---|---|---|---|---|---|---|
| ENSG00000259528 | AC068397,1 | 0,472107594 | 1,387134415 | 0,55384 | 0,76684904 | novel transcript, antisense to NEO1 |
| ENSG00000204837 | FGF7P3 | 0,472070194 | 1,387098457 | 0,4899 | 0,72424167 | fibroblast growth factor 7 pseudogene 3 |
| ENSG00000166889 | PATL1 | 0,472058465 | 1,387087179 | 0,14215 | 0,46897057 | PAT1 homolog 1, processing body mRNA decay factor |
| ENSG00000059573 | ALDH18A1 | 0,472045449 | 1,387074665 | 0,12164 | 0,44093327 | aldehyde dehydrogenase 18 family member A1 |
| ENSG00000135018 | UBQLN1 | 0,472034365 | 1,387064008 | 0,18037 | 0,50407792 | ubiquilin 1 |
| ENSG00000114439 | BBX | 0,472018338 | 1,3870486 | 0,20086 | 0,52178339 | BBX high mobility group box domain containing |
| ENSG00000086589 | RBM22 | 0,472004577 | 1,38703537 | 0,19024 | 0,51331106 | RNA binding motif protein 22 |
| ENSG00000145354 | CISD2 | 0,471950393 | 1,386983277 | 0,28034 | 0,57784691 | CDGSH iron sulfur domain 2 |
| ENSG00000182575 | NXPH3 | 0,471920094 | 1,386954148 | 0,25914 | 0,56294411 | neurexophilin 3 |
| ENSG00000165322 | ARHGAP12 | 0,471542253 | 1,386590954 | 0,24038 | 0,54950872 | Rho GTPase activating protein 12 |
| ENSG00000002587 | HS3ST1 | 0,47129725 | 1,386355498 | 0,06725 | 0,34325886 | heparan sulfate-glucosamine 3-sulfotransferase 1 |
| ENSG00000107731 | UNC5B | 0,471278341 | 1,386337327 | 0,1477 | 0,47507961 | unc-5 netrin receptor B |
| ENSG00000187889 | FYB2 | 0,471244083 | 1,386304408 | 0,29926 | 0,59123205 | FYN binding protein 2 |
| ENSG00000203644 | AC083799,1 | 0,471211524 | 1,386273122 | 0,40413 | 0,66426144 | novel transcript, sense intronic to TMCC1 |
| ENSG00000244389 | RN7SL242P | 0,471192762 | 1,386255094 | 0,47918 | 0,71679575 | RNA, 7SL, cytoplasmic 242, pseudogene |
| ENSG00000157426 | AASDH | 0,471168465 | 1,386231748 | 0,25543 | 0,55992862 | aminoadipate-semialdehyde dehydrogenase |
| ENSG00000100353 | EIF3D | 0,471136096 | 1,386200647 | 0,17909 | 0,50351388 | eukaryotic translation initiation factor 3 subunit D |
| ENSG00000115750 | TAF1B | 0,471093935 | 1,386160136 | 0,29249 | 0,58699586 | TATA-box binding protein associated factor, RNA polymerase I subunit B |
| ENSG00000126787 | DLGAP5 | 0,471066918 | 1,386134179 | 0,30257 | 0,59306846 | DLG associated protein 5 |
| ENSG00000126705 | AHDC1 | 0,471060921 | 1,386128417 | 0,08491 | 0,3782577 | AT-hook DNA binding motif containing 1 |
| ENSG00000130856 | ZNF236 | 0,471012482 | 1,386081878 | 0,25591 | 0,56048171 | zinc finger protein 236 |
| ENSG00000168264 | IRF2BP2 | 0,470884861 | 1,38595927 | 0,02181 | 0,19633144 | interferon regulatory factor 2 binding protein 2 |
| ENSG00000164164 | OTUD4 | 0,470743061 | 1,385823053 | 0,10391 | 0,41229858 | OTU deubiquitinase 4 |
| ENSG00000138430 | OLA1 | 0,470661492 | 1,385744702 | 0,20167 | 0,52214043 | Obg like ATPase 1 |
| ENSG00000074800 | ENO1 | 0,470653101 | 1,385736642 | 0,03858 | 0,26561819 | enolase 1 |
| ENSG00000196141 | SPATS2L | 0,470581517 | 1,385667886 | 0,09428 | 0,39648446 | spermatogenesis associated serine rich 2 like |
| ENSG00000129083 | COPB1 | 0,470578083 | 1,385664588 | 0,19259 | 0,51490739 | COPI coat complex subunit beta 1 |
| ENSG00000281569 | KANSL1-AS1 | 0,470575216 | 1,385661835 | 0,31647 | 0,60238875 | KANSL1 antisense RNA 1 |
| ENSG00000173156 | RHOD | 0,470538884 | 1,385626939 | 0,22019 | 0,53750545 | ras homolog family member D |
| ENSG00000023041 | ZDHHC6 | 0,470504182 | 1,385593611 | 0,29679 | 0,58995143 | zinc finger DHHC-type palmitoyltransferase 6 |
| ENSG00000274093 | AC009032,1 | 0,470484457 | 1,385574666 | 0,48161 | 0,71838485 | novel transcript, sense intronic to NFAT5 |
| ENSG00000253553 | AC090578,1 | 0,47048441 | 1,385574621 | 0,05903 | 0,32417102 | novel transcript, antisense to MMP16 |

| | | | | | | |
|---|---|---|---|---|---|---|
| ENSG00000164346 | NSA2 | 0,470394589 | 1,385488359 | 0,27999 | 0,57764051 | NSA2 ribosome biogenesis factor |
| ENSG00000276256 | AC011043,1 | 0,470375846 | 1,385470359 | 0,38868 | 0,65321475 | uncharacterized LOC389831 |
| ENSG00000230650 | AC140479,2 | 0,470329284 | 1,385425645 | 0,58515 | 0,78616171 | zinc finger protein 532 (ZNF532) pseudogene |
| ENSG00000116641 | DOCK7 | 0,470231268 | 1,385331522 | 0,13575 | 0,462686 | dedicator of cytokinesis 7 |
| ENSG00000186472 | PCLO | 0,470198812 | 1,385300358 | 0,17654 | 0,50075446 | piccolo presynaptic cytomatrix protein |
| ENSG00000171202 | TMEM126A | 0,470171831 | 1,38527445 | 0,42131 | 0,67634721 | transmembrane protein 126A |
| ENSG00000122359 | ANXA11 | 0,470141519 | 1,385245345 | 0,07753 | 0,36452608 | annexin A11 |
| ENSG00000065518 | NDUFB4 | 0,470099392 | 1,385204896 | 0,3101 | 0,59836523 | NADH:ubiquinone oxidoreductase subunit B4 |
| ENSG00000177485 | ZBTB33 | 0,470030255 | 1,385138515 | 0,36613 | 0,63617598 | zinc finger and BTB domain containing 33 |
| ENSG00000107651 | SEC23IP | 0,469924655 | 1,385037132 | 0,09035 | 0,3889575 | SEC23 interacting protein |
| ENSG00000126231 | PROZ | 0,469823313 | 1,384939844 | 0,52305 | 0,74710918 | protein Z, vitamin K dependent plasma glycoprotein |
| ENSG00000120693 | SMAD9 | 0,469799905 | 1,384917373 | 0,16571 | 0,49182669 | SMAD family member 9 |
| ENSG00000100380 | ST13 | 0,469783221 | 1,384901358 | 0,17768 | 0,50205238 | ST13 Hsp70 interacting protein |
| ENSG00000014164 | ZC3H3 | 0,469755583 | 1,384874827 | 0,17421 | 0,49867554 | zinc finger CCCH-type containing 3 |
| ENSG00000245910 | SNHG6 | 0,469663578 | 1,384786512 | 0,32826 | 0,60971379 | small nucleolar RNA host gene 6 |
| ENSG00000156599 | ZDHHC5 | 0,469577296 | 1,384703696 | 0,04729 | 0,29217708 | zinc finger DHHC-type palmitoyltransferase 5 |
| ENSG00000122034 | GTF3A | 0,469532546 | 1,384660745 | 0,28024 | 0,57784691 | general transcription factor IIIA |
| ENSG00000197381 | ADARB1 | 0,46946626 | 1,384597128 | 0,05783 | 0,32112994 | adenosine deaminase RNA specific B1 |
| ENSG00000166415 | WDR72 | 0,469364026 | 1,384499014 | 0,37628 | 0,64458342 | WD repeat domain 72 |
| ENSG00000178904 | DPY19L3 | 0,469358589 | 1,384493796 | 0,18189 | 0,50521394 | dpy-19 like C-mannosyltransferase 3 |
| ENSG00000118873 | RAB3GAP2 | 0,469353369 | 1,384488786 | 0,20042 | 0,52108722 | RAB3 GTPase activating non-catalytic protein subunit 2 |
| ENSG00000223509 | AC135983,2 | 0,469340111 | 1,384476064 | 0,2784 | 0,57651947 | WAS protein homolog associated with actin, golgi membranes and microtubules (WHAMM) pseudogene |
| ENSG00000171314 | PGAM1 | 0,46932441 | 1,384460996 | 0,21697 | 0,53505085 | phosphoglycerate mutase 1 |
| ENSG00000179218 | CALR | 0,469316065 | 1,384452988 | 0,04656 | 0,29026341 | calreticulin |
| ENSG00000120694 | HSPH1 | 0,469212891 | 1,384353983 | 0,16875 | 0,49492999 | heat shock protein family H (Hsp110) member 1 |
| ENSG00000169087 | HSPBAP1 | 0,469200995 | 1,384342568 | 0,07404 | 0,35746749 | HSPB1 associated protein 1 |
| ENSG00000236095 | AL807757,2 | 0,469160537 | 1,384303747 | 0,63544 | 0,816556 | novel transcript |
| ENSG00000149196 | HIKESHI | 0,469130445 | 1,384274873 | 0,25486 | 0,55944852 | heat shock protein nuclear import factor hikeshi |
| ENSG00000198780 | FAM169A | 0,469125624 | 1,384270248 | 0,1678 | 0,49395055 | family with sequence similarity 169 member A |
| ENSG00000258365 | AC073655,2 | 0,469105178 | 1,38425063 | 0,26426 | 0,56703954 | novel transcript, antisense to PLXNC1 |
| ENSG00000182481 | KPNA2 | 0,469078698 | 1,384225222 | 0,29458 | 0,58857672 | karyopherin subunit alpha 2 |
| ENSG00000113615 | SEC24A | 0,469054677 | 1,384202175 | 0,30516 | 0,59455047 | SEC24 homolog A, COPII coat complex component |

| | | | | | | |
|---|---|---|---|---|---|---|
| ENSG00000165775 | FUNDC2 | 0,469043168 | 1,384191133 | 0,2393 | 0,5491566 | FUN14 domain containing 2 |
| ENSG00000198791 | CNOT7 | 0,468967991 | 1,384119006 | 0,37898 | 0,64687428 | CCR4-NOT transcription complex subunit 7 |
| ENSG00000137486 | ARRB1 | 0,468860379 | 1,384015767 | 0,08197 | 0,37306719 | arrestin beta 1 |
| ENSG00000175305 | CCNE2 | 0,468844031 | 1,384000084 | 0,31653 | 0,60239064 | cyclin E2 |
| ENSG00000266313 | AC026254,2 | 0,46878645 | 1,383944847 | 0,25996 | 0,56377449 | novel transcript |
| ENSG00000107745 | MICU1 | 0,468735096 | 1,383895585 | 0,1688 | 0,49492999 | mitochondrial calcium uptake 1 |
| ENSG00000113407 | TARS1 | 0,468671015 | 1,383834117 | 0,08505 | 0,37850509 | threonyl-tRNA synthetase 1 |
| ENSG00000138378 | STAT4 | 0,4685594 | 1,38372706 | 0,32305 | 0,60584543 | signal transducer and activator of transcription 4 |
| ENSG00000066855 | MTFR1 | 0,468521506 | 1,383690715 | 0,23764 | 0,54861973 | mitochondrial fission regulator 1 |
| ENSG00000170248 | PDCD6IP | 0,468475041 | 1,383646151 | 0,2622 | 0,56507558 | programmed cell death 6 interacting protein |
| ENSG00000047634 | SCML1 | 0,468458311 | 1,383630107 | 0,43667 | 0,68681257 | Scm polycomb group protein like 1 |
| ENSG00000153879 | CEBPG | 0,468443365 | 1,383615773 | 0,22188 | 0,53821014 | CCAAT enhancer binding protein gamma |
| ENSG00000266733 | TBC1D29P | 0,468416739 | 1,383590237 | 0,38034 | 0,64797034 | TBC1 domain family member 29, pseudogene |
| ENSG00000140332 | TLE3 | 0,468275124 | 1,38345443 | 0,10324 | 0,41110033 | TLE family member 3, transcriptional corepressor |
| ENSG00000221923 | ZNF880 | 0,468203418 | 1,383385671 | 0,27719 | 0,57572679 | zinc finger protein 880 |
| ENSG00000274922 | AL139384,1 | 0,468119803 | 1,383305495 | 0,40336 | 0,66399303 | novel transcript |
| ENSG00000288078 | AL133553,2 | 0,468072188 | 1,383259842 | 0,55196 | 0,76558846 | novel transcript |
| ENSG00000095383 | TBC1D2 | 0,468059495 | 1,383247672 | 0,19905 | 0,52017864 | TBC1 domain family member 2 |
| ENSG00000196517 | SLC6A9 | 0,468028403 | 1,383217861 | 0,04199 | 0,27753946 | solute carrier family 6 member 9 |
| ENSG00000231551 | AC245100,4 | 0,468022523 | 1,383212224 | 0,12402 | 0,44495264 | phosphodiesterase 4D interacting protein (PDE4DIP) pseudogene |
| ENSG00000170473 | PYM1 | 0,467920871 | 1,383114766 | 0,02162 | 0,19544769 | PYM homolog 1, exon junction complex associated factor |
| ENSG00000197498 | RPF2 | 0,467850645 | 1,383047442 | 0,09038 | 0,3889575 | ribosome production factor 2 homolog |
| ENSG00000275311 | AC139494,3 | 0,467810995 | 1,383009432 | 0,43795 | 0,68747773 | novel transcript |
| ENSG00000118402 | ELOVL4 | 0,467777311 | 1,382977141 | 0,15602 | 0,48382611 | ELOVL fatty acid elongase 4 |
| ENSG00000205339 | IPO7 | 0,467765724 | 1,382966034 | 0,23844 | 0,54877227 | importin 7 |
| ENSG00000171566 | PLRG1 | 0,467718879 | 1,382921129 | 0,34285 | 0,62033803 | pleiotropic regulator 1 |
| ENSG00000233155 | HMGA1P8 | 0,467713315 | 1,382915795 | 0,28637 | 0,58267001 | high mobility group AT-hook 1 pseudogene 8 |
| ENSG00000115541 | HSPE1 | 0,46771149 | 1,382914046 | 0,23089 | 0,54390394 | heat shock protein family E (Hsp10) member 1 |
| ENSG00000118894 | EEF2KMT | 0,467665958 | 1,382870401 | 0,22902 | 0,54263903 | eukaryotic elongation factor 2 lysine methyltransferase |
| ENSG00000103994 | ZNF106 | 0,467585296 | 1,382793086 | 0,0139 | 0,15464569 | zinc finger protein 106 |
| ENSG00000124787 | RPP40 | 0,467486465 | 1,382698363 | 0,36769 | 0,63764567 | ribonuclease P/MRP subunit p40 |
| ENSG00000166707 | ZCCHC18 | 0,467470568 | 1,382683127 | 0,02334 | 0,20398797 | zinc finger CCHC-type containing 18 |

| | | | | | | |
|---|---|---|---|---|---|---|
| ENSG00000068885 | IFT80 | 0,467458043 | 1,382671123 | 0,29077 | 0,58579789 | intraflagellar transport 80 |
| ENSG00000138069 | RAB1A | 0,467437842 | 1,382651763 | 0,30169 | 0,59283946 | RAB1A, member RAS oncogene family |
| ENSG00000259548 | AC021483,1 | 0,4674148 | 1,38262968 | 0,61865 | 0,80665241 | novel transcript |
| ENSG00000279636 | LINC00216 | 0,467395903 | 1,382611569 | 0,46905 | 0,70862072 | long intergenic non-protein coding RNA 216 |
| ENSG00000111801 | BTN3A3 | 0,467370558 | 1,38258728 | 0,12446 | 0,44546792 | butyrophilin subfamily 3 member A3 |
| ENSG00000114786 | ABHD14A-ACY | 0,467368231 | 1,38258505 | 0,36919 | 0,63906793 | ABHD14A-ACY1 readthrough |
| ENSG00000167565 | SERTAD3 | 0,467361027 | 1,382578146 | 0,08897 | 0,3857873 | SERTA domain containing 3 |
| ENSG00000229325 | ACAP2-IT1 | 0,467334581 | 1,382552803 | 0,54377 | 0,76018847 | ACAP2 intronic transcript 1 |
| ENSG00000239474 | KLHL41 | 0,467272543 | 1,382493352 | 0,31777 | 0,60320933 | kelch like family member 41 |
| ENSG00000224955 | KCTD10P1 | 0,467269915 | 1,382490834 | 0,58382 | 0,7856659 | potassium channel tetramerization domain containing 10 pseudogene 1 |
| ENSG00000140990 | NDUFB10 | 0,467226592 | 1,382449319 | 0,27601 | 0,57533851 | NADH:ubiquinone oxidoreductase subunit B10 |
| ENSG00000184900 | SUMO3 | 0,467183229 | 1,382407767 | 0,13232 | 0,45747548 | small ubiquitin like modifier 3 |
| ENSG00000115970 | THADA | 0,467153827 | 1,382379595 | 0,12184 | 0,44093327 | THADA armadillo repeat containing |
| ENSG00000100410 | PHF5A | 0,467152204 | 1,382378039 | 0,08879 | 0,3852075 | PHD finger protein 5A |
| ENSG00000148187 | MRRF | 0,467079428 | 1,382308308 | 0,11128 | 0,42416781 | mitochondrial ribosome recycling factor |
| ENSG00000264619 | MSH5-SAPCD: | 0,467057045 | 1,382286862 | 0,4655 | 0,70591843 | MSH5-SAPCD1 readthrough (NMD candidate) |
| ENSG00000283068 | IBTK | 0,466981152 | 1,382214149 | 0,24575 | 0,55359883 | inhibitor of Bruton tyrosine kinase |
| ENSG00000159885 | ZNF222 | 0,466976669 | 1,382209854 | 0,27215 | 0,57306433 | zinc finger protein 222 |
| ENSG00000129534 | MIS18BP1 | 0,466927621 | 1,382162863 | 0,33431 | 0,61415383 | MIS18 binding protein 1 |
| ENSG00000151687 | ANKAR | 0,466877973 | 1,382115299 | 0,25026 | 0,55659392 | ankyrin and armadillo repeat containing |
| ENSG00000272148 | AC013403,2 | 0,466818551 | 1,382058373 | 0,27963 | 0,57745167 | novel transcript, antisense to AGBL5 |
| ENSG00000133226 | SRRM1 | 0,466818491 | 1,382058315 | 0,0652 | 0,33804854 | serine and arginine repetitive matrix 1 |
| ENSG00000168175 | MAPK1IP1L | 0,466793246 | 1,382034132 | 0,1729 | 0,49768904 | mitogen-activated protein kinase 1 interacting protein 1 like |
| ENSG00000231615 | AL645568,1 | 0,466790196 | 1,38203121 | 0,55462 | 0,7675495 | novel pseudogene |
| ENSG00000161955 | TNFSF13 | 0,466782263 | 1,38202361 | 0,07555 | 0,36093649 | TNF superfamily member 13 |
| ENSG00000155636 | RBM45 | 0,466721174 | 1,381965092 | 0,25747 | 0,56160845 | RNA binding motif protein 45 |
| ENSG00000151553 | FAM160B1 | 0,466698203 | 1,381943088 | 0,21665 | 0,53492477 | family with sequence similarity 160 member B1 |
| ENSG00000275318 | AL136981,2 | 0,466672451 | 1,38191842 | 0,30181 | 0,59288267 | novel zinc finger protein pseudogene |
| ENSG00000132436 | FIGNL1 | 0,466636377 | 1,381883866 | 0,30854 | 0,59736238 | fidgetin like 1 |
| ENSG00000165806 | CASP7 | 0,46662017 | 1,381868343 | 0,27561 | 0,57533171 | caspase 7 |
| ENSG00000037474 | NSUN2 | 0,466592708 | 1,381842039 | 0,07697 | 0,36357031 | NOP2/Sun RNA methyltransferase 2 |

| | | | | | | |
|---|---|---|---|---|---|---|
| ENSG00000267160 | AC091152,2 | 0,466563558 | 1,381814119 | 0,53617 | 0,75504276 | novel transcript, antisense to C17orf104 |
| ENSG00000215493 | AC007731,1 | 0,46654484 | 1,381796191 | 0,51281 | 0,73941533 | kelch-like 12 (Drosophila) (KLH12) pseudogene |
| ENSG00000134369 | NAV1 | 0,466516155 | 1,381768717 | 0,26485 | 0,56748133 | neuron navigator 1 |
| ENSG00000220323 | H2BC19P | 0,466317885 | 1,381578833 | 0,33543 | 0,61509556 | H2B clustered histone 19, pseudogene |
| ENSG00000119280 | C1orf198 | 0,46615346 | 1,381421383 | 0,00542 | 0,09153353 | chromosome 1 open reading frame 198 |
| ENSG00000203326 | ZNF525 | 0,466119253 | 1,381388628 | 0,38152 | 0,64887769 | zinc finger protein 525 |
| ENSG00000128203 | ASPHD2 | 0,466079149 | 1,38135023 | 0,02339 | 0,2040793 | aspartate beta-hydroxylase domain containing 2 |
| ENSG00000061794 | MRPS35 | 0,466039413 | 1,381312183 | 0,18534 | 0,50876814 | mitochondrial ribosomal protein S35 |
| ENSG00000286191 | AL158195,1 | 0,46581668 | 1,381098944 | 0,63281 | 0,8150808 | novel transcript |
| ENSG00000203668 | CHML | 0,465811328 | 1,381093821 | 0,33975 | 0,61872249 | CHM like Rab escort protein |
| ENSG00000138074 | SLC5A6 | 0,465619827 | 1,380910508 | 0,10098 | 0,40770184 | solute carrier family 5 member 6 |
| ENSG00000106571 | GLI3 | 0,465597545 | 1,380889181 | 0,21723 | 0,53520949 | GLI family zinc finger 3 |
| ENSG00000197601 | FAR1 | 0,465298539 | 1,380603015 | 0,28563 | 0,58212254 | fatty acyl-CoA reductase 1 |
| ENSG00000166579 | NDEL1 | 0,465297001 | 1,380601543 | 0,11921 | 0,43703483 | nudE neurodevelopment protein 1 like 1 |
| ENSG00000231697 | NANOGP5 | 0,465290421 | 1,380595245 | 0,36503 | 0,63527179 | Nanog homeobox pseudogene 5 |
| ENSG00000120697 | ALG5 | 0,465241537 | 1,380548467 | 0,17468 | 0,49886732 | ALG5 dolichyl-phosphate beta-glucosyltransferase |
| ENSG00000165417 | GTF2A1 | 0,465231114 | 1,380538493 | 0,17865 | 0,50323303 | general transcription factor IIA subunit 1 |
| ENSG00000238082 | NUDCP2 | 0,465050288 | 1,380365468 | 0,55617 | 0,76831913 | nuclear distribution C pseudogene 2 |
| ENSG00000268438 | BNIP3P27 | 0,46492366 | 1,380244316 | 0,47619 | 0,71446062 | BCL2 interacting protein 3 pseudogene 27 |
| ENSG00000275294 | LINC02340 | 0,464863453 | 1,380186717 | 0,53299 | 0,7533653 | long intergenic non-protein coding RNA 2340 |
| ENSG00000137574 | TGS1 | 0,4648614 | 1,380184753 | 0,22401 | 0,53994403 | trimethylguanosine synthase 1 |
| ENSG00000064652 | SNX24 | 0,464799092 | 1,380125146 | 0,32731 | 0,60914884 | sorting nexin 24 |
| ENSG00000275401 | AL391095,3 | 0,46479839 | 1,380124474 | 0,3309 | 0,6112353 | novel transcript |
| ENSG00000257126 | FOXG1-AS1 | 0,464758539 | 1,380086351 | 0,3411 | 0,6192144 | FOXG1 antisense RNA 1 |
| ENSG00000083720 | OXCT1 | 0,464694945 | 1,380025519 | 0,25986 | 0,56377449 | 3-oxoacid CoA-transferase 1 |
| ENSG00000115866 | DARS1 | 0,464595353 | 1,379930256 | 0,25856 | 0,5625202 | aspartyl-tRNA synthetase 1 |
| ENSG00000104327 | CALB1 | 0,464579943 | 1,379915517 | 0,42022 | 0,67567123 | calbindin 1 |
| ENSG00000120742 | SERP1 | 0,464408908 | 1,379751934 | 0,12566 | 0,44769898 | stress associated endoplasmic reticulum protein 1 |
| ENSG00000147724 | FAM135B | 0,46425687 | 1,379606537 | 0,28709 | 0,58349837 | family with sequence similarity 135 member B |
| ENSG00000163527 | STT3B | 0,464219321 | 1,37957063 | 0,18882 | 0,51219588 | STT3 oligosaccharyltransferase complex catalytic subunit B |
| ENSG00000112655 | PTK7 | 0,464189017 | 1,379541653 | 0,01361 | 0,1529196 | protein tyrosine kinase 7 (inactive) |
| ENSG00000118855 | MFSD1 | 0,464116628 | 1,379472434 | 0,04365 | 0,28188832 | major facilitator superfamily domain containing 1 |
| ENSG00000211456 | SACM1L | 0,46406114 | 1,379419379 | 0,29275 | 0,58724158 | SAC1 like phosphatidylinositide phosphatase |

| | | | | | | |
|---|---|---|---|---|---|---|
| ENSG00000176853 | FAM91A1 | 0,463877223 | 1,379243539 | 0,215 | 0,53373528 | family with sequence similarity 91 member A1 |
| ENSG00000013503 | POLR3B | 0,463860294 | 1,379227355 | 0,20342 | 0,52321822 | RNA polymerase III subunit B |
| ENSG00000197579 | TOPORS | 0,463804322 | 1,379173847 | 0,30752 | 0,59664516 | TOP1 binding arginine/serine rich protein, E3 ubiquitin ligase |
| ENSG00000108064 | TFAM | 0,463733032 | 1,379105697 | 0,1732 | 0,49786688 | transcription factor A, mitochondrial |
| ENSG00000102007 | PLP2 | 0,463505186 | 1,378887912 | 0,37713 | 0,6456352 | proteolipid protein 2 |
| ENSG00000181191 | PJA1 | 0,463411512 | 1,378798383 | 0,08604 | 0,3803347 | praja ring finger ubiquitin ligase 1 |
| ENSG00000074696 | HACD3 | 0,463331332 | 1,378721756 | 0,17234 | 0,49714741 | 3-hydroxyacyl-CoA dehydratase 3 |
| ENSG00000286970 | AC010491,2 | 0,463252099 | 1,378646039 | 0,0538 | 0,31106243 | novel transcript, antisense to ANKH |
| ENSG00000259591 | AC022898,1 | 0,46312318 | 1,378522849 | 0,46786 | 0,70757147 | novel transcript, antisense to RORA |
| ENSG00000170860 | LSM3 | 0,463089997 | 1,378491142 | 0,27922 | 0,57736567 | LSM3 homolog, U6 small nuclear RNA and mRNA degradation associated |
| ENSG00000135829 | DHX9 | 0,463024864 | 1,378428909 | 0,17836 | 0,50278602 | DExH-box helicase 9 |
| ENSG00000138795 | LEF1 | 0,462992986 | 1,378398452 | 0,2324 | 0,54513458 | lymphoid enhancer binding factor 1 |
| ENSG00000282130 | NSMCE3 | 0,462986304 | 1,378392068 | 0,17295 | 0,49768904 | NSE3 homolog, SMC5-SMC6 complex component |
| ENSG00000184481 | FOXO4 | 0,462985991 | 1,378391768 | 0,1221 | 0,44150219 | forkhead box O4 |
| ENSG00000263368 | AC069366,1 | 0,462946084 | 1,37835364 | 0,63541 | 0,81655453 | chromosome 9 open reading frame 30 (C9orf30) pseudogene |
| ENSG00000085185 | BCORL1 | 0,462925848 | 1,378334307 | 0,00826 | 0,11480072 | BCL6 corepressor like 1 |
| ENSG00000137713 | PPP2R1B | 0,462818107 | 1,378231377 | 0,13572 | 0,462686 | protein phosphatase 2 scaffold subunit Abeta |
| ENSG00000186297 | GABRA5 | 0,462813303 | 1,378226788 | 0,34624 | 0,62337014 | gamma-aminobutyric acid type A receptor subunit alpha5 |
| ENSG00000285723 | AL034430,2 | 0,462794617 | 1,378208936 | 0,33233 | 0,61225146 | novel protein |
| ENSG00000273590 | SMIM11B | 0,46278148 | 1,378196386 | 0,70254 | 0,85660698 | small integral membrane protein 11B |
| ENSG00000237373 | BRWD1-IT1 | 0,462778542 | 1,37819358 | 0,55189 | 0,76553885 | BRWD1 intronic transcript 1 |
| ENSG00000074621 | SLC24A1 | 0,462734239 | 1,378151258 | 0,04941 | 0,29915481 | solute carrier family 24 member 1 |
| ENSG00000125356 | NDUFA1 | 0,462672905 | 1,378092669 | 0,16222 | 0,48939373 | NADH:ubiquinone oxidoreductase subunit A1 |
| ENSG00000274430 | AL353804,3 | 0,4626716 | 1,378091423 | 0,65687 | 0,82856771 | |
| ENSG00000110060 | PUS3 | 0,46255411 | 1,377979199 | 0,10613 | 0,41557034 | pseudouridine synthase 3 |
| ENSG00000224379 | TCF19 | 0,462527578 | 1,377953858 | 0,17371 | 0,49846579 | transcription factor 19 |
| ENSG00000138346 | DNA2 | 0,462419455 | 1,37785059 | 0,29821 | 0,59060946 | DNA replication helicase/nuclease 2 |
| ENSG00000131873 | CHSY1 | 0,462399022 | 1,377831076 | 0,0924 | 0,39283323 | chondroitin sulfate synthase 1 |
| ENSG00000112186 | CAP2 | 0,462257864 | 1,37769627 | 0,33964 | 0,61869857 | cyclase associated actin cytoskeleton regulatory protein 2 |
| ENSG00000149548 | CCDC15 | 0,462062665 | 1,377509878 | 0,22446 | 0,54033281 | coiled-coil domain containing 15 |

| | | | | | | |
|---|---|---|---|---|---|---|
| ENSG00000154813 | DPH3 | 0,461945249 | 1,377397773 | 0,34179 | 0,6197312 | diphthamide biosynthesis 3 |
| ENSG00000176597 | B3GNT5 | 0,461851043 | 1,377307834 | 0,24741 | 0,5549105 | UDP-GlcNAc:betaGal beta-1,3-N-acetylglucosaminyltransferase 5 |
| ENSG00000251595 | ABCA11P | 0,461782885 | 1,377242766 | 0,17429 | 0,49875953 | ATP binding cassette subfamily A member 11, pseudogene |
| ENSG00000128512 | DOCK4 | 0,461661146 | 1,377126555 | 0,21448 | 0,53342315 | dedicator of cytokinesis 4 |
| ENSG00000240731 | AL139287,1 | 0,461649283 | 1,377115231 | 0,35151 | 0,62674766 | novel transcript |
| ENSG00000143553 | SNAPIN | 0,461644036 | 1,377110223 | 0,36604 | 0,63617598 | SNAP associated protein |
| ENSG00000139746 | RBM26 | 0,46141807 | 1,376894546 | 0,21336 | 0,53259344 | RNA binding motif protein 26 |
| ENSG00000265282 | AC005828,4 | 0,46137775 | 1,376856066 | 0,52769 | 0,75039004 | novel transcript |
| ENSG00000113360 | DROSHA | 0,461335559 | 1,3768158 | 0,24482 | 0,55282045 | drosha ribonuclease III |
| ENSG00000288062 | AL136981,3 | 0,461281108 | 1,376763837 | 0,59483 | 0,79299796 | uncharacterized LOC101929748 |
| ENSG00000101413 | RPRD1B | 0,461246744 | 1,376731044 | 0,15816 | 0,48521512 | regulation of nuclear pre-mRNA domain containing 1B |
| ENSG00000282220 | AC110597,5 | 0,461159246 | 1,376647549 | 0,66378 | 0,8324578 | novel transcript, antisense to DSEL |
| ENSG00000263424 | AC110597,3 | 0,461159246 | 1,376647549 | 0,66378 | 0,8324578 | novel transcript, antisense to DSEL |
| ENSG00000054148 | PHPT1 | 0,461151136 | 1,37663981 | 0,37355 | 0,6422142 | phosphohistidine phosphatase 1 |
| ENSG00000123384 | LRP1 | 0,461144192 | 1,376633184 | 0,05607 | 0,31627719 | LDL receptor related protein 1 |
| ENSG00000123416 | TUBA1B | 0,461093475 | 1,376584791 | 0,28963 | 0,58522301 | tubulin alpha 1b |
| ENSG00000269374 | AC011497,2 | 0,46104986 | 1,376543174 | 0,4499 | 0,69539135 | zinc finger, AN1-type domain 5 (ZFAND5) pseudogene |
| ENSG00000144283 | PKP4 | 0,460980263 | 1,37647677 | 0,00624 | 0,09964689 | plakophilin 4 |
| ENSG00000139505 | MTMR6 | 0,460970393 | 1,376467354 | 0,30885 | 0,59767335 | myotubularin related protein 6 |
| ENSG00000164749 | HNF4G | 0,460889626 | 1,376390296 | 0,36185 | 0,63352908 | hepatocyte nuclear factor 4 gamma |
| ENSG00000235609 | AF127577,4 | 0,46086237 | 1,376364294 | 0,33745 | 0,61713877 | novel transcript |
| ENSG00000164823 | OSGIN2 | 0,460852428 | 1,376354808 | 0,30243 | 0,59297237 | oxidative stress induced growth inhibitor family member 2 |
| ENSG00000104325 | DECR1 | 0,460827718 | 1,376331235 | 0,2759 | 0,57533851 | 2,4-dienoyl-CoA reductase 1 |
| ENSG00000054118 | THRAP3 | 0,460811704 | 1,376315957 | 0,12737 | 0,45051698 | thyroid hormone receptor associated protein 3 |
| ENSG00000101639 | CEP192 | 0,46061095 | 1,376124454 | 0,22733 | 0,54168591 | centrosomal protein 192 |
| ENSG00000180573 | H2AC6 | 0,460608827 | 1,376122429 | 0,2072 | 0,52673925 | H2A clustered histone 6 |
| ENSG00000196511 | TPK1 | 0,460598402 | 1,376112485 | 0,18589 | 0,50915451 | thiamin pyrophosphokinase 1 |
| ENSG00000177628 | GBA | 0,460544844 | 1,376061399 | 0,29159 | 0,58655914 | glucosylceramidase beta |
| ENSG00000188167 | TMPPE | 0,460538829 | 1,376055662 | 0,05446 | 0,31226071 | transmembrane protein with metallophosphoesterase domain |
| ENSG00000152208 | GRID2 | 0,460439217 | 1,375960655 | 0,17085 | 0,49641416 | glutamate ionotropic receptor delta type subunit 2 |
| ENSG00000143878 | RHOB | 0,460422152 | 1,375944379 | 0,04149 | 0,27611222 | ras homolog family member B |

| | | | | | | |
|---|---|---|---|---|---|---|
| ENSG00000104497 | SNX16 | 0,460419575 | 1,375941921 | 0,30386 | 0,59400398 | sorting nexin 16 |
| ENSG00000225900 | HSPE1P13 | 0,460343753 | 1,37586961 | 0,64448 | 0,82154386 | heat shock protein family E (Hsp10) member 1 pseudogene 13 |
| ENSG00000215199 | YWHAZP6 | 0,460265337 | 1,375794828 | 0,33245 | 0,61233149 | tyrosine 3-monooxygenase/tryptophan 5-monooxygenase activation protein zeta pseudogene 6 |
| ENSG00000187231 | SESTD1 | 0,460183891 | 1,375717161 | 0,17982 | 0,50407792 | SEC14 and spectrin domain containing 1 |
| ENSG00000106052 | TAX1BP1 | 0,459767695 | 1,375320344 | 0,27718 | 0,57572679 | Tax1 binding protein 1 |
| ENSG00000183691 | NOG | 0,45973218 | 1,375286488 | 0,44216 | 0,69003826 | noggin |
| ENSG00000090060 | PAPOLA | 0,459726706 | 1,37528127 | 0,24594 | 0,55373218 | poly(A) polymerase alpha |
| ENSG00000114391 | RPL24 | 0,459661658 | 1,375219263 | 0,12437 | 0,44539699 | ribosomal protein L24 |
| ENSG00000259180 | AC012378,1 | 0,4596176 | 1,375177266 | 0,59067 | 0,78992663 | novel transcript, antisense to PRTG |
| ENSG00000236104 | ZBTB22 | 0,459594788 | 1,375155522 | 0,71952 | 0,86592255 | zinc finger and BTB domain containing 22 |
| ENSG00000102738 | MRPS31 | 0,45937896 | 1,374949813 | 0,28204 | 0,57936013 | mitochondrial ribosomal protein S31 |
| ENSG00000170412 | GPRC5C | 0,45934547 | 1,374917896 | 0,18585 | 0,50915451 | G protein-coupled receptor class C group 5 member C |
| ENSG00000138356 | AOX1 | 0,45924233 | 1,374819606 | 0,42752 | 0,68046596 | aldehyde oxidase 1 |
| ENSG00000285807 | AL353768,2 | 0,459131407 | 1,374713905 | 0,67191 | 0,83759902 | Novel transcript, antisense to C9orf3 |
| ENSG00000213025 | COX20P1 | 0,4591114 | 1,374694841 | 0,55562 | 0,76790322 | COX20 pseudogene 1 |
| ENSG00000128590 | DNAJB9 | 0,459014794 | 1,374602791 | 0,29592 | 0,58942061 | DnaJ heat shock protein family (Hsp40) member B9 |
| ENSG00000243260 | RN7SL558P | 0,458990893 | 1,374580019 | 0,59193 | 0,79090751 | RNA, 7SL, cytoplasmic 558, pseudogene |
| ENSG00000188612 | SUMO2 | 0,458988195 | 1,374577448 | 0,1238 | 0,4446254 | small ubiquitin like modifier 2 |
| ENSG00000220744 | RPL5P18 | 0,458873088 | 1,37446778 | 0,40828 | 0,66714279 | ribosomal protein L5 pseudogene 18 |
| ENSG00000163026 | WDCP | 0,458803628 | 1,374401606 | 0,27958 | 0,57745167 | WD repeat and coiled coil containing |
| ENSG00000072571 | HMMR | 0,458791615 | 1,374390162 | 0,3061 | 0,59513628 | hyaluronan mediated motility receptor |
| ENSG00000101019 | UQCC1 | 0,458734946 | 1,374336178 | 0,17139 | 0,49665377 | ubiquinol-cytochrome c reductase complex assembly factor 1 |
| ENSG00000187801 | ZFP69B | 0,458683491 | 1,374287161 | 0,19679 | 0,51798099 | ZFP69 zinc finger protein B |
| ENSG00000118496 | FBXO30 | 0,458625091 | 1,374231531 | 0,23462 | 0,54623956 | F-box protein 30 |
| ENSG00000198146 | ZNF770 | 0,458588779 | 1,374196943 | 0,22035 | 0,53750545 | zinc finger protein 770 |
| ENSG00000134001 | EIF2S1 | 0,45853426 | 1,374145014 | 0,3042 | 0,5941708 | eukaryotic translation initiation factor 2 subunit alpha |
| ENSG00000069206 | ADAM7 | 0,458463891 | 1,37407799 | 0,73158 | 0,87270478 | ADAM metallopeptidase domain 7 |
| ENSG00000128585 | MKLN1 | 0,458428968 | 1,374044729 | 0,25402 | 0,55917212 | muskelin 1 |
| ENSG00000186448 | ZNF197 | 0,458400627 | 1,374017736 | 0,44673 | 0,69321646 | zinc finger protein 197 |
| ENSG00000108344 | PSMD3 | 0,458294271 | 1,373916447 | 0,14508 | 0,47275989 | proteasome 26S subunit, non-ATPase 3 |

| | | | | | | |
|---|---|---|---|---|---|---|
| ENSG00000116251 | RPL22 | 0,458274637 | 1,373897749 | 0,1737 | 0,49846579 | ribosomal protein L22 |
| ENSG00000172893 | DHCR7 | 0,458208803 | 1,373835056 | 0,17404 | 0,49860418 | 7-dehydrocholesterol reductase |
| ENSG00000086289 | EPDR1 | 0,45819301 | 1,373820017 | 0,1211 | 0,44016175 | ependymin related 1 |
| ENSG00000141447 | OSBPL1A | 0,458191696 | 1,373818766 | 0,06378 | 0,33485717 | oxysterol binding protein like 1A |
| ENSG00000144120 | TMEM177 | 0,458103441 | 1,373734727 | 0,13549 | 0,46221178 | transmembrane protein 177 |
| ENSG00000101337 | TM9SF4 | 0,458082086 | 1,373714392 | 0,07757 | 0,3646056 | transmembrane 9 superfamily member 4 |
| ENSG00000271880 | AGAP11 | 0,458046635 | 1,373680637 | 0,70339 | 0,8569555 | ArfGAP with GTPase domain, ankyrin repeat and PH domain 11 |
| ENSG00000258510 | AC007954,1 | 0,457986522 | 1,373623401 | 0,49297 | 0,72610638 | ATP-binding cassette, sub-family G (WHITE), member 2 (ABCG2) pseudogene |
| ENSG00000084463 | WBP11 | 0,457888284 | 1,373529869 | 0,11244 | 0,42619744 | WW domain binding protein 11 |
| ENSG00000257365 | FNTB | 0,457806121 | 1,373451647 | 0,1633 | 0,48972962 | farnesyltransferase, CAAX box, beta |
| ENSG00000153207 | AHCTF1 | 0,457789604 | 1,373435923 | 0,11724 | 0,43455623 | AT-hook containing transcription factor 1 |
| ENSG00000180398 | MCFD2 | 0,457756593 | 1,373404497 | 0,1547 | 0,48266733 | multiple coagulation factor deficiency 2, ER cargo receptor complex subunit |
| ENSG00000198018 | ENTPD7 | 0,457648011 | 1,373301134 | 0,17548 | 0,49954066 | ectonucleoside triphosphate diphosphohydrolase 7 |
| ENSG00000278516 | LENG1 | 0,457547049 | 1,373205032 | 0,43245 | 0,68411566 | leukocyte receptor cluster member 1 |
| ENSG00000101856 | PGRMC1 | 0,45748488 | 1,373145859 | 0,36064 | 0,63299251 | progesterone receptor membrane component 1 |
| ENSG00000166441 | RPL27A | 0,457466296 | 1,37312817 | 0,16272 | 0,48945635 | ribosomal protein L27a |
| ENSG00000177150 | FAM210A | 0,457239101 | 1,372911948 | 0,15367 | 0,48204779 | family with sequence similarity 210 member A |
| ENSG00000197062 | ZSCAN26 | 0,457229279 | 1,372902601 | 0,33682 | 0,61683075 | zinc finger and SCAN domain containing 26 |
| ENSG00000147124 | ZNF41 | 0,457153037 | 1,372830049 | 0,27865 | 0,57683444 | zinc finger protein 41 |
| ENSG00000051523 | CYBA | 0,457124646 | 1,372803033 | 0,32622 | 0,6085098 | cytochrome b-245 alpha chain |
| ENSG00000185621 | LMLN | 0,45706004 | 1,372741558 | 0,22072 | 0,53767462 | leishmanolysin like peptidase |
| ENSG00000273064 | AC017083,1 | 0,457003001 | 1,372687287 | 0,3272 | 0,60902638 | novel transcript |
| ENSG00000189292 | ALKAL2 | 0,456961622 | 1,372647916 | 0,31529 | 0,60158701 | ALK and LTK ligand 2 |
| ENSG00000114480 | GBE1 | 0,456906169 | 1,372595156 | 0,2447 | 0,55280101 | 1,4-alpha-glucan branching enzyme 1 |
| ENSG00000257285 | AL132780,1 | 0,456888264 | 1,372578122 | 0,29655 | 0,58979902 | novel transcript, antisense to HAUS4 |
| ENSG00000068097 | HEATR6 | 0,456845742 | 1,372537667 | 0,26094 | 0,56405761 | HEAT repeat containing 6 |
| ENSG00000242615 | AC022415,1 | 0,456759021 | 1,372455166 | 0,18092 | 0,50450233 | ribosomal protein L17 (RPL17) pseudogene |
| ENSG00000125648 | SLC25A23 | 0,456670723 | 1,372371169 | 0,11282 | 0,42686158 | solute carrier family 25 member 23 |
| ENSG00000281327 | LINC01338 | 0,456670474 | 1,372370932 | 0,59041 | 0,78974912 | long intergenic non-protein coding RNA 1338 |
| ENSG00000143222 | UFC1 | 0,45657504 | 1,372280153 | 0,18932 | 0,51254557 | ubiquitin-fold modifier conjugating enzyme 1 |

| | | | | | | |
|---|---|---|---|---|---|---|
| ENSG00000230896 | AL604028,1 | 0,456544576 | 1,372251176 | 0,26772 | 0,56999323 | novel transcript |
| ENSG00000235090 | RPL7L1P3 | 0,456511768 | 1,372219971 | 0,49985 | 0,73058737 | ribosomal protein L7 like 1 pseudogene 3 |
| ENSG00000255036 | STRA6LP | 0,456510429 | 1,372218697 | 0,20297 | 0,52282418 | STRA6 like, pseudogene |
| ENSG00000105171 | POP4 | 0,456498411 | 1,372207266 | 0,30023 | 0,59164636 | POP4 homolog, ribonuclease P/MRP subunit |
| ENSG00000023330 | ALAS1 | 0,456466571 | 1,372176982 | 0,21626 | 0,53480193 | 5'-aminolevulinate synthase 1 |
| ENSG00000115464 | USP34 | 0,456327127 | 1,37204436 | 0,3062 | 0,59517372 | ubiquitin specific peptidase 34 |
| ENSG00000005339 | CREBBP | 0,456290269 | 1,372009308 | 0,14715 | 0,47471707 | CREB binding protein |
| ENSG00000106305 | AIMP2 | 0,456157729 | 1,371883268 | 0,18932 | 0,51254557 | aminoacyl tRNA synthetase complex interacting multifunctional protein 2 |
| ENSG00000136003 | ISCU | 0,456141738 | 1,371868062 | 0,14005 | 0,46694389 | iron-sulfur cluster assembly enzyme |
| ENSG00000162601 | MYSM1 | 0,456111714 | 1,371839512 | 0,23038 | 0,54355692 | Myb like, SWIRM and MPN domains 1 |
| ENSG00000175206 | NPPA | 0,456008514 | 1,371741384 | 0,6567 | 0,82855263 | natriuretic peptide A |
| ENSG00000278000 | AC139100,2 | 0,455961096 | 1,371696299 | 0,2833 | 0,58059722 | novel transcript |
| ENSG00000169604 | ANTXR1 | 0,455835244 | 1,371576645 | 0,20245 | 0,52259159 | ANTXR cell adhesion molecule 1 |
| ENSG00000124459 | ZNF45 | 0,455818356 | 1,37156059 | 0,06772 | 0,34423169 | zinc finger protein 45 |
| ENSG00000277791 | PSMB3 | 0,455798699 | 1,371541902 | 0,35906 | 0,63201238 | proteasome 20S subunit beta 3 |
| ENSG00000258818 | RNASE4 | 0,45578837 | 1,371532083 | 0,38952 | 0,65388973 | ribonuclease A family member 4 |
| ENSG00000151748 | SAV1 | 0,4557686 | 1,371513288 | 0,05361 | 0,31071932 | salvador family WW domain containing protein 1 |
| ENSG00000261812 | TUBB8P7 | 0,45575407 | 1,371499475 | 0,51424 | 0,74051791 | tubulin beta 8 class VIII pseudogene 7 |
| ENSG00000185515 | BRCC3 | 0,455748403 | 1,371494088 | 0,23043 | 0,54355692 | BRCA1/BRCA2-containing complex subunit 3 |
| ENSG00000145907 | G3BP1 | 0,455666664 | 1,371416385 | 0,07472 | 0,35882064 | G3BP stress granule assembly factor 1 |
| ENSG00000143412 | ANXA9 | 0,455650521 | 1,37140104 | 0,20922 | 0,52826608 | annexin A9 |
| ENSG00000158435 | CNOT11 | 0,455599761 | 1,371352789 | 0,13602 | 0,46307657 | CCR4-NOT transcription complex subunit 11 |
| ENSG00000035687 | ADSS2 | 0,455471682 | 1,371231049 | 0,084 | 0,37651068 | adenylosuccinate synthase 2 |
| ENSG00000155755 | TMEM237 | 0,455461263 | 1,371221146 | 0,19025 | 0,51331106 | transmembrane protein 237 |
| ENSG00000228594 | FNDC10 | 0,455361957 | 1,371126764 | 0,16973 | 0,49597212 | fibronectin type III domain containing 10 |
| ENSG00000135926 | TMBIM1 | 0,455227256 | 1,37099875 | 0,16188 | 0,48914394 | transmembrane BAX inhibitor motif containing 1 |
| ENSG00000168000 | BSCL2 | 0,45516951 | 1,370943875 | 0,29722 | 0,59031059 | BSCL2 lipid droplet biogenesis associated, seipin |
| ENSG00000056736 | IL17RB | 0,455143423 | 1,370919085 | 0,2759 | 0,57533851 | interleukin 17 receptor B |
| ENSG00000261469 | AC020978,4 | 0,455010059 | 1,370792362 | 0,73582 | 0,8750728 | novel transcript, sense intronic to ESRP2 |
| ENSG00000274024 | AL590282,1 | 0,454869653 | 1,370658961 | 0,46336 | 0,70419128 | tigger transposable element derived 7 (TIGD7) pseudogene |
| ENSG00000152520 | PAN3 | 0,454827869 | 1,370619264 | 0,1195 | 0,43770393 | poly(A) specific ribonuclease subunit PAN3 |

| | | | | | | |
|---|---|---|---|---|---|---|
| ENSG00000139985 | ADAM21 | 0,454825921 | 1,370617413 | 0,42837 | 0,68103938 | ADAM metallopeptidase domain 21 |
| ENSG00000196466 | ZNF799 | 0,454798448 | 1,370591313 | 0,09056 | 0,38931088 | zinc finger protein 799 |
| ENSG00000103018 | CYB5B | 0,454734233 | 1,370530308 | 0,21938 | 0,53676938 | cytochrome b5 type B |
| ENSG00000137845 | ADAM10 | 0,454727122 | 1,370523553 | 0,18501 | 0,50863955 | ADAM metallopeptidase domain 10 |
| ENSG00000257666 | CBX3P5 | 0,454713703 | 1,370510806 | 0,57511 | 0,78009066 | chromobox 3 pseudogene 5 |
| ENSG00000082175 | PGR | 0,454707045 | 1,370504481 | 0,28278 | 0,58009943 | progesterone receptor |
| ENSG00000169313 | P2RY12 | 0,45470107 | 1,370498804 | 0,36426 | 0,63495751 | purinergic receptor P2Y12 |
| ENSG00000169764 | UGP2 | 0,454653012 | 1,370453153 | 0,32495 | 0,60750217 | UDP-glucose pyrophosphorylase 2 |
| ENSG00000134644 | PUM1 | 0,454639083 | 1,370439921 | 0,21655 | 0,53490925 | pumilio RNA binding family member 1 |
| ENSG00000100823 | APEX1 | 0,454559477 | 1,370364304 | 0,11744 | 0,43477245 | apurinic/apyrimidinic endodeoxyribonuclease 1 |
| ENSG00000254590 | AP003396,2 | 0,454532216 | 1,37033841 | 0,61755 | 0,80595997 | hippocampus abundant transcript 1 (HIAT1) pseudogene |
| ENSG00000115998 | C2orf42 | 0,454216152 | 1,370038231 | 0,21092 | 0,52972021 | chromosome 2 open reading frame 42 |
| ENSG00000099974 | DDTL | 0,454201285 | 1,370024112 | 0,67747 | 0,84135792 | D-dopachrome tautomerase like |
| ENSG00000196372 | ASB13 | 0,454054098 | 1,369884347 | 0,14066 | 0,46769861 | ankyrin repeat and SOCS box containing 13 |
| ENSG00000160219 | GAB3 | 0,454034204 | 1,369865457 | 0,62793 | 0,81175174 | GRB2 associated binding protein 3 |
| ENSG00000104765 | BNIP3L | 0,45398122 | 1,369815149 | 0,26574 | 0,56866772 | BCL2 interacting protein 3 like |
| ENSG00000255419 | MSH5-SAPCD: | 0,453969938 | 1,369804437 | 0,47941 | 0,71689864 | MSH5-SAPCD1 readthrough (NMD candidate) |
| ENSG00000249767 | ENPP7P10 | 0,453967767 | 1,369802375 | 0,4927 | 0,72588649 | ectonucleotide pyrophosphatase/phosphodiesterase 7 pseudogene 10 |
| ENSG00000086189 | DIMT1 | 0,45395028 | 1,369785772 | 0,0947 | 0,39724142 | DIMT1 rRNA methyltransferase and ribosome maturation factor |
| ENSG00000185149 | NPY2R | 0,453938436 | 1,369774527 | 0,59773 | 0,79474081 | neuropeptide Y receptor Y2 |
| ENSG00000147533 | GOLGA7 | 0,453863524 | 1,369703403 | 0,36664 | 0,63646696 | golgin A7 |
| ENSG00000164849 | GPR146 | 0,453832786 | 1,369674221 | 0,41025 | 0,66821634 | G protein-coupled receptor 146 |
| ENSG00000095596 | CYP26A1 | 0,453772431 | 1,369616921 | 0,65106 | 0,82521872 | cytochrome P450 family 26 subfamily A member 1 |
| ENSG00000197566 | ZNF624 | 0,453705631 | 1,369553506 | 0,24438 | 0,55253387 | zinc finger protein 624 |
| ENSG00000132321 | IQCA1 | 0,453688786 | 1,369537515 | 0,17468 | 0,49886732 | IQ motif containing with AAA domain 1 |
| ENSG00000137492 | THAP12 | 0,453677592 | 1,369526889 | 0,16479 | 0,49092529 | THAP domain containing 12 |
| ENSG00000174891 | RSRC1 | 0,453559056 | 1,36941437 | 0,26895 | 0,57085484 | arginine and serine rich coiled-coil 1 |
| ENSG00000198876 | DCAF12 | 0,453517111 | 1,369374556 | 0,04265 | 0,27826437 | DDB1 and CUL4 associated factor 12 |
| ENSG00000168036 | CTNNB1 | 0,453446412 | 1,369307451 | 0,05321 | 0,30971589 | catenin beta 1 |
| ENSG00000119328 | ABITRAM | 0,453358285 | 1,369223809 | 0,2377 | 0,54861973 | actin binding transcription modulator |
| ENSG00000083845 | RPS5 | 0,453273479 | 1,369143325 | 0,21103 | 0,52983259 | ribosomal protein S5 |

| | | | | | | |
|---|---|---|---|---|---|---|
| ENSG00000197748 | CFAP43 | 0,453112693 | 1,368990744 | 0,17299 | 0,49772429 | cilia and flagella associated protein 43 |
| ENSG00000164684 | ZNF704 | 0,453067185 | 1,368947562 | 0,19941 | 0,52039716 | zinc finger protein 704 |
| ENSG00000133107 | TRPC4 | 0,453054771 | 1,368935783 | 0,34036 | 0,61897407 | transient receptor potential cation channel subfamily C member 4 |
| ENSG00000115524 | SF3B1 | 0,453053715 | 1,368934781 | 0,20927 | 0,52826608 | splicing factor 3b subunit 1 |
| ENSG00000100697 | DICER1 | 0,452925932 | 1,368813536 | 0,15062 | 0,47885749 | dicer 1, ribonuclease III |
| ENSG00000168488 | ATXN2L | 0,452920723 | 1,368808595 | 0,12141 | 0,44078088 | ataxin 2 like |
| ENSG00000184203 | PPP1R2 | 0,452797505 | 1,368691692 | 0,07181 | 0,35363591 | protein phosphatase 1 regulatory inhibitor subunit 2 |
| ENSG00000204178 | MACO1 | 0,452734183 | 1,368631619 | 0,25629 | 0,56078582 | macoilin 1 |
| ENSG00000102978 | POLR2C | 0,452731164 | 1,368628755 | 0,20165 | 0,52214043 | RNA polymerase II subunit C |
| ENSG00000106723 | SPIN1 | 0,452654774 | 1,368556289 | 0,1853 | 0,50876814 | spindlin 1 |
| ENSG00000285070 | ADIPOR2 | 0,452540737 | 1,368448116 | 0,15465 | 0,48266733 | adiponectin receptor 2 |
| ENSG00000006831 | ADIPOR2 | 0,452540737 | 1,368448116 | 0,15465 | 0,48266733 | adiponectin receptor 2 |
| ENSG00000259040 | BLOC1S5-TXN | 0,452512702 | 1,368421525 | 0,77615 | 0,89650232 | BLOC1S5-TXNDC5 readthrough (NMD candidate) |
| ENSG00000155545 | MIER3 | 0,452512036 | 1,368420892 | 0,12544 | 0,44729629 | MIER family member 3 |
| ENSG00000275618 | BFAR | 0,452394868 | 1,368309762 | 0,16779 | 0,49395055 | bifunctional apoptosis regulator |
| ENSG00000122952 | ZWINT | 0,452366035 | 1,368282415 | 0,13243 | 0,45752148 | ZW10 interacting kinetochore protein |
| ENSG00000163645 | ERICH6 | 0,452323022 | 1,368241621 | 0,60037 | 0,79583847 | glutamate rich 6 |
| ENSG00000238061 | AL356273,2 | 0,452224869 | 1,368148537 | 0,43595 | 0,68647216 | proteasome (prosome, macropain) 26S subunit, non-ATPase, 7 (Mov34 homolog) (PSMD7) pseudogene |
| ENSG00000231066 | NPM1P9 | 0,452119819 | 1,368048919 | 0,30706 | 0,59617071 | nucleophosmin 1 pseudogene 9 |
| ENSG00000229298 | TUBB8P1 | 0,452057247 | 1,367989585 | 0,34987 | 0,62563749 | tubulin beta 8 class VIII pseudogene 1 |
| ENSG00000280870 | MIR325HG | 0,451973511 | 1,367910188 | 0,16966 | 0,49589723 | MIR325 host gene |
| ENSG00000055917 | PUM2 | 0,451861504 | 1,367803991 | 0,23667 | 0,54763386 | pumilio RNA binding family member 2 |
| ENSG00000162688 | AGL | 0,451673936 | 1,367626172 | 0,27352 | 0,57362145 | amylo-alpha-1, 6-glucosidase, 4-alpha-glucanotransferase |
| ENSG00000278463 | H2AC4 | 0,451671001 | 1,367623389 | 0,16604 | 0,49204668 | H2A clustered histone 4 |
| ENSG00000239617 | AC073610,1 | 0,451632177 | 1,367586586 | 0,7052 | 0,85797994 | ribosomal protein L32 (RPL32) pseudogene |
| ENSG00000125319 | HROB | 0,451558099 | 1,367516366 | 0,13154 | 0,45658336 | homologous recombination factor with OB-fold |
| ENSG00000080561 | MID2 | 0,451478777 | 1,36744118 | 0,3117 | 0,59912688 | midline 2 |
| ENSG00000100154 | TTC28 | 0,451461805 | 1,367425093 | 0,08848 | 0,38476379 | tetratricopeptide repeat domain 28 |
| ENSG00000213463 | SYNJ2BP | 0,451407208 | 1,367373345 | 0,01713 | 0,17198199 | synaptojanin 2 binding protein |
| ENSG00000053438 | NNAT | 0,451361219 | 1,367329758 | 0,08153 | 0,37260031 | neuronatin |
| ENSG00000284850 | DCP1B | 0,451340904 | 1,367310504 | 0,33985 | 0,61872249 | decapping mRNA 1B |

| | | | | | | |
|---|---|---|---|---|---|---|
| ENSG00000151065 | DCP1B | 0,451340904 | 1,367310504 | 0,33985 | 0,61872249 | decapping mRNA 1B |
| ENSG00000204764 | RANBP17 | 0,45128016 | 1,367252935 | 0,35385 | 0,62865377 | RAN binding protein 17 |
| ENSG00000227456 | LINC00310 | 0,451148016 | 1,367127708 | 0,11135 | 0,42416781 | long intergenic non-protein coding RNA 310 |
| ENSG00000260548 | AL035425,2 | 0,451103691 | 1,367085705 | 0,62191 | 0,80819595 | insulin receptor substrate 4 |
| ENSG00000119698 | PPP4R4 | 0,45109303 | 1,367075603 | 0,26875 | 0,57078754 | protein phosphatase 4 regulatory subunit 4 |
| ENSG00000023318 | ERP44 | 0,45104708 | 1,367032063 | 0,22877 | 0,54259385 | endoplasmic reticulum protein 44 |
| ENSG00000056291 | NPFFR2 | 0,451023596 | 1,36700981 | 0,52817 | 0,75077432 | neuropeptide FF receptor 2 |
| ENSG00000157077 | ZFYVE9 | 0,450946385 | 1,366936652 | 0,12001 | 0,43868155 | zinc finger FYVE-type containing 9 |
| ENSG00000124193 | SRSF6 | 0,450806083 | 1,366803723 | 0,2252 | 0,54058261 | serine and arginine rich splicing factor 6 |
| ENSG00000010818 | HIVEP2 | 0,450733955 | 1,366735391 | 0,24286 | 0,55182091 | HIVEP zinc finger 2 |
| ENSG00000043093 | DCUN1D1 | 0,450605584 | 1,366613784 | 0,28135 | 0,578832 | defective in cullin neddylation 1 domain containing 1 |
| ENSG00000179361 | ARID3B | 0,450558088 | 1,366568794 | 0,12252 | 0,44227785 | AT-rich interaction domain 3B |
| ENSG00000175110 | MRPS22 | 0,450556337 | 1,366567136 | 0,1175 | 0,43477245 | mitochondrial ribosomal protein S22 |
| ENSG00000153201 | RANBP2 | 0,450550867 | 1,366561955 | 0,20004 | 0,5209612 | RAN binding protein 2 |
| ENSG00000152749 | GPR180 | 0,450379883 | 1,366400003 | 0,35838 | 0,63185282 | G protein-coupled receptor 180 |
| ENSG00000285976 | AL135905,2 | 0,450313357 | 1,366336996 | 0,25388 | 0,55896983 | novel protein |
| ENSG00000234026 | AL157834,2 | 0,450256474 | 1,366283125 | 0,42261 | 0,67675106 | novel transcript, antisense to C10orf129 |
| ENSG00000145087 | STXBP5L | 0,450152815 | 1,36618496 | 0,23368 | 0,54563244 | syntaxin binding protein 5 like |
| ENSG00000044574 | HSPA5 | 0,450028184 | 1,366066943 | 0,0724 | 0,35526408 | heat shock protein family A (Hsp70) member 5 |
| ENSG00000277423 | AC069234,5 | 0,450025644 | 1,366064538 | 0,4726 | 0,71198726 | novel transcript |
| ENSG00000278211 | TRIM69 | 0,450008376 | 1,366048188 | 0,41978 | 0,67542613 | tripartite motif containing 69 |
| ENSG00000272158 | AL139022,2 | 0,449977447 | 1,366018903 | 0,54156 | 0,75882662 | novel transcript, antisense to CHURC1-FNTB |
| ENSG00000196418 | ZNF124 | 0,449924658 | 1,36596892 | 0,09101 | 0,39033853 | zinc finger protein 124 |
| ENSG00000141627 | DYM | 0,449907952 | 1,365953103 | 0,19092 | 0,51394498 | dymeclin |
| ENSG00000260743 | AC007823,1 | 0,449859364 | 1,365907099 | 0,32387 | 0,60659203 | novel transcript |
| ENSG00000270964 | AC016355,1 | 0,449814701 | 1,365864814 | 0,21643 | 0,53484961 | novel transcript |
| ENSG00000186767 | SPIN4 | 0,44972975 | 1,365784389 | 0,14451 | 0,47209141 | spindlin family member 4 |
| ENSG00000069493 | CLEC2D | 0,449694347 | 1,365750874 | 0,33448 | 0,61426654 | C-type lectin domain family 2 member D |
| ENSG00000143621 | ILF2 | 0,449413646 | 1,36548517 | 0,13811 | 0,46543358 | interleukin enhancer binding factor 2 |
| ENSG00000110721 | CHKA | 0,44935085 | 1,365425736 | 0,02209 | 0,19768196 | choline kinase alpha |
| ENSG00000180182 | MED14 | 0,449321661 | 1,365398111 | 0,24313 | 0,55199251 | mediator complex subunit 14 |
| ENSG00000030066 | NUP160 | 0,44929798 | 1,365375699 | 0,22712 | 0,54168591 | nucleoporin 160 |
| ENSG00000123607 | TTC21B | 0,449292829 | 1,365370824 | 0,12519 | 0,44690314 | tetratricopeptide repeat domain 21B |

| | | | | | | |
|---|---|---|---|---|---|---|
| ENSG00000115825 | PRKD3 | 0,449256705 | 1,365336637 | 0,0664 | 0,34113137 | protein kinase D3 |
| ENSG00000182952 | HMGN4 | 0,449213953 | 1,365296178 | 0,26462 | 0,56738489 | high mobility group nucleosomal binding domain 4 |
| ENSG00000137601 | NEK1 | 0,449195121 | 1,365278356 | 0,24337 | 0,55210404 | NIMA related kinase 1 |
| ENSG00000175193 | PARL | 0,449150855 | 1,365236466 | 0,15006 | 0,47815167 | presenilin associated rhomboid like |
| ENSG00000106028 | SSBP1 | 0,449127634 | 1,365214492 | 0,3191 | 0,60409862 | single stranded DNA binding protein 1 |
| ENSG00000115540 | MOB4 | 0,449077408 | 1,365166964 | 0,26294 | 0,56587311 | MOB family member 4, phocein |
| ENSG00000174748 | RPL15 | 0,449011684 | 1,365104774 | 0,19855 | 0,51980353 | ribosomal protein L15 |
| ENSG00000103769 | RAB11A | 0,44898625 | 1,365080708 | 0,25771 | 0,56191608 | RAB11A, member RAS oncogene family |
| ENSG00000283201 | AC092329,3 | 0,44891735 | 1,365015516 | 0,68597 | 0,84654659 | zinc finger protein 724 |
| ENSG00000005812 | FBXL3 | 0,44890654 | 1,365005288 | 0,33973 | 0,61872249 | F-box and leucine rich repeat protein 3 |
| ENSG00000174197 | MGA | 0,44875874 | 1,364865455 | 0,10206 | 0,40934973 | MAX dimerization protein MGA |
| ENSG00000121579 | NAA50 | 0,448726951 | 1,36483538 | 0,07097 | 0,35173843 | N-alpha-acetyltransferase 50, NatE catalytic subunit |
| ENSG00000083093 | PALB2 | 0,448580675 | 1,364697006 | 0,12861 | 0,45191041 | partner and localizer of BRCA2 |
| ENSG00000143398 | PIP5K1A | 0,44835247 | 1,364481156 | 0,00996 | 0,12783227 | phosphatidylinositol-4-phosphate 5-kinase type 1 alpha |
| ENSG00000164219 | PGGT1B | 0,448343003 | 1,364472202 | 0,26046 | 0,56388142 | protein geranylgeranyltransferase type I subunit beta |
| ENSG00000159873 | CCDC117 | 0,4483297 | 1,36445962 | 0,03904 | 0,26748219 | coiled-coil domain containing 117 |
| ENSG00000132549 | VPS13B | 0,448324195 | 1,364454413 | 0,23418 | 0,54600096 | vacuolar protein sorting 13 homolog B |
| ENSG00000065427 | KARS1 | 0,448293558 | 1,364425438 | 0,19873 | 0,51984196 | lysyl-tRNA synthetase 1 |
| ENSG00000272097 | AL024498,1 | 0,448197071 | 1,364334189 | 0,48062 | 0,71770193 | novel transcript |
| ENSG00000144746 | ARL6IP5 | 0,448099039 | 1,364241485 | 0,25487 | 0,55944852 | ADP ribosylation factor like GTPase 6 interacting protein 5 |
| ENSG00000016864 | GLT8D1 | 0,448092888 | 1,364235669 | 0,23821 | 0,54870569 | glycosyltransferase 8 domain containing 1 |
| ENSG00000128000 | ZNF780B | 0,448049286 | 1,364194438 | 0,47488 | 0,71364583 | zinc finger protein 780B |
| ENSG00000136448 | NMT1 | 0,447982168 | 1,364130974 | 0,04564 | 0,2883604 | N-myristoyltransferase 1 |
| ENSG00000151849 | CENPJ | 0,447833686 | 1,363990585 | 0,26107 | 0,56416552 | centromere protein J |
| ENSG00000150051 | MKX | 0,447832004 | 1,363988994 | 0,58997 | 0,78936321 | mohawk homeobox |
| ENSG00000197619 | ZNF615 | 0,447792433 | 1,363951583 | 0,21699 | 0,53505085 | zinc finger protein 615 |
| ENSG00000165637 | VDAC2 | 0,447684818 | 1,363849846 | 0,17212 | 0,49710979 | voltage dependent anion channel 2 |
| ENSG00000233639 | PANTR1 | 0,447596299 | 1,363766167 | 0,42173 | 0,67643128 | POU3F3 adjacent non-coding transcript 1 |
| ENSG00000173273 | TNKS | 0,447510479 | 1,363685044 | 0,2144 | 0,53334744 | tankyrase |
| ENSG00000102043 | MTMR8 | 0,447256198 | 1,36344471 | 0,34843 | 0,62472936 | myotubularin related protein 8 |
| ENSG00000011258 | MBTD1 | 0,447172922 | 1,363366011 | 0,24096 | 0,55003907 | mbt domain containing 1 |
| ENSG00000140396 | NCOA2 | 0,44713117 | 1,363326555 | 0,18907 | 0,51243448 | nuclear receptor coactivator 2 |
| ENSG00000121058 | COIL | 0,447086156 | 1,363284018 | 0,14413 | 0,47172279 | coilin |

| | | | | | | |
|---|---|---|---|---|---|---|
| ENSG00000116754 | SRSF11 | 0,447078974 | 1,363277232 | 0,27525 | 0,57491355 | serine and arginine rich splicing factor 11 |
| ENSG00000081386 | ZNF510 | 0,447010748 | 1,363212763 | 0,34741 | 0,62411645 | zinc finger protein 510 |
| ENSG00000172809 | RPL38 | 0,446746853 | 1,36296343 | 0,18086 | 0,50443353 | ribosomal protein L38 |
| ENSG00000138796 | HADH | 0,4466559 | 1,362877506 | 0,07194 | 0,3539609 | hydroxyacyl-CoA dehydrogenase |
| ENSG00000285542 | AC013717,1 | 0,446598846 | 1,36282361 | 0,51106 | 0,73835122 | novel protein, PPM1B-SLC3A1 readthrough |
| ENSG00000278742 | LRRC37A | 0,44657246 | 1,362798684 | 0,17016 | 0,49606609 | leucine rich repeat containing 37A |
| ENSG00000184445 | KNTC1 | 0,446571367 | 1,362797652 | 0,27942 | 0,57742338 | kinetochore associated 1 |
| ENSG00000013563 | DNASE1L1 | 0,446477183 | 1,362708687 | 0,08242 | 0,37394502 | deoxyribonuclease 1 like 1 |
| ENSG00000140299 | BNIP2 | 0,446367688 | 1,362605267 | 0,16393 | 0,4902029 | BCL2 interacting protein 2 |
| ENSG00000142945 | KIF2C | 0,446352294 | 1,362590728 | 0,23666 | 0,54763386 | kinesin family member 2C |
| ENSG00000278615 | C11orf98 | 0,446178001 | 1,362426122 | 0,24754 | 0,5550162 | chromosome 11 open reading frame 98 |
| ENSG00000126457 | PRMT1 | 0,446101796 | 1,362354158 | 0,28139 | 0,578832 | protein arginine methyltransferase 1 |
| ENSG00000243678 | NME2 | 0,44604268 | 1,362298335 | 0,17127 | 0,49653881 | NME/NM23 nucleoside diphosphate kinase 2 |
| ENSG00000167491 | GATAD2A | 0,445999684 | 1,362257736 | 0,04759 | 0,29334767 | GATA zinc finger domain containing 2A |
| ENSG00000270110 | AL353593,2 | 0,445996766 | 1,362254981 | 0,39137 | 0,65510797 | novel transcript, antisense to OBSCN |
| ENSG00000036054 | TBC1D23 | 0,445829733 | 1,362097271 | 0,1391 | 0,46648276 | TBC1 domain family member 23 |
| ENSG00000273877 | AC236972,3 | 0,445812412 | 1,362080917 | 0,60851 | 0,80122454 | uncharacterized LOC105373373 |
| ENSG00000237088 | AL163193,1 | 0,445795932 | 1,362065358 | 0,46822 | 0,70786237 | ribosomal protein L7a (RPL7A) pseudogene |
| ENSG00000228016 | RAPGEF4-AS1 | 0,445671912 | 1,361948275 | 0,54401 | 0,76034748 | RAPGEF4 antisense RNA 1 |
| ENSG00000198075 | SULT1C4 | 0,445650534 | 1,361928093 | 0,1184 | 0,43614969 | sulfotransferase family 1C member 4 |
| ENSG00000178860 | MSC | 0,445639724 | 1,361917889 | 0,41488 | 0,67170152 | musculin |
| ENSG00000110330 | BIRC2 | 0,445573738 | 1,361855598 | 0,34033 | 0,61897407 | baculoviral IAP repeat containing 2 |
| ENSG00000259494 | MRPL46 | 0,445516678 | 1,361801737 | 0,21685 | 0,53505085 | mitochondrial ribosomal protein L46 |
| ENSG00000270392 | PFN1P2 | 0,44549784 | 1,361783955 | 0,31659 | 0,60242329 | profilin 1 pseudogene 2 |
| ENSG00000171208 | NETO2 | 0,445486247 | 1,361773012 | 0,04563 | 0,2883604 | neuropilin and tolloid like 2 |
| ENSG00000071127 | WDR1 | 0,445470178 | 1,361757844 | 0,22661 | 0,54144 | WD repeat domain 1 |
| ENSG00000107290 | SETX | 0,445415238 | 1,361705987 | 0,30206 | 0,59294633 | senataxin |
| ENSG00000162976 | SLC66A3 | 0,445408991 | 1,361700091 | 0,17039 | 0,49641416 | solute carrier family 66 member 3 |
| ENSG00000138674 | SEC31A | 0,445274191 | 1,361572865 | 0,1602 | 0,48737351 | SEC31 homolog A, COPII coat complex component |
| ENSG00000139343 | SNRPF | 0,445236885 | 1,361537657 | 0,30158 | 0,59277108 | small nuclear ribonucleoprotein polypeptide F |
| ENSG00000223528 | AL359094,1 | 0,445179168 | 1,361483188 | 0,32993 | 0,61087344 | novel transcript |
| ENSG00000138459 | SLC35A5 | 0,44506824 | 1,361378508 | 0,3627 | 0,63381706 | solute carrier family 35 member A5 |
| ENSG00000154059 | IMPACT | 0,445030365 | 1,361342768 | 0,29422 | 0,5885627 | impact RWD domain protein |

| | | | | | | |
|---|---|---|---|---|---|---|
| ENSG00000112249 | ASCC3 | 0,444895376 | 1,361215397 | 0,19902 | 0,52017853 | activating signal cointegrator 1 complex subunit 3 |
| ENSG00000075618 | FSCN1 | 0,444831738 | 1,361155355 | 0,04232 | 0,27814302 | fascin actin-bundling protein 1 |
| ENSG00000104442 | ARMC1 | 0,444784165 | 1,361110472 | 0,37047 | 0,6399389 | armadillo repeat containing 1 |
| ENSG00000160211 | G6PD | 0,444741716 | 1,361070424 | 0,1589 | 0,48627932 | glucose-6-phosphate dehydrogenase |
| ENSG00000148600 | CDHR1 | 0,444705081 | 1,361035862 | 0,32544 | 0,60794826 | cadherin related family member 1 |
| ENSG00000086598 | TMED2 | 0,444669204 | 1,361002016 | 0,24011 | 0,54943734 | transmembrane p24 trafficking protein 2 |
| ENSG00000051825 | MPHOSPH9 | 0,444617452 | 1,360953196 | 0,24084 | 0,55001001 | M-phase phosphoprotein 9 |
| ENSG00000233175 | AC008105,1 | 0,444446565 | 1,360792001 | 0,4507 | 0,69592991 | novel transcript, antisense to FMNL1 |
| ENSG00000287729 | AF241728,2 | 0,444440234 | 1,360786029 | 0,62889 | 0,81237816 | novel transcript |
| ENSG00000183850 | ZNF730 | 0,444377003 | 1,360726389 | 0,4029 | 0,66398107 | zinc finger protein 730 |
| ENSG00000247728 | AC091057,2 | 0,444326281 | 1,36067855 | 0,25352 | 0,55857654 | novel transcript, antisense to ARHGAP11B |
| ENSG00000204681 | GABBR1 | 0,444296199 | 1,360650178 | 0,3203 | 0,6052338 | gamma-aminobutyric acid type B receptor subunit 1 |
| ENSG00000130711 | PRDM12 | 0,444288713 | 1,360643118 | 0,57474 | 0,77988303 | PR/SET domain 12 |
| ENSG00000145949 | MYLK4 | 0,444255322 | 1,360611626 | 0,32715 | 0,60902638 | myosin light chain kinase family member 4 |
| ENSG00000143933 | CALM2 | 0,444056404 | 1,360424039 | 0,32122 | 0,60573903 | calmodulin 2 |
| ENSG00000107020 | PLGRKT | 0,444027849 | 1,360397112 | 0,42436 | 0,67819918 | plasminogen receptor with a C-terminal lysine |
| ENSG00000101773 | RBBP8 | 0,443964577 | 1,360337451 | 0,30099 | 0,59244013 | RB binding protein 8, endonuclease |
| ENSG00000145723 | GIN1 | 0,443955814 | 1,360329188 | 0,29861 | 0,59089801 | gypsy retrotransposon integrase 1 |
| ENSG00000137876 | RSL24D1 | 0,443945207 | 1,360319187 | 0,25009 | 0,55657567 | ribosomal L24 domain containing 1 |
| ENSG00000095564 | BTAF1 | 0,443928588 | 1,360303517 | 0,22658 | 0,54144 | B-TFIID TATA-box binding protein associated factor 1 |
| ENSG00000205981 | DNAJC19 | 0,443918116 | 1,360293644 | 0,31527 | 0,60158701 | DnaJ heat shock protein family (Hsp40) member C19 |
| ENSG00000165832 | TRUB1 | 0,443910503 | 1,360286465 | 0,26017 | 0,56382871 | TruB pseudouridine synthase family member 1 |
| ENSG00000172059 | KLF11 | 0,443905573 | 1,360281817 | 0,09339 | 0,39491488 | Kruppel like factor 11 |
| ENSG00000090905 | TNRC6A | 0,443899195 | 1,360275803 | 0,25324 | 0,55836184 | trinucleotide repeat containing adaptor 6A |
| ENSG00000203791 | EEF1AKMT2 | 0,443898851 | 1,360275479 | 0,16329 | 0,48972962 | EEF1A lysine methyltransferase 2 |
| ENSG00000135521 | LTV1 | 0,443874065 | 1,360252109 | 0,11535 | 0,43106723 | LTV1 ribosome biogenesis factor |
| ENSG00000197933 | ZNF823 | 0,443683817 | 1,360072744 | 0,26884 | 0,57085484 | zinc finger protein 823 |
| ENSG00000242697 | RPL5P12 | 0,443565437 | 1,359961148 | 0,3435 | 0,62091871 | ribosomal protein L5 pseudogene 12 |
| ENSG00000135945 | REV1 | 0,443560224 | 1,359956234 | 0,16377 | 0,49000347 | REV1 DNA directed polymerase |
| ENSG00000119414 | PPP6C | 0,443504758 | 1,359903951 | 0,2188 | 0,53631957 | protein phosphatase 6 catalytic subunit |
| ENSG00000143924 | EML4 | 0,443439773 | 1,359842697 | 0,02116 | 0,19303449 | EMAP like 4 |
| ENSG00000132591 | ERAL1 | 0,443305551 | 1,359716189 | 0,12794 | 0,45145929 | Era like 12S mitochondrial rRNA chaperone 1 |
| ENSG00000118492 | ADGB | 0,44325479 | 1,359668348 | 0,48812 | 0,72309725 | androglobin |

| | | | | | | |
|---|---|---|---|---|---|---|
| ENSG00000288504 | YBX2 | 0,443244414 | 1,359658569 | 0,393 | 0,65608324 | Y-box binding protein 2 |
| ENSG00000006047 | YBX2 | 0,443244414 | 1,359658569 | 0,393 | 0,65608324 | Y-box binding protein 2 |
| ENSG00000115020 | PIKFYVE | 0,443232145 | 1,359647006 | 0,16363 | 0,49000031 | phosphoinositide kinase, FYVE-type zinc finger containing |
| ENSG00000288620 | AC092032,2 | 0,443192589 | 1,359609728 | 0,65721 | 0,82867637 | ring finger protein 216 pseudogene 1 |
| ENSG00000112640 | PPP2R5D | 0,443165722 | 1,359584408 | 0,11749 | 0,43477245 | protein phosphatase 2 regulatory subunit B'delta |
| ENSG00000264343 | NOTCH2NLA | 0,443042648 | 1,359468429 | 0,19339 | 0,51546367 | notch 2 N-terminal like A |
| ENSG00000138759 | FRAS1 | 0,442991481 | 1,359420215 | 0,10157 | 0,40840817 | Fraser extracellular matrix complex subunit 1 |
| ENSG00000115233 | PSMD14 | 0,442877874 | 1,35931317 | 0,25283 | 0,55796133 | proteasome 26S subunit, non-ATPase 14 |
| ENSG00000276023 | DUSP14 | 0,442854927 | 1,359291549 | 0,60383 | 0,79770754 | dual specificity phosphatase 14 |
| ENSG00000111252 | SH2B3 | 0,442809484 | 1,359248734 | 0,01818 | 0,17825548 | SH2B adaptor protein 3 |
| ENSG00000171940 | ZNF217 | 0,442759085 | 1,359201251 | 0,03611 | 0,25729602 | zinc finger protein 217 |
| ENSG00000125944 | HNRNPR | 0,442746173 | 1,359189086 | 0,15665 | 0,48425071 | heterogeneous nuclear ribonucleoprotein R |
| ENSG00000005189 | REXO5 | 0,4426773 | 1,359124201 | 0,31499 | 0,60145932 | RNA exonuclease 5 |
| ENSG00000242715 | CCDC169 | 0,44264545 | 1,359094196 | 0,38952 | 0,65388973 | coiled-coil domain containing 169 |
| ENSG00000102221 | JADE3 | 0,442634491 | 1,359083872 | 0,15739 | 0,48457045 | jade family PHD finger 3 |
| ENSG00000282826 | FRG1CP | 0,442628781 | 1,359078494 | 0,03835 | 0,26508383 | FSHD region gene 1 family member C, pseudogene |
| ENSG00000052126 | PLEKHA5 | 0,44258984 | 1,35904181 | 0,12141 | 0,44078088 | pleckstrin homology domain containing A5 |
| ENSG00000140307 | GTF2A2 | 0,442488589 | 1,358946433 | 0,31946 | 0,60429866 | general transcription factor IIA subunit 2 |
| ENSG00000167191 | GPRC5B | 0,442461707 | 1,358921111 | 0,1893 | 0,51254557 | G protein-coupled receptor class C group 5 member B |
| ENSG00000169131 | ZNF354A | 0,442445363 | 1,358905716 | 0,07112 | 0,35204529 | zinc finger protein 354A |
| ENSG00000254061 | AC091144,2 | 0,442331819 | 1,358798772 | 0,58449 | 0,78572744 | RNA binding motif protein 4 (RBM4) pseudogene |
| ENSG00000143363 | PRUNE1 | 0,442327783 | 1,35879497 | 0,10596 | 0,4154675 | prune exopolyphosphatase 1 |
| ENSG00000160695 | VPS11 | 0,442282944 | 1,358752739 | 0,27214 | 0,57306433 | VPS11 core subunit of CORVET and HOPS complexes |
| ENSG00000151136 | BTBD11 | 0,442246848 | 1,358718744 | 0,28365 | 0,58072192 | BTB domain containing 11 |
| ENSG00000197275 | RAD54B | 0,442194436 | 1,358669383 | 0,33845 | 0,61774255 | RAD54 homolog B |
| ENSG00000185420 | SMYD3 | 0,44213731 | 1,358615586 | 0,15356 | 0,48191866 | SET and MYND domain containing 3 |
| ENSG00000165516 | KLHDC2 | 0,442106223 | 1,358586311 | 0,19379 | 0,51547251 | kelch domain containing 2 |
| ENSG00000276463 | TAF9 | 0,442073173 | 1,358555188 | 0,20986 | 0,52877648 | TATA-box binding protein associated factor 9 |
| ENSG00000280591 | AC237191,1 | 0,442068528 | 1,358550814 | 0,3539 | 0,62865377 | novel transcript |
| ENSG00000281099 | AC244167,1 | 0,442068528 | 1,358550814 | 0,3539 | 0,62865377 | novel transcript |
| ENSG00000164754 | RAD21 | 0,441996378 | 1,358482874 | 0,27354 | 0,57362145 | RAD21 cohesin complex component |
| ENSG00000119392 | GLE1 | 0,441919245 | 1,358410245 | 0,14398 | 0,47146567 | GLE1 RNA export mediator |
| ENSG00000184384 | MAML2 | 0,441915418 | 1,358406642 | 0,28932 | 0,58494545 | mastermind like transcriptional coactivator 2 |

| | | | | | | |
|---|---|---|---|---|---|---|
| ENSG00000115808 | STRN | 0,441900819 | 1,358392895 | 0,32254 | 0,60577115 | striatin |
| ENSG00000141431 | ASXL3 | 0,441851671 | 1,358346621 | 0,25506 | 0,55954928 | ASXL transcriptional regulator 3 |
| ENSG00000247626 | MARS2 | 0,441845376 | 1,358340694 | 0,14739 | 0,47490365 | methionyl-tRNA synthetase 2, mitochondrial |
| ENSG00000224891 | ARL14EPP1 | 0,441837825 | 1,358333584 | 0,39479 | 0,65769097 | ARL14EP pseudogene 1 |
| ENSG00000161057 | PSMC2 | 0,441774231 | 1,35827371 | 0,14312 | 0,47072472 | proteasome 26S subunit, ATPase 2 |
| ENSG00000123545 | NDUFAF4 | 0,44163082 | 1,358138697 | 0,30955 | 0,59799482 | NADH:ubiquinone oxidoreductase complex assembly factor 4 |
| ENSG00000154645 | CHODL | 0,441587019 | 1,358097465 | 0,32895 | 0,60998638 | chondrolectin |
| ENSG00000010017 | RANBP9 | 0,441483966 | 1,358000458 | 0,24391 | 0,55250543 | RAN binding protein 9 |
| ENSG00000105887 | MTPN | 0,441401916 | 1,357923227 | 0,07104 | 0,35196374 | myotrophin |
| ENSG00000140848 | CPNE2 | 0,441366678 | 1,357890059 | 0,31447 | 0,6011918 | copine 2 |
| ENSG00000153827 | TRIP12 | 0,44129689 | 1,357824376 | 0,29235 | 0,58692263 | thyroid hormone receptor interactor 12 |
| ENSG00000272081 | AC008972,2 | 0,441261494 | 1,357791062 | 0,4727 | 0,71203455 | novel transcript |
| ENSG00000215795 | AL390728,2 | 0,441117305 | 1,357655366 | 0,4115 | 0,66919678 | peroxisomal biogenesis factor 5 (PEX5) pseudogene |
| ENSG00000119729 | RHOQ | 0,441113832 | 1,357652097 | 0,2141 | 0,53334744 | ras homolog family member Q |
| ENSG00000204231 | RXRB | 0,44100115 | 1,357546062 | 0,23912 | 0,5491566 | retinoid X receptor beta |
| ENSG00000115661 | STK16 | 0,440943477 | 1,357491794 | 0,19391 | 0,51547251 | serine/threonine kinase 16 |
| ENSG00000126749 | EMG1 | 0,440866173 | 1,357419057 | 0,10279 | 0,41062092 | EMG1 N1-specific pseudouridine methyltransferase |
| ENSG00000030419 | IKZF2 | 0,440762496 | 1,357321512 | 0,31508 | 0,60145932 | IKAROS family zinc finger 2 |
| ENSG00000255580 | AP000462,1 | 0,440759378 | 1,357318578 | 0,60995 | 0,80202886 | novel transcript |
| ENSG00000172244 | C5orf34 | 0,440756918 | 1,357316264 | 0,25358 | 0,55857654 | chromosome 5 open reading frame 34 |
| ENSG00000132386 | SERPINF1 | 0,440741315 | 1,357301585 | 0,3014 | 0,59258306 | serpin family F member 1 |
| ENSG00000047617 | ANO2 | 0,440695654 | 1,357258627 | 0,29008 | 0,58531219 | anoctamin 2 |
| ENSG00000198000 | NOL8 | 0,440692766 | 1,35725591 | 0,23677 | 0,54767193 | nucleolar protein 8 |
| ENSG00000162981 | LRATD1 | 0,440595148 | 1,357164077 | 0,14956 | 0,47748715 | LRAT domain containing 1 |
| ENSG00000128039 | SRD5A3 | 0,440540685 | 1,357112844 | 0,04448 | 0,28449899 | steroid 5 alpha-reductase 3 |
| ENSG00000230797 | YY2 | 0,440512745 | 1,357086561 | 0,3305 | 0,6112353 | YY2 transcription factor |
| ENSG00000152380 | FAM151B | 0,440388605 | 1,356969792 | 0,26338 | 0,56608528 | family with sequence similarity 151 member B |
| ENSG00000135414 | GDF11 | 0,440363744 | 1,356946409 | 0,11536 | 0,43106723 | growth differentiation factor 11 |
| ENSG00000160094 | ZNF362 | 0,440362681 | 1,356945409 | 0,00781 | 0,1124014 | zinc finger protein 362 |
| ENSG00000185716 | MOSMO | 0,44030156 | 1,356887922 | 0,19661 | 0,51780838 | modulator of smoothened |
| ENSG00000223973 | PAFAH1B1P2 | 0,440282312 | 1,356869819 | 0,59693 | 0,79434997 | platelet activating factor acetylhydrolase 1b regulatory subunit 1 pseudogene 2 |

| | | | | | | |
|---|---|---|---|---|---|---|
| ENSG00000179476 | C14orf28 | 0,440245256 | 1,356834967 | 0,35014 | 0,62575079 | chromosome 14 open reading frame 28 |
| ENSG00000165678 | GHITM | 0,440132047 | 1,3567285 | 0,22184 | 0,53821014 | growth hormone inducible transmembrane protein |
| ENSG00000230989 | HSBP1 | 0,440105803 | 1,356703821 | 0,3526 | 0,62750667 | heat shock factor binding protein 1 |
| ENSG00000272931 | AC099568,2 | 0,44007888 | 1,356678502 | 0,26345 | 0,56613578 | novel transcript |
| ENSG00000173145 | NOC3L | 0,440046184 | 1,356647756 | 0,30312 | 0,59345427 | NOC3 like DNA replication regulator |
| ENSG00000167981 | ZNF597 | 0,440015975 | 1,356619349 | 0,33078 | 0,6112353 | zinc finger protein 597 |
| ENSG00000187555 | USP7 | 0,439820236 | 1,356435301 | 0,17917 | 0,50364038 | ubiquitin specific peptidase 7 |
| ENSG00000267414 | SETBP1-DT | 0,439805947 | 1,356421867 | 0,36557 | 0,6357945 | SETBP1 divergent transcript |
| ENSG00000008405 | CRY1 | 0,439771023 | 1,356389032 | 0,10598 | 0,4154675 | cryptochrome circadian regulator 1 |
| ENSG00000042088 | TDP1 | 0,439709503 | 1,356331193 | 0,20464 | 0,52443222 | tyrosyl-DNA phosphodiesterase 1 |
| ENSG00000234469 | CLDN34 | 0,439671115 | 1,356295103 | 0,54722 | 0,76270053 | claudin 34 |
| ENSG00000088812 | ATRN | 0,439593498 | 1,356222137 | 0,18463 | 0,50835796 | attractin |
| ENSG00000077147 | TM9SF3 | 0,439454805 | 1,356091763 | 0,09418 | 0,39648446 | transmembrane 9 superfamily member 3 |
| ENSG00000159228 | CBR1 | 0,439429914 | 1,356068367 | 0,33322 | 0,61316259 | carbonyl reductase 1 |
| ENSG00000213250 | RBMS2P1 | 0,439385294 | 1,356026426 | 0,61334 | 0,8035281 | RNA binding motif single stranded interacting protein 2 pseudogene 1 |
| ENSG00000244693 | CTAGE8 | 0,439367247 | 1,356009464 | 0,78903 | 0,90223158 | CTAGE family member 8 |
| ENSG00000139641 | ESYT1 | 0,439366667 | 1,356008919 | 0,05616 | 0,31656215 | extended synaptotagmin 1 |
| ENSG00000162961 | DPY30 | 0,439352374 | 1,355995484 | 0,20305 | 0,52296754 | dpy-30 histone methyltransferase complex regulatory subunit |
| ENSG00000116698 | SMG7 | 0,439339698 | 1,35598357 | 0,04528 | 0,2876168 | SMG7 nonsense mediated mRNA decay factor |
| ENSG00000126218 | F10 | 0,439322661 | 1,355967557 | 0,40377 | 0,66409141 | coagulation factor X |
| ENSG00000129596 | CDO1 | 0,439308045 | 1,35595382 | 0,36024 | 0,63261878 | cysteine dioxygenase type 1 |
| ENSG00000021776 | AQR | 0,439024796 | 1,355687627 | 0,21192 | 0,53076389 | aquarius intron-binding spliceosomal factor |
| ENSG00000196396 | PTPN1 | 0,438990138 | 1,35565506 | 0,06029 | 0,32601555 | protein tyrosine phosphatase non-receptor type 1 |
| ENSG00000120526 | NUDCD1 | 0,438969024 | 1,355635219 | 0,20829 | 0,52766387 | NudC domain containing 1 |
| ENSG00000138029 | HADHB | 0,438959929 | 1,355626674 | 0,23303 | 0,54557342 | hydroxyacyl-CoA dehydrogenase trifunctional multienzyme complex subunit beta |
| ENSG00000253540 | FAM86HP | 0,438899649 | 1,355570033 | 0,23481 | 0,5463188 | family with sequence similarity 86 member H, pseudogene |
| ENSG00000072401 | UBE2D1 | 0,438778924 | 1,355456603 | 0,34552 | 0,6227624 | ubiquitin conjugating enzyme E2 D1 |
| ENSG00000226007 | BX005266,2 | 0,438569008 | 1,355259395 | 0,63988 | 0,81929791 | novel transcript |
| ENSG00000125166 | GOT2 | 0,438533708 | 1,355226235 | 0,24751 | 0,55499795 | glutamic-oxaloacetic transaminase 2 |
| ENSG00000177311 | ZBTB38 | 0,43850284 | 1,355197238 | 0,37014 | 0,63956649 | zinc finger and BTB domain containing 38 |

| | | | | | | |
|---|---|---|---|---|---|---|
| ENSG00000274364 | AL110115,1 | 0,43848243 | 1,355178066 | 0,59087 | 0,79004365 | novel transcript, sense intronic to HM13 |
| ENSG00000046647 | GEMIN8 | 0,438351141 | 1,355054747 | 0,09767 | 0,40273014 | gem nuclear organelle associated protein 8 |
| ENSG00000111913 | RIPOR2 | 0,438325033 | 1,355030226 | 0,44715 | 0,69341308 | RHO family interacting cell polarization regulator 2 |
| ENSG00000139083 | ETV6 | 0,438203283 | 1,354915879 | 0,09799 | 0,40323803 | ETS variant transcription factor 6 |
| ENSG00000105058 | FAM32A | 0,437953568 | 1,354681378 | 0,21261 | 0,53153879 | family with sequence similarity 32 member A |
| ENSG00000116473 | RAP1A | 0,437900436 | 1,354631488 | 0,28997 | 0,58531219 | RAP1A, member of RAS oncogene family |
| ENSG00000172292 | CERS6 | 0,437861464 | 1,354594895 | 0,25886 | 0,56260889 | ceramide synthase 6 |
| ENSG00000213281 | NRAS | 0,437852446 | 1,354586427 | 0,28275 | 0,58009943 | NRAS proto-oncogene, GTPase |
| ENSG00000155850 | SLC26A2 | 0,437498015 | 1,354253683 | 0,08597 | 0,38026553 | solute carrier family 26 member 2 |
| ENSG00000126804 | ZBTB1 | 0,43736628 | 1,35413003 | 0,10957 | 0,42167157 | zinc finger and BTB domain containing 1 |
| ENSG00000087077 | TRIP6 | 0,437183001 | 1,353958013 | 0,16624 | 0,49212363 | thyroid hormone receptor interactor 6 |
| ENSG00000173744 | AGFG1 | 0,437097767 | 1,353878024 | 0,2835 | 0,58065155 | ArfGAP with FG repeats 1 |
| ENSG00000107625 | DDX50 | 0,437096793 | 1,35387711 | 0,21508 | 0,53378006 | DExD-box helicase 50 |
| ENSG00000161860 | SYCE2 | 0,437064505 | 1,35384681 | 0,28585 | 0,58229607 | synaptonemal complex central element protein 2 |
| ENSG00000168994 | PXDC1 | 0,437057437 | 1,353840177 | 0,3579 | 0,63185282 | PX domain containing 1 |
| ENSG00000232450 | AL133517,1 | 0,43704264 | 1,353826292 | 0,21672 | 0,53494609 | ribosomal protein S2 (RPS2) pseudogene |
| ENSG00000157827 | FMNL2 | 0,436998788 | 1,353785142 | 0,20163 | 0,52214043 | formin like 2 |
| ENSG00000282019 | GPI | 0,436747616 | 1,353549469 | 0,20284 | 0,52276926 | glucose-6-phosphate isomerase |
| ENSG00000148143 | ZNF462 | 0,436670057 | 1,353476705 | 0,12288 | 0,44269063 | zinc finger protein 462 |
| ENSG00000177465 | ACOT4 | 0,436602387 | 1,353413221 | 0,20278 | 0,52269757 | acyl-CoA thioesterase 4 |
| ENSG00000136827 | TOR1A | 0,436583269 | 1,353395286 | 0,1095 | 0,42148306 | torsin family 1 member A |
| ENSG00000142669 | SH3BGRL3 | 0,436406867 | 1,353229814 | 0,15589 | 0,4836223 | SH3 domain binding glutamate rich protein like 3 |
| ENSG00000225611 | LINC02158 | 0,436275586 | 1,353106679 | 0,40967 | 0,6680455 | long intergenic non-protein coding RNA 2158 |
| ENSG00000235945 | AC002543,1 | 0,436235446 | 1,353069033 | 0,35244 | 0,62742225 | novel pseudogene |
| ENSG00000286161 | AC104066,4 | 0,436152582 | 1,352991318 | 0,40533 | 0,66502482 | novel transcript |
| ENSG00000100664 | EIF5 | 0,436118192 | 1,352959067 | 0,26087 | 0,56404117 | eukaryotic translation initiation factor 5 |
| ENSG00000232906 | AC092685,1 | 0,436065236 | 1,352909406 | 0,42625 | 0,67975292 | novel pseudogene containing a Zinc finger, C2H2 type domain |
| ENSG00000122824 | NUDT10 | 0,436064659 | 1,352908865 | 0,15114 | 0,47955917 | nudix hydrolase 10 |
| ENSG00000139292 | LGR5 | 0,43602598 | 1,352872593 | 0,49441 | 0,72678186 | leucine rich repeat containing G protein-coupled receptor 5 |
| ENSG00000218428 | NIP7P3 | 0,435989387 | 1,352838279 | 0,54571 | 0,76136766 | NIP7 pseudogene 3 |
| ENSG00000086666 | ZFAND6 | 0,435960533 | 1,352811223 | 0,32472 | 0,60731076 | zinc finger AN1-type containing 6 |

| | | | | | | |
|---|---|---|---|---|---|---|
| ENSG00000128829 | EIF2AK4 | 0,435876903 | 1,352732805 | 0,14136 | 0,46803321 | eukaryotic translation initiation factor 2 alpha kinase 4 |
| ENSG00000143493 | INTS7 | 0,435828678 | 1,352687588 | 0,19813 | 0,51977271 | integrator complex subunit 7 |
| ENSG00000255310 | AF131215,5 | 0,435821824 | 1,352681162 | 0,07746 | 0,36441467 | novel transcript |
| ENSG00000116030 | SUMO1 | 0,435680477 | 1,352548641 | 0,22633 | 0,54125836 | small ubiquitin like modifier 1 |
| ENSG00000170653 | ATF7 | 0,435538462 | 1,352415505 | 0,1652 | 0,49131595 | activating transcription factor 7 |
| ENSG00000242073 | AC211429,1 | 0,435368793 | 1,352256463 | 0,28284 | 0,58009943 | postmeiotic segregation increased 2-like 3 (PMS2L3) pseudogene |
| ENSG00000118454 | ANKRD13C | 0,435364521 | 1,352252459 | 0,12073 | 0,43955366 | ankyrin repeat domain 13C |
| ENSG00000121691 | CAT | 0,435267034 | 1,352161086 | 0,37224 | 0,64122758 | catalase |
| ENSG00000196597 | ZNF782 | 0,435259903 | 1,352154403 | 0,3573 | 0,63122452 | zinc finger protein 782 |
| ENSG00000184371 | CSF1 | 0,435201278 | 1,352099458 | 0,02734 | 0,22320635 | colony stimulating factor 1 |
| ENSG00000280303 | ERICD | 0,435140507 | 1,352042505 | 0,24062 | 0,54983486 | E2F1-regulated inhibitor of cell death |
| ENSG00000196504 | PRPF40A | 0,43512998 | 1,352032639 | 0,17693 | 0,5012673 | pre-mRNA processing factor 40 homolog A |
| ENSG00000167258 | CDK12 | 0,435083153 | 1,351988756 | 0,17998 | 0,50407792 | cyclin dependent kinase 12 |
| ENSG00000152782 | PANK1 | 0,435078742 | 1,351984622 | 0,12823 | 0,45171208 | pantothenate kinase 1 |
| ENSG00000197622 | CDC42SE1 | 0,434986435 | 1,351898122 | 0,09463 | 0,39709067 | CDC42 small effector 1 |
| ENSG00000117625 | RCOR3 | 0,434977845 | 1,351890072 | 0,1607 | 0,48807078 | REST corepressor 3 |
| ENSG00000008018 | PSMB1 | 0,434973119 | 1,351885644 | 0,14437 | 0,4719799 | proteasome 20S subunit beta 1 |
| ENSG00000066279 | ASPM | 0,434968191 | 1,351881026 | 0,33674 | 0,61675087 | assembly factor for spindle microtubules |
| ENSG00000150764 | DIXDC1 | 0,434945442 | 1,351859709 | 0,16083 | 0,48815237 | DIX domain containing 1 |
| ENSG00000037897 | METTL1 | 0,434936735 | 1,35185155 | 0,06822 | 0,34531713 | methyltransferase like 1 |
| ENSG00000260314 | MRC1 | 0,434716053 | 1,35164478 | 0,66356 | 0,8324578 | mannose receptor C-type 1 |
| ENSG00000165916 | PSMC3 | 0,434708356 | 1,351637569 | 0,11478 | 0,43021597 | proteasome 26S subunit, ATPase 3 |
| ENSG00000279047 | AC244517,5 | 0,434653089 | 1,351585791 | 0,79651 | 0,90597933 | novel transcript, antisense to PCDHB1 and PCDHB2 |
| ENSG00000138942 | RNF185 | 0,434632727 | 1,351566715 | 0,20881 | 0,52791772 | ring finger protein 185 |
| ENSG00000156469 | MTERF3 | 0,434596613 | 1,351532882 | 0,26214 | 0,56503489 | mitochondrial transcription termination factor 3 |
| ENSG00000185973 | TMLHE | 0,434590901 | 1,351527532 | 0,37792 | 0,64628549 | trimethyllysine hydroxylase, epsilon |
| ENSG00000123965 | PMS2P5 | 0,434580541 | 1,351517826 | 0,3001 | 0,59160746 | PMS1 homolog 2, mismatch repair system component pseudogene 5 |
| ENSG00000204792 | LINC01291 | 0,434520454 | 1,351461538 | 0,38212 | 0,64916957 | long intergenic non-protein coding RNA 1291 |
| ENSG00000132855 | ANGPTL3 | 0,434436563 | 1,351382954 | 0,20664 | 0,52622584 | angiopoietin like 3 |
| ENSG00000119927 | GPAM | 0,434418588 | 1,351366117 | 0,29265 | 0,58718198 | glycerol-3-phosphate acyltransferase, mitochondrial |
| ENSG00000117143 | UAP1 | 0,434413878 | 1,351361705 | 0,06511 | 0,33804854 | UDP-N-acetylglucosamine pyrophosphorylase 1 |

| | | | | | | |
|---|---|---|---|---|---|---|
| ENSG00000139163 | ETNK1 | 0,434310625 | 1,351264993 | 0,2509 | 0,55707572 | ethanolamine kinase 1 |
| ENSG00000165671 | NSD1 | 0,43427259 | 1,351229368 | 0,10594 | 0,4154675 | nuclear receptor binding SET domain protein 1 |
| ENSG00000224950 | AL390066,1 | 0,434218881 | 1,351179065 | 0,42229 | 0,67663581 | novel transcript |
| ENSG00000260317 | AC009812,4 | 0,434162987 | 1,351126718 | 0,51042 | 0,73798863 | novel transcript |
| ENSG00000163481 | RNF25 | 0,434076429 | 1,351045657 | 0,14074 | 0,46776292 | ring finger protein 25 |
| ENSG00000131374 | TBC1D5 | 0,434018567 | 1,350991471 | 0,28842 | 0,58455147 | TBC1 domain family member 5 |
| ENSG00000131375 | CAPN7 | 0,434016231 | 1,350989284 | 0,27281 | 0,57322782 | calpain 7 |
| ENSG00000272463 | AL357054,4 | 0,433975343 | 1,350950996 | 0,5071 | 0,73538734 | novel transcript |
| ENSG00000157191 | NECAP2 | 0,433953304 | 1,350930358 | 0,00732 | 0,10869257 | NECAP endocytosis associated 2 |
| ENSG00000147394 | ZNF185 | 0,43393572 | 1,350913893 | 0,23952 | 0,5491566 | zinc finger protein 185 with LIM domain |
| ENSG00000174912 | METTL15P1 | 0,433846875 | 1,350830702 | 0,33131 | 0,61140009 | methyltransferase like 15 pseudogene 1 |
| ENSG00000283085 | TPBG | 0,433825781 | 1,350810952 | 0,03956 | 0,26919247 | trophoblast glycoprotein |
| ENSG00000164169 | PRMT9 | 0,433771357 | 1,350759995 | 0,23602 | 0,5471053 | protein arginine methyltransferase 9 |
| ENSG00000006453 | BAIAP2L1 | 0,433711332 | 1,350703796 | 0,32375 | 0,60647553 | BAR/IMD domain containing adaptor protein 2 like 1 |
| ENSG00000152413 | HOMER1 | 0,433709824 | 1,350702384 | 0,2586 | 0,56256516 | homer scaffold protein 1 |
| ENSG00000171421 | MRPL36 | 0,433514977 | 1,350519974 | 0,36777 | 0,63775064 | mitochondrial ribosomal protein L36 |
| ENSG00000198356 | GET3 | 0,43347232 | 1,350480043 | 0,28344 | 0,58065155 | guided entry of tail-anchored proteins factor 3, ATPase |
| ENSG00000165138 | ANKS6 | 0,433434779 | 1,350444901 | 0,07852 | 0,36610837 | ankyrin repeat and sterile alpha motif domain containing 6 |
| ENSG00000142207 | URB1 | 0,433376788 | 1,35039062 | 0,10518 | 0,41409464 | URB1 ribosome biogenesis homolog |
| ENSG00000158623 | COPG2 | 0,433343203 | 1,350359184 | 0,19465 | 0,51562158 | COPI coat complex subunit gamma 2 |
| ENSG00000179958 | DCTPP1 | 0,433323631 | 1,350340864 | 0,18058 | 0,50407792 | dCTP pyrophosphatase 1 |
| ENSG00000274944 | AL139260,3 | 0,433255586 | 1,350277177 | 0,13516 | 0,46165567 | novel protein |
| ENSG00000145604 | SKP2 | 0,433246981 | 1,350269123 | 0,08279 | 0,37466043 | S-phase kinase associated protein 2 |
| ENSG00000082516 | GEMIN5 | 0,433196475 | 1,350221854 | 0,15728 | 0,48443207 | gem nuclear organelle associated protein 5 |
| ENSG00000138279 | ANXA7 | 0,433141436 | 1,350170344 | 0,22614 | 0,54121239 | annexin A7 |
| ENSG00000159377 | PSMB4 | 0,433118971 | 1,35014932 | 0,06255 | 0,33112961 | proteasome 20S subunit beta 4 |
| ENSG00000109320 | NFKB1 | 0,432978196 | 1,350017581 | 0,25009 | 0,55657567 | nuclear factor kappa B subunit 1 |
| ENSG00000105875 | WDR91 | 0,432904777 | 1,349948881 | 0,09876 | 0,40461924 | WD repeat domain 91 |
| ENSG00000120662 | MTRF1 | 0,432884119 | 1,349929551 | 0,23339 | 0,54563244 | mitochondrial translation release factor 1 |
| ENSG00000224415 | AC007683,1 | 0,432861678 | 1,349908553 | 0,61381 | 0,8035281 | ribosomal protein L19 (RPL19) pseudogene |
| ENSG00000282463 | AC216130,3 | 0,432861678 | 1,349908553 | 0,61381 | 0,8035281 | ribosomal protein L19 (RPL19) pseudogene |
| ENSG00000057657 | PRDM1 | 0,432850317 | 1,349897923 | 0,40779 | 0,66684711 | PR/SET domain 1 |

| | | | | | | |
|---|---|---|---|---|---|---|
| ENSG00000288227 | AC012435,4 | 0,432846234 | 1,349894102 | 0,67176 | 0,8375243 | coiled-coil-helix-coiled-coil-helix domain-containing protein 7 (CHCHD7) pseudogene |
| ENSG00000074219 | TEAD2 | 0,43284106 | 1,349889262 | 0,05415 | 0,31168784 | TEA domain transcription factor 2 |
| ENSG00000254932 | AP001007,1 | 0,432795585 | 1,349846713 | 0,43606 | 0,6865394 | novel transcript, antisense to PKNOX2 |
| ENSG00000197050 | ZNF420 | 0,432789093 | 1,349840638 | 0,31142 | 0,59890747 | zinc finger protein 420 |
| ENSG00000138363 | ATIC | 0,432775959 | 1,34982835 | 0,20493 | 0,52453221 | 5-aminoimidazole-4-carboxamide ribonucleotide formyltransferase/IMP cyclohydrolase |
| ENSG00000214867 | SRSF9P1 | 0,432750911 | 1,349804914 | 0,72318 | 0,86784071 | serine and arginine rich splicing factor 9 pseudogene 1 |
| ENSG00000112029 | FBXO5 | 0,432677518 | 1,349736248 | 0,38465 | 0,65052153 | F-box protein 5 |
| ENSG00000163389 | POGLUT1 | 0,432632436 | 1,349694072 | 0,36986 | 0,63943131 | protein O-glucosyltransferase 1 |
| ENSG00000143457 | GOLPH3L | 0,432569941 | 1,349635607 | 0,14709 | 0,47471707 | golgi phosphoprotein 3 like |
| ENSG00000256274 | TAS2R64P | 0,432524687 | 1,349593273 | 0,36298 | 0,63395677 | taste 2 receptor member 64 pseudogene |
| ENSG00000263083 | TAS2R64P | 0,432524687 | 1,349593273 | 0,36298 | 0,63395677 | taste 2 receptor member 64 pseudogene |
| ENSG00000186141 | POLR3C | 0,432521605 | 1,34959039 | 0,11719 | 0,43451125 | RNA polymerase III subunit C |
| ENSG00000161381 | PLXDC1 | 0,43246673 | 1,349539057 | 0,30585 | 0,59512079 | plexin domain containing 1 |
| ENSG00000089157 | RPLP0 | 0,432442463 | 1,349516357 | 0,27408 | 0,57412149 | ribosomal protein lateral stalk subunit P0 |
| ENSG00000139146 | SINHCAF | 0,432373582 | 1,349451926 | 0,31937 | 0,60429866 | SIN3-HDAC complex associated factor |
| ENSG00000112578 | BYSL | 0,432303033 | 1,349385938 | 0,27572 | 0,57533851 | bystin like |
| ENSG00000108774 | RAB5C | 0,432276297 | 1,349360932 | 0,23036 | 0,54355692 | RAB5C, member RAS oncogene family |
| ENSG00000125445 | MRPS7 | 0,432220709 | 1,349308941 | 0,1924 | 0,514872 | mitochondrial ribosomal protein S7 |
| ENSG00000284901 | RUVBL1 | 0,432217339 | 1,349305789 | 0,09649 | 0,40027716 | RuvB like AAA ATPase 1 |
| ENSG00000198755 | RPL10A | 0,432208139 | 1,349297185 | 0,14014 | 0,46710628 | ribosomal protein L10a |
| ENSG00000150779 | TIMM8B | 0,432165887 | 1,349257669 | 0,27671 | 0,57549599 | translocase of inner mitochondrial membrane 8 homolog B |
| ENSG00000112972 | HMGCS1 | 0,43213136 | 1,349225378 | 0,26648 | 0,56917012 | 3-hydroxy-3-methylglutaryl-CoA synthase 1 |
| ENSG00000169188 | APEX2 | 0,432087781 | 1,349184623 | 0,04586 | 0,28898991 | apurinic/apyrimidinic endodeoxyribonuclease 2 |
| ENSG00000250107 | CACNA1G-AS1 | 0,431902579 | 1,349011436 | 0,65855 | 0,82926071 | CACNA1G antisense RNA 1 |
| ENSG00000119787 | ATL2 | 0,431846366 | 1,348958874 | 0,06024 | 0,32601555 | atlastin GTPase 2 |
| ENSG00000134758 | RNF138 | 0,431765702 | 1,348883454 | 0,37432 | 0,64275601 | ring finger protein 138 |
| ENSG00000006327 | TNFRSF12A | 0,431734886 | 1,348854642 | 0,29122 | 0,58623355 | TNF receptor superfamily member 12A |
| ENSG00000188993 | LRRC66 | 0,431664228 | 1,348788581 | 0,4579 | 0,70103159 | leucine rich repeat containing 66 |
| ENSG00000260404 | AC110079,1 | 0,431602241 | 1,348730631 | 0,25465 | 0,55944852 | general transcription factor IIi (GTF2I) pseudogene |
| ENSG00000136932 | TRMO | 0,431593462 | 1,348722423 | 0,1372 | 0,4642578 | tRNA methyltransferase O |
| ENSG00000144320 | LNPK | 0,431536884 | 1,348669532 | 0,23348 | 0,54563244 | lunapark, ER junction formation factor |

| | | | | | | |
|---|---|---|---|---|---|---|
| ENSG00000236993 | GAPDHP21 | 0,431489081 | 1,348624845 | 0,37477 | 0,6431311 | glyceraldehyde 3 phosphate dehydrogenase pseudogene 21 |
| ENSG00000183475 | ASB7 | 0,431447493 | 1,348585969 | 0,17736 | 0,50187931 | ankyrin repeat and SOCS box containing 7 |
| ENSG00000277936 | MRPL45 | 0,431432381 | 1,348571843 | 0,16605 | 0,49204668 | mitochondrial ribosomal protein L45 |
| ENSG00000167720 | SRR | 0,431394401 | 1,348536342 | 0,40339 | 0,66399303 | serine racemase |
| ENSG00000227376 | FTH1P16 | 0,431392347 | 1,348534421 | 0,57007 | 0,77703861 | ferritin heavy chain 1 pseudogene 16 |
| ENSG00000163848 | ZNF148 | 0,43137558 | 1,348518749 | 0,24482 | 0,55282045 | zinc finger protein 148 |
| ENSG00000167107 | ACSF2 | 0,431290737 | 1,348439447 | 0,07439 | 0,35809404 | acyl-CoA synthetase family member 2 |
| ENSG00000071537 | SEL1L | 0,43110809 | 1,348268743 | 0,23734 | 0,54832106 | SEL1L adaptor subunit of ERAD E3 ubiquitin ligase |
| ENSG00000196632 | WNK3 | 0,431060784 | 1,348224534 | 0,25872 | 0,56260205 | WNK lysine deficient protein kinase 3 |
| ENSG00000168137 | SETD5 | 0,431040629 | 1,348205699 | 0,12679 | 0,44955467 | SET domain containing 5 |
| ENSG00000160741 | CRTC2 | 0,431005659 | 1,34817302 | 0,0306 | 0,23611505 | CREB regulated transcription coactivator 2 |
| ENSG00000136928 | GABBR2 | 0,430981879 | 1,348150798 | 0,30204 | 0,59294633 | gamma-aminobutyric acid type B receptor subunit 2 |
| ENSG00000205592 | MUC19 | 0,430939094 | 1,348110817 | 0,6067 | 0,79981537 | mucin 19, oligomeric |
| ENSG00000106823 | ECM2 | 0,430914789 | 1,348088106 | 0,11006 | 0,4224508 | extracellular matrix protein 2 |
| ENSG00000112624 | BICRAL | 0,430898717 | 1,348073088 | 0,40263 | 0,66396567 | BRD4 interacting chromatin remodeling complex associated protein like |
| ENSG00000153936 | HS2ST1 | 0,43085715 | 1,348034248 | 0,18562 | 0,50901954 | heparan sulfate 2-O-sulfotransferase 1 |
| ENSG00000158480 | SPATA2 | 0,430733203 | 1,347918439 | 0,16286 | 0,48945635 | spermatogenesis associated 2 |
| ENSG00000137770 | CTDSPL2 | 0,430672109 | 1,347861359 | 0,11988 | 0,43850846 | CTD small phosphatase like 2 |
| ENSG00000107959 | PITRM1 | 0,430534461 | 1,347732766 | 0,26041 | 0,56383537 | pitrilysin metallopeptidase 1 |
| ENSG00000147100 | SLC16A2 | 0,430517891 | 1,347717286 | 0,03098 | 0,23719337 | solute carrier family 16 member 2 |
| ENSG00000174417 | TRHR | 0,430473162 | 1,347675503 | 0,54378 | 0,76018847 | thyrotropin releasing hormone receptor |
| ENSG00000225859 | DDX39B | 0,430456276 | 1,347659729 | 0,13048 | 0,45477153 | DExD-box helicase 39B |
| ENSG00000116406 | EDEM3 | 0,43045016 | 1,347654016 | 0,14984 | 0,47782859 | ER degradation enhancing alpha-mannosidase like protein 3 |
| ENSG00000213977 | TAX1BP3 | 0,430394879 | 1,347602377 | 0,14368 | 0,47123322 | Tax1 binding protein 3 |
| ENSG00000254093 | PINX1 | 0,430371203 | 1,347580262 | 0,03768 | 0,26266577 | PIN2 (TERF1) interacting telomerase inhibitor 1 |
| ENSG00000226984 | AL035410,1 | 0,430337598 | 1,347548874 | 0,49634 | 0,72831792 | ribosomal protein L26 (RPL26) pseudogene |
| ENSG00000130803 | ZNF317 | 0,430189715 | 1,34741075 | 0,11972 | 0,43825427 | zinc finger protein 317 |
| ENSG00000213085 | CFAP45 | 0,430108637 | 1,34733503 | 0,48658 | 0,72196345 | cilia and flagella associated protein 45 |
| ENSG00000189007 | ADAT2 | 0,430043453 | 1,347274155 | 0,15933 | 0,4867988 | adenosine deaminase tRNA specific 2 |
| ENSG00000188725 | SMIM15 | 0,430033755 | 1,347265099 | 0,31691 | 0,60268749 | small integral membrane protein 15 |

| | | | | | | |
|---|---|---|---|---|---|---|
| ENSG00000169020 | ATP5ME | 0,429922421 | 1,347161133 | 0,24835 | 0,55546558 | ATP synthase membrane subunit e |
| ENSG00000090432 | MUL1 | 0,429884552 | 1,347125772 | 0,09897 | 0,40492885 | mitochondrial E3 ubiquitin protein ligase 1 |
| ENSG00000231934 | FAM242A | 0,429824523 | 1,347069721 | 0,33699 | 0,61684812 | family with sequence similarity 242 member A |
| ENSG00000131558 | EXOC4 | 0,429811876 | 1,347057912 | 0,21669 | 0,53494609 | exocyst complex component 4 |
| ENSG00000113231 | PDE8B | 0,429802587 | 1,347049239 | 0,289 | 0,58469988 | phosphodiesterase 8B |
| ENSG00000132952 | USPL1 | 0,429735992 | 1,34698706 | 0,2392 | 0,5491566 | ubiquitin specific peptidase like 1 |
| ENSG00000169860 | P2RY1 | 0,429722982 | 1,346974914 | 0,16756 | 0,49376742 | purinergic receptor P2Y1 |
| ENSG00000226056 | MTND4P32 | 0,429531921 | 1,346796541 | 0,64921 | 0,82436914 | MT-ND4 pseudogene 32 |
| ENSG00000148688 | RPP30 | 0,429349421 | 1,346626183 | 0,25816 | 0,56221375 | ribonuclease P/MRP subunit p30 |
| ENSG00000008118 | CAMK1G | 0,429308349 | 1,346587846 | 0,21728 | 0,53520949 | calcium/calmodulin dependent protein kinase IG |
| ENSG00000070193 | FGF10 | 0,429156126 | 1,346445771 | 0,49454 | 0,72685419 | fibroblast growth factor 10 |
| ENSG00000214093 | AL096701,1 | 0,429095411 | 1,346389108 | 0,59045 | 0,78974912 | ribosomal protein S18 (RPS18) pseudogene |
| ENSG00000198382 | UVRAG | 0,4290922 | 1,346386111 | 0,02598 | 0,21717729 | UV radiation resistance associated |
| ENSG00000132485 | ZRANB2 | 0,429063887 | 1,346359689 | 0,07729 | 0,36400794 | zinc finger RANBP2-type containing 2 |
| ENSG00000167182 | SP2 | 0,429052306 | 1,346348881 | 0,13024 | 0,45467139 | Sp2 transcription factor |
| ENSG00000125246 | CLYBL | 0,429013661 | 1,346312818 | 0,2341 | 0,54593875 | citramalyl-CoA lyase |
| ENSG00000198899 | MT-ATP6 | 0,429012438 | 1,346311677 | 0,23081 | 0,54385714 | mitochondrially encoded ATP synthase membrane subunit 6 |
| ENSG00000172269 | DPAGT1 | 0,428996073 | 1,346296405 | 0,09147 | 0,39107762 | dolichyl-phosphate N-acetylglucosaminephosphotransferase 1 |
| ENSG00000149485 | FADS1 | 0,428993576 | 1,346294074 | 0,18003 | 0,50407792 | fatty acid desaturase 1 |
| ENSG00000184319 | RPL23AP82 | 0,428961446 | 1,346264092 | 0,35882 | 0,63195238 | ribosomal protein L23a pseudogene 82 |
| ENSG00000152495 | CAMK4 | 0,428895535 | 1,346202587 | 0,3412 | 0,61934696 | calcium/calmodulin dependent protein kinase IV |
| ENSG00000105011 | ASF1B | 0,428821061 | 1,346133097 | 0,13051 | 0,45477153 | anti-silencing function 1B histone chaperone |
| ENSG00000288210 | ASF1B | 0,428821061 | 1,346133097 | 0,13051 | 0,45477153 | anti-silencing function 1B histone chaperone |
| ENSG00000266079 | SNORA59B | 0,428796993 | 1,34611064 | 0,63072 | 0,81354995 | small nucleolar RNA, H/ACA box 59B |
| ENSG00000172345 | STARD5 | 0,428763424 | 1,346079319 | 0,20895 | 0,52800078 | StAR related lipid transfer domain containing 5 |
| ENSG00000229425 | AJ009632,2 | 0,428757443 | 1,346073738 | 0,52982 | 0,75150213 | novel transcript |
| ENSG00000267187 | AC026898,1 | 0,42875629 | 1,346072662 | 0,32908 | 0,61010243 | sal-like 2 (Drosophila), pseudogene |
| ENSG00000278311 | GGNBP2 | 0,428592134 | 1,345919509 | 0,17605 | 0,50011988 | gametogenetin binding protein 2 |
| ENSG00000105926 | MPP6 | 0,428516584 | 1,345849029 | 0,17559 | 0,49954066 | membrane palmitoylated protein 6 |
| ENSG00000213533 | STIMATE | 0,428498419 | 1,345832083 | 0,06004 | 0,32549025 | STIM activating enhancer |
| ENSG00000177688 | SUMO4 | 0,428494606 | 1,345828527 | 0,31724 | 0,6027392 | small ubiquitin like modifier 4 |

| | | | | | | |
|---|---|---|---|---|---|---|
| ENSG00000198176 | TFDP1 | 0,428490944 | 1,34582511 | 0,01002 | 0,12834202 | transcription factor Dp-1 |
| ENSG00000137672 | TRPC6 | 0,428430344 | 1,345768581 | 0,60133 | 0,79647801 | transient receptor potential cation channel subfamily C member 6 |
| ENSG00000100796 | PPP4R3A | 0,428327933 | 1,345673053 | 0,17422 | 0,49867554 | protein phosphatase 4 regulatory subunit 3A |
| ENSG00000101811 | CSTF2 | 0,428259833 | 1,345609534 | 0,20808 | 0,52756478 | cleavage stimulation factor subunit 2 |
| ENSG00000198482 | ZNF808 | 0,428254038 | 1,345604129 | 0,27903 | 0,57715243 | zinc finger protein 808 |
| ENSG00000168040 | FADD | 0,428159229 | 1,345515704 | 0,07535 | 0,36046509 | Fas associated via death domain |
| ENSG00000134698 | AGO4 | 0,428140155 | 1,345497915 | 0,06751 | 0,34374257 | argonaute RISC component 4 |
| ENSG00000163785 | RYK | 0,428083691 | 1,345445256 | 0,13351 | 0,4594073 | receptor like tyrosine kinase |
| ENSG00000262904 | TMPOP2 | 0,428042633 | 1,345406966 | 0,19066 | 0,51369186 | thymopoietin pseudogene 2 |
| ENSG00000053770 | AP5M1 | 0,427998795 | 1,345366085 | 0,30343 | 0,59369944 | adaptor related protein complex 5 subunit mu 1 |
| ENSG00000163743 | RCHY1 | 0,427934951 | 1,345306549 | 0,40754 | 0,66665981 | ring finger and CHY zinc finger domain containing 1 |
| ENSG00000215482 | CALM2P3 | 0,427910773 | 1,345284004 | 0,55631 | 0,76843593 | calmodulin 2 pseudogene 3 |
| ENSG00000143442 | POGZ | 0,427716199 | 1,345102579 | 0,20829 | 0,52766387 | pogo transposable element derived with ZNF domain |
| ENSG00000228446 | ANKRD49P1 | 0,427690409 | 1,345078534 | 0,42578 | 0,67925635 | ANKRD49 pseudogene 1 |
| ENSG00000152454 | ZNF256 | 0,427640798 | 1,345032281 | 0,21274 | 0,53177822 | zinc finger protein 256 |
| ENSG00000086712 | TXLNG | 0,42761069 | 1,345004211 | 0,2783 | 0,57645922 | taxilin gamma |
| ENSG00000234358 | AC003080,1 | 0,427592672 | 1,344987413 | 0,44411 | 0,6916877 | ribosomal protein L8 (RPL8) pseudogene |
| ENSG00000075142 | SRI | 0,427557131 | 1,34495428 | 0,33368 | 0,61369834 | sorcin |
| ENSG00000177076 | ACER2 | 0,427523166 | 1,344922617 | 0,19847 | 0,51980353 | alkaline ceramidase 2 |
| ENSG00000006744 | ELAC2 | 0,427515311 | 1,344915294 | 0,13129 | 0,45625325 | elaC ribonuclease Z 2 |
| ENSG00000230658 | KLHL7-DT | 0,42749793 | 1,344899091 | 0,20851 | 0,52778377 | KLHL7 divergent transcript |
| ENSG00000104221 | BRF2 | 0,427385214 | 1,34479402 | 0,12885 | 0,45214857 | BRF2 RNA polymerase III transcription initiation factor subunit |
| ENSG00000109084 | TMEM97 | 0,427240339 | 1,344658982 | 0,15414 | 0,48239691 | transmembrane protein 97 |
| ENSG00000112237 | CCNC | 0,427114086 | 1,344541314 | 0,18753 | 0,51081411 | cyclin C |
| ENSG00000113460 | BRIX1 | 0,426772335 | 1,344222852 | 0,14931 | 0,47695763 | biogenesis of ribosomes BRX1 |
| ENSG00000053254 | FOXN3 | 0,426755053 | 1,344206749 | 0,07528 | 0,36016867 | forkhead box N3 |
| ENSG00000230370 | RPL23AP52 | 0,426749174 | 1,344201272 | 0,50802 | 0,73612961 | ribosomal protein L23a pseudogene 52 |
| ENSG00000091317 | CMTM6 | 0,426722648 | 1,344176557 | 0,2886 | 0,58455903 | CKLF like MARVEL transmembrane domain containing 6 |
| ENSG00000018510 | AGPS | 0,426719499 | 1,344173623 | 0,27434 | 0,57422483 | alkylglycerone phosphate synthase |
| ENSG00000196739 | COL27A1 | 0,426702699 | 1,34415797 | 0,45186 | 0,69682854 | collagen type XXVII alpha 1 chain |
| ENSG00000270344 | POC1B-AS1 | 0,426685098 | 1,344141572 | 0,22205 | 0,53855235 | POC1B antisense RNA 1 |

| | | | | | | |
|---|---|---|---|---|---|---|
| ENSG00000234729 | AGER | 0,426674576 | 1,344131769 | 0,35004 | 0,62564488 | advanced glycosylation end-product specific receptor |
| ENSG00000254634 | SMG1P6 | 0,426666654 | 1,344124387 | 0,23761 | 0,54861973 | SMG1 pseudogene 6 |
| ENSG00000288478 | NIPA1 | 0,426610524 | 1,344072094 | 0,2868 | 0,58325473 | NIPA magnesium transporter 1 |
| ENSG00000135597 | REPS1 | 0,426607152 | 1,344068952 | 0,17254 | 0,49744497 | RALBP1 associated Eps domain containing 1 |
| ENSG00000142875 | PRKACB | 0,426585106 | 1,344048414 | 0,22057 | 0,53759541 | protein kinase cAMP-activated catalytic subunit beta |
| ENSG00000137288 | UQCC2 | 0,426563977 | 1,34402873 | 0,2006 | 0,52141065 | ubiquinol-cytochrome c reductase complex assembly factor 2 |
| ENSG00000232040 | ZBED9 | 0,426471738 | 1,343942802 | 0,2397 | 0,5491566 | zinc finger BED-type containing 9 |
| ENSG00000143919 | CAMKMT | 0,426382896 | 1,343860044 | 0,20741 | 0,5268543 | calmodulin-lysine N-methyltransferase |
| ENSG00000196843 | ARID5A | 0,426300697 | 1,343783478 | 0,22748 | 0,54168591 | AT-rich interaction domain 5A |
| ENSG00000180329 | CCDC43 | 0,426200834 | 1,343690465 | 0,16016 | 0,48737351 | coiled-coil domain containing 43 |
| ENSG00000163938 | GNL3 | 0,426172615 | 1,343664182 | 0,04449 | 0,28449899 | G protein nucleolar 3 |
| ENSG00000279145 | AC011912,1 | 0,42608119 | 1,343579036 | 0,07175 | 0,35351167 | TEC |
| ENSG00000277406 | SEC22B4P | 0,42601202 | 1,343514619 | 0,36118 | 0,63317043 | SEC22 homolog B4, pseudogene |
| ENSG00000124151 | NCOA3 | 0,426000387 | 1,343503787 | 0,26926 | 0,57096334 | nuclear receptor coactivator 3 |
| ENSG00000274137 | MYOM2 | 0,425951531 | 1,34345829 | 0,175 | 0,49911246 | myomesin 2 |
| ENSG00000146414 | SHPRH | 0,425894326 | 1,34340502 | 0,21621 | 0,53480193 | SNF2 histone linker PHD RING helicase |
| ENSG00000215796 | AL512637,1 | 0,425841037 | 1,343355401 | 0,57494 | 0,78003515 | primase polypeptide 1 49kDa (PRIM1) pseudogene |
| ENSG00000132383 | RPA1 | 0,425680118 | 1,34320557 | 0,21877 | 0,53630871 | replication protein A1 |
| ENSG00000176092 | CRYBG2 | 0,425626641 | 1,343155781 | 0,4072 | 0,66659618 | crystallin beta-gamma domain containing 2 |
| ENSG00000254527 | ENPP7P12 | 0,425568052 | 1,343101237 | 0,3014 | 0,59258306 | ectonucleotide pyrophosphatase/phosphodiesterase 7 pseudogene 12 |
| ENSG00000224570 | AC097063,1 | 0,425567122 | 1,34310037 | 0,65324 | 0,82629807 | UDP-N-acetyl-alpha-D-galactosamine:polypeptide N-acetylgalactosaminyltransferase 11 (GalNAc-T11) (GALNT11) pseudogene |
| ENSG00000180992 | MRPL14 | 0,425505699 | 1,343043189 | 0,24266 | 0,55158827 | mitochondrial ribosomal protein L14 |
| ENSG00000118292 | C1orf54 | 0,425147298 | 1,342709585 | 0,09335 | 0,39489949 | chromosome 1 open reading frame 54 |
| ENSG00000267179 | AC008770,2 | 0,42510611 | 1,342671252 | 0,38643 | 0,65185547 | novel protein |
| ENSG00000130508 | PXDN | 0,425032246 | 1,342602511 | 0,11928 | 0,43713778 | peroxidasin |
| ENSG00000276174 | AC087683,2 | 0,4250153 | 1,342586741 | 0,61604 | 0,80475871 | novel transcript, sense intronic to PIK3C3 |
| ENSG00000198833 | UBE2J1 | 0,424997329 | 1,342570017 | 0,25718 | 0,56155289 | ubiquitin conjugating enzyme E2 J1 |
| ENSG00000138629 | UBL7 | 0,424986336 | 1,342559787 | 0,04595 | 0,28908951 | ubiquitin like 7 |
| ENSG00000288408 | UBL7 | 0,424986336 | 1,342559787 | 0,04595 | 0,28908951 | ubiquitin like 7 |

| | | | | | | |
|---|---|---|---|---|---|---|
| ENSG00000165280 | VCP | 0,424985624 | 1,342559124 | 0,10373 | 0,41194268 | valosin containing protein |
| ENSG00000284812 | AC270289,1 | 0,424878768 | 1,342459689 | 0,35017 | 0,62576449 | regulatory solute carrier protein, family 1, member 1 (RSC1A1) pseudogene |
| ENSG00000166986 | MARS1 | 0,42485441 | 1,342437024 | 0,1048 | 0,41347671 | methionyl-tRNA synthetase 1 |
| ENSG00000224470 | ATXN1L | 0,424813342 | 1,34239881 | 0,05895 | 0,3241112 | ataxin 1 like |
| ENSG00000183060 | LYSMD4 | 0,424744846 | 1,342335077 | 0,11689 | 0,43414659 | LysM domain containing 4 |
| ENSG00000171885 | AQP4 | 0,424668513 | 1,342264056 | 0,55933 | 0,77045525 | aquaporin 4 |
| ENSG00000171490 | RSL1D1 | 0,424634851 | 1,342232738 | 0,09254 | 0,39322559 | ribosomal L1 domain containing 1 |
| ENSG00000278238 | AL359513,1 | 0,424601422 | 1,342201637 | 0,54017 | 0,75814808 | novel transcript |
| ENSG00000088035 | ALG6 | 0,42456766 | 1,342170227 | 0,35458 | 0,62915556 | ALG6 alpha-1,3-glucosyltransferase |
| ENSG00000136942 | RPL35 | 0,424510953 | 1,342117473 | 0,13917 | 0,46648276 | ribosomal protein L35 |
| ENSG00000174485 | DENND4A | 0,424354485 | 1,34197192 | 0,37212 | 0,64114037 | DENN domain containing 4A |
| ENSG00000132819 | RBM38 | 0,42428712 | 1,341909261 | 0,12163 | 0,44093327 | RNA binding motif protein 38 |
| ENSG00000224097 | AC021148,1 | 0,424202404 | 1,341830465 | 0,61043 | 0,80224337 | WD repeat domain 5 (WDR5) pseudogene |
| ENSG00000162704 | ARPC5 | 0,424132302 | 1,341765266 | 0,09453 | 0,3969039 | actin related protein 2/3 complex subunit 5 |
| ENSG00000173530 | TNFRSF10D | 0,424055492 | 1,341693831 | 0,21999 | 0,53740161 | TNF receptor superfamily member 10d |
| ENSG00000114978 | MOB1A | 0,423814546 | 1,341469772 | 0,15148 | 0,47963227 | MOB kinase activator 1A |
| ENSG00000147471 | PLPBP | 0,423706129 | 1,341368966 | 0,33095 | 0,6112353 | pyridoxal phosphate binding protein |
| ENSG00000282124 | AC015849,5 | 0,423666074 | 1,341331725 | 0,25757 | 0,56171514 | leucine rich repeat containing 37 member A3 (LRRC37A3) pseudogene |
| ENSG00000127481 | UBR4 | 0,423609708 | 1,34127932 | 0,12653 | 0,44918536 | ubiquitin protein ligase E3 component n-recognin 4 |
| ENSG00000146143 | PRIM2 | 0,423455195 | 1,341135676 | 0,27374 | 0,57377252 | DNA primase subunit 2 |
| ENSG00000166913 | YWHAB | 0,423454313 | 1,341134856 | 0,23025 | 0,5434313 | tyrosine 3-monooxygenase/tryptophan 5-monooxygenase activation protein beta |
| ENSG00000251050 | RBX1P2 | 0,423327373 | 1,341016857 | 0,64978 | 0,82452434 | RBX1 pseudogene 2 |
| ENSG00000159346 | ADIPOR1 | 0,423294234 | 1,340986054 | 0,15486 | 0,48268107 | adiponectin receptor 1 |
| ENSG00000196839 | ADA | 0,423247051 | 1,340942198 | 0,44404 | 0,6916877 | adenosine deaminase |
| ENSG00000140943 | MBTPS1 | 0,4231928 | 1,340891774 | 0,16237 | 0,48945635 | membrane bound transcription factor peptidase, site 1 |
| ENSG00000284889 | PSME2 | 0,423158041 | 1,340859469 | 0,13205 | 0,45711259 | proteasome activator subunit 2 |
| ENSG00000100911 | PSME2 | 0,423158041 | 1,340859469 | 0,13205 | 0,45711259 | proteasome activator subunit 2 |
| ENSG00000062194 | GPBP1 | 0,423118431 | 1,340822655 | 0,2769 | 0,57553852 | GC-rich promoter binding protein 1 |
| ENSG00000250538 | AC104407,1 | 0,422986109 | 1,340699683 | 0,64141 | 0,82012969 | novel transcript |
| ENSG00000163947 | ARHGEF3 | 0,422894827 | 1,340614857 | 0,24027 | 0,54948131 | Rho guanine nucleotide exchange factor 3 |

| | | | | | | |
|---|---|---|---|---|---|---|
| ENSG00000085063 | CD59 | 0,422849592 | 1,340572823 | 0,2031 | 0,52299224 | CD59 molecule (CD59 blood group) |
| ENSG00000125834 | STK35 | 0,422772177 | 1,34050089 | 0,01631 | 0,16740197 | serine/threonine kinase 35 |
| ENSG00000147874 | HAUS6 | 0,42268642 | 1,34042121 | 0,23571 | 0,5471053 | HAUS augmin like complex subunit 6 |
| ENSG00000196821 | ILRUN | 0,422661991 | 1,340398513 | 0,04069 | 0,27268567 | inflammation and lipid regulator with UBA-like and NBR1-like domains |
| ENSG00000102038 | SMARCA1 | 0,422479951 | 1,340229391 | 0,15716 | 0,48443207 | SWI/SNF related, matrix associated, actin dependent regulator of chromatin, subfamily a, member 1 |
| ENSG00000102531 | FNDC3A | 0,422387618 | 1,340143619 | 0,30909 | 0,59774312 | fibronectin type III domain containing 3A |
| ENSG00000163431 | LMOD1 | 0,422221104 | 1,33998895 | 0,44878 | 0,69455511 | leiomodin 1 |
| ENSG00000286755 | AC135893,1 | 0,422149218 | 1,339922183 | 0,57629 | 0,78066251 | novel transcript, antisense to LMLN |
| ENSG00000272797 | AC092954,1 | 0,422123046 | 1,339897876 | 0,62521 | 0,81024216 | novel transcript |
| ENSG00000131480 | AOC2 | 0,422116852 | 1,339892123 | 0,30289 | 0,59331081 | amine oxidase copper containing 2 |
| ENSG00000069345 | DNAJA2 | 0,422036262 | 1,339817278 | 0,20808 | 0,52756478 | DnaJ heat shock protein family (Hsp40) member A2 |
| ENSG00000275600 | PIGW | 0,422016325 | 1,339798763 | 0,38533 | 0,65107577 | phosphatidylinositol glycan anchor biosynthesis class W |
| ENSG00000231864 | AL807752,3 | 0,421991453 | 1,339775665 | 0,78258 | 0,89903403 | novel transcript |
| ENSG00000232953 | HSPA8P18 | 0,421980099 | 1,339765121 | 0,65274 | 0,826091 | heat shock protein family A (Hsp70) member 8 pseudogene 18 |
| ENSG00000258636 | AL121821,2 | 0,421932839 | 1,339721234 | 0,37002 | 0,63943131 | novel transcript |
| ENSG00000155903 | RASA2 | 0,421881067 | 1,339673158 | 0,24523 | 0,55295131 | RAS p21 protein activator 2 |
| ENSG00000158186 | MRAS | 0,421794539 | 1,339592811 | 0,17333 | 0,4980379 | muscle RAS oncogene homolog |
| ENSG00000187049 | TMEM216 | 0,421685712 | 1,339491765 | 0,22252 | 0,53866376 | transmembrane protein 216 |
| ENSG00000226479 | TMEM185B | 0,421654909 | 1,339463166 | 0,04907 | 0,29818993 | transmembrane protein 185B |
| ENSG00000170873 | MTSS1 | 0,421644506 | 1,339453508 | 0,20364 | 0,52347929 | MTSS I-BAR domain containing 1 |
| ENSG00000119977 | TCTN3 | 0,421531721 | 1,339348798 | 0,22611 | 0,54121239 | tectonic family member 3 |
| ENSG00000104059 | FAM189A1 | 0,421514195 | 1,339332527 | 0,4268 | 0,68015803 | family with sequence similarity 189 member A1 |
| ENSG00000143727 | ACP1 | 0,421485442 | 1,339305834 | 0,33988 | 0,61872249 | acid phosphatase 1 |
| ENSG00000118407 | FILIP1 | 0,421453553 | 1,339276231 | 0,41458 | 0,67128723 | filamin A interacting protein 1 |
| ENSG00000188992 | LIPI | 0,421427653 | 1,339252188 | 0,61627 | 0,80488598 | lipase I |
| ENSG00000185433 | LINC00158 | 0,421359603 | 1,339189018 | 0,69111 | 0,84931694 | long intergenic non-protein coding RNA 158 |
| ENSG00000265069 | AP002409,1 | 0,421349564 | 1,3391797 | 0,43764 | 0,68737508 | novel transcript, antisense to LAMA1 |
| ENSG00000113658 | SMAD5 | 0,421287806 | 1,339122375 | 0,10819 | 0,41884147 | SMAD family member 5 |
| ENSG00000242661 | RPS3AP43 | 0,421214371 | 1,339054213 | 0,61197 | 0,80341001 | RPS3A pseudogene 43 |
| ENSG00000142188 | TMEM50B | 0,421212571 | 1,339052542 | 0,14154 | 0,46809977 | transmembrane protein 50B |

| | | | | | | |
|---|---|---|---|---|---|---|
| ENSG00000049883 | PTCD2 | 0,421164372 | 1,339007807 | 0,09171 | 0,39166375 | pentatricopeptide repeat domain 2 |
| ENSG00000130811 | EIF3G | 0,421109992 | 1,338957336 | 0,14671 | 0,47461701 | eukaryotic translation initiation factor 3 subunit G |
| ENSG00000164631 | ZNF12 | 0,421109421 | 1,338956807 | 0,13857 | 0,46578766 | zinc finger protein 12 |
| ENSG00000083097 | DOP1A | 0,421045301 | 1,338897298 | 0,37162 | 0,64073742 | DOP1 leucine zipper like protein A |
| ENSG00000263006 | ROCK1P1 | 0,421029811 | 1,338882922 | 0,37819 | 0,64646341 | Rho associated coiled-coil containing protein kinase 1 pseudogene 1 |
| ENSG00000198498 | TMA16 | 0,420998067 | 1,338853463 | 0,27009 | 0,57186187 | translation machinery associated 16 homolog |
| ENSG00000137337 | MDC1 | 0,420818692 | 1,338687009 | 0,1147 | 0,43014087 | mediator of DNA damage checkpoint 1 |
| ENSG00000156508 | EEF1A1 | 0,420768284 | 1,338640236 | 0,16488 | 0,49095733 | eukaryotic translation elongation factor 1 alpha 1 |
| ENSG00000110422 | HIPK3 | 0,420759865 | 1,338632425 | 0,1658 | 0,49189706 | homeodomain interacting protein kinase 3 |
| ENSG00000168876 | ANKRD49 | 0,420746454 | 1,338619981 | 0,41067 | 0,66856387 | ankyrin repeat domain 49 |
| ENSG00000148840 | PPRC1 | 0,420648676 | 1,338529259 | 0,03419 | 0,24972789 | PPARG related coactivator 1 |
| ENSG00000196230 | TUBB | 0,420586443 | 1,338471522 | 0,13528 | 0,46178224 | tubulin beta class I |
| ENSG00000104671 | DCTN6 | 0,420572241 | 1,338458345 | 0,2708 | 0,57239492 | dynactin subunit 6 |
| ENSG00000074603 | DPP8 | 0,420520272 | 1,338410132 | 0,19806 | 0,51977271 | dipeptidyl peptidase 8 |
| ENSG00000213700 | RPL17P50 | 0,420466039 | 1,33835982 | 0,59714 | 0,79434997 | ribosomal protein L17 pseudogene 50 |
| ENSG00000174444 | RPL4 | 0,420455543 | 1,338350083 | 0,1301 | 0,45452604 | ribosomal protein L4 |
| ENSG00000107938 | EDRF1 | 0,420443677 | 1,338339075 | 0,1901 | 0,51331106 | erythroid differentiation regulatory factor 1 |
| ENSG00000120278 | PLEKHG1 | 0,42043617 | 1,338332112 | 0,15621 | 0,4840074 | pleckstrin homology and RhoGEF domain containing G1 |
| ENSG00000106692 | FKTN | 0,420395265 | 1,338294166 | 0,224 | 0,53994403 | fukutin |
| ENSG00000255717 | SNHG1 | 0,420392539 | 1,338291638 | 0,22868 | 0,54252903 | small nucleolar RNA host gene 1 |
| ENSG00000213516 | RBMXL1 | 0,420228661 | 1,338139628 | 0,2318 | 0,54454192 | RBMX like 1 |
| ENSG00000154174 | TOMM70 | 0,420195156 | 1,338108551 | 0,2379 | 0,54864017 | translocase of outer mitochondrial membrane 70 |
| ENSG00000145919 | BOD1 | 0,420138373 | 1,338055885 | 0,15094 | 0,47923194 | biorientation of chromosomes in cell division 1 |
| ENSG00000104979 | C19orf53 | 0,42013606 | 1,33805374 | 0,34352 | 0,62091871 | chromosome 19 open reading frame 53 |
| ENSG00000047621 | C12orf4 | 0,420081946 | 1,338003553 | 0,14114 | 0,4679289 | chromosome 12 open reading frame 4 |
| ENSG00000219395 | HSPA8P15 | 0,420034636 | 1,337959676 | 0,45703 | 0,70056368 | heat shock protein family A (Hsp70) member 8 pseudogene 15 |
| ENSG00000152942 | RAD17 | 0,420014244 | 1,337940764 | 0,38382 | 0,65045825 | RAD17 checkpoint clamp loader component |
| ENSG00000087586 | AURKA | 0,419943604 | 1,337875255 | 0,2328 | 0,54542294 | aurora kinase A |
| ENSG00000275837 | SMARCB1 | 0,41990342 | 1,337837991 | 0,19359 | 0,51547251 | SWI/SNF related, matrix associated, actin dependent regulator of chromatin, subfamily b, member 1 |
| ENSG00000233684 | LINC01865 | 0,419902084 | 1,337836753 | 0,61877 | 0,80665241 | long intergenic non-protein coding RNA 1865 |

| | | | | | | |
|---|---|---|---|---|---|---|
| ENSG00000288114 | GTF2H1 | 0,419855353 | 1,337793419 | 0,30322 | 0,59353249 | general transcription factor IIH subunit 1 |
| ENSG00000158571 | PFKFB1 | 0,419833172 | 1,33777285 | 0,26191 | 0,56477027 | 6-phosphofructo-2-kinase/fructose-2,6-biphosphatase 1 |
| ENSG00000286416 | AL442067,3 | 0,419800176 | 1,337742255 | 0,22918 | 0,54273374 | novel transcript |
| ENSG00000162975 | KCNF1 | 0,419791105 | 1,337733844 | 0,45479 | 0,69892531 | potassium voltage-gated channel modifier subfamily F member 1 |
| ENSG00000187918 | OR51I2 | 0,419729774 | 1,337676976 | 0,62966 | 0,81315007 | olfactory receptor family 51 subfamily I member 2 |
| ENSG00000125304 | TM9SF2 | 0,41955034 | 1,337510613 | 0,12453 | 0,44559634 | transmembrane 9 superfamily member 2 |
| ENSG00000196937 | FAM3C | 0,419517707 | 1,33748036 | 0,14985 | 0,47782859 | FAM3 metabolism regulating signaling molecule C |
| ENSG00000232493 | RPL12P11 | 0,419491685 | 1,337456236 | 0,51459 | 0,74090558 | ribosomal protein L12 pseudogene 11 |
| ENSG00000068784 | SRBD1 | 0,419403555 | 1,337374538 | 0,21966 | 0,53706236 | S1 RNA binding domain 1 |
| ENSG00000113369 | ARRDC3 | 0,419192713 | 1,337179102 | 0,36946 | 0,63917123 | arrestin domain containing 3 |
| ENSG00000163161 | ERCC3 | 0,419116337 | 1,337108314 | 0,13971 | 0,4668467 | ERCC excision repair 3, TFIIH core complex helicase subunit |
| ENSG00000065882 | TBC1D1 | 0,419074306 | 1,33706936 | 0,15701 | 0,48438698 | TBC1 domain family member 1 |
| ENSG00000145012 | LPP | 0,418927142 | 1,336932976 | 0,01039 | 0,13135977 | LIM domain containing preferred translocation partner in lipoma |
| ENSG00000049860 | HEXB | 0,418856575 | 1,336867584 | 0,0904 | 0,3889575 | hexosaminidase subunit beta |
| ENSG00000197779 | ZNF81 | 0,418850045 | 1,336861534 | 0,32691 | 0,60889526 | zinc finger protein 81 |
| ENSG00000113851 | CRBN | 0,418836936 | 1,336849387 | 0,2527 | 0,55796133 | cereblon |
| ENSG00000122863 | CHST3 | 0,418770515 | 1,33678784 | 0,16351 | 0,48978436 | carbohydrate sulfotransferase 3 |
| ENSG00000165283 | STOML2 | 0,418757086 | 1,336775396 | 0,0853 | 0,3785936 | stomatin like 2 |
| ENSG00000087338 | GMCL1 | 0,418680292 | 1,336704243 | 0,09791 | 0,40323803 | germ cell-less 1, spermatogenesis associated |
| ENSG00000177853 | ZNF518A | 0,418675945 | 1,336700215 | 0,35307 | 0,62780446 | zinc finger protein 518A |
| ENSG00000152127 | MGAT5 | 0,418605959 | 1,336635372 | 0,15987 | 0,48710442 | alpha-1,6-mannosylglycoprotein 6-beta-N-acetylglucosaminyltransferase |
| ENSG00000086848 | ALG9 | 0,418367676 | 1,336414625 | 0,19703 | 0,51824743 | ALG9 alpha-1,2-mannosyltransferase |
| ENSG00000100219 | XBP1 | 0,418359498 | 1,336407049 | 0,05651 | 0,31717392 | X-box binding protein 1 |
| ENSG00000219392 | ZNF602P | 0,418175553 | 1,336236667 | 0,46608 | 0,70640556 | zinc finger protein 602, pseudogene |
| ENSG00000115616 | SLC9A2 | 0,418140347 | 1,336204059 | 0,54396 | 0,76034748 | solute carrier family 9 member A2 |
| ENSG00000115207 | GTF3C2 | 0,418110151 | 1,336176092 | 0,1483 | 0,47557349 | general transcription factor IIIC subunit 2 |
| ENSG00000285310 | AC005183,2 | 0,417993984 | 1,336068507 | 0,3662 | 0,63617598 | SDA1 domain containing 1 (SDAD1) pseudogene |
| ENSG00000271500 | AC005183,1 | 0,417993984 | 1,336068507 | 0,3662 | 0,63617598 | SDA1 domain containing 1 (SDAD1) pseudogene |
| ENSG00000132670 | PTPRA | 0,417990899 | 1,33606565 | 0,17282 | 0,4976586 | protein tyrosine phosphatase receptor type A |
| ENSG00000110108 | TMEM109 | 0,417989227 | 1,336064102 | 0,12364 | 0,44437757 | transmembrane protein 109 |

| | | | | | | |
|---|---|---|---|---|---|---|
| ENSG00000164134 | NAA15 | 0,417809386 | 1,335897563 | 0,2256 | 0,54087803 | N-alpha-acetyltransferase 15, NatA auxiliary subunit |
| ENSG00000213160 | KLHL23 | 0,417807218 | 1,335895556 | 0,24627 | 0,55395309 | kelch like family member 23 |
| ENSG00000163608 | NEPRO | 0,417783544 | 1,335873634 | 0,27513 | 0,57491355 | nucleolus and neural progenitor protein |
| ENSG00000108468 | CBX1 | 0,417757824 | 1,335849819 | 0,17075 | 0,49641416 | chromobox 1 |
| ENSG00000131115 | ZNF227 | 0,417735662 | 1,335829299 | 0,27322 | 0,57344244 | zinc finger protein 227 |
| ENSG00000213390 | ARHGAP19 | 0,417599534 | 1,33570326 | 0,18857 | 0,5119341 | Rho GTPase activating protein 19 |
| ENSG00000285509 | TBCEL-TECTA | 0,417527955 | 1,335636991 | 0,49794 | 0,72933516 | TBCEL-TECTA readthrough |
| ENSG00000270846 | MYG1P1 | 0,417491986 | 1,335603691 | 0,52042 | 0,7449397 | MYG1 pseudogene 1 |
| ENSG00000275601 | AC011330,2 | 0,417336238 | 1,335459512 | 0,47485 | 0,71364583 | novel transcript, sense intronic to CATSPER2 |
| ENSG00000162616 | DNAJB4 | 0,417238785 | 1,335369306 | 0,10442 | 0,41285329 | DnaJ heat shock protein family (Hsp40) member B4 |
| ENSG00000188342 | GTF2F2 | 0,417213041 | 1,335345478 | 0,27057 | 0,5723494 | general transcription factor IIF subunit 2 |
| ENSG00000283855 | NPIPB13 | 0,417115654 | 1,33525534 | 0,83497 | 0,92456226 | nuclear pore complex interacting protein family, member B13 |
| ENSG00000006576 | PHTF2 | 0,416968192 | 1,335118867 | 0,32708 | 0,60902638 | putative homeodomain transcription factor 2 |
| ENSG00000115008 | IL1A | 0,416936212 | 1,335089272 | 0,33312 | 0,61307256 | interleukin 1 alpha |
| ENSG00000125977 | EIF2S2 | 0,416930171 | 1,335083681 | 0,06861 | 0,34614406 | eukaryotic translation initiation factor 2 subunit beta |
| ENSG00000186638 | KIF24 | 0,416845302 | 1,335005145 | 0,23919 | 0,5491566 | kinesin family member 24 |
| ENSG00000136870 | ZNF189 | 0,41674545 | 1,33491275 | 0,22972 | 0,54324957 | zinc finger protein 189 |
| ENSG00000136451 | VEZF1 | 0,416734671 | 1,334902776 | 0,36986 | 0,63943131 | vascular endothelial zinc finger 1 |
| ENSG00000111530 | CAND1 | 0,416703139 | 1,3348736 | 0,21217 | 0,5308403 | cullin associated and neddylation dissociated 1 |
| ENSG00000224733 | AL034349,1 | 0,416694598 | 1,334865698 | 0,65992 | 0,83018671 | novel transcript, antisense to PTPRK |
| ENSG00000189050 | RNFT1 | 0,416661856 | 1,334835403 | 0,08471 | 0,37788298 | ring finger protein, transmembrane 1 |
| ENSG00000177888 | ZBTB41 | 0,416616919 | 1,334793827 | 0,2983 | 0,59073816 | zinc finger and BTB domain containing 41 |
| ENSG00000143437 | ARNT | 0,41647953 | 1,334666719 | 0,05938 | 0,32456197 | aryl hydrocarbon receptor nuclear translocator |
| ENSG00000127589 | TUBBP1 | 0,416477375 | 1,334664725 | 0,33842 | 0,61774255 | tubulin beta pseudogene 1 |
| ENSG00000101166 | PRELID3B | 0,416474952 | 1,334662484 | 0,42074 | 0,67603599 | PRELI domain containing 3B |
| ENSG00000269431 | BNIP3P8 | 0,416466904 | 1,334655038 | 0,67865 | 0,84182956 | BCL2 interacting protein 3 pseudogene 8 |
| ENSG00000140367 | UBE2Q2 | 0,416272085 | 1,334474822 | 0,23343 | 0,54563244 | ubiquitin conjugating enzyme E2 Q2 |
| ENSG00000261522 | AL845331,2 | 0,416175828 | 1,334385788 | 0,66343 | 0,83242946 | methylenetetrahydrofolate dehydrogenase (NADP+ dependent) 1-like (MTHFD1L) pseudogene |
| ENSG00000205534 | SMG1P2 | 0,416164668 | 1,334375465 | 0,78599 | 0,90065346 | SMG1 pseudogene 2 |
| ENSG00000213793 | ZNF888 | 0,416163906 | 1,334374761 | 0,35847 | 0,63185282 | zinc finger protein 888 |
| ENSG00000285372 | TNKS | 0,416080946 | 1,334298032 | 0,03142 | 0,23928335 | tankyrase |

| | | | | | | |
|---|---|---|---|---|---|---|
| ENSG00000262370 | AC108134,3 | 0,415997137 | 1,334220522 | 0,31556 | 0,60172912 | novel transcript |
| ENSG00000229743 | LINC01159 | 0,415984243 | 1,334208597 | 0,36166 | 0,63352908 | long intergenic non-protein coding RNA 1159 |
| ENSG00000120509 | PDZD11 | 0,415954298 | 1,334180905 | 0,27033 | 0,57209672 | PDZ domain containing 11 |
| ENSG00000110344 | UBE4A | 0,415948386 | 1,334175437 | 0,32493 | 0,60750217 | ubiquitination factor E4A |
| ENSG00000261315 | LARP4P | 0,415912622 | 1,334142363 | 0,45366 | 0,69783825 | LARP4 pseudogene |
| ENSG00000184117 | NIPSNAP1 | 0,41564016 | 1,333890427 | 0,23486 | 0,54635524 | nipsnap homolog 1 |
| ENSG00000273183 | AC093726,2 | 0,415519642 | 1,333779003 | 0,48577 | 0,72125477 | novel transcript, antisense to PAXIP1 |
| ENSG00000110911 | SLC11A2 | 0,41546535 | 1,33372881 | 0,08067 | 0,37077071 | solute carrier family 11 member 2 |
| ENSG00000114988 | LMAN2L | 0,415434324 | 1,333700127 | 0,1204 | 0,43926479 | lectin, mannose binding 2 like |
| ENSG00000213214 | ARHGEF35 | 0,415228706 | 1,333510057 | 0,43976 | 0,68850239 | Rho guanine nucleotide exchange factor 35 |
| ENSG00000230825 | AC005532,1 | 0,415189039 | 1,333473393 | 0,43247 | 0,68411566 | novel transcript |
| ENSG00000171867 | PRNP | 0,415128256 | 1,333417213 | 0,39203 | 0,65540679 | prion protein |
| ENSG00000115840 | SLC25A12 | 0,415056012 | 1,333350443 | 0,38369 | 0,65041894 | solute carrier family 25 member 12 |
| ENSG00000062370 | ZNF112 | 0,415047193 | 1,333342292 | 0,36636 | 0,63632065 | zinc finger protein 112 |
| ENSG00000113966 | ARL6 | 0,414937648 | 1,333241054 | 0,37705 | 0,64554654 | ADP ribosylation factor like GTPase 6 |
| ENSG00000165152 | PGAP4 | 0,414829819 | 1,33314141 | 0,04681 | 0,29102032 | post-GPI attachment to proteins GalNAc transferase 4 |
| ENSG00000260063 | AL512408,1 | 0,414824976 | 1,333136934 | 0,20107 | 0,52181085 | novel transcript, antisense to ARID1A |
| ENSG00000197905 | TEAD4 | 0,414808015 | 1,333121262 | 0,12475 | 0,44598591 | TEA domain transcription factor 4 |
| ENSG00000144445 | KANSL1L | 0,414757775 | 1,333074838 | 0,24022 | 0,54948131 | KAT8 regulatory NSL complex subunit 1 like |
| ENSG00000116237 | ICMT | 0,414711236 | 1,333031836 | 0,05383 | 0,31110058 | isoprenylcysteine carboxyl methyltransferase |
| ENSG00000053371 | AKR7A2 | 0,414675863 | 1,332999152 | 0,20313 | 0,52299224 | aldo-keto reductase family 7 member A2 |
| ENSG00000251484 | AL356488,1 | 0,41465929 | 1,332983839 | 0,3172 | 0,6027392 | poly(rC) binding protein 1 (PCBP1) pseudogene |
| ENSG00000241625 | RN7SL18P | 0,414617922 | 1,332945618 | 0,74755 | 0,88100509 | RNA, 7SL, cytoplasmic 18, pseudogene |
| ENSG00000243696 | AC006254,1 | 0,414612878 | 1,332940958 | 0,2513 | 0,55720663 | novel MUSTN1-ITIH4 readthrough |
| ENSG00000048544 | MRPS10 | 0,414609043 | 1,332937415 | 0,19481 | 0,5158697 | mitochondrial ribosomal protein S10 |
| ENSG00000288121 | MDN1 | 0,414604817 | 1,33293351 | 0,05413 | 0,31168784 | midasin AAA ATPase 1 |
| ENSG00000112159 | MDN1 | 0,414604817 | 1,33293351 | 0,05413 | 0,31168784 | midasin AAA ATPase 1 |
| ENSG00000248222 | SLIT3-AS1 | 0,41459506 | 1,332924495 | 0,68386 | 0,84517371 | SLIT3 antisense RNA 1 |
| ENSG00000121775 | TMEM39B | 0,414542714 | 1,332876133 | 0,09794 | 0,40323803 | transmembrane protein 39B |
| ENSG00000229154 | KCNQ5-AS1 | 0,41453035 | 1,33286471 | 0,69059 | 0,84900677 | KCNQ5 antisense RNA 1 |
| ENSG00000174099 | MSRB3 | 0,414339659 | 1,332688548 | 0,02899 | 0,22977124 | methionine sulfoxide reductase B3 |
| ENSG00000086062 | B4GALT1 | 0,41431978 | 1,332670184 | 0,24112 | 0,5502162 | beta-1,4-galactosyltransferase 1 |
| ENSG00000153790 | C7orf31 | 0,41418227 | 1,332543168 | 0,29494 | 0,58868965 | chromosome 7 open reading frame 31 |

| | | | | | | |
|---|---|---|---|---|---|---|
| ENSG00000184110 | EIF3C | 0,414157967 | 1,33252072 | 0,16589 | 0,49195088 | eukaryotic translation initiation factor 3 subunit C |
| ENSG00000229097 | CALM2P2 | 0,41405403 | 1,332424724 | 0,38239 | 0,64928247 | calmodulin 2 pseudogene 2 |
| ENSG00000235447 | TRAPPC13P1 | 0,413886778 | 1,332270265 | 0,35029 | 0,62578708 | trafficking protein particle complex 13 pseudogene 1 |
| ENSG00000258940 | AL132639,2 | 0,413835116 | 1,332222558 | 0,51636 | 0,74198903 | novel transcript, antisense to CTAGE5 |
| ENSG00000272777 | AC019131,2 | 0,413553466 | 1,331962501 | 0,43617 | 0,68658672 | novel transcript |
| ENSG00000243749 | TMEM35B | 0,413525022 | 1,33193624 | 0,37205 | 0,64114037 | transmembrane protein 35B |
| ENSG00000091009 | RBM27 | 0,413510052 | 1,331922419 | 0,3026 | 0,59306846 | RNA binding motif protein 27 |
| ENSG00000066422 | ZBTB11 | 0,413439706 | 1,331857476 | 0,35153 | 0,62674766 | zinc finger and BTB domain containing 11 |
| ENSG00000144736 | SHQ1 | 0,413409647 | 1,331829727 | 0,06783 | 0,34433909 | SHQ1, H/ACA ribonucleoprotein assembly factor |
| ENSG00000173163 | COMMD1 | 0,413371737 | 1,33179473 | 0,204 | 0,5236955 | copper metabolism domain containing 1 |
| ENSG00000119636 | BBOF1 | 0,413370367 | 1,331793466 | 0,3279 | 0,60964743 | basal body orientation factor 1 |
| ENSG00000130182 | ZSCAN10 | 0,413363258 | 1,331786903 | 0,65459 | 0,82719457 | zinc finger and SCAN domain containing 10 |
| ENSG00000149346 | SLX4IP | 0,413352902 | 1,331777343 | 0,20376 | 0,52348797 | SLX4 interacting protein |
| ENSG00000124615 | MOCS1 | 0,413268201 | 1,331699157 | 0,1683 | 0,49453836 | molybdenum cofactor synthesis 1 |
| ENSG00000231940 | RPS7P3 | 0,413265071 | 1,331696267 | 0,56404 | 0,77260898 | ribosomal protein S7 pseudogene 3 |
| ENSG00000174599 | TRAM1L1 | 0,41325884 | 1,331690516 | 0,39594 | 0,65863599 | translocation associated membrane protein 1 like 1 |
| ENSG00000288580 | AC023296,1 | 0,413245455 | 1,331678161 | 0,5602 | 0,7707835 | novel transcript |
| ENSG00000139372 | TDG | 0,413216889 | 1,331651794 | 0,19305 | 0,51505487 | thymine DNA glycosylase |
| ENSG00000120053 | GOT1 | 0,413191327 | 1,331628199 | 0,07614 | 0,36238326 | glutamic-oxaloacetic transaminase 1 |
| ENSG00000120137 | PANK3 | 0,413141802 | 1,331582488 | 0,23227 | 0,54499308 | pantothenate kinase 3 |
| ENSG00000079739 | PGM1 | 0,413087664 | 1,33153252 | 0,1588 | 0,48624756 | phosphoglucomutase 1 |
| ENSG00000149451 | ADAM33 | 0,41305575 | 1,331503066 | 0,27735 | 0,57586315 | ADAM metallopeptidase domain 33 |
| ENSG00000160584 | SIK3 | 0,413007506 | 1,33145854 | 0,15475 | 0,48266733 | SIK family kinase 3 |
| ENSG00000269896 | AL513477,1 | 0,41300034 | 1,331451927 | 0,19215 | 0,51468724 | small nuclear ribonucleoprotein N (SNRPN) pseudogene |
| ENSG00000232485 | RPL37A-DT | 0,412971805 | 1,331425593 | 0,5722 | 0,77846037 | RPL37A divergent transcript |
| ENSG00000164465 | DCBLD1 | 0,412917009 | 1,331375023 | 0,17798 | 0,50234382 | discoidin, CUB and LCCL domain containing 1 |
| ENSG00000224398 | CSNK2B | 0,412915094 | 1,331373256 | 0,33106 | 0,6112353 | casein kinase 2 beta |
| ENSG00000096717 | SIRT1 | 0,412732599 | 1,331204854 | 0,11434 | 0,42986344 | sirtuin 1 |
| ENSG00000285588 | AL160254,1 | 0,412691811 | 1,331167218 | 0,70136 | 0,85602538 | novel transcript, antisense to PCDH9 |
| ENSG00000234361 | AL391863,2 | 0,412666771 | 1,331144115 | 0,37908 | 0,64697338 | novel transcript |
| ENSG00000120658 | ENOX1 | 0,412514133 | 1,331003286 | 0,20553 | 0,52513499 | ecto-NOX disulfide-thiol exchanger 1 |
| ENSG00000258568 | RHOQP1 | 0,412511294 | 1,331000667 | 0,15657 | 0,48425071 | ras homolog family member Q pseudogene 1 |
| ENSG00000148484 | RSU1 | 0,412416357 | 1,330913083 | 0,10568 | 0,41518746 | Ras suppressor protein 1 |

| | | | | | | |
|---|---|---|---|---|---|---|
| ENSG00000100614 | PPM1A | 0,412371175 | 1,330871402 | 0,25108 | 0,55719428 | protein phosphatase, Mg2+/Mn2+ dependent 1A |
| ENSG00000115461 | IGFBP5 | 0,412364469 | 1,330865216 | 0,31794 | 0,60344598 | insulin like growth factor binding protein 5 |
| ENSG00000113732 | ATP6V0E1 | 0,412327123 | 1,330830766 | 0,14588 | 0,47358318 | ATPase H+ transporting V0 subunit e1 |
| ENSG00000048028 | USP28 | 0,412278758 | 1,330786151 | 0,21557 | 0,53437517 | ubiquitin specific peptidase 28 |
| ENSG00000158406 | H4C8 | 0,412258872 | 1,330767808 | 0,28269 | 0,58009943 | H4 clustered histone 8 |
| ENSG00000135870 | RC3H1 | 0,412238541 | 1,330749054 | 0,20252 | 0,52259159 | ring finger and CCCH-type domains 1 |
| ENSG00000164045 | CDC25A | 0,412169206 | 1,330685101 | 0,10656 | 0,41635243 | cell division cycle 25A |
| ENSG00000071189 | SNX13 | 0,41207637 | 1,330599475 | 0,23124 | 0,54417127 | sorting nexin 13 |
| ENSG00000237976 | RFX5-AS1 | 0,412002885 | 1,330531702 | 0,48115 | 0,71820356 | RFX5 antisense RNA 1 |
| ENSG00000204779 | FOXD4L5 | 0,41199349 | 1,330523037 | 0,24679 | 0,55449305 | forkhead box D4 like 5 |
| ENSG00000150459 | SAP18 | 0,411977797 | 1,330508564 | 0,23252 | 0,54526182 | Sin3A associated protein 18 |
| ENSG00000088832 | FKBP1A | 0,411835399 | 1,330377246 | 0,04477 | 0,28589646 | FKBP prolyl isomerase 1A |
| ENSG00000165511 | ZNF22-AS1 | 0,411810202 | 1,330354011 | 0,29119 | 0,58623007 | ZNF22 antisense RNA 1 |
| ENSG00000150991 | UBC | 0,411789017 | 1,330334476 | 0,34066 | 0,61908972 | ubiquitin C |
| ENSG00000256115 | LINC02443 | 0,411769826 | 1,330316779 | 0,36988 | 0,63943131 | long intergenic non-protein coding RNA 2443 |
| ENSG00000142528 | ZNF473 | 0,411733146 | 1,330282957 | 0,30237 | 0,59297237 | zinc finger protein 473 |
| ENSG00000115827 | DCAF17 | 0,411724925 | 1,330275377 | 0,24623 | 0,55395309 | DDB1 and CUL4 associated factor 17 |
| ENSG00000228474 | OST4 | 0,411665495 | 1,330220579 | 0,3435 | 0,62091871 | oligosaccharyltransferase complex subunit 4, non-catalytic |
| ENSG00000102309 | PIN4 | 0,411640774 | 1,330197786 | 0,29349 | 0,5879488 | peptidylprolyl cis/trans isomerase, NIMA-interacting 4 |
| ENSG00000184205 | TSPYL2 | 0,41150202 | 1,330069857 | 0,11265 | 0,42641188 | TSPY like 2 |
| ENSG00000232199 | RPL3P8 | 0,411500719 | 1,330068658 | 0,73868 | 0,87593493 | ribosomal protein L3 pseudogene 8 |
| ENSG00000121060 | TRIM25 | 0,411480541 | 1,330050055 | 0,2846 | 0,58147083 | tripartite motif containing 25 |
| ENSG00000169946 | ZFPM2 | 0,41137801 | 1,329955533 | 0,21442 | 0,53335795 | zinc finger protein, FOG family member 2 |
| ENSG00000165115 | KIF27 | 0,411350715 | 1,329930371 | 0,30784 | 0,59671558 | kinesin family member 27 |
| ENSG00000144036 | EXOC6B | 0,411321408 | 1,329903355 | 0,25557 | 0,56004204 | exocyst complex component 6B |
| ENSG00000138617 | PARP16 | 0,41129108 | 1,329875399 | 0,36185 | 0,63352908 | poly(ADP-ribose) polymerase family member 16 |
| ENSG00000183605 | SFXN4 | 0,411127563 | 1,329724677 | 0,11031 | 0,42293017 | sideroflexin 4 |
| ENSG00000105983 | LMBR1 | 0,411074936 | 1,329676172 | 0,1904 | 0,51350457 | limb development membrane protein 1 |
| ENSG00000099995 | SF3A1 | 0,411074143 | 1,329675441 | 0,14093 | 0,46791293 | splicing factor 3a subunit 1 |
| ENSG00000088325 | TPX2 | 0,410990367 | 1,32959823 | 0,1568 | 0,48425071 | TPX2 microtubule nucleation factor |
| ENSG00000150995 | ITPR1 | 0,410947607 | 1,329558823 | 0,1548 | 0,48266733 | inositol 1,4,5-trisphosphate receptor type 1 |
| ENSG00000091732 | ZC3HC1 | 0,410853867 | 1,329472437 | 0,0478 | 0,29355314 | zinc finger C3HC-type containing 1 |
| ENSG00000134324 | LPIN1 | 0,410813708 | 1,32943543 | 0,07982 | 0,36873474 | lipin 1 |

| | | | | | | |
|---|---|---|---|---|---|---|
| ENSG00000165300 | SLITRK5 | 0,410804855 | 1,329427272 | 0,19129 | 0,51431444 | SLIT and NTRK like family member 5 |
| ENSG00000033178 | UBA6 | 0,410757767 | 1,329383882 | 0,31174 | 0,59912688 | ubiquitin like modifier activating enzyme 6 |
| ENSG00000144647 | POMGNT2 | 0,410736858 | 1,329364615 | 0,11304 | 0,42730497 | protein O-linked mannose N-acetylglucosaminyltransferase 2 (beta 1,4-) |
| ENSG00000144580 | CNOT9 | 0,410729706 | 1,329358025 | 0,13997 | 0,46694389 | CCR4-NOT transcription complex subunit 9 |
| ENSG00000203724 | C1orf53 | 0,410662703 | 1,329296287 | 0,42792 | 0,68085046 | chromosome 1 open reading frame 53 |
| ENSG00000147130 | ZMYM3 | 0,410636387 | 1,32927204 | 0,07828 | 0,36569619 | zinc finger MYM-type containing 3 |
| ENSG00000267001 | AC006538,2 | 0,410608128 | 1,329246003 | 0,40974 | 0,6680455 | novel protein |
| ENSG00000282091 | AC139256,1 | 0,41059609 | 1,329234912 | 0,79159 | 0,90310057 | phospholipase A2 group X (PLA2G10) pseudogene |
| ENSG00000001461 | NIPAL3 | 0,410577056 | 1,329217374 | 0,21212 | 0,5308403 | NIPA like domain containing 3 |
| ENSG00000153989 | NUS1 | 0,410558743 | 1,329200502 | 0,14698 | 0,47471707 | NUS1 dehydrodolichyl diphosphate synthase subunit |
| ENSG00000136478 | TEX2 | 0,410493766 | 1,329140638 | 0,16502 | 0,49125639 | testis expressed 2 |
| ENSG00000228657 | RPL23AP58 | 0,410439023 | 1,329090205 | 0,53588 | 0,75497727 | ribosomal protein L23a pseudogene 58 |
| ENSG00000239626 | RPSAP41 | 0,410149576 | 1,328823577 | 0,57355 | 0,77901952 | ribosomal protein SA pseudogene 41 |
| ENSG00000115306 | SPTBN1 | 0,410136654 | 1,328811675 | 0,26188 | 0,56477027 | spectrin beta, non-erythrocytic 1 |
| ENSG00000134056 | MRPS36 | 0,410133005 | 1,328808314 | 0,4633 | 0,70416724 | mitochondrial ribosomal protein S36 |
| ENSG00000109171 | SLAIN2 | 0,410104474 | 1,328782035 | 0,21306 | 0,53205351 | SLAIN motif family member 2 |
| ENSG00000101040 | ZMYND8 | 0,409939868 | 1,328630435 | 0,29587 | 0,58942061 | zinc finger MYND-type containing 8 |
| ENSG00000121621 | KIF18A | 0,409916518 | 1,328608932 | 0,43624 | 0,68660209 | kinesin family member 18A |
| ENSG00000135250 | SRPK2 | 0,409880375 | 1,328575647 | 0,20756 | 0,52710325 | SRSF protein kinase 2 |
| ENSG00000254936 | AF131215,4 | 0,409843633 | 1,328541812 | 0,55041 | 0,76458107 | novel transcript |
| ENSG00000103051 | COG4 | 0,409841946 | 1,328540258 | 0,2152 | 0,53387837 | component of oligomeric golgi complex 4 |
| ENSG00000117262 | GPR89A | 0,409806145 | 1,32850729 | 0,24476 | 0,55280101 | G protein-coupled receptor 89A |
| ENSG00000285178 | ZFP1 | 0,409752055 | 1,328457483 | 0,43825 | 0,68763409 | ZFP1 zinc finger protein |
| ENSG00000184517 | ZFP1 | 0,409752055 | 1,328457483 | 0,43825 | 0,68763409 | ZFP1 zinc finger protein |
| ENSG00000049449 | RCN1 | 0,409750555 | 1,328456101 | 0,15468 | 0,48266733 | reticulocalbin 1 |
| ENSG00000133703 | KRAS | 0,409656046 | 1,328369079 | 0,35686 | 0,63098227 | KRAS proto-oncogene, GTPase |
| ENSG00000166794 | PPIB | 0,409568843 | 1,328288789 | 0,15051 | 0,47863728 | peptidylprolyl isomerase B |
| ENSG00000234685 | NUS1P2 | 0,409534666 | 1,328257322 | 0,48137 | 0,71822668 | NUS1 pseudogene 2 |
| ENSG00000163382 | NAXE | 0,409516661 | 1,328240746 | 0,08434 | 0,37722352 | NAD(P)HX epimerase |
| ENSG00000139496 | NUP58 | 0,409361591 | 1,328097986 | 0,20171 | 0,52214043 | nucleoporin 58 |
| ENSG00000261251 | Z97055,2 | 0,409214272 | 1,327962376 | 0,3855 | 0,65114724 | novel transcript, antisense to EFCAB6 |
| ENSG00000253236 | LINC02143 | 0,40921408 | 1,327962199 | 0,72199 | 0,86726498 | long intergenic non-protein coding RNA 2143 |

| | | | | | | |
|---|---|---|---|---|---|---|
| ENSG00000111647 | UHRF1BP1L | 0,409138848 | 1,327892951 | 0,36237 | 0,63379594 | UHRF1 binding protein 1 like |
| ENSG00000152455 | SUV39H2 | 0,409096456 | 1,327853933 | 0,17349 | 0,49809931 | suppressor of variegation 3-9 homolog 2 |
| ENSG00000110768 | GTF2H1 | 0,409019998 | 1,327783563 | 0,2224 | 0,53859118 | general transcription factor IIH subunit 1 |
| ENSG00000186369 | LINC00643 | 0,408933962 | 1,327704382 | 0,39181 | 0,65534953 | long intergenic non-protein coding RNA 643 |
| ENSG00000115561 | CHMP3 | 0,408928919 | 1,327699741 | 0,24784 | 0,55528615 | charged multivesicular body protein 3 |
| ENSG00000198646 | NCOA6 | 0,408911494 | 1,327683706 | 0,16018 | 0,48737351 | nuclear receptor coactivator 6 |
| ENSG00000065328 | MCM10 | 0,408907132 | 1,327679691 | 0,23841 | 0,54875544 | minichromosome maintenance 10 replication initiation factor |
| ENSG00000071967 | CYBRD1 | 0,408809537 | 1,32758988 | 0,14719 | 0,47471707 | cytochrome b reductase 1 |
| ENSG00000110092 | CCND1 | 0,408750978 | 1,327535994 | 0,10843 | 0,41905863 | cyclin D1 |
| ENSG00000116521 | SCAMP3 | 0,408702102 | 1,32749102 | 0,23391 | 0,54568606 | secretory carrier membrane protein 3 |
| ENSG00000090013 | BLVRB | 0,40869814 | 1,327487374 | 0,3013 | 0,59258306 | biliverdin reductase B |
| ENSG00000132646 | PCNA | 0,408665317 | 1,327457173 | 0,25901 | 0,56278415 | proliferating cell nuclear antigen |
| ENSG00000136813 | ECPAS | 0,408662018 | 1,327454137 | 0,23916 | 0,5491566 | Ecm29 proteasome adaptor and scaffold |
| ENSG00000011007 | ELOA | 0,408627466 | 1,327422346 | 0,16749 | 0,49362211 | elongin A |
| ENSG00000134802 | SLC43A3 | 0,408589877 | 1,327387761 | 0,23924 | 0,5491566 | solute carrier family 43 member 3 |
| ENSG00000224455 | VPS52 | 0,408586224 | 1,3273844 | 0,61382 | 0,8035281 | VPS52 subunit of GARP complex |
| ENSG00000236014 | VPS52 | 0,408586224 | 1,3273844 | 0,61382 | 0,8035281 | VPS52 subunit of GARP complex |
| ENSG00000228894 | HCG18 | 0,408585185 | 1,327383444 | 0,30816 | 0,59696396 | HLA complex group 18 |
| ENSG00000224619 | HCG18 | 0,408585185 | 1,327383444 | 0,30816 | 0,59696396 | HLA complex group 18 |
| ENSG00000111877 | MCM9 | 0,408567765 | 1,327367416 | 0,19728 | 0,51854293 | minichromosome maintenance 9 homologous recombination repair factor |
| ENSG00000221823 | PPP3R1 | 0,408546847 | 1,327348171 | 0,04293 | 0,27922774 | protein phosphatase 3 regulatory subunit B, alpha |
| ENSG00000116266 | STXBP3 | 0,4085099 | 1,327314178 | 0,31807 | 0,6034788 | syntaxin binding protein 3 |
| ENSG00000189056 | RELN | 0,408486997 | 1,327293107 | 0,4097 | 0,6680455 | reelin |
| ENSG00000100526 | CDKN3 | 0,408381773 | 1,327196303 | 0,39691 | 0,65942678 | cyclin dependent kinase inhibitor 3 |
| ENSG00000166454 | ATMIN | 0,40816573 | 1,326997571 | 0,22576 | 0,5409527 | ATM interactor |
| ENSG00000145632 | PLK2 | 0,408076056 | 1,326915091 | 0,11355 | 0,42829885 | polo like kinase 2 |
| ENSG00000130309 | COLGALT1 | 0,408012938 | 1,32685704 | 0,02766 | 0,22412082 | collagen beta(1-O)galactosyltransferase 1 |
| ENSG00000223516 | AFF2-IT1 | 0,40797736 | 1,326824319 | 0,67964 | 0,84263958 | AFF2 intronic transcript 1 |
| ENSG00000078043 | PIAS2 | 0,407974997 | 1,326822146 | 0,08103 | 0,37153397 | protein inhibitor of activated STAT 2 |
| ENSG00000121766 | ZCCHC17 | 0,40792375 | 1,326775016 | 0,13957 | 0,46677981 | zinc finger CCHC-type containing 17 |
| ENSG00000107672 | NSMCE4A | 0,407901771 | 1,326754803 | 0,21993 | 0,53732404 | NSE4 homolog A, SMC5-SMC6 complex component |

| | | | | | | |
|---|---|---|---|---|---|---|
| ENSG00000111729 | CLEC4A | 0,407875944 | 1,326731051 | 0,62234 | 0,80848353 | C-type lectin domain family 4 member A |
| ENSG00000189042 | ZNF567 | 0,407870541 | 1,326726083 | 0,20468 | 0,52443222 | zinc finger protein 567 |
| ENSG00000240053 | LY6G5B | 0,407790271 | 1,326652267 | 0,3575 | 0,63133849 | lymphocyte antigen 6 family member G5B |
| ENSG00000112282 | MED23 | 0,407776261 | 1,326639384 | 0,26965 | 0,57135887 | mediator complex subunit 23 |
| ENSG00000041880 | PARP3 | 0,407726319 | 1,326593461 | 0,1872 | 0,5105374 | poly(ADP-ribose) polymerase family member 3 |
| ENSG00000114354 | TFG | 0,407702104 | 1,326571194 | 0,23949 | 0,5491566 | trafficking from ER to golgi regulator |
| ENSG00000182150 | ERCC6L2 | 0,40765345 | 1,326526457 | 0,27079 | 0,57239492 | ERCC excision repair 6 like 2 |
| ENSG00000171824 | EXOSC10 | 0,407604609 | 1,326481549 | 0,13352 | 0,4594073 | exosome component 10 |
| ENSG00000013374 | NUB1 | 0,407590015 | 1,326468131 | 0,05489 | 0,31349896 | negative regulator of ubiquitin like proteins 1 |
| ENSG00000101367 | MAPRE1 | 0,407584511 | 1,326463071 | 0,10799 | 0,41856292 | microtubule associated protein RP/EB family member 1 |
| ENSG00000115307 | AUP1 | 0,407584413 | 1,326462981 | 0,17905 | 0,50344261 | AUP1 lipid droplet regulating VLDL assembly factor |
| ENSG00000258710 | LINC01193 | 0,407520694 | 1,326404397 | 0,64207 | 0,8205913 | long intergenic non-protein coding RNA 1193 |
| ENSG00000287697 | Z99127,3 | 0,407449165 | 1,326338635 | 0,22187 | 0,53821014 | novel transcript, antisense to CACYBP |
| ENSG00000164743 | C8orf48 | 0,407352547 | 1,326249813 | 0,33068 | 0,6112353 | chromosome 8 open reading frame 48 |
| ENSG00000140374 | ETFA | 0,407296089 | 1,326197912 | 0,30015 | 0,59160746 | electron transfer flavoprotein subunit alpha |
| ENSG00000164032 | H2AZ1 | 0,407294008 | 1,326195999 | 0,24607 | 0,55380958 | H2A,Z variant histone 1 |
| ENSG00000257056 | LINC02282 | 0,407285666 | 1,326188331 | 0,42751 | 0,68046596 | long intergenic non-protein coding RNA 2282 |
| ENSG00000160803 | UBQLN4 | 0,407260907 | 1,326165571 | 0,08946 | 0,38678501 | ubiquilin 4 |
| ENSG00000137414 | FAM8A1 | 0,407249793 | 1,326155356 | 0,29645 | 0,58978642 | family with sequence similarity 8 member A1 |
| ENSG00000160746 | ANO10 | 0,407118647 | 1,326034808 | 0,26435 | 0,5671001 | anoctamin 10 |
| ENSG00000189332 | SLC25A51P4 | 0,40704077 | 1,325963231 | 0,32684 | 0,60888005 | SLC25A51 pseudogene 4 |
| ENSG00000104147 | OIP5 | 0,406957009 | 1,32588625 | 0,29887 | 0,59104323 | Opa interacting protein 5 |
| ENSG00000111678 | C12orf57 | 0,40692763 | 1,32585925 | 0,13639 | 0,46359081 | chromosome 12 open reading frame 57 |
| ENSG00000256294 | ZNF225 | 0,406922461 | 1,325854499 | 0,23475 | 0,5463041 | zinc finger protein 225 |
| ENSG00000166228 | PCBD1 | 0,406870087 | 1,325806368 | 0,45146 | 0,69647916 | pterin-4 alpha-carbinolamine dehydratase 1 |
| ENSG00000183579 | ZNRF3 | 0,406866976 | 1,325803509 | 0,14868 | 0,47609532 | zinc and ring finger 3 |
| ENSG00000168116 | KIAA1586 | 0,406809799 | 1,325750966 | 0,36381 | 0,63461346 | KIAA1586 |
| ENSG00000259275 | AC087477,2 | 0,406792977 | 1,325735507 | 0,58498 | 0,78606599 | novel transcript |
| ENSG00000147274 | RBMX | 0,406689249 | 1,325640192 | 0,23467 | 0,54623956 | RNA binding motif protein X-linked |
| ENSG00000162819 | BROX | 0,406648329 | 1,325602592 | 0,30227 | 0,59294633 | BRO1 domain and CAAX motif containing |
| ENSG00000187762 | HSPE1P11 | 0,406607164 | 1,325564769 | 0,46127 | 0,70278869 | heat shock protein family E (Hsp10) member 1 pseudogene 11 |
| ENSG00000254905 | AP001318,1 | 0,406584121 | 1,325543597 | 0,48919 | 0,72376431 | novel transcript, antisense to TIRAP |

| | | | | | | |
|---|---|---|---|---|---|---|
| ENSG00000164675 | IQUB | 0,406570321 | 1,325530917 | 0,31315 | 0,59991314 | IQ motif and ubiquitin domain containing |
| ENSG00000095261 | PSMD5 | 0,406566733 | 1,325527621 | 0,13335 | 0,4594073 | proteasome 26S subunit, non-ATPase 5 |
| ENSG00000102763 | VWA8 | 0,40645085 | 1,325421154 | 0,32433 | 0,60692416 | von Willebrand factor A domain containing 8 |
| ENSG00000254450 | ALG9-IT1 | 0,406444174 | 1,32541502 | 0,51267 | 0,73941533 | ALG9 intronic transcript 1 |
| ENSG00000174446 | SNAPC5 | 0,406429471 | 1,325401513 | 0,14186 | 0,46833429 | small nuclear RNA activating complex polypeptide 5 |
| ENSG00000117475 | BLZF1 | 0,406403661 | 1,325377801 | 0,18618 | 0,50956086 | basic leucine zipper nuclear factor 1 |
| ENSG00000174705 | SH3PXD2B | 0,406362351 | 1,325339851 | 0,17494 | 0,49911246 | SH3 and PX domains 2B |
| ENSG00000140382 | HMG20A | 0,40633819 | 1,325317656 | 0,12107 | 0,44016175 | high mobility group 20A |
| ENSG00000104835 | SARS2 | 0,40630815 | 1,32529006 | 0,24529 | 0,55295131 | seryl-tRNA synthetase 2, mitochondrial |
| ENSG00000001036 | FUCA2 | 0,406266364 | 1,325251675 | 0,24858 | 0,55589057 | alpha-L-fucosidase 2 |
| ENSG00000031691 | CENPQ | 0,406252036 | 1,325238513 | 0,40023 | 0,66242349 | centromere protein Q |
| ENSG00000223842 | AL360093,1 | 0,406202093 | 1,325192637 | 0,49624 | 0,72831792 | novel transcript |
| ENSG00000118007 | STAG1 | 0,406191987 | 1,325183354 | 0,24161 | 0,55039378 | stromal antigen 1 |
| ENSG00000161016 | RPL8 | 0,406175232 | 1,325167964 | 0,14797 | 0,4751727 | ribosomal protein L8 |
| ENSG00000108578 | BLMH | 0,406131846 | 1,325128114 | 0,11103 | 0,42382951 | bleomycin hydrolase |
| ENSG00000177694 | NAALADL2 | 0,406090579 | 1,325090209 | 0,162 | 0,48916554 | N-acetylated alpha-linked acidic dipeptidase like 2 |
| ENSG00000155660 | PDIA4 | 0,405881944 | 1,324898596 | 0,11516 | 0,43077792 | protein disulfide isomerase family A member 4 |
| ENSG00000224941 | TCF19 | 0,405842853 | 1,324862697 | 0,52337 | 0,74733773 | transcription factor 19 |
| ENSG00000224472 | TCF19 | 0,405842853 | 1,324862697 | 0,52337 | 0,74733773 | transcription factor 19 |
| ENSG00000172869 | DMXL1 | 0,405836479 | 1,324856844 | 0,33853 | 0,61774255 | Dmx like 1 |
| ENSG00000133731 | IMPA1 | 0,40558389 | 1,324624907 | 0,26839 | 0,57051739 | inositol monophosphatase 1 |
| ENSG00000204359 | CFB | 0,405548436 | 1,324592355 | 0,71242 | 0,86157193 | complement factor B |
| ENSG00000120333 | MRPS14 | 0,405535408 | 1,324580393 | 0,24769 | 0,55509241 | mitochondrial ribosomal protein S14 |
| ENSG00000132141 | CCT6B | 0,405511028 | 1,324558009 | 0,05096 | 0,3037579 | chaperonin containing TCP1 subunit 6B |
| ENSG00000104356 | POP1 | 0,40544286 | 1,324495425 | 0,23859 | 0,54889276 | POP1 homolog, ribonuclease P/MRP subunit |
| ENSG00000232097 | AC079781,3 | 0,405437866 | 1,32449084 | 0,67718 | 0,84122122 | novel pseudogene |
| ENSG00000152332 | UHMK1 | 0,405386252 | 1,324443456 | 0,231 | 0,54393842 | U2AF homology motif kinase 1 |
| ENSG00000012983 | MAP4K5 | 0,405332455 | 1,324394069 | 0,21439 | 0,53334744 | mitogen-activated protein kinase kinase kinase kinase 5 |
| ENSG00000181555 | SETD2 | 0,405322589 | 1,324385012 | 0,22459 | 0,54036191 | SET domain containing 2, histone lysine methyltransferase |
| ENSG00000157020 | SEC13 | 0,405319111 | 1,324381819 | 0,1219 | 0,44093327 | SEC13 homolog, nuclear pore and COPII coat complex component |
| ENSG00000186298 | PPP1CC | 0,405297976 | 1,324362417 | 0,23119 | 0,54414957 | protein phosphatase 1 catalytic subunit gamma |
| ENSG00000164347 | GFM2 | 0,405197269 | 1,324269974 | 0,29489 | 0,58868965 | GTP dependent ribosome recycling factor mitochondrial 2 |

| | | | | | | |
|---|---|---|---|---|---|---|
| ENSG00000250479 | CHCHD10 | 0,405166136 | 1,324241397 | 0,16842 | 0,49467529 | coiled-coil-helix-coiled-coil-helix domain containing 10 |
| ENSG00000267526 | AC005702,2 | 0,405128178 | 1,324206556 | 0,51323 | 0,73963507 | UDP-N-acetyl-alpha-D-galactosamine:polypeptide N-acetylgalactosaminyltransferase 1 (GalNAc-T1) (GALNT1) pseudogene |
| ENSG00000089177 | KIF16B | 0,405110391 | 1,324190229 | 0,12081 | 0,43979929 | kinesin family member 16B |
| ENSG00000136026 | CKAP4 | 0,405049626 | 1,324134457 | 0,02443 | 0,2089849 | cytoskeleton associated protein 4 |
| ENSG00000132305 | IMMT | 0,40503674 | 1,324122631 | 0,3013 | 0,59258306 | inner membrane mitochondrial protein |
| ENSG00000114030 | KPNA1 | 0,404990167 | 1,324079886 | 0,30181 | 0,59288267 | karyopherin subunit alpha 1 |
| ENSG00000225568 | AC093155,1 | 0,404952016 | 1,324044872 | 0,46011 | 0,70210603 | pseudogene similar to part of SUMO-1 activating enzyme subunit 2 (UBA2) |
| ENSG00000177406 | NINJ2-AS1 | 0,404896206 | 1,323993653 | 0,24517 | 0,55295131 | NINJ2 antisense RNA 1 |
| ENSG00000273314 | AC005229,4 | 0,404862509 | 1,323962729 | 0,10067 | 0,4073354 | novel transcript, antisense to CUL1 |
| ENSG00000072182 | ASIC4 | 0,404837433 | 1,323939716 | 0,36098 | 0,63311211 | acid sensing ion channel subunit family member 4 |
| ENSG00000135698 | MPHOSPH6 | 0,404736195 | 1,323846815 | 0,21042 | 0,5290538 | M-phase phosphoprotein 6 |
| ENSG00000261052 | SULT1A3 | 0,404730877 | 1,323841936 | 0,2265 | 0,54144 | sulfotransferase family 1A member 3 |
| ENSG00000164244 | PRRC1 | 0,404647099 | 1,323765061 | 0,08531 | 0,3785936 | proline rich coiled-coil 1 |
| ENSG00000112763 | BTN2A1 | 0,404622155 | 1,323742174 | 0,08336 | 0,37550713 | butyrophilin subfamily 2 member A1 |
| ENSG00000114315 | HES1 | 0,404617863 | 1,323738236 | 0,17635 | 0,50050439 | hes family bHLH transcription factor 1 |
| ENSG00000105976 | MET | 0,404480025 | 1,323611769 | 0,40433 | 0,6642832 | MET proto-oncogene, receptor tyrosine kinase |
| ENSG00000133678 | TMEM254 | 0,404387345 | 1,323526742 | 0,33076 | 0,6112353 | transmembrane protein 254 |
| ENSG00000155097 | ATP6V1C1 | 0,404358055 | 1,323499872 | 0,21049 | 0,5290538 | ATPase H+ transporting V1 subunit C1 |
| ENSG00000196357 | ZNF565 | 0,404272475 | 1,323421365 | 0,23999 | 0,54936679 | zinc finger protein 565 |
| ENSG00000119318 | RAD23B | 0,404271802 | 1,323420747 | 0,18145 | 0,50490902 | RAD23 homolog B, nucleotide excision repair protein |
| ENSG00000246705 | H2AJ | 0,40425488 | 1,323405225 | 0,23167 | 0,54440987 | H2A,J histone |
| ENSG00000107223 | EDF1 | 0,404065499 | 1,323231514 | 0,17771 | 0,50205238 | endothelial differentiation related factor 1 |
| ENSG00000111371 | SLC38A1 | 0,403916043 | 1,323094441 | 0,24629 | 0,55395309 | solute carrier family 38 member 1 |
| ENSG00000278268 | MOAP1 | 0,403882624 | 1,323063793 | 0,29206 | 0,5868753 | modulator of apoptosis 1 |
| ENSG00000213920 | MDP1 | 0,403872117 | 1,323054157 | 0,25189 | 0,55773568 | magnesium dependent phosphatase 1 |
| ENSG00000285200 | MDP1 | 0,403872117 | 1,323054157 | 0,25189 | 0,55773568 | magnesium dependent phosphatase 1 |
| ENSG00000124160 | NCOA5 | 0,403826499 | 1,323012322 | 0,18557 | 0,50901954 | nuclear receptor coactivator 5 |
| ENSG00000273017 | AP000240,1 | 0,403754937 | 1,322946699 | 0,59878 | 0,79523922 | novel transcript, sense intronic BACH1 |
| ENSG00000113312 | TTC1 | 0,403705604 | 1,322901461 | 0,22406 | 0,53995756 | tetratricopeptide repeat domain 1 |
| ENSG00000154920 | EME1 | 0,403699168 | 1,32289556 | 0,22129 | 0,53805234 | essential meiotic structure-specific endonuclease 1 |

| | | | | | | |
|---|---|---|---|---|---|---|
| ENSG00000187098 | MITF | 0,403629327 | 1,32283152 | 0,37405 | 0,64254935 | melanocyte inducing transcription factor |
| ENSG00000214719 | AC005562,1 | 0,403411283 | 1,322631607 | 0,11208 | 0,42549415 | novel transcript |
| ENSG00000275902 | AC139530,3 | 0,403315819 | 1,32254409 | 0,79636 | 0,90591272 | novel transcript, antisense to HGS |
| ENSG00000237512 | UNC5B-AS1 | 0,403313603 | 1,322542059 | 0,63733 | 0,81763794 | UNC5B antisense RNA 1 |
| ENSG00000135775 | COG2 | 0,403284845 | 1,322515696 | 0,12622 | 0,44888955 | component of oligomeric golgi complex 2 |
| ENSG00000162843 | WDR64 | 0,403217527 | 1,322453987 | 0,62594 | 0,81058216 | WD repeat domain 64 |
| ENSG00000126953 | TIMM8A | 0,40301573 | 1,322269022 | 0,12439 | 0,44539699 | translocase of inner mitochondrial membrane 8A |
| ENSG00000197008 | ZNF138 | 0,402898796 | 1,322161853 | 0,39516 | 0,65806932 | zinc finger protein 138 |
| ENSG00000114956 | DGUOK | 0,402818354 | 1,322088134 | 0,22216 | 0,53859118 | deoxyguanosine kinase |
| ENSG00000255910 | AC024901,1 | 0,402771654 | 1,322045339 | 0,64163 | 0,82033064 | novel transcript |
| ENSG00000146574 | CCZ1B | 0,402758346 | 1,322033144 | 0,33044 | 0,6112353 | CCZ1 homolog B, vacuolar protein trafficking and biogenesis associated |
| ENSG00000006652 | IFRD1 | 0,40273503 | 1,322011778 | 0,2326 | 0,54529383 | interferon related developmental regulator 1 |
| ENSG00000105186 | ANKRD27 | 0,40271298 | 1,321991573 | 0,20004 | 0,5209612 | ankyrin repeat domain 27 |
| ENSG00000132967 | HMGB1P5 | 0,402698822 | 1,321978599 | 0,17555 | 0,49954066 | high mobility group box 1 pseudogene 5 |
| ENSG00000120889 | TNFRSF10B | 0,402652081 | 1,32193577 | 0,21913 | 0,53664318 | TNF receptor superfamily member 10b |
| ENSG00000113575 | PPP2CA | 0,402611358 | 1,321898456 | 0,22422 | 0,54010975 | protein phosphatase 2 catalytic subunit alpha |
| ENSG00000035928 | RFC1 | 0,402231802 | 1,321550726 | 0,19844 | 0,51980353 | replication factor C subunit 1 |
| ENSG00000066739 | ATG2B | 0,402176682 | 1,321500236 | 0,23073 | 0,54374992 | autophagy related 2B |
| ENSG00000144867 | SRPRB | 0,402148888 | 1,321474776 | 0,04727 | 0,29217708 | SRP receptor subunit beta |
| ENSG00000025434 | NR1H3 | 0,402098415 | 1,321428545 | 0,21744 | 0,53536715 | nuclear receptor subfamily 1 group H member 3 |
| ENSG00000091157 | WDR7 | 0,402043334 | 1,321378095 | 0,29768 | 0,59036937 | WD repeat domain 7 |
| ENSG00000198793 | MTOR | 0,402022668 | 1,321359167 | 0,18896 | 0,51241304 | mechanistic target of rapamycin kinase |
| ENSG00000231409 | AC018868,1 | 0,402021944 | 1,321358504 | 0,57587 | 0,78043323 | ribosomal protein L36a (RPL36A) pseudogene |
| ENSG00000109536 | FRG1 | 0,401944806 | 1,321287855 | 0,38463 | 0,65052153 | FSHD region gene 1 |
| ENSG00000283153 | FRG1 | 0,401944806 | 1,321287855 | 0,38463 | 0,65052153 | FSHD region gene 1 |
| ENSG00000283630 | FRG1 | 0,401944806 | 1,321287855 | 0,38463 | 0,65052153 | FSHD region gene 1 |
| ENSG00000271452 | AC005034,5 | 0,401940104 | 1,321283549 | 0,356 | 0,63030618 | novel transcript, sense intronic to MRPL19 |
| ENSG00000148835 | TAF5 | 0,401873285 | 1,321222354 | 0,17598 | 0,49999144 | TATA-box binding protein associated factor 5 |
| ENSG00000165533 | TTC8 | 0,401859516 | 1,321209745 | 0,08587 | 0,38009386 | tetratricopeptide repeat domain 8 |
| ENSG00000225192 | ZNF33BP1 | 0,4016589 | 1,321026036 | 0,51877 | 0,74360666 | zinc finger protein 33B pseudogene 1 |
| ENSG00000137040 | RANBP6 | 0,401583841 | 1,320957308 | 0,28497 | 0,58171017 | RAN binding protein 6 |
| ENSG00000170917 | NUDT6 | 0,401506633 | 1,320886617 | 0,16109 | 0,48841163 | nudix hydrolase 6 |

| | | | | | | |
|---|---|---|---|---|---|---|
| ENSG00000112144 | CILK1 | 0,401506449 | 1,320886448 | 0,24977 | 0,55657567 | ciliogenesis associated kinase 1 |
| ENSG00000280832 | GSEC | 0,401484114 | 1,320866 | 0,20472 | 0,52443222 | G-quadruplex forming sequence containing lncRNA |
| ENSG00000138336 | TET1 | 0,401443881 | 1,320829164 | 0,30443 | 0,59430086 | tet methylcytosine dioxygenase 1 |
| ENSG00000076641 | PAG1 | 0,401279351 | 1,320678541 | 0,24185 | 0,55068342 | phosphoprotein membrane anchor with glycosphingolipid microdomains 1 |
| ENSG00000094916 | CBX5 | 0,401279045 | 1,320678261 | 0,10834 | 0,41899991 | chromobox 5 |
| ENSG00000127720 | METTL25 | 0,401210464 | 1,320615481 | 0,11975 | 0,43830376 | methyltransferase like 25 |
| ENSG00000156831 | NSMCE2 | 0,401174421 | 1,320582489 | 0,26164 | 0,56452671 | NSE2 (MMS21) homolog, SMC5-SMC6 complex SUMO ligase |
| ENSG00000146215 | CRIP3 | 0,401163561 | 1,320572548 | 0,56491 | 0,77334048 | cysteine rich protein 3 |
| ENSG00000263001 | GTF2I | 0,401150725 | 1,320560799 | 0,22387 | 0,53971809 | general transcription factor IIi |
| ENSG00000141219 | C17orf80 | 0,401122702 | 1,320535148 | 0,39682 | 0,65936612 | chromosome 17 open reading frame 80 |
| ENSG00000272323 | AC026801,2 | 0,401096525 | 1,320511188 | 0,47963 | 0,71706545 | novel transcript, antisense to TTC23L |
| ENSG00000178409 | BEND3 | 0,401028911 | 1,320449302 | 0,13359 | 0,45946836 | BEN domain containing 3 |
| ENSG00000153234 | NR4A2 | 0,400976175 | 1,320401035 | 0,25938 | 0,56317998 | nuclear receptor subfamily 4 group A member 2 |
| ENSG00000131747 | TOP2A | 0,400788288 | 1,320229087 | 0,33483 | 0,61454298 | DNA topoisomerase II alpha |
| ENSG00000173812 | EIF1 | 0,400739427 | 1,320184374 | 0,13137 | 0,45639138 | eukaryotic translation initiation factor 1 |
| ENSG00000198727 | MT-CYB | 0,400737416 | 1,320182534 | 0,30825 | 0,59696396 | mitochondrially encoded cytochrome b |
| ENSG00000173065 | FAM222B | 0,400693455 | 1,320142306 | 0,15172 | 0,48005297 | family with sequence similarity 222 member B |
| ENSG00000278200 | LINC01971 | 0,40065353 | 1,320105773 | 0,42821 | 0,68094175 | long intergenic non-protein coding RNA 1971 |
| ENSG00000163788 | SNRK | 0,400578427 | 1,320037053 | 0,22726 | 0,54168591 | SNF related kinase |
| ENSG00000225037 | EIF1AX-AS1 | 0,400423048 | 1,319894892 | 0,45671 | 0,70037869 | EIF1AX antisense RNA 1 |
| ENSG00000118702 | GHRH | 0,40035319 | 1,319830983 | 0,59785 | 0,79484486 | growth hormone releasing hormone |
| ENSG00000260907 | AC015818,2 | 0,400309694 | 1,319791191 | 0,62638 | 0,81088056 | novel transcript |
| ENSG00000168575 | SLC20A2 | 0,400230655 | 1,319718888 | 0,01605 | 0,16606232 | solute carrier family 20 member 2 |
| ENSG00000088876 | ZNF343 | 0,400192335 | 1,319683834 | 0,1624 | 0,48945635 | zinc finger protein 343 |
| ENSG00000096872 | IFT74 | 0,400079035 | 1,319580199 | 0,33466 | 0,61439648 | intraflagellar transport 74 |
| ENSG00000278530 | CHMP1B2P | 0,400040423 | 1,319544882 | 0,41759 | 0,67413959 | charged multivesicular body protein 1B2, pseudogene |
| ENSG00000134779 | TPGS2 | 0,400025321 | 1,31953107 | 0,28038 | 0,57784691 | tubulin polyglutamylase complex subunit 2 |
| ENSG00000086991 | NOX4 | 0,399888664 | 1,319406086 | 0,34347 | 0,62091871 | NADPH oxidase 4 |
| ENSG00000175048 | ZDHHC14 | 0,399887474 | 1,319404997 | 0,17719 | 0,50187296 | zinc finger DHHC-type palmitoyltransferase 14 |
| ENSG00000124562 | SNRPC | 0,399843494 | 1,319364776 | 0,22217 | 0,53859118 | small nuclear ribonucleoprotein polypeptide C |
| ENSG00000233560 | KRT8P39 | 0,399762668 | 1,319290862 | 0,25264 | 0,55796133 | keratin 8 pseudogene 39 |
| ENSG00000118600 | RXYLT1 | 0,399568694 | 1,319113492 | 0,0228 | 0,20153127 | ribitol xylosyltransferase 1 |

| | | | | | | |
|---|---|---|---|---|---|---|
| ENSG00000176386 | CDC26 | 0,399539172 | 1,319086498 | 0,19833 | 0,51980353 | cell division cycle 26 |
| ENSG00000196387 | ZNF140 | 0,399469154 | 1,319022482 | 0,25147 | 0,55731053 | zinc finger protein 140 |
| ENSG00000198218 | QRICH1 | 0,399459889 | 1,319014011 | 0,17495 | 0,49911246 | glutamine rich 1 |
| ENSG00000100325 | ASCC2 | 0,399431632 | 1,318988176 | 0,21541 | 0,5341454 | activating signal cointegrator 1 complex subunit 2 |
| ENSG00000138413 | IDH1 | 0,399357623 | 1,318920515 | 0,32868 | 0,60992716 | isocitrate dehydrogenase (NADP(+)) 1 |
| ENSG00000172466 | ZNF24 | 0,399276219 | 1,318846097 | 0,28817 | 0,58428658 | zinc finger protein 24 |
| ENSG00000170142 | UBE2E1 | 0,399257042 | 1,318828567 | 0,34898 | 0,62518535 | ubiquitin conjugating enzyme E2 E1 |
| ENSG00000275867 | KANSL1 | 0,399211376 | 1,318786822 | 0,18962 | 0,51284737 | KAT8 regulatory NSL complex subunit 1 |
| ENSG00000258787 | SEPHS1P2 | 0,398939739 | 1,318538538 | 0,66987 | 0,83624601 | selenophosphate synthetase 1 pseudogene 2 |
| ENSG00000078061 | ARAF | 0,398892438 | 1,318495309 | 0,17524 | 0,49934416 | A-Raf proto-oncogene, serine/threonine kinase |
| ENSG00000116741 | RGS2 | 0,398880437 | 1,318484341 | 0,26812 | 0,57022339 | regulator of G protein signaling 2 |
| ENSG00000271101 | AC005181,1 | 0,398826455 | 1,318435007 | 0,56942 | 0,77665073 | SERPINE1 mRNA binding protein 1 (SERBP1) pseudogene |
| ENSG00000095059 | DHPS | 0,398824815 | 1,318433509 | 0,27102 | 0,57248407 | deoxyhypusine synthase |
| ENSG00000111605 | CPSF6 | 0,398730776 | 1,318347572 | 0,2491 | 0,55619875 | cleavage and polyadenylation specific factor 6 |
| ENSG00000154305 | MIA3 | 0,398727654 | 1,318344719 | 0,24651 | 0,5541223 | MIA SH3 domain ER export factor 3 |
| ENSG00000169446 | MMGT1 | 0,398706954 | 1,318325804 | 0,32201 | 0,60577115 | membrane magnesium transporter 1 |
| ENSG00000123737 | EXOSC9 | 0,398675141 | 1,318296734 | 0,19561 | 0,51643293 | exosome component 9 |
| ENSG00000111684 | LPCAT3 | 0,398532523 | 1,31816642 | 0,0697 | 0,34826564 | lysophosphatidylcholine acyltransferase 3 |
| ENSG00000136643 | RPS6KC1 | 0,398526868 | 1,318161252 | 0,24172 | 0,55047794 | ribosomal protein S6 kinase C1 |
| ENSG00000229519 | AL591866,1 | 0,398523153 | 1,318157858 | 0,52622 | 0,7497214 | ribosomal protein L34 (RPL34) pseudogene |
| ENSG00000107077 | KDM4C | 0,398504382 | 1,318140707 | 0,16834 | 0,4945419 | lysine demethylase 4C |
| ENSG00000204954 | C12orf73 | 0,398395717 | 1,318041428 | 0,19156 | 0,51435195 | chromosome 12 open reading frame 73 |
| ENSG00000253868 | FER1L6-AS2 | 0,398378375 | 1,318025585 | 0,63013 | 0,813276 | FER1L6 antisense RNA 2 |
| ENSG00000004487 | KDM1A | 0,398287943 | 1,31794297 | 0,12535 | 0,44716492 | lysine demethylase 1A |
| ENSG00000257835 | AC073591,1 | 0,398231858 | 1,317891735 | 0,50683 | 0,7352001 | novel transcript |
| ENSG00000267302 | RNFT1-DT | 0,398183722 | 1,317847764 | 0,20523 | 0,52486901 | RNFT1 divergent transcript |
| ENSG00000143515 | ATP8B2 | 0,398158711 | 1,317824918 | 0,02899 | 0,22977124 | ATPase phospholipid transporting 8B2 |
| ENSG00000166902 | MRPL16 | 0,398080922 | 1,317753863 | 0,17177 | 0,49710979 | mitochondrial ribosomal protein L16 |
| ENSG00000076258 | FMO4 | 0,397939725 | 1,317624901 | 0,59521 | 0,79324077 | flavin containing dimethylaniline monoxygenase 4 |
| ENSG00000151062 | CACNA2D4 | 0,397917777 | 1,317604855 | 0,54606 | 0,76166626 | calcium voltage-gated channel auxiliary subunit alpha2delta 4 |
| ENSG00000284953 | CACNA2D4 | 0,397917777 | 1,317604855 | 0,54606 | 0,76166626 | calcium voltage-gated channel auxiliary subunit alpha2delta 4 |

| | | | | | | |
|---|---|---|---|---|---|---|
| ENSG00000184076 | UQCR10 | 0,397915634 | 1,317602899 | 0,33417 | 0,61398711 | ubiquinol-cytochrome c reductase, complex III subunit X |
| ENSG00000089902 | RCOR1 | 0,397796545 | 1,31749414 | 0,17318 | 0,49786688 | REST corepressor 1 |
| ENSG00000171217 | CLDN20 | 0,3977552 | 1,317456384 | 0,44666 | 0,69321646 | claudin 20 |
| ENSG00000251550 | AC091133,5 | 0,397657821 | 1,317367461 | 0,28368 | 0,58072192 | COX assembly mitochondrial protein homolog (S, cerevisiae) (CMC1) pseudogene |
| ENSG00000132182 | NUP210 | 0,397591793 | 1,31730717 | 0,2167 | 0,53494609 | nucleoporin 210 |
| ENSG00000068912 | ERLEC1 | 0,397562999 | 1,317280879 | 0,20108 | 0,52181085 | endoplasmic reticulum lectin 1 |
| ENSG00000250546 | AC079160,1 | 0,397489075 | 1,317213383 | 0,60424 | 0,79780609 | novel transcript |
| ENSG00000286429 | BX004807,1 | 0,397393754 | 1,317126356 | 0,59274 | 0,79133235 | novel transcript, antisense to ITGB3BP |
| ENSG00000226856 | THORLNC | 0,397378032 | 1,317112002 | 0,5791 | 0,78270914 | testis associated oncogenic lncRNA |
| ENSG00000220694 | AL096711,1 | 0,39732626 | 1,317064738 | 0,69895 | 0,8544493 | ribosomal protein L17 (RPL17) pseudogene |
| ENSG00000275160 | AL354718,1 | 0,397324043 | 1,317062713 | 0,56897 | 0,77629764 | ribosomal protein L7a (RPL7A) pseudogene |
| ENSG00000113643 | RARS1 | 0,397320574 | 1,317059547 | 0,2292 | 0,54273374 | arginyl-tRNA synthetase 1 |
| ENSG00000161547 | SRSF2 | 0,397284521 | 1,317026634 | 0,11963 | 0,43810637 | serine and arginine rich splicing factor 2 |
| ENSG00000143106 | PSMA5 | 0,39728313 | 1,317025364 | 0,09392 | 0,39593973 | proteasome 20S subunit alpha 5 |
| ENSG00000131495 | NDUFA2 | 0,39725508 | 1,316999758 | 0,28903 | 0,58469988 | NADH:ubiquinone oxidoreductase subunit A2 |
| ENSG00000145700 | ANKRD31 | 0,397190928 | 1,316941196 | 0,35171 | 0,62692823 | ankyrin repeat domain 31 |
| ENSG00000170448 | NFXL1 | 0,397127259 | 1,316883078 | 0,13918 | 0,46648276 | nuclear transcription factor, X-box binding like 1 |
| ENSG00000254721 | AP000879,1 | 0,397057549 | 1,316819449 | 0,84664 | 0,930363 | novel transcript |
| ENSG00000278765 | AC004477,2 | 0,397037298 | 1,316800965 | 0,48091 | 0,71795527 | novel transcript |
| ENSG00000219355 | RPL31P52 | 0,397011749 | 1,316777645 | 0,61774 | 0,80609311 | ribosomal protein L31 pseudogene 52 |
| ENSG00000102387 | TAF7L | 0,39693734 | 1,316709732 | 0,44227 | 0,69006952 | TATA-box binding protein associated factor 7 like |
| ENSG00000136238 | RAC1 | 0,396753133 | 1,316541623 | 0,25812 | 0,56220487 | Rac family small GTPase 1 |
| ENSG00000164284 | GRPEL2 | 0,396740683 | 1,316530261 | 0,17814 | 0,50236177 | GrpE like 2, mitochondrial |
| ENSG00000141027 | NCOR1 | 0,396707612 | 1,316500082 | 0,23088 | 0,54390394 | nuclear receptor corepressor 1 |
| ENSG00000260947 | AL356489,2 | 0,396627061 | 1,31642658 | 0,35531 | 0,62970025 | novel transcript |
| ENSG00000224324 | THAP5P1 | 0,396550158 | 1,316356409 | 0,60167 | 0,79674064 | THAP domain containing 5 pseudogene 1 |
| ENSG00000165359 | INTS6L | 0,396538495 | 1,316345768 | 0,29582 | 0,58942061 | integrator complex subunit 6 like |
| ENSG00000198951 | NAGA | 0,396493876 | 1,316305057 | 0,03699 | 0,26078413 | alpha-N-acetylgalactosaminidase |
| ENSG00000055044 | NOP58 | 0,3963389 | 1,316163665 | 0,17326 | 0,49800753 | NOP58 ribonucleoprotein |
| ENSG00000156928 | MALSU1 | 0,396335207 | 1,316160297 | 0,28766 | 0,58395529 | mitochondrial assembly of ribosomal large subunit 1 |
| ENSG00000119227 | PIGZ | 0,396333583 | 1,316158815 | 0,36407 | 0,63490528 | phosphatidylinositol glycan anchor biosynthesis class Z |
| ENSG00000169692 | AGPAT2 | 0,396325217 | 1,316151182 | 0,17218 | 0,49710979 | 1-acylglycerol-3-phosphate O-acyltransferase 2 |

| | | | | | | |
|---|---|---|---|---|---|---|
| ENSG00000280906 | AC225099,1 | 0,396309002 | 1,31613639 | 0,33076 | 0,6112353 | novel transcript |
| ENSG00000102100 | SLC35A2 | 0,396303432 | 1,316131309 | 0,06319 | 0,332907 | solute carrier family 35 member A2 |
| ENSG00000221983 | UBA52 | 0,396152724 | 1,315993829 | 0,16049 | 0,48780354 | ubiquitin A-52 residue ribosomal protein fusion product 1 |
| ENSG00000123552 | USP45 | 0,395972787 | 1,315829705 | 0,32795 | 0,60970725 | ubiquitin specific peptidase 45 |
| ENSG00000023839 | ABCC2 | 0,395955277 | 1,315813735 | 0,28144 | 0,57889018 | ATP binding cassette subfamily C member 2 |
| ENSG00000116001 | TIA1 | 0,395934814 | 1,315795071 | 0,38233 | 0,64928247 | TIA1 cytotoxic granule associated RNA binding protein |
| ENSG00000116127 | ALMS1 | 0,395923458 | 1,315784714 | 0,35926 | 0,6321355 | ALMS1 centrosome and basal body associated protein |
| ENSG00000272338 | AC067838,1 | 0,395871245 | 1,315737095 | 0,58179 | 0,78454692 | novel transcript |
| ENSG00000132676 | DAP3 | 0,39578656 | 1,315659865 | 0,08469 | 0,377881 | death associated protein 3 |
| ENSG00000276840 | PMS2P10 | 0,395681527 | 1,315564084 | 0,73468 | 0,87461741 | PMS1 homolog 2, mismatch repair system component pseudogene 10 |
| ENSG00000250615 | AC008581,1 | 0,39556904 | 1,315461513 | 0,73114 | 0,87229858 | novel transcript |
| ENSG00000139618 | BRCA2 | 0,39551536 | 1,315412569 | 0,36327 | 0,63408839 | BRCA2 DNA repair associated |
| ENSG00000147383 | NSDHL | 0,395421076 | 1,315326606 | 0,1881 | 0,51128963 | NAD(P) dependent steroid dehydrogenase-like |
| ENSG00000164109 | MAD2L1 | 0,395138368 | 1,315068882 | 0,24315 | 0,55199251 | mitotic arrest deficient 2 like 1 |
| ENSG00000282958 | HNRNPR | 0,395077347 | 1,31501326 | 0,14788 | 0,4751727 | heterogeneous nuclear ribonucleoprotein R |
| ENSG00000123505 | AMD1 | 0,394941279 | 1,31488924 | 0,04951 | 0,29942963 | adenosylmethionine decarboxylase 1 |
| ENSG00000229915 | AC016999,1 | 0,394921632 | 1,314871334 | 0,71207 | 0,8615121 | novel transcript |
| ENSG00000179363 | TMEM31 | 0,394906388 | 1,314857441 | 0,51705 | 0,74231073 | transmembrane protein 31 |
| ENSG00000082438 | COBLL1 | 0,394898242 | 1,314850016 | 0,3138 | 0,600508 | cordon-bleu WH2 repeat protein like 1 |
| ENSG00000118004 | COLEC11 | 0,394875979 | 1,314829726 | 0,64529 | 0,82203909 | collectin subfamily member 11 |
| ENSG00000175155 | YPEL2 | 0,394818391 | 1,314777243 | 0,1119 | 0,42531464 | yippee like 2 |
| ENSG00000158427 | TMSB15B | 0,394722178 | 1,314689564 | 0,09381 | 0,39569427 | thymosin beta 15B |
| ENSG00000175215 | CTDSP2 | 0,394467874 | 1,314457844 | 0,09477 | 0,39724142 | CTD small phosphatase 2 |
| ENSG00000178538 | CA8 | 0,394463461 | 1,314453823 | 0,21929 | 0,53676938 | carbonic anhydrase 8 |
| ENSG00000138399 | FASTKD1 | 0,39445933 | 1,31445006 | 0,18972 | 0,512949 | FAST kinase domains 1 |
| ENSG00000243302 | AC018638,4 | 0,394336087 | 1,314337776 | 0,43949 | 0,68837688 | novel pseudogene |
| ENSG00000123106 | CCDC91 | 0,394325778 | 1,314328385 | 0,19505 | 0,51588752 | coiled-coil domain containing 91 |
| ENSG00000122591 | FAM126A | 0,394303026 | 1,314307658 | 0,17417 | 0,49867554 | family with sequence similarity 126 member A |
| ENSG00000266910 | AC008507,1 | 0,394288392 | 1,314294326 | 0,54595 | 0,7616329 | TAF9 RNA polymerase II, TATA box binding protein (TBP)-associated factor, 32kDa (TAF9) pseudogene |
| ENSG00000100591 | AHSA1 | 0,394239025 | 1,314249354 | 0,1414 | 0,46803321 | activator of HSP90 ATPase activity 1 |
| ENSG00000229427 | ANKRD26P4 | 0,394171463 | 1,314187808 | 0,44588 | 0,69284796 | ankyrin repeat domain 26 pseudogene 4 |

| | | | | | | |
|---|---|---|---|---|---|---|
| ENSG00000164338 | UTP15 | 0,394131224 | 1,314151153 | 0,17656 | 0,50077376 | UTP15 small subunit processome component |
| ENSG00000263990 | AC004253,1 | 0,394006225 | 1,314037297 | 0,53292 | 0,75333268 | novel transcript |
| ENSG00000132434 | LANCL2 | 0,39398746 | 1,314020206 | 0,19273 | 0,51490739 | LanC like 2 |
| ENSG00000132664 | POLR3F | 0,393884579 | 1,313926504 | 0,27515 | 0,57491355 | RNA polymerase III subunit F |
| ENSG00000277149 | TYW1B | 0,39385837 | 1,313902634 | 0,22607 | 0,54121239 | tRNA-yW synthesizing protein 1 homolog B |
| ENSG00000128915 | ICE2 | 0,393824915 | 1,313872166 | 0,18056 | 0,50407792 | interactor of little elongation complex ELL subunit 2 |
| ENSG00000176105 | YES1 | 0,393708849 | 1,313766468 | 0,12894 | 0,45224009 | YES proto-oncogene 1, Src family tyrosine kinase |
| ENSG00000086619 | ERO1B | 0,393673089 | 1,313733904 | 0,06158 | 0,32867767 | endoplasmic reticulum oxidoreductase 1 beta |
| ENSG00000086758 | HUWE1 | 0,393576432 | 1,313645891 | 0,14755 | 0,47504057 | HECT, UBA and WWE domain containing E3 ubiquitin protein ligase 1 |
| ENSG00000116120 | FARSB | 0,393549325 | 1,313621208 | 0,13406 | 0,46018397 | phenylalanyl-tRNA synthetase subunit beta |
| ENSG00000237438 | CECR7 | 0,393519537 | 1,313594086 | 0,21572 | 0,53440695 | cat eye syndrome chromosome region, candidate 7 |
| ENSG00000112851 | ERBIN | 0,393446605 | 1,313527682 | 0,39167 | 0,65533345 | erbb2 interacting protein |
| ENSG00000276766 | NDUFA3 | 0,393432909 | 1,313515213 | 0,30603 | 0,59512079 | NADH:ubiquinone oxidoreductase subunit A3 |
| ENSG00000278365 | NDUFA3 | 0,393432909 | 1,313515213 | 0,30603 | 0,59512079 | NADH:ubiquinone oxidoreductase subunit A3 |
| ENSG00000277722 | NDUFA3 | 0,393432909 | 1,313515213 | 0,30603 | 0,59512079 | NADH:ubiquinone oxidoreductase subunit A3 |
| ENSG00000276220 | NDUFA3 | 0,393432909 | 1,313515213 | 0,30603 | 0,59512079 | NADH:ubiquinone oxidoreductase subunit A3 |
| ENSG00000273642 | NDUFA3 | 0,393432909 | 1,313515213 | 0,30603 | 0,59512079 | NADH:ubiquinone oxidoreductase subunit A3 |
| ENSG00000274359 | NDUFA3 | 0,393432909 | 1,313515213 | 0,30603 | 0,59512079 | NADH:ubiquinone oxidoreductase subunit A3 |
| ENSG00000275724 | NDUFA3 | 0,393432909 | 1,313515213 | 0,30603 | 0,59512079 | NADH:ubiquinone oxidoreductase subunit A3 |
| ENSG00000275605 | NDUFA3 | 0,393432909 | 1,313515213 | 0,30603 | 0,59512079 | NADH:ubiquinone oxidoreductase subunit A3 |
| ENSG00000110693 | SOX6 | 0,393326012 | 1,313417891 | 0,15857 | 0,48579688 | SRY-box transcription factor 6 |
| ENSG00000277369 | AC010654,1 | 0,393189631 | 1,313293737 | 0,2665 | 0,56917012 | novel transcript |
| ENSG00000160208 | RRP1B | 0,393186901 | 1,313291251 | 0,1015 | 0,40833508 | ribosomal RNA processing 1B |
| ENSG00000071626 | DAZAP1 | 0,393181223 | 1,313286083 | 0,02862 | 0,22866679 | DAZ associated protein 1 |
| ENSG00000123595 | RAB9A | 0,393178739 | 1,313283821 | 0,38909 | 0,65363435 | RAB9A, member RAS oncogene family |
| ENSG00000113296 | THBS4 | 0,393100928 | 1,313212992 | 0,13015 | 0,45454796 | thrombospondin 4 |
| ENSG00000269600 | AC016629,2 | 0,393097573 | 1,313209938 | 0,46943 | 0,70891774 | novel transcript, antisense to RPL23AP79 |
| ENSG00000198346 | ZNF813 | 0,393083931 | 1,313197521 | 0,1362 | 0,463307 | zinc finger protein 813 |
| ENSG00000163605 | PPP4R2 | 0,393025023 | 1,313143901 | 0,13095 | 0,45558936 | protein phosphatase 4 regulatory subunit 2 |
| ENSG00000005108 | THSD7A | 0,392976997 | 1,313100189 | 0,45679 | 0,7004577 | thrombospondin type 1 domain containing 7A |
| ENSG00000138078 | PREPL | 0,392935396 | 1,313062325 | 0,31948 | 0,60429866 | prolyl endopeptidase like |
| ENSG00000157450 | RNF111 | 0,392935109 | 1,313062064 | 0,27681 | 0,57549599 | ring finger protein 111 |

| | | | | | | |
|---|---|---|---|---|---|---|
| ENSG00000278608 | MAMDC2 | 0,392932838 | 1,313059997 | 0,7998 | 0,90761856 | MAM domain containing 2 |
| ENSG00000185619 | PCGF3 | 0,392827992 | 1,312964575 | 0,20296 | 0,52282418 | polycomb group ring finger 3 |
| ENSG00000128524 | ATP6V1F | 0,392802076 | 1,31294099 | 0,18956 | 0,51280953 | ATPase H+ transporting V1 subunit F |
| ENSG00000087116 | ADAMTS2 | 0,392773687 | 1,312915154 | 0,25143 | 0,55731053 | ADAM metallopeptidase with thrombospondin type 1 motif 2 |
| ENSG00000182873 | PRKCZ-AS1 | 0,392759789 | 1,312902507 | 0,0896 | 0,38705587 | PRKCZ antisense RNA 1 |
| ENSG00000198740 | ZNF652 | 0,392643115 | 1,312796334 | 0,14727 | 0,47476862 | zinc finger protein 652 |
| ENSG00000273449 | AC093788,1 | 0,392633085 | 1,312787206 | 0,38308 | 0,64987648 | novel transcript |
| ENSG00000116221 | MRPL37 | 0,392453462 | 1,312623769 | 0,18153 | 0,50490902 | mitochondrial ribosomal protein L37 |
| ENSG00000267767 | LINC01801 | 0,392449727 | 1,31262037 | 0,48296 | 0,71925139 | long intergenic non-protein coding RNA 1801 |
| ENSG00000040275 | SPDL1 | 0,392396594 | 1,312572029 | 0,20354 | 0,52337667 | spindle apparatus coiled-coil protein 1 |
| ENSG00000110075 | PPP6R3 | 0,3922902 | 1,312475234 | 0,27776 | 0,57618023 | protein phosphatase 6 regulatory subunit 3 |
| ENSG00000183273 | CCDC60 | 0,392207978 | 1,312400436 | 0,58198 | 0,78454692 | coiled-coil domain containing 60 |
| ENSG00000077721 | UBE2A | 0,39213385 | 1,312333005 | 0,21349 | 0,53275614 | ubiquitin conjugating enzyme E2 A |
| ENSG00000064651 | SLC12A2 | 0,392075094 | 1,312279558 | 0,20676 | 0,52628775 | solute carrier family 12 member 2 |
| ENSG00000228828 | TLK2P2 | 0,392044336 | 1,312251581 | 0,39774 | 0,66032779 | tousled like kinase 2 pseudogene 2 |
| ENSG00000152763 | DNAI4 | 0,391933586 | 1,312150849 | 0,24826 | 0,55542983 | dynein axonemal intermediate chain 4 |
| ENSG00000163032 | VSNL1 | 0,391888694 | 1,312110019 | 0,21677 | 0,53497633 | visinin like 1 |
| ENSG00000189410 | SH2D5 | 0,391841235 | 1,312066857 | 0,3658 | 0,63598241 | SH2 domain containing 5 |
| ENSG00000197045 | GMFB | 0,391801614 | 1,312030824 | 0,30546 | 0,59497466 | glia maturation factor beta |
| ENSG00000127184 | COX7C | 0,391756371 | 1,311989679 | 0,45682 | 0,7004577 | cytochrome c oxidase subunit 7C |
| ENSG00000196510 | ANAPC7 | 0,391693377 | 1,311932394 | 0,28496 | 0,58171017 | anaphase promoting complex subunit 7 |
| ENSG00000256625 | AC092747,2 | 0,391691269 | 1,311930477 | 0,52451 | 0,74828894 | liprin-beta-1-like (PPFIBP1) pseudogene |
| ENSG00000069509 | FUNDC1 | 0,391658829 | 1,311900978 | 0,28885 | 0,58464073 | FUN14 domain containing 1 |
| ENSG00000141378 | PTRH2 | 0,391479923 | 1,311738301 | 0,2315 | 0,54440987 | peptidyl-tRNA hydrolase 2 |
| ENSG00000280650 | KCNIP4-IT1 | 0,391381613 | 1,311648918 | 0,35641 | 0,63043562 | KCNIP4 intronic transcript 1 |
| ENSG00000198939 | ZFP2 | 0,391359834 | 1,311629117 | 0,20866 | 0,52786574 | ZFP2 zinc finger protein |
| ENSG00000141449 | GREB1L | 0,391341715 | 1,311612644 | 0,38311 | 0,64987805 | GREB1 like retinoic acid receptor coactivator |
| ENSG00000276187 | ERMARD | 0,391264482 | 1,311542431 | 0,22624 | 0,54121239 | ER membrane associated RNA degradation |
| ENSG00000078668 | VDAC3 | 0,391254521 | 1,311533375 | 0,30289 | 0,59331081 | voltage dependent anion channel 3 |
| ENSG00000102897 | LYRM1 | 0,391253277 | 1,311532244 | 0,39119 | 0,65497307 | LYR motif containing 1 |
| ENSG00000116353 | MECR | 0,391186959 | 1,311471957 | 0,10892 | 0,42023628 | mitochondrial trans-2-enoyl-CoA reductase |
| ENSG00000137154 | RPS6 | 0,391169666 | 1,311456237 | 0,0854 | 0,37882499 | ribosomal protein S6 |

| | | | | | | |
|---|---|---|---|---|---|---|
| ENSG00000250569 | NTAN1P2 | 0,391085435 | 1,31137967 | 0,49103 | 0,72492348 | N-terminal asparagine amidase pseudogene 2 |
| ENSG00000198242 | RPL23A | 0,391051847 | 1,31134914 | 0,11929 | 0,43713778 | ribosomal protein L23a |
| ENSG00000128262 | POM121L9P | 0,390997449 | 1,311299695 | 0,51284 | 0,73941617 | POM121 transmembrane nucleoporin like 9, pseudogene |
| ENSG00000164989 | CCDC171 | 0,390979204 | 1,311283112 | 0,31482 | 0,601382 | coiled-coil domain containing 171 |
| ENSG00000243775 | OSTCP1 | 0,390865491 | 1,311179761 | 0,43227 | 0,68411566 | oligosaccharyltransferase complex subunit pseudogene 1 |
| ENSG00000101343 | CRNKL1 | 0,390834061 | 1,311151197 | 0,24431 | 0,55253387 | crooked neck pre-mRNA splicing factor 1 |
| ENSG00000272540 | AL662797,1 | 0,390728717 | 1,311055461 | 0,40991 | 0,66816149 | novel transcript, antisense to TUBB |
| ENSG00000145782 | ATG12 | 0,390716095 | 1,311043991 | 0,20297 | 0,52282418 | autophagy related 12 |
| ENSG00000138286 | FAM149B1 | 0,390649164 | 1,310983169 | 0,22241 | 0,53859118 | family with sequence similarity 149 member B1 |
| ENSG00000104133 | SPG11 | 0,39062222 | 1,310958686 | 0,24248 | 0,55137204 | SPG11 vesicle trafficking associated, spatacsin |
| ENSG00000188211 | NCR3LG1 | 0,39062151 | 1,31095804 | 0,45905 | 0,70162976 | natural killer cell cytotoxicity receptor 3 ligand 1 |
| ENSG00000115556 | PLCD4 | 0,390512587 | 1,310859067 | 0,22542 | 0,5407493 | phospholipase C delta 4 |
| ENSG00000173950 | XXYLT1 | 0,390235141 | 1,310606999 | 0,04661 | 0,29026543 | xyloside xylosyltransferase 1 |
| ENSG00000093144 | ECHDC1 | 0,39016524 | 1,310543499 | 0,18342 | 0,50701115 | ethylmalonyl-CoA decarboxylase 1 |
| ENSG00000116062 | MSH6 | 0,390049465 | 1,310438334 | 0,0991 | 0,40529643 | mutS homolog 6 |
| ENSG00000135632 | SMYD5 | 0,390020159 | 1,310411715 | 0,04823 | 0,29503004 | SMYD family member 5 |
| ENSG00000058866 | DGKG | 0,389974708 | 1,310370432 | 0,45095 | 0,69604051 | diacylglycerol kinase gamma |
| ENSG00000145868 | FBXO38 | 0,389933996 | 1,310333454 | 0,18694 | 0,51044154 | F-box protein 38 |
| ENSG00000131097 | HIGD1B | 0,389906322 | 1,310308319 | 0,70039 | 0,85531965 | HIG1 hypoxia inducible domain family member 1B |
| ENSG00000143970 | ASXL2 | 0,389877091 | 1,310281771 | 0,24108 | 0,55015592 | ASXL transcriptional regulator 2 |
| ENSG00000143971 | ETAA1 | 0,389524855 | 1,309961903 | 0,30885 | 0,59767335 | ETAA1 activator of ATR kinase |
| ENSG00000181924 | COA4 | 0,389523507 | 1,309960679 | 0,22115 | 0,53805234 | cytochrome c oxidase assembly factor 4 homolog |
| ENSG00000196262 | PPIA | 0,389522716 | 1,309959961 | 0,21642 | 0,53484961 | peptidylprolyl isomerase A |
| ENSG00000164051 | CCDC51 | 0,389458833 | 1,309901957 | 0,10787 | 0,41847368 | coiled-coil domain containing 51 |
| ENSG00000276434 | AL136221,1 | 0,389434355 | 1,309879732 | 0,63034 | 0,8133533 | novel transcript, sense intronic to GRTP1 |
| ENSG00000109519 | GRPEL1 | 0,389425384 | 1,309871587 | 0,25223 | 0,55796133 | GrpE like 1, mitochondrial |
| ENSG00000104131 | EIF3J | 0,389321516 | 1,309777285 | 0,12164 | 0,44093327 | eukaryotic translation initiation factor 3 subunit J |
| ENSG00000138660 | AP1AR | 0,389289091 | 1,309747847 | 0,22131 | 0,53805234 | adaptor related protein complex 1 associated regulatory protein |
| ENSG00000223849 | AL354893,1 | 0,389246701 | 1,309709364 | 0,64015 | 0,81935089 | ribosomal protein S26 (RPS26) pseudogene |
| ENSG00000144821 | MYH15 | 0,38913368 | 1,309606766 | 0,26082 | 0,56400166 | myosin heavy chain 15 |
| ENSG00000067829 | IDH3G | 0,389091195 | 1,3095682 | 0,3316 | 0,61165554 | isocitrate dehydrogenase (NAD(+)) 3 non-catalytic subunit gamma |

| | | | | | | |
|---|---|---|---|---|---|---|
| ENSG00000242261 | AC018635,1 | 0,389048836 | 1,309529751 | 0,40938 | 0,66795288 | aurora kinase A (AURKA) pseudogene |
| ENSG00000103227 | LMF1 | 0,388909642 | 1,30940341 | 0,15251 | 0,48061452 | lipase maturation factor 1 |
| ENSG00000092621 | PHGDH | 0,388890514 | 1,30938605 | 0,08625 | 0,38072744 | phosphoglycerate dehydrogenase |
| ENSG00000071994 | PDCD2 | 0,388884438 | 1,309380535 | 0,23072 | 0,54374992 | programmed cell death 2 |
| ENSG00000087095 | NLK | 0,388873816 | 1,309370895 | 0,38701 | 0,65213944 | nemo like kinase |
| ENSG00000215223 | CYP51A1P3 | 0,388847836 | 1,309347316 | 0,48174 | 0,71853923 | cytochrome P450 family 51 subfamily A member 1 pseudogene 3 |
| ENSG00000118308 | IRAG2 | 0,388830449 | 1,309331537 | 0,61358 | 0,8035281 | inositol 1,4,5-triphosphate receptor associated 2 |
| ENSG00000100764 | PSMC1 | 0,388779741 | 1,309285517 | 0,32269 | 0,60584543 | proteasome 26S subunit, ATPase 1 |
| ENSG00000255100 | AP003119,2 | 0,388734382 | 1,309244353 | 0,46344 | 0,70426631 | novel transcript |
| ENSG00000270977 | AC015849,5 | 0,388728394 | 1,309238919 | 0,4155 | 0,6723843 | leucine rich repeat containing 37 member A3 (LRRC37A3) pseudogene |
| ENSG00000261409 | AL035425,3 | 0,3885396 | 1,309067601 | 0,47012 | 0,70949774 | novel transcript, |
| ENSG00000163349 | HIPK1 | 0,388494272 | 1,309026471 | 0,23016 | 0,54335232 | homeodomain interacting protein kinase 1 |
| ENSG00000230623 | AC104461,1 | 0,388442285 | 1,308979302 | 0,55226 | 0,76568017 | novel transcript, antisense to ZNF281 |
| ENSG00000006451 | RALA | 0,388152238 | 1,308716165 | 0,23521 | 0,54667863 | RAS like proto-oncogene A |
| ENSG00000256771 | ZNF253 | 0,388138717 | 1,308703899 | 0,31455 | 0,60122508 | zinc finger protein 253 |
| ENSG00000123119 | NECAB1 | 0,388138595 | 1,308703789 | 0,50192 | 0,73152688 | N-terminal EF-hand calcium binding protein 1 |
| ENSG00000166439 | RNF169 | 0,388038029 | 1,308612565 | 0,16565 | 0,49182669 | ring finger protein 169 |
| ENSG00000166167 | BTRC | 0,387942001 | 1,308525465 | 0,27701 | 0,575636 | beta-transducin repeat containing E3 ubiquitin protein ligase |
| ENSG00000100462 | PRMT5 | 0,387875618 | 1,308465257 | 0,09237 | 0,39278463 | protein arginine methyltransferase 5 |
| ENSG00000101843 | PSMD10 | 0,387841562 | 1,30843437 | 0,29887 | 0,59104323 | proteasome 26S subunit, non-ATPase 10 |
| ENSG00000012963 | UBR7 | 0,38778291 | 1,308381178 | 0,35422 | 0,62892485 | ubiquitin protein ligase E3 component n-recognin 7 |
| ENSG00000114784 | EIF1B | 0,38777773 | 1,30837648 | 0,38236 | 0,64928247 | eukaryotic translation initiation factor 1B |
| ENSG00000136527 | TRA2B | 0,387725049 | 1,308328705 | 0,21602 | 0,53473714 | transformer 2 beta homolog |
| ENSG00000137807 | KIF23 | 0,387590415 | 1,308206615 | 0,34728 | 0,62403275 | kinesin family member 23 |
| ENSG00000168826 | ZBTB49 | 0,387555099 | 1,308174592 | 0,19451 | 0,51562158 | zinc finger and BTB domain containing 49 |
| ENSG00000079332 | SAR1A | 0,387449194 | 1,308078565 | 0,11476 | 0,43021597 | secretion associated Ras related GTPase 1A |
| ENSG00000164610 | RP9 | 0,387357126 | 1,30799509 | 0,27456 | 0,5745501 | RP9 pre-mRNA splicing factor |
| ENSG00000255389 | Z97989,1 | 0,387305095 | 1,307947919 | 0,34128 | 0,61939402 | chromosome 6 open reading frame 3 |
| ENSG00000109861 | CTSC | 0,387270886 | 1,307916905 | 0,19887 | 0,52005179 | cathepsin C |
| ENSG00000253420 | AC103853,1 | 0,387239554 | 1,307888501 | 0,53922 | 0,75759854 | novel transcript |

| | | | | | | |
|---|---|---|---|---|---|---|
| ENSG00000129696 | TTI2 | 0,387164693 | 1,307820636 | 0,1011 | 0,40780603 | TELO2 interacting protein 2 |
| ENSG00000174695 | TMEM167A | 0,387151332 | 1,307808525 | 0,10695 | 0,41675015 | transmembrane protein 167A |
| ENSG00000149658 | YTHDF1 | 0,38713384 | 1,307792668 | 0,1879 | 0,51107563 | YTH N6-methyladenosine RNA binding protein 1 |
| ENSG00000135372 | NAT10 | 0,387109466 | 1,307770573 | 0,10156 | 0,40840817 | N-acetyltransferase 10 |
| ENSG00000165730 | STOX1 | 0,387009555 | 1,307680009 | 0,41637 | 0,67315741 | storkhead box 1 |
| ENSG00000254132 | MTND6P3 | 0,386993457 | 1,307665418 | 0,21581 | 0,53440695 | MT-ND6 pseudogene 3 |
| ENSG00000213593 | TMX2 | 0,386975971 | 1,307649569 | 0,17596 | 0,49997447 | thioredoxin related transmembrane protein 2 |
| ENSG00000287215 | AC231759,2 | 0,386907988 | 1,307587951 | 0,42015 | 0,67563085 | novel transcript |
| ENSG00000114948 | ADAM23 | 0,3868115 | 1,307500502 | 0,25873 | 0,56260205 | ADAM metallopeptidase domain 23 |
| ENSG00000205861 | PCOTH | 0,38679762 | 1,307487922 | 0,48767 | 0,72265932 | Pro-X-Gly collagen triple helix like repeat containing |
| ENSG00000168615 | ADAM9 | 0,386740826 | 1,307436452 | 0,04272 | 0,27853965 | ADAM metallopeptidase domain 9 |
| ENSG00000143507 | DUSP10 | 0,386692493 | 1,307392651 | 0,30639 | 0,5953441 | dual specificity phosphatase 10 |
| ENSG00000143033 | MTF2 | 0,38662279 | 1,307329486 | 0,2139 | 0,53303219 | metal response element binding transcription factor 2 |
| ENSG00000102900 | NUP93 | 0,386590152 | 1,307299911 | 0,16344 | 0,48977932 | nucleoporin 93 |
| ENSG00000188242 | PP7080 | 0,386529638 | 1,307245078 | 0,17633 | 0,50050439 | uncharacterized LOC25845 |
| ENSG00000124177 | CHD6 | 0,386384637 | 1,307113697 | 0,23972 | 0,5491566 | chromodomain helicase DNA binding protein 6 |
| ENSG00000135404 | CD63 | 0,386329135 | 1,307063412 | 0,11819 | 0,43595569 | CD63 molecule |
| ENSG00000174106 | LEMD3 | 0,386327336 | 1,307061782 | 0,17108 | 0,49641416 | LEM domain containing 3 |
| ENSG00000213096 | ZNF254 | 0,386304311 | 1,307040922 | 0,407 | 0,66640966 | zinc finger protein 254 |
| ENSG00000196290 | NIF3L1 | 0,386208958 | 1,306954538 | 0,21757 | 0,53547632 | NGG1 interacting factor 3 like 1 |
| ENSG00000180917 | CMTR2 | 0,386201386 | 1,306947678 | 0,33598 | 0,61593858 | cap methyltransferase 2 |
| ENSG00000240344 | PPIL3 | 0,386197995 | 1,306944606 | 0,1537 | 0,48209239 | peptidylprolyl isomerase like 3 |
| ENSG00000164180 | TMEM161B | 0,386194047 | 1,30694103 | 0,38931 | 0,65372147 | transmembrane protein 161B |
| ENSG00000109775 | UFSP2 | 0,38610859 | 1,306863617 | 0,46324 | 0,70415312 | UFM1 specific peptidase 2 |
| ENSG00000058063 | ATP11B | 0,386081849 | 1,306839393 | 0,23368 | 0,54563244 | ATPase phospholipid transporting 11B (putative) |
| ENSG00000138071 | ACTR2 | 0,386036319 | 1,306798152 | 0,23817 | 0,54870569 | actin related protein 2 |
| ENSG00000160683 | CXCR5 | 0,385982974 | 1,306749832 | 0,42831 | 0,68103938 | C-X-C motif chemokine receptor 5 |
| ENSG00000272689 | AC004832,4 | 0,385896388 | 1,306671407 | 0,59832 | 0,79497317 | novel transcript, sense intonic to |
| ENSG00000259531 | AC022306,1 | 0,385891756 | 1,306667213 | 0,49001 | 0,72429221 | signal peptide peptidase-like 2A (SPPL2A) pseudogene |
| ENSG00000010803 | SCMH1 | 0,38587847 | 1,306655179 | 0,17867 | 0,50323303 | Scm polycomb group protein homolog 1 |
| ENSG00000262412 | AL160291,1 | 0,385784084 | 1,306569696 | 0,67587 | 0,84049293 | novel transcript |
| ENSG00000162639 | HENMT1 | 0,385742554 | 1,306532085 | 0,12806 | 0,45163061 | HEN methyltransferase 1 |
| ENSG00000169967 | MAP3K2 | 0,385729357 | 1,306520134 | 0,3446 | 0,62200601 | mitogen-activated protein kinase kinase kinase 2 |

| | | | | | | |
|---|---|---|---|---|---|---|
| ENSG00000177426 | TGIF1 | 0,385723668 | 1,306514982 | 0,20971 | 0,52867318 | TGFB induced factor homeobox 1 |
| ENSG00000234405 | Z69733,1 | 0,385716025 | 1,30650806 | 0,34693 | 0,62365773 | novel transcript |
| ENSG00000187735 | TCEA1 | 0,385634441 | 1,306434179 | 0,30784 | 0,59671558 | transcription elongation factor A1 |
| ENSG00000129646 | QRICH2 | 0,385631908 | 1,306431886 | 0,05432 | 0,3118853 | glutamine rich 2 |
| ENSG00000182782 | HCAR2 | 0,385619488 | 1,306420639 | 0,68212 | 0,84411059 | hydroxycarboxylic acid receptor 2 |
| ENSG00000154124 | OTULIN | 0,385591648 | 1,306395429 | 0,12949 | 0,45306297 | OTU deubiquitinase with linear linkage specificity |
| ENSG00000276966 | H4C5 | 0,385551442 | 1,306359022 | 0,31125 | 0,59885232 | H4 clustered histone 5 |
| ENSG00000147459 | DOCK5 | 0,385550084 | 1,306357792 | 0,07783 | 0,36497315 | dedicator of cytokinesis 5 |
| ENSG00000091039 | OSBPL8 | 0,385518136 | 1,306328864 | 0,22983 | 0,54324957 | oxysterol binding protein like 8 |
| ENSG00000230724 | LINC01001 | 0,385489532 | 1,306302964 | 0,37588 | 0,64417233 | long intergenic non-protein coding RNA 1001 |
| ENSG00000143622 | RIT1 | 0,385440575 | 1,306258636 | 0,26741 | 0,56973003 | Ras like without CAAX 1 |
| ENSG00000075568 | TMEM131 | 0,385436438 | 1,30625489 | 0,16042 | 0,48768584 | transmembrane protein 131 |
| ENSG00000120156 | TEK | 0,385405194 | 1,306226601 | 0,40512 | 0,66482913 | TEK receptor tyrosine kinase |
| ENSG00000243477 | NAA80 | 0,385400461 | 1,306222316 | 0,18039 | 0,50407792 | N-alpha-acetyltransferase 80, NatH catalytic subunit |
| ENSG00000122435 | TRMT13 | 0,385395822 | 1,306218116 | 0,30667 | 0,59571748 | tRNA methyltransferase 13 homolog |
| ENSG00000215452 | ZNF663P | 0,385366247 | 1,306191339 | 0,31146 | 0,59892475 | zinc finger protein 663, pseudogene |
| ENSG00000272812 | AC004908,2 | 0,385356851 | 1,306182832 | 0,49795 | 0,72933516 | novel transcript, sense intronic to ZNF596 |
| ENSG00000073792 | IGF2BP2 | 0,385267116 | 1,30610159 | 0,10787 | 0,41847368 | insulin like growth factor 2 mRNA binding protein 2 |
| ENSG00000163939 | PBRM1 | 0,38514416 | 1,30599028 | 0,25018 | 0,55657567 | polybromo 1 |
| ENSG00000179387 | ELMOD2 | 0,385138131 | 1,305984823 | 0,29231 | 0,58692263 | ELMO domain containing 2 |
| ENSG00000234886 | MTND5P26 | 0,385107286 | 1,305956901 | 0,5545 | 0,76745801 | MT-ND5 pseudogene 26 |
| ENSG00000230316 | FEZF1-AS1 | 0,385032829 | 1,305889502 | 0,44836 | 0,69432421 | FEZF1 antisense RNA 1 |
| ENSG00000275793 | RIMBP3 | 0,384947497 | 1,305812265 | 0,45871 | 0,70133296 | RIMS binding protein 3 |
| ENSG00000139826 | ABHD13 | 0,38478569 | 1,305665818 | 0,18981 | 0,51299724 | abhydrolase domain containing 13 |
| ENSG00000286947 | AC022467,2 | 0,38478494 | 1,305665139 | 0,63664 | 0,81709016 | novel transcript, antisense to FBN1 |
| ENSG00000286408 | AC113410,4 | 0,384776897 | 1,305657861 | 0,27603 | 0,57533851 | novel transcript |
| ENSG00000269904 | MAP2K4P1 | 0,384760439 | 1,305642966 | 0,66314 | 0,83242946 | mitogen-activated protein kinase kinase 4 pseudogene 1 |
| ENSG00000172062 | SMN1 | 0,384757475 | 1,305640283 | 0,37896 | 0,64687428 | survival of motor neuron 1, telomeric |
| ENSG00000074319 | TSG101 | 0,384642739 | 1,305536451 | 0,19913 | 0,52017864 | tumor susceptibility 101 |
| ENSG00000004142 | POLDIP2 | 0,384620033 | 1,305515905 | 0,17662 | 0,50089354 | DNA polymerase delta interacting protein 2 |
| ENSG00000198162 | MAN1A2 | 0,384514194 | 1,305420132 | 0,26337 | 0,56608528 | mannosidase alpha class 1A member 2 |
| ENSG00000163399 | ATP1A1 | 0,384489113 | 1,305397438 | 0,04362 | 0,28187766 | ATPase Na+/K+ transporting subunit alpha 1 |
| ENSG00000214274 | ANG | 0,384415391 | 1,305330734 | 0,25823 | 0,5622185 | angiogenin |

| | | | | | | |
|---|---|---|---|---|---|---|
| ENSG00000128791 | TWSG1 | 0,38439219 | 1,305309742 | 0,19083 | 0,51394498 | twisted gastrulation BMP signaling modulator 1 |
| ENSG00000142453 | CARM1 | 0,38436958 | 1,305289285 | 0,13906 | 0,46647959 | coactivator associated arginine methyltransferase 1 |
| ENSG00000259370 | AC103740,1 | 0,384365139 | 1,305285267 | 0,19861 | 0,51980353 | uncharacterized LOC105370854 |
| ENSG00000051108 | HERPUD1 | 0,384341064 | 1,305263485 | 0,15379 | 0,48214138 | homocysteine inducible ER protein with ubiquitin like domain 1 |
| ENSG00000179195 | ZNF664 | 0,384330881 | 1,305254272 | 0,35656 | 0,63063438 | zinc finger protein 664 |
| ENSG00000244926 | ALKBH3-AS1 | 0,384323078 | 1,305247213 | 0,29542 | 0,58915878 | ALKBH3 antisense RNA 1 |
| ENSG00000095794 | CREM | 0,38420458 | 1,305140009 | 0,31469 | 0,60130206 | cAMP responsive element modulator |
| ENSG00000112062 | MAPK14 | 0,384168382 | 1,305107263 | 0,23966 | 0,5491566 | mitogen-activated protein kinase 14 |
| ENSG00000112218 | GPR63 | 0,384163033 | 1,305102423 | 0,32024 | 0,60518204 | G protein-coupled receptor 63 |
| ENSG00000130312 | MRPL34 | 0,384097183 | 1,305042855 | 0,15512 | 0,48296753 | mitochondrial ribosomal protein L34 |
| ENSG00000259746 | HSPE1P3 | 0,384084322 | 1,305031221 | 0,66986 | 0,83624601 | heat shock protein family E (Hsp10) member 1 pseudogene 3 |
| ENSG00000244218 | RN7SL81P | 0,383907747 | 1,304871506 | 0,58192 | 0,78454692 | RNA, 7SL, cytoplasmic 81, pseudogene |
| ENSG00000154473 | BUB3 | 0,38390448 | 1,30486855 | 0,24099 | 0,55003907 | BUB3 mitotic checkpoint protein |
| ENSG00000158019 | BABAM2 | 0,383836342 | 1,304806923 | 0,22622 | 0,54121239 | BRISC and BRCA1 A complex member 2 |
| ENSG00000260585 | AL359851,1 | 0,383814011 | 1,304786726 | 0,54203 | 0,75894841 | novel transcript |
| ENSG00000120756 | PLS1 | 0,383716234 | 1,3046983 | 0,42467 | 0,67846394 | plastin 1 |
| ENSG00000080709 | KCNN2 | 0,383681165 | 1,304666586 | 0,18793 | 0,51107563 | potassium calcium-activated channel subfamily N member 2 |
| ENSG00000117523 | PRRC2C | 0,383643489 | 1,304632514 | 0,0911 | 0,39033853 | proline rich coiled-coil 2C |
| ENSG00000128594 | LRRC4 | 0,383623087 | 1,304614065 | 0,08002 | 0,36922825 | leucine rich repeat containing 4 |
| ENSG00000169599 | NFU1 | 0,383471955 | 1,304477405 | 0,29879 | 0,59100642 | NFU1 iron-sulfur cluster scaffold |
| ENSG00000184831 | APOO | 0,383462982 | 1,304469292 | 0,46299 | 0,70412893 | apolipoprotein O |
| ENSG00000165782 | PIP4P1 | 0,383226512 | 1,304255495 | 0,10278 | 0,41062092 | phosphatidylinositol-4,5-bisphosphate 4-phosphatase 1 |
| ENSG00000175793 | SFN | 0,383178451 | 1,304212047 | 0,63363 | 0,81549489 | stratifin |
| ENSG00000198081 | ZBTB14 | 0,38302774 | 1,30407581 | 0,34678 | 0,62352276 | zinc finger and BTB domain containing 14 |
| ENSG00000178662 | CSRNP3 | 0,382998764 | 1,304049618 | 0,31702 | 0,60268749 | cysteine and serine rich nuclear protein 3 |
| ENSG00000131981 | LGALS3 | 0,382976931 | 1,304029883 | 0,38729 | 0,65227155 | galectin 3 |
| ENSG00000281385 | AP2A2 | 0,382959104 | 1,30401377 | 0,34043 | 0,61897407 | adaptor related protein complex 2 subunit alpha 2 |
| ENSG00000235974 | VN2R19P | 0,382906328 | 1,303966068 | 0,59929 | 0,79538547 | vomeronasal 2 receptor 19, pseudogene |
| ENSG00000124571 | XPO5 | 0,382861323 | 1,303925391 | 0,21697 | 0,53505085 | exportin 5 |
| ENSG00000286905 | AC108488,2 | 0,382829008 | 1,303896185 | 0,28154 | 0,57891473 | novel protein |

| | | | | | | |
|---|---|---|---|---|---|---|
| ENSG00000170385 | SLC30A1 | 0,382701879 | 1,303781292 | 0,23374 | 0,54563244 | solute carrier family 30 member 1 |
| ENSG00000271254 | AC240274,1 | 0,382691645 | 1,303772043 | 0,10501 | 0,41375455 | neuroblastoma breakpoint family member 1 |
| ENSG00000156970 | BUB1B | 0,382600709 | 1,303689866 | 0,27128 | 0,57271346 | BUB1 mitotic checkpoint serine/threonine kinase B |
| ENSG00000010319 | SEMA3G | 0,382584201 | 1,303674949 | 0,5809 | 0,78399513 | semaphorin 3G |
| ENSG00000136631 | VPS45 | 0,382556328 | 1,303649762 | 0,40603 | 0,66571491 | vacuolar protein sorting 45 homolog |
| ENSG00000170801 | HTRA3 | 0,382553049 | 1,303646799 | 0,30404 | 0,59411013 | HtrA serine peptidase 3 |
| ENSG00000264235 | AP005329,1 | 0,382468935 | 1,303570794 | 0,39206 | 0,65540679 | novel transcript, antisense to MYL12A |
| ENSG00000212127 | TAS2R14 | 0,382438623 | 1,303543405 | 0,4034 | 0,66399303 | taste 2 receptor member 14 |
| ENSG00000260777 | AC061975,1 | 0,382428566 | 1,303534319 | 0,31945 | 0,60429866 | novel transcript |
| ENSG00000144057 | ST6GAL2 | 0,382364997 | 1,303476883 | 0,15725 | 0,48443207 | ST6 beta-galactoside alpha-2,6-sialyltransferase 2 |
| ENSG00000125676 | THOC2 | 0,382318526 | 1,303434897 | 0,28542 | 0,58187605 | THO complex 2 |
| ENSG00000143633 | C1orf131 | 0,382270984 | 1,303391945 | 0,2607 | 0,56400103 | chromosome 1 open reading frame 131 |
| ENSG00000114737 | CISH | 0,382263827 | 1,303385479 | 0,12155 | 0,44093327 | cytokine inducible SH2 containing protein |
| ENSG00000258504 | AL157871,1 | 0,382222569 | 1,303348205 | 0,34656 | 0,62350287 | novel transcript |
| ENSG00000082146 | STRADB | 0,382208382 | 1,303335389 | 0,32146 | 0,60577115 | STE20 related adaptor beta |
| ENSG00000144848 | ATG3 | 0,382195034 | 1,30332333 | 0,18991 | 0,51315877 | autophagy related 3 |
| ENSG00000197496 | SLC2A10 | 0,382161789 | 1,303293298 | 0,07914 | 0,36738337 | solute carrier family 2 member 10 |
| ENSG00000152990 | ADGRA3 | 0,382154236 | 1,303286474 | 0,12566 | 0,44769898 | adhesion G protein-coupled receptor A3 |
| ENSG00000100485 | SOS2 | 0,382140768 | 1,303274307 | 0,30722 | 0,59633879 | SOS Ras/Rho guanine nucleotide exchange factor 2 |
| ENSG00000060762 | MPC1 | 0,382118331 | 1,303254039 | 0,52573 | 0,74937147 | mitochondrial pyruvate carrier 1 |
| ENSG00000146918 | NCAPG2 | 0,382098526 | 1,303236149 | 0,26371 | 0,56629809 | non-SMC condensin II complex subunit G2 |
| ENSG00000173402 | DAG1 | 0,382046037 | 1,303188734 | 0,20468 | 0,52443222 | dystroglycan 1 |
| ENSG00000113734 | BNIP1 | 0,382026858 | 1,303171409 | 0,37177 | 0,6408454 | BCL2 interacting protein 1 |
| ENSG00000131788 | PIAS3 | 0,382024153 | 1,303168966 | 0,08183 | 0,37301152 | protein inhibitor of activated STAT 3 |
| ENSG00000223519 | KIF28P | 0,381963954 | 1,30311459 | 0,59945 | 0,79551789 | kinesin family member 28, pseudogene |
| ENSG00000062598 | ELMO2 | 0,381953613 | 1,30310525 | 0,31009 | 0,59836523 | engulfment and cell motility 2 |
| ENSG00000163714 | U2SURP | 0,381894198 | 1,303051584 | 0,18467 | 0,50835796 | U2 snRNP associated SURP domain containing |
| ENSG00000145386 | CCNA2 | 0,381892541 | 1,303050089 | 0,26652 | 0,56917012 | cyclin A2 |
| ENSG00000241420 | RN7SL505P | 0,381739909 | 1,302912238 | 0,74959 | 0,8822079 | RNA, 7SL, cytoplasmic 505, pseudogene |
| ENSG00000079134 | THOC1 | 0,381630496 | 1,302813429 | 0,36234 | 0,63379594 | THO complex 1 |
| ENSG00000102699 | PARP4 | 0,381615658 | 1,30280003 | 0,37086 | 0,64013451 | poly(ADP-ribose) polymerase family member 4 |
| ENSG00000260494 | AC002310,3 | 0,381552038 | 1,30274258 | 0,61314 | 0,8035281 | ribosomal protein L7 (RPL7) pseudogene |
| ENSG00000141076 | UTP4 | 0,38153917 | 1,30273096 | 0,27645 | 0,57546799 | UTP4 small subunit processome component |

| | | | | | | |
|---|---|---|---|---|---|---|
| ENSG00000287411 | AL031259,1 | 0,381416375 | 1,302620083 | 0,634 | 0,81549489 | novel transcript |
| ENSG00000111052 | LIN7A | 0,381296174 | 1,302511558 | 0,53224 | 0,75291701 | lin-7 homolog A, crumbs cell polarity complex component |
| ENSG00000168802 | CHTF8 | 0,381165229 | 1,302393342 | 0,15688 | 0,48425071 | chromosome transmission fidelity factor 8 |
| ENSG00000288325 | GPS2 | 0,381148261 | 1,302378024 | 0,17257 | 0,49744497 | G protein pathway suppressor 2 |
| ENSG00000132522 | GPS2 | 0,381148261 | 1,302378024 | 0,17257 | 0,49744497 | G protein pathway suppressor 2 |
| ENSG00000064726 | BTBD1 | 0,381110391 | 1,302343838 | 0,28303 | 0,58033644 | BTB domain containing 1 |
| ENSG00000187553 | CYP26C1 | 0,381082701 | 1,302318842 | 0,63509 | 0,81637102 | cytochrome P450 family 26 subfamily C member 1 |
| ENSG00000146433 | TMEM181 | 0,381063739 | 1,302301724 | 0,11162 | 0,42472977 | transmembrane protein 181 |
| ENSG00000177613 | CSTF2T | 0,38104707 | 1,302286678 | 0,2223 | 0,53859118 | cleavage stimulation factor subunit 2 tau variant |
| ENSG00000217094 | PPIAP31 | 0,380804028 | 1,302067308 | 0,61232 | 0,8035281 | peptidylprolyl isomerase A pseudogene 31 |
| ENSG00000004766 | VPS50 | 0,380772411 | 1,302038773 | 0,21789 | 0,53550245 | VPS50 subunit of EARP/GARPII complex |
| ENSG00000198945 | L3MBTL3 | 0,380693863 | 1,301967886 | 0,18367 | 0,50744379 | L3MBTL histone methyl-lysine binding protein 3 |
| ENSG00000263309 | KRT23 | 0,380680247 | 1,301955598 | 0,68098 | 0,84352125 | keratin 23 |
| ENSG00000189180 | ZNF33A | 0,380637465 | 1,30191699 | 0,36156 | 0,63347405 | zinc finger protein 33A |
| ENSG00000259347 | AC087482,1 | 0,380633677 | 1,301913572 | 0,42946 | 0,68224982 | novel transcript, antisense to SMAD3 |
| ENSG00000099290 | WASHC2A | 0,380626814 | 1,301907378 | 0,15195 | 0,48039146 | WASH complex subunit 2A |
| ENSG00000163946 | TASOR | 0,380620013 | 1,301901241 | 0,31135 | 0,59890747 | transcription activation suppressor |
| ENSG00000196172 | ZNF681 | 0,380606429 | 1,301888983 | 0,03828 | 0,264906 | zinc finger protein 681 |
| ENSG00000287299 | AC012459,1 | 0,380482391 | 1,301777056 | 0,17674 | 0,50110131 | novel transcript, antisense to SPATS2L |
| ENSG00000196505 | GDAP2 | 0,380471239 | 1,301766992 | 0,29021 | 0,58540802 | ganglioside induced differentiation associated protein 2 |
| ENSG00000117118 | SDHB | 0,380291501 | 1,301604822 | 0,35638 | 0,63043562 | succinate dehydrogenase complex iron sulfur subunit B |
| ENSG00000166949 | SMAD3 | 0,380227341 | 1,301546938 | 0,01261 | 0,14714633 | SMAD family member 3 |
| ENSG00000140750 | ARHGAP17 | 0,380209891 | 1,301531196 | 0,09336 | 0,39489949 | Rho GTPase activating protein 17 |
| ENSG00000175161 | CADM2 | 0,380162567 | 1,301488502 | 0,39915 | 0,66153677 | cell adhesion molecule 2 |
| ENSG00000185238 | PRMT3 | 0,379986135 | 1,301329349 | 0,24605 | 0,55380958 | protein arginine methyltransferase 3 |
| ENSG00000023572 | GLRX2 | 0,379978647 | 1,301322595 | 0,39894 | 0,66140951 | glutaredoxin 2 |
| ENSG00000198040 | ZNF84 | 0,37993608 | 1,3012842 | 0,37782 | 0,64626684 | zinc finger protein 84 |
| ENSG00000083817 | ZNF416 | 0,37968539 | 1,301058101 | 0,3362 | 0,61623735 | zinc finger protein 416 |
| ENSG00000155252 | PI4K2A | 0,379640081 | 1,301017241 | 0,07845 | 0,36604454 | phosphatidylinositol 4-kinase type 2 alpha |
| ENSG00000153815 | CMIP | 0,379610131 | 1,300990232 | 0,07224 | 0,35480344 | c-Maf inducing protein |
| ENSG00000070614 | NDST1 | 0,37953848 | 1,300925621 | 0,02882 | 0,2296248 | N-deacetylase and N-sulfotransferase 1 |
| ENSG00000116138 | DNAJC16 | 0,379532556 | 1,300920279 | 0,1083 | 0,4189697 | DnaJ heat shock protein family (Hsp40) member C16 |
| ENSG00000111206 | FOXM1 | 0,37952224 | 1,300910977 | 0,27737 | 0,57586315 | forkhead box M1 |

| | | | | | | |
|---|---|---|---|---|---|---|
| ENSG00000121064 | SCPEP1 | 0,379506347 | 1,300896646 | 0,05053 | 0,30240543 | serine carboxypeptidase 1 |
| ENSG00000132388 | UBE2G1 | 0,379489169 | 1,300881157 | 0,15541 | 0,48334025 | ubiquitin conjugating enzyme E2 G1 |
| ENSG00000136044 | APPL2 | 0,379454122 | 1,300849555 | 0,28995 | 0,58531219 | adaptor protein, phosphotyrosine interacting with PH domain and leucine zipper 2 |
| ENSG00000101577 | LPIN2 | 0,379362518 | 1,30076696 | 0,23329 | 0,54562736 | lipin 2 |
| ENSG00000169252 | ADRB2 | 0,379341836 | 1,300748312 | 0,72965 | 0,8715818 | adrenoceptor beta 2 |
| ENSG00000138160 | KIF11 | 0,379317313 | 1,300726203 | 0,231 | 0,54393842 | kinesin family member 11 |
| ENSG00000236502 | SIX3-AS1 | 0,37929404 | 1,30070522 | 0,55512 | 0,76778644 | SIX3 antisense RNA 1 |
| ENSG00000171121 | KCNMB3 | 0,379227801 | 1,300645502 | 0,27949 | 0,57745127 | potassium calcium-activated channel subfamily M regulatory beta subunit 3 |
| ENSG00000066827 | ZFAT | 0,379208036 | 1,300627683 | 0,28574 | 0,58215967 | zinc finger and AT-hook domain containing |
| ENSG00000187054 | TMPRSS11A | 0,379119195 | 1,300547593 | 0,60516 | 0,79831516 | transmembrane serine protease 11A |
| ENSG00000244321 | LINC01326 | 0,379049908 | 1,300485134 | 0,52125 | 0,74558236 | long intergenic non-protein coding RNA 1326 |
| ENSG00000147862 | NFIB | 0,379003228 | 1,300443056 | 0,47733 | 0,71526298 | nuclear factor I B |
| ENSG00000168300 | PCMTD1 | 0,378880748 | 1,300332657 | 0,35181 | 0,62697369 | protein-L-isoaspartate (D-aspartate) O-methyltransferase domain containing 1 |
| ENSG00000087495 | PHACTR3 | 0,378876468 | 1,3003288 | 0,32158 | 0,60577115 | phosphatase and actin regulator 3 |
| ENSG00000080986 | NDC80 | 0,378863012 | 1,300316672 | 0,2476 | 0,55506866 | NDC80 kinetochore complex component |
| ENSG00000164985 | PSIP1 | 0,378811697 | 1,300270422 | 0,30891 | 0,59770756 | PC4 and SFRS1 interacting protein 1 |
| ENSG00000273192 | AL671710,1 | 0,378794466 | 1,300254892 | 0,2643 | 0,56706657 | novel transcript, antisense to ALG12 |
| ENSG00000112739 | PRPF4B | 0,378786671 | 1,300247867 | 0,26854 | 0,57061705 | pre-mRNA processing factor 4B |
| ENSG00000096384 | HSP90AB1 | 0,37878252 | 1,300244125 | 0,08816 | 0,38442787 | heat shock protein 90 alpha family class B member 1 |
| ENSG00000162851 | TFB2M | 0,378697559 | 1,300167556 | 0,1135 | 0,42823843 | transcription factor B2, mitochondrial |
| ENSG00000229989 | MIR181A1HG | 0,378662216 | 1,300135705 | 0,2415 | 0,55037176 | MIR181A1 host gene |
| ENSG00000275197 | AC092794,2 | 0,378631183 | 1,300107739 | 0,68848 | 0,84821636 | novel transcript, antisense to KRAS |
| ENSG00000103591 | AAGAB | 0,378522068 | 1,300009412 | 0,19535 | 0,51622695 | alpha and gamma adaptin binding protein |
| ENSG00000070770 | CSNK2A2 | 0,378440429 | 1,299935849 | 0,236 | 0,5471053 | casein kinase 2 alpha 2 |
| ENSG00000135049 | AGTPBP1 | 0,378422438 | 1,299919638 | 0,31825 | 0,60349082 | ATP/GTP binding protein 1 |
| ENSG00000128908 | INO80 | 0,3783588 | 1,2998623 | 0,20191 | 0,52223654 | INO80 complex ATPase subunit |
| ENSG00000272142 | LYRM4-AS1 | 0,378349837 | 1,299854224 | 0,23225 | 0,54499308 | LYRM4 antisense RNA 1 |
| ENSG00000152484 | USP12 | 0,378336285 | 1,299842014 | 0,3255 | 0,60795521 | ubiquitin specific peptidase 12 |
| ENSG00000164934 | DCAF13 | 0,378286963 | 1,299797576 | 0,22846 | 0,54241083 | DDB1 and CUL4 associated factor 13 |
| ENSG00000138166 | DUSP5 | 0,378245566 | 1,299760281 | 0,18306 | 0,50652733 | dual specificity phosphatase 5 |

| | | | | | | |
|---|---|---|---|---|---|---|
| ENSG00000128944 | KNSTRN | 0,378206554 | 1,299725134 | 0,34999 | 0,62563749 | kinetochore localized astrin (SPAG5) binding protein |
| ENSG00000056586 | RC3H2 | 0,378100462 | 1,299629559 | 0,24802 | 0,55542983 | ring finger and CCCH-type domains 2 |
| ENSG00000259969 | AL049838,1 | 0,378093525 | 1,299623309 | 0,54524 | 0,7609742 | novel transcript, overlapping C14orf37 |
| ENSG00000287030 | AC107032,3 | 0,378088819 | 1,29961907 | 0,63775 | 0,8179135 | novel transcript |
| ENSG00000101558 | VAPA | 0,378013605 | 1,299551317 | 0,27182 | 0,57305058 | VAMP associated protein A |
| ENSG00000214194 | SMIM30 | 0,37796162 | 1,299504492 | 0,45018 | 0,69562819 | small integral membrane protein 30 |
| ENSG00000167842 | MIS12 | 0,377919883 | 1,299466897 | 0,2603 | 0,56382982 | MIS12 kinetochore complex component |
| ENSG00000164087 | POC1A | 0,377881461 | 1,29943229 | 0,40828 | 0,66714279 | POC1 centriolar protein A |
| ENSG00000153560 | UBP1 | 0,377764313 | 1,29932678 | 0,2261 | 0,54121239 | upstream binding protein 1 |
| ENSG00000105369 | CD79A | 0,377756405 | 1,299319658 | 0,69955 | 0,8548625 | CD79a molecule |
| ENSG00000114805 | PLCH1 | 0,377710577 | 1,299278384 | 0,29623 | 0,58961035 | phospholipase C eta 1 |
| ENSG00000166575 | TMEM135 | 0,377613585 | 1,299191037 | 0,42675 | 0,68012899 | transmembrane protein 135 |
| ENSG00000134247 | PTGFRN | 0,37755632 | 1,29913947 | 0,11939 | 0,43734366 | prostaglandin F2 receptor inhibitor |
| ENSG00000254180 | AC004083,1 | 0,377530367 | 1,299116099 | 0,70608 | 0,85841728 | novel transcript |
| ENSG00000253554 | LINC01414 | 0,377514623 | 1,299101922 | 0,40921 | 0,66785537 | long intergenic non-protein coding RNA 1414 |
| ENSG00000134014 | ELP3 | 0,377491411 | 1,299081021 | 0,14983 | 0,47782859 | elongator acetyltransferase complex subunit 3 |
| ENSG00000163170 | BOLA3 | 0,377362493 | 1,298964941 | 0,21238 | 0,53120227 | bolA family member 3 |
| ENSG00000181163 | NPM1 | 0,377261914 | 1,298874386 | 0,08925 | 0,38636433 | nucleophosmin 1 |
| ENSG00000144559 | TAMM41 | 0,377199181 | 1,298817907 | 0,17767 | 0,50205238 | TAM41 mitochondrial translocator assembly and maintenance homolog |
| ENSG00000146757 | ZNF92 | 0,377176641 | 1,298797615 | 0,44691 | 0,6933405 | zinc finger protein 92 |
| ENSG00000131791 | PRKAB2 | 0,377166292 | 1,298788299 | 0,36508 | 0,63531881 | protein kinase AMP-activated non-catalytic subunit beta 2 |
| ENSG00000113812 | ACTR8 | 0,377136408 | 1,298761395 | 0,32251 | 0,60577115 | actin related protein 8 |
| ENSG00000225419 | RPL21P3 | 0,377135349 | 1,298760442 | 0,7193 | 0,86579286 | ribosomal protein L21 pseudogene 3 |
| ENSG00000186462 | NAP1L2 | 0,3770689 | 1,298700624 | 0,06763 | 0,34407703 | nucleosome assembly protein 1 like 2 |
| ENSG00000151665 | PIGF | 0,377050582 | 1,298684135 | 0,38417 | 0,65052153 | phosphatidylinositol glycan anchor biosynthesis class F |
| ENSG00000281005 | LINC00921 | 0,376990664 | 1,298630199 | 0,49699 | 0,72862056 | long intergenic non-protein coding RNA 921 |
| ENSG00000238132 | CASC4P1 | 0,376950906 | 1,298594412 | 0,69582 | 0,85278034 | cancer susceptibility 4 pseudogene 1 |
| ENSG00000122068 | FYTTD1 | 0,376928206 | 1,298573979 | 0,27918 | 0,57733908 | forty-two-three domain containing 1 |
| ENSG00000114520 | SNX4 | 0,37681845 | 1,298475191 | 0,30242 | 0,59297237 | sorting nexin 4 |
| ENSG00000168890 | TMEM150A | 0,376723829 | 1,298390032 | 0,20895 | 0,52800078 | transmembrane protein 150A |
| ENSG00000156787 | TBC1D31 | 0,376719473 | 1,298386112 | 0,27997 | 0,57764051 | TBC1 domain family member 31 |
| ENSG00000111912 | NCOA7 | 0,376632425 | 1,298307774 | 0,33104 | 0,6112353 | nuclear receptor coactivator 7 |

| | | | | | | |
|---|---|---|---|---|---|---|
| ENSG00000287201 | AC009682,1 | 0,376588502 | 1,298268247 | 0,49408 | 0,72659022 | novel transcript, antisense to ASAP1 |
| ENSG00000029364 | SLC39A9 | 0,376561698 | 1,298244126 | 0,21433 | 0,53334744 | solute carrier family 39 member 9 |
| ENSG00000229520 | LINC00404 | 0,376535897 | 1,298220908 | 0,31798 | 0,60347876 | long intergenic non-protein coding RNA 404 |
| ENSG00000048991 | R3HDM1 | 0,376496424 | 1,298185389 | 0,21891 | 0,53639845 | R3H domain containing 1 |
| ENSG00000138085 | ATRAID | 0,376458098 | 1,298150903 | 0,19271 | 0,51490739 | all-trans retinoic acid induced differentiation factor |
| ENSG00000198105 | ZNF248 | 0,376370156 | 1,298071774 | 0,3022 | 0,59294633 | zinc finger protein 248 |
| ENSG00000261231 | LINC02133 | 0,376354946 | 1,298058089 | 0,43175 | 0,68379763 | long intergenic non-protein coding RNA 2133 |
| ENSG00000189403 | HMGB1 | 0,376311622 | 1,298019108 | 0,33512 | 0,61480301 | high mobility group box 1 |
| ENSG00000136450 | SRSF1 | 0,376288395 | 1,297998211 | 0,16006 | 0,48727386 | serine and arginine rich splicing factor 1 |
| ENSG00000154945 | ANKRD40 | 0,376257264 | 1,297970203 | 0,05745 | 0,31984771 | ankyrin repeat domain 40 |
| ENSG00000287166 | AC104561,4 | 0,376234706 | 1,297949908 | 0,31202 | 0,59925736 | novel transcript |
| ENSG00000187790 | FANCM | 0,376228501 | 1,297944325 | 0,36769 | 0,63764567 | FA complementation group M |
| ENSG00000278467 | AC138393,3 | 0,376184906 | 1,297905105 | 0,27624 | 0,57538841 | novel transcript |
| ENSG00000261804 | AC007342,4 | 0,376184861 | 1,297905065 | 0,26165 | 0,56452671 | novel transcript |
| ENSG00000143486 | EIF2D | 0,375989639 | 1,297729447 | 0,2632 | 0,56606892 | eukaryotic translation initiation factor 2D |
| ENSG00000163660 | CCNL1 | 0,375975362 | 1,297716605 | 0,3019 | 0,59294633 | cyclin L1 |
| ENSG00000258827 | AL079304,1 | 0,375946886 | 1,297690991 | 0,60054 | 0,79587862 | novel transcript, sense intronic to SLC25A21 |
| ENSG00000185721 | DRG1 | 0,375926658 | 1,297672796 | 0,27955 | 0,57745167 | developmentally regulated GTP binding protein 1 |
| ENSG00000148672 | GLUD1 | 0,375909393 | 1,297657267 | 0,27448 | 0,57443122 | glutamate dehydrogenase 1 |
| ENSG00000206384 | COL6A6 | 0,375901732 | 1,297650376 | 0,36129 | 0,63328947 | collagen type VI alpha 6 chain |
| ENSG00000137509 | PRCP | 0,375896539 | 1,297645705 | 0,2678 | 0,56999323 | prolylcarboxypeptidase |
| ENSG00000135974 | C2orf49 | 0,37584925 | 1,297603171 | 0,12123 | 0,44047594 | chromosome 2 open reading frame 49 |
| ENSG00000082153 | BZW1 | 0,375834837 | 1,297590208 | 0,38804 | 0,65289784 | basic leucine zipper and W2 domains 1 |
| ENSG00000172172 | MRPL13 | 0,375834647 | 1,297590036 | 0,16462 | 0,49065086 | mitochondrial ribosomal protein L13 |
| ENSG00000136522 | MRPL47 | 0,375790571 | 1,297550395 | 0,29788 | 0,59037822 | mitochondrial ribosomal protein L47 |
| ENSG00000129187 | DCTD | 0,375680878 | 1,297451741 | 0,16097 | 0,48832443 | dCMP deaminase |
| ENSG00000125459 | MSTO1 | 0,375615661 | 1,297393091 | 0,16627 | 0,49212787 | misato mitochondrial distribution and morphology regulator 1 |
| ENSG00000099282 | TSPAN15 | 0,375559205 | 1,297342322 | 0,27403 | 0,57412006 | tetraspanin 15 |
| ENSG00000067334 | DNTTIP2 | 0,375511363 | 1,297299301 | 0,14338 | 0,47083258 | deoxynucleotidyltransferase terminal interacting protein 2 |
| ENSG00000164944 | VIRMA | 0,375419287 | 1,297216507 | 0,25088 | 0,55707572 | vir like m6A methyltransferase associated |
| ENSG00000131845 | ZNF304 | 0,37539971 | 1,297198904 | 0,25957 | 0,56330128 | zinc finger protein 304 |
| ENSG00000121594 | CD80 | 0,375359872 | 1,297163084 | 0,73228 | 0,87295003 | CD80 molecule |

| | | | | | | |
|---|---|---|---|---|---|---|
| ENSG00000115365 | LANCL1 | 0,375325622 | 1,297132289 | 0,42987 | 0,68262051 | LanC like 1 |
| ENSG00000121892 | PDS5A | 0,375317811 | 1,297125267 | 0,34832 | 0,62468287 | PDS5 cohesin associated factor A |
| ENSG00000231205 | ZNF826P | 0,375258128 | 1,297071607 | 0,12638 | 0,44903997 | zinc finger protein 826, pseudogene |
| ENSG00000133103 | COG6 | 0,37522533 | 1,29704212 | 0,43176 | 0,68379763 | component of oligomeric golgi complex 6 |
| ENSG00000102144 | PGK1 | 0,375143806 | 1,296968829 | 0,24125 | 0,55027101 | phosphoglycerate kinase 1 |
| ENSG00000107201 | DDX58 | 0,375142965 | 1,296968072 | 0,35447 | 0,62908147 | DExD/H-box helicase 58 |
| ENSG00000121067 | SPOP | 0,375130848 | 1,296957179 | 0,10537 | 0,41455967 | speckle type BTB/POZ protein |
| ENSG00000138653 | NDST4 | 0,375082608 | 1,296913813 | 0,47585 | 0,71433501 | N-deacetylase and N-sulfotransferase 4 |
| ENSG00000110321 | EIF4G2 | 0,375071179 | 1,296903539 | 0,23768 | 0,54861973 | eukaryotic translation initiation factor 4 gamma 2 |
| ENSG00000138814 | PPP3CA | 0,374963941 | 1,296807142 | 0,17499 | 0,49911246 | protein phosphatase 3 catalytic subunit alpha |
| ENSG00000197728 | RPS26 | 0,374942715 | 1,296788062 | 0,16775 | 0,49395055 | ribosomal protein S26 |
| ENSG00000233820 | AL589843,2 | 0,374853447 | 1,296707825 | 0,47858 | 0,71633759 | makorin, ring finger protein, 2 (MKRN2) pseudogene |
| ENSG00000141756 | FKBP10 | 0,374820422 | 1,296678142 | 0,0457 | 0,28842796 | FKBP prolyl isomerase 10 |
| ENSG00000280627 | SURF1 | 0,374581435 | 1,296463361 | 0,28926 | 0,58493899 | SURF1 cytochrome c oxidase assembly factor |
| ENSG00000148290 | SURF1 | 0,374581435 | 1,296463361 | 0,28926 | 0,58493899 | SURF1 cytochrome c oxidase assembly factor |
| ENSG00000228793 | AL033523,1 | 0,374575735 | 1,296458239 | 0,20649 | 0,52609885 | novel transcript |
| ENSG00000106049 | HIBADH | 0,374493059 | 1,296383945 | 0,13775 | 0,46481656 | 3-hydroxyisobutyrate dehydrogenase |
| ENSG00000256083 | AC090673,1 | 0,374439848 | 1,296336131 | 0,58144 | 0,78430794 | novel transcript, antisense to HMGA2 |
| ENSG00000152795 | HNRNPDL | 0,374396527 | 1,296297205 | 0,28063 | 0,57806909 | heterogeneous nuclear ribonucleoprotein D like |
| ENSG00000225094 | SETP20 | 0,37418781 | 1,296109682 | 0,57674 | 0,7810482 | SET pseudogene 20 |
| ENSG00000233314 | PPP1R11 | 0,374156375 | 1,296081442 | 0,14775 | 0,47507961 | protein phosphatase 1 regulatory inhibitor subunit 11 |
| ENSG00000234058 | PPP1R11 | 0,374156375 | 1,296081442 | 0,14775 | 0,47507961 | protein phosphatase 1 regulatory inhibitor subunit 11 |
| ENSG00000227720 | PPP1R11 | 0,374156375 | 1,296081442 | 0,14775 | 0,47507961 | protein phosphatase 1 regulatory inhibitor subunit 11 |
| ENSG00000237403 | PPP1R11 | 0,374156375 | 1,296081442 | 0,14775 | 0,47507961 | protein phosphatase 1 regulatory inhibitor subunit 11 |
| ENSG00000164305 | CASP3 | 0,3741366 | 1,296063676 | 0,40744 | 0,66661764 | caspase 3 |
| ENSG00000182916 | TCEAL7 | 0,374083729 | 1,296016179 | 0,42622 | 0,67974949 | transcription elongation factor A like 7 |
| ENSG00000277534 | AC007996,1 | 0,374068137 | 1,296002173 | 0,38626 | 0,65168435 | novel transcript, sense intronic to KCTD1 |
| ENSG00000251354 | AC024451,1 | 0,374030997 | 1,295968809 | 0,64612 | 0,82278641 | ribosomal protein S4, X-linked (RPS4X) pseudogene |
| ENSG00000187068 | C3orf70 | 0,373983205 | 1,295925879 | 0,15792 | 0,48497123 | chromosome 3 open reading frame 70 |
| ENSG00000282986 | EIF3K | 0,373945717 | 1,295892205 | 0,2325 | 0,54526182 | eukaryotic translation initiation factor 3 subunit K |
| ENSG00000153885 | KCTD15 | 0,3738951 | 1,29584674 | 0,12804 | 0,4516239 | potassium channel tetramerization domain containing 15 |
| ENSG00000099797 | TECR | 0,373810517 | 1,295770768 | 0,05952 | 0,32482368 | trans-2,3-enoyl-CoA reductase |
| ENSG00000187118 | CMC1 | 0,373664801 | 1,295639899 | 0,35949 | 0,63230976 | C-X9-C motif containing 1 |

| | | | | | | |
|---|---|---|---|---|---|---|
| ENSG00000147421 | HMBOX1 | 0,373635357 | 1,295613456 | 0,23685 | 0,54768226 | homeobox containing 1 |
| ENSG00000031003 | FAM13B | 0,373597878 | 1,295579798 | 0,25272 | 0,55796133 | family with sequence similarity 13 member B |
| ENSG00000084112 | SSH1 | 0,373580908 | 1,295564559 | 0,03435 | 0,25061039 | slingshot protein phosphatase 1 |
| ENSG00000237704 | AP004289,2 | 0,37357153 | 1,295556137 | 0,56974 | 0,77693992 | male-specific lethal 3-like 1 (Drosophila) (MSL3L1) pseudogene |
| ENSG00000101150 | TPD52L2 | 0,373549231 | 1,295536113 | 0,10949 | 0,42148306 | TPD52 like 2 |
| ENSG00000101474 | APMAP | 0,373474291 | 1,295468819 | 0,20268 | 0,52269757 | adipocyte plasma membrane associated protein |
| ENSG00000198918 | RPL39 | 0,373463435 | 1,29545907 | 0,29567 | 0,58930611 | ribosomal protein L39 |
| ENSG00000278472 | AC009268,2 | 0,37345877 | 1,295454882 | 0,57716 | 0,78134056 | novel transcript, sense intronic to AVEN |
| ENSG00000197153 | H3C12 | 0,373415821 | 1,295416316 | 0,4397 | 0,68844641 | H3 clustered histone 12 |
| ENSG00000224363 | FP700111,1 | 0,373343865 | 1,295351708 | 0,45065 | 0,69592349 | novel transcript |
| ENSG00000169184 | MN1 | 0,373283331 | 1,295297358 | 0,2946 | 0,58857672 | MN1 proto-oncogene, transcriptional regulator |
| ENSG00000186908 | ZDHHC17 | 0,373254021 | 1,295271042 | 0,44111 | 0,68942758 | zinc finger DHHC-type palmitoyltransferase 17 |
| ENSG00000146457 | WTAP | 0,373147601 | 1,2951755 | 0,20233 | 0,52249745 | WT1 associated protein |
| ENSG00000104907 | TRMT1 | 0,373122752 | 1,295153192 | 0,10315 | 0,41083928 | tRNA methyltransferase 1 |
| ENSG00000163606 | CD200R1 | 0,373103597 | 1,295135996 | 0,70271 | 0,85665929 | CD200 receptor 1 |
| ENSG00000165905 | LARGE2 | 0,373079203 | 1,295114098 | 0,53472 | 0,7541464 | LARGE xylosyl- and glucuronyltransferase 2 |
| ENSG00000114316 | USP4 | 0,373067276 | 1,295103391 | 0,03205 | 0,24164117 | ubiquitin specific peptidase 4 |
| ENSG00000117528 | ABCD3 | 0,372976077 | 1,295021524 | 0,4075 | 0,66665981 | ATP binding cassette subfamily D member 3 |
| ENSG00000287543 | AC013486,1 | 0,372933581 | 1,294983379 | 0,39964 | 0,66184353 | novel transcript, antisense to PML |
| ENSG00000182985 | CADM1 | 0,372865128 | 1,294921936 | 0,25119 | 0,55719428 | cell adhesion molecule 1 |
| ENSG00000166340 | TPP1 | 0,3728117 | 1,294873981 | 0,08847 | 0,38476379 | tripeptidyl peptidase 1 |
| ENSG00000117114 | ADGRL2 | 0,372717912 | 1,294789805 | 0,43164 | 0,68379763 | adhesion G protein-coupled receptor L2 |
| ENSG00000232354 | VIPR1-AS1 | 0,372629292 | 1,294710274 | 0,74227 | 0,87813769 | VIPR1 antisense RNA 1 |
| ENSG00000286943 | AP001992,2 | 0,372595659 | 1,29468009 | 0,34208 | 0,61993771 | novel transcript, antisense to XRRA1 |
| ENSG00000136895 | GARNL3 | 0,372557022 | 1,294645419 | 0,20452 | 0,52439728 | GTPase activating Rap/RanGAP domain like 3 |
| ENSG00000072736 | NFATC3 | 0,372523262 | 1,294615123 | 0,14724 | 0,47471707 | nuclear factor of activated T cells 3 |
| ENSG00000066697 | MSANTD3 | 0,372435054 | 1,294535971 | 0,27662 | 0,575468 | Myb/SANT DNA binding domain containing 3 |
| ENSG00000256206 | AC018523,2 | 0,372239803 | 1,294360784 | 0,76782 | 0,89249843 | novel protein |
| ENSG00000011105 | TSPAN9 | 0,372068995 | 1,294207547 | 0,05532 | 0,31461308 | tetraspanin 9 |
| ENSG00000168806 | LCMT2 | 0,372052476 | 1,294192728 | 0,12187 | 0,44093327 | leucine carboxyl methyltransferase 2 |
| ENSG00000275549 | STPG3-AS1 | 0,371968202 | 1,294117131 | 0,70231 | 0,85651838 | STPG3 antisense RNA 1 |
| ENSG00000075239 | ACAT1 | 0,371946897 | 1,29409802 | 0,37502 | 0,64346511 | acetyl-CoA acetyltransferase 1 |

| | | | | | | |
|---|---|---|---|---|---|---|
| ENSG00000187522 | HSPA14 | 0,371891341 | 1,294048187 | 0,2165 | 0,53490925 | heat shock protein family A (Hsp70) member 14 |
| ENSG00000270580 | PKD1P6-NPIPI | 0,371873958 | 1,294032596 | 0,38917 | 0,65363446 | PKD1P6-NPIPP1 readthrough |
| ENSG00000255983 | AC007848,1 | 0,371862404 | 1,294022232 | 0,59323 | 0,79162636 | novel transcript |
| ENSG00000112977 | DAP | 0,371860794 | 1,294020788 | 0,26109 | 0,56416552 | death associated protein |
| ENSG00000146731 | CCT6A | 0,371803038 | 1,293968985 | 0,21402 | 0,53327234 | chaperonin containing TCP1 subunit 6A |
| ENSG00000134256 | CD101 | 0,371675821 | 1,293854888 | 0,24492 | 0,55288356 | CD101 molecule |
| ENSG00000288548 | AL035653,1 | 0,371648466 | 1,293830355 | 0,42848 | 0,68117835 | novel transcript |
| ENSG00000261405 | AC136944,3 | 0,371614855 | 1,293800213 | 0,66962 | 0,83623395 | CHK2 checkpoint homolog (S, pombe) (CHEK2) pseudogene |
| ENSG00000134597 | RBMX2 | 0,371541604 | 1,293734523 | 0,31459 | 0,60123933 | RNA binding motif protein X-linked 2 |
| ENSG00000234893 | HCG18 | 0,371516526 | 1,293712035 | 0,31656 | 0,60241661 | HLA complex group 18 |
| ENSG00000251022 | THAP9-AS1 | 0,371357843 | 1,293569747 | 0,13803 | 0,46527914 | THAP9 antisense RNA 1 |
| ENSG00000007341 | ST7L | 0,371353096 | 1,29356549 | 0,21893 | 0,53639845 | suppression of tumorigenicity 7 like |
| ENSG00000145416 | MARCHF1 | 0,371209093 | 1,293436379 | 0,45544 | 0,69942891 | membrane associated ring-CH-type finger 1 |
| ENSG00000140395 | WDR61 | 0,371195412 | 1,293424113 | 0,24763 | 0,55506866 | WD repeat domain 61 |
| ENSG00000080815 | PSEN1 | 0,371112761 | 1,293350016 | 0,29836 | 0,59076323 | presenilin 1 |
| ENSG00000079101 | CLUL1 | 0,371073902 | 1,293315181 | 0,42643 | 0,67987951 | clusterin like 1 |
| ENSG00000166482 | MFAP4 | 0,371073354 | 1,293314689 | 0,13951 | 0,46674345 | microfibril associated protein 4 |
| ENSG00000186866 | POFUT2 | 0,371041423 | 1,293286064 | 0,03974 | 0,26952004 | protein O-fucosyltransferase 2 |
| ENSG00000111790 | FGFR1OP2 | 0,370858223 | 1,293121848 | 0,30041 | 0,59179349 | FGFR1 oncogene partner 2 |
| ENSG00000259316 | AC087632,2 | 0,370785435 | 1,293056608 | 0,48672 | 0,72212344 | novel protein |
| ENSG00000197969 | VPS13A | 0,370783526 | 1,293054897 | 0,39552 | 0,6582534 | vacuolar protein sorting 13 homolog A |
| ENSG00000286328 | AC079052,1 | 0,37077714 | 1,293049173 | 0,56422 | 0,77278544 | novel transcript |
| ENSG00000258983 | AL162171,2 | 0,370661047 | 1,292945126 | 0,5128 | 0,73941533 | novel transcript, antisense to PTPN21 |
| ENSG00000143379 | SETDB1 | 0,37063505 | 1,292921828 | 0,20433 | 0,52425916 | SET domain bifurcated histone lysine methyltransferase 1 |
| ENSG00000100196 | KDELR3 | 0,370552593 | 1,292847933 | 0,27199 | 0,573058 | KDEL endoplasmic reticulum protein retention receptor 3 |
| ENSG00000030582 | GRN | 0,370456591 | 1,292761905 | 0,07881 | 0,36667857 | granulin precursor |
| ENSG00000132122 | SPATA6 | 0,370444175 | 1,292750779 | 0,32508 | 0,60753927 | spermatogenesis associated 6 |
| ENSG00000121210 | TMEM131L | 0,37040811 | 1,292718464 | 0,31154 | 0,59903482 | transmembrane 131 like |
| ENSG00000270733 | AL355877,1 | 0,370405655 | 1,292716263 | 0,75938 | 0,88779217 | mitochondrial ribosomal protein S6 (MRPS6) pseudogene |
| ENSG00000104853 | CLPTM1 | 0,370372999 | 1,292687003 | 0,15747 | 0,48457045 | CLPTM1 regulator of GABA type A receptor forward trafficking |
| ENSG00000165699 | TSC1 | 0,370284204 | 1,292607443 | 0,12936 | 0,45283544 | TSC complex subunit 1 |
| ENSG00000115677 | HDLBP | 0,370249986 | 1,292576785 | 0,24241 | 0,55136766 | high density lipoprotein binding protein |

| | | | | | | |
|---|---|---|---|---|---|---|
| ENSG00000076043 | REXO2 | 0,370134221 | 1,29247307 | 0,07731 | 0,36406606 | RNA exonuclease 2 |
| ENSG00000120837 | NFYB | 0,370133069 | 1,292472038 | 0,26156 | 0,56452671 | nuclear transcription factor Y subunit beta |
| ENSG00000114491 | UMPS | 0,370107223 | 1,292448884 | 0,18115 | 0,50470865 | uridine monophosphate synthetase |
| ENSG00000272109 | AC008906,2 | 0,370077927 | 1,292422639 | 0,67002 | 0,83640106 | novel transcript, antisense to ERAP1 |
| ENSG00000163558 | PRKCI | 0,369941527 | 1,292300452 | 0,13276 | 0,45830601 | protein kinase C iota |
| ENSG00000021826 | CPS1 | 0,369876045 | 1,292241797 | 0,33211 | 0,61206889 | carbamoyl-phosphate synthase 1 |
| ENSG00000242372 | EIF6 | 0,36980643 | 1,292179444 | 0,26025 | 0,56382982 | eukaryotic translation initiation factor 6 |
| ENSG00000159131 | GART | 0,369768266 | 1,292145262 | 0,20877 | 0,52791772 | phosphoribosylglycinamide formyltransferase, phosphoribosylglycinamide synthetase, phosphoribosylaminoimidazole synthetase |
| ENSG00000010072 | SPRTN | 0,369674075 | 1,292060903 | 0,19916 | 0,52017864 | SprT-like N-terminal domain |
| ENSG00000048828 | FAM120A | 0,369428085 | 1,291840616 | 0,22413 | 0,54002595 | family with sequence similarity 120A |
| ENSG00000261101 | AC234775,2 | 0,369340175 | 1,291761901 | 0,34173 | 0,6197312 | novel transcript, overlapping to ARMCX3 |
| ENSG00000237135 | DDX10P1 | 0,369144859 | 1,29158703 | 0,57808 | 0,78191189 | DEAD-box helicase 10 pseudogene 1 |
| ENSG00000100207 | TCF20 | 0,368927954 | 1,291392858 | 0,14327 | 0,47082074 | transcription factor 20 |
| ENSG00000266412 | NCOA4 | 0,36892321 | 1,291388612 | 0,2763 | 0,57542877 | nuclear receptor coactivator 4 |
| ENSG00000125454 | SLC25A19 | 0,36888594 | 1,291355251 | 0,11673 | 0,43375876 | solute carrier family 25 member 19 |
| ENSG00000181788 | SIAH2 | 0,368856244 | 1,29132867 | 0,1379 | 0,46501478 | siah E3 ubiquitin protein ligase 2 |
| ENSG00000210144 | MT-TY | 0,368836601 | 1,291311088 | 0,60882 | 0,8014808 | mitochondrially encoded tRNA-Tyr (UAU/C) |
| ENSG00000178982 | EIF3K | 0,368768672 | 1,291250289 | 0,23271 | 0,54542294 | eukaryotic translation initiation factor 3 subunit K |
| ENSG00000181381 | DDX60L | 0,368748533 | 1,291232264 | 0,33853 | 0,61774255 | DExD/H-box 60 like |
| ENSG00000006007 | GDE1 | 0,368471783 | 1,290984593 | 0,32116 | 0,60573903 | glycerophosphodiester phosphodiesterase 1 |
| ENSG00000120688 | WBP4 | 0,368437812 | 1,290954195 | 0,2248 | 0,54036191 | WW domain binding protein 4 |
| ENSG00000215717 | TMEM167B | 0,368427051 | 1,290944566 | 0,35578 | 0,63013888 | transmembrane protein 167B |
| ENSG00000169519 | METTL15 | 0,368300641 | 1,290831457 | 0,27982 | 0,57750968 | methyltransferase like 15 |
| ENSG00000102710 | SUPT20H | 0,368286342 | 1,290818663 | 0,3722 | 0,64119785 | SPT20 homolog, SAGA complex component |
| ENSG00000198417 | MT1F | 0,368031904 | 1,290591031 | 0,46535 | 0,70583978 | metallothionein 1F |
| ENSG00000164144 | ARFIP1 | 0,368011599 | 1,290572867 | 0,20395 | 0,52366423 | ADP ribosylation factor interacting protein 1 |
| ENSG00000115421 | PAPOLG | 0,367966212 | 1,290532265 | 0,26692 | 0,56935824 | poly(A) polymerase gamma |
| ENSG00000040341 | STAU2 | 0,36779252 | 1,290376902 | 0,38711 | 0,65213944 | staufen double-stranded RNA binding protein 2 |
| ENSG00000186704 | DTX2P1 | 0,367733331 | 1,290323964 | 0,52615 | 0,7497214 | DTX2 pseudogene 1 |
| ENSG00000131013 | PPIL4 | 0,367700018 | 1,290294169 | 0,23607 | 0,54714939 | peptidylprolyl isomerase like 4 |
| ENSG00000159110 | IFNAR2 | 0,367699926 | 1,290294087 | 0,19455 | 0,51562158 | interferon alpha and beta receptor subunit 2 |

| | | | | | | |
|---|---|---|---|---|---|---|
| ENSG00000115145 | STAM2 | 0,367646774 | 1,290246551 | 0,4175 | 0,67403924 | signal transducing adaptor molecule 2 |
| ENSG00000138073 | PREB | 0,36763852 | 1,290239169 | 0,12611 | 0,44869905 | prolactin regulatory element binding |
| ENSG00000106624 | AEBP1 | 0,367553419 | 1,290163063 | 0,19934 | 0,52034382 | AE binding protein 1 |
| ENSG00000162384 | CZIB | 0,367540353 | 1,290151379 | 0,20891 | 0,52800078 | CXXC motif containing zinc binding protein |
| ENSG00000183580 | FBXL7 | 0,367490131 | 1,290106468 | 0,12145 | 0,44080285 | F-box and leucine rich repeat protein 7 |
| ENSG00000198315 | ZKSCAN8 | 0,367459061 | 1,290078684 | 0,3346 | 0,61433133 | zinc finger with KRAB and SCAN domains 8 |
| ENSG00000275769 | AC068792,1 | 0,367251167 | 1,289892796 | 0,56381 | 0,77259134 | novel transcript, sense intronic to DENND5B |
| ENSG00000116199 | FAM20B | 0,367200687 | 1,289847663 | 0,06647 | 0,34124715 | FAM20B glycosaminoglycan xylosylkinase |
| ENSG00000288534 | AP001931,2 | 0,367179365 | 1,289828601 | 0,28138 | 0,578832 | TMX2-CTNND1 readthrough (NMD candidate) |
| ENSG00000169116 | PARM1 | 0,367033045 | 1,289697791 | 0,4172 | 0,6738387 | prostate androgen-regulated mucin-like protein 1 |
| ENSG00000228730 | LINC02013 | 0,367000833 | 1,289668995 | 0,49493 | 0,72730301 | long intergenic non-protein coding RNA 2013 |
| ENSG00000147419 | CCDC25 | 0,366945909 | 1,289619898 | 0,2961 | 0,58955259 | coiled-coil domain containing 25 |
| ENSG00000085999 | RAD54L | 0,366901346 | 1,289580064 | 0,1262 | 0,44887364 | RAD54 like |
| ENSG00000249006 | AL136360,1 | 0,366865081 | 1,289547648 | 0,61086 | 0,80265373 | pseudogene similar to part of HBS1-like |
| ENSG00000166261 | ZNF202 | 0,36681609 | 1,289503859 | 0,2415 | 0,55037176 | zinc finger protein 202 |
| ENSG00000276996 | SLC6A3 | 0,366747351 | 1,28944242 | 0,56343 | 0,77245642 | solute carrier family 6 member 3 |
| ENSG00000078403 | MLLT10 | 0,366723246 | 1,289420876 | 0,28138 | 0,578832 | MLLT10 histone lysine methyltransferase DOT1L cofactor |
| ENSG00000273420 | AC008738,7 | 0,366640407 | 1,28934684 | 0,72974 | 0,8715818 | novel transcript |
| ENSG00000138101 | DTNB | 0,366570336 | 1,289284218 | 0,05316 | 0,30971127 | dystrobrevin beta |
| ENSG00000117650 | NEK2 | 0,366417053 | 1,289147242 | 0,24517 | 0,55295131 | NIMA related kinase 2 |
| ENSG00000170322 | NFRKB | 0,366383985 | 1,289117694 | 0,24986 | 0,55657567 | nuclear factor related to kappaB binding protein |
| ENSG00000118655 | DCLRE1B | 0,366362589 | 1,289098576 | 0,24657 | 0,55423069 | DNA cross-link repair 1B |
| ENSG00000127989 | MTERF1 | 0,366293097 | 1,289036483 | 0,3286 | 0,60992716 | mitochondrial transcription termination factor 1 |
| ENSG00000128789 | PSMG2 | 0,366269626 | 1,289015512 | 0,34893 | 0,62514443 | proteasome assembly chaperone 2 |
| ENSG00000172469 | MANEA | 0,366229657 | 1,288979802 | 0,55138 | 0,76514664 | mannosidase endo-alpha |
| ENSG00000188501 | LCTL | 0,366173431 | 1,288929567 | 0,11421 | 0,42953156 | lactase like |
| ENSG00000182580 | EPHB3 | 0,366145083 | 1,288904241 | 0,08794 | 0,38386734 | EPH receptor B3 |
| ENSG00000167986 | DDB1 | 0,366122599 | 1,288884154 | 0,19087 | 0,51394498 | damage specific DNA binding protein 1 |
| ENSG00000269235 | ZNF350-AS1 | 0,366121263 | 1,288882961 | 0,31474 | 0,60135222 | ZNF350 antisense RNA 1 |
| ENSG00000118200 | CAMSAP2 | 0,366069746 | 1,288836937 | 0,33651 | 0,61658848 | calmodulin regulated spectrin associated protein family member 2 |
| ENSG00000043514 | TRIT1 | 0,36601491 | 1,28878795 | 0,17133 | 0,49656842 | tRNA isopentenyltransferase 1 |
| ENSG00000101084 | RAB5IF | 0,366009828 | 1,28878341 | 0,24407 | 0,55253387 | RAB5 interacting factor |

| | | | | | | |
|---|---|---|---|---|---|---|
| ENSG00000224578 | HNRNPA1P48 | 0,365979171 | 1,288756024 | 0,43967 | 0,68844641 | heterogeneous nuclear ribonucleoprotein A1 pseudogene 48 |
| ENSG00000241685 | ARPC1A | 0,365970357 | 1,28874815 | 0,17407 | 0,49860418 | actin related protein 2/3 complex subunit 1A |
| ENSG00000162927 | PUS10 | 0,365938543 | 1,288719731 | 0,12458 | 0,44559634 | pseudouridine synthase 10 |
| ENSG00000149532 | CPSF7 | 0,365902485 | 1,288687522 | 0,19988 | 0,5208744 | cleavage and polyadenylation specific factor 7 |
| ENSG00000241684 | ADAMTS9-AS2 | 0,365894983 | 1,288680821 | 0,44291 | 0,69042854 | ADAMTS9 antisense RNA 2 |
| ENSG00000089234 | BRAP | 0,365889985 | 1,288676357 | 0,25135 | 0,55724078 | BRCA1 associated protein |
| ENSG00000171793 | CTPS1 | 0,365878952 | 1,288666502 | 0,0497 | 0,30003295 | CTP synthase 1 |
| ENSG00000140988 | RPS2 | 0,365785394 | 1,288582935 | 0,17553 | 0,49954066 | ribosomal protein S2 |
| ENSG00000124549 | BTN2A3P | 0,365729172 | 1,28853272 | 0,09835 | 0,40379437 | butyrophilin subfamily 2 member A3, pseudogene |
| ENSG00000132170 | PPARG | 0,365661631 | 1,288472397 | 0,57467 | 0,77982798 | peroxisome proliferator activated receptor gamma |
| ENSG00000152822 | GRM1 | 0,365641292 | 1,288454233 | 0,43102 | 0,68331372 | glutamate metabotropic receptor 1 |
| ENSG00000281886 | TGM4 | 0,365627889 | 1,288442262 | 0,58911 | 0,78855713 | transglutaminase 4 |
| ENSG00000163810 | TGM4 | 0,365627889 | 1,288442262 | 0,58911 | 0,78855713 | transglutaminase 4 |
| ENSG00000162599 | NFIA | 0,36560329 | 1,288420294 | 0,5418 | 0,75882662 | nuclear factor I A |
| ENSG00000213467 | HMGB1P37 | 0,365584345 | 1,288403375 | 0,69922 | 0,8547152 | high mobility group box 1 pseudogene 37 |
| ENSG00000111237 | VPS29 | 0,365578813 | 1,288398435 | 0,39134 | 0,65509868 | VPS29 retromer complex component |
| ENSG00000138433 | CIR1 | 0,365559102 | 1,288380832 | 0,25238 | 0,55796133 | corepressor interacting with RBPJ, CIR1 |
| ENSG00000143155 | TIPRL | 0,3655512 | 1,288373775 | 0,31983 | 0,60470232 | TOR signaling pathway regulator |
| ENSG00000168066 | SF1 | 0,365548572 | 1,288371428 | 0,13096 | 0,45558936 | splicing factor 1 |
| ENSG00000155974 | GRIP1 | 0,365542543 | 1,288366044 | 0,3622 | 0,63365406 | glutamate receptor interacting protein 1 |
| ENSG00000110851 | PRDM4 | 0,365529486 | 1,288354384 | 0,15871 | 0,48612005 | PR/SET domain 4 |
| ENSG00000206450 | HLA-B | 0,365448913 | 1,288282432 | 0,45625 | 0,70003941 | major histocompatibility complex, class I, B |
| ENSG00000276475 | AC016769,4 | 0,36531004 | 1,28815843 | 0,44753 | 0,69342177 | BCL2/adenovirus E1B 19kDa interacting protein 3 (BNIP3) pseudogene |
| ENSG00000196591 | HDAC2 | 0,365302927 | 1,288152079 | 0,23151 | 0,54440987 | histone deacetylase 2 |
| ENSG00000156976 | EIF4A2 | 0,365296034 | 1,288145924 | 0,1726 | 0,49744497 | eukaryotic translation initiation factor 4A2 |
| ENSG00000272084 | AL137127,1 | 0,365219516 | 1,288077604 | 0,227 | 0,54168591 | novel transcript |
| ENSG00000281690 | ADAMTS12 | 0,365114273 | 1,287983644 | 0,445 | 0,69205935 | ADAM metallopeptidase with thrombospondin type 1 motif 12 |
| ENSG00000279247 | AK6 | 0,365075291 | 1,287948843 | 0,4097 | 0,6680455 | adenylate kinase 6 |
| ENSG00000135241 | PNPLA8 | 0,364929387 | 1,287818595 | 0,41973 | 0,67539312 | patatin like phospholipase domain containing 8 |
| ENSG00000166598 | HSP90B1 | 0,364760632 | 1,287667966 | 0,12872 | 0,45196697 | heat shock protein 90 beta family member 1 |

| | | | | | | |
|---|---|---|---|---|---|---|
| ENSG00000138593 | SECISBP2L | 0,364728358 | 1,28763916 | 0,32448 | 0,6070377 | SECIS binding protein 2 like |
| ENSG00000106554 | CHCHD3 | 0,364629226 | 1,287550685 | 0,19557 | 0,51643293 | coiled-coil-helix-coiled-coil-helix domain containing 3 |
| ENSG00000127955 | GNAI1 | 0,364567421 | 1,287495528 | 0,2276 | 0,54168591 | G protein subunit alpha i1 |
| ENSG00000081154 | PCNP | 0,364556509 | 1,28748579 | 0,2909 | 0,58590649 | PEST proteolytic signal containing nuclear protein |
| ENSG00000144597 | EAF1 | 0,364545665 | 1,287476113 | 0,29957 | 0,59151326 | ELL associated factor 1 |
| ENSG00000164687 | FABP5 | 0,364482411 | 1,287419665 | 0,4904 | 0,72460356 | fatty acid binding protein 5 |
| ENSG00000136381 | IREB2 | 0,364474249 | 1,287412382 | 0,12017 | 0,43872715 | iron responsive element binding protein 2 |
| ENSG00000138758 | SEPTIN11 | 0,364449863 | 1,287390621 | 0,24826 | 0,55542983 | septin 11 |
| ENSG00000109670 | FBXW7 | 0,364442742 | 1,287384266 | 0,23165 | 0,54440987 | F-box and WD repeat domain containing 7 |
| ENSG00000116830 | TTF2 | 0,3643692 | 1,287318642 | 0,08094 | 0,37135332 | transcription termination factor 2 |
| ENSG00000108984 | MAP2K6 | 0,364316619 | 1,287271726 | 0,21973 | 0,53713164 | mitogen-activated protein kinase kinase 6 |
| ENSG00000226599 | AL450344,2 | 0,364292651 | 1,28725034 | 0,70012 | 0,85521681 | novel transcript |
| ENSG00000155016 | CYP2U1 | 0,364144173 | 1,287117866 | 0,26326 | 0,56608528 | cytochrome P450 family 2 subfamily U member 1 |
| ENSG00000131023 | LATS1 | 0,364128832 | 1,28710418 | 0,30845 | 0,59731659 | large tumor suppressor kinase 1 |
| ENSG00000114062 | UBE3A | 0,364082469 | 1,287062818 | 0,22852 | 0,5424199 | ubiquitin protein ligase E3A |
| ENSG00000272699 | AC007620,3 | 0,364016104 | 1,287003614 | 0,56855 | 0,77613996 | novel transcript, antisense to GNB4 |
| ENSG00000106588 | PSMA2 | 0,363822339 | 1,286830771 | 0,13894 | 0,46633113 | proteasome 20S subunit alpha 2 |
| ENSG00000229920 | RPS4XP5 | 0,363777248 | 1,286790552 | 0,59271 | 0,79133235 | ribosomal protein S4X pseudogene 5 |
| ENSG00000139445 | FOXN4 | 0,363763125 | 1,286777955 | 0,30561 | 0,59510377 | forkhead box N4 |
| ENSG00000156795 | NTAQ1 | 0,363757942 | 1,286773332 | 0,32033 | 0,6052338 | N-terminal glutamine amidase 1 |
| ENSG00000018408 | WWTR1 | 0,363741497 | 1,286758664 | 0,25774 | 0,56191608 | WW domain containing transcription regulator 1 |
| ENSG00000109332 | UBE2D3 | 0,363658894 | 1,286684992 | 0,24606 | 0,55380958 | ubiquitin conjugating enzyme E2 D3 |
| ENSG00000198862 | LTN1 | 0,363627565 | 1,286657051 | 0,36608 | 0,63617598 | listerin E3 ubiquitin protein ligase 1 |
| ENSG00000117598 | PLPPR5 | 0,363588827 | 1,286622503 | 0,61018 | 0,80220904 | phospholipid phosphatase related 5 |
| ENSG00000104312 | RIPK2 | 0,363574649 | 1,286609859 | 0,30624 | 0,59517372 | receptor interacting serine/threonine kinase 2 |
| ENSG00000229990 | AC022400,2 | 0,363543419 | 1,286582009 | 0,58724 | 0,78717516 | novel transcript, antisense to CAMK2G |
| ENSG00000145041 | DCAF1 | 0,363534145 | 1,286573738 | 0,32359 | 0,60633847 | DDB1 and CUL4 associated factor 1 |
| ENSG00000278709 | NKILA | 0,363516114 | 1,286557658 | 0,35116 | 0,62664963 | NF-kappaB interacting lncRNA |
| ENSG00000151348 | EXT2 | 0,363319301 | 1,286382157 | 0,15306 | 0,48120112 | exostosin glycosyltransferase 2 |
| ENSG00000272196 | H2AC19 | 0,363286985 | 1,286353343 | 0,47383 | 0,71294959 | H2A clustered histone 19 |
| ENSG00000236287 | ZBED5 | 0,363279391 | 1,286346572 | 0,26874 | 0,57078754 | zinc finger BED-type containing 5 |
| ENSG00000064763 | FAR2 | 0,363160651 | 1,286240705 | 0,30935 | 0,59780895 | fatty acyl-CoA reductase 2 |
| ENSG00000156515 | HK1 | 0,363007428 | 1,286104105 | 0,19367 | 0,51547251 | hexokinase 1 |

| | | | | | | |
|---|---|---|---|---|---|---|
| ENSG00000172586 | CHCHD1 | 0,362990623 | 1,286089124 | 0,19508 | 0,51588752 | coiled-coil-helix-coiled-coil-helix domain containing 1 |
| ENSG00000164296 | TIGD6 | 0,362931613 | 1,286036521 | 0,32104 | 0,6057178 | tigger transposable element derived 6 |
| ENSG00000012779 | ALOX5 | 0,362823366 | 1,285940032 | 0,74897 | 0,88190158 | arachidonate 5-lipoxygenase |
| ENSG00000144802 | NFKBIZ | 0,362811882 | 1,285929795 | 0,14467 | 0,47224655 | NFKB inhibitor zeta |
| ENSG00000100304 | TTLL12 | 0,362683417 | 1,285815295 | 0,10661 | 0,416363 | tubulin tyrosine ligase like 12 |
| ENSG00000198924 | DCLRE1A | 0,362640246 | 1,285776819 | 0,45251 | 0,69702004 | DNA cross-link repair 1A |
| ENSG00000076685 | NT5C2 | 0,362581898 | 1,285724818 | 0,27963 | 0,57745167 | 5'-nucleotidase, cytosolic II |
| ENSG00000116761 | CTH | 0,362375276 | 1,285540691 | 0,3112 | 0,59885232 | cystathionine gamma-lyase |
| ENSG00000148444 | COMMD3 | 0,362367568 | 1,285533822 | 0,43372 | 0,68475933 | COMM domain containing 3 |
| ENSG00000285392 | AC270276,1 | 0,36228216 | 1,28545772 | 0,50623 | 0,73467375 | novel transcript |
| ENSG00000005810 | MYCBP2 | 0,362165196 | 1,285353509 | 0,19163 | 0,51436129 | MYC binding protein 2 |
| ENSG00000184007 | PTP4A2 | 0,362140682 | 1,285331669 | 0,17006 | 0,49606609 | protein tyrosine phosphatase 4A2 |
| ENSG00000138686 | BBS7 | 0,361918904 | 1,285134096 | 0,36815 | 0,6380264 | Bardet-Biedl syndrome 7 |
| ENSG00000073711 | PPP2R3A | 0,361817831 | 1,285044065 | 0,26994 | 0,57171292 | protein phosphatase 2 regulatory subunit B''alpha |
| ENSG00000132466 | ANKRD17 | 0,361784382 | 1,285014271 | 0,26583 | 0,56872241 | ankyrin repeat domain 17 |
| ENSG00000214188 | ST7-OT4 | 0,361777023 | 1,285007716 | 0,27333 | 0,57344244 | ST7 overlapping transcript 4 |
| ENSG00000169714 | CNBP | 0,361620004 | 1,284867867 | 0,12891 | 0,45222045 | CCHC-type zinc finger nucleic acid binding protein |
| ENSG00000039319 | ZFYVE16 | 0,361511324 | 1,28477108 | 0,12045 | 0,43934584 | zinc finger FYVE-type containing 16 |
| ENSG00000162735 | PEX19 | 0,361509541 | 1,284769492 | 0,1765 | 0,50072935 | peroxisomal biogenesis factor 19 |
| ENSG00000213889 | PPM1N | 0,361475292 | 1,284738993 | 0,51592 | 0,7419128 | protein phosphatase, Mg2+/Mn2+ dependent 1N (putative) |
| ENSG00000110841 | PPFIBP1 | 0,361431291 | 1,28469981 | 0,04742 | 0,29249206 | PPFIA binding protein 1 |
| ENSG00000168283 | BMI1 | 0,361395683 | 1,284668102 | 0,25454 | 0,55944852 | BMI1 proto-oncogene, polycomb ring finger |
| ENSG00000119509 | INVS | 0,361284488 | 1,284569091 | 0,2716 | 0,57291418 | inversin |
| ENSG00000244038 | DDOST | 0,361195964 | 1,284490272 | 0,07567 | 0,36117248 | dolichyl-diphosphooligosaccharide--protein glycosyltransferase non-catalytic subunit |
| ENSG00000262111 | TAS2R30 | 0,361041003 | 1,284352311 | 0,54947 | 0,76407481 | taste 2 receptor member 30 |
| ENSG00000256188 | TAS2R30 | 0,361041003 | 1,284352311 | 0,54947 | 0,76407481 | taste 2 receptor member 30 |
| ENSG00000111962 | UST | 0,361037049 | 1,284348791 | 0,16514 | 0,49131595 | uronyl 2-sulfotransferase |
| ENSG00000272434 | AC137630,3 | 0,361015158 | 1,284329303 | 0,62596 | 0,81058216 | novel transcript, antisense to IMPDH2 and QRICH1 |
| ENSG00000205571 | SMN2 | 0,361009087 | 1,284323898 | 0,3136 | 0,60035326 | survival of motor neuron 2, centromeric |
| ENSG00000148459 | PDSS1 | 0,360971964 | 1,284290851 | 0,27662 | 0,575468 | decaprenyl diphosphate synthase subunit 1 |
| ENSG00000253317 | AC078906,1 | 0,360950689 | 1,284271912 | 0,66795 | 0,83525939 | novel transcript |

| | | | | | | |
|---|---|---|---|---|---|---|
| ENSG00000042445 | RETSAT | 0,36092708 | 1,284250895 | 0,13505 | 0,46159318 | retinol saturase |
| ENSG00000129317 | PUS7L | 0,360873393 | 1,284203105 | 0,3223 | 0,60577115 | pseudouridine synthase 7 like |
| ENSG00000005893 | LAMP2 | 0,360666323 | 1,284018797 | 0,24089 | 0,55001001 | lysosomal associated membrane protein 2 |
| ENSG00000277728 | AC097641,2 | 0,360628165 | 1,283984837 | 0,63855 | 0,81860204 | novel transcript, antisense to COG1 |
| ENSG00000176165 | FOXG1 | 0,360574152 | 1,283936766 | 0,47288 | 0,71217607 | forkhead box G1 |
| ENSG00000166359 | WDR88 | 0,360572569 | 1,283935358 | 0,6157 | 0,80456457 | WD repeat domain 88 |
| ENSG00000101138 | CSTF1 | 0,360449422 | 1,283825767 | 0,11617 | 0,43268574 | cleavage stimulation factor subunit 1 |
| ENSG00000185495 | AC138393,1 | 0,360288299 | 1,283682395 | 0,35298 | 0,62780446 | septin 7 (SEPT7) pseudogene |
| ENSG00000134265 | NAPG | 0,360248506 | 1,283646989 | 0,29128 | 0,5862741 | NSF attachment protein gamma |
| ENSG00000168653 | NDUFS5 | 0,360221426 | 1,283622894 | 0,24384 | 0,55243616 | NADH:ubiquinone oxidoreductase subunit S5 |
| ENSG00000266976 | AC079466,1 | 0,360111903 | 1,283525451 | 0,41391 | 0,67094348 | novel transcript |
| ENSG00000137955 | RABGGTB | 0,360110335 | 1,283524055 | 0,10268 | 0,41043823 | Rab geranylgeranyltransferase subunit beta |
| ENSG00000103852 | TTC23 | 0,360099122 | 1,28351408 | 0,20666 | 0,52622584 | tetratricopeptide repeat domain 23 |
| ENSG00000226194 | LINC02519 | 0,360082759 | 1,283499522 | 0,55541 | 0,76788267 | long intergenic non-protein coding RNA 2519 |
| ENSG00000188958 | UTS2B | 0,360000268 | 1,283426136 | 0,17809 | 0,50236177 | urotensin 2B |
| ENSG00000069329 | VPS35 | 0,359995341 | 1,283421753 | 0,40037 | 0,66250081 | VPS35 retromer complex component |
| ENSG00000232827 | LINC01189 | 0,35998703 | 1,28341436 | 0,72201 | 0,86726498 | long intergenic non-protein coding RNA 1189 |
| ENSG00000198265 | HELZ | 0,359968545 | 1,283397916 | 0,24561 | 0,55352055 | helicase with zinc finger |
| ENSG00000075891 | PAX2 | 0,359967904 | 1,283397345 | 0,74386 | 0,87876657 | paired box 2 |
| ENSG00000118260 | CREB1 | 0,359879043 | 1,283318299 | 0,27161 | 0,57291418 | cAMP responsive element binding protein 1 |
| ENSG00000119684 | MLH3 | 0,359865367 | 1,283306133 | 0,323 | 0,60584543 | mutL homolog 3 |
| ENSG00000271105 | SCML2P2 | 0,35984535 | 1,283288327 | 0,64835 | 0,82383528 | SCML2 pseudogene 2 |
| ENSG00000144857 | BOC | 0,359812896 | 1,28325946 | 0,04212 | 0,27783166 | BOC cell adhesion associated, oncogene regulated |
| ENSG00000205352 | PRR13 | 0,359806635 | 1,283253891 | 0,11231 | 0,42601117 | proline rich 13 |
| ENSG00000265217 | AC090358,1 | 0,359729175 | 1,283184993 | 0,445 | 0,69205935 | novel transcript |
| ENSG00000230624 | DDX39B | 0,359727022 | 1,283183078 | 0,31705 | 0,60268749 | DExD-box helicase 39B |
| ENSG00000227474 | RPL6P24 | 0,359675785 | 1,283137508 | 0,66104 | 0,83105713 | ribosomal protein L6 pseudogene 24 |
| ENSG00000288586 | AL357874,3 | 0,359670987 | 1,28313324 | 0,58155 | 0,78438328 | novel transcript |
| ENSG00000215077 | BRD2 | 0,3596502 | 1,283114752 | 0,12829 | 0,45171208 | bromodomain containing 2 |
| ENSG00000103044 | HAS3 | 0,35964351 | 1,283108802 | 0,27008 | 0,57186187 | hyaluronan synthase 3 |
| ENSG00000065526 | SPEN | 0,359593498 | 1,283064323 | 0,15603 | 0,48382611 | spen family transcriptional repressor |
| ENSG00000275714 | H3C1 | 0,359591846 | 1,283062854 | 0,32618 | 0,6085098 | H3 clustered histone 1 |
| ENSG00000122483 | CCDC18 | 0,359502812 | 1,282983674 | 0,34059 | 0,61904269 | coiled-coil domain containing 18 |

| | | | | | | |
|---|---|---|---|---|---|---|
| ENSG00000221869 | CEBPD | 0,35949727 | 1,282978745 | 0,68122 | 0,84367915 | CCAAT enhancer binding protein delta |
| ENSG00000234846 | NEU1 | 0,359490538 | 1,282972759 | 0,67082 | 0,83695077 | neuraminidase 1 |
| ENSG00000224844 | AC107079,1 | 0,359414672 | 1,282905294 | 0,62558 | 0,81053676 | novel transcript |
| ENSG00000185651 | UBE2L3 | 0,359318548 | 1,282819819 | 0,24532 | 0,55295131 | ubiquitin conjugating enzyme E2 L3 |
| ENSG00000182704 | TSKU | 0,35929984 | 1,282803185 | 0,03157 | 0,23991209 | tsukushi, small leucine rich proteoglycan |
| ENSG00000166783 | MARF1 | 0,359289729 | 1,282794194 | 0,34014 | 0,61889407 | meiosis regulator and mRNA stability factor 1 |
| ENSG00000164818 | DNAAF5 | 0,359252041 | 1,282760684 | 0,07526 | 0,36012885 | dynein axonemal assembly factor 5 |
| ENSG00000278553 | Metazoa_SRP | 0,359245129 | 1,282754537 | 0,74599 | 0,88016533 | Metazoan signal recognition particle RNA |
| ENSG00000214654 | B3GNT10 | 0,359087932 | 1,282614775 | 0,31739 | 0,60281557 | UDP-GlcNAc:betaGal beta-1,3-N-acetylglucosaminyltransferase 10 (putative) |
| ENSG00000151474 | FRMD4A | 0,359012212 | 1,282547459 | 0,30048 | 0,59184847 | FERM domain containing 4A |
| ENSG00000167515 | TRAPPC2L | 0,358990698 | 1,282528334 | 0,26119 | 0,56416552 | trafficking protein particle complex 2 like |
| ENSG00000108559 | NUP88 | 0,358963029 | 1,282503736 | 0,19815 | 0,51977271 | nucleoporin 88 |
| ENSG00000152443 | ZNF776 | 0,358926447 | 1,282471217 | 0,28282 | 0,58009943 | zinc finger protein 776 |
| ENSG00000205581 | HMGN1 | 0,358910502 | 1,282457043 | 0,27546 | 0,57518625 | high mobility group nucleosome binding domain 1 |
| ENSG00000185272 | RBM11 | 0,358906051 | 1,282453086 | 0,56052 | 0,77087723 | RNA binding motif protein 11 |
| ENSG00000187210 | GCNT1 | 0,358882115 | 1,282431809 | 0,47195 | 0,71123477 | glucosaminyl (N-acetyl) transferase 1 |
| ENSG00000101596 | SMCHD1 | 0,35887893 | 1,282428978 | 0,28535 | 0,58187605 | structural maintenance of chromosomes flexible hinge domain containing 1 |
| ENSG00000146242 | TPBG | 0,358824695 | 1,282380769 | 0,22461 | 0,54036191 | trophoblast glycoprotein |
| ENSG00000118217 | ATF6 | 0,358809022 | 1,282366837 | 0,25279 | 0,55796133 | activating transcription factor 6 |
| ENSG00000100316 | RPL3 | 0,358805485 | 1,282363693 | 0,18187 | 0,50521394 | ribosomal protein L3 |
| ENSG00000270504 | AL391422,4 | 0,358741847 | 1,282307129 | 0,49682 | 0,72858829 | novel transcript, antisense to PXDC1 |
| ENSG00000124091 | GCNT7 | 0,358739934 | 1,282305428 | 0,45813 | 0,70114321 | glucosaminyl (N-acetyl) transferase family member 7 |
| ENSG00000198874 | TYW1 | 0,358724572 | 1,282291775 | 0,18328 | 0,50677286 | tRNA-yW synthesizing protein 1 homolog |
| ENSG00000135940 | COX5B | 0,358582153 | 1,282165196 | 0,22501 | 0,5404013 | cytochrome c oxidase subunit 5B |
| ENSG00000122481 | RWDD3 | 0,358565816 | 1,282150677 | 0,45184 | 0,69682854 | RWD domain containing 3 |
| ENSG00000107949 | BCCIP | 0,358532004 | 1,282120628 | 0,22472 | 0,54036191 | BRCA2 and CDKN1A interacting protein |
| ENSG00000178295 | GEN1 | 0,358453763 | 1,282051097 | 0,36315 | 0,63407828 | GEN1 Holliday junction 5' flap endonuclease |
| ENSG00000175600 | SUGCT | 0,358450256 | 1,282047981 | 0,26527 | 0,56807781 | succinyl-CoA:glutarate-CoA transferase |
| ENSG00000242795 | AC007849,1 | 0,358296841 | 1,281911656 | 0,67553 | 0,8403281 | novel transcript |
| ENSG00000187607 | ZNF286A | 0,358173697 | 1,281802241 | 0,20261 | 0,52269757 | zinc finger protein 286A |
| ENSG00000121741 | ZMYM2 | 0,358115246 | 1,281750309 | 0,27135 | 0,57274077 | zinc finger MYM-type containing 2 |

| | | | | | | |
|---|---|---|---|---|---|---|
| ENSG00000170871 | KIAA0232 | 0,358096934 | 1,28173404 | 0,33899 | 0,61800397 | KIAA0232 |
| ENSG00000256537 | SMIM10L1 | 0,358080059 | 1,281719048 | 0,3272 | 0,60902638 | small integral membrane protein 10 like 1 |
| ENSG00000117505 | DR1 | 0,35806888 | 1,281709117 | 0,09816 | 0,40362152 | down-regulator of transcription 1 |
| ENSG00000270554 | AC069431,2 | 0,358060429 | 1,281701609 | 0,56687 | 0,77482787 | eukaryotic translation initiation factor 3, subunit E, pseudogene |
| ENSG00000154957 | ZNF18 | 0,358059476 | 1,281700762 | 0,21618 | 0,53480193 | zinc finger protein 18 |
| ENSG00000217330 | SSXP10 | 0,35800707 | 1,281654205 | 0,66305 | 0,83242715 | SSX family pseudogene 10 |
| ENSG00000166479 | TMX3 | 0,357994833 | 1,281643334 | 0,45208 | 0,69691985 | thioredoxin related transmembrane protein 3 |
| ENSG00000222404 | RN7SKP281 | 0,357938333 | 1,281593142 | 0,61151 | 0,80324115 | RN7SK pseudogene 281 |
| ENSG00000119013 | NDUFB3 | 0,35793751 | 1,281592411 | 0,27054 | 0,5723494 | NADH:ubiquinone oxidoreductase subunit B3 |
| ENSG00000177173 | NAP1L4P1 | 0,357901064 | 1,281560035 | 0,35176 | 0,62692823 | nucleosome assembly protein 1 like 4 pseudogene 1 |
| ENSG00000225697 | SLC26A6 | 0,357881518 | 1,281542672 | 0,10174 | 0,40892507 | solute carrier family 26 member 6 |
| ENSG00000278677 | H2AC17 | 0,357808044 | 1,281477408 | 0,51963 | 0,74404725 | H2A clustered histone 17 |
| ENSG00000005302 | MSL3 | 0,357773901 | 1,28144708 | 0,28825 | 0,58433599 | MSL complex subunit 3 |
| ENSG00000111640 | GAPDH | 0,357718473 | 1,281397848 | 0,2271 | 0,54168591 | glyceraldehyde-3-phosphate dehydrogenase |
| ENSG00000272573 | MUSTN1 | 0,357716587 | 1,281396173 | 0,75452 | 0,8850265 | musculoskeletal, embryonic nuclear protein 1 |
| ENSG00000164597 | COG5 | 0,35762232 | 1,281312448 | 0,41255 | 0,6702125 | component of oligomeric golgi complex 5 |
| ENSG00000167658 | EEF2 | 0,357512878 | 1,281215252 | 0,17106 | 0,49641416 | eukaryotic translation elongation factor 2 |
| ENSG00000104412 | EMC2 | 0,357482414 | 1,281188198 | 0,14153 | 0,46809977 | ER membrane protein complex subunit 2 |
| ENSG00000259439 | LINC01833 | 0,357427259 | 1,281139218 | 0,44648 | 0,69314849 | long intergenic non-protein coding RNA 1833 |
| ENSG00000090520 | DNAJB11 | 0,357419518 | 1,281132345 | 0,12528 | 0,44701864 | DnaJ heat shock protein family (Hsp40) member B11 |
| ENSG00000151148 | UBE3B | 0,357361512 | 1,281080836 | 0,14311 | 0,47072472 | ubiquitin protein ligase E3B |
| ENSG00000160766 | GBAP1 | 0,357297568 | 1,281024056 | 0,27972 | 0,57750542 | glucosylceramidase beta pseudogene 1 |
| ENSG00000236802 | B3GALT4 | 0,357282077 | 1,281010301 | 0,71223 | 0,8615121 | beta-1,3-galactosyltransferase 4 |
| ENSG00000226936 | B3GALT4 | 0,357282077 | 1,281010301 | 0,71223 | 0,8615121 | beta-1,3-galactosyltransferase 4 |
| ENSG00000235155 | B3GALT4 | 0,357282077 | 1,281010301 | 0,71223 | 0,8615121 | beta-1,3-galactosyltransferase 4 |
| ENSG00000198586 | TLK1 | 0,357227585 | 1,280961917 | 0,32423 | 0,60681275 | tousled like kinase 1 |
| ENSG00000254979 | AP000781,2 | 0,3572026 | 1,280939733 | 0,7994 | 0,90744841 | novel protein |
| ENSG00000230333 | AC004160,1 | 0,357170508 | 1,28091124 | 0,45244 | 0,69702004 | novel transcript, antisense to THSD7A |
| ENSG00000135655 | USP15 | 0,357145513 | 1,280889048 | 0,36272 | 0,63381706 | ubiquitin specific peptidase 15 |
| ENSG00000158711 | ELK4 | 0,357058059 | 1,280811404 | 0,06834 | 0,34555414 | ETS transcription factor ELK4 |
| ENSG00000127325 | BEST3 | 0,356927651 | 1,280695635 | 0,71204 | 0,8615121 | bestrophin 3 |
| ENSG00000271971 | AC120053,1 | 0,356766225 | 1,280552343 | 0,56896 | 0,77629764 | novel transcript, antisense to FAM92A1 |

| | | | | | | |
|---|---|---|---|---|---|---|
| ENSG00000126903 | SLC10A3 | 0,35670371 | 1,280496855 | 0,13218 | 0,45719971 | solute carrier family 10 member 3 |
| ENSG00000115163 | CENPA | 0,356590861 | 1,280396698 | 0,30801 | 0,59691305 | centromere protein A |
| ENSG00000184924 | PTRHD1 | 0,356544935 | 1,280355938 | 0,42289 | 0,67683766 | peptidyl-tRNA hydrolase domain containing 1 |
| ENSG00000144468 | RHBDD1 | 0,356539828 | 1,280351407 | 0,31314 | 0,59991314 | rhomboid domain containing 1 |
| ENSG00000185883 | ATP6V0C | 0,356480926 | 1,280299134 | 0,14968 | 0,47775345 | ATPase H+ transporting V0 subunit c |
| ENSG00000273271 | AP000254,2 | 0,356476452 | 1,280295163 | 0,58447 | 0,78572744 | novel transcript, antisense to SOD1 |
| ENSG00000284254 | HNRNPH1 | 0,356475761 | 1,28029455 | 0,25874 | 0,56260205 | heterogeneous nuclear ribonucleoprotein H1 |
| ENSG00000135624 | CCT7 | 0,356445621 | 1,280267803 | 0,12222 | 0,44171717 | chaperonin containing TCP1 subunit 7 |
| ENSG00000100109 | TFIP11 | 0,35640446 | 1,280231277 | 0,20727 | 0,52674983 | tuftelin interacting protein 11 |
| ENSG00000257086 | AP001453,2 | 0,356354226 | 1,280186701 | 0,53339 | 0,75351438 | novel transcript |
| ENSG00000226520 | KIRREL1-IT1 | 0,356316717 | 1,280153417 | 0,52248 | 0,74667423 | KIRREL1 intronic transcript 1 |
| ENSG00000149136 | SSRP1 | 0,356224337 | 1,280071448 | 0,19986 | 0,52086852 | structure specific recognition protein 1 |
| ENSG00000101844 | ATG4A | 0,356118815 | 1,279977823 | 0,4401 | 0,68865364 | autophagy related 4A cysteine peptidase |
| ENSG00000153147 | SMARCA5 | 0,356047663 | 1,279914699 | 0,38983 | 0,65396251 | SWI/SNF related, matrix associated, actin dependent regulator of chromatin, subfamily a, member 5 |
| ENSG00000127831 | VIL1 | 0,355985499 | 1,279859549 | 0,38259 | 0,6493677 | villin 1 |
| ENSG00000142892 | PIGK | 0,355921099 | 1,279802419 | 0,41989 | 0,67551392 | phosphatidylinositol glycan anchor biosynthesis class K |
| ENSG00000182004 | SNRPE | 0,355917355 | 1,279799098 | 0,41607 | 0,67297235 | small nuclear ribonucleoprotein polypeptide E |
| ENSG00000115474 | KCNJ13 | 0,355833071 | 1,279724333 | 0,43714 | 0,68711291 | potassium inwardly rectifying channel subfamily J member 13 |
| ENSG00000166908 | PIP4K2C | 0,355812535 | 1,279706117 | 0,12493 | 0,44652417 | phosphatidylinositol-5-phosphate 4-kinase type 2 gamma |
| ENSG00000123684 | LPGAT1 | 0,355724495 | 1,279628025 | 0,29439 | 0,58857672 | lysophosphatidylglycerol acyltransferase 1 |
| ENSG00000181315 | ZNF322 | 0,355705284 | 1,279610986 | 0,31722 | 0,6027392 | zinc finger protein 322 |
| ENSG00000119408 | NEK6 | 0,355692976 | 1,279600069 | 0,25833 | 0,56226755 | NIMA related kinase 6 |
| ENSG00000177868 | SVBP | 0,355665485 | 1,279575686 | 0,38429 | 0,65052153 | small vasohibin binding protein |
| ENSG00000166451 | CENPN | 0,355504252 | 1,279432692 | 0,34282 | 0,62033803 | centromere protein N |
| ENSG00000254858 | MPV17L2 | 0,355503161 | 1,279431724 | 0,16153 | 0,48906157 | MPV17 mitochondrial inner membrane protein like 2 |
| ENSG00000163029 | SMC6 | 0,355360096 | 1,279304856 | 0,25355 | 0,55857654 | structural maintenance of chromosomes 6 |
| ENSG00000070367 | EXOC5 | 0,355339558 | 1,279286644 | 0,39589 | 0,65863599 | exocyst complex component 5 |
| ENSG00000272419 | LINC01145 | 0,355318243 | 1,279267743 | 0,26791 | 0,56999323 | long intergenic non-protein coding RNA 1145 |
| ENSG00000132010 | ZNF20 | 0,355303212 | 1,279254415 | 0,27363 | 0,57364891 | zinc finger protein 20 |
| ENSG00000167996 | FTH1 | 0,355302657 | 1,279253923 | 0,33842 | 0,61774255 | ferritin heavy chain 1 |
| ENSG00000286855 | AC074024,1 | 0,355264599 | 1,279220176 | 0,65129 | 0,82534198 | novel transcript |

| | | | | | | |
|---|---|---|---|---|---|---|
| ENSG00000237335 | PFDN6 | 0,355236762 | 1,279195494 | 0,25418 | 0,55925777 | prefoldin subunit 6 |
| ENSG00000224782 | PFDN6 | 0,355236762 | 1,279195494 | 0,25418 | 0,55925777 | prefoldin subunit 6 |
| ENSG00000265798 | AC138207,6 | 0,355111969 | 1,279084848 | 0,4065 | 0,66597268 | leucine rich repeat containing 37B (LRRC37B) pseudogene |
| ENSG00000280529 | TAF4 | 0,355006239 | 1,278991113 | 0,39078 | 0,65464997 | TATA-box binding protein associated factor 4 |
| ENSG00000145050 | MANF | 0,354991216 | 1,278977794 | 0,25806 | 0,56217022 | mesencephalic astrocyte derived neurotrophic factor |
| ENSG00000065491 | TBC1D22B | 0,354946372 | 1,278938039 | 0,22217 | 0,53859118 | TBC1 domain family member 22B |
| ENSG00000087191 | PSMC5 | 0,354942358 | 1,278934482 | 0,25118 | 0,55719428 | proteasome 26S subunit, ATPase 5 |
| ENSG00000152193 | OBI1 | 0,354916927 | 1,278911937 | 0,21561 | 0,53440695 | ORC ubiquitin ligase 1 |
| ENSG00000170606 | HSPA4 | 0,354853186 | 1,278855434 | 0,17512 | 0,49924804 | heat shock protein family A (Hsp70) member 4 |
| ENSG00000108179 | PPIF | 0,35485181 | 1,278854213 | 0,11501 | 0,43048326 | peptidylprolyl isomerase F |
| ENSG00000277053 | GTF2IP1 | 0,354785947 | 1,278795832 | 0,39674 | 0,65932701 | general transcription factor IIi pseudogene 1 |
| ENSG00000281540 | RCC2 | 0,354770564 | 1,278782197 | 0,06248 | 0,33103715 | regulator of chromosome condensation 2 |
| ENSG00000179051 | RCC2 | 0,354770564 | 1,278782197 | 0,06248 | 0,33103715 | regulator of chromosome condensation 2 |
| ENSG00000255445 | AP003072,2 | 0,354731779 | 1,278747819 | 0,51792 | 0,74291729 | aminoacyl tRNA synthetase complex-interacting multifunctional protein 2 (AIMP2) pseudogene |
| ENSG00000134308 | YWHAQ | 0,354718797 | 1,278736312 | 0,30674 | 0,59579909 | tyrosine 3-monooxygenase/tryptophan 5-monooxygenase activation protein theta |
| ENSG00000113597 | TRAPPC13 | 0,354661668 | 1,278685676 | 0,3812 | 0,64856923 | trafficking protein particle complex 13 |
| ENSG00000215817 | ZC3H11B | 0,354658751 | 1,278683091 | 0,52875 | 0,75099628 | zinc finger CCCH-type containing 11B |
| ENSG00000119986 | AVPI1 | 0,354468227 | 1,278514238 | 0,39724 | 0,65973966 | arginine vasopressin induced 1 |
| ENSG00000138092 | CENPO | 0,354448595 | 1,27849684 | 0,27827 | 0,57645922 | centromere protein O |
| ENSG00000164022 | AIMP1 | 0,354431953 | 1,278482092 | 0,24059 | 0,54983486 | aminoacyl tRNA synthetase complex interacting multifunctional protein 1 |
| ENSG00000102935 | ZNF423 | 0,354362848 | 1,278420854 | 0,12045 | 0,43934584 | zinc finger protein 423 |
| ENSG00000100603 | SNW1 | 0,354357521 | 1,278416134 | 0,18231 | 0,50576743 | SNW domain containing 1 |
| ENSG00000085365 | SCAMP1 | 0,354284141 | 1,278351111 | 0,37757 | 0,64604267 | secretory carrier membrane protein 1 |
| ENSG00000273473 | BX649601,1 | 0,354213521 | 1,278288538 | 0,6451 | 0,82193166 | novel transcript |
| ENSG00000155959 | VBP1 | 0,354091492 | 1,27818042 | 0,34933 | 0,62549288 | VHL binding protein 1 |
| ENSG00000235109 | ZSCAN31 | 0,354049744 | 1,278143433 | 0,44015 | 0,68866128 | zinc finger and SCAN domain containing 31 |
| ENSG00000176407 | KCMF1 | 0,353977013 | 1,278078999 | 0,0998 | 0,40626368 | potassium channel modulatory factor 1 |
| ENSG00000092140 | G2E3 | 0,353969221 | 1,278072096 | 0,34574 | 0,62283284 | G2/M-phase specific E3 ubiquitin protein ligase |
| ENSG00000204272 | NBDY | 0,353967209 | 1,278070314 | 0,23158 | 0,54440987 | negative regulator of P-body association |
| ENSG00000169856 | ONECUT1 | 0,353894671 | 1,278006055 | 0,12264 | 0,44239528 | one cut homeobox 1 |

| | | | | | | |
|---|---|---|---|---|---|---|
| ENSG00000067445 | TRO | 0,353823997 | 1,27794345 | 0,06052 | 0,32669488 | trophinin |
| ENSG00000164002 | EXO5 | 0,353822056 | 1,277941731 | 0,06778 | 0,34426294 | exonuclease 5 |
| ENSG00000116918 | TSNAX | 0,353537886 | 1,277690037 | 0,09701 | 0,40141222 | translin associated factor X |
| ENSG00000214425 | LRRC37A4P | 0,3535204 | 1,277674551 | 0,36908 | 0,63899712 | leucine rich repeat containing 37 member A4, pseudogene |
| ENSG00000149260 | CAPN5 | 0,353363846 | 1,277535911 | 0,30236 | 0,59297237 | calpain 5 |
| ENSG00000182149 | IST1 | 0,353344518 | 1,277518797 | 0,08122 | 0,3719701 | IST1 factor associated with ESCRT-III |
| ENSG00000112312 | GMNN | 0,353161091 | 1,277356381 | 0,21581 | 0,53440695 | geminin DNA replication inhibitor |
| ENSG00000159128 | IFNGR2 | 0,353149038 | 1,277345709 | 0,32823 | 0,60971379 | interferon gamma receptor 2 |
| ENSG00000123146 | ADGRE5 | 0,353140132 | 1,277337824 | 0,40558 | 0,66528281 | adhesion G protein-coupled receptor E5 |
| ENSG00000130348 | QRSL1 | 0,352998969 | 1,277212846 | 0,21845 | 0,53578053 | glutaminyl-tRNA amidotransferase subunit QRSL1 |
| ENSG00000168538 | TRAPPC11 | 0,352890142 | 1,277116506 | 0,38934 | 0,65372387 | trafficking protein particle complex 11 |
| ENSG00000153896 | ZNF599 | 0,352886384 | 1,27711318 | 0,16392 | 0,4902029 | zinc finger protein 599 |
| ENSG00000109270 | LAMTOR3 | 0,352852578 | 1,277083253 | 0,31032 | 0,59842274 | late endosomal/lysosomal adaptor, MAPK and MTOR activator 3 |
| ENSG00000166398 | GARRE1 | 0,352831455 | 1,277064556 | 0,16199 | 0,48916554 | granule associated Rac and RHOG effector 1 |
| ENSG00000111231 | GPN3 | 0,352710867 | 1,276957816 | 0,17752 | 0,50196932 | GPN-loop GTPase 3 |
| ENSG00000286437 | AF111169,4 | 0,352666626 | 1,276918658 | 0,58433 | 0,78572744 | novel transcript, antisense to ANGEL1 |
| ENSG00000115109 | EPB41L5 | 0,35256676 | 1,276830271 | 0,32016 | 0,60511306 | erythrocyte membrane protein band 4,1 like 5 |
| ENSG00000142731 | PLK4 | 0,352497582 | 1,276769047 | 0,38773 | 0,65272082 | polo like kinase 4 |
| ENSG00000133134 | BEX2 | 0,352494015 | 1,27676589 | 0,28085 | 0,57822547 | brain expressed X-linked 2 |
| ENSG00000196670 | ZFP62 | 0,35245403 | 1,276730505 | 0,44768 | 0,69362056 | ZFP62 zinc finger protein |
| ENSG00000237827 | RPS15AP29 | 0,352412109 | 1,276693407 | 0,6046 | 0,79801151 | ribosomal protein S15a pseudogene 29 |
| ENSG00000205918 | PDPK2P | 0,352337336 | 1,27662724 | 0,18907 | 0,51243448 | 3-phosphoinositide dependent protein kinase 2, pseudogene |
| ENSG00000197323 | TRIM33 | 0,352239866 | 1,276540992 | 0,30763 | 0,59671558 | tripartite motif containing 33 |
| ENSG00000196862 | RGPD4 | 0,352221876 | 1,276525074 | 0,30239 | 0,59297237 | RANBP2 like and GRIP domain containing 4 |
| ENSG00000251455 | AC092611,1 | 0,352215224 | 1,276519188 | 0,71904 | 0,8656088 | novel transcript, antisense to PET112L |
| ENSG00000102543 | CDADC1 | 0,3522026 | 1,276508018 | 0,31103 | 0,59883009 | cytidine and dCMP deaminase domain containing 1 |
| ENSG00000139651 | ZNF740 | 0,352198444 | 1,276504341 | 0,04155 | 0,27616337 | zinc finger protein 740 |
| ENSG00000071243 | ING3 | 0,352185831 | 1,276493181 | 0,44743 | 0,69342177 | inhibitor of growth family member 3 |
| ENSG00000140263 | SORD | 0,35215738 | 1,276468008 | 0,15977 | 0,4870975 | sorbitol dehydrogenase |
| ENSG00000229152 | ANKRD10-IT1 | 0,352107448 | 1,27642383 | 0,48966 | 0,72400256 | ANKRD10 intronic transcript 1 |
| ENSG00000104450 | SPAG1 | 0,352030679 | 1,27635591 | 0,33169 | 0,61166253 | sperm associated antigen 1 |

| | | | | | | |
|---|---|---|---|---|---|---|
| ENSG00000276393 | AC105429,2 | 0,351992695 | 1,276322306 | 0,69037 | 0,84895522 | nuclear receptor subfamily 3, group C, member 1 (glucocorticoid receptor) (NR3C1) pseudogene |
| ENSG00000244513 | AC109587,1 | 0,35194487 | 1,276279997 | 0,35532 | 0,62970025 | novel transcript |
| ENSG00000147475 | ERLIN2 | 0,351897837 | 1,27623839 | 0,22027 | 0,53750545 | ER lipid raft associated 2 |
| ENSG00000185722 | ANKFY1 | 0,351856876 | 1,276202156 | 0,30744 | 0,59658791 | ankyrin repeat and FYVE domain containing 1 |
| ENSG00000122729 | ACO1 | 0,351843871 | 1,276190652 | 0,18923 | 0,51249536 | aconitase 1 |
| ENSG00000179031 | NSA2P3 | 0,351791095 | 1,276143968 | 0,798 | 0,90685415 | NSA2 pseudogene 3 |
| ENSG00000153113 | CAST | 0,351715609 | 1,276077198 | 0,4034 | 0,66399303 | calpastatin |
| ENSG00000115091 | ACTR3 | 0,351698691 | 1,276062233 | 0,32406 | 0,60668177 | actin related protein 3 |
| ENSG00000146859 | TMEM140 | 0,351666793 | 1,276034021 | 0,25319 | 0,55832664 | transmembrane protein 140 |
| ENSG00000134480 | CCNH | 0,351619455 | 1,275992151 | 0,40891 | 0,66773124 | cyclin H |
| ENSG00000068383 | INPP5A | 0,351541847 | 1,275923512 | 0,19838 | 0,51980353 | inositol polyphosphate-5-phosphatase A |
| ENSG00000214216 | IQCJ | 0,351446716 | 1,275839381 | 0,81047 | 0,91251177 | IQ motif containing J |
| ENSG00000104522 | GFUS | 0,351405057 | 1,275802541 | 0,33342 | 0,6133607 | GDP-L-fucose synthase |
| ENSG00000196743 | GM2A | 0,351331617 | 1,275737598 | 0,14429 | 0,47189017 | GM2 ganglioside activator |
| ENSG00000164117 | FBXO8 | 0,351249318 | 1,275664825 | 0,43319 | 0,68455289 | F-box protein 8 |
| ENSG00000134352 | IL6ST | 0,351209722 | 1,275629814 | 0,45249 | 0,69702004 | interleukin 6 signal transducer |
| ENSG00000108424 | KPNB1 | 0,351188064 | 1,275610665 | 0,09653 | 0,40027716 | karyopherin subunit beta 1 |
| ENSG00000139112 | GABARAPL1 | 0,351161571 | 1,27558724 | 0,19184 | 0,51440231 | GABA type A receptor associated protein like 1 |
| ENSG00000148690 | FRA10AC1 | 0,351089618 | 1,275523623 | 0,23793 | 0,54864017 | FRA10A associated CGG repeat 1 |
| ENSG00000171943 | SRGAP2C | 0,350988273 | 1,275434024 | 0,26475 | 0,56748133 | SLIT-ROBO Rho GTPase activating protein 2C |
| ENSG00000235018 | AL137077,1 | 0,350916914 | 1,27537094 | 0,63991 | 0,81929791 | ribosomal protein L17 (RPL17) pseudogene |
| ENSG00000065978 | YBX1 | 0,350880959 | 1,275339155 | 0,32303 | 0,60584543 | Y-box binding protein 1 |
| ENSG00000255031 | AP002807,1 | 0,350822217 | 1,275287228 | 0,61566 | 0,80454806 | novel transcript, antisense to CHKA |
| ENSG00000154518 | ATP5MC3 | 0,350752399 | 1,275225514 | 0,19286 | 0,51493916 | ATP synthase membrane subunit c locus 3 |
| ENSG00000102471 | NDFIP2 | 0,350631289 | 1,275118466 | 0,42143 | 0,67643128 | Nedd4 family interacting protein 2 |
| ENSG00000102893 | PHKB | 0,350599557 | 1,275090421 | 0,41928 | 0,67515889 | phosphorylase kinase regulatory subunit beta |
| ENSG00000153822 | KCNJ16 | 0,350534111 | 1,27503258 | 0,60372 | 0,79770754 | potassium inwardly rectifying channel subfamily J member 16 |
| ENSG00000204604 | ZNF468 | 0,350449344 | 1,274957666 | 0,21334 | 0,53259344 | zinc finger protein 468 |
| ENSG00000262911 | DNAJC28 | 0,350412231 | 1,274924868 | 0,43868 | 0,6878469 | DnaJ heat shock protein family (Hsp40) member C28 |
| ENSG00000282947 | HNRNPL | 0,350399564 | 1,274913674 | 0,21501 | 0,53373528 | heterogeneous nuclear ribonucleoprotein L |
| ENSG00000104824 | HNRNPL | 0,350399564 | 1,274913674 | 0,21501 | 0,53373528 | heterogeneous nuclear ribonucleoprotein L |

| | | | | | | |
|---|---|---|---|---|---|---|
| ENSG00000177463 | NR2C2 | 0,350393894 | 1,274908664 | 0,27267 | 0,57315967 | nuclear receptor subfamily 2 group C member 2 |
| ENSG00000230456 | DDR1 | 0,350371808 | 1,274889146 | 0,6786 | 0,84182956 | discoidin domain receptor tyrosine kinase 1 |
| ENSG00000102393 | GLA | 0,350349367 | 1,274869316 | 0,1442 | 0,47175088 | galactosidase alpha |
| ENSG00000236449 | AC010894,2 | 0,35032912 | 1,274851424 | 0,43429 | 0,68505296 | novel transcript, antisense to WIPF1 and CHRNA1 |
| ENSG00000108960 | MMD | 0,350236226 | 1,27476934 | 0,22559 | 0,54087803 | monocyte to macrophage differentiation associated |
| ENSG00000087470 | DNM1L | 0,350220398 | 1,274755354 | 0,3157 | 0,60176284 | dynamin 1 like |
| ENSG00000141576 | RNF157 | 0,350210797 | 1,274746871 | 0,15676 | 0,48425071 | ring finger protein 157 |
| ENSG00000186792 | HYAL3 | 0,350202062 | 1,274739153 | 0,32849 | 0,60990374 | hyaluronidase 3 |
| ENSG00000172262 | ZNF131 | 0,350186702 | 1,274725581 | 0,22122 | 0,53805234 | zinc finger protein 131 |
| ENSG00000062485 | CS | 0,350179563 | 1,274719273 | 0,27678 | 0,57549599 | citrate synthase |
| ENSG00000182220 | ATP6AP2 | 0,350063347 | 1,274616593 | 0,37012 | 0,63956649 | ATPase H+ transporting accessory protein 2 |
| ENSG00000288323 | FAM86DP | 0,349959047 | 1,274524447 | 0,05419 | 0,31168784 | family with sequence similarity 86 member D, pseudogene |
| ENSG00000072135 | PTPN18 | 0,34988295 | 1,274457223 | 0,0922 | 0,39260493 | protein tyrosine phosphatase non-receptor type 18 |
| ENSG00000256349 | AP002748,5 | 0,349872235 | 1,274447757 | 0,90578 | 0,95975315 | novel protein |
| ENSG00000229666 | MAST4-AS1 | 0,349736878 | 1,274328191 | 0,57482 | 0,77995956 | MAST4 antisense RNA 1 |
| ENSG00000076321 | KLHL20 | 0,349709708 | 1,274304192 | 0,27928 | 0,57742338 | kelch like family member 20 |
| ENSG00000213839 | TMX2P1 | 0,349612728 | 1,274218534 | 0,30466 | 0,59445922 | thioredoxin related transmembrane protein 2 pseudogene 1 |
| ENSG00000106460 | TMEM106B | 0,349594642 | 1,274202561 | 0,41235 | 0,67001327 | transmembrane protein 106B |
| ENSG00000109452 | INPP4B | 0,349587279 | 1,274196058 | 0,50904 | 0,7370222 | inositol polyphosphate-4-phosphatase type II B |
| ENSG00000278275 | AC103705,1 | 0,349584177 | 1,274193318 | 0,56593 | 0,77413633 | protein tyrosine phosphatase type IVA, member 1 (PTP4A1) pseudogene |
| ENSG00000227917 | AL449043,1 | 0,349551867 | 1,274164782 | 0,76868 | 0,89277929 | novel transcript |
| ENSG00000126838 | PZP | 0,349468435 | 1,274091098 | 0,37376 | 0,6423379 | PZP alpha-2-macroglobulin like |
| ENSG00000136866 | ZFP37 | 0,349384833 | 1,274017269 | 0,40823 | 0,66714279 | ZFP37 zinc finger protein |
| ENSG00000143612 | C1orf43 | 0,349256386 | 1,273903844 | 0,22538 | 0,5407493 | chromosome 1 open reading frame 43 |
| ENSG00000236515 | ZBTB9 | 0,349252095 | 1,273900056 | 0,13889 | 0,46622186 | zinc finger and BTB domain containing 9 |
| ENSG00000264937 | AC100830,2 | 0,349128757 | 1,273791153 | 0,64708 | 0,82343226 | novel transcript, antisense to ornithine decarboxylase antizyme 2 OAZ2 |
| ENSG00000151458 | ANKRD50 | 0,349094089 | 1,273760544 | 0,32181 | 0,60577115 | ankyrin repeat domain 50 |
| ENSG00000165948 | IFI27L1 | 0,349028142 | 1,273702321 | 0,39435 | 0,65725123 | interferon alpha inducible protein 27 like 1 |
| ENSG00000082512 | TRAF5 | 0,34896824 | 1,273649437 | 0,0983 | 0,40379437 | TNF receptor associated factor 5 |
| ENSG00000240804 | NPM1P28 | 0,348928666 | 1,2736145 | 0,62669 | 0,81110476 | nucleophosmin 1 pseudogene 28 |

| | | | | | | |
|---|---|---|---|---|---|---|
| ENSG00000184227 | ACOT1 | 0,34892752 | 1,273613488 | 0,5632 | 0,77230901 | acyl-CoA thioesterase 1 |
| ENSG00000172594 | SMPDL3A | 0,348822387 | 1,27352068 | 0,39936 | 0,66169316 | sphingomyelin phosphodiesterase acid like 3A |
| ENSG00000082014 | SMARCD3 | 0,348778889 | 1,273482283 | 0,07094 | 0,35169607 | SWI/SNF related, matrix associated, actin dependent regulator of chromatin, subfamily d, member 3 |
| ENSG00000127337 | YEATS4 | 0,348760025 | 1,273465632 | 0,28381 | 0,58078109 | YEATS domain containing 4 |
| ENSG00000173598 | NUDT4 | 0,348740954 | 1,273448798 | 0,07929 | 0,36776284 | nudix hydrolase 4 |
| ENSG00000110711 | AIP | 0,348662881 | 1,273379886 | 0,10009 | 0,40666505 | aryl hydrocarbon receptor interacting protein |
| ENSG00000147164 | SNX12 | 0,348556041 | 1,273285588 | 0,23588 | 0,5471053 | sorting nexin 12 |
| ENSG00000113319 | RASGRF2 | 0,348486132 | 1,27322389 | 0,37542 | 0,64381159 | Ras protein specific guanine nucleotide releasing factor 2 |
| ENSG00000151292 | CSNK1G3 | 0,348354102 | 1,273107375 | 0,16941 | 0,49554671 | casein kinase 1 gamma 3 |
| ENSG00000174944 | P2RY14 | 0,348298528 | 1,273058334 | 0,52642 | 0,74978877 | purinergic receptor P2Y14 |
| ENSG00000111181 | SLC6A12 | 0,348274465 | 1,273037101 | 0,70713 | 0,85902171 | solute carrier family 6 member 12 |
| ENSG00000204970 | PCDHA1 | 0,348211622 | 1,272981649 | 0,74402 | 0,87882457 | protocadherin alpha 1 |
| ENSG00000173456 | RNF26 | 0,348155095 | 1,272931772 | 0,08576 | 0,37982002 | ring finger protein 26 |
| ENSG00000169439 | SDC2 | 0,348026609 | 1,272818411 | 0,04879 | 0,29705113 | syndecan 2 |
| ENSG00000183978 | COA3 | 0,348008401 | 1,272802347 | 0,43756 | 0,68735017 | cytochrome c oxidase assembly factor 3 |
| ENSG00000127990 | SGCE | 0,348006416 | 1,272800595 | 0,20548 | 0,52508239 | sarcoglycan epsilon |
| ENSG00000132694 | ARHGEF11 | 0,347996017 | 1,272791421 | 0,10506 | 0,4138867 | Rho guanine nucleotide exchange factor 11 |
| ENSG00000142227 | EMP3 | 0,347930999 | 1,272734062 | 0,21778 | 0,53550245 | epithelial membrane protein 3 |
| ENSG00000115295 | CLIP4 | 0,347841667 | 1,272655256 | 0,19322 | 0,51520939 | CAP-Gly domain containing linker protein family member 4 |
| ENSG00000163607 | GTPBP8 | 0,347831141 | 1,272645971 | 0,33895 | 0,61800397 | GTP binding protein 8 (putative) |
| ENSG00000273455 | AC072039,2 | 0,347734553 | 1,272560771 | 0,66909 | 0,83595057 | novel transcript, antisense to PPP2R3A |
| ENSG00000125812 | GZF1 | 0,34771773 | 1,272545931 | 0,41664 | 0,6734221 | GDNF inducible zinc finger protein 1 |
| ENSG00000224114 | AL591846,1 | 0,347664194 | 1,27249871 | 0,4805 | 0,71764212 | ribosomal protein S14 (RPS14) pseudogene |
| ENSG00000130940 | CASZ1 | 0,347658546 | 1,272493728 | 0,31707 | 0,60268749 | castor zinc finger 1 |
| ENSG00000165458 | INPPL1 | 0,347636473 | 1,27247426 | 0,0832 | 0,37547217 | inositol polyphosphate phosphatase like 1 |
| ENSG00000213927 | CCL27 | 0,347616122 | 1,27245631 | 0,63326 | 0,81541029 | C-C motif chemokine ligand 27 |
| ENSG00000106682 | EIF4H | 0,347539431 | 1,27238867 | 0,17019 | 0,49606609 | eukaryotic translation initiation factor 4H |
| ENSG00000224370 | AC093392,2 | 0,347460282 | 1,272318867 | 0,5526 | 0,76595895 | calmodulin 3 (phosphorylase kinase, delta) (CALM3) pseudogene |
| ENSG00000076944 | STXBP2 | 0,347446371 | 1,272306599 | 0,62797 | 0,81175174 | syntaxin binding protein 2 |
| ENSG00000125450 | NUP85 | 0,347423472 | 1,272286404 | 0,14313 | 0,47072472 | nucleoporin 85 |
| ENSG00000146085 | MMUT | 0,347409315 | 1,272273919 | 0,34956 | 0,62563749 | methylmalonyl-CoA mutase |

| | | | | | | |
|---|---|---|---|---|---|---|
| ENSG00000172661 | WASHC2C | 0,347359185 | 1,272229712 | 0,18966 | 0,51284737 | WASH complex subunit 2C |
| ENSG00000272654 | AL358472,2 | 0,347345516 | 1,272217658 | 0,30212 | 0,59294633 | novel transcript |
| ENSG00000086232 | EIF2AK1 | 0,347339982 | 1,272212778 | 0,25599 | 0,56057876 | eukaryotic translation initiation factor 2 alpha kinase 1 |
| ENSG00000225400 | RAB28P5 | 0,347334975 | 1,272208363 | 0,44965 | 0,69525704 | RAB28, member RAS oncogene family pseudogene 5 |
| ENSG00000260469 | INSYN1-AS1 | 0,347302329 | 1,272179575 | 0,37127 | 0,64045979 | INSYN1 antisense RNA 1 |
| ENSG00000116885 | OSCP1 | 0,347260035 | 1,27214228 | 0,28512 | 0,58171017 | organic solute carrier partner 1 |
| ENSG00000270277 | AC009948,1 | 0,347249117 | 1,272132653 | 0,50584 | 0,73418581 | novel transcript, antisense to titin |
| ENSG00000169288 | MRPL1 | 0,347187721 | 1,272078516 | 0,26929 | 0,57098657 | mitochondrial ribosomal protein L1 |
| ENSG00000112651 | MRPL2 | 0,346936647 | 1,271857155 | 0,11637 | 0,43288152 | mitochondrial ribosomal protein L2 |
| ENSG00000181038 | METTL23 | 0,346843836 | 1,271775337 | 0,26892 | 0,57085484 | methyltransferase like 23 |
| ENSG00000118246 | FASTKD2 | 0,346654984 | 1,271608869 | 0,24366 | 0,55235189 | FAST kinase domains 2 |
| ENSG00000171792 | RHNO1 | 0,346617964 | 1,27157624 | 0,306 | 0,59512079 | RAD9-HUS1-RAD1 interacting nuclear orphan 1 |
| ENSG00000224418 | STK24-AS1 | 0,346546728 | 1,271513455 | 0,50383 | 0,73294771 | STK24 antisense RNA 1 |
| ENSG00000077232 | DNAJC10 | 0,346537022 | 1,2715049 | 0,22346 | 0,53953208 | DnaJ heat shock protein family (Hsp40) member C10 |
| ENSG00000218730 | AL117342,1 | 0,346327061 | 1,271319867 | 0,71015 | 0,86036733 | A kinase (PRKA) interacting protein 1 (AKIP1) pseudogene |
| ENSG00000158813 | EDA | 0,346280655 | 1,271278974 | 0,37792 | 0,64628549 | ectodysplasin A |
| ENSG00000164329 | TENT2 | 0,346279869 | 1,271278282 | 0,29589 | 0,58942061 | terminal nucleotidyltransferase 2 |
| ENSG00000131966 | ACTR10 | 0,346189072 | 1,271198275 | 0,29109 | 0,58613153 | actin related protein 10 |
| ENSG00000117500 | TMED5 | 0,346183709 | 1,27119355 | 0,33792 | 0,61755283 | transmembrane p24 trafficking protein 5 |
| ENSG00000093217 | XYLB | 0,346166897 | 1,271178736 | 0,08755 | 0,38313766 | xylulokinase |
| ENSG00000249459 | ZNF286B | 0,346105667 | 1,271124786 | 0,05524 | 0,31443527 | zinc finger protein 286B (pseudogene) |
| ENSG00000132341 | RAN | 0,34610446 | 1,271123723 | 0,25955 | 0,56330128 | RAN, member RAS oncogene family |
| ENSG00000254592 | AC123788,1 | 0,346100859 | 1,271120551 | 0,71038 | 0,86051357 | novel transcript |
| ENSG00000288559 | AC016394,2 | 0,346066313 | 1,271090113 | 0,26802 | 0,5700759 | novel transcript |
| ENSG00000015479 | MATR3 | 0,346058258 | 1,271083017 | 0,29528 | 0,58901329 | matrin 3 |
| ENSG00000111012 | CYP27B1 | 0,346040748 | 1,271067589 | 0,36329 | 0,63408839 | cytochrome P450 family 27 subfamily B member 1 |
| ENSG00000272602 | ZNF595 | 0,34600691 | 1,271037778 | 0,12989 | 0,45410275 | zinc finger protein 595 |
| ENSG00000105261 | OVOL3 | 0,345904056 | 1,270947165 | 0,65986 | 0,83018671 | ovo like zinc finger 3 |
| ENSG00000100297 | MCM5 | 0,345794131 | 1,270850329 | 0,18807 | 0,51128963 | minichromosome maintenance complex component 5 |
| ENSG00000100084 | HIRA | 0,345753107 | 1,270814193 | 0,20125 | 0,52181085 | histone cell cycle regulator |
| ENSG00000287581 | AC016549,1 | 0,345711679 | 1,270777701 | 0,26978 | 0,5714611 | novel transcript, antisense to TRIO |
| ENSG00000166887 | VPS39 | 0,345710776 | 1,270776905 | 0,19995 | 0,52092803 | VPS39 subunit of HOPS complex |
| ENSG00000132912 | DCTN4 | 0,345695279 | 1,270763255 | 0,37337 | 0,64204597 | dynactin subunit 4 |

| | | | | | | |
|---|---|---|---|---|---|---|
| ENSG00000172817 | CYP7B1 | 0,345674686 | 1,270745116 | 0,3322 | 0,61216934 | cytochrome P450 family 7 subfamily B member 1 |
| ENSG00000119844 | AFTPH | 0,345586857 | 1,270667757 | 0,40322 | 0,66399303 | aftiphilin |
| ENSG00000240489 | SETP14 | 0,345500932 | 1,270592081 | 0,6592 | 0,82978434 | SET pseudogene 14 |
| ENSG00000246859 | STARD4-AS1 | 0,34548228 | 1,270575653 | 0,31989 | 0,60473824 | STARD4 antisense RNA 1 |
| ENSG00000112110 | MRPL18 | 0,345421348 | 1,270521992 | 0,30222 | 0,59294633 | mitochondrial ribosomal protein L18 |
| ENSG00000138182 | KIF20B | 0,345401942 | 1,270504902 | 0,35077 | 0,62618267 | kinesin family member 20B |
| ENSG00000137309 | HMGA1 | 0,345250164 | 1,270371247 | 0,1987 | 0,51984196 | high mobility group AT-hook 1 |
| ENSG00000106993 | CDC37L1 | 0,345080086 | 1,270221493 | 0,30288 | 0,59331081 | cell division cycle 37 like 1 |
| ENSG00000198551 | ZNF627 | 0,345055342 | 1,270199706 | 0,30139 | 0,59258306 | zinc finger protein 627 |
| ENSG00000101444 | AHCY | 0,345054552 | 1,270199011 | 0,3873 | 0,65227155 | adenosylhomocysteinase |
| ENSG00000181904 | C5orf24 | 0,345051313 | 1,270196159 | 0,18719 | 0,5105374 | chromosome 5 open reading frame 24 |
| ENSG00000017260 | ATP2C1 | 0,345030936 | 1,270178219 | 0,25828 | 0,56224798 | ATPase secretory pathway Ca2+ transporting 1 |
| ENSG00000104093 | DMXL2 | 0,344918708 | 1,270079415 | 0,3425 | 0,62018255 | Dmx like 2 |
| ENSG00000103160 | HSDL1 | 0,344893963 | 1,270057631 | 0,17559 | 0,49954066 | hydroxysteroid dehydrogenase like 1 |
| ENSG00000101363 | MANBAL | 0,344793917 | 1,26996956 | 0,29555 | 0,58929511 | mannosidase beta like |
| ENSG00000102317 | RBM3 | 0,344775005 | 1,269952912 | 0,15625 | 0,4840074 | RNA binding motif protein 3 |
| ENSG00000110756 | HPS5 | 0,344701674 | 1,269888363 | 0,14814 | 0,47527573 | HPS5 biogenesis of lysosomal organelles complex 2 subunit 2 |
| ENSG00000168275 | COA6 | 0,344636057 | 1,269830607 | 0,37385 | 0,6423379 | cytochrome c oxidase assembly factor 6 |
| ENSG00000010244 | ZNF207 | 0,344623484 | 1,269819541 | 0,21296 | 0,53198758 | zinc finger protein 207 |
| ENSG00000074054 | CLASP1 | 0,344610538 | 1,269808145 | 0,22649 | 0,54144 | cytoplasmic linker associated protein 1 |
| ENSG00000134684 | YARS1 | 0,34460738 | 1,269805366 | 0,12144 | 0,44080285 | tyrosyl-tRNA synthetase 1 |
| ENSG00000229058 | AGER | 0,344592571 | 1,269792332 | 0,497 | 0,72862056 | advanced glycosylation end-product specific receptor |
| ENSG00000237405 | AGER | 0,344592571 | 1,269792332 | 0,497 | 0,72862056 | advanced glycosylation end-product specific receptor |
| ENSG00000206320 | AGER | 0,344592571 | 1,269792332 | 0,497 | 0,72862056 | advanced glycosylation end-product specific receptor |
| ENSG00000171135 | JAGN1 | 0,344575713 | 1,269777494 | 0,27483 | 0,57463533 | jagunal homolog 1 |
| ENSG00000138777 | PPA2 | 0,344462039 | 1,269677449 | 0,25056 | 0,55690541 | inorganic pyrophosphatase 2 |
| ENSG00000100916 | BRMS1L | 0,344446261 | 1,269663563 | 0,33815 | 0,6177255 | BRMS1 like transcriptional repressor |
| ENSG00000124098 | FAM210B | 0,344422939 | 1,269643039 | 0,32649 | 0,60866572 | family with sequence similarity 210 member B |
| ENSG00000198836 | OPA1 | 0,344411088 | 1,269632609 | 0,25023 | 0,55657567 | OPA1 mitochondrial dynamin like GTPase |
| ENSG00000249492 | AC114956,2 | 0,344254145 | 1,269494501 | 0,34156 | 0,61963031 | novel transcript |
| ENSG00000099957 | P2RX6 | 0,344176716 | 1,269426369 | 0,44669 | 0,69321646 | purinergic receptor P2X 6 |
| ENSG00000273802 | H2BC8 | 0,344165211 | 1,269416245 | 0,56435 | 0,77292062 | H2B clustered histone 8 |

| | | | | | | |
|---|---|---|---|---|---|---|
| ENSG00000271964 | AC090948,1 | 0,344144234 | 1,269397789 | 0,53061 | 0,75212511 | novel transcript, antisense to RFTN1 |
| ENSG00000127423 | AUNIP | 0,344111198 | 1,269368721 | 0,16098 | 0,48832443 | aurora kinase A and ninein interacting protein |
| ENSG00000172671 | ZFAND4 | 0,344099053 | 1,269358035 | 0,16931 | 0,49548445 | zinc finger AN1-type containing 4 |
| ENSG00000165219 | GAPVD1 | 0,343995773 | 1,269267167 | 0,16331 | 0,48972962 | GTPase activating protein and VPS9 domains 1 |
| ENSG00000170759 | KIF5B | 0,343928738 | 1,269208192 | 0,33979 | 0,61872249 | kinesin family member 5B |
| ENSG00000259366 | AC108449,2 | 0,343917937 | 1,26919869 | 0,40927 | 0,66785781 | novel transcript, sense intronic to HMBOX1 |
| ENSG00000151718 | WWC2 | 0,343808454 | 1,269102377 | 0,09715 | 0,40152057 | WW and C2 domain containing 2 |
| ENSG00000136709 | WDR33 | 0,343783964 | 1,269080834 | 0,23622 | 0,54725688 | WD repeat domain 33 |
| ENSG00000165271 | NOL6 | 0,343780597 | 1,269077872 | 0,08682 | 0,3816108 | nucleolar protein 6 |
| ENSG00000065135 | GNAI3 | 0,34377934 | 1,269076766 | 0,23161 | 0,54440987 | G protein subunit alpha i3 |
| ENSG00000271147 | ARMCX5-GPR | 0,343635774 | 1,268950483 | 0,19358 | 0,51547251 | ARMCX5-GPRASP2 readthrough |
| ENSG00000174946 | GPR171 | 0,343604773 | 1,268923216 | 0,47798 | 0,71567419 | G protein-coupled receptor 171 |
| ENSG00000119953 | SMNDC1 | 0,343259731 | 1,26861977 | 0,27182 | 0,57305058 | survival motor neuron domain containing 1 |
| ENSG00000168724 | DNAJC21 | 0,343223788 | 1,268588165 | 0,23969 | 0,5491566 | DnaJ heat shock protein family (Hsp40) member C21 |
| ENSG00000272529 | AC090948,3 | 0,343178914 | 1,268548707 | 0,50541 | 0,73393676 | novel transcript, antisense to RFTN1 |
| ENSG00000268089 | GABRQ | 0,343173115 | 1,268543608 | 0,3591 | 0,63201238 | gamma-aminobutyric acid type A receptor subunit theta |
| ENSG00000100220 | RTCB | 0,343171011 | 1,268541758 | 0,11024 | 0,42280938 | RNA 2',3'-cyclic phosphate and 5'-OH ligase |
| ENSG00000253719 | ATXN7L3B | 0,34307936 | 1,268461173 | 0,16452 | 0,49065086 | ataxin 7 like 3B |
| ENSG00000160213 | CSTB | 0,343066691 | 1,268450035 | 0,23003 | 0,54335232 | cystatin B |
| ENSG00000134769 | DTNA | 0,343060519 | 1,268444607 | 0,41597 | 0,67284446 | dystrobrevin alpha |
| ENSG00000272398 | CD24 | 0,342880888 | 1,268286682 | 0,34189 | 0,6197312 | CD24 molecule |
| ENSG00000171649 | ZIK1 | 0,342865734 | 1,268273361 | 0,23293 | 0,54548847 | zinc finger protein interacting with K protein 1 |
| ENSG00000265808 | SEC22B | 0,342797112 | 1,268213036 | 0,29862 | 0,59089801 | SEC22 homolog B, vesicle trafficking protein |
| ENSG00000168958 | MFF | 0,342787533 | 1,268204616 | 0,37179 | 0,6408454 | mitochondrial fission factor |
| ENSG00000113971 | NPHP3 | 0,342733298 | 1,268156942 | 0,13408 | 0,46020578 | nephrocystin 3 |
| ENSG00000037749 | MFAP3 | 0,342718508 | 1,268143941 | 0,25273 | 0,55796133 | microfibril associated protein 3 |
| ENSG00000108510 | MED13 | 0,342650036 | 1,268083755 | 0,39913 | 0,66153677 | mediator complex subunit 13 |
| ENSG00000167380 | ZNF226 | 0,342610235 | 1,268048771 | 0,31242 | 0,59934192 | zinc finger protein 226 |
| ENSG00000206828 | RNVU1-30 | 0,342603027 | 1,268042436 | 0,66229 | 0,83183445 | RNA, variant U1 small nuclear 30 |
| ENSG00000258708 | SLC25A21-AS1 | 0,342590873 | 1,268031753 | 0,30814 | 0,59696396 | SLC25A21 antisense RNA 1 |
| ENSG00000148450 | MSRB2 | 0,342545068 | 1,267991495 | 0,28798 | 0,5840701 | methionine sulfoxide reductase B2 |
| ENSG00000107719 | PALD1 | 0,34253996 | 1,267987005 | 0,2388 | 0,54892264 | phosphatase domain containing paladin 1 |
| ENSG00000076003 | MCM6 | 0,342465401 | 1,267921476 | 0,18547 | 0,50893545 | minichromosome maintenance complex component 6 |

| | | | | | | |
|---|---|---|---|---|---|---|
| ENSG00000177674 | AGTRAP | 0,342437875 | 1,267897286 | 0,19368 | 0,51547251 | angiotensin II receptor associated protein |
| ENSG00000172409 | CLP1 | 0,342373282 | 1,267840519 | 0,26661 | 0,5691939 | cleavage factor polyribonucleotide kinase subunit 1 |
| ENSG00000229524 | BAG6 | 0,342269115 | 1,267748981 | 0,26775 | 0,56999323 | BAG cochaperone 6 |
| ENSG00000082068 | WDR70 | 0,342149607 | 1,26764397 | 0,15355 | 0,48191866 | WD repeat domain 70 |
| ENSG00000116906 | GNPAT | 0,342127982 | 1,267624968 | 0,29324 | 0,58756756 | glyceronephosphate O-acyltransferase |
| ENSG00000125772 | GPCPD1 | 0,342103803 | 1,267603724 | 0,35597 | 0,63029851 | glycerophosphocholine phosphodiesterase 1 |
| ENSG00000257083 | AC073530,1 | 0,342087921 | 1,267589769 | 0,58214 | 0,78454692 | novel transcript |
| ENSG00000274383 | AC103691,1 | 0,341984548 | 1,267498946 | 0,52114 | 0,74558236 | novel transcript, antisense to ANKDD1A |
| ENSG00000114812 | VIPR1 | 0,341965291 | 1,267482028 | 0,62466 | 0,80990161 | vasoactive intestinal peptide receptor 1 |
| ENSG00000008324 | SS18L2 | 0,34193113 | 1,267452016 | 0,27411 | 0,57412149 | SS18 like 2 |
| ENSG00000223646 | AC002463,1 | 0,34191626 | 1,267438952 | 0,69503 | 0,85206252 | novel transcript |
| ENSG00000118965 | WDR35 | 0,341822158 | 1,267356284 | 0,36957 | 0,63929758 | WD repeat domain 35 |
| ENSG00000224751 | SHMT1P1 | 0,341753547 | 1,267296014 | 0,44688 | 0,6933405 | serine hydroxymethyltransferase 1 (soluble) pseudogene 1 |
| ENSG00000145414 | NAF1 | 0,341701542 | 1,267250332 | 0,05364 | 0,31073157 | nuclear assembly factor 1 ribonucleoprotein |
| ENSG00000272037 | AP002907,1 | 0,341666124 | 1,267219221 | 0,46373 | 0,70449472 | novel transcript, antisense to UBR5 |
| ENSG00000225548 | LINC01980 | 0,341615496 | 1,267174752 | 0,31945 | 0,60429866 | long intergenic non-protein coding RNA 1980 |
| ENSG00000137942 | FNBP1L | 0,341611869 | 1,267171566 | 0,3843 | 0,65052153 | formin binding protein 1 like |
| ENSG00000067560 | RHOA | 0,341590707 | 1,26715298 | 0,18533 | 0,50876814 | ras homolog family member A |
| ENSG00000273797 | AL133445,2 | 0,341507514 | 1,267079911 | 0,67082 | 0,83695077 | novel transcript, sense intronic to KIAA0247 |
| ENSG00000135473 | PAN2 | 0,341430632 | 1,26701239 | 0,3373 | 0,61706507 | poly(A) specific ribonuclease subunit PAN2 |
| ENSG00000218313 | AL139274,1 | 0,341294957 | 1,266893242 | 0,52617 | 0,7497214 | novel TWIST neighbor (TWISTNB) pseudogene |
| ENSG00000272750 | AL592148,3 | 0,341229153 | 1,266835458 | 0,62982 | 0,8132682 | novel transcript, antisense to MIA3 |
| ENSG00000166199 | ALKBH3 | 0,341125337 | 1,2667443 | 0,28701 | 0,58346069 | alkB homolog 3, alpha-ketoglutarate dependent dioxygenase |
| ENSG00000160014 | CALM3 | 0,341049193 | 1,266677444 | 0,0907 | 0,38975298 | calmodulin 3 |
| ENSG00000136940 | PDCL | 0,341034066 | 1,266664163 | 0,14642 | 0,47433542 | phosducin like |
| ENSG00000101079 | NDRG3 | 0,340950555 | 1,266590844 | 0,25714 | 0,56155289 | NDRG family member 3 |
| ENSG00000092203 | TOX4 | 0,340669247 | 1,266343898 | 0,16255 | 0,48945635 | TOX high mobility group box family member 4 |
| ENSG00000169826 | CSGALNACT2 | 0,340663889 | 1,266339195 | 0,31209 | 0,59925736 | chondroitin sulfate N-acetylgalactosaminyltransferase 2 |
| ENSG00000170748 | RBMXL2 | 0,340603525 | 1,266286211 | 0,69208 | 0,85016274 | RBMX like 2 |
| ENSG00000215712 | TMEM242 | 0,340577802 | 1,266263633 | 0,31221 | 0,59925736 | transmembrane protein 242 |

| | | | | | | |
|---|---|---|---|---|---|---|
| ENSG00000271086 | NAMA | 0,34053881 | 1,26622941 | 0,45499 | 0,69913628 | non-protein coding RNA, associated with MAP kinase pathway and growth arrest |
| ENSG00000257591 | ZNF625 | 0,340515541 | 1,266208988 | 0,1817 | 0,50497494 | zinc finger protein 625 |
| ENSG00000257376 | AC079601,2 | 0,340435559 | 1,266138792 | 0,66243 | 0,83191849 | fragile X mental retardation, autosomal homolog 1 (FXR1) pseudogene |
| ENSG00000105865 | DUS4L | 0,340425029 | 1,266129551 | 0,39898 | 0,66140951 | dihydrouridine synthase 4 like |
| ENSG00000284103 | DUS4L | 0,340425029 | 1,266129551 | 0,39898 | 0,66140951 | dihydrouridine synthase 4 like |
| ENSG00000102921 | N4BP1 | 0,340397092 | 1,266105033 | 0,10838 | 0,4190049 | NEDD4 binding protein 1 |
| ENSG00000272990 | AC084036,1 | 0,340173361 | 1,265908702 | 0,71643 | 0,86426583 | novel transcript, antisense to KCNAB1 |
| ENSG00000231125 | AF129075,1 | 0,340171871 | 1,265907395 | 0,57179 | 0,77829419 | novel transcript, sense intronic to CCT8 |
| ENSG00000169740 | ZNF32 | 0,340148004 | 1,265886452 | 0,40828 | 0,66714279 | zinc finger protein 32 |
| ENSG00000270557 | AC013731,1 | 0,340047943 | 1,265798658 | 0,50632 | 0,73476122 | novel transcript |
| ENSG00000231162 | COX11P1 | 0,33982556 | 1,265603557 | 0,69902 | 0,85450694 | COX11 pseudogene 1 |
| ENSG00000272916 | AC022400,5 | 0,339674922 | 1,265471417 | 0,11394 | 0,42901302 | novel transcript |
| ENSG00000089057 | SLC23A2 | 0,339541221 | 1,265354145 | 0,24821 | 0,55542983 | solute carrier family 23 member 2 |
| ENSG00000105176 | URI1 | 0,339527644 | 1,265342237 | 0,21185 | 0,53070055 | URI1 prefoldin like chaperone |
| ENSG00000148158 | SNX30 | 0,339523371 | 1,26533849 | 0,24002 | 0,54936679 | sorting nexin family member 30 |
| ENSG00000178053 | MLF1 | 0,33948056 | 1,265300942 | 0,43223 | 0,68411566 | myeloid leukemia factor 1 |
| ENSG00000236576 | AC241520,1 | 0,339458388 | 1,265281497 | 0,61234 | 0,8035281 | zinc finger protein pseudogene |
| ENSG00000135272 | MDFIC | 0,339407865 | 1,265237187 | 0,13497 | 0,46159318 | MyoD family inhibitor domain containing |
| ENSG00000117115 | PADI2 | 0,339400181 | 1,265230449 | 0,51574 | 0,74180801 | peptidyl arginine deiminase 2 |
| ENSG00000234332 | BCAS2P2 | 0,339378583 | 1,265211507 | 0,53555 | 0,75478533 | BCAS2 pseudogene 2 |
| ENSG00000139734 | DIAPH3 | 0,339307512 | 1,265149182 | 0,14147 | 0,46809363 | diaphanous related formin 3 |
| ENSG00000196605 | ZNF846 | 0,339122267 | 1,264986745 | 0,12026 | 0,43881019 | zinc finger protein 846 |
| ENSG00000026508 | CD44 | 0,33910431 | 1,264970999 | 0,53821 | 0,75658794 | CD44 molecule (Indian blood group) |
| ENSG00000114107 | CEP70 | 0,339046499 | 1,264920311 | 0,38482 | 0,65062567 | centrosomal protein 70 |
| ENSG00000198554 | WDHD1 | 0,338991818 | 1,264872369 | 0,28424 | 0,58107964 | WD repeat and HMG-box DNA binding protein 1 |
| ENSG00000166557 | TMED3 | 0,338921395 | 1,264810627 | 0,21766 | 0,53547632 | transmembrane p24 trafficking protein 3 |
| ENSG00000143952 | VPS54 | 0,338845751 | 1,264744312 | 0,24433 | 0,55253387 | VPS54 subunit of GARP complex |
| ENSG00000176542 | USF3 | 0,33882147 | 1,264723026 | 0,31832 | 0,60349082 | upstream transcription factor family member 3 |
| ENSG00000132300 | PTCD3 | 0,338818861 | 1,264720739 | 0,26287 | 0,56582568 | pentatricopeptide repeat domain 3 |
| ENSG00000096060 | FKBP5 | 0,338807777 | 1,264711023 | 0,35455 | 0,62915556 | FKBP prolyl isomerase 5 |
| ENSG00000128242 | GAL3ST1 | 0,338675428 | 1,264595007 | 0,27625 | 0,57538841 | galactose-3-O-sulfotransferase 1 |

| | | | | | | |
|---|---|---|---|---|---|---|
| ENSG00000109180 | OCIAD1 | 0,338670557 | 1,264590737 | 0,26512 | 0,56788357 | OCIA domain containing 1 |
| ENSG00000119812 | FAM98A | 0,338663781 | 1,264584797 | 0,33662 | 0,61666883 | family with sequence similarity 98 member A |
| ENSG00000111046 | MYF6 | 0,338620332 | 1,264546713 | 0,65314 | 0,82628153 | myogenic factor 6 |
| ENSG00000163082 | SGPP2 | 0,33861868 | 1,264545265 | 0,40961 | 0,6680455 | sphingosine-1-phosphate phosphatase 2 |
| ENSG00000253079 | NRON | 0,338523292 | 1,264461659 | 0,58445 | 0,78572744 | non-coding repressor of NFAT |
| ENSG00000277140 | MARF1 | 0,338521617 | 1,264460191 | 0,31616 | 0,60224605 | meiosis regulator and mRNA stability factor 1 |
| ENSG00000276541 | TAS2R14 | 0,33843493 | 1,264384216 | 0,7482 | 0,8814896 | taste 2 receptor member 14 |
| ENSG00000217128 | FNIP1 | 0,338421416 | 1,264372372 | 0,09428 | 0,39648446 | folliculin interacting protein 1 |
| ENSG00000265683 | SYPL1P2 | 0,338336477 | 1,264297934 | 0,5692 | 0,77646605 | synaptophysin like 1 pseudogene 2 |
| ENSG00000100393 | EP300 | 0,338330373 | 1,264292584 | 0,15438 | 0,48266733 | E1A binding protein p300 |
| ENSG00000160714 | UBE2Q1 | 0,338204889 | 1,264182623 | 0,07631 | 0,36243358 | ubiquitin conjugating enzyme E2 Q1 |
| ENSG00000254781 | GVINP2 | 0,338198036 | 1,264176618 | 0,64976 | 0,82452434 | GTPase, very large interferon inducible pseudogene 2 |
| ENSG00000116922 | C1orf109 | 0,3381928 | 1,264172029 | 0,1598 | 0,4870975 | chromosome 1 open reading frame 109 |
| ENSG00000169189 | NSMCE1 | 0,338105159 | 1,264095236 | 0,17069 | 0,49641416 | NSE1 homolog, SMC5-SMC6 complex component |
| ENSG00000072756 | TRNT1 | 0,33808877 | 1,264080876 | 0,16808 | 0,49417662 | tRNA nucleotidyl transferase 1 |
| ENSG00000102898 | NUTF2 | 0,338076105 | 1,264069779 | 0,15978 | 0,4870975 | nuclear transport factor 2 |
| ENSG00000281183 | NPTN-IT1 | 0,338068681 | 1,264063274 | 0,11862 | 0,43646336 | NPTN intronic transcript 1 |
| ENSG00000102053 | ZC3H12B | 0,338054072 | 1,264050474 | 0,12265 | 0,44239528 | zinc finger CCCH-type containing 12B |
| ENSG00000160783 | PMF1 | 0,338040027 | 1,264038169 | 0,16235 | 0,48945635 | polyamine modulated factor 1 |
| ENSG00000265142 | MIR133A1HG | 0,337932273 | 1,263943761 | 0,5696 | 0,7768187 | MIR133A1 host gene |
| ENSG00000274726 | ARHGEF10 | 0,337778138 | 1,263808732 | 0,17875 | 0,50325871 | Rho guanine nucleotide exchange factor 10 |
| ENSG00000152977 | ZIC1 | 0,337660433 | 1,263705625 | 0,18816 | 0,51129824 | Zic family member 1 |
| ENSG00000201558 | RNVU1-6 | 0,337617787 | 1,263668271 | 0,72389 | 0,86817061 | RNA, variant U1 small nuclear 6 |
| ENSG00000092020 | PPP2R3C | 0,337550678 | 1,263609491 | 0,46586 | 0,70619029 | protein phosphatase 2 regulatory subunit B''gamma |
| ENSG00000178386 | ZNF223 | 0,337548896 | 1,26360793 | 0,37056 | 0,63997937 | zinc finger protein 223 |
| ENSG00000175806 | MSRA | 0,337525438 | 1,263587384 | 0,49355 | 0,7263798 | methionine sulfoxide reductase A |
| ENSG00000255154 | HTD2 | 0,337500495 | 1,263565538 | 0,31073 | 0,5987267 | hydroxyacyl-thioester dehydratase type 2 |
| ENSG00000159063 | ALG8 | 0,337458397 | 1,263528667 | 0,20065 | 0,5214727 | ALG8 alpha-1,3-glucosyltransferase |
| ENSG00000272886 | DCP1A | 0,337448752 | 1,26352022 | 0,27527 | 0,57491355 | decapping mRNA 1A |
| ENSG00000109572 | CLCN3 | 0,337379008 | 1,26345914 | 0,37849 | 0,64679625 | chloride voltage-gated channel 3 |
| ENSG00000120885 | CLU | 0,337355128 | 1,263438226 | 0,27057 | 0,5723494 | clusterin |
| ENSG00000111880 | RNGTT | 0,337351721 | 1,263435243 | 0,22478 | 0,54036191 | RNA guanylyltransferase and 5'-phosphatase |
| ENSG00000158161 | EYA3 | 0,337334621 | 1,263420268 | 0,0917 | 0,39166375 | EYA transcriptional coactivator and phosphatase 3 |

| | | | | | | |
|---|---|---|---|---|---|---|
| ENSG00000079335 | CDC14A | 0,337323448 | 1,263410483 | 0,41016 | 0,66821634 | cell division cycle 14A |
| ENSG00000241112 | RPL29P14 | 0,33732101 | 1,263408348 | 0,57958 | 0,78292908 | ribosomal protein L29 pseudogene 14 |
| ENSG00000205213 | LGR4 | 0,337302351 | 1,263392008 | 0,13342 | 0,4594073 | leucine rich repeat containing G protein-coupled receptor 4 |
| ENSG00000141401 | IMPA2 | 0,337219417 | 1,263319384 | 0,27858 | 0,5767806 | inositol monophosphatase 2 |
| ENSG00000182810 | DDX28 | 0,33711593 | 1,263228767 | 0,12156 | 0,44093327 | DEAD-box helicase 28 |
| ENSG00000117984 | CTSD | 0,337084647 | 1,263201375 | 0,07471 | 0,35882064 | cathepsin D |
| ENSG00000178917 | ZNF852 | 0,337061043 | 1,263180708 | 0,18794 | 0,51107563 | zinc finger protein 852 |
| ENSG00000281626 | ZNF852 | 0,337061043 | 1,263180708 | 0,18794 | 0,51107563 | zinc finger protein 852 |
| ENSG00000147155 | EBP | 0,337049772 | 1,26317084 | 0,14343 | 0,47083258 | EBP cholestenol delta-isomerase |
| ENSG00000166833 | NAV2 | 0,337047627 | 1,263168962 | 0,40865 | 0,66743928 | neuron navigator 2 |
| ENSG00000066926 | FECH | 0,337002747 | 1,263129667 | 0,21256 | 0,53151956 | ferrochelatase |
| ENSG00000243431 | RPL5P30 | 0,336993384 | 1,263121469 | 0,3872 | 0,65225036 | ribosomal protein L5 pseudogene 30 |
| ENSG00000126756 | UXT | 0,336934913 | 1,263070277 | 0,40199 | 0,66341002 | ubiquitously expressed prefoldin like chaperone |
| ENSG00000213047 | DENND1B | 0,336931473 | 1,263067265 | 0,49057 | 0,72473187 | DENN domain containing 1B |
| ENSG00000284024 | AC069544,2 | 0,336822698 | 1,262972038 | 0,06568 | 0,33945082 | zinc finger and SCAN domain containing 29 |
| ENSG00000284898 | AL136419,1 | 0,336806279 | 1,262957664 | 0,59578 | 0,79349831 | novel transcript |
| ENSG00000285467 | AL136419,1 | 0,336806279 | 1,262957664 | 0,59578 | 0,79349831 | novel transcript |
| ENSG00000263158 | TAS2R63P | 0,336800438 | 1,26295255 | 0,62091 | 0,80776776 | taste 2 receptor member 63 pseudogene |
| ENSG00000256019 | TAS2R63P | 0,336800438 | 1,26295255 | 0,62091 | 0,80776776 | taste 2 receptor member 63 pseudogene |
| ENSG00000171806 | METTL18 | 0,336706336 | 1,262870175 | 0,14822 | 0,47545432 | methyltransferase like 18 |
| ENSG00000283293 | RN7SK | 0,336616575 | 1,262791605 | 0,33425 | 0,6140941 | RNA component of 7SK nuclear ribonucleoprotein |
| ENSG00000225376 | TMEM246-AS | 0,336568299 | 1,26274935 | 0,57342 | 0,77892286 | TMEM246 antisense RNA 1 |
| ENSG00000177119 | ANO6 | 0,336355921 | 1,262563475 | 0,34285 | 0,62033803 | anoctamin 6 |
| ENSG00000119685 | TTLL5 | 0,336235595 | 1,262458177 | 0,28031 | 0,57784691 | tubulin tyrosine ligase like 5 |
| ENSG00000286502 | AL645768,1 | 0,336194836 | 1,262422511 | 0,48338 | 0,71965182 | novel transcript, antisense to COL5A1 |
| ENSG00000141664 | ZCCHC2 | 0,336193165 | 1,262421048 | 0,18456 | 0,50835796 | zinc finger CCHC-type containing 2 |
| ENSG00000169684 | CHRNA5 | 0,336050049 | 1,262295822 | 0,24127 | 0,55027101 | cholinergic receptor nicotinic alpha 5 subunit |
| ENSG00000288130 | TNRC6A | 0,335982834 | 1,262237013 | 0,28288 | 0,58011711 | trinucleotide repeat containing adaptor 6A |
| ENSG00000172775 | PSME3IP1 | 0,33594941 | 1,26220777 | 0,18094 | 0,50450233 | proteasome activator subunit 3 interacting protein 1 |
| ENSG00000260091 | AC093752,3 | 0,335903614 | 1,262167704 | 0,57208 | 0,77840742 | novel transcript |
| ENSG00000229404 | LINC00858 | 0,335801586 | 1,262078447 | 0,47204 | 0,71132328 | long intergenic non-protein coding RNA 858 |
| ENSG00000148773 | MKI67 | 0,335733602 | 1,262018975 | 0,3899 | 0,65400705 | marker of proliferation Ki-67 |

| | | | | | | |
|---|---|---|---|---|---|---|
| ENSG00000086102 | NFX1 | 0,335732265 | 1,262017805 | 0,27015 | 0,57192517 | nuclear transcription factor, X-box binding 1 |
| ENSG00000198431 | TXNRD1 | 0,335731233 | 1,262016903 | 0,15268 | 0,48061452 | thioredoxin reductase 1 |
| ENSG00000065183 | WDR3 | 0,335600062 | 1,261902164 | 0,11016 | 0,42269499 | WD repeat domain 3 |
| ENSG00000152422 | XRCC4 | 0,335586392 | 1,261890207 | 0,30324 | 0,59353249 | X-ray repair cross complementing 4 |
| ENSG00000215105 | TTC3P1 | 0,335583291 | 1,261887495 | 0,4397 | 0,68844641 | tetratricopeptide repeat domain 3 pseudogene 1 |
| ENSG00000131653 | TRAF7 | 0,335563457 | 1,261870147 | 0,13899 | 0,4664494 | TNF receptor associated factor 7 |
| ENSG00000258476 | LINC02207 | 0,335553176 | 1,261861154 | 0,43839 | 0,68769293 | long intergenic non-protein coding RNA 2207 |
| ENSG00000116903 | EXOC8 | 0,335522248 | 1,261834104 | 0,16145 | 0,48896201 | exocyst complex component 8 |
| ENSG00000083290 | ULK2 | 0,33546843 | 1,261787033 | 0,22419 | 0,54007888 | unc-51 like autophagy activating kinase 2 |
| ENSG00000172167 | MTBP | 0,335379726 | 1,261709454 | 0,27313 | 0,57344244 | MDM2 binding protein |
| ENSG00000249558 | RCC2P4 | 0,335353103 | 1,261686171 | 0,62069 | 0,80776776 | regulator of chromosome condensation 2 pseudogene 4 |
| ENSG00000176986 | SEC24C | 0,335297626 | 1,261637656 | 0,24461 | 0,55271519 | SEC24 homolog C, COPII coat complex component |
| ENSG00000186260 | MRTFB | 0,335272348 | 1,261615551 | 0,34039 | 0,61897407 | myocardin related transcription factor B |
| ENSG00000080546 | SESN1 | 0,335153419 | 1,261511553 | 0,33106 | 0,6112353 | sestrin 1 |
| ENSG00000156052 | GNAQ | 0,335145509 | 1,261504637 | 0,20974 | 0,52867318 | G protein subunit alpha q |
| ENSG00000139546 | TARBP2 | 0,335102277 | 1,261466835 | 0,19521 | 0,51603779 | TARBP2 subunit of RISC loading complex |
| ENSG00000197142 | ACSL5 | 0,335096813 | 1,261462057 | 0,5027 | 0,73209345 | acyl-CoA synthetase long chain family member 5 |
| ENSG00000248559 | AC109454,2 | 0,335060335 | 1,261430162 | 0,45267 | 0,69712152 | novel transcript |
| ENSG00000109184 | DCUN1D4 | 0,334916305 | 1,261304234 | 0,44257 | 0,69016964 | defective in cullin neddylation 1 domain containing 4 |
| ENSG00000183530 | PRR14L | 0,334546043 | 1,260980568 | 0,23995 | 0,54936679 | proline rich 14 like |
| ENSG00000109618 | SEPSECS | 0,334520034 | 1,260957835 | 0,46641 | 0,70665143 | Sep (O-phosphoserine) tRNA:Sec (selenocysteine) tRNA synthase |
| ENSG00000243649 | CFB | 0,334509056 | 1,26094824 | 0,48345 | 0,71971559 | complement factor B |
| ENSG00000273564 | FAM189A1 | 0,334502698 | 1,260942682 | 0,28217 | 0,57949345 | family with sequence similarity 189 member A1 |
| ENSG00000075151 | EIF4G3 | 0,334486275 | 1,260928329 | 0,20922 | 0,52826608 | eukaryotic translation initiation factor 4 gamma 3 |
| ENSG00000124380 | SNRNP27 | 0,334486266 | 1,260928321 | 0,31389 | 0,60059594 | small nuclear ribonucleoprotein U4/U6,U5 subunit 27 |
| ENSG00000099901 | RANBP1 | 0,334483544 | 1,260925942 | 0,16702 | 0,49293221 | RAN binding protein 1 |
| ENSG00000107518 | ATRNL1 | 0,334445614 | 1,260892791 | 0,29517 | 0,58886617 | attractin like 1 |
| ENSG00000284869 | EEFSEC | 0,334359899 | 1,26081788 | 0,2402 | 0,54948131 | eukaryotic elongation factor, selenocysteine-tRNA specific |
| ENSG00000251141 | MRPS30-DT | 0,334277004 | 1,260745438 | 0,60545 | 0,79865565 | MRPS30 divergent transcript |
| ENSG00000270170 | NCBP2AS2 | 0,334276805 | 1,260745264 | 0,34949 | 0,62561317 | NCBP2 antisense 2 (head to head) |
| ENSG00000117602 | RCAN3 | 0,334239176 | 1,26071238 | 0,19895 | 0,52006724 | RCAN family member 3 |
| ENSG00000128708 | HAT1 | 0,334175134 | 1,260656418 | 0,37628 | 0,64458342 | histone acetyltransferase 1 |

| | | | | | | |
|---|---|---|---|---|---|---|
| ENSG00000129055 | ANAPC13 | 0,334022054 | 1,260522661 | 0,27967 | 0,57745167 | anaphase promoting complex subunit 13 |
| ENSG00000233589 | AL138789,1 | 0,333988805 | 1,26049361 | 0,56249 | 0,77186635 | novel transcript |
| ENSG00000255153 | TOLLIP-AS1 | 0,333981779 | 1,260487471 | 0,5244 | 0,74828894 | TOLLIP antisense RNA 1 (head to head) |
| ENSG00000164919 | COX6C | 0,333928325 | 1,26044077 | 0,36889 | 0,63870499 | cytochrome c oxidase subunit 6C |
| ENSG00000147526 | TACC1 | 0,333897209 | 1,260413585 | 0,41768 | 0,67421607 | transforming acidic coiled-coil containing protein 1 |
| ENSG00000278505 | C17orf78 | 0,333863328 | 1,260383985 | 0,76124 | 0,88879633 | chromosome 17 open reading frame 78 |
| ENSG00000083535 | PIBF1 | 0,333860858 | 1,260381827 | 0,38914 | 0,65363435 | progesterone immunomodulatory binding factor 1 |
| ENSG00000044115 | CTNNA1 | 0,333852567 | 1,260374583 | 0,3017 | 0,59283946 | catenin alpha 1 |
| ENSG00000175548 | ALG10B | 0,333599076 | 1,260153147 | 0,28683 | 0,58326438 | ALG10 alpha-1,2-glucosyltransferase B |
| ENSG00000112183 | RBM24 | 0,33356981 | 1,260127584 | 0,44502 | 0,69205935 | RNA binding motif protein 24 |
| ENSG00000122140 | MRPS2 | 0,333520601 | 1,260084603 | 0,14414 | 0,47172279 | mitochondrial ribosomal protein S2 |
| ENSG00000040633 | PHF23 | 0,33351312 | 1,26007807 | 0,01746 | 0,17420662 | PHD finger protein 23 |
| ENSG00000164715 | LMTK2 | 0,33348849 | 1,260056558 | 0,23525 | 0,54672923 | lemur tyrosine kinase 2 |
| ENSG00000136937 | NCBP1 | 0,333441454 | 1,260015476 | 0,21853 | 0,53584458 | nuclear cap binding protein subunit 1 |
| ENSG00000280639 | LINC02204 | 0,333413386 | 1,259990963 | 0,68827 | 0,84815887 | long intergenic non-protein coding RNA 2204 |
| ENSG00000229273 | BX664615,1 | 0,333324509 | 1,259913343 | 0,61725 | 0,8057177 | novel pseudogene |
| ENSG00000198276 | UCKL1 | 0,333302014 | 1,259893699 | 0,17057 | 0,49641416 | uridine-cytidine kinase 1 like 1 |
| ENSG00000085832 | EPS15 | 0,333262837 | 1,259859486 | 0,29721 | 0,59031059 | epidermal growth factor receptor pathway substrate 15 |
| ENSG00000118420 | UBE3D | 0,333154788 | 1,259765134 | 0,12007 | 0,43868155 | ubiquitin protein ligase E3D |
| ENSG00000114331 | ACAP2 | 0,333097245 | 1,259714889 | 0,11284 | 0,42691181 | ArfGAP with coiled-coil, ankyrin repeat and PH domains 2 |
| ENSG00000216937 | CCDC7 | 0,333034321 | 1,259659946 | 0,43798 | 0,68747773 | coiled-coil domain containing 7 |
| ENSG00000176387 | HSD11B2 | 0,333002383 | 1,259632061 | 0,40837 | 0,66721014 | hydroxysteroid 11-beta dehydrogenase 2 |
| ENSG00000278644 | PCGF2 | 0,332981493 | 1,259613821 | 0,17562 | 0,49954066 | polycomb group ring finger 2 |
| ENSG00000102081 | FMR1 | 0,332973116 | 1,259606507 | 0,24966 | 0,55657567 | FMRP translational regulator 1 |
| ENSG00000176102 | CSTF3 | 0,332953028 | 1,259588969 | 0,36723 | 0,63724986 | cleavage stimulation factor subunit 3 |
| ENSG00000206053 | JPT2 | 0,332933233 | 1,259571686 | 0,17016 | 0,49606609 | Jupiter microtubule associated homolog 2 |
| ENSG00000175061 | SNHG29 | 0,33285875 | 1,25950666 | 0,26915 | 0,57086439 | small nucleolar RNA host gene 29 |
| ENSG00000110958 | PTGES3 | 0,332731693 | 1,259395741 | 0,26714 | 0,56958284 | prostaglandin E synthase 3 |
| ENSG00000155897 | ADCY8 | 0,332684649 | 1,259354674 | 0,49733 | 0,72869621 | adenylate cyclase 8 |
| ENSG00000261542 | AC011978,2 | 0,332606462 | 1,259286425 | 0,28542 | 0,58187605 | novel transcript |
| ENSG00000106484 | MEST | 0,332577675 | 1,259261298 | 0,45506 | 0,69914733 | mesoderm specific transcript |
| ENSG00000158158 | CNNM4 | 0,332546715 | 1,259234275 | 0,22761 | 0,54168591 | cyclin and CBS domain divalent metal cation transport mediator 4 |

| | | | | | | |
|---|---|---|---|---|---|---|
| ENSG00000287888 | AC087463,3 | 0,332522761 | 1,259213367 | 0,63007 | 0,8132682 | novel transcript |
| ENSG00000288448 | AC087463,4 | 0,332522761 | 1,259213367 | 0,63007 | 0,8132682 | novel transcript |
| ENSG00000182512 | GLRX5 | 0,332477519 | 1,25917388 | 0,25747 | 0,56160845 | glutaredoxin 5 |
| ENSG00000007202 | KIAA0100 | 0,332443668 | 1,259144335 | 0,21159 | 0,53038627 | KIAA0100 |
| ENSG00000256664 | AC025423,2 | 0,332441353 | 1,259142314 | 0,82992 | 0,92193085 | ribosomal L24 domain containing 1 (RSL24D1) pseudogene |
| ENSG00000126216 | TUBGCP3 | 0,33225597 | 1,258980528 | 0,34993 | 0,62563749 | tubulin gamma complex associated protein 3 |
| ENSG00000146247 | PHIP | 0,332250684 | 1,258975915 | 0,26972 | 0,57140742 | pleckstrin homology domain interacting protein |
| ENSG00000131400 | NAPSA | 0,332190965 | 1,258923803 | 0,36478 | 0,6350493 | napsin A aspartic peptidase |
| ENSG00000178234 | GALNT11 | 0,332189609 | 1,258922619 | 0,20075 | 0,52160854 | polypeptide N-acetylgalactosaminyltransferase 11 |
| ENSG00000119912 | IDE | 0,332150703 | 1,258888669 | 0,10039 | 0,4067769 | insulin degrading enzyme |
| ENSG00000112339 | HBS1L | 0,332149563 | 1,258887675 | 0,30077 | 0,59208755 | HBS1 like translational GTPase |
| ENSG00000161905 | ALOX15 | 0,332130891 | 1,258871382 | 0,52951 | 0,7512527 | arachidonate 15-lipoxygenase |
| ENSG00000183148 | ANKRD20A2P | 0,332103111 | 1,258847141 | 0,77078 | 0,89382253 | ankyrin repeat domain 20 family member A2, pseudogene |
| ENSG00000144015 | TRIM43 | 0,33196568 | 1,25872723 | 0,66581 | 0,83366537 | tripartite motif containing 43 |
| ENSG00000286456 | AC016769,5 | 0,331908391 | 1,258677247 | 0,23896 | 0,54898114 | novel transcript |
| ENSG00000112541 | PDE10A | 0,331902395 | 1,258672016 | 0,30746 | 0,59658969 | phosphodiesterase 10A |
| ENSG00000256589 | ENPP7P5 | 0,331871982 | 1,258645482 | 0,66182 | 0,83156405 | ectonucleotide pyrophosphatase/phosphodiesterase 7 pseudogene 5 |
| ENSG00000173692 | PSMD1 | 0,331867196 | 1,258641307 | 0,17288 | 0,49768904 | proteasome 26S subunit, non-ATPase 1 |
| ENSG00000169251 | NMD3 | 0,331839064 | 1,258616764 | 0,3505 | 0,62601861 | NMD3 ribosome export adaptor |
| ENSG00000166037 | CEP57 | 0,331823822 | 1,258603467 | 0,35064 | 0,62606342 | centrosomal protein 57 |
| ENSG00000274392 | AC093724,1 | 0,331815924 | 1,258596577 | 0,68709 | 0,8473827 | neurobeachin (NBEA) pseudogene |
| ENSG00000141034 | GID4 | 0,33180319 | 1,258585468 | 0,13852 | 0,46578766 | GID complex subunit 4 homolog |
| ENSG00000117697 | NSL1 | 0,331592824 | 1,258401961 | 0,16346 | 0,48977932 | NSL1 component of MIS12 kinetochore complex |
| ENSG00000102024 | PLS3 | 0,331423824 | 1,258254558 | 0,46937 | 0,70890278 | plastin 3 |
| ENSG00000122674 | CCZ1 | 0,331347024 | 1,258187578 | 0,28709 | 0,58349837 | CCZ1 homolog, vacuolar protein trafficking and biogenesis associated |
| ENSG00000148358 | GPR107 | 0,331344775 | 1,258185617 | 0,26649 | 0,56917012 | G protein-coupled receptor 107 |
| ENSG00000100075 | SLC25A1 | 0,331287167 | 1,258135378 | 0,15141 | 0,47956749 | solute carrier family 25 member 1 |
| ENSG00000185009 | AP3M1 | 0,331223097 | 1,258079505 | 0,34381 | 0,62120234 | adaptor related protein complex 3 subunit mu 1 |
| ENSG00000182010 | RTKN2 | 0,331196486 | 1,2580563 | 0,49772 | 0,72908464 | rhotekin 2 |
| ENSG00000097033 | SH3GLB1 | 0,331049617 | 1,257928234 | 0,37286 | 0,64172884 | SH3 domain containing GRB2 like, endophilin B1 |
| ENSG00000089289 | IGBP1 | 0,331042318 | 1,25792187 | 0,27127 | 0,57271346 | immunoglobulin binding protein 1 |

| | | | | | | |
|---|---|---|---|---|---|---|
| ENSG00000259814 | AC102941,1 | 0,330964029 | 1,257853609 | 0,68837 | 0,84817878 | novel transcript |
| ENSG00000230988 | RPL23AP11 | 0,330941262 | 1,257833759 | 0,69676 | 0,85309963 | ribosomal protein L23a pseudogene 11 |
| ENSG00000205060 | SLC35B4 | 0,330920733 | 1,257815861 | 0,22233 | 0,53859118 | solute carrier family 35 member B4 |
| ENSG00000108588 | CCDC47 | 0,330875808 | 1,257776693 | 0,29608 | 0,58954675 | coiled-coil domain containing 47 |
| ENSG00000221962 | TMEM14EP | 0,330868946 | 1,257770711 | 0,37503 | 0,64346511 | transmembrane protein 14E, pseudogene |
| ENSG00000159459 | UBR1 | 0,330860697 | 1,257763519 | 0,37406 | 0,64254935 | ubiquitin protein ligase E3 component n-recognin 1 |
| ENSG00000067113 | PLPP1 | 0,330828845 | 1,257735751 | 0,13399 | 0,4601411 | phospholipid phosphatase 1 |
| ENSG00000140650 | PMM2 | 0,330791653 | 1,257703327 | 0,19416 | 0,51562158 | phosphomannomutase 2 |
| ENSG00000119950 | MXI1 | 0,330675839 | 1,257602368 | 0,34467 | 0,62202834 | MAX interactor 1, dimerization protein |
| ENSG00000104517 | UBR5 | 0,330632167 | 1,257564299 | 0,25609 | 0,56065266 | ubiquitin protein ligase E3 component n-recognin 5 |
| ENSG00000230734 | RPL10P3 | 0,330607427 | 1,257542735 | 0,60263 | 0,79710677 | ribosomal protein L10 pseudogene 3 |
| ENSG00000235700 | CYCSP52 | 0,330602489 | 1,25753843 | 0,70867 | 0,85960075 | CYCS pseudogene 52 |
| ENSG00000241537 | AC134050,1 | 0,330593222 | 1,257530352 | 0,62662 | 0,81105911 | U2-associated SR140 protein (SR140) pseudogene |
| ENSG00000259165 | DDX18P1 | 0,330563598 | 1,257504531 | 0,45329 | 0,69750019 | DEAD-box helicase 18 pseudogene 1 |
| ENSG00000229992 | HMGB3P9 | 0,330549737 | 1,257492449 | 0,54447 | 0,76055278 | high mobility group box 3 pseudogene 9 |
| ENSG00000206559 | ZCWPW2 | 0,330515045 | 1,257462211 | 0,46716 | 0,70714082 | zinc finger CW-type and PWWP domain containing 2 |
| ENSG00000244045 | TMEM199 | 0,330504798 | 1,25745328 | 0,09008 | 0,38819633 | transmembrane protein 199 |
| ENSG00000224680 | PLA2G12AP1 | 0,330428938 | 1,257387162 | 0,51611 | 0,74198903 | phospholipase A2 group XIIA pseudogene 1 |
| ENSG00000236254 | MTND4P14 | 0,330379654 | 1,257344208 | 0,65307 | 0,8262455 | MT-ND4 pseudogene 14 |
| ENSG00000025796 | SEC63 | 0,330223779 | 1,257208367 | 0,41777 | 0,6742688 | SEC63 homolog, protein translocation regulator |
| ENSG00000131503 | ANKHD1 | 0,330189357 | 1,257178371 | 0,3138 | 0,600508 | ankyrin repeat and KH domain containing 1 |
| ENSG00000183207 | RUVBL2 | 0,330083594 | 1,257086212 | 0,10776 | 0,41834811 | RuvB like AAA ATPase 2 |
| ENSG00000119682 | AREL1 | 0,330019276 | 1,25703017 | 0,14158 | 0,46809977 | apoptosis resistant E3 ubiquitin protein ligase 1 |
| ENSG00000123213 | NLN | 0,33000946 | 1,257021617 | 0,3194 | 0,60429866 | neurolysin |
| ENSG00000260089 | ADAM3B | 0,329926543 | 1,256949373 | 0,47793 | 0,71563665 | ADAM metallopeptidase domain 3B (pseudogene) |
| ENSG00000135249 | RINT1 | 0,329834898 | 1,25686953 | 0,23767 | 0,54861973 | RAD50 interactor 1 |
| ENSG00000113638 | TTC33 | 0,329696257 | 1,256748752 | 0,28202 | 0,57936013 | tetratricopeptide repeat domain 33 |
| ENSG00000281977 | AC244131,3 | 0,329640626 | 1,256700293 | 0,67538 | 0,84021377 | taste receptor, type 2, member 7 pseudogene |
| ENSG00000256657 | AC244131,1 | 0,329640626 | 1,256700293 | 0,67538 | 0,84021377 | taste receptor, type 2, member 7 pseudogene |
| ENSG00000144233 | AMMECR1L | 0,329630875 | 1,256691799 | 0,11522 | 0,43084701 | AMMECR1 like |
| ENSG00000214226 | C17orf67 | 0,329506626 | 1,256583573 | 0,30926 | 0,59776375 | chromosome 17 open reading frame 67 |
| ENSG00000013016 | EHD3 | 0,329504837 | 1,256582016 | 0,20976 | 0,52867318 | EH domain containing 3 |
| ENSG00000140598 | EFL1 | 0,32936106 | 1,256456793 | 0,41634 | 0,67315741 | elongation factor like GTPase 1 |

| | | | | | | |
|---|---|---|---|---|---|---|
| ENSG00000149084 | HSD17B12 | 0,32934123 | 1,256439522 | 0,22081 | 0,53776578 | hydroxysteroid 17-beta dehydrogenase 12 |
| ENSG00000179899 | PHC1P1 | 0,329262952 | 1,256371352 | 0,5159 | 0,7419128 | polyhomeotic homolog 1 pseudogene 1 |
| ENSG00000235795 | AC093157,2 | 0,329153363 | 1,25627592 | 0,75733 | 0,88668471 | novel transcript |
| ENSG00000142252 | GEMIN7 | 0,329144067 | 1,256267825 | 0,42438 | 0,67819918 | gem nuclear organelle associated protein 7 |
| ENSG00000030110 | BAK1 | 0,329082428 | 1,256214152 | 0,0737 | 0,35703856 | BCL2 antagonist/killer 1 |
| ENSG00000170325 | PRDM10 | 0,329068305 | 1,256201855 | 0,30621 | 0,59517372 | PR/SET domain 10 |
| ENSG00000287129 | AC097500,1 | 0,329021705 | 1,256161279 | 0,6618 | 0,83156405 | novel transcript |
| ENSG00000112874 | NUDT12 | 0,328920657 | 1,2560733 | 0,4539 | 0,69794076 | nudix hydrolase 12 |
| ENSG00000085872 | CHERP | 0,328872702 | 1,256031549 | 0,13137 | 0,45639138 | calcium homeostasis endoplasmic reticulum protein |
| ENSG00000221866 | PLXNA4 | 0,328862366 | 1,25602255 | 0,47973 | 0,71714401 | plexin A4 |
| ENSG00000124225 | PMEPA1 | 0,328839852 | 1,25600295 | 0,12868 | 0,45191041 | prostate transmembrane protein, androgen induced 1 |
| ENSG00000180113 | TDRD6 | 0,328821684 | 1,255987133 | 0,43533 | 0,68601186 | tudor domain containing 6 |
| ENSG00000198919 | DZIP3 | 0,328808785 | 1,255975903 | 0,44472 | 0,69205935 | DAZ interacting zinc finger protein 3 |
| ENSG00000114933 | INO80D | 0,328801882 | 1,255969893 | 0,39968 | 0,66186365 | INO80 complex subunit D |
| ENSG00000283118 | AC009951,5 | 0,328760085 | 1,255933506 | 0,732 | 0,8728363 | novel transcript, antisense to ZEB2 |
| ENSG00000136830 | NIBAN2 | 0,328735547 | 1,255912145 | 0,3033 | 0,59355788 | niban apoptosis regulator 2 |
| ENSG00000175606 | TMEM70 | 0,328707878 | 1,255888059 | 0,30915 | 0,59774312 | transmembrane protein 70 |
| ENSG00000242220 | TCP10L | 0,328686139 | 1,255869134 | 0,38535 | 0,65107577 | t-complex 10 like |
| ENSG00000251614 | HSPA8P19 | 0,328621108 | 1,255812527 | 0,63863 | 0,81863386 | heat shock protein family A (Hsp70) member pseudogene 19 |
| ENSG00000233967 | AL359715,1 | 0,328565689 | 1,255764287 | 0,22573 | 0,5409527 | novel transcript |
| ENSG00000120008 | WDR11 | 0,328564217 | 1,255763006 | 0,25116 | 0,55719428 | WD repeat domain 11 |
| ENSG00000250848 | AC021087,2 | 0,328490904 | 1,255699193 | 0,67188 | 0,83759902 | protein phosphatase 4, regulatory subunit 2 (PPP4R2) pseudogene |
| ENSG00000167608 | TMC4 | 0,328471792 | 1,255682559 | 0,74036 | 0,87691301 | transmembrane channel like 4 |
| ENSG00000014641 | MDH1 | 0,328265518 | 1,255503037 | 0,33935 | 0,61834932 | malate dehydrogenase 1 |
| ENSG00000178999 | AURKB | 0,328241395 | 1,255482044 | 0,17062 | 0,49641416 | aurora kinase B |
| ENSG00000257267 | ZNF271P | 0,328218101 | 1,255461773 | 0,36999 | 0,63943131 | zinc finger protein 271, pseudogene |
| ENSG00000127920 | GNG11 | 0,328169359 | 1,255419358 | 0,5092 | 0,73706413 | G protein subunit gamma 11 |
| ENSG00000204850 | AC011484,1 | 0,32811997 | 1,25537638 | 0,86024 | 0,93716999 | novel transcript, antisense to PPP5D1 |
| ENSG00000167434 | CA4 | 0,328110141 | 1,255367827 | 0,57175 | 0,77829419 | carbonic anhydrase 4 |
| ENSG00000283510 | INO80D | 0,328095833 | 1,255355377 | 0,39978 | 0,66193009 | INO80 complex subunit D |
| ENSG00000140743 | CDR2 | 0,328078197 | 1,255340031 | 0,19679 | 0,51798099 | cerebellar degeneration related protein 2 |

| | | | | | | |
|---|---|---|---|---|---|---|
| ENSG00000263238 | CTSO | 0,328077018 | 1,255339006 | 0,17218 | 0,49710979 | cathepsin O |
| ENSG00000131051 | RBM39 | 0,327975246 | 1,255250453 | 0,25941 | 0,56317998 | RNA binding motif protein 39 |
| ENSG00000104756 | KCTD9 | 0,327956309 | 1,255233977 | 0,28015 | 0,57784691 | potassium channel tetramerization domain containing 9 |
| ENSG00000168438 | CDC40 | 0,327927429 | 1,25520885 | 0,22285 | 0,53904413 | cell division cycle 40 |
| ENSG00000224356 | AL356966,1 | 0,327892131 | 1,255178139 | 0,43025 | 0,68294374 | novel transcript |
| ENSG00000222724 | RNU2-63P | 0,327786077 | 1,255085874 | 0,48053 | 0,71764212 | RNA, U2 small nuclear 63, pseudogene |
| ENSG00000149547 | EI24 | 0,327735418 | 1,255041802 | 0,17731 | 0,50187931 | EI24 autophagy associated transmembrane protein |
| ENSG00000132970 | WASF3 | 0,327718838 | 1,25502738 | 0,3657 | 0,6359214 | WASP family member 3 |
| ENSG00000144730 | IL17RD | 0,327678636 | 1,254992408 | 0,22453 | 0,54035379 | interleukin 17 receptor D |
| ENSG00000136045 | PWP1 | 0,327584112 | 1,254910184 | 0,29181 | 0,5867146 | PWP1 homolog, endonuclein |
| ENSG00000234370 | GTF2H4 | 0,327569822 | 1,254897755 | 0,71781 | 0,8648887 | general transcription factor IIH subunit 4 |
| ENSG00000174448 | STARD6 | 0,327521795 | 1,25485598 | 0,66512 | 0,83332469 | StAR related lipid transfer domain containing 6 |
| ENSG00000170779 | CDCA4 | 0,327429501 | 1,254775705 | 0,14852 | 0,47603542 | cell division cycle associated 4 |
| ENSG00000002016 | RAD52 | 0,327363834 | 1,254718592 | 0,17375 | 0,49846579 | RAD52 homolog, DNA repair protein |
| ENSG00000267560 | AC027514,2 | 0,327352181 | 1,254708458 | 0,57559 | 0,78031972 | novel transcript, antisense to KIAA1468 |
| ENSG00000058799 | YIPF1 | 0,327349669 | 1,254706273 | 0,21235 | 0,53117701 | Yip1 domain family member 1 |
| ENSG00000196367 | TRRAP | 0,327323421 | 1,254683446 | 0,12862 | 0,45191041 | transformation/transcription domain associated protein |
| ENSG00000237071 | TRIM27 | 0,327286245 | 1,254651115 | 0,22443 | 0,54033281 | tripartite motif containing 27 |
| ENSG00000102931 | ARL2BP | 0,327273152 | 1,254639728 | 0,42099 | 0,67619901 | ADP ribosylation factor like GTPase 2 binding protein |
| ENSG00000276855 | AC015922,3 | 0,327266785 | 1,254634192 | 0,55446 | 0,76745801 | novel transcript |
| ENSG00000140564 | FURIN | 0,327219471 | 1,254593046 | 0,05441 | 0,3122594 | furin, paired basic amino acid cleaving enzyme |
| ENSG00000143256 | PFDN2 | 0,327200857 | 1,254576859 | 0,24928 | 0,55642837 | prefoldin subunit 2 |
| ENSG00000232070 | TMEM253 | 0,327160655 | 1,254541899 | 0,45022 | 0,69562948 | transmembrane protein 253 |
| ENSG00000102096 | PIM2 | 0,327142852 | 1,254526419 | 0,1718 | 0,49710979 | Pim-2 proto-oncogene, serine/threonine kinase |
| ENSG00000072364 | AFF4 | 0,327108728 | 1,254496746 | 0,33512 | 0,61480301 | AF4/FMR2 family member 4 |
| ENSG00000007402 | CACNA2D2 | 0,327071428 | 1,254464311 | 0,14478 | 0,4723264 | calcium voltage-gated channel auxiliary subunit alpha2delta 2 |
| ENSG00000217512 | AL356776,1 | 0,327025009 | 1,254423949 | 0,62737 | 0,8114294 | nucleoporin 155kDa pseudogene |
| ENSG00000249167 | SEMA6A-AS2 | 0,327002079 | 1,254404012 | 0,51874 | 0,74360005 | SEMA6A antisense RNA 2 |
| ENSG00000185519 | FAM131C | 0,326998017 | 1,25440048 | 0,56937 | 0,77665073 | family with sequence similarity 131 member C |
| ENSG00000100804 | PSMB5 | 0,326877448 | 1,254295652 | 0,27717 | 0,57572679 | proteasome 20S subunit beta 5 |
| ENSG00000188846 | RPL14 | 0,326752251 | 1,254186809 | 0,18745 | 0,51074735 | ribosomal protein L14 |
| ENSG00000066322 | ELOVL1 | 0,326711934 | 1,254151761 | 0,24353 | 0,55229687 | ELOVL fatty acid elongase 1 |

| | | | | | | |
|---|---|---|---|---|---|---|
| ENSG00000115657 | ABCB6 | 0,326690212 | 1,254132877 | 0,22383 | 0,53967355 | ATP binding cassette subfamily B member 6 (Langereis blood group) |
| ENSG00000085719 | CPNE3 | 0,326688397 | 1,254131299 | 0,49834 | 0,7296 | copine 3 |
| ENSG00000140876 | NUDT7 | 0,326687679 | 1,254130676 | 0,51721 | 0,74250326 | nudix hydrolase 7 |
| ENSG00000137575 | SDCBP | 0,326665519 | 1,254111412 | 0,47131 | 0,7106605 | syndecan binding protein |
| ENSG00000135750 | KCNK1 | 0,32662813 | 1,254078911 | 0,49386 | 0,72653694 | potassium two pore domain channel subfamily K member 1 |
| ENSG00000128591 | FLNC | 0,326590215 | 1,254045953 | 0,15693 | 0,48425071 | filamin C |
| ENSG00000127328 | RAB3IP | 0,32658323 | 1,254039881 | 0,24257 | 0,551495 | RAB3A interacting protein |
| ENSG00000110048 | OSBP | 0,326564573 | 1,254023664 | 0,21426 | 0,53334744 | oxysterol binding protein |
| ENSG00000126226 | PCID2 | 0,326523911 | 1,25398832 | 0,22262 | 0,53875487 | PCI domain containing 2 |
| ENSG00000168795 | ZBTB5 | 0,326509953 | 1,253976188 | 0,30587 | 0,59512079 | zinc finger and BTB domain containing 5 |
| ENSG00000145439 | CBR4 | 0,326438343 | 1,253913947 | 0,41111 | 0,6689497 | carbonyl reductase 4 |
| ENSG00000103507 | BCKDK | 0,326334481 | 1,253823679 | 0,16059 | 0,48791491 | branched chain keto acid dehydrogenase kinase |
| ENSG00000288445 | HPS5 | 0,326282882 | 1,253778836 | 0,1614 | 0,48892403 | HPS5 biogenesis of lysosomal organelles complex 2 subunit 2 |
| ENSG00000066084 | DIP2B | 0,326282826 | 1,253778787 | 0,24953 | 0,55657567 | disco interacting protein 2 homolog B |
| ENSG00000113649 | TCERG1 | 0,326235167 | 1,25373737 | 0,19398 | 0,51551719 | transcription elongation regulator 1 |
| ENSG00000272316 | AL021368,2 | 0,32613211 | 1,253647814 | 0,47911 | 0,71677199 | novel transcript |
| ENSG00000106125 | MINDY4 | 0,326088849 | 1,253610222 | 0,24118 | 0,55024202 | MINDY lysine 48 deubiquitinase 4 |
| ENSG00000248487 | ABHD14A | 0,325980581 | 1,253516148 | 0,19566 | 0,51643293 | abhydrolase domain containing 14A |
| ENSG00000155096 | AZIN1 | 0,325939599 | 1,25348054 | 0,24962 | 0,55657567 | antizyme inhibitor 1 |
| ENSG00000078369 | GNB1 | 0,325908404 | 1,253453437 | 0,31024 | 0,59841998 | G protein subunit beta 1 |
| ENSG00000019582 | CD74 | 0,325824739 | 1,253380748 | 0,50911 | 0,73703253 | CD74 molecule |
| ENSG00000197903 | H2BC12 | 0,325803466 | 1,253362267 | 0,57341 | 0,77892286 | H2B clustered histone 12 |
| ENSG00000104324 | CPQ | 0,325570011 | 1,253159466 | 0,42523 | 0,67890168 | carboxypeptidase Q |
| ENSG00000205268 | PDE7A | 0,325498349 | 1,25309722 | 0,16647 | 0,49245911 | phosphodiesterase 7A |
| ENSG00000156802 | ATAD2 | 0,325471974 | 1,253074312 | 0,35158 | 0,62675995 | ATPase family AAA domain containing 2 |
| ENSG00000100644 | HIF1A | 0,325436472 | 1,253043476 | 0,277 | 0,575636 | hypoxia inducible factor 1 subunit alpha |
| ENSG00000271366 | AC002128,2 | 0,325415239 | 1,253025034 | 0,64713 | 0,82343226 | novel transcript |
| ENSG00000169629 | RGPD8 | 0,325323989 | 1,252945784 | 0,40253 | 0,66388949 | RANBP2 like and GRIP domain containing 8 |
| ENSG00000136986 | DERL1 | 0,32529076 | 1,252916925 | 0,42057 | 0,67595875 | derlin 1 |
| ENSG00000144224 | UBXN4 | 0,325227885 | 1,252862322 | 0,23501 | 0,5465505 | UBX domain protein 4 |

| | | | | | | |
|---|---|---|---|---|---|---|
| ENSG00000164190 | NIPBL | 0,325210734 | 1,252847428 | 0,25251 | 0,55796133 | NIPBL cohesin loading factor |
| ENSG00000204856 | FAM216A | 0,325204106 | 1,252841672 | 0,16142 | 0,48892403 | family with sequence similarity 216 member A |
| ENSG00000186051 | TAL2 | 0,325016984 | 1,252679186 | 0,74089 | 0,87717177 | TAL bHLH transcription factor 2 |
| ENSG00000060237 | WNK1 | 0,324959945 | 1,25262966 | 0,19312 | 0,51517146 | WNK lysine deficient protein kinase 1 |
| ENSG00000023516 | AKAP11 | 0,324904487 | 1,252581509 | 0,36812 | 0,6380264 | A-kinase anchoring protein 11 |
| ENSG00000163812 | ZDHHC3 | 0,324859934 | 1,252542828 | 0,07412 | 0,35756831 | zinc finger DHHC-type palmitoyltransferase 3 |
| ENSG00000119929 | CUTC | 0,324850466 | 1,252534608 | 0,2856 | 0,58212096 | cutC copper transporter |
| ENSG00000159199 | ATP5MC1 | 0,324832197 | 1,252518747 | 0,32826 | 0,60971379 | ATP synthase membrane subunit c locus 1 |
| ENSG00000073584 | SMARCE1 | 0,324801313 | 1,252491934 | 0,35313 | 0,6278392 | SWI/SNF related, matrix associated, actin dependent regulator of chromatin, subfamily e, member 1 |
| ENSG00000277440 | AC012676,4 | 0,324701442 | 1,252405234 | 0,76099 | 0,88874409 | novel transcript, sense intronic with DNAJA3 |
| ENSG00000273056 | AL354694,1 | 0,324683134 | 1,25238934 | 0,75536 | 0,88541196 | novel transcript |
| ENSG00000287458 | AL033519,5 | 0,324597883 | 1,252315336 | 0,36092 | 0,63311211 | novel transcript, antisense to FKBP5 |
| ENSG00000213693 | SEC14L1P1 | 0,324461474 | 1,252196934 | 0,40571 | 0,66544818 | SEC14 like 1 pseudogene 1 |
| ENSG00000271857 | AL096865,1 | 0,324321614 | 1,252075548 | 0,72234 | 0,86734497 | novel transcript, antisense to RUNX2 |
| ENSG00000233797 | UFL1-AS1 | 0,324254498 | 1,252017301 | 0,56511 | 0,773443 | UFL1 antisense RNA 1 |
| ENSG00000163320 | CGGBP1 | 0,324122776 | 1,251902994 | 0,39987 | 0,66197344 | CGG triplet repeat binding protein 1 |
| ENSG00000155970 | MICU3 | 0,324116467 | 1,251897519 | 0,33705 | 0,61684812 | mitochondrial calcium uptake family member 3 |
| ENSG00000229688 | CRPPA-AS1 | 0,32385239 | 1,251668387 | 0,67469 | 0,83975921 | CRPPA antisense RNA 1 |
| ENSG00000165983 | PTER | 0,323836044 | 1,251654206 | 0,33674 | 0,61675087 | phosphotriesterase related |
| ENSG00000255776 | VDAC2P2 | 0,32381472 | 1,251635706 | 0,66429 | 0,83267806 | VDAC2 pseudogene 2 |
| ENSG00000103429 | BFAR | 0,323810848 | 1,251632346 | 0,35356 | 0,62829557 | bifunctional apoptosis regulator |
| ENSG00000135999 | EPC2 | 0,323782031 | 1,251607346 | 0,31292 | 0,59987747 | enhancer of polycomb homolog 2 |
| ENSG00000253250 | C8orf88 | 0,323750077 | 1,251579625 | 0,54511 | 0,7609742 | chromosome 8 open reading frame 88 |
| ENSG00000167797 | CDK2AP2 | 0,323727564 | 1,251560094 | 0,18582 | 0,50915451 | cyclin dependent kinase 2 associated protein 2 |
| ENSG00000151247 | EIF4E | 0,323624121 | 1,251470359 | 0,18568 | 0,50907679 | eukaryotic translation initiation factor 4E |
| ENSG00000179912 | R3HDM2 | 0,323593321 | 1,251443641 | 0,1152 | 0,43084185 | R3H domain containing 2 |
| ENSG00000063601 | MTMR1 | 0,323576518 | 1,251429066 | 0,30705 | 0,59617071 | myotubularin related protein 1 |
| ENSG00000070731 | ST6GALNAC2 | 0,323573092 | 1,251426094 | 0,61927 | 0,80699858 | ST6 N-acetylgalactosaminide alpha-2,6-sialyltransferase 2 |
| ENSG00000107758 | PPP3CB | 0,32357259 | 1,251425659 | 0,29239 | 0,58694246 | protein phosphatase 3 catalytic subunit beta |
| ENSG00000153933 | DGKE | 0,323563022 | 1,251417359 | 0,14098 | 0,46791293 | diacylglycerol kinase epsilon |
| ENSG00000186184 | POLR1D | 0,323506276 | 1,251368138 | 0,21468 | 0,53362255 | RNA polymerase I and III subunit D |
| ENSG00000141580 | WDR45B | 0,323482266 | 1,251347312 | 0,35534 | 0,62970025 | WD repeat domain 45B |

| | | | | | | |
|---|---|---|---|---|---|---|
| ENSG00000114388 | NPRL2 | 0,323476826 | 1,251342594 | 0,39741 | 0,65990329 | NPR2 like, GATOR1 complex subunit |
| ENSG00000276421 | AC012314,11 | 0,323404008 | 1,251279435 | 0,72057 | 0,86628014 | pre-mRNA processing factor 31 |
| ENSG00000274894 | AC012314,6 | 0,323404008 | 1,251279435 | 0,72057 | 0,86628014 | pre-mRNA processing factor 31 |
| ENSG00000274651 | AC012314,5 | 0,323404008 | 1,251279435 | 0,72057 | 0,86628014 | pre-mRNA processing factor 31 |
| ENSG00000275117 | AC012314,7 | 0,323404008 | 1,251279435 | 0,72057 | 0,86628014 | pre-mRNA processing factor 31 |
| ENSG00000277953 | AC012314,14 | 0,323404008 | 1,251279435 | 0,72057 | 0,86628014 | pre-mRNA processing factor 31 |
| ENSG00000274144 | AC012314,3 | 0,323404008 | 1,251279435 | 0,72057 | 0,86628014 | pre-mRNA processing factor 31 |
| ENSG00000277707 | AC012314,13 | 0,323404008 | 1,251279435 | 0,72057 | 0,86628014 | pre-mRNA processing factor 31 |
| ENSG00000277154 | AC012314,12 | 0,323404008 | 1,251279435 | 0,72057 | 0,86628014 | pre-mRNA processing factor 31 |
| ENSG00000198830 | HMGN2 | 0,323290802 | 1,251181254 | 0,34686 | 0,62361524 | high mobility group nucleosomal binding domain 2 |
| ENSG00000167384 | ZNF180 | 0,323173396 | 1,251079437 | 0,38026 | 0,64794085 | zinc finger protein 180 |
| ENSG00000140386 | SCAPER | 0,323166735 | 1,251073661 | 0,40587 | 0,66553796 | S-phase cyclin A associated protein in the ER |
| ENSG00000189046 | ALKBH2 | 0,323105363 | 1,251020441 | 0,31509 | 0,60145932 | alkB homolog 2, alpha-ketoglutarate dependent dioxygenase |
| ENSG00000276336 | SCARF1 | 0,323066317 | 1,250986583 | 0,79546 | 0,90529628 | scavenger receptor class F member 1 |
| ENSG00000171989 | LDHAL6B | 0,322998491 | 1,250927771 | 0,52629 | 0,74973856 | lactate dehydrogenase A like 6B |
| ENSG00000116747 | RO60 | 0,322958965 | 1,2508935 | 0,38532 | 0,65107577 | Ro60, Y RNA binding protein |
| ENSG00000230946 | HNRNPA1P68 | 0,32294133 | 1,250878209 | 0,44431 | 0,69172919 | heterogeneous nuclear ribonucleoprotein A1 pseudogene 68 |
| ENSG00000107554 | DNMBP | 0,322923613 | 1,250862848 | 0,24781 | 0,55526645 | dynamin binding protein |
| ENSG00000230010 | AL049646,1 | 0,322908039 | 1,250849346 | 0,36744 | 0,6374214 | novel transcript |
| ENSG00000283780 | MAML1 | 0,322838061 | 1,250788674 | 0,13691 | 0,4641716 | mastermind like transcriptional coactivator 1 |
| ENSG00000096401 | CDC5L | 0,322701041 | 1,250669886 | 0,27149 | 0,57285347 | cell division cycle 5 like |
| ENSG00000146282 | RARS2 | 0,322661386 | 1,250635509 | 0,17217 | 0,49710979 | arginyl-tRNA synthetase 2, mitochondrial |
| ENSG00000147854 | UHRF2 | 0,32253166 | 1,250523059 | 0,23951 | 0,5491566 | ubiquitin like with PHD and ring finger domains 2 |
| ENSG00000120458 | MSANTD2 | 0,322519564 | 1,250512574 | 0,23674 | 0,54767193 | Myb/SANT DNA binding domain containing 2 |
| ENSG00000143013 | LMO4 | 0,322389902 | 1,250400189 | 0,32902 | 0,61006602 | LIM domain only 4 |
| ENSG00000173041 | ZNF680 | 0,322361742 | 1,250375783 | 0,44625 | 0,69299062 | zinc finger protein 680 |
| ENSG00000111361 | EIF2B1 | 0,32227501 | 1,250300615 | 0,2816 | 0,57891473 | eukaryotic translation initiation factor 2B subunit alpha |
| ENSG00000178952 | TUFM | 0,322204041 | 1,250239112 | 0,19385 | 0,51547251 | Tu translation elongation factor, mitochondrial |
| ENSG00000151500 | THYN1 | 0,322162364 | 1,250202995 | 0,43011 | 0,68280659 | thymocyte nuclear protein 1 |
| ENSG00000134371 | CDC73 | 0,322158385 | 1,250199547 | 0,23215 | 0,54496653 | cell division cycle 73 |
| ENSG00000116005 | PCYOX1 | 0,322085915 | 1,250136748 | 0,10121 | 0,40785299 | prenylcysteine oxidase 1 |

| | | | | | | |
|---|---|---|---|---|---|---|
| ENSG00000106025 | TSPAN12 | 0,322020004 | 1,250079636 | 0,11601 | 0,4324056 | tetraspanin 12 |
| ENSG00000170881 | RNF139 | 0,321934958 | 1,250005946 | 0,43341 | 0,68473723 | ring finger protein 139 |
| ENSG00000131771 | PPP1R1B | 0,321933119 | 1,250004353 | 0,55547 | 0,76788267 | protein phosphatase 1 regulatory inhibitor subunit 1B |
| ENSG00000144048 | DUSP11 | 0,321933097 | 1,250004334 | 0,1328 | 0,45830601 | dual specificity phosphatase 11 |
| ENSG00000128016 | ZFP36 | 0,321906426 | 1,249981226 | 0,22666 | 0,54144 | ZFP36 ring finger protein |
| ENSG00000009335 | UBE3C | 0,321902325 | 1,249977672 | 0,17575 | 0,49972993 | ubiquitin protein ligase E3C |
| ENSG00000231119 | AL031666,1 | 0,321797354 | 1,249886727 | 0,37797 | 0,64631202 | novel transcript, antisense to ZMYND8 |
| ENSG00000261996 | AC004706,1 | 0,321796484 | 1,249885973 | 0,53132 | 0,75260265 | novel transcript |
| ENSG00000178852 | EFCAB13 | 0,321771682 | 1,249864486 | 0,37432 | 0,64275601 | EF-hand calcium binding domain 13 |
| ENSG00000288310 | AL133216,7 | 0,321759778 | 1,249854173 | 0,5363 | 0,75512187 | general transcription factor IIi (GTF2I) pseudogene |
| ENSG00000165006 | UBAP1 | 0,321730219 | 1,249828565 | 0,12456 | 0,44559634 | ubiquitin associated protein 1 |
| ENSG00000124145 | SDC4 | 0,32168725 | 1,249791341 | 0,16865 | 0,49492999 | syndecan 4 |
| ENSG00000155508 | CNOT8 | 0,321618889 | 1,249732122 | 0,22007 | 0,53749754 | CCR4-NOT transcription complex subunit 8 |
| ENSG00000224043 | CCNT2-AS1 | 0,321420048 | 1,249559889 | 0,13236 | 0,45750519 | CCNT2 antisense RNA 1 |
| ENSG00000146729 | NIPSNAP2 | 0,321368477 | 1,249515222 | 0,28593 | 0,58229607 | nipsnap homolog 2 |
| ENSG00000127995 | CASD1 | 0,321280223 | 1,249438788 | 0,45606 | 0,69998075 | CAS1 domain containing 1 |
| ENSG00000235750 | KIAA0040 | 0,32115845 | 1,249333332 | 0,60344 | 0,79757929 | KIAA0040 |
| ENSG00000255639 | AC005833,1 | 0,321117828 | 1,249298155 | 0,36854 | 0,63822157 | novel protein |
| ENSG00000178966 | RMI1 | 0,321101375 | 1,249283907 | 0,4282 | 0,68094175 | RecQ mediated genome instability 1 |
| ENSG00000134697 | GNL2 | 0,321087804 | 1,249272156 | 0,09495 | 0,39751728 | G protein nucleolar 2 |
| ENSG00000134809 | TIMM10 | 0,321083069 | 1,249268055 | 0,28951 | 0,58522301 | translocase of inner mitochondrial membrane 10 |
| ENSG00000051341 | POLQ | 0,321075349 | 1,24926137 | 0,24742 | 0,5549105 | DNA polymerase theta |
| ENSG00000169490 | TM2D2 | 0,321056644 | 1,249245173 | 0,15626 | 0,4840074 | TM2 domain containing 2 |
| ENSG00000167695 | TLCD3A | 0,320914667 | 1,24912224 | 0,14995 | 0,47794655 | TLC domain containing 3A |
| ENSG00000117481 | NSUN4 | 0,320884214 | 1,249095873 | 0,04934 | 0,29886777 | NOP2/Sun RNA methyltransferase 4 |
| ENSG00000175854 | SWI5 | 0,320837271 | 1,249055231 | 0,31195 | 0,59925736 | SWI5 homologous recombination repair protein |
| ENSG00000253304 | TMEM200B | 0,320786942 | 1,249011657 | 0,37583 | 0,64415653 | transmembrane protein 200B |
| ENSG00000169607 | CKAP2L | 0,320760383 | 1,248988664 | 0,49147 | 0,72510994 | cytoskeleton associated protein 2 like |
| ENSG00000131871 | SELENOS | 0,320639713 | 1,248884201 | 0,30799 | 0,59691026 | selenoprotein S |
| ENSG00000275956 | AC104446,2 | 0,320635854 | 1,24888086 | 0,46585 | 0,70619029 | zinc finger protein 610 (ZNF610) pseudogene |
| ENSG00000129071 | MBD4 | 0,320635341 | 1,248880416 | 0,28822 | 0,58430378 | methyl-CpG binding domain 4, DNA glycosylase |
| ENSG00000244217 | RPS4XP10 | 0,32052138 | 1,248781769 | 0,66542 | 0,83351136 | ribosomal protein S4X pseudogene 10 |
| ENSG00000118193 | KIF14 | 0,320465229 | 1,248733165 | 0,38175 | 0,64905556 | kinesin family member 14 |

| | | | | | | |
|---|---|---|---|---|---|---|
| ENSG00000186814 | ZSCAN30 | 0,320435305 | 1,248707265 | 0,27322 | 0,57344244 | zinc finger and SCAN domain containing 30 |
| ENSG00000100347 | SAMM50 | 0,320338432 | 1,248623421 | 0,22621 | 0,54121239 | SAMM50 sorting and assembly machinery component |
| ENSG00000226957 | AL358075,1 | 0,320335515 | 1,248620896 | 0,62087 | 0,80776776 | novel transcript |
| ENSG00000164830 | OXR1 | 0,320328532 | 1,248614852 | 0,40713 | 0,66657364 | oxidation resistance 1 |
| ENSG00000099810 | MTAP | 0,320327492 | 1,248613953 | 0,05905 | 0,32417102 | methylthioadenosine phosphorylase |
| ENSG00000285090 | AC002074,1 | 0,320285918 | 1,248577971 | 0,71923 | 0,86574114 | COL1A2 antisense RNA 1 |
| ENSG00000196715 | VKORC1L1 | 0,32018327 | 1,248489139 | 0,21201 | 0,53079538 | vitamin K epoxide reductase complex subunit 1 like 1 |
| ENSG00000253738 | OTUD6B-AS1 | 0,320146735 | 1,248457521 | 0,4219 | 0,67643128 | OTUD6B antisense RNA 1 (head to head) |
| ENSG00000174238 | PITPNA | 0,319993801 | 1,248325185 | 0,24282 | 0,55182091 | phosphatidylinositol transfer protein alpha |
| ENSG00000276266 | FP325318,1 | 0,319927288 | 1,248267634 | 0,50948 | 0,73730159 | novel transcript |
| ENSG00000281379 | SEPTIN14P19 | 0,319896617 | 1,248241097 | 0,74574 | 0,88009222 | septin 14 pseudogene 19 |
| ENSG00000110442 | COMMD9 | 0,319857447 | 1,248207207 | 0,15687 | 0,48425071 | COMM domain containing 9 |
| ENSG00000198730 | CTR9 | 0,319822242 | 1,248176749 | 0,36867 | 0,63840086 | CTR9 homolog, Paf1/RNA polymerase II complex component |
| ENSG00000052795 | FNIP2 | 0,319818179 | 1,248173233 | 0,33756 | 0,61721828 | folliculin interacting protein 2 |
| ENSG00000111652 | COPS7A | 0,319656237 | 1,248033134 | 0,09793 | 0,40323803 | COP9 signalosome subunit 7A |
| ENSG00000287477 | AL356495,1 | 0,319535996 | 1,247929122 | 0,44626 | 0,69299062 | novel transcript, antisense to TRPC4 |
| ENSG00000186501 | TMEM222 | 0,319331527 | 1,247752269 | 0,21478 | 0,53373102 | transmembrane protein 222 |
| ENSG00000066651 | TRMT11 | 0,319290778 | 1,247717026 | 0,32805 | 0,60971379 | tRNA methyltransferase 11 homolog |
| ENSG00000165891 | E2F7 | 0,319281045 | 1,247708609 | 0,25614 | 0,56068598 | E2F transcription factor 7 |
| ENSG00000136153 | LMO7 | 0,319158967 | 1,247603034 | 0,44601 | 0,69284796 | LIM domain 7 |
| ENSG00000263412 | NFE2L1-DT | 0,319136187 | 1,247583335 | 0,25288 | 0,55798781 | NFE2L1 divergent transcript |
| ENSG00000113595 | TRIM23 | 0,319131488 | 1,247579272 | 0,35708 | 0,63118328 | tripartite motif containing 23 |
| ENSG00000087494 | PTHLH | 0,319118487 | 1,247568029 | 0,63379 | 0,81549489 | parathyroid hormone like hormone |
| ENSG00000213371 | NAP1L1P3 | 0,319097783 | 1,247550126 | 0,78224 | 0,8988906 | nucleosome assembly protein 1 like 1 pseudogene 3 |
| ENSG00000106804 | C5 | 0,319097587 | 1,247549956 | 0,47771 | 0,71545897 | complement C5 |
| ENSG00000230262 | LINC02603 | 0,319026667 | 1,24748863 | 0,51686 | 0,74218125 | long intergenic non-protein coding RNA 2603 |
| ENSG00000218690 | H2AC10P | 0,318994207 | 1,247460563 | 0,62943 | 0,81292831 | H2A clustered histone 10, pseudogene |
| ENSG00000068796 | KIF2A | 0,318977692 | 1,247446283 | 0,37889 | 0,64679625 | kinesin family member 2A |
| ENSG00000090863 | GLG1 | 0,318976716 | 1,247445439 | 0,11186 | 0,42531464 | golgi glycoprotein 1 |
| ENSG00000262470 | TVP23CP2 | 0,318965782 | 1,247435985 | 0,56306 | 0,77224059 | TVP23C pseudogene 2 |
| ENSG00000100731 | PCNX1 | 0,318911988 | 1,247389472 | 0,27829 | 0,57645922 | pecanex 1 |
| ENSG00000155363 | MOV10 | 0,318822195 | 1,247311838 | 0,12419 | 0,44504227 | Mov10 RISC complex RNA helicase |

| | | | | | | |
|---|---|---|---|---|---|---|
| ENSG00000157653 | C9orf43 | 0,318787647 | 1,247281969 | 0,53392 | 0,75377522 | chromosome 9 open reading frame 43 |
| ENSG00000140931 | CMTM3 | 0,318785618 | 1,247280215 | 0,29728 | 0,59031059 | CKLF like MARVEL transmembrane domain containing 3 |
| ENSG00000162407 | PLPP3 | 0,318663722 | 1,247174834 | 0,39936 | 0,66169316 | phospholipid phosphatase 3 |
| ENSG00000186675 | MAGEE2 | 0,318606006 | 1,24712494 | 0,52951 | 0,7512527 | MAGE family member E2 |
| ENSG00000138138 | ATAD1 | 0,318574581 | 1,247097776 | 0,34189 | 0,6197312 | ATPase family AAA domain containing 1 |
| ENSG00000283024 | ATAD1 | 0,318574581 | 1,247097776 | 0,34189 | 0,6197312 | ATPase family AAA domain containing 1 |
| ENSG00000171467 | ZNF318 | 0,318493004 | 1,247027261 | 0,10735 | 0,41771747 | zinc finger protein 318 |
| ENSG00000138463 | SLC49A4 | 0,318447346 | 1,246987796 | 0,24326 | 0,55199251 | solute carrier family 49 member 4 |
| ENSG00000233210 | GPANK1 | 0,318439967 | 1,246981418 | 0,7456 | 0,87996245 | G-patch domain and ankyrin repeats 1 |
| ENSG00000138411 | HECW2 | 0,318247335 | 1,246814929 | 0,2398 | 0,54926442 | HECT, C2 and WW domain containing E3 ubiquitin protein ligase 2 |
| ENSG00000229470 | GNL1 | 0,318192015 | 1,246767122 | 0,74531 | 0,87977252 | G protein nucleolar 1 (putative) |
| ENSG00000100030 | MAPK1 | 0,318106884 | 1,246693554 | 0,3498 | 0,62563749 | mitogen-activated protein kinase 1 |
| ENSG00000283236 | AC074141,1 | 0,317994484 | 1,246596428 | 0,33403 | 0,61387439 | novel zinc finger protein pseudogene |
| ENSG00000123136 | DDX39A | 0,317890245 | 1,246506361 | 0,2736 | 0,57364891 | DExD-box helicase 39A |
| ENSG00000186106 | ANKRD46 | 0,317805031 | 1,246432737 | 0,31561 | 0,60172912 | ankyrin repeat domain 46 |
| ENSG00000230736 | AL021937,1 | 0,317755189 | 1,246389676 | 0,50904 | 0,7370222 | novel transcript |
| ENSG00000248816 | AC079226,2 | 0,317750609 | 1,246385719 | 0,65785 | 0,82887634 | novel transcript |
| ENSG00000179152 | TCAIM | 0,317627466 | 1,246279337 | 0,4185 | 0,67477592 | T cell activation inhibitor, mitochondrial |
| ENSG00000125484 | GTF3C4 | 0,31760504 | 1,246259965 | 0,18981 | 0,51299724 | general transcription factor IIIC subunit 4 |
| ENSG00000062650 | WAPL | 0,317582794 | 1,246240748 | 0,24796 | 0,55537155 | WAPL cohesin release factor |
| ENSG00000006062 | MAP3K14 | 0,317319338 | 1,246013188 | 0,17079 | 0,49641416 | mitogen-activated protein kinase kinase kinase 14 |
| ENSG00000227256 | MIS18A-AS1 | 0,31728437 | 1,245982988 | 0,70677 | 0,85889796 | MIS18A antisense RNA 1 |
| ENSG00000139218 | SCAF11 | 0,317259759 | 1,245961732 | 0,19135 | 0,51435195 | SR-related CTD associated factor 11 |
| ENSG00000106799 | TGFBR1 | 0,317199879 | 1,245910019 | 0,27604 | 0,57533851 | transforming growth factor beta receptor 1 |
| ENSG00000186073 | CDIN1 | 0,317152558 | 1,245869153 | 0,13578 | 0,462686 | CDAN1 interacting nuclease 1 |
| ENSG00000111669 | TPI1 | 0,317060867 | 1,245789974 | 0,08701 | 0,38214042 | triosephosphate isomerase 1 |
| ENSG00000188215 | DCUN1D3 | 0,317046488 | 1,245777558 | 0,15045 | 0,47857814 | defective in cullin neddylation 1 domain containing 3 |
| ENSG00000150540 | HNMT | 0,316979213 | 1,245719466 | 0,30356 | 0,59380161 | histamine N-methyltransferase |
| ENSG00000286299 | AL121938,1 | 0,316811816 | 1,245574933 | 0,57504 | 0,78004262 | novel transcript, antisense to HDDC2 |
| ENSG00000143207 | COP1 | 0,316775792 | 1,245543832 | 0,19261 | 0,51490739 | COP1 E3 ubiquitin ligase |
| ENSG00000063761 | ADCK1 | 0,316752391 | 1,245523629 | 0,15512 | 0,48296753 | aarF domain containing kinase 1 |
| ENSG00000111554 | MDM1 | 0,316682958 | 1,245463686 | 0,42191 | 0,67643128 | Mdm1 nuclear protein |

| | | | | | | |
|---|---|---|---|---|---|---|
| ENSG00000175520 | UBQLN3 | 0,316679303 | 1,245460531 | 0,64772 | 0,82380454 | ubiquilin 3 |
| ENSG00000138823 | MTTP | 0,316650899 | 1,245436011 | 0,63711 | 0,81747939 | microsomal triglyceride transfer protein |
| ENSG00000006740 | ARHGAP44 | 0,316636979 | 1,245423994 | 0,1193 | 0,43713778 | Rho GTPase activating protein 44 |
| ENSG00000259915 | AC017071,1 | 0,316517054 | 1,245320471 | 0,65809 | 0,82897081 | novel transcript |
| ENSG00000164924 | YWHAZ | 0,31620096 | 1,245047652 | 0,38434 | 0,65052153 | tyrosine 3-monooxygenase/tryptophan 5-monooxygenase activation protein zeta |
| ENSG00000125447 | GGA3 | 0,316194166 | 1,245041789 | 0,13336 | 0,4594073 | golgi associated, gamma adaptin ear containing, ARF binding protein 3 |
| ENSG00000288140 | AC006144,3 | 0,316133897 | 1,244989778 | 0,73189 | 0,87276602 | novel transcript |
| ENSG00000287379 | AC006144,2 | 0,316133897 | 1,244989778 | 0,73189 | 0,87276602 | novel transcript |
| ENSG00000185787 | MORF4L1 | 0,316121555 | 1,244979127 | 0,39599 | 0,65863599 | mortality factor 4 like 1 |
| ENSG00000167858 | TEKT1 | 0,315965574 | 1,244844531 | 0,63621 | 0,81694486 | tektin 1 |
| ENSG00000112079 | STK38 | 0,315946052 | 1,244827686 | 0,14543 | 0,47300165 | serine/threonine kinase 38 |
| ENSG00000197935 | ZNF311 | 0,315823717 | 1,244722133 | 0,7615 | 0,88879633 | zinc finger protein 311 |
| ENSG00000112941 | TENT4A | 0,315789921 | 1,244692975 | 0,05724 | 0,31908628 | terminal nucleotidyltransferase 4A |
| ENSG00000100888 | CHD8 | 0,315788665 | 1,244691892 | 0,19115 | 0,51406315 | chromodomain helicase DNA binding protein 8 |
| ENSG00000154114 | TBCEL | 0,315765003 | 1,244671478 | 0,22826 | 0,54221266 | tubulin folding cofactor E like |
| ENSG00000134258 | VTCN1 | 0,315723525 | 1,244635693 | 0,57402 | 0,77929088 | V-set domain containing T cell activation inhibitor 1 |
| ENSG00000151461 | UPF2 | 0,315704313 | 1,244619119 | 0,3287 | 0,60992716 | UPF2 regulator of nonsense mediated mRNA decay |
| ENSG00000112992 | NNT | 0,31566452 | 1,24458479 | 0,24726 | 0,55483158 | nicotinamide nucleotide transhydrogenase |
| ENSG00000166529 | ZSCAN21 | 0,315658335 | 1,244579454 | 0,24283 | 0,55182091 | zinc finger and SCAN domain containing 21 |
| ENSG00000176533 | GNG7 | 0,315643859 | 1,244566966 | 0,21993 | 0,53732404 | G protein subunit gamma 7 |
| ENSG00000144228 | SPOPL | 0,315631513 | 1,244556315 | 0,42241 | 0,67663831 | speckle type BTB/POZ protein like |
| ENSG00000135002 | RFK | 0,315586898 | 1,244517829 | 0,38773 | 0,65272082 | riboflavin kinase |
| ENSG00000269502 | DMRTC1 | 0,315479041 | 1,244424791 | 0,70418 | 0,85726326 | DMRT like family C1 |
| ENSG00000261893 | CLK2 | 0,31540745 | 1,24436304 | 0,22044 | 0,53750545 | CDC like kinase 2 |
| ENSG00000146802 | TMEM168 | 0,315384232 | 1,244343014 | 0,26364 | 0,56629809 | transmembrane protein 168 |
| ENSG00000183134 | PTGDR2 | 0,315327247 | 1,244293865 | 0,53071 | 0,75212511 | prostaglandin D2 receptor 2 |
| ENSG00000125633 | CCDC93 | 0,315294582 | 1,244265693 | 0,3912 | 0,65497307 | coiled-coil domain containing 93 |
| ENSG00000182670 | TTC3 | 0,315238005 | 1,244216898 | 0,25325 | 0,55836184 | tetratricopeptide repeat domain 3 |
| ENSG00000206418 | RAB12 | 0,315220158 | 1,244201507 | 0,38982 | 0,65396251 | RAB12, member RAS oncogene family |
| ENSG00000186566 | GPATCH8 | 0,315147806 | 1,244139111 | 0,31725 | 0,6027392 | G-patch domain containing 8 |
| ENSG00000272335 | AC093297,2 | 0,315105745 | 1,244102839 | 0,14058 | 0,46757653 | novel transcript |

| | | | | | | |
|---|---|---|---|---|---|---|
| ENSG00000108100 | CCNY | 0,315033055 | 1,244040156 | 0,16267 | 0,48945635 | cyclin Y |
| ENSG00000117425 | PTCH2 | 0,315017562 | 1,244026797 | 0,31235 | 0,59932999 | patched 2 |
| ENSG00000107929 | LARP4B | 0,314996289 | 1,244008454 | 0,0552 | 0,31442055 | La ribonucleoprotein 4B |
| ENSG00000164331 | ANKRA2 | 0,314928682 | 1,243950158 | 0,21972 | 0,53713164 | ankyrin repeat family A member 2 |
| ENSG00000148498 | PARD3 | 0,31491273 | 1,243936404 | 0,28207 | 0,57938416 | par-3 family cell polarity regulator |
| ENSG00000167617 | CDC42EP5 | 0,314912192 | 1,24393594 | 0,90927 | 0,96104995 | CDC42 effector protein 5 |
| ENSG00000064313 | TAF2 | 0,314873732 | 1,243902779 | 0,28495 | 0,58171017 | TATA-box binding protein associated factor 2 |
| ENSG00000131669 | NINJ1 | 0,314854477 | 1,243886177 | 0,28434 | 0,58119223 | ninjurin 1 |
| ENSG00000185158 | LRRC37B | 0,314825644 | 1,243861318 | 0,35343 | 0,62818386 | leucine rich repeat containing 37B |
| ENSG00000001561 | ENPP4 | 0,314784401 | 1,24382576 | 0,58279 | 0,78477395 | ectonucleotide pyrophosphatase/phosphodiesterase 4 |
| ENSG00000065060 | UHRF1BP1 | 0,314751118 | 1,243797065 | 0,29992 | 0,59160746 | UHRF1 binding protein 1 |
| ENSG00000084676 | NCOA1 | 0,314702595 | 1,243755232 | 0,42984 | 0,68261024 | nuclear receptor coactivator 1 |
| ENSG00000139842 | CUL4A | 0,314689694 | 1,24374411 | 0,12665 | 0,44928017 | cullin 4A |
| ENSG00000178691 | SUZ12 | 0,314633076 | 1,243695301 | 0,21723 | 0,53520949 | SUZ12 polycomb repressive complex 2 subunit |
| ENSG00000257859 | CASC18 | 0,314540636 | 1,243615615 | 0,7104 | 0,86051357 | cancer susceptibility 18 |
| ENSG00000146701 | MDH2 | 0,31448981 | 1,243571803 | 0,13684 | 0,4641716 | malate dehydrogenase 2 |
| ENSG00000121749 | TBC1D15 | 0,314480267 | 1,243563577 | 0,24576 | 0,55359883 | TBC1 domain family member 15 |
| ENSG00000109436 | TBC1D9 | 0,314456746 | 1,243543302 | 0,23786 | 0,54864017 | TBC1 domain family member 9 |
| ENSG00000133704 | IPO8 | 0,314432679 | 1,243522558 | 0,23815 | 0,54870569 | importin 8 |
| ENSG00000269486 | ERVK9-11 | 0,314262616 | 1,243375982 | 0,43675 | 0,68681257 | endogenous retrovirus group K9, member 11 |
| ENSG00000282930 | ERVK9-11 | 0,314262616 | 1,243375982 | 0,43675 | 0,68681257 | endogenous retrovirus group K9, member 11 |
| ENSG00000167257 | RNF214 | 0,314196785 | 1,243319247 | 0,28078 | 0,57814942 | ring finger protein 214 |
| ENSG00000267496 | FAM215A | 0,314151064 | 1,243279845 | 0,74258 | 0,87827554 | family with sequence similarity 215 member A |
| ENSG00000151693 | ASAP2 | 0,314126589 | 1,243258754 | 0,33854 | 0,61774255 | ArfGAP with SH3 domain, ankyrin repeat and PH domain 2 |
| ENSG00000101052 | IFT52 | 0,314005054 | 1,243154024 | 0,35152 | 0,62674766 | intraflagellar transport 52 |
| ENSG00000174574 | AKIRIN1 | 0,31398131 | 1,243133564 | 0,08405 | 0,37661338 | akirin 1 |
| ENSG00000258289 | CHURC1 | 0,313964938 | 1,243119457 | 0,38461 | 0,65052153 | churchill domain containing 1 |
| ENSG00000163312 | HELQ | 0,313885264 | 1,243050806 | 0,25744 | 0,56160845 | helicase, POLQ like |
| ENSG00000116649 | SRM | 0,313742639 | 1,242927925 | 0,17084 | 0,49641416 | spermidine synthase |
| ENSG00000284879 | AC133644,2 | 0,313721295 | 1,242909536 | 0,41296 | 0,67055663 | novel transcript |
| ENSG00000254682 | AP002387,1 | 0,313569506 | 1,242778774 | 0,34504 | 0,62228057 | novel transcript |
| ENSG00000092036 | HAUS4 | 0,313539199 | 1,242752667 | 0,16569 | 0,49182669 | HAUS augmin like complex subunit 4 |

| | | | | | | |
|---|---|---|---|---|---|---|
| ENSG00000231831 | MTHFD1P1 | 0,313488468 | 1,242708967 | 0,58053 | 0,78361129 | methylenetetrahydrofolate dehydrogenase (NADP+ dependent) 1 pseudogene 1 |
| ENSG00000103257 | SLC7A5 | 0,313460612 | 1,242684973 | 0,11892 | 0,43696075 | solute carrier family 7 member 5 |
| ENSG00000151470 | C4orf33 | 0,31344353 | 1,242670259 | 0,42126 | 0,67634721 | chromosome 4 open reading frame 33 |
| ENSG00000230993 | RPL12P15 | 0,313289936 | 1,242537967 | 0,62521 | 0,81024216 | ribosomal protein L12 pseudogene 15 |
| ENSG00000153922 | CHD1 | 0,313285914 | 1,242534504 | 0,3162 | 0,60226167 | chromodomain helicase DNA binding protein 1 |
| ENSG00000125630 | POLR1B | 0,313228958 | 1,24248545 | 0,14928 | 0,47695763 | RNA polymerase I subunit B |
| ENSG00000163728 | TTC14 | 0,313220622 | 1,242478272 | 0,3619 | 0,63352908 | tetratricopeptide repeat domain 14 |
| ENSG00000205629 | LCMT1 | 0,313215673 | 1,242474009 | 0,30464 | 0,59445922 | leucine carboxyl methyltransferase 1 |
| ENSG00000224551 | HMGB3P21 | 0,313145651 | 1,242413707 | 0,67735 | 0,84132177 | high mobility group box 3 pseudogene 21 |
| ENSG00000188428 | BLOC1S5 | 0,313088281 | 1,242364302 | 0,25383 | 0,55891 | biogenesis of lysosomal organelles complex 1 subunit 5 |
| ENSG00000169045 | HNRNPH1 | 0,312952592 | 1,24224746 | 0,27829 | 0,57645922 | heterogeneous nuclear ribonucleoprotein H1 |
| ENSG00000232912 | RERE-AS1 | 0,312799784 | 1,24211589 | 0,54545 | 0,76113525 | RERE antisense RNA 1 |
| ENSG00000260329 | AC007541,1 | 0,312665576 | 1,242000346 | 0,4184 | 0,67471324 | novel transcript, antisense to C12orf23 |
| ENSG00000141040 | ZNF287 | 0,312637872 | 1,241976496 | 0,36728 | 0,63729309 | zinc finger protein 287 |
| ENSG00000269911 | FAM226B | 0,312607372 | 1,24195024 | 0,28135 | 0,578832 | family with sequence similarity 226 member B |
| ENSG00000113558 | SKP1 | 0,312589694 | 1,241935022 | 0,25856 | 0,5625202 | S-phase kinase associated protein 1 |
| ENSG00000143390 | RFX5 | 0,312564244 | 1,241913114 | 0,2851 | 0,58171017 | regulatory factor X5 |
| ENSG00000132824 | SERINC3 | 0,312555631 | 1,2419057 | 0,17727 | 0,50187931 | serine incorporator 3 |
| ENSG00000215301 | DDX3X | 0,312452184 | 1,241816653 | 0,31575 | 0,60181942 | DEAD-box helicase 3 X-linked |
| ENSG00000143702 | CEP170 | 0,31245037 | 1,241815092 | 0,3928 | 0,6559444 | centrosomal protein 170 |
| ENSG00000159917 | ZNF235 | 0,312374829 | 1,24175007 | 0,29039 | 0,58561852 | zinc finger protein 235 |
| ENSG00000051180 | RAD51 | 0,312309295 | 1,241693666 | 0,10144 | 0,40833508 | RAD51 recombinase |
| ENSG00000164574 | GALNT10 | 0,31229783 | 1,241683798 | 0,34764 | 0,62423931 | polypeptide N-acetylgalactosaminyltransferase 10 |
| ENSG00000112242 | E2F3 | 0,312266872 | 1,241657154 | 0,11233 | 0,42601117 | E2F transcription factor 3 |
| ENSG00000146112 | PPP1R18 | 0,312228534 | 1,241624159 | 0,28337 | 0,58063542 | protein phosphatase 1 regulatory subunit 18 |
| ENSG00000161203 | AP2M1 | 0,312106218 | 1,241518895 | 0,27228 | 0,57315552 | adaptor related protein complex 2 subunit mu 1 |
| ENSG00000176714 | CCDC121 | 0,312046239 | 1,24146728 | 0,45977 | 0,70184297 | coiled-coil domain containing 121 |
| ENSG00000122778 | KIAA1549 | 0,312039739 | 1,241461687 | 0,32406 | 0,60668177 | KIAA1549 |
| ENSG00000221937 | TAS2R40 | 0,312017373 | 1,241442441 | 0,78641 | 0,90080813 | taste 2 receptor member 40 |
| ENSG00000271853 | AL162258,1 | 0,312009205 | 1,241435412 | 0,70605 | 0,85841728 | novel transcript |
| ENSG00000061676 | NCKAP1 | 0,311893696 | 1,241336021 | 0,33237 | 0,61227899 | NCK associated protein 1 |
| ENSG00000117054 | ACADM | 0,311839465 | 1,241289361 | 0,25715 | 0,56155289 | acyl-CoA dehydrogenase medium chain |

| | | | | | | |
|---|---|---|---|---|---|---|
| ENSG00000260196 | AC124798,1 | 0,311833404 | 1,241284145 | 0,57059 | 0,77748547 | antisense to KCNJ11 and overlapping to a novel gene |
| ENSG00000136158 | SPRY2 | 0,311798694 | 1,241254282 | 0,477 | 0,71492678 | sprouty RTK signaling antagonist 2 |
| ENSG00000102595 | UGGT2 | 0,311780947 | 1,241239013 | 0,31302 | 0,59987747 | UDP-glucose glycoprotein glucosyltransferase 2 |
| ENSG00000284938 | FAM180B | 0,311680202 | 1,241152339 | 0,71953 | 0,86592255 | family with sequence similarity 180 member B |
| ENSG00000023734 | STRAP | 0,311670005 | 1,241143566 | 0,15862 | 0,48587478 | serine/threonine kinase receptor associated protein |
| ENSG00000055609 | KMT2C | 0,311663898 | 1,241138312 | 0,31101 | 0,59883009 | lysine methyltransferase 2C |
| ENSG00000215154 | AC141586,1 | 0,311616702 | 1,241097711 | 0,08359 | 0,37573324 | potassium channel tetramerisation domain containing 5 (KCTD5) pseudogene |
| ENSG00000114745 | GORASP1 | 0,311603397 | 1,241086265 | 0,07155 | 0,35342971 | golgi reassembly stacking protein 1 |
| ENSG00000285387 | NDUFS3 | 0,311543884 | 1,24103507 | 0,28821 | 0,58430378 | NADH:ubiquinone oxidoreductase core subunit S3 |
| ENSG00000254317 | AC022973,5 | 0,311533116 | 1,241025807 | 0,69961 | 0,85488711 | novel transcript |
| ENSG00000182827 | ACBD3 | 0,311500982 | 1,240998165 | 0,18109 | 0,50470865 | acyl-CoA binding domain containing 3 |
| ENSG00000184009 | ACTG1 | 0,311408305 | 1,240918447 | 0,29937 | 0,59128796 | actin gamma 1 |
| ENSG00000166363 | OR10A5 | 0,311290785 | 1,240817368 | 0,63958 | 0,81913513 | olfactory receptor family 10 subfamily A member 5 |
| ENSG00000121005 | CRISPLD1 | 0,311262766 | 1,24079327 | 0,44622 | 0,69299062 | cysteine rich secretory protein LCCL domain containing 1 |
| ENSG00000172315 | TP53RK | 0,311235311 | 1,240769658 | 0,26499 | 0,56769728 | TP53 regulating kinase |
| ENSG00000228513 | AC023271,2 | 0,311132645 | 1,240681363 | 0,65709 | 0,82862827 | zinc finger protein 14 (ZNF14) pseudogene |
| ENSG00000126970 | ZC4H2 | 0,311107167 | 1,240659454 | 0,40432 | 0,6642832 | zinc finger C4H2-type containing |
| ENSG00000137764 | MAP2K5 | 0,311011944 | 1,240577568 | 0,26364 | 0,56629809 | mitogen-activated protein kinase kinase 5 |
| ENSG00000122299 | ZC3H7A | 0,311008728 | 1,240574803 | 0,25517 | 0,55966322 | zinc finger CCCH-type containing 7A |
| ENSG00000104331 | BPNT2 | 0,310925027 | 1,24050283 | 0,25007 | 0,55657567 | 3'(2'), 5'-bisphosphate nucleotidase 2 |
| ENSG00000204659 | CBY3 | 0,310883729 | 1,240467321 | 0,69011 | 0,8488605 | chibby family member 3 |
| ENSG00000283718 | CBY3 | 0,310883729 | 1,240467321 | 0,69011 | 0,8488605 | chibby family member 3 |
| ENSG00000196233 | LCOR | 0,310876386 | 1,240461007 | 0,38957 | 0,6539298 | ligand dependent nuclear receptor corepressor |
| ENSG00000111860 | CEP85L | 0,310848272 | 1,240436834 | 0,46535 | 0,70583978 | centrosomal protein 85 like |
| ENSG00000234264 | DEPDC1-AS1 | 0,31080154 | 1,240396654 | 0,72817 | 0,87073219 | DEPDC1 antisense RNA 1 |
| ENSG00000072110 | ACTN1 | 0,310740873 | 1,240344496 | 0,05576 | 0,31564067 | actinin alpha 1 |
| ENSG00000197024 | ZNF398 | 0,310716465 | 1,240323511 | 0,22339 | 0,53953208 | zinc finger protein 398 |
| ENSG00000158615 | PPP1R15B | 0,310710172 | 1,240318101 | 0,27307 | 0,57344244 | protein phosphatase 1 regulatory subunit 15B |
| ENSG00000168827 | GFM1 | 0,310679227 | 1,240291497 | 0,17563 | 0,49954066 | G elongation factor mitochondrial 1 |
| ENSG00000280780 | JAKMIP2-AS1 | 0,310663127 | 1,240277656 | 0,52385 | 0,74775411 | JAKMIP2 antisense RNA 1 |
| ENSG00000174564 | IL20RB | 0,310626708 | 1,240246347 | 0,41407 | 0,67096878 | interleukin 20 receptor subunit beta |
| ENSG00000155324 | GRAMD2B | 0,310585651 | 1,240211052 | 0,4069 | 0,6662711 | GRAM domain containing 2B |

| | | | | | | |
|---|---|---|---|---|---|---|
| ENSG00000164406 | LEAP2 | 0,3105379 | 1,240170004 | 0,33305 | 0,61306511 | liver enriched antimicrobial peptide 2 |
| ENSG00000113368 | LMNB1 | 0,310359293 | 1,240016479 | 0,23374 | 0,54563244 | lamin B1 |
| ENSG00000204116 | CHIC1 | 0,310132184 | 1,239821291 | 0,43121 | 0,68350322 | cysteine rich hydrophobic domain 1 |
| ENSG00000126767 | ELK1 | 0,309995212 | 1,239703586 | 0,08835 | 0,38462876 | ETS transcription factor ELK1 |
| ENSG00000203812 | H2AC18 | 0,309981078 | 1,23969144 | 0,45803 | 0,7010995 | H2A clustered histone 18 |
| ENSG00000115267 | IFIH1 | 0,309880405 | 1,239604936 | 0,61014 | 0,80220238 | interferon induced with helicase C domain 1 |
| ENSG00000200312 | RN7SKP255 | 0,309870015 | 1,239596009 | 0,61605 | 0,80475871 | RN7SK pseudogene 255 |
| ENSG00000154144 | TBRG1 | 0,309826868 | 1,239558936 | 0,26364 | 0,56629809 | transforming growth factor beta regulator 1 |
| ENSG00000162600 | OMA1 | 0,30978488 | 1,239522861 | 0,16508 | 0,49125883 | OMA1 zinc metallopeptidase |
| ENSG00000144747 | TMF1 | 0,309670574 | 1,239424656 | 0,35404 | 0,62877376 | TATA element modulatory factor 1 |
| ENSG00000120727 | PAIP2 | 0,309657178 | 1,239413148 | 0,39661 | 0,65923689 | poly(A) binding protein interacting protein 2 |
| ENSG00000111142 | METAP2 | 0,30964755 | 1,239404877 | 0,24037 | 0,54950872 | methionyl aminopeptidase 2 |
| ENSG00000151422 | FER | 0,309483327 | 1,239263803 | 0,46763 | 0,70749135 | FER tyrosine kinase |
| ENSG00000133943 | DGLUCY | 0,309413568 | 1,239203882 | 0,21284 | 0,53189271 | D-glutamate cyclase |
| ENSG00000104738 | MCM4 | 0,309410348 | 1,239201115 | 0,21004 | 0,52897377 | minichromosome maintenance complex component 4 |
| ENSG00000198130 | HIBCH | 0,309220562 | 1,239038109 | 0,34354 | 0,62091871 | 3-hydroxyisobutyryl-CoA hydrolase |
| ENSG00000166164 | BRD7 | 0,3092069 | 1,239026377 | 0,27684 | 0,57549599 | bromodomain containing 7 |
| ENSG00000173085 | COQ2 | 0,309183898 | 1,239006622 | 0,46657 | 0,7067394 | coenzyme Q2, polyprenyltransferase |
| ENSG00000282641 | AC183084,1 | 0,309035918 | 1,238879541 | 0,63232 | 0,8146065 | aminopeptidase puromycin sensitive (NPEPPS) pseudogene |
| ENSG00000094914 | AAAS | 0,308999276 | 1,238848076 | 0,15224 | 0,48061452 | aladin WD repeat nucleoporin |
| ENSG00000273792 | AC022898,2 | 0,3088437 | 1,23871449 | 0,63784 | 0,81798655 | novel transcript, sense intronic to RORA |
| ENSG00000147162 | OGT | 0,308834195 | 1,238706328 | 0,34995 | 0,62563749 | O-linked N-acetylglucosamine (GlcNAc) transferase |
| ENSG00000278601 | AL158163,2 | 0,308539443 | 1,238453279 | 0,61216 | 0,8035281 | novel transcript, antisense to BBIP1 |
| ENSG00000124226 | RNF114 | 0,308515112 | 1,238432392 | 0,18784 | 0,51107563 | ring finger protein 114 |
| ENSG00000121988 | ZRANB3 | 0,30846827 | 1,238392183 | 0,2441 | 0,55253387 | zinc finger RANBP2-type containing 3 |
| ENSG00000162779 | AXDND1 | 0,308446638 | 1,238373614 | 0,62433 | 0,80968303 | axonemal dynein light chain domain containing 1 |
| ENSG00000248476 | BACH1-IT1 | 0,308405209 | 1,238338053 | 0,4229 | 0,67683766 | BACH1 intronic transcript 1 |
| ENSG00000120539 | MASTL | 0,308401438 | 1,238334816 | 0,34479 | 0,62213186 | microtubule associated serine/threonine kinase like |
| ENSG00000164587 | RPS14 | 0,30838876 | 1,238323934 | 0,22981 | 0,54324957 | ribosomal protein S14 |
| ENSG00000196419 | XRCC6 | 0,308376204 | 1,238313157 | 0,19446 | 0,51562158 | X-ray repair cross complementing 6 |
| ENSG00000177788 | AL162595,1 | 0,308365538 | 1,238304002 | 0,17007 | 0,49606609 | novel transcript |
| ENSG00000158856 | DMTN | 0,308360294 | 1,238299501 | 0,44002 | 0,6885781 | dematin actin binding protein |

| | | | | | | |
|---|---|---|---|---|---|---|
| ENSG00000271664 | AC004890,3 | 0,308237125 | 1,238193787 | 0,61527 | 0,80431041 | SECIS binding protein 2, pseudogene |
| ENSG00000186479 | RGS7BP | 0,308166125 | 1,238132852 | 0,55008 | 0,76444876 | regulator of G protein signaling 7 binding protein |
| ENSG00000088367 | EPB41L1 | 0,308126282 | 1,238098659 | 0,24364 | 0,55235189 | erythrocyte membrane protein band 4,1 like 1 |
| ENSG00000165716 | DIPK1B | 0,308072508 | 1,238052512 | 0,14572 | 0,47317517 | divergent protein kinase domain 1B |
| ENSG00000035862 | TIMP2 | 0,308006703 | 1,237996042 | 0,24786 | 0,55528615 | TIMP metallopeptidase inhibitor 2 |
| ENSG00000156011 | PSD3 | 0,307981796 | 1,23797467 | 0,40375 | 0,66409141 | pleckstrin and Sec7 domain containing 3 |
| ENSG00000119514 | GALNT12 | 0,307928933 | 1,237929309 | 0,47445 | 0,71319993 | polypeptide N-acetylgalactosaminyltransferase 12 |
| ENSG00000143375 | CGN | 0,30789481 | 1,23790003 | 0,34801 | 0,62441612 | cingulin |
| ENSG00000225808 | DNAJC19P5 | 0,307879828 | 1,237887174 | 0,67171 | 0,8375243 | DnaJ heat shock protein family (Hsp40) member C19 pseudogene 5 |
| ENSG00000142149 | HUNK | 0,307846756 | 1,237858798 | 0,1837 | 0,50744379 | hormonally up-regulated Neu-associated kinase |
| ENSG00000036672 | USP2 | 0,307840066 | 1,237853058 | 0,20017 | 0,52106859 | ubiquitin specific peptidase 2 |
| ENSG00000109576 | AADAT | 0,307814346 | 1,237830989 | 0,42793 | 0,68085046 | aminoadipate aminotransferase |
| ENSG00000111727 | HCFC2 | 0,307773753 | 1,237796161 | 0,43262 | 0,68418465 | host cell factor C2 |
| ENSG00000134755 | DSC2 | 0,307754427 | 1,237779581 | 0,40487 | 0,66481282 | desmocollin 2 |
| ENSG00000187498 | COL4A1 | 0,307586499 | 1,237635513 | 0,44425 | 0,69172919 | collagen type IV alpha 1 chain |
| ENSG00000171848 | RRM2 | 0,30751929 | 1,237577857 | 0,20106 | 0,52181085 | ribonucleotide reductase regulatory subunit M2 |
| ENSG00000134323 | MYCN | 0,307511116 | 1,237570846 | 0,34404 | 0,621447 | MYCN proto-oncogene, bHLH transcription factor |
| ENSG00000253454 | NDUFA5P2 | 0,307480123 | 1,23754426 | 0,7651 | 0,8907696 | NADH:ubiquinone oxidoreductase subunit A5 pseudogene 2 |
| ENSG00000261736 | AC002551,1 | 0,307413054 | 1,237486729 | 0,76087 | 0,88871285 | novel transcript |
| ENSG00000175518 | UBQLNL | 0,307366456 | 1,23744676 | 0,59436 | 0,79263193 | ubiquilin like |
| ENSG00000259322 | AC090607,1 | 0,307338611 | 1,237422877 | 0,60424 | 0,79780609 | novel transcript, antisense to WDR61 |
| ENSG00000159348 | CYB5R1 | 0,307331781 | 1,237417019 | 0,26637 | 0,56917012 | cytochrome b5 reductase 1 |
| ENSG00000115204 | MPV17 | 0,307252703 | 1,237349194 | 0,23353 | 0,54563244 | mitochondrial inner membrane protein MPV17 |
| ENSG00000131043 | AAR2 | 0,307218512 | 1,23731987 | 0,23801 | 0,54864017 | AAR2 splicing factor |
| ENSG00000197894 | ADH5 | 0,307216823 | 1,237318421 | 0,40163 | 0,66306607 | alcohol dehydrogenase 5 (class III), chi polypeptide |
| ENSG00000162521 | RBBP4 | 0,30720768 | 1,23731058 | 0,23469 | 0,54623956 | RB binding protein 4, chromatin remodeling factor |
| ENSG00000152219 | ARL14EP | 0,307103444 | 1,237221187 | 0,35667 | 0,63070014 | ADP ribosylation factor like GTPase 14 effector protein |
| ENSG00000163512 | AZI2 | 0,306944337 | 1,237084748 | 0,25265 | 0,55796133 | 5-azacytidine induced 2 |
| ENSG00000236554 | ASNSP3 | 0,306867507 | 1,237018869 | 0,71696 | 0,86453146 | asparagine synthetase pseudogene 3 |
| ENSG00000007001 | UPP2 | 0,306760463 | 1,236927089 | 0,59866 | 0,7951852 | uridine phosphorylase 2 |

| | | | | | | |
|---|---|---|---|---|---|---|
| ENSG00000181929 | PRKAG1 | 0,306710436 | 1,236884198 | 0,43133 | 0,68358509 | protein kinase AMP-activated non-catalytic subunit gamma 1 |
| ENSG00000196693 | ZNF33B | 0,306699289 | 1,236874641 | 0,40459 | 0,66451829 | zinc finger protein 33B |
| ENSG00000196865 | NHLRC2 | 0,306686334 | 1,236863534 | 0,19494 | 0,51588752 | NHL repeat containing 2 |
| ENSG00000172171 | TEFM | 0,306576183 | 1,236769102 | 0,30984 | 0,59807209 | transcription elongation factor, mitochondrial |
| ENSG00000271361 | HTATSF1P2 | 0,306564433 | 1,23675903 | 0,60857 | 0,80122454 | HIV-1 Tat specific factor 1 pseudogene 2 |
| ENSG00000198967 | OR10Z1 | 0,306476859 | 1,236683959 | 0,77349 | 0,89545981 | olfactory receptor family 10 subfamily Z member 1 |
| ENSG00000160868 | CYP3A4 | 0,306354425 | 1,236579012 | 0,6567 | 0,82855263 | cytochrome P450 family 3 subfamily A member 4 |
| ENSG00000228878 | SEPTIN7-DT | 0,306267847 | 1,236504806 | 0,36467 | 0,63504684 | SEPTIN7 divergent transcript |
| ENSG00000087460 | GNAS | 0,306243633 | 1,236484053 | 0,17793 | 0,50233162 | GNAS complex locus |
| ENSG00000246596 | AC139795,1 | 0,306209374 | 1,236454691 | 0,25109 | 0,55719428 | SUMO interacting motifs containing 1 (SIMC1) pseudogene |
| ENSG00000163050 | COQ8A | 0,306144736 | 1,236399294 | 0,22524 | 0,54058261 | coenzyme Q8A |
| ENSG00000074266 | EED | 0,306143678 | 1,236398388 | 0,16317 | 0,48972962 | embryonic ectoderm development |
| ENSG00000138785 | INTS12 | 0,306130319 | 1,236386939 | 0,50066 | 0,73097006 | integrator complex subunit 12 |
| ENSG00000161277 | THAP8 | 0,306102883 | 1,236363427 | 0,3624 | 0,63379594 | THAP domain containing 8 |
| ENSG00000181938 | GINS3 | 0,306076718 | 1,236341004 | 0,29651 | 0,58979902 | GINS complex subunit 3 |
| ENSG00000197044 | ZNF441 | 0,306039858 | 1,236309416 | 0,51193 | 0,73893491 | zinc finger protein 441 |
| ENSG00000138381 | ASNSD1 | 0,306011829 | 1,236285398 | 0,20665 | 0,52622584 | asparagine synthetase domain containing 1 |
| ENSG00000284946 | AC068831,6 | 0,305993589 | 1,236269768 | 0,31297 | 0,59987747 | novel protein |
| ENSG00000165724 | ZMYND19 | 0,305984495 | 1,236261975 | 0,11746 | 0,43477245 | zinc finger MYND-type containing 19 |
| ENSG00000274191 | AC026333,4 | 0,305959359 | 1,236240435 | 0,53188 | 0,75280487 | novel transcript |
| ENSG00000136824 | SMC2 | 0,305897922 | 1,236187791 | 0,38394 | 0,65047768 | structural maintenance of chromosomes 2 |
| ENSG00000111875 | ASF1A | 0,305875147 | 1,236168277 | 0,5175 | 0,74270246 | anti-silencing function 1A histone chaperone |
| ENSG00000232978 | AL161909,2 | 0,305862938 | 1,236157815 | 0,72035 | 0,86628014 | novel transcript |
| ENSG00000259705 | AC084757,3 | 0,305843564 | 1,236141215 | 0,47741 | 0,71533146 | novel transcript |
| ENSG00000272370 | AC092354,2 | 0,3058107 | 1,236113057 | 0,63306 | 0,81529025 | novel transcript |
| ENSG00000134717 | BTF3L4 | 0,305724247 | 1,236038985 | 0,34607 | 0,62327403 | basic transcription factor 3 like 4 |
| ENSG00000189144 | ZNF573 | 0,305724208 | 1,236038951 | 0,38586 | 0,65129675 | zinc finger protein 573 |
| ENSG00000149050 | ZNF214 | 0,305675104 | 1,235996882 | 0,57189 | 0,77838663 | zinc finger protein 214 |
| ENSG00000139168 | ZCRB1 | 0,305389046 | 1,235751832 | 0,23954 | 0,5491566 | zinc finger CCHC-type and RNA binding motif containing 1 |
| ENSG00000257953 | AC083805,2 | 0,305384541 | 1,235747974 | 0,58557 | 0,78640335 | novel transcript, antisense to CTDSP2 |
| ENSG00000257446 | ZNF878 | 0,305358735 | 1,23572587 | 0,40192 | 0,66338822 | zinc finger protein 878 |
| ENSG00000286077 | AL158141,1 | 0,305325406 | 1,235697323 | 0,80609 | 0,91078302 | novel transcript, antisense to EDA |

| | | | | | | |
|---|---|---|---|---|---|---|
| ENSG00000164488 | DACT2 | 0,305325346 | 1,235697271 | 0,59574 | 0,79349831 | dishevelled binding antagonist of beta catenin 2 |
| ENSG00000036257 | CUL3 | 0,305246881 | 1,235630066 | 0,2668 | 0,56923396 | cullin 3 |
| ENSG00000164707 | SLC13A4 | 0,305234282 | 1,235619276 | 0,19489 | 0,51588752 | solute carrier family 13 member 4 |
| ENSG00000175792 | RUVBL1 | 0,305225183 | 1,235611482 | 0,22705 | 0,54168591 | RuvB like AAA ATPase 1 |
| ENSG00000147905 | ZCCHC7 | 0,305194066 | 1,235584832 | 0,43189 | 0,68387935 | zinc finger CCHC-type containing 7 |
| ENSG00000142794 | NBPF3 | 0,30518899 | 1,235580485 | 0,13989 | 0,46689302 | NBPF member 3 |
| ENSG00000132825 | PPP1R3D | 0,305089175 | 1,235495003 | 0,31833 | 0,60349082 | protein phosphatase 1 regulatory subunit 3D |
| ENSG00000183513 | COA5 | 0,305065918 | 1,235475086 | 0,42018 | 0,6756366 | cytochrome c oxidase assembly factor 5 |
| ENSG00000256269 | HMBS | 0,305042019 | 1,23545462 | 0,17422 | 0,49867554 | hydroxymethylbilane synthase |
| ENSG00000005882 | PDK2 | 0,30491924 | 1,235349482 | 0,38736 | 0,65229648 | pyruvate dehydrogenase kinase 2 |
| ENSG00000227502 | MROCKI | 0,304883966 | 1,235319278 | 0,41736 | 0,67399507 | MARCKS cis regulating lncRNA promoter of cytokines and inflammation |
| ENSG00000110917 | MLEC | 0,3048527 | 1,235292507 | 0,16665 | 0,49255702 | malectin |
| ENSG00000128609 | NDUFA5 | 0,304847217 | 1,235287812 | 0,40307 | 0,66399303 | NADH:ubiquinone oxidoreductase subunit A5 |
| ENSG00000163535 | SGO2 | 0,304828061 | 1,23527141 | 0,37928 | 0,64710751 | shugoshin 2 |
| ENSG00000228252 | COL6A4P2 | 0,304784107 | 1,235233776 | 0,47404 | 0,71294959 | collagen type VI alpha 4 pseudogene 2 |
| ENSG00000164751 | PEX2 | 0,304760161 | 1,235213273 | 0,31114 | 0,59885232 | peroxisomal biogenesis factor 2 |
| ENSG00000197841 | ZNF181 | 0,304710747 | 1,235170967 | 0,37914 | 0,64699394 | zinc finger protein 181 |
| ENSG00000119396 | RAB14 | 0,304641491 | 1,235111674 | 0,37297 | 0,64180957 | RAB14, member RAS oncogene family |
| ENSG00000141425 | RPRD1A | 0,304577727 | 1,235057086 | 0,32365 | 0,60636122 | regulation of nuclear pre-mRNA domain containing 1A |
| ENSG00000213762 | ZNF134 | 0,304558934 | 1,235040998 | 0,21121 | 0,53012655 | zinc finger protein 134 |
| ENSG00000177156 | TALDO1 | 0,304436399 | 1,234936105 | 0,31249 | 0,59939093 | transaldolase 1 |
| ENSG00000225891 | DHDDS-AS1 | 0,304417844 | 1,234920222 | 0,53627 | 0,75512187 | DHDDS antisense RNA 1 |
| ENSG00000116574 | RHOU | 0,304341175 | 1,234854597 | 0,58042 | 0,78359603 | ras homolog family member U |
| ENSG00000284984 | RHOU | 0,304341175 | 1,234854597 | 0,58042 | 0,78359603 | ras homolog family member U |
| ENSG00000005194 | CIAPIN1 | 0,304333583 | 1,234848098 | 0,22818 | 0,54221266 | cytokine induced apoptosis inhibitor 1 |
| ENSG00000168811 | IL12A | 0,304272294 | 1,23479564 | 0,59802 | 0,79484486 | interleukin 12A |
| ENSG00000117010 | ZNF684 | 0,304224902 | 1,234755078 | 0,20157 | 0,52214043 | zinc finger protein 684 |
| ENSG00000214160 | ALG3 | 0,304148596 | 1,234689772 | 0,23619 | 0,54725688 | ALG3 alpha-1,3- mannosyltransferase |
| ENSG00000231104 | AC022395,1 | 0,304125495 | 1,234670002 | 0,46736 | 0,70725568 | novel transcript, antisense to RAB11FIP2 |
| ENSG00000140015 | KCNH5 | 0,304062711 | 1,234616272 | 0,64433 | 0,82151477 | potassium voltage-gated channel subfamily H member 5 |
| ENSG00000269887 | AL391001,1 | 0,304033305 | 1,234591107 | 0,61757 | 0,80595997 | novel transcript, sense intronic to PBX1 |
| ENSG00000005889 | ZFX | 0,303980915 | 1,234546276 | 0,32342 | 0,60621131 | zinc finger protein X-linked |

| | | | | | | |
|---|---|---|---|---|---|---|
| ENSG00000229308 | AC010737,1 | 0,303863278 | 1,234445615 | 0,73364 | 0,87368351 | novel transcript |
| ENSG00000238222 | MKRN4P | 0,303819516 | 1,23440817 | 0,74788 | 0,88128464 | makorin ring finger protein 4, pseudogene |
| ENSG00000166747 | AP1G1 | 0,303814812 | 1,234404145 | 0,2879 | 0,5840701 | adaptor related protein complex 1 subunit gamma 1 |
| ENSG00000214992 | AKAP17BP | 0,303704885 | 1,234310093 | 0,44714 | 0,69341308 | A-kinase anchoring protein 17B, pseudogene |
| ENSG00000187783 | TMEM72 | 0,303665965 | 1,234276795 | 0,53775 | 0,75631707 | transmembrane protein 72 |
| ENSG00000253352 | TUG1 | 0,303560997 | 1,234186995 | 0,14139 | 0,46803321 | taurine up-regulated 1 |
| ENSG00000228380 | MYL6BP1 | 0,303441159 | 1,23408448 | 0,76793 | 0,89249843 | myosin light chain 6B pseudogene 1 |
| ENSG00000124783 | SSR1 | 0,303422176 | 1,234068243 | 0,23584 | 0,5471053 | signal sequence receptor subunit 1 |
| ENSG00000105677 | TMEM147 | 0,303389789 | 1,234040539 | 0,36247 | 0,63379594 | transmembrane protein 147 |
| ENSG00000187824 | TMEM220 | 0,303371879 | 1,23402522 | 0,20542 | 0,52503195 | transmembrane protein 220 |
| ENSG00000109920 | FNBP4 | 0,303288197 | 1,233953644 | 0,26384 | 0,56634699 | formin binding protein 4 |
| ENSG00000285182 | FNBP4 | 0,303288197 | 1,233953644 | 0,26384 | 0,56634699 | formin binding protein 4 |
| ENSG00000144357 | UBR3 | 0,303282803 | 1,23394903 | 0,48145 | 0,71826037 | ubiquitin protein ligase E3 component n-recognin 3 |
| ENSG00000119943 | PYROXD2 | 0,303248734 | 1,233919891 | 0,16937 | 0,49554671 | pyridine nucleotide-disulphide oxidoreductase domain 2 |
| ENSG00000163902 | RPN1 | 0,303120819 | 1,233810491 | 0,13946 | 0,46666406 | ribophorin I |
| ENSG00000114904 | NEK4 | 0,303111479 | 1,233802503 | 0,35459 | 0,62915556 | NIMA related kinase 4 |
| ENSG00000152404 | CWF19L2 | 0,302977092 | 1,23368758 | 0,30006 | 0,59160746 | CWF19 like cell cycle control factor 2 |
| ENSG00000061987 | MON2 | 0,302932781 | 1,233649689 | 0,28894 | 0,58464073 | MON2 homolog, regulator of endosome-to-Golgi trafficking |
| ENSG00000171659 | GPR34 | 0,302846393 | 1,233575821 | 0,44987 | 0,69539135 | G protein-coupled receptor 34 |
| ENSG00000196411 | EPHB4 | 0,302791249 | 1,233528672 | 0,14786 | 0,4751727 | EPH receptor B4 |
| ENSG00000140153 | WDR20 | 0,302696993 | 1,233448084 | 0,26827 | 0,5704222 | WD repeat domain 20 |
| ENSG00000121871 | SLITRK3 | 0,302606232 | 1,233370489 | 0,58724 | 0,78717516 | SLIT and NTRK like family member 3 |
| ENSG00000168874 | ATOH8 | 0,302595864 | 1,233361626 | 0,52274 | 0,74689824 | atonal bHLH transcription factor 8 |
| ENSG00000203485 | INF2 | 0,302452681 | 1,233239224 | 0,37086 | 0,64013451 | inverted formin 2 |
| ENSG00000229456 | RLIMP1 | 0,302361374 | 1,233161175 | 0,63742 | 0,81771796 | ring finger protein, LIM domain interacting pseudogene 1 |
| ENSG00000214517 | PPME1 | 0,302234575 | 1,233052797 | 0,28762 | 0,58395529 | protein phosphatase methylesterase 1 |
| ENSG00000103343 | ZNF174 | 0,302233002 | 1,233051453 | 0,3421 | 0,61993771 | zinc finger protein 174 |
| ENSG00000066629 | EML1 | 0,302220632 | 1,23304088 | 0,39034 | 0,65441492 | EMAP like 1 |
| ENSG00000218757 | AL121952,1 | 0,302219921 | 1,233040273 | 0,52126 | 0,74558236 | 60S ribosomal protein L17 (RPL17) pseudogene |
| ENSG00000106780 | MEGF9 | 0,302191828 | 1,233016262 | 0,28155 | 0,57891473 | multiple EGF like domains 9 |
| ENSG00000198886 | MT-ND4 | 0,302099818 | 1,232937628 | 0,37018 | 0,63960227 | mitochondrially encoded NADH:ubiquinone oxidoreductase core subunit 4 |

| | | | | | | |
|---|---|---|---|---|---|---|
| ENSG00000182473 | EXOC7 | 0,302097373 | 1,232935538 | 0,29292 | 0,58737078 | exocyst complex component 7 |
| ENSG00000236063 | ABHD16A | 0,302097346 | 1,232935515 | 0,44501 | 0,69205935 | abhydrolase domain containing 16A, phospholipase |
| ENSG00000231488 | ABHD16A | 0,302097346 | 1,232935515 | 0,44501 | 0,69205935 | abhydrolase domain containing 16A, phospholipase |
| ENSG00000235676 | ABHD16A | 0,302097346 | 1,232935515 | 0,44501 | 0,69205935 | abhydrolase domain containing 16A, phospholipase |
| ENSG00000197647 | ZNF433 | 0,302060084 | 1,232903671 | 0,40181 | 0,66324659 | zinc finger protein 433 |
| ENSG00000171097 | KYAT1 | 0,301897267 | 1,232764539 | 0,36421 | 0,634951 | kynurenine aminotransferase 1 |
| ENSG00000163053 | SLC16A14 | 0,301891678 | 1,232759762 | 0,52437 | 0,74828894 | solute carrier family 16 member 14 |
| ENSG00000176208 | ATAD5 | 0,301890585 | 1,232758829 | 0,29991 | 0,59160746 | ATPase family AAA domain containing 5 |
| ENSG00000171316 | CHD7 | 0,301873346 | 1,232744098 | 0,16348 | 0,48977932 | chromodomain helicase DNA binding protein 7 |
| ENSG00000232811 | AL360270,1 | 0,301777591 | 1,232662281 | 0,48527 | 0,72089329 | novel transcript |
| ENSG00000196247 | ZNF107 | 0,301732078 | 1,232623395 | 0,42302 | 0,67691679 | zinc finger protein 107 |
| ENSG00000172738 | TMEM217 | 0,301639793 | 1,23254455 | 0,60356 | 0,79765228 | transmembrane protein 217 |
| ENSG00000157315 | TMED6 | 0,301609642 | 1,232518791 | 0,53424 | 0,75405583 | transmembrane p24 trafficking protein 6 |
| ENSG00000272054 | AC007390,1 | 0,301585821 | 1,232498441 | 0,5421 | 0,75894841 | novel transcript |
| ENSG00000131470 | PSMC3IP | 0,301575349 | 1,232489494 | 0,31239 | 0,59934192 | PSMC3 interacting protein |
| ENSG00000103671 | TRIP4 | 0,301433825 | 1,232368597 | 0,31887 | 0,60399503 | thyroid hormone receptor interactor 4 |
| ENSG00000134551 | PRH2 | 0,301404543 | 1,232343583 | 0,34574 | 0,62283284 | proline rich protein HaeIII subfamily 2 |
| ENSG00000196368 | NUDT11 | 0,301354369 | 1,232300726 | 0,27752 | 0,57589302 | nudix hydrolase 11 |
| ENSG00000151806 | GUF1 | 0,301352006 | 1,232298708 | 0,295 | 0,58869206 | GUF1 homolog, GTPase |
| ENSG00000152242 | C18orf25 | 0,301342781 | 1,232290828 | 0,36347 | 0,63428574 | chromosome 18 open reading frame 25 |
| ENSG00000285390 | ATP5IF1 | 0,301262703 | 1,232222431 | 0,23941 | 0,5491566 | ATP synthase inhibitory factor subunit 1 |
| ENSG00000130770 | ATP5IF1 | 0,301262703 | 1,232222431 | 0,23941 | 0,5491566 | ATP synthase inhibitory factor subunit 1 |
| ENSG00000284484 | AC025287,4 | 0,301123412 | 1,232103466 | 0,53565 | 0,75485128 | novel protein similar to cytoplasmic polyadenylated homeobox like CPHXL |
| ENSG00000259349 | AC011921,1 | 0,301081853 | 1,232067974 | 0,5042 | 0,73315506 | novel transcript, antisense to APPBP2 |
| ENSG00000196284 | SUPT3H | 0,301055078 | 1,232045109 | 0,47075 | 0,71022965 | SPT3 homolog, SAGA and STAGA complex component |
| ENSG00000136925 | TSTD2 | 0,301026566 | 1,23202076 | 0,39802 | 0,66065114 | thiosulfate sulfurtransferase like domain containing 2 |
| ENSG00000188001 | TPRG1 | 0,301007553 | 1,232004523 | 0,57261 | 0,77858457 | tumor protein p63 regulated 1 |
| ENSG00000230084 | AC006059,1 | 0,300752973 | 1,231787141 | 0,58258 | 0,7847455 | novel transcript |
| ENSG00000106105 | GARS1 | 0,300715451 | 1,231755105 | 0,10167 | 0,40871061 | glycyl-tRNA synthetase 1 |
| ENSG00000115268 | RPS15 | 0,30067147 | 1,231717556 | 0,23822 | 0,54870569 | ribosomal protein S15 |
| ENSG00000250131 | AC078881,1 | 0,300656975 | 1,23170518 | 0,53468 | 0,7541464 | novel transcript |
| ENSG00000226887 | ERVMER34-1 | 0,300601731 | 1,231658016 | 0,54424 | 0,76045619 | endogenous retrovirus group MER34 member 1, envelope |

| | | | | | | |
|---|---|---|---|---|---|---|
| ENSG00000228315 | GUSBP11 | 0,300553883 | 1,231617168 | 0,14446 | 0,47208992 | GUSB pseudogene 11 |
| ENSG00000093000 | NUP50 | 0,300529635 | 1,231596468 | 0,18751 | 0,51081411 | nucleoporin 50 |
| ENSG00000213139 | CRYGS | 0,300526109 | 1,231593458 | 0,45738 | 0,70074793 | crystallin gamma S |
| ENSG00000103540 | CCP110 | 0,300454308 | 1,231532165 | 0,43443 | 0,68505666 | centriolar coiled-coil protein 110 |
| ENSG00000078177 | N4BP2 | 0,300445419 | 1,231524577 | 0,29797 | 0,59037822 | NEDD4 binding protein 2 |
| ENSG00000241131 | LINC02032 | 0,300370115 | 1,231460297 | 0,50175 | 0,73147015 | long intergenic non-protein coding RNA 2032 |
| ENSG00000172053 | QARS1 | 0,300335423 | 1,231430685 | 0,28456 | 0,58143647 | glutaminyl-tRNA synthetase 1 |
| ENSG00000103404 | USP31 | 0,300322653 | 1,231419784 | 0,38318 | 0,649885 | ubiquitin specific peptidase 31 |
| ENSG00000081087 | OSTM1 | 0,300309395 | 1,231408468 | 0,12398 | 0,44485003 | osteoclastogenesis associated transmembrane protein 1 |
| ENSG00000256712 | AC134349,1 | 0,300289849 | 1,231391785 | 0,66381 | 0,8324578 | novel transcript |
| ENSG00000262531 | AC134349,4 | 0,300289849 | 1,231391785 | 0,66381 | 0,8324578 | novel transcript |
| ENSG00000282546 | AC245078,4 | 0,300289849 | 1,231391785 | 0,66381 | 0,8324578 | novel transcript |
| ENSG00000092199 | HNRNPC | 0,300278618 | 1,231382199 | 0,17522 | 0,49934416 | heterogeneous nuclear ribonucleoprotein C |
| ENSG00000105393 | BABAM1 | 0,300245962 | 1,231354327 | 0,07293 | 0,35631792 | BRISC and BRCA1 A complex member 1 |
| ENSG00000170089 | AC106795,1 | 0,300220042 | 1,231332204 | 0,21916 | 0,53664318 | THO complex 3 (THOC3) pseudogene |
| ENSG00000132475 | H3-3B | 0,300212154 | 1,231325471 | 0,2334 | 0,54563244 | H3,3 histone B |
| ENSG00000165030 | NFIL3 | 0,300191488 | 1,231307833 | 0,44858 | 0,6944286 | nuclear factor, interleukin 3 regulated |
| ENSG00000161958 | FGF11 | 0,300166968 | 1,231286906 | 0,27518 | 0,57491355 | fibroblast growth factor 11 |
| ENSG00000166503 | HDGFL3 | 0,300143216 | 1,231266634 | 0,20272 | 0,52269757 | HDGF like 3 |
| ENSG00000015171 | ZMYND11 | 0,300090295 | 1,23122147 | 0,42408 | 0,67782838 | zinc finger MYND-type containing 11 |
| ENSG00000125885 | MCM8 | 0,300005097 | 1,231148763 | 0,11264 | 0,42641188 | minichromosome maintenance 8 homologous recombination repair factor |
| ENSG00000234651 | BAG6 | 0,299844084 | 1,231011367 | 0,21458 | 0,53352305 | BAG cochaperone 6 |
| ENSG00000272438 | AL645608,6 | 0,29972177 | 1,230907005 | 0,76005 | 0,88816166 | novel transcript |
| ENSG00000250979 | AC022905,1 | 0,299710843 | 1,230897682 | 0,59737 | 0,79443333 | transient receptor potential cation channel, subfamily A, member 1 (TRPA1) pseudogene |
| ENSG00000282638 | AC243742,3 | 0,299691271 | 1,230880983 | 0,60082 | 0,79598946 | novel transcript |
| ENSG00000280206 | AC026401,3 | 0,299691271 | 1,230880983 | 0,60082 | 0,79598946 | novel transcript |
| ENSG00000286535 | AC139019,1 | 0,299636043 | 1,230833865 | 0,36391 | 0,63474657 | novel transcript, antisense to ASAP1 |
| ENSG00000117724 | CENPF | 0,299575624 | 1,230782319 | 0,4151 | 0,67198389 | centromere protein F |
| ENSG00000005471 | ABCB4 | 0,299474432 | 1,230695994 | 0,48689 | 0,72215091 | ATP binding cassette subfamily B member 4 |
| ENSG00000080345 | RIF1 | 0,299458721 | 1,230682591 | 0,31647 | 0,60238875 | replication timing regulatory factor 1 |
| ENSG00000159086 | PAXBP1 | 0,299426724 | 1,230655297 | 0,38856 | 0,65310114 | PAX3 and PAX7 binding protein 1 |

| | | | | | | |
|---|---|---|---|---|---|---|
| ENSG00000228817 | BACH1-IT2 | 0,299408379 | 1,230639649 | 0,52073 | 0,74526832 | BACH1 intronic transcript 2 |
| ENSG00000186148 | AC013268,1 | 0,299385452 | 1,230620092 | 0,7356 | 0,8750728 | two-pore calcium channel 3 (TPCN3) pseudogene |
| ENSG00000237887 | RPL23AP32 | 0,299327249 | 1,230570446 | 0,5681 | 0,77582773 | ribosomal protein L23a pseudogene 32 |
| ENSG00000101751 | POLI | 0,299075209 | 1,230355482 | 0,38335 | 0,65009654 | DNA polymerase iota |
| ENSG00000077348 | EXOSC5 | 0,299074781 | 1,230355118 | 0,17722 | 0,50187296 | exosome component 5 |
| ENSG00000138376 | BARD1 | 0,299070452 | 1,230351426 | 0,29896 | 0,5910969 | BRCA1 associated RING domain 1 |
| ENSG00000163935 | SFMBT1 | 0,298768538 | 1,230093976 | 0,12685 | 0,44957921 | Scm like with four mbt domains 1 |
| ENSG00000166860 | ZBTB39 | 0,298752378 | 1,230080198 | 0,16486 | 0,49095733 | zinc finger and BTB domain containing 39 |
| ENSG00000125449 | ARMC7 | 0,298743025 | 1,230072223 | 0,09976 | 0,40626368 | armadillo repeat containing 7 |
| ENSG00000225484 | NUTM2B-AS1 | 0,298580782 | 1,229933899 | 0,3339 | 0,61382024 | NUTM2B antisense RNA 1 |
| ENSG00000167978 | SRRM2 | 0,298483687 | 1,229851126 | 0,14255 | 0,46989963 | serine/arginine repetitive matrix 2 |
| ENSG00000040933 | INPP4A | 0,298469404 | 1,22983895 | 0,32712 | 0,60902638 | inositol polyphosphate-4-phosphatase type I A |
| ENSG00000112290 | WASF1 | 0,298455712 | 1,229827278 | 0,42965 | 0,68237934 | WASP family member 1 |
| ENSG00000075292 | ZNF638 | 0,29841298 | 1,229790852 | 0,27085 | 0,57239492 | zinc finger protein 638 |
| ENSG00000213608 | SLC25A14P1 | 0,298304399 | 1,229698298 | 0,60944 | 0,80172326 | solute carrier family 25 member 14 pseudogene 1 |
| ENSG00000137710 | RDX | 0,298235014 | 1,229639159 | 0,36741 | 0,63739285 | radixin |
| ENSG00000105948 | TTC26 | 0,298208649 | 1,229616687 | 0,42962 | 0,6823653 | tetratricopeptide repeat domain 26 |
| ENSG00000180776 | ZDHHC20 | 0,29819453 | 1,229604653 | 0,16621 | 0,49212363 | zinc finger DHHC-type palmitoyltransferase 20 |
| ENSG00000125734 | GPR108 | 0,298164729 | 1,229579255 | 0,34671 | 0,62350287 | G protein-coupled receptor 108 |
| ENSG00000138738 | PRDM5 | 0,298141363 | 1,229559341 | 0,36101 | 0,63311211 | PR/SET domain 5 |
| ENSG00000120915 | EPHX2 | 0,298110757 | 1,229533256 | 0,42813 | 0,68094175 | epoxide hydrolase 2 |
| ENSG00000232692 | AP001596,1 | 0,298013509 | 1,229450379 | 0,59812 | 0,79484486 | novel transcript |
| ENSG00000164978 | NUDT2 | 0,297949804 | 1,229396092 | 0,39889 | 0,66138533 | nudix hydrolase 2 |
| ENSG00000251669 | FAM86EP | 0,297884879 | 1,229340767 | 0,3478 | 0,62427624 | family with sequence similarity 86 member E, pseudogene |
| ENSG00000043355 | ZIC2 | 0,297882181 | 1,229338468 | 0,38293 | 0,64965769 | Zic family member 2 |
| ENSG00000170364 | SETMAR | 0,297873434 | 1,229331015 | 0,31366 | 0,60035326 | SET domain and mariner transposase fusion gene |
| ENSG00000080819 | CPOX | 0,297776305 | 1,229248253 | 0,1944 | 0,51562158 | coproporphyrinogen oxidase |
| ENSG00000108946 | PRKAR1A | 0,297710614 | 1,229192283 | 0,30934 | 0,59780895 | protein kinase cAMP-dependent type I regulatory subunit alpha |
| ENSG00000213551 | DNAJC9 | 0,297710173 | 1,229191907 | 0,28201 | 0,57936013 | DnaJ heat shock protein family (Hsp40) member C9 |
| ENSG00000259396 | AC087721,1 | 0,297674759 | 1,229161734 | 0,71591 | 0,86378773 | novel transcript, sense intronic to NUSAP1 |
| ENSG00000261512 | AC092368,3 | 0,297599873 | 1,229097934 | 0,60075 | 0,79598946 | novel transcript |
| ENSG00000115875 | SRSF7 | 0,297580857 | 1,229081733 | 0,19449 | 0,51562158 | serine and arginine rich splicing factor 7 |

| | | | | | | |
|---|---|---|---|---|---|---|
| ENSG00000204380 | PKP4-AS1 | 0,297515356 | 1,229025932 | 0,42236 | 0,67663581 | PKP4 antisense RNA 1 |
| ENSG00000065675 | PRKCQ | 0,297503366 | 1,229015717 | 0,58947 | 0,78887837 | protein kinase C theta |
| ENSG00000242992 | FTH1P4 | 0,297501437 | 1,229014074 | 0,75975 | 0,88801075 | ferritin heavy chain 1 pseudogene 4 |
| ENSG00000258944 | AC004846,2 | 0,29748465 | 1,228999774 | 0,59055 | 0,78980722 | novel transcript, antisense to PAPLN |
| ENSG00000251676 | SNHG27 | 0,29745892 | 1,228977855 | 0,50245 | 0,73208008 | small nucleolar RNA host gene 27 |
| ENSG00000119689 | DLST | 0,297427718 | 1,228951276 | 0,20978 | 0,52867318 | dihydrolipoamide S-succinyltransferase |
| ENSG00000081721 | DUSP12 | 0,297406306 | 1,228933036 | 0,37884 | 0,64679625 | dual specificity phosphatase 12 |
| ENSG00000126746 | ZNF384 | 0,297369555 | 1,22890173 | 0,20628 | 0,5259882 | zinc finger protein 384 |
| ENSG00000103202 | NME4 | 0,297314886 | 1,228855164 | 0,18652 | 0,509961 | NME/NM23 nucleoside diphosphate kinase 4 |
| ENSG00000108829 | LRRC59 | 0,297242262 | 1,228793306 | 0,276 | 0,57533851 | leucine rich repeat containing 59 |
| ENSG00000198920 | KIAA0753 | 0,297125222 | 1,228693623 | 0,46979 | 0,70922292 | KIAA0753 |
| ENSG00000080822 | CLDND1 | 0,297106792 | 1,228677927 | 0,36323 | 0,63407828 | claudin domain containing 1 |
| ENSG00000182021 | AL591379,1 | 0,297013087 | 1,228598126 | 0,49887 | 0,72996654 | methylenetetrahydrofolate dehydrogenase (NADP+ dependent) 1 like pseudogene |
| ENSG00000122545 | SEPTIN7 | 0,296979121 | 1,2285692 | 0,36067 | 0,63299251 | septin 7 |
| ENSG00000242247 | ARFGAP3 | 0,296970935 | 1,228562229 | 0,49661 | 0,72841371 | ADP ribosylation factor GTPase activating protein 3 |
| ENSG00000197472 | ZNF695 | 0,296937686 | 1,228533915 | 0,30807 | 0,59696396 | zinc finger protein 695 |
| ENSG00000159899 | NPR2 | 0,296926457 | 1,228524353 | 0,15156 | 0,47972295 | natriuretic peptide receptor 2 |
| ENSG00000124541 | RRP36 | 0,296906516 | 1,228507373 | 0,09507 | 0,39768318 | ribosomal RNA processing 36 |
| ENSG00000020426 | MNAT1 | 0,296905386 | 1,228506411 | 0,2589 | 0,56264327 | MNAT1 component of CDK activating kinase |
| ENSG00000131504 | DIAPH1 | 0,296891758 | 1,228494806 | 0,16258 | 0,48945635 | diaphanous related formin 1 |
| ENSG00000256196 | AP003721,1 | 0,296875851 | 1,228481261 | 0,2115 | 0,53032236 | novel transcript |
| ENSG00000175267 | VWA3A | 0,296842034 | 1,228452465 | 0,55378 | 0,7667945 | von Willebrand factor A domain containing 3A |
| ENSG00000259866 | AC007599,1 | 0,296828967 | 1,228441338 | 0,64662 | 0,82303578 | Rho GTPase activating protein 20 (ARHGAP20) pseudogene |
| ENSG00000271967 | AL583856,2 | 0,296708599 | 1,228338851 | 0,78435 | 0,89962715 | novel transcript |
| ENSG00000249363 | AC011411,1 | 0,296695633 | 1,228327811 | 0,6619 | 0,83159381 | transient receptor potential cation channel family (TRPC7) pseudogene |
| ENSG00000114473 | IQCG | 0,296650877 | 1,228289706 | 0,30472 | 0,59446033 | IQ motif containing G |
| ENSG00000070831 | CDC42 | 0,296639749 | 1,228280232 | 0,09432 | 0,39648446 | cell division cycle 42 |
| ENSG00000137522 | RNF121 | 0,296629734 | 1,228271705 | 0,27807 | 0,57635142 | ring finger protein 121 |
| ENSG00000091972 | CD200 | 0,296573337 | 1,228223692 | 0,43381 | 0,68475933 | CD200 molecule |
| ENSG00000142627 | EPHA2 | 0,296542569 | 1,228197498 | 0,29727 | 0,59031059 | EPH receptor A2 |
| ENSG00000136169 | SETDB2 | 0,296511659 | 1,228171184 | 0,24226 | 0,55112431 | SET domain bifurcated histone lysine methyltransferase 2 |

| | | | | | | |
|---|---|---|---|---|---|---|
| ENSG00000244716 | BX679664,3 | 0,296487623 | 1,228150722 | 0,80443 | 0,90996631 | ribosomal protein L17 (RPL17) pseudogene |
| ENSG00000156587 | UBE2L6 | 0,296369079 | 1,228049811 | 0,36793 | 0,63791537 | ubiquitin conjugating enzyme E2 L6 |
| ENSG00000183604 | SMG1P5 | 0,29635346 | 1,228036516 | 0,25317 | 0,55832664 | SMG1 pseudogene 5 |
| ENSG00000158050 | DUSP2 | 0,296315246 | 1,228003988 | 0,62694 | 0,81116451 | dual specificity phosphatase 2 |
| ENSG00000156411 | ATP5MPL | 0,296306552 | 1,227996588 | 0,33492 | 0,61467789 | ATP synthase membrane subunit 6,8PL |
| ENSG00000144649 | GASK1A | 0,296232956 | 1,227933946 | 0,49007 | 0,72433291 | golgi associated kinase 1A |
| ENSG00000272677 | AC124016,1 | 0,295959135 | 1,227700909 | 0,43176 | 0,68379763 | novel transcript, antisense to HNRNPD |
| ENSG00000115685 | PPP1R7 | 0,295868432 | 1,227623724 | 0,27389 | 0,57387352 | protein phosphatase 1 regulatory subunit 7 |
| ENSG00000109381 | ELF2 | 0,29585647 | 1,227613546 | 0,32388 | 0,60659203 | E74 like ETS transcription factor 2 |
| ENSG00000130347 | RTN4IP1 | 0,295838651 | 1,227598383 | 0,30413 | 0,59411013 | reticulon 4 interacting protein 1 |
| ENSG00000270264 | NDUFB8P2 | 0,295815131 | 1,22757837 | 0,68958 | 0,84865812 | NADH:ubiquinone oxidoreductase subunit B8 pseudogene 2 |
| ENSG00000176393 | RNPEP | 0,295704365 | 1,227484124 | 0,13621 | 0,463307 | arginyl aminopeptidase |
| ENSG00000259001 | AL355075,4 | 0,295694627 | 1,227475839 | 0,69334 | 0,85096709 | ribonuclease P RNA component H1 |
| ENSG00000171604 | CXXC5 | 0,29568736 | 1,227469656 | 0,17072 | 0,49641416 | CXXC finger protein 5 |
| ENSG00000241863 | RPP21 | 0,295686932 | 1,227469292 | 0,32637 | 0,60866572 | ribonuclease P/MRP subunit p21 |
| ENSG00000270772 | PDHA1P1 | 0,295669134 | 1,227454148 | 0,70312 | 0,8568579 | PDHA1 pseudogene 1 |
| ENSG00000234329 | AL604028,2 | 0,295596018 | 1,227391943 | 0,44193 | 0,68994635 | ribosomal protein L7a (RPL7A) pseudogene |
| ENSG00000267645 | AC105052,3 | 0,295593223 | 1,227389565 | 0,87889 | 0,94615035 | novel protein, POLR2J2-UPK3BL readthrough |
| ENSG00000140694 | PARN | 0,295571781 | 1,227371323 | 0,20449 | 0,52439728 | poly(A)-specific ribonuclease |
| ENSG00000101057 | MYBL2 | 0,295553932 | 1,227356138 | 0,22954 | 0,54307963 | MYB proto-oncogene like 2 |
| ENSG00000174953 | DHX36 | 0,29550626 | 1,227315583 | 0,30904 | 0,59774312 | DEAH-box helicase 36 |
| ENSG00000012174 | MBTPS2 | 0,295483066 | 1,227295851 | 0,18224 | 0,50571799 | membrane bound transcription factor peptidase, site 2 |
| ENSG00000108375 | RNF43 | 0,29543359 | 1,227253763 | 0,61184 | 0,80337676 | ring finger protein 43 |
| ENSG00000076382 | SPAG5 | 0,295384413 | 1,227211931 | 0,27129 | 0,57271346 | sperm associated antigen 5 |
| ENSG00000237654 | AP003025,1 | 0,295345936 | 1,227179201 | 0,7636 | 0,88983737 | novel transcript |
| ENSG00000138764 | CCNG2 | 0,295311716 | 1,227150093 | 0,29005 | 0,58531219 | cyclin G2 |
| ENSG00000164402 | SEPTIN8 | 0,295276522 | 1,227120158 | 0,31815 | 0,60349082 | septin 8 |
| ENSG00000111725 | PRKAB1 | 0,295254792 | 1,227101674 | 0,32123 | 0,60573903 | protein kinase AMP-activated non-catalytic subunit beta 1 |
| ENSG00000158373 | H2BC5 | 0,295246145 | 1,22709432 | 0,59926 | 0,79538547 | H2B clustered histone 5 |
| ENSG00000094975 | SUCO | 0,295123457 | 1,226989971 | 0,45242 | 0,69702004 | SUN domain containing ossification factor |
| ENSG00000111731 | C2CD5 | 0,295114542 | 1,226982389 | 0,40431 | 0,6642832 | C2 calcium dependent domain containing 5 |
| ENSG00000005156 | LIG3 | 0,295026901 | 1,226907854 | 0,1559 | 0,4836223 | DNA ligase 3 |

| | | | | | | |
|---|---|---|---|---|---|---|
| ENSG00000273562 | NAP1L4 | 0,29499619 | 1,226881737 | 0,31514 | 0,60150855 | nucleosome assembly protein 1 like 4 |
| ENSG00000186432 | KPNA4 | 0,294919131 | 1,226816207 | 0,13453 | 0,46072811 | karyopherin subunit alpha 4 |
| ENSG00000188243 | COMMD6 | 0,294888326 | 1,226790012 | 0,53082 | 0,75219176 | COMM domain containing 6 |
| ENSG00000127838 | PNKD | 0,294730831 | 1,226656094 | 0,17217 | 0,49710979 | PNKD metallo-beta-lactamase domain containing |
| ENSG00000168014 | C2CD3 | 0,294662982 | 1,226598407 | 0,23853 | 0,5488078 | C2 domain containing 3 centriole elongation regulator |
| ENSG00000240288 | GHRLOS | 0,294662064 | 1,226597626 | 0,24603 | 0,55380958 | ghrelin opposite strand/antisense RNA |
| ENSG00000121053 | EPX | 0,294637047 | 1,226576357 | 0,77375 | 0,89570857 | eosinophil peroxidase |
| ENSG00000138050 | THUMPD2 | 0,294625601 | 1,226566625 | 0,3149 | 0,60144815 | THUMP domain containing 2 |
| ENSG00000011376 | LARS2 | 0,294594854 | 1,226540485 | 0,1596 | 0,48693054 | leucyl-tRNA synthetase 2, mitochondrial |
| ENSG00000213523 | SRA1 | 0,294549536 | 1,226501957 | 0,20857 | 0,52778377 | steroid receptor RNA activator 1 |
| ENSG00000273226 | AL391834,2 | 0,294481698 | 1,226444286 | 0,43153 | 0,68377875 | novel transcript, antisense to RPS6 |
| ENSG00000069248 | NUP133 | 0,294400552 | 1,226375305 | 0,35969 | 0,63242662 | nucleoporin 133 |
| ENSG00000239884 | RN7SL608P | 0,294398976 | 1,226373966 | 0,68761 | 0,8478032 | RNA, 7SL, cytoplasmic 608, pseudogene |
| ENSG00000106399 | RPA3 | 0,294348672 | 1,226331205 | 0,39524 | 0,6581785 | replication protein A3 |
| ENSG00000284770 | TBCE | 0,294312759 | 1,226300678 | 0,12936 | 0,45283544 | tubulin folding cofactor E |
| ENSG00000065154 | OAT | 0,294260968 | 1,226256656 | 0,39648 | 0,6591566 | ornithine aminotransferase |
| ENSG00000145569 | OTULINL | 0,294239112 | 1,226238079 | 0,33986 | 0,61872249 | OTU deubiquitinase with linear linkage specificity like |
| ENSG00000173465 | ZNRD2 | 0,294226369 | 1,226227248 | 0,30537 | 0,59483988 | zinc ribbon domain containing 2 |
| ENSG00000109586 | GALNT7 | 0,29421406 | 1,226216787 | 0,52946 | 0,7512527 | polypeptide N-acetylgalactosaminyltransferase 7 |
| ENSG00000219481 | NBPF1 | 0,29417826 | 1,226186359 | 0,22313 | 0,53927144 | NBPF member 1 |
| ENSG00000179399 | GPC5 | 0,294177662 | 1,226185851 | 0,70245 | 0,85653047 | glypican 5 |
| ENSG00000078304 | PPP2R5C | 0,294161161 | 1,226171826 | 0,27405 | 0,57412006 | protein phosphatase 2 regulatory subunit B'gamma |
| ENSG00000188610 | FAM72B | 0,294130509 | 1,226145775 | 0,44364 | 0,69133101 | family with sequence similarity 72 member B |
| ENSG00000153574 | RPIA | 0,294090011 | 1,226111356 | 0,09822 | 0,40371781 | ribose 5-phosphate isomerase A |
| ENSG00000187984 | ANKRD19P | 0,29398642 | 1,226023319 | 0,49269 | 0,72588649 | ankyrin repeat domain 19, pseudogene |
| ENSG00000049618 | ARID1B | 0,293841638 | 1,225900288 | 0,1924 | 0,514872 | AT-rich interaction domain 1B |
| ENSG00000137502 | RAB30 | 0,29383202 | 1,225892115 | 0,41804 | 0,67432365 | RAB30, member RAS oncogene family |
| ENSG00000271270 | TMCC1-AS1 | 0,293816244 | 1,22587871 | 0,45995 | 0,70201575 | TMCC1 antisense RNA 1 (head to head) |
| ENSG00000219384 | AL138880,1 | 0,293717458 | 1,225794773 | 0,7018 | 0,85629987 | pseudogene similar to part of poly(rC) binding protein 3 (PCBP3) |
| ENSG00000084674 | APOB | 0,293696062 | 1,225776594 | 0,64894 | 0,82427495 | apolipoprotein B |
| ENSG00000175197 | DDIT3 | 0,293677719 | 1,225761009 | 0,41483 | 0,67165882 | DNA damage inducible transcript 3 |
| ENSG00000287012 | AC108667,2 | 0,293625995 | 1,225717064 | 0,60767 | 0,80041268 | novel transcript |

| | | | | | | |
|---|---|---|---|---|---|---|
| ENSG00000277221 | LRRC37A2 | 0,293618211 | 1,22571045 | 0,33027 | 0,61114402 | leucine rich repeat containing 37 member A2 |
| ENSG00000139990 | DCAF5 | 0,293606552 | 1,225700545 | 0,3486 | 0,62475089 | DDB1 and CUL4 associated factor 5 |
| ENSG00000177034 | MTX3 | 0,293578675 | 1,225676861 | 0,18645 | 0,50994868 | metaxin 3 |
| ENSG00000160703 | NLRX1 | 0,293577999 | 1,225676286 | 0,2527 | 0,55796133 | NLR family member X1 |
| ENSG00000249637 | AC008438,1 | 0,293576623 | 1,225675118 | 0,50024 | 0,73078774 | novel transcript |
| ENSG00000146360 | GPR6 | 0,293524827 | 1,225631113 | 0,7203 | 0,86628014 | G protein-coupled receptor 6 |
| ENSG00000186019 | AC021092,1 | 0,293523265 | 1,225629786 | 0,0639 | 0,33507507 | novel transcript, antisense to ZNF225 and ZNF224 |
| ENSG00000151445 | VIPAS39 | 0,293491846 | 1,225603096 | 0,26004 | 0,56382871 | VPS33B interacting protein, apical-basolateral polarity regulator, spe-39 homolog |
| ENSG00000123297 | TSFM | 0,293470939 | 1,225585334 | 0,16142 | 0,48892403 | Ts translation elongation factor, mitochondrial |
| ENSG00000179023 | KLHDC7A | 0,293389836 | 1,225516438 | 0,64278 | 0,82094129 | kelch domain containing 7A |
| ENSG00000168843 | FSTL5 | 0,293383274 | 1,225510864 | 0,53837 | 0,7567407 | follistatin like 5 |
| ENSG00000204571 | KRTAP5-11 | 0,293365584 | 1,225495837 | 0,60591 | 0,79907418 | keratin associated protein 5-11 |
| ENSG00000187764 | SEMA4D | 0,29331213 | 1,225450432 | 0,30278 | 0,59321687 | semaphorin 4D |
| ENSG00000116667 | C1orf21 | 0,29330979 | 1,225448444 | 0,29922 | 0,59123205 | chromosome 1 open reading frame 21 |
| ENSG00000068878 | PSME4 | 0,293303114 | 1,225442773 | 0,33442 | 0,61423173 | proteasome activator subunit 4 |
| ENSG00000114503 | NCBP2 | 0,293198308 | 1,225353753 | 0,41426 | 0,67104294 | nuclear cap binding protein subunit 2 |
| ENSG00000278771 | RN7SL3 | 0,293136759 | 1,225301478 | 0,23016 | 0,54335232 | RNA component of signal recognition particle 7SL3 |
| ENSG00000170471 | RALGAPB | 0,29308171 | 1,225254725 | 0,25544 | 0,55992862 | Ral GTPase activating protein non-catalytic subunit beta |
| ENSG00000143643 | TTC13 | 0,292978494 | 1,225167068 | 0,48262 | 0,719011 | tetratricopeptide repeat domain 13 |
| ENSG00000078747 | ITCH | 0,29292649 | 1,225122906 | 0,38723 | 0,65225328 | itchy E3 ubiquitin protein ligase |
| ENSG00000241030 | RPL29P24 | 0,292832283 | 1,225042909 | 0,68697 | 0,84727142 | ribosomal protein L29 pseudogene 24 |
| ENSG00000259065 | AC005520,2 | 0,292678384 | 1,224912235 | 0,6076 | 0,80036065 | novel transcript, antisense to C14orf43 |
| ENSG00000224994 | TRIM39 | 0,29260802 | 1,224852495 | 0,19116 | 0,51406315 | tripartite motif containing 39 |
| ENSG00000229929 | TRIM39 | 0,29260802 | 1,224852495 | 0,19116 | 0,51406315 | tripartite motif containing 39 |
| ENSG00000232839 | TRIM39 | 0,29260802 | 1,224852495 | 0,19116 | 0,51406315 | tripartite motif containing 39 |
| ENSG00000204599 | TRIM39 | 0,29260802 | 1,224852495 | 0,19116 | 0,51406315 | tripartite motif containing 39 |
| ENSG00000230308 | TRIM39 | 0,29260802 | 1,224852495 | 0,19116 | 0,51406315 | tripartite motif containing 39 |
| ENSG00000168894 | RNF181 | 0,292523649 | 1,224780865 | 0,33049 | 0,6112353 | ring finger protein 181 |
| ENSG00000227197 | AC009365,3 | 0,29246813 | 1,224733733 | 0,6257 | 0,81057207 | novel transcript |
| ENSG00000115318 | LOXL3 | 0,292402875 | 1,224678338 | 0,23572 | 0,5471053 | lysyl oxidase like 3 |
| ENSG00000259959 | AC107068,1 | 0,292389361 | 1,224666866 | 0,21476 | 0,53372266 | novel transcript |
| ENSG00000108515 | ENO3 | 0,292279413 | 1,224573538 | 0,42179 | 0,67643128 | enolase 3 |

| | | | | | | |
|---|---|---|---|---|---|---|
| ENSG00000174010 | KLHL15 | 0,292216973 | 1,224520539 | 0,46994 | 0,7093087 | kelch like family member 15 |
| ENSG00000213341 | CHUK | 0,292206532 | 1,224511677 | 0,28232 | 0,5797164 | component of inhibitor of nuclear factor kappa B kinase complex |
| ENSG00000171456 | ASXL1 | 0,292201234 | 1,224507181 | 0,14756 | 0,47504057 | ASXL transcriptional regulator 1 |
| ENSG00000135315 | CEP162 | 0,292072191 | 1,224397658 | 0,49397 | 0,72657717 | centrosomal protein 162 |
| ENSG00000287580 | AC073349,4 | 0,292021654 | 1,224354769 | 0,53554 | 0,75478533 | novel transcript, antisense to ZNF138 |
| ENSG00000225673 | AC104164,1 | 0,291883109 | 1,224237198 | 0,72925 | 0,87157404 | transmembrane emp24 domain trafficking protein 2 (TMED2) pseudogene |
| ENSG00000123219 | CENPK | 0,291859914 | 1,224217515 | 0,33229 | 0,61221238 | centromere protein K |
| ENSG00000165591 | FAAH2 | 0,291836703 | 1,224197819 | 0,50266 | 0,73209345 | fatty acid amide hydrolase 2 |
| ENSG00000118363 | SPCS2 | 0,291830785 | 1,224192797 | 0,1874 | 0,51074619 | signal peptidase complex subunit 2 |
| ENSG00000115216 | NRBP1 | 0,291780938 | 1,224150501 | 0,25777 | 0,56191608 | nuclear receptor binding protein 1 |
| ENSG00000168883 | USP39 | 0,291686851 | 1,224070669 | 0,14044 | 0,46735399 | ubiquitin specific peptidase 39 |
| ENSG00000205730 | ITPRIPL2 | 0,291658672 | 1,22404676 | 0,38685 | 0,6521009 | ITPRIP like 2 |
| ENSG00000065911 | MTHFD2 | 0,291626555 | 1,224019511 | 0,25736 | 0,56160845 | methylenetetrahydrofolate dehydrogenase (NADP+ dependent) 2, methenyltetrahydrofolate cyclohydrolase |
| ENSG00000278915 | AC005609,3 | 0,291624562 | 1,22401782 | 0,46068 | 0,70247195 | novel transcript, antisense to PCDHA12 |
| ENSG00000077713 | SLC25A43 | 0,291589902 | 1,223988414 | 0,14133 | 0,46803321 | solute carrier family 25 member 43 |
| ENSG00000228000 | RPL7AP65 | 0,291525769 | 1,223934005 | 0,63618 | 0,81694486 | ribosomal protein L7a pseudogene 65 |
| ENSG00000284212 | RPL7AP65 | 0,291525769 | 1,223934005 | 0,63618 | 0,81694486 | ribosomal protein L7a pseudogene 65 |
| ENSG00000188529 | SRSF10 | 0,291498866 | 1,223911181 | 0,34505 | 0,62228057 | serine and arginine rich splicing factor 10 |
| ENSG00000234728 | SNHG32 | 0,291458234 | 1,223876712 | 0,36644 | 0,6363423 | small nucleolar RNA host gene 32 |
| ENSG00000206380 | SNHG32 | 0,291458234 | 1,223876712 | 0,36644 | 0,6363423 | small nucleolar RNA host gene 32 |
| ENSG00000214174 | AMZ2P1 | 0,291450522 | 1,22387017 | 0,37887 | 0,64679625 | archaelysin family metallopeptidase 2 pseudogene 1 |
| ENSG00000276340 | ZNF623 | 0,291394436 | 1,223822591 | 0,44256 | 0,69016964 | zinc finger protein 623 |
| ENSG00000155100 | OTUD6B | 0,291338666 | 1,223775283 | 0,43288 | 0,68436596 | OTU deubiquitinase 6B |
| ENSG00000009844 | VTA1 | 0,291303643 | 1,223745575 | 0,37473 | 0,6431311 | vesicle trafficking 1 |
| ENSG00000213050 | TPM3P1 | 0,291272335 | 1,223719019 | 0,69226 | 0,85023519 | tropomyosin 3 pseudogene 1 |
| ENSG00000156697 | UTP14A | 0,291261821 | 1,223710101 | 0,1374 | 0,46439773 | UTP14A small subunit processome component |
| ENSG00000169105 | CHST14 | 0,291237745 | 1,223689679 | 0,15567 | 0,48345635 | carbohydrate sulfotransferase 14 |
| ENSG00000183562 | AC131971,1 | 0,291183727 | 1,223643862 | 0,32748 | 0,60929711 | novel transcript, antisense to NAP1L4 |
| ENSG00000282438 | AC001228,2 | 0,291183727 | 1,223643862 | 0,32748 | 0,60929711 | novel transcript, antisense to NAP1L4 |
| ENSG00000112592 | TBP | 0,291179358 | 1,223640156 | 0,14704 | 0,47471707 | TATA-box binding protein |

| | | | | | | |
|---|---|---|---|---|---|---|
| ENSG00000114993 | RTKN | 0,291177192 | 1,22363832 | 0,18599 | 0,50932713 | rhotekin |
| ENSG00000145293 | ENOPH1 | 0,291142489 | 1,223608886 | 0,38833 | 0,6529693 | enolase-phosphatase 1 |
| ENSG00000249456 | AL731577,2 | 0,291136064 | 1,223603437 | 0,26924 | 0,57096273 | novel transcript, sense overlapping ZRANB1 |
| ENSG00000258843 | AL133485,2 | 0,291101794 | 1,223574372 | 0,66165 | 0,831498 | novel transcript, antisense to NIN |
| ENSG00000164949 | GEM | 0,291071468 | 1,223548652 | 0,4654 | 0,70588034 | GTP binding protein overexpressed in skeletal muscle |
| ENSG00000223390 | AL445685,1 | 0,291031915 | 1,223515107 | 0,81274 | 0,91386772 | novel transcript |
| ENSG00000117713 | ARID1A | 0,29099023 | 1,223479756 | 0,13037 | 0,45477153 | AT-rich interaction domain 1A |
| ENSG00000169018 | FEM1B | 0,290955082 | 1,223449949 | 0,21019 | 0,52904355 | fem-1 homolog B |
| ENSG00000085760 | MTIF2 | 0,290952975 | 1,223448162 | 0,34721 | 0,62400742 | mitochondrial translational initiation factor 2 |
| ENSG00000227769 | AC072062,1 | 0,290895398 | 1,223399336 | 0,59878 | 0,79523922 | novel transcript, antisense to ABCA12 |
| ENSG00000184635 | ZNF93 | 0,290822756 | 1,223337737 | 0,3759 | 0,64417233 | zinc finger protein 93 |
| ENSG00000177707 | NECTIN3 | 0,29079457 | 1,223313837 | 0,50583 | 0,73418581 | nectin cell adhesion molecule 3 |
| ENSG00000163536 | SERPINI1 | 0,290610146 | 1,223157468 | 0,53783 | 0,75635915 | serpin family I member 1 |
| ENSG00000158747 | NBL1 | 0,290589959 | 1,223140353 | 0,45843 | 0,70119611 | NBL1, DAN family BMP antagonist |
| ENSG00000257594 | GALNT4 | 0,290558256 | 1,223113474 | 0,31468 | 0,60130206 | polypeptide N-acetylgalactosaminyltransferase 4 |
| ENSG00000175334 | BANF1 | 0,290530604 | 1,223090031 | 0,2213 | 0,53805234 | BAF nuclear assembly factor 1 |
| ENSG00000110218 | PANX1 | 0,290494913 | 1,223059774 | 0,34904 | 0,62519057 | pannexin 1 |
| ENSG00000213754 | KPNA4P1 | 0,290483676 | 1,223050247 | 0,59413 | 0,79243905 | KPNA4 pseudogene 1 |
| ENSG00000286563 | AC078983,1 | 0,290344193 | 1,222932005 | 0,63102 | 0,81390227 | novel transcript |
| ENSG00000259103 | AF107885,2 | 0,290319027 | 1,222910673 | 0,59571 | 0,79349831 | novel transcript, antisense to TTLL5 |
| ENSG00000168003 | SLC3A2 | 0,290317158 | 1,222909089 | 0,23092 | 0,54391581 | solute carrier family 3 member 2 |
| ENSG00000277873 | AC131159,2 | 0,290248706 | 1,222851067 | 0,55326 | 0,76630282 | novel transcript, antisense to RFC5 |
| ENSG00000129103 | SUMF2 | 0,290108424 | 1,222732168 | 0,09173 | 0,39166375 | sulfatase modifying factor 2 |
| ENSG00000156253 | RWDD2B | 0,290049646 | 1,222682352 | 0,19582 | 0,51663034 | RWD domain containing 2B |
| ENSG00000270956 | AC009948,2 | 0,290039568 | 1,222673811 | 0,70228 | 0,85651838 | novel transcript, antisense to TTN |
| ENSG00000201901 | RN7SKP48 | 0,290005798 | 1,222645191 | 0,78706 | 0,90114442 | RN7SK pseudogene 48 |
| ENSG00000127081 | ZNF484 | 0,28998833 | 1,222630388 | 0,57399 | 0,77928773 | zinc finger protein 484 |
| ENSG00000086061 | DNAJA1 | 0,289930144 | 1,222581078 | 0,28644 | 0,58273069 | DnaJ heat shock protein family (Hsp40) member A1 |
| ENSG00000183833 | CFAP91 | 0,289904455 | 1,222559309 | 0,49125 | 0,72507957 | cilia and flagella associated protein 91 |
| ENSG00000104626 | ERI1 | 0,289835511 | 1,222500886 | 0,48119 | 0,71820653 | exoribonuclease 1 |
| ENSG00000100023 | PPIL2 | 0,28967672 | 1,222366339 | 0,35461 | 0,62915556 | peptidylprolyl isomerase like 2 |
| ENSG00000248967 | AC022493,2 | 0,289560798 | 1,222268124 | 0,73555 | 0,8750728 | S-phase kinase-associated protein 1 (SKP1) pseudogene |
| ENSG00000237686 | AL109615,3 | 0,28953879 | 1,222249478 | 0,69709 | 0,85330122 | novel transcript, antisense to C6orf223 |

| | | | | | | |
|---|---|---|---|---|---|---|
| ENSG00000241829 | RPL21P54 | 0,289477733 | 1,222197752 | 0,76805 | 0,89257351 | ribosomal protein L21 pseudogene 54 |
| ENSG00000169180 | XPO6 | 0,289421036 | 1,222149722 | 0,17956 | 0,50389568 | exportin 6 |
| ENSG00000124721 | DNAH8 | 0,28942051 | 1,222149276 | 0,60305 | 0,7971742 | dynein axonemal heavy chain 8 |
| ENSG00000141391 | PRELID3A | 0,289376992 | 1,222112411 | 0,19256 | 0,51490739 | PRELI domain containing 3A |
| ENSG00000273345 | AC104109,4 | 0,28925228 | 1,222006772 | 0,62863 | 0,81215334 | novel transcript |
| ENSG00000156136 | DCK | 0,289223894 | 1,221982728 | 0,48049 | 0,71764212 | deoxycytidine kinase |
| ENSG00000074657 | ZNF532 | 0,289039198 | 1,221826298 | 0,37718 | 0,6456352 | zinc finger protein 532 |
| ENSG00000001626 | CFTR | 0,289025575 | 1,221814761 | 0,54964 | 0,76419314 | CF transmembrane conductance regulator |
| ENSG00000174963 | ZIC4 | 0,28899292 | 1,221787106 | 0,37625 | 0,64458342 | Zic family member 4 |
| ENSG00000258439 | AC007956,2 | 0,288922517 | 1,221727485 | 0,70519 | 0,85797994 | ribosomal protein S2 (RPS2) pseudogene |
| ENSG00000185532 | PRKG1 | 0,288840906 | 1,221658375 | 0,59446 | 0,79269155 | protein kinase cGMP-dependent 1 |
| ENSG00000225423 | TNPO1P1 | 0,288673036 | 1,221516233 | 0,67589 | 0,84049293 | transportin 1 pseudogene 1 |
| ENSG00000187079 | TEAD1 | 0,288550137 | 1,22141218 | 0,34451 | 0,62198289 | TEA domain transcription factor 1 |
| ENSG00000150527 | MIA2 | 0,288376681 | 1,221265338 | 0,08917 | 0,38617197 | MIA SH3 domain ER export factor 2 |
| ENSG00000006695 | COX10 | 0,288337819 | 1,221232441 | 0,35055 | 0,62606342 | cytochrome c oxidase assembly factor heme A:farnesyltransferase COX10 |
| ENSG00000258634 | AL160006,1 | 0,288244402 | 1,221153367 | 0,12661 | 0,4492748 | novel transcript, antisense to ALX3 |
| ENSG00000257957 | QRSL1P3 | 0,28821609 | 1,221129402 | 0,68844 | 0,84821636 | QRSL1 pseudogene 3 |
| ENSG00000022277 | RTF2 | 0,288164053 | 1,221085358 | 0,35715 | 0,63118399 | replication termination factor 2 |
| ENSG00000267680 | ZNF224 | 0,287969613 | 1,220920797 | 0,4819 | 0,71869839 | zinc finger protein 224 |
| ENSG00000115289 | PCGF1 | 0,287858627 | 1,220826875 | 0,22513 | 0,54058261 | polycomb group ring finger 1 |
| ENSG00000174327 | SLC16A13 | 0,287822335 | 1,220796165 | 0,26658 | 0,5691939 | solute carrier family 16 member 13 |
| ENSG00000186272 | ZNF17 | 0,287726388 | 1,220714978 | 0,38971 | 0,65396251 | zinc finger protein 17 |
| ENSG00000169330 | MINAR1 | 0,287602825 | 1,220610431 | 0,50375 | 0,73293685 | membrane integral NOTCH2 associated receptor 1 |
| ENSG00000112200 | ZNF451 | 0,287458658 | 1,220488463 | 0,34849 | 0,62474431 | zinc finger protein 451 |
| ENSG00000103196 | CRISPLD2 | 0,287395552 | 1,220435078 | 0,62124 | 0,80787022 | cysteine rich secretory protein LCCL domain containing 2 |
| ENSG00000156482 | RPL30 | 0,287375879 | 1,220418436 | 0,22522 | 0,54058261 | ribosomal protein L30 |
| ENSG00000189060 | H1-0 | 0,287371922 | 1,220415089 | 0,20591 | 0,5253263 | H1,0 linker histone |
| ENSG00000132801 | ZSWIM3 | 0,287290465 | 1,220346184 | 0,35976 | 0,63246012 | zinc finger SWIM-type containing 3 |
| ENSG00000255553 | LINC02733 | 0,287265495 | 1,220325062 | 0,55068 | 0,76476271 | long intergenic non-protein coding RNA 2733 |
| ENSG00000024048 | UBR2 | 0,287242868 | 1,220305924 | 0,35836 | 0,63185282 | ubiquitin protein ligase E3 component n-recognin 2 |
| ENSG00000173473 | SMARCC1 | 0,287225005 | 1,220290814 | 0,20348 | 0,52332405 | SWI/SNF related, matrix associated, actin dependent regulator of chromatin subfamily c member 1 |

| | | | | | | |
|---|---|---|---|---|---|---|
| ENSG00000122042 | UBL3 | 0,287204214 | 1,220273228 | 0,36106 | 0,63315528 | ubiquitin like 3 |
| ENSG00000119231 | SENP5 | 0,287150589 | 1,220227871 | 0,2441 | 0,55253387 | SUMO specific peptidase 5 |
| ENSG00000260912 | AL158206,1 | 0,28701232 | 1,22011093 | 0,38398 | 0,65049725 | novel transcript, overlapping ACER2 |
| ENSG00000100426 | ZBED4 | 0,287002801 | 1,220102879 | 0,24529 | 0,55295131 | zinc finger BED-type containing 4 |
| ENSG00000278903 | CU633906,2 | 0,28699024 | 1,220092256 | 0,40782 | 0,66684711 | novel transcript |
| ENSG00000104472 | CHRAC1 | 0,286987852 | 1,220090236 | 0,14796 | 0,4751727 | chromatin accessibility complex subunit 1 |
| ENSG00000276500 | BMS1P14 | 0,286920605 | 1,220033367 | 0,60282 | 0,7971742 | BMS1 pseudogene 14 |
| ENSG00000162409 | PRKAA2 | 0,286916524 | 1,220029916 | 0,4416 | 0,68981532 | protein kinase AMP-activated catalytic subunit alpha 2 |
| ENSG00000070087 | PFN2 | 0,286873986 | 1,219993944 | 0,43321 | 0,68455289 | profilin 2 |
| ENSG00000273343 | AC007663,4 | 0,286802075 | 1,219933135 | 0,58833 | 0,78811087 | novel transcript, sense intronic to DGCR6L |
| ENSG00000270574 | AC010680,2 | 0,286779528 | 1,219914069 | 0,50792 | 0,73602279 | novel transcript, antisense to titin |
| ENSG00000243943 | ZNF512 | 0,286714037 | 1,219858692 | 0,35853 | 0,63185282 | zinc finger protein 512 |
| ENSG00000274194 | MBOAT7 | 0,286710475 | 1,219855681 | 0,90417 | 0,95888764 | membrane bound O-acyltransferase domain containing 7 |
| ENSG00000277923 | MBOAT7 | 0,286710475 | 1,219855681 | 0,90417 | 0,95888764 | membrane bound O-acyltransferase domain containing 7 |
| ENSG00000278519 | MBOAT7 | 0,286710475 | 1,219855681 | 0,90417 | 0,95888764 | membrane bound O-acyltransferase domain containing 7 |
| ENSG00000277733 | MBOAT7 | 0,286710475 | 1,219855681 | 0,90417 | 0,95888764 | membrane bound O-acyltransferase domain containing 7 |
| ENSG00000277025 | MBOAT7 | 0,286710475 | 1,219855681 | 0,90417 | 0,95888764 | membrane bound O-acyltransferase domain containing 7 |
| ENSG00000278322 | MBOAT7 | 0,286710475 | 1,219855681 | 0,90417 | 0,95888764 | membrane bound O-acyltransferase domain containing 7 |
| ENSG00000122026 | RPL21 | 0,28670699 | 1,219852734 | 0,28808 | 0,58415113 | ribosomal protein L21 |
| ENSG00000184611 | KCNH7 | 0,2866904 | 1,219838707 | 0,53212 | 0,7528508 | potassium voltage-gated channel subfamily H member 7 |
| ENSG00000234741 | GAS5 | 0,286622978 | 1,219781701 | 0,10809 | 0,41875186 | growth arrest specific 5 |
| ENSG00000140022 | STON2 | 0,286622196 | 1,21978104 | 0,39849 | 0,66104262 | stonin 2 |
| ENSG00000269535 | AC010320,4 | 0,286583523 | 1,219748343 | 0,587 | 0,78704355 | novel transcript, antisense to ZNF480 and ZNF610 |
| ENSG00000145191 | EIF2B5 | 0,286572373 | 1,219738916 | 0,3597 | 0,63242662 | eukaryotic translation initiation factor 2B subunit epsilon |
| ENSG00000143162 | CREG1 | 0,286557085 | 1,219725991 | 0,26952 | 0,57130325 | cellular repressor of E1A stimulated genes 1 |
| ENSG00000188523 | CFAP77 | 0,286486101 | 1,219665978 | 0,5877 | 0,78764925 | cilia and flagella associated protein 77 |
| ENSG00000131721 | RHOXF2 | 0,286484302 | 1,219664458 | 0,51651 | 0,74198903 | Rhox homeobox family member 2 |
| ENSG00000266839 | AC008088,1 | 0,286427731 | 1,219616633 | 0,75354 | 0,88453179 | novel transcript |
| ENSG00000287022 | AC079841,2 | 0,286416981 | 1,219607545 | 0,5382 | 0,75658794 | novel transcript, antisense to STXBP5L |
| ENSG00000138669 | PRKG2 | 0,286380301 | 1,219576538 | 0,58019 | 0,78348519 | protein kinase cGMP-dependent 2 |
| ENSG00000253263 | AP003354,1 | 0,286362033 | 1,219561095 | 0,65983 | 0,83018671 | novel transcript |
| ENSG00000264112 | AC015813,1 | 0,286338101 | 1,219540865 | 0,23468 | 0,54623956 | novel transcript |
| ENSG00000278053 | DDX52 | 0,286319622 | 1,219525244 | 0,1041 | 0,41245634 | DExD-box helicase 52 |

| | | | | | | |
|---|---|---|---|---|---|---|
| ENSG00000132603 | NIP7 | 0,286242246 | 1,219459839 | 0,1402 | 0,46710628 | nucleolar pre-rRNA processing protein NIP7 |
| ENSG00000144043 | TEX261 | 0,286160349 | 1,219390616 | 0,34785 | 0,62428208 | testis expressed 261 |
| ENSG00000233436 | BTBD18 | 0,286146832 | 1,219379192 | 0,29002 | 0,58531219 | BTB domain containing 18 |
| ENSG00000143977 | SNRPG | 0,286133914 | 1,219368273 | 0,38239 | 0,64928247 | small nuclear ribonucleoprotein polypeptide G |
| ENSG00000133835 | HSD17B4 | 0,286102446 | 1,219341676 | 0,1808 | 0,50437478 | hydroxysteroid 17-beta dehydrogenase 4 |
| ENSG00000108061 | SHOC2 | 0,286063321 | 1,219308609 | 0,40739 | 0,66661764 | SHOC2 leucine rich repeat scaffold protein |
| ENSG00000124224 | PPP4R1L | 0,286033751 | 1,219283618 | 0,36818 | 0,6380264 | protein phosphatase 4 regulatory subunit 1 like (pseudogene) |
| ENSG00000130939 | UBE4B | 0,28583724 | 1,21911755 | 0,15009 | 0,47818779 | ubiquitination factor E4B |
| ENSG00000109911 | ELP4 | 0,285782999 | 1,219071716 | 0,39945 | 0,66176242 | elongator acetyltransferase complex subunit 4 |
| ENSG00000196338 | NLGN3 | 0,28577515 | 1,219065083 | 0,29871 | 0,59094252 | neuroligin 3 |
| ENSG00000253688 | AC026904,2 | 0,285758499 | 1,219051013 | 0,75882 | 0,88742567 | novel transcript |
| ENSG00000285780 | AC109129,1 | 0,285752158 | 1,219045655 | 0,76589 | 0,89114666 | Novel transcript, antisense to EPHA3 |
| ENSG00000113555 | PCDH12 | 0,285666066 | 1,218972911 | 0,29868 | 0,59094252 | protocadherin 12 |
| ENSG00000257027 | AC010186,3 | 0,285662585 | 1,21896997 | 0,46251 | 0,70358709 | novel transcript |
| ENSG00000274641 | H2BC17 | 0,285626555 | 1,218939528 | 0,60131 | 0,79647801 | H2B clustered histone 17 |
| ENSG00000166133 | RPUSD2 | 0,285553793 | 1,218878052 | 0,13905 | 0,46647959 | RNA pseudouridine synthase domain containing 2 |
| ENSG00000083223 | TUT7 | 0,285549132 | 1,218874115 | 0,41068 | 0,66856387 | terminal uridylyl transferase 7 |
| ENSG00000221978 | CCNL2 | 0,285359431 | 1,218713854 | 0,27596 | 0,57533851 | cyclin L2 |
| ENSG00000198231 | DDX42 | 0,285349275 | 1,218705275 | 0,36265 | 0,63381553 | DEAD-box helicase 42 |
| ENSG00000169902 | TPST1 | 0,285336144 | 1,218694183 | 0,22229 | 0,53859118 | tyrosylprotein sulfotransferase 1 |
| ENSG00000106538 | RARRES2 | 0,285267035 | 1,218635806 | 0,54405 | 0,76034748 | retinoic acid receptor responder 2 |
| ENSG00000235286 | AP000925,1 | 0,285223429 | 1,218598972 | 0,63376 | 0,81549489 | keratin 8 (KRT8) pseudogene |
| ENSG00000116016 | EPAS1 | 0,285217626 | 1,218594071 | 0,57134 | 0,77809338 | endothelial PAS domain protein 1 |
| ENSG00000254349 | MIR2052HG | 0,285112168 | 1,218504997 | 0,57145 | 0,77809338 | MIR2052 host gene |
| ENSG00000163348 | PYGO2 | 0,285105859 | 1,218499669 | 0,17173 | 0,49710979 | pygopus family PHD finger 2 |
| ENSG00000141452 | RMC1 | 0,285061786 | 1,218462446 | 0,51766 | 0,74272749 | regulator of MON1-CCZ1 |
| ENSG00000285035 | AC091057,6 | 0,285058122 | 1,21845935 | 0,69609 | 0,85287022 | Serine/threonine-protein kinase ULK4 (ULK4), pseudogene |
| ENSG00000272023 | AC010240,3 | 0,28501736 | 1,218424925 | 0,73699 | 0,87539732 | novel transcript |
| ENSG00000286388 | AC026748,7 | 0,285012653 | 1,21842095 | 0,16974 | 0,49597212 | novel transcript |
| ENSG00000261147 | AC091167,2 | 0,284949625 | 1,218367721 | 0,61451 | 0,80382182 | novel protein |
| ENSG00000273373 | AL355488,1 | 0,284947186 | 1,218365661 | 0,4576 | 0,70082333 | novel transcript, antisense to SLC16A4 |
| ENSG00000106609 | TMEM248 | 0,284893818 | 1,218320592 | 0,16276 | 0,48945635 | transmembrane protein 248 |

| | | | | | | |
|---|---|---|---|---|---|---|
| ENSG00000163808 | KIF15 | 0,284873792 | 1,218303681 | 0,45019 | 0,69562819 | kinesin family member 15 |
| ENSG00000280610 | KIF15 | 0,284873792 | 1,218303681 | 0,45019 | 0,69562819 | kinesin family member 15 |
| ENSG00000186010 | NDUFA13 | 0,284814451 | 1,21825357 | 0,37025 | 0,63967954 | NADH:ubiquinone oxidoreductase subunit A13 |
| ENSG00000234383 | CTBP2P8 | 0,284693678 | 1,218151591 | 0,6327 | 0,81497431 | CTBP2 pseudogene 8 |
| ENSG00000257335 | MGAM | 0,284692089 | 1,218150249 | 0,67466 | 0,83975921 | maltase-glucoamylase |
| ENSG00000116455 | WDR77 | 0,284684908 | 1,218144185 | 0,12219 | 0,44167604 | WD repeat domain 77 |
| ENSG00000254088 | SLC2A3P4 | 0,284658342 | 1,218121755 | 0,70257 | 0,8566072 | solute carrier family 2 member 3 pseudogene 4 |
| ENSG00000084234 | APLP2 | 0,284658297 | 1,218121717 | 0,03832 | 0,264906 | amyloid beta precursor like protein 2 |
| ENSG00000179262 | RAD23A | 0,284630397 | 1,21809816 | 0,17334 | 0,4980379 | RAD23 homolog A, nucleotide excision repair protein |
| ENSG00000107372 | ZFAND5 | 0,284607792 | 1,218079074 | 0,38424 | 0,65052153 | zinc finger AN1-type containing 5 |
| ENSG00000189269 | DRICH1 | 0,284512522 | 1,217998639 | 0,69079 | 0,84906639 | aspartate rich 1 |
| ENSG00000261520 | DLGAP1-AS5 | 0,284502149 | 1,217989883 | 0,55124 | 0,76505157 | DLGAP1 antisense RNA 5 |
| ENSG00000283267 | FAM237B | 0,28447501 | 1,217966971 | 0,69368 | 0,8512419 | family with sequence similarity 237 member B |
| ENSG00000143416 | SELENBP1 | 0,284334234 | 1,217848129 | 0,41684 | 0,67362915 | selenium binding protein 1 |
| ENSG00000227171 | RNF39 | 0,284210872 | 1,217743998 | 0,68172 | 0,84383965 | ring finger protein 39 |
| ENSG00000186301 | MST1P2 | 0,284207079 | 1,217740796 | 0,49631 | 0,72831792 | macrophage stimulating 1 pseudogene 2 |
| ENSG00000240771 | ARHGEF25 | 0,28415457 | 1,217696475 | 0,24029 | 0,54948131 | Rho guanine nucleotide exchange factor 25 |
| ENSG00000173852 | DPY19L1 | 0,2841097 | 1,217658604 | 0,31805 | 0,6034788 | dpy-19 like C-mannosyltransferase 1 |
| ENSG00000226759 | DAB1-AS1 | 0,284104943 | 1,217654589 | 0,59973 | 0,79552021 | DAB1 antisense RNA 1 |
| ENSG00000261796 | ISY1-RAB43 | 0,283967556 | 1,217538638 | 0,62199 | 0,80819595 | ISY1-RAB43 readthrough |
| ENSG00000132196 | HSD17B7 | 0,283864609 | 1,21745176 | 0,26016 | 0,56382871 | hydroxysteroid 17-beta dehydrogenase 7 |
| ENSG00000227449 | FGF7P6 | 0,283834672 | 1,217426498 | 0,71312 | 0,86197885 | fibroblast growth factor 7 pseudogene 6 |
| ENSG00000204070 | SYS1 | 0,283804929 | 1,2174014 | 0,34906 | 0,62519057 | SYS1 golgi trafficking protein |
| ENSG00000236676 | AC117944,1 | 0,283696999 | 1,217310327 | 0,81327 | 0,91426124 | novel transcript |
| ENSG00000065243 | PKN2 | 0,283644904 | 1,217266371 | 0,2579 | 0,56199743 | protein kinase N2 |
| ENSG00000166881 | NEMP1 | 0,283622987 | 1,217247879 | 0,32667 | 0,60877026 | nuclear envelope integral membrane protein 1 |
| ENSG00000271550 | BNIP3P11 | 0,283620841 | 1,217246068 | 0,78574 | 0,90048284 | BCL2 interacting protein 3 pseudogene 11 |
| ENSG00000188021 | UBQLN2 | 0,283596831 | 1,217225811 | 0,31497 | 0,60145932 | ubiquilin 2 |
| ENSG00000109606 | DHX15 | 0,283582117 | 1,217213397 | 0,14721 | 0,47471707 | DEAH-box helicase 15 |
| ENSG00000143512 | HHIPL2 | 0,283570667 | 1,217203736 | 0,64442 | 0,82154386 | HHIP like 2 |
| ENSG00000059758 | CDK17 | 0,283554545 | 1,217190134 | 0,47172 | 0,71108681 | cyclin dependent kinase 17 |
| ENSG00000136854 | STXBP1 | 0,283514753 | 1,217156562 | 0,17221 | 0,49710979 | syntaxin binding protein 1 |
| ENSG00000237296 | SMG1P1 | 0,283475365 | 1,217123332 | 0,42391 | 0,67773865 | SMG1 pseudogene 1 |

| | | | | | | |
|---|---|---|---|---|---|---|
| ENSG00000136933 | RABEPK | 0,283463694 | 1,217113486 | 0,22608 | 0,54121239 | Rab9 effector protein with kelch motifs |
| ENSG00000124496 | TRERF1 | 0,283337667 | 1,21700717 | 0,33063 | 0,6112353 | transcriptional regulating factor 1 |
| ENSG00000135953 | MFSD9 | 0,283283212 | 1,216961234 | 0,20474 | 0,52443315 | major facilitator superfamily domain containing 9 |
| ENSG00000136718 | IMP4 | 0,28326585 | 1,216946588 | 0,14869 | 0,47609532 | IMP U3 small nucleolar ribonucleoprotein 4 |
| ENSG00000119725 | ZNF410 | 0,283211453 | 1,216900704 | 0,32955 | 0,61056241 | zinc finger protein 410 |
| ENSG00000245864 | MEF2C-AS2 | 0,283172261 | 1,216867647 | 0,60075 | 0,79598946 | MEF2C antisense RNA 2 |
| ENSG00000124209 | RAB22A | 0,283138894 | 1,216839503 | 0,3603 | 0,63261878 | RAB22A, member RAS oncogene family |
| ENSG00000116679 | IVNS1ABP | 0,283081171 | 1,216790818 | 0,30297 | 0,593345 | influenza virus NS1A binding protein |
| ENSG00000100568 | VTI1B | 0,282931341 | 1,216664455 | 0,34772 | 0,62423931 | vesicle transport through interaction with t-SNAREs 1B |
| ENSG00000172954 | LCLAT1 | 0,28289456 | 1,216633437 | 0,20389 | 0,52362952 | lysocardiolipin acyltransferase 1 |
| ENSG00000158483 | FAM86C1P | 0,282886754 | 1,216626854 | 0,15578 | 0,48358141 | family with sequence similarity 86 member C1, pseudogene |
| ENSG00000242125 | SNHG3 | 0,28276871 | 1,216527312 | 0,29513 | 0,58883222 | small nucleolar RNA host gene 3 |
| ENSG00000198763 | MT-ND2 | 0,282762077 | 1,216521719 | 0,34823 | 0,62460194 | mitochondrially encoded NADH:ubiquinone oxidoreductase core subunit 2 |
| ENSG00000287979 | AC253572,1 | 0,282535642 | 1,216330797 | 0,27967 | 0,57745167 | novel transcript |
| ENSG00000165195 | PIGA | 0,282486672 | 1,216289512 | 0,10785 | 0,41847368 | phosphatidylinositol glycan anchor biosynthesis class A |
| ENSG00000067596 | DHX8 | 0,282461041 | 1,216267903 | 0,24626 | 0,55395309 | DEAH-box helicase 8 |
| ENSG00000271425 | NBPF10 | 0,282321394 | 1,216150179 | 0,30599 | 0,59512079 | NBPF member 10 |
| ENSG00000180376 | CCDC66 | 0,282292284 | 1,21612564 | 0,29949 | 0,59139511 | coiled-coil domain containing 66 |
| ENSG00000270039 | AC025165,5 | 0,282281927 | 1,21611691 | 0,63386 | 0,81549489 | novel transcript |
| ENSG00000107643 | MAPK8 | 0,282270645 | 1,2161074 | 0,44085 | 0,68924957 | mitogen-activated protein kinase 8 |
| ENSG00000115993 | TRAK2 | 0,282268664 | 1,216105729 | 0,47954 | 0,71700404 | trafficking kinesin protein 2 |
| ENSG00000180626 | ZNF594 | 0,282247105 | 1,216087557 | 0,40628 | 0,66584658 | zinc finger protein 594 |
| ENSG00000175216 | CKAP5 | 0,282202328 | 1,216049813 | 0,39078 | 0,65464997 | cytoskeleton associated protein 5 |
| ENSG00000164808 | SPIDR | 0,282146914 | 1,216003106 | 0,20649 | 0,52609885 | scaffold protein involved in DNA repair |
| ENSG00000076108 | BAZ2A | 0,282140813 | 1,215997964 | 0,21518 | 0,53387837 | bromodomain adjacent to zinc finger domain 2A |
| ENSG00000183891 | TTC32 | 0,282087479 | 1,215953011 | 0,39535 | 0,65820274 | tetratricopeptide repeat domain 32 |
| ENSG00000162344 | FGF19 | 0,281992832 | 1,215873242 | 0,52619 | 0,7497214 | fibroblast growth factor 19 |
| ENSG00000168610 | STAT3 | 0,281883395 | 1,215781014 | 0,13577 | 0,462686 | signal transducer and activator of transcription 3 |
| ENSG00000145332 | KLHL8 | 0,281832717 | 1,215738308 | 0,24037 | 0,54950872 | kelch like family member 8 |
| ENSG00000233426 | EIF3FP3 | 0,281734961 | 1,215655933 | 0,65616 | 0,82829288 | eukaryotic translation initiation factor 3 subunit F pseudogene 3 |

| | | | | | | |
|---|---|---|---|---|---|---|
| ENSG00000259411 | HNRNPA1P45 | 0,281554556 | 1,215503928 | 0,74875 | 0,88178784 | heterogeneous nuclear ribonucleoprotein A1 pseudogene 45 |
| ENSG00000270231 | NBPF8 | 0,281436131 | 1,215404157 | 0,17872 | 0,50323303 | NBPF member 8 |
| ENSG00000048707 | VPS13D | 0,281322673 | 1,215308578 | 0,35839 | 0,63185282 | vacuolar protein sorting 13 homolog D |
| ENSG00000228368 | AC106873,1 | 0,281295467 | 1,21528566 | 0,75695 | 0,88650931 | novel transcript, antisense to ST7 |
| ENSG00000080189 | SLC35C2 | 0,281275512 | 1,21526885 | 0,11075 | 0,42337179 | solute carrier family 35 member C2 |
| ENSG00000198818 | SFT2D1 | 0,281246631 | 1,215244522 | 0,23985 | 0,54931011 | SFT2 domain containing 1 |
| ENSG00000087263 | OGFOD1 | 0,281233203 | 1,215233211 | 0,11796 | 0,43562473 | 2-oxoglutarate and iron dependent oxygenase domain containing 1 |
| ENSG00000145715 | RASA1 | 0,281225624 | 1,215226827 | 0,36134 | 0,63331577 | RAS p21 protein activator 1 |
| ENSG00000196335 | STK31 | 0,281200352 | 1,21520554 | 0,43162 | 0,68379763 | serine/threonine kinase 31 |
| ENSG00000179409 | GEMIN4 | 0,280908231 | 1,214959507 | 0,09186 | 0,39178722 | gem nuclear organelle associated protein 4 |
| ENSG00000114127 | XRN1 | 0,280907079 | 1,214958536 | 0,3867 | 0,65199475 | 5'-3' exoribonuclease 1 |
| ENSG00000141646 | SMAD4 | 0,280837555 | 1,214899988 | 0,25481 | 0,55944852 | SMAD family member 4 |
| ENSG00000154814 | OXNAD1 | 0,280774112 | 1,214846564 | 0,16695 | 0,49293221 | oxidoreductase NAD binding domain containing 1 |
| ENSG00000284727 | AC116562,3 | 0,280678468 | 1,214766028 | 0,31905 | 0,60403146 | RNA binding protein, fox-1 homolog (C, elegans) 1 (RBFOX1) pseudogene |
| ENSG00000161800 | RACGAP1 | 0,280658985 | 1,214749623 | 0,43559 | 0,68618266 | Rac GTPase activating protein 1 |
| ENSG00000206489 | PPP1R10 | 0,280637355 | 1,214731411 | 0,27092 | 0,57244549 | protein phosphatase 1 regulatory subunit 10 |
| ENSG00000231887 | PRH1 | 0,280579794 | 1,214682946 | 0,60298 | 0,7971742 | proline rich protein HaeIII subfamily 1 |
| ENSG00000164023 | SGMS2 | 0,280570007 | 1,214674706 | 0,52756 | 0,75037219 | sphingomyelin synthase 2 |
| ENSG00000166086 | JAM3 | 0,280541756 | 1,214650921 | 0,19401 | 0,51552252 | junctional adhesion molecule 3 |
| ENSG00000188647 | PTAR1 | 0,28053659 | 1,214646571 | 0,51696 | 0,74226247 | protein prenyltransferase alpha subunit repeat containing 1 |
| ENSG00000023287 | RB1CC1 | 0,28053016 | 1,214641158 | 0,38975 | 0,65396251 | RB1 inducible coiled-coil 1 |
| ENSG00000245105 | A2M-AS1 | 0,280459635 | 1,214581782 | 0,46107 | 0,70278639 | A2M antisense RNA 1 |
| ENSG00000164091 | WDR82 | 0,280432208 | 1,214558692 | 0,27978 | 0,5775083 | WD repeat domain 82 |
| ENSG00000144810 | COL8A1 | 0,28042925 | 1,214556202 | 0,36993 | 0,63943131 | collagen type VIII alpha 1 chain |
| ENSG00000124635 | H2BC11 | 0,280383233 | 1,214517462 | 0,66408 | 0,83250729 | H2B clustered histone 11 |
| ENSG00000272576 | AC027271,1 | 0,280309169 | 1,214455114 | 0,54959 | 0,7641582 | novel transcript |
| ENSG00000198815 | FOXJ3 | 0,280235888 | 1,214393428 | 0,12662 | 0,4492748 | forkhead box J3 |
| ENSG00000116560 | SFPQ | 0,280076753 | 1,214259483 | 0,25942 | 0,56317998 | splicing factor proline and glutamine rich |
| ENSG00000273570 | AC007351,1 | 0,280017712 | 1,214209791 | 0,81691 | 0,91580038 | |

| | | | | | | |
|---|---|---|---|---|---|---|
| ENSG00000126602 | TRAP1 | 0,279961188 | 1,21416222 | 0,37683 | 0,64537182 | TNF receptor associated protein 1 |
| ENSG00000241837 | ATP5PO | 0,279875505 | 1,214090112 | 0,35866 | 0,63185282 | ATP synthase peripheral stalk subunit OSCP |
| ENSG00000056277 | ZNF280C | 0,279854232 | 1,21407221 | 0,25334 | 0,55839162 | zinc finger protein 280C |
| ENSG00000130741 | EIF2S3 | 0,279763533 | 1,213995887 | 0,25766 | 0,5618698 | eukaryotic translation initiation factor 2 subunit gamma |
| ENSG00000154122 | ANKH | 0,279706538 | 1,213947927 | 0,2375 | 0,54850427 | ANKH inorganic pyrophosphate transport regulator |
| ENSG00000287063 | AL356489,3 | 0,279654568 | 1,213904198 | 0,53882 | 0,75721606 | novel transcript |
| ENSG00000131089 | ARHGEF9 | 0,279588112 | 1,213848282 | 0,27666 | 0,57549599 | Cdc42 guanine nucleotide exchange factor 9 |
| ENSG00000167635 | ZNF146 | 0,279502644 | 1,213776374 | 0,44094 | 0,68930383 | zinc finger protein 146 |
| ENSG00000103423 | DNAJA3 | 0,279439054 | 1,213722875 | 0,2179 | 0,53550245 | DnaJ heat shock protein family (Hsp40) member A3 |
| ENSG00000167220 | HDHD2 | 0,279389344 | 1,213681056 | 0,38816 | 0,6529525 | haloacid dehalogenase like hydrolase domain containing 2 |
| ENSG00000131508 | UBE2D2 | 0,279385292 | 1,213677646 | 0,32434 | 0,60692416 | ubiquitin conjugating enzyme E2 D2 |
| ENSG00000183576 | SETD3 | 0,279384453 | 1,213676941 | 0,14372 | 0,47123322 | SET domain containing 3, actin histidine methyltransferase |
| ENSG00000230006 | ANKRD36BP2 | 0,279330092 | 1,213631211 | 0,37969 | 0,64745085 | ankyrin repeat domain 36B pseudogene 2 |
| ENSG00000142192 | APP | 0,279313384 | 1,213617155 | 0,12276 | 0,44256265 | amyloid beta precursor protein |
| ENSG00000119401 | TRIM32 | 0,279231826 | 1,213548549 | 0,19195 | 0,5145257 | tripartite motif containing 32 |
| ENSG00000167193 | CRK | 0,279201566 | 1,213523096 | 0,21046 | 0,5290538 | CRK proto-oncogene, adaptor protein |
| ENSG00000132356 | PRKAA1 | 0,279107702 | 1,213444144 | 0,05224 | 0,30693571 | protein kinase AMP-activated catalytic subunit alpha 1 |
| ENSG00000260804 | LINC01963 | 0,279100996 | 1,213438504 | 0,52449 | 0,74828894 | long intergenic non-protein coding RNA 1963 |
| ENSG00000130150 | MOSPD2 | 0,279088666 | 1,213428133 | 0,35983 | 0,63249219 | motile sperm domain containing 2 |
| ENSG00000121454 | LHX4 | 0,27890513 | 1,213273774 | 0,26574 | 0,56866772 | LIM homeobox 4 |
| ENSG00000249131 | PSD2-AS1 | 0,27879965 | 1,213185071 | 0,76715 | 0,89196675 | PSD2 antisense RNA 1 |
| ENSG00000204634 | TBC1D8 | 0,27878618 | 1,213173744 | 0,20386 | 0,52361295 | TBC1 domain family member 8 |
| ENSG00000138867 | GUCD1 | 0,278721665 | 1,213119494 | 0,24764 | 0,55506866 | guanylyl cyclase domain containing 1 |
| ENSG00000170502 | NUDT9 | 0,278710363 | 1,213109991 | 0,23061 | 0,54367096 | nudix hydrolase 9 |
| ENSG00000072134 | EPN2 | 0,278704223 | 1,213104828 | 0,32498 | 0,60752552 | epsin 2 |
| ENSG00000092201 | SUPT16H | 0,278699293 | 1,213100682 | 0,16674 | 0,49257009 | SPT16 homolog, facilitates chromatin remodeling subunit |
| ENSG00000204334 | ERICH2 | 0,278685338 | 1,213088948 | 0,80897 | 0,91201277 | glutamate rich 2 |
| ENSG00000196730 | DAPK1 | 0,278680644 | 1,213085001 | 0,32086 | 0,60558211 | death associated protein kinase 1 |
| ENSG00000237765 | FAM200B | 0,278526324 | 1,212955249 | 0,518 | 0,7429275 | family with sequence similarity 200 member B |
| ENSG00000178226 | PRSS36 | 0,278356686 | 1,212812632 | 0,59832 | 0,79497317 | serine protease 36 |
| ENSG00000147650 | LRP12 | 0,278345185 | 1,212802964 | 0,12242 | 0,4421901 | LDL receptor related protein 12 |
| ENSG00000101158 | NELFCD | 0,278179098 | 1,212663351 | 0,33472 | 0,61442925 | negative elongation factor complex member C/D |
| ENSG00000283709 | FAM238C | 0,278157949 | 1,212645575 | 0,54919 | 0,76404333 | family with sequence similarity 238 member C |

| | | | | | | |
|---|---|---|---|---|---|---|
| ENSG00000112782 | CLIC5 | 0,278152202 | 1,212640744 | 0,77611 | 0,89650232 | chloride intracellular channel 5 |
| ENSG00000204147 | ASAH2B | 0,278064462 | 1,212566997 | 0,40353 | 0,66399439 | N-acylsphingosine amidohydrolase 2B |
| ENSG00000175928 | LRRN1 | 0,277985517 | 1,212500647 | 0,19777 | 0,51938324 | leucine rich repeat neuronal 1 |
| ENSG00000143742 | SRP9 | 0,277960511 | 1,212479631 | 0,45232 | 0,69702004 | signal recognition particle 9 |
| ENSG00000259033 | AL356804,1 | 0,277934207 | 1,212457525 | 0,66566 | 0,83358761 | novel transcript |
| ENSG00000140157 | NIPA2 | 0,277894677 | 1,212424304 | 0,1849 | 0,50844776 | NIPA magnesium transporter 2 |
| ENSG00000184677 | ZBTB40 | 0,277796303 | 1,212341634 | 0,26298 | 0,56587311 | zinc finger and BTB domain containing 40 |
| ENSG00000142396 | ERVK3-1 | 0,277788664 | 1,212335215 | 0,36965 | 0,63934748 | endogenous retrovirus group K3 member 1 |
| ENSG00000131171 | SH3BGRL | 0,277777031 | 1,212325439 | 0,50962 | 0,73740498 | SH3 domain binding glutamate rich protein like |
| ENSG00000105821 | DNAJC2 | 0,277762317 | 1,212313075 | 0,13982 | 0,46689302 | DnaJ heat shock protein family (Hsp40) member C2 |
| ENSG00000126945 | HNRNPH2 | 0,277683901 | 1,212247183 | 0,24088 | 0,55001001 | heterogeneous nuclear ribonucleoprotein H2 |
| ENSG00000180008 | SOCS4 | 0,277653142 | 1,212221337 | 0,4856 | 0,72107911 | suppressor of cytokine signaling 4 |
| ENSG00000230995 | PPP1R10 | 0,277613168 | 1,21218775 | 0,27247 | 0,57315552 | protein phosphatase 1 regulatory subunit 10 |
| ENSG00000227804 | PPP1R10 | 0,277613168 | 1,21218775 | 0,27247 | 0,57315552 | protein phosphatase 1 regulatory subunit 10 |
| ENSG00000238104 | PPP1R10 | 0,277613168 | 1,21218775 | 0,27247 | 0,57315552 | protein phosphatase 1 regulatory subunit 10 |
| ENSG00000283847 | HBP1 | 0,277579798 | 1,212159712 | 0,33904 | 0,61802993 | HMG-box transcription factor 1 |
| ENSG00000257738 | AC009248,1 | 0,277555552 | 1,212139341 | 0,75496 | 0,88516173 | carbonyl reductase 3 (CBR3) pseudogene |
| ENSG00000281761 | AL078605,3 | 0,277501185 | 1,212093662 | 0,72741 | 0,87021076 | novel transcript |
| ENSG00000271234 | AL078605,1 | 0,277501185 | 1,212093662 | 0,72741 | 0,87021076 | novel transcript |
| ENSG00000197712 | FAM114A1 | 0,277488489 | 1,212082996 | 0,34451 | 0,62198289 | family with sequence similarity 114 member A1 |
| ENSG00000278165 | DUSP8 | 0,277364584 | 1,211978902 | 0,86649 | 0,94074101 | dual specificity phosphatase 8 |
| ENSG00000134313 | KIDINS220 | 0,277348824 | 1,211965661 | 0,45566 | 0,69957443 | kinase D interacting substrate 220 |
| ENSG00000112308 | C6orf62 | 0,277307455 | 1,211930909 | 0,35505 | 0,62946175 | chromosome 6 open reading frame 62 |
| ENSG00000214184 | GCC2-AS1 | 0,277269076 | 1,211898669 | 0,6311 | 0,81392891 | GCC2 antisense RNA 1 |
| ENSG00000255565 | AC073651,1 | 0,277258574 | 1,211889848 | 0,66721 | 0,83474892 | novel transcript |
| ENSG00000082701 | GSK3B | 0,277252802 | 1,211884999 | 0,29789 | 0,59037822 | glycogen synthase kinase 3 beta |
| ENSG00000227141 | AL160286,1 | 0,277243572 | 1,211877246 | 0,62168 | 0,80812645 | centaurin, gamma 2 (CENTG2) pseudogene |
| ENSG00000163875 | MEAF6 | 0,277171985 | 1,211817114 | 0,39268 | 0,65593262 | MYST/Esa1 associated factor 6 |
| ENSG00000163960 | UBXN7 | 0,277169892 | 1,211815355 | 0,19142 | 0,51435195 | UBX domain protein 7 |
| ENSG00000162631 | NTNG1 | 0,277110397 | 1,211765383 | 0,45851 | 0,70126479 | netrin G1 |
| ENSG00000078618 | NRDC | 0,277060675 | 1,211723621 | 0,1498 | 0,47782859 | nardilysin convertase |
| ENSG00000179299 | NSUN7 | 0,276983097 | 1,211658464 | 0,53826 | 0,75662205 | NOP2/Sun RNA methyltransferase family member 7 |
| ENSG00000274561 | AC005332,3 | 0,27696392 | 1,211642358 | 0,53006 | 0,75176608 | novel transcript |

| | | | | | | |
|---|---|---|---|---|---|---|
| ENSG00000260648 | AC020658,3 | 0,276853332 | 1,211549485 | 0,51255 | 0,73941327 | novel transcript, sense intronic to PAK6 |
| ENSG00000150787 | PTS | 0,276850154 | 1,211546816 | 0,40931 | 0,66788818 | 6-pyruvoyltetrahydropterin synthase |
| ENSG00000155229 | MMS19 | 0,276831883 | 1,211531473 | 0,27065 | 0,57239075 | MMS19 homolog, cytosolic iron-sulfur assembly component |
| ENSG00000168944 | CEP120 | 0,276776152 | 1,211484672 | 0,42393 | 0,67773865 | centrosomal protein 120 |
| ENSG00000147894 | C9orf72 | 0,276766164 | 1,211476285 | 0,53812 | 0,75658794 | C9orf72-SMCR8 complex subunit |
| ENSG00000005436 | GCFC2 | 0,276714218 | 1,211432665 | 0,38315 | 0,64987805 | GC-rich sequence DNA-binding factor 2 |
| ENSG00000229721 | AC104115,1 | 0,276661781 | 1,211388635 | 0,72717 | 0,87016629 | ribosomal protein S8 (RPS8) pseudogene |
| ENSG00000275191 | AC007497,1 | 0,276648632 | 1,211377594 | 0,51963 | 0,74404725 | novel transcript, sense intronic to RPGRIP1L |
| ENSG00000277778 | PGM5P2 | 0,276609509 | 1,211344744 | 0,43048 | 0,68295004 | phosphoglucomutase 5 pseudogene 2 |
| ENSG00000100038 | TOP3B | 0,276597173 | 1,211334386 | 0,3564 | 0,63043562 | DNA topoisomerase III beta |
| ENSG00000157617 | C2CD2 | 0,276539056 | 1,21128559 | 0,18039 | 0,50407792 | C2 calcium dependent domain containing 2 |
| ENSG00000276462 | BX255923,2 | 0,276503523 | 1,211255757 | 0,67533 | 0,84021377 | novel transcript |
| ENSG00000143157 | POGK | 0,276472919 | 1,211230063 | 0,39085 | 0,65464997 | pogo transposable element derived with KRAB domain |
| ENSG00000182923 | CEP63 | 0,276336862 | 1,21111584 | 0,31242 | 0,59934192 | centrosomal protein 63 |
| ENSG00000196204 | RNF216P1 | 0,276311972 | 1,211094945 | 0,18241 | 0,50594375 | ring finger protein 216 pseudogene 1 |
| ENSG00000100201 | DDX17 | 0,276297836 | 1,211083079 | 0,25207 | 0,5579463 | DEAD-box helicase 17 |
| ENSG00000229056 | HECW2-AS1 | 0,276253381 | 1,211045762 | 0,7422 | 0,87810032 | HECW2 antisense RNA 1 |
| ENSG00000270068 | AC125494,3 | 0,276237608 | 1,211032521 | 0,77218 | 0,89466114 | novel transcript, antisense to ZNF384 |
| ENSG00000125386 | FAM193A | 0,276200431 | 1,211001314 | 0,28383 | 0,58078109 | family with sequence similarity 193 member A |
| ENSG00000084693 | AGBL5 | 0,276155608 | 1,210963691 | 0,19711 | 0,51837936 | ATP/GTP binding protein like 5 |
| ENSG00000230312 | SLC25A53P1 | 0,276136482 | 1,210947637 | 0,54756 | 0,76293749 | SLC25A53 pseudogene 1 |
| ENSG00000198934 | MAGEE1 | 0,27612237 | 1,210935792 | 0,15655 | 0,48425071 | MAGE family member E1 |
| ENSG00000176171 | BNIP3 | 0,276108916 | 1,210924499 | 0,21812 | 0,53569444 | BCL2 interacting protein 3 |
| ENSG00000151893 | CACUL1 | 0,276030679 | 1,210858833 | 0,22605 | 0,54121239 | CDK2 associated cullin domain 1 |
| ENSG00000285728 | AC098484,4 | 0,275993489 | 1,210827619 | 0,29406 | 0,58837921 | novel transcript, antisense to PPIH |
| ENSG00000276718 | AC005840,4 | 0,275940454 | 1,210783109 | 0,53214 | 0,7528508 | novel transcript, antisense to VAMP1 |
| ENSG00000115946 | PNO1 | 0,275904619 | 1,210753035 | 0,14161 | 0,46809977 | partner of NOB1 homolog |
| ENSG00000198721 | ECI2 | 0,275869935 | 1,210723927 | 0,43742 | 0,68717331 | enoyl-CoA delta isomerase 2 |
| ENSG00000104047 | DTWD1 | 0,275866172 | 1,210720769 | 0,35708 | 0,63118328 | DTW domain containing 1 |
| ENSG00000235291 | PPP1R10 | 0,275763141 | 1,210634308 | 0,27431 | 0,57420712 | protein phosphatase 1 regulatory subunit 10 |
| ENSG00000231737 | PPP1R10 | 0,275763141 | 1,210634308 | 0,27431 | 0,57420712 | protein phosphatase 1 regulatory subunit 10 |
| ENSG00000104343 | UBE2W | 0,275632859 | 1,210524987 | 0,42149 | 0,67643128 | ubiquitin conjugating enzyme E2 W |

| | | | | | | |
|---|---|---|---|---|---|---|
| ENSG00000259287 | AC010809,1 | 0,275572157 | 1,210474055 | 0,49522 | 0,72754843 | novel transcript, antisense to RYR3 |
| ENSG00000100104 | SRRD | 0,275540442 | 1,210447445 | 0,15086 | 0,47912084 | SRR1 domain containing |
| ENSG00000103342 | GSPT1 | 0,275446511 | 1,210368638 | 0,29315 | 0,58745839 | G1 to S phase transition 1 |
| ENSG00000092148 | HECTD1 | 0,275369567 | 1,210304086 | 0,24644 | 0,55411366 | HECT domain E3 ubiquitin protein ligase 1 |
| ENSG00000100348 | TXN2 | 0,275356984 | 1,21029353 | 0,32219 | 0,60577115 | thioredoxin 2 |
| ENSG00000131398 | KCNC3 | 0,275354833 | 1,210291726 | 0,30126 | 0,59258306 | potassium voltage-gated channel subfamily C member 3 |
| ENSG00000198538 | ZNF28 | 0,275313863 | 1,210257356 | 0,27717 | 0,57572679 | zinc finger protein 28 |
| ENSG00000214534 | ZNF705E | 0,275292746 | 1,210239642 | 0,60304 | 0,7971742 | zinc finger protein 705E |
| ENSG00000170631 | ZNF16 | 0,275227574 | 1,210184971 | 0,29573 | 0,5893128 | zinc finger protein 16 |
| ENSG00000286643 | AP000248,1 | 0,275196072 | 1,210158546 | 0,6167 | 0,8052709 | novel transcript, antisense to TIAM1 |
| ENSG00000112679 | DUSP22 | 0,275161001 | 1,210129129 | 0,06727 | 0,34325886 | dual specificity phosphatase 22 |
| ENSG00000155754 | C2CD6 | 0,27510152 | 1,210079238 | 0,66001 | 0,83025282 | C2 calcium dependent domain containing 6 |
| ENSG00000188801 | ZNF322P1 | 0,274903368 | 1,209913046 | 0,76384 | 0,8899284 | zinc finger protein 322 pseudogene 1 |
| ENSG00000198642 | KLHL9 | 0,274900619 | 1,20991074 | 0,48717 | 0,72231397 | kelch like family member 9 |
| ENSG00000227345 | PARG | 0,274857961 | 1,209874966 | 0,35536 | 0,62970025 | poly(ADP-ribose) glycohydrolase |
| ENSG00000189419 | SPATA41 | 0,274818839 | 1,209842158 | 0,566 | 0,77413633 | spermatogenesis associated 41 |
| ENSG00000008838 | MED24 | 0,274790066 | 1,20981803 | 0,27263 | 0,57315967 | mediator complex subunit 24 |
| ENSG00000275723 | NAPEPLD | 0,274712908 | 1,209753328 | 0,29385 | 0,58832005 | N-acyl phosphatidylethanolamine phospholipase D |
| ENSG00000111602 | TIMELESS | 0,274675956 | 1,209722343 | 0,2208 | 0,53776578 | timeless circadian regulator |
| ENSG00000284697 | AC006511,5 | 0,274551581 | 1,209618057 | 0,72493 | 0,86868749 | novel transcript |
| ENSG00000138081 | FBXO11 | 0,274538823 | 1,20960736 | 0,30001 | 0,59160746 | F-box protein 11 |
| ENSG00000070961 | ATP2B1 | 0,274507829 | 1,209581374 | 0,32284 | 0,60584543 | ATPase plasma membrane Ca2+ transporting 1 |
| ENSG00000198700 | IPO9 | 0,274473671 | 1,209552736 | 0,20373 | 0,52348472 | importin 9 |
| ENSG00000147251 | DOCK11 | 0,274448618 | 1,209531731 | 0,52111 | 0,74558236 | dedicator of cytokinesis 11 |
| ENSG00000148730 | EIF4EBP2 | 0,27441508 | 1,209503614 | 0,25684 | 0,56132991 | eukaryotic translation initiation factor 4E binding protein 2 |
| ENSG00000105726 | ATP13A1 | 0,274344911 | 1,209444788 | 0,17757 | 0,50200419 | ATPase 13A1 |
| ENSG00000138587 | MNS1 | 0,274339037 | 1,209439864 | 0,42001 | 0,67552612 | meiosis specific nuclear structural 1 |
| ENSG00000245384 | CXXC4-AS1 | 0,274301835 | 1,209408677 | 0,5363 | 0,75512187 | CXXC4 antisense RNA 1 |
| ENSG00000198718 | TOGARAM1 | 0,274259963 | 1,209373577 | 0,35057 | 0,62606342 | TOG array regulator of axonemal microtubules 1 |
| ENSG00000062725 | APPBP2 | 0,274204636 | 1,209327198 | 0,42235 | 0,67663581 | amyloid beta precursor protein binding protein 2 |
| ENSG00000137819 | PAQR5 | 0,274162695 | 1,209292042 | 0,43152 | 0,68377875 | progestin and adipoQ receptor family member 5 |
| ENSG00000218586 | AC006971,1 | 0,274136404 | 1,209270005 | 0,77498 | 0,8960241 | Rho GTPase activating protein 5 (ARHGAP5) pseudogene |
| ENSG00000066933 | MYO9A | 0,27397436 | 1,209134186 | 0,45413 | 0,6981789 | myosin IXA |

| | | | | | | |
|---|---|---|---|---|---|---|
| ENSG00000101447 | FAM83D | 0,273950538 | 1,209114222 | 0,27189 | 0,57305058 | family with sequence similarity 83 member D |
| ENSG00000082996 | RNF13 | 0,273943222 | 1,209108089 | 0,51849 | 0,74331541 | ring finger protein 13 |
| ENSG00000077935 | SMC1B | 0,273886698 | 1,209060718 | 0,60584 | 0,79901957 | structural maintenance of chromosomes 1B |
| ENSG00000175390 | EIF3F | 0,273885257 | 1,209059511 | 0,32293 | 0,60584543 | eukaryotic translation initiation factor 3 subunit F |
| ENSG00000116039 | ATP6V1B1 | 0,273863523 | 1,209041297 | 0,72005 | 0,86623639 | ATPase H+ transporting V1 subunit B1 |
| ENSG00000053900 | ANAPC4 | 0,273855918 | 1,209034923 | 0,35271 | 0,62754548 | anaphase promoting complex subunit 4 |
| ENSG00000240729 | AC146507,2 | 0,273855418 | 1,209034504 | 0,59603 | 0,79365122 | ribosomal protein L21 (RPL21) pseudogene |
| ENSG00000135211 | TMEM60 | 0,273797632 | 1,208986079 | 0,37847 | 0,64679625 | transmembrane protein 60 |
| ENSG00000052802 | MSMO1 | 0,273771674 | 1,208964326 | 0,32242 | 0,60577115 | methylsterol monooxygenase 1 |
| ENSG00000131475 | VPS25 | 0,273696842 | 1,208901619 | 0,16314 | 0,48972962 | vacuolar protein sorting 25 homolog |
| ENSG00000271976 | AC012467,2 | 0,273648464 | 1,208861081 | 0,43944 | 0,68834432 | novel transcript, antisense to IL17RB |
| ENSG00000136271 | DDX56 | 0,273579767 | 1,20880352 | 0,21947 | 0,53687235 | DEAD-box helicase 56 |
| ENSG00000205765 | C5orf51 | 0,273221958 | 1,208503757 | 0,3409 | 0,6192089 | chromosome 5 open reading frame 51 |
| ENSG00000185437 | SH3BGR | 0,273209045 | 1,20849294 | 0,40291 | 0,66398107 | SH3 domain binding glutamate rich protein |
| ENSG00000135093 | USP30 | 0,273110006 | 1,208409982 | 0,51754 | 0,74270246 | ubiquitin specific peptidase 30 |
| ENSG00000112234 | FBXL4 | 0,273088584 | 1,208392038 | 0,35623 | 0,63035942 | F-box and leucine rich repeat protein 4 |
| ENSG00000142961 | MOB3C | 0,27299462 | 1,208313338 | 0,22414 | 0,54002595 | MOB kinase activator 3C |
| ENSG00000060140 | STYK1 | 0,272920782 | 1,208251497 | 0,65624 | 0,82829288 | serine/threonine/tyrosine kinase 1 |
| ENSG00000146122 | DAAM2 | 0,272915197 | 1,208246819 | 0,57516 | 0,78011735 | dishevelled associated activator of morphogenesis 2 |
| ENSG00000167323 | STIM1 | 0,272856239 | 1,208197444 | 0,14297 | 0,47060958 | stromal interaction molecule 1 |
| ENSG00000003393 | ALS2 | 0,272843577 | 1,20818684 | 0,29856 | 0,59089801 | alsin Rho guanine nucleotide exchange factor ALS2 |
| ENSG00000169621 | APLF | 0,272759278 | 1,208116245 | 0,51351 | 0,73984584 | aprataxin and PNKP like factor |
| ENSG00000123472 | ATPAF1 | 0,272739791 | 1,208099927 | 0,33002 | 0,61091362 | ATP synthase mitochondrial F1 complex assembly factor 1 |
| ENSG00000137815 | RTF1 | 0,272674036 | 1,208044866 | 0,2993 | 0,59123205 | RTF1 homolog, Paf1/RNA polymerase II complex component |
| ENSG00000166321 | NUDT13 | 0,272656429 | 1,208030123 | 0,56268 | 0,77194048 | nudix hydrolase 13 |
| ENSG00000282854 | ASAP3 | 0,272641417 | 1,208017552 | 0,26789 | 0,56999323 | ArfGAP with SH3 domain, ankyrin repeat and PH domain 3 |
| ENSG00000088280 | ASAP3 | 0,272641417 | 1,208017552 | 0,26789 | 0,56999323 | ArfGAP with SH3 domain, ankyrin repeat and PH domain 3 |
| ENSG00000276358 | PLEKHM1 | 0,272625044 | 1,208003843 | 0,60077 | 0,79598946 | pleckstrin homology and RUN domain containing M1 |
| ENSG00000157741 | UBN2 | 0,272490476 | 1,207891171 | 0,27127 | 0,57271346 | ubinuclein 2 |
| ENSG00000106610 | STAG3L4 | 0,27239602 | 1,207812091 | 0,42742 | 0,68046596 | stromal antigen 3-like 4 (pseudogene) |
| ENSG00000164506 | STXBP5 | 0,272370239 | 1,207790507 | 0,52832 | 0,7507973 | syntaxin binding protein 5 |

| | | | | | | |
|---|---|---|---|---|---|---|
| ENSG00000113845 | TIMMDC1 | 0,272354292 | 1,207777157 | 0,27537 | 0,57503857 | translocase of inner mitochondrial membrane domain containing 1 |
| ENSG00000017797 | RALBP1 | 0,272217147 | 1,207662349 | 0,14064 | 0,46767265 | ralA binding protein 1 |
| ENSG00000109103 | UNC119 | 0,272182879 | 1,207633664 | 0,33662 | 0,61666883 | unc-119 lipid binding chaperone |
| ENSG00000135446 | CDK4 | 0,272167219 | 1,207620556 | 0,14544 | 0,47300165 | cyclin dependent kinase 4 |
| ENSG00000286849 | AC009152,6 | 0,272142463 | 1,207599834 | 0,6781 | 0,84159488 | novel transcript, antisense to XYLT1 |
| ENSG00000131143 | COX4I1 | 0,272043924 | 1,207517355 | 0,22884 | 0,54261107 | cytochrome c oxidase subunit 4I1 |
| ENSG00000154582 | ELOC | 0,272023466 | 1,207500232 | 0,30501 | 0,59446033 | elongin C |
| ENSG00000274425 | AC114271,1 | 0,271988338 | 1,207470831 | 0,21448 | 0,53342315 | novel transcript, antisense to ICAM3 |
| ENSG00000120306 | CYSTM1 | 0,271936826 | 1,207427718 | 0,63909 | 0,81893766 | cysteine rich transmembrane module containing 1 |
| ENSG00000180884 | ZNF792 | 0,271912367 | 1,207407248 | 0,40432 | 0,6642832 | zinc finger protein 792 |
| ENSG00000263400 | TMEM220-AS | 0,271902974 | 1,207399387 | 0,6338 | 0,81549489 | TMEM220 antisense RNA 1 |
| ENSG00000163874 | ZC3H12A | 0,271885743 | 1,207384967 | 0,40131 | 0,6630277 | zinc finger CCCH-type containing 12A |
| ENSG00000088247 | KHSRP | 0,271882414 | 1,20738218 | 0,16303 | 0,48967932 | KH-type splicing regulatory protein |
| ENSG00000284319 | AC275454,2 | 0,271880584 | 1,207380649 | 0,61263 | 0,8035281 | novel transcript, antisense to SLC25A24 |
| ENSG00000260879 | AL359258,2 | 0,271880584 | 1,207380649 | 0,61263 | 0,8035281 | novel transcript, antisense to SLC25A24 |
| ENSG00000281324 | LRP6 | 0,271746593 | 1,207268518 | 0,31644 | 0,60238875 | LDL receptor related protein 6 |
| ENSG00000070018 | LRP6 | 0,271746593 | 1,207268518 | 0,31644 | 0,60238875 | LDL receptor related protein 6 |
| ENSG00000111837 | MAK | 0,27170199 | 1,207231194 | 0,34562 | 0,62278421 | male germ cell associated kinase |
| ENSG00000018869 | ZNF582 | 0,271655509 | 1,2071923 | 0,45748 | 0,70075609 | zinc finger protein 582 |
| ENSG00000168310 | IRF2 | 0,27163794 | 1,207177599 | 0,48838 | 0,72325321 | interferon regulatory factor 2 |
| ENSG00000258982 | AL133523,1 | 0,271566478 | 1,207117805 | 0,56226 | 0,77176502 | novel transcript, antisense to YY1 |
| ENSG00000103043 | VAC14 | 0,27156542 | 1,207116919 | 0,22891 | 0,54262488 | VAC14 component of PIKFYVE complex |
| ENSG00000175087 | PDIK1L | 0,271522422 | 1,207080943 | 0,3921 | 0,65542687 | PDLIM1 interacting kinase 1 like |
| ENSG00000089327 | FXYD5 | 0,271502415 | 1,207064203 | 0,70985 | 0,86017148 | FXYD domain containing ion transport regulator 5 |
| ENSG00000114767 | RRP9 | 0,271464662 | 1,207032618 | 0,32231 | 0,60577115 | ribosomal RNA processing 9, U3 small nucleolar RNA binding protein |
| ENSG00000109534 | GAR1 | 0,271262617 | 1,206863588 | 0,42652 | 0,67990398 | GAR1 ribonucleoprotein |
| ENSG00000110871 | COQ5 | 0,271242462 | 1,206846727 | 0,18208 | 0,5053975 | coenzyme Q5, methyltransferase |
| ENSG00000111665 | CDCA3 | 0,271225825 | 1,20683281 | 0,28795 | 0,5840701 | cell division cycle associated 3 |
| ENSG00000206549 | AC109583,1 | 0,271162338 | 1,206779704 | 0,60198 | 0,79680798 | Probable threonine protease PRSS50 |
| ENSG00000166435 | XRRA1 | 0,271161523 | 1,206779022 | 0,28427 | 0,58108664 | X-ray radiation resistance associated 1 |
| ENSG00000224080 | UBE2FP1 | 0,271130773 | 1,206753301 | 0,70063 | 0,85539553 | UBE2F pseudogene 1 |

| | | | | | | |
|---|---|---|---|---|---|---|
| ENSG00000169981 | ZNF35 | 0,271079066 | 1,206710051 | 0,24222 | 0,55112431 | zinc finger protein 35 |
| ENSG00000281306 | ZNF35 | 0,271079066 | 1,206710051 | 0,24222 | 0,55112431 | zinc finger protein 35 |
| ENSG00000081059 | TCF7 | 0,270936517 | 1,206590825 | 0,09999 | 0,40655436 | transcription factor 7 |
| ENSG00000165972 | CCDC38 | 0,270886093 | 1,206548654 | 0,56169 | 0,77136826 | coiled-coil domain containing 38 |
| ENSG00000089916 | GPATCH2L | 0,270860598 | 1,206527332 | 0,28046 | 0,57795109 | G-patch domain containing 2 like |
| ENSG00000239207 | GAPDHP39 | 0,27084157 | 1,206511419 | 0,71604 | 0,86386517 | glyceraldehyde 3 phosphate dehydrogenase pseudogene 39 |
| ENSG00000047230 | CTPS2 | 0,270798378 | 1,206475298 | 0,43377 | 0,68475933 | CTP synthase 2 |
| ENSG00000249087 | ZNF436-AS1 | 0,27079463 | 1,206472164 | 0,43301 | 0,68446573 | ZNF436 antisense RNA 1 |
| ENSG00000283106 | ZNF436-AS1 | 0,27079463 | 1,206472164 | 0,43301 | 0,68446573 | ZNF436 antisense RNA 1 |
| ENSG00000134452 | FBH1 | 0,270763947 | 1,206446506 | 0,36613 | 0,63617598 | F-box DNA helicase 1 |
| ENSG00000183172 | SMDT1 | 0,270757011 | 1,206440706 | 0,36479 | 0,6350493 | single-pass membrane protein with aspartate rich tail 1 |
| ENSG00000129559 | NEDD8 | 0,270749246 | 1,206434211 | 0,23129 | 0,54422787 | NEDD8 ubiquitin like modifier |
| ENSG00000260423 | LINC02367 | 0,270737582 | 1,206424458 | 0,41526 | 0,67212646 | long intergenic non-protein coding RNA 2367 |
| ENSG00000037042 | TUBG2 | 0,270602623 | 1,206311607 | 0,4181 | 0,67437072 | tubulin gamma 2 |
| ENSG00000218682 | AC064847,1 | 0,270601707 | 1,206310841 | 0,77848 | 0,89745058 | enolase 2 (gamma, neuronal) (ENO2) pseudogene |
| ENSG00000126858 | RHOT1 | 0,270549947 | 1,206267562 | 0,2944 | 0,58857672 | ras homolog family member T1 |
| ENSG00000266472 | MRPS21 | 0,270537649 | 1,20625728 | 0,28971 | 0,58522301 | mitochondrial ribosomal protein S21 |
| ENSG00000261754 | AC008555,1 | 0,270443959 | 1,206178947 | 0,60483 | 0,79819795 | novel transcript |
| ENSG00000236364 | AL358115,1 | 0,270413437 | 1,206153429 | 0,3467 | 0,62350287 | novel transcript, antisense to UCK2 |
| ENSG00000143093 | STRIP1 | 0,270412183 | 1,20615238 | 0,12862 | 0,45191041 | striatin interacting protein 1 |
| ENSG00000259116 | AL049869,2 | 0,270369869 | 1,206117004 | 0,71765 | 0,86483656 | novel transcript, antisense to HSPA2 & ZBTB1 |
| ENSG00000155876 | RRAGA | 0,270288036 | 1,206048593 | 0,33065 | 0,6112353 | Ras related GTP binding A |
| ENSG00000162777 | DENND2D | 0,27026305 | 1,206027706 | 0,64849 | 0,82396865 | DENN domain containing 2D |
| ENSG00000137547 | MRPL15 | 0,270260447 | 1,206025529 | 0,20674 | 0,52628269 | mitochondrial ribosomal protein L15 |
| ENSG00000137409 | MTCH1 | 0,270259727 | 1,206024927 | 0,2476 | 0,55506866 | mitochondrial carrier 1 |
| ENSG00000242689 | CNTF | 0,270219662 | 1,205991436 | 0,59709 | 0,79434997 | ciliary neurotrophic factor |
| ENSG00000249915 | PDCD6 | 0,270184354 | 1,205961921 | 0,2492 | 0,55635288 | programmed cell death 6 |
| ENSG00000113810 | SMC4 | 0,270173796 | 1,205953096 | 0,46642 | 0,70665143 | structural maintenance of chromosomes 4 |
| ENSG00000198860 | TSEN15 | 0,270122788 | 1,205910459 | 0,36379 | 0,63461346 | tRNA splicing endonuclease subunit 15 |
| ENSG00000086570 | FAT2 | 0,270109566 | 1,205899407 | 0,39259 | 0,65593262 | FAT atypical cadherin 2 |
| ENSG00000183439 | TRIM61 | 0,270041511 | 1,205842523 | 0,4821 | 0,71880798 | tripartite motif containing 61 |
| ENSG00000250324 | MRPL22P1 | 0,270023424 | 1,205827406 | 0,66556 | 0,83358761 | mitochondrial ribosomal protein L22 pseudogene 1 |

| | | | | | | |
|---|---|---|---|---|---|---|
| ENSG00000089876 | DHX32 | 0,270006447 | 1,205813216 | 0,46983 | 0,70922292 | DEAH-box helicase 32 (putative) |
| ENSG00000286788 | AC004687,3 | 0,269995307 | 1,205803905 | 0,74835 | 0,88150915 | novel transcript, antisense to LPO |
| ENSG00000287619 | AL021308,1 | 0,269946413 | 1,20576304 | 0,47503 | 0,71364583 | novel transcript |
| ENSG00000243701 | DUBR | 0,269755842 | 1,205603777 | 0,43244 | 0,68411566 | DPPA2 upstream binding RNA |
| ENSG00000128654 | MTX2 | 0,269728063 | 1,205580563 | 0,13609 | 0,46320633 | metaxin 2 |
| ENSG00000164609 | SLU7 | 0,269615442 | 1,205486456 | 0,32033 | 0,6052338 | SLU7 homolog, splicing factor |
| ENSG00000164983 | TMEM65 | 0,269607589 | 1,205479895 | 0,26941 | 0,57119597 | transmembrane protein 65 |
| ENSG00000134440 | NARS1 | 0,269522988 | 1,205409206 | 0,3717 | 0,64079608 | asparaginyl-tRNA synthetase 1 |
| ENSG00000140391 | TSPAN3 | 0,269519399 | 1,205406207 | 0,3699 | 0,63943131 | tetraspanin 3 |
| ENSG00000100330 | MTMR3 | 0,269493303 | 1,205384404 | 0,28992 | 0,58531219 | myotubularin related protein 3 |
| ENSG00000140525 | FANCI | 0,269403772 | 1,205309602 | 0,41922 | 0,67514348 | FA complementation group I |
| ENSG00000034677 | RNF19A | 0,269347153 | 1,2052623 | 0,38509 | 0,65088045 | ring finger protein 19A, RBR E3 ubiquitin protein ligase |
| ENSG00000165689 | ENTR1 | 0,269177056 | 1,205120206 | 0,19149 | 0,51435195 | endosome associated trafficking regulator 1 |
| ENSG00000152760 | TCTEX1D1 | 0,269139646 | 1,205088957 | 0,71365 | 0,86220659 | Tctex1 domain containing 1 |
| ENSG00000014919 | COX15 | 0,269117244 | 1,205070244 | 0,22693 | 0,54168591 | cytochrome c oxidase assembly homolog COX15 |
| ENSG00000227712 | AL359915,1 | 0,269106695 | 1,205061432 | 0,71881 | 0,86552951 | novel transcript |
| ENSG00000273494 | PANK4 | 0,269041754 | 1,205007189 | 0,19282 | 0,51493916 | pantothenate kinase 4 (inactive) |
| ENSG00000183671 | GPR1 | 0,26904141 | 1,205006902 | 0,78624 | 0,90080813 | G protein-coupled receptor 1 |
| ENSG00000136750 | GAD2 | 0,269015582 | 1,204985329 | 0,63513 | 0,81638093 | glutamate decarboxylase 2 |
| ENSG00000103415 | HMOX2 | 0,268975356 | 1,204951732 | 0,41079 | 0,66856387 | heme oxygenase 2 |
| ENSG00000182903 | ZNF721 | 0,268853507 | 1,204849967 | 0,42182 | 0,67643128 | zinc finger protein 721 |
| ENSG00000278198 | AC011043,2 | 0,268842515 | 1,204840787 | 0,7764 | 0,89655595 | |
| ENSG00000273496 | AC011841,1 | 0,268842515 | 1,204840787 | 0,7764 | 0,89655595 | |
| ENSG00000008256 | CYTH3 | 0,268825975 | 1,204826974 | 0,36207 | 0,63354671 | cytohesin 3 |
| ENSG00000239653 | PSMD6-AS2 | 0,268698932 | 1,204720882 | 0,29744 | 0,59034513 | PSMD6 antisense RNA 2 |
| ENSG00000230202 | AL450405,1 | 0,26868233 | 1,204707019 | 0,75077 | 0,88302317 | ribosomal protein L29 (RPL29) pseudogene |
| ENSG00000142864 | SERBP1 | 0,268647678 | 1,204678084 | 0,16902 | 0,49510117 | SERPINE1 mRNA binding protein 1 |
| ENSG00000118058 | KMT2A | 0,268590177 | 1,20463007 | 0,34484 | 0,62213186 | lysine methyltransferase 2A |
| ENSG00000284689 | OR7E84P | 0,268584954 | 1,204625709 | 0,74259 | 0,87827554 | olfactory receptor family 7 subfamily E member 84 pseudogene |
| ENSG00000216866 | RPS2P55 | 0,268583714 | 1,204624673 | 0,7876 | 0,90122495 | ribosomal protein S2 pseudogene 55 |
| ENSG00000229893 | TAX1BP1-AS1 | 0,268467447 | 1,204527597 | 0,73457 | 0,87452837 | TAX1BP1 antisense RNA 1 |
| ENSG00000134504 | KCTD1 | 0,268406747 | 1,204476919 | 0,13451 | 0,46072811 | potassium channel tetramerization domain containing 1 |

| | | | | | | |
|---|---|---|---|---|---|---|
| ENSG00000056097 | ZFR | 0,268294693 | 1,20438337 | 0,42264 | 0,67675846 | zinc finger RNA binding protein |
| ENSG00000225171 | DUTP6 | 0,268286382 | 1,204376432 | 0,52343 | 0,74736255 | deoxyuridine triphosphatase pseudogene 6 |
| ENSG00000009724 | MASP2 | 0,268274182 | 1,204366248 | 0,23603 | 0,5471053 | mannan binding lectin serine peptidase 2 |
| ENSG00000257803 | PIGAP1 | 0,268235924 | 1,20433431 | 0,53107 | 0,75245598 | phosphatidylinositol glycan anchor biosynthesis class A pseudogene 1 |
| ENSG00000103275 | UBE2I | 0,268204453 | 1,204308039 | 0,19468 | 0,51562158 | ubiquitin conjugating enzyme E2 I |
| ENSG00000213304 | AC008481,2 | 0,268181873 | 1,20428919 | 0,70643 | 0,85870246 | ribosomal protein L18a (RPL18A) pseudogene |
| ENSG00000242100 | RPL9P32 | 0,268180439 | 1,204287993 | 0,59715 | 0,79434997 | ribosomal protein L9 pseudogene 32 |
| ENSG00000272910 | AC090425,3 | 0,268179027 | 1,204286815 | 0,7285 | 0,87102838 | novel transcript, antisense to ACTL6A |
| ENSG00000105656 | ELL | 0,268142559 | 1,204256373 | 0,17453 | 0,49884018 | elongation factor for RNA polymerase II |
| ENSG00000142186 | SCYL1 | 0,268136875 | 1,204251629 | 0,16637 | 0,49232464 | SCY1 like pseudokinase 1 |
| ENSG00000218336 | TENM3 | 0,267961083 | 1,2041049 | 0,42895 | 0,68169807 | teneurin transmembrane protein 3 |
| ENSG00000088298 | EDEM2 | 0,267933761 | 1,204082097 | 0,45027 | 0,69566873 | ER degradation enhancing alpha-mannosidase like protein 2 |
| ENSG00000188994 | ZNF292 | 0,267913951 | 1,204065563 | 0,47086 | 0,71034754 | zinc finger protein 292 |
| ENSG00000223572 | CKMT1A | 0,26790922 | 1,204061615 | 0,58286 | 0,78477395 | creatine kinase, mitochondrial 1A |
| ENSG00000119760 | SUPT7L | 0,267892603 | 1,204047746 | 0,18722 | 0,5105374 | SPT7 like, STAGA complex subunit gamma |
| ENSG00000178913 | TAF7 | 0,26784801 | 1,20401053 | 0,20448 | 0,52439728 | TATA-box binding protein associated factor 7 |
| ENSG00000284128 | AP000356,3 | 0,267828775 | 1,203994477 | 0,62137 | 0,80787859 | breakpoint cluster region pseudogene 3 |
| ENSG00000273772 | SMN2 | 0,267706153 | 1,203892148 | 0,32241 | 0,60577115 | survival of motor neuron 2, centromeric |
| ENSG00000156966 | B3GNT7 | 0,267622299 | 1,203822177 | 0,59619 | 0,79379314 | UDP-GlcNAc:betaGal beta-1,3-N-acetylglucosaminyltransferase 7 |
| ENSG00000277475 | AC213203,2 | 0,267582247 | 1,203788757 | 0,74015 | 0,87681378 | |
| ENSG00000204469 | PRRC2A | 0,267536344 | 1,203750456 | 0,51636 | 0,74198903 | proline rich coiled-coil 2A |
| ENSG00000156925 | ZIC3 | 0,267514436 | 1,203732176 | 0,40505 | 0,66481282 | Zic family member 3 |
| ENSG00000008952 | SEC62 | 0,267482325 | 1,203705384 | 0,42637 | 0,67986793 | SEC62 homolog, preprotein translocation factor |
| ENSG00000281994 | NPIPA1 | 0,267412199 | 1,203646877 | 0,46502 | 0,70556892 | nuclear pore complex interacting protein family member A1 |
| ENSG00000258539 | AC068896,1 | 0,267340828 | 1,203587333 | 0,38702 | 0,65213944 | novel transcript, METTL10-FAM53B readthrough |
| ENSG00000093167 | LRRFIP2 | 0,267335342 | 1,203582756 | 0,37276 | 0,64166433 | LRR binding FLII interacting protein 2 |
| ENSG00000148335 | NTMT1 | 0,267272924 | 1,203530684 | 0,31538 | 0,60163243 | N-terminal Xaa-Pro-Lys N-methyltransferase 1 |
| ENSG00000085840 | ORC1 | 0,267165372 | 1,203440965 | 0,24504 | 0,55295131 | origin recognition complex subunit 1 |
| ENSG00000273181 | AC131235,3 | 0,267142211 | 1,203421645 | 0,74972 | 0,8822333 | novel transcript |

| | | | | | | |
|---|---|---|---|---|---|---|
| ENSG00000083520 | DIS3 | 0,267120358 | 1,203403417 | 0,32416 | 0,60675074 | DIS3 homolog, exosome endoribonuclease and 3'-5' exoribonuclease |
| ENSG00000074201 | CLNS1A | 0,267007174 | 1,203309009 | 0,31167 | 0,59911644 | chloride nucleotide-sensitive channel 1A |
| ENSG00000160445 | ZER1 | 0,266938254 | 1,203251527 | 0,25196 | 0,55779669 | zyg-11 related cell cycle regulator |
| ENSG00000229867 | STEAP3-AS1 | 0,266709989 | 1,203061162 | 0,78344 | 0,89918294 | STEAP3 antisense RNA 1 |
| ENSG00000234607 | AL355994,4 | 0,266702508 | 1,203054924 | 0,72533 | 0,86898797 | chromosome 16 open reading frame 52 (C16orf52) pseudogene |
| ENSG00000147548 | NSD3 | 0,266612391 | 1,202979778 | 0,41161 | 0,66926602 | nuclear receptor binding SET domain protein 3 |
| ENSG00000163625 | WDFY3 | 0,266593457 | 1,20296399 | 0,38001 | 0,64776602 | WD repeat and FYVE domain containing 3 |
| ENSG00000083457 | ITGAE | 0,266574663 | 1,202948319 | 0,52484 | 0,74859526 | integrin subunit alpha E |
| ENSG00000072803 | FBXW11 | 0,266457374 | 1,202850525 | 0,30798 | 0,59691026 | F-box and WD repeat domain containing 11 |
| ENSG00000169016 | E2F6 | 0,266338899 | 1,20275175 | 0,20239 | 0,52256429 | E2F transcription factor 6 |
| ENSG00000198363 | ASPH | 0,26627966 | 1,202702365 | 0,15565 | 0,48345635 | aspartate beta-hydroxylase |
| ENSG00000277840 | AC026368,1 | 0,266269936 | 1,202694258 | 0,69377 | 0,8512419 | novel transcript, sense intronic to FBXW8 |
| ENSG00000123643 | SLC36A1 | 0,266256585 | 1,202683128 | 0,28771 | 0,58395529 | solute carrier family 36 member 1 |
| ENSG00000170100 | ZNF778 | 0,266249409 | 1,202677147 | 0,08681 | 0,3816108 | zinc finger protein 778 |
| ENSG00000135577 | NMBR | 0,266246541 | 1,202674755 | 0,755 | 0,88516173 | neuromedin B receptor |
| ENSG00000272921 | AC005832,4 | 0,266185267 | 1,202623677 | 0,60261 | 0,79710677 | novel protein |
| ENSG00000185340 | GAS2L1 | 0,266130081 | 1,202577675 | 0,38689 | 0,65212432 | growth arrest specific 2 like 1 |
| ENSG00000286362 | AP005717,2 | 0,266100356 | 1,202552897 | 0,59139 | 0,79058842 | novel transcript, antisense to DSCC1 |
| ENSG00000204628 | RACK1 | 0,26608188 | 1,202537496 | 0,24725 | 0,55483158 | receptor for activated C kinase 1 |
| ENSG00000167721 | TSR1 | 0,266065192 | 1,202523587 | 0,16218 | 0,48939373 | TSR1 ribosome maturation factor |
| ENSG00000249084 | AC027627,1 | 0,266035012 | 1,202498432 | 0,55729 | 0,76904341 | novel transcript |
| ENSG00000166128 | RAB8B | 0,266013236 | 1,202480281 | 0,60993 | 0,80202886 | RAB8B, member RAS oncogene family |
| ENSG00000232956 | SNHG15 | 0,265939552 | 1,202418867 | 0,19501 | 0,51588752 | small nucleolar RNA host gene 15 |
| ENSG00000147118 | ZNF182 | 0,265938629 | 1,202418098 | 0,45262 | 0,69708742 | zinc finger protein 182 |
| ENSG00000162148 | PPP1R32 | 0,265894401 | 1,202381237 | 0,51052 | 0,73802717 | protein phosphatase 1 regulatory subunit 32 |
| ENSG00000285804 | AC025774,1 | 0,265885052 | 1,202373445 | 0,47771 | 0,71545897 | novel transcript |
| ENSG00000168906 | MAT2A | 0,265762207 | 1,202271067 | 0,20129 | 0,52181085 | methionine adenosyltransferase 2A |
| ENSG00000164086 | DUSP7 | 0,265761111 | 1,202270154 | 0,32287 | 0,60584543 | dual specificity phosphatase 7 |
| ENSG00000101452 | DHX35 | 0,265742345 | 1,202254516 | 0,21954 | 0,53694774 | DEAH-box helicase 35 |
| ENSG00000121570 | DPPA4 | 0,265727225 | 1,202241916 | 0,26946 | 0,57124604 | developmental pluripotency associated 4 |
| ENSG00000248593 | DSTNP2 | 0,265619799 | 1,202152398 | 0,29223 | 0,58692263 | DSTN pseudogene 2 |

| | | | | | | |
|---|---|---|---|---|---|---|
| ENSG00000127946 | HIP1 | 0,265605529 | 1,202140507 | 0,12283 | 0,44269063 | huntingtin interacting protein 1 |
| ENSG00000172977 | KAT5 | 0,265587855 | 1,20212578 | 0,40233 | 0,66359858 | lysine acetyltransferase 5 |
| ENSG00000006459 | KDM7A | 0,265534652 | 1,20208145 | 0,38376 | 0,65041894 | lysine demethylase 7A |
| ENSG00000237916 | RPL37P12 | 0,265397671 | 1,201967319 | 0,69062 | 0,84900677 | ribosomal protein L37 pseudogene 12 |
| ENSG00000197702 | PARVA | 0,265319052 | 1,201901821 | 0,21885 | 0,53639331 | parvin alpha |
| ENSG00000148842 | CNNM2 | 0,265301155 | 1,201886911 | 0,14547 | 0,47300165 | cyclin and CBS domain divalent metal cation transport mediator 2 |
| ENSG00000172331 | BPGM | 0,265272672 | 1,201863182 | 0,43648 | 0,68678001 | bisphosphoglycerate mutase |
| ENSG00000010292 | NCAPD2 | 0,265261021 | 1,201853477 | 0,24236 | 0,55130426 | non-SMC condensin I complex subunit D2 |
| ENSG00000145428 | RNF175 | 0,265259667 | 1,201852348 | 0,49209 | 0,72541367 | ring finger protein 175 |
| ENSG00000198521 | ZNF43 | 0,26525887 | 1,201851685 | 0,4695 | 0,70898874 | zinc finger protein 43 |
| ENSG00000277135 | AC012409,3 | 0,26525477 | 1,201848269 | 0,83958 | 0,92668389 | novel transcript |
| ENSG00000226826 | HLA-DPB1 | 0,265185994 | 1,201790976 | 0,48883 | 0,72350189 | major histocompatibility complex, class II, DP beta 1 |
| ENSG00000237710 | HLA-DPB1 | 0,265185994 | 1,201790976 | 0,48883 | 0,72350189 | major histocompatibility complex, class II, DP beta 1 |
| ENSG00000251537 | AC005324,2 | 0,265130354 | 1,201744627 | 0,55102 | 0,76487238 | novel tripartite motif-containing 16 (TRIM16) and CMT1A duplicated region transcript 1 (CDRT1) protein |
| ENSG00000111859 | NEDD9 | 0,265077958 | 1,201700983 | 0,55453 | 0,76746279 | neural precursor cell expressed, developmentally down-regulated 9 |
| ENSG00000261118 | AC092123,1 | 0,26504817 | 1,201676171 | 0,64817 | 0,82383528 | novel transcript, antisense to SPG7 |
| ENSG00000272752 | STAG3L5P-PV | 0,265022413 | 1,201654717 | 0,55297 | 0,76620716 | STAG3L5P-PVRIG2P-PILRB readthrough |
| ENSG00000106459 | NRF1 | 0,264964848 | 1,201606772 | 0,35866 | 0,63185282 | nuclear respiratory factor 1 |
| ENSG00000108387 | SEPTIN4 | 0,264957704 | 1,201600822 | 0,53283 | 0,75333268 | septin 4 |
| ENSG00000215021 | PHB2 | 0,264955832 | 1,201599262 | 0,23304 | 0,54557342 | prohibitin 2 |
| ENSG00000152433 | ZNF547 | 0,264793748 | 1,201464272 | 0,47507 | 0,71364583 | zinc finger protein 547 |
| ENSG00000140948 | ZCCHC14 | 0,264725582 | 1,201407505 | 0,37357 | 0,64221436 | zinc finger CCHC-type containing 14 |
| ENSG00000213246 | SUPT4H1 | 0,264682712 | 1,201371806 | 0,21425 | 0,53334744 | SPT4 homolog, DSIF elongation factor subunit |
| ENSG00000281195 | AC007878,1 | 0,264582761 | 1,201288577 | 0,51532 | 0,74139472 | novel transcript |
| ENSG00000124313 | IQSEC2 | 0,264565436 | 1,201274151 | 0,41136 | 0,66904284 | IQ motif and Sec7 domain ArfGEF 2 |
| ENSG00000004779 | NDUFAB1 | 0,264516395 | 1,201233317 | 0,4759 | 0,71433501 | NADH:ubiquinone oxidoreductase subunit AB1 |
| ENSG00000240592 | EGFL8 | 0,264435637 | 1,201166077 | 0,70727 | 0,85910429 | EGF like domain multiple 8 |
| ENSG00000164252 | AGGF1 | 0,26441469 | 1,201148637 | 0,44078 | 0,68917797 | angiogenic factor with G-patch and FHA domains 1 |
| ENSG00000124641 | MED20 | 0,26431818 | 1,201068289 | 0,28198 | 0,57936013 | mediator complex subunit 20 |
| ENSG00000274471 | HERC2P7 | 0,264288237 | 1,201043361 | 0,13666 | 0,46406381 | hect domain and RLD 2 pseudogene 7 |

| | | | | | | |
|---|---|---|---|---|---|---|
| ENSG00000230561 | CCDC192 | 0,26422013 | 1,200986663 | 0,5566 | 0,76850571 | coiled-coil domain containing 192 |
| ENSG00000171970 | ZNF57 | 0,264217885 | 1,200984794 | 0,49483 | 0,72723033 | zinc finger protein 57 |
| ENSG00000101310 | SEC23B | 0,264188192 | 1,200960076 | 0,27827 | 0,57645922 | SEC23 homolog B, COPII coat complex component |
| ENSG00000066185 | ZMYND12 | 0,264177852 | 1,200951469 | 0,77406 | 0,89572556 | zinc finger MYND-type containing 12 |
| ENSG00000011454 | RABGAP1 | 0,264098795 | 1,20088566 | 0,37989 | 0,64769466 | RAB GTPase activating protein 1 |
| ENSG00000197256 | KANK2 | 0,264097554 | 1,200884628 | 0,62034 | 0,80773684 | KN motif and ankyrin repeat domains 2 |
| ENSG00000154511 | DIPK1A | 0,264095033 | 1,200882529 | 0,28476 | 0,58163496 | divergent protein kinase domain 1A |
| ENSG00000267606 | AC006116,8 | 0,264090087 | 1,200878413 | 0,73593 | 0,8750728 | novel transcript |
| ENSG00000237194 | SNAI1P1 | 0,264062432 | 1,200855393 | 0,75598 | 0,88573119 | snail family zinc finger 1 pseudogene 1 |
| ENSG00000214211 | CTAGE14P | 0,264042779 | 1,200839035 | 0,63913 | 0,81893766 | CTAGE family member 14, pseudogene |
| ENSG00000231192 | OR5H1 | 0,263999555 | 1,200803057 | 0,58466 | 0,78582465 | olfactory receptor family 5 subfamily H member 1 |
| ENSG00000072506 | HSD17B10 | 0,263973025 | 1,200780976 | 0,18039 | 0,50407792 | hydroxysteroid 17-beta dehydrogenase 10 |
| ENSG00000281128 | PTENP1-AS | 0,263951342 | 1,200762929 | 0,41703 | 0,6738387 | PTENP1 antisense RNA |
| ENSG00000125952 | MAX | 0,263923359 | 1,200739639 | 0,22744 | 0,54168591 | MYC associated factor X |
| ENSG00000287202 | AC008119,1 | 0,263885362 | 1,200708014 | 0,38659 | 0,65194712 | novel transcript, antisense to SART3 |
| ENSG00000213079 | SCAF8 | 0,263728804 | 1,200577724 | 0,33072 | 0,6112353 | SR-related CTD associated factor 8 |
| ENSG00000113300 | CNOT6 | 0,263627057 | 1,200493055 | 0,4759 | 0,71433501 | CCR4-NOT transcription complex subunit 6 |
| ENSG00000116539 | ASH1L | 0,263621277 | 1,200488245 | 0,36367 | 0,63452784 | ASH1 like histone lysine methyltransferase |
| ENSG00000182362 | YBEY | 0,26352424 | 1,200407501 | 0,44234 | 0,69006979 | ybeY metalloendoribonuclease |
| ENSG00000138792 | ENPEP | 0,263485415 | 1,200375197 | 0,60012 | 0,79579686 | glutamyl aminopeptidase |
| ENSG00000165821 | SALL2 | 0,263431442 | 1,200330291 | 0,23311 | 0,54557342 | spalt like transcription factor 2 |
| ENSG00000075785 | RAB7A | 0,263295883 | 1,20021751 | 0,14027 | 0,46714558 | RAB7A, member RAS oncogene family |
| ENSG00000156853 | ZNF689 | 0,263287845 | 1,200210823 | 0,28237 | 0,57972352 | zinc finger protein 689 |
| ENSG00000287879 | AC114341,2 | 0,263236238 | 1,200167891 | 0,43897 | 0,68799204 | novel transcript |
| ENSG00000129810 | SGO1 | 0,263231023 | 1,200163553 | 0,33731 | 0,61706507 | shugoshin 1 |
| ENSG00000158352 | SHROOM4 | 0,263204532 | 1,200141515 | 0,18857 | 0,5119341 | shroom family member 4 |
| ENSG00000164742 | ADCY1 | 0,263198776 | 1,200136727 | 0,18181 | 0,50517706 | adenylate cyclase 1 |
| ENSG00000166595 | CIAO2B | 0,263155905 | 1,200101064 | 0,4912 | 0,7250491 | cytosolic iron-sulfur assembly component 2B |
| ENSG00000205808 | PLPP6 | 0,263105171 | 1,200058862 | 0,29173 | 0,58660475 | phospholipid phosphatase 6 |
| ENSG00000127980 | PEX1 | 0,263072965 | 1,200032073 | 0,35876 | 0,63191767 | peroxisomal biogenesis factor 1 |
| ENSG00000188582 | PAQR9 | 0,263057199 | 1,200018959 | 0,50146 | 0,73139105 | progestin and adipoQ receptor family member 9 |
| ENSG00000261008 | LINC01572 | 0,263042944 | 1,200007101 | 0,32287 | 0,60584543 | long intergenic non-protein coding RNA 1572 |
| ENSG00000224003 | YES1P1 | 0,262994702 | 1,199966976 | 0,64793 | 0,82381635 | YES1 pseudogene 1 |

| | | | | | | |
|---|---|---|---|---|---|---|
| ENSG00000162687 | KCNT2 | 0,262914135 | 1,199899965 | 0,51509 | 0,74133616 | potassium sodium-activated channel subfamily T member 2 |
| ENSG00000165209 | STRBP | 0,262887554 | 1,199877858 | 0,42188 | 0,67643128 | spermatid perinuclear RNA binding protein |
| ENSG00000253649 | PRSS51 | 0,262876732 | 1,199868858 | 0,59516 | 0,79320995 | serine protease 51 |
| ENSG00000178567 | EPM2AIP1 | 0,262858243 | 1,199853481 | 0,50545 | 0,7339567 | EPM2A interacting protein 1 |
| ENSG00000102226 | USP11 | 0,262836958 | 1,199835779 | 0,26787 | 0,56999323 | ubiquitin specific peptidase 11 |
| ENSG00000162746 | FCRLB | 0,262805871 | 1,199809926 | 0,56514 | 0,773443 | Fc receptor like B |
| ENSG00000249572 | AC034231,1 | 0,262794535 | 1,199800498 | 0,66561 | 0,83358761 | novel transcript |
| ENSG00000267216 | ZNF8-ERVK3-1 | 0,262707461 | 1,199728086 | 0,27641 | 0,57542877 | ZNF8-ERVK3-1 readthrough |
| ENSG00000100722 | ZC3H14 | 0,262548432 | 1,199595846 | 0,32939 | 0,61044057 | zinc finger CCCH-type containing 14 |
| ENSG00000168152 | THAP9 | 0,262454805 | 1,199517998 | 0,51685 | 0,74218125 | THAP domain containing 9 |
| ENSG00000152223 | EPG5 | 0,262361962 | 1,199440807 | 0,36249 | 0,63379594 | ectopic P-granules autophagy protein 5 homolog |
| ENSG00000101294 | HM13 | 0,262348067 | 1,199429256 | 0,11088 | 0,42363726 | histocompatibility minor 13 |
| ENSG00000149313 | AASDHPPT | 0,262346301 | 1,199427787 | 0,40061 | 0,66264525 | aminoadipate-semialdehyde dehydrogenase-phosphopantetheinyl transferase |
| ENSG00000141858 | SAMD1 | 0,262257131 | 1,199353655 | 0,25183 | 0,55773568 | sterile alpha motif domain containing 1 |
| ENSG00000128050 | PAICS | 0,26222181 | 1,199324292 | 0,20197 | 0,52230208 | phosphoribosylaminoimidazole carboxylase and phosphoribosylaminoimidazolesuccinocarboxamide synthase |
| ENSG00000174527 | MYO1H | 0,262131629 | 1,199249327 | 0,47669 | 0,71481671 | myosin IH |
| ENSG00000265552 | AC104961,1 | 0,262118778 | 1,199238644 | 0,74053 | 0,87708444 | novel transcript |
| ENSG00000232632 | GABBR1 | 0,262080108 | 1,1992065 | 0,34599 | 0,6231701 | gamma-aminobutyric acid type B receptor subunit 1 |
| ENSG00000122692 | SMU1 | 0,262075993 | 1,19920308 | 0,23469 | 0,54623956 | SMU1 DNA replication regulator and spliceosomal factor |
| ENSG00000140600 | SH3GL3 | 0,261967111 | 1,199112578 | 0,30114 | 0,59252799 | SH3 domain containing GRB2 like 3, endophilin A3 |
| ENSG00000198373 | WWP2 | 0,261944603 | 1,19909387 | 0,27641 | 0,57542877 | WW domain containing E3 ubiquitin protein ligase 2 |
| ENSG00000175768 | TOMM5 | 0,261925243 | 1,199077778 | 0,27462 | 0,57463533 | translocase of outer mitochondrial membrane 5 |
| ENSG00000140284 | SLC27A2 | 0,261911838 | 1,199066638 | 0,6751 | 0,84004757 | solute carrier family 27 member 2 |
| ENSG00000186603 | HPDL | 0,261799258 | 1,198973073 | 0,36421 | 0,634951 | 4-hydroxyphenylpyruvate dioxygenase like |
| ENSG00000226784 | PGAM4 | 0,261684428 | 1,198877645 | 0,79905 | 0,90735534 | phosphoglycerate mutase family member 4 |
| ENSG00000182973 | CNOT10 | 0,261669133 | 1,198864935 | 0,25363 | 0,55863744 | CCR4-NOT transcription complex subunit 10 |
| ENSG00000178057 | NDUFAF3 | 0,261656375 | 1,198854333 | 0,31192 | 0,59925736 | NADH:ubiquinone oxidoreductase complex assembly factor 3 |
| ENSG00000276345 | AC004556,3 | 0,261642557 | 1,198842851 | 0,19388 | 0,51547251 | 39S ribosomal protein L23, mitochondrial |

| | | | | | | |
|---|---|---|---|---|---|---|
| ENSG00000141568 | FOXK2 | 0,261628753 | 1,19883138 | 0,20953 | 0,52856218 | forkhead box K2 |
| ENSG00000102547 | CAB39L | 0,261597969 | 1,1988058 | 0,54168 | 0,75882662 | calcium binding protein 39 like |
| ENSG00000261535 | AC005086,2 | 0,261553852 | 1,198769142 | 0,70524 | 0,85799582 | novel transcript, antisense to CUX1 |
| ENSG00000038210 | PI4K2B | 0,26149653 | 1,198721513 | 0,20627 | 0,5259882 | phosphatidylinositol 4-kinase type 2 beta |
| ENSG00000184014 | DENND5A | 0,261494072 | 1,19871947 | 0,33064 | 0,6112353 | DENN domain containing 5A |
| ENSG00000175895 | PLEKHF2 | 0,261471054 | 1,198700345 | 0,42059 | 0,67595875 | pleckstrin homology and FYVE domain containing 2 |
| ENSG00000213676 | ATF6B | 0,261455211 | 1,198687181 | 0,2742 | 0,57420712 | activating transcription factor 6 beta |
| ENSG00000122882 | ECD | 0,261426308 | 1,198663167 | 0,10651 | 0,41625157 | ecdysoneless cell cycle regulator |
| ENSG00000103222 | ABCC1 | 0,26133683 | 1,198588826 | 0,35699 | 0,63110115 | ATP binding cassette subfamily C member 1 |
| ENSG00000100902 | PSMA6 | 0,261184697 | 1,198462441 | 0,18322 | 0,50663141 | proteasome 20S subunit alpha 6 |
| ENSG00000267886 | AC074135,1 | 0,261164753 | 1,198445874 | 0,47501 | 0,71364583 | novel transcript |
| ENSG00000277972 | CISD3 | 0,261121136 | 1,198409641 | 0,57935 | 0,78270914 | CDGSH iron sulfur domain 3 |
| ENSG00000105258 | POLR2I | 0,261089528 | 1,198383386 | 0,27935 | 0,57742338 | RNA polymerase II subunit I |
| ENSG00000173451 | THAP2 | 0,261015906 | 1,198322233 | 0,61357 | 0,8035281 | THAP domain containing 2 |
| ENSG00000112118 | MCM3 | 0,260976377 | 1,1982894 | 0,18253 | 0,5059851 | minichromosome maintenance complex component 3 |
| ENSG00000173221 | GLRX | 0,260897764 | 1,198224107 | 0,30947 | 0,59793596 | glutaredoxin |
| ENSG00000134897 | BIVM | 0,260815596 | 1,198155864 | 0,32591 | 0,60844548 | basic, immunoglobulin-like variable motif containing |
| ENSG00000149577 | SIDT2 | 0,260686532 | 1,198048682 | 0,20813 | 0,52756478 | SID1 transmembrane family member 2 |
| ENSG00000254274 | TDGF1P5 | 0,260542484 | 1,197929066 | 0,66838 | 0,83536064 | teratocarcinoma-derived growth factor 1 pseudogene 5 |
| ENSG00000113522 | RAD50 | 0,260507983 | 1,19790042 | 0,48729 | 0,72237072 | RAD50 double strand break repair protein |
| ENSG00000273033 | LINC02035 | 0,260502329 | 1,197895725 | 0,2897 | 0,58522301 | long intergenic non-protein coding RNA 2035 |
| ENSG00000100519 | PSMC6 | 0,260358897 | 1,197776637 | 0,28029 | 0,57784691 | proteasome 26S subunit, ATPase 6 |
| ENSG00000254004 | ZNF260 | 0,260284697 | 1,197715035 | 0,42186 | 0,67643128 | zinc finger protein 260 |
| ENSG00000164385 | LINC01600 | 0,260259821 | 1,197694383 | 0,56134 | 0,77136826 | long intergenic non-protein coding RNA 1600 |
| ENSG00000184162 | NR2C2AP | 0,260075977 | 1,19754177 | 0,24507 | 0,55295131 | nuclear receptor 2C2 associated protein |
| ENSG00000175224 | ATG13 | 0,259973907 | 1,197457047 | 0,14118 | 0,4679289 | autophagy related 13 |
| ENSG00000175455 | CCDC14 | 0,259939478 | 1,197428471 | 0,59202 | 0,79093233 | coiled-coil domain containing 14 |
| ENSG00000105866 | SP4 | 0,259931551 | 1,197421891 | 0,35851 | 0,63185282 | Sp4 transcription factor |
| ENSG00000215271 | HOMEZ | 0,259917991 | 1,197410637 | 0,22236 | 0,53859118 | homeobox and leucine zipper encoding |
| ENSG00000058272 | PPP1R12A | 0,259907723 | 1,197402115 | 0,33815 | 0,6177255 | protein phosphatase 1 regulatory subunit 12A |
| ENSG00000135968 | GCC2 | 0,259863823 | 1,197365679 | 0,41349 | 0,67078587 | GRIP and coiled-coil domain containing 2 |
| ENSG00000136875 | PRPF4 | 0,259851839 | 1,197355733 | 0,30944 | 0,59790134 | pre-mRNA processing factor 4 |
| ENSG00000101972 | STAG2 | 0,259850699 | 1,197354787 | 0,47178 | 0,71113267 | stromal antigen 2 |

| | | | | | | |
|---|---|---|---|---|---|---|
| ENSG00000255302 | EID1 | 0,259836618 | 1,197343101 | 0,30499 | 0,59446033 | EP300 interacting inhibitor of differentiation 1 |
| ENSG00000229727 | AC013460,1 | 0,259826991 | 1,197335111 | 0,52866 | 0,75098717 | novel transcript |
| ENSG00000135930 | EIF4E2 | 0,25978679 | 1,197301747 | 0,25425 | 0,55925777 | eukaryotic translation initiation factor 4E family member 2 |
| ENSG00000120948 | TARDBP | 0,259776395 | 1,19729312 | 0,17991 | 0,50407792 | TAR DNA binding protein |
| ENSG00000119718 | EIF2B2 | 0,259760787 | 1,197280167 | 0,31211 | 0,59925736 | eukaryotic translation initiation factor 2B subunit beta |
| ENSG00000197889 | MEIG1 | 0,259694193 | 1,197224903 | 0,58395 | 0,78572744 | meiosis/spermiogenesis associated 1 |
| ENSG00000152078 | TLCD4 | 0,259482811 | 1,197049499 | 0,5662 | 0,77421829 | TLC domain containing 4 |
| ENSG00000136100 | VPS36 | 0,259228906 | 1,196838845 | 0,06888 | 0,34681313 | vacuolar protein sorting 36 homolog |
| ENSG00000279457 | WASH9P | 0,258942957 | 1,196601649 | 0,34199 | 0,61986127 | WAS protein family homolog 9, pseudogene |
| ENSG00000250001 | AC027335,1 | 0,25890305 | 1,196568551 | 0,70978 | 0,86015146 | novel transcript |
| ENSG00000118690 | ARMC2 | 0,25884127 | 1,196517311 | 0,49044 | 0,72461398 | armadillo repeat containing 2 |
| ENSG00000142867 | BCL10 | 0,258723578 | 1,196419706 | 0,37964 | 0,64740658 | BCL10 immune signaling adaptor |
| ENSG00000171953 | ATPAF2 | 0,258702287 | 1,196402049 | 0,16189 | 0,48914394 | ATP synthase mitochondrial F1 complex assembly factor 2 |
| ENSG00000007923 | DNAJC11 | 0,258676414 | 1,196380594 | 0,17989 | 0,50407792 | DnaJ heat shock protein family (Hsp40) member C11 |
| ENSG00000112304 | ACOT13 | 0,258665416 | 1,196371474 | 0,23392 | 0,54568606 | acyl-CoA thioesterase 13 |
| ENSG00000114648 | KLHL18 | 0,258625415 | 1,196338302 | 0,33497 | 0,61467789 | kelch like family member 18 |
| ENSG00000143882 | ATP6V1C2 | 0,258582627 | 1,196302821 | 0,47903 | 0,71671228 | ATPase H+ transporting V1 subunit C2 |
| ENSG00000170293 | CMTM8 | 0,258575694 | 1,196297073 | 0,64918 | 0,82436914 | CKLF like MARVEL transmembrane domain containing 8 |
| ENSG00000253710 | ALG11 | 0,258527048 | 1,196256736 | 0,18668 | 0,51013726 | ALG11 alpha-1,2-mannosyltransferase |
| ENSG00000143258 | USP21 | 0,258457144 | 1,196198774 | 0,18036 | 0,50407792 | ubiquitin specific peptidase 21 |
| ENSG00000105993 | DNAJB6 | 0,258454342 | 1,196196451 | 0,27475 | 0,57463533 | DnaJ heat shock protein family (Hsp40) member B6 |
| ENSG00000279078 | SND1-IT1 | 0,25844482 | 1,196188556 | 0,27351 | 0,57362145 | SND1 intronic transcript 1 |
| ENSG00000119801 | YPEL5 | 0,25843082 | 1,196176948 | 0,42564 | 0,67920713 | yippee like 5 |
| ENSG00000171634 | BPTF | 0,25842851 | 1,196175033 | 0,34263 | 0,62025394 | bromodomain PHD finger transcription factor |
| ENSG00000123427 | EEF1AKMT3 | 0,258318984 | 1,196084225 | 0,49684 | 0,72858829 | EEF1A lysine methyltransferase 3 |
| ENSG00000112081 | SRSF3 | 0,258263572 | 1,196038286 | 0,19722 | 0,5184837 | serine and arginine rich splicing factor 3 |
| ENSG00000248100 | AC087257,1 | 0,258246917 | 1,196024479 | 0,53039 | 0,75200826 | novel transcript |
| ENSG00000287205 | AL451054,4 | 0,258194924 | 1,195981376 | 0,79399 | 0,90426294 | novel transcript |
| ENSG00000143393 | PI4KB | 0,258126294 | 1,195924484 | 0,19854 | 0,51980353 | phosphatidylinositol 4-kinase beta |
| ENSG00000065457 | ADAT1 | 0,25799348 | 1,195814392 | 0,26716 | 0,56958284 | adenosine deaminase tRNA specific 1 |
| ENSG00000143811 | PYCR2 | 0,257951046 | 1,195779221 | 0,21659 | 0,53490925 | pyrroline-5-carboxylate reductase 2 |
| ENSG00000102225 | CDK16 | 0,257845831 | 1,195692016 | 0,3037 | 0,59385655 | cyclin dependent kinase 16 |
| ENSG00000157625 | TAB3 | 0,257787065 | 1,195643312 | 0,33266 | 0,61256564 | TGF-beta activated kinase 1 (MAP3K7) binding protein 3 |

| | | | | | | |
|---|---|---|---|---|---|---|
| ENSG00000161654 | LSM12 | 0,257752659 | 1,195614798 | 0,19314 | 0,51517146 | LSM12 homolog |
| ENSG00000283312 | AC017104,3 | 0,257715742 | 1,195584205 | 0,76312 | 0,88980026 | GTF2I repeat domain containing 2(GTF2IRD2) pseudogene |
| ENSG00000145779 | TNFAIP8 | 0,257691807 | 1,195564369 | 0,31689 | 0,60268749 | TNF alpha induced protein 8 |
| ENSG00000177200 | CHD9 | 0,257660207 | 1,195538182 | 0,49255 | 0,72575639 | chromodomain helicase DNA binding protein 9 |
| ENSG00000127947 | PTPN12 | 0,257638898 | 1,195520524 | 0,48997 | 0,72429221 | protein tyrosine phosphatase non-receptor type 12 |
| ENSG00000213995 | NAXD | 0,257632495 | 1,195515218 | 0,218 | 0,53566935 | NAD(P)HX dehydratase |
| ENSG00000145362 | ANK2 | 0,257575095 | 1,195467654 | 0,43362 | 0,68473723 | ankyrin 2 |
| ENSG00000251396 | LINC01301 | 0,257535557 | 1,195434892 | 0,45333 | 0,69750019 | long intergenic non-protein coding RNA 1301 |
| ENSG00000047932 | GOPC | 0,257530713 | 1,195430878 | 0,49085 | 0,72482729 | golgi associated PDZ and coiled-coil motif containing |
| ENSG00000254585 | MAGEL2 | 0,257364575 | 1,195293222 | 0,28746 | 0,58386207 | MAGE family member L2 |
| ENSG00000241322 | CDRT1 | 0,257348324 | 1,195279758 | 0,49141 | 0,72510994 | CMT1A duplicated region transcript 1 |
| ENSG00000243304 | AC008494,1 | 0,257347476 | 1,195279056 | 0,7436 | 0,87867571 | ribosomal protein S2 (RPS2) pseudogene |
| ENSG00000122507 | BBS9 | 0,257341201 | 1,195273857 | 0,50185 | 0,73149732 | Bardet-Biedl syndrome 9 |
| ENSG00000272498 | AC090948,2 | 0,257335826 | 1,195269404 | 0,5108 | 0,73828855 | novel transcript, antisense to RFTN1 |
| ENSG00000065613 | SLK | 0,257316552 | 1,195253435 | 0,27038 | 0,57217543 | STE20 like kinase |
| ENSG00000054983 | GALC | 0,257253557 | 1,195201246 | 0,38725 | 0,65225328 | galactosylceramidase |
| ENSG00000244041 | LINC01011 | 0,25721717 | 1,195171102 | 0,39226 | 0,6556164 | long intergenic non-protein coding RNA 1011 |
| ENSG00000129235 | TXNDC17 | 0,257082673 | 1,195059685 | 0,49739 | 0,72870789 | thioredoxin domain containing 17 |
| ENSG00000237757 | EEF1A1P30 | 0,256997229 | 1,194988909 | 0,60686 | 0,79994037 | eukaryotic translation elongation factor 1 alpha 1 pseudogene 30 |
| ENSG00000071794 | HLTF | 0,256924704 | 1,194928838 | 0,54957 | 0,7641582 | helicase like transcription factor |
| ENSG00000164209 | SLC25A46 | 0,256906784 | 1,194913996 | 0,40267 | 0,66396567 | solute carrier family 25 member 46 |
| ENSG00000103742 | IGDCC4 | 0,256868961 | 1,194882669 | 0,32258 | 0,60577617 | immunoglobulin superfamily DCC subclass member 4 |
| ENSG00000196704 | AMZ2 | 0,256830388 | 1,194850722 | 0,32708 | 0,60902638 | archaelysin family metallopeptidase 2 |
| ENSG00000029725 | RABEP1 | 0,256805779 | 1,194830342 | 0,408 | 0,66699097 | rabaptin, RAB GTPase binding effector protein 1 |
| ENSG00000152683 | SLC30A6 | 0,2567203 | 1,194759551 | 0,12996 | 0,45419678 | solute carrier family 30 member 6 |
| ENSG00000135821 | GLUL | 0,256645329 | 1,194697465 | 0,28695 | 0,58337791 | glutamate-ammonia ligase |
| ENSG00000282491 | AC243807,6 | 0,256624599 | 1,194680299 | 0,61275 | 0,8035281 | novel transcript |
| ENSG00000109079 | TNFAIP1 | 0,256614075 | 1,194671584 | 0,38708 | 0,65213944 | TNF alpha induced protein 1 |
| ENSG00000157349 | DDX19B | 0,25658143 | 1,194644552 | 0,4166 | 0,67338802 | DEAD-box helicase 19B |
| ENSG00000023171 | GRAMD1B | 0,256566371 | 1,194632082 | 0,18731 | 0,51056538 | GRAM domain containing 1B |
| ENSG00000182195 | LDOC1 | 0,256455398 | 1,194540193 | 0,23774 | 0,54863436 | LDOC1 regulator of NFKB signaling |

| | | | | | | |
|---|---|---|---|---|---|---|
| ENSG00000259505 | AC009268,1 | 0,256429251 | 1,194518544 | 0,74928 | 0,88204547 | Sjogren syndrome antigen B (autoantigen La) (SSB) pseudogene |
| ENSG00000225987 | PBX2 | 0,256414337 | 1,194506196 | 0,40233 | 0,66359858 | PBX homeobox 2 |
| ENSG00000125629 | INSIG2 | 0,256400394 | 1,194494651 | 0,50893 | 0,7370222 | insulin induced gene 2 |
| ENSG00000131148 | EMC8 | 0,256396248 | 1,194491219 | 0,28376 | 0,5807739 | ER membrane protein complex subunit 8 |
| ENSG00000261341 | AC010325,1 | 0,256347387 | 1,194450764 | 0,75358 | 0,88453179 | novel protein |
| ENSG00000144381 | HSPD1 | 0,256338226 | 1,19444318 | 0,27008 | 0,57186187 | heat shock protein family D (Hsp60) member 1 |
| ENSG00000137269 | LRRC1 | 0,256290093 | 1,19440333 | 0,38189 | 0,64905556 | leucine rich repeat containing 1 |
| ENSG00000047410 | TPR | 0,256273063 | 1,194389231 | 0,31076 | 0,5987267 | translocated promoter region, nuclear basket protein |
| ENSG00000160207 | HSF2BP | 0,256084273 | 1,194232945 | 0,40825 | 0,66714279 | heat shock transcription factor 2 binding protein |
| ENSG00000107242 | PIP5K1B | 0,255983692 | 1,194149688 | 0,66391 | 0,83247324 | phosphatidylinositol-4-phosphate 5-kinase type 1 beta |
| ENSG00000102057 | KCND1 | 0,255896651 | 1,194077645 | 0,50755 | 0,73579864 | potassium voltage-gated channel subfamily D member 1 |
| ENSG00000162745 | OLFML2B | 0,255882799 | 1,19406618 | 0,56248 | 0,77186635 | olfactomedin like 2B |
| ENSG00000170959 | DCDC1 | 0,255873617 | 1,194058581 | 0,29757 | 0,59036937 | doublecortin domain containing 1 |
| ENSG00000185710 | SMG1P4 | 0,255858723 | 1,194046254 | 0,55314 | 0,76626672 | SMG1 pseudogene 4 |
| ENSG00000176624 | MEX3C | 0,255803153 | 1,194000262 | 0,20126 | 0,52181085 | mex-3 RNA binding family member C |
| ENSG00000232859 | LYRM9 | 0,255681284 | 1,193899405 | 0,43907 | 0,68808849 | LYR motif containing 9 |
| ENSG00000143891 | GALM | 0,25565081 | 1,193874187 | 0,32735 | 0,60914884 | galactose mutarotase |
| ENSG00000234219 | CDCA4P4 | 0,255441926 | 1,193701342 | 0,51988 | 0,74432381 | cell division cycle associated 4 pseudogene 4 |
| ENSG00000167985 | SDHAF2 | 0,255362561 | 1,193635676 | 0,40215 | 0,66350593 | succinate dehydrogenase complex assembly factor 2 |
| ENSG00000108389 | MTMR4 | 0,255331586 | 1,193610049 | 0,38375 | 0,65041894 | myotubularin related protein 4 |
| ENSG00000111981 | ULBP1 | 0,255250006 | 1,193542555 | 0,48284 | 0,7191814 | UL16 binding protein 1 |
| ENSG00000139517 | LNX2 | 0,255204383 | 1,193504812 | 0,24393 | 0,55250543 | ligand of numb-protein X 2 |
| ENSG00000271121 | NUDT4P2 | 0,255138783 | 1,193450544 | 0,31345 | 0,6002104 | nudix hydrolase 4 pseudogene 2 |
| ENSG00000285081 | AC004593,1 | 0,255093873 | 1,193413393 | 0,66805 | 0,83531052 | Novel transcript, sense intronic to CPVL |
| ENSG00000111834 | RSPH4A | 0,255061132 | 1,19338631 | 0,51208 | 0,73906991 | radial spoke head component 4A |
| ENSG00000169398 | PTK2 | 0,255052674 | 1,193379313 | 0,20165 | 0,52214043 | protein tyrosine kinase 2 |
| ENSG00000108733 | PEX12 | 0,25501157 | 1,193345314 | 0,4354 | 0,68601186 | peroxisomal biogenesis factor 12 |
| ENSG00000125675 | GRIA3 | 0,254770631 | 1,193146034 | 0,41075 | 0,66856387 | glutamate ionotropic receptor AMPA type subunit 3 |
| ENSG00000187699 | C2orf88 | 0,254752086 | 1,193130697 | 0,49332 | 0,72631 | chromosome 2 open reading frame 88 |
| ENSG00000059769 | DNAJC25 | 0,254737222 | 1,193118404 | 0,33821 | 0,6177255 | DnaJ heat shock protein family (Hsp40) member C25 |
| ENSG00000248200 | AC093770,1 | 0,254711112 | 1,193096811 | 0,80276 | 0,9091124 | ribosomal protein L17 (RPL17) pseudogene |
| ENSG00000048052 | HDAC9 | 0,254688626 | 1,193078215 | 0,50074 | 0,73097006 | histone deacetylase 9 |

| | | | | | | |
|---|---|---|---|---|---|---|
| ENSG00000178150 | ZNF114 | 0,254636858 | 1,193035406 | 0,19021 | 0,51331106 | zinc finger protein 114 |
| ENSG00000162402 | USP24 | 0,254526801 | 1,192944397 | 0,30622 | 0,59517372 | ubiquitin specific peptidase 24 |
| ENSG00000185013 | NT5C1B | 0,254365111 | 1,192810705 | 0,67959 | 0,84261173 | 5'-nucleotidase, cytosolic IB |
| ENSG00000088538 | DOCK3 | 0,254128145 | 1,1926148 | 0,27612 | 0,57533851 | dedicator of cytokinesis 3 |
| ENSG00000285210 | LINC02795 | 0,254125804 | 1,192612864 | 0,6184 | 0,80662722 | long intergenic non-protein coding RNA 2795 |
| ENSG00000272758 | AC083798,2 | 0,254087448 | 1,192581158 | 0,42486 | 0,67857395 | novel transcript, antisense to KPNA1 |
| ENSG00000137818 | RPLP1 | 0,254049941 | 1,192550153 | 0,33896 | 0,61800397 | ribosomal protein lateral stalk subunit P1 |
| ENSG00000260000 | AL133338,1 | 0,253998517 | 1,192507646 | 0,40126 | 0,6630277 | novel transcript, antisense to ASCC3 |
| ENSG00000223905 | AC103592,1 | 0,253941138 | 1,192460219 | 0,8202 | 0,91725087 | peroxiredoxin 5 (PRDX5) pseudogene |
| ENSG00000263883 | EEF1DP7 | 0,253938271 | 1,192457849 | 0,70709 | 0,85902171 | eukaryotic translation elongation factor 1 delta pseudogene 7 |
| ENSG00000168734 | PKIG | 0,253913797 | 1,192437621 | 0,4009 | 0,6628292 | cAMP-dependent protein kinase inhibitor gamma |
| ENSG00000128272 | ATF4 | 0,253895025 | 1,192422105 | 0,12995 | 0,45419678 | activating transcription factor 4 |
| ENSG00000117614 | SYF2 | 0,253841152 | 1,192377578 | 0,22127 | 0,53805234 | SYF2 pre-mRNA splicing factor |
| ENSG00000148985 | PGAP2 | 0,25382041 | 1,192360435 | 0,37045 | 0,6399389 | post-GPI attachment to proteins 2 |
| ENSG00000123908 | AGO2 | 0,253816482 | 1,192357189 | 0,3724 | 0,64134584 | argonaute RISC catalytic component 2 |
| ENSG00000196154 | S100A4 | 0,253801569 | 1,192344863 | 0,61344 | 0,8035281 | S100 calcium binding protein A4 |
| ENSG00000174384 | PMS2P6 | 0,253756794 | 1,192307859 | 0,62413 | 0,80958576 | PMS1 homolog 2, mismatch repair system component pseudogene 6 |
| ENSG00000141639 | MAPK4 | 0,253743247 | 1,192296663 | 0,38561 | 0,65122192 | mitogen-activated protein kinase 4 |
| ENSG00000174720 | LARP7 | 0,253717104 | 1,192275058 | 0,38681 | 0,65210024 | La ribonucleoprotein 7, transcriptional regulator |
| ENSG00000154429 | CCSAP | 0,253684328 | 1,192247971 | 0,22242 | 0,53859118 | centriole, cilia and spindle associated protein |
| ENSG00000113384 | GOLPH3 | 0,253662755 | 1,192230144 | 0,16306 | 0,48971747 | golgi phosphoprotein 3 |
| ENSG00000205856 | C22orf42 | 0,253610246 | 1,192186751 | 0,53997 | 0,75807575 | chromosome 22 open reading frame 42 |
| ENSG00000129691 | ASH2L | 0,25360377 | 1,1921814 | 0,25712 | 0,56155289 | ASH2 like, histone lysine methyltransferase complex subunit |
| ENSG00000254315 | AC023632,5 | 0,253591147 | 1,192170969 | 0,80853 | 0,91193281 | novel transcript |
| ENSG00000133739 | LRRCC1 | 0,253581768 | 1,192163219 | 0,57742 | 0,78140182 | leucine rich repeat and coiled-coil centrosomal protein 1 |
| ENSG00000164941 | INTS8 | 0,253574835 | 1,192157489 | 0,2624 | 0,56534125 | integrator complex subunit 8 |
| ENSG00000285516 | NAALAD2 | 0,253543496 | 1,192131593 | 0,56267 | 0,77194048 | N-acetylated alpha-linked acidic dipeptidase 2 |
| ENSG00000077616 | NAALAD2 | 0,253543496 | 1,192131593 | 0,56267 | 0,77194048 | N-acetylated alpha-linked acidic dipeptidase 2 |
| ENSG00000287252 | AL009179,1 | 0,253522855 | 1,192114537 | 0,79105 | 0,90292452 | novel transcript |
| ENSG00000261063 | AC009139,2 | 0,253467219 | 1,192068565 | 0,71772 | 0,86483656 | novel transcript, antisense to MON1B & SYCE1L |

| | | | | | | |
|---|---|---|---|---|---|---|
| ENSG00000229300 | SNHG32 | 0,253411301 | 1,192022362 | 0,42093 | 0,67616334 | small nucleolar RNA host gene 32 |
| ENSG00000164306 | PRIMPOL | 0,253347523 | 1,191969667 | 0,2549 | 0,5594539 | primase and DNA directed polymerase |
| ENSG00000144635 | DYNC1LI1 | 0,253299436 | 1,191929937 | 0,28324 | 0,58053323 | dynein cytoplasmic 1 light intermediate chain 1 |
| ENSG00000066379 | POLR1H | 0,25323031 | 1,191872829 | 0,5697 | 0,77690984 | RNA polymerase I subunit H |
| ENSG00000181016 | LSMEM1 | 0,253165673 | 1,191819431 | 0,72759 | 0,87031567 | leucine rich single-pass membrane protein 1 |
| ENSG00000144579 | CTDSP1 | 0,253147335 | 1,191804281 | 0,22759 | 0,54168591 | CTD small phosphatase 1 |
| ENSG00000091656 | ZFHX4 | 0,253090131 | 1,191757026 | 0,5176 | 0,74271521 | zinc finger homeobox 4 |
| ENSG00000198663 | C6orf89 | 0,253070851 | 1,1917411 | 0,39791 | 0,66050333 | chromosome 6 open reading frame 89 |
| ENSG00000156467 | UQCRB | 0,252992648 | 1,191676501 | 0,39356 | 0,65651919 | ubiquinol-cytochrome c reductase binding protein |
| ENSG00000175040 | CHST2 | 0,252966546 | 1,191654942 | 0,55884 | 0,77025265 | carbohydrate sulfotransferase 2 |
| ENSG00000283022 | HYDIN | 0,252957532 | 1,191647496 | 0,45253 | 0,69702004 | HYDIN axonemal central pair apparatus protein |
| ENSG00000157423 | HYDIN | 0,252957532 | 1,191647496 | 0,45253 | 0,69702004 | HYDIN axonemal central pair apparatus protein |
| ENSG00000270019 | AC110769,2 | 0,252796517 | 1,191514507 | 0,3441 | 0,62149652 | novel transcript |
| ENSG00000285639 | AL133259,1 | 0,252750042 | 1,191476125 | 0,75859 | 0,88727479 | novel transcript |
| ENSG00000214706 | IFRD2 | 0,252678251 | 1,191416836 | 0,33533 | 0,61497665 | interferon related developmental regulator 2 |
| ENSG00000242737 | AC012170,1 | 0,252565225 | 1,1913235 | 0,757 | 0,88651662 | ribosomal protein S20 (RPS20) pseudogene |
| ENSG00000227761 | BAG6 | 0,252454386 | 1,191231976 | 0,37888 | 0,64679625 | BAG cochaperone 6 |
| ENSG00000250508 | LINC02701 | 0,252436206 | 1,191216965 | 0,62133 | 0,80787022 | long intergenic non-protein coding RNA 2701 |
| ENSG00000143815 | LBR | 0,252424577 | 1,191207363 | 0,38586 | 0,65129675 | lamin B receptor |
| ENSG00000172733 | PURG | 0,252306222 | 1,191109644 | 0,35569 | 0,63008438 | purine rich element binding protein G |
| ENSG00000280858 | RPL7A | 0,252240061 | 1,191055021 | 0,21366 | 0,53275614 | ribosomal protein L7a |
| ENSG00000148303 | RPL7A | 0,252240061 | 1,191055021 | 0,21366 | 0,53275614 | ribosomal protein L7a |
| ENSG00000179902 | C1orf194 | 0,252238997 | 1,191054143 | 0,61586 | 0,80469924 | chromosome 1 open reading frame 194 |
| ENSG00000104897 | SF3A2 | 0,252217719 | 1,191036577 | 0,24329 | 0,55199251 | splicing factor 3a subunit 2 |
| ENSG00000101654 | RNMT | 0,252202979 | 1,191024408 | 0,18005 | 0,50407792 | RNA guanine-7 methyltransferase |
| ENSG00000258890 | CEP95 | 0,252180556 | 1,191005897 | 0,27086 | 0,57239492 | centrosomal protein 95 |
| ENSG00000149089 | APIP | 0,252180298 | 1,191005684 | 0,36735 | 0,63733285 | APAF1 interacting protein |
| ENSG00000131732 | ZCCHC9 | 0,252165989 | 1,190993871 | 0,12796 | 0,45145929 | zinc finger CCHC-type containing 9 |
| ENSG00000119203 | CPSF3 | 0,252049557 | 1,190897757 | 0,19447 | 0,51562158 | cleavage and polyadenylation specific factor 3 |
| ENSG00000119640 | ACYP1 | 0,252029913 | 1,190881541 | 0,63072 | 0,81354995 | acylphosphatase 1 |
| ENSG00000124181 | PLCG1 | 0,25196423 | 1,190827324 | 0,22563 | 0,54087803 | phospholipase C gamma 1 |
| ENSG00000269925 | Z98884,2 | 0,251824221 | 1,190711764 | 0,59218 | 0,79101863 | novel transcript, sense intronic to VAMP3 |

| | | | | | | |
|---|---|---|---|---|---|---|
| ENSG00000265967 | AC100830,3 | 0,251821493 | 1,190709512 | 0,84129 | 0,92775849 | novel transcript, antisense to ornithine decarboxylase antizyme 2 OAZ2 |
| ENSG00000287302 | AL133477,2 | 0,251705079 | 1,190613435 | 0,80848 | 0,91193281 | novel transcript |
| ENSG00000287244 | AL020997,5 | 0,251682314 | 1,190594647 | 0,73701 | 0,87539732 | novel transcript |
| ENSG00000006607 | FARP2 | 0,251658013 | 1,190574593 | 0,2285 | 0,5424199 | FERM, ARH/RhoGEF and pleckstrin domain protein 2 |
| ENSG00000108256 | NUFIP2 | 0,251657789 | 1,190574408 | 0,33703 | 0,61684812 | nuclear FMR1 interacting protein 2 |
| ENSG00000251303 | CAB39P1 | 0,251656325 | 1,1905732 | 0,7093 | 0,85982913 | calcium binding protein 39 pseudogene 1 |
| ENSG00000197961 | ZNF121 | 0,251577236 | 1,190507934 | 0,15189 | 0,48031011 | zinc finger protein 121 |
| ENSG00000132256 | TRIM5 | 0,251465591 | 1,190415809 | 0,28411 | 0,58097626 | tripartite motif containing 5 |
| ENSG00000226950 | DANCR | 0,251442365 | 1,190396645 | 0,29771 | 0,59036937 | differentiation antagonizing non-protein coding RNA |
| ENSG00000250869 | AC087359,1 | 0,251427366 | 1,190384269 | 0,68927 | 0,84861676 | tigger transposable element derived 1 (TIGD1) pseudogene |
| ENSG00000176871 | WSB2 | 0,251367164 | 1,190334596 | 0,40208 | 0,66346196 | WD repeat and SOCS box containing 2 |
| ENSG00000012048 | BRCA1 | 0,251331936 | 1,190305531 | 0,39543 | 0,6582534 | BRCA1 DNA repair associated |
| ENSG00000120265 | PCMT1 | 0,251319428 | 1,190295211 | 0,44979 | 0,69538334 | protein-L-isoaspartate (D-aspartate) O-methyltransferase |
| ENSG00000010270 | STARD3NL | 0,251319376 | 1,190295168 | 0,48537 | 0,72097949 | STARD3 N-terminal like |
| ENSG00000132622 | HSPA12B | 0,251273439 | 1,190257268 | 0,4704 | 0,70985356 | heat shock protein family A (Hsp70) member 12B |
| ENSG00000259170 | AC090985,2 | 0,25122709 | 1,19021903 | 0,74231 | 0,87815402 | novel transcript, antisense to ST8SIA2 |
| ENSG00000288353 | ARHGAP17 | 0,251199215 | 1,190196034 | 0,15716 | 0,48443207 | Rho GTPase activating protein 17 |
| ENSG00000186020 | ZNF529 | 0,251183421 | 1,190183004 | 0,19398 | 0,51551719 | zinc finger protein 529 |
| ENSG00000132109 | TRIM21 | 0,251058909 | 1,190080289 | 0,40923 | 0,66785537 | tripartite motif containing 21 |
| ENSG00000254701 | AC138866,1 | 0,251024094 | 1,190051571 | 0,33928 | 0,61831023 | glucuronidase, beta (GUSB) pseudogene |
| ENSG00000133858 | ZFC3H1 | 0,25097464 | 1,190010778 | 0,35227 | 0,62740642 | zinc finger C3H1-type containing |
| ENSG00000117758 | STX12 | 0,250907501 | 1,189955399 | 0,50531 | 0,73393676 | syntaxin 12 |
| ENSG00000213639 | PPP1CB | 0,250865326 | 1,189920613 | 0,49055 | 0,72473187 | protein phosphatase 1 catalytic subunit beta |
| ENSG00000110367 | DDX6 | 0,250848741 | 1,189906934 | 0,30814 | 0,59696396 | DEAD-box helicase 6 |
| ENSG00000164591 | MYOZ3 | 0,250841725 | 1,189901147 | 0,51798 | 0,7429275 | myozenin 3 |
| ENSG00000100906 | NFKBIA | 0,250789823 | 1,189858341 | 0,30968 | 0,59807209 | NFKB inhibitor alpha |
| ENSG00000197409 | H3C4 | 0,250780491 | 1,189850644 | 0,40454 | 0,66449258 | H3 clustered histone 4 |
| ENSG00000103353 | UBFD1 | 0,250572945 | 1,189679485 | 0,17538 | 0,49944679 | ubiquitin family domain containing 1 |
| ENSG00000205659 | LIN52 | 0,250559536 | 1,189668428 | 0,12382 | 0,4446254 | lin-52 DREAM MuvB core complex component |
| ENSG00000232995 | RGS5 | 0,250472621 | 1,189596759 | 0,57851 | 0,78223356 | regulator of G protein signaling 5 |
| ENSG00000107669 | ATE1 | 0,250460222 | 1,189586534 | 0,48584 | 0,72131258 | arginyltransferase 1 |

| | | | | | | |
|---|---|---|---|---|---|---|
| ENSG00000133794 | ARNTL | 0,250445554 | 1,18957444 | 0,51303 | 0,7394947 | aryl hydrocarbon receptor nuclear translocator like |
| ENSG00000213640 | EEF1DP4 | 0,250399379 | 1,189536367 | 0,71286 | 0,86184574 | eukaryotic translation elongation factor 1 delta pseudogene 4 |
| ENSG00000100346 | CACNA1I | 0,250373643 | 1,189515147 | 0,68549 | 0,84625716 | calcium voltage-gated channel subunit alpha1 I |
| ENSG00000150938 | CRIM1 | 0,250200986 | 1,189372799 | 0,22254 | 0,53866376 | cysteine rich transmembrane BMP regulator 1 |
| ENSG00000139697 | SBNO1 | 0,250176896 | 1,189352939 | 0,18413 | 0,50785017 | strawberry notch homolog 1 |
| ENSG00000187866 | FAM122A | 0,250158844 | 1,189338056 | 0,4474 | 0,69342177 | family with sequence similarity 122A |
| ENSG00000156603 | MED19 | 0,249950151 | 1,189166026 | 0,524 | 0,74789361 | mediator complex subunit 19 |
| ENSG00000266076 | AC004805,1 | 0,249921822 | 1,189142675 | 0,80792 | 0,91168773 | novel transcript |
| ENSG00000171790 | SLFNL1 | 0,249859745 | 1,189091509 | 0,26373 | 0,56629809 | schlafen like 1 |
| ENSG00000110011 | DNAJC4 | 0,249817195 | 1,189056439 | 0,28748 | 0,58386917 | DnaJ heat shock protein family (Hsp40) member C4 |
| ENSG00000158796 | DEDD | 0,249794681 | 1,189037883 | 0,37516 | 0,64350562 | death effector domain containing |
| ENSG00000007866 | TEAD3 | 0,249773044 | 1,189020051 | 0,18157 | 0,50490902 | TEA domain transcription factor 3 |
| ENSG00000100982 | PCIF1 | 0,249472576 | 1,188772441 | 0,31875 | 0,60383175 | PDX1 C-terminal inhibiting factor 1 |
| ENSG00000189057 | FAM111B | 0,249363088 | 1,188682227 | 0,61328 | 0,8035281 | family with sequence similarity 111 member B |
| ENSG00000172915 | NBEA | 0,249327706 | 1,188653075 | 0,42533 | 0,67893062 | neurobeachin |
| ENSG00000007384 | RHBDF1 | 0,249212646 | 1,188558279 | 0,39406 | 0,65702282 | rhomboid 5 homolog 1 |
| ENSG00000146021 | KLHL3 | 0,249191513 | 1,188540869 | 0,26532 | 0,56810437 | kelch like family member 3 |
| ENSG00000285139 | KF459542,2 | 0,249114547 | 1,188477464 | 0,80395 | 0,909789 | novel transcript, antisense to KBTBD4 |
| ENSG00000231880 | KF459542,1 | 0,249114547 | 1,188477464 | 0,80395 | 0,909789 | novel transcript, antisense to KBTBD4 |
| ENSG00000046651 | OFD1 | 0,249073833 | 1,188443925 | 0,45174 | 0,69675201 | OFD1 centriole and centriolar satellite protein |
| ENSG00000176761 | ZNF285B | 0,248987252 | 1,188372604 | 0,65004 | 0,8246264 | zinc finger protein 285B (pseudogene) |
| ENSG00000172071 | EIF2AK3 | 0,248958102 | 1,188348593 | 0,38389 | 0,65045825 | eukaryotic translation initiation factor 2 alpha kinase 3 |
| ENSG00000133195 | SLC39A11 | 0,248927052 | 1,188323018 | 0,23094 | 0,54392721 | solute carrier family 39 member 11 |
| ENSG00000106346 | USP42 | 0,248898471 | 1,188299476 | 0,25472 | 0,55944852 | ubiquitin specific peptidase 42 |
| ENSG00000188611 | ASAH2 | 0,248792907 | 1,18821253 | 0,35111 | 0,62663934 | N-acylsphingosine amidohydrolase 2 |
| ENSG00000228253 | MT-ATP8 | 0,248740379 | 1,188169268 | 0,4044 | 0,66435702 | mitochondrially encoded ATP synthase membrane subunit 8 |
| ENSG00000063046 | EIF4B | 0,248728383 | 1,188159388 | 0,13457 | 0,46080955 | eukaryotic translation initiation factor 4B |
| ENSG00000229036 | VDAC1P8 | 0,248632761 | 1,18808064 | 0,44573 | 0,69282012 | voltage dependent anion channel 1 pseudogene 8 |
| ENSG00000237080 | EHMT2-AS1 | 0,248601953 | 1,188055269 | 0,52524 | 0,74895722 | EHMT2 and SLC44A4 antisense RNA 1 |
| ENSG00000121417 | ZNF211 | 0,248583813 | 1,188040331 | 0,38027 | 0,64794085 | zinc finger protein 211 |
| ENSG00000171603 | CLSTN1 | 0,248482931 | 1,187957259 | 0,19133 | 0,51435195 | calsyntenin 1 |

| | | | | | | |
|---|---|---|---|---|---|---|
| ENSG00000197406 | DIO3 | 0,248384925 | 1,187876561 | 0,69152 | 0,849672 | iodothyronine deiodinase 3 |
| ENSG00000085788 | DDHD2 | 0,248322485 | 1,18782515 | 0,3172 | 0,6027392 | DDHD domain containing 2 |
| ENSG00000112759 | SLC29A1 | 0,248273831 | 1,187785092 | 0,3548 | 0,62936991 | solute carrier family 29 member 1 (Augustine blood group) |
| ENSG00000124172 | ATP5F1E | 0,248024072 | 1,187579481 | 0,51899 | 0,7437068 | ATP synthase F1 subunit epsilon |
| ENSG00000108599 | AKAP10 | 0,248020713 | 1,187576716 | 0,33289 | 0,61286447 | A-kinase anchoring protein 10 |
| ENSG00000126003 | PLAGL2 | 0,248018741 | 1,187575093 | 0,16334 | 0,48976376 | PLAG1 like zinc finger 2 |
| ENSG00000076513 | ANKRD13A | 0,247963617 | 1,187529718 | 0,21981 | 0,53722186 | ankyrin repeat domain 13A |
| ENSG00000162769 | FLVCR1 | 0,247817131 | 1,187409146 | 0,13768 | 0,46480929 | FLVCR heme transporter 1 |
| ENSG00000233776 | LINC01251 | 0,247793907 | 1,187390032 | 0,70896 | 0,85977658 | long intergenic non-protein coding RNA 1251 |
| ENSG00000284773 | AC114490,3 | 0,247784034 | 1,187381906 | 0,48654 | 0,72196345 | novel protein |
| ENSG00000237990 | CNTN4-AS1 | 0,247736525 | 1,187342806 | 0,78961 | 0,90258817 | CNTN4 antisense RNA 1 |
| ENSG00000147145 | LPAR4 | 0,247616934 | 1,187244385 | 0,52062 | 0,74515898 | lysophosphatidic acid receptor 4 |
| ENSG00000212907 | MT-ND4L | 0,247570847 | 1,18720646 | 0,45841 | 0,70119611 | mitochondrially encoded NADH:ubiquinone oxidoreductase core subunit 4L |
| ENSG00000153406 | NMRAL1 | 0,247523791 | 1,187167738 | 0,34717 | 0,62397921 | NmrA like redox sensor 1 |
| ENSG00000233073 | GRM3-AS1 | 0,247426607 | 1,187087769 | 0,67762 | 0,84135792 | GRM3 antisense RNA 1 |
| ENSG00000141446 | ESCO1 | 0,2474074 | 1,187071966 | 0,3045 | 0,59430086 | establishment of sister chromatid cohesion N-acetyltransferase 1 |
| ENSG00000166262 | FAM227B | 0,247377569 | 1,187047421 | 0,49987 | 0,73058737 | family with sequence similarity 227 member B |
| ENSG00000187987 | ZSCAN23 | 0,24736317 | 1,187035573 | 0,50866 | 0,73679246 | zinc finger and SCAN domain containing 23 |
| ENSG00000131174 | COX7B | 0,247301609 | 1,186984922 | 0,40345 | 0,66399363 | cytochrome c oxidase subunit 7B |
| ENSG00000214198 | TTC41P | 0,247301187 | 1,186984575 | 0,5154 | 0,74146765 | tetratricopeptide repeat domain 41, pseudogene |
| ENSG00000132471 | WBP2 | 0,24729458 | 1,186979139 | 0,21079 | 0,52944479 | WW domain binding protein 2 |
| ENSG00000203761 | MSTO2P | 0,247288895 | 1,186974461 | 0,52753 | 0,75037219 | misato family member 2, pseudogene |
| ENSG00000121774 | KHDRBS1 | 0,247198312 | 1,186899937 | 0,28283 | 0,58009943 | KH RNA binding domain containing, signal transduction associated 1 |
| ENSG00000035499 | DEPDC1B | 0,247111316 | 1,186828368 | 0,50169 | 0,73146914 | DEP domain containing 1B |
| ENSG00000114770 | ABCC5 | 0,246961706 | 1,186705298 | 0,3387 | 0,61789765 | ATP binding cassette subfamily C member 5 |
| ENSG00000215114 | UBXN2B | 0,246908128 | 1,186661228 | 0,52399 | 0,74789361 | UBX domain protein 2B |
| ENSG00000281763 | DHX36 | 0,246894206 | 1,186649777 | 0,53218 | 0,75287063 | DEAH-box helicase 36 |
| ENSG00000170540 | ARL6IP1 | 0,246803317 | 1,18657502 | 0,42163 | 0,67643128 | ADP ribosylation factor like GTPase 6 interacting protein 1 |
| ENSG00000159202 | UBE2Z | 0,246800718 | 1,186572883 | 0,07628 | 0,36242671 | ubiquitin conjugating enzyme E2 Z |
| ENSG00000100883 | SRP54 | 0,24679331 | 1,18656679 | 0,23018 | 0,54335232 | signal recognition particle 54 |

| | | | | | | |
|---|---|---|---|---|---|---|
| ENSG00000260092 | AC009163,3 | 0,246722285 | 1,186508376 | 0,7188 | 0,86552951 | TMEM231-CHST5 readthrough protein |
| ENSG00000265496 | AC100778,3 | 0,246603736 | 1,186410882 | 0,80015 | 0,90795227 | microRNA 1539 |
| ENSG00000075131 | TIPIN | 0,246583844 | 1,186394524 | 0,19554 | 0,51643293 | TIMELESS interacting protein |
| ENSG00000105402 | NAPA | 0,24654933 | 1,186366142 | 0,32215 | 0,60577115 | NSF attachment protein alpha |
| ENSG00000240857 | RDH14 | 0,246513611 | 1,18633677 | 0,3497 | 0,62563749 | retinol dehydrogenase 14 |
| ENSG00000174437 | ATP2A2 | 0,246422261 | 1,186261655 | 0,16247 | 0,48945635 | ATPase sarcoplasmic/endoplasmic reticulum Ca2+ transporting 2 |
| ENSG00000164603 | BMT2 | 0,24640118 | 1,186244321 | 0,55794 | 0,76961995 | base methyltransferase of 25S rRNA 2 homolog |
| ENSG00000104447 | TRPS1 | 0,246351585 | 1,186203542 | 0,54095 | 0,75863636 | transcriptional repressor GATA binding 1 |
| ENSG00000205111 | CDKL4 | 0,246268205 | 1,186134988 | 0,77187 | 0,89437474 | cyclin dependent kinase like 4 |
| ENSG00000165215 | CLDN3 | 0,246224805 | 1,186099307 | 0,68777 | 0,84787163 | claudin 3 |
| ENSG00000099942 | CRKL | 0,246222548 | 1,186097451 | 0,18167 | 0,50497494 | CRK like proto-oncogene, adaptor protein |
| ENSG00000106070 | GRB10 | 0,245901351 | 1,185833411 | 0,43386 | 0,68477203 | growth factor receptor bound protein 10 |
| ENSG00000083123 | BCKDHB | 0,245852468 | 1,185793233 | 0,2392 | 0,5491566 | branched chain keto acid dehydrogenase E1 subunit beta |
| ENSG00000070010 | UFD1 | 0,2457347 | 1,185696439 | 0,17334 | 0,4980379 | ubiquitin recognition factor in ER associated degradation 1 |
| ENSG00000153006 | SREK1IP1 | 0,245695036 | 1,185663842 | 0,452 | 0,69691985 | SREK1 interacting protein 1 |
| ENSG00000103876 | FAH | 0,245684698 | 1,185655345 | 0,40203 | 0,66341002 | fumarylacetoacetate hydrolase |
| ENSG00000171723 | GPHN | 0,245630732 | 1,185610995 | 0,4918 | 0,72530727 | gephyrin |
| ENSG00000228753 | EIF4BP2 | 0,245605557 | 1,185590306 | 0,81013 | 0,91247714 | eukaryotic translation initiation factor 4B pseudogene 2 |
| ENSG00000169021 | UQCRFS1 | 0,245548474 | 1,185543397 | 0,3366 | 0,61666883 | ubiquinol-cytochrome c reductase, Rieske iron-sulfur polypeptide 1 |
| ENSG00000225449 | RAB6C-AS1 | 0,245545461 | 1,185540921 | 0,80737 | 0,91131389 | RAB6C antisense RNA 1 |
| ENSG00000198205 | ZXDA | 0,245534168 | 1,185531641 | 0,42754 | 0,68046596 | zinc finger X-linked duplicated A |
| ENSG00000163879 | DNALI1 | 0,24553054 | 1,18552866 | 0,44178 | 0,68993396 | dynein axonemal light intermediate chain 1 |
| ENSG00000236187 | GJA6P | 0,245494574 | 1,185499106 | 0,77523 | 0,89624737 | gap junction protein alpha 6, pseudogene |
| ENSG00000274677 | AC040169,3 | 0,245442505 | 1,18545632 | 0,67635 | 0,84075837 | novel transcript, sense intronic to MBTPS1 |
| ENSG00000268442 | AC073534,1 | 0,245432303 | 1,185447936 | 0,58729 | 0,78721215 | hepatitis A virus cellular receptor 1 (HAVCR1) pseudogene |
| ENSG00000104723 | TUSC3 | 0,245389833 | 1,18541304 | 0,43691 | 0,68701656 | tumor suppressor candidate 3 |
| ENSG00000278488 | NAPRT | 0,245349533 | 1,185379927 | 0,6361 | 0,81694486 | nicotinate phosphoribosyltransferase |
| ENSG00000155957 | TMBIM4 | 0,245272489 | 1,185316626 | 0,34633 | 0,62342969 | transmembrane BAX inhibitor motif containing 4 |
| ENSG00000154710 | RABGEF1 | 0,245261338 | 1,185307465 | 0,45931 | 0,7017037 | RAB guanine nucleotide exchange factor 1 |
| ENSG00000061273 | HDAC7 | 0,24521536 | 1,18526969 | 0,26093 | 0,56405761 | histone deacetylase 7 |
| ENSG00000230433 | AL031770,1 | 0,245205085 | 1,185261249 | 0,71255 | 0,86169874 | novel transcript |

| | | | | | | |
|---|---|---|---|---|---|---|
| ENSG00000107186 | MPDZ | 0,245140483 | 1,185208175 | 0,49727 | 0,72868655 | multiple PDZ domain crumbs cell polarity complex component |
| ENSG00000089006 | SNX5 | 0,245054663 | 1,185137674 | 0,2604 | 0,56383537 | sorting nexin 5 |
| ENSG00000274336 | CDIP1 | 0,245052864 | 1,185136196 | 0,45086 | 0,69594229 | cell death inducing p53 target 1 |
| ENSG00000099194 | SCD | 0,245034347 | 1,185120986 | 0,41908 | 0,67510996 | stearoyl-CoA desaturase |
| ENSG00000259100 | COILP1 | 0,245017226 | 1,185106921 | 0,66742 | 0,83493154 | coilin pseudogene 1 |
| ENSG00000198843 | SELENOT | 0,244998705 | 1,185091707 | 0,57569 | 0,78042244 | selenoprotein T |
| ENSG00000179008 | C14orf39 | 0,244975507 | 1,185072652 | 0,6261 | 0,81067751 | chromosome 14 open reading frame 39 |
| ENSG00000282834 | AP000787,3 | 0,244867611 | 1,184984026 | 0,76366 | 0,88986704 | ribosomal protein S16 (RPS16) pseudogene |
| ENSG00000164161 | HHIP | 0,244861552 | 1,184979049 | 0,60299 | 0,7971742 | hedgehog interacting protein |
| ENSG00000197451 | HNRNPAB | 0,244834067 | 1,184956474 | 0,20571 | 0,52518691 | heterogeneous nuclear ribonucleoprotein A/B |
| ENSG00000162543 | UBXN10 | 0,244808205 | 1,184935233 | 0,53655 | 0,75531624 | UBX domain protein 10 |
| ENSG00000066230 | SLC9A3 | 0,244789088 | 1,184919531 | 0,42297 | 0,67690977 | solute carrier family 9 member A3 |
| ENSG00000050344 | NFE2L3 | 0,244713619 | 1,184857549 | 0,44064 | 0,68910892 | nuclear factor, erythroid 2 like 3 |
| ENSG00000213757 | RPS3AP20 | 0,244646055 | 1,184802061 | 0,65404 | 0,82680983 | RPS3A pseudogene 20 |
| ENSG00000184708 | EIF4ENIF1 | 0,244469557 | 1,184657122 | 0,26644 | 0,56917012 | eukaryotic translation initiation factor 4E nuclear import factor 1 |
| ENSG00000258525 | AL049830,3 | 0,244422282 | 1,184618303 | 0,46902 | 0,70862072 | novel transcript, antisense to COCH |
| ENSG00000140262 | TCF12 | 0,244309984 | 1,184526097 | 0,59042 | 0,78974912 | transcription factor 12 |
| ENSG00000273820 | USP27X | 0,244264631 | 1,184488861 | 0,33754 | 0,61721828 | ubiquitin specific peptidase 27 X-linked |
| ENSG00000090686 | USP48 | 0,244234137 | 1,184463825 | 0,32977 | 0,61069811 | ubiquitin specific peptidase 48 |
| ENSG00000284792 | PTEN | 0,24409409 | 1,18434885 | 0,31531 | 0,60159235 | phosphatase and tensin homolog |
| ENSG00000272267 | AC021242,3 | 0,244090856 | 1,184346196 | 0,61628 | 0,80488598 | novel transcript, antisense to TNKS |
| ENSG00000236065 | AL020995,1 | 0,244086156 | 1,184342337 | 0,59998 | 0,79568184 | novel transcript |
| ENSG00000116977 | LGALS8 | 0,244043939 | 1,184307681 | 0,54221 | 0,75894841 | galectin 8 |
| ENSG00000288598 | AL354833,1 | 0,243563623 | 1,183913455 | 0,50441 | 0,7332113 | uncharacterized LOC105370174 |
| ENSG00000158985 | CDC42SE2 | 0,243401639 | 1,183780534 | 0,4928 | 0,72597062 | CDC42 small effector 2 |
| ENSG00000175893 | ZDHHC21 | 0,243326909 | 1,183719217 | 0,5165 | 0,74198903 | zinc finger DHHC-type palmitoyltransferase 21 |
| ENSG00000234160 | AL513165,1 | 0,243321638 | 1,183714893 | 0,67757 | 0,84135792 | novel transcript |
| ENSG00000268731 | AC010615,3 | 0,24331303 | 1,18370783 | 0,77406 | 0,89572556 | iron-sulfur cluster assembly 1 homolog (S, cerevisiae) (ISCA1) pseudogene |
| ENSG00000141642 | ELAC1 | 0,24330721 | 1,183703055 | 0,34663 | 0,62350287 | elaC ribonuclease Z 1 |
| ENSG00000073464 | CLCN4 | 0,243298469 | 1,183695883 | 0,30197 | 0,59294633 | chloride voltage-gated channel 4 |

| | | | | | | |
|---|---|---|---|---|---|---|
| ENSG00000174405 | LIG4 | 0,243288215 | 1,18368747 | 0,49171 | 0,72526787 | DNA ligase 4 |
| ENSG00000214018 | RRM2P3 | 0,2432778 | 1,183678925 | 0,70661 | 0,85885395 | ribonucleotide reductase M2 polypeptide pseudogene 3 |
| ENSG00000224287 | MSL3P1 | 0,242990092 | 1,183442894 | 0,60186 | 0,79680798 | MSL complex subunit 3 pseudogene 1 |
| ENSG00000178694 | NSUN3 | 0,242977586 | 1,183432635 | 0,26151 | 0,56452671 | NOP2/Sun RNA methyltransferase 3 |
| ENSG00000275428 | AC024940,6 | 0,242902662 | 1,183371178 | 0,64233 | 0,82066205 | ovostatin 2 |
| ENSG00000155506 | LARP1 | 0,242755659 | 1,183250605 | 0,24979 | 0,55657567 | La ribonucleoprotein 1, translational regulator |
| ENSG00000143549 | TPM3 | 0,242729869 | 1,183229452 | 0,21516 | 0,53387837 | tropomyosin 3 |
| ENSG00000100014 | SPECC1L | 0,242541304 | 1,18307481 | 0,35867 | 0,63185282 | sperm antigen with calponin homology and coiled-coil domains 1 like |
| ENSG00000136636 | KCTD3 | 0,242487478 | 1,183030672 | 0,11503 | 0,43048326 | potassium channel tetramerization domain containing 3 |
| ENSG00000272741 | AC069257,3 | 0,24241496 | 1,182971207 | 0,75817 | 0,88722916 | novel protein |
| ENSG00000113716 | HMGXB3 | 0,242408644 | 1,182966028 | 0,13098 | 0,45558936 | HMG-box containing 3 |
| ENSG00000173588 | CEP83 | 0,242372665 | 1,182936527 | 0,27748 | 0,57588377 | centrosomal protein 83 |
| ENSG00000244187 | TMEM141 | 0,242329481 | 1,182901119 | 0,19862 | 0,51980353 | transmembrane protein 141 |
| ENSG00000261701 | HPR | 0,242262504 | 1,182846204 | 0,73822 | 0,87571125 | haptoglobin-related protein |
| ENSG00000259075 | POC1B-GALNT | 0,242228099 | 1,182817996 | 0,46351 | 0,70434395 | POC1B-GALNT4 readthrough |
| ENSG00000180210 | F2 | 0,242175497 | 1,182774871 | 0,7697 | 0,89301171 | coagulation factor II, thrombin |
| ENSG00000157657 | ZNF618 | 0,242069981 | 1,182688367 | 0,22149 | 0,53805234 | zinc finger protein 618 |
| ENSG00000156097 | GPR61 | 0,242036645 | 1,18266104 | 0,54088 | 0,75863464 | G protein-coupled receptor 61 |
| ENSG00000267515 | AP001029,3 | 0,241971776 | 1,182607864 | 0,70407 | 0,85725494 | high mobility group box 3 (HMGB3) pseudogene |
| ENSG00000122687 | MRM2 | 0,241958745 | 1,182597183 | 0,17535 | 0,49944679 | mitochondrial rRNA methyltransferase 2 |
| ENSG00000132478 | UNK | 0,241938332 | 1,18258045 | 0,27878 | 0,57698038 | unk zinc finger |
| ENSG00000188641 | DPYD | 0,241907862 | 1,182555474 | 0,3529 | 0,6277271 | dihydropyrimidine dehydrogenase |
| ENSG00000139180 | NDUFA9 | 0,241904642 | 1,182552834 | 0,14031 | 0,46718058 | NADH:ubiquinone oxidoreductase subunit A9 |
| ENSG00000249855 | EEF1A1P19 | 0,241868405 | 1,182523132 | 0,52121 | 0,74558236 | eukaryotic translation elongation factor 1 alpha 1 pseudogene 19 |
| ENSG00000006757 | PNPLA4 | 0,24181609 | 1,182480252 | 0,54213 | 0,75894841 | patatin like phospholipase domain containing 4 |
| ENSG00000232739 | AL607028,1 | 0,241722819 | 1,182403806 | 0,66478 | 0,83304443 | novel transcript |
| ENSG00000271862 | AC104118,1 | 0,24166756 | 1,182358518 | 0,543 | 0,75965603 | novel transcript |
| ENSG00000273707 | CDKN1C | 0,241639238 | 1,182335307 | 0,42286 | 0,67683766 | cyclin dependent kinase inhibitor 1C |
| ENSG00000248685 | LINC02484 | 0,241609352 | 1,182310815 | 0,31561 | 0,60172912 | long intergenic non-protein coding RNA 2484 |
| ENSG00000180574 | EIF2S3B | 0,24145817 | 1,182186925 | 0,43536 | 0,68601186 | eukaryotic translation initiation factor 2 subunit gamma B |
| ENSG00000238083 | LRRC37A2 | 0,241381788 | 1,182124337 | 0,31443 | 0,6011918 | leucine rich repeat containing 37 member A2 |

| | | | | | | |
|---|---|---|---|---|---|---|
| ENSG00000278673 | AC136352,5 | 0,241323583 | 1,182076646 | 0,80732 | 0,91131389 | |
| ENSG00000115816 | CEBPZ | 0,24128709 | 1,182046745 | 0,27912 | 0,57725738 | CCAAT enhancer binding protein zeta |
| ENSG00000272986 | AC009570,1 | 0,241270664 | 1,182033288 | 0,66879 | 0,83569335 | novel transcript, antisense to RUFY3 |
| ENSG00000165502 | RPL36AL | 0,241234595 | 1,182003735 | 0,43052 | 0,68295004 | ribosomal protein L36a like |
| ENSG00000269951 | AC090181,2 | 0,241173516 | 1,181953694 | 0,43605 | 0,6865394 | novel transcript, sense intronic to TSPAN3 |
| ENSG00000197934 | CYYR1-AS1 | 0,24109946 | 1,181893024 | 0,54771 | 0,76306829 | CYYR1 antisense RNA 1 |
| ENSG00000134744 | TUT4 | 0,241084277 | 1,181880586 | 0,29145 | 0,58649877 | terminal uridylyl transferase 4 |
| ENSG00000106462 | EZH2 | 0,241037222 | 1,181842038 | 0,38068 | 0,64823548 | enhancer of zeste 2 polycomb repressive complex 2 subunit |
| ENSG00000213619 | NDUFS3 | 0,241034928 | 1,181840159 | 0,30017 | 0,59160746 | NADH:ubiquinone oxidoreductase core subunit S3 |
| ENSG00000100225 | FBXO7 | 0,241030991 | 1,181836934 | 0,19529 | 0,51611615 | F-box protein 7 |
| ENSG00000276482 | SLC12A7 | 0,240913553 | 1,181740735 | 0,29834 | 0,59076323 | solute carrier family 12 member 7 |
| ENSG00000175324 | LSM1 | 0,240679063 | 1,181548674 | 0,50568 | 0,73418581 | LSM1 homolog, mRNA degradation associated |
| ENSG00000231969 | MMADHC-DT | 0,240646323 | 1,181521861 | 0,63058 | 0,81354995 | MMADHC divergent transcript |
| ENSG00000177084 | POLE | 0,240581186 | 1,181468517 | 0,19891 | 0,52005179 | DNA polymerase epsilon, catalytic subunit |
| ENSG00000183340 | JRKL | 0,240569146 | 1,181458657 | 0,49037 | 0,72458682 | JRK like |
| ENSG00000228748 | AL450306,1 | 0,240559205 | 1,181450516 | 0,52601 | 0,74968838 | novel transcript |
| ENSG00000104427 | ZC2HC1A | 0,240532231 | 1,181428427 | 0,64985 | 0,82453893 | zinc finger C2HC-type containing 1A |
| ENSG00000084092 | NOA1 | 0,240460084 | 1,181369347 | 0,41886 | 0,67490937 | nitric oxide associated 1 |
| ENSG00000239917 | RPS10P16 | 0,2403271 | 1,181260457 | 0,7812 | 0,898729 | ribosomal protein S10 pseudogene 16 |
| ENSG00000099250 | NRP1 | 0,240324251 | 1,181258124 | 0,42598 | 0,67940091 | neuropilin 1 |
| ENSG00000135205 | CCDC146 | 0,240265364 | 1,181209909 | 0,40495 | 0,66481282 | coiled-coil domain containing 146 |
| ENSG00000259985 | AC017100,1 | 0,240181282 | 1,181141069 | 0,58699 | 0,78704355 | novel transcript, antisense to B4GALT6 |
| ENSG00000157869 | RAB28 | 0,24002914 | 1,181016516 | 0,28024 | 0,57784691 | RAB28, member RAS oncogene family |
| ENSG00000196787 | H2AC11 | 0,239902896 | 1,180913175 | 0,52147 | 0,74580669 | H2A clustered histone 11 |
| ENSG00000182185 | RAD51B | 0,239897 | 1,180908349 | 0,38694 | 0,65213944 | RAD51 paralog B |
| ENSG00000270130 | AC068790,7 | 0,239798433 | 1,18082767 | 0,69096 | 0,84920479 | novel transcript, sense intronic to CCDC92 |
| ENSG00000273355 | AP000894,4 | 0,239791544 | 1,180822031 | 0,72496 | 0,86868749 | novel transcript |
| ENSG00000109220 | CHIC2 | 0,239727139 | 1,180769318 | 0,36433 | 0,6349692 | cysteine rich hydrophobic domain 2 |
| ENSG00000054392 | HHAT | 0,239683999 | 1,180734011 | 0,38576 | 0,65123178 | hedgehog acyltransferase |
| ENSG00000280680 | HHAT | 0,239683999 | 1,180734011 | 0,38576 | 0,65123178 | hedgehog acyltransferase |
| ENSG00000142599 | RERE | 0,239590222 | 1,180657264 | 0,15069 | 0,47887343 | arginine-glutamic acid dipeptide repeats |
| ENSG00000286162 | AL162253,2 | 0,239449724 | 1,18054229 | 0,53467 | 0,7541464 | novel transcript |

| | | | | | | |
|---|---|---|---|---|---|---|
| ENSG00000185565 | LSAMP | 0,23940379 | 1,180504704 | 0,57066 | 0,77748547 | limbic system associated membrane protein |
| ENSG00000213385 | AC105052,2 | 0,23933808 | 1,180450936 | 0,74407 | 0,87882457 | ribosomal protein L7a (RPL7A) pseudogene |
| ENSG00000186153 | WWOX | 0,239206103 | 1,180342955 | 0,29478 | 0,58867017 | WW domain containing oxidoreductase |
| ENSG00000163463 | KRTCAP2 | 0,239136652 | 1,180286134 | 0,18029 | 0,50407792 | keratinocyte associated protein 2 |
| ENSG00000272379 | AL008729,2 | 0,239063493 | 1,180226284 | 0,59032 | 0,78972365 | novel transcript |
| ENSG00000287373 | AL122014,1 | 0,239055043 | 1,180219371 | 0,79345 | 0,90390791 | novel transcript, antisense to DCDC1 |
| ENSG00000278372 | MYO19 | 0,239053698 | 1,180218271 | 0,42947 | 0,68224982 | myosin XIX |
| ENSG00000047249 | ATP6V1H | 0,239042375 | 1,180209008 | 0,32179 | 0,60577115 | ATPase H+ transporting V1 subunit H |
| ENSG00000279094 | LINC01670 | 0,238983292 | 1,180160676 | 0,52741 | 0,75037219 | long intergenic non-protein coding RNA 1670 |
| ENSG00000111615 | KRR1 | 0,23891764 | 1,180106972 | 0,3721 | 0,64114037 | KRR1 small subunit processome component homolog |
| ENSG00000147588 | PMP2 | 0,238916051 | 1,180105672 | 0,80674 | 0,91120963 | peripheral myelin protein 2 |
| ENSG00000284832 | POLR2A | 0,238813353 | 1,180021669 | 0,13414 | 0,46034978 | RNA polymerase II subunit A |
| ENSG00000047188 | YTHDC2 | 0,238722927 | 1,17994771 | 0,33626 | 0,61623735 | YTH domain containing 2 |
| ENSG00000280164 | CU638689,5 | 0,238705011 | 1,179933057 | 0,47385 | 0,71294959 | novel transcript |
| ENSG00000259081 | AF111169,3 | 0,238608969 | 1,17985451 | 0,61245 | 0,8035281 | novel transcript |
| ENSG00000103365 | GGA2 | 0,238598684 | 1,179846099 | 0,22517 | 0,54058261 | golgi associated, gamma adaptin ear containing, ARF binding protein 2 |
| ENSG00000283765 | AC131160,1 | 0,238553528 | 1,17980917 | 0,53306 | 0,75338858 | presenilin associated rhomboid like |
| ENSG00000100227 | POLDIP3 | 0,238367438 | 1,179656999 | 0,14089 | 0,46791293 | DNA polymerase delta interacting protein 3 |
| ENSG00000264207 | AC239868,1 | 0,23835716 | 1,179648595 | 0,36835 | 0,63803074 | novel transcript, antisense HIST2H4A |
| ENSG00000257698 | GIHCG | 0,238182159 | 1,179505511 | 0,55507 | 0,76775076 | GIHCG inhibitor of miR-200b/200a/429 expression |
| ENSG00000214866 | DCDC2C | 0,238137787 | 1,179469234 | 0,73432 | 0,87430159 | doublecortin domain containing 2C |
| ENSG00000114850 | SSR3 | 0,238133236 | 1,179465513 | 0,107 | 0,41677702 | signal sequence receptor subunit 3 |
| ENSG00000230869 | AGAP10P | 0,238103242 | 1,179440992 | 0,80592 | 0,91077382 | ArfGAP with GTPase domain, ankyrin repeat and PH domain 10, pseudogene |
| ENSG00000132661 | NXT1 | 0,237984367 | 1,179343813 | 0,25553 | 0,56000169 | nuclear transport factor 2 like export factor 1 |
| ENSG00000197299 | BLM | 0,237921762 | 1,179292637 | 0,3092 | 0,59774312 | BLM RecQ like helicase |
| ENSG00000269226 | TMSB15B | 0,237913026 | 1,179285496 | 0,43275 | 0,68421938 | thymosin beta 15B |
| ENSG00000173320 | STOX2 | 0,237889338 | 1,179266133 | 0,46315 | 0,70415312 | storkhead box 2 |
| ENSG00000223584 | TVP23CP1 | 0,237862931 | 1,179244548 | 0,62595 | 0,81058216 | TVP23C pseudogene 1 |
| ENSG00000100600 | LGMN | 0,237862899 | 1,179244522 | 0,28486 | 0,5816767 | legumain |
| ENSG00000116141 | MARK1 | 0,23779942 | 1,179192636 | 0,42816 | 0,68094175 | microtubule affinity regulating kinase 1 |
| ENSG00000113569 | NUP155 | 0,237788051 | 1,179183344 | 0,14499 | 0,47263441 | nucleoporin 155 |

| | | | | | | |
|---|---|---|---|---|---|---|
| ENSG00000241945 | PWP2 | 0,237700444 | 1,17911174 | 0,14721 | 0,47471707 | PWP2 small subunit processome component |
| ENSG00000186231 | KLHL32 | 0,237675968 | 1,179091736 | 0,54775 | 0,76307321 | kelch like family member 32 |
| ENSG00000255150 | EID3 | 0,237585347 | 1,179017675 | 0,35486 | 0,62936991 | EP300 interacting inhibitor of differentiation 3 |
| ENSG00000064933 | PMS1 | 0,237537095 | 1,178978243 | 0,43126 | 0,68350322 | PMS1 homolog 1, mismatch repair system component |
| ENSG00000112305 | SMAP1 | 0,237355424 | 1,17882979 | 0,35841 | 0,63185282 | small ArfGAP 1 |
| ENSG00000187626 | ZKSCAN4 | 0,237337462 | 1,178815113 | 0,45398 | 0,69799269 | zinc finger with KRAB and SCAN domains 4 |
| ENSG00000078687 | TNRC6C | 0,23732268 | 1,178803035 | 0,32942 | 0,61044693 | trinucleotide repeat containing adaptor 6C |
| ENSG00000168710 | AHCYL1 | 0,237250496 | 1,178744056 | 0,40823 | 0,66714279 | adenosylhomocysteinase like 1 |
| ENSG00000198908 | BHLHB9 | 0,237244739 | 1,178739352 | 0,41999 | 0,67552612 | basic helix-loop-helix family member b9 |
| ENSG00000267203 | SNRPGP4 | 0,237241272 | 1,17873652 | 0,8129 | 0,91395885 | small nuclear ribonucleoprotein polypeptide G pseudogene 4 |
| ENSG00000170242 | USP47 | 0,23707877 | 1,178603757 | 0,44845 | 0,69432421 | ubiquitin specific peptidase 47 |
| ENSG00000126267 | COX6B1 | 0,237038699 | 1,178571022 | 0,36458 | 0,63504684 | cytochrome c oxidase subunit 6B1 |
| ENSG00000212866 | HSPA1B | 0,237023507 | 1,178558611 | 0,50505 | 0,73378796 | heat shock protein family A (Hsp70) member 1B |
| ENSG00000227230 | AL606534,1 | 0,236986059 | 1,178528019 | 0,6517 | 0,82550101 | novel transcript |
| ENSG00000139405 | RITA1 | 0,236949238 | 1,178497941 | 0,28166 | 0,57900853 | RBPJ interacting and tubulin associated 1 |
| ENSG00000183048 | SLC25A10 | 0,236821122 | 1,178393291 | 0,2469 | 0,55457136 | solute carrier family 25 member 10 |
| ENSG00000178750 | STX19 | 0,236807752 | 1,17838237 | 0,56368 | 0,77255679 | syntaxin 19 |
| ENSG00000096092 | TMEM14A | 0,23679002 | 1,178367888 | 0,43955 | 0,68843579 | transmembrane protein 14A |
| ENSG00000179889 | PDXDC1 | 0,236760649 | 1,178343898 | 0,48446 | 0,72028271 | pyridoxal dependent decarboxylase domain containing 1 |
| ENSG00000166900 | STX3 | 0,236568579 | 1,178187033 | 0,57278 | 0,77868876 | syntaxin 3 |
| ENSG00000155886 | SLC24A2 | 0,236469811 | 1,178106376 | 0,71856 | 0,86541453 | solute carrier family 24 member 2 |
| ENSG00000286071 | AL136171,2 | 0,236371386 | 1,178026004 | 0,55492 | 0,76763832 | disrupted in schizophrenia 2 |
| ENSG00000129007 | CALML4 | 0,236359382 | 1,178016203 | 0,4777 | 0,71545897 | calmodulin like 4 |
| ENSG00000114999 | TTL | 0,236175082 | 1,177865724 | 0,41203 | 0,66982472 | tubulin tyrosine ligase |
| ENSG00000162923 | WDR26 | 0,236147608 | 1,177843293 | 0,30275 | 0,59321687 | WD repeat domain 26 |
| ENSG00000109066 | TMEM104 | 0,236052654 | 1,177765774 | 0,25413 | 0,55925777 | transmembrane protein 104 |
| ENSG00000067533 | RRP15 | 0,236040112 | 1,177755535 | 0,18013 | 0,50407792 | ribosomal RNA processing 15 homolog |
| ENSG00000171444 | MCC | 0,236038302 | 1,177754058 | 0,44229 | 0,69006952 | MCC regulator of WNT signaling pathway |
| ENSG00000167460 | TPM4 | 0,235989914 | 1,177714556 | 0,2795 | 0,57745127 | tropomyosin 4 |
| ENSG00000114867 | EIF4G1 | 0,235958996 | 1,177689317 | 0,19138 | 0,51435195 | eukaryotic translation initiation factor 4 gamma 1 |
| ENSG00000138688 | KIAA1109 | 0,235886345 | 1,177630013 | 0,51666 | 0,74206471 | KIAA1109 |
| ENSG00000277837 | AC090340,1 | 0,235885602 | 1,177629406 | 0,78765 | 0,90122495 | novel transcript |

| | | | | | | |
|---|---|---|---|---|---|---|
| ENSG00000159267 | HLCS | 0,235865484 | 1,177612985 | 0,43719 | 0,68711446 | holocarboxylase synthetase |
| ENSG00000276021 | WDR81 | 0,235851407 | 1,177601494 | 0,4312 | 0,68350322 | WD repeat domain 81 |
| ENSG00000166411 | IDH3A | 0,235850955 | 1,177601125 | 0,30003 | 0,59160746 | isocitrate dehydrogenase (NAD(+)) 3 catalytic subunit alpha |
| ENSG00000158169 | FANCC | 0,235847638 | 1,177598418 | 0,18158 | 0,50490902 | FA complementation group C |
| ENSG00000145743 | FBXL17 | 0,235838394 | 1,177590872 | 0,31894 | 0,60399503 | F-box and leucine rich repeat protein 17 |
| ENSG00000115282 | TTC31 | 0,235797376 | 1,177557392 | 0,41306 | 0,67067821 | tetratricopeptide repeat domain 31 |
| ENSG00000213149 | CNN2P9 | 0,235784367 | 1,177546774 | 0,77985 | 0,8981559 | calponin 2 pseudogene 9 |
| ENSG00000198960 | ARMCX6 | 0,235719812 | 1,177494084 | 0,32688 | 0,60888005 | armadillo repeat containing X-linked 6 |
| ENSG00000275580 | AC022306,2 | 0,235680304 | 1,17746184 | 0,77628 | 0,89654301 | novel transcript, sense intronic to GALK2 |
| ENSG00000272848 | AL135910,1 | 0,23567495 | 1,17745747 | 0,52905 | 0,75111082 | novel transcript |
| ENSG00000114544 | SLC41A3 | 0,235655541 | 1,177441629 | 0,34654 | 0,62350287 | solute carrier family 41 member 3 |
| ENSG00000071282 | LMCD1 | 0,235634475 | 1,177424437 | 0,4538 | 0,69789417 | LIM and cysteine rich domains 1 |
| ENSG00000101266 | CSNK2A1 | 0,235549569 | 1,177355145 | 0,16919 | 0,49530908 | casein kinase 2 alpha 1 |
| ENSG00000072849 | DERL2 | 0,235503611 | 1,177317639 | 0,44696 | 0,69334376 | derlin 2 |
| ENSG00000244342 | LINC00698 | 0,235449565 | 1,177273536 | 0,73785 | 0,87544492 | long intergenic non-protein coding RNA 698 |
| ENSG00000203943 | SAMD13 | 0,235357899 | 1,177198737 | 0,60206 | 0,79687124 | sterile alpha motif domain containing 13 |
| ENSG00000266893 | AC005394,1 | 0,235221571 | 1,177087502 | 0,77794 | 0,89728573 | novel transcript |
| ENSG00000232388 | SMIM26 | 0,23520033 | 1,177070172 | 0,46361 | 0,70441254 | small integral membrane protein 26 |
| ENSG00000152818 | UTRN | 0,235085968 | 1,176976869 | 0,3373 | 0,61706507 | utrophin |
| ENSG00000172765 | TMCC1 | 0,235031713 | 1,176932608 | 0,29568 | 0,58930611 | transmembrane and coiled-coil domain family 1 |
| ENSG00000270959 | LPP-AS2 | 0,234990134 | 1,176898689 | 0,38183 | 0,64905556 | LPP antisense RNA 2 |
| ENSG00000275349 | SMN1 | 0,234971838 | 1,176883763 | 0,62289 | 0,80880395 | survival of motor neuron 1, telomeric |
| ENSG00000237649 | KIFC1 | 0,234905078 | 1,176829306 | 0,36943 | 0,63917123 | kinesin family member C1 |
| ENSG00000164828 | SUN1 | 0,234817965 | 1,176758248 | 0,39306 | 0,65610193 | Sad1 and UNC84 domain containing 1 |
| ENSG00000229222 | KRT18P4 | 0,234805561 | 1,176748131 | 0,65604 | 0,82824314 | keratin 18 pseudogene 4 |
| ENSG00000259381 | AC090695,1 | 0,234803192 | 1,176746199 | 0,80947 | 0,91213228 | WD repeat domain 12 (WDR12) pseudogene |
| ENSG00000134283 | PPHLN1 | 0,234798888 | 1,176742688 | 0,14949 | 0,47729587 | periphilin 1 |
| ENSG00000215305 | VPS16 | 0,23469347 | 1,176656706 | 0,21916 | 0,53664318 | VPS16 core subunit of CORVET and HOPS complexes |
| ENSG00000119402 | FBXW2 | 0,23461732 | 1,1765946 | 0,23766 | 0,54861973 | F-box and WD repeat domain containing 2 |
| ENSG00000267481 | AC011477,1 | 0,23459206 | 1,176574 | 0,46531 | 0,70583978 | novel transcript, sense intronic to ZNF506 |
| ENSG00000187608 | ISG15 | 0,234436163 | 1,176446866 | 0,49304 | 0,72615908 | ISG15 ubiquitin like modifier |
| ENSG00000160200 | CBS | 0,234434556 | 1,176445555 | 0,35042 | 0,62595491 | cystathionine beta-synthase |

| | | | | | | |
|---|---|---|---|---|---|---|
| ENSG00000284916 | PSME1 | 0,234398732 | 1,176416343 | 0,50085 | 0,73097006 | proteasome activator subunit 1 |
| ENSG00000092010 | PSME1 | 0,234398732 | 1,176416343 | 0,50085 | 0,73097006 | proteasome activator subunit 1 |
| ENSG00000136021 | SCYL2 | 0,234342246 | 1,176370284 | 0,59083 | 0,79001985 | SCY1 like pseudokinase 2 |
| ENSG00000173482 | PTPRM | 0,234319249 | 1,176351532 | 0,3585 | 0,63185282 | protein tyrosine phosphatase receptor type M |
| ENSG00000163444 | TMEM183A | 0,23428706 | 1,176325286 | 0,18998 | 0,51329223 | transmembrane protein 183A |
| ENSG00000124702 | KLHDC3 | 0,234179991 | 1,176237989 | 0,19363 | 0,51547251 | kelch domain containing 3 |
| ENSG00000283761 | AC118553,2 | 0,234156951 | 1,176219205 | 0,45805 | 0,7010995 | novel protein |
| ENSG00000282233 | AL450306,2 | 0,234133671 | 1,176200225 | 0,53401 | 0,75385628 | novel transcript |
| ENSG00000271533 | Z83843,1 | 0,23413033 | 1,176197501 | 0,52699 | 0,75012908 | novel transcript, sense intronic FTX |
| ENSG00000142611 | PRDM16 | 0,234064534 | 1,17614386 | 0,57886 | 0,78256244 | PR/SET domain 16 |
| ENSG00000122986 | HVCN1 | 0,233997528 | 1,176089235 | 0,35878 | 0,63191767 | hydrogen voltage gated channel 1 |
| ENSG00000225689 | AL008633,1 | 0,233993586 | 1,176086022 | 0,50363 | 0,73287346 | novel transcript |
| ENSG00000149554 | CHEK1 | 0,233935168 | 1,1760384 | 0,4335 | 0,68473723 | checkpoint kinase 1 |
| ENSG00000266340 | AC138207,7 | 0,233899499 | 1,176009324 | 0,39235 | 0,65568918 | novel transcript, antisense RNF135 |
| ENSG00000115526 | CHST10 | 0,233814604 | 1,175940125 | 0,29681 | 0,58995143 | carbohydrate sulfotransferase 10 |
| ENSG00000237938 | AL450998,2 | 0,233734437 | 1,175874782 | 0,62596 | 0,81058216 | novel transcript, antisense to SLC25A34 and PLEKHM2 |
| ENSG00000112996 | MRPS30 | 0,233706847 | 1,175852295 | 0,24958 | 0,55657567 | mitochondrial ribosomal protein S30 |
| ENSG00000153246 | PLA2R1 | 0,233677916 | 1,175828715 | 0,45813 | 0,70114321 | phospholipase A2 receptor 1 |
| ENSG00000150776 | NKAPD1 | 0,233564419 | 1,175736217 | 0,34853 | 0,62475089 | NKAP domain containing 1 |
| ENSG00000139350 | NEDD1 | 0,233496088 | 1,175680531 | 0,58147 | 0,78431121 | NEDD1 gamma-tubulin ring complex targeting factor |
| ENSG00000142230 | SAE1 | 0,233450467 | 1,175643354 | 0,23016 | 0,54335232 | SUMO1 activating enzyme subunit 1 |
| ENSG00000186976 | EFCAB6 | 0,233377114 | 1,175583581 | 0,51622 | 0,74198903 | EF-hand calcium binding domain 6 |
| ENSG00000187747 | OR52B6 | 0,233374475 | 1,17558143 | 0,75067 | 0,8829851 | olfactory receptor family 52 subfamily B member 6 |
| ENSG00000285027 | ALG1L13P | 0,233309655 | 1,175528612 | 0,671 | 0,83695245 | ALG1 like 13, pseudogene |
| ENSG00000253981 | ALG1L13P | 0,233309655 | 1,175528612 | 0,671 | 0,83695245 | ALG1 like 13, pseudogene |
| ENSG00000284807 | DHRS4 | 0,2331903 | 1,175431365 | 0,29469 | 0,58857672 | dehydrogenase/reductase 4 |
| ENSG00000157326 | DHRS4 | 0,2331903 | 1,175431365 | 0,29469 | 0,58857672 | dehydrogenase/reductase 4 |
| ENSG00000134490 | TMEM241 | 0,23318773 | 1,175429271 | 0,27281 | 0,57322782 | transmembrane protein 241 |
| ENSG00000106853 | PTGR1 | 0,233181049 | 1,175423827 | 0,22127 | 0,53805234 | prostaglandin reductase 1 |
| ENSG00000256745 | ARPC3P3 | 0,233068165 | 1,17533186 | 0,70493 | 0,857835 | actin related protein 2/3 complex subunit 3 pseudogene 3 |
| ENSG00000019995 | ZRANB1 | 0,233047544 | 1,17531506 | 0,22028 | 0,53750545 | zinc finger RANBP2-type containing 1 |
| ENSG00000143748 | NVL | 0,233022361 | 1,175294545 | 0,24627 | 0,55395309 | nuclear VCP like |

| | | | | | | |
|---|---|---|---|---|---|---|
| ENSG00000101365 | IDH3B | 0,2330053 | 1,175280646 | 0,21372 | 0,53279085 | isocitrate dehydrogenase (NAD(+)) 3 non-catalytic subunit beta |
| ENSG00000285262 | MCPH1 | 0,232950968 | 1,175236386 | 0,46136 | 0,70278869 | microcephalin 1 |
| ENSG00000104695 | PPP2CB | 0,232883658 | 1,175181556 | 0,46379 | 0,70453935 | protein phosphatase 2 catalytic subunit beta |
| ENSG00000148606 | POLR3A | 0,23287927 | 1,175177981 | 0,19598 | 0,51680811 | RNA polymerase III subunit A |
| ENSG00000188124 | OR2AG2 | 0,232770085 | 1,175089045 | 0,61025 | 0,80221019 | olfactory receptor family 2 subfamily AG member 2 |
| ENSG00000196652 | ZKSCAN5 | 0,232752364 | 1,175074612 | 0,37277 | 0,64166433 | zinc finger with KRAB and SCAN domains 5 |
| ENSG00000213462 | ERV3-1 | 0,232711591 | 1,175041403 | 0,48238 | 0,71892541 | endogenous retrovirus group 3 member 1, envelope |
| ENSG00000116793 | PHTF1 | 0,232688713 | 1,175022769 | 0,43392 | 0,68480274 | putative homeodomain transcription factor 1 |
| ENSG00000172731 | LRRC20 | 0,232675924 | 1,175012353 | 0,47695 | 0,71488724 | leucine rich repeat containing 20 |
| ENSG00000170633 | RNF34 | 0,232609846 | 1,174958536 | 0,36194 | 0,63352908 | ring finger protein 34 |
| ENSG00000066557 | LRRC40 | 0,232576293 | 1,174931211 | 0,5107 | 0,73822353 | leucine rich repeat containing 40 |
| ENSG00000226869 | LHFPL3-AS1 | 0,232490098 | 1,174861015 | 0,81896 | 0,91689679 | LHFPL3 antisense RNA 1 |
| ENSG00000149273 | RPS3 | 0,232399956 | 1,174787611 | 0,17893 | 0,5033932 | ribosomal protein S3 |
| ENSG00000143537 | ADAM15 | 0,232312654 | 1,174716523 | 0,38375 | 0,65041894 | ADAM metallopeptidase domain 15 |
| ENSG00000268573 | AC011815,1 | 0,232282243 | 1,174691761 | 0,46208 | 0,70341975 | novel transcript |
| ENSG00000112981 | NME5 | 0,23227183 | 1,174683282 | 0,58568 | 0,78640335 | NME/NM23 family member 5 |
| ENSG00000166507 | NDST2 | 0,232167185 | 1,17459808 | 0,35831 | 0,63185282 | N-deacetylase and N-sulfotransferase 2 |
| ENSG00000229863 | AC097358,1 | 0,232155451 | 1,174588527 | 0,61725 | 0,8057177 | ribosomal protein L19 (RPL19) pseudogene |
| ENSG00000258758 | AL137100,1 | 0,232144137 | 1,174579316 | 0,80607 | 0,91078302 | zinc finger, CCHC domain containing 7 (ZCCHC7) pseudogene |
| ENSG00000204010 | IFIT1B | 0,232086532 | 1,174532417 | 0,68779 | 0,84787163 | interferon induced protein with tetratricopeptide repeats 1B |
| ENSG00000069943 | PIGB | 0,232050805 | 1,174503331 | 0,4734 | 0,71264815 | phosphatidylinositol glycan anchor biosynthesis class B |
| ENSG00000187676 | B3GLCT | 0,231762893 | 1,174268964 | 0,64108 | 0,81993426 | beta 3-glucosyltransferase |
| ENSG00000139645 | ANKRD52 | 0,231683216 | 1,174204113 | 0,11704 | 0,43438325 | ankyrin repeat domain 52 |
| ENSG00000177352 | CCDC71 | 0,23160617 | 1,174141408 | 0,27187 | 0,57305058 | coiled-coil domain containing 71 |
| ENSG00000198026 | ZNF335 | 0,231577213 | 1,174117841 | 0,36293 | 0,63395677 | zinc finger protein 335 |
| ENSG00000261431 | AL023803,1 | 0,231509813 | 1,17406299 | 0,83331 | 0,92389797 | novel transcript, antisense to DHX35 |
| ENSG00000121903 | ZSCAN20 | 0,2315011 | 1,174055899 | 0,28877 | 0,58464073 | zinc finger and SCAN domain containing 20 |
| ENSG00000137338 | PGBD1 | 0,231419209 | 1,173989259 | 0,51134 | 0,73853921 | piggyBac transposable element derived 1 |
| ENSG00000269481 | AC010319,4 | 0,23141291 | 1,173984133 | 0,6737 | 0,83919798 | novel transcript, antisense FAM125A |
| ENSG00000001084 | GCLC | 0,231402129 | 1,173975361 | 0,155 | 0,48292767 | glutamate-cysteine ligase catalytic subunit |

| | | | | | | |
|---|---|---|---|---|---|---|
| ENSG00000144199 | FAHD2B | 0,231371151 | 1,173950153 | 0,50992 | 0,73755734 | fumarylacetoacetate hydrolase domain containing 2B |
| ENSG00000265480 | KRT18P55 | 0,231201396 | 1,173812028 | 0,76924 | 0,89297049 | keratin 18 pseudogene 55 |
| ENSG00000120738 | EGR1 | 0,231114518 | 1,173741344 | 0,70335 | 0,8569555 | early growth response 1 |
| ENSG00000104408 | EIF3E | 0,231111916 | 1,173739227 | 0,23103 | 0,54395664 | eukaryotic translation initiation factor 3 subunit E |
| ENSG00000128917 | DLL4 | 0,23107542 | 1,173709535 | 0,68367 | 0,84505352 | delta like canonical Notch ligand 4 |
| ENSG00000108506 | INTS2 | 0,231026592 | 1,173669812 | 0,27694 | 0,57557548 | integrator complex subunit 2 |
| ENSG00000232479 | KRT8P52 | 0,23101499 | 1,173660373 | 0,69619 | 0,85287022 | keratin 8 pseudogene 52 |
| ENSG00000100802 | C14orf93 | 0,23092772 | 1,173589379 | 0,33895 | 0,61800397 | chromosome 14 open reading frame 93 |
| ENSG00000163697 | APBB2 | 0,230895735 | 1,173563361 | 0,5471 | 0,76264866 | amyloid beta precursor protein binding family B member 2 |
| ENSG00000172301 | COPRS | 0,230864713 | 1,173538126 | 0,36151 | 0,63347405 | coordinator of PRMT5 and differentiation stimulator |
| ENSG00000182359 | KBTBD3 | 0,230785763 | 1,173473907 | 0,65151 | 0,82541155 | kelch repeat and BTB domain containing 3 |
| ENSG00000165934 | CPSF2 | 0,230776353 | 1,173466253 | 0,14974 | 0,47782859 | cleavage and polyadenylation specific factor 2 |
| ENSG00000205922 | ONECUT3 | 0,230760249 | 1,173453154 | 0,78091 | 0,8985082 | one cut homeobox 3 |
| ENSG00000281649 | EBLN3P | 0,230731376 | 1,17342967 | 0,42131 | 0,67634721 | endogenous Bornavirus like nucleoprotein 3, pseudogene |
| ENSG00000087088 | BAX | 0,2307225 | 1,17342245 | 0,2261 | 0,54121239 | BCL2 associated X, apoptosis regulator |
| ENSG00000099875 | MKNK2 | 0,230639698 | 1,173355105 | 0,32928 | 0,61035569 | MAPK interacting serine/threonine kinase 2 |
| ENSG00000168916 | ZNF608 | 0,230501297 | 1,173242548 | 0,39931 | 0,66167983 | zinc finger protein 608 |
| ENSG00000135316 | SYNCRIP | 0,230482672 | 1,173227402 | 0,46939 | 0,70890278 | synaptotagmin binding cytoplasmic RNA interacting protein |
| ENSG00000165525 | NEMF | 0,230387308 | 1,173149853 | 0,34137 | 0,61943649 | nuclear export mediator factor |
| ENSG00000260171 | AC008269,2 | 0,230349002 | 1,173118704 | 0,73581 | 0,8750728 | novel transcript |
| ENSG00000165240 | ATP7A | 0,23031969 | 1,173094869 | 0,47188 | 0,71121104 | ATPase copper transporting alpha |
| ENSG00000229205 | LINC00200 | 0,230194418 | 1,172993011 | 0,76427 | 0,89015718 | long intergenic non-protein coding RNA 200 |
| ENSG00000079785 | DDX1 | 0,230127375 | 1,172938503 | 0,3288 | 0,60992716 | DEAD-box helicase 1 |
| ENSG00000115423 | DNAH6 | 0,230019864 | 1,172851098 | 0,59341 | 0,7917651 | dynein axonemal heavy chain 6 |
| ENSG00000198547 | C20orf203 | 0,230014587 | 1,172846808 | 0,80265 | 0,90908968 | chromosome 20 open reading frame 203 |
| ENSG00000109654 | TRIM2 | 0,229992024 | 1,172828465 | 0,59525 | 0,79324077 | tripartite motif containing 2 |
| ENSG00000164088 | PPM1M | 0,229889565 | 1,172745175 | 0,29461 | 0,58857672 | protein phosphatase, Mg2+/Mn2+ dependent 1M |
| ENSG00000108479 | GALK1 | 0,229721697 | 1,172608725 | 0,28523 | 0,5817483 | galactokinase 1 |
| ENSG00000272994 | AC012360,3 | 0,229676067 | 1,172571639 | 0,55839 | 0,7700003 | novel transcript |
| ENSG00000170315 | UBB | 0,229565947 | 1,17248214 | 0,38858 | 0,65310114 | ubiquitin B |
| ENSG00000011260 | UTP18 | 0,229518545 | 1,172443617 | 0,1813 | 0,50482416 | UTP18 small subunit processome component |
| ENSG00000139644 | TMBIM6 | 0,229498767 | 1,172427544 | 0,46877 | 0,70833955 | transmembrane BAX inhibitor motif containing 6 |

| | | | | | | |
|---|---|---|---|---|---|---|
| ENSG00000285121 | MTCH2 | 0,229465217 | 1,17240028 | 0,36999 | 0,63943131 | mitochondrial carrier 2 |
| ENSG00000109919 | MTCH2 | 0,229465217 | 1,17240028 | 0,36999 | 0,63943131 | mitochondrial carrier 2 |
| ENSG00000172939 | OXSR1 | 0,229408215 | 1,172353958 | 0,29033 | 0,58553923 | oxidative stress responsive kinase 1 |
| ENSG00000105732 | ZNF574 | 0,229339758 | 1,17229833 | 0,29235 | 0,58692263 | zinc finger protein 574 |
| ENSG00000143641 | GALNT2 | 0,229322637 | 1,172284418 | 0,29922 | 0,59123205 | polypeptide N-acetylgalactosaminyltransferase 2 |
| ENSG00000261366 | MANEA-DT | 0,229297457 | 1,172263958 | 0,61786 | 0,80610038 | MANEA divergent transcript |
| ENSG00000197415 | VEPH1 | 0,229284609 | 1,172253518 | 0,58266 | 0,7847455 | ventricular zone expressed PH domain containing 1 |
| ENSG00000228021 | AL158835,1 | 0,229172242 | 1,172162219 | 0,73497 | 0,87477884 | novel transcript |
| ENSG00000162994 | CLHC1 | 0,228998084 | 1,172020728 | 0,43793 | 0,68747773 | clathrin heavy chain linker domain containing 1 |
| ENSG00000133393 | CEP20 | 0,228987718 | 1,172012306 | 0,38401 | 0,65051083 | centrosomal protein 20 |
| ENSG00000185664 | PMEL | 0,228893735 | 1,171935959 | 0,70476 | 0,85775118 | premelanosome protein |
| ENSG00000159840 | ZYX | 0,228792148 | 1,17185344 | 0,43048 | 0,68295004 | zyxin |
| ENSG00000285443 | ZYX | 0,228792148 | 1,17185344 | 0,43048 | 0,68295004 | zyxin |
| ENSG00000218418 | AL591135,1 | 0,22873367 | 1,171805942 | 0,70839 | 0,85937051 | Rho GTPase activating protein 21 (ARHGAP21) pseudogene |
| ENSG00000178177 | LCORL | 0,228691938 | 1,171772046 | 0,556 | 0,76827287 | ligand dependent nuclear receptor corepressor like |
| ENSG00000179295 | PTPN11 | 0,228656618 | 1,171743359 | 0,34673 | 0,62350287 | protein tyrosine phosphatase non-receptor type 11 |
| ENSG00000164118 | CEP44 | 0,228635709 | 1,171726377 | 0,2779 | 0,5762948 | centrosomal protein 44 |
| ENSG00000140264 | SERF2 | 0,22860912 | 1,171704782 | 0,27655 | 0,575468 | small EDRK-rich factor 2 |
| ENSG00000006125 | AP2B1 | 0,228522365 | 1,171634325 | 0,43728 | 0,68715611 | adaptor related protein complex 2 subunit beta 1 |
| ENSG00000005483 | KMT2E | 0,228514506 | 1,171627942 | 0,47553 | 0,71413261 | lysine methyltransferase 2E (inactive) |
| ENSG00000198925 | ATG9A | 0,228435987 | 1,171564178 | 0,28367 | 0,58072192 | autophagy related 9A |
| ENSG00000136273 | HUS1 | 0,228391245 | 1,171527845 | 0,1319 | 0,45708171 | HUS1 checkpoint clamp component |
| ENSG00000132406 | TMEM128 | 0,228370927 | 1,171511347 | 0,38343 | 0,650151 | transmembrane protein 128 |
| ENSG00000104763 | ASAH1 | 0,228357109 | 1,171500126 | 0,53967 | 0,75796312 | N-acylsphingosine amidohydrolase 1 |
| ENSG00000004455 | AK2 | 0,22835325 | 1,171496992 | 0,1652 | 0,49131595 | adenylate kinase 2 |
| ENSG00000120438 | TCP1 | 0,228294833 | 1,171449557 | 0,2363 | 0,54725688 | t-complex 1 |
| ENSG00000173559 | NABP1 | 0,22826422 | 1,1714247 | 0,61964 | 0,80726991 | nucleic acid binding protein 1 |
| ENSG00000100271 | TTLL1 | 0,228199432 | 1,171372095 | 0,44385 | 0,69154673 | tubulin tyrosine ligase like 1 |
| ENSG00000131389 | SLC6A6 | 0,228178848 | 1,171355382 | 0,45417 | 0,69818495 | solute carrier family 6 member 6 |
| ENSG00000108883 | EFTUD2 | 0,228151371 | 1,171333074 | 0,28395 | 0,58089675 | elongation factor Tu GTP binding domain containing 2 |
| ENSG00000213930 | GALT | 0,228125105 | 1,171311749 | 0,24385 | 0,55243616 | galactose-1-phosphate uridylyltransferase |
| ENSG00000167325 | RRM1 | 0,228018401 | 1,17122512 | 0,41025 | 0,66821634 | ribonucleotide reductase catalytic subunit M1 |
| ENSG00000138385 | SSB | 0,227980462 | 1,17119432 | 0,5109 | 0,73829835 | small RNA binding exonuclease protection factor La |

| | | | | | | |
|---|---|---|---|---|---|---|
| ENSG00000076604 | TRAF4 | 0,227971533 | 1,171187071 | 0,29002 | 0,58531219 | TNF receptor associated factor 4 |
| ENSG00000197124 | ZNF682 | 0,227910256 | 1,171137327 | 0,39929 | 0,66167983 | zinc finger protein 682 |
| ENSG00000156502 | SUPV3L1 | 0,22779546 | 1,171044143 | 0,23328 | 0,54562736 | Suv3 like RNA helicase |
| ENSG00000189308 | LIN54 | 0,227735658 | 1,170995603 | 0,38815 | 0,6529525 | lin-54 DREAM MuvB core complex component |
| ENSG00000153814 | JAZF1 | 0,227680663 | 1,170950966 | 0,65173 | 0,82550688 | JAZF zinc finger 1 |
| ENSG00000204581 | ACOXL-AS1 | 0,2276641 | 1,170937522 | 0,51278 | 0,73941533 | ACOXL antisense RNA 1 |
| ENSG00000120868 | APAF1 | 0,227651131 | 1,170926996 | 0,15267 | 0,48061452 | apoptotic peptidase activating factor 1 |
| ENSG00000204574 | ABCF1 | 0,227644374 | 1,170921512 | 0,53609 | 0,75500755 | ATP binding cassette subfamily F member 1 |
| ENSG00000258315 | C17orf49 | 0,227609495 | 1,170893204 | 0,26161 | 0,56452671 | chromosome 17 open reading frame 49 |
| ENSG00000157540 | DYRK1A | 0,227505361 | 1,170808692 | 0,24569 | 0,55358893 | dual specificity tyrosine phosphorylation regulated kinase 1A |
| ENSG00000273723 | SUGT1-DT | 0,227478473 | 1,170786871 | 0,71347 | 0,86217438 | SUGT1 divergent transcript |
| ENSG00000260708 | AL118516,1 | 0,22740464 | 1,170726955 | 0,59193 | 0,79090751 | novel transcript, antisense to TBC1D22A |
| ENSG00000147065 | MSN | 0,227389781 | 1,170714898 | 0,39178 | 0,65534953 | moesin |
| ENSG00000104375 | STK3 | 0,227385885 | 1,170711736 | 0,20034 | 0,52107152 | serine/threonine kinase 3 |
| ENSG00000111481 | COPZ1 | 0,227336969 | 1,170672043 | 0,34355 | 0,62091871 | COPI coat complex subunit zeta 1 |
| ENSG00000101457 | DNTTIP1 | 0,227315529 | 1,170654645 | 0,39651 | 0,6591566 | deoxynucleotidyltransferase terminal interacting protein 1 |
| ENSG00000123374 | CDK2 | 0,227302224 | 1,170643849 | 0,29495 | 0,58868965 | cyclin dependent kinase 2 |
| ENSG00000197548 | ATG7 | 0,22726371 | 1,170612598 | 0,33369 | 0,61369834 | autophagy related 7 |
| ENSG00000182489 | XKRX | 0,22719888 | 1,170559996 | 0,71409 | 0,8623286 | XK related X-linked |
| ENSG00000144306 | SCRN3 | 0,227151376 | 1,170521453 | 0,33046 | 0,6112353 | secernin 3 |
| ENSG00000251660 | AC007036,3 | 0,227124274 | 1,170499464 | 0,74419 | 0,87889181 | novel transcript |
| ENSG00000287354 | AL034351,4 | 0,227059185 | 1,170446657 | 0,76788 | 0,89249843 | novel transcript, antisense to HHAT |
| ENSG00000287371 | AL034351,5 | 0,227059185 | 1,170446657 | 0,76788 | 0,89249843 | novel transcript, antisense to HHAT |
| ENSG00000133874 | RNF122 | 0,226976712 | 1,170379749 | 0,42304 | 0,67691679 | ring finger protein 122 |
| ENSG00000226259 | GTF2H2B | 0,226854081 | 1,17028027 | 0,35025 | 0,62578708 | general transcription factor IIH subunit 2B (pseudogene) |
| ENSG00000256751 | PLBD1-AS1 | 0,2267168 | 1,170168916 | 0,60893 | 0,80155195 | PLBD1 antisense RNA 1 |
| ENSG00000072818 | ACAP1 | 0,226667621 | 1,170129027 | 0,66603 | 0,83386604 | ArfGAP with coiled-coil, ankyrin repeat and PH domains 1 |
| ENSG00000135447 | PPP1R1A | 0,226617416 | 1,170088308 | 0,3811 | 0,64852799 | protein phosphatase 1 regulatory inhibitor subunit 1A |
| ENSG00000004478 | FKBP4 | 0,226568824 | 1,170048898 | 0,19068 | 0,51369186 | FKBP prolyl isomerase 4 |
| ENSG00000145365 | TIFA | 0,226536181 | 1,170022424 | 0,58395 | 0,78572744 | TRAF interacting protein with forkhead associated domain |
| ENSG00000109163 | GNRHR | 0,226522581 | 1,170011396 | 0,62803 | 0,81175174 | gonadotropin releasing hormone receptor |
| ENSG00000065268 | WDR18 | 0,226512498 | 1,170003218 | 0,22551 | 0,54081943 | WD repeat domain 18 |

| | | | | | | |
|---|---|---|---|---|---|---|
| ENSG00000250909 | AC138956,1 | 0,226492017 | 1,169986609 | 0,76331 | 0,88980026 | novel transcript |
| ENSG00000182518 | FAM104B | 0,226410985 | 1,169920895 | 0,46674 | 0,70676695 | family with sequence similarity 104 member B |
| ENSG00000157916 | RER1 | 0,226409973 | 1,169920075 | 0,22732 | 0,54168591 | retention in endoplasmic reticulum sorting receptor 1 |
| ENSG00000264215 | AC126365,1 | 0,226355548 | 1,169875941 | 0,70287 | 0,85674436 | novel transcript |
| ENSG00000138780 | GSTCD | 0,226335498 | 1,169859683 | 0,34242 | 0,62008204 | glutathione S-transferase C-terminal domain containing |
| ENSG00000276127 | AL589935,2 | 0,226328068 | 1,169853658 | 0,76899 | 0,89287402 | coiled-coil domain containing 112 (CCDC112) pseudogene |
| ENSG00000146066 | HIGD2A | 0,226316204 | 1,169844038 | 0,35909 | 0,63201238 | HIG1 hypoxia inducible domain family member 2A |
| ENSG00000101353 | MROH8 | 0,226172844 | 1,169727796 | 0,38105 | 0,64852799 | maestro heat like repeat family member 8 |
| ENSG00000102974 | CTCF | 0,225895982 | 1,16950334 | 0,31947 | 0,60429866 | CCCTC-binding factor |
| ENSG00000178397 | FAM220A | 0,225839752 | 1,169457759 | 0,47742 | 0,71533146 | family with sequence similarity 220 member A |
| ENSG00000187815 | ZFP69 | 0,225831866 | 1,169451366 | 0,47425 | 0,71305872 | ZFP69 zinc finger protein |
| ENSG00000258957 | AL359317,2 | 0,225806892 | 1,169431123 | 0,70167 | 0,85618008 | novel transcript |
| ENSG00000283777 | CANX | 0,225806233 | 1,169430588 | 0,45119 | 0,69626785 | calnexin |
| ENSG00000025800 | KPNA6 | 0,225804443 | 1,169429138 | 0,14488 | 0,47244928 | karyopherin subunit alpha 6 |
| ENSG00000130706 | ADRM1 | 0,225764437 | 1,16939671 | 0,37099 | 0,64024107 | adhesion regulating molecule 1 |
| ENSG00000115839 | RAB3GAP1 | 0,225754184 | 1,169388399 | 0,29871 | 0,59094252 | RAB3 GTPase activating protein catalytic subunit 1 |
| ENSG00000203965 | EFCAB7 | 0,225690214 | 1,169336548 | 0,5898 | 0,78917756 | EF-hand calcium binding domain 7 |
| ENSG00000278829 | AC099811,5 | 0,225594083 | 1,169258635 | 0,56829 | 0,7759239 | novel transcript, sense intronic to STAT5B |
| ENSG00000103061 | SLC7A6OS | 0,225576545 | 1,169244421 | 0,39688 | 0,65942065 | solute carrier family 7 member 6 opposite strand |
| ENSG00000178307 | TMEM11 | 0,225556753 | 1,16922838 | 0,27409 | 0,57412149 | transmembrane protein 11 |
| ENSG00000134109 | EDEM1 | 0,225454139 | 1,16914522 | 0,39313 | 0,65617519 | ER degradation enhancing alpha-mannosidase like protein 1 |
| ENSG00000206432 | TMEM200C | 0,225368259 | 1,169075626 | 0,37644 | 0,64481513 | transmembrane protein 200C |
| ENSG00000117899 | MESD | 0,225347011 | 1,169058408 | 0,14998 | 0,47799251 | mesoderm development LRP chaperone |
| ENSG00000145241 | CENPC | 0,225308208 | 1,169026965 | 0,31262 | 0,59951563 | centromere protein C |
| ENSG00000197714 | ZNF460 | 0,225301033 | 1,169021152 | 0,41134 | 0,66904284 | zinc finger protein 460 |
| ENSG00000155868 | MED7 | 0,225293467 | 1,16901502 | 0,57818 | 0,78191189 | mediator complex subunit 7 |
| ENSG00000133789 | SWAP70 | 0,225259421 | 1,168987434 | 0,25908 | 0,56284741 | switching B cell complex subunit SWAP70 |
| ENSG00000122390 | NAA60 | 0,225173334 | 1,168917681 | 0,28079 | 0,57814942 | N-alpha-acetyltransferase 60, NatF catalytic subunit |
| ENSG00000160345 | C9orf116 | 0,225110726 | 1,168866955 | 0,59219 | 0,79101863 | chromosome 9 open reading frame 116 |
| ENSG00000163040 | CCDC74A | 0,225097085 | 1,168855903 | 0,50153 | 0,73145255 | coiled-coil domain containing 74A |
| ENSG00000136051 | WASHC4 | 0,225076106 | 1,168838907 | 0,23502 | 0,5465505 | WASH complex subunit 4 |
| ENSG00000271699 | SNX29P2 | 0,225038957 | 1,168808809 | 0,88519 | 0,94923177 | sorting nexin 29 pseudogene 2 |

| | | | | | | |
|---|---|---|---|---|---|---|
| ENSG00000126822 | PLEKHG3 | 0,225017893 | 1,168791744 | 0,3662 | 0,63617598 | pleckstrin homology and RhoGEF domain containing G3 |
| ENSG00000230393 | AC092667,1 | 0,2249724 | 1,168754889 | 0,5938 | 0,7920712 | novel transcript |
| ENSG00000149531 | FRG1BP | 0,224893365 | 1,168690863 | 0,63496 | 0,8162492 | FSHD region gene 1 family member B, pseudogene |
| ENSG00000166348 | USP54 | 0,22488167 | 1,16868139 | 0,34085 | 0,61918628 | ubiquitin specific peptidase 54 |
| ENSG00000196914 | ARHGEF12 | 0,224872007 | 1,168673562 | 0,25731 | 0,56160845 | Rho guanine nucleotide exchange factor 12 |
| ENSG00000254433 | AP001001,1 | 0,224835686 | 1,16864414 | 0,62333 | 0,8091109 | novel transcript |
| ENSG00000122574 | WIPF3 | 0,224667644 | 1,168508026 | 0,49807 | 0,72937175 | WAS/WASL interacting protein family member 3 |
| ENSG00000165527 | ARF6 | 0,22455469 | 1,168416543 | 0,38215 | 0,64916957 | ADP ribosylation factor 6 |
| ENSG00000197930 | ERO1A | 0,224509249 | 1,168379741 | 0,17222 | 0,49710979 | endoplasmic reticulum oxidoreductase 1 alpha |
| ENSG00000137404 | NRM | 0,224424114 | 1,168310796 | 0,32214 | 0,60577115 | nurim |
| ENSG00000228867 | NRM | 0,224424114 | 1,168310796 | 0,32214 | 0,60577115 | nurim |
| ENSG00000235773 | NRM | 0,224424114 | 1,168310796 | 0,32214 | 0,60577115 | nurim |
| ENSG00000234809 | NRM | 0,224424114 | 1,168310796 | 0,32214 | 0,60577115 | nurim |
| ENSG00000236843 | NRM | 0,224424114 | 1,168310796 | 0,32214 | 0,60577115 | nurim |
| ENSG00000228854 | NRM | 0,224424114 | 1,168310796 | 0,32214 | 0,60577115 | nurim |
| ENSG00000206484 | NRM | 0,224424114 | 1,168310796 | 0,32214 | 0,60577115 | nurim |
| ENSG00000118557 | PMFBP1 | 0,224422598 | 1,168309569 | 0,52896 | 0,75111082 | polyamine modulated factor 1 binding protein 1 |
| ENSG00000149792 | MRPL49 | 0,224402713 | 1,168293466 | 0,22446 | 0,54033281 | mitochondrial ribosomal protein L49 |
| ENSG00000173171 | MTX1 | 0,224394596 | 1,168286893 | 0,2438 | 0,55243616 | metaxin 1 |
| ENSG00000114686 | MRPL3 | 0,224352102 | 1,168252482 | 0,47501 | 0,71364583 | mitochondrial ribosomal protein L3 |
| ENSG00000101189 | MRGBP | 0,224343674 | 1,168245657 | 0,20184 | 0,52217102 | MRG domain binding protein |
| ENSG00000229530 | AL136097,1 | 0,224240381 | 1,168162017 | 0,81243 | 0,91379618 | novel pseudogene |
| ENSG00000185989 | RASA3 | 0,224233559 | 1,168156493 | 0,53162 | 0,75280487 | RAS p21 protein activator 3 |
| ENSG00000280477 | RASA3 | 0,224233559 | 1,168156493 | 0,53162 | 0,75280487 | RAS p21 protein activator 3 |
| ENSG00000122218 | COPA | 0,224065385 | 1,16802033 | 0,22861 | 0,54249621 | COPI coat complex subunit alpha |
| ENSG00000277104 | TADA2A | 0,224023611 | 1,16798651 | 0,50172 | 0,73146914 | transcriptional adaptor 2A |
| ENSG00000138685 | FGF2 | 0,223989938 | 1,167959249 | 0,5211 | 0,74558236 | fibroblast growth factor 2 |
| ENSG00000231770 | TMEM44-AS1 | 0,223966947 | 1,167940636 | 0,57493 | 0,78003515 | TMEM44 antisense RNA 1 |
| ENSG00000177646 | ACAD9 | 0,22371357 | 1,167735531 | 0,34325 | 0,62075265 | acyl-CoA dehydrogenase family member 9 |
| ENSG00000170791 | CHCHD7 | 0,223695808 | 1,167721155 | 0,2542 | 0,55925777 | coiled-coil-helix-coiled-coil-helix domain containing 7 |
| ENSG00000196981 | WDR5B | 0,223564287 | 1,167614707 | 0,41024 | 0,66821634 | WD repeat domain 5B |
| ENSG00000198380 | GFPT1 | 0,223464427 | 1,16753389 | 0,30396 | 0,59407983 | glutamine--fructose-6-phosphate transaminase 1 |
| ENSG00000272047 | GTF2H5 | 0,223396035 | 1,167478543 | 0,42813 | 0,68094175 | general transcription factor IIH subunit 5 |

| | | | | | | |
|---|---|---|---|---|---|---|
| ENSG00000277157 | H4C4 | 0,2233704 | 1,167457798 | 0,49645 | 0,72831792 | H4 clustered histone 4 |
| ENSG00000285316 | AC275401,5 | 0,223264095 | 1,167371777 | 0,73734 | 0,87539732 | |
| ENSG00000259357 | AL590133,2 | 0,223204794 | 1,167323795 | 0,61527 | 0,80431041 | novel transcript, antisense to CERS2 |
| ENSG00000183690 | EFHC2 | 0,223199708 | 1,167319679 | 0,75517 | 0,88526692 | EF-hand domain containing 2 |
| ENSG00000221164 | SNORA11F | 0,22319775 | 1,167318095 | 0,71966 | 0,86592885 | small nucleolar RNA, H/ACA box 11F |
| ENSG00000268471 | MIR4453HG | 0,223145325 | 1,167275678 | 0,47615 | 0,71446062 | MIR4453 host gene |
| ENSG00000089123 | TASP1 | 0,223124585 | 1,167258897 | 0,2874 | 0,58382745 | taspase 1 |
| ENSG00000065970 | FOXJ2 | 0,223038895 | 1,167189568 | 0,18489 | 0,50844776 | forkhead box J2 |
| ENSG00000197372 | ZNF675 | 0,222935793 | 1,167106159 | 0,4305 | 0,68295004 | zinc finger protein 675 |
| ENSG00000287957 | AC007695,1 | 0,222891478 | 1,167070309 | 0,43153 | 0,68377875 | novel transcript, antisense to RIC8B |
| ENSG00000131263 | RLIM | 0,222880135 | 1,167061134 | 0,36886 | 0,63868683 | ring finger protein, LIM domain interacting |
| ENSG00000258648 | UBE2CP1 | 0,222853559 | 1,167039635 | 0,64354 | 0,82109591 | ubiquitin conjugating enzyme E2 C pseudogene 1 |
| ENSG00000271137 | AC006254,3 | 0,222839882 | 1,167028571 | 0,72597 | 0,86949831 | AKT interacting protein, pseudogene |
| ENSG00000271778 | AC080013,2 | 0,222833697 | 1,167023568 | 0,67982 | 0,8427192 | novel transcript, sense intronic to MFSD1 |
| ENSG00000181610 | MRPS23 | 0,222806536 | 1,167001598 | 0,13713 | 0,4642578 | mitochondrial ribosomal protein S23 |
| ENSG00000231806 | PCAT7 | 0,222805066 | 1,167000408 | 0,34005 | 0,61888023 | prostate cancer associated transcript 7 |
| ENSG00000227057 | WDR46 | 0,222763255 | 1,166966588 | 0,78307 | 0,89918294 | WD repeat domain 46 |
| ENSG00000240996 | AL157400,4 | 0,222762069 | 1,166965628 | 0,74902 | 0,88190158 | novel transcript |
| ENSG00000103657 | HERC1 | 0,222534475 | 1,166781547 | 0,43996 | 0,6885781 | HECT and RLD domain containing E3 ubiquitin protein ligase family member 1 |
| ENSG00000168769 | TET2 | 0,22251586 | 1,166766492 | 0,61334 | 0,8035281 | tet methylcytosine dioxygenase 2 |
| ENSG00000228063 | LYPLAL1-DT | 0,222412675 | 1,166683045 | 0,70751 | 0,85915509 | LYPLAL1 divergent transcript |
| ENSG00000140993 | TIGD7 | 0,222388849 | 1,166663778 | 0,53842 | 0,75676698 | tigger transposable element derived 7 |
| ENSG00000228261 | ITPRIP-AS1 | 0,222241168 | 1,166544359 | 0,82513 | 0,91900665 | ITPRIP antisense RNA 1 |
| ENSG00000090861 | AARS1 | 0,222220375 | 1,166527546 | 0,23446 | 0,5462109 | alanyl-tRNA synthetase 1 |
| ENSG00000156256 | USP16 | 0,222177062 | 1,166492524 | 0,50686 | 0,73521263 | ubiquitin specific peptidase 16 |
| ENSG00000134287 | ARF3 | 0,222134376 | 1,166458011 | 0,25881 | 0,56260423 | ADP ribosylation factor 3 |
| ENSG00000163374 | YY1AP1 | 0,22212215 | 1,166448127 | 0,24005 | 0,54938115 | YY1 associated protein 1 |
| ENSG00000081052 | COL4A4 | 0,222054791 | 1,166393666 | 0,58259 | 0,7847455 | collagen type IV alpha 4 chain |
| ENSG00000228133 | AC099684,1 | 0,222002868 | 1,166351688 | 0,80743 | 0,91133754 | novel transcript, antisense to RTN4RL1 |
| ENSG00000104643 | MTMR9 | 0,221964994 | 1,166321069 | 0,33133 | 0,61140339 | myotubularin related protein 9 |
| ENSG00000164197 | RNF180 | 0,221946076 | 1,166305776 | 0,58653 | 0,78686419 | ring finger protein 180 |
| ENSG00000272732 | AC004982,1 | 0,221945484 | 1,166305297 | 0,75475 | 0,88510422 | novel transcript |

| | | | | | | |
|---|---|---|---|---|---|---|
| ENSG00000138180 | CEP55 | 0,221945343 | 1,166305183 | 0,61453 | 0,80382182 | centrosomal protein 55 |
| ENSG00000260025 | CRIM1-DT | 0,221927606 | 1,166290844 | 0,82828 | 0,92076203 | CRIM1 divergent transcript |
| ENSG00000075415 | SLC25A3 | 0,221906209 | 1,166273547 | 0,23887 | 0,54892264 | solute carrier family 25 member 3 |
| ENSG00000105738 | SIPA1L3 | 0,221883449 | 1,166255147 | 0,40919 | 0,66785537 | signal induced proliferation associated 1 like 3 |
| ENSG00000185630 | PBX1 | 0,221860022 | 1,16623621 | 0,44153 | 0,68977736 | PBX homeobox 1 |
| ENSG00000126777 | KTN1 | 0,221731606 | 1,166132406 | 0,37063 | 0,63997937 | kinectin 1 |
| ENSG00000166170 | BAG5 | 0,221707764 | 1,166113135 | 0,4839 | 0,71999322 | BAG cochaperone 5 |
| ENSG00000141543 | EIF4A3 | 0,221700358 | 1,166107149 | 0,31048 | 0,59844173 | eukaryotic translation initiation factor 4A3 |
| ENSG00000198736 | MSRB1 | 0,221694576 | 1,166102475 | 0,53069 | 0,75212511 | methionine sulfoxide reductase B1 |
| ENSG00000196712 | NF1 | 0,221694064 | 1,166102062 | 0,48721 | 0,72231397 | neurofibromin 1 |
| ENSG00000168209 | DDIT4 | 0,22149866 | 1,165944131 | 0,36657 | 0,63642135 | DNA damage inducible transcript 4 |
| ENSG00000120963 | ZNF706 | 0,22148442 | 1,165932622 | 0,49448 | 0,72681059 | zinc finger protein 706 |
| ENSG00000227288 | ARID3BP1 | 0,221428707 | 1,165887599 | 0,80301 | 0,90922086 | ARID3B pseudogene 1 |
| ENSG00000179222 | MAGED1 | 0,221398344 | 1,165863062 | 0,39185 | 0,65534953 | MAGE family member D1 |
| ENSG00000171155 | C1GALT1C1 | 0,221359407 | 1,165831596 | 0,52901 | 0,75111082 | C1GALT1 specific chaperone 1 |
| ENSG00000288368 | C1GALT1C1 | 0,221359407 | 1,165831596 | 0,52901 | 0,75111082 | C1GALT1 specific chaperone 1 |
| ENSG00000115255 | REEP6 | 0,2213334 | 1,165810581 | 0,51567 | 0,7417843 | receptor accessory protein 6 |
| ENSG00000228697 | AL023755,1 | 0,221263176 | 1,165753836 | 0,77056 | 0,89367734 | novel transcript |
| ENSG00000177054 | ZDHHC13 | 0,22126307 | 1,16575375 | 0,51202 | 0,73902604 | zinc finger DHHC-type palmitoyltransferase 13 |
| ENSG00000104044 | OCA2 | 0,221220891 | 1,165719668 | 0,72723 | 0,87016629 | OCA2 melanosomal transmembrane protein |
| ENSG00000134686 | PHC2 | 0,221184143 | 1,165689975 | 0,42146 | 0,67643128 | polyhomeotic homolog 2 |
| ENSG00000261652 | C15orf65 | 0,221154419 | 1,165665958 | 0,78156 | 0,89877377 | chromosome 15 open reading frame 65 |
| ENSG00000011405 | PIK3C2A | 0,221112667 | 1,165632224 | 0,55532 | 0,76788267 | phosphatidylinositol-4-phosphate 3-kinase catalytic subunit type 2 alpha |
| ENSG00000204623 | ZNRD1ASP | 0,221091507 | 1,165615128 | 0,56384 | 0,77259134 | zinc ribbon domain containing 1 antisense, pseudogene |
| ENSG00000236051 | MYCBP2-AS1 | 0,221072072 | 1,165599426 | 0,69974 | 0,85496872 | MYCBP2 antisense RNA 1 |
| ENSG00000147586 | MRPS28 | 0,221020478 | 1,165557742 | 0,5398 | 0,75803648 | mitochondrial ribosomal protein S28 |
| ENSG00000248668 | OXCT1-AS1 | 0,221011303 | 1,165550329 | 0,63216 | 0,81450663 | OXCT1 antisense RNA 1 |
| ENSG00000172164 | SNTB1 | 0,220990865 | 1,165533818 | 0,41949 | 0,6752986 | syntrophin beta 1 |
| ENSG00000276814 | AC004801,6 | 0,220915121 | 1,165472627 | 0,6473 | 0,82349878 | novel transcript |
| ENSG00000119906 | SLF2 | 0,220898149 | 1,165458916 | 0,49985 | 0,73058737 | SMC5-SMC6 complex localization factor 2 |
| ENSG00000111707 | SUDS3 | 0,22065757 | 1,165264585 | 0,20656 | 0,5262225 | SDS3 homolog, SIN3A corepressor complex component |
| ENSG00000285973 | AC063960,2 | 0,220608536 | 1,165224981 | 0,54905 | 0,76394298 | novel transcript, antisense to TACC2 |

| | | | | | | |
|---|---|---|---|---|---|---|
| ENSG00000169230 | PRELID1 | 0,220557388 | 1,165183671 | 0,28196 | 0,57936013 | PRELI domain containing 1 |
| ENSG00000115942 | ORC2 | 0,220538522 | 1,165168433 | 0,19558 | 0,51643293 | origin recognition complex subunit 2 |
| ENSG00000119523 | ALG2 | 0,220534617 | 1,16516528 | 0,26581 | 0,56871984 | ALG2 alpha-1,3/1,6-mannosyltransferase |
| ENSG00000271553 | AC018638,7 | 0,22052104 | 1,165154314 | 0,5956 | 0,79342232 | novel transcript |
| ENSG00000224185 | SNX18P9 | 0,220494664 | 1,165133013 | 0,68907 | 0,84854717 | sorting nexin 18 pseudogene 9 |
| ENSG00000279712 | FP325330,3 | 0,220262998 | 1,164945932 | 0,66201 | 0,83168744 | novel transcript |
| ENSG00000182154 | MRPL41 | 0,220204409 | 1,164898624 | 0,25117 | 0,55719428 | mitochondrial ribosomal protein L41 |
| ENSG00000164040 | PGRMC2 | 0,220132314 | 1,164840413 | 0,54487 | 0,76084582 | progesterone receptor membrane component 2 |
| ENSG00000131620 | ANO1 | 0,220112922 | 1,164824756 | 0,69675 | 0,85309963 | anoctamin 1 |
| ENSG00000155287 | SLC25A28 | 0,220089452 | 1,164805806 | 0,53481 | 0,75419058 | solute carrier family 25 member 28 |
| ENSG00000147912 | FBXO10 | 0,220057064 | 1,164779657 | 0,27423 | 0,57420712 | F-box protein 10 |
| ENSG00000260924 | LINC01311 | 0,220031723 | 1,164759198 | 0,48037 | 0,71764212 | long intergenic non-protein coding RNA 1311 |
| ENSG00000156973 | PDE6D | 0,219999678 | 1,164733326 | 0,53686 | 0,75562287 | phosphodiesterase 6D |
| ENSG00000120451 | SNX19 | 0,219942757 | 1,164687374 | 0,215 | 0,53373528 | sorting nexin 19 |
| ENSG00000169989 | TIGD4 | 0,219860727 | 1,164621152 | 0,62134 | 0,80787022 | tigger transposable element derived 4 |
| ENSG00000130402 | ACTN4 | 0,219561409 | 1,164379552 | 0,41823 | 0,67451754 | actinin alpha 4 |
| ENSG00000139278 | GLIPR1 | 0,2195248 | 1,164350006 | 0,66613 | 0,83391259 | GLI pathogenesis related 1 |
| ENSG00000242616 | GNG10 | 0,219483249 | 1,164316472 | 0,67644 | 0,8407877 | G protein subunit gamma 10 |
| ENSG00000137124 | ALDH1B1 | 0,219466763 | 1,164303167 | 0,20131 | 0,52181085 | aldehyde dehydrogenase 1 family member B1 |
| ENSG00000116985 | BMP8B | 0,219366393 | 1,164222167 | 0,44188 | 0,68994635 | bone morphogenetic protein 8b |
| ENSG00000149256 | TENM4 | 0,219281522 | 1,164153681 | 0,41157 | 0,6692379 | teneurin transmembrane protein 4 |
| ENSG00000187013 | LINC02875 | 0,219281062 | 1,16415331 | 0,80712 | 0,91125307 | long intergenic non-protein coding RNA 2875 |
| ENSG00000105708 | ZNF14 | 0,219107443 | 1,164013219 | 0,54631 | 0,76190511 | zinc finger protein 14 |
| ENSG00000162650 | ATXN7L2 | 0,219064554 | 1,163978616 | 0,32215 | 0,60577115 | ataxin 7 like 2 |
| ENSG00000107882 | SUFU | 0,218953336 | 1,163888887 | 0,23465 | 0,54623956 | SUFU negative regulator of hedgehog signaling |
| ENSG00000143127 | ITGA10 | 0,218924687 | 1,163865775 | 0,81684 | 0,91577924 | integrin subunit alpha 10 |
| ENSG00000075945 | KIFAP3 | 0,218907019 | 1,163851523 | 0,49885 | 0,72996654 | kinesin associated protein 3 |
| ENSG00000085465 | OVGP1 | 0,218851981 | 1,163807123 | 0,56852 | 0,77613996 | oviductal glycoprotein 1 |
| ENSG00000147059 | SPIN2A | 0,218761408 | 1,163734061 | 0,44648 | 0,69314849 | spindlin family member 2A |
| ENSG00000197121 | PGAP1 | 0,218735763 | 1,163713375 | 0,5422 | 0,75894841 | post-GPI attachment to proteins inositol deacylase 1 |
| ENSG00000133816 | MICAL2 | 0,218715823 | 1,16369729 | 0,59919 | 0,79538547 | microtubule associated monooxygenase, calponin and LIM domain containing 2 |
| ENSG00000170185 | USP38 | 0,218632042 | 1,163629714 | 0,54405 | 0,76034748 | ubiquitin specific peptidase 38 |

| | | | | | | |
|---|---|---|---|---|---|---|
| ENSG00000171617 | ENC1 | 0,21856133 | 1,163572682 | 0,53638 | 0,75518515 | ectodermal-neural cortex 1 |
| ENSG00000105483 | CARD8 | 0,218480746 | 1,16350769 | 0,44703 | 0,69339238 | caspase recruitment domain family member 8 |
| ENSG00000163961 | RNF168 | 0,218451399 | 1,163484022 | 0,24832 | 0,55546558 | ring finger protein 168 |
| ENSG00000280071 | GATD3B | 0,218347865 | 1,163400529 | 0,23949 | 0,5491566 | glutamine amidotransferase like class 1 domain containing 3B |
| ENSG00000145982 | FARS2 | 0,218289169 | 1,163353197 | 0,20494 | 0,52453221 | phenylalanyl-tRNA synthetase 2, mitochondrial |
| ENSG00000179151 | EDC3 | 0,218167497 | 1,163255088 | 0,20248 | 0,52259159 | enhancer of mRNA decapping 3 |
| ENSG00000102870 | ZNF629 | 0,218131313 | 1,163225913 | 0,36255 | 0,63379594 | zinc finger protein 629 |
| ENSG00000148337 | CIZ1 | 0,218105127 | 1,163204799 | 0,31853 | 0,6036094 | CDKN1A interacting zinc finger protein 1 |
| ENSG00000218596 | AL162578,1 | 0,218092032 | 1,163194241 | 0,76859 | 0,89277929 | ribosomal protein S15a (RPS15A) pseudogene |
| ENSG00000235724 | AC009299,2 | 0,218024791 | 1,163140029 | 0,76695 | 0,89180676 | novel transcript, antisense to TANK |
| ENSG00000130943 | PKDREJ | 0,217908336 | 1,163046143 | 0,6574 | 0,82872268 | polycystin family receptor for egg jelly |
| ENSG00000118816 | CCNI | 0,217907304 | 1,163045311 | 0,29675 | 0,58995143 | cyclin I |
| ENSG00000231555 | HSPA1B | 0,217720765 | 1,16289494 | 0,64357 | 0,82109675 | heat shock protein family A (Hsp70) member 1B |
| ENSG00000196369 | SRGAP2B | 0,217582812 | 1,162783748 | 0,39124 | 0,65500675 | SLIT-ROBO Rho GTPase activating protein 2B |
| ENSG00000268705 | BNIP3P26 | 0,217575437 | 1,162777803 | 0,72179 | 0,86716786 | BCL2 interacting protein 3 pseudogene 26 |
| ENSG00000186364 | NUDT17 | 0,217564857 | 1,162769277 | 0,49372 | 0,72643895 | nudix hydrolase 17 |
| ENSG00000130775 | THEMIS2 | 0,217539996 | 1,162749239 | 0,49709 | 0,72865531 | thymocyte selection associated family member 2 |
| ENSG00000242588 | AC108010,1 | 0,21749728 | 1,162714812 | 0,47544 | 0,71404521 | novel transcript |
| ENSG00000106244 | PDAP1 | 0,217461414 | 1,162685907 | 0,42533 | 0,67893062 | PDGFA associated protein 1 |
| ENSG00000106537 | TSPAN13 | 0,217401646 | 1,162637741 | 0,58426 | 0,78572744 | tetraspanin 13 |
| ENSG00000205978 | NYNRIN | 0,217369289 | 1,162611665 | 0,31078 | 0,5987267 | NYN domain and retroviral integrase containing |
| ENSG00000170236 | USP50 | 0,217327646 | 1,162578107 | 0,55556 | 0,76790322 | ubiquitin specific peptidase 50 |
| ENSG00000125827 | TMX4 | 0,21732052 | 1,162572365 | 0,3189 | 0,60399503 | thioredoxin related transmembrane protein 4 |
| ENSG00000113712 | CSNK1A1 | 0,217308838 | 1,162562951 | 0,39464 | 0,65756576 | casein kinase 1 alpha 1 |
| ENSG00000273542 | H4C12 | 0,217234713 | 1,16250322 | 0,68417 | 0,84539665 | H4 clustered histone 12 |
| ENSG00000286207 | AC012443,2 | 0,217116229 | 1,162407752 | 0,50015 | 0,73078774 | novel transcript, antisense to FMNL2 |
| ENSG00000272240 | AC004908,1 | 0,217085972 | 1,162383373 | 0,66262 | 0,83204447 | novel transcript, antisense to ZNF596 |
| ENSG00000140632 | GLYR1 | 0,217029057 | 1,162337518 | 0,27267 | 0,57315967 | glyoxylate reductase 1 homolog |
| ENSG00000178449 | COX14 | 0,21698399 | 1,162301209 | 0,4655 | 0,70591843 | cytochrome c oxidase assembly factor COX14 |
| ENSG00000168002 | POLR2G | 0,216952315 | 1,162275691 | 0,49349 | 0,72636471 | RNA polymerase II subunit G |
| ENSG00000125818 | PSMF1 | 0,216925308 | 1,162253933 | 0,30445 | 0,59430086 | proteasome inhibitor subunit 1 |
| ENSG00000285248 | IPO4 | 0,216892939 | 1,162227857 | 0,32293 | 0,60584543 | importin 4 |

| | | | | | | |
|---|---|---|---|---|---|---|
| ENSG00000196497 | IPO4 | 0,216892939 | 1,162227857 | 0,32293 | 0,60584543 | importin 4 |
| ENSG00000187601 | MAGEH1 | 0,216781501 | 1,162138086 | 0,38267 | 0,64939435 | MAGE family member H1 |
| ENSG00000283828 | AL137002,2 | 0,216776652 | 1,16213418 | 0,78825 | 0,90156082 | WAS/WASL-interacting protein family member 2-like |
| ENSG00000180189 | HMGB1P14 | 0,216638282 | 1,162022724 | 0,6011 | 0,79627587 | high mobility group box 1 pseudogene 14 |
| ENSG00000275221 | H2AC15 | 0,216604236 | 1,161995302 | 0,61337 | 0,8035281 | H2A clustered histone 15 |
| ENSG00000177692 | DNAJC28 | 0,216574532 | 1,161971377 | 0,67315 | 0,83865474 | DnaJ heat shock protein family (Hsp40) member C28 |
| ENSG00000105341 | DMAC2 | 0,216507906 | 1,161917717 | 0,20872 | 0,52791772 | distal membrane arm assembly complex 2 |
| ENSG00000138162 | TACC2 | 0,216420022 | 1,161846939 | 0,23303 | 0,54557342 | transforming acidic coiled-coil containing protein 2 |
| ENSG00000230124 | ACBD6 | 0,216419851 | 1,161846801 | 0,22154 | 0,53805234 | acyl-CoA binding domain containing 6 |
| ENSG00000226853 | AC010894,1 | 0,216413548 | 1,161841726 | 0,67821 | 0,84161088 | novel transcript |
| ENSG00000103995 | CEP152 | 0,216359692 | 1,161798355 | 0,44209 | 0,69003109 | centrosomal protein 152 |
| ENSG00000136869 | TLR4 | 0,216346482 | 1,161787717 | 0,75859 | 0,88727479 | toll like receptor 4 |
| ENSG00000039123 | MTREX | 0,216345774 | 1,161787147 | 0,2089 | 0,52800078 | Mtr4 exosome RNA helicase |
| ENSG00000198046 | ZNF667 | 0,216276988 | 1,161731755 | 0,69885 | 0,85437411 | zinc finger protein 667 |
| ENSG00000237816 | RPL21P109 | 0,216239286 | 1,161701396 | 0,8202 | 0,91725087 | ribosomal protein L21 pseudogene 109 |
| ENSG00000202198 | 7SK | 0,21617751 | 1,161651653 | 0,76376 | 0,88986791 | 7SK RNA |
| ENSG00000240445 | FOXO3B | 0,216086813 | 1,161578627 | 0,27275 | 0,57322782 | forkhead box O3B |
| ENSG00000147485 | PXDNL | 0,216053179 | 1,161551547 | 0,58688 | 0,78703464 | peroxidasin like |
| ENSG00000180901 | KCTD2 | 0,216027127 | 1,161530572 | 0,21152 | 0,53032236 | potassium channel tetramerization domain containing 2 |
| ENSG00000270314 | AC007161,2 | 0,216026224 | 1,161529845 | 0,74003 | 0,87675329 | 5-azacytidine induced 2 (AZI2) pseudogene |
| ENSG00000199691 | RN7SKP173 | 0,215860837 | 1,161396698 | 0,79731 | 0,90661857 | RN7SK pseudogene 173 |
| ENSG00000263080 | AC009121,3 | 0,215852685 | 1,161390136 | 0,69305 | 0,85087762 | novel transcript, antisense to RMI2 |
| ENSG00000279208 | CR381653,2 | 0,215793615 | 1,161342584 | 0,90748 | 0,96044232 | ankyrin repeat domain 18A (ANKRD18A) pseudogene |
| ENSG00000163872 | YEATS2 | 0,215742881 | 1,161301745 | 0,39394 | 0,65694093 | YEATS domain containing 2 |
| ENSG00000171161 | ZNF672 | 0,215676614 | 1,161248405 | 0,29431 | 0,58857672 | zinc finger protein 672 |
| ENSG00000282256 | TAS2R31 | 0,215648881 | 1,161226081 | 0,76941 | 0,89297049 | taste 2 receptor member 31 |
| ENSG00000164172 | MOCS2 | 0,215630923 | 1,161211627 | 0,4437 | 0,69139564 | molybdenum cofactor synthesis 2 |
| ENSG00000288295 | MCTS1 | 0,215620451 | 1,161203199 | 0,67453 | 0,83975333 | MCTS1 re-initiation and release factor |
| ENSG00000117395 | EBNA1BP2 | 0,215598744 | 1,161185727 | 0,2332 | 0,54558536 | EBNA1 binding protein 2 |
| ENSG00000196465 | MYL6B | 0,215562892 | 1,161156871 | 0,27732 | 0,57586315 | myosin light chain 6B |
| ENSG00000248027 | ARHGAP42-AS | 0,215560538 | 1,161154977 | 0,73672 | 0,87529332 | ARHGAP42 antisense RNA 1 |
| ENSG00000183495 | EP400 | 0,215556603 | 1,161151809 | 0,2575 | 0,56160845 | E1A binding protein p400 |
| ENSG00000254900 | AC091053,2 | 0,215545267 | 1,161142686 | 0,85303 | 0,9332295 | novel transcript |

| | | | | | | |
|---|---|---|---|---|---|---|
| ENSG00000260410 | AC040169,2 | 0,215534457 | 1,161133986 | 0,77017 | 0,89333117 | polo-like kinase 1 (PLK1) pseudogene |
| ENSG00000179104 | TMTC2 | 0,21551483 | 1,161118189 | 0,59263 | 0,79130579 | transmembrane O-mannosyltransferase targeting cadherins 2 |
| ENSG00000146950 | SHROOM2 | 0,215488289 | 1,161096828 | 0,37703 | 0,64554193 | shroom family member 2 |
| ENSG00000087502 | ERGIC2 | 0,215456171 | 1,16107098 | 0,35506 | 0,62946175 | ERGIC and golgi 2 |
| ENSG00000163069 | SGCB | 0,215453251 | 1,16106863 | 0,37767 | 0,64613367 | sarcoglycan beta |
| ENSG00000105085 | MED26 | 0,215402221 | 1,161027562 | 0,33696 | 0,61684812 | mediator complex subunit 26 |
| ENSG00000225675 | LINC01771 | 0,215335114 | 1,160973558 | 0,85025 | 0,93185051 | long intergenic non-protein coding RNA 1771 |
| ENSG00000182944 | EWSR1 | 0,215292407 | 1,160939191 | 0,37207 | 0,64114037 | EWS RNA binding protein 1 |
| ENSG00000064199 | SPA17 | 0,215234887 | 1,160892905 | 0,55484 | 0,76763832 | sperm autoantigenic protein 17 |
| ENSG00000214070 | MANEALP1 | 0,21517656 | 1,160845972 | 0,78239 | 0,89892226 | MANEAL pseudogene 1 |
| ENSG00000223462 | AL353801,1 | 0,215120467 | 1,160800839 | 0,82488 | 0,91900665 | novel transcript |
| ENSG00000047644 | WWC3 | 0,215103837 | 1,160787458 | 0,21216 | 0,5308403 | WWC family member 3 |
| ENSG00000267058 | AC006213,2 | 0,215016509 | 1,160717197 | 0,53023 | 0,75184556 | novel transcript |
| ENSG00000282859 | AC003992,1 | 0,214970939 | 1,160680534 | 0,87531 | 0,94485247 | novel transcript, sense intronic to FOXP2 |
| ENSG00000184863 | RBM33 | 0,214869648 | 1,160599046 | 0,2896 | 0,58522301 | RNA binding motif protein 33 |
| ENSG00000124406 | ATP8A1 | 0,214795872 | 1,160539697 | 0,58827 | 0,78810304 | ATPase phospholipid transporting 8A1 |
| ENSG00000240356 | RPL23AP7 | 0,214757025 | 1,160508448 | 0,63141 | 0,81403245 | ribosomal protein L23a pseudogene 7 |
| ENSG00000214113 | LYRM4 | 0,214655388 | 1,160426694 | 0,41058 | 0,66856387 | LYR motif containing 4 |
| ENSG00000213300 | HNRNPA3P6 | 0,214575667 | 1,160362572 | 0,90769 | 0,96044232 | heterogeneous nuclear ribonucleoprotein A3 pseudogene 6 |
| ENSG00000077549 | CAPZB | 0,214508928 | 1,160308895 | 0,26542 | 0,56823546 | capping actin protein of muscle Z-line subunit beta |
| ENSG00000272077 | AC124045,1 | 0,214412549 | 1,160231383 | 0,57776 | 0,78160743 | novel transcript |
| ENSG00000281102 | AC092046,2 | 0,214412549 | 1,160231383 | 0,57776 | 0,78160743 | novel transcript |
| ENSG00000284900 | KBTBD4 | 0,214401833 | 1,160222765 | 0,48737 | 0,72237072 | kelch repeat and BTB domain containing 4 |
| ENSG00000123444 | KBTBD4 | 0,214401833 | 1,160222765 | 0,48737 | 0,72237072 | kelch repeat and BTB domain containing 4 |
| ENSG00000181192 | DHTKD1 | 0,214367503 | 1,160195157 | 0,24909 | 0,55619875 | dehydrogenase E1 and transketolase domain containing 1 |
| ENSG00000256683 | ZNF350 | 0,214248505 | 1,160099465 | 0,48431 | 0,72019446 | zinc finger protein 350 |
| ENSG00000152767 | FARP1 | 0,214175158 | 1,160040487 | 0,17522 | 0,49934416 | FERM, ARH/RhoGEF and pleckstrin domain protein 1 |
| ENSG00000175387 | SMAD2 | 0,21405338 | 1,159942572 | 0,21034 | 0,5290538 | SMAD family member 2 |
| ENSG00000038532 | CLEC16A | 0,214040644 | 1,159932332 | 0,27467 | 0,57463533 | C-type lectin domain containing 16A |
| ENSG00000171843 | MLLT3 | 0,214025215 | 1,159919927 | 0,60421 | 0,79780609 | MLLT3 super elongation complex subunit |
| ENSG00000267500 | ZNF887P | 0,213970056 | 1,15987558 | 0,66958 | 0,83623395 | zinc finger protein 887, pseudogene |

| | | | | | | |
|---|---|---|---|---|---|---|
| ENSG00000196923 | PDLIM7 | 0,213941462 | 1,159852592 | 0,3183 | 0,60349082 | PDZ and LIM domain 7 |
| ENSG00000138794 | CASP6 | 0,213842697 | 1,159773193 | 0,31977 | 0,60470232 | caspase 6 |
| ENSG00000129460 | NGDN | 0,213838024 | 1,159769436 | 0,22498 | 0,54037642 | neuroguidin |
| ENSG00000259305 | ZHX1-C8orf76 | 0,21382575 | 1,15975957 | 0,64192 | 0,8205913 | ZHX1-C8orf76 readthrough |
| ENSG00000167747 | C19orf48 | 0,21373566 | 1,159687149 | 0,39952 | 0,66176242 | chromosome 19 open reading frame 48 |
| ENSG00000099954 | CECR2 | 0,213720994 | 1,15967536 | 0,31083 | 0,59878622 | CECR2 histone acetyl-lysine reader |
| ENSG00000178974 | FBXO34 | 0,213691155 | 1,159651375 | 0,52686 | 0,75003528 | F-box protein 34 |
| ENSG00000150712 | MTMR12 | 0,213572887 | 1,159556315 | 0,17903 | 0,50343511 | myotubularin related protein 12 |
| ENSG00000258056 | AC009779,2 | 0,213558832 | 1,159545018 | 0,3773 | 0,64577365 | novel transcript, antisense to CD63 |
| ENSG00000287630 | AC011370,1 | 0,213512314 | 1,159507631 | 0,55763 | 0,76931947 | novel transcript, antisense to JAKMIP2 |
| ENSG00000204406 | MBD5 | 0,213498527 | 1,15949655 | 0,38709 | 0,65213944 | methyl-CpG binding domain protein 5 |
| ENSG00000116786 | PLEKHM2 | 0,213496656 | 1,159495046 | 0,34235 | 0,62005694 | pleckstrin homology and RUN domain containing M2 |
| ENSG00000106771 | TMEM245 | 0,213464732 | 1,159469389 | 0,40736 | 0,66661764 | transmembrane protein 245 |
| ENSG00000198824 | CHAMP1 | 0,213427373 | 1,159439364 | 0,2093 | 0,52826608 | chromosome alignment maintaining phosphoprotein 1 |
| ENSG00000224715 | Z82186,1 | 0,213356166 | 1,15938214 | 0,77737 | 0,8969723 | novel transcript |
| ENSG00000205937 | RNPS1 | 0,213140226 | 1,159208618 | 0,24518 | 0,55295131 | RNA binding protein with serine rich domain 1 |
| ENSG00000138767 | CNOT6L | 0,213094336 | 1,159171746 | 0,57242 | 0,77849297 | CCR4-NOT transcription complex subunit 6 like |
| ENSG00000270020 | AC009108,3 | 0,213089444 | 1,159167816 | 0,62791 | 0,81175174 | novel transcript |
| ENSG00000286846 | Z93943,1 | 0,21305998 | 1,159144142 | 0,66782 | 0,83521885 | novel transcript |
| ENSG00000165312 | OTUD1 | 0,213013906 | 1,159107125 | 0,39022 | 0,65427905 | OTU deubiquitinase 1 |
| ENSG00000244604 | AC025518,1 | 0,212902436 | 1,159017569 | 0,78753 | 0,90121071 | ribosomal protein S26 (RPS26) pseudogene |
| ENSG00000188352 | FOCAD | 0,212783314 | 1,158921874 | 0,52499 | 0,74873866 | focadhesin |
| ENSG00000166446 | CDYL2 | 0,212727567 | 1,158877093 | 0,49845 | 0,72960863 | chromodomain Y like 2 |
| ENSG00000011485 | PPP5C | 0,21271811 | 1,158869497 | 0,42503 | 0,67871429 | protein phosphatase 5 catalytic subunit |
| ENSG00000174886 | NDUFA11 | 0,21268704 | 1,15884454 | 0,40501 | 0,66481282 | NADH:ubiquinone oxidoreductase subunit A11 |
| ENSG00000260733 | AC046158,1 | 0,212685168 | 1,158843036 | 0,76128 | 0,88879633 | novel transcript, antisense to WWOX |
| ENSG00000187123 | LYPD6 | 0,212684296 | 1,158842335 | 0,6397 | 0,81918091 | LY6/PLAUR domain containing 6 |
| ENSG00000042753 | AP2S1 | 0,212620891 | 1,158791406 | 0,31219 | 0,59925736 | adaptor related protein complex 2 subunit sigma 1 |
| ENSG00000259161 | NPM1P20 | 0,212551548 | 1,158735711 | 0,72692 | 0,87006668 | nucleophosmin 1 pseudogene 20 |
| ENSG00000273117 | AC144652,1 | 0,212530889 | 1,158719118 | 0,48938 | 0,72387044 | novel transcript |
| ENSG00000196876 | SCN8A | 0,212522478 | 1,158712362 | 0,42786 | 0,68085041 | sodium voltage-gated channel alpha subunit 8 |
| ENSG00000151917 | BEND6 | 0,212445363 | 1,158650429 | 0,53158 | 0,75280487 | BEN domain containing 6 |
| ENSG00000119862 | LGALSL | 0,21234088 | 1,15856652 | 0,68869 | 0,84822863 | galectin like |

| | | | | | | |
|---|---|---|---|---|---|---|
| ENSG00000154639 | CXADR | 0,212329601 | 1,158557462 | 0,56005 | 0,77076855 | CXADR Ig-like cell adhesion molecule |
| ENSG00000276550 | HERC2P2 | 0,212255104 | 1,158497639 | 0,43264 | 0,68418465 | hect domain and RLD 2 pseudogene 2 |
| ENSG00000135677 | GNS | 0,21221321 | 1,158463998 | 0,27143 | 0,57285347 | glucosamine (N-acetyl)-6-sulfatase |
| ENSG00000134444 | RELCH | 0,21219081 | 1,158446011 | 0,45086 | 0,69594229 | RAB11 binding and LisH domain, coiled-coil and HEAT repeat containing |
| ENSG00000102796 | DHRS12 | 0,212147141 | 1,158410947 | 0,45363 | 0,69783624 | dehydrogenase/reductase 12 |
| ENSG00000147896 | IFNK | 0,212094069 | 1,158368334 | 0,68969 | 0,84865812 | interferon kappa |
| ENSG00000105372 | RPS19 | 0,212053374 | 1,158335659 | 0,3932 | 0,65625452 | ribosomal protein S19 |
| ENSG00000186577 | SMIM29 | 0,212033452 | 1,158319664 | 0,42819 | 0,68094175 | small integral membrane protein 29 |
| ENSG00000266642 | AC024267,6 | 0,211947605 | 1,158250741 | 0,89016 | 0,95161642 | novel transcript, antisense to DHRS13 |
| ENSG00000156639 | ZFAND3 | 0,211924947 | 1,15823255 | 0,33191 | 0,61191629 | zinc finger AN1-type containing 3 |
| ENSG00000227232 | WASH7P | 0,211844227 | 1,158167748 | 0,41745 | 0,67403924 | WASP family homolog 7, pseudogene |
| ENSG00000138760 | SCARB2 | 0,211717719 | 1,158066194 | 0,45591 | 0,69987751 | scavenger receptor class B member 2 |
| ENSG00000114857 | NKTR | 0,211361219 | 1,157780063 | 0,51671 | 0,7420923 | natural killer cell triggering receptor |
| ENSG00000149970 | CNKSR2 | 0,211356798 | 1,157776516 | 0,48561 | 0,72107911 | connector enhancer of kinase suppressor of Ras 2 |
| ENSG00000283903 | FGF11 | 0,211305811 | 1,157735598 | 0,44452 | 0,69185302 | fibroblast growth factor 11 |
| ENSG00000262410 | AC024361,1 | 0,211241893 | 1,157684307 | 0,62046 | 0,80776776 | novel transcript, sense intronic FN3K |
| ENSG00000280987 | MATR3 | 0,211209747 | 1,157658512 | 0,79942 | 0,90744841 | matrin 3 |
| ENSG00000111716 | LDHB | 0,211148389 | 1,157609278 | 0,31612 | 0,60224605 | lactate dehydrogenase B |
| ENSG00000170291 | ELP5 | 0,211134438 | 1,157598083 | 0,3044 | 0,59430086 | elongator acetyltransferase complex subunit 5 |
| ENSG00000288485 | ELP5 | 0,211134438 | 1,157598083 | 0,3044 | 0,59430086 | elongator acetyltransferase complex subunit 5 |
| ENSG00000271795 | AC011337,1 | 0,211093141 | 1,157564947 | 0,62222 | 0,80838627 | novel transcript, antisense to FAT2 |
| ENSG00000141506 | PIK3R5 | 0,21100984 | 1,157498111 | 0,8171 | 0,91592225 | phosphoinositide-3-kinase regulatory subunit 5 |
| ENSG00000101182 | PSMA7 | 0,211002814 | 1,157492475 | 0,23885 | 0,54892264 | proteasome 20S subunit alpha 7 |
| ENSG00000196313 | POM121 | 0,210964564 | 1,157461787 | 0,27485 | 0,57463533 | POM121 transmembrane nucleoporin |
| ENSG00000267501 | AC104365,2 | 0,210891545 | 1,157403206 | 0,80736 | 0,91131389 | novel transcript |
| ENSG00000259318 | HMGN1P1 | 0,210882284 | 1,157395776 | 0,73084 | 0,87205836 | high mobility group nucleosome binding domain 1 pseudogene 1 |
| ENSG00000287151 | C2orf27A | 0,210822093 | 1,157347489 | 0,60915 | 0,80161331 | chromosome 2 open reading frame 27A |
| ENSG00000135441 | BLOC1S1 | 0,210531998 | 1,157114795 | 0,61345 | 0,8035281 | biogenesis of lysosomal organelles complex 1 subunit 1 |
| ENSG00000159915 | ZNF233 | 0,210507522 | 1,157095163 | 0,49886 | 0,72996654 | zinc finger protein 233 |
| ENSG00000118096 | IFT46 | 0,210468171 | 1,157063604 | 0,35124 | 0,626706 | intraflagellar transport 46 |
| ENSG00000183723 | CMTM4 | 0,210446342 | 1,157046096 | 0,32155 | 0,60577115 | CKLF like MARVEL transmembrane domain containing 4 |

| | | | | | | |
|---|---|---|---|---|---|---|
| ENSG00000063241 | ISOC2 | 0,210397735 | 1,157007114 | 0,35373 | 0,62855856 | isochorismatase domain containing 2 |
| ENSG00000197054 | ZNF763 | 0,21033264 | 1,156954911 | 0,51047 | 0,73800292 | zinc finger protein 763 |
| ENSG00000092978 | GPATCH2 | 0,210317831 | 1,156943035 | 0,34108 | 0,6192144 | G-patch domain containing 2 |
| ENSG00000263181 | CYP2D7 | 0,210304171 | 1,156932081 | 0,75919 | 0,8876058 | cytochrome P450 family 2 subfamily D member 7 (gene/pseudogene) |
| ENSG00000148123 | PLPPR1 | 0,21024568 | 1,156885176 | 0,62576 | 0,81058216 | phospholipid phosphatase related 1 |
| ENSG00000281877 | RPS6KA1 | 0,210226483 | 1,156869782 | 0,40162 | 0,66306607 | ribosomal protein S6 kinase A1 |
| ENSG00000117676 | RPS6KA1 | 0,210226483 | 1,156869782 | 0,40162 | 0,66306607 | ribosomal protein S6 kinase A1 |
| ENSG00000285000 | AC008581,2 | 0,210217967 | 1,156862954 | 0,2733 | 0,57344244 | novel protein, AGGF1-ZBED3-AS1 readthrough |
| ENSG00000100897 | DCAF11 | 0,210142215 | 1,156802211 | 0,54629 | 0,76190511 | DDB1 and CUL4 associated factor 11 |
| ENSG00000185219 | ZNF445 | 0,210120485 | 1,156784787 | 0,53333 | 0,75351438 | zinc finger protein 445 |
| ENSG00000115226 | FNDC4 | 0,210084085 | 1,156755602 | 0,44895 | 0,69460209 | fibronectin type III domain containing 4 |
| ENSG00000155975 | VPS37A | 0,21008367 | 1,156755269 | 0,42066 | 0,67598168 | VPS37A subunit of ESCRT-I |
| ENSG00000151692 | RNF144A | 0,210055589 | 1,156732754 | 0,36575 | 0,63597214 | ring finger protein 144A |
| ENSG00000174851 | YIF1A | 0,210045655 | 1,156724789 | 0,41344 | 0,67078587 | Yip1 interacting factor homolog A, membrane trafficking protein |
| ENSG00000092470 | WDR76 | 0,21000888 | 1,156695304 | 0,43846 | 0,68769465 | WD repeat domain 76 |
| ENSG00000176209 | SMIM19 | 0,209950924 | 1,156648838 | 0,35825 | 0,63185282 | small integral membrane protein 19 |
| ENSG00000244184 | AC091153,3 | 0,209940122 | 1,156640177 | 0,71272 | 0,86178349 | novel transcript |
| ENSG00000215319 | EIF5P1 | 0,209868784 | 1,156582986 | 0,6901 | 0,8488605 | eukaryotic translation initiation factor 5 pseudogene 1 |
| ENSG00000107371 | EXOSC3 | 0,20984216 | 1,156561641 | 0,195 | 0,51588752 | exosome component 3 |
| ENSG00000148843 | PDCD11 | 0,209764751 | 1,156499587 | 0,34639 | 0,62342969 | programmed cell death 11 |
| ENSG00000271265 | AL355297,2 | 0,209750844 | 1,156488439 | 0,79813 | 0,90685415 | novel transcript |
| ENSG00000137216 | TMEM63B | 0,20974063 | 1,156480251 | 0,35743 | 0,63132419 | transmembrane protein 63B |
| ENSG00000253966 | AC008514,2 | 0,209668167 | 1,156422166 | 0,75108 | 0,88327589 | tubulin, epsilon 1 (TUBE1) pseudogene |
| ENSG00000108523 | RNF167 | 0,209655883 | 1,156412319 | 0,32923 | 0,61033361 | ring finger protein 167 |
| ENSG00000115211 | EIF2B4 | 0,209439028 | 1,156238509 | 0,35434 | 0,62899743 | eukaryotic translation initiation factor 2B subunit delta |
| ENSG00000134077 | THUMPD3 | 0,209385328 | 1,156195472 | 0,38459 | 0,65052153 | THUMP domain containing 3 |
| ENSG00000138757 | G3BP2 | 0,209365222 | 1,156179359 | 0,23155 | 0,54440987 | G3BP stress granule assembly factor 2 |
| ENSG00000214776 | AC092821,1 | 0,209324243 | 1,156146519 | 0,4681 | 0,70771758 | alpha-2-macroglobulin like 1 pseudogene |
| ENSG00000118707 | TGIF2 | 0,209293896 | 1,1561222 | 0,28616 | 0,58259176 | TGFB induced factor homeobox 2 |
| ENSG00000134057 | CCNB1 | 0,209241725 | 1,156080393 | 0,30779 | 0,59671558 | cyclin B1 |
| ENSG00000073614 | KDM5A | 0,209216885 | 1,156060488 | 0,27552 | 0,57522081 | lysine demethylase 5A |

| | | | | | | |
|---|---|---|---|---|---|---|
| ENSG00000222011 | FAM185A | 0,209167782 | 1,156021141 | 0,46981 | 0,70922292 | family with sequence similarity 185 member A |
| ENSG00000075975 | MKRN2 | 0,209150244 | 1,156007088 | 0,48821 | 0,72315487 | makorin ring finger protein 2 |
| ENSG00000054116 | TRAPPC3 | 0,209084491 | 1,155954403 | 0,32545 | 0,60794826 | trafficking protein particle complex 3 |
| ENSG00000234474 | MIR3663HG | 0,20894941 | 1,155846174 | 0,81898 | 0,91689679 | MIR3663 host gene |
| ENSG00000148719 | DNAJB12 | 0,208885156 | 1,155794698 | 0,33801 | 0,6176778 | DnaJ heat shock protein family (Hsp40) member B12 |
| ENSG00000121073 | SLC35B1 | 0,208875541 | 1,155786995 | 0,29224 | 0,58692263 | solute carrier family 35 member B1 |
| ENSG00000185658 | BRWD1 | 0,208870363 | 1,155782846 | 0,46593 | 0,70625067 | bromodomain and WD repeat domain containing 1 |
| ENSG00000089159 | PXN | 0,208745037 | 1,155682448 | 0,31751 | 0,60289482 | paxillin |
| ENSG00000120733 | KDM3B | 0,208726074 | 1,155667258 | 0,36073 | 0,63305669 | lysine demethylase 3B |
| ENSG00000187510 | PLEKHG7 | 0,208659335 | 1,155613798 | 0,80711 | 0,91125307 | pleckstrin homology and RhoGEF domain containing G7 |
| ENSG00000070718 | AP3M2 | 0,208460763 | 1,155454751 | 0,39394 | 0,65694093 | adaptor related protein complex 3 subunit mu 2 |
| ENSG00000259940 | AC109449,1 | 0,2084502 | 1,155446291 | 0,67071 | 0,83689479 | novel transcript, antisense to JMJD5 |
| ENSG00000285077 | ARHGAP11B | 0,208396599 | 1,155403363 | 0,67361 | 0,83912754 | Rho GTPase activating protein 11B |
| ENSG00000286623 | AC005041,4 | 0,208369378 | 1,155381563 | 0,68299 | 0,8447391 | novel transcript, antisense to C2orf81 |
| ENSG00000123609 | NMI | 0,208300735 | 1,155326591 | 0,6127 | 0,8035281 | N-myc and STAT interactor |
| ENSG00000174611 | KY | 0,208208968 | 1,155253106 | 0,62236 | 0,80848353 | kyphoscoliosis peptidase |
| ENSG00000237461 | AL359710,1 | 0,20819257 | 1,155239975 | 0,65553 | 0,82781192 | novel transcript |
| ENSG00000255843 | AP000593,3 | 0,208142282 | 1,155199707 | 0,78939 | 0,90249403 | novel transcript |
| ENSG00000156030 | MIDEAS | 0,208120232 | 1,155182052 | 0,44414 | 0,6916877 | mitotic deacetylase associated SANT domain protein |
| ENSG00000104549 | SQLE | 0,20806449 | 1,155137419 | 0,29687 | 0,58996365 | squalene epoxidase |
| ENSG00000160209 | PDXK | 0,207804049 | 1,154928909 | 0,29586 | 0,58942061 | pyridoxal kinase |
| ENSG00000084774 | CAD | 0,207759283 | 1,154893072 | 0,29792 | 0,59037822 | carbamoyl-phosphate synthetase 2, aspartate transcarbamylase, and dihydroorotase |
| ENSG00000168487 | BMP1 | 0,207705303 | 1,154849861 | 0,27836 | 0,57651947 | bone morphogenetic protein 1 |
| ENSG00000288661 | AL451106,1 | 0,207633287 | 1,154792215 | 0,8664 | 0,94074101 | novel protein |
| ENSG00000136699 | SMPD4 | 0,20750647 | 1,15469071 | 0,29461 | 0,58857672 | sphingomyelin phosphodiesterase 4 |
| ENSG00000263846 | CIAPIN1P | 0,207462592 | 1,154655592 | 0,80231 | 0,90900383 | cytokine induced apoptosis inhibitor 1 pseudogene |
| ENSG00000132153 | DHX30 | 0,207373085 | 1,154583957 | 0,22145 | 0,53805234 | DExH-box helicase 30 |
| ENSG00000111653 | ING4 | 0,207355001 | 1,154569485 | 0,50032 | 0,73081513 | inhibitor of growth family member 4 |
| ENSG00000010327 | STAB1 | 0,207292876 | 1,154519768 | 0,78364 | 0,89918294 | stabilin 1 |
| ENSG00000250624 | MTCYBP43 | 0,207223186 | 1,154464001 | 0,6282 | 0,81178582 | MT-CYB pseudogene 43 |
| ENSG00000284980 | AGPAT5 | 0,207218679 | 1,154460394 | 0,5715 | 0,77809338 | 1-acylglycerol-3-phosphate O-acyltransferase 5 |
| ENSG00000155189 | AGPAT5 | 0,207218679 | 1,154460394 | 0,5715 | 0,77809338 | 1-acylglycerol-3-phosphate O-acyltransferase 5 |

| | | | | | | |
|---|---|---|---|---|---|---|
| ENSG00000152492 | CCDC50 | 0,207129892 | 1,154389348 | 0,51268 | 0,73941533 | coiled-coil domain containing 50 |
| ENSG00000285853 | AL137802,3 | 0,207114246 | 1,154376828 | 0,59853 | 0,7951852 | novel transcript |
| ENSG00000135503 | ACVR1B | 0,207062668 | 1,154335558 | 0,36113 | 0,63316311 | activin A receptor type 1B |
| ENSG00000204977 | TRIM13 | 0,206941239 | 1,154238404 | 0,45821 | 0,70115011 | tripartite motif containing 13 |
| ENSG00000273136 | NBPF26 | 0,206921464 | 1,154222584 | 0,20574 | 0,52518691 | NBPF member 26 |
| ENSG00000173548 | SNX33 | 0,206873396 | 1,154184128 | 0,21289 | 0,53196825 | sorting nexin 33 |
| ENSG00000164168 | TMEM184C | 0,206842299 | 1,154159249 | 0,35723 | 0,63120669 | transmembrane protein 184C |
| ENSG00000266900 | AC027514,1 | 0,206758396 | 1,154092128 | 0,6677 | 0,83516218 | novel transcript, sense intronic to KIAA1468 |
| ENSG00000134627 | PIWIL4 | 0,20672617 | 1,15406635 | 0,74318 | 0,87840582 | piwi like RNA-mediated gene silencing 4 |
| ENSG00000132680 | KHDC4 | 0,206617074 | 1,153979083 | 0,42335 | 0,67712191 | KH domain containing 4, pre-mRNA splicing factor |
| ENSG00000137700 | SLC37A4 | 0,206610986 | 1,153974214 | 0,48479 | 0,72052523 | solute carrier family 37 member 4 |
| ENSG00000135362 | PRR5L | 0,206610404 | 1,153973748 | 0,53235 | 0,75291763 | proline rich 5 like |
| ENSG00000124588 | NQO2 | 0,206558111 | 1,153931921 | 0,42318 | 0,67696528 | N-ribosyldihydronicotinamide:quinone reductase 2 |
| ENSG00000225806 | AL121917,1 | 0,206546912 | 1,153922963 | 0,79319 | 0,90371578 | novel transcript, antisense to GNAS |
| ENSG00000284282 | POLR3E | 0,206455953 | 1,153850213 | 0,71167 | 0,86129697 | RNA polymerase III subunit E |
| ENSG00000134291 | TMEM106C | 0,206441224 | 1,153838433 | 0,67517 | 0,84009781 | transmembrane protein 106C |
| ENSG00000170234 | PWWP2A | 0,206404008 | 1,153808669 | 0,41638 | 0,67315741 | PWWP domain containing 2A |
| ENSG00000205707 | ETFRF1 | 0,206392185 | 1,153799213 | 0,64826 | 0,82383528 | electron transfer flavoprotein regulatory factor 1 |
| ENSG00000117139 | KDM5B | 0,206385608 | 1,153793953 | 0,35763 | 0,63145383 | lysine demethylase 5B |
| ENSG00000179091 | CYC1 | 0,206323551 | 1,153744325 | 0,33308 | 0,61307023 | cytochrome c1 |
| ENSG00000144724 | PTPRG | 0,206237366 | 1,153675403 | 0,61381 | 0,8035281 | protein tyrosine phosphatase receptor type G |
| ENSG00000233757 | AC092835,1 | 0,206201695 | 1,153646879 | 0,27592 | 0,57533851 | novel C2H2 type zinc finger protein |
| ENSG00000184661 | CDCA2 | 0,206168083 | 1,153620001 | 0,32503 | 0,60753927 | cell division cycle associated 2 |
| ENSG00000048162 | NOP16 | 0,206104025 | 1,15356878 | 0,2347 | 0,54623956 | NOP16 nucleolar protein |
| ENSG00000250151 | ARPC4-TTLL3 | 0,206068382 | 1,15354028 | 0,79193 | 0,90336706 | ARPC4-TTLL3 readthrough |
| ENSG00000107338 | SHB | 0,206006166 | 1,153490535 | 0,57115 | 0,77790703 | SH2 domain containing adaptor protein B |
| ENSG00000154930 | ACSS1 | 0,205989513 | 1,153477221 | 0,75347 | 0,88453179 | acyl-CoA synthetase short chain family member 1 |
| ENSG00000120159 | CAAP1 | 0,205965889 | 1,153458332 | 0,49029 | 0,72451286 | caspase activity and apoptosis inhibitor 1 |
| ENSG00000158423 | RIBC1 | 0,205948643 | 1,153444544 | 0,66913 | 0,83595685 | RIB43A domain with coiled-coils 1 |
| ENSG00000089101 | CFAP61 | 0,205918856 | 1,15342073 | 0,66833 | 0,83536064 | cilia and flagella associated protein 61 |
| ENSG00000288054 | AL360178,2 | 0,205915729 | 1,15341823 | 0,76338 | 0,88980026 | novel transcript, antisense to PDE7B |
| ENSG00000147955 | SIGMAR1 | 0,205914894 | 1,153417562 | 0,31047 | 0,59844173 | sigma non-opioid intracellular receptor 1 |
| ENSG00000127125 | PPCS | 0,205914488 | 1,153417238 | 0,51113 | 0,738352 | phosphopantothenoylcysteine synthetase |

| | | | | | | |
|---|---|---|---|---|---|---|
| ENSG00000151611 | MMAA | 0,205853215 | 1,153368251 | 0,54856 | 0,76372599 | metabolism of cobalamin associated A |
| ENSG00000196792 | STRN3 | 0,205823423 | 1,153344434 | 0,35717 | 0,63118399 | striatin 3 |
| ENSG00000163597 | SNHG16 | 0,205822933 | 1,153344043 | 0,30136 | 0,59258306 | small nucleolar RNA host gene 16 |
| ENSG00000131828 | PDHA1 | 0,205705239 | 1,153249958 | 0,21039 | 0,5290538 | pyruvate dehydrogenase E1 subunit alpha 1 |
| ENSG00000119778 | ATAD2B | 0,205620731 | 1,153182406 | 0,3891 | 0,65363435 | ATPase family AAA domain containing 2B |
| ENSG00000136936 | XPA | 0,205594757 | 1,153161644 | 0,43513 | 0,68584256 | XPA, DNA damage recognition and repair factor |
| ENSG00000155961 | RAB39B | 0,205565224 | 1,153138039 | 0,67479 | 0,83976962 | RAB39B, member RAS oncogene family |
| ENSG00000124422 | USP22 | 0,205509864 | 1,15309379 | 0,23439 | 0,5461771 | ubiquitin specific peptidase 22 |
| ENSG00000145912 | NHP2 | 0,205362992 | 1,152976407 | 0,4033 | 0,66399303 | NHP2 ribonucleoprotein |
| ENSG00000008083 | JARID2 | 0,205333968 | 1,152953211 | 0,36199 | 0,63352908 | jumonji and AT-rich interaction domain containing 2 |
| ENSG00000125971 | DYNLRB1 | 0,205327329 | 1,152947906 | 0,47401 | 0,71294959 | dynein light chain roadblock-type 1 |
| ENSG00000176225 | RTTN | 0,205306775 | 1,15293148 | 0,37127 | 0,64045979 | rotatin |
| ENSG00000186732 | MPPED1 | 0,20528583 | 1,152914743 | 0,78108 | 0,89863435 | metallophosphoesterase domain containing 1 |
| ENSG00000082898 | XPO1 | 0,205174454 | 1,152825741 | 0,34938 | 0,62549486 | exportin 1 |
| ENSG00000234704 | BRD2 | 0,205170585 | 1,152822649 | 0,25708 | 0,56155289 | bromodomain containing 2 |
| ENSG00000168495 | POLR3D | 0,205164767 | 1,152818 | 0,21782 | 0,53550245 | RNA polymerase III subunit D |
| ENSG00000111802 | TDP2 | 0,205150123 | 1,152806299 | 0,43411 | 0,68493098 | tyrosyl-DNA phosphodiesterase 2 |
| ENSG00000206142 | FAM230H | 0,205149732 | 1,152805986 | 0,72752 | 0,87027254 | family with sequence similarity 230 member H |
| ENSG00000181915 | ADO | 0,205063548 | 1,152737121 | 0,43091 | 0,68325486 | 2-aminoethanethiol dioxygenase |
| ENSG00000213178 | RPL22P1 | 0,20485759 | 1,15257257 | 0,31854 | 0,6036094 | ribosomal protein L22 pseudogene 1 |
| ENSG00000099246 | RAB18 | 0,204747601 | 1,152484702 | 0,58491 | 0,78605624 | RAB18, member RAS oncogene family |
| ENSG00000116209 | TMEM59 | 0,204674652 | 1,152426429 | 0,38193 | 0,64905556 | transmembrane protein 59 |
| ENSG00000183666 | GUSBP1 | 0,204580197 | 1,152350981 | 0,5005 | 0,73086101 | GUSB pseudogene 1 |
| ENSG00000228506 | AL513550,1 | 0,204543524 | 1,152321689 | 0,26074 | 0,56400103 | novel transcript |
| ENSG00000116273 | PHF13 | 0,204503745 | 1,152289916 | 0,22703 | 0,54168591 | PHD finger protein 13 |
| ENSG00000180481 | GLIPR1L2 | 0,204448843 | 1,152246067 | 0,48039 | 0,71764212 | GLIPR1 like 2 |
| ENSG00000235501 | AC105942,1 | 0,204378493 | 1,152189881 | 0,5381 | 0,75658794 | novel transcript |
| ENSG00000143228 | NUF2 | 0,20437155 | 1,152184337 | 0,56283 | 0,77200886 | NUF2 component of NDC80 kinetochore complex |
| ENSG00000140905 | GCSH | 0,204259558 | 1,152094899 | 0,50431 | 0,7332113 | glycine cleavage system protein H |
| ENSG00000224934 | AL391684,1 | 0,204179059 | 1,152030617 | 0,31588 | 0,60199254 | novel transcript |
| ENSG00000136874 | STX17 | 0,20417648 | 1,152028558 | 0,42744 | 0,68046596 | syntaxin 17 |
| ENSG00000204351 | SKIV2L | 0,204160312 | 1,152015647 | 0,37873 | 0,64679625 | Ski2 like RNA helicase |
| ENSG00000106615 | RHEB | 0,204041595 | 1,151920853 | 0,28512 | 0,58171017 | Ras homolog, mTORC1 binding |

| | | | | | | |
|---|---|---|---|---|---|---|
| ENSG00000074582 | BCS1L | 0,204006851 | 1,151893112 | 0,42813 | 0,68094175 | BCS1 homolog, ubiquinol-cytochrome c reductase complex chaperone |
| ENSG00000187837 | H1-2 | 0,20389486 | 1,151803699 | 0,54714 | 0,76264866 | H1,2 linker histone, cluster member |
| ENSG00000166938 | DIS3L | 0,203648704 | 1,151607192 | 0,29623 | 0,58961035 | DIS3 like exosome 3'-5' exoribonuclease |
| ENSG00000141424 | SLC39A6 | 0,203619864 | 1,151584171 | 0,62419 | 0,80963125 | solute carrier family 39 member 6 |
| ENSG00000136943 | CTSV | 0,20349662 | 1,1514858 | 0,40783 | 0,66684711 | cathepsin V |
| ENSG00000179403 | VWA1 | 0,203488153 | 1,151479042 | 0,60163 | 0,79671505 | von Willebrand factor A domain containing 1 |
| ENSG00000102384 | CENPI | 0,203411873 | 1,151418161 | 0,51701 | 0,74230144 | centromere protein I |
| ENSG00000236514 | AL135791,1 | 0,203152725 | 1,151211353 | 0,78305 | 0,89918294 | novel transcript |
| ENSG00000139192 | TAPBPL | 0,203137421 | 1,151199141 | 0,70932 | 0,85982913 | TAP binding protein like |
| ENSG00000252355 | RN7SKP287 | 0,203095112 | 1,151165381 | 0,81471 | 0,91470265 | RN7SK pseudogene 287 |
| ENSG00000260064 | AC092332,1 | 0,203073652 | 1,151148258 | 0,76656 | 0,89139706 | novel transcript, sense intronic to CDYL2 |
| ENSG00000122012 | SV2C | 0,20300307 | 1,151091941 | 0,55708 | 0,76893697 | synaptic vesicle glycoprotein 2C |
| ENSG00000243979 | AC087752,1 | 0,202998562 | 1,151088344 | 0,81963 | 0,91708488 | ribosomal protein L17 (RPL17) pseudogene |
| ENSG00000138190 | EXOC6 | 0,202893139 | 1,151004233 | 0,64062 | 0,81954047 | exocyst complex component 6 |
| ENSG00000264057 | AC103810,1 | 0,202872485 | 1,150987755 | 0,74192 | 0,87795766 | bromodomain PHD finger transcription factor (BPTF) pseudogene |
| ENSG00000144452 | ABCA12 | 0,202749049 | 1,150889282 | 0,64202 | 0,8205913 | ATP binding cassette subfamily A member 12 |
| ENSG00000179271 | GADD45GIP1 | 0,202652166 | 1,150811997 | 0,33335 | 0,61333573 | GADD45G interacting protein 1 |
| ENSG00000281404 | LINC01176 | 0,202510086 | 1,150698668 | 0,7004 | 0,85531965 | long intergenic non-protein coding RNA 1176 |
| ENSG00000255152 | MSH5-SAPCD | 0,202343213 | 1,150565577 | 0,8423 | 0,92821072 | MSH5-SAPCD1 readthrough (NMD candidate) |
| ENSG00000076201 | PTPN23 | 0,202332219 | 1,150556809 | 0,28558 | 0,58210705 | protein tyrosine phosphatase non-receptor type 23 |
| ENSG00000120029 | ARMH3 | 0,202295348 | 1,150527405 | 0,44632 | 0,69300607 | armadillo like helical domain containing 3 |
| ENSG00000270820 | AC016727,1 | 0,202083112 | 1,150358162 | 0,56388 | 0,77259134 | novel transcript, antisense to XPO1 |
| ENSG00000165156 | ZHX1 | 0,202034585 | 1,150319469 | 0,60241 | 0,7970763 | zinc fingers and homeoboxes 1 |
| ENSG00000167920 | TMEM99 | 0,202030837 | 1,150316481 | 0,43433 | 0,68505666 | transmembrane protein 99 (putative) |
| ENSG00000229414 | KCNQ1-AS1 | 0,201959952 | 1,150259962 | 0,81845 | 0,91665476 | KCNQ1 antisense RNA 1 |
| ENSG00000217643 | PTGES3P2 | 0,201916735 | 1,150225506 | 0,84342 | 0,92850266 | prostaglandin E synthase 3 pseudogene 2 |
| ENSG00000137822 | TUBGCP4 | 0,201904498 | 1,15021575 | 0,41721 | 0,6738387 | tubulin gamma complex associated protein 4 |
| ENSG00000189369 | GSPT2 | 0,201874764 | 1,150192044 | 0,49632 | 0,72831792 | G1 to S phase transition 2 |
| ENSG00000119772 | DNMT3A | 0,201745805 | 1,150089236 | 0,34672 | 0,62350287 | DNA methyltransferase 3 alpha |
| ENSG00000111670 | GNPTAB | 0,201701701 | 1,150054078 | 0,61306 | 0,8035281 | N-acetylglucosamine-1-phosphate transferase subunits alpha and beta |

| | | | | | | |
|---|---|---|---|---|---|---|
| ENSG00000120910 | PPP3CC | 0,201681992 | 1,150038367 | 0,50514 | 0,73378796 | protein phosphatase 3 catalytic subunit gamma |
| ENSG00000117385 | P3H1 | 0,201639225 | 1,150004275 | 0,52348 | 0,74738502 | prolyl 3-hydroxylase 1 |
| ENSG00000172115 | CYCS | 0,201591289 | 1,149966066 | 0,31123 | 0,59885232 | cytochrome c, somatic |
| ENSG00000172785 | CBWD1 | 0,201549454 | 1,14993272 | 0,2056 | 0,52513499 | COBW domain containing 1 |
| ENSG00000224093 | BCAR3-AS1 | 0,201430853 | 1,149838189 | 0,68456 | 0,84564064 | BCAR3 antisense RNA 1 |
| ENSG00000242516 | LINC00960 | 0,201342464 | 1,149767745 | 0,64235 | 0,82066205 | long intergenic non-protein coding RNA 960 |
| ENSG00000276903 | H2AC16 | 0,201311154 | 1,149742792 | 0,50494 | 0,73369183 | H2A clustered histone 16 |
| ENSG00000271576 | AL359504,1 | 0,201222478 | 1,149672125 | 0,60393 | 0,79773702 | novel transcript |
| ENSG00000147180 | ZNF711 | 0,201174876 | 1,149634193 | 0,57766 | 0,7815762 | zinc finger protein 711 |
| ENSG00000163867 | ZMYM6 | 0,201136898 | 1,149603929 | 0,38118 | 0,64856923 | zinc finger MYM-type containing 6 |
| ENSG00000103494 | RPGRIP1L | 0,201119414 | 1,149589997 | 0,37379 | 0,6423379 | RPGRIP1 like |
| ENSG00000272086 | AC025181,2 | 0,200848469 | 1,149374119 | 0,50129 | 0,7312466 | novel transcript |
| ENSG00000102265 | TIMP1 | 0,200842919 | 1,149369698 | 0,47678 | 0,71481671 | TIMP metallopeptidase inhibitor 1 |
| ENSG00000198900 | TOP1 | 0,200768807 | 1,149310655 | 0,39169 | 0,65533345 | DNA topoisomerase I |
| ENSG00000135334 | AKIRIN2 | 0,200667589 | 1,149230024 | 0,46314 | 0,70415312 | akirin 2 |
| ENSG00000116586 | LAMTOR2 | 0,200466108 | 1,149069538 | 0,57685 | 0,7810839 | late endosomal/lysosomal adaptor, MAPK and MTOR activator 2 |
| ENSG00000171960 | PPIH | 0,200451952 | 1,149058263 | 0,30235 | 0,59297237 | peptidylprolyl isomerase H |
| ENSG00000075073 | TACR2 | 0,200407288 | 1,14902269 | 0,72866 | 0,87118284 | tachykinin receptor 2 |
| ENSG00000261188 | Z95115,1 | 0,200407256 | 1,149022665 | 0,40806 | 0,66701349 | novel transcript, antisense to TFIP11 |
| ENSG00000124374 | PAIP2B | 0,200370464 | 1,148993363 | 0,44841 | 0,69432421 | poly(A) binding protein interacting protein 2B |
| ENSG00000171551 | ECEL1 | 0,200357678 | 1,14898318 | 0,60705 | 0,80003503 | endothelin converting enzyme like 1 |
| ENSG00000092208 | GEMIN2 | 0,200256895 | 1,148902917 | 0,35659 | 0,63063438 | gem nuclear organelle associated protein 2 |
| ENSG00000205940 | HSP90AB2P | 0,200249889 | 1,148897338 | 0,54683 | 0,76240845 | heat shock protein 90 alpha family class B member 2, pseudogene |
| ENSG00000189372 | AL034369,1 | 0,200208424 | 1,148864318 | 0,84761 | 0,93078523 | pancreatic lipase-related protein 2 (PNLIPRP2) pseudogene |
| ENSG00000197170 | PSMD12 | 0,200133876 | 1,148804954 | 0,2438 | 0,55243616 | proteasome 26S subunit, non-ATPase 12 |
| ENSG00000276524 | AC010999,2 | 0,200123163 | 1,148796423 | 0,74647 | 0,88046587 | novel transcript |
| ENSG00000197780 | TAF13 | 0,20011164 | 1,148787248 | 0,49772 | 0,72908464 | TATA-box binding protein associated factor 13 |
| ENSG00000286129 | AC245060,7 | 0,200049432 | 1,148737714 | 0,60569 | 0,79885773 | novel transcript |
| ENSG00000064419 | TNPO3 | 0,199907113 | 1,1486244 | 0,33548 | 0,61510039 | transportin 3 |
| ENSG00000156304 | SCAF4 | 0,199776624 | 1,148520513 | 0,25856 | 0,5625202 | SR-related CTD associated factor 4 |

| | | | | | | |
|---|---|---|---|---|---|---|
| ENSG00000180747 | SMG1P3 | 0,199770703 | 1,148515799 | 0,46629 | 0,70661104 | SMG1 pseudogene 3 |
| ENSG00000250241 | AC105383,1 | 0,199737238 | 1,148489158 | 0,51242 | 0,73930522 | novel transcript |
| ENSG00000108106 | UBE2S | 0,199635319 | 1,148408027 | 0,2383 | 0,54875544 | ubiquitin conjugating enzyme E2 S |
| ENSG00000237877 | LINC01473 | 0,199616877 | 1,148393346 | 0,54206 | 0,75894841 | long intergenic non-protein coding RNA 1473 |
| ENSG00000233851 | LATS2-AS1 | 0,199505016 | 1,148304308 | 0,71491 | 0,8628412 | LATS2 antisense RNA 1 |
| ENSG00000183793 | NPIPA5 | 0,199378228 | 1,148203396 | 0,77021 | 0,89334263 | nuclear pore complex interacting protein family member A5 |
| ENSG00000198894 | CIPC | 0,199327211 | 1,148162794 | 0,40069 | 0,66268564 | CLOCK interacting pacemaker |
| ENSG00000286518 | AL354743,1 | 0,199240558 | 1,148093833 | 0,71911 | 0,86563516 | novel transcript, antisense to PRDM16 |
| ENSG00000114416 | FXR1 | 0,1991734 | 1,148040391 | 0,35343 | 0,62818386 | FMR1 autosomal homolog 1 |
| ENSG00000255417 | MTCO2P15 | 0,199167734 | 1,148035882 | 0,82289 | 0,9181175 | MT-CO2 pseudogene 15 |
| ENSG00000124875 | CXCL6 | 0,199166524 | 1,148034919 | 0,82567 | 0,91917221 | C-X-C motif chemokine ligand 6 |
| ENSG00000164414 | SLC35A1 | 0,199162728 | 1,148031898 | 0,69476 | 0,8520376 | solute carrier family 35 member A1 |
| ENSG00000118418 | HMGN3 | 0,199149033 | 1,148021001 | 0,27565 | 0,57533851 | high mobility group nucleosomal binding domain 3 |
| ENSG00000084090 | STARD7 | 0,199122119 | 1,147999583 | 0,35864 | 0,63185282 | StAR related lipid transfer domain containing 7 |
| ENSG00000269526 | ERVV-1 | 0,199120244 | 1,147998092 | 0,81587 | 0,91534523 | endogenous retrovirus group V member 1, envelope |
| ENSG00000136982 | DSCC1 | 0,199118297 | 1,147996543 | 0,47994 | 0,71730153 | DNA replication and sister chromatid cohesion 1 |
| ENSG00000033050 | ABCF2 | 0,19910426 | 1,147985373 | 0,28158 | 0,57891473 | ATP binding cassette subfamily F member 2 |
| ENSG00000173418 | NAA20 | 0,199062751 | 1,147952344 | 0,39512 | 0,65804585 | N-alpha-acetyltransferase 20, NatB catalytic subunit |
| ENSG00000148396 | SEC16A | 0,199060067 | 1,147950208 | 0,27382 | 0,57384741 | SEC16 homolog A, endoplasmic reticulum export factor |
| ENSG00000091490 | SEL1L3 | 0,199003376 | 1,1479051 | 0,741 | 0,87724557 | SEL1L family member 3 |
| ENSG00000169508 | GPR183 | 0,198796531 | 1,147740532 | 0,63952 | 0,81913513 | G protein-coupled receptor 183 |
| ENSG00000258441 | LINC00641 | 0,198785004 | 1,147731361 | 0,53128 | 0,75258369 | long intergenic non-protein coding RNA 641 |
| ENSG00000135960 | EDAR | 0,19875396 | 1,147706665 | 0,74676 | 0,88057664 | ectodysplasin A receptor |
| ENSG00000076248 | UNG | 0,198735498 | 1,147691978 | 0,22958 | 0,54311366 | uracil DNA glycosylase |
| ENSG00000143776 | CDC42BPA | 0,198697487 | 1,14766174 | 0,44577 | 0,69284743 | CDC42 binding protein kinase alpha |
| ENSG00000109738 | GLRB | 0,198635787 | 1,147612658 | 0,6402 | 0,81935089 | glycine receptor beta |
| ENSG00000215571 | GRK6P1 | 0,19853878 | 1,147535496 | 0,6951 | 0,85206252 | G protein-coupled receptor kinase 6 pseudogene 1 |
| ENSG00000276334 | AL133243,2 | 0,198537936 | 1,147534825 | 0,49143 | 0,72510994 | novel transcript, sense intronic to BIRC6 |
| ENSG00000235651 | G3BP1P1 | 0,198516097 | 1,147517453 | 0,60382 | 0,79770754 | G3BP1 pseudogene 1 |
| ENSG00000271709 | AC017033,1 | 0,198400348 | 1,147425391 | 0,74402 | 0,87882457 | novel transcript |
| ENSG00000213186 | TRIM59 | 0,19834515 | 1,147381491 | 0,52518 | 0,74892434 | tripartite motif containing 59 |
| ENSG00000137038 | DMAC1 | 0,198242107 | 1,147299543 | 0,31159 | 0,59905757 | distal membrane arm assembly complex 1 |

| | | | | | | |
|---|---|---|---|---|---|---|
| ENSG00000171819 | ANGPTL7 | 0,198203477 | 1,147268823 | 0,55602 | 0,76827287 | angiopoietin like 7 |
| ENSG00000100412 | ACO2 | 0,198178367 | 1,147248855 | 0,27429 | 0,57420712 | aconitase 2 |
| ENSG00000125740 | FOSB | 0,198139093 | 1,147217624 | 0,79268 | 0,90363236 | FosB proto-oncogene, AP-1 transcription factor subunit |
| ENSG00000138175 | ARL3 | 0,198067908 | 1,14716102 | 0,49078 | 0,72480683 | ADP ribosylation factor like GTPase 3 |
| ENSG00000266027 | MSH5-SAPCD: | 0,198018857 | 1,147122018 | 0,87698 | 0,94549531 | MSH5-SAPCD1 readthrough (NMD candidate) |
| ENSG00000100580 | TMED8 | 0,198002682 | 1,147109156 | 0,44791 | 0,69394017 | transmembrane p24 trafficking protein family member 8 |
| ENSG00000113318 | MSH3 | 0,197992691 | 1,147101213 | 0,20977 | 0,52867318 | mutS homolog 3 |
| ENSG00000137207 | YIPF3 | 0,197925059 | 1,147047439 | 0,35488 | 0,62936991 | Yip1 domain family member 3 |
| ENSG00000241058 | NSUN6 | 0,19788695 | 1,14701714 | 0,55785 | 0,76954098 | NOP2/Sun RNA methyltransferase 6 |
| ENSG00000250250 | CTD-2350J17, | 0,197870471 | 1,147004039 | 0,81908 | 0,91689679 | uncharacterized LOC101929472 |
| ENSG00000157954 | WIPI2 | 0,197800704 | 1,146948572 | 0,28778 | 0,5840168 | WD repeat domain, phosphoinositide interacting 2 |
| ENSG00000258302 | AC025034,1 | 0,197713461 | 1,146879216 | 0,51507 | 0,74133616 | novel transcript, antisense to ATP2B1 |
| ENSG00000260536 | AL035071,2 | 0,197489679 | 1,146701332 | 0,81556 | 0,91511005 | novel transcript, antisense to MAPRE1 |
| ENSG00000229894 | GK3P | 0,197379043 | 1,146613399 | 0,72537 | 0,86899186 | glycerol kinase 3 pseudogene |
| ENSG00000272807 | AC007038,1 | 0,197275763 | 1,146531318 | 0,67452 | 0,83975333 | novel transcript, antisense to KANSL1L |
| ENSG00000099910 | KLHL22 | 0,19712666 | 1,14641283 | 0,35507 | 0,62946175 | kelch like family member 22 |
| ENSG00000204256 | BRD2 | 0,197077899 | 1,146374083 | 0,27581 | 0,57533851 | bromodomain containing 2 |
| ENSG00000260874 | AC106820,5 | 0,197045523 | 1,146348357 | 0,80736 | 0,91131389 | novel transcript |
| ENSG00000278878 | CU638689,1 | 0,196880607 | 1,146217324 | 0,60347 | 0,79758282 | novel transcript |
| ENSG00000198420 | TCAF1 | 0,19686357 | 1,146203789 | 0,38509 | 0,65088045 | TRPM8 channel associated factor 1 |
| ENSG00000184304 | PRKD1 | 0,196741365 | 1,146106703 | 0,55108 | 0,76487698 | protein kinase D1 |
| ENSG00000271754 | AL355802,3 | 0,196720149 | 1,146089848 | 0,69612 | 0,85287022 | novel transcript, antisense to POLR1C |
| ENSG00000213228 | RPL12P38 | 0,196698922 | 1,146072985 | 0,80275 | 0,9091124 | ribosomal protein L12 pseudogene 38 |
| ENSG00000165392 | WRN | 0,196695051 | 1,14606991 | 0,28372 | 0,58072192 | WRN RecQ like helicase |
| ENSG00000173914 | RBM4B | 0,196573016 | 1,14597297 | 0,27967 | 0,57745167 | RNA binding motif protein 4B |
| ENSG00000149179 | C11orf49 | 0,19655071 | 1,145955252 | 0,4015 | 0,6630277 | chromosome 11 open reading frame 49 |
| ENSG00000180638 | SLC47A2 | 0,196541001 | 1,14594754 | 0,69029 | 0,84893069 | solute carrier family 47 member 2 |
| ENSG00000089693 | MLF2 | 0,196495267 | 1,145911214 | 0,30855 | 0,59736238 | myeloid leukemia factor 2 |
| ENSG00000168038 | ULK4 | 0,196452639 | 1,145877355 | 0,45689 | 0,7004577 | unc-51 like kinase 4 |
| ENSG00000236519 | LINC01424 | 0,196403543 | 1,145838361 | 0,64385 | 0,82128583 | long intergenic non-protein coding RNA 1424 |
| ENSG00000130638 | ATXN10 | 0,196377083 | 1,145817346 | 0,50901 | 0,7370222 | ataxin 10 |
| ENSG00000152256 | PDK1 | 0,196330893 | 1,145780661 | 0,36251 | 0,63379594 | pyruvate dehydrogenase kinase 1 |
| ENSG00000115107 | STEAP3 | 0,196309235 | 1,145763461 | 0,6333 | 0,81541029 | STEAP3 metalloreductase |

| | | | | | | |
|---|---|---|---|---|---|---|
| ENSG00000272489 | AL132656,3 | 0,19624783 | 1,145714695 | 0,75989 | 0,88801075 | novel transcript |
| ENSG00000115317 | HTRA2 | 0,196244625 | 1,14571215 | 0,3586 | 0,63185282 | HtrA serine peptidase 2 |
| ENSG00000228405 | RNF5 | 0,196109479 | 1,145604829 | 0,34311 | 0,62060539 | ring finger protein 5 |
| ENSG00000279486 | OR2AG1 | 0,195990817 | 1,145510606 | 0,55487 | 0,76763832 | olfactory receptor family 2 subfamily AG member 1 |
| ENSG00000236772 | AL034550,1 | 0,195944902 | 1,145474151 | 0,76518 | 0,89082001 | novel transcript, antisense to NOL4L |
| ENSG00000274487 | NPEPPSP1 | 0,195860623 | 1,145407237 | 0,59949 | 0,79552021 | NPEPPS pseudogene 1 |
| ENSG00000103363 | ELOB | 0,195858198 | 1,145405311 | 0,53247 | 0,75300764 | elongin B |
| ENSG00000260634 | AC012508,2 | 0,19585065 | 1,145399318 | 0,79148 | 0,90310057 | novel transcript |
| ENSG00000185825 | BCAP31 | 0,195849421 | 1,145398343 | 0,26703 | 0,56950942 | B cell receptor associated protein 31 |
| ENSG00000138031 | ADCY3 | 0,195737772 | 1,145309705 | 0,31031 | 0,59842274 | adenylate cyclase 3 |
| ENSG00000145592 | RPL37 | 0,195633493 | 1,145226924 | 0,27788 | 0,5762948 | ribosomal protein L37 |
| ENSG00000072121 | ZFYVE26 | 0,195580147 | 1,145184578 | 0,16446 | 0,49065086 | zinc finger FYVE-type containing 26 |
| ENSG00000225472 | AL136366,1 | 0,195557592 | 1,145166674 | 0,78681 | 0,90104289 | novel transcript |
| ENSG00000288407 | CCDC33 | 0,195376137 | 1,14502265 | 0,62818 | 0,81178582 | coiled-coil domain containing 33 |
| ENSG00000140481 | CCDC33 | 0,195376137 | 1,14502265 | 0,62818 | 0,81178582 | coiled-coil domain containing 33 |
| ENSG00000241839 | PLEKHO2 | 0,195277022 | 1,144943988 | 0,44872 | 0,69454448 | pleckstrin homology domain containing O2 |
| ENSG00000165494 | PCF11 | 0,195266333 | 1,144935505 | 0,44981 | 0,69538334 | PCF11 cleavage and polyadenylation factor subunit |
| ENSG00000112294 | ALDH5A1 | 0,195260666 | 1,144931008 | 0,46431 | 0,70493977 | aldehyde dehydrogenase 5 family member A1 |
| ENSG00000169181 | GSG1L | 0,195260136 | 1,144930587 | 0,83105 | 0,92270607 | GSG1 like |
| ENSG00000219451 | RPL23P8 | 0,195220796 | 1,144899367 | 0,83775 | 0,92596606 | ribosomal protein L23 pseudogene 8 |
| ENSG00000117593 | DARS2 | 0,195188216 | 1,144873513 | 0,3233 | 0,60606867 | aspartyl-tRNA synthetase 2, mitochondrial |
| ENSG00000120256 | LRP11 | 0,195160978 | 1,144851898 | 0,42491 | 0,67857395 | LDL receptor related protein 11 |
| ENSG00000127452 | FBXL12 | 0,195096891 | 1,144801042 | 0,32883 | 0,60992716 | F-box and leucine rich repeat protein 12 |
| ENSG00000155366 | RHOC | 0,195081533 | 1,144788855 | 0,39715 | 0,65966023 | ras homolog family member C |
| ENSG00000170426 | SDR9C7 | 0,195048255 | 1,14476245 | 0,77423 | 0,89572556 | short chain dehydrogenase/reductase family 9C member 7 |
| ENSG00000221909 | FAM200A | 0,195034355 | 1,144751421 | 0,54903 | 0,76394298 | family with sequence similarity 200 member A |
| ENSG00000261326 | LINC01355 | 0,195003081 | 1,144726606 | 0,63021 | 0,81330068 | long intergenic non-protein coding RNA 1355 |
| ENSG00000283124 | LINC01355 | 0,195003081 | 1,144726606 | 0,63021 | 0,81330068 | long intergenic non-protein coding RNA 1355 |
| ENSG00000165819 | METTL3 | 0,19496575 | 1,144696985 | 0,33259 | 0,61252597 | methyltransferase like 3 |
| ENSG00000105829 | BET1 | 0,194925426 | 1,144664991 | 0,38663 | 0,6519582 | Bet1 golgi vesicular membrane trafficking protein |
| ENSG00000285996 | AC006116,9 | 0,194813362 | 1,14457608 | 0,54803 | 0,76335169 | novel transcript, antisense to ZNF582 |
| ENSG00000089248 | ERP29 | 0,194527059 | 1,144348961 | 0,42307 | 0,67691679 | endoplasmic reticulum protein 29 |
| ENSG00000177663 | IL17RA | 0,19448294 | 1,144313966 | 0,59772 | 0,79474081 | interleukin 17 receptor A |

| | | | | | | |
|---|---|---|---|---|---|---|
| ENSG00000147121 | KRBOX4 | 0,194467523 | 1,144301738 | 0,27772 | 0,57617849 | KRAB box domain containing 4 |
| ENSG00000261770 | AC006504,1 | 0,194427513 | 1,144270004 | 0,3809 | 0,6484556 | novel transcript |
| ENSG00000285778 | AL591463,1 | 0,194341458 | 1,144201752 | 0,75987 | 0,88801075 | novel transcript |
| ENSG00000173660 | UQCRH | 0,194341112 | 1,144201477 | 0,29048 | 0,58565836 | ubiquinol-cytochrome c reductase hinge protein |
| ENSG00000270361 | AL451085,1 | 0,194299357 | 1,144168362 | 0,82266 | 0,91804426 | novel transcript |
| ENSG00000115548 | KDM3A | 0,194296039 | 1,14416573 | 0,49967 | 0,73058737 | lysine demethylase 3A |
| ENSG00000198964 | SGMS1 | 0,194248525 | 1,144128048 | 0,49725 | 0,72868655 | sphingomyelin synthase 1 |
| ENSG00000008988 | RPS20 | 0,194226456 | 1,144110547 | 0,49853 | 0,72969953 | ribosomal protein S20 |
| ENSG00000236526 | AL035448,1 | 0,194067993 | 1,143984887 | 0,47941 | 0,71689864 | novel transcript |
| ENSG00000153130 | SCOC | 0,194052007 | 1,143972211 | 0,62008 | 0,80755243 | short coiled-coil protein |
| ENSG00000165406 | MARCHF8 | 0,194050315 | 1,143970869 | 0,43322 | 0,68455289 | membrane associated ring-CH-type finger 8 |
| ENSG00000288258 | BX470111,1 | 0,19401762 | 1,143944945 | 0,87868 | 0,94608807 | novel protein |
| ENSG00000236268 | LINC01361 | 0,193970318 | 1,143907438 | 0,61554 | 0,80447376 | long intergenic non-protein coding RNA 1361 |
| ENSG00000111215 | PRR4 | 0,193920531 | 1,143867963 | 0,76042 | 0,8884526 | proline rich 4 |
| ENSG00000120832 | MTERF2 | 0,193716736 | 1,143706391 | 0,39501 | 0,65794526 | mitochondrial transcription termination factor 2 |
| ENSG00000223305 | RN7SKP30 | 0,193712515 | 1,143703045 | 0,7883 | 0,90157293 | RN7SK pseudogene 30 |
| ENSG00000237788 | AL162615,1 | 0,19368377 | 1,143680258 | 0,67629 | 0,84075837 | cytokine-like nuclear factor n-pac (N-PAC) pseudogene |
| ENSG00000228716 | DHFR | 0,19367431 | 1,143672758 | 0,33503 | 0,61471383 | dihydrofolate reductase |
| ENSG00000287188 | AC068989,1 | 0,193649782 | 1,143653315 | 0,76538 | 0,89085946 | novel transcript, antisense to ANXA10 |
| ENSG00000203760 | CENPW | 0,193643304 | 1,143648179 | 0,32818 | 0,60971379 | centromere protein W |
| ENSG00000183520 | UTP11 | 0,193614971 | 1,14362572 | 0,31 | 0,59828567 | UTP11 small subunit processome component |
| ENSG00000259849 | VENTXP1 | 0,193568106 | 1,14358857 | 0,65259 | 0,826091 | VENT homeobox pseudogene 1 |
| ENSG00000164867 | NOS3 | 0,193559568 | 1,143581802 | 0,62472 | 0,80993965 | nitric oxide synthase 3 |
| ENSG00000259212 | AC103739,1 | 0,193487178 | 1,143524422 | 0,76278 | 0,88972756 | novel transcript, antisense to CRTC3 |
| ENSG00000267115 | AC022148,2 | 0,193435016 | 1,143483078 | 0,81756 | 0,91621442 | novel transcript |
| ENSG00000108406 | DHX40 | 0,193434941 | 1,143483018 | 0,65463 | 0,82719457 | DEAH-box helicase 40 |
| ENSG00000215218 | UBE2QL1 | 0,193424649 | 1,143474861 | 0,61981 | 0,80730752 | ubiquitin conjugating enzyme E2 Q family like 1 |
| ENSG00000170365 | SMAD1 | 0,193379256 | 1,143438883 | 0,39777 | 0,66034494 | SMAD family member 1 |
| ENSG00000174483 | BBS1 | 0,193195704 | 1,143293415 | 0,43393 | 0,68480274 | Bardet-Biedl syndrome 1 |
| ENSG00000164694 | FNDC1 | 0,193163645 | 1,143268009 | 0,76307 | 0,88980026 | fibronectin type III domain containing 1 |
| ENSG00000196550 | FAM72A | 0,193120889 | 1,143234127 | 0,48323 | 0,71961312 | family with sequence similarity 72 member A |
| ENSG00000155592 | ZKSCAN2 | 0,193076096 | 1,143198633 | 0,47684 | 0,71481671 | zinc finger with KRAB and SCAN domains 2 |
| ENSG00000108669 | CYTH1 | 0,193030306 | 1,143162349 | 0,51736 | 0,74268179 | cytohesin 1 |

| | | | | | | |
|---|---|---|---|---|---|---|
| ENSG00000164054 | SHISA5 | 0,193025168 | 1,143158278 | 0,39249 | 0,65581327 | shisa family member 5 |
| ENSG00000057252 | SOAT1 | 0,192941099 | 1,143091665 | 0,28474 | 0,58161933 | sterol O-acyltransferase 1 |
| ENSG00000217442 | SYCE3 | 0,192899688 | 1,143058855 | 0,81138 | 0,91319213 | synaptonemal complex central element protein 3 |
| ENSG00000047849 | MAP4 | 0,192639151 | 1,142852448 | 0,41748 | 0,67403924 | microtubule associated protein 4 |
| ENSG00000136231 | IGF2BP3 | 0,192553085 | 1,142784271 | 0,51223 | 0,73918504 | insulin like growth factor 2 mRNA binding protein 3 |
| ENSG00000114346 | ECT2 | 0,192473095 | 1,142720911 | 0,4562 | 0,70002015 | epithelial cell transforming 2 |
| ENSG00000250770 | AC005865,1 | 0,19242328 | 1,142681455 | 0,69952 | 0,8548625 | tetraspanin 11 (TSPAN11) pseudogene |
| ENSG00000198546 | ZNF511 | 0,19230862 | 1,142590642 | 0,4911 | 0,72493845 | zinc finger protein 511 |
| ENSG00000135299 | ANKRD6 | 0,192279784 | 1,142567805 | 0,51339 | 0,73975613 | ankyrin repeat domain 6 |
| ENSG00000138658 | ZGRF1 | 0,192272395 | 1,142561954 | 0,40363 | 0,66399439 | zinc finger GRF-type containing 1 |
| ENSG00000163166 | IWS1 | 0,192243237 | 1,142538862 | 0,33407 | 0,61387439 | interacts with SUPT6H, CTD assembly factor 1 |
| ENSG00000204049 | AL391421,1 | 0,192169096 | 1,142480147 | 0,70511 | 0,85794356 | novel transcript, antisense to DLG5 |
| ENSG00000282690 | AL391421,2 | 0,192169096 | 1,142480147 | 0,70511 | 0,85794356 | novel transcript, antisense to DLG5 |
| ENSG00000285815 | GET1-SH3BGR | 0,192131888 | 1,142450682 | 0,85754 | 0,93569939 | GET1-SH3BGR readthrough |
| ENSG00000114902 | SPCS1 | 0,192006954 | 1,142351753 | 0,27384 | 0,57384741 | signal peptidase complex subunit 1 |
| ENSG00000265817 | FSBP | 0,191878936 | 1,142250391 | 0,47059 | 0,71006214 | fibrinogen silencer binding protein |
| ENSG00000283812 | GPR22 | 0,191857575 | 1,142233478 | 0,73502 | 0,87477884 | G protein-coupled receptor 22 |
| ENSG00000172209 | GPR22 | 0,191857575 | 1,142233478 | 0,73502 | 0,87477884 | G protein-coupled receptor 22 |
| ENSG00000266338 | NBPF15 | 0,191800171 | 1,14218803 | 0,29436 | 0,58857672 | NBPF member 15 |
| ENSG00000161558 | TMEM143 | 0,191784123 | 1,142175325 | 0,53543 | 0,75468833 | transmembrane protein 143 |
| ENSG00000160131 | VMA21 | 0,191777464 | 1,142170054 | 0,36271 | 0,63381706 | vacuolar ATPase assembly factor VMA21 |
| ENSG00000078142 | PIK3C3 | 0,19177537 | 1,142168396 | 0,41945 | 0,67527436 | phosphatidylinositol 3-kinase catalytic subunit type 3 |
| ENSG00000173406 | DAB1 | 0,191533217 | 1,141976701 | 0,62265 | 0,80866078 | DAB adaptor protein 1 |
| ENSG00000079805 | DNM2 | 0,191483622 | 1,141937444 | 0,22751 | 0,54168591 | dynamin 2 |
| ENSG00000106245 | BUD31 | 0,191454768 | 1,141914606 | 0,42556 | 0,67914804 | BUD31 homolog |
| ENSG00000116670 | MAD2L2 | 0,191440199 | 1,141903075 | 0,34166 | 0,6197312 | mitotic arrest deficient 2 like 2 |
| ENSG00000287817 | AL096701,4 | 0,191395714 | 1,141867865 | 0,65791 | 0,82891212 | novel transcript, antisense to EIF4ENIF1 |
| ENSG00000102178 | UBL4A | 0,191336985 | 1,141821383 | 0,4142 | 0,67104294 | ubiquitin like 4A |
| ENSG00000243279 | PRAF2 | 0,191332568 | 1,141817887 | 0,5333 | 0,75351438 | PRA1 domain family member 2 |
| ENSG00000242797 | GLYCTK-AS1 | 0,191264399 | 1,141763936 | 0,68111 | 0,84363978 | GLYCTK antisense RNA 1 |
| ENSG00000145088 | EAF2 | 0,191250075 | 1,1417526 | 0,70278 | 0,85666784 | ELL associated factor 2 |
| ENSG00000286613 | AC006213,7 | 0,191208646 | 1,141719813 | 0,70775 | 0,85918434 | novel transcript, sense intronic to ZNF404 |
| ENSG00000116898 | MRPS15 | 0,191195001 | 1,141709015 | 0,33494 | 0,61467789 | mitochondrial ribosomal protein S15 |

| | | | | | | |
|---|---|---|---|---|---|---|
| ENSG00000272150 | NBPF25P | 0,191191559 | 1,141706291 | 0,34402 | 0,621447 | NBPF member 25, pseudogene |
| ENSG00000181991 | MRPS11 | 0,191168715 | 1,141688213 | 0,34857 | 0,62475089 | mitochondrial ribosomal protein S11 |
| ENSG00000029639 | TFB1M | 0,19114253 | 1,141667492 | 0,58131 | 0,78419712 | transcription factor B1, mitochondrial |
| ENSG00000247796 | AC008966,1 | 0,191133803 | 1,141660586 | 0,56075 | 0,77105099 | novel transcript, antisense to MOCS2 |
| ENSG00000100744 | GSKIP | 0,19113185 | 1,14165904 | 0,73485 | 0,87477884 | GSK3B interacting protein |
| ENSG00000102119 | EMD | 0,191126587 | 1,141654875 | 0,34993 | 0,62563749 | emerin |
| ENSG00000143164 | DCAF6 | 0,191122032 | 1,141651271 | 0,38114 | 0,64854231 | DDB1 and CUL4 associated factor 6 |
| ENSG00000119487 | MAPKAP1 | 0,191110509 | 1,141642153 | 0,36283 | 0,63388743 | MAPK associated protein 1 |
| ENSG00000144619 | CNTN4 | 0,191101915 | 1,141635352 | 0,44815 | 0,69411284 | contactin 4 |
| ENSG00000114573 | ATP6V1A | 0,191073406 | 1,141612792 | 0,38245 | 0,64928247 | ATPase H+ transporting V1 subunit A |
| ENSG00000185760 | KCNQ5 | 0,191029145 | 1,141577769 | 0,81043 | 0,91251177 | potassium voltage-gated channel subfamily Q member 5 |
| ENSG00000140534 | TICRR | 0,190977142 | 1,14153662 | 0,30917 | 0,59774312 | TOPBP1 interacting checkpoint and replication regulator |
| ENSG00000263290 | SCAMP3 | 0,190894797 | 1,141471467 | 0,36834 | 0,63803074 | secretory carrier membrane protein 3 |
| ENSG00000156804 | FBXO32 | 0,190877569 | 1,141457836 | 0,60266 | 0,7971128 | F-box protein 32 |
| ENSG00000063978 | RNF4 | 0,19083444 | 1,141423713 | 0,32675 | 0,60880597 | ring finger protein 4 |
| ENSG00000164104 | HMGB2 | 0,19080803 | 1,141402818 | 0,38803 | 0,65289784 | high mobility group box 2 |
| ENSG00000254263 | AC022973,4 | 0,190791257 | 1,141389548 | 0,75202 | 0,88386445 | novel transcript |
| ENSG00000069188 | SDK2 | 0,190677622 | 1,141299649 | 0,36994 | 0,63943131 | sidekick cell adhesion molecule 2 |
| ENSG00000155792 | DEPTOR | 0,190626271 | 1,141259026 | 0,70957 | 0,85993974 | DEP domain containing MTOR interacting protein |
| ENSG00000255837 | TAS2R20 | 0,190471306 | 1,141136447 | 0,72962 | 0,8715818 | taste 2 receptor member 20 |
| ENSG00000273092 | TAS2R20 | 0,190471306 | 1,141136447 | 0,72962 | 0,8715818 | taste 2 receptor member 20 |
| ENSG00000106733 | NMRK1 | 0,190439337 | 1,14111116 | 0,41366 | 0,67085666 | nicotinamide riboside kinase 1 |
| ENSG00000108947 | EFNB3 | 0,1904372 | 1,141109469 | 0,51514 | 0,74136036 | ephrin B3 |
| ENSG00000222041 | CYTOR | 0,190402353 | 1,141081908 | 0,41551 | 0,6723843 | cytoskeleton regulator RNA |
| ENSG00000267165 | CHMP1B-AS1 | 0,190231128 | 1,140946487 | 0,86195 | 0,93834335 | CHMP1B antisense RNA 1 |
| ENSG00000162877 | PM20D1 | 0,190047354 | 1,14080116 | 0,67352 | 0,8390573 | peptidase M20 domain containing 1 |
| ENSG00000176946 | THAP4 | 0,190040423 | 1,140795679 | 0,26393 | 0,56649862 | THAP domain containing 4 |
| ENSG00000163781 | TOPBP1 | 0,190002916 | 1,140766022 | 0,26612 | 0,56908449 | DNA topoisomerase II binding protein 1 |
| ENSG00000186376 | ZNF75D | 0,189948026 | 1,14072262 | 0,35539 | 0,62971233 | zinc finger protein 75D |
| ENSG00000238113 | LINC01410 | 0,189943066 | 1,140718698 | 0,51529 | 0,74138689 | long intergenic non-protein coding RNA 1410 |
| ENSG00000143171 | RXRG | 0,189895397 | 1,140681008 | 0,74266 | 0,87827554 | retinoid X receptor gamma |
| ENSG00000172014 | ANKRD20A4P | 0,189841941 | 1,140638743 | 0,74712 | 0,88064941 | ankyrin repeat domain 20 family member A4, pseudogene |
| ENSG00000286315 | AC087175,1 | 0,189824354 | 1,140624838 | 0,71774 | 0,86483656 | novel transcript |

| | | | | | | |
|---|---|---|---|---|---|---|
| ENSG00000117419 | ERI3 | 0,189824093 | 1,140624631 | 0,3848 | 0,65062567 | ERI1 exoribonuclease family member 3 |
| ENSG00000100354 | TNRC6B | 0,189810821 | 1,140614138 | 0,46219 | 0,70347763 | trinucleotide repeat containing adaptor 6B |
| ENSG00000271580 | AL583832,1 | 0,189804354 | 1,140609025 | 0,71453 | 0,8626416 | glycogenin 2 (GYG2) pseudogene |
| ENSG00000081791 | DELE1 | 0,189789063 | 1,140596936 | 0,4748 | 0,71364583 | DAP3 binding cell death enhancer 1 |
| ENSG00000081019 | RSBN1 | 0,189772672 | 1,140583978 | 0,49328 | 0,72631 | round spermatid basic protein 1 |
| ENSG00000136710 | CCDC115 | 0,189674238 | 1,140506159 | 0,33241 | 0,61230666 | coiled-coil domain containing 115 |
| ENSG00000241878 | PISD | 0,189671818 | 1,140504246 | 0,46024 | 0,70210714 | phosphatidylserine decarboxylase |
| ENSG00000075884 | ARHGAP15 | 0,189494552 | 1,14036412 | 0,74587 | 0,88010204 | Rho GTPase activating protein 15 |
| ENSG00000176293 | ZNF135 | 0,189456149 | 1,140333764 | 0,5329 | 0,75333268 | zinc finger protein 135 |
| ENSG00000135476 | ESPL1 | 0,189425398 | 1,140309459 | 0,39548 | 0,6582534 | extra spindle pole bodies like 1, separase |
| ENSG00000008869 | HEATR5B | 0,189370906 | 1,140266389 | 0,61449 | 0,80382182 | HEAT repeat containing 5B |
| ENSG00000261716 | H2BC20P | 0,189344157 | 1,140245247 | 0,55162 | 0,76535107 | H2B clustered histone 20, pseudogene |
| ENSG00000275797 | AC008746,4 | 0,189281606 | 1,14019581 | 0,69661 | 0,85309963 | novel transcript |
| ENSG00000267838 | AC245884,8 | 0,189281606 | 1,14019581 | 0,69661 | 0,85309963 | novel transcript |
| ENSG00000175564 | UCP3 | 0,189169336 | 1,140107085 | 0,43262 | 0,68418465 | uncoupling protein 3 |
| ENSG00000165630 | PRPF18 | 0,189164018 | 1,140102882 | 0,55088 | 0,76486471 | pre-mRNA processing factor 18 |
| ENSG00000125835 | SNRPB | 0,189139258 | 1,140083316 | 0,29402 | 0,58837921 | small nuclear ribonucleoprotein polypeptides B and B1 |
| ENSG00000118762 | PKD2 | 0,18911053 | 1,140060614 | 0,31735 | 0,60276767 | polycystin 2, transient receptor potential cation channel |
| ENSG00000159140 | SON | 0,189028143 | 1,139995511 | 0,29425 | 0,58857672 | SON DNA and RNA binding protein |
| ENSG00000196352 | CD55 | 0,188992542 | 1,139967379 | 0,68659 | 0,84702526 | CD55 molecule (Cromer blood group) |
| ENSG00000230913 | NPM1P51 | 0,18893898 | 1,139925058 | 0,61545 | 0,80439941 | nucleophosmin 1 pseudogene 51 |
| ENSG00000110429 | FBXO3 | 0,188711342 | 1,139745207 | 0,47094 | 0,71043734 | F-box protein 3 |
| ENSG00000224895 | VPS26BP1 | 0,18868538 | 1,139724697 | 0,75541 | 0,88543251 | VPS26B pseudogene 1 |
| ENSG00000167840 | ZNF232 | 0,188677582 | 1,139718536 | 0,29816 | 0,59055479 | zinc finger protein 232 |
| ENSG00000037637 | FBXO42 | 0,188656521 | 1,139701898 | 0,25048 | 0,55682585 | F-box protein 42 |
| ENSG00000164548 | TRA2A | 0,188643106 | 1,139691301 | 0,3564 | 0,63043562 | transformer 2 alpha homolog |
| ENSG00000141198 | TOM1L1 | 0,188584181 | 1,139644752 | 0,63824 | 0,81832361 | target of myb1 like 1 membrane trafficking protein |
| ENSG00000268129 | AC026304,1 | 0,188544961 | 1,139613772 | 0,69221 | 0,85023519 | novel transcript |
| ENSG00000272375 | AC026979,3 | 0,188493548 | 1,13957316 | 0,74653 | 0,88046587 | novel transcript |
| ENSG00000167815 | PRDX2 | 0,188364242 | 1,139471027 | 0,53203 | 0,75285051 | peroxiredoxin 2 |
| ENSG00000089053 | ANAPC5 | 0,188352584 | 1,139461819 | 0,28971 | 0,58522301 | anaphase promoting complex subunit 5 |
| ENSG00000197696 | NMB | 0,188333389 | 1,139446659 | 0,51355 | 0,73987113 | neuromedin B |
| ENSG00000134594 | RAB33A | 0,188233763 | 1,139367977 | 0,66721 | 0,83474892 | RAB33A, member RAS oncogene family |

| | | | | | | |
|---|---|---|---|---|---|---|
| ENSG00000135776 | ABCB10 | 0,188214285 | 1,139352594 | 0,27892 | 0,57700804 | ATP binding cassette subfamily B member 10 |
| ENSG00000285407 | AL033530,1 | 0,188188903 | 1,139332549 | 0,6671 | 0,83471429 | novel transcript |
| ENSG00000143498 | TAF1A | 0,18811952 | 1,139277757 | 0,27521 | 0,57491355 | TATA-box binding protein associated factor, RNA polymerase I subunit A |
| ENSG00000196683 | TOMM7 | 0,18805081 | 1,139223498 | 0,47659 | 0,71481671 | translocase of outer mitochondrial membrane 7 |
| ENSG00000153993 | SEMA3D | 0,188040264 | 1,139215171 | 0,61514 | 0,80421986 | semaphorin 3D |
| ENSG00000073536 | NLE1 | 0,187975501 | 1,139164033 | 0,35754 | 0,63137382 | notchless homolog 1 |
| ENSG00000013523 | ANGEL1 | 0,187925908 | 1,139124874 | 0,3955 | 0,6582534 | angel homolog 1 |
| ENSG00000187479 | C11orf96 | 0,187918123 | 1,139118727 | 0,80213 | 0,90897727 | chromosome 11 open reading frame 96 |
| ENSG00000185946 | RNPC3 | 0,187844169 | 1,139060336 | 0,59301 | 0,79148028 | RNA binding region (RNP1, RRM) containing 3 |
| ENSG00000145358 | DDIT4L | 0,187726671 | 1,138967571 | 0,77165 | 0,89423898 | DNA damage inducible transcript 4 like |
| ENSG00000243107 | AC000120,1 | 0,187726355 | 1,138967322 | 0,56897 | 0,77629764 | novel transcript |
| ENSG00000002834 | LASP1 | 0,187611167 | 1,138876388 | 0,31681 | 0,60268695 | LIM and SH3 protein 1 |
| ENSG00000248734 | AC008906,1 | 0,187567172 | 1,138841658 | 0,64751 | 0,82364696 | novel transcript |
| ENSG00000168852 | TPTE2P5 | 0,187519943 | 1,138804377 | 0,68585 | 0,84654659 | transmembrane phosphoinositide 3-phosphatase and tensin homolog 2 pseudogene 5 |
| ENSG00000267745 | AC060766,7 | 0,187506655 | 1,138793888 | 0,73655 | 0,87529332 | novel transcript |
| ENSG00000102804 | TSC22D1 | 0,187396388 | 1,138706851 | 0,56336 | 0,7724391 | TSC22 domain family member 1 |
| ENSG00000259583 | AC015712,2 | 0,187391626 | 1,138703093 | 0,87051 | 0,94289917 | novel transcript, antisense to LRRK1 and ALDH1A3 |
| ENSG00000143669 | LYST | 0,18727273 | 1,138609254 | 0,65485 | 0,82733348 | lysosomal trafficking regulator |
| ENSG00000225460 | AL590233,1 | 0,187257005 | 1,138596843 | 0,83612 | 0,92493625 | ubiquitin-conjugating enzyme E2 pseudogene |
| ENSG00000204434 | POTEKP | 0,187199217 | 1,138551237 | 0,78701 | 0,90114442 | POTE ankyrin domain family member K, pseudogene |
| ENSG00000214455 | RCN1P2 | 0,187178583 | 1,138534953 | 0,56141 | 0,77136826 | reticulocalbin 1 pseudogene 2 |
| ENSG00000229267 | SNHG31 | 0,187166188 | 1,138525171 | 0,44278 | 0,69040116 | small nucleolar RNA host gene 31 |
| ENSG00000128607 | KLHDC10 | 0,187146628 | 1,138509736 | 0,47622 | 0,71446062 | kelch domain containing 10 |
| ENSG00000153310 | CYRIB | 0,18713914 | 1,138503826 | 0,32809 | 0,60971379 | CYFIP related Rac1 interactor B |
| ENSG00000135537 | AFG1L | 0,186903346 | 1,138317765 | 0,46215 | 0,70344988 | AFG1 like ATPase |
| ENSG00000261487 | AC135048,1 | 0,186900246 | 1,138315318 | 0,67017 | 0,83650134 | novel transcript |
| ENSG00000143799 | PARP1 | 0,18687086 | 1,138292133 | 0,4628 | 0,70391423 | poly(ADP-ribose) polymerase 1 |
| ENSG00000215421 | ZNF407 | 0,186828941 | 1,138259059 | 0,57613 | 0,78053116 | zinc finger protein 407 |
| ENSG00000083750 | RRAGB | 0,186808867 | 1,138243221 | 0,49102 | 0,72492348 | Ras related GTP binding B |
| ENSG00000286128 | AC022413,2 | 0,186670097 | 1,138133741 | 0,75918 | 0,8876058 | novel transcript |
| ENSG00000099991 | CABIN1 | 0,186508416 | 1,138006199 | 0,2702 | 0,57192517 | calcineurin binding protein 1 |

| | | | | | | |
|---|---|---|---|---|---|---|
| ENSG00000183935 | HTR7P1 | 0,186442202 | 1,13795397 | 0,38012 | 0,64784628 | 5-hydroxytryptamine receptor 7 pseudogene 1 |
| ENSG00000176261 | ZBTB8OS | 0,186398094 | 1,137919179 | 0,29483 | 0,58868965 | zinc finger and BTB domain containing 8 opposite strand |
| ENSG00000188811 | NHLRC3 | 0,186326553 | 1,137862753 | 0,48291 | 0,719207 | NHL repeat containing 3 |
| ENSG00000188554 | NBR1 | 0,18629123 | 1,137834894 | 0,26503 | 0,56773732 | NBR1 autophagy cargo receptor |
| ENSG00000287814 | AC011407,1 | 0,186285223 | 1,137830156 | 0,81235 | 0,91377341 | novel transcript |
| ENSG00000102189 | EEA1 | 0,186199943 | 1,137762899 | 0,61736 | 0,8057555 | early endosome antigen 1 |
| ENSG00000249609 | AC080188,1 | 0,186104294 | 1,137687469 | 0,80314 | 0,90931794 | novel transcript |
| ENSG00000141084 | RANBP10 | 0,186036252 | 1,137633814 | 0,36033 | 0,63263188 | RAN binding protein 10 |
| ENSG00000135845 | PIGC | 0,185996651 | 1,137602587 | 0,31031 | 0,59842274 | phosphatidylinositol glycan anchor biosynthesis class C |
| ENSG00000117707 | PROX1 | 0,185990498 | 1,137597735 | 0,61865 | 0,80665241 | prospero homeobox 1 |
| ENSG00000134759 | ELP2 | 0,185983048 | 1,137591861 | 0,36323 | 0,63407828 | elongator acetyltransferase complex subunit 2 |
| ENSG00000037280 | FLT4 | 0,185925111 | 1,137546177 | 0,53392 | 0,75377522 | fms related receptor tyrosine kinase 4 |
| ENSG00000270462 | AC005034,2 | 0,185900828 | 1,137527031 | 0,72367 | 0,86802997 | novel transcript, antisense to C2orf3 |
| ENSG00000196131 | VN1R2 | 0,185859149 | 1,137494168 | 0,67807 | 0,84159488 | vomeronasal 1 receptor 2 |
| ENSG00000264448 | MIR378D2HG | 0,185849764 | 1,137486769 | 0,83273 | 0,92348349 | MIR378D2 host gene |
| ENSG00000116717 | GADD45A | 0,185801968 | 1,137449085 | 0,48878 | 0,72350189 | growth arrest and DNA damage inducible alpha |
| ENSG00000137210 | TMEM14B | 0,185779493 | 1,137431365 | 0,51441 | 0,74069047 | transmembrane protein 14B |
| ENSG00000272841 | MAP3K4-AS1 | 0,185698772 | 1,137367726 | 0,44431 | 0,69172919 | MAP3K4 antisense RNA 1 |
| ENSG00000151575 | TEX9 | 0,185545972 | 1,137247271 | 0,61124 | 0,80300957 | testis expressed 9 |
| ENSG00000259151 | CAP2P1 | 0,185361117 | 1,137101562 | 0,77464 | 0,89586241 | cyclase associated actin cytoskeleton regulatory protein 2 pseudogene 1 |
| ENSG00000278023 | RDM1 | 0,185305965 | 1,137058093 | 0,57934 | 0,78270914 | RAD52 motif containing 1 |
| ENSG00000164404 | GDF9 | 0,185201342 | 1,136975638 | 0,72969 | 0,8715818 | growth differentiation factor 9 |
| ENSG00000117360 | PRPF3 | 0,185156411 | 1,136940229 | 0,27617 | 0,57535709 | pre-mRNA processing factor 3 |
| ENSG00000100376 | FAM118A | 0,185123655 | 1,136914415 | 0,24936 | 0,55642837 | family with sequence similarity 118 member A |
| ENSG00000236106 | AC010729,1 | 0,185118441 | 1,136910306 | 0,68616 | 0,84663586 | novel transcript |
| ENSG00000079616 | KIF22 | 0,185091932 | 1,136889416 | 0,34427 | 0,6217351 | kinesin family member 22 |
| ENSG00000010256 | UQCRC1 | 0,185042855 | 1,136850742 | 0,40607 | 0,66573389 | ubiquinol-cytochrome c reductase core protein 1 |
| ENSG00000197879 | MYO1C | 0,184916815 | 1,136751427 | 0,32255 | 0,60577115 | myosin IC |
| ENSG00000267191 | AC006213,3 | 0,18491016 | 1,136746183 | 0,67892 | 0,84203421 | novel transcript, antisense to ZNF45 |
| ENSG00000152620 | NADK2 | 0,184906914 | 1,136743625 | 0,41952 | 0,67530041 | NAD kinase 2, mitochondrial |
| ENSG00000272144 | AC025171,4 | 0,184777501 | 1,136641662 | 0,64027 | 0,81936319 | novel transcript |
| ENSG00000152284 | TCF7L1 | 0,184770805 | 1,136636386 | 0,35467 | 0,62922462 | transcription factor 7 like 1 |

| | | | | | | |
|---|---|---|---|---|---|---|
| ENSG00000146410 | MTFR2 | 0,184752708 | 1,136622129 | 0,55968 | 0,7706142 | mitochondrial fission regulator 2 |
| ENSG00000169925 | BRD3 | 0,184690826 | 1,136573376 | 0,27683 | 0,57549599 | bromodomain containing 3 |
| ENSG00000227382 | EIF4A2P2 | 0,184657575 | 1,13654718 | 0,74983 | 0,88232127 | eukaryotic translation initiation factor 4A2 pseudogene 2 |
| ENSG00000090487 | SPG21 | 0,184624322 | 1,136520985 | 0,32256 | 0,60577115 | SPG21 abhydrolase domain containing, maspardin |
| ENSG00000129295 | LRRC6 | 0,184454052 | 1,136386858 | 0,51951 | 0,74402056 | leucine rich repeat containing 6 |
| ENSG00000167232 | ZNF91 | 0,184443304 | 1,136378392 | 0,59219 | 0,79101863 | zinc finger protein 91 |
| ENSG00000152926 | ZNF117 | 0,184403853 | 1,136347318 | 0,56373 | 0,77255679 | zinc finger protein 117 |
| ENSG00000233231 | HNRNPA1P49 | 0,18436992 | 1,136320591 | 0,69305 | 0,85087762 | heterogeneous nuclear ribonucleoprotein A1 pseudogene 49 |
| ENSG00000128694 | OSGEPL1 | 0,184307085 | 1,1362711 | 0,52754 | 0,75037219 | O-sialoglycoprotein endopeptidase like 1 |
| ENSG00000110931 | CAMKK2 | 0,184234824 | 1,136214189 | 0,48912 | 0,72372012 | calcium/calmodulin dependent protein kinase kinase 2 |
| ENSG00000198873 | GRK5 | 0,184215691 | 1,13619912 | 0,44212 | 0,69003826 | G protein-coupled receptor kinase 5 |
| ENSG00000277292 | PIP4K2B | 0,184189597 | 1,13617857 | 0,4222 | 0,67663014 | phosphatidylinositol-5-phosphate 4-kinase type 2 beta |
| ENSG00000276293 | PIP4K2B | 0,184189597 | 1,13617857 | 0,4222 | 0,67663014 | phosphatidylinositol-5-phosphate 4-kinase type 2 beta |
| ENSG00000068323 | TFE3 | 0,184142481 | 1,136141465 | 0,28534 | 0,58187605 | transcription factor binding to IGHM enhancer 3 |
| ENSG00000264736 | BDP1P | 0,184075693 | 1,136088869 | 0,71657 | 0,86437402 | B double prime 1 pseudogene |
| ENSG00000275764 | AC092747,4 | 0,184047593 | 1,136066742 | 0,56513 | 0,773443 | novel transcript |
| ENSG00000172296 | SPTLC3 | 0,184038555 | 1,136059624 | 0,49951 | 0,73058737 | serine palmitoyltransferase long chain base subunit 3 |
| ENSG00000108433 | GOSR2 | 0,184037343 | 1,136058671 | 0,27553 | 0,57522081 | golgi SNAP receptor complex member 2 |
| ENSG00000116668 | SWT1 | 0,183982168 | 1,136015223 | 0,63001 | 0,8132682 | SWT1 RNA endoribonuclease homolog |
| ENSG00000237984 | PTENP1 | 0,183969694 | 1,136005401 | 0,73593 | 0,8750728 | phosphatase and tensin homolog pseudogene 1 |
| ENSG00000127952 | STYXL1 | 0,183960793 | 1,135998393 | 0,39508 | 0,65801556 | serine/threonine/tyrosine interacting like 1 |
| ENSG00000166912 | MTMR10 | 0,183912878 | 1,135960664 | 0,63457 | 0,81595759 | myotubularin related protein 10 |
| ENSG00000273058 | AL359921,2 | 0,183863759 | 1,135921989 | 0,74219 | 0,87810032 | novel transcript |
| ENSG00000287620 | AC092053,4 | 0,183863366 | 1,13592168 | 0,6664 | 0,83411217 | novel transcript |
| ENSG00000100483 | VCPKMT | 0,183784832 | 1,135859846 | 0,42065 | 0,67598168 | valosin containing protein lysine methyltransferase |
| ENSG00000261762 | AC027228,2 | 0,183590277 | 1,135706681 | 0,59439 | 0,79263193 | novel transcript, antisense to CHRNA5 |
| ENSG00000111269 | CREBL2 | 0,183580592 | 1,135699056 | 0,35065 | 0,62606342 | cAMP responsive element binding protein like 2 |
| ENSG00000102181 | CD99L2 | 0,183540061 | 1,13566715 | 0,31728 | 0,60276767 | CD99 molecule like 2 |
| ENSG00000219163 | HMGB1P20 | 0,183466843 | 1,135609516 | 0,80221 | 0,90899209 | high mobility group box 1 pseudogene 20 |
| ENSG00000173486 | FKBP2 | 0,183445309 | 1,135592565 | 0,44725 | 0,69342177 | FKBP prolyl isomerase 2 |
| ENSG00000244229 | RPL26P35 | 0,183428384 | 1,135579243 | 0,79639 | 0,90591272 | ribosomal protein L26 pseudogene 35 |
| ENSG00000276107 | AC037198,1 | 0,183422844 | 1,135574883 | 0,7315 | 0,87265295 | novel transcript, sense intronic to THBS1 |

| | | | | | | |
|---|---|---|---|---|---|---|
| ENSG00000109680 | TBC1D19 | 0,183404924 | 1,135560777 | 0,55719 | 0,76897426 | TBC1 domain family member 19 |
| ENSG00000101493 | ZNF516 | 0,183372473 | 1,135535235 | 0,3395 | 0,61852579 | zinc finger protein 516 |
| ENSG00000283913 | AL512662,1 | 0,18332291 | 1,135496225 | 0,64993 | 0,82459691 | novel transcript |
| ENSG00000105655 | ISYNA1 | 0,183285469 | 1,135466757 | 0,42089 | 0,67615615 | inositol-3-phosphate synthase 1 |
| ENSG00000136404 | TM6SF1 | 0,183081235 | 1,135306027 | 0,81623 | 0,91550954 | transmembrane 6 superfamily member 1 |
| ENSG00000198783 | ZNF830 | 0,183038702 | 1,135272557 | 0,53578 | 0,75487955 | zinc finger protein 830 |
| ENSG00000288612 | AL133351,4 | 0,18303546 | 1,135270006 | 0,75588 | 0,8857133 | novel transcript |
| ENSG00000242103 | AC079841,1 | 0,18290319 | 1,135165926 | 0,8175 | 0,91619921 | actin related protein 2/3 complex, subunit 1A, 41kDa (ARPC1A) pseudogene |
| ENSG00000140992 | PDPK1 | 0,182816507 | 1,135097722 | 0,35993 | 0,63249219 | 3-phosphoinositide dependent protein kinase 1 |
| ENSG00000249863 | AC021106,1 | 0,182787254 | 1,135074707 | 0,70716 | 0,85902171 | synaptotagmin binding, cytoplasmic RNA interacting protein (SYNCRIP) pseudogene |
| ENSG00000230225 | MTND5P14 | 0,182726269 | 1,135026727 | 0,8478 | 0,9309317 | MT-ND5 pseudogene 14 |
| ENSG00000188916 | INSYN2A | 0,182712276 | 1,135015718 | 0,53777 | 0,75631707 | inhibitory synaptic factor 2A |
| ENSG00000128534 | LSM8 | 0,182640181 | 1,134958999 | 0,51653 | 0,74198903 | LSM8 homolog, U6 small nuclear RNA associated |
| ENSG00000165169 | DYNLT3 | 0,182435645 | 1,134798104 | 0,61882 | 0,80665241 | dynein light chain Tctex-type 3 |
| ENSG00000244625 | MIATNB | 0,182248095 | 1,13465059 | 0,49311 | 0,72619695 | MIAT neighbor |
| ENSG00000258968 | AL049830,4 | 0,182244171 | 1,134647503 | 0,83256 | 0,92339761 | intraflagellar transport 20 homolog (Chlamydomonas) (IFT20) pseudogene |
| ENSG00000165813 | CCDC186 | 0,182176211 | 1,134594056 | 0,55809 | 0,76973191 | coiled-coil domain containing 186 |
| ENSG00000264772 | AC016876,2 | 0,182130143 | 1,134557826 | 0,42069 | 0,67598913 | novel transcript |
| ENSG00000116580 | GON4L | 0,182127067 | 1,134555408 | 0,30382 | 0,59399806 | gon-4 like |
| ENSG00000128989 | ARPP19 | 0,182122016 | 1,134551436 | 0,48782 | 0,72279694 | cAMP regulated phosphoprotein 19 |
| ENSG00000109674 | NEIL3 | 0,181901473 | 1,134378011 | 0,55023 | 0,76449335 | nei like DNA glycosylase 3 |
| ENSG00000039139 | DNAH5 | 0,181875197 | 1,134357351 | 0,6448 | 0,82169377 | dynein axonemal heavy chain 5 |
| ENSG00000115042 | FAHD2A | 0,181734906 | 1,134247049 | 0,48874 | 0,72350189 | fumarylacetoacetate hydrolase domain containing 2A |
| ENSG00000186660 | ZFP91 | 0,181719113 | 1,134234632 | 0,47115 | 0,71052357 | ZFP91 zinc finger protein, atypical E3 ubiquitin ligase |
| ENSG00000131067 | GGT7 | 0,181636163 | 1,134169419 | 0,32313 | 0,60590706 | gamma-glutamyltransferase 7 |
| ENSG00000117682 | DHDDS | 0,181517429 | 1,134076082 | 0,25616 | 0,56068598 | dehydrodolichyl diphosphate synthase subunit |
| ENSG00000132768 | DPH2 | 0,181500983 | 1,134063153 | 0,46702 | 0,70706711 | diphthamide biosynthesis 2 |
| ENSG00000108528 | SLC25A11 | 0,181345119 | 1,13394064 | 0,25351 | 0,55857654 | solute carrier family 25 member 11 |
| ENSG00000100811 | YY1 | 0,181291926 | 1,133898832 | 0,36178 | 0,63352908 | YY1 transcription factor |
| ENSG00000271966 | AC021321,1 | 0,181283085 | 1,133891883 | 0,57276 | 0,77868876 | novel transcript |

| | | | | | | |
|---|---|---|---|---|---|---|
| ENSG00000170836 | PPM1D | 0,181257623 | 1,133871871 | 0,43255 | 0,68418465 | protein phosphatase, Mg2+/Mn2+ dependent 1D |
| ENSG00000173334 | TRIB1 | 0,181240542 | 1,133858447 | 0,52716 | 0,75025748 | tribbles pseudokinase 1 |
| ENSG00000143797 | MBOAT2 | 0,18118002 | 1,133810881 | 0,588 | 0,78793103 | membrane bound O-acyltransferase domain containing 2 |
| ENSG00000198952 | SMG5 | 0,181160284 | 1,133795371 | 0,32351 | 0,60631901 | SMG5 nonsense mediated mRNA decay factor |
| ENSG00000172239 | PAIP1 | 0,181150537 | 1,133787711 | 0,51666 | 0,74206471 | poly(A) binding protein interacting protein 1 |
| ENSG00000230069 | LRRC37A15P | 0,18113045 | 1,133771925 | 0,82105 | 0,91738668 | leucine rich repeat containing 37 member A15, pseudogene |
| ENSG00000110318 | CEP126 | 0,181129596 | 1,133771254 | 0,35497 | 0,6294354 | centrosomal protein 126 |
| ENSG00000119969 | HELLS | 0,181051281 | 1,13370971 | 0,37702 | 0,64554193 | helicase, lymphoid specific |
| ENSG00000144010 | TRIM43B | 0,181042748 | 1,133703004 | 0,88397 | 0,94847506 | tripartite motif containing 43B |
| ENSG00000260439 | LMF1-AS1 | 0,180976394 | 1,133650863 | 0,74128 | 0,8774838 | LMF1 antisense RNA 1 |
| ENSG00000163807 | KIAA1143 | 0,180969515 | 1,133645458 | 0,69281 | 0,85072778 | KIAA1143 |
| ENSG00000110315 | RNF141 | 0,180791241 | 1,133505382 | 0,43796 | 0,68747773 | ring finger protein 141 |
| ENSG00000166685 | COG1 | 0,180640448 | 1,133386912 | 0,4239 | 0,67773865 | component of oligomeric golgi complex 1 |
| ENSG00000175166 | PSMD2 | 0,180618732 | 1,133369852 | 0,49587 | 0,72815206 | proteasome 26S subunit, non-ATPase 2 |
| ENSG00000253168 | AC124069,1 | 0,180566101 | 1,133328506 | 0,75254 | 0,88402345 | novel transcript |
| ENSG00000159055 | MIS18A | 0,180336597 | 1,13314823 | 0,21501 | 0,53373528 | MIS18 kinetochore protein A |
| ENSG00000087086 | FTL | 0,180321909 | 1,133136694 | 0,51686 | 0,74218125 | ferritin light chain |
| ENSG00000250045 | CBX3P3 | 0,180302124 | 1,133121155 | 0,82719 | 0,92010219 | chromobox 3 pseudogene 3 |
| ENSG00000186591 | UBE2H | 0,180152064 | 1,133003301 | 0,34747 | 0,62411645 | ubiquitin conjugating enzyme E2 H |
| ENSG00000125482 | TTF1 | 0,180144273 | 1,132997182 | 0,45386 | 0,6979134 | transcription termination factor 1 |
| ENSG00000235758 | AGPAT1 | 0,180035666 | 1,132911893 | 0,44753 | 0,69342177 | 1-acylglycerol-3-phosphate O-acyltransferase 1 |
| ENSG00000206324 | AGPAT1 | 0,180035666 | 1,132911893 | 0,44753 | 0,69342177 | 1-acylglycerol-3-phosphate O-acyltransferase 1 |
| ENSG00000226467 | AGPAT1 | 0,180035666 | 1,132911893 | 0,44753 | 0,69342177 | 1-acylglycerol-3-phosphate O-acyltransferase 1 |
| ENSG00000178338 | ZNF354B | 0,17999948 | 1,132883477 | 0,33875 | 0,61792351 | zinc finger protein 354B |
| ENSG00000226650 | KIF4B | 0,179998822 | 1,13288296 | 0,69432 | 0,85164591 | kinesin family member 4B |
| ENSG00000184574 | LPAR5 | 0,179995151 | 1,132880078 | 0,85497 | 0,93417381 | lysophosphatidic acid receptor 5 |
| ENSG00000203685 | STUM | 0,179930128 | 1,132829019 | 0,68955 | 0,84865812 | stum, mechanosensory transduction mediator homolog |
| ENSG00000132792 | CTNNBL1 | 0,179856617 | 1,132771298 | 0,46965 | 0,70914554 | catenin beta like 1 |
| ENSG00000244451 | RPL34P21 | 0,179819996 | 1,132742545 | 0,88684 | 0,95005038 | ribosomal protein L34 pseudogene 21 |
| ENSG00000110427 | KIAA1549L | 0,179815179 | 1,132738763 | 0,58811 | 0,78803625 | KIAA1549 like |
| ENSG00000170396 | ZNF804A | 0,179648107 | 1,132607593 | 0,72113 | 0,86669144 | zinc finger protein 804A |
| ENSG00000163467 | TSACC | 0,179647003 | 1,132606726 | 0,7475 | 0,88098027 | TSSK6 activating cochaperone |

| | | | | | | |
|---|---|---|---|---|---|---|
| ENSG00000196152 | ZNF79 | 0,179595532 | 1,132566319 | 0,41125 | 0,66903348 | zinc finger protein 79 |
| ENSG00000154734 | ADAMTS1 | 0,179586702 | 1,132559388 | 0,45835 | 0,70119611 | ADAM metallopeptidase with thrombospondin type 1 motif 1 |
| ENSG00000134262 | AP4B1 | 0,179550365 | 1,132530862 | 0,37558 | 0,64393288 | adaptor related protein complex 4 subunit beta 1 |
| ENSG00000152291 | TGOLN2 | 0,179529365 | 1,132514377 | 0,37818 | 0,64646341 | trans-golgi network protein 2 |
| ENSG00000151923 | TIAL1 | 0,179520971 | 1,132507787 | 0,22044 | 0,53750545 | TIA1 cytotoxic granule associated RNA binding protein like 1 |
| ENSG00000281189 | GHET1 | 0,17947392 | 1,132470853 | 0,48679 | 0,72215091 | gastric carcinoma proliferation enhancing transcript 1 |
| ENSG00000168297 | PXK | 0,17924044 | 1,132287593 | 0,55305 | 0,7662484 | PX domain containing serine/threonine kinase like |
| ENSG00000224905 | AP001347,1 | 0,179177281 | 1,132238025 | 0,59797 | 0,79484486 | novel transcript, antisense to LIPI |
| ENSG00000134982 | APC | 0,179116048 | 1,13218997 | 0,64873 | 0,82411802 | APC regulator of WNT signaling pathway |
| ENSG00000164011 | ZNF691 | 0,179040327 | 1,132130547 | 0,4736 | 0,71283511 | zinc finger protein 691 |
| ENSG00000110395 | CBL | 0,178981564 | 1,132084435 | 0,45501 | 0,69913628 | Cbl proto-oncogene |
| ENSG00000278183 | ABCC1 | 0,178849576 | 1,131980869 | 0,34045 | 0,61897407 | ATP binding cassette subfamily C member 1 |
| ENSG00000258012 | AC026111,1 | 0,178807685 | 1,131948 | 0,748 | 0,88136669 | methionine adenosyltransferase II, alpha (MAT2A) pseudogene |
| ENSG00000259330 | INAFM2 | 0,178505022 | 1,131710554 | 0,59881 | 0,79524043 | InaF motif containing 2 |
| ENSG00000125945 | ZNF436 | 0,178393232 | 1,131622864 | 0,55546 | 0,76788267 | zinc finger protein 436 |
| ENSG00000283009 | ZNF436 | 0,178393232 | 1,131622864 | 0,55546 | 0,76788267 | zinc finger protein 436 |
| ENSG00000267940 | AC022762,2 | 0,178345407 | 1,131585352 | 0,61103 | 0,80279899 | novel transcript |
| ENSG00000198786 | MT-ND5 | 0,178322034 | 1,131567019 | 0,34002 | 0,61888023 | mitochondrially encoded NADH:ubiquinone oxidoreductase core subunit 5 |
| ENSG00000260209 | AP000842,3 | 0,178275955 | 1,131530879 | 0,86702 | 0,94116774 | novel transcript |
| ENSG00000241278 | ENPP7P4 | 0,178219577 | 1,131486661 | 0,71089 | 0,86079022 | ectonucleotide pyrophosphatase/phosphodiesterase 7 pseudogene 4 |
| ENSG00000275236 | AC009120,5 | 0,178199985 | 1,131471295 | 0,72246 | 0,86734497 | novel transcript |
| ENSG00000278229 | RPS17 | 0,178140572 | 1,1314247 | 0,53206 | 0,75285051 | ribosomal protein S17 |
| ENSG00000162804 | SNED1 | 0,178108606 | 1,131399631 | 0,6213 | 0,80787022 | sushi, nidogen and EGF like domains 1 |
| ENSG00000213492 | NT5C3AP1 | 0,178061549 | 1,131362729 | 0,8589 | 0,93646304 | NT5C3A pseudogene 1 |
| ENSG00000246363 | LINC02458 | 0,178019837 | 1,131330018 | 0,76634 | 0,89138648 | long intergenic non-protein coding RNA 2458 |
| ENSG00000067177 | PHKA1 | 0,178008805 | 1,131321368 | 0,65042 | 0,82492554 | phosphorylase kinase regulatory subunit alpha 1 |
| ENSG00000075426 | FOSL2 | 0,177989059 | 1,131305884 | 0,68319 | 0,84490651 | FOS like 2, AP-1 transcription factor subunit |
| ENSG00000223687 | ZNF311 | 0,177868732 | 1,131211532 | 0,76741 | 0,89211771 | zinc finger protein 311 |

| | | | | | | |
|---|---|---|---|---|---|---|
| ENSG00000269937 | AC093525,7 | 0,17762142 | 1,131017632 | 0,36929 | 0,63912679 | novel transcript, antisense to PDPK1 |
| ENSG00000144554 | FANCD2 | 0,177598453 | 1,130999627 | 0,37267 | 0,64161496 | FA complementation group D2 |
| ENSG00000164182 | NDUFAF2 | 0,1775699 | 1,130977242 | 0,4108 | 0,66856387 | NADH:ubiquinone oxidoreductase complex assembly factor 2 |
| ENSG00000163882 | POLR2H | 0,177556637 | 1,130966846 | 0,40721 | 0,66659618 | RNA polymerase II, I and III subunit H |
| ENSG00000137074 | APTX | 0,177530987 | 1,130946738 | 0,29913 | 0,59122003 | aprataxin |
| ENSG00000172748 | ZNF596 | 0,177518361 | 1,13093684 | 0,50654 | 0,73497152 | zinc finger protein 596 |
| ENSG00000239570 | SETP11 | 0,177515046 | 1,130934241 | 0,75726 | 0,88663582 | SET pseudogene 11 |
| ENSG00000188629 | ZNF177 | 0,17748702 | 1,130912272 | 0,66155 | 0,83148228 | zinc finger protein 177 |
| ENSG00000132002 | DNAJB1 | 0,177404881 | 1,130847886 | 0,35617 | 0,63035942 | DnaJ heat shock protein family (Hsp40) member B1 |
| ENSG00000135469 | COQ10A | 0,177298329 | 1,130764369 | 0,49576 | 0,72803885 | coenzyme Q10A |
| ENSG00000130821 | SLC6A8 | 0,17718065 | 1,130672137 | 0,41422 | 0,67104294 | solute carrier family 6 member 8 |
| ENSG00000112033 | PPARD | 0,177136303 | 1,130637383 | 0,48942 | 0,72387251 | peroxisome proliferator activated receptor delta |
| ENSG00000077254 | USP33 | 0,177102364 | 1,130610785 | 0,42337 | 0,67712191 | ubiquitin specific peptidase 33 |
| ENSG00000285777 | AL139142,2 | 0,176993864 | 1,130525759 | 0,82511 | 0,91900665 | novel protein |
| ENSG00000039560 | RAI14 | 0,176951987 | 1,130492943 | 0,32985 | 0,61078709 | retinoic acid induced 14 |
| ENSG00000239388 | ASB14 | 0,176830511 | 1,130397759 | 0,55681 | 0,76863826 | ankyrin repeat and SOCS box containing 14 |
| ENSG00000132424 | PNISR | 0,176793203 | 1,130368528 | 0,62047 | 0,80776776 | PNN interacting serine and arginine rich protein |
| ENSG00000125970 | RALY | 0,176751854 | 1,13033613 | 0,46256 | 0,70358709 | RALY heterogeneous nuclear ribonucleoprotein |
| ENSG00000112796 | ENPP5 | 0,176649266 | 1,130255757 | 0,73929 | 0,8763893 | ectonucleotide pyrophosphatase/phosphodiesterase family member 5 |
| ENSG00000139668 | WDFY2 | 0,176574566 | 1,130197236 | 0,46066 | 0,70247195 | WD repeat and FYVE domain containing 2 |
| ENSG00000247498 | GPRC5D-AS1 | 0,176529112 | 1,130161628 | 0,7057 | 0,85825904 | GPRC5D and HEBP1 antisense RNA 1 |
| ENSG00000028528 | SNX1 | 0,176503434 | 1,130141513 | 0,26459 | 0,56738489 | sorting nexin 1 |
| ENSG00000267006 | AC008507,2 | 0,17638344 | 1,130047519 | 0,74693 | 0,88059941 | novel transcript |
| ENSG00000172766 | NAA16 | 0,176363228 | 1,130031687 | 0,41993 | 0,67551392 | N-alpha-acetyltransferase 16, NatA auxiliary subunit |
| ENSG00000223650 | UHRF2P1 | 0,176210952 | 1,129912419 | 0,69435 | 0,85164591 | ubiquitin like with PHD and ring finger domains 2 pseudogene 1 |
| ENSG00000055483 | USP36 | 0,176191826 | 1,12989744 | 0,30515 | 0,59455047 | ubiquitin specific peptidase 36 |
| ENSG00000157764 | BRAF | 0,176108342 | 1,129832058 | 0,42857 | 0,68124412 | B-Raf proto-oncogene, serine/threonine kinase |
| ENSG00000166341 | DCHS1 | 0,176055041 | 1,129790317 | 0,41439 | 0,67106867 | dachsous cadherin-related 1 |
| ENSG00000260685 | AC027104,1 | 0,175977562 | 1,129729644 | 0,85438 | 0,93397962 | RWD domain containing 1 (RWDD1) pseudogene |
| ENSG00000105397 | TYK2 | 0,175928105 | 1,129690917 | 0,39241 | 0,6557477 | tyrosine kinase 2 |

| | | | | | | |
|---|---|---|---|---|---|---|
| ENSG00000102524 | TNFSF13B | 0,175925128 | 1,129688585 | 0,77794 | 0,89728573 | TNF superfamily member 13b |
| ENSG00000270528 | FAM133EP | 0,175919293 | 1,129684016 | 0,83878 | 0,92638462 | family with sequence similarity 133 member E, pseudogene |
| ENSG00000180370 | PAK2 | 0,175846147 | 1,129626741 | 0,50886 | 0,7369661 | p21 (RAC1) activated kinase 2 |
| ENSG00000108055 | SMC3 | 0,175836056 | 1,129618841 | 0,59373 | 0,79201823 | structural maintenance of chromosomes 3 |
| ENSG00000047578 | KATNIP | 0,17583111 | 1,129614968 | 0,39195 | 0,65536998 | katanin interacting protein |
| ENSG00000101442 | ACTR5 | 0,175802537 | 1,129592595 | 0,47826 | 0,71600883 | actin related protein 5 |
| ENSG00000282800 | MIR22HG | 0,17574827 | 1,129550107 | 0,63758 | 0,81781506 | MIR22 host gene |
| ENSG00000186594 | MIR22HG | 0,17574827 | 1,129550107 | 0,63758 | 0,81781506 | MIR22 host gene |
| ENSG00000163719 | MTMR14 | 0,17572492 | 1,129531825 | 0,32456 | 0,60709243 | myotubularin related protein 14 |
| ENSG00000285246 | NEDD8 | 0,175656032 | 1,129477892 | 0,41589 | 0,67283255 | NEDD8 ubiquitin like modifier |
| ENSG00000261799 | AC007406,4 | 0,175621876 | 1,129451152 | 0,51688 | 0,74218125 | novel transcript |
| ENSG00000171865 | RNASEH1 | 0,175612518 | 1,129443825 | 0,31009 | 0,59836523 | ribonuclease H1 |
| ENSG00000028203 | VEZT | 0,175582556 | 1,129420369 | 0,52907 | 0,75111082 | vezatin, adherens junctions transmembrane protein |
| ENSG00000276016 | ABR | 0,175499163 | 1,129355086 | 0,64645 | 0,82293404 | ABR activator of RhoGEF and GTPase |
| ENSG00000129353 | SLC44A2 | 0,175497081 | 1,129353457 | 0,4223 | 0,67663581 | solute carrier family 44 member 2 |
| ENSG00000266028 | SRGAP2 | 0,175474453 | 1,129335744 | 0,536 | 0,75500755 | SLIT-ROBO Rho GTPase activating protein 2 |
| ENSG00000216813 | AL358178,1 | 0,175456473 | 1,129321669 | 0,77613 | 0,89650232 | 40S ribosomal protein S12 (RPS12) pseudogene |
| ENSG00000265763 | ZNF488 | 0,175432015 | 1,129302523 | 0,71975 | 0,86599113 | zinc finger protein 488 |
| ENSG00000220721 | OR1F12 | 0,175406719 | 1,129282723 | 0,80145 | 0,90860375 | olfactory receptor family 1 subfamily F member 12 |
| ENSG00000198857 | HSD3BP5 | 0,175387435 | 1,129267628 | 0,72546 | 0,86906859 | hydroxy-delta-5-steroid dehydrogenase, 3 beta, pseudogene 5 |
| ENSG00000105700 | KXD1 | 0,175376441 | 1,129259023 | 0,3693 | 0,63912679 | KxDL motif containing 1 |
| ENSG00000159352 | PSMD4 | 0,175331106 | 1,129223538 | 0,23798 | 0,54864017 | proteasome 26S subunit, non-ATPase 4 |
| ENSG00000148660 | CAMK2G | 0,175318907 | 1,129213989 | 0,30477 | 0,59446033 | calcium/calmodulin dependent protein kinase II gamma |
| ENSG00000111247 | RAD51AP1 | 0,175251199 | 1,129160994 | 0,45839 | 0,70119611 | RAD51 associated protein 1 |
| ENSG00000170927 | PKHD1 | 0,175215794 | 1,129133285 | 0,73379 | 0,87382179 | PKHD1 ciliary IPT domain containing fibrocystin/polyductin |
| ENSG00000254616 | AP001775,1 | 0,175169644 | 1,129097165 | 0,8248 | 0,91900665 | Zinc finger protein 628 (ZNF628) pseudogene |
| ENSG00000175063 | UBE2C | 0,175123182 | 1,129060803 | 0,35801 | 0,63185282 | ubiquitin conjugating enzyme E2 C |
| ENSG00000127603 | MACF1 | 0,175118923 | 1,12905747 | 0,43446 | 0,68505666 | microtubule actin crosslinking factor 1 |
| ENSG00000241973 | PI4KA | 0,174881176 | 1,128871424 | 0,40756 | 0,66665981 | phosphatidylinositol 4-kinase alpha |
| ENSG00000181744 | DIPK2A | 0,174795321 | 1,128804247 | 0,44712 | 0,69341308 | divergent protein kinase domain 2A |
| ENSG00000228278 | ORM2 | 0,174757233 | 1,128774446 | 0,86388 | 0,93914712 | orosomucoid 2 |

| | | | | | | |
|---|---|---|---|---|---|---|
| ENSG00000070061 | ELP1 | 0,174710588 | 1,128737952 | 0,43934 | 0,68826702 | elongator complex protein 1 |
| ENSG00000117155 | SSX2IP | 0,174587286 | 1,128641487 | 0,59273 | 0,79133235 | SSX family member 2 interacting protein |
| ENSG00000089505 | CMTM1 | 0,174556461 | 1,128617372 | 0,64788 | 0,82380454 | CKLF like MARVEL transmembrane domain containing 1 |
| ENSG00000117632 | STMN1 | 0,174555756 | 1,128616821 | 0,43656 | 0,68681257 | stathmin 1 |
| ENSG00000139173 | TMEM117 | 0,17449049 | 1,128565764 | 0,71595 | 0,86379502 | transmembrane protein 117 |
| ENSG00000184939 | ZFP90 | 0,174436512 | 1,12852354 | 0,49596 | 0,72824205 | ZFP90 zinc finger protein |
| ENSG00000260604 | AL590004,3 | 0,174404985 | 1,128498878 | 0,67209 | 0,8377111 | novel transcript |
| ENSG00000213397 | HAUS7 | 0,174399875 | 1,128494881 | 0,57175 | 0,77829419 | HAUS augmin like complex subunit 7 |
| ENSG00000133997 | MED6 | 0,174379119 | 1,128478646 | 0,25285 | 0,55796133 | mediator complex subunit 6 |
| ENSG00000270808 | AC022400,3 | 0,174353584 | 1,128458673 | 0,84268 | 0,92821072 | ataxin 1-like (ATXN1L) pseudogene |
| ENSG00000205531 | NAP1L4 | 0,174262157 | 1,128387162 | 0,43419 | 0,68500084 | nucleosome assembly protein 1 like 4 |
| ENSG00000203999 | LINC01270 | 0,174139247 | 1,128291033 | 0,81394 | 0,9144566 | long intergenic non-protein coding RNA 1270 |
| ENSG00000196866 | H2AC7 | 0,17412451 | 1,128279508 | 0,634 | 0,81549489 | H2A clustered histone 7 |
| ENSG00000206353 | SKIV2L | 0,174078896 | 1,128243836 | 0,53182 | 0,75280487 | Ski2 like RNA helicase |
| ENSG00000228896 | SKIV2L | 0,174078896 | 1,128243836 | 0,53182 | 0,75280487 | Ski2 like RNA helicase |
| ENSG00000232616 | SKIV2L | 0,174078896 | 1,128243836 | 0,53182 | 0,75280487 | Ski2 like RNA helicase |
| ENSG00000197620 | EOLA1 | 0,174045218 | 1,128217498 | 0,50762 | 0,73585078 | endothelium and lymphocyte associated ASCH domain 1 |
| ENSG00000167315 | ACAA2 | 0,174009694 | 1,128189718 | 0,3346 | 0,61433133 | acetyl-CoA acyltransferase 2 |
| ENSG00000100991 | TRPC4AP | 0,17399707 | 1,128179846 | 0,32968 | 0,61069033 | transient receptor potential cation channel subfamily C member 4 associated protein |
| ENSG00000146872 | TLK2 | 0,173971405 | 1,128159776 | 0,36782 | 0,63779082 | tousled like kinase 2 |
| ENSG00000115468 | EFHD1 | 0,17392982 | 1,128127258 | 0,6818 | 0,84389903 | EF-hand domain family member D1 |
| ENSG00000230453 | ANKRD18B | 0,173908237 | 1,128110381 | 0,78674 | 0,9010162 | ankyrin repeat domain 18B |
| ENSG00000282174 | AC006518,7 | 0,173837509 | 1,128055077 | 0,82628 | 0,91953499 | taste receptor, type 2 pseudogene |
| ENSG00000272718 | AC006518,3 | 0,173837509 | 1,128055077 | 0,82628 | 0,91953499 | taste receptor, type 2 pseudogene |
| ENSG00000256651 | AC006518,1 | 0,173837509 | 1,128055077 | 0,82628 | 0,91953499 | taste receptor, type 2 pseudogene |
| ENSG00000188050 | RNF133 | 0,173829532 | 1,12804884 | 0,85242 | 0,93292666 | ring finger protein 133 |
| ENSG00000153157 | SYCP2L | 0,173817709 | 1,128039596 | 0,79151 | 0,90310057 | synaptonemal complex protein 2 like |
| ENSG00000159256 | MORC3 | 0,173721419 | 1,127964309 | 0,50538 | 0,73393676 | MORC family CW-type zinc finger 3 |
| ENSG00000067369 | TP53BP1 | 0,173659672 | 1,127916033 | 0,46905 | 0,70862072 | tumor protein p53 binding protein 1 |
| ENSG00000137656 | BUD13 | 0,173569645 | 1,127845652 | 0,3503 | 0,62578708 | BUD13 homolog |
| ENSG00000217930 | PAM16 | 0,17356756 | 1,127844022 | 0,58097 | 0,78401175 | presequence translocase associated motor 16 |
| ENSG00000144366 | GULP1 | 0,173539225 | 1,12782187 | 0,53191 | 0,75280487 | GULP PTB domain containing engulfment adaptor 1 |

| | | | | | | |
|---|---|---|---|---|---|---|
| ENSG00000163214 | DHX57 | 0,173487633 | 1,127781539 | 0,37297 | 0,64180957 | DExH-box helicase 57 |
| ENSG00000135823 | STX6 | 0,173403019 | 1,127715397 | 0,53495 | 0,75428051 | syntaxin 6 |
| ENSG00000198237 | GUSBP13 | 0,173123966 | 1,12749729 | 0,72786 | 0,87053185 | GUSB pseudogene 13 |
| ENSG00000163530 | DPPA2 | 0,1730852 | 1,127466994 | 0,8126 | 0,91386173 | developmental pluripotency associated 2 |
| ENSG00000259585 | RBM17P4 | 0,17305694 | 1,127444909 | 0,78045 | 0,89830719 | RNA binding motif protein 17 pseudogene 4 |
| ENSG00000130640 | TUBGCP2 | 0,173052527 | 1,12744146 | 0,40151 | 0,6630277 | tubulin gamma complex associated protein 2 |
| ENSG00000285571 | AL513548,4 | 0,172963463 | 1,12737186 | 0,5866 | 0,78690792 | novel transcript |
| ENSG00000097021 | ACOT7 | 0,172951147 | 1,127362236 | 0,49485 | 0,72723033 | acyl-CoA thioesterase 7 |
| ENSG00000110900 | TSPAN11 | 0,172926398 | 1,127342897 | 0,50489 | 0,73368838 | tetraspanin 11 |
| ENSG00000254772 | EEF1G | 0,172923043 | 1,127340275 | 0,32528 | 0,60778059 | eukaryotic translation elongation factor 1 gamma |
| ENSG00000114861 | FOXP1 | 0,172889821 | 1,127314316 | 0,51527 | 0,74138689 | forkhead box P1 |
| ENSG00000169744 | LDB2 | 0,172865013 | 1,127294931 | 0,75167 | 0,88359932 | LIM domain binding 2 |
| ENSG00000183067 | IGSF5 | 0,172850485 | 1,127283579 | 0,80664 | 0,91118635 | immunoglobulin superfamily member 5 |
| ENSG00000165650 | PDZD8 | 0,172810806 | 1,127252575 | 0,32129 | 0,60577115 | PDZ domain containing 8 |
| ENSG00000157796 | WDR19 | 0,172799299 | 1,127243584 | 0,54358 | 0,76018238 | WD repeat domain 19 |
| ENSG00000125149 | C16orf70 | 0,172786429 | 1,127233528 | 0,31134 | 0,59890747 | chromosome 16 open reading frame 70 |
| ENSG00000185261 | KIAA0825 | 0,172726932 | 1,127187042 | 0,65448 | 0,8271533 | KIAA0825 |
| ENSG00000282489 | AC016482,1 | 0,172686905 | 1,127155769 | 0,84401 | 0,92886643 | ribosomal protein L18a (RPL18A) pseudogene |
| ENSG00000276810 | AC244093,2 | 0,172686905 | 1,127155769 | 0,84401 | 0,92886643 | ribosomal protein L18a (RPL18A) pseudogene |
| ENSG00000234177 | LINC01114 | 0,172439726 | 1,126962669 | 0,76982 | 0,89306835 | long intergenic non-protein coding RNA 1114 |
| ENSG00000133065 | SLC41A1 | 0,172222409 | 1,126792924 | 0,37746 | 0,64594766 | solute carrier family 41 member 1 |
| ENSG00000274997 | H2AC12 | 0,172022781 | 1,126637018 | 0,60054 | 0,79587862 | H2A clustered histone 12 |
| ENSG00000164073 | MFSD8 | 0,17201888 | 1,126633972 | 0,51653 | 0,74198903 | major facilitator superfamily domain containing 8 |
| ENSG00000231427 | LINC01445 | 0,172008854 | 1,126626143 | 0,55323 | 0,76630282 | long intergenic non-protein coding RNA 1445 |
| ENSG00000177479 | ARIH2 | 0,171705572 | 1,126389329 | 0,30972 | 0,59807209 | ariadne RBR E3 ubiquitin protein ligase 2 |
| ENSG00000100280 | AP1B1 | 0,171673924 | 1,12636462 | 0,43219 | 0,68411566 | adaptor related protein complex 1 subunit beta 1 |
| ENSG00000144909 | OSBPL11 | 0,171635388 | 1,126334534 | 0,27512 | 0,57491355 | oxysterol binding protein like 11 |
| ENSG00000152778 | IFIT5 | 0,17161905 | 1,126321779 | 0,66369 | 0,8324578 | interferon induced protein with tetratricopeptide repeats 5 |
| ENSG00000171735 | CAMTA1 | 0,171592771 | 1,126301263 | 0,5279 | 0,75063016 | calmodulin binding transcription activator 1 |
| ENSG00000276030 | AC073534,2 | 0,171564522 | 1,12627921 | 0,76056 | 0,88850235 | novel transcript |
| ENSG00000237724 | HSPA1A | 0,171502062 | 1,126230449 | 0,66221 | 0,83183445 | heat shock protein family A (Hsp70) member 1A |
| ENSG00000123411 | IKZF4 | 0,17148361 | 1,126216045 | 0,44255 | 0,69016964 | IKAROS family zinc finger 4 |

| | | | | | | |
|---|---|---|---|---|---|---|
| ENSG00000152457 | DCLRE1C | 0,171279111 | 1,126056417 | 0,46121 | 0,70278639 | DNA cross-link repair 1C |
| ENSG00000251259 | AC004069,1 | 0,171250196 | 1,126033849 | 0,84445 | 0,92906179 | novel transcript |
| ENSG00000162817 | C1orf115 | 0,171243845 | 1,126028892 | 0,64692 | 0,82331024 | chromosome 1 open reading frame 115 |
| ENSG00000100554 | ATP6V1D | 0,17124272 | 1,126028014 | 0,38332 | 0,65008116 | ATPase H+ transporting V1 subunit D |
| ENSG00000286843 | AL160159,1 | 0,171219166 | 1,12600963 | 0,80651 | 0,9111068 | novel transcript, antisense to GPC6 |
| ENSG00000257818 | C1GALT1P1 | 0,171193791 | 1,125989826 | 0,80835 | 0,91190957 | C1GALT1 pseudogene 1 |
| ENSG00000001630 | CYP51A1 | 0,171134052 | 1,125943201 | 0,36381 | 0,63461346 | cytochrome P450 family 51 subfamily A member 1 |
| ENSG00000213997 | PGAM1P7 | 0,171062898 | 1,125887671 | 0,80686 | 0,91120963 | phosphoglycerate mutase 1 pseudogene 7 |
| ENSG00000205643 | CDPF1 | 0,171013145 | 1,125848844 | 0,59434 | 0,79263193 | cysteine rich DPF motif domain containing 1 |
| ENSG00000237973 | MTCO1P12 | 0,170986684 | 1,125828195 | 0,35153 | 0,62674766 | MT-CO1 pseudogene 12 |
| ENSG00000122644 | ARL4A | 0,170900995 | 1,125761328 | 0,6767 | 0,8409179 | ADP ribosylation factor like GTPase 4A |
| ENSG00000088992 | TESC | 0,170884963 | 1,125748818 | 0,82463 | 0,91900665 | tescalcin |
| ENSG00000153044 | CENPH | 0,170842178 | 1,125715433 | 0,42359 | 0,67732256 | centromere protein H |
| ENSG00000228863 | AL121985,1 | 0,170740998 | 1,125636487 | 0,71796 | 0,86489865 | novel transcript, antisense to CD48 |
| ENSG00000106809 | OGN | 0,170667892 | 1,125579448 | 0,73996 | 0,8767413 | osteoglycin |
| ENSG00000204584 | AC027801,1 | 0,170662902 | 1,125575555 | 0,72478 | 0,86866311 | novel transcript, antisense to TAC4 |
| ENSG00000115758 | ODC1 | 0,17061655 | 1,125539392 | 0,6203 | 0,80773684 | ornithine decarboxylase 1 |
| ENSG00000000457 | SCYL3 | 0,170584429 | 1,125514333 | 0,49134 | 0,72510994 | SCY1 like pseudokinase 3 |
| ENSG00000181467 | RAP2B | 0,170500567 | 1,12544891 | 0,29173 | 0,58660475 | RAP2B, member of RAS oncogene family |
| ENSG00000250950 | AC093752,2 | 0,170437937 | 1,125400053 | 0,65321 | 0,82629393 | novel transcript |
| ENSG00000164885 | CDK5 | 0,170422752 | 1,125388208 | 0,41876 | 0,67486451 | cyclin dependent kinase 5 |
| ENSG00000275056 | AC020663,3 | 0,170411186 | 1,125379186 | 0,70601 | 0,85841728 | novel transcript, sense intronic to GLYR1 |
| ENSG00000147457 | CHMP7 | 0,170358049 | 1,125337737 | 0,28865 | 0,58455903 | charged multivesicular body protein 7 |
| ENSG00000108591 | DRG2 | 0,17032608 | 1,1253128 | 0,50573 | 0,73418581 | developmentally regulated GTP binding protein 2 |
| ENSG00000214875 | MED28P1 | 0,170325478 | 1,125312331 | 0,81481 | 0,91470265 | mediator complex subunit 28 pseudogene 1 |
| ENSG00000274624 | AC007671,1 | 0,170262422 | 1,125263148 | 0,79795 | 0,90685415 | novel transcript, sense intronic to ST8SIA1 |
| ENSG00000089220 | PEBP1 | 0,170235897 | 1,12524246 | 0,39873 | 0,66126752 | phosphatidylethanolamine binding protein 1 |
| ENSG00000003756 | RBM5 | 0,170118468 | 1,125150873 | 0,32665 | 0,60877026 | RNA binding motif protein 5 |
| ENSG00000168291 | PDHB | 0,170107332 | 1,125142189 | 0,3071 | 0,59617071 | pyruvate dehydrogenase E1 subunit beta |
| ENSG00000196968 | FUT11 | 0,17008855 | 1,125127541 | 0,40525 | 0,6649751 | fucosyltransferase 11 |
| ENSG00000185697 | MYBL1 | 0,170055177 | 1,125101514 | 0,55279 | 0,7660829 | MYB proto-oncogene like 1 |
| ENSG00000134748 | PRPF38A | 0,170045617 | 1,125094059 | 0,34182 | 0,6197312 | pre-mRNA processing factor 38A |
| ENSG00000214765 | SEPTIN7P2 | 0,170013174 | 1,125068758 | 0,3643 | 0,6349692 | septin 7 pseudogene 2 |

| | | | | | | |
|---|---|---|---|---|---|---|
| ENSG00000204519 | ZNF551 | 0,169995043 | 1,125054619 | 0,36259 | 0,63379594 | zinc finger protein 551 |
| ENSG00000162517 | PEF1 | 0,169975776 | 1,125039594 | 0,42314 | 0,67694038 | penta-EF-hand domain containing 1 |
| ENSG00000025293 | PHF20 | 0,169970168 | 1,125035221 | 0,30473 | 0,59446033 | PHD finger protein 20 |
| ENSG00000248735 | R3HDM2P1 | 0,169815165 | 1,124914354 | 0,67794 | 0,84156256 | R3H domain containing 2 pseudogene 1 |
| ENSG00000179331 | RAB39A | 0,169524009 | 1,124687353 | 0,47002 | 0,70939245 | RAB39A, member RAS oncogene family |
| ENSG00000217648 | AL136116,3 | 0,1695034 | 1,124671287 | 0,75713 | 0,88663582 | pseudogene similar to presenilin associated, rhomboid-like PARL |
| ENSG00000280641 | CDH4 | 0,169485404 | 1,124657259 | 0,72337 | 0,86789521 | cadherin 4 |
| ENSG00000287715 | AC026120,3 | 0,169396562 | 1,124588003 | 0,71231 | 0,86152011 | novel transcript |
| ENSG00000205250 | E2F4 | 0,169394593 | 1,124586468 | 0,42488 | 0,67857395 | E2F transcription factor 4 |
| ENSG00000075624 | ACTB | 0,169301544 | 1,124513938 | 0,49196 | 0,72531636 | actin beta |
| ENSG00000115271 | GCA | 0,169236432 | 1,124463188 | 0,72549 | 0,86906859 | grancalcin |
| ENSG00000134815 | DHX34 | 0,169233432 | 1,12446085 | 0,35901 | 0,63201238 | DExH-box helicase 34 |
| ENSG00000100442 | FKBP3 | 0,16921755 | 1,124448471 | 0,5694 | 0,77665073 | FKBP prolyl isomerase 3 |
| ENSG00000116652 | DLEU2L | 0,169204209 | 1,124438073 | 0,71753 | 0,86483656 | deleted in lymphocytic leukemia 2 like |
| ENSG00000122565 | CBX3 | 0,169170882 | 1,124412098 | 0,46727 | 0,70722674 | chromobox 3 |
| ENSG00000135541 | AHI1 | 0,169089767 | 1,12434888 | 0,64577 | 0,82244907 | Abelson helper integration site 1 |
| ENSG00000152954 | NRSN1 | 0,169075061 | 1,12433742 | 0,71417 | 0,86238618 | neurensin 1 |
| ENSG00000276291 | FRG1HP | 0,169059407 | 1,12432522 | 0,34764 | 0,62423931 | FSHD region gene 1 family member H, pseudogene |
| ENSG00000258131 | AC007656,2 | 0,16899679 | 1,124276423 | 0,88515 | 0,94922997 | novel transcript |
| ENSG00000104728 | ARHGEF10 | 0,168955145 | 1,124243969 | 0,43938 | 0,68828216 | Rho guanine nucleotide exchange factor 10 |
| ENSG00000162642 | C1orf52 | 0,168946949 | 1,124237582 | 0,43965 | 0,68844641 | chromosome 1 open reading frame 52 |
| ENSG00000180479 | ZNF571 | 0,168923851 | 1,124219583 | 0,63175 | 0,81431541 | zinc finger protein 571 |
| ENSG00000285465 | TM9SF1 | 0,168914252 | 1,124212103 | 0,37917 | 0,64699394 | transmembrane 9 superfamily member 1 |
| ENSG00000100926 | TM9SF1 | 0,168914252 | 1,124212103 | 0,37917 | 0,64699394 | transmembrane 9 superfamily member 1 |
| ENSG00000241163 | LINC00877 | 0,16889999 | 1,124200989 | 0,67813 | 0,84159488 | long intergenic non-protein coding RNA 877 |
| ENSG00000235888 | AF064858,1 | 0,168744802 | 1,124080068 | 0,63493 | 0,8162492 | novel transcript |
| ENSG00000188171 | ZNF626 | 0,168740309 | 1,124076567 | 0,38992 | 0,65400705 | zinc finger protein 626 |
| ENSG00000162522 | KIAA1522 | 0,168638371 | 1,123997145 | 0,40146 | 0,6630277 | KIAA1522 |
| ENSG00000130176 | CNN1 | 0,168613026 | 1,123977399 | 0,75361 | 0,88453179 | calponin 1 |
| ENSG00000138400 | MDH1B | 0,16859085 | 1,123960122 | 0,67644 | 0,8407877 | malate dehydrogenase 1B |
| ENSG00000286665 | AP000648,9 | 0,168576144 | 1,123948665 | 0,83616 | 0,92493625 | novel transcript |
| ENSG00000288058 | AP000648,10 | 0,168576144 | 1,123948665 | 0,83616 | 0,92493625 | novel transcript |

| | | | | | | |
|---|---|---|---|---|---|---|
| ENSG00000008311 | AASS | 0,168574733 | 1,123947566 | 0,41931 | 0,67517108 | aminoadipate-semialdehyde synthase |
| ENSG00000164078 | MST1R | 0,168561158 | 1,12393699 | 0,70405 | 0,85725494 | macrophage stimulating 1 receptor |
| ENSG00000141510 | TP53 | 0,168329554 | 1,123756573 | 0,4814 | 0,71822668 | tumor protein p53 |
| ENSG00000115944 | COX7A2L | 0,168181207 | 1,123641027 | 0,38404 | 0,65052153 | cytochrome c oxidase subunit 7A2 like |
| ENSG00000131408 | NR1H2 | 0,168124632 | 1,123596964 | 0,46423 | 0,7048936 | nuclear receptor subfamily 1 group H member 2 |
| ENSG00000140497 | SCAMP2 | 0,168017482 | 1,123513517 | 0,36761 | 0,63758887 | secretory carrier membrane protein 2 |
| ENSG00000285612 | AC004974,1 | 0,167975235 | 1,123480617 | 0,80556 | 0,91065997 | novel transcript, antisense to SIPA1L1 |
| ENSG00000186017 | ZNF566 | 0,167903127 | 1,123424466 | 0,37958 | 0,64735373 | zinc finger protein 566 |
| ENSG00000186952 | TMEM232 | 0,167889488 | 1,123413845 | 0,61929 | 0,80699858 | transmembrane protein 232 |
| ENSG00000158042 | MRPL17 | 0,167883211 | 1,123408957 | 0,36456 | 0,63504684 | mitochondrial ribosomal protein L17 |
| ENSG00000228363 | AC015971,1 | 0,167817839 | 1,123358054 | 0,62241 | 0,8084993 | novel transcript |
| ENSG00000153914 | SREK1 | 0,167776952 | 1,123326217 | 0,54739 | 0,76285014 | splicing regulatory glutamic acid and lysine rich protein 1 |
| ENSG00000241253 | CFB | 0,167668491 | 1,12324177 | 0,80943 | 0,91213228 | complement factor B |
| ENSG00000164879 | CA3 | 0,167642999 | 1,123221923 | 0,74007 | 0,87675792 | carbonic anhydrase 3 |
| ENSG00000105671 | DDX49 | 0,167636909 | 1,123217181 | 0,49661 | 0,72841371 | DEAD-box helicase 49 |
| ENSG00000184979 | USP18 | 0,167517238 | 1,123124014 | 0,64219 | 0,82066205 | ubiquitin specific peptidase 18 |
| ENSG00000154781 | CCDC174 | 0,167458208 | 1,123078061 | 0,45318 | 0,69750019 | coiled-coil domain containing 174 |
| ENSG00000155906 | RMND1 | 0,167342985 | 1,122988368 | 0,53789 | 0,75640122 | required for meiotic nuclear division 1 homolog |
| ENSG00000278351 | AC009248,2 | 0,167213877 | 1,122887876 | 0,87383 | 0,94405019 | novel transcript, sense intronic to ANO6 |
| ENSG00000259924 | AC011939,1 | 0,167185219 | 1,122865571 | 0,81505 | 0,91490386 | DnaJ (Hsp40) homolog, subfamily B, member 14 pseudogene |
| ENSG00000108395 | TRIM37 | 0,167180089 | 1,122861578 | 0,44288 | 0,69042854 | tripartite motif containing 37 |
| ENSG00000141698 | NT5C3B | 0,167130107 | 1,122822678 | 0,45941 | 0,7017037 | 5'-nucleotidase, cytosolic IIIB |
| ENSG00000175395 | ZNF25 | 0,167125995 | 1,122819477 | 0,73033 | 0,87185115 | zinc finger protein 25 |
| ENSG00000287329 | FP326651,1 | 0,167023593 | 1,122739783 | 0,67872 | 0,84182956 | novel transcript |
| ENSG00000087274 | ADD1 | 0,167011935 | 1,12273071 | 0,43173 | 0,68379763 | adducin 1 |
| ENSG00000183155 | RABIF | 0,166979687 | 1,122705614 | 0,65651 | 0,82841463 | RAB interacting factor |
| ENSG00000271789 | AL080317,1 | 0,166972654 | 1,122700142 | 0,68254 | 0,84443323 | novel transcript |
| ENSG00000186318 | BACE1 | 0,166899275 | 1,12264304 | 0,46359 | 0,70441254 | beta-secretase 1 |
| ENSG00000168092 | PAFAH1B2 | 0,16686083 | 1,122613123 | 0,31436 | 0,60115498 | platelet activating factor acetylhydrolase 1b catalytic subunit 2 |
| ENSG00000197724 | PHF2 | 0,166842949 | 1,12259921 | 0,5285 | 0,75090627 | PHD finger protein 2 |
| ENSG00000229311 | BX088651,2 | 0,166741975 | 1,122520642 | 0,74554 | 0,87996245 | novel transcript |

| | | | | | | |
|---|---|---|---|---|---|---|
| ENSG00000234917 | AC098484,2 | 0,166668305 | 1,122463323 | 0,75593 | 0,8857133 | novel transcript |
| ENSG00000187266 | EPOR | 0,166601771 | 1,122411559 | 0,67158 | 0,8374118 | erythropoietin receptor |
| ENSG00000278259 | MYO19 | 0,166582111 | 1,122396263 | 0,5159 | 0,7419128 | myosin XIX |
| ENSG00000213996 | TM6SF2 | 0,166550989 | 1,122372051 | 0,71807 | 0,86497602 | transmembrane 6 superfamily member 2 |
| ENSG00000285758 | AC036214,4 | 0,166491778 | 1,122325988 | 0,72527 | 0,86894646 | novel transcript, antisense to MRPS28 |
| ENSG00000022840 | RNF10 | 0,166487566 | 1,122322711 | 0,31051 | 0,5984598 | ring finger protein 10 |
| ENSG00000163913 | IFT122 | 0,166484273 | 1,122320149 | 0,5394 | 0,75771283 | intraflagellar transport 122 |
| ENSG00000278458 | KANSL1 | 0,166458434 | 1,122300048 | 0,73861 | 0,87591843 | KAT8 regulatory NSL complex subunit 1 |
| ENSG00000180817 | PPA1 | 0,166445196 | 1,12228975 | 0,54971 | 0,76425279 | inorganic pyrophosphatase 1 |
| ENSG00000106927 | AMBP | 0,166417734 | 1,122268388 | 0,68437 | 0,84554424 | alpha-1-microglobulin/bikunin precursor |
| ENSG00000080644 | CHRNA3 | 0,166399386 | 1,122254115 | 0,67986 | 0,8427192 | cholinergic receptor nicotinic alpha 3 subunit |
| ENSG00000272008 | AL139274,2 | 0,166321047 | 1,122193178 | 0,47342 | 0,71264815 | novel transcript, antisense to ZNF292 |
| ENSG00000260274 | AC068338,2 | 0,166320009 | 1,12219237 | 0,67201 | 0,83765677 | novel transcript |
| ENSG00000167081 | PBX3 | 0,16626753 | 1,122151551 | 0,61322 | 0,8035281 | PBX homeobox 3 |
| ENSG00000111276 | CDKN1B | 0,166249279 | 1,122137355 | 0,32254 | 0,60577115 | cyclin dependent kinase inhibitor 1B |
| ENSG00000224072 | VEZTP1 | 0,166246831 | 1,12213545 | 0,74407 | 0,87882457 | VEZT pseudogene 1 |
| ENSG00000237651 | C2orf74 | 0,166238196 | 1,122128734 | 0,66527 | 0,83339621 | chromosome 2 open reading frame 74 |
| ENSG00000170832 | USP32 | 0,166237887 | 1,122128494 | 0,67149 | 0,83734318 | ubiquitin specific peptidase 32 |
| ENSG00000124688 | MAD2L1BP | 0,166199615 | 1,122098727 | 0,47271 | 0,71203455 | MAD2L1 binding protein |
| ENSG00000042317 | SPATA7 | 0,166084199 | 1,122008962 | 0,68918 | 0,8485712 | spermatogenesis associated 7 |
| ENSG00000157152 | SYN2 | 0,166046964 | 1,121980004 | 0,68441 | 0,84555824 | synapsin II |
| ENSG00000172348 | RCAN2 | 0,166044924 | 1,121978417 | 0,69619 | 0,85287022 | regulator of calcineurin 2 |
| ENSG00000087111 | PIGS | 0,165946777 | 1,121902092 | 0,35349 | 0,62819991 | phosphatidylinositol glycan anchor biosynthesis class S |
| ENSG00000118137 | APOA1 | 0,165931675 | 1,121890348 | 0,81124 | 0,91307458 | apolipoprotein A1 |
| ENSG00000121104 | FAM117A | 0,16592879 | 1,121888104 | 0,5981 | 0,79484486 | family with sequence similarity 117 member A |
| ENSG00000171861 | MRM3 | 0,165882907 | 1,121852424 | 0,42078 | 0,67606003 | mitochondrial rRNA methyltransferase 3 |
| ENSG00000161835 | TAMALIN | 0,165811417 | 1,121796834 | 0,70741 | 0,85910429 | trafficking regulator and scaffold protein tamalin |
| ENSG00000170903 | MSANTD4 | 0,165770704 | 1,121765178 | 0,59966 | 0,79552021 | Myb/SANT DNA binding domain containing 4 with coiled-coils |
| ENSG00000258044 | AC073569,1 | 0,165760153 | 1,121756973 | 0,81885 | 0,91689247 | novel transcript, antisense to PPP1R12A |
| ENSG00000255084 | AP003110,1 | 0,165706673 | 1,121715391 | 0,86572 | 0,94036783 | novel transcript |
| ENSG00000041802 | LSG1 | 0,165677242 | 1,121692509 | 0,42888 | 0,68161494 | large 60S subunit nuclear export GTPase 1 |
| ENSG00000212743 | LINC02656 | 0,165669226 | 1,121686276 | 0,82904 | 0,92120896 | long intergenic non-protein coding RNA 2656 |

| | | | | | | |
|---|---|---|---|---|---|---|
| ENSG00000260855 | AL591848,3 | 0,165665546 | 1,121683415 | 0,57247 | 0,77849297 | novel transcript |
| ENSG00000277072 | STAG3L2 | 0,165626786 | 1,12165328 | 0,44956 | 0,69519764 | stromal antigen 3-like 2 (pseudogene) |
| ENSG00000162430 | SELENON | 0,1655564 | 1,121598558 | 0,3922 | 0,65555739 | selenoprotein N |
| ENSG00000242094 | FOXP1-IT1 | 0,165523067 | 1,121572644 | 0,67387 | 0,8393266 | FOXP1 intronic transcript 1 |
| ENSG00000148229 | POLE3 | 0,16551133 | 1,12156352 | 0,40831 | 0,66714392 | DNA polymerase epsilon 3, accessory subunit |
| ENSG00000115504 | EHBP1 | 0,16546544 | 1,121527846 | 0,6192 | 0,80699858 | EH domain binding protein 1 |
| ENSG00000283101 | BMS1P21 | 0,165373388 | 1,121456288 | 0,82219 | 0,91790528 | BMS1 pseudogene 21 |
| ENSG00000184507 | NUTM1 | 0,165339055 | 1,1214296 | 0,76195 | 0,88909725 | NUT midline carcinoma family member 1 |
| ENSG00000285972 | CERNA2 | 0,165204411 | 1,121324944 | 0,64469 | 0,82162908 | competing endogenous lncRNA 2 for microRNA let-7b |
| ENSG00000165309 | ARMC3 | 0,165170688 | 1,121298733 | 0,58627 | 0,78667706 | armadillo repeat containing 3 |
| ENSG00000120699 | EXOSC8 | 0,16509962 | 1,121243499 | 0,31401 | 0,60070179 | exosome component 8 |
| ENSG00000253676 | TAGLN2P1 | 0,165056462 | 1,121209958 | 0,84269 | 0,92821072 | transgelin 2 pseudogene 1 |
| ENSG00000287408 | AC027811,1 | 0,164941396 | 1,121120536 | 0,73048 | 0,87199414 | novel transcript, antisense to TMED3 |
| ENSG00000172493 | AFF1 | 0,164919609 | 1,121103605 | 0,61031 | 0,80221019 | AF4/FMR2 family member 1 |
| ENSG00000088682 | COQ9 | 0,164749677 | 1,120971561 | 0,67005 | 0,8364011 | coenzyme Q9 |
| ENSG00000120324 | PCDHB10 | 0,164747253 | 1,120969677 | 0,6016 | 0,79671505 | protocadherin beta 10 |
| ENSG00000285449 | H2BU1 | 0,164661558 | 1,120903094 | 0,92315 | 0,96667029 | H2B,U histone 1 |
| ENSG00000119616 | FCF1 | 0,164628902 | 1,120877722 | 0,34662 | 0,62350287 | FCF1 rRNA-processing protein |
| ENSG00000280169 | AL136526,1 | 0,164613056 | 1,120865411 | 0,76319 | 0,88980026 | novel transcript, antisense to GGACT |
| ENSG00000121390 | PSPC1 | 0,164560339 | 1,120824455 | 0,43925 | 0,68816791 | paraspeckle component 1 |
| ENSG00000090447 | TFAP4 | 0,164560287 | 1,120824415 | 0,5657 | 0,77393796 | transcription factor AP-4 |
| ENSG00000260257 | AL035071,1 | 0,164496022 | 1,120774489 | 0,41859 | 0,67477592 | novel transcript |
| ENSG00000137203 | TFAP2A | 0,164427968 | 1,120721621 | 0,61852 | 0,80665241 | transcription factor AP-2 alpha |
| ENSG00000198890 | PRMT6 | 0,164341839 | 1,120654716 | 0,36915 | 0,63903096 | protein arginine methyltransferase 6 |
| ENSG00000184083 | FAM120C | 0,164261256 | 1,120592122 | 0,48287 | 0,71919726 | family with sequence similarity 120C |
| ENSG00000136811 | ODF2 | 0,164257397 | 1,120589125 | 0,58631 | 0,78668524 | outer dense fiber of sperm tails 2 |
| ENSG00000131652 | THOC6 | 0,164229232 | 1,120567249 | 0,3845 | 0,65052153 | THO complex 6 |
| ENSG00000250326 | AC104596,1 | 0,164127771 | 1,120488445 | 0,59509 | 0,79320995 | novel transcript |
| ENSG00000149600 | COMMD7 | 0,164109392 | 1,12047417 | 0,59514 | 0,79320995 | COMM domain containing 7 |
| ENSG00000168246 | UBTD2 | 0,163988301 | 1,120380129 | 0,60995 | 0,80202886 | ubiquitin domain containing 2 |
| ENSG00000185122 | HSF1 | 0,163986904 | 1,120379044 | 0,34217 | 0,61998187 | heat shock transcription factor 1 |
| ENSG00000284774 | HSF1 | 0,163986904 | 1,120379044 | 0,34217 | 0,61998187 | heat shock transcription factor 1 |
| ENSG00000260231 | KDM7A-DT | 0,163904733 | 1,120315232 | 0,57071 | 0,77748547 | KDM7A divergent transcript |

| | | | | | | |
|---|---|---|---|---|---|---|
| ENSG00000141564 | RPTOR | 0,16376766 | 1,120208795 | 0,32779 | 0,60960585 | regulatory associated protein of MTOR complex 1 |
| ENSG00000110172 | CHORDC1 | 0,163669456 | 1,120132545 | 0,6386 | 0,8186299 | cysteine and histidine rich domain containing 1 |
| ENSG00000288467 | AL513166,2 | 0,163532465 | 1,120026188 | 0,80539 | 0,91057421 | novel transcript |
| ENSG00000287001 | AC010624,5 | 0,163518739 | 1,120015532 | 0,62313 | 0,80892518 | novel transcript |
| ENSG00000128408 | RIBC2 | 0,163500576 | 1,120001432 | 0,6018 | 0,79680798 | RIB43A domain with coiled-coils 2 |
| ENSG00000174469 | CNTNAP2 | 0,163485624 | 1,119989824 | 0,80258 | 0,90906425 | contactin associated protein 2 |
| ENSG00000177700 | POLR2L | 0,163474843 | 1,119981454 | 0,46555 | 0,70595153 | RNA polymerase II, I and III subunit L |
| ENSG00000179157 | RPS2P28 | 0,163287336 | 1,1198359 | 0,82117 | 0,91738668 | ribosomal protein S2 pseudogene 28 |
| ENSG00000280145 | CU638689,4 | 0,163194272 | 1,119763665 | 0,62281 | 0,80873034 | novel transcript |
| ENSG00000135052 | GOLM1 | 0,163091276 | 1,119683727 | 0,51378 | 0,74004886 | golgi membrane protein 1 |
| ENSG00000038358 | EDC4 | 0,163006346 | 1,119617813 | 0,50409 | 0,73311005 | enhancer of mRNA decapping 4 |
| ENSG00000263753 | LINC00667 | 0,163002044 | 1,119614475 | 0,60297 | 0,7971742 | long intergenic non-protein coding RNA 667 |
| ENSG00000269886 | AC022382,1 | 0,162974989 | 1,119593479 | 0,85071 | 0,93206349 | novel transcript, antisense to TTLL3 |
| ENSG00000267430 | AC036176,2 | 0,162925832 | 1,119555332 | 0,7874 | 0,90121071 | proteasome (prosome, macropain) 26S subunit, ATPase, 5 (PSMC5) pseudogene |
| ENSG00000166272 | WBP1L | 0,16287095 | 1,119512743 | 0,48771 | 0,72268512 | WW domain binding protein 1 like |
| ENSG00000225950 | NTF4 | 0,162835987 | 1,119485612 | 0,81961 | 0,91708488 | neurotrophin 4 |
| ENSG00000165194 | PCDH19 | 0,162828167 | 1,119479544 | 0,4833 | 0,71964105 | protocadherin 19 |
| ENSG00000100714 | MTHFD1 | 0,162778288 | 1,119440841 | 0,40554 | 0,66526837 | methylenetetrahydrofolate dehydrogenase, cyclohydrolase and formyltetrahydrofolate synthetase 1 |
| ENSG00000164344 | KLKB1 | 0,162676992 | 1,119362244 | 0,81375 | 0,9144566 | kallikrein B1 |
| ENSG00000100360 | IFT27 | 0,162672672 | 1,119358892 | 0,49219 | 0,72544199 | intraflagellar transport 27 |
| ENSG00000180340 | FZD2 | 0,162646792 | 1,119338813 | 0,49983 | 0,73058737 | frizzled class receptor 2 |
| ENSG00000109790 | KLHL5 | 0,162630761 | 1,119326375 | 0,36012 | 0,63261458 | kelch like family member 5 |
| ENSG00000273249 | BX649632,1 | 0,162606217 | 1,119307333 | 0,83578 | 0,92493625 | novel transcript, antisense to WDR5 |
| ENSG00000034239 | EFCAB1 | 0,162596251 | 1,1192996 | 0,6649 | 0,83314543 | EF-hand calcium binding domain 1 |
| ENSG00000161594 | KLHL10 | 0,162525532 | 1,119244735 | 0,84092 | 0,92751395 | kelch like family member 10 |
| ENSG00000100335 | MIEF1 | 0,16243954 | 1,119178024 | 0,3541 | 0,62877376 | mitochondrial elongation factor 1 |
| ENSG00000170899 | GSTA4 | 0,162342814 | 1,119102991 | 0,54431 | 0,76051055 | glutathione S-transferase alpha 4 |
| ENSG00000110906 | KCTD10 | 0,162297975 | 1,11906821 | 0,4603 | 0,70217029 | potassium channel tetramerization domain containing 10 |
| ENSG00000273084 | AC092171,5 | 0,162290739 | 1,119062598 | 0,6833 | 0,84495012 | novel transcript |
| ENSG00000066777 | ARFGEF1 | 0,162088955 | 1,118906089 | 0,49696 | 0,72862056 | ADP ribosylation factor guanine nucleotide exchange factor 1 |

| | | | | | | |
|---|---|---|---|---|---|---|
| ENSG00000171530 | TBCA | 0,162081147 | 1,118900034 | 0,55315 | 0,76626672 | tubulin folding cofactor A |
| ENSG00000166477 | LEO1 | 0,162068234 | 1,118890019 | 0,57373 | 0,77921182 | LEO1 homolog, Paf1/RNA polymerase II complex component |
| ENSG00000108825 | PTGES3L-AAR | 0,162045967 | 1,11887275 | 0,77012 | 0,89333117 | PTGES3L-AARSD1 readthrough |
| ENSG00000158850 | B4GALT3 | 0,162032581 | 1,118862369 | 0,37742 | 0,64593913 | beta-1,4-galactosyltransferase 3 |
| ENSG00000125534 | PPDPF | 0,162025191 | 1,118856638 | 0,55862 | 0,77025265 | pancreatic progenitor cell differentiation and proliferation factor |
| ENSG00000230565 | ZNF32-AS2 | 0,162020761 | 1,118853201 | 0,71205 | 0,8615121 | ZNF32 antisense RNA 2 |
| ENSG00000179954 | SSC5D | 0,162019729 | 1,118852401 | 0,66403 | 0,83250729 | scavenger receptor cysteine rich family member with 5 domains |
| ENSG00000256250 | AC073912,1 | 0,162009581 | 1,118844532 | 0,75071 | 0,88299301 | novel transcript |
| ENSG00000107771 | CCSER2 | 0,161998926 | 1,118836268 | 0,48161 | 0,71838485 | coiled-coil serine rich protein 2 |
| ENSG00000153066 | TXNDC11 | 0,161972324 | 1,118815638 | 0,39286 | 0,65599749 | thioredoxin domain containing 11 |
| ENSG00000184508 | HDDC3 | 0,1619564 | 1,118803289 | 0,45959 | 0,7017275 | HD domain containing 3 |
| ENSG00000286619 | AC119673,2 | 0,161953296 | 1,118800882 | 0,51769 | 0,74272749 | novel transcript, antisense to PM20D1 |
| ENSG00000273847 | AC139494,2 | 0,161792142 | 1,118675915 | 0,70677 | 0,85889796 | novel transcript |
| ENSG00000107263 | RAPGEF1 | 0,161684554 | 1,118592494 | 0,44023 | 0,68866128 | Rap guanine nucleotide exchange factor 1 |
| ENSG00000114841 | DNAH1 | 0,16162093 | 1,118543164 | 0,57746 | 0,78141582 | dynein axonemal heavy chain 1 |
| ENSG00000141337 | ARSG | 0,16154422 | 1,118483691 | 0,5289 | 0,75110319 | arylsulfatase G |
| ENSG00000285856 | AL353704,1 | 0,161469992 | 1,118426145 | 0,77624 | 0,89654301 | novel transcript, antisense to ARHGEF7 |
| ENSG00000143740 | SNAP47 | 0,161464361 | 1,11842178 | 0,45521 | 0,69925766 | synaptosome associated protein 47 |
| ENSG00000155926 | SLA | 0,1613407 | 1,118325918 | 0,88068 | 0,94687991 | Src like adaptor |
| ENSG00000286753 | AC091978,1 | 0,161117331 | 1,118152784 | 0,68168 | 0,84383965 | novel transcript, antisense to ICE1 |
| ENSG00000101115 | SALL4 | 0,16102075 | 1,118077932 | 0,64627 | 0,82289551 | spalt like transcription factor 4 |
| ENSG00000123342 | MMP19 | 0,160917065 | 1,11799758 | 0,67438 | 0,83968055 | matrix metallopeptidase 19 |
| ENSG00000106976 | DNM1 | 0,160851003 | 1,117946387 | 0,61375 | 0,8035281 | dynamin 1 |
| ENSG00000144659 | SLC25A38 | 0,160837598 | 1,117936 | 0,41413 | 0,67099447 | solute carrier family 25 member 38 |
| ENSG00000113272 | THG1L | 0,160798039 | 1,117905346 | 0,42094 | 0,67616334 | tRNA-histidine guanylyltransferase 1 like |
| ENSG00000152642 | GPD1L | 0,160765028 | 1,117879767 | 0,74364 | 0,87867571 | glycerol-3-phosphate dehydrogenase 1 like |
| ENSG00000261740 | BOLA2-SMG1 | 0,160730384 | 1,117852923 | 0,32206 | 0,60577115 | BOLA2-SMG1P6 readthrough |
| ENSG00000170043 | TRAPPC1 | 0,160701215 | 1,117830322 | 0,49805 | 0,72937175 | trafficking protein particle complex 1 |
| ENSG00000270558 | AC025449,1 | 0,160493274 | 1,117669216 | 0,69719 | 0,85330122 | chromobox homolog 3 (CBX3) pseudogene |
| ENSG00000011677 | GABRA3 | 0,160462957 | 1,11764573 | 0,66937 | 0,83617862 | gamma-aminobutyric acid type A receptor subunit alpha3 |

| | | | | | | |
|---|---|---|---|---|---|---|
| ENSG00000277209 | RPPH1 | 0,160455019 | 1,117639581 | 0,75197 | 0,88386445 | ribonuclease P RNA component H1 |
| ENSG00000128626 | MRPS12 | 0,160383299 | 1,117584021 | 0,53168 | 0,75280487 | mitochondrial ribosomal protein S12 |
| ENSG00000162909 | CAPN2 | 0,160378712 | 1,117580468 | 0,45964 | 0,701759 | calpain 2 |
| ENSG00000163682 | RPL9 | 0,160329549 | 1,117542385 | 0,40626 | 0,66584658 | ribosomal protein L9 |
| ENSG00000078674 | PCM1 | 0,16023691 | 1,117470627 | 0,52406 | 0,74793732 | pericentriolar material 1 |
| ENSG00000269713 | NBPF9 | 0,160164545 | 1,117414576 | 0,25067 | 0,55690541 | NBPF member 9 |
| ENSG00000286532 | PARTICL | 0,160138516 | 1,117394416 | 0,53989 | 0,75807575 | promoter of MAT2A antisense radiation-induced circulating long non-coding RNA |
| ENSG00000104964 | TLE5 | 0,160033781 | 1,1173133 | 0,70385 | 0,85724036 | TLE family member 5, transcriptional modulator |
| ENSG00000090975 | PITPNM2 | 0,160030112 | 1,117310458 | 0,52857 | 0,75090627 | phosphatidylinositol transfer protein membrane associated 2 |
| ENSG00000095787 | WAC | 0,159966089 | 1,117260876 | 0,27261 | 0,57315967 | WW domain containing adaptor with coiled-coil |
| ENSG00000136783 | NIPSNAP3A | 0,159958767 | 1,117255206 | 0,7328 | 0,87320251 | nipsnap homolog 3A |
| ENSG00000124155 | PIGT | 0,159941847 | 1,117242103 | 0,41863 | 0,67477592 | phosphatidylinositol glycan anchor biosynthesis class T |
| ENSG00000188010 | MORN2 | 0,159931782 | 1,117234309 | 0,61843 | 0,80663169 | MORN repeat containing 2 |
| ENSG00000049541 | RFC2 | 0,159885912 | 1,117198786 | 0,36013 | 0,63261458 | replication factor C subunit 2 |
| ENSG00000181631 | P2RY13 | 0,159844738 | 1,117166903 | 0,71466 | 0,86265094 | purinergic receptor P2Y13 |
| ENSG00000178125 | PPP1R42 | 0,159773276 | 1,117111567 | 0,79101 | 0,90292452 | protein phosphatase 1 regulatory subunit 42 |
| ENSG00000156990 | RPUSD3 | 0,159734615 | 1,117081631 | 0,35176 | 0,62692823 | RNA pseudouridine synthase D3 |
| ENSG00000181097 | LINC02878 | 0,159665676 | 1,117028252 | 0,86816 | 0,94176112 | long intergenic non-protein coding RNA 2878 |
| ENSG00000107537 | PHYH | 0,159578544 | 1,116960791 | 0,6279 | 0,81175174 | phytanoyl-CoA 2-hydroxylase |
| ENSG00000283757 | AL031686,1 | 0,159540298 | 1,116931181 | 0,74956 | 0,8822079 | novel transcript |
| ENSG00000153786 | ZDHHC7 | 0,159481954 | 1,116886012 | 0,36275 | 0,63382442 | zinc finger DHHC-type palmitoyltransferase 7 |
| ENSG00000096070 | BRPF3 | 0,159349297 | 1,116783318 | 0,30779 | 0,59671558 | bromodomain and PHD finger containing 3 |
| ENSG00000179603 | GRM8 | 0,159312926 | 1,116755164 | 0,56113 | 0,77125979 | glutamate metabotropic receptor 8 |
| ENSG00000275464 | FP565260,1 | 0,159239615 | 1,116698418 | 0,69377 | 0,8512419 | periodic tryptophan protein 2 homolog |
| ENSG00000174844 | DNAH12 | 0,15923547 | 1,116695209 | 0,73173 | 0,87274707 | dynein axonemal heavy chain 12 |
| ENSG00000275476 | AC009318,4 | 0,159215197 | 1,116679517 | 0,82816 | 0,9207216 | novel transcript, antisense to FAR2 |
| ENSG00000235376 | RPEL1 | 0,159203837 | 1,116670724 | 0,83939 | 0,92655196 | ribulose-5-phosphate-3-epimerase like 1 |
| ENSG00000117408 | IPO13 | 0,159128955 | 1,116612766 | 0,51955 | 0,74404402 | importin 13 |
| ENSG00000102218 | RP2 | 0,159126375 | 1,116610769 | 0,68447 | 0,84559757 | RP2 activator of ARL3 GTPase |
| ENSG00000180773 | SLC36A4 | 0,159096222 | 1,116587432 | 0,60662 | 0,79981537 | solute carrier family 36 member 4 |
| ENSG00000154328 | NEIL2 | 0,159090228 | 1,116582792 | 0,38556 | 0,65117983 | nei like DNA glycosylase 2 |

| | | | | | | |
|---|---|---|---|---|---|---|
| ENSG00000151229 | SLC2A13 | 0,159039765 | 1,116543737 | 0,44807 | 0,69410917 | solute carrier family 2 member 13 |
| ENSG00000105856 | HBP1 | 0,15903171 | 1,116537503 | 0,6024 | 0,7970763 | HMG-box transcription factor 1 |
| ENSG00000242671 | LINC02010 | 0,158885181 | 1,116424106 | 0,78995 | 0,90273003 | long intergenic non-protein coding RNA 2010 |
| ENSG00000255921 | AC026310,1 | 0,158696645 | 1,116278218 | 0,83904 | 0,92640383 | novel transcript, antisense to BCAT1 |
| ENSG00000255517 | AP002748,4 | 0,158647744 | 1,116240381 | 0,58936 | 0,78883783 | novel transcript |
| ENSG00000104904 | OAZ1 | 0,158592981 | 1,116198011 | 0,318 | 0,6034788 | ornithine decarboxylase antizyme 1 |
| ENSG00000032742 | IFT88 | 0,158555184 | 1,116168768 | 0,5233 | 0,7473071 | intraflagellar transport 88 |
| ENSG00000123124 | WWP1 | 0,158391288 | 1,116041974 | 0,54712 | 0,76264866 | WW domain containing E3 ubiquitin protein ligase 1 |
| ENSG00000109685 | NSD2 | 0,158383004 | 1,116035566 | 0,49499 | 0,72732671 | nuclear receptor binding SET domain protein 2 |
| ENSG00000120217 | CD274 | 0,158350203 | 1,116010193 | 0,8453 | 0,92946127 | CD274 molecule |
| ENSG00000198814 | GK | 0,158345466 | 1,116006528 | 0,6493 | 0,82439462 | glycerol kinase |
| ENSG00000167130 | DOLPP1 | 0,15834068 | 1,116002826 | 0,51182 | 0,7388352 | dolichyldiphosphatase 1 |
| ENSG00000216775 | AL109918,1 | 0,158335637 | 1,115998925 | 0,64789 | 0,82380454 | heterogeneous nuclear ribonucleoprotein A/B (HNRNPAB) pseudogene |
| ENSG00000186132 | C2orf76 | 0,158196605 | 1,115891382 | 0,67992 | 0,8427192 | chromosome 2 open reading frame 76 |
| ENSG00000198258 | UBL5 | 0,15810167 | 1,115817954 | 0,57383 | 0,77921182 | ubiquitin like 5 |
| ENSG00000188013 | MEIS3P2 | 0,158098598 | 1,115815578 | 0,68364 | 0,84505352 | Meis homeobox 3 pseudogene 2 |
| ENSG00000135451 | TROAP | 0,158048407 | 1,115776759 | 0,54057 | 0,75839186 | trophinin associated protein |
| ENSG00000150756 | ATPSCKMT | 0,157998539 | 1,115738193 | 0,53534 | 0,75460305 | ATP synthase c subunit lysine N-methyltransferase |
| ENSG00000284238 | TOP3A | 0,157979857 | 1,115723744 | 0,65814 | 0,82898078 | DNA topoisomerase III alpha |
| ENSG00000197226 | TBC1D9B | 0,157933588 | 1,115687963 | 0,35583 | 0,63013888 | TBC1 domain family member 9B |
| ENSG00000149798 | CDC42EP2 | 0,157843813 | 1,115618538 | 0,76619 | 0,89136081 | CDC42 effector protein 2 |
| ENSG00000160336 | ZNF761 | 0,157836695 | 1,115613034 | 0,46969 | 0,70916811 | zinc finger protein 761 |
| ENSG00000250041 | AC069360,1 | 0,157752806 | 1,115548166 | 0,75669 | 0,88633961 | novel transcript |
| ENSG00000173801 | JUP | 0,157723038 | 1,115525148 | 0,54406 | 0,76034748 | junction plakoglobin |
| ENSG00000135185 | TMEM243 | 0,157706896 | 1,115512667 | 0,5724 | 0,77849297 | transmembrane protein 243 |
| ENSG00000235437 | LINC01278 | 0,157703797 | 1,115510271 | 0,46245 | 0,70356854 | long intergenic non-protein coding RNA 1278 |
| ENSG00000206511 | GABBR1 | 0,157646345 | 1,115465849 | 0,63892 | 0,81889694 | gamma-aminobutyric acid type B receptor subunit 1 |
| ENSG00000288107 | AL731769,2 | 0,157643301 | 1,115463495 | 0,53571 | 0,75485762 | novel transcript, antisense to FRG2Band SYCE1 |
| ENSG00000198839 | ZNF277 | 0,15764046 | 1,115461299 | 0,55976 | 0,77066501 | zinc finger protein 277 |
| ENSG00000103248 | MTHFSD | 0,157553369 | 1,115393964 | 0,52525 | 0,74895722 | methenyltetrahydrofolate synthetase domain containing |
| ENSG00000112367 | FIG4 | 0,157480018 | 1,115337255 | 0,57288 | 0,77868876 | FIG4 phosphoinositide 5-phosphatase |
| ENSG00000286621 | AC064843,1 | 0,157305137 | 1,115202065 | 0,7098 | 0,86015146 | novel transcript, antisense to DAPL1 |

| | | | | | | |
|---|---|---|---|---|---|---|
| ENSG00000125875 | TBC1D20 | 0,157273564 | 1,115177658 | 0,3904 | 0,65441492 | TBC1 domain family member 20 |
| ENSG00000226306 | NPY6R | 0,157260797 | 1,11516779 | 0,81101 | 0,91288837 | neuropeptide Y receptor Y6 (pseudogene) |
| ENSG00000184436 | THAP7 | 0,157216171 | 1,115133296 | 0,42125 | 0,67634721 | THAP domain containing 7 |
| ENSG00000111271 | ACAD10 | 0,157192105 | 1,115114695 | 0,50804 | 0,73612961 | acyl-CoA dehydrogenase family member 10 |
| ENSG00000179038 | AP001885,1 | 0,157063281 | 1,115015126 | 0,77144 | 0,8940638 | tetratricopeptide repeat domain 3 (TTC3) pseudogene |
| ENSG00000238058 | AL355574,1 | 0,157058251 | 1,115011238 | 0,75351 | 0,88453179 | novel transcript |
| ENSG00000135686 | KLHL36 | 0,156850194 | 1,11485045 | 0,45903 | 0,70162976 | kelch like family member 36 |
| ENSG00000088356 | PDRG1 | 0,156823373 | 1,114829723 | 0,5447 | 0,76079293 | p53 and DNA damage regulated 1 |
| ENSG00000214783 | POLR2J4 | 0,156772169 | 1,114790157 | 0,58613 | 0,78664449 | RNA polymerase II subunit J4, pseudogene |
| ENSG00000104973 | MED25 | 0,156758806 | 1,114779831 | 0,48702 | 0,72215091 | mediator complex subunit 25 |
| ENSG00000271011 | AC010680,3 | 0,15674852 | 1,114771883 | 0,78928 | 0,9024333 | novel transcript, antisense to TTN |
| ENSG00000143252 | SDHC | 0,156745274 | 1,114769375 | 0,32352 | 0,60631901 | succinate dehydrogenase complex subunit C |
| ENSG00000142784 | WDTC1 | 0,156701724 | 1,114735724 | 0,33212 | 0,61206889 | WD and tetratricopeptide repeats 1 |
| ENSG00000198561 | CTNND1 | 0,156692285 | 1,114728431 | 0,24447 | 0,55253387 | catenin delta 1 |
| ENSG00000267370 | AC008752,3 | 0,156681964 | 1,114720456 | 0,78946 | 0,90251229 | zinc finger protein 420 (ZNF420) pseudogene |
| ENSG00000198087 | CD2AP | 0,156632044 | 1,114681886 | 0,49345 | 0,72634871 | CD2 associated protein |
| ENSG00000173193 | PARP14 | 0,156616458 | 1,114669843 | 0,58409 | 0,78572744 | poly(ADP-ribose) polymerase family member 14 |
| ENSG00000164626 | KCNK5 | 0,156614511 | 1,114668339 | 0,71963 | 0,86592885 | potassium two pore domain channel subfamily K member 5 |
| ENSG00000133961 | NUMB | 0,156477055 | 1,114562141 | 0,3049 | 0,59446033 | NUMB endocytic adaptor protein |
| ENSG00000103485 | QPRT | 0,156404038 | 1,114505733 | 0,64664 | 0,82303578 | quinolinate phosphoribosyltransferase |
| ENSG00000165028 | NIPSNAP3B | 0,156353683 | 1,114466834 | 0,6725 | 0,83803462 | nipsnap homolog 3B |
| ENSG00000170464 | DNAJC18 | 0,1563535 | 1,114466693 | 0,829 | 0,92119722 | DnaJ heat shock protein family (Hsp40) member C18 |
| ENSG00000254717 | GLYATL1P2 | 0,156331672 | 1,11444983 | 0,75805 | 0,88719949 | glycine-N-acyltransferase like 1 pseudogene 2 |
| ENSG00000068120 | COASY | 0,156289648 | 1,114417369 | 0,41373 | 0,67085666 | Coenzyme A synthase |
| ENSG00000181982 | CCDC149 | 0,156258524 | 1,114393327 | 0,54667 | 0,76233291 | coiled-coil domain containing 149 |
| ENSG00000205133 | TRIQK | 0,156200904 | 1,114348819 | 0,68041 | 0,84309883 | triple QxxK/R motif containing |
| ENSG00000215424 | MCM3AP-AS1 | 0,156183848 | 1,114335646 | 0,64905 | 0,82434007 | MCM3AP antisense RNA 1 |
| ENSG00000147416 | ATP6V1B2 | 0,15607511 | 1,11425166 | 0,49771 | 0,72908464 | ATPase H+ transporting V1 subunit B2 |
| ENSG00000175354 | PTPN2 | 0,156066715 | 1,114245176 | 0,32392 | 0,60659203 | protein tyrosine phosphatase non-receptor type 2 |
| ENSG00000130055 | GDPD2 | 0,156047789 | 1,114230559 | 0,44041 | 0,6888598 | glycerophosphodiester phosphodiesterase domain containing 2 |
| ENSG00000100216 | TOMM22 | 0,155989829 | 1,114185796 | 0,32479 | 0,60732633 | translocase of outer mitochondrial membrane 22 |

| | | | | | | |
|---|---|---|---|---|---|---|
| ENSG00000262435 | WASH8P | 0,155945961 | 1,114151917 | 0,48702 | 0,72215091 | WAS protein family homolog 8, pseudogene |
| ENSG00000226210 | WASH8P | 0,155945961 | 1,114151917 | 0,48702 | 0,72215091 | WAS protein family homolog 8, pseudogene |
| ENSG00000136950 | ARPC5L | 0,155909857 | 1,114124036 | 0,36655 | 0,63641349 | actin related protein 2/3 complex subunit 5 like |
| ENSG00000206206 | DAXX | 0,155871012 | 1,114094038 | 0,35152 | 0,62674766 | death domain associated protein |
| ENSG00000227046 | DAXX | 0,155871012 | 1,114094038 | 0,35152 | 0,62674766 | death domain associated protein |
| ENSG00000198848 | CES1 | 0,155813802 | 1,114049859 | 0,88681 | 0,95005038 | carboxylesterase 1 |
| ENSG00000170027 | YWHAG | 0,155797938 | 1,114037609 | 0,31507 | 0,60145932 | tyrosine 3-monooxygenase/tryptophan 5-monooxygenase activation protein gamma |
| ENSG00000099284 | MACROH2A2 | 0,155713539 | 1,113972439 | 0,34846 | 0,62472936 | macroH2A,2 histone |
| ENSG00000166851 | PLK1 | 0,155664405 | 1,113934501 | 0,4135 | 0,67078587 | polo like kinase 1 |
| ENSG00000111752 | PHC1 | 0,155375697 | 1,113711605 | 0,58847 | 0,78818673 | polyhomeotic homolog 1 |
| ENSG00000124212 | PTGIS | 0,155361364 | 1,113700542 | 0,65119 | 0,8253103 | prostaglandin I2 synthase |
| ENSG00000084207 | GSTP1 | 0,155355974 | 1,113696381 | 0,59811 | 0,79484486 | glutathione S-transferase pi 1 |
| ENSG00000125753 | VASP | 0,155329164 | 1,113675685 | 0,45911 | 0,70168704 | vasodilator stimulated phosphoprotein |
| ENSG00000163734 | CXCL3 | 0,155311005 | 1,113661667 | 0,79938 | 0,90744841 | C-X-C motif chemokine ligand 3 |
| ENSG00000226266 | AC009961,1 | 0,155266054 | 1,113626969 | 0,72248 | 0,86734497 | novel transcript |
| ENSG00000214135 | AC132008,2 | 0,155226613 | 1,113596524 | 0,58115 | 0,78406473 | succinate dehydrogenase complex, subunit A, flavoprotein (Fp) (SDHA) pseudogene |
| ENSG00000104267 | CA2 | 0,155222279 | 1,113593179 | 0,70794 | 0,85926733 | carbonic anhydrase 2 |
| ENSG00000286608 | AC010201,3 | 0,155147265 | 1,113535278 | 0,80909 | 0,91204664 | novel transcript |
| ENSG00000187778 | MCRS1 | 0,155116178 | 1,113511284 | 0,44294 | 0,69043269 | microspherule protein 1 |
| ENSG00000132405 | TBC1D14 | 0,155018077 | 1,113435569 | 0,47393 | 0,71294959 | TBC1 domain family member 14 |
| ENSG00000125872 | LRRN4 | 0,154814966 | 1,113278825 | 0,79259 | 0,90361513 | leucine rich repeat neuronal 4 |
| ENSG00000109927 | TECTA | 0,154807188 | 1,113272823 | 0,59354 | 0,79186889 | tectorin alpha |
| ENSG00000075089 | ACTR6 | 0,154713752 | 1,113200724 | 0,57813 | 0,78191189 | actin related protein 6 |
| ENSG00000117906 | RCN2 | 0,154707725 | 1,113196073 | 0,42686 | 0,6801859 | reticulocalbin 2 |
| ENSG00000160172 | FAM86C2P | 0,15469076 | 1,113182983 | 0,53012 | 0,75179571 | family with sequence similarity 86 member C2, pseudogene |
| ENSG00000009780 | FAM76A | 0,154659913 | 1,113159182 | 0,47381 | 0,71294959 | family with sequence similarity 76 member A |
| ENSG00000050748 | MAPK9 | 0,15458543 | 1,113101714 | 0,46754 | 0,70739549 | mitogen-activated protein kinase 9 |
| ENSG00000110002 | VWA5A | 0,154572741 | 1,113091924 | 0,6346 | 0,81596468 | von Willebrand factor A domain containing 5A |
| ENSG00000139364 | TMEM132B | 0,154525967 | 1,113055836 | 0,75898 | 0,88747312 | transmembrane protein 132B |
| ENSG00000136861 | CDK5RAP2 | 0,15446559 | 1,113009256 | 0,47106 | 0,71051384 | CDK5 regulatory subunit associated protein 2 |

| | | | | | | |
|---|---|---|---|---|---|---|
| ENSG00000080824 | HSP90AA1 | 0,154419742 | 1,112973886 | 0,47601 | 0,71443502 | heat shock protein 90 alpha family class A member 1 |
| ENSG00000251579 | AC055733,3 | 0,1543514 | 1,112921164 | 0,79129 | 0,90300322 | pseudogene similar to part of makorin ring finger protein 1 |
| ENSG00000258744 | AL132800,1 | 0,154262707 | 1,112852746 | 0,66571 | 0,83361269 | novel transcript, antisense to CMA1 |
| ENSG00000050405 | LIMA1 | 0,15421303 | 1,112814428 | 0,71637 | 0,86423072 | LIM domain and actin binding 1 |
| ENSG00000166797 | CIAO2A | 0,154097174 | 1,112725066 | 0,47716 | 0,71510081 | cytosolic iron-sulfur assembly component 2A |
| ENSG00000119541 | VPS4B | 0,154024974 | 1,112669381 | 0,30016 | 0,59160746 | vacuolar protein sorting 4 homolog B |
| ENSG00000275052 | PPP4R3B | 0,15394258 | 1,112605837 | 0,45703 | 0,70056368 | protein phosphatase 4 regulatory subunit 3B |
| ENSG00000103042 | SLC38A7 | 0,153908776 | 1,112579768 | 0,36449 | 0,63501401 | solute carrier family 38 member 7 |
| ENSG00000106003 | LFNG | 0,153825889 | 1,112515849 | 0,65143 | 0,82538041 | LFNG O-fucosylpeptide 3-beta-N-acetylglucosaminyltransferase |
| ENSG00000197756 | RPL37A | 0,153796008 | 1,112492807 | 0,38455 | 0,65052153 | ribosomal protein L37a |
| ENSG00000122085 | MTERF4 | 0,153705518 | 1,11242303 | 0,49193 | 0,72531636 | mitochondrial transcription termination factor 4 |
| ENSG00000132205 | EMILIN2 | 0,15367008 | 1,112395705 | 0,61038 | 0,80221019 | elastin microfibril interfacer 2 |
| ENSG00000275582 | AL031670,1 | 0,153541842 | 1,112296831 | 0,60381 | 0,79770754 | novel transcript, antisense to PANK2 |
| ENSG00000028116 | VRK2 | 0,153501778 | 1,112265943 | 0,61403 | 0,80367735 | VRK serine/threonine kinase 2 |
| ENSG00000132275 | RRP8 | 0,153498525 | 1,112263435 | 0,45085 | 0,69594229 | ribosomal RNA processing 8 |
| ENSG00000268810 | AC007193,1 | 0,153490744 | 1,112257437 | 0,92022 | 0,96530017 | novel transcript, antisense to PPP5C |
| ENSG00000059691 | GATB | 0,15347593 | 1,112246015 | 0,38609 | 0,65152373 | glutamyl-tRNA amidotransferase subunit B |
| ENSG00000188177 | ZC3H6 | 0,153360818 | 1,112157273 | 0,47913 | 0,71677199 | zinc finger CCCH-type containing 6 |
| ENSG00000256464 | YWHABP2 | 0,153231482 | 1,112057574 | 0,83233 | 0,92328639 | tyrosine 3-monooxygenase/tryptophan 5-monooxygenase activation protein beta pseudogene 2 |
| ENSG00000013392 | RWDD2A | 0,153196291 | 1,112030449 | 0,64021 | 0,81935089 | RWD domain containing 2A |
| ENSG00000178440 | TIMM23B-AG. | 0,153188627 | 1,112024541 | 0,65211 | 0,82576571 | TIMM23B-AGAP6 readthrough (NMD candidate) |
| ENSG00000169100 | SLC25A6 | 0,153166532 | 1,11200751 | 0,41967 | 0,67536133 | solute carrier family 25 member 6 |
| ENSG00000196371 | FUT4 | 0,153091959 | 1,111950033 | 0,57002 | 0,77703861 | fucosyltransferase 4 |
| ENSG00000228316 | MTATP6P19 | 0,153034587 | 1,111905814 | 0,87041 | 0,942849 | MT-ATP6 pseudogene 19 |
| ENSG00000130202 | NECTIN2 | 0,152942647 | 1,111834957 | 0,46405 | 0,70478475 | nectin cell adhesion molecule 2 |
| ENSG00000266498 | AC055811,3 | 0,152871119 | 1,111779834 | 0,80676 | 0,91120963 | novel transcript |
| ENSG00000253305 | PCDHGB6 | 0,15280242 | 1,111726894 | 0,62557 | 0,81053676 | protocadherin gamma subfamily B, 6 |
| ENSG00000273763 | AC007318,1 | 0,152802188 | 1,111726715 | 0,70048 | 0,85531965 | marker of proliferation Ki-67 (MKI67) pseudogene |
| ENSG00000198961 | PJA2 | 0,152666266 | 1,11162198 | 0,73738 | 0,87539732 | praja ring finger ubiquitin ligase 2 |
| ENSG00000162236 | STX5 | 0,152554363 | 1,11153576 | 0,37993 | 0,64769466 | syntaxin 5 |
| ENSG00000090889 | KIF4A | 0,152504171 | 1,11149709 | 0,50437 | 0,7332113 | kinesin family member 4A |

| | | | | | | |
|---|---|---|---|---|---|---|
| ENSG00000125037 | EMC3 | 0,15242167 | 1,11143353 | 0,38665 | 0,65196019 | ER membrane protein complex subunit 3 |
| ENSG00000274428 | RNVU1-25 | 0,152417511 | 1,111430326 | 0,78985 | 0,90272027 | RNA, variant U1 small nuclear 25 |
| ENSG00000072133 | RPS6KA6 | 0,152315112 | 1,111351443 | 0,52877 | 0,75099628 | ribosomal protein S6 kinase A6 |
| ENSG00000234684 | SDCBP2-AS1 | 0,152267936 | 1,111315102 | 0,55565 | 0,76790322 | SDCBP2 antisense RNA 1 |
| ENSG00000140612 | SEC11A | 0,15220631 | 1,111267632 | 0,41223 | 0,66996291 | SEC11 homolog A, signal peptidase complex subunit |
| ENSG00000259332 | ST20-MTHFS | 0,152181571 | 1,111248576 | 0,7022 | 0,85651838 | ST20-MTHFS readthrough |
| ENSG00000250433 | CLSTN2-AS1 | 0,152159704 | 1,111231734 | 0,80984 | 0,91237449 | CLSTN2 antisense RNA 1 |
| ENSG00000137106 | GRHPR | 0,152139114 | 1,111215875 | 0,35798 | 0,63185282 | glyoxylate and hydroxypyruvate reductase |
| ENSG00000118620 | ZNF430 | 0,152121597 | 1,111202382 | 0,4278 | 0,68079193 | zinc finger protein 430 |
| ENSG00000286342 | AC073210,3 | 0,152107104 | 1,111191219 | 0,79874 | 0,9071596 | novel transcript |
| ENSG00000244005 | NFS1 | 0,152074225 | 1,111165895 | 0,49338 | 0,72634709 | NFS1 cysteine desulfurase |
| ENSG00000105202 | FBL | 0,151973966 | 1,111088679 | 0,49023 | 0,72445532 | fibrillarin |
| ENSG00000182568 | SATB1 | 0,151939605 | 1,111062216 | 0,5026 | 0,73209345 | SATB homeobox 1 |
| ENSG00000214050 | FBXO16 | 0,151931379 | 1,111055881 | 0,62746 | 0,8114733 | F-box protein 16 |
| ENSG00000163950 | SLBP | 0,151825499 | 1,110974343 | 0,41073 | 0,66856387 | stem-loop binding protein |
| ENSG00000142655 | PEX14 | 0,151659726 | 1,110846694 | 0,44204 | 0,68998374 | peroxisomal biogenesis factor 14 |
| ENSG00000274081 | PUF60 | 0,151580763 | 1,110785895 | 0,59178 | 0,79088049 | poly(U) binding splicing factor 60 |
| ENSG00000105643 | ARRDC2 | 0,151510453 | 1,110731762 | 0,82525 | 0,91907311 | arrestin domain containing 2 |
| ENSG00000197863 | ZNF790 | 0,151509277 | 1,110730856 | 0,46584 | 0,70619029 | zinc finger protein 790 |
| ENSG00000130560 | UBAC1 | 0,151436514 | 1,110674838 | 0,37002 | 0,63943131 | UBA domain containing 1 |
| ENSG00000100243 | CYB5R3 | 0,151370504 | 1,11062402 | 0,38278 | 0,64947958 | cytochrome b5 reductase 3 |
| ENSG00000141867 | BRD4 | 0,151347329 | 1,11060618 | 0,40508 | 0,66481282 | bromodomain containing 4 |
| ENSG00000275066 | SYNRG | 0,151346975 | 1,110605907 | 0,54549 | 0,76113525 | synergin gamma |
| ENSG00000001631 | KRIT1 | 0,151299626 | 1,110569458 | 0,49184 | 0,72530727 | KRIT1 ankyrin repeat containing |
| ENSG00000112182 | BACH2 | 0,151273231 | 1,11054914 | 0,47173 | 0,71108681 | BTB domain and CNC homolog 2 |
| ENSG00000287467 | AC037459,4 | 0,151245868 | 1,110528077 | 0,8994 | 0,95714792 | novel transcript, sense intronic to SORBS3 |
| ENSG00000234719 | NPIPB2 | 0,151239028 | 1,110522811 | 0,75719 | 0,88663582 | nuclear pore complex interacting protein family member B2 |
| ENSG00000125089 | SH3TC1 | 0,15122708 | 1,110513614 | 0,77824 | 0,89735311 | SH3 domain and tetratricopeptide repeats 1 |
| ENSG00000283580 | AC098484,3 | 0,151155661 | 1,110458641 | 0,65828 | 0,8290823 | novel protein |
| ENSG00000198887 | SMC5 | 0,151148617 | 1,11045322 | 0,47818 | 0,71592551 | structural maintenance of chromosomes 5 |
| ENSG00000079156 | OSBPL6 | 0,151072371 | 1,110394534 | 0,70462 | 0,85764572 | oxysterol binding protein like 6 |
| ENSG00000189227 | C15orf61 | 0,151033 | 1,110364232 | 0,60729 | 0,80012503 | chromosome 15 open reading frame 61 |

| | | | | | | |
|---|---|---|---|---|---|---|
| ENSG00000101391 | CDK5RAP1 | 0,151018062 | 1,110352735 | 0,45283 | 0,69717623 | CDK5 regulatory subunit associated protein 1 |
| ENSG00000115129 | TP53I3 | 0,151004125 | 1,110342008 | 0,5603 | 0,7707835 | tumor protein p53 inducible protein 3 |
| ENSG00000254635 | WAC-AS1 | 0,150929171 | 1,110284323 | 0,46241 | 0,70356854 | WAC antisense RNA 1 (head to head) |
| ENSG00000257354 | AC048341,1 | 0,150789702 | 1,110176994 | 0,71769 | 0,86483656 | novel transcript |
| ENSG00000250033 | SLC7A11-AS1 | 0,150777116 | 1,110167309 | 0,77743 | 0,89700087 | SLC7A11 antisense RNA 1 |
| ENSG00000272638 | AC006027,1 | 0,150682504 | 1,110094506 | 0,87084 | 0,94302816 | novel transcript, antisense to NOD1 |
| ENSG00000129351 | ILF3 | 0,150643625 | 1,110064591 | 0,48079 | 0,71788144 | interleukin enhancer binding factor 3 |
| ENSG00000163472 | TMEM79 | 0,150595106 | 1,11002726 | 0,54373 | 0,76018847 | transmembrane protein 79 |
| ENSG00000227331 | RPL7AP22 | 0,15058876 | 1,110022377 | 0,73031 | 0,87185115 | ribosomal protein L7a pseudogene 22 |
| ENSG00000144227 | NXPH2 | 0,150553318 | 1,109995107 | 0,76338 | 0,88980026 | neurexophilin 2 |
| ENSG00000166508 | MCM7 | 0,15048043 | 1,10993903 | 0,41005 | 0,66821634 | minichromosome maintenance complex component 7 |
| ENSG00000281954 | AC208069,1 | 0,150174518 | 1,109703701 | 0,96162 | 0,98401207 | novel transcript, antisense to DNAH10 |
| ENSG00000235374 | SSR4P1 | 0,150089098 | 1,109637999 | 0,65202 | 0,82568422 | signal sequence receptor subunit 4 pseudogene 1 |
| ENSG00000067182 | TNFRSF1A | 0,150087553 | 1,109636811 | 0,5582 | 0,76982396 | TNF receptor superfamily member 1A |
| ENSG00000234674 | TCF19 | 0,150070146 | 1,109623422 | 0,67594 | 0,84049293 | transcription factor 19 |
| ENSG00000206455 | TCF19 | 0,150070146 | 1,109623422 | 0,67594 | 0,84049293 | transcription factor 19 |
| ENSG00000198792 | TMEM184B | 0,150061948 | 1,109617117 | 0,39101 | 0,65473229 | transmembrane protein 184B |
| ENSG00000228409 | CCT6P1 | 0,150002725 | 1,109571568 | 0,63341 | 0,81543903 | chaperonin containing TCP1 subunit 6 pseudogene 1 |
| ENSG00000274047 | SYNRG | 0,149981409 | 1,109555174 | 0,56019 | 0,7707835 | synergin gamma |
| ENSG00000242265 | PEG10 | 0,149974237 | 1,109549658 | 0,57628 | 0,78066251 | paternally expressed 10 |
| ENSG00000249258 | AC079193,2 | 0,149930816 | 1,109516264 | 0,77418 | 0,89572556 | novel transcript |
| ENSG00000287601 | BX323046,2 | 0,149897209 | 1,109490419 | 0,78265 | 0,89905848 | novel transcript |
| ENSG00000132716 | DCAF8 | 0,149874286 | 1,10947279 | 0,3566 | 0,63063438 | DDB1 and CUL4 associated factor 8 |
| ENSG00000159720 | ATP6V0D1 | 0,149832813 | 1,109440896 | 0,47314 | 0,71242761 | ATPase H+ transporting V0 subunit d1 |
| ENSG00000148019 | CEP78 | 0,149662094 | 1,10930962 | 0,40481 | 0,66475559 | centrosomal protein 78 |
| ENSG00000275045 | GTF2H2 | 0,149633084 | 1,109287315 | 0,62086 | 0,80776776 | general transcription factor IIH subunit 2 |
| ENSG00000268105 | AC124856,1 | 0,149580163 | 1,109246625 | 0,56159 | 0,77136826 | ribosomal protein L5 (RPL5) pseudogene |
| ENSG00000095321 | CRAT | 0,149535712 | 1,109212448 | 0,61407 | 0,80367735 | carnitine O-acetyltransferase |
| ENSG00000047056 | WDR37 | 0,149465577 | 1,109158526 | 0,50711 | 0,73538734 | WD repeat domain 37 |
| ENSG00000176783 | RUFY1 | 0,149361617 | 1,109078604 | 0,52909 | 0,75111125 | RUN and FYVE domain containing 1 |
| ENSG00000152061 | RABGAP1L | 0,149349704 | 1,109069446 | 0,43795 | 0,68747773 | RAB GTPase activating protein 1 like |
| ENSG00000165244 | ZNF367 | 0,149343595 | 1,109064749 | 0,55213 | 0,76565817 | zinc finger protein 367 |
| ENSG00000197265 | GTF2E2 | 0,149271609 | 1,109009411 | 0,44236 | 0,69006979 | general transcription factor IIE subunit 2 |

| | | | | | | |
|---|---|---|---|---|---|---|
| ENSG00000113552 | GNPDA1 | 0,149252493 | 1,108994717 | 0,63373 | 0,81549489 | glucosamine-6-phosphate deaminase 1 |
| ENSG00000224281 | SLC25A5-AS1 | 0,14924863 | 1,108991747 | 0,65858 | 0,82926071 | SLC25A5 antisense RNA 1 |
| ENSG00000059728 | MXD1 | 0,149186735 | 1,10894417 | 0,58457 | 0,78577262 | MAX dimerization protein 1 |
| ENSG00000254507 | DEFB131E | 0,149132925 | 1,108902809 | 0,82075 | 0,9173395 | defensin beta 131E (pseudogene) |
| ENSG00000285431 | DEFB131E | 0,149132925 | 1,108902809 | 0,82075 | 0,9173395 | defensin beta 131E (pseudogene) |
| ENSG00000006042 | TMEM98 | 0,149122694 | 1,108894946 | 0,47325 | 0,7125462 | transmembrane protein 98 |
| ENSG00000073050 | XRCC1 | 0,149075763 | 1,108858873 | 0,56467 | 0,77306193 | X-ray repair cross complementing 1 |
| ENSG00000163482 | STK36 | 0,14897763 | 1,108783451 | 0,5803 | 0,78353223 | serine/threonine kinase 36 |
| ENSG00000243232 | PCDHAC2 | 0,148967737 | 1,108775848 | 0,59514 | 0,79320995 | protocadherin alpha subfamily C, 2 |
| ENSG00000175970 | UNC119B | 0,148883238 | 1,108710908 | 0,39164 | 0,65533345 | unc-119 lipid binding chaperone B |
| ENSG00000260194 | AC007496,1 | 0,148878827 | 1,108707519 | 0,81649 | 0,91556164 | novel transcript, antisense to FTO |
| ENSG00000287222 | AC234917,3 | 0,148821733 | 1,108663643 | 0,48357 | 0,71980944 | novel transcript, antisense to PUF60 |
| ENSG00000103047 | TANGO6 | 0,148690071 | 1,10856247 | 0,53384 | 0,75373779 | transport and golgi organization 6 homolog |
| ENSG00000171435 | KSR2 | 0,148629851 | 1,108516197 | 0,63802 | 0,81814424 | kinase suppressor of ras 2 |
| ENSG00000163479 | SSR2 | 0,148620058 | 1,108508673 | 0,34805 | 0,62442306 | signal sequence receptor subunit 2 |
| ENSG00000118518 | RNF146 | 0,14854372 | 1,10845002 | 0,54288 | 0,75963423 | ring finger protein 146 |
| ENSG00000197872 | CYRIA | 0,148510821 | 1,108424743 | 0,75293 | 0,88426071 | CYFIP related Rac1 interactor A |
| ENSG00000120685 | PROSER1 | 0,148509305 | 1,108423578 | 0,51472 | 0,74102666 | proline and serine rich 1 |
| ENSG00000110066 | KMT5B | 0,148404158 | 1,108342797 | 0,61533 | 0,80434562 | lysine methyltransferase 5B |
| ENSG00000176029 | C11orf16 | 0,148399696 | 1,108339368 | 0,7766 | 0,89658055 | chromosome 11 open reading frame 16 |
| ENSG00000234206 | AL121957,1 | 0,148379729 | 1,10832403 | 0,81547 | 0,91505837 | novel transcript |
| ENSG00000154001 | PPP2R5E | 0,148299626 | 1,108262494 | 0,34792 | 0,62435416 | protein phosphatase 2 regulatory subunit B'epsilon |
| ENSG00000140835 | CHST4 | 0,148119321 | 1,108123994 | 0,82776 | 0,92048005 | carbohydrate sulfotransferase 4 |
| ENSG00000068654 | POLR1A | 0,148113908 | 1,108119836 | 0,43469 | 0,68529786 | RNA polymerase I subunit A |
| ENSG00000197183 | NOL4L | 0,147966634 | 1,108006722 | 0,54546 | 0,76113525 | nucleolar protein 4 like |
| ENSG00000228444 | AL137244,1 | 0,147950313 | 1,107994188 | 0,86946 | 0,9425782 | novel transcript, sense intronic to LMO7 |
| ENSG00000213020 | ZNF611 | 0,14786919 | 1,107931886 | 0,48274 | 0,71910922 | zinc finger protein 611 |
| ENSG00000064666 | CNN2 | 0,147767163 | 1,107853537 | 0,46717 | 0,70714082 | calponin 2 |
| ENSG00000249019 | AC108471,1 | 0,147765986 | 1,107852633 | 0,81693 | 0,91580038 | ribosomal protein S7 (RPS7) pseudogene |
| ENSG00000065665 | SEC61A2 | 0,147687774 | 1,107792575 | 0,68297 | 0,8447391 | SEC61 translocon subunit alpha 2 |
| ENSG00000101901 | ALG13 | 0,14768468 | 1,107790199 | 0,55355 | 0,76655124 | ALG13 UDP-N-acetylglucosaminyltransferase subunit |
| ENSG00000257103 | LSM14A | 0,147604514 | 1,107728645 | 0,57201 | 0,77840347 | LSM14A mRNA processing body assembly factor |
| ENSG00000256779 | AP002383,3 | 0,147521194 | 1,107664672 | 0,87043 | 0,942849 | serine/arginine-rich splicing factor 8 (SRSF8) pseudogene |

| | | | | | | |
|---|---|---|---|---|---|---|
| ENSG00000179796 | LRRC3B | 0,147503592 | 1,107651158 | 0,75302 | 0,88429223 | leucine rich repeat containing 3B |
| ENSG00000162377 | COA7 | 0,147334223 | 1,107521129 | 0,4656 | 0,70598256 | cytochrome c oxidase assembly factor 7 (putative) |
| ENSG00000253875 | AC013643,2 | 0,147328683 | 1,107516877 | 0,66607 | 0,83387778 | novel transcript |
| ENSG00000259158 | ADAM20P1 | 0,147219004 | 1,107432682 | 0,70774 | 0,85918434 | ADAM metallopeptidase domain 20 pseudogene 1 |
| ENSG00000151502 | VPS26B | 0,14716517 | 1,10739136 | 0,41536 | 0,67224051 | VPS26, retromer complex component B |
| ENSG00000106628 | POLD2 | 0,147094352 | 1,107337001 | 0,48395 | 0,71999322 | DNA polymerase delta 2, accessory subunit |
| ENSG00000266777 | SH3GL1P1 | 0,147090436 | 1,107333996 | 0,62854 | 0,81209878 | SH3 domain containing GRB2 like 1, endophilin A2 pseudogene 1 |
| ENSG00000277687 | AL139407,1 | 0,147040727 | 1,107295843 | 0,6178 | 0,80610038 | novel transcript, sense intronic to CACUL1 |
| ENSG00000160007 | ARHGAP35 | 0,147020524 | 1,107280337 | 0,4427 | 0,69033311 | Rho GTPase activating protein 35 |
| ENSG00000260260 | SNHG19 | 0,146956756 | 1,107231395 | 0,73206 | 0,87285234 | small nucleolar RNA host gene 19 |
| ENSG00000283208 | AC001226,2 | 0,146935274 | 1,107214909 | 0,69503 | 0,85206252 | novel protein |
| ENSG00000065600 | PACC1 | 0,146884163 | 1,107175683 | 0,68304 | 0,84475757 | proton activated chloride channel 1 |
| ENSG00000235643 | LINC01647 | 0,14666911 | 1,107010657 | 0,77964 | 0,89815441 | long intergenic non-protein coding RNA 1647 |
| ENSG00000188002 | AC026412,1 | 0,146553095 | 1,106921639 | 0,72969 | 0,8715818 | programmed cell death 6 (PDCD6) pseudogene |
| ENSG00000073578 | SDHA | 0,146477791 | 1,106863863 | 0,36978 | 0,63943131 | succinate dehydrogenase complex flavoprotein subunit A |
| ENSG00000272604 | AC073073,2 | 0,14646338 | 1,106852807 | 0,74578 | 0,88010204 | novel transcript, antisense to EFCAB10 |
| ENSG00000226982 | CENPCP1 | 0,14617514 | 1,106631688 | 0,7877 | 0,90122495 | centromere protein C pseudogene 1 |
| ENSG00000182636 | NDN | 0,146096439 | 1,106571321 | 0,69956 | 0,8548625 | necdin, MAGE family member |
| ENSG00000182093 | GET1 | 0,146087183 | 1,106564221 | 0,58289 | 0,78477395 | guided entry of tail-anchored proteins factor 1 |
| ENSG00000130255 | RPL36 | 0,145961884 | 1,10646812 | 0,43661 | 0,68681257 | ribosomal protein L36 |
| ENSG00000278574 | ZNHIT3 | 0,14596088 | 1,10646735 | 0,55087 | 0,76486471 | zinc finger HIT-type containing 3 |
| ENSG00000169567 | HINT1 | 0,145939838 | 1,106451212 | 0,51935 | 0,74394224 | histidine triad nucleotide binding protein 1 |
| ENSG00000163635 | ATXN7 | 0,145883537 | 1,106408034 | 0,43043 | 0,68295004 | ataxin 7 |
| ENSG00000065357 | DGKA | 0,145874991 | 1,10640148 | 0,72897 | 0,87140581 | diacylglycerol kinase alpha |
| ENSG00000254880 | PKNOX2-AS1 | 0,145869409 | 1,106397199 | 0,89187 | 0,95246557 | PKNOX2 antisense RNA 1 (head to head) |
| ENSG00000176896 | TCEANC | 0,145808552 | 1,106350529 | 0,59297 | 0,79148028 | transcription elongation factor A N-terminal and central domain containing |
| ENSG00000163884 | KLF15 | 0,145805577 | 1,106348248 | 0,71378 | 0,86224977 | Kruppel like factor 15 |
| ENSG00000064393 | HIPK2 | 0,14578386 | 1,106331594 | 0,57151 | 0,77809338 | homeodomain interacting protein kinase 2 |
| ENSG00000125780 | TGM3 | 0,145744609 | 1,106301494 | 0,91838 | 0,96446185 | transglutaminase 3 |
| ENSG00000255040 | MORF4L1P3 | 0,1457435 | 1,106300644 | 0,81465 | 0,91470265 | mortality factor 4 like 1 pseudogene 3 |
| ENSG00000168646 | AXIN2 | 0,145736358 | 1,106295167 | 0,5792 | 0,78270914 | axin 2 |

| | | | | | | |
|---|---|---|---|---|---|---|
| ENSG00000134152 | KATNBL1 | 0,145674346 | 1,106247616 | 0,58017 | 0,78348519 | katanin regulatory subunit B1 like 1 |
| ENSG00000233072 | RPS15AP6 | 0,145605342 | 1,106194706 | 0,8213 | 0,91740344 | ribosomal protein S15a pseudogene 6 |
| ENSG00000183643 | C15orf32 | 0,145546303 | 1,106149438 | 0,83873 | 0,92638462 | chromosome 15 putative open reading frame 32 |
| ENSG00000276523 | AC025287,3 | 0,14552699 | 1,10613463 | 0,7823 | 0,8988906 | novel transcript, antisense to GABARAPL2 |
| ENSG00000089060 | SLC8B1 | 0,145453579 | 1,106078346 | 0,58793 | 0,78787891 | solute carrier family 8 member B1 |
| ENSG00000183665 | TRMT12 | 0,145336954 | 1,105988937 | 0,45101 | 0,69608679 | tRNA methyltransferase 12 homolog |
| ENSG00000259956 | RBM15B | 0,145324531 | 1,105979413 | 0,40542 | 0,66513295 | RNA binding motif protein 15B |
| ENSG00000284454 | NPIPB3 | 0,145236828 | 1,105912181 | 0,43856 | 0,68776759 | nuclear pore complex interacting protein family member B3 |
| ENSG00000157800 | SLC37A3 | 0,145210298 | 1,105891844 | 0,40151 | 0,6630277 | solute carrier family 37 member 3 |
| ENSG00000113916 | BCL6 | 0,145165911 | 1,105857821 | 0,702 | 0,85646941 | BCL6 transcription repressor |
| ENSG00000226410 | AC104058,1 | 0,145159465 | 1,105852879 | 0,87951 | 0,94632406 | ribosomal protein L29 (RPL29) pseudogene |
| ENSG00000174579 | MSL2 | 0,145153279 | 1,105848138 | 0,39337 | 0,65634348 | MSL complex subunit 2 |
| ENSG00000249693 | THEGL | 0,145113064 | 1,105817313 | 0,7144 | 0,86255727 | theg spermatid protein like |
| ENSG00000288191 | MISP3 | 0,145090092 | 1,105799705 | 0,74317 | 0,87840582 | MISP family member 3 |
| ENSG00000141854 | MISP3 | 0,145090092 | 1,105799705 | 0,74317 | 0,87840582 | MISP family member 3 |
| ENSG00000174606 | ANGEL2 | 0,145082923 | 1,10579421 | 0,43885 | 0,6878469 | angel homolog 2 |
| ENSG00000138439 | FAM117B | 0,145035706 | 1,10575802 | 0,48752 | 0,72250947 | family with sequence similarity 117 member B |
| ENSG00000161048 | NAPEPLD | 0,145031038 | 1,105754442 | 0,40506 | 0,66481282 | N-acyl phosphatidylethanolamine phospholipase D |
| ENSG00000197417 | SHPK | 0,145003492 | 1,10573333 | 0,3794 | 0,64720307 | sedoheptulokinase |
| ENSG00000125733 | TRIP10 | 0,14498247 | 1,105717218 | 0,43273 | 0,68421938 | thyroid hormone receptor interactor 10 |
| ENSG00000215472 | RPL17-C18orf | 0,144842491 | 1,10560994 | 0,82706 | 0,92010136 | RPL17-C18orf32 readthrough |
| ENSG00000130816 | DNMT1 | 0,144802022 | 1,105578926 | 0,44406 | 0,6916877 | DNA methyltransferase 1 |
| ENSG00000277150 | F8A3 | 0,144660712 | 1,105470642 | 0,63448 | 0,81588861 | coagulation factor VIII associated 3 |
| ENSG00000171388 | APLN | 0,144627563 | 1,105445242 | 0,7054 | 0,85804888 | apelin |
| ENSG00000274429 | DLG5 | 0,144495976 | 1,10534442 | 0,74181 | 0,87793115 | discs large MAGUK scaffold protein 5 |
| ENSG00000163012 | ZSWIM2 | 0,14444754 | 1,10530731 | 0,86289 | 0,9388179 | zinc finger SWIM-type containing 2 |
| ENSG00000185100 | ADSS1 | 0,144443797 | 1,105304443 | 0,65099 | 0,82521872 | adenylosuccinate synthase 1 |
| ENSG00000186714 | CCDC73 | 0,144420202 | 1,105286366 | 0,88339 | 0,94837933 | coiled-coil domain containing 73 |
| ENSG00000276302 | AL021997,2 | 0,144109163 | 1,105048096 | 0,7932 | 0,90371578 | novel protein |
| ENSG00000152439 | ZNF773 | 0,143845965 | 1,104846515 | 0,58585 | 0,78648151 | zinc finger protein 773 |
| ENSG00000184588 | PDE4B | 0,143771766 | 1,104789694 | 0,7707 | 0,89376602 | phosphodiesterase 4B |
| ENSG00000224846 | NQO2-AS1 | 0,143621193 | 1,104674393 | 0,87565 | 0,94497474 | NQO2 antisense RNA 1 |

| | | | | | | |
|---|---|---|---|---|---|---|
| ENSG00000250506 | CDK3 | 0,143601749 | 1,104659505 | 0,66059 | 0,8307262 | cyclin dependent kinase 3 |
| ENSG00000167216 | KATNAL2 | 0,143571515 | 1,104636355 | 0,65357 | 0,82638628 | katanin catalytic subunit A1 like 2 |
| ENSG00000285747 | AL133485,3 | 0,143566062 | 1,10463218 | 0,80932 | 0,91213228 | novel transcript |
| ENSG00000152582 | SPEF2 | 0,143513686 | 1,104592078 | 0,60221 | 0,79696185 | sperm flagellar 2 |
| ENSG00000164405 | UQCRQ | 0,143482253 | 1,104568012 | 0,66108 | 0,83105713 | ubiquinol-cytochrome c reductase complex III subunit VII |
| ENSG00000224023 | EDRF1-DT | 0,143381576 | 1,104490933 | 0,71068 | 0,860669 | EDRF1 divergent transcript |
| ENSG00000148291 | SURF2 | 0,143370263 | 1,104482273 | 0,57236 | 0,77849297 | surfeit 2 |
| ENSG00000281024 | SURF2 | 0,143370263 | 1,104482273 | 0,57236 | 0,77849297 | surfeit 2 |
| ENSG00000275055 | AC011468,5 | 0,143346652 | 1,104464197 | 0,64804 | 0,82383528 | novel transcript, antisense to ZNF432 |
| ENSG00000166402 | TUB | 0,143341107 | 1,104459952 | 0,54944 | 0,76407481 | TUB bipartite transcription factor |
| ENSG00000131153 | GINS2 | 0,14331048 | 1,104436505 | 0,40201 | 0,66341002 | GINS complex subunit 2 |
| ENSG00000164850 | GPER1 | 0,143271489 | 1,104406657 | 0,69845 | 0,85403406 | G protein-coupled estrogen receptor 1 |
| ENSG00000064490 | RFXANK | 0,14326331 | 1,104400395 | 0,48468 | 0,72039842 | regulatory factor X associated ankyrin containing protein |
| ENSG00000079950 | STX7 | 0,143217803 | 1,10436556 | 0,72703 | 0,87013394 | syntaxin 7 |
| ENSG00000105355 | PLIN3 | 0,143176727 | 1,104334117 | 0,40278 | 0,66396567 | perilipin 3 |
| ENSG00000163904 | SENP2 | 0,142890861 | 1,104115318 | 0,37461 | 0,6431311 | SUMO specific peptidase 2 |
| ENSG00000170852 | KBTBD2 | 0,142872157 | 1,104101004 | 0,47332 | 0,71260733 | kelch repeat and BTB domain containing 2 |
| ENSG00000139921 | TMX1 | 0,142779661 | 1,104030218 | 0,49604 | 0,72828309 | thioredoxin related transmembrane protein 1 |
| ENSG00000231738 | TSPAN19 | 0,142743683 | 1,104002686 | 0,75363 | 0,88453179 | tetraspanin 19 |
| ENSG00000001629 | ANKIB1 | 0,142740829 | 1,104000503 | 0,64809 | 0,82383528 | ankyrin repeat and IBR domain containing 1 |
| ENSG00000166987 | MBD6 | 0,142716561 | 1,103981932 | 0,48658 | 0,72196345 | methyl-CpG binding domain protein 6 |
| ENSG00000133265 | HSPBP1 | 0,142653476 | 1,103933659 | 0,4743 | 0,71307604 | HSPA (Hsp70) binding protein 1 |
| ENSG00000261069 | AC124312,3 | 0,142611273 | 1,103901366 | 0,62792 | 0,81175174 | novel transcript, SNORD host |
| ENSG00000103479 | RBL2 | 0,142574995 | 1,103873608 | 0,66498 | 0,83318628 | RB transcriptional corepressor like 2 |
| ENSG00000111087 | GLI1 | 0,142526883 | 1,103836796 | 0,81184 | 0,91334888 | GLI family zinc finger 1 |
| ENSG00000184786 | TCTE3 | 0,142524169 | 1,103834719 | 0,65157 | 0,82541975 | t-complex-associated-testis-expressed 3 |
| ENSG00000115956 | PLEK | 0,142402756 | 1,103741828 | 0,85012 | 0,93177313 | pleckstrin |
| ENSG00000185668 | POU3F1 | 0,142388496 | 1,103730918 | 0,69071 | 0,84906099 | POU class 3 homeobox 1 |
| ENSG00000101098 | RIMS4 | 0,142317266 | 1,103676425 | 0,60737 | 0,8001725 | regulating synaptic membrane exocytosis 4 |
| ENSG00000197555 | SIPA1L1 | 0,142275953 | 1,103644821 | 0,64889 | 0,82425307 | signal induced proliferation associated 1 like 1 |
| ENSG00000196961 | AP2A1 | 0,142195204 | 1,10358305 | 0,37581 | 0,64415653 | adaptor related protein complex 2 subunit alpha 1 |
| ENSG00000271732 | AL137798,1 | 0,142164875 | 1,103559851 | 0,90492 | 0,9593272 | novel transcript |
| ENSG00000277276 | OTUB2 | 0,142164018 | 1,103559195 | 0,89942 | 0,95714792 | OTU deubiquitinase, ubiquitin aldehyde binding 2 |

| | | | | | | |
|---|---|---|---|---|---|---|
| ENSG00000133243 | BTBD2 | 0,142102413 | 1,103512073 | 0,4943 | 0,72675894 | BTB domain containing 2 |
| ENSG00000029993 | HMGB3 | 0,142072428 | 1,103489137 | 0,51944 | 0,74396498 | high mobility group box 3 |
| ENSG00000234742 | AC144530,1 | 0,142022595 | 1,103451022 | 0,79936 | 0,90744841 | ribosomal protein L17 (RPL17) pseudogene |
| ENSG00000119650 | IFT43 | 0,141971976 | 1,103412306 | 0,5891 | 0,78855713 | intraflagellar transport 43 |
| ENSG00000224914 | LINC00863 | 0,14184917 | 1,103318385 | 0,41878 | 0,67486451 | long intergenic non-protein coding RNA 863 |
| ENSG00000170153 | RNF150 | 0,141807825 | 1,103286766 | 0,68031 | 0,84309883 | ring finger protein 150 |
| ENSG00000273489 | AC008264,2 | 0,141770767 | 1,103258427 | 0,56241 | 0,77186635 | novel transcript, antisense to MKLN1 |
| ENSG00000246731 | MGC16275 | 0,141724332 | 1,103222918 | 0,71503 | 0,86295089 | uncharacterized protein MGC16275 |
| ENSG00000104915 | STX10 | 0,141719884 | 1,103219516 | 0,43038 | 0,68295004 | syntaxin 10 |
| ENSG00000267454 | ZNF582-AS1 | 0,141578662 | 1,10311153 | 0,66836 | 0,83536064 | ZNF582 antisense RNA 1 (head to head) |
| ENSG00000137313 | TRIM26 | 0,141493429 | 1,103046362 | 0,77938 | 0,89794855 | tripartite motif containing 26 |
| ENSG00000124795 | DEK | 0,14142925 | 1,102997293 | 0,5857 | 0,78640335 | DEK proto-oncogene |
| ENSG00000189091 | SF3B3 | 0,141385698 | 1,102963996 | 0,37885 | 0,64679625 | splicing factor 3b subunit 3 |
| ENSG00000272667 | AC012306,2 | 0,141314512 | 1,102909574 | 0,65692 | 0,82856771 | novel transcript |
| ENSG00000081320 | STK17B | 0,141300992 | 1,102899239 | 0,45008 | 0,69557689 | serine/threonine kinase 17b |
| ENSG00000125656 | CLPP | 0,141227816 | 1,102843299 | 0,48205 | 0,71880798 | caseinolytic mitochondrial matrix peptidase proteolytic subunit |
| ENSG00000267172 | AC022031,1 | 0,141221955 | 1,102838819 | 0,90866 | 0,9609322 | novel transcript |
| ENSG00000196453 | ZNF777 | 0,141211774 | 1,102831036 | 0,51526 | 0,74138689 | zinc finger protein 777 |
| ENSG00000156486 | KCNS2 | 0,141175578 | 1,102803368 | 0,74152 | 0,87769924 | potassium voltage-gated channel modifier subfamily S member 2 |
| ENSG00000065548 | ZC3H15 | 0,141015454 | 1,102680975 | 0,46314 | 0,70415312 | zinc finger CCCH-type containing 15 |
| ENSG00000232850 | PTGES2-AS1 | 0,140943484 | 1,102625968 | 0,58791 | 0,78787891 | PTGES2 antisense RNA 1 (head to head) |
| ENSG00000286771 | AC097658,3 | 0,140908411 | 1,102599163 | 0,79605 | 0,90571355 | novel transcript, antisense to GAB1 |
| ENSG00000247400 | DNAJC3-DT | 0,140867565 | 1,102567946 | 0,6258 | 0,81058216 | DNAJC3 divergent transcript |
| ENSG00000183856 | IQGAP3 | 0,140778739 | 1,102500063 | 0,58217 | 0,78454692 | IQ motif containing GTPase activating protein 3 |
| ENSG00000137513 | NARS2 | 0,140559591 | 1,102332605 | 0,34743 | 0,62411645 | asparaginyl-tRNA synthetase 2, mitochondrial |
| ENSG00000213799 | ZNF845 | 0,140371683 | 1,102189037 | 0,48487 | 0,72052946 | zinc finger protein 845 |
| ENSG00000089818 | NECAP1 | 0,140366951 | 1,102185422 | 0,56387 | 0,77259134 | NECAP endocytosis associated 1 |
| ENSG00000254051 | AC011853,1 | 0,140320972 | 1,102150296 | 0,88066 | 0,94687991 | NADH dehydrogenase [ubiquinone] flavoprotein 2, mitochondrial (NDUFV2) pseudogene |
| ENSG00000233367 | AL133480,1 | 0,140268017 | 1,102109842 | 0,74474 | 0,87921382 | novel transcript |
| ENSG00000088305 | DNMT3B | 0,140257644 | 1,102101917 | 0,71239 | 0,86156979 | DNA methyltransferase 3 beta |

| | | | | | | |
|---|---|---|---|---|---|---|
| ENSG00000139865 | TTC6 | 0,140220461 | 1,102073513 | 0,82238 | 0,91794576 | tetratricopeptide repeat domain 6 |
| ENSG00000102786 | INTS6 | 0,140200654 | 1,102058383 | 0,41439 | 0,67106867 | integrator complex subunit 6 |
| ENSG00000183283 | DAZAP2 | 0,140137509 | 1,102010148 | 0,69021 | 0,84887944 | DAZ associated protein 2 |
| ENSG00000181830 | SLC35C1 | 0,140119964 | 1,101996746 | 0,57968 | 0,78299396 | solute carrier family 35 member C1 |
| ENSG00000100814 | CCNB1IP1 | 0,140021431 | 1,101921485 | 0,56449 | 0,77300039 | cyclin B1 interacting protein 1 |
| ENSG00000225791 | TRAM2-AS1 | 0,139972527 | 1,101884133 | 0,48402 | 0,72002429 | TRAM2 antisense RNA 1 (head to head) |
| ENSG00000131080 | EDA2R | 0,139817488 | 1,101765726 | 0,80138 | 0,90860375 | ectodysplasin A2 receptor |
| ENSG00000145002 | FAM86B2 | 0,1397843 | 1,10174038 | 0,89054 | 0,95184844 | family with sequence similarity 86 member B2 |
| ENSG00000138771 | SHROOM3 | 0,139734496 | 1,101702348 | 0,53499 | 0,75428901 | shroom family member 3 |
| ENSG00000118579 | MED28 | 0,139722393 | 1,101693105 | 0,36296 | 0,63395677 | mediator complex subunit 28 |
| ENSG00000110492 | MDK | 0,139719308 | 1,101690749 | 0,64765 | 0,82375544 | midkine |
| ENSG00000230769 | Z98048,1 | 0,139658843 | 1,101644577 | 0,82022 | 0,91725087 | riboflavin kinase (RFK) pseudogene |
| ENSG00000026652 | AGPAT4 | 0,139647038 | 1,101635563 | 0,52663 | 0,74985411 | 1-acylglycerol-3-phosphate O-acyltransferase 4 |
| ENSG00000145721 | LIX1 | 0,139615836 | 1,101611737 | 0,76948 | 0,89297049 | limb and CNS expressed 1 |
| ENSG00000253961 | AC009563,1 | 0,139593637 | 1,101594786 | 0,85851 | 0,93621468 | novel transcript |
| ENSG00000164967 | RPP25L | 0,139530953 | 1,101546925 | 0,6219 | 0,80819595 | ribonuclease P/MRP subunit p25 like |
| ENSG00000147649 | MTDH | 0,139503496 | 1,10152596 | 0,41294 | 0,67055663 | metadherin |
| ENSG00000187953 | PMS2CL | 0,139465138 | 1,101496673 | 0,56809 | 0,77582773 | PMS2 C-terminal like pseudogene |
| ENSG00000274646 | RPS9 | 0,139439276 | 1,101476928 | 0,86154 | 0,93811282 | ribosomal protein S9 |
| ENSG00000151657 | KIN | 0,139371771 | 1,10142539 | 0,40029 | 0,66242349 | Kin17 DNA and RNA binding protein |
| ENSG00000241749 | RPSAP52 | 0,13931267 | 1,101380271 | 0,64033 | 0,81936319 | ribosomal protein SA pseudogene 52 |
| ENSG00000234377 | OBI1-AS1 | 0,13927045 | 1,101348039 | 0,84728 | 0,93060479 | OBI1 antisense RNA 1 |
| ENSG00000145882 | PCYOX1L | 0,139201781 | 1,101295619 | 0,61619 | 0,80487059 | prenylcysteine oxidase 1 like |
| ENSG00000232112 | TMA7 | 0,139148084 | 1,101254629 | 0,58723 | 0,78717516 | translation machinery associated 7 homolog |
| ENSG00000171302 | CANT1 | 0,139139655 | 1,101248196 | 0,40784 | 0,66684711 | calcium activated nucleotidase 1 |
| ENSG00000135605 | TEC | 0,139117598 | 1,101231359 | 0,78752 | 0,90121071 | tec protein tyrosine kinase |
| ENSG00000286053 | ASDURF | 0,139109577 | 1,101225237 | 0,83324 | 0,92386058 | ASNSD1 upstream reading frame |
| ENSG00000286512 | AL031768,2 | 0,139015607 | 1,10115351 | 0,74443 | 0,87904799 | novel transcript |
| ENSG00000023892 | DEF6 | 0,1388944 | 1,101061001 | 0,74427 | 0,87895391 | DEF6 guanine nucleotide exchange factor |
| ENSG00000169855 | ROBO1 | 0,138890349 | 1,10105791 | 0,72598 | 0,86949831 | roundabout guidance receptor 1 |
| ENSG00000265043 | AC068418,2 | 0,138780158 | 1,100973816 | 0,80949 | 0,91213228 | novel transcript |
| ENSG00000286104 | AC016629,3 | 0,138749676 | 1,100950554 | 0,6069 | 0,79996159 | Novel transcript |
| ENSG00000234773 | AC012618,3 | 0,138699106 | 1,100911964 | 0,61168 | 0,80334296 | novel zinc finger protein pseudogene |

| | | | | | | |
|---|---|---|---|---|---|---|
| ENSG00000197122 | SRC | 0,138642992 | 1,100869144 | 0,51183 | 0,7388352 | SRC proto-oncogene, non-receptor tyrosine kinase |
| ENSG00000274020 | LINC01138 | 0,138534039 | 1,100786009 | 0,52108 | 0,74558236 | long intergenic non-protein coding RNA 1138 |
| ENSG00000232960 | CSNK2B | 0,13852426 | 1,100778548 | 0,77933 | 0,89794855 | casein kinase 2 beta |
| ENSG00000112146 | FBXO9 | 0,138453667 | 1,100724687 | 0,40328 | 0,66399303 | F-box protein 9 |
| ENSG00000136319 | TTC5 | 0,138448392 | 1,100720662 | 0,468 | 0,70769773 | tetratricopeptide repeat domain 5 |
| ENSG00000173811 | CCDC13-AS1 | 0,138416688 | 1,100696473 | 0,81399 | 0,91447468 | CCDC13 antisense RNA 1 |
| ENSG00000185065 | AC000068,1 | 0,13838317 | 1,100670901 | 0,63682 | 0,81728501 | novel transcript, antisense to C22orf39 |
| ENSG00000263028 | TAS2R19 | 0,138302249 | 1,100609166 | 0,74961 | 0,8822079 | taste 2 receptor member 19 |
| ENSG00000212124 | TAS2R19 | 0,138302249 | 1,100609166 | 0,74961 | 0,8822079 | taste 2 receptor member 19 |
| ENSG00000066117 | SMARCD1 | 0,138222028 | 1,100547968 | 0,44956 | 0,69519764 | SWI/SNF related, matrix associated, actin dependent regulator of chromatin, subfamily d, member 1 |
| ENSG00000140443 | IGF1R | 0,138094472 | 1,100450668 | 0,3811 | 0,64852799 | insulin like growth factor 1 receptor |
| ENSG00000224264 | VARS1 | 0,137948073 | 1,100339004 | 0,76858 | 0,89277929 | valyl-tRNA synthetase 1 |
| ENSG00000231116 | VARS1 | 0,137948073 | 1,100339004 | 0,76858 | 0,89277929 | valyl-tRNA synthetase 1 |
| ENSG00000285497 | USP17L7 | 0,137860172 | 1,100271964 | 0,79316 | 0,90371578 | ubiquitin specific peptidase 17 like family member 7 |
| ENSG00000271427 | AL358072,1 | 0,137790234 | 1,100218627 | 0,88908 | 0,9510223 | novel transcript |
| ENSG00000259644 | AC022613,3 | 0,137754354 | 1,100191265 | 0,63592 | 0,81684802 | novel transcript |
| ENSG00000232344 | AC087163,1 | 0,137702358 | 1,100151614 | 0,79045 | 0,90286944 | ribosomal protein S2 (RPS2) pseudogene |
| ENSG00000282706 | LINC01043 | 0,137681738 | 1,10013589 | 0,83411 | 0,92405734 | long intergenic non-protein coding RNA 1043 |
| ENSG00000156110 | ADK | 0,137647484 | 1,10010977 | 0,605 | 0,79821396 | adenosine kinase |
| ENSG00000090621 | PABPC4 | 0,137612973 | 1,100083454 | 0,3523 | 0,62740642 | poly(A) binding protein cytoplasmic 4 |
| ENSG00000227953 | LINC01341 | 0,137603361 | 1,100076125 | 0,64858 | 0,82403409 | long intergenic non-protein coding RNA 1341 |
| ENSG00000117543 | DPH5 | 0,137561439 | 1,100044159 | 0,4842 | 0,72005601 | diphthamide biosynthesis 5 |
| ENSG00000136682 | CBWD2 | 0,13744212 | 1,099953183 | 0,40064 | 0,66264525 | COBW domain containing 2 |
| ENSG00000177951 | BET1L | 0,137433713 | 1,099946773 | 0,42696 | 0,68019102 | Bet1 golgi vesicular membrane trafficking protein like |
| ENSG00000185306 | C12orf56 | 0,137386416 | 1,099910713 | 0,78775 | 0,90123584 | chromosome 12 open reading frame 56 |
| ENSG00000143543 | JTB | 0,137302258 | 1,099846553 | 0,53462 | 0,7541464 | jumping translocation breakpoint |
| ENSG00000077380 | DYNC1I2 | 0,137232532 | 1,099793398 | 0,61561 | 0,80451901 | dynein cytoplasmic 1 intermediate chain 2 |
| ENSG00000266916 | ZNF793-AS1 | 0,137223611 | 1,099786598 | 0,72229 | 0,86734497 | ZNF793 antisense RNA 1 (head to head) |
| ENSG00000163171 | CDC42EP3 | 0,137183903 | 1,099756328 | 0,71069 | 0,860669 | CDC42 effector protein 3 |
| ENSG00000164307 | ERAP1 | 0,137152854 | 1,09973266 | 0,68468 | 0,84571019 | endoplasmic reticulum aminopeptidase 1 |
| ENSG00000287843 | AL161628,1 | 0,137145229 | 1,099726848 | 0,82161 | 0,91753858 | novel transcript |
| ENSG00000143178 | TBX19 | 0,137020486 | 1,099631764 | 0,75909 | 0,88755659 | T-box transcription factor 19 |

| | | | | | | |
|---|---|---|---|---|---|---|
| ENSG00000181638 | ZFP41 | 0,136947834 | 1,099576389 | 0,40268 | 0,66396567 | ZFP41 zinc finger protein |
| ENSG00000285082 | AL160272,2 | 0,136943984 | 1,099573455 | 0,85287 | 0,93314746 | novel protein |
| ENSG00000213542 | AC007000,1 | 0,136937377 | 1,099568419 | 0,79582 | 0,9055252 | ribosomal protein L7a (RPL7A) pseudogene |
| ENSG00000117477 | CCDC181 | 0,136767081 | 1,099438634 | 0,60442 | 0,79794393 | coiled-coil domain containing 181 |
| ENSG00000285999 | AC025442,2 | 0,136742744 | 1,099420088 | 0,70491 | 0,857835 | novel transcipt |
| ENSG00000211445 | GPX3 | 0,136698827 | 1,099386621 | 0,66233 | 0,83183445 | glutathione peroxidase 3 |
| ENSG00000197971 | MBP | 0,136693196 | 1,09938233 | 0,55912 | 0,77044204 | myelin basic protein |
| ENSG00000274512 | TBC1D3L | 0,136629894 | 1,099334092 | 0,69326 | 0,85094387 | TBC1 domain family member 3L |
| ENSG00000151632 | AKR1C2 | 0,136628918 | 1,099333349 | 0,65716 | 0,82867637 | aldo-keto reductase family 1 member C2 |
| ENSG00000148399 | DPH7 | 0,136547996 | 1,099271687 | 0,53949 | 0,75774769 | diphthamide biosynthesis 7 |
| ENSG00000179094 | PER1 | 0,13649236 | 1,099229296 | 0,52226 | 0,74651799 | period circadian regulator 1 |
| ENSG00000010165 | EEF1AKNMT | 0,136490185 | 1,099227639 | 0,37798 | 0,64631202 | eEF1A lysine and N-terminal methyltransferase |
| ENSG00000127948 | POR | 0,136449927 | 1,099196966 | 0,55714 | 0,76897426 | cytochrome p450 oxidoreductase |
| ENSG00000136521 | NDUFB5 | 0,136408989 | 1,099165776 | 0,40638 | 0,66593291 | NADH:ubiquinone oxidoreductase subunit B5 |
| ENSG00000269556 | TMEM185A | 0,136363291 | 1,09913096 | 0,51275 | 0,73941533 | transmembrane protein 185A |
| ENSG00000106635 | BCL7B | 0,136329965 | 1,09910557 | 0,46409 | 0,70479912 | BAF chromatin remodeling complex subunit BCL7B |
| ENSG00000129422 | MTUS1 | 0,136255145 | 1,099048571 | 0,75322 | 0,88441977 | microtubule associated scaffold protein 1 |
| ENSG00000235244 | DANT2 | 0,136212003 | 1,099015705 | 0,63845 | 0,81851383 | DXZ4 associated non-coding transcript 2, distal |
| ENSG00000169762 | TAPT1 | 0,136154371 | 1,098971803 | 0,63539 | 0,81655453 | transmembrane anterior posterior transformation 1 |
| ENSG00000119965 | C10orf88 | 0,13615006 | 1,098968519 | 0,64313 | 0,82098995 | chromosome 10 open reading frame 88 |
| ENSG00000232864 | NUCKS1P1 | 0,136138723 | 1,098959884 | 0,82296 | 0,91812142 | nuclear casein kinase and cyclin dependent kinase substrate 1 pseudogene 1 |
| ENSG00000179523 | EIF3J-DT | 0,135940621 | 1,098808992 | 0,49013 | 0,72437872 | EIF3J divergent transcript |
| ENSG00000146409 | SLC18B1 | 0,135904272 | 1,098781307 | 0,5451 | 0,7609742 | solute carrier family 18 member B1 |
| ENSG00000230266 | XXYLT1-AS2 | 0,135825943 | 1,098721652 | 0,83569 | 0,92493625 | XXYLT1 antisense RNA 2 |
| ENSG00000078053 | AMPH | 0,135685515 | 1,09861471 | 0,67108 | 0,83700963 | amphiphysin |
| ENSG00000180694 | TMEM64 | 0,13564861 | 1,098586608 | 0,63812 | 0,81820518 | transmembrane protein 64 |
| ENSG00000102302 | FGD1 | 0,13562681 | 1,098570007 | 0,54195 | 0,75894841 | FYVE, RhoGEF and PH domain containing 1 |
| ENSG00000137714 | FDX1 | 0,135544356 | 1,098507223 | 0,50291 | 0,73217708 | ferredoxin 1 |
| ENSG00000268750 | AC010522,1 | 0,135542852 | 1,098506078 | 0,72248 | 0,86734497 | novel protein |
| ENSG00000162775 | RBM15 | 0,135447376 | 1,098433382 | 0,40452 | 0,66449258 | RNA binding motif protein 15 |
| ENSG00000256533 | AP002373,2 | 0,13542133 | 1,098413552 | 0,87765 | 0,94566951 | CCR4 carbon catabolite repression 4-like (S, cerevisiae) (CCRN4L) pseudogene |

| | | | | | | |
|---|---|---|---|---|---|---|
| ENSG00000239219 | AC008040,1 | 0,135329942 | 1,098343975 | 0,74777 | 0,88118711 | novel transcript, antisense to SEC62 |
| ENSG00000149480 | MTA2 | 0,135255803 | 1,098287533 | 0,44882 | 0,69458723 | metastasis associated 1 family member 2 |
| ENSG00000286459 | AP006287,3 | 0,135155806 | 1,09821141 | 0,83039 | 0,92226505 | novel transcript, antisense to GAL3ST3 |
| ENSG00000113648 | MACROH2A1 | 0,135056544 | 1,098135853 | 0,54524 | 0,7609742 | macroH2A,1 histone |
| ENSG00000160752 | FDPS | 0,135004127 | 1,098095955 | 0,40391 | 0,66409141 | farnesyl diphosphate synthase |
| ENSG00000174206 | C12orf66 | 0,134960166 | 1,098062495 | 0,64822 | 0,82383528 | chromosome 12 open reading frame 66 |
| ENSG00000235613 | NSRP1P1 | 0,134953671 | 1,098057551 | 0,72952 | 0,8715818 | nuclear speckle splicing regulatory protein 1 pseudogene 1 |
| ENSG00000285733 | AL031315,1 | 0,134843608 | 1,097973784 | 0,7531 | 0,88435474 | novel protein |
| ENSG00000151835 | SACS | 0,134651724 | 1,097827759 | 0,74715 | 0,88065286 | sacsin molecular chaperone |
| ENSG00000130332 | LSM7 | 0,134580585 | 1,097773627 | 0,48648 | 0,72196345 | LSM7 homolog, U6 small nuclear RNA and mRNA degradation associated |
| ENSG00000234772 | LINC00412 | 0,13456719 | 1,097763434 | 0,86733 | 0,94137971 | long intergenic non-protein coding RNA 412 |
| ENSG00000096155 | BAG6 | 0,134546854 | 1,097747961 | 0,62387 | 0,80952593 | BAG cochaperone 6 |
| ENSG00000139977 | NAA30 | 0,134544021 | 1,097745804 | 0,45984 | 0,7019161 | N-alpha-acetyltransferase 30, NatC catalytic subunit |
| ENSG00000092439 | TRPM7 | 0,13453078 | 1,09773573 | 0,41678 | 0,67361437 | transient receptor potential cation channel subfamily M member 7 |
| ENSG00000137161 | CNPY3 | 0,134440058 | 1,097666702 | 0,45209 | 0,69691985 | canopy FGF signaling regulator 3 |
| ENSG00000285084 | SLC35G5 | 0,134390714 | 1,09762916 | 0,82713 | 0,92010219 | solute carrier family 35 member G5 |
| ENSG00000173262 | SLC2A14 | 0,134352477 | 1,097600069 | 0,76332 | 0,88980026 | solute carrier family 2 member 14 |
| ENSG00000169733 | RFNG | 0,134346171 | 1,097595271 | 0,66618 | 0,83391259 | RFNG O-fucosylpeptide 3-beta-N-acetylglucosaminyltransferase |
| ENSG00000182979 | MTA1 | 0,134327233 | 1,097580863 | 0,42195 | 0,67643128 | metastasis associated 1 |
| ENSG00000080200 | CRYBG3 | 0,13431553 | 1,09757196 | 0,68587 | 0,84654659 | crystallin beta-gamma domain containing 3 |
| ENSG00000141441 | GAREM1 | 0,134162667 | 1,097455671 | 0,71399 | 0,86227395 | GRB2 associated regulator of MAPK1 subtype 1 |
| ENSG00000187688 | TRPV2 | 0,13405283 | 1,097372122 | 0,88355 | 0,94837933 | transient receptor potential cation channel subfamily V member 2 |
| ENSG00000103356 | EARS2 | 0,134007421 | 1,097337582 | 0,43124 | 0,68350322 | glutamyl-tRNA synthetase 2, mitochondrial |
| ENSG00000047346 | FAM214A | 0,133996811 | 1,097329512 | 0,73518 | 0,87480865 | family with sequence similarity 214 member A |
| ENSG00000140682 | TGFB1I1 | 0,133828179 | 1,097201256 | 0,73976 | 0,87659348 | transforming growth factor beta 1 induced transcript 1 |
| ENSG00000185278 | ZBTB37 | 0,133663288 | 1,09707586 | 0,51574 | 0,74180801 | zinc finger and BTB domain containing 37 |
| ENSG00000251474 | RPL32P3 | 0,13366124 | 1,097074302 | 0,4956 | 0,7278408 | ribosomal protein L32 pseudogene 3 |
| ENSG00000105778 | AVL9 | 0,13360161 | 1,097028959 | 0,44545 | 0,69255042 | AVL9 cell migration associated |
| ENSG00000144674 | GOLGA4 | 0,133552292 | 1,096991458 | 0,66955 | 0,83623395 | golgin A4 |

| | | | | | | |
|---|---|---|---|---|---|---|
| ENSG00000176476 | SGF29 | 0,133524449 | 1,096970287 | 0,59544 | 0,79336828 | SAGA complex associated factor 29 |
| ENSG00000136444 | RSAD1 | 0,133522814 | 1,096969044 | 0,50785 | 0,73602012 | radical S-adenosyl methionine domain containing 1 |
| ENSG00000105538 | RASIP1 | 0,133376947 | 1,096858138 | 0,71793 | 0,86489865 | Ras interacting protein 1 |
| ENSG00000100221 | JOSD1 | 0,133323145 | 1,096817234 | 0,43376 | 0,68475933 | Josephin domain containing 1 |
| ENSG00000165102 | HGSNAT | 0,133293453 | 1,09679466 | 0,50416 | 0,73315506 | heparan-alpha-glucosaminide N-acetyltransferase |
| ENSG00000145375 | SPATA5 | 0,133211579 | 1,096732418 | 0,34964 | 0,62563749 | spermatogenesis associated 5 |
| ENSG00000130768 | SMPDL3B | 0,1331549 | 1,096689332 | 0,84872 | 0,93107027 | sphingomyelin phosphodiesterase acid like 3B |
| ENSG00000145390 | USP53 | 0,132916724 | 1,096508294 | 0,74184 | 0,87793115 | ubiquitin specific peptidase 53 |
| ENSG00000288146 | NOC4L | 0,132880762 | 1,096480961 | 0,5862 | 0,78664449 | nucleolar complex associated 4 homolog |
| ENSG00000184967 | NOC4L | 0,132880762 | 1,096480961 | 0,5862 | 0,78664449 | nucleolar complex associated 4 homolog |
| ENSG00000204463 | BAG6 | 0,132835774 | 1,09644677 | 0,71512 | 0,86301991 | BAG cochaperone 6 |
| ENSG00000080802 | CNOT4 | 0,132698636 | 1,09634255 | 0,48096 | 0,71798792 | CCR4-NOT transcription complex subunit 4 |
| ENSG00000168779 | SHOX2 | 0,132696953 | 1,096341271 | 0,64315 | 0,82098995 | short stature homeobox 2 |
| ENSG00000177106 | EPS8L2 | 0,13268494 | 1,096332143 | 0,79797 | 0,90685415 | EPS8 like 2 |
| ENSG00000185627 | PSMD13 | 0,132632257 | 1,096292108 | 0,40625 | 0,66584658 | proteasome 26S subunit, non-ATPase 13 |
| ENSG00000055163 | CYFIP2 | 0,132600701 | 1,09626813 | 0,7306 | 0,87202167 | cytoplasmic FMR1 interacting protein 2 |
| ENSG00000270800 | RPS10-NUDT3 | 0,132562694 | 1,096239249 | 0,56321 | 0,77230901 | RPS10-NUDT3 readthrough |
| ENSG00000168026 | TTC21A | 0,132448875 | 1,096152766 | 0,61179 | 0,80337676 | tetratricopeptide repeat domain 21A |
| ENSG00000135365 | PHF21A | 0,132349216 | 1,096077049 | 0,53169 | 0,75280487 | PHD finger protein 21A |
| ENSG00000213024 | NUP62 | 0,132318597 | 1,096053786 | 0,42954 | 0,68228592 | nucleoporin 62 |
| ENSG00000264924 | AC090772,1 | 0,132279508 | 1,09602409 | 0,77233 | 0,89476608 | novel transcript, antisense to TTC39C |
| ENSG00000286786 | AC116158,3 | 0,132239387 | 1,09599361 | 0,57304 | 0,77878023 | novel transcript, antisense to RASGRP1 |
| ENSG00000071553 | ATP6AP1 | 0,132239297 | 1,095993541 | 0,47411 | 0,71294959 | ATPase H+ transporting accessory protein 1 |
| ENSG00000143499 | SMYD2 | 0,132084664 | 1,095876076 | 0,61906 | 0,80689825 | SET and MYND domain containing 2 |
| ENSG00000288038 | AL445588,1 | 0,132074322 | 1,09586822 | 0,8612 | 0,93791776 | novel transcript |
| ENSG00000224802 | TUBB4BP2 | 0,13204127 | 1,095843114 | 0,90851 | 0,96083985 | TUBB4B pseudogene 2 |
| ENSG00000086504 | MRPL28 | 0,131984021 | 1,095799629 | 0,53447 | 0,75405583 | mitochondrial ribosomal protein L28 |
| ENSG00000145979 | TBC1D7 | 0,131719047 | 1,095598387 | 0,57936 | 0,78270914 | TBC1 domain family member 7 |
| ENSG00000110844 | PRPF40B | 0,131601373 | 1,095509028 | 0,4254 | 0,67896955 | pre-mRNA processing factor 40 homolog B |
| ENSG00000286904 | AC093675,1 | 0,131597396 | 1,095506008 | 0,71311 | 0,86197885 | novel transcript, antisense to TMEM87B |
| ENSG00000006715 | VPS41 | 0,131576029 | 1,095489783 | 0,45776 | 0,70096886 | VPS41 subunit of HOPS complex |
| ENSG00000115137 | DNAJC27 | 0,13141345 | 1,095366338 | 0,64781 | 0,82380454 | DnaJ heat shock protein family (Hsp40) member C27 |
| ENSG00000268322 | BNIP3P25 | 0,131409374 | 1,095363243 | 0,79319 | 0,90371578 | BCL2 interacting protein 3 pseudogene 25 |

| | | | | | | |
|---|---|---|---|---|---|---|
| ENSG00000269981 | AL627309,7 | 0,131384945 | 1,095344696 | 0,85736 | 0,93562394 | novel pseudogene |
| ENSG00000126391 | FRMD8 | 0,131329242 | 1,095302405 | 0,66747 | 0,83496253 | FERM domain containing 8 |
| ENSG00000187742 | SECISBP2 | 0,131326045 | 1,095299978 | 0,53918 | 0,75757364 | SECIS binding protein 2 |
| ENSG00000204673 | AKT1S1 | 0,13127507 | 1,095261278 | 0,58674 | 0,78699486 | AKT1 substrate 1 |
| ENSG00000070269 | TMEM260 | 0,13122734 | 1,095225043 | 0,71042 | 0,86051357 | transmembrane protein 260 |
| ENSG00000215022 | AL008729,1 | 0,131205029 | 1,095208105 | 0,69761 | 0,8534853 | novel transcript, antisense to PHACTR1 |
| ENSG00000134453 | RBM17 | 0,131200982 | 1,095205033 | 0,45831 | 0,70119611 | RNA binding motif protein 17 |
| ENSG00000160058 | BSDC1 | 0,131158397 | 1,095172706 | 0,41264 | 0,6702832 | BSD domain containing 1 |
| ENSG00000135164 | DMTF1 | 0,131152973 | 1,095168588 | 0,58662 | 0,78690792 | cyclin D binding myb like transcription factor 1 |
| ENSG00000015676 | NUDCD3 | 0,131117786 | 1,095141878 | 0,42973 | 0,68246875 | NudC domain containing 3 |
| ENSG00000130985 | UBA1 | 0,131038659 | 1,095081815 | 0,48611 | 0,72163783 | ubiquitin like modifier activating enzyme 1 |
| ENSG00000287584 | AC004837,4 | 0,130963357 | 1,095024658 | 0,76087 | 0,88871285 | novel transcript |
| ENSG00000144560 | VGLL4 | 0,13084968 | 1,094938379 | 0,46446 | 0,70506031 | vestigial like family member 4 |
| ENSG00000114098 | ARMC8 | 0,130817133 | 1,094913678 | 0,52992 | 0,7516047 | armadillo repeat containing 8 |
| ENSG00000144028 | SNRNP200 | 0,130757792 | 1,094868642 | 0,51332 | 0,73972098 | small nuclear ribonucleoprotein U5 subunit 200 |
| ENSG00000156983 | BRPF1 | 0,130725176 | 1,094843891 | 0,50835 | 0,73649214 | bromodomain and PHD finger containing 1 |
| ENSG00000198142 | SOWAHC | 0,130707835 | 1,094830731 | 0,62812 | 0,81178582 | sosondowah ankyrin repeat domain family member C |
| ENSG00000141252 | VPS53 | 0,130690702 | 1,094817728 | 0,73953 | 0,87648643 | VPS53 subunit of GARP complex |
| ENSG00000119537 | KDSR | 0,130639693 | 1,09477902 | 0,38135 | 0,64874863 | 3-ketodihydrosphingosine reductase |
| ENSG00000288455 | SEMA7A | 0,130600724 | 1,09474945 | 0,68895 | 0,84847234 | semaphorin 7A (John Milton Hagen blood group) |
| ENSG00000138623 | SEMA7A | 0,130600724 | 1,09474945 | 0,68895 | 0,84847234 | semaphorin 7A (John Milton Hagen blood group) |
| ENSG00000275763 | ZNF516-DT | 0,130486059 | 1,094662442 | 0,82074 | 0,9173395 | ZNF516 divergent transcript |
| ENSG00000245694 | CRNDE | 0,130449048 | 1,09463436 | 0,68538 | 0,84616325 | colorectal neoplasia differentially expressed |
| ENSG00000173230 | GOLGB1 | 0,130424401 | 1,094615659 | 0,64911 | 0,82436914 | golgin B1 |
| ENSG00000144426 | NBEAL1 | 0,130309967 | 1,094528839 | 0,62081 | 0,80776776 | neurobeachin like 1 |
| ENSG00000167554 | ZNF610 | 0,130308785 | 1,094527942 | 0,56163 | 0,77136826 | zinc finger protein 610 |
| ENSG00000225163 | LINC00618 | 0,130277708 | 1,094504365 | 0,82447 | 0,91900665 | long intergenic non-protein coding RNA 618 |
| ENSG00000100596 | SPTLC2 | 0,130237762 | 1,09447406 | 0,54175 | 0,75882662 | serine palmitoyltransferase long chain base subunit 2 |
| ENSG00000011009 | LYPLA2 | 0,13015239 | 1,094409297 | 0,56395 | 0,77259188 | lysophospholipase 2 |
| ENSG00000182134 | TDRKH | 0,130074205 | 1,094349988 | 0,69825 | 0,85390309 | tudor and KH domain containing |
| ENSG00000256340 | ABCC6P1 | 0,130015888 | 1,094305752 | 0,8875 | 0,95041376 | ATP binding cassette subfamily C member 6 pseudogene 1 |
| ENSG00000128951 | DUT | 0,129931958 | 1,094242092 | 0,68607 | 0,84660425 | deoxyuridine triphosphatase |

| | | | | | | |
|---|---|---|---|---|---|---|
| ENSG00000286220 | AC007546,3 | 0,129889631 | 1,094209989 | 0,63978 | 0,81925013 | purinergic receptor P2X, ligand-gated ion channel, 7 (P2RX7) pseudogene |
| ENSG00000186417 | GLDN | 0,129756252 | 1,094108833 | 0,77908 | 0,89781842 | gliomedin |
| ENSG00000196776 | CD47 | 0,129725807 | 1,094085744 | 0,65069 | 0,82499358 | CD47 molecule |
| ENSG00000167080 | B4GALNT2 | 0,129649982 | 1,094028243 | 0,71292 | 0,86184574 | beta-1,4-N-acetyl-galactosaminyltransferase 2 |
| ENSG00000111737 | RAB35 | 0,129647318 | 1,094026222 | 0,55993 | 0,77076855 | RAB35, member RAS oncogene family |
| ENSG00000112167 | SAYSD1 | 0,129557708 | 1,093958271 | 0,64403 | 0,82138283 | SAYSVFN motif domain containing 1 |
| ENSG00000234129 | AC073529,1 | 0,129525297 | 1,093933695 | 0,79225 | 0,90351559 | novel transcript |
| ENSG00000282961 | PRNCR1 | 0,129490297 | 1,093907156 | 0,636 | 0,81690517 | prostate cancer associated non-coding RNA 1 |
| ENSG00000286365 | AP000553,9 | 0,129481956 | 1,093900832 | 0,75281 | 0,88421955 | novel transcript, antisense to YPEL1 |
| ENSG00000261173 | AC018845,3 | 0,129468236 | 1,093890429 | 0,88111 | 0,947073 | novel transcript, antisense to DNAJA2 |
| ENSG00000184381 | PLA2G6 | 0,129464082 | 1,093887279 | 0,53578 | 0,75487955 | phospholipase A2 group VI |
| ENSG00000265095 | FTLP12 | 0,129457774 | 1,093882496 | 0,90138 | 0,95798991 | ferritin light chain pseudogene 12 |
| ENSG00000140577 | CRTC3 | 0,129426146 | 1,093858515 | 0,54047 | 0,75829455 | CREB regulated transcription coactivator 3 |
| ENSG00000146700 | SSC4D | 0,129374689 | 1,093819501 | 0,79802 | 0,90685415 | scavenger receptor cysteine rich family member with 4 domains |
| ENSG00000183444 | OR7E38P | 0,12926023 | 1,093732724 | 0,78779 | 0,90125214 | olfactory receptor family 7 subfamily E member 38 pseudogene |
| ENSG00000103550 | KNOP1 | 0,129258206 | 1,09373119 | 0,48139 | 0,71822668 | lysine rich nucleolar protein 1 |
| ENSG00000151466 | SCLT1 | 0,129244802 | 1,093721029 | 0,48237 | 0,71892541 | sodium channel and clathrin linker 1 |
| ENSG00000113240 | CLK4 | 0,129104699 | 1,09361482 | 0,53438 | 0,75405583 | CDC like kinase 4 |
| ENSG00000165914 | TTC7B | 0,129026478 | 1,093555528 | 0,38894 | 0,65351197 | tetratricopeptide repeat domain 7B |
| ENSG00000253741 | LNCOC1 | 0,128967676 | 1,093510957 | 0,78371 | 0,89918294 | lncRNA associated with ovarian cancer 1 |
| ENSG00000164129 | NPY5R | 0,128927335 | 1,093480381 | 0,84203 | 0,92818855 | neuropeptide Y receptor Y5 |
| ENSG00000134046 | MBD2 | 0,12884858 | 1,09342069 | 0,46913 | 0,7086228 | methyl-CpG binding domain protein 2 |
| ENSG00000147481 | SNTG1 | 0,128843882 | 1,093417129 | 0,90056 | 0,95772302 | syntrophin gamma 1 |
| ENSG00000236044 | FABP5P2 | 0,12883652 | 1,09341155 | 0,91138 | 0,96169573 | fatty acid binding protein 5 pseudogene 2 |
| ENSG00000188559 | RALGAPA2 | 0,12874355 | 1,093341091 | 0,65553 | 0,82781192 | Ral GTPase activating protein catalytic subunit alpha 2 |
| ENSG00000230131 | LINC02641 | 0,128722561 | 1,093325184 | 0,90664 | 0,96009372 | long intergenic non-protein coding RNA 2641 |
| ENSG00000088833 | NSFL1C | 0,128696279 | 1,093305267 | 0,45251 | 0,69702004 | NSFL1 cofactor |
| ENSG00000015568 | RGPD5 | 0,128633301 | 1,093257542 | 0,8193 | 0,91699219 | RANBP2 like and GRIP domain containing 5 |
| ENSG00000128487 | SPECC1 | 0,128629031 | 1,093254306 | 0,40355 | 0,66399439 | sperm antigen with calponin homology and coiled-coil domains 1 |

| | | | | | | |
|---|---|---|---|---|---|---|
| ENSG00000226144 | RPS27AP3 | 0,128502935 | 1,093158757 | 0,86163 | 0,93811282 | RPS27A pseudogene 3 |
| ENSG00000224184 | MIR3681HG | 0,128438837 | 1,093110189 | 0,7897 | 0,90258817 | MIR3681 host gene |
| ENSG00000166971 | AKTIP | 0,128434828 | 1,093107151 | 0,69734 | 0,85330122 | AKT interacting protein |
| ENSG00000115241 | PPM1G | 0,128398231 | 1,093079423 | 0,50826 | 0,73640172 | protein phosphatase, Mg2+/Mn2+ dependent 1G |
| ENSG00000174989 | FBXW8 | 0,128369743 | 1,093057839 | 0,42219 | 0,67663014 | F-box and WD repeat domain containing 8 |
| ENSG00000271009 | AC116667,1 | 0,128358003 | 1,093048944 | 0,85776 | 0,93584401 | novel transcript |
| ENSG00000011198 | ABHD5 | 0,128199979 | 1,092929225 | 0,63699 | 0,81742435 | abhydrolase domain containing 5, lysophosphatidic acid acyltransferase |
| ENSG00000185842 | DNAH14 | 0,128197212 | 1,092927129 | 0,5036 | 0,732867 | dynein axonemal heavy chain 14 |
| ENSG00000286951 | AL356113,1 | 0,128177423 | 1,092912137 | 0,79062 | 0,90286944 | novel transcript, antisense to KIAA1217 |
| ENSG00000244968 | LIFR-AS1 | 0,128029009 | 1,092799713 | 0,644 | 0,82138283 | LIFR antisense RNA 1 |
| ENSG00000205726 | ITSN1 | 0,127805991 | 1,092630796 | 0,54014 | 0,75814808 | intersectin 1 |
| ENSG00000262899 | AC004232,1 | 0,127736831 | 1,092578418 | 0,95622 | 0,98189243 | novel transcript |
| ENSG00000149100 | EIF3M | 0,127716372 | 1,092562925 | 0,49155 | 0,72511953 | eukaryotic translation initiation factor 3 subunit M |
| ENSG00000281697 | AP003392,10 | 0,127678569 | 1,092534296 | 0,85413 | 0,93380868 | potassium channel tetramerisation domain containing 6 (KCTD6) pseudogene |
| ENSG00000254909 | AP003392,2 | 0,127678569 | 1,092534296 | 0,85413 | 0,93380868 | potassium channel tetramerisation domain containing 6 (KCTD6) pseudogene |
| ENSG00000272173 | U47924,2 | 0,127651 | 1,092513419 | 0,62585 | 0,81058216 | novel transcript, antisense to C12orf57 |
| ENSG00000133895 | MEN1 | 0,127619662 | 1,092489688 | 0,47138 | 0,71072395 | menin 1 |
| ENSG00000130876 | SLC7A10 | 0,127613183 | 1,092484782 | 0,7688 | 0,89277929 | solute carrier family 7 member 10 |
| ENSG00000141522 | ARHGDIA | 0,127586401 | 1,092464501 | 0,61375 | 0,8035281 | Rho GDP dissociation inhibitor alpha |
| ENSG00000238105 | GOLGA2P5 | 0,127493502 | 1,092394157 | 0,73978 | 0,87659348 | GOLGA2 pseudogene 5 |
| ENSG00000178719 | GRINA | 0,127446033 | 1,092358214 | 0,53488 | 0,75424675 | glutamate ionotropic receptor NMDA type subunit associated protein 1 |
| ENSG00000272163 | AF106564,1 | 0,127443151 | 1,092356032 | 0,59586 | 0,79353726 | novel transcript, antisense to NEFM |
| ENSG00000128973 | CLN6 | 0,127369303 | 1,092300119 | 0,5986 | 0,7951852 | CLN6 transmembrane ER protein |
| ENSG00000207205 | RNVU1-15 | 0,127224812 | 1,092190726 | 0,8068 | 0,91120963 | RNA, variant U1 small nuclear 15 |
| ENSG00000287893 | AC083864,5 | 0,12722368 | 1,092189869 | 0,74374 | 0,87871466 | novel transcript |
| ENSG00000134899 | ERCC5 | 0,127071924 | 1,092074988 | 0,57699 | 0,78124459 | ERCC excision repair 5, endonuclease |
| ENSG00000166169 | POLL | 0,126875799 | 1,091926538 | 0,53341 | 0,75351438 | DNA polymerase lambda |
| ENSG00000106789 | CORO2A | 0,126854295 | 1,091910263 | 0,79235 | 0,90358057 | coronin 2A |

| | | | | | | |
|---|---|---|---|---|---|---|
| ENSG00000256159 | AC022080,2 | 0,126818486 | 1,091883161 | 0,84994 | 0,93170188 | mitochondrial ribosomal protein L30 (MRPL30) pseudogene |
| ENSG00000288283 | RFX1 | 0,126787895 | 1,091860009 | 0,50971 | 0,73740498 | regulatory factor X1 |
| ENSG00000132005 | RFX1 | 0,126787895 | 1,091860009 | 0,50971 | 0,73740498 | regulatory factor X1 |
| ENSG00000164588 | HCN1 | 0,126784188 | 1,091857203 | 0,82213 | 0,91790528 | hyperpolarization activated cyclic nucleotide gated potassium channel 1 |
| ENSG00000259113 | AL118556,1 | 0,126747957 | 1,091829783 | 0,86284 | 0,9388179 | novel transcript |
| ENSG00000181072 | CHRM2 | 0,126529411 | 1,0916644 | 0,78884 | 0,90208756 | cholinergic receptor muscarinic 2 |
| ENSG00000136935 | GOLGA1 | 0,126478284 | 1,091625714 | 0,66618 | 0,83391259 | golgin A1 |
| ENSG00000259155 | STK16P1 | 0,126439251 | 1,09159618 | 0,80544 | 0,91059395 | serine/threonine kinase 16 pseudogene 1 |
| ENSG00000166669 | ATF7IP2 | 0,126429063 | 1,091588471 | 0,6397 | 0,81918091 | activating transcription factor 7 interacting protein 2 |
| ENSG00000187109 | NAP1L1 | 0,126381174 | 1,091552238 | 0,40768 | 0,66682187 | nucleosome assembly protein 1 like 1 |
| ENSG00000111011 | RSRC2 | 0,126368933 | 1,091542976 | 0,59026 | 0,78971843 | arginine and serine rich coiled-coil 2 |
| ENSG00000136463 | TACO1 | 0,126350617 | 1,091529118 | 0,48001 | 0,71732148 | translational activator of cytochrome c oxidase I |
| ENSG00000231845 | HMGB3P14 | 0,126309474 | 1,09149799 | 0,84853 | 0,93094401 | high mobility group box 3 pseudogene 14 |
| ENSG00000214223 | HNRNPA1P10 | 0,126157351 | 1,091382905 | 0,73067 | 0,87205836 | heterogeneous nuclear ribonucleoprotein A1 pseudogene 10 |
| ENSG00000175183 | CSRP2 | 0,126151449 | 1,09137844 | 0,7226 | 0,86745114 | cysteine and glycine rich protein 2 |
| ENSG00000224831 | TMEM183B | 0,126093552 | 1,091334642 | 0,8899 | 0,95152291 | transmembrane protein 183B |
| ENSG00000251615 | AC104825,1 | 0,12599426 | 1,091259535 | 0,69615 | 0,85287022 | novel transcript |
| ENSG00000134882 | UBAC2 | 0,125827956 | 1,09113375 | 0,52208 | 0,7463592 | UBA domain containing 2 |
| ENSG00000254463 | PPIAP41 | 0,12582637 | 1,09113255 | 0,8037 | 0,90969283 | peptidylprolyl isomerase A pseudogene 41 |
| ENSG00000276726 | DNAJA3 | 0,125819448 | 1,091127315 | 0,509 | 0,7370222 | DnaJ heat shock protein family (Hsp40) member A3 |
| ENSG00000128739 | SNRPN | 0,125693453 | 1,091032028 | 0,60249 | 0,79710174 | small nuclear ribonucleoprotein polypeptide N |
| ENSG00000109062 | SLC9A3R1 | 0,12564269 | 1,090993639 | 0,63494 | 0,8162492 | SLC9A3 regulator 1 |
| ENSG00000197603 | CPLANE1 | 0,125618758 | 1,090975541 | 0,6119 | 0,80337676 | ciliogenesis and planar polarity effector 1 |
| ENSG00000173715 | C11orf80 | 0,125561454 | 1,090932209 | 0,56536 | 0,77366076 | chromosome 11 open reading frame 80 |
| ENSG00000113068 | PFDN1 | 0,125488626 | 1,090877139 | 0,55157 | 0,76535107 | prefoldin subunit 1 |
| ENSG00000166233 | ARIH1 | 0,125459583 | 1,090855179 | 0,41626 | 0,67307944 | ariadne RBR E3 ubiquitin protein ligase 1 |
| ENSG00000166311 | SMPD1 | 0,12544649 | 1,090845279 | 0,57733 | 0,78136026 | sphingomyelin phosphodiesterase 1 |
| ENSG00000259096 | SHLD2P2 | 0,125328661 | 1,09075619 | 0,80485 | 0,91011425 | shieldin complex subunit 2 pseudogene 2 |
| ENSG00000269907 | PTMAP11 | 0,125258161 | 1,090702889 | 0,84422 | 0,92888169 | prothymosin alpha pseudogene 11 |
| ENSG00000005448 | WDR54 | 0,125224172 | 1,090677193 | 0,68532 | 0,84616069 | WD repeat domain 54 |

| | | | | | | |
|---|---|---|---|---|---|---|
| ENSG00000231468 | PRDX3P2 | 0,125223983 | 1,090677051 | 0,90105 | 0,95787401 | peroxiredoxin 3 pseudogene 2 |
| ENSG00000139323 | POC1B | 0,125213156 | 1,090668866 | 0,57245 | 0,77849297 | POC1 centriolar protein B |
| ENSG00000106772 | PRUNE2 | 0,125200954 | 1,090659641 | 0,7794 | 0,89794855 | prune homolog 2 with BCH domain |
| ENSG00000270292 | AC079880,2 | 0,125200722 | 1,090659466 | 0,79534 | 0,90519396 | tubulin, alpha 4a (TUBA4A) pseudogene |
| ENSG00000198963 | RORB | 0,125020156 | 1,090522969 | 0,77589 | 0,89635267 | RAR related orphan receptor B |
| ENSG00000269189 | AL445223,1 | 0,125012342 | 1,090517061 | 0,81905 | 0,91689679 | novel transcript, sense intronic to FARP1 |
| ENSG00000259659 | AC009996,1 | 0,124974771 | 1,090488663 | 0,79074 | 0,90287625 | novel transcript |
| ENSG00000099817 | POLR2E | 0,124945815 | 1,090466776 | 0,4373 | 0,68715611 | RNA polymerase II, I and III subunit E |
| ENSG00000141480 | ARRB2 | 0,12491458 | 1,090443167 | 0,60277 | 0,7971742 | arrestin beta 2 |
| ENSG00000245849 | RAD51-AS1 | 0,124887591 | 1,090422768 | 0,60888 | 0,80151558 | RAD51 antisense RNA 1 |
| ENSG00000120594 | PLXDC2 | 0,124834785 | 1,090382857 | 0,68966 | 0,84865812 | plexin domain containing 2 |
| ENSG00000162695 | SLC30A7 | 0,124692049 | 1,090274983 | 0,46234 | 0,70356854 | solute carrier family 30 member 7 |
| ENSG00000065054 | SLC9A3R2 | 0,124684449 | 1,090269239 | 0,75045 | 0,88291013 | SLC9A3 regulator 2 |
| ENSG00000257303 | AC073896,2 | 0,124655631 | 1,090247461 | 0,75172 | 0,88361939 | novel transcript, antisense to CNPY2 and CS |
| ENSG00000165259 | HDX | 0,124634069 | 1,090231167 | 0,75205 | 0,88386445 | highly divergent homeobox |
| ENSG00000211452 | DIO1 | 0,124510568 | 1,090137842 | 0,83205 | 0,92312766 | iodothyronine deiodinase 1 |
| ENSG00000273013 | AC117490,2 | 0,12449725 | 1,090127779 | 0,86132 | 0,9380166 | novel transcript, antisense to RNF168 |
| ENSG00000272430 | LINC02637 | 0,124447383 | 1,090090099 | 0,89429 | 0,95389664 | long intergenic non-protein coding RNA 2637 |
| ENSG00000146425 | DYNLT1 | 0,124396029 | 1,090051297 | 0,60706 | 0,80003503 | dynein light chain Tctex-type 1 |
| ENSG00000258461 | AC012651,1 | 0,12433185 | 1,090002806 | 0,90302 | 0,95846794 | novel transcript |
| ENSG00000227671 | AL390728,4 | 0,124315617 | 1,089990542 | 0,66343 | 0,83242946 | zinc finger protein pseudogene |
| ENSG00000258732 | AC025884,1 | 0,124286136 | 1,089968269 | 0,68625 | 0,84663586 | arginine-glutamic acid dipeptide (RE) repeats (RERE) pseudogene |
| ENSG00000099256 | PRTFDC1 | 0,124251543 | 1,089942134 | 0,76508 | 0,8907696 | phosphoribosyl transferase domain containing 1 |
| ENSG00000287466 | AL355531,1 | 0,124118432 | 1,089841575 | 0,80993 | 0,91240493 | novel transcript, antisense to MGMT |
| ENSG00000258405 | ZNF578 | 0,124040909 | 1,089783014 | 0,66799 | 0,83527385 | zinc finger protein 578 |
| ENSG00000094631 | HDAC6 | 0,123935159 | 1,089703135 | 0,56037 | 0,77078479 | histone deacetylase 6 |
| ENSG00000229800 | ATP8A2P2 | 0,123898608 | 1,089675528 | 0,84636 | 0,93016887 | ATPase phospholipid transporting 8A2 pseudogene 2 |
| ENSG00000162814 | SPATA17 | 0,123887611 | 1,089667221 | 0,67786 | 0,84154221 | spermatogenesis associated 17 |
| ENSG00000104957 | CCDC130 | 0,123799963 | 1,089601023 | 0,53511 | 0,7543529 | coiled-coil domain containing 130 |
| ENSG00000226741 | LINC02554 | 0,123793781 | 1,089596354 | 0,85536 | 0,93435351 | long intergenic non-protein coding RNA 2554 |
| ENSG00000120251 | GRIA2 | 0,123753922 | 1,089566251 | 0,84697 | 0,93041089 | glutamate ionotropic receptor AMPA type subunit 2 |
| ENSG00000044446 | PHKA2 | 0,123722647 | 1,089542631 | 0,57677 | 0,7810482 | phosphorylase kinase regulatory subunit alpha 2 |

| | | | | | | |
|---|---|---|---|---|---|---|
| ENSG00000137200 | CMTR1 | 0,123659207 | 1,089494721 | 0,40287 | 0,66398107 | cap methyltransferase 1 |
| ENSG00000153015 | CWC27 | 0,123635262 | 1,089476639 | 0,63207 | 0,81450663 | CWC27 spliceosome associated cyclophilin |
| ENSG00000102554 | KLF5 | 0,123612687 | 1,089459591 | 0,73763 | 0,87540065 | Kruppel like factor 5 |
| ENSG00000119335 | SET | 0,123523645 | 1,089392353 | 0,40292 | 0,66398107 | SET nuclear proto-oncogene |
| ENSG00000260876 | LINC01229 | 0,123483847 | 1,089362301 | 0,74665 | 0,88057211 | long intergenic non-protein coding RNA 1229 |
| ENSG00000055950 | MRPL43 | 0,123444555 | 1,089332633 | 0,66081 | 0,83092106 | mitochondrial ribosomal protein L43 |
| ENSG00000068745 | IP6K2 | 0,123443732 | 1,089332012 | 0,47496 | 0,71364583 | inositol hexakisphosphate kinase 2 |
| ENSG00000091986 | CCDC80 | 0,123412218 | 1,089308217 | 0,75468 | 0,88505717 | coiled-coil domain containing 80 |
| ENSG00000171552 | BCL2L1 | 0,123384548 | 1,089287325 | 0,47729 | 0,71524229 | BCL2 like 1 |
| ENSG00000162664 | ZNF326 | 0,123308663 | 1,08923003 | 0,47342 | 0,71264815 | zinc finger protein 326 |
| ENSG00000228728 | PPIAP91 | 0,123213182 | 1,089157944 | 0,87177 | 0,94335287 | peptidylprolyl isomerase A pseudogene 91 |
| ENSG00000109323 | MANBA | 0,123140538 | 1,089103104 | 0,75442 | 0,8849381 | mannosidase beta |
| ENSG00000227051 | C14orf132 | 0,123093666 | 1,08906772 | 0,56308 | 0,77224059 | chromosome 14 open reading frame 132 |
| ENSG00000254582 | PSMA2P1 | 0,123025867 | 1,089016541 | 0,85185 | 0,93259393 | proteasome subunit alpha 2 pseudogene 1 |
| ENSG00000172943 | PHF8 | 0,123010325 | 1,089004809 | 0,54487 | 0,76084582 | PHD finger protein 8 |
| ENSG00000204271 | SPIN3 | 0,122823123 | 1,088863511 | 0,65546 | 0,82781124 | spindlin family member 3 |
| ENSG00000168781 | PPIP5K1 | 0,122818012 | 1,088859653 | 0,54682 | 0,76240845 | diphosphoinositol pentakisphosphate kinase 1 |
| ENSG00000106636 | YKT6 | 0,122815787 | 1,088857974 | 0,47195 | 0,71123477 | YKT6 v-SNARE homolog |
| ENSG00000204160 | ZDHHC18 | 0,122793589 | 1,08884122 | 0,43276 | 0,68421938 | zinc finger DHHC-type palmitoyltransferase 18 |
| ENSG00000188706 | ZDHHC9 | 0,122783133 | 1,088833329 | 0,65639 | 0,82832937 | zinc finger DHHC-type palmitoyltransferase 9 |
| ENSG00000274266 | SNORA73A | 0,122737188 | 1,088798653 | 0,66982 | 0,83624601 | small nucleolar RNA, H/ACA box 73A |
| ENSG00000226752 | CUTALP | 0,122672348 | 1,088749721 | 0,55108 | 0,76487698 | cutA divalent cation tolerance like, pseudogene |
| ENSG00000223668 | EEF1A1P24 | 0,122664063 | 1,088743468 | 0,8635 | 0,93904284 | eukaryotic translation elongation factor 1 alpha 1 pseudogene 24 |
| ENSG00000152102 | FAM168B | 0,122623613 | 1,088712942 | 0,56182 | 0,77145675 | family with sequence similarity 168 member B |
| ENSG00000082805 | ERC1 | 0,122576237 | 1,088677191 | 0,53378 | 0,75368609 | ELKS/RAB6-interacting/CAST family member 1 |
| ENSG00000248550 | OTX2-AS1 | 0,122567808 | 1,08867083 | 0,81548 | 0,91505837 | OTX2 antisense RNA 1 (head to head) |
| ENSG00000057294 | PKP2 | 0,122539864 | 1,088649744 | 0,78638 | 0,90080813 | plakophilin 2 |
| ENSG00000168939 | SPRY3 | 0,122461826 | 1,088590859 | 0,66934 | 0,83617811 | sprouty RTK signaling antagonist 3 |
| ENSG00000132612 | VPS4A | 0,122441693 | 1,088575667 | 0,44186 | 0,68994635 | vacuolar protein sorting 4 homolog A |
| ENSG00000100445 | SDR39U1 | 0,122421757 | 1,088560624 | 0,6252 | 0,81024216 | short chain dehydrogenase/reductase family 39U member 1 |
| ENSG00000229927 | RHEBP1 | 0,122406847 | 1,088549375 | 0,91656 | 0,9636258 | RHEB pseudogene 1 |

| | | | | | | |
|---|---|---|---|---|---|---|
| ENSG00000276571 | AC002550,2 | 0,122390534 | 1,088537066 | 0,8082 | 0,9117763 | novel transcript, antisense to C16orf62 |
| ENSG00000075218 | GTSE1 | 0,122249524 | 1,088430677 | 0,54042 | 0,75829455 | G2 and S-phase expressed 1 |
| ENSG00000185745 | IFIT1 | 0,122233387 | 1,088418502 | 0,64011 | 0,81935089 | interferon induced protein with tetratricopeptide repeats 1 |
| ENSG00000263465 | SRSF8 | 0,122230442 | 1,088416281 | 0,65384 | 0,82668269 | serine and arginine rich splicing factor 8 |
| ENSG00000152932 | RAB3C | 0,122223587 | 1,088411109 | 0,80135 | 0,90860375 | RAB3C, member RAS oncogene family |
| ENSG00000214077 | GNAQP1 | 0,122222493 | 1,088410284 | 0,82568 | 0,91917221 | G protein subunit alpha q pseudogene 1 |
| ENSG00000187239 | FNBP1 | 0,122197766 | 1,088391629 | 0,61054 | 0,80235359 | formin binding protein 1 |
| ENSG00000227518 | MIR1302-9HG | 0,122174703 | 1,08837423 | 0,46791 | 0,70761432 | MIR1302-9 host gene |
| ENSG00000227455 | AL450345,1 | 0,121971074 | 1,088220623 | 0,88074 | 0,94687991 | novel transcript |
| ENSG00000244509 | APOBEC3C | 0,121918347 | 1,088180852 | 0,73016 | 0,87180309 | apolipoprotein B mRNA editing enzyme catalytic subunit 3C |
| ENSG00000175938 | ORAI3 | 0,121856268 | 1,088134028 | 0,65927 | 0,82981653 | ORAI calcium release-activated calcium modulator 3 |
| ENSG00000141298 | SSH2 | 0,121800191 | 1,088091734 | 0,50661 | 0,73503941 | slingshot protein phosphatase 2 |
| ENSG00000143436 | MRPL9 | 0,121668102 | 1,087992115 | 0,49095 | 0,7249177 | mitochondrial ribosomal protein L9 |
| ENSG00000011638 | TMEM159 | 0,121605314 | 1,087944766 | 0,80411 | 0,909789 | transmembrane protein 159 |
| ENSG00000153707 | PTPRD | 0,121556938 | 1,087908286 | 0,81763 | 0,91621442 | protein tyrosine phosphatase receptor type D |
| ENSG00000244515 | KRT18P34 | 0,12154706 | 1,087900837 | 0,82324 | 0,91829029 | keratin 18 pseudogene 34 |
| ENSG00000164062 | APEH | 0,121495103 | 1,087861658 | 0,54275 | 0,75956237 | acylaminoacyl-peptide hydrolase |
| ENSG00000179604 | CDC42EP4 | 0,121467443 | 1,087840802 | 0,58953 | 0,78890126 | CDC42 effector protein 4 |
| ENSG00000109133 | TMEM33 | 0,121465882 | 1,087839625 | 0,4022 | 0,66353215 | transmembrane protein 33 |
| ENSG00000134864 | GGACT | 0,121463026 | 1,087837471 | 0,6265 | 0,81097268 | gamma-glutamylamine cyclotransferase |
| ENSG00000230359 | TPI1P2 | 0,121459937 | 1,087835142 | 0,77466 | 0,89586241 | triosephosphate isomerase 1 pseudogene 2 |
| ENSG00000097007 | ABL1 | 0,121408171 | 1,087796109 | 0,5077 | 0,73588248 | ABL proto-oncogene 1, non-receptor tyrosine kinase |
| ENSG00000103978 | TMEM87A | 0,121381892 | 1,087776295 | 0,50068 | 0,73097006 | transmembrane protein 87A |
| ENSG00000108773 | KAT2A | 0,121331553 | 1,087738341 | 0,60252 | 0,79710677 | lysine acetyltransferase 2A |
| ENSG00000204131 | NHSL2 | 0,121327318 | 1,087735148 | 0,79911 | 0,90737821 | NHS like 2 |
| ENSG00000273611 | ZNHIT3 | 0,121319674 | 1,087729384 | 0,65731 | 0,82868311 | zinc finger HIT-type containing 3 |
| ENSG00000131941 | RHPN2 | 0,121208207 | 1,087645347 | 0,65321 | 0,82629393 | rhophilin Rho GTPase binding protein 2 |
| ENSG00000287839 | AL353807,5 | 0,121170154 | 1,087616659 | 0,73756 | 0,87540065 | novel transcript |
| ENSG00000167930 | FAM234A | 0,121168907 | 1,087615718 | 0,43721 | 0,68711446 | family with sequence similarity 234 member A |
| ENSG00000225930 | LINC02249 | 0,121168721 | 1,087615578 | 0,69058 | 0,84900677 | long intergenic non-protein coding RNA 2249 |
| ENSG00000102606 | ARHGEF7 | 0,121130903 | 1,087587069 | 0,58197 | 0,78454692 | Rho guanine nucleotide exchange factor 7 |

| | | | | | | |
|---|---|---|---|---|---|---|
| ENSG00000273080 | AC009309,1 | 0,121111598 | 1,087572515 | 0,84295 | 0,9282764 | novel transcript, antisense to IMMT |
| ENSG00000201772 | SNORA5C | 0,120949616 | 1,087450413 | 0,83624 | 0,92493625 | small nucleolar RNA, H/ACA box 5C |
| ENSG00000149636 | DSN1 | 0,120631286 | 1,087210494 | 0,59158 | 0,79071076 | DSN1 component of MIS12 kinetochore complex |
| ENSG00000175322 | ZNF519 | 0,120499604 | 1,087111263 | 0,54903 | 0,76394298 | zinc finger protein 519 |
| ENSG00000137054 | POLR1E | 0,120496326 | 1,087108793 | 0,52541 | 0,74903593 | RNA polymerase I subunit E |
| ENSG00000160075 | SSU72 | 0,120449259 | 1,087073327 | 0,47564 | 0,71419272 | SSU72 homolog, RNA polymerase II CTD phosphatase |
| ENSG00000141349 | G6PC3 | 0,120414834 | 1,087047388 | 0,47402 | 0,71294959 | glucose-6-phosphatase catalytic subunit 3 |
| ENSG00000105568 | PPP2R1A | 0,12039654 | 1,087033604 | 0,48249 | 0,71900202 | protein phosphatase 2 scaffold subunit Aalpha |
| ENSG00000287064 | AL606500,1 | 0,120388849 | 1,08702781 | 0,73089 | 0,87206482 | novel transcript |
| ENSG00000132155 | RAF1 | 0,12035024 | 1,086998719 | 0,49361 | 0,72640121 | Raf-1 proto-oncogene, serine/threonine kinase |
| ENSG00000182544 | MFSD5 | 0,120326319 | 1,086980695 | 0,61304 | 0,8035281 | major facilitator superfamily domain containing 5 |
| ENSG00000257596 | SCAT2 | 0,12029029 | 1,086953551 | 0,7465 | 0,88046587 | S-phase cancer associated transcript 2 |
| ENSG00000157703 | SVOPL | 0,120227918 | 1,086906559 | 0,85285 | 0,93314746 | SVOP like |
| ENSG00000231105 | ECE1-AS1 | 0,120225817 | 1,086904977 | 0,83479 | 0,9244867 | ECE1 antisense RNA 1 |
| ENSG00000225193 | RPS12P26 | 0,120206916 | 1,086890737 | 0,87822 | 0,9458146 | ribosomal protein S12 pseudogene 26 |
| ENSG00000168807 | SNTB2 | 0,120199708 | 1,086885306 | 0,47072 | 0,71021824 | syntrophin beta 2 |
| ENSG00000119888 | EPCAM | 0,120123517 | 1,086827908 | 0,71094 | 0,86079022 | epithelial cell adhesion molecule |
| ENSG00000269559 | AC093677,2 | 0,120043961 | 1,086767978 | 0,74971 | 0,8822333 | novel transcript, antisense to MTHFD2L |
| ENSG00000168062 | BATF2 | 0,119991283 | 1,086728296 | 0,92134 | 0,96592951 | basic leucine zipper ATF-like transcription factor 2 |
| ENSG00000271383 | NBPF19 | 0,119961613 | 1,086705947 | 0,59284 | 0,79142743 | NBPF member 19 |
| ENSG00000137563 | GGH | 0,119930241 | 1,086682317 | 0,72971 | 0,8715818 | gamma-glutamyl hydrolase |
| ENSG00000203811 | H3C14 | 0,119909115 | 1,086666404 | 0,63127 | 0,81395758 | H3 clustered histone 14 |
| ENSG00000203852 | H3C15 | 0,119909115 | 1,086666404 | 0,63127 | 0,81395758 | H3 clustered histone 15 |
| ENSG00000135148 | TRAFD1 | 0,119881504 | 1,086645607 | 0,56806 | 0,77582773 | TRAF-type zinc finger domain containing 1 |
| ENSG00000115568 | ZNF142 | 0,119808919 | 1,086590937 | 0,45728 | 0,70073404 | zinc finger protein 142 |
| ENSG00000111540 | RAB5B | 0,11975281 | 1,086548679 | 0,41098 | 0,66881669 | RAB5B, member RAS oncogene family |
| ENSG00000258515 | AL355075,2 | 0,119676558 | 1,086491251 | 0,70804 | 0,85927396 | novel transcript, antisense to OSGEP |
| ENSG00000186517 | ARHGAP30 | 0,119603944 | 1,086436568 | 0,63029 | 0,81332648 | Rho GTPase activating protein 30 |
| ENSG00000204740 | MALRD1 | 0,119580997 | 1,086419287 | 0,79648 | 0,90597933 | MAM and LDL receptor class A domain containing 1 |
| ENSG00000152503 | TRIM36 | 0,119565557 | 1,08640766 | 0,81513 | 0,91495549 | tripartite motif containing 36 |
| ENSG00000048140 | TSPAN17 | 0,11952178 | 1,086374695 | 0,43439 | 0,68505666 | tetraspanin 17 |
| ENSG00000048405 | ZNF800 | 0,119480242 | 1,086343416 | 0,68099 | 0,84352125 | zinc finger protein 800 |
| ENSG00000223675 | AC093117,1 | 0,119341407 | 1,086238879 | 0,87397 | 0,94410944 | novel transcript |

| | | | | | | |
|---|---|---|---|---|---|---|
| ENSG00000103544 | VPS35L | 0,119294928 | 1,086203885 | 0,43577 | 0,68634331 | VPS35 endosomal protein sorting factor like |
| ENSG00000167114 | SLC27A4 | 0,119287085 | 1,086197979 | 0,73029 | 0,87185115 | solute carrier family 27 member 4 |
| ENSG00000236296 | GUSBP5 | 0,119271494 | 1,086186241 | 0,81291 | 0,91395885 | GUSB pseudogene 5 |
| ENSG00000075391 | RASAL2 | 0,119100826 | 1,086057755 | 0,65547 | 0,82781124 | RAS protein activator like 2 |
| ENSG00000134297 | PLEKHA8P1 | 0,118893551 | 1,08590173 | 0,71443 | 0,86255792 | pleckstrin homology domain containing A8 pseudogene 1 |
| ENSG00000014138 | POLA2 | 0,118831418 | 1,085854965 | 0,57017 | 0,77710133 | DNA polymerase alpha 2, accessory subunit |
| ENSG00000186104 | CYP2R1 | 0,118777825 | 1,085814628 | 0,76544 | 0,89085946 | cytochrome P450 family 2 subfamily R member 1 |
| ENSG00000164930 | FZD6 | 0,118762267 | 1,085802918 | 0,69714 | 0,85330122 | frizzled class receptor 6 |
| ENSG00000080845 | DLGAP4 | 0,118697307 | 1,085754029 | 0,48652 | 0,72196345 | DLG associated protein 4 |
| ENSG00000267405 | AC005180,1 | 0,118688356 | 1,085747293 | 0,84217 | 0,92821072 | novel transcript |
| ENSG00000139318 | DUSP6 | 0,118640486 | 1,085711267 | 0,773 | 0,89517309 | dual specificity phosphatase 6 |
| ENSG00000224501 | HSPA1B | 0,118619977 | 1,085695834 | 0,68373 | 0,84509131 | heat shock protein family A (Hsp70) member 1B |
| ENSG00000114739 | ACVR2B | 0,118602029 | 1,085682327 | 0,44592 | 0,69284796 | activin A receptor type 2B |
| ENSG00000257869 | AL139023,1 | 0,118588253 | 1,08567196 | 0,91955 | 0,96506391 | novel transcript |
| ENSG00000272275 | AC092687,3 | 0,118514625 | 1,085616554 | 0,87974 | 0,94649956 | novel transcript |
| ENSG00000177483 | RBM44 | 0,118496952 | 1,085603255 | 0,80283 | 0,90915255 | RNA binding motif protein 44 |
| ENSG00000111641 | NOP2 | 0,118479265 | 1,085589946 | 0,51486 | 0,74115184 | NOP2 nucleolar protein |
| ENSG00000198901 | PRC1 | 0,118402509 | 1,085532191 | 0,60517 | 0,79831516 | protein regulator of cytokinesis 1 |
| ENSG00000166483 | WEE1 | 0,118389856 | 1,08552267 | 0,45651 | 0,70015414 | WEE1 G2 checkpoint kinase |
| ENSG00000213625 | LEPROT | 0,118321778 | 1,085471448 | 0,45062 | 0,69592349 | leptin receptor overlapping transcript |
| ENSG00000236390 | AC092800,1 | 0,118252337 | 1,085419203 | 0,85098 | 0,93218292 | novel transcript |
| ENSG00000167964 | RAB26 | 0,11824157 | 1,085411101 | 0,74196 | 0,87796092 | RAB26, member RAS oncogene family |
| ENSG00000230046 | BIRC6-AS1 | 0,11817824 | 1,085363456 | 0,88163 | 0,94737759 | BIRC6 antisense RNA 1 |
| ENSG00000282012 | AP000357,2 | 0,118097872 | 1,085302996 | 0,86686 | 0,94109529 | novel transcript |
| ENSG00000175711 | B3GNTL1 | 0,117967939 | 1,085205255 | 0,58286 | 0,78477395 | UDP-GlcNAc:betaGal beta-1,3-N-acetylglucosaminyltransferase like 1 |
| ENSG00000172888 | ZNF621 | 0,117917878 | 1,085167599 | 0,45052 | 0,69587159 | zinc finger protein 621 |
| ENSG00000282419 | TEX13D | 0,117872695 | 1,085133615 | 0,87156 | 0,94322889 | TEX13 family member D |
| ENSG00000091640 | SPAG7 | 0,117862037 | 1,085125598 | 0,53349 | 0,75351438 | sperm associated antigen 7 |
| ENSG00000171570 | RAB4B-EGLN2 | 0,11784585 | 1,085113423 | 0,91657 | 0,9636258 | RAB4B-EGLN2 readthrough (NMD candidate) |
| ENSG00000187944 | C2orf66 | 0,117726225 | 1,085023452 | 0,8785 | 0,94600145 | chromosome 2 open reading frame 66 |
| ENSG00000105676 | ARMC6 | 0,117723783 | 1,085021615 | 0,52934 | 0,75112358 | armadillo repeat containing 6 |
| ENSG00000229496 | DDX39B | 0,117545946 | 1,084887875 | 0,4833 | 0,71964105 | DExD-box helicase 39B |

| | | | | | | |
|---|---|---|---|---|---|---|
| ENSG00000148308 | GTF3C5 | 0,117534374 | 1,084879173 | 0,45873 | 0,70133296 | general transcription factor IIIC subunit 5 |
| ENSG00000249839 | AC011330,1 | 0,117440143 | 1,084808316 | 0,91596 | 0,96339511 | histidine acid phosphatase domain containing 2A (HISPPD2A) pseudogene |
| ENSG00000177731 | FLII | 0,117403075 | 1,084780444 | 0,60191 | 0,79680798 | FLII actin remodeling protein |
| ENSG00000284571 | FLII | 0,117403075 | 1,084780444 | 0,60191 | 0,79680798 | FLII actin remodeling protein |
| ENSG00000269416 | LINC01224 | 0,117339572 | 1,084732696 | 0,50382 | 0,73294771 | long intergenic non-protein coding RNA 1224 |
| ENSG00000112406 | HECA | 0,117289042 | 1,084694704 | 0,55052 | 0,76469497 | hdc homolog, cell cycle regulator |
| ENSG00000255441 | SIGLEC10-AS1 | 0,117256555 | 1,084670279 | 0,78542 | 0,90030008 | SIGLEC10 antisense RNA 1 |
| ENSG00000267920 | SNX6P1 | 0,117255945 | 1,08466982 | 0,84146 | 0,92787857 | sorting nexin 6 pseudogene 1 |
| ENSG00000175203 | DCTN2 | 0,117156237 | 1,084594859 | 0,5602 | 0,7707835 | dynactin subunit 2 |
| ENSG00000158716 | DUSP23 | 0,11712843 | 1,084573954 | 0,75239 | 0,88396524 | dual specificity phosphatase 23 |
| ENSG00000106144 | CASP2 | 0,117123016 | 1,084569884 | 0,45954 | 0,7017275 | caspase 2 |
| ENSG00000255471 | AP001528,1 | 0,117105663 | 1,084556839 | 0,84127 | 0,92775849 | novel transcript |
| ENSG00000225868 | AC016582,1 | 0,117024977 | 1,084496185 | 0,8019 | 0,90880281 | WD repeat domain 87 (WDR87) pseudogene |
| ENSG00000258777 | HIF1A-AS1 | 0,116922508 | 1,084419159 | 0,84981 | 0,93163876 | HIF1A antisense RNA 1 |
| ENSG00000197302 | ZNF720 | 0,116913688 | 1,08441253 | 0,6507 | 0,82499358 | zinc finger protein 720 |
| ENSG00000134330 | IAH1 | 0,116901277 | 1,084403201 | 0,60474 | 0,79812088 | isoamyl acetate hydrolyzing esterase 1 (putative) |
| ENSG00000235967 | COX5BP3 | 0,116861179 | 1,084373061 | 0,90001 | 0,95740913 | cytochrome c oxidase subunit 5B pseudogene 3 |
| ENSG00000272211 | AC114760,2 | 0,116816174 | 1,084339235 | 0,81047 | 0,91251177 | novel transcript, antisense to STK17B |
| ENSG00000150773 | PIH1D2 | 0,1166293 | 1,084198789 | 0,78277 | 0,89910316 | PIH1 domain containing 2 |
| ENSG00000285898 | AC007317,1 | 0,116556005 | 1,084143708 | 0,77455 | 0,89586241 | Novel transcript |
| ENSG00000105755 | ETHE1 | 0,116526043 | 1,084121192 | 0,70825 | 0,85927396 | ETHE1 persulfide dioxygenase |
| ENSG00000226963 | AC078883,2 | 0,11640359 | 1,084029178 | 0,87956 | 0,94634942 | novel transcript |
| ENSG00000089154 | GCN1 | 0,116378629 | 1,084010424 | 0,43716 | 0,68711291 | GCN1 activator of EIF2AK4 |
| ENSG00000105072 | C19orf44 | 0,116218379 | 1,083890022 | 0,51893 | 0,7437068 | chromosome 19 open reading frame 44 |
| ENSG00000163754 | GYG1 | 0,116186027 | 1,083865716 | 0,73083 | 0,87205836 | glycogenin 1 |
| ENSG00000135040 | NAA35 | 0,116165191 | 1,083850062 | 0,59302 | 0,79148028 | N-alpha-acetyltransferase 35, NatC auxiliary subunit |
| ENSG00000033867 | SLC4A7 | 0,116106004 | 1,083805598 | 0,69134 | 0,84952103 | solute carrier family 4 member 7 |
| ENSG00000124596 | OARD1 | 0,116024893 | 1,083744666 | 0,705 | 0,8578842 | O-acyl-ADP-ribose deacylase 1 |
| ENSG00000168522 | FNTA | 0,115887788 | 1,083641679 | 0,52656 | 0,7497927 | farnesyltransferase, CAAX box, alpha |
| ENSG00000132581 | SDF2 | 0,115820549 | 1,083591175 | 0,45203 | 0,69691985 | stromal cell derived factor 2 |
| ENSG00000080371 | RAB21 | 0,115746997 | 1,083535932 | 0,47418 | 0,7129821 | RAB21, member RAS oncogene family |
| ENSG00000133313 | CNDP2 | 0,115729253 | 1,083522606 | 0,52199 | 0,7463592 | carnosine dipeptidase 2 |

| | | | | | | |
|---|---|---|---|---|---|---|
| ENSG00000142546 | NOSIP | 0,115712193 | 1,083509793 | 0,58332 | 0,78514291 | nitric oxide synthase interacting protein |
| ENSG00000160183 | TMPRSS3 | 0,115679036 | 1,083484892 | 0,91104 | 0,9616927 | transmembrane serine protease 3 |
| ENSG00000213366 | GSTM2 | 0,115654693 | 1,083466609 | 0,4792 | 0,71679575 | glutathione S-transferase mu 2 |
| ENSG00000258984 | UBE2F-SCLY | 0,115638047 | 1,083454108 | 0,88818 | 0,95069428 | UBE2F-SCLY readthrough (NMD candidate) |
| ENSG00000085733 | CTTN | 0,115480293 | 1,083335643 | 0,54109 | 0,75868526 | cortactin |
| ENSG00000288401 | CTTN | 0,115480293 | 1,083335643 | 0,54109 | 0,75868526 | cortactin |
| ENSG00000251463 | FKBP4P1 | 0,115443266 | 1,083307839 | 0,90328 | 0,95856163 | FKBP prolyl isomerase 4 pseudogene 1 |
| ENSG00000287934 | AP001885,3 | 0,115410038 | 1,083282889 | 0,74201 | 0,87798329 | novel transcript, antisense to KDM2A |
| ENSG00000287031 | AC093307,1 | 0,115355765 | 1,083242137 | 0,90028 | 0,95760242 | novel transcript |
| ENSG00000120896 | SORBS3 | 0,115276999 | 1,083182998 | 0,59592 | 0,793576 | sorbin and SH3 domain containing 3 |
| ENSG00000265728 | AP001099,1 | 0,115181307 | 1,083111154 | 0,78404 | 0,89941845 | novel transcript |
| ENSG00000275368 | TCTE3 | 0,115105 | 1,083053868 | 0,71486 | 0,86282308 | t-complex-associated-testis-expressed 3 |
| ENSG00000250012 | AC079848,1 | 0,11500498 | 1,082978784 | 0,81305 | 0,91408807 | novel transcript |
| ENSG00000101850 | GPR143 | 0,114973806 | 1,082955383 | 0,7166 | 0,86437402 | G protein-coupled receptor 143 |
| ENSG00000111832 | RWDD1 | 0,11493475 | 1,082926066 | 0,56144 | 0,77136826 | RWD domain containing 1 |
| ENSG00000178761 | FAM219B | 0,114898823 | 1,082899098 | 0,65744 | 0,82872324 | family with sequence similarity 219 member B |
| ENSG00000153140 | CETN3 | 0,114777572 | 1,08280809 | 0,60669 | 0,79981537 | centrin 3 |
| ENSG00000258136 | AC007622,2 | 0,114625775 | 1,082694166 | 0,71089 | 0,86079022 | novel transcript, antisense to PRDM4 |
| ENSG00000137992 | DBT | 0,114606221 | 1,082679491 | 0,44149 | 0,68977736 | dihydrolipoamide branched chain transacylase E2 |
| ENSG00000272223 | AL136304,1 | 0,114490407 | 1,082592581 | 0,82731 | 0,92015788 | novel transcript, antisense to RRP36 |
| ENSG00000185689 | C6orf201 | 0,114426404 | 1,082544555 | 0,84821 | 0,93094401 | chromosome 6 open reading frame 201 |
| ENSG00000256315 | AP000462,2 | 0,114322666 | 1,082466717 | 0,81924 | 0,91696956 | novel transcript |
| ENSG00000196663 | TECPR2 | 0,11419437 | 1,082370459 | 0,64351 | 0,82109591 | tectonin beta-propeller repeat containing 2 |
| ENSG00000214736 | TOMM6 | 0,114142591 | 1,082331613 | 0,53408 | 0,75391449 | translocase of outer mitochondrial membrane 6 |
| ENSG00000183763 | TRAIP | 0,114074786 | 1,082280746 | 0,64354 | 0,82109591 | TRAF interacting protein |
| ENSG00000107104 | KANK1 | 0,114032451 | 1,082248988 | 0,68939 | 0,84865116 | KN motif and ankyrin repeat domains 1 |
| ENSG00000057935 | MTA3 | 0,113905745 | 1,082153942 | 0,5781 | 0,78191189 | metastasis associated 1 family member 3 |
| ENSG00000275645 | AC068338,3 | 0,113902078 | 1,082151192 | 0,85331 | 0,93334646 | novel transcript, antisense to NEIL1 |
| ENSG00000269893 | SNHG8 | 0,113857622 | 1,082117847 | 0,61974 | 0,80726991 | small nucleolar RNA host gene 8 |
| ENSG00000084652 | TXLNA | 0,113822681 | 1,082091638 | 0,54174 | 0,75882662 | taxilin alpha |
| ENSG00000160551 | TAOK1 | 0,113809206 | 1,082081531 | 0,60774 | 0,80046176 | TAO kinase 1 |
| ENSG00000222489 | SNORA79B | 0,113713267 | 1,082009576 | 0,79955 | 0,90748957 | small nucleolar RNA, H/ACA box 79B |
| ENSG00000168159 | RNF187 | 0,113678097 | 1,081983199 | 0,6268 | 0,81113811 | ring finger protein 187 |

| | | | | | | |
|---|---|---|---|---|---|---|
| ENSG00000285426 | RNF187 | 0,113678097 | 1,081983199 | 0,6268 | 0,81113811 | ring finger protein 187 |
| ENSG00000066044 | ELAVL1 | 0,113675162 | 1,081980998 | 0,42455 | 0,67841584 | ELAV like RNA binding protein 1 |
| ENSG00000072864 | NDE1 | 0,113665192 | 1,08197352 | 0,77586 | 0,89635267 | nudE neurodevelopment protein 1 |
| ENSG00000182307 | C8orf33 | 0,113564204 | 1,081897785 | 0,64037 | 0,81936319 | chromosome 8 open reading frame 33 |
| ENSG00000224117 | PTPN2P2 | 0,11346951 | 1,081826775 | 0,88151 | 0,94729476 | PTPN2 pseudogene 2 |
| ENSG00000167196 | FBXO22 | 0,113426005 | 1,081794153 | 0,51321 | 0,73963507 | F-box protein 22 |
| ENSG00000274684 | NMRAL1 | 0,113386197 | 1,081764303 | 0,71786 | 0,86489865 | NmrA like redox sensor 1 |
| ENSG00000203875 | SNHG5 | 0,113254942 | 1,081665891 | 0,53057 | 0,75212511 | small nucleolar RNA host gene 5 |
| ENSG00000253539 | AC018442,2 | 0,113247212 | 1,081660095 | 0,88742 | 0,95041376 | novel transcript |
| ENSG00000258716 | AL139193,2 | 0,113232434 | 1,081649015 | 0,8137 | 0,9144566 | novel transcript, antisense to TTC7B |
| ENSG00000108349 | CASC3 | 0,113196783 | 1,081622286 | 0,57838 | 0,78213794 | CASC3 exon junction complex subunit |
| ENSG00000100028 | SNRPD3 | 0,113159082 | 1,081594021 | 0,49374 | 0,72643895 | small nuclear ribonucleoprotein D3 polypeptide |
| ENSG00000183258 | DDX41 | 0,113018943 | 1,081488963 | 0,50517 | 0,73378796 | DEAD-box helicase 41 |
| ENSG00000127527 | EPS15L1 | 0,112975276 | 1,08145623 | 0,58557 | 0,78640335 | epidermal growth factor receptor pathway substrate 15 like 1 |
| ENSG00000007062 | PROM1 | 0,112958475 | 1,081443636 | 0,71991 | 0,86614912 | prominin 1 |
| ENSG00000120327 | PCDHB14 | 0,11288941 | 1,081391866 | 0,63612 | 0,81694486 | protocadherin beta 14 |
| ENSG00000147439 | BIN3 | 0,112886347 | 1,08138957 | 0,57328 | 0,77886693 | bridging integrator 3 |
| ENSG00000163159 | VPS72 | 0,11286456 | 1,081373239 | 0,45036 | 0,69574391 | vacuolar protein sorting 72 homolog |
| ENSG00000230470 | AL078645,1 | 0,1127229 | 1,081267064 | 0,8884 | 0,95069428 | novel transcript |
| ENSG00000137177 | KIF13A | 0,11261521 | 1,081186355 | 0,64018 | 0,81935089 | kinesin family member 13A |
| ENSG00000146707 | POMZP3 | 0,112614062 | 1,081185495 | 0,56824 | 0,77590889 | POM121 and ZP3 fusion |
| ENSG00000155111 | CDK19 | 0,112613662 | 1,081185195 | 0,65915 | 0,82977637 | cyclin dependent kinase 19 |
| ENSG00000278774 | U2 | 0,112585352 | 1,081163979 | 0,97427 | 0,98933043 | U2 spliceosomal RNA |
| ENSG00000108654 | DDX5 | 0,112385675 | 1,081014351 | 0,55013 | 0,76444876 | DEAD-box helicase 5 |
| ENSG00000144278 | GALNT13 | 0,112346395 | 1,080984918 | 0,66358 | 0,8324578 | polypeptide N-acetylgalactosaminyltransferase 13 |
| ENSG00000044459 | CNTLN | 0,112276832 | 1,080932797 | 0,643 | 0,82098995 | centlein |
| ENSG00000250312 | ZNF718 | 0,112239441 | 1,080904783 | 0,44732 | 0,69342177 | zinc finger protein 718 |
| ENSG00000137145 | DENND4C | 0,112221746 | 1,080891525 | 0,71864 | 0,86547418 | DENN domain containing 4C |
| ENSG00000277203 | F8A1 | 0,112139426 | 1,080829852 | 0,65671 | 0,82855263 | coagulation factor VIII associated 1 |
| ENSG00000147454 | SLC25A37 | 0,112100141 | 1,08080042 | 0,5992 | 0,79538547 | solute carrier family 25 member 37 |
| ENSG00000205269 | TMEM170B | 0,112008435 | 1,080731721 | 0,68796 | 0,84804486 | transmembrane protein 170B |
| ENSG00000196267 | ZNF836 | 0,11191808 | 1,080664037 | 0,74639 | 0,88045757 | zinc finger protein 836 |

| | | | | | | |
|---|---|---|---|---|---|---|
| ENSG00000198185 | ZNF334 | 0,111900052 | 1,080650533 | 0,75602 | 0,88573899 | zinc finger protein 334 |
| ENSG00000150636 | CCDC102B | 0,111844177 | 1,080608681 | 0,71846 | 0,86533732 | coiled-coil domain containing 102B |
| ENSG00000127314 | RAP1B | 0,11174683 | 1,080535769 | 0,56911 | 0,77637061 | RAP1B, member of RAS oncogene family |
| ENSG00000246263 | UBR5-AS1 | 0,111707916 | 1,080506624 | 0,7268 | 0,86997018 | UBR5 antisense RNA 1 |
| ENSG00000228852 | AC092802,2 | 0,111651786 | 1,080464585 | 0,76604 | 0,89122725 | novel transcript, antisense to RWDD3 |
| ENSG00000234788 | HSPA8P3 | 0,111638378 | 1,080454544 | 0,85893 | 0,93646304 | heat shock protein family A (Hsp70) member 8 pseudogene 3 |
| ENSG00000146556 | WASH2P | 0,111636797 | 1,08045336 | 0,48072 | 0,71781176 | WASP family homolog 2, pseudogene |
| ENSG00000104691 | UBXN8 | 0,111607593 | 1,080431489 | 0,73634 | 0,87525505 | UBX domain protein 8 |
| ENSG00000117122 | MFAP2 | 0,111595812 | 1,080422667 | 0,69058 | 0,84900677 | microfibril associated protein 2 |
| ENSG00000226310 | AL022157,1 | 0,111573056 | 1,080405625 | 0,82119 | 0,91738668 | novel transcript |
| ENSG00000166780 | BMERB1 | 0,111553586 | 1,080391044 | 0,71767 | 0,86483656 | bMERB domain containing 1 |
| ENSG00000075539 | FRYL | 0,111473158 | 1,080330815 | 0,76429 | 0,89015718 | FRY like transcription coactivator |
| ENSG00000243364 | EFNA4 | 0,111446224 | 1,080310647 | 0,65618 | 0,82829288 | ephrin A4 |
| ENSG00000287360 | AC006296,3 | 0,111280444 | 1,080186516 | 0,87942 | 0,94631723 | novel transcript |
| ENSG00000273891 | AL731566,2 | 0,111274169 | 1,080181817 | 0,66411 | 0,83250729 | novel transcript, antisense to NSMCE4A |
| ENSG00000053747 | LAMA3 | 0,111221046 | 1,080142044 | 0,77686 | 0,89660077 | laminin subunit alpha 3 |
| ENSG00000264577 | AC010761,1 | 0,111205087 | 1,080130095 | 0,59555 | 0,79339314 | novel transcript, antisense to RPL23A |
| ENSG00000265261 | AC027575,1 | 0,111180241 | 1,080111493 | 0,86464 | 0,93973773 | exportin, tRNA (nuclear export receptor for tRNAs) (XPOT) pseudogene |
| ENSG00000135269 | TES | 0,111122963 | 1,080068612 | 0,69648 | 0,85303733 | testin LIM domain protein |
| ENSG00000087448 | KLHL42 | 0,111022711 | 1,079993561 | 0,65978 | 0,83018671 | kelch like family member 42 |
| ENSG00000196646 | ZNF136 | 0,110989895 | 1,079968996 | 0,57883 | 0,78256244 | zinc finger protein 136 |
| ENSG00000125846 | ZNF133 | 0,11096056 | 1,079947036 | 0,68154 | 0,84375324 | zinc finger protein 133 |
| ENSG00000111786 | SRSF9 | 0,110955796 | 1,07994347 | 0,50569 | 0,73418581 | serine and arginine rich splicing factor 9 |
| ENSG00000147041 | SYTL5 | 0,110887737 | 1,079892525 | 0,88924 | 0,95110607 | synaptotagmin like 5 |
| ENSG00000154978 | VOPP1 | 0,110785812 | 1,079816235 | 0,50257 | 0,73209345 | VOPP1 WW domain binding protein |
| ENSG00000132207 | SLX1A | 0,110785099 | 1,079815701 | 0,59496 | 0,79311029 | SLX1 homolog A, structure-specific endonuclease subunit |
| ENSG00000067225 | PKM | 0,110690434 | 1,079744849 | 0,51473 | 0,74102666 | pyruvate kinase M1/2 |
| ENSG00000145555 | MYO10 | 0,110681637 | 1,079738265 | 0,66308 | 0,83242715 | myosin X |
| ENSG00000105662 | CRTC1 | 0,110522556 | 1,079619213 | 0,57177 | 0,77829419 | CREB regulated transcription coactivator 1 |
| ENSG00000009413 | REV3L | 0,110503143 | 1,079604685 | 0,55937 | 0,77045525 | REV3 like, DNA directed polymerase zeta catalytic subunit |
| ENSG00000205414 | AC007608,1 | 0,11047833 | 1,079586117 | 0,83575 | 0,92493625 | novel transcript, antisense to NKD1 |

| | | | | | | |
|---|---|---|---|---|---|---|
| ENSG00000280374 | AC019080,5 | 0,110415976 | 1,079539458 | 0,85234 | 0,93290976 | novel transcript |
| ENSG00000232938 | RPL23AP87 | 0,110332784 | 1,079477209 | 0,77567 | 0,89630112 | ribosomal protein L23a pseudogene 87 |
| ENSG00000010626 | LRRC23 | 0,110245983 | 1,079412263 | 0,62616 | 0,81072048 | leucine rich repeat containing 23 |
| ENSG00000110987 | BCL7A | 0,110185725 | 1,07936718 | 0,64645 | 0,82293404 | BAF chromatin remodeling complex subunit BCL7A |
| ENSG00000279897 | BIRC6-AS2 | 0,110034435 | 1,079253997 | 0,84044 | 0,92722187 | BIRC6 antisense RNA 2 |
| ENSG00000099953 | MMP11 | 0,110024251 | 1,079246378 | 0,78267 | 0,89905848 | matrix metallopeptidase 11 |
| ENSG00000261490 | AC005674,1 | 0,109745978 | 1,079038228 | 0,77116 | 0,89404837 | novel transcript, overlapping WDR1 |
| ENSG00000288382 | AC256223,4 | 0,109726421 | 1,079023601 | 0,88815 | 0,95069428 | |
| ENSG00000157833 | GAREM2 | 0,109610091 | 1,078936599 | 0,68564 | 0,84637134 | GRB2 associated regulator of MAPK1 subtype 2 |
| ENSG00000231074 | HCG18 | 0,109600926 | 1,078929745 | 0,53916 | 0,75757364 | HLA complex group 18 |
| ENSG00000204822 | MRPL53 | 0,109480979 | 1,078840045 | 0,61608 | 0,80476728 | mitochondrial ribosomal protein L53 |
| ENSG00000140471 | LINS1 | 0,109452581 | 1,078818809 | 0,55189 | 0,76553885 | lines homolog 1 |
| ENSG00000137070 | IL11RA | 0,109298015 | 1,078703235 | 0,69017 | 0,84887944 | interleukin 11 receptor subunit alpha |
| ENSG00000118849 | RARRES1 | 0,109282019 | 1,078691274 | 0,82461 | 0,91900665 | retinoic acid receptor responder 1 |
| ENSG00000085224 | ATRX | 0,109280511 | 1,078690147 | 0,71396 | 0,86227395 | ATRX chromatin remodeler |
| ENSG00000275478 | AL671879,2 | 0,109216428 | 1,078642234 | 0,80945 | 0,91213228 | novel transcript |
| ENSG00000259299 | AC061965,2 | 0,109165948 | 1,078604492 | 0,88284 | 0,94822558 | DDB1 and CUL4 associated factor 6 (DCAF6) pseudogene |
| ENSG00000247092 | SNHG10 | 0,109090259 | 1,078547907 | 0,60898 | 0,80157606 | small nucleolar RNA host gene 10 |
| ENSG00000134575 | ACP2 | 0,109062569 | 1,078527206 | 0,55308 | 0,7662484 | acid phosphatase 2, lysosomal |
| ENSG00000249624 | AP000295,1 | 0,109053504 | 1,078520429 | 0,90547 | 0,95961188 | novel protein |
| ENSG00000152348 | ATG10 | 0,108967702 | 1,078456288 | 0,60213 | 0,7969257 | autophagy related 10 |
| ENSG00000184887 | BTBD6 | 0,108915061 | 1,078416938 | 0,60085 | 0,79599359 | BTB domain containing 6 |
| ENSG00000072062 | PRKACA | 0,108910786 | 1,078413742 | 0,57074 | 0,77748547 | protein kinase cAMP-activated catalytic subunit alpha |
| ENSG00000138231 | DBR1 | 0,108746283 | 1,078290783 | 0,53707 | 0,75574932 | debranching RNA lariats 1 |
| ENSG00000282932 | PTPRD | 0,108690041 | 1,078248748 | 0,7916 | 0,90310057 | protein tyrosine phosphatase receptor type D |
| ENSG00000151240 | DIP2C | 0,108642516 | 1,078213229 | 0,59991 | 0,79564899 | disco interacting protein 2 homolog C |
| ENSG00000132376 | INPP5K | 0,108609262 | 1,078188377 | 0,61512 | 0,80421986 | inositol polyphosphate-5-phosphatase K |
| ENSG00000142507 | PSMB6 | 0,108580242 | 1,078166689 | 0,53806 | 0,75658794 | proteasome 20S subunit beta 6 |
| ENSG00000255508 | AP002990,1 | 0,108507985 | 1,078112691 | 0,66957 | 0,83623395 | novel protein |
| ENSG00000188234 | AGAP4 | 0,108465698 | 1,078081091 | 0,6483 | 0,82383528 | ArfGAP with GTPase domain, ankyrin repeat and PH domain 4 |
| ENSG00000077684 | JADE1 | 0,108449286 | 1,078068826 | 0,56695 | 0,77482793 | jade family PHD finger 1 |
| ENSG00000228677 | TTC3-AS1 | 0,108430972 | 1,078055141 | 0,83602 | 0,92493625 | TTC3 antisense RNA 1 |

| | | | | | | |
|---|---|---|---|---|---|---|
| ENSG00000145861 | C1QTNF2 | 0,108369819 | 1,078009446 | 0,79418 | 0,9043512 | C1q and TNF related 2 |
| ENSG00000243570 | CFB | 0,108326789 | 1,077977293 | 0,87698 | 0,94549531 | complement factor B |
| ENSG00000242335 | CFB | 0,108326789 | 1,077977293 | 0,87698 | 0,94549531 | complement factor B |
| ENSG00000226919 | AL365184,1 | 0,108223728 | 1,077900289 | 0,88876 | 0,95089681 | novel transcript, antisense to RGS7 |
| ENSG00000218305 | CDC14C | 0,108188316 | 1,077873831 | 0,83985 | 0,92680022 | cell division cycle 14C |
| ENSG00000110925 | CSRNP2 | 0,108157004 | 1,077850438 | 0,55064 | 0,76475342 | cysteine and serine rich nuclear protein 2 |
| ENSG00000223764 | LINC02593 | 0,108133658 | 1,077832996 | 0,75527 | 0,88533971 | long intergenic non-protein coding RNA 2593 |
| ENSG00000160218 | TRAPPC10 | 0,108019393 | 1,077747632 | 0,49525 | 0,7275528 | trafficking protein particle complex 10 |
| ENSG00000164292 | RHOBTB3 | 0,108011533 | 1,077741761 | 0,75244 | 0,88398552 | Rho related BTB domain containing 3 |
| ENSG00000105287 | PRKD2 | 0,107976847 | 1,077715849 | 0,5451 | 0,7609742 | protein kinase D2 |
| ENSG00000130558 | OLFM1 | 0,107917061 | 1,077671189 | 0,81779 | 0,91632405 | olfactomedin 1 |
| ENSG00000166130 | IKBIP | 0,107909948 | 1,077665876 | 0,61694 | 0,80547324 | IKBKB interacting protein |
| ENSG00000082196 | C1QTNF3 | 0,107819135 | 1,077598042 | 0,71357 | 0,86217438 | C1q and TNF related 3 |
| ENSG00000166224 | SGPL1 | 0,107799401 | 1,077583302 | 0,55925 | 0,77045525 | sphingosine-1-phosphate lyase 1 |
| ENSG00000198060 | MARCHF5 | 0,107656656 | 1,077476688 | 0,58876 | 0,78830778 | membrane associated ring-CH-type finger 5 |
| ENSG00000075234 | TTC38 | 0,107637948 | 1,077462716 | 0,61629 | 0,80488598 | tetratricopeptide repeat domain 38 |
| ENSG00000162869 | PPP1R21 | 0,107617368 | 1,077447346 | 0,78186 | 0,8987738 | protein phosphatase 1 regulatory subunit 21 |
| ENSG00000260032 | NORAD | 0,107588449 | 1,077425749 | 0,62269 | 0,80866078 | non-coding RNA activated by DNA damage |
| ENSG00000124831 | LRRFIP1 | 0,107578522 | 1,077418335 | 0,60388 | 0,79770754 | LRR binding FLII interacting protein 1 |
| ENSG00000166704 | ZNF606 | 0,107554392 | 1,077400315 | 0,62194 | 0,80819595 | zinc finger protein 606 |
| ENSG00000142541 | RPL13A | 0,107396092 | 1,077282103 | 0,63909 | 0,81893766 | ribosomal protein L13a |
| ENSG00000266369 | AC090774,2 | 0,107260407 | 1,07718079 | 0,84854 | 0,93094401 | novel transcript, antisense to CCDC144NL |
| ENSG00000272391 | POM121C | 0,107212185 | 1,077144786 | 0,56364 | 0,77255679 | POM121 transmembrane nucleoporin C |
| ENSG00000101439 | CST3 | 0,107125706 | 1,077080221 | 0,71047 | 0,86051357 | cystatin C |
| ENSG00000132423 | COQ3 | 0,107060849 | 1,077031802 | 0,64951 | 0,82442734 | coenzyme Q3, methyltransferase |
| ENSG00000160294 | MCM3AP | 0,106829879 | 1,076859387 | 0,50741 | 0,73566394 | minichromosome maintenance complex component 3 associated protein |
| ENSG00000260306 | AC092375,2 | 0,106781874 | 1,076823555 | 0,83192 | 0,92312766 | novel transcript |
| ENSG00000149923 | PPP4C | 0,106703366 | 1,076764959 | 0,53695 | 0,75565547 | protein phosphatase 4 catalytic subunit |
| ENSG00000077097 | TOP2B | 0,106669068 | 1,07673936 | 0,74002 | 0,87675329 | DNA topoisomerase II beta |
| ENSG00000172379 | ARNT2 | 0,106650966 | 1,07672585 | 0,63616 | 0,81694486 | aryl hydrocarbon receptor nuclear translocator 2 |
| ENSG00000144524 | COPS7B | 0,106572124 | 1,07666701 | 0,58645 | 0,78678775 | COP9 signalosome subunit 7B |
| ENSG00000111581 | NUP107 | 0,106568981 | 1,076664664 | 0,54225 | 0,75895614 | nucleoporin 107 |

| | | | | | | |
|---|---|---|---|---|---|---|
| ENSG00000219891 | ZSCAN12P1 | 0,106437325 | 1,076566415 | 0,83617 | 0,92493625 | zinc finger and SCAN domain containing 12 pseudogene 1 |
| ENSG00000072310 | SREBF1 | 0,10632946 | 1,076485928 | 0,62589 | 0,81058216 | sterol regulatory element binding transcription factor 1 |
| ENSG00000111266 | DUSP16 | 0,106276187 | 1,076446178 | 0,61429 | 0,80370125 | dual specificity phosphatase 16 |
| ENSG00000280962 | DUSP16 | 0,106276187 | 1,076446178 | 0,61429 | 0,80370125 | dual specificity phosphatase 16 |
| ENSG00000283415 | AC087280,2 | 0,106254827 | 1,076430241 | 0,7929 | 0,90370819 | novel transcript |
| ENSG00000102103 | PQBP1 | 0,106184203 | 1,076377548 | 0,58642 | 0,78678775 | polyglutamine binding protein 1 |
| ENSG00000160563 | MED27 | 0,106146436 | 1,07634937 | 0,57725 | 0,78134056 | mediator complex subunit 27 |
| ENSG00000129195 | PIMREG | 0,106108689 | 1,076321209 | 0,73508 | 0,87480251 | PICALM interacting mitotic regulator |
| ENSG00000113140 | SPARC | 0,106106471 | 1,076319554 | 0,65228 | 0,82578944 | secreted protein acidic and cysteine rich |
| ENSG00000174744 | BRMS1 | 0,106084357 | 1,076303056 | 0,66251 | 0,83197976 | BRMS1 transcriptional repressor and anoikis regulator |
| ENSG00000214447 | FAM187A | 0,106078006 | 1,076298318 | 0,82437 | 0,91900665 | family with sequence similarity 187 member A |
| ENSG00000181026 | AEN | 0,106058287 | 1,076283607 | 0,54167 | 0,75882662 | apoptosis enhancing nuclease |
| ENSG00000137996 | RTCA | 0,105865614 | 1,076139879 | 0,50096 | 0,73099828 | RNA 3'-terminal phosphate cyclase |
| ENSG00000272205 | AL451050,2 | 0,105646052 | 1,075976115 | 0,86279 | 0,9388179 | novel transcript |
| ENSG00000186026 | ZNF284 | 0,105619585 | 1,075956375 | 0,64118 | 0,81994133 | zinc finger protein 284 |
| ENSG00000141026 | MED9 | 0,105590196 | 1,075934457 | 0,65685 | 0,82856771 | mediator complex subunit 9 |
| ENSG00000259224 | SLC35G6 | 0,105539289 | 1,075896493 | 0,89845 | 0,95636487 | solute carrier family 35 member G6 |
| ENSG00000284515 | SLC35G6 | 0,105539289 | 1,075896493 | 0,89845 | 0,95636487 | solute carrier family 35 member G6 |
| ENSG00000021300 | PLEKHB1 | 0,105533356 | 1,075892068 | 0,6582 | 0,82902424 | pleckstrin homology domain containing B1 |
| ENSG00000135720 | DYNC1LI2 | 0,105511472 | 1,075875748 | 0,72972 | 0,8715818 | dynein cytoplasmic 1 light intermediate chain 2 |
| ENSG00000101109 | STK4 | 0,105416884 | 1,075805212 | 0,48523 | 0,72086542 | serine/threonine kinase 4 |
| ENSG00000256436 | TAS2R31 | 0,105402243 | 1,075794295 | 0,79775 | 0,90685415 | taste 2 receptor member 31 |
| ENSG00000263097 | TAS2R31 | 0,105402243 | 1,075794295 | 0,79775 | 0,90685415 | taste 2 receptor member 31 |
| ENSG00000158458 | NRG2 | 0,105386367 | 1,075782456 | 0,82718 | 0,92010219 | neuregulin 2 |
| ENSG00000259499 | AC090519,1 | 0,105338396 | 1,075746686 | 0,88035 | 0,94683782 | novel transcript |
| ENSG00000143674 | MAP3K21 | 0,105265709 | 1,075692488 | 0,84175 | 0,92804468 | mitogen-activated protein kinase kinase kinase 21 |
| ENSG00000268066 | FMR1-AS1 | 0,105254764 | 1,075684327 | 0,87999 | 0,94664997 | FMR1 antisense RNA 1 |
| ENSG00000234420 | ZNF37BP | 0,105210515 | 1,075651335 | 0,56661 | 0,77458849 | zinc finger protein 37B, pseudogene |
| ENSG00000143889 | HNRNPLL | 0,105208933 | 1,075650156 | 0,49985 | 0,73058737 | heterogeneous nuclear ribonucleoprotein L like |
| ENSG00000229320 | KRT8P12 | 0,105189024 | 1,075635313 | 0,74066 | 0,87717177 | keratin 8 pseudogene 12 |
| ENSG00000188848 | BEND4 | 0,105186308 | 1,075633287 | 0,86205 | 0,93836958 | BEN domain containing 4 |
| ENSG00000230685 | CLIC1 | 0,105144351 | 1,075602006 | 0,6961 | 0,85287022 | chloride intracellular channel 1 |
| ENSG00000233614 | DDX11L10 | 0,1051201 | 1,075583926 | 0,83365 | 0,92396169 | DEAD/H-box helicase 11 like 10 (pseudogene) |

| | | | | | | |
|---|---|---|---|---|---|---|
| ENSG00000232907 | DLGAP4-AS1 | 0,104949937 | 1,07545707 | 0,78435 | 0,89962715 | DLGAP4 antisense RNA 1 |
| ENSG00000125247 | TMTC4 | 0,104908833 | 1,07542643 | 0,57138 | 0,77809338 | transmembrane O-mannosyltransferase targeting cadherins 4 |
| ENSG00000177963 | RIC8A | 0,104874534 | 1,075400863 | 0,53164 | 0,75280487 | RIC8 guanine nucleotide exchange factor A |
| ENSG00000275127 | SNORD116-22 | 0,104762486 | 1,075317344 | 0,86644 | 0,94074101 | small nucleolar RNA, C/D box 116-22 |
| ENSG00000212128 | TAS2R13 | 0,104742027 | 1,075302095 | 0,87042 | 0,942849 | taste 2 receptor member 13 |
| ENSG00000277254 | TAS2R13 | 0,104742027 | 1,075302095 | 0,87042 | 0,942849 | taste 2 receptor member 13 |
| ENSG00000273457 | TAS2R13 | 0,104742027 | 1,075302095 | 0,87042 | 0,942849 | taste 2 receptor member 13 |
| ENSG00000104980 | TIMM44 | 0,104741763 | 1,075301898 | 0,5369 | 0,75562287 | translocase of inner mitochondrial membrane 44 |
| ENSG00000163681 | SLMAP | 0,104676644 | 1,075253363 | 0,5341 | 0,75391449 | sarcolemma associated protein |
| ENSG00000006530 | AGK | 0,104661864 | 1,075242348 | 0,57574 | 0,78043323 | acylglycerol kinase |
| ENSG00000172830 | SSH3 | 0,104499565 | 1,075121393 | 0,80081 | 0,90830749 | slingshot protein phosphatase 3 |
| ENSG00000143324 | XPR1 | 0,104427057 | 1,07506736 | 0,67231 | 0,83787466 | xenotropic and polytropic retrovirus receptor 1 |
| ENSG00000149300 | C11orf52 | 0,104383541 | 1,075034933 | 0,9052 | 0,95950958 | chromosome 11 open reading frame 52 |
| ENSG00000117305 | HMGCL | 0,104358784 | 1,075016485 | 0,5908 | 0,79001985 | 3-hydroxy-3-methylglutaryl-CoA lyase |
| ENSG00000067606 | PRKCZ | 0,104346472 | 1,075007312 | 0,71392 | 0,86227395 | protein kinase C zeta |
| ENSG00000185085 | INTS5 | 0,104335439 | 1,07499909 | 0,54817 | 0,76347743 | integrator complex subunit 5 |
| ENSG00000174775 | HRAS | 0,10419522 | 1,074894614 | 0,78242 | 0,89892226 | HRas proto-oncogene, GTPase |
| ENSG00000105705 | SUGP1 | 0,104157668 | 1,074866635 | 0,60876 | 0,80142967 | SURP and G-patch domain containing 1 |
| ENSG00000147133 | TAF1 | 0,104103847 | 1,074826537 | 0,60299 | 0,7971742 | TATA-box binding protein associated factor 1 |
| ENSG00000118162 | KPTN | 0,103978237 | 1,07473296 | 0,62053 | 0,80776776 | kaptin, actin binding protein |
| ENSG00000164253 | WDR41 | 0,103748218 | 1,074561622 | 0,53206 | 0,75285051 | WD repeat domain 41 |
| ENSG00000266053 | NDUFV2-AS1 | 0,103728142 | 1,074546668 | 0,78528 | 0,90017138 | NDUFV2 antisense RNA 1 |
| ENSG00000186222 | BLOC1S4 | 0,10370109 | 1,07452652 | 0,63172 | 0,81431541 | biogenesis of lysosomal organelles complex 1 subunit 4 |
| ENSG00000175073 | VCPIP1 | 0,103682237 | 1,074512478 | 0,64458 | 0,82154386 | valosin containing protein interacting protein 1 |
| ENSG00000158301 | GPRASP2 | 0,103589512 | 1,074443419 | 0,54777 | 0,76307321 | G protein-coupled receptor associated sorting protein 2 |
| ENSG00000137135 | ARHGEF39 | 0,103352257 | 1,074266739 | 0,71359 | 0,86217438 | Rho guanine nucleotide exchange factor 39 |
| ENSG00000125266 | EFNB2 | 0,103232194 | 1,074177341 | 0,78139 | 0,89877377 | ephrin B2 |
| ENSG00000258101 | AC010173,1 | 0,10320759 | 1,074159022 | 0,89015 | 0,95161642 | novel transcript, antisense to TUBA1C |
| ENSG00000125877 | ITPA | 0,103204965 | 1,074157067 | 0,62243 | 0,8084993 | inosine triphosphatase |
| ENSG00000260277 | AC009019,1 | 0,103172125 | 1,074132617 | 0,91256 | 0,96205682 | novel transcript, antisense to C16orf52 and VWA3A |
| ENSG00000140995 | DEF8 | 0,103116651 | 1,074091316 | 0,53981 | 0,75803648 | differentially expressed in FDCP 8 homolog |

| | | | | | | |
|---|---|---|---|---|---|---|
| ENSG00000243716 | NPIPB5 | 0,103098801 | 1,074078026 | 0,53845 | 0,7567699 | nuclear pore complex interacting protein family member B5 |
| ENSG00000107521 | HPS1 | 0,103039831 | 1,074034124 | 0,68968 | 0,84865812 | HPS1 biogenesis of lysosomal organelles complex 3 subunit 1 |
| ENSG00000282226 | AC093614,2 | 0,103027513 | 1,074024953 | 0,81029 | 0,91251177 | novel transcript |
| ENSG00000174948 | GPR149 | 0,102971258 | 1,073983075 | 0,82135 | 0,91742056 | G protein-coupled receptor 149 |
| ENSG00000197989 | SNHG12 | 0,102970598 | 1,073982584 | 0,66337 | 0,83242946 | small nucleolar RNA host gene 12 |
| ENSG00000165959 | CLMN | 0,102947114 | 1,073965102 | 0,7023 | 0,85651838 | calmin |
| ENSG00000131323 | TRAF3 | 0,102840858 | 1,073886006 | 0,58592 | 0,78652174 | TNF receptor associated factor 3 |
| ENSG00000100890 | PRORP | 0,102838909 | 1,073884556 | 0,53231 | 0,75291763 | protein only RNase P catalytic subunit |
| ENSG00000100266 | PACSIN2 | 0,102827933 | 1,073876386 | 0,60856 | 0,80122454 | protein kinase C and casein kinase substrate in neurons 2 |
| ENSG00000268201 | AC020915,1 | 0,102774985 | 1,073836974 | 0,81124 | 0,91307458 | novel transcript, sense intronic to ZNF544 |
| ENSG00000034510 | TMSB10 | 0,102711135 | 1,073789449 | 0,66699 | 0,83465859 | thymosin beta 10 |
| ENSG00000165476 | REEP3 | 0,102701469 | 1,073782255 | 0,6696 | 0,83623395 | receptor accessory protein 3 |
| ENSG00000229887 | HNRNPA1P6 | 0,102690335 | 1,073773968 | 0,85937 | 0,93683711 | heterogeneous nuclear ribonucleoprotein A1 pseudogene 6 |
| ENSG00000196405 | EVL | 0,102662457 | 1,07375322 | 0,59162 | 0,79071076 | Enah/Vasp-like |
| ENSG00000068724 | TTC7A | 0,102617487 | 1,07371975 | 0,64422 | 0,82151477 | tetratricopeptide repeat domain 7A |
| ENSG00000228809 | AL034428,1 | 0,102528493 | 1,073653519 | 0,86963 | 0,9425949 | novel transcript, antisense to SNRPB2 |
| ENSG00000154928 | EPHB1 | 0,102461899 | 1,07360396 | 0,70076 | 0,85546985 | EPH receptor B1 |
| ENSG00000133316 | WDR74 | 0,102439392 | 1,073587212 | 0,63958 | 0,81913513 | WD repeat domain 74 |
| ENSG00000272703 | AP005137,2 | 0,102320771 | 1,073498943 | 0,93506 | 0,9721963 | novel transcript |
| ENSG00000114547 | ROPN1B | 0,102311365 | 1,073491944 | 0,81585 | 0,91534523 | rhophilin associated tail protein 1B |
| ENSG00000117569 | PTBP2 | 0,102282681 | 1,073470601 | 0,73548 | 0,87505874 | polypyrimidine tract binding protein 2 |
| ENSG00000168028 | RPSA | 0,102194023 | 1,073404635 | 0,62388 | 0,80952593 | ribosomal protein SA |
| ENSG00000165572 | KBTBD6 | 0,102183027 | 1,073396454 | 0,55015 | 0,76444876 | kelch repeat and BTB domain containing 6 |
| ENSG00000178573 | MAF | 0,102035296 | 1,073286544 | 0,82811 | 0,9207216 | MAF bZIP transcription factor |
| ENSG00000143942 | CHAC2 | 0,101998745 | 1,073259353 | 0,73655 | 0,87529332 | ChaC glutathione specific gamma-glutamylcyclotransferase 2 |
| ENSG00000223431 | MTND6P21 | 0,101927428 | 1,073206299 | 0,91828 | 0,96446185 | MT-ND6 pseudogene 21 |
| ENSG00000231652 | AL590428,1 | 0,10192634 | 1,07320549 | 0,86534 | 0,94013317 | novel transcript |
| ENSG00000186665 | C17orf58 | 0,101902154 | 1,073187498 | 0,66016 | 0,83029149 | chromosome 17 open reading frame 58 |
| ENSG00000168298 | H1-4 | 0,101833148 | 1,073136168 | 0,72038 | 0,86628014 | H1,4 linker histone, cluster member |

| | | | | | | |
|---|---|---|---|---|---|---|
| ENSG00000255395 | AP001972,4 | 0,101809735 | 1,073118753 | 0,83335 | 0,923909 | novel transcript, antisense to SLCO2B1 |
| ENSG00000224775 | BRAFP1 | 0,10179872 | 1,073110559 | 0,87365 | 0,94405019 | BRAF pseudogene 1 |
| ENSG00000149295 | DRD2 | 0,101731801 | 1,073060784 | 0,86984 | 0,94261122 | dopamine receptor D2 |
| ENSG00000132394 | EEFSEC | 0,101556766 | 1,072930603 | 0,87945 | 0,94631723 | eukaryotic elongation factor, selenocysteine-tRNA specific |
| ENSG00000103522 | IL21R | 0,101552946 | 1,072927762 | 0,8556 | 0,93454951 | interleukin 21 receptor |
| ENSG00000198742 | SMURF1 | 0,101464128 | 1,07286171 | 0,90372 | 0,95873972 | SMAD specific E3 ubiquitin protein ligase 1 |
| ENSG00000217624 | YWHAZP10 | 0,101456357 | 1,072855932 | 0,8394 | 0,92655196 | YWHAZ pseudogene 10 |
| ENSG00000278768 | BACE1-AS | 0,101441979 | 1,07284524 | 0,55413 | 0,76713437 | BACE1 antisense RNA |
| ENSG00000221988 | PPT2 | 0,101433017 | 1,072838575 | 0,75237 | 0,88396524 | palmitoyl-protein thioesterase 2 |
| ENSG00000073670 | ADAM11 | 0,101258258 | 1,072708626 | 0,74954 | 0,8822079 | ADAM metallopeptidase domain 11 |
| ENSG00000264575 | LINC00526 | 0,101229095 | 1,072686942 | 0,72351 | 0,86795614 | long intergenic non-protein coding RNA 526 |
| ENSG00000149292 | TTC12 | 0,10120119 | 1,072666194 | 0,70806 | 0,85927396 | tetratricopeptide repeat domain 12 |
| ENSG00000268058 | BNIP3P40 | 0,101190678 | 1,072658378 | 0,84851 | 0,93094401 | BCL2 interacting protein 3 pseudogene 40 |
| ENSG00000259802 | AC012640,2 | 0,101189719 | 1,072657666 | 0,74395 | 0,87882457 | novel transcript, antisense to MARCH6 |
| ENSG00000233384 | CNIH3-AS2 | 0,101088926 | 1,072582728 | 0,77403 | 0,89572556 | CNIH3 antisense RNA 2 |
| ENSG00000105722 | ERF | 0,101073687 | 1,072571398 | 0,72293 | 0,86768626 | ETS2 repressor factor |
| ENSG00000175115 | PACS1 | 0,101020498 | 1,072531855 | 0,60488 | 0,79819795 | phosphofurin acidic cluster sorting protein 1 |
| ENSG00000236618 | PITPNA-AS1 | 0,100968878 | 1,07249348 | 0,74822 | 0,8814896 | PITPNA antisense RNA 1 |
| ENSG00000090674 | MCOLN1 | 0,10077861 | 1,072352045 | 0,6359 | 0,81684802 | mucolipin 1 |
| ENSG00000155761 | SPAG17 | 0,100651798 | 1,072257791 | 0,7925 | 0,90361513 | sperm associated antigen 17 |
| ENSG00000259709 | AC023906,4 | 0,100651383 | 1,072257482 | 0,84422 | 0,92888169 | novel transcript, antisense to GNB5 |
| ENSG00000285360 | LONRF1 | 0,100635698 | 1,072245825 | 0,74869 | 0,8817805 | LON peptidase N-terminal domain and ring finger 1 |
| ENSG00000235381 | AL596202,1 | 0,100514486 | 1,072155741 | 0,67175 | 0,8375243 | novel transcript |
| ENSG00000095380 | NANS | 0,100463631 | 1,072117948 | 0,75143 | 0,88347546 | N-acetylneuraminate synthase |
| ENSG00000223770 | CACNA2D1-AS | 0,100449066 | 1,072107124 | 0,83348 | 0,92396169 | CACNA2D1 antisense RNA 1 |
| ENSG00000170876 | TMEM43 | 0,100019202 | 1,071787728 | 0,61011 | 0,80219814 | transmembrane protein 43 |
| ENSG00000258693 | AL133368,2 | 0,099964241 | 1,071746898 | 0,91238 | 0,96205682 | novel transcript |
| ENSG00000107282 | APBA1 | 0,099926814 | 1,071719095 | 0,67706 | 0,84113215 | amyloid beta precursor protein binding family A member 1 |
| ENSG00000231519 | AC007285,1 | 0,099924076 | 1,071717061 | 0,86818 | 0,94176112 | novel transcript |
| ENSG00000261668 | AC093591,2 | 0,099913273 | 1,071709035 | 0,81791 | 0,9163876 | novel transcript, antisense HSPA4L |
| ENSG00000132846 | ZBED3 | 0,099873776 | 1,071679695 | 0,57009 | 0,7770391 | zinc finger BED-type containing 3 |
| ENSG00000270629 | NBPF14 | 0,099838887 | 1,071653779 | 0,68416 | 0,84539665 | NBPF member 14 |
| ENSG00000174373 | RALGAPA1 | 0,099798298 | 1,071623629 | 0,69863 | 0,85413915 | Ral GTPase activating protein catalytic subunit alpha 1 |

| | | | | | | |
|---|---|---|---|---|---|---|
| ENSG00000261141 | AC092718,5 | 0,099680832 | 1,07153638 | 0,85472 | 0,93416235 | novel transcript, antisense to C16orf46 |
| ENSG00000141012 | GALNS | 0,099626266 | 1,071495853 | 0,61601 | 0,80475871 | galactosamine (N-acetyl)-6-sulfatase |
| ENSG00000235872 | AC078777,1 | 0,099573907 | 1,071456966 | 0,87912 | 0,94623795 | novel transcript |
| ENSG00000150471 | ADGRL3 | 0,099535947 | 1,071428774 | 0,7445 | 0,87904799 | adhesion G protein-coupled receptor L3 |
| ENSG00000198408 | OGA | 0,099446405 | 1,071362278 | 0,63877 | 0,81877064 | O-GlcNAcase |
| ENSG00000172590 | MRPL52 | 0,099432159 | 1,071351699 | 0,56358 | 0,77255679 | mitochondrial ribosomal protein L52 |
| ENSG00000109956 | B3GAT1 | 0,09940656 | 1,071332689 | 0,59545 | 0,79336828 | beta-1,3-glucuronyltransferase 1 |
| ENSG00000261613 | AC093525,6 | 0,099253252 | 1,071218849 | 0,66565 | 0,83358761 | novel transcript, antisense to PDPK1 |
| ENSG00000272968 | RBAK-RBAKDI | 0,099214989 | 1,071190439 | 0,87697 | 0,94549531 | RBAK-RBAKDN readthrough |
| ENSG00000159322 | ADPGK | 0,099145531 | 1,071138868 | 0,61292 | 0,8035281 | ADP dependent glucokinase |
| ENSG00000093183 | SEC22C | 0,099128968 | 1,071126571 | 0,60157 | 0,79671037 | SEC22 homolog C, vesicle trafficking protein |
| ENSG00000143314 | MRPL24 | 0,099064315 | 1,071078571 | 0,63183 | 0,81438644 | mitochondrial ribosomal protein L24 |
| ENSG00000100147 | CCDC134 | 0,099035934 | 1,0710575 | 0,50981 | 0,73746722 | coiled-coil domain containing 134 |
| ENSG00000286522 | H3C2 | 0,098998989 | 1,071030072 | 0,69798 | 0,85378759 | H3 clustered histone 2 |
| ENSG00000189376 | C8orf76 | 0,098921328 | 1,07097242 | 0,67957 | 0,84261173 | chromosome 8 open reading frame 76 |
| ENSG00000154640 | BTG3 | 0,098839446 | 1,070911638 | 0,8148 | 0,91470265 | BTG anti-proliferation factor 3 |
| ENSG00000119705 | SLIRP | 0,098781676 | 1,070868755 | 0,55139 | 0,76514664 | SRA stem-loop interacting RNA binding protein |
| ENSG00000253194 | AL137009,1 | 0,098767509 | 1,07085824 | 0,87086 | 0,94302816 | novel transcript, antisense to FAM184A |
| ENSG00000237522 | NONOP2 | 0,098721582 | 1,070824151 | 0,81269 | 0,91386173 | non-POU domain containing, octamer-binding pseudogene 2 |
| ENSG00000213339 | QTRT1 | 0,098599609 | 1,070733622 | 0,6586 | 0,82926071 | queuine tRNA-ribosyltransferase catalytic subunit 1 |
| ENSG00000092108 | SCFD1 | 0,098587852 | 1,070724896 | 0,54113 | 0,75868526 | sec1 family domain containing 1 |
| ENSG00000253284 | AC092828,1 | 0,09857573 | 1,070715899 | 0,76324 | 0,88980026 | novel transcript |
| ENSG00000212916 | MAP10 | 0,098566579 | 1,070709108 | 0,78001 | 0,89820865 | microtubule associated protein 10 |
| ENSG00000119638 | NEK9 | 0,098507301 | 1,070665115 | 0,64458 | 0,82154386 | NIMA related kinase 9 |
| ENSG00000186806 | VSIG10L | 0,098473153 | 1,070639773 | 0,78356 | 0,89918294 | V-set and immunoglobulin domain containing 10 like |
| ENSG00000268469 | BNIP3P38 | 0,098436807 | 1,070612801 | 0,86738 | 0,94137971 | BCL2 interacting protein 3 pseudogene 38 |
| ENSG00000270179 | AP002840,2 | 0,098429394 | 1,070607299 | 0,84812 | 0,93094401 | novel transcript |
| ENSG00000159023 | EPB41 | 0,098426648 | 1,070605262 | 0,56771 | 0,77563329 | erythrocyte membrane protein band 4,1 |
| ENSG00000197885 | NKIRAS1 | 0,098415013 | 1,070596627 | 0,60215 | 0,7969257 | NFKB inhibitor interacting Ras like 1 |
| ENSG00000272514 | CFAP206 | 0,098319749 | 1,070525936 | 0,7922 | 0,90351559 | cilia and flagella associated protein 206 |
| ENSG00000151690 | MFSD6 | 0,098319202 | 1,07052553 | 0,80151 | 0,90860375 | major facilitator superfamily domain containing 6 |
| ENSG00000260141 | AC145350,2 | 0,098306769 | 1,070516305 | 0,7757 | 0,89630112 | novel transcript |

| | | | | | | |
|---|---|---|---|---|---|---|
| ENSG00000179277 | MEIS3P1 | 0,098305835 | 1,070515611 | 0,77142 | 0,8940638 | Meis homeobox 3 pseudogene 1 |
| ENSG00000257643 | AC084824,2 | 0,098214088 | 1,070447535 | 0,91577 | 0,9633821 | mitochondrial assembly of ribosomal large subunit 1 (MALSU1) pseudogene |
| ENSG00000187741 | FANCA | 0,0981961 | 1,070434188 | 0,57721 | 0,78134056 | FA complementation group A |
| ENSG00000156958 | GALK2 | 0,098185764 | 1,07042652 | 0,62878 | 0,81227675 | galactokinase 2 |
| ENSG00000221886 | ZBED8 | 0,098182789 | 1,070424313 | 0,6697 | 0,83623395 | zinc finger BED-type containing 8 |
| ENSG00000259456 | ADNP-AS1 | 0,098091918 | 1,070356892 | 0,76322 | 0,88980026 | ADNP antisense RNA 1 |
| ENSG00000177556 | ATOX1 | 0,097999672 | 1,070288455 | 0,61407 | 0,80367735 | antioxidant 1 copper chaperone |
| ENSG00000183032 | SLC25A21 | 0,097960154 | 1,070259138 | 0,72566 | 0,8692267 | solute carrier family 25 member 21 |
| ENSG00000243008 | AL359538,1 | 0,097913902 | 1,070224827 | 0,8866 | 0,94987292 | novel transcript |
| ENSG00000136758 | YME1L1 | 0,09788573 | 1,070203928 | 0,4727 | 0,71203455 | YME1 like 1 ATPase |
| ENSG00000137872 | SEMA6D | 0,097870908 | 1,070192933 | 0,83464 | 0,92444381 | semaphorin 6D |
| ENSG00000237100 | C6orf136 | 0,097856835 | 1,070182495 | 0,77684 | 0,89660077 | chromosome 6 open reading frame 136 |
| ENSG00000233641 | C6orf136 | 0,097856835 | 1,070182495 | 0,77684 | 0,89660077 | chromosome 6 open reading frame 136 |
| ENSG00000224120 | C6orf136 | 0,097856835 | 1,070182495 | 0,77684 | 0,89660077 | chromosome 6 open reading frame 136 |
| ENSG00000237012 | C6orf136 | 0,097856835 | 1,070182495 | 0,77684 | 0,89660077 | chromosome 6 open reading frame 136 |
| ENSG00000281396 | AL122023,2 | 0,097837685 | 1,070168289 | 0,90632 | 0,95993807 | novel transcript, sense intronic UNC79 |
| ENSG00000278396 | AL122023,1 | 0,097837685 | 1,070168289 | 0,90632 | 0,95993807 | novel transcript, sense intronic UNC79 |
| ENSG00000128641 | MYO1B | 0,097636061 | 1,070018738 | 0,80171 | 0,90870853 | myosin IB |
| ENSG00000108469 | RECQL5 | 0,097455613 | 1,069884912 | 0,63219 | 0,81450663 | RecQ like helicase 5 |
| ENSG00000160221 | GATD3A | 0,09730171 | 1,069770785 | 0,6395 | 0,81913513 | glutamine amidotransferase like class 1 domain containing 3A |
| ENSG00000109906 | ZBTB16 | 0,097293982 | 1,069765055 | 0,81446 | 0,91470265 | zinc finger and BTB domain containing 16 |
| ENSG00000267648 | AC060766,5 | 0,097132164 | 1,069645073 | 0,88051 | 0,94687991 | DTW domain containing 2 (DTWD2) pseudogene |
| ENSG00000164604 | GPR85 | 0,097092616 | 1,069615751 | 0,85329 | 0,93334646 | G protein-coupled receptor 85 |
| ENSG00000151914 | DST | 0,097073731 | 1,06960175 | 0,78374 | 0,89918294 | dystonin |
| ENSG00000177511 | ST8SIA3 | 0,096959077 | 1,06951675 | 0,83144 | 0,9229059 | ST8 alpha-N-acetyl-neuraminide alpha-2,8-sialyltransferase 3 |
| ENSG00000129933 | MAU2 | 0,096951494 | 1,069511128 | 0,61293 | 0,8035281 | MAU2 sister chromatid cohesion factor |
| ENSG00000175573 | C11orf68 | 0,096803691 | 1,069401563 | 0,61365 | 0,8035281 | chromosome 11 open reading frame 68 |
| ENSG00000189157 | FAM47E | 0,096708375 | 1,069330913 | 0,663 | 0,83240968 | family with sequence similarity 47 member E |
| ENSG00000139620 | KANSL2 | 0,096699445 | 1,069324293 | 0,60237 | 0,7970763 | KAT8 regulatory NSL complex subunit 2 |
| ENSG00000105583 | WDR83OS | 0,096666241 | 1,069299683 | 0,65704 | 0,82860793 | WD repeat domain 83 opposite strand |

| | | | | | | |
|---|---|---|---|---|---|---|
| ENSG00000271737 | AC008608,2 | 0,096621252 | 1,069266338 | 0,89736 | 0,95578297 | novel transcript |
| ENSG00000012232 | EXTL3 | 0,096616541 | 1,069262847 | 0,54945 | 0,76407481 | exostosin like glycosyltransferase 3 |
| ENSG00000288275 | DNAJC18 | 0,096595578 | 1,06924731 | 0,89109 | 0,95208954 | DnaJ heat shock protein family (Hsp40) member C18 |
| ENSG00000266907 | AC006116,1 | 0,096549104 | 1,069212867 | 0,8828 | 0,94822558 | glycerol-3-phosphate dehydrogenase 2 (mitochondrial), pseudogene |
| ENSG00000198954 | KIFBP | 0,09650089 | 1,069177135 | 0,72984 | 0,87166544 | kinesin family binding protein |
| ENSG00000224668 | IPO8P1 | 0,096414793 | 1,069113331 | 0,92577 | 0,96768961 | importin 8 pseudogene 1 |
| ENSG00000170776 | AKAP13 | 0,09630358 | 1,069030919 | 0,59953 | 0,79552021 | A-kinase anchoring protein 13 |
| ENSG00000161040 | FBXL13 | 0,096182102 | 1,068940908 | 0,85311 | 0,93323752 | F-box and leucine rich repeat protein 13 |
| ENSG00000170892 | TSEN34 | 0,096172901 | 1,068934091 | 0,8168 | 0,91576125 | tRNA splicing endonuclease subunit 34 |
| ENSG00000276935 | MBOAT7 | 0,096087678 | 1,068870948 | 0,71219 | 0,8615121 | membrane bound O-acyltransferase domain containing 7 |
| ENSG00000101974 | ATP11C | 0,096055251 | 1,068846924 | 0,70825 | 0,85927396 | ATPase phospholipid transporting 11C |
| ENSG00000107779 | BMPR1A | 0,096011924 | 1,068814825 | 0,77857 | 0,89750716 | bone morphogenetic protein receptor type 1A |
| ENSG00000182057 | OGFRP1 | 0,09594736 | 1,068766994 | 0,78702 | 0,90114442 | opioid growth factor receptor pseudogene 1 |
| ENSG00000120860 | WASHC3 | 0,0959372 | 1,068759468 | 0,83881 | 0,92638462 | WASH complex subunit 3 |
| ENSG00000215012 | RTL10 | 0,095900766 | 1,068732477 | 0,56601 | 0,77413633 | retrotransposon Gag like 10 |
| ENSG00000105438 | KDELR1 | 0,095889119 | 1,068723849 | 0,55446 | 0,76745801 | KDEL endoplasmic reticulum protein retention receptor 1 |
| ENSG00000270562 | AC097634,1 | 0,09582999 | 1,068680048 | 0,8812 | 0,9471287 | novel transcript |
| ENSG00000173933 | RBM4 | 0,095817048 | 1,068670461 | 0,5566 | 0,76850571 | RNA binding motif protein 4 |
| ENSG00000168672 | LRATD2 | 0,095800596 | 1,068658275 | 0,77979 | 0,8981559 | LRAT domain containing 2 |
| ENSG00000165507 | DEPP1 | 0,095767278 | 1,068633595 | 0,7827 | 0,89905956 | DEPP1 autophagy regulator |
| ENSG00000115392 | FANCL | 0,09570385 | 1,068586614 | 0,64574 | 0,82244907 | FA complementation group L |
| ENSG00000144134 | RABL2A | 0,095694281 | 1,068579527 | 0,61434 | 0,8037333 | RAB, member of RAS oncogene family like 2A |
| ENSG00000260385 | AL356961,1 | 0,095642608 | 1,068541254 | 0,90979 | 0,96110658 | novel transcript |
| ENSG00000178568 | ERBB4 | 0,095447984 | 1,068397114 | 0,85411 | 0,93380868 | erb-b2 receptor tyrosine kinase 4 |
| ENSG00000006071 | ABCC8 | 0,095406215 | 1,068366183 | 0,76827 | 0,89274589 | ATP binding cassette subfamily C member 8 |
| ENSG00000141385 | AFG3L2 | 0,095222489 | 1,068230136 | 0,55665 | 0,76853996 | AFG3 like matrix AAA peptidase subunit 2 |
| ENSG00000173894 | CBX2 | 0,095217465 | 1,068226416 | 0,47578 | 0,71433501 | chromobox 2 |
| ENSG00000185104 | FAF1 | 0,095205259 | 1,068217377 | 0,59967 | 0,79552021 | Fas associated factor 1 |
| ENSG00000122966 | CIT | 0,095176976 | 1,068196436 | 0,67191 | 0,83759902 | citron rho-interacting serine/threonine kinase |
| ENSG00000132330 | SCLY | 0,095153313 | 1,068178916 | 0,60411 | 0,79780609 | selenocysteine lyase |
| ENSG00000258435 | AC048337,1 | 0,095109346 | 1,068146363 | 0,94085 | 0,97486264 | novel transcript |
| ENSG00000120942 | UBIAD1 | 0,095056763 | 1,068107432 | 0,66447 | 0,83277805 | UbiA prenyltransferase domain containing 1 |

| | | | | | | |
|---|---|---|---|---|---|---|
| ENSG00000267327 | AC009271,1 | 0,095043071 | 1,068097295 | 0,83696 | 0,92549623 | novel transcript |
| ENSG00000165185 | KIAA1958 | 0,095035392 | 1,06809161 | 0,59135 | 0,79058842 | KIAA1958 |
| ENSG00000112379 | ARFGEF3 | 0,095006533 | 1,068070245 | 0,82101 | 0,91738668 | ARFGEF family member 3 |
| ENSG00000116604 | MEF2D | 0,095002344 | 1,068067144 | 0,6856 | 0,84635545 | myocyte enhancer factor 2D |
| ENSG00000287106 | AC106743,1 | 0,094950084 | 1,068028455 | 0,83493 | 0,92456089 | novel transcript, antisense to WWOX |
| ENSG00000090612 | ZNF268 | 0,094945322 | 1,068024929 | 0,52742 | 0,75037219 | zinc finger protein 268 |
| ENSG00000221843 | C2orf16 | 0,094915513 | 1,068002862 | 0,67254 | 0,83803462 | chromosome 2 open reading frame 16 |
| ENSG00000177600 | RPLP2 | 0,094838424 | 1,067945795 | 0,60936 | 0,80165403 | ribosomal protein lateral stalk subunit P2 |
| ENSG00000124733 | MEA1 | 0,094765868 | 1,067892088 | 0,59098 | 0,79015638 | male-enhanced antigen 1 |
| ENSG00000072210 | ALDH3A2 | 0,094762101 | 1,0678893 | 0,69079 | 0,84906639 | aldehyde dehydrogenase 3 family member A2 |
| ENSG00000277424 | HMOX2 | 0,094752423 | 1,067882136 | 0,75771 | 0,88698721 | heme oxygenase 2 |
| ENSG00000136883 | KIF12 | 0,094738153 | 1,067871573 | 0,90608 | 0,95982458 | kinesin family member 12 |
| ENSG00000031823 | RANBP3 | 0,094676093 | 1,067825638 | 0,61076 | 0,80255859 | RAN binding protein 3 |
| ENSG00000186446 | ZNF501 | 0,094618538 | 1,067783039 | 0,79109 | 0,90292452 | zinc finger protein 501 |
| ENSG00000058600 | POLR3E | 0,094546447 | 1,067729684 | 0,60929 | 0,80164829 | RNA polymerase III subunit E |
| ENSG00000174227 | PIGG | 0,094371682 | 1,067600349 | 0,621 | 0,80782102 | phosphatidylinositol glycan anchor biosynthesis class G |
| ENSG00000179918 | SEPHS2 | 0,09421613 | 1,067485246 | 0,5445 | 0,76055278 | selenophosphate synthetase 2 |
| ENSG00000281856 | AC106017,2 | 0,09419859 | 1,067472268 | 0,90604 | 0,95982149 | novel transcript |
| ENSG00000152234 | ATP5F1A | 0,094161845 | 1,06744508 | 0,58328 | 0,78513431 | ATP synthase F1 subunit alpha |
| ENSG00000049759 | NEDD4L | 0,094128451 | 1,067420372 | 0,66158 | 0,83148232 | NEDD4 like E3 ubiquitin protein ligase |
| ENSG00000285884 | AL022345,4 | 0,094097743 | 1,067397652 | 0,90782 | 0,96044232 | novel transcript |
| ENSG00000286686 | AL136040,2 | 0,094052735 | 1,067364353 | 0,80717 | 0,91127337 | novel transcript |
| ENSG00000120322 | PCDHB8 | 0,094004912 | 1,067328972 | 0,81952 | 0,91708488 | protocadherin beta 8 |
| ENSG00000197385 | ZNF860 | 0,093914093 | 1,067261785 | 0,91061 | 0,96143161 | zinc finger protein 860 |
| ENSG00000026025 | VIM | 0,093908708 | 1,067257801 | 0,74908 | 0,88190158 | vimentin |
| ENSG00000105877 | DNAH11 | 0,093818015 | 1,067190711 | 0,75398 | 0,88467924 | dynein axonemal heavy chain 11 |
| ENSG00000138083 | SIX3 | 0,093766071 | 1,067152288 | 0,85821 | 0,93612837 | SIX homeobox 3 |
| ENSG00000183598 | H3C13 | 0,0936671 | 1,067079082 | 0,64478 | 0,82169377 | H3 clustered histone 13 |
| ENSG00000118307 | CFAP94 | 0,093598113 | 1,067028058 | 0,84801 | 0,9309317 | cilia and flagella associated protein 94 |
| ENSG00000164114 | MAP9 | 0,093580609 | 1,067015112 | 0,80657 | 0,91114466 | microtubule associated protein 9 |
| ENSG00000239523 | MYLK-AS1 | 0,093493277 | 1,066950523 | 0,62146 | 0,80788604 | MYLK antisense RNA 1 |
| ENSG00000153291 | SLC25A27 | 0,093385171 | 1,066870576 | 0,73725 | 0,87539732 | solute carrier family 25 member 27 |
| ENSG00000095397 | WHRN | 0,093351874 | 1,066845954 | 0,7306 | 0,87202167 | whirlin |

| | | | | | | |
|---|---|---|---|---|---|---|
| ENSG00000131943 | C19orf12 | 0,093332727 | 1,066831795 | 0,59725 | 0,79440664 | chromosome 19 open reading frame 12 |
| ENSG00000277662 | AL354696,1 | 0,093279361 | 1,066792333 | 0,90028 | 0,95760242 | novel transcript, sense intronic to MTRF1 |
| ENSG00000226281 | AL031123,1 | 0,093229981 | 1,06675582 | 0,91956 | 0,96506391 | novel transcript |
| ENSG00000234925 | ATP5PDP4 | 0,093157633 | 1,066702325 | 0,87041 | 0,942849 | ATP synthase peripheral stalk subunit d pseudogene 4 |
| ENSG00000134278 | SPIRE1 | 0,093124651 | 1,066677939 | 0,74692 | 0,88059941 | spire type actin nucleation factor 1 |
| ENSG00000214189 | ZNF788P | 0,093083433 | 1,066647465 | 0,67688 | 0,84109542 | zinc finger family member 788, pseudogene |
| ENSG00000184220 | CMSS1 | 0,093069707 | 1,066637317 | 0,64717 | 0,82344958 | cms1 ribosomal small subunit homolog |
| ENSG00000071051 | NCK2 | 0,093054538 | 1,066626101 | 0,68863 | 0,84822736 | NCK adaptor protein 2 |
| ENSG00000154451 | GBP5 | 0,093016744 | 1,06659816 | 0,8448 | 0,92922901 | guanylate binding protein 5 |
| ENSG00000267272 | LINC01140 | 0,093010014 | 1,066593184 | 0,86307 | 0,93891554 | long intergenic non-protein coding RNA 1140 |
| ENSG00000130826 | DKC1 | 0,093001169 | 1,066586645 | 0,64251 | 0,82079227 | dyskerin pseudouridine synthase 1 |
| ENSG00000197016 | ZNF470 | 0,092973408 | 1,066566122 | 0,61634 | 0,80492045 | zinc finger protein 470 |
| ENSG00000142687 | KIAA0319L | 0,09297086 | 1,066564238 | 0,53341 | 0,75351438 | KIAA0319 like |
| ENSG00000157837 | SPPL3 | 0,092934643 | 1,066537464 | 0,5413 | 0,75881984 | signal peptide peptidase like 3 |
| ENSG00000284862 | CCDC39 | 0,09292344 | 1,066529181 | 0,77898 | 0,89781842 | coiled-coil domain containing 39 |
| ENSG00000188690 | UROS | 0,092832873 | 1,066462231 | 0,63428 | 0,81566722 | uroporphyrinogen III synthase |
| ENSG00000214078 | CPNE1 | 0,092726273 | 1,066383433 | 0,61034 | 0,80221019 | copine 1 |
| ENSG00000124507 | PACSIN1 | 0,092576275 | 1,066272566 | 0,88084 | 0,94688557 | protein kinase C and casein kinase substrate in neurons 1 |
| ENSG00000276111 | SDCCAG8 | 0,092516961 | 1,066228729 | 0,81978 | 0,91716029 | SHH signaling and ciliogenesis regulator SDCCAG8 |
| ENSG00000158411 | MITD1 | 0,092503222 | 1,066218575 | 0,63002 | 0,8132682 | microtubule interacting and trafficking domain containing 1 |
| ENSG00000182224 | CYB5D1 | 0,092413913 | 1,066152574 | 0,67754 | 0,84135792 | cytochrome b5 domain containing 1 |
| ENSG00000177683 | THAP5 | 0,092367436 | 1,066118228 | 0,58902 | 0,7885434 | THAP domain containing 5 |
| ENSG00000122557 | HERPUD2 | 0,092310294 | 1,066076002 | 0,61265 | 0,8035281 | HERPUD family member 2 |
| ENSG00000005020 | SKAP2 | 0,092288029 | 1,06605955 | 0,82212 | 0,91790528 | src kinase associated phosphoprotein 2 |
| ENSG00000279072 | AC118758,2 | 0,092275101 | 1,066049997 | 0,83387 | 0,92404752 | novel piRNA host transcript |
| ENSG00000135063 | FAM189A2 | 0,092272095 | 1,066047775 | 0,86796 | 0,94175463 | family with sequence similarity 189 member A2 |
| ENSG00000180198 | RCC1 | 0,092216494 | 1,066006692 | 0,62773 | 0,81173104 | regulator of chromosome condensation 1 |
| ENSG00000170545 | SMAGP | 0,092207081 | 1,065999736 | 0,83146 | 0,9229059 | small cell adhesion glycoprotein |
| ENSG00000281661 | ZNF501 | 0,092152368 | 1,06595931 | 0,79517 | 0,90519297 | zinc finger protein 501 |
| ENSG00000260282 | EIF4EBP2P2 | 0,092102776 | 1,065922668 | 0,91594 | 0,96339511 | eukaryotic translation initiation factor 4E binding protein 2 pseudogene 2 |
| ENSG00000198826 | ARHGAP11A | 0,092101806 | 1,065921952 | 0,78083 | 0,89846287 | Rho GTPase activating protein 11A |

| | | | | | | |
|---|---|---|---|---|---|---|
| ENSG00000224584 | UBE2V1P13 | 0,092082802 | 1,065907911 | 0,88043 | 0,94687991 | ubiquitin conjugating enzyme E2 V1 pseudogene 13 |
| ENSG00000287218 | AL359962,3 | 0,092073366 | 1,06590094 | 0,89829 | 0,95630226 | novel transcript, antisense to CD247 |
| ENSG00000185129 | PURA | 0,092033401 | 1,065871413 | 0,72233 | 0,86734497 | purine rich element binding protein A |
| ENSG00000107862 | GBF1 | 0,09199353 | 1,065841956 | 0,54886 | 0,76394298 | golgi brefeldin A resistant guanine nucleotide exchange factor 1 |
| ENSG00000164933 | SLC25A32 | 0,091869028 | 1,06574998 | 0,6169 | 0,80545869 | solute carrier family 25 member 32 |
| ENSG00000269376 | AL356740,3 | 0,0917698 | 1,065676681 | 0,88529 | 0,94924301 | novel transcript |
| ENSG00000232671 | ZNF687-AS1 | 0,091675194 | 1,0656068 | 0,81529 | 0,91502073 | ZNF687 antisense RNA 1 |
| ENSG00000269940 | AL049840,4 | 0,091653179 | 1,06559054 | 0,75268 | 0,88411623 | novel transcript, sense intronic to KLC1 |
| ENSG00000160917 | CPSF4 | 0,09161325 | 1,065561048 | 0,64373 | 0,82122916 | cleavage and polyadenylation specific factor 4 |
| ENSG00000175505 | CLCF1 | 0,091585706 | 1,065540704 | 0,82095 | 0,91738668 | cardiotrophin like cytokine factor 1 |
| ENSG00000198670 | LPA | 0,091582945 | 1,065538665 | 0,86881 | 0,94211995 | lipoprotein(a) |
| ENSG00000286760 | AL022069,3 | 0,091555442 | 1,065518352 | 0,76126 | 0,88879633 | novel transcript |
| ENSG00000280548 | FBL | 0,091545326 | 1,065510881 | 0,66109 | 0,83105713 | fibrillarin |
| ENSG00000184787 | UBE2G2 | 0,091515232 | 1,065488655 | 0,57392 | 0,77926229 | ubiquitin conjugating enzyme E2 G2 |
| ENSG00000103888 | CEMIP | 0,091485261 | 1,06546652 | 0,80084 | 0,90830749 | cell migration inducing hyaluronidase 1 |
| ENSG00000233012 | HDAC1P2 | 0,0913342 | 1,065354964 | 0,82747 | 0,92023045 | histone deacetylase 1 pseudogene 2 |
| ENSG00000078549 | ADCYAP1R1 | 0,091246004 | 1,065289838 | 0,90112 | 0,95787401 | ADCYAP receptor type I |
| ENSG00000196470 | SIAH1 | 0,091222388 | 1,0652724 | 0,56442 | 0,7729862 | siah E3 ubiquitin protein ligase 1 |
| ENSG00000085982 | USP40 | 0,091216048 | 1,065267719 | 0,62735 | 0,8114294 | ubiquitin specific peptidase 40 |
| ENSG00000072778 | ACADVL | 0,09110018 | 1,065182166 | 0,60927 | 0,80164829 | acyl-CoA dehydrogenase very long chain |
| ENSG00000165526 | RPUSD4 | 0,09107804 | 1,06516582 | 0,64985 | 0,82453893 | RNA pseudouridine synthase D4 |
| ENSG00000173120 | KDM2A | 0,091070473 | 1,065160234 | 0,54137 | 0,75881984 | lysine demethylase 2A |
| ENSG00000181754 | AMIGO1 | 0,091007024 | 1,065113389 | 0,60424 | 0,79780609 | adhesion molecule with Ig like domain 1 |
| ENSG00000285222 | GCNT2 | 0,090998635 | 1,065107196 | 0,81642 | 0,91554153 | glucosaminyl (N-acetyl) transferase 2 (I blood group) |
| ENSG00000183955 | KMT5A | 0,090866697 | 1,065009793 | 0,71326 | 0,86208742 | lysine methyltransferase 5A |
| ENSG00000135457 | TFCP2 | 0,090854787 | 1,065001002 | 0,69634 | 0,85299392 | transcription factor CP2 |
| ENSG00000076351 | SLC46A1 | 0,090788713 | 1,064952226 | 0,80399 | 0,909789 | solute carrier family 46 member 1 |
| ENSG00000204619 | PPP1R11 | 0,090726974 | 1,064906654 | 0,68816 | 0,84809542 | protein phosphatase 1 regulatory inhibitor subunit 11 |
| ENSG00000108826 | MRPL27 | 0,090679921 | 1,064871923 | 0,60289 | 0,7971742 | mitochondrial ribosomal protein L27 |
| ENSG00000119471 | HSDL2 | 0,090642009 | 1,06484394 | 0,84685 | 0,93041089 | hydroxysteroid dehydrogenase like 2 |
| ENSG00000233369 | GTF2IP4 | 0,090599545 | 1,064812598 | 0,68265 | 0,84454346 | general transcription factor IIi pseudogene 4 |
| ENSG00000068001 | HYAL2 | 0,090515391 | 1,064750488 | 0,6337 | 0,81549489 | hyaluronidase 2 |

| | | | | | | |
|---|---|---|---|---|---|---|
| ENSG00000176148 | TCP11L1 | 0,090511299 | 1,064747468 | 0,57004 | 0,77703861 | t-complex 11 like 1 |
| ENSG00000143443 | C1orf56 | 0,09050657 | 1,064743978 | 0,71347 | 0,86217438 | chromosome 1 open reading frame 56 |
| ENSG00000102078 | SLC25A14 | 0,090474697 | 1,064720455 | 0,78018 | 0,89822236 | solute carrier family 25 member 14 |
| ENSG00000223974 | BNIP3P42 | 0,090396036 | 1,064662404 | 0,87919 | 0,94623795 | BCL2 interacting protein 3 pseudogene 42 |
| ENSG00000162063 | CCNF | 0,090394272 | 1,064661102 | 0,64876 | 0,82412601 | cyclin F |
| ENSG00000249359 | AC093274,1 | 0,090157091 | 1,064486085 | 0,89726 | 0,955727 | novel transcript |
| ENSG00000092094 | OSGEP | 0,090108806 | 1,064450459 | 0,58106 | 0,78406342 | O-sialoglycoprotein endopeptidase |
| ENSG00000103653 | CSK | 0,090061484 | 1,064415544 | 0,67872 | 0,84182956 | C-terminal Src kinase |
| ENSG00000179632 | MAF1 | 0,090054703 | 1,064410541 | 0,62086 | 0,80776776 | MAF1 homolog, negative regulator of RNA polymerase III |
| ENSG00000132128 | LRRC41 | 0,090048557 | 1,064406006 | 0,52805 | 0,75070014 | leucine rich repeat containing 41 |
| ENSG00000070366 | SMG6 | 0,090020766 | 1,064385503 | 0,65695 | 0,82856771 | SMG6 nonsense mediated mRNA decay factor |
| ENSG00000164258 | NDUFS4 | 0,090001981 | 1,064371644 | 0,79807 | 0,90685415 | NADH:ubiquinone oxidoreductase subunit S4 |
| ENSG00000162374 | ELAVL4 | 0,089833213 | 1,06424714 | 0,86632 | 0,94074101 | ELAV like RNA binding protein 4 |
| ENSG00000143199 | ADCY10 | 0,089792224 | 1,064216903 | 0,87722 | 0,94559723 | adenylate cyclase 10 |
| ENSG00000174740 | PABPC5 | 0,089709775 | 1,064156086 | 0,77917 | 0,89786635 | poly(A) binding protein cytoplasmic 5 |
| ENSG00000115525 | ST3GAL5 | 0,089684682 | 1,064137578 | 0,70662 | 0,85885395 | ST3 beta-galactoside alpha-2,3-sialyltransferase 5 |
| ENSG00000172965 | MIR4435-2HG | 0,089684084 | 1,064137136 | 0,64822 | 0,82383528 | MIR4435-2 host gene |
| ENSG00000105750 | ZNF85 | 0,089635167 | 1,064101056 | 0,88614 | 0,94956686 | zinc finger protein 85 |
| ENSG00000160877 | NACC1 | 0,089609993 | 1,064082488 | 0,57289 | 0,77868876 | nucleus accumbens associated 1 |
| ENSG00000128881 | TTBK2 | 0,089600419 | 1,064075427 | 0,73713 | 0,87539732 | tau tubulin kinase 2 |
| ENSG00000206283 | PFDN6 | 0,08958268 | 1,064062343 | 0,78515 | 0,90010552 | prefoldin subunit 6 |
| ENSG00000140455 | USP3 | 0,089492965 | 1,063996175 | 0,75397 | 0,88467924 | ubiquitin specific peptidase 3 |
| ENSG00000136770 | DNAJC1 | 0,089366071 | 1,063902594 | 0,78804 | 0,90149492 | DnaJ heat shock protein family (Hsp40) member C1 |
| ENSG00000279278 | AC245060,6 | 0,089282072 | 1,063840652 | 0,75214 | 0,88389154 | novel transcript |
| ENSG00000137411 | VARS2 | 0,089074338 | 1,06368748 | 0,84463 | 0,92917984 | valyl-tRNA synthetase 2, mitochondrial |
| ENSG00000283154 | IQCJ-SCHIP1 | 0,089068837 | 1,063683424 | 0,70116 | 0,85581073 | IQCJ-SCHIP1 readthrough |
| ENSG00000256628 | ZBTB11-AS1 | 0,089058689 | 1,063675942 | 0,72399 | 0,8682296 | ZBTB11 antisense RNA 1 |
| ENSG00000287268 | AL050350,1 | 0,08887171 | 1,063538095 | 0,87312 | 0,94385071 | novel transcript, antisense to METTL24 |
| ENSG00000283403 | AC005394,2 | 0,088827706 | 1,063505656 | 0,80701 | 0,91125307 | novel transcript |
| ENSG00000274626 | RPS9 | 0,088790994 | 1,063478593 | 0,6188 | 0,80665241 | ribosomal protein S9 |
| ENSG00000277359 | RPS9 | 0,088790994 | 1,063478593 | 0,6188 | 0,80665241 | ribosomal protein S9 |
| ENSG00000274005 | RPS9 | 0,088790994 | 1,063478593 | 0,6188 | 0,80665241 | ribosomal protein S9 |
| ENSG00000278081 | RPS9 | 0,088790994 | 1,063478593 | 0,6188 | 0,80665241 | ribosomal protein S9 |

| | | | | | | |
|---|---|---|---|---|---|---|
| ENSG00000275323 | RPS9 | 0,088790994 | 1,063478593 | 0,6188 | 0,80665241 | ribosomal protein S9 |
| ENSG00000277079 | RPS9 | 0,088790994 | 1,063478593 | 0,6188 | 0,80665241 | ribosomal protein S9 |
| ENSG00000136270 | TBRG4 | 0,088782879 | 1,063472612 | 0,62087 | 0,80776776 | transforming growth factor beta regulator 4 |
| ENSG00000178381 | ZFAND2A | 0,088781624 | 1,063471686 | 0,6835 | 0,84495012 | zinc finger AN1-type containing 2A |
| ENSG00000149311 | ATM | 0,088758788 | 1,063454853 | 0,63324 | 0,81541029 | ATM serine/threonine kinase |
| ENSG00000198604 | BAZ1A | 0,088757466 | 1,063453879 | 0,68866 | 0,84822736 | bromodomain adjacent to zinc finger domain 1A |
| ENSG00000259397 | AC021231,2 | 0,08874963 | 1,063448103 | 0,93964 | 0,9743009 | U2 small nuclear RNA auxiliary factor 1 (U2AF1) pseudogene |
| ENSG00000175344 | CHRNA7 | 0,088696947 | 1,06340927 | 0,76879 | 0,89277929 | cholinergic receptor nicotinic alpha 7 subunit |
| ENSG00000262209 | PCDHGB3 | 0,088607276 | 1,063343175 | 0,82944 | 0,92157796 | protocadherin gamma subfamily B, 3 |
| ENSG00000182628 | SKA2 | 0,088576143 | 1,063320229 | 0,6097 | 0,80189841 | spindle and kinetochore associated complex subunit 2 |
| ENSG00000135317 | SNX14 | 0,088352304 | 1,063155263 | 0,65543 | 0,82781124 | sorting nexin 14 |
| ENSG00000287385 | AL121820,3 | 0,088095878 | 1,062966314 | 0,82237 | 0,91794576 | uncharacterized LOC107984016 |
| ENSG00000271824 | SMIM32 | 0,087922424 | 1,062838522 | 0,89421 | 0,95389664 | small integral membrane protein 32 |
| ENSG00000176933 | TOB2P1 | 0,087864944 | 1,062796178 | 0,87441 | 0,9443876 | transducer of ERBB2, 2 pseudogene 1 |
| ENSG00000009830 | POMT2 | 0,087844928 | 1,062781432 | 0,68037 | 0,84309883 | protein O-mannosyltransferase 2 |
| ENSG00000119397 | CNTRL | 0,087745501 | 1,06270819 | 0,76791 | 0,89249843 | centriolin |
| ENSG00000206490 | ABCF1 | 0,087655407 | 1,062641828 | 0,77924 | 0,89791639 | ATP binding cassette subfamily F member 1 |
| ENSG00000174791 | RIN1 | 0,087648503 | 1,062636743 | 0,77494 | 0,8960241 | Ras and Rab interactor 1 |
| ENSG00000154040 | CABYR | 0,087637874 | 1,062628914 | 0,70767 | 0,85918434 | calcium binding tyrosine phosphorylation regulated |
| ENSG00000288045 | AL352979,4 | 0,087541013 | 1,062557573 | 0,81478 | 0,91470265 | novel transcript, antisense to FERMT2 |
| ENSG00000238795 | SCARNA12 | 0,087521851 | 1,06254346 | 0,80872 | 0,91198969 | small Cajal body-specific RNA 12 |
| ENSG00000267784 | AC010680,1 | 0,087500082 | 1,062527427 | 0,83823 | 0,92618329 | novel transcript, antisense to titin |
| ENSG00000162415 | ZSWIM5 | 0,087350106 | 1,062416977 | 0,76527 | 0,89082001 | zinc finger SWIM-type containing 5 |
| ENSG00000012822 | CALCOCO1 | 0,087350003 | 1,062416901 | 0,68523 | 0,84608974 | calcium binding and coiled-coil domain 1 |
| ENSG00000279669 | AC079801,1 | 0,087330291 | 1,062402386 | 0,91421 | 0,96287665 | novel transcript |
| ENSG00000132254 | ARFIP2 | 0,087326807 | 1,062399819 | 0,65102 | 0,82521872 | ADP ribosylation factor interacting protein 2 |
| ENSG00000164038 | SLC9B2 | 0,087161146 | 1,062277834 | 0,76898 | 0,89287402 | solute carrier family 9 member B2 |
| ENSG00000076242 | MLH1 | 0,087109691 | 1,062239948 | 0,56739 | 0,77531691 | mutL homolog 1 |
| ENSG00000275003 | DDT | 0,086968713 | 1,062136153 | 0,75613 | 0,88583532 | D-dopachrome tautomerase |
| ENSG00000239306 | RBM14 | 0,086896137 | 1,062082722 | 0,58244 | 0,7847455 | RNA binding motif protein 14 |
| ENSG00000157538 | VPS26C | 0,086879958 | 1,062070811 | 0,59966 | 0,79552021 | VPS26 endosomal protein sorting factor C |
| ENSG00000267313 | KC6 | 0,086557748 | 1,061833636 | 0,78448 | 0,89962715 | keratoconus gene 6 |

| | | | | | | |
|---|---|---|---|---|---|---|
| ENSG00000111639 | MRPL51 | 0,086518488 | 1,061804741 | 0,6336 | 0,81549489 | mitochondrial ribosomal protein L51 |
| ENSG00000283703 | VSIG10L2 | 0,086508584 | 1,061797452 | 0,91874 | 0,9646736 | V-set and immunoglobulin domain containing 10 like 2 |
| ENSG00000130684 | ZNF337 | 0,086508528 | 1,06179741 | 0,84689 | 0,93041089 | zinc finger protein 337 |
| ENSG00000197818 | SLC9A8 | 0,086447192 | 1,06175227 | 0,6817 | 0,84383965 | solute carrier family 9 member A8 |
| ENSG00000223959 | AFG3L1P | 0,086417275 | 1,061730252 | 0,63024 | 0,81330068 | AFG3 like matrix AAA peptidase subunit 1, pseudogene |
| ENSG00000148584 | A1CF | 0,086387283 | 1,06170818 | 0,84311 | 0,92838808 | APOBEC1 complementation factor |
| ENSG00000142798 | HSPG2 | 0,086355509 | 1,061684797 | 0,59467 | 0,7928588 | heparan sulfate proteoglycan 2 |
| ENSG00000151498 | ACAD8 | 0,086325091 | 1,061662412 | 0,57891 | 0,78259415 | acyl-CoA dehydrogenase family member 8 |
| ENSG00000287670 | AC108479,1 | 0,086300033 | 1,061643973 | 0,87659 | 0,94547358 | novel transcript |
| ENSG00000272305 | AC096887,1 | 0,086276171 | 1,061626414 | 0,93929 | 0,97411824 | novel transcript |
| ENSG00000148541 | FAM13C | 0,086254967 | 1,061610811 | 0,86812 | 0,94176112 | family with sequence similarity 13 member C |
| ENSG00000148090 | AUH | 0,0861833 | 1,061558075 | 0,70591 | 0,85838918 | AU RNA binding methylglutaconyl-CoA hydratase |
| ENSG00000101945 | SUV39H1 | 0,086151037 | 1,061534336 | 0,68242 | 0,84436596 | suppressor of variegation 3-9 homolog 1 |
| ENSG00000105669 | COPE | 0,086146748 | 1,06153118 | 0,65034 | 0,82485623 | COPI coat complex subunit epsilon |
| ENSG00000233993 | DTD1-AS1 | 0,086134692 | 1,06152231 | 0,84932 | 0,93142453 | DTD1 antisense RNA 1 |
| ENSG00000196268 | ZNF493 | 0,086085463 | 1,061486088 | 0,72181 | 0,86716786 | zinc finger protein 493 |
| ENSG00000142684 | ZNF593 | 0,086082032 | 1,061483563 | 0,78147 | 0,89877377 | zinc finger protein 593 |
| ENSG00000118495 | PLAGL1 | 0,086067628 | 1,061472965 | 0,84793 | 0,9309317 | PLAG1 like zinc finger 1 |
| ENSG00000100629 | CEP128 | 0,086051065 | 1,06146078 | 0,78807 | 0,90150029 | centrosomal protein 128 |
| ENSG00000143570 | SLC39A1 | 0,086043825 | 1,061455453 | 0,68419 | 0,84539665 | solute carrier family 39 member 1 |
| ENSG00000146963 | LUC7L2 | 0,085991873 | 1,061417229 | 0,58724 | 0,78717516 | LUC7 like 2, pre-mRNA splicing factor |
| ENSG00000256594 | AC010186,2 | 0,085956949 | 1,061391536 | 0,77821 | 0,89735311 | C-type lectin domain family 2 (CLEC2) pseudogene |
| ENSG00000278619 | MRM1 | 0,085937091 | 1,061376926 | 0,89134 | 0,95208954 | mitochondrial rRNA methyltransferase 1 |
| ENSG00000137185 | ZSCAN9 | 0,085907417 | 1,061355096 | 0,74241 | 0,87819795 | zinc finger and SCAN domain containing 9 |
| ENSG00000171307 | ZDHHC16 | 0,085793179 | 1,061271057 | 0,65261 | 0,826091 | zinc finger DHHC-type palmitoyltransferase 16 |
| ENSG00000259630 | FABP5P9 | 0,085776825 | 1,061259027 | 0,88586 | 0,94944442 | fatty acid binding protein 5 pseudogene 9 |
| ENSG00000229544 | NKX1-2 | 0,085765466 | 1,061250671 | 0,92062 | 0,9656015 | NK1 homeobox 2 |
| ENSG00000250651 | PABPC1P7 | 0,085699208 | 1,061201933 | 0,93295 | 0,97143736 | poly(A) binding protein cytoplasmic 1 pseudogene 7 |
| ENSG00000235692 | PFDN6 | 0,085696035 | 1,061199599 | 0,75779 | 0,88699366 | prefoldin subunit 6 |
| ENSG00000123131 | PRDX4 | 0,085674748 | 1,061183941 | 0,67953 | 0,84260739 | peroxiredoxin 4 |
| ENSG00000104341 | LAPTM4B | 0,085648165 | 1,061164387 | 0,70788 | 0,85923505 | lysosomal protein transmembrane 4 beta |
| ENSG00000188599 | NPIPP1 | 0,085641935 | 1,061159805 | 0,69674 | 0,85309963 | nuclear pore complex interacting protein pseudogene 1 |
| ENSG00000164938 | TP53INP1 | 0,08561199 | 1,06113778 | 0,74465 | 0,87914393 | tumor protein p53 inducible nuclear protein 1 |

| | | | | | | |
|---|---|---|---|---|---|---|
| ENSG00000240764 | PCDHGC5 | 0,085541048 | 1,061085602 | 0,72651 | 0,86978598 | protocadherin gamma subfamily C, 5 |
| ENSG00000132938 | MTUS2 | 0,085383858 | 1,060969996 | 0,83613 | 0,92493625 | microtubule associated scaffold protein 2 |
| ENSG00000280661 | ZNF660 | 0,085343886 | 1,060940601 | 0,88607 | 0,94954542 | zinc finger protein 660 |
| ENSG00000265413 | AP001094,2 | 0,085258896 | 1,060878102 | 0,90141 | 0,95798991 | novel transcript, sense intronic to PTPRM |
| ENSG00000226686 | LINC01535 | 0,085250218 | 1,060871721 | 0,8949 | 0,95417507 | long intergenic non-protein coding RNA 1535 |
| ENSG00000287965 | AC018557,3 | 0,085223559 | 1,060852117 | 0,88867 | 0,9508882 | novel transcript, antisense to DYNC1LI2 |
| ENSG00000171262 | FAM98B | 0,085142048 | 1,060792182 | 0,79041 | 0,90286944 | family with sequence similarity 98 member B |
| ENSG00000074803 | SLC12A1 | 0,084983317 | 1,060675476 | 0,87386 | 0,94405019 | solute carrier family 12 member 1 |
| ENSG00000119599 | DCAF4 | 0,084937921 | 1,060642101 | 0,77285 | 0,89507232 | DDB1 and CUL4 associated factor 4 |
| ENSG00000151779 | NBAS | 0,084848316 | 1,060576227 | 0,78515 | 0,90010552 | NBAS subunit of NRZ tethering complex |
| ENSG00000173436 | MICOS10 | 0,084769024 | 1,060517938 | 0,64785 | 0,82380454 | mitochondrial contact site and cristae organizing system subunit 10 |
| ENSG00000204805 | AL391987,2 | 0,084767214 | 1,060516608 | 0,91096 | 0,96168891 | |
| ENSG00000105339 | DENND3 | 0,084689383 | 1,060459397 | 0,70244 | 0,85653047 | DENN domain containing 3 |
| ENSG00000110536 | PTPMT1 | 0,084671852 | 1,06044651 | 0,80462 | 0,90996631 | protein tyrosine phosphatase mitochondrial 1 |
| ENSG00000285206 | PTPMT1 | 0,084671852 | 1,06044651 | 0,80462 | 0,90996631 | protein tyrosine phosphatase mitochondrial 1 |
| ENSG00000118564 | FBXL5 | 0,084618831 | 1,060407538 | 0,76978 | 0,89306695 | F-box and leucine rich repeat protein 5 |
| ENSG00000259732 | AC092757,2 | 0,084617461 | 1,060406531 | 0,89032 | 0,95170091 | novel transcript, sense overlapping to CCNB2 |
| ENSG00000275441 | AC020765,2 | 0,084554363 | 1,060360154 | 0,9302 | 0,97010144 | novel transcript, antisense to CCDC101 |
| ENSG00000284872 | AC270289,3 | 0,08454944 | 1,060356536 | 0,87652 | 0,94547358 | tRNA methyltransferase 9B (putative) |
| ENSG00000232862 | AL138787,1 | 0,084486778 | 1,060310481 | 0,91582 | 0,9633821 | implantation-associated protein (DKFZp564K142) pseudogene |
| ENSG00000136848 | DAB2IP | 0,084431143 | 1,060269593 | 0,61364 | 0,8035281 | DAB2 interacting protein |
| ENSG00000245205 | EEF1A1P4 | 0,08432431 | 1,060191082 | 0,87758 | 0,94566578 | eukaryotic translation elongation factor 1 alpha 1 pseudogene 4 |
| ENSG00000167874 | TMEM88 | 0,084231232 | 1,060122684 | 0,84181 | 0,92804468 | transmembrane protein 88 |
| ENSG00000108797 | CNTNAP1 | 0,084227495 | 1,060119938 | 0,80755 | 0,91143945 | contactin associated protein 1 |
| ENSG00000175130 | MARCKSL1 | 0,084126454 | 1,060045693 | 0,60492 | 0,79821396 | MARCKS like 1 |
| ENSG00000170456 | DENND5B | 0,084109132 | 1,060032966 | 0,75833 | 0,88727479 | DENN domain containing 5B |
| ENSG00000225648 | SBDSP1 | 0,084066829 | 1,060001884 | 0,72487 | 0,86868749 | SBDS pseudogene 1 |
| ENSG00000014216 | CAPN1 | 0,084024652 | 1,059970895 | 0,63555 | 0,81657648 | calpain 1 |
| ENSG00000241228 | RPL12P31 | 0,083963935 | 1,059926287 | 0,90828 | 0,96078264 | ribosomal protein L12 pseudogene 31 |
| ENSG00000175463 | TBC1D10C | 0,083928409 | 1,059900187 | 0,81936 | 0,91699877 | TBC1 domain family member 10C |

| | | | | | | |
|---|---|---|---|---|---|---|
| ENSG00000273014 | AC018645,3 | 0,08391814 | 1,059892642 | 0,9349 | 0,9721963 | novel transcript |
| ENSG00000247363 | AC090061,1 | 0,083846122 | 1,059839735 | 0,83574 | 0,92493625 | novel transcript |
| ENSG00000261158 | AC109597,2 | 0,083794556 | 1,059801853 | 0,92565 | 0,96768961 | novel transcript, antisense to CPPED1 |
| ENSG00000135932 | CAB39 | 0,083776219 | 1,059788383 | 0,67754 | 0,84135792 | calcium binding protein 39 |
| ENSG00000140511 | HAPLN3 | 0,083771628 | 1,059785011 | 0,84223 | 0,92821072 | hyaluronan and proteoglycan link protein 3 |
| ENSG00000095587 | TLL2 | 0,083725224 | 1,059750924 | 0,87157 | 0,94322889 | tolloid like 2 |
| ENSG00000159082 | SYNJ1 | 0,083714717 | 1,059743206 | 0,82057 | 0,91732362 | synaptojanin 1 |
| ENSG00000254388 | DUTP2 | 0,083666512 | 1,059707797 | 0,85621 | 0,9349542 | deoxyuridine triphosphatase pseudogene 2 |
| ENSG00000277801 | AL138478,1 | 0,083650764 | 1,05969623 | 0,85401 | 0,93380868 | novel transcript |
| ENSG00000284634 | AC092821,3 | 0,083611364 | 1,05966729 | 0,73232 | 0,87295647 | novel transcript |
| ENSG00000148834 | GSTO1 | 0,083588062 | 1,059650174 | 0,79787 | 0,90685415 | glutathione S-transferase omega 1 |
| ENSG00000139910 | NOVA1 | 0,083348683 | 1,059474367 | 0,83595 | 0,92493625 | NOVA alternative splicing regulator 1 |
| ENSG00000162493 | PDPN | 0,083314171 | 1,059449023 | 0,86649 | 0,94074101 | podoplanin |
| ENSG00000164187 | LMBRD2 | 0,083254854 | 1,059405463 | 0,82844 | 0,92081006 | LMBR1 domain containing 2 |
| ENSG00000168496 | FEN1 | 0,083217624 | 1,059378125 | 0,65561 | 0,82787332 | flap structure-specific endonuclease 1 |
| ENSG00000115364 | MRPL19 | 0,083073322 | 1,059272169 | 0,72272 | 0,86756424 | mitochondrial ribosomal protein L19 |
| ENSG00000147140 | NONO | 0,083033293 | 1,059242778 | 0,61486 | 0,80404192 | non-POU domain containing octamer binding |
| ENSG00000130717 | UCK1 | 0,083018094 | 1,059231619 | 0,60453 | 0,79797731 | uridine-cytidine kinase 1 |
| ENSG00000213672 | NCKIPSD | 0,082998589 | 1,059217299 | 0,72951 | 0,8715818 | NCK interacting protein with SH3 domain |
| ENSG00000286545 | AC092645,2 | 0,082941282 | 1,059175225 | 0,75547 | 0,88547359 | novel transcript, antisense to ZC3H8 |
| ENSG00000107874 | CUEDC2 | 0,082888287 | 1,059136319 | 0,67871 | 0,84182956 | CUE domain containing 2 |
| ENSG00000157933 | SKI | 0,082779693 | 1,059056599 | 0,75027 | 0,8827292 | SKI proto-oncogene |
| ENSG00000284526 | AC015802,6 | 0,08274542 | 1,05903144 | 0,78833 | 0,90157293 | novel protein |
| ENSG00000281582 | AP000350,16 | 0,082694839 | 1,058994311 | 0,79002 | 0,90273405 | novel transcript |
| ENSG00000273295 | AP000350,6 | 0,082694839 | 1,058994311 | 0,79002 | 0,90273405 | novel transcript |
| ENSG00000104856 | RELB | 0,082685933 | 1,058987774 | 0,76148 | 0,88879633 | RELB proto-oncogene, NF-kB subunit |
| ENSG00000164304 | CAGE1 | 0,082590488 | 1,058917716 | 0,91019 | 0,9613425 | cancer antigen 1 |
| ENSG00000109339 | MAPK10 | 0,082573423 | 1,05890519 | 0,76916 | 0,89297049 | mitogen-activated protein kinase 10 |
| ENSG00000262919 | CCNQ | 0,082533905 | 1,058876186 | 0,68683 | 0,84724466 | cyclin Q |
| ENSG00000259378 | DCAF13P3 | 0,082361064 | 1,058749336 | 0,90113 | 0,95787401 | DDB1 and CUL4 associated factor 13 pseudogene 3 |
| ENSG00000163539 | CLASP2 | 0,082346324 | 1,058738518 | 0,77817 | 0,89735311 | cytoplasmic linker associated protein 2 |
| ENSG00000122335 | SERAC1 | 0,082308306 | 1,058710619 | 0,60443 | 0,79794393 | serine active site containing 1 |
| ENSG00000166847 | DCTN5 | 0,082252878 | 1,058669944 | 0,64836 | 0,82383528 | dynactin subunit 5 |

| | | | | | | |
|---|---|---|---|---|---|---|
| ENSG00000169410 | PTPN9 | 0,082139939 | 1,058587071 | 0,65163 | 0,82544945 | protein tyrosine phosphatase non-receptor type 9 |
| ENSG00000122783 | CYREN | 0,082120281 | 1,058572647 | 0,68457 | 0,84564064 | cell cycle regulator of NHEJ |
| ENSG00000091947 | TMEM101 | 0,082089409 | 1,058549995 | 0,72496 | 0,86868749 | transmembrane protein 101 |
| ENSG00000173599 | PC | 0,082046991 | 1,058518872 | 0,74305 | 0,87840582 | pyruvate carboxylase |
| ENSG00000270728 | AL035413,1 | 0,082027297 | 1,058504423 | 0,84956 | 0,93150269 | synaptonemal complex protein 3 (SYCP3) pseudogene |
| ENSG00000167987 | VPS37C | 0,081936412 | 1,058437743 | 0,73593 | 0,8750728 | VPS37C subunit of ESCRT-I |
| ENSG00000275825 | AC139494,4 | 0,081847139 | 1,058372249 | 0,88458 | 0,94883769 | novel transcript |
| ENSG00000184478 | OR56A3 | 0,081819355 | 1,058351866 | 0,92425 | 0,96705923 | olfactory receptor family 56 subfamily A member 3 |
| ENSG00000283269 | AC005524,1 | 0,081614356 | 1,058201491 | 0,86363 | 0,93904284 | SHC SH2-domain binding protein 1 (SHCBP1) pseudogene |
| ENSG00000276603 | AL109614,1 | 0,081483011 | 1,058105156 | 0,85803 | 0,93610959 | novel transcript, sense intronic to BLCAP |
| ENSG00000223459 | TCAF1P1 | 0,081458987 | 1,058087536 | 0,73872 | 0,87593493 | TRPM8 channel associated factor 1 pseudogene 1 |
| ENSG00000198841 | KTI12 | 0,081437974 | 1,058072125 | 0,72701 | 0,87013394 | KTI12 chromatin associated homolog |
| ENSG00000281406 | BLACAT1 | 0,081210825 | 1,057905547 | 0,85064 | 0,93206349 | bladder cancer associated transcript 1 |
| ENSG00000108961 | RANGRF | 0,081157522 | 1,057866462 | 0,69723 | 0,85330122 | RAN guanine nucleotide release factor |
| ENSG00000232627 | AC021876,1 | 0,081032983 | 1,057775147 | 0,91336 | 0,96249114 | ribosomal protein L31 (RPL31) pseudogene |
| ENSG00000110435 | PDHX | 0,080771991 | 1,057583806 | 0,67036 | 0,83656528 | pyruvate dehydrogenase complex component X |
| ENSG00000172992 | DCAKD | 0,080687977 | 1,05752222 | 0,74539 | 0,87982881 | dephospho-CoA kinase domain containing |
| ENSG00000075275 | CELSR1 | 0,080680956 | 1,057517074 | 0,75489 | 0,88513155 | cadherin EGF LAG seven-pass G-type receptor 1 |
| ENSG00000124383 | MPHOSPH10 | 0,080614046 | 1,057468029 | 0,71682 | 0,8644737 | M-phase phosphoprotein 10 |
| ENSG00000048649 | RSF1 | 0,080609571 | 1,057464749 | 0,69929 | 0,85475604 | remodeling and spacing factor 1 |
| ENSG00000020577 | SAMD4A | 0,080551829 | 1,057422426 | 0,70739 | 0,85910429 | sterile alpha motif domain containing 4A |
| ENSG00000129292 | PHF20L1 | 0,080533033 | 1,057408649 | 0,70848 | 0,85940638 | PHD finger protein 20 like 1 |
| ENSG00000285828 | AC254945,4 | 0,080034304 | 1,057043174 | 0,88834 | 0,95069428 | novel transcript, antisense to ARHGAP11A |
| ENSG00000285948 | AC123768,4 | 0,080034304 | 1,057043174 | 0,88834 | 0,95069428 | novel transcript, antisense to ARHGAP11A |
| ENSG00000277909 | SPC25 | 0,080020437 | 1,057033014 | 0,83651 | 0,92514377 | SPC25 component of NDC80 kinetochore complex |
| ENSG00000264449 | AP001021,1 | 0,079962287 | 1,05699041 | 0,78824 | 0,90156082 | novel transcript, antisense to TMEM200C |
| ENSG00000162542 | TMCO4 | 0,079942105 | 1,056975624 | 0,91598 | 0,96339511 | transmembrane and coiled-coil domains 4 |
| ENSG00000099968 | BCL2L13 | 0,079781479 | 1,056857949 | 0,63836 | 0,81843416 | BCL2 like 13 |
| ENSG00000264707 | L3MBTL4-AS1 | 0,079712895 | 1,056807709 | 0,89439 | 0,95392565 | L3MBTL4 antisense RNA 1 |
| ENSG00000075856 | SART3 | 0,079489878 | 1,056644356 | 0,67775 | 0,84147813 | spliceosome associated factor 3, U4/U6 recycling protein |
| ENSG00000179630 | LACC1 | 0,079429674 | 1,056600263 | 0,76647 | 0,89139706 | laccase domain containing 1 |
| ENSG00000237853 | NFIA-AS1 | 0,07937956 | 1,056563561 | 0,91621 | 0,96347905 | NFIA antisense RNA 1 |
| ENSG00000227466 | PDE4B-AS1 | 0,079330641 | 1,056527736 | 0,93159 | 0,9707979 | PDE4B antisense RNA 1 |

| | | | | | | |
|---|---|---|---|---|---|---|
| ENSG00000168952 | STXBP6 | 0,079265096 | 1,056479736 | 0,86156 | 0,93811282 | syntaxin binding protein 6 |
| ENSG00000234750 | AC012618,2 | 0,079264218 | 1,056479094 | 0,87476 | 0,94461928 | ribosomal protein S2 (RPS2) pseudogene |
| ENSG00000238286 | SLC35E1P1 | 0,079242025 | 1,056462842 | 0,89208 | 0,95255977 | solute carrier family 35 member E1 pseudogene 1 |
| ENSG00000285250 | MSRA | 0,079207182 | 1,056437327 | 0,75587 | 0,8857133 | methionine sulfoxide reductase A |
| ENSG00000182986 | ZNF320 | 0,079157765 | 1,056401141 | 0,69431 | 0,85164591 | zinc finger protein 320 |
| ENSG00000130413 | STK33 | 0,078939606 | 1,056241408 | 0,73859 | 0,87591843 | serine/threonine kinase 33 |
| ENSG00000269906 | AL606834,1 | 0,078936844 | 1,056239386 | 0,88433 | 0,9486819 | novel transcript, sense intronic to SAV1 |
| ENSG00000284126 | SMURF1 | 0,078868034 | 1,056189009 | 0,68272 | 0,84458531 | SMAD specific E3 ubiquitin protein ligase 1 |
| ENSG00000166526 | ZNF3 | 0,078865995 | 1,056187517 | 0,65478 | 0,8272805 | zinc finger protein 3 |
| ENSG00000230392 | AC004835,1 | 0,078865653 | 1,056187266 | 0,74989 | 0,88236188 | novel transcript |
| ENSG00000256525 | POLG2 | 0,078843284 | 1,05617089 | 0,68697 | 0,84727142 | DNA polymerase gamma 2, accessory subunit |
| ENSG00000079691 | CARMIL1 | 0,078783382 | 1,056127038 | 0,80857 | 0,91193281 | capping protein regulator and myosin 1 linker 1 |
| ENSG00000154080 | CHST9 | 0,078723671 | 1,056083327 | 0,88367 | 0,94839496 | carbohydrate sulfotransferase 9 |
| ENSG00000168476 | REEP4 | 0,078699675 | 1,056065762 | 0,76938 | 0,89297049 | receptor accessory protein 4 |
| ENSG00000137142 | IGFBPL1 | 0,07862804 | 1,056013326 | 0,819 | 0,91689679 | insulin like growth factor binding protein like 1 |
| ENSG00000152409 | JMY | 0,078600176 | 1,05599293 | 0,75297 | 0,88427886 | junction mediating and regulatory protein, p53 cofactor |
| ENSG00000110497 | AMBRA1 | 0,078338699 | 1,055801557 | 0,58572 | 0,78640335 | autophagy and beclin 1 regulator 1 |
| ENSG00000142632 | ARHGEF19 | 0,078307447 | 1,055778687 | 0,83431 | 0,92416267 | Rho guanine nucleotide exchange factor 19 |
| ENSG00000287146 | AC000085,1 | 0,078305938 | 1,055777582 | 0,89364 | 0,95359241 | novel transcript |
| ENSG00000183117 | CSMD1 | 0,07828909 | 1,055765252 | 0,89135 | 0,95208954 | CUB and Sushi multiple domains 1 |
| ENSG00000176857 | GJA1P1 | 0,078245092 | 1,055733056 | 0,84887 | 0,93118549 | gap junction protein alpha 1 pseudogene 1 |
| ENSG00000235106 | BRD3OS | 0,078087293 | 1,055617588 | 0,71341 | 0,86217438 | BRD3 opposite strand |
| ENSG00000124299 | PEPD | 0,078081159 | 1,055613099 | 0,75696 | 0,88650931 | peptidase D |
| ENSG00000113163 | CERT1 | 0,078075307 | 1,055608818 | 0,62708 | 0,81124422 | ceramide transporter 1 |
| ENSG00000157445 | CACNA2D3 | 0,078073213 | 1,055607285 | 0,81461 | 0,91470265 | calcium voltage-gated channel auxiliary subunit alpha2delta 3 |
| ENSG00000214182 | PTMAP5 | 0,078071922 | 1,055606341 | 0,73934 | 0,8763893 | prothymosin alpha pseudogene 5 |
| ENSG00000183354 | KIAA2026 | 0,078034306 | 1,055578818 | 0,72639 | 0,86975255 | KIAA2026 |
| ENSG00000249158 | PCDHA11 | 0,077940744 | 1,055510364 | 0,77112 | 0,89404837 | protocadherin alpha 11 |
| ENSG00000172139 | SLC9C1 | 0,077852625 | 1,055445896 | 0,91654 | 0,9636258 | solute carrier family 9 member C1 |
| ENSG00000285044 | SLC9C1 | 0,077852625 | 1,055445896 | 0,91654 | 0,9636258 | solute carrier family 9 member C1 |
| ENSG00000286659 | Z99716,2 | 0,077767078 | 1,055383313 | 0,92009 | 0,96530017 | novel transcript, antisense to POLDIP3 |
| ENSG00000281089 | ATG16L1 | 0,077702302 | 1,055335928 | 0,73174 | 0,87274707 | autophagy related 16 like 1 |

| | | | | | | |
|---|---|---|---|---|---|---|
| ENSG00000085978 | ATG16L1 | 0,077702302 | 1,055335928 | 0,73174 | 0,87274707 | autophagy related 16 like 1 |
| ENSG00000100350 | FOXRED2 | 0,077682927 | 1,055321755 | 0,65218 | 0,82578944 | FAD dependent oxidoreductase domain containing 2 |
| ENSG00000136840 | ST6GALNAC4 | 0,077643536 | 1,055292941 | 0,74678 | 0,88057664 | ST6 N-acetylgalactosaminide alpha-2,6-sialyltransferase 4 |
| ENSG00000243317 | STMP1 | 0,077631248 | 1,055283953 | 0,6502 | 0,82472422 | short transmembrane mitochondrial protein 1 |
| ENSG00000178802 | MPI | 0,077539387 | 1,055216762 | 0,64723 | 0,82349145 | mannose phosphate isomerase |
| ENSG00000226367 | ST7-AS2 | 0,077447101 | 1,055149264 | 0,84912 | 0,93130502 | ST7 antisense RNA 2 |
| ENSG00000243710 | CFAP57 | 0,077389675 | 1,055107265 | 0,88001 | 0,94664997 | cilia and flagella associated protein 57 |
| ENSG00000171160 | MORN4 | 0,077351053 | 1,05507902 | 0,67194 | 0,83760327 | MORN repeat containing 4 |
| ENSG00000227638 | HNRNPA1P14 | 0,077304087 | 1,055044673 | 0,82369 | 0,91860411 | heterogeneous nuclear ribonucleoprotein A1 pseudogene 14 |
| ENSG00000275199 | AKT3 | 0,077105238 | 1,054899265 | 0,86703 | 0,94116774 | AKT serine/threonine kinase 3 |
| ENSG00000170584 | NUDCD2 | 0,077057658 | 1,054864475 | 0,70643 | 0,85870246 | NudC domain containing 2 |
| ENSG00000183798 | EMILIN3 | 0,077027537 | 1,054842451 | 0,8254 | 0,91911153 | elastin microfibril interfacer 3 |
| ENSG00000286177 | AC011462,5 | 0,076754023 | 1,054642487 | 0,7542 | 0,88479657 | novel transcript, antisense to CCDC97 and HNRNPUL1 |
| ENSG00000108010 | GLRX3 | 0,076665075 | 1,054577466 | 0,67762 | 0,84135792 | glutaredoxin 3 |
| ENSG00000227322 | RXRB | 0,076629363 | 1,054551362 | 0,70682 | 0,85889796 | retinoid X receptor beta |
| ENSG00000103067 | ESRP2 | 0,076606293 | 1,054534499 | 0,84595 | 0,92992612 | epithelial splicing regulatory protein 2 |
| ENSG00000077235 | GTF3C1 | 0,076600043 | 1,05452993 | 0,62794 | 0,81175174 | general transcription factor IIIC subunit 1 |
| ENSG00000232830 | AC110994,1 | 0,076559838 | 1,054500543 | 0,95073 | 0,97934603 | family with sequence similarity 136, member A (FAM136A) pseudogene |
| ENSG00000183426 | NPIPA1 | 0,076539354 | 1,054485571 | 0,72236 | 0,86734497 | nuclear pore complex interacting protein family member A1 |
| ENSG00000270681 | AC095055,1 | 0,07651919 | 1,054470833 | 0,90657 | 0,96009372 | novel transcript, antisense to SH3D19 |
| ENSG00000266967 | AARSD1 | 0,076498732 | 1,05445588 | 0,69728 | 0,85330122 | alanyl-tRNA synthetase domain containing 1 |
| ENSG00000277036 | AC139494,6 | 0,076493094 | 1,054451759 | 0,87659 | 0,94547358 | novel transcript |
| ENSG00000173786 | CNP | 0,07643363 | 1,054408299 | 0,6412 | 0,81994133 | 2',3'-cyclic nucleotide 3' phosphodiesterase |
| ENSG00000166135 | HIF1AN | 0,076089807 | 1,054157042 | 0,58874 | 0,78830778 | hypoxia inducible factor 1 subunit alpha inhibitor |
| ENSG00000281016 | AC092574,2 | 0,07606094 | 1,05413595 | 0,89899 | 0,95683014 | novel transcript, antisense to ZNF721 |
| ENSG00000100321 | SYNGR1 | 0,076056738 | 1,054132879 | 0,7549 | 0,88513155 | synaptogyrin 1 |
| ENSG00000100372 | SLC25A17 | 0,075940966 | 1,054048292 | 0,71359 | 0,86217438 | solute carrier family 25 member 17 |
| ENSG00000145901 | TNIP1 | 0,075924893 | 1,054036548 | 0,78323 | 0,89918294 | TNFAIP3 interacting protein 1 |
| ENSG00000124614 | RPS10 | 0,075771898 | 1,053924776 | 0,7371 | 0,87539732 | ribosomal protein S10 |
| ENSG00000100100 | PIK3IP1 | 0,075673485 | 1,053852885 | 0,78005 | 0,89820865 | phosphoinositide-3-kinase interacting protein 1 |

| | | | | | | |
|---|---|---|---|---|---|---|
| ENSG00000221829 | FANCG | 0,075483169 | 1,053713874 | 0,74193 | 0,87795766 | FA complementation group G |
| ENSG00000183090 | FREM3 | 0,07546704 | 1,053702093 | 0,9133 | 0,96246104 | FRAS1 related extracellular matrix 3 |
| ENSG00000237344 | PBX2 | 0,075463757 | 1,053699695 | 0,64647 | 0,82293404 | PBX homeobox 2 |
| ENSG00000213046 | EEF1A1P32 | 0,075463192 | 1,053699283 | 0,89972 | 0,95730568 | eukaryotic translation elongation factor 1 alpha 1 pseudogene 32 |
| ENSG00000225073 | DDX39B | 0,075417295 | 1,053665761 | 0,68125 | 0,84367915 | DExD-box helicase 39B |
| ENSG00000051009 | FAM160A2 | 0,075338458 | 1,053608185 | 0,70028 | 0,85526044 | family with sequence similarity 160 member A2 |
| ENSG00000170214 | ADRA1B | 0,075333889 | 1,053604848 | 0,92231 | 0,9663588 | adrenoceptor alpha 1B |
| ENSG00000268912 | AC012313,5 | 0,075308258 | 1,05358613 | 0,74297 | 0,87840582 | novel transcript |
| ENSG00000115648 | MLPH | 0,075305636 | 1,053584215 | 0,95197 | 0,97966441 | melanophilin |
| ENSG00000062282 | DGAT2 | 0,075271932 | 1,053559601 | 0,8919 | 0,95246557 | diacylglycerol O-acyltransferase 2 |
| ENSG00000254258 | AC087354,1 | 0,075256227 | 1,053548132 | 0,80359 | 0,90964529 | novel transcript |
| ENSG00000165416 | SUGT1 | 0,075163703 | 1,053480568 | 0,71463 | 0,86265094 | SGT1 homolog, MIS12 kinetochore complex assembly cochaperone |
| ENSG00000224096 | HCG25 | 0,075014343 | 1,053371509 | 0,90174 | 0,95802163 | HLA complex group 25 |
| ENSG00000233772 | HCG25 | 0,075014343 | 1,053371509 | 0,90174 | 0,95802163 | HLA complex group 25 |
| ENSG00000231037 | HCG25 | 0,075014343 | 1,053371509 | 0,90174 | 0,95802163 | HLA complex group 25 |
| ENSG00000010278 | CD9 | 0,074998897 | 1,053360231 | 0,88286 | 0,94822558 | CD9 molecule |
| ENSG00000105364 | MRPL4 | 0,074991785 | 1,053355038 | 0,70401 | 0,85725494 | mitochondrial ribosomal protein L4 |
| ENSG00000116183 | PAPPA2 | 0,074889558 | 1,053280401 | 0,84802 | 0,9309317 | pappalysin 2 |
| ENSG00000104043 | ATP8B4 | 0,07482245 | 1,053231408 | 0,90563 | 0,95974361 | ATPase phospholipid transporting 8B4 (putative) |
| ENSG00000168434 | COG7 | 0,074740454 | 1,05317155 | 0,74672 | 0,88057664 | component of oligomeric golgi complex 7 |
| ENSG00000111850 | SMIM8 | 0,074515194 | 1,053007122 | 0,78932 | 0,9024443 | small integral membrane protein 8 |
| ENSG00000250116 | RHOQ-AS1 | 0,074504121 | 1,05299904 | 0,8519 | 0,93260477 | RHOQ antisense RNA 1 |
| ENSG00000169118 | CSNK1G1 | 0,074477475 | 1,052979592 | 0,68153 | 0,84375324 | casein kinase 1 gamma 1 |
| ENSG00000171724 | VAT1L | 0,074452471 | 1,052961342 | 0,86288 | 0,9388179 | vesicle amine transport 1 like |
| ENSG00000286892 | AC084212,1 | 0,074416721 | 1,05293525 | 0,80794 | 0,91168773 | novel transcript |
| ENSG00000269894 | AC018809,1 | 0,074402024 | 1,052924524 | 0,90631 | 0,95993807 | novel transcript, sense intronic to CRELD1 |
| ENSG00000123091 | RNF11 | 0,07438428 | 1,052911574 | 0,72807 | 0,87069932 | ring finger protein 11 |
| ENSG00000232721 | AC239800,2 | 0,074215186 | 1,052788172 | 0,83597 | 0,92493625 | novel transcript |
| ENSG00000230426 | LINC01036 | 0,074213819 | 1,052787175 | 0,90376 | 0,95873972 | long intergenic non-protein coding RNA 1036 |
| ENSG00000071889 | FAM3A | 0,074212367 | 1,052786115 | 0,6953 | 0,85222093 | FAM3 metabolism regulating signaling molecule A |
| ENSG00000131351 | HAUS8 | 0,073994912 | 1,052627442 | 0,73085 | 0,87205836 | HAUS augmin like complex subunit 8 |

| | | | | | | |
|---|---|---|---|---|---|---|
| ENSG00000269843 | AC008537,2 | 0,073979479 | 1,052616182 | 0,91206 | 0,96198423 | novel transcript |
| ENSG00000004059 | ARF5 | 0,073899708 | 1,052557981 | 0,70062 | 0,85539553 | ADP ribosylation factor 5 |
| ENSG00000284981 | UPK3BL2 | 0,073897853 | 1,052556628 | 0,81983 | 0,91718681 | uroplakin 3B like 2 |
| ENSG00000146250 | PRSS35 | 0,073861075 | 1,052529796 | 0,82383 | 0,91865473 | serine protease 35 |
| ENSG00000268555 | AC123912,4 | 0,07378796 | 1,052476455 | 0,87861 | 0,94605315 | novel transcript |
| ENSG00000109424 | UCP1 | 0,073751475 | 1,05244984 | 0,92699 | 0,96837518 | uncoupling protein 1 |
| ENSG00000166578 | IQCD | 0,073749506 | 1,052448403 | 0,81806 | 0,91652411 | IQ motif containing D |
| ENSG00000100422 | CERK | 0,073746793 | 1,052446424 | 0,59919 | 0,79538547 | ceramide kinase |
| ENSG00000152192 | POU4F1 | 0,073601444 | 1,052340397 | 0,90309 | 0,95850777 | POU class 4 homeobox 1 |
| ENSG00000277027 | RMRP | 0,073557289 | 1,05230819 | 0,89637 | 0,95523334 | RNA component of mitochondrial RNA processing endoribonuclease |
| ENSG00000064042 | LIMCH1 | 0,073399352 | 1,052192996 | 0,88977 | 0,95146267 | LIM and calponin homology domains 1 |
| ENSG00000239883 | PARGP1 | 0,073332827 | 1,052144479 | 0,6814 | 0,84375324 | poly(ADP-ribose) glycohydrolase pseudogene 1 |
| ENSG00000177728 | TMEM94 | 0,073309545 | 1,0521275 | 0,64866 | 0,82406756 | transmembrane protein 94 |
| ENSG00000158109 | TPRG1L | 0,073302242 | 1,052122174 | 0,67598 | 0,84051682 | tumor protein p63 regulated 1 like |
| ENSG00000156671 | SAMD8 | 0,073283851 | 1,052108762 | 0,74698 | 0,88059941 | sterile alpha motif domain containing 8 |
| ENSG00000108679 | LGALS3BP | 0,073279855 | 1,052105848 | 0,7959 | 0,90557616 | galectin 3 binding protein |
| ENSG00000249249 | AC010226,1 | 0,073207706 | 1,052053233 | 0,80671 | 0,91120963 | novel transcript |
| ENSG00000015532 | XYLT2 | 0,073090331 | 1,051967644 | 0,73109 | 0,87227623 | xylosyltransferase 2 |
| ENSG00000260455 | NBAT1 | 0,073076583 | 1,051957619 | 0,78072 | 0,89846287 | neuroblastoma associated transcript 1 |
| ENSG00000232284 | GNG12-AS1 | 0,073031282 | 1,051924588 | 0,87723 | 0,94559723 | GNG12, DIRAS3 and WLS antisense RNA 1 |
| ENSG00000225434 | LINC01504 | 0,072993821 | 1,051897274 | 0,8538 | 0,93377737 | long intergenic non-protein coding RNA 1504 |
| ENSG00000278550 | SLC43A2 | 0,072977414 | 1,051885312 | 0,77796 | 0,89728573 | solute carrier family 43 member 2 |
| ENSG00000183773 | AIFM3 | 0,072931233 | 1,051851641 | 0,90191 | 0,95804366 | apoptosis inducing factor mitochondria associated 3 |
| ENSG00000134851 | TMEM165 | 0,07292112 | 1,051844267 | 0,65201 | 0,82568422 | transmembrane protein 165 |
| ENSG00000164683 | HEY1 | 0,072879171 | 1,051813684 | 0,86144 | 0,93807534 | hes related family bHLH transcription factor with YRPW motif 1 |
| ENSG00000105401 | CDC37 | 0,072816442 | 1,051767951 | 0,65752 | 0,82875603 | cell division cycle 37, HSP90 cochaperone |
| ENSG00000259467 | NDUFAF4P1 | 0,072809416 | 1,051762829 | 0,94985 | 0,97899157 | NADH:ubiquinone oxidoreductase complex assembly factor 4 pseudogene 1 |
| ENSG00000011275 | RNF216 | 0,072750527 | 1,051719899 | 0,68187 | 0,84394068 | ring finger protein 216 |
| ENSG00000196747 | H2AC13 | 0,072685538 | 1,051672523 | 0,72789 | 0,87053185 | H2A clustered histone 13 |
| ENSG00000078967 | UBE2D4 | 0,072576215 | 1,051592834 | 0,7144 | 0,86255727 | ubiquitin conjugating enzyme E2 D4 (putative) |

| | | | | | | |
|---|---|---|---|---|---|---|
| ENSG00000231707 | PABPC1P1 | 0,072495771 | 1,051534199 | 0,85394 | 0,93380868 | poly(A) binding protein cytoplasmic 1 pseudogene 1 |
| ENSG00000050130 | JKAMP | 0,072487745 | 1,051528349 | 0,6935 | 0,85109296 | JNK1/MAPK8 associated membrane protein |
| ENSG00000158863 | FAM160B2 | 0,072364627 | 1,051438616 | 0,66901 | 0,83591711 | family with sequence similarity 160 member B2 |
| ENSG00000070423 | RNF126 | 0,072282359 | 1,051378661 | 0,72096 | 0,86659896 | ring finger protein 126 |
| ENSG00000215354 | FAM90A24P | 0,072249733 | 1,051354885 | 0,91568 | 0,9633821 | family with sequence similarity 90 member A24, pseudogene |
| ENSG00000110063 | DCPS | 0,072226987 | 1,051338309 | 0,78471 | 0,89974349 | decapping enzyme, scavenger |
| ENSG00000125257 | ABCC4 | 0,072220685 | 1,051333716 | 0,83 | 0,921946 | ATP binding cassette subfamily C member 4 |
| ENSG00000282381 | AC104073,4 | 0,072074125 | 1,05122692 | 0,81333 | 0,91428978 | novel transcript |
| ENSG00000204959 | ARHGEF34P | 0,072070968 | 1,051224619 | 0,94612 | 0,97704781 | Rho guanine nucleotide exchange factor 34, pseudogene |
| ENSG00000084070 | SMAP2 | 0,071921366 | 1,051115617 | 0,76796 | 0,89250304 | small ArfGAP2 |
| ENSG00000247934 | AC022364,1 | 0,071908905 | 1,051106537 | 0,82552 | 0,91915045 | novel transcript, antisense to CCDC91 |
| ENSG00000147576 | ADHFE1 | 0,071850169 | 1,051063745 | 0,86369 | 0,93906863 | alcohol dehydrogenase iron containing 1 |
| ENSG00000124201 | ZNFX1 | 0,071833687 | 1,051051737 | 0,72766 | 0,87036169 | zinc finger NFX1-type containing 1 |
| ENSG00000212694 | LINC01089 | 0,071826786 | 1,05104671 | 0,79281 | 0,90366929 | long intergenic non-protein coding RNA 1089 |
| ENSG00000144535 | DIS3L2 | 0,071822129 | 1,051043317 | 0,6423 | 0,82066205 | DIS3 like 3'-5' exoribonuclease 2 |
| ENSG00000162959 | MEMO1 | 0,071787886 | 1,051018371 | 0,65275 | 0,826091 | mediator of cell motility 1 |
| ENSG00000127311 | HELB | 0,07176013 | 1,05099815 | 0,8423 | 0,92821072 | DNA helicase B |
| ENSG00000172500 | FIBP | 0,071619175 | 1,05089547 | 0,7091 | 0,85978251 | FGF1 intracellular binding protein |
| ENSG00000187398 | LUZP2 | 0,071595324 | 1,050878097 | 0,87497 | 0,94463878 | leucine zipper protein 2 |
| ENSG00000136040 | PLXNC1 | 0,071587611 | 1,050872478 | 0,81058 | 0,91251177 | plexin C1 |
| ENSG00000128656 | CHN1 | 0,071551113 | 1,050845893 | 0,81276 | 0,91386772 | chimerin 1 |
| ENSG00000168385 | SEPTIN2 | 0,071550776 | 1,050845647 | 0,72359 | 0,86800739 | septin 2 |
| ENSG00000181585 | TMIE | 0,071389734 | 1,050728353 | 0,90268 | 0,95838699 | transmembrane inner ear |
| ENSG00000283018 | MRPS12 | 0,071327568 | 1,050683078 | 0,79509 | 0,90513847 | mitochondrial ribosomal protein S12 |
| ENSG00000276734 | KIF5C | 0,071291079 | 1,050656504 | 0,76457 | 0,89036868 | kinesin family member 5C |
| ENSG00000182901 | RGS7 | 0,071252852 | 1,050628665 | 0,83499 | 0,92456226 | regulator of G protein signaling 7 |
| ENSG00000118900 | UBN1 | 0,071242511 | 1,050621134 | 0,68535 | 0,84616325 | ubinuclein 1 |
| ENSG00000101160 | CTSZ | 0,071237098 | 1,050617192 | 0,6685 | 0,83543142 | cathepsin Z |
| ENSG00000255072 | PIGY | 0,071212672 | 1,050599405 | 0,70341 | 0,8569555 | phosphatidylinositol glycan anchor biosynthesis class Y |
| ENSG00000145337 | PYURF | 0,071212672 | 1,050599405 | 0,70341 | 0,8569555 | PIGY upstream reading frame |
| ENSG00000162512 | SDC3 | 0,071136575 | 1,050543991 | 0,81664 | 0,91570045 | syndecan 3 |
| ENSG00000250267 | AC021237,1 | 0,071072642 | 1,050497437 | 0,87666 | 0,94547358 | novel transcript |

| | | | | | | |
|---|---|---|---|---|---|---|
| ENSG00000237491 | LINC01409 | 0,071016609 | 1,050456638 | 0,81381 | 0,9144566 | long intergenic non-protein coding RNA 1409 |
| ENSG00000101162 | TUBB1 | 0,070851655 | 1,050336538 | 0,83277 | 0,92348349 | tubulin beta 1 class VI |
| ENSG00000259768 | AC004943,2 | 0,070801726 | 1,050300188 | 0,80935 | 0,91213228 | novel transcript, antisense to ZFHX3 |
| ENSG00000249965 | CDC42P4 | 0,070748897 | 1,050261729 | 0,93937 | 0,97413085 | cell division cycle 42 pseudogene 4 |
| ENSG00000183535 | COL18A1-AS1 | 0,070722541 | 1,050242542 | 0,91131 | 0,9616927 | COL18A1 antisense RNA 1 |
| ENSG00000183889 | AC138969,1 | 0,070675094 | 1,050208002 | 0,70907 | 0,85978194 | novel member of the nuclear pore complex interacting protein NPIP gene family |
| ENSG00000164845 | FAM86FP | 0,070628348 | 1,050173975 | 0,92258 | 0,96643081 | family with sequence similarity 86 member F, pseudogene |
| ENSG00000177971 | IMP3 | 0,070620295 | 1,050168112 | 0,74897 | 0,88190158 | IMP U3 small nucleolar ribonucleoprotein 3 |
| ENSG00000228681 | AC008072,1 | 0,07060249 | 1,050155152 | 0,85119 | 0,93226967 | ribosomal protein S27a (RPS27A) pseudogene |
| ENSG00000274605 | PCCA-DT | 0,070537612 | 1,050107927 | 0,825 | 0,91900665 | PCCA divergent transcript |
| ENSG00000157343 | ARMC12 | 0,070502313 | 1,050082234 | 0,87563 | 0,94497474 | armadillo repeat containing 12 |
| ENSG00000144481 | TRPM8 | 0,07048423 | 1,050069072 | 0,90144 | 0,95798991 | transient receptor potential cation channel subfamily M member 8 |
| ENSG00000049656 | CLPTM1L | 0,070457134 | 1,05004935 | 0,63522 | 0,81645836 | CLPTM1 like |
| ENSG00000260101 | AC008074,2 | 0,070377807 | 1,049991615 | 0,91443 | 0,96299769 | novel transcript |
| ENSG00000138709 | LARP1B | 0,070362744 | 1,049980652 | 0,76047 | 0,88845438 | La ribonucleoprotein 1B |
| ENSG00000176358 | TAC4 | 0,070360997 | 1,049979381 | 0,91159 | 0,96182676 | tachykinin precursor 4 |
| ENSG00000176658 | MYO1D | 0,070356564 | 1,049976155 | 0,83904 | 0,92640383 | myosin ID |
| ENSG00000184270 | H2AC21 | 0,070349256 | 1,049970836 | 0,70586 | 0,85838918 | H2A clustered histone 21 |
| ENSG00000178971 | CTC1 | 0,070336305 | 1,04996141 | 0,71359 | 0,86217438 | CST telomere replication complex component 1 |
| ENSG00000271926 | AC008972,1 | 0,070208643 | 1,049868505 | 0,94341 | 0,97611183 | novel transcript |
| ENSG00000171720 | HDAC3 | 0,070154874 | 1,049829377 | 0,6873 | 0,8475632 | histone deacetylase 3 |
| ENSG00000010361 | FUZ | 0,07003865 | 1,049744806 | 0,78193 | 0,8987738 | fuzzy planar cell polarity protein |
| ENSG00000008277 | ADAM22 | 0,069927687 | 1,04966407 | 0,84207 | 0,92818855 | ADAM metallopeptidase domain 22 |
| ENSG00000267796 | LIN37 | 0,069893882 | 1,049639474 | 0,71582 | 0,86371705 | lin-37 DREAM MuvB core complex component |
| ENSG00000144045 | DQX1 | 0,06988703 | 1,049634489 | 0,92979 | 0,96989348 | DEAQ-box RNA dependent ATPase 1 |
| ENSG00000157014 | TATDN2 | 0,069880484 | 1,049629727 | 0,71797 | 0,86489865 | TatD DNase domain containing 2 |
| ENSG00000169783 | LINGO1 | 0,069866997 | 1,049619914 | 0,74288 | 0,87840582 | leucine rich repeat and Ig domain containing 1 |
| ENSG00000217767 | NDUFAB1P1 | 0,069780038 | 1,04955665 | 0,94029 | 0,97464787 | NADH:ubiquinone oxidoreductase subunit AB1 pseudogene 1 |
| ENSG00000162734 | PEA15 | 0,069773553 | 1,049551932 | 0,85241 | 0,93292666 | proliferation and apoptosis adaptor protein 15 |
| ENSG00000203667 | COX20 | 0,069546119 | 1,049386488 | 0,72626 | 0,86965491 | cytochrome c oxidase assembly factor COX20 |

| | | | | | | |
|---|---|---|---|---|---|---|
| ENSG00000138185 | ENTPD1 | 0,069529135 | 1,049374134 | 0,72656 | 0,86978598 | ectonucleoside triphosphate diphosphohydrolase 1 |
| ENSG00000285309 | AL136295,18 | 0,069519659 | 1,049367242 | 0,85709 | 0,9354716 | novel transcript |
| ENSG00000278784 | AL136295,7 | 0,069519659 | 1,049367242 | 0,85709 | 0,9354716 | novel transcript |
| ENSG00000162924 | REL | 0,069517944 | 1,049365995 | 0,7322 | 0,87293156 | REL proto-oncogene, NF-kB subunit |
| ENSG00000136048 | DRAM1 | 0,069428341 | 1,049300823 | 0,83556 | 0,92493625 | DNA damage regulated autophagy modulator 1 |
| ENSG00000253816 | GUSBP14 | 0,069313461 | 1,049217271 | 0,89109 | 0,95208954 | GUSB pseudogene 14 |
| ENSG00000141127 | PRPSAP2 | 0,069282239 | 1,049194565 | 0,67756 | 0,84135792 | phosphoribosyl pyrophosphate synthetase associated protein 2 |
| ENSG00000284738 | AL358472,5 | 0,069248641 | 1,049170131 | 0,88568 | 0,94943038 | novel transcript, antisense to DENND4B |
| ENSG00000139428 | MMAB | 0,069243323 | 1,049166264 | 0,66668 | 0,8343869 | metabolism of cobalamin associated B |
| ENSG00000142166 | IFNAR1 | 0,069137347 | 1,049089198 | 0,76968 | 0,89301171 | interferon alpha and beta receptor subunit 1 |
| ENSG00000229994 | RPL5P4 | 0,069111101 | 1,049070113 | 0,89037 | 0,95170091 | ribosomal protein L5 pseudogene 4 |
| ENSG00000275485 | AL512652,1 | 0,069083892 | 1,049050327 | 0,93321 | 0,97152506 | novel transcript, sense intronic to N6AMT2 |
| ENSG00000233325 | MIPEPP3 | 0,068945044 | 1,048949369 | 0,91048 | 0,96136765 | mitochondrial intermediate peptidase pseudogene 3 |
| ENSG00000056998 | GYG2 | 0,068873969 | 1,048897694 | 0,68599 | 0,84654659 | glycogenin 2 |
| ENSG00000166171 | DPCD | 0,06887295 | 1,048896953 | 0,76128 | 0,88879633 | deleted in primary ciliary dyskinesia homolog (mouse) |
| ENSG00000225164 | PRRC2A | 0,068804522 | 1,048847204 | 0,67812 | 0,84159488 | proline rich coiled-coil 2A |
| ENSG00000213689 | TREX1 | 0,068793502 | 1,048839193 | 0,82165 | 0,91753858 | three prime repair exonuclease 1 |
| ENSG00000259581 | TYRO3P | 0,068773365 | 1,048824553 | 0,84422 | 0,92888169 | TYRO3P protein tyrosine kinase pseudogene |
| ENSG00000082212 | ME2 | 0,06874098 | 1,04880101 | 0,67481 | 0,83976962 | malic enzyme 2 |
| ENSG00000287591 | AL360169,3 | 0,068704832 | 1,048774732 | 0,81477 | 0,91470265 | novel transcript |
| ENSG00000246477 | AF131216,1 | 0,068652402 | 1,048736618 | 0,79891 | 0,90723937 | uncharacterized LOC101929290 |
| ENSG00000285340 | AC270277,5 | 0,068652402 | 1,048736618 | 0,79891 | 0,90723937 | uncharacterized LOC101929290 |
| ENSG00000167264 | DUS2 | 0,068590623 | 1,04869171 | 0,73628 | 0,87522991 | dihydrouridine synthase 2 |
| ENSG00000145687 | SSBP2 | 0,068572265 | 1,048678366 | 0,79099 | 0,90292452 | single stranded DNA binding protein 2 |
| ENSG00000147099 | HDAC8 | 0,068143387 | 1,048366666 | 0,70321 | 0,8568579 | histone deacetylase 8 |
| ENSG00000281601 | ZNF780B | 0,068105064 | 1,048338818 | 0,89066 | 0,9519419 | zinc finger protein 780B |
| ENSG00000104805 | NUCB1 | 0,068053331 | 1,048301227 | 0,73518 | 0,87480865 | nucleobindin 1 |
| ENSG00000154767 | XPC | 0,067887743 | 1,048180913 | 0,74451 | 0,87904799 | XPC complex subunit, DNA damage recognition and repair factor |
| ENSG00000180185 | FAHD1 | 0,06784265 | 1,048148151 | 0,72343 | 0,86793272 | fumarylacetoacetate hydrolase domain containing 1 |
| ENSG00000130204 | TOMM40 | 0,067842349 | 1,048147932 | 0,73771 | 0,87540065 | translocase of outer mitochondrial membrane 40 |
| ENSG00000205293 | LINC01602 | 0,067743649 | 1,048076227 | 0,92951 | 0,96983055 | long intergenic non-protein coding RNA 1602 |

| | | | | | | |
|---|---|---|---|---|---|---|
| ENSG00000013583 | HEBP1 | 0,067732097 | 1,048067835 | 0,77139 | 0,8940638 | heme binding protein 1 |
| ENSG00000184408 | KCND2 | 0,067707117 | 1,048049688 | 0,79038 | 0,90286944 | potassium voltage-gated channel subfamily D member 2 |
| ENSG00000154175 | ABI3BP | 0,067627578 | 1,047991909 | 0,83696 | 0,92549623 | ABI family member 3 binding protein |
| ENSG00000278191 | CARS1 | 0,067622737 | 1,047988392 | 0,65726 | 0,82868311 | cysteinyl-tRNA synthetase 1 |
| ENSG00000159461 | AMFR | 0,06758 | 1,047957348 | 0,74801 | 0,88136669 | autocrine motility factor receptor |
| ENSG00000223561 | AC005165,1 | 0,06752421 | 1,047916824 | 0,92423 | 0,96705923 | novel transcript |
| ENSG00000185485 | SDHAP1 | 0,067487272 | 1,047889993 | 0,69733 | 0,85330122 | succinate dehydrogenase complex flavoprotein subunit A pseudogene 1 |
| ENSG00000083168 | KAT6A | 0,067482771 | 1,047886724 | 0,71899 | 0,8656088 | lysine acetyltransferase 6A |
| ENSG00000235522 | AC010978,1 | 0,067463081 | 1,047872423 | 0,8853 | 0,94924301 | novel transcript |
| ENSG00000260802 | SERTM2 | 0,067400914 | 1,04782727 | 0,92728 | 0,9685704 | serine rich and transmembrane domain containing 2 |
| ENSG00000288475 | CASP8AP2 | 0,06739919 | 1,047826017 | 0,8596 | 0,93687112 | caspase 8 associated protein 2 |
| ENSG00000117868 | ESYT2 | 0,067380018 | 1,047812093 | 0,78593 | 0,90062293 | extended synaptotagmin 2 |
| ENSG00000137075 | RNF38 | 0,067172811 | 1,047661612 | 0,86374 | 0,93907743 | ring finger protein 38 |
| ENSG00000287130 | AC068616,3 | 0,067139394 | 1,047637346 | 0,85269 | 0,93310506 | novel transcript |
| ENSG00000258917 | ZMYND19P1 | 0,067080819 | 1,047594811 | 0,93537 | 0,97222198 | zinc finger MYND-type containing 19 pseudogene 1 |
| ENSG00000140718 | FTO | 0,067069125 | 1,04758632 | 0,70713 | 0,85902171 | FTO alpha-ketoglutarate dependent dioxygenase |
| ENSG00000170161 | AL512625,1 | 0,067059464 | 1,047579304 | 0,82061 | 0,91732362 | uncharacterized LOC554249 |
| ENSG00000126947 | ARMCX1 | 0,067030932 | 1,047558587 | 0,7402 | 0,87681509 | armadillo repeat containing X-linked 1 |
| ENSG00000172530 | BANP | 0,066886589 | 1,047453783 | 0,71674 | 0,86441315 | BTG3 associated nuclear protein |
| ENSG00000174796 | THAP6 | 0,066876575 | 1,047446513 | 0,72611 | 0,86954182 | THAP domain containing 6 |
| ENSG00000119185 | ITGB1BP1 | 0,066870004 | 1,047441742 | 0,7204 | 0,86628014 | integrin subunit beta 1 binding protein 1 |
| ENSG00000102910 | LONP2 | 0,066749514 | 1,047354266 | 0,68415 | 0,84539665 | lon peptidase 2, peroxisomal |
| ENSG00000270874 | RPAP2P1 | 0,066697764 | 1,047316697 | 0,92333 | 0,96667848 | RNA polymerase II associated protein 2 pseudogene 1 |
| ENSG00000183648 | NDUFB1 | 0,066662839 | 1,047291344 | 0,75068 | 0,8829851 | NADH:ubiquinone oxidoreductase subunit B1 |
| ENSG00000165733 | BMS1 | 0,066461731 | 1,047145364 | 0,6967 | 0,85309963 | BMS1 ribosome biogenesis factor |
| ENSG00000165097 | KDM1B | 0,066447562 | 1,04713508 | 0,69453 | 0,85179889 | lysine demethylase 1B |
| ENSG00000267102 | AC060766,1 | 0,066301803 | 1,047029291 | 0,87516 | 0,9447268 | RAB32, member RAS oncogene family (RAB32) pseudogene |
| ENSG00000251294 | AC091885,2 | 0,066294664 | 1,04702411 | 0,91707 | 0,9640264 | novel transcript |
| ENSG00000174233 | ADCY6 | 0,066284624 | 1,047016823 | 0,8203 | 0,9172552 | adenylate cyclase 6 |
| ENSG00000167526 | RPL13 | 0,066185833 | 1,04694513 | 0,72935 | 0,8715818 | ribosomal protein L13 |
| ENSG00000165121 | AL353743,1 | 0,066155259 | 1,046922943 | 0,8209 | 0,91738668 | kinesin family member 27 (KIF27) pseudogene |
| ENSG00000261644 | AC007728,2 | 0,066085199 | 1,046872103 | 0,93599 | 0,97249457 | novel transcript, antisense to CYLD and NOD2 |

| | | | | | | |
|---|---|---|---|---|---|---|
| ENSG00000122970 | IFT81 | 0,066076699 | 1,046865935 | 0,79787 | 0,90685415 | intraflagellar transport 81 |
| ENSG00000114353 | GNAI2 | 0,066050461 | 1,046846897 | 0,70925 | 0,85981894 | G protein subunit alpha i2 |
| ENSG00000152217 | SETBP1 | 0,06604886 | 1,046845735 | 0,79913 | 0,90737821 | SET binding protein 1 |
| ENSG00000117226 | GBP3 | 0,066040267 | 1,0468395 | 0,91387 | 0,96272082 | guanylate binding protein 3 |
| ENSG00000267346 | EIF5AP3 | 0,065987003 | 1,046800852 | 0,94581 | 0,97689657 | eukaryotic translation initiation factor 5A pseudogene 3 |
| ENSG00000226055 | PAICSP1 | 0,065948621 | 1,046773002 | 0,89097 | 0,95208039 | phosphoribosylaminoimidazole carboxylase, phosphoribosylaminoimidazole succinocarboxamide synthetase pseudogene 1 |
| ENSG00000105618 | PRPF31 | 0,065927626 | 1,046757769 | 0,76867 | 0,89277929 | pre-mRNA processing factor 31 |
| ENSG00000196724 | ZNF418 | 0,065892554 | 1,046732323 | 0,7844 | 0,89962715 | zinc finger protein 418 |
| ENSG00000225951 | ODF2-AS1 | 0,065865463 | 1,046712668 | 0,9205 | 0,96551423 | ODF2 antisense RNA 1 |
| ENSG00000167601 | AXL | 0,065778908 | 1,046649871 | 0,87269 | 0,94374455 | AXL receptor tyrosine kinase |
| ENSG00000286360 | AC107081,3 | 0,065746567 | 1,046626409 | 0,82844 | 0,92081006 | novel transcript, antisense to CCT4 |
| ENSG00000253174 | AC009630,2 | 0,065737059 | 1,046619511 | 0,96722 | 0,98623935 | novel transcript, antisense to GINS4 |
| ENSG00000281131 | SCHLAP1 | 0,065693512 | 1,04658792 | 0,91881 | 0,96471905 | SWI/SNF complex antagonist associated with prostate cancer 1 |
| ENSG00000072415 | MPP5 | 0,065663975 | 1,046566493 | 0,81751 | 0,91619921 | membrane palmitoylated protein 5 |
| ENSG00000187323 | DCC | 0,065639169 | 1,046548498 | 0,85043 | 0,93198471 | DCC netrin 1 receptor |
| ENSG00000245059 | AC092718,1 | 0,065595676 | 1,046516949 | 0,92258 | 0,96643081 | novel transcript |
| ENSG00000099940 | SNAP29 | 0,065566186 | 1,046495557 | 0,70843 | 0,85937893 | synaptosome associated protein 29 |
| ENSG00000173064 | HECTD4 | 0,065467817 | 1,046424205 | 0,78522 | 0,90014 | HECT domain E3 ubiquitin protein ligase 4 |
| ENSG00000129625 | REEP5 | 0,065389869 | 1,046367668 | 0,84066 | 0,92739891 | receptor accessory protein 5 |
| ENSG00000230982 | DSTNP1 | 0,065339467 | 1,046331113 | 0,87919 | 0,94623795 | DSTN pseudogene 1 |
| ENSG00000282684 | ZC3H3 | 0,065223648 | 1,046247117 | 0,87117 | 0,94321347 | zinc finger CCCH-type containing 3 |
| ENSG00000101782 | RIOK3 | 0,065136772 | 1,046184117 | 0,74685 | 0,88058971 | RIO kinase 3 |
| ENSG00000267879 | AC011483,1 | 0,065120916 | 1,046172619 | 0,91805 | 0,9644207 | novel transcript, antisense to KLK6, KLK7, KLK8 and KLK9 |
| ENSG00000229419 | RALGAPA1P1 | 0,065106239 | 1,046161976 | 0,82416 | 0,91883662 | RALGAPA1 pseudogene 1 |
| ENSG00000050165 | DKK3 | 0,065094307 | 1,046153323 | 0,77122 | 0,8940638 | dickkopf WNT signaling pathway inhibitor 3 |
| ENSG00000135047 | CTSL | 0,064984462 | 1,046073673 | 0,70916 | 0,859809 | cathepsin L |
| ENSG00000144061 | NPHP1 | 0,064978008 | 1,046068993 | 0,73745 | 0,87540065 | nephrocystin 1 |
| ENSG00000075711 | DLG1 | 0,064964324 | 1,046059072 | 0,70711 | 0,85902171 | discs large MAGUK scaffold protein 1 |
| ENSG00000217334 | AL355615,1 | 0,064821943 | 1,04595584 | 0,91342 | 0,96249683 | pseudogene similar to part of heat shock 90kD protein 1, beta HSPCB) |

| | | | | | | |
|---|---|---|---|---|---|---|
| ENSG00000215375 | MYL5 | 0,064814235 | 1,045950252 | 0,76956 | 0,89301171 | myosin light chain 5 |
| ENSG00000077585 | GPR137B | 0,064810295 | 1,045947395 | 0,77825 | 0,89735311 | G protein-coupled receptor 137B |
| ENSG00000148925 | BTBD10 | 0,064744202 | 1,04589948 | 0,87064 | 0,94300286 | BTB domain containing 10 |
| ENSG00000124762 | CDKN1A | 0,064695537 | 1,045864199 | 0,88226 | 0,94783281 | cyclin dependent kinase inhibitor 1A |
| ENSG00000111450 | STX2 | 0,06468995 | 1,045860149 | 0,69326 | 0,85094387 | syntaxin 2 |
| ENSG00000091651 | ORC6 | 0,06466391 | 1,045841272 | 0,7102 | 0,86037536 | origin recognition complex subunit 6 |
| ENSG00000116337 | AMPD2 | 0,064530243 | 1,045744378 | 0,78334 | 0,89918294 | adenosine monophosphate deaminase 2 |
| ENSG00000145495 | MARCHF6 | 0,064441847 | 1,045680306 | 0,77134 | 0,8940638 | membrane associated ring-CH-type finger 6 |
| ENSG00000157193 | LRP8 | 0,064420563 | 1,04566488 | 0,79458 | 0,90473839 | LDL receptor related protein 8 |
| ENSG00000139687 | RB1 | 0,064345392 | 1,045610397 | 0,89331 | 0,95343413 | RB transcriptional corepressor 1 |
| ENSG00000229955 | Z98749,1 | 0,064264544 | 1,045551803 | 0,8864 | 0,9498079 | novel transcript, sense intronic to CSNK1E |
| ENSG00000133740 | E2F5 | 0,064128582 | 1,045453273 | 0,75692 | 0,88650931 | E2F transcription factor 5 |
| ENSG00000115275 | MOGS | 0,064122328 | 1,045448742 | 0,78031 | 0,89823499 | mannosyl-oligosaccharide glucosidase |
| ENSG00000113273 | ARSB | 0,064084114 | 1,04542105 | 0,74088 | 0,87717177 | arylsulfatase B |
| ENSG00000135390 | ATP5MC2 | 0,064007949 | 1,04536586 | 0,7659 | 0,89114666 | ATP synthase membrane subunit c locus 2 |
| ENSG00000160679 | CHTOP | 0,063977089 | 1,045343499 | 0,66106 | 0,83105713 | chromatin target of PRMT1 |
| ENSG00000274492 | AL121768,1 | 0,063893782 | 1,045283139 | 0,85071 | 0,93206349 | novel transcript |
| ENSG00000136010 | ALDH1L2 | 0,063870924 | 1,045266577 | 0,76343 | 0,88981244 | aldehyde dehydrogenase 1 family member L2 |
| ENSG00000135778 | NTPCR | 0,063846146 | 1,045248625 | 0,71153 | 0,86116229 | nucleoside-triphosphatase, cancer-related |
| ENSG00000112701 | SENP6 | 0,063841498 | 1,045245258 | 0,68223 | 0,84420893 | SUMO specific peptidase 6 |
| ENSG00000141560 | FN3KRP | 0,063819495 | 1,045229317 | 0,7619 | 0,88909725 | fructosamine 3 kinase related protein |
| ENSG00000251301 | LINC02384 | 0,06367312 | 1,045123273 | 0,89502 | 0,95419386 | long intergenic non-protein coding RNA 2384 |
| ENSG00000225527 | AL450384,1 | 0,063619173 | 1,045084194 | 0,92955 | 0,96983055 | novel transcript |
| ENSG00000187257 | RSBN1L | 0,063596843 | 1,045068018 | 0,71116 | 0,86096523 | round spermatid basic protein 1 like |
| ENSG00000099203 | TMED1 | 0,063589355 | 1,045062594 | 0,79007 | 0,90275167 | transmembrane p24 trafficking protein 1 |
| ENSG00000013441 | CLK1 | 0,063569677 | 1,04504834 | 0,79724 | 0,90659204 | CDC like kinase 1 |
| ENSG00000101311 | FERMT1 | 0,063493279 | 1,044993001 | 0,94466 | 0,97650311 | fermitin family member 1 |
| ENSG00000204387 | SNHG32 | 0,063459808 | 1,044968757 | 0,87371 | 0,94405019 | small nucleolar RNA host gene 32 |
| ENSG00000187080 | OR2AK2 | 0,063436856 | 1,044952133 | 0,9329 | 0,97143711 | olfactory receptor family 2 subfamily AK member 2 |
| ENSG00000144868 | TMEM108 | 0,063421668 | 1,044941132 | 0,85497 | 0,93417381 | transmembrane protein 108 |
| ENSG00000288010 | AL035587,3 | 0,063305061 | 1,044856677 | 0,83195 | 0,92312766 | novel transcript |
| ENSG00000236888 | RPS20P5 | 0,063295906 | 1,044850046 | 0,95062 | 0,97934603 | ribosomal protein S20 pseudogene 5 |
| ENSG00000040531 | CTNS | 0,063193761 | 1,044776072 | 0,77552 | 0,89630112 | cystinosin, lysosomal cystine transporter |

| | | | | | | |
|---|---|---|---|---|---|---|
| ENSG00000162913 | OBSCN-AS1 | 0,063180497 | 1,044766467 | 0,83625 | 0,92493625 | OBSCN antisense RNA 1 |
| ENSG00000102054 | RBBP7 | 0,063163759 | 1,044754346 | 0,81391 | 0,9144566 | RB binding protein 7, chromatin remodeling factor |
| ENSG00000064225 | ST3GAL6 | 0,063070282 | 1,044686654 | 0,82892 | 0,92114387 | ST3 beta-galactoside alpha-2,3-sialyltransferase 6 |
| ENSG00000285427 | SOD2-OT1 | 0,063012772 | 1,044645011 | 0,76965 | 0,89301171 | SOD2 overlapping transcript 1 |
| ENSG00000087157 | PGS1 | 0,062907018 | 1,044568438 | 0,68811 | 0,84808917 | phosphatidylglycerophosphate synthase 1 |
| ENSG00000143502 | SUSD4 | 0,062693493 | 1,044413849 | 0,92139 | 0,96594467 | sushi domain containing 4 |
| ENSG00000259706 | HSP90B2P | 0,062631339 | 1,044368854 | 0,85631 | 0,93503131 | heat shock protein 90 beta family member 2, pseudogene |
| ENSG00000271133 | AC004130,2 | 0,062595955 | 1,04434324 | 0,92865 | 0,96934915 | novel transcript, antisense to ITGB8 |
| ENSG00000139675 | HNRNPA1L2 | 0,062500638 | 1,044274244 | 0,82445 | 0,91900665 | heterogeneous nuclear ribonucleoprotein A1 like 2 |
| ENSG00000129003 | VPS13C | 0,062485957 | 1,044263618 | 0,86714 | 0,94124126 | vacuolar protein sorting 13 homolog C |
| ENSG00000032389 | EIPR1 | 0,062459332 | 1,044244346 | 0,74636 | 0,88045757 | EARP complex and GARP complex interacting protein 1 |
| ENSG00000171100 | MTM1 | 0,062397113 | 1,044199311 | 0,85596 | 0,93482725 | myotubularin 1 |
| ENSG00000162298 | SYVN1 | 0,062384217 | 1,044189978 | 0,71999 | 0,86621694 | synoviolin 1 |
| ENSG00000244691 | RPL10AP1 | 0,062373073 | 1,044181912 | 0,89985 | 0,95730568 | ribosomal protein L10a pseudogene 1 |
| ENSG00000199683 | RN7SKP185 | 0,062362267 | 1,044174091 | 0,92325 | 0,96667029 | RN7SK pseudogene 185 |
| ENSG00000175662 | TOM1L2 | 0,062303331 | 1,044131436 | 0,76192 | 0,88909725 | target of myb1 like 2 membrane trafficking protein |
| ENSG00000251348 | HSPD1P11 | 0,062192638 | 1,044051327 | 0,90834 | 0,96080099 | heat shock protein family D (Hsp60) member 1 pseudogene 11 |
| ENSG00000235079 | ZRANB2-AS1 | 0,062180272 | 1,044042378 | 0,90178 | 0,95802163 | ZRANB2 antisense RNA 1 |
| ENSG00000178988 | MRFAP1L1 | 0,062141247 | 1,044014137 | 0,68594 | 0,84654659 | Morf4 family associated protein 1 like 1 |
| ENSG00000273592 | MBOAT7 | 0,062031133 | 1,043934455 | 0,85098 | 0,93218292 | membrane bound O-acyltransferase domain containing 7 |
| ENSG00000198855 | FICD | 0,061994045 | 1,043907618 | 0,84833 | 0,93094401 | FIC domain protein adenylyltransferase |
| ENSG00000185808 | PIGP | 0,061882206 | 1,043826697 | 0,79869 | 0,90713957 | phosphatidylinositol glycan anchor biosynthesis class P |
| ENSG00000149657 | LSM14B | 0,061824088 | 1,043784648 | 0,782 | 0,8987738 | LSM family member 14B |
| ENSG00000155008 | APOOL | 0,061714695 | 1,043705505 | 0,70814 | 0,85927396 | apolipoprotein O like |
| ENSG00000188636 | RTL6 | 0,061635875 | 1,043648485 | 0,73692 | 0,87539732 | retrotransposon Gag like 6 |
| ENSG00000170634 | ACYP2 | 0,061485764 | 1,043539901 | 0,82376 | 0,91861952 | acylphosphatase 2 |
| ENSG00000167186 | COQ7 | 0,061433652 | 1,043502207 | 0,7936 | 0,9039896 | coenzyme Q7, hydroxylase |
| ENSG00000248216 | KCTD9P5 | 0,061414396 | 1,04348828 | 0,91959 | 0,96506391 | potassium channel tetramerization domain containing 9 pseudogene 5 |
| ENSG00000051596 | THOC3 | 0,061356064 | 1,043446089 | 0,7738 | 0,89571194 | THO complex 3 |
| ENSG00000085382 | HACE1 | 0,061350568 | 1,043442114 | 0,68145 | 0,84375324 | HECT domain and ankyrin repeat containing E3 ubiquitin protein ligase 1 |

| | | | | | | |
|---|---|---|---|---|---|---|
| ENSG00000229905 | AL669831,2 | 0,061322197 | 1,043421595 | 0,90592 | 0,95980391 | novel transcript |
| ENSG00000120533 | ENY2 | 0,061257859 | 1,043375064 | 0,75724 | 0,88663582 | ENY2 transcription and export complex 2 subunit |
| ENSG00000143545 | RAB13 | 0,061196556 | 1,04333073 | 0,76117 | 0,88879633 | RAB13, member RAS oncogene family |
| ENSG00000236397 | DDX11L2 | 0,061172003 | 1,043312974 | 0,81369 | 0,9144566 | DEAD/H-box helicase 11 like 2 (pseudogene) |
| ENSG00000285043 | AC093512,2 | 0,061168966 | 1,043310777 | 0,81381 | 0,9144566 | uncharaterized LOC112694756 |
| ENSG00000157106 | SMG1 | 0,060997596 | 1,043186855 | 0,69979 | 0,85497086 | SMG1 nonsense mediated mRNA decay associated PI3K related kinase |
| ENSG00000275581 | SERF1A | 0,060733354 | 1,042995804 | 0,92767 | 0,96872012 | small EDRK-rich factor 1A |
| ENSG00000224669 | AC016769,2 | 0,060673255 | 1,042952356 | 0,89347 | 0,9534926 | glycine cleavage system protein H (aminomethyl carrier) (GCSH) pseudogene |
| ENSG00000260337 | AC091544,4 | 0,060416853 | 1,042767015 | 0,95419 | 0,9807577 | novel transcript |
| ENSG00000091542 | ALKBH5 | 0,060386326 | 1,042744951 | 0,71302 | 0,86193361 | alkB homolog 5, RNA demethylase |
| ENSG00000261087 | ZNNT1 | 0,06033734 | 1,042709546 | 0,82266 | 0,91804426 | ZNF706 neighboring transcript 1 |
| ENSG00000165449 | SLC16A9 | 0,060313508 | 1,042692321 | 0,87223 | 0,94371063 | solute carrier family 16 member 9 |
| ENSG00000151092 | NGLY1 | 0,06031071 | 1,042690299 | 0,71146 | 0,86114994 | N-glycanase 1 |
| ENSG00000174137 | FAM53A | 0,060304373 | 1,042685719 | 0,92622 | 0,96789681 | family with sequence similarity 53 member A |
| ENSG00000101247 | NDUFAF5 | 0,060243749 | 1,042641905 | 0,74846 | 0,88159265 | NADH:ubiquinone oxidoreductase complex assembly factor 5 |
| ENSG00000257386 | AC025257,1 | 0,060117812 | 1,042550893 | 0,79472 | 0,90486565 | novel transcript, antisense to ATXN7L3B |
| ENSG00000266173 | STRADA | 0,060093114 | 1,042533046 | 0,77113 | 0,89404837 | STE20 related adaptor alpha |
| ENSG00000119979 | DENND10 | 0,060013403 | 1,042475446 | 0,72925 | 0,87157404 | DENN domain containing 10 |
| ENSG00000089094 | KDM2B | 0,059990465 | 1,042458871 | 0,78555 | 0,90037696 | lysine demethylase 2B |
| ENSG00000184613 | NELL2 | 0,059942129 | 1,042423945 | 0,84809 | 0,93093915 | neural EGFL like 2 |
| ENSG00000227304 | AC067942,1 | 0,059893802 | 1,042389027 | 0,9037 | 0,95873972 | ribosomal protein S6 (RPS6) pseudogene |
| ENSG00000065029 | ZNF76 | 0,059847313 | 1,042355438 | 0,80462 | 0,90996631 | zinc finger protein 76 |
| ENSG00000148950 | IMMP1L | 0,059816594 | 1,042333243 | 0,8086 | 0,91193281 | inner mitochondrial membrane peptidase subunit 1 |
| ENSG00000101193 | GID8 | 0,05980197 | 1,042322678 | 0,70703 | 0,85902171 | GID complex subunit 8 homolog |
| ENSG00000137275 | RIPK1 | 0,05943171 | 1,042055205 | 0,79066 | 0,90286944 | receptor interacting serine/threonine kinase 1 |
| ENSG00000008130 | NADK | 0,059385458 | 1,042021799 | 0,71906 | 0,8656088 | NAD kinase |
| ENSG00000283031 | AC009242,1 | 0,05935289 | 1,041998276 | 0,86975 | 0,9425949 | novel transcript |
| ENSG00000177885 | GRB2 | 0,059313181 | 1,041969596 | 0,77209 | 0,89459692 | growth factor receptor bound protein 2 |
| ENSG00000229833 | PET100 | 0,059221401 | 1,041903311 | 0,8051 | 0,9103203 | PET100 cytochrome c oxidase chaperone |
| ENSG00000234718 | AC007161,1 | 0,059184924 | 1,041876968 | 0,93269 | 0,97136054 | ribosomal protein L23a (RPL23A) pseudogene |

| | | | | | | |
|---|---|---|---|---|---|---|
| ENSG00000101290 | CDS2 | 0,059171115 | 1,041866995 | 0,82145 | 0,91749859 | CDP-diacylglycerol synthase 2 |
| ENSG00000069020 | MAST4 | 0,059103951 | 1,041818493 | 0,77998 | 0,89820865 | microtubule associated serine/threonine kinase family member 4 |
| ENSG00000136425 | CIB2 | 0,05909535 | 1,041812282 | 0,79611 | 0,90574377 | calcium and integrin binding family member 2 |
| ENSG00000114982 | KANSL3 | 0,059005806 | 1,041747622 | 0,74591 | 0,88010501 | KAT8 regulatory NSL complex subunit 3 |
| ENSG00000178631 | ACTG1P1 | 0,058989514 | 1,041735857 | 0,92921 | 0,96975046 | actin gamma 1 pseudogene 1 |
| ENSG00000272235 | AL590438,1 | 0,058888067 | 1,041662607 | 0,89601 | 0,9549591 | novel transcript |
| ENSG00000139132 | FGD4 | 0,058881628 | 1,041657958 | 0,86208 | 0,93836958 | FYVE, RhoGEF and PH domain containing 4 |
| ENSG00000205763 | RP9P | 0,058753796 | 1,041565665 | 0,80819 | 0,9117763 | RP9 pseudogene |
| ENSG00000245571 | FAM111A-DT | 0,058720962 | 1,04154196 | 0,86572 | 0,94036783 | FAM111A divergent transcript |
| ENSG00000231562 | AL512593,1 | 0,058664279 | 1,041501039 | 0,86164 | 0,93811282 | ribosomal protein L7a (RPL7A) pseudogene |
| ENSG00000163138 | PACRGL | 0,058660551 | 1,041498348 | 0,73017 | 0,87180309 | parkin coregulated like |
| ENSG00000282813 | ARMC10 | 0,058462921 | 1,041355687 | 0,91302 | 0,96234107 | armadillo repeat containing 10 |
| ENSG00000241772 | AC092620,1 | 0,058442568 | 1,041340995 | 0,94677 | 0,97731612 | novel transcript |
| ENSG00000166869 | CHP2 | 0,058397533 | 1,041308489 | 0,88859 | 0,95084729 | calcineurin like EF-hand protein 2 |
| ENSG00000121879 | PIK3CA | 0,05837782 | 1,041294261 | 0,79782 | 0,90685415 | phosphatidylinositol-4,5-bisphosphate 3-kinase catalytic subunit alpha |
| ENSG00000226205 | RPS3AP27 | 0,058336955 | 1,041264766 | 0,94053 | 0,97474573 | RPS3A pseudogene 27 |
| ENSG00000170703 | TTLL6 | 0,05826638 | 1,04121383 | 0,94799 | 0,97814651 | tubulin tyrosine ligase like 6 |
| ENSG00000140526 | ABHD2 | 0,05822837 | 1,041186398 | 0,81851 | 0,91665476 | abhydrolase domain containing 2, acylglycerol lipase |
| ENSG00000165175 | MID1IP1 | 0,058058182 | 1,041063581 | 0,73723 | 0,87539732 | MID1 interacting protein 1 |
| ENSG00000046653 | GPM6B | 0,057977478 | 1,041005346 | 0,90905 | 0,96102407 | glycoprotein M6B |
| ENSG00000138443 | ABI2 | 0,057935517 | 1,040975069 | 0,78007 | 0,89820865 | abl interactor 2 |
| ENSG00000239465 | AC090543,1 | 0,057820041 | 1,040891751 | 0,94407 | 0,97642036 | ribosomal protein L21 (RPL21) pseudogene |
| ENSG00000126460 | PRRG2 | 0,057730218 | 1,040826946 | 0,92304 | 0,96662141 | proline rich and Gla domain 2 |
| ENSG00000160767 | FAM189B | 0,057453915 | 1,040627628 | 0,82269 | 0,91804426 | family with sequence similarity 189 member B |
| ENSG00000144152 | FBLN7 | 0,057437678 | 1,040615916 | 0,8196 | 0,91708488 | fibulin 7 |
| ENSG00000267397 | AC090229,1 | 0,057316597 | 1,040528584 | 0,89763 | 0,95581835 | novel transcript |
| ENSG00000286667 | AC010641,2 | 0,057315625 | 1,040527882 | 0,8624 | 0,93853472 | novel transcript |
| ENSG00000180537 | RNF182 | 0,05729218 | 1,040510974 | 0,85727 | 0,93557143 | ring finger protein 182 |
| ENSG00000281026 | N4BP2L2-IT2 | 0,057273467 | 1,040497477 | 0,81765 | 0,91621442 | N4BPL2 intronic transcript 2 |
| ENSG00000150551 | LYPD1 | 0,057187974 | 1,04043582 | 0,91808 | 0,9644207 | LY6/PLAUR domain containing 1 |
| ENSG00000113811 | SELENOK | 0,057115003 | 1,040383196 | 0,80481 | 0,91010357 | selenoprotein K |

| | | | | | | |
|---|---|---|---|---|---|---|
| ENSG00000259515 | AL117209,1 | 0,05710999 | 1,040379581 | 0,90771 | 0,96044232 | novel transcript, antisense to CDC42BPB |
| ENSG00000111897 | SERINC1 | 0,057107874 | 1,040378055 | 0,80132 | 0,90860375 | serine incorporator 1 |
| ENSG00000253558 | AC024568,1 | 0,057052332 | 1,040338003 | 0,88751 | 0,95041376 | heat shock protein 90kDa alpha (cytosolic), class B member 1 (HSP90AB1) pseudogene |
| ENSG00000136802 | LRRC8A | 0,05701842 | 1,040313549 | 0,73527 | 0,87483589 | leucine rich repeat containing 8 VRAC subunit A |
| ENSG00000088766 | CRLS1 | 0,056947404 | 1,040262341 | 0,76935 | 0,89297049 | cardiolipin synthase 1 |
| ENSG00000174529 | TMEM81 | 0,056895044 | 1,040224587 | 0,81039 | 0,91251177 | transmembrane protein 81 |
| ENSG00000130311 | DDA1 | 0,056880708 | 1,040214251 | 0,76819 | 0,89269797 | DET1 and DDB1 associated 1 |
| ENSG00000173821 | RNF213 | 0,056877187 | 1,040211712 | 0,72298 | 0,86768626 | ring finger protein 213 |
| ENSG00000287837 | AL590064,1 | 0,056843283 | 1,040187267 | 0,8887 | 0,9508882 | novel transcript |
| ENSG00000287918 | AF274853,1 | 0,056819867 | 1,040170384 | 0,90183 | 0,95803509 | novel transcript, antisense to VMA21 |
| ENSG00000235183 | SRP14P3 | 0,05678457 | 1,040144936 | 0,93125 | 0,97061998 | signal recognition particle 14 pseudogene 3 |
| ENSG00000105699 | LSR | 0,056771806 | 1,040135733 | 0,92201 | 0,96618214 | lipolysis stimulated lipoprotein receptor |
| ENSG00000114455 | HHLA2 | 0,056698583 | 1,040082943 | 0,88501 | 0,94911792 | HERV-H LTR-associating 2 |
| ENSG00000125841 | NRSN2 | 0,05656239 | 1,039984762 | 0,78174 | 0,89877377 | neurensin 2 |
| ENSG00000164120 | HPGD | 0,056491805 | 1,039933881 | 0,95877 | 0,98291816 | 15-hydroxyprostaglandin dehydrogenase |
| ENSG00000170468 | RIOX1 | 0,056420899 | 1,039882771 | 0,81964 | 0,91708488 | ribosomal oxygenase 1 |
| ENSG00000139719 | VPS33A | 0,05635191 | 1,039833046 | 0,74465 | 0,87914393 | VPS33A core subunit of CORVET and HOPS complexes |
| ENSG00000180116 | C12orf40 | 0,056344992 | 1,03982806 | 0,93593 | 0,97249457 | chromosome 12 open reading frame 40 |
| ENSG00000213398 | LCAT | 0,056327132 | 1,039815188 | 0,86947 | 0,9425782 | lecithin-cholesterol acyltransferase |
| ENSG00000176182 | MYPOP | 0,056287292 | 1,039786473 | 0,84682 | 0,93041089 | Myb related transcription factor, partner of profilin |
| ENSG00000158552 | ZFAND2B | 0,056160678 | 1,039695223 | 0,7815 | 0,89877377 | zinc finger AN1-type containing 2B |
| ENSG00000159423 | ALDH4A1 | 0,056120558 | 1,03966631 | 0,77616 | 0,89650232 | aldehyde dehydrogenase 4 family member A1 |
| ENSG00000164953 | TMEM67 | 0,056056121 | 1,039619876 | 0,80402 | 0,909789 | transmembrane protein 67 |
| ENSG00000228487 | AL450263,1 | 0,056035277 | 1,039604855 | 0,91914 | 0,96485808 | novel transcript, antisense to RALGPS1 |
| ENSG00000266904 | LINC00663 | 0,056018121 | 1,039592493 | 0,84118 | 0,92771379 | long intergenic non-protein coding RNA 663 |
| ENSG00000284841 | H2AW | 0,055985317 | 1,039568855 | 0,85132 | 0,93233257 | H2A,W histone |
| ENSG00000181218 | H2AW | 0,055985317 | 1,039568855 | 0,85132 | 0,93233257 | H2A,W histone |
| ENSG00000005249 | PRKAR2B | 0,055978172 | 1,039563706 | 0,86825 | 0,94176198 | protein kinase cAMP-dependent type II regulatory subunit beta |
| ENSG00000284096 | PRKAR2B | 0,055978172 | 1,039563706 | 0,86825 | 0,94176198 | protein kinase cAMP-dependent type II regulatory subunit beta |
| ENSG00000143469 | SYT14 | 0,055943932 | 1,039539034 | 0,87947 | 0,94631723 | synaptotagmin 14 |

| | | | | | | |
|---|---|---|---|---|---|---|
| ENSG00000167394 | ZNF668 | 0,055752996 | 1,039401464 | 0,7557 | 0,88566553 | zinc finger protein 668 |
| ENSG00000287724 | AC096536,3 | 0,055600286 | 1,039291449 | 0,93819 | 0,97361277 | novel transcript, antisense to DHCR24 |
| ENSG00000184986 | TMEM121 | 0,05558889 | 1,039283239 | 0,91425 | 0,96287922 | transmembrane protein 121 |
| ENSG00000167863 | ATP5PD | 0,055484606 | 1,039208118 | 0,7588 | 0,88742567 | ATP synthase peripheral stalk subunit d |
| ENSG00000282493 | AC100810,2 | 0,055390479 | 1,039140318 | 0,93832 | 0,97361277 | DNA fragmentation factor, 45kDa, alpha polypeptide (DFFA) pseudogene |
| ENSG00000282021 | AC100810,3 | 0,055390479 | 1,039140318 | 0,93832 | 0,97361277 | novel transcript |
| ENSG00000119655 | NPC2 | 0,055343267 | 1,039106313 | 0,8943 | 0,95389664 | NPC intracellular cholesterol transporter 2 |
| ENSG00000235944 | ZNF815P | 0,055302135 | 1,039076688 | 0,84748 | 0,93071557 | zinc finger protein 815, pseudogene |
| ENSG00000132669 | RIN2 | 0,055295647 | 1,039072015 | 0,8168 | 0,91576125 | Ras and Rab interactor 2 |
| ENSG00000241546 | AC092910,2 | 0,0552869 | 1,039065715 | 0,93288 | 0,97143711 | riboflavin kinase (RFK) pseudogene |
| ENSG00000288040 | AC097713,2 | 0,055218627 | 1,039016544 | 0,85354 | 0,93356371 | novel transcript |
| ENSG00000238142 | BX284668,5 | 0,055208019 | 1,039008905 | 0,84207 | 0,92818855 | novel transcript |
| ENSG00000239881 | RPS27P25 | 0,05516003 | 1,038974344 | 0,9383 | 0,97361277 | ribosomal protein S27 pseudogene 25 |
| ENSG00000058085 | LAMC2 | 0,055154263 | 1,038970191 | 0,89248 | 0,95285354 | laminin subunit gamma 2 |
| ENSG00000257642 | KCCAT198 | 0,055078088 | 1,038915335 | 0,92003 | 0,96530017 | renal clear cell carcinoma-associated transcript 198 |
| ENSG00000205323 | SARNP | 0,054978889 | 1,038843902 | 0,75057 | 0,88297665 | SAP domain containing ribonucleoprotein |
| ENSG00000081177 | EXD2 | 0,054862639 | 1,038760197 | 0,82244 | 0,91798143 | exonuclease 3'-5' domain containing 2 |
| ENSG00000288049 | AC010889,2 | 0,054834159 | 1,038739691 | 0,91677 | 0,96378593 | novel transcript, antisense to KDM5D |
| ENSG00000250749 | AC122714,1 | 0,054831913 | 1,038738074 | 0,93119 | 0,97060346 | proteolipid protein 2 (colonic epithelium-enriched) (PLP2) pseudogene |
| ENSG00000046604 | DSG2 | 0,054789924 | 1,038707842 | 0,89332 | 0,95343413 | desmoglein 2 |
| ENSG00000180423 | HARBI1 | 0,054782734 | 1,038702666 | 0,81175 | 0,91334888 | harbinger transposase derived 1 |
| ENSG00000237943 | PRKCQ-AS1 | 0,054718383 | 1,038656335 | 0,91463 | 0,96309662 | PRKCQ antisense RNA 1 |
| ENSG00000253958 | CLDN23 | 0,054698479 | 1,038642006 | 0,91805 | 0,9644207 | claudin 23 |
| ENSG00000140265 | ZSCAN29 | 0,054697092 | 1,038641007 | 0,71225 | 0,8615121 | zinc finger and SCAN domain containing 29 |
| ENSG00000285715 | AC063960,1 | 0,054673771 | 1,038624218 | 0,90494 | 0,9593272 | novel transcript, antisense to TACC2 |
| ENSG00000238228 | OR7E7P | 0,054634287 | 1,038595794 | 0,91116 | 0,9616927 | olfactory receptor family 7 subfamily E member 7 pseudogene |
| ENSG00000067208 | EVI5 | 0,054612479 | 1,038580094 | 0,71719 | 0,86477199 | ecotropic viral integration site 5 |
| ENSG00000277290 | AC136475,10 | 0,054552545 | 1,038536949 | 0,94636 | 0,97714436 | signal recognition particle 19kDa (SRP19) pseudogene |
| ENSG00000288531 | AL669831,7 | 0,054504318 | 1,038502233 | 0,8994 | 0,95714792 | novel transcript |
| ENSG00000224531 | SMIM13 | 0,054470418 | 1,038477831 | 0,8656 | 0,94030154 | small integral membrane protein 13 |

| | | | | | | |
|---|---|---|---|---|---|---|
| ENSG00000245498 | AP000866,1 | 0,054384139 | 1,038415727 | 0,78084 | 0,89846287 | novel transcript, antisense to C11orf61 |
| ENSG00000263648 | AC005244,1 | 0,054215777 | 1,038294552 | 0,95153 | 0,97948342 | dynein, light chain, Tctex-type 1 (DYNLT1) pseudogene |
| ENSG00000287626 | AL021978,1 | 0,054214654 | 1,038293743 | 0,92475 | 0,9672926 | novel transcript, antisense to DTNBP1 |
| ENSG00000248079 | DPH6-DT | 0,054106202 | 1,038215695 | 0,94486 | 0,97662603 | DPH6 divergent transcript |
| ENSG00000205423 | CNEP1R1 | 0,054003515 | 1,0381418 | 0,75825 | 0,88727479 | CTD nuclear envelope phosphatase 1 regulatory subunit 1 |
| ENSG00000137100 | DCTN3 | 0,053965162 | 1,038114202 | 0,82399 | 0,91876332 | dynactin subunit 3 |
| ENSG00000236015 | AC011290,2 | 0,053952146 | 1,038104836 | 0,85847 | 0,93621468 | thioredoxin domain containing 1 (TXNDC1) pseudogene |
| ENSG00000254726 | MEX3A | 0,053896118 | 1,038064521 | 0,75434 | 0,88487815 | mex-3 RNA binding family member A |
| ENSG00000240898 | AC132942,1 | 0,053867481 | 1,038043917 | 0,93317 | 0,97152506 | ribosomal protein L21 (RPL21)pseudogene |
| ENSG00000281630 | MUC20 | 0,053838555 | 1,038023104 | 0,96815 | 0,98639465 | mucin 20, cell surface associated |
| ENSG00000157216 | SSBP3 | 0,053797227 | 1,037993368 | 0,74681 | 0,88057664 | single stranded DNA binding protein 3 |
| ENSG00000183826 | BTBD9 | 0,053686175 | 1,037913472 | 0,83858 | 0,92627183 | BTB domain containing 9 |
| ENSG00000070214 | SLC44A1 | 0,053573593 | 1,037832481 | 0,85674 | 0,93535509 | solute carrier family 44 member 1 |
| ENSG00000151612 | ZNF827 | 0,053546144 | 1,037812735 | 0,80074 | 0,90828402 | zinc finger protein 827 |
| ENSG00000235066 | AC009303,2 | 0,053464302 | 1,037753863 | 0,92657 | 0,96811087 | novel transcript, antisense to CCDC93 |
| ENSG00000101421 | CHMP4B | 0,053442479 | 1,037738165 | 0,83696 | 0,92549623 | charged multivesicular body protein 4B |
| ENSG00000185043 | CIB1 | 0,053289954 | 1,037628459 | 0,89473 | 0,9541449 | calcium and integrin binding 1 |
| ENSG00000060138 | YBX3 | 0,053188902 | 1,037555782 | 0,87452 | 0,9444767 | Y-box binding protein 3 |
| ENSG00000172260 | NEGR1 | 0,053120799 | 1,037506805 | 0,85752 | 0,93569939 | neuronal growth regulator 1 |
| ENSG00000231307 | RPS3P2 | 0,05310894 | 1,037498276 | 0,93549 | 0,9722573 | ribosomal protein S3 pseudogene 2 |
| ENSG00000246560 | UBE2D3-AS1 | 0,053058665 | 1,037462122 | 0,89492 | 0,95417507 | UBE2D3 antisense RNA 1 |
| ENSG00000132698 | RAB25 | 0,053023277 | 1,037436675 | 0,90663 | 0,96009372 | RAB25, member RAS oncogene family |
| ENSG00000260948 | AL390195,2 | 0,052995434 | 1,037416653 | 0,84607 | 0,92995515 | novel transcript, sense overlapping WDR77 & OVGP1 |
| ENSG00000103148 | NPRL3 | 0,052914655 | 1,037358568 | 0,80477 | 0,91009327 | NPR3 like, GATOR1 complex subunit |
| ENSG00000103145 | HCFC1R1 | 0,052793194 | 1,037271236 | 0,81644 | 0,91554153 | host cell factor C1 regulator 1 |
| ENSG00000075213 | SEMA3A | 0,052779456 | 1,037261359 | 0,92912 | 0,96969273 | semaphorin 3A |
| ENSG00000253878 | AC087752,3 | 0,052767028 | 1,037252424 | 0,91362 | 0,96257531 | novel transcript |
| ENSG00000276216 | AC245014,3 | 0,052702072 | 1,037205723 | 0,8399 | 0,92681214 | novel transcript |
| ENSG00000213197 | NUDCP1 | 0,052675442 | 1,037186578 | 0,89409 | 0,95389664 | nuclear distribution C pseudogene 1 |
| ENSG00000139508 | SLC46A3 | 0,05265533 | 1,037172119 | 0,94021 | 0,97463843 | solute carrier family 46 member 3 |
| ENSG00000165443 | PHYHIPL | 0,05264426 | 1,037164161 | 0,92024 | 0,96530017 | phytanoyl-CoA 2-hydroxylase interacting protein like |
| ENSG00000136159 | NUDT15 | 0,052577329 | 1,037116045 | 0,81018 | 0,91249823 | nudix hydrolase 15 |
| ENSG00000135749 | PCNX2 | 0,052537092 | 1,03708712 | 0,84243 | 0,92821072 | pecanex 2 |

| | | | | | | |
|---|---|---|---|---|---|---|
| ENSG00000130037 | KCNA5 | 0,05244384 | 1,037020087 | 0,95461 | 0,98100885 | potassium voltage-gated channel subfamily A member 5 |
| ENSG00000147119 | CHST7 | 0,052408034 | 1,03699435 | 0,89353 | 0,95351752 | carbohydrate sulfotransferase 7 |
| ENSG00000178537 | SLC25A20 | 0,052387919 | 1,036979892 | 0,84292 | 0,9282764 | solute carrier family 25 member 20 |
| ENSG00000130023 | ERMARD | 0,052383437 | 1,036976671 | 0,91072 | 0,96150464 | ER membrane associated RNA degradation |
| ENSG00000281794 | MUC20-OT1 | 0,052339664 | 1,036945208 | 0,84436 | 0,92899739 | MUC20 overlapping transcript |
| ENSG00000278032 | LY6E | 0,052280367 | 1,036902588 | 0,90372 | 0,95873972 | lymphocyte antigen 6 family member E |
| ENSG00000204186 | ZDBF2 | 0,052249535 | 1,036880429 | 0,89117 | 0,95208954 | zinc finger DBF-type containing 2 |
| ENSG00000278817 | AC007325,4 | 0,05224784 | 1,036879211 | 0,93484 | 0,9721963 | protein DGCR6 |
| ENSG00000055332 | EIF2AK2 | 0,052119858 | 1,036787233 | 0,73915 | 0,87625511 | eukaryotic translation initiation factor 2 alpha kinase 2 |
| ENSG00000188931 | CFAP126 | 0,052087241 | 1,036763793 | 0,8893 | 0,95110855 | cilia and flagella associated protein 126 |
| ENSG00000155666 | KDM8 | 0,052076486 | 1,036756064 | 0,78141 | 0,89877377 | lysine demethylase 8 |
| ENSG00000132635 | PCED1A | 0,052055388 | 1,036740903 | 0,82693 | 0,91999458 | PC-esterase domain containing 1A |
| ENSG00000197535 | MYO5A | 0,051991982 | 1,036695339 | 0,86356 | 0,93904284 | myosin VA |
| ENSG00000148429 | USP6NL | 0,051867068 | 1,036605583 | 0,75641 | 0,88605383 | USP6 N-terminal like |
| ENSG00000226849 | AL109811,1 | 0,051863827 | 1,036603254 | 0,89575 | 0,95482399 | novel transcript |
| ENSG00000159259 | CHAF1B | 0,05182496 | 1,036575327 | 0,78194 | 0,8987738 | chromatin assembly factor 1 subunit B |
| ENSG00000186645 | SPDYE17 | 0,051814059 | 1,036567495 | 0,93658 | 0,9727206 | speedy/RINGO cell cycle regulator family member E17 |
| ENSG00000267228 | AC012254,2 | 0,051797249 | 1,036555417 | 0,93715 | 0,97305801 | novel transcript |
| ENSG00000101997 | CCDC22 | 0,051796409 | 1,036554813 | 0,77693 | 0,89665303 | coiled-coil domain containing 22 |
| ENSG00000171189 | GRIK1 | 0,051785272 | 1,036546812 | 0,91186 | 0,96184653 | glutamate ionotropic receptor kainate type subunit 1 |
| ENSG00000104960 | PTOV1 | 0,051771882 | 1,036537191 | 0,74364 | 0,87867571 | PTOV1 extended AT-hook containing adaptor protein |
| ENSG00000151729 | SLC25A4 | 0,05176328 | 1,036531011 | 0,84564 | 0,92978831 | solute carrier family 25 member 4 |
| ENSG00000213558 | HMGN2P32 | 0,051736613 | 1,036511852 | 0,93618 | 0,97254199 | high mobility group nucleosomal binding domain 2 pseudogene 32 |
| ENSG00000197375 | SLC22A5 | 0,051629051 | 1,036434576 | 0,81613 | 0,91549169 | solute carrier family 22 member 5 |
| ENSG00000214988 | RPL7AP26 | 0,051575229 | 1,036395912 | 0,95771 | 0,98257839 | ribosomal protein L7a pseudogene 26 |
| ENSG00000136436 | CALCOCO2 | 0,051525514 | 1,036360198 | 0,82803 | 0,92070145 | calcium binding and coiled-coil domain 2 |
| ENSG00000064601 | CTSA | 0,05151016 | 1,036349169 | 0,83182 | 0,92312748 | cathepsin A |
| ENSG00000105647 | PIK3R2 | 0,05150844 | 1,036347933 | 0,7829 | 0,89917991 | phosphoinositide-3-kinase regulatory subunit 2 |
| ENSG00000112855 | HARS2 | 0,05149335 | 1,036337093 | 0,74366 | 0,87867571 | histidyl-tRNA synthetase 2, mitochondrial |
| ENSG00000169857 | AVEN | 0,051442755 | 1,03630075 | 0,83831 | 0,92622127 | apoptosis and caspase activation inhibitor |
| ENSG00000089723 | OTUB2 | 0,051404005 | 1,036272916 | 0,87488 | 0,94463878 | OTU deubiquitinase, ubiquitin aldehyde binding 2 |
| ENSG00000031698 | SARS1 | 0,051290735 | 1,036191558 | 0,75229 | 0,88394752 | seryl-tRNA synthetase 1 |

| | | | | | | |
|---|---|---|---|---|---|---|
| ENSG00000273026 | AL358472,3 | 0,05120681 | 1,036131282 | 0,91009 | 0,96128046 | novel transcript, antisense to SLC39A1 |
| ENSG00000271973 | AC141002,1 | 0,051194077 | 1,036122137 | 0,94395 | 0,97635191 | novel transcript |
| ENSG00000248890 | HHIP-AS1 | 0,051142443 | 1,036085055 | 0,90457 | 0,95918368 | HHIP antisense RNA 1 |
| ENSG00000025423 | HSD17B6 | 0,051121375 | 1,036069925 | 0,89929 | 0,95711577 | hydroxysteroid 17-beta dehydrogenase 6 |
| ENSG00000246528 | AC079089,1 | 0,051102598 | 1,036056441 | 0,89625 | 0,95515151 | novel transcript, antisense to SLCO5A1 |
| ENSG00000132323 | ILKAP | 0,051099948 | 1,036054537 | 0,73705 | 0,87539732 | ILK associated serine/threonine phosphatase |
| ENSG00000229956 | ZRANB2-AS2 | 0,05107751 | 1,036038424 | 0,84351 | 0,92855259 | ZRANB2 antisense RNA 2 (head to head) |
| ENSG00000020922 | MRE11 | 0,051059576 | 1,036025546 | 0,81035 | 0,91251177 | MRE11 homolog, double strand break repair nuclease |
| ENSG00000286847 | AC007036,5 | 0,051038611 | 1,03601049 | 0,91488 | 0,9632313 | novel transcript, antisense to PLEKHA8 |
| ENSG00000197020 | ZNF100 | 0,051012383 | 1,035991656 | 0,79575 | 0,90548349 | zinc finger protein 100 |
| ENSG00000144136 | SLC20A1 | 0,051005637 | 1,035986812 | 0,75837 | 0,88727479 | solute carrier family 20 member 1 |
| ENSG00000231769 | AL035701,1 | 0,050893341 | 1,035906175 | 0,95395 | 0,98062659 | novel transcript |
| ENSG00000241120 | HMGN1P8 | 0,050873417 | 1,03589187 | 0,93223 | 0,97114134 | high mobility group nucleosome binding domain 1 pseudogene 8 |
| ENSG00000156521 | TYSND1 | 0,050853135 | 1,035877307 | 0,8373 | 0,92554257 | trypsin domain containing 1 |
| ENSG00000161642 | ZNF385A | 0,050848127 | 1,035873711 | 0,88328 | 0,94837933 | zinc finger protein 385A |
| ENSG00000105357 | MYH14 | 0,050846787 | 1,035872749 | 0,94859 | 0,97847634 | myosin heavy chain 14 |
| ENSG00000220370 | AL078595,1 | 0,050810547 | 1,035846729 | 0,86322 | 0,93891554 | ribosomal protein (RPL7A) pseudogene |
| ENSG00000229558 | SACS-AS1 | 0,050790569 | 1,035832385 | 0,9089 | 0,96099548 | SACS antisense RNA 1 |
| ENSG00000186047 | DLEU7 | 0,050784513 | 1,035828036 | 0,84605 | 0,92995515 | deleted in lymphocytic leukemia 7 |
| ENSG00000187627 | RGPD1 | 0,050589323 | 1,035687903 | 0,891 | 0,95208039 | RANBP2 like and GRIP domain containing 1 |
| ENSG00000104613 | INTS10 | 0,050566413 | 1,035671456 | 0,8152 | 0,91495565 | integrator complex subunit 10 |
| ENSG00000174365 | SNHG11 | 0,050505849 | 1,03562798 | 0,83804 | 0,92611649 | small nucleolar RNA host gene 11 |
| ENSG00000076924 | XAB2 | 0,050488615 | 1,035615609 | 0,78149 | 0,89877377 | XPA binding protein 2 |
| ENSG00000282022 | AC006581,2 | 0,050398499 | 1,035550922 | 0,96344 | 0,98448919 | novel transcript, antisense to PHC1 and A2ML1 |
| ENSG00000231638 | LUARIS | 0,050394373 | 1,035547961 | 0,91968 | 0,96507001 | lncRNA upregulator of antiviral response interferon signaling |
| ENSG00000143420 | ENSA | 0,050314374 | 1,03549054 | 0,74909 | 0,88190158 | endosulfine alpha |
| ENSG00000169223 | LMAN2 | 0,050293096 | 1,035475268 | 0,76643 | 0,89138648 | lectin, mannose binding 2 |
| ENSG00000288025 | AC093849,4 | 0,050276616 | 1,03546344 | 0,86322 | 0,93891554 | novel transcript, antisense to SCRG1 |
| ENSG00000282503 | AC243686,1 | 0,050214955 | 1,035419185 | 0,91766 | 0,96426294 | novel transcript |
| ENSG00000259690 | AC107980,1 | 0,050214955 | 1,035419185 | 0,91766 | 0,96426294 | novel transcript |
| ENSG00000171587 | DSCAM | 0,050200853 | 1,035409064 | 0,91453 | 0,96303629 | DS cell adhesion molecule |

| | | | | | | |
|---|---|---|---|---|---|---|
| ENSG00000275746 | KLF13 | 0,050164567 | 1,035383022 | 0,85105 | 0,93222304 | Kruppel like factor 13 |
| ENSG00000131149 | GSE1 | 0,050042236 | 1,035295233 | 0,81391 | 0,9144566 | Gse1 coiled-coil protein |
| ENSG00000235070 | AC062015,1 | 0,049909175 | 1,03519975 | 0,94446 | 0,97650311 | novel transcript |
| ENSG00000005469 | CROT | 0,049848941 | 1,035156531 | 0,90695 | 0,96017342 | carnitine O-octanoyltransferase |
| ENSG00000143067 | ZNF697 | 0,049825722 | 1,035139871 | 0,81591 | 0,91535843 | zinc finger protein 697 |
| ENSG00000254910 | AC136475,2 | 0,049659723 | 1,035020773 | 0,95147 | 0,97948342 | novel transcript |
| ENSG00000126249 | PDCD2L | 0,049654556 | 1,035017066 | 0,84472 | 0,92922421 | programmed cell death 2 like |
| ENSG00000101868 | POLA1 | 0,049627482 | 1,034997643 | 0,83161 | 0,92300125 | DNA polymerase alpha 1, catalytic subunit |
| ENSG00000211455 | STK38L | 0,049522407 | 1,034922264 | 0,81447 | 0,91470265 | serine/threonine kinase 38 like |
| ENSG00000272696 | AL359091,3 | 0,0494036 | 1,034837041 | 0,91896 | 0,9647667 | novel transcript, antisense to TRUB2 |
| ENSG00000137198 | GMPR | 0,049376027 | 1,034817263 | 0,95564 | 0,98169255 | guanosine monophosphate reductase |
| ENSG00000287336 | AL137157,1 | 0,049347539 | 1,03479683 | 0,94464 | 0,97650311 | novel transcript, antisense to DNM3 |
| ENSG00000101464 | PIGU | 0,049291195 | 1,034756417 | 0,83149 | 0,92291031 | phosphatidylinositol glycan anchor biosynthesis class U |
| ENSG00000204934 | ATP6V0E2-AS | 0,049224356 | 1,034708478 | 0,84708 | 0,93048578 | ATP6V0E2 antisense RNA 1 |
| ENSG00000166925 | TSC22D4 | 0,049159302 | 1,034661822 | 0,81339 | 0,91431467 | TSC22 domain family member 4 |
| ENSG00000011295 | TTC19 | 0,049083023 | 1,034607118 | 0,77405 | 0,89572556 | tetratricopeptide repeat domain 19 |
| ENSG00000167977 | KCTD5 | 0,049011895 | 1,034556111 | 0,80287 | 0,90916096 | potassium channel tetramerization domain containing 5 |
| ENSG00000176473 | WDR25 | 0,048889846 | 1,034468594 | 0,85007 | 0,93176425 | WD repeat domain 25 |
| ENSG00000100336 | APOL4 | 0,048876772 | 1,034459219 | 0,90461 | 0,95918368 | apolipoprotein L4 |
| ENSG00000258545 | RHOXF1-AS1 | 0,048828937 | 1,034424921 | 0,90235 | 0,95825434 | RHOXF1 antisense RNA 1 |
| ENSG00000237190 | CDKN2AIPNL | 0,048800297 | 1,034404386 | 0,79772 | 0,90685415 | CDKN2A interacting protein N-terminal like |
| ENSG00000167283 | ATP5MG | 0,048793272 | 1,034399349 | 0,78702 | 0,90114442 | ATP synthase membrane subunit g |
| ENSG00000197362 | ZNF786 | 0,048782973 | 1,034391965 | 0,81532 | 0,91502219 | zinc finger protein 786 |
| ENSG00000238035 | AC138035,1 | 0,04873932 | 1,034360666 | 0,87371 | 0,94405019 | novel transcript |
| ENSG00000272885 | AC092574,1 | 0,048644165 | 1,034292446 | 0,92832 | 0,96914921 | novel transcript |
| ENSG00000229020 | AKR7A2P1 | 0,04856716 | 1,034237241 | 0,92887 | 0,96947219 | AKR7A2 pseudogene 1 |
| ENSG00000280751 | AKR7A2P1 | 0,04856716 | 1,034237241 | 0,92887 | 0,96947219 | AKR7A2 pseudogene 1 |
| ENSG00000285339 | VPS28 | 0,048465216 | 1,034164162 | 0,8139 | 0,9144566 | VPS28 subunit of ESCRT-I |
| ENSG00000161981 | SNRNP25 | 0,048417623 | 1,034130047 | 0,81889 | 0,91689679 | small nuclear ribonucleoprotein U11/U12 subunit 25 |
| ENSG00000236474 | GCNT1P1 | 0,048388452 | 1,034109138 | 0,94964 | 0,97895198 | glucosaminyl (N-acetyl) transferase 1 pseudogene 1 |
| ENSG00000076650 | GPATCH1 | 0,048321118 | 1,034060874 | 0,80586 | 0,91077382 | G-patch domain containing 1 |
| ENSG00000213465 | ARL2 | 0,048316752 | 1,034057745 | 0,85284 | 0,93314746 | ADP ribosylation factor like GTPase 2 |
| ENSG00000139154 | AEBP2 | 0,048305041 | 1,034049351 | 0,84501 | 0,92930882 | AE binding protein 2 |

| | | | | | | |
|---|---|---|---|---|---|---|
| ENSG00000280802 | ITPK1-AS1 | 0,048173822 | 1,033955304 | 0,96196 | 0,9840176 | ITPK1 antisense RNA 1 |
| ENSG00000280873 | DGKD | 0,048168752 | 1,03395167 | 0,85221 | 0,93280134 | diacylglycerol kinase delta |
| ENSG00000256616 | AP002414,2 | 0,048078316 | 1,033886859 | 0,86456 | 0,93968603 | G protein-coupled receptor 125 (GPR125) pseudogene |
| ENSG00000130164 | LDLR | 0,047963419 | 1,033804523 | 0,81064 | 0,91254003 | low density lipoprotein receptor |
| ENSG00000231268 | AGER | 0,047874966 | 1,033741141 | 0,93419 | 0,97206381 | advanced glycosylation end-product specific receptor |
| ENSG00000230514 | AGER | 0,047874966 | 1,033741141 | 0,93419 | 0,97206381 | advanced glycosylation end-product specific receptor |
| ENSG00000137814 | HAUS2 | 0,047843122 | 1,033718324 | 0,76733 | 0,89210064 | HAUS augmin like complex subunit 2 |
| ENSG00000256588 | AC027544,2 | 0,04774756 | 1,033649854 | 0,93939 | 0,97413085 | novel transcript |
| ENSG00000102128 | RAB40AL | 0,047712825 | 1,033624968 | 0,92617 | 0,96789681 | RAB40A like |
| ENSG00000271888 | AL136162,1 | 0,047580144 | 1,033529913 | 0,92505 | 0,96744841 | novel transcript |
| ENSG00000177791 | MYOZ1 | 0,047498784 | 1,033471629 | 0,94975 | 0,97899157 | myozenin 1 |
| ENSG00000272483 | AC022001,3 | 0,047368424 | 1,03337825 | 0,91793 | 0,9644207 | novel transcript, antisense to VGLL4 |
| ENSG00000198898 | CAPZA2 | 0,047343458 | 1,033360367 | 0,84837 | 0,93094401 | capping actin protein of muscle Z-line subunit alpha 2 |
| ENSG00000081026 | MAGI3 | 0,047339539 | 1,03335756 | 0,8397 | 0,92673681 | membrane associated guanylate kinase, WW and PDZ domain containing 3 |
| ENSG00000171116 | HSFX1 | 0,047325312 | 1,03334737 | 0,87133 | 0,94321633 | heat shock transcription factor family, X-linked 1 |
| ENSG00000067141 | NEO1 | 0,047316447 | 1,03334102 | 0,86322 | 0,93891554 | neogenin 1 |
| ENSG00000151353 | TMEM18 | 0,047286107 | 1,033319289 | 0,80248 | 0,90903011 | transmembrane protein 18 |
| ENSG00000197324 | LRP10 | 0,047260513 | 1,033300958 | 0,87588 | 0,94507291 | LDL receptor related protein 10 |
| ENSG00000269889 | AC078802,1 | 0,047223597 | 1,033274518 | 0,94433 | 0,97650311 | novel transcript, antisense to ACTRT3 |
| ENSG00000240048 | DDX50P2 | 0,047078669 | 1,033170724 | 0,91118 | 0,9616927 | DEAD-box helicase 50 pseudogene 2 |
| ENSG00000126768 | TIMM17B | 0,046908057 | 1,033048549 | 0,84947 | 0,93147902 | translocase of inner mitochondrial membrane 17B |
| ENSG00000272221 | AL645933,2 | 0,046859493 | 1,033013776 | 0,93632 | 0,97254199 | novel transcript |
| ENSG00000138075 | ABCG5 | 0,0468343 | 1,032995736 | 0,95359 | 0,98051225 | ATP binding cassette subfamily G member 5 |
| ENSG00000119508 | NR4A3 | 0,046833884 | 1,032995439 | 0,90274 | 0,95839144 | nuclear receptor subfamily 4 group A member 3 |
| ENSG00000256552 | AC092745,2 | 0,046820329 | 1,032985733 | 0,92176 | 0,96611557 | ribosomal protein S24 (RPS24) pseudogene |
| ENSG00000282379 | DTX2 | 0,046785015 | 1,032960448 | 0,93498 | 0,9721963 | deltex E3 ubiquitin ligase 2 |
| ENSG00000264932 | AC115989,1 | 0,04667483 | 1,03288156 | 0,9483 | 0,97829358 | novel transcript, antisense to AL3A2 |
| ENSG00000273680 | AC009318,2 | 0,046640401 | 1,032856911 | 0,92084 | 0,96576056 | novel transcript, antisense to FAR2 |
| ENSG00000161202 | DVL3 | 0,046510089 | 1,032763622 | 0,82221 | 0,91790528 | dishevelled segment polarity protein 3 |
| ENSG00000144681 | STAC | 0,046283782 | 1,032601631 | 0,94501 | 0,97666034 | SH3 and cysteine rich domain |
| ENSG00000271643 | AC112220,2 | 0,04625049 | 1,032577802 | 0,89412 | 0,95389664 | novel transcript |
| ENSG00000019505 | SYT13 | 0,046179891 | 1,032527275 | 0,83402 | 0,92405734 | synaptotagmin 13 |

| | | | | | | |
|---|---|---|---|---|---|---|
| ENSG00000242199 | AC022336,1 | 0,04613808 | 1,032497351 | 0,95053 | 0,97934063 | ribosomal protein S26 (RPS26) pseudogene |
| ENSG00000163357 | DCST1 | 0,046130074 | 1,032491621 | 0,95396 | 0,98062659 | DC-STAMP domain containing 1 |
| ENSG00000056558 | TRAF1 | 0,046085678 | 1,032459849 | 0,85275 | 0,93313847 | TNF receptor associated factor 1 |
| ENSG00000167766 | ZNF83 | 0,046046655 | 1,032431923 | 0,81321 | 0,9142243 | zinc finger protein 83 |
| ENSG00000158077 | NLRP14 | 0,045978857 | 1,032383405 | 0,89747 | 0,95579119 | NLR family pyrin domain containing 14 |
| ENSG00000096093 | EFHC1 | 0,045932563 | 1,032350278 | 0,85028 | 0,93185051 | EF-hand domain containing 1 |
| ENSG00000120925 | RNF170 | 0,045859022 | 1,032297656 | 0,81914 | 0,91691915 | ring finger protein 170 |
| ENSG00000147202 | DIAPH2 | 0,04585135 | 1,032292167 | 0,90465 | 0,95918368 | diaphanous related formin 2 |
| ENSG00000204220 | PFDN6 | 0,045849958 | 1,032291171 | 0,84039 | 0,92722187 | prefoldin subunit 6 |
| ENSG00000160404 | TOR2A | 0,04556697 | 1,032088704 | 0,8194 | 0,91699877 | torsin family 2 member A |
| ENSG00000174669 | SLC29A2 | 0,045565986 | 1,032088 | 0,91147 | 0,96175738 | solute carrier family 29 member 2 |
| ENSG00000155962 | CLIC2 | 0,045558788 | 1,03208285 | 0,90267 | 0,95838699 | chloride intracellular channel 2 |
| ENSG00000123992 | DNPEP | 0,045550816 | 1,032077148 | 0,85462 | 0,93416235 | aspartyl aminopeptidase |
| ENSG00000241975 | ELOCP19 | 0,045504991 | 1,032044366 | 0,91328 | 0,96246104 | elongin C pseudogene 19 |
| ENSG00000149476 | TKFC | 0,045460982 | 1,032012884 | 0,81412 | 0,91457884 | triokinase and FMN cyclase |
| ENSG00000172531 | PPP1CA | 0,045419985 | 1,031983558 | 0,83105 | 0,92270607 | protein phosphatase 1 catalytic subunit alpha |
| ENSG00000261114 | AC012181,1 | 0,045282315 | 1,031885085 | 0,94577 | 0,97689657 | novel transcript, sense intronic to HERPUD1 |
| ENSG00000110514 | MADD | 0,045168517 | 1,031803694 | 0,79811 | 0,90685415 | MAP kinase activating death domain |
| ENSG00000139971 | ARMH4 | 0,045121866 | 1,03177033 | 0,85906 | 0,93657672 | armadillo like helical domain containing 4 |
| ENSG00000204348 | DXO | 0,045031143 | 1,03170545 | 0,90204 | 0,95811717 | decapping exoribonuclease |
| ENSG00000287009 | AC021037,1 | 0,044974097 | 1,031664656 | 0,8735 | 0,94398015 | novel transcript, antisense to ZNF518A |
| ENSG00000273015 | AC008124,1 | 0,04482342 | 1,031556913 | 0,85309 | 0,93323752 | novel transcript |
| ENSG00000122203 | KIAA1191 | 0,044770048 | 1,031518752 | 0,75873 | 0,88740301 | KIAA1191 |
| ENSG00000248196 | AC092658,1 | 0,044762764 | 1,031513544 | 0,9187 | 0,9646736 | uracil-DNA glycosylase (UNG) pseudogene |
| ENSG00000240184 | PCDHGC3 | 0,044755626 | 1,03150844 | 0,85014 | 0,93177313 | protocadherin gamma subfamily C, 3 |
| ENSG00000267143 | AP001120,2 | 0,044740458 | 1,031497595 | 0,95627 | 0,98189243 | novel transcript |
| ENSG00000168005 | SPINDOC | 0,044721735 | 1,031484209 | 0,83251 | 0,92337841 | spindlin interactor and repressor of chromatin binding |
| ENSG00000243335 | KCTD7 | 0,044564363 | 1,031371699 | 0,84272 | 0,92821072 | potassium channel tetramerization domain containing 7 |
| ENSG00000248843 | AL645949,2 | 0,044513332 | 1,031335217 | 0,95554 | 0,98167649 | ribosomal protein L7a (RPL7A) pseudogene |
| ENSG00000165792 | METTL17 | 0,04449027 | 1,031318731 | 0,83893 | 0,92640383 | methyltransferase like 17 |
| ENSG00000233613 | DCUN1D2-AS | 0,044480188 | 1,031311524 | 0,9408 | 0,97486264 | DCUN1D2 antisense RNA |
| ENSG00000146038 | DCDC2 | 0,04446956 | 1,031303927 | 0,93743 | 0,97320741 | doublecortin domain containing 2 |

| | | | | | | |
|---|---|---|---|---|---|---|
| ENSG00000156564 | LRFN2 | 0,044413952 | 1,031264177 | 0,95501 | 0,98123965 | leucine rich repeat and fibronectin type III domain containing 2 |
| ENSG00000185442 | FAM174B | 0,044349615 | 1,031218188 | 0,87587 | 0,94507291 | family with sequence similarity 174 member B |
| ENSG00000168528 | SERINC2 | 0,044197827 | 1,031109698 | 0,85376 | 0,93377502 | serine incorporator 2 |
| ENSG00000127483 | HP1BP3 | 0,044171616 | 1,031090965 | 0,74329 | 0,87846196 | heterochromatin protein 1 binding protein 3 |
| ENSG00000167562 | ZNF701 | 0,044169805 | 1,031089671 | 0,80615 | 0,91079954 | zinc finger protein 701 |
| ENSG00000176601 | MAP3K19 | 0,044156597 | 1,031080231 | 0,95165 | 0,97948342 | mitogen-activated protein kinase kinase kinase 19 |
| ENSG00000109390 | NDUFC1 | 0,043914664 | 1,030907339 | 0,82397 | 0,91876332 | NADH:ubiquinone oxidoreductase subunit C1 |
| ENSG00000280670 | CCDC163 | 0,043881868 | 1,030883904 | 0,9016 | 0,95801185 | coiled-coil domain containing 163 |
| ENSG00000229686 | SNORD56 | 0,043794413 | 1,030821414 | 0,96125 | 0,98401207 | small nucleolar RNA, C/D box 56 |
| ENSG00000015475 | BID | 0,043709985 | 1,030761091 | 0,89668 | 0,95542881 | BH3 interacting domain death agonist |
| ENSG00000261117 | AC009486,1 | 0,043661983 | 1,030726796 | 0,94158 | 0,97505054 | novel transcript |
| ENSG00000277775 | H3C7 | 0,04358187 | 1,030669561 | 0,88382 | 0,94846868 | H3 clustered histone 7 |
| ENSG00000205189 | ZBTB10 | 0,043544912 | 1,030643158 | 0,85264 | 0,93309251 | zinc finger and BTB domain containing 10 |
| ENSG00000107902 | LHPP | 0,043516047 | 1,030622538 | 0,85178 | 0,93254608 | phospholysine phosphohistidine inorganic pyrophosphate phosphatase |
| ENSG00000080298 | RFX3 | 0,043487124 | 1,030601876 | 0,91833 | 0,96446185 | regulatory factor X3 |
| ENSG00000005961 | ITGA2B | 0,04326196 | 1,030441041 | 0,9305 | 0,97023398 | integrin subunit alpha 2b |
| ENSG00000179163 | FUCA1 | 0,043213274 | 1,030406268 | 0,91626 | 0,96348118 | alpha-L-fucosidase 1 |
| ENSG00000184616 | SPDYE12P | 0,043120359 | 1,030339908 | 0,93744 | 0,97320741 | speedy/RINGO cell cycle regulator family member E12, pseudogene |
| ENSG00000131378 | RFTN1 | 0,042935807 | 1,030208114 | 0,91582 | 0,9633821 | raftlin, lipid raft linker 1 |
| ENSG00000171574 | ZNF584 | 0,042800843 | 1,030111742 | 0,82545 | 0,91911153 | zinc finger protein 584 |
| ENSG00000180233 | ZNRF2 | 0,042661524 | 1,03001227 | 0,90087 | 0,95787401 | zinc and ring finger 2 |
| ENSG00000274803 | BDP1 | 0,042624578 | 1,029985893 | 0,88477 | 0,94893347 | B double prime 1, subunit of RNA polymerase III transcription initiation factor IIIB |
| ENSG00000179994 | SPDYE7P | 0,042579133 | 1,029953449 | 0,96712 | 0,98623935 | speedy/RINGO cell cycle regulator family member E7, pseudogene |
| ENSG00000132382 | MYBBP1A | 0,042578972 | 1,029953334 | 0,85171 | 0,93251117 | MYB binding protein 1a |
| ENSG00000163155 | LYSMD1 | 0,042437295 | 1,029852194 | 0,83708 | 0,92549623 | LysM domain containing 1 |
| ENSG00000229954 | MTND2P2 | 0,042398319 | 1,029824372 | 0,95653 | 0,98197334 | MT-ND2 pseudogene 2 |
| ENSG00000278622 | TSEN34 | 0,042325188 | 1,029772171 | 0,87576 | 0,94505344 | tRNA splicing endonuclease subunit 34 |
| ENSG00000246350 | AL049543,1 | 0,042207667 | 1,02968829 | 0,98649 | 0,99423097 | novel transcript |

| | | | | | | |
|---|---|---|---|---|---|---|
| ENSG00000227290 | LINC01364 | 0,042027021 | 1,029559367 | 0,95045 | 0,97932267 | long intergenic non-protein coding RNA 1364 |
| ENSG00000165949 | IFI27 | 0,04201978 | 1,029554199 | 0,91713 | 0,9640264 | interferon alpha inducible protein 27 |
| ENSG00000287875 | AC068193,1 | 0,042014153 | 1,029550183 | 0,9451 | 0,97666034 | novel transcript, antisense to VRK2 |
| ENSG00000133466 | C1QTNF6 | 0,041943047 | 1,029499441 | 0,83973 | 0,92673905 | C1q and TNF related 6 |
| ENSG00000114405 | C3orf14 | 0,041904364 | 1,029471838 | 0,94108 | 0,97489099 | chromosome 3 open reading frame 14 |
| ENSG00000278002 | AL627171,1 | 0,041901737 | 1,029469963 | 0,93476 | 0,9721963 | novel transcript |
| ENSG00000183624 | HMCES | 0,041834414 | 1,029421924 | 0,7817 | 0,89877377 | 5-hydroxymethylcytosine binding, ES cell specific |
| ENSG00000167965 | MLST8 | 0,041774265 | 1,029379006 | 0,84597 | 0,92992612 | MTOR associated protein, LST8 homolog |
| ENSG00000228288 | PCAT6 | 0,041755491 | 1,029365611 | 0,90271 | 0,95838699 | prostate cancer associated transcript 6 |
| ENSG00000248049 | UBA6-AS1 | 0,04169931 | 1,029325526 | 0,84179 | 0,92804468 | UBA6 antisense RNA 1 (head to head) |
| ENSG00000271410 | AC020728,1 | 0,041686443 | 1,029316346 | 0,9235 | 0,96671006 | protein serine kinase H1 (PSKH1) pseudogene |
| ENSG00000247228 | AC009060,1 | 0,041618393 | 1,029267796 | 0,92369 | 0,96675951 | novel transcript, antisense to PDPR & CLEC18C |
| ENSG00000110651 | CD81 | 0,041612573 | 1,029263644 | 0,82544 | 0,91911153 | CD81 molecule |
| ENSG00000266397 | AP000919,2 | 0,041544447 | 1,029215042 | 0,96269 | 0,98432594 | novel transcript, antisense to LPIN2 |
| ENSG00000176049 | JAKMIP2 | 0,041413097 | 1,029121341 | 0,87244 | 0,94374455 | janus kinase and microtubule interacting protein 2 |
| ENSG00000254064 | AC105206,2 | 0,041318421 | 1,029053808 | 0,94273 | 0,9756583 | novel transcript |
| ENSG00000277948 | AC007435,1 | 0,041218155 | 1,028982292 | 0,95314 | 0,98025619 | |
| ENSG00000198168 | SVIP | 0,041201892 | 1,028970693 | 0,90719 | 0,96034215 | small VCP interacting protein |
| ENSG00000176095 | IP6K1 | 0,041171003 | 1,028948662 | 0,8136 | 0,9144566 | inositol hexakisphosphate kinase 1 |
| ENSG00000232000 | CLCN3P1 | 0,041128152 | 1,028918101 | 0,87663 | 0,94547358 | chloride voltage-gated channel 3 pseudogene 1 |
| ENSG00000188493 | C19orf54 | 0,041084889 | 1,028887246 | 0,86081 | 0,93761195 | chromosome 19 open reading frame 54 |
| ENSG00000261584 | AL513548,1 | 0,041052879 | 1,028864418 | 0,88654 | 0,94987292 | novel transcript |
| ENSG00000280026 | AC005618,3 | 0,04093864 | 1,028782952 | 0,94955 | 0,97888896 | novel transcript, antisense to PCDHGB1 |
| ENSG00000257842 | LINC02588 | 0,040834946 | 1,02870901 | 0,94815 | 0,9782417 | long intergenic non-protein coding RNA 2588 |
| ENSG00000166206 | GABRB3 | 0,040759789 | 1,028655421 | 0,90135 | 0,95798991 | gamma-aminobutyric acid type A receptor subunit beta3 |
| ENSG00000198015 | MRPL42 | 0,04061779 | 1,028554179 | 0,85289 | 0,93314746 | mitochondrial ribosomal protein L42 |
| ENSG00000286984 | AC090578,3 | 0,040617706 | 1,028554119 | 0,95587 | 0,98173232 | novel transcript |
| ENSG00000132600 | PRMT7 | 0,040422469 | 1,028414937 | 0,815 | 0,91488358 | protein arginine methyltransferase 7 |
| ENSG00000166532 | RIMKLB | 0,040350064 | 1,028363325 | 0,77525 | 0,89624737 | ribosomal modification protein rimK like family member B |
| ENSG00000143322 | ABL2 | 0,040336174 | 1,028353423 | 0,79346 | 0,90390791 | ABL proto-oncogene 2, non-receptor tyrosine kinase |
| ENSG00000226457 | RPL22P3 | 0,040213249 | 1,028265806 | 0,94381 | 0,97627205 | ribosomal protein L22 pseudogene 3 |
| ENSG00000116095 | PLEKHA3 | 0,040208564 | 1,028262467 | 0,80457 | 0,90996631 | pleckstrin homology domain containing A3 |
| ENSG00000198892 | SHISA4 | 0,040197256 | 1,028254408 | 0,89496 | 0,95417507 | shisa family member 4 |

| | | | | | | |
|---|---|---|---|---|---|---|
| ENSG00000136114 | THSD1 | 0,04013809 | 1,028212239 | 0,91127 | 0,9616927 | thrombospondin type 1 domain containing 1 |
| ENSG00000196072 | BLOC1S2 | 0,040105808 | 1,028189232 | 0,83649 | 0,92514377 | biogenesis of lysosomal organelles complex 1 subunit 2 |
| ENSG00000167118 | URM1 | 0,039934145 | 1,028066897 | 0,86288 | 0,9388179 | ubiquitin related modifier 1 |
| ENSG00000242889 | RN7SL449P | 0,039856887 | 1,028011844 | 0,93308 | 0,97150994 | RNA, 7SL, cytoplasmic 449, pseudogene |
| ENSG00000102901 | CENPT | 0,039848473 | 1,028005849 | 0,8442 | 0,92888169 | centromere protein T |
| ENSG00000153487 | ING1 | 0,03979892 | 1,02797054 | 0,82172 | 0,91757875 | inhibitor of growth family member 1 |
| ENSG00000203778 | FAM229B | 0,039747518 | 1,027933915 | 0,8939 | 0,95376978 | family with sequence similarity 229 member B |
| ENSG00000282675 | AC245078,5 | 0,039721031 | 1,027915043 | 0,90281 | 0,95842435 | taste receptor, type 2, member 18 pseudogene |
| ENSG00000166596 | CFAP52 | 0,039715998 | 1,027911457 | 0,92683 | 0,96825947 | cilia and flagella associated protein 52 |
| ENSG00000130724 | CHMP2A | 0,039683107 | 1,027888022 | 0,84489 | 0,92925254 | charged multivesicular body protein 2A |
| ENSG00000237170 | RPS7P15 | 0,039652151 | 1,027865967 | 0,96111 | 0,98401207 | ribosomal protein S7 pseudogene 15 |
| ENSG00000167333 | TRIM68 | 0,039609057 | 1,027835265 | 0,82802 | 0,92070145 | tripartite motif containing 68 |
| ENSG00000127080 | IPPK | 0,039575602 | 1,02781143 | 0,85837 | 0,9361457 | inositol-pentakisphosphate 2-kinase |
| ENSG00000110660 | SLC35F2 | 0,03954275 | 1,027788026 | 0,93511 | 0,9721963 | solute carrier family 35 member F2 |
| ENSG00000057757 | PITHD1 | 0,039452303 | 1,027723593 | 0,85527 | 0,93431435 | PITH domain containing 1 |
| ENSG00000174899 | SLC66A1L | 0,039437724 | 1,027713208 | 0,95587 | 0,98173232 | solute carrier family 66 member 1 like |
| ENSG00000242299 | AC073861,1 | 0,039392454 | 1,02768096 | 0,94131 | 0,97501891 | ribosomal protein S18 (RPS18) pseudogene |
| ENSG00000145388 | METTL14 | 0,039359213 | 1,027657281 | 0,79049 | 0,90286944 | methyltransferase like 14 |
| ENSG00000173575 | CHD2 | 0,039354171 | 1,02765369 | 0,85739 | 0,93562394 | chromodomain helicase DNA binding protein 2 |
| ENSG00000085433 | WDR47 | 0,039349597 | 1,027650432 | 0,9036 | 0,95873972 | WD repeat domain 47 |
| ENSG00000265451 | AC012447,1 | 0,039336423 | 1,027641048 | 0,96129 | 0,98401207 | novel transcript, antisense to CLASP1 |
| ENSG00000102030 | NAA10 | 0,039312253 | 1,027623832 | 0,83365 | 0,92396169 | N-alpha-acetyltransferase 10, NatA catalytic subunit |
| ENSG00000197150 | ABCB8 | 0,03930634 | 1,027619619 | 0,83978 | 0,92674902 | ATP binding cassette subfamily B member 8 |
| ENSG00000185033 | SEMA4B | 0,039210414 | 1,027551294 | 0,85853 | 0,93621468 | semaphorin 4B |
| ENSG00000157184 | CPT2 | 0,039063899 | 1,027446945 | 0,82194 | 0,91779154 | carnitine palmitoyltransferase 2 |
| ENSG00000133665 | DYDC2 | 0,039033757 | 1,027425479 | 0,93239 | 0,97120063 | DPY30 domain containing 2 |
| ENSG00000246662 | LINC00535 | 0,038996719 | 1,027399103 | 0,91415 | 0,96285318 | long intergenic non-protein coding RNA 535 |
| ENSG00000204217 | BMPR2 | 0,03894478 | 1,027362116 | 0,88105 | 0,94704 | bone morphogenetic protein receptor type 2 |
| ENSG00000254877 | AP001636,2 | 0,038843045 | 1,027289671 | 0,95566 | 0,98169255 | cAMP-dependent protein kinase type II-alpha regulatory subunit (PRKAR2A) pseudogene |
| ENSG00000112983 | BRD8 | 0,038840253 | 1,027287683 | 0,82498 | 0,91900665 | bromodomain containing 8 |
| ENSG00000274950 | RPS9 | 0,038772948 | 1,027239759 | 0,84162 | 0,92798669 | ribosomal protein S9 |
| ENSG00000257167 | TMPO-AS1 | 0,038608268 | 1,027122509 | 0,82728 | 0,92015725 | TMPO antisense RNA 1 |

| | | | | | | |
|---|---|---|---|---|---|---|
| ENSG00000262587 | AC133552,2 | 0,038456091 | 1,027014173 | 0,94774 | 0,97799866 | leucine carboxyl methyltransferase 1 (LCMT1) pseudogene |
| ENSG00000278535 | DHRS11 | 0,038420886 | 1,026989112 | 0,89383 | 0,95373049 | dehydrogenase/reductase 11 |
| ENSG00000233932 | CTXN2 | 0,03840936 | 1,026980907 | 0,95788 | 0,98267656 | cortexin 2 |
| ENSG00000150433 | TMEM218 | 0,038259318 | 1,026874105 | 0,90264 | 0,95838699 | transmembrane protein 218 |
| ENSG00000268119 | AC010615,2 | 0,038155732 | 1,026800378 | 0,88765 | 0,95052778 | novel transcript |
| ENSG00000213569 | GTF3C6P2 | 0,037928737 | 1,026638833 | 0,9787 | 0,99131549 | GTF3C6 pseudogene 2 |
| ENSG00000285893 | AL358777,4 | 0,037630757 | 1,026426809 | 0,95208 | 0,97971456 | novel transcript, antisense to C6orf52 |
| ENSG00000165282 | PIGO | 0,037474803 | 1,026315859 | 0,86233 | 0,93853472 | phosphatidylinositol glycan anchor biosynthesis class O |
| ENSG00000180914 | OXTR | 0,037472188 | 1,026313999 | 0,903 | 0,95846794 | oxytocin receptor |
| ENSG00000138193 | PLCE1 | 0,037334263 | 1,026215886 | 0,85858 | 0,93622982 | phospholipase C epsilon 1 |
| ENSG00000254266 | PKIA-AS1 | 0,037309853 | 1,026198522 | 0,91949 | 0,96506391 | PKIA antisense RNA 1 |
| ENSG00000267293 | AC012569,1 | 0,037229881 | 1,026141639 | 0,92746 | 0,96860856 | U3 small nucleolar RNA-associated protein 18 homolog, pseudogene |
| ENSG00000236035 | NSA2P1 | 0,037090845 | 1,026042752 | 0,96224 | 0,98415736 | NSA2 pseudogene 1 |
| ENSG00000235489 | DBF4P1 | 0,037000684 | 1,025978632 | 0,93784 | 0,97351346 | DBF4 zinc finger pseudogene 1 |
| ENSG00000107614 | TRDMT1 | 0,036725978 | 1,025783292 | 0,9029 | 0,95846794 | tRNA aspartic acid methyltransferase 1 |
| ENSG00000149806 | FAU | 0,036710787 | 1,025772491 | 0,84166 | 0,92799135 | FAU ubiquitin like and ribosomal protein S30 fusion |
| ENSG00000047579 | DTNBP1 | 0,036671678 | 1,025744685 | 0,86073 | 0,93759071 | dystrobrevin binding protein 1 |
| ENSG00000158793 | NIT1 | 0,036670363 | 1,025743749 | 0,84947 | 0,93147902 | nitrilase 1 |
| ENSG00000260034 | LCMT1-AS2 | 0,036648123 | 1,025727937 | 0,86493 | 0,93998058 | LCMT1 antisense RNA 2 |
| ENSG00000225748 | PRRC2A | 0,036581652 | 1,025680679 | 0,83101 | 0,92270607 | proline rich coiled-coil 2A |
| ENSG00000179799 | OR7E22P | 0,036467267 | 1,02559936 | 0,98079 | 0,99205474 | olfactory receptor family 7 subfamily E member 22 pseudogene |
| ENSG00000171475 | WIPF2 | 0,036368314 | 1,025529017 | 0,81162 | 0,91334888 | WAS/WASL interacting protein family member 2 |
| ENSG00000197361 | FBXL22 | 0,036346987 | 1,025513858 | 0,93646 | 0,97265426 | F-box and leucine rich repeat protein 22 |
| ENSG00000230105 | AL354793,1 | 0,036278358 | 1,025465075 | 0,94113 | 0,97490097 | novel transcript |
| ENSG00000236583 | AP004290,1 | 0,036141232 | 1,025367611 | 0,92736 | 0,96860856 | insulin-like growth factor 2 mRNA binding protein 2 (IGF2BP2) pseudogene |
| ENSG00000267769 | AC011498,6 | 0,035914949 | 1,025206797 | 0,96581 | 0,98552708 | novel transcript, antisense to UBX6 |
| ENSG00000117640 | MTFR1L | 0,035882638 | 1,025183836 | 0,85829 | 0,93612837 | mitochondrial fission regulator 1 like |
| ENSG00000286140 | DERPC | 0,035793242 | 1,025120313 | 0,87802 | 0,94581036 | DERPC proline and glycine rich nuclear protein |
| ENSG00000236782 | AL391650,1 | 0,035782503 | 1,025112683 | 0,9398 | 0,97442764 | novel protein |

| | | | | | | |
|---|---|---|---|---|---|---|
| ENSG00000216990 | HSPD1P10 | 0,035776094 | 1,025108129 | 0,96548 | 0,9854196 | heat shock protein family D (Hsp60) member 1 pseudogene 10 |
| ENSG00000275214 | IFI27 | 0,035747553 | 1,025087849 | 0,93088 | 0,97037753 | interferon alpha inducible protein 27 |
| ENSG00000230154 | AIDAP1 | 0,035684252 | 1,025042873 | 0,96097 | 0,98401207 | AIDA pseudogene 1 |
| ENSG00000167377 | ZNF23 | 0,035682658 | 1,02504174 | 0,87684 | 0,94549531 | zinc finger protein 23 |
| ENSG00000166821 | PEX11A | 0,035472667 | 1,024892551 | 0,87977 | 0,94649956 | peroxisomal biogenesis factor 11 alpha |
| ENSG00000275031 | METRNL | 0,035458321 | 1,02488236 | 0,92638 | 0,96799343 | meteorin like, glial cell differentiation regulator |
| ENSG00000204172 | AGAP9 | 0,035451778 | 1,024877712 | 0,85495 | 0,93417381 | ArfGAP with GTPase domain, ankyrin repeat and PH domain 9 |
| ENSG00000263041 | AL513324,1 | 0,035397356 | 1,024839052 | 0,97119 | 0,98773415 | novel transcript |
| ENSG00000260219 | CD2BP2-DT | 0,035349631 | 1,02480515 | 0,92835 | 0,96914921 | CD2BP2 divergent transcript |
| ENSG00000259024 | TVP23C-CDRT | 0,035086681 | 1,024618383 | 0,92685 | 0,96825947 | TVP23C-CDRT4 readthrough |
| ENSG00000196696 | PDXDC2P-NPI | 0,035055995 | 1,02459659 | 0,89247 | 0,95285354 | nuclear pore complex-interacting protein |
| ENSG00000205611 | LINC01597 | 0,035025219 | 1,024574733 | 0,9271 | 0,96842124 | long intergenic non-protein coding RNA 1597 |
| ENSG00000267641 | BNIP3P16 | 0,034941456 | 1,024515248 | 0,95281 | 0,98009667 | BCL2 interacting protein 3 pseudogene 16 |
| ENSG00000147050 | KDM6A | 0,034901959 | 1,0244872 | 0,85948 | 0,93687112 | lysine demethylase 6A |
| ENSG00000267466 | AC021683,1 | 0,034890771 | 1,024479255 | 0,93401 | 0,97201193 | novel transcript |
| ENSG00000206535 | LNP1 | 0,034815868 | 1,024426066 | 0,88686 | 0,95005038 | leukemia NUP98 fusion partner 1 |
| ENSG00000175147 | TMEM51-AS1 | 0,03474048 | 1,024372537 | 0,94462 | 0,97650311 | TMEM51 antisense RNA 1 |
| ENSG00000248445 | SEMA6A-AS1 | 0,034600454 | 1,024273117 | 0,94952 | 0,97888896 | SEMA6A antisense RNA 1 |
| ENSG00000105771 | SMG9 | 0,034597904 | 1,024271306 | 0,85386 | 0,93380868 | SMG9 nonsense mediated mRNA decay factor |
| ENSG00000288076 | AC018398,2 | 0,034578115 | 1,024257257 | 0,93434 | 0,97214033 | novel transcript, antisense to MCPH1 |
| ENSG00000102401 | ARMCX3 | 0,034544558 | 1,024233434 | 0,86158 | 0,93811282 | armadillo repeat containing X-linked 3 |
| ENSG00000107362 | ABHD17B | 0,034537584 | 1,024228482 | 0,83845 | 0,92626651 | abhydrolase domain containing 17B, depalmitoylase |
| ENSG00000177640 | CASC2 | 0,034525568 | 1,024219952 | 0,90603 | 0,95982149 | cancer susceptibility 2 |
| ENSG00000114744 | COMMD2 | 0,034488799 | 1,024193849 | 0,82008 | 0,91723726 | COMM domain containing 2 |
| ENSG00000106633 | GCK | 0,034486854 | 1,024192468 | 0,93286 | 0,97143711 | glucokinase |
| ENSG00000117411 | B4GALT2 | 0,034482857 | 1,02418963 | 0,82882 | 0,92107649 | beta-1,4-galactosyltransferase 2 |
| ENSG00000287857 | AL592546,3 | 0,034433386 | 1,02415451 | 0,92332 | 0,96667848 | novel transcript, antisense to NHLRC2 |
| ENSG00000091409 | ITGA6 | 0,034389863 | 1,024123615 | 0,9246 | 0,96724903 | integrin subunit alpha 6 |
| ENSG00000158882 | TOMM40L | 0,034385973 | 1,024120853 | 0,83617 | 0,92493625 | translocase of outer mitochondrial membrane 40 like |
| ENSG00000175643 | RMI2 | 0,034341515 | 1,024089294 | 0,87173 | 0,94334603 | RecQ mediated genome instability 2 |
| ENSG00000285595 | AC105114,2 | 0,034332138 | 1,024082638 | 0,90699 | 0,96017342 | novel transcript, antisense to ZNF521 |

| | | | | | | |
|---|---|---|---|---|---|---|
| ENSG00000286735 | AC126182,3 | 0,034319635 | 1,024073763 | 0,96184 | 0,98401207 | novel transcript, antisense to FSTL1 |
| ENSG00000162073 | PAQR4 | 0,034298989 | 1,024059108 | 0,86941 | 0,9425782 | progestin and adipoQ receptor family member 4 |
| ENSG00000266708 | AP001793,1 | 0,034252796 | 1,024026319 | 0,96161 | 0,98401207 | novel transcript, antisense to RAB12 |
| ENSG00000213903 | LTB4R | 0,034223067 | 1,024005218 | 0,91742 | 0,96419315 | leukotriene B4 receptor |
| ENSG00000125124 | BBS2 | 0,034191116 | 1,02398254 | 0,82409 | 0,91880473 | Bardet-Biedl syndrome 2 |
| ENSG00000007080 | CCDC124 | 0,034151744 | 1,023954595 | 0,83363 | 0,92396169 | coiled-coil domain containing 124 |
| ENSG00000285276 | AC270301,7 | 0,034104762 | 1,02392125 | 0,95194 | 0,97966441 | novel transcript |
| ENSG00000259090 | SEPTIN7P1 | 0,034012947 | 1,023856089 | 0,95928 | 0,98321703 | septin 7 pseudogene 1 |
| ENSG00000134775 | FHOD3 | 0,033965605 | 1,023822491 | 0,89811 | 0,95618956 | formin homology 2 domain containing 3 |
| ENSG00000115963 | RND3 | 0,03392582 | 1,023794258 | 0,94224 | 0,97544115 | Rho family GTPase 3 |
| ENSG00000236204 | LINC01376 | 0,033841833 | 1,023734659 | 0,91576 | 0,9633821 | long intergenic non-protein coding RNA 1376 |
| ENSG00000272491 | AL109659,2 | 0,033824233 | 1,02372217 | 0,9363 | 0,97254199 | novel transcript, antisense to SLC5A9 |
| ENSG00000274474 | YWHAE | 0,033789237 | 1,023697338 | 0,87891 | 0,94615035 | tyrosine 3-monooxygenase/tryptophan 5-monooxygenase activation protein epsilon |
| ENSG00000198399 | ITSN2 | 0,033715677 | 1,023645143 | 0,92592 | 0,96772813 | intersectin 2 |
| ENSG00000225345 | SNX18P3 | 0,033711819 | 1,023642406 | 0,93073 | 0,97029435 | sorting nexin 18 pseudogene 3 |
| ENSG00000225234 | TRAPPC12-AS | 0,033668406 | 1,023611603 | 0,92466 | 0,96725125 | TRAPPC12 antisense RNA 1 |
| ENSG00000273594 | EEF1D | 0,033658812 | 1,023604796 | 0,8701 | 0,94277661 | eukaryotic translation elongation factor 1 delta |
| ENSG00000197951 | ZNF71 | 0,033648615 | 1,023597561 | 0,84267 | 0,92821072 | zinc finger protein 71 |
| ENSG00000244482 | LILRA6 | 0,03364526 | 1,023595181 | 0,97473 | 0,98939607 | leukocyte immunoglobulin like receptor A6 |
| ENSG00000177830 | CHID1 | 0,033597485 | 1,023561285 | 0,84263 | 0,92821072 | chitinase domain containing 1 |
| ENSG00000184903 | IMMP2L | 0,033544283 | 1,02352354 | 0,92105 | 0,96580009 | inner mitochondrial membrane peptidase subunit 2 |
| ENSG00000135617 | PRADC1 | 0,033532133 | 1,02351492 | 0,89654 | 0,95534526 | protease associated domain containing 1 |
| ENSG00000151093 | OXSM | 0,033407991 | 1,023426852 | 0,92287 | 0,96662141 | 3-oxoacyl-ACP synthase, mitochondrial |
| ENSG00000237945 | LINC00649 | 0,033390762 | 1,02341463 | 0,90686 | 0,96017342 | long intergenic non-protein coding RNA 649 |
| ENSG00000168268 | NT5DC2 | 0,033389522 | 1,02341375 | 0,88353 | 0,94837933 | 5'-nucleotidase domain containing 2 |
| ENSG00000218574 | HNRNPA1P37 | 0,033324182 | 1,023367401 | 0,9703 | 0,98749329 | heterogeneous nuclear ribonucleoprotein A1 pseudogene 37 |
| ENSG00000252699 | SNORA21B | 0,033307282 | 1,023355413 | 0,9649 | 0,98525685 | small nucleolar RNA, H/ACA box 21B |
| ENSG00000133706 | LARS1 | 0,033292763 | 1,023345114 | 0,84746 | 0,93071557 | leucyl-tRNA synthetase 1 |
| ENSG00000089048 | ESF1 | 0,033240833 | 1,023308279 | 0,88749 | 0,95041376 | ESF1 nucleolar pre-rRNA processing protein homolog |
| ENSG00000180229 | HERC2P3 | 0,033231909 | 1,023301949 | 0,82465 | 0,91900665 | hect domain and RLD 2 pseudogene 3 |
| ENSG00000169976 | SF3B5 | 0,033223055 | 1,023295669 | 0,87277 | 0,94374571 | splicing factor 3b subunit 5 |

| | | | | | | |
|---|---|---|---|---|---|---|
| ENSG00000143337 | TOR1AIP1 | 0,03321673 | 1,023291183 | 0,9053 | 0,95953664 | torsin 1A interacting protein 1 |
| ENSG00000273243 | Z82243,1 | 0,033124972 | 1,023226102 | 0,94271 | 0,9756583 | novel transcript |
| ENSG00000115073 | ACTR1B | 0,033123982 | 1,0232254 | 0,86936 | 0,94257563 | actin related protein 1B |
| ENSG00000164327 | RICTOR | 0,033123253 | 1,023224882 | 0,81845 | 0,91665476 | RPTOR independent companion of MTOR complex 2 |
| ENSG00000149782 | PLCB3 | 0,033095779 | 1,023205397 | 0,875 | 0,94463878 | phospholipase C beta 3 |
| ENSG00000274891 | TRAPPC12 | 0,032730238 | 1,022946177 | 0,8808 | 0,94688182 | trafficking protein particle complex 12 |
| ENSG00000147364 | FBXO25 | 0,032704025 | 1,022927591 | 0,90926 | 0,96104995 | F-box protein 25 |
| ENSG00000274423 | SEC22B2P | 0,032690713 | 1,022918151 | 0,98264 | 0,99247242 | SEC22 homolog B2, pseudogene |
| ENSG00000276742 | AL731566,3 | 0,032680769 | 1,022911101 | 0,97113 | 0,98773415 | novel transcript, antisense to NSMCE4A |
| ENSG00000165943 | MOAP1 | 0,032680191 | 1,022910691 | 0,91258 | 0,96205682 | modulator of apoptosis 1 |
| ENSG00000269845 | CCNYL6 | 0,032520483 | 1,02279746 | 0,92203 | 0,96618214 | cyclin Y like 6 (pseudogene) |
| ENSG00000279112 | BRI3BPP1 | 0,032520483 | 1,02279746 | 0,92203 | 0,96618214 | BRI3 binding protein pseudogene 1 |
| ENSG00000138621 | PPCDC | 0,032513432 | 1,022792461 | 0,85585 | 0,9347432 | phosphopantothenoylcysteine decarboxylase |
| ENSG00000188486 | H2AX | 0,032425545 | 1,022730156 | 0,87678 | 0,94549531 | H2A,X variant histone |
| ENSG00000172725 | CORO1B | 0,032399132 | 1,022711432 | 0,8569 | 0,93541811 | coronin 1B |
| ENSG00000166548 | TK2 | 0,032282957 | 1,02262908 | 0,84338 | 0,92850187 | thymidine kinase 2 |
| ENSG00000272977 | AL008721,2 | 0,032281305 | 1,022627909 | 0,88909 | 0,9510223 | novel transcript |
| ENSG00000003987 | MTMR7 | 0,032249487 | 1,022605355 | 0,94427 | 0,97649967 | myotubularin related protein 7 |
| ENSG00000240225 | ZNF542P | 0,03221624 | 1,02258179 | 0,883 | 0,94823653 | zinc finger protein 542, pseudogene |
| ENSG00000105698 | USF2 | 0,032101466 | 1,022500441 | 0,87137 | 0,94321633 | upstream transcription factor 2, c-fos interacting |
| ENSG00000253716 | MINCR | 0,031980201 | 1,022414499 | 0,92536 | 0,96760657 | MYC-induced long non-coding RNA |
| ENSG00000165661 | QSOX2 | 0,031942146 | 1,022387531 | 0,84457 | 0,92915822 | quiescin sulfhydryl oxidase 2 |
| ENSG00000143315 | PIGM | 0,031844529 | 1,022318355 | 0,86547 | 0,94024247 | phosphatidylinositol glycan anchor biosynthesis class M |
| ENSG00000109472 | CPE | 0,031699069 | 1,022215285 | 0,91313 | 0,96242609 | carboxypeptidase E |
| ENSG00000134326 | CMPK2 | 0,031688308 | 1,022207661 | 0,93598 | 0,97249457 | cytidine/uridine monophosphate kinase 2 |
| ENSG00000234740 | AL162386,2 | 0,031654898 | 1,022183988 | 0,95573 | 0,9817293 | novel transcript |
| ENSG00000271524 | BNIP3P17 | 0,031600262 | 1,022145278 | 0,90966 | 0,96105018 | BCL2 interacting protein 3 pseudogene 17 |
| ENSG00000272033 | AL136984,1 | 0,031561638 | 1,022117913 | 0,90533 | 0,95953664 | novel transcript, sense intronic to POU2F1 |
| ENSG00000119522 | DENND1A | 0,031558575 | 1,022115743 | 0,89214 | 0,95257296 | DENN domain containing 1A |
| ENSG00000269902 | AC234772,1 | 0,031538094 | 1,022101233 | 0,97322 | 0,98872813 | novel transcript |
| ENSG00000128185 | DGCR6L | 0,031260768 | 1,021904776 | 0,88909 | 0,9510223 | DiGeorge syndrome critical region gene 6 like |
| ENSG00000168286 | THAP11 | 0,031247632 | 1,021895471 | 0,85479 | 0,93417381 | THAP domain containing 11 |

| | | | | | | |
|---|---|---|---|---|---|---|
| ENSG00000249626 | AC024560,2 | 0,031149885 | 1,021826237 | 0,94104 | 0,97489099 | protein phosphatase 4, regulatory subunit 2 (PPP4R2) pseudogene |
| ENSG00000286732 | AC091181,2 | 0,031122497 | 1,021806839 | 0,9656 | 0,98542381 | novel transcript, antisense to MSI2 |
| ENSG00000085415 | SEH1L | 0,031017972 | 1,021732811 | 0,87746 | 0,94566578 | SEH1 like nucleoporin |
| ENSG00000078018 | MAP2 | 0,031013755 | 1,021729824 | 0,93006 | 0,97006737 | microtubule associated protein 2 |
| ENSG00000248180 | GAPDHP60 | 0,030964288 | 1,021694792 | 0,94869 | 0,97853939 | glyceraldehyde-3-phosphate dehydrogenase pseudogene 60 |
| ENSG00000108924 | HLF | 0,030933383 | 1,021672905 | 0,95696 | 0,98218743 | HLF transcription factor, PAR bZIP family member |
| ENSG00000164695 | CHMP4C | 0,030906897 | 1,021654149 | 0,95541 | 0,98158195 | charged multivesicular body protein 4C |
| ENSG00000184402 | SS18L1 | 0,030882135 | 1,021636614 | 0,86307 | 0,93891554 | SS18L1 subunit of BAF chromatin remodeling complex |
| ENSG00000023228 | NDUFS1 | 0,030876795 | 1,021632833 | 0,92326 | 0,96667029 | NADH:ubiquinone oxidoreductase core subunit S1 |
| ENSG00000234545 | FAM133B | 0,030706742 | 1,021512417 | 0,8408 | 0,92747732 | family with sequence similarity 133 member B |
| ENSG00000164542 | KIAA0895 | 0,030580101 | 1,021422752 | 0,91601 | 0,96339511 | KIAA0895 |
| ENSG00000101306 | MYLK2 | 0,030558494 | 1,021407455 | 0,93158 | 0,9707979 | myosin light chain kinase 2 |
| ENSG00000197021 | EOLA2 | 0,030549954 | 1,021401409 | 0,88883 | 0,95091757 | endothelium and lymphocyte associated ASCH domain 2 |
| ENSG00000152133 | GPATCH11 | 0,030539657 | 1,021394119 | 0,91473 | 0,96317264 | G-patch domain containing 11 |
| ENSG00000171858 | RPS21 | 0,030515412 | 1,021376954 | 0,88983 | 0,9514844 | ribosomal protein S21 |
| ENSG00000123415 | SMUG1 | 0,030512715 | 1,021375044 | 0,87313 | 0,94385071 | single-strand-selective monofunctional uracil-DNA glycosylase 1 |
| ENSG00000247137 | AP000873,2 | 0,030468145 | 1,021343491 | 0,89092 | 0,95208039 | novel transcript |
| ENSG00000182968 | SOX1 | 0,030393734 | 1,021290814 | 0,88303 | 0,94823653 | SRY-box transcription factor 1 |
| ENSG00000251323 | LINC02728 | 0,030311845 | 1,021232846 | 0,91568 | 0,9633821 | long intergenic non-protein coding RNA 2728 |
| ENSG00000143147 | GPR161 | 0,030134234 | 1,021107129 | 0,87636 | 0,94545974 | G protein-coupled receptor 161 |
| ENSG00000259248 | USP3-AS1 | 0,030115956 | 1,021094193 | 0,92096 | 0,96576077 | USP3 antisense RNA 1 |
| ENSG00000253180 | AC104986,1 | 0,030097285 | 1,021080978 | 0,96289 | 0,98432594 | ubiquitin-conjugating enzyme E2 variant 1 (UBE2V1) pseudogene |
| ENSG00000100246 | DNAL4 | 0,029964963 | 1,02098733 | 0,89307 | 0,95327499 | dynein axonemal light chain 4 |
| ENSG00000268889 | AC008750,6 | 0,029804211 | 1,020873573 | 0,95161 | 0,97948342 | novel transcript |
| ENSG00000238287 | AL603839,3 | 0,029719565 | 1,020813678 | 0,96103 | 0,98401207 | novel transcript |
| ENSG00000184669 | OR7E14P | 0,029699337 | 1,020799365 | 0,93731 | 0,97313558 | olfactory receptor family 7 subfamily E member 14 pseudogene |
| ENSG00000152076 | CCDC74B | 0,029652691 | 1,020766361 | 0,9312 | 0,97060346 | coiled-coil domain containing 74B |
| ENSG00000268364 | SMC5-AS1 | 0,029605318 | 1,020732843 | 0,9414 | 0,97504246 | SMC5 antisense RNA 1 (head to head) |

| | | | | | | |
|---|---|---|---|---|---|---|
| ENSG00000167513 | CDT1 | 0,029552997 | 1,020695826 | 0,88363 | 0,94839496 | chromatin licensing and DNA replication factor 1 |
| ENSG00000249465 | RBMXP4 | 0,029552702 | 1,020695617 | 0,98131 | 0,99236691 | RBMX pseudogene 4 |
| ENSG00000189195 | BTBD8 | 0,0292751 | 1,020499235 | 0,95452 | 0,98095268 | BTB domain containing 8 |
| ENSG00000279968 | CCDC28A-AS1 | 0,029174389 | 1,020427999 | 0,94227 | 0,975443 | CCDC28A antisense RNA 1 |
| ENSG00000223984 | HNRNPRP1 | 0,029096536 | 1,020372934 | 0,94221 | 0,97544115 | heterogeneous nuclear ribonucleoprotein R pseudogene 1 |
| ENSG00000005486 | RHBDD2 | 0,029071966 | 1,020355556 | 0,89982 | 0,95730568 | rhomboid domain containing 2 |
| ENSG00000007255 | TRAPPC6A | 0,029030621 | 1,020326315 | 0,91034 | 0,9613584 | trafficking protein particle complex 6A |
| ENSG00000286163 | AC002377,1 | 0,029021751 | 1,020320042 | 0,97185 | 0,98800339 | novel transcript |
| ENSG00000123352 | SPATS2 | 0,029005909 | 1,020308838 | 0,8883 | 0,95069428 | spermatogenesis associated serine rich 2 |
| ENSG00000159884 | CCDC107 | 0,028998462 | 1,020303571 | 0,93512 | 0,9721963 | coiled-coil domain containing 107 |
| ENSG00000258759 | AL049779,2 | 0,028986403 | 1,020295043 | 0,97746 | 0,99059321 | DnaJ (Hsp40) homolog, subfamily C, member 9 (DNAJC9) pseudogene |
| ENSG00000100226 | GTPBP1 | 0,028902418 | 1,020235649 | 0,86801 | 0,94175526 | GTP binding protein 1 |
| ENSG00000241360 | PDXP | 0,028829064 | 1,020183777 | 0,89806 | 0,95616327 | pyridoxal phosphatase |
| ENSG00000174236 | REP15 | 0,028787088 | 1,020154094 | 0,96857 | 0,98658951 | RAB15 effector protein |
| ENSG00000127463 | EMC1 | 0,028732039 | 1,020115169 | 0,89339 | 0,95343413 | ER membrane protein complex subunit 1 |
| ENSG00000161653 | NAGS | 0,028729998 | 1,020113726 | 0,92821 | 0,96910984 | N-acetylglutamate synthase |
| ENSG00000164080 | RAD54L2 | 0,028634245 | 1,020046023 | 0,88538 | 0,94929432 | RAD54 like 2 |
| ENSG00000129214 | SHBG | 0,028621983 | 1,020037353 | 0,93159 | 0,9707979 | sex hormone binding globulin |
| ENSG00000158715 | SLC45A3 | 0,028478391 | 1,019935833 | 0,95151 | 0,97948342 | solute carrier family 45 member 3 |
| ENSG00000249577 | AC010424,1 | 0,028407478 | 1,019885701 | 0,98128 | 0,99236691 | ARP3 actin-related protein 3 homolog (yeast) pseudogene |
| ENSG00000140836 | ZFHX3 | 0,028357407 | 1,019850305 | 0,93873 | 0,97378588 | zinc finger homeobox 3 |
| ENSG00000167074 | TEF | 0,028310723 | 1,019817304 | 0,91244 | 0,96205682 | TEF transcription factor, PAR bZIP family member |
| ENSG00000078070 | MCCC1 | 0,028244137 | 1,019770237 | 0,86111 | 0,93786377 | methylcrotonoyl-CoA carboxylase 1 |
| ENSG00000278705 | H4C2 | 0,028171787 | 1,019719098 | 0,87825 | 0,9458146 | H4 clustered histone 2 |
| ENSG00000163145 | C1QTNF7 | 0,028113674 | 1,019678023 | 0,94702 | 0,9774744 | C1q and TNF related 7 |
| ENSG00000169057 | MECP2 | 0,028089028 | 1,019660604 | 0,84794 | 0,9309317 | methyl-CpG binding protein 2 |
| ENSG00000100116 | GCAT | 0,028085901 | 1,019658394 | 0,90237 | 0,95825434 | glycine C-acetyltransferase |
| ENSG00000262612 | TAS2R43 | 0,028085292 | 1,019657964 | 0,95306 | 0,98024505 | taste 2 receptor member 43 |
| ENSG00000060642 | PIGV | 0,028067314 | 1,019645257 | 0,89209 | 0,95255977 | phosphatidylinositol glycan anchor biosynthesis class V |
| ENSG00000223642 | AC008277,1 | 0,028048127 | 1,019631696 | 0,95804 | 0,98276081 | novel transcript, antisense to BAZ2B |
| ENSG00000127884 | ECHS1 | 0,027947131 | 1,019560319 | 0,87732 | 0,94563566 | enoyl-CoA hydratase, short chain 1 |
| ENSG00000114383 | TUSC2 | 0,027928788 | 1,019547356 | 0,88064 | 0,94687991 | tumor suppressor 2, mitochondrial calcium regulator |

| | | | | | | |
|---|---|---|---|---|---|---|
| ENSG00000225934 | FAM133FP | 0,02792525 | 1,019544856 | 0,96322 | 0,98440677 | family with sequence similarity 133 member F, pseudogene |
| ENSG00000173366 | AC097637,1 | 0,027778952 | 1,019441473 | 0,98529 | 0,99382561 | novel twinfilin, actin-binding protein, homolog 2 (Drosophila) (TWF2) and toll-like receptor 9 (TLR9) protein |
| ENSG00000101624 | CEP76 | 0,027713828 | 1,019395456 | 0,90725 | 0,96034215 | centrosomal protein 76 |
| ENSG00000125352 | RNF113A | 0,027712233 | 1,019394329 | 0,91489 | 0,9632313 | ring finger protein 113A |
| ENSG00000167272 | POP5 | 0,027603002 | 1,01931715 | 0,88571 | 0,94943038 | POP5 homolog, ribonuclease P/MRP subunit |
| ENSG00000106992 | AK1 | 0,027555699 | 1,01928373 | 0,9025 | 0,95832496 | adenylate kinase 1 |
| ENSG00000153237 | CCDC148 | 0,02748641 | 1,019234777 | 0,9326 | 0,9713043 | coiled-coil domain containing 148 |
| ENSG00000133019 | CHRM3 | 0,027454565 | 1,01921228 | 0,92133 | 0,96592951 | cholinergic receptor muscarinic 3 |
| ENSG00000274270 | AL137060,3 | 0,027203853 | 1,019035176 | 0,9699 | 0,98728333 | novel transcript, sense intronic to DLEU1 |
| ENSG00000177239 | MAN1B1 | 0,027164023 | 1,019007043 | 0,89756 | 0,95581155 | mannosidase alpha class 1B member 1 |
| ENSG00000111726 | CMAS | 0,027123668 | 1,01897854 | 0,88544 | 0,94932414 | cytidine monophosphate N-acetylneuraminic acid synthetase |
| ENSG00000170345 | FOS | 0,027112005 | 1,018970302 | 0,92976 | 0,96989348 | Fos proto-oncogene, AP-1 transcription factor subunit |
| ENSG00000188672 | RHCE | 0,027035695 | 1,018916406 | 0,96394 | 0,98461596 | Rh blood group CcEe antigens |
| ENSG00000184261 | KCNK12 | 0,027007743 | 1,018896665 | 0,93398 | 0,97201193 | potassium two pore domain channel subfamily K member 12 |
| ENSG00000246214 | AC024588,1 | 0,026952234 | 1,018857463 | 0,9244 | 0,96714797 | novel transcript, antisense to FAM134B |
| ENSG00000141971 | MVB12A | 0,026856736 | 1,018790023 | 0,90905 | 0,96102407 | multivesicular body subunit 12A |
| ENSG00000164535 | DAGLB | 0,026807133 | 1,018754995 | 0,87147 | 0,94322889 | diacylglycerol lipase beta |
| ENSG00000145730 | PAM | 0,026703482 | 1,018681805 | 0,93462 | 0,9721963 | peptidylglycine alpha-amidating monooxygenase |
| ENSG00000286327 | AL035427,2 | 0,026644947 | 1,018640475 | 0,93629 | 0,97254199 | novel transcript |
| ENSG00000234881 | PIGFP2 | 0,026396855 | 1,01846532 | 0,97657 | 0,9901566 | phosphatidylinositol glycan anchor biosynthesis class F pseudogene 2 |
| ENSG00000270265 | AC104819,3 | 0,026388622 | 1,018459508 | 0,96783 | 0,98624732 | novel transcript |
| ENSG00000235865 | GSN-AS1 | 0,026269295 | 1,018375273 | 0,96205 | 0,984073 | GSN antisense RNA 1 |
| ENSG00000243368 | MCCC1-AS1 | 0,026233582 | 1,018350064 | 0,92554 | 0,96765626 | MCCC1 antisense RNA 1 |
| ENSG00000188786 | MTF1 | 0,026213625 | 1,018335977 | 0,87273 | 0,94374455 | metal regulatory transcription factor 1 |
| ENSG00000236324 | SNX9-AS1 | 0,026204873 | 1,018329799 | 0,98096 | 0,99215365 | SNX9 antisense RNA 1 |
| ENSG00000243406 | MRPS31P5 | 0,026175118 | 1,018308797 | 0,88319 | 0,94832116 | mitochondrial ribosomal protein S31 pseudogene 5 |
| ENSG00000172404 | DNAJB7 | 0,026168818 | 1,01830435 | 0,93886 | 0,97388366 | DnaJ heat shock protein family (Hsp40) member B7 |
| ENSG00000181322 | NME9 | 0,026160853 | 1,018298728 | 0,91749 | 0,96423084 | NME/NM23 family member 9 |

| | | | | | | |
|---|---|---|---|---|---|---|
| ENSG00000136531 | SCN2A | 0,026158682 | 1,018297196 | 0,9655 | 0,9854196 | sodium voltage-gated channel alpha subunit 2 |
| ENSG00000267147 | LINC01842 | 0,026133334 | 1,018279305 | 0,97203 | 0,98800339 | long intergenic non-protein coding RNA 1842 |
| ENSG00000203780 | FANK1 | 0,02607013 | 1,018234696 | 0,9246 | 0,96724903 | fibronectin type III and ankyrin repeat domains 1 |
| ENSG00000157680 | DGKI | 0,026067326 | 1,018232716 | 0,93243 | 0,97120063 | diacylglycerol kinase iota |
| ENSG00000248228 | SLIT2-IT1 | 0,02604101 | 1,018214143 | 0,95665 | 0,98202499 | SLIT2 intronic transcript 1 |
| ENSG00000180720 | CHRM4 | 0,02600712 | 1,018190225 | 0,95259 | 0,97993161 | cholinergic receptor muscarinic 4 |
| ENSG00000198597 | ZNF536 | 0,025961263 | 1,018157862 | 0,95262 | 0,97993161 | zinc finger protein 536 |
| ENSG00000272462 | U91328,1 | 0,025939438 | 1,018142459 | 0,88455 | 0,94883769 | novel transcript |
| ENSG00000184207 | PGP | 0,025806049 | 1,018048328 | 0,89469 | 0,95413182 | phosphoglycolate phosphatase |
| ENSG00000167528 | ZNF641 | 0,025663115 | 1,01794747 | 0,94285 | 0,97567445 | zinc finger protein 641 |
| ENSG00000271917 | AL357568,1 | 0,02564481 | 1,017934554 | 0,96773 | 0,98623935 | novel transcript |
| ENSG00000107951 | MTPAP | 0,025602238 | 1,017904517 | 0,85716 | 0,93548553 | mitochondrial poly(A) polymerase |
| ENSG00000100099 | HPS4 | 0,025521733 | 1,017847718 | 0,87134 | 0,94321633 | HPS4 biogenesis of lysosomal organelles complex 3 subunit 2 |
| ENSG00000115446 | UNC50 | 0,025510775 | 1,017839986 | 0,89838 | 0,95636187 | unc-50 inner nuclear membrane RNA binding protein |
| ENSG00000118507 | AKAP7 | 0,02548463 | 1,017821541 | 0,91203 | 0,96198423 | A-kinase anchoring protein 7 |
| ENSG00000110717 | NDUFS8 | 0,025402435 | 1,017763554 | 0,89086 | 0,95207499 | NADH:ubiquinone oxidoreductase core subunit S8 |
| ENSG00000158163 | DZIP1L | 0,025360787 | 1,017734173 | 0,9124 | 0,96205682 | DAZ interacting zinc finger protein 1 like |
| ENSG00000117461 | PIK3R3 | 0,02535036 | 1,017726818 | 0,90459 | 0,95918368 | phosphoinositide-3-kinase regulatory subunit 3 |
| ENSG00000165669 | FAM204A | 0,025326233 | 1,017709799 | 0,89468 | 0,95413182 | family with sequence similarity 204 member A |
| ENSG00000169246 | NPIPB3 | 0,025322277 | 1,017707008 | 0,92299 | 0,96662141 | nuclear pore complex interacting protein family member B3 |
| ENSG00000225731 | PDE9A-AS1 | 0,025289144 | 1,017683635 | 0,96061 | 0,98383172 | PDE9A antisense RNA 1 |
| ENSG00000136816 | TOR1B | 0,025176168 | 1,017603944 | 0,92437 | 0,967147 | torsin family 1 member B |
| ENSG00000231584 | FAHD2CP | 0,025119995 | 1,017564324 | 0,9258 | 0,96768961 | fumarylacetoacetate hydrolase domain containing 2C, pseudogene |
| ENSG00000239900 | ADSL | 0,025036264 | 1,017505268 | 0,89424 | 0,95389664 | adenylosuccinate lyase |
| ENSG00000122257 | RBBP6 | 0,024891529 | 1,017403194 | 0,86717 | 0,94124126 | RB binding protein 6, ubiquitin ligase |
| ENSG00000188219 | POTEE | 0,024824739 | 1,017356095 | 0,94488 | 0,97662603 | POTE ankyrin domain family member E |
| ENSG00000083799 | CYLD | 0,02479048 | 1,017331936 | 0,91183 | 0,96184653 | CYLD lysine 63 deubiquitinase |
| ENSG00000240497 | AC092919,1 | 0,024627313 | 1,017216884 | 0,9759 | 0,98986897 | novel transcript |
| ENSG00000163249 | CCNYL1 | 0,024603323 | 1,017199969 | 0,91186 | 0,96184653 | cyclin Y like 1 |
| ENSG00000126016 | AMOT | 0,024542828 | 1,017157317 | 0,93226 | 0,97114134 | angiomotin |

| | | | | | | |
|---|---|---|---|---|---|---|
| ENSG00000239407 | Z68871,1 | 0,024499389 | 1,017126691 | 0,9254 | 0,96761679 | novel transcript |
| ENSG00000234373 | SNX18P7 | 0,024392659 | 1,017051447 | 0,97822 | 0,99099142 | sorting nexin 18 pseudogene 7 |
| ENSG00000259118 | AL139022,1 | 0,024384938 | 1,017046004 | 0,95443 | 0,98093594 | novel transcript, antisense to MAX |
| ENSG00000282717 | AC243651,2 | 0,02433442 | 1,017010391 | 0,96309 | 0,98432594 | novel transcript, antisense to PDXDC1 |
| ENSG00000275910 | AC138932,4 | 0,02433442 | 1,017010391 | 0,96309 | 0,98432594 | novel transcript, antisense to PDXDC1 |
| ENSG00000276547 | PCDHGB5 | 0,024320155 | 1,017000336 | 0,95813 | 0,98276081 | protocadherin gamma subfamily B, 5 |
| ENSG00000153551 | CMTM7 | 0,024267023 | 1,016962881 | 0,93461 | 0,9721963 | CKLF like MARVEL transmembrane domain containing 7 |
| ENSG00000137817 | PARP6 | 0,024208915 | 1,016921922 | 0,8912 | 0,95208954 | poly(ADP-ribose) polymerase family member 6 |
| ENSG00000229127 | KANSL1L-AS1 | 0,024080089 | 1,016831119 | 0,92936 | 0,96977202 | KANSL1L antisense RNA 1 |
| ENSG00000227630 | LINC01132 | 0,02406581 | 1,016821056 | 0,97412 | 0,98928341 | long intergenic non-protein coding RNA 1132 |
| ENSG00000278434 | AC023830,3 | 0,024022297 | 1,016790388 | 0,94245 | 0,97556707 | novel transcript, sense intronic to CDIP1 |
| ENSG00000264668 | AC138696,1 | 0,02401478 | 1,01678509 | 0,9729 | 0,98865206 | novel protein |
| ENSG00000100092 | SH3BP1 | 0,023904428 | 1,016707319 | 0,92162 | 0,96607272 | SH3 domain binding protein 1 |
| ENSG00000168259 | DNAJC7 | 0,023885079 | 1,016693683 | 0,88468 | 0,94887389 | DnaJ heat shock protein family (Hsp40) member C7 |
| ENSG00000250371 | AC007370,2 | 0,023851076 | 1,016669721 | 0,95814 | 0,98276081 | novel transcript |
| ENSG00000271318 | MOB4P2 | 0,023847592 | 1,016667266 | 0,98186 | 0,99242247 | MOB4 pseudogene 2 |
| ENSG00000163568 | AIM2 | 0,02384032 | 1,016662141 | 0,95725 | 0,98231979 | absent in melanoma 2 |
| ENSG00000114796 | KLHL24 | 0,023835574 | 1,016658797 | 0,93363 | 0,97168925 | kelch like family member 24 |
| ENSG00000263786 | AC022211,1 | 0,023765108 | 1,016609141 | 0,94593 | 0,97689657 | novel transcript, sense intronic to HN1 |
| ENSG00000285593 | AC055872,2 | 0,023670594 | 1,016542543 | 0,96683 | 0,98609266 | novel transcript |
| ENSG00000119703 | ZC2HC1C | 0,023662314 | 1,016536709 | 0,9294 | 0,96977202 | zinc finger C2HC-type containing 1C |
| ENSG00000141644 | MBD1 | 0,023514502 | 1,016432564 | 0,88388 | 0,94847506 | methyl-CpG binding domain protein 1 |
| ENSG00000265298 | AC132812,1 | 0,023491567 | 1,016416406 | 0,96846 | 0,98654786 | small nuclear ribonucleoprotein 200kDa (U5) (SNRNP200) pseudogene |
| ENSG00000164758 | MED30 | 0,023448372 | 1,016385974 | 0,92749 | 0,96860856 | mediator complex subunit 30 |
| ENSG00000260029 | AC007608,2 | 0,023368413 | 1,016329644 | 0,95856 | 0,98291816 | novel transcript, antisense to NKD1 |
| ENSG00000157483 | MYO1E | 0,023269493 | 1,016259961 | 0,9435 | 0,97617306 | myosin IE |
| ENSG00000166510 | CCDC68 | 0,023262764 | 1,016255221 | 0,97572 | 0,9897621 | coiled-coil domain containing 68 |
| ENSG00000166263 | STXBP4 | 0,023259848 | 1,016253167 | 0,89507 | 0,95421556 | syntaxin binding protein 4 |
| ENSG00000124920 | MYRF | 0,023219758 | 1,016224927 | 0,91134 | 0,9616927 | myelin regulatory factor |
| ENSG00000100578 | KIAA0586 | 0,023043231 | 1,01610059 | 0,88806 | 0,95069428 | KIAA0586 |
| ENSG00000169964 | TMEM42 | 0,022941226 | 1,01602875 | 0,92531 | 0,96758768 | transmembrane protein 42 |
| ENSG00000280516 | TMEM42 | 0,022941226 | 1,01602875 | 0,92531 | 0,96758768 | transmembrane protein 42 |

| | | | | | | |
|---|---|---|---|---|---|---|
| ENSG00000067066 | SP100 | 0,022872584 | 1,01598041 | 0,92079 | 0,96574531 | SP100 nuclear antigen |
| ENSG00000163319 | MRPS18C | 0,022829566 | 1,015950116 | 0,90954 | 0,96105018 | mitochondrial ribosomal protein S18C |
| ENSG00000151948 | GLT1D1 | 0,022766944 | 1,015906018 | 0,91131 | 0,9616927 | glycosyltransferase 1 domain containing 1 |
| ENSG00000186174 | BCL9L | 0,022721873 | 1,015874281 | 0,90751 | 0,96044232 | BCL9 like |
| ENSG00000230158 | MTND1P28 | 0,022711182 | 1,015866753 | 0,9816 | 0,99239184 | MT-ND1 pseudogene 28 |
| ENSG00000188997 | KCTD21 | 0,022645129 | 1,015820243 | 0,9302 | 0,97010144 | potassium channel tetramerization domain containing 21 |
| ENSG00000250240 | AC008840,1 | 0,022490893 | 1,015711649 | 0,94095 | 0,97489099 | novel transcript |
| ENSG00000117410 | ATP6V0B | 0,022480707 | 1,015704478 | 0,89573 | 0,95482399 | ATPase H+ transporting V0 subunit b |
| ENSG00000121671 | CRY2 | 0,022472275 | 1,015698541 | 0,89337 | 0,95343413 | cryptochrome circadian regulator 2 |
| ENSG00000125826 | RBCK1 | 0,022465938 | 1,01569408 | 0,91004 | 0,96126119 | RANBP2-type and C3HC4-type zinc finger containing 1 |
| ENSG00000232936 | AL157400,2 | 0,022429613 | 1,015668506 | 0,95826 | 0,98276958 | novel transcript |
| ENSG00000181104 | F2R | 0,022333796 | 1,015601053 | 0,90159 | 0,95801185 | coagulation factor II thrombin receptor |
| ENSG00000054611 | TBC1D22A | 0,022307597 | 1,01558261 | 0,91814 | 0,96444601 | TBC1 domain family member 22A |
| ENSG00000218896 | TUBB8P2 | 0,022277106 | 1,015561146 | 0,97198 | 0,98800339 | tubulin beta 8 class VIII pseudogene 2 |
| ENSG00000146535 | GNA12 | 0,02227379 | 1,015558812 | 0,90957 | 0,96105018 | G protein subunit alpha 12 |
| ENSG00000044090 | CUL7 | 0,022256588 | 1,015546703 | 0,91919 | 0,96485808 | cullin 7 |
| ENSG00000286964 | AL136169,1 | 0,022149243 | 1,015471143 | 0,96055 | 0,98383172 | novel transcript |
| ENSG00000272273 | IER3-AS1 | 0,022097057 | 1,015434411 | 0,98101 | 0,99215365 | IER3 antisense RNA 1 |
| ENSG00000137834 | SMAD6 | 0,022091442 | 1,01543046 | 0,96359 | 0,98450096 | SMAD family member 6 |
| ENSG00000261474 | AC026471,4 | 0,022059955 | 1,015408298 | 0,97425 | 0,98933043 | novel transcript |
| ENSG00000175697 | GPR156 | 0,022041514 | 1,015395318 | 0,96041 | 0,98381824 | G protein-coupled receptor 156 |
| ENSG00000273144 | TAS2R15P | 0,022013619 | 1,015375686 | 0,95705 | 0,98218743 | taste 2 receptor member 15 pseudogene |
| ENSG00000276217 | TAS2R15P | 0,022013619 | 1,015375686 | 0,95705 | 0,98218743 | taste 2 receptor member 15 pseudogene |
| ENSG00000212125 | TAS2R15P | 0,022013619 | 1,015375686 | 0,95705 | 0,98218743 | taste 2 receptor member 15 pseudogene |
| ENSG00000160932 | LY6E | 0,021981245 | 1,015352901 | 0,96769 | 0,98623935 | lymphocyte antigen 6 family member E |
| ENSG00000101216 | GMEB2 | 0,021955963 | 1,015335108 | 0,89495 | 0,95417507 | glucocorticoid modulatory element binding protein 2 |
| ENSG00000115902 | SLC1A4 | 0,021938042 | 1,015322496 | 0,92581 | 0,96768961 | solute carrier family 1 member 4 |
| ENSG00000154016 | GRAP | 0,021830423 | 1,01524676 | 0,95826 | 0,98276958 | GRB2 related adaptor protein |
| ENSG00000278733 | AC022079,2 | 0,021770997 | 1,015204941 | 0,983 | 0,9926386 | novel transcript, antisense to CCDC91 |
| ENSG00000179029 | TMEM107 | 0,021751447 | 1,015191185 | 0,91409 | 0,96285224 | transmembrane protein 107 |
| ENSG00000198648 | STK39 | 0,021659405 | 1,015126419 | 0,93727 | 0,97312989 | serine/threonine kinase 39 |
| ENSG00000135486 | HNRNPA1 | 0,021646035 | 1,015117012 | 0,90327 | 0,95856163 | heterogeneous nuclear ribonucleoprotein A1 |
| ENSG00000186007 | LEMD1 | 0,021615803 | 1,01509574 | 0,94416 | 0,97642036 | LEM domain containing 1 |

| | | | | | | |
|---|---|---|---|---|---|---|
| ENSG00000255443 | CD44-AS1 | 0,021555797 | 1,01505352 | 0,97096 | 0,98771801 | CD44 antisense RNA 1 |
| ENSG00000269985 | AL021328,1 | 0,021477175 | 1,014998204 | 0,95858 | 0,98291816 | novel transcript, antisense to FARS2 |
| ENSG00000006025 | OSBPL7 | 0,021432081 | 1,014966479 | 0,94014 | 0,97460197 | oxysterol binding protein like 7 |
| ENSG00000263499 | AC007431,1 | 0,021375155 | 1,014926431 | 0,95384 | 0,98057921 | novel transcript, antisense to MSI2 |
| ENSG00000198580 | AC073343,1 | 0,02122955 | 1,014824004 | 0,97988 | 0,99172545 | zinc finger protein pseudogene |
| ENSG00000109881 | CCDC34 | 0,021191779 | 1,014797436 | 0,92344 | 0,96667848 | coiled-coil domain containing 34 |
| ENSG00000204308 | RNF5 | 0,021187064 | 1,014794119 | 0,92515 | 0,96749513 | ring finger protein 5 |
| ENSG00000136908 | DPM2 | 0,021179387 | 1,014788719 | 0,9004 | 0,95760678 | dolichyl-phosphate mannosyltransferase subunit 2, regulatory |
| ENSG00000169627 | BOLA2B | 0,021125983 | 1,014751156 | 0,93708 | 0,97304999 | bolA family member 2B |
| ENSG00000235032 | BMP7-AS1 | 0,020933507 | 1,014615782 | 0,98154 | 0,99239184 | BMP7 antisense RNA 1 |
| ENSG00000272115 | AC233992,3 | 0,020843295 | 1,01455234 | 0,9794 | 0,99154627 | novel transcript, antisense to SLC39A4 |
| ENSG00000284888 | AC233992,5 | 0,020843295 | 1,01455234 | 0,9794 | 0,99154627 | novel transcript, antisense to SLC39A4 |
| ENSG00000101104 | PABPC1L | 0,020809099 | 1,014528293 | 0,93227 | 0,97114134 | poly(A) binding protein cytoplasmic 1 like |
| ENSG00000133661 | SFTPD | 0,020654686 | 1,014419712 | 0,95971 | 0,98338726 | surfactant protein D |
| ENSG00000107130 | NCS1 | 0,020602078 | 1,014382722 | 0,91766 | 0,96426294 | neuronal calcium sensor 1 |
| ENSG00000172465 | TCEAL1 | 0,020518683 | 1,014324087 | 0,95129 | 0,97948342 | transcription elongation factor A like 1 |
| ENSG00000101255 | TRIB3 | 0,020369393 | 1,014219131 | 0,93063 | 0,97025341 | tribbles pseudokinase 3 |
| ENSG00000186907 | RTN4RL2 | 0,020359929 | 1,014212478 | 0,96164 | 0,98401207 | reticulon 4 receptor like 2 |
| ENSG00000245534 | RORA-AS1 | 0,020319622 | 1,014184142 | 0,95959 | 0,98338726 | RORA antisense RNA 1 |
| ENSG00000184719 | RNLS | 0,020164956 | 1,014075421 | 0,93427 | 0,97210301 | renalase, FAD dependent amine oxidase |
| ENSG00000286808 | AC092338,3 | 0,02015253 | 1,014066687 | 0,95154 | 0,97948342 | novel transcript, antisense to POLR3E |
| ENSG00000251620 | STPG2-AS1 | 0,020110444 | 1,014037106 | 0,94056 | 0,97474573 | STPG2 antisense RNA 1 |
| ENSG00000236088 | COX10-AS1 | 0,020021889 | 1,013974864 | 0,9271 | 0,96842124 | COX10 antisense RNA 1 |
| ENSG00000258102 | MAP1LC3B2 | 0,019830686 | 1,013840489 | 0,97459 | 0,98933402 | microtubule associated protein 1 light chain 3 beta 2 |
| ENSG00000283589 | GOLGA8N | 0,01978866 | 1,013810956 | 0,93894 | 0,97393487 | golgin A8 family member N |
| ENSG00000132906 | CASP9 | 0,0197194 | 1,013762287 | 0,91236 | 0,96205682 | caspase 9 |
| ENSG00000273983 | H3C8 | 0,019679593 | 1,013734316 | 0,9445 | 0,97650311 | H3 clustered histone 8 |
| ENSG00000236991 | EDRF1-AS1 | 0,019674926 | 1,013731036 | 0,95897 | 0,98304779 | EDRF1 antisense RNA 1 |
| ENSG00000157890 | MEGF11 | 0,019623877 | 1,013695166 | 0,93196 | 0,97097872 | multiple EGF like domains 11 |
| ENSG00000237336 | AL445489,1 | 0,019564052 | 1,013653131 | 0,97334 | 0,98881219 | novel transcript |
| ENSG00000271919 | AC005828,7 | 0,019557079 | 1,013648233 | 0,97028 | 0,98749329 | density-regulated protein (DENR) pseudogene |
| ENSG00000196227 | FAM217B | 0,019553094 | 1,013645433 | 0,92821 | 0,96910984 | family with sequence similarity 217 member B |

| | | | | | | |
|---|---|---|---|---|---|---|
| ENSG00000287513 | AL133390,1 | 0,019537688 | 1,013634608 | 0,98056 | 0,99198429 | novel transcript |
| ENSG00000285918 | AC092376,3 | 0,019471687 | 1,013588237 | 0,97073 | 0,98765168 | novel transcript, antisense to WWOX |
| ENSG00000132437 | DDC | 0,019449233 | 1,013572462 | 0,973 | 0,9886748 | dopa decarboxylase |
| ENSG00000267248 | AC025048,2 | 0,019392285 | 1,013532454 | 0,93419 | 0,97206381 | novel transcript |
| ENSG00000137731 | FXYD2 | 0,019366611 | 1,013514417 | 0,97482 | 0,98941297 | FXYD domain containing ion transport regulator 2 |
| ENSG00000120802 | TMPO | 0,019359089 | 1,013509133 | 0,91116 | 0,9616927 | thymopoietin |
| ENSG00000177432 | NAP1L5 | 0,019308929 | 1,013473895 | 0,95169 | 0,97948419 | nucleosome assembly protein 1 like 5 |
| ENSG00000137101 | CD72 | 0,019281353 | 1,013454524 | 0,95132 | 0,97948342 | CD72 molecule |
| ENSG00000280759 | AP2A2 | 0,01923703 | 1,013423389 | 0,92488 | 0,9673913 | adaptor related protein complex 2 subunit alpha 2 |
| ENSG00000038219 | BOD1L1 | 0,019150893 | 1,013362883 | 0,93908 | 0,97400466 | biorientation of chromosomes in cell division 1 like 1 |
| ENSG00000186918 | ZNF395 | 0,019131274 | 1,013349103 | 0,93259 | 0,9713043 | zinc finger protein 395 |
| ENSG00000149403 | GRIK4 | 0,019091014 | 1,013320825 | 0,95963 | 0,98338726 | glutamate ionotropic receptor kainate type subunit 4 |
| ENSG00000287021 | AC097462,3 | 0,018992511 | 1,013251641 | 0,96774 | 0,98623935 | novel transcript, antisense to INTU |
| ENSG00000253200 | AC037459,2 | 0,01894757 | 1,013220078 | 0,92591 | 0,96772813 | novel transcript, antisense to KIAA1967 |
| ENSG00000161912 | ADCY10P1 | 0,01892412 | 1,013203609 | 0,95148 | 0,97948342 | ADCY10 pseudogene 1 |
| ENSG00000177182 | CLVS1 | 0,018912307 | 1,013195312 | 0,96633 | 0,98579843 | clavesin 1 |
| ENSG00000274129 | TSEN34 | 0,018896107 | 1,013183935 | 0,93901 | 0,97396769 | tRNA splicing endonuclease subunit 34 |
| ENSG00000275086 | TFPT | 0,018830777 | 1,013138056 | 0,96192 | 0,98401207 | TCF3 fusion partner |
| ENSG00000278161 | TFPT | 0,018830777 | 1,013138056 | 0,96192 | 0,98401207 | TCF3 fusion partner |
| ENSG00000276022 | TFPT | 0,018830777 | 1,013138056 | 0,96192 | 0,98401207 | TCF3 fusion partner |
| ENSG00000276323 | TFPT | 0,018830777 | 1,013138056 | 0,96192 | 0,98401207 | TCF3 fusion partner |
| ENSG00000276296 | TFPT | 0,018830777 | 1,013138056 | 0,96192 | 0,98401207 | TCF3 fusion partner |
| ENSG00000274073 | TFPT | 0,018830777 | 1,013138056 | 0,96192 | 0,98401207 | TCF3 fusion partner |
| ENSG00000276263 | TFPT | 0,018830777 | 1,013138056 | 0,96192 | 0,98401207 | TCF3 fusion partner |
| ENSG00000273833 | TFPT | 0,018830777 | 1,013138056 | 0,96192 | 0,98401207 | TCF3 fusion partner |
| ENSG00000260035 | AC051619,7 | 0,018745413 | 1,01307811 | 0,9617 | 0,98401207 | novel transcript |
| ENSG00000227252 | AC012063,1 | 0,018582534 | 1,012963741 | 0,94669 | 0,97731612 | novel transcript, antisense to COPS8 |
| ENSG00000158669 | GPAT4 | 0,018569937 | 1,012954897 | 0,90496 | 0,9593272 | glycerol-3-phosphate acyltransferase 4 |
| ENSG00000091844 | RGS17 | 0,018485853 | 1,01289586 | 0,95971 | 0,98338726 | regulator of G protein signaling 17 |
| ENSG00000256134 | EGLN3P1 | 0,018289472 | 1,012757994 | 0,97411 | 0,98928341 | egl-9 family hypoxia-inducible factor 3 pseudogene 1 |
| ENSG00000185359 | HGS | 0,018173267 | 1,012676422 | 0,90948 | 0,96105018 | hepatocyte growth factor-regulated tyrosine kinase substrate |
| ENSG00000242398 | RN7SL800P | 0,018171679 | 1,012675307 | 0,97716 | 0,99046478 | RNA, 7SL, cytoplasmic 800, pseudogene |

| | | | | | | |
|---|---|---|---|---|---|---|
| ENSG00000228794 | LINC01128 | 0,018132931 | 1,01264811 | 0,93311 | 0,97150994 | long intergenic non-protein coding RNA 1128 |
| ENSG00000259712 | AC023906,5 | 0,018063624 | 1,012599462 | 0,96766 | 0,98623935 | novel transcript, antisense to MAPK6 |
| ENSG00000117020 | AKT3 | 0,018043368 | 1,012585246 | 0,915 | 0,9632313 | AKT serine/threonine kinase 3 |
| ENSG00000226754 | AL606760,1 | 0,018041241 | 1,012583753 | 0,96952 | 0,98718244 | novel transcript, antisense to MAGOH |
| ENSG00000286110 | AC090983,2 | 0,018006777 | 1,012559564 | 0,96227 | 0,98415736 | novel transcript |
| ENSG00000235121 | AL645504,1 | 0,017953297 | 1,012522029 | 0,96285 | 0,98432594 | novel transcript |
| ENSG00000280868 | LINC00242 | 0,017864053 | 1,012459398 | 0,96357 | 0,98450096 | long intergenic non-protein coding RNA 242 |
| ENSG00000188807 | TMEM201 | 0,017855384 | 1,012453314 | 0,90964 | 0,96105018 | transmembrane protein 201 |
| ENSG00000197857 | ZNF44 | 0,017811671 | 1,012422637 | 0,92112 | 0,96582196 | zinc finger protein 44 |
| ENSG00000261645 | DISC1FP1 | 0,017743622 | 1,012374884 | 0,98007 | 0,9917697 | DISC1 fusion partner 1 |
| ENSG00000205108 | FAM205A | 0,017671808 | 1,012324492 | 0,97675 | 0,99026923 | family with sequence similarity 205 member A |
| ENSG00000284640 | AL590434,1 | 0,017624326 | 1,012291175 | 0,97609 | 0,98994851 | novel transcript |
| ENSG00000197019 | SERTAD1 | 0,017611485 | 1,012282165 | 0,96078 | 0,9839024 | SERTA domain containing 1 |
| ENSG00000127957 | PMS2P3 | 0,017587721 | 1,012265491 | 0,92496 | 0,96744548 | PMS1 homolog 2, mismatch repair system component pseudogene 3 |
| ENSG00000092964 | DPYSL2 | 0,017505994 | 1,012208149 | 0,93629 | 0,97254199 | dihydropyrimidinase like 2 |
| ENSG00000011021 | CLCN6 | 0,017504683 | 1,012207229 | 0,92243 | 0,96637639 | chloride voltage-gated channel 6 |
| ENSG00000128833 | MYO5C | 0,017108325 | 1,011929179 | 0,96974 | 0,98725952 | myosin VC |
| ENSG00000258485 | SRMP2 | 0,017058912 | 1,011894521 | 0,97299 | 0,9886748 | SRM pseudogene 2 |
| ENSG00000174903 | RAB1B | 0,017052049 | 1,011889707 | 0,9363 | 0,97254199 | RAB1B, member RAS oncogene family |
| ENSG00000198728 | LDB1 | 0,017048606 | 1,011887292 | 0,91377 | 0,96270153 | LIM domain binding 1 |
| ENSG00000274828 | AC068473,5 | 0,017023199 | 1,011869472 | 0,95863 | 0,98291816 | novel transcript |
| ENSG00000282330 | AC068473,8 | 0,017023199 | 1,011869472 | 0,95863 | 0,98291816 | novel transcript |
| ENSG00000136147 | PHF11 | 0,016991317 | 1,011847111 | 0,9297 | 0,96987987 | PHD finger protein 11 |
| ENSG00000170035 | UBE2E3 | 0,016826909 | 1,011731809 | 0,95065 | 0,97934603 | ubiquitin conjugating enzyme E2 E3 |
| ENSG00000196363 | WDR5 | 0,016826612 | 1,0117316 | 0,92416 | 0,96703664 | WD repeat domain 5 |
| ENSG00000175137 | SH3BP5L | 0,01680455 | 1,011716129 | 0,92568 | 0,96768961 | SH3 binding domain protein 5 like |
| ENSG00000286340 | AL356776,2 | 0,016803682 | 1,01171552 | 0,9453 | 0,97666034 | novel transcript, antisense to PHIP |
| ENSG00000236998 | AL354794,2 | 0,016612843 | 1,0115817 | 0,97539 | 0,98956832 | adaptor-related protein complex 3, mu 2 subunit (AP3M2) pseudogene |
| ENSG00000276564 | AC130650,2 | 0,01647267 | 1,011483419 | 0,96054 | 0,98383172 | novel transcript |
| ENSG00000100625 | SIX4 | 0,01624843 | 1,011326215 | 0,97489 | 0,98941297 | SIX homeobox 4 |
| ENSG00000172432 | GTPBP2 | 0,016182074 | 1,011279701 | 0,92805 | 0,96901646 | GTP binding protein 2 |

| | | | | | | |
|---|---|---|---|---|---|---|
| ENSG00000269900 | RMRP | 0,016170108 | 1,011271313 | 0,98187 | 0,99242247 | RNA component of mitochondrial RNA processing endoribonuclease |
| ENSG00000163923 | RPL39L | 0,01615022 | 1,011257372 | 0,94746 | 0,97781662 | ribosomal protein L39 like |
| ENSG00000198715 | GLMP | 0,016099613 | 1,0112219 | 0,96257 | 0,98430469 | glycosylated lysosomal membrane protein |
| ENSG00000288510 | AC148477,19 | 0,016080718 | 1,011208656 | 0,98236 | 0,99244329 | novel transcript |
| ENSG00000256542 | AC148477,2 | 0,016080718 | 1,011208656 | 0,98236 | 0,99244329 | novel transcript |
| ENSG00000143862 | ARL8A | 0,015921208 | 1,011096859 | 0,95123 | 0,97948342 | ADP ribosylation factor like GTPase 8A |
| ENSG00000162910 | MRPL55 | 0,015650664 | 1,010907269 | 0,94009 | 0,97457888 | mitochondrial ribosomal protein L55 |
| ENSG00000062716 | VMP1 | 0,015527036 | 1,010820645 | 0,95476 | 0,98110426 | vacuole membrane protein 1 |
| ENSG00000119737 | GPR75 | 0,015333349 | 1,010684948 | 0,96688 | 0,98611004 | G protein-coupled receptor 75 |
| ENSG00000183808 | RBM12B | 0,015291936 | 1,010655937 | 0,9224 | 0,96637639 | RNA binding motif protein 12B |
| ENSG00000273987 | AC121761,2 | 0,015258802 | 1,010632726 | 0,98742 | 0,99447743 | novel transcript, sense intronic to CAPS2 |
| ENSG00000126773 | PCNX4 | 0,01522304 | 1,010607674 | 0,9238 | 0,96681926 | pecanex 4 |
| ENSG00000184224 | C11orf72 | 0,015092445 | 1,010516196 | 0,96275 | 0,98432594 | chromosome 11 putative open reading frame 72 |
| ENSG00000148426 | PROSER2 | 0,015069736 | 1,01050029 | 0,98387 | 0,99300402 | proline and serine rich 2 |
| ENSG00000282034 | AC106886,5 | 0,015026835 | 1,010470241 | 0,93489 | 0,9721963 | novel transcript |
| ENSG00000101752 | MIB1 | 0,014939855 | 1,010409323 | 0,93336 | 0,97160226 | mindbomb E3 ubiquitin protein ligase 1 |
| ENSG00000233806 | LINC01237 | 0,014935289 | 1,010406124 | 0,97936 | 0,99154627 | long intergenic non-protein coding RNA 1237 |
| ENSG00000117448 | AKR1A1 | 0,014811075 | 1,010319134 | 0,94673 | 0,97731612 | aldo-keto reductase family 1 member A1 |
| ENSG00000186283 | TOR3A | 0,014750511 | 1,010276722 | 0,93555 | 0,97228877 | torsin family 3 member A |
| ENSG00000139537 | CCDC65 | 0,014727412 | 1,010260546 | 0,96493 | 0,98525685 | coiled-coil domain containing 65 |
| ENSG00000282872 | C1orf232 | 0,014639206 | 1,010198781 | 0,9821 | 0,99244329 | chromosome 1 open reading frame 230 |
| ENSG00000125821 | DTD1 | 0,014554032 | 1,010139142 | 0,93801 | 0,97361277 | D-aminoacyl-tRNA deacylase 1 |
| ENSG00000166405 | RIC3 | 0,014534124 | 1,010125203 | 0,96764 | 0,98623935 | RIC3 acetylcholine receptor chaperone |
| ENSG00000197586 | ENTPD6 | 0,01449404 | 1,010097138 | 0,93159 | 0,9707979 | ectonucleoside triphosphate diphosphohydrolase 6 |
| ENSG00000131507 | NDFIP1 | 0,014478658 | 1,010086369 | 0,94505 | 0,97666034 | Nedd4 family interacting protein 1 |
| ENSG00000101350 | KIF3B | 0,014425712 | 1,0100493 | 0,93948 | 0,97416733 | kinesin family member 3B |
| ENSG00000163377 | TAFA4 | 0,014424276 | 1,010048295 | 0,98405 | 0,99309929 | TAFA chemokine like family member 4 |
| ENSG00000267257 | AC105105,1 | 0,014362054 | 1,010004733 | 0,98595 | 0,99413637 | novel transcript, antisense to ALPK2 |
| ENSG00000106477 | CEP41 | 0,01433583 | 1,009986374 | 0,93192 | 0,97097872 | centrosomal protein 41 |
| ENSG00000169371 | SNUPN | 0,014309291 | 1,009967796 | 0,96086 | 0,98393555 | snurportin 1 |
| ENSG00000128245 | YWHAH | 0,014141256 | 1,009850168 | 0,93815 | 0,97361277 | tyrosine 3-monooxygenase/tryptophan 5-monooxygenase activation protein eta |

| | | | | | | |
|---|---|---|---|---|---|---|
| ENSG00000103966 | EHD4 | 0,014010687 | 1,009758777 | 0,95356 | 0,98051225 | EH domain containing 4 |
| ENSG00000258521 | AL157871,2 | 0,014008837 | 1,009757483 | 0,98812 | 0,99471957 | novel transcript, antisense to WARS |
| ENSG00000066135 | KDM4A | 0,014005633 | 1,00975524 | 0,94335 | 0,97609158 | lysine demethylase 4A |
| ENSG00000131269 | ABCB7 | 0,013824255 | 1,0096283 | 0,94158 | 0,97505054 | ATP binding cassette subfamily B member 7 |
| ENSG00000273156 | AC124016,2 | 0,013748805 | 1,0095755 | 0,97107 | 0,98773415 | novel transcript |
| ENSG00000108839 | ALOX12 | 0,013721972 | 1,009556723 | 0,97824 | 0,99099142 | arachidonate 12-lipoxygenase, 12S type |
| ENSG00000077157 | PPP1R12B | 0,013717945 | 1,009553905 | 0,93407 | 0,97204024 | protein phosphatase 1 regulatory subunit 12B |
| ENSG00000167969 | ECI1 | 0,013704015 | 1,009544157 | 0,94912 | 0,97866986 | enoyl-CoA delta isomerase 1 |
| ENSG00000243224 | AC006252,1 | 0,013689377 | 1,009533914 | 0,97935 | 0,99154627 | novel transcript, antisense to TWF2 |
| ENSG00000183496 | MEX3B | 0,013667815 | 1,009518826 | 0,92849 | 0,969217 | mex-3 RNA binding family member B |
| ENSG00000100211 | CBY1 | 0,013665018 | 1,009516869 | 0,94029 | 0,97464787 | chibby family member 1, beta catenin antagonist |
| ENSG00000136286 | MYO1G | 0,01365124 | 1,009507228 | 0,98069 | 0,99204224 | myosin IG |
| ENSG00000088448 | ANKRD10 | 0,013354237 | 1,009299425 | 0,94259 | 0,97560255 | ankyrin repeat domain 10 |
| ENSG00000177374 | HIC1 | 0,013342063 | 1,009290908 | 0,98022 | 0,99188844 | HIC ZBTB transcriptional repressor 1 |
| ENSG00000176383 | B3GNT4 | 0,013298052 | 1,009260119 | 0,96973 | 0,98725952 | UDP-GlcNAc:betaGal beta-1,3-N-acetylglucosaminyltransferase 4 |
| ENSG00000132781 | MUTYH | 0,0132856 | 1,009251408 | 0,95803 | 0,98276081 | mutY DNA glycosylase |
| ENSG00000165264 | NDUFB6 | 0,013171995 | 1,009171938 | 0,94464 | 0,97650311 | NADH:ubiquinone oxidoreductase subunit B6 |
| ENSG00000181449 | SOX2 | 0,013125874 | 1,009139677 | 0,96937 | 0,98709824 | SRY-box transcription factor 2 |
| ENSG00000090776 | EFNB1 | 0,012932551 | 1,00900446 | 0,96552 | 0,9854196 | ephrin B1 |
| ENSG00000288640 | AC005192,1 | 0,012886997 | 1,0089726 | 0,98456 | 0,99331785 | novel protein |
| ENSG00000272606 | AC015982,2 | 0,012861064 | 1,008954464 | 0,97998 | 0,9917697 | novel transcript |
| ENSG00000236144 | TMEM147-AS | 0,012816071 | 1,008922998 | 0,94946 | 0,97888896 | TMEM147 antisense RNA 1 |
| ENSG00000095539 | SEMA4G | 0,012806651 | 1,008916411 | 0,94571 | 0,97689657 | semaphorin 4G |
| ENSG00000232228 | RPL36AP16 | 0,012793089 | 1,008906926 | 0,98865 | 0,99502165 | ribosomal protein L36a pseudogene 16 |
| ENSG00000249790 | AC092490,1 | 0,012613729 | 1,008781504 | 0,96983 | 0,98728333 | novel transcript |
| ENSG00000171163 | ZNF692 | 0,012596496 | 1,008769454 | 0,9545 | 0,98095268 | zinc finger protein 692 |
| ENSG00000145990 | GFOD1 | 0,012538473 | 1,008728884 | 0,9761 | 0,98994851 | glucose-fructose oxidoreductase domain containing 1 |
| ENSG00000200087 | SNORA73B | 0,012459264 | 1,008673502 | 0,95368 | 0,9805199 | small nucleolar RNA, H/ACA box 73B |
| ENSG00000271639 | PPFIA1P1 | 0,012433246 | 1,008655312 | 0,98302 | 0,9926386 | PPFIA1 pseudogene 1 |
| ENSG00000228335 | AC073063,1 | 0,012395629 | 1,008629013 | 0,98203 | 0,99244329 | cytokine induced protein 29 kDa (CIP29) pseudogene |
| ENSG00000198799 | LRIG2 | 0,01231868 | 1,008575217 | 0,94594 | 0,97689657 | leucine rich repeats and immunoglobulin like domains 2 |
| ENSG00000105248 | YJU2 | 0,012295915 | 1,008559302 | 0,95976 | 0,98338726 | YJU2 splicing factor homolog |

| | | | | | | |
|---|---|---|---|---|---|---|
| ENSG00000196814 | MVB12B | 0,012178238 | 1,00847704 | 0,9576 | 0,98257839 | multivesicular body subunit 12B |
| ENSG00000136717 | BIN1 | 0,012085265 | 1,008412052 | 0,96296 | 0,98432594 | bridging integrator 1 |
| ENSG00000068394 | GPKOW | 0,012044467 | 1,008383535 | 0,94139 | 0,97504246 | G-patch domain and KOW motifs |
| ENSG00000288055 | AC107373,3 | 0,012011851 | 1,008360738 | 0,98245 | 0,99246154 | novel transcript |
| ENSG00000160318 | CLDND2 | 0,011911849 | 1,008290845 | 0,98776 | 0,99464284 | claudin domain containing 2 |
| ENSG00000251629 | LINC02241 | 0,011884458 | 1,008271702 | 0,98259 | 0,99246154 | long intergenic non-protein coding RNA 2241 |
| ENSG00000124116 | WFDC3 | 0,011728716 | 1,008162863 | 0,98963 | 0,99572035 | WAP four-disulfide core domain 3 |
| ENSG00000273302 | AC016747,3 | 0,011663729 | 1,00811745 | 0,97571 | 0,9897621 | novel transcript |
| ENSG00000271382 | AC012464,3 | 0,011600918 | 1,008073561 | 0,98818 | 0,99471957 | novel transcript |
| ENSG00000169314 | C22orf15 | 0,011568219 | 1,008050713 | 0,98533 | 0,99383022 | chromosome 22 open reading frame 15 |
| ENSG00000175189 | INHBC | 0,01154667 | 1,008035655 | 0,98333 | 0,99281596 | inhibin subunit beta C |
| ENSG00000186350 | RXRA | 0,011470473 | 1,007982417 | 0,96449 | 0,98495288 | retinoid X receptor alpha |
| ENSG00000278119 | GOLGA8J | 0,011412788 | 1,007942114 | 0,98236 | 0,99244329 | golgin A8 family member J |
| ENSG00000254346 | MTND1P6 | 0,011172171 | 1,007774021 | 0,98659 | 0,99423972 | MT-ND1 pseudogene 6 |
| ENSG00000250031 | AC009927,1 | 0,011160719 | 1,007766021 | 0,98034 | 0,99195802 | protein tyrosine phosphatase, non-receptor type 11 (PTPN11) pseudogene |
| ENSG00000153132 | CLGN | 0,011128592 | 1,00774358 | 0,97588 | 0,98986897 | calmegin |
| ENSG00000138032 | PPM1B | 0,011096357 | 1,007721064 | 0,95058 | 0,97934063 | protein phosphatase, Mg2+/Mn2+ dependent 1B |
| ENSG00000126581 | BECN1 | 0,011028612 | 1,007673745 | 0,94743 | 0,97781662 | beclin 1 |
| ENSG00000267338 | AP000365,1 | 0,010999624 | 1,007653498 | 0,98833 | 0,99477886 | novel transcript |
| ENSG00000287608 | AC022137,3 | 0,010992866 | 1,007648778 | 0,97529 | 0,98956405 | novel transcript, antisense to ZNF761and ZNF813 |
| ENSG00000172046 | USP19 | 0,010958255 | 1,007624604 | 0,94769 | 0,97798027 | ubiquitin specific peptidase 19 |
| ENSG00000237978 | KCNMB2-AS1 | 0,010956946 | 1,00762369 | 0,9725 | 0,98835697 | KCNMB2 antisense RNA 1 |
| ENSG00000196388 | INCA1 | 0,010870196 | 1,007563103 | 0,97619 | 0,98994851 | inhibitor of CDK, cyclin A1 interacting protein 1 |
| ENSG00000071054 | MAP4K4 | 0,010863113 | 1,007558156 | 0,96259 | 0,98430469 | mitogen-activated protein kinase kinase kinase kinase 4 |
| ENSG00000198440 | ZNF583 | 0,010830852 | 1,007535625 | 0,95898 | 0,98304779 | zinc finger protein 583 |
| ENSG00000187838 | PLSCR3 | 0,010799472 | 1,007513711 | 0,96386 | 0,98461596 | phospholipid scramblase 3 |
| ENSG00000284009 | PLSCR3 | 0,010799472 | 1,007513711 | 0,96386 | 0,98461596 | phospholipid scramblase 3 |
| ENSG00000271141 | AC010680,4 | 0,010769219 | 1,007492584 | 0,98344 | 0,99281596 | novel transcript |
| ENSG00000248905 | FMN1 | 0,010768174 | 1,007491854 | 0,98727 | 0,99446724 | formin 1 |
| ENSG00000248713 | C4orf54 | 0,010752819 | 1,007481131 | 0,98734 | 0,99446724 | chromosome 4 open reading frame 54 |
| ENSG00000198821 | CD247 | 0,010635586 | 1,007399267 | 0,9851 | 0,9937454 | CD247 molecule |
| ENSG00000229692 | SOS1-IT1 | 0,010537868 | 1,007331035 | 0,97309 | 0,9886748 | SOS1 intronic transcript 1 |

| | | | | | | |
|---|---|---|---|---|---|---|
| ENSG00000250366 | TUNAR | 0,010509132 | 1,007310971 | 0,97054 | 0,98754526 | TCL1 upstream neural differentiation-associated RNA |
| ENSG00000135643 | KCNMB4 | 0,010439698 | 1,007262492 | 0,96247 | 0,98429303 | potassium calcium-activated channel subfamily M regulatory beta subunit 4 |
| ENSG00000164733 | CTSB | 0,010420566 | 1,007249135 | 0,96488 | 0,98525685 | cathepsin B |
| ENSG00000287220 | AL590138,1 | 0,010268096 | 1,00714269 | 0,98622 | 0,99415471 | novel transcript, antisense to PLXNA2 |
| ENSG00000084072 | PPIE | 0,010243645 | 1,007125621 | 0,95034 | 0,97927894 | peptidylprolyl isomerase E |
| ENSG00000171033 | PKIA | 0,010229196 | 1,007115534 | 0,98351 | 0,99281596 | cAMP-dependent protein kinase inhibitor alpha |
| ENSG00000233471 | KRT18P62 | 0,010210529 | 1,007102504 | 0,98977 | 0,99572035 | keratin 18 pseudogene 62 |
| ENSG00000119878 | CRIPT | 0,010203462 | 1,00709757 | 0,95055 | 0,97934063 | CXXC repeat containing interactor of PDZ3 domain |
| ENSG00000205220 | PSMB10 | 0,010159171 | 1,007066653 | 0,9734 | 0,98882674 | proteasome 20S subunit beta 10 |
| ENSG00000235760 | MSH2-OT1 | 0,010142176 | 1,00705479 | 0,98444 | 0,99330857 | MSH2 overlapping transcript 1 |
| ENSG00000196214 | ZNF766 | 0,01011141 | 1,007033313 | 0,94626 | 0,97711344 | zinc finger protein 766 |
| ENSG00000079841 | RIMS1 | 0,010105751 | 1,007029364 | 0,97868 | 0,99131549 | regulating synaptic membrane exocytosis 1 |
| ENSG00000112812 | PRSS16 | 0,010089669 | 1,007018138 | 0,983 | 0,9926386 | serine protease 16 |
| ENSG00000170525 | PFKFB3 | 0,010035689 | 1,00698046 | 0,95965 | 0,98338726 | 6-phosphofructo-2-kinase/fructose-2,6-biphosphatase 3 |
| ENSG00000117308 | GALE | 0,01002167 | 1,006970675 | 0,95877 | 0,98291816 | UDP-galactose-4-epimerase |
| ENSG00000197217 | ENTPD4 | 0,009941816 | 1,00691494 | 0,9544 | 0,98093594 | ectonucleoside triphosphate diphosphohydrolase 4 |
| ENSG00000228327 | AL669831,1 | 0,009883922 | 1,006874534 | 0,96244 | 0,98429217 | general transcription factor IIi (GTF2I) pseudogene |
| ENSG00000092421 | SEMA6A | 0,009880319 | 1,00687202 | 0,97647 | 0,9901323 | semaphorin 6A |
| ENSG00000259321 | AL136295,2 | 0,009873899 | 1,006867539 | 0,98328 | 0,99281596 | novel transcript |
| ENSG00000285298 | AL136295,16 | 0,009873899 | 1,006867539 | 0,98328 | 0,99281596 | novel transcript |
| ENSG00000113209 | PCDHB5 | 0,00979021 | 1,006809134 | 0,96554 | 0,9854196 | protocadherin beta 5 |
| ENSG00000169031 | COL4A3 | 0,009733366 | 1,006769465 | 0,98556 | 0,9940276 | collagen type IV alpha 3 chain |
| ENSG00000280787 | HERC2P2 | 0,009713973 | 1,006755932 | 0,9975 | 0,99910705 | hect domain and RLD 2 pseudogene 2 |
| ENSG00000136485 | DCAF7 | 0,009597326 | 1,006674536 | 0,96181 | 0,98401207 | DDB1 and CUL4 associated factor 7 |
| ENSG00000106443 | PHF14 | 0,009564318 | 1,006651503 | 0,95956 | 0,98338726 | PHD finger protein 14 |
| ENSG00000117335 | CD46 | 0,009528796 | 1,006626718 | 0,95778 | 0,98261326 | CD46 molecule |
| ENSG00000229794 | MTCYBP32 | 0,009471594 | 1,006586807 | 0,98984 | 0,99573375 | MT-CYB pseudogene 32 |
| ENSG00000137274 | BPHL | 0,009463581 | 1,006581216 | 0,95003 | 0,97913156 | biphenyl hydrolase like |
| ENSG00000233276 | GPX1 | 0,009458702 | 1,006577812 | 0,96775 | 0,98623935 | glutathione peroxidase 1 |
| ENSG00000178149 | DALRD3 | 0,009258142 | 1,00643789 | 0,96595 | 0,98557781 | DALR anticodon binding domain containing 3 |
| ENSG00000171877 | FRMD5 | 0,009256942 | 1,006437052 | 0,97277 | 0,98855428 | FERM domain containing 5 |
| ENSG00000204842 | ATXN2 | 0,009067011 | 1,006304564 | 0,96388 | 0,98461596 | ataxin 2 |

| | | | | | | |
|---|---|---|---|---|---|---|
| ENSG00000136247 | ZDHHC4 | 0,008930358 | 1,00620925 | 0,97457 | 0,98933402 | zinc finger DHHC-type palmitoyltransferase 4 |
| ENSG00000280594 | BTG3-AS1 | 0,008859748 | 1,006160005 | 0,97695 | 0,99033775 | BTG3 antisense RNA 1 |
| ENSG00000281317 | BTG3-AS1 | 0,008859748 | 1,006160005 | 0,97695 | 0,99033775 | BTG3 antisense RNA 1 |
| ENSG00000169689 | CENPX | 0,008803262 | 1,006120611 | 0,96944 | 0,98713404 | centromere protein X |
| ENSG00000234539 | ATF6B | 0,008766166 | 1,006094741 | 0,9631 | 0,98432594 | activating transcription factor 6 beta |
| ENSG00000169302 | STK32A | 0,008760096 | 1,006090508 | 0,98457 | 0,99331785 | serine/threonine kinase 32A |
| ENSG00000110876 | SELPLG | 0,008740334 | 1,006076727 | 0,98795 | 0,99471733 | selectin P ligand |
| ENSG00000226124 | FTCDNL1 | 0,008712777 | 1,00605751 | 0,9919 | 0,99651645 | formiminotransferase cyclodeaminase N-terminal like |
| ENSG00000251045 | SLC25A48-AS1 | 0,008612898 | 1,005987862 | 0,99236 | 0,99669518 | SLC25A48 antisense RNA 1 |
| ENSG00000136243 | NUP42 | 0,008560382 | 1,005951243 | 0,97179 | 0,98800339 | nucleoporin 42 |
| ENSG00000099917 | MED15 | 0,008557389 | 1,005949157 | 0,9635 | 0,98448919 | mediator complex subunit 15 |
| ENSG00000214026 | MRPL23 | 0,0085001 | 1,005909211 | 0,96993 | 0,98728333 | mitochondrial ribosomal protein L23 |
| ENSG00000288141 | MRPL23 | 0,0085001 | 1,005909211 | 0,96993 | 0,98728333 | mitochondrial ribosomal protein L23 |
| ENSG00000260161 | MTCYBP28 | 0,008471007 | 1,005888927 | 0,99228 | 0,99668951 | MT-CYB pseudogene 28 |
| ENSG00000234281 | LANCL1-AS1 | 0,008315961 | 1,00578083 | 0,98671 | 0,99425859 | LANCL1 antisense RNA 1 |
| ENSG00000255524 | NPIPB8 | 0,008306063 | 1,005773929 | 0,98904 | 0,99524652 | nuclear pore complex interacting protein family member B8 |
| ENSG00000282917 | AC107068,2 | 0,008305143 | 1,005773288 | 0,98259 | 0,99246154 | novel transcript, antisense to CORIN and NFXL1 |
| ENSG00000226833 | AC092164,1 | 0,008255006 | 1,005738335 | 0,97304 | 0,9886748 | novel transcript |
| ENSG00000116288 | PARK7 | 0,008244437 | 1,005730968 | 0,96515 | 0,98539831 | Parkinsonism associated deglycase |
| ENSG00000151743 | AMN1 | 0,008149062 | 1,005664483 | 0,97416 | 0,98928341 | antagonist of mitotic exit network 1 homolog |
| ENSG00000214063 | TSPAN4 | 0,008093087 | 1,005625464 | 0,97696 | 0,99033775 | tetraspanin 4 |
| ENSG00000110171 | TRIM3 | 0,008021547 | 1,005575599 | 0,97006 | 0,9873786 | tripartite motif containing 3 |
| ENSG00000170271 | FAXDC2 | 0,007991629 | 1,005554746 | 0,97442 | 0,98933402 | fatty acid hydroxylase domain containing 2 |
| ENSG00000257923 | CUX1 | 0,007923039 | 1,00550694 | 0,96927 | 0,98707447 | cut like homeobox 1 |
| ENSG00000142552 | RCN3 | 0,007832955 | 1,005444156 | 0,97759 | 0,99069536 | reticulocalbin 3 |
| ENSG00000196476 | C20orf96 | 0,00778977 | 1,005414061 | 0,97974 | 0,99170352 | chromosome 20 open reading frame 96 |
| ENSG00000067900 | ROCK1 | 0,00777909 | 1,005406618 | 0,96542 | 0,9854196 | Rho associated coiled-coil containing protein kinase 1 |
| ENSG00000261502 | AC040174,1 | 0,007748265 | 1,005385136 | 0,9924 | 0,99670212 | novel transcript |
| ENSG00000130958 | SLC35D2 | 0,007731678 | 1,005373577 | 0,96859 | 0,98658951 | solute carrier family 35 member D2 |
| ENSG00000248616 | ANKRD49P3 | 0,007714213 | 1,005361406 | 0,99314 | 0,99690338 | ANKRD49 pseudogene 3 |
| ENSG00000175416 | CLTB | 0,007677555 | 1,005335861 | 0,97618 | 0,98994851 | clathrin light chain B |
| ENSG00000184378 | ACTRT3 | 0,007622276 | 1,005297341 | 0,98515 | 0,99375271 | actin related protein T3 |

| | | | | | | |
|---|---|---|---|---|---|---|
| ENSG00000243660 | ZNF487 | 0,00755757 | 1,005252253 | 0,97135 | 0,98775885 | zinc finger protein 487 |
| ENSG00000164309 | CMYA5 | 0,007539051 | 1,00523935 | 0,98411 | 0,99310064 | cardiomyopathy associated 5 |
| ENSG00000277701 | AC159540,2 | 0,007522857 | 1,005228066 | 0,98145 | 0,99239184 | novel transcript |
| ENSG00000257060 | AC091078,1 | 0,007520166 | 1,005226191 | 0,96733 | 0,98623935 | novel transcript |
| ENSG00000127578 | WFIKKN1 | 0,007437017 | 1,005168257 | 0,9881 | 0,99471957 | WAP, follistatin/kazal, immunoglobulin, kunitz and netrin domain containing 1 |
| ENSG00000104687 | GSR | 0,007416979 | 1,005154296 | 0,96931 | 0,98708123 | glutathione-disulfide reductase |
| ENSG00000101871 | MID1 | 0,00738854 | 1,005134482 | 0,98246 | 0,99246154 | midline 1 |
| ENSG00000112130 | RNF8 | 0,007305204 | 1,005076423 | 0,96363 | 0,9845058 | ring finger protein 8 |
| ENSG00000182700 | IGIP | 0,00728295 | 1,00506092 | 0,98689 | 0,99437646 | IgA inducing protein |
| ENSG00000188321 | ZNF559 | 0,007250057 | 1,005038005 | 0,96847 | 0,98654786 | zinc finger protein 559 |
| ENSG00000116478 | HDAC1 | 0,007193821 | 1,004998829 | 0,97394 | 0,98924139 | histone deacetylase 1 |
| ENSG00000244754 | N4BP2L2 | 0,007184142 | 1,004992087 | 0,9671 | 0,98623935 | NEDD4 binding protein 2 like 2 |
| ENSG00000168393 | DTYMK | 0,007149793 | 1,004968159 | 0,96649 | 0,98588576 | deoxythymidylate kinase |
| ENSG00000165113 | GKAP1 | 0,007148544 | 1,004967289 | 0,98212 | 0,99244329 | G kinase anchoring protein 1 |
| ENSG00000274089 | PRIMA1 | 0,007111596 | 1,004941552 | 0,98654 | 0,99423097 | proline rich membrane anchor 1 |
| ENSG00000175785 | PRIMA1 | 0,007111596 | 1,004941552 | 0,98654 | 0,99423097 | proline rich membrane anchor 1 |
| ENSG00000282537 | TAS2R43 | 0,007026618 | 1,004882361 | 0,98607 | 0,99413637 | taste 2 receptor member 43 |
| ENSG00000261936 | TAS2R45 | 0,007026618 | 1,004882361 | 0,98607 | 0,99413637 | taste 2 receptor member 45 |
| ENSG00000177984 | LCN15 | 0,007022135 | 1,004879238 | 0,99098 | 0,99631601 | lipocalin 15 |
| ENSG00000116675 | DNAJC6 | 0,006880882 | 1,004780856 | 0,98596 | 0,99413637 | DnaJ heat shock protein family (Hsp40) member C6 |
| ENSG00000160796 | NBEAL2 | 0,006865144 | 1,004769895 | 0,98356 | 0,99283272 | neurobeachin like 2 |
| ENSG00000007047 | MARK4 | 0,006863252 | 1,004768577 | 0,97209 | 0,98800339 | microtubule affinity regulating kinase 4 |
| ENSG00000230847 | OCLNP1 | 0,006797599 | 1,004722854 | 0,99333 | 0,99693507 | OCLN pseudogene 1 |
| ENSG00000049167 | ERCC8 | 0,006758095 | 1,004695343 | 0,97094 | 0,98771801 | ERCC excision repair 8, CSA ubiquitin ligase complex subunit |
| ENSG00000172456 | FGGY | 0,006682794 | 1,004642905 | 0,98064 | 0,9920229 | FGGY carbohydrate kinase domain containing |
| ENSG00000266100 | AC007431,2 | 0,006627905 | 1,004604683 | 0,99346 | 0,99694625 | novel transcript, antisense to MSI2 |
| ENSG00000276180 | H4C9 | 0,006612885 | 1,004594224 | 0,97523 | 0,98956405 | H4 clustered histone 9 |
| ENSG00000113742 | CPEB4 | 0,00655547 | 1,004554245 | 0,9794 | 0,99154627 | cytoplasmic polyadenylation element binding protein 4 |
| ENSG00000234432 | AC092171,3 | 0,006516299 | 1,00452697 | 0,98583 | 0,99413637 | uncharacterized LOC100129484 |
| ENSG00000215126 | CBWD6 | 0,006504286 | 1,004518606 | 0,98177 | 0,99242247 | COBW domain containing 6 |
| ENSG00000104112 | SCG3 | 0,006415791 | 1,004456991 | 0,98948 | 0,99562246 | secretogranin III |

| | | | | | | |
|---|---|---|---|---|---|---|
| ENSG00000281151 | KCNQ2 | 0,006259374 | 1,004348093 | 0,98881 | 0,9950859 | potassium voltage-gated channel subfamily Q member 2 |
| ENSG00000261159 | AC112484,3 | 0,006174715 | 1,004289159 | 0,9863 | 0,99417149 | novel transcript |
| ENSG00000231084 | RPL22P24 | 0,00612453 | 1,004254224 | 0,99198 | 0,99651645 | ribosomal protein L22 pseudogene 24 |
| ENSG00000229962 | AP000221,1 | 0,006082883 | 1,004225234 | 0,99508 | 0,99799225 | novel transcript |
| ENSG00000163412 | EIF4E3 | 0,006056861 | 1,004207121 | 0,98612 | 0,99413637 | eukaryotic translation initiation factor 4E family member 3 |
| ENSG00000158321 | AUTS2 | 0,006036765 | 1,004193133 | 0,97979 | 0,99170352 | activator of transcription and developmental regulator AUTS2 |
| ENSG00000286123 | AL049828,2 | 0,006033084 | 1,004190571 | 0,99551 | 0,99812418 | novel transcript |
| ENSG00000229488 | PRRT1 | 0,005990066 | 1,004160629 | 0,98971 | 0,99572035 | proline rich transmembrane protein 1 |
| ENSG00000082458 | DLG3 | 0,005769927 | 1,004007417 | 0,98183 | 0,99242247 | discs large MAGUK scaffold protein 3 |
| ENSG00000159792 | PSKH1 | 0,005734105 | 1,003982488 | 0,97926 | 0,99154627 | protein serine kinase H1 |
| ENSG00000141504 | SAT2 | 0,005711259 | 1,003966589 | 0,97438 | 0,98933402 | spermidine/spermine N1-acetyltransferase family member 2 |
| ENSG00000196604 | POTEF | 0,005556162 | 1,003858663 | 0,97454 | 0,98933402 | POTE ankyrin domain family member F |
| ENSG00000158246 | TENT5B | 0,005524263 | 1,003836468 | 0,98704 | 0,99446724 | terminal nucleotidyltransferase 5B |
| ENSG00000276464 | POMZP3 | 0,005522055 | 1,003834931 | 0,97822 | 0,99099142 | POM121 and ZP3 fusion |
| ENSG00000153086 | ACMSD | 0,005517114 | 1,003831493 | 0,99255 | 0,99670754 | aminocarboxymuconate semialdehyde decarboxylase |
| ENSG00000113749 | HRH2 | 0,005440223 | 1,003777994 | 0,98569 | 0,99412365 | histamine receptor H2 |
| ENSG00000189212 | DPY19L2P1 | 0,005391169 | 1,003743864 | 0,99033 | 0,99596348 | DPY19L2 pseudogene 1 |
| ENSG00000154277 | UCHL1 | 0,005352643 | 1,003717061 | 0,98173 | 0,99242247 | ubiquitin C-terminal hydrolase L1 |
| ENSG00000229214 | LINC00242 | 0,005253431 | 1,003648039 | 0,9869 | 0,99437646 | long intergenic non-protein coding RNA 242 |
| ENSG00000145996 | CDKAL1 | 0,005197309 | 1,003608997 | 0,97535 | 0,98956832 | CDK5 regulatory subunit associated protein 1 like 1 |
| ENSG00000099949 | LZTR1 | 0,005146161 | 1,003573417 | 0,97727 | 0,99051207 | leucine zipper like transcription regulator 1 |
| ENSG00000041988 | THAP3 | 0,005135236 | 1,003565817 | 0,98041 | 0,99195802 | THAP domain containing 3 |
| ENSG00000238825 | RNVU1-2 | 0,005117489 | 1,003553472 | 0,99636 | 0,99859001 | RNA, variant U1 small nuclear 2 |
| ENSG00000178896 | EXOSC4 | 0,005083178 | 1,003529605 | 0,98 | 0,9917697 | exosome component 4 |
| ENSG00000134376 | CRB1 | 0,005023158 | 1,003487857 | 0,99182 | 0,99651645 | crumbs cell polarity complex component 1 |
| ENSG00000100583 | SAMD15 | 0,004958788 | 1,003443084 | 0,98523 | 0,99379729 | sterile alpha motif domain containing 15 |
| ENSG00000151365 | THRSP | 0,004879132 | 1,003387682 | 0,99293 | 0,99678701 | thyroid hormone responsive |
| ENSG00000009954 | BAZ1B | 0,004876547 | 1,003385884 | 0,98202 | 0,99244329 | bromodomain adjacent to zinc finger domain 1B |
| ENSG00000229622 | MTND5P2 | 0,004772685 | 1,003313651 | 0,99518 | 0,99805801 | MT-ND5 pseudogene 2 |
| ENSG00000171503 | ETFDH | 0,004720925 | 1,003277656 | 0,98454 | 0,99331785 | electron transfer flavoprotein dehydrogenase |
| ENSG00000285007 | REC8 | 0,004532769 | 1,003146817 | 0,9837 | 0,99290113 | REC8 meiotic recombination protein |

| | | | | | | |
|---|---|---|---|---|---|---|
| ENSG00000100918 | REC8 | 0,004532769 | 1,003146817 | 0,9837 | 0,99290113 | REC8 meiotic recombination protein |
| ENSG00000183255 | PTTG1IP | 0,004481221 | 1,003110975 | 0,97816 | 0,99099142 | PTTG1 interacting protein |
| ENSG00000159733 | ZFYVE28 | 0,004291627 | 1,002979158 | 0,99451 | 0,99763557 | zinc finger FYVE-type containing 28 |
| ENSG00000012171 | SEMA3B | 0,004262315 | 1,00295878 | 0,99295 | 0,99678701 | semaphorin 3B |
| ENSG00000001497 | LAS1L | 0,004052675 | 1,00281305 | 0,98343 | 0,99281596 | LAS1 like ribosome biogenesis factor |
| ENSG00000272817 | AL359198,1 | 0,004028984 | 1,002796582 | 0,99274 | 0,99671114 | novel transcript, antisense to FGFBP3 |
| ENSG00000131187 | F12 | 0,003962481 | 1,002750358 | 0,99612 | 0,99850694 | coagulation factor XII |
| ENSG00000111254 | AKAP3 | 0,003865804 | 1,002683165 | 0,98815 | 0,99471957 | A-kinase anchoring protein 3 |
| ENSG00000288490 | GALNT9 | 0,00373147 | 1,002589806 | 0,99673 | 0,9987577 | polypeptide N-acetylgalactosaminyltransferase 9 |
| ENSG00000231817 | LINC01198 | 0,003703481 | 1,002570355 | 0,98979 | 0,99572035 | long intergenic non-protein coding RNA 1198 |
| ENSG00000285993 | AC018931,1 | 0,003615411 | 1,002509155 | 0,99474 | 0,99772791 | novel transcript, antisense to RNF165 |
| ENSG00000227863 | AC002383,1 | 0,003613396 | 1,002507755 | 0,99333 | 0,99693507 | novel transcript |
| ENSG00000133401 | PDZD2 | 0,003520563 | 1,002443248 | 0,99356 | 0,99696919 | PDZ domain containing 2 |
| ENSG00000157570 | TSPAN18 | 0,003485325 | 1,002418763 | 0,98476 | 0,99343186 | tetraspanin 18 |
| ENSG00000125912 | NCLN | 0,003371022 | 1,002339346 | 0,99121 | 0,99634744 | nicalin |
| ENSG00000102349 | KLF8 | 0,003355031 | 1,002328236 | 0,9918 | 0,99651645 | Kruppel like factor 8 |
| ENSG00000126860 | EVI2A | 0,003145373 | 1,002182584 | 0,99255 | 0,99670754 | ecotropic viral integration site 2A |
| ENSG00000286807 | AC093639,2 | 0,003093118 | 1,002146286 | 0,99649 | 0,99859001 | novel transcript |
| ENSG00000183206 | POTEC | 0,003081353 | 1,002138114 | 0,99309 | 0,99688937 | POTE ankyrin domain family member C |
| ENSG00000207375 | SNORD116-23 | 0,003069697 | 1,002130017 | 0,99728 | 0,998989 | small nucleolar RNA, C/D box 116-23 |
| ENSG00000080603 | SRCAP | 0,002888 | 1,002003814 | 0,98829 | 0,99477886 | Snf2 related CREBBP activator protein |
| ENSG00000164096 | C4orf3 | 0,002664798 | 1,001848804 | 0,98763 | 0,99461882 | chromosome 4 open reading frame 3 |
| ENSG00000197238 | H4C11 | 0,002612053 | 1,001812177 | 0,99393 | 0,99726631 | H4 clustered histone 11 |
| ENSG00000054654 | SYNE2 | 0,002581212 | 1,001790761 | 0,99274 | 0,99671114 | spectrin repeat containing nuclear envelope protein 2 |
| ENSG00000134318 | ROCK2 | 0,002573605 | 1,001785479 | 0,99153 | 0,99649731 | Rho associated coiled-coil containing protein kinase 2 |
| ENSG00000165506 | DNAAF2 | 0,002485582 | 1,001724359 | 0,9925 | 0,99670754 | dynein axonemal assembly factor 2 |
| ENSG00000242999 | RN7SL239P | 0,002414155 | 1,001674765 | 0,9977 | 0,99923163 | RNA, 7SL, cytoplasmic 239, pseudogene |
| ENSG00000263219 | RYKP1 | 0,002394861 | 1,00166137 | 0,99542 | 0,99812418 | RYK pseudogene 1 |
| ENSG00000162971 | TYW5 | 0,002355943 | 1,001634349 | 0,98933 | 0,99549932 | tRNA-yW synthesizing protein 5 |
| ENSG00000248932 | AC046134,2 | 0,002316841 | 1,001607202 | 0,98991 | 0,99574641 | novel transcript, antisense to RBP2, RBP1 & NMNAT3 |
| ENSG00000135919 | SERPINE2 | 0,002303903 | 1,00159822 | 0,98993 | 0,99574641 | serpin family E member 2 |
| ENSG00000140280 | LYSMD2 | 0,002205736 | 1,001530069 | 0,99648 | 0,99859001 | LysM domain containing 2 |
| ENSG00000164053 | ATRIP | 0,001986705 | 1,001378027 | 0,99103 | 0,99632521 | ATR interacting protein |

| | | | | | | |
|---|---|---|---|---|---|---|
| ENSG00000237094 | AL732372,2 | 0,001929983 | 1,001338657 | 0,99204 | 0,99651645 | pseudogene similar to part of septin 14 SEPT14 |
| ENSG00000139540 | SLC39A5 | 0,001909343 | 1,001324332 | 0,99719 | 0,99897141 | solute carrier family 39 member 5 |
| ENSG00000039523 | RIPOR1 | 0,001886091 | 1,001308193 | 0,99218 | 0,99661925 | RHO family interacting cell polarization regulator 1 |
| ENSG00000109686 | SH3D19 | 0,001711965 | 1,001187348 | 0,99641 | 0,99859001 | SH3 domain containing 19 |
| ENSG00000186205 | MTARC1 | 0,001699035 | 1,001178375 | 0,99356 | 0,99696919 | mitochondrial amidoxime reducing component 1 |
| ENSG00000113761 | ZNF346 | 0,001651921 | 1,00114568 | 0,99197 | 0,99651645 | zinc finger protein 346 |
| ENSG00000253596 | AC103952,1 | 0,001557995 | 1,001080503 | 0,99842 | 0,99956168 | novel transcript |
| ENSG00000282084 | UBE2Q2P2 | 0,001525272 | 1,001057797 | 0,99404 | 0,99731406 | ubiquitin conjugating enzyme E2 Q2 pseudogene 2 |
| ENSG00000263956 | NBPF11 | 0,001453069 | 1,001007698 | 0,99277 | 0,99671114 | NBPF member 11 |
| ENSG00000147316 | MCPH1 | 0,001425917 | 1,000988859 | 0,99627 | 0,99858544 | microcephalin 1 |
| ENSG00000065413 | ANKRD44 | 0,00132621 | 1,000919682 | 0,99618 | 0,99852746 | ankyrin repeat domain 44 |
| ENSG00000100068 | LRP5L | 0,001304507 | 1,000904624 | 0,99649 | 0,99859001 | LDL receptor related protein 5 like |
| ENSG00000282933 | RHOXF1P3 | 0,001225776 | 1,000850004 | 0,99825 | 0,99947928 | Rhox homeobox family member 1 pseudogene 3 |
| ENSG00000020256 | ZFP64 | 0,001132844 | 1,000785536 | 0,99465 | 0,99769817 | ZFP64 zinc finger protein |
| ENSG00000034713 | GABARAPL2 | 0,001085085 | 1,000752406 | 0,99558 | 0,99812418 | GABA type A receptor associated protein like 2 |
| ENSG00000260464 | AL049796,1 | 0,001035963 | 1,000718333 | 0,99827 | 0,99947928 | novel transcript |
| ENSG00000126107 | HECTD3 | 0,000981976 | 1,000680885 | 0,9958 | 0,9982573 | HECT domain E3 ubiquitin protein ligase 3 |
| ENSG00000232818 | RPS2P32 | 0,000904579 | 1,000627203 | 0,99813 | 0,99945242 | ribosomal protein S2 pseudogene 32 |
| ENSG00000261764 | KRT18P18 | 0,00089261 | 1,000618902 | 0,99877 | 0,99970131 | keratin 18 pseudogene 18 |
| ENSG00000116497 | S100PBP | 0,000787299 | 1,000545863 | 0,99557 | 0,99812418 | S100P binding protein |
| ENSG00000286895 | AC092745,5 | 0,000752443 | 1,00052169 | 0,99859 | 0,99960886 | novel transcript |
| ENSG00000241599 | AL627309,4 | 0,000715881 | 1,000496334 | 0,99912 | 0,99978149 | novel transcript |
| ENSG00000139579 | NABP2 | 0,000610389 | 1,000423179 | 0,99726 | 0,998989 | nucleic acid binding protein 2 |
| ENSG00000286311 | AL034550,3 | 0,000463597 | 1,000321393 | 0,9993 | 0,99987765 | novel transcript, antisense to NOL4L |
| ENSG00000226225 | RPS18 | 0,000387196 | 1,00026842 | 0,99903 | 0,999764 | ribosomal protein S18 |
| ENSG00000273350 | AC004832,5 | 0,000350096 | 1,000242697 | 0,99949 | 0,99994283 | novel transcript, sense intronic to SEC14L2 |
| ENSG00000275932 | DUSP14 | 0,000318827 | 1,000221019 | 0,99934 | 0,99987765 | dual specificity phosphatase 14 |
| ENSG00000180104 | EXOC3 | 0,000311364 | 1,000215844 | 0,99857 | 0,99960886 | exocyst complex component 3 |
| ENSG00000185920 | PTCH1 | 0,000275637 | 1,000191076 | 0,99931 | 0,99987765 | patched 1 |
| ENSG00000213871 | TAF9BP1 | 0,000275111 | 1,000190711 | 0,99958 | 0,99994283 | TATA-box binding protein associated factor 9b pseudogene 1 |
| ENSG00000105576 | TNPO2 | 0,000234134 | 1,000162302 | 0,99892 | 0,99973974 | transportin 2 |
| ENSG00000283483 | AC068205,4 | 0,000163101 | 1,000113059 | 0,99971 | 0,99994283 | prohibitin (PHB) pseudogene |

| | | | | | | |
|---|---|---|---|---|---|---|
| ENSG00000120616 | EPC1 | 8,53E-05 | 1,000059158 | 0,99957 | 0,99994283 | enhancer of polycomb homolog 1 |
| ENSG00000257259 | LINC02388 | 7,38E-05 | 1,000051125 | 0,9998 | 0,99994283 | long intergenic non-protein coding RNA 2388 |
| ENSG00000284237 | LINC02767 | 4,05E-05 | 1,000028107 | 0,99996 | 0,99999442 | long intergenic non-protein coding RNA 2767 |
| ENSG00000269040 | BNIP3P24 | -4,57E-06 | 0,999996831 | 0,99999 | 0,99999442 | BCL2 interacting protein 3 pseudogene 24 |
| ENSG00000280428 | BNIP3P24 | -4,57E-06 | 0,999996831 | 0,99999 | 0,99999442 | BCL2 interacting protein 3 pseudogene 24 |
| ENSG00000006704 | GTF2IRD1 | -7,44E-05 | 0,999948407 | 0,99974 | 0,99994283 | GTF2I repeat domain containing 1 |
| ENSG00000168397 | ATG4B | -0,000107022 | 0,999925821 | 0,99952 | 0,99994283 | autophagy related 4B cysteine peptidase |
| ENSG00000253660 | AC008464,1 | -0,000136013 | 0,999905728 | 0,99987 | 0,99998166 | novel transcript |
| ENSG00000213967 | ZNF726 | -0,00016158 | 0,999888007 | 0,99973 | 0,99994283 | zinc finger protein 726 |
| ENSG00000285125 | AL136295,13 | -0,000227806 | 0,999842109 | 0,99977 | 0,99994283 | novel protein |
| ENSG00000259522 | AL136295,4 | -0,000227806 | 0,999842109 | 0,99977 | 0,99994283 | novel protein |
| ENSG00000134824 | FADS2 | -0,000235714 | 0,999836629 | 0,99861 | 0,99960886 | fatty acid desaturase 2 |
| ENSG00000006837 | CDKL3 | -0,00041667 | 0,999711228 | 0,99858 | 0,99960886 | cyclin dependent kinase like 3 |
| ENSG00000273402 | AC004908,3 | -0,000430994 | 0,999701302 | 0,99966 | 0,99994283 | novel transcript, antisense to ZNF596 |
| ENSG00000166046 | TCP11L2 | -0,000460427 | 0,999680907 | 0,99882 | 0,99970766 | t-complex 11 like 2 |
| ENSG00000102034 | ELF4 | -0,000474139 | 0,999671406 | 0,99905 | 0,999764 | E74 like ETS transcription factor 4 |
| ENSG00000143653 | SCCPDH | -0,000504156 | 0,999650607 | 0,99777 | 0,99926914 | saccharopine dehydrogenase (putative) |
| ENSG00000149187 | CELF1 | -0,000567638 | 0,999606621 | 0,99712 | 0,99894911 | CUGBP Elav-like family member 1 |
| ENSG00000278730 | AC005332,6 | -0,000685064 | 0,999525262 | 0,99813 | 0,99945242 | novel transcript |
| ENSG00000241889 | AC079944,2 | -0,000753241 | 0,999478029 | 0,99897 | 0,99975574 | dullard homolog (Xenopus laevis) (DULLARD) pseudogene |
| ENSG00000143624 | INTS3 | -0,000850495 | 0,999410656 | 0,99825 | 0,99947928 | integrator complex subunit 3 |
| ENSG00000226953 | NCKAP5-AS2 | -0,000856884 | 0,99940623 | 0,99803 | 0,99945242 | NCKAP5 antisense RNA 2 |
| ENSG00000267709 | AC024592,2 | -0,000866511 | 0,999399561 | 0,99888 | 0,99973508 | novel transcript, antisense to FUT3 |
| ENSG00000114019 | AMOTL2 | -0,000869518 | 0,999397477 | 0,9975 | 0,99910705 | angiomotin like 2 |
| ENSG00000256070 | AC016956,1 | -0,000898699 | 0,999377263 | 0,99914 | 0,99978149 | suppression of tumorigenicity 13 (colon carcinoma) (Hsp70 interacting protein) (ST13) pseudogene |
| ENSG00000204839 | MROH6 | -0,001017035 | 0,999295294 | 0,99866 | 0,9996198 | maestro heat like repeat family member 6 |
| ENSG00000172458 | IL17D | -0,001114687 | 0,999227656 | 0,99757 | 0,99913485 | interleukin 17D |
| ENSG00000259408 | AC010809,2 | -0,001212011 | 0,999160251 | 0,99686 | 0,99883674 | novel transcript, antisense to RYR3 |
| ENSG00000261308 | FIGNL2 | -0,001240092 | 0,999140803 | 0,99781 | 0,99927254 | fidgetin like 2 |
| ENSG00000198832 | SELENOM | -0,001251093 | 0,999133184 | 0,99711 | 0,99894911 | selenoprotein M |
| ENSG00000284610 | AC107918,4 | -0,001424169 | 0,999013329 | 0,99656 | 0,99862114 | RNA binding protein, fox-1 homolog (C, elegans) 1 (RBFOX1) pseudogene |

| | | | | | | |
|---|---|---|---|---|---|---|
| ENSG00000169093 | ASMTL | -0,001469843 | 0,998981702 | 0,99532 | 0,99809315 | acetylserotonin O-methyltransferase like |
| ENSG00000137959 | IFI44L | -0,001626562 | 0,998873188 | 0,99831 | 0,99948165 | interferon induced protein 44 like |
| ENSG00000274363 | AC087742,1 | -0,001821616 | 0,998738149 | 0,99739 | 0,99905887 | novel transcript |
| ENSG00000273762 | VN1R76P | -0,001830635 | 0,998731905 | 0,9969 | 0,99883674 | vomeronasal 1 receptor 76 pseudogene |
| ENSG00000225921 | NOL7 | -0,001845489 | 0,998721622 | 0,99172 | 0,99651645 | nucleolar protein 7 |
| ENSG00000274987 | AC092794,1 | -0,001854313 | 0,998715514 | 0,99813 | 0,99945242 | novel transcript, antisense to KRAS |
| ENSG00000182446 | NPLOC4 | -0,00187125 | 0,998703789 | 0,9916 | 0,99649731 | NPL4 homolog, ubiquitin recognition factor |
| ENSG00000257221 | AC007569,1 | -0,001988292 | 0,99862277 | 0,99607 | 0,99849121 | novel transcript, antisense to SELPLG |
| ENSG00000163565 | IFI16 | -0,002067469 | 0,998567966 | 0,99713 | 0,99894911 | interferon gamma inducible protein 16 |
| ENSG00000112877 | CEP72 | -0,002093875 | 0,998549689 | 0,99326 | 0,99693507 | centrosomal protein 72 |
| ENSG00000105982 | RNF32 | -0,002117836 | 0,998533105 | 0,99427 | 0,99750488 | ring finger protein 32 |
| ENSG00000166455 | C16orf46 | -0,002273561 | 0,998425329 | 0,99691 | 0,99883674 | chromosome 16 open reading frame 46 |
| ENSG00000168263 | KCNV2 | -0,002285418 | 0,998417123 | 0,99643 | 0,99859001 | potassium voltage-gated channel modifier subfamily V member 2 |
| ENSG00000227500 | SCAMP4 | -0,002375654 | 0,998354677 | 0,99202 | 0,99651645 | secretory carrier membrane protein 4 |
| ENSG00000133606 | MKRN1 | -0,002514191 | 0,998258813 | 0,99264 | 0,99671114 | makorin ring finger protein 1 |
| ENSG00000182504 | CEP97 | -0,002612914 | 0,998190505 | 0,98763 | 0,99461882 | centrosomal protein 97 |
| ENSG00000235194 | PPP1R3E | -0,002625411 | 0,998181858 | 0,99026 | 0,99596348 | protein phosphatase 1 regulatory subunit 3E |
| ENSG00000165029 | ABCA1 | -0,002631961 | 0,998177326 | 0,9933 | 0,99693507 | ATP binding cassette subfamily A member 1 |
| ENSG00000164099 | PRSS12 | -0,002755028 | 0,998092183 | 0,9949 | 0,99785291 | serine protease 12 |
| ENSG00000145244 | CORIN | -0,002836393 | 0,998035894 | 0,99579 | 0,9982573 | corin, serine peptidase |
| ENSG00000107679 | PLEKHA1 | -0,002860503 | 0,998019215 | 0,99195 | 0,99651645 | pleckstrin homology domain containing A1 |
| ENSG00000128283 | CDC42EP1 | -0,002883072 | 0,998003602 | 0,99247 | 0,99670754 | CDC42 effector protein 1 |
| ENSG00000224837 | GCSHP5 | -0,003171219 | 0,997804293 | 0,99468 | 0,99769817 | glycine cleavage system protein H pseudogene 5 |
| ENSG00000151176 | PLBD2 | -0,003256217 | 0,997745507 | 0,98793 | 0,99471733 | phospholipase B domain containing 2 |
| ENSG00000236353 | PBX2 | -0,003301582 | 0,997714134 | 0,99521 | 0,99806005 | PBX homeobox 2 |
| ENSG00000131368 | MRPS25 | -0,003327242 | 0,997696389 | 0,98275 | 0,99251385 | mitochondrial ribosomal protein S25 |
| ENSG00000253390 | AC104561,1 | -0,003374543 | 0,997663678 | 0,99558 | 0,99812418 | novel transcript |
| ENSG00000258359 | PCNPP1 | -0,003465251 | 0,997600954 | 0,99529 | 0,99809315 | PEST containing nuclear protein pseudogene 1 |
| ENSG00000268516 | AC020915,2 | -0,003627621 | 0,997488683 | 0,98577 | 0,99413307 | novel transcript, antisense to ZNF544 |
| ENSG00000135090 | TAOK3 | -0,003673496 | 0,997456965 | 0,99123 | 0,99634744 | TAO kinase 3 |
| ENSG00000198121 | LPAR1 | -0,003739712 | 0,997411186 | 0,99539 | 0,99812418 | lysophosphatidic acid receptor 1 |
| ENSG00000175581 | MRPL48 | -0,003745386 | 0,997407263 | 0,98348 | 0,99281596 | mitochondrial ribosomal protein L48 |

| | | | | | | |
|---|---|---|---|---|---|---|
| ENSG00000274210 | RNVU1-27 | -0,003801569 | 0,997368422 | 0,9888 | 0,9950859 | RNA, variant U1 small nuclear 27 |
| ENSG00000127083 | OMD | -0,003858959 | 0,997328748 | 0,99281 | 0,99671845 | osteomodulin |
| ENSG00000142621 | FHAD1 | -0,004031863 | 0,997209227 | 0,98719 | 0,99446724 | forkhead associated phosphopeptide binding domain 1 |
| ENSG00000123353 | ORMDL2 | -0,004128893 | 0,997142161 | 0,98714 | 0,99446724 | ORMDL sphingolipid biosynthesis regulator 2 |
| ENSG00000286504 | AC092745,4 | -0,004148516 | 0,997128598 | 0,99144 | 0,99645159 | novel transcript |
| ENSG00000286655 | AL158011,1 | -0,004276123 | 0,997040405 | 0,99346 | 0,99694625 | novel transcript, antisense to TSEN15 |
| ENSG00000228775 | WEE2-AS1 | -0,004299616 | 0,99702417 | 0,99089 | 0,99629071 | WEE2 antisense RNA 1 |
| ENSG00000234456 | MAGI2-AS3 | -0,004400994 | 0,996954111 | 0,98642 | 0,99421624 | MAGI2 antisense RNA 3 |
| ENSG00000281106 | TMEM272 | -0,004409114 | 0,996948501 | 0,9944 | 0,99756272 | transmembrane protein 272 |
| ENSG00000175348 | TMEM9B | -0,004424041 | 0,996938186 | 0,98774 | 0,99464284 | TMEM9 domain family member B |
| ENSG00000273828 | AL133227,1 | -0,00443292 | 0,99693205 | 0,98613 | 0,99413637 | novel transcript, antisense to ELMO2 |
| ENSG00000011422 | PLAUR | -0,004464007 | 0,996910569 | 0,98801 | 0,99471957 | plasminogen activator, urokinase receptor |
| ENSG00000197343 | ZNF655 | -0,004465058 | 0,996909842 | 0,97511 | 0,98950949 | zinc finger protein 655 |
| ENSG00000247311 | AC010255,1 | -0,004512763 | 0,996876878 | 0,99184 | 0,99651645 | novel transcript |
| ENSG00000274675 | GTF2H2C_2 | -0,004547815 | 0,996852658 | 0,98727 | 0,99446724 | GTF2H2 family member C, copy 2 |
| ENSG00000065989 | PDE4A | -0,004548852 | 0,996851942 | 0,99048 | 0,99596348 | phosphodiesterase 4A |
| ENSG00000213741 | RPS29 | -0,004565571 | 0,99684039 | 0,98133 | 0,99236691 | ribosomal protein S29 |
| ENSG00000175764 | TTLL11 | -0,004596986 | 0,996818683 | 0,98894 | 0,99518765 | tubulin tyrosine ligase like 11 |
| ENSG00000130725 | UBE2M | -0,004608596 | 0,996810662 | 0,98081 | 0,99205474 | ubiquitin conjugating enzyme E2 M |
| ENSG00000143384 | MCL1 | -0,004637926 | 0,996790397 | 0,97406 | 0,98928341 | MCL1 apoptosis regulator, BCL2 family member |
| ENSG00000273899 | NOL12 | -0,004664494 | 0,99677204 | 0,98436 | 0,99330857 | nucleolar protein 12 |
| ENSG00000259475 | AC036108,1 | -0,004700367 | 0,996747256 | 0,99467 | 0,99769817 | novel transcript, antisense to IGF1R and SYNM |
| ENSG00000169738 | DCXR | -0,004702313 | 0,996745911 | 0,98577 | 0,99413307 | dicarbonyl and L-xylulose reductase |
| ENSG00000111224 | PARP11 | -0,004704173 | 0,996744626 | 0,981 | 0,99215365 | poly(ADP-ribose) polymerase family member 11 |
| ENSG00000167770 | OTUB1 | -0,004735771 | 0,996722796 | 0,97859 | 0,99131203 | OTU deubiquitinase, ubiquitin aldehyde binding 1 |
| ENSG00000251186 | AC105345,2 | -0,004885801 | 0,996619149 | 0,9956 | 0,99812418 | novel transcript |
| ENSG00000239463 | HLA-DMA | -0,005003963 | 0,996537525 | 0,9905 | 0,99596348 | major histocompatibility complex, class II, DM alpha |
| ENSG00000243215 | HLA-DMA | -0,005003963 | 0,996537525 | 0,9905 | 0,99596348 | major histocompatibility complex, class II, DM alpha |
| ENSG00000243189 | HLA-DMA | -0,005003963 | 0,996537525 | 0,9905 | 0,99596348 | major histocompatibility complex, class II, DM alpha |
| ENSG00000242685 | HLA-DMA | -0,005003963 | 0,996537525 | 0,9905 | 0,99596348 | major histocompatibility complex, class II, DM alpha |
| ENSG00000243719 | HLA-DMA | -0,005003963 | 0,996537525 | 0,9905 | 0,99596348 | major histocompatibility complex, class II, DM alpha |
| ENSG00000129028 | THAP10 | -0,005011348 | 0,996532424 | 0,9863 | 0,99417149 | THAP domain containing 10 |
| ENSG00000248206 | AC074194,1 | -0,005090091 | 0,996478034 | 0,98663 | 0,99424613 | novel transcript |

| | | | | | | |
|---|---|---|---|---|---|---|
| ENSG00000232630 | PRPS1P2 | -0,005130799 | 0,996449918 | 0,99099 | 0,99631601 | phosphoribosyl pyrophosphate synthetase 1 pseudogene 2 |
| ENSG00000105939 | ZC3HAV1 | -0,005221014 | 0,996387609 | 0,98867 | 0,99502165 | zinc finger CCCH-type containing, antiviral 1 |
| ENSG00000231607 | DLEU2 | -0,00537853 | 0,996278828 | 0,98222 | 0,99244329 | deleted in lymphocytic leukemia 2 |
| ENSG00000287122 | AC026495,2 | -0,005401021 | 0,996263296 | 0,98813 | 0,99471957 | novel transcript |
| ENSG00000165868 | HSPA12A | -0,005417285 | 0,996252065 | 0,98347 | 0,99281596 | heat shock protein family A (Hsp70) member 12A |
| ENSG00000137462 | TLR2 | -0,005454123 | 0,996226627 | 0,99117 | 0,99634744 | toll like receptor 2 |
| ENSG00000270194 | AC097359,2 | -0,005473297 | 0,996213387 | 0,98164 | 0,99239184 | novel transcript, antisense to GOLGA4 |
| ENSG00000185905 | C16orf54 | -0,005486228 | 0,996204458 | 0,99335 | 0,99693507 | chromosome 16 open reading frame 54 |
| ENSG00000087589 | CASS4 | -0,005495076 | 0,996198348 | 0,99271 | 0,99671114 | Cas scaffold protein family member 4 |
| ENSG00000262333 | HNRNPA1P16 | -0,005623469 | 0,996109695 | 0,98336 | 0,99281596 | heterogeneous nuclear ribonucleoprotein A1 pseudogene 16 |
| ENSG00000179134 | SAMD4B | -0,005647964 | 0,996092783 | 0,97452 | 0,98933402 | sterile alpha motif domain containing 4B |
| ENSG00000250299 | MRPS31P4 | -0,005903793 | 0,995916164 | 0,99134 | 0,99641558 | mitochondrial ribosomal protein S31 pseudogene 4 |
| ENSG00000174501 | ANKRD36C | -0,006030766 | 0,995828516 | 0,98254 | 0,99246154 | ankyrin repeat domain 36C |
| ENSG00000166246 | C16orf71 | -0,00608826 | 0,995788832 | 0,98635 | 0,99418036 | chromosome 16 open reading frame 71 |
| ENSG00000285870 | AC012673,1 | -0,006110854 | 0,995773237 | 0,99381 | 0,99718342 | novel transcript |
| ENSG00000285827 | AP001267,5 | -0,006189027 | 0,995719282 | 0,98273 | 0,99251385 | novel protein |
| ENSG00000174109 | C16orf91 | -0,00623478 | 0,995687704 | 0,97804 | 0,99099142 | chromosome 16 open reading frame 91 |
| ENSG00000272254 | AC022893,3 | -0,006249266 | 0,995677707 | 0,99157 | 0,99649731 | novel transcript, antisense to TERF1 |
| ENSG00000086300 | SNX10 | -0,006252498 | 0,995675477 | 0,99163 | 0,99649731 | sorting nexin 10 |
| ENSG00000171791 | BCL2 | -0,006348352 | 0,995609325 | 0,98397 | 0,99306837 | BCL2 apoptosis regulator |
| ENSG00000245958 | AC093752,1 | -0,006412567 | 0,995565011 | 0,97951 | 0,99159589 | septin 4 (sept4) pseudogene |
| ENSG00000237337 | ELAVL4-AS1 | -0,006419172 | 0,995560453 | 0,99433 | 0,99752551 | ELAVL4 antisense RNA 1 |
| ENSG00000173545 | ZNF622 | -0,0066024 | 0,995434021 | 0,97034 | 0,98749329 | zinc finger protein 622 |
| ENSG00000259964 | THSD4-AS1 | -0,006626498 | 0,995417394 | 0,99188 | 0,99651645 | THSD4 antisense RNA 1 |
| ENSG00000187189 | TSPYL4 | -0,006726729 | 0,99534824 | 0,96827 | 0,98646784 | TSPY like 4 |
| ENSG00000261611 | AC010547,4 | -0,006800459 | 0,995297373 | 0,98794 | 0,99471733 | novel transcript |
| ENSG00000248408 | LINC02469 | -0,006866356 | 0,995251913 | 0,99144 | 0,99645159 | long intergenic non-protein coding RNA 2469 |
| ENSG00000184898 | RBM43 | -0,006942203 | 0,995199591 | 0,97111 | 0,98773415 | RNA binding motif protein 43 |
| ENSG00000260565 | ERVK13-1 | -0,007007569 | 0,995154501 | 0,97898 | 0,99141951 | endogenous retrovirus group K13 member 1 |
| ENSG00000205086 | C2orf91 | -0,007032557 | 0,995137265 | 0,98979 | 0,99572035 | chromosome 2 putative open reading frame 91 |
| ENSG00000143473 | KCNH1 | -0,007047714 | 0,99512681 | 0,97813 | 0,99099142 | potassium voltage-gated channel subfamily H member 1 |
| ENSG00000236490 | TAPBP | -0,007049196 | 0,995125788 | 0,99339 | 0,99694012 | TAP binding protein |

| | | | | | | |
|---|---|---|---|---|---|---|
| ENSG00000287710 | AC067852,5 | -0,007059145 | 0,995118925 | 0,98444 | 0,99330857 | novel transcript, antisense to PSMC3IP |
| ENSG00000228057 | SEC63P1 | -0,007061673 | 0,995117181 | 0,99007 | 0,99582133 | SEC63 homolog, protein translocation regulator pseudogene 1 |
| ENSG00000170296 | GABARAP | -0,00731195 | 0,994944565 | 0,96255 | 0,98430469 | GABA type A receptor-associated protein |
| ENSG00000165915 | SLC39A13 | -0,007345983 | 0,994921094 | 0,97785 | 0,99088131 | solute carrier family 39 member 13 |
| ENSG00000115234 | SNX17 | -0,007483389 | 0,99482634 | 0,96148 | 0,98401207 | sorting nexin 17 |
| ENSG00000157110 | RBPMS | -0,007515559 | 0,994804157 | 0,97048 | 0,98751893 | RNA binding protein, mRNA processing factor |
| ENSG00000180879 | SSR4 | -0,00758994 | 0,994752869 | 0,96676 | 0,98609023 | signal sequence receptor subunit 4 |
| ENSG00000124222 | STX16 | -0,007606447 | 0,994741487 | 0,96567 | 0,98544814 | syntaxin 16 |
| ENSG00000130175 | PRKCSH | -0,007619508 | 0,994732482 | 0,97452 | 0,98933402 | protein kinase C substrate 80K-H |
| ENSG00000231441 | DST-AS1 | -0,007677141 | 0,994692745 | 0,99396 | 0,99726631 | DST antisense RNA 1 |
| ENSG00000169231 | THBS3 | -0,007746956 | 0,994644611 | 0,96762 | 0,98623935 | thrombospondin 3 |
| ENSG00000241111 | PRICKLE2-AS1 | -0,00778468 | 0,994618603 | 0,98739 | 0,99447743 | PRICKLE2 antisense RNA 1 |
| ENSG00000168658 | VWA3B | -0,008038533 | 0,994443608 | 0,98728 | 0,99446724 | von Willebrand factor A domain containing 3B |
| ENSG00000133872 | SARAF | -0,008098484 | 0,994402285 | 0,97081 | 0,98765476 | store-operated calcium entry associated regulatory factor |
| ENSG00000162889 | MAPKAPK2 | -0,008120602 | 0,99438704 | 0,96173 | 0,98401207 | MAPK activated protein kinase 2 |
| ENSG00000275026 | GXYLT1P4 | -0,00812367 | 0,994384925 | 0,99236 | 0,99669518 | GXYLT1 pseudogene 4 |
| ENSG00000159658 | EFCAB14 | -0,008292377 | 0,99426865 | 0,95219 | 0,97974576 | EF-hand calcium binding domain 14 |
| ENSG00000188730 | VWC2 | -0,008294685 | 0,994267059 | 0,98706 | 0,99446724 | von Willebrand factor C domain containing 2 |
| ENSG00000149269 | PAK1 | -0,008340038 | 0,994235803 | 0,96419 | 0,98478217 | p21 (RAC1) activated kinase 1 |
| ENSG00000144711 | IQSEC1 | -0,00842696 | 0,994175902 | 0,96829 | 0,98646784 | IQ motif and Sec7 domain ArfGEF 1 |
| ENSG00000164611 | PTTG1 | -0,008603478 | 0,99405427 | 0,96306 | 0,98432594 | PTTG1 regulator of sister chromatid separation, securin |
| ENSG00000281709 | ZNF197 | -0,008649077 | 0,994022852 | 0,95487 | 0,98116612 | zinc finger protein 197 |
| ENSG00000180357 | ZNF609 | -0,008695132 | 0,99399112 | 0,95876 | 0,98291816 | zinc finger protein 609 |
| ENSG00000184047 | DIABLO | -0,008765854 | 0,993942395 | 0,95878 | 0,98291816 | diablo IAP-binding mitochondrial protein |
| ENSG00000062582 | MRPS24 | -0,008772147 | 0,993938059 | 0,96553 | 0,9854196 | mitochondrial ribosomal protein S24 |
| ENSG00000122515 | ZMIZ2 | -0,008864578 | 0,993874381 | 0,96778 | 0,98623935 | zinc finger MIZ-type containing 2 |
| ENSG00000178502 | KLHL11 | -0,008955707 | 0,993811604 | 0,94917 | 0,97868075 | kelch like family member 11 |
| ENSG00000158006 | PAFAH2 | -0,008967877 | 0,993803221 | 0,96561 | 0,98542381 | platelet activating factor acetylhydrolase 2 |
| ENSG00000130377 | ACSBG2 | -0,009008947 | 0,993774931 | 0,9821 | 0,99244329 | acyl-CoA synthetase bubblegum family member 2 |
| ENSG00000116863 | ADPRS | -0,00917662 | 0,993659439 | 0,96398 | 0,98461596 | ADP-ribosylserine hydrolase |
| ENSG00000232533 | AC093673,1 | -0,009189129 | 0,993650823 | 0,97489 | 0,98941297 | novel transcript |
| ENSG00000262227 | AC004771,2 | -0,009194786 | 0,993646927 | 0,98608 | 0,99413637 | novel transcript, antisense to INCA1 |

| | | | | | | |
|---|---|---|---|---|---|---|
| ENSG00000134248 | LAMTOR5 | -0,009240796 | 0,993615238 | 0,96537 | 0,9854196 | late endosomal/lysosomal adaptor, MAPK and MTOR activator 5 |
| ENSG00000178028 | DMAP1 | -0,009253685 | 0,993606361 | 0,95828 | 0,98276958 | DNA methyltransferase 1 associated protein 1 |
| ENSG00000174460 | ZCCHC12 | -0,009285375 | 0,993584536 | 0,98161 | 0,99239184 | zinc finger CCHC-type containing 12 |
| ENSG00000287169 | AC107050,1 | -0,009322027 | 0,993559294 | 0,97687 | 0,99033775 | novel transcript, antisense to FAT1 |
| ENSG00000132639 | SNAP25 | -0,009322872 | 0,993558712 | 0,98188 | 0,99242247 | synaptosome associated protein 25 |
| ENSG00000272905 | AC018648,1 | -0,009378831 | 0,993520175 | 0,9926 | 0,99671114 | novel transcript |
| ENSG00000138468 | SENP7 | -0,009451524 | 0,993470115 | 0,95923 | 0,98320317 | SUMO specific peptidase 7 |
| ENSG00000049239 | H6PD | -0,009457105 | 0,993466272 | 0,96528 | 0,9854196 | hexose-6-phosphate dehydrogenase/glucose 1-dehydrogenase |
| ENSG00000169213 | RAB3B | -0,009465504 | 0,993460489 | 0,98044 | 0,99195802 | RAB3B, member RAS oncogene family |
| ENSG00000273576 | AC009283,1 | -0,009548563 | 0,993403295 | 0,97883 | 0,99136324 | novel transcript |
| ENSG00000188316 | ENO4 | -0,009603683 | 0,993365341 | 0,98161 | 0,99239184 | enolase 4 |
| ENSG00000136881 | BAAT | -0,009621399 | 0,993353143 | 0,97978 | 0,99170352 | bile acid-CoA:amino acid N-acyltransferase |
| ENSG00000106479 | ZNF862 | -0,009720312 | 0,99328504 | 0,9581 | 0,98276081 | zinc finger protein 862 |
| ENSG00000090372 | STRN4 | -0,009776461 | 0,993246383 | 0,96188 | 0,98401207 | striatin 4 |
| ENSG00000233473 | RAD1P2 | -0,009797082 | 0,993232186 | 0,9908 | 0,99623172 | RAD1 pseudogene 2 |
| ENSG00000164543 | STK17A | -0,009818106 | 0,993217712 | 0,98148 | 0,99239184 | serine/threonine kinase 17a |
| ENSG00000273138 | AC005021,1 | -0,009822844 | 0,99321445 | 0,98621 | 0,99415471 | novel transcript, antisense to PON2 |
| ENSG00000119431 | HDHD3 | -0,00985571 | 0,993191823 | 0,97245 | 0,98833499 | haloacid dehalogenase like hydrolase domain containing 3 |
| ENSG00000161204 | ABCF3 | -0,009902316 | 0,993159739 | 0,95632 | 0,98190071 | ATP binding cassette subfamily F member 3 |
| ENSG00000157103 | SLC6A1 | -0,009971555 | 0,993112076 | 0,9882 | 0,99471957 | solute carrier family 6 member 1 |
| ENSG00000253133 | AC009630,1 | -0,009977823 | 0,993107761 | 0,97075 | 0,98765168 | novel transcript, antisense to AGPAT6 and GINS4 |
| ENSG00000169919 | GUSB | -0,009988768 | 0,993100227 | 0,96775 | 0,98623935 | glucuronidase beta |
| ENSG00000281912 | LINC01144 | -0,010199961 | 0,99295486 | 0,97207 | 0,98800339 | long intergenic non-protein coding RNA 1144 |
| ENSG00000286873 | AC012306,3 | -0,010264375 | 0,992910527 | 0,96553 | 0,9854196 | novel transcript |
| ENSG00000154027 | AK5 | -0,010384007 | 0,992828196 | 0,98076 | 0,99205474 | adenylate kinase 5 |
| ENSG00000233122 | CTAGE7P | -0,010448955 | 0,992783501 | 0,97552 | 0,98962884 | CTAGE family member 7, pseudogene |
| ENSG00000164366 | CCDC127 | -0,010533243 | 0,9927255 | 0,9497 | 0,97897619 | coiled-coil domain containing 127 |
| ENSG00000285967 | NIPBL-DT | -0,010645262 | 0,992648423 | 0,95496 | 0,98122749 | NIPBL divergent transcript |
| ENSG00000147536 | GINS4 | -0,010779835 | 0,992555834 | 0,95601 | 0,98181191 | GINS complex subunit 4 |
| ENSG00000226882 | GNL1 | -0,010852621 | 0,992505759 | 0,98589 | 0,99413637 | G protein nucleolar 1 (putative) |
| ENSG00000286921 | AC004522,4 | -0,010948352 | 0,992439903 | 0,97767 | 0,9907329 | novel transcript, antisense to AZGP1 |

| | | | | | | |
|---|---|---|---|---|---|---|
| ENSG00000273993 | PTPRK | -0,011025153 | 0,992387073 | 0,97201 | 0,98800339 | protein tyrosine phosphatase receptor type K |
| ENSG00000285873 | AL445253,1 | -0,01107957 | 0,992349641 | 0,98668 | 0,99425859 | novel transcript |
| ENSG00000175756 | AURKAIP1 | -0,011138211 | 0,992309307 | 0,96284 | 0,98432594 | aurora kinase A interacting protein 1 |
| ENSG00000146197 | SCUBE3 | -0,011175816 | 0,992283441 | 0,94379 | 0,97627205 | signal peptide, CUB domain and EGF like domain containing 3 |
| ENSG00000163832 | ELP6 | -0,011206518 | 0,992262325 | 0,95976 | 0,98338726 | elongator acetyltransferase complex subunit 6 |
| ENSG00000004838 | ZMYND10 | -0,011208846 | 0,992260723 | 0,97885 | 0,99136324 | zinc finger MYND-type containing 10 |
| ENSG00000261460 | AC009690,2 | -0,011221692 | 0,992251889 | 0,9655 | 0,9854196 | novel transcript, antisense to HEXA and CELF6 |
| ENSG00000143158 | MPC2 | -0,011248185 | 0,992233667 | 0,96432 | 0,9848531 | mitochondrial pyruvate carrier 2 |
| ENSG00000157429 | ZNF19 | -0,01131674 | 0,992186519 | 0,95248 | 0,97986142 | zinc finger protein 19 |
| ENSG00000286584 | AC023494,1 | -0,011389273 | 0,992136637 | 0,97673 | 0,99026923 | novel transcript, antisense to COX17 |
| ENSG00000286473 | AC133485,7 | -0,011511879 | 0,992052325 | 0,9805 | 0,99195802 | novel transcript |
| ENSG00000254692 | AL136295,1 | -0,011718099 | 0,99191053 | 0,99121 | 0,99634744 | novel protein |
| ENSG00000285502 | AL136295,20 | -0,011718099 | 0,99191053 | 0,99121 | 0,99634744 | novel protein |
| ENSG00000204130 | RUFY2 | -0,011725489 | 0,991905449 | 0,96586 | 0,98553345 | RUN and FYVE domain containing 2 |
| ENSG00000287164 | AC112254,1 | -0,011760421 | 0,991881432 | 0,98967 | 0,99572035 | novel transcript, antisense to AFAP1 |
| ENSG00000189067 | LITAF | -0,011817331 | 0,991842306 | 0,97885 | 0,99136324 | lipopolysaccharide induced TNF factor |
| ENSG00000171320 | ESCO2 | -0,011939483 | 0,991758332 | 0,95075 | 0,97934603 | establishment of sister chromatid cohesion N-acetyltransferase 2 |
| ENSG00000124171 | PARD6B | -0,011939968 | 0,991757998 | 0,97035 | 0,98749329 | par-6 family cell polarity regulator beta |
| ENSG00000049323 | LTBP1 | -0,011996269 | 0,991719295 | 0,97954 | 0,99159589 | latent transforming growth factor beta binding protein 1 |
| ENSG00000196741 | LINC01560 | -0,012149998 | 0,991613627 | 0,97656 | 0,9901566 | long intergenic non-protein coding RNA 1560 |
| ENSG00000173889 | PHC3 | -0,012375881 | 0,991458381 | 0,93477 | 0,9721963 | polyhomeotic homolog 3 |
| ENSG00000163930 | BAP1 | -0,012378866 | 0,99145633 | 0,93465 | 0,9721963 | BRCA1 associated protein 1 |
| ENSG00000106415 | GLCCI1 | -0,012506407 | 0,991368685 | 0,9713 | 0,98775203 | glucocorticoid induced 1 |
| ENSG00000121022 | COPS5 | -0,012674284 | 0,991253333 | 0,94539 | 0,9766664 | COP9 signalosome subunit 5 |
| ENSG00000224713 | AC025165,1 | -0,012687831 | 0,991244025 | 0,9646 | 0,98503092 | novel transcript, antisense to ARHGEF25 |
| ENSG00000205559 | CHKB-DT | -0,012857526 | 0,991127438 | 0,94888 | 0,97862975 | CHKB divergent transcript |
| ENSG00000286204 | AC025475,2 | -0,012919922 | 0,991084573 | 0,97542 | 0,98956832 | novel transcript |
| ENSG00000198690 | FAN1 | -0,013010843 | 0,991022115 | 0,94839 | 0,97834225 | FANCD2 and FANCI associated nuclease 1 |
| ENSG00000253276 | CCDC71L | -0,013023445 | 0,991013459 | 0,94585 | 0,97689657 | coiled-coil domain containing 71 like |
| ENSG00000186310 | NAP1L3 | -0,01305928 | 0,990988843 | 0,94642 | 0,97715566 | nucleosome assembly protein 1 like 3 |
| ENSG00000172534 | HCFC1 | -0,013150672 | 0,990926067 | 0,94979 | 0,97899157 | host cell factor C1 |

| | | | | | | |
|---|---|---|---|---|---|---|
| ENSG00000213190 | MLLT11 | -0,013224846 | 0,990875122 | 0,95371 | 0,9805199 | MLLT11 transcription factor 7 cofactor |
| ENSG00000114450 | GNB4 | -0,013241332 | 0,990863799 | 0,96343 | 0,98448919 | G protein subunit beta 4 |
| ENSG00000249207 | AC079921,1 | -0,013378901 | 0,990769319 | 0,96973 | 0,98725952 | novel transcript |
| ENSG00000171425 | ZNF581 | -0,013380201 | 0,990768426 | 0,95672 | 0,98205915 | zinc finger protein 581 |
| ENSG00000109046 | WSB1 | -0,01339226 | 0,990760145 | 0,95625 | 0,98189243 | WD repeat and SOCS box containing 1 |
| ENSG00000250406 | AC109811,1 | -0,013401481 | 0,990753813 | 0,98855 | 0,99496822 | novel transcript |
| ENSG00000054267 | ARID4B | -0,013422638 | 0,990739283 | 0,9417 | 0,97505821 | AT-rich interaction domain 4B |
| ENSG00000266993 | AL050343,2 | -0,013492755 | 0,990691133 | 0,97124 | 0,98775203 | novel transcript, antisense to NRD1 |
| ENSG00000109917 | ZPR1 | -0,013497127 | 0,990688131 | 0,94532 | 0,97666034 | ZPR1 zinc finger |
| ENSG00000223697 | AF230666,1 | -0,013549525 | 0,99065215 | 0,97918 | 0,99154627 | novel transcript, antisense to PHF20L1 |
| ENSG00000271204 | AC016831,4 | -0,013594746 | 0,990621099 | 0,97492 | 0,98941297 | novel transcript |
| ENSG00000101191 | DIDO1 | -0,013684563 | 0,990559428 | 0,94495 | 0,97666034 | death inducer-obliterator 1 |
| ENSG00000261609 | GAN | -0,013685688 | 0,990558656 | 0,94412 | 0,97642036 | gigaxonin |
| ENSG00000146083 | RNF44 | -0,01370962 | 0,990542224 | 0,96218 | 0,98413404 | ring finger protein 44 |
| ENSG00000137692 | DCUN1D5 | -0,013776927 | 0,990496013 | 0,93018 | 0,97010144 | defective in cullin neddylation 1 domain containing 5 |
| ENSG00000107611 | CUBN | -0,013783782 | 0,990491307 | 0,97615 | 0,98994851 | cubilin |
| ENSG00000225439 | BOLA3-AS1 | -0,013841869 | 0,990451427 | 0,95008 | 0,97915169 | BOLA3 divergent transcript |
| ENSG00000235347 | Z84478,1 | -0,013843267 | 0,990450468 | 0,97492 | 0,98941297 | tRNA nucleotidyl transferase, CCA-adding, 1 (TRNT1) pseudogene |
| ENSG00000187048 | CYP4A11 | -0,013852378 | 0,990444213 | 0,98446 | 0,99330857 | cytochrome P450 family 4 subfamily A member 11 |
| ENSG00000151023 | ENKUR | -0,013881948 | 0,990423912 | 0,97167 | 0,98794066 | enkurin, TRPC channel interacting protein |
| ENSG00000213297 | ZNF625-ZNF2( | -0,013886597 | 0,990420721 | 0,95581 | 0,98173232 | ZNF625-ZNF20 readthrough (NMD candidate) |
| ENSG00000143774 | GUK1 | -0,013923737 | 0,990395224 | 0,95105 | 0,97947047 | guanylate kinase 1 |
| ENSG00000169155 | ZBTB43 | -0,013942925 | 0,990382052 | 0,92747 | 0,96860856 | zinc finger and BTB domain containing 43 |
| ENSG00000225733 | FGD5-AS1 | -0,013952369 | 0,990375569 | 0,92385 | 0,96681926 | FGD5 antisense RNA 1 |
| ENSG00000239665 | AL157392,3 | -0,014037294 | 0,990317272 | 0,94912 | 0,97866986 | novel transcript |
| ENSG00000165899 | OTOGL | -0,014057011 | 0,990303738 | 0,97942 | 0,99154627 | otogelin like |
| ENSG00000110888 | CAPRIN2 | -0,014162787 | 0,990231132 | 0,94106 | 0,97489099 | caprin family member 2 |
| ENSG00000183114 | FAM43B | -0,014295428 | 0,990140095 | 0,97738 | 0,99055154 | family with sequence similarity 43 member B |
| ENSG00000250318 | AC003072,1 | -0,01434501 | 0,990106067 | 0,97369 | 0,98906507 | ribosomal protein L13a (RPL13A) pseudogene |
| ENSG00000226977 | HMGN1P24 | -0,014398657 | 0,990069251 | 0,98729 | 0,99446724 | high mobility group nucleosome binding domain 1 pseudogene 24 |
| ENSG00000092200 | RPGRIP1 | -0,014451113 | 0,990033252 | 0,98733 | 0,99446724 | RPGR interacting protein 1 |

| | | | | | | |
|---|---|---|---|---|---|---|
| ENSG00000064995 | TAF11 | -0,014547115 | 0,989967374 | 0,93499 | 0,9721963 | TATA-box binding protein associated factor 11 |
| ENSG00000197102 | DYNC1H1 | -0,014606164 | 0,989926856 | 0,94077 | 0,97486264 | dynein cytoplasmic 1 heavy chain 1 |
| ENSG00000140474 | ULK3 | -0,014713918 | 0,989852921 | 0,94454 | 0,97650311 | unc-51 like kinase 3 |
| ENSG00000070610 | GBA2 | -0,014793602 | 0,989798251 | 0,9425 | 0,97556707 | glucosylceramidase beta 2 |
| ENSG00000272506 | AL357078,2 | -0,014936547 | 0,989700185 | 0,98601 | 0,99413637 | novel transcript |
| ENSG00000144395 | CCDC150 | -0,015004811 | 0,989653356 | 0,95128 | 0,97948342 | coiled-coil domain containing 150 |
| ENSG00000227471 | AKR1B15 | -0,015006406 | 0,989652262 | 0,94676 | 0,97731612 | aldo-keto reductase family 1 member B15 |
| ENSG00000267259 | ERVE-1 | -0,015017427 | 0,989644702 | 0,97142 | 0,98777425 | endogenous retrovirus group E member 1 |
| ENSG00000139624 | CERS5 | -0,015058975 | 0,989616202 | 0,92635 | 0,96798905 | ceramide synthase 5 |
| ENSG00000174125 | TLR1 | -0,01506606 | 0,989611342 | 0,98407 | 0,99309929 | toll like receptor 1 |
| ENSG00000226632 | UBE2V1P1 | -0,015169289 | 0,989540534 | 0,9838 | 0,9929677 | ubiquitin conjugating enzyme E2 V1 pseudogene 1 |
| ENSG00000270270 | BNIP3P10 | -0,015392406 | 0,989387512 | 0,96777 | 0,98623935 | BCL2 interacting protein 3 pseudogene 10 |
| ENSG00000124659 | TBCC | -0,015394797 | 0,989385872 | 0,94321 | 0,97601141 | tubulin folding cofactor C |
| ENSG00000285805 | AC012378,2 | -0,015414161 | 0,989372592 | 0,97701 | 0,99035539 | novel transcript, antisense to PYGO1 |
| ENSG00000129158 | SERGEF | -0,015539659 | 0,989286532 | 0,95157 | 0,97948342 | secretion regulating guanine nucleotide exchange factor |
| ENSG00000140873 | ADAMTS18 | -0,015583208 | 0,989256669 | 0,97645 | 0,9901323 | ADAM metallopeptidase with thrombospondin type 1 motif 18 |
| ENSG00000288066 | AC097448,1 | -0,015594265 | 0,989249088 | 0,94931 | 0,97879465 | novel transcript, antisense to ARL5A |
| ENSG00000174516 | PELI3 | -0,015661776 | 0,989202797 | 0,95765 | 0,98257839 | pellino E3 ubiquitin protein ligase family member 3 |
| ENSG00000237972 | TUBG1P | -0,015828325 | 0,989088607 | 0,97512 | 0,98950949 | tubulin gamma 1 pseudogene |
| ENSG00000175634 | RPS6KB2 | -0,015884556 | 0,989050056 | 0,93671 | 0,97279937 | ribosomal protein S6 kinase B2 |
| ENSG00000277283 | AC004812,2 | -0,015891939 | 0,989044995 | 0,94983 | 0,97899157 | novel transcript, antisense to RAB35 |
| ENSG00000225880 | LINC00115 | -0,016034189 | 0,98894748 | 0,95996 | 0,98353421 | long intergenic non-protein coding RNA 115 |
| ENSG00000228076 | AL591719,1 | -0,016077461 | 0,988917818 | 0,98226 | 0,99244329 | chromosome 20 open reading frame 147 (C20orf147) pseudogene |
| ENSG00000166801 | FAM111A | -0,016114079 | 0,988892718 | 0,9688 | 0,98677126 | family with sequence similarity 111 member A |
| ENSG00000267534 | S1PR2 | -0,01613551 | 0,988878028 | 0,96801 | 0,98637425 | sphingosine-1-phosphate receptor 2 |
| ENSG00000157601 | MX1 | -0,016426153 | 0,988678831 | 0,96613 | 0,98562284 | MX dynamin like GTPase 1 |
| ENSG00000285257 | AL157893,1 | -0,01661847 | 0,988547045 | 0,98217 | 0,99244329 | synaptotagmin 15 (SYT15) pseudogene |
| ENSG00000183864 | TOB2 | -0,016622298 | 0,988544422 | 0,94414 | 0,97642036 | transducer of ERBB2, 2 |
| ENSG00000241764 | AC002467,1 | -0,016647469 | 0,988527174 | 0,96598 | 0,98557781 | novel transcript, antisense to CBLL1 |
| ENSG00000141748 | ARL5C | -0,016667233 | 0,988513632 | 0,97504 | 0,98950522 | ADP ribosylation factor like GTPase 5C |
| ENSG00000264859 | DSG2-AS1 | -0,016694728 | 0,988494793 | 0,97801 | 0,99099142 | DSG2 antisense RNA 1 |

| | | | | | | |
|---|---|---|---|---|---|---|
| ENSG00000128463 | EMC4 | -0,016749865 | 0,988457016 | 0,92938 | 0,96977202 | ER membrane protein complex subunit 4 |
| ENSG00000273807 | AC244489,1 | -0,016776954 | 0,988438456 | 0,9731 | 0,9886748 | novel member of the nuclear pore complex interacting protein NPIP gene family |
| ENSG00000011523 | CEP68 | -0,016789258 | 0,988430026 | 0,90916 | 0,96104995 | centrosomal protein 68 |
| ENSG00000237927 | AL078604,2 | -0,016806487 | 0,988418222 | 0,98475 | 0,99343186 | novel transcript |
| ENSG00000169220 | RGS14 | -0,016864234 | 0,988378659 | 0,94847 | 0,97838613 | regulator of G protein signaling 14 |
| ENSG00000108352 | RAPGEFL1 | -0,016965711 | 0,988309141 | 0,95313 | 0,98025619 | Rap guanine nucleotide exchange factor like 1 |
| ENSG00000288631 | ELOA3 | -0,017108489 | 0,988211336 | 0,96679 | 0,98609023 | elongin A3 |
| ENSG00000111667 | USP5 | -0,017117855 | 0,988204921 | 0,92014 | 0,96530017 | ubiquitin specific peptidase 5 |
| ENSG00000142661 | MYOM3 | -0,017437452 | 0,98798603 | 0,97414 | 0,98928341 | myomesin 3 |
| ENSG00000165125 | TRPV6 | -0,017771056 | 0,987757598 | 0,98155 | 0,99239184 | transient receptor potential cation channel subfamily V member 6 |
| ENSG00000232043 | AL133230,1 | -0,017957075 | 0,987630246 | 0,96517 | 0,98539831 | novel transcript, antisense to PTPN1 |
| ENSG00000239779 | WBP1 | -0,017971812 | 0,987620158 | 0,93533 | 0,97222198 | WW domain binding protein 1 |
| ENSG00000197746 | PSAP | -0,018114391 | 0,987522558 | 0,93353 | 0,97162107 | prosaposin |
| ENSG00000186212 | SOWAHB | -0,018125129 | 0,987515208 | 0,98284 | 0,99256924 | sosondowah ankyrin repeat domain family member B |
| ENSG00000175931 | UBE2O | -0,018167979 | 0,987485878 | 0,93992 | 0,97444851 | ubiquitin conjugating enzyme E2 O |
| ENSG00000281931 | SCG5 | -0,018296046 | 0,987398223 | 0,97144 | 0,98777425 | secretogranin V |
| ENSG00000126259 | KIRREL2 | -0,018336115 | 0,9873708 | 0,96972 | 0,98725952 | kirre like nephrin family adhesion molecule 2 |
| ENSG00000261195 | AC027130,1 | -0,018412065 | 0,987318822 | 0,97605 | 0,98994851 | novel transcript, antisense to IQCK |
| ENSG00000152910 | CNTNAP4 | -0,018455116 | 0,98728936 | 0,97083 | 0,98765476 | contactin associated protein family member 4 |
| ENSG00000263310 | SALL3 | -0,018529994 | 0,987238119 | 0,98048 | 0,99195802 | spalt like transcription factor 3 |
| ENSG00000230660 | HCG18 | -0,018673193 | 0,987140133 | 0,97387 | 0,98920952 | HLA complex group 18 |
| ENSG00000244371 | PFN1P8 | -0,018769761 | 0,98707406 | 0,94526 | 0,97666034 | profilin 1 pseudogene 8 |
| ENSG00000273715 | AC138827,1 | -0,018831814 | 0,987031605 | 0,96898 | 0,98688154 | NLR family, apoptosis inhibitory protein (NAIP) pseudogene |
| ENSG00000232186 | TERF1P7 | -0,018844266 | 0,987023086 | 0,97527 | 0,98956405 | TERF1 pseudogene 7 |
| ENSG00000227417 | AC114402,1 | -0,018930343 | 0,986964198 | 0,97065 | 0,98761994 | SNF7 domain containing 2 (SNF7DC20) pseudogene |
| ENSG00000227159 | DDX11L16 | -0,018933297 | 0,986962177 | 0,97988 | 0,99172545 | DEAD/H-box helicase 11 like 16 (pseudogene) |
| ENSG00000270589 | AL158163,1 | -0,0189708 | 0,986936521 | 0,95643 | 0,98194036 | novel transcript, antisense to BBIP1 |
| ENSG00000259039 | AL161804,1 | -0,018992913 | 0,986921394 | 0,97978 | 0,99170352 | novel transcript, antisense to SLC35F4 |
| ENSG00000187243 | MAGED4B | -0,019086724 | 0,986857222 | 0,91935 | 0,96495401 | MAGE family member D4B |
| ENSG00000167792 | NDUFV1 | -0,019190099 | 0,986786512 | 0,90842 | 0,96080514 | NADH:ubiquinone oxidoreductase core subunit V1 |

| | | | | | | |
|---|---|---|---|---|---|---|
| ENSG00000139197 | PEX5 | -0,019295398 | 0,986714491 | 0,94168 | 0,97505821 | peroxisomal biogenesis factor 5 |
| ENSG00000148218 | ALAD | -0,019448886 | 0,98660952 | 0,92026 | 0,96530017 | aminolevulinate dehydratase |
| ENSG00000148120 | AOPEP | -0,019504026 | 0,986571813 | 0,93044 | 0,97020757 | aminopeptidase O (putative) |
| ENSG00000168961 | LGALS9 | -0,019521747 | 0,986559695 | 0,963 | 0,98432594 | galectin 9 |
| ENSG00000078487 | ZCWPW1 | -0,01953648 | 0,986549619 | 0,93941 | 0,97413085 | zinc finger CW-type and PWWP domain containing 1 |
| ENSG00000172361 | CFAP53 | -0,019548511 | 0,986541393 | 0,96759 | 0,98623935 | cilia and flagella associated protein 53 |
| ENSG00000198911 | SREBF2 | -0,019556859 | 0,986535684 | 0,91522 | 0,9633004 | sterol regulatory element binding transcription factor 2 |
| ENSG00000085721 | RRN3 | -0,019564578 | 0,986530406 | 0,96005 | 0,98353421 | RRN3 homolog, RNA polymerase I transcription factor |
| ENSG00000186230 | ZNF749 | -0,019721848 | 0,986422869 | 0,90936 | 0,96105018 | zinc finger protein 749 |
| ENSG00000247950 | SEC24B-AS1 | -0,019775122 | 0,986386444 | 0,95915 | 0,98315781 | SEC24B antisense RNA 1 |
| ENSG00000255240 | AP001636,3 | -0,019838384 | 0,986343192 | 0,9611 | 0,98401207 | novel transcript |
| ENSG00000051620 | HEBP2 | -0,019895334 | 0,986304257 | 0,90125 | 0,95796772 | heme binding protein 2 |
| ENSG00000067992 | PDK3 | -0,019958185 | 0,98626129 | 0,92771 | 0,96873207 | pyruvate dehydrogenase kinase 3 |
| ENSG00000005421 | PON1 | -0,020073485 | 0,986182471 | 0,96079 | 0,9839024 | paraoxonase 1 |
| ENSG00000236178 | VARS2 | -0,020467342 | 0,985913279 | 0,91278 | 0,96223361 | valyl-tRNA synthetase 2, mitochondrial |
| ENSG00000282092 | MED16 | -0,020471262 | 0,9859106 | 0,94276 | 0,97566033 | mediator complex subunit 16 |
| ENSG00000187736 | NHEJ1 | -0,020493725 | 0,98589525 | 0,9436 | 0,97618492 | non-homologous end joining factor 1 |
| ENSG00000228742 | LINC02577 | -0,020565544 | 0,985846172 | 0,96749 | 0,98623935 | long intergenic non-protein coding RNA 2577 |
| ENSG00000165802 | NSMF | -0,020577432 | 0,985838049 | 0,90903 | 0,96102407 | NMDA receptor synaptonuclear signaling and neuronal migration factor |
| ENSG00000143156 | NME7 | -0,020682473 | 0,985766274 | 0,90238 | 0,95825434 | NME/NM23 family member 7 |
| ENSG00000158987 | RAPGEF6 | -0,020761585 | 0,985712219 | 0,91117 | 0,9616927 | Rap guanine nucleotide exchange factor 6 |
| ENSG00000251456 | AC113398,2 | -0,020774607 | 0,985703322 | 0,9615 | 0,98401207 | novel transcript |
| ENSG00000232578 | AC093311,1 | -0,02081805 | 0,98567364 | 0,97733 | 0,99052937 | FAST kinase domains 1 (FASTKD1) pseudogene |
| ENSG00000087085 | ACHE | -0,020832589 | 0,985663707 | 0,95705 | 0,98218743 | acetylcholinesterase (Cartwright blood group) |
| ENSG00000259343 | TMC3-AS1 | -0,021010104 | 0,985542435 | 0,94793 | 0,97811704 | TMC3 antisense RNA 1 |
| ENSG00000135900 | MRPL44 | -0,021173544 | 0,985430791 | 0,90082 | 0,95787401 | mitochondrial ribosomal protein L44 |
| ENSG00000265218 | AC103810,2 | -0,021190813 | 0,985418995 | 0,95962 | 0,98338726 | novel transcript, antisense to LRRC37A3 |
| ENSG00000145491 | ROPN1L | -0,021217287 | 0,985400913 | 0,96421 | 0,98478217 | rhophilin associated tail protein 1 like |
| ENSG00000262879 | AC005670,3 | -0,021261197 | 0,985370921 | 0,89186 | 0,95246557 | novel transcript |
| ENSG00000236992 | RPL12P12 | -0,021356331 | 0,985305947 | 0,96722 | 0,98623935 | ribosomal protein L12 pseudogene 12 |
| ENSG00000148677 | ANKRD1 | -0,02135825 | 0,985304636 | 0,97898 | 0,99141951 | ankyrin repeat domain 1 |
| ENSG00000187713 | TMEM203 | -0,02135834 | 0,985304575 | 0,92993 | 0,9699953 | transmembrane protein 203 |

| | | | | | | |
|---|---|---|---|---|---|---|
| ENSG00000125434 | SLC25A35 | -0,021433853 | 0,985253004 | 0,95526 | 0,98146656 | solute carrier family 25 member 35 |
| ENSG00000116171 | SCP2 | -0,021434177 | 0,985252782 | 0,9174 | 0,96419315 | sterol carrier protein 2 |
| ENSG00000267278 | MAP3K14-AS1 | -0,021612716 | 0,985130861 | 0,94894 | 0,97862975 | MAP3K14 antisense RNA 1 |
| ENSG00000105514 | RAB3D | -0,021625663 | 0,98512202 | 0,91022 | 0,9613425 | RAB3D, member RAS oncogene family |
| ENSG00000215425 | DDX39B | -0,02165013 | 0,985105313 | 0,90001 | 0,95740913 | DExD-box helicase 39B |
| ENSG00000173728 | C1orf100 | -0,021650433 | 0,985105106 | 0,98047 | 0,99195802 | chromosome 1 open reading frame 100 |
| ENSG00000196381 | ZNF781 | -0,021667833 | 0,985093225 | 0,96043 | 0,98381824 | zinc finger protein 781 |
| ENSG00000165140 | FBP1 | -0,021703044 | 0,985069183 | 0,98253 | 0,99246154 | fructose-bisphosphatase 1 |
| ENSG00000254017 | IGHEP2 | -0,021781126 | 0,98501587 | 0,97912 | 0,99152159 | immunoglobulin heavy constant epsilon P2 (pseudogene) |
| ENSG00000137266 | SLC22A23 | -0,021797069 | 0,985004985 | 0,92098 | 0,96576077 | solute carrier family 22 member 23 |
| ENSG00000178096 | BOLA1 | -0,021988548 | 0,984874261 | 0,94167 | 0,97505821 | bolA family member 1 |
| ENSG00000153012 | LGI2 | -0,022195544 | 0,984732963 | 0,96212 | 0,98410918 | leucine rich repeat LGI family member 2 |
| ENSG00000103266 | STUB1 | -0,022376259 | 0,984609621 | 0,92833 | 0,96914921 | STIP1 homology and U-box containing protein 1 |
| ENSG00000270706 | PRMT1P1 | -0,022415895 | 0,98458257 | 0,95382 | 0,98057921 | protein arginine methyltransferase 1 pseudogene 1 |
| ENSG00000227799 | CDPF1P1 | -0,022451202 | 0,984558475 | 0,94035 | 0,97466991 | CDPF1 pseudogene 1 |
| ENSG00000275594 | ADAM32 | -0,022467249 | 0,984547523 | 0,94948 | 0,97888896 | ADAM metallopeptidase domain 32 |
| ENSG00000236745 | YRDCP2 | -0,022604505 | 0,984453859 | 0,96725 | 0,98623935 | yrdC N(6)-threonylcarbamoyltransferase domain containing pseudogene 2 |
| ENSG00000013810 | TACC3 | -0,022609008 | 0,984450787 | 0,89279 | 0,9531296 | transforming acidic coiled-coil containing protein 3 |
| ENSG00000271991 | AC013400,1 | -0,022621301 | 0,984442399 | 0,9635 | 0,98448919 | novel transcript |
| ENSG00000103502 | CDIPT | -0,022772396 | 0,984339302 | 0,92114 | 0,96582196 | CDP-diacylglycerol--inositol 3-phosphatidyltransferase |
| ENSG00000253726 | AC013562,1 | -0,022820962 | 0,984306167 | 0,9619 | 0,98401207 | novel transcript |
| ENSG00000288432 | AC270091,1 | -0,022857398 | 0,984281307 | 0,98777 | 0,99464284 | |
| ENSG00000185900 | POMK | -0,022889668 | 0,984259292 | 0,90697 | 0,96017342 | protein O-mannose kinase |
| ENSG00000251013 | GAPDHP62 | -0,022911077 | 0,984244686 | 0,96367 | 0,98451153 | glyceraldehyde 3 phosphate dehydrogenase pseudogene 62 |
| ENSG00000224046 | AC005076,1 | -0,022926376 | 0,984234249 | 0,94824 | 0,97829358 | novel transcript, antisense to DMTF1 |
| ENSG00000198910 | L1CAM | -0,022943588 | 0,984222506 | 0,9502 | 0,97920193 | L1 cell adhesion molecule |
| ENSG00000102390 | PBDC1 | -0,022987186 | 0,984192764 | 0,91615 | 0,96347905 | polysaccharide biosynthesis domain containing 1 |
| ENSG00000257851 | HNRNPA3P10 | -0,023002704 | 0,984182178 | 0,971 | 0,98771801 | heterogeneous nuclear ribonucleoprotein A3 pseudogene 10 |
| ENSG00000198455 | ZXDB | -0,023025943 | 0,984166324 | 0,92338 | 0,96667848 | zinc finger X-linked duplicated B |
| ENSG00000176697 | BDNF | -0,023098481 | 0,984116842 | 0,94528 | 0,97666034 | brain derived neurotrophic factor |

| | | | | | | |
|---|---|---|---|---|---|---|
| ENSG00000103121 | CMC2 | -0,023212927 | 0,984038777 | 0,90844 | 0,96080514 | C-X9-C motif containing 2 |
| ENSG00000239969 | AC091390,3 | -0,023219659 | 0,984034185 | 0,96054 | 0,98383172 | uncharacterized LOC100289561 |
| ENSG00000106404 | CLDN15 | -0,02343708 | 0,983885898 | 0,93114 | 0,97060346 | claudin 15 |
| ENSG00000253567 | AC025871,1 | -0,023455598 | 0,983873269 | 0,97634 | 0,99006347 | novel transcript |
| ENSG00000104812 | GYS1 | -0,02346121 | 0,983869442 | 0,89761 | 0,95581835 | glycogen synthase 1 |
| ENSG00000285618 | AC074132,1 | -0,023517842 | 0,983830821 | 0,97206 | 0,98800339 | novel transcript, antisense to ADGRV1 |
| ENSG00000150403 | TMCO3 | -0,023593226 | 0,983779416 | 0,89419 | 0,95389664 | transmembrane and coiled-coil domains 3 |
| ENSG00000105270 | CLIP3 | -0,023683543 | 0,98371783 | 0,92192 | 0,96616853 | CAP-Gly domain containing linker protein 3 |
| ENSG00000168490 | PHYHIP | -0,023736668 | 0,983681607 | 0,95329 | 0,98030368 | phytanoyl-CoA 2-hydroxylase interacting protein |
| ENSG00000196993 | NPIPB9 | -0,023807943 | 0,98363301 | 0,96043 | 0,98381824 | nuclear pore complex interacting protein family member B9 |
| ENSG00000265241 | RBM8A | -0,023895836 | 0,983573086 | 0,87254 | 0,94374455 | RNA binding motif protein 8A |
| ENSG00000183647 | ZNF530 | -0,024189027 | 0,983373221 | 0,92395 | 0,96689592 | zinc finger protein 530 |
| ENSG00000254034 | INTS9-AS1 | -0,02421621 | 0,983354692 | 0,94148 | 0,97505054 | INTS9 antisense RNA 1 |
| ENSG00000227317 | DDAH2 | -0,024267947 | 0,983319429 | 0,95046 | 0,97932267 | dimethylarginine dimethylaminohydrolase 2 |
| ENSG00000245573 | BDNF-AS | -0,024271336 | 0,983317119 | 0,91061 | 0,96143161 | BDNF antisense RNA |
| ENSG00000241343 | RPL36A | -0,024506885 | 0,983156585 | 0,8809 | 0,94692171 | ribosomal protein L36a |
| ENSG00000171055 | FEZ2 | -0,024616365 | 0,98308198 | 0,9007 | 0,95781447 | fasciculation and elongation protein zeta 2 |
| ENSG00000111300 | NAA25 | -0,024779803 | 0,982970617 | 0,91453 | 0,96303629 | N-alpha-acetyltransferase 25, NatB auxiliary subunit |
| ENSG00000257494 | AC004217,1 | -0,024806032 | 0,982952746 | 0,98226 | 0,99244329 | novel transcript |
| ENSG00000121966 | CXCR4 | -0,024831667 | 0,98293528 | 0,94541 | 0,9766664 | C-X-C motif chemokine receptor 4 |
| ENSG00000261215 | AL162231,4 | -0,024947031 | 0,982856684 | 0,96151 | 0,98401207 | novel transcript (novel gene-CCL27 readthrough) |
| ENSG00000154380 | ENAH | -0,024952458 | 0,982852987 | 0,89116 | 0,95208954 | ENAH actin regulator |
| ENSG00000224740 | FLOT1 | -0,025070495 | 0,982772576 | 0,94644 | 0,97715566 | flotillin 1 |
| ENSG00000253869 | PIGFP1 | -0,025156758 | 0,982713815 | 0,97868 | 0,99131549 | phosphatidylinositol glycan anchor biosynthesis class F pseudogene 1 |
| ENSG00000169926 | KLF13 | -0,025311585 | 0,982608358 | 0,90384 | 0,95878714 | Kruppel like factor 13 |
| ENSG00000102683 | SGCG | -0,025360902 | 0,982574769 | 0,96496 | 0,98525685 | sarcoglycan gamma |
| ENSG00000235217 | TSPY26P | -0,02542665 | 0,982529991 | 0,88257 | 0,94809798 | testis specific protein Y-linked 26, pseudogene |
| ENSG00000228492 | RAB11FIP1P1 | -0,025430356 | 0,982527467 | 0,9433 | 0,97607277 | RAB11 family interacting protein 1 pseudogene 1 |
| ENSG00000135838 | NPL | -0,025579151 | 0,982426137 | 0,96837 | 0,9865184 | N-acetylneuraminate pyruvate lyase |
| ENSG00000121101 | TEX14 | -0,025674006 | 0,982361546 | 0,95765 | 0,98257839 | testis expressed 14, intercellular bridge forming factor |
| ENSG00000258477 | PPIAP6 | -0,02568531 | 0,982353849 | 0,9744 | 0,98933402 | peptidylprolyl isomerase A pseudogene 6 |

| | | | | | | |
|---|---|---|---|---|---|---|
| ENSG00000281665 | KIAA1143 | -0,025723094 | 0,982328122 | 0,93066 | 0,97025341 | KIAA1143 |
| ENSG00000197448 | GSTK1 | -0,025804451 | 0,982272728 | 0,91321 | 0,96246104 | glutathione S-transferase kappa 1 |
| ENSG00000172974 | VDAC2P5 | -0,025813074 | 0,982266856 | 0,98205 | 0,99244329 | VDAC2 pseudogene 5 |
| ENSG00000104365 | IKBKB | -0,025914512 | 0,982197795 | 0,885 | 0,94911792 | inhibitor of nuclear factor kappa B kinase subunit beta |
| ENSG00000178498 | DTX3 | -0,025932169 | 0,982185773 | 0,89699 | 0,95561124 | deltex E3 ubiquitin ligase 3 |
| ENSG00000223705 | NSUN5P1 | -0,026065476 | 0,982095023 | 0,91327 | 0,96246104 | NSUN5 pseudogene 1 |
| ENSG00000270062 | AL606834,2 | -0,026079829 | 0,982085252 | 0,9732 | 0,98872813 | novel transcript, sense intronic to NIN |
| ENSG00000270427 | NRBF2P5 | -0,026147312 | 0,982039315 | 0,94442 | 0,97650311 | nuclear receptor binding factor 2 pseudogene 5 |
| ENSG00000054965 | FAM168A | -0,026174036 | 0,982021124 | 0,92321 | 0,96667029 | family with sequence similarity 168 member A |
| ENSG00000108828 | VAT1 | -0,02623886 | 0,981977001 | 0,90153 | 0,95801185 | vesicle amine transport 1 |
| ENSG00000270813 | NANOGNBP3 | -0,026354398 | 0,981898362 | 0,96898 | 0,98688154 | NANOGNB pseudogene 3 |
| ENSG00000272232 | RN7SL726P | -0,026442521 | 0,981838388 | 0,97048 | 0,98751893 | RNA, 7SL, cytoplasmic 726, pseudogene |
| ENSG00000142208 | AKT1 | -0,026446052 | 0,981835985 | 0,88819 | 0,95069428 | AKT serine/threonine kinase 1 |
| ENSG00000120071 | KANSL1 | -0,026508982 | 0,981793158 | 0,87935 | 0,94630103 | KAT8 regulatory NSL complex subunit 1 |
| ENSG00000234807 | LINC01135 | -0,026511567 | 0,981791399 | 0,96919 | 0,98702411 | long intergenic non-protein coding RNA 1135 |
| ENSG00000171928 | TVP23B | -0,026527913 | 0,981780275 | 0,88601 | 0,94953808 | trans-golgi network vesicle protein 23 homolog B |
| ENSG00000203876 | ADD3-AS1 | -0,026584771 | 0,981741583 | 0,94157 | 0,97505054 | ADD3 antisense RNA 1 |
| ENSG00000287042 | AC090945,1 | -0,026873146 | 0,981545366 | 0,96665 | 0,9860143 | novel transcript |
| ENSG00000239291 | AC002558,1 | -0,026879832 | 0,981540817 | 0,95241 | 0,9798257 | ribosomal protein S2 (RPS2) pseudogene |
| ENSG00000170921 | TANC2 | -0,026926023 | 0,981509392 | 0,92235 | 0,96637446 | tetratricopeptide repeat, ankyrin repeat and coiled-coil containing 2 |
| ENSG00000204310 | AGPAT1 | -0,026932724 | 0,981504832 | 0,92133 | 0,96592951 | 1-acylglycerol-3-phosphate O-acyltransferase 1 |
| ENSG00000262599 | DENND11 | -0,027190698 | 0,981329341 | 0,94173 | 0,97505821 | DENN domain containing 11 |
| ENSG00000224810 | AL355482,1 | -0,027190927 | 0,981329185 | 0,91838 | 0,96446185 | novel transcript (LOC149157) |
| ENSG00000178075 | GRAMD1C | -0,027242292 | 0,981294248 | 0,94815 | 0,9782417 | GRAM domain containing 1C |
| ENSG00000242600 | MBL1P | -0,027511775 | 0,981110967 | 0,95329 | 0,98030368 | mannose binding lectin 1, pseudogene |
| ENSG00000278486 | YRDCP1 | -0,027544515 | 0,981088703 | 0,96813 | 0,98639465 | yrdC N(6)-threonylcarbamoyltransferase domain containing pseudogene 1 |
| ENSG00000145216 | FIP1L1 | -0,027642355 | 0,98102217 | 0,89967 | 0,95730568 | factor interacting with PAPOLA and CPSF1 |
| ENSG00000242485 | MRPL20 | -0,027694137 | 0,980986959 | 0,89037 | 0,95170091 | mitochondrial ribosomal protein L20 |
| ENSG00000147996 | CBWD5 | -0,027835299 | 0,980890978 | 0,91894 | 0,9647667 | COBW domain containing 5 |
| ENSG00000287553 | AC090617,10 | -0,0278766 | 0,980862898 | 0,96351 | 0,98448919 | novel transcript, antisense to DPH1 |

| | | | | | | |
|---|---|---|---|---|---|---|
| ENSG00000102886 | GDPD3 | -0,027908525 | 0,980841193 | 0,93705 | 0,97304927 | glycerophosphodiester phosphodiesterase domain containing 3 |
| ENSG00000100003 | SEC14L2 | -0,027957449 | 0,980807931 | 0,90097 | 0,95787401 | SEC14 like lipid binding 2 |
| ENSG00000260816 | AC027279,1 | -0,027996961 | 0,98078107 | 0,95124 | 0,97948342 | novel transcript, sense intronic to WWOX |
| ENSG00000175221 | MED16 | -0,028027003 | 0,980760647 | 0,90175 | 0,95802163 | mediator complex subunit 16 |
| ENSG00000003509 | NDUFAF7 | -0,028104097 | 0,980708239 | 0,86253 | 0,93863824 | NADH:ubiquinone oxidoreductase complex assembly factor 7 |
| ENSG00000162413 | KLHL21 | -0,028116218 | 0,9807 | 0,91806 | 0,9644207 | kelch like family member 21 |
| ENSG00000107021 | TBC1D13 | -0,028303645 | 0,9805726 | 0,87417 | 0,94423986 | TBC1 domain family member 13 |
| ENSG00000135083 | CCNJL | -0,028318775 | 0,980562317 | 0,93353 | 0,97162107 | cyclin J like |
| ENSG00000163154 | TNFAIP8L2 | -0,028327067 | 0,980556681 | 0,96325 | 0,98440677 | TNF alpha induced protein 8 like 2 |
| ENSG00000223724 | RAD17P2 | -0,028408891 | 0,980501069 | 0,96443 | 0,98493011 | RAD17 pseudogene 2 |
| ENSG00000186684 | CYP27C1 | -0,028502118 | 0,980437712 | 0,93606 | 0,97253068 | cytochrome P450 family 27 subfamily C member 1 |
| ENSG00000249065 | PCNAP1 | -0,028523653 | 0,980423076 | 0,97808 | 0,99099142 | proliferating cell nuclear antigen pseudogene 1 |
| ENSG00000285818 | AL591893,1 | -0,028537003 | 0,980414004 | 0,96149 | 0,98401207 | putative UPF0607 protein ENSP00000383783 |
| ENSG00000287616 | AL357522,1 | -0,028565915 | 0,980394357 | 0,97193 | 0,98800339 | novel transcript |
| ENSG00000130005 | GAMT | -0,028610006 | 0,980364395 | 0,92613 | 0,96789681 | guanidinoacetate N-methyltransferase |
| ENSG00000146676 | PURB | -0,028701388 | 0,980302299 | 0,88187 | 0,94752399 | purine rich element binding protein B |
| ENSG00000287658 | AC109642,1 | -0,028736582 | 0,980278385 | 0,91501 | 0,9632313 | novel transcript |
| ENSG00000172375 | C2CD2L | -0,028738407 | 0,980277145 | 0,91437 | 0,96296851 | C2CD2 like |
| ENSG00000137474 | MYO7A | -0,02879459 | 0,980238971 | 0,93455 | 0,9721963 | myosin VIIA |
| ENSG00000182533 | CAV3 | -0,028818238 | 0,980222903 | 0,97131 | 0,98775203 | caveolin 3 |
| ENSG00000228838 | AL355483,1 | -0,028863359 | 0,980192247 | 0,96608 | 0,98562284 | novel transcript |
| ENSG00000079974 | RABL2B | -0,02889482 | 0,980170872 | 0,8701 | 0,94277661 | RAB, member of RAS oncogene family like 2B |
| ENSG00000005513 | SOX8 | -0,028900972 | 0,980166692 | 0,94908 | 0,97866986 | SRY-box transcription factor 8 |
| ENSG00000073008 | PVR | -0,028927278 | 0,980148821 | 0,87712 | 0,94557577 | PVR cell adhesion molecule |
| ENSG00000220785 | MTMR9LP | -0,029026374 | 0,980081498 | 0,94592 | 0,97689657 | myotubularin related protein 9 like, pseudogene |
| ENSG00000108641 | B9D1 | -0,029069405 | 0,980052266 | 0,88317 | 0,94832116 | B9 domain containing 1 |
| ENSG00000106089 | STX1A | -0,029101679 | 0,980030342 | 0,90929 | 0,96104995 | syntaxin 1A |
| ENSG00000217488 | AL356057,1 | -0,029161686 | 0,979989579 | 0,95327 | 0,98030368 | LSM14A, SCD6 homolog A (S, cerevisiae) (LSM14A) pseudogene |
| ENSG00000188738 | FSIP2 | -0,029235666 | 0,979939328 | 0,93197 | 0,97097872 | fibrous sheath interacting protein 2 |
| ENSG00000270100 | AC012065,3 | -0,029243815 | 0,979933793 | 0,97721 | 0,99048622 | novel transcript |

| | | | | | | |
|---|---|---|---|---|---|---|
| ENSG00000196177 | ACADSB | -0,029283239 | 0,979907015 | 0,84377 | 0,92878539 | acyl-CoA dehydrogenase short/branched chain |
| ENSG00000128311 | TST | -0,02937059 | 0,979847686 | 0,89616 | 0,95508888 | thiosulfate sulfurtransferase |
| ENSG00000269102 | AC010320,3 | -0,029396849 | 0,979829852 | 0,97118 | 0,98773415 | novel transcript, antisense to ZNF766 and ZNF480 |
| ENSG00000272024 | AC064807,4 | -0,029416475 | 0,979816523 | 0,97041 | 0,98751545 | novel transcript |
| ENSG00000277991 | FP236241,1 | -0,029425879 | 0,979810136 | 0,9409 | 0,97488728 | uncharacterized LOC102723360 |
| ENSG00000099251 | HSD17B7P2 | -0,029558849 | 0,979719833 | 0,90964 | 0,96105018 | hydroxysteroid 17-beta dehydrogenase 7 pseudogene 2 |
| ENSG00000288374 | HSD17B7P2 | -0,029558849 | 0,979719833 | 0,90964 | 0,96105018 | hydroxysteroid 17-beta dehydrogenase 7 pseudogene 2 |
| ENSG00000100416 | TRMU | -0,029597164 | 0,979693814 | 0,85408 | 0,93380868 | tRNA 5-methylaminomethyl-2-thiouridylate methyltransferase |
| ENSG00000168078 | PBK | -0,029676827 | 0,979639719 | 0,94284 | 0,97567445 | PDZ binding kinase |
| ENSG00000233927 | RPS28 | -0,029709583 | 0,979617477 | 0,91255 | 0,96205682 | ribosomal protein S28 |
| ENSG00000134184 | GSTM1 | -0,029725834 | 0,979606442 | 0,93716 | 0,97305801 | glutathione S-transferase mu 1 |
| ENSG00000101473 | ACOT8 | -0,029754549 | 0,979586944 | 0,87934 | 0,94630103 | acyl-CoA thioesterase 8 |
| ENSG00000259171 | AL163636,1 | -0,029784422 | 0,979566661 | 0,98996 | 0,99574948 | novel protein, ANG-RNASE4 readthrough |
| ENSG00000231050 | AL109917,1 | -0,029815331 | 0,979545674 | 0,96395 | 0,98461596 | novel transcript |
| ENSG00000127774 | EMC6 | -0,030006531 | 0,979415864 | 0,876 | 0,94516507 | ER membrane protein complex subunit 6 |
| ENSG00000132199 | ENOSF1 | -0,030016711 | 0,979408953 | 0,87398 | 0,94410944 | enolase superfamily member 1 |
| ENSG00000255435 | AP001267,3 | -0,030094531 | 0,979356124 | 0,90144 | 0,95798991 | novel transcript, antisense to MLL and TTC36 |
| ENSG00000233860 | SHROOM3-AS | -0,030122668 | 0,979337024 | 0,87922 | 0,94623795 | SHROOM3 antisense RNA 1 |
| ENSG00000158528 | PPP1R9A | -0,030144465 | 0,979322228 | 0,91347 | 0,96249683 | protein phosphatase 1 regulatory subunit 9A |
| ENSG00000175104 | TRAF6 | -0,030207847 | 0,979279204 | 0,88197 | 0,94759691 | TNF receptor associated factor 6 |
| ENSG00000264044 | AC005726,2 | -0,030233438 | 0,979261834 | 0,94831 | 0,97829358 | novel transcript, antisense to KIAA0100 |
| ENSG00000285710 | AC012184,4 | -0,030270714 | 0,979236532 | 0,96916 | 0,98702411 | novel transcript, antisense to DDX19A |
| ENSG00000171126 | KCNG3 | -0,030413347 | 0,979139724 | 0,96612 | 0,98562284 | potassium voltage-gated channel modifier subfamily G member 3 |
| ENSG00000273008 | AC010864,1 | -0,030426204 | 0,979130998 | 0,93512 | 0,9721963 | novel transcript |
| ENSG00000215049 | PRDX2P1 | -0,030666984 | 0,978967599 | 0,96143 | 0,98401207 | peroxiredoxin 2 pseudogene 1 |
| ENSG00000218189 | POM121L14P | -0,030724323 | 0,978928691 | 0,96262 | 0,98430614 | POM121 transmembrane nucleoporin like 14, pseudogene |
| ENSG00000214413 | BBIP1 | -0,030740071 | 0,978918006 | 0,86637 | 0,94074101 | BBSome interacting protein 1 |
| ENSG00000182858 | ALG12 | -0,030818207 | 0,978864989 | 0,88799 | 0,95069428 | ALG12 alpha-1,6-mannosyltransferase |
| ENSG00000285041 | AL035412,1 | -0,030849864 | 0,978843509 | 0,97025 | 0,98749329 | novel transcript |
| ENSG00000147234 | FRMPD3 | -0,030860352 | 0,978836394 | 0,95123 | 0,97948342 | FERM and PDZ domain containing 3 |
| ENSG00000130584 | ZBTB46 | -0,030901809 | 0,978808266 | 0,89439 | 0,95392565 | zinc finger and BTB domain containing 46 |

| | | | | | | |
|---|---|---|---|---|---|---|
| ENSG00000126062 | TMEM115 | -0,030961225 | 0,978767956 | 0,87655 | 0,94547358 | transmembrane protein 115 |
| ENSG00000272154 | AC244517,2 | -0,031018946 | 0,978728797 | 0,95637 | 0,98191931 | novel transcript, antisense to PCDHB3, PCDHB4, PCDHB5, PCDHB6 and PCDHB7 |
| ENSG00000236915 | CLCA4-AS1 | -0,031090364 | 0,978680348 | 0,95558 | 0,98167846 | CLCA4 antisense RNA 1 |
| ENSG00000224078 | SNHG14 | -0,031240808 | 0,978578297 | 0,87236 | 0,94374455 | small nucleolar RNA host gene 14 |
| ENSG00000103932 | RPAP1 | -0,031282513 | 0,978550009 | 0,87076 | 0,94302816 | RNA polymerase II associated protein 1 |
| ENSG00000160310 | PRMT2 | -0,031311439 | 0,978530389 | 0,88525 | 0,94924301 | protein arginine methyltransferase 2 |
| ENSG00000278828 | H3C10 | -0,031350422 | 0,978503949 | 0,88727 | 0,950382 | H3 clustered histone 10 |
| ENSG00000233665 | AC060234,2 | -0,031524928 | 0,978385597 | 0,93524 | 0,97222198 | novel transcript |
| ENSG00000226314 | ZNF192P1 | -0,031623864 | 0,978318505 | 0,93483 | 0,9721963 | zinc finger protein 192 pseudogene 1 |
| ENSG00000224870 | MRPL20-AS1 | -0,031760588 | 0,978225794 | 0,87478 | 0,94461928 | MRPL20 antisense RNA 1 |
| ENSG00000240219 | AL512306,2 | -0,031824461 | 0,978182486 | 0,92401 | 0,96692212 | novel transcript |
| ENSG00000087206 | UIMC1 | -0,031853468 | 0,978162819 | 0,84845 | 0,93094401 | ubiquitin interaction motif containing 1 |
| ENSG00000150625 | GPM6A | -0,031880858 | 0,978144248 | 0,9324 | 0,97120063 | glycoprotein M6A |
| ENSG00000267002 | AC060780,1 | -0,032068514 | 0,978017026 | 0,85418 | 0,93380868 | novel transcript |
| ENSG00000245970 | AP003352,1 | -0,032083238 | 0,978007044 | 0,91969 | 0,96507001 | novel transcript |
| ENSG00000177425 | PAWR | -0,03216273 | 0,977953158 | 0,91382 | 0,96272082 | pro-apoptotic WT1 regulator |
| ENSG00000235748 | SEPTIN14P12 | -0,032163192 | 0,977952845 | 0,94529 | 0,97666034 | septin 14 pseudogene 12 |
| ENSG00000277656 | GSTT1 | -0,032188834 | 0,977935463 | 0,87136 | 0,94321633 | glutathione S-transferase theta 1 |
| ENSG00000146007 | ZMAT2 | -0,032244296 | 0,977897869 | 0,8547 | 0,93416235 | zinc finger matrin-type 2 |
| ENSG00000231466 | AL022324,2 | -0,032256372 | 0,977889684 | 0,95477 | 0,98110426 | PHD finger protein 10 (PHF10) pseudogene |
| ENSG00000170581 | STAT2 | -0,032312026 | 0,977851961 | 0,87809 | 0,9458146 | signal transducer and activator of transcription 2 |
| ENSG00000183150 | GPR19 | -0,032377161 | 0,977807813 | 0,93836 | 0,97361475 | G protein-coupled receptor 19 |
| ENSG00000161542 | PRPSAP1 | -0,032439551 | 0,977765529 | 0,90433 | 0,95902207 | phosphoribosyl pyrophosphate synthetase associated protein 1 |
| ENSG00000231148 | HMGB1P7 | -0,032453665 | 0,977755963 | 0,96741 | 0,98623935 | high mobility group box 1 pseudogene 7 |
| ENSG00000064655 | EYA2 | -0,032591885 | 0,977662292 | 0,92501 | 0,96744841 | EYA transcriptional coactivator and phosphatase 2 |
| ENSG00000275272 | YTHDC1 | -0,032694265 | 0,977592915 | 0,85665 | 0,93532903 | YTH domain containing 1 |
| ENSG00000180878 | C11orf42 | -0,032700263 | 0,977588851 | 0,9509 | 0,9793462 | chromosome 11 open reading frame 42 |
| ENSG00000197956 | S100A6 | -0,032713974 | 0,97757956 | 0,93297 | 0,97143736 | S100 calcium binding protein A6 |
| ENSG00000132849 | PATJ | -0,032735783 | 0,977564782 | 0,88706 | 0,95021901 | PATJ crumbs cell polarity complex component |
| ENSG00000233270 | SNRPEP4 | -0,03279371 | 0,977525532 | 0,97531 | 0,98956405 | SNRPE pseudogene 4 |
| ENSG00000130165 | ELOF1 | -0,033042561 | 0,977356933 | 0,87876 | 0,94612705 | elongation factor 1 homolog |

| | | | | | | |
|---|---|---|---|---|---|---|
| ENSG00000250198 | LINC02199 | -0,033068448 | 0,977339396 | 0,95289 | 0,9801387 | long intergenic non-protein coding RNA 2199 |
| ENSG00000196159 | FAT4 | -0,033091042 | 0,97732409 | 0,90746 | 0,96044232 | FAT atypical cadherin 4 |
| ENSG00000260735 | AC139256,1 | -0,033144488 | 0,977287885 | 0,88609 | 0,94954542 | phospholipase A2 group X (PLA2G10) pseudogene |
| ENSG00000186105 | LRRC70 | -0,033430521 | 0,977094144 | 0,96003 | 0,98353421 | leucine rich repeat containing 70 |
| ENSG00000146281 | PM20D2 | -0,03357533 | 0,976996074 | 0,87294 | 0,94378579 | peptidase M20 domain containing 2 |
| ENSG00000057019 | DCBLD2 | -0,033610816 | 0,976972043 | 0,87323 | 0,94386143 | discoidin, CUB and LCCL domain containing 2 |
| ENSG00000101745 | ANKRD12 | -0,033929318 | 0,976756382 | 0,90474 | 0,95924013 | ankyrin repeat domain 12 |
| ENSG00000110031 | LPXN | -0,033969722 | 0,976729028 | 0,90298 | 0,95846794 | leupaxin |
| ENSG00000133985 | TTC9 | -0,034010455 | 0,976701451 | 0,89697 | 0,95561124 | tetratricopeptide repeat domain 9 |
| ENSG00000229820 | CR391992,1 | -0,034025066 | 0,976691559 | 0,97342 | 0,98882674 | DnaJ (Hsp40) homolog, subfamily C member 8 (DNAJC8) pseudogene |
| ENSG00000084623 | EIF3I | -0,034120703 | 0,976626816 | 0,89163 | 0,95235303 | eukaryotic translation initiation factor 3 subunit I |
| ENSG00000143429 | LSP1P4 | -0,034153427 | 0,976604664 | 0,92341 | 0,96667848 | LSP1 pseudogene 4 |
| ENSG00000136807 | CDK9 | -0,034220704 | 0,976559123 | 0,83528 | 0,92477092 | cyclin dependent kinase 9 |
| ENSG00000221838 | AP4M1 | -0,034349711 | 0,976471803 | 0,88877 | 0,95089681 | adaptor related protein complex 4 subunit mu 1 |
| ENSG00000286237 | ARMCX5-GPR | -0,034354587 | 0,976468502 | 0,88397 | 0,94847506 | ARMCX5-GPRASP2 readthrough |
| ENSG00000250462 | LRRC37BP1 | -0,034442261 | 0,976409163 | 0,8426 | 0,92821072 | leucine rich repeat containing 37B pseudogene 1 |
| ENSG00000278270 | RPS9 | -0,034552867 | 0,976334308 | 0,92621 | 0,96789681 | ribosomal protein S9 |
| ENSG00000243176 | AC092944,1 | -0,034636763 | 0,976277534 | 0,87762 | 0,94566951 | novel transcript |
| ENSG00000170113 | NIPA1 | -0,034720094 | 0,976221145 | 0,90983 | 0,96111314 | NIPA magnesium transporter 1 |
| ENSG00000255345 | AP002957,1 | -0,034743976 | 0,976204985 | 0,93921 | 0,9741012 | novel transcript |
| ENSG00000174804 | FZD4 | -0,034788254 | 0,976175025 | 0,92176 | 0,96611557 | frizzled class receptor 4 |
| ENSG00000228459 | LINC01546 | -0,034925993 | 0,976081831 | 0,96582 | 0,98552708 | long intergenic non-protein coding RNA 1546 |
| ENSG00000196187 | TMEM63A | -0,034977433 | 0,976047028 | 0,86958 | 0,9425949 | transmembrane protein 63A |
| ENSG00000172426 | RSPH9 | -0,034994824 | 0,976035263 | 0,88551 | 0,94936604 | radial spoke head component 9 |
| ENSG00000286819 | Z98745,1 | -0,035040105 | 0,976004629 | 0,93349 | 0,97162107 | novel transcript |
| ENSG00000155760 | FZD7 | -0,035065198 | 0,975987653 | 0,92549 | 0,9676347 | frizzled class receptor 7 |
| ENSG00000104872 | PIH1D1 | -0,035092911 | 0,975968905 | 0,90867 | 0,9609322 | PIH1 domain containing 1 |
| ENSG00000168273 | SMIM4 | -0,035141488 | 0,975936044 | 0,86839 | 0,9418797 | small integral membrane protein 4 |
| ENSG00000285979 | AC009090,6 | -0,035256544 | 0,975858215 | 0,88648 | 0,94985626 | novel transcript, antisense to FAM192A |
| ENSG00000254712 | AC087280,1 | -0,035332756 | 0,975806666 | 0,95768 | 0,98257839 | phosphorylated adaptor for RNA export (PHAX) pseudogene |
| ENSG00000287117 | AC105916,1 | -0,035398756 | 0,975762026 | 0,89206 | 0,95255977 | novel transcript |

| | | | | | | |
|---|---|---|---|---|---|---|
| ENSG00000260400 | AL513534,2 | -0,035636331 | 0,975601357 | 0,88266 | 0,94815391 | novel transcript, overlapping to TET1 |
| ENSG00000231345 | BEND3P1 | -0,035696503 | 0,975560667 | 0,9293 | 0,96977202 | BEN domain containing 3 pseudogene 1 |
| ENSG00000155330 | C16orf87 | -0,035720761 | 0,975544263 | 0,84408 | 0,92888169 | chromosome 16 open reading frame 87 |
| ENSG00000233435 | AGGF1P2 | -0,035778191 | 0,975505431 | 0,94052 | 0,97474573 | angiogenic factor with G-patch and FHA domains 1 pseudogene 2 |
| ENSG00000177519 | RPRM | -0,035810585 | 0,975483527 | 0,93337 | 0,97160226 | reprimo, TP53 dependent G2 arrest mediator homolog |
| ENSG00000133114 | GPALPP1 | -0,035981737 | 0,975367808 | 0,85704 | 0,9354716 | GPALPP motifs containing 1 |
| ENSG00000246877 | DNM1P35 | -0,036045535 | 0,975324677 | 0,94872 | 0,97853939 | dynamin 1 pseudogene 35 |
| ENSG00000226609 | AL390067,1 | -0,036087742 | 0,975296144 | 0,95867 | 0,98291816 | novel transcript |
| ENSG00000140557 | ST8SIA2 | -0,036123201 | 0,975272173 | 0,87813 | 0,9458146 | ST8 alpha-N-acetyl-neuraminide alpha-2,8-sialyltransferase 2 |
| ENSG00000133687 | TMTC1 | -0,036144191 | 0,975257984 | 0,90926 | 0,96104995 | transmembrane O-mannosyltransferase targeting cadherins 1 |
| ENSG00000233348 | BAG6 | -0,036307839 | 0,975147365 | 0,9129 | 0,96228253 | BAG cochaperone 6 |
| ENSG00000196498 | NCOR2 | -0,036346155 | 0,975121466 | 0,91099 | 0,96168891 | nuclear receptor corepressor 2 |
| ENSG00000127914 | AKAP9 | -0,03644013 | 0,975057951 | 0,89593 | 0,95491162 | A-kinase anchoring protein 9 |
| ENSG00000269929 | MIRLET7A1HG | -0,036476831 | 0,975033146 | 0,86504 | 0,93999031 | miRlet-7a-1/let-7f-1/let-7d cluster host gene |
| ENSG00000196810 | CTBP1-DT | -0,036603126 | 0,974947794 | 0,87411 | 0,94420896 | CTBP1 divergent transcript |
| ENSG00000162300 | ZFPL1 | -0,036704463 | 0,974879315 | 0,83093 | 0,92270607 | zinc finger protein like 1 |
| ENSG00000161999 | JMJD8 | -0,036767365 | 0,974836811 | 0,87262 | 0,94374455 | jumonji domain containing 8 |
| ENSG00000002933 | TMEM176A | -0,036800137 | 0,974814667 | 0,95414 | 0,9807577 | transmembrane protein 176A |
| ENSG00000257883 | AC125603,2 | -0,036929006 | 0,974727596 | 0,96275 | 0,98432594 | novel transcript |
| ENSG00000187951 | AC091057,1 | -0,036982933 | 0,974691161 | 0,91047 | 0,96136765 | OTU deubiquitinase 7A pseudogene |
| ENSG00000176623 | RMDN1 | -0,036986407 | 0,974688814 | 0,8197 | 0,91711385 | regulator of microtubule dynamics 1 |
| ENSG00000120784 | ZFP30 | -0,037054668 | 0,974642698 | 0,80592 | 0,91077382 | ZFP30 zinc finger protein |
| ENSG00000237513 | AC007384,1 | -0,037066688 | 0,974634578 | 0,91178 | 0,96184653 | novel transcript |
| ENSG00000259967 | AC087761,1 | -0,037189576 | 0,974551562 | 0,97209 | 0,98800339 | novel transcript |
| ENSG00000239002 | SCARNA10 | -0,037237363 | 0,974519282 | 0,91556 | 0,9633821 | small Cajal body-specific RNA 10 |
| ENSG00000198752 | CDC42BPB | -0,03725629 | 0,974506498 | 0,87786 | 0,94568249 | CDC42 binding protein kinase beta |
| ENSG00000213753 | CENPBD1P1 | -0,037415255 | 0,974399127 | 0,81634 | 0,91550954 | CENPB DNA-binding domains containing 1 pseudogene 1 |
| ENSG00000175567 | UCP2 | -0,037481574 | 0,974354336 | 0,90201 | 0,95811717 | uncoupling protein 2 |
| ENSG00000288071 | AL591135,2 | -0,037505315 | 0,974338302 | 0,96733 | 0,98623935 | novel transcript |
| ENSG00000234650 | PCCA-AS1 | -0,037521816 | 0,974327158 | 0,9536 | 0,98051225 | PCCA antisense RNA 1 |

| | | | | | | |
|---|---|---|---|---|---|---|
| ENSG00000240522 | RPL7AP10 | -0,037535599 | 0,974317849 | 0,92667 | 0,96815248 | ribosomal protein L7a pseudogene 10 |
| ENSG00000253667 | AC100821,1 | -0,037542843 | 0,974312957 | 0,9304 | 0,97019767 | ESF1, nucleolar pre-rRNA processing protein, homolog (S, cerevisiae) (ESF1) pseudogene |
| ENSG00000258670 | AL049874,3 | -0,037736081 | 0,974182464 | 0,92274 | 0,96656551 | novel transcript, antisense to C14orf39 |
| ENSG00000236018 | AC004898,1 | -0,037853728 | 0,974103026 | 0,90996 | 0,96121516 | UBX domain protein 2B (UBXN2B) pseudogene |
| ENSG00000283738 | AC241640,1 | -0,037857553 | 0,974100443 | 0,88202 | 0,94761312 | |
| ENSG00000032444 | PNPLA6 | -0,037963039 | 0,974029222 | 0,87198 | 0,94350937 | patatin like phospholipase domain containing 6 |
| ENSG00000259605 | AC074212,1 | -0,038152662 | 0,973901208 | 0,90582 | 0,95975315 | novel transcript, antisense to SIX5 |
| ENSG00000235941 | HSPA1A | -0,038236896 | 0,973844346 | 0,94098 | 0,97489099 | heat shock protein family A (Hsp70) member 1A |
| ENSG00000229337 | AC079305,2 | -0,038254261 | 0,973832625 | 0,90789 | 0,96044232 | novel transcript |
| ENSG00000154743 | TSEN2 | -0,038318254 | 0,97378943 | 0,81197 | 0,9134209 | tRNA splicing endonuclease subunit 2 |
| ENSG00000197037 | ZSCAN25 | -0,038538654 | 0,973640676 | 0,8336 | 0,92396169 | zinc finger and SCAN domain containing 25 |
| ENSG00000287315 | AL670729,3 | -0,038542192 | 0,973638288 | 0,89858 | 0,95646662 | novel transcript, antisense to OBSCN |
| ENSG00000100503 | NIN | -0,038574445 | 0,973616522 | 0,89121 | 0,95208954 | ninein |
| ENSG00000224596 | ZMIZ1-AS1 | -0,038651206 | 0,97356472 | 0,95224 | 0,97974576 | ZMIZ1 antisense RNA 1 |
| ENSG00000237162 | RPS20P9 | -0,038678017 | 0,973546627 | 0,96983 | 0,98728333 | ribosomal protein S20 pseudogene 9 |
| ENSG00000022556 | NLRP2 | -0,038686433 | 0,973540949 | 0,92093 | 0,96576077 | NLR family pyrin domain containing 2 |
| ENSG00000169379 | ARL13B | -0,0386941 | 0,973535774 | 0,84093 | 0,92751395 | ADP ribosylation factor like GTPase 13B |
| ENSG00000228049 | POLR2J2 | -0,038750596 | 0,973497652 | 0,9136 | 0,96257531 | RNA polymerase II subunit J2 |
| ENSG00000012061 | ERCC1 | -0,038807623 | 0,973459172 | 0,83407 | 0,92405734 | ERCC excision repair 1, endonuclease non-catalytic subunit |
| ENSG00000178301 | AQP11 | -0,03882119 | 0,973450018 | 0,91582 | 0,9633821 | aquaporin 11 |
| ENSG00000166963 | MAP1A | -0,038828865 | 0,973444839 | 0,85807 | 0,93611043 | microtubule associated protein 1A |
| ENSG00000198885 | ITPRIPL1 | -0,038857073 | 0,973425806 | 0,90693 | 0,96017342 | ITPRIP like 1 |
| ENSG00000173209 | AHSA2P | -0,038875176 | 0,973413592 | 0,85756 | 0,93569939 | activator of HSP90 ATPase homolog 2, pseudogene |
| ENSG00000189367 | KIAA0408 | -0,03905465 | 0,973292505 | 0,93546 | 0,9722573 | KIAA0408 |
| ENSG00000267582 | AC020916,2 | -0,039075431 | 0,973278485 | 0,95617 | 0,98189243 | novel transcript, antisense to ZSWIM4 |
| ENSG00000288519 | AC020916,6 | -0,039075431 | 0,973278485 | 0,95617 | 0,98189243 | novel transcript, antisense to ZSWIM4 |
| ENSG00000116514 | RNF19B | -0,039128754 | 0,973242513 | 0,87779 | 0,94568249 | ring finger protein 19B |
| ENSG00000184602 | SNN | -0,039161532 | 0,973220401 | 0,87633 | 0,94545974 | stannin |
| ENSG00000170417 | TMEM182 | -0,039245134 | 0,973164006 | 0,84938 | 0,93144541 | transmembrane protein 182 |
| ENSG00000270421 | AC022022,2 | -0,039290545 | 0,973133374 | 0,95903 | 0,98306356 | leucine rich repeat containing 40 (LRRC40) pseudogene |
| ENSG00000166183 | ASPG | -0,039317084 | 0,973115474 | 0,97167 | 0,98794066 | asparaginase |
| ENSG00000236809 | SNX25P1 | -0,039702493 | 0,972855546 | 0,96804 | 0,98637425 | sorting nexin 25 pseudogene 1 |

| | | | | | | |
|---|---|---|---|---|---|---|
| ENSG00000144406 | UNC80 | -0,039703638 | 0,972854773 | 0,86975 | 0,9425949 | unc-80 homolog, NALCN channel complex subunit |
| ENSG00000225572 | DOCK4-AS1 | -0,039724751 | 0,972840536 | 0,93323 | 0,97152506 | DOCK4 antisense RNA 1 |
| ENSG00000102977 | ACD | -0,039925505 | 0,972705173 | 0,84476 | 0,92922421 | ACD shelterin complex subunit and telomerase recruitment factor |
| ENSG00000236708 | SDK1-AS1 | -0,039940448 | 0,972695098 | 0,95226 | 0,97974576 | SDK1 antisense RNA 1 |
| ENSG00000197858 | GPAA1 | -0,039989302 | 0,97266216 | 0,84292 | 0,9282764 | glycosylphosphatidylinositol anchor attachment 1 |
| ENSG00000248540 | AC010931,1 | -0,040003688 | 0,972652461 | 0,92969 | 0,96987987 | novel transcript |
| ENSG00000144840 | RABL3 | -0,040049938 | 0,97262128 | 0,84477 | 0,92922421 | RAB, member of RAS oncogene family like 3 |
| ENSG00000162062 | TEDC2 | -0,040070718 | 0,972607271 | 0,86973 | 0,9425949 | tubulin epsilon and delta complex 2 |
| ENSG00000276488 | AC008735,4 | -0,040130714 | 0,972566825 | 0,94884 | 0,97862581 | novel transcript, antisense to ZNF865 |
| ENSG00000277111 | PLEKHM1 | -0,040227274 | 0,972501733 | 0,82233 | 0,91794576 | pleckstrin homology and RUN domain containing M1 |
| ENSG00000183092 | BEGAIN | -0,040251873 | 0,972485151 | 0,94533 | 0,97666034 | brain enriched guanylate kinase associated |
| ENSG00000175727 | MLXIP | -0,040295057 | 0,972456043 | 0,85493 | 0,93417381 | MLX interacting protein |
| ENSG00000255794 | RMST | -0,040320737 | 0,972438733 | 0,91752 | 0,96423084 | rhabdomyosarcoma 2 associated transcript |
| ENSG00000181852 | RNF41 | -0,04032339 | 0,972436944 | 0,83618 | 0,92493625 | ring finger protein 41 |
| ENSG00000090266 | NDUFB2 | -0,040324809 | 0,972435988 | 0,85999 | 0,93704372 | NADH:ubiquinone oxidoreductase subunit B2 |
| ENSG00000140830 | TXNL4B | -0,040342991 | 0,972423733 | 0,8547 | 0,93416235 | thioredoxin like 4B |
| ENSG00000234084 | AL049552,1 | -0,040382527 | 0,972397084 | 0,94694 | 0,97745567 | novel transcript |
| ENSG00000226254 | PTP4A1P3 | -0,040391283 | 0,972391183 | 0,95652 | 0,98197334 | PTP4A1 pseudogene 3 |
| ENSG00000286463 | AP002755,1 | -0,040472757 | 0,97233627 | 0,93866 | 0,9737443 | novel transcript, antisense to ZBTB16 |
| ENSG00000099849 | RASSF7 | -0,040598365 | 0,972251617 | 0,92005 | 0,96530017 | Ras association domain family member 7 |
| ENSG00000282086 | PKD1P6 | -0,040619309 | 0,972237503 | 0,82629 | 0,91953499 | polycystin 1, transient receptor potential channel interacting pseudogene 6 |
| ENSG00000023909 | GCLM | -0,04091047 | 0,972041309 | 0,86394 | 0,93915148 | glutamate-cysteine ligase modifier subunit |
| ENSG00000143373 | ZNF687 | -0,0410371 | 0,971955993 | 0,80366 | 0,90968333 | zinc finger protein 687 |
| ENSG00000285293 | TBILA | -0,041118799 | 0,971900953 | 0,95085 | 0,9793462 | TGF-beta induced lncRNA |
| ENSG00000261488 | TBILA | -0,041118799 | 0,971900953 | 0,95085 | 0,9793462 | TGF-beta induced lncRNA |
| ENSG00000159685 | CHCHD6 | -0,041140771 | 0,971886151 | 0,8733 | 0,94386143 | coiled-coil-helix-coiled-coil-helix domain containing 6 |
| ENSG00000099800 | TIMM13 | -0,041455663 | 0,971674045 | 0,83837 | 0,92625269 | translocase of inner mitochondrial membrane 13 |
| ENSG00000281917 | SLC16A1 | -0,041480881 | 0,97165706 | 0,87159 | 0,94322889 | solute carrier family 16 member 1 |
| ENSG00000248477 | NAIPP3 | -0,041567918 | 0,971598442 | 0,90754 | 0,96044232 | NAIP pseudogene 3 |
| ENSG00000225989 | ABCF1 | -0,041603928 | 0,971574191 | 0,87671 | 0,94549417 | ATP binding cassette subfamily F member 1 |
| ENSG00000158008 | EXTL1 | -0,04161963 | 0,971563617 | 0,89376 | 0,95368671 | exostosin like glycosyltransferase 1 |

| | | | | | | |
|---|---|---|---|---|---|---|
| ENSG00000072071 | ADGRL1 | -0,041975096 | 0,971324263 | 0,90883 | 0,96099548 | adhesion G protein-coupled receptor L1 |
| ENSG00000288324 | ADGRL1 | -0,041975096 | 0,971324263 | 0,90883 | 0,96099548 | adhesion G protein-coupled receptor L1 |
| ENSG00000159363 | ATP13A2 | -0,042089264 | 0,9712474 | 0,83134 | 0,9229059 | ATPase cation transporting 13A2 |
| ENSG00000285049 | CPSF1 | -0,042114933 | 0,971230119 | 0,80976 | 0,91232038 | cleavage and polyadenylation specific factor 1 |
| ENSG00000071894 | CPSF1 | -0,042114933 | 0,971230119 | 0,80976 | 0,91232038 | cleavage and polyadenylation specific factor 1 |
| ENSG00000274554 | AC083806,2 | -0,042130167 | 0,971219864 | 0,87695 | 0,94549531 | novel transcript, antisense to FBXW8 |
| ENSG00000177025 | C19orf18 | -0,042135127 | 0,971216524 | 0,94125 | 0,97499833 | chromosome 19 open reading frame 18 |
| ENSG00000225830 | ERCC6 | -0,042182856 | 0,971184394 | 0,85969 | 0,93692606 | ERCC excision repair 6, chromatin remodeling factor |
| ENSG00000247556 | OIP5-AS1 | -0,042222328 | 0,971157823 | 0,81626 | 0,91550954 | OIP5 antisense RNA 1 |
| ENSG00000284391 | AL139398,1 | -0,042223101 | 0,971157302 | 0,93081 | 0,97034563 | uncharacterized LOC105373244 |
| ENSG00000282405 | AC019257,7 | -0,04224849 | 0,971140212 | 0,90505 | 0,95938369 | novel transcript, antisense to ARHGEF10 |
| ENSG00000237298 | TTN-AS1 | -0,042293646 | 0,971109816 | 0,88059 | 0,94687991 | TTN antisense RNA 1 |
| ENSG00000197608 | ZNF841 | -0,042313144 | 0,971096692 | 0,88841 | 0,95069428 | zinc finger protein 841 |
| ENSG00000230424 | EMC1-AS1 | -0,042325511 | 0,971088367 | 0,94577 | 0,97689657 | EMC1 antisense RNA 1 |
| ENSG00000196593 | ANKRD20A19 | -0,04237157 | 0,971057365 | 0,91282 | 0,96223725 | ankyrin repeat domain 20 family member A19, pseudogene |
| ENSG00000160688 | FLAD1 | -0,042395018 | 0,971041583 | 0,83598 | 0,92493625 | flavin adenine dinucleotide synthetase 1 |
| ENSG00000236409 | NRADDP | -0,04243598 | 0,971014012 | 0,94249 | 0,97556707 | neurotrophin receptor associated death domain, pseudogene |
| ENSG00000043143 | JADE2 | -0,042525272 | 0,970953916 | 0,91856 | 0,96459263 | jade family PHD finger 2 |
| ENSG00000103381 | CPPED1 | -0,042541424 | 0,970943045 | 0,85817 | 0,93612837 | calcineurin like phosphoesterase domain containing 1 |
| ENSG00000145916 | RMND5B | -0,042657893 | 0,970864664 | 0,80601 | 0,91078302 | required for meiotic nuclear division 5 homolog B |
| ENSG00000102104 | RS1 | -0,04270111 | 0,970835582 | 0,94083 | 0,97486264 | retinoschisin 1 |
| ENSG00000254449 | SF3A3P2 | -0,042703756 | 0,970833801 | 0,93806 | 0,97361277 | splicing factor 3a, subunit 3 pseudogene 2 |
| ENSG00000286358 | AC096647,1 | -0,042860502 | 0,970728328 | 0,83824 | 0,92618329 | novel transcript |
| ENSG00000143294 | PRCC | -0,042952713 | 0,970666284 | 0,8639 | 0,93914712 | proline rich mitotic checkpoint control factor |
| ENSG00000139974 | SLC38A6 | -0,043078262 | 0,970581817 | 0,87086 | 0,94302816 | solute carrier family 38 member 6 |
| ENSG00000234889 | KPNB1P1 | -0,043100597 | 0,970566791 | 0,98006 | 0,9917697 | KPNB1 pseudogene 1 |
| ENSG00000204685 | STARD7-AS1 | -0,043144353 | 0,970537355 | 0,8401 | 0,92699467 | STARD7 antisense RNA 1 |
| ENSG00000123977 | DAW1 | -0,043172624 | 0,970518337 | 0,95235 | 0,97980306 | dynein assembly factor with WD repeats 1 |
| ENSG00000271743 | AF287957,1 | -0,043175524 | 0,970516385 | 0,92301 | 0,96662141 | novel transcript |
| ENSG00000285267 | AF287957,2 | -0,043175524 | 0,970516385 | 0,92301 | 0,96662141 | novel transcript |
| ENSG00000169756 | LIMS1 | -0,04322485 | 0,970483204 | 0,77468 | 0,89586241 | LIM zinc finger domain containing 1 |

| | | | | | | |
|---|---|---|---|---|---|---|
| ENSG00000274292 | AC084018,2 | -0,043312721 | 0,970424096 | 0,85456 | 0,93413847 | novel transcript |
| ENSG00000272505 | AC104964,3 | -0,04331537 | 0,970422314 | 0,94508 | 0,97666034 | novel transcript |
| ENSG00000206656 | SNORD116-17 | -0,043325586 | 0,970415443 | 0,95023 | 0,97920193 | small nucleolar RNA, C/D box 116-17 |
| ENSG00000207460 | SNORD116-19 | -0,043325586 | 0,970415443 | 0,95023 | 0,97920193 | small nucleolar RNA, C/D box 116-19 |
| ENSG00000257802 | MRS2P2 | -0,043341279 | 0,970404887 | 0,93521 | 0,97222198 | MRS2 pseudogene 2 |
| ENSG00000153208 | MERTK | -0,043342575 | 0,970404015 | 0,91238 | 0,96205682 | MER proto-oncogene, tyrosine kinase |
| ENSG00000112531 | QKI | -0,0434782 | 0,970312794 | 0,90058 | 0,95772302 | QKI, KH domain containing RNA binding |
| ENSG00000142549 | IGLON5 | -0,043503436 | 0,970295821 | 0,85761 | 0,93572248 | IgLON family member 5 |
| ENSG00000083896 | YTHDC1 | -0,043511891 | 0,970290134 | 0,80625 | 0,91085216 | YTH domain containing 1 |
| ENSG00000240808 | AC126389,1 | -0,043528991 | 0,970278634 | 0,95418 | 0,9807577 | ribosomal protein L34 (RPL34) pseudogene |
| ENSG00000091831 | ESR1 | -0,043578114 | 0,970245597 | 0,91497 | 0,9632313 | estrogen receptor 1 |
| ENSG00000253734 | LINC01289 | -0,043595731 | 0,970233749 | 0,94528 | 0,97666034 | long intergenic non-protein coding RNA 1289 |
| ENSG00000226472 | AC008013,1 | -0,043622871 | 0,970215497 | 0,91232 | 0,96205682 | tetraspanin family pseudogene |
| ENSG00000142082 | SIRT3 | -0,043673048 | 0,970181754 | 0,86197 | 0,93834335 | sirtuin 3 |
| ENSG00000234602 | MCIDAS | -0,043790487 | 0,970102782 | 0,91872 | 0,9646736 | multiciliate differentiation and DNA synthesis associated cell cycle protein |
| ENSG00000260007 | AC107871,1 | -0,043803523 | 0,970094016 | 0,84824 | 0,93094401 | novel protein |
| ENSG00000178445 | GLDC | -0,043858366 | 0,970057139 | 0,8975 | 0,95579119 | glycine decarboxylase |
| ENSG00000167900 | TK1 | -0,043897464 | 0,97003085 | 0,87756 | 0,94566578 | thymidine kinase 1 |
| ENSG00000005801 | ZNF195 | -0,043956009 | 0,969991487 | 0,77565 | 0,89630112 | zinc finger protein 195 |
| ENSG00000100647 | SUSD6 | -0,044079111 | 0,969908723 | 0,85118 | 0,93226967 | sushi domain containing 6 |
| ENSG00000141985 | SH3GL1 | -0,044082805 | 0,96990624 | 0,82736 | 0,92017947 | SH3 domain containing GRB2 like 1, endophilin A2 |
| ENSG00000133639 | BTG1 | -0,044169833 | 0,969847734 | 0,85068 | 0,93206349 | BTG anti-proliferation factor 1 |
| ENSG00000233554 | B4GALT1-AS1 | -0,044171029 | 0,96984693 | 0,94464 | 0,97650311 | B4GALT1 antisense RNA 1 |
| ENSG00000259571 | BLID | -0,044192528 | 0,969832478 | 0,95697 | 0,98218743 | BH3-like motif containing, cell death inducer |
| ENSG00000010810 | FYN | -0,04428278 | 0,969771808 | 0,82521 | 0,91906552 | FYN proto-oncogene, Src family tyrosine kinase |
| ENSG00000010539 | ZNF200 | -0,044382881 | 0,969704523 | 0,84897 | 0,93125441 | zinc finger protein 200 |
| ENSG00000286145 | AC068282,1 | -0,044439185 | 0,96966668 | 0,88809 | 0,95069428 | novel transcript, antisense to MAP3K2 |
| ENSG00000107863 | ARHGAP21 | -0,044515624 | 0,969615305 | 0,90804 | 0,96056328 | Rho GTPase activating protein 21 |
| ENSG00000102984 | ZNF821 | -0,044585679 | 0,969568223 | 0,82453 | 0,91900665 | zinc finger protein 821 |
| ENSG00000106344 | RBM28 | -0,044640338 | 0,96953149 | 0,78635 | 0,90080813 | RNA binding motif protein 28 |
| ENSG00000086015 | MAST2 | -0,044667492 | 0,969513242 | 0,84336 | 0,92850187 | microtubule associated serine/threonine kinase 2 |
| ENSG00000163322 | ABRAXAS1 | -0,044733023 | 0,969469205 | 0,88301 | 0,94823653 | abraxas 1, BRCA1 A complex subunit |

| | | | | | | |
|---|---|---|---|---|---|---|
| ENSG00000287340 | AC007222,2 | -0,044735329 | 0,969467655 | 0,95305 | 0,98024505 | novel transcript, antisense to RBFOX1 |
| ENSG00000143494 | VASH2 | -0,044784476 | 0,96943463 | 0,88799 | 0,95069428 | vasohibin 2 |
| ENSG00000116584 | ARHGEF2 | -0,04485087 | 0,969390017 | 0,78075 | 0,89846287 | Rho/Rac guanine nucleotide exchange factor 2 |
| ENSG00000188785 | ZNF548 | -0,044883796 | 0,969367893 | 0,79254 | 0,90361513 | zinc finger protein 548 |
| ENSG00000122678 | POLM | -0,044884298 | 0,969367556 | 0,82051 | 0,91732362 | DNA polymerase mu |
| ENSG00000143376 | SNX27 | -0,044887654 | 0,969365301 | 0,78965 | 0,90258817 | sorting nexin 27 |
| ENSG00000025039 | RRAGD | -0,044890021 | 0,96936371 | 0,86239 | 0,93853472 | Ras related GTP binding D |
| ENSG00000167468 | GPX4 | -0,04489947 | 0,969357361 | 0,81051 | 0,91251177 | glutathione peroxidase 4 |
| ENSG00000273209 | AC069148,1 | -0,044908949 | 0,969350993 | 0,92875 | 0,96941592 | novel transcript |
| ENSG00000155256 | ZFYVE27 | -0,044972199 | 0,969308495 | 0,78357 | 0,89918294 | zinc finger FYVE-type containing 27 |
| ENSG00000197063 | MAFG | -0,044987554 | 0,969298179 | 0,87663 | 0,94547358 | MAF bZIP transcription factor G |
| ENSG00000118976 | OTUD4P1 | -0,045102605 | 0,969220883 | 0,91718 | 0,9640264 | OTUD4 pseudogene 1 |
| ENSG00000144852 | NR1I2 | -0,045152748 | 0,969187197 | 0,91293 | 0,96228253 | nuclear receptor subfamily 1 group I member 2 |
| ENSG00000270095 | AC068790,6 | -0,045212846 | 0,969146825 | 0,94905 | 0,97866986 | novel transcript, sense intronic to CCDC92 |
| ENSG00000235078 | AC231981,1 | -0,04530037 | 0,969088031 | 0,93472 | 0,9721963 | novel transcript, antisense to ADI1 |
| ENSG00000165898 | ISCA2 | -0,045343323 | 0,969059179 | 0,84258 | 0,92821072 | iron-sulfur cluster assembly 2 |
| ENSG00000087008 | ACOX3 | -0,045368417 | 0,969042324 | 0,81867 | 0,91672491 | acyl-CoA oxidase 3, pristanoyl |
| ENSG00000039600 | SOX30 | -0,045589372 | 0,968893922 | 0,91415 | 0,96285318 | SRY-box transcription factor 30 |
| ENSG00000175832 | ETV4 | -0,045631915 | 0,968865351 | 0,89578 | 0,95482399 | ETS variant transcription factor 4 |
| ENSG00000287794 | AC119150,2 | -0,045679917 | 0,968833115 | 0,95856 | 0,98291816 | novel ancient vomeronasal 1 receptor pseudogene |
| ENSG00000079462 | PAFAH1B3 | -0,045811905 | 0,968744484 | 0,81438 | 0,91470265 | platelet activating factor acetylhydrolase 1b catalytic subunit 3 |
| ENSG00000257913 | DDN-AS1 | -0,045855132 | 0,968715458 | 0,89748 | 0,95579119 | DDN and PRKAG1 antisense RNA 1 |
| ENSG00000134222 | PSRC1 | -0,045927421 | 0,96866692 | 0,7967 | 0,90605632 | proline and serine rich coiled-coil 1 |
| ENSG00000157036 | EXOG | -0,045950336 | 0,968651534 | 0,79533 | 0,90519396 | exo/endonuclease G |
| ENSG00000148468 | FAM171A1 | -0,046026676 | 0,96860028 | 0,808 | 0,91168773 | family with sequence similarity 171 member A1 |
| ENSG00000235233 | MICA | -0,046052609 | 0,968582869 | 0,92023 | 0,96530017 | MHC class I polypeptide-related sequence A |
| ENSG00000023608 | SNAPC1 | -0,046208064 | 0,968478506 | 0,83026 | 0,92216235 | small nuclear RNA activating complex polypeptide 1 |
| ENSG00000277171 | AC010975,1 | -0,046208873 | 0,968477963 | 0,95826 | 0,98276958 | novel zinc finger protein pseudogene |
| ENSG00000120709 | FAM53C | -0,04641921 | 0,968336775 | 0,81574 | 0,91527054 | family with sequence similarity 53 member C |
| ENSG00000122008 | POLK | -0,046427227 | 0,968331394 | 0,78201 | 0,8987738 | DNA polymerase kappa |
| ENSG00000171497 | PPID | -0,046431593 | 0,968328463 | 0,77402 | 0,89572556 | peptidylprolyl isomerase D |
| ENSG00000261476 | AC009163,4 | -0,046487433 | 0,968290985 | 0,93155 | 0,9707979 | novel transcript |

| | | | | | | |
|---|---|---|---|---|---|---|
| ENSG00000165985 | C1QL3 | -0,046522534 | 0,968267426 | 0,94373 | 0,9762643 | complement C1q like 3 |
| ENSG00000213760 | ATP6V1G2 | -0,046580716 | 0,968228378 | 0,94895 | 0,97862975 | ATPase H+ transporting V1 subunit G2 |
| ENSG00000167699 | GLOD4 | -0,046618126 | 0,968203271 | 0,83044 | 0,92228969 | glyoxalase domain containing 4 |
| ENSG00000173295 | FAM86B3P | -0,046665843 | 0,968171248 | 0,82485 | 0,91900665 | family with sequence similarity 86 member B3, pseudogene |
| ENSG00000236137 | AL445231,1 | -0,046683141 | 0,96815964 | 0,9184 | 0,96446185 | novel transcript |
| ENSG00000229241 | PNPT1P1 | -0,046708255 | 0,968142787 | 0,93824 | 0,97361277 | polyribonucleotide nucleotidyltransferase 1 pseudogene 1 |
| ENSG00000110921 | MVK | -0,046778429 | 0,968095697 | 0,78736 | 0,90121071 | mevalonate kinase |
| ENSG00000132463 | GRSF1 | -0,04687099 | 0,968033587 | 0,83385 | 0,92404752 | G-rich RNA sequence binding factor 1 |
| ENSG00000264596 | AP000897,1 | -0,046894173 | 0,968018032 | 0,928 | 0,96899998 | novel transcript, sense intronic to PTPRM |
| ENSG00000138380 | CARF | -0,046936574 | 0,967989582 | 0,81833 | 0,91663479 | calcium responsive transcription factor |
| ENSG00000167632 | TRAPPC9 | -0,046949949 | 0,967980608 | 0,78074 | 0,89846287 | trafficking protein particle complex 9 |
| ENSG00000285048 | IRF9 | -0,046963332 | 0,967971629 | 0,86321 | 0,93891554 | interferon regulatory factor 9 |
| ENSG00000240230 | COX19 | -0,047009894 | 0,967940389 | 0,79405 | 0,90426294 | cytochrome c oxidase assembly factor COX19 |
| ENSG00000196083 | IL1RAP | -0,047041438 | 0,967919225 | 0,91162 | 0,96182676 | interleukin 1 receptor accessory protein |
| ENSG00000119632 | IFI27L2 | -0,047150928 | 0,96784577 | 0,90781 | 0,96044232 | interferon alpha inducible protein 27 like 2 |
| ENSG00000282308 | DPRXP3 | -0,047196866 | 0,967814953 | 0,93674 | 0,97279937 | divergent-paired related homeobox pseudogene 3 |
| ENSG00000196150 | ZNF250 | -0,047211276 | 0,967805286 | 0,78571 | 0,90048075 | zinc finger protein 250 |
| ENSG00000237840 | FAM21FP | -0,047304756 | 0,967742578 | 0,92568 | 0,96768961 | family with sequence similarity 21 member F, pseudogene |
| ENSG00000233937 | CTC-338M12,4 | -0,047436157 | 0,967654441 | 0,89746 | 0,95579119 | uncharacterized LOC101928649 |
| ENSG00000283196 | LSP1P5 | -0,047453203 | 0,967643007 | 0,90689 | 0,96017342 | LSP1 pseudogene 5 |
| ENSG00000105127 | AKAP8 | -0,047480089 | 0,967624974 | 0,78988 | 0,90272027 | A-kinase anchoring protein 8 |
| ENSG00000285731 | AC090644,1 | -0,047492661 | 0,967616543 | 0,94354 | 0,97617591 | Novel transcript, antisense to TBC1D5 |
| ENSG00000100473 | COCH | -0,047675925 | 0,967493635 | 0,91179 | 0,96184653 | cochlin |
| ENSG00000108799 | EZH1 | -0,047728911 | 0,967458102 | 0,75377 | 0,88462383 | enhancer of zeste 1 polycomb repressive complex 2 subunit |
| ENSG00000136731 | UGGT1 | -0,04774701 | 0,967445966 | 0,83092 | 0,92270607 | UDP-glucose glycoprotein glucosyltransferase 1 |
| ENSG00000262202 | AC007952,4 | -0,047904237 | 0,967340538 | 0,87145 | 0,94322889 | novel transcript |
| ENSG00000160055 | TMEM234 | -0,048109263 | 0,967203076 | 0,7922 | 0,90351559 | transmembrane protein 234 |
| ENSG00000284991 | AC275401,4 | -0,048185731 | 0,967151811 | 0,96066 | 0,98384009 | ubiquitin specific peptidase 17-like 2 ( USP17L2) pseudogene |
| ENSG00000082269 | FAM135A | -0,048194275 | 0,967146084 | 0,75516 | 0,88526692 | family with sequence similarity 135 member A |
| ENSG00000128298 | BAIAP2L2 | -0,048229931 | 0,967122181 | 0,9478 | 0,97802717 | BAR/IMD domain containing adaptor protein 2 like 2 |

| | | | | | | |
|---|---|---|---|---|---|---|
| ENSG00000267009 | AC007780,1 | -0,048294775 | 0,967078713 | 0,94733 | 0,97775136 | novel transcript |
| ENSG00000149231 | CCDC82 | -0,048323958 | 0,967059151 | 0,85508 | 0,93419851 | coiled-coil domain containing 82 |
| ENSG00000272159 | AC087623,3 | -0,048351284 | 0,967040835 | 0,93865 | 0,9737443 | novel transcript, antisense to LETM2 |
| ENSG00000141499 | WRAP53 | -0,04838298 | 0,967019589 | 0,82296 | 0,91812142 | WD repeat containing antisense to TP53 |
| ENSG00000278545 | MARCHF8 | -0,048396459 | 0,967010555 | 0,85881 | 0,93641306 | membrane associated ring-CH-type finger 8 |
| ENSG00000117281 | CD160 | -0,048493325 | 0,966945629 | 0,87826 | 0,9458146 | CD160 molecule |
| ENSG00000127580 | WDR24 | -0,048499472 | 0,966941509 | 0,84486 | 0,92925254 | WD repeat domain 24 |
| ENSG00000225446 | LINC01763 | -0,048518925 | 0,966928472 | 0,94632 | 0,97713881 | long intergenic non-protein coding RNA 1763 |
| ENSG00000286068 | AC010974,2 | -0,048625112 | 0,966857305 | 0,94043 | 0,97472523 | novel transcript, antisense to NCKAP5 |
| ENSG00000068308 | OTUD5 | -0,048649448 | 0,966840995 | 0,82032 | 0,9172552 | OTU deubiquitinase 5 |
| ENSG00000223482 | NUTM2A-AS1 | -0,048777004 | 0,966755516 | 0,80185 | 0,90880281 | NUTM2A antisense RNA 1 |
| ENSG00000042429 | MED17 | -0,048783691 | 0,966751035 | 0,79283 | 0,90366929 | mediator complex subunit 17 |
| ENSG00000270069 | MIR222HG | -0,048791129 | 0,966746051 | 0,8942 | 0,95389664 | miR222/221 cluster host gene |
| ENSG00000286878 | AC100805,1 | -0,048822676 | 0,966724911 | 0,95087 | 0,9793462 | novel transcript |
| ENSG00000113391 | FAM172A | -0,048889013 | 0,966680461 | 0,80379 | 0,90974015 | family with sequence similarity 172 member A |
| ENSG00000163516 | ANKZF1 | -0,048901601 | 0,966672027 | 0,80086 | 0,90830749 | ankyrin repeat and zinc finger peptidyl tRNA hydrolase 1 |
| ENSG00000223969 | AC002456,1 | -0,049079354 | 0,966552931 | 0,89085 | 0,95207499 | novel transcript |
| ENSG00000180035 | ZNF48 | -0,049104465 | 0,966536108 | 0,78316 | 0,89918294 | zinc finger protein 48 |
| ENSG00000118997 | DNAH7 | -0,049452462 | 0,966302995 | 0,86797 | 0,94175463 | dynein axonemal heavy chain 7 |
| ENSG00000131979 | GCH1 | -0,049456725 | 0,96630014 | 0,92092 | 0,96576077 | GTP cyclohydrolase 1 |
| ENSG00000105229 | PIAS4 | -0,049525465 | 0,9662541 | 0,81244 | 0,91379618 | protein inhibitor of activated STAT 4 |
| ENSG00000282535 | AC092192,1 | -0,049525621 | 0,966253995 | 0,93924 | 0,9741012 | novel transcript |
| ENSG00000226800 | CACTIN-AS1 | -0,049627618 | 0,966185684 | 0,96726 | 0,98623935 | CACTIN antisense RNA 1 |
| ENSG00000164010 | ERMAP | -0,04976592 | 0,966093067 | 0,77969 | 0,8981559 | erythroblast membrane associated protein (Scianna blood group) |
| ENSG00000251246 | AL691442,1 | -0,049860075 | 0,966030018 | 0,96772 | 0,98623935 | novel ephrin-A4 (EFNA4) and ephrin-A3 (EFNA3) protein |
| ENSG00000159496 | RGL4 | -0,049952815 | 0,965967922 | 0,90104 | 0,95787401 | ral guanine nucleotide dissociation stimulator like 4 |
| ENSG00000074590 | NUAK1 | -0,050242552 | 0,965773945 | 0,8336 | 0,92396169 | NUAK family kinase 1 |
| ENSG00000164068 | RNF123 | -0,050252712 | 0,965767144 | 0,82341 | 0,9183689 | ring finger protein 123 |
| ENSG00000068024 | HDAC4 | -0,050365114 | 0,965691903 | 0,77812 | 0,89735311 | histone deacetylase 4 |
| ENSG00000175274 | TP53I11 | -0,050420292 | 0,965654969 | 0,9021 | 0,95812378 | tumor protein p53 inducible protein 11 |
| ENSG00000272485 | AL392183,1 | -0,050469718 | 0,965621887 | 0,96062 | 0,98383172 | novel transcript |
| ENSG00000226191 | CLK3P2 | -0,050488902 | 0,965609047 | 0,96288 | 0,98432594 | CDC like kinase 3 pseudogene 2 |

| | | | | | | |
|---|---|---|---|---|---|---|
| ENSG00000105220 | GPI | -0,050491947 | 0,965607009 | 0,77393 | 0,89572556 | glucose-6-phosphate isomerase |
| ENSG00000164663 | USP49 | -0,050584101 | 0,965545331 | 0,84956 | 0,93150269 | ubiquitin specific peptidase 49 |
| ENSG00000133398 | MED10 | -0,050728514 | 0,965448686 | 0,82215 | 0,91790528 | mediator complex subunit 10 |
| ENSG00000186111 | PIP5K1C | -0,050896572 | 0,965336228 | 0,81519 | 0,91495565 | phosphatidylinositol-4-phosphate 5-kinase type 1 gamma |
| ENSG00000180015 | AC093909,1 | -0,050911766 | 0,965326061 | 0,93463 | 0,9721963 | mitochondrial translational release factor 1-like (MTRF1L) pseudogene |
| ENSG00000286444 | AL035661,2 | -0,050926545 | 0,965316173 | 0,94147 | 0,97505054 | novel transcript, antisense to APMAP |
| ENSG00000170638 | TRABD | -0,050951766 | 0,965299298 | 0,82648 | 0,91967458 | TraB domain containing |
| ENSG00000169905 | TOR1AIP2 | -0,050971218 | 0,965286282 | 0,79237 | 0,90358057 | torsin 1A interacting protein 2 |
| ENSG00000084733 | RAB10 | -0,050977393 | 0,965282151 | 0,75483 | 0,88512447 | RAB10, member RAS oncogene family |
| ENSG00000260793 | AC003102,1 | -0,051007235 | 0,965262184 | 0,88128 | 0,94715141 | novel transcript, antisense to UBTF |
| ENSG00000126067 | PSMB2 | -0,051022688 | 0,965251845 | 0,83666 | 0,92527363 | proteasome 20S subunit beta 2 |
| ENSG00000263826 | AC112907,3 | -0,051040176 | 0,965240144 | 0,83805 | 0,92611649 | novel transcript, antisense to eukaryotic translation initiation factor 4A, isoform 2 EIF4A2 |
| ENSG00000166619 | BLCAP | -0,051129288 | 0,965180525 | 0,82836 | 0,92081006 | BLCAP apoptosis inducing factor |
| ENSG00000065057 | NTHL1 | -0,05115964 | 0,96516022 | 0,81419 | 0,91457884 | nth like DNA glycosylase 1 |
| ENSG00000205084 | TMEM231 | -0,051214359 | 0,965123614 | 0,78632 | 0,90080813 | transmembrane protein 231 |
| ENSG00000139973 | SYT16 | -0,051428444 | 0,964980408 | 0,91164 | 0,96182676 | synaptotagmin 16 |
| ENSG00000140688 | RUSF1 | -0,051449252 | 0,96496649 | 0,78035 | 0,89823499 | RUS family member 1 |
| ENSG00000287348 | AC099535,2 | -0,051483034 | 0,964943894 | 0,92278 | 0,96656995 | novel transcript, antisense to SLC4A7 |
| ENSG00000277005 | AC006449,4 | -0,051694558 | 0,964802427 | 0,90789 | 0,96044232 | |
| ENSG00000117362 | APH1A | -0,051789589 | 0,964738878 | 0,75947 | 0,88782455 | aph-1 homolog A, gamma-secretase subunit |
| ENSG00000166136 | NDUFB8 | -0,051832058 | 0,964710479 | 0,77459 | 0,89586241 | NADH:ubiquinone oxidoreductase subunit B8 |
| ENSG00000176108 | CHMP6 | -0,051849771 | 0,964698634 | 0,79141 | 0,90306634 | charged multivesicular body protein 6 |
| ENSG00000275155 | AC027348,1 | -0,051889867 | 0,964671823 | 0,85153 | 0,9324506 | novel transcript, sense intronic to ZNF423 |
| ENSG00000267281 | ATF7-NPFF | -0,051914282 | 0,964655498 | 0,89669 | 0,95542881 | ATF7-NPFF readthrough |
| ENSG00000256028 | AC026362,1 | -0,051917882 | 0,964653091 | 0,82374 | 0,91861952 | novel transcript |
| ENSG00000217897 | HSPE1P8 | -0,051949468 | 0,964631971 | 0,9399 | 0,97444851 | heat shock protein family E (Hsp10) member 1 pseudogene 8 |
| ENSG00000197747 | S100A10 | -0,052000467 | 0,964597872 | 0,89972 | 0,95730568 | S100 calcium binding protein A10 |
| ENSG00000144589 | STK11IP | -0,052089544 | 0,964538317 | 0,81837 | 0,91664258 | serine/threonine kinase 11 interacting protein |
| ENSG00000227236 | AL583844,1 | -0,052093069 | 0,96453596 | 0,9097 | 0,96105018 | solute carrier family 39 (zinc transporter), member 14 pseudogene (SLC39A14) |

| | | | | | | |
|---|---|---|---|---|---|---|
| ENSG00000206621 | SNORD116-14 | -0,05210196 | 0,964530016 | 0,95222 | 0,97974576 | small nucleolar RNA, C/D box 116-14 |
| ENSG00000181211 | HECW1-IT1 | -0,052102287 | 0,964529797 | 0,92956 | 0,96983055 | HECW1 intronic transcript 1 |
| ENSG00000213613 | RPL11P3 | -0,052128186 | 0,964512482 | 0,9153 | 0,9633004 | ribosomal protein L11 pseudogene 3 |
| ENSG00000283120 | RPL11P3 | -0,052128186 | 0,964512482 | 0,9153 | 0,9633004 | ribosomal protein L11 pseudogene 3 |
| ENSG00000268043 | NBPF12 | -0,052149523 | 0,964498217 | 0,74336 | 0,87846501 | NBPF member 12 |
| ENSG00000275389 | AC068790,9 | -0,052155113 | 0,96449448 | 0,92219 | 0,96631215 | novel transcript |
| ENSG00000177917 | ARL6IP6 | -0,052237455 | 0,964439433 | 0,77591 | 0,89635267 | ADP ribosylation factor like GTPase 6 interacting protein 6 |
| ENSG00000162231 | NXF1 | -0,052252879 | 0,964429122 | 0,76725 | 0,8920483 | nuclear RNA export factor 1 |
| ENSG00000285706 | AL807776,1 | -0,05235023 | 0,964364046 | 0,94447 | 0,97650311 | Novel transcript |
| ENSG00000261693 | AC134682,1 | -0,052385645 | 0,964340373 | 0,95582 | 0,98173232 | novel transcript, antisense to TSNARE1 |
| ENSG00000226148 | SLC25A39P1 | -0,05243462 | 0,964307638 | 0,89301 | 0,95325328 | solute carrier family 25 member 39 pseudogene 1 |
| ENSG00000186451 | SPATA12 | -0,052474567 | 0,964280937 | 0,85529 | 0,93431435 | spermatogenesis associated 12 |
| ENSG00000122965 | RBM19 | -0,052488826 | 0,964271407 | 0,83228 | 0,92327049 | RNA binding motif protein 19 |
| ENSG00000257472 | AL355112,1 | -0,052542631 | 0,964235445 | 0,95602 | 0,98181191 | novel transcript |
| ENSG00000135740 | SLC9A5 | -0,052597281 | 0,96419892 | 0,89781 | 0,95593639 | solute carrier family 9 member A5 |
| ENSG00000275619 | GON7 | -0,052643992 | 0,964167702 | 0,97259 | 0,98841317 | GON7 subunit of KEOPS complex |
| ENSG00000162881 | OXER1 | -0,052745319 | 0,964099987 | 0,93534 | 0,97222198 | oxoeicosanoid receptor 1 |
| ENSG00000198342 | ZNF442 | -0,052806384 | 0,96405918 | 0,87381 | 0,94405019 | zinc finger protein 442 |
| ENSG00000273437 | AC108673,3 | -0,05283586 | 0,964039483 | 0,91042 | 0,96136765 | novel transcript |
| ENSG00000120253 | NUP43 | -0,052850013 | 0,964030026 | 0,82457 | 0,91900665 | nucleoporin 43 |
| ENSG00000116396 | KCNC4 | -0,052939054 | 0,963970529 | 0,87503 | 0,94463878 | potassium voltage-gated channel subfamily C member 4 |
| ENSG00000255449 | AP002812,5 | -0,052981065 | 0,963942459 | 0,91509 | 0,96323337 | novel transcript, antisense to AAMDC |
| ENSG00000274712 | AC005332,4 | -0,052985223 | 0,963939681 | 0,79119 | 0,90297979 | bromodomain PHD finger transcription factor (BPTF) pseudogene |
| ENSG00000100162 | CENPM | -0,053027903 | 0,963911165 | 0,81849 | 0,91665476 | centromere protein M |
| ENSG00000100592 | DAAM1 | -0,05303093 | 0,963909142 | 0,86739 | 0,94137971 | dishevelled associated activator of morphogenesis 1 |
| ENSG00000103021 | CCDC113 | -0,05310933 | 0,963856762 | 0,85148 | 0,9324506 | coiled-coil domain containing 113 |
| ENSG00000114650 | SCAP | -0,053168001 | 0,963817565 | 0,78657 | 0,90089706 | SREBF chaperone |
| ENSG00000133624 | ZNF767P | -0,05333698 | 0,963704682 | 0,78167 | 0,89877377 | zinc finger family member 767, pseudogene |
| ENSG00000146416 | AIG1 | -0,053526844 | 0,963577863 | 0,81456 | 0,91470265 | androgen induced 1 |
| ENSG00000177169 | ULK1 | -0,053546333 | 0,963564847 | 0,83394 | 0,92405734 | unc-51 like autophagy activating kinase 1 |
| ENSG00000172828 | CES3 | -0,053619525 | 0,963515964 | 0,83898 | 0,92640383 | carboxylesterase 3 |
| ENSG00000223427 | RBM7P1 | -0,053671341 | 0,963481359 | 0,93984 | 0,97443135 | RBM7 pseudogene 1 |

| | | | | | | |
|---|---|---|---|---|---|---|
| ENSG00000139726 | DENR | -0,053705803 | 0,963458344 | 0,78658 | 0,90089706 | density regulated re-initiation and release factor |
| ENSG00000137078 | SIT1 | -0,053734543 | 0,963439151 | 0,9162 | 0,96347905 | signaling threshold regulating transmembrane adaptor 1 |
| ENSG00000258424 | AL512791,1 | -0,053739345 | 0,963435945 | 0,9317 | 0,97087386 | novel transcript, antisense to CALM1 |
| ENSG00000138398 | PPIG | -0,053854268 | 0,963359201 | 0,80164 | 0,90870853 | peptidylprolyl isomerase G |
| ENSG00000183684 | ALYREF | -0,053897234 | 0,963330512 | 0,80462 | 0,90996631 | Aly/REF export factor |
| ENSG00000070081 | NUCB2 | -0,053922745 | 0,963313477 | 0,75944 | 0,88781725 | nucleobindin 2 |
| ENSG00000263612 | AF186192,3 | -0,054018995 | 0,963249212 | 0,95076 | 0,97934603 | novel transcript |
| ENSG00000184557 | SOCS3 | -0,054026592 | 0,963244139 | 0,86063 | 0,93752064 | suppressor of cytokine signaling 3 |
| ENSG00000116213 | WRAP73 | -0,054189052 | 0,963135676 | 0,76076 | 0,88869869 | WD repeat containing, antisense to TP73 |
| ENSG00000091436 | MAP3K20 | -0,054214909 | 0,963118414 | 0,84849 | 0,93094401 | mitogen-activated protein kinase kinase kinase 20 |
| ENSG00000272843 | AC211476,2 | -0,05428346 | 0,963072651 | 0,91246 | 0,96205682 | novel transcript, antisense to POM121 |
| ENSG00000070540 | WIPI1 | -0,054486737 | 0,962936963 | 0,89126 | 0,95208954 | WD repeat domain, phosphoinositide interacting 1 |
| ENSG00000114268 | PFKFB4 | -0,054590646 | 0,962867611 | 0,80617 | 0,91079954 | 6-phosphofructo-2-kinase/fructose-2,6-biphosphatase 4 |
| ENSG00000272321 | AP003355,2 | -0,054639374 | 0,96283509 | 0,9426 | 0,97560255 | novel transcript, antisense to STK3 |
| ENSG00000261556 | SMG1P7 | -0,05468741 | 0,962803032 | 0,79108 | 0,90292452 | SMG1 pseudogene 7 |
| ENSG00000137869 | CYP19A1 | -0,05470579 | 0,962790765 | 0,91534 | 0,96331022 | cytochrome P450 family 19 subfamily A member 1 |
| ENSG00000232645 | LINC01431 | -0,054763893 | 0,962751991 | 0,92467 | 0,96725125 | long intergenic non-protein coding RNA 1431 |
| ENSG00000264895 | AC006141,1 | -0,05479388 | 0,96273198 | 0,81266 | 0,91386173 | novel transcript, sense intronic to MBTD1 |
| ENSG00000188710 | QRFP | -0,054813017 | 0,96271921 | 0,87781 | 0,94568249 | pyroglutamylated RFamide peptide |
| ENSG00000135108 | FBXO21 | -0,054825341 | 0,962710986 | 0,76085 | 0,88871285 | F-box protein 21 |
| ENSG00000284260 | RUFY1 | -0,054856909 | 0,962689921 | 0,82231 | 0,91794576 | RUN and FYVE domain containing 1 |
| ENSG00000229802 | SLC39A7 | -0,054860371 | 0,962687611 | 0,82484 | 0,91900665 | solute carrier family 39 member 7 |
| ENSG00000206288 | SLC39A7 | -0,054860371 | 0,962687611 | 0,82484 | 0,91900665 | solute carrier family 39 member 7 |
| ENSG00000227402 | SLC39A7 | -0,054860371 | 0,962687611 | 0,82484 | 0,91900665 | solute carrier family 39 member 7 |
| ENSG00000224399 | SLC39A7 | -0,054860371 | 0,962687611 | 0,82484 | 0,91900665 | solute carrier family 39 member 7 |
| ENSG00000226614 | SLC39A7 | -0,054860371 | 0,962687611 | 0,82484 | 0,91900665 | solute carrier family 39 member 7 |
| ENSG00000149091 | DGKZ | -0,054921685 | 0,962646698 | 0,79965 | 0,90755972 | diacylglycerol kinase zeta |
| ENSG00000136504 | KAT7 | -0,054927197 | 0,96264302 | 0,81433 | 0,9146754 | lysine acetyltransferase 7 |
| ENSG00000258443 | AC005225,1 | -0,054942803 | 0,962632607 | 0,96806 | 0,98637425 | novel transcript, sense intronic to HEATR4 |
| ENSG00000213983 | AP1G2 | -0,055026074 | 0,962577046 | 0,86589 | 0,94044334 | adaptor related protein complex 1 subunit gamma 2 |
| ENSG00000013306 | SLC25A39 | -0,055052847 | 0,962559183 | 0,82812 | 0,9207216 | solute carrier family 25 member 39 |
| ENSG00000255966 | AC006064,3 | -0,055116366 | 0,962516804 | 0,94763 | 0,97795803 | novel transcript |
| ENSG00000271895 | AL109811,2 | -0,05523499 | 0,962437666 | 0,82059 | 0,91732362 | novel transcript, antisense to MASP2 |

| | | | | | | |
|---|---|---|---|---|---|---|
| ENSG00000139998 | RAB15 | -0,055246613 | 0,962429912 | 0,81615 | 0,91549169 | RAB15, member RAS oncogene family |
| ENSG00000179136 | LINC00670 | -0,055250742 | 0,962427158 | 0,92582 | 0,96768961 | long intergenic non-protein coding RNA 670 |
| ENSG00000246090 | AP002026,1 | -0,05540456 | 0,962324551 | 0,87904 | 0,94623795 | novel transcript, antisense to ADH4, ADH6 & ADH1A |
| ENSG00000231305 | AC112484,1 | -0,055405121 | 0,962324176 | 0,86857 | 0,9419405 | novel transcript |
| ENSG00000183831 | ANKRD45 | -0,055435458 | 0,962303941 | 0,76337 | 0,88980026 | ankyrin repeat domain 45 |
| ENSG00000187416 | LHFPL3 | -0,055439872 | 0,962300997 | 0,91044 | 0,96136765 | LHFPL tetraspan subfamily member 3 |
| ENSG00000184441 | AP001062,1 | -0,055475014 | 0,962277557 | 0,85521 | 0,93430161 | novel transcript, antisense to C21orf2 |
| ENSG00000132359 | RAP1GAP2 | -0,055605573 | 0,962190478 | 0,8634 | 0,93900203 | RAP1 GTPase activating protein 2 |
| ENSG00000278741 | ABR | -0,055682067 | 0,962139462 | 0,87249 | 0,94374455 | ABR activator of RhoGEF and GTPase |
| ENSG00000178821 | TMEM52 | -0,055706848 | 0,962122935 | 0,90759 | 0,96044232 | transmembrane protein 52 |
| ENSG00000105784 | RUNDC3B | -0,055978546 | 0,96194176 | 0,88076 | 0,94687991 | RUN domain containing 3B |
| ENSG00000283192 | AC007383,6 | -0,056027163 | 0,961909344 | 0,86849 | 0,94191673 | novel transcript |
| ENSG00000227946 | AC007383,1 | -0,056027163 | 0,961909344 | 0,86849 | 0,94191673 | novel transcript |
| ENSG00000156050 | FAM161B | -0,056072415 | 0,961879172 | 0,82612 | 0,91953132 | FAM161 centrosomal protein B |
| ENSG00000262803 | AL354943,1 | -0,056197744 | 0,961795617 | 0,94362 | 0,97618492 | transmembrane protein 192 (TMEM192) pseudogene |
| ENSG00000157593 | SLC35B2 | -0,056210211 | 0,961787305 | 0,80962 | 0,91223535 | solute carrier family 35 member B2 |
| ENSG00000285634 | AL583827,1 | -0,056274744 | 0,961744284 | 0,86514 | 0,93999031 | Novel transcript |
| ENSG00000068028 | RASSF1 | -0,056283485 | 0,961738458 | 0,79936 | 0,90744841 | Ras association domain family member 1 |
| ENSG00000272870 | SAP30-DT | -0,056303513 | 0,961725106 | 0,83623 | 0,92493625 | SAP30 divergent transcript |
| ENSG00000102755 | FLT1 | -0,056325402 | 0,961710515 | 0,93283 | 0,97143711 | fms related receptor tyrosine kinase 1 |
| ENSG00000241852 | C8orf58 | -0,05640903 | 0,961654769 | 0,81184 | 0,91334888 | chromosome 8 open reading frame 58 |
| ENSG00000249850 | KRT18P31 | -0,05644516 | 0,961630687 | 0,88939 | 0,95110855 | keratin 18 pseudogene 31 |
| ENSG00000260927 | AC009107,2 | -0,056549974 | 0,961560825 | 0,83571 | 0,92493625 | uncharacterized LOC101927556 |
| ENSG00000184992 | BRI3BP | -0,056604295 | 0,961524621 | 0,81039 | 0,91251177 | BRI3 binding protein |
| ENSG00000227212 | PFN1P6 | -0,056701377 | 0,96145992 | 0,9183 | 0,96446185 | profilin 1 pseudogene 6 |
| ENSG00000181619 | GPR135 | -0,056705441 | 0,961457212 | 0,82996 | 0,92194059 | G protein-coupled receptor 135 |
| ENSG00000213287 | AP001102,1 | -0,056983339 | 0,96127203 | 0,93596 | 0,97249457 | translocase of outer mitochondrial membrane 22 homolog (yeast) (TOMM22) pseudogene |
| ENSG00000148400 | NOTCH1 | -0,057069847 | 0,961214391 | 0,75838 | 0,88727479 | notch receptor 1 |
| ENSG00000100983 | GSS | -0,057077619 | 0,961209212 | 0,76133 | 0,88879633 | glutathione synthetase |
| ENSG00000283103 | AC010642,2 | -0,05708245 | 0,961205994 | 0,75132 | 0,88347546 | novel transcript, antisense to ERVK3-1 |
| ENSG00000165105 | RASEF | -0,057153121 | 0,96115891 | 0,81203 | 0,91344855 | RAS and EF-hand domain containing |
| ENSG00000166407 | LMO1 | -0,057381971 | 0,961006457 | 0,88348 | 0,94837933 | LIM domain only 1 |

| | | | | | | |
|---|---|---|---|---|---|---|
| ENSG00000196182 | STK40 | -0,057662129 | 0,960819856 | 0,68812 | 0,84808917 | serine/threonine kinase 40 |
| ENSG00000204611 | ZNF616 | -0,057828768 | 0,960708883 | 0,78259 | 0,89903403 | zinc finger protein 616 |
| ENSG00000114395 | CYB561D2 | -0,057865662 | 0,960684315 | 0,80434 | 0,90993834 | cytochrome b561 family member D2 |
| ENSG00000222046 | DCDC2B | -0,057958697 | 0,960622365 | 0,8467 | 0,930363 | doublecortin domain containing 2B |
| ENSG00000153558 | FBXL2 | -0,057976205 | 0,960610707 | 0,76702 | 0,89185843 | F-box and leucine rich repeat protein 2 |
| ENSG00000231977 | AL096828,1 | -0,058022172 | 0,960580101 | 0,93659 | 0,9727206 | novel transcript |
| ENSG00000163798 | SLC4A1AP | -0,058097015 | 0,96053027 | 0,77909 | 0,89781842 | solute carrier family 4 member 1 adaptor protein |
| ENSG00000172831 | CES2 | -0,058187565 | 0,960469985 | 0,79055 | 0,90286944 | carboxylesterase 2 |
| ENSG00000280770 | AC216281,1 | -0,058291577 | 0,960400742 | 0,91918 | 0,96485808 | novel transcript sense intronic to TRAPPC12 |
| ENSG00000185986 | SDHAP3 | -0,058411106 | 0,960321174 | 0,7795 | 0,89802258 | succinate dehydrogenase complex flavoprotein subunit A pseudogene 3 |
| ENSG00000277511 | AC116407,2 | -0,058690914 | 0,960134941 | 0,94481 | 0,97662366 | novel transcript |
| ENSG00000235253 | AC010240,2 | -0,058872174 | 0,960014317 | 0,84673 | 0,930363 | eukaryotic translation elongation factor 1 alpha 1 (EEF1A1) pseudogene |
| ENSG00000161940 | BCL6B | -0,058931972 | 0,959974526 | 0,88071 | 0,94687991 | BCL6B transcription repressor |
| ENSG00000268858 | AL118506,1 | -0,058995485 | 0,959932265 | 0,83434 | 0,92416267 | uncharacterized LOC112268269 |
| ENSG00000113657 | DPYSL3 | -0,059007234 | 0,959924448 | 0,809 | 0,91201277 | dihydropyrimidinase like 3 |
| ENSG00000273702 | AC091271,1 | -0,059037846 | 0,95990408 | 0,82669 | 0,91979905 | novel transcript |
| ENSG00000261526 | AC012615,1 | -0,059088508 | 0,959870372 | 0,84499 | 0,92930882 | novel transcript |
| ENSG00000197694 | SPTAN1 | -0,059145251 | 0,95983262 | 0,82585 | 0,91929949 | spectrin alpha, non-erythrocytic 1 |
| ENSG00000245975 | AC090515,2 | -0,059225314 | 0,959779355 | 0,84727 | 0,93060479 | novel transcript |
| ENSG00000153944 | MSI2 | -0,059333578 | 0,959707333 | 0,74703 | 0,88059941 | musashi RNA binding protein 2 |
| ENSG00000232795 | SCAND3P1 | -0,059422486 | 0,959648191 | 0,94196 | 0,97522303 | SCAN domain containing 3 pseudogene 1 |
| ENSG00000234608 | MAPKAPK5-A! | -0,059495804 | 0,959599423 | 0,80803 | 0,91168773 | MAPKAPK5 antisense RNA 1 |
| ENSG00000225590 | VPS52 | -0,059510378 | 0,95958973 | 0,9383 | 0,97361277 | VPS52 subunit of GARP complex |
| ENSG00000228425 | VPS52 | -0,059510378 | 0,95958973 | 0,9383 | 0,97361277 | VPS52 subunit of GARP complex |
| ENSG00000116857 | TMEM9 | -0,059527873 | 0,959578093 | 0,7301 | 0,87180309 | transmembrane protein 9 |
| ENSG00000264016 | AC015908,3 | -0,059643009 | 0,959501516 | 0,9197 | 0,96507001 | novel transcript |
| ENSG00000060339 | CCAR1 | -0,059659386 | 0,959490624 | 0,69177 | 0,84985941 | cell division cycle and apoptosis regulator 1 |
| ENSG00000277033 | B3GNTL1 | -0,059857712 | 0,959358733 | 0,88568 | 0,94943038 | UDP-GlcNAc:betaGal beta-1,3-N-acetylglucosaminyltransferase like 1 |
| ENSG00000166484 | MAPK7 | -0,05990562 | 0,959326876 | 0,74646 | 0,88046587 | mitogen-activated protein kinase 7 |
| ENSG00000108219 | TSPAN14 | -0,059985002 | 0,959274092 | 0,7757 | 0,89630112 | tetraspanin 14 |

| | | | | | | |
|---|---|---|---|---|---|---|
| ENSG00000182872 | RBM10 | -0,060047288 | 0,959232678 | 0,78346 | 0,89918294 | RNA binding motif protein 10 |
| ENSG00000183346 | CABCOCO1 | -0,060079403 | 0,959211325 | 0,93217 | 0,97114134 | ciliary associated calcium binding coiled-coil 1 |
| ENSG00000115539 | PDCL3 | -0,060201894 | 0,959129887 | 0,80858 | 0,91193281 | phosducin like 3 |
| ENSG00000256879 | AC129102,1 | -0,060227285 | 0,959113007 | 0,97465 | 0,98935656 | novel transcript |
| ENSG00000133142 | TCEAL4 | -0,06027955 | 0,959078262 | 0,76848 | 0,89277929 | transcription elongation factor A like 4 |
| ENSG00000198156 | NPIPB6 | -0,060563413 | 0,958889573 | 0,91029 | 0,9613425 | nuclear pore complex interacting protein family member B6 |
| ENSG00000230772 | VN1R108P | -0,060571452 | 0,95888423 | 0,9571 | 0,98220263 | vomeronasal 1 receptor 108 pseudogene |
| ENSG00000175097 | RAG2 | -0,060629202 | 0,958845847 | 0,92318 | 0,96667029 | recombination activating 2 |
| ENSG00000111775 | COX6A1 | -0,060775432 | 0,958748664 | 0,74913 | 0,88190373 | cytochrome c oxidase subunit 6A1 |
| ENSG00000145247 | OCIAD2 | -0,060808078 | 0,958726969 | 0,89885 | 0,95672027 | OCIA domain containing 2 |
| ENSG00000286977 | Z83745,1 | -0,060810992 | 0,958725033 | 0,85543 | 0,93439957 | novel transcript |
| ENSG00000073111 | MCM2 | -0,060929948 | 0,958645986 | 0,76245 | 0,88952703 | minichromosome maintenance complex component 2 |
| ENSG00000120875 | DUSP4 | -0,06099153 | 0,958605067 | 0,84412 | 0,92888169 | dual specificity phosphatase 4 |
| ENSG00000184357 | H1-5 | -0,061005714 | 0,958595642 | 0,8115 | 0,9132701 | H1,5 linker histone, cluster member |
| ENSG00000163354 | DCST2 | -0,06107209 | 0,958551539 | 0,91347 | 0,96249683 | DC-STAMP domain containing 2 |
| ENSG00000156298 | TSPAN7 | -0,061086735 | 0,958541809 | 0,88008 | 0,94664997 | tetraspanin 7 |
| ENSG00000159069 | FBXW5 | -0,06119963 | 0,958466803 | 0,71045 | 0,86051357 | F-box and WD repeat domain containing 5 |
| ENSG00000130024 | PHF10 | -0,061228219 | 0,95844781 | 0,80568 | 0,91072757 | PHD finger protein 10 |
| ENSG00000081479 | LRP2 | -0,061242475 | 0,958438339 | 0,88005 | 0,94664997 | LDL receptor related protein 2 |
| ENSG00000151150 | ANK3 | -0,06134915 | 0,958367474 | 0,85671 | 0,93535509 | ankyrin 3 |
| ENSG00000132361 | CLUH | -0,06142679 | 0,958315899 | 0,79528 | 0,90519396 | clustered mitochondria homolog |
| ENSG00000070413 | DGCR2 | -0,061453588 | 0,958298099 | 0,76639 | 0,89138648 | DiGeorge syndrome critical region gene 2 |
| ENSG00000167566 | NCKAP5L | -0,061472771 | 0,958285357 | 0,74586 | 0,88010204 | NCK associated protein 5 like |
| ENSG00000276644 | DACH1 | -0,061588702 | 0,958208355 | 0,88915 | 0,95104435 | dachshund family transcription factor 1 |
| ENSG00000242888 | AL133368,1 | -0,061708772 | 0,95812861 | 0,86984 | 0,94261122 | ribosomal protein S2 (RPS2) pseudogene |
| ENSG00000206149 | HERC2P9 | -0,061732467 | 0,958112874 | 0,72444 | 0,86847556 | hect domain and RLD 2 pseudogene 9 |
| ENSG00000274397 | AL357153,4 | -0,061740098 | 0,958107806 | 0,89311 | 0,9532892 | ADAM metallopeptidase domain 20 (ADAM20) pseudogene |
| ENSG00000259668 | AC066613,1 | -0,06179936 | 0,958068451 | 0,87213 | 0,94363878 | novel transcript, antisense to DMXL2 |
| ENSG00000130193 | THEM6 | -0,061816995 | 0,95805674 | 0,86809 | 0,94176112 | thioesterase superfamily member 6 |
| ENSG00000260641 | AC114811,2 | -0,061825074 | 0,958051375 | 0,88242 | 0,94796839 | novel transcript, antisense TSPAN5 |
| ENSG00000262728 | AC123768,2 | -0,061869444 | 0,95802191 | 0,91087 | 0,96163156 | novel transcript, antisense to GOLGA8N |

| | | | | | | |
|---|---|---|---|---|---|---|
| ENSG00000274818 | AC004825,2 | -0,061886342 | 0,958010689 | 0,92303 | 0,96662141 | novel transcript |
| ENSG00000284406 | AC004492,3 | -0,061941868 | 0,957973818 | 0,88941 | 0,95110855 | novel transcript, antisense to HBP1 |
| ENSG00000272072 | AC004492,1 | -0,061941868 | 0,957973818 | 0,88941 | 0,95110855 | novel transcript, antisense to HBP1 |
| ENSG00000259948 | AC124068,1 | -0,06199963 | 0,957935464 | 0,96638 | 0,98580842 | VAMP (vesicle-associated membrane protein)-associated protein A, 33kDa (VAPA) pseudogene |
| ENSG00000182013 | PNMA8A | -0,062007011 | 0,957930563 | 0,77411 | 0,89572556 | PNMA family member 8A |
| ENSG00000168743 | NPNT | -0,062007793 | 0,957930044 | 0,95732 | 0,98235923 | nephronectin |
| ENSG00000287406 | AC098656,1 | -0,062078706 | 0,95788296 | 0,8948 | 0,95417507 | novel transcript |
| ENSG00000234947 | STK19 | -0,062273833 | 0,957753413 | 0,78241 | 0,89892226 | serine/threonine kinase 19 |
| ENSG00000158691 | ZSCAN12 | -0,062282608 | 0,957747587 | 0,68988 | 0,84877222 | zinc finger and SCAN domain containing 12 |
| ENSG00000033100 | CHPF2 | -0,062296063 | 0,957738656 | 0,78325 | 0,89918294 | chondroitin polymerizing factor 2 |
| ENSG00000136819 | C9orf78 | -0,062323881 | 0,957720189 | 0,79069 | 0,90286944 | chromosome 9 open reading frame 78 |
| ENSG00000008056 | SYN1 | -0,062352426 | 0,95770124 | 0,86514 | 0,93999031 | synapsin I |
| ENSG00000225589 | MDC1 | -0,062405836 | 0,957665785 | 0,85959 | 0,93687112 | mediator of DNA damage checkpoint 1 |
| ENSG00000234012 | MDC1 | -0,062405836 | 0,957665785 | 0,85959 | 0,93687112 | mediator of DNA damage checkpoint 1 |
| ENSG00000231135 | MDC1 | -0,062405836 | 0,957665785 | 0,85959 | 0,93687112 | mediator of DNA damage checkpoint 1 |
| ENSG00000235660 | LINC00345 | -0,062409972 | 0,957663039 | 0,86403 | 0,93918171 | long intergenic non-protein coding RNA 345 |
| ENSG00000246863 | GPR176-DT | -0,06254447 | 0,957573764 | 0,90033 | 0,95760242 | GPR176-DT |
| ENSG00000105137 | SYDE1 | -0,062563534 | 0,95756111 | 0,79404 | 0,90426294 | synapse defective Rho GTPase homolog 1 |
| ENSG00000258096 | AC025031,2 | -0,062573205 | 0,957554691 | 0,94621 | 0,97710458 | novel transcript, antisense to SLC38A2 |
| ENSG00000164713 | BRI3 | -0,062610411 | 0,957529997 | 0,75892 | 0,88743316 | brain protein I3 |
| ENSG00000132554 | RGS22 | -0,062888548 | 0,957345413 | 0,91776 | 0,96433167 | regulator of G protein signaling 22 |
| ENSG00000215630 | GUSBP9 | -0,06301968 | 0,9572584 | 0,76223 | 0,88933378 | GUSB pseudogene 9 |
| ENSG00000020129 | NCDN | -0,063034445 | 0,957248603 | 0,79378 | 0,90411641 | neurochondrin |
| ENSG00000261915 | AC026954,2 | -0,063104055 | 0,957202417 | 0,95166 | 0,97948342 | novel protein |
| ENSG00000288456 | AC026954,6 | -0,063104055 | 0,957202417 | 0,95166 | 0,97948342 | novel protein |
| ENSG00000166323 | C11orf65 | -0,063267726 | 0,957093831 | 0,87784 | 0,94568249 | chromosome 11 open reading frame 65 |
| ENSG00000276276 | ARL17B | -0,063280132 | 0,9570856 | 0,84524 | 0,92946127 | ADP ribosylation factor like GTPase 17B |
| ENSG00000118960 | HS1BP3 | -0,063359306 | 0,957033078 | 0,76196 | 0,88909725 | HCLS1 binding protein 3 |
| ENSG00000087884 | AAMDC | -0,063364867 | 0,957029388 | 0,80533 | 0,91055339 | adipogenesis associated Mth938 domain containing |
| ENSG00000260095 | AC106820,3 | -0,063397212 | 0,957007932 | 0,93574 | 0,97240996 | novel transcript, antisense to CCNF |
| ENSG00000198931 | APRT | -0,063427617 | 0,956987763 | 0,78204 | 0,89877579 | adenine phosphoribosyltransferase |
| ENSG00000166856 | GPR182 | -0,063510939 | 0,956932495 | 0,85206 | 0,9327167 | G protein-coupled receptor 182 |

| | | | | | | |
|---|---|---|---|---|---|---|
| ENSG00000154162 | CDH12 | -0,06357891 | 0,956887411 | 0,8925 | 0,95285354 | cadherin 12 |
| ENSG00000233124 | LINC00456 | -0,063620447 | 0,956859862 | 0,93447 | 0,9721963 | long intergenic non-protein coding RNA 456 |
| ENSG00000249244 | AC110373,1 | -0,063629513 | 0,956853848 | 0,93754 | 0,97327398 | glycerol kinase 2 pseudogene |
| ENSG00000274523 | RCC1L | -0,063759779 | 0,956767455 | 0,747 | 0,88059941 | RCC1 like |
| ENSG00000215018 | COL28A1 | -0,063771629 | 0,956759596 | 0,8262 | 0,91953499 | collagen type XXVIII alpha 1 chain |
| ENSG00000177383 | MAGEF1 | -0,063786598 | 0,956749669 | 0,73496 | 0,87477884 | MAGE family member F1 |
| ENSG00000274776 | AC090241,3 | -0,063828932 | 0,956721595 | 0,91028 | 0,9613425 | novel transcript |
| ENSG00000259153 | AC004816,1 | -0,064258979 | 0,956436452 | 0,90319 | 0,95854863 | novel transcript |
| ENSG00000172803 | SNX32 | -0,064264173 | 0,956433009 | 0,84042 | 0,92722187 | sorting nexin 32 |
| ENSG00000135766 | EGLN1 | -0,064267863 | 0,956430563 | 0,71746 | 0,86483656 | egl-9 family hypoxia inducible factor 1 |
| ENSG00000262243 | CES1 | -0,064268602 | 0,956430073 | 0,90939 | 0,96105018 | carboxylesterase 1 |
| ENSG00000183741 | CBX6 | -0,064274135 | 0,956426405 | 0,73023 | 0,87183859 | chromobox 6 |
| ENSG00000231370 | PRRC2A | -0,064335394 | 0,956385794 | 0,89706 | 0,95564404 | proline rich coiled-coil 2A |
| ENSG00000273142 | LINC02604 | -0,064338638 | 0,956383644 | 0,84962 | 0,93152542 | long intergenic non-protein coding RNA 2604 |
| ENSG00000278831 | AL513190,1 | -0,064380626 | 0,95635581 | 0,89681 | 0,95548853 | novel transcript, antisense to FAM53B |
| ENSG00000246082 | NUDT16P1 | -0,064396845 | 0,956345058 | 0,89781 | 0,95593639 | nudix hydrolase 16 pseudogene 1 |
| ENSG00000116688 | MFN2 | -0,06440603 | 0,95633897 | 0,73262 | 0,87313685 | mitofusin 2 |
| ENSG00000238056 | PRRT1 | -0,064424123 | 0,956326976 | 0,89288 | 0,95314522 | proline rich transmembrane protein 1 |
| ENSG00000235956 | PRRT1 | -0,064424123 | 0,956326976 | 0,89288 | 0,95314522 | proline rich transmembrane protein 1 |
| ENSG00000165370 | GPR101 | -0,064480907 | 0,956289336 | 0,88743 | 0,95041376 | G protein-coupled receptor 101 |
| ENSG00000180155 | LYNX1 | -0,064544294 | 0,956247321 | 0,80694 | 0,91123384 | Ly6/neurotoxin 1 |
| ENSG00000070476 | ZXDC | -0,064571018 | 0,956229608 | 0,73844 | 0,87583043 | ZXD family zinc finger C |
| ENSG00000205560 | CPT1B | -0,064635564 | 0,956186828 | 0,83722 | 0,92549623 | carnitine palmitoyltransferase 1B |
| ENSG00000157911 | PEX10 | -0,064714621 | 0,956134432 | 0,76274 | 0,88972756 | peroxisomal biogenesis factor 10 |
| ENSG00000168701 | TMEM208 | -0,064831292 | 0,956057112 | 0,7368 | 0,87532747 | transmembrane protein 208 |
| ENSG00000140543 | DET1 | -0,064895421 | 0,956014615 | 0,71204 | 0,8615121 | DET1 partner of COP1 E3 ubiquitin ligase |
| ENSG00000277781 | MROH6 | -0,064980423 | 0,955958289 | 0,89961 | 0,95730568 | maestro heat like repeat family member 6 |
| ENSG00000114541 | FRMD4B | -0,065055191 | 0,955908748 | 0,87483 | 0,94462576 | FERM domain containing 4B |
| ENSG00000107140 | TESK1 | -0,065130505 | 0,955858848 | 0,76543 | 0,89085946 | testis associated actin remodelling kinase 1 |
| ENSG00000224081 | SLC44A3-AS1 | -0,065148395 | 0,955846995 | 0,88347 | 0,94837933 | SLC44A3 antisense RNA 1 |
| ENSG00000163075 | CFAP221 | -0,065282762 | 0,955757975 | 0,75329 | 0,88446342 | cilia and flagella associated protein 221 |
| ENSG00000223403 | MEG9 | -0,065307184 | 0,955741796 | 0,94392 | 0,97635124 | maternally expressed 9 |
| ENSG00000138668 | HNRNPD | -0,065329874 | 0,955726765 | 0,7383 | 0,87571125 | heterogeneous nuclear ribonucleoprotein D |

| | | | | | | |
|---|---|---|---|---|---|---|
| ENSG00000155275 | TRMT44 | -0,065353992 | 0,955710787 | 0,77542 | 0,89630112 | tRNA methyltransferase 44 homolog |
| ENSG00000213509 | PPIAP16 | -0,065358264 | 0,955707958 | 0,95196 | 0,97966441 | peptidylprolyl isomerase A pseudogene 16 |
| ENSG00000124570 | SERPINB6 | -0,065369322 | 0,955700632 | 0,79125 | 0,90299428 | serpin family B member 6 |
| ENSG00000135976 | ANKRD36 | -0,065441474 | 0,955652837 | 0,73257 | 0,87310563 | ankyrin repeat domain 36 |
| ENSG00000217555 | CKLF | -0,065487087 | 0,955622623 | 0,78546 | 0,90031115 | chemokine like factor |
| ENSG00000050820 | BCAR1 | -0,065495789 | 0,955616859 | 0,7515 | 0,88347546 | BCAR1 scaffold protein, Cas family member |
| ENSG00000276934 | AC009704,2 | -0,065611172 | 0,955540434 | 0,92368 | 0,96675951 | novel transcript, sense intronic to CNDP2 |
| ENSG00000169203 | NPIPB12 | -0,065685838 | 0,955490982 | 0,70731 | 0,85910429 | nuclear pore complex interacting protein family member B12 |
| ENSG00000106829 | TLE4 | -0,065904008 | 0,9553465 | 0,87113 | 0,943209 | TLE family member 4, transcriptional corepressor |
| ENSG00000236463 | LINC00427 | -0,065913918 | 0,955339937 | 0,94073 | 0,97486264 | long intergenic non-protein coding RNA 427 |
| ENSG00000275498 | PDXDC1 | -0,066002904 | 0,955281014 | 0,80203 | 0,90890569 | pyridoxal dependent decarboxylase domain containing 1 |
| ENSG00000280241 | AC079298,3 | -0,066028299 | 0,955264198 | 0,90787 | 0,96044232 | novel transcript, antisense to DCHS2 |
| ENSG00000283917 | AC104657,1 | -0,066028299 | 0,955264198 | 0,90787 | 0,96044232 | novel transcript, antisense to DCHS2 |
| ENSG00000224952 | PBX2 | -0,066183497 | 0,955161441 | 0,84531 | 0,92946127 | PBX homeobox 2 |
| ENSG00000139190 | VAMP1 | -0,066194242 | 0,955154327 | 0,77534 | 0,89624737 | vesicle associated membrane protein 1 |
| ENSG00000132470 | ITGB4 | -0,066256355 | 0,955113206 | 0,93697 | 0,97300612 | integrin subunit beta 4 |
| ENSG00000185475 | TMEM179B | -0,066264555 | 0,955107777 | 0,76887 | 0,89281519 | transmembrane protein 179B |
| ENSG00000141279 | NPEPPS | -0,06629481 | 0,955087748 | 0,66331 | 0,83242946 | aminopeptidase puromycin sensitive |
| ENSG00000264273 | AC107982,2 | -0,066420339 | 0,955004649 | 0,91719 | 0,9640264 | novel transcript, antisense to PRPSAP2 |
| ENSG00000247572 | CKMT2-AS1 | -0,0664264 | 0,955000636 | 0,78433 | 0,89962715 | CKMT2 antisense RNA 1 |
| ENSG00000250027 | AC022272,1 | -0,066438478 | 0,954992641 | 0,93829 | 0,97361277 | novel transcript |
| ENSG00000273024 | INTS4P2 | -0,066449797 | 0,954985149 | 0,87752 | 0,94566578 | integrator complex subunit 4 pseudogene 2 |
| ENSG00000184361 | SPATA32 | -0,066534181 | 0,954929293 | 0,9171 | 0,9640264 | spermatogenesis associated 32 |
| ENSG00000108861 | DUSP3 | -0,066663082 | 0,954843977 | 0,7897 | 0,90258817 | dual specificity phosphatase 3 |
| ENSG00000147535 | PLPP5 | -0,066772152 | 0,954771791 | 0,76129 | 0,88879633 | phospholipid phosphatase 5 |
| ENSG00000265625 | AC104564,5 | -0,066839873 | 0,954726975 | 0,87691 | 0,94549531 | novel transcript, sense intronic to SSH2 |
| ENSG00000282866 | CFL1P1 | -0,066975261 | 0,954637384 | 0,90215 | 0,95812378 | cofilin 1 pseudogene 1 |
| ENSG00000223820 | CFL1P1 | -0,066975261 | 0,954637384 | 0,90215 | 0,95812378 | cofilin 1 pseudogene 1 |
| ENSG00000120725 | SIL1 | -0,067054764 | 0,954584778 | 0,70227 | 0,85651838 | SIL1 nucleotide exchange factor |
| ENSG00000272953 | AC092171,4 | -0,067110562 | 0,954547859 | 0,84523 | 0,92946127 | novel transcript |
| ENSG00000274421 | AL162390,1 | -0,067135344 | 0,954531462 | 0,84578 | 0,92982715 | novel transcript |
| ENSG00000280734 | LINC01232 | -0,067136332 | 0,954530809 | 0,74418 | 0,87889181 | long intergenic non-protein coding RNA 1232 |

| | | | | | | |
|---|---|---|---|---|---|---|
| ENSG00000263513 | FAM72C | -0,067219832 | 0,954475564 | 0,803 | 0,90922086 | family with sequence similarity 72 member C |
| ENSG00000198523 | PLN | -0,06727985 | 0,954435858 | 0,91503 | 0,9632313 | phospholamban |
| ENSG00000250644 | AC068580,4 | -0,067309665 | 0,954416133 | 0,83297 | 0,92363433 | novel protein |
| ENSG00000162373 | BEND5 | -0,067354902 | 0,954386207 | 0,75149 | 0,88347546 | BEN domain containing 5 |
| ENSG00000287644 | AC080112,4 | -0,067394954 | 0,954359712 | 0,95146 | 0,97948342 | novel transcript, antisense to RARA |
| ENSG00000242337 | INHCAP | -0,067449667 | 0,954323519 | 0,9284 | 0,96916354 | inhibitor of carbonic anhydrase pseudogene |
| ENSG00000220920 | AL023807,1 | -0,067503592 | 0,954287849 | 0,9386 | 0,9737443 | ribosomal protein S12 (RPS12) pseudogene |
| ENSG00000174799 | CEP135 | -0,067549847 | 0,954257254 | 0,67792 | 0,84156256 | centrosomal protein 135 |
| ENSG00000288551 | AL023806,5 | -0,067757534 | 0,954119891 | 0,86858 | 0,9419405 | novel transcript, antisense to SHPRH |
| ENSG00000116685 | KIAA2013 | -0,067856746 | 0,954054279 | 0,77987 | 0,8981559 | KIAA2013 |
| ENSG00000171311 | EXOSC1 | -0,067914284 | 0,954016231 | 0,79076 | 0,90287625 | exosome component 1 |
| ENSG00000257191 | AC090709,1 | -0,067916293 | 0,954014902 | 0,92444 | 0,96715259 | novel transcript, antisense to SYT1 |
| ENSG00000286850 | AL358232,2 | -0,067982922 | 0,953970843 | 0,92149 | 0,9660069 | novel transcript |
| ENSG00000213434 | VTI1BP2 | -0,06801055 | 0,953952575 | 0,91221 | 0,96205682 | vesicle transport through interaction with t-SNAREs 1B pseudogene 2 |
| ENSG00000257122 | RRN3P3 | -0,068036478 | 0,953935431 | 0,79225 | 0,90351559 | RRN3 pseudogene 3 |
| ENSG00000187446 | CHP1 | -0,068037556 | 0,953934718 | 0,80191 | 0,90880281 | calcineurin like EF-hand protein 1 |
| ENSG00000160818 | GPATCH4 | -0,068166722 | 0,953849315 | 0,7578 | 0,88699366 | G-patch domain containing 4 |
| ENSG00000280018 | CU634019,5 | -0,068281023 | 0,953773746 | 0,78644 | 0,90080813 | novel transcript |
| ENSG00000106638 | TBL2 | -0,068304499 | 0,953758226 | 0,79016 | 0,9027893 | transducin beta like 2 |
| ENSG00000257135 | ODC1-DT | -0,068320988 | 0,953747326 | 0,78675 | 0,9010162 | ODC1 divergent transcript |
| ENSG00000278338 | VWA8-AS1 | -0,068406589 | 0,953690738 | 0,93539 | 0,97222198 | VWA8 antisense RNA 1 (head to head) |
| ENSG00000011143 | MKS1 | -0,068473801 | 0,953646309 | 0,69617 | 0,85287022 | MKS transition zone complex subunit 1 |
| ENSG00000227066 | Z98257,1 | -0,068474831 | 0,953645627 | 0,91182 | 0,96184653 | novel transcript |
| ENSG00000253057 | RN7SKP57 | -0,068492637 | 0,953633858 | 0,93565 | 0,97235376 | RN7SK pseudogene 57 |
| ENSG00000063322 | MED29 | -0,06862623 | 0,953545555 | 0,765 | 0,89075368 | mediator complex subunit 29 |
| ENSG00000149499 | EML3 | -0,068661751 | 0,953522079 | 0,8491 | 0,93130502 | EMAP like 3 |
| ENSG00000143224 | PPOX | -0,068835387 | 0,953407324 | 0,76535 | 0,89085946 | protoporphyrinogen oxidase |
| ENSG00000168993 | CPLX1 | -0,068868992 | 0,953385116 | 0,92384 | 0,96681926 | complexin 1 |
| ENSG00000270893 | AC008277,2 | -0,068872521 | 0,953382784 | 0,90686 | 0,96017342 | elongation factor Tu GTP binding domain containing 1 (EFTUD1) pseudogene |
| ENSG00000160191 | PDE9A | -0,068896447 | 0,953366973 | 0,73273 | 0,87318352 | phosphodiesterase 9A |
| ENSG00000109854 | HTATIP2 | -0,068918089 | 0,953352672 | 0,8894 | 0,95110855 | HIV-1 Tat interactive protein 2 |

| | | | | | | |
|---|---|---|---|---|---|---|
| ENSG00000278911 | AC015819,3 | -0,068970026 | 0,953318351 | 0,94573 | 0,97689657 | novel transcript |
| ENSG00000219665 | ZNF433-AS1 | -0,068976242 | 0,953314244 | 0,6938 | 0,8512419 | ZNF433 and ZNF878 antisense RNA 1 |
| ENSG00000182173 | TSEN54 | -0,069011217 | 0,953291133 | 0,75762 | 0,88691095 | tRNA splicing endonuclease subunit 54 |
| ENSG00000133460 | SLC2A11 | -0,069025512 | 0,953281688 | 0,83224 | 0,92325856 | solute carrier family 2 member 11 |
| ENSG00000070778 | PTPN21 | -0,069095693 | 0,953235316 | 0,74481 | 0,87925125 | protein tyrosine phosphatase non-receptor type 21 |
| ENSG00000134884 | ARGLU1 | -0,069164463 | 0,953189878 | 0,64921 | 0,82436914 | arginine and glutamate rich 1 |
| ENSG00000182141 | ZNF708 | -0,069176725 | 0,953181777 | 0,73242 | 0,87300686 | zinc finger protein 708 |
| ENSG00000100029 | PES1 | -0,069329798 | 0,953080648 | 0,76945 | 0,89297049 | pescadillo ribosomal biogenesis factor 1 |
| ENSG00000168679 | SLC16A4 | -0,069355728 | 0,953063517 | 0,86331 | 0,9389486 | solute carrier family 16 member 4 |
| ENSG00000143842 | SOX13 | -0,069390063 | 0,953040836 | 0,87326 | 0,94386143 | SRY-box transcription factor 13 |
| ENSG00000169169 | CPT1C | -0,069391982 | 0,953039568 | 0,79656 | 0,90599703 | carnitine palmitoyltransferase 1C |
| ENSG00000265287 | AC005726,4 | -0,069421942 | 0,953019777 | 0,91538 | 0,96331365 | novel transcript, antisense to novel protein |
| ENSG00000165632 | TAF3 | -0,069650847 | 0,952868578 | 0,78559 | 0,90038248 | TATA-box binding protein associated factor 3 |
| ENSG00000134186 | PRPF38B | -0,069762903 | 0,95279457 | 0,78503 | 0,90004085 | pre-mRNA processing factor 38B |
| ENSG00000079999 | KEAP1 | -0,06979095 | 0,952776048 | 0,67785 | 0,84154221 | kelch like ECH associated protein 1 |
| ENSG00000288589 | AC226119,3 | -0,069805101 | 0,952766702 | 0,85829 | 0,93612837 | novel transcript |
| ENSG00000142102 | PGGHG | -0,069809514 | 0,952763788 | 0,83939 | 0,92655196 | protein-glucosylgalactosylhydroxylysine glucosidase |
| ENSG00000235833 | AC017099,1 | -0,069853542 | 0,952734712 | 0,86797 | 0,94175463 | coiled-coil domain containing 144C (CCDC144C) pseudogene |
| ENSG00000272686 | WASL-DT | -0,069895722 | 0,952706857 | 0,77828 | 0,89735311 | WASL divergent transcript |
| ENSG00000187720 | THSD4 | -0,070025417 | 0,952621215 | 0,86508 | 0,93999031 | thrombospondin type 1 domain containing 4 |
| ENSG00000231556 | RSL24D1P3 | -0,070060036 | 0,952598356 | 0,93188 | 0,97097872 | ribosomal L24 domain containing 1 pseudogene 3 |
| ENSG00000162496 | DHRS3 | -0,070107281 | 0,952567161 | 0,89727 | 0,955727 | dehydrogenase/reductase 3 |
| ENSG00000275185 | AC130324,3 | -0,070180791 | 0,952518626 | 0,87314 | 0,94385071 | novel transcript, antisense to TEFM |
| ENSG00000147789 | ZNF7 | -0,070306222 | 0,952435816 | 0,77977 | 0,8981559 | zinc finger protein 7 |
| ENSG00000076356 | PLXNA2 | -0,070356097 | 0,95240289 | 0,89099 | 0,95208039 | plexin A2 |
| ENSG00000165349 | SLC7A3 | -0,070555194 | 0,952271464 | 0,82125 | 0,91738668 | solute carrier family 7 member 3 |
| ENSG00000240695 | AC117382,1 | -0,070664649 | 0,952199219 | 0,88152 | 0,94729476 | DnaJ (Hsp40) homolog, subfamily C, member 8 (DNAJC8) pseudogene |
| ENSG00000064999 | ANKS1A | -0,070752689 | 0,952141113 | 0,68938 | 0,84865116 | ankyrin repeat and sterile alpha motif domain containing 1A |
| ENSG00000136305 | CIDEB | -0,070952903 | 0,952008986 | 0,9156 | 0,9633821 | cell death inducing DFFA like effector b |
| ENSG00000285199 | CIDEB | -0,070952903 | 0,952008986 | 0,9156 | 0,9633821 | cell death inducing DFFA like effector b |

| | | | | | | |
|---|---|---|---|---|---|---|
| ENSG00000263327 | TAPT1-AS1 | -0,070966187 | 0,952000221 | 0,78188 | 0,8987738 | TAPT1 antisense RNA 1 (head to head) |
| ENSG00000273445 | AC133644,1 | -0,070971509 | 0,951996709 | 0,89723 | 0,955727 | novel transcript |
| ENSG00000157379 | DHRS1 | -0,0709736 | 0,951995329 | 0,73518 | 0,87480865 | dehydrogenase/reductase 1 |
| ENSG00000250130 | GAPDHP43 | -0,071000506 | 0,951977575 | 0,8919 | 0,95246557 | glyceraldehyde 3 phosphate dehydrogenase pseudogene 43 |
| ENSG00000088727 | KIF9 | -0,071032446 | 0,951956499 | 0,79101 | 0,90292452 | kinesin family member 9 |
| ENSG00000155744 | FAM126B | -0,071066573 | 0,951933981 | 0,61868 | 0,80665241 | family with sequence similarity 126 member B |
| ENSG00000280081 | LINC01667 | -0,071140612 | 0,951885129 | 0,84188 | 0,92809021 | long intergenic non-protein coding RNA 1667 |
| ENSG00000162670 | BRINP3 | -0,071141715 | 0,951884401 | 0,9016 | 0,95801185 | BMP/retinoic acid inducible neural specific 3 |
| ENSG00000288453 | DCAF15 | -0,071146632 | 0,951881156 | 0,7248 | 0,86866311 | DDB1 and CUL4 associated factor 15 |
| ENSG00000132017 | DCAF15 | -0,071146632 | 0,951881156 | 0,7248 | 0,86866311 | DDB1 and CUL4 associated factor 15 |
| ENSG00000138594 | TMOD3 | -0,071194226 | 0,951849755 | 0,80604 | 0,91078302 | tropomodulin 3 |
| ENSG00000254273 | AC018620,1 | -0,071201028 | 0,951845267 | 0,92753 | 0,96861617 | chromatin accessibility complex protein 1 (CHRAC1) pseudogene |
| ENSG00000163704 | PRRT3 | -0,071270603 | 0,951799365 | 0,81253 | 0,91386173 | proline rich transmembrane protein 3 |
| ENSG00000178026 | LRRC75B | -0,071352031 | 0,951745645 | 0,70873 | 0,85960107 | leucine rich repeat containing 75B |
| ENSG00000184428 | TOP1MT | -0,071397542 | 0,951715622 | 0,81002 | 0,91245196 | DNA topoisomerase I mitochondrial |
| ENSG00000226088 | HHLA3-AS1 | -0,071409624 | 0,951707652 | 0,93592 | 0,97249457 | HHLA3 antisense RNA 1 |
| ENSG00000234215 | CT66 | -0,07149553 | 0,951650984 | 0,91892 | 0,9647667 | cancer/testis associated transcript 66 |
| ENSG00000143753 | DEGS1 | -0,071677996 | 0,951530631 | 0,74068 | 0,87717177 | delta 4-desaturase, sphingolipid 1 |
| ENSG00000130766 | SESN2 | -0,071790014 | 0,951456752 | 0,75395 | 0,88467924 | sestrin 2 |
| ENSG00000285069 | SESN2 | -0,071790014 | 0,951456752 | 0,75395 | 0,88467924 | sestrin 2 |
| ENSG00000204316 | MRPL38 | -0,071852815 | 0,951415336 | 0,72329 | 0,86784071 | mitochondrial ribosomal protein L38 |
| ENSG00000134243 | SORT1 | -0,071867898 | 0,951405389 | 0,80332 | 0,90946649 | sortilin 1 |
| ENSG00000063177 | RPL18 | -0,071890965 | 0,951390177 | 0,77495 | 0,8960241 | ribosomal protein L18 |
| ENSG00000206527 | HACD2 | -0,071922119 | 0,951369633 | 0,82151 | 0,91749859 | 3-hydroxyacyl-CoA dehydratase 2 |
| ENSG00000176826 | FKBP9P1 | -0,07194959 | 0,951351518 | 0,88414 | 0,9485968 | FKBP prolyl isomerase 9 pseudogene 1 |
| ENSG00000280663 | PCMTD2 | -0,072122411 | 0,951237561 | 0,70214 | 0,85651838 | protein-L-isoaspartate (D-aspartate) O-methyltransferase domain containing 2 |
| ENSG00000115274 | INO80B | -0,072355598 | 0,951083823 | 0,70293 | 0,8567871 | INO80 complex subunit B |
| ENSG00000154874 | CCDC144B | -0,072539059 | 0,950962885 | 0,73253 | 0,87309896 | coiled-coil domain containing 144B (pseudogene) |
| ENSG00000142303 | ADAMTS10 | -0,07260116 | 0,950921952 | 0,70388 | 0,85724036 | ADAM metallopeptidase with thrombospondin type 1 motif 10 |

| | | | | | | |
|---|---|---|---|---|---|---|
| ENSG00000276724 | AC123768,3 | -0,07273726 | 0,950832249 | 0,88008 | 0,94664997 | novel transcript |
| ENSG00000078237 | TIGAR | -0,072868366 | 0,950745845 | 0,73221 | 0,87293156 | TP53 induced glycolysis regulatory phosphatase |
| ENSG00000275894 | AL021578,1 | -0,072906746 | 0,950720552 | 0,91482 | 0,9632249 | novel transcript, sense intronic to SDC4 |
| ENSG00000123739 | PLA2G12A | -0,072918788 | 0,950712617 | 0,73641 | 0,87529332 | phospholipase A2 group XIIA |
| ENSG00000285700 | AC104260,3 | -0,073000708 | 0,950658635 | 0,93506 | 0,9721963 | novel transcript |
| ENSG00000288021 | AL450338,2 | -0,073051553 | 0,950625131 | 0,88334 | 0,94837933 | novel transcript |
| ENSG00000286411 | AC002458,1 | -0,073143213 | 0,950564736 | 0,78918 | 0,90236217 | novel transcript, antisense to CDK14 |
| ENSG00000154263 | ABCA10 | -0,073208919 | 0,950521445 | 0,85994 | 0,93704372 | ATP binding cassette subfamily A member 10 |
| ENSG00000237036 | ZEB1-AS1 | -0,073230978 | 0,950506911 | 0,69405 | 0,85147285 | ZEB1 antisense RNA 1 |
| ENSG00000287242 | AC073593,2 | -0,07323538 | 0,950504011 | 0,85583 | 0,9347432 | novel transcript, antisense to SCARB1 |
| ENSG00000101349 | PAK5 | -0,073285159 | 0,950471215 | 0,86261 | 0,93868982 | p21 (RAC1) activated kinase 5 |
| ENSG00000254873 | AP001267,1 | -0,073288173 | 0,95046923 | 0,93025 | 0,97011853 | novel transcript, antisense to ATP5L and UBE4A |
| ENSG00000251675 | AC010260,1 | -0,073390078 | 0,950402095 | 0,93858 | 0,9737443 | novel transcript |
| ENSG00000135392 | DNAJC14 | -0,073426781 | 0,950377917 | 0,74316 | 0,87840582 | DnaJ heat shock protein family (Hsp40) member C14 |
| ENSG00000261758 | AC117382,2 | -0,073508688 | 0,950323962 | 0,78748 | 0,90121071 | novel transcript, antisense to STAG1 |
| ENSG00000219188 | CACYBPP3 | -0,073568347 | 0,950284665 | 0,93527 | 0,97222198 | calcyclin binding protein pseudogene 3 |
| ENSG00000162999 | DUSP19 | -0,073673655 | 0,950215302 | 0,77422 | 0,89572556 | dual specificity phosphatase 19 |
| ENSG00000158156 | XKR8 | -0,073685811 | 0,950207296 | 0,68591 | 0,84654659 | XK related 8 |
| ENSG00000007168 | PAFAH1B1 | -0,073720219 | 0,950184634 | 0,68936 | 0,84865116 | platelet activating factor acetylhydrolase 1b regulatory subunit 1 |
| ENSG00000169710 | FASN | -0,073757035 | 0,950160387 | 0,77982 | 0,8981559 | fatty acid synthase |
| ENSG00000141551 | CSNK1D | -0,073780808 | 0,95014473 | 0,73849 | 0,87584394 | casein kinase 1 delta |
| ENSG00000278916 | CEP83-DT | -0,07386349 | 0,950090278 | 0,79818 | 0,90685415 | CEP83 divergent transcript |
| ENSG00000166734 | GOLM2 | -0,073954888 | 0,950030089 | 0,72381 | 0,86812308 | golgi membrane protein 2 |
| ENSG00000106261 | ZKSCAN1 | -0,073971441 | 0,950019189 | 0,66578 | 0,8336647 | zinc finger with KRAB and SCAN domains 1 |
| ENSG00000224786 | CETN4P | -0,07398583 | 0,950009714 | 0,84091 | 0,92751395 | centrin 4, pseudogene |
| ENSG00000277978 | AC010542,5 | -0,074055875 | 0,94996359 | 0,90604 | 0,95982149 | novel transcript, antisense to CKLF |
| ENSG00000162627 | SNX7 | -0,074148847 | 0,949902374 | 0,84805 | 0,9309317 | sorting nexin 7 |
| ENSG00000135931 | ARMC9 | -0,074237465 | 0,949844028 | 0,6964 | 0,85299392 | armadillo repeat containing 9 |
| ENSG00000223979 | SMCR2 | -0,07434475 | 0,949773395 | 0,9223 | 0,9663588 | Smith-Magenis syndrome chromosome region, candidate 2 |
| ENSG00000099899 | TRMT2A | -0,074461174 | 0,949696753 | 0,69254 | 0,85051331 | tRNA methyltransferase 2 homolog A |
| ENSG00000152904 | GGPS1 | -0,074548837 | 0,949639048 | 0,63662 | 0,81709016 | geranylgeranyl diphosphate synthase 1 |

| | | | | | | |
|---|---|---|---|---|---|---|
| ENSG00000162997 | PRORSD1P | -0,074709693 | 0,949533172 | 0,77415 | 0,89572556 | prolyl-tRNA synthetase associated domain containing 1, pseudogene |
| ENSG00000203865 | ATP1A1-AS1 | -0,07487315 | 0,949425597 | 0,71203 | 0,8615121 | ATP1A1 antisense RNA 1 |
| ENSG00000105223 | PLD3 | -0,074893618 | 0,949412126 | 0,69227 | 0,85023519 | phospholipase D family member 3 |
| ENSG00000272711 | AC019069,1 | -0,074961673 | 0,949367341 | 0,7974 | 0,90666364 | novel transcript |
| ENSG00000121940 | CLCC1 | -0,075041107 | 0,949315072 | 0,69783 | 0,85363988 | chloride channel CLIC like 1 |
| ENSG00000100300 | TSPO | -0,075170932 | 0,949229648 | 0,85074 | 0,93206349 | translocator protein |
| ENSG00000232160 | RAP2C-AS1 | -0,07520904 | 0,949204575 | 0,73077 | 0,87205836 | RAP2C antisense RNA 1 |
| ENSG00000275074 | NUDT18 | -0,075245313 | 0,94918071 | 0,8362 | 0,92493625 | nudix hydrolase 18 |
| ENSG00000154548 | SRSF12 | -0,075265676 | 0,949167314 | 0,71695 | 0,86453146 | serine and arginine rich splicing factor 12 |
| ENSG00000278513 | AC091046,2 | -0,075294475 | 0,949148366 | 0,92189 | 0,96616853 | zinc finger protein 551 (ZNF551) pseudogene |
| ENSG00000172650 | AGAP5 | -0,075432947 | 0,94905727 | 0,78483 | 0,89984877 | ArfGAP with GTPase domain, ankyrin repeat and PH domain 5 |
| ENSG00000196189 | SEMA4A | -0,075441624 | 0,949051562 | 0,8391 | 0,92640383 | semaphorin 4A |
| ENSG00000103351 | CLUAP1 | -0,075550494 | 0,948979946 | 0,66703 | 0,83466874 | clusterin associated protein 1 |
| ENSG00000285958 | AC073174,1 | -0,075565369 | 0,948970162 | 0,84783 | 0,9309317 | novel transcript |
| ENSG00000240370 | RPL13P5 | -0,075648094 | 0,948915749 | 0,77117 | 0,89404837 | ribosomal protein L13 pseudogene 5 |
| ENSG00000178605 | GTPBP6 | -0,075764631 | 0,948839101 | 0,7195 | 0,86592255 | GTP binding protein 6 (putative) |
| ENSG00000120899 | PTK2B | -0,07586625 | 0,94877227 | 0,6926 | 0,85054353 | protein tyrosine kinase 2 beta |
| ENSG00000244757 | AL137024,1 | -0,075925459 | 0,948733333 | 0,93194 | 0,97097872 | novel transcript |
| ENSG00000261485 | PAN3-AS1 | -0,076617247 | 0,948278514 | 0,78066 | 0,89846287 | PAN3 antisense RNA 1 |
| ENSG00000168569 | TMEM223 | -0,076617765 | 0,948278174 | 0,7041 | 0,85725494 | transmembrane protein 223 |
| ENSG00000173567 | ADGRF3 | -0,076656563 | 0,948252672 | 0,86868 | 0,94200797 | adhesion G protein-coupled receptor F3 |
| ENSG00000133027 | PEMT | -0,076674399 | 0,948240949 | 0,74294 | 0,87840582 | phosphatidylethanolamine N-methyltransferase |
| ENSG00000175229 | GAL3ST3 | -0,076756024 | 0,9481873 | 0,80041 | 0,90809698 | galactose-3-O-sulfotransferase 3 |
| ENSG00000283741 | SMCR8 | -0,076842872 | 0,948130223 | 0,96 | 0,98353421 | SMCR8-C9orf72 complex subunit |
| ENSG00000182836 | PLCXD3 | -0,076953025 | 0,948057834 | 0,91588 | 0,96339511 | phosphatidylinositol specific phospholipase C X domain containing 3 |
| ENSG00000143771 | CNIH4 | -0,076954288 | 0,948057004 | 0,69295 | 0,85086877 | cornichon family AMPA receptor auxiliary protein 4 |
| ENSG00000178229 | ZNF543 | -0,076982877 | 0,948038217 | 0,69045 | 0,84900677 | zinc finger protein 543 |
| ENSG00000129355 | CDKN2D | -0,077068361 | 0,947982045 | 0,79709 | 0,90645712 | cyclin dependent kinase inhibitor 2D |
| ENSG00000170558 | CDH2 | -0,077076642 | 0,947976603 | 0,82591 | 0,91932995 | cadherin 2 |
| ENSG00000246084 | LINC02325 | -0,077189898 | 0,947902187 | 0,87439 | 0,9443876 | long intergenic non-protein coding RNA 2325 |

| | | | | | | |
|---|---|---|---|---|---|---|
| ENSG00000213123 | TCTEX1D2 | -0,077289491 | 0,947836753 | 0,70547 | 0,85808932 | Tctex1 domain containing 2 |
| ENSG00000287004 | AC007686,5 | -0,077522725 | 0,947683533 | 0,92549 | 0,9676347 | novel transcript |
| ENSG00000110786 | PTPN5 | -0,077523975 | 0,947682712 | 0,87856 | 0,94602899 | protein tyrosine phosphatase non-receptor type 5 |
| ENSG00000282364 | AC004980,6 | -0,07753805 | 0,947673466 | 0,87882 | 0,94612705 | pseudogene similar to part of RB-associated KRAB zinc finger RBAK |
| ENSG00000214243 | AC004980,2 | -0,07753805 | 0,947673466 | 0,87882 | 0,94612705 | pseudogene similar to part of RB-associated KRAB zinc finger RBAK |
| ENSG00000169136 | ATF5 | -0,077580769 | 0,947645406 | 0,76855 | 0,89277929 | activating transcription factor 5 |
| ENSG00000196547 | MAN2A2 | -0,077612635 | 0,947624475 | 0,75814 | 0,88722916 | mannosidase alpha class 2A member 2 |
| ENSG00000265751 | AC015878,1 | -0,077719994 | 0,947553959 | 0,9376 | 0,97330214 | novel transcript, antisense to GREB1L |
| ENSG00000105793 | GTPBP10 | -0,077790572 | 0,947507605 | 0,65756 | 0,82877 | GTP binding protein 10 |
| ENSG00000105607 | GCDH | -0,077798068 | 0,947502682 | 0,71834 | 0,86522478 | glutaryl-CoA dehydrogenase |
| ENSG00000106330 | MOSPD3 | -0,077949164 | 0,947403453 | 0,7395 | 0,87648588 | motile sperm domain containing 3 |
| ENSG00000240429 | LRRFIP1P1 | -0,077968834 | 0,947390537 | 0,83789 | 0,92608361 | LRR binding FLII interacting protein 1 pseudogene 1 |
| ENSG00000010318 | PHF7 | -0,078023201 | 0,947354835 | 0,7782 | 0,89735311 | PHD finger protein 7 |
| ENSG00000108950 | FAM20A | -0,078084549 | 0,947314552 | 0,77574 | 0,89631083 | FAM20A golgi associated secretory pathway pseudokinase |
| ENSG00000131730 | CKMT2 | -0,078108761 | 0,947298654 | 0,77777 | 0,89717046 | creatine kinase, mitochondrial 2 |
| ENSG00000228925 | AC016722,2 | -0,078376787 | 0,94712268 | 0,81616 | 0,91549169 | novel transcript, antisense to MCFD2 |
| ENSG00000240204 | SMKR1 | -0,078404187 | 0,947104692 | 0,82317 | 0,91828654 | small lysine rich protein 1 |
| ENSG00000206491 | PRR3 | -0,078440705 | 0,947080719 | 0,87725 | 0,94559723 | proline rich 3 |
| ENSG00000277215 | SPANXA2-OT1 | -0,078504836 | 0,947038619 | 0,76877 | 0,89277929 | SPANXA2 overlapping transcript 1 |
| ENSG00000183617 | MRPL54 | -0,078523756 | 0,9470262 | 0,70417 | 0,85726326 | mitochondrial ribosomal protein L54 |
| ENSG00000158467 | AHCYL2 | -0,078557695 | 0,947003921 | 0,7235 | 0,86795614 | adenosylhomocysteinase like 2 |
| ENSG00000254986 | DPP3 | -0,078853634 | 0,946809683 | 0,67834 | 0,84169189 | dipeptidyl peptidase 3 |
| ENSG00000168348 | INSM2 | -0,078888955 | 0,946786503 | 0,84395 | 0,92886643 | INSM transcriptional repressor 2 |
| ENSG00000108465 | CDK5RAP3 | -0,079006529 | 0,946709347 | 0,69822 | 0,85390309 | CDK5 regulatory subunit associated protein 3 |
| ENSG00000130234 | ACE2 | -0,07904173 | 0,946686248 | 0,7936 | 0,9039896 | angiotensin I converting enzyme 2 |
| ENSG00000273791 | AC007204,1 | -0,07906069 | 0,946673807 | 0,88596 | 0,94951344 | BCL2/adenovirus E1B 19kDa interacting protein 3 (BNIP3) pseudogene |
| ENSG00000273045 | C2orf15 | -0,079076329 | 0,946663545 | 0,86015 | 0,93714201 | chromosome 2 open reading frame 15 |
| ENSG00000234327 | ZNF232-AS1 | -0,079239242 | 0,946556651 | 0,72065 | 0,86633316 | ZNF232 antisense RNA 1 |
| ENSG00000134375 | TIMM17A | -0,079255211 | 0,946546173 | 0,78146 | 0,89877377 | translocase of inner mitochondrial membrane 17A |
| ENSG00000122779 | TRIM24 | -0,07934765 | 0,946485526 | 0,75227 | 0,88394752 | tripartite motif containing 24 |

| | | | | | | |
|---|---|---|---|---|---|---|
| ENSG00000272663 | AC093635,1 | -0,079347911 | 0,946485355 | 0,87185 | 0,9434099 | novel transcript |
| ENSG00000081014 | AP4E1 | -0,079372613 | 0,94646915 | 0,68494 | 0,84583571 | adaptor related protein complex 4 subunit epsilon 1 |
| ENSG00000254911 | SCARNA9 | -0,079398716 | 0,946452025 | 0,75684 | 0,88648088 | small Cajal body-specific RNA 9 |
| ENSG00000250365 | AL139353,2 | -0,079416739 | 0,946440202 | 0,84019 | 0,92705697 | novel transcript, antisense to C14orf126 |
| ENSG00000138778 | CENPE | -0,079656813 | 0,946282721 | 0,82559 | 0,91917221 | centromere protein E |
| ENSG00000128191 | DGCR8 | -0,079667911 | 0,946275442 | 0,66547 | 0,83353389 | DGCR8 microprocessor complex subunit |
| ENSG00000174177 | CTU2 | -0,079729612 | 0,946234972 | 0,73599 | 0,87510758 | cytosolic thiouridylase subunit 2 |
| ENSG00000274528 | AC090970,2 | -0,079810162 | 0,946182143 | 0,86325 | 0,93891554 | novel transcript |
| ENSG00000142530 | FAM71E1 | -0,079833882 | 0,946166586 | 0,83567 | 0,92493625 | family with sequence similarity 71 member E1 |
| ENSG00000277151 | AL138820,1 | -0,079860021 | 0,946149444 | 0,80025 | 0,90801903 | novel transcript |
| ENSG00000224138 | AC000123,1 | -0,079943876 | 0,946094451 | 0,91648 | 0,9636258 | novel transcript |
| ENSG00000106603 | COA1 | -0,079960244 | 0,946083718 | 0,66474 | 0,83302909 | cytochrome c oxidase assembly factor 1 homolog |
| ENSG00000104889 | RNASEH2A | -0,079963348 | 0,946081682 | 0,649 | 0,82431124 | ribonuclease H2 subunit A |
| ENSG00000147601 | TERF1 | -0,079975129 | 0,946073956 | 0,66333 | 0,83242946 | telomeric repeat binding factor 1 |
| ENSG00000213614 | HEXA | -0,080003208 | 0,946055543 | 0,69542 | 0,85232854 | hexosaminidase subunit alpha |
| ENSG00000240265 | AC099328,1 | -0,080020622 | 0,946044124 | 0,90741 | 0,96044232 | H1 histone family, member O, oocyte-specific (H1FOO) pseudogene |
| ENSG00000249825 | THBS4-AS1 | -0,080042471 | 0,946029796 | 0,85712 | 0,9354716 | THBS4 antisense RNA 1 |
| ENSG00000223508 | RPL23AP53 | -0,080069351 | 0,946012171 | 0,79272 | 0,90364189 | ribosomal protein L23a pseudogene 53 |
| ENSG00000260229 | PPIAP51 | -0,080107812 | 0,945986951 | 0,84655 | 0,93031425 | peptidylprolyl isomerase A pseudogene 51 |
| ENSG00000270177 | AC104109,2 | -0,080179829 | 0,94593973 | 0,83858 | 0,92627183 | novel transcript |
| ENSG00000111671 | SPSB2 | -0,08020995 | 0,94591998 | 0,7642 | 0,89012684 | splA/ryanodine receptor domain and SOCS box containing 2 |
| ENSG00000225756 | DBH-AS1 | -0,080327376 | 0,945842992 | 0,83284 | 0,92352564 | DBH antisense RNA 1 |
| ENSG00000126453 | BCL2L12 | -0,080353841 | 0,945825641 | 0,79479 | 0,90488348 | BCL2 like 12 |
| ENSG00000178950 | GAK | -0,080676584 | 0,945614076 | 0,67921 | 0,84232543 | cyclin G associated kinase |
| ENSG00000197006 | METTL9 | -0,080770539 | 0,945552495 | 0,65246 | 0,8259831 | methyltransferase like 9 |
| ENSG00000100307 | CBX7 | -0,080772867 | 0,945550969 | 0,78463 | 0,89968774 | chromobox 7 |
| ENSG00000249673 | NOP14-AS1 | -0,08080809 | 0,945527884 | 0,63696 | 0,81742435 | NOP14 antisense RNA 1 |
| ENSG00000171962 | DRC3 | -0,081005181 | 0,945398722 | 0,7514 | 0,88347546 | dynein regulatory complex subunit 3 |
| ENSG00000182208 | MOB2 | -0,081029781 | 0,945382601 | 0,73223 | 0,87293156 | MOB kinase activator 2 |
| ENSG00000161956 | SENP3 | -0,081096605 | 0,945338813 | 0,72812 | 0,87073013 | SUMO specific peptidase 3 |
| ENSG00000275164 | MUC4 | -0,081155819 | 0,945300013 | 0,947 | 0,9774744 | mucin 4, cell surface associated |

| | | | | | | |
|---|---|---|---|---|---|---|
| ENSG00000143845 | ETNK2 | -0,081189398 | 0,945278012 | 0,79178 | 0,90327669 | ethanolamine kinase 2 |
| ENSG00000241015 | TPM3P9 | -0,081276604 | 0,945220875 | 0,66436 | 0,83269974 | tropomyosin 3 pseudogene 9 |
| ENSG00000151778 | SERP2 | -0,081326896 | 0,945187925 | 0,84289 | 0,9282764 | stress associated endoplasmic reticulum protein family member 2 |
| ENSG00000145833 | DDX46 | -0,08136418 | 0,945163499 | 0,70241 | 0,85653047 | DEAD-box helicase 46 |
| ENSG00000288390 | UBL7-AS1 | -0,08137772 | 0,945154628 | 0,66394 | 0,83247324 | UBL7 antisense RNA 1 (head to head) |
| ENSG00000255492 | H3P33 | -0,081412724 | 0,945131696 | 0,91798 | 0,9644207 | H3 histone pseudogene 33 |
| ENSG00000197081 | IGF2R | -0,081421059 | 0,945126236 | 0,75135 | 0,88347546 | insulin like growth factor 2 receptor |
| ENSG00000278307 | TAS2R19 | -0,081547124 | 0,945043653 | 0,83078 | 0,92262505 | taste 2 receptor member 19 |
| ENSG00000163633 | C4orf36 | -0,08155836 | 0,945036293 | 0,89582 | 0,95483699 | chromosome 4 open reading frame 36 |
| ENSG00000223694 | ADH5P3 | -0,081572126 | 0,945027276 | 0,90493 | 0,9593272 | ADH5 pseudogene 3 |
| ENSG00000204706 | MAMDC2-AS1 | -0,081619444 | 0,944996281 | 0,87273 | 0,94374455 | MAMDC2 antisense RNA 1 |
| ENSG00000146278 | PNRC1 | -0,081637932 | 0,944984171 | 0,74101 | 0,87724557 | proline rich nuclear receptor coactivator 1 |
| ENSG00000180777 | ANKRD30B | -0,08166634 | 0,944965563 | 0,82986 | 0,92189428 | ankyrin repeat domain 30B |
| ENSG00000184640 | SEPTIN9 | -0,081693737 | 0,944947618 | 0,78229 | 0,8988906 | septin 9 |
| ENSG00000286257 | AL137100,3 | -0,081792717 | 0,94488279 | 0,84527 | 0,92946127 | novel transcript |
| ENSG00000180425 | C11orf71 | -0,081802901 | 0,94487612 | 0,81696 | 0,9158035 | chromosome 11 open reading frame 71 |
| ENSG00000213923 | CSNK1E | -0,081817853 | 0,944866328 | 0,63005 | 0,8132682 | casein kinase 1 epsilon |
| ENSG00000230799 | RPS2P16 | -0,081829384 | 0,944858775 | 0,87129 | 0,94321633 | ribosomal protein S2 pseudogene 16 |
| ENSG00000274958 | ITPK1 | -0,081943183 | 0,944784248 | 0,77838 | 0,89744206 | inositol-tetrakisphosphate 1-kinase |
| ENSG00000259458 | MGC15885 | -0,081983061 | 0,944758133 | 0,89719 | 0,955727 | uncharacterized protein MGC15885 |
| ENSG00000232282 | MTND1P32 | -0,081984067 | 0,944757475 | 0,88181 | 0,94750352 | MT-ND1 pseudogene 32 |
| ENSG00000107937 | GTPBP4 | -0,082011837 | 0,944739289 | 0,74608 | 0,88023223 | GTP binding protein 4 |
| ENSG00000158828 | PINK1 | -0,082073701 | 0,944698779 | 0,75406 | 0,88473546 | PTEN induced kinase 1 |
| ENSG00000127922 | SEM1 | -0,082137011 | 0,944657324 | 0,68922 | 0,84859173 | SEM1 26S proteasome complex subunit |
| ENSG00000285868 | AC008676,3 | -0,08214098 | 0,944654725 | 0,84143 | 0,92787857 | Novel protein |
| ENSG00000063244 | U2AF2 | -0,082174259 | 0,944632935 | 0,69948 | 0,8548625 | U2 small nuclear RNA auxiliary factor 2 |
| ENSG00000272432 | AL031432,3 | -0,082255713 | 0,944579602 | 0,90888 | 0,96099548 | novel transcript |
| ENSG00000232648 | AC107214,1 | -0,082275274 | 0,944566796 | 0,77466 | 0,89586241 | novel transcript |
| ENSG00000157693 | TMEM268 | -0,082450937 | 0,944451792 | 0,74333 | 0,87846407 | transmembrane protein 268 |
| ENSG00000204947 | ZNF425 | -0,082527508 | 0,944401666 | 0,67485 | 0,83977504 | zinc finger protein 425 |
| ENSG00000251287 | ALG1L2 | -0,082538269 | 0,944394622 | 0,8385 | 0,92627183 | ALG1 chitobiosyldiphosphodolichol beta-mannosyltransferase like 2 |

| | | | | | | |
|---|---|---|---|---|---|---|
| ENSG00000228723 | SRGAP3-AS2 | -0,082863568 | 0,944181703 | 0,87771 | 0,94568249 | SRGAP3 antisense RNA 2 |
| ENSG00000134030 | CTIF | -0,08289223 | 0,944162946 | 0,77461 | 0,89586241 | cap binding complex dependent translation initiation factor |
| ENSG00000260910 | LINC00565 | -0,082931473 | 0,944137264 | 0,86199 | 0,93834335 | long intergenic non-protein coding RNA 565 |
| ENSG00000196081 | ZNF724 | -0,082975278 | 0,944108597 | 0,65777 | 0,82881843 | zinc finger protein 724 |
| ENSG00000144792 | ZNF660 | -0,082984964 | 0,944102258 | 0,87814 | 0,9458146 | zinc finger protein 660 |
| ENSG00000210140 | MT-TC | -0,083072903 | 0,944044713 | 0,9306 | 0,97025341 | mitochondrially encoded tRNA-Cys (UGU/C) |
| ENSG00000162738 | VANGL2 | -0,083083919 | 0,944037505 | 0,58505 | 0,7860906 | VANGL planar cell polarity protein 2 |
| ENSG00000061936 | SFSWAP | -0,083099388 | 0,944027382 | 0,70319 | 0,8568579 | splicing factor SWAP |
| ENSG00000201428 | RN7SKP71 | -0,083125924 | 0,944010019 | 0,88292 | 0,94823653 | RN7SK pseudogene 71 |
| ENSG00000107105 | ELAVL2 | -0,083333318 | 0,943874323 | 0,85001 | 0,93174363 | ELAV like RNA binding protein 2 |
| ENSG00000227068 | AL162584,1 | -0,083399093 | 0,943831291 | 0,91819 | 0,96445665 | novel transcript |
| ENSG00000128266 | GNAZ | -0,083489094 | 0,943772413 | 0,62819 | 0,81178582 | G protein subunit alpha z |
| ENSG00000241007 | SEPTIN7P6 | -0,083489438 | 0,943772188 | 0,90881 | 0,96099548 | septin 7 pseudogene 6 |
| ENSG00000196458 | ZNF605 | -0,083528457 | 0,943746663 | 0,6601 | 0,83029149 | zinc finger protein 605 |
| ENSG00000164048 | ZNF589 | -0,083713098 | 0,943625886 | 0,71738 | 0,86483656 | zinc finger protein 589 |
| ENSG00000170889 | RPS9 | -0,083741935 | 0,943607025 | 0,79621 | 0,90581597 | ribosomal protein S9 |
| ENSG00000241553 | ARPC4 | -0,083754679 | 0,94359869 | 0,65936 | 0,82984259 | actin related protein 2/3 complex subunit 4 |
| ENSG00000227017 | AC007036,1 | -0,083806591 | 0,943564738 | 0,94655 | 0,97724021 | novel transcript |
| ENSG00000287719 | AP001531,1 | -0,083831446 | 0,943548482 | 0,82005 | 0,91723726 | novel transcript |
| ENSG00000100949 | RABGGTA | -0,084163965 | 0,943331033 | 0,77569 | 0,89630112 | Rab geranylgeranyltransferase subunit alpha |
| ENSG00000285193 | RABGGTA | -0,084163965 | 0,943331033 | 0,77569 | 0,89630112 | Rab geranylgeranyltransferase subunit alpha |
| ENSG00000172428 | COPS9 | -0,084441104 | 0,943149839 | 0,68194 | 0,84396124 | COP9 signalosome subunit 9 |
| ENSG00000233143 | DIRC3-AS1 | -0,084451707 | 0,943142907 | 0,87561 | 0,94497474 | DIRC3 antisense RNA 1 |
| ENSG00000286924 | AL139815,1 | -0,084758811 | 0,942942163 | 0,92646 | 0,96803855 | novel transcript, antisense to ARMC3and MSRB2 |
| ENSG00000228554 | AC004837,2 | -0,084813904 | 0,942906155 | 0,90723 | 0,96034215 | novel transcript |
| ENSG00000213928 | IRF9 | -0,084815649 | 0,942905015 | 0,76527 | 0,89082001 | interferon regulatory factor 9 |
| ENSG00000153162 | BMP6 | -0,084868664 | 0,942870366 | 0,80495 | 0,91018722 | bone morphogenetic protein 6 |
| ENSG00000146263 | MMS22L | -0,084924532 | 0,942833855 | 0,65357 | 0,82638628 | MMS22 like, DNA repair protein |
| ENSG00000270900 | NUDT19P4 | -0,085144462 | 0,942690136 | 0,92971 | 0,96987987 | NUDT19 pseudogene 4 |
| ENSG00000225569 | CCT4P2 | -0,085203126 | 0,942651805 | 0,87638 | 0,94545974 | chaperonin containing TCP1 subunit 4 pseudogene 2 |
| ENSG00000110583 | NAA40 | -0,08523889 | 0,942628437 | 0,62131 | 0,80787022 | N-alpha-acetyltransferase 40, NatD catalytic subunit |
| ENSG00000176809 | LRRC37A3 | -0,085310197 | 0,942581848 | 0,6274 | 0,8114294 | leucine rich repeat containing 37 member A3 |

| | | | | | | |
|---|---|---|---|---|---|---|
| ENSG00000105552 | BCAT2 | -0,085489742 | 0,942464549 | 0,798 | 0,90685415 | branched chain amino acid transaminase 2 |
| ENSG00000110400 | NECTIN1 | -0,085496975 | 0,942459825 | 0,71016 | 0,86036733 | nectin cell adhesion molecule 1 |
| ENSG00000106069 | CHN2 | -0,085665268 | 0,942349891 | 0,82253 | 0,9180405 | chimerin 2 |
| ENSG00000116205 | TCEANC2 | -0,085734223 | 0,942304852 | 0,72521 | 0,86894646 | transcription elongation factor A N-terminal and central domain containing 2 |
| ENSG00000117242 | PINK1-AS | -0,085745593 | 0,942297425 | 0,72717 | 0,87016629 | PINK1 antisense RNA |
| ENSG00000213003 | BTF3P12 | -0,085841665 | 0,942234678 | 0,88585 | 0,94944442 | basic transcription factor 3 pseudogene 12 |
| ENSG00000269662 | BNIP3P39 | -0,085848236 | 0,942230386 | 0,8807 | 0,94687991 | BCL2 interacting protein 3 pseudogene 39 |
| ENSG00000167548 | KMT2D | -0,085967777 | 0,942152317 | 0,75967 | 0,88798255 | lysine methyltransferase 2D |
| ENSG00000225855 | RUSC1-AS1 | -0,086132101 | 0,942045011 | 0,77 | 0,89324517 | RUSC1 antisense RNA 1 |
| ENSG00000033030 | ZCCHC8 | -0,086247835 | 0,941969443 | 0,70044 | 0,85531965 | zinc finger CCHC-type containing 8 |
| ENSG00000140259 | MFAP1 | -0,086262395 | 0,941959936 | 0,64198 | 0,8205913 | microfibril associated protein 1 |
| ENSG00000109016 | DHRS7B | -0,086371286 | 0,941888842 | 0,75389 | 0,88467924 | dehydrogenase/reductase 7B |
| ENSG00000109501 | WFS1 | -0,086397048 | 0,941872023 | 0,67585 | 0,84049293 | wolframin ER transmembrane glycoprotein |
| ENSG00000287430 | AL136090,2 | -0,086579481 | 0,941752928 | 0,94182 | 0,97511532 | novel transcript |
| ENSG00000227268 | KLLN | -0,086644593 | 0,941710426 | 0,78729 | 0,90121071 | killin, p53 regulated DNA replication inhibitor |
| ENSG00000177108 | ZDHHC22 | -0,086678502 | 0,941688292 | 0,89131 | 0,95208954 | zinc finger DHHC-type palmitoyltransferase 22 |
| ENSG00000196586 | MYO6 | -0,086693414 | 0,941678559 | 0,77246 | 0,89482448 | myosin VI |
| ENSG00000133275 | CSNK1G2 | -0,086787049 | 0,941617443 | 0,67985 | 0,8427192 | casein kinase 1 gamma 2 |
| ENSG00000131779 | PEX11B | -0,086790985 | 0,941614874 | 0,71669 | 0,86439754 | peroxisomal biogenesis factor 11 beta |
| ENSG00000053372 | MRTO4 | -0,086849007 | 0,941577005 | 0,76632 | 0,89138648 | MRT4 homolog, ribosome maturation factor |
| ENSG00000069535 | MAOB | -0,08702717 | 0,941460734 | 0,80268 | 0,90909092 | monoamine oxidase B |
| ENSG00000160097 | FNDC5 | -0,087053712 | 0,941443414 | 0,75371 | 0,88459403 | fibronectin type III domain containing 5 |
| ENSG00000262119 | AL079343,1 | -0,087105551 | 0,941409586 | 0,84623 | 0,93006588 | novel transcript, sense intronic to NOVA1 |
| ENSG00000104290 | FZD3 | -0,087289815 | 0,941289355 | 0,74559 | 0,87996245 | frizzled class receptor 3 |
| ENSG00000236011 | GPANK1 | -0,08735384 | 0,941247583 | 0,81184 | 0,91334888 | G-patch domain and ankyrin repeats 1 |
| ENSG00000100197 | CYP2D6 | -0,087377445 | 0,941232182 | 0,8163 | 0,91550954 | cytochrome P450 family 2 subfamily D member 6 |
| ENSG00000229847 | EMX2OS | -0,087468119 | 0,941173028 | 0,836 | 0,92493625 | EMX2 opposite strand/antisense RNA |
| ENSG00000271749 | AC005828,6 | -0,087473967 | 0,941169212 | 0,92166 | 0,96608203 | mitochondrially encoded cytochrome c oxidase III (MT-CO3) pseudogene |
| ENSG00000010295 | IFFO1 | -0,087545829 | 0,941122333 | 0,72194 | 0,86726498 | intermediate filament family orphan 1 |
| ENSG00000071462 | BUD23 | -0,087588849 | 0,94109427 | 0,69596 | 0,85283771 | BUD23 rRNA methyltransferase and ribosome maturation factor |

| | | | | | | |
|---|---|---|---|---|---|---|
| ENSG00000197180 | CH17-340M24 | -0,087786668 | 0,940965238 | 0,77802 | 0,89732098 | uncharacterized protein BC009467 |
| ENSG00000179562 | GCC1 | -0,087889894 | 0,940897914 | 0,68761 | 0,8478032 | GRIP and coiled-coil domain containing 1 |
| ENSG00000226721 | EEF1DP2 | -0,087963003 | 0,940850235 | 0,88017 | 0,94667622 | eukaryotic translation elongation factor 1 delta pseudogene 2 |
| ENSG00000103264 | FBXO31 | -0,088010993 | 0,940818939 | 0,71401 | 0,86227395 | F-box protein 31 |
| ENSG00000276651 | AC007950,2 | -0,088091616 | 0,940766364 | 0,83469 | 0,92444381 | novel transcript |
| ENSG00000127220 | ABHD8 | -0,08822175 | 0,940681508 | 0,73772 | 0,87540065 | abhydrolase domain containing 8 |
| ENSG00000270011 | ZNF559-ZNF1 | -0,088235147 | 0,940672773 | 0,73824 | 0,87571125 | ZNF559-ZNF177 readthrough |
| ENSG00000166123 | GPT2 | -0,0883359 | 0,940607082 | 0,5476 | 0,76294624 | glutamic--pyruvic transaminase 2 |
| ENSG00000168421 | RHOH | -0,088420031 | 0,940552232 | 0,86355 | 0,93904284 | ras homolog family member H |
| ENSG00000122705 | CLTA | -0,08844247 | 0,940537603 | 0,77351 | 0,89545981 | clathrin light chain A |
| ENSG00000126709 | IFI6 | -0,08864349 | 0,940406562 | 0,86897 | 0,94225628 | interferon alpha inducible protein 6 |
| ENSG00000161847 | RAVER1 | -0,088656654 | 0,94039798 | 0,68914 | 0,8485712 | ribonucleoprotein, PTB binding 1 |
| ENSG00000163738 | MTHFD2L | -0,088727352 | 0,940351898 | 0,81458 | 0,91470265 | methylenetetrahydrofolate dehydrogenase (NADP+ dependent) 2 like |
| ENSG00000263002 | ZNF234 | -0,088755582 | 0,940333498 | 0,68385 | 0,84517371 | zinc finger protein 234 |
| ENSG00000156689 | GLYATL2 | -0,08881957 | 0,940291792 | 0,8758 | 0,94506005 | glycine-N-acyltransferase like 2 |
| ENSG00000185432 | METTL7A | -0,088851234 | 0,940271155 | 0,86605 | 0,9405388 | methyltransferase like 7A |
| ENSG00000223575 | RBMX2P3 | -0,088885355 | 0,940248917 | 0,90906 | 0,96102407 | RBMX2 pseudogene 3 |
| ENSG00000238039 | AC244197,2 | -0,088912512 | 0,940231218 | 0,82828 | 0,92076203 | novel transcript |
| ENSG00000112576 | CCND3 | -0,089000366 | 0,940173964 | 0,64092 | 0,81980451 | cyclin D3 |
| ENSG00000162755 | KLHDC9 | -0,089042925 | 0,94014623 | 0,78753 | 0,90121071 | kelch domain containing 9 |
| ENSG00000274553 | AC138035,3 | -0,089101713 | 0,940107921 | 0,77684 | 0,89660077 | capicua transcriptional repressor (CIC) pseudogene |
| ENSG00000131797 | CLUHP3 | -0,089157577 | 0,940071519 | 0,68202 | 0,84401479 | clustered mitochondria homolog pseudogene 3 |
| ENSG00000287662 | AC109824,1 | -0,089208723 | 0,940038192 | 0,93472 | 0,9721963 | novel transcript |
| ENSG00000184857 | TMEM186 | -0,089277933 | 0,939993097 | 0,6935 | 0,85109296 | transmembrane protein 186 |
| ENSG00000145740 | SLC30A5 | -0,089360472 | 0,93993932 | 0,7245 | 0,86850994 | solute carrier family 30 member 5 |
| ENSG00000100650 | SRSF5 | -0,089465161 | 0,939871116 | 0,67086 | 0,83695077 | serine and arginine rich splicing factor 5 |
| ENSG00000143303 | RRNAD1 | -0,089529237 | 0,939829373 | 0,65922 | 0,82978434 | ribosomal RNA adenine dimethylase domain containing 1 |
| ENSG00000259209 | AC004943,1 | -0,089736647 | 0,939694267 | 0,80173 | 0,90870853 | novel transcript, antisense to ZFHX3 |
| ENSG00000257954 | AC125611,2 | -0,089846677 | 0,939622603 | 0,87843 | 0,94596522 | FGFR1 oncogene partner 2 (FGFR1OP2) pseudogene |
| ENSG00000070501 | POLB | -0,089875017 | 0,939604145 | 0,73825 | 0,87571125 | DNA polymerase beta |
| ENSG00000162572 | SCNN1D | -0,090020075 | 0,939509676 | 0,83528 | 0,92477092 | sodium channel epithelial 1 subunit delta |

| | | | | | | |
|---|---|---|---|---|---|---|
| ENSG00000254706 | AL512785,2 | -0,090137394 | 0,939433279 | 0,81731 | 0,91606335 | novel protein |
| ENSG00000170949 | ZNF160 | -0,090237444 | 0,939368132 | 0,59162 | 0,79071076 | zinc finger protein 160 |
| ENSG00000232169 | ABCF1 | -0,090237661 | 0,939367991 | 0,73935 | 0,8763893 | ATP binding cassette subfamily F member 1 |
| ENSG00000184368 | MAP7D2 | -0,090255272 | 0,939356523 | 0,7481 | 0,88143479 | MAP7 domain containing 2 |
| ENSG00000168256 | NKIRAS2 | -0,090275714 | 0,939343214 | 0,65587 | 0,82812732 | NFKB inhibitor interacting Ras like 2 |
| ENSG00000106290 | TAF6 | -0,090309638 | 0,939321126 | 0,66774 | 0,83516218 | TATA-box binding protein associated factor 6 |
| ENSG00000080839 | RBL1 | -0,090319006 | 0,939315026 | 0,62684 | 0,81115024 | RB transcriptional corepressor like 1 |
| ENSG00000088826 | SMOX | -0,090354358 | 0,93929201 | 0,719 | 0,8656088 | spermine oxidase |
| ENSG00000166922 | SCG5 | -0,090379338 | 0,939275746 | 0,82301 | 0,91813876 | secretogranin V |
| ENSG00000138613 | APH1B | -0,090380017 | 0,939275304 | 0,80692 | 0,91123384 | aph-1 homolog B, gamma-secretase subunit |
| ENSG00000064961 | HMG20B | -0,090381357 | 0,939274432 | 0,70025 | 0,85526044 | high mobility group 20B |
| ENSG00000065615 | CYB5R4 | -0,090425734 | 0,939245541 | 0,54854 | 0,76372599 | cytochrome b5 reductase 4 |
| ENSG00000224494 | HNRNPA3P14 | -0,090442569 | 0,939234581 | 0,91908 | 0,96485227 | heterogeneous nuclear ribonucleoprotein A3 pseudogene 14 |
| ENSG00000240891 | PLCXD2 | -0,090448204 | 0,939230912 | 0,83242 | 0,92331399 | phosphatidylinositol specific phospholipase C X domain containing 2 |
| ENSG00000183011 | NAA38 | -0,090559029 | 0,939158765 | 0,64287 | 0,82098907 | N-alpha-acetyltransferase 38, NatC auxiliary subunit |
| ENSG00000263393 | AC011825,2 | -0,090642544 | 0,9391044 | 0,88886 | 0,95091757 | novel transcript, antisense to RNF138 |
| ENSG00000198689 | SLC9A6 | -0,090728415 | 0,939048505 | 0,62015 | 0,807595 | solute carrier family 9 member A6 |
| ENSG00000129038 | LOXL1 | -0,090878708 | 0,938950685 | 0,82426 | 0,91891548 | lysyl oxidase like 1 |
| ENSG00000178980 | SELENOW | -0,090951536 | 0,938903287 | 0,73503 | 0,87477884 | selenoprotein W |
| ENSG00000224598 | RPS5P2 | -0,09109081 | 0,938812652 | 0,92355 | 0,96672273 | RPS5 pseudogene 2 |
| ENSG00000234352 | AC009264,1 | -0,091142107 | 0,938779272 | 0,84716 | 0,93053893 | novel transcript, antisense to CHRM2 |
| ENSG00000010322 | NISCH | -0,091241423 | 0,938714648 | 0,50701 | 0,735349 | nischarin |
| ENSG00000134215 | VAV3 | -0,091370421 | 0,938630717 | 0,80089 | 0,90831319 | vav guanine nucleotide exchange factor 3 |
| ENSG00000108854 | SMURF2 | -0,091462046 | 0,938571107 | 0,60065 | 0,79594503 | SMAD specific E3 ubiquitin protein ligase 2 |
| ENSG00000243147 | MRPL33 | -0,091463399 | 0,938570227 | 0,63383 | 0,81549489 | mitochondrial ribosomal protein L33 |
| ENSG00000007952 | NOX1 | -0,091520735 | 0,938532927 | 0,90967 | 0,96105018 | NADPH oxidase 1 |
| ENSG00000136193 | SCRN1 | -0,091638358 | 0,938456411 | 0,79839 | 0,90701149 | secernin 1 |
| ENSG00000178700 | DHFR2 | -0,091675853 | 0,938432021 | 0,59928 | 0,79538547 | dihydrofolate reductase 2 |
| ENSG00000287216 | AC106865,2 | -0,091821957 | 0,93833699 | 0,92932 | 0,96977202 | novel transcript, antisense to KIAA0922 |
| ENSG00000283930 | AL117339,4 | -0,091822693 | 0,938336511 | 0,67937 | 0,84248953 | novel transcript |
| ENSG00000099795 | NDUFB7 | -0,091997832 | 0,938222606 | 0,73352 | 0,87368351 | NADH:ubiquinone oxidoreductase subunit B7 |

| | | | | | | |
|---|---|---|---|---|---|---|
| ENSG00000281530 | AC004461,2 | -0,092149574 | 0,93812393 | 0,81058 | 0,91251177 | DiGeorge syndrome critical region gene 12 (non-protein coding) |
| ENSG00000213413 | PVRIG | -0,092200306 | 0,938090941 | 0,85684 | 0,93541397 | PVR related immunoglobulin domain containing |
| ENSG00000269883 | AC007375,3 | -0,092205958 | 0,938087266 | 0,91995 | 0,96530017 | novel transcript, antisense to ZDHHC22 |
| ENSG00000265478 | AC107982,3 | -0,09223113 | 0,938070899 | 0,7912 | 0,90297979 | novel transcript |
| ENSG00000196526 | AFAP1 | -0,092378308 | 0,937975205 | 0,75359 | 0,88453179 | actin filament associated protein 1 |
| ENSG00000262766 | AC135050,5 | -0,092507469 | 0,937891235 | 0,83721 | 0,92549623 | novel transcript |
| ENSG00000114626 | ABTB1 | -0,09254085 | 0,937869534 | 0,82747 | 0,92023045 | ankyrin repeat and BTB domain containing 1 |
| ENSG00000273397 | NDUFA6 | -0,092547438 | 0,937865251 | 0,90105 | 0,95787401 | NADH:ubiquinone oxidoreductase subunit A6 |
| ENSG00000277365 | NDUFA6 | -0,092547438 | 0,937865251 | 0,90105 | 0,95787401 | NADH:ubiquinone oxidoreductase subunit A6 |
| ENSG00000143590 | EFNA3 | -0,092607621 | 0,937826128 | 0,67378 | 0,83926014 | ephrin A3 |
| ENSG00000261390 | MAFTRR | -0,092625325 | 0,93781462 | 0,81995 | 0,91720659 | MAF transcriptional regulator RNA |
| ENSG00000288400 | EP400P1 | -0,092637328 | 0,937806818 | 0,86755 | 0,94147331 | EP400 pseudogene 1 |
| ENSG00000160051 | IQCC | -0,092751313 | 0,937732726 | 0,62379 | 0,80952593 | IQ motif containing C |
| ENSG00000215859 | PDZK1P1 | -0,092768711 | 0,937721418 | 0,80548 | 0,91060982 | PDZ domain containing 1 pseudogene 1 |
| ENSG00000100167 | SEPTIN3 | -0,092774707 | 0,93771752 | 0,70538 | 0,85804888 | septin 3 |
| ENSG00000254226 | LINC01933 | -0,092980488 | 0,937583777 | 0,91238 | 0,96205682 | long intergenic non-protein coding RNA 1933 |
| ENSG00000106263 | EIF3B | -0,093030751 | 0,937551112 | 0,73894 | 0,87611723 | eukaryotic translation initiation factor 3 subunit B |
| ENSG00000196781 | TLE1 | -0,093102192 | 0,937504687 | 0,68472 | 0,84571718 | TLE family member 1, transcriptional corepressor |
| ENSG00000154760 | SLFN13 | -0,093122543 | 0,937491462 | 0,85247 | 0,93294074 | schlafen family member 13 |
| ENSG00000184156 | KCNQ3 | -0,093138583 | 0,937481039 | 0,87095 | 0,94305486 | potassium voltage-gated channel subfamily Q member 3 |
| ENSG00000163521 | GLB1L | -0,093250178 | 0,937408526 | 0,67309 | 0,83863326 | galactosidase beta 1 like |
| ENSG00000198570 | RD3 | -0,093271945 | 0,937394383 | 0,78012 | 0,89822236 | retinal degeneration 3, GUCY2D regulator |
| ENSG00000146151 | HMGCLL1 | -0,09334684 | 0,937345721 | 0,82766 | 0,92040131 | 3-hydroxymethyl-3-methylglutaryl-CoA lyase like 1 |
| ENSG00000222000 | AC092675,1 | -0,093388703 | 0,937318522 | 0,89818 | 0,95621754 | novel transcript, antisense to CNGA3 |
| ENSG00000108784 | NAGLU | -0,093446365 | 0,93728106 | 0,64451 | 0,82154386 | N-acetyl-alpha-glucosaminidase |
| ENSG00000170482 | SLC23A1 | -0,093460099 | 0,937272137 | 0,85096 | 0,93218292 | solute carrier family 23 member 1 |
| ENSG00000223752 | ATAT1 | -0,093486638 | 0,937254896 | 0,68326 | 0,84495012 | alpha tubulin acetyltransferase 1 |
| ENSG00000248885 | AC118465,1 | -0,093516952 | 0,937235203 | 0,90653 | 0,96009372 | septin family pseudogene |
| ENSG00000177757 | FAM87B | -0,09359731 | 0,937183 | 0,81826 | 0,91663479 | family with sequence similarity 87 member B |
| ENSG00000128335 | APOL2 | -0,093703148 | 0,93711425 | 0,64369 | 0,82121082 | apolipoprotein L2 |
| ENSG00000281894 | ZKSCAN7 | -0,093717612 | 0,937104855 | 0,6998 | 0,85497086 | zinc finger with KRAB and SCAN domains 7 |
| ENSG00000103152 | MPG | -0,093751176 | 0,937083053 | 0,76289 | 0,889794 | N-methylpurine DNA glycosylase |

| | | | | | | |
|---|---|---|---|---|---|---|
| ENSG00000254389 | RHPN1-AS1 | -0,093904931 | 0,936983189 | 0,71942 | 0,86589115 | RHPN1 antisense RNA 1 (head to head) |
| ENSG00000174684 | B4GAT1 | -0,093913715 | 0,936977484 | 0,63392 | 0,81549489 | beta-1,4-glucuronyltransferase 1 |
| ENSG00000163877 | SNIP1 | -0,093941411 | 0,936959497 | 0,73903 | 0,87619277 | Smad nuclear interacting protein 1 |
| ENSG00000226348 | VN2R10P | -0,093979477 | 0,936934775 | 0,85111 | 0,9322574 | vomeronasal 2 receptor 10, pseudogene |
| ENSG00000177990 | DPY19L2 | -0,094043094 | 0,936893461 | 0,75319 | 0,88441977 | dpy-19 like 2 |
| ENSG00000141759 | TXNL4A | -0,094081307 | 0,936868646 | 0,65068 | 0,82499358 | thioredoxin like 4A |
| ENSG00000142675 | CNKSR1 | -0,094120648 | 0,936843098 | 0,85495 | 0,93417381 | connector enhancer of kinase suppressor of Ras 1 |
| ENSG00000275430 | MUC20 | -0,094193663 | 0,936795686 | 0,86048 | 0,93739825 | mucin 20, cell surface associated |
| ENSG00000276495 | APBA2 | -0,094233086 | 0,936770087 | 0,52139 | 0,74572351 | amyloid beta precursor protein binding family A member 2 |
| ENSG00000214439 | FAM185BP | -0,094312517 | 0,936718513 | 0,81197 | 0,9134209 | family with sequence similarity 185 member B, pseudogene |
| ENSG00000284196 | AL158064,1 | -0,094390574 | 0,936667833 | 0,69307 | 0,85087762 | novel transcript |
| ENSG00000269984 | AC078795,1 | -0,094398316 | 0,936662806 | 0,62662 | 0,81105911 | novel transcript, antisense to MYNN |
| ENSG00000170145 | SIK2 | -0,094415233 | 0,936651823 | 0,62801 | 0,81175174 | salt inducible kinase 2 |
| ENSG00000259244 | AC048382,2 | -0,094625044 | 0,936515616 | 0,88334 | 0,94837933 | novel transcript |
| ENSG00000179115 | FARSA | -0,094632651 | 0,936510678 | 0,71001 | 0,86028876 | phenylalanyl-tRNA synthetase subunit alpha |
| ENSG00000244119 | PDCL3P4 | -0,094722363 | 0,936452444 | 0,88383 | 0,94846868 | PDCL3 pseudogene 4 |
| ENSG00000248529 | AC026746,1 | -0,094738363 | 0,936442058 | 0,92666 | 0,96815248 | novel transcript |
| ENSG00000275851 | MLLT6 | -0,094927666 | 0,936319191 | 0,79035 | 0,90286944 | MLLT6, PHD finger containing |
| ENSG00000111412 | SPRING1 | -0,095112328 | 0,936199352 | 0,67667 | 0,8409179 | SREBF pathway regulator in golgi 1 |
| ENSG00000232527 | LINC02802 | -0,095156764 | 0,936170517 | 0,77017 | 0,89333117 | long intergenic non-protein coding RNA 2802 |
| ENSG00000170579 | DLGAP1 | -0,095178706 | 0,936156279 | 0,73611 | 0,87517709 | DLG associated protein 1 |
| ENSG00000251410 | AC007106,1 | -0,095216563 | 0,936131714 | 0,78715 | 0,90117606 | novel transcript |
| ENSG00000277450 | AC002094,4 | -0,095232803 | 0,936121176 | 0,87151 | 0,94322889 | novel transcript, sense intronic to transcript readthrough between TMEM199 and SARM1 |
| ENSG00000286244 | AC073046,3 | -0,095239276 | 0,936116976 | 0,88304 | 0,94823653 | novel transcript, antisense to STAMBP |
| ENSG00000227583 | RPS3AP37 | -0,095310716 | 0,936070622 | 0,86018 | 0,93714201 | RPS3A pseudogene 37 |
| ENSG00000139880 | CDH24 | -0,095431485 | 0,935992267 | 0,66959 | 0,83623395 | cadherin 24 |
| ENSG00000196967 | ZNF585A | -0,095466541 | 0,935969523 | 0,66564 | 0,83358761 | zinc finger protein 585A |
| ENSG00000184984 | CHRM5 | -0,095745162 | 0,935788781 | 0,85616 | 0,9349542 | cholinergic receptor muscarinic 5 |
| ENSG00000288044 | AL096870,12 | -0,095747223 | 0,935787444 | 0,83814 | 0,92614673 | novel transcript, antisense to RABGGTAand DHRS1 |
| ENSG00000287436 | AL096870,11 | -0,095747223 | 0,935787444 | 0,83814 | 0,92614673 | novel transcript, antisense to RABGGTAand DHRS1 |
| ENSG00000178202 | POGLUT3 | -0,095779089 | 0,935766775 | 0,73313 | 0,87350302 | protein O-glucosyltransferase 3 |

| | | | | | | |
|---|---|---|---|---|---|---|
| ENSG00000236021 | AL591043,2 | -0,095844693 | 0,935724224 | 0,87477 | 0,94461928 | novel transcript |
| ENSG00000140464 | PML | -0,095903186 | 0,935686287 | 0,72428 | 0,86840844 | PML nuclear body scaffold |
| ENSG00000179909 | ZNF154 | -0,096097248 | 0,935560432 | 0,59693 | 0,79434997 | zinc finger protein 154 |
| ENSG00000232377 | AC016910,1 | -0,096269703 | 0,935448605 | 0,90922 | 0,96104995 | novel transcript |
| ENSG00000223887 | PRR3 | -0,096271143 | 0,935447671 | 0,81184 | 0,91334888 | proline rich 3 |
| ENSG00000272701 | MESTIT1 | -0,096311318 | 0,935421622 | 0,82339 | 0,9183689 | MEST intronic transcript 1, antisense RNA |
| ENSG00000007392 | LUC7L | -0,096423213 | 0,935349074 | 0,63664 | 0,81709016 | LUC7 like |
| ENSG00000122741 | DCAF10 | -0,096508915 | 0,935293512 | 0,60015 | 0,79579686 | DDB1 and CUL4 associated factor 10 |
| ENSG00000164221 | CCDC112 | -0,096680158 | 0,935182503 | 0,7663 | 0,89138648 | coiled-coil domain containing 112 |
| ENSG00000137802 | MAPKBP1 | -0,096699882 | 0,935169717 | 0,59295 | 0,79148028 | mitogen-activated protein kinase binding protein 1 |
| ENSG00000121989 | ACVR2A | -0,096749953 | 0,935137261 | 0,75965 | 0,88798255 | activin A receptor type 2A |
| ENSG00000126882 | FAM78A | -0,096752712 | 0,935135473 | 0,82511 | 0,91900665 | family with sequence similarity 78 member A |
| ENSG00000258343 | AC090001,1 | -0,096813071 | 0,935096349 | 0,90659 | 0,96009372 | novel transcript |
| ENSG00000105825 | TFPI2 | -0,096836701 | 0,935081034 | 0,85499 | 0,93417381 | tissue factor pathway inhibitor 2 |
| ENSG00000229593 | SUCLA2P3 | -0,096863587 | 0,935063608 | 0,87493 | 0,94463878 | SUCLA2 pseudogene 3 |
| ENSG00000179532 | DNHD1 | -0,096903078 | 0,935038013 | 0,60959 | 0,80181076 | dynein heavy chain domain 1 |
| ENSG00000072518 | MARK2 | -0,097055574 | 0,934939182 | 0,62384 | 0,80952593 | microtubule affinity regulating kinase 2 |
| ENSG00000286874 | AC072054,1 | -0,097113959 | 0,934901347 | 0,92192 | 0,96616853 | novel transcript |
| ENSG00000230568 | SF3A3P1 | -0,097160985 | 0,934870873 | 0,91623 | 0,96347905 | splicing factor 3a, subunit 3 pseudogene 1 |
| ENSG00000065485 | PDIA5 | -0,097190567 | 0,934851704 | 0,77238 | 0,89477953 | protein disulfide isomerase family A member 5 |
| ENSG00000167535 | CACNB3 | -0,097219324 | 0,93483307 | 0,58209 | 0,78454692 | calcium voltage-gated channel auxiliary subunit beta 3 |
| ENSG00000188976 | NOC2L | -0,097453277 | 0,934681486 | 0,70205 | 0,85648523 | NOC2 like nucleolar associated transcriptional repressor |
| ENSG00000230399 | RBBP8P1 | -0,097466235 | 0,934673091 | 0,87351 | 0,94398015 | RBBP8 pseudogene 1 |
| ENSG00000261369 | AC007533,1 | -0,097494288 | 0,934654917 | 0,88557 | 0,94939105 | novel transcript, antisense to ITFG1 |
| ENSG00000104142 | VPS18 | -0,097650451 | 0,934553751 | 0,62489 | 0,81001052 | VPS18 core subunit of CORVET and HOPS complexes |
| ENSG00000234969 | AARSD1P1 | -0,097655651 | 0,934550383 | 0,84156 | 0,92795437 | AARSD1 pseudogene 1 |
| ENSG00000187091 | PLCD1 | -0,097708404 | 0,934516211 | 0,64164 | 0,82033064 | phospholipase C delta 1 |
| ENSG00000227122 | PRRT1 | -0,097773372 | 0,934474129 | 0,82849 | 0,92081006 | proline rich transmembrane protein 1 |
| ENSG00000225141 | PRRT1 | -0,097773372 | 0,934474129 | 0,82849 | 0,92081006 | proline rich transmembrane protein 1 |
| ENSG00000126775 | ATG14 | -0,09783597 | 0,934433583 | 0,72133 | 0,86689814 | autophagy related 14 |
| ENSG00000226469 | ADAM1B | -0,097936533 | 0,934368451 | 0,75205 | 0,88386445 | ADAM metallopeptidase domain 1B (pseudogene) |
| ENSG00000188895 | MSL1 | -0,097961353 | 0,934352376 | 0,64228 | 0,82066205 | MSL complex subunit 1 |
| ENSG00000273989 | AC022079,1 | -0,098005821 | 0,934323577 | 0,92363 | 0,96675951 | novel transcript, sense intronic to CCDC91 |

| | | | | | | |
|---|---|---|---|---|---|---|
| ENSG00000166068 | SPRED1 | -0,098058394 | 0,93428953 | 0,71327 | 0,86208742 | sprouty related EVH1 domain containing 1 |
| ENSG00000198732 | SMOC1 | -0,098119408 | 0,934250018 | 0,83373 | 0,92400213 | SPARC related modular calcium binding 1 |
| ENSG00000161526 | SAP30BP | -0,098177027 | 0,934212707 | 0,6428 | 0,82094129 | SAP30 binding protein |
| ENSG00000129911 | KLF16 | -0,098246656 | 0,934167619 | 0,80921 | 0,91206591 | Kruppel like factor 16 |
| ENSG00000286310 | AC008443,6 | -0,098363157 | 0,934092186 | 0,90544 | 0,95961188 | novel transcript |
| ENSG00000282907 | Z98883,1 | -0,098390724 | 0,934074338 | 0,90372 | 0,95873972 | novel transcript |
| ENSG00000134940 | ACRV1 | -0,098512077 | 0,933995771 | 0,79218 | 0,90351559 | acrosomal vesicle protein 1 |
| ENSG00000171766 | GATM | -0,098551871 | 0,933970009 | 0,79326 | 0,90374812 | glycine amidinotransferase |
| ENSG00000197568 | HHLA3 | -0,098591574 | 0,933944307 | 0,74262 | 0,87827554 | HERV-H LTR-associating 3 |
| ENSG00000267717 | SRSF10P1 | -0,098606977 | 0,933934335 | 0,92018 | 0,96530017 | serine and arginine rich splicing factor 10 pseudogene 1 |
| ENSG00000154721 | JAM2 | -0,098629808 | 0,933919556 | 0,76563 | 0,89093624 | junctional adhesion molecule 2 |
| ENSG00000285229 | LINCR-0001 | -0,098795629 | 0,933812218 | 0,83203 | 0,92312766 | uncharacterized LINCR-0001 |
| ENSG00000175175 | PPM1E | -0,098801953 | 0,933808125 | 0,58856 | 0,78826488 | protein phosphatase, Mg2+/Mn2+ dependent 1E |
| ENSG00000175213 | ZNF408 | -0,098840566 | 0,933783133 | 0,70275 | 0,85666784 | zinc finger protein 408 |
| ENSG00000236039 | AC019117,1 | -0,098873863 | 0,933761581 | 0,88466 | 0,94887389 | novel transcript |
| ENSG00000259943 | AL050341,2 | -0,099048281 | 0,933648699 | 0,7078 | 0,85921019 | novel transcript, antisense to ZMPSTE24 |
| ENSG00000168237 | GLYCTK | -0,099049146 | 0,933648139 | 0,58498 | 0,78606599 | glycerate kinase |
| ENSG00000008294 | SPAG9 | -0,099102874 | 0,933613369 | 0,65842 | 0,82918448 | sperm associated antigen 9 |
| ENSG00000150627 | WDR17 | -0,099229352 | 0,933531525 | 0,68903 | 0,84853604 | WD repeat domain 17 |
| ENSG00000100564 | PIGH | -0,099352461 | 0,933451868 | 0,57337 | 0,77892286 | phosphatidylinositol glycan anchor biosynthesis class H |
| ENSG00000255311 | DLG2-AS2 | -0,099395111 | 0,933424273 | 0,9032 | 0,95854863 | DLG2 antisense RNA 2 |
| ENSG00000105278 | ZFR2 | -0,099478905 | 0,933370059 | 0,85508 | 0,93419851 | zinc finger RNA binding protein 2 |
| ENSG00000114125 | RNF7 | -0,099520594 | 0,933343089 | 0,66974 | 0,83623395 | ring finger protein 7 |
| ENSG00000283549 | AC007161,3 | -0,099569523 | 0,933311435 | 0,85827 | 0,93612837 | novel transcript |
| ENSG00000268565 | AC005339,1 | -0,099606287 | 0,933287652 | 0,92742 | 0,96860856 | novel transcript, antisense to TNFAIP8L1 |
| ENSG00000277361 | OCA2 | -0,099672947 | 0,93324453 | 0,88709 | 0,95021901 | OCA2 melanosomal transmembrane protein |
| ENSG00000154898 | CCDC144CP | -0,099755139 | 0,933191364 | 0,65008 | 0,8246401 | coiled-coil domain containing 144C, pseudogene |
| ENSG00000224707 | E2F3-IT1 | -0,099813542 | 0,933153587 | 0,8665 | 0,94074101 | E2F3 intronic transcript 1 |
| ENSG00000171056 | SOX7 | -0,09982525 | 0,933146015 | 0,88823 | 0,95069428 | SRY-box transcription factor 7 |
| ENSG00000059378 | PARP12 | -0,099856203 | 0,933125994 | 0,74764 | 0,8810788 | poly(ADP-ribose) polymerase family member 12 |
| ENSG00000180139 | ACTA2-AS1 | -0,099872202 | 0,933115646 | 0,84819 | 0,93094401 | ACTA2 antisense RNA 1 |
| ENSG00000138131 | LOXL4 | -0,099888815 | 0,933104901 | 0,7328 | 0,87320251 | lysyl oxidase like 4 |
| ENSG00000248871 | TNFSF12-TNF! | -0,099970078 | 0,933052343 | 0,95656 | 0,98197334 | TNFSF12-TNFSF13 readthrough |

| | | | | | | |
|---|---|---|---|---|---|---|
| ENSG00000104969 | SGTA | -0,100044137 | 0,933004447 | 0,66685 | 0,83455372 | small glutamine rich tetratricopeptide repeat containing alpha |
| ENSG00000259612 | EEF1A1P22 | -0,100051435 | 0,932999728 | 0,8117 | 0,91334888 | eukaryotic translation elongation factor 1 alpha 1 pseudogene 22 |
| ENSG00000173480 | ZNF417 | -0,100052524 | 0,932999024 | 0,5545 | 0,76745801 | zinc finger protein 417 |
| ENSG00000162086 | ZNF75A | -0,100108433 | 0,932962867 | 0,54373 | 0,76018847 | zinc finger protein 75a |
| ENSG00000196972 | SMIM10L2B | -0,100125194 | 0,932952028 | 0,70541 | 0,85804888 | small integral membrane protein 10 like 2B |
| ENSG00000275748 | ARL17B | -0,100224093 | 0,932888075 | 0,6372 | 0,81750458 | ADP ribosylation factor like GTPase 17B |
| ENSG00000270903 | HNRNPA3P9 | -0,100251771 | 0,932870178 | 0,84855 | 0,93094401 | heterogeneous nuclear ribonucleoprotein A3 pseudogene 9 |
| ENSG00000280055 | TMEM75 | -0,10027147 | 0,932857441 | 0,78337 | 0,89918294 | transmembrane protein 75 |
| ENSG00000242971 | RN7SL233P | -0,10030781 | 0,932833943 | 0,92507 | 0,96744841 | RNA, 7SL, cytoplasmic 233, pseudogene |
| ENSG00000248423 | AC100850,1 | -0,100368461 | 0,932794727 | 0,84265 | 0,92821072 | clathrin, light chain (Lca) (CLTA) pseudogene |
| ENSG00000288063 | AC099050,1 | -0,100394759 | 0,932777724 | 0,82027 | 0,9172552 | novel transcript, antisense to RAD54L2and TEX264 |
| ENSG00000235548 | HEBP2P1 | -0,100577145 | 0,932659809 | 0,86196 | 0,93834335 | HEBP2 pseudogene 1 |
| ENSG00000173875 | ZNF791 | -0,100593315 | 0,932649356 | 0,61223 | 0,8035281 | zinc finger protein 791 |
| ENSG00000262848 | AC136624,2 | -0,100632911 | 0,932623759 | 0,87291 | 0,94378579 | novel transcript, antisense to NOMO3 |
| ENSG00000282332 | AC244489,4 | -0,100632911 | 0,932623759 | 0,87291 | 0,94378579 | novel transcript, antisense to NOMO3 |
| ENSG00000283979 | RFLNB | -0,100655921 | 0,932608885 | 0,86767 | 0,94153368 | refilin B |
| ENSG00000111642 | CHD4 | -0,100695361 | 0,932583389 | 0,58169 | 0,78453349 | chromodomain helicase DNA binding protein 4 |
| ENSG00000170835 | CEL | -0,100876578 | 0,932466255 | 0,73354 | 0,87368351 | carboxyl ester lipase |
| ENSG00000078900 | TP73 | -0,100920857 | 0,932437636 | 0,87265 | 0,94374455 | tumor protein p73 |
| ENSG00000169972 | PUSL1 | -0,10114769 | 0,932291042 | 0,7835 | 0,89918294 | pseudouridine synthase like 1 |
| ENSG00000226647 | AL365356,1 | -0,101162035 | 0,932281772 | 0,86525 | 0,94006758 | novel transcript |
| ENSG00000262877 | AC110285,2 | -0,101236697 | 0,932233526 | 0,76408 | 0,89012684 | novel transcript |
| ENSG00000177337 | DLGAP1-AS1 | -0,101313865 | 0,932183664 | 0,70045 | 0,85531965 | DLGAP1 antisense RNA 1 |
| ENSG00000253767 | PCDHGA8 | -0,101319251 | 0,932180184 | 0,78289 | 0,89917991 | protocadherin gamma subfamily A, 8 |
| ENSG00000146834 | MEPCE | -0,101427282 | 0,932110383 | 0,71269 | 0,86178349 | methylphosphate capping enzyme |
| ENSG00000006468 | ETV1 | -0,101498029 | 0,932064675 | 0,85156 | 0,9324506 | ETS variant transcription factor 1 |
| ENSG00000270157 | AC004918,3 | -0,10175883 | 0,931896198 | 0,73058 | 0,87202167 | novel transcript |
| ENSG00000006194 | ZNF263 | -0,10176597 | 0,931891586 | 0,60193 | 0,79680798 | zinc finger protein 263 |
| ENSG00000226360 | RPL10AP6 | -0,10182648 | 0,931852501 | 0,88182 | 0,94750352 | ribosomal protein L10a pseudogene 6 |
| ENSG00000147123 | NDUFB11 | -0,101862367 | 0,931829321 | 0,6569 | 0,82856771 | NADH:ubiquinone oxidoreductase subunit B11 |

| | | | | | | |
|---|---|---|---|---|---|---|
| ENSG00000104435 | STMN2 | -0,102092494 | 0,931680695 | 0,82856 | 0,92085329 | stathmin 2 |
| ENSG00000231548 | OR55B1P | -0,102184878 | 0,931621037 | 0,80576 | 0,91073695 | olfactory receptor family 55 subfamily B member 1 pseudogene |
| ENSG00000230606 | AC092683,1 | -0,102222451 | 0,931596774 | 0,6245 | 0,80980546 | novel transcript |
| ENSG00000283706 | PRSS50 | -0,102243292 | 0,931583316 | 0,90354 | 0,95873972 | serine protease 50 |
| ENSG00000149357 | LAMTOR1 | -0,102354862 | 0,931511276 | 0,54929 | 0,76407481 | late endosomal/lysosomal adaptor, MAPK and MTOR activator 1 |
| ENSG00000115486 | GGCX | -0,102443327 | 0,931454158 | 0,72026 | 0,86628014 | gamma-glutamyl carboxylase |
| ENSG00000251609 | SETP12 | -0,102568079 | 0,931373617 | 0,87086 | 0,94302816 | SET pseudogene 12 |
| ENSG00000225998 | LSM2 | -0,102628627 | 0,93133453 | 0,80414 | 0,909789 | LSM2 homolog, U6 small nuclear RNA and mRNA degradation associated |
| ENSG00000224979 | LSM2 | -0,102628627 | 0,93133453 | 0,80414 | 0,909789 | LSM2 homolog, U6 small nuclear RNA and mRNA degradation associated |
| ENSG00000231502 | LSM2 | -0,102628627 | 0,93133453 | 0,80414 | 0,909789 | LSM2 homolog, U6 small nuclear RNA and mRNA degradation associated |
| ENSG00000172850 | LSM2 | -0,102628627 | 0,93133453 | 0,80414 | 0,909789 | LSM2 homolog, U6 small nuclear RNA and mRNA degradation associated |
| ENSG00000271936 | AC012073,1 | -0,102713187 | 0,931279944 | 0,65308 | 0,8262455 | novel transcript, antisense to ADCY3 |
| ENSG00000160201 | U2AF1 | -0,102810581 | 0,931217076 | 0,80052 | 0,90812402 | U2 small nuclear RNA auxiliary factor 1 |
| ENSG00000100359 | SGSM3 | -0,10284217 | 0,931196687 | 0,63298 | 0,81526144 | small G protein signaling modulator 3 |
| ENSG00000204219 | TCEA3 | -0,102850294 | 0,931191443 | 0,80129 | 0,90860375 | transcription elongation factor A3 |
| ENSG00000282941 | TCEA3 | -0,102850294 | 0,931191443 | 0,80129 | 0,90860375 | transcription elongation factor A3 |
| ENSG00000177030 | DEAF1 | -0,102891407 | 0,931164907 | 0,52723 | 0,75029458 | DEAF1 transcription factor |
| ENSG00000265656 | AC106037,3 | -0,102970925 | 0,931113585 | 0,88806 | 0,95069428 | novel transcript, antisense to ABHD3 |
| ENSG00000265194 | AL359922,2 | -0,10300428 | 0,931092058 | 0,73653 | 0,87529332 | novel transcript, antisense MTAP |
| ENSG00000259191 | AC027237,1 | -0,103083139 | 0,931041164 | 0,82358 | 0,91852182 | fumarylacetoacetate hydrolase domain containing 1 (FAHD1) pseudogene |
| ENSG00000137812 | KNL1 | -0,103087163 | 0,931038568 | 0,5682 | 0,77590889 | kinetochore scaffold 1 |
| ENSG00000112232 | KHDRBS2 | -0,103097792 | 0,931031708 | 0,66823 | 0,83536064 | KH RNA binding domain containing, signal transduction associated 2 |
| ENSG00000230815 | AL807757,1 | -0,103212619 | 0,930957609 | 0,89187 | 0,95246557 | novel transcript |
| ENSG00000139178 | C1RL | -0,103269518 | 0,930920893 | 0,84988 | 0,93166725 | complement C1r subcomponent like |
| ENSG00000277273 | CDK7 | -0,103341976 | 0,930874139 | 0,83708 | 0,92549623 | cyclin dependent kinase 7 |

| | | | | | | |
|---|---|---|---|---|---|---|
| ENSG00000003989 | SLC7A2 | -0,103475475 | 0,930788006 | 0,84337 | 0,92850187 | solute carrier family 7 member 2 |
| ENSG00000270839 | AC091393,1 | -0,103496808 | 0,930774242 | 0,88129 | 0,94715141 | nuclear transcription factor Y, beta (NFYB) pseudogene |
| ENSG00000170632 | ARMC10 | -0,103636445 | 0,930684158 | 0,66832 | 0,83536064 | armadillo repeat containing 10 |
| ENSG00000268520 | AC008750,4 | -0,10369576 | 0,930645894 | 0,89984 | 0,95730568 | novel transcript, sense intronic to ETFB |
| ENSG00000237711 | MTCO3P11 | -0,103737717 | 0,93061883 | 0,89547 | 0,95460904 | MT-CO3 pseudogene 11 |
| ENSG00000115459 | ELMOD3 | -0,103742653 | 0,930615646 | 0,51503 | 0,74133616 | ELMO domain containing 3 |
| ENSG00000109111 | SUPT6H | -0,103747351 | 0,930612615 | 0,66908 | 0,83595057 | SPT6 homolog, histone chaperone and transcription elongation factor |
| ENSG00000119596 | YLPM1 | -0,103799656 | 0,930578876 | 0,6425 | 0,82079227 | YLP motif containing 1 |
| ENSG00000131016 | AKAP12 | -0,103835056 | 0,930556042 | 0,66409 | 0,83250729 | A-kinase anchoring protein 12 |
| ENSG00000231961 | AL035461,1 | -0,103989583 | 0,930456376 | 0,90354 | 0,95873972 | pseudogene similar to part of cytochrome c oxidase I |
| ENSG00000112425 | EPM2A | -0,103997441 | 0,930451308 | 0,62112 | 0,80787022 | EPM2A glucan phosphatase, laforin |
| ENSG00000156858 | PRR14 | -0,104047825 | 0,930418814 | 0,62869 | 0,81220195 | proline rich 14 |
| ENSG00000286424 | AL157996,2 | -0,10408201 | 0,930396768 | 0,85957 | 0,93687112 | novel transcript |
| ENSG00000223866 | AC002486,1 | -0,104118049 | 0,930373527 | 0,85972 | 0,93692606 | pseudogene similar to part of pituitary tumor-transforming 1 PTTG1 |
| ENSG00000140553 | UNC45A | -0,104239315 | 0,930295327 | 0,69162 | 0,84971913 | unc-45 myosin chaperone A |
| ENSG00000124532 | MRS2 | -0,10427419 | 0,930272839 | 0,55946 | 0,77045624 | magnesium transporter MRS2 |
| ENSG00000137494 | ANKRD42 | -0,104429474 | 0,930172714 | 0,6241 | 0,80958576 | ankyrin repeat domain 42 |
| ENSG00000206315 | PBX2 | -0,104521517 | 0,930113372 | 0,87536 | 0,9448685 | PBX homeobox 2 |
| ENSG00000130511 | SSBP4 | -0,104555774 | 0,930091287 | 0,7037 | 0,85720409 | single stranded DNA binding protein 4 |
| ENSG00000131370 | SH3BP5 | -0,104594639 | 0,930066231 | 0,76202 | 0,88913841 | SH3 domain binding protein 5 |
| ENSG00000164077 | MON1A | -0,10465123 | 0,930029749 | 0,63536 | 0,81655453 | MON1 homolog A, secretory trafficking associated |
| ENSG00000165861 | ZFYVE1 | -0,104663236 | 0,930022009 | 0,60262 | 0,79710677 | zinc finger FYVE-type containing 1 |
| ENSG00000226801 | OSTCP8 | -0,104754365 | 0,929963266 | 0,90582 | 0,95975315 | oligosaccharyltransferase complex subunit pseudogene 8 |
| ENSG00000141736 | ERBB2 | -0,104755811 | 0,929962334 | 0,68624 | 0,84663586 | erb-b2 receptor tyrosine kinase 2 |
| ENSG00000105131 | EPHX3 | -0,104801518 | 0,929932872 | 0,8038 | 0,90974015 | epoxide hydrolase 3 |
| ENSG00000286409 | AL391839,3 | -0,104806617 | 0,929929585 | 0,91392 | 0,96272082 | novel transcript |
| ENSG00000138172 | CALHM2 | -0,104944154 | 0,929840935 | 0,75781 | 0,88699366 | calcium homeostasis modulator family member 2 |
| ENSG00000262814 | MRPL12 | -0,10500176 | 0,929803809 | 0,63188 | 0,81440143 | mitochondrial ribosomal protein L12 |
| ENSG00000040199 | PHLPP2 | -0,105121232 | 0,929726813 | 0,52754 | 0,75037219 | PH domain and leucine rich repeat protein phosphatase 2 |
| ENSG00000126106 | TMEM53 | -0,105131988 | 0,929719882 | 0,60948 | 0,80173647 | transmembrane protein 53 |

| | | | | | | |
|---|---|---|---|---|---|---|
| ENSG00000159713 | TPPP3 | -0,105214456 | 0,929666738 | 0,82075 | 0,9173395 | tubulin polymerization promoting protein family member 3 |
| ENSG00000287232 | AC079848,2 | -0,105226347 | 0,929659076 | 0,67477 | 0,83976962 | novel transcript |
| ENSG00000281103 | TRG-AS1 | -0,105235552 | 0,929653144 | 0,83917 | 0,92642589 | T cell receptor gamma locus antisense RNA 1 |
| ENSG00000269958 | AL049840,5 | -0,105398459 | 0,929548175 | 0,74966 | 0,8822333 | novel transcript, sense intronic to KLC1 |
| ENSG00000104490 | NCALD | -0,105464421 | 0,929505675 | 0,70739 | 0,85910429 | neurocalcin delta |
| ENSG00000280682 | HYOU1 | -0,105596096 | 0,929420843 | 0,72454 | 0,86851584 | hypoxia up-regulated 1 |
| ENSG00000130540 | SULT4A1 | -0,105631826 | 0,929397825 | 0,70923 | 0,85981894 | sulfotransferase family 4A member 1 |
| ENSG00000236194 | AC099811,1 | -0,105663543 | 0,929377393 | 0,81363 | 0,9144566 | novel transcript, sense intronic to STAT5B |
| ENSG00000257467 | PPFIA2-AS1 | -0,105773876 | 0,92930632 | 0,86581 | 0,94042433 | PPFIA2 antisense RNA 1 |
| ENSG00000196767 | POU3F4 | -0,10592004 | 0,929212173 | 0,87504 | 0,94463878 | POU class 3 homeobox 4 |
| ENSG00000083838 | ZNF446 | -0,105979214 | 0,929174061 | 0,62638 | 0,81088056 | zinc finger protein 446 |
| ENSG00000169727 | GPS1 | -0,106000195 | 0,929160548 | 0,6248 | 0,8100082 | G protein pathway suppressor 1 |
| ENSG00000161082 | CELF5 | -0,106052885 | 0,929126615 | 0,73012 | 0,87180309 | CUGBP Elav-like family member 5 |
| ENSG00000236811 | GAPDHP2 | -0,106066529 | 0,929117828 | 0,87125 | 0,94321633 | glyceraldehyde-3-phosphate dehydrogenase pseudogene 2 |
| ENSG00000074071 | MRPS34 | -0,106109333 | 0,929090261 | 0,6679 | 0,83525939 | mitochondrial ribosomal protein S34 |
| ENSG00000265688 | MAFG-DT | -0,106121144 | 0,929082655 | 0,66389 | 0,83247324 | MAFG divergent transcript |
| ENSG00000237493 | AC034102,1 | -0,106132421 | 0,929075393 | 0,85981 | 0,93699188 | RAB13 member RAS oncogene family pseudogene |
| ENSG00000062822 | POLD1 | -0,106187742 | 0,929039768 | 0,67407 | 0,83946548 | DNA polymerase delta 1, catalytic subunit |
| ENSG00000149428 | HYOU1 | -0,106251233 | 0,928998883 | 0,72362 | 0,86800739 | hypoxia up-regulated 1 |
| ENSG00000169718 | DUS1L | -0,106511967 | 0,928831003 | 0,61467 | 0,80393884 | dihydrouridine synthase 1 like |
| ENSG00000235065 | RPL24P2 | -0,106540814 | 0,928812431 | 0,90374 | 0,95873972 | RPL24 pseudogene 2 |
| ENSG00000235772 | AP001625,2 | -0,106717414 | 0,928698742 | 0,88016 | 0,94667622 | novel transcript |
| ENSG00000125848 | FLRT3 | -0,106833732 | 0,928623869 | 0,80864 | 0,91193418 | fibronectin leucine rich transmembrane protein 3 |
| ENSG00000247240 | UBL7-AS1 | -0,106850534 | 0,928613054 | 0,57197 | 0,77838663 | UBL7 antisense RNA 1 (head to head) |
| ENSG00000160953 | PWWP3A | -0,106888558 | 0,928588579 | 0,54135 | 0,75881984 | PWWP domain containing 3A, DNA repair factor |
| ENSG00000150630 | VEGFC | -0,106905873 | 0,928577434 | 0,7635 | 0,88982985 | vascular endothelial growth factor C |
| ENSG00000262402 | MCUR1P1 | -0,107032423 | 0,928495985 | 0,82659 | 0,91973568 | MCUR1 pseudogene 1 |
| ENSG00000154832 | CXXC1 | -0,107123309 | 0,928437494 | 0,61273 | 0,8035281 | CXXC finger protein 1 |
| ENSG00000096746 | HNRNPH3 | -0,107123466 | 0,928437393 | 0,64328 | 0,82098995 | heterogeneous nuclear ribonucleoprotein H3 |
| ENSG00000286694 | AL161935,3 | -0,107191198 | 0,928393806 | 0,81727 | 0,91606335 | novel transcript |
| ENSG00000081307 | UBA5 | -0,107227931 | 0,928370168 | 0,49286 | 0,72601934 | ubiquitin like modifier activating enzyme 5 |

| | | | | | | |
|---|---|---|---|---|---|---|
| ENSG00000285156 | GS1-24F4,2 | -0,107259602 | 0,928349788 | 0,79566 | 0,90544794 | uncharacterized LOC100652791 |
| ENSG00000168016 | TRANK1 | -0,107312578 | 0,928315699 | 0,56564 | 0,77390317 | tetratricopeptide repeat and ankyrin repeat containing 1 |
| ENSG00000269924 | AC024451,4 | -0,107382153 | 0,928270932 | 0,82863 | 0,92089347 | novel transcript, sense intronic to KIAA0146 |
| ENSG00000101152 | DNAJC5 | -0,107641307 | 0,9281042 | 0,64534 | 0,82203909 | DnaJ heat shock protein family (Hsp40) member C5 |
| ENSG00000258701 | LINC00638 | -0,107724987 | 0,928050369 | 0,81054 | 0,91251177 | long intergenic non-protein coding RNA 638 |
| ENSG00000162378 | ZYG11B | -0,107746604 | 0,928036463 | 0,49429 | 0,72675894 | zyg-11 family member B, cell cycle regulator |
| ENSG00000172794 | RAB37 | -0,107753142 | 0,928032258 | 0,89984 | 0,95730568 | RAB37, member RAS oncogene family |
| ENSG00000285218 | AC026316,4 | -0,107754225 | 0,928031561 | 0,95367 | 0,9805199 | novel protein |
| ENSG00000260689 | HNRNPA3P11 | -0,107974358 | 0,927889969 | 0,82114 | 0,91738668 | heterogeneous nuclear ribonucleoprotein A3 pseudogene 11 |
| ENSG00000213089 | PDCL3P5 | -0,107995596 | 0,927876309 | 0,72326 | 0,86784071 | PDCL3 pseudogene 5 |
| ENSG00000254884 | PRR13P2 | -0,108007845 | 0,927868431 | 0,89002 | 0,95158772 | proline rich 13 pseudogene 2 |
| ENSG00000236149 | ABCF1 | -0,108088876 | 0,927816318 | 0,69509 | 0,85206252 | ATP binding cassette subfamily F member 1 |
| ENSG00000088808 | PPP1R13B | -0,108123054 | 0,927794337 | 0,68491 | 0,84583571 | protein phosphatase 1 regulatory subunit 13B |
| ENSG00000168959 | GRM5 | -0,108197038 | 0,92774676 | 0,80224 | 0,90899209 | glutamate metabotropic receptor 5 |
| ENSG00000108262 | GIT1 | -0,108223896 | 0,927729489 | 0,55476 | 0,76762759 | GIT ArfGAP 1 |
| ENSG00000180998 | GPR137C | -0,108230196 | 0,927725437 | 0,58049 | 0,78359603 | G protein-coupled receptor 137C |
| ENSG00000271869 | AC026979,2 | -0,108230261 | 0,927725396 | 0,82124 | 0,91738668 | novel transcript |
| ENSG00000163703 | CRELD1 | -0,108259993 | 0,927706276 | 0,58187 | 0,78454692 | cysteine rich with EGF like domains 1 |
| ENSG00000162337 | LRP5 | -0,10827762 | 0,927694942 | 0,47555 | 0,71413261 | LDL receptor related protein 5 |
| ENSG00000004534 | RBM6 | -0,108299647 | 0,927680777 | 0,54697 | 0,76255801 | RNA binding motif protein 6 |
| ENSG00000176788 | BASP1 | -0,108320587 | 0,927667313 | 0,67693 | 0,84109542 | brain abundant membrane attached signal protein 1 |
| ENSG00000112773 | TENT5A | -0,108418406 | 0,927604416 | 0,84692 | 0,93041089 | terminal nucleotidyltransferase 5A |
| ENSG00000177370 | TIMM22 | -0,108475256 | 0,927567865 | 0,73007 | 0,87180309 | translocase of inner mitochondrial membrane 22 |
| ENSG00000288442 | AC256231,3 | -0,108502515 | 0,927550339 | 0,90525 | 0,95952631 | |
| ENSG00000258366 | RTEL1 | -0,108635384 | 0,927464917 | 0,64424 | 0,82151477 | regulator of telomere elongation helicase 1 |
| ENSG00000232656 | IDI2-AS1 | -0,108658858 | 0,927449827 | 0,70351 | 0,857007 | IDI2 antisense RNA 1 |
| ENSG00000254413 | CHKB-CPT1B | -0,108672859 | 0,927440826 | 0,65288 | 0,8261381 | CHKB-CPT1B readthrough (NMD candidate) |
| ENSG00000287104 | AC097382,3 | -0,108675523 | 0,927439113 | 0,77329 | 0,89536254 | novel transcript, antisense to TBC1D14 |
| ENSG00000267673 | FDX2 | -0,108691762 | 0,927428674 | 0,64946 | 0,82442734 | ferredoxin 2 |
| ENSG00000207274 | SNORA70I | -0,108701321 | 0,927422529 | 0,84874 | 0,93107027 | small nucleolar RNA, H/ACA box 70I |
| ENSG00000139287 | TPH2 | -0,108760755 | 0,927384324 | 0,76187 | 0,88909725 | tryptophan hydroxylase 2 |
| ENSG00000286221 | AC009070,1 | -0,108835188 | 0,927336478 | 0,91926 | 0,96489517 | novel protein |

| | | | | | | |
|---|---|---|---|---|---|---|
| ENSG00000115604 | IL18R1 | -0,108864517 | 0,927317626 | 0,85693 | 0,93541811 | interleukin 18 receptor 1 |
| ENSG00000256806 | C17orf100 | -0,108995392 | 0,927233508 | 0,66816 | 0,83536064 | chromosome 17 open reading frame 100 |
| ENSG00000110074 | FOXRED1 | -0,109057631 | 0,927193507 | 0,52831 | 0,7507973 | FAD dependent oxidoreductase domain containing 1 |
| ENSG00000229018 | PMS2P7 | -0,109199154 | 0,927102557 | 0,77315 | 0,89523792 | PMS1 homolog 2, mismatch repair system component pseudogene 7 |
| ENSG00000137831 | UACA | -0,10933442 | 0,927015637 | 0,74639 | 0,88045757 | uveal autoantigen with coiled-coil domains and ankyrin repeats |
| ENSG00000285791 | AC009879,3 | -0,109357178 | 0,927001014 | 0,93351 | 0,97162107 | novel transcript |
| ENSG00000227543 | SPAG5-AS1 | -0,109463008 | 0,926933016 | 0,70744 | 0,85910429 | SPAG5 antisense RNA 1 |
| ENSG00000286813 | AC015871,6 | -0,109473907 | 0,926926013 | 0,76375 | 0,88986791 | novel transcript, sense intronic to ST20-MTHFSand MTHFS |
| ENSG00000106268 | NUDT1 | -0,109562003 | 0,926869413 | 0,60608 | 0,79925497 | nudix hydrolase 1 |
| ENSG00000100997 | ABHD12 | -0,109637002 | 0,926821231 | 0,54673 | 0,76237709 | abhydrolase domain containing 12, lysophospholipase |
| ENSG00000097096 | SYDE2 | -0,109736255 | 0,926757471 | 0,7432 | 0,87840582 | synapse defective Rho GTPase homolog 2 |
| ENSG00000154997 | SEPTIN14 | -0,109795273 | 0,926719559 | 0,76876 | 0,89277929 | septin 14 |
| ENSG00000288060 | AL355338,1 | -0,109866443 | 0,926673845 | 0,80302 | 0,90922086 | novel transcript |
| ENSG00000287720 | AC023480,1 | -0,110074976 | 0,926539909 | 0,78402 | 0,89941845 | novel transcript, antisense to SUMF1 |
| ENSG00000099308 | MAST3 | -0,110139406 | 0,926498531 | 0,55948 | 0,77045624 | microtubule associated serine/threonine kinase 3 |
| ENSG00000162222 | TTC9C | -0,110153603 | 0,926489414 | 0,6931 | 0,85087762 | tetratricopeptide repeat domain 9C |
| ENSG00000232119 | MCTS1 | -0,110379489 | 0,926344363 | 0,70381 | 0,85724036 | MCTS1 re-initiation and release factor |
| ENSG00000277969 | AC006449,6 | -0,110485967 | 0,926275996 | 0,83908 | 0,92640383 | novel transcript |
| ENSG00000282494 | AC006449,9 | -0,110485967 | 0,926275996 | 0,83908 | 0,92640383 | novel transcript |
| ENSG00000183963 | SMTN | -0,110486384 | 0,926275729 | 0,76865 | 0,89277929 | smoothelin |
| ENSG00000239329 | HLA-DMB | -0,110496815 | 0,926269032 | 0,90679 | 0,96017342 | major histocompatibility complex, class II, DM beta |
| ENSG00000182389 | CACNB4 | -0,110533343 | 0,926245579 | 0,59327 | 0,79162636 | calcium voltage-gated channel auxiliary subunit beta 4 |
| ENSG00000239519 | CADM2-AS1 | -0,11054691 | 0,926236869 | 0,82504 | 0,91900665 | CADM2 antisense RNA 1 |
| ENSG00000090238 | YPEL3 | -0,110682962 | 0,926149525 | 0,73793 | 0,87548397 | yippee like 3 |
| ENSG00000260077 | AC104794,3 | -0,110694942 | 0,926141834 | 0,70398 | 0,85725494 | novel transcript |
| ENSG00000251661 | AC136475,1 | -0,110794352 | 0,926078021 | 0,72429 | 0,86840844 | novel transcript, antisense to IFITM3 |
| ENSG00000138835 | RGS3 | -0,110890624 | 0,926016225 | 0,65974 | 0,83018671 | regulator of G protein signaling 3 |
| ENSG00000224552 | ABHD16A | -0,11093604 | 0,925987074 | 0,83222 | 0,92325856 | abhydrolase domain containing 16A, phospholipase |
| ENSG00000287300 | AL080275,1 | -0,110951356 | 0,925977244 | 0,85873 | 0,93635424 | novel transcript |
| ENSG00000255330 | AL096711,2 | -0,111074527 | 0,925898191 | 0,66431 | 0,83267806 | novel protein |
| ENSG00000168394 | TAP1 | -0,111146301 | 0,925852129 | 0,90618 | 0,95989899 | transporter 1, ATP binding cassette subfamily B member |

| | | | | | | |
|---|---|---|---|---|---|---|
| ENSG00000173992 | CCS | -0,111190407 | 0,925823824 | 0,65356 | 0,82638628 | copper chaperone for superoxide dismutase |
| ENSG00000237357 | BX088651,4 | -0,1112442 | 0,925789304 | 0,74907 | 0,88190158 | novel transcript |
| ENSG00000073803 | MAP3K13 | -0,111330379 | 0,925734004 | 0,77129 | 0,8940638 | mitogen-activated protein kinase kinase kinase 13 |
| ENSG00000100815 | TRIP11 | -0,111369362 | 0,92570899 | 0,52008 | 0,74456236 | thyroid hormone receptor interactor 11 |
| ENSG00000148824 | MTG1 | -0,111451652 | 0,92565619 | 0,62133 | 0,80787022 | mitochondrial ribosome associated GTPase 1 |
| ENSG00000254757 | AC127526,1 | -0,111838985 | 0,925407704 | 0,87479 | 0,94461928 | novel transcript |
| ENSG00000285447 | ZNF883 | -0,111898343 | 0,92536963 | 0,97446 | 0,98933402 | zinc finger protein 883 |
| ENSG00000160539 | PLPP7 | -0,111980313 | 0,925317055 | 0,72618 | 0,86959327 | phospholipid phosphatase 7 (inactive) |
| ENSG00000266466 | AC233702,7 | -0,111981785 | 0,92531611 | 0,87656 | 0,94547358 | novel transcript, antisense to C17orf51 |
| ENSG00000259584 | AC021755,2 | -0,11204997 | 0,925272379 | 0,89005 | 0,95158772 | novel transcript |
| ENSG00000105953 | OGDH | -0,112094451 | 0,925243852 | 0,59045 | 0,78974912 | oxoglutarate dehydrogenase |
| ENSG00000179242 | CDH4 | -0,112125883 | 0,925223693 | 0,78178 | 0,89877377 | cadherin 4 |
| ENSG00000283692 | Z96074,1 | -0,112195598 | 0,925178986 | 0,77455 | 0,89586241 | novel transcript |
| ENSG00000248458 | AL139147,1 | -0,112354509 | 0,925077084 | 0,73316 | 0,87350302 | novel transcript, antisense to SGIP1 |
| ENSG00000159692 | CTBP1 | -0,112380368 | 0,925060502 | 0,45781 | 0,70096886 | C-terminal binding protein 1 |
| ENSG00000100239 | PPP6R2 | -0,112452728 | 0,925014106 | 0,5422 | 0,75894841 | protein phosphatase 6 regulatory subunit 2 |
| ENSG00000156162 | DPY19L4 | -0,112458913 | 0,925010141 | 0,4655 | 0,70591843 | dpy-19 like 4 |
| ENSG00000164649 | CDCA7L | -0,112690588 | 0,92486161 | 0,70821 | 0,85927396 | cell division cycle associated 7 like |
| ENSG00000137133 | HINT2 | -0,112702474 | 0,92485399 | 0,60807 | 0,8008277 | histidine triad nucleotide binding protein 2 |
| ENSG00000280893 | AC009133,6 | -0,112871031 | 0,924745941 | 0,7101 | 0,86036543 | novel transcript |
| ENSG00000109466 | KLHL2 | -0,112888106 | 0,924734997 | 0,61037 | 0,80221019 | kelch like family member 2 |
| ENSG00000105323 | HNRNPUL1 | -0,112890776 | 0,924733286 | 0,65418 | 0,82689051 | heterogeneous nuclear ribonucleoprotein U like 1 |
| ENSG00000250496 | ABT1P1 | -0,113068032 | 0,924619676 | 0,85395 | 0,93380868 | activator of basal transcription 1 pseudogene 1 |
| ENSG00000236545 | AP001619,2 | -0,113089949 | 0,924605629 | 0,86937 | 0,94257563 | novel transcript |
| ENSG00000139908 | TSSK4 | -0,113124002 | 0,924583805 | 0,72725 | 0,87016629 | testis specific serine kinase 4 |
| ENSG00000285140 | TSSK4 | -0,113124002 | 0,924583805 | 0,72725 | 0,87016629 | testis specific serine kinase 4 |
| ENSG00000065802 | ASB1 | -0,113357723 | 0,924434032 | 0,6042 | 0,79780609 | ankyrin repeat and SOCS box containing 1 |
| ENSG00000198837 | DENND4B | -0,113370802 | 0,924425652 | 0,60358 | 0,79765228 | DENN domain containing 4B |
| ENSG00000089685 | BIRC5 | -0,11338808 | 0,92441458 | 0,5593 | 0,77045525 | baculoviral IAP repeat containing 5 |
| ENSG00000177725 | AC105206,1 | -0,113433912 | 0,924385214 | 0,78196 | 0,8987738 | novel transcript, sense overlapping BMP1 |
| ENSG00000284990 | USP17L2 | -0,113472031 | 0,92436079 | 0,80942 | 0,91213228 | ubiquitin specific peptidase 17 like family member 2 |
| ENSG00000273451 | AL031666,3 | -0,113536938 | 0,924319204 | 0,73318 | 0,87350302 | novel transcript, sense intronic to ZMYND8 |
| ENSG00000263916 | AC100778,2 | -0,113885096 | 0,92409617 | 0,87016 | 0,94277661 | novel transcript, sense intronic to C18orf32 |

| | | | | | | |
|---|---|---|---|---|---|---|
| ENSG00000133884 | DPF2 | -0,113994311 | 0,924026216 | 0,64939 | 0,82442734 | double PHD fingers 2 |
| ENSG00000248752 | AC116362,1 | -0,114053057 | 0,923988591 | 0,79937 | 0,90744841 | novel transcript |
| ENSG00000288637 | AC013477,1 | -0,114072911 | 0,923975875 | 0,83196 | 0,92312766 | novel protein |
| ENSG00000260278 | AC098818,2 | -0,114343068 | 0,923802869 | 0,87331 | 0,94386143 | novel transcript, antisense to BMP2K |
| ENSG00000248235 | AC037459,1 | -0,114409799 | 0,92376014 | 0,83584 | 0,92493625 | novel protein |
| ENSG00000256060 | TRAPPC2B | -0,114535394 | 0,923679725 | 0,62782 | 0,81175174 | trafficking protein particle complex 2B |
| ENSG00000272610 | MAGI1-IT1 | -0,114575597 | 0,923653986 | 0,79193 | 0,90336706 | MAGI1 intronic transcript 1 |
| ENSG00000105968 | H2AZ2 | -0,11458051 | 0,92365084 | 0,40918 | 0,66785537 | H2A,Z variant histone 2 |
| ENSG00000163162 | RNF149 | -0,114611359 | 0,92363109 | 0,59597 | 0,79361445 | ring finger protein 149 |
| ENSG00000258813 | AL442663,3 | -0,114688228 | 0,923581879 | 0,72298 | 0,86768626 | novel transcript, antisense to Rbm25 |
| ENSG00000219755 | AL137784,1 | -0,114737466 | 0,923550358 | 0,91253 | 0,96205682 | 40S ribososmal protein 3 (RPS3) pseudogene |
| ENSG00000126803 | HSPA2 | -0,11491519 | 0,923436594 | 0,81031 | 0,91251177 | heat shock protein family A (Hsp70) member 2 |
| ENSG00000231177 | LINC00852 | -0,115169142 | 0,923274059 | 0,6969 | 0,85317851 | long intergenic non-protein coding RNA 852 |
| ENSG00000090097 | PCBP4 | -0,115224748 | 0,923238474 | 0,6445 | 0,82154386 | poly(rC) binding protein 4 |
| ENSG00000213514 | AL731556,1 | -0,115296184 | 0,923192761 | 0,83616 | 0,92493625 | novel zinc finger pseudogene (KIAA1629) |
| ENSG00000224032 | EPB41L4A-AS1 | -0,115545723 | 0,923033092 | 0,54331 | 0,75999594 | EPB41L4A antisense RNA 1 |
| ENSG00000013561 | RNF14 | -0,115628183 | 0,922980336 | 0,56109 | 0,77125979 | ring finger protein 14 |
| ENSG00000239835 | AC092910,1 | -0,115701142 | 0,922933661 | 0,80781 | 0,91168773 | presenilin associated, rhomboid-like (PARL) pseudogene |
| ENSG00000206476 | VARS2 | -0,115706415 | 0,922930288 | 0,612 | 0,80341001 | valyl-tRNA synthetase 2, mitochondrial |
| ENSG00000263590 | LINC02591 | -0,115765424 | 0,922892539 | 0,93631 | 0,97254199 | long intergenic non-protein coding RNA 2591 |
| ENSG00000187229 | AC100800,1 | -0,115854302 | 0,922835685 | 0,7906 | 0,90286944 | novel transcript, antisense to PSD3 |
| ENSG00000197345 | MRPL21 | -0,115952336 | 0,922772979 | 0,60922 | 0,80163067 | mitochondrial ribosomal protein L21 |
| ENSG00000215256 | DHRS4-AS1 | -0,116007891 | 0,922737446 | 0,58436 | 0,78572744 | DHRS4 antisense RNA 1 |
| ENSG00000153774 | CFDP1 | -0,116072428 | 0,922696169 | 0,59497 | 0,79311029 | craniofacial development protein 1 |
| ENSG00000287532 | AL353689,3 | -0,116117462 | 0,922667367 | 0,8624 | 0,93853472 | novel transcript, antisense to CDC42BPA |
| ENSG00000086544 | ITPKC | -0,116129543 | 0,922659641 | 0,56623 | 0,77421829 | inositol-trisphosphate 3-kinase C |
| ENSG00000140350 | ANP32A | -0,116187426 | 0,922622624 | 0,67553 | 0,8403281 | acidic nuclear phosphoprotein 32 family member A |
| ENSG00000213137 | ARF1P2 | -0,116274546 | 0,922566911 | 0,8582 | 0,93612837 | ADP ribosylation factor 1 pseudogene 2 |
| ENSG00000172320 | OR5A1 | -0,116282377 | 0,922561903 | 0,70374 | 0,85721029 | olfactory receptor family 5 subfamily A member 1 |
| ENSG00000224490 | TTC21B-AS1 | -0,116283788 | 0,922561 | 0,73271 | 0,87318352 | TTC21B antisense RNA 1 |
| ENSG00000141556 | TBCD | -0,116313055 | 0,922542286 | 0,53719 | 0,75583332 | tubulin folding cofactor D |
| ENSG00000288174 | SLC2A4 | -0,116333215 | 0,922529394 | 0,87745 | 0,94566578 | solute carrier family 2 member 4 |
| ENSG00000156049 | GNA14 | -0,116363172 | 0,922510238 | 0,84802 | 0,9309317 | G protein subunit alpha 14 |

| | | | | | | |
|---|---|---|---|---|---|---|
| ENSG00000127804 | METTL16 | -0,116385884 | 0,922495716 | 0,60908 | 0,80161331 | methyltransferase like 16 |
| ENSG00000254489 | MPPED2-AS1 | -0,116421026 | 0,922473245 | 0,87084 | 0,94302816 | MPPED2 antisense RNA 1 |
| ENSG00000238196 | NOTCH4 | -0,116537146 | 0,922399 | 0,79251 | 0,90361513 | notch receptor 4 |
| ENSG00000187514 | PTMA | -0,116594246 | 0,922362494 | 0,60984 | 0,80199287 | prothymosin alpha |
| ENSG00000253591 | KCNIP1-AS1 | -0,116599842 | 0,922358915 | 0,89036 | 0,95170091 | KCNIP1 antisense RNA 1 |
| ENSG00000137449 | CPEB2 | -0,116625381 | 0,922342588 | 0,78821 | 0,90156082 | cytoplasmic polyadenylation element binding protein 2 |
| ENSG00000106701 | FSD1L | -0,116904415 | 0,922164213 | 0,78991 | 0,90272027 | fibronectin type III and SPRY domain containing 1 like |
| ENSG00000232046 | LINC01798 | -0,116985853 | 0,92211216 | 0,87013 | 0,94277661 | long intergenic non-protein coding RNA 1798 |
| ENSG00000224109 | CENPVL3 | -0,117153958 | 0,92200472 | 0,58172 | 0,78454045 | centromere protein V like 3 |
| ENSG00000254907 | AC087276,2 | -0,117216787 | 0,921964568 | 0,85063 | 0,93206349 | novel transcript, antisense to API5 and TTC17 |
| ENSG00000048540 | LMO3 | -0,117268441 | 0,921931559 | 0,77112 | 0,89404837 | LIM domain only 3 |
| ENSG00000257752 | AC091516,1 | -0,117564647 | 0,921742293 | 0,83404 | 0,92405734 | transmembrane protein 38A (TMEM38A) |
| ENSG00000278206 | AL031320,2 | -0,117630976 | 0,921699916 | 0,67816 | 0,84159488 | novel transcript |
| ENSG00000224289 | IFIT6P | -0,117880121 | 0,921540757 | 0,92996 | 0,9699953 | interferon induced protein with tetratricopeptide repeats 6, pseudogene |
| ENSG00000120129 | DUSP1 | -0,117986589 | 0,921472752 | 0,64308 | 0,82098995 | dual specificity phosphatase 1 |
| ENSG00000138670 | RASGEF1B | -0,117988067 | 0,921471808 | 0,73191 | 0,87276602 | RasGEF domain family member 1B |
| ENSG00000253978 | CTB-178M22, | -0,118018733 | 0,921452221 | 0,8701 | 0,94277661 | uncharacterized LOC101927862 |
| ENSG00000155367 | PPM1J | -0,118070328 | 0,921419268 | 0,77879 | 0,89768807 | protein phosphatase, Mg2+/Mn2+ dependent 1J |
| ENSG00000099904 | ZDHHC8 | -0,118144346 | 0,921371996 | 0,55182 | 0,76553885 | zinc finger DHHC-type palmitoyltransferase 8 |
| ENSG00000141699 | RETREG3 | -0,118270962 | 0,921291136 | 0,55355 | 0,76655124 | reticulophagy regulator family member 3 |
| ENSG00000003096 | KLHL13 | -0,118331437 | 0,921252519 | 0,69449 | 0,85178253 | kelch like family member 13 |
| ENSG00000143947 | RPS27A | -0,118363198 | 0,921232237 | 0,56402 | 0,77260898 | ribosomal protein S27a |
| ENSG00000274695 | AC108704,1 | -0,118455688 | 0,92117318 | 0,87311 | 0,94385071 | novel transcript |
| ENSG00000235236 | AC137630,2 | -0,118514842 | 0,92113541 | 0,77129 | 0,8940638 | novel transcript |
| ENSG00000261360 | AC010491,1 | -0,11859679 | 0,921083089 | 0,76277 | 0,88972756 | novel transcript, antisense to FAM105B |
| ENSG00000203872 | C6orf163 | -0,11870301 | 0,921015276 | 0,6307 | 0,81354995 | chromosome 6 open reading frame 163 |
| ENSG00000176101 | SSNA1 | -0,11877801 | 0,920967397 | 0,58517 | 0,78616171 | SS nuclear autoantigen 1 |
| ENSG00000224843 | LINC00240 | -0,118779772 | 0,920966273 | 0,76317 | 0,88980026 | long intergenic non-protein coding RNA 240 |
| ENSG00000131626 | PPFIA1 | -0,11885992 | 0,92091511 | 0,58879 | 0,788309 | PTPRF interacting protein alpha 1 |
| ENSG00000284648 | AC097493,3 | -0,118933229 | 0,920868316 | 0,90765 | 0,96044232 | RNA binding protein, fox-1 homolog (C, elegans) 1 (RBFOX1) pseudogene |
| ENSG00000287032 | AC018647,3 | -0,118988882 | 0,920832794 | 0,75413 | 0,88474264 | novel transcript |

| | | | | | | |
|---|---|---|---|---|---|---|
| ENSG00000229474 | PATL2 | -0,119060298 | 0,920787212 | 0,87267 | 0,94374455 | PAT1 homolog 2 |
| ENSG00000172183 | ISG20 | -0,11906872 | 0,920781836 | 0,78584 | 0,90056507 | interferon stimulated exonuclease gene 20 |
| ENSG00000230002 | ALMS1-IT1 | -0,119102478 | 0,920760291 | 0,72744 | 0,87021418 | ALMS1 intronic transcript 1 |
| ENSG00000198513 | ATL1 | -0,119171241 | 0,920716406 | 0,75631 | 0,88597527 | atlastin GTPase 1 |
| ENSG00000177733 | HNRNPA0 | -0,119184209 | 0,92070813 | 0,53906 | 0,75752257 | heterogeneous nuclear ribonucleoprotein A0 |
| ENSG00000285793 | ANAPC1P2 | -0,119287856 | 0,920641986 | 0,65452 | 0,82716884 | ANAPC1 pseudogene 2 |
| ENSG00000115649 | CNPPD1 | -0,119373076 | 0,920587606 | 0,47245 | 0,71184073 | cyclin Pas1/PHO80 domain containing 1 |
| ENSG00000231473 | RB1-DT | -0,11941243 | 0,920562494 | 0,71811 | 0,86499068 | RB1 divergent transcript |
| ENSG00000268144 | NIFKP6 | -0,119627545 | 0,920425243 | 0,90838 | 0,96080514 | nucleolar protein interacting with the FHA domain of MKI67 pseudogene 6 |
| ENSG00000084764 | MAPRE3 | -0,119847184 | 0,920285126 | 0,5907 | 0,78992663 | microtubule associated protein RP/EB family member 3 |
| ENSG00000251369 | ZNF550 | -0,119912632 | 0,920243378 | 0,63394 | 0,81549489 | zinc finger protein 550 |
| ENSG00000271151 | AC016737,1 | -0,120026949 | 0,920170462 | 0,78311 | 0,89918294 | novel transcript |
| ENSG00000102890 | ELMO3 | -0,120049848 | 0,920155857 | 0,75283 | 0,88421955 | engulfment and cell motility 3 |
| ENSG00000135298 | ADGRB3 | -0,120058777 | 0,920150162 | 0,74824 | 0,8814896 | adhesion G protein-coupled receptor B3 |
| ENSG00000269570 | AP001350,1 | -0,12014665 | 0,920094118 | 0,72039 | 0,86628014 | novel transcript, antisense to ZFP91 |
| ENSG00000243620 | AC092957,1 | -0,120315457 | 0,919986466 | 0,85998 | 0,93704372 | novel transcript |
| ENSG00000260452 | TPRKBP2 | -0,120413038 | 0,919924242 | 0,91132 | 0,9616927 | TP53RK binding protein pseudogene 2 |
| ENSG00000257176 | AC009318,1 | -0,120509618 | 0,919862661 | 0,59622 | 0,79379314 | novel transcript, antisense to FAR2 |
| ENSG00000167291 | TBC1D16 | -0,120528755 | 0,919850459 | 0,55985 | 0,77074768 | TBC1 domain family member 16 |
| ENSG00000163637 | PRICKLE2 | -0,120606267 | 0,919801039 | 0,65592 | 0,82815262 | prickle planar cell polarity protein 2 |
| ENSG00000159445 | THEM4 | -0,120619991 | 0,919792289 | 0,59701 | 0,79434997 | thioesterase superfamily member 4 |
| ENSG00000170469 | SPATA24 | -0,120735682 | 0,919718534 | 0,70321 | 0,8568579 | spermatogenesis associated 24 |
| ENSG00000204859 | ZBTB48 | -0,120788431 | 0,919684907 | 0,46673 | 0,70676695 | zinc finger and BTB domain containing 48 |
| ENSG00000148300 | REXO4 | -0,120803971 | 0,919675 | 0,57226 | 0,77846037 | REX4 homolog, 3'-5' exonuclease |
| ENSG00000280706 | REXO4 | -0,120803971 | 0,919675 | 0,57226 | 0,77846037 | REX4 homolog, 3'-5' exonuclease |
| ENSG00000130844 | ZNF331 | -0,120847312 | 0,919647372 | 0,58451 | 0,78572744 | zinc finger protein 331 |
| ENSG00000259556 | AC090971,3 | -0,12101815 | 0,919538478 | 0,8636 | 0,93904284 | Leo1, Paf1/RNA polymerase II complex component, homolog (S, cerevisiae) (LEO1) pseudogene |
| ENSG00000110811 | P3H3 | -0,121054065 | 0,919515587 | 0,78225 | 0,8988906 | prolyl 3-hydroxylase 3 |
| ENSG00000170382 | LRRN2 | -0,121073481 | 0,919503212 | 0,7067 | 0,85889796 | leucine rich repeat neuronal 2 |
| ENSG00000287577 | AC009533,4 | -0,121083487 | 0,919496835 | 0,90296 | 0,95846794 | novel transcript |
| ENSG00000286787 | AL110114,1 | -0,121190268 | 0,919428781 | 0,68077 | 0,84338282 | novel transcript, antisense to PSMF1 |

| | | | | | | |
|---|---|---|---|---|---|---|
| ENSG00000172175 | MALT1 | -0,121199221 | 0,919423075 | 0,49098 | 0,72492257 | MALT1 paracaspase |
| ENSG00000283360 | AL592295,3 | -0,12120033 | 0,919422368 | 0,9003 | 0,95760242 | novel transcript |
| ENSG00000227973 | PIN4P1 | -0,121215452 | 0,919412731 | 0,86377 | 0,93907743 | peptidylprolyl cis/trans isomerase, NIMA-interacting 4 pseudogene 1 |
| ENSG00000196502 | SULT1A1 | -0,121217941 | 0,919411145 | 0,83466 | 0,92444381 | sulfotransferase family 1A member 1 |
| ENSG00000285036 | DHRS4L1 | -0,12124084 | 0,919396551 | 0,90581 | 0,95975315 | dehydrogenase/reductase 4 like 1 |
| ENSG00000225766 | DHRS4L1 | -0,12124084 | 0,919396551 | 0,90581 | 0,95975315 | dehydrogenase/reductase 4 like 1 |
| ENSG00000087903 | RFX2 | -0,121242152 | 0,919395715 | 0,5165 | 0,74198903 | regulatory factor X2 |
| ENSG00000088854 | C20orf194 | -0,121311517 | 0,919351512 | 0,5704 | 0,77733579 | chromosome 20 open reading frame 194 |
| ENSG00000283530 | MGAT4C | -0,121338407 | 0,919334376 | 0,89006 | 0,95158772 | MGAT4 family member C |
| ENSG00000133318 | RTN3 | -0,121474685 | 0,91924754 | 0,48914 | 0,72372012 | reticulon 3 |
| ENSG00000186575 | NF2 | -0,121603779 | 0,919165288 | 0,63539 | 0,81655453 | neurofibromin 2 |
| ENSG00000272405 | AL365181,3 | -0,12165481 | 0,919132776 | 0,79773 | 0,90685415 | novel transcript, antisense to BCAN |
| ENSG00000002586 | CD99 | -0,121777155 | 0,919054834 | 0,60508 | 0,79827042 | CD99 molecule (Xg blood group) |
| ENSG00000182108 | DEXI | -0,121863032 | 0,919000128 | 0,54893 | 0,76394298 | Dexi homolog |
| ENSG00000078902 | TOLLIP | -0,121912121 | 0,918968859 | 0,56751 | 0,7754074 | toll interacting protein |
| ENSG00000283413 | AC008703,1 | -0,121928011 | 0,918958738 | 0,87753 | 0,94566578 | novel transcript |
| ENSG00000164106 | SCRG1 | -0,121973021 | 0,918930068 | 0,76569 | 0,89097507 | stimulator of chondrogenesis 1 |
| ENSG00000110324 | IL10RA | -0,122027652 | 0,918895271 | 0,82934 | 0,9215027 | interleukin 10 receptor subunit alpha |
| ENSG00000214106 | PAXIP1-AS2 | -0,122087109 | 0,918857402 | 0,67389 | 0,8393266 | PAXIP1 antisense RNA 2 |
| ENSG00000176454 | LPCAT4 | -0,122263281 | 0,918745204 | 0,55357 | 0,76655124 | lysophosphatidylcholine acyltransferase 4 |
| ENSG00000115355 | CCDC88A | -0,122272507 | 0,918739328 | 0,48695 | 0,72215091 | coiled-coil domain containing 88A |
| ENSG00000162836 | ACP6 | -0,122332814 | 0,918700925 | 0,54204 | 0,75894841 | acid phosphatase 6, lysophosphatidic |
| ENSG00000165186 | PTCHD1 | -0,122336449 | 0,91869861 | 0,83189 | 0,92312766 | patched domain containing 1 |
| ENSG00000229124 | VIM-AS1 | -0,122361252 | 0,918682816 | 0,65641 | 0,82832937 | VIM antisense RNA 1 |
| ENSG00000179456 | ZBTB18 | -0,122506191 | 0,918590526 | 0,63144 | 0,81403245 | zinc finger and BTB domain containing 18 |
| ENSG00000167088 | SNRPD1 | -0,122531048 | 0,918574699 | 0,58969 | 0,78906453 | small nuclear ribonucleoprotein D1 polypeptide |
| ENSG00000011451 | WIZ | -0,122581813 | 0,918542377 | 0,44091 | 0,68929518 | WIZ zinc finger |
| ENSG00000252481 | SCARNA13 | -0,122620895 | 0,918517494 | 0,65327 | 0,82629807 | small Cajal body-specific RNA 13 |
| ENSG00000227028 | SLC8A1-AS1 | -0,122677378 | 0,918481534 | 0,71199 | 0,8615121 | SLC8A1 antisense RNA 1 |
| ENSG00000152117 | SMPD4BP | -0,122842209 | 0,918376602 | 0,53345 | 0,75351438 | sphingomyelin phosphodiesterase 4B, pseudogene |
| ENSG00000125538 | IL1B | -0,122843106 | 0,918376031 | 0,80801 | 0,91168773 | interleukin 1 beta |
| ENSG00000112039 | FANCE | -0,122929222 | 0,918321214 | 0,65076 | 0,82501561 | FA complementation group E |

| | | | | | | |
|---|---|---|---|---|---|---|
| ENSG00000233621 | LINC01137 | -0,12316757 | 0,91816951 | 0,83006 | 0,92197668 | long intergenic non-protein coding RNA 1137 |
| ENSG00000264169 | RN7SL665P | -0,123169883 | 0,918168038 | 0,91393 | 0,96272082 | RNA, 7SL, cytoplasmic 665, pseudogene |
| ENSG00000132467 | UTP3 | -0,123194039 | 0,918152664 | 0,5924 | 0,79117578 | UTP3 small subunit processome component |
| ENSG00000137463 | MGARP | -0,123203293 | 0,918146776 | 0,90776 | 0,96044232 | mitochondria localized glutamic acid rich protein |
| ENSG00000239650 | GUSBP4 | -0,123206085 | 0,918144998 | 0,88807 | 0,95069428 | GUSB pseudogene 4 |
| ENSG00000130147 | SH3BP4 | -0,123356754 | 0,918049116 | 0,61343 | 0,8035281 | SH3 domain binding protein 4 |
| ENSG00000161011 | SQSTM1 | -0,123376686 | 0,918036433 | 0,64319 | 0,82098995 | sequestosome 1 |
| ENSG00000105204 | DYRK1B | -0,123468139 | 0,91797824 | 0,57284 | 0,77868876 | dual specificity tyrosine phosphorylation regulated kinase 1B |
| ENSG00000281320 | DYRK1B | -0,123468139 | 0,91797824 | 0,57284 | 0,77868876 | dual specificity tyrosine phosphorylation regulated kinase 1B |
| ENSG00000279855 | AC005618,2 | -0,123576744 | 0,917909138 | 0,93038 | 0,97019767 | novel transcript, antisense to protocadherin gamma cluster |
| ENSG00000198707 | CEP290 | -0,123640677 | 0,917868462 | 0,53286 | 0,75333268 | centrosomal protein 290 |
| ENSG00000285650 | AL157827,2 | -0,123740426 | 0,917805001 | 0,66977 | 0,83623597 | Novel transcript, antisense to CENPP |
| ENSG00000164669 | INTS4P1 | -0,123759111 | 0,917793115 | 0,82087 | 0,91738668 | integrator complex subunit 4 pseudogene 1 |
| ENSG00000084444 | FAM234B | -0,12393818 | 0,917679204 | 0,49235 | 0,72556208 | family with sequence similarity 234 member B |
| ENSG00000124466 | LYPD3 | -0,124176751 | 0,917527465 | 0,77864 | 0,89755385 | LY6/PLAUR domain containing 3 |
| ENSG00000172461 | FUT9 | -0,12422398 | 0,917497429 | 0,75853 | 0,88727479 | fucosyltransferase 9 |
| ENSG00000232341 | RPL4P2 | -0,124322486 | 0,917434785 | 0,82717 | 0,92010219 | ribosomal protein L4 pseudogene 2 |
| ENSG00000113763 | UNC5A | -0,124406261 | 0,917381513 | 0,77768 | 0,89714267 | unc-5 netrin receptor A |
| ENSG00000090470 | PDCD7 | -0,124443726 | 0,91735769 | 0,50203 | 0,73163635 | programmed cell death 7 |
| ENSG00000187066 | TMEM262 | -0,124471821 | 0,917339825 | 0,74206 | 0,87800726 | transmembrane protein 262 |
| ENSG00000164975 | SNAPC3 | -0,124591972 | 0,91726343 | 0,50113 | 0,73116951 | small nuclear RNA activating complex polypeptide 3 |
| ENSG00000133169 | BEX1 | -0,124641886 | 0,917231696 | 0,61487 | 0,80404192 | brain expressed X-linked 1 |
| ENSG00000109771 | LRP2BP | -0,124818451 | 0,917119447 | 0,71396 | 0,86227395 | LRP2 binding protein |
| ENSG00000256746 | MADD-AS1 | -0,12482201 | 0,917117184 | 0,87543 | 0,94491504 | MADD antisense RNA 1 |
| ENSG00000121236 | TRIM6 | -0,1248342 | 0,917109435 | 0,58839 | 0,78811087 | tripartite motif containing 6 |
| ENSG00000196576 | PLXNB2 | -0,124972037 | 0,917021817 | 0,57399 | 0,77928773 | plexin B2 |
| ENSG00000173627 | APOBEC4 | -0,125064144 | 0,916963273 | 0,76264 | 0,88967304 | apolipoprotein B mRNA editing enzyme catalytic polypeptide like 4 |
| ENSG00000181284 | TMEM102 | -0,125072123 | 0,916958201 | 0,8696 | 0,9425949 | transmembrane protein 102 |
| ENSG00000284228 | TMEM102 | -0,125072123 | 0,916958201 | 0,8696 | 0,9425949 | transmembrane protein 102 |
| ENSG00000106133 | NSUN5P2 | -0,125079973 | 0,916953212 | 0,61292 | 0,8035281 | NSUN5 pseudogene 2 |

| | | | | | | |
|---|---|---|---|---|---|---|
| ENSG00000253955 | AC008663,2 | -0,125204396 | 0,916874134 | 0,82406 | 0,91879719 | novel transcript |
| ENSG00000197128 | ZNF772 | -0,125242079 | 0,916850186 | 0,60983 | 0,80199287 | zinc finger protein 772 |
| ENSG00000151116 | UEVLD | -0,125263841 | 0,916836356 | 0,51762 | 0,74271521 | UEV and lactate/malate dehyrogenase domains |
| ENSG00000234617 | SNRK-AS1 | -0,125335943 | 0,916790536 | 0,67732 | 0,84132177 | SNRK antisense RNA 1 |
| ENSG00000218823 | PAPOLB | -0,125369061 | 0,916769491 | 0,80899 | 0,91201277 | poly(A) polymerase beta |
| ENSG00000133805 | AMPD3 | -0,125372235 | 0,916767474 | 0,83517 | 0,92472031 | adenosine monophosphate deaminase 3 |
| ENSG00000270739 | AC090953,1 | -0,125536404 | 0,916663158 | 0,87294 | 0,94378579 | THAP domain containing 5, pseudogene |
| ENSG00000237310 | GS1-124K5,4 | -0,125617866 | 0,9166114 | 0,7331 | 0,87350302 | uncharacterized LOC100289098 |
| ENSG00000271810 | AL603832,3 | -0,125652083 | 0,916589661 | 0,93845 | 0,97367474 | novel protein, PPM1J-RHOC readthrough |
| ENSG00000271013 | LRRC37A9P | -0,125684518 | 0,916569054 | 0,70309 | 0,8568579 | leucine rich repeat containing 37 member A9, pseudogene |
| ENSG00000117525 | F3 | -0,125731212 | 0,916539389 | 0,75339 | 0,88453179 | coagulation factor III, tissue factor |
| ENSG00000234062 | AL390879,1 | -0,125822302 | 0,916481522 | 0,62674 | 0,81113561 | transmembrane 9 superfamily member 2 (TM9SF2) pseudogene |
| ENSG00000239677 | PDZRN3-AS1 | -0,125825115 | 0,916479735 | 0,87093 | 0,94305486 | PDZRN3 antisense RNA 1 |
| ENSG00000275506 | H3P25 | -0,125882447 | 0,916443315 | 0,91807 | 0,9644207 | H3 histone pseudogene 25 |
| ENSG00000184492 | FOXD4L1 | -0,125946026 | 0,916402928 | 0,82979 | 0,92186898 | forkhead box D4 like 1 |
| ENSG00000141956 | PRDM15 | -0,125958978 | 0,916394702 | 0,46697 | 0,70703542 | PR/SET domain 15 |
| ENSG00000164746 | C7orf57 | -0,126234616 | 0,916219634 | 0,87914 | 0,94623795 | chromosome 7 open reading frame 57 |
| ENSG00000272901 | SMDT1 | -0,126272363 | 0,916195662 | 0,74108 | 0,87729166 | single-pass membrane protein with aspartate rich tail 1 |
| ENSG00000272406 | AC108102,1 | -0,126273961 | 0,916194647 | 0,90189 | 0,95804366 | novel transcript |
| ENSG00000277600 | CNOT3 | -0,126317535 | 0,916166976 | 0,82678 | 0,91986424 | CCR4-NOT transcription complex subunit 3 |
| ENSG00000227541 | SFR1P1 | -0,126337684 | 0,91615418 | 0,84799 | 0,9309317 | SFR1 pseudogene 1 |
| ENSG00000205038 | PKHD1L1 | -0,12637378 | 0,916131258 | 0,75249 | 0,88400878 | PKHD1 like 1 |
| ENSG00000152556 | PFKM | -0,126402645 | 0,916112929 | 0,59308 | 0,79148977 | phosphofructokinase, muscle |
| ENSG00000158062 | UBXN11 | -0,126447369 | 0,91608453 | 0,51892 | 0,7437068 | UBX domain protein 11 |
| ENSG00000272625 | AP000919,3 | -0,126467364 | 0,916071833 | 0,86318 | 0,93891554 | novel transcript, antisense to LPIN2 |
| ENSG00000263272 | AC004148,1 | -0,126573619 | 0,916004367 | 0,63757 | 0,81781506 | novel transcript, antisense to RPAIN |
| ENSG00000273177 | AC092954,2 | -0,126716121 | 0,915913893 | 0,90251 | 0,95832496 | novel transcript |
| ENSG00000131446 | MGAT1 | -0,126731124 | 0,915904368 | 0,63987 | 0,81929791 | alpha-1,3-mannosyl-glycoprotein 2-beta-N-acetylglucosaminyltransferase |
| ENSG00000264434 | AC110603,1 | -0,126733567 | 0,915902817 | 0,84696 | 0,93041089 | novel transcript, antisense to ZNF521 |
| ENSG00000112659 | CUL9 | -0,126873882 | 0,915813742 | 0,47591 | 0,71433501 | cullin 9 |

| | | | | | | |
|---|---|---|---|---|---|---|
| ENSG00000275976 | AC211476,3 | -0,126915013 | 0,915787633 | 0,80875 | 0,91198969 | speedy/RINGO cell cycle regulator family member E1 (SPDYE1) pseudogene |
| ENSG00000108592 | FTSJ3 | -0,127018123 | 0,915722183 | 0,59527 | 0,79324077 | FtsJ RNA 2'-O-methyltransferase 3 |
| ENSG00000227600 | PPT2 | -0,127097784 | 0,915671621 | 0,80142 | 0,90860375 | palmitoyl-protein thioesterase 2 |
| ENSG00000169372 | CRADD | -0,12710452 | 0,915667346 | 0,66144 | 0,83138572 | CASP2 and RIPK1 domain containing adaptor with death domain |
| ENSG00000249141 | AL159163,1 | -0,127124578 | 0,915654616 | 0,84397 | 0,92886643 | novel protein |
| ENSG00000251596 | HADHAP1 | -0,127137331 | 0,915646522 | 0,71661 | 0,86437402 | HADHA pseudogene 1 |
| ENSG00000143627 | PKLR | -0,127176094 | 0,91562192 | 0,92251 | 0,96642782 | pyruvate kinase L/R |
| ENSG00000164466 | SFXN1 | -0,127281938 | 0,915554747 | 0,40002 | 0,66216538 | sideroflexin 1 |
| ENSG00000175664 | TEX26 | -0,127282789 | 0,915554207 | 0,90463 | 0,95918368 | testis expressed 26 |
| ENSG00000234280 | TSBP1 | -0,127288159 | 0,915550799 | 0,93065 | 0,97025341 | testis expressed basic protein 1 |
| ENSG00000182796 | TMEM198B | -0,127429887 | 0,915460862 | 0,65987 | 0,83018671 | transmembrane protein 198B (pseudogene) |
| ENSG00000143321 | HDGF | -0,127478534 | 0,915429993 | 0,61686 | 0,80544374 | heparin binding growth factor |
| ENSG00000277147 | LINC00869 | -0,127592709 | 0,915357549 | 0,62569 | 0,81057207 | long intergenic non-protein coding RNA 869 |
| ENSG00000092841 | MYL6 | -0,127614961 | 0,915343431 | 0,58873 | 0,78830778 | myosin light chain 6 |
| ENSG00000287535 | AC009630,4 | -0,12769123 | 0,915295041 | 0,85832 | 0,93612837 | novel transcript |
| ENSG00000236986 | AL157938,2 | -0,127770269 | 0,915244898 | 0,76437 | 0,89017297 | novel transcript, antisense to NUP214 |
| ENSG00000132964 | CDK8 | -0,127920953 | 0,915149309 | 0,52821 | 0,75077432 | cyclin dependent kinase 8 |
| ENSG00000151552 | QDPR | -0,127934802 | 0,915140524 | 0,42694 | 0,68019102 | quinoid dihydropteridine reductase |
| ENSG00000100258 | LMF2 | -0,127941056 | 0,915136557 | 0,61463 | 0,80391491 | lipase maturation factor 2 |
| ENSG00000284656 | AL645937,4 | -0,128009542 | 0,915093116 | 0,87566 | 0,94497474 | novel transcript |
| ENSG00000087365 | SF3B2 | -0,12809586 | 0,915038366 | 0,5597 | 0,7706142 | splicing factor 3b subunit 2 |
| ENSG00000259362 | AC046168,1 | -0,128191213 | 0,91497789 | 0,83116 | 0,92276172 | novel transcript |
| ENSG00000104129 | DNAJC17 | -0,128207961 | 0,914967268 | 0,58426 | 0,78572744 | DnaJ heat shock protein family (Hsp40) member C17 |
| ENSG00000066405 | CLDN18 | -0,128236246 | 0,91494933 | 0,72476 | 0,86866311 | claudin 18 |
| ENSG00000072682 | P4HA2 | -0,128238217 | 0,91494808 | 0,59856 | 0,7951852 | prolyl 4-hydroxylase subunit alpha 2 |
| ENSG00000248975 | AL133372,2 | -0,128560273 | 0,914743857 | 0,89494 | 0,95417507 | novel transcript |
| ENSG00000136828 | RALGPS1 | -0,128624769 | 0,914702964 | 0,64327 | 0,82098995 | Ral GEF with PH domain and SH3 binding motif 1 |
| ENSG00000140950 | MEAK7 | -0,128780556 | 0,914604197 | 0,51811 | 0,74303983 | MTOR associated protein, eak-7 homolog |
| ENSG00000078725 | BRINP1 | -0,128980579 | 0,9144774 | 0,80234 | 0,90900383 | BMP/retinoic acid inducible neural specific 1 |
| ENSG00000166848 | TERF2IP | -0,128990469 | 0,914471131 | 0,57227 | 0,77846037 | TERF2 interacting protein |
| ENSG00000117616 | RSRP1 | -0,128999685 | 0,914465289 | 0,40179 | 0,66324659 | arginine and serine rich protein 1 |

| | | | | | | |
|---|---|---|---|---|---|---|
| ENSG00000156170 | NDUFAF6 | -0,129002829 | 0,914463296 | 0,65733 | 0,82868311 | NADH:ubiquinone oxidoreductase complex assembly factor 6 |
| ENSG00000275888 | AC132872,3 | -0,129073491 | 0,914418508 | 0,84263 | 0,92821072 | novel transcript, antisense to CSNK1D |
| ENSG00000273133 | AC116651,1 | -0,129097416 | 0,914403344 | 0,73831 | 0,87571125 | novel transcript, antisense to CC2D2A |
| ENSG00000168216 | LMBRD1 | -0,129190769 | 0,914344177 | 0,53458 | 0,75412503 | LMBR1 domain containing 1 |
| ENSG00000213222 | TOMM40P4 | -0,129259404 | 0,914300679 | 0,76876 | 0,89277929 | TOMM40 pseudogene 4 |
| ENSG00000281848 | TOMM40P4 | -0,129259404 | 0,914300679 | 0,76876 | 0,89277929 | TOMM40 pseudogene 4 |
| ENSG00000232684 | ATP11A-AS1 | -0,129308787 | 0,914269383 | 0,77654 | 0,89657313 | ATP11A antisense RNA 1 |
| ENSG00000130396 | AFDN | -0,129337739 | 0,914251036 | 0,52538 | 0,74902159 | afadin, adherens junction formation factor |
| ENSG00000200693 | U3 | -0,129348188 | 0,914244414 | 0,88901 | 0,9510223 | Small nucleolar RNA U3 |
| ENSG00000188831 | DPPA3P2 | -0,129545966 | 0,914119089 | 0,88269 | 0,94815391 | developmental pluripotency associated 3 pseudogene 2 |
| ENSG00000139970 | RTN1 | -0,129647622 | 0,914054681 | 0,7024 | 0,85653047 | reticulon 1 |
| ENSG00000170681 | CAVIN4 | -0,129652024 | 0,914051892 | 0,52921 | 0,75112358 | caveolae associated protein 4 |
| ENSG00000238109 | RNF14P3 | -0,129744536 | 0,91399328 | 0,7293 | 0,87157671 | RNF14 pseudogene 3 |
| ENSG00000232150 | ST13P4 | -0,129757296 | 0,913985197 | 0,86812 | 0,94176112 | ST13, Hsp70 interacting protein pseudogene 4 |
| ENSG00000143552 | NUP210L | -0,129819562 | 0,91394575 | 0,76019 | 0,88825236 | nucleoporin 210 like |
| ENSG00000256061 | DNAAF4 | -0,12982896 | 0,913939797 | 0,60032 | 0,79581466 | dynein axonemal assembly factor 4 |
| ENSG00000268895 | A1BG-AS1 | -0,129845339 | 0,913929421 | 0,59363 | 0,79192667 | A1BG antisense RNA 1 |
| ENSG00000273362 | AP000547,2 | -0,1298467 | 0,913928558 | 0,76225 | 0,88933378 | novel transcript |
| ENSG00000274104 | AC020910,5 | -0,129851844 | 0,9139253 | 0,86398 | 0,9391623 | novel transcript |
| ENSG00000099622 | CIRBP | -0,129880644 | 0,913907055 | 0,56031 | 0,7707835 | cold inducible RNA binding protein |
| ENSG00000160271 | RALGDS | -0,129929563 | 0,913876068 | 0,51024 | 0,7378597 | ral guanine nucleotide dissociation stimulator |
| ENSG00000196376 | SLC35F1 | -0,130076774 | 0,913782822 | 0,65185 | 0,82561449 | solute carrier family 35 member F1 |
| ENSG00000228061 | AC131571,1 | -0,130116523 | 0,913757645 | 0,71886 | 0,86556176 | novel transcript, antisense to ELP4 |
| ENSG00000090530 | P3H2 | -0,130210294 | 0,913698255 | 0,82046 | 0,91732362 | prolyl 3-hydroxylase 2 |
| ENSG00000171298 | GAA | -0,130233278 | 0,913683699 | 0,55886 | 0,77025265 | glucosidase alpha, acid |
| ENSG00000223773 | CD99P1 | -0,130260587 | 0,913666404 | 0,56499 | 0,77338195 | CD99 molecule pseudogene 1 |
| ENSG00000111664 | GNB3 | -0,130293252 | 0,913645718 | 0,64711 | 0,82343226 | G protein subunit beta 3 |
| ENSG00000272779 | AC245060,4 | -0,130391337 | 0,913583603 | 0,59865 | 0,7951852 | BMS1, ribosome biogenesis factor (BSM1) pseudogene |
| ENSG00000277342 | AC048344,4 | -0,130433976 | 0,913556603 | 0,85418 | 0,93380868 | novel transcript, sense intronic to BICD1 |
| ENSG00000218283 | MORF4L1P1 | -0,130438582 | 0,913553686 | 0,7997 | 0,90758156 | mortality factor 4 like 1 pseudogene 1 |
| ENSG00000250539 | KRT8P33 | -0,130466388 | 0,913536079 | 0,70469 | 0,85769579 | keratin 8 pseudogene 33 |
| ENSG00000197646 | PDCD1LG2 | -0,13054382 | 0,913487049 | 0,88068 | 0,94687991 | programmed cell death 1 ligand 2 |

| | | | | | | |
|---|---|---|---|---|---|---|
| ENSG00000123329 | ARHGAP9 | -0,130661173 | 0,913412746 | 0,79979 | 0,90761856 | Rho GTPase activating protein 9 |
| ENSG00000158258 | CLSTN2 | -0,130883246 | 0,913272156 | 0,70019 | 0,85526044 | calsyntenin 2 |
| ENSG00000261096 | AC073476,2 | -0,130891394 | 0,913266998 | 0,66318 | 0,83242946 | novel transcript |
| ENSG00000287447 | AC099501,1 | -0,130907643 | 0,913256712 | 0,8298 | 0,92186898 | novel transcript, antisense to FAM172A |
| ENSG00000275005 | AL354950,1 | -0,131107969 | 0,91312991 | 0,90107 | 0,95787401 | novel transcript, antisense to EBF3 |
| ENSG00000130477 | UNC13A | -0,131139882 | 0,913109712 | 0,76551 | 0,8908718 | unc-13 homolog A |
| ENSG00000065618 | COL17A1 | -0,131179846 | 0,913084418 | 0,77249 | 0,89482448 | collagen type XVII alpha 1 chain |
| ENSG00000254556 | AF131215,2 | -0,131225536 | 0,913055501 | 0,72144 | 0,86698039 | novel transcript |
| ENSG00000113070 | HBEGF | -0,131258348 | 0,913034735 | 0,68802 | 0,8480754 | heparin binding EGF like growth factor |
| ENSG00000171757 | LRRC34 | -0,131296599 | 0,913010528 | 0,6522 | 0,82578944 | leucine rich repeat containing 34 |
| ENSG00000188158 | NHS | -0,131373336 | 0,912961966 | 0,77937 | 0,89794855 | NHS actin remodeling regulator |
| ENSG00000261646 | AC093849,1 | -0,13194114 | 0,912602721 | 0,84982 | 0,93163876 | novel transcript |
| ENSG00000237037 | NDUFA6-DT | -0,131977346 | 0,912579818 | 0,65545 | 0,82781124 | NDUFA6 divergent transcript |
| ENSG00000237645 | AC092967,1 | -0,132147648 | 0,9124721 | 0,86136 | 0,93801973 | chromosome 14 open reading frame 83 (C14orf83) pseudogene |
| ENSG00000027847 | B4GALT7 | -0,132152197 | 0,912469223 | 0,49642 | 0,72831792 | beta-1,4-galactosyltransferase 7 |
| ENSG00000227666 | CYCSP24 | -0,132191209 | 0,912444549 | 0,90584 | 0,95975315 | CYCS pseudogene 24 |
| ENSG00000127054 | INTS11 | -0,13222855 | 0,912420932 | 0,58318 | 0,7850707 | integrator complex subunit 11 |
| ENSG00000254672 | AL136146,1 | -0,132249057 | 0,912407963 | 0,73189 | 0,87276602 | WEE1 homolog (S, pombe) (WEE1) pseudogene |
| ENSG00000135902 | CHRND | -0,132339121 | 0,912351006 | 0,78228 | 0,8988906 | cholinergic receptor nicotinic delta subunit |
| ENSG00000229948 | B3GALNT1P1 | -0,13242976 | 0,912293688 | 0,86012 | 0,93714201 | beta-1,3-N-acetylgalactosaminyltransferase 1 pseudogene 1 |
| ENSG00000113108 | APBB3 | -0,132449776 | 0,912281031 | 0,6144 | 0,80376648 | amyloid beta precursor protein binding family B member 3 |
| ENSG00000115239 | ASB3 | -0,132495531 | 0,912252098 | 0,47681 | 0,71481671 | ankyrin repeat and SOCS box containing 3 |
| ENSG00000071564 | TCF3 | -0,132575418 | 0,912201585 | 0,46805 | 0,70769773 | transcription factor 3 |
| ENSG00000140854 | KATNB1 | -0,132648833 | 0,912155167 | 0,45464 | 0,69879733 | katanin regulatory subunit B1 |
| ENSG00000142039 | CCDC97 | -0,132667874 | 0,912143128 | 0,61546 | 0,80439941 | coiled-coil domain containing 97 |
| ENSG00000165046 | LETM2 | -0,132782852 | 0,912070436 | 0,64168 | 0,82033064 | leucine zipper and EF-hand containing transmembrane protein 2 |
| ENSG00000229944 | EIF4EP2 | -0,132848056 | 0,912029215 | 0,91582 | 0,9633821 | eukaryotic translation initiation factor 4E pseudogene 2 |
| ENSG00000267342 | AC087289,2 | -0,132879478 | 0,912009351 | 0,80229 | 0,90900383 | novel transcript, antisense to MRPL38 |
| ENSG00000102879 | CORO1A | -0,13301428 | 0,911924139 | 0,47157 | 0,71092925 | coronin 1A |

| | | | | | | |
|---|---|---|---|---|---|---|
| ENSG00000271396 | AC091167,4 | -0,133115882 | 0,911859919 | 0,77344 | 0,89545683 | MKI67 (FHA domain) interacting nucleolar phosphoprotein (MKI67IP) pseudogene |
| ENSG00000237065 | NANOGP4 | -0,133185518 | 0,911815907 | 0,80218 | 0,90899209 | Nanog homeobox pseudogene 4 |
| ENSG00000203397 | SHISA5P2 | -0,133282914 | 0,911754352 | 0,86612 | 0,94058212 | SHISA5 pseudogene 2 |
| ENSG00000216977 | RPL21P65 | -0,133386753 | 0,91168873 | 0,8257 | 0,91917221 | ribosomal protein L21 pseudogene 65 |
| ENSG00000156873 | PHKG2 | -0,133407212 | 0,911675802 | 0,58408 | 0,78572744 | phosphorylase kinase catalytic subunit gamma 2 |
| ENSG00000273604 | EPOP | -0,133416298 | 0,91167006 | 0,69715 | 0,85330122 | elongin BC and polycomb repressive complex 2 associated protein |
| ENSG00000271918 | AC034236,2 | -0,133449018 | 0,911649384 | 0,66972 | 0,83623395 | novel transcript, antisense to COMMD10 |
| ENSG00000134780 | DAGLA | -0,133518646 | 0,911605386 | 0,42161 | 0,67643128 | diacylglycerol lipase alpha |
| ENSG00000122547 | EEPD1 | -0,133520279 | 0,911604355 | 0,54048 | 0,75829455 | endonuclease/exonuclease/phosphatase family domain containing 1 |
| ENSG00000174243 | DDX23 | -0,133535748 | 0,91159458 | 0,52171 | 0,7460228 | DEAD-box helicase 23 |
| ENSG00000249212 | ATP1B1P1 | -0,133546485 | 0,911587796 | 0,78826 | 0,90156082 | ATPase Na+/K+ transporting subunit beta 1 pseudogene 1 |
| ENSG00000217447 | AL021331,1 | -0,133621154 | 0,911540616 | 0,88817 | 0,95069428 | MAD2 mitotic arrest deficient-like 1 (yeast) (MAD2L1) pseudogene |
| ENSG00000231731 | AC010976,1 | -0,133715068 | 0,91148128 | 0,61515 | 0,80421986 | novel transcript, antisense to IWS1 |
| ENSG00000167397 | VKORC1 | -0,133731278 | 0,911471039 | 0,62362 | 0,80937074 | vitamin K epoxide reductase complex subunit 1 |
| ENSG00000286268 | AF196969,1 | -0,133787554 | 0,911435486 | 0,82328 | 0,9182945 | novel protein |
| ENSG00000178187 | ZNF454 | -0,133801617 | 0,911426601 | 0,83203 | 0,92312766 | zinc finger protein 454 |
| ENSG00000252690 | AC105339,2 | -0,134264182 | 0,911134421 | 0,6663 | 0,83402227 | TEC |
| ENSG00000130520 | LSM4 | -0,13426461 | 0,911134151 | 0,5554 | 0,76788267 | LSM4 homolog, U6 small nuclear RNA and mRNA degradation associated |
| ENSG00000238184 | CD81-AS1 | -0,134318177 | 0,911100321 | 0,83387 | 0,92404752 | CD81 antisense RNA 1 |
| ENSG00000206331 | PRRT1 | -0,134353491 | 0,911078019 | 0,76654 | 0,89139706 | proline rich transmembrane protein 1 |
| ENSG00000160410 | SHKBP1 | -0,134785386 | 0,910805314 | 0,53188 | 0,75280487 | SH3KBP1 binding protein 1 |
| ENSG00000103168 | TAF1C | -0,134790352 | 0,910802179 | 0,5276 | 0,75037219 | TATA-box binding protein associated factor, RNA polymerase I subunit C |
| ENSG00000206573 | THUMPD3-AS | -0,134893677 | 0,91073695 | 0,40115 | 0,6630277 | THUMPD3 antisense RNA 1 |
| ENSG00000104883 | PEX11G | -0,134996966 | 0,910671749 | 0,70759 | 0,85916455 | peroxisomal biogenesis factor 11 gamma |
| ENSG00000133640 | LRRIQ1 | -0,13510179 | 0,910605583 | 0,68961 | 0,84865812 | leucine rich repeats and IQ motif containing 1 |
| ENSG00000100056 | ESS2 | -0,135194623 | 0,91054699 | 0,50484 | 0,73365359 | ess-2 splicing factor homolog |
| ENSG00000233885 | YEATS2-AS1 | -0,135253388 | 0,910509902 | 0,62457 | 0,80982446 | YEATS2 antisense RNA 1 |

| | | | | | | |
|---|---|---|---|---|---|---|
| ENSG00000073331 | ALPK1 | -0,135286932 | 0,910488732 | 0,68766 | 0,84782493 | alpha kinase 1 |
| ENSG00000273162 | AL133215,2 | -0,135287719 | 0,910488235 | 0,65188 | 0,82561636 | novel transcript |
| ENSG00000237325 | AP000563,1 | -0,135317762 | 0,910469275 | 0,77172 | 0,894281 | chromosome 6 open reading frame 203 (C6orf203) pseudogene |
| ENSG00000278016 | TIGD5 | -0,135361363 | 0,91044176 | 0,73776 | 0,87540065 | tigger transposable element derived 5 |
| ENSG00000267413 | LINC01901 | -0,135389401 | 0,910424066 | 0,80858 | 0,91193281 | long intergenic non-protein coding RNA 1901 |
| ENSG00000240849 | PEDS1 | -0,135431478 | 0,910397513 | 0,67088 | 0,83695077 | plasmanylethanolamine desaturase 1 |
| ENSG00000185813 | PCYT2 | -0,135439574 | 0,910392404 | 0,47561 | 0,71417866 | phosphate cytidylyltransferase 2, ethanolamine |
| ENSG00000156042 | CFAP70 | -0,135525898 | 0,910337933 | 0,68353 | 0,84495012 | cilia and flagella associated protein 70 |
| ENSG00000183161 | FANCF | -0,135638239 | 0,910267048 | 0,4863 | 0,72181021 | FA complementation group F |
| ENSG00000204194 | RPL12P1 | -0,135705034 | 0,910224905 | 0,83715 | 0,92549623 | ribosomal protein L12 pseudogene 1 |
| ENSG00000232733 | RPL12P1 | -0,135705034 | 0,910224905 | 0,83715 | 0,92549623 | ribosomal protein L12 pseudogene 1 |
| ENSG00000231169 | EEF1B2P1 | -0,135793777 | 0,910168917 | 0,84383 | 0,92880828 | eukaryotic translation elongation factor 1 beta 2 pseudogene 1 |
| ENSG00000258920 | FOXN3-AS1 | -0,135862896 | 0,910125312 | 0,7291 | 0,87149657 | FOXN3 antisense RNA 1 |
| ENSG00000074211 | PPP2R2C | -0,135956695 | 0,910066141 | 0,81266 | 0,91386173 | protein phosphatase 2 regulatory subunit Bgamma |
| ENSG00000175376 | EIF1AD | -0,135973651 | 0,910055445 | 0,6486 | 0,82403409 | eukaryotic translation initiation factor 1A domain containing |
| ENSG00000171860 | C3AR1 | -0,136109366 | 0,909969839 | 0,83271 | 0,92348349 | complement C3a receptor 1 |
| ENSG00000261618 | LINC02605 | -0,13611864 | 0,90996399 | 0,74291 | 0,87840582 | long intergenic non-protein coding RNA 2605 |
| ENSG00000231952 | DPY19L1P2 | -0,136142804 | 0,909948749 | 0,84255 | 0,92821072 | DPY19L1 pseudogene 2 |
| ENSG00000273329 | AC078846,1 | -0,136290043 | 0,909855886 | 0,59227 | 0,79106659 | novel transcript |
| ENSG00000102383 | ZDHHC15 | -0,136360123 | 0,90981169 | 0,4481 | 0,69411284 | zinc finger DHHC-type palmitoyltransferase 15 |
| ENSG00000277946 | RN7SL478P | -0,136400194 | 0,90978642 | 0,82121 | 0,91738668 | RNA, 7SL, cytoplasmic 478, pseudogene |
| ENSG00000250933 | GAPDHP66 | -0,136426532 | 0,909769811 | 0,84921 | 0,93136769 | glyceraldehyde-3-phosphate dehydrogenase pseudogene 66 |
| ENSG00000114126 | TFDP2 | -0,136437467 | 0,909762916 | 0,66014 | 0,83029149 | transcription factor Dp-2 |
| ENSG00000187942 | LDLRAD2 | -0,136459065 | 0,909749296 | 0,83418 | 0,92405867 | low density lipoprotein receptor class A domain containing 2 |
| ENSG00000160408 | ST6GALNAC6 | -0,136470854 | 0,909741862 | 0,59431 | 0,79263193 | ST6 N-acetylgalactosaminide alpha-2,6-sialyltransferase 6 |
| ENSG00000242082 | SLC5A4-AS1 | -0,136535543 | 0,909701071 | 0,74072 | 0,87717177 | SLC5A4 antisense RNA 1 |
| ENSG00000225442 | MPRIP-AS1 | -0,136598916 | 0,909661112 | 0,76463 | 0,89039679 | MPRIP antisense RNA 1 |
| ENSG00000204802 | AL590399,1 | -0,136607341 | 0,9096558 | 0,60731 | 0,80012503 | novel transcript |

| | | | | | | |
|---|---|---|---|---|---|---|
| ENSG00000118503 | TNFAIP3 | -0,136671604 | 0,909615281 | 0,76656 | 0,89139706 | TNF alpha induced protein 3 |
| ENSG00000228544 | CCDC183-AS1 | -0,136750431 | 0,909565582 | 0,71966 | 0,86592885 | CCDC183 antisense RNA 1 |
| ENSG00000163072 | NOSTRIN | -0,136861274 | 0,909495703 | 0,72736 | 0,87021076 | nitric oxide synthase trafficking |
| ENSG00000219438 | TAFA5 | -0,136878141 | 0,909485069 | 0,69678 | 0,85309963 | TAFA chemokine like family member 5 |
| ENSG00000232587 | EEF1A1P3 | -0,136940059 | 0,909446036 | 0,72644 | 0,86975399 | eukaryotic translation elongation factor 1 alpha 1 pseudogene 3 |
| ENSG00000187630 | DHRS4L2 | -0,136956613 | 0,909435601 | 0,65639 | 0,82832937 | dehydrogenase/reductase 4 like 2 |
| ENSG00000112208 | BAG2 | -0,136957044 | 0,90943533 | 0,69853 | 0,85408806 | BAG cochaperone 2 |
| ENSG00000258033 | SNX5P2 | -0,136966384 | 0,909429442 | 0,84754 | 0,93074315 | sorting nexin 5 pseudogene 2 |
| ENSG00000243646 | IL10RB | -0,136990779 | 0,909414064 | 0,61701 | 0,80553112 | interleukin 10 receptor subunit beta |
| ENSG00000179397 | CATSPERE | -0,137079859 | 0,909357914 | 0,70112 | 0,85579724 | catsper channel auxiliary subunit epsilon |
| ENSG00000258909 | AC091078,2 | -0,137145463 | 0,909316563 | 0,569 | 0,77629764 | novel transcript |
| ENSG00000272288 | AL451165,2 | -0,137171443 | 0,909300188 | 0,7176 | 0,86483656 | novel transcript, antisense to C6orf106 |
| ENSG00000049768 | FOXP3 | -0,137196666 | 0,909284291 | 0,7845 | 0,89962715 | forkhead box P3 |
| ENSG00000247708 | STX18-AS1 | -0,137209577 | 0,909276154 | 0,59536 | 0,79331764 | STX18 antisense RNA 1 (head to head) |
| ENSG00000273943 | CNOT3 | -0,137232304 | 0,90926183 | 0,73672 | 0,87529332 | CCR4-NOT transcription complex subunit 3 |
| ENSG00000274176 | CNOT3 | -0,137232304 | 0,90926183 | 0,73672 | 0,87529332 | CCR4-NOT transcription complex subunit 3 |
| ENSG00000274941 | CNOT3 | -0,137232304 | 0,90926183 | 0,73672 | 0,87529332 | CCR4-NOT transcription complex subunit 3 |
| ENSG00000276082 | CNOT3 | -0,137232304 | 0,90926183 | 0,73672 | 0,87529332 | CCR4-NOT transcription complex subunit 3 |
| ENSG00000275979 | CNOT3 | -0,137232304 | 0,90926183 | 0,73672 | 0,87529332 | CCR4-NOT transcription complex subunit 3 |
| ENSG00000277615 | CNOT3 | -0,137232304 | 0,90926183 | 0,73672 | 0,87529332 | CCR4-NOT transcription complex subunit 3 |
| ENSG00000164411 | GJB7 | -0,137251944 | 0,909249452 | 0,72094 | 0,86659896 | gap junction protein beta 7 |
| ENSG00000214376 | VSTM5 | -0,137266292 | 0,909240409 | 0,70564 | 0,85821762 | V-set and transmembrane domain containing 5 |
| ENSG00000122733 | PHF24 | -0,137281735 | 0,909230676 | 0,78178 | 0,89877377 | PHD finger protein 24 |
| ENSG00000177842 | ZNF620 | -0,137401157 | 0,909155416 | 0,56634 | 0,77432588 | zinc finger protein 620 |
| ENSG00000286448 | AP006222,2 | -0,137452836 | 0,90912285 | 0,90411 | 0,95888764 | novel transcript |
| ENSG00000122756 | CNTFR | -0,137462054 | 0,909117041 | 0,69743 | 0,8533426 | ciliary neurotrophic factor receptor |
| ENSG00000266805 | AP005432,1 | -0,137473911 | 0,909109569 | 0,77027 | 0,89337052 | novel transcript, sense intronic to RALBP1 |
| ENSG00000229046 | HMGN1P2 | -0,137572156 | 0,909047663 | 0,84113 | 0,92769747 | high mobility group nucleosome binding domain 1 pseudogene 2 |
| ENSG00000171044 | XKR6 | -0,137681108 | 0,908979014 | 0,65283 | 0,82611554 | XK related 6 |
| ENSG00000169084 | DHRSX | -0,137694378 | 0,908970654 | 0,54162 | 0,75882662 | dehydrogenase/reductase X-linked |
| ENSG00000244161 | FLNB-AS1 | -0,137703156 | 0,908965123 | 0,73981 | 0,87659348 | FLNB antisense RNA 1 |

| | | | | | | |
|---|---|---|---|---|---|---|
| ENSG00000272425 | AC009902,3 | -0,137812926 | 0,908895965 | 0,8427 | 0,92821072 | novel transcript |
| ENSG00000234995 | RPS3AP28 | -0,137904041 | 0,908838564 | 0,84296 | 0,9282764 | RPS3A pseudogene 28 |
| ENSG00000171763 | SPATA5L1 | -0,138091526 | 0,908720464 | 0,59204 | 0,79093233 | spermatogenesis associated 5 like 1 |
| ENSG00000105723 | GSK3A | -0,13813978 | 0,908690071 | 0,48484 | 0,72052946 | glycogen synthase kinase 3 alpha |
| ENSG00000110696 | C11orf58 | -0,13823181 | 0,908632107 | 0,58693 | 0,7870367 | chromosome 11 open reading frame 58 |
| ENSG00000227542 | AC092614,1 | -0,138288036 | 0,908596696 | 0,63593 | 0,81684802 | novel transcript |
| ENSG00000153060 | TEKT5 | -0,138330559 | 0,908569916 | 0,86556 | 0,94029773 | tektin 5 |
| ENSG00000250049 | AC114316,1 | -0,138330692 | 0,908569832 | 0,82537 | 0,91911153 | novel transcript |
| ENSG00000173674 | EIF1AX | -0,138352439 | 0,908556136 | 0,73739 | 0,87539732 | eukaryotic translation initiation factor 1A X-linked |
| ENSG00000189241 | TSPYL1 | -0,138458981 | 0,908489042 | 0,56674 | 0,77469399 | TSPY like 1 |
| ENSG00000264176 | MAGOH2P | -0,138465681 | 0,908484823 | 0,86217 | 0,93843207 | mago homolog 2, pseudogene |
| ENSG00000282689 | ULK4P2 | -0,138486147 | 0,908471936 | 0,87252 | 0,94374455 | ULK4 pseudogene 2 |
| ENSG00000260128 | ULK4P2 | -0,138486147 | 0,908471936 | 0,87252 | 0,94374455 | ULK4 pseudogene 2 |
| ENSG00000203739 | PRDX6-AS1 | -0,138571656 | 0,908418092 | 0,70812 | 0,85927396 | PRDX6 antisense RNA 1 |
| ENSG00000005981 | ASB4 | -0,138616142 | 0,908390081 | 0,80708 | 0,91125307 | ankyrin repeat and SOCS box containing 4 |
| ENSG00000276728 | AC142472,1 | -0,138870812 | 0,908229743 | 0,53913 | 0,75757364 | novel transcript |
| ENSG00000258230 | AC063950,1 | -0,138998954 | 0,908149076 | 0,87244 | 0,94374455 | suppression of tumorigenicity 13 (colon carcinoma) (Hsp70 interacting protein) (ST13) pseudogene |
| ENSG00000171612 | SLC25A33 | -0,139014862 | 0,908139062 | 0,65695 | 0,82856771 | solute carrier family 25 member 33 |
| ENSG00000079263 | SP140 | -0,139224435 | 0,908007151 | 0,86692 | 0,94112274 | SP140 nuclear body protein |
| ENSG00000181264 | TLCD5 | -0,139247864 | 0,907992406 | 0,60417 | 0,79780609 | TLC domain containing 5 |
| ENSG00000163885 | CFAP100 | -0,139262241 | 0,907983357 | 0,74831 | 0,88150078 | cilia and flagella associated protein 100 |
| ENSG00000160293 | VAV2 | -0,139280656 | 0,907971767 | 0,46724 | 0,70721963 | vav guanine nucleotide exchange factor 2 |
| ENSG00000286480 | AC009501,2 | -0,139688899 | 0,907714872 | 0,79733 | 0,90661857 | novel transcript, antisense to WDPCP |
| ENSG00000213368 | ST13P11 | -0,139702936 | 0,907706041 | 0,85621 | 0,9349542 | ST13, Hsp70 interacting protein pseudogene 11 |
| ENSG00000272465 | AL031768,1 | -0,139762581 | 0,907668515 | 0,84743 | 0,93071557 | novel transcript |
| ENSG00000104660 | LEPROTL1 | -0,139784035 | 0,907655017 | 0,77759 | 0,89712515 | leptin receptor overlapping transcript like 1 |
| ENSG00000154545 | MAGED4 | -0,139785579 | 0,907654046 | 0,53114 | 0,75245598 | MAGE family member D4 |
| ENSG00000234810 | AL603840,1 | -0,139804428 | 0,907642187 | 0,57533 | 0,7801789 | novel transcript |
| ENSG00000225518 | LINC01703 | -0,139866148 | 0,907603358 | 0,83406 | 0,92405734 | long intergenic non-protein coding RNA 1703 |
| ENSG00000070159 | PTPN3 | -0,140112822 | 0,907448188 | 0,65633 | 0,82832937 | protein tyrosine phosphatase non-receptor type 3 |
| ENSG00000104067 | TJP1 | -0,14021731 | 0,907382468 | 0,62824 | 0,81179986 | tight junction protein 1 |
| ENSG00000176641 | RNF152 | -0,140255127 | 0,907358683 | 0,64332 | 0,82100089 | ring finger protein 152 |

| | | | | | | |
|---|---|---|---|---|---|---|
| ENSG00000276763 | AC069499,2 | -0,140267494 | 0,907350906 | 0,78398 | 0,89941845 | ribosomal protein L10 (RPL10) pseudogene |
| ENSG00000176230 | OR4K17 | -0,140314358 | 0,907321432 | 0,56096 | 0,7712266 | olfactory receptor family 4 subfamily K member 17 |
| ENSG00000164494 | PDSS2 | -0,140474403 | 0,907220784 | 0,58111 | 0,78406473 | decaprenyl diphosphate synthase subunit 2 |
| ENSG00000186056 | MATN1-AS1 | -0,140490325 | 0,907210771 | 0,80794 | 0,91168773 | MATN1 antisense RNA 1 |
| ENSG00000268107 | AC003005,1 | -0,140522751 | 0,907190381 | 0,82322 | 0,91829029 | novel transcript |
| ENSG00000250635 | CXXC5-AS1 | -0,140666043 | 0,907100281 | 0,81988 | 0,91720659 | CXXC5 antisense RNA 1 |
| ENSG00000232928 | DDX3P1 | -0,140789249 | 0,907022819 | 0,76551 | 0,8908718 | DEAD-box helicase 3 pseudogene 1 |
| ENSG00000238103 | RPL9P7 | -0,140792854 | 0,907020552 | 0,90091 | 0,95787401 | ribosomal protein L9 pseudogene 7 |
| ENSG00000275885 | PRPF31 | -0,140816513 | 0,907005678 | 0,70192 | 0,85640523 | pre-mRNA processing factor 31 |
| ENSG00000122378 | PRXL2A | -0,140891626 | 0,906958456 | 0,49289 | 0,72602358 | peroxiredoxin like 2A |
| ENSG00000276709 | AC245049,1 | -0,140958658 | 0,906916317 | 0,46239 | 0,70356854 | Pyridoxal-dependent decarboxylase domain-containing protein 1 |
| ENSG00000132837 | DMGDH | -0,14096717 | 0,906910967 | 0,7515 | 0,88347546 | dimethylglycine dehydrogenase |
| ENSG00000137496 | IL18BP | -0,141039118 | 0,906865739 | 0,54723 | 0,76270053 | interleukin 18 binding protein |
| ENSG00000204394 | VARS1 | -0,141054191 | 0,906856265 | 0,58557 | 0,78640335 | valyl-tRNA synthetase 1 |
| ENSG00000189306 | RRP7A | -0,141110331 | 0,906820976 | 0,61541 | 0,80439941 | ribosomal RNA processing 7 homolog A |
| ENSG00000104228 | TRIM35 | -0,141115513 | 0,906817719 | 0,45976 | 0,70184297 | tripartite motif containing 35 |
| ENSG00000175866 | BAIAP2 | -0,141181911 | 0,906775985 | 0,46063 | 0,7024702 | BAR/IMD domain containing adaptor protein 2 |
| ENSG00000113361 | CDH6 | -0,14126212 | 0,906725573 | 0,77667 | 0,89660077 | cadherin 6 |
| ENSG00000100124 | ANKRD54 | -0,141286415 | 0,906710304 | 0,49015 | 0,72437872 | ankyrin repeat domain 54 |
| ENSG00000112514 | CUTA | -0,141336745 | 0,906678673 | 0,63954 | 0,81913513 | cutA divalent cation tolerance homolog |
| ENSG00000146576 | C7orf26 | -0,141339386 | 0,906677013 | 0,4643 | 0,70493977 | chromosome 7 open reading frame 26 |
| ENSG00000109445 | ZNF330 | -0,14138026 | 0,906651326 | 0,54364 | 0,76018442 | zinc finger protein 330 |
| ENSG00000198055 | GRK6 | -0,141380266 | 0,906651322 | 0,48602 | 0,72154217 | G protein-coupled receptor kinase 6 |
| ENSG00000141665 | FBXO15 | -0,141535033 | 0,906554065 | 0,53082 | 0,75219176 | F-box protein 15 |
| ENSG00000180628 | PCGF5 | -0,141594656 | 0,9065166 | 0,68917 | 0,8485712 | polycomb group ring finger 5 |
| ENSG00000267274 | AC008770,3 | -0,141856749 | 0,906351929 | 0,73719 | 0,87539732 | novel transcript, sense overlapping with ZNF700 |
| ENSG00000184465 | WDR27 | -0,14192428 | 0,906309505 | 0,53446 | 0,75405583 | WD repeat domain 27 |
| ENSG00000187605 | TET3 | -0,141945548 | 0,906296144 | 0,5856 | 0,78640335 | tet methylcytosine dioxygenase 3 |
| ENSG00000247596 | TWF2 | -0,142020437 | 0,9062491 | 0,56588 | 0,77410921 | twinfilin actin binding protein 2 |
| ENSG00000204084 | INPP5B | -0,142048543 | 0,906231446 | 0,62843 | 0,81201392 | inositol polyphosphate-5-phosphatase B |
| ENSG00000174453 | VWC2L | -0,142052746 | 0,906228805 | 0,79526 | 0,90519396 | von Willebrand factor C domain containing 2 like |
| ENSG00000170260 | ZNF212 | -0,142052935 | 0,906228686 | 0,48057 | 0,71766872 | zinc finger protein 212 |

| | | | | | | |
|---|---|---|---|---|---|---|
| ENSG00000105254 | TBCB | -0,142107874 | 0,906194177 | 0,5212 | 0,74558236 | tubulin folding cofactor B |
| ENSG00000234367 | PFN1P3 | -0,142341809 | 0,906047249 | 0,87352 | 0,94398015 | profilin 1 pseudogene 3 |
| ENSG00000272572 | AL138762,1 | -0,142372728 | 0,906027831 | 0,77425 | 0,89572556 | novel transcript, antisense to FAM178A |
| ENSG00000175265 | GOLGA8A | -0,142442772 | 0,905983844 | 0,56146 | 0,77136826 | golgin A8 family member A |
| ENSG00000131697 | NPHP4 | -0,142506481 | 0,905943836 | 0,49209 | 0,72541367 | nephrocystin 4 |
| ENSG00000266962 | AC067852,2 | -0,142530633 | 0,90592867 | 0,4821 | 0,71880798 | novel transcript, antisense to HSD17B1 |
| ENSG00000267904 | AC024075,1 | -0,142559887 | 0,9059103 | 0,70535 | 0,85804888 | novel transcript, sense intronic C19orf42 |
| ENSG00000198198 | SZT2 | -0,14258637 | 0,905893671 | 0,37862 | 0,64679625 | SZT2 subunit of KICSTOR complex |
| ENSG00000102858 | MGRN1 | -0,142641199 | 0,905859244 | 0,45802 | 0,7010995 | mahogunin ring finger 1 |
| ENSG00000064607 | SUGP2 | -0,142745397 | 0,905793821 | 0,29677 | 0,58995143 | SURP and G-patch domain containing 2 |
| ENSG00000243004 | AC005062,1 | -0,142803385 | 0,905757414 | 0,85641 | 0,93509883 | novel transcript, sense overlapping to MACC1 |
| ENSG00000185252 | ZNF74 | -0,142805907 | 0,905755831 | 0,40304 | 0,66399303 | zinc finger protein 74 |
| ENSG00000105221 | AKT2 | -0,142916605 | 0,905686335 | 0,41526 | 0,67212646 | AKT serine/threonine kinase 2 |
| ENSG00000197355 | UAP1L1 | -0,142956017 | 0,905661593 | 0,86486 | 0,93993509 | UDP-N-acetylglucosamine pyrophosphorylase 1 like 1 |
| ENSG00000080224 | EPHA6 | -0,14310066 | 0,905570797 | 0,61936 | 0,80705741 | EPH receptor A6 |
| ENSG00000253372 | AC016405,1 | -0,143105274 | 0,905567901 | 0,83853 | 0,92627183 | novel transcript |
| ENSG00000231464 | AC024937,2 | -0,143125319 | 0,905555319 | 0,89678 | 0,95548853 | protein phosphatase 4, regulatory subunit 2 (PPP4R2) pseudogene |
| ENSG00000231784 | DBIL5P | -0,143326506 | 0,905429046 | 0,61173 | 0,80334296 | diazepam binding inhibitor-like 5, pseudogene |
| ENSG00000213950 | RPS10P2 | -0,143402668 | 0,905381249 | 0,87818 | 0,9458146 | ribosomal protein S10 pseudogene 2 |
| ENSG00000131437 | KIF3A | -0,143411521 | 0,905375693 | 0,43537 | 0,68601186 | kinesin family member 3A |
| ENSG00000175550 | DRAP1 | -0,143472556 | 0,90533739 | 0,45043 | 0,69580372 | DR1 associated protein 1 |
| ENSG00000237750 | KCNH7-AS1 | -0,143482163 | 0,905331362 | 0,82496 | 0,91900665 | KCNH7 antisense RNA 1 |
| ENSG00000239282 | CASTOR1 | -0,143563579 | 0,905280272 | 0,69675 | 0,85309963 | cytosolic arginine sensor for mTORC1 subunit 1 |
| ENSG00000107014 | RLN2 | -0,1436169 | 0,905246814 | 0,88422 | 0,9485968 | relaxin 2 |
| ENSG00000185187 | SIGIRR | -0,143635871 | 0,905234911 | 0,59928 | 0,79538547 | single Ig and TIR domain containing |
| ENSG00000180385 | EMC3-AS1 | -0,143655745 | 0,905222441 | 0,55695 | 0,76879045 | EMC3 antisense RNA 1 |
| ENSG00000272620 | AFAP1-AS1 | -0,143660668 | 0,905219352 | 0,66225 | 0,83183445 | AFAP1 antisense RNA 1 |
| ENSG00000105792 | CFAP69 | -0,14378498 | 0,905141355 | 0,60971 | 0,80189841 | cilia and flagella associated protein 69 |
| ENSG00000123240 | OPTN | -0,143855048 | 0,905097396 | 0,65147 | 0,82540452 | optineurin |
| ENSG00000275326 | NOSTRIN | -0,143864394 | 0,905091533 | 0,68148 | 0,84375324 | nitric oxide synthase trafficking |
| ENSG00000074755 | ZZEF1 | -0,143975122 | 0,905022069 | 0,58938 | 0,78883783 | zinc finger ZZ-type and EF-hand domain containing 1 |
| ENSG00000088888 | MAVS | -0,144152201 | 0,904910992 | 0,6213 | 0,80787022 | mitochondrial antiviral signaling protein |

| | | | | | | |
|---|---|---|---|---|---|---|
| ENSG00000175691 | ZNF77 | -0,144190116 | 0,904887211 | 0,5382 | 0,75658794 | zinc finger protein 77 |
| ENSG00000231688 | RPL21P43 | -0,1442266 | 0,904864327 | 0,82962 | 0,92174649 | ribosomal protein L21 pseudogene 43 |
| ENSG00000180834 | MAP6D1 | -0,144299809 | 0,904818411 | 0,71742 | 0,86483656 | MAP6 domain containing 1 |
| ENSG00000130414 | NDUFA10 | -0,144337116 | 0,904795014 | 0,40353 | 0,66399439 | NADH:ubiquinone oxidoreductase subunit A10 |
| ENSG00000147166 | ITGB1BP2 | -0,144337948 | 0,904794492 | 0,60025 | 0,79579686 | integrin subunit beta 1 binding protein 2 |
| ENSG00000261094 | AC007066,2 | -0,144345194 | 0,904789947 | 0,60386 | 0,79770754 | novel transcript |
| ENSG00000149289 | ZC3H12C | -0,144400216 | 0,904755441 | 0,72248 | 0,86734497 | zinc finger CCCH-type containing 12C |
| ENSG00000119383 | PTPA | -0,144421183 | 0,904742292 | 0,49324 | 0,72631 | protein phosphatase 2 phosphatase activator |
| ENSG00000284294 | AC007326,5 | -0,14454021 | 0,904667651 | 0,49306 | 0,72615908 | novel transcript |
| ENSG00000005238 | FAM214B | -0,144646157 | 0,904601217 | 0,51644 | 0,74198903 | family with sequence similarity 214 member B |
| ENSG00000138375 | SMARCAL1 | -0,144672087 | 0,904584959 | 0,53949 | 0,75774769 | SWI/SNF related, matrix associated, actin dependent regulator of chromatin, subfamily a like 1 |
| ENSG00000255320 | AP000759,1 | -0,144691173 | 0,904572992 | 0,86934 | 0,94257563 | novel transcript |
| ENSG00000225972 | MTND1P23 | -0,144807576 | 0,90450001 | 0,72481 | 0,86866311 | MT-ND1 pseudogene 23 |
| ENSG00000269895 | AP000654,1 | -0,144990399 | 0,904385396 | 0,75433 | 0,88487815 | novel transcript |
| ENSG00000189159 | JPT1 | -0,145006843 | 0,904375088 | 0,42334 | 0,67712191 | Jupiter microtubule associated homolog 1 |
| ENSG00000249286 | AMD1P3 | -0,145316026 | 0,904181293 | 0,81789 | 0,9163876 | adenosylmethionine decarboxylase 1 pseudogene 3 |
| ENSG00000225329 | LHFPL3-AS2 | -0,145349727 | 0,904160171 | 0,83718 | 0,92549623 | LHFPL3 antisense RNA 2 |
| ENSG00000164291 | ARSK | -0,145425294 | 0,904112814 | 0,52321 | 0,74726158 | arylsulfatase family member K |
| ENSG00000135506 | OS9 | -0,145457505 | 0,904092628 | 0,54705 | 0,76263922 | OS9 endoplasmic reticulum lectin |
| ENSG00000273574 | LENG9 | -0,145489744 | 0,904072425 | 0,88416 | 0,9485968 | leukocyte receptor cluster member 9 |
| ENSG00000173542 | MOB1B | -0,145507324 | 0,904061409 | 0,65157 | 0,82541975 | MOB kinase activator 1B |
| ENSG00000239412 | RPL21P71 | -0,145577815 | 0,904017236 | 0,86503 | 0,93999031 | ribosomal protein L21 pseudogene 71 |
| ENSG00000129151 | BBOX1 | -0,145637577 | 0,903979789 | 0,8457 | 0,92981009 | gamma-butyrobetaine hydroxylase 1 |
| ENSG00000234076 | TPRG1-AS1 | -0,145656125 | 0,903968167 | 0,92161 | 0,96607272 | TPRG1 antisense RNA 1 |
| ENSG00000100413 | POLR3H | -0,145672765 | 0,903957741 | 0,49998 | 0,73071355 | RNA polymerase III subunit H |
| ENSG00000272656 | AC024933,1 | -0,145684688 | 0,90395027 | 0,89453 | 0,95403341 | novel transcript, antisense to MRPS22 |
| ENSG00000104903 | LYL1 | -0,145725609 | 0,903924631 | 0,76304 | 0,88980026 | LYL1 basic helix-loop-helix family member |
| ENSG00000172757 | CFL1 | -0,145729679 | 0,903922081 | 0,43917 | 0,68811742 | cofilin 1 |
| ENSG00000278175 | GLIDR | -0,145772342 | 0,903895351 | 0,53368 | 0,75366502 | glioblastoma down-regulated RNA |
| ENSG00000125457 | MIF4GD | -0,14599367 | 0,903756692 | 0,5321 | 0,7528508 | MIF4G domain containing |
| ENSG00000155530 | LRGUK | -0,146003059 | 0,903750811 | 0,68098 | 0,84352125 | leucine rich repeats and guanylate kinase domain containing |

| | | | | | | |
|---|---|---|---|---|---|---|
| ENSG00000168778 | TCTN2 | -0,146107503 | 0,903685386 | 0,55892 | 0,77025265 | tectonic family member 2 |
| ENSG00000115947 | ORC4 | -0,146450634 | 0,903470478 | 0,38808 | 0,65291326 | origin recognition complex subunit 4 |
| ENSG00000232472 | EEF1B2P3 | -0,146524957 | 0,903423936 | 0,71797 | 0,86489865 | eukaryotic translation elongation factor 1 beta 2 pseudogene 3 |
| ENSG00000256139 | AC007637,1 | -0,146580664 | 0,903389053 | 0,75835 | 0,88727479 | novel transcript |
| ENSG00000105426 | PTPRS | -0,14674372 | 0,903286956 | 0,3583 | 0,63185282 | protein tyrosine phosphatase receptor type S |
| ENSG00000259657 | PIGHP1 | -0,146755762 | 0,903279416 | 0,80712 | 0,91125307 | phosphatidylinositol glycan anchor biosynthesis class H pseudogene 1 |
| ENSG00000147400 | CETN2 | -0,146772713 | 0,903268803 | 0,45386 | 0,6979134 | centrin 2 |
| ENSG00000288031 | AC093838,2 | -0,146791684 | 0,903256925 | 0,77653 | 0,89657313 | novel pseudogene |
| ENSG00000278383 | AL031673,1 | -0,146861838 | 0,903213004 | 0,76967 | 0,89301171 | novel transcript, antisense to ZNF337 |
| ENSG00000130513 | GDF15 | -0,146943345 | 0,903161977 | 0,8324 | 0,92331399 | growth differentiation factor 15 |
| ENSG00000277399 | GPR179 | -0,146967947 | 0,903146576 | 0,72149 | 0,86698039 | G protein-coupled receptor 179 |
| ENSG00000262671 | AC004918,1 | -0,146996789 | 0,90312852 | 0,78371 | 0,89918294 | novel transcript, antisense to AGK |
| ENSG00000244701 | AC004918,1 | -0,146996789 | 0,90312852 | 0,78371 | 0,89918294 | novel transcript, antisense to AGK |
| ENSG00000227896 | AL731569,1 | -0,14702121 | 0,903113233 | 0,55224 | 0,76568017 | novel transcript |
| ENSG00000288427 | CLSTN3 | -0,147032589 | 0,903106109 | 0,57674 | 0,7810482 | calsyntenin 3 |
| ENSG00000139182 | CLSTN3 | -0,147032589 | 0,903106109 | 0,57674 | 0,7810482 | calsyntenin 3 |
| ENSG00000123191 | ATP7B | -0,14706299 | 0,903087079 | 0,44134 | 0,68969561 | ATPase copper transporting beta |
| ENSG00000259345 | AC013652,1 | -0,147098072 | 0,903065119 | 0,69489 | 0,85206252 | novel transcript |
| ENSG00000248213 | CICP16 | -0,147201966 | 0,903000088 | 0,67568 | 0,84047234 | capicua transcriptional repressor pseudogene 16 |
| ENSG00000275131 | AC241952,1 | -0,147300437 | 0,902938456 | 0,59137 | 0,79058842 | phosphodiesterase 4D interacting protein (PDE4DIP) pseudogene |
| ENSG00000021488 | SLC7A9 | -0,147351643 | 0,902906409 | 0,76901 | 0,89287402 | solute carrier family 7 member 9 |
| ENSG00000204946 | ZNF783 | -0,147379579 | 0,902888925 | 0,62486 | 0,81001052 | zinc finger family member 783 |
| ENSG00000172732 | MUS81 | -0,147447834 | 0,90284621 | 0,43506 | 0,68577552 | MUS81 structure-specific endonuclease subunit |
| ENSG00000284845 | AL136295,10 | -0,14753808 | 0,902789735 | 0,68052 | 0,84316527 | novel transcript |
| ENSG00000259529 | AL136295,5 | -0,14753808 | 0,902789735 | 0,68052 | 0,84316527 | novel transcript |
| ENSG00000102981 | PARD6A | -0,147585294 | 0,902760191 | 0,63366 | 0,81549489 | par-6 family cell polarity regulator alpha |
| ENSG00000240616 | RPS6P25 | -0,147602924 | 0,902749159 | 0,77273 | 0,89496764 | ribosomal protein S6 pseudogene 25 |
| ENSG00000280852 | AC025048,6 | -0,147619929 | 0,902738518 | 0,77432 | 0,89573411 | adaptor-related protein complex 1, sigma 2 subunit (AP1S2) pseudogene |
| ENSG00000267080 | ASB16-AS1 | -0,147635853 | 0,902728554 | 0,57926 | 0,78270914 | ASB16 antisense RNA 1 |

| | | | | | | |
|---|---|---|---|---|---|---|
| ENSG00000267649 | AC010327,5 | -0,147724252 | 0,902673243 | 0,75625 | 0,88593758 | novel transcript |
| ENSG00000114670 | NEK11 | -0,147792696 | 0,902630419 | 0,57769 | 0,7815762 | NIMA related kinase 11 |
| ENSG00000183166 | CALN1 | -0,147822845 | 0,902611557 | 0,82542 | 0,91911153 | calneuron 1 |
| ENSG00000254865 | AC100763,1 | -0,147892383 | 0,902568052 | 0,82004 | 0,91723726 | novel transcript |
| ENSG00000260253 | AC084262,1 | -0,147941772 | 0,902537154 | 0,80737 | 0,91131389 | novel transcript, overlapping to DUSP4 |
| ENSG00000143194 | MAEL | -0,148057599 | 0,902464696 | 0,78642 | 0,90080813 | maelstrom spermatogenic transposon silencer |
| ENSG00000120318 | ARAP3 | -0,14809746 | 0,902439762 | 0,53186 | 0,75280487 | ArfGAP with RhoGAP domain, ankyrin repeat and PH domain 3 |
| ENSG00000168765 | GSTM4 | -0,148178804 | 0,902388881 | 0,39093 | 0,65464997 | glutathione S-transferase mu 4 |
| ENSG00000228686 | AL590723,1 | -0,148233832 | 0,902354462 | 0,80583 | 0,91077382 | novel transcript |
| ENSG00000242294 | STAG3L5P | -0,148236873 | 0,90235256 | 0,38842 | 0,6529693 | stromal antigen 3-like 5 pseudogene |
| ENSG00000088543 | C3orf18 | -0,148357483 | 0,902277126 | 0,53378 | 0,75368609 | chromosome 3 open reading frame 18 |
| ENSG00000286705 | AC079140,6 | -0,148421261 | 0,902237239 | 0,68431 | 0,84550748 | novel transcript |
| ENSG00000220205 | VAMP2 | -0,148487942 | 0,902195539 | 0,39434 | 0,65725123 | vesicle associated membrane protein 2 |
| ENSG00000198734 | F5 | -0,148544368 | 0,902160254 | 0,65524 | 0,82770864 | coagulation factor V |
| ENSG00000160959 | LRRC14 | -0,148546498 | 0,902158922 | 0,40732 | 0,66661761 | leucine rich repeat containing 14 |
| ENSG00000126950 | TMEM35A | -0,148563592 | 0,902148232 | 0,63316 | 0,81538829 | transmembrane protein 35A |
| ENSG00000226674 | TEX41 | -0,148610876 | 0,902118665 | 0,84353 | 0,92855259 | testis expressed 41 |
| ENSG00000250731 | TPM3P6 | -0,148740958 | 0,902037328 | 0,82085 | 0,91738668 | tropomyosin 3 pseudogene 6 |
| ENSG00000260368 | AC027373,1 | -0,148749633 | 0,902031905 | 0,68138 | 0,84375324 | novel transcript, overlapping YWHAZ |
| ENSG00000224320 | HLA-A | -0,148831338 | 0,901980821 | 0,45929 | 0,7017037 | major histocompatibility complex, class I, A |
| ENSG00000198933 | TBKBP1 | -0,149100586 | 0,901812501 | 0,63055 | 0,81354995 | TBK1 binding protein 1 |
| ENSG00000283649 | ZDBF2 | -0,149102667 | 0,9018112 | 0,67258 | 0,83803462 | zinc finger DBF-type containing 2 |
| ENSG00000283000 | LINC02635 | -0,149152762 | 0,901779887 | 0,8423 | 0,92821072 | long intergenic non-protein coding RNA 2635 |
| ENSG00000198354 | DCAF12L2 | -0,149239865 | 0,901725443 | 0,59463 | 0,79284456 | DDB1 and CUL4 associated factor 12 like 2 |
| ENSG00000233924 | RPSAP13 | -0,14928101 | 0,901699727 | 0,79366 | 0,90402496 | ribosomal protein SA pseudogene 13 |
| ENSG00000269469 | AC010619,1 | -0,14935963 | 0,90165059 | 0,87437 | 0,9443876 | novel protein |
| ENSG00000069998 | HDHD5 | -0,149376449 | 0,901640078 | 0,58215 | 0,78454692 | haloacid dehalogenase like hydrolase domain containing 5 |
| ENSG00000229215 | HLA-A | -0,149430673 | 0,901606191 | 0,45688 | 0,7004577 | major histocompatibility complex, class I, A |
| ENSG00000169592 | INO80E | -0,149489755 | 0,901569269 | 0,44885 | 0,69459126 | INO80 complex subunit E |
| ENSG00000261120 | AL133299,1 | -0,149495274 | 0,90156582 | 0,80133 | 0,90860375 | novel transcript, antisense to LRRC9 |
| ENSG00000176845 | METRNL | -0,149566327 | 0,901521419 | 0,72926 | 0,87157404 | meteorin like, glial cell differentiation regulator |
| ENSG00000074410 | CA12 | -0,149589874 | 0,901506705 | 0,75479 | 0,88512083 | carbonic anhydrase 12 |

| | | | | | | |
|---|---|---|---|---|---|---|
| ENSG00000183778 | B3GALT5 | -0,149609045 | 0,901494725 | 0,50338 | 0,73266852 | beta-1,3-galactosyltransferase 5 |
| ENSG00000245614 | DDX11-AS1 | -0,149791379 | 0,901380798 | 0,54018 | 0,75814808 | DDX11 antisense RNA 1 |
| ENSG00000180011 | ZADH2 | -0,150028179 | 0,90123286 | 0,52625 | 0,74972979 | zinc binding alcohol dehydrogenase domain containing 2 |
| ENSG00000147872 | PLIN2 | -0,150081471 | 0,901199569 | 0,6581 | 0,82897081 | perilipin 2 |
| ENSG00000118785 | SPP1 | -0,150081479 | 0,901199564 | 0,8264 | 0,91961778 | secreted phosphoprotein 1 |
| ENSG00000143355 | LHX9 | -0,15017136 | 0,901143421 | 0,70146 | 0,8560724 | LIM homeobox 9 |
| ENSG00000253598 | SLC10A5 | -0,150416546 | 0,900990284 | 0,69311 | 0,85087762 | solute carrier family 10 member 5 |
| ENSG00000234171 | RNASEH1-AS1 | -0,150418632 | 0,900988981 | 0,47613 | 0,71446062 | RNASEH1 antisense RNA 1 |
| ENSG00000078808 | SDF4 | -0,150456157 | 0,900965547 | 0,57519 | 0,7801176 | stromal cell derived factor 4 |
| ENSG00000276997 | AL513314,2 | -0,15050606 | 0,900934383 | 0,60321 | 0,79730504 | novel transcript |
| ENSG00000275575 | PTP4A3 | -0,150546199 | 0,900909317 | 0,5236 | 0,74747909 | protein tyrosine phosphatase 4A3 |
| ENSG00000162191 | UBXN1 | -0,150640888 | 0,900850189 | 0,50772 | 0,73588248 | UBX domain protein 1 |
| ENSG00000069966 | GNB5 | -0,150663439 | 0,900836108 | 0,58665 | 0,78690792 | G protein subunit beta 5 |
| ENSG00000103855 | CD276 | -0,150752984 | 0,900780197 | 0,56111 | 0,77125979 | CD276 molecule |
| ENSG00000150990 | DHX37 | -0,151035514 | 0,90060381 | 0,57196 | 0,77838663 | DEAH-box helicase 37 |
| ENSG00000228875 | CSNK2B | -0,151099527 | 0,900563851 | 0,74887 | 0,88189246 | casein kinase 2 beta |
| ENSG00000166188 | ZNF319 | -0,15113067 | 0,900544411 | 0,56463 | 0,77306193 | zinc finger protein 319 |
| ENSG00000198673 | TAFA2 | -0,151166211 | 0,900522226 | 0,71063 | 0,86066702 | TAFA chemokine like family member 2 |
| ENSG00000271963 | AC026786,2 | -0,151209172 | 0,90049541 | 0,88581 | 0,94944442 | novel transcript, antisense to ZNF891 |
| ENSG00000273598 | AC211486,3 | -0,151429461 | 0,900357922 | 0,53352 | 0,75351438 | postmeiotic segregation increased 2 pseudogene 6 |
| ENSG00000248249 | AC110794,1 | -0,1516375 | 0,900228098 | 0,84323 | 0,92847654 | Rac GTPase activating protein 1 (RACGAP1) pseudogene |
| ENSG00000285501 | AGBL2 | -0,151684834 | 0,900198563 | 0,81007 | 0,91245196 | ATP/GTP binding protein like 2 |
| ENSG00000165923 | AGBL2 | -0,151684834 | 0,900198563 | 0,81007 | 0,91245196 | ATP/GTP binding protein like 2 |
| ENSG00000136014 | USP44 | -0,151701616 | 0,900188091 | 0,73568 | 0,8750728 | ubiquitin specific peptidase 44 |
| ENSG00000213433 | RPLP1P6 | -0,151724405 | 0,900173872 | 0,79524 | 0,90519396 | ribosomal protein lateral stalk subunit P1 pseudogene 6 |
| ENSG00000281015 | LINC01596 | -0,151748207 | 0,900159021 | 0,8881 | 0,95069428 | long intergenic non-protein coding RNA 1596 |
| ENSG00000277561 | GOLGA8IP | -0,151838393 | 0,900102751 | 0,7559 | 0,8857133 | golgin A8 family member I, pseudogene |
| ENSG00000215187 | FAM166B | -0,152071181 | 0,899957526 | 0,78059 | 0,8984342 | family with sequence similarity 166 member B |
| ENSG00000250510 | GPR162 | -0,152250665 | 0,89984557 | 0,58836 | 0,78811087 | G protein-coupled receptor 162 |
| ENSG00000227248 | FAM155A-IT1 | -0,152279241 | 0,899827746 | 0,73397 | 0,87400155 | FAM155A intronic transcript 1 |
| ENSG00000275345 | AC091167,5 | -0,152318944 | 0,899802984 | 0,81172 | 0,91334888 | novel transcript |
| ENSG00000271828 | AC008937,3 | -0,152448944 | 0,899721907 | 0,6132 | 0,8035281 | novel transcript, antisense to MIER3 |
| ENSG00000265750 | AC090772,3 | -0,152470211 | 0,899708644 | 0,88736 | 0,95041376 | novel transcript |

| | | | | | | |
|---|---|---|---|---|---|---|
| ENSG00000276662 | AL713922,2 | -0,152579363 | 0,899640576 | 0,77654 | 0,89657313 | DEAD (Asp-Glu-Ala-Asp) box polypeptide 20 (DDX20) pseudogene |
| ENSG00000166278 | C2 | -0,152593904 | 0,899631509 | 0,9151 | 0,96323337 | complement C2 |
| ENSG00000139291 | TMEM19 | -0,152602518 | 0,899626137 | 0,52204 | 0,7463592 | transmembrane protein 19 |
| ENSG00000241743 | XACT | -0,152790448 | 0,899508957 | 0,74321 | 0,87840582 | X active specific transcript |
| ENSG00000139636 | LMBR1L | -0,152829696 | 0,899484486 | 0,42598 | 0,67940091 | limb development membrane protein 1 like |
| ENSG00000102781 | KATNAL1 | -0,152964676 | 0,899400333 | 0,48931 | 0,7238264 | katanin catalytic subunit A1 like 1 |
| ENSG00000132604 | TERF2 | -0,153023341 | 0,899363761 | 0,4783 | 0,71602854 | telomeric repeat binding factor 2 |
| ENSG00000197980 | LEKR1 | -0,153121012 | 0,899302876 | 0,72835 | 0,87088765 | leucine, glutamate and lysine rich 1 |
| ENSG00000258561 | AL359232,1 | -0,153227208 | 0,899236682 | 0,7707 | 0,89376602 | novel transcript |
| ENSG00000186281 | GPAT2 | -0,153237566 | 0,899230226 | 0,65605 | 0,82824314 | glycerol-3-phosphate acyltransferase 2, mitochondrial |
| ENSG00000169891 | REPS2 | -0,153303834 | 0,899188921 | 0,79101 | 0,90292452 | RALBP1 associated Eps domain containing 2 |
| ENSG00000166326 | TRIM44 | -0,153364348 | 0,899151206 | 0,30939 | 0,59786185 | tripartite motif containing 44 |
| ENSG00000077782 | FGFR1 | -0,153724006 | 0,898927079 | 0,60684 | 0,79994037 | fibroblast growth factor receptor 1 |
| ENSG00000204624 | DISP3 | -0,153957554 | 0,898781569 | 0,49256 | 0,72575639 | dispatched RND transporter family member 3 |
| ENSG00000244558 | KCNK15-AS1 | -0,153964577 | 0,898777195 | 0,88395 | 0,94847506 | KCNK15 and WISP2 antisense RNA 1 |
| ENSG00000089737 | DDX24 | -0,153979766 | 0,898767732 | 0,57729 | 0,78134056 | DEAD-box helicase 24 |
| ENSG00000177854 | TMEM187 | -0,153983508 | 0,8987654 | 0,63259 | 0,81490582 | transmembrane protein 187 |
| ENSG00000285103 | AL451123,1 | -0,154031132 | 0,898735733 | 0,60043 | 0,79584102 | novel transcript |
| ENSG00000174007 | CEP19 | -0,154155557 | 0,898658225 | 0,56615 | 0,77421829 | centrosomal protein 19 |
| ENSG00000274577 | LHX1 | -0,154428324 | 0,898488333 | 0,76908 | 0,8929156 | LIM homeobox 1 |
| ENSG00000259932 | AC051619,6 | -0,154428514 | 0,898488215 | 0,63786 | 0,81798655 | novel transcript |
| ENSG00000249055 | TBCAP3 | -0,154464695 | 0,898465682 | 0,83802 | 0,92611649 | tubulin folding cofactor A pseudogene 3 |
| ENSG00000108587 | GOSR1 | -0,154476688 | 0,898458214 | 0,52302 | 0,74710918 | golgi SNAP receptor complex member 1 |
| ENSG00000275035 | CCDC92 | -0,154542694 | 0,898417109 | 0,49495 | 0,72730301 | coiled-coil domain containing 92 |
| ENSG00000172123 | SLFN12 | -0,154556003 | 0,89840882 | 0,64067 | 0,81956256 | schlafen family member 12 |
| ENSG00000152430 | BOLL | -0,154563021 | 0,89840445 | 0,86081 | 0,93761195 | boule homolog, RNA binding protein |
| ENSG00000241061 | RPL5P1 | -0,154605844 | 0,898377784 | 0,74705 | 0,88059941 | ribosomal protein L5 pseudogene 1 |
| ENSG00000110047 | EHD1 | -0,154625423 | 0,898365592 | 0,39144 | 0,65510995 | EH domain containing 1 |
| ENSG00000251552 | COQ10BP2 | -0,154664164 | 0,898341468 | 0,87747 | 0,94566578 | coenzyme Q10B pseudogene 2 |
| ENSG00000110881 | ASIC1 | -0,154758202 | 0,898282914 | 0,46319 | 0,70415312 | acid sensing ion channel subunit 1 |
| ENSG00000103832 | GOLGA8UP | -0,1547731 | 0,898273638 | 0,8532 | 0,93330335 | golgin A8 family member U, pseudogene |
| ENSG00000273951 | AL031667,3 | -0,154776319 | 0,898271633 | 0,78768 | 0,90122495 | novel transcript, sense intronic to CHD6 |

| | | | | | | |
|---|---|---|---|---|---|---|
| ENSG00000226508 | LINC01918 | -0,154788628 | 0,89826397 | 0,61729 | 0,8057177 | long intergenic non-protein coding RNA 1918 |
| ENSG00000183780 | SLC35F3 | -0,15480841 | 0,898251653 | 0,70639 | 0,85870246 | solute carrier family 35 member F3 |
| ENSG00000257501 | PAFAH1B2P2 | -0,154900722 | 0,898194179 | 0,77428 | 0,89572556 | PAFAH1B2 pseudogene 2 |
| ENSG00000112293 | GPLD1 | -0,155032812 | 0,898111946 | 0,59903 | 0,79538547 | glycosylphosphatidylinositol specific phospholipase D1 |
| ENSG00000286379 | AL591845,1 | -0,155129926 | 0,898051492 | 0,80573 | 0,91073695 | novel transcript |
| ENSG00000111364 | DDX55 | -0,15513557 | 0,898047979 | 0,53733 | 0,75591439 | DEAD-box helicase 55 |
| ENSG00000154556 | SORBS2 | -0,155208806 | 0,898002393 | 0,64816 | 0,82383528 | sorbin and SH3 domain containing 2 |
| ENSG00000152270 | PDE3B | -0,155281419 | 0,897957196 | 0,51229 | 0,73922987 | phosphodiesterase 3B |
| ENSG00000259335 | HNRNPMP1 | -0,155424624 | 0,897868067 | 0,75411 | 0,88474264 | heterogeneous nuclear ribonucleoprotein M pseudogene 1 |
| ENSG00000254815 | LMNTD2-AS1 | -0,155540318 | 0,897796067 | 0,84338 | 0,92850187 | LMNTD2 antisense RNA 1 |
| ENSG00000240591 | AL096701,3 | -0,15554747 | 0,897791616 | 0,66959 | 0,83623395 | novel transcript |
| ENSG00000266236 | NARF-IT1 | -0,155610641 | 0,897752306 | 0,79671 | 0,90605632 | NARF intronic transcript 1 |
| ENSG00000232265 | LINC02805 | -0,155615782 | 0,897749107 | 0,79802 | 0,90685415 | long intergenic non-protein coding RNA 2805 |
| ENSG00000149182 | ARFGAP2 | -0,155664657 | 0,897718694 | 0,51588 | 0,7419128 | ADP ribosylation factor GTPase activating protein 2 |
| ENSG00000082641 | NFE2L1 | -0,155726106 | 0,897680458 | 0,56789 | 0,77572201 | nuclear factor, erythroid 2 like 1 |
| ENSG00000281165 | SLC2A6 | -0,155769732 | 0,897653314 | 0,56533 | 0,77365864 | solute carrier family 2 member 6 |
| ENSG00000171453 | POLR1C | -0,155840982 | 0,897608982 | 0,48956 | 0,72395897 | RNA polymerase I and III subunit C |
| ENSG00000278637 | H4C1 | -0,155986987 | 0,897518146 | 0,45944 | 0,7017037 | H4 clustered histone 1 |
| ENSG00000248124 | RRN3P1 | -0,156075496 | 0,897463086 | 0,37147 | 0,64063729 | RRN3 pseudogene 1 |
| ENSG00000164484 | TMEM200A | -0,15610079 | 0,897447351 | 0,64386 | 0,82128583 | transmembrane protein 200A |
| ENSG00000170906 | NDUFA3 | -0,156104033 | 0,897445333 | 0,73352 | 0,87368351 | NADH:ubiquinone oxidoreductase subunit A3 |
| ENSG00000130338 | TULP4 | -0,156116077 | 0,897437841 | 0,38452 | 0,65052153 | TUB like protein 4 |
| ENSG00000260093 | AC034111,1 | -0,156120455 | 0,897435118 | 0,73871 | 0,87593493 | novel transcript, antisense to MSRA |
| ENSG00000284966 | AL138689,2 | -0,156257473 | 0,897349889 | 0,73073 | 0,87205836 | novel transcript, antisense to EFNB2 |
| ENSG00000104976 | SNAPC2 | -0,156290462 | 0,89732937 | 0,56165 | 0,77136826 | small nuclear RNA activating complex polypeptide 2 |
| ENSG00000228929 | RPS13P2 | -0,156502356 | 0,897197586 | 0,84672 | 0,930363 | ribosomal protein S13 pseudogene 2 |
| ENSG00000171984 | SHLD1 | -0,156538971 | 0,897174816 | 0,63889 | 0,81888518 | shieldin complex subunit 1 |
| ENSG00000074047 | GLI2 | -0,156565504 | 0,897158316 | 0,51323 | 0,73963507 | GLI family zinc finger 2 |
| ENSG00000100150 | DEPDC5 | -0,156640687 | 0,897111563 | 0,48626 | 0,72178885 | DEP domain containing 5, GATOR1 subcomplex subunit |
| ENSG00000167065 | DUSP18 | -0,156641103 | 0,897111305 | 0,49736 | 0,72869621 | dual specificity phosphatase 18 |
| ENSG00000007372 | PAX6 | -0,156641504 | 0,897111055 | 0,72316 | 0,86784071 | paired box 6 |

| | | | | | | |
|---|---|---|---|---|---|---|
| ENSG00000244471 | AL137058,1 | -0,156683469 | 0,897084961 | 0,78079 | 0,89846287 | transmembrane phosphatase with tensin homology (TPTE) pseudogene |
| ENSG00000113621 | TXNDC15 | -0,156752743 | 0,897041886 | 0,46095 | 0,70273497 | thioredoxin domain containing 15 |
| ENSG00000231257 | ATAT1 | -0,156809086 | 0,897006854 | 0,61776 | 0,80609311 | alpha tubulin acetyltransferase 1 |
| ENSG00000235658 | ATAT1 | -0,156809086 | 0,897006854 | 0,61776 | 0,80609311 | alpha tubulin acetyltransferase 1 |
| ENSG00000265753 | RN7SL444P | -0,156910274 | 0,896943942 | 0,86363 | 0,93904284 | RNA, 7SL, cytoplasmic 444, pseudogene |
| ENSG00000277114 | CNOT3 | -0,156998169 | 0,896889298 | 0,59356 | 0,79186889 | CCR4-NOT transcription complex subunit 3 |
| ENSG00000258644 | SYNJ2BP-COX | -0,15706188 | 0,896849691 | 0,72221 | 0,86734497 | SYNJ2BP-COX16 readthrough |
| ENSG00000123636 | BAZ2B | -0,157262929 | 0,896724718 | 0,34466 | 0,62202834 | bromodomain adjacent to zinc finger domain 2B |
| ENSG00000121542 | SEC22A | -0,157281757 | 0,896713015 | 0,47112 | 0,71051384 | SEC22 homolog A, vesicle trafficking protein |
| ENSG00000251209 | LINC00923 | -0,157370036 | 0,896658147 | 0,3462 | 0,62335555 | long intergenic non-protein coding RNA 923 |
| ENSG00000260484 | AC131902,1 | -0,157420285 | 0,896626917 | 0,84966 | 0,93153852 | novel transcript |
| ENSG00000269069 | AC007842,1 | -0,157438185 | 0,896615792 | 0,58552 | 0,78640335 | protein associated with topoisomerase II homolog 1 (yeast) (PATL1) pseudogene |
| ENSG00000281233 | AC007842,3 | -0,157438185 | 0,896615792 | 0,58552 | 0,78640335 | protein associated with topoisomerase II homolog 1 (yeast) (PATL1) pseudogene |
| ENSG00000100652 | SLC10A1 | -0,157596737 | 0,89651726 | 0,65673 | 0,82855263 | solute carrier family 10 member 1 |
| ENSG00000232027 | AL671986,1 | -0,157682887 | 0,896463726 | 0,82011 | 0,91723726 | proteasome (prosome, macropain) 26S subunit, non-ATPase, 7 (Mov34 homolog) (PSMD7) pseudogene |
| ENSG00000131094 | C1QL1 | -0,157769795 | 0,896409724 | 0,7589 | 0,88743316 | complement C1q like 1 |
| ENSG00000278133 | AC135050,6 | -0,157927637 | 0,896311656 | 0,62195 | 0,80819595 | novel transcript, sense intronic to KAT8 |
| ENSG00000126351 | THRA | -0,157944473 | 0,896301196 | 0,56623 | 0,77421829 | thyroid hormone receptor alpha |
| ENSG00000172037 | LAMB2 | -0,158004901 | 0,896263655 | 0,48698 | 0,72215091 | laminin subunit beta 2 |
| ENSG00000272256 | AC044849,1 | -0,158043018 | 0,896239975 | 0,8964 | 0,95523599 | novel transcript, antisense to TMEM66 |
| ENSG00000260398 | AC068700,1 | -0,158115417 | 0,896195 | 0,70232 | 0,85651838 | novel transcript, overlapping to PKIA |
| ENSG00000235481 | UBE2R2-AS1 | -0,158225846 | 0,896126405 | 0,60744 | 0,80021902 | UBE2R2 antisense RNA 1 |
| ENSG00000197959 | DNM3 | -0,158277986 | 0,896094019 | 0,64376 | 0,82123365 | dynamin 3 |
| ENSG00000168032 | ENTPD3 | -0,158346686 | 0,896051348 | 0,8025 | 0,90903011 | ectonucleoside triphosphate diphosphohydrolase 3 |
| ENSG00000241219 | AC078785,2 | -0,158482596 | 0,895966939 | 0,81089 | 0,91279122 | novel transcript |
| ENSG00000111203 | ITFG2 | -0,158622513 | 0,89588005 | 0,40147 | 0,6630277 | integrin alpha FG-GAP repeat containing 2 |
| ENSG00000171119 | NRTN | -0,158635393 | 0,895872052 | 0,81541 | 0,9150533 | neurturin |
| ENSG00000135951 | TSGA10 | -0,1587447 | 0,895804178 | 0,51335 | 0,73972767 | testis specific 10 |

| | | | | | | |
|---|---|---|---|---|---|---|
| ENSG00000113532 | ST8SIA4 | -0,15895673 | 0,895672533 | 0,72674 | 0,86994325 | ST8 alpha-N-acetyl-neuraminide alpha-2,8-sialyltransferase 4 |
| ENSG00000162032 | SPSB3 | -0,158962261 | 0,8956691 | 0,44166 | 0,68986044 | splA/ryanodine receptor domain and SOCS box containing 3 |
| ENSG00000168612 | ZSWIM1 | -0,158966347 | 0,895666562 | 0,61508 | 0,80421986 | zinc finger SWIM-type containing 1 |
| ENSG00000100813 | ACIN1 | -0,159115011 | 0,895574272 | 0,40409 | 0,66423756 | apoptotic chromatin condensation inducer 1 |
| ENSG00000236709 | DAPK1-IT1 | -0,159180469 | 0,895533639 | 0,83169 | 0,9230208 | DAPK1 intronic transcript 1 |
| ENSG00000123500 | COL10A1 | -0,159236205 | 0,895499042 | 0,61418 | 0,80370125 | collagen type X alpha 1 chain |
| ENSG00000060656 | PTPRU | -0,159249378 | 0,895490866 | 0,67043 | 0,83657765 | protein tyrosine phosphatase receptor type U |
| ENSG00000242071 | RPL7AP6 | -0,159295342 | 0,895462336 | 0,78747 | 0,90121071 | ribosomal protein L7a pseudogene 6 |
| ENSG00000133083 | DCLK1 | -0,159326937 | 0,895442726 | 0,63704 | 0,81744914 | doublecortin like kinase 1 |
| ENSG00000169306 | IL1RAPL1 | -0,159347305 | 0,895430084 | 0,65097 | 0,82521872 | interleukin 1 receptor accessory protein like 1 |
| ENSG00000240970 | RPL23AP64 | -0,159379731 | 0,895409959 | 0,79307 | 0,90371578 | ribosomal protein L23a pseudogene 64 |
| ENSG00000281913 | RPL23AP64 | -0,159379731 | 0,895409959 | 0,79307 | 0,90371578 | ribosomal protein L23a pseudogene 64 |
| ENSG00000157856 | DRC1 | -0,159398953 | 0,895398028 | 0,69594 | 0,85283771 | dynein regulatory complex subunit 1 |
| ENSG00000273486 | AC096992,2 | -0,159406812 | 0,895393151 | 0,59678 | 0,79431386 | novel transcript, antisense to SLC35G2 |
| ENSG00000126215 | XRCC3 | -0,159467103 | 0,895355733 | 0,40729 | 0,66661761 | X-ray repair cross complementing 3 |
| ENSG00000154217 | PITPNC1 | -0,159645276 | 0,895245163 | 0,3788 | 0,64679625 | phosphatidylinositol transfer protein cytoplasmic 1 |
| ENSG00000182919 | C11orf54 | -0,159691008 | 0,895216785 | 0,34097 | 0,6192144 | chromosome 11 open reading frame 54 |
| ENSG00000164904 | ALDH7A1 | -0,159752813 | 0,895178435 | 0,30467 | 0,59445922 | aldehyde dehydrogenase 7 family member A1 |
| ENSG00000148803 | FUOM | -0,159799112 | 0,895149708 | 0,65002 | 0,8246264 | fucose mutarotase |
| ENSG00000257702 | LBX2-AS1 | -0,159801852 | 0,895148007 | 0,85419 | 0,93380868 | LBX2 antisense RNA 1 |
| ENSG00000074842 | MYDGF | -0,159888348 | 0,895094341 | 0,45525 | 0,69928454 | myeloid derived growth factor |
| ENSG00000101977 | MCF2 | -0,159894552 | 0,895090492 | 0,75134 | 0,88347546 | MCF,2 cell line derived transforming sequence |
| ENSG00000101335 | MYL9 | -0,160045885 | 0,894996605 | 0,61922 | 0,80699858 | myosin light chain 9 |
| ENSG00000197077 | KIAA1671 | -0,160054138 | 0,894991485 | 0,51613 | 0,74198903 | KIAA1671 |
| ENSG00000164081 | TEX264 | -0,160098408 | 0,894964022 | 0,4135 | 0,67078587 | testis expressed 264, ER-phagy receptor |
| ENSG00000224019 | RPL21P32 | -0,160161744 | 0,894924733 | 0,82047 | 0,91732362 | ribosomal protein L21 pseudogene 32 |
| ENSG00000235313 | HM13-IT1 | -0,160221854 | 0,894887447 | 0,81466 | 0,91470265 | HM13 intronic transcript 1 |
| ENSG00000272328 | AC007128,2 | -0,160254234 | 0,894867362 | 0,72882 | 0,87130946 | novel transcript, antisense to ICA1 |
| ENSG00000272138 | LINC01607 | -0,160390654 | 0,894782748 | 0,76133 | 0,88879633 | long intergenic non-protein coding RNA 1607 |
| ENSG00000147676 | MAL2 | -0,160579754 | 0,894665473 | 0,72675 | 0,86994325 | mal, T cell differentiation protein 2 |
| ENSG00000287221 | AL589794,2 | -0,160591884 | 0,894657951 | 0,70686 | 0,85889796 | novel transcript, antisense to ASAH2B |

| | | | | | | |
|---|---|---|---|---|---|---|
| ENSG00000158966 | CACHD1 | -0,160599934 | 0,894652959 | 0,49959 | 0,73058737 | cache domain containing 1 |
| ENSG00000137821 | LRRC49 | -0,160653638 | 0,894619656 | 0,51621 | 0,74198903 | leucine rich repeat containing 49 |
| ENSG00000261534 | AL596244,1 | -0,160665314 | 0,894612416 | 0,66466 | 0,83296965 | novel transcript, overlapping TTLL11 |
| ENSG00000100490 | CDKL1 | -0,160711077 | 0,894584039 | 0,49617 | 0,72831792 | cyclin dependent kinase like 1 |
| ENSG00000131899 | LLGL1 | -0,161102771 | 0,894341191 | 0,76357 | 0,88983737 | LLGL scribble cell polarity complex component 1 |
| ENSG00000188566 | NDOR1 | -0,161111751 | 0,894335624 | 0,47973 | 0,71714401 | NADPH dependent diflavin oxidoreductase 1 |
| ENSG00000226060 | TRIM26 | -0,161160315 | 0,894305519 | 0,68501 | 0,84585686 | tripartite motif containing 26 |
| ENSG00000228881 | TRIM26 | -0,161160315 | 0,894305519 | 0,68501 | 0,84585686 | tripartite motif containing 26 |
| ENSG00000104529 | EEF1D | -0,161205158 | 0,894277722 | 0,5507 | 0,76476271 | eukaryotic translation elongation factor 1 delta |
| ENSG00000176371 | ZSCAN2 | -0,161217576 | 0,894270025 | 0,49231 | 0,72555473 | zinc finger and SCAN domain containing 2 |
| ENSG00000235560 | AC002310,1 | -0,161327675 | 0,894201781 | 0,6208 | 0,80776776 | novel transcript, antisense to ZNF747 |
| ENSG00000185298 | CCDC137 | -0,161425835 | 0,894140942 | 0,45669 | 0,70037869 | coiled-coil domain containing 137 |
| ENSG00000269891 | ARHGAP19-SL | -0,161468697 | 0,894114378 | 0,49178 | 0,72530727 | ARHGAP19-SLIT1 readthrough (NMD candidate) |
| ENSG00000197653 | DNAH10 | -0,161512227 | 0,894087401 | 0,57496 | 0,78003515 | dynein axonemal heavy chain 10 |
| ENSG00000256087 | ZNF432 | -0,161527574 | 0,89407789 | 0,47443 | 0,71319993 | zinc finger protein 432 |
| ENSG00000281935 | DNAH10 | -0,161547536 | 0,894065519 | 0,69021 | 0,84887944 | dynein axonemal heavy chain 10 |
| ENSG00000137776 | SLTM | -0,161697461 | 0,893972612 | 0,43567 | 0,68623199 | SAFB like transcription modulator |
| ENSG00000133030 | MPRIP | -0,161706441 | 0,893967048 | 0,43583 | 0,68637175 | myosin phosphatase Rho interacting protein |
| ENSG00000214293 | APTR | -0,161817635 | 0,893898149 | 0,41993 | 0,67551392 | Alu-mediated CDKN1A/p21 transcriptional regulator |
| ENSG00000233157 | Z97055,1 | -0,161833875 | 0,893888087 | 0,8183 | 0,91663479 | ribosomal protein L4 (RPL4) pseudogene |
| ENSG00000230715 | AC018638,2 | -0,161869683 | 0,893865901 | 0,59935 | 0,79542234 | novel pseudogene |
| ENSG00000130382 | MLLT1 | -0,161969088 | 0,893804313 | 0,34995 | 0,62563749 | MLLT1 super elongation complex subunit |
| ENSG00000219409 | AL590704,1 | -0,162245117 | 0,893633319 | 0,81264 | 0,91386173 | rhomboid domain containing 1 (RHBDD1) pseudogene |
| ENSG00000168675 | LDLRAD4 | -0,16224996 | 0,893630319 | 0,50276 | 0,73210009 | low density lipoprotein receptor class A domain containing 4 |
| ENSG00000136720 | HS6ST1 | -0,162252365 | 0,89362883 | 0,39852 | 0,66104262 | heparan sulfate 6-O-sulfotransferase 1 |
| ENSG00000185291 | IL3RA | -0,162336683 | 0,893576603 | 0,63661 | 0,81709016 | interleukin 3 receptor subunit alpha |
| ENSG00000162676 | GFI1 | -0,162354512 | 0,89356556 | 0,77697 | 0,89665717 | growth factor independent 1 transcriptional repressor |
| ENSG00000258057 | BCDIN3D-AS1 | -0,162365444 | 0,893558789 | 0,64029 | 0,81936319 | BCDIN3D antisense RNA 1 |
| ENSG00000164951 | PDP1 | -0,162369552 | 0,893556245 | 0,64753 | 0,82364696 | pyruvate dehyrogenase phosphatase catalytic subunit 1 |
| ENSG00000145545 | SRD5A1 | -0,162450843 | 0,893505898 | 0,43673 | 0,68681257 | steroid 5 alpha-reductase 1 |
| ENSG00000148082 | SHC3 | -0,162451516 | 0,893505481 | 0,56129 | 0,77136826 | SHC adaptor protein 3 |
| ENSG00000129925 | PGAP6 | -0,162533416 | 0,893454759 | 0,48204 | 0,71880798 | post-glycosylphosphatidylinositol attachment to proteins 6 |

| | | | | | | |
|---|---|---|---|---|---|---|
| ENSG00000241547 | ACTG1P20 | -0,162560023 | 0,893438282 | 0,83795 | 0,92611649 | actin gamma 1 pseudogene 20 |
| ENSG00000234585 | CCT6P3 | -0,162617836 | 0,89340248 | 0,40132 | 0,6630277 | chaperonin containing TCP1 subunit 6 pseudogene 3 |
| ENSG00000242735 | RPSAP26 | -0,162622622 | 0,893399516 | 0,88835 | 0,95069428 | ribosomal protein SA pseudogene 26 |
| ENSG00000156453 | PCDH1 | -0,162699101 | 0,893352157 | 0,69034 | 0,84895522 | protocadherin 1 |
| ENSG00000038002 | AGA | -0,16284744 | 0,893260306 | 0,54338 | 0,76001687 | aspartylglucosaminidase |
| ENSG00000148297 | MED22 | -0,162864698 | 0,893249621 | 0,5732 | 0,77881262 | mediator complex subunit 22 |
| ENSG00000130349 | MTRES1 | -0,162869763 | 0,893246485 | 0,44076 | 0,68917395 | mitochondrial transcription rescue factor 1 |
| ENSG00000286190 | AC055839,2 | -0,162872038 | 0,893245077 | 0,4971 | 0,72865531 | uncharacterized LOC728392 |
| ENSG00000247679 | AC139795,2 | -0,162911243 | 0,893220803 | 0,44585 | 0,69284796 | novel transcript |
| ENSG00000055070 | SZRD1 | -0,162934904 | 0,893206154 | 0,50849 | 0,7365844 | SUZ RNA binding domain containing 1 |
| ENSG00000139613 | SMARCC2 | -0,162964463 | 0,893187854 | 0,4101 | 0,66821634 | SWI/SNF related, matrix associated, actin dependent regulator of chromatin subfamily c member 2 |
| ENSG00000167552 | TUBA1A | -0,162996893 | 0,893167776 | 0,57929 | 0,78270914 | tubulin alpha 1a |
| ENSG00000278970 | HEIH | -0,16303795 | 0,893142358 | 0,43925 | 0,68816791 | hepatocellular carcinoma up-regulated EZH2-associated long non-coding RNA |
| ENSG00000185972 | CCIN | -0,163058131 | 0,893129865 | 0,86441 | 0,93955564 | calicin |
| ENSG00000198695 | MT-ND6 | -0,163139512 | 0,893079485 | 0,51626 | 0,74198903 | mitochondrially encoded NADH:ubiquinone oxidoreductase core subunit 6 |
| ENSG00000102805 | CLN5 | -0,163234097 | 0,893020936 | 0,30335 | 0,59362941 | CLN5 intracellular trafficking protein |
| ENSG00000198131 | ZNF544 | -0,163296543 | 0,892982283 | 0,39376 | 0,65672538 | zinc finger protein 544 |
| ENSG00000183137 | CEP57L1 | -0,16351131 | 0,892849359 | 0,49449 | 0,72681059 | centrosomal protein 57 like 1 |
| ENSG00000286182 | AC112172,2 | -0,163675639 | 0,892747665 | 0,599 | 0,79538547 | novel transcript |
| ENSG00000167774 | AC010323,1 | -0,163732497 | 0,892712482 | 0,80567 | 0,91072757 | novel transcript |
| ENSG00000130559 | CAMSAP1 | -0,163810515 | 0,892664207 | 0,32708 | 0,60902638 | calmodulin regulated spectrin associated protein 1 |
| ENSG00000170846 | AC093323,1 | -0,163815781 | 0,892660949 | 0,35048 | 0,62601861 | Putative MORF4 family-associated protein 1-like protein UPP |
| ENSG00000176732 | PFN4 | -0,163836119 | 0,892648365 | 0,7271 | 0,87016629 | profilin family member 4 |
| ENSG00000173905 | GOLIM4 | -0,163886233 | 0,892617358 | 0,55611 | 0,76831913 | golgi integral membrane protein 4 |
| ENSG00000273002 | AL355388,1 | -0,163894675 | 0,892612135 | 0,69275 | 0,85069507 | uncharacterized LOC107985209 |
| ENSG00000204086 | RPA4 | -0,163925529 | 0,892593045 | 0,85221 | 0,93280134 | replication protein A4 |
| ENSG00000278099 | RNVU1-2A | -0,16397524 | 0,89256229 | 0,82261 | 0,91804426 | RNA, variant U1 small nuclear 2A |
| ENSG00000273757 | RNVU1-2A | -0,16397524 | 0,89256229 | 0,82261 | 0,91804426 | RNA, variant U1 small nuclear 2A |
| ENSG00000245275 | SAP30L-AS1 | -0,16398813 | 0,892554315 | 0,57817 | 0,78191189 | SAP30L antisense RNA 1 (head to head) |

| | | | | | | |
|---|---|---|---|---|---|---|
| ENSG00000272426 | BX284668,6 | -0,163995373 | 0,892549834 | 0,78124 | 0,89873504 | novel transcript |
| ENSG00000198169 | ZNF251 | -0,164114444 | 0,892476172 | 0,62693 | 0,81116451 | zinc finger protein 251 |
| ENSG00000185650 | ZFP36L1 | -0,16420088 | 0,892422702 | 0,38058 | 0,64818855 | ZFP36 ring finger protein like 1 |
| ENSG00000288512 | C1R | -0,164237139 | 0,892400273 | 0,54757 | 0,76293749 | complement C1r |
| ENSG00000127191 | TRAF2 | -0,16431022 | 0,892355069 | 0,38822 | 0,6529693 | TNF receptor associated factor 2 |
| ENSG00000235989 | MORC2-AS1 | -0,164398861 | 0,892300243 | 0,75231 | 0,88394752 | MORC2 antisense RNA 1 |
| ENSG00000156384 | SFR1 | -0,164421875 | 0,89228601 | 0,39478 | 0,65769097 | SWI5 dependent homologous recombination repair protein 1 |
| ENSG00000176396 | EID2 | -0,164523889 | 0,892222917 | 0,46524 | 0,70581615 | EP300 interacting inhibitor of differentiation 2 |
| ENSG00000135454 | B4GALNT1 | -0,164642655 | 0,892149471 | 0,55783 | 0,76954098 | beta-1,4-N-acetyl-galactosaminyltransferase 1 |
| ENSG00000122512 | PMS2 | -0,164785347 | 0,892061236 | 0,41003 | 0,66821634 | PMS1 homolog 2, mismatch repair system component |
| ENSG00000151967 | SCHIP1 | -0,164803993 | 0,892049706 | 0,68289 | 0,84472467 | schwannomin interacting protein 1 |
| ENSG00000184612 | RPL7P26 | -0,164839042 | 0,892028035 | 0,77382 | 0,89571194 | ribosomal protein L7 pseudogene 26 |
| ENSG00000223768 | LINC00205 | -0,16490331 | 0,891988299 | 0,50008 | 0,73078606 | long intergenic non-protein coding RNA 205 |
| ENSG00000125991 | ERGIC3 | -0,164934471 | 0,891969033 | 0,50579 | 0,73418581 | ERGIC and golgi 3 |
| ENSG00000091879 | ANGPT2 | -0,165063412 | 0,891889316 | 0,72324 | 0,86784071 | angiopoietin 2 |
| ENSG00000257195 | HNRNPA1P50 | -0,165145487 | 0,891838578 | 0,79051 | 0,90286944 | heterogeneous nuclear ribonucleoprotein A1 pseudogene 50 |
| ENSG00000266094 | RASSF5 | -0,165286613 | 0,891751342 | 0,68487 | 0,84582677 | Ras association domain family member 5 |
| ENSG00000162928 | PEX13 | -0,165311764 | 0,891735796 | 0,53494 | 0,75428051 | peroxisomal biogenesis factor 13 |
| ENSG00000213380 | COG8 | -0,165474441 | 0,89163525 | 0,56089 | 0,77115738 | component of oligomeric golgi complex 8 |
| ENSG00000107890 | ANKRD26 | -0,165645338 | 0,891529636 | 0,40972 | 0,6680455 | ankyrin repeat domain 26 |
| ENSG00000152402 | GUCY1A2 | -0,165648591 | 0,891527626 | 0,65303 | 0,8262455 | guanylate cyclase 1 soluble subunit alpha 2 |
| ENSG00000134007 | ADAM20 | -0,165715743 | 0,89148613 | 0,66324 | 0,83242946 | ADAM metallopeptidase domain 20 |
| ENSG00000255561 | FDXACB1 | -0,165884085 | 0,891382112 | 0,39089 | 0,65464997 | ferredoxin-fold anticodon binding domain containing 1 |
| ENSG00000189377 | CXCL17 | -0,165953479 | 0,891339237 | 0,87555 | 0,94497474 | C-X-C motif chemokine ligand 17 |
| ENSG00000237161 | AC068446,1 | -0,165963651 | 0,891332953 | 0,8814 | 0,94723445 | p21 protein (Cdc42/Rac)-activated kinase 2 (PAK2) pseudogene |
| ENSG00000173124 | ACSM6 | -0,165972203 | 0,891327669 | 0,58742 | 0,78734837 | acyl-CoA synthetase medium chain family member 6 |
| ENSG00000226465 | AL390198,1 | -0,166095467 | 0,891251517 | 0,77088 | 0,89390008 | novel transcript |
| ENSG00000286387 | AC007432,1 | -0,16618455 | 0,891196487 | 0,76558 | 0,89091205 | novel transcript |
| ENSG00000182791 | CCDC87 | -0,166314165 | 0,891116423 | 0,64728 | 0,82349878 | coiled-coil domain containing 87 |
| ENSG00000287878 | AC022382,2 | -0,166332758 | 0,891104939 | 0,64633 | 0,82293278 | novel transcript, antisense to CPNE9and MTMR14 |

| | | | | | | |
|---|---|---|---|---|---|---|
| ENSG00000156140 | ADAMTS3 | -0,16642894 | 0,891045532 | 0,70769 | 0,85918434 | ADAM metallopeptidase with thrombospondin type 1 motif 3 |
| ENSG00000214827 | MTCP1 | -0,166458364 | 0,89102736 | 0,66134 | 0,83129468 | mature T cell proliferation 1 |
| ENSG00000123810 | B9D2 | -0,166469623 | 0,891020406 | 0,60454 | 0,79797731 | B9 domain containing 2 |
| ENSG00000282390 | AL392083,1 | -0,166472124 | 0,891018861 | 0,78172 | 0,89877377 | novel transcript, sense intronic to ADARB2 |
| ENSG00000221944 | TIGD1 | -0,166515635 | 0,890991988 | 0,33792 | 0,61755283 | tigger transposable element derived 1 |
| ENSG00000286529 | AC090398,2 | -0,166719239 | 0,890866254 | 0,61119 | 0,80297503 | novel transcript |
| ENSG00000140319 | SRP14 | -0,166754093 | 0,890844732 | 0,53561 | 0,75482321 | signal recognition particle 14 |
| ENSG00000230445 | LRRC37A6P | -0,166789015 | 0,890823168 | 0,54363 | 0,76018442 | leucine rich repeat containing 37 member A6, pseudogene |
| ENSG00000136378 | ADAMTS7 | -0,166855158 | 0,890782328 | 0,52982 | 0,75150213 | ADAM metallopeptidase with thrombospondin type 1 motif 7 |
| ENSG00000130734 | ATG4D | -0,167118772 | 0,890619576 | 0,43043 | 0,68295004 | autophagy related 4D cysteine peptidase |
| ENSG00000104901 | DKKL1 | -0,167151428 | 0,890599417 | 0,60635 | 0,79949515 | dickkopf like acrosomal protein 1 |
| ENSG00000254447 | OR7E11P | -0,167191394 | 0,890574745 | 0,86588 | 0,94044334 | olfactory receptor family 7 subfamily E member 11 pseudogene |
| ENSG00000262001 | DLGAP1-AS2 | -0,167196316 | 0,890571707 | 0,579 | 0,78267043 | DLGAP1 antisense RNA 2 |
| ENSG00000285628 | AC126755,5 | -0,167329685 | 0,890489383 | 0,67099 | 0,83695245 | novel transcript |
| ENSG00000232063 | AL691447,2 | -0,167345492 | 0,890479626 | 0,5151 | 0,74133616 | novel transcript |
| ENSG00000262829 | AC242308,2 | -0,167414388 | 0,890437102 | 0,59969 | 0,79552021 | novel taste receptor, type 2 family pseudogene |
| ENSG00000260024 | MRPS21P7 | -0,167433683 | 0,890425193 | 0,83304 | 0,92367721 | mitochondrial ribosomal protein S21 pseudogene 7 |
| ENSG00000119403 | PHF19 | -0,16744909 | 0,890415684 | 0,44484 | 0,69205935 | PHD finger protein 19 |
| ENSG00000050393 | MCUR1 | -0,167449328 | 0,890415537 | 0,51931 | 0,74394224 | mitochondrial calcium uniporter regulator 1 |
| ENSG00000166803 | PCLAF | -0,167508075 | 0,89037928 | 0,3978 | 0,66036173 | PCNA clamp associated factor |
| ENSG00000287419 | AL121929,3 | -0,167580405 | 0,890334641 | 0,76925 | 0,89297049 | novel transcript, antisense to NEURL1 |
| ENSG00000235726 | AC010148,1 | -0,167636194 | 0,890300213 | 0,70625 | 0,85859137 | novel transcript |
| ENSG00000011114 | BTBD7 | -0,167640523 | 0,890297541 | 0,58293 | 0,78477395 | BTB domain containing 7 |
| ENSG00000144021 | CIAO1 | -0,167704809 | 0,890257871 | 0,3594 | 0,63226238 | cytosolic iron-sulfur assembly component 1 |
| ENSG00000266932 | AC006116,2 | -0,16773276 | 0,890240623 | 0,77703 | 0,89668942 | NADH dehydrogenase (ubiquinone) complex I, assembly factor 1, pseudogene |
| ENSG00000137720 | C11orf1 | -0,167843431 | 0,890172334 | 0,55599 | 0,76827287 | chromosome 11 open reading frame 1 |
| ENSG00000273432 | AC004951,4 | -0,167847612 | 0,890169755 | 0,69062 | 0,84900677 | novel transcript, POLR2J4 pseudogene-RASA4 pseudogene readthrough |
| ENSG00000241597 | AC022121,1 | -0,167897216 | 0,890139149 | 0,7764 | 0,89655595 | ribosomal protein S9 (RPS9) pseudogene |

| | | | | | | |
|---|---|---|---|---|---|---|
| ENSG00000168785 | TSPAN5 | -0,168062344 | 0,89003727 | 0,57505 | 0,78004262 | tetraspanin 5 |
| ENSG00000215845 | TSTD1 | -0,168075128 | 0,890029384 | 0,79829 | 0,9069401 | thiosulfate sulfurtransferase like domain containing 1 |
| ENSG00000180354 | MTURN | -0,168091552 | 0,890019252 | 0,53937 | 0,75771283 | maturin, neural progenitor differentiation regulator homolog |
| ENSG00000232373 | MTCYBP3 | -0,168151733 | 0,889982126 | 0,83882 | 0,92638462 | MT-CYB pseudogene 3 |
| ENSG00000060971 | ACAA1 | -0,168226605 | 0,889935939 | 0,3628 | 0,63387459 | acetyl-CoA acyltransferase 1 |
| ENSG00000168389 | MFSD2A | -0,168228199 | 0,889934956 | 0,48186 | 0,71868571 | major facilitator superfamily domain containing 2A |
| ENSG00000248415 | GAPDHP61 | -0,168240567 | 0,889927327 | 0,68718 | 0,84745675 | glyceraldehyde 3 phosphate dehydrogenase pseudogene 61 |
| ENSG00000261527 | AC026464,5 | -0,168302815 | 0,88988893 | 0,8034 | 0,90947335 | transmembrane protein 111 (TMEM111) pseudogene |
| ENSG00000146205 | ANO7 | -0,168319581 | 0,889878588 | 0,69487 | 0,85206252 | anoctamin 7 |
| ENSG00000169914 | OTUD3 | -0,168347222 | 0,889861539 | 0,52305 | 0,74710918 | OTU deubiquitinase 3 |
| ENSG00000087152 | ATXN7L3 | -0,168369905 | 0,889847549 | 0,48734 | 0,72237072 | ataxin 7 like 3 |
| ENSG00000138107 | ACTR1A | -0,168411747 | 0,889821741 | 0,38075 | 0,64832095 | actin related protein 1A |
| ENSG00000274964 | AC026356,1 | -0,168647828 | 0,889676143 | 0,63222 | 0,81450663 | novel transcript, sense intronic to BICD1 |
| ENSG00000260339 | HEXA-AS1 | -0,168681014 | 0,889655679 | 0,71487 | 0,86282308 | HEXA antisense RNA 1 |
| ENSG00000255559 | ZNF252P-AS1 | -0,168716817 | 0,8896336 | 0,70107 | 0,85578362 | ZNF252P antisense RNA 1 |
| ENSG00000163900 | TMEM41A | -0,168792774 | 0,889586763 | 0,49641 | 0,72831792 | transmembrane protein 41A |
| ENSG00000229619 | MBNL1-AS1 | -0,168865726 | 0,889541781 | 0,68825 | 0,84815887 | MBNL1 antisense RNA 1 |
| ENSG00000240024 | LINC00888 | -0,169001744 | 0,889457918 | 0,40389 | 0,66409141 | long intergenic non-protein coding RNA 888 |
| ENSG00000092847 | AGO1 | -0,169165522 | 0,889356951 | 0,39852 | 0,66104262 | argonaute RISC component 1 |
| ENSG00000125170 | DOK4 | -0,169244303 | 0,889308387 | 0,52868 | 0,75098717 | docking protein 4 |
| ENSG00000151640 | DPYSL4 | -0,169353785 | 0,889240902 | 0,29053 | 0,58565836 | dihydropyrimidinase like 4 |
| ENSG00000233183 | AL138889,1 | -0,169544968 | 0,88912307 | 0,46116 | 0,70278639 | novel transcript |
| ENSG00000272858 | Z93930,3 | -0,169617606 | 0,889078305 | 0,77735 | 0,8969723 | novel transcript |
| ENSG00000164651 | SP8 | -0,169778066 | 0,888979425 | 0,7526 | 0,88406666 | Sp8 transcription factor |
| ENSG00000158201 | ABHD3 | -0,169812416 | 0,888958259 | 0,61257 | 0,8035281 | abhydrolase domain containing 3, phospholipase |
| ENSG00000274240 | AC136352,1 | -0,169899969 | 0,888904312 | 0,82049 | 0,91732362 | p21 protein (Cdc42/Rac)-activated kinase 2 (PAK2) pseudogene |
| ENSG00000214300 | SPDYE3 | -0,169977613 | 0,888856474 | 0,67866 | 0,84182956 | speedy/RINGO cell cycle regulator family member E3 |
| ENSG00000234636 | MED14OS | -0,170169063 | 0,888738528 | 0,65626 | 0,82829288 | MED14 opposite strand |
| ENSG00000231713 | AF064860,2 | -0,170175701 | 0,888734439 | 0,79058 | 0,90286944 | novel transcript |
| ENSG00000262074 | SNORD3B-2 | -0,170185638 | 0,888728317 | 0,64623 | 0,82287969 | small nucleolar RNA, C/D box 3B-2 |

| | | | | | | |
|---|---|---|---|---|---|---|
| ENSG00000252010 | SCARNA5 | -0,170189427 | 0,888725983 | 0,68349 | 0,84495012 | small Cajal body-specific RNA 5 |
| ENSG00000280652 | SCARNA5 | -0,170189427 | 0,888725983 | 0,68349 | 0,84495012 | small Cajal body-specific RNA 5 |
| ENSG00000233339 | AC018878,1 | -0,170372829 | 0,888613012 | 0,79284 | 0,90366929 | novel transcript |
| ENSG00000196782 | MAML3 | -0,170514534 | 0,888525734 | 0,31225 | 0,59925736 | mastermind like transcriptional coactivator 3 |
| ENSG00000144712 | CAND2 | -0,170603604 | 0,888470879 | 0,44195 | 0,68994635 | cullin associated and neddylation dissociated 2 (putative) |
| ENSG00000287969 | AL355474,1 | -0,170913523 | 0,888280039 | 0,86105 | 0,9378327 | novel transcript, antisense to FAM13C |
| ENSG00000230649 | AC024084,1 | -0,171107561 | 0,888160576 | 0,79202 | 0,90343182 | novel transcript |
| ENSG00000272325 | NUDT3 | -0,171110838 | 0,888158559 | 0,40329 | 0,66399303 | nudix hydrolase 3 |
| ENSG00000099381 | SETD1A | -0,171187906 | 0,888111115 | 0,53019 | 0,75182721 | SET domain containing 1A, histone lysine methyltransferase |
| ENSG00000206344 | HCG27 | -0,171230735 | 0,88808475 | 0,87912 | 0,94623795 | HLA complex group 27 |
| ENSG00000277130 | AC073569,3 | -0,171347385 | 0,888012946 | 0,55916 | 0,77044645 | novel transcript, antisense to PPP1R12A |
| ENSG00000139597 | N4BP2L1 | -0,171665031 | 0,887817449 | 0,64036 | 0,81936319 | NEDD4 binding protein 2 like 1 |
| ENSG00000153048 | CARHSP1 | -0,171719028 | 0,887784221 | 0,26911 | 0,57085484 | calcium regulated heat stable protein 1 |
| ENSG00000077150 | NFKB2 | -0,171724742 | 0,887780705 | 0,4915 | 0,72510994 | nuclear factor kappa B subunit 2 |
| ENSG00000185803 | SLC52A2 | -0,171811719 | 0,887727184 | 0,57682 | 0,78107792 | solute carrier family 52 member 2 |
| ENSG00000224376 | AC017104,1 | -0,171825094 | 0,887718954 | 0,74448 | 0,87904799 | novel transcript |
| ENSG00000131477 | RAMP2 | -0,171957198 | 0,887637671 | 0,62185 | 0,80819595 | receptor activity modifying protein 2 |
| ENSG00000257243 | AC020612,1 | -0,171987365 | 0,887619111 | 0,76388 | 0,88993069 | Protein DJ-1 (PARK7) pseudogene |
| ENSG00000223960 | CHROMR | -0,172031186 | 0,88759215 | 0,36322 | 0,63407828 | cholesterol induced regulator of metabolism RNA |
| ENSG00000082293 | COL19A1 | -0,172050035 | 0,887580554 | 0,75217 | 0,88389568 | collagen type XIX alpha 1 chain |
| ENSG00000186834 | HEXIM1 | -0,172141368 | 0,887524365 | 0,55887 | 0,77025265 | HEXIM P-TEFb complex subunit 1 |
| ENSG00000251513 | LIX1-AS1 | -0,172144488 | 0,887522446 | 0,81863 | 0,91672281 | LIX1 and RIOK2 antisense RNA 1 |
| ENSG00000169218 | RSPO1 | -0,172304838 | 0,887423807 | 0,76008 | 0,88816392 | R-spondin 1 |
| ENSG00000285847 | AL713852,1 | -0,172348376 | 0,887397027 | 0,61291 | 0,8035281 | novel transcript |
| ENSG00000152527 | PLEKHH2 | -0,172385987 | 0,887373892 | 0,5361 | 0,75500755 | pleckstrin homology, MyTH4 and FERM domain containing H2 |
| ENSG00000196975 | ANXA4 | -0,172546417 | 0,88727522 | 0,58572 | 0,78640335 | annexin A4 |
| ENSG00000261266 | AC008870,3 | -0,172547332 | 0,887274658 | 0,88342 | 0,94837933 | novel transcript |
| ENSG00000135097 | MSI1 | -0,172671241 | 0,887198455 | 0,36978 | 0,63943131 | musashi RNA binding protein 1 |
| ENSG00000170324 | FRMPD2 | -0,172754021 | 0,88714755 | 0,79457 | 0,90473839 | FERM and PDZ domain containing 2 |
| ENSG00000196312 | MFSD14C | -0,172870021 | 0,887076222 | 0,42476 | 0,67855129 | major facilitator superfamily domain containing 14C |
| ENSG00000181007 | ZFP82 | -0,172927162 | 0,887041088 | 0,39501 | 0,65794526 | ZFP82 zinc finger protein |

| | | | | | | |
|---|---|---|---|---|---|---|
| ENSG00000124249 | KCNK15 | -0,172937009 | 0,887035034 | 0,80683 | 0,91120963 | potassium two pore domain channel subfamily K member 15 |
| ENSG00000226498 | RPSAP21 | -0,172943207 | 0,887031223 | 0,78159 | 0,89877377 | ribosomal protein SA pseudogene 21 |
| ENSG00000131848 | ZSCAN5A | -0,173023114 | 0,886982095 | 0,44104 | 0,68939121 | zinc finger and SCAN domain containing 5A |
| ENSG00000218510 | LINC00339 | -0,173058973 | 0,886960048 | 0,46456 | 0,7051 | long intergenic non-protein coding RNA 339 |
| ENSG00000100395 | L3MBTL2 | -0,173084913 | 0,886944101 | 0,5418 | 0,75882662 | L3MBTL histone methyl-lysine binding protein 2 |
| ENSG00000213648 | SULT1A4 | -0,173085451 | 0,88694377 | 0,75817 | 0,88722916 | sulfotransferase family 1A member 4 |
| ENSG00000261504 | LINC01686 | -0,173218698 | 0,886861856 | 0,74263 | 0,87827554 | long intergenic non-protein coding RNA 1686 |
| ENSG00000166436 | TRIM66 | -0,173270848 | 0,886829799 | 0,41132 | 0,66904284 | tripartite motif containing 66 |
| ENSG00000224141 | MIR548XHG | -0,173299809 | 0,886811996 | 0,70761 | 0,85916455 | MIR548X host gene |
| ENSG00000287044 | AL139106,1 | -0,173332366 | 0,886791984 | 0,6577 | 0,82881843 | novel transcript, antisense to LACE1 |
| ENSG00000143367 | TUFT1 | -0,173394396 | 0,886753856 | 0,53338 | 0,75351438 | tuftelin 1 |
| ENSG00000161395 | PGAP3 | -0,173476301 | 0,886703515 | 0,28799 | 0,5840701 | post-GPI attachment to proteins phospholipase 3 |
| ENSG00000013573 | DDX11 | -0,173490213 | 0,886694964 | 0,40919 | 0,66785537 | DEAD/H-box helicase 11 |
| ENSG00000161013 | MGAT4B | -0,173538398 | 0,88666535 | 0,4595 | 0,7017037 | alpha-1,3-mannosyl-glycoprotein 4-beta-N-acetylglucosaminyltransferase B |
| ENSG00000284501 | MGAT4B | -0,173538398 | 0,88666535 | 0,4595 | 0,7017037 | alpha-1,3-mannosyl-glycoprotein 4-beta-N-acetylglucosaminyltransferase B |
| ENSG00000180891 | CUEDC1 | -0,173571744 | 0,886644856 | 0,50083 | 0,73097006 | CUE domain containing 1 |
| ENSG00000155393 | HEATR3 | -0,173624959 | 0,886612152 | 0,60188 | 0,79680798 | HEAT repeat containing 3 |
| ENSG00000186523 | FAM86B1 | -0,173647237 | 0,886598461 | 0,47685 | 0,71481671 | family with sequence similarity 86 member B1 |
| ENSG00000233321 | LINC02669 | -0,17375401 | 0,886532847 | 0,7308 | 0,87205836 | long intergenic non-protein coding RNA 2669 |
| ENSG00000239305 | RNF103 | -0,173844976 | 0,88647695 | 0,44801 | 0,69404938 | ring finger protein 103 |
| ENSG00000234261 | AL138720,1 | -0,173848614 | 0,886474714 | 0,47985 | 0,71724282 | novel transcript |
| ENSG00000109065 | NAT9 | -0,173913617 | 0,886434774 | 0,35968 | 0,63242662 | N-acetyltransferase 9 (putative) |
| ENSG00000118523 | CCN2 | -0,173928544 | 0,886425602 | 0,62034 | 0,80773684 | cellular communication network factor 2 |
| ENSG00000171853 | TRAPPC12 | -0,173982837 | 0,886392244 | 0,44483 | 0,69205935 | trafficking protein particle complex 12 |
| ENSG00000165181 | SHOC1 | -0,174124707 | 0,886305084 | 0,70902 | 0,85978194 | shortage in chiasmata 1 |
| ENSG00000286403 | AC010378,1 | -0,174321403 | 0,886184253 | 0,44933 | 0,69502656 | novel transcript |
| ENSG00000172922 | RNASEH2C | -0,174323658 | 0,886182868 | 0,4667 | 0,70676695 | ribonuclease H2 subunit C |
| ENSG00000248925 | AC021087,1 | -0,174446972 | 0,886107125 | 0,77661 | 0,89658055 | novel transcript |
| ENSG00000159882 | ZNF230 | -0,174458024 | 0,886100337 | 0,55409 | 0,76712518 | zinc finger protein 230 |
| ENSG00000286527 | AL161665,2 | -0,174528001 | 0,886057358 | 0,85061 | 0,93206349 | novel transcript |

| | | | | | | |
|---|---|---|---|---|---|---|
| ENSG00000075790 | BCAP29 | -0,174569869 | 0,886031645 | 0,44607 | 0,69284796 | B cell receptor associated protein 29 |
| ENSG00000121957 | GPSM2 | -0,174627196 | 0,885996438 | 0,41432 | 0,67106864 | G protein signaling modulator 2 |
| ENSG00000131484 | AC091132,1 | -0,174809338 | 0,885884587 | 0,70733 | 0,85910429 | novel transcript |
| ENSG00000123685 | BATF3 | -0,174841039 | 0,885865122 | 0,83474 | 0,92446289 | basic leucine zipper ATF-like transcription factor 3 |
| ENSG00000136999 | CCN3 | -0,17502874 | 0,885749874 | 0,837 | 0,92549623 | cellular communication network factor 3 |
| ENSG00000226798 | AL138749,1 | -0,175145679 | 0,885678081 | 0,87779 | 0,94568249 | putatitive novel transcript |
| ENSG00000188827 | SLX4 | -0,175185522 | 0,885653622 | 0,49514 | 0,72746032 | SLX4 structure-specific endonuclease subunit |
| ENSG00000275265 | AC127002,2 | -0,175189074 | 0,885651441 | 0,87375 | 0,94405019 | novel transcript |
| ENSG00000155827 | RNF20 | -0,175231975 | 0,885625105 | 0,36009 | 0,63261458 | ring finger protein 20 |
| ENSG00000127022 | CANX | -0,175269837 | 0,885601864 | 0,37771 | 0,64615983 | calnexin |
| ENSG00000160190 | SLC37A1 | -0,175349568 | 0,885552922 | 0,67024 | 0,83650134 | solute carrier family 37 member 1 |
| ENSG00000007376 | RPUSD1 | -0,17536781 | 0,885541724 | 0,51632 | 0,74198903 | RNA pseudouridine synthase domain containing 1 |
| ENSG00000146067 | FAM193B | -0,175459211 | 0,885485623 | 0,40364 | 0,66399439 | family with sequence similarity 193 member B |
| ENSG00000155849 | ELMO1 | -0,175632265 | 0,885379414 | 0,38962 | 0,65396251 | engulfment and cell motility 1 |
| ENSG00000163393 | SLC22A15 | -0,175663851 | 0,88536003 | 0,66075 | 0,83089245 | solute carrier family 22 member 15 |
| ENSG00000148153 | INIP | -0,175742777 | 0,885311596 | 0,24464 | 0,55271519 | INTS3 and NABP interacting protein |
| ENSG00000173511 | VEGFB | -0,175757046 | 0,88530284 | 0,66041 | 0,83053271 | vascular endothelial growth factor B |
| ENSG00000140545 | MFGE8 | -0,175780843 | 0,885288237 | 0,50127 | 0,7312466 | milk fat globule EGF and factor V/VIII domain containing |
| ENSG00000285704 | AC004765,1 | -0,175910268 | 0,885208821 | 0,61783 | 0,80610038 | novel transcript, antisense to WNK1 |
| ENSG00000104774 | MAN2B1 | -0,176030304 | 0,885135172 | 0,45309 | 0,69746212 | mannosidase alpha class 2B member 1 |
| ENSG00000285569 | AC099344,3 | -0,176080619 | 0,885104303 | 0,6342 | 0,81563073 | novel transcript |
| ENSG00000177000 | MTHFR | -0,17613691 | 0,885069769 | 0,55063 | 0,76475342 | methylenetetrahydrofolate reductase |
| ENSG00000177822 | TENM3-AS1 | -0,176234394 | 0,885009966 | 0,61329 | 0,8035281 | TENM3 antisense RNA 1 |
| ENSG00000235652 | FBXO30-DT | -0,176245512 | 0,885003146 | 0,3706 | 0,63997937 | FBXO30 divergent transcript |
| ENSG00000249035 | CLMAT3 | -0,176261703 | 0,884993213 | 0,75804 | 0,88719949 | colorectal liver metastasis associated transcript 3 |
| ENSG00000169618 | PROKR1 | -0,176408551 | 0,884903137 | 0,74488 | 0,87929941 | prokineticin receptor 1 |
| ENSG00000169554 | ZEB2 | -0,176412394 | 0,884900779 | 0,56372 | 0,77255679 | zinc finger E-box binding homeobox 2 |
| ENSG00000270116 | AP001429,1 | -0,176460402 | 0,884871334 | 0,57314 | 0,77878548 | novel transcript, sense intronic to TTC3 |
| ENSG00000140365 | COMMD4 | -0,176626117 | 0,884769699 | 0,52279 | 0,74692337 | COMM domain containing 4 |
| ENSG00000275367 | AC092111,1 | -0,176663113 | 0,88474701 | 0,44414 | 0,6916877 | novel transcript |
| ENSG00000248971 | KRT8P46 | -0,176814873 | 0,884653947 | 0,58284 | 0,78477395 | keratin 8 pseudogene 46 |
| ENSG00000158234 | FAIM | -0,176963049 | 0,884563091 | 0,4091 | 0,66785537 | Fas apoptotic inhibitory molecule |
| ENSG00000183154 | AC138356,1 | -0,177221756 | 0,884404483 | 0,79629 | 0,90587222 | novel transcript, antisense to ERLIN2 |

| | | | | | | |
|---|---|---|---|---|---|---|
| ENSG00000213066 | CEP43 | -0,177295575 | 0,884359232 | 0,52974 | 0,75147469 | centrosomal protein 43 |
| ENSG00000259420 | AC046168,2 | -0,17733896 | 0,884332638 | 0,86758 | 0,94147581 | novel transcript |
| ENSG00000228649 | SNHG26 | -0,177442506 | 0,884269169 | 0,73609 | 0,87517709 | small nucleolar RNA host gene 26 |
| ENSG00000122417 | ODF2L | -0,177446583 | 0,88426667 | 0,31717 | 0,6027392 | outer dense fiber of sperm tails 2 like |
| ENSG00000055211 | GINM1 | -0,177495513 | 0,88423668 | 0,47274 | 0,71204009 | glycoprotein integral membrane 1 |
| ENSG00000198039 | ZNF273 | -0,177511285 | 0,884227014 | 0,4203 | 0,67570188 | zinc finger protein 273 |
| ENSG00000232792 | FTH1P25 | -0,177555017 | 0,88420021 | 0,80334 | 0,90946649 | ferritin heavy chain 1 pseudogene 25 |
| ENSG00000117016 | RIMS3 | -0,17755614 | 0,884199523 | 0,46326 | 0,70415312 | regulating synaptic membrane exocytosis 3 |
| ENSG00000164251 | F2RL1 | -0,177586408 | 0,884180972 | 0,73364 | 0,87368351 | F2R like trypsin receptor 1 |
| ENSG00000175513 | TSGA10IP | -0,177609243 | 0,884166977 | 0,58105 | 0,78406342 | testis specific 10 interacting protein |
| ENSG00000057468 | MSH4 | -0,177684351 | 0,884120948 | 0,84591 | 0,92992612 | mutS homolog 4 |
| ENSG00000245768 | AC106793,1 | -0,177713794 | 0,884102904 | 0,76872 | 0,89277929 | novel transcript |
| ENSG00000160229 | ZNF66 | -0,177855607 | 0,884016004 | 0,351 | 0,62648918 | zinc finger protein 66 |
| ENSG00000287690 | AL023775,2 | -0,177865686 | 0,884009828 | 0,78141 | 0,89877377 | novel transcript, antisense to RPS6KA2 |
| ENSG00000083814 | ZNF671 | -0,177884658 | 0,883998203 | 0,40393 | 0,66409141 | zinc finger protein 671 |
| ENSG00000232775 | BMS1P22 | -0,178051844 | 0,883895767 | 0,7748 | 0,89592738 | BMS1 pseudogene 22 |
| ENSG00000166432 | ZMAT1 | -0,178176532 | 0,883819378 | 0,6758 | 0,84049293 | zinc finger matrin-type 1 |
| ENSG00000135048 | CEMIP2 | -0,178222148 | 0,883791433 | 0,60315 | 0,79726763 | cell migration inducing hyaluronidase 2 |
| ENSG00000226533 | BTBD9-AS1 | -0,178278604 | 0,883756849 | 0,84609 | 0,92995515 | BTBD9 antisense RNA 1 |
| ENSG00000139116 | KIF21A | -0,17828409 | 0,883753489 | 0,64054 | 0,81950381 | kinesin family member 21A |
| ENSG00000214773 | AC112512,1 | -0,178350479 | 0,883712822 | 0,60707 | 0,80003503 | novel transcript, sense intronic SMARCC1 |
| ENSG00000261839 | AL358933,1 | -0,178392783 | 0,883686909 | 0,65105 | 0,82521872 | novel transcript, overlapping to ZNF435 |
| ENSG00000081377 | CDC14B | -0,178455778 | 0,883648323 | 0,62059 | 0,80776776 | cell division cycle 14B |
| ENSG00000279277 | AC012020,1 | -0,178549228 | 0,883591088 | 0,64599 | 0,82269302 | novel transcript |
| ENSG00000074370 | ATP2A3 | -0,178707375 | 0,883494235 | 0,62919 | 0,81265687 | ATPase sarcoplasmic/endoplasmic reticulum Ca2+ transporting 3 |
| ENSG00000261115 | TMEM178B | -0,178712906 | 0,883490848 | 0,35267 | 0,62754548 | transmembrane protein 178B |
| ENSG00000213963 | AC019080,1 | -0,178727932 | 0,883481646 | 0,50098 | 0,73099828 | novel transcript |
| ENSG00000147853 | AK3 | -0,178797214 | 0,88343922 | 0,42014 | 0,67563085 | adenylate kinase 3 |
| ENSG00000237586 | AC090192,1 | -0,178806776 | 0,883433364 | 0,88444 | 0,94876126 | SRY (sex determining region Y)-box 2 (SOX2) pseudogene |
| ENSG00000100577 | GSTZ1 | -0,178839794 | 0,883413146 | 0,37114 | 0,64038503 | glutathione S-transferase zeta 1 |
| ENSG00000169241 | SLC50A1 | -0,178865499 | 0,883397406 | 0,69636 | 0,85299392 | solute carrier family 50 member 1 |
| ENSG00000184840 | TMED9 | -0,178969234 | 0,883333889 | 0,44845 | 0,69432421 | transmembrane p24 trafficking protein 9 |

| | | | | | | |
|---|---|---|---|---|---|---|
| ENSG00000106341 | PPP1R17 | -0,179038812 | 0,883291289 | 0,73738 | 0,87539732 | protein phosphatase 1 regulatory subunit 17 |
| ENSG00000257252 | AC124947,1 | -0,179095412 | 0,883256636 | 0,82286 | 0,9181175 | novel transcript, antisense to NUDT4 |
| ENSG00000183111 | ARHGEF37 | -0,179161508 | 0,883216171 | 0,55341 | 0,76647245 | Rho guanine nucleotide exchange factor 37 |
| ENSG00000272980 | Z94721,2 | -0,179400693 | 0,883069755 | 0,44445 | 0,69185302 | novel transcript |
| ENSG00000137968 | SLC44A5 | -0,179438597 | 0,883046554 | 0,64103 | 0,81991652 | solute carrier family 44 member 5 |
| ENSG00000176953 | NFATC2IP | -0,179471514 | 0,883026406 | 0,47494 | 0,71364583 | nuclear factor of activated T cells 2 interacting protein |
| ENSG00000103355 | PRSS33 | -0,179548348 | 0,88297938 | 0,74674 | 0,88057664 | serine protease 33 |
| ENSG00000272601 | AC010655,3 | -0,179649569 | 0,882917431 | 0,50404 | 0,73311005 | EF-hand calcium binding domain 3 (EFCAB3) pseudogene |
| ENSG00000003147 | ICA1 | -0,179711617 | 0,882879459 | 0,62539 | 0,81042981 | islet cell autoantigen 1 |
| ENSG00000033170 | FUT8 | -0,17973617 | 0,882864434 | 0,40846 | 0,66723037 | fucosyltransferase 8 |
| ENSG00000132872 | SYT4 | -0,179748905 | 0,882856641 | 0,70059 | 0,85539553 | synaptotagmin 4 |
| ENSG00000163686 | ABHD6 | -0,179937371 | 0,882741316 | 0,50964 | 0,73740498 | abhydrolase domain containing 6, acylglycerol lipase |
| ENSG00000259774 | AC243562,3 | -0,180083504 | 0,882651906 | 0,59473 | 0,79290095 | novel transcript |
| ENSG00000139718 | SETD1B | -0,180205519 | 0,88257726 | 0,44552 | 0,69256445 | SET domain containing 1B, histone lysine methyltransferase |
| ENSG00000272733 | AP000345,2 | -0,180275367 | 0,882534531 | 0,80812 | 0,91176063 | novel transcript |
| ENSG00000117480 | FAAH | -0,1802754 | 0,882534511 | 0,57052 | 0,77743553 | fatty acid amide hydrolase |
| ENSG00000257511 | AC084824,1 | -0,180414354 | 0,882449513 | 0,58954 | 0,78890126 | nucleosome assembly protein 1-like 1 (NAP1L1) pseudogene |
| ENSG00000204120 | GIGYF2 | -0,180455949 | 0,882424071 | 0,39925 | 0,66166661 | GRB10 interacting GYF protein 2 |
| ENSG00000257391 | AC126763,1 | -0,180531048 | 0,882378138 | 0,67453 | 0,83975333 | microRNA 3180-4 |
| ENSG00000257246 | PHBP19 | -0,180561489 | 0,88235952 | 0,76096 | 0,88874219 | prohibitin pseudogene 19 |
| ENSG00000261669 | AC008731,1 | -0,180569733 | 0,882354478 | 0,78449 | 0,89962715 | novel transcript, antisense to TNRC6A |
| ENSG00000226330 | AL606489,1 | -0,180583947 | 0,882345785 | 0,8194 | 0,91699877 | novel transcript |
| ENSG00000285072 | AC270272,2 | -0,18061422 | 0,88232727 | 0,77644 | 0,89656608 | |
| ENSG00000189319 | FAM53B | -0,180706848 | 0,882270622 | 0,3256 | 0,60804921 | family with sequence similarity 53 member B |
| ENSG00000122642 | FKBP9 | -0,18071021 | 0,882268566 | 0,32859 | 0,60992716 | FKBP prolyl isomerase 9 |
| ENSG00000225691 | HLA-C | -0,180904775 | 0,88214959 | 0,66492 | 0,83314543 | major histocompatibility complex, class I, C |
| ENSG00000276005 | AC138749,8 | -0,180978314 | 0,882104624 | 0,5422 | 0,75894841 | hect domain and RLD 2 (HERC2) pseudogene |
| ENSG00000231905 | SETP10 | -0,180982105 | 0,882102306 | 0,74078 | 0,87717177 | SET pseudogene 10 |
| ENSG00000111679 | PTPN6 | -0,181266926 | 0,881928176 | 0,73525 | 0,87483589 | protein tyrosine phosphatase non-receptor type 6 |
| ENSG00000279206 | AC004943,3 | -0,181279497 | 0,881920492 | 0,73966 | 0,87653492 | novel transcript |
| ENSG00000253711 | AC023866,1 | -0,181375643 | 0,88186172 | 0,85686 | 0,93541397 | novel transcript |

| | | | | | | |
|---|---|---|---|---|---|---|
| ENSG00000158864 | NDUFS2 | -0,181433963 | 0,881826072 | 0,45251 | 0,69702004 | NADH:ubiquinone oxidoreductase core subunit S2 |
| ENSG00000236914 | LINC01852 | -0,181558599 | 0,881749893 | 0,49893 | 0,73001923 | long intergenic non-protein coding RNA 1852 |
| ENSG00000225649 | AC064875,1 | -0,181711119 | 0,88165668 | 0,79305 | 0,90371578 | novel transcript |
| ENSG00000138449 | SLC40A1 | -0,181754424 | 0,881630217 | 0,64785 | 0,82380454 | solute carrier family 40 member 1 |
| ENSG00000267529 | AP005131,4 | -0,18177533 | 0,881617441 | 0,76938 | 0,89297049 | novel transcript, antisense C18orf1 |
| ENSG00000228691 | NEU1 | -0,181798935 | 0,881603016 | 0,56876 | 0,77623532 | neuraminidase 1 |
| ENSG00000206062 | AL121823,1 | -0,181812664 | 0,881594627 | 0,83143 | 0,9229059 | novel transcript |
| ENSG00000183020 | AP2A2 | -0,181868659 | 0,88156041 | 0,48237 | 0,71892541 | adaptor related protein complex 2 subunit alpha 2 |
| ENSG00000185262 | UBALD2 | -0,181878104 | 0,881554639 | 0,41956 | 0,67530234 | UBA like domain containing 2 |
| ENSG00000002919 | SNX11 | -0,18189584 | 0,881543801 | 0,41943 | 0,67527436 | sorting nexin 11 |
| ENSG00000242457 | RBBP4P2 | -0,181954424 | 0,881508005 | 0,68837 | 0,84817878 | RBBP4 pseudogene 2 |
| ENSG00000287011 | AC003077,1 | -0,181999953 | 0,881480187 | 0,68126 | 0,84367915 | novel transcript, antisense to DOCK4 |
| ENSG00000225187 | AC073283,1 | -0,182023523 | 0,881465786 | 0,82165 | 0,91753858 | novel transcript |
| ENSG00000183977 | PP2D1 | -0,182084866 | 0,881428307 | 0,7137 | 0,8622254 | protein phosphatase 2C like domain containing 1 |
| ENSG00000263247 | PRR4 | -0,18221095 | 0,881351278 | 0,64638 | 0,82293278 | proline rich 4 |
| ENSG00000287158 | AC117464,1 | -0,182222802 | 0,881344038 | 0,80897 | 0,91201277 | novel transcript |
| ENSG00000288369 | RBP5 | -0,182335499 | 0,881275193 | 0,77763 | 0,89712515 | retinol binding protein 5 |
| ENSG00000139194 | RBP5 | -0,182335499 | 0,881275193 | 0,77763 | 0,89712515 | retinol binding protein 5 |
| ENSG00000163870 | TPRA1 | -0,182382649 | 0,881246392 | 0,46784 | 0,70757147 | transmembrane protein adipocyte associated 1 |
| ENSG00000214753 | HNRNPUL2 | -0,182624202 | 0,881098856 | 0,46673 | 0,70676695 | heterogeneous nuclear ribonucleoprotein U like 2 |
| ENSG00000256069 | A2MP1 | -0,182782674 | 0,881002077 | 0,83484 | 0,92450066 | alpha-2-macroglobulin pseudogene 1 |
| ENSG00000233382 | NKAPP1 | -0,182852435 | 0,880959478 | 0,59761 | 0,79470049 | NFKB activating protein pseudogene 1 |
| ENSG00000276888 | AL512624,2 | -0,183007607 | 0,880864729 | 0,85936 | 0,93683711 | novel transcript, antisense to POTEM |
| ENSG00000182054 | IDH2 | -0,183024606 | 0,88085435 | 0,22484 | 0,5403718 | isocitrate dehydrogenase (NADP(+)) 2 |
| ENSG00000272574 | AL596325,2 | -0,183075829 | 0,880823076 | 0,80147 | 0,90860375 | novel transcript, sense intronic to UAP1 |
| ENSG00000126254 | RBM42 | -0,183098917 | 0,88080898 | 0,50421 | 0,73315506 | RNA binding motif protein 42 |
| ENSG00000232838 | PET117 | -0,183174643 | 0,880762749 | 0,30217 | 0,59294633 | PET117 cytochrome c oxidase chaperone |
| ENSG00000119899 | SLC17A5 | -0,18321119 | 0,880740437 | 0,30863 | 0,5974479 | solute carrier family 17 member 5 |
| ENSG00000105136 | ZNF419 | -0,183252544 | 0,880715192 | 0,25643 | 0,56092974 | zinc finger protein 419 |
| ENSG00000148384 | INPP5E | -0,183268313 | 0,880705565 | 0,40623 | 0,66584658 | inositol polyphosphate-5-phosphatase E |
| ENSG00000225087 | AL583808,1 | -0,183422007 | 0,880611746 | 0,58757 | 0,78750866 | novel transcript |
| ENSG00000113205 | PCDHB3 | -0,183477314 | 0,880577988 | 0,38197 | 0,64905556 | protocadherin beta 3 |
| ENSG00000050030 | NEXMIF | -0,18368509 | 0,880451177 | 0,62563 | 0,81055893 | neurite extension and migration factor |

| | | | | | | |
|---|---|---|---|---|---|---|
| ENSG00000075413 | MARK3 | -0,183873404 | 0,880336259 | 0,46161 | 0,70290608 | microtubule affinity regulating kinase 3 |
| ENSG00000100842 | EFS | -0,1839506 | 0,880289156 | 0,43263 | 0,68418465 | embryonal Fyn-associated substrate |
| ENSG00000104219 | ZDHHC2 | -0,183978056 | 0,880272403 | 0,70797 | 0,85927265 | zinc finger DHHC-type palmitoyltransferase 2 |
| ENSG00000228128 | DDAH2 | -0,184011639 | 0,880251912 | 0,54897 | 0,76394298 | dimethylarginine dimethylaminohydrolase 2 |
| ENSG00000225635 | DDAH2 | -0,184011639 | 0,880251912 | 0,54897 | 0,76394298 | dimethylarginine dimethylaminohydrolase 2 |
| ENSG00000233076 | DDAH2 | -0,184011639 | 0,880251912 | 0,54897 | 0,76394298 | dimethylarginine dimethylaminohydrolase 2 |
| ENSG00000113211 | PCDHB6 | -0,184033786 | 0,8802384 | 0,5733 | 0,77886693 | protocadherin beta 6 |
| ENSG00000272562 | AL512343,2 | -0,184283143 | 0,880086271 | 0,66408 | 0,83250729 | novel transcript, antisense to H3F3A |
| ENSG00000287229 | AC009731,1 | -0,184383487 | 0,88002506 | 0,62591 | 0,81058216 | novel transcript, antisense to RBM19 |
| ENSG00000260583 | AP000223,1 | -0,184445786 | 0,87998706 | 0,73568 | 0,8750728 | novel transcript, antisense to MRPL39 |
| ENSG00000130731 | METTL26 | -0,184478955 | 0,879966828 | 0,49356 | 0,7263798 | methyltransferase like 26 |
| ENSG00000239887 | C1orf226 | -0,184525856 | 0,879938222 | 0,45533 | 0,6993739 | chromosome 1 open reading frame 226 |
| ENSG00000276754 | AC006111,3 | -0,184525942 | 0,879938169 | 0,83098 | 0,92270607 | novel transcript, antisense to TRAP1 |
| ENSG00000272556 | GTF2IP13 | -0,18481097 | 0,87976434 | 0,49213 | 0,72542601 | general transcription factor IIi pseudogene 13 |
| ENSG00000249635 | AC109361,1 | -0,184911573 | 0,879702994 | 0,7916 | 0,90310057 | novel transcript |
| ENSG00000277870 | FAM230A | -0,184940412 | 0,879685409 | 0,73327 | 0,87353852 | family with sequence similarity 230 member A |
| ENSG00000276509 | AC239799,1 | -0,185115757 | 0,879578499 | 0,68348 | 0,84495012 | novel transcript |
| ENSG00000232913 | PLCE1-AS2 | -0,185151862 | 0,879556487 | 0,80425 | 0,90987458 | PLCE1 antisense RNA 2 |
| ENSG00000173581 | CCDC106 | -0,185166604 | 0,879547499 | 0,50187 | 0,73149732 | coiled-coil domain containing 106 |
| ENSG00000107819 | SFXN3 | -0,18517277 | 0,87954374 | 0,48465 | 0,72038194 | sideroflexin 3 |
| ENSG00000272969 | AC024243,1 | -0,185407044 | 0,879400926 | 0,77144 | 0,8940638 | novel transcript, antisense to clock |
| ENSG00000125520 | SLC2A4RG | -0,185477284 | 0,879358112 | 0,45737 | 0,70074793 | SLC2A4 regulator |
| ENSG00000267422 | AC016582,2 | -0,185487146 | 0,8793521 | 0,729 | 0,87140602 | increased sodium tolerance 1 homolog (yeast) (IST1) pseudogene |
| ENSG00000224307 | AL161785,1 | -0,185624856 | 0,879268167 | 0,80796 | 0,91168773 | novel transcript |
| ENSG00000144029 | MRPS5 | -0,185643169 | 0,879257006 | 0,39882 | 0,66135164 | mitochondrial ribosomal protein S5 |
| ENSG00000228192 | AL512353,1 | -0,185807677 | 0,879156752 | 0,57378 | 0,77921182 | novel transcript, antisense to ERMAP |
| ENSG00000235026 | DPP10-AS1 | -0,185838739 | 0,879137823 | 0,79087 | 0,90292452 | DPP10 antisense RNA 1 |
| ENSG00000089163 | SIRT4 | -0,185899618 | 0,879100726 | 0,7082 | 0,85927396 | sirtuin 4 |
| ENSG00000269293 | ZSCAN16-AS1 | -0,185972512 | 0,879056309 | 0,48695 | 0,72215091 | ZSCAN16 antisense RNA 1 |
| ENSG00000270175 | AC023509,3 | -0,18600707 | 0,879035253 | 0,58416 | 0,78572744 | novel transcript, antisense to TARBP2 |
| ENSG00000164877 | MICALL2 | -0,186086819 | 0,878986663 | 0,56037 | 0,77078479 | MICAL like 2 |
| ENSG00000184897 | H1-10 | -0,186114441 | 0,878969834 | 0,43054 | 0,68295004 | H1,10 linker histone |

| | | | | | | |
|---|---|---|---|---|---|---|
| ENSG00000100105 | PATZ1 | -0,186119501 | 0,878966752 | 0,28766 | 0,58395529 | POZ/BTB and AT hook containing zinc finger 1 |
| ENSG00000135324 | MRAP2 | -0,186121298 | 0,878965657 | 0,65687 | 0,82856771 | melanocortin 2 receptor accessory protein 2 |
| ENSG00000232682 | AL592430,1 | -0,18616159 | 0,878941109 | 0,75722 | 0,88663582 | novel transcript, antisense to ANK3 |
| ENSG00000273247 | AC097376,3 | -0,18633066 | 0,878838112 | 0,43204 | 0,68401174 | novel transcript, antisense to RAB33B |
| ENSG00000172403 | SYNPO2 | -0,18640092 | 0,878795313 | 0,80883 | 0,9120028 | synaptopodin 2 |
| ENSG00000241907 | RPS20P4 | -0,186535558 | 0,878713304 | 0,82119 | 0,91738668 | ribosomal protein S20 pseudogene 4 |
| ENSG00000166452 | AKIP1 | -0,186599277 | 0,878674495 | 0,65071 | 0,82499358 | A-kinase interacting protein 1 |
| ENSG00000270276 | H4C15 | -0,186638624 | 0,878650531 | 0,61371 | 0,8035281 | H4 clustered histone 15 |
| ENSG00000272103 | AC026741,1 | -0,186673242 | 0,878629448 | 0,54151 | 0,75882662 | novel transcript |
| ENSG00000183023 | SLC8A1 | -0,186726135 | 0,878597235 | 0,56864 | 0,77619389 | solute carrier family 8 member A1 |
| ENSG00000223599 | AL513523,1 | -0,186750648 | 0,878582307 | 0,65906 | 0,82969668 | adipose differentiation-related protein (ADFP) pseudogene |
| ENSG00000262025 | AL513523,5 | -0,186750648 | 0,878582307 | 0,65906 | 0,82969668 | adipose differentiation-related protein (ADFP) pseudogene |
| ENSG00000236360 | RRAS2P1 | -0,186759687 | 0,878576803 | 0,84267 | 0,92821072 | RRAS2 pseudogene 1 |
| ENSG00000260190 | AL807752,5 | -0,186760238 | 0,878576467 | 0,73618 | 0,87519603 | novel transcript, overlapping FBXW5 |
| ENSG00000134905 | CARS2 | -0,187026186 | 0,878414524 | 0,29895 | 0,5910969 | cysteinyl-tRNA synthetase 2, mitochondrial |
| ENSG00000122550 | KLHL7 | -0,187135642 | 0,878347882 | 0,46372 | 0,70449472 | kelch like family member 7 |
| ENSG00000128731 | HERC2 | -0,187315396 | 0,87823845 | 0,41884 | 0,67490937 | HECT and RLD domain containing E3 ubiquitin protein ligase 2 |
| ENSG00000198858 | R3HDM4 | -0,187333482 | 0,87822744 | 0,38497 | 0,65080233 | R3H domain containing 4 |
| ENSG00000251580 | LINC02482 | -0,187379776 | 0,87819926 | 0,53534 | 0,75460305 | long intergenic non-protein coding RNA 2482 |
| ENSG00000176945 | MUC20 | -0,187507877 | 0,878121286 | 0,57303 | 0,77878023 | mucin 20, cell surface associated |
| ENSG00000231443 | AC124944,1 | -0,187544159 | 0,878099202 | 0,71458 | 0,86265094 | protein phosphatase 4, regulatory subunit 2 (PPP4R2) pseudogene |
| ENSG00000065000 | AP3D1 | -0,187582995 | 0,878075565 | 0,38039 | 0,64797239 | adaptor related protein complex 3 subunit delta 1 |
| ENSG00000158445 | KCNB1 | -0,187624076 | 0,878050562 | 0,41888 | 0,67490937 | potassium voltage-gated channel subfamily B member 1 |
| ENSG00000131591 | C1orf159 | -0,187639284 | 0,878041306 | 0,27364 | 0,57364891 | chromosome 1 open reading frame 159 |
| ENSG00000248449 | PCDHGB8P | -0,187695937 | 0,878006827 | 0,53357 | 0,75353661 | protocadherin gamma subfamily B, 8 pseudogene |
| ENSG00000159433 | STARD9 | -0,187737076 | 0,877981791 | 0,42167 | 0,67643128 | StAR related lipid transfer domain containing 9 |
| ENSG00000274211 | SOCS7 | -0,187807488 | 0,877938941 | 0,42181 | 0,67643128 | suppressor of cytokine signaling 7 |
| ENSG00000141448 | GATA6 | -0,187850075 | 0,877913026 | 0,76641 | 0,89138648 | GATA binding protein 6 |
| ENSG00000285132 | AC270285,3 | -0,188362356 | 0,877601346 | 0,50911 | 0,73703253 | cathepsin B |

| | | | | | | |
|---|---|---|---|---|---|---|
| ENSG00000100218 | RSPH14 | -0,188458904 | 0,877542618 | 0,64744 | 0,82360839 | radial spoke head 14 homolog |
| ENSG00000276143 | AC138749,7 | -0,18849331 | 0,87752169 | 0,86594 | 0,94045351 | family with sequence similarity 208, member A (FAM208A) pseudogene |
| ENSG00000277809 | AC012354,8 | -0,188505942 | 0,877514006 | 0,74083 | 0,87717177 | |
| ENSG00000246695 | RASSF8-AS1 | -0,188659373 | 0,877420687 | 0,42935 | 0,68217673 | RASSF8 antisense RNA 1 |
| ENSG00000274334 | VASN | -0,18867538 | 0,877410952 | 0,70089 | 0,85559159 | vasorin |
| ENSG00000280634 | THRIL | -0,188678001 | 0,877409358 | 0,79502 | 0,90508829 | TNF and HNRNPL related immunoregulatory long non-coding RNA |
| ENSG00000287735 | AL355075,6 | -0,188832429 | 0,877315444 | 0,65499 | 0,82746625 | novel transcript, antisense to CCNB1IP1 |
| ENSG00000219712 | AL357054,1 | -0,188838667 | 0,877311651 | 0,825 | 0,91900665 | novel serine/threonine kinase pseudogene |
| ENSG00000184313 | MROH7 | -0,188866736 | 0,877294582 | 0,81421 | 0,91457884 | maestro heat like repeat family member 7 |
| ENSG00000145348 | TBCK | -0,188894005 | 0,877278 | 0,45869 | 0,70133296 | TBC1 domain containing kinase |
| ENSG00000255248 | MIR100HG | -0,189021833 | 0,877200274 | 0,64925 | 0,82436914 | mir-100-let-7a-2-mir-125b-1 cluster host gene |
| ENSG00000214756 | CSKMT | -0,18908303 | 0,877163065 | 0,41124 | 0,66903348 | citrate synthase lysine methyltransferase |
| ENSG00000269956 | MKNK1-AS1 | -0,189145024 | 0,877125373 | 0,6543 | 0,82697207 | MKNK1 antisense RNA 1 |
| ENSG00000018189 | RUFY3 | -0,189274293 | 0,877046784 | 0,44609 | 0,69284796 | RUN and FYVE domain containing 3 |
| ENSG00000129245 | FXR2 | -0,189329833 | 0,877013021 | 0,43885 | 0,6878469 | FMR1 autosomal homolog 2 |
| ENSG00000254044 | AC034154,1 | -0,189343476 | 0,877004727 | 0,58202 | 0,78454692 | novel transcript, sense overlapping to STPG2 |
| ENSG00000204113 | BMP2KL | -0,189424199 | 0,876955657 | 0,74584 | 0,88010204 | BMP2 inducible kinase like (pseudogene) |
| ENSG00000246820 | RIC3-DT | -0,189477238 | 0,876923418 | 0,70004 | 0,85515696 | RIC3 divergent transcript |
| ENSG00000285739 | AC010157,2 | -0,189492582 | 0,876914091 | 0,73406 | 0,87407106 | novel transcript, antisense to NRG3 |
| ENSG00000261033 | AC005730,2 | -0,189497242 | 0,876911259 | 0,73705 | 0,87539732 | novel transcript, antisense to SPECC1 |
| ENSG00000176153 | GPX2 | -0,189593559 | 0,876852716 | 0,6685 | 0,83543142 | glutathione peroxidase 2 |
| ENSG00000283393 | AC068205,3 | -0,189597296 | 0,876850445 | 0,76767 | 0,89238586 | ribosomal protein L23a (RPL23A) pseudogene |
| ENSG00000197136 | PCNX3 | -0,189637982 | 0,876825717 | 0,49675 | 0,72853804 | pecanex 3 |
| ENSG00000229696 | KARS1P1 | -0,189691416 | 0,876793242 | 0,81633 | 0,91550954 | lysyl-tRNA synthetase 1 pseudogene 1 |
| ENSG00000286953 | AC122697,1 | -0,189732916 | 0,876768021 | 0,83919 | 0,92642589 | novel transcript |
| ENSG00000230143 | FLOT1 | -0,189881948 | 0,876677455 | 0,72728 | 0,87016629 | flotillin 1 |
| ENSG00000100994 | PYGB | -0,189894198 | 0,876670011 | 0,23471 | 0,54623956 | glycogen phosphorylase B |
| ENSG00000214140 | PRCD | -0,189907496 | 0,87666193 | 0,80605 | 0,91078302 | photoreceptor disc component |
| ENSG00000277767 | AL442128,2 | -0,189989211 | 0,876612277 | 0,55726 | 0,76903027 | novel transcript |
| ENSG00000178104 | PDE4DIP | -0,190003436 | 0,876603634 | 0,31272 | 0,59962409 | phosphodiesterase 4D interacting protein |
| ENSG00000103091 | WDR59 | -0,190031729 | 0,876586443 | 0,44659 | 0,69321646 | WD repeat domain 59 |

| | | | | | | |
|---|---|---|---|---|---|---|
| ENSG00000246523 | FZD4-DT | -0,19016425 | 0,876505926 | 0,74297 | 0,87840582 | FZD4 divergent transcript |
| ENSG00000238227 | TMEM250 | -0,190245026 | 0,876456852 | 0,56369 | 0,77255679 | transmembrane protein 250 |
| ENSG00000227375 | DLG1-AS1 | -0,190273342 | 0,87643965 | 0,73776 | 0,87540065 | DLG1 antisense RNA 1 |
| ENSG00000151838 | CCDC175 | -0,190280679 | 0,876435192 | 0,77901 | 0,89781842 | coiled-coil domain containing 175 |
| ENSG00000271122 | AC018647,2 | -0,190446446 | 0,876334495 | 0,27234 | 0,57315552 | novel transcript |
| ENSG00000173281 | PPP1R3B | -0,190544499 | 0,876274937 | 0,59731 | 0,79440664 | protein phosphatase 1 regulatory subunit 3B |
| ENSG00000128228 | SDF2L1 | -0,190809281 | 0,876114126 | 0,55871 | 0,77025265 | stromal cell derived factor 2 like 1 |
| ENSG00000261761 | LINC02616 | -0,190833485 | 0,876099428 | 0,79868 | 0,90713957 | long intergenic non-protein coding RNA 2616 |
| ENSG00000123159 | GIPC1 | -0,190894009 | 0,876062674 | 0,48455 | 0,72031474 | GIPC PDZ domain containing family member 1 |
| ENSG00000285702 | AC211476,5 | -0,191063103 | 0,87596 | 0,37038 | 0,63986978 | novel transcript |
| ENSG00000286945 | AL513478,4 | -0,191149337 | 0,875907643 | 0,9004 | 0,95760678 | novel transcript |
| ENSG00000245648 | KLRK1-AS1 | -0,191200296 | 0,875876705 | 0,52364 | 0,7474875 | KLRK1 antisense RNA 1 |
| ENSG00000270673 | YTHDF3-AS1 | -0,19125353 | 0,875844386 | 0,68663 | 0,84703257 | YTHDF3 antisense RNA 1 (head to head) |
| ENSG00000229816 | DDX50P1 | -0,19125524 | 0,875843348 | 0,51142 | 0,73853921 | DEAD-box helicase 50 pseudogene 1 |
| ENSG00000162482 | AKR7A3 | -0,191337448 | 0,875793442 | 0,56467 | 0,77306193 | aldo-keto reductase family 7 member A3 |
| ENSG00000188394 | GPR21 | -0,191470833 | 0,875712474 | 0,56986 | 0,777027 | G protein-coupled receptor 21 |
| ENSG00000116350 | SRSF4 | -0,191492819 | 0,875699128 | 0,34154 | 0,61963031 | serine and arginine rich splicing factor 4 |
| ENSG00000224007 | USP12P2 | -0,191765658 | 0,875533534 | 0,7741 | 0,89572556 | USP12 pseudogene 2 |
| ENSG00000174473 | GALNTL6 | -0,19181447 | 0,875503912 | 0,70141 | 0,85604483 | polypeptide N-acetylgalactosaminyltransferase like 6 |
| ENSG00000272942 | AL022324,3 | -0,191936389 | 0,875429928 | 0,84651 | 0,93030659 | novel transcript |
| ENSG00000286859 | AL158214,2 | -0,192023045 | 0,875377347 | 0,65477 | 0,8272805 | novel transcript, antisense to COG2 |
| ENSG00000272950 | AC093799,1 | -0,192051585 | 0,87536003 | 0,81862 | 0,91672281 | novel transcript, antisense to BAIAP2L1 |
| ENSG00000228495 | LINC01013 | -0,192057738 | 0,875356297 | 0,78713 | 0,90117606 | long intergenic non-protein coding RNA 1013 |
| ENSG00000169217 | CD2BP2 | -0,192150842 | 0,875299807 | 0,37806 | 0,64636276 | CD2 cytoplasmic tail binding protein 2 |
| ENSG00000185049 | NELFA | -0,192234145 | 0,875249268 | 0,39063 | 0,65464817 | negative elongation factor complex member A |
| ENSG00000188921 | HACD4 | -0,192359175 | 0,875173418 | 0,38132 | 0,64873107 | 3-hydroxyacyl-CoA dehydratase 4 |
| ENSG00000163866 | SMIM12 | -0,192407293 | 0,87514423 | 0,36218 | 0,63365406 | small integral membrane protein 12 |
| ENSG00000285312 | C6orf52 | -0,19241719 | 0,875138226 | 0,7244 | 0,8684669 | chromosome 6 open reading frame 52 |
| ENSG00000137434 | C6orf52 | -0,19241719 | 0,875138226 | 0,7244 | 0,8684669 | chromosome 6 open reading frame 52 |
| ENSG00000146350 | TBC1D32 | -0,192562931 | 0,875049824 | 0,28257 | 0,57997723 | TBC1 domain family member 32 |
| ENSG00000273240 | AC013468,1 | -0,192635264 | 0,875005952 | 0,79317 | 0,90371578 | novel transcript |
| ENSG00000133612 | AGAP3 | -0,192758141 | 0,87493143 | 0,52885 | 0,75107579 | ArfGAP with GTPase domain, ankyrin repeat and PH domain 3 |

| | | | | | | |
|---|---|---|---|---|---|---|
| ENSG00000281909 | HERC2P7 | -0,1928207 | 0,874893491 | 0,80908 | 0,91204664 | hect domain and RLD 2 pseudogene 7 |
| ENSG00000271958 | AC021860,1 | -0,192940247 | 0,874820997 | 0,80448 | 0,90996631 | novel transcript |
| ENSG00000236032 | OR5H14 | -0,192991036 | 0,874790201 | 0,71262 | 0,86173807 | olfactory receptor family 5 subfamily H member 14 |
| ENSG00000271109 | AC008555,4 | -0,193093979 | 0,874727782 | 0,29768 | 0,59036937 | novel transcript |
| ENSG00000173040 | EVC2 | -0,193154019 | 0,87469138 | 0,57505 | 0,78004262 | EvC ciliary complex subunit 2 |
| ENSG00000115155 | OTOF | -0,193178249 | 0,874676689 | 0,81994 | 0,91720659 | otoferlin |
| ENSG00000164576 | SAP30L | -0,193179988 | 0,874675635 | 0,38685 | 0,6521009 | SAP30 like |
| ENSG00000213055 | EEF1B2P7 | -0,193193178 | 0,874667638 | 0,69099 | 0,84920479 | eukaryotic translation elongation factor 1 beta 2 pseudogene 7 |
| ENSG00000158604 | TMED4 | -0,193205558 | 0,874660133 | 0,34315 | 0,62060539 | transmembrane p24 trafficking protein 4 |
| ENSG00000100027 | YPEL1 | -0,193224914 | 0,874648398 | 0,3966 | 0,65923689 | yippee like 1 |
| ENSG00000205838 | TTC23L | -0,193273124 | 0,87461917 | 0,6609 | 0,83097133 | tetratricopeptide repeat domain 23 like |
| ENSG00000104689 | TNFRSF10A | -0,19346964 | 0,874500043 | 0,68039 | 0,84309883 | TNF receptor superfamily member 10a |
| ENSG00000272735 | AC007881,3 | -0,193531803 | 0,874462363 | 0,81393 | 0,9144566 | novel transcript |
| ENSG00000161217 | PCYT1A | -0,193751353 | 0,874329297 | 0,54368 | 0,76018847 | phosphate cytidylyltransferase 1, choline, alpha |
| ENSG00000269825 | AC022150,4 | -0,193923673 | 0,874224871 | 0,31692 | 0,60268749 | novel transcript, sense intronic to ZNF83 |
| ENSG00000163110 | PDLIM5 | -0,193927544 | 0,874222525 | 0,49357 | 0,7263798 | PDZ and LIM domain 5 |
| ENSG00000272371 | AL591167,1 | -0,193996814 | 0,874180551 | 0,69379 | 0,8512419 | novel transcript |
| ENSG00000146592 | CREB5 | -0,193997637 | 0,874180052 | 0,51752 | 0,74270246 | cAMP responsive element binding protein 5 |
| ENSG00000218226 | TATDN2P2 | -0,194020362 | 0,874166283 | 0,4964 | 0,72831792 | TatD DNase domain containing 2 pseudogene 2 |
| ENSG00000234040 | RPL10P12 | -0,194151046 | 0,874087101 | 0,81597 | 0,91538504 | ribosomal protein L10 pseudogene 12 |
| ENSG00000170619 | COMMD5 | -0,194198243 | 0,874058506 | 0,3375 | 0,61718742 | COMM domain containing 5 |
| ENSG00000215158 | AC138409,2 | -0,194232281 | 0,874037885 | 0,39206 | 0,65540679 | Putative beta-glucuronidase-like protein FLJ75287 pseudogene |
| ENSG00000274437 | DERL3 | -0,194239559 | 0,874033475 | 0,63411 | 0,81555826 | derlin 3 |
| ENSG00000105516 | DBP | -0,194267275 | 0,874016684 | 0,56822 | 0,77590889 | D-box binding PAR bZIP transcription factor |
| ENSG00000272140 | AC022400,4 | -0,194299016 | 0,873997455 | 0,54043 | 0,75829455 | novel transcript |
| ENSG00000277128 | AL589743,3 | -0,194345716 | 0,873969164 | 0,65018 | 0,82472422 | novel transcript |
| ENSG00000164105 | SAP30 | -0,194359611 | 0,873960747 | 0,40118 | 0,6630277 | Sin3A associated protein 30 |
| ENSG00000157985 | AGAP1 | -0,1943684 | 0,873955423 | 0,29499 | 0,58869206 | ArfGAP with GTPase domain, ankyrin repeat and PH domain 1 |
| ENSG00000270108 | AL049840,6 | -0,194493511 | 0,873879636 | 0,56187 | 0,77147224 | novel transcript, sense intronic to KLC1 |
| ENSG00000153930 | ANKFN1 | -0,194531862 | 0,873856406 | 0,69767 | 0,85352639 | ankyrin repeat and fibronectin type III domain containing 1 |

| | | | | | | |
|---|---|---|---|---|---|---|
| ENSG00000238141 | BRWD1-AS1 | -0,194609234 | 0,873809542 | 0,77182 | 0,89435615 | BRWD1 antisense RNA 1 |
| ENSG00000265713 | AC023389,2 | -0,194678616 | 0,87376752 | 0,67636 | 0,84075837 | RNA terminal phosphate cyclase-like 1 (RCL1) pseudogene |
| ENSG00000214676 | RPL9P16 | -0,194815335 | 0,873684721 | 0,73807 | 0,87561417 | ribosomal protein L9 pseudogene 16 |
| ENSG00000198853 | RUSC2 | -0,1949403 | 0,873609046 | 0,45762 | 0,70082333 | RUN and SH3 domain containing 2 |
| ENSG00000217120 | Z98755,1 | -0,195089034 | 0,873518986 | 0,57811 | 0,78191189 | ClpX caseinolytic protease X homolog (E, coli) (CLPX) pseudogene |
| ENSG00000119865 | CNRIP1 | -0,195125401 | 0,873496967 | 0,64432 | 0,82151477 | cannabinoid receptor interacting protein 1 |
| ENSG00000204560 | DHX16 | -0,195143196 | 0,873486193 | 0,55263 | 0,76596429 | DEAH-box helicase 16 |
| ENSG00000230188 | AL158156,1 | -0,195315512 | 0,873381869 | 0,80053 | 0,90812402 | ribosomal protein L21 (RPL21) pseudogene |
| ENSG00000239246 | AC008026,1 | -0,195379668 | 0,873343031 | 0,8651 | 0,93999031 | ribosomal protein S10 (RPS10) pseudogene |
| ENSG00000172336 | POP7 | -0,195493472 | 0,873274143 | 0,52856 | 0,75090627 | POP7 homolog, ribonuclease P/MRP subunit |
| ENSG00000125755 | SYMPK | -0,195674821 | 0,873164378 | 0,41866 | 0,67477592 | symplekin |
| ENSG00000237054 | PRMT5-AS1 | -0,195707014 | 0,873144893 | 0,6212 | 0,80787022 | PRMT5 antisense RNA 1 |
| ENSG00000148704 | VAX1 | -0,19573573 | 0,873127514 | 0,55493 | 0,76763832 | ventral anterior homeobox 1 |
| ENSG00000229221 | HNRNPA1P66 | -0,195848651 | 0,873059177 | 0,80341 | 0,90947335 | heterogeneous nuclear ribonucleoprotein A1 pseudogene 66 |
| ENSG00000197565 | COL4A6 | -0,195852366 | 0,873056929 | 0,67409 | 0,83946548 | collagen type IV alpha 6 chain |
| ENSG00000241884 | AC114401,1 | -0,195870216 | 0,873046126 | 0,76942 | 0,89297049 | novel transcript |
| ENSG00000182831 | C16orf72 | -0,195884903 | 0,873037239 | 0,29495 | 0,58868965 | chromosome 16 open reading frame 72 |
| ENSG00000248401 | AC114781,1 | -0,195892285 | 0,873032771 | 0,78017 | 0,89822236 | aminomethyltransferase (AMT) pseudogene |
| ENSG00000121057 | AKAP1 | -0,195907509 | 0,873023559 | 0,34315 | 0,62060539 | A-kinase anchoring protein 1 |
| ENSG00000130726 | TRIM28 | -0,195932309 | 0,873008552 | 0,44838 | 0,69432421 | tripartite motif containing 28 |
| ENSG00000253320 | MAILR | -0,195934432 | 0,873007267 | 0,29077 | 0,58579789 | macrophage interferon regulatory lncRNA |
| ENSG00000133424 | LARGE1 | -0,1959509 | 0,872997302 | 0,59384 | 0,79208467 | LARGE xylosyl- and glucuronyltransferase 1 |
| ENSG00000275050 | ZNF66 | -0,1961929 | 0,872850876 | 0,55325 | 0,76630282 | zinc finger protein 66 |
| ENSG00000160888 | IER2 | -0,196285165 | 0,872795056 | 0,65807 | 0,82897081 | immediate early response 2 |
| ENSG00000282015 | AC004099,2 | -0,196586523 | 0,872612761 | 0,8092 | 0,91206591 | ribosomal protein L24 (RPL24) pseudogene |
| ENSG00000275839 | AC243654,2 | -0,196586523 | 0,872612761 | 0,8092 | 0,91206591 | ribosomal protein L24 (RPL24) pseudogene |
| ENSG00000197013 | ZNF429 | -0,196648108 | 0,872575512 | 0,47752 | 0,71544156 | zinc finger protein 429 |
| ENSG00000241316 | SUCLG2-AS1 | -0,196687883 | 0,872551456 | 0,33739 | 0,61706766 | SUCLG2 antisense RNA 1 (head to head) |
| ENSG00000183751 | TBL3 | -0,196737082 | 0,872521701 | 0,4564 | 0,70003941 | transducin beta like 3 |
| ENSG00000254141 | AC103853,2 | -0,196915264 | 0,872413945 | 0,7505 | 0,88293152 | novel transcript |
| ENSG00000166140 | ZFYVE19 | -0,196943412 | 0,872396924 | 0,44019 | 0,68866128 | zinc finger FYVE-type containing 19 |

| | | | | | | |
|---|---|---|---|---|---|---|
| ENSG00000158941 | CCAR2 | -0,196963271 | 0,872384915 | 0,35118 | 0,62664963 | cell cycle and apoptosis regulator 2 |
| ENSG00000183979 | NPB | -0,19701673 | 0,87235259 | 0,77312 | 0,89523715 | neuropeptide B |
| ENSG00000270953 | AC007938,3 | -0,197155576 | 0,872268638 | 0,56791 | 0,77572201 | novel transcript, antisense to COPG2 |
| ENSG00000126432 | PRDX5 | -0,197301118 | 0,872180646 | 0,50651 | 0,73497152 | peroxiredoxin 5 |
| ENSG00000271998 | AC034229,3 | -0,197450665 | 0,872090242 | 0,87383 | 0,94405019 | novel transcript |
| ENSG00000223813 | AC007255,1 | -0,197456904 | 0,87208647 | 0,67461 | 0,83975921 | novel transcript, antisense to CHN2 |
| ENSG00000100324 | TAB1 | -0,197470402 | 0,872078312 | 0,2012 | 0,52181085 | TGF-beta activated kinase 1 (MAP3K7) binding protein 1 |
| ENSG00000267855 | NDUFA7 | -0,197580593 | 0,872011706 | 0,53376 | 0,75368609 | NADH:ubiquinone oxidoreductase subunit A7 |
| ENSG00000283879 | MTND6P35 | -0,197634114 | 0,871979357 | 0,85484 | 0,93417381 | MT-ND6 pseudogene 35 |
| ENSG00000287975 | AC113143,2 | -0,197677263 | 0,871953277 | 0,71385 | 0,86226331 | novel transcript |
| ENSG00000069431 | ABCC9 | -0,197870607 | 0,87183643 | 0,54584 | 0,76151932 | ATP binding cassette subfamily C member 9 |
| ENSG00000151876 | FBXO4 | -0,197875396 | 0,871833536 | 0,3506 | 0,62606342 | F-box protein 4 |
| ENSG00000267199 | AP001029,2 | -0,198188615 | 0,871644275 | 0,49826 | 0,72956698 | novel transcript, antisense SPIRE1 |
| ENSG00000177599 | ZNF491 | -0,198233028 | 0,871617442 | 0,29538 | 0,58912175 | zinc finger protein 491 |
| ENSG00000235258 | NDUFB4P6 | -0,198233307 | 0,871617274 | 0,77734 | 0,8969723 | NADH:ubiquinone oxidoreductase subunit B4 pseudogene 6 |
| ENSG00000184911 | DMRTC1B | -0,19831261 | 0,871569363 | 0,59875 | 0,79523922 | DMRT like family C1B |
| ENSG00000235919 | ASH1L-AS1 | -0,19847215 | 0,871472986 | 0,48228 | 0,71892541 | ASH1L antisense RNA 1 |
| ENSG00000272888 | CHASERR | -0,19850695 | 0,871451966 | 0,20988 | 0,52877648 | CHD2 adjacent suppressive regulatory RNA |
| ENSG00000253230 | MIR124-1HG | -0,198532289 | 0,871436659 | 0,63001 | 0,8132682 | MIR124-1 host gene |
| ENSG00000198917 | SPOUT1 | -0,198818704 | 0,871263673 | 0,31297 | 0,59987747 | SPOUT domain containing methyltransferase 1 |
| ENSG00000257727 | CNPY2 | -0,198906645 | 0,871210565 | 0,52432 | 0,74827564 | canopy FGF signaling regulator 2 |
| ENSG00000267808 | AC018755,1 | -0,198961761 | 0,871177283 | 0,79991 | 0,90771269 | novel transcript |
| ENSG00000236047 | RPL13AP12 | -0,199028499 | 0,871136984 | 0,632 | 0,81449147 | ribosomal protein L13a pseudogene 12 |
| ENSG00000261072 | AC084783,1 | -0,199030235 | 0,871135936 | 0,58597 | 0,78652771 | hydroxyacyl-CoA dehydrogenase (HADH) pseudogene |
| ENSG00000235316 | DUSP8P5 | -0,199041209 | 0,871129309 | 0,54115 | 0,75868526 | dual specificity phosphatase 8 pseudogene 5 |
| ENSG00000076928 | ARHGEF1 | -0,199066017 | 0,871114329 | 0,5044 | 0,7332113 | Rho guanine nucleotide exchange factor 1 |
| ENSG00000144362 | PHOSPHO2 | -0,199070918 | 0,87111137 | 0,51085 | 0,73828855 | phosphatase, orphan 2 |
| ENSG00000224295 | OLFM5P | -0,199395006 | 0,870915705 | 0,7629 | 0,889794 | olfactomedin family member 5, pseudogene |
| ENSG00000248434 | AC107463,1 | -0,199421386 | 0,87089978 | 0,57588 | 0,78043323 | novel transcript |
| ENSG00000228412 | LNC-LBCS | -0,199468248 | 0,870871492 | 0,47353 | 0,71276528 | lncRNA bladder and prostate cancer suppressor, hnRNPK interacting |
| ENSG00000172932 | ANKRD13D | -0,199494835 | 0,870855443 | 0,33695 | 0,61684812 | ankyrin repeat domain 13D |

| | | | | | | |
|---|---|---|---|---|---|---|
| ENSG00000271401 | AC010680,5 | -0,199564224 | 0,870813559 | 0,84577 | 0,92982715 | novel transcript |
| ENSG00000170144 | HNRNPA3 | -0,199698218 | 0,870732684 | 0,36165 | 0,63352908 | heterogeneous nuclear ribonucleoprotein A3 |
| ENSG00000270919 | AC108451,2 | -0,199807138 | 0,870666948 | 0,76414 | 0,89012684 | novel transcript, antisense to ARNT2 |
| ENSG00000234632 | NECAP1P2 | -0,199867635 | 0,870630438 | 0,78947 | 0,90251229 | NECAP endocytosis associated 1 pseudogene 2 |
| ENSG00000182484 | WASH6P | -0,200132528 | 0,870470597 | 0,28311 | 0,58042135 | WASP family homolog 6, pseudogene |
| ENSG00000261242 | AL136038,3 | -0,200178924 | 0,870442604 | 0,72248 | 0,86734497 | novel transcript, antisense to PPP2R5E |
| ENSG00000018236 | CNTN1 | -0,200308172 | 0,870364626 | 0,54449 | 0,76055278 | contactin 1 |
| ENSG00000243667 | WDR92 | -0,20047211 | 0,87026573 | 0,435 | 0,68571197 | WD repeat domain 92 |
| ENSG00000130714 | POMT1 | -0,200478952 | 0,870261602 | 0,27203 | 0,57306433 | protein O-mannosyltransferase 1 |
| ENSG00000163352 | LENEP | -0,200512149 | 0,870241578 | 0,83736 | 0,92557879 | lens epithelial protein |
| ENSG00000280518 | AC244260,1 | -0,200542862 | 0,870223052 | 0,7926 | 0,90361513 | NLR family, apoptosis inhibitory protein pseudogene |
| ENSG00000188095 | MESP2 | -0,200648981 | 0,870159044 | 0,69683 | 0,85312597 | mesoderm posterior bHLH transcription factor 2 |
| ENSG00000188199 | NUTM2B | -0,200799511 | 0,870068257 | 0,5221 | 0,7463592 | NUT family member 2B |
| ENSG00000183091 | NEB | -0,200963749 | 0,869969212 | 0,474 | 0,71294959 | nebulin |
| ENSG00000189366 | ALG1L | -0,201056409 | 0,869913339 | 0,76319 | 0,88980026 | ALG1 chitobiosyldiphosphodolichol beta-mannosyltransferase like |
| ENSG00000272865 | AL591686,2 | -0,20109388 | 0,869890745 | 0,86751 | 0,94146404 | novel transcript, antisense to PLD5 |
| ENSG00000270098 | LDHAL6EP | -0,201139006 | 0,869863536 | 0,81363 | 0,9144566 | lactate dehydrogenase A like 6E, pseudogene |
| ENSG00000145349 | CAMK2D | -0,201180263 | 0,869838661 | 0,39817 | 0,66078008 | calcium/calmodulin dependent protein kinase II delta |
| ENSG00000125551 | PLGLB2 | -0,201428405 | 0,869689062 | 0,45226 | 0,69698561 | plasminogen like B2 |
| ENSG00000103642 | LACTB | -0,201450274 | 0,869675879 | 0,27874 | 0,57694485 | lactamase beta |
| ENSG00000279623 | AL359697,1 | -0,201531217 | 0,869627087 | 0,78612 | 0,90077646 | novel transcript, sense intronic to DNAJC1 |
| ENSG00000111490 | TBC1D30 | -0,201607651 | 0,869581015 | 0,69813 | 0,85390204 | TBC1 domain family member 30 |
| ENSG00000198912 | C1orf174 | -0,201730257 | 0,869507118 | 0,53303 | 0,75337932 | chromosome 1 open reading frame 174 |
| ENSG00000151694 | ADAM17 | -0,201777698 | 0,869478526 | 0,22724 | 0,54168591 | ADAM metallopeptidase domain 17 |
| ENSG00000162426 | SLC45A1 | -0,202003286 | 0,86934258 | 0,55887 | 0,77025265 | solute carrier family 45 member 1 |
| ENSG00000022976 | ZNF839 | -0,202005207 | 0,869341422 | 0,50698 | 0,73534235 | zinc finger protein 839 |
| ENSG00000179454 | KLHL28 | -0,202138685 | 0,869260995 | 0,30111 | 0,59252799 | kelch like family member 28 |
| ENSG00000259736 | CRTC3-AS1 | -0,202151415 | 0,869253325 | 0,5887 | 0,78830778 | CRTC3 antisense RNA 1 |
| ENSG00000255618 | LINC02440 | -0,202206379 | 0,869220208 | 0,64636 | 0,82293278 | long intergenic non-protein coding RNA 2440 |
| ENSG00000163702 | IL17RC | -0,202225894 | 0,869208451 | 0,56897 | 0,77629764 | interleukin 17 receptor C |
| ENSG00000100612 | DHRS7 | -0,202333215 | 0,869143794 | 0,50017 | 0,73078774 | dehydrogenase/reductase 7 |
| ENSG00000286074 | AC110795,1 | -0,202544254 | 0,869016663 | 0,77979 | 0,8981559 | novel transcript |

| | | | | | | |
|---|---|---|---|---|---|---|
| ENSG00000176022 | B3GALT6 | -0,202547386 | 0,869014777 | 0,52655 | 0,7497927 | beta-1,3-galactosyltransferase 6 |
| ENSG00000077463 | SIRT6 | -0,202606324 | 0,868979276 | 0,37399 | 0,64249775 | sirtuin 6 |
| ENSG00000272468 | AL021807,1 | -0,202607544 | 0,868978541 | 0,72147 | 0,86698039 | novel transcript |
| ENSG00000273204 | AC104506,1 | -0,202627268 | 0,868966661 | 0,73621 | 0,87519603 | novel transcript, antisense to EXTL2 |
| ENSG00000254400 | AC091564,2 | -0,202670347 | 0,868940714 | 0,91528 | 0,9633004 | novel transcript |
| ENSG00000248015 | AC005329,1 | -0,202718782 | 0,868911542 | 0,64968 | 0,82449863 | novel transcript, antisense to NDUFS7 |
| ENSG00000254469 | AP002495,1 | -0,202723706 | 0,868908576 | 0,46004 | 0,70205579 | XRCC1 N-terminal domain containing 1-like |
| ENSG00000276545 | PCDHGB9P | -0,202774778 | 0,868877817 | 0,528 | 0,75065864 | protocadherin gamma subfamily B, 9 pseudogene |
| ENSG00000273472 | AC096733,2 | -0,202819821 | 0,868850689 | 0,61429 | 0,80370125 | novel transcript |
| ENSG00000255321 | AC068389,3 | -0,202829379 | 0,868844933 | 0,7574 | 0,88672876 | novel transcript |
| ENSG00000249196 | TMEM132D-A | -0,202942842 | 0,868776604 | 0,79954 | 0,90748957 | TMEM132D antisense RNA 1 |
| ENSG00000102882 | MAPK3 | -0,202963542 | 0,868764139 | 0,43667 | 0,68681257 | mitogen-activated protein kinase 3 |
| ENSG00000105063 | PPP6R1 | -0,203201118 | 0,868621087 | 0,43791 | 0,68747773 | protein phosphatase 6 regulatory subunit 1 |
| ENSG00000114115 | RBP1 | -0,203323607 | 0,868547342 | 0,42747 | 0,68046596 | retinol binding protein 1 |
| ENSG00000118298 | CA14 | -0,203389766 | 0,868507513 | 0,6511 | 0,82522584 | carbonic anhydrase 14 |
| ENSG00000162714 | ZNF496 | -0,20347348 | 0,868457118 | 0,40583 | 0,66553796 | zinc finger protein 496 |
| ENSG00000153904 | DDAH1 | -0,203629347 | 0,868363296 | 0,46374 | 0,70449472 | dimethylarginine dimethylaminohydrolase 1 |
| ENSG00000177189 | RPS6KA3 | -0,203663404 | 0,868342798 | 0,54354 | 0,76016422 | ribosomal protein S6 kinase A3 |
| ENSG00000284161 | EEF2K | -0,2036832 | 0,868330883 | 0,61499 | 0,80416871 | eukaryotic elongation factor 2 kinase |
| ENSG00000116833 | NR5A2 | -0,203777245 | 0,868274281 | 0,85167 | 0,93249922 | nuclear receptor subfamily 5 group A member 2 |
| ENSG00000119720 | NRDE2 | -0,20378902 | 0,868267194 | 0,36689 | 0,6368162 | NRDE-2, necessary for RNA interference, domain containing |
| ENSG00000204022 | LIPJ | -0,203830715 | 0,868242101 | 0,78704 | 0,90114442 | lipase family member J |
| ENSG00000282228 | PAM16 | -0,203846316 | 0,868232712 | 0,66083 | 0,83092106 | presequence translocase associated motor 16 |
| ENSG00000213277 | MARCKSL1P1 | -0,203857025 | 0,868226267 | 0,81151 | 0,9132701 | MARCKS like 1 pseudogene 1 |
| ENSG00000269091 | AC010624,3 | -0,204062103 | 0,868102858 | 0,58279 | 0,78477395 | novel transcript, antisense to ZNF473 |
| ENSG00000166352 | IFTAP | -0,204107718 | 0,868075411 | 0,29755 | 0,59036937 | intraflagellar transport associated protein |
| ENSG00000165025 | SYK | -0,204273871 | 0,867975442 | 0,67468 | 0,83975921 | spleen associated tyrosine kinase |
| ENSG00000117222 | RBBP5 | -0,204366601 | 0,867919654 | 0,46322 | 0,70415312 | RB binding protein 5, histone lysine methyltransferase complex subunit |
| ENSG00000278834 | AC073508,3 | -0,204447678 | 0,86787088 | 0,46081 | 0,70258879 | novel transcript, antisense to SMARCE1 |
| ENSG00000266820 | KPNA2P3 | -0,204449011 | 0,867870077 | 0,78349 | 0,89918294 | karyopherin subunit alpha 2 pseudogene 3 |
| ENSG00000132780 | NASP | -0,204489322 | 0,867845829 | 0,39844 | 0,66102743 | nuclear autoantigenic sperm protein |

| | | | | | | |
|---|---|---|---|---|---|---|
| ENSG00000278177 | AL354811,1 | -0,204510875 | 0,867832863 | 0,69157 | 0,84969293 | novel transcript, antisense to GPC6 |
| ENSG00000288307 | CTDNEP1 | -0,204673436 | 0,867735083 | 0,49972 | 0,73058737 | CTD nuclear envelope phosphatase 1 |
| ENSG00000175826 | CTDNEP1 | -0,204673436 | 0,867735083 | 0,49972 | 0,73058737 | CTD nuclear envelope phosphatase 1 |
| ENSG00000150510 | FAM124A | -0,204692982 | 0,867723326 | 0,40773 | 0,66684304 | family with sequence similarity 124 member A |
| ENSG00000228995 | MTND1P9 | -0,204734014 | 0,867698648 | 0,6497 | 0,82449863 | MT-ND1 pseudogene 9 |
| ENSG00000273656 | AC139494,1 | -0,204858779 | 0,867623612 | 0,30825 | 0,59696396 | glucuronidase, beta (GUSB) pseudogene |
| ENSG00000203722 | RAET1G | -0,204935211 | 0,867577648 | 0,7903 | 0,90286944 | retinoic acid early transcript 1G |
| ENSG00000232126 | HLA-B | -0,204992584 | 0,867543146 | 0,84931 | 0,93142453 | major histocompatibility complex, class I, B |
| ENSG00000259905 | PWRN1 | -0,205157446 | 0,867444015 | 0,42704 | 0,68023201 | Prader-Willi region non-protein coding RNA 1 |
| ENSG00000186625 | KATNA1 | -0,205315128 | 0,867349211 | 0,41113 | 0,6689497 | katanin catalytic subunit A1 |
| ENSG00000216906 | AL355312,1 | -0,205482501 | 0,867248593 | 0,78969 | 0,90258817 | multimerin 2 (MMRN2) pseudogene |
| ENSG00000197826 | CFAP299 | -0,205488 | 0,867245287 | 0,7317 | 0,87274707 | cilia and flagella associated protein 299 |
| ENSG00000099960 | SLC7A4 | -0,205509102 | 0,867232602 | 0,83842 | 0,92626651 | solute carrier family 7 member 4 |
| ENSG00000205138 | SDHAF1 | -0,205552219 | 0,867206684 | 0,47257 | 0,71197113 | succinate dehydrogenase complex assembly factor 1 |
| ENSG00000149328 | GLB1L2 | -0,20570349 | 0,867115759 | 0,51031 | 0,73792002 | galactosidase beta 1 like 2 |
| ENSG00000146670 | CDCA5 | -0,205801258 | 0,867056999 | 0,40087 | 0,66282495 | cell division cycle associated 5 |
| ENSG00000145014 | TMEM44 | -0,205856039 | 0,867024076 | 0,42267 | 0,67678159 | transmembrane protein 44 |
| ENSG00000256862 | PARP11-AS1 | -0,206017928 | 0,866926791 | 0,61589 | 0,80470415 | PARP11 antisense RNA 1 |
| ENSG00000235493 | LINC01967 | -0,206184498 | 0,866826703 | 0,50915 | 0,73704572 | long intergenic non-protein coding RNA 1967 |
| ENSG00000005007 | UPF1 | -0,206194877 | 0,866820467 | 0,28935 | 0,58495488 | UPF1 RNA helicase and ATPase |
| ENSG00000219200 | RNASEK | -0,20639344 | 0,866701172 | 0,50341 | 0,73266852 | ribonuclease K |
| ENSG00000163440 | PDCL2 | -0,206430997 | 0,86667861 | 0,83137 | 0,9229059 | phosducin like 2 |
| ENSG00000286649 | AC006566,2 | -0,206442831 | 0,8666715 | 0,78031 | 0,89823499 | novel transcript, antisense to PTPRM |
| ENSG00000148331 | ASB6 | -0,20646075 | 0,866660736 | 0,39291 | 0,65604611 | ankyrin repeat and SOCS box containing 6 |
| ENSG00000281919 | SLC45A2 | -0,206645005 | 0,866550057 | 0,86976 | 0,9425949 | solute carrier family 45 member 2 |
| ENSG00000274403 | AC090510,2 | -0,206733459 | 0,866496929 | 0,55308 | 0,7662484 | novel transcript, antisense to CDAN1 |
| ENSG00000250220 | AC053527,2 | -0,206786937 | 0,86646481 | 0,72295 | 0,86768626 | novel transcript |
| ENSG00000100425 | BRD1 | -0,206808546 | 0,866451832 | 0,33634 | 0,61635233 | bromodomain containing 1 |
| ENSG00000182541 | LIMK2 | -0,206832652 | 0,866437355 | 0,39742 | 0,65990329 | LIM domain kinase 2 |
| ENSG00000288579 | AC025472,1 | -0,206858649 | 0,866421742 | 0,77474 | 0,89588588 | tigger transposable element derived 1 (TIGD1) pseudogene |
| ENSG00000237631 | AL161454,1 | -0,206862527 | 0,866419412 | 0,83967 | 0,92673661 | parkinson disease 7 (PARK7) pseudogene |
| ENSG00000136367 | ZFHX2 | -0,207108305 | 0,866271822 | 0,34613 | 0,62331777 | zinc finger homeobox 2 |

| | | | | | | |
|---|---|---|---|---|---|---|
| ENSG00000223756 | TSSC2 | -0,207252584 | 0,866185193 | 0,37573 | 0,64406408 | tumor suppressing subtransferable candidate 2 (pseudogene) |
| ENSG00000214367 | HAUS3 | -0,207339926 | 0,866132755 | 0,40575 | 0,66548388 | HAUS augmin like complex subunit 3 |
| ENSG00000277182 | AC006449,5 | -0,207344742 | 0,866129864 | 0,57309 | 0,77878277 | novel transcript, antisense to PCGF2 |
| ENSG00000282749 | AC006449,10 | -0,207344742 | 0,866129864 | 0,57309 | 0,77878277 | novel transcript, antisense to PCGF2 |
| ENSG00000282988 | AL031777,2 | -0,207372214 | 0,866113371 | 0,51274 | 0,73941533 | novel protein |
| ENSG00000227477 | STK4-AS1 | -0,207385692 | 0,86610528 | 0,85988 | 0,9370297 | STK4 antisense RNA 1 (head to head) |
| ENSG00000002330 | BAD | -0,207491574 | 0,866041717 | 0,20392 | 0,52363163 | BCL2 associated agonist of cell death |
| ENSG00000146733 | PSPH | -0,207572246 | 0,865993291 | 0,519 | 0,7437068 | phosphoserine phosphatase |
| ENSG00000274614 | AL390754,1 | -0,207732065 | 0,865897364 | 0,69941 | 0,85482742 | spalt-like transcription factor 4 (SALL4) pseudogene |
| ENSG00000135436 | FAM186B | -0,20783494 | 0,865835621 | 0,54443 | 0,76055278 | family with sequence similarity 186 member B |
| ENSG00000250980 | NSA2P6 | -0,20790517 | 0,865793473 | 0,73726 | 0,87539732 | NSA2 pseudogene 6 |
| ENSG00000197479 | PCDHB11 | -0,208173043 | 0,865632731 | 0,37516 | 0,64350562 | protocadherin beta 11 |
| ENSG00000109255 | NMU | -0,208222063 | 0,865603319 | 0,65342 | 0,82638159 | neuromedin U |
| ENSG00000099331 | MYO9B | -0,208285615 | 0,86556519 | 0,38254 | 0,64932251 | myosin IXB |
| ENSG00000272094 | AC091614,1 | -0,208295067 | 0,865559519 | 0,5602 | 0,7707835 | novel transcript |
| ENSG00000224609 | HSD52 | -0,208345801 | 0,865529081 | 0,4954 | 0,7276636 | uncharacterized LOC729467 |
| ENSG00000282100 | HSP90AB4P | -0,208418349 | 0,865485558 | 0,65461 | 0,82719457 | heat shock protein 90 alpha family class B member 4, pseudogene |
| ENSG00000101890 | GUCY2F | -0,208478252 | 0,865449622 | 0,72207 | 0,8672968 | guanylate cyclase 2F, retinal |
| ENSG00000176563 | CNTD1 | -0,208532971 | 0,865416798 | 0,61599 | 0,80475871 | cyclin N-terminal domain containing 1 |
| ENSG00000112137 | PHACTR1 | -0,208581811 | 0,865387501 | 0,42651 | 0,67990398 | phosphatase and actin regulator 1 |
| ENSG00000225264 | ZNRF2P2 | -0,208788133 | 0,865263749 | 0,52922 | 0,75112358 | zinc and ring finger 2 pseudogene 2 |
| ENSG00000272619 | AC093673,2 | -0,208855064 | 0,865223608 | 0,72888 | 0,87134251 | novel transcript |
| ENSG00000286282 | AC021733,4 | -0,209402169 | 0,864895557 | 0,81535 | 0,91502219 | novel transcript, antisense to NOV |
| ENSG00000183311 | TUBB | -0,209407149 | 0,864892571 | 0,45126 | 0,69631879 | tubulin beta class I |
| ENSG00000280195 | AC245140,2 | -0,209424697 | 0,864882051 | 0,54071 | 0,75849891 | novel transcript, antisense to RPL10 |
| ENSG00000187164 | SHTN1 | -0,209624004 | 0,864762577 | 0,66024 | 0,83036507 | shootin 1 |
| ENSG00000207442 | SNORD116-6 | -0,209677066 | 0,864730772 | 0,84227 | 0,92821072 | small nucleolar RNA, C/D box 116-6 |
| ENSG00000233668 | AL353662,1 | -0,209931326 | 0,864578385 | 0,76376 | 0,88986791 | ribosomal protein S8 (RPS8) pseudogene |
| ENSG00000239705 | AL354710,2 | -0,209951294 | 0,864566419 | 0,80899 | 0,91201277 | novel transcript |
| ENSG00000233626 | AC092814,2 | -0,210024908 | 0,864522305 | 0,8336 | 0,92396169 | protein (peptidyl-prolyl cis\\/trans isomerase) NIMA-interacting, 4 (parvulin) (PIN4) pseudogene |

| | | | | | | |
|---|---|---|---|---|---|---|
| ENSG00000184785 | SMIM10 | -0,210050276 | 0,864507104 | 0,6897 | 0,84865812 | small integral membrane protein 10 |
| ENSG00000178607 | ERN1 | -0,210066284 | 0,864497511 | 0,43875 | 0,6878469 | endoplasmic reticulum to nucleus signaling 1 |
| ENSG00000160961 | ZNF333 | -0,210190897 | 0,864422844 | 0,29876 | 0,59098155 | zinc finger protein 333 |
| ENSG00000256124 | LINC01152 | -0,210207421 | 0,864412943 | 0,4582 | 0,70115011 | long intergenic non-protein coding RNA 1152 |
| ENSG00000155858 | LSM11 | -0,210242412 | 0,864391978 | 0,46243 | 0,70356854 | LSM11, U7 small nuclear RNA associated |
| ENSG00000279168 | AC105052,4 | -0,210275202 | 0,864372332 | 0,54242 | 0,75915391 | novel transcript, sense overlapping POLR2J2, UPK3BL and RASA4 |
| ENSG00000285839 | AL445685,3 | -0,210375254 | 0,864312389 | 0,95209 | 0,97971456 | novel transcript |
| ENSG00000178460 | MCMDC2 | -0,210376734 | 0,864311502 | 0,45189 | 0,69683967 | minichromosome maintenance domain containing 2 |
| ENSG00000259536 | AC091045,1 | -0,21051554 | 0,864228348 | 0,75556 | 0,88554206 | novel transcript, antisense to C15orf57 |
| ENSG00000113141 | IK | -0,210528685 | 0,864220474 | 0,34076 | 0,61914946 | IK cytokine |
| ENSG00000276836 | ARHGAP27 | -0,210587403 | 0,864185301 | 0,72951 | 0,8715818 | Rho GTPase activating protein 27 |
| ENSG00000267284 | AC022031,2 | -0,21075219 | 0,864086598 | 0,3802 | 0,64789138 | novel transcript |
| ENSG00000108651 | UTP6 | -0,210916615 | 0,863988123 | 0,45724 | 0,70073404 | UTP6 small subunit processome component |
| ENSG00000288396 | AC213033,3 | -0,210924856 | 0,863983188 | 0,56538 | 0,77366076 | |
| ENSG00000238251 | NSA2P7 | -0,210935196 | 0,863976995 | 0,67991 | 0,8427192 | NSA2 pseudogene 7 |
| ENSG00000118473 | SGIP1 | -0,210945799 | 0,863970646 | 0,63809 | 0,81820499 | SH3GL interacting endocytic adaptor 1 |
| ENSG00000269938 | AC068790,2 | -0,211001637 | 0,863937207 | 0,56886 | 0,77629764 | novel transcript, sense intronic to CCDC92 |
| ENSG00000273311 | DGCR11 | -0,211070972 | 0,863895688 | 0,49074 | 0,72478928 | DiGeorge syndrome critical region gene 11 |
| ENSG00000185055 | EFCAB10 | -0,211083468 | 0,863888205 | 0,61881 | 0,80665241 | EF-hand calcium binding domain 10 |
| ENSG00000168172 | HOOK3 | -0,211114672 | 0,86386952 | 0,3715 | 0,64066064 | hook microtubule tethering protein 3 |
| ENSG00000247828 | TMEM161B-A | -0,211120397 | 0,863866092 | 0,35725 | 0,63120669 | TMEM161B antisense RNA 1 |
| ENSG00000179455 | MKRN3 | -0,211144039 | 0,863851936 | 0,34261 | 0,62025394 | makorin ring finger protein 3 |
| ENSG00000023902 | PLEKHO1 | -0,211167184 | 0,863838077 | 0,31842 | 0,60358233 | pleckstrin homology domain containing O1 |
| ENSG00000258035 | AC123567,2 | -0,211173444 | 0,863834329 | 0,66792 | 0,83525939 | novel transcript, antisense to PLXNC1 |
| ENSG00000256817 | TPT1P12 | -0,211205427 | 0,863815179 | 0,79561 | 0,90543123 | tumor protein, translationally-controlled 1 pseudogene 12 |
| ENSG00000108312 | UBTF | -0,211209666 | 0,863812641 | 0,46746 | 0,70731446 | upstream binding transcription factor |
| ENSG00000265185 | SNORD3B-1 | -0,21132036 | 0,863746365 | 0,59148 | 0,79063841 | small nucleolar RNA, C/D box 3B-1 |
| ENSG00000228501 | RPL15P18 | -0,211409054 | 0,863693266 | 0,79407 | 0,90426294 | ribosomal protein L15 pseudogene 18 |
| ENSG00000146094 | DOK3 | -0,211587039 | 0,863586719 | 0,62263 | 0,80866078 | docking protein 3 |
| ENSG00000104941 | RSPH6A | -0,211599442 | 0,863579294 | 0,76493 | 0,89071863 | radial spoke head 6 homolog A |
| ENSG00000253377 | AC068672,2 | -0,211604713 | 0,863576139 | 0,78097 | 0,89853519 | novel transcript |
| ENSG00000282953 | MUC20-OT1 | -0,211631841 | 0,863559901 | 0,5946 | 0,79283643 | MUC20 overlapping transcript |

| | | | | | | |
|---|---|---|---|---|---|---|
| ENSG00000215790 | SLC35E2A | -0,211916315 | 0,863389639 | 0,38571 | 0,65122858 | solute carrier family 35 member E2A |
| ENSG00000233723 | LINC01122 | -0,211937157 | 0,863377166 | 0,51756 | 0,74270246 | long intergenic non-protein coding RNA 1122 |
| ENSG00000160781 | PAQR6 | -0,212096048 | 0,863282083 | 0,61159 | 0,80328691 | progestin and adipoQ receptor family member 6 |
| ENSG00000272100 | AL513218,1 | -0,212146326 | 0,863251998 | 0,75106 | 0,88327589 | novel transcript, antisense to CC2D1B |
| ENSG00000133059 | DSTYK | -0,212148197 | 0,863250879 | 0,45864 | 0,70133296 | dual serine/threonine and tyrosine protein kinase |
| ENSG00000214655 | ZSWIM8 | -0,212149443 | 0,863250133 | 0,34373 | 0,62112663 | zinc finger SWIM-type containing 8 |
| ENSG00000213067 | AC092447,1 | -0,212367861 | 0,863119451 | 0,74898 | 0,88190158 | insulin-like growth factor 2 mRNA binding protein 3 (IGF2BP3) pseudogene |
| ENSG00000256482 | AC023796,2 | -0,212433092 | 0,863080426 | 0,79948 | 0,90748078 | novel transcript, antisense to BCAT1 |
| ENSG00000158869 | FCER1G | -0,212561671 | 0,863003508 | 0,88419 | 0,9485968 | Fc fragment of IgE receptor Ig |
| ENSG00000127616 | SMARCA4 | -0,21260192 | 0,862979432 | 0,337 | 0,61684812 | SWI/SNF related, matrix associated, actin dependent regulator of chromatin, subfamily a, member 4 |
| ENSG00000107864 | CPEB3 | -0,212647594 | 0,862952111 | 0,49276 | 0,72594839 | cytoplasmic polyadenylation element binding protein 3 |
| ENSG00000225920 | RIMKLBP2 | -0,212780343 | 0,862872711 | 0,72113 | 0,86669144 | ribosomal modification protein rimK like family member B pseudogene 2 |
| ENSG00000241429 | EEF1A1P25 | -0,212940764 | 0,862776769 | 0,72418 | 0,86838632 | eukaryotic translation elongation factor 1 alpha 1 pseudogene 25 |
| ENSG00000167962 | ZNF598 | -0,212960333 | 0,862765066 | 0,21634 | 0,53480193 | zinc finger protein 598, E3 ubiquitin ligase |
| ENSG00000261329 | AC016597,1 | -0,212989236 | 0,862747782 | 0,52762 | 0,75037219 | novel transcript, antisense to a novel protein |
| ENSG00000255545 | B3GAT1-DT | -0,213016868 | 0,862731257 | 0,62393 | 0,80955151 | B3GAT1 diverget transcript |
| ENSG00000287508 | AC009560,4 | -0,213201411 | 0,862620908 | 0,78454 | 0,89962715 | novel transcript, antisense to RORA |
| ENSG00000141994 | DUS3L | -0,21325144 | 0,862590995 | 0,48416 | 0,72005601 | dihydrouridine synthase 3 like |
| ENSG00000269044 | AC024075,2 | -0,213393283 | 0,862506191 | 0,43244 | 0,68411566 | novel transcript, sense intronic C19orf42 |
| ENSG00000273125 | LINC01990 | -0,213450564 | 0,862471946 | 0,80167 | 0,90870853 | long intergenic non-protein coding RNA 1990 |
| ENSG00000106400 | ZNHIT1 | -0,213555774 | 0,862409052 | 0,46617 | 0,70650474 | zinc finger HIT-type containing 1 |
| ENSG00000283839 | AC096667,1 | -0,213582137 | 0,862393293 | 0,69813 | 0,85390204 | proline-rich protein 18-like |
| ENSG00000164066 | INTU | -0,213588249 | 0,862389639 | 0,25082 | 0,55702488 | inturned planar cell polarity protein |
| ENSG00000250497 | AC007126,1 | -0,21361553 | 0,862373332 | 0,35993 | 0,63249219 | novel transcript |
| ENSG00000101144 | BMP7 | -0,213708825 | 0,862317567 | 0,60912 | 0,80161331 | bone morphogenetic protein 7 |
| ENSG00000160285 | LSS | -0,213712048 | 0,86231564 | 0,36295 | 0,63395677 | lanosterol synthase |
| ENSG00000154655 | L3MBTL4 | -0,213750696 | 0,86229254 | 0,6364 | 0,81701009 | L3MBTL histone methyl-lysine binding protein 4 |
| ENSG00000090263 | MRPS33 | -0,21390668 | 0,862199314 | 0,37623 | 0,64458342 | mitochondrial ribosomal protein S33 |
| ENSG00000100461 | RBM23 | -0,213977904 | 0,862156749 | 0,408 | 0,66699097 | RNA binding motif protein 23 |

| | | | | | | |
|---|---|---|---|---|---|---|
| ENSG00000256282 | AC112694,1 | -0,214023634 | 0,862129422 | 0,68144 | 0,84375324 | mitochondrial carrier homolog 1 (C, elegans) (MTCH1) |
| ENSG00000122484 | RPAP2 | -0,214071833 | 0,862100619 | 0,2697 | 0,57140742 | RNA polymerase II associated protein 2 |
| ENSG00000230366 | DSCR9 | -0,214243492 | 0,861998048 | 0,62076 | 0,80776776 | Down syndrome critical region 9 |
| ENSG00000117834 | SLC5A9 | -0,21424674 | 0,861996108 | 0,54147 | 0,75882662 | solute carrier family 5 member 9 |
| ENSG00000110200 | ANAPC15 | -0,214260392 | 0,86198795 | 0,33821 | 0,6177255 | anaphase promoting complex subunit 15 |
| ENSG00000213445 | SIPA1 | -0,214344445 | 0,861937732 | 0,58613 | 0,78664449 | signal-induced proliferation-associated 1 |
| ENSG00000090924 | PLEKHG2 | -0,214415276 | 0,861895415 | 0,41337 | 0,67078587 | pleckstrin homology and RhoGEF domain containing G2 |
| ENSG00000287789 | AL158136,1 | -0,214425654 | 0,861889215 | 0,57444 | 0,77959212 | novel transcript, antisense to BACH2 |
| ENSG00000213918 | DNASE1 | -0,214431111 | 0,861885955 | 0,42082 | 0,6760947 | deoxyribonuclease 1 |
| ENSG00000232036 | AL078645,2 | -0,214434412 | 0,861883983 | 0,72607 | 0,86953997 | novel pseudogene |
| ENSG00000105447 | GRWD1 | -0,214471522 | 0,861861813 | 0,36814 | 0,6380264 | glutamate rich WD repeat containing 1 |
| ENSG00000188306 | LRRIQ4 | -0,214529402 | 0,861827237 | 0,72068 | 0,86633947 | leucine rich repeats and IQ motif containing 4 |
| ENSG00000257390 | AC023055,1 | -0,214622083 | 0,861771873 | 0,67733 | 0,84132177 | novel protein |
| ENSG00000065534 | MYLK | -0,214649947 | 0,861755229 | 0,47717 | 0,71510081 | myosin light chain kinase |
| ENSG00000180730 | SHISA2 | -0,214655108 | 0,861752146 | 0,75459 | 0,8850495 | shisa family member 2 |
| ENSG00000248283 | CCNL2P1 | -0,214736706 | 0,861703407 | 0,821 | 0,91738668 | cyclin L2 pseudogene 1 |
| ENSG00000198355 | PIM3 | -0,215196278 | 0,861428955 | 0,55234 | 0,76574901 | Pim-3 proto-oncogene, serine/threonine kinase |
| ENSG00000149743 | TRPT1 | -0,215221853 | 0,861413684 | 0,38421 | 0,65052153 | tRNA phosphotransferase 1 |
| ENSG00000182648 | LINC01006 | -0,215229221 | 0,861409285 | 0,32849 | 0,60990374 | long intergenic non-protein coding RNA 1006 |
| ENSG00000236988 | CTDSPL2P1 | -0,215244337 | 0,861400259 | 0,77233 | 0,89476608 | CTDSPL2 pseudogene 1 |
| ENSG00000183734 | ASCL2 | -0,215247376 | 0,861398445 | 0,6194 | 0,80706028 | achaete-scute family bHLH transcription factor 2 |
| ENSG00000260551 | PWRN2 | -0,215382216 | 0,861317939 | 0,56606 | 0,77416801 | Prader-Willi region non-protein coding RNA 2 |
| ENSG00000124120 | TTPAL | -0,215510385 | 0,861241423 | 0,34138 | 0,61943649 | alpha tocopherol transfer protein like |
| ENSG00000230551 | AC021078,1 | -0,215571703 | 0,861204819 | 0,30722 | 0,59633879 | novel transcript |
| ENSG00000143450 | OAZ3 | -0,215693266 | 0,861132256 | 0,53142 | 0,75269971 | ornithine decarboxylase antizyme 3 |
| ENSG00000225083 | GRTP1-AS1 | -0,215784244 | 0,861077954 | 0,70155 | 0,85609945 | GRTP1 antisense RNA 1 |
| ENSG00000287951 | AC026402,2 | -0,215842679 | 0,861043077 | 0,85197 | 0,93265377 | novel transcript |
| ENSG00000079482 | OPHN1 | -0,216052921 | 0,860917607 | 0,35536 | 0,62970025 | oligophrenin 1 |
| ENSG00000063169 | BICRA | -0,216078459 | 0,860902368 | 0,41223 | 0,66996291 | BRD4 interacting chromatin remodeling complex associated protein |
| ENSG00000115762 | PLEKHB2 | -0,216280912 | 0,860781566 | 0,17381 | 0,49846579 | pleckstrin homology domain containing B2 |
| ENSG00000125657 | TNFSF9 | -0,216342223 | 0,860744986 | 0,66172 | 0,83154613 | TNF superfamily member 9 |
| ENSG00000231165 | TRBV26OR9-2 | -0,216443016 | 0,860684853 | 0,74165 | 0,87781553 | T cell receptor beta variable 26/OR9-2 (pseudogene) |

| | | | | | | |
|---|---|---|---|---|---|---|
| ENSG00000228729 | AL627308,1 | -0,216455553 | 0,860677373 | 0,74084 | 0,87717177 | cathepsin B (CTSB) pseudogene |
| ENSG00000188825 | LINC00910 | -0,216533876 | 0,860630649 | 0,55614 | 0,76831913 | long intergenic non-protein coding RNA 910 |
| ENSG00000130810 | PPAN | -0,216550442 | 0,860620767 | 0,48271 | 0,71909811 | peter pan homolog |
| ENSG00000196873 | CBWD3 | -0,216589274 | 0,860597602 | 0,3776 | 0,64605678 | COBW domain containing 3 |
| ENSG00000113456 | RAD1 | -0,21660079 | 0,860590733 | 0,33853 | 0,61774255 | RAD1 checkpoint DNA exonuclease |
| ENSG00000105518 | TMEM205 | -0,216624379 | 0,860576662 | 0,47682 | 0,71481671 | transmembrane protein 205 |
| ENSG00000270915 | H3P8 | -0,216676738 | 0,86054543 | 0,83276 | 0,92348349 | H3 histone pseudogene 8 |
| ENSG00000286588 | AC104841,1 | -0,216716581 | 0,860521664 | 0,48878 | 0,72350189 | novel transcript, antisense to SEPT2 |
| ENSG00000257169 | AC125612,1 | -0,216726446 | 0,86051578 | 0,76092 | 0,88874162 | small nuclear ribonucleoprotein D1 polypeptide 16kDa (SNRPD1) pseudogene |
| ENSG00000102385 | DRP2 | -0,216766596 | 0,860491832 | 0,56244 | 0,77186635 | dystrophin related protein 2 |
| ENSG00000237001 | WASF3-AS1 | -0,216796595 | 0,86047394 | 0,64603 | 0,82269956 | WASF3 antisense RNA 1 |
| ENSG00000285848 | AC025280,3 | -0,216805603 | 0,860468567 | 0,67248 | 0,83803462 | novel transcript, antisense to CRISPLD2 |
| ENSG00000166965 | RCCD1 | -0,216840137 | 0,86044797 | 0,37245 | 0,64138554 | RCC1 domain containing 1 |
| ENSG00000103111 | MON1B | -0,2169521 | 0,860381196 | 0,21506 | 0,53378006 | MON1 homolog B, secretory trafficking associated |
| ENSG00000142634 | EFHD2 | -0,21697088 | 0,860369997 | 0,43673 | 0,68681257 | EF-hand domain family member D2 |
| ENSG00000287160 | Z82195,3 | -0,217122742 | 0,860279436 | 0,37194 | 0,64101991 | novel transcript |
| ENSG00000166741 | NNMT | -0,21731777 | 0,860163149 | 0,69092 | 0,84918509 | nicotinamide N-methyltransferase |
| ENSG00000224429 | LINC00539 | -0,217374046 | 0,860129597 | 0,44893 | 0,69460209 | long intergenic non-protein coding RNA 539 |
| ENSG00000108666 | C17orf75 | -0,217393978 | 0,860117713 | 0,44 | 0,6885781 | chromosome 17 open reading frame 75 |
| ENSG00000227359 | AC017074,1 | -0,217447249 | 0,860085954 | 0,83168 | 0,9230208 | novel transcript |
| ENSG00000171105 | INSR | -0,217495337 | 0,860057287 | 0,2433 | 0,55199251 | insulin receptor |
| ENSG00000113269 | RNF130 | -0,217531805 | 0,860035547 | 0,12707 | 0,45009139 | ring finger protein 130 |
| ENSG00000285991 | AL355312,5 | -0,217588439 | 0,860001786 | 0,64813 | 0,82383528 | novel transcript |
| ENSG00000179335 | CLK3 | -0,217630509 | 0,859976708 | 0,33517 | 0,61485832 | CDC like kinase 3 |
| ENSG00000121406 | ZNF549 | -0,217679471 | 0,859947523 | 0,35806 | 0,63185282 | zinc finger protein 549 |
| ENSG00000269043 | AC008554,1 | -0,217746163 | 0,859907771 | 0,76151 | 0,88879633 | novel transcript |
| ENSG00000286629 | AC005803,1 | -0,217863458 | 0,85983786 | 0,80241 | 0,90900521 | novel transcript, antisense to ERN1 |
| ENSG00000280807 | TUBGCP5 | -0,217890893 | 0,859821509 | 0,75464 | 0,88505036 | tubulin gamma complex associated protein 5 |
| ENSG00000166840 | GLYATL1 | -0,217916389 | 0,859806314 | 0,66538 | 0,8334992 | glycine-N-acyltransferase like 1 |
| ENSG00000270923 | TAS2R6P | -0,217945259 | 0,859789109 | 0,76253 | 0,88957921 | taste 2 receptor member 6 pseudogene |
| ENSG00000230992 | FAM201B | -0,217947694 | 0,859787658 | 0,68393 | 0,84523058 | family with sequence similarity 201 member B |
| ENSG00000267309 | AC092295,2 | -0,218109825 | 0,85969104 | 0,51963 | 0,74404725 | novel transcript, antisense to ZNF566 |

| | | | | | | |
|---|---|---|---|---|---|---|
| ENSG00000068354 | TBC1D25 | -0,218188382 | 0,85964423 | 0,35503 | 0,62946175 | TBC1 domain family member 25 |
| ENSG00000287842 | AL161714,1 | -0,218295963 | 0,859580129 | 0,77311 | 0,89523715 | novel transcript, sense intronic to ENOX1 |
| ENSG00000140521 | POLG | -0,218301284 | 0,859576959 | 0,36262 | 0,63379972 | DNA polymerase gamma, catalytic subunit |
| ENSG00000139438 | FAM222A | -0,218304922 | 0,859574791 | 0,43082 | 0,68325486 | family with sequence similarity 222 member A |
| ENSG00000259592 | PRELID1P4 | -0,218400897 | 0,85951761 | 0,85296 | 0,93318855 | PRELID1 pseudogene 4 |
| ENSG00000276131 | AC009118,2 | -0,218431155 | 0,859499583 | 0,77906 | 0,89781842 | novel transcript, antisense to GINS3 |
| ENSG00000180974 | OR52E4 | -0,218441803 | 0,859493239 | 0,72641 | 0,86975255 | olfactory receptor family 52 subfamily E member 4 |
| ENSG00000131165 | CHMP1A | -0,218454416 | 0,859485725 | 0,39683 | 0,65936612 | charged multivesicular body protein 1A |
| ENSG00000285241 | PCK2 | -0,218471914 | 0,859475301 | 0,54998 | 0,7643663 | phosphoenolpyruvate carboxykinase 2, mitochondrial |
| ENSG00000100889 | PCK2 | -0,218471914 | 0,859475301 | 0,54998 | 0,7643663 | phosphoenolpyruvate carboxykinase 2, mitochondrial |
| ENSG00000107341 | UBE2R2 | -0,218500666 | 0,859458172 | 0,45805 | 0,7010995 | ubiquitin conjugating enzyme E2 R2 |
| ENSG00000144824 | PHLDB2 | -0,2185286 | 0,859441531 | 0,66284 | 0,83225294 | pleckstrin homology like domain family B member 2 |
| ENSG00000197822 | OCLN | -0,218691491 | 0,8593445 | 0,62412 | 0,80958576 | occludin |
| ENSG00000156535 | CD109 | -0,218823944 | 0,859265607 | 0,55932 | 0,77045525 | CD109 molecule |
| ENSG00000183779 | ZNF703 | -0,218879384 | 0,859232588 | 0,45729 | 0,70073404 | zinc finger protein 703 |
| ENSG00000162139 | NEU3 | -0,218897792 | 0,859221625 | 0,36351 | 0,63433098 | neuraminidase 3 |
| ENSG00000182366 | FAM87A | -0,218963352 | 0,85918258 | 0,60807 | 0,8008277 | family with sequence similarity 87 member A |
| ENSG00000162825 | NBPF20 | -0,219180823 | 0,859053077 | 0,36797 | 0,63794202 | NBPF member 20 |
| ENSG00000255052 | FAM66D | -0,219205113 | 0,859038614 | 0,47674 | 0,71481671 | family with sequence similarity 66 member D |
| ENSG00000159625 | DRC7 | -0,219261583 | 0,85900499 | 0,67431 | 0,83962967 | dynein regulatory complex subunit 7 |
| ENSG00000205649 | HTN3 | -0,219286354 | 0,858990241 | 0,78349 | 0,89918294 | histatin 3 |
| ENSG00000282967 | HTN3 | -0,219286354 | 0,858990241 | 0,78349 | 0,89918294 | histatin 3 |
| ENSG00000196588 | MRTFA | -0,219366742 | 0,858942379 | 0,28515 | 0,58171017 | myocardin related transcription factor A |
| ENSG00000288075 | AC108519,1 | -0,21940173 | 0,858921548 | 0,29245 | 0,58699586 | novel transcript |
| ENSG00000213131 | YWHAZP4 | -0,219403057 | 0,858920758 | 0,61397 | 0,80365116 | tyrosine 3-monooxygenase/tryptophan 5-monooxygenase activation protein zeta pseudogene 4 |
| ENSG00000260978 | AC126407,1 | -0,219439817 | 0,858898873 | 0,56508 | 0,773443 | MKRN3 antisense RNA 1 |
| ENSG00000179085 | DPM3 | -0,219585619 | 0,858812075 | 0,41682 | 0,67362915 | dolichyl-phosphate mannosyltransferase subunit 3, regulatory |
| ENSG00000067601 | PMS2P4 | -0,21980252 | 0,858682967 | 0,341 | 0,6192144 | PMS1 homolog 2, mismatch repair system component pseudogene 4 |
| ENSG00000144229 | THSD7B | -0,219829473 | 0,858666925 | 0,72224 | 0,86734497 | thrombospondin type 1 domain containing 7B |
| ENSG00000141002 | TCF25 | -0,219918234 | 0,858614098 | 0,4329 | 0,68436943 | transcription factor 25 |

| | | | | | | |
|---|---|---|---|---|---|---|
| ENSG00000141570 | CBX8 | -0,219944075 | 0,858598718 | 0,4058 | 0,66552827 | chromobox 8 |
| ENSG00000104381 | GDAP1 | -0,220046739 | 0,858537622 | 0,23934 | 0,5491566 | ganglioside induced differentiation associated protein 1 |
| ENSG00000262668 | AJ003147,2 | -0,220245068 | 0,858419606 | 0,56162 | 0,77136826 | olfactory receptor family 1 subfamily F member 1 |
| ENSG00000286670 | AL450467,1 | -0,220337638 | 0,858364527 | 0,73009 | 0,87180309 | novel transcript, antisense to VLDLR |
| ENSG00000149582 | TMEM25 | -0,220636011 | 0,858187022 | 0,22345 | 0,53953208 | transmembrane protein 25 |
| ENSG00000269139 | AC010336,5 | -0,220661082 | 0,858172109 | 0,69694 | 0,85319224 | novel transcript, antisense to CTXN1 and TIMM44 |
| ENSG00000214717 | ZBED1 | -0,22071303 | 0,858141209 | 0,32675 | 0,60880597 | zinc finger BED-type containing 1 |
| ENSG00000239789 | MRPS17 | -0,220842431 | 0,858064242 | 0,40955 | 0,66801281 | mitochondrial ribosomal protein S17 |
| ENSG00000255495 | AC145124,1 | -0,220857986 | 0,858054991 | 0,38675 | 0,65203769 | novel transcript |
| ENSG00000234806 | RPL26P29 | -0,220929408 | 0,858012512 | 0,69141 | 0,8495691 | ribosomal protein L26 pseudogene 29 |
| ENSG00000271933 | AL603756,1 | -0,220979873 | 0,8579825 | 0,49904 | 0,73002372 | novel transcript, antisense to GHITM |
| ENSG00000131781 | FMO5 | -0,22105894 | 0,857935479 | 0,55103 | 0,76487238 | flavin containing dimethylaniline monoxygenase 5 |
| ENSG00000146090 | RASGEF1C | -0,22114103 | 0,857886664 | 0,56003 | 0,77076855 | RasGEF domain family member 1C |
| ENSG00000135587 | SMPD2 | -0,221143669 | 0,857885095 | 0,44736 | 0,69342177 | sphingomyelin phosphodiesterase 2 |
| ENSG00000203709 | MIR29B2CHG | -0,221155256 | 0,857878205 | 0,20517 | 0,52486901 | MIR29B2 and MIR29C host gene |
| ENSG00000254485 | AC026191,1 | -0,221155681 | 0,857877952 | 0,82818 | 0,9207216 | novel transcript, antisense to THUMPD3 |
| ENSG00000145888 | GLRA1 | -0,221248585 | 0,85782271 | 0,71381 | 0,86224977 | glycine receptor alpha 1 |
| ENSG00000276681 | LENG8 | -0,221284032 | 0,857801633 | 0,28617 | 0,58259176 | leukocyte receptor cluster member 8 |
| ENSG00000235554 | SRSF6P2 | -0,221328842 | 0,85777499 | 0,83416 | 0,92405867 | SRSF6 pseudogene 2 |
| ENSG00000250062 | MAPK10-AS1 | -0,22136061 | 0,857756103 | 0,70687 | 0,85889796 | MAPK10 antisense RNA 1 |
| ENSG00000178947 | SMIM10L2A | -0,221396314 | 0,857734875 | 0,45396 | 0,69799145 | small integral membrane protein 10 like 2A |
| ENSG00000282727 | AC091132,3 | -0,221422008 | 0,857719599 | 0,33762 | 0,61728862 | TEC |
| ENSG00000255112 | CHMP1B | -0,221691743 | 0,85755925 | 0,45614 | 0,69998075 | charged multivesicular body protein 1B |
| ENSG00000284691 | AC073111,3 | -0,221700751 | 0,857553895 | 0,41369 | 0,67085666 | novel zinc finger protein |
| ENSG00000176410 | DNAJC30 | -0,221824354 | 0,857480427 | 0,4043 | 0,6642832 | DnaJ heat shock protein family (Hsp40) member C30 |
| ENSG00000253300 | AC018541,1 | -0,221850938 | 0,857464627 | 0,73956 | 0,87648643 | novel transcript |
| ENSG00000272583 | AL592494,2 | -0,221914522 | 0,857426837 | 0,67527 | 0,8401859 | novel transcript |
| ENSG00000286729 | AC137695,3 | -0,221990379 | 0,857381754 | 0,59664 | 0,79419995 | novel transcript, antisense to COPG1 |
| ENSG00000157870 | PRXL2B | -0,2221554 | 0,857283689 | 0,43163 | 0,68379763 | peroxiredoxin like 2B |
| ENSG00000164171 | ITGA2 | -0,222162553 | 0,857279439 | 0,63901 | 0,81893471 | integrin subunit alpha 2 |
| ENSG00000092853 | CLSPN | -0,222505942 | 0,857075414 | 0,41935 | 0,67519089 | claspin |
| ENSG00000132259 | CNGA4 | -0,2225316 | 0,857060171 | 0,78736 | 0,90121071 | cyclic nucleotide gated channel subunit alpha 4 |
| ENSG00000242748 | RPL23AP81 | -0,222544706 | 0,857052385 | 0,78903 | 0,90223158 | ribosomal protein L23a pseudogene 81 |

| | | | | | | |
|---|---|---|---|---|---|---|
| ENSG00000118898 | PPL | -0,222613032 | 0,857011796 | 0,5541 | 0,76712518 | periplakin |
| ENSG00000248858 | FLJ46284 | -0,22263489 | 0,856998812 | 0,65938 | 0,82984259 | uncharacterized LOC441369 |
| ENSG00000288393 | AP003086,3 | -0,22269747 | 0,856961639 | 0,57006 | 0,77703861 | zinc finger protein 75C, pseudogene |
| ENSG00000171681 | ATF7IP | -0,222699944 | 0,85696017 | 0,15988 | 0,48710442 | activating transcription factor 7 interacting protein |
| ENSG00000211454 | AKR7L | -0,22271866 | 0,856949052 | 0,51747 | 0,74270246 | aldo-keto reductase family 7 like (gene/pseudogene) |
| ENSG00000099804 | CDC34 | -0,222903135 | 0,856839483 | 0,27551 | 0,57522081 | cell division cycle 34, ubiqiutin conjugating enzyme |
| ENSG00000272822 | AC073610,2 | -0,222922908 | 0,856827739 | 0,70309 | 0,8568579 | novel protein |
| ENSG00000214711 | CAPN14 | -0,22295458 | 0,856808929 | 0,76361 | 0,88983737 | calpain 14 |
| ENSG00000129194 | SOX15 | -0,222961814 | 0,856804633 | 0,63357 | 0,81549489 | SRY-box transcription factor 15 |
| ENSG00000253298 | LINC01845 | -0,223025006 | 0,856767104 | 0,7349 | 0,87477884 | long intergenic non-protein coding RNA 1845 |
| ENSG00000282770 | DLG5-AS1 | -0,223119974 | 0,856710708 | 0,4838 | 0,71996444 | DLG5 antisense RNA 1 |
| ENSG00000254779 | EGLN1P1 | -0,223138619 | 0,856699636 | 0,88369 | 0,94839496 | egl-9 family hypoxia inducible factor 1 pseudogene 1 |
| ENSG00000106948 | AKNA | -0,223228852 | 0,856646056 | 0,2518 | 0,55773568 | AT-hook transcription factor |
| ENSG00000254685 | FPGT | -0,223229749 | 0,856645524 | 0,71104 | 0,86087272 | fucose-1-phosphate guanylyltransferase |
| ENSG00000282418 | AC092811,1 | -0,223255536 | 0,856630211 | 0,75213 | 0,88389154 | novel transcript |
| ENSG00000115828 | QPCT | -0,223353595 | 0,856571989 | 0,7483 | 0,88150078 | glutaminyl-peptide cyclotransferase |
| ENSG00000168872 | DDX19A | -0,223513373 | 0,856477129 | 0,4428 | 0,69040116 | DEAD-box helicase 19A |
| ENSG00000172955 | ADH6 | -0,223600832 | 0,85642521 | 0,48562 | 0,72107911 | alcohol dehydrogenase 6 (class V) |
| ENSG00000223476 | VN1R42P | -0,223711551 | 0,856359486 | 0,79818 | 0,90685415 | vomeronasal 1 receptor 42 pseudogene |
| ENSG00000254901 | BORCS8 | -0,223740956 | 0,856342032 | 0,40361 | 0,66399439 | BLOC-1 related complex subunit 8 |
| ENSG00000240682 | ISY1 | -0,223928097 | 0,856230957 | 0,44513 | 0,69216024 | ISY1 splicing factor homolog |
| ENSG00000148362 | PAXX | -0,224052277 | 0,85615726 | 0,29145 | 0,58649877 | PAXX non-homologous end joining factor |
| ENSG00000164764 | SBSPON | -0,224235416 | 0,856048585 | 0,5846 | 0,78577493 | somatomedin B and thrombospondin type 1 domain containing |
| ENSG00000228451 | SDAD1P1 | -0,224289476 | 0,856016508 | 0,58117 | 0,78406473 | SDA1 domain containing 1 pseudogene 1 |
| ENSG00000100284 | TOM1 | -0,224549307 | 0,855862352 | 0,45712 | 0,70067038 | target of myb1 membrane trafficking protein |
| ENSG00000111783 | RFX4 | -0,224571802 | 0,855849007 | 0,66649 | 0,83418752 | regulatory factor X4 |
| ENSG00000267565 | AC011477,2 | -0,2246069 | 0,855828187 | 0,50205 | 0,73163635 | novel transcript, antisense to ZNF253 |
| ENSG00000254510 | AP001107,5 | -0,224625416 | 0,855817203 | 0,50341 | 0,73266852 | novel transcript |
| ENSG00000188223 | AD000671,1 | -0,2246879 | 0,855780138 | 0,84837 | 0,93094401 | novel protein |
| ENSG00000227379 | PPIAP2 | -0,224689608 | 0,855779125 | 0,74022 | 0,87681509 | peptidylprolyl isomerase A pseudogene 2 |
| ENSG00000259498 | TPM1-AS | -0,224894856 | 0,855657384 | 0,64519 | 0,82200417 | TPM1 antisense RNA |
| ENSG00000173269 | MMRN2 | -0,224915302 | 0,855645257 | 0,55036 | 0,76458107 | multimerin 2 |

| | | | | | | |
|---|---|---|---|---|---|---|
| ENSG00000273616 | NDUFA6-DT | -0,225022075 | 0,855581934 | 0,64831 | 0,82383528 | NDUFA6 divergent transcript |
| ENSG00000213699 | SLC35F6 | -0,225034874 | 0,855574344 | 0,40133 | 0,6630277 | solute carrier family 35 member F6 |
| ENSG00000140326 | CDAN1 | -0,225044799 | 0,855568458 | 0,31425 | 0,60102501 | codanin 1 |
| ENSG00000267279 | AC090409,1 | -0,225065878 | 0,855555958 | 0,79569 | 0,90545311 | novel transcript |
| ENSG00000287357 | AL512484,1 | -0,225076296 | 0,855549779 | 0,57096 | 0,77772342 | novel transcript |
| ENSG00000249894 | AC024581,1 | -0,225208114 | 0,855471612 | 0,56172 | 0,77136826 | novel transcript |
| ENSG00000287133 | AC007279,1 | -0,225275233 | 0,855431814 | 0,62919 | 0,81265687 | novel transcript, antisense to MPP4and TMEM237 |
| ENSG00000119688 | ABCD4 | -0,225363768 | 0,855379319 | 0,31805 | 0,6034788 | ATP binding cassette subfamily D member 4 |
| ENSG00000133983 | COX16 | -0,225382315 | 0,855368322 | 0,38447 | 0,65052153 | cytochrome c oxidase assembly factor COX16 |
| ENSG00000282825 | CTIF | -0,225530973 | 0,855280188 | 0,34767 | 0,62423931 | cap binding complex dependent translation initiation factor |
| ENSG00000235703 | LINC00894 | -0,225540834 | 0,855274343 | 0,50041 | 0,73083859 | long intergenic non-protein coding RNA 894 |
| ENSG00000127526 | SLC35E1 | -0,225702432 | 0,855178547 | 0,43994 | 0,6885781 | solute carrier family 35 member E1 |
| ENSG00000166173 | LARP6 | -0,225794113 | 0,855124204 | 0,43959 | 0,68844641 | La ribonucleoprotein 6, translational regulator |
| ENSG00000288095 | AC005618,4 | -0,225835274 | 0,855099807 | 0,71745 | 0,86483656 | novel transcript |
| ENSG00000127561 | SYNGR3 | -0,225918926 | 0,855050227 | 0,52906 | 0,75111082 | synaptogyrin 3 |
| ENSG00000254788 | CKLF-CMTM1 | -0,225960071 | 0,855025842 | 0,51291 | 0,73946844 | CKLF-CMTM1 readthrough |
| ENSG00000272320 | AL445309,1 | -0,226044241 | 0,854975959 | 0,40774 | 0,66684304 | novel transcript, antisense to SLC22A23 |
| ENSG00000110455 | ACCS | -0,226183404 | 0,854893492 | 0,37944 | 0,6472232 | 1-aminocyclopropane-1-carboxylate synthase homolog (inactive) |
| ENSG00000175356 | SCUBE2 | -0,226190085 | 0,854889533 | 0,47315 | 0,71242761 | signal peptide, CUB domain and EGF like domain containing 2 |
| ENSG00000011028 | MRC2 | -0,22619643 | 0,854885773 | 0,41069 | 0,66856387 | mannose receptor C type 2 |
| ENSG00000139354 | GAS2L3 | -0,226221246 | 0,854871068 | 0,26834 | 0,57047423 | growth arrest specific 2 like 3 |
| ENSG00000178201 | VN1R1 | -0,226255898 | 0,854850535 | 0,48924 | 0,72376431 | vomeronasal 1 receptor 1 |
| ENSG00000100403 | ZC3H7B | -0,226364074 | 0,854786439 | 0,36485 | 0,63505392 | zinc finger CCCH-type containing 7B |
| ENSG00000181444 | ZNF467 | -0,226485126 | 0,85471472 | 0,49876 | 0,72990941 | zinc finger protein 467 |
| ENSG00000267698 | AC002116,2 | -0,226502508 | 0,854704422 | 0,67943 | 0,84252145 | uncharacterized LOC101927572 |
| ENSG00000268174 | BNIP3P22 | -0,226506758 | 0,854701904 | 0,63926 | 0,81902269 | BCL2 interacting protein 3 pseudogene 22 |
| ENSG00000214253 | FIS1 | -0,226543001 | 0,854680433 | 0,31322 | 0,59997051 | fission, mitochondrial 1 |
| ENSG00000130119 | GNL3L | -0,226588201 | 0,854653656 | 0,46655 | 0,7067394 | G protein nucleolar 3 like |
| ENSG00000197776 | KLHDC1 | -0,226618829 | 0,854635512 | 0,42288 | 0,67683766 | kelch domain containing 1 |
| ENSG00000221571 | RNU6ATAC35 | -0,226735985 | 0,854566113 | 0,82562 | 0,91917221 | RNA, U6atac small nuclear 35, pseudogene |

| | | | | | | |
|---|---|---|---|---|---|---|
| ENSG00000249115 | HAUS5 | -0,22676616 | 0,854548239 | 0,36443 | 0,63501401 | HAUS augmin like complex subunit 5 |
| ENSG00000266794 | RN7SL7P | -0,226869118 | 0,854487257 | 0,57726 | 0,78134056 | RNA, 7SL, cytoplasmic 7, pseudogene |
| ENSG00000140009 | ESR2 | -0,22687418 | 0,854484259 | 0,39534 | 0,65820274 | estrogen receptor 2 |
| ENSG00000100441 | KHNYN | -0,226991009 | 0,854415066 | 0,35619 | 0,63035942 | KH and NYN domain containing |
| ENSG00000158292 | GPR153 | -0,227028689 | 0,85439275 | 0,38607 | 0,65152373 | G protein-coupled receptor 153 |
| ENSG00000197562 | RAB40C | -0,227043154 | 0,854384184 | 0,27211 | 0,57306433 | RAB40C, member RAS oncogene family |
| ENSG00000279571 | AL162426,1 | -0,227054843 | 0,854377262 | 0,66731 | 0,83484024 | novel transcript |
| ENSG00000288016 | AL162493,1 | -0,227064967 | 0,854371266 | 0,75461 | 0,8850495 | novel transcript |
| ENSG00000177042 | TMEM80 | -0,227117944 | 0,854339894 | 0,32588 | 0,60842402 | transmembrane protein 80 |
| ENSG00000168884 | TNIP2 | -0,227157633 | 0,854316391 | 0,34729 | 0,62403275 | TNFAIP3 interacting protein 2 |
| ENSG00000213199 | ASIC3 | -0,227208733 | 0,854286131 | 0,59228 | 0,79106659 | acid sensing ion channel subunit 3 |
| ENSG00000101236 | RNF24 | -0,227322239 | 0,854218922 | 0,40921 | 0,66785537 | ring finger protein 24 |
| ENSG00000065883 | CDK13 | -0,22741812 | 0,854162153 | 0,25867 | 0,56260205 | cyclin dependent kinase 13 |
| ENSG00000248585 | AC084024,2 | -0,227441105 | 0,854148544 | 0,8341 | 0,92405734 | origin recognition complex, subunit 3-like (yeast) (ORC3L) pseudogene |
| ENSG00000274227 | AC073575,2 | -0,227493948 | 0,854117259 | 0,81815 | 0,91658434 | novel transcript, antisense to ERP29 |
| ENSG00000283669 | AC112487,2 | -0,227593231 | 0,854058483 | 0,8183 | 0,91663479 | novel pseudogene |
| ENSG00000180592 | SKIDA1 | -0,227701923 | 0,853994141 | 0,47118 | 0,71053521 | SKI/DACH domain containing 1 |
| ENSG00000171714 | ANO5 | -0,227721557 | 0,853982519 | 0,58206 | 0,78454692 | anoctamin 5 |
| ENSG00000162104 | ADCY9 | -0,227871625 | 0,853893693 | 0,45636 | 0,70003941 | adenylate cyclase 9 |
| ENSG00000107175 | CREB3 | -0,227874642 | 0,853891907 | 0,35946 | 0,63230932 | cAMP responsive element binding protein 3 |
| ENSG00000248019 | FAM13A-AS1 | -0,227928634 | 0,853859952 | 0,42124 | 0,67634721 | FAM13A antisense RNA 1 |
| ENSG00000164056 | SPRY1 | -0,228018139 | 0,85380698 | 0,21935 | 0,53676938 | sprouty RTK signaling antagonist 1 |
| ENSG00000112539 | C6orf118 | -0,228112629 | 0,853751061 | 0,20542 | 0,52503195 | chromosome 6 open reading frame 118 |
| ENSG00000115687 | PASK | -0,228383361 | 0,853590863 | 0,29164 | 0,58655914 | PAS domain containing serine/threonine kinase |
| ENSG00000227110 | LMCD1-AS1 | -0,228386254 | 0,853589151 | 0,41229 | 0,6699869 | LMCD1 antisense RNA 1 |
| ENSG00000198520 | ARMH1 | -0,228549023 | 0,853492853 | 0,52703 | 0,75012908 | armadillo like helical domain containing 1 |
| ENSG00000137824 | RMDN3 | -0,228773722 | 0,853359932 | 0,37425 | 0,64275387 | regulator of microtubule dynamics 3 |
| ENSG00000264985 | AC124804,1 | -0,228859364 | 0,853309276 | 0,56272 | 0,77194048 | novel transcript, antisense to SDK2 |
| ENSG00000175066 | GK5 | -0,228945312 | 0,853258442 | 0,39082 | 0,65464997 | glycerol kinase 5 |
| ENSG00000259767 | AC022558,1 | -0,229006275 | 0,853222387 | 0,77563 | 0,89630112 | novel transcript, sense intronic to BTBD1 |
| ENSG00000150054 | MPP7 | -0,229033884 | 0,853206059 | 0,37867 | 0,64679625 | membrane palmitoylated protein 7 |
| ENSG00000269821 | KCNQ1OT1 | -0,229051238 | 0,853195796 | 0,37318 | 0,64189779 | KCNQ1 opposite strand/antisense transcript 1 |

| | | | | | | |
|---|---|---|---|---|---|---|
| ENSG00000177710 | SLC35G5 | -0,229099026 | 0,853167535 | 0,8723 | 0,94374455 | solute carrier family 35 member G5 |
| ENSG00000267800 | AC012254,4 | -0,229198534 | 0,853108691 | 0,84298 | 0,9282764 | SUMO2 SMT3 suppressor of mif two 3 homolog 2 (S, cerevisiae), pseudogene |
| ENSG00000255206 | AC011853,2 | -0,229230485 | 0,853089797 | 0,73562 | 0,8750728 | novel transcript |
| ENSG00000254427 | LINC02696 | -0,229379536 | 0,853001665 | 0,65268 | 0,826091 | long intergenic non-protein coding RNA 2696 |
| ENSG00000237734 | AL163192,1 | -0,229393786 | 0,852993241 | 0,73576 | 0,8750728 | novel transcript |
| ENSG00000059588 | TARBP1 | -0,229462343 | 0,852952707 | 0,49841 | 0,72960863 | TAR (HIV-1) RNA binding protein 1 |
| ENSG00000005059 | MCUB | -0,229652337 | 0,852840386 | 0,52655 | 0,7497927 | mitochondrial calcium uniporter dominant negative subunit beta |
| ENSG00000172890 | NADSYN1 | -0,22965736 | 0,852837416 | 0,19973 | 0,52077515 | NAD synthetase 1 |
| ENSG00000271815 | AC008897,3 | -0,229693775 | 0,852815891 | 0,58414 | 0,78572744 | novel transcript |
| ENSG00000287167 | AL133313,1 | -0,229961109 | 0,852657877 | 0,73236 | 0,87296547 | novel transcript, antisense to RAD51B |
| ENSG00000279529 | AC008764,8 | -0,229979758 | 0,852646855 | 0,41186 | 0,66963061 | novel transcript, antisense to MED26 |
| ENSG00000269980 | AC068768,1 | -0,230154832 | 0,852543391 | 0,77251 | 0,89482448 | novel transcript, antisense to CDK2AP1 |
| ENSG00000227794 | RPS18 | -0,230227344 | 0,852500542 | 0,7021 | 0,85651838 | ribosomal protein S18 |
| ENSG00000063854 | HAGH | -0,23048877 | 0,852346077 | 0,35695 | 0,63108111 | hydroxyacylglutathione hydrolase |
| ENSG00000276449 | AC004076,2 | -0,230492064 | 0,852344131 | 0,35062 | 0,62606342 | novel transcript |
| ENSG00000269335 | IKBKG | -0,230624209 | 0,852266063 | 0,33093 | 0,6112353 | inhibitor of nuclear factor kappa B kinase regulatory subunit gamma |
| ENSG00000126861 | OMG | -0,230745615 | 0,852194346 | 0,48515 | 0,72079261 | oligodendrocyte myelin glycoprotein |
| ENSG00000154822 | PLCL2 | -0,230886274 | 0,852111263 | 0,56031 | 0,7707835 | phospholipase C like 2 |
| ENSG00000284017 | PLCL2 | -0,230886274 | 0,852111263 | 0,56031 | 0,7707835 | phospholipase C like 2 |
| ENSG00000271714 | AC010501,2 | -0,230957189 | 0,852069379 | 0,56166 | 0,77136826 | novel transcript, antisense to FAM169A |
| ENSG00000108423 | TUBD1 | -0,230973832 | 0,85205955 | 0,54932 | 0,76407481 | tubulin delta 1 |
| ENSG00000226359 | ACTG1P24 | -0,231013331 | 0,852036222 | 0,69832 | 0,85390309 | actin gamma 1 pseudogene 24 |
| ENSG00000229176 | SAPCD1 | -0,231223904 | 0,851911869 | 0,73713 | 0,87539732 | suppressor APC domain containing 1 |
| ENSG00000092330 | TINF2 | -0,231228407 | 0,85190921 | 0,3183 | 0,60349082 | TERF1 interacting nuclear factor 2 |
| ENSG00000160305 | DIP2A | -0,231264426 | 0,851887941 | 0,29188 | 0,58674733 | disco interacting protein 2 homolog A |
| ENSG00000108852 | MPP2 | -0,231338961 | 0,851843931 | 0,39043 | 0,65443234 | membrane palmitoylated protein 2 |
| ENSG00000168131 | OR2B2 | -0,23135123 | 0,851836687 | 0,79862 | 0,90712651 | olfactory receptor family 2 subfamily B member 2 |
| ENSG00000128254 | C22orf24 | -0,231374343 | 0,85182304 | 0,47848 | 0,71622997 | chromosome 22 open reading frame 24 |
| ENSG00000287007 | AC087683,3 | -0,231536434 | 0,85172734 | 0,80028 | 0,90802741 | novel transcript |
| ENSG00000224525 | AL591686,1 | -0,231658839 | 0,851655079 | 0,51082 | 0,73828855 | novel transcript, antisense to PLD5 |

| | | | | | | |
|---|---|---|---|---|---|---|
| ENSG00000230269 | LINC02525 | -0,231707847 | 0,851626149 | 0,67858 | 0,84182956 | long intergenic non-protein coding RNA 2525 |
| ENSG00000251152 | AC025539,1 | -0,231721322 | 0,851618194 | 0,74448 | 0,87904799 | novel transcript |
| ENSG00000261168 | AL592424,1 | -0,231834792 | 0,851551216 | 0,59905 | 0,79538547 | novel transcript, sense overlapping SEMA6C |
| ENSG00000054523 | KIF1B | -0,231881544 | 0,851523621 | 0,43357 | 0,68473723 | kinesin family member 1B |
| ENSG00000249471 | ZNF324B | -0,232009022 | 0,851448383 | 0,34024 | 0,61897407 | zinc finger protein 324B |
| ENSG00000281013 | NDUFA6 | -0,232087586 | 0,851402018 | 0,49808 | 0,72937175 | NADH:ubiquinone oxidoreductase subunit A6 |
| ENSG00000254539 | AC239804,1 | -0,232195312 | 0,851338446 | 0,69416 | 0,851567 | novel transcript |
| ENSG00000261924 | AC127496,1 | -0,23229779 | 0,851277975 | 0,59731 | 0,79440664 | novel transcript, antisense to RPTOR |
| ENSG00000240038 | AMY2B | -0,232343431 | 0,851251045 | 0,52853 | 0,75090627 | amylase alpha 2B |
| ENSG00000231365 | WARS2-AS1 | -0,232562262 | 0,851121935 | 0,36531 | 0,6354528 | WARS2 antisense RNA 1 |
| ENSG00000143409 | MINDY1 | -0,232613295 | 0,851091828 | 0,42581 | 0,67925635 | MINDY lysine 48 deubiquitinase 1 |
| ENSG00000119917 | IFIT3 | -0,232628515 | 0,85108285 | 0,56783 | 0,77568344 | interferon induced protein with tetratricopeptide repeats 3 |
| ENSG00000229587 | AL158825,2 | -0,23285364 | 0,850950053 | 0,73325 | 0,87353852 | novel transcript |
| ENSG00000243508 | DNAJB6P7 | -0,232886845 | 0,850930468 | 0,57611 | 0,78053116 | DNAJB6 pseudogene 7 |
| ENSG00000196155 | PLEKHG4 | -0,232908374 | 0,85091777 | 0,58113 | 0,78406473 | pleckstrin homology and RhoGEF domain containing G4 |
| ENSG00000075240 | GRAMD4 | -0,233189897 | 0,850751741 | 0,42174 | 0,67643128 | GRAM domain containing 4 |
| ENSG00000224055 | GAPDHP55 | -0,233243759 | 0,850719979 | 0,77583 | 0,89635267 | glyceraldehyde 3 phosphate dehydrogenase pseudogene 55 |
| ENSG00000262874 | C19orf84 | -0,233448897 | 0,850599023 | 0,7188 | 0,86552951 | chromosome 19 open reading frame 84 |
| ENSG00000133422 | MORC2 | -0,233462827 | 0,85059081 | 0,39194 | 0,65536998 | MORC family CW-type zinc finger 2 |
| ENSG00000224411 | HSP90AA2P | -0,233595797 | 0,850512417 | 0,56695 | 0,77482793 | heat shock protein 90 alpha family class A member 2, pseudogene |
| ENSG00000268649 | AL132655,2 | -0,233864829 | 0,850353829 | 0,68865 | 0,84822736 | novel transcript |
| ENSG00000245317 | AC008393,1 | -0,23410223 | 0,850213911 | 0,62435 | 0,80968303 | novel transcript |
| ENSG00000284274 | AC008393,3 | -0,23410223 | 0,850213911 | 0,62435 | 0,80968303 | novel transcript |
| ENSG00000136379 | ABHD17C | -0,234180104 | 0,850168019 | 0,56153 | 0,77136826 | abhydrolase domain containing 17C, depalmitoylase |
| ENSG00000235547 | NDUFB11P1 | -0,234285541 | 0,850105889 | 0,7902 | 0,9027921 | NADH:ubiquinone oxidoreductase subunit B11 pseudogene 1 |
| ENSG00000181274 | FRAT2 | -0,234323601 | 0,850083462 | 0,54011 | 0,75814808 | FRAT regulator of WNT signaling pathway 2 |
| ENSG00000149809 | TM7SF2 | -0,234361282 | 0,85006126 | 0,37921 | 0,6470219 | transmembrane 7 superfamily member 2 |
| ENSG00000180447 | GAS1 | -0,234381636 | 0,850049267 | 0,53471 | 0,7541464 | growth arrest specific 1 |
| ENSG00000104884 | ERCC2 | -0,234443885 | 0,85001259 | 0,42762 | 0,68054957 | ERCC excision repair 2, TFIIH core complex helicase subunit |

| | | | | | | |
|---|---|---|---|---|---|---|
| ENSG00000006712 | PAF1 | -0,23453274 | 0,84996024 | 0,23452 | 0,54623956 | PAF1 homolog, Paf1/RNA polymerase II complex component |
| ENSG00000130479 | MAP1S | -0,234547458 | 0,849951569 | 0,3127 | 0,59962409 | microtubule associated protein 1S |
| ENSG00000260744 | AC007494,2 | -0,234633219 | 0,849901045 | 0,69774 | 0,85357135 | novel transcript, antisense to ITFG1 |
| ENSG00000130066 | SAT1 | -0,234718715 | 0,84985068 | 0,57411 | 0,77933181 | spermidine/spermine N1-acetyltransferase 1 |
| ENSG00000282670 | AC254944,4 | -0,234768587 | 0,849821302 | 0,82287 | 0,9181175 | novel transcript |
| ENSG00000269930 | AC091057,3 | -0,234768587 | 0,849821302 | 0,82287 | 0,9181175 | novel transcript |
| ENSG00000286762 | AL096870,9 | -0,234917546 | 0,849733562 | 0,70271 | 0,85665929 | novel transcript, antisense to RABGGTA |
| ENSG00000286825 | AL096870,10 | -0,234917546 | 0,849733562 | 0,70271 | 0,85665929 | novel transcript, antisense to RABGGTA |
| ENSG00000267520 | AC010733,1 | -0,235035147 | 0,849664299 | 0,38437 | 0,65052153 | novel transcript |
| ENSG00000241549 | GUSBP2 | -0,235053172 | 0,849653684 | 0,27806 | 0,57635142 | GUSB pseudogene 2 |
| ENSG00000197191 | CYSRT1 | -0,235154671 | 0,84959391 | 0,70918 | 0,859809 | cysteine rich tail 1 |
| ENSG00000278638 | AC244093,5 | -0,235428706 | 0,849432547 | 0,78023 | 0,89823499 | novel transcript, antisense AATF |
| ENSG00000169814 | BTD | -0,235460338 | 0,849413923 | 0,44018 | 0,68866128 | biotinidase |
| ENSG00000273597 | AL355377,1 | -0,235501396 | 0,84938975 | 0,71563 | 0,86355924 | SPATA31 subfamily D, member 1 (SPATA31D1) pseudogene |
| ENSG00000238134 | EHMT2 | -0,235786572 | 0,849221869 | 0,35729 | 0,63122452 | euchromatic histone lysine methyltransferase 2 |
| ENSG00000135842 | NIBAN1 | -0,235812567 | 0,849206567 | 0,6067 | 0,79981537 | niban apoptosis regulator 1 |
| ENSG00000254934 | LINC00678 | -0,235864557 | 0,849175965 | 0,67817 | 0,84159488 | long intergenic non-protein coding RNA 678 |
| ENSG00000110237 | ARHGEF17 | -0,235876403 | 0,849168993 | 0,42251 | 0,67663831 | Rho guanine nucleotide exchange factor 17 |
| ENSG00000145309 | CABS1 | -0,235916274 | 0,849145525 | 0,77342 | 0,89545683 | calcium binding protein, spermatid associated 1 |
| ENSG00000213669 | AL137074,1 | -0,235919636 | 0,849143546 | 0,79932 | 0,90744841 | ribosomal protein L21 (RPL21) pseudogene |
| ENSG00000254895 | MTCO3P15 | -0,236030129 | 0,849078514 | 0,78363 | 0,89918294 | MT-CO3 pseudogene 15 |
| ENSG00000261136 | AC023908,3 | -0,236074601 | 0,849052341 | 0,34765 | 0,62423931 | novel transcript, antisense to FSIP1 |
| ENSG00000021355 | SERPINB1 | -0,236124011 | 0,849023263 | 0,26376 | 0,56631093 | serpin family B member 1 |
| ENSG00000161677 | JOSD2 | -0,236131903 | 0,849018619 | 0,59698 | 0,79434997 | Josephin domain containing 2 |
| ENSG00000198064 | NPIPB13 | -0,236289639 | 0,848925797 | 0,48258 | 0,71901075 | nuclear pore complex interacting protein family, member B13 |
| ENSG00000271780 | AL118558,3 | -0,236326204 | 0,848904281 | 0,59638 | 0,79392867 | novel transcript |
| ENSG00000196924 | FLNA | -0,236333174 | 0,84890018 | 0,2683 | 0,5704222 | filamin A |
| ENSG00000261292 | AC110491,1 | -0,236399116 | 0,84886138 | 0,54011 | 0,75814808 | novel transcript |
| ENSG00000247775 | SNCA-AS1 | -0,236419374 | 0,84884946 | 0,69487 | 0,85206252 | SNCA antisense RNA 1 |
| ENSG00000205155 | PSENEN | -0,236941619 | 0,848542239 | 0,25518 | 0,55966322 | presenilin enhancer, gamma-secretase subunit |

| | | | | | | |
|---|---|---|---|---|---|---|
| ENSG00000285119 | FAM86B3P | -0,237017593 | 0,848497554 | 0,43001 | 0,68268477 | family with sequence similarity 86 member B3, pseudogene |
| ENSG00000107295 | SH3GL2 | -0,237055951 | 0,848474995 | 0,64789 | 0,82380454 | SH3 domain containing GRB2 like 2, endophilin A1 |
| ENSG00000246451 | AL049840,2 | -0,237159468 | 0,848414117 | 0,2093 | 0,52826608 | novel transcript, antisense to KLC1 |
| ENSG00000100626 | GALNT16 | -0,237344176 | 0,848305502 | 0,33703 | 0,61684812 | polypeptide N-acetylgalactosaminyltransferase 16 |
| ENSG00000182179 | UBA7 | -0,237352243 | 0,848300759 | 0,60727 | 0,80012503 | ubiquitin like modifier activating enzyme 7 |
| ENSG00000233693 | PBX1-AS1 | -0,237411499 | 0,848265917 | 0,65271 | 0,826091 | PBX1 antisense RNA 1 |
| ENSG00000281501 | SEPSECS-AS1 | -0,237439536 | 0,848249432 | 0,29999 | 0,59160746 | SEPSECS antisense RNA 1 (head to head) |
| ENSG00000268472 | AP002884,3 | -0,237592217 | 0,848159666 | 0,81733 | 0,91606335 | novel transcript, antisense to C11orf34 |
| ENSG00000256977 | LIMS3 | -0,237630291 | 0,848137283 | 0,56734 | 0,77527465 | LIM zinc finger domain containing 3 |
| ENSG00000225929 | AC000036,1 | -0,237690413 | 0,848101939 | 0,76739 | 0,89211771 | novel transcript |
| ENSG00000269578 | AC008764,7 | -0,237714941 | 0,84808752 | 0,75162 | 0,88357423 | novel transcript, sense intronic MED26 |
| ENSG00000106683 | LIMK1 | -0,237760055 | 0,848061 | 0,32304 | 0,60584543 | LIM domain kinase 1 |
| ENSG00000087087 | SRRT | -0,237767077 | 0,848056872 | 0,31445 | 0,6011918 | serrate, RNA effector molecule |
| ENSG00000223843 | EFCAB6-AS1 | -0,237819875 | 0,848025837 | 0,33123 | 0,61131843 | EFCAB6 antisense RNA 1 |
| ENSG00000103245 | CIAO3 | -0,237908873 | 0,847973525 | 0,38266 | 0,64939435 | cytosolic iron-sulfur assembly component 3 |
| ENSG00000230773 | AC092650,1 | -0,237910086 | 0,847972812 | 0,67849 | 0,84182956 | novel transcript |
| ENSG00000185963 | BICD2 | -0,237939991 | 0,847955235 | 0,25821 | 0,5622185 | BICD cargo adaptor 2 |
| ENSG00000158457 | TSPAN33 | -0,237961239 | 0,847942746 | 0,53508 | 0,75434242 | tetraspanin 33 |
| ENSG00000261286 | ATP2C2-AS1 | -0,237979176 | 0,847932203 | 0,6002 | 0,79579686 | ATP2C2 antisense RNA 1 |
| ENSG00000064115 | TM7SF3 | -0,238056728 | 0,847886624 | 0,2887 | 0,58455903 | transmembrane 7 superfamily member 3 |
| ENSG00000111452 | ADGRD1 | -0,238163773 | 0,847823715 | 0,56752 | 0,7754074 | adhesion G protein-coupled receptor D1 |
| ENSG00000213064 | SFT2D2 | -0,238168785 | 0,84782077 | 0,35889 | 0,63196414 | SFT2 domain containing 2 |
| ENSG00000162174 | ASRGL1 | -0,238170721 | 0,847819632 | 0,26117 | 0,56416552 | asparaginase and isoaspartyl peptidase 1 |
| ENSG00000084710 | EFR3B | -0,238596132 | 0,847569671 | 0,29453 | 0,58857672 | EFR3 homolog B |
| ENSG00000286550 | AL591002,1 | -0,238792242 | 0,847454465 | 0,62085 | 0,80776776 | novel transcript, antisense to ANKS1A |
| ENSG00000177666 | PNPLA2 | -0,238816397 | 0,847440277 | 0,51102 | 0,73835122 | patatin like phospholipase domain containing 2 |
| ENSG00000088256 | GNA11 | -0,238844828 | 0,847423576 | 0,46744 | 0,70731446 | G protein subunit alpha 11 |
| ENSG00000167311 | ART5 | -0,238940206 | 0,847367554 | 0,65223 | 0,82578944 | ADP-ribosyltransferase 5 |
| ENSG00000260456 | C16orf95 | -0,239005235 | 0,84732936 | 0,39406 | 0,65702282 | chromosome 16 open reading frame 95 |
| ENSG00000141294 | LRRC46 | -0,239184189 | 0,847224263 | 0,40058 | 0,66264525 | leucine rich repeat containing 46 |
| ENSG00000181798 | LINC00471 | -0,239473817 | 0,847054196 | 0,6765 | 0,84080793 | long intergenic non-protein coding RNA 471 |
| ENSG00000174996 | KLC2 | -0,239484294 | 0,847048044 | 0,25794 | 0,56199743 | kinesin light chain 2 |

| | | | | | | |
|---|---|---|---|---|---|---|
| ENSG00000136280 | CCM2 | -0,239643405 | 0,846954631 | 0,24192 | 0,55071239 | CCM2 scaffold protein |
| ENSG00000138696 | BMPR1B | -0,239682942 | 0,84693142 | 0,54893 | 0,76394298 | bone morphogenetic protein receptor type 1B |
| ENSG00000231329 | AL031772,1 | -0,239839021 | 0,846839799 | 0,57603 | 0,78050563 | novel transcript |
| ENSG00000279068 | AC244517,6 | -0,239849049 | 0,846833913 | 0,71213 | 0,8615121 | novel transcript |
| ENSG00000234036 | TXNP6 | -0,239896202 | 0,846806235 | 0,72391 | 0,86817061 | thioredoxin pseudogene 6 |
| ENSG00000123395 | ATG101 | -0,239957369 | 0,846770334 | 0,45217 | 0,69691985 | autophagy related 101 |
| ENSG00000232931 | LINC00342 | -0,24003082 | 0,846727223 | 0,30303 | 0,59337005 | long intergenic non-protein coding RNA 342 |
| ENSG00000059377 | TBXAS1 | -0,240195168 | 0,846630772 | 0,51887 | 0,7437068 | thromboxane A synthase 1 |
| ENSG00000167302 | TEPSIN | -0,240213903 | 0,846619778 | 0,31617 | 0,60224605 | TEPSIN adaptor related protein complex 4 accessory protein |
| ENSG00000076864 | RAP1GAP | -0,240229055 | 0,846610886 | 0,46081 | 0,70258879 | RAP1 GTPase activating protein |
| ENSG00000288581 | AL137139,3 | -0,240322488 | 0,846556059 | 0,70877 | 0,85960107 | novel transcript, antisense to CLYBL |
| ENSG00000285456 | LTB4R | -0,24032343 | 0,846555506 | 0,55015 | 0,76444876 | leukotriene B4 receptor |
| ENSG00000089091 | DZANK1 | -0,240621372 | 0,846380696 | 0,29056 | 0,58565836 | double zinc ribbon and ankyrin repeat domains 1 |
| ENSG00000260083 | MIR762HG | -0,24072182 | 0,846321768 | 0,34096 | 0,6192144 | MIR762 host gene |
| ENSG00000243970 | PPIEL | -0,240767095 | 0,846295209 | 0,41369 | 0,67085666 | peptidylprolyl isomerase E like pseudogene |
| ENSG00000169228 | RAB24 | -0,240935495 | 0,846196431 | 0,41285 | 0,6705041 | RAB24, member RAS oncogene family |
| ENSG00000169249 | ZRSR2 | -0,24119822 | 0,846042346 | 0,3662 | 0,63617598 | zinc finger CCCH-type, RNA binding motif and serine/arginine rich 2 |
| ENSG00000186399 | GOLGA8R | -0,241249557 | 0,846012241 | 0,27891 | 0,57700804 | golgin A8 family member R |
| ENSG00000234835 | PHBP13 | -0,241341534 | 0,845958306 | 0,73749 | 0,87540065 | prohibitin pseudogene 13 |
| ENSG00000083099 | LYRM2 | -0,241343813 | 0,84595697 | 0,35409 | 0,62877376 | LYR motif containing 2 |
| ENSG00000139625 | MAP3K12 | -0,241363348 | 0,845945515 | 0,3288 | 0,60992716 | mitogen-activated protein kinase kinase kinase 12 |
| ENSG00000204442 | FAM155A | -0,241470281 | 0,845882816 | 0,46605 | 0,70640325 | family with sequence similarity 155 member A |
| ENSG00000244627 | TPTEP2 | -0,241548591 | 0,845836902 | 0,39598 | 0,65863599 | TPTE pseudogene 2 |
| ENSG00000165325 | DEUP1 | -0,241795286 | 0,84569228 | 0,67636 | 0,84075837 | deuterosome assembly protein 1 |
| ENSG00000239569 | KMT2E-AS1 | -0,241823622 | 0,84567567 | 0,45142 | 0,69646078 | KMT2E antisense RNA 1 |
| ENSG00000157368 | IL34 | -0,241868164 | 0,845649561 | 0,76325 | 0,88980026 | interleukin 34 |
| ENSG00000269386 | RAB11B-AS1 | -0,241895724 | 0,845633406 | 0,42947 | 0,68224982 | RAB11B antisense RNA 1 |
| ENSG00000214087 | ARL16 | -0,242048161 | 0,84554406 | 0,39038 | 0,65441492 | ADP ribosylation factor like GTPase 16 |
| ENSG00000147408 | CSGALNACT1 | -0,242059876 | 0,845537194 | 0,72419 | 0,86838632 | chondroitin sulfate N-acetylgalactosaminyltransferase 1 |
| ENSG00000233021 | AL669841,1 | -0,242136009 | 0,845492576 | 0,78749 | 0,90121071 | novel transcript |
| ENSG00000132589 | FLOT2 | -0,242192211 | 0,845459639 | 0,08876 | 0,38517778 | flotillin 2 |

| | | | | | | |
|---|---|---|---|---|---|---|
| ENSG00000220506 | AL136310,1 | -0,242204583 | 0,845452389 | 0,75978 | 0,88801075 | novel pseudogene |
| ENSG00000135679 | MDM2 | -0,242238795 | 0,84543234 | 0,32064 | 0,60553919 | MDM2 proto-oncogene |
| ENSG00000161692 | DBF4B | -0,242244549 | 0,845428968 | 0,34238 | 0,62005694 | DBF4 zinc finger B |
| ENSG00000250903 | GMDS-DT | -0,242249352 | 0,845426153 | 0,24711 | 0,55483158 | GMDS divergent transcript |
| ENSG00000254827 | SLC22A18AS | -0,242258178 | 0,845420981 | 0,62998 | 0,8132682 | solute carrier family 22 member 18 antisense |
| ENSG00000167615 | LENG8 | -0,242304776 | 0,845393675 | 0,29355 | 0,58800349 | leukocyte receptor cluster member 8 |
| ENSG00000185829 | ARL17A | -0,242423797 | 0,845323934 | 0,35929 | 0,63214783 | ADP ribosylation factor like GTPase 17A |
| ENSG00000160785 | SLC25A44 | -0,242482796 | 0,845289365 | 0,41014 | 0,66821634 | solute carrier family 25 member 44 |
| ENSG00000138442 | WDR12 | -0,242543407 | 0,845253853 | 0,50276 | 0,73210009 | WD repeat domain 12 |
| ENSG00000204371 | EHMT2 | -0,242587056 | 0,84522828 | 0,3388 | 0,61792351 | euchromatic histone lysine methyltransferase 2 |
| ENSG00000092445 | TYRO3 | -0,242616596 | 0,845210974 | 0,43994 | 0,6885781 | TYRO3 protein tyrosine kinase |
| ENSG00000021762 | OSBPL5 | -0,242635112 | 0,845200126 | 0,33057 | 0,6112353 | oxysterol binding protein like 5 |
| ENSG00000181222 | POLR2A | -0,242755648 | 0,845129514 | 0,60424 | 0,79780609 | RNA polymerase II subunit A |
| ENSG00000179820 | MYADM | -0,242756513 | 0,845129007 | 0,55272 | 0,76604561 | myeloid associated differentiation marker |
| ENSG00000234494 | SP2-AS1 | -0,242768892 | 0,845121755 | 0,44225 | 0,69006952 | SP2 antisense RNA 1 |
| ENSG00000187902 | SHISA7 | -0,242846626 | 0,84507622 | 0,53288 | 0,75333268 | shisa family member 7 |
| ENSG00000270269 | IMMP1LP1 | -0,243102969 | 0,844926077 | 0,82152 | 0,91749859 | inner mitochondrial membrane peptidase subunit 1 pseudogene 1 |
| ENSG00000121680 | PEX16 | -0,243111605 | 0,84492102 | 0,41741 | 0,67403924 | peroxisomal biogenesis factor 16 |
| ENSG00000179010 | MRFAP1 | -0,243135 | 0,844907319 | 0,3438 | 0,62120234 | Morf4 family associated protein 1 |
| ENSG00000154917 | RAB6B | -0,243146121 | 0,844900806 | 0,4162 | 0,67305137 | RAB6B, member RAS oncogene family |
| ENSG00000232453 | LINC02777 | -0,243229774 | 0,844851816 | 0,69589 | 0,85282958 | long intergenic non-protein coding RNA 2777 |
| ENSG00000286676 | ACTBP4 | -0,243524983 | 0,844678958 | 0,7167 | 0,86439754 | ACTB pseudogene 4 |
| ENSG00000196834 | POTEI | -0,243537694 | 0,844671515 | 0,29783 | 0,59037822 | POTE ankyrin domain family member I |
| ENSG00000187672 | ERC2 | -0,243572771 | 0,844650979 | 0,17003 | 0,49606609 | ELKS/RAB6-interacting/CAST family member 2 |
| ENSG00000179774 | ATOH7 | -0,243598911 | 0,844635675 | 0,64167 | 0,82033064 | atonal bHLH transcription factor 7 |
| ENSG00000283078 | AL137077,2 | -0,24374604 | 0,844549542 | 0,46566 | 0,70604054 | novel transcript |
| ENSG00000063660 | GPC1 | -0,24374867 | 0,844548002 | 0,38444 | 0,65052153 | glypican 1 |
| ENSG00000162076 | FLYWCH2 | -0,243794691 | 0,844521062 | 0,36698 | 0,63692813 | FLYWCH family member 2 |
| ENSG00000074356 | NCBP3 | -0,243810849 | 0,844511604 | 0,30675 | 0,59579909 | nuclear cap binding subunit 3 |
| ENSG00000257242 | LINC01619 | -0,243914299 | 0,844451049 | 0,73207 | 0,87285234 | long intergenic non-protein coding RNA 1619 |
| ENSG00000254531 | FLJ20021 | -0,243954911 | 0,844427278 | 0,62307 | 0,80892518 | uncharacterized LOC90024 |
| ENSG00000270074 | AC087203,3 | -0,243958614 | 0,84442511 | 0,42645 | 0,67987951 | novel transcript |

| | | | | | | |
|---|---|---|---|---|---|---|
| ENSG00000130733 | YIPF2 | -0,244076494 | 0,844356117 | 0,32827 | 0,60971379 | Yip1 domain family member 2 |
| ENSG00000147684 | NDUFB9 | -0,244181195 | 0,844294842 | 0,34314 | 0,62060539 | NADH:ubiquinone oxidoreductase subunit B9 |
| ENSG00000148343 | MIGA2 | -0,24423587 | 0,844262846 | 0,24734 | 0,5549105 | mitoguardin 2 |
| ENSG00000198453 | ZNF568 | -0,244241692 | 0,844259438 | 0,30696 | 0,5960776 | zinc finger protein 568 |
| ENSG00000185414 | MRPL30 | -0,24440107 | 0,844166176 | 0,44376 | 0,69144937 | mitochondrial ribosomal protein L30 |
| ENSG00000197429 | IPP | -0,244428374 | 0,8441502 | 0,22906 | 0,54263903 | intracisternal A particle-promoted polypeptide |
| ENSG00000214870 | AC004540,1 | -0,244432419 | 0,844147833 | 0,49844 | 0,72960863 | novel transcript |
| ENSG00000281538 | AC254562,3 | -0,244439618 | 0,844143621 | 0,75859 | 0,88727479 | novel transcript |
| ENSG00000282969 | AC235563,1 | -0,244439618 | 0,844143621 | 0,75859 | 0,88727479 | novel transcript |
| ENSG00000282874 | AC254847,1 | -0,244439618 | 0,844143621 | 0,75859 | 0,88727479 | novel transcript |
| ENSG00000240964 | RN7SL751P | -0,244442205 | 0,844142107 | 0,77255 | 0,89483773 | RNA, 7SL, cytoplasmic 751, pseudogene |
| ENSG00000263324 | HCN3 | -0,244483406 | 0,844118 | 0,53684 | 0,75562287 | hyperpolarization activated cyclic nucleotide gated potassium channel 3 |
| ENSG00000168772 | CXXC4 | -0,244665432 | 0,844011504 | 0,60288 | 0,7971742 | CXXC finger protein 4 |
| ENSG00000285400 | MCPH1-AS1 | -0,24482341 | 0,843919088 | 0,3617 | 0,63352908 | MCPH1 antisense RNA 1 |
| ENSG00000196544 | BORCS6 | -0,244909275 | 0,843868862 | 0,48696 | 0,72215091 | BLOC-1 related complex subunit 6 |
| ENSG00000182768 | NGRN | -0,244999419 | 0,843816136 | 0,37172 | 0,64079608 | neugrin, neurite outgrowth associated |
| ENSG00000159588 | CCDC17 | -0,245013528 | 0,843807884 | 0,39865 | 0,66118205 | coiled-coil domain containing 17 |
| ENSG00000270911 | AC114878,2 | -0,245073859 | 0,843772598 | 0,80044 | 0,90809799 | CTAGE family, member 5 (CTAGE5) pseudogene |
| ENSG00000183655 | KLHL25 | -0,245127029 | 0,843741501 | 0,20294 | 0,52282418 | kelch like family member 25 |
| ENSG00000156398 | SFXN2 | -0,245155758 | 0,8437247 | 0,43642 | 0,68676106 | sideroflexin 2 |
| ENSG00000270343 | UNGP3 | -0,24524783 | 0,843670856 | 0,63144 | 0,81403245 | uracil-DNA glycosylase pseudogene 3 |
| ENSG00000108509 | CAMTA2 | -0,245360158 | 0,84360517 | 0,2007 | 0,5215633 | calmodulin binding transcription activator 2 |
| ENSG00000271882 | AP001330,4 | -0,245415688 | 0,8435727 | 0,70411 | 0,85725494 | novel transcript |
| ENSG00000244244 | RPS3AP42 | -0,245525387 | 0,843508559 | 0,77991 | 0,89817238 | RPS3A pseudogene 42 |
| ENSG00000280537 | AC068946,1 | -0,245594018 | 0,843468433 | 0,7389 | 0,87610803 | novel protein, SLC23A3-NHEJ1 readthrough |
| ENSG00000287439 | AC092078,3 | -0,245622943 | 0,843451523 | 0,62123 | 0,80787022 | novel transcript |
| ENSG00000197208 | SLC22A4 | -0,245661721 | 0,843428852 | 0,66688 | 0,83455977 | solute carrier family 22 member 4 |
| ENSG00000171466 | ZNF562 | -0,245713962 | 0,843398311 | 0,20338 | 0,52321822 | zinc finger protein 562 |
| ENSG00000249717 | PARM1-AS1 | -0,245763649 | 0,843369265 | 0,46472 | 0,70529302 | PARM1 antisense RNA 1 |
| ENSG00000214629 | RPSAP6 | -0,245843925 | 0,843322338 | 0,75989 | 0,88801075 | ribosomal protein SA pseudogene 6 |
| ENSG00000285813 | AP000813,1 | -0,245894429 | 0,843292817 | 0,48541 | 0,72098266 | novel transcript, antisense to MSANTD4 |
| ENSG00000196208 | GREB1 | -0,245894676 | 0,843292672 | 0,49369 | 0,7264386 | growth regulating estrogen receptor binding 1 |

| | | | | | | |
|---|---|---|---|---|---|---|
| ENSG00000281677 | AC005393,2 | -0,245971311 | 0,843247879 | 0,71728 | 0,8648058 | novel transcript |
| ENSG00000276445 | AC005393,1 | -0,245971311 | 0,843247879 | 0,71728 | 0,8648058 | novel transcript |
| ENSG00000286033 | CR381670,2 | -0,246263823 | 0,843076924 | 0,68119 | 0,84367915 | novel transcript |
| ENSG00000108255 | CRYBA1 | -0,246407011 | 0,842993253 | 0,75088 | 0,88311592 | crystallin beta A1 |
| ENSG00000275381 | AC019206,1 | -0,246438589 | 0,842974801 | 0,658 | 0,82895829 | Zinc finger protein 607 (ZNF607) pseudogene |
| ENSG00000189143 | CLDN4 | -0,246583188 | 0,842890315 | 0,63004 | 0,8132682 | claudin 4 |
| ENSG00000133101 | CCNA1 | -0,246603875 | 0,842878229 | 0,58291 | 0,78477395 | cyclin A1 |
| ENSG00000187559 | FOXD4L3 | -0,246685249 | 0,842830688 | 0,81671 | 0,91574173 | forkhead box D4 like 3 |
| ENSG00000126012 | KDM5C | -0,246750792 | 0,842792399 | 0,38556 | 0,65117983 | lysine demethylase 5C |
| ENSG00000183765 | CHEK2 | -0,246920227 | 0,842693425 | 0,47395 | 0,71294959 | checkpoint kinase 2 |
| ENSG00000147036 | LANCL3 | -0,246935124 | 0,842684723 | 0,7376 | 0,87540065 | LanC like 3 |
| ENSG00000258274 | AC012085,2 | -0,246972185 | 0,842663076 | 0,59817 | 0,79488132 | novel transcript |
| ENSG00000237025 | AL121601,2 | -0,247204332 | 0,842527492 | 0,63713 | 0,81747939 | par-6 partitioning defective 6 homolog beta (C, elegans) (PARD6B) pseudogene |
| ENSG00000109832 | DDX25 | -0,247251921 | 0,842499701 | 0,44428 | 0,69172919 | DEAD-box helicase 25 |
| ENSG00000228463 | AP006222,1 | -0,247385914 | 0,842421456 | 0,15952 | 0,48693054 | ribosomal protein L23a (RPL23A) pseudogene |
| ENSG00000137500 | CCDC90B | -0,24741501 | 0,842404466 | 0,15785 | 0,48490111 | coiled-coil domain containing 90B |
| ENSG00000186889 | TMEM17 | -0,247428905 | 0,842396353 | 0,23941 | 0,5491566 | transmembrane protein 17 |
| ENSG00000135517 | MIP | -0,247571838 | 0,842312898 | 0,7201 | 0,86623639 | major intrinsic protein of lens fiber |
| ENSG00000270218 | AC013553,2 | -0,247613977 | 0,842288295 | 0,78423 | 0,89959305 | transmembrane protein 126A (TMEM126A) pseudogene |
| ENSG00000125458 | NT5C | -0,247743167 | 0,842212873 | 0,26695 | 0,56937393 | 5', 3'-nucleotidase, cytosolic |
| ENSG00000224986 | PPP1R8P1 | -0,247791247 | 0,842184806 | 0,58825 | 0,78810304 | protein phosphatase 1 regulatory subunit 8 pseudogene 1 |
| ENSG00000225968 | ELFN1 | -0,247816043 | 0,842170331 | 0,32262 | 0,60581965 | extracellular leucine rich repeat and fibronectin type III domain containing 1 |
| ENSG00000284868 | DHRS1 | -0,24786179 | 0,842143627 | 0,33287 | 0,61286447 | dehydrogenase/reductase 1 |
| ENSG00000176927 | EFCAB5 | -0,248064953 | 0,842025043 | 0,7059 | 0,85838918 | EF-hand calcium binding domain 5 |
| ENSG00000261186 | AC093642,2 | -0,24807236 | 0,84202072 | 0,65351 | 0,82638628 | novel transcript |
| ENSG00000100288 | CHKB | -0,248072918 | 0,842020394 | 0,22036 | 0,53750545 | choline kinase beta |
| ENSG00000066248 | NGEF | -0,248123952 | 0,841990609 | 0,54282 | 0,75960352 | neuronal guanine nucleotide exchange factor |
| ENSG00000162604 | TM2D1 | -0,248262791 | 0,841909583 | 0,39962 | 0,66184353 | TM2 domain containing 1 |
| ENSG00000101448 | EPPIN | -0,248279318 | 0,841899939 | 0,57262 | 0,77858457 | epididymal peptidase inhibitor |
| ENSG00000285872 | AC007240,3 | -0,24829341 | 0,841891715 | 0,68754 | 0,8478032 | novel transcript |
| ENSG00000197892 | KIF13B | -0,248426521 | 0,841814041 | 0,21602 | 0,53473714 | kinesin family member 13B |

| | | | | | | |
|---|---|---|---|---|---|---|
| ENSG00000164620 | RELL2 | -0,248571202 | 0,841729624 | 0,27282 | 0,57322782 | RELT like 2 |
| ENSG00000105559 | PLEKHA4 | -0,248638549 | 0,841690332 | 0,6356 | 0,81657648 | pleckstrin homology domain containing A4 |
| ENSG00000152669 | CCNO | -0,248692281 | 0,841658984 | 0,66768 | 0,83516218 | cyclin O |
| ENSG00000269038 | AP001462,1 | -0,248955085 | 0,84150568 | 0,72161 | 0,8670761 | novel transcript |
| ENSG00000254238 | AC100782,1 | -0,249029225 | 0,841462436 | 0,7035 | 0,857007 | novel transcript |
| ENSG00000260371 | AC026464,3 | -0,24906602 | 0,841440976 | 0,90967 | 0,96105018 | novel protein |
| ENSG00000101384 | JAG1 | -0,249112352 | 0,841413953 | 0,51108 | 0,73835122 | jagged canonical Notch ligand 1 |
| ENSG00000118263 | KLF7 | -0,249115582 | 0,84141207 | 0,42691 | 0,68018708 | Kruppel like factor 7 |
| ENSG00000242979 | RPS15AP18 | -0,249213265 | 0,841355101 | 0,8088 | 0,9120028 | ribosomal protein S15a pseudogene 18 |
| ENSG00000201600 | RN7SKP124 | -0,249291371 | 0,841309552 | 0,63106 | 0,81391092 | RN7SK pseudogene 124 |
| ENSG00000219470 | AL355802,1 | -0,249413967 | 0,841238062 | 0,58271 | 0,78477395 | 40S ribosomal protein S2 (RPS2) pseudogene |
| ENSG00000226318 | RPS3AP38 | -0,249437413 | 0,841224391 | 0,57042 | 0,77733701 | RPS3A pseudogene 38 |
| ENSG00000226393 | IFNA20P | -0,249464442 | 0,841208631 | 0,76194 | 0,88909725 | interferon alpha 20, pseudogene |
| ENSG00000137843 | PAK6 | -0,249508135 | 0,841183155 | 0,54781 | 0,76308811 | p21 (RAC1) activated kinase 6 |
| ENSG00000085231 | AK6 | -0,249521248 | 0,841175509 | 0,49369 | 0,7264386 | adenylate kinase 6 |
| ENSG00000077327 | SPAG6 | -0,24955014 | 0,841158663 | 0,61372 | 0,8035281 | sperm associated antigen 6 |
| ENSG00000162419 | GMEB1 | -0,249557615 | 0,841154305 | 0,48907 | 0,7236969 | glucocorticoid modulatory element binding protein 1 |
| ENSG00000274963 | RN7SL600P | -0,249586886 | 0,841137239 | 0,71128 | 0,86101138 | RNA, 7SL, cytoplasmic 600, pseudogene |
| ENSG00000218739 | CEBPZOS | -0,249713893 | 0,841063194 | 0,50733 | 0,73559605 | CEBPZ opposite strand |
| ENSG00000180304 | OAZ2 | -0,249777011 | 0,841026398 | 0,35839 | 0,63185282 | ornithine decarboxylase antizyme 2 |
| ENSG00000271590 | AC108463,3 | -0,249824544 | 0,840998689 | 0,5721 | 0,77840742 | novel transcript |
| ENSG00000141959 | PFKL | -0,250116932 | 0,840828262 | 0,34147 | 0,61956597 | phosphofructokinase, liver type |
| ENSG00000144741 | SLC25A26 | -0,250174052 | 0,840794973 | 0,51154 | 0,73860667 | solute carrier family 25 member 26 |
| ENSG00000227486 | AL050309,1 | -0,25026547 | 0,840741696 | 0,27245 | 0,57315552 | novel transcript |
| ENSG00000187372 | PCDHB13 | -0,25028146 | 0,840732378 | 0,19847 | 0,51980353 | protocadherin beta 13 |
| ENSG00000213305 | HNRNPCP6 | -0,25038068 | 0,84067456 | 0,6697 | 0,83623395 | heterogeneous nuclear ribonucleoprotein C pseudogene 6 |
| ENSG00000287757 | AC244636,3 | -0,250408982 | 0,840658068 | 0,31013 | 0,5983747 | novel transcript, antisense to ZNF630 |
| ENSG00000239521 | CASTOR3 | -0,250646929 | 0,840519428 | 0,29681 | 0,58995143 | CASTOR family member 3 |
| ENSG00000151746 | BICD1 | -0,250688255 | 0,840495351 | 0,49415 | 0,72666176 | BICD cargo adaptor 1 |
| ENSG00000224763 | FDPSP8 | -0,250768711 | 0,84044848 | 0,76334 | 0,88980026 | farnesyl diphosphate synthase pseudogene 8 |
| ENSG00000273167 | AL359736,1 | -0,250927476 | 0,840355996 | 0,52684 | 0,75003528 | novel transcript |
| ENSG00000146826 | MAP11 | -0,250931427 | 0,840353694 | 0,39188 | 0,65534953 | microtubule associated protein 11 |
| ENSG00000244055 | AC007566,1 | -0,250959928 | 0,840337093 | 0,40137 | 0,6630277 | novel transcript, antisense to ERVW-1 and PEX1 |

| | | | | | | |
|---|---|---|---|---|---|---|
| ENSG00000184515 | BEX5 | -0,251008934 | 0,840308549 | 0,42706 | 0,68023201 | brain expressed X-linked 5 |
| ENSG00000248592 | STIMATE-MU! | -0,25108441 | 0,840264588 | 0,38453 | 0,65052153 | STIMATE-MUSTN1 readthrough |
| ENSG00000287558 | AL109933,5 | -0,251418098 | 0,840070262 | 0,45836 | 0,70119611 | novel transcript, antisense to PLG |
| ENSG00000139220 | PPFIA2 | -0,251449933 | 0,840051725 | 0,55822 | 0,76982396 | PTPRF interacting protein alpha 2 |
| ENSG00000130045 | NXNL2 | -0,251708905 | 0,839900944 | 0,43879 | 0,6878469 | nucleoredoxin like 2 |
| ENSG00000126870 | DYNC2I1 | -0,251737921 | 0,839884052 | 0,22475 | 0,54036191 | dynein 2 intermediate chain 1 |
| ENSG00000286730 | AL365295,2 | -0,251896551 | 0,839791708 | 0,45378 | 0,69789417 | novel transcript |
| ENSG00000273799 | AC146335,1 | -0,251969683 | 0,839749139 | 0,49109 | 0,72493845 | glucuronidase, beta (GUSB) pseudogene |
| ENSG00000272143 | FGF14-AS2 | -0,252288218 | 0,83956375 | 0,31859 | 0,60360964 | FGF14 antisense RNA 2 |
| ENSG00000263841 | RN7SL44P | -0,252308199 | 0,839552122 | 0,64433 | 0,82151477 | RNA, 7SL, cytoplasmic 44, pseudogene |
| ENSG00000253121 | AC120042,2 | -0,252521829 | 0,839427813 | 0,76032 | 0,88836884 | novel transcript |
| ENSG00000102780 | DGKH | -0,25255781 | 0,839406878 | 0,51368 | 0,73998037 | diacylglycerol kinase eta |
| ENSG00000011132 | APBA3 | -0,252661414 | 0,8393466 | 0,48224 | 0,71892541 | amyloid beta precursor protein binding family A member 3 |
| ENSG00000130299 | GTPBP3 | -0,252713867 | 0,839316084 | 0,52698 | 0,75012908 | GTP binding protein 3, mitochondrial |
| ENSG00000130669 | PAK4 | -0,252714218 | 0,83931588 | 0,3281 | 0,60971379 | p21 (RAC1) activated kinase 4 |
| ENSG00000278722 | AL445288,1 | -0,252743314 | 0,839298953 | 0,85153 | 0,9324506 | novel transcript, antisense to PCDH17 |
| ENSG00000236882 | LINC01554 | -0,252756707 | 0,839291162 | 0,60497 | 0,79821396 | long intergenic non-protein coding RNA 1554 |
| ENSG00000085552 | IGSF9 | -0,252805889 | 0,83926255 | 0,33 | 0,61091085 | immunoglobulin superfamily member 9 |
| ENSG00000100605 | ITPK1 | -0,25282298 | 0,839252608 | 0,43839 | 0,68769293 | inositol-tetrakisphosphate 1-kinase |
| ENSG00000147144 | CCDC120 | -0,252896151 | 0,839210043 | 0,36029 | 0,63261878 | coiled-coil domain containing 120 |
| ENSG00000278416 | PMS2P2 | -0,25295173 | 0,839177714 | 0,38835 | 0,6529693 | PMS1 homolog 2, mismatch repair system component pseudogene 2 |
| ENSG00000276416 | UNC79 | -0,25302646 | 0,839134247 | 0,5111 | 0,73835122 | unc-79 homolog, NALCN channel complex subunit |
| ENSG00000255159 | AC026894,1 | -0,25321188 | 0,839026405 | 0,65275 | 0,826091 | novel transcript |
| ENSG00000134138 | MEIS2 | -0,253270615 | 0,838992248 | 0,41344 | 0,67078587 | Meis homeobox 2 |
| ENSG00000129197 | RPAIN | -0,253319244 | 0,838963968 | 0,2348 | 0,5463188 | RPA interacting protein |
| ENSG00000085514 | PILRA | -0,253431124 | 0,838898909 | 0,5524 | 0,76579928 | paired immunoglobin like type 2 receptor alpha |
| ENSG00000272800 | AC021851,1 | -0,253469408 | 0,838876648 | 0,56372 | 0,77255679 | novel transcript |
| ENSG00000237480 | AL355306,2 | -0,25354326 | 0,838833707 | 0,54566 | 0,76134666 | novel transcript |
| ENSG00000235695 | HIGD2AP2 | -0,253589694 | 0,838806709 | 0,69732 | 0,85330122 | HIGD2A pseudogene 2 |
| ENSG00000013288 | MAN2B2 | -0,253633186 | 0,838781422 | 0,20988 | 0,52877648 | mannosidase alpha class 2B member 2 |
| ENSG00000196911 | KPNA5 | -0,253699206 | 0,838743039 | 0,27758 | 0,57597292 | karyopherin subunit alpha 5 |
| ENSG00000251379 | AC099550,1 | -0,253719958 | 0,838730975 | 0,32088 | 0,60558211 | novel transcript |

| | | | | | | |
|---|---|---|---|---|---|---|
| ENSG00000150667 | FSIP1 | -0,253725942 | 0,838727496 | 0,25461 | 0,55944852 | fibrous sheath interacting protein 1 |
| ENSG00000120440 | TTLL2 | -0,253865449 | 0,838646396 | 0,65746 | 0,82872324 | tubulin tyrosine ligase like 2 |
| ENSG00000233821 | ENOX1-AS1 | -0,25391878 | 0,838615395 | 0,64059 | 0,8195307 | ENOX1 antisense RNA 1 |
| ENSG00000103460 | TOX3 | -0,253933433 | 0,838606878 | 0,43634 | 0,68667195 | TOX high mobility group box family member 3 |
| ENSG00000083812 | ZNF324 | -0,254025935 | 0,83855311 | 0,21848 | 0,53578053 | zinc finger protein 324 |
| ENSG00000101180 | HRH3 | -0,25412073 | 0,838498013 | 0,55159 | 0,76535107 | histamine receptor H3 |
| ENSG00000108342 | CSF3 | -0,254150115 | 0,838480934 | 0,83893 | 0,92640383 | colony stimulating factor 3 |
| ENSG00000213150 | AL512430,1 | -0,254252669 | 0,838421333 | 0,74615 | 0,8802836 | novel pseudogene |
| ENSG00000282367 | PTDSS2 | -0,254310039 | 0,838387993 | 0,45725 | 0,70073404 | phosphatidylserine synthase 2 |
| ENSG00000232631 | MTND2P23 | -0,254395446 | 0,838338362 | 0,64674 | 0,82312376 | MT-ND2 pseudogene 23 |
| ENSG00000133216 | EPHB2 | -0,25452069 | 0,838265587 | 0,19465 | 0,51562158 | EPH receptor B2 |
| ENSG00000114302 | PRKAR2A | -0,254537564 | 0,838255783 | 0,4649 | 0,70550186 | protein kinase cAMP-dependent type II regulatory subunit alpha |
| ENSG00000253948 | VPS13B-DT | -0,25453889 | 0,838255012 | 0,34794 | 0,62435764 | VPS13B divergent transcript |
| ENSG00000179981 | TSHZ1 | -0,254590219 | 0,838225189 | 0,44898 | 0,69460209 | teashirt zinc finger homeobox 1 |
| ENSG00000141562 | NARF | -0,254620072 | 0,838207844 | 0,21243 | 0,53128603 | nuclear prelamin A recognition factor |
| ENSG00000168282 | MGAT2 | -0,254790399 | 0,83810889 | 0,37874 | 0,64679625 | alpha-1,6-mannosyl-glycoprotein 2-beta-N-acetylglucosaminyltransferase |
| ENSG00000103174 | NAGPA | -0,254864827 | 0,838065653 | 0,48869 | 0,72350189 | N-acetylglucosamine-1-phosphodiester alpha-N-acetylglucosaminidase |
| ENSG00000232045 | EHMT2 | -0,254927158 | 0,838029446 | 0,34523 | 0,6224039 | euchromatic histone lysine methyltransferase 2 |
| ENSG00000244227 | LRRC77P | -0,254944456 | 0,838019398 | 0,61187 | 0,80337676 | leucine rich repeat containing 77, pseudogene |
| ENSG00000205702 | CYP2D7 | -0,255032445 | 0,837968289 | 0,80844 | 0,91193281 | cytochrome P450 family 2 subfamily D member 7 (gene/pseudogene) |
| ENSG00000146477 | SLC22A3 | -0,255055416 | 0,837954947 | 0,73755 | 0,87540065 | solute carrier family 22 member 3 |
| ENSG00000224769 | MUC20P1 | -0,255094734 | 0,837932111 | 0,83948 | 0,9266077 | mucin 20, cell surface associated pseudogene 1 |
| ENSG00000184305 | CCSER1 | -0,255172065 | 0,837887197 | 0,58439 | 0,78572744 | coiled-coil serine rich protein 1 |
| ENSG00000162992 | NEUROD1 | -0,255245072 | 0,837844797 | 0,5958 | 0,79349831 | neuronal differentiation 1 |
| ENSG00000123307 | NEUROD4 | -0,255300187 | 0,83781279 | 0,4881 | 0,72309725 | neuronal differentiation 4 |
| ENSG00000288565 | AC068057,3 | -0,255345824 | 0,837786287 | 0,63715 | 0,81747939 | novel transcript |
| ENSG00000225928 | CACYBPP1 | -0,255566441 | 0,837658183 | 0,74379 | 0,87871595 | calcyclin binding protein pseudogene 1 |
| ENSG00000251621 | AC009487,2 | -0,255686738 | 0,837588339 | 0,76602 | 0,89122725 | novel transcript |
| ENSG00000148334 | PTGES2 | -0,255712552 | 0,837573352 | 0,34867 | 0,62475089 | prostaglandin E synthase 2 |

| | | | | | | |
|---|---|---|---|---|---|---|
| ENSG00000234031 | RPS3AP44 | -0,255907657 | 0,837460089 | 0,63366 | 0,81549489 | RPS3A pseudogene 44 |
| ENSG00000228709 | LINC02575 | -0,255963405 | 0,837427729 | 0,61268 | 0,8035281 | long intergenic non-protein coding RNA 2575 |
| ENSG00000181220 | ZNF746 | -0,255989432 | 0,837412622 | 0,29593 | 0,58942061 | zinc finger protein 746 |
| ENSG00000117643 | MAN1C1 | -0,256120657 | 0,837336456 | 0,56983 | 0,77701569 | mannosidase alpha class 1C member 1 |
| ENSG00000173327 | MAP3K11 | -0,256247572 | 0,837262797 | 0,47932 | 0,71687041 | mitogen-activated protein kinase kinase kinase 11 |
| ENSG00000108848 | LUC7L3 | -0,256283558 | 0,837241914 | 0,15451 | 0,48266733 | LUC7 like 3 pre-mRNA splicing factor |
| ENSG00000145692 | BHMT | -0,256785067 | 0,836950922 | 0,62775 | 0,81173104 | betaine--homocysteine S-methyltransferase |
| ENSG00000264801 | ERVFRD-3 | -0,256808695 | 0,836937215 | 0,65849 | 0,82923456 | endogenous retrovirus group FRD member 3 |
| ENSG00000071575 | TRIB2 | -0,256827721 | 0,836926178 | 0,37718 | 0,6456352 | tribbles pseudokinase 2 |
| ENSG00000269846 | AL136172,1 | -0,256907873 | 0,836879682 | 0,55489 | 0,76763832 | novel transcript, antisense to RBL1 |
| ENSG00000222012 | AC005481,1 | -0,25697499 | 0,83684075 | 0,47888 | 0,7165637 | novel transcript, antisense to PTPRN2 |
| ENSG00000184617 | ZNF840P | -0,257325592 | 0,836637407 | 0,61354 | 0,8035281 | zinc finger protein 840, pseudogene |
| ENSG00000228981 | AC097658,1 | -0,257371857 | 0,836610577 | 0,54294 | 0,75965603 | ribosomal protein S2 (RPS2) pseudogene |
| ENSG00000198959 | TGM2 | -0,257399494 | 0,836594551 | 0,76435 | 0,89017297 | transglutaminase 2 |
| ENSG00000285277 | AC254813,1 | -0,257494164 | 0,836539655 | 0,80259 | 0,90906425 | family with sequence similarity 115, member C (FAM115C) pseudogene |
| ENSG00000214745 | GOT2P6 | -0,257500529 | 0,836535964 | 0,6423 | 0,82066205 | GOT2 pseudogene 6 |
| ENSG00000123454 | DBH | -0,257614533 | 0,836469863 | 0,64962 | 0,82449863 | dopamine beta-hydroxylase |
| ENSG00000130054 | FAM155B | -0,257688263 | 0,836427115 | 0,47508 | 0,71364583 | family with sequence similarity 155 member B |
| ENSG00000162408 | NOL9 | -0,257814428 | 0,836353972 | 0,41552 | 0,6723843 | nucleolar protein 9 |
| ENSG00000162227 | TAF6L | -0,257852074 | 0,836332148 | 0,3626 | 0,63379594 | TATA-box binding protein associated factor 6 like |
| ENSG00000284852 | YPEL5P2 | -0,257890531 | 0,836309855 | 0,7642 | 0,89012684 | YPEL5 pseudogene 2 |
| ENSG00000271557 | YPEL5P2 | -0,257890531 | 0,836309855 | 0,7642 | 0,89012684 | YPEL5 pseudogene 2 |
| ENSG00000235278 | ZNF652P1 | -0,257905541 | 0,836301154 | 0,81418 | 0,91457884 | zinc finger protein 652 pseudogene 1 |
| ENSG00000258718 | LINC02311 | -0,258001253 | 0,836245674 | 0,40843 | 0,66723037 | long intergenic non-protein coding RNA 2311 |
| ENSG00000179564 | LSMEM2 | -0,258045245 | 0,836220174 | 0,54419 | 0,76041947 | leucine rich single-pass membrane protein 2 |
| ENSG00000213612 | FAM220CP | -0,258076347 | 0,836202147 | 0,79301 | 0,90371578 | family with sequence similarity 220 member C, pseudogene |
| ENSG00000267150 | AC006557,1 | -0,258111734 | 0,836181637 | 0,42674 | 0,68012899 | novel transcript, antisense ZNF519 |
| ENSG00000160145 | KALRN | -0,258127662 | 0,836172405 | 0,27306 | 0,57344244 | kalirin RhoGEF kinase |
| ENSG00000166401 | SERPINB8 | -0,258202576 | 0,836128987 | 0,43883 | 0,6878469 | serpin family B member 8 |
| ENSG00000180190 | TDRP | -0,25822042 | 0,836118645 | 0,24044 | 0,54955956 | testis development related protein |
| ENSG00000248724 | NPHP3-AS1 | -0,258388565 | 0,836021202 | 0,69318 | 0,85092062 | NPHP3 antisense RNA 1 |

| | | | | | | |
|---|---|---|---|---|---|---|
| ENSG00000241511 | RPS15AP24 | -0,258465121 | 0,83597684 | 0,83366 | 0,92396169 | ribosomal protein S15a pseudogene 24 |
| ENSG00000100077 | GRK3 | -0,258521297 | 0,835944289 | 0,28204 | 0,57936013 | G protein-coupled receptor kinase 3 |
| ENSG00000237872 | POU5F1P4 | -0,258537753 | 0,835934754 | 0,73004 | 0,87180309 | POU class 5 homeobox 1 pseudogene 4 |
| ENSG00000134668 | SPOCD1 | -0,258576695 | 0,83591219 | 0,72227 | 0,86734497 | SPOC domain containing 1 |
| ENSG00000196096 | SPAG16-DT | -0,258654757 | 0,835866961 | 0,31612 | 0,60224605 | SPAG16 divergent transcript |
| ENSG00000287961 | AC020911,3 | -0,258676937 | 0,835854111 | 0,73942 | 0,87642873 | novel transcript, antisense to AP1M1 |
| ENSG00000205542 | TMSB4X | -0,258741409 | 0,835816758 | 0,41361 | 0,67085666 | thymosin beta 4 X-linked |
| ENSG00000089280 | FUS | -0,258757616 | 0,835807369 | 0,33083 | 0,6112353 | FUS RNA binding protein |
| ENSG00000239300 | AC080162,1 | -0,259031641 | 0,835648631 | 0,79847 | 0,90704887 | novel transcript |
| ENSG00000214960 | CRPPA | -0,259086959 | 0,83561659 | 0,31137 | 0,59890747 | CDP-L-ribitol pyrophosphorylase A |
| ENSG00000118160 | SLC8A2 | -0,259096131 | 0,835611278 | 0,65279 | 0,82610447 | solute carrier family 8 member A2 |
| ENSG00000188729 | OSTN | -0,259172878 | 0,835566827 | 0,69496 | 0,85206252 | osteocrin |
| ENSG00000283859 | NLGN2 | -0,259204301 | 0,835548628 | 0,43627 | 0,68660209 | neuroligin 2 |
| ENSG00000169992 | NLGN2 | -0,259204301 | 0,835548628 | 0,43627 | 0,68660209 | neuroligin 2 |
| ENSG00000198668 | CALM1 | -0,259205051 | 0,835548193 | 0,309 | 0,59774312 | calmodulin 1 |
| ENSG00000272853 | AC069544,1 | -0,259348539 | 0,835465096 | 0,63635 | 0,81700605 | novel transcript |
| ENSG00000173457 | PPP1R14B | -0,259385505 | 0,835443689 | 0,38466 | 0,65052153 | protein phosphatase 1 regulatory inhibitor subunit 14B |
| ENSG00000179750 | APOBEC3B | -0,25939245 | 0,835439667 | 0,59251 | 0,79121909 | apolipoprotein B mRNA editing enzyme catalytic subunit 3B |
| ENSG00000176903 | PNMA1 | -0,25943216 | 0,835416672 | 0,24964 | 0,55657567 | PNMA family member 1 |
| ENSG00000270012 | AC232271,1 | -0,259443492 | 0,83541011 | 0,28727 | 0,58369609 | novel transcript |
| ENSG00000251405 | AC016571,1 | -0,259520365 | 0,835365597 | 0,5871 | 0,78713886 | novel transcript, sense overlapping ADAM19 |
| ENSG00000124486 | USP9X | -0,259585109 | 0,835328109 | 0,38546 | 0,65112398 | ubiquitin specific peptidase 9 X-linked |
| ENSG00000180098 | TRNAU1AP | -0,259715142 | 0,835252822 | 0,40387 | 0,66409141 | tRNA selenocysteine 1 associated protein 1 |
| ENSG00000100139 | MICALL1 | -0,259715411 | 0,835252667 | 0,33624 | 0,61623735 | MICAL like 1 |
| ENSG00000230461 | PROX1-AS1 | -0,259771557 | 0,835220162 | 0,545 | 0,7609742 | PROX1 antisense RNA 1 |
| ENSG00000183474 | GTF2H2C | -0,259908891 | 0,835140659 | 0,30766 | 0,59671558 | GTF2H2 family member C |
| ENSG00000186185 | KIF18B | -0,259926073 | 0,835130712 | 0,33881 | 0,61792351 | kinesin family member 18B |
| ENSG00000170629 | DPY19L2P2 | -0,260045557 | 0,83506155 | 0,23518 | 0,54667863 | DPY19L2 pseudogene 2 |
| ENSG00000175198 | PCCA | -0,260148785 | 0,835001801 | 0,30375 | 0,59391804 | propionyl-CoA carboxylase subunit alpha |
| ENSG00000214401 | KANSL1-AS1 | -0,260243871 | 0,834946769 | 0,53612 | 0,75500755 | KANSL1 antisense RNA 1 |
| ENSG00000282799 | KANSL1-AS1 | -0,260243871 | 0,834946769 | 0,53612 | 0,75500755 | KANSL1 antisense RNA 1 |
| ENSG00000284976 | BX255925,3 | -0,260418792 | 0,834845541 | 0,41342 | 0,67078587 | novel RING finger protein |

| | | | | | | |
|---|---|---|---|---|---|---|
| ENSG00000120334 | CENPL | -0,260480402 | 0,83480989 | 0,38633 | 0,651759 | centromere protein L |
| ENSG00000259826 | AC072061,1 | -0,260742898 | 0,834658012 | 0,40225 | 0,66358835 | novel transcript, antisense to CDC2L5 |
| ENSG00000228741 | AL445985,1 | -0,260853571 | 0,834593986 | 0,3439 | 0,62131246 | novel transcript |
| ENSG00000175220 | ARHGAP1 | -0,260914346 | 0,834558828 | 0,4345 | 0,6850888 | Rho GTPase activating protein 1 |
| ENSG00000225140 | AL358216,1 | -0,261015211 | 0,834500482 | 0,59987 | 0,79564899 | novel transcript |
| ENSG00000169427 | KCNK9 | -0,261068292 | 0,834469779 | 0,74084 | 0,87717177 | potassium two pore domain channel subfamily K member 9 |
| ENSG00000245937 | LINC01184 | -0,261075168 | 0,834465802 | 0,28594 | 0,58229607 | long intergenic non-protein coding RNA 1184 |
| ENSG00000274768 | CISD3 | -0,261136587 | 0,834430278 | 0,56026 | 0,7707835 | CDGSH iron sulfur domain 3 |
| ENSG00000261596 | AC005632,3 | -0,261236286 | 0,834372616 | 0,29089 | 0,58590649 | novel transcript, sense intronic to METTL9 |
| ENSG00000163001 | CFAP36 | -0,261346189 | 0,834309057 | 0,29504 | 0,58874428 | cilia and flagella associated protein 36 |
| ENSG00000286890 | AC055840,1 | -0,261530767 | 0,834202322 | 0,41387 | 0,67092036 | novel transcript, antisense to FHOD3 |
| ENSG00000198400 | NTRK1 | -0,261580166 | 0,834173759 | 0,5886 | 0,78828113 | neurotrophic receptor tyrosine kinase 1 |
| ENSG00000196584 | XRCC2 | -0,262046598 | 0,833904109 | 0,32409 | 0,60671008 | X-ray repair cross complementing 2 |
| ENSG00000236342 | ABCF1 | -0,262078453 | 0,833885696 | 0,77157 | 0,89417611 | ATP binding cassette subfamily F member 1 |
| ENSG00000179526 | SHARPIN | -0,262167961 | 0,833833962 | 0,27587 | 0,57533851 | SHANK associated RH domain interactor |
| ENSG00000227199 | ST7-AS1 | -0,262246676 | 0,833788468 | 0,41135 | 0,66904284 | ST7 antisense RNA 1 |
| ENSG00000275212 | LINC02826 | -0,262710465 | 0,83352047 | 0,44422 | 0,69172919 | long intergenic non-protein coding RNA 2826 |
| ENSG00000179041 | RRS1 | -0,262799838 | 0,833468836 | 0,31246 | 0,59937766 | ribosome biogenesis regulator 1 homolog |
| ENSG00000132510 | KDM6B | -0,263135074 | 0,833275187 | 0,3898 | 0,65396251 | lysine demethylase 6B |
| ENSG00000242276 | RPL5P3 | -0,263162499 | 0,833259347 | 0,70783 | 0,8592125 | ribosomal protein L5 pseudogene 3 |
| ENSG00000152154 | TMEM178A | -0,263258721 | 0,833203774 | 0,30513 | 0,59455047 | transmembrane protein 178A |
| ENSG00000100296 | THOC5 | -0,263316092 | 0,833170641 | 0,42507 | 0,67871429 | THO complex 5 |
| ENSG00000174917 | MICOS13 | -0,263418913 | 0,833111263 | 0,25933 | 0,56317998 | mitochondrial contact site and cristae organizing system subunit 13 |
| ENSG00000241782 | AP002812,1 | -0,263463637 | 0,833085436 | 0,72606 | 0,86953997 | ribosomal protein L21 (RPL21) pseudogene |
| ENSG00000070814 | TCOF1 | -0,263619363 | 0,832995517 | 0,27115 | 0,57263095 | treacle ribosome biogenesis factor 1 |
| ENSG00000187134 | AKR1C1 | -0,263830737 | 0,832873481 | 0,4061 | 0,66574548 | aldo-keto reductase family 1 member C1 |
| ENSG00000124067 | SLC12A4 | -0,263936483 | 0,832812436 | 0,34262 | 0,62025394 | solute carrier family 12 member 4 |
| ENSG00000286941 | AC244250,4 | -0,263962188 | 0,832797598 | 0,64129 | 0,82001731 | novel transcript |
| ENSG00000233030 | AC243772,2 | -0,26400762 | 0,832771372 | 0,55563 | 0,76790322 | novel transcript, antisense to FCGR1A |
| ENSG00000150347 | ARID5B | -0,264115656 | 0,832709013 | 0,54189 | 0,75891031 | AT-rich interaction domain 5B |
| ENSG00000021574 | SPAST | -0,264122734 | 0,832704927 | 0,26906 | 0,57085484 | spastin |

| | | | | | | |
|---|---|---|---|---|---|---|
| ENSG00000167925 | GHDC | -0,264168622 | 0,832678442 | 0,57965 | 0,7829899 | GH3 domain containing |
| ENSG00000186715 | MST1L | -0,26431279 | 0,832595237 | 0,51971 | 0,74412419 | macrophage stimulating 1 like |
| ENSG00000236759 | EHMT2 | -0,264424122 | 0,832530988 | 0,24429 | 0,55253387 | euchromatic histone lysine methyltransferase 2 |
| ENSG00000268333 | AL132655,1 | -0,264529997 | 0,832469893 | 0,68026 | 0,84309883 | novel transcript |
| ENSG00000146555 | SDK1 | -0,264536662 | 0,832466048 | 0,11824 | 0,43604372 | sidekick cell adhesion molecule 1 |
| ENSG00000233237 | LINC00472 | -0,264539224 | 0,83246457 | 0,31037 | 0,59842274 | long intergenic non-protein coding RNA 472 |
| ENSG00000181090 | EHMT1 | -0,264624625 | 0,832415293 | 0,1962 | 0,51720037 | euchromatic histone lysine methyltransferase 1 |
| ENSG00000160396 | HIPK4 | -0,264690657 | 0,832377194 | 0,74936 | 0,88210735 | homeodomain interacting protein kinase 4 |
| ENSG00000198556 | ZNF789 | -0,264708342 | 0,832366991 | 0,29292 | 0,58737078 | zinc finger protein 789 |
| ENSG00000272170 | AL355385,1 | -0,264722335 | 0,832358917 | 0,70556 | 0,85816192 | novel transcript, antisense to KLC4 |
| ENSG00000221926 | TRIM16 | -0,264726974 | 0,832356241 | 0,45283 | 0,69717623 | tripartite motif containing 16 |
| ENSG00000177410 | ZFAS1 | -0,264803376 | 0,832312163 | 0,37216 | 0,64116374 | ZNFX1 antisense RNA 1 |
| ENSG00000206357 | NELFE | -0,264850867 | 0,832284764 | 0,33109 | 0,6112353 | negative elongation factor complex member E |
| ENSG00000231044 | NELFE | -0,264850867 | 0,832284764 | 0,33109 | 0,6112353 | negative elongation factor complex member E |
| ENSG00000229363 | NELFE | -0,264850867 | 0,832284764 | 0,33109 | 0,6112353 | negative elongation factor complex member E |
| ENSG00000233801 | NELFE | -0,264850867 | 0,832284764 | 0,33109 | 0,6112353 | negative elongation factor complex member E |
| ENSG00000206268 | NELFE | -0,264850867 | 0,832284764 | 0,33109 | 0,6112353 | negative elongation factor complex member E |
| ENSG00000189337 | KAZN | -0,264882879 | 0,832266297 | 0,58012 | 0,78347322 | kazrin, periplakin interacting protein |
| ENSG00000163617 | CCDC191 | -0,264936737 | 0,832235228 | 0,31866 | 0,60371722 | coiled-coil domain containing 191 |
| ENSG00000239524 | RPL32P34 | -0,265203098 | 0,832081589 | 0,74235 | 0,87815637 | ribosomal protein L32 pseudogene 34 |
| ENSG00000226648 | PLCG1-AS1 | -0,265290164 | 0,832031374 | 0,37332 | 0,64204597 | PLCG1 antisense RNA 1 |
| ENSG00000138036 | DYNC2LI1 | -0,265336966 | 0,832004383 | 0,18247 | 0,50597301 | dynein cytoplasmic 2 light intermediate chain 1 |
| ENSG00000268670 | AC016586,1 | -0,265502186 | 0,831909106 | 0,64969 | 0,82449863 | novel transcript, sense intronic to ZBTB7A |
| ENSG00000265107 | GJA5 | -0,265507385 | 0,831906108 | 0,67093 | 0,83695245 | gap junction protein alpha 5 |
| ENSG00000237693 | IRGM | -0,265511224 | 0,831903895 | 0,76843 | 0,89277929 | immunity related GTPase M |
| ENSG00000186088 | GSAP | -0,265512281 | 0,831903285 | 0,57908 | 0,78270914 | gamma-secretase activating protein |
| ENSG00000137221 | TJAP1 | -0,265723423 | 0,831781543 | 0,34647 | 0,62342969 | tight junction associated protein 1 |
| ENSG00000110104 | CCDC86 | -0,265729512 | 0,831778032 | 0,51118 | 0,73838863 | coiled-coil domain containing 86 |
| ENSG00000172197 | MBOAT1 | -0,265766355 | 0,831756791 | 0,51776 | 0,74276913 | membrane bound O-acyltransferase domain containing 1 |
| ENSG00000121864 | ZNF639 | -0,265816973 | 0,831727609 | 0,28449 | 0,58137706 | zinc finger protein 639 |
| ENSG00000234136 | AC055764,1 | -0,265923461 | 0,831666219 | 0,76509 | 0,8907696 | novel transcript |
| ENSG00000006016 | CRLF1 | -0,266237322 | 0,831485308 | 0,59186 | 0,79090751 | cytokine receptor like factor 1 |
| ENSG00000180953 | ST20 | -0,266300618 | 0,831448829 | 0,27967 | 0,57745167 | suppressor of tumorigenicity 20 |

| | | | | | | |
|---|---|---|---|---|---|---|
| ENSG00000084628 | NKAIN1 | -0,266313315 | 0,831441512 | 0,26289 | 0,56582568 | sodium/potassium transporting ATPase interacting 1 |
| ENSG00000272914 | AL359532,1 | -0,266502514 | 0,831332482 | 0,75066 | 0,8829851 | novel transcript, sense intronic to ZNF438 |
| ENSG00000268621 | IGFL2-AS1 | -0,266625504 | 0,831261613 | 0,7359 | 0,8750728 | IGFL2 antisense RNA 1 |
| ENSG00000197536 | IRF1-AS1 | -0,266646818 | 0,831249333 | 0,2946 | 0,58857672 | IRF1 antisense RNA 1 |
| ENSG00000271119 | AC026412,3 | -0,266652078 | 0,831246302 | 0,50159 | 0,73146091 | novel transcript |
| ENSG00000103249 | CLCN7 | -0,266684348 | 0,831227709 | 0,38013 | 0,64784628 | chloride voltage-gated channel 7 |
| ENSG00000156345 | CDK20 | -0,266752895 | 0,831188216 | 0,22497 | 0,54037642 | cyclin dependent kinase 20 |
| ENSG00000278062 | ZNF251 | -0,266792557 | 0,831165365 | 0,62699 | 0,81119765 | zinc finger protein 251 |
| ENSG00000120158 | RCL1 | -0,266831736 | 0,831142793 | 0,4042 | 0,6642832 | RNA terminal phosphate cyclase like 1 |
| ENSG00000175077 | RTP1 | -0,266931333 | 0,831085417 | 0,65227 | 0,82578944 | receptor transporter protein 1 |
| ENSG00000120162 | MOB3B | -0,266938507 | 0,831081284 | 0,32955 | 0,61056241 | MOB kinase activator 3B |
| ENSG00000260037 | AC104938,1 | -0,267061885 | 0,831010215 | 0,67022 | 0,83650134 | novel transcript, antisense to MYO9A |
| ENSG00000196428 | TSC22D2 | -0,267074361 | 0,831003028 | 0,35303 | 0,62780446 | TSC22 domain family member 2 |
| ENSG00000153187 | HNRNPU | -0,267141785 | 0,830964192 | 0,24958 | 0,55657567 | heterogeneous nuclear ribonucleoprotein U |
| ENSG00000248431 | AC021134,1 | -0,267237585 | 0,830909015 | 0,6626 | 0,83204447 | novel transcript |
| ENSG00000177873 | ZNF619 | -0,26725668 | 0,830898017 | 0,45917 | 0,7017037 | zinc finger protein 619 |
| ENSG00000204176 | SYT15 | -0,267294003 | 0,830876522 | 0,24001 | 0,54936679 | synaptotagmin 15 |
| ENSG00000284534 | AC092634,7 | -0,267319397 | 0,830861897 | 0,74866 | 0,8817805 | novel transcript |
| ENSG00000127993 | RBM48 | -0,267458032 | 0,83078206 | 0,34342 | 0,62091871 | RNA binding motif protein 48 |
| ENSG00000287185 | AC041005,1 | -0,267464899 | 0,830778106 | 0,73178 | 0,87276602 | novel transcript |
| ENSG00000112096 | SOD2 | -0,267557048 | 0,830725043 | 0,2603 | 0,56382982 | superoxide dismutase 2 |
| ENSG00000188152 | NUTM2G | -0,267645806 | 0,830673937 | 0,50058 | 0,73094504 | NUT family member 2G |
| ENSG00000267758 | AC099811,4 | -0,267718212 | 0,830632248 | 0,6495 | 0,82442734 | novel transcript |
| ENSG00000255480 | AL353699,1 | -0,26780755 | 0,830580813 | 0,6638 | 0,8324578 | novel transcript |
| ENSG00000256399 | AC079866,1 | -0,267923824 | 0,830513875 | 0,63559 | 0,81657648 | CTAGE family, member 5 (CTAGE5) pseudogene |
| ENSG00000266402 | SNHG25 | -0,267944506 | 0,830501969 | 0,56253 | 0,77186635 | small nucleolar RNA host gene 25 |
| ENSG00000230516 | AL589642,2 | -0,267994071 | 0,830473437 | 0,75981 | 0,88801075 | novel transcript |
| ENSG00000205309 | NT5M | -0,268022806 | 0,830456896 | 0,3521 | 0,62724407 | 5',3'-nucleotidase, mitochondrial |
| ENSG00000173376 | NDNF | -0,268130475 | 0,830394921 | 0,65514 | 0,82762048 | neuron derived neurotrophic factor |
| ENSG00000155093 | PTPRN2 | -0,268173328 | 0,830370256 | 0,33639 | 0,61640718 | protein tyrosine phosphatase receptor type N2 |
| ENSG00000213904 | LIPE-AS1 | -0,268248711 | 0,830326869 | 0,37472 | 0,6431311 | LIPE antisense RNA 1 |
| ENSG00000143479 | DYRK3 | -0,268330251 | 0,830279941 | 0,47245 | 0,71184073 | dual specificity tyrosine phosphorylation regulated kinase 3 |

| | | | | | | |
|---|---|---|---|---|---|---|
| ENSG00000166546 | BEAN1 | -0,268479682 | 0,830193947 | 0,51525 | 0,74138689 | brain expressed associated with NEDD4 1 |
| ENSG00000145029 | NICN1 | -0,268532585 | 0,830163504 | 0,37064 | 0,63997937 | nicolin 1 |
| ENSG00000256420 | OSBPL9P4 | -0,268785479 | 0,830017996 | 0,63328 | 0,81541029 | oxysterol binding protein like 9 pseudogene 4 |
| ENSG00000276432 | RDM1 | -0,268791292 | 0,830014651 | 0,50318 | 0,73248965 | RAD52 motif containing 1 |
| ENSG00000226200 | SGMS1-AS1 | -0,268886879 | 0,82995966 | 0,24575 | 0,55359883 | SGMS1 antisense RNA 1 |
| ENSG00000243015 | RN7SL737P | -0,268959833 | 0,829917692 | 0,83113 | 0,92276009 | RNA, 7SL, cytoplasmic 737, pseudogene |
| ENSG00000114554 | PLXNA1 | -0,26900103 | 0,829893993 | 0,11191 | 0,42531464 | plexin A1 |
| ENSG00000159164 | SV2A | -0,269012459 | 0,829887419 | 0,1409 | 0,46791293 | synaptic vesicle glycoprotein 2A |
| ENSG00000242950 | ERVW-1 | -0,269035205 | 0,829874335 | 0,29024 | 0,58540802 | endogenous retrovirus group W member 1, envelope |
| ENSG00000223944 | AL353604,1 | -0,269048731 | 0,829866554 | 0,6877 | 0,84783397 | novel transcript |
| ENSG00000229043 | ZFAND2A-DT | -0,269067538 | 0,829855736 | 0,3449 | 0,62213186 | ZFAND2A divergent transcript |
| ENSG00000122694 | GLIPR2 | -0,269330898 | 0,829704262 | 0,31493 | 0,60144815 | GLI pathogenesis related 2 |
| ENSG00000154485 | MMP21 | -0,269339193 | 0,829699492 | 0,47663 | 0,71481671 | matrix metallopeptidase 21 |
| ENSG00000262319 | AC007952,6 | -0,269405108 | 0,829661584 | 0,63337 | 0,81542987 | novel transcript, antisense to GRAPL |
| ENSG00000225553 | PHF1 | -0,26944568 | 0,829638253 | 0,31203 | 0,59925736 | PHD finger protein 1 |
| ENSG00000231671 | LINC01307 | -0,269528251 | 0,82959077 | 0,76137 | 0,88879633 | long intergenic non-protein coding RNA 1307 |
| ENSG00000125967 | NECAB3 | -0,269654156 | 0,829518375 | 0,34635 | 0,62342969 | N-terminal EF-hand calcium binding protein 3 |
| ENSG00000282417 | HERC2P9 | -0,269719077 | 0,829481048 | 0,2606 | 0,56400103 | hect domain and RLD 2 pseudogene 9 |
| ENSG00000240083 | RPS3AP22 | -0,269751588 | 0,829462355 | 0,69933 | 0,85477558 | RPS3A pseudogene 22 |
| ENSG00000206284 | WDR46 | -0,269874237 | 0,829391843 | 0,61183 | 0,80337676 | WD repeat domain 46 |
| ENSG00000198003 | CCDC151 | -0,269891747 | 0,829381776 | 0,42994 | 0,68268477 | coiled-coil domain containing 151 |
| ENSG00000036828 | CASR | -0,269914499 | 0,829368697 | 0,17244 | 0,49737713 | calcium sensing receptor |
| ENSG00000261334 | AL353803,4 | -0,270030398 | 0,829302072 | 0,72655 | 0,86978598 | novel transcript |
| ENSG00000168913 | ENHO | -0,270221761 | 0,829192079 | 0,25219 | 0,55796133 | energy homeostasis associated |
| ENSG00000277941 | Z99774,4 | -0,270227242 | 0,829188928 | 0,81906 | 0,91689679 | |
| ENSG00000254778 | AL662828,1 | -0,270233688 | 0,829185224 | 0,43087 | 0,68325486 | palmitoyl-protein thioesterase 2 (PPT2) and EGF-like-domain, multiple 8 (EGFL8) readthrough |
| ENSG00000228213 | NLGN1-AS1 | -0,27027497 | 0,829161497 | 0,62855 | 0,81209878 | NLGN1 antisense RNA 1 |
| ENSG00000141873 | SLC39A3 | -0,270348307 | 0,829119349 | 0,29757 | 0,59036937 | solute carrier family 39 member 3 |
| ENSG00000129244 | ATP1B2 | -0,270375195 | 0,829103897 | 0,33983 | 0,61872249 | ATPase Na+/K+ transporting subunit beta 2 |
| ENSG00000150394 | CDH8 | -0,270513984 | 0,82902414 | 0,22376 | 0,53967355 | cadherin 8 |
| ENSG00000285982 | AC012213,5 | -0,27064986 | 0,828946064 | 0,79253 | 0,90361513 | novel protein |

| | | | | | | |
|---|---|---|---|---|---|---|
| ENSG00000180228 | PRKRA | -0,270650736 | 0,828945561 | 0,29901 | 0,59115664 | protein activator of interferon induced protein kinase EIF2AK2 |
| ENSG00000267426 | AC087289,3 | -0,270697097 | 0,828918923 | 0,75587 | 0,8857133 | novel protein |
| ENSG00000130699 | TAF4 | -0,270723751 | 0,828903609 | 0,25148 | 0,55731053 | TATA-box binding protein associated factor 4 |
| ENSG00000272078 | AL139423,1 | -0,270728679 | 0,828900777 | 0,52116 | 0,74558236 | novel transcript, antisense to CASZ1 |
| ENSG00000007545 | CRAMP1 | -0,270919111 | 0,828791372 | 0,1168 | 0,43387136 | cramped chromatin regulator homolog 1 |
| ENSG00000175470 | PPP2R2D | -0,270978039 | 0,82875752 | 0,30979 | 0,59807209 | protein phosphatase 2 regulatory subunit Bdelta |
| ENSG00000141503 | MINK1 | -0,27102725 | 0,828729251 | 0,28417 | 0,5810616 | misshapen like kinase 1 |
| ENSG00000285601 | AC084262,2 | -0,271033159 | 0,828725857 | 0,55507 | 0,76775076 | novel transcript |
| ENSG00000247675 | LRP4-AS1 | -0,271094719 | 0,828690496 | 0,6116 | 0,80328691 | LRP4 antisense RNA 1 |
| ENSG00000225226 | LINC01795 | -0,271124059 | 0,828673643 | 0,45513 | 0,69921882 | long intergenic non-protein coding RNA 1795 |
| ENSG00000046889 | PREX2 | -0,271156347 | 0,828655097 | 0,50969 | 0,73740498 | phosphatidylinositol-3,4,5-trisphosphate dependent Rac exchange factor 2 |
| ENSG00000266405 | CBX3P2 | -0,271235043 | 0,828609897 | 0,45277 | 0,69717623 | chromobox 3 pseudogene 2 |
| ENSG00000150593 | PDCD4 | -0,271252089 | 0,828600107 | 0,29841 | 0,59076323 | programmed cell death 4 |
| ENSG00000255277 | ABCC6P2 | -0,271535658 | 0,828437257 | 0,7114 | 0,86111501 | ATP binding cassette subfamily C member 6 pseudogene 2 |
| ENSG00000232454 | AL138752,1 | -0,271544416 | 0,828432228 | 0,79885 | 0,90723937 | mitochondrial ribosomal protein S10 (MRPS10) pseudogene |
| ENSG00000123453 | SARDH | -0,271619599 | 0,828389057 | 0,23054 | 0,54361941 | sarcosine dehydrogenase |
| ENSG00000100294 | MCAT | -0,27166983 | 0,828360215 | 0,32632 | 0,60863796 | malonyl-CoA-acyl carrier protein transacylase |
| ENSG00000184566 | AC132216,1 | -0,271796157 | 0,828287685 | 0,63214 | 0,81450663 | novel transcript, antisense to LMO2 |
| ENSG00000170445 | HARS1 | -0,27183093 | 0,828267721 | 0,26532 | 0,56810437 | histidyl-tRNA synthetase 1 |
| ENSG00000244073 | RPS4XP6 | -0,271876419 | 0,828241606 | 0,58421 | 0,78572744 | ribosomal protein S4X pseudogene 6 |
| ENSG00000139352 | ASCL1 | -0,271964682 | 0,828190935 | 0,19929 | 0,52034238 | achaete-scute family bHLH transcription factor 1 |
| ENSG00000228625 | AL451047,1 | -0,272112801 | 0,828105911 | 0,71695 | 0,86453146 | novel transcript |
| ENSG00000224888 | AC138028,2 | -0,272129172 | 0,828096514 | 0,71119 | 0,86096523 | novel transcript, antisense to FAM38A |
| ENSG00000261098 | AP000766,1 | -0,272169683 | 0,828073262 | 0,31365 | 0,60035326 | novel transcript |
| ENSG00000144815 | NXPE3 | -0,272254215 | 0,828024743 | 0,09982 | 0,40626368 | neurexophilin and PC-esterase domain family member 3 |
| ENSG00000118094 | TREH | -0,272254622 | 0,82802451 | 0,57583 | 0,78043323 | trehalase |
| ENSG00000285970 | AC106820,6 | -0,27229839 | 0,82799939 | 0,73354 | 0,87368351 | novel transcript, antisense to CCNF |
| ENSG00000233644 | ARHGEF7-IT1 | -0,272348644 | 0,827970549 | 0,70942 | 0,85983746 | ARHGEF7 intronic transcript 1 |
| ENSG00000236670 | KRT18P5 | -0,272472476 | 0,827899484 | 0,55938 | 0,77045525 | keratin 18 pseudogene 5 |
| ENSG00000115325 | DOK1 | -0,272605776 | 0,827822992 | 0,44876 | 0,69455511 | docking protein 1 |

| | | | | | | |
|---|---|---|---|---|---|---|
| ENSG00000269609 | RPARP-AS1 | -0,272632201 | 0,827807829 | 0,43236 | 0,68411566 | RPARP antisense RNA 1 |
| ENSG00000168924 | LETM1 | -0,273137583 | 0,827517896 | 0,30065 | 0,59200188 | leucine zipper and EF-hand containing transmembrane protein 1 |
| ENSG00000235408 | SNORA71B | -0,273337145 | 0,827403437 | 0,49952 | 0,73058737 | small nucleolar RNA, H/ACA box 71B |
| ENSG00000273108 | AL121929,2 | -0,273475157 | 0,827324289 | 0,67127 | 0,83720965 | novel transcript, antisense to SH3PXD2A |
| ENSG00000153832 | FBXO36 | -0,273565747 | 0,827272341 | 0,25384 | 0,55891 | F-box protein 36 |
| ENSG00000224051 | CPTP | -0,273574538 | 0,8272673 | 0,30497 | 0,59446033 | ceramide-1-phosphate transfer protein |
| ENSG00000072501 | SMC1A | -0,273607222 | 0,827248559 | 0,24565 | 0,55357646 | structural maintenance of chromosomes 1A |
| ENSG00000280795 | MPRIPP1 | -0,27361129 | 0,827246226 | 0,53445 | 0,75405583 | myosin phosphatase Rho interacting protein pseudogene 1 |
| ENSG00000214820 | MPRIPP1 | -0,27361129 | 0,827246226 | 0,53445 | 0,75405583 | myosin phosphatase Rho interacting protein pseudogene 1 |
| ENSG00000255655 | AC007570,1 | -0,27378858 | 0,827144574 | 0,70905 | 0,85978194 | novel transcript, antisense to MYO1H |
| ENSG00000259336 | AC021231,1 | -0,273856858 | 0,827105429 | 0,6505 | 0,82492577 | novel transcript |
| ENSG00000280733 | AC093642,2 | -0,273914524 | 0,827072369 | 0,61261 | 0,8035281 | novel transcript |
| ENSG00000281213 | AC093642,2 | -0,273914524 | 0,827072369 | 0,61261 | 0,8035281 | novel transcript |
| ENSG00000277218 | AL139123,1 | -0,274013587 | 0,82701558 | 0,7158 | 0,86371705 | novel transcript |
| ENSG00000070886 | EPHA8 | -0,274277466 | 0,826864327 | 0,78637 | 0,90080813 | EPH receptor A8 |
| ENSG00000182196 | ARL6IP4 | -0,274319108 | 0,82684046 | 0,35348 | 0,62819991 | ADP ribosylation factor like GTPase 6 interacting protein 4 |
| ENSG00000105146 | AURKC | -0,274354739 | 0,82682004 | 0,58683 | 0,78701992 | aurora kinase C |
| ENSG00000168077 | SCARA3 | -0,274654396 | 0,826648322 | 0,37476 | 0,6431311 | scavenger receptor class A member 3 |
| ENSG00000235875 | ARHGEF7-AS2 | -0,274671535 | 0,826638501 | 0,50445 | 0,73323943 | ARHGEF7 antisense RNA 2 |
| ENSG00000218521 | AL589655,1 | -0,274744773 | 0,826596539 | 0,68943 | 0,84865812 | pseudogene similar to an integrin Alpha-E precursor fragment |
| ENSG00000108187 | PBLD | -0,274760085 | 0,826587765 | 0,17688 | 0,50123064 | phenazine biosynthesis like protein domain containing |
| ENSG00000215914 | MMP23A | -0,2748788 | 0,826519751 | 0,85161 | 0,93247452 | matrix metallopeptidase 23A (pseudogene) |
| ENSG00000265790 | RNASEH1P1 | -0,274930814 | 0,826489953 | 0,47417 | 0,7129821 | ribonuclease H1 pseudogene 1 |
| ENSG00000178184 | PARD6G | -0,275139812 | 0,82637023 | 0,48116 | 0,71820356 | par-6 family cell polarity regulator gamma |
| ENSG00000287574 | AC004938,2 | -0,275509079 | 0,826158743 | 0,56581 | 0,77406022 | novel transcript |
| ENSG00000162341 | TPCN2 | -0,275523825 | 0,826150299 | 0,27612 | 0,57533851 | two pore segment channel 2 |
| ENSG00000123144 | TRIR | -0,275667154 | 0,826068226 | 0,353 | 0,62780446 | telomerase RNA component interacting RNase |
| ENSG00000167671 | UBXN6 | -0,275715557 | 0,826040511 | 0,16193 | 0,48914394 | UBX domain protein 6 |
| ENSG00000189283 | FHIT | -0,275801052 | 0,825991561 | 0,32669 | 0,60877026 | fragile histidine triad diadenosine triphosphatase |
| ENSG00000197140 | ADAM32 | -0,275901619 | 0,825933985 | 0,28484 | 0,5816767 | ADAM metallopeptidase domain 32 |
| ENSG00000227780 | ZBTB22 | -0,276047151 | 0,825850673 | 0,32209 | 0,60577115 | zinc finger and BTB domain containing 22 |

| | | | | | | |
|---|---|---|---|---|---|---|
| ENSG00000229253 | ZBTB22 | -0,276047151 | 0,825850673 | 0,32209 | 0,60577115 | zinc finger and BTB domain containing 22 |
| ENSG00000272894 | AC004982,2 | -0,2761731 | 0,825778579 | 0,26119 | 0,56416552 | novel transcript |
| ENSG00000176749 | CDK5R1 | -0,276226111 | 0,825748236 | 0,37334 | 0,64204597 | cyclin dependent kinase 5 regulatory subunit 1 |
| ENSG00000257621 | PSMA3-AS1 | -0,276241104 | 0,825739656 | 0,16966 | 0,49589723 | PSMA3 antisense RNA 1 |
| ENSG00000128573 | FOXP2 | -0,276363817 | 0,825669422 | 0,60194 | 0,79680798 | forkhead box P2 |
| ENSG00000183570 | PCBP3 | -0,276459489 | 0,82561467 | 0,381 | 0,64851669 | poly(rC) binding protein 3 |
| ENSG00000104946 | TBC1D17 | -0,276480336 | 0,82560274 | 0,40139 | 0,6630277 | TBC1 domain family member 17 |
| ENSG00000250517 | LDHAL6CP | -0,276486647 | 0,825599129 | 0,64325 | 0,82098995 | lactate dehydrogenase A like 6C, pseudogene |
| ENSG00000134115 | CNTN6 | -0,276534808 | 0,825571568 | 0,37465 | 0,6431311 | contactin 6 |
| ENSG00000114656 | CFAP92 | -0,276539772 | 0,825568727 | 0,36451 | 0,63501401 | cilia and flagella associated protein 92 (putative) |
| ENSG00000124159 | MATN4 | -0,276552982 | 0,825561169 | 0,79393 | 0,90424974 | matrilin 4 |
| ENSG00000268416 | AC010329,1 | -0,276622635 | 0,825521312 | 0,25483 | 0,55944852 | novel transcript |
| ENSG00000280030 | AC010329,4 | -0,276622635 | 0,825521312 | 0,25483 | 0,55944852 | novel transcript |
| ENSG00000115257 | PCSK4 | -0,277039912 | 0,825282577 | 0,43524 | 0,68594493 | proprotein convertase subtilisin/kexin type 4 |
| ENSG00000214562 | NUTM2D | -0,27708776 | 0,825255206 | 0,18658 | 0,509961 | NUT family member 2D |
| ENSG00000250860 | AC093225,1 | -0,277121393 | 0,825235968 | 0,72959 | 0,8715818 | novel zinc finger protein pseudogene |
| ENSG00000287029 | AL162713,2 | -0,277179177 | 0,825202915 | 0,63556 | 0,81657648 | novel transcript, antisense to ENOX1 |
| ENSG00000080823 | MOK | -0,277256964 | 0,825158423 | 0,1926 | 0,51490739 | MOK protein kinase |
| ENSG00000280604 | AJ239328,1 | -0,277317502 | 0,825123799 | 0,51054 | 0,73802717 | PCBP3 overlapping transcript 1 |
| ENSG00000177669 | MBOAT4 | -0,277417403 | 0,825066664 | 0,71092 | 0,86079022 | membrane bound O-acyltransferase domain containing 4 |
| ENSG00000174428 | GTF2IRD2B | -0,277733164 | 0,824886103 | 0,24586 | 0,55372423 | GTF2I repeat domain containing 2B |
| ENSG00000283402 | AC087821,2 | -0,277801624 | 0,82484696 | 0,53435 | 0,75405583 | mitochondrially encoded cytochrome b (MT-CYB) pseudogene |
| ENSG00000225313 | AL513327,1 | -0,27785493 | 0,824816484 | 0,27659 | 0,575468 | novel transcript |
| ENSG00000170049 | KCNAB3 | -0,277878836 | 0,824802816 | 0,39344 | 0,65641419 | potassium voltage-gated channel subfamily A regulatory beta subunit 3 |
| ENSG00000181722 | ZBTB20 | -0,277957488 | 0,824757851 | 0,37934 | 0,64718149 | zinc finger and BTB domain containing 20 |
| ENSG00000253764 | AC019257,1 | -0,278168754 | 0,824637084 | 0,38034 | 0,64797034 | novel transcript |
| ENSG00000286552 | AL390839,1 | -0,278239314 | 0,824596753 | 0,70816 | 0,85927396 | novel transcript |
| ENSG00000226864 | ATE1-AS1 | -0,278292504 | 0,824566352 | 0,55058 | 0,7647513 | ATE1 antisense RNA 1 |
| ENSG00000273540 | AGBL1 | -0,278586199 | 0,824398509 | 0,67135 | 0,837283 | ATP/GTP binding protein like 1 |
| ENSG00000181541 | MAB21L2 | -0,278719034 | 0,824322607 | 0,38192 | 0,64905556 | mab-21 like 2 |
| ENSG00000131127 | ZNF141 | -0,279217283 | 0,824037968 | 0,36135 | 0,63331577 | zinc finger protein 141 |

| | | | | | | |
|---|---|---|---|---|---|---|
| ENSG00000189339 | SLC35E2B | -0,279238948 | 0,824025593 | 0,33695 | 0,61684812 | solute carrier family 35 member E2B |
| ENSG00000101412 | E2F1 | -0,279324684 | 0,823976625 | 0,26335 | 0,56608528 | E2F transcription factor 1 |
| ENSG00000269997 | AC068790,3 | -0,279418015 | 0,823923322 | 0,5277 | 0,75039004 | novel transcript, sense intronic to CCDC92 |
| ENSG00000225849 | MKRN7P | -0,279440127 | 0,823910694 | 0,49237 | 0,72556208 | makorin ring finger protein 7, pseudogene |
| ENSG00000101400 | SNTA1 | -0,279482787 | 0,823886331 | 0,34487 | 0,62213186 | syntrophin alpha 1 |
| ENSG00000280975 | CENATAC | -0,279543285 | 0,823851783 | 0,29907 | 0,59117691 | centrosomal AT-AC splicing factor |
| ENSG00000268403 | AC132192,2 | -0,279634953 | 0,823799438 | 0,2811 | 0,57848427 | novel transcript, antisense to ZNF143 |
| ENSG00000133131 | MORC4 | -0,279682943 | 0,823772036 | 0,37158 | 0,64073742 | MORC family CW-type zinc finger 4 |
| ENSG00000284308 | C2orf81 | -0,279856665 | 0,823672847 | 0,43075 | 0,68324475 | chromosome 2 open reading frame 81 |
| ENSG00000255920 | CCND2-AS1 | -0,279917472 | 0,823638131 | 0,56042 | 0,77082225 | CCND2 antisense RNA 1 |
| ENSG00000218233 | NEPNP | -0,280008159 | 0,82358636 | 0,78768 | 0,90122495 | nephrocan, pseudogene |
| ENSG00000223883 | LINC01707 | -0,28009521 | 0,823536667 | 0,7138 | 0,86224977 | long intergenic non-protein coding RNA 1707 |
| ENSG00000261324 | AC010168,2 | -0,280111205 | 0,823527536 | 0,33213 | 0,61206889 | novel transcript, overlapping HIST4H4 |
| ENSG00000269176 | AP001160,3 | -0,280254898 | 0,823445517 | 0,40276 | 0,66396567 | novel transcript, antisense to TAF6L |
| ENSG00000280495 | TRAPPC4 | -0,280326895 | 0,823404424 | 0,36377 | 0,63461346 | trafficking protein particle complex 4 |
| ENSG00000128965 | CHAC1 | -0,280346654 | 0,823393147 | 0,35443 | 0,62908147 | ChaC glutathione specific gamma-glutamylcyclotransferase 1 |
| ENSG00000112619 | PRPH2 | -0,28038417 | 0,823371736 | 0,61788 | 0,80610038 | peripherin 2 |
| ENSG00000174945 | AMZ1 | -0,280435653 | 0,823342354 | 0,51167 | 0,73874688 | archaelysin family metallopeptidase 1 |
| ENSG00000239577 | RN7SL388P | -0,280444125 | 0,823337519 | 0,77295 | 0,89514572 | RNA, 7SL, cytoplasmic 388, pseudogene |
| ENSG00000160049 | DFFA | -0,280508406 | 0,823300835 | 0,46684 | 0,70686749 | DNA fragmentation factor subunit alpha |
| ENSG00000204650 | LINC02210 | -0,280527117 | 0,823290157 | 0,24226 | 0,55112431 | long intergenic non-protein coding RNA 2210 |
| ENSG00000032219 | ARID4A | -0,280608782 | 0,823243555 | 0,18015 | 0,50407792 | AT-rich interaction domain 4A |
| ENSG00000235173 | HGH1 | -0,280617698 | 0,823238468 | 0,30825 | 0,59696396 | HGH1 homolog |
| ENSG00000106367 | AP1S1 | -0,280725219 | 0,823177116 | 0,13643 | 0,46362513 | adaptor related protein complex 1 subunit sigma 1 |
| ENSG00000146352 | CLVS2 | -0,280736934 | 0,823170432 | 0,50326 | 0,73256439 | clavesin 2 |
| ENSG00000090273 | NUDC | -0,280747813 | 0,823164224 | 0,32965 | 0,61066538 | nuclear distribution C, dynein complex regulator |
| ENSG00000233716 | MTFR2P1 | -0,280774259 | 0,823149135 | 0,54751 | 0,76293749 | MTFR2 pseudogene 1 |
| ENSG00000163701 | IL17RE | -0,280778473 | 0,82314673 | 0,58574 | 0,78640335 | interleukin 17 receptor E |
| ENSG00000223896 | CCNJP2 | -0,280857987 | 0,823101364 | 0,59764 | 0,79470049 | cyclin J pseudogene 2 |
| ENSG00000204177 | BMS1P1 | -0,280970563 | 0,823037139 | 0,18629 | 0,50961491 | BMS1 pseudogene 1 |
| ENSG00000144827 | ABHD10 | -0,280977209 | 0,823033347 | 0,40315 | 0,66399303 | abhydrolase domain containing 10, depalmitoylase |
| ENSG00000167487 | KLHL26 | -0,28100273 | 0,823018788 | 0,32035 | 0,60523419 | kelch like family member 26 |

| | | | | | | |
|---|---|---|---|---|---|---|
| ENSG00000068976 | PYGM | -0,281011067 | 0,823014032 | 0,66374 | 0,8324578 | glycogen phosphorylase, muscle associated |
| ENSG00000259577 | CERNA1 | -0,28106634 | 0,822982501 | 0,56834 | 0,77592553 | competing endogenous lncRNA 1 for miR-4707-5p and miR-4767 |
| ENSG00000082482 | KCNK2 | -0,281122573 | 0,822950423 | 0,19402 | 0,51552252 | potassium two pore domain channel subfamily K member 2 |
| ENSG00000151338 | MIPOL1 | -0,281375793 | 0,822805993 | 0,24636 | 0,55403311 | mirror-image polydactyly 1 |
| ENSG00000063180 | CA11 | -0,281380934 | 0,822803061 | 0,44821 | 0,69418084 | carbonic anhydrase 11 |
| ENSG00000260677 | AL445531,1 | -0,281493616 | 0,822738798 | 0,69227 | 0,85023519 | novel transcript |
| ENSG00000124140 | SLC12A5 | -0,281680201 | 0,8226324 | 0,29646 | 0,58978642 | solute carrier family 12 member 5 |
| ENSG00000166793 | YPEL4 | -0,281709304 | 0,822615805 | 0,29464 | 0,58857672 | yippee like 4 |
| ENSG00000237773 | AC073332,1 | -0,281961635 | 0,82247194 | 0,77771 | 0,89714267 | novel transcript |
| ENSG00000136878 | USP20 | -0,282141785 | 0,822369244 | 0,14432 | 0,47189017 | ubiquitin specific peptidase 20 |
| ENSG00000076555 | ACACB | -0,282210335 | 0,82233017 | 0,2003 | 0,52107152 | acetyl-CoA carboxylase beta |
| ENSG00000274565 | AC080038,1 | -0,282240477 | 0,822312989 | 0,47592 | 0,71433501 | novel transcript |
| ENSG00000133812 | SBF2 | -0,282312663 | 0,822271845 | 0,27217 | 0,57306433 | SET binding factor 2 |
| ENSG00000067836 | ROGDI | -0,282445093 | 0,82219637 | 0,16459 | 0,49065086 | rogdi atypical leucine zipper |
| ENSG00000273419 | AC004877,1 | -0,282447091 | 0,822195231 | 0,47888 | 0,7165637 | novel transcript, antisense to ZNF862 |
| ENSG00000288187 | CU633906,5 | -0,282525551 | 0,822150518 | 0,43828 | 0,6876359 | novel transcript |
| ENSG00000241769 | LINC00893 | -0,2825564 | 0,822132938 | 0,33404 | 0,61387439 | long intergenic non-protein coding RNA 893 |
| ENSG00000064547 | LPAR2 | -0,28257853 | 0,822120327 | 0,23047 | 0,54360012 | lysophosphatidic acid receptor 2 |
| ENSG00000225075 | AL603832,1 | -0,282695771 | 0,82205352 | 0,67575 | 0,84049293 | novel transcript |
| ENSG00000166295 | ANAPC16 | -0,282858638 | 0,821960723 | 0,35855 | 0,63185282 | anaphase promoting complex subunit 16 |
| ENSG00000154240 | CEP112 | -0,282905463 | 0,821934045 | 0,17233 | 0,49714741 | centrosomal protein 112 |
| ENSG00000235672 | AC090286,3 | -0,282916701 | 0,821927642 | 0,68866 | 0,84822736 | RAN binding protein 1 (RANBP1) pseudogene |
| ENSG00000085998 | POMGNT1 | -0,28315006 | 0,821794705 | 0,29708 | 0,59016006 | protein O-linked mannose N-acetylglucosaminyltransferase 1 (beta 1,2-) |
| ENSG00000231908 | IDH1-AS1 | -0,283272943 | 0,821724711 | 0,55041 | 0,76458107 | IDH1 antisense RNA 1 |
| ENSG00000184702 | SEPTIN5 | -0,283316428 | 0,821699943 | 0,3707 | 0,63998958 | septin 5 |
| ENSG00000175773 | ZBTB44-DT | -0,283706088 | 0,821478039 | 0,73421 | 0,87420968 | ZBTB44 divergent transcript |
| ENSG00000231651 | DLG3-AS1 | -0,283895582 | 0,821370147 | 0,67806 | 0,84159488 | DLG3 antisense RNA 1 |
| ENSG00000151881 | TMEM267 | -0,283922907 | 0,82135459 | 0,31385 | 0,60056652 | transmembrane protein 267 |
| ENSG00000117691 | NENF | -0,284070404 | 0,821270622 | 0,1722 | 0,49710979 | neudesin neurotrophic factor |
| ENSG00000224255 | PDCL3P6 | -0,284216292 | 0,821187578 | 0,75346 | 0,88453179 | PDCL3 pseudogene 6 |

| | | | | | | |
|---|---|---|---|---|---|---|
| ENSG00000135482 | ZC3H10 | -0,284311102 | 0,821133613 | 0,13352 | 0,4594073 | zinc finger CCCH-type containing 10 |
| ENSG00000163964 | PIGX | -0,284316707 | 0,821130423 | 0,3037 | 0,59385655 | phosphatidylinositol glycan anchor biosynthesis class X |
| ENSG00000206395 | DDAH2 | -0,284324769 | 0,821125834 | 0,48137 | 0,71822668 | dimethylarginine dimethylaminohydrolase 2 |
| ENSG00000226634 | DDAH2 | -0,284324769 | 0,821125834 | 0,48137 | 0,71822668 | dimethylarginine dimethylaminohydrolase 2 |
| ENSG00000231340 | ACTG1P10 | -0,284377298 | 0,821095937 | 0,45379 | 0,69789417 | actin gamma 1 pseudogene 10 |
| ENSG00000139410 | SDSL | -0,284555273 | 0,82099465 | 0,31853 | 0,6036094 | serine dehydratase like |
| ENSG00000260615 | RPL23AP97 | -0,284689023 | 0,820918541 | 0,78451 | 0,89962715 | ribosomal protein L23a pseudogene 97 |
| ENSG00000125744 | RTN2 | -0,284699325 | 0,820912679 | 0,24722 | 0,55483158 | reticulon 2 |
| ENSG00000154153 | RETREG1 | -0,28475482 | 0,820881102 | 0,3827 | 0,64939435 | reticulophagy regulator 1 |
| ENSG00000269210 | AC019171,1 | -0,284758075 | 0,82087925 | 0,80993 | 0,91240493 | novel transcript, antisense to ARHGEF33 |
| ENSG00000117174 | ZNHIT6 | -0,284833924 | 0,820836094 | 0,234 | 0,54581446 | zinc finger HIT-type containing 6 |
| ENSG00000187624 | C17orf97 | -0,284852521 | 0,820825513 | 0,34844 | 0,62472936 | chromosome 17 open reading frame 97 |
| ENSG00000255397 | NASPP1 | -0,284885972 | 0,820806482 | 0,68599 | 0,84654659 | nuclear autoantigenic sperm protein pseudogene 1 |
| ENSG00000154447 | SH3RF1 | -0,285113408 | 0,820677094 | 0,29513 | 0,58883222 | SH3 domain containing ring finger 1 |
| ENSG00000267083 | KRT18P61 | -0,285115961 | 0,820675642 | 0,77909 | 0,89781842 | keratin 18 pseudogene 61 |
| ENSG00000276802 | HERC2 | -0,285172076 | 0,820643721 | 0,39988 | 0,66197344 | HECT and RLD domain containing E3 ubiquitin protein ligase 2 |
| ENSG00000263724 | DLGAP1-AS3 | -0,285203244 | 0,820625992 | 0,54823 | 0,76347743 | DLGAP1 antisense RNA 3 |
| ENSG00000232593 | KANTR | -0,285224634 | 0,820613826 | 0,18981 | 0,51299724 | KDM5C adjacent transcript |
| ENSG00000138495 | COX17 | -0,285228746 | 0,820611487 | 0,30495 | 0,59446033 | cytochrome c oxidase copper chaperone COX17 |
| ENSG00000273447 | AC004067,1 | -0,285252916 | 0,820597739 | 0,43799 | 0,68747773 | novel transcript, antisense to CASP6 |
| ENSG00000187550 | SBK2 | -0,28531688 | 0,820561357 | 0,70606 | 0,85841728 | SH3 domain binding kinase family member 2 |
| ENSG00000135452 | TSPAN31 | -0,285347517 | 0,820543932 | 0,34072 | 0,61914946 | tetraspanin 31 |
| ENSG00000287825 | AL035587,2 | -0,28553027 | 0,820439996 | 0,5273 | 0,75031847 | novel transcript, antisense to CNPY3 |
| ENSG00000143554 | SLC27A3 | -0,285550273 | 0,820428621 | 0,46149 | 0,70284173 | solute carrier family 27 member 3 |
| ENSG00000170458 | CD14 | -0,28558929 | 0,820406434 | 0,71277 | 0,86181743 | CD14 molecule |
| ENSG00000132932 | ATP8A2 | -0,285599757 | 0,820400481 | 0,50261 | 0,73209345 | ATPase phospholipid transporting 8A2 |
| ENSG00000182087 | TMEM259 | -0,28572234 | 0,820330776 | 0,26643 | 0,56917012 | transmembrane protein 259 |
| ENSG00000170264 | FAM161A | -0,286100025 | 0,820116049 | 0,1514 | 0,47956749 | FAM161 centrosomal protein A |
| ENSG00000105649 | RAB3A | -0,286107525 | 0,820111785 | 0,44546 | 0,69255042 | RAB3A, member RAS oncogene family |
| ENSG00000229358 | DPY19L1P1 | -0,286163443 | 0,820079999 | 0,36595 | 0,63616426 | DPY19L1 pseudogene 1 |
| ENSG00000181350 | LRRC75A | -0,286192128 | 0,820063693 | 0,35965 | 0,63242662 | leucine rich repeat containing 75A |
| ENSG00000137094 | DNAJB5 | -0,286198082 | 0,820060309 | 0,3204 | 0,60528007 | DnaJ heat shock protein family (Hsp40) member B5 |

| | | | | | | |
|---|---|---|---|---|---|---|
| ENSG00000122122 | SASH3 | -0,28632667 | 0,81998722 | 0,7144 | 0,86255727 | SAM and SH3 domain containing 3 |
| ENSG00000109819 | PPARGC1A | -0,286338525 | 0,819980482 | 0,49558 | 0,7278408 | PPARG coactivator 1 alpha |
| ENSG00000236751 | LINC01186 | -0,286355381 | 0,819970902 | 0,75887 | 0,88743316 | long intergenic non-protein coding RNA 1186 |
| ENSG00000168303 | MPLKIP | -0,286471337 | 0,819904999 | 0,28975 | 0,58525534 | M-phase specific PLK1 interacting protein |
| ENSG00000284070 | AP000356,2 | -0,286508482 | 0,81988389 | 0,31355 | 0,60030528 | novel transcript |
| ENSG00000257218 | GATC | -0,286642162 | 0,819807923 | 0,34996 | 0,62563749 | glutamyl-tRNA amidotransferase subunit C |
| ENSG00000174915 | PTDSS2 | -0,286671165 | 0,819791442 | 0,31989 | 0,60473824 | phosphatidylserine synthase 2 |
| ENSG00000228223 | HCG11 | -0,286682226 | 0,819785157 | 0,40048 | 0,66257606 | HLA complex group 11 |
| ENSG00000167642 | SPINT2 | -0,286813527 | 0,819710551 | 0,47676 | 0,71481671 | serine peptidase inhibitor, Kunitz type 2 |
| ENSG00000126214 | KLC1 | -0,28690825 | 0,819656733 | 0,12255 | 0,44232452 | kinesin light chain 1 |
| ENSG00000253873 | PCDHGA11 | -0,286919204 | 0,81965051 | 0,36525 | 0,63543797 | protocadherin gamma subfamily A, 11 |
| ENSG00000187187 | ZNF546 | -0,286979544 | 0,819616229 | 0,66279 | 0,83222119 | zinc finger protein 546 |
| ENSG00000100095 | SEZ6L | -0,286982275 | 0,819614677 | 0,53249 | 0,75300764 | seizure related 6 homolog like |
| ENSG00000248769 | AC139495,1 | -0,287106014 | 0,819544383 | 0,40277 | 0,66396567 | POM121 membrane glycoprotein-like 1 (POM121L1) pseudogene |
| ENSG00000055732 | MCOLN3 | -0,287111991 | 0,819540987 | 0,78302 | 0,89918294 | mucolipin 3 |
| ENSG00000278632 | TBC1D3B | -0,287191236 | 0,819495972 | 0,62488 | 0,81001052 | TBC1 domain family member 3B |
| ENSG00000250850 | AL161781,2 | -0,287234408 | 0,819471449 | 0,77849 | 0,89745058 | novel transcript, antisense to PAX5 |
| ENSG00000160298 | C21orf58 | -0,287330093 | 0,819417101 | 0,17669 | 0,50102357 | chromosome 21 open reading frame 58 |
| ENSG00000181378 | CFAP65 | -0,28737102 | 0,819393856 | 0,43556 | 0,68618266 | cilia and flagella associated protein 65 |
| ENSG00000235422 | KATNBL1P3 | -0,287457394 | 0,8193448 | 0,79851 | 0,90704887 | katanin regulatory subunit B1 like 1 pseudogene 3 |
| ENSG00000255008 | LINC02739 | -0,287613872 | 0,819255937 | 0,65722 | 0,82867637 | long intergenic non-protein coding RNA 2739 |
| ENSG00000079435 | LIPE | -0,287628207 | 0,819247797 | 0,27797 | 0,57629868 | lipase E, hormone sensitive type |
| ENSG00000228212 | OFD1P17 | -0,287633233 | 0,819244943 | 0,44196 | 0,68994635 | OFD1 pseudogene 17 |
| ENSG00000167131 | CCDC103 | -0,287703163 | 0,819205233 | 0,60914 | 0,80161331 | coiled-coil domain containing 103 |
| ENSG00000228430 | AL162726,3 | -0,287860011 | 0,819116175 | 0,78357 | 0,89918294 | serine palmitoyltransferase long chain base subunit 1 (SPTLC1) pseudogene |
| ENSG00000269950 | AP000962,1 | -0,287903047 | 0,819091741 | 0,58694 | 0,7870367 | novel transcript |
| ENSG00000285980 | AP003025,2 | -0,28791933 | 0,819082496 | 0,74871 | 0,8817805 | novel transcript |
| ENSG00000233602 | ERI3-IT1 | -0,287928478 | 0,819077303 | 0,68032 | 0,84309883 | ERI3 intronic transcript 1 |
| ENSG00000102466 | FGF14 | -0,287970075 | 0,819053687 | 0,62915 | 0,81265687 | fibroblast growth factor 14 |
| ENSG00000228300 | FAM174C | -0,288088937 | 0,818986209 | 0,45549 | 0,69942891 | family with sequence similarity 174 member C |
| ENSG00000111666 | CHPT1 | -0,288152943 | 0,818949875 | 0,13669 | 0,46406381 | choline phosphotransferase 1 |

| | | | | | | |
|---|---|---|---|---|---|---|
| ENSG00000273076 | AL021707,7 | -0,288345233 | 0,818840728 | 0,80437 | 0,90994462 | novel transcript, antisense to SUN2 |
| ENSG00000164440 | TXLNB | -0,288552344 | 0,818723185 | 0,62558 | 0,81053676 | taxilin beta |
| ENSG00000215097 | AL731733,1 | -0,288999673 | 0,818469367 | 0,5594 | 0,77045525 | dual specificity phosphatase 8 pseudogene 2 |
| ENSG00000255769 | GOLGA2P10 | -0,289181698 | 0,818366107 | 0,22309 | 0,53927144 | GOLGA2 pseudogene 10 |
| ENSG00000220201 | ZGLP1 | -0,289196147 | 0,818357911 | 0,3908 | 0,65464997 | zinc finger GATA like protein 1 |
| ENSG00000223953 | C1QTNF5 | -0,289222687 | 0,818342857 | 0,56048 | 0,77086257 | C1q and TNF related 5 |
| ENSG00000269707 | AC018730,1 | -0,289305609 | 0,818295822 | 0,28722 | 0,58364712 | novel transcript, sense overlapping POU3F3 |
| ENSG00000153266 | FEZF2 | -0,289554875 | 0,818154451 | 0,61063 | 0,80242603 | FEZ family zinc finger 2 |
| ENSG00000260916 | CCPG1 | -0,289562754 | 0,818149983 | 0,26642 | 0,56917012 | cell cycle progression 1 |
| ENSG00000283985 | C17orf97 | -0,289603723 | 0,818126749 | 0,34031 | 0,61897407 | chromosome 17 open reading frame 97 |
| ENSG00000269794 | AC010642,1 | -0,289818557 | 0,81800493 | 0,58185 | 0,78454692 | zinc finger protein family pseudogene |
| ENSG00000279726 | AC005609,5 | -0,289964077 | 0,817922424 | 0,46444 | 0,70506031 | novel transcript, antisense to PCDHA5, PCDHA4, PCDHA3, PCDHA2 and PCDHA1 |
| ENSG00000048471 | SNX29 | -0,289981202 | 0,817912715 | 0,212 | 0,53079538 | sorting nexin 29 |
| ENSG00000110076 | NRXN2 | -0,290027283 | 0,817886591 | 0,31251 | 0,59939093 | neurexin 2 |
| ENSG00000223949 | ROR1-AS1 | -0,290076369 | 0,817858764 | 0,32231 | 0,60577115 | ROR1 antisense RNA 1 |
| ENSG00000177398 | UMODL1 | -0,290097496 | 0,817846787 | 0,72789 | 0,87053185 | uromodulin like 1 |
| ENSG00000184166 | OR1D2 | -0,290146763 | 0,817818859 | 0,27975 | 0,5775083 | olfactory receptor family 1 subfamily D member 2 |
| ENSG00000151366 | NDUFC2 | -0,290273191 | 0,817747194 | 0,3026 | 0,59306846 | NADH:ubiquinone oxidoreductase subunit C2 |
| ENSG00000267549 | AC006116,7 | -0,290300221 | 0,817731873 | 0,39581 | 0,6585697 | novel transcript, antisense to ZSCAN5A |
| ENSG00000165521 | EML5 | -0,290412531 | 0,817668217 | 0,30507 | 0,59453935 | EMAP like 5 |
| ENSG00000253771 | TPTE2P1 | -0,29049483 | 0,817621574 | 0,37804 | 0,64636276 | transmembrane phosphoinositide 3-phosphatase and tensin homolog 2 pseudogene 1 |
| ENSG00000271913 | AL035530,2 | -0,290888987 | 0,817398223 | 0,23212 | 0,54496653 | novel transcript, antisense to TAGAP |
| ENSG00000129566 | TEP1 | -0,290977 | 0,817348359 | 0,33547 | 0,61510039 | telomerase associated protein 1 |
| ENSG00000224452 | RSL24D1P6 | -0,291239928 | 0,817199413 | 0,78334 | 0,89918294 | ribosomal L24 domain containing 1 pseudogene 6 |
| ENSG00000272341 | AL137003,1 | -0,291404497 | 0,8171062 | 0,79263 | 0,90361513 | novel transcript |
| ENSG00000206181 | ELOA2 | -0,291407649 | 0,817104414 | 0,64925 | 0,82436914 | elongin A2 |
| ENSG00000162490 | DRAXIN | -0,291427027 | 0,817093439 | 0,4179 | 0,6742866 | dorsal inhibitory axon guidance protein |
| ENSG00000157999 | ANKRD61 | -0,291430194 | 0,817091646 | 0,32479 | 0,60732633 | ankyrin repeat domain 61 |
| ENSG00000076053 | RBM7 | -0,291518993 | 0,817041355 | 0,38387 | 0,65045825 | RNA binding motif protein 7 |
| ENSG00000275917 | CHRFAM7A | -0,29155682 | 0,817019932 | 0,37512 | 0,64350562 | CHRNA7 (exons 5-10) and FAM7A (exons A-E) fusion |
| ENSG00000135245 | HILPDA | -0,291667063 | 0,816957502 | 0,31132 | 0,59890747 | hypoxia inducible lipid droplet associated |

| | | | | | | |
|---|---|---|---|---|---|---|
| ENSG00000267064 | UXT-AS1 | -0,292017388 | 0,816759147 | 0,48563 | 0,72107911 | UXT antisense RNA 1 |
| ENSG00000258571 | PTTG4P | -0,292029787 | 0,816752128 | 0,62195 | 0,80819595 | pituitary tumor-transforming 4 pseudogene |
| ENSG00000248810 | LINC02432 | -0,292076509 | 0,816725677 | 0,55559 | 0,76790322 | long intergenic non-protein coding RNA 2432 |
| ENSG00000224687 | RASAL2-AS1 | -0,292115651 | 0,816703519 | 0,25749 | 0,56160845 | RASAL2 antisense RNA 1 |
| ENSG00000166886 | NAB2 | -0,292167631 | 0,816674094 | 0,25916 | 0,56295427 | NGFI-A binding protein 2 |
| ENSG00000279762 | AC005899,8 | -0,292215791 | 0,816646832 | 0,4342 | 0,68500084 | novel transcript, antisense to PSMD11 |
| ENSG00000256690 | AP001160,1 | -0,292218991 | 0,816645021 | 0,61989 | 0,80737281 | novel transcript |
| ENSG00000281691 | RBM5-AS1 | -0,292228618 | 0,816639571 | 0,24857 | 0,55589057 | RBM5 antisense RNA 1 |
| ENSG00000214046 | SMIM7 | -0,292319372 | 0,816588201 | 0,18685 | 0,51035306 | small integral membrane protein 7 |
| ENSG00000224543 | SNRPGP15 | -0,292369941 | 0,816559579 | 0,63399 | 0,81549489 | small nuclear ribonucleoprotein polypeptide G pseudogene 15 |
| ENSG00000073910 | FRY | -0,292408723 | 0,816537629 | 0,43541 | 0,68601186 | FRY microtubule binding protein |
| ENSG00000106258 | CYP3A5 | -0,292526703 | 0,816470857 | 0,24325 | 0,55199251 | cytochrome P450 family 3 subfamily A member 5 |
| ENSG00000284948 | AC107959,4 | -0,292606925 | 0,816425458 | 0,72177 | 0,86716786 | novel transcript |
| ENSG00000207263 | SNORD116-16 | -0,292615043 | 0,816420864 | 0,77548 | 0,89630112 | small nucleolar RNA, C/D box 116-16 |
| ENSG00000236698 | EIF1AXP1 | -0,292640783 | 0,816406298 | 0,50716 | 0,7354206 | EIF1AX pseudogene 1 |
| ENSG00000130723 | PRRC2B | -0,292651583 | 0,816400186 | 0,20336 | 0,52321822 | proline rich coiled-coil 2B |
| ENSG00000258754 | LINC01579 | -0,29266733 | 0,816391276 | 0,4102 | 0,66821634 | long intergenic non-protein coding RNA 1579 |
| ENSG00000239503 | MARK2P8 | -0,292676184 | 0,816386265 | 0,43364 | 0,68473723 | microtubule affinity regulating kinase 2 pseudogene 8 |
| ENSG00000241697 | TMEFF1 | -0,292724879 | 0,81635871 | 0,36418 | 0,634951 | transmembrane protein with EGF like and two follistatin like domains 1 |
| ENSG00000224464 | PGAM1P6 | -0,292833827 | 0,816297064 | 0,66874 | 0,83569335 | phosphoglycerate mutase 1 pseudogene 6 |
| ENSG00000288217 | PEX5 | -0,292955757 | 0,816228077 | 0,22546 | 0,5407493 | peroxisomal biogenesis factor 5 |
| ENSG00000225411 | CR786580,1 | -0,293029912 | 0,816186124 | 0,48562 | 0,72107911 | novel transcript |
| ENSG00000260070 | AC006960,3 | -0,29309367 | 0,816150054 | 0,58022 | 0,78349398 | novel transcript, antisense to KIAA0895 |
| ENSG00000286872 | AC024270,4 | -0,293129011 | 0,816130062 | 0,53192 | 0,75280487 | novel transcript, antisense to TM6SF1 |
| ENSG00000115694 | STK25 | -0,293169124 | 0,81610737 | 0,21481 | 0,53373102 | serine/threonine kinase 25 |
| ENSG00000249898 | MCPH1-AS1 | -0,293223834 | 0,816076422 | 0,2147 | 0,53362255 | MCPH1 antisense RNA 1 |
| ENSG00000272871 | AL159169,2 | -0,293372111 | 0,815992552 | 0,45213 | 0,69691985 | novel transcript |
| ENSG00000159387 | IRX6 | -0,293374467 | 0,81599122 | 0,71761 | 0,86483656 | iroquois homeobox 6 |
| ENSG00000287180 | AC106818,2 | -0,293400508 | 0,815976491 | 0,60731 | 0,80012503 | novel transcript, antisense to FAM172A |
| ENSG00000173662 | TAS1R1 | -0,293480493 | 0,815931253 | 0,69511 | 0,85206252 | taste 1 receptor member 1 |
| ENSG00000187634 | SAMD11 | -0,293500583 | 0,815919891 | 0,64834 | 0,82383528 | sterile alpha motif domain containing 11 |

| | | | | | | |
|---|---|---|---|---|---|---|
| ENSG00000137941 | TTLL7 | -0,293601947 | 0,815862567 | 0,20591 | 0,5253263 | tubulin tyrosine ligase like 7 |
| ENSG00000276870 | PLA2G10 | -0,293754675 | 0,815776201 | 0,69822 | 0,85390309 | phospholipase A2 group X |
| ENSG00000144115 | THNSL2 | -0,29380356 | 0,81574856 | 0,18196 | 0,50521773 | threonine synthase like 2 |
| ENSG00000134321 | RSAD2 | -0,293846234 | 0,815724431 | 0,5084 | 0,73653319 | radical S-adenosyl methionine domain containing 2 |
| ENSG00000112667 | DNPH1 | -0,293914279 | 0,815685958 | 0,36932 | 0,63912679 | 2'-deoxynucleoside 5'-phosphate N-hydrolase 1 |
| ENSG00000260160 | AC011468,1 | -0,293946937 | 0,815667493 | 0,64406 | 0,82138445 | novel transcript |
| ENSG00000186666 | BCDIN3D | -0,294060963 | 0,815603028 | 0,30966 | 0,59807209 | BCDIN3 domain containing RNA methyltransferase |
| ENSG00000206346 | DXO | -0,29407896 | 0,815592854 | 0,41779 | 0,6742688 | decapping exoribonuclease |
| ENSG00000206480 | FLOT1 | -0,29407934 | 0,815592639 | 0,56397 | 0,77259188 | flotillin 1 |
| ENSG00000206379 | FLOT1 | -0,29407934 | 0,815592639 | 0,56397 | 0,77259188 | flotillin 1 |
| ENSG00000276272 | AC024884,2 | -0,294370913 | 0,815427822 | 0,47873 | 0,71645045 | novel transcript |
| ENSG00000186642 | PDE2A | -0,294400881 | 0,815410884 | 0,52513 | 0,74889591 | phosphodiesterase 2A |
| ENSG00000119661 | DNAL1 | -0,294583057 | 0,815307924 | 0,28371 | 0,58072192 | dynein axonemal light chain 1 |
| ENSG00000232970 | POLHP1 | -0,294735385 | 0,815221844 | 0,52975 | 0,75147469 | DNA polymerase eta pseudogene 1 |
| ENSG00000100865 | CINP | -0,294754458 | 0,815211067 | 0,38786 | 0,65289784 | cyclin dependent kinase 2 interacting protein |
| ENSG00000108557 | RAI1 | -0,294831913 | 0,815167301 | 0,30167 | 0,59283946 | retinoic acid induced 1 |
| ENSG00000182165 | TP53TG1 | -0,294997558 | 0,815073712 | 0,30919 | 0,59774312 | TP53 target 1 |
| ENSG00000214761 | HNRNPA1P15 | -0,295020093 | 0,81506098 | 0,62404 | 0,80957813 | heterogeneous nuclear ribonucleoprotein A1 pseudogene 15 |
| ENSG00000243960 | AL390195,1 | -0,295050594 | 0,815043749 | 0,54335 | 0,76000878 | novel transcript |
| ENSG00000260391 | AC022336,2 | -0,295309876 | 0,814897282 | 0,78164 | 0,89877377 | novel transcript, overlapping to KALRN |
| ENSG00000258824 | AL122035,1 | -0,295329986 | 0,814885923 | 0,63053 | 0,81354995 | novel transcript |
| ENSG00000088881 | EBF4 | -0,295366309 | 0,814865407 | 0,27326 | 0,57344244 | EBF family member 4 |
| ENSG00000253159 | PCDHGA12 | -0,29540383 | 0,814844215 | 0,25987 | 0,56377449 | protocadherin gamma subfamily A, 12 |
| ENSG00000171823 | FBXL14 | -0,295510224 | 0,814784125 | 0,298 | 0,59038354 | F-box and leucine rich repeat protein 14 |
| ENSG00000072163 | LIMS2 | -0,295528599 | 0,814773747 | 0,59999 | 0,79568184 | LIM zinc finger domain containing 2 |
| ENSG00000105854 | PON2 | -0,295766444 | 0,814639433 | 0,38913 | 0,65363435 | paraoxonase 2 |
| ENSG00000170011 | MYRIP | -0,295830624 | 0,814603194 | 0,37088 | 0,64013451 | myosin VIIA and Rab interacting protein |
| ENSG00000248710 | AC079594,1 | -0,29585415 | 0,81458991 | 0,45468 | 0,69880158 | TRIM59 and iFT80 readthrough |
| ENSG00000242083 | RPL7AP31 | -0,295895369 | 0,814566637 | 0,80065 | 0,90821843 | ribosomal protein L7a pseudogene 31 |
| ENSG00000229299 | AL121845,1 | -0,296092554 | 0,814455311 | 0,61172 | 0,80334296 | novel transcript |
| ENSG00000278759 | TBCD | -0,296101536 | 0,814450241 | 0,2556 | 0,56006988 | tubulin folding cofactor D |
| ENSG00000134253 | TRIM45 | -0,296135442 | 0,8144311 | 0,22733 | 0,54168591 | tripartite motif containing 45 |

| | | | | | | |
|---|---|---|---|---|---|---|
| ENSG00000260528 | FAM157C | -0,296159602 | 0,814417461 | 0,46784 | 0,70757147 | family with sequence similarity 157 member C |
| ENSG00000228198 | OR2M3 | -0,296201879 | 0,814393596 | 0,38172 | 0,64905556 | olfactory receptor family 2 subfamily M member 3 |
| ENSG00000167771 | RCOR2 | -0,296231582 | 0,814376829 | 0,37994 | 0,64769466 | REST corepressor 2 |
| ENSG00000254806 | SYS1-DBNDD2 | -0,296368031 | 0,814299809 | 0,70897 | 0,85977658 | SYS1-DBNDD2 readthrough (NMD candidate) |
| ENSG00000241560 | ZBTB20-AS1 | -0,296371882 | 0,814297635 | 0,4318 | 0,6838061 | ZBTB20 antisense RNA 1 |
| ENSG00000251011 | TMEM108-AS | -0,296407712 | 0,814277412 | 0,45608 | 0,69998075 | TMEM108 antisense RNA 1 |
| ENSG00000285825 | CCDC15-DT | -0,296459716 | 0,814248061 | 0,44687 | 0,6933405 | CCDC15 divergent transcript |
| ENSG00000174306 | ZHX3 | -0,296618236 | 0,814158598 | 0,21377 | 0,53281812 | zinc fingers and homeoboxes 3 |
| ENSG00000131748 | STARD3 | -0,296655782 | 0,81413741 | 0,35992 | 0,63249219 | StAR related lipid transfer domain containing 3 |
| ENSG00000139547 | RDH16 | -0,296679632 | 0,814123951 | 0,33622 | 0,61623735 | retinol dehydrogenase 16 |
| ENSG00000236863 | RPL23AP23 | -0,296703624 | 0,814110413 | 0,65773 | 0,82881843 | ribosomal protein L23a pseudogene 23 |
| ENSG00000116260 | QSOX1 | -0,29683689 | 0,814035214 | 0,38085 | 0,64840997 | quiescin sulfhydryl oxidase 1 |
| ENSG00000265579 | AC023301,1 | -0,297035541 | 0,813923134 | 0,87705 | 0,94553042 | novel transcript , sense intronic to NETO1 |
| ENSG00000173638 | SLC19A1 | -0,297122579 | 0,813874031 | 0,25587 | 0,56047793 | solute carrier family 19 member 1 |
| ENSG00000101198 | NKAIN4 | -0,297134535 | 0,813867286 | 0,24369 | 0,55235189 | sodium/potassium transporting ATPase interacting 4 |
| ENSG00000184060 | ADAP2 | -0,29715278 | 0,813856994 | 0,42251 | 0,67663831 | ArfGAP with dual PH domains 2 |
| ENSG00000164287 | CDC20B | -0,29721299 | 0,813823029 | 0,58531 | 0,78627961 | cell division cycle 20B |
| ENSG00000256937 | KRT17P8 | -0,297298215 | 0,813774955 | 0,63426 | 0,81566722 | keratin 17 pseudogene 8 |
| ENSG00000225092 | AL355336,1 | -0,297420659 | 0,813705891 | 0,45547 | 0,69942891 | novel transcript |
| ENSG00000179542 | SLITRK4 | -0,297531205 | 0,813643544 | 0,44495 | 0,69205935 | SLIT and NTRK like family member 4 |
| ENSG00000229185 | OR2H2 | -0,297586192 | 0,813612533 | 0,69988 | 0,85503778 | olfactory receptor family 2 subfamily H member 2 |
| ENSG00000148344 | PTGES | -0,297589733 | 0,813610536 | 0,55102 | 0,76487238 | prostaglandin E synthase |
| ENSG00000169836 | TACR3 | -0,297633737 | 0,81358572 | 0,36178 | 0,63352908 | tachykinin receptor 3 |
| ENSG00000240399 | AC004801,2 | -0,297685778 | 0,813556373 | 0,65608 | 0,82824314 | ribosomal protein L37 (RPL37) pseudogene |
| ENSG00000282458 | WASH5P | -0,297691426 | 0,813553188 | 0,21984 | 0,53722186 | WASP family homolog 5, pseudogene |
| ENSG00000089351 | GRAMD1A | -0,297715851 | 0,813539415 | 0,34644 | 0,62342969 | GRAM domain containing 1A |
| ENSG00000152082 | MZT2B | -0,297981729 | 0,813389499 | 0,24448 | 0,55253387 | mitotic spindle organizing protein 2B |
| ENSG00000281289 | LSS | -0,298185436 | 0,813274658 | 0,25198 | 0,55779669 | lanosterol synthase |
| ENSG00000267506 | AC021683,2 | -0,298195083 | 0,813269219 | 0,21052 | 0,5290538 | novel transcript |
| ENSG00000168234 | TTC39C | -0,29824778 | 0,813239514 | 0,33984 | 0,61872249 | tetratricopeptide repeat domain 39C |
| ENSG00000245060 | LINC00847 | -0,298261871 | 0,813231571 | 0,14843 | 0,47586237 | long intergenic non-protein coding RNA 847 |
| ENSG00000228407 | AL139156,2 | -0,298306012 | 0,813206689 | 0,73777 | 0,87540065 | pseudogene similar to part of alcohol dehydrogenase 5 (class III), chi polypeptidem (ADH5) |

| | | | | | | |
|---|---|---|---|---|---|---|
| ENSG00000148634 | HERC4 | -0,298343119 | 0,813185774 | 0,34588 | 0,62300518 | HECT and RLD domain containing E3 ubiquitin protein ligase 4 |
| ENSG00000120645 | IQSEC3 | -0,298369045 | 0,81317116 | 0,49651 | 0,728373 | IQ motif and Sec7 domain ArfGEF 3 |
| ENSG00000261537 | AC027279,2 | -0,298659562 | 0,813007428 | 0,67588 | 0,84049293 | novel transcript, sense intronic to WWOX |
| ENSG00000099330 | OCEL1 | -0,298834675 | 0,812908752 | 0,43445 | 0,68505666 | occludin/ELL domain containing 1 |
| ENSG00000237126 | AC073254,1 | -0,29888943 | 0,812877899 | 0,54009 | 0,75814808 | novel transcript |
| ENSG00000234043 | NUDT9P1 | -0,298949844 | 0,812843861 | 0,77629 | 0,89654301 | nudix hydrolase 9 pseudogene 1 |
| ENSG00000099937 | SERPIND1 | -0,29903679 | 0,812794875 | 0,50246 | 0,73208008 | serpin family D member 1 |
| ENSG00000185909 | KLHDC8B | -0,299102525 | 0,812757842 | 0,25277 | 0,55796133 | kelch domain containing 8B |
| ENSG00000140932 | CMTM2 | -0,299218953 | 0,812692253 | 0,66227 | 0,83183445 | CKLF like MARVEL transmembrane domain containing 2 |
| ENSG00000262652 | AC124283,3 | -0,299340912 | 0,812623554 | 0,57753 | 0,78147914 | novel transcript, sense intronic FOXK2 |
| ENSG00000078124 | ACER3 | -0,299394244 | 0,812593515 | 0,288 | 0,5840701 | alkaline ceramidase 3 |
| ENSG00000103449 | SALL1 | -0,299533783 | 0,812514924 | 0,4684 | 0,7080558 | spalt like transcription factor 1 |
| ENSG00000128872 | TMOD2 | -0,299915273 | 0,8123001 | 0,34224 | 0,62000209 | tropomodulin 2 |
| ENSG00000188191 | PRKAR1B | -0,30011867 | 0,812185587 | 0,44548 | 0,69255042 | protein kinase cAMP-dependent type I regulatory subunit beta |
| ENSG00000103056 | SMPD3 | -0,300198214 | 0,812140808 | 0,45213 | 0,69691985 | sphingomyelin phosphodiesterase 3 |
| ENSG00000184792 | OSBP2 | -0,300206042 | 0,812136401 | 0,18062 | 0,50411652 | oxysterol binding protein 2 |
| ENSG00000213337 | ANKRD39 | -0,300325139 | 0,81206936 | 0,08397 | 0,37651068 | ankyrin repeat domain 39 |
| ENSG00000223691 | AC068044,1 | -0,300415546 | 0,812018473 | 0,7693 | 0,89297049 | Mdm2, transformed 3T3 cell double minute 2, p53 binding protein (mouse) (MDM2) pseudogene |
| ENSG00000250132 | AC004803,1 | -0,300489415 | 0,811976898 | 0,1524 | 0,48061452 | novel transcript |
| ENSG00000174514 | MFSD4A | -0,300511746 | 0,811964329 | 0,32641 | 0,60866572 | major facilitator superfamily domain containing 4A |
| ENSG00000287403 | AC016245,2 | -0,300526345 | 0,811956113 | 0,41998 | 0,67552612 | novel transcript, antisense to RPTOR |
| ENSG00000287725 | AP003071,5 | -0,300532844 | 0,811952455 | 0,33315 | 0,61307256 | novel protein, TPCN2 - SMIM38 readthrough |
| ENSG00000224635 | AL391095,1 | -0,300648823 | 0,811887185 | 0,27376 | 0,57377712 | novel transcript |
| ENSG00000225125 | RANP4 | -0,300745416 | 0,811832828 | 0,64209 | 0,8205913 | RAN pseudogene 4 |
| ENSG00000188613 | NANOS1 | -0,300748919 | 0,811830857 | 0,31589 | 0,60199254 | nanos C2HC-type zinc finger 1 |
| ENSG00000258739 | ENO1P2 | -0,300851014 | 0,811773408 | 0,44154 | 0,68977736 | enolase 1 pseudogene 2 |
| ENSG00000255408 | PCDHA3 | -0,300891151 | 0,811750824 | 0,33297 | 0,61297335 | protocadherin alpha 3 |
| ENSG00000283162 | AL390726,5 | -0,300993804 | 0,811693067 | 0,63061 | 0,81354995 | STRA6-like (Stra6l) pseudogene |
| ENSG00000111911 | HINT3 | -0,301182233 | 0,811587059 | 0,07587 | 0,36178429 | histidine triad nucleotide binding protein 3 |
| ENSG00000106608 | URGCP | -0,301342249 | 0,811497047 | 0,16598 | 0,49197859 | upregulator of cell proliferation |

| | | | | | | |
|---|---|---|---|---|---|---|
| ENSG00000274721 | AC091825,1 | -0,301536942 | 0,811387542 | 0,81454 | 0,91470265 | |
| ENSG00000230866 | LINC02558 | -0,301554269 | 0,811377798 | 0,20905 | 0,52809624 | long intergenic non-protein coding RNA 2558 |
| ENSG00000231942 | HNRNPA1P36 | -0,301554495 | 0,811377671 | 0,70593 | 0,85838918 | heterogeneous nuclear ribonucleoprotein A1 pseudogene 36 |
| ENSG00000134207 | SYT6 | -0,301691401 | 0,811300678 | 0,50988 | 0,73752989 | synaptotagmin 6 |
| ENSG00000232901 | CYCSP10 | -0,301862255 | 0,811204604 | 0,52591 | 0,74959092 | CYCS pseudogene 10 |
| ENSG00000262703 | AC009121,2 | -0,30187883 | 0,811195284 | 0,31562 | 0,60172912 | novel transcript, antisense to RMI2 |
| ENSG00000236542 | MED28P7 | -0,302037527 | 0,811106057 | 0,70481 | 0,85777377 | mediator complex subunit 28 pseudogene 7 |
| ENSG00000138646 | HERC5 | -0,302068987 | 0,81108837 | 0,40999 | 0,66821634 | HECT and RLD domain containing E3 ubiquitin protein ligase 5 |
| ENSG00000281388 | SCARNA6 | -0,302118021 | 0,811060803 | 0,38193 | 0,64905556 | small Cajal body-specific RNA 6 |
| ENSG00000251791 | SCARNA6 | -0,302118021 | 0,811060803 | 0,38193 | 0,64905556 | small Cajal body-specific RNA 6 |
| ENSG00000254416 | LINC02732 | -0,30231027 | 0,810952731 | 0,45283 | 0,69717623 | long intergenic non-protein coding RNA 2732 |
| ENSG00000110025 | SNX15 | -0,302416125 | 0,810893231 | 0,16106 | 0,48841163 | sorting nexin 15 |
| ENSG00000166148 | AVPR1A | -0,302661712 | 0,810755206 | 0,7108 | 0,86076616 | arginine vasopressin receptor 1A |
| ENSG00000285644 | AC108448,3 | -0,302815625 | 0,810668716 | 0,18426 | 0,50802406 | novel transcript, antisense to CARS |
| ENSG00000253015 | RN7SKP64 | -0,302839161 | 0,810655491 | 0,70424 | 0,85730406 | RN7SK pseudogene 64 |
| ENSG00000099783 | HNRNPM | -0,303041841 | 0,810541612 | 0,25123 | 0,55719434 | heterogeneous nuclear ribonucleoprotein M |
| ENSG00000159388 | BTG2 | -0,30307092 | 0,810525275 | 0,21778 | 0,53550245 | BTG anti-proliferation factor 2 |
| ENSG00000087253 | LPCAT2 | -0,303101535 | 0,810508075 | 0,42569 | 0,67923925 | lysophosphatidylcholine acyltransferase 2 |
| ENSG00000287629 | AC006059,5 | -0,303219207 | 0,81044197 | 0,45686 | 0,7004577 | novel transcript, antisense to ZBTB47 |
| ENSG00000183775 | KCTD16 | -0,303755582 | 0,810140714 | 0,47862 | 0,71634089 | potassium channel tetramerization domain containing 16 |
| ENSG00000164615 | CAMLG | -0,303796534 | 0,810117718 | 0,24319 | 0,55199251 | calcium modulating ligand |
| ENSG00000263165 | AC087292,1 | -0,303871693 | 0,810075515 | 0,42535 | 0,67893062 | novel transcript, antisense to ANKFY1 |
| ENSG00000282534 | IMPDH1P5 | -0,304031273 | 0,809985916 | 0,55953 | 0,77045624 | inosine monophosphate dehydrogenase 1 pseudogene 5 |
| ENSG00000213513 | IMPDH1P5 | -0,304031273 | 0,809985916 | 0,55953 | 0,77045624 | inosine monophosphate dehydrogenase 1 pseudogene 5 |
| ENSG00000040608 | RTN4R | -0,304245175 | 0,809865831 | 0,20307 | 0,52296754 | reticulon 4 receptor |
| ENSG00000198722 | UNC13B | -0,304261407 | 0,80985672 | 0,15925 | 0,4867709 | unc-13 homolog B |
| ENSG00000224063 | AC007319,1 | -0,304305473 | 0,809831984 | 0,61275 | 0,8035281 | novel transcript, antisense to TFPI and CALCRL |
| ENSG00000130159 | ECSIT | -0,304306836 | 0,809831218 | 0,23274 | 0,54542294 | ECSIT signaling integrator |
| ENSG00000131018 | SYNE1 | -0,304433102 | 0,809760344 | 0,24935 | 0,55642837 | spectrin repeat containing nuclear envelope protein 1 |
| ENSG00000212747 | RTL8B | -0,304819274 | 0,809543622 | 0,41313 | 0,67067821 | retrotransposon Gag like 8B |
| ENSG00000185236 | RAB11B | -0,305227536 | 0,809314565 | 0,31481 | 0,601382 | RAB11B, member RAS oncogene family |

| | | | | | | |
|---|---|---|---|---|---|---|
| ENSG00000186187 | ZNRF1 | -0,30538366 | 0,809226988 | 0,34972 | 0,62563749 | zinc and ring finger 1 |
| ENSG00000124104 | SNX21 | -0,305687109 | 0,809056797 | 0,14153 | 0,46809977 | sorting nexin family member 21 |
| ENSG00000178814 | OPLAH | -0,305688684 | 0,809055914 | 0,38385 | 0,65045825 | 5-oxoprolinase, ATP-hydrolysing |
| ENSG00000237188 | AC242426,2 | -0,305779474 | 0,809005001 | 0,65797 | 0,82895662 | novel transcript, antisense to FMO5 |
| ENSG00000251414 | AC138956,2 | -0,305837191 | 0,808972636 | 0,61379 | 0,8035281 | novel transcript |
| ENSG00000275482 | EPHB6 | -0,305843329 | 0,808969195 | 0,18907 | 0,51243448 | EPH receptor B6 |
| ENSG00000287378 | AC129803,1 | -0,306047762 | 0,80885457 | 0,60694 | 0,79997116 | novel transcript, antisense to CMSS1 |
| ENSG00000175309 | PHYKPL | -0,306190971 | 0,808774283 | 0,11295 | 0,42702832 | 5-phosphohydroxy-L-lysine phospho-lyase |
| ENSG00000135956 | TMEM127 | -0,306310685 | 0,808707174 | 0,28364 | 0,58072192 | transmembrane protein 127 |
| ENSG00000092850 | TEKT2 | -0,30635787 | 0,808680725 | 0,23533 | 0,54677606 | tektin 2 |
| ENSG00000173258 | ZNF483 | -0,306373344 | 0,808672051 | 0,12827 | 0,45171208 | zinc finger protein 483 |
| ENSG00000132004 | FBXW9 | -0,306506113 | 0,808597634 | 0,20716 | 0,52670822 | F-box and WD repeat domain containing 9 |
| ENSG00000260643 | AC092718,3 | -0,306552853 | 0,808571438 | 0,40878 | 0,6675559 | novel protein |
| ENSG00000226033 | STK19 | -0,306586136 | 0,808552784 | 0,43405 | 0,6848964 | serine/threonine kinase 19 |
| ENSG00000236250 | STK19 | -0,306586136 | 0,808552784 | 0,43405 | 0,6848964 | serine/threonine kinase 19 |
| ENSG00000048740 | CELF2 | -0,30665765 | 0,808512705 | 0,43323 | 0,68455289 | CUGBP Elav-like family member 2 |
| ENSG00000113119 | TMCO6 | -0,307103463 | 0,808262902 | 0,41046 | 0,66848416 | transmembrane and coiled-coil domains 6 |
| ENSG00000100836 | PABPN1 | -0,30719872 | 0,808209537 | 0,30974 | 0,59807209 | poly(A) binding protein nuclear 1 |
| ENSG00000269019 | HOMER3-AS1 | -0,307527396 | 0,80802543 | 0,32295 | 0,60584543 | HOMER3 antisense RNA 1 |
| ENSG00000244476 | ERVFRD-1 | -0,307575265 | 0,80799862 | 0,35173 | 0,62692823 | endogenous retrovirus group FRD member 1, envelope |
| ENSG00000083067 | TRPM3 | -0,307580679 | 0,807995588 | 0,23143 | 0,54440536 | transient receptor potential cation channel subfamily M member 3 |
| ENSG00000176974 | SHMT1 | -0,307619649 | 0,807973763 | 0,35748 | 0,63133849 | serine hydroxymethyltransferase 1 |
| ENSG00000106278 | PTPRZ1 | -0,307778771 | 0,807884652 | 0,51216 | 0,73911262 | protein tyrosine phosphatase receptor type Z1 |
| ENSG00000186166 | CENATAC | -0,308018903 | 0,807750194 | 0,28437 | 0,58120847 | centrosomal AT-AC splicing factor |
| ENSG00000275438 | AL031008,1 | -0,308119353 | 0,807693955 | 0,67909 | 0,84221681 | novel transcript |
| ENSG00000253973 | AC079296,1 | -0,308295267 | 0,807595475 | 0,4361 | 0,68655459 | novel transcript |
| ENSG00000286828 | AC008883,3 | -0,308295937 | 0,8075951 | 0,2656 | 0,56848959 | novel transcript |
| ENSG00000183389 | OR56A4 | -0,308419936 | 0,807525691 | 0,52117 | 0,74558236 | olfactory receptor family 56 subfamily A member 4 |
| ENSG00000180919 | OR56B4 | -0,308733875 | 0,807349988 | 0,77531 | 0,89624737 | olfactory receptor family 56 subfamily B member 4 |
| ENSG00000137806 | NDUFAF1 | -0,308916044 | 0,80724805 | 0,27948 | 0,57745127 | NADH:ubiquinone oxidoreductase complex assembly factor 1 |
| ENSG00000124535 | WRNIP1 | -0,308917246 | 0,807247377 | 0,31329 | 0,60005442 | WRN helicase interacting protein 1 |

| | | | | | | |
|---|---|---|---|---|---|---|
| ENSG00000203950 | RTL8A | -0,309028452 | 0,807185156 | 0,23173 | 0,54442759 | retrotransposon Gag like 8A |
| ENSG00000214941 | ZSWIM7 | -0,309114811 | 0,807136839 | 0,27582 | 0,57533851 | zinc finger SWIM-type containing 7 |
| ENSG00000071859 | FAM50A | -0,30927435 | 0,807047588 | 0,24044 | 0,54955956 | family with sequence similarity 50 member A |
| ENSG00000221947 | XKR9 | -0,309312154 | 0,80702644 | 0,28386 | 0,5807907 | XK related 9 |
| ENSG00000169059 | VCX3A | -0,309323968 | 0,807019832 | 0,69508 | 0,85206252 | variable charge X-linked 3A |
| ENSG00000128923 | MINDY2 | -0,309364733 | 0,806997029 | 0,1471 | 0,47471707 | MINDY lysine 48 deubiquitinase 2 |
| ENSG00000105486 | LIG1 | -0,309387292 | 0,80698441 | 0,27632 | 0,57542877 | DNA ligase 1 |
| ENSG00000273319 | AC058791,1 | -0,309400769 | 0,806976872 | 0,54522 | 0,7609742 | novel transcript |
| ENSG00000182180 | MRPS16 | -0,309491553 | 0,806926093 | 0,28159 | 0,57891473 | mitochondrial ribosomal protein S16 |
| ENSG00000169752 | NRG4 | -0,309601207 | 0,806864764 | 0,28859 | 0,58455903 | neuregulin 4 |
| ENSG00000274317 | LINC02334 | -0,309614133 | 0,806857535 | 0,22959 | 0,54311366 | long intergenic non-protein coding RNA 2334 |
| ENSG00000276794 | AC108108,1 | -0,309766229 | 0,806772476 | 0,24742 | 0,5549105 | glucuronidase, beta (GUSB) pseudogene |
| ENSG00000173757 | STAT5B | -0,309823829 | 0,806740266 | 0,15976 | 0,4870975 | signal transducer and activator of transcription 5B |
| ENSG00000163684 | RPP14 | -0,309871398 | 0,806713667 | 0,29801 | 0,59038354 | ribonuclease P/MRP subunit p14 |
| ENSG00000266537 | SPDYE22P | -0,310036863 | 0,806621149 | 0,61027 | 0,80221019 | speedy/RINGO cell cycle regulator family member E22, pseudogene |
| ENSG00000219926 | OR7E104P | -0,310041743 | 0,80661842 | 0,64489 | 0,82173349 | olfactory receptor family 7 subfamily E member 104 pseudogene |
| ENSG00000207174 | SNORD116-15 | -0,310226563 | 0,806515093 | 0,50999 | 0,73758064 | small nucleolar RNA, C/D box 116-15 |
| ENSG00000287156 | AL138479,2 | -0,310256798 | 0,806498191 | 0,75588 | 0,8857133 | novel transcript, antisense to BMP4 |
| ENSG00000251664 | PCDHA12 | -0,310301356 | 0,806473282 | 0,20275 | 0,52269757 | protocadherin alpha 12 |
| ENSG00000249464 | LINC01091 | -0,310388256 | 0,806424706 | 0,41381 | 0,67085666 | long intergenic non-protein coding RNA 1091 |
| ENSG00000111886 | GABRR2 | -0,310467668 | 0,806380318 | 0,57592 | 0,78043323 | gamma-aminobutyric acid type A receptor subunit rho2 |
| ENSG00000286723 | AL021368,5 | -0,310472551 | 0,806377589 | 0,51381 | 0,74005591 | novel transcript |
| ENSG00000259920 | AC007938,1 | -0,310497985 | 0,806363373 | 0,64053 | 0,81950381 | novel transcript |
| ENSG00000270480 | AC073413,1 | -0,310587169 | 0,806313527 | 0,58398 | 0,78572744 | PQ loop repeat containing 1 (PQLC1) pseudogene |
| ENSG00000148288 | GBGT1 | -0,310597704 | 0,806307639 | 0,32292 | 0,60584543 | globoside alpha-1,3-N-acetylgalactosaminyltransferase 1 (FORS blood group) |
| ENSG00000252757 | RN7SKP96 | -0,31063069 | 0,806289204 | 0,6881 | 0,84808917 | RN7SK pseudogene 96 |
| ENSG00000151117 | TMEM86A | -0,310637354 | 0,80628548 | 0,26547 | 0,56829304 | transmembrane protein 86A |
| ENSG00000184906 | AMYH020208 | -0,310704339 | 0,806248044 | 0,71121 | 0,86096523 | methylenetetrahydrofolate dehydrogenase (NADP+ dependent) 1 like pseudogene |

| | | | | | | |
|---|---|---|---|---|---|---|
| ENSG00000180867 | PDIA3P1 | -0,310797741 | 0,806195848 | 0,36513 | 0,63537609 | protein disulfide isomerase family A member 3 pseudogene 1 |
| ENSG00000287650 | AL390071,1 | -0,310869405 | 0,806155802 | 0,39826 | 0,66085176 | novel transcript, antisense to NBEA |
| ENSG00000125901 | MRPS26 | -0,31096626 | 0,806101683 | 0,23438 | 0,5461771 | mitochondrial ribosomal protein S26 |
| ENSG00000276533 | AC018926,2 | -0,311035483 | 0,806063006 | 0,72581 | 0,86937392 | novel transcript, sense intronic to RAB27A |
| ENSG00000160870 | CYP3A7 | -0,311116756 | 0,806017599 | 0,64484 | 0,82171358 | cytochrome P450 family 3 subfamily A member 7 |
| ENSG00000176700 | SCAND2P | -0,311125799 | 0,806012546 | 0,29254 | 0,58705154 | SCAN domain containing 2 pseudogene |
| ENSG00000154914 | USP43 | -0,311260181 | 0,805937473 | 0,34863 | 0,62475089 | ubiquitin specific peptidase 43 |
| ENSG00000276672 | AL161891,1 | -0,311260247 | 0,805937436 | 0,36184 | 0,63352908 | novel transcript, sense intronic to RFC3 |
| ENSG00000138028 | CGREF1 | -0,311265268 | 0,805934631 | 0,59954 | 0,79552021 | cell growth regulator with EF-hand domain 1 |
| ENSG00000206503 | HLA-A | -0,311284492 | 0,805923892 | 0,24313 | 0,55199251 | major histocompatibility complex, class I, A |
| ENSG00000236274 | AC004865,1 | -0,311292944 | 0,805919171 | 0,63628 | 0,81699776 | zinc finger protein pseudogene |
| ENSG00000158717 | RNF166 | -0,311298707 | 0,805915951 | 0,28755 | 0,58395529 | ring finger protein 166 |
| ENSG00000173862 | AC008080,1 | -0,311327733 | 0,805899737 | 0,80094 | 0,90833055 | novel transcript |
| ENSG00000196557 | CACNA1H | -0,311431966 | 0,805841514 | 0,41723 | 0,6738387 | calcium voltage-gated channel subunit alpha1 H |
| ENSG00000140104 | CLBA1 | -0,311443262 | 0,805835204 | 0,17317 | 0,49786688 | clathrin binding box of aftiphilin containing 1 |
| ENSG00000272374 | Z97832,2 | -0,3114889 | 0,805809713 | 0,21529 | 0,53398925 | novel transcript, antisense to SCUBE3 |
| ENSG00000172081 | MOB3A | -0,31162836 | 0,805731822 | 0,2609 | 0,56405761 | MOB kinase activator 3A |
| ENSG00000285799 | AL645929,2 | -0,311741686 | 0,805668533 | 0,50905 | 0,7370222 | MHC class I polypeptide-related sequence F pseudogene |
| ENSG00000245552 | AP000787,1 | -0,312028729 | 0,80550825 | 0,17742 | 0,50191142 | novel transcript, antisense to SESN3 |
| ENSG00000226010 | AL355852,1 | -0,312039614 | 0,805502173 | 0,57592 | 0,78043323 | zinc finger protein pseudogene |
| ENSG00000256229 | ZNF486 | -0,312207395 | 0,805408501 | 0,2301 | 0,54335232 | zinc finger protein 486 |
| ENSG00000137103 | TMEM8B | -0,31229053 | 0,805362091 | 0,16775 | 0,49395055 | transmembrane protein 8B |
| ENSG00000276968 | AL158196,1 | -0,312480296 | 0,805256164 | 0,61349 | 0,8035281 | novel transcript, antisense to NUDT15 |
| ENSG00000107331 | ABCA2 | -0,312487543 | 0,805252119 | 0,20114 | 0,52181085 | ATP binding cassette subfamily A member 2 |
| ENSG00000235413 | KRT18P63 | -0,312558948 | 0,805212265 | 0,49152 | 0,72510994 | keratin 18 pseudogene 63 |
| ENSG00000274611 | TBC1D3 | -0,312601282 | 0,805188637 | 0,19953 | 0,52047306 | TBC1 domain family member 3 |
| ENSG00000167106 | FAM102A | -0,312661754 | 0,805154888 | 0,30038 | 0,59177961 | family with sequence similarity 102 member A |
| ENSG00000212464 | SNORA12 | -0,312731772 | 0,805115812 | 0,37434 | 0,64275601 | small nucleolar RNA, H/ACA box 12 |
| ENSG00000268947 | AC002128,1 | -0,312811262 | 0,805071453 | 0,29976 | 0,59160746 | novel transcript |
| ENSG00000231028 | LINC00271 | -0,312816949 | 0,805068279 | 0,18593 | 0,50921804 | long intergenic non-protein coding RNA 271 |
| ENSG00000144451 | SPAG16 | -0,313050438 | 0,804937996 | 0,07488 | 0,35909576 | sperm associated antigen 16 |
| ENSG00000214694 | ARHGEF33 | -0,313156818 | 0,804878644 | 0,50315 | 0,73248308 | Rho guanine nucleotide exchange factor 33 |

| | | | | | | |
|---|---|---|---|---|---|---|
| ENSG00000164674 | SYTL3 | -0,313405455 | 0,804739942 | 0,52274 | 0,74689824 | synaptotagmin like 3 |
| ENSG00000226746 | SMCR5 | -0,313421885 | 0,804730777 | 0,35191 | 0,62699036 | Smith-Magenis syndrome chromosome region, candidate 5 |
| ENSG00000244607 | CCDC13 | -0,313525045 | 0,804673237 | 0,42949 | 0,68224982 | coiled-coil domain containing 13 |
| ENSG00000243854 | RN7SL67P | -0,313639889 | 0,804609184 | 0,51395 | 0,74022017 | RNA, 7SL, cytoplasmic 67, pseudogene |
| ENSG00000100628 | ASB2 | -0,313746498 | 0,804549729 | 0,79789 | 0,90685415 | ankyrin repeat and SOCS box containing 2 |
| ENSG00000276938 | AC073135,4 | -0,313891715 | 0,80446875 | 0,64309 | 0,82098995 | family with sequence similarity 157, member A |
| ENSG00000196436 | NPIPB15 | -0,313906056 | 0,804460753 | 0,23725 | 0,54819454 | nuclear pore complex interacting protein family member B15 |
| ENSG00000276863 | AC124319,2 | -0,314095659 | 0,804355036 | 0,70022 | 0,85526044 | novel transcript, antisense to RNF213 |
| ENSG00000125851 | PCSK2 | -0,314127415 | 0,804337331 | 0,36465 | 0,63504684 | proprotein convertase subtilisin/kexin type 2 |
| ENSG00000167775 | CD320 | -0,314437524 | 0,804164456 | 0,17679 | 0,50110131 | CD320 molecule |
| ENSG00000229759 | MRPS18AP1 | -0,314480904 | 0,804140277 | 0,5981 | 0,79484486 | mitochondrial ribosomal protein S18A pseudogene 1 |
| ENSG00000184990 | SIVA1 | -0,314520023 | 0,804118472 | 0,22866 | 0,54252903 | SIVA1 apoptosis inducing factor |
| ENSG00000213886 | UBD | -0,314640369 | 0,804051398 | 0,87317 | 0,94385148 | ubiquitin D |
| ENSG00000287624 | AL590666,5 | -0,314655047 | 0,804043217 | 0,53771 | 0,75631707 | novel transcript, antisense to PRCC |
| ENSG00000136535 | TBR1 | -0,314657569 | 0,804041811 | 0,46396 | 0,70468629 | T-box brain transcription factor 1 |
| ENSG00000279170 | TSTD3 | -0,314676269 | 0,80403139 | 0,27347 | 0,57362145 | thiosulfate sulfurtransferase like domain containing 3 |
| ENSG00000168453 | HR | -0,314680023 | 0,804029298 | 0,66819 | 0,83536064 | HR lysine demethylase and nuclear receptor corepressor |
| ENSG00000143401 | ANP32E | -0,31471026 | 0,804012446 | 0,24137 | 0,55033747 | acidic nuclear phosphoprotein 32 family member E |
| ENSG00000114446 | IFT57 | -0,314719617 | 0,804007232 | 0,32193 | 0,60577115 | intraflagellar transport 57 |
| ENSG00000179869 | ABCA13 | -0,314766489 | 0,803981111 | 0,32927 | 0,61035569 | ATP binding cassette subfamily A member 13 |
| ENSG00000235659 | AL391987,4 | -0,31499247 | 0,803855186 | 0,70943 | 0,85983746 | novel transcript |
| ENSG00000137726 | FXYD6 | -0,31503245 | 0,80383291 | 0,24369 | 0,55235189 | FXYD domain containing ion transport regulator 6 |
| ENSG00000261697 | AC010531,4 | -0,315172551 | 0,803754853 | 0,60566 | 0,7988563 | novel transcript, sense intronic to C16orf95 |
| ENSG00000215475 | SIAH3 | -0,315209871 | 0,803734062 | 0,50764 | 0,7358546 | siah E3 ubiquitin protein ligase family member 3 |
| ENSG00000166800 | LDHAL6A | -0,315215231 | 0,803731076 | 0,39333 | 0,6563386 | lactate dehydrogenase A like 6A |
| ENSG00000104177 | MYEF2 | -0,315376891 | 0,803641019 | 0,07832 | 0,36577348 | myelin expression factor 2 |
| ENSG00000270402 | AC011458,1 | -0,315410943 | 0,803622051 | 0,73674 | 0,87529332 | ankyrin repeat domain 26 (ANKRD26) pseudogene |
| ENSG00000130158 | DOCK6 | -0,315625524 | 0,803502533 | 0,44607 | 0,69284796 | dedicator of cytokinesis 6 |
| ENSG00000124593 | AL365205,1 | -0,31571864 | 0,803450673 | 0,2921 | 0,58691588 | novel protein |
| ENSG00000175414 | ARL10 | -0,315722147 | 0,80344872 | 0,20808 | 0,52756478 | ADP ribosylation factor like GTPase 10 |
| ENSG00000283683 | MYOCOS | -0,315928428 | 0,803333849 | 0,42404 | 0,6778073 | myocilin opposite strand |
| ENSG00000142449 | FBN3 | -0,315933074 | 0,803331262 | 0,31352 | 0,60029566 | fibrillin 3 |

| | | | | | | |
|---|---|---|---|---|---|---|
| ENSG00000241345 | AC004690,2 | -0,316018893 | 0,803283477 | 0,66775 | 0,83516218 | novel transcript |
| ENSG00000132749 | TESMIN | -0,316059614 | 0,803260804 | 0,38571 | 0,65122858 | testis expressed metallothionein like protein |
| ENSG00000165555 | NOXRED1 | -0,316159341 | 0,803205281 | 0,42662 | 0,68002571 | NADP dependent oxidoreductase domain containing 1 |
| ENSG00000099385 | BCL7C | -0,316190546 | 0,803187908 | 0,22436 | 0,54029213 | BAF chromatin remodeling complex subunit BCL7C |
| ENSG00000176695 | OR4F17 | -0,316275798 | 0,803140447 | 0,29166 | 0,58655914 | olfactory receptor family 4 subfamily F member 17 |
| ENSG00000261783 | AC009054,2 | -0,316291357 | 0,803131785 | 0,71287 | 0,86184574 | novel transcript, sense intronic to CFDP1 |
| ENSG00000125351 | UPF3B | -0,316328305 | 0,803111217 | 0,13228 | 0,4573773 | UPF3B regulator of nonsense mediated mRNA decay |
| ENSG00000234798 | DXO | -0,316372113 | 0,80308683 | 0,38523 | 0,65101234 | decapping exoribonuclease |
| ENSG00000224313 | DXO | -0,316372113 | 0,80308683 | 0,38523 | 0,65101234 | decapping exoribonuclease |
| ENSG00000225682 | DXO | -0,316372113 | 0,80308683 | 0,38523 | 0,65101234 | decapping exoribonuclease |
| ENSG00000267419 | ZNF56 | -0,316395192 | 0,803073984 | 0,50012 | 0,73078774 | zinc finger protein 56 |
| ENSG00000264673 | AC098850,1 | -0,316786696 | 0,802856083 | 0,69229 | 0,85023519 | novel transcript |
| ENSG00000286092 | AC117503,5 | -0,316841355 | 0,802825666 | 0,38615 | 0,65158326 | novel transcript, antisense to ATP6V0A2 |
| ENSG00000115977 | AAK1 | -0,316961931 | 0,802758571 | 0,20834 | 0,52769445 | AP2 associated kinase 1 |
| ENSG00000260018 | AC040169,1 | -0,31697746 | 0,802749931 | 0,49442 | 0,72678186 | novel transcript, antisense to MBTPS1 |
| ENSG00000135540 | NHSL1 | -0,317025904 | 0,802722975 | 0,283 | 0,58030723 | NHS like 1 |
| ENSG00000132535 | DLG4 | -0,31711342 | 0,802674282 | 0,15693 | 0,48425071 | discs large MAGUK scaffold protein 4 |
| ENSG00000275621 | LRRC37A3 | -0,317186218 | 0,802633781 | 0,63962 | 0,81915869 | leucine rich repeat containing 37 member A3 |
| ENSG00000146243 | IRAK1BP1 | -0,317317436 | 0,802560782 | 0,08564 | 0,3794467 | interleukin 1 receptor associated kinase 1 binding protein 1 |
| ENSG00000084731 | KIF3C | -0,317589826 | 0,802409268 | 0,24942 | 0,55652928 | kinesin family member 3C |
| ENSG00000139233 | LLPH | -0,317619745 | 0,802392627 | 0,28261 | 0,58002653 | LLP homolog, long-term synaptic facilitation factor |
| ENSG00000139269 | INHBE | -0,317639512 | 0,802381633 | 0,66448 | 0,83277805 | inhibin subunit beta E |
| ENSG00000250500 | AC017007,4 | -0,317644708 | 0,802378744 | 0,72327 | 0,86784071 | eukaryotic translation initiation factor 2, subunit 3 gamma, 52kDa (EIF2S3) pseudogene |
| ENSG00000177212 | OR2T33 | -0,31768982 | 0,802353654 | 0,56375 | 0,77255679 | olfactory receptor family 2 subfamily T member 33 |
| ENSG00000281021 | AL078581,4 | -0,317692984 | 0,802351895 | 0,71171 | 0,86130402 | novel transcript |
| ENSG00000111728 | ST8SIA1 | -0,317845913 | 0,802266848 | 0,16261 | 0,48945635 | ST8 alpha-N-acetyl-neuraminide alpha-2,8-sialyltransferase 1 |
| ENSG00000114021 | NIT2 | -0,317987505 | 0,802188114 | 0,22332 | 0,53953208 | nitrilase family member 2 |
| ENSG00000171241 | SHCBP1 | -0,318242631 | 0,802046268 | 0,17455 | 0,49884018 | SHC binding and spindle associated 1 |
| ENSG00000259453 | AC087286,4 | -0,318325344 | 0,802000286 | 0,51636 | 0,74198903 | novel transcript, sense intronic to AKAP13 |
| ENSG00000229017 | LINC01277 | -0,318454328 | 0,801928586 | 0,75845 | 0,88727479 | long intergenic non-protein coding RNA 1277 |

| | | | | | | |
|---|---|---|---|---|---|---|
| ENSG00000275467 | AC083806,3 | -0,318831589 | 0,801718912 | 0,76049 | 0,88845438 | novel transcript |
| ENSG00000101746 | NOL4 | -0,318920665 | 0,801669413 | 0,54171 | 0,75882662 | nucleolar protein 4 |
| ENSG00000173210 | ABLIM3 | -0,318953597 | 0,801651113 | 0,36198 | 0,63352908 | actin binding LIM protein family member 3 |
| ENSG00000285570 | AL590666,4 | -0,319028119 | 0,801609705 | 0,63115 | 0,81395758 | Novel transcript, antisense to CRABP2 |
| ENSG00000235280 | MCF2L-AS1 | -0,319035229 | 0,801605755 | 0,51544 | 0,74148081 | MCF2L antisense RNA 1 |
| ENSG00000173141 | MRPL57 | -0,319291714 | 0,801463256 | 0,24977 | 0,55657567 | mitochondrial ribosomal protein L57 |
| ENSG00000276740 | AL445649,1 | -0,319333557 | 0,801440012 | 0,48816 | 0,72311306 | novel transcript, sense intronic to FAM155A |
| ENSG00000127837 | AAMP | -0,319773341 | 0,801195742 | 0,26299 | 0,56587311 | angio associated migratory cell protein |
| ENSG00000231079 | KIF5C-AS1 | -0,320144739 | 0,800989514 | 0,24993 | 0,55657567 | KIF5C antisense RNA 1 |
| ENSG00000119782 | FKBP1B | -0,320148464 | 0,800987446 | 0,3022 | 0,59294633 | FKBP prolyl isomerase 1B |
| ENSG00000119333 | DYNC2I2 | -0,320203262 | 0,800957023 | 0,19934 | 0,52034382 | dynein 2 intermediate chain 2 |
| ENSG00000089050 | RBBP9 | -0,320312635 | 0,800896303 | 0,20893 | 0,52800078 | RB binding protein 9, serine hydrolase |
| ENSG00000144791 | LIMD1 | -0,320365881 | 0,800866744 | 0,43233 | 0,68411566 | LIM domains containing 1 |
| ENSG00000198937 | CCDC167 | -0,320450323 | 0,80081987 | 0,14263 | 0,47004319 | coiled-coil domain containing 167 |
| ENSG00000125492 | BARHL1 | -0,320457031 | 0,800816147 | 0,71894 | 0,8656088 | BarH like homeobox 1 |
| ENSG00000274315 | AC009318,3 | -0,320507126 | 0,800788341 | 0,37698 | 0,64554193 | novel transcript, antisense to FAR2 |
| ENSG00000183044 | ABAT | -0,320570575 | 0,800753123 | 0,34366 | 0,62107724 | 4-aminobutyrate aminotransferase |
| ENSG00000287189 | AL121956,7 | -0,320667603 | 0,80069927 | 0,18744 | 0,51074735 | novel transcript |
| ENSG00000144550 | CPNE9 | -0,320739233 | 0,800659516 | 0,48798 | 0,72295768 | copine family member 9 |
| ENSG00000197860 | SGTB | -0,320808402 | 0,80062113 | 0,11438 | 0,42986344 | small glutamine rich tetratricopeptide repeat containing beta |
| ENSG00000108963 | DPH1 | -0,3208178 | 0,800615915 | 0,35906 | 0,63201238 | diphthamide biosynthesis 1 |
| ENSG00000273686 | B2M | -0,32084967 | 0,800598229 | 0,6443 | 0,82151477 | beta-2-microglobulin |
| ENSG00000233647 | NENFP1 | -0,321137325 | 0,800438616 | 0,63334 | 0,81542248 | neudesin neurotrophic factor pseudogene 1 |
| ENSG00000243069 | ARHGEF26-AS | -0,321153946 | 0,800429394 | 0,20166 | 0,52214043 | ARHGEF26 antisense RNA 1 |
| ENSG00000229808 | AL391825,1 | -0,321169796 | 0,8004206 | 0,56997 | 0,77703861 | proteasome (prosome, macropain) activator subunit 3 (PA28 gamma; Ki) pseudogene |
| ENSG00000224514 | LINC00620 | -0,321375441 | 0,800306515 | 0,69743 | 0,8533426 | long intergenic non-protein coding RNA 620 |
| ENSG00000174840 | PDE12 | -0,321422615 | 0,800280347 | 0,1573 | 0,48443207 | phosphodiesterase 12 |
| ENSG00000223532 | HLA-B | -0,321488547 | 0,800243774 | 0,49506 | 0,72738312 | major histocompatibility complex, class I, B |
| ENSG00000228157 | AC007952,2 | -0,321516691 | 0,800228164 | 0,44855 | 0,6944286 | novel transcript |
| ENSG00000149557 | FEZ1 | -0,321538579 | 0,800216022 | 0,27371 | 0,5737685 | fasciculation and elongation protein zeta 1 |
| ENSG00000141519 | CCDC40 | -0,321560365 | 0,800203939 | 0,16324 | 0,48972962 | coiled-coil domain containing 40 |

| | | | | | | |
|---|---|---|---|---|---|---|
| ENSG00000277507 | AP000351,2 | -0,321711629 | 0,800120043 | 0,63999 | 0,81935089 | D-dopachrome tautomerase (DDT) pseudogene |
| ENSG00000226756 | AC007365,1 | -0,321760425 | 0,800092981 | 0,69193 | 0,85002709 | novel transcript |
| ENSG00000223591 | CENPVL1 | -0,321987138 | 0,79996726 | 0,3965 | 0,6591566 | centromere protein V like 1 |
| ENSG00000283093 | CENPVL2 | -0,321987138 | 0,79996726 | 0,3965 | 0,6591566 | centromere protein V like 2 |
| ENSG00000267127 | AC090360,1 | -0,322024065 | 0,799946784 | 0,32094 | 0,6056608 | novel protein |
| ENSG00000214544 | GTF2IRD2P1 | -0,32203146 | 0,799942684 | 0,58395 | 0,78572744 | GTF2I repeat domain containing 2 pseudogene 1 |
| ENSG00000111788 | AC009533,1 | -0,322052902 | 0,799930795 | 0,27385 | 0,57384741 | DEAD/H (Asp-Glu-Ala-Asp/His) box polypeptide like pseudogene |
| ENSG00000054967 | RELT | -0,322329682 | 0,799777344 | 0,16477 | 0,49092529 | RELT TNF receptor |
| ENSG00000248275 | TRIM52-AS1 | -0,322363473 | 0,799758612 | 0,1723 | 0,49714741 | TRIM52 antisense RNA 1 (head to head) |
| ENSG00000101199 | ARFGAP1 | -0,32251928 | 0,799672245 | 0,25234 | 0,55796133 | ADP ribosylation factor GTPase activating protein 1 |
| ENSG00000279177 | CTBP2P9 | -0,322583947 | 0,799636401 | 0,5041 | 0,73311005 | CTBP2 pseudogene 9 |
| ENSG00000280330 | CTBP2P10 | -0,322583947 | 0,799636401 | 0,5041 | 0,73311005 | CTBP2 pseudogene 10 |
| ENSG00000251034 | AC087854,1 | -0,322622333 | 0,799615126 | 0,19446 | 0,51562158 | novel transcript |
| ENSG00000151532 | VTI1A | -0,322678432 | 0,799584033 | 0,28635 | 0,58267001 | vesicle transport through interaction with t-SNAREs 1A |
| ENSG00000099866 | MADCAM1 | -0,322843018 | 0,79949282 | 0,57405 | 0,77929088 | mucosal vascular addressin cell adhesion molecule 1 |
| ENSG00000229891 | LINC01315 | -0,322854822 | 0,799486279 | 0,29567 | 0,58930611 | long intergenic non-protein coding RNA 1315 |
| ENSG00000167778 | SPRYD3 | -0,323219224 | 0,799284366 | 0,22843 | 0,54241083 | SPRY domain containing 3 |
| ENSG00000231690 | LINC00574 | -0,323229682 | 0,799278572 | 0,66524 | 0,83339621 | long intergenic non-protein coding RNA 574 |
| ENSG00000111186 | WNT5B | -0,323273639 | 0,79925422 | 0,54982 | 0,76429591 | Wnt family member 5B |
| ENSG00000270638 | AL023806,1 | -0,323406721 | 0,799180496 | 0,29221 | 0,58692263 | novel transcript |
| ENSG00000172354 | GNB2 | -0,32365358 | 0,79904376 | 0,10998 | 0,42231571 | G protein subunit beta 2 |
| ENSG00000100031 | GGT1 | -0,323679681 | 0,799029304 | 0,51911 | 0,74382271 | gamma-glutamyltransferase 1 |
| ENSG00000229536 | LINC02572 | -0,323754932 | 0,798987628 | 0,499 | 0,73002372 | long intergenic non-protein coding RNA 2572 |
| ENSG00000239880 | RALBP1P1 | -0,323832317 | 0,798944772 | 0,7615 | 0,88879633 | RALBP1 pseudogene 1 |
| ENSG00000132016 | C19orf57 | -0,323844099 | 0,798938247 | 0,14537 | 0,47300165 | chromosome 19 open reading frame 57 |
| ENSG00000288152 | C19orf57 | -0,323844099 | 0,798938247 | 0,14537 | 0,47300165 | chromosome 19 open reading frame 57 |
| ENSG00000184056 | VPS33B | -0,323846178 | 0,798937095 | 0,26156 | 0,56452671 | VPS33B late endosome and lysosome associated |
| ENSG00000247324 | AC010547,1 | -0,323860727 | 0,798929039 | 0,3417 | 0,6197312 | novel transcript, antisense to ZNF23 |
| ENSG00000141622 | RNF165 | -0,324033722 | 0,798833244 | 0,26242 | 0,5653453 | ring finger protein 165 |
| ENSG00000167740 | CYB5D2 | -0,324101072 | 0,798795953 | 0,18656 | 0,509961 | cytochrome b5 domain containing 2 |
| ENSG00000261971 | MMP25-AS1 | -0,324118159 | 0,798786492 | 0,39427 | 0,65724328 | MMP25 antisense RNA 1 |
| ENSG00000255875 | PABPN1P2 | -0,32415728 | 0,798764832 | 0,58241 | 0,7847455 | PABPN1 pseudogene 2 |

| | | | | | | |
|---|---|---|---|---|---|---|
| ENSG00000102468 | HTR2A | -0,324167436 | 0,798759209 | 0,61139 | 0,80316168 | 5-hydroxytryptamine receptor 2A |
| ENSG00000270823 | AC007938,2 | -0,324278258 | 0,798697854 | 0,31624 | 0,60226233 | novel transcript, antisense to MEST |
| ENSG00000130305 | NSUN5 | -0,324286235 | 0,798693438 | 0,23414 | 0,54597007 | NOP2/Sun RNA methyltransferase 5 |
| ENSG00000283463 | HSFX4 | -0,32433887 | 0,798664298 | 0,67708 | 0,84113215 | heat shock transcription factor family, X-linked member 4 |
| ENSG00000117394 | SLC2A1 | -0,324486384 | 0,79858264 | 0,3353 | 0,61497665 | solute carrier family 2 member 1 |
| ENSG00000226677 | IGBP1P1 | -0,324591466 | 0,798524476 | 0,67872 | 0,84182956 | immunoglobulin (CD79A) binding protein 1 pseudogene 1 |
| ENSG00000246067 | RAB30-DT | -0,324650238 | 0,798491946 | 0,29311 | 0,58745839 | RAB30 divergent transcript |
| ENSG00000188051 | TMEM221 | -0,324654207 | 0,79848975 | 0,50233 | 0,73197662 | transmembrane protein 221 |
| ENSG00000282805 | AC023315,3 | -0,324688977 | 0,798470506 | 0,71751 | 0,86483656 | novel transcript, sense intronic AATF |
| ENSG00000277589 | AC244093,4 | -0,324688977 | 0,798470506 | 0,71751 | 0,86483656 | novel transcript, sense intronic AATF |
| ENSG00000253392 | AC119403,1 | -0,324821302 | 0,798397273 | 0,61235 | 0,8035281 | novel transcript, antisense to ZNF57 |
| ENSG00000164463 | CREBRF | -0,324907582 | 0,798349526 | 0,28452 | 0,58138883 | CREB3 regulatory factor |
| ENSG00000259728 | LINC00933 | -0,325266177 | 0,798151114 | 0,64265 | 0,82090182 | long intergenic non-protein coding RNA 933 |
| ENSG00000277047 | AL390964,1 | -0,325314182 | 0,798124556 | 0,62516 | 0,81024216 | novel transcript |
| ENSG00000129493 | HEATR5A | -0,325325603 | 0,798118238 | 0,09935 | 0,40586731 | HEAT repeat containing 5A |
| ENSG00000180881 | CAPS2 | -0,325346548 | 0,798106651 | 0,36944 | 0,63917123 | calcyphosine 2 |
| ENSG00000244468 | AC093001,1 | -0,32538754 | 0,798083975 | 0,65137 | 0,82534198 | novel transcript |
| ENSG00000231057 | AC096637,2 | -0,325416216 | 0,798068111 | 0,33026 | 0,61114402 | novel transcript |
| ENSG00000170004 | CHD3 | -0,325419182 | 0,79806647 | 0,34178 | 0,6197312 | chromodomain helicase DNA binding protein 3 |
| ENSG00000126856 | PRDM7 | -0,325467292 | 0,798039858 | 0,45991 | 0,70199338 | PR/SET domain 7 |
| ENSG00000175485 | OR52W1 | -0,325632714 | 0,797948358 | 0,59719 | 0,79436765 | olfactory receptor family 52 subfamily W member 1 |
| ENSG00000175575 | PAAF1 | -0,325721561 | 0,797899219 | 0,24452 | 0,55258874 | proteasomal ATPase associated factor 1 |
| ENSG00000183506 | PI4KAP2 | -0,325972558 | 0,797760414 | 0,21133 | 0,53032107 | phosphatidylinositol 4-kinase alpha pseudogene 2 |
| ENSG00000277769 | RBM22P4 | -0,326047936 | 0,797718734 | 0,56173 | 0,77136826 | RNA binding motif protein 22 pseudogene 4 |
| ENSG00000176681 | LRRC37A | -0,326054628 | 0,797715033 | 0,68851 | 0,84821636 | leucine rich repeat containing 37A |
| ENSG00000131725 | WDR44 | -0,326054894 | 0,797714887 | 0,31983 | 0,60470232 | WD repeat domain 44 |
| ENSG00000286688 | AP002748,6 | -0,326276088 | 0,79759259 | 0,50786 | 0,73602012 | novel transcript, antisense to BBS1 |
| ENSG00000255358 | NDUFAF2P2 | -0,326397045 | 0,797525722 | 0,64111 | 0,81993567 | NDUFAF2 pseudogene 2 |
| ENSG00000186469 | GNG2 | -0,326416886 | 0,797514754 | 0,48256 | 0,71901075 | G protein subunit gamma 2 |
| ENSG00000236197 | PPP1R9A-AS1 | -0,326478798 | 0,79748053 | 0,35308 | 0,62780446 | PPP1R9A antisense RNA 1 |
| ENSG00000177051 | FBXO46 | -0,326782166 | 0,797312855 | 0,2081 | 0,52756478 | F-box protein 46 |
| ENSG00000221116 | SNORD110 | -0,326788298 | 0,797309466 | 0,67426 | 0,83959502 | small nucleolar RNA, C/D box 110 |
| ENSG00000271943 | AC098614,2 | -0,327127618 | 0,797121962 | 0,54136 | 0,75881984 | tropomyosin 4 (TPM4) pseudogene |

| | | | | | | |
|---|---|---|---|---|---|---|
| ENSG00000177542 | SLC25A22 | -0,327166625 | 0,79710041 | 0,27217 | 0,57306433 | solute carrier family 25 member 22 |
| ENSG00000157778 | PSMG3 | -0,327284525 | 0,797035272 | 0,2443 | 0,55253387 | proteasome assembly chaperone 3 |
| ENSG00000250197 | HMGN1P15 | -0,327320123 | 0,797015605 | 0,68293 | 0,8447391 | high mobility group nucleosome binding domain 1 pseudogene 15 |
| ENSG00000025770 | NCAPH2 | -0,327552802 | 0,796887072 | 0,20201 | 0,52230208 | non-SMC condensin II complex subunit H2 |
| ENSG00000015153 | YAF2 | -0,327638774 | 0,796839586 | 0,21839 | 0,53578053 | YY1 associated factor 2 |
| ENSG00000282532 | AC016482,2 | -0,327732346 | 0,796787905 | 0,57537 | 0,7801789 | ribosomal protein S2 (RPS2) pseudogene |
| ENSG00000274308 | AC244093,1 | -0,327732346 | 0,796787905 | 0,57537 | 0,7801789 | ribosomal protein S2 (RPS2) pseudogene |
| ENSG00000272661 | AC021097,1 | -0,327957532 | 0,796663547 | 0,81925 | 0,91696956 | novel transcript, antisense to SMARCD3 |
| ENSG00000007174 | DNAH9 | -0,328014757 | 0,796631948 | 0,14823 | 0,47545606 | dynein axonemal heavy chain 9 |
| ENSG00000231395 | ARL4AP4 | -0,328092205 | 0,796589183 | 0,61487 | 0,80404192 | ADP ribosylation factor like GTPase 4A pseudogene 4 |
| ENSG00000249140 | PRDX2P3 | -0,328363614 | 0,796439338 | 0,56664 | 0,77458849 | peroxiredoxin 2 pseudogene 3 |
| ENSG00000284556 | PRDX2P3 | -0,328363614 | 0,796439338 | 0,56664 | 0,77458849 | peroxiredoxin 2 pseudogene 3 |
| ENSG00000261823 | AC084782,2 | -0,328418728 | 0,796408913 | 0,54726 | 0,76270174 | novel transcript |
| ENSG00000143751 | SDE2 | -0,328523956 | 0,796350826 | 0,27132 | 0,57272978 | SDE2 telomere maintenance homolog |
| ENSG00000254480 | LINC02749 | -0,328624036 | 0,796295585 | 0,65195 | 0,82566892 | long intergenic non-protein coding RNA 2749 |
| ENSG00000267690 | LDLRAD4-AS1 | -0,328950592 | 0,796115363 | 0,65332 | 0,82632712 | LDLRAD4 antisense RNA 1 |
| ENSG00000215374 | FAM66B | -0,329112637 | 0,796025947 | 0,27518 | 0,57491355 | family with sequence similarity 66 member B |
| ENSG00000230185 | C9orf147 | -0,329188819 | 0,795983914 | 0,17341 | 0,49804988 | chromosome 9 open reading frame 147 |
| ENSG00000277957 | SENP3-EIF4A1 | -0,329276078 | 0,795935772 | 0,2229 | 0,53906657 | SENP3-EIF4A1 readthrough (NMD candidate) |
| ENSG00000159761 | C16orf86 | -0,329626549 | 0,79574244 | 0,41702 | 0,6738387 | chromosome 16 open reading frame 86 |
| ENSG00000105404 | RABAC1 | -0,329932831 | 0,795573523 | 0,29054 | 0,58565836 | Rab acceptor 1 |
| ENSG00000139187 | KLRG1 | -0,329996502 | 0,795538413 | 0,21605 | 0,53474184 | killer cell lectin like receptor G1 |
| ENSG00000079102 | RUNX1T1 | -0,330090271 | 0,795486708 | 0,2328 | 0,54542294 | RUNX1 partner transcriptional co-repressor 1 |
| ENSG00000130956 | HABP4 | -0,330520777 | 0,795249367 | 0,168 | 0,49417064 | hyaluronan binding protein 4 |
| ENSG00000090316 | MAEA | -0,330646767 | 0,795179921 | 0,3499 | 0,62563749 | macrophage erythroblast attacher, E3 ubiquitin ligase |
| ENSG00000245281 | AC124242,1 | -0,33076486 | 0,795114833 | 0,3395 | 0,61852579 | novel transcript |
| ENSG00000285152 | RNF31 | -0,330842339 | 0,795072133 | 0,38603 | 0,65149512 | ring finger protein 31 |
| ENSG00000259172 | AC023024,1 | -0,330887207 | 0,795047407 | 0,49724 | 0,72868655 | novel transcript, antisense to SNRPA1 and PCSK6 |
| ENSG00000169635 | HIC2 | -0,330985609 | 0,794993181 | 0,2359 | 0,5471053 | HIC ZBTB transcriptional repressor 2 |
| ENSG00000283529 | AL035685,1 | -0,331138069 | 0,794909173 | 0,57418 | 0,77939135 | somatomedin B and thrombospondin type 1 domain containing (SBSPON) pseudogene |
| ENSG00000282355 | CARS1-AS1 | -0,331150652 | 0,79490224 | 0,62708 | 0,81124422 | CARS1 antisense RNA 1 |

| | | | | | | |
|---|---|---|---|---|---|---|
| ENSG00000236295 | AC007557,3 | -0,331200033 | 0,794875032 | 0,72107 | 0,86669144 | novel transcript |
| ENSG00000247271 | ZBED5-AS1 | -0,331262075 | 0,79484085 | 0,21941 | 0,53676938 | ZBED5 antisense RNA 1 |
| ENSG00000163508 | EOMES | -0,331384765 | 0,794773258 | 0,52718 | 0,75025748 | eomesodermin |
| ENSG00000261221 | ZNF865 | -0,331411581 | 0,794758485 | 0,25231 | 0,55796133 | zinc finger protein 865 |
| ENSG00000274602 | PI4KAP1 | -0,331429526 | 0,7947486 | 0,27652 | 0,575468 | phosphatidylinositol 4-kinase alpha pseudogene 1 |
| ENSG00000114166 | KAT2B | -0,331446723 | 0,794739126 | 0,39768 | 0,66026872 | lysine acetyltransferase 2B |
| ENSG00000137504 | CREBZF | -0,331448 | 0,794738423 | 0,23191 | 0,5446505 | CREB/ATF bZIP transcription factor |
| ENSG00000285841 | AC005692,3 | -0,331551926 | 0,794681175 | 0,29698 | 0,59008525 | novel transcript |
| ENSG00000258520 | AL359317,1 | -0,331554967 | 0,7946795 | 0,60125 | 0,79644593 | novel transcript |
| ENSG00000272788 | AP000864,1 | -0,331570058 | 0,794671187 | 0,57667 | 0,7810482 | novel transcript, sense intronic to SOGA2 |
| ENSG00000278356 | AC005911,1 | -0,331586063 | 0,794662372 | 0,4132 | 0,67067821 | novel transcript |
| ENSG00000171295 | ZNF440 | -0,331711309 | 0,794593387 | 0,20498 | 0,52453221 | zinc finger protein 440 |
| ENSG00000225470 | JPX | -0,331853375 | 0,794515145 | 0,14576 | 0,47326216 | JPX transcript, XIST activator |
| ENSG00000131019 | ULBP3 | -0,332098508 | 0,794380157 | 0,15381 | 0,48214138 | UL16 binding protein 3 |
| ENSG00000198711 | SSBP3-AS1 | -0,332266772 | 0,794287513 | 0,32878 | 0,60992716 | SSBP3 antisense RNA 1 |
| ENSG00000274145 | RILP | -0,332369381 | 0,794231023 | 0,58229 | 0,78466391 | Rab interacting lysosomal protein |
| ENSG00000135406 | PRPH | -0,332515272 | 0,794150711 | 0,73787 | 0,87544492 | peripherin |
| ENSG00000106246 | PTCD1 | -0,332725397 | 0,794035054 | 0,33049 | 0,6112353 | pentatricopeptide repeat domain 1 |
| ENSG00000166313 | APBB1 | -0,332765528 | 0,794012966 | 0,15214 | 0,48061452 | amyloid beta precursor protein binding family B member 1 |
| ENSG00000223639 | CLIC1 | -0,332847108 | 0,793968069 | 0,27489 | 0,57463533 | chloride intracellular channel 1 |
| ENSG00000226248 | CLIC1 | -0,332847108 | 0,793968069 | 0,27489 | 0,57463533 | chloride intracellular channel 1 |
| ENSG00000226417 | CLIC1 | -0,332847108 | 0,793968069 | 0,27489 | 0,57463533 | chloride intracellular channel 1 |
| ENSG00000213719 | CLIC1 | -0,332847108 | 0,793968069 | 0,27489 | 0,57463533 | chloride intracellular channel 1 |
| ENSG00000267713 | AC012433,1 | -0,332869364 | 0,79395582 | 0,76527 | 0,89082001 | NADH dehydrogenase pseudogene |
| ENSG00000182963 | GJC1 | -0,332904723 | 0,793936361 | 0,30587 | 0,59512079 | gap junction protein gamma 1 |
| ENSG00000144026 | ZNF514 | -0,332964751 | 0,793903328 | 0,09516 | 0,39774693 | zinc finger protein 514 |
| ENSG00000248268 | AC010275,1 | -0,333000364 | 0,793883731 | 0,53755 | 0,7561567 | novel transcript |
| ENSG00000272146 | ARF4-AS1 | -0,333044564 | 0,793859409 | 0,4944 | 0,72678186 | ARF4 antisense RNA 1 |
| ENSG00000260558 | AC018557,2 | -0,333255309 | 0,793743452 | 0,47992 | 0,71730153 | novel transcript |
| ENSG00000249859 | PVT1 | -0,33350446 | 0,793606386 | 0,3707 | 0,63998958 | Pvt1 oncogene |
| ENSG00000286974 | AL662884,5 | -0,333596789 | 0,793555599 | 0,32267 | 0,60584543 | novel transcript, antisense to TNXB |
| ENSG00000165801 | ARHGEF40 | -0,333824588 | 0,793430308 | 0,35257 | 0,62748267 | Rho guanine nucleotide exchange factor 40 |
| ENSG00000225300 | AL591623,1 | -0,333861219 | 0,793410162 | 0,42642 | 0,67987951 | novel transcript |

| | | | | | | |
|---|---|---|---|---|---|---|
| ENSG00000256120 | SOX5-AS1 | -0,333893082 | 0,793392639 | 0,31662 | 0,60243844 | SOX5 antisense RNA 1 |
| ENSG00000237489 | C10orf143 | -0,334049143 | 0,79330682 | 0,24037 | 0,54950872 | chromosome 10 open reading frame 143 |
| ENSG00000253116 | AC027698,1 | -0,334220248 | 0,793212739 | 0,80033 | 0,90804014 | novel transcript |
| ENSG00000249245 | NCOA4P3 | -0,334248612 | 0,793197144 | 0,51139 | 0,73853921 | nuclear receptor coactivator 4 pseudogene 3 |
| ENSG00000140406 | TLNRD1 | -0,334286231 | 0,793176461 | 0,31899 | 0,60399503 | talin rod domain containing 1 |
| ENSG00000110887 | DAO | -0,334354622 | 0,793138862 | 0,74728 | 0,88076162 | D-amino acid oxidase |
| ENSG00000286961 | AC091938,1 | -0,334543101 | 0,79303525 | 0,56445 | 0,77299069 | novel transcript, antisense to CDH12 |
| ENSG00000162614 | NEXN | -0,334649691 | 0,792976661 | 0,18587 | 0,50915451 | nexilin F-actin binding protein |
| ENSG00000271327 | AC010201,2 | -0,33466614 | 0,792967619 | 0,43917 | 0,68811742 | novel transcript |
| ENSG00000078804 | TP53INP2 | -0,334744886 | 0,792924338 | 0,21951 | 0,53691892 | tumor protein p53 inducible nuclear protein 2 |
| ENSG00000176915 | ANKLE2 | -0,33479173 | 0,792898593 | 0,2488 | 0,55591283 | ankyrin repeat and LEM domain containing 2 |
| ENSG00000228624 | HDAC2-AS2 | -0,334833507 | 0,792875632 | 0,16585 | 0,49195088 | HDAC2 and HS3ST5 antisense RNA 2 |
| ENSG00000001460 | STPG1 | -0,335378179 | 0,792576348 | 0,31397 | 0,60070179 | sperm tail PG-rich repeat containing 1 |
| ENSG00000267249 | AP005482,2 | -0,335401844 | 0,792563347 | 0,60551 | 0,79868878 | novel transcript, sense intronic CEP76 |
| ENSG00000125779 | PANK2 | -0,335525969 | 0,792495161 | 0,29202 | 0,5868753 | pantothenate kinase 2 |
| ENSG00000222020 | HDAC4-AS1 | -0,335722923 | 0,792386978 | 0,62101 | 0,80782102 | HDAC4 antisense RNA 1 |
| ENSG00000114026 | OGG1 | -0,335733253 | 0,792381305 | 0,08312 | 0,37523979 | 8-oxoguanine DNA glycosylase |
| ENSG00000237605 | DYRK3-AS1 | -0,335765397 | 0,79236365 | 0,79852 | 0,90704887 | DYRK3 antisense RNA 1 |
| ENSG00000140675 | SLC5A2 | -0,335769346 | 0,792361481 | 0,57438 | 0,77954257 | solute carrier family 5 member 2 |
| ENSG00000185418 | TARS3 | -0,335899659 | 0,792289914 | 0,04629 | 0,28930497 | threonyl-tRNA synthetase 3 |
| ENSG00000132334 | PTPRE | -0,335903265 | 0,792287933 | 0,3847 | 0,65054085 | protein tyrosine phosphatase receptor type E |
| ENSG00000266803 | AC127540,1 | -0,335964904 | 0,792254083 | 0,67457 | 0,83975729 | novel transcript |
| ENSG00000166337 | TAF10 | -0,335997796 | 0,792236021 | 0,2546 | 0,55944852 | TATA-box binding protein associated factor 10 |
| ENSG00000100142 | POLR2F | -0,336133459 | 0,792161527 | 0,36199 | 0,63352908 | RNA polymerase II, I and III subunit F |
| ENSG00000080493 | SLC4A4 | -0,336140999 | 0,792157387 | 0,53995 | 0,75807575 | solute carrier family 4 member 4 |
| ENSG00000284975 | AC275401,3 | -0,336172322 | 0,792140189 | 0,74024 | 0,87681509 | ubiquitin specific peptidase 17-like 2 (USP17L2) pseudogene |
| ENSG00000237217 | AC008074,1 | -0,336210783 | 0,792119071 | 0,70389 | 0,85724036 | novel pseudogene |
| ENSG00000173080 | RXFP4 | -0,336245869 | 0,792099807 | 0,55084 | 0,76486471 | relaxin family peptide/INSL5 receptor 4 |
| ENSG00000172548 | NIPAL4 | -0,336260976 | 0,792091513 | 0,39357 | 0,65651919 | NIPA like domain containing 4 |
| ENSG00000285584 | AL158800,1 | -0,336269151 | 0,792087024 | 0,72375 | 0,86808717 | novel transcript |
| ENSG00000225706 | PTPRD-AS1 | -0,336311004 | 0,792064046 | 0,32441 | 0,60694578 | PTPRD antisense RNA 1 |

| | | | | | | |
|---|---|---|---|---|---|---|
| ENSG00000260782 | AC007225,1 | -0,336486002 | 0,791967975 | 0,45075 | 0,69592991 | nuclear factor (erythroid-derived 2)-like 3 (NFE2L3) pseudogene |
| ENSG00000251442 | LINC01094 | -0,336617025 | 0,791896053 | 0,39141 | 0,65510995 | long intergenic non-protein coding RNA 1094 |
| ENSG00000236086 | HMGN2P28 | -0,336816941 | 0,791786327 | 0,64506 | 0,82191409 | high mobility group nucleosomal binding domain 2 pseudogene 28 |
| ENSG00000175106 | TVP23C | -0,336840035 | 0,791773652 | 0,16167 | 0,48914394 | trans-golgi network vesicle protein 23 homolog C |
| ENSG00000213777 | AC011487,1 | -0,336887649 | 0,791747521 | 0,677 | 0,84111349 | vomeronasal 1 receptor (VN1R) pseudogene |
| ENSG00000109118 | PHF12 | -0,336927077 | 0,791725884 | 0,26908 | 0,57085484 | PHD finger protein 12 |
| ENSG00000237257 | MTND2P8 | -0,336957885 | 0,791708977 | 0,72009 | 0,86623639 | MT-ND2 pseudogene 8 |
| ENSG00000119574 | ZBTB45 | -0,33702469 | 0,791672317 | 0,27749 | 0,57588377 | zinc finger and BTB domain containing 45 |
| ENSG00000284959 | AC007262,2 | -0,337035339 | 0,791666473 | 0,2457 | 0,55358893 | novel transcript, antisense to TSHR |
| ENSG00000287933 | AL357141,1 | -0,337068547 | 0,791648252 | 0,14551 | 0,47304491 | novel transcript, antisense to FRK |
| ENSG00000235910 | APOA1-AS | -0,337089775 | 0,791636603 | 0,53014 | 0,75179571 | APOA1 antisense RNA |
| ENSG00000164236 | ANKRD33B | -0,337128393 | 0,791615413 | 0,41812 | 0,67437072 | ankyrin repeat domain 33B |
| ENSG00000102230 | PCYT1B | -0,337279879 | 0,791532296 | 0,4919 | 0,72531636 | phosphate cytidylyltransferase 1, choline, beta |
| ENSG00000152092 | ASTN1 | -0,337453258 | 0,791437177 | 0,30878 | 0,59767335 | astrotactin 1 |
| ENSG00000163449 | TMEM169 | -0,337458698 | 0,791434193 | 0,20097 | 0,52181085 | transmembrane protein 169 |
| ENSG00000233016 | SNHG7 | -0,337473192 | 0,791426242 | 0,3608 | 0,63310436 | small nucleolar RNA host gene 7 |
| ENSG00000104859 | CLASRP | -0,337523417 | 0,79139869 | 0,09504 | 0,39768318 | CLK4 associating serine/arginine rich protein |
| ENSG00000201793 | RN7SKP9 | -0,337566971 | 0,791374799 | 0,60904 | 0,80161331 | RN7SK pseudogene 9 |
| ENSG00000149716 | LTO1 | -0,337649641 | 0,791329452 | 0,17113 | 0,49641416 | LTO1 maturation factor of ABCE1 |
| ENSG00000099994 | SUSD2 | -0,338048853 | 0,791110512 | 0,5068 | 0,7352001 | sushi domain containing 2 |
| ENSG00000101347 | SAMHD1 | -0,33805508 | 0,791107097 | 0,08572 | 0,37967355 | SAM and HD domain containing deoxynucleoside triphosphate triphosphohydrolase 1 |
| ENSG00000168280 | KIF5C | -0,338145416 | 0,791057563 | 0,20971 | 0,52867318 | kinesin family member 5C |
| ENSG00000237399 | PITRM1-AS1 | -0,338159231 | 0,791049988 | 0,43908 | 0,68808849 | PITRM1 antisense RNA 1 |
| ENSG00000270228 | AC079880,1 | -0,338166318 | 0,791046102 | 0,46115 | 0,70278639 | HEAT repeat containing 2 (HEATR2) pseudogene |
| ENSG00000267348 | GEMIN7-AS1 | -0,338199756 | 0,791027768 | 0,4616 | 0,70290608 | GEMIN7 antisense RNA 1 |
| ENSG00000159784 | FAM131B | -0,338204468 | 0,791025184 | 0,07139 | 0,35295176 | family with sequence similarity 131 member B |
| ENSG00000231533 | AL391840,1 | -0,338230765 | 0,791010766 | 0,438 | 0,68747773 | novel transcript |
| ENSG00000137878 | GCOM1 | -0,338375713 | 0,790931296 | 0,61421 | 0,80370125 | GRINL1A complex locus 1 |
| ENSG00000165071 | TMEM71 | -0,338539808 | 0,79084134 | 0,65973 | 0,83018671 | transmembrane protein 71 |
| ENSG00000140043 | PTGR2 | -0,338672628 | 0,790768535 | 0,18379 | 0,50744379 | prostaglandin reductase 2 |

| | | | | | | |
|---|---|---|---|---|---|---|
| ENSG00000146842 | TMEM209 | -0,338673275 | 0,790768181 | 0,28885 | 0,58464073 | transmembrane protein 209 |
| ENSG00000136856 | SLC2A8 | -0,338945952 | 0,790618735 | 0,14801 | 0,4751727 | solute carrier family 2 member 8 |
| ENSG00000153789 | CIBAR2 | -0,339054784 | 0,790559096 | 0,53931 | 0,75767367 | CBY1 interacting BAR domain containing 2 |
| ENSG00000178921 | PFAS | -0,339073181 | 0,790549015 | 0,2725 | 0,57315552 | phosphoribosylformylglycinamidine synthase |
| ENSG00000274322 | AL136531,2 | -0,339200257 | 0,790479384 | 0,77533 | 0,89624737 | novel protein |
| ENSG00000105135 | ILVBL | -0,339263917 | 0,790444505 | 0,23676 | 0,54767193 | ilvB acetolactate synthase like |
| ENSG00000186532 | SMYD4 | -0,339314366 | 0,790416865 | 0,32975 | 0,61069811 | SET and MYND domain containing 4 |
| ENSG00000206282 | RGL2 | -0,339344942 | 0,790400113 | 0,32784 | 0,60960585 | ral guanine nucleotide dissociation stimulator like 2 |
| ENSG00000138606 | SHF | -0,339504373 | 0,790312771 | 0,36837 | 0,63803074 | Src homology 2 domain containing F |
| ENSG00000176973 | FAM89B | -0,339595457 | 0,790262877 | 0,22879 | 0,54259385 | family with sequence similarity 89 member B |
| ENSG00000286601 | AC007563,3 | -0,339715839 | 0,790196938 | 0,70945 | 0,85983746 | novel transcript |
| ENSG00000228242 | XPC-AS1 | -0,339771047 | 0,7901667 | 0,1972 | 0,5184837 | XPC antisense RNA 1 |
| ENSG00000204990 | AC005692,1 | -0,339979004 | 0,79005281 | 0,46005 | 0,70205579 | novel transcript |
| ENSG00000103496 | STX4 | -0,340057529 | 0,790009809 | 0,31456 | 0,60122508 | syntaxin 4 |
| ENSG00000246982 | Z84485,1 | -0,340100792 | 0,789986118 | 0,37231 | 0,64126849 | novel transcript, antisense to BRPF3 |
| ENSG00000278769 | AC090510,3 | -0,340117574 | 0,789976929 | 0,74175 | 0,8778969 | novel transcript, antisense to CDAN1 |
| ENSG00000188712 | OR13D3P | -0,34032978 | 0,78986074 | 0,75747 | 0,88677814 | olfactory receptor family 13 subfamily D member 3 pseudogene |
| ENSG00000278668 | AC005899,7 | -0,340355604 | 0,789846602 | 0,57291 | 0,77868876 | novel transcript |
| ENSG00000237977 | EIF4HP2 | -0,340442298 | 0,78979914 | 0,50426 | 0,73317978 | eukaryotic translation initiation factor 4H pseudogene 2 |
| ENSG00000176771 | NCKAP5 | -0,340451884 | 0,789793892 | 0,44962 | 0,69525077 | NCK associated protein 5 |
| ENSG00000126464 | PRR12 | -0,340822868 | 0,789590826 | 0,28768 | 0,58395529 | proline rich 12 |
| ENSG00000261794 | GOLGA8H | -0,340879753 | 0,789559693 | 0,43245 | 0,68411566 | golgin A8 family member H |
| ENSG00000142197 | DOP1B | -0,340885031 | 0,789556804 | 0,05943 | 0,32465136 | DOP1 leucine zipper like protein B |
| ENSG00000188042 | ARL4C | -0,341099241 | 0,78943958 | 0,26005 | 0,56382871 | ADP ribosylation factor like GTPase 4C |
| ENSG00000215067 | ALOX12-AS1 | -0,341150249 | 0,78941167 | 0,20673 | 0,52628269 | ALOX12 antisense RNA 1 |
| ENSG00000214942 | AC113167,1 | -0,341229016 | 0,789368571 | 0,5722 | 0,77846037 | novel transcript |
| ENSG00000234478 | ACBD3-AS1 | -0,341322501 | 0,789317423 | 0,68613 | 0,84663586 | ACBD3 antisense RNA 1 |
| ENSG00000120937 | NPPB | -0,341359998 | 0,789296907 | 0,78326 | 0,89918294 | natriuretic peptide B |
| ENSG00000114631 | PODXL2 | -0,341398815 | 0,789275671 | 0,09642 | 0,4002147 | podocalyxin like 2 |
| ENSG00000226651 | CLIC1 | -0,341547006 | 0,789194602 | 0,26796 | 0,57001388 | chloride intracellular channel 1 |
| ENSG00000206394 | CLIC1 | -0,341547006 | 0,789194602 | 0,26796 | 0,57001388 | chloride intracellular channel 1 |
| ENSG00000123989 | CHPF | -0,341573099 | 0,789180329 | 0,24347 | 0,55221671 | chondroitin polymerizing factor |

| | | | | | | |
|---|---|---|---|---|---|---|
| ENSG00000241535 | CBX5P1 | -0,341746695 | 0,789085374 | 0,55251 | 0,76590732 | chromobox 5 pseudogene 1 |
| ENSG00000135924 | DNAJB2 | -0,341801648 | 0,789055318 | 0,17224 | 0,49710979 | DnaJ heat shock protein family (Hsp40) member B2 |
| ENSG00000259904 | ACTG1P15 | -0,341839206 | 0,789034777 | 0,76969 | 0,89301171 | actin gamma 1 pseudogene 15 |
| ENSG00000136104 | RNASEH2B | -0,341855178 | 0,789026042 | 0,16101 | 0,48836951 | ribonuclease H2 subunit B |
| ENSG00000277548 | AC018926,3 | -0,341915821 | 0,788992876 | 0,62053 | 0,80776776 | novel transcript, sense intronic to CCPG1 |
| ENSG00000181523 | SGSH | -0,34205874 | 0,788914719 | 0,27641 | 0,57542877 | N-sulfoglucosamine sulfohydrolase |
| ENSG00000197670 | AL157838,1 | -0,342194326 | 0,78884058 | 0,45162 | 0,69664148 | novel transcript, antisense to ZNF217 |
| ENSG00000171223 | JUNB | -0,342209459 | 0,788832305 | 0,29607 | 0,58954675 | JunB proto-oncogene, AP-1 transcription factor subunit |
| ENSG00000225205 | AC078883,1 | -0,342222279 | 0,788825296 | 0,55296 | 0,76620716 | novel transcript |
| ENSG00000178467 | P4HTM | -0,342538552 | 0,788652385 | 0,12151 | 0,44093327 | prolyl 4-hydroxylase, transmembrane |
| ENSG00000170743 | SYT9 | -0,342581367 | 0,788628981 | 0,38838 | 0,6529693 | synaptotagmin 9 |
| ENSG00000173113 | TRMT112 | -0,34259848 | 0,788619626 | 0,29926 | 0,59123205 | tRNA methyltransferase subunit 11-2 |
| ENSG00000162441 | LZIC | -0,342658416 | 0,788586864 | 0,1975 | 0,518964 | leucine zipper and CTNNBIP1 domain containing |
| ENSG00000086205 | FOLH1 | -0,342689983 | 0,788569609 | 0,4003 | 0,66242349 | folate hydrolase 1 |
| ENSG00000242602 | RPL35AP13 | -0,342741047 | 0,788541699 | 0,67424 | 0,83959502 | ribosomal protein L35a pseudogene 13 |
| ENSG00000113594 | LIFR | -0,342770185 | 0,788525773 | 0,43475 | 0,68535313 | LIF receptor subunit alpha |
| ENSG00000163344 | PMVK | -0,342790225 | 0,78851482 | 0,27228 | 0,57315552 | phosphomevalonate kinase |
| ENSG00000214022 | REPIN1 | -0,342924493 | 0,788441438 | 0,27261 | 0,57315967 | replication initiator 1 |
| ENSG00000285696 | AP002433,2 | -0,342956576 | 0,788423905 | 0,48442 | 0,72028271 | novel transcript, antisense to ACAT1 |
| ENSG00000002746 | HECW1 | -0,342993307 | 0,788403832 | 0,34271 | 0,62029585 | HECT, C2 and WW domain containing E3 ubiquitin protein ligase 1 |
| ENSG00000258535 | AL079307,1 | -0,343094656 | 0,788348448 | 0,33373 | 0,61370377 | novel transcript |
| ENSG00000272941 | AC083862,1 | -0,343126592 | 0,788330997 | 0,33247 | 0,61233149 | novel transcript, antisense to C7orf49 |
| ENSG00000275552 | AC243965,2 | -0,343193385 | 0,788294501 | 0,59628 | 0,79382929 | novel transcript, antisense ABHD4 |
| ENSG00000068615 | REEP1 | -0,343208935 | 0,788286004 | 0,38802 | 0,65289784 | receptor accessory protein 1 |
| ENSG00000287399 | AC011503,4 | -0,343251437 | 0,788262781 | 0,49639 | 0,72831792 | novel transcript |
| ENSG00000130544 | ZNF557 | -0,343274145 | 0,788250374 | 0,29002 | 0,58531219 | zinc finger protein 557 |
| ENSG00000134198 | TSPAN2 | -0,34329531 | 0,788238811 | 0,47759 | 0,71545897 | tetraspanin 2 |
| ENSG00000102312 | PORCN | -0,343324314 | 0,788222964 | 0,18908 | 0,51243448 | porcupine O-acyltransferase |
| ENSG00000284620 | AF228730,5 | -0,343402058 | 0,788180489 | 0,26209 | 0,56501119 | RNA binding protein, fox-1 homolog (C, elegans) 1 (RBFOX1) pseudogene |
| ENSG00000141295 | SCRN2 | -0,343404378 | 0,788179221 | 0,27903 | 0,57715243 | secernin 2 |

| | | | | | | |
|---|---|---|---|---|---|---|
| ENSG00000249646 | OR7E94P | -0,343722455 | 0,788005468 | 0,54903 | 0,76394298 | olfactory receptor family 7 subfamily E member 94 pseudogene |
| ENSG00000177303 | CASKIN2 | -0,343802083 | 0,787961976 | 0,10799 | 0,41856292 | CASK interacting protein 2 |
| ENSG00000275565 | ALOX5 | -0,34385948 | 0,787930627 | 0,62143 | 0,80788604 | arachidonate 5-lipoxygenase |
| ENSG00000225555 | AP000320,1 | -0,343923081 | 0,787895892 | 0,71291 | 0,86184574 | novel transcript |
| ENSG00000185000 | DGAT1 | -0,344043006 | 0,787830401 | 0,34867 | 0,62475089 | diacylglycerol O-acyltransferase 1 |
| ENSG00000285482 | DGAT1 | -0,344043006 | 0,787830401 | 0,34867 | 0,62475089 | diacylglycerol O-acyltransferase 1 |
| ENSG00000270071 | AP001172,1 | -0,344154433 | 0,787769555 | 0,44245 | 0,69016149 | novel transcript |
| ENSG00000151164 | RAD9B | -0,344258122 | 0,787712939 | 0,3981 | 0,66073349 | RAD9 checkpoint clamp component B |
| ENSG00000265972 | TXNIP | -0,344290157 | 0,787695448 | 0,30023 | 0,59164636 | thioredoxin interacting protein |
| ENSG00000182534 | MXRA7 | -0,344291261 | 0,787694845 | 0,14332 | 0,47083258 | matrix remodeling associated 7 |
| ENSG00000169255 | B3GALNT1 | -0,344315294 | 0,787681723 | 0,13318 | 0,45912454 | beta-1,3-N-acetylgalactosaminyltransferase 1 (globoside blood group) |
| ENSG00000274615 | AC233968,1 | -0,344356426 | 0,787659266 | 0,08199 | 0,37306719 | aminopeptidase puromycin sensitive (NPEPPS) pseudogene |
| ENSG00000255836 | AC131206,1 | -0,344478493 | 0,787592625 | 0,49381 | 0,7265016 | M-phase phosphoprotein 8 (MPHOSPH8) pseudogene |
| ENSG00000182824 | FAM230E | -0,344518807 | 0,787570617 | 0,6776 | 0,84135792 | family with sequence similarity 230 member E |
| ENSG00000278019 | AC245690,1 | -0,344576396 | 0,78753918 | 0,73121 | 0,87233924 | glucuronidase, beta (GUSB) pseudogene |
| ENSG00000284915 | TINF2 | -0,34470446 | 0,787469276 | 0,18588 | 0,50915451 | TERF1 interacting nuclear factor 2 |
| ENSG00000183960 | KCNH8 | -0,344771864 | 0,787432485 | 0,41215 | 0,66994473 | potassium voltage-gated channel subfamily H member 8 |
| ENSG00000186635 | ARAP1 | -0,344969258 | 0,787324753 | 0,39952 | 0,66176242 | ArfGAP with RhoGAP domain, ankyrin repeat and PH domain 1 |
| ENSG00000023191 | RNH1 | -0,345010733 | 0,787302119 | 0,23795 | 0,54864017 | ribonuclease/angiogenin inhibitor 1 |
| ENSG00000260639 | AC025678,1 | -0,345037548 | 0,787287486 | 0,62222 | 0,80838627 | mitochondrially encoded cytochrome c oxidase III (MT-CO3) pseudogene |
| ENSG00000285641 | AL358472,6 | -0,345342279 | 0,78712121 | 0,38855 | 0,65310114 | novel protein |
| ENSG00000272217 | AL645940,1 | -0,345410864 | 0,787083792 | 0,60632 | 0,79949515 | novel transcript |
| ENSG00000206195 | DUXAP8 | -0,345477662 | 0,78704735 | 0,43247 | 0,68411566 | double homeobox A pseudogene 8 |
| ENSG00000129204 | USP6 | -0,345514463 | 0,787027274 | 0,19692 | 0,51817451 | ubiquitin specific peptidase 6 |
| ENSG00000257545 | AC079385,1 | -0,345564488 | 0,786999984 | 0,49441 | 0,72678186 | novel transcript, antisense to POLR3B, RFX4 and RIC8B, |
| ENSG00000287678 | AL031778,1 | -0,345675778 | 0,786939277 | 0,38064 | 0,6482096 | novel transcript, antisense to NFYA |
| ENSG00000224795 | NTM-AS1 | -0,345695176 | 0,786928696 | 0,5611 | 0,77125979 | NTM antisense RNA 1 |
| ENSG00000113073 | SLC4A9 | -0,345785254 | 0,786879564 | 0,65574 | 0,82800391 | solute carrier family 4 member 9 |

| | | | | | | |
|---|---|---|---|---|---|---|
| ENSG00000241168 | AC128685,1 | -0,345800875 | 0,786871044 | 0,26431 | 0,56706657 | novel transcript |
| ENSG00000272070 | AC005618,1 | -0,345826881 | 0,78685686 | 0,25475 | 0,55944852 | novel transcript |
| ENSG00000225311 | AL024474,2 | -0,345850463 | 0,786843999 | 0,67397 | 0,83939184 | novel transcript, antisense to UTRN |
| ENSG00000232204 | TET1P1 | -0,345980174 | 0,786773257 | 0,36948 | 0,63917123 | tet methylcytosine dioxygenase 1 pseudogene 1 |
| ENSG00000186118 | TEX38 | -0,34611083 | 0,786702008 | 0,6231 | 0,80892518 | testis expressed 38 |
| ENSG00000287884 | AC005594,1 | -0,346202574 | 0,786651981 | 0,49822 | 0,72954885 | novel transcript, antisense to DPP9 |
| ENSG00000125355 | TMEM255A | -0,34634529 | 0,786574167 | 0,52051 | 0,74503377 | transmembrane protein 255A |
| ENSG00000226622 | AC092155,1 | -0,34636845 | 0,78656154 | 0,27828 | 0,57645922 | novel transcript |
| ENSG00000068971 | PPP2R5B | -0,346371825 | 0,7865597 | 0,29162 | 0,58655914 | protein phosphatase 2 regulatory subunit B'beta |
| ENSG00000259426 | AC027237,3 | -0,346397641 | 0,786545625 | 0,46083 | 0,70258879 | novel transcript, antisense to PAQR5 |
| ENSG00000002726 | AOC1 | -0,346531127 | 0,786472853 | 0,78866 | 0,90191888 | amine oxidase copper containing 1 |
| ENSG00000159399 | HK2 | -0,346633807 | 0,78641688 | 0,12858 | 0,45191041 | hexokinase 2 |
| ENSG00000281010 | AC009065,9 | -0,346724654 | 0,78636736 | 0,72421 | 0,86838632 | |
| ENSG00000256092 | SBNO1-AS1 | -0,346761448 | 0,786347305 | 0,25188 | 0,55773568 | SBNO1 antisense RNA 1 |
| ENSG00000234196 | ZBTB12 | -0,34676355 | 0,78634616 | 0,51842 | 0,74326177 | zinc finger and BTB domain containing 12 |
| ENSG00000262349 | GIHCG | -0,346801156 | 0,786325663 | 0,36177 | 0,63352908 | GIHCG inhibitor of miR-200b/200a/429 expression |
| ENSG00000127419 | TMEM175 | -0,34681372 | 0,786318815 | 0,21828 | 0,535736 | transmembrane protein 175 |
| ENSG00000255141 | HNRNPA1P76 | -0,346876016 | 0,786284862 | 0,63386 | 0,81549489 | heterogeneous nuclear ribonucleoprotein A1 pseudogene 76 |
| ENSG00000267642 | AC091198,1 | -0,346990353 | 0,786222549 | 0,74376 | 0,87871466 | novel transcript |
| ENSG00000224152 | AC009506,1 | -0,347154156 | 0,786133287 | 0,19823 | 0,51980353 | novel transcript, antisense to BAZ2B |
| ENSG00000227398 | KIF9-AS1 | -0,34717336 | 0,786122823 | 0,29975 | 0,59160746 | KIF9 antisense RNA 1 |
| ENSG00000217733 | CCT7P1 | -0,347310927 | 0,786047866 | 0,61927 | 0,80699858 | chaperonin containing TCP1 subunit 7 pseudogene 1 |
| ENSG00000041515 | MYO16 | -0,347367704 | 0,786016932 | 0,24696 | 0,55461513 | myosin XVI |
| ENSG00000286242 | AC018797,3 | -0,347595845 | 0,785892645 | 0,33671 | 0,61675087 | novel transcript, antisense to UBE2D3 |
| ENSG00000186838 | SELENOV | -0,347635113 | 0,785871254 | 0,56461 | 0,77306193 | selenoprotein V |
| ENSG00000160856 | FCRL3 | -0,347658835 | 0,785858332 | 0,68832 | 0,84817878 | Fc receptor like 3 |
| ENSG00000187051 | RPS19BP1 | -0,34778198 | 0,785791257 | 0,30755 | 0,59664516 | ribosomal protein S19 binding protein 1 |
| ENSG00000237716 | AL136309,2 | -0,347826292 | 0,785767122 | 0,48387 | 0,71999322 | novel transcript |
| ENSG00000235683 | AC018442,1 | -0,347849968 | 0,785754227 | 0,60499 | 0,79821396 | leucine zipper-EF-hand containing transmembrane protein 1 (LETM1) pseudogene |
| ENSG00000272631 | AC067750,1 | -0,347894251 | 0,785730108 | 0,12769 | 0,45102957 | novel transcript, antisense to BMPR1A |
| ENSG00000219133 | AL592114,1 | -0,347931918 | 0,785709594 | 0,57849 | 0,78223356 | ribosomal protein L21 (RPL21) pseudogene |

| | | | | | | |
|---|---|---|---|---|---|---|
| ENSG00000244151 | AC010973,2 | -0,347965113 | 0,785691516 | 0,44072 | 0,68917395 | novel transcript |
| ENSG00000285901 | AC008012,1 | -0,348069242 | 0,785634809 | 0,20663 | 0,52622584 | novel transcript |
| ENSG00000242445 | RPL7AP11 | -0,348082442 | 0,785627621 | 0,51814 | 0,74305171 | ribosomal protein L7a pseudogene 11 |
| ENSG00000243323 | PTPRVP | -0,348155264 | 0,785587967 | 0,55996 | 0,77076855 | protein tyrosine phosphatase receptor type V, pseudogene |
| ENSG00000285377 | EMC9 | -0,348225973 | 0,785549464 | 0,20218 | 0,52245781 | ER membrane protein complex subunit 9 |
| ENSG00000100908 | EMC9 | -0,348225973 | 0,785549464 | 0,20218 | 0,52245781 | ER membrane protein complex subunit 9 |
| ENSG00000280722 | AL008628,2 | -0,348231189 | 0,785546624 | 0,85097 | 0,93218292 | novel transcript, overlapping to FAM120B |
| ENSG00000237934 | AL590729,1 | -0,348394782 | 0,785457553 | 0,66009 | 0,83029149 | novel transcript |
| ENSG00000131711 | MAP1B | -0,348496343 | 0,785402261 | 0,2229 | 0,53906657 | microtubule associated protein 1B |
| ENSG00000286934 | AC027251,1 | -0,348577756 | 0,785357941 | 0,7948 | 0,90488348 | novel transcript, antisense to CSMD1 |
| ENSG00000156265 | MAP3K7CL | -0,348724856 | 0,785277869 | 0,68231 | 0,84426936 | MAP3K7 C-terminal like |
| ENSG00000273893 | AL133520,1 | -0,348764718 | 0,785256172 | 0,34037 | 0,61897407 | novel transcript, sense intronic to SLC13A3 |
| ENSG00000228599 | RPL7P52 | -0,348927816 | 0,785167403 | 0,68338 | 0,84495012 | ribosomal protein L7 pseudogene 52 |
| ENSG00000129654 | FOXJ1 | -0,348992678 | 0,785132103 | 0,42835 | 0,68103938 | forkhead box J1 |
| ENSG00000162620 | LRRIQ3 | -0,349216317 | 0,785010406 | 0,64323 | 0,82098995 | leucine rich repeats and IQ motif containing 3 |
| ENSG00000248356 | AC093864,1 | -0,349305373 | 0,784961949 | 0,73577 | 0,8750728 | novel transcript |
| ENSG00000125651 | GTF2F1 | -0,34936511 | 0,784929448 | 0,15044 | 0,47857814 | general transcription factor IIF subunit 1 |
| ENSG00000113758 | DBN1 | -0,349436519 | 0,784890597 | 0,19305 | 0,51505487 | drebrin 1 |
| ENSG00000188153 | COL4A5 | -0,349521989 | 0,784844099 | 0,26366 | 0,56629809 | collagen type IV alpha 5 chain |
| ENSG00000162929 | KIAA1841 | -0,349724897 | 0,784733722 | 0,12024 | 0,43879231 | KIAA1841 |
| ENSG00000104164 | BLOC1S6 | -0,349773836 | 0,784707103 | 0,18051 | 0,50407792 | biogenesis of lysosomal organelles complex 1 subunit 6 |
| ENSG00000101049 | SGK2 | -0,349854091 | 0,784663452 | 0,51305 | 0,7394947 | serum/glucocorticoid regulated kinase 2 |
| ENSG00000240342 | RPS2P5 | -0,34991391 | 0,784630918 | 0,40387 | 0,66409141 | ribosomal protein S2 pseudogene 5 |
| ENSG00000277795 | AC244517,3 | -0,349989878 | 0,784589603 | 0,65985 | 0,83018671 | novel zinc finger pseudogene |
| ENSG00000281309 | SURF6 | -0,350043286 | 0,784560558 | 0,30205 | 0,59294633 | surfeit 6 |
| ENSG00000148296 | SURF6 | -0,350043286 | 0,784560558 | 0,30205 | 0,59294633 | surfeit 6 |
| ENSG00000260442 | ATP2A1-AS1 | -0,350057788 | 0,784552671 | 0,18269 | 0,50615118 | ATP2A1 antisense RNA 1 |
| ENSG00000250127 | LINC02108 | -0,350333148 | 0,784402942 | 0,34556 | 0,62278421 | long intergenic non-protein coding RNA 2108 |
| ENSG00000283982 | AL445623,2 | -0,35035699 | 0,784389979 | 0,36002 | 0,63260537 | novel transcript |
| ENSG00000261175 | LINC02188 | -0,350422 | 0,784354634 | 0,63941 | 0,81910693 | long intergenic non-protein coding RNA 2188 |
| ENSG00000261616 | AC036108,3 | -0,350483535 | 0,78432118 | 0,30783 | 0,59671558 | novel transcript, antisense to TTC23 |
| ENSG00000205476 | CCDC85C | -0,350560011 | 0,784279605 | 0,249 | 0,55616532 | coiled-coil domain containing 85C |

| | | | | | | |
|---|---|---|---|---|---|---|
| ENSG00000168621 | GDNF | -0,350563717 | 0,78427759 | 0,64696 | 0,82332882 | glial cell derived neurotrophic factor |
| ENSG00000266074 | BAHCC1 | -0,350702071 | 0,784202382 | 0,46529 | 0,70583978 | BAH domain and coiled-coil containing 1 |
| ENSG00000271746 | AL031848,2 | -0,350713356 | 0,784196248 | 0,52969 | 0,75147239 | novel transcript, antisense to ACOT7 |
| ENSG00000234690 | EPCAM-DT | -0,350737166 | 0,784183306 | 0,47938 | 0,71689864 | EPCAM divergent transcript |
| ENSG00000287238 | AP002954,2 | -0,350915723 | 0,784086256 | 0,24212 | 0,55105388 | novel transcript |
| ENSG00000221994 | ZNF630 | -0,350938476 | 0,78407389 | 0,29365 | 0,58809888 | zinc finger protein 630 |
| ENSG00000174326 | SLC16A11 | -0,350986686 | 0,78404769 | 0,60919 | 0,8016224 | solute carrier family 16 member 11 |
| ENSG00000287807 | AL390242,1 | -0,350993445 | 0,784044016 | 0,57457 | 0,7797355 | novel transcript |
| ENSG00000103037 | SETD6 | -0,351030875 | 0,784023675 | 0,26627 | 0,56917012 | SET domain containing 6, protein lysine methyltransferase |
| ENSG00000182511 | FES | -0,351035132 | 0,784021362 | 0,30211 | 0,59294633 | FES proto-oncogene, tyrosine kinase |
| ENSG00000155729 | KCTD18 | -0,351098274 | 0,783987048 | 0,19862 | 0,51980353 | potassium channel tetramerization domain containing 18 |
| ENSG00000141404 | GNAL | -0,351110524 | 0,783980391 | 0,20853 | 0,52778377 | G protein subunit alpha L |
| ENSG00000258743 | LINC02301 | -0,351114772 | 0,783978083 | 0,55352 | 0,76655124 | long intergenic non-protein coding RNA 2301 |
| ENSG00000215184 | RPS12P16 | -0,351182303 | 0,783941387 | 0,41801 | 0,67432365 | ribosomal protein S12 pseudogene 16 |
| ENSG00000182308 | DCAF4L1 | -0,351536744 | 0,783748812 | 0,45117 | 0,69626785 | DDB1 and CUL4 associated factor 4 like 1 |
| ENSG00000143786 | CNIH3 | -0,351566836 | 0,783732464 | 0,40106 | 0,66295784 | cornichon family AMPA receptor auxiliary protein 3 |
| ENSG00000281500 | SLC37A4 | -0,351653552 | 0,783685358 | 0,44308 | 0,69057052 | solute carrier family 37 member 4 |
| ENSG00000153391 | INO80C | -0,35165872 | 0,783682551 | 0,24367 | 0,55235189 | INO80 complex subunit C |
| ENSG00000228423 | AL357552,2 | -0,351736076 | 0,783640532 | 0,63636 | 0,81700605 | novel transcript |
| ENSG00000164008 | C1orf50 | -0,351838934 | 0,783584663 | 0,04008 | 0,27058857 | chromosome 1 open reading frame 50 |
| ENSG00000242882 | RPL5P11 | -0,351857491 | 0,783574584 | 0,74137 | 0,8775607 | ribosomal protein L5 pseudogene 11 |
| ENSG00000274810 | NPHP3-ACAD1 | -0,351898642 | 0,783552234 | 0,46873 | 0,70833955 | NPHP3-ACAD11 readthrough (NMD candidate) |
| ENSG00000273301 | AC016717,2 | -0,351954963 | 0,783521646 | 0,57722 | 0,78134056 | novel transcript |
| ENSG00000176390 | CRLF3 | -0,351982102 | 0,783506907 | 0,2313 | 0,54422787 | cytokine receptor like factor 3 |
| ENSG00000243230 | AC011005,4 | -0,352303269 | 0,783332505 | 0,54141 | 0,75881984 | novel transcript |
| ENSG00000181785 | OR5AS1 | -0,35239001 | 0,783285409 | 0,21034 | 0,5290538 | olfactory receptor family 5 subfamily AS member 1 |
| ENSG00000121851 | POLR3GL | -0,352413728 | 0,783272532 | 0,14341 | 0,47083258 | RNA polymerase III subunit GL |
| ENSG00000267059 | AC005943,1 | -0,352463313 | 0,783245612 | 0,28387 | 0,5807907 | novel transcript, readthrough between UQCR11 and MBD3 |
| ENSG00000066427 | ATXN3 | -0,352472045 | 0,783240871 | 0,23004 | 0,54335232 | ataxin 3 |
| ENSG00000184451 | CCR10 | -0,352531075 | 0,783208824 | 0,52684 | 0,75003528 | C-C motif chemokine receptor 10 |
| ENSG00000168243 | GNG4 | -0,352578574 | 0,783183038 | 0,11325 | 0,42788135 | G protein subunit gamma 4 |
| ENSG00000259404 | EFL1P1 | -0,352657238 | 0,783140336 | 0,47663 | 0,71481671 | elongation factor like GTPase 1 pseudogene 1 |
| ENSG00000259594 | AC023034,1 | -0,352804299 | 0,783060511 | 0,53502 | 0,75429628 | novel transcript |

| | | | | | | |
|---|---|---|---|---|---|---|
| ENSG00000142959 | BEST4 | -0,352870897 | 0,783024364 | 0,5444 | 0,76055278 | bestrophin 4 |
| ENSG00000145020 | AMT | -0,352972746 | 0,782969087 | 0,27611 | 0,57533851 | aminomethyltransferase |
| ENSG00000081665 | ZNF506 | -0,353096837 | 0,782901744 | 0,18967 | 0,51284737 | zinc finger protein 506 |
| ENSG00000104870 | FCGRT | -0,353228773 | 0,78283015 | 0,36946 | 0,63917123 | Fc fragment of IgG receptor and transporter |
| ENSG00000184828 | ZBTB7C | -0,353394574 | 0,782740189 | 0,50269 | 0,73209345 | zinc finger and BTB domain containing 7C |
| ENSG00000287600 | AC122719,3 | -0,353576376 | 0,782641558 | 0,21037 | 0,5290538 | novel transcript, antisense to TRIP13 |
| ENSG00000117600 | PLPPR4 | -0,353590302 | 0,782634003 | 0,62331 | 0,8091109 | phospholipid phosphatase related 4 |
| ENSG00000205771 | CATSPER2P1 | -0,353648679 | 0,782602335 | 0,38817 | 0,6529525 | cation channel sperm associated 2 pseudogene 1 |
| ENSG00000272655 | POLR2J4 | -0,353672007 | 0,782589681 | 0,16578 | 0,49189706 | RNA polymerase II subunit J4, pseudogene |
| ENSG00000287937 | AC079753,2 | -0,353730502 | 0,782557951 | 0,51897 | 0,7437068 | novel transcript |
| ENSG00000259485 | LINC02253 | -0,353748216 | 0,782548343 | 0,30011 | 0,59160746 | long intergenic non-protein coding RNA 2253 |
| ENSG00000213540 | TXN2P1 | -0,353860107 | 0,782487653 | 0,48238 | 0,71892541 | TXN2 pseudogene 1 |
| ENSG00000074181 | NOTCH3 | -0,353918694 | 0,782455878 | 0,24436 | 0,55253387 | notch receptor 3 |
| ENSG00000182871 | COL18A1 | -0,353942239 | 0,782443108 | 0,58198 | 0,78454692 | collagen type XVIII alpha 1 chain |
| ENSG00000075886 | TUBA3D | -0,353977935 | 0,782423748 | 0,4665 | 0,7067229 | tubulin alpha 3d |
| ENSG00000276910 | GTF2H2 | -0,353980351 | 0,782422438 | 0,38348 | 0,65020746 | general transcription factor IIH subunit 2 |
| ENSG00000150455 | TIRAP | -0,354003486 | 0,782409891 | 0,31598 | 0,60212384 | TIR domain containing adaptor protein |
| ENSG00000106351 | AGFG2 | -0,354128204 | 0,782342257 | 0,0834 | 0,37550713 | ArfGAP with FG repeats 2 |
| ENSG00000153037 | SRP19 | -0,35423039 | 0,782286845 | 0,21773 | 0,53550245 | signal recognition particle 19 |
| ENSG00000142871 | CCN1 | -0,354363188 | 0,78221484 | 0,23169 | 0,54440999 | cellular communication network factor 1 |
| ENSG00000158417 | EIF5B | -0,354398744 | 0,782195562 | 0,13066 | 0,45504746 | eukaryotic translation initiation factor 5B |
| ENSG00000120328 | PCDHB12 | -0,354505246 | 0,782137821 | 0,07026 | 0,34990935 | protocadherin beta 12 |
| ENSG00000234353 | AP000346,2 | -0,354521345 | 0,782129094 | 0,71879 | 0,86552951 | immunoglobulin lambda-like polypeptide 1 (IGLL1) pseudogene |
| ENSG00000213599 | SLX1A-SULT1/ | -0,354626586 | 0,782072042 | 0,11079 | 0,42340354 | SLX1A-SULT1A3 readthrough (NMD candidate) |
| ENSG00000223692 | DIP2A-IT1 | -0,35462997 | 0,782070207 | 0,65906 | 0,82969668 | DIP2A intronic transcript 1 |
| ENSG00000116151 | MORN1 | -0,354653483 | 0,782057461 | 0,14694 | 0,47471707 | MORN repeat containing 1 |
| ENSG00000286192 | AC069288,1 | -0,354825789 | 0,781964062 | 0,47105 | 0,71051384 | Novel protein |
| ENSG00000254565 | MTND2P26 | -0,354935658 | 0,781904514 | 0,6619 | 0,83159381 | MT-ND2 pseudogene 26 |
| ENSG00000285485 | NFATC4 | -0,35500669 | 0,781866017 | 0,4248 | 0,67855129 | nuclear factor of activated T cells 4 |
| ENSG00000100968 | NFATC4 | -0,35500669 | 0,781866017 | 0,4248 | 0,67855129 | nuclear factor of activated T cells 4 |
| ENSG00000246851 | AL157938,3 | -0,355180172 | 0,781772005 | 0,28505 | 0,58171017 | novel transcript |
| ENSG00000143951 | WDPCP | -0,355237886 | 0,781740731 | 0,20998 | 0,52887688 | WD repeat containing planar cell polarity effector |

| | | | | | | |
|---|---|---|---|---|---|---|
| ENSG00000141582 | CBX4 | -0,355351025 | 0,781679428 | 0,44049 | 0,68895352 | chromobox 4 |
| ENSG00000287646 | AC021701,1 | -0,355409067 | 0,78164798 | 0,44143 | 0,68976733 | novel transcript, antisense to RTTN |
| ENSG00000198358 | AL354714,1 | -0,355424334 | 0,781639709 | 0,2572 | 0,56156298 | antisense to ITLN2 |
| ENSG00000165886 | UBTD1 | -0,355430636 | 0,781636294 | 0,39716 | 0,65966023 | ubiquitin domain containing 1 |
| ENSG00000269910 | AL049840,3 | -0,355520749 | 0,781587474 | 0,46442 | 0,70506031 | novel transcript, antisense to KLC1 |
| ENSG00000244257 | PKD1P1 | -0,355531917 | 0,781581424 | 0,63003 | 0,8132682 | polycystin 1, transient receptor potential channel interacting pseudogene 1 |
| ENSG00000120329 | SLC25A2 | -0,355555583 | 0,781568603 | 0,41381 | 0,67085666 | solute carrier family 25 member 2 |
| ENSG00000286544 | AL359502,2 | -0,355658867 | 0,781512651 | 0,67475 | 0,83976962 | novel transcript |
| ENSG00000115310 | RTN4 | -0,355732685 | 0,781472665 | 0,07806 | 0,36538634 | reticulon 4 |
| ENSG00000227014 | FKBP14-AS1 | -0,355746409 | 0,781465231 | 0,35269 | 0,62754548 | FKBP14 antisense RNA 1 |
| ENSG00000237248 | LINC00987 | -0,35575535 | 0,781460388 | 0,41437 | 0,67106867 | long intergenic non-protein coding RNA 987 |
| ENSG00000144567 | RETREG2 | -0,355815378 | 0,781427873 | 0,20495 | 0,52453221 | reticulophagy regulator family member 2 |
| ENSG00000178922 | HYI | -0,355851656 | 0,781408224 | 0,13215 | 0,45719971 | hydroxypyruvate isomerase (putative) |
| ENSG00000175426 | PCSK1 | -0,355868857 | 0,781398907 | 0,34636 | 0,62342969 | proprotein convertase subtilisin/kexin type 1 |
| ENSG00000101331 | CCM2L | -0,355892577 | 0,78138606 | 0,67039 | 0,83656528 | CCM2 like scaffold protein |
| ENSG00000278145 | C17orf78 | -0,355956541 | 0,781351417 | 0,55894 | 0,77025265 | chromosome 17 open reading frame 78 |
| ENSG00000285567 | AC074051,3 | -0,356126374 | 0,781259442 | 0,38252 | 0,64932239 | novel transcript |
| ENSG00000173208 | ABCD2 | -0,35615794 | 0,781242349 | 0,49664 | 0,72841371 | ATP binding cassette subfamily D member 2 |
| ENSG00000181800 | CELF2-AS1 | -0,356188258 | 0,781225931 | 0,47688 | 0,71482093 | CELF2 antisense RNA 1 |
| ENSG00000196296 | ATP2A1 | -0,356238868 | 0,781198526 | 0,17466 | 0,49886732 | ATPase sarcoplasmic/endoplasmic reticulum Ca2+ transporting 1 |
| ENSG00000228328 | PGK1P1 | -0,356301699 | 0,781164505 | 0,51841 | 0,74326177 | phosphoglycerate kinase 1 pseudogene 1 |
| ENSG00000175967 | LINC02880 | -0,356336913 | 0,781145438 | 0,63761 | 0,81781528 | long intergenic non-protein coding RNA 2880 |
| ENSG00000285865 | AC010285,3 | -0,356396138 | 0,781113371 | 0,27185 | 0,57305058 | Novel transcript, antisense to RNF130 |
| ENSG00000173988 | LRRC63 | -0,35644318 | 0,781087902 | 0,46038 | 0,7022453 | leucine rich repeat containing 63 |
| ENSG00000267653 | AC002546,1 | -0,35654194 | 0,781034434 | 0,60401 | 0,79779897 | novel transcript |
| ENSG00000230826 | AL390774,2 | -0,356604165 | 0,781000748 | 0,3682 | 0,6380264 | novel transcript |
| ENSG00000286626 | AC090469,1 | -0,3568189 | 0,78088451 | 0,47956 | 0,71700404 | novel transcript, antisense to CAPRIN1 |
| ENSG00000033627 | ATP6V0A1 | -0,356825108 | 0,78088115 | 0,12088 | 0,43993759 | ATPase H+ transporting V0 subunit a1 |
| ENSG00000255622 | PCDHB17P | -0,356910256 | 0,780835064 | 0,29312 | 0,58745839 | protocadherin beta 17 pseudogene |
| ENSG00000260428 | SCX | -0,356932853 | 0,780822833 | 0,64575 | 0,82244907 | scleraxis bHLH transcription factor |
| ENSG00000259276 | AC087286,1 | -0,356977567 | 0,780798633 | 0,64296 | 0,82098995 | novel transcript, antisense to AKAP13 |

| | | | | | | |
|---|---|---|---|---|---|---|
| ENSG00000189077 | TMEM120A | -0,356978206 | 0,780798287 | 0,23159 | 0,54440987 | transmembrane protein 120A |
| ENSG00000099992 | TBC1D10A | -0,35711295 | 0,780725366 | 0,24436 | 0,55253387 | TBC1 domain family member 10A |
| ENSG00000159212 | CLIC6 | -0,357175765 | 0,780691375 | 0,62218 | 0,80838627 | chloride intracellular channel 6 |
| ENSG00000240991 | RPL23AP67 | -0,357225284 | 0,780664579 | 0,61579 | 0,80464855 | ribosomal protein L23a pseudogene 67 |
| ENSG00000269514 | AC024257,3 | -0,35723066 | 0,78066167 | 0,16446 | 0,49065086 | novel transcript, antisense to OR10AD1 |
| ENSG00000137727 | ARHGAP20 | -0,357254267 | 0,780648896 | 0,6869 | 0,84725905 | Rho GTPase activating protein 20 |
| ENSG00000249242 | TMEM150C | -0,357281709 | 0,780634047 | 0,10625 | 0,4157803 | transmembrane protein 150C |
| ENSG00000226257 | STK19 | -0,357686576 | 0,780415006 | 0,32191 | 0,60577115 | serine/threonine kinase 19 |
| ENSG00000132950 | ZMYM5 | -0,35775925 | 0,780375694 | 0,19415 | 0,51562158 | zinc finger MYM-type containing 5 |
| ENSG00000231527 | FAM27C | -0,357870978 | 0,780315262 | 0,25963 | 0,5633887 | family with sequence similarity 27 member C |
| ENSG00000165475 | CRYL1 | -0,358012148 | 0,78023891 | 0,24085 | 0,55001001 | crystallin lambda 1 |
| ENSG00000214357 | NEURL1B | -0,358174245 | 0,78015125 | 0,27097 | 0,57248186 | neuralized E3 ubiquitin protein ligase 1B |
| ENSG00000119900 | OGFRL1 | -0,358177566 | 0,780149454 | 0,40279 | 0,66396567 | opioid growth factor receptor like 1 |
| ENSG00000164889 | SLC4A2 | -0,358235631 | 0,780118055 | 0,3496 | 0,62563749 | solute carrier family 4 member 2 |
| ENSG00000254505 | CHMP4A | -0,358292154 | 0,780087492 | 0,23627 | 0,54725688 | charged multivesicular body protein 4A |
| ENSG00000275911 | NDE1 | -0,358434725 | 0,780010405 | 0,34522 | 0,6224039 | nudE neurodevelopment protein 1 |
| ENSG00000102967 | DHODH | -0,358451946 | 0,780001095 | 0,29395 | 0,58836931 | dihydroorotate dehydrogenase (quinone) |
| ENSG00000153802 | TMPRSS11D | -0,358474368 | 0,779988972 | 0,53429 | 0,75405583 | transmembrane serine protease 11D |
| ENSG00000229431 | MED8-AS1 | -0,358486408 | 0,779982463 | 0,67226 | 0,8378522 | MED8 antisense RNA 1 |
| ENSG00000258725 | PRC1-AS1 | -0,358536359 | 0,779955458 | 0,19427 | 0,51562158 | PRC1 antisense RNA 1 |
| ENSG00000287306 | AC016821,1 | -0,358552246 | 0,779946869 | 0,1796 | 0,50392705 | novel transcript |
| ENSG00000287495 | AL356489,4 | -0,358560224 | 0,779942556 | 0,61261 | 0,8035281 | novel transcript |
| ENSG00000263160 | TMEM50B | -0,358796641 | 0,779814756 | 0,41917 | 0,67514348 | transmembrane protein 50B |
| ENSG00000248265 | FLJ12825 | -0,358830301 | 0,779796562 | 0,45272 | 0,6971565 | uncharacterized LOC440101 |
| ENSG00000287525 | AL627308,3 | -0,358899427 | 0,779759199 | 0,43862 | 0,68782243 | novel transcript, antisense to CDC42BPA |
| ENSG00000165566 | AMER2 | -0,358914118 | 0,779751259 | 0,40343 | 0,66399363 | APC membrane recruitment protein 2 |
| ENSG00000171811 | CFAP46 | -0,359009612 | 0,779699648 | 0,17794 | 0,50233162 | cilia and flagella associated protein 46 |
| ENSG00000099812 | MISP | -0,359018974 | 0,779694588 | 0,68078 | 0,84338282 | mitotic spindle positioning |
| ENSG00000225963 | AC009950,1 | -0,359177509 | 0,779608914 | 0,1504 | 0,47857814 | novel transcript, antisense to SP110 |
| ENSG00000213785 | AKR1B1P3 | -0,359276661 | 0,779555335 | 0,67256 | 0,83803462 | aldo-keto reductase family 1 member B1 pseudogene 3 |
| ENSG00000203647 | AC106827,1 | -0,359415724 | 0,779480197 | 0,61942 | 0,80706028 | novel pseudogene |
| ENSG00000113504 | SLC12A7 | -0,359539683 | 0,779413226 | 0,23875 | 0,54892264 | solute carrier family 12 member 7 |
| ENSG00000218537 | MIF-AS1 | -0,35974348 | 0,779303132 | 0,40651 | 0,66597268 | MIF antisense RNA 1 |

| | | | | | | |
|---|---|---|---|---|---|---|
| ENSG00000281335 | MIF-AS1 | -0,35974348 | 0,779303132 | 0,40651 | 0,66597268 | MIF antisense RNA 1 |
| ENSG00000224616 | RTCA-AS1 | -0,35984183 | 0,779250008 | 0,20148 | 0,52214043 | RTCA antisense RNA 1 |
| ENSG00000184307 | ZDHHC23 | -0,359852056 | 0,779244485 | 0,49863 | 0,72979594 | zinc finger DHHC-type palmitoyltransferase 23 |
| ENSG00000225618 | ZNRD1ASP | -0,359945811 | 0,779193846 | 0,40047 | 0,66257606 | zinc ribbon domain containing 1 antisense, pseudogene |
| ENSG00000236598 | ZNRD1ASP | -0,359945811 | 0,779193846 | 0,40047 | 0,66257606 | zinc ribbon domain containing 1 antisense, pseudogene |
| ENSG00000176406 | RIMS2 | -0,359976595 | 0,77917722 | 0,05398 | 0,31167721 | regulating synaptic membrane exocytosis 2 |
| ENSG00000173157 | ADAMTS20 | -0,360107039 | 0,779106773 | 0,339 | 0,61800397 | ADAM metallopeptidase with thrombospondin type 1 motif 20 |
| ENSG00000214199 | EEF1A1P12 | -0,360258667 | 0,779024892 | 0,65409 | 0,82680983 | eukaryotic translation elongation factor 1 alpha 1 pseudogene 12 |
| ENSG00000197283 | SYNGAP1 | -0,360339059 | 0,778981484 | 0,29472 | 0,58860102 | synaptic Ras GTPase activating protein 1 |
| ENSG00000223963 | THAP12P8 | -0,360623811 | 0,778827747 | 0,64349 | 0,82109591 | THAP domain containing 12 pseudogene 8 |
| ENSG00000146677 | AC004453,1 | -0,36068428 | 0,778795104 | 0,61712 | 0,80563454 | ribosomal protein L32 (RPL32) pseudogene |
| ENSG00000130313 | PGLS | -0,360899339 | 0,77867902 | 0,25563 | 0,5600812 | 6-phosphogluconolactonase |
| ENSG00000260266 | PPIAP46 | -0,361003789 | 0,778622646 | 0,35151 | 0,62674766 | peptidylprolyl isomerase A pseudogene 46 |
| ENSG00000288314 | PPIAP46 | -0,361003789 | 0,778622646 | 0,35151 | 0,62674766 | peptidylprolyl isomerase A pseudogene 46 |
| ENSG00000172057 | ORMDL3 | -0,361036323 | 0,778605088 | 0,18199 | 0,50524085 | ORMDL sphingolipid biosynthesis regulator 3 |
| ENSG00000228393 | LINC01004 | -0,361083872 | 0,778579427 | 0,23533 | 0,54677606 | long intergenic non-protein coding RNA 1004 |
| ENSG00000273238 | TMEM271 | -0,36114005 | 0,778549109 | 0,5979 | 0,79484486 | transmembrane protein 271 |
| ENSG00000224728 | IMPDH1P8 | -0,361327838 | 0,778447776 | 0,43344 | 0,68473723 | inosine monophosphate dehydrogenase 1 pseudogene 8 |
| ENSG00000142065 | ZFP14 | -0,361395475 | 0,778411282 | 0,1916 | 0,51435702 | ZFP14 zinc finger protein |
| ENSG00000233189 | RPL12P29 | -0,361508938 | 0,778350065 | 0,56282 | 0,77200886 | ribosomal protein L12 pseudogene 29 |
| ENSG00000259495 | AC016705,2 | -0,361688582 | 0,778253151 | 0,10604 | 0,41557034 | novel transcript |
| ENSG00000260664 | AC004158,1 | -0,36181655 | 0,778184123 | 0,36208 | 0,63354671 | novel transcript |
| ENSG00000283380 | AL807742,1 | -0,362006824 | 0,778081496 | 0,61307 | 0,8035281 | novel transcript |
| ENSG00000233597 | AC133435,1 | -0,362071173 | 0,778046792 | 0,57075 | 0,77748547 | ribosomal protein L19 (RPL19) pseudogene |
| ENSG00000228736 | RGL2 | -0,362088134 | 0,778037645 | 0,32137 | 0,60577115 | ral guanine nucleotide dissociation stimulator like 2 |
| ENSG00000224841 | RGL2 | -0,362088134 | 0,778037645 | 0,32137 | 0,60577115 | ral guanine nucleotide dissociation stimulator like 2 |
| ENSG00000237825 | RGL2 | -0,362088134 | 0,778037645 | 0,32137 | 0,60577115 | ral guanine nucleotide dissociation stimulator like 2 |
| ENSG00000213626 | LBH | -0,362092713 | 0,778035176 | 0,32248 | 0,60577115 | LBH regulator of WNT signaling pathway |
| ENSG00000236779 | AL512306,1 | -0,362125802 | 0,778017331 | 0,52728 | 0,75031847 | PEF protein with a long N-terminal hydrophobic domain (peflin) (PEF) pseudogene |
| ENSG00000277868 | CCDC125 | -0,362162382 | 0,777997605 | 0,42237 | 0,67663581 | coiled-coil domain containing 125 |

| | | | | | | |
|---|---|---|---|---|---|---|
| ENSG00000260493 | AC011773,4 | -0,362212152 | 0,777970765 | 0,537 | 0,75567847 | novel transcript, antisense to E2F5 |
| ENSG00000113838 | TBCCD1 | -0,362224437 | 0,777964141 | 0,29519 | 0,58886617 | TBCC domain containing 1 |
| ENSG00000130590 | SAMD10 | -0,362248358 | 0,777951242 | 0,19019 | 0,51331106 | sterile alpha motif domain containing 10 |
| ENSG00000213315 | AL122020,1 | -0,362544494 | 0,777791571 | 0,65051 | 0,82492577 | ribosomal protein S18 (RPS18) pseudogene |
| ENSG00000139737 | SLAIN1 | -0,362634771 | 0,777742903 | 0,06492 | 0,33778346 | SLAIN motif family member 1 |
| ENSG00000076984 | MAP2K7 | -0,362661742 | 0,777728363 | 0,21575 | 0,53440695 | mitogen-activated protein kinase kinase 7 |
| ENSG00000261824 | LINC00662 | -0,362681996 | 0,777717444 | 0,09241 | 0,39283968 | long intergenic non-protein coding RNA 662 |
| ENSG00000149016 | TUT1 | -0,362697366 | 0,777709159 | 0,17015 | 0,49606609 | terminal uridylyl transferase 1, U6 snRNA-specific |
| ENSG00000256980 | KHDC1L | -0,362867684 | 0,777617351 | 0,68332 | 0,84495012 | KH domain containing 1 like |
| ENSG00000253369 | AC022034,3 | -0,362956419 | 0,777569524 | 0,36179 | 0,63352908 | novel transcript |
| ENSG00000146856 | AGBL3 | -0,363032651 | 0,777528439 | 0,2102 | 0,52904355 | ATP/GTP binding protein like 3 |
| ENSG00000140853 | NLRC5 | -0,363190656 | 0,777443288 | 0,6639 | 0,83247324 | NLR family CARD domain containing 5 |
| ENSG00000230943 | LINC02541 | -0,363191638 | 0,777442759 | 0,64442 | 0,82154386 | long intergenic non-protein coding RNA 2541 |
| ENSG00000261970 | AL391840,2 | -0,363218625 | 0,777428216 | 0,70438 | 0,85743518 | solute carrier family 19, member 3 (SLC19A3) pseudogene |
| ENSG00000116176 | TPSG1 | -0,363593648 | 0,777226153 | 0,74307 | 0,87840582 | tryptase gamma 1 |
| ENSG00000240622 | RPL7P15 | -0,363716853 | 0,777159781 | 0,54435 | 0,76052802 | ribosomal protein L7 pseudogene 15 |
| ENSG00000243414 | TICAM2 | -0,363773995 | 0,777129 | 0,76145 | 0,88879633 | toll like receptor adaptor molecule 2 |
| ENSG00000263126 | AC040162,3 | -0,363888835 | 0,777067142 | 0,40943 | 0,66795766 | novel transcript |
| ENSG00000248152 | ANK2-AS1 | -0,363995351 | 0,777009773 | 0,45257 | 0,69704777 | ANK2 antisense RNA 1 |
| ENSG00000196655 | TRAPPC4 | -0,36433916 | 0,776824625 | 0,27233 | 0,57315552 | trafficking protein particle complex 4 |
| ENSG00000284753 | EEF1AKMT4 | -0,364395208 | 0,776794447 | 0,45718 | 0,7007187 | EEF1A lysine methyltransferase 4 |
| ENSG00000258311 | AC009779,3 | -0,364535476 | 0,776718926 | 0,07818 | 0,36540845 | novel protein |
| ENSG00000282143 | AL137798,2 | -0,364730464 | 0,776613955 | 0,91679 | 0,96378593 | novel transcript |
| ENSG00000260942 | CAPN10-DT | -0,364886417 | 0,776530008 | 0,23461 | 0,54623956 | CAPN10 divergent transcript |
| ENSG00000042286 | AIFM2 | -0,365136792 | 0,776395256 | 0,1465 | 0,47433542 | apoptosis inducing factor mitochondria associated 2 |
| ENSG00000255224 | AC109322,1 | -0,36548709 | 0,776206764 | 0,27798 | 0,57629868 | TEC |
| ENSG00000127540 | UQCR11 | -0,365621168 | 0,77613463 | 0,29167 | 0,58655914 | ubiquinol-cytochrome c reductase, complex III subunit XI |
| ENSG00000208892 | SNORA49 | -0,36569127 | 0,776096918 | 0,29562 | 0,58930611 | small nucleolar RNA, H/ACA box 49 |
| ENSG00000104885 | DOT1L | -0,365729072 | 0,776076582 | 0,22377 | 0,53967355 | DOT1 like histone lysine methyltransferase |
| ENSG00000277581 | AL023803,3 | -0,365763846 | 0,776057876 | 0,62058 | 0,80776776 | novel transcript, sense intronic to DHX35 |
| ENSG00000107018 | RLN1 | -0,365804935 | 0,776035774 | 0,56319 | 0,77230901 | relaxin 1 |
| ENSG00000257150 | PGAM1P5 | -0,365812962 | 0,776031456 | 0,55941 | 0,77045525 | phosphoglycerate mutase 1 pseudogene 5 |
| ENSG00000220008 | LINGO3 | -0,365954773 | 0,775955179 | 0,71231 | 0,86152011 | leucine rich repeat and Ig domain containing 3 |

| | | | | | | |
|---|---|---|---|---|---|---|
| ENSG00000239704 | CDRT4 | -0,3659832 | 0,77593989 | 0,54822 | 0,76347743 | CMT1A duplicated region transcript 4 |
| ENSG00000263551 | AP000829,1 | -0,366001054 | 0,775930288 | 0,39022 | 0,65427905 | novel transcript |
| ENSG00000204315 | FKBPL | -0,366211945 | 0,775816871 | 0,69424 | 0,85162906 | FKBP prolyl isomerase like |
| ENSG00000232164 | LINC01873 | -0,366386254 | 0,775723142 | 0,2384 | 0,54875544 | long intergenic non-protein coding RNA 1873 |
| ENSG00000187961 | KLHL17 | -0,366465357 | 0,77568061 | 0,31208 | 0,59925736 | kelch like family member 17 |
| ENSG00000227660 | UST-AS1 | -0,366592197 | 0,775612416 | 0,63386 | 0,81549489 | UST antisense RNA 1 |
| ENSG00000263843 | AC022211,2 | -0,366598469 | 0,775609044 | 0,40315 | 0,66399303 | novel transcript, antisense to SLC25A19 |
| ENSG00000225721 | AL592166,1 | -0,366613674 | 0,77560087 | 0,50373 | 0,73293685 | novel transcript |
| ENSG00000103226 | NOMO3 | -0,366779851 | 0,775511537 | 0,38653 | 0,65194712 | NODAL modulator 3 |
| ENSG00000109089 | CDR2L | -0,366828472 | 0,775485402 | 0,24029 | 0,54948131 | cerebellar degeneration related protein 2 like |
| ENSG00000198585 | NUDT16 | -0,366955594 | 0,775417073 | 0,05911 | 0,32424653 | nudix hydrolase 16 |
| ENSG00000181240 | SLC25A41 | -0,3670346 | 0,77537461 | 0,48002 | 0,71732148 | solute carrier family 25 member 41 |
| ENSG00000255555 | AP000857,2 | -0,367109158 | 0,77533454 | 0,68948 | 0,84865812 | novel transcript |
| ENSG00000181666 | ZNF875 | -0,367201899 | 0,775284701 | 0,23033 | 0,54355692 | zinc finger protein 875 |
| ENSG00000132692 | BCAN | -0,367304298 | 0,775229675 | 0,53975 | 0,75802973 | brevican |
| ENSG00000100941 | PNN | -0,367351191 | 0,775204478 | 0,07765 | 0,36468824 | pinin, desmosome associated protein |
| ENSG00000283122 | HYMAI | -0,367505862 | 0,775121373 | 0,31057 | 0,59852059 | hydatidiform mole associated and imprinted |
| ENSG00000099840 | IZUMO4 | -0,367592258 | 0,775074955 | 0,41074 | 0,66856387 | IZUMO family member 4 |
| ENSG00000196118 | CCDC189 | -0,367593947 | 0,775074048 | 0,24128 | 0,55027101 | coiled-coil domain containing 189 |
| ENSG00000284707 | AC079781,5 | -0,367623078 | 0,775058398 | 0,19201 | 0,5145257 | novel transcript |
| ENSG00000249621 | AC113349,1 | -0,367697476 | 0,77501843 | 0,64533 | 0,82203909 | novel transcript |
| ENSG00000163362 | INAVA | -0,367749856 | 0,774990292 | 0,29856 | 0,59089801 | innate immunity activator |
| ENSG00000287322 | AC068189,2 | -0,367801733 | 0,774962425 | 0,36568 | 0,6359214 | novel transcript, antisense to NDUFAF6 |
| ENSG00000171159 | C9orf16 | -0,367871076 | 0,774925177 | 0,04163 | 0,27639057 | chromosome 9 open reading frame 16 |
| ENSG00000129514 | FOXA1 | -0,368001618 | 0,774855062 | 0,6869 | 0,84725905 | forkhead box A1 |
| ENSG00000007516 | BAIAP3 | -0,368240966 | 0,774726522 | 0,3825 | 0,64932112 | BAI1 associated protein 3 |
| ENSG00000287094 | AL049552,2 | -0,368244387 | 0,774724684 | 0,53729 | 0,7559015 | novel transcript, antisense to AHI1 |
| ENSG00000119242 | CCDC92 | -0,36833756 | 0,774674652 | 0,05798 | 0,321265 | coiled-coil domain containing 92 |
| ENSG00000267871 | ZNF460-AS1 | -0,368452144 | 0,774613128 | 0,30885 | 0,59767335 | ZNF460 antisense RNA 1 |
| ENSG00000101161 | PRPF6 | -0,368482299 | 0,774596937 | 0,1582 | 0,48525382 | pre-mRNA processing factor 6 |
| ENSG00000105379 | ETFB | -0,368576868 | 0,774546163 | 0,22135 | 0,53805234 | electron transfer flavoprotein subunit beta |
| ENSG00000245522 | LINC02709 | -0,368669547 | 0,774496408 | 0,53594 | 0,75499048 | long intergenic non-protein coding RNA 2709 |
| ENSG00000115084 | SLC35F5 | -0,368758587 | 0,774448609 | 0,2675 | 0,56973003 | solute carrier family 35 member F5 |

| | | | | | | |
|---|---|---|---|---|---|---|
| ENSG00000179950 | PUF60 | -0,36893989 | 0,77435129 | 0,34907 | 0,62519057 | poly(U) binding splicing factor 60 |
| ENSG00000112787 | FBRSL1 | -0,369068024 | 0,774282519 | 0,33306 | 0,61306511 | fibrosin like 1 |
| ENSG00000211584 | SLC48A1 | -0,369079792 | 0,774276203 | 0,14526 | 0,47289051 | solute carrier family 48 member 1 |
| ENSG00000230844 | ZNF674-AS1 | -0,369084917 | 0,774273453 | 0,11371 | 0,42855874 | ZNF674 antisense RNA 1 (head to head) |
| ENSG00000259726 | CSPG4P11 | -0,369122061 | 0,774253518 | 0,29748 | 0,59034513 | chondroitin sulfate proteoglycan 4 pseudogene 11 |
| ENSG00000276128 | AL591441,1 | -0,369179559 | 0,774222662 | 0,44704 | 0,69339238 | glucoside xylosyltransferase 1 pseudogene 5 |
| ENSG00000269934 | AL353593,1 | -0,369472264 | 0,774065597 | 0,23995 | 0,54936679 | novel transcript, antisense to OBSCN |
| ENSG00000224727 | FCF1P7 | -0,369502664 | 0,774049287 | 0,72527 | 0,86894646 | FCF1 pseudogene 7 |
| ENSG00000160446 | ZDHHC12 | -0,369518065 | 0,774041023 | 0,37338 | 0,64204597 | zinc finger DHHC-type palmitoyltransferase 12 |
| ENSG00000224451 | ATP5PBP1 | -0,369529299 | 0,774034996 | 0,54743 | 0,76285787 | ATP synthase peripheral stalk-membrane subunit b pseudogene 1 |
| ENSG00000258752 | AL357093,2 | -0,369724697 | 0,773930168 | 0,4293 | 0,68217673 | novel transcript, antisense to FOXN3 |
| ENSG00000275494 | AC133552,5 | -0,369742952 | 0,773920375 | 0,37065 | 0,63997937 | novel transcript |
| ENSG00000275029 | HMGB1P24 | -0,369952726 | 0,773807852 | 0,42575 | 0,67925635 | high mobility group box 1 pseudogene 24 |
| ENSG00000282695 | HMGB1P24 | -0,369952726 | 0,773807852 | 0,42575 | 0,67925635 | high mobility group box 1 pseudogene 24 |
| ENSG00000275282 | CFL1P5 | -0,370089739 | 0,773734367 | 0,64953 | 0,82442734 | cofilin 1 pseudogene 5 |
| ENSG00000213830 | CFL1P5 | -0,370089739 | 0,773734367 | 0,64953 | 0,82442734 | cofilin 1 pseudogene 5 |
| ENSG00000103005 | USB1 | -0,370101284 | 0,773728176 | 0,21994 | 0,53732404 | U6 snRNA biogenesis phosphodiesterase 1 |
| ENSG00000145022 | TCTA | -0,3701266 | 0,773714599 | 0,24413 | 0,55253387 | T cell leukemia translocation altered |
| ENSG00000278621 | AC037198,2 | -0,370219071 | 0,773665008 | 0,50535 | 0,73393676 | novel transcript, antisense to THBS1 |
| ENSG00000102158 | MAGT1 | -0,370248835 | 0,773649047 | 0,35585 | 0,63013888 | magnesium transporter 1 |
| ENSG00000139890 | REM2 | -0,370302329 | 0,773620361 | 0,18688 | 0,51036936 | RRAD and GEM like GTPase 2 |
| ENSG00000184113 | CLDN5 | -0,370335558 | 0,773602543 | 0,40508 | 0,66481282 | claudin 5 |
| ENSG00000230201 | ATP6V0CP1 | -0,370457795 | 0,773537 | 0,74319 | 0,87840582 | ATPase H+ transporting V0 subunit c pseudogene 1 |
| ENSG00000250056 | LINC01018 | -0,370518187 | 0,77350462 | 0,51405 | 0,74032076 | long intergenic non-protein coding RNA 1018 |
| ENSG00000271698 | AC233992,2 | -0,370575858 | 0,7734737 | 0,52929 | 0,75112358 | novel transcript |
| ENSG00000285047 | AC233992,6 | -0,370575858 | 0,7734737 | 0,52929 | 0,75112358 | novel transcript |
| ENSG00000207166 | SNORA68 | -0,370793249 | 0,773357158 | 0,48924 | 0,72376431 | small nucleolar RNA, H/ACA box 68 |
| ENSG00000176978 | DPP7 | -0,370846897 | 0,773328401 | 0,36305 | 0,63401731 | dipeptidyl peptidase 7 |
| ENSG00000258738 | AL121603,2 | -0,370902116 | 0,773298802 | 0,16022 | 0,48737351 | novel transcript, antisense to BAZ1A |
| ENSG00000174282 | ZBTB4 | -0,370924518 | 0,773286795 | 0,03945 | 0,2687757 | zinc finger and BTB domain containing 4 |
| ENSG00000283868 | ZBTB4 | -0,370924518 | 0,773286795 | 0,03945 | 0,2687757 | zinc finger and BTB domain containing 4 |
| ENSG00000227825 | SLC9A7P1 | -0,371118151 | 0,773183014 | 0,42129 | 0,67634721 | solute carrier family 9 member 7 pseudogene 1 |

| | | | | | | |
|---|---|---|---|---|---|---|
| ENSG00000177697 | CD151 | -0,371227234 | 0,773124556 | 0,21703 | 0,53505085 | CD151 molecule (Raph blood group) |
| ENSG00000274180 | NATD1 | -0,371608128 | 0,772920466 | 0,31093 | 0,59883009 | N-acetyltransferase domain containing 1 |
| ENSG00000145506 | NKD2 | -0,371767201 | 0,772835248 | 0,66353 | 0,8324578 | NKD inhibitor of WNT signaling pathway 2 |
| ENSG00000139899 | CBLN3 | -0,371782854 | 0,772826862 | 0,2868 | 0,58325473 | cerebellin 3 precursor |
| ENSG00000287623 | AC037193,1 | -0,371968039 | 0,772727668 | 0,22745 | 0,54168591 | novel transcript, antisense to SRGAP3 |
| ENSG00000267811 | AP001160,2 | -0,372163745 | 0,772622852 | 0,24365 | 0,55235189 | novel transcript, antisense to TAF6L |
| ENSG00000135828 | RNASEL | -0,372211998 | 0,772597011 | 0,12051 | 0,43944057 | ribonuclease L |
| ENSG00000203799 | CCDC162P | -0,372392218 | 0,772500506 | 0,45139 | 0,69645146 | coiled-coil domain containing 162, pseudogene |
| ENSG00000259088 | AL137779,2 | -0,372454597 | 0,772467105 | 0,55472 | 0,76760293 | novel transcript, antisense to PPP2R5C |
| ENSG00000263873 | THY1-AS1 | -0,372493166 | 0,772446454 | 0,52764 | 0,75037219 | THY1 antisense RNA 1 |
| ENSG00000132793 | LPIN3 | -0,372501158 | 0,772442175 | 0,33785 | 0,61751351 | lipin 3 |
| ENSG00000227528 | DIAPH3-AS1 | -0,372595836 | 0,772391485 | 0,64277 | 0,82094129 | DIAPH3 antisense RNA 1 |
| ENSG00000269737 | AL691432,1 | -0,372597019 | 0,772390851 | 0,14381 | 0,47134131 | ribosomal protein S7 (RPS7) pseudogene |
| ENSG00000149932 | TMEM219 | -0,372601841 | 0,77238827 | 0,28004 | 0,57770942 | transmembrane protein 219 |
| ENSG00000259228 | HNRNPA1P62 | -0,372814302 | 0,772274531 | 0,57206 | 0,77840742 | heterogeneous nuclear ribonucleoprotein A1 pseudogene 62 |
| ENSG00000160352 | ZNF714 | -0,372849397 | 0,772255745 | 0,2078 | 0,52736343 | zinc finger protein 714 |
| ENSG00000184206 | GOLGA6L4 | -0,372909672 | 0,772223481 | 0,1918 | 0,51436129 | golgin A6 family like 4 |
| ENSG00000260285 | AL133367,1 | -0,373148245 | 0,772095792 | 0,16278 | 0,48945635 | novel transcript, antisense to TRMT61A |
| ENSG00000197119 | SLC25A29 | -0,373187797 | 0,772074625 | 0,20009 | 0,5209612 | solute carrier family 25 member 29 |
| ENSG00000229236 | TTTY10 | -0,373198954 | 0,772068654 | 0,17837 | 0,50278602 | testis-specific transcript, Y-linked 10 |
| ENSG00000267568 | AC016168,2 | -0,373237476 | 0,772048039 | 0,73963 | 0,87653492 | novel transcript |
| ENSG00000178115 | GOLGA8Q | -0,373300356 | 0,77201439 | 0,66712 | 0,83471429 | golgin A8 family member Q |
| ENSG00000175283 | DOLK | -0,373466202 | 0,771925648 | 0,31141 | 0,59890747 | dolichol kinase |
| ENSG00000137166 | FOXP4 | -0,3736083 | 0,771849621 | 0,18513 | 0,50872634 | forkhead box P4 |
| ENSG00000256222 | MTRNR2L3 | -0,373706608 | 0,771797027 | 0,47893 | 0,71659623 | MT-RNR2 like 3 |
| ENSG00000163590 | PPM1L | -0,373804952 | 0,771744418 | 0,13689 | 0,4641716 | protein phosphatase, Mg2+/Mn2+ dependent 1L |
| ENSG00000111644 | ACRBP | -0,37408847 | 0,77159277 | 0,50391 | 0,73297531 | acrosin binding protein |
| ENSG00000269973 | AC010969,2 | -0,374310175 | 0,771474205 | 0,25436 | 0,55940142 | novel transcript |
| ENSG00000161091 | MFSD12 | -0,374726534 | 0,771251591 | 0,24141 | 0,55033747 | major facilitator superfamily domain containing 12 |
| ENSG00000280623 | PCAT14 | -0,374888561 | 0,771164978 | 0,1368 | 0,4641716 | prostate cancer associated transcript 14 |
| ENSG00000249867 | LINC02742 | -0,374911036 | 0,771152965 | 0,41319 | 0,67067821 | long intergenic non-protein coding RNA 2742 |
| ENSG00000165323 | FAT3 | -0,374995633 | 0,771107747 | 0,25424 | 0,55925777 | FAT atypical cadherin 3 |

| | | | | | | |
|---|---|---|---|---|---|---|
| ENSG00000286269 | AC092661,1 | -0,375146048 | 0,771027355 | 0,21788 | 0,53550245 | novel transcript |
| ENSG00000081692 | JMJD4 | -0,375204728 | 0,770995996 | 0,18628 | 0,50961491 | jumonji domain containing 4 |
| ENSG00000264475 | AP005062,1 | -0,375219425 | 0,770988141 | 0,51036 | 0,7379578 | novel transcript, antisense to LAMA1 |
| ENSG00000183762 | KREMEN1 | -0,375251012 | 0,770971261 | 0,43673 | 0,68681257 | kringle containing transmembrane protein 1 |
| ENSG00000256984 | AC008013,2 | -0,375452397 | 0,770863649 | 0,47243 | 0,71184073 | mitochondrially encoded cytochrome c oxidase I (MT-CO1) pseudogene |
| ENSG00000153767 | GTF2E1 | -0,375671814 | 0,770746419 | 0,34518 | 0,6224039 | general transcription factor IIE subunit 1 |
| ENSG00000120314 | WDR55 | -0,375677542 | 0,770743359 | 0,23297 | 0,54555709 | WD repeat domain 55 |
| ENSG00000273010 | AL360270,3 | -0,375702951 | 0,770729784 | 0,57535 | 0,7801789 | novel transcript, antisense to LRIF1 |
| ENSG00000243055 | GK-AS1 | -0,375712925 | 0,770724456 | 0,65471 | 0,82726721 | GK antisense RNA 1 |
| ENSG00000279006 | AL022323,1 | -0,375978636 | 0,770582519 | 0,25903 | 0,5628026 | novel transcript |
| ENSG00000153093 | ACOXL | -0,375994356 | 0,770574123 | 0,40657 | 0,66597268 | acyl-CoA oxidase like |
| ENSG00000198753 | PLXNB3 | -0,376176741 | 0,770476714 | 0,45881 | 0,70137313 | plexin B3 |
| ENSG00000256053 | COA8 | -0,376383569 | 0,770366264 | 0,04768 | 0,29344099 | cytochrome c oxidase assembly factor 8 |
| ENSG00000242575 | TUBAP13 | -0,376402078 | 0,770356381 | 0,56002 | 0,77076855 | tubulin alpha pseudogene 13 |
| ENSG00000245556 | SCAMP1-AS1 | -0,376516675 | 0,770295192 | 0,07738 | 0,36426829 | SCAMP1 antisense RNA 1 |
| ENSG00000103260 | METRN | -0,376573303 | 0,770264957 | 0,40063 | 0,66264525 | meteorin, glial cell differentiation regulator |
| ENSG00000287555 | AC023424,2 | -0,376608952 | 0,770245924 | 0,26344 | 0,56613578 | novel transcript, antisense to FBXW7 |
| ENSG00000285940 | AC009899,1 | -0,376790466 | 0,770149021 | 0,45061 | 0,69592349 | novel transcript, antisense to CELF4 |
| ENSG00000225727 | AL139184,1 | -0,377021898 | 0,770025486 | 0,63552 | 0,81657648 | filamin A interacting protein 1-like (FILIP1L) pseudogene |
| ENSG00000240163 | AC087385,1 | -0,377023553 | 0,770024603 | 0,64402 | 0,82138283 | ribosomal protein L36a (RPL36A) pseudogene |
| ENSG00000197532 | OR6Y1 | -0,377031324 | 0,770020455 | 0,22822 | 0,54221266 | olfactory receptor family 6 subfamily Y member 1 |
| ENSG00000149541 | B3GAT3 | -0,377110158 | 0,76997838 | 0,14805 | 0,47519813 | beta-1,3-glucuronyltransferase 3 |
| ENSG00000234290 | AC116366,1 | -0,377209055 | 0,7699256 | 0,36524 | 0,63543797 | novel transcript |
| ENSG00000116691 | MIIP | -0,377265347 | 0,769895559 | 0,2656 | 0,56848959 | migration and invasion inhibitory protein |
| ENSG00000232626 | AC099336,1 | -0,377376771 | 0,769836099 | 0,57585 | 0,78043323 | FK506 binding protein 8, 38kDa (FKBP8) pseudogene |
| ENSG00000213328 | AC104117,1 | -0,377460507 | 0,769791418 | 0,4602 | 0,70210714 | DCN1, defective in cullin neddylation 1, domain containing 3 (S, cerevisiae) (DCUN1D3) pseudogene |
| ENSG00000235008 | AL023775,1 | -0,377471225 | 0,7697857 | 0,45868 | 0,70133296 | IGF-II mRNA-binding protein 3 (KOC1) pseudogene |
| ENSG00000167113 | COQ4 | -0,377513581 | 0,7697631 | 0,26083 | 0,56400166 | coenzyme Q4 |
| ENSG00000267681 | AC135721,1 | -0,377611764 | 0,769710715 | 0,42222 | 0,67663014 | high mobility group nucleosome binding domain 1 (HMGN1) pseudogene |
| ENSG00000244274 | DBNDD2 | -0,377637726 | 0,769696864 | 0,18839 | 0,51166676 | dysbindin domain containing 2 |

| | | | | | | |
|---|---|---|---|---|---|---|
| ENSG00000090661 | CERS4 | -0,377648874 | 0,769690916 | 0,26138 | 0,56439768 | ceramide synthase 4 |
| ENSG00000280708 | AC006359,3 | -0,377725693 | 0,769649934 | 0,46889 | 0,70848626 | novel transcript |
| ENSG00000284287 | AC270107,7 | -0,378070493 | 0,769466012 | 0,17169 | 0,49704622 | |
| ENSG00000284036 | AC270107,3 | -0,378070493 | 0,769466012 | 0,17169 | 0,49704622 | |
| ENSG00000284230 | AC270107,6 | -0,378070493 | 0,769466012 | 0,17169 | 0,49704622 | |
| ENSG00000283963 | AC270107,1 | -0,378070493 | 0,769466012 | 0,17169 | 0,49704622 | |
| ENSG00000284151 | AC270107,5 | -0,378070493 | 0,769466012 | 0,17169 | 0,49704622 | |
| ENSG00000283989 | AC270107,2 | -0,378070493 | 0,769466012 | 0,17169 | 0,49704622 | |
| ENSG00000160602 | NEK8 | -0,378677413 | 0,769142377 | 0,24592 | 0,55372423 | NIMA related kinase 8 |
| ENSG00000186998 | EMID1 | -0,37879035 | 0,769082169 | 0,31368 | 0,60035326 | EMI domain containing 1 |
| ENSG00000272523 | LINC01023 | -0,378897241 | 0,769025189 | 0,48155 | 0,71837579 | long intergenic non-protein coding RNA 1023 |
| ENSG00000237031 | AC008067,1 | -0,378916601 | 0,76901487 | 0,69385 | 0,85125839 | novel transcript |
| ENSG00000286669 | AC008532,1 | -0,378943769 | 0,769000388 | 0,65346 | 0,82638628 | novel transcript |
| ENSG00000258422 | AL160191,1 | -0,378964932 | 0,768989107 | 0,25261 | 0,55796133 | novel transcript, antisense to SLC8A3 |
| ENSG00000250686 | KHDRBS2-OT1 | -0,378998064 | 0,768971447 | 0,51179 | 0,7388352 | KHDRBS2 3'UTR overlapping transcript 1 |
| ENSG00000250182 | EEF1A1P13 | -0,379164989 | 0,76888248 | 0,58262 | 0,7847455 | eukaryotic translation elongation factor 1 alpha 1 pseudogene 13 |
| ENSG00000125844 | RRBP1 | -0,379237491 | 0,768843841 | 0,32882 | 0,60992716 | ribosome binding protein 1 |
| ENSG00000198625 | MDM4 | -0,379398226 | 0,768758187 | 0,2377 | 0,54861973 | MDM4 regulator of p53 |
| ENSG00000101224 | CDC25B | -0,379515103 | 0,76869591 | 0,30684 | 0,59592193 | cell division cycle 25B |
| ENSG00000275111 | ZNF2 | -0,37953761 | 0,768683918 | 0,25707 | 0,56155289 | zinc finger protein 2 |
| ENSG00000164136 | IL15 | -0,379693975 | 0,768600609 | 0,31946 | 0,60429866 | interleukin 15 |
| ENSG00000157470 | FAM81A | -0,380015594 | 0,768429284 | 0,41116 | 0,668955 | family with sequence similarity 81 member A |
| ENSG00000285933 | AP003498,2 | -0,380067237 | 0,768401778 | 0,32693 | 0,60889526 | novel transcript |
| ENSG00000232466 | H3P31 | -0,380126536 | 0,768370196 | 0,623 | 0,80887719 | H3 histone pseudogene 31 |
| ENSG00000125255 | SLC10A2 | -0,380164441 | 0,768350008 | 0,60224 | 0,79696356 | solute carrier family 10 member 2 |
| ENSG00000137642 | SORL1 | -0,380183565 | 0,768339823 | 0,29657 | 0,58979902 | sortilin related receptor 1 |
| ENSG00000156509 | FBXO43 | -0,380323246 | 0,768265436 | 0,39669 | 0,65929197 | F-box protein 43 |
| ENSG00000184811 | TRARG1 | -0,380435204 | 0,768205819 | 0,44862 | 0,6944286 | trafficking regulator of GLUT4 (SLC2A4) 1 |
| ENSG00000105325 | FZR1 | -0,380489984 | 0,76817665 | 0,15666 | 0,48425071 | fizzy and cell division cycle 20 related 1 |
| ENSG00000264247 | LINC00909 | -0,380524688 | 0,768158172 | 0,28854 | 0,58455903 | long intergenic non-protein coding RNA 909 |
| ENSG00000286293 | AP005901,5 | -0,380574682 | 0,768131553 | 0,1825 | 0,5059851 | novel transcript |
| ENSG00000198879 | SFMBT2 | -0,380669037 | 0,768081317 | 0,34664 | 0,62350287 | Scm like with four mbt domains 2 |

| | | | | | | |
|---|---|---|---|---|---|---|
| ENSG00000249803 | AC112178,1 | -0,380719235 | 0,768054593 | 0,31833 | 0,60349082 | novel transcript |
| ENSG00000028310 | BRD9 | -0,380841032 | 0,767989753 | 0,15822 | 0,48525382 | bromodomain containing 9 |
| ENSG00000286039 | AC093849,2 | -0,380872976 | 0,767972749 | 0,64327 | 0,82098995 | novel transcript |
| ENSG00000232677 | LINC00665 | -0,380873995 | 0,767972207 | 0,32392 | 0,60659203 | long intergenic non-protein coding RNA 665 |
| ENSG00000099617 | EFNA2 | -0,380876859 | 0,767970682 | 0,34924 | 0,62541064 | ephrin A2 |
| ENSG00000212864 | RNF208 | -0,381121329 | 0,767840558 | 0,37541 | 0,64381159 | ring finger protein 208 |
| ENSG00000174938 | SEZ6L2 | -0,381167892 | 0,767815776 | 0,28074 | 0,57812368 | seizure related 6 homolog like 2 |
| ENSG00000185823 | NPAP1 | -0,381302989 | 0,76774388 | 0,20468 | 0,52443222 | nuclear pore associated protein 1 |
| ENSG00000231849 | UBE2V2P4 | -0,381407513 | 0,767688258 | 0,59254 | 0,79121945 | ubiquitin conjugating enzyme E2 V2 pseudogene 4 |
| ENSG00000230590 | FTX | -0,381694931 | 0,767535332 | 0,1933 | 0,51532949 | FTX transcript, XIST regulator |
| ENSG00000196302 | GUSBP15 | -0,381809914 | 0,767474162 | 0,27682 | 0,57549599 | GUSB pseudogene 15 |
| ENSG00000187686 | KRT18P59 | -0,38195902 | 0,767394846 | 0,3845 | 0,65052153 | keratin 18 pseudogene 59 |
| ENSG00000285531 | Z83840,1 | -0,381962939 | 0,767392761 | 0,53666 | 0,75543626 | LLP homolog, long-term synaptic facilitation ( LLPH) pseudogene |
| ENSG00000146540 | C7orf50 | -0,382081438 | 0,767329732 | 0,15251 | 0,48061452 | chromosome 7 open reading frame 50 |
| ENSG00000033327 | GAB2 | -0,382117132 | 0,767310748 | 0,08602 | 0,38030214 | GRB2 associated binding protein 2 |
| ENSG00000261240 | AC009065,6 | -0,382181361 | 0,767276588 | 0,4467 | 0,69321646 | novel transcript, antisense to PKD1 |
| ENSG00000269967 | AL136115,2 | -0,382227488 | 0,767252056 | 0,48614 | 0,72164423 | novel transcript |
| ENSG00000250764 | AC025178,1 | -0,382239672 | 0,767245576 | 0,60378 | 0,79770754 | novel transcript |
| ENSG00000253330 | AC024451,2 | -0,38224654 | 0,767241924 | 0,3121 | 0,59925736 | NSFL1 (p97) cofactor (p47) (NSFL1C) pseudogene |
| ENSG00000182606 | TRAK1 | -0,382361399 | 0,767180843 | 0,01757 | 0,1746915 | trafficking kinesin protein 1 |
| ENSG00000251364 | AC107884,1 | -0,382586874 | 0,767060952 | 0,22715 | 0,54168591 | novel transcript, antisense to OLFML1 |
| ENSG00000286316 | Z93020,1 | -0,382605361 | 0,767051122 | 0,61096 | 0,80275524 | novel transcript, antisense to RANBP9 |
| ENSG00000136877 | FPGS | -0,382825159 | 0,76693427 | 0,19919 | 0,52019001 | folylpolyglutamate synthase |
| ENSG00000254786 | AP001636,1 | -0,382858332 | 0,766916635 | 0,64228 | 0,82066205 | Glycine N-acyltransferase-like protein 1 (GLYATL1) pseudogene |
| ENSG00000223922 | ASS1P2 | -0,382882912 | 0,766903569 | 0,53234 | 0,75291763 | argininosuccinate synthetase 1 pseudogene 2 |
| ENSG00000288043 | AC006511,6 | -0,382931533 | 0,766877723 | 0,62053 | 0,80776776 | novel transcript |
| ENSG00000162194 | LBHD1 | -0,382970008 | 0,766857272 | 0,33314 | 0,61307256 | LBH domain containing 1 |
| ENSG00000273212 | AC000068,2 | -0,383009675 | 0,766836187 | 0,48999 | 0,72429221 | novel transcript, antisense to UFD1L |
| ENSG00000156313 | RPGR | -0,383144346 | 0,766764609 | 0,20786 | 0,52746553 | retinitis pigmentosa GTPase regulator |
| ENSG00000127445 | PIN1 | -0,383228498 | 0,766719885 | 0,19844 | 0,51980353 | peptidylprolyl cis/trans isomerase, NIMA-interacting 1 |
| ENSG00000196616 | ADH1B | -0,383341586 | 0,766659787 | 0,46666 | 0,70676695 | alcohol dehydrogenase 1B (class I), beta polypeptide |

| | | | | | | |
|---|---|---|---|---|---|---|
| ENSG00000067221 | STOML1 | -0,383347987 | 0,766656385 | 0,19115 | 0,51406315 | stomatin like 1 |
| ENSG00000237515 | SHISA9 | -0,383446408 | 0,766604085 | 0,19807 | 0,51977271 | shisa family member 9 |
| ENSG00000152253 | SPC25 | -0,383454175 | 0,766599958 | 0,42395 | 0,67773865 | SPC25 component of NDC80 kinetochore complex |
| ENSG00000203930 | LINC00632 | -0,38348212 | 0,766585109 | 0,13026 | 0,45467139 | long intergenic non-protein coding RNA 632 |
| ENSG00000142686 | C1orf216 | -0,383729562 | 0,766453641 | 0,28859 | 0,58455903 | chromosome 1 open reading frame 216 |
| ENSG00000287191 | AC055807,1 | -0,383849988 | 0,766389666 | 0,5293 | 0,75112358 | novel transcript, antisense to IGF1R |
| ENSG00000184434 | LRRC19 | -0,383861047 | 0,766383791 | 0,31748 | 0,60289058 | leucine rich repeat containing 19 |
| ENSG00000105928 | GSDME | -0,383944001 | 0,766339725 | 0,16569 | 0,49182669 | gasdermin E |
| ENSG00000260986 | AC126335,1 | -0,384207839 | 0,766199591 | 0,62403 | 0,80957813 | novel transcript |
| ENSG00000285296 | AL096870,5 | -0,384230738 | 0,76618743 | 0,75288 | 0,88423768 | novel transcript |
| ENSG00000100418 | DESI1 | -0,384383636 | 0,766106233 | 0,19439 | 0,51562158 | desumoylating isopeptidase 1 |
| ENSG00000241962 | AC079447,1 | -0,384529054 | 0,766029016 | 0,44735 | 0,69342177 | novel protein, C2orf15-MRPL30-novel lincRNA readthrough |
| ENSG00000101342 | TLDC2 | -0,384633525 | 0,765973547 | 0,56831 | 0,7759239 | TBC/LysM-associated domain containing 2 |
| ENSG00000281813 | KAT6B | -0,384640299 | 0,765969951 | 0,07864 | 0,3663904 | lysine acetyltransferase 6B |
| ENSG00000196912 | ANKRD36B | -0,384726154 | 0,765924369 | 0,06481 | 0,33757437 | ankyrin repeat domain 36B |
| ENSG00000164082 | GRM2 | -0,384841151 | 0,76586332 | 0,19784 | 0,51942408 | glutamate metabotropic receptor 2 |
| ENSG00000272829 | AC002470,1 | -0,384873345 | 0,76584623 | 0,5997 | 0,79552021 | novel transcript |
| ENSG00000139835 | GRTP1 | -0,385113019 | 0,765719011 | 0,45179 | 0,69679392 | growth hormone regulated TBC protein 1 |
| ENSG00000136279 | DBNL | -0,385256294 | 0,765642971 | 0,1562 | 0,4840074 | drebrin like |
| ENSG00000078589 | P2RY10 | -0,385271322 | 0,765634995 | 0,32315 | 0,60590706 | P2Y receptor family member 10 |
| ENSG00000259511 | UBE2Q2L | -0,385384692 | 0,765574832 | 0,39913 | 0,66153677 | ubiquitin conjugating enzyme E2 Q2 like |
| ENSG00000270614 | AC011451,2 | -0,385415449 | 0,765558511 | 0,28485 | 0,5816767 | UBX domain protein 2A (UBXN2A) pseudogene |
| ENSG00000242267 | SKINT1L | -0,385455914 | 0,765537039 | 0,40669 | 0,66608043 | Skint1 like (pseudogene) |
| ENSG00000245148 | ARAP1-AS2 | -0,385507112 | 0,765509872 | 0,30435 | 0,59430086 | ARAP1 antisense RNA 2 |
| ENSG00000261790 | AC005606,2 | -0,385596448 | 0,765462471 | 0,32179 | 0,60577115 | novel transcript |
| ENSG00000272106 | AL691432,2 | -0,385596939 | 0,76546221 | 0,07384 | 0,35713141 | artifact |
| ENSG00000154370 | TRIM11 | -0,385769434 | 0,765370694 | 0,23865 | 0,54890467 | tripartite motif containing 11 |
| ENSG00000256897 | AC018410,1 | -0,385816967 | 0,765345477 | 0,43619 | 0,68658672 | novel transcript |
| ENSG00000287643 | AC004825,3 | -0,385967069 | 0,765265853 | 0,21837 | 0,53578053 | novel transcript, antisense to PCNX1 |
| ENSG00000183654 | MARCHF11 | -0,385996281 | 0,765250357 | 0,61482 | 0,80404192 | membrane associated ring-CH-type finger 11 |
| ENSG00000274378 | AC005702,3 | -0,386117295 | 0,765186171 | 0,59738 | 0,79443333 | double C2-like domains, beta (DOC2B) pseudogene |
| ENSG00000162951 | LRRTM1 | -0,386271675 | 0,765104294 | 0,33843 | 0,61774255 | leucine rich repeat transmembrane neuronal 1 |
| ENSG00000105373 | NOP53 | -0,386355004 | 0,765060103 | 0,17676 | 0,50110131 | NOP53 ribosome biogenesis factor |

| | | | | | | |
|---|---|---|---|---|---|---|
| ENSG00000260267 | AC026471,1 | -0,386357491 | 0,765058785 | 0,23255 | 0,54529383 | novel transcript, antisense to ARMC5 |
| ENSG00000100364 | KIAA0930 | -0,386497443 | 0,764984572 | 0,15642 | 0,48425071 | KIAA0930 |
| ENSG00000156650 | KAT6B | -0,386786474 | 0,76483133 | 0,14167 | 0,46821296 | lysine acetyltransferase 6B |
| ENSG00000265800 | AC022211,3 | -0,386836356 | 0,764804885 | 0,50457 | 0,73328552 | novel transcript, sense intronic to HN1 |
| ENSG00000103126 | AXIN1 | -0,386853907 | 0,764795581 | 0,22242 | 0,53859118 | axin 1 |
| ENSG00000276701 | MIF | -0,386993673 | 0,764721492 | 0,25464 | 0,55944852 | macrophage migration inhibitory factor |
| ENSG00000111325 | OGFOD2 | -0,387006217 | 0,764714843 | 0,21626 | 0,53480193 | 2-oxoglutarate and iron dependent oxygenase domain containing 2 |
| ENSG00000167941 | SOST | -0,38705999 | 0,764686341 | 0,69329 | 0,85095173 | sclerostin |
| ENSG00000079459 | FDFT1 | -0,38725282 | 0,764584141 | 0,29406 | 0,58837921 | farnesyl-diphosphate farnesyltransferase 1 |
| ENSG00000228703 | AL355310,2 | -0,387290442 | 0,764564202 | 0,63173 | 0,81431541 | novel transcript |
| ENSG00000231999 | LRRC8C-DT | -0,387321416 | 0,764547788 | 0,37418 | 0,64270809 | LRRC8C divergent transcript |
| ENSG00000232762 | AL355483,2 | -0,387434612 | 0,764487802 | 0,73738 | 0,87539732 | novel transcript |
| ENSG00000118596 | SLC16A7 | -0,387730453 | 0,764331051 | 0,21293 | 0,53198758 | solute carrier family 16 member 7 |
| ENSG00000286817 | AC105339,6 | -0,387745233 | 0,764323221 | 0,47922 | 0,71679575 | novel transcript |
| ENSG00000111674 | ENO2 | -0,387804663 | 0,764291737 | 0,27891 | 0,57700804 | enolase 2 |
| ENSG00000265514 | LINC01892 | -0,388008635 | 0,764183687 | 0,24494 | 0,55288356 | long intergenic non-protein coding RNA 1892 |
| ENSG00000282260 | AC019257,6 | -0,38818308 | 0,76409129 | 0,61975 | 0,80726991 | novel transcript |
| ENSG00000270988 | AC019257,2 | -0,38818308 | 0,76409129 | 0,61975 | 0,80726991 | novel transcript |
| ENSG00000235016 | SEMA3F-AS1 | -0,388275981 | 0,764042089 | 0,20005 | 0,5209612 | SEMA3F antisense RNA 1 |
| ENSG00000274070 | CASTOR2 | -0,388463167 | 0,763942963 | 0,19363 | 0,51547251 | cytosolic arginine sensor for mTORC1 subunit 2 |
| ENSG00000286747 | AL627095,2 | -0,388551976 | 0,763895938 | 0,49407 | 0,72659022 | novel transcript |
| ENSG00000232855 | AF165147,1 | -0,388689496 | 0,763823125 | 0,28853 | 0,58455903 | novel transcript |
| ENSG00000178105 | DDX10 | -0,388699488 | 0,763817835 | 0,31305 | 0,59987747 | DEAD-box helicase 10 |
| ENSG00000137218 | FRS3 | -0,388874136 | 0,763725375 | 0,25794 | 0,56199743 | fibroblast growth factor receptor substrate 3 |
| ENSG00000128602 | SMO | -0,388977683 | 0,763670562 | 0,18359 | 0,50731682 | smoothened, frizzled class receptor |
| ENSG00000224940 | PRRT4 | -0,389012272 | 0,763652253 | 0,68477 | 0,84574397 | proline rich transmembrane protein 4 |
| ENSG00000213714 | FAM209B | -0,389111629 | 0,763599663 | 0,68061 | 0,84323972 | family with sequence similarity 209 member B |
| ENSG00000258232 | AC125611,3 | -0,389118393 | 0,763596083 | 0,30329 | 0,59355788 | novel transcript, antisense to TUBA1C |
| ENSG00000185482 | STAC3 | -0,389262269 | 0,763519935 | 0,35446 | 0,62908147 | SH3 and cysteine rich domain 3 |
| ENSG00000126934 | MAP2K2 | -0,389271564 | 0,763515016 | 0,27019 | 0,57192517 | mitogen-activated protein kinase kinase 2 |
| ENSG00000276791 | AC092117,1 | -0,389557995 | 0,763363444 | 0,30922 | 0,59774312 | novel transcript |

| | | | | | | |
|---|---|---|---|---|---|---|
| ENSG00000206561 | COLQ | -0,389592579 | 0,763345145 | 0,31225 | 0,59925736 | collagen like tail subunit of asymmetric acetylcholinesterase |
| ENSG00000197462 | AC003975,1 | -0,38961843 | 0,763331467 | 0,33704 | 0,61684812 | novel transcript |
| ENSG00000259073 | FOXN3-AS2 | -0,389643617 | 0,76331814 | 0,69377 | 0,8512419 | FOXN3 antisense RNA 2 |
| ENSG00000285151 | AC140134,1 | -0,389833881 | 0,76321748 | 0,66345 | 0,83242946 | novel transcript, antisense to SMN2 |
| ENSG00000285204 | AC139834,1 | -0,389833881 | 0,76321748 | 0,66345 | 0,83242946 | novel transcript, antisense to SMN1 |
| ENSG00000275709 | AC090527,3 | -0,390015378 | 0,76312147 | 0,60041 | 0,79584102 | novel transcript |
| ENSG00000251503 | CENPS-CORT | -0,390068814 | 0,763093205 | 0,34562 | 0,62278421 | CENPS-CORT readthrough |
| ENSG00000168395 | ING5 | -0,390159509 | 0,763045235 | 0,2567 | 0,56112699 | inhibitor of growth family member 5 |
| ENSG00000146285 | SCML4 | -0,390221492 | 0,763012453 | 0,45049 | 0,69586245 | Scm polycomb group protein like 4 |
| ENSG00000156860 | FBRS | -0,390401083 | 0,762917476 | 0,23017 | 0,54335232 | fibrosin |
| ENSG00000242028 | HYPK | -0,390415725 | 0,762909734 | 0,2675 | 0,56973003 | huntingtin interacting protein K |
| ENSG00000258900 | HNRNPCP1 | -0,390461089 | 0,762885745 | 0,35838 | 0,63185282 | heterogeneous nuclear ribonucleoprotein C pseudogene 1 |
| ENSG00000229839 | AC018462,1 | -0,390550931 | 0,762838239 | 0,65992 | 0,83018671 | novel transcript |
| ENSG00000104880 | ARHGEF18 | -0,390636464 | 0,762793014 | 0,09499 | 0,39751728 | Rho/Rac guanine nucleotide exchange factor 18 |
| ENSG00000273599 | AL731571,1 | -0,390770128 | 0,762722345 | 0,28065 | 0,5780801 | novel transcript, antisense to CTBP2 |
| ENSG00000273344 | PAXIP1-AS1 | -0,390991917 | 0,762605099 | 0,08206 | 0,37329007 | PAXIP1 antisense RNA 1 (head to head) |
| ENSG00000281022 | MED22 | -0,391284612 | 0,762450397 | 0,22896 | 0,54262488 | mediator complex subunit 22 |
| ENSG00000288125 | AC023300,4 | -0,391397023 | 0,762390991 | 0,59709 | 0,79434997 | novel transcript, antisense to CCDC33 |
| ENSG00000277749 | AC023300,2 | -0,391397023 | 0,762390991 | 0,59709 | 0,79434997 | novel transcript, antisense to CCDC33 |
| ENSG00000232077 | LINC01031 | -0,391511299 | 0,762330605 | 0,5826 | 0,7847455 | long intergenic non-protein coding RNA 1031 |
| ENSG00000102904 | TSNAXIP1 | -0,391658643 | 0,762252751 | 0,23655 | 0,54760364 | translin associated factor X interacting protein 1 |
| ENSG00000226352 | PSPC1-AS2 | -0,391726248 | 0,762217033 | 0,60853 | 0,80122454 | PSPC1 antisense RNA 2 |
| ENSG00000130021 | PUDP | -0,391877087 | 0,762137344 | 0,32359 | 0,60633847 | pseudouridine 5'-phosphatase |
| ENSG00000099999 | RNF215 | -0,391999723 | 0,762072561 | 0,16939 | 0,49554671 | ring finger protein 215 |
| ENSG00000185670 | ZBTB3 | -0,392030339 | 0,762056389 | 0,2691 | 0,57085484 | zinc finger and BTB domain containing 3 |
| ENSG00000230230 | TRIM26 | -0,392253358 | 0,761938596 | 0,31036 | 0,59842274 | tripartite motif containing 26 |
| ENSG00000285879 | AC018628,2 | -0,392304227 | 0,761911731 | 0,20322 | 0,52311966 | novel transcript, antisense to MED13 |
| ENSG00000225650 | EIF2S2P5 | -0,392314203 | 0,761906463 | 0,32227 | 0,60577115 | eukaryotic translation initiation factor 2 subunit 2 beta pseudogene 5 |
| ENSG00000273973 | AC025162,2 | -0,39249359 | 0,761811731 | 0,61334 | 0,8035281 | novel transcript, sense intronic to DGKA |
| ENSG00000136059 | VILL | -0,39250445 | 0,761805997 | 0,13217 | 0,45719971 | villin like |
| ENSG00000261553 | AL137782,1 | -0,392633805 | 0,761737695 | 0,3155 | 0,60172912 | novel transcript |

| | | | | | | |
|---|---|---|---|---|---|---|
| ENSG00000179292 | TMEM151A | -0,392739506 | 0,761681887 | 0,55796 | 0,76961995 | transmembrane protein 151A |
| ENSG00000232633 | AC079465,1 | -0,392757751 | 0,761672255 | 0,47604 | 0,71443502 | novel transcript |
| ENSG00000158089 | GALNT14 | -0,392825744 | 0,761636359 | 0,55295 | 0,76620716 | polypeptide N-acetylgalactosaminyltransferase 14 |
| ENSG00000228166 | MTND1P11 | -0,39283451 | 0,761631731 | 0,51146 | 0,73856651 | MT-ND1 pseudogene 11 |
| ENSG00000069974 | RAB27A | -0,392885725 | 0,761604694 | 0,33458 | 0,61433133 | RAB27A, member RAS oncogene family |
| ENSG00000235802 | HCFC1-AS1 | -0,392919292 | 0,761586974 | 0,5795 | 0,78285648 | HCFC1 antisense RNA 1 |
| ENSG00000196337 | CGB7 | -0,392928142 | 0,761582302 | 0,58679 | 0,78701992 | chorionic gonadotropin subunit beta 7 |
| ENSG00000146938 | NLGN4X | -0,393008034 | 0,761540129 | 0,2901 | 0,58531219 | neuroligin 4 X-linked |
| ENSG00000154102 | C16orf74 | -0,393009251 | 0,761539487 | 0,657 | 0,82858357 | chromosome 16 open reading frame 74 |
| ENSG00000273691 | AC087284,1 | -0,393117215 | 0,761482499 | 0,48415 | 0,72005601 | novel transcript, sense intronic to ZNF710 |
| ENSG00000272944 | AC079834,1 | -0,393216648 | 0,761430018 | 0,26865 | 0,57070906 | novel transcript |
| ENSG00000272145 | NFYC-AS1 | -0,393308645 | 0,761381465 | 0,17795 | 0,50233162 | NFYC antisense RNA 1 |
| ENSG00000278839 | SERF1B | -0,393323595 | 0,761373575 | 0,22138 | 0,53805234 | small EDRK-rich factor 1B |
| ENSG00000235298 | AL354733,3 | -0,393574786 | 0,761241022 | 0,28568 | 0,58214567 | novel transcript |
| ENSG00000135596 | MICAL1 | -0,393764136 | 0,761141118 | 0,02836 | 0,22760667 | microtubule associated monooxygenase, calponin and LIM domain containing 1 |
| ENSG00000184497 | TMEM255B | -0,393791413 | 0,761126727 | 0,40096 | 0,66290164 | transmembrane protein 255B |
| ENSG00000273825 | FP700111,2 | -0,394123398 | 0,760951601 | 0,78743 | 0,90121071 | otopetrin 1 (OTOP1) pseudogene |
| ENSG00000253925 | AC011369,1 | -0,394151372 | 0,760936847 | 0,52613 | 0,7497214 | novel transcript |
| ENSG00000103942 | HOMER2 | -0,394163557 | 0,76093042 | 0,02048 | 0,18979312 | homer scaffold protein 2 |
| ENSG00000213872 | AC092798,1 | -0,39422361 | 0,760898746 | 0,45947 | 0,7017037 | ribosomal protein L32 (RPL32) pseudogene |
| ENSG00000145016 | RUBCN | -0,394464697 | 0,760771604 | 0,26269 | 0,56575908 | rubicon autophagy regulator |
| ENSG00000267213 | AC007773,1 | -0,394502979 | 0,760751417 | 0,41959 | 0,67530409 | novel transcript, antisense to DPY19L3 |
| ENSG00000286045 | AC005034,6 | -0,394562119 | 0,760720232 | 0,18138 | 0,50490902 | novel transcript, antisense to GCFC2 |
| ENSG00000244731 | C4A | -0,39476871 | 0,760611307 | 0,55746 | 0,76915913 | complement C4A (Rodgers blood group) |
| ENSG00000267077 | AC020663,2 | -0,394940675 | 0,76052065 | 0,73362 | 0,87368351 | novel transcript, antisense to novel protein |
| ENSG00000015285 | WAS | -0,394988055 | 0,760495673 | 0,40231 | 0,66359858 | WASP actin nucleation promoting factor |
| ENSG00000160193 | WDR4 | -0,395073479 | 0,760450644 | 0,22989 | 0,54326776 | WD repeat domain 4 |
| ENSG00000178585 | CTNNBIP1 | -0,395110565 | 0,760431097 | 0,11527 | 0,43096576 | catenin beta interacting protein 1 |
| ENSG00000157851 | DPYSL5 | -0,395115559 | 0,760428464 | 0,01977 | 0,18665735 | dihydropyrimidinase like 5 |
| ENSG00000260729 | AC009690,1 | -0,395219797 | 0,760373524 | 0,46419 | 0,70487632 | novel protein |
| ENSG00000270986 | HMGB1P51 | -0,395321185 | 0,760320089 | 0,3524 | 0,62742225 | high mobility group box 1 pseudogene 51 |
| ENSG00000143319 | ISG20L2 | -0,395399292 | 0,760278927 | 0,30031 | 0,59172892 | interferon stimulated exonuclease gene 20 like 2 |

| | | | | | | |
|---|---|---|---|---|---|---|
| ENSG00000072958 | AP1M1 | -0,395455847 | 0,760249123 | 0,17263 | 0,49744497 | adaptor related protein complex 1 subunit mu 1 |
| ENSG00000144455 | SUMF1 | -0,395480199 | 0,760236291 | 0,30354 | 0,59380161 | sulfatase modifying factor 1 |
| ENSG00000276168 | RN7SL1 | -0,395516112 | 0,760217366 | 0,15727 | 0,48443207 | RNA component of signal recognition particle 7SL1 |
| ENSG00000107404 | DVL1 | -0,395636934 | 0,760153703 | 0,16948 | 0,49562276 | dishevelled segment polarity protein 1 |
| ENSG00000269961 | AC010359,2 | -0,39566999 | 0,760136286 | 0,57599 | 0,78048142 | novel transcript |
| ENSG00000112658 | SRF | -0,395766392 | 0,760085495 | 0,28054 | 0,57800643 | serum response factor |
| ENSG00000226471 | Z93930,2 | -0,395990588 | 0,759967386 | 0,48538 | 0,72097949 | novel transcript |
| ENSG00000166855 | CLPX | -0,396178182 | 0,759868574 | 0,23818 | 0,54870569 | caseinolytic mitochondrial matrix peptidase chaperone subunit X |
| ENSG00000151320 | AKAP6 | -0,396285724 | 0,759811933 | 0,25303 | 0,55809014 | A-kinase anchoring protein 6 |
| ENSG00000099956 | SMARCB1 | -0,396298379 | 0,759805269 | 0,18095 | 0,50450233 | SWI/SNF related, matrix associated, actin dependent regulator of chromatin, subfamily b, member 1 |
| ENSG00000204344 | STK19 | -0,396685382 | 0,759601478 | 0,41147 | 0,66919097 | serine/threonine kinase 19 |
| ENSG00000188522 | FAM83G | -0,396782945 | 0,759550111 | 0,51137 | 0,73853921 | family with sequence similarity 83 member G |
| ENSG00000247315 | ZCCHC3 | -0,396883291 | 0,759497283 | 0,29658 | 0,58979902 | zinc finger CCHC-type containing 3 |
| ENSG00000108001 | EBF3 | -0,39706064 | 0,759403925 | 0,34268 | 0,62029024 | EBF transcription factor 3 |
| ENSG00000180422 | LINC00304 | -0,39713261 | 0,759366042 | 0,65537 | 0,82781124 | long intergenic non-protein coding RNA 304 |
| ENSG00000176472 | ZNF575 | -0,397182186 | 0,759339948 | 0,36039 | 0,63265655 | zinc finger protein 575 |
| ENSG00000271993 | AC126118,1 | -0,397336247 | 0,759258865 | 0,38842 | 0,6529693 | novel transcript, antisense to LRRFIP2 |
| ENSG00000132718 | SYT11 | -0,397414711 | 0,759217572 | 0,11359 | 0,42836024 | synaptotagmin 11 |
| ENSG00000239840 | RPL23AP72 | -0,397770533 | 0,759030344 | 0,63124 | 0,81395758 | ribosomal protein L23a pseudogene 72 |
| ENSG00000250687 | NAIPP1 | -0,398245576 | 0,758780456 | 0,23423 | 0,54602618 | NAIP pseudogene 1 |
| ENSG00000119139 | TJP2 | -0,398418465 | 0,758689531 | 0,15023 | 0,47830235 | tight junction protein 2 |
| ENSG00000237628 | MTCO2P19 | -0,398512813 | 0,758639916 | 0,6631 | 0,83242715 | MT-CO2 pseudogene 19 |
| ENSG00000149503 | INCENP | -0,398616432 | 0,758585431 | 0,12556 | 0,44748549 | inner centromere protein |
| ENSG00000177294 | FBXO39 | -0,398837605 | 0,758469144 | 0,71153 | 0,86116229 | F-box protein 39 |
| ENSG00000286158 | AL034548,2 | -0,399067711 | 0,75834818 | 0,44922 | 0,69489321 | novel transcript |
| ENSG00000271522 | AC016831,5 | -0,399247421 | 0,758253722 | 0,45536 | 0,69938057 | novel transcript |
| ENSG00000243746 | EEF1A1P10 | -0,399297895 | 0,758227194 | 0,60023 | 0,79579686 | eukaryotic translation elongation factor 1 alpha 1 pseudogene 10 |
| ENSG00000287925 | AL353576,1 | -0,399381438 | 0,758183288 | 0,56871 | 0,77620121 | novel transcript, antisense to C1QL3 |
| ENSG00000164621 | SMAD5-AS1 | -0,399565884 | 0,758086362 | 0,34667 | 0,62350287 | SMAD5 antisense RNA 1 |
| ENSG00000226742 | HSBP1L1 | -0,399576044 | 0,758081023 | 0,1956 | 0,51643293 | heat shock factor binding protein 1 like 1 |

| | | | | | | |
|---|---|---|---|---|---|---|
| ENSG00000266602 | AP005262,1 | -0,399742947 | 0,757993327 | 0,19642 | 0,51750057 | novel transcript |
| ENSG00000286376 | AC006041,2 | -0,399743864 | 0,757992845 | 0,5967 | 0,79424321 | novel transcript |
| ENSG00000251322 | SHANK3 | -0,399808301 | 0,757958991 | 0,2181 | 0,53569444 | SH3 and multiple ankyrin repeat domains 3 |
| ENSG00000287115 | AC009951,6 | -0,399925577 | 0,757897379 | 0,60754 | 0,80031274 | novel transcript |
| ENSG00000166839 | ANKDD1A | -0,399930488 | 0,757894799 | 0,24207 | 0,55097978 | ankyrin repeat and death domain containing 1A |
| ENSG00000226103 | C6orf47 | -0,399964763 | 0,757876793 | 0,45935 | 0,7017037 | chromosome 6 open reading frame 47 |
| ENSG00000203623 | C6orf47 | -0,399964763 | 0,757876793 | 0,45935 | 0,7017037 | chromosome 6 open reading frame 47 |
| ENSG00000226531 | C6orf47 | -0,399964763 | 0,757876793 | 0,45935 | 0,7017037 | chromosome 6 open reading frame 47 |
| ENSG00000286886 | AJ011931,2 | -0,40015567 | 0,757776513 | 0,5549 | 0,76763832 | novel transcript, antisense to PCBP3 |
| ENSG00000106541 | AGR2 | -0,400159097 | 0,757774713 | 0,58628 | 0,78667706 | anterior gradient 2, protein disulphide isomerase family member |
| ENSG00000068489 | PRR11 | -0,400580192 | 0,757553566 | 0,22664 | 0,54144 | proline rich 11 |
| ENSG00000219027 | RPS3AP2 | -0,400778869 | 0,757449248 | 0,62341 | 0,8091647 | RPS3A pseudogene 2 |
| ENSG00000072786 | STK10 | -0,400827412 | 0,757423762 | 0,35952 | 0,63231401 | serine/threonine kinase 10 |
| ENSG00000133028 | SCO1 | -0,400989266 | 0,757338793 | 0,28494 | 0,58171017 | synthesis of cytochrome C oxidase 1 |
| ENSG00000169551 | CT55 | -0,401062875 | 0,757300153 | 0,69004 | 0,8488605 | cancer/testis antigen 55 |
| ENSG00000185379 | RAD51D | -0,40106733 | 0,757297814 | 0,17483 | 0,49908125 | RAD51 paralog D |
| ENSG00000263585 | AC145207,4 | -0,401073162 | 0,757294753 | 0,88582 | 0,94944442 | novel transcript, antisense PYCR1 |
| ENSG00000243244 | STON1 | -0,401168566 | 0,757244676 | 0,19633 | 0,51733996 | stonin 1 |
| ENSG00000275198 | AL512791,2 | -0,40116897 | 0,757244463 | 0,45747 | 0,70075609 | novel transcript |
| ENSG00000257509 | AC073487,1 | -0,401391951 | 0,757127434 | 0,25111 | 0,55719428 | novel transcript, sense intronic to SARNP |
| ENSG00000286583 | AC105101,2 | -0,401552993 | 0,757042924 | 0,69501 | 0,85206252 | novel transcript, antisense to ZBTB7C |
| ENSG00000136492 | BRIP1 | -0,401771072 | 0,756928497 | 0,22041 | 0,53750545 | BRCA1 interacting protein C-terminal helicase 1 |
| ENSG00000285957 | AC019257,9 | -0,401907319 | 0,756857017 | 0,46022 | 0,70210714 | novel transcript, antisense to ARHGEF10 |
| ENSG00000287695 | AC008625,1 | -0,401910594 | 0,756855299 | 0,43713 | 0,68711291 | novel transcript, sense intronic to GALNT10 |
| ENSG00000111816 | FRK | -0,401947311 | 0,756836036 | 0,184 | 0,50777976 | fyn related Src family tyrosine kinase |
| ENSG00000254003 | SRI-AS1 | -0,402064695 | 0,75677446 | 0,57607 | 0,78052092 | SRI antisense RNA 1 |
| ENSG00000233680 | HNRNPA1P27 | -0,402106364 | 0,756752602 | 0,61249 | 0,8035281 | heterogeneous nuclear ribonucleoprotein A1 pseudogene 27 |
| ENSG00000261441 | AC124068,2 | -0,402137137 | 0,756736461 | 0,50126 | 0,7312466 | novel transcript, antisense to POLG |
| ENSG00000267383 | AC011447,3 | -0,40216475 | 0,756721977 | 0,17047 | 0,49641416 | novel transcript, antisense to ZNF486 and ZNF90 |
| ENSG00000151322 | NPAS3 | -0,402286787 | 0,756657969 | 0,27526 | 0,57491355 | neuronal PAS domain protein 3 |
| ENSG00000269110 | AC010636,2 | -0,402577292 | 0,756505621 | 0,65421 | 0,82689795 | novel transcript |

| | | | | | | |
|---|---|---|---|---|---|---|
| ENSG00000270878 | GCATP1 | -0,402587744 | 0,756500141 | 0,26911 | 0,57085484 | glycine C-acetyltransferase pseudogene 1 |
| ENSG00000081923 | ATP8B1 | -0,402638881 | 0,756473327 | 0,24445 | 0,55253387 | ATPase phospholipid transporting 8B1 |
| ENSG00000138798 | EGF | -0,402678066 | 0,75645278 | 0,42534 | 0,67893062 | epidermal growth factor |
| ENSG00000213234 | ST13P10 | -0,402720602 | 0,756430478 | 0,69724 | 0,85330122 | ST13, Hsp70 interacting protein pseudogene 10 |
| ENSG00000228843 | AL133410,2 | -0,402807451 | 0,756384943 | 0,68987 | 0,84877222 | novel transcript |
| ENSG00000273004 | AL078644,1 | -0,402866049 | 0,756354221 | 0,29224 | 0,58692263 | novel transcript |
| ENSG00000288568 | AC092720,3 | -0,402993408 | 0,756287454 | 0,5201 | 0,74456236 | novel transcript |
| ENSG00000282171 | LINC02210 | -0,403013963 | 0,756276679 | 0,1887 | 0,5120698 | long intergenic non-protein coding RNA 2210 |
| ENSG00000275700 | AATF | -0,403153282 | 0,75620365 | 0,17879 | 0,50328925 | apoptosis antagonizing transcription factor |
| ENSG00000141569 | TRIM65 | -0,403163863 | 0,756198104 | 0,34998 | 0,62563749 | tripartite motif containing 65 |
| ENSG00000285581 | AC091196,1 | -0,403168815 | 0,756195508 | 0,48875 | 0,72350189 | novel transcript |
| ENSG00000230487 | PSMG3-AS1 | -0,40319461 | 0,756181988 | 0,06767 | 0,34407703 | PSMG3 antisense RNA 1 (head to head) |
| ENSG00000113212 | PCDHB7 | -0,403234739 | 0,756160955 | 0,08451 | 0,37735388 | protocadherin beta 7 |
| ENSG00000137343 | ATAT1 | -0,403343615 | 0,756103891 | 0,23921 | 0,5491566 | alpha tubulin acetyltransferase 1 |
| ENSG00000176720 | BOK | -0,403443257 | 0,756051672 | 0,22617 | 0,54121239 | BCL2 family apoptosis regulator BOK |
| ENSG00000035115 | SH3YL1 | -0,403558923 | 0,755991059 | 0,10765 | 0,41827415 | SH3 and SYLF domain containing 1 |
| ENSG00000276916 | AL442125,2 | -0,403830149 | 0,755848946 | 0,62632 | 0,81088056 | novel transcript, sense intronic |
| ENSG00000236611 | LINC02556 | -0,403873021 | 0,755826485 | 0,65045 | 0,82492554 | long intergenic non-protein coding RNA 2556 |
| ENSG00000267128 | RNF157-AS1 | -0,403926984 | 0,755798215 | 0,48652 | 0,72196345 | RNF157 antisense RNA 1 |
| ENSG00000235904 | RBMS3-AS3 | -0,404008678 | 0,755755418 | 0,30484 | 0,59446033 | RBMS3 antisense RNA 3 |
| ENSG00000243144 | AC079760,2 | -0,404091242 | 0,755712168 | 0,6135 | 0,8035281 | novel transcript |
| ENSG00000145242 | EPHA5 | -0,404119877 | 0,755697169 | 0,49613 | 0,72831792 | EPH receptor A5 |
| ENSG00000267497 | NFE2L3P1 | -0,404150042 | 0,755681368 | 0,51602 | 0,74198903 | nuclear factor, erythroid 2 like 3 pseudogene 1 |
| ENSG00000198840 | MT-ND3 | -0,404303208 | 0,755601145 | 0,18288 | 0,50636151 | mitochondrially encoded NADH:ubiquinone oxidoreductase core subunit 3 |
| ENSG00000158805 | ZNF276 | -0,40435493 | 0,755574056 | 0,14873 | 0,47609532 | zinc finger protein 276 |
| ENSG00000179965 | ZNF771 | -0,404375056 | 0,755563516 | 0,26988 | 0,57162985 | zinc finger protein 771 |
| ENSG00000236290 | EEF1GP7 | -0,404391887 | 0,755554701 | 0,43581 | 0,68637175 | eukaryotic translation elongation factor 1 gamma pseudogene 7 |
| ENSG00000277421 | ZNF280B | -0,404397233 | 0,755551901 | 0,38938 | 0,65372387 | zinc finger protein 280B |
| ENSG00000228436 | AL139260,1 | -0,404471811 | 0,755512845 | 0,41227 | 0,6699869 | novel transcript |
| ENSG00000275342 | PRAG1 | -0,404532337 | 0,755481149 | 0,16556 | 0,49182669 | PEAK1 related, kinase-activating pseudokinase 1 |
| ENSG00000196756 | SNHG17 | -0,404649317 | 0,755419894 | 0,31448 | 0,6011918 | small nucleolar RNA host gene 17 |

| | | | | | | |
|---|---|---|---|---|---|---|
| ENSG00000188938 | FAM120AOS | -0,404736799 | 0,755374089 | 0,206 | 0,52551102 | family with sequence similarity 120A opposite strand |
| ENSG00000284796 | DCAF11 | -0,404799126 | 0,755341456 | 0,32867 | 0,60992716 | DDB1 and CUL4 associated factor 11 |
| ENSG00000063245 | EPN1 | -0,404919417 | 0,755278478 | 0,22066 | 0,53767067 | epsin 1 |
| ENSG00000250906 | AC131953,1 | -0,405005999 | 0,755233152 | 0,54411 | 0,76035025 | novel transcript |
| ENSG00000234504 | AL353801,2 | -0,405044557 | 0,755212968 | 0,63939 | 0,81910693 | novel transcript |
| ENSG00000085662 | AKR1B1 | -0,405125574 | 0,755170559 | 0,22362 | 0,53960526 | aldo-keto reductase family 1 member B |
| ENSG00000140450 | ARRDC4 | -0,405153743 | 0,755155814 | 0,14302 | 0,47061408 | arrestin domain containing 4 |
| ENSG00000168412 | MTNR1A | -0,405189889 | 0,755136894 | 0,55606 | 0,76828102 | melatonin receptor 1A |
| ENSG00000140199 | SLC12A6 | -0,405246491 | 0,755107268 | 0,14667 | 0,47455277 | solute carrier family 12 member 6 |
| ENSG00000172717 | FAM71D | -0,405321255 | 0,755068138 | 0,61146 | 0,80322383 | family with sequence similarity 71 member D |
| ENSG00000068781 | STON1-GTF2A | -0,405345504 | 0,755055446 | 0,52449 | 0,74828894 | STON1-GTF2A1L readthrough |
| ENSG00000259269 | AC109630,1 | -0,405421766 | 0,755015535 | 0,5844 | 0,78572744 | novel transcript |
| ENSG00000189099 | PRSS48 | -0,405545378 | 0,754950847 | 0,56547 | 0,7737418 | serine protease 48 |
| ENSG00000232915 | RNF14P1 | -0,405699249 | 0,754870331 | 0,66822 | 0,83536064 | RNF14 pseudogene 1 |
| ENSG00000165837 | ERICH6B | -0,406022382 | 0,754701276 | 0,46768 | 0,70751769 | glutamate rich 6B |
| ENSG00000255350 | MTCO1P15 | -0,4062257 | 0,754594924 | 0,43223 | 0,68411566 | MT-CO1 pseudogene 15 |
| ENSG00000255557 | AP001266,2 | -0,406282155 | 0,754565396 | 0,62752 | 0,81150424 | novel transcript |
| ENSG00000231172 | AC007099,1 | -0,406304805 | 0,75455355 | 0,11163 | 0,42472977 | novel transcript, antisense to EVA1A |
| ENSG00000234860 | AL162394,1 | -0,406438117 | 0,754483828 | 0,66825 | 0,83536064 | novel transcript |
| ENSG00000228109 | MELTF-AS1 | -0,406476818 | 0,754463589 | 0,17262 | 0,49744497 | MELTF antisense RNA 1 |
| ENSG00000249307 | LINC01088 | -0,406517778 | 0,754442169 | 0,16442 | 0,49065086 | long intergenic non-protein coding RNA 1088 |
| ENSG00000079313 | REXO1 | -0,406626467 | 0,754385333 | 0,19496 | 0,51588752 | RNA exonuclease 1 homolog |
| ENSG00000145494 | NDUFS6 | -0,406666379 | 0,754364464 | 0,19197 | 0,5145257 | NADH:ubiquinone oxidoreductase subunit S6 |
| ENSG00000249846 | LINC02021 | -0,4067143 | 0,754339407 | 0,15361 | 0,48191866 | long intergenic non-protein coding RNA 2021 |
| ENSG00000242802 | AP5Z1 | -0,40694752 | 0,754217474 | 0,18723 | 0,5105374 | adaptor related protein complex 5 subunit zeta 1 |
| ENSG00000130475 | FCHO1 | -0,407009443 | 0,754185102 | 0,16142 | 0,48892403 | FCH and mu domain containing endocytic adaptor 1 |
| ENSG00000198870 | STKLD1 | -0,407223554 | 0,754073181 | 0,35432 | 0,62899743 | serine/threonine kinase like domain containing 1 |
| ENSG00000182397 | DNM1P46 | -0,407444723 | 0,753957588 | 0,29345 | 0,58790464 | dynamin 1 pseudogene 46 |
| ENSG00000263394 | AC007952,7 | -0,407472108 | 0,753943277 | 0,59555 | 0,79339314 | novel transcript |
| ENSG00000224597 | SVIL-AS1 | -0,407477241 | 0,753940595 | 0,28689 | 0,58334004 | SVIL antisense RNA 1 |
| ENSG00000162813 | BPNT1 | -0,407500191 | 0,753928601 | 0,27297 | 0,573359 | 3'(2'), 5'-bisphosphate nucleotidase 1 |
| ENSG00000197921 | HES5 | -0,407609455 | 0,753871504 | 0,51608 | 0,74198903 | hes family bHLH transcription factor 5 |
| ENSG00000288011 | AC024597,1 | -0,407664909 | 0,753842528 | 0,39493 | 0,65788813 | novel transcript, antisense to ZNF365 |

| | | | | | | |
|---|---|---|---|---|---|---|
| ENSG00000248367 | AC008610,1 | -0,407780166 | 0,753782306 | 0,53347 | 0,75351438 | novel transcript |
| ENSG00000110169 | HPX | -0,407830712 | 0,753755897 | 0,35843 | 0,63185282 | hemopexin |
| ENSG00000213935 | RPL22P16 | -0,40785905 | 0,753741091 | 0,58548 | 0,78640335 | ribosomal protein L22 pseudogene 16 |
| ENSG00000237211 | SETP4 | -0,407963142 | 0,75368671 | 0,65134 | 0,82534198 | SET pseudogene 4 |
| ENSG00000259575 | AC012404,2 | -0,408068199 | 0,753631828 | 0,64114 | 0,81993567 | novel transcript, antisense to RORA |
| ENSG00000276657 | PYCR3 | -0,408085501 | 0,75362279 | 0,31859 | 0,60360964 | pyrroline-5-carboxylate reductase 3 |
| ENSG00000157259 | GATAD1 | -0,408166738 | 0,753580355 | 0,11915 | 0,43696826 | GATA zinc finger domain containing 1 |
| ENSG00000187105 | HEATR4 | -0,408304565 | 0,753508366 | 0,51129 | 0,73851043 | HEAT repeat containing 4 |
| ENSG00000244556 | ODCP | -0,408379457 | 0,753469252 | 0,31629 | 0,60227795 | ornithine decarboxylase pseudogene |
| ENSG00000168255 | POLR2J3 | -0,408490337 | 0,753411345 | 0,23375 | 0,54563244 | RNA polymerase II subunit J3 |
| ENSG00000287998 | AC104232,3 | -0,408637456 | 0,75333452 | 0,11992 | 0,43853551 | novel transcript |
| ENSG00000126522 | ASL | -0,408677589 | 0,753313564 | 0,22799 | 0,54213739 | argininosuccinate lyase |
| ENSG00000242498 | ARPIN | -0,408695378 | 0,753304275 | 0,1795 | 0,50387897 | actin related protein 2/3 complex inhibitor |
| ENSG00000231752 | EMBP1 | -0,40871283 | 0,753295162 | 0,07032 | 0,34995627 | embigin pseudogene 1 |
| ENSG00000204657 | OR2H2 | -0,408723811 | 0,753289429 | 0,45497 | 0,69913628 | olfactory receptor family 2 subfamily H member 2 |
| ENSG00000135773 | CAPN9 | -0,408727656 | 0,753287421 | 0,2774 | 0,57586315 | calpain 9 |
| ENSG00000282954 | FBXO17 | -0,408848627 | 0,75322426 | 0,20031 | 0,52107152 | F-box protein 17 |
| ENSG00000269190 | FBXO17 | -0,408848627 | 0,75322426 | 0,20031 | 0,52107152 | F-box protein 17 |
| ENSG00000227006 | AL136988,2 | -0,408953451 | 0,753169534 | 0,52934 | 0,75112358 | novel transcript |
| ENSG00000067191 | CACNB1 | -0,409071215 | 0,753108057 | 0,17018 | 0,49606609 | calcium voltage-gated channel auxiliary subunit beta 1 |
| ENSG00000272895 | AC009303,3 | -0,409094461 | 0,753095922 | 0,57147 | 0,77809338 | novel transcript, antisense to CCDC93 |
| ENSG00000185774 | KCNIP4 | -0,409267273 | 0,753005719 | 0,53655 | 0,75531624 | potassium voltage-gated channel interacting protein 4 |
| ENSG00000132763 | MMACHC | -0,409377874 | 0,752947993 | 0,29988 | 0,59160746 | metabolism of cobalamin associated C |
| ENSG00000287609 | AL589787,2 | -0,409563603 | 0,752851067 | 0,19265 | 0,51490739 | novel transcript |
| ENSG00000236901 | MIR600HG | -0,409707407 | 0,752776029 | 0,1386 | 0,46578766 | MIR600 host gene |
| ENSG00000158555 | GDPD5 | -0,40978921 | 0,752733346 | 0,15247 | 0,48061452 | glycerophosphodiester phosphodiesterase domain containing 5 |
| ENSG00000178252 | WDR6 | -0,409971224 | 0,752638386 | 0,11495 | 0,43039322 | WD repeat domain 6 |
| ENSG00000241489 | AC244197,3 | -0,409991521 | 0,752627797 | 0,18291 | 0,50638964 | novel protein |
| ENSG00000139797 | RNF113B | -0,410003938 | 0,75262132 | 0,31934 | 0,60429866 | ring finger protein 113B |
| ENSG00000233098 | CCDC144NL-A | -0,410057755 | 0,752593245 | 0,11718 | 0,43451125 | CCDC144NL antisense RNA 1 |
| ENSG00000101197 | BIRC7 | -0,410147277 | 0,752546546 | 0,72819 | 0,87073219 | baculoviral IAP repeat containing 7 |
| ENSG00000270207 | AC077690,1 | -0,410195446 | 0,752521421 | 0,19911 | 0,52017864 | novel transcript, antisense to GRM7 |

| | | | | | | |
|---|---|---|---|---|---|---|
| ENSG00000127666 | TICAM1 | -0,410235199 | 0,752500685 | 0,48685 | 0,72215091 | toll like receptor adaptor molecule 1 |
| ENSG00000154479 | CCDC173 | -0,41045437 | 0,752386376 | 0,1352 | 0,46165567 | coiled-coil domain containing 173 |
| ENSG00000276371 | SINHCAF | -0,410719235 | 0,752248258 | 0,42167 | 0,67643128 | SIN3-HDAC complex associated factor |
| ENSG00000251867 | AC009812,1 | -0,410764878 | 0,752224459 | 0,29276 | 0,58724158 | novel transcript, antisense to ZBTB10 |
| ENSG00000278728 | CNTNAP2 | -0,410915076 | 0,752146149 | 0,38177 | 0,64905556 | contactin associated protein 2 |
| ENSG00000083828 | ZNF586 | -0,410954545 | 0,752125572 | 0,14351 | 0,47092552 | zinc finger protein 586 |
| ENSG00000092931 | MFSD11 | -0,411017632 | 0,752092684 | 0,27048 | 0,57232537 | major facilitator superfamily domain containing 11 |
| ENSG00000286500 | AL512643,2 | -0,411164768 | 0,752015984 | 0,46769 | 0,70751769 | novel transcript, antisense to FREM1 |
| ENSG00000222036 | POTEM | -0,41118718 | 0,752004302 | 0,34522 | 0,6224039 | POTE ankyrin domain family member M |
| ENSG00000119673 | ACOT2 | -0,411277544 | 0,751957201 | 0,30411 | 0,59411013 | acyl-CoA thioesterase 2 |
| ENSG00000225364 | ATP6V0E1P1 | -0,411479806 | 0,751851786 | 0,64208 | 0,8205913 | ATPase H+ transporting V0 subunit e1 pseudogene 1 |
| ENSG00000260412 | AL353746,1 | -0,411493961 | 0,751844409 | 0,50075 | 0,73097006 | novel transcript, overlapping LINGO2 |
| ENSG00000050426 | LETMD1 | -0,411500009 | 0,751841258 | 0,2631 | 0,56589635 | LETM1 domain containing 1 |
| ENSG00000285904 | AC006452,1 | -0,411680881 | 0,751747005 | 0,55217 | 0,76565817 | novel transcript |
| ENSG00000185904 | LINC00839 | -0,411688633 | 0,751742965 | 0,40379 | 0,66409141 | long intergenic non-protein coding RNA 839 |
| ENSG00000248174 | LINC02268 | -0,411696863 | 0,751738677 | 0,15723 | 0,48443207 | long intergenic non-protein coding RNA 2268 |
| ENSG00000122386 | ZNF205 | -0,411704998 | 0,751734438 | 0,09608 | 0,39928792 | zinc finger protein 205 |
| ENSG00000198624 | CCDC69 | -0,4118172 | 0,751675976 | 0,23782 | 0,54864017 | coiled-coil domain containing 69 |
| ENSG00000124228 | DDX27 | -0,411852645 | 0,751657508 | 0,25458 | 0,55944852 | DEAD-box helicase 27 |
| ENSG00000243819 | RN7SL832P | -0,411995204 | 0,751583238 | 0,32397 | 0,60664834 | RNA, 7SL, cytoplasmic 832, pseudogene |
| ENSG00000267198 | AC091132,4 | -0,412220298 | 0,751465982 | 0,33965 | 0,61869857 | novel transcript |
| ENSG00000244230 | RN7SL151P | -0,412375694 | 0,751385044 | 0,14375 | 0,47127006 | RNA, 7SL, cytoplasmic 151, pseudogene |
| ENSG00000181404 | WASHC1 | -0,412473453 | 0,751334132 | 0,20715 | 0,52670822 | WASH complex subunit 1 |
| ENSG00000272168 | CASC15 | -0,412525116 | 0,751307226 | 0,07449 | 0,35841642 | cancer susceptibility 15 |
| ENSG00000260572 | AC069224,1 | -0,412615523 | 0,751260147 | 0,29597 | 0,58942061 | novel transcript, antisense to PPM1L |
| ENSG00000184277 | TM2D3 | -0,412746647 | 0,751191869 | 0,1547 | 0,48266733 | TM2 domain containing 3 |
| ENSG00000160948 | VPS28 | -0,412845588 | 0,751140354 | 0,06369 | 0,33458308 | VPS28 subunit of ESCRT-I |
| ENSG00000244124 | ATP1B3-AS1 | -0,412989469 | 0,751065446 | 0,45078 | 0,69592991 | ATP1B3 antisense RNA 1 |
| ENSG00000228549 | BX284668,2 | -0,413017374 | 0,751050919 | 0,10051 | 0,40713164 | novel transcript |
| ENSG00000114757 | PEX5L | -0,413024798 | 0,751047054 | 0,12101 | 0,44006069 | peroxisomal biogenesis factor 5 like |
| ENSG00000243679 | AC018638,5 | -0,413025836 | 0,751046513 | 0,12429 | 0,44533764 | novel pseudogene |
| ENSG00000215386 | MIR99AHG | -0,413037651 | 0,751040363 | 0,24227 | 0,55112431 | mir-99a-let-7c cluster host gene |
| ENSG00000180178 | FAR2P1 | -0,413071194 | 0,751022901 | 0,1986 | 0,51980353 | fatty acyl-CoA reductase 2 pseudogene 1 |

| | | | | | | |
|---|---|---|---|---|---|---|
| ENSG00000164089 | ETNPPL | -0,413098693 | 0,751008586 | 0,63768 | 0,81786886 | ethanolamine-phosphate phospho-lyase |
| ENSG00000132740 | IGHMBP2 | -0,413204806 | 0,75095335 | 0,20187 | 0,52217657 | immunoglobulin mu DNA binding protein 2 |
| ENSG00000116991 | SIPA1L2 | -0,413233276 | 0,750938531 | 0,21369 | 0,53276331 | signal induced proliferation associated 1 like 2 |
| ENSG00000134183 | GNAT2 | -0,413235742 | 0,750937248 | 0,41014 | 0,66821634 | G protein subunit alpha transducin 2 |
| ENSG00000196345 | ZKSCAN7 | -0,413301598 | 0,750902969 | 0,06051 | 0,32669488 | zinc finger with KRAB and SCAN domains 7 |
| ENSG00000179141 | MTUS2-AS1 | -0,41330625 | 0,750900548 | 0,57878 | 0,78253224 | MTUS2 antisense RNA 1 |
| ENSG00000259291 | ZNF710-AS1 | -0,413417855 | 0,750842462 | 0,0397 | 0,26947142 | ZNF710 antisense RNA 1 |
| ENSG00000249014 | HMGN2P4 | -0,41360103 | 0,750747136 | 0,39275 | 0,6559444 | high mobility group nucleosomal binding domain 2 pseudogene 4 |
| ENSG00000178690 | DYNAP | -0,413669395 | 0,75071156 | 0,46003 | 0,70205579 | dynactin associated protein |
| ENSG00000164946 | FREM1 | -0,413728954 | 0,750680569 | 0,4336 | 0,68473723 | FRAS1 related extracellular matrix 1 |
| ENSG00000215283 | HMGB3P24 | -0,413774696 | 0,750656769 | 0,48695 | 0,72215091 | high mobility group box 3 pseudogene 24 |
| ENSG00000287126 | AC108488,3 | -0,413827949 | 0,750629061 | 0,18099 | 0,50451145 | novel transcript |
| ENSG00000287683 | AL591519,1 | -0,413900607 | 0,750591258 | 0,64075 | 0,81963066 | novel transcript, antisense to EPHA7 |
| ENSG00000159921 | GNE | -0,413920099 | 0,750581117 | 0,29052 | 0,58565836 | glucosamine (UDP-N-acetyl)-2-epimerase/N-acetylmannosamine kinase |
| ENSG00000259358 | DEPDC1P1 | -0,413953504 | 0,750563738 | 0,41925 | 0,67514348 | DEP domain containing 1 pseudogene 1 |
| ENSG00000162433 | AK4 | -0,414105769 | 0,750484526 | 0,40869 | 0,66745626 | adenylate kinase 4 |
| ENSG00000236132 | AL022331,1 | -0,414132073 | 0,750470843 | 0,70679 | 0,85889796 | ribosomal protein L17 (RPL17) pseudogene |
| ENSG00000282923 | AC093567,2 | -0,414155074 | 0,750458878 | 0,61415 | 0,80370125 | novel transcript, antisense to CTIF |
| ENSG00000267764 | AC093567,1 | -0,414155074 | 0,750458878 | 0,61415 | 0,80370125 | novel transcript, antisense to CTIF |
| ENSG00000171502 | COL24A1 | -0,414156175 | 0,750458306 | 0,16455 | 0,49065086 | collagen type XXIV alpha 1 chain |
| ENSG00000151651 | ADAM8 | -0,414312298 | 0,750377098 | 0,29278 | 0,58724158 | ADAM metallopeptidase domain 8 |
| ENSG00000139915 | MDGA2 | -0,414362141 | 0,750351174 | 0,31045 | 0,59844173 | MAM domain containing glycosylphosphatidylinositol anchor 2 |
| ENSG00000220804 | LINC01881 | -0,41436335 | 0,750350545 | 0,18057 | 0,50407792 | long intergenic non-protein coding RNA 1881 |
| ENSG00000274616 | CNOT3 | -0,414391388 | 0,750335963 | 0,43796 | 0,68747773 | CCR4-NOT transcription complex subunit 3 |
| ENSG00000229766 | AL021396,1 | -0,414673821 | 0,750189086 | 0,44002 | 0,6885781 | novel transcript, antisense to TMX4 |
| ENSG00000226567 | LINC00606 | -0,414688447 | 0,75018148 | 0,53941 | 0,75771283 | long intergenic non-protein coding RNA 606 |
| ENSG00000231550 | PTCHD3P2 | -0,414720347 | 0,750164893 | 0,2316 | 0,54440987 | patched domain containing 3 pseudogene 2 |
| ENSG00000147799 | ARHGAP39 | -0,41489582 | 0,750073657 | 0,21706 | 0,53505085 | Rho GTPase activating protein 39 |
| ENSG00000226702 | MIR217HG | -0,414906541 | 0,750068083 | 0,5691 | 0,77637061 | MIR217 host gene |
| ENSG00000279070 | AC073263,2 | -0,41524336 | 0,749892989 | 0,5986 | 0,7951852 | novel transcript |

| | | | | | | |
|---|---|---|---|---|---|---|
| ENSG00000215208 | KRT18P60 | -0,415270786 | 0,749878734 | 0,62505 | 0,81017716 | keratin 18 pseudogene 60 |
| ENSG00000172007 | RAB33B | -0,415341107 | 0,749842183 | 0,13176 | 0,4570124 | RAB33B, member RAS oncogene family |
| ENSG00000270773 | AC137695,2 | -0,415362312 | 0,749831162 | 0,54932 | 0,76407481 | MAP/microtubule affinity-regulating kinase 2, pseudogene |
| ENSG00000230585 | PHBP12 | -0,41567903 | 0,749666568 | 0,37183 | 0,64086851 | prohibitin pseudogene 12 |
| ENSG00000162302 | RPS6KA4 | -0,415712969 | 0,749648932 | 0,22044 | 0,53750545 | ribosomal protein S6 kinase A4 |
| ENSG00000268205 | AC005261,1 | -0,415885086 | 0,749559503 | 0,15847 | 0,485647 | novel transcript |
| ENSG00000236814 | AC046176,1 | -0,415892553 | 0,749555623 | 0,61804 | 0,80622686 | proteasome (prosome, macropain) 26S subunit, ATPase, 6 (PSMC6) pseudogene |
| ENSG00000171310 | CHST11 | -0,416055147 | 0,749471152 | 0,20479 | 0,52445677 | carbohydrate sulfotransferase 11 |
| ENSG00000246089 | AC016065,1 | -0,416141972 | 0,749426048 | 0,34576 | 0,62283284 | novel transcript |
| ENSG00000114378 | HYAL1 | -0,416268001 | 0,749360583 | 0,19747 | 0,51895349 | hyaluronidase 1 |
| ENSG00000169047 | IRS1 | -0,416319573 | 0,749333797 | 0,11256 | 0,42638197 | insulin receptor substrate 1 |
| ENSG00000147437 | GNRH1 | -0,416330474 | 0,749328135 | 0,13368 | 0,45957971 | gonadotropin releasing hormone 1 |
| ENSG00000274718 | AL136964,1 | -0,416486256 | 0,749247227 | 0,4671 | 0,70712115 | novel transcript |
| ENSG00000178636 | AC092656,1 | -0,416666992 | 0,74915337 | 0,45957 | 0,7017275 | mitochondrial ribosomal protein L42 (MRPL42) pseudogene |
| ENSG00000228299 | HLA-C | -0,416689864 | 0,749141493 | 0,54096 | 0,75863636 | major histocompatibility complex, class I, C |
| ENSG00000251136 | AF117829,1 | -0,416712734 | 0,749129617 | 0,18149 | 0,50490902 | novel transcript |
| ENSG00000244480 | AC005154,2 | -0,416740992 | 0,749114944 | 0,31415 | 0,60088745 | pseudogene similar golgi autoantigen, golgin subfamily a, 8A GOLGA8A |
| ENSG00000159403 | C1R | -0,417015927 | 0,748972199 | 0,25656 | 0,56097949 | complement C1r |
| ENSG00000239754 | CFB | -0,417125971 | 0,748915072 | 0,43846 | 0,68769465 | complement factor B |
| ENSG00000168936 | TMEM129 | -0,417269543 | 0,748840546 | 0,07472 | 0,35882064 | transmembrane protein 129, E3 ubiquitin ligase |
| ENSG00000272690 | LINC02018 | -0,417298324 | 0,748825607 | 0,193 | 0,51505487 | long intergenic non-protein coding RNA 2018 |
| ENSG00000282485 | LINC00235 | -0,417343376 | 0,748802223 | 0,34977 | 0,62563749 | long intergenic non-protein coding RNA 235 |
| ENSG00000277142 | LINC00235 | -0,417343376 | 0,748802223 | 0,34977 | 0,62563749 | long intergenic non-protein coding RNA 235 |
| ENSG00000264522 | OTUD7B | -0,417413697 | 0,748765726 | 0,23369 | 0,54563244 | OTU deubiquitinase 7B |
| ENSG00000260269 | AC105036,3 | -0,417470313 | 0,748736342 | 0,57006 | 0,77703861 | novel transcript |
| ENSG00000105053 | VRK3 | -0,417513695 | 0,748713828 | 0,19147 | 0,51435195 | VRK serine/threonine kinase 3 |
| ENSG00000268070 | AC006539,2 | -0,417513891 | 0,748713726 | 0,26711 | 0,56958284 | BCL2/adenovirus E1B 19kDa interacting protein 3 (BNIP3) pseudogene |
| ENSG00000240673 | AC006539,1 | -0,417639656 | 0,748648461 | 0,55675 | 0,76859106 | ribosomal protein S8 (RPS8) pseudogene |
| ENSG00000140983 | RHOT2 | -0,417703103 | 0,748615537 | 0,12517 | 0,44690314 | ras homolog family member T2 |

| | | | | | | |
|---|---|---|---|---|---|---|
| ENSG00000181588 | MEX3D | -0,417727535 | 0,74860286 | 0,22147 | 0,53805234 | mex-3 RNA binding family member D |
| ENSG00000267391 | MIR122HG | -0,417874933 | 0,74852638 | 0,60148 | 0,7966316 | MIR122 host gene |
| ENSG00000121964 | GTDC1 | -0,418037011 | 0,748442292 | 0,34809 | 0,62442761 | glycosyltransferase like domain containing 1 |
| ENSG00000124126 | PREX1 | -0,418060018 | 0,748430357 | 0,36447 | 0,63501401 | phosphatidylinositol-3,4,5-trisphosphate dependent Rac exchange factor 1 |
| ENSG00000261056 | AC079416,2 | -0,418110137 | 0,748404357 | 0,24948 | 0,55657567 | thyroid hormone receptor interactor 13 (TRIP13) pseudogene |
| ENSG00000270492 | AC137695,1 | -0,418199436 | 0,748358035 | 0,4953 | 0,72758381 | MAP/microtubule affinity-regulating kinase 2, pseudogene |
| ENSG00000161533 | ACOX1 | -0,418264484 | 0,748324293 | 0,07833 | 0,36577348 | acyl-CoA oxidase 1 |
| ENSG00000140057 | AK7 | -0,418281027 | 0,748315713 | 0,25147 | 0,55731053 | adenylate kinase 7 |
| ENSG00000282880 | B3GALNT2 | -0,418288521 | 0,748311825 | 0,16869 | 0,49492999 | beta-1,3-N-acetylgalactosaminyltransferase 2 |
| ENSG00000206199 | ANKUB1 | -0,418289494 | 0,748311321 | 0,60715 | 0,80010281 | ankyrin repeat and ubiquitin domain containing 1 |
| ENSG00000165698 | SPACA9 | -0,418457858 | 0,748223997 | 0,15241 | 0,48061452 | sperm acrosome associated 9 |
| ENSG00000231889 | TRAF3IP2-AS1 | -0,418643608 | 0,748127668 | 0,21041 | 0,5290538 | TRAF3IP2 antisense RNA 1 |
| ENSG00000076067 | RBMS2 | -0,418771527 | 0,748061337 | 0,27565 | 0,57533851 | RNA binding motif single stranded interacting protein 2 |
| ENSG00000160753 | RUSC1 | -0,418781575 | 0,748056127 | 0,12932 | 0,45283544 | RUN and SH3 domain containing 1 |
| ENSG00000223745 | CCDC18-AS1 | -0,418850449 | 0,748020415 | 0,23019 | 0,54335232 | CCDC18 antisense RNA 1 |
| ENSG00000162066 | AMDHD2 | -0,418906021 | 0,747991603 | 0,23756 | 0,54860019 | amidohydrolase domain containing 2 |
| ENSG00000268628 | AL121761,1 | -0,418913361 | 0,747987797 | 0,23747 | 0,54849149 | novel transcript |
| ENSG00000173020 | GRK2 | -0,418936619 | 0,747975739 | 0,16808 | 0,49417662 | G protein-coupled receptor kinase 2 |
| ENSG00000164691 | TAGAP | -0,419069135 | 0,747907038 | 0,42101 | 0,67619901 | T cell activation RhoGTPase activating protein |
| ENSG00000163817 | SLC6A20 | -0,419120104 | 0,747880616 | 0,31315 | 0,59991314 | solute carrier family 6 member 20 |
| ENSG00000188859 | FAM78B | -0,419125152 | 0,747877999 | 0,27875 | 0,57694485 | family with sequence similarity 78 member B |
| ENSG00000228208 | LINC02869 | -0,419472134 | 0,747698149 | 0,68194 | 0,84396124 | long intergenic non-protein coding RNA 2869 |
| ENSG00000287828 | AL354743,2 | -0,419485077 | 0,747691441 | 0,5565 | 0,76843732 | novel transcript, antisense to PRDM16 |
| ENSG00000188542 | DUSP28 | -0,419551948 | 0,747656785 | 0,25824 | 0,5622185 | dual specificity phosphatase 28 |
| ENSG00000170390 | DCLK2 | -0,419658942 | 0,747601339 | 0,1281 | 0,45166627 | doublecortin like kinase 2 |
| ENSG00000237406 | NDUFA9P1 | -0,419869083 | 0,747492452 | 0,36711 | 0,63709691 | NADH:ubiquinone oxidoreductase subunit A9 pseudogene 1 |
| ENSG00000197070 | ARRDC1 | -0,419906971 | 0,747472822 | 0,0638 | 0,33487879 | arrestin domain containing 1 |
| ENSG00000261336 | EIF4BP5 | -0,420120426 | 0,747362237 | 0,42705 | 0,68023201 | eukaryotic translation initiation factor 4B pseudogene 5 |
| ENSG00000236826 | LSM2 | -0,420169587 | 0,747336771 | 0,55101 | 0,76487238 | LSM2 homolog, U6 small nuclear RNA and mRNA degradation associated |

| | | | | | | |
|---|---|---|---|---|---|---|
| ENSG00000232729 | AC211433,1 | -0,420192257 | 0,747325027 | 0,13013 | 0,45454796 | novel transcript, antisense to GTF2I |
| ENSG00000240207 | AC080013,1 | -0,420296483 | 0,747271039 | 0,05888 | 0,3241112 | novel transcript, antisense to RARRES1 |
| ENSG00000054793 | ATP9A | -0,42068569 | 0,74706947 | 0,20587 | 0,52532321 | ATPase phospholipid transporting 9A (putative) |
| ENSG00000186812 | ZNF397 | -0,420741946 | 0,747040339 | 0,14347 | 0,47083258 | zinc finger protein 397 |
| ENSG00000225715 | PSMA1P1 | -0,420746188 | 0,747038143 | 0,58241 | 0,7847455 | PSMA1 pseudogene 1 |
| ENSG00000206680 | SNORD21 | -0,420888705 | 0,74696435 | 0,63637 | 0,81700605 | small nucleolar RNA, C/D box 21 |
| ENSG00000198252 | STYX | -0,421007435 | 0,746902879 | 0,28202 | 0,57936013 | serine/threonine/tyrosine interacting protein |
| ENSG00000188372 | ZP3 | -0,421173813 | 0,746816748 | 0,0949 | 0,39743157 | zona pellucida glycoprotein 3 |
| ENSG00000233266 | HMGB1P31 | -0,421182706 | 0,746812145 | 0,45332 | 0,69750019 | high mobility group box 1 pseudogene 31 |
| ENSG00000082781 | ITGB5 | -0,421257566 | 0,746773394 | 0,07608 | 0,36217039 | integrin subunit beta 5 |
| ENSG00000167646 | DNAAF3 | -0,421286653 | 0,746758338 | 0,25099 | 0,55719428 | dynein axonemal assembly factor 3 |
| ENSG00000151303 | AL136982,1 | -0,421293623 | 0,746754731 | 0,04948 | 0,29935562 | novel transcript |
| ENSG00000273300 | AC000068,3 | -0,421442811 | 0,746677513 | 0,76349 | 0,88982985 | novel transcript, antisense to UFD1L |
| ENSG00000149599 | DUSP15 | -0,421499296 | 0,74664828 | 0,29808 | 0,59043587 | dual specificity phosphatase 15 |
| ENSG00000279064 | FP236315,1 | -0,421659551 | 0,746565346 | 0,56005 | 0,77076855 | novel transcript |
| ENSG00000261832 | AC138894,1 | -0,421701637 | 0,746543568 | 0,44642 | 0,69311695 | novel protein |
| ENSG00000181031 | RPH3AL | -0,421740484 | 0,746523466 | 0,39093 | 0,65464997 | rabphilin 3A like (without C2 domains) |
| ENSG00000235890 | TSPEAR-AS1 | -0,421818832 | 0,746482926 | 0,16619 | 0,49212363 | TSPEAR antisense RNA 1 |
| ENSG00000236327 | AL831737,1 | -0,421917521 | 0,746431864 | 0,65838 | 0,82917969 | ribosomal protein L30 (RPL30) pseudogene |
| ENSG00000197948 | FCHSD1 | -0,422059944 | 0,74635818 | 0,07422 | 0,35761188 | FCH and double SH3 domains 1 |
| ENSG00000284137 | LLGL1 | -0,422185009 | 0,746293482 | 0,30072 | 0,59206269 | LLGL scribble cell polarity complex component 1 |
| ENSG00000272899 | ATP6V1FNB | -0,422210642 | 0,746280223 | 0,30418 | 0,5941708 | ATP6V1F neighbor |
| ENSG00000108788 | MLX | -0,422364385 | 0,746200698 | 0,24382 | 0,55243616 | MAX dimerization protein MLX |
| ENSG00000259699 | HMGB1P8 | -0,422435831 | 0,746163745 | 0,36837 | 0,63803074 | high mobility group box 1 pseudogene 8 |
| ENSG00000204852 | TCTN1 | -0,422718974 | 0,746017317 | 0,1012 | 0,40785299 | tectonic family member 1 |
| ENSG00000204065 | TCEAL5 | -0,422984452 | 0,745880052 | 0,1113 | 0,42416781 | transcription elongation factor A like 5 |
| ENSG00000231663 | COA6-AS1 | -0,423014063 | 0,745864743 | 0,38157 | 0,64892048 | COA6 antisense RNA 1 |
| ENSG00000196998 | WDR45 | -0,423033758 | 0,745854561 | 0,30977 | 0,59807209 | WD repeat domain 45 |
| ENSG00000287066 | AP000962,3 | -0,423083037 | 0,745829084 | 0,35994 | 0,63249219 | novel transcript |
| ENSG00000007541 | PIGQ | -0,423136343 | 0,745801528 | 0,22264 | 0,53875487 | phosphatidylinositol glycan anchor biosynthesis class Q |
| ENSG00000214946 | TBC1D26 | -0,423155503 | 0,745791623 | 0,50551 | 0,73401376 | TBC1 domain family member 26 |
| ENSG00000089820 | ARHGAP4 | -0,423277064 | 0,745728785 | 0,21897 | 0,53641439 | Rho GTPase activating protein 4 |
| ENSG00000144231 | POLR2D | -0,423438114 | 0,745645543 | 0,20883 | 0,52791772 | RNA polymerase II subunit D |

| | | | | | | |
|---|---|---|---|---|---|---|
| ENSG00000151789 | ZNF385D | -0,423443709 | 0,745642652 | 0,38101 | 0,64851669 | zinc finger protein 385D |
| ENSG00000205336 | ADGRG1 | -0,42344811 | 0,745640377 | 0,17108 | 0,49641416 | adhesion G protein-coupled receptor G1 |
| ENSG00000257433 | AC004241,1 | -0,423646128 | 0,74553804 | 0,30172 | 0,59283946 | novel transcript, antisense to ENDOU and RAPGEF3 |
| ENSG00000284776 | AL121900,1 | -0,423661346 | 0,745530176 | 0,13617 | 0,463307 | novel protein |
| ENSG00000149294 | NCAM1 | -0,423674239 | 0,745523514 | 0,28763 | 0,58395529 | neural cell adhesion molecule 1 |
| ENSG00000260848 | AC092114,1 | -0,423674584 | 0,745523336 | 0,55471 | 0,76760293 | novel transcript |
| ENSG00000267696 | ERVK-28 | -0,423699144 | 0,745510644 | 0,43364 | 0,68473723 | endogenous retrovirus group K member 28 |
| ENSG00000134461 | ANKRD16 | -0,423836521 | 0,745439658 | 0,17402 | 0,49860418 | ankyrin repeat domain 16 |
| ENSG00000223901 | AP001469,1 | -0,424037743 | 0,745335694 | 0,40866 | 0,66743928 | novel transcript |
| ENSG00000115353 | TACR1 | -0,424258242 | 0,745221787 | 0,37548 | 0,6438135 | tachykinin receptor 1 |
| ENSG00000164627 | KIF6 | -0,424347416 | 0,745175726 | 0,35241 | 0,62742225 | kinesin family member 6 |
| ENSG00000140961 | OSGIN1 | -0,424442081 | 0,745126831 | 0,19669 | 0,51792163 | oxidative stress induced growth inhibitor 1 |
| ENSG00000089116 | LHX5 | -0,424516683 | 0,745088302 | 0,3626 | 0,63379594 | LIM homeobox 5 |
| ENSG00000199883 | RN7SKP90 | -0,424565798 | 0,745062936 | 0,51923 | 0,74388669 | RN7SK pseudogene 90 |
| ENSG00000254760 | AC008750,1 | -0,424578851 | 0,745056195 | 0,49127 | 0,72507957 | novel transcript, antisense to SIGLEC10 |
| ENSG00000186493 | C5orf38 | -0,424620377 | 0,74503475 | 0,48338 | 0,71965182 | chromosome 5 open reading frame 38 |
| ENSG00000261546 | AC135782,3 | -0,424627799 | 0,745030918 | 0,3799 | 0,64769466 | novel transcript, antisense to ACSF3 |
| ENSG00000167034 | NKX3-1 | -0,424812339 | 0,744935624 | 0,58492 | 0,78605624 | NK3 homeobox 1 |
| ENSG00000286485 | Z80897,1 | -0,424829165 | 0,744926936 | 0,29262 | 0,58716506 | novel transcript, antisense to ANKRD54and IFT27 |
| ENSG00000231530 | AL157932,1 | -0,424922661 | 0,744878662 | 0,60384 | 0,79770754 | novel transcript |
| ENSG00000272979 | AC093388,1 | -0,425310991 | 0,74467819 | 0,60935 | 0,80165403 | novel transcript, antisense to MFSD6 |
| ENSG00000121207 | LRAT | -0,42531963 | 0,744673731 | 0,45742 | 0,70074793 | lecithin retinol acyltransferase |
| ENSG00000167645 | YIF1B | -0,425354648 | 0,744655655 | 0,20123 | 0,52181085 | Yip1 interacting factor homolog B, membrane trafficking protein |
| ENSG00000235902 | AC108472,1 | -0,425477803 | 0,744592091 | 0,47537 | 0,71396741 | novel transcript |
| ENSG00000271537 | AC009220,3 | -0,425514624 | 0,744573087 | 0,5094 | 0,73725396 | FCF1 small subunit (SSU) processome component homolog (S, cerevisiae), pseudogene |
| ENSG00000255135 | AP002360,1 | -0,425567331 | 0,744545886 | 0,17265 | 0,49744497 | novel transcript |
| ENSG00000233527 | ZNF529-AS1 | -0,425594456 | 0,744531887 | 0,08674 | 0,38151737 | ZNF529 antisense RNA 1 |
| ENSG00000184709 | LRRC26 | -0,42571183 | 0,744471317 | 0,73447 | 0,87444606 | leucine rich repeat containing 26 |
| ENSG00000250333 | AC093725,2 | -0,425897539 | 0,744375491 | 0,20929 | 0,52826608 | novel transcript |
| ENSG00000188818 | ZDHHC11 | -0,426143269 | 0,744248715 | 0,11011 | 0,42253609 | zinc finger DHHC-type containing 11 |
| ENSG00000213212 | NCLP1 | -0,426277914 | 0,744179259 | 0,39322 | 0,65625452 | nucleolin pseudogene 1 |

| | | | | | | |
|---|---|---|---|---|---|---|
| ENSG00000197993 | KEL | -0,426595207 | 0,744015609 | 0,68994 | 0,84879823 | Kell metallo-endopeptidase (Kell blood group) |
| ENSG00000288091 | AC062022,2 | -0,426991346 | 0,743811343 | 0,08599 | 0,38027313 | novel transcript, antisense to SLC4A10 |
| ENSG00000225017 | AP004289,1 | -0,426991839 | 0,743811089 | 0,42731 | 0,68043324 | aconitase 2, mitochondrial (ACO2) pseudogene |
| ENSG00000239382 | ALKBH6 | -0,427009467 | 0,743802 | 0,21746 | 0,53536715 | alkB homolog 6 |
| ENSG00000287214 | AC021321,2 | -0,427120047 | 0,743744991 | 0,28091 | 0,57822547 | novel transcript, sense intronic to ARFGEF1 |
| ENSG00000285095 | AC025887,2 | -0,427248808 | 0,743678615 | 0,51818 | 0,74306841 | novel transcript |
| ENSG00000104432 | IL7 | -0,427477322 | 0,74356083 | 0,34767 | 0,62423931 | interleukin 7 |
| ENSG00000281526 | ZNF546 | -0,427584674 | 0,743505503 | 0,5492 | 0,76404333 | zinc finger protein 546 |
| ENSG00000287474 | AC110614,1 | -0,427600205 | 0,743497499 | 0,17015 | 0,49606609 | novel transcript |
| ENSG00000281950 | SPDYE16 | -0,427606985 | 0,743494005 | 0,15333 | 0,48157724 | speedy/RINGO cell cycle regulator family member E16 |
| ENSG00000157551 | KCNJ15 | -0,427613537 | 0,743490629 | 0,2011 | 0,52181085 | potassium inwardly rectifying channel subfamily J member 15 |
| ENSG00000165076 | PRSS37 | -0,427619228 | 0,743487696 | 0,61388 | 0,80357704 | serine protease 37 |
| ENSG00000177201 | OR2T12 | -0,427647079 | 0,743473343 | 0,35247 | 0,62742225 | olfactory receptor family 2 subfamily T member 12 |
| ENSG00000204044 | SLC12A5-AS1 | -0,427911103 | 0,743337294 | 0,37787 | 0,64628549 | SLC12A5 and MMP9 antisense RNA 1 |
| ENSG00000182993 | C12orf60 | -0,427932764 | 0,743326134 | 0,3053 | 0,59473483 | chromosome 12 open reading frame 60 |
| ENSG00000205090 | TMEM240 | -0,428186839 | 0,743195237 | 0,18578 | 0,50915451 | transmembrane protein 240 |
| ENSG00000186235 | LINC02610 | -0,428240898 | 0,74316739 | 0,29384 | 0,58832005 | long intergenic non-protein coding RNA 2610 |
| ENSG00000224420 | ADM5 | -0,428331994 | 0,743120465 | 0,42934 | 0,68217673 | adrenomedullin 5 (putative) |
| ENSG00000236776 | RPL21P23 | -0,428385794 | 0,743092754 | 0,70406 | 0,85725494 | ribosomal protein L21 pseudogene 23 |
| ENSG00000287005 | AC090018,2 | -0,428426864 | 0,7430716 | 0,41205 | 0,66983104 | novel transcript, antisense to XXYLT1 |
| ENSG00000124713 | GNMT | -0,42845249 | 0,743058401 | 0,45945 | 0,7017037 | glycine N-methyltransferase |
| ENSG00000267023 | LRRC37A16P | -0,42846333 | 0,743052818 | 0,2085 | 0,52778377 | leucine rich repeat containing 37 member A16, pseudogene |
| ENSG00000234982 | AL441963,1 | -0,428496259 | 0,743035858 | 0,57579 | 0,78043323 | ribosomal protein L7a (RPL7A) pseudogene |
| ENSG00000286994 | AC069243,1 | -0,428716638 | 0,742922365 | 0,26168 | 0,56453855 | novel transcript, sense intronic to GOLIM4 |
| ENSG00000160789 | LMNA | -0,428741183 | 0,742909725 | 0,36446 | 0,63501401 | lamin A/C |
| ENSG00000136295 | TTYH3 | -0,428838998 | 0,742859357 | 0,11559 | 0,43159073 | tweety family member 3 |
| ENSG00000185739 | SRL | -0,428885905 | 0,742835205 | 0,3509 | 0,62636083 | sarcalumenin |
| ENSG00000250634 | LINC01182 | -0,429290629 | 0,742626844 | 0,3206 | 0,60553826 | long intergenic non-protein coding RNA 1182 |
| ENSG00000219085 | NPM1P37 | -0,429293149 | 0,742625547 | 0,39362 | 0,65656332 | nucleophosmin 1 pseudogene 37 |
| ENSG00000160050 | CCDC28B | -0,429337955 | 0,742602483 | 0,10416 | 0,41252273 | coiled-coil domain containing 28B |
| ENSG00000224208 | ZCRB1P1 | -0,429451562 | 0,742544008 | 0,54408 | 0,76034748 | ZCRB1 pseudogene 1 |

| | | | | | | |
|---|---|---|---|---|---|---|
| ENSG00000165752 | STK32C | -0,429509712 | 0,74251408 | 0,12858 | 0,45191041 | serine/threonine kinase 32C |
| ENSG00000285693 | AP002381,2 | -0,429653427 | 0,742440118 | 0,33084 | 0,6112353 | Novel transcript |
| ENSG00000215515 | IFIT1P1 | -0,429662398 | 0,742435501 | 0,56805 | 0,77582773 | interferon induced protein with tetratricopeptide repeats 1 pseudogene 1 |
| ENSG00000250358 | LINC02200 | -0,429705525 | 0,742413307 | 0,4615 | 0,70284173 | long intergenic non-protein coding RNA 2200 |
| ENSG00000256564 | AC007552,2 | -0,429773224 | 0,74237847 | 0,67223 | 0,83784792 | novel transcript |
| ENSG00000221916 | C19orf73 | -0,429795335 | 0,742367092 | 0,3418 | 0,6197312 | chromosome 19 open reading frame 73 |
| ENSG00000240853 | RN7SL328P | -0,429937337 | 0,742294026 | 0,64347 | 0,82109591 | RNA, 7SL, cytoplasmic 328, pseudogene |
| ENSG00000160199 | PKNOX1 | -0,429946226 | 0,742289452 | 0,04606 | 0,28930497 | PBX/knotted 1 homeobox 1 |
| ENSG00000283125 | AC022726,2 | -0,430046016 | 0,742238111 | 0,69758 | 0,8534853 | novel transcript, antisense to BCL2 |
| ENSG00000091262 | ABCC6 | -0,43061978 | 0,741942979 | 0,61201 | 0,80341001 | ATP binding cassette subfamily C member 6 |
| ENSG00000283771 | TMEM256 | -0,430853242 | 0,741822925 | 0,26307 | 0,56588348 | transmembrane protein 256 |
| ENSG00000205544 | TMEM256 | -0,430853242 | 0,741822925 | 0,26307 | 0,56588348 | transmembrane protein 256 |
| ENSG00000260012 | AC007494,1 | -0,430904351 | 0,741796646 | 0,59959 | 0,79552021 | proteasome (prosome, macropain) subunit, beta type, 5 (PSMB5) pseudogene |
| ENSG00000228663 | PSMD10P1 | -0,430905388 | 0,741796112 | 0,19504 | 0,51588752 | proteasome 26S subunit, non-ATPase, 10 pseudogene 1 |
| ENSG00000166592 | RRAD | -0,430951008 | 0,741772656 | 0,70311 | 0,8568579 | RRAD, Ras related glycolysis inhibitor and calcium channel regulator |
| ENSG00000268926 | AL354861,3 | -0,431051527 | 0,741720975 | 0,65064 | 0,82499358 | novel transcript |
| ENSG00000228889 | UBAC2-AS1 | -0,431097971 | 0,741697098 | 0,1511 | 0,47950432 | UBAC2 antisense RNA 1 |
| ENSG00000238197 | PAXBP1-AS1 | -0,43129377 | 0,741596443 | 0,13521 | 0,46165567 | PAXBP1 antisense RNA 1 |
| ENSG00000231043 | AC007238,1 | -0,431484024 | 0,741498652 | 0,59031 | 0,78972365 | IK cytokine, down-regulator of HLA II (IK) pseudogene |
| ENSG00000236007 | MTCO1P46 | -0,431515306 | 0,741482575 | 0,44284 | 0,69042854 | MT-CO1 pseudogene 46 |
| ENSG00000205790 | DPP9-AS1 | -0,431681451 | 0,741397188 | 0,3788 | 0,64679625 | DPP9 antisense RNA 1 |
| ENSG00000231793 | DOC2GP | -0,43172432 | 0,741375159 | 0,35602 | 0,63030618 | double C2 domain gamma, pseudogene |
| ENSG00000105696 | TMEM59L | -0,431863539 | 0,74130362 | 0,34781 | 0,62427624 | transmembrane protein 59 like |
| ENSG00000151490 | PTPRO | -0,43191456 | 0,741277404 | 0,29304 | 0,58745839 | protein tyrosine phosphatase receptor type O |
| ENSG00000163517 | HDAC11 | -0,431926613 | 0,741271211 | 0,10697 | 0,41675015 | histone deacetylase 11 |
| ENSG00000146223 | RPL7L1 | -0,431933584 | 0,741267629 | 0,26673 | 0,56923396 | ribosomal protein L7 like 1 |
| ENSG00000133056 | PIK3C2B | -0,432017483 | 0,741224522 | 0,16438 | 0,49065086 | phosphatidylinositol-4-phosphate 3-kinase catalytic subunit type 2 beta |
| ENSG00000248677 | LINC02102 | -0,432092468 | 0,741185998 | 0,59292 | 0,79148028 | long intergenic non-protein coding RNA 2102 |
| ENSG00000242732 | RTL5 | -0,432124543 | 0,741169519 | 0,15587 | 0,4836223 | retrotransposon Gag like 5 |

| | | | | | | |
|---|---|---|---|---|---|---|
| ENSG00000273199 | AP000692,2 | -0,432282635 | 0,741088306 | 0,40314 | 0,66399303 | novel transcript, antisense to MORC3 |
| ENSG00000228577 | AC010731,1 | -0,432335464 | 0,741061169 | 0,6349 | 0,8162492 | novel transcript |
| ENSG00000276691 | AC004466,3 | -0,432469388 | 0,74099238 | 0,54506 | 0,7609742 | novel transcript, antisense to HDAC7 |
| ENSG00000205740 | AL359878,1 | -0,432485098 | 0,740984311 | 0,23224 | 0,54499308 | uncharacterized LOC107984285 |
| ENSG00000204323 | SMIM5 | -0,432500374 | 0,740976465 | 0,38923 | 0,65363949 | small integral membrane protein 5 |
| ENSG00000228196 | PTPN2P1 | -0,432580497 | 0,740935315 | 0,57912 | 0,78270914 | PTPN2 pseudogene 1 |
| ENSG00000275496 | CU633906,1 | -0,432606105 | 0,740922163 | 0,53603 | 0,75500755 | uncharacterized LOC102724701 |
| ENSG00000169062 | UPF3A | -0,432647311 | 0,740901001 | 0,08438 | 0,37731853 | UPF3A regulator of nonsense mediated mRNA decay |
| ENSG00000260691 | ANKRD20A1 | -0,432682839 | 0,740882756 | 0,51368 | 0,73998037 | ankyrin repeat domain 20 family member A1 |
| ENSG00000184281 | TSSC4 | -0,432710397 | 0,740868604 | 0,17932 | 0,50368273 | tumor suppressing subtransferable candidate 4 |
| ENSG00000268362 | AC092279,1 | -0,432830017 | 0,740807178 | 0,13648 | 0,46372738 | novel transcript |
| ENSG00000267040 | AC027097,1 | -0,432899555 | 0,740771472 | 0,13814 | 0,46543358 | novel transcript, antisense to ATP8B1 |
| ENSG00000180336 | MEIOC | -0,433017069 | 0,740711135 | 0,29621 | 0,58961035 | meiosis specific with coiled-coil domain |
| ENSG00000213891 | RPL3P6 | -0,433069007 | 0,74068447 | 0,2082 | 0,52760275 | ribosomal protein L3 pseudogene 6 |
| ENSG00000189409 | MMP23B | -0,433234373 | 0,740599575 | 0,56075 | 0,77105099 | matrix metallopeptidase 23B |
| ENSG00000249028 | AC004672,1 | -0,433301066 | 0,74056534 | 0,49395 | 0,72657717 | novel transcript |
| ENSG00000126243 | LRFN3 | -0,433329715 | 0,740550633 | 0,20171 | 0,52214043 | leucine rich repeat and fibronectin type III domain containing 3 |
| ENSG00000284952 | AC104472,3 | -0,433453301 | 0,740487198 | 0,3814 | 0,64874863 | novel transcript |
| ENSG00000262516 | AC007151,1 | -0,433572892 | 0,740425819 | 0,42751 | 0,68046596 | developmental pluripotency associated 3 (DPPA3) pseudogene |
| ENSG00000253452 | AC124290,1 | -0,433625389 | 0,740398876 | 0,3216 | 0,60577115 | novel transcript |
| ENSG00000167799 | NUDT8 | -0,433694119 | 0,740363605 | 0,26778 | 0,56999323 | nudix hydrolase 8 |
| ENSG00000112852 | PCDHB2 | -0,433749681 | 0,740335092 | 0,07458 | 0,35861227 | protocadherin beta 2 |
| ENSG00000237854 | LINC00674 | -0,433839706 | 0,740288896 | 0,25542 | 0,55992862 | long intergenic non-protein coding RNA 674 |
| ENSG00000087266 | SH3BP2 | -0,433880014 | 0,740268213 | 0,20668 | 0,52622584 | SH3 domain binding protein 2 |
| ENSG00000173114 | LRRN3 | -0,43388548 | 0,740265408 | 0,3297 | 0,61069033 | leucine rich repeat neuronal 3 |
| ENSG00000173531 | MST1 | -0,434030324 | 0,740191091 | 0,12379 | 0,4446254 | macrophage stimulating 1 |
| ENSG00000138079 | SLC3A1 | -0,43406778 | 0,740171874 | 0,35266 | 0,62754548 | solute carrier family 3 member 1 |
| ENSG00000157017 | GHRL | -0,43407963 | 0,740165794 | 0,28892 | 0,58464073 | ghrelin and obestatin prepropeptide |
| ENSG00000258592 | AL391152,1 | -0,434082275 | 0,740164437 | 0,33095 | 0,6112353 | novel transcript |
| ENSG00000173714 | WFIKKN2 | -0,434100725 | 0,740154971 | 0,49897 | 0,73002372 | WAP, follistatin/kazal, immunoglobulin, kunitz and netrin domain containing 2 |

| | | | | | | |
|---|---|---|---|---|---|---|
| ENSG00000148339 | SLC25A25 | -0,434166116 | 0,740121425 | 0,02572 | 0,21626994 | solute carrier family 25 member 25 |
| ENSG00000236671 | PRKG1-AS1 | -0,434377358 | 0,740013063 | 0,12892 | 0,45222045 | PRKG1 antisense RNA 1 |
| ENSG00000204524 | ZNF805 | -0,434484145 | 0,739958289 | 0,19507 | 0,51588752 | zinc finger protein 805 |
| ENSG00000204237 | OXLD1 | -0,434630431 | 0,739883263 | 0,17008 | 0,49606609 | oxidoreductase like domain containing 1 |
| ENSG00000173705 | SUSD5 | -0,434630765 | 0,739883092 | 0,16237 | 0,48945635 | sushi domain containing 5 |
| ENSG00000267651 | AC015961,1 | -0,434808081 | 0,739792161 | 0,29041 | 0,58561852 | novel transcript |
| ENSG00000170122 | FOXD4 | -0,434824857 | 0,739783559 | 0,47048 | 0,70993321 | forkhead box D4 |
| ENSG00000075461 | CACNG4 | -0,435046486 | 0,739669921 | 0,05277 | 0,30840541 | calcium voltage-gated channel auxiliary subunit gamma 4 |
| ENSG00000092098 | RNF31 | -0,435114041 | 0,739635286 | 0,27981 | 0,57750968 | ring finger protein 31 |
| ENSG00000105419 | MEIS3 | -0,4351405 | 0,739621722 | 0,13191 | 0,45708171 | Meis homeobox 3 |
| ENSG00000272030 | AL162258,2 | -0,43514739 | 0,739618189 | 0,43181 | 0,6838061 | novel transcript, antisense to S100A1 and CHTOP |
| ENSG00000132613 | MTSS2 | -0,43548322 | 0,739446041 | 0,20736 | 0,52677688 | MTSS I-BAR domain containing 2 |
| ENSG00000126091 | ST3GAL3 | -0,435494766 | 0,739440123 | 0,1798 | 0,50407792 | ST3 beta-galactoside alpha-2,3-sialyltransferase 3 |
| ENSG00000106689 | LHX2 | -0,435534581 | 0,739419717 | 0,38985 | 0,65396251 | LIM homeobox 2 |
| ENSG00000273568 | AC131009,3 | -0,435622805 | 0,739374501 | 0,68621 | 0,84663586 | novel transcript, sense intronic to PUS1 |
| ENSG00000170265 | ZNF282 | -0,435755204 | 0,73930665 | 0,17526 | 0,49934416 | zinc finger protein 282 |
| ENSG00000205236 | AC105052,1 | -0,435998683 | 0,73918189 | 0,50027 | 0,73078774 | novel protein, UPK3BL-RASA4 readthrough |
| ENSG00000172346 | CSDC2 | -0,436012483 | 0,73917482 | 0,1626 | 0,48945635 | cold shock domain containing C2 |
| ENSG00000259417 | CTXND1 | -0,4360138 | 0,739174145 | 0,41157 | 0,6692379 | cortexin domain containing 1 |
| ENSG00000230005 | SNAP47-AS1 | -0,436029079 | 0,739166317 | 0,35739 | 0,63130076 | SNAP47 antisense RNA 1 |
| ENSG00000154134 | ROBO3 | -0,436083328 | 0,739138523 | 0,26154 | 0,56452671 | roundabout guidance receptor 3 |
| ENSG00000230806 | SRGAP2-AS1 | -0,43611336 | 0,739123137 | 0,27707 | 0,57569532 | SRGAP2 antisense RNA 1 |
| ENSG00000075914 | EXOSC7 | -0,436132365 | 0,7391134 | 0,1745 | 0,49884018 | exosome component 7 |
| ENSG00000278981 | AC079298,1 | -0,436301112 | 0,739026953 | 0,59197 | 0,79090751 | novel transcript, antisense to DCHS2 |
| ENSG00000284467 | AC079298,5 | -0,436301112 | 0,739026953 | 0,59197 | 0,79090751 | novel transcript, antisense to DCHS2 |
| ENSG00000287258 | AL512283,2 | -0,436325105 | 0,739014663 | 0,19088 | 0,51394498 | novel transcript, antisense to PKIB |
| ENSG00000187566 | NHLRC1 | -0,436358698 | 0,738997455 | 0,44023 | 0,68866128 | NHL repeat containing E3 ubiquitin protein ligase 1 |
| ENSG00000231825 | PRRC2A | -0,436374182 | 0,738989524 | 0,19319 | 0,51517146 | proline rich coiled-coil 2A |
| ENSG00000206427 | PRRC2A | -0,436374182 | 0,738989524 | 0,19319 | 0,51517146 | proline rich coiled-coil 2A |
| ENSG00000197483 | ZNF628 | -0,436488681 | 0,738930877 | 0,22754 | 0,54168591 | zinc finger protein 628 |
| ENSG00000112139 | MDGA1 | -0,436525829 | 0,73891185 | 0,11201 | 0,42546126 | MAM domain containing glycosylphosphatidylinositol anchor 1 |
| ENSG00000287959 | AC021236,1 | -0,437075488 | 0,738630383 | 0,26773 | 0,56999323 | novel transcript, antisense to PRKDC |

| | | | | | | |
|---|---|---|---|---|---|---|
| ENSG00000169991 | IFFO2 | -0,437159905 | 0,738587164 | 0,09285 | 0,39343521 | intermediate filament family orphan 2 |
| ENSG00000163661 | PTX3 | -0,437212016 | 0,738560486 | 0,32104 | 0,6057178 | pentraxin 3 |
| ENSG00000011243 | AKAP8L | -0,437385158 | 0,738471855 | 0,10205 | 0,40934973 | A-kinase anchoring protein 8 like |
| ENSG00000270021 | AC026691,1 | -0,437399966 | 0,738464275 | 0,39531 | 0,65820274 | novel transcript, antisense to H2AFY |
| ENSG00000242318 | AC058823,1 | -0,437443704 | 0,738441887 | 0,70757 | 0,85916455 | ribosomal protein L21 (RPL21) pseudogene |
| ENSG00000206337 | HCP5 | -0,43745908 | 0,738434017 | 0,5229 | 0,7470043 | HLA complex P5 |
| ENSG00000117245 | KIF17 | -0,437502554 | 0,738411766 | 0,17989 | 0,50407792 | kinesin family member 17 |
| ENSG00000093072 | ADA2 | -0,437583053 | 0,738370565 | 0,10526 | 0,41429627 | adenosine deaminase 2 |
| ENSG00000228906 | AL353804,1 | -0,437630023 | 0,738346526 | 0,25881 | 0,56260423 | novel transcript |
| ENSG00000276398 | AC018659,5 | -0,437676803 | 0,738322586 | 0,46195 | 0,70326917 | |
| ENSG00000165269 | AQP7 | -0,437810211 | 0,738254315 | 0,4597 | 0,70182254 | aquaporin 7 |
| ENSG00000150672 | DLG2 | -0,437816009 | 0,738251348 | 0,1061 | 0,41557034 | discs large MAGUK scaffold protein 2 |
| ENSG00000188295 | ZNF669 | -0,437862198 | 0,738227713 | 0,24326 | 0,55199251 | zinc finger protein 669 |
| ENSG00000232295 | AL589935,1 | -0,43824503 | 0,738031844 | 0,32645 | 0,60866572 | novel transcript, antisense OGFRL1 |
| ENSG00000287764 | AL136038,5 | -0,438250885 | 0,738028849 | 0,41788 | 0,6742866 | novel transcript |
| ENSG00000274015 | AL136038,4 | -0,438250885 | 0,738028849 | 0,41788 | 0,6742866 | novel transcript |
| ENSG00000258951 | KRT18P7 | -0,438310003 | 0,737998606 | 0,40396 | 0,66410381 | keratin 18 pseudogene 7 |
| ENSG00000260971 | AC119674,1 | -0,438332422 | 0,737987138 | 0,32615 | 0,6085098 | novel transcript |
| ENSG00000228203 | GRASLND | -0,438337078 | 0,737984756 | 0,25635 | 0,560877 | glycosaminoglycan regulatory associated long non-coding RNA |
| ENSG00000111358 | GTF2H3 | -0,438347019 | 0,737979671 | 0,22355 | 0,53953208 | general transcription factor IIH subunit 3 |
| ENSG00000183722 | LHFPL6 | -0,438505211 | 0,737898756 | 0,18742 | 0,51074735 | LHFPL tetraspan subfamily member 6 |
| ENSG00000229853 | AL034418,1 | -0,438521953 | 0,737890193 | 0,59251 | 0,79121909 | ribosomal protein S3A (RPS3A) pseudogene |
| ENSG00000100106 | TRIOBP | -0,438543796 | 0,737879021 | 0,04116 | 0,27485265 | TRIO and F-actin binding protein |
| ENSG00000257599 | OVCH1-AS1 | -0,438593198 | 0,737853755 | 0,23317 | 0,54558536 | OVCH1 antisense RNA 1 |
| ENSG00000272056 | AC013472,2 | -0,438603255 | 0,737848611 | 0,25944 | 0,56317998 | novel transcript, antisense to AGBL5 |
| ENSG00000133247 | KMT5C | -0,43861428 | 0,737842973 | 0,30781 | 0,59671558 | lysine methyltransferase 5C |
| ENSG00000249947 | XBP1P1 | -0,439010236 | 0,737640495 | 0,41749 | 0,67403924 | X-box binding protein 1 pseudogene 1 |
| ENSG00000227684 | CROCCP4 | -0,439013064 | 0,737639049 | 0,63573 | 0,81671084 | CROCC pseudogene 4 |
| ENSG00000174871 | CNIH2 | -0,439224318 | 0,737531044 | 0,07995 | 0,36910486 | cornichon family AMPA receptor auxiliary protein 2 |
| ENSG00000281357 | ARRDC3-AS1 | -0,439457649 | 0,737411771 | 0,07624 | 0,36242671 | ARRDC3 antisense RNA 1 |
| ENSG00000226015 | CCT8P1 | -0,439480358 | 0,737400164 | 0,4872 | 0,72231397 | chaperonin containing TCP1 subunit 8 pseudogene 1 |
| ENSG00000274853 | MRM1 | -0,43952366 | 0,737378031 | 0,44991 | 0,69539135 | mitochondrial rRNA methyltransferase 1 |

| | | | | | | |
|---|---|---|---|---|---|---|
| ENSG00000204152 | TIMM23B | -0,439544991 | 0,737367129 | 0,16863 | 0,49492999 | translocase of inner mitochondrial membrane 23 homolog B |
| ENSG00000124766 | SOX4 | -0,439554979 | 0,737362024 | 0,10186 | 0,40926385 | SRY-box transcription factor 4 |
| ENSG00000197380 | DACT3 | -0,439601295 | 0,737338352 | 0,4685 | 0,70812799 | dishevelled binding antagonist of beta catenin 3 |
| ENSG00000254377 | MIR124-2HG | -0,439608887 | 0,737334472 | 0,02124 | 0,19328293 | MIR124-2 host gene |
| ENSG00000160111 | CPAMD8 | -0,439757373 | 0,737258588 | 0,20404 | 0,52370308 | C3 and PZP like alpha-2-macroglobulin domain containing 8 |
| ENSG00000279873 | LINC01126 | -0,439819094 | 0,737227047 | 0,21804 | 0,5356729 | long intergenic non-protein coding RNA 1126 |
| ENSG00000135314 | KHDC1 | -0,439837685 | 0,737217547 | 0,01686 | 0,17057452 | KH domain containing 1 |
| ENSG00000076554 | TPD52 | -0,439841647 | 0,737215522 | 0,32879 | 0,60992716 | tumor protein D52 |
| ENSG00000215217 | C5orf49 | -0,439918634 | 0,737176183 | 0,25436 | 0,55940142 | chromosome 5 open reading frame 49 |
| ENSG00000163485 | ADORA1 | -0,439920637 | 0,73717516 | 0,31898 | 0,60399503 | adenosine A1 receptor |
| ENSG00000185344 | ATP6V0A2 | -0,439921065 | 0,737174941 | 0,23107 | 0,54397035 | ATPase H+ transporting V0 subunit a2 |
| ENSG00000253519 | AC106801,1 | -0,43992684 | 0,73717199 | 0,42461 | 0,67841584 | novel transcript |
| ENSG00000099977 | DDT | -0,439971709 | 0,737149064 | 0,20779 | 0,52736343 | D-dopachrome tautomerase |
| ENSG00000145819 | ARHGAP26 | -0,440014428 | 0,737127237 | 0,0921 | 0,39241636 | Rho GTPase activating protein 26 |
| ENSG00000185040 | SPDYE16 | -0,440218971 | 0,737022735 | 0,13519 | 0,46165567 | speedy/RINGO cell cycle regulator family member E16 |
| ENSG00000258457 | AL132780,2 | -0,440252689 | 0,73700551 | 0,28129 | 0,578832 | novel transcript, antisense to C14orf93 |
| ENSG00000100302 | RASD2 | -0,440288937 | 0,736986993 | 0,72171 | 0,86715946 | RASD family member 2 |
| ENSG00000105717 | PBX4 | -0,440319999 | 0,736971126 | 0,04729 | 0,29217708 | PBX homeobox 4 |
| ENSG00000138111 | MFSD13A | -0,440342793 | 0,736959482 | 0,19461 | 0,51562158 | major facilitator superfamily domain containing 13A |
| ENSG00000172687 | ZNF738 | -0,44042355 | 0,736918231 | 0,12311 | 0,44322266 | zinc finger protein 738 |
| ENSG00000279685 | MAPT-IT1 | -0,440521941 | 0,736867975 | 0,79666 | 0,90605632 | MAPT intronic transcript 1 |
| ENSG00000259867 | AC105411,1 | -0,440522017 | 0,736867936 | 0,24791 | 0,55532274 | novel transcript, antisense DYNLRB2 |
| ENSG00000163933 | RFT1 | -0,440559634 | 0,736848723 | 0,26337 | 0,56608528 | RFT1 homolog |
| ENSG00000101958 | GLRA2 | -0,440610715 | 0,736822634 | 0,50977 | 0,73745081 | glycine receptor alpha 2 |
| ENSG00000246922 | UBAP1L | -0,440643349 | 0,736805967 | 0,09318 | 0,39449045 | ubiquitin associated protein 1 like |
| ENSG00000213057 | C1orf220 | -0,440713513 | 0,736770135 | 0,20182 | 0,52216179 | chromosome 1 putative open reading frame 220 |
| ENSG00000271855 | AC073195,1 | -0,440765971 | 0,736743345 | 0,14723 | 0,47471707 | novel transcript |
| ENSG00000115286 | NDUFS7 | -0,440796269 | 0,736727873 | 0,19361 | 0,51547251 | NADH:ubiquinone oxidoreductase core subunit S7 |
| ENSG00000286703 | AC005682,2 | -0,440855614 | 0,736697569 | 0,57543 | 0,78019408 | novel transcript |
| ENSG00000137193 | PIM1 | -0,441006422 | 0,736620564 | 0,25022 | 0,55657567 | Pim-1 proto-oncogene, serine/threonine kinase |
| ENSG00000112530 | PACRG | -0,441038714 | 0,736604076 | 0,37165 | 0,64074902 | parkin coregulated |
| ENSG00000196935 | SRGAP1 | -0,441193294 | 0,736525156 | 0,17381 | 0,49846579 | SLIT-ROBO Rho GTPase activating protein 1 |

| | | | | | | |
|---|---|---|---|---|---|---|
| ENSG00000287677 | AC025839,1 | -0,441235646 | 0,736503535 | 0,6072 | 0,80012503 | novel transcript |
| ENSG00000248092 | NNT-AS1 | -0,441332402 | 0,736454142 | 0,07481 | 0,35899472 | NNT antisense RNA 1 |
| ENSG00000127152 | BCL11B | -0,441349421 | 0,736445454 | 0,34157 | 0,61963031 | BAF chromatin remodeling complex subunit BCL11B |
| ENSG00000197785 | ATAD3A | -0,441364784 | 0,736437612 | 0,1699 | 0,49606609 | ATPase family AAA domain containing 3A |
| ENSG00000237731 | RNGTTP1 | -0,441389361 | 0,736425066 | 0,42834 | 0,68103938 | RNA guanylyltransferase and 5'-phosphatase pseudogene 1 |
| ENSG00000112357 | PEX7 | -0,441392772 | 0,736423325 | 0,26964 | 0,57135887 | peroxisomal biogenesis factor 7 |
| ENSG00000228150 | AL357140,2 | -0,441538382 | 0,736349002 | 0,64934 | 0,8244072 | novel transcript |
| ENSG00000167395 | ZNF646 | -0,441693829 | 0,736269667 | 0,1189 | 0,43696075 | zinc finger protein 646 |
| ENSG00000257529 | RPL36A-HNRN | -0,441783595 | 0,736223857 | 0,48366 | 0,71986841 | RPL36A-HNRNPH2 readthrough |
| ENSG00000275846 | AL513548,3 | -0,441832607 | 0,736198846 | 0,38829 | 0,6529693 | novel transcript |
| ENSG00000254604 | AP002336,2 | -0,442001766 | 0,73611253 | 0,51918 | 0,74385663 | novel transcript |
| ENSG00000288343 | AP002336,6 | -0,442001766 | 0,73611253 | 0,51918 | 0,74385663 | novel transcript |
| ENSG00000272912 | AL356608,1 | -0,442048008 | 0,736088936 | 0,54997 | 0,7643663 | novel transcript |
| ENSG00000183287 | CCBE1 | -0,442125092 | 0,736049608 | 0,48835 | 0,72325321 | collagen and calcium binding EGF domains 1 |
| ENSG00000233542 | AL391845,2 | -0,44216562 | 0,736028931 | 0,5799 | 0,78324364 | novel transcript |
| ENSG00000235740 | PHACTR2-AS1 | -0,442176368 | 0,736023447 | 0,3907 | 0,65464997 | PHACTR2 antisense RNA 1 |
| ENSG00000108551 | RASD1 | -0,442221048 | 0,736000653 | 0,13044 | 0,45477153 | ras related dexamethasone induced 1 |
| ENSG00000283897 | AC011416,3 | -0,442258899 | 0,735981344 | 0,46672 | 0,70676695 | novel transcript |
| ENSG00000223039 | RN7SKP268 | -0,442477132 | 0,735870022 | 0,56135 | 0,77136826 | RN7SK pseudogene 268 |
| ENSG00000282788 | AC243791,4 | -0,442556303 | 0,73582964 | 0,11073 | 0,42337179 | novel transcript |
| ENSG00000099365 | STX1B | -0,442575992 | 0,735819599 | 0,09606 | 0,39928497 | syntaxin 1B |
| ENSG00000156475 | PPP2R2B | -0,442859309 | 0,735675112 | 0,25689 | 0,56139699 | protein phosphatase 2 regulatory subunit Bbeta |
| ENSG00000161180 | CCDC116 | -0,44291319 | 0,735647637 | 0,31159 | 0,59905757 | coiled-coil domain containing 116 |
| ENSG00000173285 | OR10K1 | -0,442947778 | 0,735630001 | 0,15619 | 0,4840074 | olfactory receptor family 10 subfamily K member 1 |
| ENSG00000246889 | AP000487,1 | -0,443184041 | 0,73550954 | 0,20222 | 0,5224664 | novel transcript |
| ENSG00000288166 | AP000487,3 | -0,443184041 | 0,73550954 | 0,20222 | 0,5224664 | novel transcript |
| ENSG00000130032 | PRRG3 | -0,443202737 | 0,735500008 | 0,1644 | 0,49065086 | proline rich and Gla domain 3 |
| ENSG00000171148 | TADA3 | -0,443248096 | 0,735476884 | 0,10774 | 0,41834811 | transcriptional adaptor 3 |
| ENSG00000213406 | ANXA2P1 | -0,443286445 | 0,735457335 | 0,44255 | 0,69016964 | annexin A2 pseudogene 1 |
| ENSG00000232542 | DPYD-IT1 | -0,443310548 | 0,735445047 | 0,58046 | 0,78359603 | DPYD intronic transcript 1 |
| ENSG00000256802 | AC022613,1 | -0,443444451 | 0,735376791 | 0,47383 | 0,71294959 | novel transcript, antisense to TJP1 |
| ENSG00000278616 | BEND3P3 | -0,443446537 | 0,735375727 | 0,16673 | 0,49257009 | BEN domain containing 3 pseudogene 3 |

| | | | | | | |
|---|---|---|---|---|---|---|
| ENSG00000204843 | DCTN1 | -0,443610879 | 0,735291963 | 0,10788 | 0,41847368 | dynactin subunit 1 |
| ENSG00000258947 | TUBB3 | -0,443737823 | 0,735227267 | 0,07423 | 0,35761188 | tubulin beta 3 class III |
| ENSG00000138600 | SPPL2A | -0,443792504 | 0,735199401 | 0,14095 | 0,46791293 | signal peptide peptidase like 2A |
| ENSG00000239922 | AC092958,1 | -0,443838779 | 0,735175819 | 0,24581 | 0,55366148 | novel transcript |
| ENSG00000287480 | AL158832,2 | -0,443859125 | 0,735165452 | 0,42247 | 0,67663831 | novel transcript |
| ENSG00000257497 | AC121761,1 | -0,444042735 | 0,735071894 | 0,27156 | 0,57288834 | novel transcript, antisense to GLIPR1 |
| ENSG00000172367 | PDZD3 | -0,444052255 | 0,735067043 | 0,64737 | 0,8235478 | PDZ domain containing 3 |
| ENSG00000157823 | AP3S2 | -0,444179406 | 0,735002261 | 0,16931 | 0,49548445 | adaptor related protein complex 3 subunit sigma 2 |
| ENSG00000130518 | IQCN | -0,444349311 | 0,734915706 | 0,09802 | 0,4033099 | IQ motif containing N |
| ENSG00000227627 | AL080276,2 | -0,444495206 | 0,73484139 | 0,16771 | 0,49395055 | novel transcript |
| ENSG00000133138 | TBC1D8B | -0,44452701 | 0,734825191 | 0,15565 | 0,48345635 | TBC1 domain family member 8B |
| ENSG00000110274 | CEP164 | -0,444639605 | 0,734767844 | 0,11 | 0,42231571 | centrosomal protein 164 |
| ENSG00000263766 | AC025682,1 | -0,44470216 | 0,734735985 | 0,2689 | 0,57085484 | novel transcript, antisense to NPEPPS |
| ENSG00000197847 | SLC22A20P | -0,444729339 | 0,734722143 | 0,19886 | 0,52005179 | solute carrier family 22 member 20, pseudogene |
| ENSG00000219433 | BTBD10P2 | -0,444788036 | 0,734692252 | 0,62087 | 0,80776776 | BTB domain containing 10 pseudogene 2 |
| ENSG00000167595 | PROSER3 | -0,445116813 | 0,734524841 | 0,18509 | 0,50871188 | proline and serine rich 3 |
| ENSG00000089022 | MAPKAPK5 | -0,445213591 | 0,734475569 | 0,18678 | 0,51023628 | MAPK activated protein kinase 5 |
| ENSG00000112562 | SMOC2 | -0,445376914 | 0,734392427 | 0,56194 | 0,77147224 | SPARC related modular calcium binding 2 |
| ENSG00000229659 | RPL26P6 | -0,445430948 | 0,734364921 | 0,52647 | 0,7497927 | ribosomal protein L26 pseudogene 6 |
| ENSG00000170379 | TCAF2 | -0,445568019 | 0,734295152 | 0,2247 | 0,54036191 | TRPM8 channel associated factor 2 |
| ENSG00000248588 | AC008517,1 | -0,44576721 | 0,734193776 | 0,3501 | 0,62570584 | novel transcript |
| ENSG00000185838 | GNB1L | -0,445863143 | 0,734144957 | 0,10404 | 0,41241152 | G protein subunit beta 1 like |
| ENSG00000227080 | AC012441,1 | -0,44587493 | 0,734138959 | 0,41792 | 0,6742866 | novel pseudogene |
| ENSG00000235954 | TTC28-AS1 | -0,445885424 | 0,734133619 | 0,1079 | 0,41847368 | TTC28 antisense RNA 1 |
| ENSG00000276136 | AC016957,2 | -0,445922803 | 0,734114599 | 0,33868 | 0,61789765 | novel transcript |
| ENSG00000139572 | GPR84 | -0,44612365 | 0,734012405 | 0,40197 | 0,66341002 | G protein-coupled receptor 84 |
| ENSG00000115520 | COQ10B | -0,446135248 | 0,734006504 | 0,125 | 0,44663415 | coenzyme Q10B |
| ENSG00000161298 | ZNF382 | -0,446238458 | 0,733953995 | 0,12867 | 0,45191041 | zinc finger protein 382 |
| ENSG00000258344 | AC078778,1 | -0,446364697 | 0,733889775 | 0,38019 | 0,64789138 | novel transcript, sense overlapping to COPZ1 & HNRNPA1 |
| ENSG00000286918 | AL035411,3 | -0,446418343 | 0,733862487 | 0,09088 | 0,39016581 | novel transcript, antisense to OMA1 |
| ENSG00000214700 | C12orf71 | -0,446440287 | 0,733851324 | 0,66597 | 0,83383368 | chromosome 12 open reading frame 71 |
| ENSG00000119866 | BCL11A | -0,446480414 | 0,733830913 | 0,2974 | 0,59033838 | BAF chromatin remodeling complex subunit BCL11A |
| ENSG00000227333 | EHMT2 | -0,44650506 | 0,733818377 | 0,07898 | 0,3669053 | euchromatic histone lysine methyltransferase 2 |

| | | | | | | |
|---|---|---|---|---|---|---|
| ENSG00000251432 | LINC02615 | -0,446511886 | 0,733814905 | 0,27352 | 0,57362145 | long intergenic non-protein coding RNA 2615 |
| ENSG00000271200 | AC099791,2 | -0,44664079 | 0,733749342 | 0,49541 | 0,7276636 | novel transcript |
| ENSG00000229422 | AL512625,2 | -0,446954716 | 0,733589698 | 0,26959 | 0,57135887 | novel transcript |
| ENSG00000230937 | MIR205HG | -0,446964762 | 0,73358459 | 0,79132 | 0,90300322 | MIR205 host gene |
| ENSG00000085831 | TTC39A | -0,44700931 | 0,733561938 | 0,35254 | 0,62747731 | tetratricopeptide repeat domain 39A |
| ENSG00000263961 | RHEX | -0,447077518 | 0,733527258 | 0,72869 | 0,87118284 | regulator of hemoglobinization and erythroid cell expansion |
| ENSG00000182586 | LINC00334 | -0,447094581 | 0,733518582 | 0,65768 | 0,82881843 | long intergenic non-protein coding RNA 334 |
| ENSG00000163626 | COX18 | -0,447355708 | 0,733385828 | 0,19007 | 0,51331106 | cytochrome c oxidase assembly factor COX18 |
| ENSG00000242163 | AL121769,1 | -0,447469796 | 0,733327834 | 0,6264 | 0,81088056 | ribosomal protein S24 (RPS24) pseudogene |
| ENSG00000230321 | MTCO2P27 | -0,447500058 | 0,733312452 | 0,5276 | 0,75037219 | MT-CO2 pseudogene 27 |
| ENSG00000136052 | SLC41A2 | -0,44758788 | 0,733267814 | 0,1885 | 0,51186769 | solute carrier family 41 member 2 |
| ENSG00000228477 | AL663070,1 | -0,447646937 | 0,733237798 | 0,26305 | 0,56588348 | ribosomal protein S2 (RPS2) pseudogene |
| ENSG00000250816 | DCAF13P2 | -0,447735352 | 0,733192863 | 0,64279 | 0,82094129 | DDB1 and CUL4 associated factor 13 pseudogene 2 |
| ENSG00000275576 | AL049539,1 | -0,447852035 | 0,733133566 | 0,566 | 0,77413633 | novel transcript, sense intronic to TM9SF4 |
| ENSG00000242753 | AL445430,2 | -0,447861105 | 0,733128957 | 0,33731 | 0,61706507 | novel transcript |
| ENSG00000225351 | ZNF311 | -0,447939998 | 0,733088867 | 0,54526 | 0,7609742 | zinc finger protein 311 |
| ENSG00000235178 | ZNF311 | -0,447939998 | 0,733088867 | 0,54526 | 0,7609742 | zinc finger protein 311 |
| ENSG00000235589 | ZNF311 | -0,447939998 | 0,733088867 | 0,54526 | 0,7609742 | zinc finger protein 311 |
| ENSG00000110013 | SIAE | -0,448056948 | 0,733029443 | 0,10817 | 0,41884147 | sialic acid acetylesterase |
| ENSG00000276167 | TAS2R50 | -0,448064918 | 0,733025394 | 0,46136 | 0,70278869 | taste 2 receptor member 50 |
| ENSG00000273431 | TAS2R50 | -0,448064918 | 0,733025394 | 0,46136 | 0,70278869 | taste 2 receptor member 50 |
| ENSG00000212126 | TAS2R50 | -0,448064918 | 0,733025394 | 0,46136 | 0,70278869 | taste 2 receptor member 50 |
| ENSG00000284925 | GCSAM | -0,448214557 | 0,732949367 | 0,38213 | 0,64916957 | germinal center associated signaling and motility |
| ENSG00000174500 | GCSAM | -0,448214557 | 0,732949367 | 0,38213 | 0,64916957 | germinal center associated signaling and motility |
| ENSG00000146966 | DENND2A | -0,448223567 | 0,732944789 | 0,18388 | 0,50755749 | DENN domain containing 2A |
| ENSG00000285809 | AC093295,1 | -0,448227046 | 0,732943022 | 0,5807 | 0,78380423 | Novel transcript |
| ENSG00000235363 | SNRPGP10 | -0,448330655 | 0,732890386 | 0,5687 | 0,77620121 | small nuclear ribonucleoprotein polypeptide G pseudogene 10 |
| ENSG00000143630 | HCN3 | -0,448445304 | 0,732832147 | 0,04509 | 0,2872181 | hyperpolarization activated cyclic nucleotide gated potassium channel 3 |
| ENSG00000272769 | AC097532,2 | -0,448626065 | 0,732740333 | 0,19826 | 0,51980353 | novel transcript |
| ENSG00000196422 | PPP1R26 | -0,448645792 | 0,732730314 | 0,03587 | 0,25662951 | protein phosphatase 1 regulatory subunit 26 |

| | | | | | | |
|---|---|---|---|---|---|---|
| ENSG00000283025 | AC027045,3 | -0,44881929 | 0,732642201 | 0,59715 | 0,79434997 | novel transcript, antisense to novel protein |
| ENSG00000102241 | HTATSF1 | -0,449074505 | 0,732512607 | 0,10741 | 0,41771747 | HIV-1 Tat specific factor 1 |
| ENSG00000213221 | DNLZ | -0,449107349 | 0,732495931 | 0,21337 | 0,53259344 | DNL-type zinc finger |
| ENSG00000281836 | AC136352,12 | -0,449120328 | 0,732489342 | 0,11388 | 0,42900168 | novel transcript |
| ENSG00000106348 | IMPDH1 | -0,449201093 | 0,732448337 | 0,07975 | 0,36858935 | inosine monophosphate dehydrogenase 1 |
| ENSG00000173540 | GMPPB | -0,449223641 | 0,732436889 | 0,08707 | 0,3821708 | GDP-mannose pyrophosphorylase B |
| ENSG00000255276 | RRM1-AS1 | -0,449236211 | 0,732430508 | 0,68286 | 0,84472218 | RRM1 antisense RNA 1 |
| ENSG00000260882 | AC026458,1 | -0,449272411 | 0,732412129 | 0,6296 | 0,81311163 | novel pseudogene |
| ENSG00000133026 | MYH10 | -0,449501426 | 0,732295875 | 0,03661 | 0,25940941 | myosin heavy chain 10 |
| ENSG00000286658 | AC005153,1 | -0,449504915 | 0,732294104 | 0,18699 | 0,5105217 | novel transcript, antisense to GRB10 |
| ENSG00000108602 | ALDH3A1 | -0,449525902 | 0,732283451 | 0,53294 | 0,75333268 | aldehyde dehydrogenase 3 family member A1 |
| ENSG00000169955 | ZNF747 | -0,449526942 | 0,732282923 | 0,33719 | 0,61706507 | zinc finger protein 747 |
| ENSG00000213204 | AL049697,1 | -0,449550769 | 0,73227083 | 0,34108 | 0,6192144 | novel transcript, C6orf165-SLC35A1 readthrough |
| ENSG00000285529 | AC016816,1 | -0,449581287 | 0,732255339 | 0,4287 | 0,68136875 | novel transcript |
| ENSG00000203879 | GDI1 | -0,449629044 | 0,7322311 | 0,14389 | 0,47144689 | GDP dissociation inhibitor 1 |
| ENSG00000270300 | PHACTR2P1 | -0,449845975 | 0,732121006 | 0,29277 | 0,58724158 | phosphatase and actin regulator 2 pseudogene 1 |
| ENSG00000267277 | AC024575,1 | -0,449877508 | 0,732105005 | 0,58944 | 0,78887672 | novel transcript, antisense to SWSAP1 |
| ENSG00000228857 | AC104653,1 | -0,450049014 | 0,732017978 | 0,71464 | 0,86265094 | novel transcript |
| ENSG00000280907 | ZNF197-AS1 | -0,450150686 | 0,731966392 | 0,47763 | 0,71545897 | ZNF197 antisense RNA 1 |
| ENSG00000233509 | ZNF197-AS1 | -0,450150686 | 0,731966392 | 0,47763 | 0,71545897 | ZNF197 antisense RNA 1 |
| ENSG00000178773 | CPNE7 | -0,45024038 | 0,731920886 | 0,65892 | 0,82963066 | copine 7 |
| ENSG00000197852 | INKA2 | -0,450346931 | 0,731866831 | 0,10889 | 0,42014849 | inka box actin regulator 2 |
| ENSG00000286577 | AC117528,1 | -0,450426757 | 0,731826338 | 0,67696 | 0,84109786 | novel transcript, antisense to KIAA0825 |
| ENSG00000179588 | ZFPM1 | -0,450470581 | 0,731804108 | 0,2946 | 0,58857672 | zinc finger protein, FOG family member 1 |
| ENSG00000130779 | CLIP1 | -0,450548994 | 0,731764334 | 0,07354 | 0,35672172 | CAP-Gly domain containing linker protein 1 |
| ENSG00000256085 | AC130404,1 | -0,450556435 | 0,73176056 | 0,55867 | 0,77025265 | novel transcript |
| ENSG00000182674 | KCNB2 | -0,450591639 | 0,731742704 | 0,08777 | 0,38337966 | potassium voltage-gated channel subfamily B member 2 |
| ENSG00000135709 | KIAA0513 | -0,450733871 | 0,731670567 | 0,22048 | 0,53750545 | KIAA0513 |
| ENSG00000259677 | AC027176,2 | -0,450792902 | 0,731640629 | 0,40803 | 0,66700475 | Arginine/serine-rich coiled-coil protein (RSRC2) pseudogene |
| ENSG00000127329 | PTPRB | -0,450844802 | 0,73161431 | 0,09969 | 0,40615323 | protein tyrosine phosphatase receptor type B |
| ENSG00000177807 | KCNJ10 | -0,450895577 | 0,731588561 | 0,34227 | 0,62000209 | potassium inwardly rectifying channel subfamily J member 10 |

| | | | | | | |
|---|---|---|---|---|---|---|
| ENSG00000225964 | NRIR | -0,450987812 | 0,73154179 | 0,33732 | 0,61706507 | negative regulator of interferon response |
| ENSG00000105143 | SLC1A6 | -0,451018967 | 0,731525993 | 0,26328 | 0,56608528 | solute carrier family 1 member 6 |
| ENSG00000237883 | DGUOK-AS1 | -0,451103854 | 0,731482952 | 0,18821 | 0,51132417 | DGUOK antisense RNA 1 |
| ENSG00000235071 | AC073909,1 | -0,451181254 | 0,731443709 | 0,60627 | 0,79946546 | heat shock protein pseudogene |
| ENSG00000236816 | ANKRD20A7P | -0,451348456 | 0,731358943 | 0,32963 | 0,61066538 | ankyrin repeat domain 20 family member A7, pseudogene |
| ENSG00000249437 | NAIP | -0,451355811 | 0,731355214 | 0,0721 | 0,35433046 | NLR family apoptosis inhibitory protein |
| ENSG00000213853 | EMP2 | -0,451539834 | 0,731261932 | 0,25956 | 0,56330128 | epithelial membrane protein 2 |
| ENSG00000189420 | ZFP92 | -0,451665682 | 0,731198146 | 0,68342 | 0,84495012 | ZFP92 zinc finger protein |
| ENSG00000280993 | MUC20-OT1 | -0,451687818 | 0,731186927 | 0,25184 | 0,55773568 | MUC20 overlapping transcript |
| ENSG00000161671 | EMC10 | -0,451887817 | 0,731085571 | 0,16966 | 0,49589723 | ER membrane protein complex subunit 10 |
| ENSG00000258034 | AC012157,2 | -0,45202029 | 0,731018443 | 0,55556 | 0,76790322 | novel transcript, antisense to MED13L |
| ENSG00000229728 | AL136531,1 | -0,452066075 | 0,730995244 | 0,33282 | 0,61282727 | novel transcript |
| ENSG00000245112 | SMARCA5-AS1 | -0,452152109 | 0,730951653 | 0,1416 | 0,46809977 | SMARCA5 antisense RNA 1 |
| ENSG00000180543 | TSPYL5 | -0,452434539 | 0,730808572 | 0,19373 | 0,51547251 | TSPY like 5 |
| ENSG00000212283 | SNORD89 | -0,452516247 | 0,730767183 | 0,21192 | 0,53076389 | small nucleolar RNA, C/D box 89 |
| ENSG00000187135 | VSTM2B | -0,452574128 | 0,730737865 | 0,51091 | 0,73829835 | V-set and transmembrane domain containing 2B |
| ENSG00000160447 | PKN3 | -0,452739262 | 0,730654228 | 0,16021 | 0,48737351 | protein kinase N3 |
| ENSG00000260672 | AC100827,4 | -0,452869962 | 0,730588038 | 0,4141 | 0,67098035 | novel transcript |
| ENSG00000168386 | FILIP1L | -0,452890044 | 0,730577869 | 0,22977 | 0,54324957 | filamin A interacting protein 1 like |
| ENSG00000177096 | PHETA2 | -0,453104114 | 0,730469472 | 0,13399 | 0,4601411 | PH domain containing endocytic trafficking adaptor 2 |
| ENSG00000172771 | EFCAB12 | -0,453132354 | 0,730455173 | 0,302 | 0,59294633 | EF-hand calcium binding domain 12 |
| ENSG00000267640 | AC016582,3 | -0,453283401 | 0,730378701 | 0,11912 | 0,43696075 | novel zinc finger protein pseudogene |
| ENSG00000187003 | ACTL7A | -0,453357594 | 0,730341141 | 0,72195 | 0,86726498 | actin like 7A |
| ENSG00000143869 | GDF7 | -0,453556268 | 0,730240572 | 0,09749 | 0,40230362 | growth differentiation factor 7 |
| ENSG00000226429 | AL590399,2 | -0,453564177 | 0,730236569 | 0,49974 | 0,73058737 | novel pseudogene |
| ENSG00000253505 | AC103957,1 | -0,453566828 | 0,730235227 | 0,12322 | 0,44350459 | novel transcript |
| ENSG00000166377 | ATP9B | -0,453571748 | 0,730232737 | 0,13105 | 0,45576197 | ATPase phospholipid transporting 9B (putative) |
| ENSG00000008226 | DLEC1 | -0,453612836 | 0,73021194 | 0,07955 | 0,3682969 | DLEC1 cilia and flagella associated protein |
| ENSG00000230513 | THAP7-AS1 | -0,453712302 | 0,730161597 | 0,21141 | 0,53032236 | THAP7 antisense RNA 1 |
| ENSG00000264204 | AGAP7P | -0,453758436 | 0,730138249 | 0,26671 | 0,56923396 | ArfGAP with GTPase domain, ankyrin repeat and PH domain 7, pseudogene |
| ENSG00000109944 | JHY | -0,45389781 | 0,730067716 | 0,22747 | 0,54168591 | junctional cadherin complex regulator |
| ENSG00000248489 | LINC02062 | -0,453916846 | 0,730058083 | 0,22268 | 0,53876885 | long intergenic non-protein coding RNA 2062 |

| | | | | | | |
|---|---|---|---|---|---|---|
| ENSG00000235351 | AC114730,3 | -0,453942455 | 0,730045124 | 0,63651 | 0,81706957 | novel transcript |
| ENSG00000173653 | RCE1 | -0,453956642 | 0,730037945 | 0,18116 | 0,50470865 | Ras converting CAAX endopeptidase 1 |
| ENSG00000224731 | AC016716,1 | -0,454065622 | 0,729982801 | 0,65338 | 0,82636171 | novel transcript |
| ENSG00000231534 | FBXO36-IT1 | -0,454068942 | 0,729981121 | 0,58076 | 0,78384297 | FBXO36 intronic transcript 1 |
| ENSG00000176728 | TTTY14 | -0,454072028 | 0,72997956 | 0,32611 | 0,6085098 | testis-specific transcript, Y-linked 14 |
| ENSG00000153404 | PLEKHG4B | -0,454147194 | 0,729941528 | 0,06826 | 0,34544445 | pleckstrin homology and RhoGEF domain containing G4B |
| ENSG00000148341 | SH3GLB2 | -0,454182618 | 0,729923605 | 0,07291 | 0,35631792 | SH3 domain containing GRB2 like, endophilin B2 |
| ENSG00000282183 | GOLGA2P10 | -0,45419015 | 0,729919794 | 0,21536 | 0,53409502 | GOLGA2 pseudogene 10 |
| ENSG00000224402 | OR6D1P | -0,454280687 | 0,729873989 | 0,12123 | 0,44047594 | olfactory receptor family 6 subfamily D member 1 pseudogene |
| ENSG00000162645 | GBP2 | -0,454280891 | 0,729873886 | 0,51938 | 0,74395272 | guanylate binding protein 2 |
| ENSG00000109072 | VTN | -0,454374277 | 0,729826643 | 0,43648 | 0,68678001 | vitronectin |
| ENSG00000266835 | GAPLINC | -0,454415322 | 0,729805879 | 0,56194 | 0,77147224 | gastric adenocarcinoma associated, positive CD44 regulator, long intergenic non-coding RNA |
| ENSG00000105701 | FKBP8 | -0,454422627 | 0,729802184 | 0,19594 | 0,51680811 | FKBP prolyl isomerase 8 |
| ENSG00000187147 | RNF220 | -0,454471803 | 0,729777308 | 0,02782 | 0,22494202 | ring finger protein 220 |
| ENSG00000226121 | AHCTF1P1 | -0,454553036 | 0,729736218 | 0,3694 | 0,63917123 | AT-hook containing transcription factor 1 pseudogene 1 |
| ENSG00000205209 | SCGB2B2 | -0,454716049 | 0,729653768 | 0,61806 | 0,80622686 | secretoglobin family 2B member 2 |
| ENSG00000187595 | ZNF385C | -0,454815526 | 0,729603459 | 0,21033 | 0,5290538 | zinc finger protein 385C |
| ENSG00000124557 | BTN1A1 | -0,454859023 | 0,729581462 | 0,5858 | 0,78644815 | butyrophilin subfamily 1 member A1 |
| ENSG00000204314 | PRRT1 | -0,454860718 | 0,729580605 | 0,52167 | 0,74600315 | proline rich transmembrane protein 1 |
| ENSG00000236973 | GAPDHP51 | -0,454891192 | 0,729565194 | 0,60837 | 0,80115227 | glyceraldehyde 3 phosphate dehydrogenase pseudogene 51 |
| ENSG00000260647 | AC127537,1 | -0,454894758 | 0,72956339 | 0,46807 | 0,70769773 | novel transcript, antisense to TOP3A |
| ENSG00000284449 | AC127537,5 | -0,454894758 | 0,72956339 | 0,46807 | 0,70769773 | novel transcript, antisense to TOP3A |
| ENSG00000174032 | SLC25A30 | -0,454895148 | 0,729563193 | 0,28612 | 0,58258764 | solute carrier family 25 member 30 |
| ENSG00000271797 | AC008494,3 | -0,454895334 | 0,729563099 | 0,41571 | 0,67261802 | novel transcript, antisense to PGGT1B |
| ENSG00000234616 | JRK | -0,454955328 | 0,729532761 | 0,1066 | 0,416363 | Jrk helix-turn-helix protein |
| ENSG00000273320 | AC007032,1 | -0,454972118 | 0,729524271 | 0,78631 | 0,90080813 | novel transcript, antisense to NAMPT |
| ENSG00000096150 | RPS18 | -0,455029939 | 0,729495033 | 0,47928 | 0,71685748 | ribosomal protein S18 |
| ENSG00000019102 | VSIG2 | -0,455040846 | 0,729489518 | 0,45466 | 0,69879733 | V-set and immunoglobulin domain containing 2 |
| ENSG00000285701 | AL357140,4 | -0,455127265 | 0,729445822 | 0,44003 | 0,6885781 | novel transcript, antisense to CTNNBIP1 |
| ENSG00000101940 | WDR13 | -0,455159585 | 0,729429481 | 0,10984 | 0,42203441 | WD repeat domain 13 |

| | | | | | | |
|---|---|---|---|---|---|---|
| ENSG00000142694 | EVA1B | -0,455452398 | 0,72928145 | 0,23036 | 0,54355692 | eva-1 homolog B |
| ENSG00000166822 | TMEM170A | -0,455472294 | 0,729271392 | 0,17643 | 0,50064995 | transmembrane protein 170A |
| ENSG00000065609 | SNAP91 | -0,455493254 | 0,729260797 | 0,26855 | 0,57061705 | synaptosome associated protein 91 |
| ENSG00000119669 | IRF2BPL | -0,455575623 | 0,729219162 | 0,25611 | 0,56066081 | interferon regulatory factor 2 binding protein like |
| ENSG00000262503 | AC027763,1 | -0,455866246 | 0,729072279 | 0,67143 | 0,83732104 | Spi-C transcription factor (Spi-1/PU,1 related) (SPIC) pseudogene |
| ENSG00000156374 | PCGF6 | -0,455962411 | 0,729023684 | 0,22739 | 0,54168591 | polycomb group ring finger 6 |
| ENSG00000233579 | KRT8P15 | -0,456064789 | 0,728971952 | 0,30134 | 0,59258306 | keratin 8 pseudogene 15 |
| ENSG00000224029 | AL450344,1 | -0,456079709 | 0,728964413 | 0,53479 | 0,75419058 | novel transcript |
| ENSG00000103495 | MAZ | -0,456189834 | 0,728908771 | 0,06306 | 0,33249564 | MYC associated zinc finger protein |
| ENSG00000230221 | AL157384,1 | -0,456316341 | 0,728844857 | 0,22738 | 0,54168591 | ATM interactor (ATMIN) pseudogene |
| ENSG00000227372 | TP73-AS1 | -0,456544041 | 0,728729833 | 0,12288 | 0,44269063 | TP73 antisense RNA 1 |
| ENSG00000173991 | TCAP | -0,457129889 | 0,728433971 | 0,58325 | 0,78512624 | titin-cap |
| ENSG00000105520 | PLPPR2 | -0,457139558 | 0,728429089 | 0,07346 | 0,35662549 | phospholipid phosphatase related 2 |
| ENSG00000229353 | TNXB | -0,457188903 | 0,728404175 | 0,63125 | 0,81395758 | tenascin XB |
| ENSG00000160113 | NR2F6 | -0,457219831 | 0,72838856 | 0,1982 | 0,51980353 | nuclear receptor subfamily 2 group F member 6 |
| ENSG00000181045 | SLC26A11 | -0,457220072 | 0,728388438 | 0,19517 | 0,51599597 | solute carrier family 26 member 11 |
| ENSG00000160957 | RECQL4 | -0,457318479 | 0,728338757 | 0,1498 | 0,47782859 | RecQ like helicase 4 |
| ENSG00000233044 | AL049820,1 | -0,457334117 | 0,728330862 | 0,50223 | 0,73186474 | novel transcript |
| ENSG00000203727 | SAMD5 | -0,457401765 | 0,728296711 | 0,17868 | 0,50323303 | sterile alpha motif domain containing 5 |
| ENSG00000212195 | U3 | -0,457459094 | 0,728267771 | 0,3123 | 0,59927258 | Small nucleolar RNA U3 |
| ENSG00000251079 | BMS1P2 | -0,457530819 | 0,728231565 | 0,13945 | 0,46666406 | BMS1 pseudogene 2 |
| ENSG00000270605 | AL353622,1 | -0,457559597 | 0,728217039 | 0,84064 | 0,92739891 | novel transcript |
| ENSG00000256663 | AC112777,1 | -0,457586759 | 0,728203329 | 0,13991 | 0,46689302 | ubiquitin-like with PHD and ring finger domains 1 (UHRF1) pseudogene |
| ENSG00000205758 | CRYZL1 | -0,457587296 | 0,728203058 | 0,08071 | 0,37077071 | crystallin zeta like 1 |
| ENSG00000205464 | ATP6AP1L | -0,457649726 | 0,728171547 | 0,38434 | 0,65052153 | ATPase H+ transporting accessory protein 1 like |
| ENSG00000267270 | PARD6G-AS1 | -0,457780646 | 0,728105471 | 0,09712 | 0,40152057 | PARD6G antisense RNA 1 |
| ENSG00000136099 | PCDH8 | -0,45794795 | 0,72802104 | 0,59833 | 0,79497317 | protocadherin 8 |
| ENSG00000105255 | FSD1 | -0,458051982 | 0,727968544 | 0,03461 | 0,25152438 | fibronectin type III and SPRY domain containing 1 |
| ENSG00000163975 | MELTF | -0,458179743 | 0,727904081 | 0,34313 | 0,62060539 | melanotransferrin |
| ENSG00000231890 | DARS-AS1 | -0,458268839 | 0,727859129 | 0,20536 | 0,52503195 | DARS antisense RNA 1 |
| ENSG00000254165 | AC090739,1 | -0,458284401 | 0,727851278 | 0,28856 | 0,58455903 | novel transcript |

| | | | | | | |
|---|---|---|---|---|---|---|
| ENSG00000170667 | RASA4B | -0,45830823 | 0,727839256 | 0,15066 | 0,47887343 | RAS p21 protein activator 4B |
| ENSG00000108840 | HDAC5 | -0,458406047 | 0,727789909 | 0,16506 | 0,49125639 | histone deacetylase 5 |
| ENSG00000237818 | RPS3AP29 | -0,458479526 | 0,727752842 | 0,3865 | 0,65193563 | RPS3A pseudogene 29 |
| ENSG00000226029 | LINC01772 | -0,458485897 | 0,727749628 | 0,21706 | 0,53505085 | long intergenic non-protein coding RNA 1772 |
| ENSG00000274204 | AL138689,1 | -0,458496407 | 0,727744327 | 0,36101 | 0,63311211 | novel transcript, sense intronic to EFNB2 |
| ENSG00000250838 | AC091133,3 | -0,458562598 | 0,727710939 | 0,2565 | 0,56092974 | novel transcript |
| ENSG00000143850 | PLEKHA6 | -0,458784681 | 0,727598926 | 0,14118 | 0,4679289 | pleckstrin homology domain containing A6 |
| ENSG00000120664 | SPART-AS1 | -0,459144554 | 0,727417453 | 0,40321 | 0,66399303 | SPART antisense RNA 1 |
| ENSG00000184922 | FMNL1 | -0,459156616 | 0,727411371 | 0,41284 | 0,6705041 | formin like 1 |
| ENSG00000233907 | LINC01761 | -0,459162885 | 0,727408211 | 0,15944 | 0,48684894 | long intergenic non-protein coding RNA 1761 |
| ENSG00000146833 | TRIM4 | -0,459451711 | 0,727262598 | 0,20705 | 0,52670822 | tripartite motif containing 4 |
| ENSG00000008283 | CYB561 | -0,459857127 | 0,727058257 | 0,19205 | 0,5145257 | cytochrome b561 |
| ENSG00000160256 | FAM207A | -0,459879259 | 0,727047104 | 0,24821 | 0,55542983 | family with sequence similarity 207 member A |
| ENSG00000182841 | RRP7BP | -0,459957186 | 0,727007833 | 0,1569 | 0,48425071 | ribosomal RNA processing 7 homolog B, pseudogene |
| ENSG00000276615 | KEL | -0,460059319 | 0,726956368 | 0,34784 | 0,62428208 | Kell metallo-endopeptidase (Kell blood group) |
| ENSG00000230911 | PPIHP1 | -0,460105368 | 0,726933165 | 0,33225 | 0,61221107 | peptidylprolyl isomerase H pseudogene 1 |
| ENSG00000287081 | AC104656,1 | -0,460168459 | 0,726901376 | 0,5365 | 0,75531624 | novel transcript, antisense to LDB2 |
| ENSG00000237989 | LINC01679 | -0,460277706 | 0,726846334 | 0,60028 | 0,79579686 | long intergenic non-protein coding RNA 1679 |
| ENSG00000270429 | KNOP1P2 | -0,460458718 | 0,726755143 | 0,373 | 0,64180957 | lysine rich nucleolar protein 1 pseudogene 2 |
| ENSG00000197587 | DMBX1 | -0,46051989 | 0,726724329 | 0,68953 | 0,84865812 | diencephalon/mesencephalon homeobox 1 |
| ENSG00000143126 | CELSR2 | -0,460520536 | 0,726724003 | 0,02426 | 0,20811806 | cadherin EGF LAG seven-pass G-type receptor 2 |
| ENSG00000261451 | AC104964,2 | -0,46053519 | 0,726716622 | 0,20367 | 0,52347929 | novel transcript, overlapping to MSRA |
| ENSG00000233705 | SLC26A4-AS1 | -0,460648949 | 0,726659321 | 0,16193 | 0,48914394 | SLC26A4 antisense RNA 1 |
| ENSG00000225151 | GOLGA2P7 | -0,46071281 | 0,726627156 | 0,07398 | 0,35740873 | GOLGA2 pseudogene 7 |
| ENSG00000259407 | AC021739,2 | -0,460766107 | 0,726600313 | 0,2181 | 0,53569444 | novel transcript, antisense to AKAP13 |
| ENSG00000164512 | ANKRD55 | -0,460784345 | 0,726591128 | 0,47296 | 0,71225062 | ankyrin repeat domain 55 |
| ENSG00000039650 | PNKP | -0,460902915 | 0,726531415 | 0,19464 | 0,51562158 | polynucleotide kinase 3'-phosphatase |
| ENSG00000156113 | KCNMA1 | -0,461166918 | 0,726398476 | 0,18288 | 0,50636151 | potassium calcium-activated channel subfamily M alpha 1 |
| ENSG00000206144 | MAPK6P1 | -0,461311307 | 0,72632578 | 0,18166 | 0,50497494 | mitogen-activated protein kinase 6 pseudogene 1 |
| ENSG00000267498 | AC007786,1 | -0,461351579 | 0,726305506 | 0,51648 | 0,74198903 | uncharacterized LOC105372352 |
| ENSG00000210825 | SNORA40 | -0,461378938 | 0,726291732 | 0,64021 | 0,81935089 | small nucleolar RNA, H/ACA box 40 |
| ENSG00000136206 | SPDYE1 | -0,46140946 | 0,726276367 | 0,22121 | 0,53805234 | speedy/RINGO cell cycle regulator family member E1 |
| ENSG00000136842 | TMOD1 | -0,46142523 | 0,726268428 | 0,55531 | 0,76788267 | tropomodulin 1 |

| | | | | | | |
|---|---|---|---|---|---|---|
| ENSG00000239332 | LINC01119 | -0,461451463 | 0,726255222 | 0,52153 | 0,74585386 | long intergenic non-protein coding RNA 1119 |
| ENSG00000254454 | RCC2P6 | -0,46161748 | 0,726171653 | 0,23863 | 0,54890467 | regulator of chromosome condensation 2 pseudogene 6 |
| ENSG00000228818 | AL359918,1 | -0,46164262 | 0,726158999 | 0,33092 | 0,6112353 | proteasome 26S non-ATPase subunit 2 (PSMD2) pseudogene |
| ENSG00000238140 | AC104170,1 | -0,461696266 | 0,726131998 | 0,5553 | 0,76788267 | novel transcript |
| ENSG00000127951 | FGL2 | -0,46172698 | 0,726116539 | 0,16803 | 0,49417662 | fibrinogen like 2 |
| ENSG00000225057 | AC012485,1 | -0,461894795 | 0,726032082 | 0,37614 | 0,64448883 | novel transcript |
| ENSG00000226658 | RPL23AP30 | -0,461972341 | 0,725993059 | 0,62269 | 0,80866078 | ribosomal protein L23a pseudogene 30 |
| ENSG00000232234 | AL355499,1 | -0,461988752 | 0,7259848 | 0,73762 | 0,87540065 | novel transcript |
| ENSG00000107185 | RGP1 | -0,462055075 | 0,725951426 | 0,05985 | 0,32535617 | RGP1 homolog, RAB6A GEF complex partner 1 |
| ENSG00000260918 | AC107398,3 | -0,462082922 | 0,725937414 | 0,50048 | 0,73086101 | novel transcript, overlapping GABRB1 |
| ENSG00000186897 | C1QL4 | -0,462092327 | 0,725932681 | 0,45224 | 0,69698561 | complement C1q like 4 |
| ENSG00000137285 | TUBB2B | -0,462168395 | 0,725894407 | 0,07366 | 0,35701531 | tubulin beta 2B class IIb |
| ENSG00000264538 | SUZ12P1 | -0,462362135 | 0,725796933 | 0,16031 | 0,48753847 | SUZ12 pseudogene 1 |
| ENSG00000262788 | UTP4 | -0,462452309 | 0,725751569 | 0,4993 | 0,73036634 | UTP4 small subunit processome component |
| ENSG00000125895 | TMEM74B | -0,462495182 | 0,725730002 | 0,33831 | 0,61774255 | transmembrane protein 74B |
| ENSG00000249731 | AC126768,2 | -0,462496616 | 0,725729281 | 0,1318 | 0,45708114 | novel transcript |
| ENSG00000233230 | AC079807,1 | -0,46267678 | 0,725638657 | 0,43654 | 0,68681257 | novel transcript |
| ENSG00000240579 | AC006529,1 | -0,462769387 | 0,72559208 | 0,52016 | 0,74461374 | pseudogene similar to part of CD2 (cytoplasmic tail) binding protein 2 (CD2BP2) |
| ENSG00000285637 | AL161449,2 | -0,462786764 | 0,72558334 | 0,52753 | 0,75037219 | Novel transcript |
| ENSG00000260230 | FRRS1L | -0,462839829 | 0,725556652 | 0,27817 | 0,57645922 | ferric chelate reductase 1 like |
| ENSG00000214353 | VAC14-AS1 | -0,46287369 | 0,725539623 | 0,32354 | 0,60631901 | VAC14 antisense RNA 1 |
| ENSG00000239697 | TNFSF12 | -0,462878422 | 0,725537243 | 0,20761 | 0,52713166 | TNF superfamily member 12 |
| ENSG00000281344 | HELLPAR | -0,462897566 | 0,725527616 | 0,13652 | 0,46380898 | HELLP associated long non-coding RNA |
| ENSG00000242622 | AC092910,3 | -0,462952504 | 0,725499988 | 0,1345 | 0,46072811 | novel transcript |
| ENSG00000196123 | KIAA0895L | -0,463065319 | 0,725443258 | 0,02534 | 0,21419157 | KIAA0895 like |
| ENSG00000181894 | ZNF329 | -0,463237142 | 0,725356864 | 0,20251 | 0,52259159 | zinc finger protein 329 |
| ENSG00000160570 | DEDD2 | -0,463245359 | 0,725352733 | 0,20736 | 0,52677688 | death effector domain containing 2 |
| ENSG00000117152 | RGS4 | -0,463377138 | 0,72528648 | 0,34351 | 0,62091871 | regulator of G protein signaling 4 |
| ENSG00000085644 | ZNF213 | -0,463401717 | 0,725274124 | 0,10333 | 0,41114168 | zinc finger protein 213 |
| ENSG00000167508 | MVD | -0,463463482 | 0,725243074 | 0,25945 | 0,56317998 | mevalonate diphosphate decarboxylase |
| ENSG00000229146 | SNX18P4 | -0,463876487 | 0,725035486 | 0,67967 | 0,84263958 | sorting nexin 18 pseudogene 4 |

| | | | | | | |
|---|---|---|---|---|---|---|
| ENSG00000214029 | ZNF891 | -0,463879889 | 0,725033776 | 0,17052 | 0,49641416 | zinc finger protein 891 |
| ENSG00000260570 | AC133550,2 | -0,463899697 | 0,725023822 | 0,19148 | 0,51435195 | novel transcript, antisense to ATXN2L |
| ENSG00000165688 | PMPCA | -0,463918565 | 0,72501434 | 0,20123 | 0,52181085 | peptidase, mitochondrial processing subunit alpha |
| ENSG00000248711 | THUMPD3P1 | -0,464000062 | 0,724973385 | 0,60954 | 0,80179036 | THUMP domain containing 3 pseudogene 1 |
| ENSG00000287281 | AC012101,2 | -0,464105348 | 0,724920479 | 0,26754 | 0,56973003 | novel transcript |
| ENSG00000181741 | FDX1P1 | -0,464129866 | 0,72490816 | 0,67652 | 0,84080793 | ferredoxin 1 pseudogene 1 |
| ENSG00000143195 | ILDR2 | -0,46427376 | 0,724835861 | 0,16453 | 0,49065086 | immunoglobulin like domain containing receptor 2 |
| ENSG00000169075 | BRD7P3 | -0,464277834 | 0,724833815 | 0,40924 | 0,66785537 | bromodomain containing 7 pseudogene 3 |
| ENSG00000183615 | FAM167B | -0,464291085 | 0,724827157 | 0,40496 | 0,66481282 | family with sequence similarity 167 member B |
| ENSG00000157350 | ST3GAL2 | -0,464359774 | 0,724792648 | 0,23666 | 0,54763386 | ST3 beta-galactoside alpha-2,3-sialyltransferase 2 |
| ENSG00000185324 | CDK10 | -0,464393416 | 0,724775747 | 0,16053 | 0,48780354 | cyclin dependent kinase 10 |
| ENSG00000130751 | NPAS1 | -0,464405583 | 0,724769634 | 0,41721 | 0,6738387 | neuronal PAS domain protein 1 |
| ENSG00000272948 | AP001412,1 | -0,464441413 | 0,724751634 | 0,38472 | 0,65054348 | novel transcript, antisense to DSCR3 |
| ENSG00000263004 | AC007114,1 | -0,464580696 | 0,724681668 | 0,28958 | 0,58522301 | novel transcript |
| ENSG00000283045 | AC103703,1 | -0,464665619 | 0,724639011 | 0,43407 | 0,6848964 | novel transcript, sense intronic to GPATCH8 |
| ENSG00000168546 | GFRA2 | -0,464667115 | 0,72463826 | 0,20964 | 0,5286431 | GDNF family receptor alpha 2 |
| ENSG00000147852 | VLDLR | -0,46472026 | 0,724611566 | 0,15616 | 0,4840074 | very low density lipoprotein receptor |
| ENSG00000161888 | SPC24 | -0,465213206 | 0,724364021 | 0,20172 | 0,52214043 | SPC24 component of NDC80 kinetochore complex |
| ENSG00000281245 | STKLD1 | -0,465269509 | 0,724335752 | 0,30727 | 0,59637875 | serine/threonine kinase like domain containing 1 |
| ENSG00000260959 | AC087463,2 | -0,465304854 | 0,724318006 | 0,54866 | 0,76381464 | novel transcript |
| ENSG00000248527 | MTATP6P1 | -0,46532574 | 0,72430752 | 0,1406 | 0,46760662 | MT-ATP6 pseudogene 1 |
| ENSG00000091181 | IL5RA | -0,465416322 | 0,724262045 | 0,5072 | 0,73543706 | interleukin 5 receptor subunit alpha |
| ENSG00000183662 | TAFA1 | -0,465536301 | 0,724201815 | 0,34697 | 0,62369436 | TAFA chemokine like family member 1 |
| ENSG00000100083 | GGA1 | -0,465644857 | 0,724147325 | 0,14028 | 0,46714558 | golgi associated, gamma adaptin ear containing, ARF binding protein 1 |
| ENSG00000226232 | NPIPB14P | -0,465656705 | 0,724141378 | 0,60263 | 0,79710677 | nuclear pore complex interacting protein family member B14, pseudogene |
| ENSG00000115896 | PLCL1 | -0,465668733 | 0,724135341 | 0,19655 | 0,51769283 | phospholipase C like 1 (inactive) |
| ENSG00000136490 | LIMD2 | -0,465717426 | 0,7241109 | 0,08937 | 0,38668036 | LIM domain containing 2 |
| ENSG00000285922 | AC104453,1 | -0,46590071 | 0,724018913 | 0,41398 | 0,67096878 | novel transcript |
| ENSG00000261243 | AC022165,1 | -0,465923523 | 0,724007465 | 0,50133 | 0,73127574 | novel transcript, antisense to KIAA1609 |
| ENSG00000127554 | GFER | -0,465961079 | 0,723988618 | 0,14672 | 0,47461701 | growth factor, augmenter of liver regeneration |
| ENSG00000099326 | MZF1 | -0,465970647 | 0,723983816 | 0,05793 | 0,32123433 | myeloid zinc finger 1 |

| | | | | | | |
|---|---|---|---|---|---|---|
| ENSG00000251247 | ZNF345 | -0,466319628 | 0,723808709 | 0,23801 | 0,54864017 | zinc finger protein 345 |
| ENSG00000251623 | AC139491,5 | -0,466478961 | 0,723728775 | 0,598 | 0,79484486 | centrosomal protein 192 (CEP192), pseudogene |
| ENSG00000110436 | SLC1A2 | -0,466841086 | 0,723547138 | 0,20047 | 0,52110549 | solute carrier family 1 member 2 |
| ENSG00000273727 | U1 | -0,46689599 | 0,723519603 | 0,43841 | 0,68769465 | U1 spliceosomal RNA |
| ENSG00000286040 | AL158058,2 | -0,467035341 | 0,723449721 | 0,62557 | 0,81053676 | novel transcript |
| ENSG00000287663 | AL122020,2 | -0,467052358 | 0,723441187 | 0,35812 | 0,63185282 | novel transcript, antisense to TTC7B |
| ENSG00000251229 | AL645924,2 | -0,467066559 | 0,723434066 | 0,29771 | 0,59036937 | ribosomal protein S3A (RPS3A) pseudogene |
| ENSG00000234032 | VARS2 | -0,467084588 | 0,723425026 | 0,18934 | 0,51255421 | valyl-tRNA synthetase 2, mitochondrial |
| ENSG00000013619 | MAMLD1 | -0,467519676 | 0,723206888 | 0,20534 | 0,52503195 | mastermind like domain containing 1 |
| ENSG00000121905 | HPCA | -0,467661148 | 0,723135973 | 0,21825 | 0,53571161 | hippocalcin |
| ENSG00000213015 | ZNF580 | -0,467667302 | 0,723132889 | 0,14674 | 0,47461781 | zinc finger protein 580 |
| ENSG00000185753 | CXorf38 | -0,467685718 | 0,723123658 | 0,15957 | 0,48693054 | chromosome X open reading frame 38 |
| ENSG00000285677 | AL355388,2 | -0,467723373 | 0,723104785 | 0,41309 | 0,67067821 | Novel transcript, antisense to SSR2 |
| ENSG00000120160 | EQTN | -0,467740317 | 0,723096292 | 0,41311 | 0,67067821 | equatorin |
| ENSG00000275437 | AL121832,3 | -0,467795722 | 0,723068523 | 0,2743 | 0,57420712 | novel transcript, sense intronic to CABLES2 |
| ENSG00000187535 | IFT140 | -0,467856578 | 0,723038023 | 0,06831 | 0,34555414 | intraflagellar transport 140 |
| ENSG00000281332 | LINC00997 | -0,468022787 | 0,722954728 | 0,07111 | 0,35204529 | long intergenic non-protein coding RNA 997 |
| ENSG00000110697 | PITPNM1 | -0,46848389 | 0,7227237 | 0,15308 | 0,48120112 | phosphatidylinositol transfer protein membrane associated 1 |
| ENSG00000215498 | FAM230B | -0,468619832 | 0,722655603 | 0,56463 | 0,77306193 | family with sequence similarity 230 member B |
| ENSG00000226803 | ZNF451-AS1 | -0,468688916 | 0,722620999 | 0,23448 | 0,54621919 | ZNF451 regulatory antisense RNA 1 |
| ENSG00000146909 | NOM1 | -0,468708796 | 0,722611041 | 0,21751 | 0,53541589 | nucleolar protein with MIF4G domain 1 |
| ENSG00000276945 | AC008498,1 | -0,468931349 | 0,722499579 | 0,48866 | 0,72350189 | protocadherin 19 (PCDH19) pseudogene |
| ENSG00000135094 | SDS | -0,468957479 | 0,722486493 | 0,32329 | 0,60606867 | serine dehydratase |
| ENSG00000258727 | AL135999,1 | -0,469002266 | 0,722464064 | 0,24871 | 0,55591283 | novel transcript, antisense to AP1G2 |
| ENSG00000128268 | MGAT3 | -0,469132743 | 0,722398728 | 0,18757 | 0,51082796 | beta-1,4-mannosyl-glycoprotein 4-beta-N-acetylglucosaminyltransferase |
| ENSG00000131116 | ZNF428 | -0,469353706 | 0,722288094 | 0,12794 | 0,45145929 | zinc finger protein 428 |
| ENSG00000177191 | B3GNT8 | -0,469376949 | 0,722276457 | 0,26022 | 0,56382982 | UDP-GlcNAc:betaGal beta-1,3-N-acetylglucosaminyltransferase 8 |
| ENSG00000179935 | LINC00652 | -0,469421767 | 0,72225402 | 0,16257 | 0,48945635 | long intergenic non-protein coding RNA 652 |
| ENSG00000269054 | AC012313,6 | -0,469524542 | 0,722202569 | 0,11035 | 0,42293369 | novel transcript, antisense to ZNF497 |
| ENSG00000272383 | AC006270,2 | -0,469528575 | 0,722200551 | 0,67039 | 0,83656528 | zinc finger protein 329 (ZNF329) pseudogene |

| | | | | | | |
|---|---|---|---|---|---|---|
| ENSG00000108239 | TBC1D12 | -0,469612263 | 0,722158658 | 0,06883 | 0,34677117 | TBC1 domain family member 12 |
| ENSG00000078319 | PMS2P1 | -0,469665027 | 0,722132247 | 0,13924 | 0,46655993 | PMS1 homolog 2, mismatch repair system component pseudogene 1 |
| ENSG00000232439 | RPL18AP7 | -0,46990053 | 0,722014377 | 0,3603 | 0,63261878 | ribosomal protein L18a pseudogene 7 |
| ENSG00000280336 | AC244517,12 | -0,469975705 | 0,721976756 | 0,42863 | 0,6812979 | novel transcript |
| ENSG00000287693 | AC091691,3 | -0,469987039 | 0,721971084 | 0,4671 | 0,70712115 | novel transcript |
| ENSG00000186868 | MAPT | -0,470152903 | 0,721888085 | 0,38701 | 0,65213944 | microtubule associated protein tau |
| ENSG00000286233 | AC087885,1 | -0,470216522 | 0,721856252 | 0,51752 | 0,74270246 | novel transcript, antisense to SRRM4 |
| ENSG00000173846 | PLK3 | -0,470282142 | 0,72182342 | 0,09652 | 0,40027716 | polo like kinase 3 |
| ENSG00000286826 | AL136129,1 | -0,470363333 | 0,721782799 | 0,40299 | 0,66399303 | novel transcript |
| ENSG00000270060 | AC090589,3 | -0,470554756 | 0,721687036 | 0,40456 | 0,66449258 | novel transcript |
| ENSG00000072609 | CHFR | -0,470616885 | 0,721655958 | 0,14391 | 0,47144689 | checkpoint with forkhead and ring finger domains |
| ENSG00000168060 | NAALADL1 | -0,470627269 | 0,721650763 | 0,50082 | 0,73097006 | N-acetylated alpha-linked acidic dipeptidase like 1 |
| ENSG00000235370 | DNM1P51 | -0,470704664 | 0,72161205 | 0,23042 | 0,54355692 | dynamin 1 pseudogene 51 |
| ENSG00000285627 | AC005343,4 | -0,470830793 | 0,721548966 | 0,62431 | 0,80968303 | novel transcript, antisense to ADIPOR2 |
| ENSG00000187758 | ADH1A | -0,470990543 | 0,721469073 | 0,42423 | 0,67803888 | alcohol dehydrogenase 1A (class I), alpha polypeptide |
| ENSG00000267493 | CIRBP-AS1 | -0,470994705 | 0,721466991 | 0,26708 | 0,5695616 | CIRBP antisense RNA 1 |
| ENSG00000284431 | AL022238,3 | -0,471047962 | 0,721440359 | 0,44175 | 0,68993396 | novel protein |
| ENSG00000178385 | PLEKHM3 | -0,471134426 | 0,721397123 | 0,21551 | 0,53430995 | pleckstrin homology domain containing M3 |
| ENSG00000241651 | AC099340,1 | -0,471188264 | 0,721370202 | 0,62453 | 0,80980546 | ribosomal protein L15 (RPL15) pseudogene |
| ENSG00000139531 | SUOX | -0,471269212 | 0,721329728 | 0,19549 | 0,51643293 | sulfite oxidase |
| ENSG00000108018 | SORCS1 | -0,47143373 | 0,721247476 | 0,26894 | 0,57085484 | sortilin related VPS10 domain containing receptor 1 |
| ENSG00000237689 | AC007064,2 | -0,471558375 | 0,721185165 | 0,44178 | 0,68993396 | novel transcript |
| ENSG00000155621 | C9orf85 | -0,471568101 | 0,721180303 | 0,21311 | 0,53213032 | chromosome 9 open reading frame 85 |
| ENSG00000184216 | IRAK1 | -0,471569181 | 0,721179763 | 0,01141 | 0,13878661 | interleukin 1 receptor associated kinase 1 |
| ENSG00000278129 | ZNF8 | -0,471675599 | 0,721126568 | 0,20145 | 0,52211246 | zinc finger protein 8 |
| ENSG00000234751 | AP002381,1 | -0,471791733 | 0,721068522 | 0,53117 | 0,75245876 | ribosomal protein L15 (RPL15) pseudogene |
| ENSG00000144834 | TAGLN3 | -0,471815917 | 0,721056434 | 0,14282 | 0,47034016 | transgelin 3 |
| ENSG00000133315 | MACROD1 | -0,471878095 | 0,721025358 | 0,17381 | 0,49846579 | mono-ADP ribosylhydrolase 1 |
| ENSG00000276710 | CSPG4P10 | -0,471929287 | 0,720999774 | 0,23385 | 0,54565913 | chondroitin sulfate proteoglycan 4 pseudogene 10 |
| ENSG00000259972 | AC009120,2 | -0,471980198 | 0,720974332 | 0,13496 | 0,46159318 | novel transcript, antisense to PSMD7 |
| ENSG00000233928 | AL591501,1 | -0,472072336 | 0,720928288 | 0,11192 | 0,42531464 | novel transcript |
| ENSG00000260097 | SPDYE6 | -0,472166284 | 0,720881343 | 0,19157 | 0,51435195 | speedy/RINGO cell cycle regulator family member E6 |

| | | | | | | |
|---|---|---|---|---|---|---|
| ENSG00000132561 | MATN2 | -0,472295851 | 0,720816604 | 0,3854 | 0,65110417 | matrilin 2 |
| ENSG00000161904 | LEMD2 | -0,472369887 | 0,720779614 | 0,13247 | 0,45754597 | LEM domain nuclear envelope protein 2 |
| ENSG00000283100 | SIRT2 | -0,472498881 | 0,720715171 | 0,15988 | 0,48710442 | sirtuin 2 |
| ENSG00000226180 | AC010536,1 | -0,472566021 | 0,720681631 | 0,15261 | 0,48061452 | novel transcript |
| ENSG00000130528 | HRC | -0,47260024 | 0,720664537 | 0,30446 | 0,59430086 | histidine rich calcium binding protein |
| ENSG00000206412 | GNL1 | -0,472987445 | 0,720471144 | 0,46144 | 0,7028392 | G protein nucleolar 1 (putative) |
| ENSG00000231616 | AL354733,2 | -0,473031572 | 0,720449108 | 0,21391 | 0,53303219 | novel transcript, antisense to KIF27 |
| ENSG00000143858 | SYT2 | -0,473034116 | 0,720447838 | 0,26126 | 0,56427754 | synaptotagmin 2 |
| ENSG00000100242 | SUN2 | -0,473077161 | 0,720426342 | 0,19773 | 0,51937526 | Sad1 and UNC84 domain containing 2 |
| ENSG00000137171 | KLC4 | -0,473251888 | 0,720339096 | 0,04998 | 0,30089233 | kinesin light chain 4 |
| ENSG00000288068 | AC027088,5 | -0,473265553 | 0,720332273 | 0,50386 | 0,73294771 | novel transcript, sense intronic to NOX5 |
| ENSG00000286108 | AL359916,1 | -0,473447771 | 0,720241298 | 0,31502 | 0,60145932 | novel transcript, antisense to STK35 |
| ENSG00000105519 | CAPS | -0,473675623 | 0,720127555 | 0,22727 | 0,54168591 | calcyphosine |
| ENSG00000185608 | MRPL40 | -0,473693591 | 0,720118586 | 0,06577 | 0,33961566 | mitochondrial ribosomal protein L40 |
| ENSG00000108639 | SYNGR2 | -0,473976204 | 0,719977534 | 0,18058 | 0,50407792 | synaptogyrin 2 |
| ENSG00000213023 | SYT3 | -0,474018237 | 0,719956558 | 0,19177 | 0,51436129 | synaptotagmin 3 |
| ENSG00000246777 | AC044802,1 | -0,474335231 | 0,719798384 | 0,38209 | 0,64916957 | novel transcript |
| ENSG00000224448 | AC005020,1 | -0,47439736 | 0,719767387 | 0,51422 | 0,74051791 | ADP-ribosylation factor 6 (ARF6) pseudogene |
| ENSG00000286651 | PRSS43P | -0,474470933 | 0,719730682 | 0,52678 | 0,75002373 | serine protease 43, pseudogene |
| ENSG00000225828 | FAM229A | -0,474685178 | 0,719623808 | 0,11557 | 0,43158502 | family with sequence similarity 229 member A |
| ENSG00000170092 | SPDYE5 | -0,474705828 | 0,719613508 | 0,14891 | 0,47638416 | speedy/RINGO cell cycle regulator family member E5 |
| ENSG00000233087 | RAB6D | -0,474769694 | 0,719581652 | 0,57179 | 0,77829419 | RAB6D, member RAS oncogene family |
| ENSG00000170946 | DNAJC24 | -0,474855578 | 0,719538817 | 0,14879 | 0,47616437 | DnaJ heat shock protein family (Hsp40) member C24 |
| ENSG00000188451 | SRP72P2 | -0,474871523 | 0,719530864 | 0,21462 | 0,53356177 | signal recognition particle 72 pseudogene 2 |
| ENSG00000287061 | AC124248,2 | -0,474992158 | 0,719470701 | 0,27676 | 0,57549599 | novel transcript, antisense to BLM |
| ENSG00000287766 | AC006130,3 | -0,47508281 | 0,719425494 | 0,49342 | 0,72634709 | novel transcript |
| ENSG00000129682 | FGF13 | -0,475087811 | 0,719423 | 0,12471 | 0,44595911 | fibroblast growth factor 13 |
| ENSG00000204514 | ZNF814 | -0,475102095 | 0,719415877 | 0,1708 | 0,49641416 | zinc finger protein 814 |
| ENSG00000171224 | FAM241B | -0,475103979 | 0,719414938 | 0,16836 | 0,4945419 | family with sequence similarity 241 member B |
| ENSG00000150687 | PRSS23 | -0,475191406 | 0,719371343 | 0,29551 | 0,58925888 | serine protease 23 |
| ENSG00000213578 | CPLX3 | -0,475236181 | 0,719349017 | 0,58838 | 0,78811087 | complexin 3 |
| ENSG00000178217 | SH2D4B | -0,475262766 | 0,719335761 | 0,33588 | 0,61578286 | SH2 domain containing 4B |
| ENSG00000207191 | SNORD116-5 | -0,475270278 | 0,719332016 | 0,50089 | 0,73097006 | small nucleolar RNA, C/D box 116-5 |

| | | | | | | |
|---|---|---|---|---|---|---|
| ENSG00000207133 | SNORD116-7 | -0,475270278 | 0,719332016 | 0,50089 | 0,73097006 | small nucleolar RNA, C/D box 116-7 |
| ENSG00000234072 | AC074117,1 | -0,475510089 | 0,719212455 | 0,1591 | 0,48655958 | novel transcript, antisense to GTF3C2 and EIF2B4 |
| ENSG00000145284 | SCD5 | -0,475617602 | 0,71915886 | 0,03114 | 0,23785663 | stearoyl-CoA desaturase 5 |
| ENSG00000140488 | CELF6 | -0,47563277 | 0,719151299 | 0,35981 | 0,63249219 | CUGBP Elav-like family member 6 |
| ENSG00000231993 | EP300-AS1 | -0,475676198 | 0,719129652 | 0,50187 | 0,73149732 | EP300 antisense RNA 1 |
| ENSG00000225393 | BX571846,1 | -0,475751889 | 0,719091923 | 0,26029 | 0,56382982 | novel transcript |
| ENSG00000250746 | AC105250,1 | -0,475855916 | 0,719040074 | 0,56898 | 0,77629764 | heat shock 70kDa protein 5 (glucose-regulated protein, 78kDa) (HSPA5) pseudogene |
| ENSG00000150477 | KIAA1328 | -0,476181819 | 0,718877663 | 0,10464 | 0,41313952 | KIAA1328 |
| ENSG00000234929 | AC018685,2 | -0,476193061 | 0,718872061 | 0,81766 | 0,91621442 | novel transcript |
| ENSG00000285672 | AL160396,2 | -0,476341485 | 0,718798107 | 0,36257 | 0,63379594 | novel transcript |
| ENSG00000244052 | RPL5P24 | -0,476362506 | 0,718787634 | 0,64536 | 0,82203909 | ribosomal protein L5 pseudogene 24 |
| ENSG00000124357 | NAGK | -0,476502625 | 0,718717827 | 0,09835 | 0,40379437 | N-acetylglucosamine kinase |
| ENSG00000206299 | TAP2 | -0,476517373 | 0,71871048 | 0,73907 | 0,87620747 | transporter 2, ATP binding cassette subfamily B member |
| ENSG00000112493 | TAPBP | -0,476634869 | 0,718651949 | 0,45065 | 0,69592349 | TAP binding protein |
| ENSG00000282393 | AC016588,2 | -0,476839484 | 0,718550031 | 0,13708 | 0,4642578 | novel transcript |
| ENSG00000127249 | ATP13A4 | -0,476980426 | 0,718479836 | 0,21294 | 0,53198758 | ATPase 13A4 |
| ENSG00000204175 | GPRIN2 | -0,477131762 | 0,718404473 | 0,27338 | 0,57349735 | G protein regulated inducer of neurite outgrowth 2 |
| ENSG00000243156 | MICAL3 | -0,477308904 | 0,718316269 | 0,13428 | 0,46056769 | microtubule associated monooxygenase, calponin and LIM domain containing 3 |
| ENSG00000230071 | RPL4P6 | -0,477318747 | 0,718311368 | 0,25789 | 0,56199743 | ribosomal protein L4 pseudogene 6 |
| ENSG00000169499 | PLEKHA2 | -0,477343303 | 0,718299142 | 0,37235 | 0,64128645 | pleckstrin homology domain containing A2 |
| ENSG00000184608 | FAM167A-AS1 | -0,477725811 | 0,718108721 | 0,48635 | 0,72184113 | FAM167A antisense RNA 1 |
| ENSG00000225076 | WWC3-AS1 | -0,477779053 | 0,71808222 | 0,60839 | 0,80115227 | WWC3 antisense RNA 1 |
| ENSG00000286827 | AC072026,2 | -0,477797412 | 0,718073083 | 0,10358 | 0,41166534 | novel transcript, antisense to STXBP5L |
| ENSG00000205683 | DPF3 | -0,478118469 | 0,7179133 | 0,20424 | 0,52406249 | double PHD fingers 3 |
| ENSG00000186094 | AGBL4 | -0,478466799 | 0,717739986 | 0,39083 | 0,65464997 | ATP/GTP binding protein like 4 |
| ENSG00000258682 | AL132989,1 | -0,478521375 | 0,717712835 | 0,30519 | 0,59456936 | novel transcript, sense overlapping PSMA3 |
| ENSG00000276255 | LINC02809 | -0,478580027 | 0,717683657 | 0,24474 | 0,55280101 | long intergenic non-protein coding RNA 2809 |
| ENSG00000141068 | KSR1 | -0,478632239 | 0,717657684 | 0,03746 | 0,2618019 | kinase suppressor of ras 1 |
| ENSG00000095739 | BAMBI | -0,478633345 | 0,717657133 | 0,39173 | 0,65534953 | BMP and activin membrane bound inhibitor |
| ENSG00000267646 | AC008543,5 | -0,478708408 | 0,717619795 | 0,53051 | 0,75210231 | novel transcript, antisense to ZNF491 |
| ENSG00000159714 | ZDHHC1 | -0,478762505 | 0,717592887 | 0,23422 | 0,54602618 | zinc finger DHHC-type containing 1 |

| | | | | | | |
|---|---|---|---|---|---|---|
| ENSG00000261426 | AC144833,1 | -0,478816644 | 0,717565959 | 0,46975 | 0,70921406 | novel transcript, sense intronic to GABRG3 |
| ENSG00000123130 | ACOT9 | -0,478826995 | 0,71756081 | 0,22285 | 0,53904413 | acyl-CoA thioesterase 9 |
| ENSG00000281705 | CENATAC-DT | -0,478858906 | 0,717544939 | 0,17898 | 0,50339563 | CENATAC divergent transcript |
| ENSG00000255121 | CENATAC-DT | -0,478858906 | 0,717544939 | 0,17898 | 0,50339563 | CENATAC divergent transcript |
| ENSG00000261471 | AC092145,1 | -0,478862829 | 0,717542988 | 0,43324 | 0,68455289 | novel transcript, antisense to COTL1 |
| ENSG00000261684 | AC018362,1 | -0,479197762 | 0,717376424 | 0,30513 | 0,59455047 | novel transcript, antisense to SNAP23 |
| ENSG00000204695 | OR14J1 | -0,479309222 | 0,717321003 | 0,09682 | 0,40094776 | olfactory receptor family 14 subfamily J member 1 |
| ENSG00000284512 | AC092718,8 | -0,479502451 | 0,717224934 | 0,28072 | 0,57812368 | novel protein |
| ENSG00000230583 | GTF2IRD1P1 | -0,479619584 | 0,717166705 | 0,39436 | 0,65725123 | GTF2I repeat domain containing 1 pseudogene 1 |
| ENSG00000185864 | NPIPB4 | -0,479810497 | 0,717071808 | 0,13629 | 0,46342304 | nuclear pore complex interacting protein family member B4 |
| ENSG00000130702 | LAMA5 | -0,479942927 | 0,717005988 | 0,33772 | 0,6174316 | laminin subunit alpha 5 |
| ENSG00000272455 | MRPL20-DT | -0,480047277 | 0,716954129 | 0,22986 | 0,54324957 | MRPL20 divergent transcript |
| ENSG00000286218 | AC005040,2 | -0,480067279 | 0,716944189 | 0,50026 | 0,73078774 | novel transcript |
| ENSG00000285541 | AC005548,1 | -0,480158509 | 0,716898854 | 0,23403 | 0,54584395 | novel transcript, antisense to SHISA6 |
| ENSG00000143793 | C1orf35 | -0,480287503 | 0,716834757 | 0,16249 | 0,48945635 | chromosome 1 open reading frame 35 |
| ENSG00000259398 | AC090970,1 | -0,480402754 | 0,716777495 | 0,55751 | 0,76919576 | novel transcript, intronic to MAPK6 |
| ENSG00000267429 | AC006116,6 | -0,480458676 | 0,716749711 | 0,29059 | 0,58565836 | novel transcript, sense intronic to ZSCAN5A |
| ENSG00000232653 | GOLGA8N | -0,480480002 | 0,716739117 | 0,28733 | 0,58372981 | golgin A8 family member N |
| ENSG00000257337 | AC068888,1 | -0,480606957 | 0,716676048 | 0,22134 | 0,53805234 | novel transcript, antisense to TENC1 & EIF4B |
| ENSG00000103066 | PLA2G15 | -0,480713016 | 0,716623363 | 0,08924 | 0,38636433 | phospholipase A2 group XV |
| ENSG00000175749 | EIF3KP1 | -0,480729197 | 0,716615326 | 0,48006 | 0,71732148 | eukaryotic translation initiation factor 3 subunit K pseudogene 1 |
| ENSG00000278022 | AC118658,1 | -0,480766197 | 0,716596947 | 0,39 | 0,65410436 | novel transcript, sense intronic to IGF1R |
| ENSG00000185504 | FAAP100 | -0,480936011 | 0,716512604 | 0,15857 | 0,48579688 | FA core complex associated protein 100 |
| ENSG00000187775 | DNAH17 | -0,481043512 | 0,716459216 | 0,20772 | 0,52735573 | dynein axonemal heavy chain 17 |
| ENSG00000287967 | AC004882,3 | -0,481272687 | 0,716345415 | 0,47456 | 0,71332138 | novel transcript, antisense to UQCR10 |
| ENSG00000139370 | SLC15A4 | -0,481314267 | 0,716324769 | 0,16113 | 0,48841573 | solute carrier family 15 member 4 |
| ENSG00000143297 | FCRL5 | -0,481531698 | 0,716216818 | 0,33532 | 0,61497665 | Fc receptor like 5 |
| ENSG00000274653 | AC106782,5 | -0,481655101 | 0,716155558 | 0,48556 | 0,72107911 | novel transcript, antisense to TBC1D10B |
| ENSG00000104524 | PYCR3 | -0,481716518 | 0,716125072 | 0,20714 | 0,52670822 | pyrroline-5-carboxylate reductase 3 |
| ENSG00000130635 | COL5A1 | -0,48179934 | 0,716083962 | 0,41559 | 0,67246712 | collagen type V alpha 1 chain |
| ENSG00000272501 | AL662844,4 | -0,481831389 | 0,716068054 | 0,1426 | 0,47000875 | novel transcript, antisense to HCG27 |

| | | | | | | |
|---|---|---|---|---|---|---|
| ENSG00000274843 | MUC20 | -0,481860251 | 0,716053729 | 0,70835 | 0,85935158 | mucin 20, cell surface associated |
| ENSG00000257935 | LHX5-AS1 | -0,481937468 | 0,716015405 | 0,25648 | 0,56092974 | LHX5 antisense RNA 1 |
| ENSG00000129657 | SEC14L1 | -0,482036575 | 0,715966219 | 0,12176 | 0,44093327 | SEC14 like lipid binding 1 |
| ENSG00000160216 | AGPAT3 | -0,482062633 | 0,715953288 | 0,08342 | 0,37550713 | 1-acylglycerol-3-phosphate O-acyltransferase 3 |
| ENSG00000256596 | AC026358,1 | -0,482149429 | 0,715910216 | 0,34559 | 0,62278421 | novel transcript |
| ENSG00000237646 | HLCS-IT1 | -0,48216276 | 0,7159036 | 0,56998 | 0,77703861 | HLCS intronic transcript 1 |
| ENSG00000234203 | AC004771,1 | -0,482203044 | 0,715883611 | 0,63497 | 0,8162492 | novel transcript, antisense to CAMTA2 |
| ENSG00000241890 | RPL13P4 | -0,482313634 | 0,715828736 | 0,55882 | 0,77025265 | ribosomal protein L13 pseudogene 4 |
| ENSG00000196417 | ZNF765 | -0,482337516 | 0,715816887 | 0,14558 | 0,47304491 | zinc finger protein 765 |
| ENSG00000235140 | AC026391,1 | -0,482471288 | 0,715750517 | 0,45613 | 0,69998075 | uncharacterized LOC105378318 |
| ENSG00000249715 | FER1L5 | -0,4824794 | 0,715746492 | 0,22713 | 0,54168591 | fer-1 like family member 5 |
| ENSG00000176834 | VSIG10 | -0,482517752 | 0,715727465 | 0,0229 | 0,20200592 | V-set and immunoglobulin domain containing 10 |
| ENSG00000053918 | KCNQ1 | -0,482521944 | 0,715725386 | 0,38169 | 0,64905556 | potassium voltage-gated channel subfamily Q member 1 |
| ENSG00000108306 | FBXL20 | -0,482537883 | 0,715717478 | 0,21732 | 0,53520949 | F-box and leucine rich repeat protein 20 |
| ENSG00000239857 | GET4 | -0,482840368 | 0,715567432 | 0,16791 | 0,49407144 | guided entry of tail-anchored proteins factor 4 |
| ENSG00000171827 | ZNF570 | -0,482924481 | 0,715525714 | 0,15806 | 0,48506908 | zinc finger protein 570 |
| ENSG00000160179 | ABCG1 | -0,48294951 | 0,7155133 | 0,20031 | 0,52107152 | ATP binding cassette subfamily G member 1 |
| ENSG00000188878 | FBF1 | -0,483230282 | 0,715374063 | 0,06142 | 0,32865897 | Fas binding factor 1 |
| ENSG00000259562 | AC090607,2 | -0,4832473 | 0,715365625 | 0,33527 | 0,61497665 | zinc finger CCHC-type and RNA binding motif 1 (ZCRB1) pseudogene |
| ENSG00000197584 | KCNMB2 | -0,483275471 | 0,715351656 | 0,46877 | 0,70833955 | potassium calcium-activated channel subfamily M regulatory beta subunit 2 |
| ENSG00000177675 | CD163L1 | -0,48348964 | 0,71524547 | 0,34717 | 0,62397921 | CD163 molecule like 1 |
| ENSG00000265590 | CFAP298-TCP: | -0,483689445 | 0,715146419 | 0,13034 | 0,45477153 | CFAP298-TCP10L readthrough |
| ENSG00000253213 | USP12P1 | -0,483966153 | 0,715009268 | 0,44996 | 0,69542581 | USP12 pseudogene 1 |
| ENSG00000160188 | RSPH1 | -0,484047511 | 0,714968948 | 0,17736 | 0,50187931 | radial spoke head component 1 |
| ENSG00000275097 | AC024940,5 | -0,484281734 | 0,714852881 | 0,54982 | 0,76429591 | novel transcript, overlapping 3'UTR of FAM60A |
| ENSG00000166682 | TMPRSS5 | -0,484452548 | 0,714768248 | 0,16118 | 0,48849437 | transmembrane serine protease 5 |
| ENSG00000212123 | PRR22 | -0,484549748 | 0,714720093 | 0,23277 | 0,54542294 | proline rich 22 |
| ENSG00000260630 | SNAI3-AS1 | -0,484575625 | 0,714707273 | 0,16922 | 0,49530908 | SNAI3 antisense RNA 1 |
| ENSG00000198794 | SCAMP5 | -0,484763849 | 0,714614034 | 0,14503 | 0,47264896 | secretory carrier membrane protein 5 |
| ENSG00000130224 | LRCH2 | -0,484884988 | 0,714554032 | 0,23853 | 0,5488078 | leucine rich repeats and calponin homology domain containing 2 |

| | | | | | | |
|---|---|---|---|---|---|---|
| ENSG00000187726 | DNAJB13 | -0,485011513 | 0,714491368 | 0,51301 | 0,7394947 | DnaJ heat shock protein family (Hsp40) member B13 |
| ENSG00000235816 | PRELID1P3 | -0,485226919 | 0,714384697 | 0,3927 | 0,65593262 | PRELID1 pseudogene 3 |
| ENSG00000143226 | FCGR2A | -0,485290044 | 0,71435344 | 0,52814 | 0,75077432 | Fc fragment of IgG receptor IIa |
| ENSG00000165060 | FXN | -0,485356451 | 0,714320559 | 0,22325 | 0,53944353 | frataxin |
| ENSG00000274507 | AL121772,2 | -0,485491173 | 0,714253857 | 0,59926 | 0,79538547 | novel transcript, antisense to PYGB |
| ENSG00000198496 | NBR2 | -0,485612196 | 0,714193943 | 0,17525 | 0,49934416 | neighbor of BRCA1 lncRNA 2 |
| ENSG00000184489 | PTP4A3 | -0,485723056 | 0,714139065 | 0,15387 | 0,48214138 | protein tyrosine phosphatase 4A3 |
| ENSG00000228801 | AC064807,1 | -0,48582231 | 0,714089936 | 0,21674 | 0,53494609 | novel transcript |
| ENSG00000154359 | LONRF1 | -0,485855813 | 0,714073353 | 0,07809 | 0,36538634 | LON peptidase N-terminal domain and ring finger 1 |
| ENSG00000111676 | ATN1 | -0,485914233 | 0,714044438 | 0,1571 | 0,48443207 | atrophin 1 |
| ENSG00000230950 | MTND2P21 | -0,486126918 | 0,71393918 | 0,51255 | 0,73941327 | MT-ND2 pseudogene 21 |
| ENSG00000104886 | PLEKHJ1 | -0,486174747 | 0,713915511 | 0,12953 | 0,4530895 | pleckstrin homology domain containing J1 |
| ENSG00000288033 | AC137894,4 | -0,486186937 | 0,713909479 | 0,15238 | 0,48061452 | novel transcript |
| ENSG00000250608 | AC010210,1 | -0,48626551 | 0,713870599 | 0,0549 | 0,31350641 | novel transcript |
| ENSG00000217801 | AL390719,1 | -0,486783552 | 0,713614308 | 0,29626 | 0,58961815 | protein tyrosine phosphatase family pseudogene |
| ENSG00000186470 | BTN3A2 | -0,486965325 | 0,713524402 | 0,35923 | 0,6321355 | butyrophilin subfamily 3 member A2 |
| ENSG00000237846 | AL773545,3 | -0,486973307 | 0,713520455 | 0,39732 | 0,65983856 | novel pseudogene |
| ENSG00000231187 | AL356056,2 | -0,487018428 | 0,713498139 | 0,1185 | 0,43617111 | uncharacterized LOC102724593 |
| ENSG00000284099 | SQSTM1 | -0,487034189 | 0,713490344 | 0,17296 | 0,49768904 | sequestosome 1 |
| ENSG00000287578 | AL034348,1 | -0,48705601 | 0,713479553 | 0,46017 | 0,70210714 | novel transcript |
| ENSG00000261172 | AC133919,2 | -0,487078114 | 0,713468621 | 0,65407 | 0,82680983 | novel transcript |
| ENSG00000149929 | HIRIP3 | -0,487119038 | 0,713448383 | 0,0891 | 0,38611566 | HIRA interacting protein 3 |
| ENSG00000231006 | RPL7P32 | -0,487197608 | 0,71340953 | 0,49197 | 0,72531636 | ribosomal protein L7 pseudogene 32 |
| ENSG00000287727 | AL139423,2 | -0,487394756 | 0,713312047 | 0,22825 | 0,54221266 | novel transcript, antisense to PEX14 |
| ENSG00000284606 | AC105233,4 | -0,487485534 | 0,713267165 | 0,05141 | 0,30462001 | novel transcript |
| ENSG00000223930 | AC117453,1 | -0,487530188 | 0,713245088 | 0,43519 | 0,68590519 | novel transcript, antisense to KCNMB2 |
| ENSG00000266578 | AP001025,1 | -0,487822904 | 0,713100389 | 0,57065 | 0,77748547 | novel transcript |
| ENSG00000283008 | CYP2D7 | -0,487860038 | 0,713082035 | 0,22802 | 0,54214771 | cytochrome P450 family 2 subfamily D member 7 (gene/pseudogene) |
| ENSG00000260279 | AC137932,1 | -0,48801892 | 0,713003508 | 0,09916 | 0,4054333 | novel transcript, antisense to ANKRD11 |
| ENSG00000250571 | GLI4 | -0,488053905 | 0,712986218 | 0,18708 | 0,5105374 | GLI family zinc finger 4 |
| ENSG00000177576 | C18orf32 | -0,488462434 | 0,71278435 | 0,13189 | 0,45708171 | chromosome 18 open reading frame 32 |
| ENSG00000224143 | EHMT2 | -0,488493256 | 0,712769122 | 0,22808 | 0,54220742 | euchromatic histone lysine methyltransferase 2 |

| | | | | | | |
|---|---|---|---|---|---|---|
| ENSG00000163293 | NIPAL1 | -0,488539034 | 0,712746506 | 0,38587 | 0,65129675 | NIPA like domain containing 1 |
| ENSG00000276071 | AC074138,1 | -0,488662124 | 0,712685697 | 0,22742 | 0,54168591 | novel transcript |
| ENSG00000119922 | IFIT2 | -0,489048803 | 0,712494705 | 0,31565 | 0,601741 | interferon induced protein with tetratricopeptide repeats 2 |
| ENSG00000236222 | WDR46 | -0,489130239 | 0,712454487 | 0,3857 | 0,65122858 | WD repeat domain 46 |
| ENSG00000148057 | IDNK | -0,489357491 | 0,712342271 | 0,06901 | 0,34717336 | IDNK gluconokinase |
| ENSG00000163959 | SLC51A | -0,48942037 | 0,712311225 | 0,27481 | 0,57463533 | solute carrier family 51 subunit alpha |
| ENSG00000270607 | AC009549,1 | -0,489461114 | 0,712291108 | 0,43358 | 0,68473723 | novel transcript |
| ENSG00000241532 | AGGF1P3 | -0,489547836 | 0,712248293 | 0,62688 | 0,81116451 | angiogenic factor with G-patch and FHA domains 1 pseudogene 3 |
| ENSG00000169570 | DTWD2 | -0,489570207 | 0,712237248 | 0,18645 | 0,50994868 | DTW domain containing 2 |
| ENSG00000251497 | PITPNM2-AS1 | -0,489676266 | 0,712184891 | 0,47132 | 0,7106605 | PITPNM2 antisense RNA 1 |
| ENSG00000231327 | LINC01816 | -0,48986022 | 0,712094088 | 0,10699 | 0,41677007 | long intergenic non-protein coding RNA 1816 |
| ENSG00000103647 | CORO2B | -0,489970302 | 0,712039755 | 0,09982 | 0,40626368 | coronin 2B |
| ENSG00000287917 | AP000944,7 | -0,489988399 | 0,712030823 | 0,35911 | 0,63201238 | novel transcript, antisense to POLA2 |
| ENSG00000112706 | IMPG1 | -0,490134435 | 0,711958752 | 0,5625 | 0,77186635 | interphotoreceptor matrix proteoglycan 1 |
| ENSG00000250331 | LINC01340 | -0,490164066 | 0,711944129 | 0,58519 | 0,78616171 | long intergenic non-protein coding RNA 1340 |
| ENSG00000287153 | AL161851,1 | -0,490214425 | 0,711919279 | 0,40956 | 0,66801281 | novel transcript, antisense to NPAS3 |
| ENSG00000083307 | GRHL2 | -0,490268439 | 0,711892625 | 0,23718 | 0,54813266 | grainyhead like transcription factor 2 |
| ENSG00000198932 | GPRASP1 | -0,49027473 | 0,711889521 | 0,19571 | 0,51646992 | G protein-coupled receptor associated sorting protein 1 |
| ENSG00000226704 | HSPA1L | -0,490527075 | 0,711765014 | 0,64294 | 0,82098995 | heat shock protein family A (Hsp70) member 1 like |
| ENSG00000223730 | Z83838,1 | -0,49061716 | 0,711720571 | 0,22626 | 0,54121239 | ribosomal protein L6 (RPL6) pseudogene |
| ENSG00000124788 | ATXN1 | -0,490703884 | 0,711677789 | 0,11045 | 0,42311032 | ataxin 1 |
| ENSG00000175445 | LPL | -0,490794633 | 0,711633024 | 0,45035 | 0,69574391 | lipoprotein lipase |
| ENSG00000244313 | AC024293,1 | -0,49083815 | 0,711611559 | 0,53323 | 0,75351438 | ribosomal protein S10 (RPS10) pseudogene |
| ENSG00000250422 | AC117529,2 | -0,490873002 | 0,711594368 | 0,58594 | 0,78652174 | tigger transposable element derived 1 (TIGD1) pseudogene |
| ENSG00000187193 | MT1X | -0,491126212 | 0,711469486 | 0,3275 | 0,60929711 | metallothionein 1X |
| ENSG00000260996 | BX255925,1 | -0,491240553 | 0,711413101 | 0,56194 | 0,77147224 | novel transcript, overlapping NRARP |
| ENSG00000275559 | AL137060,5 | -0,491255299 | 0,711405829 | 0,36472 | 0,6350493 | |
| ENSG00000185669 | SNAI3 | -0,491282293 | 0,711392518 | 0,19274 | 0,51490739 | snail family transcriptional repressor 3 |
| ENSG00000263938 | AC100832,2 | -0,491370468 | 0,711349041 | 0,52819 | 0,75077432 | novel transcript, sense intronic to PPM1E |
| ENSG00000286927 | AC103955,1 | -0,491427406 | 0,711320967 | 0,09633 | 0,39998624 | novel transcript |

| | | | | | | |
|---|---|---|---|---|---|---|
| ENSG00000159788 | RGS12 | -0,491476996 | 0,711296517 | 0,08054 | 0,37032385 | regulator of G protein signaling 12 |
| ENSG00000132530 | XAF1 | -0,491543274 | 0,71126384 | 0,1702 | 0,49606609 | XIAP associated factor 1 |
| ENSG00000151131 | C12orf45 | -0,491591097 | 0,711240264 | 0,11714 | 0,43444592 | chromosome 12 open reading frame 45 |
| ENSG00000272442 | AL353588,1 | -0,491620961 | 0,711225541 | 0,26202 | 0,56490812 | novel protein, TMEM151B-SPATS1 readtrough |
| ENSG00000188933 | USP32P1 | -0,491854132 | 0,7111106 | 0,11832 | 0,43614969 | ubiquitin specific peptidase 32 pseudogene 1 |
| ENSG00000271207 | MTCO1P22 | -0,492028825 | 0,711024499 | 0,59055 | 0,78980722 | MT-CO1 pseudogene 22 |
| ENSG00000249738 | AC008691,1 | -0,492291922 | 0,710894845 | 0,18858 | 0,5119341 | novel transcript |
| ENSG00000231134 | TCF7L1-IT1 | -0,492330637 | 0,710875768 | 0,51796 | 0,7429275 | TCF7L1 intronic transcript 1 |
| ENSG00000101846 | STS | -0,492370788 | 0,710855984 | 0,03635 | 0,25801846 | steroid sulfatase |
| ENSG00000212443 | SNORA53 | -0,492402673 | 0,710840274 | 0,05716 | 0,31890625 | small nucleolar RNA, H/ACA box 53 |
| ENSG00000260806 | AL163051,1 | -0,492433242 | 0,710825212 | 0,11567 | 0,43162108 | novel transcript, antisense to PAPOLA |
| ENSG00000210135 | MT-TN | -0,492434827 | 0,710824431 | 0,44988 | 0,69539135 | mitochondrially encoded tRNA-Asn (AAU/C) |
| ENSG00000154839 | SKA1 | -0,492542047 | 0,710771605 | 0,18535 | 0,50876814 | spindle and kinetochore associated complex subunit 1 |
| ENSG00000249228 | AC068944,1 | -0,492653449 | 0,710716723 | 0,0692 | 0,34748485 | novel transcript |
| ENSG00000101460 | MAP1LC3A | -0,492844957 | 0,710622386 | 0,21149 | 0,53032236 | microtubule associated protein 1 light chain 3 alpha |
| ENSG00000283445 | AL136985,3 | -0,493061336 | 0,710515814 | 0,12917 | 0,45253718 | novel transcript |
| ENSG00000165995 | CACNB2 | -0,493088494 | 0,710502438 | 0,03579 | 0,25662951 | calcium voltage-gated channel auxiliary subunit beta 2 |
| ENSG00000183309 | ZNF623 | -0,49313264 | 0,710480697 | 0,21428 | 0,53334744 | zinc finger protein 623 |
| ENSG00000241738 | ZNF90P1 | -0,493286055 | 0,71040515 | 0,23803 | 0,54864017 | zinc finger protein 90 pseudogene 1 |
| ENSG00000286069 | AC076968,2 | -0,493315837 | 0,710390485 | 0,24527 | 0,55295131 | novel transcript |
| ENSG00000285730 | Z94721,3 | -0,493378325 | 0,710359716 | 0,17296 | 0,49768904 | novel transcript |
| ENSG00000259797 | AC020978,1 | -0,49348179 | 0,710308773 | 0,48954 | 0,72395897 | novel transcript, intronic to PRMT7 |
| ENSG00000263164 | AC087500,2 | -0,493493701 | 0,710302909 | 0,46579 | 0,70619029 | novel transcript |
| ENSG00000138592 | USP8 | -0,493526401 | 0,71028681 | 0,14763 | 0,47507961 | ubiquitin specific peptidase 8 |
| ENSG00000218857 | AL137222,1 | -0,493533903 | 0,710283116 | 0,5175 | 0,74270246 | D4, zinc and double PHD fingers family 2 (DPF2) pseudogene |
| ENSG00000198929 | NOS1AP | -0,493647808 | 0,710227039 | 0,1079 | 0,41847368 | nitric oxide synthase 1 adaptor protein |
| ENSG00000010404 | IDS | -0,493654192 | 0,710223897 | 0,09039 | 0,3889575 | iduronate 2-sulfatase |
| ENSG00000171206 | TRIM8 | -0,493654634 | 0,710223679 | 0,06753 | 0,34379132 | tripartite motif containing 8 |
| ENSG00000112276 | BVES | -0,493676465 | 0,710212932 | 0,10017 | 0,4067769 | blood vessel epicardial substance |
| ENSG00000124602 | UNC5CL | -0,493898302 | 0,710103734 | 0,17555 | 0,49954066 | unc-5 family C-terminal like |
| ENSG00000281354 | AP000350,15 | -0,493904293 | 0,710100785 | 0,52797 | 0,75065864 | novel transcript, antisense to SLC2A11 |
| ENSG00000272973 | AP000350,5 | -0,493904293 | 0,710100785 | 0,52797 | 0,75065864 | novel transcript, antisense to SLC2A11 |

| | | | | | | |
|---|---|---|---|---|---|---|
| ENSG00000249504 | PCDHA14 | -0,494059702 | 0,710024296 | 0,85621 | 0,9349542 | protocadherin alpha 14 pseudogene |
| ENSG00000104679 | R3HCC1 | -0,494202256 | 0,709954142 | 0,07162 | 0,35351167 | R3H domain and coiled-coil containing 1 |
| ENSG00000255082 | GRM5-AS1 | -0,49425093 | 0,70993019 | 0,66232 | 0,83183445 | GRM5 antisense RNA 1 |
| ENSG00000259664 | LINC02254 | -0,494408984 | 0,709852417 | 0,12757 | 0,45080057 | long intergenic non-protein coding RNA 2254 |
| ENSG00000224747 | MTCYBP21 | -0,494433872 | 0,709840172 | 0,29293 | 0,58737078 | MT-CYB pseudogene 21 |
| ENSG00000119707 | RBM25 | -0,494621322 | 0,709747948 | 0,07559 | 0,36093649 | RNA binding motif protein 25 |
| ENSG00000178927 | CYBC1 | -0,494715415 | 0,709701659 | 0,1092 | 0,42097623 | cytochrome b-245 chaperone 1 |
| ENSG00000155719 | OTOA | -0,494778378 | 0,709670687 | 0,42116 | 0,67634721 | otoancorin |
| ENSG00000071246 | VASH1 | -0,494780402 | 0,709669691 | 0,21276 | 0,53177822 | vasohibin 1 |
| ENSG00000131188 | PRR7 | -0,494812708 | 0,7096538 | 0,27977 | 0,5775083 | proline rich 7, synaptic |
| ENSG00000257920 | LINC02409 | -0,494819928 | 0,709650249 | 0,55095 | 0,76487238 | long intergenic non-protein coding RNA 2409 |
| ENSG00000225335 | AC016027,1 | -0,494845458 | 0,709637691 | 0,15325 | 0,48147318 | novel transcript |
| ENSG00000160087 | UBE2J2 | -0,494920255 | 0,7096009 | 0,18693 | 0,51044154 | ubiquitin conjugating enzyme E2 J2 |
| ENSG00000158486 | DNAH3 | -0,494949843 | 0,709586347 | 0,50517 | 0,73378796 | dynein axonemal heavy chain 3 |
| ENSG00000256973 | AC053513,2 | -0,49501081 | 0,709556361 | 0,22018 | 0,53750545 | novel transcript |
| ENSG00000103150 | MLYCD | -0,495287879 | 0,709420105 | 0,02922 | 0,2303054 | malonyl-CoA decarboxylase |
| ENSG00000284922 | AP000812,4 | -0,495444028 | 0,709343325 | 0,04103 | 0,27431646 | leucine rich transmembrane and O-methyltransferase domain containing |
| ENSG00000233264 | AC006042,2 | -0,495486077 | 0,709322651 | 0,18123 | 0,50472253 | peptidylprolyl isomerase (cyclophilin)-like 4 (PPIL4) pseudogene |
| ENSG00000243492 | HLA-L | -0,495686168 | 0,70922428 | 0,08557 | 0,37920193 | major histocompatibility complex, class I, L (pseudogene) |
| ENSG00000260600 | AC092125,1 | -0,495871416 | 0,709133218 | 0,69051 | 0,84900677 | novel transcript |
| ENSG00000267219 | AC010504,1 | -0,495872047 | 0,709132908 | 0,73585 | 0,8750728 | novel transcript, antisense to KIAA0355 |
| ENSG00000071242 | RPS6KA2 | -0,495940146 | 0,709099436 | 0,02152 | 0,19510325 | ribosomal protein S6 kinase A2 |
| ENSG00000247853 | AC006064,2 | -0,495944724 | 0,709097186 | 0,20795 | 0,52755008 | novel transcript, antisense to CHD4 |
| ENSG00000103197 | TSC2 | -0,496106595 | 0,70901763 | 0,16979 | 0,49606538 | TSC complex subunit 2 |
| ENSG00000162585 | FAAP20 | -0,496342408 | 0,708901748 | 0,19942 | 0,52039716 | FA core complex associated protein 20 |
| ENSG00000196456 | ZNF775 | -0,496846113 | 0,708654284 | 0,08635 | 0,38089752 | zinc finger protein 775 |
| ENSG00000117595 | IRF6 | -0,496980753 | 0,708588151 | 0,37598 | 0,64426406 | interferon regulatory factor 6 |
| ENSG00000243609 | RPS2P44 | -0,497066768 | 0,708545906 | 0,35028 | 0,62578708 | ribosomal protein S2 pseudogene 44 |
| ENSG00000276846 | AC016590,3 | -0,497069157 | 0,708544732 | 0,319 | 0,60399503 | novel transcript, antisense to HKR1 |
| ENSG00000163577 | EIF5A2 | -0,497173487 | 0,708493495 | 0,1963 | 0,51733996 | eukaryotic translation initiation factor 5A2 |
| ENSG00000177981 | ASB8 | -0,497251097 | 0,708455383 | 0,06446 | 0,33652241 | ankyrin repeat and SOCS box containing 8 |

| | | | | | | |
|---|---|---|---|---|---|---|
| ENSG00000231856 | AL162377,1 | -0,49730493 | 0,708428948 | 0,14698 | 0,47471707 | novel transcript |
| ENSG00000182809 | CRIP2 | -0,497377209 | 0,708393457 | 0,04131 | 0,27529279 | cysteine rich protein 2 |
| ENSG00000250280 | AC026124,1 | -0,497391682 | 0,70838635 | 0,32777 | 0,60960585 | novel transcript |
| ENSG00000242412 | DBIL5P2 | -0,497481091 | 0,70834245 | 0,43275 | 0,68421938 | diazepam binding inhibitor-like 5 pseudogene 2 |
| ENSG00000277895 | AC135279,3 | -0,497827276 | 0,708172499 | 0,27091 | 0,57244549 | novel transcript, antisense to TBK1 |
| ENSG00000263829 | SINHCAFP1 | -0,497892307 | 0,708140578 | 0,53026 | 0,75185463 | SINHCAF pseudogene 1 |
| ENSG00000172146 | OR1A1 | -0,498026764 | 0,708074583 | 0,08032 | 0,37007763 | olfactory receptor family 1 subfamily A member 1 |
| ENSG00000283104 | SARS2 | -0,498200116 | 0,707989507 | 0,2389 | 0,54892264 | seryl-tRNA synthetase 2, mitochondrial |
| ENSG00000112297 | CRYBG1 | -0,498238888 | 0,707970481 | 0,2541 | 0,55925777 | crystallin beta-gamma domain containing 1 |
| ENSG00000230679 | ENO1-AS1 | -0,498290587 | 0,707945111 | 0,63913 | 0,81893766 | ENO1 antisense RNA 1 |
| ENSG00000159079 | CFAP298 | -0,49834197 | 0,707919897 | 0,14808 | 0,47519813 | cilia and flagella associated protein 298 |
| ENSG00000122566 | HNRNPA2B1 | -0,498355434 | 0,70791329 | 0,076 | 0,36206975 | heterogeneous nuclear ribonucleoprotein A2/B1 |
| ENSG00000178038 | ALS2CL | -0,498578217 | 0,707803982 | 0,3041 | 0,59411013 | ALS2 C-terminal like |
| ENSG00000100036 | SLC35E4 | -0,498624935 | 0,707781062 | 0,2193 | 0,53676938 | solute carrier family 35 member E4 |
| ENSG00000196116 | TDRD7 | -0,498771346 | 0,707709237 | 0,24403 | 0,55253387 | tudor domain containing 7 |
| ENSG00000268496 | AC245884,9 | -0,498906551 | 0,707642915 | 0,20377 | 0,52348797 | novel transcript, sense intronic to TTYH1 |
| ENSG00000100065 | CARD10 | -0,499081417 | 0,707557149 | 0,37747 | 0,64594766 | caspase recruitment domain family member 10 |
| ENSG00000166896 | ATP23 | -0,499111682 | 0,707542306 | 0,04722 | 0,29217708 | ATP23 metallopeptidase and ATP synthase assembly factor homolog |
| ENSG00000236540 | AC006547,1 | -0,49934023 | 0,707430227 | 0,54073 | 0,75849891 | novel transcript, antisense to C22orf25 |
| ENSG00000128346 | C22orf23 | -0,499851961 | 0,707179343 | 0,09172 | 0,39166375 | chromosome 22 open reading frame 23 |
| ENSG00000286021 | LINC02822 | -0,500113331 | 0,707051236 | 0,15407 | 0,48233496 | long intergenic non-protein coding RNA 2822 |
| ENSG00000153253 | SCN3A | -0,50016313 | 0,707026831 | 0,3231 | 0,60589758 | sodium voltage-gated channel alpha subunit 3 |
| ENSG00000122735 | DNAI1 | -0,500168257 | 0,707024318 | 0,06996 | 0,34916596 | dynein axonemal intermediate chain 1 |
| ENSG00000275397 | DHRS11 | -0,500195548 | 0,707010944 | 0,38171 | 0,64905556 | dehydrogenase/reductase 11 |
| ENSG00000079432 | CIC | -0,500256864 | 0,706980896 | 0,14635 | 0,47433542 | capicua transcriptional repressor |
| ENSG00000130988 | RGN | -0,500367046 | 0,706926904 | 0,20371 | 0,52348472 | regucalcin |
| ENSG00000227036 | LINC00511 | -0,5003913 | 0,70691502 | 0,12406 | 0,44501797 | long intergenic non-protein coding RNA 511 |
| ENSG00000253898 | LINC01419 | -0,500408352 | 0,706906664 | 0,30323 | 0,59353249 | long intergenic non-protein coding RNA 1419 |
| ENSG00000230832 | SSBL4P | -0,500460597 | 0,706881065 | 0,55198 | 0,76558846 | SSB like 4, pseudogene |
| ENSG00000229321 | AC008269,1 | -0,500477879 | 0,706872598 | 0,16561 | 0,49182669 | novel transcript |
| ENSG00000235369 | RPL36AP15 | -0,500649513 | 0,706788508 | 0,42005 | 0,67554068 | ribosomal protein L36a pseudogene 15 |
| ENSG00000285344 | TRIM17 | -0,500750385 | 0,706739091 | 0,08331 | 0,37550713 | tripartite motif containing 17 |

| | | | | | | |
|---|---|---|---|---|---|---|
| ENSG00000162931 | TRIM17 | -0,500750385 | 0,706739091 | 0,08331 | 0,37550713 | tripartite motif containing 17 |
| ENSG00000170430 | MGMT | -0,500876802 | 0,706677166 | 0,11541 | 0,43113218 | O-6-methylguanine-DNA methyltransferase |
| ENSG00000234394 | AL353608,3 | -0,500916218 | 0,706657859 | 0,4964 | 0,72831792 | uncharacterized LOC440896 |
| ENSG00000267577 | AC010327,4 | -0,501145393 | 0,706545614 | 0,47644 | 0,71465252 | novel transcript, antisense to C19orf51 and SYT5 |
| ENSG00000189114 | BLOC1S3 | -0,501159885 | 0,706538516 | 0,22676 | 0,54154849 | biogenesis of lysosomal organelles complex 1 subunit 3 |
| ENSG00000285458 | AC093827,5 | -0,501358767 | 0,706441123 | 0,09954 | 0,40609117 | novel protein |
| ENSG00000270141 | TERC | -0,501383343 | 0,70642909 | 0,18469 | 0,50835796 | telomerase RNA component |
| ENSG00000284624 | AC092902,4 | -0,501425569 | 0,706408414 | 0,0941 | 0,39622576 | novel transcript |
| ENSG00000239508 | PLCH1-AS1 | -0,501537584 | 0,706353568 | 0,16427 | 0,49065086 | PLCH1 antisense RNA 1 |
| ENSG00000233750 | CICP27 | -0,50164174 | 0,706302574 | 0,09296 | 0,39384605 | capicua transcriptional repressor pseudogene 27 |
| ENSG00000271344 | AC018638,6 | -0,50167643 | 0,706285591 | 0,29005 | 0,58531219 | novel transcript |
| ENSG00000250656 | ST3GAL1P1 | -0,501790389 | 0,706229804 | 0,30693 | 0,59605674 | ST3GAL1 pseudogene 1 |
| ENSG00000258384 | RCCD1-AS1 | -0,501812278 | 0,706219089 | 0,43446 | 0,68505666 | RCCD1 and UNC45A antisense RNA 1 |
| ENSG00000160808 | MYL3 | -0,501833833 | 0,706208537 | 0,43739 | 0,68717176 | myosin light chain 3 |
| ENSG00000059145 | UNKL | -0,501862038 | 0,706194731 | 0,16263 | 0,48945635 | unk like zinc finger |
| ENSG00000233690 | EBAG9P1 | -0,501915041 | 0,706168787 | 0,37513 | 0,64350562 | EBAG9 pseudogene 1 |
| ENSG00000196167 | COLCA1 | -0,502172392 | 0,70604283 | 0,11399 | 0,42906718 | colorectal cancer associated 1 |
| ENSG00000182578 | CSF1R | -0,502172816 | 0,706042623 | 0,24998 | 0,55657567 | colony stimulating factor 1 receptor |
| ENSG00000287343 | AL022397,1 | -0,502228413 | 0,706015414 | 0,30651 | 0,59552485 | novel transcript, antisense to SYT14 |
| ENSG00000129467 | ADCY4 | -0,502251885 | 0,706003928 | 0,32986 | 0,61078709 | adenylate cyclase 4 |
| ENSG00000275450 | PTGER4P2-CD | -0,502292314 | 0,705984144 | 0,28648 | 0,5827707 | PTGER4P2-CDK2AP2P2 readthrough, transcribed pseudogene |
| ENSG00000204564 | C6orf136 | -0,502364888 | 0,70594863 | 0,15295 | 0,4811885 | chromosome 6 open reading frame 136 |
| ENSG00000286591 | AC006197,2 | -0,502557165 | 0,70585455 | 0,21101 | 0,52983259 | novel transcript |
| ENSG00000235448 | LURAP1L-AS1 | -0,502633047 | 0,705817425 | 0,50455 | 0,73328552 | LURAP1L antisense RNA 1 |
| ENSG00000282712 | DEAF1 | -0,50284135 | 0,705715523 | 0,52632 | 0,74973856 | DEAF1 transcription factor |
| ENSG00000106128 | GHRHR | -0,502844406 | 0,705714029 | 0,52844 | 0,75090627 | growth hormone releasing hormone receptor |
| ENSG00000262246 | CORO7 | -0,502899193 | 0,705687229 | 0,45615 | 0,69998075 | coronin 7 |
| ENSG00000160226 | CFAP410 | -0,503136477 | 0,705571172 | 0,14407 | 0,47164913 | cilia and flagella associated protein 410 |
| ENSG00000071909 | MYO3B | -0,503226675 | 0,705527061 | 0,47525 | 0,71382243 | myosin IIIB |
| ENSG00000048342 | CC2D2A | -0,503351859 | 0,705465844 | 0,12 | 0,43868155 | coiled-coil and C2 domain containing 2A |
| ENSG00000286116 | AL157394,3 | -0,503490015 | 0,705398291 | 0,53711 | 0,75575829 | FAS antisense RNA 1 |
| ENSG00000286835 | AL133477,1 | -0,503518116 | 0,705384551 | 0,35246 | 0,62742225 | novel transcript, sense intronic to ZNF367 |

| | | | | | | |
|---|---|---|---|---|---|---|
| ENSG00000110801 | PSMD9 | -0,503527253 | 0,705380084 | 0,21354 | 0,53275614 | proteasome 26S subunit, non-ATPase 9 |
| ENSG00000287417 | AC006059,4 | -0,503570288 | 0,705359043 | 0,11 | 0,42231571 | novel transcript, antisense to CCDC13 |
| ENSG00000157240 | FZD1 | -0,503768009 | 0,70526238 | 0,02304 | 0,20244972 | frizzled class receptor 1 |
| ENSG00000100379 | KCTD17 | -0,504170257 | 0,705065768 | 0,03838 | 0,26521398 | potassium channel tetramerization domain containing 17 |
| ENSG00000229071 | PRRT1 | -0,504362258 | 0,704971941 | 0,24725 | 0,55483158 | proline rich transmembrane protein 1 |
| ENSG00000250091 | DNAH10OS | -0,504413688 | 0,70494681 | 0,01137 | 0,13842343 | dynein axonemal heavy chain 10 opposite strand |
| ENSG00000101003 | GINS1 | -0,504413956 | 0,704946679 | 0,1853 | 0,50876814 | GINS complex subunit 1 |
| ENSG00000148604 | RGR | -0,504542737 | 0,704883756 | 0,50944 | 0,73727892 | retinal G protein coupled receptor |
| ENSG00000124251 | TP53TG5 | -0,50455703 | 0,704876772 | 0,17885 | 0,50335172 | TP53 target 5 |
| ENSG00000278932 | CR381653,1 | -0,504642717 | 0,704834908 | 0,25179 | 0,55773568 | novel transcript |
| ENSG00000244226 | ILF2P1 | -0,504743309 | 0,704785765 | 0,28487 | 0,5816767 | interleukin enhancer binding factor 2 pseudogene 1 |
| ENSG00000261061 | AC092718,4 | -0,504890883 | 0,704713676 | 0,16747 | 0,49362211 | novel transcript |
| ENSG00000204922 | UQCC3 | -0,505110993 | 0,704606167 | 0,18917 | 0,51249536 | ubiquinol-cytochrome c reductase complex assembly factor 3 |
| ENSG00000227861 | SAPCD1 | -0,505113084 | 0,704605146 | 0,36094 | 0,63311211 | suppressor APC domain containing 1 |
| ENSG00000234951 | SAPCD1 | -0,505113084 | 0,704605146 | 0,36094 | 0,63311211 | suppressor APC domain containing 1 |
| ENSG00000227074 | SAPCD1 | -0,505113084 | 0,704605146 | 0,36094 | 0,63311211 | suppressor APC domain containing 1 |
| ENSG00000160284 | SPATC1L | -0,505127736 | 0,70459799 | 0,07548 | 0,36080119 | spermatogenesis and centriole associated 1 like |
| ENSG00000272945 | AL356299,2 | -0,505166868 | 0,704578878 | 0,55024 | 0,76449335 | novel transcript |
| ENSG00000214826 | DDX12P | -0,505199163 | 0,704563106 | 0,12009 | 0,43868155 | DEAD/H-box helicase 12, pseudogene |
| ENSG00000168228 | ZCCHC4 | -0,505210445 | 0,704557597 | 0,19382 | 0,51547251 | zinc finger CCHC-type containing 4 |
| ENSG00000267255 | AC011498,3 | -0,505311534 | 0,704508231 | 0,35229 | 0,62740642 | novel transcript, antisense to CHAF1A |
| ENSG00000165996 | HACD1 | -0,505544348 | 0,70439455 | 0,08945 | 0,38678501 | 3-hydroxyacyl-CoA dehydratase 1 |
| ENSG00000281392 | LINC00506 | -0,505608508 | 0,704363225 | 0,13052 | 0,45477153 | long intergenic non-protein coding RNA 506 |
| ENSG00000165805 | C12orf50 | -0,505774609 | 0,704282134 | 0,07403 | 0,35746749 | chromosome 12 open reading frame 50 |
| ENSG00000225737 | SKIV2L | -0,505986209 | 0,704178845 | 0,1816 | 0,50490902 | Ski2 like RNA helicase |
| ENSG00000267390 | AC036176,1 | -0,506116405 | 0,704115299 | 0,22951 | 0,54305428 | novel transcript, antisense to KDSR |
| ENSG00000273091 | AP000255,1 | -0,506127919 | 0,70410968 | 0,45418 | 0,69818495 | novel transcript |
| ENSG00000254614 | AP003068,2 | -0,506128352 | 0,704109468 | 0,24719 | 0,55483158 | novel transcript, antisense to CAPN1 |
| ENSG00000133958 | UNC79 | -0,506150546 | 0,704098637 | 0,10254 | 0,41028143 | unc-79 homolog, NALCN channel complex subunit |
| ENSG00000243986 | ENO1P3 | -0,506160689 | 0,704093686 | 0,33605 | 0,6160164 | enolase 1 pseudogene 3 |
| ENSG00000144161 | ZC3H8 | -0,506318842 | 0,704016506 | 0,18941 | 0,51258824 | zinc finger CCCH-type containing 8 |
| ENSG00000160062 | ZBTB8A | -0,506383347 | 0,703985029 | 0,20816 | 0,52756478 | zinc finger and BTB domain containing 8A |

| | | | | | | |
|---|---|---|---|---|---|---|
| ENSG00000265666 | RARA-AS1 | -0,506396598 | 0,703978562 | 0,38564 | 0,65122858 | RARA antisense RNA 1 |
| ENSG00000213859 | KCTD11 | -0,506482066 | 0,703936859 | 0,13286 | 0,45832113 | potassium channel tetramerization domain containing 11 |
| ENSG00000288399 | KCTD11 | -0,506482066 | 0,703936859 | 0,13286 | 0,45832113 | potassium channel tetramerization domain containing 11 |
| ENSG00000129991 | TNNI3 | -0,506540368 | 0,703908412 | 0,32623 | 0,6085098 | troponin I3, cardiac type |
| ENSG00000230368 | FAM41C | -0,50654426 | 0,703906513 | 0,403 | 0,66399303 | family with sequence similarity 41 member C |
| ENSG00000231247 | PPP1R18 | -0,506726507 | 0,703817598 | 0,38194 | 0,64905556 | protein phosphatase 1 regulatory subunit 18 |
| ENSG00000151376 | ME3 | -0,507021532 | 0,703673685 | 0,3037 | 0,59385655 | malic enzyme 3 |
| ENSG00000100228 | RAB36 | -0,507084466 | 0,70364299 | 0,08818 | 0,38444882 | RAB36, member RAS oncogene family |
| ENSG00000228205 | AC131235,1 | -0,507271598 | 0,703551726 | 0,45165 | 0,69665972 | ribosomal protein S3 (RPS3) pseudogene |
| ENSG00000278017 | AC064801,1 | -0,507357907 | 0,703509638 | 0,53062 | 0,75212511 | novel transcript, antisense to ZCCHC2 |
| ENSG00000103510 | KAT8 | -0,507469276 | 0,703455332 | 0,17105 | 0,49641416 | lysine acetyltransferase 8 |
| ENSG00000235125 | NFKBIL1 | -0,507599763 | 0,70339171 | 0,09422 | 0,39648446 | NFKB inhibitor like 1 |
| ENSG00000088053 | GP6 | -0,507872194 | 0,703258898 | 0,65299 | 0,8262455 | glycoprotein VI platelet |
| ENSG00000278232 | CRHR1 | -0,508079617 | 0,703157794 | 0,39552 | 0,6582534 | corticotropin releasing hormone receptor 1 |
| ENSG00000264026 | LINC02003 | -0,508236617 | 0,703081278 | 0,63012 | 0,813276 | long intergenic non-protein coding RNA 2003 |
| ENSG00000172425 | TTC36 | -0,5082763 | 0,703061939 | 0,41431 | 0,67106864 | tetratricopeptide repeat domain 36 |
| ENSG00000229980 | TOB1-AS1 | -0,508285151 | 0,703057626 | 0,27672 | 0,57549599 | TOB1 antisense RNA 1 |
| ENSG00000178951 | ZBTB7A | -0,508366826 | 0,703017825 | 0,19629 | 0,51733996 | zinc finger and BTB domain containing 7A |
| ENSG00000272457 | AC113194,1 | -0,508656451 | 0,702876706 | 0,50848 | 0,7365844 | novel transcript, antisense to RGS20 |
| ENSG00000245888 | NSMCE1-DT | -0,508693167 | 0,702858819 | 0,45831 | 0,70119611 | NSMCE1 divergent transcript |
| ENSG00000167614 | TTYH1 | -0,508772499 | 0,70282017 | 0,10552 | 0,41477267 | tweety family member 1 |
| ENSG00000237594 | AP000251,1 | -0,508778461 | 0,702817266 | 0,26484 | 0,56748133 | uncharacterized LOC150051 |
| ENSG00000131409 | LRRC4B | -0,508817184 | 0,702798402 | 0,11442 | 0,42986344 | leucine rich repeat containing 4B |
| ENSG00000100263 | RHBDD3 | -0,508881141 | 0,702767246 | 0,14407 | 0,47164913 | rhomboid domain containing 3 |
| ENSG00000077522 | ACTN2 | -0,508906096 | 0,70275509 | 0,33156 | 0,61165554 | actinin alpha 2 |
| ENSG00000263220 | AC015727,1 | -0,508906256 | 0,702755012 | 0,4963 | 0,72831792 | novel transcript, antisense to RABEP1 |
| ENSG00000184164 | CRELD2 | -0,508985571 | 0,702716378 | 0,12272 | 0,44254346 | cysteine rich with EGF like domains 2 |
| ENSG00000262096 | PCDHB19P | -0,509162082 | 0,702630408 | 0,08639 | 0,38089807 | protocadherin beta 19 pseudogene |
| ENSG00000271978 | AL359643,2 | -0,509330879 | 0,702548204 | 0,22869 | 0,54252903 | novel transcript |
| ENSG00000130881 | LRP3 | -0,509338036 | 0,702544719 | 0,08269 | 0,37450725 | LDL receptor related protein 3 |
| ENSG00000215244 | LINC02649 | -0,509344958 | 0,702541347 | 0,39979 | 0,66193009 | long intergenic non-protein coding RNA 2649 |
| ENSG00000233396 | LINC01719 | -0,509569416 | 0,702432053 | 0,03877 | 0,26656704 | long intergenic non-protein coding RNA 1719 |
| ENSG00000184985 | SORCS2 | -0,509750794 | 0,702343748 | 0,09863 | 0,40451471 | sortilin related VPS10 domain containing receptor 2 |

| | | | | | | |
|---|---|---|---|---|---|---|
| ENSG00000226765 | RPSAP33 | -0,509823315 | 0,702308443 | 0,50156 | 0,73145255 | ribosomal protein SA pseudogene 33 |
| ENSG00000266903 | AC243964,3 | -0,509833973 | 0,702303255 | 0,38981 | 0,65396251 | novel transcript, antisense to CEACAM16, CEACAM19, PVR & IGSF23 |
| ENSG00000125505 | MBOAT7 | -0,509974924 | 0,702234643 | 0,17733 | 0,50187931 | membrane bound O-acyltransferase domain containing 7 |
| ENSG00000233670 | PIRT | -0,51003522 | 0,702205295 | 0,43242 | 0,68411566 | phosphoinositide interacting regulator of transient receptor potential channels |
| ENSG00000267041 | ZNF850 | -0,510420125 | 0,702017974 | 0,15893 | 0,48628471 | zinc finger protein 850 |
| ENSG00000140538 | NTRK3 | -0,510881574 | 0,701793469 | 0,15732 | 0,48443207 | neurotrophic receptor tyrosine kinase 3 |
| ENSG00000268560 | AC010636,1 | -0,51096552 | 0,701752634 | 0,51343 | 0,73977095 | novel transcript |
| ENSG00000272831 | AC027644,3 | -0,511003133 | 0,701734339 | 0,07332 | 0,35655334 | novel transcript, antisense to RABGEF1 |
| ENSG00000180535 | BHLHA15 | -0,51106562 | 0,701703946 | 0,404 | 0,66411876 | basic helix-loop-helix family member a15 |
| ENSG00000261302 | AC106779,1 | -0,511322709 | 0,701578913 | 0,54849 | 0,76372599 | novel transcript, antisense to NUP93 |
| ENSG00000153443 | UBALD1 | -0,511333633 | 0,701573601 | 0,17928 | 0,50368273 | UBA like domain containing 1 |
| ENSG00000225973 | PIGBOS1 | -0,511403397 | 0,701539676 | 0,17113 | 0,49641416 | PIGB opposite strand 1 |
| ENSG00000185513 | L3MBTL1 | -0,511410167 | 0,701536384 | 0,05772 | 0,32084684 | L3MBTL histone methyl-lysine binding protein 1 |
| ENSG00000185737 | NRG3 | -0,511625315 | 0,701431772 | 0,05251 | 0,30766476 | neuregulin 3 |
| ENSG00000271825 | AC019080,3 | -0,511707329 | 0,701391898 | 0,47947 | 0,71694889 | novel transcript, antisense to NFE2L2 |
| ENSG00000232021 | LEF1-AS1 | -0,511812795 | 0,701340626 | 0,42194 | 0,67643128 | LEF1 antisense RNA 1 |
| ENSG00000196365 | LONP1 | -0,511977546 | 0,70126054 | 0,13416 | 0,46036439 | lon peptidase 1, mitochondrial |
| ENSG00000261253 | AC137932,2 | -0,511993834 | 0,701252622 | 0,23431 | 0,54614318 | novel transcript, antisense to ANKRD11 |
| ENSG00000257894 | AC027288,3 | -0,512034737 | 0,701232741 | 0,04246 | 0,2782038 | novel transcript |
| ENSG00000197487 | GALP | -0,512100745 | 0,701200658 | 0,41782 | 0,67428013 | galanin like peptide |
| ENSG00000225339 | AL354740,1 | -0,512169595 | 0,701167195 | 0,31913 | 0,60410856 | novel transcript, antisense to C6orf1 |
| ENSG00000281948 | CES1P1 | -0,512207409 | 0,701148817 | 0,88361 | 0,94839496 | carboxylesterase 1 pseudogene 1 |
| ENSG00000260618 | AC025917,1 | -0,512271497 | 0,701117671 | 0,45335 | 0,69750019 | novel transcript, antisense to KIAA1370 |
| ENSG00000198720 | ANKRD13B | -0,512623102 | 0,70094682 | 0,00741 | 0,10952546 | ankyrin repeat domain 13B |
| ENSG00000007520 | TSR3 | -0,512743706 | 0,700888226 | 0,13841 | 0,46568278 | TSR3 ribosome maturation factor |
| ENSG00000101181 | MTG2 | -0,512788321 | 0,700866552 | 0,11089 | 0,42363726 | mitochondrial ribosome associated GTPase 2 |
| ENSG00000215840 | AL592295,1 | -0,512871916 | 0,700825942 | 0,53995 | 0,75807575 | ribosomal protein S2 (RPS2) pseudogene |
| ENSG00000163820 | FYCO1 | -0,513097693 | 0,700716273 | 0,00252 | 0,05844485 | FYVE and coiled-coil domain autophagy adaptor 1 |
| ENSG00000226854 | AL451105,1 | -0,513108606 | 0,700710973 | 0,17068 | 0,49641416 | novel transcript |
| ENSG00000241859 | ANOS2P | -0,513209556 | 0,700661944 | 0,11054 | 0,42322615 | anosmin 2, pseudogene |
| ENSG00000234534 | CSNK1G2P1 | -0,513330333 | 0,70060329 | 0,59149 | 0,79063841 | casein kinase 1 gamma 2 pseudogene 1 |

| | | | | | | |
|---|---|---|---|---|---|---|
| ENSG00000213018 | PABPN1P1 | -0,513462243 | 0,700539234 | 0,46912 | 0,7086228 | PABPN1 pseudogene 1 |
| ENSG00000265749 | AC135178,4 | -0,513641117 | 0,700452382 | 0,16725 | 0,49324785 | novel transcript, antisense to KRBA2 and RPL26 |
| ENSG00000128563 | PRKRIP1 | -0,513716155 | 0,700415951 | 0,10667 | 0,41648041 | PRKR interacting protein 1 |
| ENSG00000259163 | AL096869,2 | -0,513737418 | 0,700405628 | 0,29109 | 0,58613153 | novel transcript, antisense to TTC7B |
| ENSG00000173928 | SWSAP1 | -0,513768142 | 0,700390712 | 0,15395 | 0,48226548 | SWIM-type zinc finger 7 associated protein 1 |
| ENSG00000205837 | LINC00487 | -0,513780064 | 0,700384925 | 0,30782 | 0,59671558 | long intergenic non-protein coding RNA 487 |
| ENSG00000213801 | ZNF321P | -0,513984968 | 0,700285457 | 0,23954 | 0,5491566 | zinc finger protein 321, pseudogene |
| ENSG00000232804 | HSPA1B | -0,514120956 | 0,700219451 | 0,07057 | 0,3507077 | heat shock protein family A (Hsp70) member 1B |
| ENSG00000276007 | AC079414,3 | -0,51412895 | 0,700215572 | 0,34105 | 0,6192144 | novel transcript, antisense WWOX |
| ENSG00000233117 | LINC00702 | -0,514329452 | 0,700118264 | 0,2674 | 0,56973003 | long intergenic non-protein coding RNA 702 |
| ENSG00000250461 | AC122718,1 | -0,514459485 | 0,700055164 | 0,15502 | 0,48293856 | family with sequence similarity 133, member B (FAM133B) pseudogene |
| ENSG00000249691 | AC026117,1 | -0,514493767 | 0,700038529 | 0,44075 | 0,68917395 | developmental pluripotency associated 4 (DPPA4) pseudogene |
| ENSG00000206562 | METTL6 | -0,514724334 | 0,69992666 | 0,14873 | 0,47609532 | methyltransferase like 6 |
| ENSG00000142751 | GPN2 | -0,514783384 | 0,699898012 | 0,15675 | 0,48425071 | GPN-loop GTPase 2 |
| ENSG00000242816 | AC092691,3 | -0,515343088 | 0,699626535 | 0,42797 | 0,68087556 | novel transcript |
| ENSG00000143198 | MGST3 | -0,515433767 | 0,699582562 | 0,02847 | 0,22807919 | microsomal glutathione S-transferase 3 |
| ENSG00000077080 | ACTL6B | -0,515439549 | 0,699579758 | 0,23013 | 0,54335232 | actin like 6B |
| ENSG00000272645 | GTF2IP20 | -0,515536437 | 0,699532777 | 0,10817 | 0,41884147 | general transcription factor IIi pseudogene 20 |
| ENSG00000087269 | NOP14 | -0,515633909 | 0,699485517 | 0,2034 | 0,52321822 | NOP14 nucleolar protein |
| ENSG00000286980 | AC020584,1 | -0,515650225 | 0,699477606 | 0,53068 | 0,75212511 | novel transcript |
| ENSG00000205710 | C17orf107 | -0,515875185 | 0,699368545 | 0,36008 | 0,63261458 | chromosome 17 open reading frame 107 |
| ENSG00000225742 | LINC02036 | -0,515875652 | 0,699368318 | 0,44108 | 0,68940607 | long intergenic non-protein coding RNA 2036 |
| ENSG00000123815 | COQ8B | -0,516324471 | 0,69915078 | 0,13364 | 0,45954133 | coenzyme Q8B |
| ENSG00000139626 | ITGB7 | -0,51644135 | 0,699094141 | 0,68464 | 0,84569463 | integrin subunit beta 7 |
| ENSG00000172264 | MACROD2 | -0,516456698 | 0,699086704 | 0,02939 | 0,23068546 | mono-ADP ribosylhydrolase 2 |
| ENSG00000121764 | HCRTR1 | -0,516558727 | 0,699037266 | 0,31412 | 0,60087803 | hypocretin receptor 1 |
| ENSG00000270184 | AC018695,4 | -0,516587311 | 0,699023416 | 0,34275 | 0,62031192 | novel transcript, antisense to COX4NB |
| ENSG00000261140 | AC093525,4 | -0,516654936 | 0,698990651 | 0,27638 | 0,57542877 | novel transcript, antisense to PDPK1 |
| ENSG00000233242 | AL512604,2 | -0,516703674 | 0,698967037 | 0,45782 | 0,70096886 | novel transcript |
| ENSG00000273032 | DGCR5 | -0,516840373 | 0,698900812 | 0,03797 | 0,26375851 | DiGeorge syndrome critical region gene 5 |
| ENSG00000273507 | AL354809,1 | -0,516987915 | 0,69882934 | 0,10675 | 0,41660052 | novel transcript |

| | | | | | | |
|---|---|---|---|---|---|---|
| ENSG00000160799 | CCDC12 | -0,516995841 | 0,698825501 | 0,15133 | 0,47956749 | coiled-coil domain containing 12 |
| ENSG00000156709 | AIFM1 | -0,517006822 | 0,698820181 | 0,20947 | 0,52856218 | apoptosis inducing factor mitochondria associated 1 |
| ENSG00000255571 | MIR9-3HG | -0,517093424 | 0,698778234 | 0,48506 | 0,72069686 | MIR9-3 host gene |
| ENSG00000284491 | THSD8 | -0,517117543 | 0,698766552 | 0,39376 | 0,65672538 | thrombospondin type 1 domain containing 8 |
| ENSG00000288032 | AC104472,5 | -0,517217033 | 0,698718366 | 0,23536 | 0,54680313 | novel transcript, sense intronic to GMPS |
| ENSG00000214283 | RAD51AP1P1 | -0,517236602 | 0,698708888 | 0,35975 | 0,63246012 | RAD51AP1 pseudogene 1 |
| ENSG00000253669 | GASAL1 | -0,517340174 | 0,698658729 | 0,54855 | 0,76372599 | growth arrest associated lncRNA 1 |
| ENSG00000235478 | LINC01664 | -0,517489097 | 0,698586614 | 0,54408 | 0,76034748 | long intergenic non-protein coding RNA 1664 |
| ENSG00000157064 | NMNAT2 | -0,517494673 | 0,698583914 | 0,26869 | 0,57073706 | nicotinamide nucleotide adenylyltransferase 2 |
| ENSG00000283020 | DUX4L37 | -0,517575341 | 0,698544854 | 0,36814 | 0,6380264 | double homeobox 4 like 37 (pseudogene) |
| ENSG00000145736 | GTF2H2 | -0,517624986 | 0,698520816 | 0,1662 | 0,49212363 | general transcription factor IIH subunit 2 |
| ENSG00000173267 | SNCG | -0,517892776 | 0,69839117 | 0,10297 | 0,41078404 | synuclein gamma |
| ENSG00000213347 | MXD3 | -0,517907603 | 0,698383993 | 0,024 | 0,20707741 | MAX dimerization protein 3 |
| ENSG00000235360 | C6orf47 | -0,517949554 | 0,698363685 | 0,08873 | 0,38517778 | chromosome 6 open reading frame 47 |
| ENSG00000235569 | MSH5 | -0,518068217 | 0,698306246 | 0,40591 | 0,66554977 | mutS homolog 5 |
| ENSG00000214290 | COLCA2 | -0,518072377 | 0,698304233 | 0,14524 | 0,47289051 | colorectal cancer associated 2 |
| ENSG00000235440 | RPS2P15 | -0,518236934 | 0,698224587 | 0,41803 | 0,67432365 | ribosomal protein S2 pseudogene 15 |
| ENSG00000224533 | TMLHE-AS1 | -0,518266386 | 0,698210333 | 0,37266 | 0,64161496 | TMLHE antisense RNA 1 |
| ENSG00000179627 | ZBTB42 | -0,518362525 | 0,698163807 | 0,25147 | 0,55731053 | zinc finger and BTB domain containing 42 |
| ENSG00000167549 | CORO6 | -0,51863136 | 0,698033722 | 0,14027 | 0,46714558 | coronin 6 |
| ENSG00000262619 | LINC00621 | -0,518961312 | 0,697874096 | 0,15268 | 0,48061452 | long intergenic non-protein coding RNA 621 |
| ENSG00000237158 | AL353718,1 | -0,519005243 | 0,697852846 | 0,32018 | 0,60511306 | pseudogene similar to part of DCN1, defective in cullin neddylation 1, domain containing 1 (S, cerevisiae) DCUN1D1 |
| ENSG00000003402 | CFLAR | -0,519079501 | 0,697816927 | 0,06841 | 0,34567487 | CASP8 and FADD like apoptosis regulator |
| ENSG00000167525 | PROCA1 | -0,519088363 | 0,697812641 | 0,0542 | 0,31168784 | protein interacting with cyclin A1 |
| ENSG00000135631 | RAB11FIP5 | -0,519113488 | 0,697800488 | 0,05465 | 0,31304183 | RAB11 family interacting protein 5 |
| ENSG00000225870 | LINC00368 | -0,519316445 | 0,697702329 | 0,52353 | 0,74740993 | long intergenic non-protein coding RNA 368 |
| ENSG00000273850 | PPP2R3B | -0,51935123 | 0,697685507 | 0,19781 | 0,51942408 | protein phosphatase 2 regulatory subunit B''beta |
| ENSG00000153485 | TMEM251 | -0,519370641 | 0,69767612 | 0,04441 | 0,28428403 | transmembrane protein 251 |
| ENSG00000204950 | LRRC10B | -0,519372965 | 0,697674996 | 0,15139 | 0,47956749 | leucine rich repeat containing 10B |
| ENSG00000147174 | GCNA | -0,519505876 | 0,697610724 | 0,08164 | 0,37277532 | germ cell nuclear acidic peptidase |
| ENSG00000179314 | WSCD1 | -0,519584511 | 0,697572702 | 0,06635 | 0,34103636 | WSC domain containing 1 |

| | | | | | | |
|---|---|---|---|---|---|---|
| ENSG00000185800 | DMWD | -0,519614641 | 0,697558133 | 0,07927 | 0,36776284 | DM1 locus, WD repeat containing |
| ENSG00000224238 | WARS2-IT1 | -0,520142905 | 0,697302759 | 0,16198 | 0,48916554 | WARS2 intronic transcript 1 |
| ENSG00000249685 | AC079921,2 | -0,520242845 | 0,697254456 | 0,42683 | 0,68017963 | novel transcript |
| ENSG00000258212 | ZNF75BP | -0,520354919 | 0,697200293 | 0,43358 | 0,68473723 | zinc finger protein 75B, pseudogene |
| ENSG00000242086 | MUC20-OT1 | -0,520467874 | 0,697145708 | 0,24119 | 0,55024202 | MUC20 overlapping transcript |
| ENSG00000269343 | ZNF587B | -0,520501946 | 0,697129244 | 0,21306 | 0,53205351 | zinc finger protein 587B |
| ENSG00000215068 | AC025171,2 | -0,520640668 | 0,697062215 | 0,11486 | 0,43031399 | novel transcript, antisense to ANXA2R |
| ENSG00000174943 | KCTD13 | -0,520642347 | 0,697061404 | 0,17361 | 0,49838936 | potassium channel tetramerization domain containing 13 |
| ENSG00000206952 | SNORA50A | -0,520683241 | 0,697041646 | 0,45331 | 0,69750019 | small nucleolar RNA, H/ACA box 50A |
| ENSG00000184194 | GPR173 | -0,520703104 | 0,697032049 | 0,08049 | 0,37018111 | G protein-coupled receptor 173 |
| ENSG00000140548 | ZNF710 | -0,520794829 | 0,696987734 | 0,00894 | 0,11991469 | zinc finger protein 710 |
| ENSG00000276900 | AC023157,2 | -0,520870224 | 0,69695131 | 0,1924 | 0,514872 | novel transcript, antisense to AMN1 |
| ENSG00000146166 | LGSN | -0,520905029 | 0,696934496 | 0,43467 | 0,68529786 | lengsin, lens protein with glutamine synthetase domain |
| ENSG00000225224 | RPS27AP12 | -0,521049816 | 0,696864556 | 0,21067 | 0,52921344 | RPS27A pseudogene 12 |
| ENSG00000273174 | AC108673,2 | -0,521056213 | 0,696861466 | 0,58684 | 0,78701992 | novel transcript, antisense to RAB43 |
| ENSG00000177736 | FXNP2 | -0,521163556 | 0,696809619 | 0,42229 | 0,67663581 | frataxin pseudogene 2 |
| ENSG00000259104 | PTCSC3 | -0,521286973 | 0,696750012 | 0,21027 | 0,5290538 | papillary thyroid carcinoma susceptibility candidate 3 |
| ENSG00000197808 | ZNF461 | -0,521348875 | 0,696720117 | 0,19003 | 0,51331106 | zinc finger protein 461 |
| ENSG00000226530 | AL158055,1 | -0,521362229 | 0,696713668 | 0,13523 | 0,46165567 | novel transcript |
| ENSG00000106785 | TRIM14 | -0,521498622 | 0,696647803 | 0,0946 | 0,39709067 | tripartite motif containing 14 |
| ENSG00000146006 | LRRTM2 | -0,52153218 | 0,696631599 | 0,2329 | 0,54547822 | leucine rich repeat transmembrane neuronal 2 |
| ENSG00000245466 | AL357153,1 | -0,521540748 | 0,696627462 | 0,50885 | 0,7369661 | novel transcript |
| ENSG00000258316 | KLF17P1 | -0,521589784 | 0,696603785 | 0,49904 | 0,73002372 | Kruppel like factor 17 pseudogene 1 |
| ENSG00000154736 | ADAMTS5 | -0,521612577 | 0,696592779 | 0,24513 | 0,55295131 | ADAM metallopeptidase with thrombospondin type 1 motif 5 |
| ENSG00000286445 | AL355334,2 | -0,521631842 | 0,696583477 | 0,24011 | 0,54943734 | novel transcript |
| ENSG00000128805 | ARHGAP22 | -0,521737119 | 0,696532647 | 0,11861 | 0,43646336 | Rho GTPase activating protein 22 |
| ENSG00000188766 | SPRED3 | -0,521882123 | 0,696462643 | 0,12247 | 0,4421901 | sprouty related EVH1 domain containing 3 |
| ENSG00000135363 | LMO2 | -0,521999285 | 0,696406085 | 0,26166 | 0,56452671 | LIM domain only 2 |
| ENSG00000285849 | AL021918,4 | -0,522107362 | 0,696353917 | 0,20809 | 0,52756478 | novel transcript, antisense to ZNF184 |
| ENSG00000156869 | FRRS1 | -0,522260685 | 0,696279916 | 0,29733 | 0,59031059 | ferric chelate reductase 1 |
| ENSG00000260232 | PWRN4 | -0,522549513 | 0,696140535 | 0,20358 | 0,52343204 | Prader-Willi region non-protein coding RNA 4 |
| ENSG00000251562 | MALAT1 | -0,5227572 | 0,696040327 | 0,02703 | 0,22193038 | metastasis associated lung adenocarcinoma transcript 1 |

| | | | | | | |
|---|---|---|---|---|---|---|
| ENSG00000167523 | SPATA33 | -0,522876374 | 0,695982833 | 0,11713 | 0,43444592 | spermatogenesis associated 33 |
| ENSG00000253641 | LINCR-0001 | -0,523078004 | 0,69588557 | 0,5357 | 0,75485762 | uncharacterized LINCR-0001 |
| ENSG00000257989 | AC078864,1 | -0,52318006 | 0,695836344 | 0,73622 | 0,87519603 | novel transcript |
| ENSG00000129480 | DTD2 | -0,523263394 | 0,695796152 | 0,11779 | 0,43532264 | D-aminoacyl-tRNA deacylase 2 |
| ENSG00000268350 | FAM156A | -0,523365455 | 0,695746931 | 0,38444 | 0,65052153 | family with sequence similarity 156 member A |
| ENSG00000234614 | C2CD4D-AS1 | -0,523427696 | 0,695716916 | 0,56699 | 0,77484339 | C2CD4D and THEM5 antisense RNA 1 |
| ENSG00000243243 | AC073130,2 | -0,523473943 | 0,695694614 | 0,44449 | 0,69185302 | novel transcript |
| ENSG00000105088 | OLFM2 | -0,523880577 | 0,695498555 | 0,0679 | 0,34439007 | olfactomedin 2 |
| ENSG00000237329 | AL356320,2 | -0,52428976 | 0,695301323 | 0,30385 | 0,59400398 | TAF9 RNA polymerase II, TATA box binding protein (TBP)-associated factor, 32kDa (TAF9) pseudogene |
| ENSG00000250073 | AP000866,2 | -0,524347514 | 0,695273489 | 0,13873 | 0,46603966 | novel transcript, antisense to ESAM |
| ENSG00000287528 | AL121749,2 | -0,524354939 | 0,695269911 | 0,41643 | 0,67315741 | novel transcript, antisense to FZD8 |
| ENSG00000206127 | GOLGA8O | -0,524699988 | 0,695103644 | 0,18628 | 0,50961491 | golgin A8 family member O |
| ENSG00000180934 | OR56A1 | -0,524874709 | 0,695019466 | 0,18459 | 0,50835796 | olfactory receptor family 56 subfamily A member 1 |
| ENSG00000175497 | DPP10 | -0,524924238 | 0,694995606 | 0,01765 | 0,1750863 | dipeptidyl peptidase like 10 |
| ENSG00000259459 | LINC02568 | -0,525006099 | 0,694956172 | 0,59326 | 0,79162636 | long intergenic non-protein coding RNA 2568 |
| ENSG00000177133 | PRDM16-DT | -0,525148487 | 0,694887586 | 0,26016 | 0,56382871 | PRDM16 divergent transcript |
| ENSG00000235128 | AC013474,1 | -0,525169956 | 0,694877245 | 0,42207 | 0,67659005 | novel pseudogene |
| ENSG00000105245 | NUMBL | -0,525176161 | 0,694874257 | 0,11571 | 0,43162108 | NUMB like endocytic adaptor protein |
| ENSG00000238045 | AC009133,1 | -0,525259008 | 0,694834355 | 0,0234 | 0,2040793 | novel transcript, antisense to C16orf53 and MVP |
| ENSG00000273007 | AC021205,3 | -0,525319725 | 0,694805113 | 0,16338 | 0,48977932 | novel transcript, antisense to FGF2 |
| ENSG00000249695 | AC026369,1 | -0,525570727 | 0,69468424 | 0,58374 | 0,78560543 | novel transcript, antisense to IQSEC3 |
| ENSG00000263137 | AC026369,1 | -0,525570727 | 0,69468424 | 0,58374 | 0,78560543 | novel transcript, antisense to IQSEC3 |
| ENSG00000073169 | SELENOO | -0,525628379 | 0,69465648 | 0,08154 | 0,37260031 | selenoprotein O |
| ENSG00000285725 | AC004967,2 | -0,525633409 | 0,694654058 | 0,13793 | 0,46505993 | novel transcript |
| ENSG00000259946 | BX005019,1 | -0,525634817 | 0,69465338 | 0,22611 | 0,54121239 | novel transcript |
| ENSG00000136872 | ALDOB | -0,525692082 | 0,694625808 | 0,23711 | 0,54809872 | aldolase, fructose-bisphosphate B |
| ENSG00000231532 | LINC01249 | -0,525763511 | 0,694591417 | 0,50516 | 0,73378796 | long intergenic non-protein coding RNA 1249 |
| ENSG00000226618 | PRRC2A | -0,525898329 | 0,694526512 | 0,56253 | 0,77186635 | proline rich coiled-coil 2A |
| ENSG00000231087 | FDPSP7 | -0,526337384 | 0,694315179 | 0,47278 | 0,71206872 | farnesyl diphosphate synthase pseudogene 7 |
| ENSG00000134285 | FKBP11 | -0,526445112 | 0,694263335 | 0,10094 | 0,40770184 | FKBP prolyl isomerase 11 |
| ENSG00000228835 | AC012123,1 | -0,526574219 | 0,694201208 | 0,3661 | 0,63617598 | novel transcript, antisense to KLHL14 |
| ENSG00000240869 | RN7SL128P | -0,52661883 | 0,694179742 | 0,28504 | 0,58171017 | RNA, 7SL, cytoplasmic 128, pseudogene |

| | | | | | | |
|---|---|---|---|---|---|---|
| ENSG00000161980 | POLR3K | -0,526745466 | 0,694118812 | 0,22148 | 0,53805234 | RNA polymerase III subunit K |
| ENSG00000227107 | AC104667,1 | -0,526793259 | 0,694095818 | 0,43695 | 0,6870169 | novel transcript |
| ENSG00000079277 | MKNK1 | -0,526892629 | 0,694048011 | 0,19444 | 0,51562158 | MAPK interacting serine/threonine kinase 1 |
| ENSG00000254305 | MRPL9P1 | -0,526923088 | 0,694033358 | 0,44161 | 0,68981532 | mitochondrial ribosomal protein L9 pseudogene 1 |
| ENSG00000080031 | PTPRH | -0,527426853 | 0,693791056 | 0,2619 | 0,56477027 | protein tyrosine phosphatase receptor type H |
| ENSG00000226581 | LINC02848 | -0,527429453 | 0,693789805 | 0,25299 | 0,55809014 | long intergenic non-protein coding RNA 2848 |
| ENSG00000122490 | SLC66A2 | -0,528093961 | 0,693470318 | 0,15453 | 0,48266733 | solute carrier family 66 member 2 |
| ENSG00000261420 | AL022069,1 | -0,528477864 | 0,69328581 | 0,39591 | 0,65863599 | novel transcript, antisense to BRP44L |
| ENSG00000187796 | CARD9 | -0,528523335 | 0,693263959 | 0,22227 | 0,53859118 | caspase recruitment domain family member 9 |
| ENSG00000242634 | RPS24P16 | -0,528545745 | 0,69325319 | 0,53591 | 0,75497727 | ribosomal protein S24 pseudogene 16 |
| ENSG00000285756 | BX890604,1 | -0,528554544 | 0,693248962 | 0,00021 | 0,01067558 | novel transcript |
| ENSG00000187045 | TMPRSS6 | -0,528631715 | 0,69321188 | 0,31454 | 0,60122508 | transmembrane serine protease 6 |
| ENSG00000288595 | AC236668,1 | -0,52867124 | 0,693192889 | 0,55526 | 0,76788267 | novel transcript |
| ENSG00000164796 | CSMD3 | -0,52870732 | 0,693175553 | 0,30301 | 0,59337005 | CUB and Sushi multiple domains 3 |
| ENSG00000165434 | PGM2L1 | -0,528790451 | 0,693135612 | 0,08102 | 0,37153397 | phosphoglucomutase 2 like 1 |
| ENSG00000006756 | ARSD | -0,528856982 | 0,693103648 | 0,11331 | 0,42797362 | arylsulfatase D |
| ENSG00000239415 | AP001469,3 | -0,529090955 | 0,692991252 | 0,03666 | 0,25954613 | novel transcript, antisense to MCM3APAS |
| ENSG00000286523 | AL022151,1 | -0,529132556 | 0,692971269 | 0,07818 | 0,36540845 | novel transcript |
| ENSG00000227740 | LINC02803 | -0,529173636 | 0,692951538 | 0,26662 | 0,5691939 | long intergenic non-protein coding RNA 2803 |
| ENSG00000184486 | POU3F2 | -0,529229372 | 0,692924767 | 0,34284 | 0,62033803 | POU class 3 homeobox 2 |
| ENSG00000272983 | AL117339,3 | -0,529307147 | 0,692887413 | 0,15653 | 0,48425071 | novel transcript |
| ENSG00000197332 | AC008543,1 | -0,529320934 | 0,692880791 | 0,13393 | 0,46007586 | novel transcript |
| ENSG00000233251 | AC007743,1 | -0,529360635 | 0,692861725 | 0,1291 | 0,4524852 | novel transcript, antisense to CCDC85A |
| ENSG00000223551 | TMSB4XP4 | -0,529363846 | 0,692860182 | 0,31225 | 0,59925736 | TMSB4X pseudogene 4 |
| ENSG00000154222 | CC2D1B | -0,529377522 | 0,692853614 | 0,13912 | 0,46648276 | coiled-coil and C2 domain containing 1B |
| ENSG00000184209 | SNRNP35 | -0,529705196 | 0,692696267 | 0,14159 | 0,46809977 | small nuclear ribonucleoprotein U11/U12 subunit 35 |
| ENSG00000169918 | OTUD7A | -0,529738716 | 0,692680173 | 0,05537 | 0,31482368 | OTU deubiquitinase 7A |
| ENSG00000140829 | DHX38 | -0,529786502 | 0,69265723 | 0,06128 | 0,32861192 | DEAH-box helicase 38 |
| ENSG00000128891 | CCDC32 | -0,52986604 | 0,692619043 | 0,15803 | 0,48506908 | coiled-coil domain containing 32 |
| ENSG00000123200 | ZC3H13 | -0,52989112 | 0,692607003 | 0,03802 | 0,26375851 | zinc finger CCCH-type containing 13 |
| ENSG00000188659 | SAXO2 | -0,530113915 | 0,692500052 | 0,11235 | 0,42601117 | stabilizer of axonemal microtubules 2 |
| ENSG00000075399 | VPS9D1 | -0,530122977 | 0,692495703 | 0,07067 | 0,35071472 | VPS9 domain containing 1 |
| ENSG00000269978 | AL359881,1 | -0,530258208 | 0,692430794 | 0,51043 | 0,73798863 | novel transcript |

| | | | | | | |
|---|---|---|---|---|---|---|
| ENSG00000287377 | AC090948,4 | -0,530346846 | 0,692388253 | 0,27208 | 0,57306433 | novel transcript, antisense to RFTN1 |
| ENSG00000283046 | HTN1 | -0,530354068 | 0,692384787 | 0,25074 | 0,55690541 | histatin 1 |
| ENSG00000126550 | HTN1 | -0,530354068 | 0,692384787 | 0,25074 | 0,55690541 | histatin 1 |
| ENSG00000231609 | AC007098,1 | -0,530374166 | 0,692375142 | 0,25193 | 0,55777361 | novel transcript, antisense to EHBP1 |
| ENSG00000271937 | AC104187,1 | -0,530569076 | 0,692281607 | 0,44523 | 0,69227211 | novel transcript, antisense to TCAIM |
| ENSG00000160469 | BRSK1 | -0,530758964 | 0,692190495 | 0,11783 | 0,43532264 | BR serine/threonine kinase 1 |
| ENSG00000263620 | AC129492,3 | -0,530785165 | 0,692177924 | 0,34693 | 0,62365773 | novel protein |
| ENSG00000105472 | CLEC11A | -0,530922956 | 0,692111818 | 0,17499 | 0,49911246 | C-type lectin domain containing 11A |
| ENSG00000144230 | GPR17 | -0,530952647 | 0,692097574 | 0,56058 | 0,7708846 | G protein-coupled receptor 17 |
| ENSG00000147027 | TMEM47 | -0,53098446 | 0,692082313 | 0,07046 | 0,35048491 | transmembrane protein 47 |
| ENSG00000272333 | KMT2B | -0,531077819 | 0,692037529 | 0,11291 | 0,42702832 | lysine methyltransferase 2B |
| ENSG00000229118 | AC068491,1 | -0,531403245 | 0,691881444 | 0,54061 | 0,75840587 | novel transcript |
| ENSG00000250303 | LINC02762 | -0,531421406 | 0,691872735 | 0,12136 | 0,44078088 | long intergenic non-protein coding RNA 2762 |
| ENSG00000174013 | FBXO45 | -0,531567215 | 0,691802813 | 0,15253 | 0,48061452 | F-box protein 45 |
| ENSG00000287287 | AL133476,1 | -0,531600606 | 0,691786802 | 0,48004 | 0,71732148 | novel transcript, antisense to RUSC2 |
| ENSG00000262420 | AC007613,1 | -0,532268056 | 0,691466826 | 0,42755 | 0,68046596 | novel transcript, antisense to TXNDC11 |
| ENSG00000260582 | TPST2P1 | -0,532319101 | 0,691442362 | 0,64202 | 0,8205913 | tyrosylprotein sulfotransferase 2 pseudogene 1 |
| ENSG00000204967 | PCDHA4 | -0,532388529 | 0,691409087 | 0,00112 | 0,03489155 | protocadherin alpha 4 |
| ENSG00000253661 | ZFHX4-AS1 | -0,532471437 | 0,691369355 | 0,19426 | 0,51562158 | ZFHX4 antisense RNA 1 |
| ENSG00000267136 | AP005131,1 | -0,532627742 | 0,691294455 | 0,18087 | 0,50443353 | novel transcript, antisense C18orf1 |
| ENSG00000233084 | RPL23AP25 | -0,532713726 | 0,691253255 | 0,77845 | 0,89745058 | ribosomal protein L23a pseudogene 25 |
| ENSG00000260343 | LINC01043 | -0,532750775 | 0,691235503 | 0,23173 | 0,54442759 | long intergenic non-protein coding RNA 1043 |
| ENSG00000260086 | AC007611,1 | -0,532764534 | 0,691228911 | 0,51631 | 0,74198903 | novel transcript |
| ENSG00000233056 | ERVH48-1 | -0,533043803 | 0,69109512 | 0,19082 | 0,51394498 | endogenous retrovirus group 48 member 1 |
| ENSG00000178935 | ZNF552 | -0,533122498 | 0,691057424 | 0,12501 | 0,44663415 | zinc finger protein 552 |
| ENSG00000149823 | VPS51 | -0,533124095 | 0,691056658 | 0,09684 | 0,40095852 | VPS51 subunit of GARP complex |
| ENSG00000198768 | APCDD1L | -0,533125753 | 0,691055864 | 0,29786 | 0,59037822 | APC down-regulated 1 like |
| ENSG00000103154 | NECAB2 | -0,533352925 | 0,690947057 | 0,25263 | 0,55796133 | N-terminal EF-hand calcium binding protein 2 |
| ENSG00000177994 | C2orf73 | -0,533358825 | 0,690944231 | 0,4684 | 0,7080558 | chromosome 2 open reading frame 73 |
| ENSG00000178162 | FAR2P2 | -0,533508665 | 0,690872473 | 0,1728 | 0,49764744 | fatty acyl-CoA reductase 2 pseudogene 2 |
| ENSG00000238169 | LINC01053 | -0,533663255 | 0,690798447 | 0,38642 | 0,65185547 | long intergenic non-protein coding RNA 1053 |
| ENSG00000185052 | SLC24A3 | -0,533666433 | 0,690796926 | 0,32464 | 0,60720224 | solute carrier family 24 member 3 |
| ENSG00000231212 | AL590399,3 | -0,533839548 | 0,690714039 | 0,36018 | 0,63261458 | novel transcript |

| | | | | | | |
|---|---|---|---|---|---|---|
| ENSG00000100151 | PICK1 | -0,533908 | 0,690681267 | 0,06153 | 0,32865897 | protein interacting with PRKCA 1 |
| ENSG00000124587 | PEX6 | -0,534072484 | 0,690602526 | 0,10049 | 0,40712841 | peroxisomal biogenesis factor 6 |
| ENSG00000236819 | LINC01563 | -0,534088023 | 0,690595087 | 0,29394 | 0,58836931 | long intergenic non-protein coding RNA 1563 |
| ENSG00000257815 | PRANCR | -0,534166769 | 0,690557394 | 0,19697 | 0,51824743 | progenitor renewal associated non-coding RNA |
| ENSG00000236154 | AL450311,1 | -0,534304036 | 0,690491693 | 0,50384 | 0,73294771 | novel transcript |
| ENSG00000167100 | SAMD14 | -0,53430892 | 0,690489356 | 0,10082 | 0,40770184 | sterile alpha motif domain containing 14 |
| ENSG00000281603 | MUC20-OT1 | -0,534431805 | 0,690430544 | 0,14475 | 0,47232606 | MUC20 overlapping transcript |
| ENSG00000286122 | AC016074,2 | -0,534456017 | 0,690418957 | 0,08194 | 0,37306719 | novel transcript |
| ENSG00000185022 | MAFF | -0,534507689 | 0,690394229 | 0,19393 | 0,51547251 | MAF bZIP transcription factor F |
| ENSG00000280616 | VPS11 | -0,53459335 | 0,690353238 | 0,2794 | 0,57742338 | VPS11 core subunit of CORVET and HOPS complexes |
| ENSG00000267317 | AC027307,2 | -0,534635113 | 0,690333254 | 0,20823 | 0,52760275 | novel transcript, antisense to APC2 |
| ENSG00000124523 | SIRT5 | -0,534688099 | 0,690307901 | 0,15626 | 0,4840074 | sirtuin 5 |
| ENSG00000106868 | SUSD1 | -0,534699518 | 0,690302437 | 0,13715 | 0,4642578 | sushi domain containing 1 |
| ENSG00000143318 | CASQ1 | -0,534915471 | 0,690199115 | 0,11982 | 0,43834197 | calsequestrin 1 |
| ENSG00000160392 | C19orf47 | -0,535366103 | 0,689983562 | 0,15507 | 0,48296753 | chromosome 19 open reading frame 47 |
| ENSG00000258789 | AL162171,1 | -0,535417989 | 0,689958748 | 0,32154 | 0,60577115 | novel transcript, antisense to PTPN21 |
| ENSG00000286581 | AL450992,3 | -0,535538464 | 0,689901134 | 0,48889 | 0,72355061 | novel transcript |
| ENSG00000138641 | HERC3 | -0,535613934 | 0,689865045 | 0,09729 | 0,40183698 | HECT and RLD domain containing E3 ubiquitin protein ligase 3 |
| ENSG00000107815 | TWNK | -0,535618811 | 0,689862713 | 0,12878 | 0,45206676 | twinkle mtDNA helicase |
| ENSG00000275512 | AC007998,4 | -0,535725401 | 0,689811746 | 0,08519 | 0,37857645 | novel transcript |
| ENSG00000187954 | CYHR1 | -0,535741563 | 0,689804018 | 0,08657 | 0,38115611 | cysteine and histidine rich 1 |
| ENSG00000110057 | UNC93B1 | -0,535891725 | 0,689732224 | 0,22442 | 0,54033281 | unc-93 homolog B1, TLR signaling regulator |
| ENSG00000186329 | TMEM212 | -0,536019444 | 0,689671166 | 0,06346 | 0,33383762 | transmembrane protein 212 |
| ENSG00000282885 | AL627171,2 | -0,536140875 | 0,689613119 | 0,07872 | 0,36645437 | novel transcript |
| ENSG00000158526 | TSR2 | -0,5361444 | 0,689611434 | 0,15245 | 0,48061452 | TSR2 ribosome maturation factor |
| ENSG00000106080 | FKBP14 | -0,536214783 | 0,689577791 | 0,20623 | 0,52594338 | FKBP prolyl isomerase 14 |
| ENSG00000160117 | ANKLE1 | -0,536319506 | 0,689527738 | 0,2588 | 0,56260423 | ankyrin repeat and LEM domain containing 1 |
| ENSG00000287014 | AC087883,2 | -0,536497335 | 0,689442751 | 0,12138 | 0,44078088 | novel transcript |
| ENSG00000172031 | EPHX4 | -0,536614039 | 0,689386982 | 0,16716 | 0,49319879 | epoxide hydrolase 4 |
| ENSG00000256988 | AC005833,2 | -0,536768093 | 0,689313372 | 0,11064 | 0,42326288 | new variant |
| ENSG00000126456 | IRF3 | -0,536969395 | 0,689217197 | 0,12413 | 0,44504227 | interferon regulatory factor 3 |
| ENSG00000125995 | ROMO1 | -0,537047524 | 0,689179874 | 0,14027 | 0,46714558 | reactive oxygen species modulator 1 |

| | | | | | | |
|---|---|---|---|---|---|---|
| ENSG00000123933 | MXD4 | -0,537165861 | 0,689123346 | 0,0794 | 0,3679672 | MAX dimerization protein 4 |
| ENSG00000121753 | ADGRB2 | -0,537227774 | 0,689093773 | 0,09175 | 0,39166375 | adhesion G protein-coupled receptor B2 |
| ENSG00000227201 | CNN2P1 | -0,537235319 | 0,689090169 | 0,43362 | 0,68473723 | calponin 2 pseudogene 1 |
| ENSG00000286273 | AC078816,1 | -0,537237976 | 0,6890889 | 0,60054 | 0,79587862 | novel transcript, antisense to GPR87 |
| ENSG00000176124 | DLEU1 | -0,537607825 | 0,688912268 | 0,18321 | 0,50663141 | deleted in lymphocytic leukemia 1 |
| ENSG00000260122 | AC068987,1 | -0,537889965 | 0,688777555 | 0,78035 | 0,89823499 | novel transcript |
| ENSG00000177570 | SAMD12 | -0,538044049 | 0,688703995 | 0,11707 | 0,43438325 | sterile alpha motif domain containing 12 |
| ENSG00000152705 | CATSPER3 | -0,53806063 | 0,68869608 | 0,4554 | 0,69940453 | cation channel sperm associated 3 |
| ENSG00000100417 | PMM1 | -0,538116763 | 0,688669284 | 0,07967 | 0,36840099 | phosphomannomutase 1 |
| ENSG00000128567 | PODXL | -0,538144272 | 0,688656153 | 0,26676 | 0,56923396 | podocalyxin like |
| ENSG00000125878 | TCF15 | -0,538234761 | 0,68861296 | 0,45751 | 0,70075729 | transcription factor 15 |
| ENSG00000179915 | NRXN1 | -0,538255266 | 0,688603173 | 0,19152 | 0,51435195 | neurexin 1 |
| ENSG00000267250 | AC011591,1 | -0,538337344 | 0,688563998 | 0,53344 | 0,75351438 | novel transcript |
| ENSG00000132821 | VSTM2L | -0,538612223 | 0,688432818 | 0,21203 | 0,53079538 | V-set and transmembrane domain containing 2 like |
| ENSG00000269964 | MEI4 | -0,538738691 | 0,688372472 | 0,4616 | 0,70290608 | meiotic double-stranded break formation protein 4 |
| ENSG00000249267 | LINC00939 | -0,538751906 | 0,688366166 | 0,04186 | 0,27719832 | long intergenic non-protein coding RNA 939 |
| ENSG00000235034 | C19orf81 | -0,538887147 | 0,68830164 | 0,26526 | 0,56807781 | chromosome 19 open reading frame 81 |
| ENSG00000286894 | AC009141,1 | -0,538914569 | 0,688288557 | 0,26687 | 0,56929757 | novel transcript, antisense to WWOX |
| ENSG00000248712 | CCDC153 | -0,539080097 | 0,688209591 | 0,39091 | 0,65464997 | coiled-coil domain containing 153 |
| ENSG00000248521 | AC092612,1 | -0,53913273 | 0,688184484 | 0,48959 | 0,72395897 | nei like 2 (E, coli) (NEIL2) pseudogene |
| ENSG00000083844 | ZNF264 | -0,53964181 | 0,687941689 | 0,168 | 0,49417064 | zinc finger protein 264 |
| ENSG00000182732 | RGS6 | -0,539709446 | 0,687909438 | 0,10682 | 0,41660052 | regulator of G protein signaling 6 |
| ENSG00000223429 | AL139156,1 | -0,539968322 | 0,687786011 | 0,34223 | 0,62000209 | transformer-2 alpha (TRA2A) pseudogene |
| ENSG00000255043 | NAV2-AS5 | -0,540145034 | 0,687701771 | 0,40167 | 0,66308566 | NAV2 antisense RNA 5 |
| ENSG00000270996 | AC005034,4 | -0,540183128 | 0,687683613 | 0,46773 | 0,70752795 | novel transcript |
| ENSG00000180245 | RRH | -0,540191183 | 0,687679773 | 0,37821 | 0,64646341 | retinal pigment epithelium-derived rhodopsin homolog |
| ENSG00000186001 | LRCH3 | -0,540259855 | 0,687647041 | 0,14131 | 0,46803321 | leucine rich repeats and calponin homology domain containing 3 |
| ENSG00000120341 | SEC16B | -0,540268551 | 0,687642896 | 0,31665 | 0,60246102 | SEC16 homolog B, endoplasmic reticulum export factor |
| ENSG00000119138 | KLF9 | -0,54034753 | 0,687605252 | 0,15219 | 0,48061452 | Kruppel like factor 9 |
| ENSG00000183230 | CTNNA3 | -0,540424622 | 0,68756851 | 0,33784 | 0,61751351 | catenin alpha 3 |
| ENSG00000197982 | C1orf122 | -0,540513336 | 0,687526232 | 0,13199 | 0,45708171 | chromosome 1 open reading frame 122 |
| ENSG00000233406 | AL162430,1 | -0,540622712 | 0,68747411 | 0,26862 | 0,57070906 | ribosomal protein S2 (RPS2) pseudogene |

| | | | | | | |
|---|---|---|---|---|---|---|
| ENSG00000188033 | ZNF490 | -0,540690349 | 0,68744188 | 0,16579 | 0,49189706 | zinc finger protein 490 |
| ENSG00000265479 | DTX2P1-UPK3 | -0,540694505 | 0,6874399 | 0,07884 | 0,36668483 | DTX2P1-UPK3BP1-PMS2P11 readthrough, transcribed pseudogene |
| ENSG00000259363 | AC090825,1 | -0,54086052 | 0,687360799 | 0,12807 | 0,45163061 | novel transcript |
| ENSG00000240698 | RPS2P39 | -0,540918172 | 0,687333331 | 0,6327 | 0,81497431 | ribosomal protein S2 pseudogene 39 |
| ENSG00000183250 | LINC01547 | -0,541021212 | 0,687284242 | 0,19754 | 0,51902178 | long intergenic non-protein coding RNA 1547 |
| ENSG00000183569 | SERHL2 | -0,541178472 | 0,68720933 | 0,25131 | 0,55720663 | serine hydrolase like 2 |
| ENSG00000182632 | CCNYL2 | -0,541455425 | 0,687077419 | 0,50711 | 0,73538734 | cyclin Y like 2 (pseudogene) |
| ENSG00000286548 | AL596214,1 | -0,541802345 | 0,68691222 | 0,12106 | 0,44016175 | novel transcript, antisense to LRRC8B |
| ENSG00000026559 | KCNG1 | -0,541832003 | 0,686898099 | 0,40721 | 0,66659618 | potassium voltage-gated channel modifier subfamily G member 1 |
| ENSG00000165879 | FRAT1 | -0,54183339 | 0,686897438 | 0,26628 | 0,56917012 | FRAT regulator of WNT signaling pathway 1 |
| ENSG00000229447 | AC114495,2 | -0,541900114 | 0,686865671 | 0,34092 | 0,6192089 | general transcription factor IIA, 2, 12kDa (GTF2A2) pseudogene |
| ENSG00000169895 | SYAP1 | -0,541964838 | 0,686834857 | 0,15358 | 0,48191866 | synapse associated protein 1 |
| ENSG00000132958 | TPTE2 | -0,542161334 | 0,686741315 | 0,51787 | 0,74287921 | transmembrane phosphoinositide 3-phosphatase and tensin homolog 2 |
| ENSG00000064692 | SNCAIP | -0,542377391 | 0,686638477 | 0,15957 | 0,48693054 | synuclein alpha interacting protein |
| ENSG00000174255 | ZNF80 | -0,54238252 | 0,686636036 | 0,42714 | 0,68031759 | zinc finger protein 80 |
| ENSG00000286883 | AC005041,5 | -0,542385766 | 0,686634491 | 0,14689 | 0,47471707 | novel transcript, antisense to CCDC142and MRPL53and MOGS |
| ENSG00000248671 | ALG1L9P | -0,542520848 | 0,686570203 | 0,08188 | 0,37305174 | ALG1 like 9, pseudogene |
| ENSG00000235421 | AC114501,2 | -0,54271232 | 0,686479089 | 0,76622 | 0,89136081 | novel pseudogene similar to part of integrator complex subunit 4 INTS4 |
| ENSG00000154803 | FLCN | -0,542797519 | 0,68643855 | 0,16864 | 0,49492999 | folliculin |
| ENSG00000225684 | FAM225B | -0,542860571 | 0,68640855 | 0,67664 | 0,8409179 | family with sequence similarity 225 member B |
| ENSG00000276241 | AC243829,2 | -0,54287238 | 0,686402932 | 0,30483 | 0,59446033 | novel transcript, antisense TBC1D3B |
| ENSG00000287847 | AC018678,1 | -0,542993173 | 0,686345463 | 0,21653 | 0,53490925 | novel transcript |
| ENSG00000198435 | NRARP | -0,543106356 | 0,68629162 | 0,09712 | 0,40152057 | NOTCH regulated ankyrin repeat protein |
| ENSG00000227730 | MTND6P5 | -0,543381656 | 0,686160672 | 0,48041 | 0,71764212 | MT-ND6 pseudogene 5 |
| ENSG00000196943 | NOP9 | -0,543567423 | 0,686072325 | 0,09376 | 0,39568176 | NOP9 nucleolar protein |
| ENSG00000285326 | NOP9 | -0,543567423 | 0,686072325 | 0,09376 | 0,39568176 | NOP9 nucleolar protein |
| ENSG00000137561 | TTPA | -0,543681641 | 0,686018011 | 0,30214 | 0,59294633 | alpha tocopherol transfer protein |

| | | | | | | |
|---|---|---|---|---|---|---|
| ENSG00000285442 | FBXL6 | -0,543774886 | 0,685973673 | 0,10309 | 0,41078404 | F-box and leucine rich repeat protein 6 |
| ENSG00000182325 | FBXL6 | -0,543774886 | 0,685973673 | 0,10309 | 0,41078404 | F-box and leucine rich repeat protein 6 |
| ENSG00000267049 | AC002398,1 | -0,543779281 | 0,685971584 | 0,39948 | 0,66176242 | novel transcript |
| ENSG00000216809 | AL589993,1 | -0,54382908 | 0,685947905 | 0,16038 | 0,48767145 | DEAD/H (Asp-Glu-Ala-Asp/His) box polypeptide 24 (DDX24) pseudogene |
| ENSG00000231160 | KLF3-AS1 | -0,543835261 | 0,685944967 | 0,10671 | 0,41658493 | KLF3 antisense RNA 1 |
| ENSG00000228696 | ARL17B | -0,543885786 | 0,685920944 | 0,20563 | 0,52513499 | ADP ribosylation factor like GTPase 17B |
| ENSG00000242259 | C22orf39 | -0,544023109 | 0,685855658 | 0,06357 | 0,33422972 | chromosome 22 open reading frame 39 |
| ENSG00000231312 | MAP4K3-DT | -0,544177485 | 0,685782272 | 0,15545 | 0,48334025 | MAP4K3 divergent transcript |
| ENSG00000254330 | AC104232,2 | -0,54439676 | 0,685678048 | 0,5483 | 0,76354047 | novel transcript |
| ENSG00000241370 | RPP21 | -0,544749949 | 0,685510206 | 0,27265 | 0,57315967 | ribonuclease P/MRP subunit p21 |
| ENSG00000255872 | AL138752,2 | -0,544802457 | 0,685485257 | 0,63646 | 0,81704929 | novel transcript |
| ENSG00000124785 | NRN1 | -0,545037869 | 0,685373412 | 0,09436 | 0,39653034 | neuritin 1 |
| ENSG00000087510 | TFAP2C | -0,545065651 | 0,685360214 | 0,192 | 0,5145257 | transcription factor AP-2 gamma |
| ENSG00000240253 | FAR2P3 | -0,545110645 | 0,685338839 | 0,12826 | 0,45171208 | fatty acyl-CoA reductase 2 pseudogene 3 |
| ENSG00000236411 | NDUFAF4P3 | -0,54519188 | 0,685300251 | 0,5162 | 0,74198903 | NADH:ubiquinone oxidoreductase complex assembly factor 4 pseudogene 3 |
| ENSG00000266717 | AC134407,1 | -0,545368623 | 0,6852163 | 0,41405 | 0,67096878 | novel transcript, antisense to BPTF |
| ENSG00000267605 | AC016590,1 | -0,545410671 | 0,68519633 | 0,28722 | 0,58364712 | novel transcript, antisense to ZNF383 |
| ENSG00000260259 | LINC02166 | -0,54565843 | 0,685078669 | 0,35072 | 0,62614142 | long intergenic non-protein coding RNA 2166 |
| ENSG00000237767 | LINC01370 | -0,545749462 | 0,685035443 | 0,47642 | 0,71465252 | long intergenic non-protein coding RNA 1370 |
| ENSG00000273579 | C22orf15 | -0,545821567 | 0,685001206 | 0,39578 | 0,6585603 | chromosome 22 open reading frame 15 |
| ENSG00000144191 | CNGA3 | -0,545958971 | 0,684935969 | 0,1713 | 0,49653881 | cyclic nucleotide gated channel subunit alpha 3 |
| ENSG00000233841 | HLA-C | -0,546185577 | 0,684828393 | 0,241 | 0,55003907 | major histocompatibility complex, class I, C |
| ENSG00000243770 | RN7SL65P | -0,546383385 | 0,684734503 | 0,48336 | 0,71965182 | RNA, 7SL, cytoplasmic 65, pseudogene |
| ENSG00000205277 | MUC12 | -0,546642654 | 0,684611459 | 0,17266 | 0,49744497 | mucin 12, cell surface associated |
| ENSG00000070495 | JMJD6 | -0,546737142 | 0,684566623 | 0,09236 | 0,39278463 | jumonji domain containing 6, arginine demethylase and lysine hydroxylase |
| ENSG00000263017 | AC036164,1 | -0,546749774 | 0,684560629 | 0,42521 | 0,67890168 | chromosome 19 open reading frame 70 (C19orf70) pseudogene |
| ENSG00000265136 | AC124283,4 | -0,546930135 | 0,684475052 | 0,44199 | 0,68994635 | ADP-ribosylation factor-like 2 binding protein (ARL2BP) pseudogene |
| ENSG00000174353 | STAG3L3 | -0,546962824 | 0,684459544 | 0,17825 | 0,50257132 | stromal antigen 3-like 3 (pseudogene) |

| | | | | | | |
|---|---|---|---|---|---|---|
| ENSG00000253549 | CA3-AS1 | -0,54696308 | 0,684459422 | 0,17591 | 0,49987596 | CA3 antisense RNA 1 |
| ENSG00000236829 | Z97634,1 | -0,546999109 | 0,684442329 | 0,53351 | 0,75351438 | ribosomal protein L23a (RPL23A) pseudogene |
| ENSG00000286245 | AC125437,3 | -0,547043377 | 0,684421328 | 0,14874 | 0,47609532 | novel transcript, antisense to ATP9B |
| ENSG00000100058 | CRYBB2P1 | -0,547109776 | 0,684389829 | 0,13942 | 0,46666406 | crystallin beta B2 pseudogene 1 |
| ENSG00000230910 | AL391807,1 | -0,547155181 | 0,68436829 | 0,26081 | 0,56400166 | novel transcript |
| ENSG00000179364 | PACS2 | -0,547177941 | 0,684357493 | 0,06242 | 0,33099652 | phosphofurin acidic cluster sorting protein 2 |
| ENSG00000287356 | AL590822,3 | -0,547214342 | 0,684340226 | 0,24464 | 0,55271519 | novel transcript, antisense to MORN1 |
| ENSG00000269837 | IPO5P1 | -0,547336896 | 0,684282095 | 0,1573 | 0,48443207 | importin 5 pseudogene 1 |
| ENSG00000182253 | SYNM | -0,54735562 | 0,684273214 | 0,08723 | 0,38241146 | synemin |
| ENSG00000282685 | AC105219,8 | -0,547413783 | 0,684245628 | 0,38505 | 0,65088045 | IQ motif and ankyrin repeat containing 1 |
| ENSG00000185008 | ROBO2 | -0,547476238 | 0,684216008 | 0,08513 | 0,37851884 | roundabout guidance receptor 2 |
| ENSG00000178752 | ERFE | -0,547560067 | 0,684176252 | 0,42819 | 0,68094175 | erythroferrone |
| ENSG00000181458 | TMEM45A | -0,547716064 | 0,684102276 | 0,10945 | 0,42142456 | transmembrane protein 45A |
| ENSG00000286773 | AL353593,3 | -0,547745601 | 0,684088271 | 0,59615 | 0,79377068 | novel transcript, sense intronic to OBSCN |
| ENSG00000118946 | PCDH17 | -0,547791987 | 0,684066276 | 0,07799 | 0,36536048 | protocadherin 17 |
| ENSG00000250519 | AP002784,1 | -0,547881484 | 0,684023841 | 0,11082 | 0,42347501 | novel transcript |
| ENSG00000271723 | MROH7-TTC4 | -0,547902543 | 0,684013857 | 0,46502 | 0,70556892 | MROH7-TTC4 readthrough (NMD candidate) |
| ENSG00000233049 | DHX16 | -0,547920294 | 0,684005441 | 0,64013 | 0,81935089 | DEAH-box helicase 16 |
| ENSG00000231377 | DHX16 | -0,547920294 | 0,684005441 | 0,64013 | 0,81935089 | DEAH-box helicase 16 |
| ENSG00000181001 | OR52N1 | -0,548281538 | 0,68383419 | 0,20544 | 0,52503195 | olfactory receptor family 52 subfamily N member 1 |
| ENSG00000166166 | TRMT61A | -0,54831334 | 0,683819117 | 0,08261 | 0,37429151 | tRNA methyltransferase 61A |
| ENSG00000066923 | STAG3 | -0,548664252 | 0,683652809 | 0,07417 | 0,35759867 | stromal antigen 3 |
| ENSG00000187531 | SIRT7 | -0,548715902 | 0,683628334 | 0,06953 | 0,34816245 | sirtuin 7 |
| ENSG00000113790 | EHHADH | -0,548737995 | 0,683617865 | 0,25716 | 0,56155289 | enoyl-CoA hydratase and 3-hydroxyacyl CoA dehydrogenase |
| ENSG00000227999 | MTND5P1 | -0,548777559 | 0,683599118 | 0,29733 | 0,59031059 | MT-ND5 pseudogene 1 |
| ENSG00000172936 | MYD88 | -0,548830708 | 0,683573935 | 0,04591 | 0,28908129 | MYD88 innate immune signal transduction adaptor |
| ENSG00000068831 | RASGRP2 | -0,548863278 | 0,683558503 | 0,22371 | 0,53967355 | RAS guanyl releasing protein 2 |
| ENSG00000171889 | MIR31HG | -0,549039322 | 0,683475097 | 0,64813 | 0,82383528 | MIR31 host gene |
| ENSG00000163531 | NFASC | -0,549121317 | 0,683436253 | 0,06483 | 0,33757437 | neurofascin |
| ENSG00000259780 | AC009065,1 | -0,549172975 | 0,683411782 | 0,7156 | 0,86355924 | novel transcript |
| ENSG00000235245 | AL360181,2 | -0,549240594 | 0,683379751 | 0,35252 | 0,62747731 | novel transcript |
| ENSG00000180458 | AC022148,1 | -0,549257602 | 0,683371695 | 0,26382 | 0,56634699 | novel transcript, antisense to ZNF793 |

| | | | | | | |
|---|---|---|---|---|---|---|
| ENSG00000229525 | AC053503,1 | -0,549340886 | 0,683332246 | 0,51263 | 0,73941533 | novel transcript, antisense to DNPEP |
| ENSG00000160993 | ALKBH4 | -0,549343449 | 0,683331032 | 0,13861 | 0,46578766 | alkB homolog 4, lysine demethylase |
| ENSG00000259240 | MIR4713HG | -0,54959216 | 0,683213241 | 0,61292 | 0,8035281 | MIR4713 host gene |
| ENSG00000204754 | LINC01951 | -0,549599104 | 0,683209952 | 0,25219 | 0,55796133 | long intergenic non-protein coding RNA 1951 |
| ENSG00000237523 | LINC00857 | -0,549626929 | 0,683196776 | 0,41877 | 0,67486451 | long intergenic non-protein coding RNA 857 |
| ENSG00000177302 | TOP3A | -0,549984739 | 0,683027353 | 0,14106 | 0,4679289 | DNA topoisomerase III alpha |
| ENSG00000198822 | GRM3 | -0,550009524 | 0,68301562 | 0,29668 | 0,58991157 | glutamate metabotropic receptor 3 |
| ENSG00000250767 | AC139783,1 | -0,550059248 | 0,682992079 | 0,52314 | 0,74719504 | fumarate hydratase (FH) pseudogene |
| ENSG00000147679 | UTP23 | -0,550154392 | 0,682947038 | 0,12453 | 0,44559634 | UTP23 small subunit processome component |
| ENSG00000241449 | AC092666,1 | -0,550229754 | 0,682911364 | 0,28659 | 0,58291115 | novel transcript |
| ENSG00000149972 | CNTN5 | -0,550262108 | 0,682896049 | 0,17942 | 0,5038121 | contactin 5 |
| ENSG00000081148 | IMPG2 | -0,550327947 | 0,682864885 | 0,11131 | 0,42416781 | interphotoreceptor matrix proteoglycan 2 |
| ENSG00000203497 | PDCD4-AS1 | -0,550545855 | 0,682761751 | 0,19954 | 0,52047306 | PDCD4 antisense RNA 1 |
| ENSG00000167522 | ANKRD11 | -0,550575292 | 0,68274782 | 0,07063 | 0,3507077 | ankyrin repeat domain 11 |
| ENSG00000181027 | FKRP | -0,550767656 | 0,682656791 | 0,15693 | 0,48425071 | fukutin related protein |
| ENSG00000271614 | ATP2B1-AS1 | -0,550960852 | 0,68256538 | 0,14799 | 0,4751727 | ATP2B1 antisense RNA 1 |
| ENSG00000264956 | AC138761,1 | -0,550994376 | 0,68254952 | 0,3961 | 0,65876705 | novel transcript |
| ENSG00000286673 | LO018686,4 | -0,551012875 | 0,682540768 | 0,11125 | 0,42416781 | novel transcript |
| ENSG00000286931 | AL136419,3 | -0,551012875 | 0,682540768 | 0,11125 | 0,42416781 | novel transcript |
| ENSG00000286814 | AC009152,5 | -0,551157563 | 0,682472319 | 0,45844 | 0,70119611 | novel transcript, antisense to XYLT1 |
| ENSG00000124701 | APOBEC2 | -0,551183601 | 0,682460002 | 0,40026 | 0,66242349 | apolipoprotein B mRNA editing enzyme catalytic subunit 2 |
| ENSG00000284207 | HMGB3P22 | -0,551220109 | 0,682442732 | 0,56692 | 0,77482793 | high mobility group box 3 pseudogene 22 |
| ENSG00000229081 | LINC01165 | -0,551360979 | 0,682376099 | 0,45194 | 0,69686926 | long intergenic non-protein coding RNA 1165 |
| ENSG00000250122 | AC122694,1 | -0,551363729 | 0,682374799 | 0,33124 | 0,61131843 | novel transcript |
| ENSG00000141552 | ANAPC11 | -0,551407638 | 0,68235403 | 0,06383 | 0,33492932 | anaphase promoting complex subunit 11 |
| ENSG00000166689 | PLEKHA7 | -0,551444434 | 0,682336627 | 0,16224 | 0,48939373 | pleckstrin homology domain containing A7 |
| ENSG00000236548 | RNF217-AS1 | -0,551730704 | 0,682201246 | 0,36334 | 0,6341399 | RNF217 antisense RNA 1 (head to head) |
| ENSG00000181195 | PENK | -0,551796776 | 0,682170004 | 0,43567 | 0,68623199 | proenkephalin |
| ENSG00000229339 | AC093414,1 | -0,551887725 | 0,682127001 | 0,41062 | 0,66856387 | ribosomal protein L7a (RPL7A) pseudogene |
| ENSG00000091622 | PITPNM3 | -0,551904578 | 0,682119032 | 0,06883 | 0,34677117 | PITPNM family member 3 |
| ENSG00000129675 | ARHGEF6 | -0,552015169 | 0,682066746 | 0,1057 | 0,41520886 | Rac/Cdc42 guanine nucleotide exchange factor 6 |
| ENSG00000247199 | AC011346,1 | -0,552030077 | 0,682059698 | 0,0963 | 0,39990052 | novel transcript, antisense to SPINK9, SPINK7 and SPINK13 |
| ENSG00000276043 | UHRF1 | -0,552106092 | 0,682023761 | 0,05664 | 0,31746133 | ubiquitin like with PHD and ring finger domains 1 |

| | | | | | | |
|---|---|---|---|---|---|---|
| ENSG00000188385 | JAKMIP3 | -0,552121813 | 0,682016329 | 0,12719 | 0,45032932 | Janus kinase and microtubule interacting protein 3 |
| ENSG00000130649 | CYP2E1 | -0,552300016 | 0,681932091 | 0,29468 | 0,58857672 | cytochrome P450 family 2 subfamily E member 1 |
| ENSG00000198785 | GRIN3A | -0,552650622 | 0,681766387 | 0,06269 | 0,33130676 | glutamate ionotropic receptor NMDA type subunit 3A |
| ENSG00000286834 | AL360020,1 | -0,552661179 | 0,681761399 | 0,13402 | 0,46018397 | novel transcript |
| ENSG00000156463 | SH3RF2 | -0,552823752 | 0,681684577 | 0,29593 | 0,58942061 | SH3 domain containing ring finger 2 |
| ENSG00000241472 | PTPRG-AS1 | -0,552916836 | 0,681640596 | 0,04018 | 0,2708918 | PTPRG antisense RNA 1 |
| ENSG00000284974 | AC270301,4 | -0,553012265 | 0,681595509 | 0,155 | 0,48292767 | novel transcript |
| ENSG00000162753 | SLC9C2 | -0,553022472 | 0,681590687 | 0,36486 | 0,63505392 | solute carrier family 9 member C2 (putative) |
| ENSG00000229191 | AL358473,1 | -0,553063858 | 0,681571134 | 0,07204 | 0,354195 | novel transcript, antisense KIF21B |
| ENSG00000100034 | PPM1F | -0,553182462 | 0,681515105 | 0,19865 | 0,51980353 | protein phosphatase, Mg2+/Mn2+ dependent 1F |
| ENSG00000164896 | FASTK | -0,553472847 | 0,681377944 | 0,15426 | 0,48258262 | Fas activated serine/threonine kinase |
| ENSG00000153395 | LPCAT1 | -0,553647124 | 0,681295638 | 0,10983 | 0,42203441 | lysophosphatidylcholine acyltransferase 1 |
| ENSG00000087299 | L2HGDH | -0,553665253 | 0,681287077 | 0,15248 | 0,48061452 | L-2-hydroxyglutarate dehydrogenase |
| ENSG00000274529 | SEBOX | -0,553745523 | 0,681249172 | 0,74256 | 0,87827554 | SEBOX homeobox |
| ENSG00000092096 | SLC22A17 | -0,55397763 | 0,681139579 | 0,17744 | 0,50191142 | solute carrier family 22 member 17 |
| ENSG00000053328 | METTL24 | -0,554030906 | 0,681114426 | 0,311 | 0,59883009 | methyltransferase like 24 |
| ENSG00000267098 | AC113137,1 | -0,554052126 | 0,681104408 | 0,12507 | 0,44678334 | novel transcript |
| ENSG00000165684 | SNAPC4 | -0,5541352 | 0,681065189 | 0,05173 | 0,30583008 | small nuclear RNA activating complex polypeptide 4 |
| ENSG00000236546 | MYCL-AS1 | -0,554179261 | 0,68104439 | 0,35931 | 0,63214783 | MYCL antisense RNA 1 |
| ENSG00000157045 | NTAN1 | -0,554447205 | 0,680917915 | 0,04804 | 0,29434174 | N-terminal asparagine amidase |
| ENSG00000233729 | NCKAP5-AS1 | -0,554519735 | 0,680883683 | 0,1551 | 0,48296753 | NCKAP5 antisense RNA 1 |
| ENSG00000147813 | NAPRT | -0,554557379 | 0,680865917 | 0,23832 | 0,54875544 | nicotinate phosphoribosyltransferase |
| ENSG00000267587 | AC090241,1 | -0,554829937 | 0,680737298 | 0,43698 | 0,68702836 | Ubiquitin carboxyl-terminal hydrolase 4, pseudogene |
| ENSG00000204815 | TTC25 | -0,554833527 | 0,680735604 | 0,03709 | 0,26086671 | tetratricopeptide repeat domain 25 |
| ENSG00000132563 | REEP2 | -0,554930702 | 0,680689754 | 0,02757 | 0,22376406 | receptor accessory protein 2 |
| ENSG00000253846 | PCDHGA10 | -0,555098505 | 0,680610586 | 0,05813 | 0,32179302 | protocadherin gamma subfamily A, 10 |
| ENSG00000272420 | AL513477,2 | -0,555105392 | 0,680607337 | 0,32005 | 0,60496053 | novel transcript, sense intronic to MORN1 |
| ENSG00000078295 | ADCY2 | -0,555413552 | 0,680461974 | 0,15139 | 0,47956749 | adenylate cyclase 2 |
| ENSG00000174749 | FAM241A | -0,55544382 | 0,680447698 | 0,10869 | 0,41972109 | family with sequence similarity 241 member A |
| ENSG00000262468 | LINC01569 | -0,555475498 | 0,680432757 | 0,15346 | 0,48185226 | long intergenic non-protein coding RNA 1569 |
| ENSG00000082397 | EPB41L3 | -0,555681608 | 0,680335554 | 0,18617 | 0,50956086 | erythrocyte membrane protein band 4,1 like 3 |
| ENSG00000274698 | AC099521,1 | -0,555952391 | 0,680207873 | 0,39071 | 0,65464997 | novel transcript |
| ENSG00000164880 | INTS1 | -0,556121763 | 0,680128021 | 0,12632 | 0,44903997 | integrator complex subunit 1 |

| | | | | | | |
|---|---|---|---|---|---|---|
| ENSG00000161265 | U2AF1L4 | -0,556193806 | 0,680094059 | 0,19596 | 0,51680811 | U2 small nuclear RNA auxiliary factor 1 like 4 |
| ENSG00000230535 | BASP1P1 | -0,556236527 | 0,68007392 | 0,51438 | 0,74068623 | BASP1 pseudogene 1 |
| ENSG00000266709 | AC005224,3 | -0,5563084 | 0,680040041 | 0,41401 | 0,67096878 | novel transcript |
| ENSG00000255145 | STX17-AS1 | -0,55639194 | 0,680000664 | 0,10177 | 0,4089368 | STX17 antisense RNA 1 |
| ENSG00000280484 | AL732314,1 | -0,556413598 | 0,679990456 | 0,52468 | 0,7484123 | novel transcript, antisense to PPP2R3B |
| ENSG00000281849 | AL732314,10 | -0,556413598 | 0,679990456 | 0,52468 | 0,7484123 | novel transcript, antisense to PPP2R3B |
| ENSG00000280593 | AL732314,3 | -0,556413598 | 0,679990456 | 0,52468 | 0,7484123 | novel transcript, antisense to PPP2R3B |
| ENSG00000272578 | AP000347,1 | -0,556782155 | 0,679816764 | 0,09405 | 0,39609058 | glucuronidase, beta (GUSB) pseudogene |
| ENSG00000250938 | AC073475,1 | -0,556933816 | 0,679745304 | 0,33738 | 0,61706766 | novel transcript, antisense to MAD2L1 |
| ENSG00000269534 | AC011466,3 | -0,5573772 | 0,679536429 | 0,47682 | 0,71481671 | novel transcript, antisense to LIG1 |
| ENSG00000236125 | USP17L4 | -0,557555508 | 0,679452448 | 0,53322 | 0,75351438 | ubiquitin specific peptidase 17 like family member 4 |
| ENSG00000232339 | NOTCH4 | -0,557621545 | 0,679421348 | 0,21638 | 0,53482962 | notch receptor 4 |
| ENSG00000162390 | ACOT11 | -0,557696047 | 0,679386263 | 0,28593 | 0,58229607 | acyl-CoA thioesterase 11 |
| ENSG00000258811 | AL583810,2 | -0,557765688 | 0,679353469 | 0,54142 | 0,75881984 | novel transcript |
| ENSG00000234390 | USP27X-AS1 | -0,557927539 | 0,679277259 | 0,13366 | 0,45954245 | USP27X antisense RNA 1 (head to head) |
| ENSG00000283341 | AC068205,2 | -0,558008449 | 0,679239164 | 0,15288 | 0,48105429 | novel transcript |
| ENSG00000277443 | MARCKS | -0,55811973 | 0,679186774 | 0,16591 | 0,49195088 | myristoylated alanine rich protein kinase C substrate |
| ENSG00000159374 | M1AP | -0,558214505 | 0,679142158 | 0,3906 | 0,65464381 | meiosis 1 associated protein |
| ENSG00000228284 | HLA-DQA1 | -0,558249467 | 0,679125699 | 0,09001 | 0,38804781 | major histocompatibility complex, class II, DQ alpha 1 |
| ENSG00000129250 | KIF1C | -0,558292951 | 0,679105231 | 0,16991 | 0,49606609 | kinesin family member 1C |
| ENSG00000236549 | RPS27AP7 | -0,558321594 | 0,679091748 | 0,3157 | 0,60176284 | RPS27A pseudogene 7 |
| ENSG00000281852 | LINC00891 | -0,558384131 | 0,679062312 | 0,06597 | 0,34005786 | long intergenic non-protein coding RNA 891 |
| ENSG00000115507 | OTX1 | -0,558415093 | 0,679047738 | 0,30178 | 0,59288267 | orthodenticle homeobox 1 |
| ENSG00000198466 | ZNF587 | -0,558444715 | 0,679033796 | 0,13501 | 0,46159318 | zinc finger protein 587 |
| ENSG00000259781 | HMGB1P6 | -0,558446608 | 0,679032905 | 0,08762 | 0,38323183 | high mobility group box 1 pseudogene 6 |
| ENSG00000223549 | MTND5P28 | -0,558521237 | 0,67899778 | 0,51623 | 0,74198903 | MT-ND5 pseudogene 28 |
| ENSG00000237212 | AL449403,1 | -0,558747061 | 0,678891505 | 0,21432 | 0,53334744 | novel transcript |
| ENSG00000205746 | PKD1P4 | -0,55874797 | 0,678891077 | 0,05471 | 0,31313954 | polycystin 1, transient receptor potential channel interacting pseudogene 4 |
| ENSG00000258399 | AL117190,1 | -0,558813415 | 0,678860282 | 0,60449 | 0,79797731 | novel transcript |
| ENSG00000227935 | AL390243,1 | -0,558913649 | 0,678813119 | 0,42851 | 0,68118413 | novel transcript |
| ENSG00000169432 | SCN9A | -0,55892244 | 0,678808982 | 0,21051 | 0,5290538 | sodium voltage-gated channel alpha subunit 9 |
| ENSG00000174171 | AC020659,1 | -0,559011866 | 0,678766907 | 0,51233 | 0,73924429 | novel transcript, antisense to SPTBN5 |

| | | | | | | |
|---|---|---|---|---|---|---|
| ENSG00000258667 | HIF1A-AS3 | -0,559055647 | 0,678746309 | 0,09268 | 0,39335125 | HIF1A antisense RNA 3 |
| ENSG00000233539 | AC011294,1 | -0,55906766 | 0,678740657 | 0,52611 | 0,7497214 | novel transcript |
| ENSG00000255198 | SNHG9 | -0,559238924 | 0,678660088 | 0,0409 | 0,27365037 | small nucleolar RNA host gene 9 |
| ENSG00000258590 | NBEAP1 | -0,559413214 | 0,678578105 | 0,25936 | 0,56317998 | neurobeachin pseudogene 1 |
| ENSG00000281484 | BTG3 | -0,559561925 | 0,678508162 | 0,35884 | 0,63195238 | BTG anti-proliferation factor 3 |
| ENSG00000140400 | MAN2C1 | -0,559831999 | 0,678381156 | 0,10226 | 0,40974995 | mannosidase alpha class 2C member 1 |
| ENSG00000168061 | SAC3D1 | -0,559898938 | 0,678349681 | 0,121 | 0,44006069 | SAC3 domain containing 1 |
| ENSG00000167363 | FN3K | -0,559909371 | 0,678344775 | 0,11742 | 0,43477245 | fructosamine 3 kinase |
| ENSG00000182776 | AC239585,1 | -0,560208964 | 0,678203924 | 0,45641 | 0,70003941 | novel transcript |
| ENSG00000179028 | AC245177,1 | -0,560208964 | 0,678203924 | 0,45641 | 0,70003941 | novel transcript |
| ENSG00000265766 | CXADRP3 | -0,560525573 | 0,678055103 | 0,50963 | 0,73740498 | CXADR pseudogene 3 |
| ENSG00000137332 | DDR1 | -0,560526655 | 0,678054595 | 0,00061 | 0,02348863 | discoidin domain receptor tyrosine kinase 1 |
| ENSG00000123064 | DDX54 | -0,560901584 | 0,677878404 | 0,16288 | 0,48945635 | DEAD-box helicase 54 |
| ENSG00000188833 | ENTPD8 | -0,561294569 | 0,677693778 | 0,64523 | 0,82202778 | ectonucleoside triphosphate diphosphohydrolase 8 |
| ENSG00000236200 | KDM4A-AS1 | -0,561364671 | 0,677660849 | 0,1909 | 0,51394498 | KDM4A antisense RNA 1 |
| ENSG00000248596 | AC139491,2 | -0,561494619 | 0,677599813 | 0,24434 | 0,55253387 | novel transcript |
| ENSG00000142949 | PTPRF | -0,561639342 | 0,677531843 | 0,01451 | 0,15751168 | protein tyrosine phosphatase receptor type F |
| ENSG00000166343 | MSS51 | -0,561682984 | 0,677511348 | 0,18816 | 0,51129824 | MSS51 mitochondrial translational activator |
| ENSG00000276230 | RNH1 | -0,561795235 | 0,677458635 | 0,1257 | 0,44776504 | ribonuclease/angiogenin inhibitor 1 |
| ENSG00000237672 | KRR1P1 | -0,562119702 | 0,677306289 | 0,20467 | 0,52443222 | KRR1 pseudogene 1 |
| ENSG00000211450 | SELENOH | -0,562487213 | 0,677133775 | 0,09755 | 0,4024644 | selenoprotein H |
| ENSG00000275099 | GGNBP2 | -0,562663081 | 0,677051236 | 0,04614 | 0,28930497 | gametogenetin binding protein 2 |
| ENSG00000233110 | AC093797,1 | -0,562749785 | 0,677010547 | 0,22765 | 0,54168591 | novel transcript, antisense to SORBS2 |
| ENSG00000233223 | AC016876,1 | -0,562881266 | 0,67694885 | 0,1414 | 0,46803321 | novel transcript, antisense to MPDU1 and CD68 |
| ENSG00000250327 | RPSAP70 | -0,562911283 | 0,676934766 | 0,39019 | 0,65427905 | ribosomal protein SA pseudogene 70 |
| ENSG00000226220 | CICP22 | -0,563000684 | 0,676892819 | 0,34727 | 0,62403275 | capicua transcriptional repressor pseudogene 22 |
| ENSG00000251155 | SEPTIN14P4 | -0,563150859 | 0,676822362 | 0,54822 | 0,76347743 | septin 14 pseudogene 4 |
| ENSG00000237641 | RPL28P2 | -0,563155535 | 0,676820169 | 0,33837 | 0,61774255 | ribosomal protein L28 pseudogene 2 |
| ENSG00000163257 | DCAF16 | -0,563157421 | 0,676819284 | 0,1592 | 0,4867709 | DDB1 and CUL4 associated factor 16 |
| ENSG00000243795 | LINC02044 | -0,563269467 | 0,676766721 | 0,23382 | 0,54565913 | long intergenic non-protein coding RNA 2044 |
| ENSG00000171303 | KCNK3 | -0,563405851 | 0,676702746 | 0,2568 | 0,56130308 | potassium two pore domain channel subfamily K member 3 |
| ENSG00000214671 | RPL6P12 | -0,563803523 | 0,676516243 | 0,59552 | 0,79339314 | ribosomal protein L6 pseudogene 12 |

| | | | | | | |
|---|---|---|---|---|---|---|
| ENSG00000273391 | AC083880,1 | -0,563860111 | 0,676489707 | 0,21959 | 0,53699318 | novel transcript, antisense to LUC7L2 |
| ENSG00000170919 | TPT1-AS1 | -0,563950211 | 0,67644746 | 0,08807 | 0,38423687 | TPT1 antisense RNA 1 |
| ENSG00000249565 | SERBP1P5 | -0,564219812 | 0,676321062 | 0,44736 | 0,69342177 | SERPINE1 mRNA binding protein 1 pseudogene 5 |
| ENSG00000272674 | PCDHB16 | -0,564475443 | 0,676201235 | 0,06908 | 0,34726337 | protocadherin beta 16 |
| ENSG00000276561 | IRF7 | -0,565091581 | 0,675912509 | 0,06954 | 0,34816245 | interferon regulatory factor 7 |
| ENSG00000255270 | NAV2-IT1 | -0,565268974 | 0,675829404 | 0,70876 | 0,85960107 | NAV2 intronic transcript 1 |
| ENSG00000223704 | LINC01422 | -0,565419251 | 0,675759011 | 0,18236 | 0,50585943 | long intergenic non-protein coding RNA 1422 |
| ENSG00000105197 | TIMM50 | -0,565611354 | 0,675669035 | 0,18243 | 0,50594375 | translocase of inner mitochondrial membrane 50 |
| ENSG00000103549 | RNF40 | -0,565857034 | 0,675553984 | 0,05273 | 0,30832283 | ring finger protein 40 |
| ENSG00000260051 | AL031600,1 | -0,565929283 | 0,675520154 | 0,41407 | 0,67096878 | novel transcript, antisense to CLCN7 |
| ENSG00000138435 | CHRNA1 | -0,566119778 | 0,675430963 | 0,40475 | 0,66473708 | cholinergic receptor nicotinic alpha 1 subunit |
| ENSG00000198746 | GPATCH3 | -0,566200461 | 0,675393191 | 0,11934 | 0,43721667 | G-patch domain containing 3 |
| ENSG00000286415 | AC017050,1 | -0,566205115 | 0,675391012 | 0,16191 | 0,48914394 | novel transcript |
| ENSG00000172009 | THOP1 | -0,566352133 | 0,675322189 | 0,108 | 0,41856292 | thimet oligopeptidase 1 |
| ENSG00000205670 | SMIM11A | -0,566359103 | 0,675318927 | 0,15045 | 0,47857814 | small integral membrane protein 11A |
| ENSG00000162065 | TBC1D24 | -0,56663432 | 0,675190111 | 0,08501 | 0,37843844 | TBC1 domain family member 24 |
| ENSG00000137261 | KIAA0319 | -0,566731032 | 0,675144851 | 0,11913 | 0,43696075 | KIAA0319 |
| ENSG00000139433 | GLTP | -0,566903379 | 0,675064202 | 0,18524 | 0,50876814 | glycolipid transfer protein |
| ENSG00000250444 | CCT5P1 | -0,566990822 | 0,675023287 | 0,40392 | 0,66409141 | chaperonin containing TCP1 subunit 5 pseudogene 1 |
| ENSG00000008517 | IL32 | -0,567130124 | 0,674958112 | 0,50098 | 0,73099828 | interleukin 32 |
| ENSG00000198324 | PHETA1 | -0,567268686 | 0,67489329 | 0,08831 | 0,38452862 | PH domain containing endocytic trafficking adaptor 1 |
| ENSG00000174576 | NPAS4 | -0,567274761 | 0,674890448 | 0,17455 | 0,49884018 | neuronal PAS domain protein 4 |
| ENSG00000100060 | MFNG | -0,567602214 | 0,674737283 | 0,28381 | 0,58078109 | MFNG O-fucosylpeptide 3-beta-N-acetylglucosaminyltransferase |
| ENSG00000284626 | AC091167,7 | -0,567623861 | 0,674727159 | 0,59186 | 0,79090751 | novel protein |
| ENSG00000283050 | GTF2IP12 | -0,567777078 | 0,674655505 | 0,14396 | 0,47144689 | general transcription factor IIi pseudogene 12 |
| ENSG00000266456 | AP001178,3 | -0,567920602 | 0,674588392 | 0,19014 | 0,51331106 | novel transcript, antisense to C18orf56 |
| ENSG00000273888 | FRMD6-AS1 | -0,568078004 | 0,674514797 | 0,30043 | 0,59179349 | FRMD6 antisense RNA 1 |
| ENSG00000133878 | DUSP26 | -0,568568495 | 0,674285512 | 0,13576 | 0,462686 | dual specificity phosphatase 26 |
| ENSG00000126759 | CFP | -0,568695169 | 0,67422631 | 0,2851 | 0,58171017 | complement factor properdin |
| ENSG00000128581 | IFT22 | -0,568802079 | 0,674176349 | 0,10233 | 0,40985316 | intraflagellar transport 22 |
| ENSG00000223878 | PPIAP53 | -0,568864814 | 0,674147033 | 0,27824 | 0,57645922 | peptidylprolyl isomerase A pseudogene 53 |
| ENSG00000244486 | SCARF2 | -0,568910804 | 0,674125543 | 0,2397 | 0,5491566 | scavenger receptor class F member 2 |

| | | | | | | |
|---|---|---|---|---|---|---|
| ENSG00000287335 | AL139351,3 | -0,56940781 | 0,673893348 | 0,24072 | 0,5500079 | novel transcript |
| ENSG00000115457 | IGFBP2 | -0,569515273 | 0,673843153 | 0,09534 | 0,39794903 | insulin like growth factor binding protein 2 |
| ENSG00000286147 | AC092445,1 | -0,569666101 | 0,673772709 | 0,12389 | 0,44467458 | novel transcript, antisense to DKK2 |
| ENSG00000159214 | CCDC24 | -0,569929626 | 0,673649648 | 0,1887 | 0,5120698 | coiled-coil domain containing 24 |
| ENSG00000099785 | MARCHF2 | -0,56993727 | 0,673646079 | 0,05796 | 0,32124805 | membrane associated ring-CH-type finger 2 |
| ENSG00000274281 | AC022929,2 | -0,570063632 | 0,673587078 | 0,0886 | 0,38491339 | novel transcript |
| ENSG00000231744 | AL669818,1 | -0,570174375 | 0,673535375 | 0,53987 | 0,75807575 | RAN binding protein 1 (RANBP1) pseudogene |
| ENSG00000233360 | PDXP-DT | -0,570175258 | 0,673534963 | 0,4593 | 0,7017037 | PDXP divergent transcript |
| ENSG00000253273 | AC022826,1 | -0,570573537 | 0,673349048 | 0,43917 | 0,68811742 | regulator of G-protein signaling 10 (RGS10) pseudogene |
| ENSG00000244004 | RPS3AP19 | -0,57061578 | 0,673329332 | 0,33887 | 0,61799811 | RPS3A pseudogene 19 |
| ENSG00000235453 | SMIM27 | -0,570684788 | 0,673297126 | 0,05786 | 0,32112994 | small integral membrane protein 27 |
| ENSG00000204060 | FOXO6 | -0,570718632 | 0,673281331 | 0,36962 | 0,63934345 | forkhead box O6 |
| ENSG00000173818 | ENDOV | -0,570852357 | 0,673218927 | 0,07765 | 0,36468824 | endonuclease V |
| ENSG00000122641 | INHBA | -0,57105286 | 0,673125371 | 0,37084 | 0,64013451 | inhibin subunit beta A |
| ENSG00000241832 | CECR3 | -0,571244809 | 0,673035818 | 0,43882 | 0,6878469 | cat eye syndrome chromosome region, candidate 3 |
| ENSG00000233051 | MICA | -0,571343792 | 0,672989643 | 0,15587 | 0,4836223 | MHC class I polypeptide-related sequence A |
| ENSG00000233418 | DHX16 | -0,571350398 | 0,672986561 | 0,61349 | 0,8035281 | DEAH-box helicase 16 |
| ENSG00000233561 | DHX16 | -0,571350398 | 0,672986561 | 0,61349 | 0,8035281 | DEAH-box helicase 16 |
| ENSG00000225578 | NCBP2-AS1 | -0,571400636 | 0,672963127 | 0,40625 | 0,66584658 | NCBP2 antisense RNA 1 |
| ENSG00000072840 | EVC | -0,571667055 | 0,672838864 | 0,22424 | 0,5401098 | EvC ciliary complex subunit 1 |
| ENSG00000117472 | TSPAN1 | -0,571703049 | 0,672822078 | 0,39816 | 0,66078008 | tetraspanin 1 |
| ENSG00000254602 | AP000662,1 | -0,571719011 | 0,672814633 | 0,07966 | 0,36840099 | novel transcript, sense overlapping CLP1 |
| ENSG00000067365 | METTL22 | -0,571814218 | 0,672770234 | 0,15139 | 0,47956749 | methyltransferase like 22 |
| ENSG00000283443 | AC018688,1 | -0,571827452 | 0,672764063 | 0,31758 | 0,60294242 | zinc finger protein 285 (ZNF285) pseudogene |
| ENSG00000242522 | KLHL6-AS1 | -0,571833252 | 0,672761358 | 0,09011 | 0,38827032 | KLHL6 antisense RNA 1 |
| ENSG00000235303 | AL023495,1 | -0,571902449 | 0,672729091 | 0,50178 | 0,73147839 | novel transcript |
| ENSG00000288674 | AL391628,1 | -0,57205636 | 0,672657326 | 0,16676 | 0,49259924 | Atypical kinase COQ8A, mitochondrial |
| ENSG00000258741 | H2AZ2P1 | -0,572111461 | 0,672631636 | 0,06304 | 0,33245086 | H2AZ2 pseudogene 1 |
| ENSG00000271830 | AC012213,4 | -0,572138472 | 0,672619042 | 0,23878 | 0,54892264 | novel transcript |
| ENSG00000242791 | AC117395,1 | -0,572245011 | 0,672569373 | 0,43776 | 0,68744895 | novel transcript |
| ENSG00000221821 | C6orf226 | -0,572249938 | 0,672567076 | 0,13489 | 0,46159318 | chromosome 6 open reading frame 226 |
| ENSG00000286741 | AC008766,1 | -0,57276139 | 0,672328686 | 0,23096 | 0,54392721 | novel transcript |
| ENSG00000141086 | CTRL | -0,57281713 | 0,67230271 | 0,36542 | 0,63559906 | chymotrypsin like |

| | | | | | | |
|---|---|---|---|---|---|---|
| ENSG00000263884 | AP000845,1 | -0,572936559 | 0,672247058 | 0,16488 | 0,49095733 | novel transcript |
| ENSG00000188596 | CFAP54 | -0,573202345 | 0,672123222 | 0,05891 | 0,3241112 | cilia and flagella associated protein 54 |
| ENSG00000162129 | CLPB | -0,573343572 | 0,67205743 | 0,16895 | 0,4950359 | caseinolytic mitochondrial matrix peptidase chaperone subunit B |
| ENSG00000267251 | AC139100,1 | -0,573432268 | 0,672016114 | 0,07771 | 0,36480631 | novel transcript, antisense to PARD6G |
| ENSG00000174004 | NRROS | -0,573469332 | 0,671998849 | 0,22857 | 0,54244516 | negative regulator of reactive oxygen species |
| ENSG00000101265 | RASSF2 | -0,573606107 | 0,671935144 | 0,18808 | 0,51128963 | Ras association domain family member 2 |
| ENSG00000206481 | MDC1 | -0,573607888 | 0,671934314 | 0,39049 | 0,65448813 | mediator of DNA damage checkpoint 1 |
| ENSG00000133619 | KRBA1 | -0,573841546 | 0,671825497 | 0,08823 | 0,38444882 | KRAB-A domain containing 1 |
| ENSG00000185950 | IRS2 | -0,574058786 | 0,671724341 | 0,26658 | 0,5691939 | insulin receptor substrate 2 |
| ENSG00000219881 | GAPDHP42 | -0,574059045 | 0,671724221 | 0,32547 | 0,60794826 | glyceraldehyde 3 phosphate dehydrogenase pseudogene 42 |
| ENSG00000232442 | MHENCR | -0,574122786 | 0,671694543 | 0,05635 | 0,31700069 | melanoma highly expressed competing endogenous lncRNA for miR-425 and miR-489 |
| ENSG00000232433 | GXYLT1P3 | -0,574359722 | 0,671584239 | 0,27438 | 0,57427267 | GXYLT1 pseudogene 3 |
| ENSG00000186481 | ANKRD20A5P | -0,574394553 | 0,671568025 | 0,05057 | 0,30253733 | ankyrin repeat domain 20 family member A5, pseudogene |
| ENSG00000162783 | IER5 | -0,574460573 | 0,671537294 | 0,11807 | 0,43562845 | immediate early response 5 |
| ENSG00000196235 | SUPT5H | -0,574471008 | 0,671532437 | 0,02286 | 0,20180274 | SPT5 homolog, DSIF elongation factor subunit |
| ENSG00000272904 | AL390726,2 | -0,574501538 | 0,671518226 | 0,25069 | 0,55690541 | novel transcript |
| ENSG00000196295 | GARS1-DT | -0,574566069 | 0,67148819 | 0,12011 | 0,43868155 | GARS1 divergent transcript |
| ENSG00000283034 | ZNF445 | -0,574573994 | 0,671484502 | 0,00587 | 0,09646421 | zinc finger protein 445 |
| ENSG00000117877 | POLR1G | -0,574751488 | 0,671401894 | 0,29796 | 0,59037822 | RNA polymerase I subunit G |
| ENSG00000058404 | CAMK2B | -0,574937823 | 0,671315183 | 0,18768 | 0,51107292 | calcium/calmodulin dependent protein kinase II beta |
| ENSG00000105497 | ZNF175 | -0,575000827 | 0,671285866 | 0,1191 | 0,43696075 | zinc finger protein 175 |
| ENSG00000288093 | AL592295,5 | -0,575007422 | 0,671282798 | 0,29613 | 0,58956107 | novel transcript |
| ENSG00000148219 | ASTN2 | -0,575290691 | 0,671151006 | 0,11138 | 0,42416781 | astrotactin 2 |
| ENSG00000160326 | SLC2A6 | -0,575454498 | 0,671074807 | 0,03942 | 0,2687016 | solute carrier family 2 member 6 |
| ENSG00000236560 | PPP1R11 | -0,575529888 | 0,67103974 | 0,21796 | 0,53560075 | protein phosphatase 1 regulatory inhibitor subunit 11 |
| ENSG00000232788 | ITGA6-AS1 | -0,575564697 | 0,671023549 | 0,48455 | 0,72031474 | ITGA6 antisense RNA 1 |
| ENSG00000167967 | E4F1 | -0,575599485 | 0,671007369 | 0,02496 | 0,21192916 | E4F transcription factor 1 |
| ENSG00000176894 | PXMP2 | -0,575758608 | 0,670933364 | 0,00371 | 0,07421558 | peroxisomal membrane protein 2 |
| ENSG00000040487 | SLC66A1 | -0,575874544 | 0,670879449 | 0,20046 | 0,52110549 | solute carrier family 66 member 1 |
| ENSG00000287559 | AL136305,1 | -0,575947146 | 0,670845689 | 0,09602 | 0,39921834 | novel transcript, antisense to RBM24 |

| | | | | | | |
|---|---|---|---|---|---|---|
| ENSG00000217612 | AL031119,1 | -0,575977436 | 0,670831604 | 0,51658 | 0,74202621 | BTB/POZ domain zinc finger protein pseudogene |
| ENSG00000112964 | GHR | -0,575999834 | 0,67082119 | 0,15654 | 0,48425071 | growth hormone receptor |
| ENSG00000105509 | HAS1 | -0,57614176 | 0,670755201 | 0,5588 | 0,77025265 | hyaluronan synthase 1 |
| ENSG00000237414 | AL365204,2 | -0,576390437 | 0,670639593 | 0,11263 | 0,42641188 | novel transcript |
| ENSG00000223884 | AC068481,1 | -0,576485044 | 0,670595616 | 0,26041 | 0,56383537 | novel transcript, antisense to RNF144A |
| ENSG00000287826 | AC090994,1 | -0,576525087 | 0,670577003 | 0,12733 | 0,45051698 | novel transcript |
| ENSG00000128928 | IVD | -0,576574562 | 0,670554007 | 0,09477 | 0,39724142 | isovaleryl-CoA dehydrogenase |
| ENSG00000167904 | TMEM68 | -0,576788596 | 0,670454533 | 0,09764 | 0,40273014 | transmembrane protein 68 |
| ENSG00000129255 | MPDU1 | -0,576857132 | 0,670422684 | 0,15424 | 0,48258262 | mannose-P-dolichol utilization defect 1 |
| ENSG00000155875 | SAXO1 | -0,576887895 | 0,670408388 | 0,38037 | 0,64797239 | stabilizer of axonemal microtubules 1 |
| ENSG00000081818 | PCDHB4 | -0,577060312 | 0,670328272 | 0,007 | 0,10655525 | protocadherin beta 4 |
| ENSG00000105612 | DNASE2 | -0,577211023 | 0,67025825 | 0,25794 | 0,56199743 | deoxyribonuclease 2, lysosomal |
| ENSG00000248266 | AC108142,1 | -0,577246462 | 0,670241786 | 0,56859 | 0,7761508 | novel transcript |
| ENSG00000124440 | HIF3A | -0,577341737 | 0,670197524 | 0,10468 | 0,41317844 | hypoxia inducible factor 3 subunit alpha |
| ENSG00000228045 | DNAJA1P6 | -0,577397602 | 0,670171573 | 0,49197 | 0,72531636 | DnaJ heat shock protein family (Hsp40) member A1 pseudogene 6 |
| ENSG00000216475 | AL024474,1 | -0,577624296 | 0,670066276 | 0,46494 | 0,7055215 | single-stranded DNA binding protein 1 (SSBP1) pseudogene |
| ENSG00000214465 | SMARCE1P6 | -0,577657484 | 0,670050862 | 0,51106 | 0,73835122 | SWI/SNF related, matrix associated, actin dependent regulator of chromatin, subfamily e, member 1 pseudogene 6 |
| ENSG00000161681 | SHANK1 | -0,578072142 | 0,669858304 | 0,18305 | 0,50652733 | SH3 and multiple ankyrin repeat domains 1 |
| ENSG00000223891 | OSER1-DT | -0,578134675 | 0,66982927 | 0,0871 | 0,3821708 | OSER1 divergent transcript |
| ENSG00000287867 | AC007402,2 | -0,578135043 | 0,669829099 | 0,14372 | 0,47123322 | novel transcript |
| ENSG00000236008 | LINC01814 | -0,578205288 | 0,669796486 | 0,04526 | 0,2876168 | long intergenic non-protein coding RNA 1814 |
| ENSG00000120210 | INSL6 | -0,578395279 | 0,669708285 | 0,37861 | 0,64679625 | insulin like 6 |
| ENSG00000244560 | AC004890,2 | -0,578461737 | 0,669677435 | 0,07356 | 0,35672172 | zinc finger protein pseudogene |
| ENSG00000235823 | OLMALINC | -0,57853704 | 0,669642482 | 0,08591 | 0,38009386 | oligodendrocyte maturation-associated long intergenic non-coding RNA |
| ENSG00000095906 | NUBP2 | -0,578711624 | 0,669561452 | 0,07592 | 0,36193676 | nucleotide binding protein 2 |
| ENSG00000129473 | BCL2L2 | -0,578960391 | 0,669446008 | 0,09577 | 0,39859038 | BCL2 like 2 |
| ENSG00000144040 | SFXN5 | -0,578971827 | 0,669440701 | 0,02764 | 0,22402593 | sideroflexin 5 |
| ENSG00000258640 | RPL21P5 | -0,578998209 | 0,66942846 | 0,4961 | 0,72831792 | ribosomal protein L21 pseudogene 5 |

| | | | | | | |
|---|---|---|---|---|---|---|
| ENSG00000120471 | TP53AIP1 | -0,579028141 | 0,669414571 | 0,43982 | 0,6885488 | tumor protein p53 regulated apoptosis inducing protein 1 |
| ENSG00000243915 | THAP12P2 | -0,579186599 | 0,66934105 | 0,43434 | 0,68505666 | THAP domain containing 12 pseudogene 2 |
| ENSG00000219410 | AC125494,1 | -0,579272981 | 0,669300974 | 0,26059 | 0,56400103 | novel transcript |
| ENSG00000230373 | GOLGA6L5P | -0,579983087 | 0,66897162 | 0,08652 | 0,38100206 | golgin A6 family like 5, pseudogene |
| ENSG00000272088 | AL512413,1 | -0,580003142 | 0,668962321 | 0,62343 | 0,8091647 | novel transcript |
| ENSG00000237484 | LINC01684 | -0,580214903 | 0,668864136 | 0,05507 | 0,31407633 | long intergenic non-protein coding RNA 1684 |
| ENSG00000278231 | AL133342,1 | -0,580244171 | 0,668850567 | 0,50359 | 0,732867 | novel transcript |
| ENSG00000269858 | EGLN2 | -0,580382648 | 0,668786371 | 0,10658 | 0,41635243 | egl-9 family hypoxia inducible factor 2 |
| ENSG00000234426 | AL139042,1 | -0,580456471 | 0,66875215 | 0,22547 | 0,5407493 | novel transcript |
| ENSG00000146221 | TCTE1 | -0,580653628 | 0,668660765 | 0,1803 | 0,50407792 | t-complex-associated-testis-expressed 1 |
| ENSG00000254995 | STX16-NPEPL1 | -0,580670281 | 0,668653047 | 0,02356 | 0,20483723 | STX16-NPEPL1 readthrough (NMD candidate) |
| ENSG00000237596 | AL138828,1 | -0,580915587 | 0,668539363 | 0,13937 | 0,46666406 | novel transcript |
| ENSG00000284057 | AP001273,2 | -0,580927044 | 0,668534054 | 0,84837 | 0,93094401 | novel protein, C11orf54-MED17 readthrough |
| ENSG00000180044 | C3orf80 | -0,580957263 | 0,668520051 | 0,41922 | 0,67514348 | chromosome 3 open reading frame 80 |
| ENSG00000227463 | AL390838,1 | -0,58119542 | 0,668409702 | 0,08365 | 0,37578481 | novel transcript |
| ENSG00000205238 | SPDYE2 | -0,581239437 | 0,668389309 | 0,08019 | 0,36972652 | speedy/RINGO cell cycle regulator family member E2 |
| ENSG00000129347 | KRI1 | -0,581334606 | 0,668345219 | 0,05683 | 0,31784869 | KRI1 homolog |
| ENSG00000234933 | CDC42P1 | -0,581334868 | 0,668345098 | 0,53438 | 0,75405583 | cell division cycle 42 pseudogene 1 |
| ENSG00000109625 | CPZ | -0,581355314 | 0,668335626 | 0,58133 | 0,78419712 | carboxypeptidase Z |
| ENSG00000255367 | AC127526,2 | -0,581470054 | 0,668282475 | 0,19551 | 0,51643293 | novel transcript |
| ENSG00000101220 | C20orf27 | -0,581498889 | 0,668269118 | 0,15747 | 0,48457045 | chromosome 20 open reading frame 27 |
| ENSG00000253517 | XRCC6P4 | -0,581551529 | 0,668244735 | 0,32251 | 0,60577115 | X-ray repair cross complementing 6 pseudogene 4 |
| ENSG00000130762 | ARHGEF16 | -0,581591406 | 0,668226265 | 0,01049 | 0,13225385 | Rho guanine nucleotide exchange factor 16 |
| ENSG00000165548 | TMEM63C | -0,581680649 | 0,668184931 | 0,22763 | 0,54168591 | transmembrane protein 63C |
| ENSG00000213592 | PPIAP42 | -0,581725404 | 0,668164202 | 0,44355 | 0,69123144 | peptidylprolyl isomerase A pseudogene 42 |
| ENSG00000175779 | LINC02694 | -0,581831829 | 0,668114915 | 0,27332 | 0,57344244 | long intergenic non-protein coding RNA 2694 |
| ENSG00000262049 | AC139530,1 | -0,581861769 | 0,66810105 | 0,14098 | 0,46791293 | novel transcript, antisense to HGS |
| ENSG00000049089 | COL9A2 | -0,582034096 | 0,668021251 | 0,32786 | 0,60960585 | collagen type IX alpha 2 chain |
| ENSG00000255874 | PRECSIT | -0,582258355 | 0,667917419 | 0,16664 | 0,49255702 | p53 regulated carcinoma associated Stat3 activating long intergenic non-protein coding transcript |
| ENSG00000224167 | LINC01357 | -0,582417117 | 0,667843922 | 0,43559 | 0,68618266 | long intergenic non-protein coding RNA 1357 |
| ENSG00000182685 | BRICD5 | -0,582461708 | 0,66782328 | 0,08318 | 0,37544257 | BRICHOS domain containing 5 |
| ENSG00000090615 | GOLGA3 | -0,582463085 | 0,667822643 | 0,11156 | 0,42463997 | golgin A3 |

| | | | | | | |
|---|---|---|---|---|---|---|
| ENSG00000267053 | AC092296,2 | -0,582543752 | 0,667785303 | 0,36117 | 0,63317043 | novel transcript |
| ENSG00000067715 | SYT1 | -0,582558401 | 0,667778523 | 0,16749 | 0,49362211 | synaptotagmin 1 |
| ENSG00000112164 | GLP1R | -0,582628609 | 0,667746026 | 0,31639 | 0,60238875 | glucagon like peptide 1 receptor |
| ENSG00000101670 | LIPG | -0,58267805 | 0,667723143 | 0,12971 | 0,45363524 | lipase G, endothelial type |
| ENSG00000159216 | RUNX1 | -0,582804667 | 0,667664544 | 0,10364 | 0,41173831 | RUNX family transcription factor 1 |
| ENSG00000261642 | AL713866,1 | -0,582878478 | 0,667630385 | 0,25094 | 0,55711287 | novel transcript |
| ENSG00000276250 | AC127024,6 | -0,582887529 | 0,667626197 | 0,17935 | 0,5036829 | novel transcript, antisense to CRLF3 |
| ENSG00000171817 | ZNF540 | -0,582891957 | 0,667624148 | 0,0723 | 0,35498117 | zinc finger protein 540 |
| ENSG00000286627 | AP005900,1 | -0,582894539 | 0,667622953 | 0,34309 | 0,62060539 | novel transcript, antisense to PTPRM |
| ENSG00000231691 | AL592114,3 | -0,582998379 | 0,667574902 | 0,46384 | 0,70455791 | novel transcript |
| ENSG00000029534 | ANK1 | -0,5830631 | 0,667544954 | 0,21909 | 0,53659265 | ankyrin 1 |
| ENSG00000250802 | ZBED3-AS1 | -0,583097299 | 0,66752913 | 0,08829 | 0,38452862 | ZBED3 antisense RNA 1 |
| ENSG00000166471 | TMEM41B | -0,583314919 | 0,667428446 | 0,13349 | 0,4594073 | transmembrane protein 41B |
| ENSG00000108474 | PIGL | -0,583328455 | 0,667422184 | 0,0821 | 0,37331809 | phosphatidylinositol glycan anchor biosynthesis class L |
| ENSG00000172324 | OR5A2 | -0,583340282 | 0,667416713 | 0,13787 | 0,46497045 | olfactory receptor family 5 subfamily A member 2 |
| ENSG00000282556 | AC068733,3 | -0,583496259 | 0,667344559 | 0,03424 | 0,24991804 | novel transcript, antisense to PRKCDBP |
| ENSG00000220305 | HNRNPH1P1 | -0,58355629 | 0,667316791 | 0,24288 | 0,55182091 | heterogeneous nuclear ribonucleoprotein H1 pseudogene 1 |
| ENSG00000214435 | AS3MT | -0,583636603 | 0,667279643 | 0,15145 | 0,47962207 | arsenite methyltransferase |
| ENSG00000054938 | CHRDL2 | -0,583656038 | 0,667270654 | 0,36653 | 0,63641349 | chordin like 2 |
| ENSG00000159176 | CSRP1 | -0,583670935 | 0,667263764 | 0,0354 | 0,25490938 | cysteine and glycine rich protein 1 |
| ENSG00000179406 | LINC00174 | -0,583704424 | 0,667248275 | 0,10022 | 0,4067769 | long intergenic non-protein coding RNA 174 |
| ENSG00000253051 | SNORA31B | -0,583731025 | 0,667235972 | 0,16134 | 0,48892403 | small nucleolar RNA, H/ACA box 31B |
| ENSG00000287733 | AC083862,2 | -0,583965561 | 0,66712751 | 0,1841 | 0,50785017 | novel transcript |
| ENSG00000237941 | KCNQ1DN | -0,584059959 | 0,66708386 | 0,30224 | 0,59294633 | KCNQ1 downstream neighbor |
| ENSG00000285971 | AC023095,1 | -0,584190818 | 0,667023355 | 0,18749 | 0,51079428 | novel transcript |
| ENSG00000176532 | PRR15 | -0,584213586 | 0,667012829 | 0,52852 | 0,75090627 | proline rich 15 |
| ENSG00000229950 | TFAP2A-AS1 | -0,584414433 | 0,666919976 | 0,53234 | 0,75291763 | TFAP2A antisense RNA 1 |
| ENSG00000278464 | AC068506,1 | -0,584612497 | 0,666828422 | 0,57074 | 0,77748547 | novel transcript |
| ENSG00000163795 | ZNF513 | -0,58480274 | 0,666740496 | 0,0872 | 0,38234284 | zinc finger protein 513 |
| ENSG00000196951 | SCOC-AS1 | -0,584972099 | 0,666662231 | 0,29863 | 0,59089801 | SCOC antisense RNA 1 |
| ENSG00000108830 | RND2 | -0,585095354 | 0,666605278 | 0,06681 | 0,34222837 | Rho family GTPase 2 |
| ENSG00000169668 | BCRP2 | -0,585473244 | 0,666430695 | 0,0777 | 0,36480631 | BCR pseudogene 2 |

| | | | | | | |
|---|---|---|---|---|---|---|
| ENSG00000261330 | PPIAP50 | -0,585655734 | 0,666346402 | 0,29841 | 0,59076323 | peptidylprolyl isomerase A pseudogene 50 |
| ENSG00000105290 | APLP1 | -0,585869182 | 0,666247823 | 0,05871 | 0,32371785 | amyloid beta precursor like protein 1 |
| ENSG00000260693 | AC026150,1 | -0,585884682 | 0,666240664 | 0,44606 | 0,69284796 | novel transcript |
| ENSG00000281960 | AC254960,1 | -0,585884682 | 0,666240664 | 0,44606 | 0,69284796 | novel transcript |
| ENSG00000249937 | LINC02223 | -0,586556436 | 0,665930519 | 0,31341 | 0,6002104 | long intergenic non-protein coding RNA 2223 |
| ENSG00000133069 | TMCC2 | -0,586606395 | 0,665907459 | 0,11447 | 0,42986344 | transmembrane and coiled-coil domain family 2 |
| ENSG00000163945 | UVSSA | -0,586617648 | 0,665902265 | 0,08828 | 0,38452862 | UV stimulated scaffold protein A |
| ENSG00000231680 | LINC02723 | -0,586623639 | 0,665899499 | 0,53097 | 0,75236575 | long intergenic non-protein coding RNA 2723 |
| ENSG00000173452 | TMEM196 | -0,586809057 | 0,665813922 | 0,14568 | 0,4731066 | transmembrane protein 196 |
| ENSG00000223776 | LGALS8-AS1 | -0,586964723 | 0,665742085 | 0,33898 | 0,61800397 | LGALS8 antisense RNA 1 |
| ENSG00000056487 | PHF21B | -0,587011015 | 0,665720724 | 0,00642 | 0,10141052 | PHD finger protein 21B |
| ENSG00000260552 | AC023043,1 | -0,587094465 | 0,665682218 | 0,63221 | 0,81450663 | novel transcript, antisense to MOCOS |
| ENSG00000234127 | TRIM26 | -0,587288723 | 0,66559259 | 0,10064 | 0,40726911 | tripartite motif containing 26 |
| ENSG00000173673 | HES3 | -0,587501134 | 0,6654946 | 0,47581 | 0,71433501 | hes family bHLH transcription factor 3 |
| ENSG00000175701 | MTLN | -0,587724838 | 0,665391417 | 0,07247 | 0,35547283 | mitoregulin |
| ENSG00000173264 | GPR137 | -0,587725495 | 0,665391114 | 0,17482 | 0,49908125 | G protein-coupled receptor 137 |
| ENSG00000180089 | TMEM86B | -0,588212029 | 0,665166756 | 0,18524 | 0,50876814 | transmembrane protein 86B |
| ENSG00000003249 | DBNDD1 | -0,588263908 | 0,665142837 | 0,02043 | 0,18959706 | dysbindin domain containing 1 |
| ENSG00000230756 | RHOQP3 | -0,588285692 | 0,665132794 | 0,58436 | 0,78572744 | ras homolog family member Q pseudogene 3 |
| ENSG00000135966 | TGFBRAP1 | -0,588342068 | 0,665106803 | 0,16745 | 0,49362211 | transforming growth factor beta receptor associated protein 1 |
| ENSG00000271380 | AL451085,2 | -0,588345798 | 0,665105083 | 0,25551 | 0,55999807 | uncharacterized LOC101928120 |
| ENSG00000284456 | LINC02076 | -0,588423081 | 0,665069456 | 0,23837 | 0,54875544 | long intergenic non-protein coding RNA 2076 |
| ENSG00000033011 | ALG1 | -0,588444177 | 0,66505973 | 0,1678 | 0,49395055 | ALG1 chitobiosyldiphosphodolichol beta-mannosyltransferase |
| ENSG00000143374 | TARS2 | -0,588594649 | 0,664990369 | 0,18846 | 0,51181123 | threonyl-tRNA synthetase 2, mitochondrial |
| ENSG00000261351 | AC116913,1 | -0,588771013 | 0,664909081 | 0,10798 | 0,41856292 | novel transcript, antisense to MAP2K1 |
| ENSG00000261840 | AC093249,6 | -0,588800735 | 0,664895383 | 0,19381 | 0,51547251 | uncharacterized LOC730183 |
| ENSG00000265366 | GLUD1P2 | -0,588811113 | 0,6648906 | 0,2852 | 0,58172571 | glutamate dehydrogenase 1 pseudogene 2 |
| ENSG00000138134 | STAMBPL1 | -0,588917043 | 0,664841782 | 0,06892 | 0,34690198 | STAM binding protein like 1 |
| ENSG00000231859 | AC079781,1 | -0,589066215 | 0,664773042 | 0,16869 | 0,49492999 | novel pseudogene |
| ENSG00000259017 | AL133163,3 | -0,589082476 | 0,66476555 | 0,31693 | 0,60268749 | novel transcript |
| ENSG00000224477 | AL109933,1 | -0,589128849 | 0,664744182 | 0,57438 | 0,77954257 | novel transcript |

| | | | | | | |
|---|---|---|---|---|---|---|
| ENSG00000130764 | LRRC47 | -0,589136974 | 0,664740439 | 0,1386 | 0,46578766 | leucine rich repeat containing 47 |
| ENSG00000151006 | PRSS53 | -0,589342807 | 0,664645605 | 0,0912 | 0,39063588 | serine protease 53 |
| ENSG00000243197 | TUSC7 | -0,589549606 | 0,66455034 | 0,18707 | 0,5105374 | tumor suppressor candidate 7 |
| ENSG00000184619 | KRBA2 | -0,589702123 | 0,66448009 | 0,10087 | 0,40770184 | KRAB-A domain containing 2 |
| ENSG00000281490 | CICP14 | -0,589759347 | 0,664453734 | 0,06418 | 0,33569685 | capicua transcriptional repressor pseudogene 14 |
| ENSG00000260898 | ADPGK-AS1 | -0,590097783 | 0,664297881 | 0,12189 | 0,44093327 | ADPGK antisense RNA 1 |
| ENSG00000286973 | AC087490,1 | -0,590213893 | 0,664244419 | 0,43246 | 0,68411566 | novel transcript |
| ENSG00000165973 | NELL1 | -0,590290118 | 0,664209325 | 0,26106 | 0,56416552 | neural EGFL like 1 |
| ENSG00000271155 | AL161729,1 | -0,590390426 | 0,664163145 | 0,20019 | 0,52107026 | novel transcript, antisense to PTCH1 |
| ENSG00000168904 | LRRC28 | -0,590411357 | 0,664153509 | 0,15078 | 0,47896194 | leucine rich repeat containing 28 |
| ENSG00000272764 | AL596094,1 | -0,590572933 | 0,664079131 | 0,46908 | 0,70862237 | novel transcript |
| ENSG00000198270 | TMEM116 | -0,590748353 | 0,663998389 | 0,12776 | 0,45114976 | transmembrane protein 116 |
| ENSG00000138769 | CDKL2 | -0,590918639 | 0,66392002 | 0,07401 | 0,35746749 | cyclin dependent kinase like 2 |
| ENSG00000146426 | TIAM2 | -0,591053427 | 0,663857994 | 0,03938 | 0,26865324 | TIAM Rac1 associated GEF 2 |
| ENSG00000077264 | PAK3 | -0,59117665 | 0,663801295 | 0,04528 | 0,2876168 | p21 (RAC1) activated kinase 3 |
| ENSG00000213409 | C1QBPP2 | -0,591235761 | 0,663774098 | 0,11697 | 0,4343753 | complement C1q binding protein pseudogene 2 |
| ENSG00000185917 | SETD4 | -0,591463836 | 0,663669171 | 0,11033 | 0,42293369 | SET domain containing 4 |
| ENSG00000282057 | AC092807,3 | -0,591581559 | 0,663615018 | 0,07736 | 0,36423167 | novel transcript |
| ENSG00000122367 | LDB3 | -0,59166939 | 0,663574618 | 0,04768 | 0,29344099 | LIM domain binding 3 |
| ENSG00000127418 | FGFRL1 | -0,591673246 | 0,663572845 | 0,00241 | 0,05654945 | fibroblast growth factor receptor like 1 |
| ENSG00000202408 | RNVU1-21 | -0,591708429 | 0,663556663 | 0,34011 | 0,61889407 | RNA, variant U1 small nuclear 21 |
| ENSG00000267044 | AC005757,1 | -0,59172742 | 0,663547928 | 0,56206 | 0,77155921 | novel transcript, antisense to PPP1R37 |
| ENSG00000135824 | RGS8 | -0,591899795 | 0,663468651 | 0,16329 | 0,48972962 | regulator of G protein signaling 8 |
| ENSG00000108370 | RGS9 | -0,591940999 | 0,663449702 | 0,28324 | 0,58053323 | regulator of G protein signaling 9 |
| ENSG00000260923 | LINC02193 | -0,592135086 | 0,663360454 | 0,04498 | 0,28673676 | long intergenic non-protein coding RNA 2193 |
| ENSG00000144407 | PTH2R | -0,592192138 | 0,663334222 | 0,31648 | 0,60238875 | parathyroid hormone 2 receptor |
| ENSG00000239482 | AC112487,1 | -0,592221418 | 0,663320759 | 0,28216 | 0,57949345 | novel transcript |
| ENSG00000137841 | PLCB2 | -0,592369572 | 0,663252645 | 0,02099 | 0,19244681 | phospholipase C beta 2 |
| ENSG00000139044 | B4GALNT3 | -0,592411885 | 0,663233192 | 0,14126 | 0,46803321 | beta-1,4-N-acetyl-galactosaminyltransferase 3 |
| ENSG00000185133 | INPP5J | -0,592883902 | 0,663016233 | 0,20369 | 0,52347929 | inositol polyphosphate-5-phosphatase J |
| ENSG00000162444 | RBP7 | -0,59291411 | 0,66300235 | 0,55084 | 0,76486471 | retinol binding protein 7 |
| ENSG00000198429 | ZNF69 | -0,592931923 | 0,662994164 | 0,01695 | 0,17088104 | zinc finger protein 69 |
| ENSG00000255129 | AP000880,1 | -0,592955472 | 0,662983342 | 0,37254 | 0,64149653 | novel transcript |

| | | | | | | |
|---|---|---|---|---|---|---|
| ENSG00000232934 | AL157786,1 | -0,593005362 | 0,662960416 | 0,40285 | 0,66398107 | novel transcript, antisense to ACSL5 |
| ENSG00000230495 | AL132765,1 | -0,593168743 | 0,662885342 | 0,55131 | 0,76511664 | ras-related pseudogene |
| ENSG00000229791 | AL356259,1 | -0,593222351 | 0,662860711 | 0,57036 | 0,77733192 | novel transcript |
| ENSG00000070808 | CAMK2A | -0,593352154 | 0,662801074 | 0,1726 | 0,49744497 | calcium/calmodulin dependent protein kinase II alpha |
| ENSG00000171533 | MAP6 | -0,593361544 | 0,66279676 | 0,02757 | 0,22376406 | microtubule associated protein 6 |
| ENSG00000285212 | AL009178,3 | -0,593824638 | 0,662584042 | 0,4908 | 0,72480683 | novel transcript |
| ENSG00000115919 | KYNU | -0,593875696 | 0,662560593 | 0,05187 | 0,30606616 | kynureninase |
| ENSG00000164970 | FAM219A | -0,593976998 | 0,662514071 | 0,0661 | 0,34035709 | family with sequence similarity 219 member A |
| ENSG00000279442 | AP000542,2 | -0,594004604 | 0,662501395 | 0,11132 | 0,42416781 | novel transcript |
| ENSG00000176401 | EID2B | -0,594057228 | 0,662477229 | 0,11476 | 0,43021597 | EP300 interacting inhibitor of differentiation 2B |
| ENSG00000197928 | ZNF677 | -0,594286787 | 0,662371825 | 0,02193 | 0,19698513 | zinc finger protein 677 |
| ENSG00000259917 | HNRNPLP2 | -0,594316707 | 0,662358089 | 0,35832 | 0,63185282 | heterogeneous nuclear ribonucleoprotein L pseudogene 2 |
| ENSG00000204335 | SP5 | -0,594760418 | 0,662154407 | 0,32227 | 0,60577115 | Sp5 transcription factor |
| ENSG00000256940 | PPP1R14B-AS | -0,594800748 | 0,662135897 | 0,21778 | 0,53550245 | PPP1R14B antisense RNA 1 |
| ENSG00000287762 | AC108050,1 | -0,594828383 | 0,662123214 | 0,41718 | 0,6738387 | novel transcript, antisense to LIMCH1 |
| ENSG00000236107 | SCN1A-AS1 | -0,594928972 | 0,66207705 | 0,31093 | 0,59883009 | SCN1A and SCN9A antisense RNA 1 |
| ENSG00000028277 | POU2F2 | -0,594979982 | 0,662053641 | 0,09881 | 0,40463341 | POU class 2 homeobox 2 |
| ENSG00000276856 | TUBGCP5 | -0,595324049 | 0,661895767 | 0,18818 | 0,5113023 | tubulin gamma complex associated protein 5 |
| ENSG00000149201 | CCDC81 | -0,595539816 | 0,661796783 | 0,05207 | 0,30670474 | coiled-coil domain containing 81 |
| ENSG00000135297 | MTO1 | -0,595550062 | 0,661792083 | 0,16267 | 0,48945635 | mitochondrial tRNA translation optimization 1 |
| ENSG00000267246 | AC091132,5 | -0,5955516 | 0,661791378 | 0,14409 | 0,47166592 | novel transcript |
| ENSG00000124257 | NEURL2 | -0,595620945 | 0,661759568 | 0,61998 | 0,80745864 | neuralized E3 ubiquitin protein ligase 2 |
| ENSG00000280512 | MUC20-OT1 | -0,595630647 | 0,661755118 | 0,09378 | 0,39568176 | MUC20 overlapping transcript |
| ENSG00000273454 | AC112503,2 | -0,595631607 | 0,661754678 | 0,5492 | 0,76404333 | novel transcript |
| ENSG00000125814 | NAPB | -0,595832386 | 0,661662588 | 0,07906 | 0,36719701 | NSF attachment protein beta |
| ENSG00000260136 | AC008915,2 | -0,59599844 | 0,661586436 | 0,29114 | 0,58616212 | novel transcript |
| ENSG00000274012 | RN7SL2 | -0,596414351 | 0,661395736 | 0,06003 | 0,32549025 | RNA component of signal recognition particle 7SL2 |
| ENSG00000161509 | GRIN2C | -0,596696806 | 0,661266258 | 0,17575 | 0,49972993 | glutamate ionotropic receptor NMDA type subunit 2C |
| ENSG00000125804 | FAM182A | -0,596738974 | 0,661246931 | 0,19442 | 0,51562158 | family with sequence similarity 182 member A |
| ENSG00000227314 | MSH5 | -0,596767164 | 0,66123401 | 0,48855 | 0,72346703 | mutS homolog 5 |
| ENSG00000267169 | AC022098,1 | -0,596906403 | 0,661170196 | 0,14283 | 0,47034016 | novel transcript, antisense to LPHN1 |
| ENSG00000288348 | AC022098,6 | -0,596906403 | 0,661170196 | 0,14283 | 0,47034016 | novel transcript, antisense to LPHN1 |
| ENSG00000228839 | PIK3IP1-DT | -0,59691652 | 0,661165559 | 0,43731 | 0,68715611 | PIK3IP1 divergent transcript |

| | | | | | | |
|---|---|---|---|---|---|---|
| ENSG00000182324 | KCNJ14 | -0,597086689 | 0,661087578 | 0,12358 | 0,4443454 | potassium inwardly rectifying channel subfamily J member 14 |
| ENSG00000133519 | ZDHHC8P1 | -0,597342574 | 0,660970334 | 0,41712 | 0,6738387 | ZDHHC8 pseudogene 1 |
| ENSG00000261079 | AC009053,2 | -0,597391417 | 0,660947957 | 0,0885 | 0,38476689 | novel transcript |
| ENSG00000159753 | CARMIL2 | -0,597544379 | 0,660877883 | 0,25711 | 0,56155289 | capping protein regulator and myosin 1 linker 2 |
| ENSG00000236289 | GACAT3 | -0,597577933 | 0,660862513 | 0,43 | 0,68268477 | gastric cancer associated transcript 3 |
| ENSG00000258099 | ATXN2-AS | -0,597629065 | 0,660839091 | 0,62608 | 0,81067751 | ATXN2 antisense RNA |
| ENSG00000235098 | ANKRD65 | -0,597693598 | 0,660809532 | 0,12917 | 0,45253718 | ankyrin repeat domain 65 |
| ENSG00000229202 | PRR3 | -0,597745845 | 0,660785601 | 0,01607 | 0,16620111 | proline rich 3 |
| ENSG00000170837 | GPR27 | -0,597814532 | 0,660754142 | 0,03071 | 0,23637289 | G protein-coupled receptor 27 |
| ENSG00000197594 | ENPP1 | -0,598010232 | 0,660664517 | 0,13508 | 0,46159318 | ectonucleotide pyrophosphatase/phosphodiesterase 1 |
| ENSG00000284337 | AC013271,1 | -0,598148971 | 0,660600986 | 0,69856 | 0,85408973 | novel transcript |
| ENSG00000254812 | AC067930,2 | -0,598277108 | 0,660542316 | 0,33779 | 0,61751351 | novel transcript |
| ENSG00000236778 | INTS6-AS1 | -0,598344523 | 0,660511451 | 0,06419 | 0,33569685 | INTS6 antisense RNA 1 |
| ENSG00000278611 | ZNF426-DT | -0,598420314 | 0,660476752 | 0,14366 | 0,47123322 | ZNF426 divergent transcript |
| ENSG00000235411 | DRG1P1 | -0,598576907 | 0,660405066 | 0,47408 | 0,71294959 | DRG1 pseudogene 1 |
| ENSG00000163046 | ANKRD30BL | -0,598609185 | 0,660390291 | 0,49717 | 0,72865711 | ankyrin repeat domain 30B like |
| ENSG00000270139 | AP001172,2 | -0,598654099 | 0,660369732 | 0,38227 | 0,64928247 | novel transcript |
| ENSG00000140945 | CDH13 | -0,598686299 | 0,660354993 | 0,15827 | 0,48529549 | cadherin 13 |
| ENSG00000273747 | AC022558,3 | -0,598686695 | 0,660354812 | 0,17472 | 0,4989163 | novel transcript, antisense to BTBD1 |
| ENSG00000268173 | AC007192,1 | -0,598712386 | 0,660343053 | 0,6761 | 0,84061748 | novel protein, readthrough between PIK3R2 and IFI30 |
| ENSG00000157895 | C12orf43 | -0,598726039 | 0,660336803 | 0,15767 | 0,48476347 | chromosome 12 open reading frame 43 |
| ENSG00000284814 | ADCY4 | -0,598740646 | 0,660330118 | 0,24427 | 0,55253387 | adenylate cyclase 4 |
| ENSG00000253363 | AC090809,1 | -0,598951564 | 0,660233586 | 0,28742 | 0,58382745 | novel transcript |
| ENSG00000254708 | MMADHCP2 | -0,598966225 | 0,660226877 | 0,27152 | 0,57285347 | MMADHC pseudogene 2 |
| ENSG00000134590 | RTL8C | -0,599193668 | 0,660122799 | 0,05355 | 0,31056376 | retrotransposon Gag like 8C |
| ENSG00000234323 | LINC01505 | -0,599209708 | 0,66011546 | 0,14369 | 0,47123322 | long intergenic non-protein coding RNA 1505 |
| ENSG00000062038 | CDH3 | -0,599271524 | 0,660087177 | 0,26903 | 0,57085484 | cadherin 3 |
| ENSG00000274679 | SPATC1L | -0,599332584 | 0,66005924 | 0,03671 | 0,25964506 | spermatogenesis and centriole associated 1 like |
| ENSG00000239212 | RPL6P7 | -0,59945173 | 0,660004731 | 0,32298 | 0,60584543 | ribosomal protein L6 pseudogene 7 |
| ENSG00000154265 | ABCA5 | -0,599491141 | 0,659986701 | 0,21215 | 0,5308403 | ATP binding cassette subfamily A member 5 |
| ENSG00000165275 | TRMT10B | -0,599539921 | 0,659964386 | 0,12058 | 0,43944057 | tRNA methyltransferase 10B |
| ENSG00000229207 | SERPINH1P1 | -0,599557957 | 0,659956135 | 0,34941 | 0,62550599 | serpin family H member 1 pseudogene 1 |

| | | | | | | |
|---|---|---|---|---|---|---|
| ENSG00000255081 | AP003168,2 | -0,599558951 | 0,659955681 | 0,48984 | 0,7241832 | novel transcript |
| ENSG00000168517 | HEXIM2 | -0,599622429 | 0,659926644 | 0,14052 | 0,46742953 | HEXIM P-TEFb complex subunit 2 |
| ENSG00000064932 | SBNO2 | -0,599871199 | 0,659812859 | 0,06606 | 0,34031663 | strawberry notch homolog 2 |
| ENSG00000251250 | AC091951,3 | -0,599911049 | 0,659794635 | 0,25585 | 0,56047793 | pseudogene similar to part of hect domain and RLD 2 HERC2 |
| ENSG00000260331 | AC079148,1 | -0,600002965 | 0,659752599 | 0,38859 | 0,65310114 | novel transcript |
| ENSG00000171084 | FAM86JP | -0,600070722 | 0,659721614 | 0,17205 | 0,49710979 | family with sequence similarity 86 member J, pseudogene |
| ENSG00000173626 | TRAPPC3L | -0,600226997 | 0,659650156 | 0,48126 | 0,71822668 | trafficking protein particle complex 3 like |
| ENSG00000029153 | ARNTL2 | -0,600239789 | 0,659644307 | 0,06082 | 0,32750733 | aryl hydrocarbon receptor nuclear translocator like 2 |
| ENSG00000176912 | TYMSOS | -0,600386996 | 0,659577004 | 0,12063 | 0,43944057 | TYMS opposite strand |
| ENSG00000286406 | AC138776,1 | -0,600532661 | 0,659510411 | 0,07507 | 0,35949119 | novel ankyrin repeat domain family pseudogene |
| ENSG00000255410 | AC091564,5 | -0,600623974 | 0,65946867 | 0,14915 | 0,47671235 | novel transcript |
| ENSG00000235288 | AC099329,1 | -0,600657503 | 0,659453343 | 0,15096 | 0,47923194 | novel transcript, antisense to CCBP2 |
| ENSG00000284194 | SCO2 | -0,600897287 | 0,659343748 | 0,22914 | 0,54270085 | synthesis of cytochrome C oxidase 2 |
| ENSG00000250582 | SMAD1-AS2 | -0,600977016 | 0,659307311 | 0,23105 | 0,54396771 | SMAD1 antisense RNA 2 |
| ENSG00000131831 | RAI2 | -0,601000705 | 0,659296485 | 0,31443 | 0,6011918 | retinoic acid induced 2 |
| ENSG00000274010 | ZFP91P1 | -0,601184549 | 0,659212475 | 0,31298 | 0,59987747 | ZFP91 pseudogene 1 |
| ENSG00000286989 | AL691432,4 | -0,601382318 | 0,659122115 | 0,29357 | 0,58800349 | novel transcript |
| ENSG00000126217 | MCF2L | -0,601439317 | 0,659096074 | 0,04531 | 0,2876168 | MCF,2 cell line derived transforming sequence like |
| ENSG00000196542 | SPTSSB | -0,601609852 | 0,65901817 | 0,30012 | 0,59160746 | serine palmitoyltransferase small subunit B |
| ENSG00000185915 | KLHL34 | -0,601622478 | 0,659012403 | 0,23986 | 0,54931011 | kelch like family member 34 |
| ENSG00000262663 | AC087222,1 | -0,601732343 | 0,658962219 | 0,10959 | 0,42172157 | novel transcript, sense intronic TBCD |
| ENSG00000196642 | RABL6 | -0,601901981 | 0,65888474 | 0,12327 | 0,44354688 | RAB, member RAS oncogene family like 6 |
| ENSG00000005206 | SPPL2B | -0,602020899 | 0,658830432 | 0,11216 | 0,4257168 | signal peptide peptidase like 2B |
| ENSG00000279851 | CR382287,1 | -0,602029448 | 0,658826528 | 0,53688 | 0,75562287 | novel transcript |
| ENSG00000275613 | AC243830,1 | -0,602174228 | 0,658760415 | 0,34951 | 0,62561612 | novel transcript |
| ENSG00000224897 | POT1-AS1 | -0,602178447 | 0,658758489 | 0,13579 | 0,462686 | POT1 antisense RNA 1 |
| ENSG00000254459 | AP002812,2 | -0,602241716 | 0,6587296 | 0,38377 | 0,65041894 | novel transcript, antisense to AAMDC |
| ENSG00000140320 | BAHD1 | -0,60235397 | 0,658678347 | 0,05642 | 0,31712073 | bromo adjacent homology domain containing 1 |
| ENSG00000286482 | AL117344,2 | -0,602386327 | 0,658663574 | 0,13695 | 0,46423633 | novel transcript |
| ENSG00000100314 | CABP7 | -0,602732775 | 0,658505422 | 0,23312 | 0,54557342 | calcium binding protein 7 |
| ENSG00000158792 | SPATA2L | -0,60279602 | 0,658476555 | 0,13091 | 0,45558936 | spermatogenesis associated 2 like |
| ENSG00000152700 | SAR1B | -0,603110895 | 0,658332855 | 0,10364 | 0,41173831 | secretion associated Ras related GTPase 1B |

| | | | | | | |
|---|---|---|---|---|---|---|
| ENSG00000229623 | METTL21AP1 | -0,603175324 | 0,658303456 | 0,31109 | 0,59885232 | methyltransferase like 21A pseudogene 1 |
| ENSG00000254630 | AP001922,2 | -0,603301597 | 0,65824584 | 0,27987 | 0,57751493 | novel transcript |
| ENSG00000261624 | NDUFB10P2 | -0,603504643 | 0,658153204 | 0,39244 | 0,65576476 | NADH:ubiquinone oxidoreductase subunit B10 pseudogene 2 |
| ENSG00000274184 | AC011815,2 | -0,603594747 | 0,6581121 | 0,19257 | 0,51490739 | novel transcript, antisense to ZSCAN30 |
| ENSG00000285651 | AL450992,2 | -0,603602539 | 0,658108546 | 0,12784 | 0,45133265 | novel transcript, antisense to THEM4 |
| ENSG00000132570 | PCBD2 | -0,603672513 | 0,658076627 | 0,05081 | 0,30326674 | pterin-4 alpha-carbinolamine dehydratase 2 |
| ENSG00000159307 | SCUBE1 | -0,603697981 | 0,65806501 | 0,35473 | 0,62930239 | signal peptide, CUB domain and EGF like domain containing 1 |
| ENSG00000116809 | ZBTB17 | -0,603812157 | 0,658012932 | 0,04061 | 0,2725433 | zinc finger and BTB domain containing 17 |
| ENSG00000120913 | PDLIM2 | -0,60390948 | 0,657968544 | 0,18483 | 0,50844776 | PDZ and LIM domain 2 |
| ENSG00000091664 | SLC17A6 | -0,604077927 | 0,657891726 | 0,22307 | 0,53927144 | solute carrier family 17 member 6 |
| ENSG00000282773 | GOLGA8Q | -0,604199777 | 0,657836163 | 0,46227 | 0,70351872 | golgin A8 family member Q |
| ENSG00000234566 | RPL7AP71 | -0,604428098 | 0,657732061 | 0,42312 | 0,67694038 | ribosomal protein L7a pseudogene 71 |
| ENSG00000278390 | AL354696,2 | -0,604462154 | 0,657716535 | 0,15115 | 0,47955917 | novel transcript, antisense to KBTBD7 and MTRF1 |
| ENSG00000131381 | RBSN | -0,604630509 | 0,657639788 | 0,07711 | 0,36384125 | rabenosyn, RAB effector |
| ENSG00000259868 | AL163952,1 | -0,60471305 | 0,657602163 | 0,42306 | 0,67691679 | novel transcript |
| ENSG00000116198 | CEP104 | -0,604821481 | 0,657552741 | 0,12758 | 0,45080057 | centrosomal protein 104 |
| ENSG00000253226 | HAUS1P3 | -0,605039535 | 0,657453363 | 0,36582 | 0,63598241 | HAUS augmin like complex subunit 1 pseudogene 3 |
| ENSG00000090006 | LTBP4 | -0,605092296 | 0,65742932 | 0,1984 | 0,51980353 | latent transforming growth factor beta binding protein 4 |
| ENSG00000011347 | SYT7 | -0,60516906 | 0,65739434 | 0,13337 | 0,4594073 | synaptotagmin 7 |
| ENSG00000256101 | AC092745,1 | -0,605217921 | 0,657372076 | 0,21374 | 0,53279778 | novel transcript |
| ENSG00000100399 | CHADL | -0,605278101 | 0,657344655 | 0,30591 | 0,59512079 | chondroadherin like |
| ENSG00000274225 | AP001065,3 | -0,605458013 | 0,657262686 | 0,08666 | 0,38138884 | novel transcript |
| ENSG00000157087 | ATP2B2 | -0,605589113 | 0,657202962 | 0,37055 | 0,63997937 | ATPase plasma membrane Ca2+ transporting 2 |
| ENSG00000288383 | AC018653,7 | -0,605613442 | 0,657191879 | 0,36832 | 0,63803074 | novel transcript |
| ENSG00000256967 | AC018653,3 | -0,605613442 | 0,657191879 | 0,36832 | 0,63803074 | novel transcript |
| ENSG00000235986 | GNL1 | -0,605624368 | 0,657186902 | 0,22153 | 0,53805234 | G protein nucleolar 1 (putative) |
| ENSG00000223355 | NOTCH4 | -0,605728741 | 0,657139359 | 0,105 | 0,41375455 | notch receptor 4 |
| ENSG00000196263 | ZNF471 | -0,605959152 | 0,657034416 | 0,08512 | 0,37850509 | zinc finger protein 471 |
| ENSG00000115425 | PECR | -0,606121236 | 0,656960604 | 0,08352 | 0,37570927 | peroxisomal trans-2-enoyl-CoA reductase |
| ENSG00000170160 | CCDC144A | -0,606758618 | 0,656670423 | 0,09905 | 0,40513672 | coiled-coil domain containing 144A |
| ENSG00000072832 | CRMP1 | -0,606983475 | 0,656568083 | 0,00079 | 0,0274628 | collapsin response mediator protein 1 |

| | | | | | | |
|---|---|---|---|---|---|---|
| ENSG00000102743 | SLC25A15 | -0,607325378 | 0,656412502 | 0,18481 | 0,50844776 | solute carrier family 25 member 15 |
| ENSG00000261210 | CLEC19A | -0,607355476 | 0,656398808 | 0,38734 | 0,65229554 | C-type lectin domain containing 19A |
| ENSG00000250920 | AC105460,1 | -0,607378774 | 0,656388208 | 0,16256 | 0,48945635 | novel transcript |
| ENSG00000092871 | RFFL | -0,607382527 | 0,6563865 | 0,08913 | 0,38617197 | ring finger and FYVE like domain containing E3 ubiquitin protein ligase |
| ENSG00000286689 | AC090946,1 | -0,607458489 | 0,656351941 | 0,14682 | 0,47471707 | novel transcript |
| ENSG00000285597 | AC051619,10 | -0,607539852 | 0,656314926 | 0,46857 | 0,70819843 | novel transcript |
| ENSG00000175730 | BAK1P1 | -0,607600536 | 0,65628732 | 0,36526 | 0,63543797 | BCL2 antagonist/killer 1 pseudogene 1 |
| ENSG00000206538 | VGLL3 | -0,607761839 | 0,656213946 | 0,08181 | 0,37301152 | vestigial like family member 3 |
| ENSG00000141456 | PELP1 | -0,608021702 | 0,656095758 | 0,14466 | 0,47224655 | proline, glutamate and leucine rich protein 1 |
| ENSG00000227848 | SUCLA2-AS1 | -0,608152645 | 0,656036211 | 0,43605 | 0,6865394 | SUCLA2 antisense RNA 1 |
| ENSG00000111249 | CUX2 | -0,608545253 | 0,655857705 | 0,01788 | 0,17623976 | cut like homeobox 2 |
| ENSG00000259120 | SMIM6 | -0,608595336 | 0,655834938 | 0,27597 | 0,57533851 | small integral membrane protein 6 |
| ENSG00000167524 | RSKR | -0,60864561 | 0,655812084 | 0,08233 | 0,37389555 | ribosomal protein S6 kinase related |
| ENSG00000278627 | AC005962,1 | -0,608710476 | 0,655782598 | 0,22448 | 0,54033492 | novel transcript, antisense to EPX |
| ENSG00000150628 | SPATA4 | -0,609443239 | 0,655449602 | 0,46183 | 0,70316 | spermatogenesis associated 4 |
| ENSG00000111275 | ALDH2 | -0,60971986 | 0,655323939 | 0,17436 | 0,49877327 | aldehyde dehydrogenase 2 family member |
| ENSG00000120049 | KCNIP2 | -0,609972896 | 0,655209011 | 0,06508 | 0,33804854 | potassium voltage-gated channel interacting protein 2 |
| ENSG00000259415 | AC012291,2 | -0,610050908 | 0,655173583 | 0,54943 | 0,76407481 | novel transcript |
| ENSG00000197580 | BCO2 | -0,61013512 | 0,65513534 | 0,14517 | 0,47289051 | beta-carotene oxygenase 2 |
| ENSG00000132879 | FBXO44 | -0,610746861 | 0,654857604 | 0,10844 | 0,41905863 | F-box protein 44 |
| ENSG00000285410 | GABPB1-IT1 | -0,610841923 | 0,654814456 | 0,08746 | 0,38296462 | GABPB1 intronic transcript |
| ENSG00000226396 | AL031727,1 | -0,61110143 | 0,654696681 | 0,19266 | 0,51490739 | ribosomal protein S14 (RPS14) pseudogene |
| ENSG00000228606 | AL139011,1 | -0,611256826 | 0,654626166 | 0,07265 | 0,35556123 | novel transcript |
| ENSG00000264558 | AC015674,1 | -0,611343687 | 0,654586754 | 0,25044 | 0,55681669 | novel transcript, antisense to KPNB1 |
| ENSG00000235397 | EPN2-AS1 | -0,611402981 | 0,654559851 | 0,29941 | 0,59132045 | EPN2 antisense RNA 1 |
| ENSG00000257711 | AC079385,2 | -0,611407095 | 0,654557984 | 0,28063 | 0,57806909 | novel transcript, antisense to RFX4 |
| ENSG00000175170 | FAM182B | -0,611493672 | 0,654518705 | 0,16427 | 0,49065086 | family with sequence similarity 182 member B |
| ENSG00000262810 | AC090617,4 | -0,611615668 | 0,654463361 | 0,71352 | 0,86217438 | novel transcript, antisense to SMG6 |
| ENSG00000236432 | MFF-DT | -0,611865527 | 0,654350025 | 0,17541 | 0,49948561 | MFF divergent transcript |
| ENSG00000226545 | AL357552,1 | -0,611975386 | 0,654300199 | 0,42041 | 0,67578401 | ribosomal protein L27 (RPL27) pseudogene |
| ENSG00000158769 | F11R | -0,611988935 | 0,654294054 | 0,28881 | 0,58464073 | F11 receptor |
| ENSG00000160716 | CHRNB2 | -0,612145561 | 0,654223024 | 0,01054 | 0,1325438 | cholinergic receptor nicotinic beta 2 subunit |

| | | | | | | |
|---|---|---|---|---|---|---|
| ENSG00000064545 | TMEM161A | -0,612441576 | 0,654088803 | 0,11173 | 0,42501069 | transmembrane protein 161A |
| ENSG00000277232 | GTSE1-DT | -0,61248511 | 0,654069066 | 0,15676 | 0,48425071 | GTSE1 divergent transcript |
| ENSG00000249353 | NPM1P27 | -0,612487304 | 0,654068072 | 0,38791 | 0,65289784 | nucleophosmin 1 pseudogene 27 |
| ENSG00000233421 | LINC01783 | -0,61294663 | 0,653859862 | 0,06371 | 0,33460472 | long intergenic non-protein coding RNA 1783 |
| ENSG00000261054 | AC036108,2 | -0,61296222 | 0,653852796 | 0,46118 | 0,70278639 | novel transcript, antisense to SYNM |
| ENSG00000127511 | SIN3B | -0,613076319 | 0,653801087 | 0,0842 | 0,37695424 | SIN3 transcription regulator family member B |
| ENSG00000275996 | SNORD27 | -0,613256517 | 0,65371943 | 0,29223 | 0,58692263 | small nucleolar RNA, C/D box 27 |
| ENSG00000280341 | AP000542,3 | -0,613379453 | 0,653663727 | 0,08211 | 0,37331809 | novel transcript |
| ENSG00000174839 | DENND6A | -0,613561517 | 0,653581242 | 0,08659 | 0,38118368 | DENN domain containing 6A |
| ENSG00000139722 | VPS37B | -0,613706937 | 0,653515366 | 0,08172 | 0,37295776 | VPS37B subunit of ESCRT-I |
| ENSG00000184208 | C22orf46 | -0,613863065 | 0,653444646 | 0,05807 | 0,32157493 | chromosome 22 open reading frame 46 |
| ENSG00000107829 | FBXW4 | -0,613984651 | 0,653389578 | 0,06744 | 0,34371006 | F-box and WD repeat domain containing 4 |
| ENSG00000144214 | LYG1 | -0,613987362 | 0,653388351 | 0,44301 | 0,69050803 | lysozyme g1 |
| ENSG00000226171 | DHX16 | -0,614331292 | 0,653232605 | 0,66323 | 0,83242946 | DEAH-box helicase 16 |
| ENSG00000284428 | AC092329,4 | -0,614390112 | 0,653205973 | 0,10437 | 0,4128259 | novel transcript |
| ENSG00000213579 | AL096829,1 | -0,614745426 | 0,653045118 | 0,35571 | 0,63008438 | ribosomal protein L29 (RPL29) pseudogene |
| ENSG00000182378 | PLCXD1 | -0,614764007 | 0,653036707 | 0,11837 | 0,43614969 | phosphatidylinositol specific phospholipase C X domain containing 1 |
| ENSG00000173153 | ESRRA | -0,614789401 | 0,653025213 | 0,12099 | 0,44006069 | estrogen related receptor alpha |
| ENSG00000255817 | AC025576,2 | -0,61485006 | 0,652997756 | 0,43084 | 0,68325486 | novel transcript |
| ENSG00000065923 | SLC9A7 | -0,614958246 | 0,652948791 | 0,02782 | 0,22494202 | solute carrier family 9 member A7 |
| ENSG00000139767 | SRRM4 | -0,614982953 | 0,652937609 | 0,03367 | 0,24788708 | serine/arginine repetitive matrix 4 |
| ENSG00000265907 | AP000919,1 | -0,615240525 | 0,652821047 | 0,21218 | 0,5308403 | novel transcript, antisense to LPIN2 |
| ENSG00000285282 | DHRS4-AS1 | -0,615313195 | 0,652788164 | 0,11447 | 0,42986344 | DHRS4 antisense RNA 1 |
| ENSG00000275202 | AL161421,1 | -0,615492014 | 0,652707258 | 0,06073 | 0,32716307 | novel transcript, antisense to FNDC3A |
| ENSG00000286997 | AC005856,1 | -0,615558668 | 0,652677102 | 0,11903 | 0,43696075 | novel transcript, antisense to BCAS3 |
| ENSG00000087995 | METTL2A | -0,615762135 | 0,65258506 | 0,11392 | 0,42901302 | methyltransferase like 2A |
| ENSG00000134917 | ADAMTS8 | -0,615794565 | 0,652570391 | 0,1782 | 0,50248137 | ADAM metallopeptidase with thrombospondin type 1 motif 8 |
| ENSG00000281060 | MUC20-OT1 | -0,61588468 | 0,652529631 | 0,05351 | 0,31052177 | MUC20 overlapping transcript |
| ENSG00000235706 | DICER1-AS1 | -0,615913476 | 0,652516607 | 0,07454 | 0,35852216 | DICER1 antisense RNA 1 |
| ENSG00000259723 | AL354993,2 | -0,61599047 | 0,652481784 | 0,27505 | 0,57491355 | novel transcript, antisense to TSHZ2 |
| ENSG00000287330 | AL353697,1 | -0,616008709 | 0,652473535 | 0,09538 | 0,39803067 | novel transcript, antisense to PCCA |

| | | | | | | |
|---|---|---|---|---|---|---|
| ENSG00000184788 | SATL1 | -0,616472041 | 0,652264022 | 0,02272 | 0,20117395 | spermidine/spermine N1-acetyl transferase like 1 |
| ENSG00000287109 | AP000224,1 | -0,616978381 | 0,652035138 | 0,15708 | 0,48443207 | novel transcript, antisense to JAM2 |
| ENSG00000140749 | IGSF6 | -0,617362505 | 0,651861554 | 0,34042 | 0,61897407 | immunoglobulin superfamily member 6 |
| ENSG00000063438 | AHRR | -0,617486786 | 0,651805402 | 0,07721 | 0,36386039 | aryl-hydrocarbon receptor repressor |
| ENSG00000223653 | AL078459,1 | -0,617746998 | 0,65168785 | 0,07721 | 0,36386039 | novel transcript |
| ENSG00000282956 | MAGI1 | -0,618225741 | 0,65147163 | 0,03587 | 0,25662951 | membrane associated guanylate kinase, WW and PDZ domain containing 1 |
| ENSG00000151276 | MAGI1 | -0,618225741 | 0,65147163 | 0,03587 | 0,25662951 | membrane associated guanylate kinase, WW and PDZ domain containing 1 |
| ENSG00000281650 | AC243922,3 | -0,618227212 | 0,651470965 | 0,07637 | 0,36251343 | glucuronidase, beta (GUSB) pseudogene |
| ENSG00000268120 | AC010336,3 | -0,618255667 | 0,651458116 | 0,07019 | 0,34975886 | novel transcript |
| ENSG00000259642 | ST20-AS1 | -0,618427068 | 0,651380724 | 0,12013 | 0,43868155 | ST20 antisense RNA 1 |
| ENSG00000276184 | UPK3B | -0,618470914 | 0,651360928 | 0,36579 | 0,63598241 | uroplakin 3B |
| ENSG00000166016 | ABTB2 | -0,618574236 | 0,651314281 | 0,13121 | 0,45610339 | ankyrin repeat and BTB domain containing 2 |
| ENSG00000251026 | LINC02163 | -0,618676421 | 0,65126815 | 0,25001 | 0,55657567 | long intergenic non-protein coding RNA 2163 |
| ENSG00000077279 | DCX | -0,618689691 | 0,65126216 | 0,09041 | 0,3889575 | doublecortin |
| ENSG00000272668 | AL590560,3 | -0,619084209 | 0,651084091 | 0,26601 | 0,56896415 | uncharacterized LOC107985216 |
| ENSG00000184111 | RPL11P4 | -0,619095627 | 0,651078938 | 0,42245 | 0,67663831 | ribosomal protein L11 pseudogene 4 |
| ENSG00000255339 | AL133352,1 | -0,619145355 | 0,651056496 | 0,07473 | 0,35882064 | NADH dehydrogenase (ubiquinone) 1 beta subcomplex, 8, 19kDa (NDUFB8) and SEC31 homolog B (S, cerevisiae) (SEC31B) readthrough |
| ENSG00000286031 | AC232271,2 | -0,619206413 | 0,651028943 | 0,40311 | 0,66399303 | novel transcript, antisense to PPP1R3F |
| ENSG00000139437 | TCHP | -0,619347256 | 0,650965389 | 0,06286 | 0,33177754 | trichoplein keratin filament binding |
| ENSG00000173825 | TIGD3 | -0,619349019 | 0,650964593 | 0,13145 | 0,45657505 | tigger transposable element derived 3 |
| ENSG00000287936 | AL391425,1 | -0,619434503 | 0,650926023 | 0,17872 | 0,50323303 | novel transcript |
| ENSG00000171703 | TCEA2 | -0,619491377 | 0,650900362 | 0,07857 | 0,36624708 | transcription elongation factor A2 |
| ENSG00000269806 | BICRA-AS1 | -0,619601331 | 0,650850756 | 0,2618 | 0,56475385 | BICRA antisense RNA 1 |
| ENSG00000206297 | TAP1 | -0,61985395 | 0,650736801 | 0,63066 | 0,81354995 | transporter 1, ATP binding cassette subfamily B member |
| ENSG00000161647 | MPP3 | -0,620109726 | 0,650621442 | 0,04824 | 0,29503004 | membrane palmitoylated protein 3 |
| ENSG00000176248 | ANAPC2 | -0,620261799 | 0,650552864 | 0,07065 | 0,3507077 | anaphase promoting complex subunit 2 |
| ENSG00000273210 | AP001437,1 | -0,620296198 | 0,650537353 | 0,28407 | 0,58097536 | novel transcript, antisense to DYRK1A |
| ENSG00000236928 | AC008267,4 | -0,620485032 | 0,65045221 | 0,3997 | 0,6618663 | GrpE-like 1, mitochondrial (E, coli) (GRPEL1) pseudogene |
| ENSG00000260440 | LINC01544 | -0,620495097 | 0,650447672 | 0,29971 | 0,59160746 | long intergenic non-protein coding RNA 1544 |

| | | | | | | |
|---|---|---|---|---|---|---|
| ENSG00000160633 | SAFB | -0,620574013 | 0,650412093 | 0,04107 | 0,27450724 | scaffold attachment factor B |
| ENSG00000117971 | CHRNB4 | -0,620631003 | 0,650386401 | 0,21416 | 0,53334744 | cholinergic receptor nicotinic beta 4 subunit |
| ENSG00000269403 | AC008750,7 | -0,620846366 | 0,650289319 | 0,75502 | 0,88516173 | novel protein |
| ENSG00000167536 | DHRS13 | -0,620917231 | 0,650257378 | 0,1303 | 0,45474562 | dehydrogenase/reductase 13 |
| ENSG00000282249 | AC254962,1 | -0,620992755 | 0,650223339 | 0,35661 | 0,63063438 | hyperpolarization activated cyclic nucleotide-gated potassium channel 2 (HCN2) pseudogene |
| ENSG00000167693 | NXN | -0,620995545 | 0,650222081 | 0,15211 | 0,48061452 | nucleoredoxin |
| ENSG00000273521 | AL162274,1 | -0,620999505 | 0,650220297 | 0,15748 | 0,48457045 | novel transcript, antisense to BNIP3 |
| ENSG00000223547 | ZNF844 | -0,621218037 | 0,650121812 | 0,14624 | 0,47428193 | zinc finger protein 844 |
| ENSG00000185633 | NDUFA4L2 | -0,62135776 | 0,650058852 | 0,16142 | 0,48892403 | NDUFA4 mitochondrial complex associated like 2 |
| ENSG00000253347 | AC040934,1 | -0,621386315 | 0,650045985 | 0,5487 | 0,76383004 | novel transcript |
| ENSG00000233290 | AL157944,1 | -0,621843367 | 0,64984008 | 0,38195 | 0,64905556 | novel transcript |
| ENSG00000115053 | NCL | -0,62196108 | 0,649787061 | 0,10829 | 0,4189697 | nucleolin |
| ENSG00000140859 | KIFC3 | -0,622114573 | 0,649717931 | 0,20151 | 0,52214043 | kinesin family member C3 |
| ENSG00000234056 | LINC00463 | -0,622150215 | 0,64970188 | 0,25468 | 0,55944852 | long intergenic non-protein coding RNA 463 |
| ENSG00000283662 | AC138904,3 | -0,622162167 | 0,649696498 | 0,13946 | 0,46666406 | novel transcript |
| ENSG00000260350 | AC012173,1 | -0,622303364 | 0,649632915 | 0,38536 | 0,65107577 | novel transcript, antisense to PMM2 |
| ENSG00000159409 | CELF3 | -0,622414164 | 0,649583025 | 0,09404 | 0,39609058 | CUGBP Elav-like family member 3 |
| ENSG00000274352 | ZNF707 | -0,622453385 | 0,649565365 | 0,14888 | 0,47633105 | zinc finger protein 707 |
| ENSG00000206440 | NFKBIL1 | -0,622712567 | 0,64944868 | 0,2217 | 0,53805234 | NFKB inhibitor like 1 |
| ENSG00000236346 | NFKBIL1 | -0,622712567 | 0,64944868 | 0,2217 | 0,53805234 | NFKB inhibitor like 1 |
| ENSG00000234530 | NFKBIL1 | -0,622712567 | 0,64944868 | 0,2217 | 0,53805234 | NFKB inhibitor like 1 |
| ENSG00000227565 | NFKBIL1 | -0,622712567 | 0,64944868 | 0,2217 | 0,53805234 | NFKB inhibitor like 1 |
| ENSG00000236196 | NFKBIL1 | -0,622712567 | 0,64944868 | 0,2217 | 0,53805234 | NFKB inhibitor like 1 |
| ENSG00000156172 | C8orf37 | -0,623202575 | 0,649228134 | 0,06554 | 0,33914981 | chromosome 8 open reading frame 37 |
| ENSG00000243207 | PPAN-P2RY11 | -0,623285581 | 0,649190781 | 0,67317 | 0,83865474 | PPAN-P2RY11 readthrough |
| ENSG00000019144 | PHLDB1 | -0,6235626 | 0,649066139 | 0,09873 | 0,40461924 | pleckstrin homology like domain family B member 1 |
| ENSG00000074527 | NTN4 | -0,623772599 | 0,648971668 | 0,04526 | 0,2876168 | netrin 4 |
| ENSG00000235436 | DPY19L2P4 | -0,623803401 | 0,648957812 | 0,05854 | 0,32311626 | DPY19L2 pseudogene 4 |
| ENSG00000226562 | CYP4F26P | -0,624111187 | 0,648819378 | 0,44026 | 0,68866748 | cytochrome P450 family 4 subfamily F member 26, pseudogene |
| ENSG00000173597 | SULT1B1 | -0,624407695 | 0,648686044 | 0,08291 | 0,37487894 | sulfotransferase family 1B member 1 |
| ENSG00000269242 | AC010422,3 | -0,624448806 | 0,648667559 | 0,39168 | 0,65533345 | novel transcript |

| | | | | | | |
|---|---|---|---|---|---|---|
| ENSG00000170037 | CNTROB | -0,624523675 | 0,648633897 | 0,09729 | 0,40183698 | centrobin, centriole duplication and spindle assembly protein |
| ENSG00000285287 | AL354855,1 | -0,624798058 | 0,648510547 | 0,44607 | 0,69284796 | novel pseudogene similar to avian transmembrane protein 180 |
| ENSG00000275348 | AC096861,1 | -0,624867316 | 0,648479415 | 0,60197 | 0,79680798 | high mobility group nucleosome binding domain 1 (HMGN1) pseudogene |
| ENSG00000164061 | BSN | -0,625184973 | 0,648336646 | 0,0392 | 0,26793262 | bassoon presynaptic cytomatrix protein |
| ENSG00000196503 | ARL9 | -0,625227353 | 0,648317602 | 0,09575 | 0,39858189 | ADP ribosylation factor like GTPase 9 |
| ENSG00000213638 | ADAT3 | -0,625400834 | 0,648239647 | 0,22704 | 0,54168591 | adenosine deaminase tRNA specific 3 |
| ENSG00000273139 | AC007663,3 | -0,62548939 | 0,648199858 | 0,33448 | 0,61426654 | novel transcript |
| ENSG00000135835 | KIAA1614 | -0,625700841 | 0,648104861 | 0,09651 | 0,40027716 | KIAA1614 |
| ENSG00000270606 | PPIAP52 | -0,625882753 | 0,648023145 | 0,14142 | 0,4680519 | peptidylprolyl isomerase A pseudogene 52 |
| ENSG00000081985 | IL12RB2 | -0,625941143 | 0,647996918 | 0,25736 | 0,56160845 | interleukin 12 receptor subunit beta 2 |
| ENSG00000223546 | LINC00630 | -0,625950544 | 0,647992696 | 0,07849 | 0,36604454 | long intergenic non-protein coding RNA 630 |
| ENSG00000273525 | SNORA54 | -0,626103387 | 0,647924049 | 0,06969 | 0,34826564 | small nucleolar RNA, H/ACA box 54 |
| ENSG00000207008 | SNORA54 | -0,626103387 | 0,647924049 | 0,06969 | 0,34826564 | small nucleolar RNA, H/ACA box 54 |
| ENSG00000135119 | RNFT2 | -0,626199755 | 0,647880771 | 0,06928 | 0,34763785 | ring finger protein, transmembrane 2 |
| ENSG00000164123 | C4orf45 | -0,626358006 | 0,647809708 | 0,47408 | 0,71294959 | chromosome 4 open reading frame 45 |
| ENSG00000285254 | AL513128,2 | -0,626610187 | 0,647696482 | 0,31494 | 0,60144815 | novel pseudogene |
| ENSG00000229212 | AC044860,1 | -0,626702631 | 0,647654981 | 0,18614 | 0,50956086 | golgin subfamily A pseudogene |
| ENSG00000239351 | NPM1P29 | -0,626774235 | 0,647622837 | 0,1688 | 0,49492999 | nucleophosmin 1 pseudogene 29 |
| ENSG00000171608 | PIK3CD | -0,626887933 | 0,6475718 | 0,09237 | 0,39278463 | phosphatidylinositol-4,5-bisphosphate 3-kinase catalytic subunit delta |
| ENSG00000279643 | TAMM41 | -0,626909938 | 0,647561923 | 0,13824 | 0,46543358 | TAM41 mitochondrial translocator assembly and maintenance homolog |
| ENSG00000186340 | THBS2 | -0,626913735 | 0,647560219 | 0,16568 | 0,49182669 | thrombospondin 2 |
| ENSG00000205583 | STAG3L1 | -0,627074291 | 0,647488157 | 0,08415 | 0,37695424 | stromal antigen 3-like 1 (pseudogene) |
| ENSG00000185271 | KLHL33 | -0,62751896 | 0,647288618 | 0,12019 | 0,43872715 | kelch like family member 33 |
| ENSG00000267065 | LINC02080 | -0,627655266 | 0,647227465 | 0,0918 | 0,39174496 | long intergenic non-protein coding RNA 2080 |
| ENSG00000164776 | PHKG1 | -0,627684148 | 0,647214508 | 0,25301 | 0,55809014 | phosphorylase kinase catalytic subunit gamma 1 |
| ENSG00000254027 | AC009902,2 | -0,627784277 | 0,64716959 | 0,3437 | 0,62110958 | novel transcript |
| ENSG00000276537 | TTYH1 | -0,627824333 | 0,647151622 | 0,0752 | 0,35996907 | tweety family member 1 |
| ENSG00000174721 | FGFBP3 | -0,627871101 | 0,647130643 | 0,05725 | 0,31908628 | fibroblast growth factor binding protein 3 |

| | | | | | | |
|---|---|---|---|---|---|---|
| ENSG00000283183 | AC116565,1 | -0,627931165 | 0,647103702 | 0,33046 | 0,6112353 | novel transcript |
| ENSG00000261949 | GFY | -0,627935646 | 0,647101692 | 0,2349 | 0,54637125 | golgi associated olfactory signaling regulator |
| ENSG00000258162 | AC069228,1 | -0,62810307 | 0,6470266 | 0,19026 | 0,51331106 | novel transcript, antisense to PPFIA2 |
| ENSG00000163428 | LRRC58 | -0,628104074 | 0,64702615 | 0,1745 | 0,49884018 | leucine rich repeat containing 58 |
| ENSG00000287972 | AC073648,7 | -0,628226948 | 0,646971045 | 0,13976 | 0,4668467 | novel transcript |
| ENSG00000136274 | NACAD | -0,62833076 | 0,646924493 | 0,06233 | 0,33087235 | NAC alpha domain containing |
| ENSG00000006118 | TMEM132A | -0,628380462 | 0,646902206 | 0,09941 | 0,40596459 | transmembrane protein 132A |
| ENSG00000227598 | Z94721,1 | -0,628383079 | 0,646901033 | 0,2492 | 0,55635288 | novel transcript |
| ENSG00000255581 | AC006511,2 | -0,62864778 | 0,646782352 | 0,33855 | 0,61774255 | heat shock 60kDa protein 1 (chaperonin) (HSPD1) pseudogene |
| ENSG00000279232 | AC008522,1 | -0,628719248 | 0,646750313 | 0,23261 | 0,54529383 | novel transcript, antisense to RGMB |
| ENSG00000234936 | AC010883,1 | -0,628787972 | 0,646719505 | 0,39889 | 0,66138533 | novel transcript |
| ENSG00000231672 | DIRC3 | -0,628893016 | 0,646672418 | 0,15937 | 0,48684738 | disrupted in renal carcinoma 3 |
| ENSG00000129993 | CBFA2T3 | -0,628895081 | 0,646671493 | 0,09704 | 0,40143759 | CBFA2/RUNX1 partner transcriptional co-repressor 3 |
| ENSG00000141696 | P3H4 | -0,628949334 | 0,646647175 | 0,1553 | 0,48334025 | prolyl 3-hydroxylase family member 4 (inactive) |
| ENSG00000275544 | BDH1 | -0,629166016 | 0,646550061 | 0,08184 | 0,37301152 | 3-hydroxybutyrate dehydrogenase 1 |
| ENSG00000246250 | AC087521,2 | -0,629197575 | 0,646535918 | 0,10772 | 0,41834811 | novel transcript |
| ENSG00000197321 | SVIL | -0,629222133 | 0,646524912 | 0,02631 | 0,21861252 | supervillin |
| ENSG00000167972 | ABCA3 | -0,629418747 | 0,646436808 | 0,00157 | 0,04253303 | ATP binding cassette subfamily A member 3 |
| ENSG00000231064 | THBS3-AS1 | -0,629433781 | 0,646430072 | 0,02197 | 0,19708179 | THBS3 antisense RNA 1 |
| ENSG00000275488 | AC023509,5 | -0,629458799 | 0,646418862 | 0,34894 | 0,62514443 | zinc finger protein 337 (ZNF337) pseudogene |
| ENSG00000087074 | PPP1R15A | -0,6295401 | 0,646382435 | 0,08151 | 0,37255227 | protein phosphatase 1 regulatory subunit 15A |
| ENSG00000257341 | AL928654,3 | -0,629559876 | 0,646373575 | 0,32181 | 0,60577115 | novel protein |
| ENSG00000264364 | DYNLL2 | -0,629781228 | 0,64627441 | 0,06941 | 0,34799331 | dynein light chain LC8-type 2 |
| ENSG00000103024 | NME3 | -0,629884317 | 0,646228231 | 0,06759 | 0,34401832 | NME/NM23 nucleoside diphosphate kinase 3 |
| ENSG00000145911 | N4BP3 | -0,629941413 | 0,646202657 | 0,12189 | 0,44093327 | NEDD4 binding protein 3 |
| ENSG00000232774 | AL355916,1 | -0,63007505 | 0,646142802 | 0,11377 | 0,42869237 | novel transcript |
| ENSG00000276045 | ORAI1 | -0,630122517 | 0,646121543 | 0,21148 | 0,53032236 | ORAI calcium release-activated calcium modulator 1 |
| ENSG00000240305 | AC091152,1 | -0,630283055 | 0,646049649 | 0,16312 | 0,48972962 | ribosomal protein L6 (RPL6) pseudogene |
| ENSG00000184923 | NUTM2A | -0,630305794 | 0,646039466 | 0,01936 | 0,18420145 | NUT family member 2A |
| ENSG00000227966 | LINC01623 | -0,630318281 | 0,646033874 | 0,65388 | 0,82669364 | long intergenic non-protein coding RNA 1623 |
| ENSG00000276061 | NDUFA3 | -0,630324628 | 0,646031032 | 0,05225 | 0,3069525 | NADH:ubiquinone oxidoreductase subunit A3 |
| ENSG00000274578 | AC006249,1 | -0,630358933 | 0,646015671 | 0,08039 | 0,37007763 | novel transcript, sense intronic to CDH2 |

| | | | | | | |
|---|---|---|---|---|---|---|
| ENSG00000169221 | TBC1D10B | -0,630876897 | 0,645783777 | 0,12544 | 0,44729629 | TBC1 domain family member 10B |
| ENSG00000129159 | KCNC1 | -0,630896249 | 0,645775114 | 0,00406 | 0,07819491 | potassium voltage-gated channel subfamily C member 1 |
| ENSG00000172663 | TMEM134 | -0,631122714 | 0,645673753 | 0,16478 | 0,49092529 | transmembrane protein 134 |
| ENSG00000223478 | AL441992,1 | -0,631156495 | 0,645658634 | 0,15471 | 0,48266733 | novel transcript |
| ENSG00000111696 | NT5DC3 | -0,631243054 | 0,645619897 | 0,28467 | 0,58151922 | 5'-nucleotidase domain containing 3 |
| ENSG00000257918 | AC079385,3 | -0,63125218 | 0,645615813 | 0,36037 | 0,63265655 | novel transcript, antisense to RFX4 |
| ENSG00000164898 | FMC1 | -0,631762599 | 0,645387437 | 0,12335 | 0,44379017 | formation of mitochondrial complex V assembly factor 1 homolog |
| ENSG00000225872 | LINC01529 | -0,631844309 | 0,645350886 | 0,06116 | 0,32813211 | long intergenic non-protein coding RNA 1529 |
| ENSG00000179938 | GOLGA8J | -0,631991502 | 0,645285046 | 0,09511 | 0,39772599 | golgin A8 family member J |
| ENSG00000179846 | NKPD1 | -0,632000409 | 0,645281062 | 0,28787 | 0,5840701 | NTPase KAP family P-loop domain containing 1 |
| ENSG00000256422 | LINC02552 | -0,632000914 | 0,645280837 | 0,06638 | 0,34113137 | long intergenic non-protein coding RNA 2552 |
| ENSG00000161664 | ASB16 | -0,632094105 | 0,645239156 | 0,13399 | 0,4601411 | ankyrin repeat and SOCS box containing 16 |
| ENSG00000100319 | ZMAT5 | -0,632101691 | 0,645235763 | 0,09263 | 0,39335125 | zinc finger matrin-type 5 |
| ENSG00000263731 | AC145207,5 | -0,632217803 | 0,645183835 | 0,125 | 0,44663415 | novel transcript |
| ENSG00000267827 | AC011468,2 | -0,632245006 | 0,645171669 | 0,37283 | 0,64172617 | novel transcript |
| ENSG00000199293 | SNORA21 | -0,632290206 | 0,645151456 | 0,2441 | 0,55253387 | small nucleolar RNA, H/ACA box 21 |
| ENSG00000008853 | RHOBTB2 | -0,632684229 | 0,644975279 | 0,09835 | 0,40379437 | Rho related BTB domain containing 2 |
| ENSG00000126005 | MMP24OS | -0,63270636 | 0,644965385 | 0,11206 | 0,42549415 | MMP24 opposite strand |
| ENSG00000266983 | AC011444,1 | -0,632755776 | 0,644943294 | 0,40725 | 0,66661466 | novel transcript, antisense to RFX2 |
| ENSG00000168758 | SEMA4C | -0,633453505 | 0,644631456 | 0,14657 | 0,47446965 | semaphorin 4C |
| ENSG00000285932 | AL133353,2 | -0,633462174 | 0,644627583 | 0,21184 | 0,53070055 | novel transcript |
| ENSG00000286412 | AC007682,1 | -0,633534618 | 0,644595214 | 0,03176 | 0,24047554 | novel transcript, antisense to NRXN1 |
| ENSG00000241984 | RPL7AP2 | -0,633658265 | 0,644539971 | 0,24408 | 0,55253387 | ribosomal protein L7a pseudogene 2 |
| ENSG00000109736 | MFSD10 | -0,633688161 | 0,644526615 | 0,07896 | 0,3669053 | major facilitator superfamily domain containing 10 |
| ENSG00000185942 | NKAIN3 | -0,633723474 | 0,644510839 | 0,01678 | 0,17000427 | sodium/potassium transporting ATPase interacting 3 |
| ENSG00000273936 | TNFRSF14 | -0,633935072 | 0,644416317 | 0,52834 | 0,7507973 | TNF receptor superfamily member 14 |
| ENSG00000157873 | TNFRSF14 | -0,633935072 | 0,644416317 | 0,52834 | 0,7507973 | TNF receptor superfamily member 14 |
| ENSG00000179743 | FLJ37453 | -0,634194672 | 0,64430037 | 0,05877 | 0,32381025 | uncharacterized LOC729614 |
| ENSG00000259257 | AC037479,1 | -0,634284243 | 0,644260369 | 0,37313 | 0,64185106 | Coiled-coil domain-containing protein 117 (CC117) pseudogene |
| ENSG00000270061 | AC068790,5 | -0,634305899 | 0,644250698 | 0,12716 | 0,45032932 | novel transcript, sense intronic to CCDC92 |
| ENSG00000237457 | LINC01351 | -0,634331628 | 0,644239209 | 0,43377 | 0,68475933 | long intergenic non-protein coding RNA 1351 |

| | | | | | | |
|---|---|---|---|---|---|---|
| ENSG00000287195 | AC131568,1 | -0,634420688 | 0,64419944 | 0,304 | 0,59410954 | novel transcript, antisense to FAM49B |
| ENSG00000215160 | OR7E122P | -0,634425619 | 0,644197238 | 0,07762 | 0,36468824 | olfactory receptor family 7 subfamily E member 122 pseudogene |
| ENSG00000261455 | LINC01003 | -0,63454575 | 0,644143599 | 0,06882 | 0,34677117 | long intergenic non-protein coding RNA 1003 |
| ENSG00000132326 | PER2 | -0,634804806 | 0,644027944 | 0,2025 | 0,52259159 | period circadian regulator 2 |
| ENSG00000126246 | IGFLR1 | -0,634895135 | 0,643987622 | 0,09528 | 0,39794903 | IGF like family receptor 1 |
| ENSG00000267505 | AC005180,2 | -0,634976177 | 0,643951448 | 0,34771 | 0,62423931 | novel transcript |
| ENSG00000172366 | MCRIP2 | -0,635222227 | 0,643841632 | 0,1269 | 0,44963011 | MAPK regulated corepressor interacting protein 2 |
| ENSG00000228175 | GEMIN8P4 | -0,635241584 | 0,643832994 | 0,18119 | 0,50470865 | gem nuclear organelle associated protein 8 pseudogene 4 |
| ENSG00000121653 | MAPK8IP1 | -0,635842472 | 0,643564891 | 0,03158 | 0,23994313 | mitogen-activated protein kinase 8 interacting protein 1 |
| ENSG00000277498 | AC107081,2 | -0,635995119 | 0,643496801 | 0,19284 | 0,51493916 | serine/arginine repetitive matrix 1 (SRRM1) pseudogene |
| ENSG00000285301 | BOP1 | -0,636116958 | 0,643442458 | 0,16321 | 0,48972962 | BOP1 ribosomal biogenesis factor |
| ENSG00000261236 | BOP1 | -0,636116958 | 0,643442458 | 0,16321 | 0,48972962 | BOP1 ribosomal biogenesis factor |
| ENSG00000214021 | TTLL3 | -0,636128363 | 0,643437372 | 0,03209 | 0,24164117 | tubulin tyrosine ligase like 3 |
| ENSG00000177551 | NHLH2 | -0,636338876 | 0,64334349 | 0,13875 | 0,46603966 | nescient helix-loop-helix 2 |
| ENSG00000259098 | AC025884,2 | -0,636406091 | 0,643313518 | 0,12738 | 0,45051698 | zinc finger, CCHC domain containing 2 (ZCCHC2) pseudogene |
| ENSG00000196778 | OR52K1 | -0,636646713 | 0,643206231 | 0,15027 | 0,47831211 | olfactory receptor family 52 subfamily K member 1 |
| ENSG00000249112 | AC112196,1 | -0,636722335 | 0,643172517 | 0,19318 | 0,51517146 | novel transcript, antisense to ZNF608 |
| ENSG00000154096 | THY1 | -0,636908284 | 0,643089623 | 0,11451 | 0,42992088 | Thy-1 cell surface antigen |
| ENSG00000227775 | AL031282,1 | -0,636976588 | 0,643059177 | 0,08276 | 0,37461566 | ribosomal protein S7 (RPS7) pseudogene |
| ENSG00000273113 | AC133528,1 | -0,637201292 | 0,642959026 | 0,36452 | 0,63501401 | novel transcript |
| ENSG00000177272 | KCNA3 | -0,637259204 | 0,642933218 | 0,15547 | 0,48334025 | potassium voltage-gated channel subfamily A member 3 |
| ENSG00000224383 | PRR29 | -0,637475396 | 0,64283688 | 0,1388 | 0,46610395 | proline rich 29 |
| ENSG00000006210 | CX3CL1 | -0,637651109 | 0,64275859 | 0,08394 | 0,37651068 | C-X3-C motif chemokine ligand 1 |
| ENSG00000249921 | AC034207,1 | -0,637876727 | 0,642658079 | 0,34807 | 0,62442306 | guanine nucleotide binding protein (G protein), beta polypeptide 2-like 1 (GNB2L1) pseudogene |
| ENSG00000203663 | OR2L2 | -0,637887581 | 0,642653244 | 0,0853 | 0,3785936 | olfactory receptor family 2 subfamily L member 2 |
| ENSG00000249274 | PDLIM1P4 | -0,637888169 | 0,642652982 | 0,5499 | 0,76436158 | PDZ and LIM domain 1 pseudogene 4 |
| ENSG00000258783 | KRT18P6 | -0,638013038 | 0,642597361 | 0,21484 | 0,53373102 | keratin 18 pseudogene 6 |
| ENSG00000270885 | RASL10B | -0,638200833 | 0,64251372 | 0,11421 | 0,42953156 | RAS like family 10 member B |
| ENSG00000230612 | AC004039,1 | -0,638221822 | 0,642504373 | 0,05649 | 0,31717392 | novel transcript, antisense to KIF3A |
| ENSG00000178977 | LINC00324 | -0,638273637 | 0,642481297 | 0,19763 | 0,51915907 | long intergenic non-protein coding RNA 324 |

| | | | | | | |
|---|---|---|---|---|---|---|
| ENSG00000066468 | FGFR2 | -0,638307168 | 0,642466365 | 0,15248 | 0,48061452 | fibroblast growth factor receptor 2 |
| ENSG00000197978 | GOLGA6L9 | -0,638343402 | 0,642450229 | 0,01057 | 0,1325438 | golgin A6 family like 9 |
| ENSG00000232079 | LINC01697 | -0,638440615 | 0,642406941 | 0,27417 | 0,5742005 | long intergenic non-protein coding RNA 1697 |
| ENSG00000246548 | LINC02288 | -0,638509447 | 0,642376292 | 0,2298 | 0,54324957 | long intergenic non-protein coding RNA 2288 |
| ENSG00000270799 | AC013467,1 | -0,638518718 | 0,642372164 | 0,30961 | 0,59806916 | Y box binding protein 1 (YBX1) pseudogene |
| ENSG00000268069 | AC004466,1 | -0,638669048 | 0,642305231 | 0,19243 | 0,514872 | novel transcript, antisense to HDAC7 |
| ENSG00000258559 | AC005519,1 | -0,638800048 | 0,642246911 | 0,10834 | 0,41899991 | novel transcript, sense overlapping ABCD4 |
| ENSG00000265511 | AC020558,2 | -0,63884043 | 0,642228935 | 0,2207 | 0,53767462 | novel transcript |
| ENSG00000140839 | CLEC18B | -0,638890982 | 0,642206431 | 0,25036 | 0,55672178 | C-type lectin domain family 18 member B |
| ENSG00000160325 | CACFD1 | -0,638893061 | 0,642205506 | 0,35687 | 0,63098227 | calcium channel flower domain containing 1 |
| ENSG00000205609 | EIF3CL | -0,63903085 | 0,642144173 | 0,30081 | 0,59211453 | eukaryotic translation initiation factor 3 subunit C like |
| ENSG00000262461 | SPDYE9 | -0,639127422 | 0,64210119 | 0,04591 | 0,28908129 | speedy/RINGO cell cycle regulator family member E9 |
| ENSG00000122971 | ACADS | -0,639296459 | 0,642025961 | 0,05645 | 0,31712073 | acyl-CoA dehydrogenase short chain |
| ENSG00000255867 | DENND5B-AS: | -0,639369776 | 0,641993335 | 0,33703 | 0,61684812 | DENND5B antisense RNA 1 |
| ENSG00000259380 | AC087473,1 | -0,639643497 | 0,641871542 | 0,07894 | 0,3669053 | novel transcript |
| ENSG00000286524 | AC092567,2 | -0,640088468 | 0,641673599 | 0,16902 | 0,49510117 | novel transcript, antisense to EHBP1 |
| ENSG00000230267 | HERC2P4 | -0,640406889 | 0,641531989 | 0,20618 | 0,52587153 | hect domain and RLD 2 pseudogene 4 |
| ENSG00000185090 | MANEAL | -0,640482306 | 0,641498454 | 0,0833 | 0,37550713 | mannosidase endo-alpha like |
| ENSG00000135973 | GPR45 | -0,640652225 | 0,641422904 | 0,37565 | 0,64399236 | G protein-coupled receptor 45 |
| ENSG00000101246 | ARFRP1 | -0,64081427 | 0,641350862 | 0,06239 | 0,33099652 | ADP ribosylation factor related protein 1 |
| ENSG00000214872 | SMTNL1 | -0,640891264 | 0,641316636 | 0,09432 | 0,39648446 | smoothelin like 1 |
| ENSG00000168056 | LTBP3 | -0,640966158 | 0,641283344 | 0,16098 | 0,48832443 | latent transforming growth factor beta binding protein 3 |
| ENSG00000161036 | LRWD1 | -0,641067845 | 0,641238146 | 0,08307 | 0,37504927 | leucine rich repeats and WD repeat domain containing 1 |
| ENSG00000287310 | AL133445,3 | -0,641128469 | 0,6412112 | 0,1474 | 0,47490365 | novel transcript |
| ENSG00000131379 | C3orf20 | -0,641210254 | 0,641174852 | 0,35724 | 0,63120669 | chromosome 3 open reading frame 20 |
| ENSG00000249592 | AC139887,2 | -0,641242354 | 0,641160586 | 0,08426 | 0,37695424 | novel transcript, antisense to PCGF3 |
| ENSG00000258445 | MAD2L1P1 | -0,641455475 | 0,641065878 | 0,38221 | 0,64922637 | mitotic arrest deficient 2 like 1 pseudogene 1 |
| ENSG00000235012 | JCADP1 | -0,641651493 | 0,640978782 | 0,3664 | 0,6363423 | JCAD pseudogene 1 |
| ENSG00000279213 | SNX18P10 | -0,641675565 | 0,640968088 | 0,37866 | 0,64679625 | sorting nexin 18 pseudogene 10 |
| ENSG00000283667 | AC009802,1 | -0,641708361 | 0,640953517 | 0,04811 | 0,29469582 | novel transcript, antisense to SERPINB8 |
| ENSG00000249876 | AC010285,2 | -0,641943612 | 0,640849009 | 0,16368 | 0,49000347 | DnaJ (Hsp40) homolog, subfamily B, member 6 (DNAJB6) pseudogene |
| ENSG00000228295 | LINC00392 | -0,642010737 | 0,640819193 | 0,07173 | 0,35351167 | long intergenic non-protein coding RNA 392 |

| | | | | | | |
|---|---|---|---|---|---|---|
| ENSG00000228727 | SAPCD1 | -0,642102704 | 0,640778344 | 0,24268 | 0,55158827 | suppressor APC domain containing 1 |
| ENSG00000178188 | SH2B1 | -0,642118441 | 0,640771355 | 0,09266 | 0,39335125 | SH2B adaptor protein 1 |
| ENSG00000270127 | AC027020,2 | -0,642200654 | 0,640734841 | 0,15589 | 0,4836223 | novel transcript |
| ENSG00000235674 | LDHAP2 | -0,642205736 | 0,640732584 | 0,37818 | 0,64646341 | lactate dehydrogenase A pseudogene 2 |
| ENSG00000123358 | NR4A1 | -0,642222161 | 0,640725289 | 0,22029 | 0,53750545 | nuclear receptor subfamily 4 group A member 1 |
| ENSG00000270641 | TSIX | -0,642305961 | 0,640688073 | 0,16281 | 0,48945635 | TSIX transcript, XIST antisense RNA |
| ENSG00000273456 | AC064836,2 | -0,642354635 | 0,640666458 | 0,20325 | 0,5231305 | novel transcript |
| ENSG00000188322 | SBK1 | -0,642371813 | 0,64065883 | 0,01887 | 0,1820996 | SH3 domain binding kinase 1 |
| ENSG00000163888 | CAMK2N2 | -0,642375144 | 0,64065735 | 0,10272 | 0,4105152 | calcium/calmodulin dependent protein kinase II inhibitor 2 |
| ENSG00000279652 | Z82217,1 | -0,642550018 | 0,640579699 | 0,20848 | 0,52778377 | novel transcript |
| ENSG00000169515 | CCDC8 | -0,642658606 | 0,640531486 | 0,06485 | 0,33759365 | coiled-coil domain containing 8 |
| ENSG00000172058 | SERF1A | -0,642717174 | 0,640505483 | 0,114 | 0,42906718 | small EDRK-rich factor 1A |
| ENSG00000185015 | CA13 | -0,642757314 | 0,640487663 | 0,14718 | 0,47471707 | carbonic anhydrase 13 |
| ENSG00000204520 | MICA | -0,642796817 | 0,640470126 | 0,20557 | 0,52513499 | MHC class I polypeptide-related sequence A |
| ENSG00000135423 | GLS2 | -0,642914379 | 0,640417937 | 0,33782 | 0,61751351 | glutaminase 2 |
| ENSG00000189423 | USP32P3 | -0,642925849 | 0,640412845 | 0,10577 | 0,41527838 | ubiquitin specific peptidase 32 pseudogene 3 |
| ENSG00000127564 | PKMYT1 | -0,643085311 | 0,640342064 | 0,09879 | 0,40461924 | protein kinase, membrane associated tyrosine/threonine 1 |
| ENSG00000286035 | AC073840,1 | -0,643254127 | 0,640267139 | 0,13351 | 0,4594073 | novel transcript |
| ENSG00000253476 | AC091185,1 | -0,6438082 | 0,640021289 | 0,11655 | 0,4332779 | novel transcript |
| ENSG00000238057 | ZEB2-AS1 | -0,643921468 | 0,639971042 | 0,07618 | 0,36240925 | ZEB2 antisense RNA 1 |
| ENSG00000106009 | BRAT1 | -0,6441537 | 0,639868034 | 0,10056 | 0,40713342 | BRCA1 associated ATM activator 1 |
| ENSG00000125843 | AP5S1 | -0,644204893 | 0,639845329 | 0,08283 | 0,37470115 | adaptor related protein complex 5 subunit sigma 1 |
| ENSG00000265254 | AC015917,2 | -0,644351881 | 0,639780142 | 0,50292 | 0,73217708 | novel transcript, antisense to SLC46A1 |
| ENSG00000128271 | ADORA2A | -0,644390661 | 0,639762945 | 0,52086 | 0,74542331 | adenosine A2a receptor |
| ENSG00000107581 | EIF3A | -0,644558204 | 0,639688652 | 0,05496 | 0,31372599 | eukaryotic translation initiation factor 3 subunit A |
| ENSG00000106605 | BLVRA | -0,644632949 | 0,639655511 | 0,00736 | 0,10888451 | biliverdin reductase A |
| ENSG00000163864 | NMNAT3 | -0,644851585 | 0,639558581 | 0,07445 | 0,35829793 | nicotinamide nucleotide adenylyltransferase 3 |
| ENSG00000206452 | HLA-C | -0,645054084 | 0,639468818 | 0,40739 | 0,66661764 | major histocompatibility complex, class I, C |
| ENSG00000172568 | FNDC9 | -0,645237495 | 0,639387527 | 0,45378 | 0,69789417 | fibronectin type III domain containing 9 |
| ENSG00000204138 | PHACTR4 | -0,645307894 | 0,639356327 | 0,15032 | 0,47842252 | phosphatase and actin regulator 4 |
| ENSG00000285413 | AP001056,2 | -0,645462671 | 0,639287739 | 0,39328 | 0,65631298 | novel transcript |
| ENSG00000234899 | SOX9-AS1 | -0,645684633 | 0,639189391 | 0,26035 | 0,56383537 | SOX9 antisense RNA 1 |
| ENSG00000130363 | RSPH3 | -0,645872919 | 0,639105976 | 0,06839 | 0,34567487 | radial spoke head 3 |

| | | | | | | |
|---|---|---|---|---|---|---|
| ENSG00000225726 | AC007000,2 | -0,645893817 | 0,639096718 | 0,1567 | 0,48425071 | pseudogene similar to part of SWI/SNF related, matrix associated, actin dependent regulator of chromatin, subfamily a, member 1 SMARCA1 |
| ENSG00000242607 | RPS3AP34 | -0,645932916 | 0,639079398 | 0,5318 | 0,75280487 | RPS3A pseudogene 34 |
| ENSG00000206376 | EHMT2 | -0,645968963 | 0,63906343 | 0,33379 | 0,61372258 | euchromatic histone lysine methyltransferase 2 |
| ENSG00000164076 | CAMKV | -0,646079169 | 0,639014614 | 0,02272 | 0,20117395 | CaM kinase like vesicle associated |
| ENSG00000100401 | RANGAP1 | -0,646284865 | 0,638923511 | 0,06393 | 0,33512301 | Ran GTPase activating protein 1 |
| ENSG00000143801 | PSEN2 | -0,646410663 | 0,638867802 | 0,07395 | 0,35740873 | presenilin 2 |
| ENSG00000259112 | NDUFC2-KCTD | -0,646678647 | 0,638749142 | 0,26735 | 0,56973003 | NDUFC2-KCTD14 readthrough |
| ENSG00000288670 | AL592295,6 | -0,64670531 | 0,638737337 | 0,14799 | 0,4751727 | novel transcript |
| ENSG00000197363 | ZNF517 | -0,646736437 | 0,638723556 | 0,08161 | 0,37277532 | zinc finger protein 517 |
| ENSG00000167112 | TRUB2 | -0,646749706 | 0,638717681 | 0,16595 | 0,49197859 | TruB pseudouridine synthase family member 2 |
| ENSG00000160972 | PPP1R16A | -0,646768368 | 0,638709419 | 0,04554 | 0,28824362 | protein phosphatase 1 regulatory subunit 16A |
| ENSG00000164074 | ABHD18 | -0,6468663 | 0,638666064 | 0,1219 | 0,44093327 | abhydrolase domain containing 18 |
| ENSG00000265908 | AC024267,5 | -0,646933873 | 0,638636151 | 0,18717 | 0,5105374 | novel transcript, antisense to PHF12 |
| ENSG00000259551 | AC048382,4 | -0,646984246 | 0,638613853 | 0,49957 | 0,73058737 | novel transcript |
| ENSG00000259694 | AC136698,1 | -0,646984246 | 0,638613853 | 0,49957 | 0,73058737 | novel transcript |
| ENSG00000127124 | HIVEP3 | -0,647282916 | 0,638481659 | 0,01487 | 0,16002963 | HIVEP zinc finger 3 |
| ENSG00000196220 | SRGAP3 | -0,647337702 | 0,638457414 | 0,00038 | 0,01694838 | SLIT-ROBO Rho GTPase activating protein 3 |
| ENSG00000233718 | MYCNOS | -0,647380164 | 0,638438623 | 0,28282 | 0,58009943 | MYCN opposite strand |
| ENSG00000130755 | GMFG | -0,647397154 | 0,638431104 | 0,3102 | 0,5983937 | glia maturation factor gamma |
| ENSG00000115009 | CCL20 | -0,647472514 | 0,638397756 | 0,4914 | 0,72510994 | C-C motif chemokine ligand 20 |
| ENSG00000107262 | BAG1 | -0,6479598 | 0,638182167 | 0,08337 | 0,37550713 | BAG cochaperone 1 |
| ENSG00000176422 | SPRYD4 | -0,648157025 | 0,638094929 | 0,05228 | 0,30698377 | SPRY domain containing 4 |
| ENSG00000130304 | SLC27A1 | -0,64828552 | 0,6380381 | 0,10716 | 0,41721821 | solute carrier family 27 member 1 |
| ENSG00000243056 | EIF4EBP3 | -0,648359371 | 0,638005439 | 0,07515 | 0,35979766 | eukaryotic translation initiation factor 4E binding protein 3 |
| ENSG00000285682 | AC005343,5 | -0,648362759 | 0,638003941 | 0,32934 | 0,61040605 | novel transcript, antisense to ADIPOR2 |
| ENSG00000101134 | DOK5 | -0,648495028 | 0,63794545 | 0,35027 | 0,62578708 | docking protein 5 |
| ENSG00000157734 | SNX22 | -0,648774655 | 0,637821814 | 0,10155 | 0,40840817 | sorting nexin 22 |
| ENSG00000285302 | CHMP4A | -0,648859105 | 0,63778448 | 0,16663 | 0,49255702 | charged multivesicular body protein 4A |
| ENSG00000070404 | FSTL3 | -0,648944388 | 0,637746779 | 0,15309 | 0,48120112 | follistatin like 3 |
| ENSG00000232412 | AL121601,1 | -0,64910986 | 0,637673636 | 0,25024 | 0,55657567 | novel transcript |
| ENSG00000149131 | SERPING1 | -0,649211274 | 0,637628812 | 0,19086 | 0,51394498 | serpin family G member 1 |

| | | | | | | |
|---|---|---|---|---|---|---|
| ENSG00000234965 | SHISA8 | -0,649328065 | 0,637577196 | 0,49832 | 0,7296 | shisa family member 8 |
| ENSG00000275740 | AC091959,1 | -0,649334656 | 0,637574283 | 0,12385 | 0,4446254 | novel readthrough transcript |
| ENSG00000248319 | LINC02275 | -0,649551692 | 0,637478375 | 0,12798 | 0,45147497 | long intergenic non-protein coding RNA 2275 |
| ENSG00000198719 | DLL1 | -0,649572757 | 0,637469067 | 0,12263 | 0,44239528 | delta like canonical Notch ligand 1 |
| ENSG00000140403 | DNAJA4 | -0,649825673 | 0,637357324 | 0,06049 | 0,32668736 | DnaJ heat shock protein family (Hsp40) member A4 |
| ENSG00000285203 | LTB4R2 | -0,649864782 | 0,637340046 | 0,13686 | 0,4641716 | leukotriene B4 receptor 2 |
| ENSG00000213906 | LTB4R2 | -0,649864782 | 0,637340046 | 0,13686 | 0,4641716 | leukotriene B4 receptor 2 |
| ENSG00000159208 | CIART | -0,650046973 | 0,637259565 | 0,21456 | 0,53351996 | circadian associated repressor of transcription |
| ENSG00000267934 | AC010300,1 | -0,650069081 | 0,637249799 | 0,12771 | 0,45102957 | novel transcript, antisense to ZNF91 |
| ENSG00000166189 | HPS6 | -0,650071566 | 0,637248702 | 0,07662 | 0,36266919 | HPS6 biogenesis of lysosomal organelles complex 2 subunit 3 |
| ENSG00000116990 | MYCL | -0,650125006 | 0,637225097 | 0,03313 | 0,24586374 | MYCL proto-oncogene, bHLH transcription factor |
| ENSG00000214796 | TUBA5P | -0,650149956 | 0,637214077 | 0,13022 | 0,45467139 | tubulin alpha 5, pseudogene |
| ENSG00000288124 | C1RL | -0,650239223 | 0,637174651 | 0,38268 | 0,64939435 | complement C1r subcomponent like |
| ENSG00000249962 | AL157400,5 | -0,650259624 | 0,637165641 | 0,23969 | 0,5491566 | pantothenate kinase 1 (PANK1) pseudogene |
| ENSG00000176160 | HSF5 | -0,650354608 | 0,637123692 | 0,53723 | 0,75585927 | heat shock transcription factor 5 |
| ENSG00000133121 | STARD13 | -0,650487711 | 0,637064914 | 0,0429 | 0,27914798 | StAR related lipid transfer domain containing 13 |
| ENSG00000061337 | LZTS1 | -0,650608973 | 0,637011369 | 0,01629 | 0,16730761 | leucine zipper tumor suppressor 1 |
| ENSG00000230337 | EXOSC10-AS1 | -0,650609934 | 0,637010945 | 0,30591 | 0,59512079 | EXOSC10 antisense RNA 1 |
| ENSG00000251365 | LINC02236 | -0,65076882 | 0,636940794 | 0,43199 | 0,68400795 | long intergenic non-protein coding RNA 2236 |
| ENSG00000234964 | FABP5P7 | -0,650863888 | 0,636898823 | 0,07899 | 0,3669327 | fatty acid binding protein 5 pseudogene 7 |
| ENSG00000243694 | LINC02027 | -0,650917261 | 0,636875262 | 0,12783 | 0,45133265 | long intergenic non-protein coding RNA 2027 |
| ENSG00000232268 | OR52I1 | -0,651252631 | 0,63672723 | 0,09679 | 0,40094776 | olfactory receptor family 52 subfamily I member 1 |
| ENSG00000259673 | IQCH-AS1 | -0,651558292 | 0,636592343 | 0,16373 | 0,49000347 | IQCH antisense RNA 1 |
| ENSG00000171951 | SCG2 | -0,651689499 | 0,63653445 | 0,20098 | 0,52181085 | secretogranin II |
| ENSG00000157542 | KCNJ6 | -0,651736203 | 0,636513844 | 0,26304 | 0,56588348 | potassium inwardly rectifying channel subfamily J member 6 |
| ENSG00000274442 | PRPF8 | -0,651754421 | 0,636505806 | 0,3384 | 0,61774255 | pre-mRNA processing factor 8 |
| ENSG00000197444 | OGDHL | -0,651807203 | 0,636482519 | 0,04517 | 0,2876168 | oxoglutarate dehydrogenase L |
| ENSG00000176236 | RPP38-DT | -0,651809324 | 0,636481584 | 0,11707 | 0,43438325 | RPP38 divergent transcript |
| ENSG00000231407 | GORAB-AS1 | -0,651950571 | 0,636419272 | 0,13199 | 0,45708171 | GORAB antisense RNA 1 |
| ENSG00000171357 | LURAP1 | -0,652037142 | 0,636381084 | 0,0804 | 0,37007763 | leucine rich adaptor protein 1 |
| ENSG00000285460 | BCAR1 | -0,652057703 | 0,636372014 | 0,11779 | 0,43532264 | BCAR1 scaffold protein, Cas family member |

| | | | | | | |
|---|---|---|---|---|---|---|
| ENSG00000230149 | AL021707,3 | -0,652059233 | 0,636371339 | 0,41336 | 0,67078587 | novel transcript, antisense to SUN2 |
| ENSG00000026297 | RNASET2 | -0,652284577 | 0,636271948 | 0,11324 | 0,42787814 | ribonuclease T2 |
| ENSG00000275223 | AL121906,2 | -0,652555154 | 0,636152627 | 0,24446 | 0,55253387 | novel transcript |
| ENSG00000104983 | CCDC61 | -0,652649351 | 0,636111092 | 0,09738 | 0,40206391 | coiled-coil domain containing 61 |
| ENSG00000169495 | HTRA4 | -0,652664443 | 0,636104438 | 0,26445 | 0,56725636 | HtrA serine peptidase 4 |
| ENSG00000189298 | ZKSCAN3 | -0,652738397 | 0,636071832 | 0,10776 | 0,41834811 | zinc finger with KRAB and SCAN domains 3 |
| ENSG00000267858 | MZF1-AS1 | -0,652819944 | 0,636035879 | 0,15592 | 0,4836223 | MZF1 antisense RNA 1 |
| ENSG00000170684 | ZNF296 | -0,652948629 | 0,635979149 | 0,15207 | 0,48055609 | zinc finger protein 296 |
| ENSG00000204428 | LY6G5C | -0,653066936 | 0,635926998 | 0,27604 | 0,57533851 | lymphocyte antigen 6 family member G5C |
| ENSG00000186716 | BCR | -0,653512346 | 0,635730695 | 0,10938 | 0,42137238 | BCR activator of RhoGEF and GTPase |
| ENSG00000241860 | AL627309,5 | -0,653514758 | 0,635729633 | 0,11196 | 0,42539842 | novel transcript |
| ENSG00000213988 | ZNF90 | -0,653539655 | 0,635718662 | 0,1447 | 0,47224655 | zinc finger protein 90 |
| ENSG00000100241 | SBF1 | -0,653604837 | 0,63568994 | 0,03547 | 0,25521132 | SET binding factor 1 |
| ENSG00000114735 | HEMK1 | -0,653776213 | 0,635614432 | 0,07331 | 0,35655334 | HemK methyltransferase family member 1 |
| ENSG00000250120 | PCDHA10 | -0,65382254 | 0,635594022 | 0,03737 | 0,26161103 | protocadherin alpha 10 |
| ENSG00000274373 | AC148476,1 | -0,653928422 | 0,635547376 | 0,21356 | 0,53275614 | novel transcript |
| ENSG00000260276 | AC022167,2 | -0,653965796 | 0,635530912 | 0,08296 | 0,37497922 | novel transcript, antisense to CARHSP1 |
| ENSG00000281915 | MUC20-OT1 | -0,654071305 | 0,635484435 | 0,1713 | 0,49653881 | MUC20 overlapping transcript |
| ENSG00000239335 | LLPH-DT | -0,654119825 | 0,635463063 | 0,36997 | 0,63943131 | LLPH divergent transcript |
| ENSG00000140398 | NEIL1 | -0,654200339 | 0,6354276 | 0,07722 | 0,36386039 | nei like DNA glycosylase 1 |
| ENSG00000228628 | ATF6B | -0,654388823 | 0,635344589 | 0,43021 | 0,68293021 | activating transcription factor 6 beta |
| ENSG00000228008 | AC105935,1 | -0,654419076 | 0,635331266 | 0,42031 | 0,67570188 | novel transcript, antisense to MON1A |
| ENSG00000176428 | VPS37D | -0,654428113 | 0,635327286 | 0,08305 | 0,37504927 | VPS37D subunit of ESCRT-I |
| ENSG00000272849 | AC084018,1 | -0,654478961 | 0,635304895 | 0,22375 | 0,53967355 | novel transcript |
| ENSG00000104967 | NOVA2 | -0,654594619 | 0,635253965 | 0,01987 | 0,18711351 | NOVA alternative splicing regulator 2 |
| ENSG00000121898 | CPXM2 | -0,654625344 | 0,635240437 | 0,11647 | 0,43301667 | carboxypeptidase X, M14 family member 2 |
| ENSG00000266865 | AC138207,8 | -0,654883567 | 0,635126747 | 0,06619 | 0,34056415 | SMAD specific E3 ubiquitin protein ligase 2 (SMURF2) pseudogene |
| ENSG00000161996 | WDR90 | -0,654939416 | 0,635102161 | 0,08187 | 0,37305174 | WD repeat domain 90 |
| ENSG00000267666 | AC004156,1 | -0,655382576 | 0,634907103 | 0,08807 | 0,38423687 | novel transcript |
| ENSG00000232044 | SILC1 | -0,655587468 | 0,63481694 | 0,0121 | 0,14307474 | sciatic injury induced lincRNA upregulator of SOX11 |
| ENSG00000273254 | AF129075,2 | -0,655633533 | 0,634796671 | 0,23369 | 0,54563244 | novel transcript, antisense to USP16 |
| ENSG00000286402 | AC090629,1 | -0,655976468 | 0,634645795 | 0,361 | 0,63311211 | novel transcript |

| | | | | | | |
|---|---|---|---|---|---|---|
| ENSG00000259834 | AL365361,1 | -0,655987564 | 0,634640914 | 0,29903 | 0,59115804 | novel transcript |
| ENSG00000285994 | AL731559,1 | -0,656138732 | 0,634574418 | 0,07692 | 0,36356213 | novel transcript, sense intronic to CAMK1D |
| ENSG00000267583 | AC007998,3 | -0,656248754 | 0,634526027 | 0,48749 | 0,72249902 | novel transcript |
| ENSG00000123178 | SPRYD7 | -0,656261093 | 0,6345206 | 0,14382 | 0,47134131 | SPRY domain containing 7 |
| ENSG00000151892 | GFRA1 | -0,656521372 | 0,634406135 | 0,09186 | 0,39178722 | GDNF family receptor alpha 1 |
| ENSG00000268992 | CDC42EP3P1 | -0,656613254 | 0,634365732 | 0,17126 | 0,49653881 | CDC42 effector protein 3 pseudogene 1 |
| ENSG00000224593 | AC092427,1 | -0,656715241 | 0,634320889 | 0,05139 | 0,30462001 | ribosomal protein L21 (RPL21) pseudogene |
| ENSG00000219626 | FAM228B | -0,656860724 | 0,634256927 | 0,04659 | 0,29026543 | family with sequence similarity 228 member B |
| ENSG00000213742 | ZNF337-AS1 | -0,656942345 | 0,634221045 | 0,18827 | 0,51144311 | ZNF337 antisense RNA 1 |
| ENSG00000267062 | AC018761,2 | -0,657057308 | 0,634170508 | 0,35621 | 0,63035942 | novel transcript, antisense to PRDX2 |
| ENSG00000246334 | PRR7-AS1 | -0,657109909 | 0,634147386 | 0,2039 | 0,52362952 | PRR7 antisense RNA 1 |
| ENSG00000188986 | NELFB | -0,657508127 | 0,633972371 | 0,06871 | 0,34648746 | negative elongation factor complex member B |
| ENSG00000160323 | ADAMTS13 | -0,65757882 | 0,633941307 | 0,21824 | 0,53571161 | ADAM metallopeptidase with thrombospondin type 1 motif 13 |
| ENSG00000286291 | AC004047,1 | -0,657621523 | 0,633922543 | 0,32895 | 0,60998638 | novel transcript |
| ENSG00000272281 | TPTE2P2 | -0,657675235 | 0,633898942 | 0,22819 | 0,54221266 | transmembrane phosphoinositide 3-phosphatase and tensin homolog 2 pseudogene 2 |
| ENSG00000255568 | BRWD1-AS2 | -0,657706115 | 0,633885374 | 0,19463 | 0,51562158 | BRWD1 antisense RNA 2 |
| ENSG00000169682 | SPNS1 | -0,657983306 | 0,633763594 | 0,16392 | 0,4902029 | sphingolipid transporter 1 (putative) |
| ENSG00000224981 | AC000367,1 | -0,658001801 | 0,63375547 | 0,20295 | 0,52282418 | tigger transposable element derived 1 (TIGD1) pseudogene |
| ENSG00000286831 | AC244472,1 | -0,658021852 | 0,633746662 | 0,13796 | 0,4651173 | novel transcript |
| ENSG00000183248 | PRR36 | -0,658412473 | 0,633575093 | 0,09531 | 0,39794903 | proline rich 36 |
| ENSG00000050438 | SLC4A8 | -0,65841863 | 0,633572389 | 0,03922 | 0,26800218 | solute carrier family 4 member 8 |
| ENSG00000187122 | SLIT1 | -0,658456559 | 0,633555733 | 0,15072 | 0,47890008 | slit guidance ligand 1 |
| ENSG00000063176 | SPHK2 | -0,658745357 | 0,63342892 | 0,13136 | 0,45639138 | sphingosine kinase 2 |
| ENSG00000116014 | KISS1R | -0,659482213 | 0,633105479 | 0,40981 | 0,6680777 | KISS1 receptor |
| ENSG00000245149 | RNF139-AS1 | -0,659510958 | 0,633092865 | 0,12815 | 0,45171208 | RNF139 antisense RNA 1 (head to head) |
| ENSG00000006432 | MAP3K9 | -0,659569939 | 0,633066984 | 0,07695 | 0,36356213 | mitogen-activated protein kinase kinase kinase 9 |
| ENSG00000225783 | MIAT | -0,659596054 | 0,633055524 | 0,10344 | 0,41135663 | myocardial infarction associated transcript |
| ENSG00000288009 | AL353135,2 | -0,659597291 | 0,633054981 | 0,11441 | 0,42986344 | novel transcript |
| ENSG00000261842 | AC073314,1 | -0,659659972 | 0,633027477 | 0,28928 | 0,5849426 | novel transcript |
| ENSG00000259818 | AL606760,3 | -0,659739945 | 0,632992388 | 0,15968 | 0,4869487 | novel transcript, antisense to C1orf123 & MAGOH |

| | | | | | | |
|---|---|---|---|---|---|---|
| ENSG00000227144 | VWA7 | -0,660233272 | 0,632775974 | 0,54312 | 0,75976694 | von Willebrand factor A domain containing 7 |
| ENSG00000268288 | AL589765,5 | -0,660362559 | 0,632719271 | 0,57545 | 0,78019408 | novel transcript, antisense to OAZ3 |
| ENSG00000186496 | ZNF396 | -0,660503952 | 0,632657263 | 0,0796 | 0,36831832 | zinc finger protein 396 |
| ENSG00000185730 | ZNF696 | -0,660526414 | 0,632647413 | 0,08305 | 0,37504927 | zinc finger protein 696 |
| ENSG00000259648 | AL132640,2 | -0,660619296 | 0,632606684 | 0,5887 | 0,78830778 | high mobility group box 1 (HMGB1) pseudogene |
| ENSG00000235060 | VDAC1P4 | -0,660735733 | 0,63255563 | 0,48276 | 0,71910922 | voltage dependent anion channel 1 pseudogene 4 |
| ENSG00000231129 | ABCF1 | -0,660772787 | 0,632539383 | 0,45134 | 0,69640448 | ATP binding cassette subfamily F member 1 |
| ENSG00000223571 | DHRSX-IT1 | -0,66105646 | 0,632415021 | 0,52236 | 0,74653683 | DHRSX intronic transcript 1 |
| ENSG00000255200 | PGAM1P8 | -0,661182378 | 0,632359826 | 0,44152 | 0,68977736 | phosphoglycerate mutase 1 pseudogene 8 |
| ENSG00000138311 | ZNF365 | -0,66121444 | 0,632345773 | 0,22495 | 0,54037642 | zinc finger protein 365 |
| ENSG00000270171 | AL359881,3 | -0,661296581 | 0,632309771 | 0,17931 | 0,50368273 | novel transcript |
| ENSG00000174279 | EVX2 | -0,661355172 | 0,632284092 | 0,67613 | 0,840628 | even-skipped homeobox 2 |
| ENSG00000226701 | RPL15P14 | -0,661656488 | 0,63215205 | 0,53777 | 0,75631707 | ribosomal protein L15 pseudogene 14 |
| ENSG00000164902 | PHAX | -0,66238805 | 0,631831579 | 0,13675 | 0,46409456 | phosphorylated adaptor for RNA export |
| ENSG00000269653 | AC011479,3 | -0,662445851 | 0,631806266 | 0,35114 | 0,62664963 | novel transcript, sense intronic to SIPA1L3 |
| ENSG00000142444 | TIMM29 | -0,662721069 | 0,631685749 | 0,10326 | 0,41111142 | translocase of inner mitochondrial membrane 29 |
| ENSG00000069869 | NEDD4 | -0,662787837 | 0,631656516 | 0,07185 | 0,35364999 | NEDD4 E3 ubiquitin protein ligase |
| ENSG00000234161 | PABPC5-AS1 | -0,662814932 | 0,631644653 | 0,22636 | 0,54126693 | PABPC5 antisense RNA 1 |
| ENSG00000008128 | CDK11A | -0,662909194 | 0,631603384 | 0,04995 | 0,30082943 | cyclin dependent kinase 11A |
| ENSG00000273485 | AL139339,2 | -0,662925366 | 0,631596304 | 0,46931 | 0,70885494 | novel transcript, antisense to CALHM2 |
| ENSG00000274026 | FAM27E3 | -0,663065604 | 0,631534912 | 0,11731 | 0,43468107 | family with sequence similarity 27 member E3 |
| ENSG00000129680 | MAP7D3 | -0,663080526 | 0,63152838 | 0,19389 | 0,51547251 | MAP7 domain containing 3 |
| ENSG00000281376 | ABALON | -0,663084068 | 0,63152683 | 0,25996 | 0,56377449 | apoptotic BCL2L1-antisense long non-coding RNA |
| ENSG00000221995 | TIAF1 | -0,663173649 | 0,631487618 | 0,10441 | 0,41285329 | TGFB1-induced anti-apoptotic factor 1 |
| ENSG00000288089 | AC011303,1 | -0,663297471 | 0,631433422 | 0,39169 | 0,65533345 | novel transcript |
| ENSG00000099822 | HCN2 | -0,663431956 | 0,631374563 | 0,04029 | 0,27141252 | hyperpolarization activated cyclic nucleotide gated potassium and sodium channel 2 |
| ENSG00000171045 | TSNARE1 | -0,663513132 | 0,631339039 | 0,08305 | 0,37504927 | t-SNARE domain containing 1 |
| ENSG00000261635 | AC103988,1 | -0,663552405 | 0,631321853 | 0,29394 | 0,58836931 | tropomyosin 3 (TPM3) pseudogene |
| ENSG00000287112 | AC068787,3 | -0,663746251 | 0,631237032 | 0,32735 | 0,60914884 | novel transcript |
| ENSG00000049540 | ELN | -0,663774806 | 0,631224538 | 0,2664 | 0,56917012 | elastin |
| ENSG00000134516 | DOCK2 | -0,664185774 | 0,631044752 | 0,26783 | 0,56999323 | dedicator of cytokinesis 2 |
| ENSG00000187186 | AL162231,1 | -0,664211117 | 0,631033667 | 0,10896 | 0,42024426 | uncharacterized LOC730098 |

| | | | | | | |
|---|---|---|---|---|---|---|
| ENSG00000160439 | RDH13 | -0,664342088 | 0,630976383 | 0,24607 | 0,55380958 | retinol dehydrogenase 13 |
| ENSG00000242282 | AC108488,1 | -0,66438209 | 0,630958888 | 0,16184 | 0,48914394 | novel transcript |
| ENSG00000226492 | CUTA | -0,66453682 | 0,630891221 | 0,18378 | 0,50744379 | cutA divalent cation tolerance homolog |
| ENSG00000134463 | ECHDC3 | -0,664661939 | 0,630836508 | 0,05326 | 0,30991339 | enoyl-CoA hydratase domain containing 3 |
| ENSG00000227948 | AC003989,1 | -0,664797782 | 0,630777112 | 0,31982 | 0,60470232 | pseudogene similar to part of argininosuccinate synthetase 1 (ASS1) |
| ENSG00000276887 | TTYH1 | -0,664844814 | 0,630756549 | 0,13846 | 0,46575418 | tweety family member 1 |
| ENSG00000277112 | ANKRD20A21 | -0,664965119 | 0,630703953 | 0,35912 | 0,63201238 | ankyrin repeat domain 20 family member A21, pseudogene |
| ENSG00000259314 | AC103739,2 | -0,665029072 | 0,630675995 | 0,35992 | 0,63249219 | novel transcript, antisense to CRTC3 |
| ENSG00000105605 | CACNG7 | -0,665153928 | 0,630621417 | 0,09154 | 0,39126619 | calcium voltage-gated channel auxiliary subunit gamma 7 |
| ENSG00000198797 | BRINP2 | -0,665183546 | 0,63060847 | 0,13227 | 0,4573773 | BMP/retinoic acid inducible neural specific 2 |
| ENSG00000285934 | AC091074,2 | -0,665195869 | 0,630603084 | 0,04131 | 0,27529279 | novel transcript |
| ENSG00000105327 | BBC3 | -0,665336233 | 0,630541734 | 0,10541 | 0,41466216 | BCL2 binding component 3 |
| ENSG00000247081 | BAALC-AS1 | -0,665613448 | 0,630420587 | 0,05381 | 0,31106243 | BAALC antisense RNA 1 |
| ENSG00000163596 | ICA1L | -0,66578132 | 0,630347235 | 0,02455 | 0,20956669 | islet cell autoantigen 1 like |
| ENSG00000198171 | DDRGK1 | -0,665923032 | 0,630285321 | 0,06815 | 0,34506926 | DDRGK domain containing 1 |
| ENSG00000168065 | SLC22A11 | -0,666005079 | 0,630249477 | 0,41954 | 0,67530041 | solute carrier family 22 member 11 |
| ENSG00000163545 | NUAK2 | -0,666206978 | 0,630161282 | 0,14659 | 0,47448174 | NUAK family kinase 2 |
| ENSG00000264529 | DNAJB6P8 | -0,666297855 | 0,630121589 | 0,29796 | 0,59037822 | DNAJB6 pseudogene 8 |
| ENSG00000287679 | AL355347,1 | -0,666323559 | 0,630110363 | 0,07452 | 0,35851426 | novel transcript, antisense to KHDRBS2 |
| ENSG00000160685 | ZBTB7B | -0,666484303 | 0,63004016 | 0,2106 | 0,52915889 | zinc finger and BTB domain containing 7B |
| ENSG00000228662 | MCCD1 | -0,666513308 | 0,630027493 | 0,27253 | 0,57315967 | mitochondrial coiled-coil domain 1 |
| ENSG00000281750 | AC213220,3 | -0,666536026 | 0,630017572 | 0,32622 | 0,6085098 | novel transcript, sense intronic to NDUFA6 |
| ENSG00000280750 | AL021878,9 | -0,666536026 | 0,630017572 | 0,32622 | 0,6085098 | novel transcript, sense intronic to NDUFA6 |
| ENSG00000281542 | AL021878,11 | -0,666536026 | 0,630017572 | 0,32622 | 0,6085098 | novel transcript, sense intronic to NDUFA6 |
| ENSG00000270083 | AL021878,4 | -0,666536026 | 0,630017572 | 0,32622 | 0,6085098 | novel transcript, sense intronic to NDUFA6 |
| ENSG00000257800 | FNBP1P1 | -0,666751619 | 0,629923431 | 0,00502 | 0,0883535 | formin binding protein 1 pseudogene 1 |
| ENSG00000162188 | GNG3 | -0,66685995 | 0,629876132 | 0,07029 | 0,34990935 | G protein subunit gamma 3 |
| ENSG00000288030 | AC066585,1 | -0,666885152 | 0,62986513 | 0,51282 | 0,73941533 | novel transcript, sense intronic to GRM7 |
| ENSG00000105556 | MIER2 | -0,666933549 | 0,629844 | 0,10641 | 0,41609301 | MIER family member 2 |
| ENSG00000211451 | GNRHR2 | -0,667070385 | 0,629784264 | 0,07378 | 0,3570536 | gonadotropin releasing hormone receptor 2 (pseudogene) |
| ENSG00000129535 | NRL | -0,667267125 | 0,629698386 | 0,0755 | 0,36080715 | neural retina leucine zipper |

| | | | | | | |
|---|---|---|---|---|---|---|
| ENSG00000262634 | SKA1 | -0,667547585 | 0,629575984 | 0,23019 | 0,54335232 | spindle and kinetochore associated complex subunit 1 |
| ENSG00000134398 | ERN2 | -0,667742578 | 0,629490898 | 0,30559 | 0,59509691 | endoplasmic reticulum to nucleus signaling 2 |
| ENSG00000214851 | LINC00612 | -0,668148788 | 0,629313681 | 0,05471 | 0,31313954 | long intergenic non-protein coding RNA 612 |
| ENSG00000267296 | CEBPA-DT | -0,668228073 | 0,629279097 | 0,16192 | 0,48914394 | CEBPA divergent transcript |
| ENSG00000198467 | TPM2 | -0,668343722 | 0,629228655 | 0,02042 | 0,18959706 | tropomyosin 2 |
| ENSG00000281416 | AC006359,4 | -0,668350051 | 0,629225895 | 0,32803 | 0,60971379 | novel transcript |
| ENSG00000256361 | AC027544,1 | -0,668571608 | 0,629129271 | 0,26077 | 0,56400103 | glycine cleavage system protein H (aminomethyl carrier) (GCSH) |
| ENSG00000274737 | AC004466,2 | -0,668692327 | 0,62907663 | 0,42029 | 0,67570188 | novel transcript, sense intronic to HDAC7 |
| ENSG00000184949 | FAM227A | -0,668705495 | 0,629070888 | 0,09475 | 0,39724142 | family with sequence similarity 227 member A |
| ENSG00000285697 | AC002059,3 | -0,668807211 | 0,629026538 | 0,31528 | 0,60158701 | chromosome 22 open reading frame 42 (C22orf42) pseudogene |
| ENSG00000286881 | AL358780,1 | -0,66890827 | 0,628982477 | 0,0978 | 0,4029082 | novel transcript, sense intronic to MLLT10 |
| ENSG00000234921 | AC006450,2 | -0,668926354 | 0,628974593 | 0,32276 | 0,60584543 | novel transcript, antisense to LHX2 |
| ENSG00000254689 | LINC02235 | -0,668987931 | 0,628947747 | 0,18977 | 0,51299724 | long intergenic non-protein coding RNA 2235 |
| ENSG00000229376 | CICP3 | -0,669009439 | 0,628938371 | 0,14561 | 0,47304491 | capicua transcriptional repressor pseudogene 3 |
| ENSG00000277429 | SERF1B | -0,669033426 | 0,628927914 | 0,38918 | 0,65363446 | small EDRK-rich factor 1B |
| ENSG00000170419 | VSTM2A | -0,669163877 | 0,628871048 | 0,28157 | 0,57891473 | V-set and transmembrane domain containing 2A |
| ENSG00000144583 | MARCHF4 | -0,669267492 | 0,628825884 | 0,09269 | 0,39335125 | membrane associated ring-CH-type finger 4 |
| ENSG00000236279 | CLEC2L | -0,669274516 | 0,628822822 | 0,31031 | 0,59842274 | C-type lectin domain family 2 member L |
| ENSG00000148798 | INA | -0,669405359 | 0,628765795 | 0,0555 | 0,31503413 | internexin neuronal intermediate filament protein alpha |
| ENSG00000179943 | FIZ1 | -0,669430485 | 0,628754844 | 0,07971 | 0,36854037 | FLT3 interacting zinc finger 1 |
| ENSG00000178171 | AMER3 | -0,669571259 | 0,628693495 | 0,07065 | 0,3507077 | APC membrane recruitment protein 3 |
| ENSG00000257522 | AL133166,1 | -0,669668115 | 0,628651289 | 0,34125 | 0,61939402 | novel transcript |
| ENSG00000134202 | GSTM3 | -0,669762948 | 0,628609967 | 0,11064 | 0,42326288 | glutathione S-transferase mu 3 |
| ENSG00000274259 | SYNGAP1-AS1 | -0,669987182 | 0,628512271 | 0,24091 | 0,55001001 | SYNGAP1 antisense RNA 1 |
| ENSG00000283647 | AL512643,1 | -0,670030291 | 0,628493491 | 0,39558 | 0,65831344 | novel transcript, antisense to FREM1 |
| ENSG00000196110 | ZNF699 | -0,670222623 | 0,62840971 | 0,12294 | 0,44282207 | zinc finger protein 699 |
| ENSG00000170537 | TMC7 | -0,670412926 | 0,628326823 | 0,07173 | 0,35351167 | transmembrane channel like 7 |
| ENSG00000232474 | NCKAP5-IT1 | -0,670475182 | 0,628299709 | 0,29156 | 0,58655792 | NCKAP5 intronic transcript 1 |
| ENSG00000125869 | LAMP5 | -0,670592948 | 0,628248424 | 0,50022 | 0,73078774 | lysosomal associated membrane protein family member 5 |
| ENSG00000238168 | IFITM3P5 | -0,670688974 | 0,628206609 | 0,10791 | 0,41847368 | IFITM3 pseudogene 5 |
| ENSG00000060069 | CTDP1 | -0,670709848 | 0,62819752 | 0,09237 | 0,39278463 | CTD phosphatase subunit 1 |

| | | | | | | |
|---|---|---|---|---|---|---|
| ENSG00000155816 | FMN2 | -0,670889535 | 0,628119283 | 0,00534 | 0,09086835 | formin 2 |
| ENSG00000130822 | PNCK | -0,671058685 | 0,628045643 | 0,21119 | 0,53012655 | pregnancy up-regulated nonubiquitous CaM kinase |
| ENSG00000125531 | FNDC11 | -0,671119516 | 0,628019162 | 0,09082 | 0,39006208 | fibronectin type III domain containing 11 |
| ENSG00000231227 | AC006450,1 | -0,671131983 | 0,628013735 | 0,40099 | 0,66290375 | novel transcript |
| ENSG00000150977 | RILPL2 | -0,671343794 | 0,627921539 | 0,08589 | 0,38009386 | Rab interacting lysosomal protein like 2 |
| ENSG00000235731 | LINC02250 | -0,671811331 | 0,62771808 | 0,06417 | 0,33569685 | long intergenic non-protein coding RNA 2250 |
| ENSG00000169951 | ZNF764 | -0,671885991 | 0,627685596 | 0,08958 | 0,38705587 | zinc finger protein 764 |
| ENSG00000272476 | AL024507,2 | -0,671898622 | 0,627680101 | 0,11516 | 0,43077792 | novel transcript, antisense to SEC63 |
| ENSG00000241258 | CRCP | -0,671962252 | 0,627652418 | 0,08985 | 0,38767975 | CGRP receptor component |
| ENSG00000198914 | POU3F3 | -0,671987611 | 0,627641385 | 0,23594 | 0,5471053 | POU class 3 homeobox 3 |
| ENSG00000187650 | VMAC | -0,672186885 | 0,627554698 | 0,15302 | 0,48120112 | vimentin type intermediate filament associated coiled-coil protein |
| ENSG00000212695 | AL583805,1 | -0,672465216 | 0,627433639 | 0,36467 | 0,63504684 | ribosomal protein 18 (RPL18) pseudogene |
| ENSG00000130035 | GALNT8 | -0,672705102 | 0,62732932 | 0,04731 | 0,29224089 | polypeptide N-acetylgalactosaminyltransferase 8 |
| ENSG00000160233 | LRRC3 | -0,672767926 | 0,627302003 | 0,02227 | 0,19831519 | leucine rich repeat containing 3 |
| ENSG00000241939 | RN7SL517P | -0,672933845 | 0,627229863 | 0,25718 | 0,56155289 | RNA, 7SL, cytoplasmic 517, pseudogene |
| ENSG00000128274 | A4GALT | -0,673018633 | 0,627193002 | 0,42917 | 0,6820074 | alpha 1,4-galactosyltransferase (P blood group) |
| ENSG00000287625 | AC022210,1 | -0,673038687 | 0,627184284 | 0,17069 | 0,49641416 | novel transcript |
| ENSG00000145934 | TENM2 | -0,673140696 | 0,627139939 | 0,04593 | 0,28908951 | teneurin transmembrane protein 2 |
| ENSG00000201183 | RNVU1-3 | -0,673164893 | 0,62712942 | 0,23513 | 0,54666157 | RNA, variant U1 small nuclear 3 |
| ENSG00000157303 | SUSD3 | -0,673430453 | 0,627013994 | 0,50435 | 0,7332113 | sushi domain containing 3 |
| ENSG00000173838 | MARCHF10 | -0,673517609 | 0,626976116 | 0,05831 | 0,32231869 | membrane associated ring-CH-type finger 10 |
| ENSG00000236255 | AC009404,1 | -0,67353564 | 0,62696828 | 0,11131 | 0,42416781 | novel transcript |
| ENSG00000204227 | RING1 | -0,673705819 | 0,626894327 | 0,07853 | 0,36610837 | ring finger protein 1 |
| ENSG00000285331 | AC090517,5 | -0,673946336 | 0,626789825 | 0,0641 | 0,33569301 | novel transcript |
| ENSG00000226332 | AL354836,1 | -0,674050499 | 0,626744572 | 0,22437 | 0,54029213 | novel transcript |
| ENSG00000185736 | ADARB2 | -0,67410993 | 0,626718754 | 0,16401 | 0,49033897 | adenosine deaminase RNA specific B2 (inactive) |
| ENSG00000110328 | GALNT18 | -0,674242585 | 0,62666113 | 0,03068 | 0,23627456 | polypeptide N-acetylgalactosaminyltransferase 18 |
| ENSG00000267368 | UPK3BL1 | -0,67436315 | 0,626608763 | 0,10823 | 0,41892484 | uroplakin 3B like 1 |
| ENSG00000278195 | SSTR3 | -0,674852154 | 0,626396409 | 0,24173 | 0,55047794 | somatostatin receptor 3 |
| ENSG00000245532 | NEAT1 | -0,675449345 | 0,626137171 | 0,04277 | 0,27875629 | nuclear paraspeckle assembly transcript 1 |
| ENSG00000284837 | AC007846,1 | -0,67547963 | 0,626124027 | 0,14876 | 0,47610555 | novel transcript |
| ENSG00000286058 | AC018865,2 | -0,675553174 | 0,62609211 | 0,02904 | 0,22988377 | novel transcript |

| | | | | | | |
|---|---|---|---|---|---|---|
| ENSG00000271938 | AC103724,4 | -0,675580571 | 0,626080221 | 0,43085 | 0,68325486 | novel transcript |
| ENSG00000203666 | EFCAB2 | -0,675593618 | 0,626074559 | 0,0407 | 0,27272229 | EF-hand calcium binding domain 2 |
| ENSG00000218018 | RBM38-AS1 | -0,675940265 | 0,625924145 | 0,30369 | 0,59385655 | RBM38 antisense RNA 1 |
| ENSG00000271849 | MAN2A1-DT | -0,676047937 | 0,625877433 | 0,35638 | 0,63043562 | MAN2A1 divergent transcript |
| ENSG00000288360 | ZSWIM4 | -0,676129384 | 0,6258421 | 0,04398 | 0,28274932 | zinc finger SWIM-type containing 4 |
| ENSG00000132003 | ZSWIM4 | -0,676129384 | 0,6258421 | 0,04398 | 0,28274932 | zinc finger SWIM-type containing 4 |
| ENSG00000151576 | QTRT2 | -0,676164043 | 0,625827065 | 0,0986 | 0,40441723 | queuine tRNA-ribosyltransferase accessory subunit 2 |
| ENSG00000269646 | AC010487,1 | -0,676261546 | 0,62578477 | 0,42793 | 0,68085046 | novel transcript |
| ENSG00000288499 | FAM20C | -0,676978315 | 0,625473941 | 0,12674 | 0,44948266 | FAM20C golgi associated secretory pathway kinase |
| ENSG00000177706 | FAM20C | -0,676978315 | 0,625473941 | 0,12674 | 0,44948266 | FAM20C golgi associated secretory pathway kinase |
| ENSG00000229048 | DUTP1 | -0,677559787 | 0,625221898 | 0,36199 | 0,63352908 | deoxyuridine triphosphatase pseudogene 1 |
| ENSG00000265984 | AC022809,1 | -0,677639042 | 0,625187552 | 0,38489 | 0,65071151 | novel transcript |
| ENSG00000275125 | PRXL2B | -0,677705848 | 0,625158602 | 0,15659 | 0,48425071 | peroxiredoxin like 2B |
| ENSG00000287078 | AL513218,2 | -0,677729002 | 0,625148569 | 0,39429 | 0,65724328 | novel transcript, antisense to CC2D1B |
| ENSG00000139631 | CSAD | -0,678114898 | 0,624981375 | 0,12924 | 0,45267249 | cysteine sulfinic acid decarboxylase |
| ENSG00000197912 | SPG7 | -0,678153001 | 0,624964869 | 0,08472 | 0,37790018 | SPG7 matrix AAA peptidase subunit, paraplegin |
| ENSG00000242880 | AC026341,1 | -0,678411696 | 0,624852814 | 0,35291 | 0,6277271 | novel transcript |
| ENSG00000178235 | SLITRK1 | -0,678416767 | 0,624850618 | 0,03965 | 0,26944441 | SLIT and NTRK like family member 1 |
| ENSG00000109756 | RAPGEF2 | -0,678478663 | 0,624823811 | 0,00461 | 0,08360185 | Rap guanine nucleotide exchange factor 2 |
| ENSG00000167912 | AC090152,1 | -0,678498436 | 0,624815247 | 0,02619 | 0,21809028 | novel transcript |
| ENSG00000248863 | AC097376,2 | -0,678561828 | 0,624787793 | 0,23251 | 0,54526182 | glucose-fructose oxidoreductase domain containing 2 (GFOD2) pseudogene |
| ENSG00000261837 | AC046158,2 | -0,678651101 | 0,624749133 | 0,3736 | 0,6422203 | novel transcript, sense intronic to WWOX |
| ENSG00000149489 | ROM1 | -0,678685674 | 0,624734161 | 0,18882 | 0,51219588 | retinal outer segment membrane protein 1 |
| ENSG00000142327 | RNPEPL1 | -0,678946796 | 0,624621097 | 0,00716 | 0,10764638 | arginyl aminopeptidase like 1 |
| ENSG00000286457 | AC009560,3 | -0,679075102 | 0,624565549 | 0,28792 | 0,5840701 | novel transcript, antisense to RORA |
| ENSG00000077800 | FKBP6 | -0,679098807 | 0,624555287 | 0,33825 | 0,61774255 | FKBP prolyl isomerase 6 |
| ENSG00000286106 | NOTCH2NLR | -0,679148979 | 0,624533568 | 0,08783 | 0,38352337 | notch 2 N-terminal like R (pseudogene) |
| ENSG00000138741 | TRPC3 | -0,679149317 | 0,624533421 | 0,06027 | 0,32601555 | transient receptor potential cation channel subfamily C member 3 |
| ENSG00000269815 | AC010463,3 | -0,679244122 | 0,624492382 | 0,13198 | 0,45708171 | novel transcript, sense intronic ABHD8 |
| ENSG00000083454 | P2RX5 | -0,679448337 | 0,624403991 | 0,25471 | 0,55944852 | purinergic receptor P2X 5 |
| ENSG00000277268 | LHX1-DT | -0,679461007 | 0,624398507 | 0,19854 | 0,51980353 | LHX1 divergent transcript |

| | | | | | | |
|---|---|---|---|---|---|---|
| ENSG00000204104 | TRAF3IP1 | -0,679462121 | 0,624398025 | 0,0122 | 0,14356185 | TRAF3 interacting protein 1 |
| ENSG00000177721 | ANXA2R | -0,679750994 | 0,624273013 | 0,26031 | 0,56382982 | annexin A2 receptor |
| ENSG00000288028 | AL049839,3 | -0,679832661 | 0,624237676 | 0,12747 | 0,45071101 | novel transcript |
| ENSG00000286789 | AL161645,2 | -0,67984213 | 0,624233579 | 0,13903 | 0,46647959 | novel transcript, sense intronic to CYP2E1 |
| ENSG00000172716 | SLFN11 | -0,679912959 | 0,624202933 | 0,15333 | 0,48157724 | schlafen family member 11 |
| ENSG00000247735 | AC120114,1 | -0,680047079 | 0,624144907 | 0,11435 | 0,42986344 | novel transcript, antisense to KCTD13 |
| ENSG00000161267 | BDH1 | -0,68019583 | 0,624080556 | 0,07537 | 0,36047139 | 3-hydroxybutyrate dehydrogenase 1 |
| ENSG00000287257 | AC111182,2 | -0,680278562 | 0,624044769 | 0,4738 | 0,71294959 | novel transcript, antisense to SEPT9 |
| ENSG00000180822 | PSMG4 | -0,680634876 | 0,623890663 | 0,06154 | 0,32865897 | proteasome assembly chaperone 4 |
| ENSG00000165171 | METTL27 | -0,680954113 | 0,623752625 | 0,2573 | 0,56160845 | methyltransferase like 27 |
| ENSG00000278576 | AL162171,3 | -0,680972186 | 0,623744811 | 0,10192 | 0,40932218 | novel transcript, sense intronic to ZC3H14 |
| ENSG00000269918 | AF131215,6 | -0,681251808 | 0,623623929 | 0,2177 | 0,53548046 | novel transcript |
| ENSG00000177946 | CENPBD1 | -0,681317331 | 0,623595607 | 0,13907 | 0,46647959 | CENPB DNA-binding domain containing 1 |
| ENSG00000134762 | DSC3 | -0,681322634 | 0,623593314 | 0,2273 | 0,54168591 | desmocollin 3 |
| ENSG00000251002 | AC244502,1 | -0,681564434 | 0,623488807 | 0,0395 | 0,2689738 | novel transcript, antisense to IG_gene_cluster |
| ENSG00000228395 | AL356481,1 | -0,681686092 | 0,623436233 | 0,18404 | 0,50781294 | novel transcript |
| ENSG00000240882 | AC063952,2 | -0,681784545 | 0,623393689 | 0,54682 | 0,76240845 | prostate tumor overexpressed 1 (PTOV1) pseudogene |
| ENSG00000167716 | WDR81 | -0,681938083 | 0,623327348 | 0,28347 | 0,58065155 | WD repeat domain 81 |
| ENSG00000250959 | GLUD1P3 | -0,682039975 | 0,623283326 | 0,03737 | 0,26161103 | glutamate dehydrogenase 1 pseudogene 3 |
| ENSG00000120332 | TNN | -0,682313488 | 0,623165173 | 0,36926 | 0,63912679 | tenascin N |
| ENSG00000099821 | POLRMT | -0,682353171 | 0,623148032 | 0,07653 | 0,36266919 | RNA polymerase mitochondrial |
| ENSG00000286001 | AC021087,5 | -0,68291986 | 0,622903309 | 0,45943 | 0,7017037 | novel protein |
| ENSG00000277606 | GUSBP1 | -0,682984964 | 0,622875199 | 0,01819 | 0,1782976 | GUSB pseudogene 1 |
| ENSG00000173214 | MFSD4B | -0,683131408 | 0,622811976 | 0,12509 | 0,44678334 | major facilitator superfamily domain containing 4B |
| ENSG00000237413 | MGC27382 | -0,683183698 | 0,622789403 | 0,13783 | 0,46490262 | uncharacterized MGC27382 |
| ENSG00000160401 | CFAP157 | -0,683194954 | 0,622784544 | 0,10464 | 0,41313952 | cilia and flagella associated protein 157 |
| ENSG00000232237 | ASCL5 | -0,683203727 | 0,622780757 | 0,25115 | 0,55719428 | achaete-scute family bHLH transcription factor 5 |
| ENSG00000287223 | Z94277,1 | -0,683218873 | 0,622774219 | 0,08344 | 0,37551246 | novel transcript |
| ENSG00000228317 | AL158070,1 | -0,683295325 | 0,622741217 | 0,31888 | 0,60399503 | novel transcript |
| ENSG00000170745 | KCNS3 | -0,683497828 | 0,622653813 | 0,1251 | 0,44678334 | potassium voltage-gated channel modifier subfamily S member 3 |
| ENSG00000260683 | AL591643,1 | -0,683583383 | 0,622616889 | 0,2642 | 0,56696403 | novel transcript |
| ENSG00000254737 | OR10G4 | -0,683868654 | 0,622493788 | 0,19854 | 0,51980353 | olfactory receptor family 10 subfamily G member 4 |

| | | | | | | |
|---|---|---|---|---|---|---|
| ENSG00000232335 | AL354864,1 | -0,683960063 | 0,622454348 | 0,2993 | 0,59123205 | novel transcript |
| ENSG00000168081 | PNOC | -0,684002857 | 0,622435885 | 0,49797 | 0,72933516 | prepronociceptin |
| ENSG00000262327 | AGK | -0,684005384 | 0,622434795 | 0,63922 | 0,81901202 | acylglycerol kinase |
| ENSG00000229180 | AC006001,3 | -0,684010771 | 0,622432471 | 0,12068 | 0,4394821 | RAB guanine nucleotide exchange factor (GEF) 1 (RABGEF1) pseudogene |
| ENSG00000286724 | AL732509,1 | -0,684111801 | 0,622388884 | 0,12575 | 0,44784483 | novel transcript, antisense to ADGRG2and PHKA2 |
| ENSG00000179431 | FJX1 | -0,684158756 | 0,622368628 | 0,0748 | 0,35899472 | four-jointed box kinase 1 |
| ENSG00000225321 | LINC01427 | -0,684230966 | 0,622337478 | 0,07323 | 0,35655334 | long intergenic non-protein coding RNA 1427 |
| ENSG00000174502 | SLC26A9 | -0,684463238 | 0,62223729 | 0,48032 | 0,71764212 | solute carrier family 26 member 9 |
| ENSG00000238243 | OR2W3 | -0,684476253 | 0,622231677 | 0,30277 | 0,59321687 | olfactory receptor family 2 subfamily W member 3 |
| ENSG00000275395 | FCGBP | -0,684496932 | 0,622222758 | 0,45503 | 0,69913628 | Fc fragment of IgG binding protein |
| ENSG00000253877 | LINC01608 | -0,684575094 | 0,622189048 | 0,06744 | 0,34371006 | long intergenic non-protein coding RNA 1608 |
| ENSG00000286746 | AL590807,1 | -0,684592678 | 0,622181465 | 0,07375 | 0,35703856 | novel transcript |
| ENSG00000064201 | TSPAN32 | -0,684774314 | 0,622103137 | 0,40991 | 0,66816149 | tetraspanin 32 |
| ENSG00000203993 | ARRDC1-AS1 | -0,685062566 | 0,621978853 | 0,07107 | 0,35204529 | ARRDC1 antisense RNA 1 |
| ENSG00000249784 | SCARNA22 | -0,685213296 | 0,621913873 | 0,07298 | 0,35631792 | small Cajal body-specific RNA 22 |
| ENSG00000215883 | CYB5RL | -0,685340377 | 0,621859093 | 0,09776 | 0,40289531 | cytochrome b5 reductase like |
| ENSG00000226009 | KCNIP2-AS1 | -0,685343085 | 0,621857926 | 0,27674 | 0,57549599 | KCNIP2 antisense RNA 1 |
| ENSG00000204983 | PRSS1 | -0,685395514 | 0,621835328 | 0,39717 | 0,65966023 | serine protease 1 |
| ENSG00000064270 | ATP2C2 | -0,685401981 | 0,62183254 | 0,06248 | 0,33103715 | ATPase secretory pathway Ca2+ transporting 2 |
| ENSG00000272588 | AC139887,4 | -0,685632327 | 0,621733264 | 0,23541 | 0,54683669 | novel transcript, antisense to PCGF3 |
| ENSG00000265778 | AC018413,1 | -0,68571934 | 0,621695767 | 0,11912 | 0,43696075 | novel transcript, antisense to ZFN156 |
| ENSG00000287277 | AL392086,3 | -0,686066103 | 0,621546355 | 0,05843 | 0,32267557 | novel transcript |
| ENSG00000165055 | METTL2B | -0,686205532 | 0,621486289 | 0,10099 | 0,40770184 | methyltransferase like 2B |
| ENSG00000216819 | TUBB2BP1 | -0,686221637 | 0,621479351 | 0,43157 | 0,68379763 | tubulin beta 2B class IIb pseudogene 1 |
| ENSG00000254618 | TMED10P1 | -0,686261772 | 0,621462062 | 0,40267 | 0,66396567 | transmembrane p24 trafficking protein 10 pseudogene 1 |
| ENSG00000262155 | LINC02175 | -0,686278702 | 0,621454769 | 0,02281 | 0,20153127 | long intergenic non-protein coding RNA 2175 |
| ENSG00000228172 | AL020996,1 | -0,686311272 | 0,62144074 | 0,10416 | 0,41252273 | novel transcript |
| ENSG00000128309 | MPST | -0,686784833 | 0,621236787 | 0,08763 | 0,38323183 | mercaptopyruvate sulfurtransferase |
| ENSG00000223834 | AL161935,1 | -0,686863471 | 0,621202925 | 0,19217 | 0,51471664 | novel transcript |
| ENSG00000275457 | AL117332,1 | -0,686864102 | 0,621202654 | 0,08613 | 0,38043908 | novel transcript, antisense to XRN2 |
| ENSG00000027644 | INSRR | -0,686945455 | 0,621167625 | 0,39676 | 0,65932701 | insulin receptor related receptor |
| ENSG00000163956 | LRPAP1 | -0,686996528 | 0,621145636 | 0,07235 | 0,35512127 | LDL receptor related protein associated protein 1 |

| | | | | | | |
|---|---|---|---|---|---|---|
| ENSG00000225798 | AC025918,1 | -0,687036495 | 0,621128428 | 0,47623 | 0,71446062 | novel transcript |
| ENSG00000183718 | TRIM52 | -0,687105049 | 0,621098914 | 0,07844 | 0,36604454 | tripartite motif containing 52 |
| ENSG00000260352 | AC092287,1 | -0,687162068 | 0,621074367 | 0,38143 | 0,64876277 | novel transcript, antisense to SMG1 |
| ENSG00000108878 | CACNG1 | -0,687171347 | 0,621070373 | 0,46385 | 0,70455791 | calcium voltage-gated channel auxiliary subunit gamma 1 |
| ENSG00000287778 | AC006230,1 | -0,687796885 | 0,620801141 | 0,21846 | 0,53578053 | novel transcript, antisense to HS3ST1 |
| ENSG00000261734 | AC116096,1 | -0,687823973 | 0,620789485 | 0,31193 | 0,59925736 | novel transcript |
| ENSG00000232778 | RPL23AP50 | -0,688016516 | 0,62070664 | 0,22055 | 0,53759099 | ribosomal protein L23a pseudogene 50 |
| ENSG00000164050 | PLXNB1 | -0,688029922 | 0,620700872 | 0,00779 | 0,11225873 | plexin B1 |
| ENSG00000287966 | AL590062,1 | -0,688072592 | 0,620682514 | 0,04725 | 0,29217708 | novel transcript |
| ENSG00000288056 | AL023806,4 | -0,688245726 | 0,620608032 | 0,02973 | 0,23215706 | novel transcript |
| ENSG00000234944 | LINC02623 | -0,688314326 | 0,620578523 | 0,5233 | 0,7473071 | long intergenic non-protein coding RNA 2623 |
| ENSG00000240240 | BX664727,3 | -0,688503238 | 0,620497267 | 0,14423 | 0,47177175 | novel transcript |
| ENSG00000186648 | CARMIL3 | -0,688696638 | 0,620414092 | 0,00064 | 0,02439336 | capping protein regulator and myosin 1 linker 3 |
| ENSG00000284765 | CARMIL3 | -0,688696638 | 0,620414092 | 0,00064 | 0,02439336 | capping protein regulator and myosin 1 linker 3 |
| ENSG00000179598 | PLD6 | -0,688801173 | 0,62036914 | 0,22315 | 0,53927144 | phospholipase D family member 6 |
| ENSG00000273687 | AC004223,4 | -0,688830748 | 0,620356422 | 0,44096 | 0,68930383 | novel transcript, antisense to RFFL |
| ENSG00000206120 | EGFEM1P | -0,688879408 | 0,620335499 | 0,09866 | 0,40453901 | EGF like and EMI domain containing 1, pseudogene |
| ENSG00000224479 | AC104162,1 | -0,688901468 | 0,620326014 | 0,36257 | 0,63379594 | novel cadherin domain containing pseudogene |
| ENSG00000285942 | AC018362,3 | -0,689221642 | 0,620188362 | 0,54974 | 0,76425279 | novel protein |
| ENSG00000264294 | SNORD55 | -0,689340876 | 0,620137107 | 0,22382 | 0,53967355 | small nucleolar RNA, C/D box 55 |
| ENSG00000181135 | ZNF707 | -0,689497173 | 0,620069927 | 0,11189 | 0,42531464 | zinc finger protein 707 |
| ENSG00000162711 | NLRP3 | -0,689542426 | 0,620050478 | 0,25218 | 0,55796133 | NLR family pyrin domain containing 3 |
| ENSG00000286093 | AC022267,1 | -0,689920578 | 0,619887974 | 0,12787 | 0,45135897 | novel transcript |
| ENSG00000266783 | AP005136,3 | -0,689986829 | 0,619859509 | 0,1374 | 0,46439773 | novel transcript, sense intronic METTL4 |
| ENSG00000230699 | AL645608,2 | -0,690248646 | 0,619747029 | 0,10615 | 0,41557034 | novel transcript |
| ENSG00000105737 | GRIK5 | -0,690304342 | 0,619723103 | 0,04402 | 0,28274932 | glutamate ionotropic receptor kainate type subunit 5 |
| ENSG00000274600 | RIMBP3B | -0,690314298 | 0,619718827 | 0,22725 | 0,54168591 | RIMS binding protein 3B |
| ENSG00000288457 | LJII02000414, | -0,690509348 | 0,619635047 | 0,1933 | 0,51532949 | |
| ENSG00000285603 | AL035693,1 | -0,690775639 | 0,619520686 | 0,46116 | 0,70278639 | novel transcript, antisense to GMDS |
| ENSG00000129946 | SHC2 | -0,690871977 | 0,619479318 | 0,15375 | 0,48214138 | SHC adaptor protein 2 |
| ENSG00000185189 | NRBP2 | -0,690902937 | 0,619466025 | 0,00104 | 0,03303848 | nuclear receptor binding protein 2 |
| ENSG00000149930 | TAOK2 | -0,690948384 | 0,619446511 | 0,04969 | 0,30003295 | TAO kinase 2 |
| ENSG00000198001 | IRAK4 | -0,691210727 | 0,61933388 | 0,09147 | 0,39107762 | interleukin 1 receptor associated kinase 4 |

| | | | | | | |
|---|---|---|---|---|---|---|
| ENSG00000259277 | AC127522,1 | -0,691319999 | 0,619286972 | 0,42059 | 0,67595875 | novel transcript |
| ENSG00000287868 | AL355377,4 | -0,691739041 | 0,619107121 | 0,09343 | 0,39496269 | novel transcript |
| ENSG00000173198 | CYSLTR1 | -0,691763437 | 0,619096652 | 0,07304 | 0,35640071 | cysteinyl leukotriene receptor 1 |
| ENSG00000257607 | AC073957,2 | -0,691772957 | 0,619092567 | 0,64319 | 0,82098995 | novel transcript, antisense to GPR146 |
| ENSG00000150782 | IL18 | -0,691886308 | 0,619043927 | 0,25279 | 0,55796133 | interleukin 18 |
| ENSG00000186300 | ZNF555 | -0,692121169 | 0,618943159 | 0,09199 | 0,39215144 | zinc finger protein 555 |
| ENSG00000234592 | AL359845,1 | -0,692710246 | 0,618690486 | 0,34769 | 0,62423931 | tight junction protein 4 (peripheral) (TJP4) pseudogene |
| ENSG00000228069 | MTCO3P29 | -0,692755693 | 0,618670997 | 0,41394 | 0,67095641 | MT-CO3 pseudogene 29 |
| ENSG00000089775 | ZBTB25 | -0,692837787 | 0,618635793 | 0,10588 | 0,4154675 | zinc finger and BTB domain containing 25 |
| ENSG00000258332 | LINC02394 | -0,692932102 | 0,618595352 | 0,2559 | 0,56048171 | long intergenic non-protein coding RNA 2394 |
| ENSG00000231023 | LINC00326 | -0,692938931 | 0,618592424 | 0,5521 | 0,76565817 | long intergenic non-protein coding RNA 326 |
| ENSG00000004468 | CD38 | -0,693133893 | 0,618508834 | 0,35305 | 0,62780446 | CD38 molecule |
| ENSG00000118804 | STBD1 | -0,693549008 | 0,618330893 | 0,13991 | 0,46689302 | starch binding domain 1 |
| ENSG00000180613 | GSX2 | -0,693750649 | 0,618244477 | 0,29788 | 0,59037822 | GS homeobox 2 |
| ENSG00000176715 | ACSF3 | -0,693815719 | 0,618216593 | 0,02847 | 0,22807919 | acyl-CoA synthetase family member 3 |
| ENSG00000149564 | ESAM | -0,69383359 | 0,618208935 | 0,42356 | 0,67732256 | endothelial cell adhesion molecule |
| ENSG00000215873 | FEN1P1 | -0,69384392 | 0,618204508 | 0,30406 | 0,59411013 | flap structure-specific endonuclease 1 pseudogene 1 |
| ENSG00000163462 | TRIM46 | -0,693940734 | 0,618163024 | 0,04799 | 0,29426648 | tripartite motif containing 46 |
| ENSG00000272902 | TBC1D8-AS1 | -0,694089265 | 0,618099385 | 0,36531 | 0,6354528 | TBC1D8 antisense RNA 1 |
| ENSG00000277831 | AL138960,1 | -0,694406753 | 0,617963378 | 0,42726 | 0,68038916 | novel transcript |
| ENSG00000188386 | PPP3R2 | -0,694627874 | 0,61786867 | 0,44391 | 0,69159506 | protein phosphatase 3 regulatory subunit B, beta |
| ENSG00000133742 | CA1 | -0,694647806 | 0,617860133 | 0,5725 | 0,77850309 | carbonic anhydrase 1 |
| ENSG00000253333 | AC146944,1 | -0,694985646 | 0,617715464 | 0,31522 | 0,60158701 | glucuronidase, beta (GUSB) pseudogene |
| ENSG00000224559 | LINC01087 | -0,695020528 | 0,617700529 | 0,26446 | 0,56725636 | long intergenic non-protein coding RNA 1087 |
| ENSG00000157510 | AFAP1L1 | -0,695156133 | 0,617642472 | 0,24172 | 0,55047794 | actin filament associated protein 1 like 1 |
| ENSG00000285871 | AL359844,1 | -0,695233992 | 0,61760914 | 0,2479 | 0,55532274 | novel transcript |
| ENSG00000102452 | NALCN | -0,695298084 | 0,617581703 | 0,20813 | 0,52756478 | sodium leak channel, non-selective |
| ENSG00000258135 | LINC02396 | -0,69537372 | 0,617549326 | 0,58047 | 0,78359603 | long intergenic non-protein coding RNA 2396 |
| ENSG00000282598 | AC138374,2 | -0,695702262 | 0,617408709 | 0,30103 | 0,59247662 | novel transcript |
| ENSG00000228116 | PPT2 | -0,695782397 | 0,617374415 | 0,3211 | 0,60573903 | palmitoyl-protein thioesterase 2 |
| ENSG00000249348 | UGDH-AS1 | -0,695866733 | 0,617338327 | 0,0674 | 0,34363554 | UGDH antisense RNA 1 |
| ENSG00000102125 | TAZ | -0,696053851 | 0,617258263 | 0,09742 | 0,40211931 | tafazzin |
| ENSG00000259070 | LINC00639 | -0,696109445 | 0,617234477 | 0,06526 | 0,33813363 | long intergenic non-protein coding RNA 639 |

| | | | | | | |
|---|---|---|---|---|---|---|
| ENSG00000101966 | XIAP | -0,696347721 | 0,617132543 | 0,10306 | 0,41078404 | X-linked inhibitor of apoptosis |
| ENSG00000230751 | AC007036,2 | -0,696595637 | 0,617026503 | 0,5999 | 0,79564899 | novel transcript |
| ENSG00000258354 | AC138932,1 | -0,696608564 | 0,617020974 | 0,55894 | 0,77025265 | microRNA 3180-1 |
| ENSG00000186687 | LYRM7 | -0,696712706 | 0,616976435 | 0,10973 | 0,42183783 | LYR motif containing 7 |
| ENSG00000278107 | AC027575,2 | -0,696809894 | 0,616934874 | 0,23966 | 0,5491566 | novel transcript |
| ENSG00000198099 | ADH4 | -0,69681446 | 0,616932921 | 0,08033 | 0,37007763 | alcohol dehydrogenase 4 (class II), pi polypeptide |
| ENSG00000285106 | AC016831,6 | -0,696951028 | 0,616874524 | 0,23013 | 0,54335232 | novel transcript |
| ENSG00000141505 | ASGR1 | -0,69697106 | 0,616865959 | 0,13165 | 0,45678344 | asialoglycoprotein receptor 1 |
| ENSG00000237022 | HLA-C | -0,697240467 | 0,616750777 | 0,44251 | 0,69016964 | major histocompatibility complex, class I, C |
| ENSG00000255467 | AP002433,1 | -0,697632386 | 0,616583255 | 0,66204 | 0,83168744 | novel transcript |
| ENSG00000233024 | NPIPA9 | -0,697799685 | 0,616511758 | 0,03605 | 0,25711935 | nuclear pore complex interacting protein family, member A9 |
| ENSG00000248994 | AC124852,1 | -0,697839212 | 0,616494867 | 0,06141 | 0,32865897 | novel transcript |
| ENSG00000101489 | CELF4 | -0,697963992 | 0,616441548 | 0,05869 | 0,32371785 | CUGBP Elav-like family member 4 |
| ENSG00000241785 | RN7SL390P | -0,697984097 | 0,616432958 | 0,33472 | 0,61442925 | RNA, 7SL, cytoplasmic 390, pseudogene |
| ENSG00000285675 | AC023385,1 | -0,697986321 | 0,616432007 | 0,40372 | 0,66409141 | novel transcript |
| ENSG00000186162 | CIDECP1 | -0,69816031 | 0,61635767 | 0,13226 | 0,4573773 | cell death inducing DFFA like effector c pseudogene 1 |
| ENSG00000238741 | SCARNA7 | -0,69835777 | 0,616273316 | 0,04555 | 0,28824362 | small Cajal body-specific RNA 7 |
| ENSG00000240086 | AC092969,1 | -0,69847744 | 0,616222199 | 0,02172 | 0,19584506 | novel transcript, antisense to EPHB1 |
| ENSG00000276308 | AC078880,4 | -0,698514955 | 0,616206175 | 0,04119 | 0,27486753 | novel transcript |
| ENSG00000282377 | AC011295,3 | -0,698816234 | 0,616077506 | 0,07047 | 0,35049863 | chondroitin sulfate proteoglycan 4 (CSPG4) pseudogene |
| ENSG00000105855 | ITGB8 | -0,698833508 | 0,616070129 | 0,00016 | 0,0090156 | integrin subunit beta 8 |
| ENSG00000286909 | AC024560,4 | -0,698886062 | 0,616047688 | 0,06709 | 0,34303817 | novel transcript |
| ENSG00000265148 | TSPOAP1-AS1 | -0,698897403 | 0,616042845 | 0,02861 | 0,22866679 | TSPOAP1, SUPT4H1 and RNF43 antisense RNA 1 |
| ENSG00000128617 | OPN1SW | -0,698934332 | 0,616027076 | 0,35606 | 0,6303154 | opsin 1, short wave sensitive |
| ENSG00000260211 | AC139426,1 | -0,698939129 | 0,616025028 | 0,25314 | 0,55829991 | hyperpolarization activated cyclic nucleotide-gated potassium channel 2 (HCN2) pseudogene |
| ENSG00000213280 | AC090114,1 | -0,699017347 | 0,61599163 | 0,50681 | 0,7352001 | ribosomal protein S10 (RPS10) pseudogene |
| ENSG00000162729 | IGSF8 | -0,699350527 | 0,615849388 | 0,06391 | 0,33507507 | immunoglobulin superfamily member 8 |
| ENSG00000236259 | PPIAP64 | -0,69942169 | 0,615819011 | 0,4474 | 0,69342177 | peptidylprolyl isomerase A pseudogene 64 |
| ENSG00000237353 | PATE4 | -0,699451586 | 0,61580625 | 0,32552 | 0,60796134 | prostate and testis expressed 4 |
| ENSG00000283674 | AC068587,4 | -0,699720202 | 0,615691603 | 0,03267 | 0,24390361 | novel transcript |
| ENSG00000230074 | AL162231,2 | -0,699848932 | 0,615636668 | 0,20726 | 0,52674983 | novel transcript |

| | | | | | | |
|---|---|---|---|---|---|---|
| ENSG00000253103 | LINC01609 | -0,700020097 | 0,615563632 | 0,07374 | 0,35703856 | long intergenic non-protein coding RNA 1609 |
| ENSG00000235162 | C12orf75 | -0,700026386 | 0,615560948 | 0,02966 | 0,23170642 | chromosome 12 open reading frame 75 |
| ENSG00000206262 | FOXL2NB | -0,700158286 | 0,615504673 | 0,2366 | 0,54760364 | FOXL2 neighbor |
| ENSG00000171443 | ZNF524 | -0,700207292 | 0,615483765 | 0,11708 | 0,43438325 | zinc finger protein 524 |
| ENSG00000165863 | C10orf82 | -0,700467946 | 0,615372575 | 0,29191 | 0,58674733 | chromosome 10 open reading frame 82 |
| ENSG00000015592 | STMN4 | -0,700618834 | 0,615308218 | 0,15462 | 0,48266733 | stathmin 4 |
| ENSG00000286478 | AC007130,1 | -0,700680232 | 0,615282032 | 0,24995 | 0,55657567 | novel transcript, antisense to HIBADH |
| ENSG00000172318 | B3GALT1 | -0,701180926 | 0,615068533 | 0,16289 | 0,48945635 | beta-1,3-galactosyltransferase 1 |
| ENSG00000265788 | NOS2P1 | -0,701713689 | 0,61484144 | 0,44722 | 0,69342177 | nitric oxide synthase 2 pseudogene 1 |
| ENSG00000274112 | SMDT1 | -0,701799087 | 0,614805047 | 0,04989 | 0,30075789 | single-pass membrane protein with aspartate rich tail 1 |
| ENSG00000272835 | SMDT1 | -0,701799087 | 0,614805047 | 0,04989 | 0,30075789 | single-pass membrane protein with aspartate rich tail 1 |
| ENSG00000276429 | PLCH2 | -0,701954107 | 0,614738989 | 0,46182 | 0,70316 | phospholipase C eta 2 |
| ENSG00000105429 | MEGF8 | -0,702140161 | 0,614659715 | 0,09297 | 0,39384605 | multiple EGF like domains 8 |
| ENSG00000232098 | AC012313,1 | -0,70227902 | 0,614600557 | 0,06147 | 0,32865897 | novel transcript |
| ENSG00000091129 | NRCAM | -0,702373003 | 0,614560521 | 0,11869 | 0,43667604 | neuronal cell adhesion molecule |
| ENSG00000245848 | CEBPA | -0,70265172 | 0,614441804 | 0,29156 | 0,58655792 | CCAAT enhancer binding protein alpha |
| ENSG00000276633 | AJ011931,1 | -0,702693132 | 0,614424167 | 0,32619 | 0,6085098 | novel transcript |
| ENSG00000236668 | AC006450,3 | -0,702737303 | 0,614405356 | 0,09922 | 0,40550949 | novel transcript, antisense to LHX2 |
| ENSG00000068903 | SIRT2 | -0,702788624 | 0,6143835 | 0,07628 | 0,36242671 | sirtuin 2 |
| ENSG00000263307 | AC007216,4 | -0,702847279 | 0,614358522 | 0,19872 | 0,51984196 | novel transcript, antisense to GSPT1 |
| ENSG00000283447 | NDUFS1 | -0,702955484 | 0,614312445 | 0,0474 | 0,29245799 | NADH:ubiquinone oxidoreductase core subunit S1 |
| ENSG00000185024 | BRF1 | -0,703104572 | 0,614248966 | 0,05161 | 0,30535791 | BRF1 RNA polymerase III transcription initiation factor subunit |
| ENSG00000230833 | RPEP3 | -0,703180183 | 0,614216774 | 0,44566 | 0,69274629 | ribulose-5-phosphate-3-epimerase pseudogene 3 |
| ENSG00000183837 | PNMA3 | -0,703336953 | 0,614150034 | 0,07718 | 0,36386039 | PNMA family member 3 |
| ENSG00000143333 | RGS16 | -0,703512037 | 0,614075506 | 0,03762 | 0,26251324 | regulator of G protein signaling 16 |
| ENSG00000285402 | AC244230,2 | -0,703551581 | 0,614058674 | 0,46414 | 0,70484551 | novel transcript |
| ENSG00000258903 | AL390816,2 | -0,704021568 | 0,613858665 | 0,23228 | 0,54499308 | novel transcript |
| ENSG00000282989 | PDCL3P2 | -0,704128124 | 0,613813327 | 0,07549 | 0,36080119 | PDCL3 pseudogene 2 |
| ENSG00000244617 | ASPRV1 | -0,704145135 | 0,61380609 | 0,06571 | 0,33945082 | aspartic peptidase retroviral like 1 |
| ENSG00000179528 | LBX2 | -0,704173416 | 0,613794058 | 0,54354 | 0,76016422 | ladybird homeobox 2 |
| ENSG00000167637 | ZNF283 | -0,704263946 | 0,613755543 | 0,10036 | 0,4067769 | zinc finger protein 283 |
| ENSG00000233930 | KRTAP5-AS1 | -0,704477166 | 0,613664841 | 0,24689 | 0,55457136 | KRTAP5-1/KRTAP5-2 antisense RNA 1 |

| | | | | | | |
|---|---|---|---|---|---|---|
| ENSG00000286139 | ARHGAP11B | -0,704514522 | 0,613648952 | 0,30824 | 0,59696396 | Rho GTPase activating protein 11B |
| ENSG00000267100 | ILF3-DT | -0,70465072 | 0,613591023 | 0,06833 | 0,34555414 | ILF3 divergent transcript |
| ENSG00000258134 | AC016954,1 | -0,704742004 | 0,6135522 | 0,25282 | 0,55796133 | WW domain binding protein 2 (WBP2) pseudogene |
| ENSG00000112511 | PHF1 | -0,70529933 | 0,613315225 | 0,26803 | 0,5700759 | PHD finger protein 1 |
| ENSG00000273729 | AC007686,3 | -0,70535273 | 0,613292524 | 0,17059 | 0,49641416 | novel transcript, antisense to IRF2BPL |
| ENSG00000287813 | AC016705,3 | -0,705377301 | 0,613282079 | 0,21012 | 0,52904355 | novel transcript, antisense to ARNT2 |
| ENSG00000278376 | AP004609,3 | -0,705547303 | 0,613209816 | 0,20443 | 0,52437248 | novel transcript |
| ENSG00000176490 | DIRAS1 | -0,70555084 | 0,613208313 | 0,05971 | 0,32521812 | DIRAS family GTPase 1 |
| ENSG00000260612 | AC009097,3 | -0,705808313 | 0,613098885 | 0,37849 | 0,64679625 | novel transcript |
| ENSG00000286274 | AC008679,1 | -0,706047785 | 0,612997126 | 0,15443 | 0,48266733 | novel transcript |
| ENSG00000235958 | UBOX5-AS1 | -0,706218429 | 0,612924624 | 0,24522 | 0,55295131 | UBOX5 antisense RNA 1 |
| ENSG00000234562 | TPMTP2 | -0,706373698 | 0,612858662 | 0,28348 | 0,58065155 | thiopurine S-methyltransferase pseudogene 2 |
| ENSG00000279765 | AC013394,1 | -0,706438735 | 0,612831035 | 0,06787 | 0,34437708 | novel protein |
| ENSG00000287777 | AL161851,2 | -0,706452735 | 0,612825088 | 0,33202 | 0,61203982 | novel transcript, antisense to NPAS3 |
| ENSG00000182183 | SHISAL2A | -0,706478043 | 0,612814338 | 0,38078 | 0,64832095 | shisa like 2A |
| ENSG00000233828 | MIR4280HG | -0,706912165 | 0,612629963 | 0,0276 | 0,22390311 | MIR4280 host gene |
| ENSG00000115350 | POLE4 | -0,707253127 | 0,612485193 | 0,04631 | 0,28930497 | DNA polymerase epsilon 4, accessory subunit |
| ENSG00000276438 | PPP2R3B | -0,707487174 | 0,612385839 | 0,55204 | 0,76563921 | protein phosphatase 2 regulatory subunit B''beta |
| ENSG00000160999 | SH2B2 | -0,707574187 | 0,612348905 | 0,04325 | 0,28025067 | SH2B adaptor protein 2 |
| ENSG00000141314 | RHBDL3 | -0,707636255 | 0,612322561 | 0,01389 | 0,15462634 | rhomboid like 3 |
| ENSG00000095585 | BLNK | -0,707786902 | 0,612258625 | 0,04083 | 0,27335275 | B cell linker |
| ENSG00000285774 | AL133444,1 | -0,708020357 | 0,612159558 | 0,39304 | 0,65610193 | novel transcript, antisense to SAMD4A |
| ENSG00000134871 | COL4A2 | -0,708102286 | 0,612124796 | 0,12684 | 0,44957921 | collagen type IV alpha 2 chain |
| ENSG00000254931 | MTATP6P15 | -0,708328035 | 0,612029019 | 0,23825 | 0,54870569 | MT-ATP6 pseudogene 15 |
| ENSG00000137571 | SLCO5A1 | -0,708384069 | 0,612005249 | 0,06728 | 0,34325886 | solute carrier organic anion transporter family member 5A1 |
| ENSG00000124608 | AARS2 | -0,708414965 | 0,611992143 | 0,06916 | 0,3474201 | alanyl-tRNA synthetase 2, mitochondrial |
| ENSG00000236924 | AL162411,1 | -0,708631 | 0,611900507 | 0,06728 | 0,34325886 | novel transcript |
| ENSG00000113249 | HAVCR1 | -0,70886586 | 0,611800902 | 0,19945 | 0,52039716 | hepatitis A virus cellular receptor 1 |
| ENSG00000127928 | GNGT1 | -0,70905068 | 0,611722531 | 0,3691 | 0,63899712 | G protein subunit gamma transducin 1 |
| ENSG00000064687 | ABCA7 | -0,709080189 | 0,611710019 | 0,12278 | 0,44259775 | ATP binding cassette subfamily A member 7 |
| ENSG00000213121 | AL590867,1 | -0,709217725 | 0,611651706 | 0,0789 | 0,36683128 | novel transcript |
| ENSG00000266965 | AC090220,1 | -0,709256561 | 0,611635241 | 0,08427 | 0,37695424 | novel transcript |

| | | | | | | |
|---|---|---|---|---|---|---|
| ENSG00000285844 | FO393414,3 | -0,709307442 | 0,61161367 | 0,48415 | 0,72005601 | novel transcript |
| ENSG00000174672 | BRSK2 | -0,70932237 | 0,611607342 | 0,0136 | 0,1529196 | BR serine/threonine kinase 2 |
| ENSG00000149634 | SPATA25 | -0,70932827 | 0,611604841 | 0,20494 | 0,52453221 | spermatogenesis associated 25 |
| ENSG00000196700 | ZNF512B | -0,709345108 | 0,611597702 | 0,02844 | 0,22805204 | zinc finger protein 512B |
| ENSG00000158113 | LRRC43 | -0,709531481 | 0,611518699 | 0,06199 | 0,32961785 | leucine rich repeat containing 43 |
| ENSG00000125746 | EML2 | -0,709631519 | 0,611476297 | 0,08704 | 0,3821708 | EMAP like 2 |
| ENSG00000165272 | AQP3 | -0,70974096 | 0,611429913 | 0,18538 | 0,50881578 | aquaporin 3 (Gill blood group) |
| ENSG00000283001 | AL450426,1 | -0,709773627 | 0,611416068 | 0,07425 | 0,35762981 | novel transcript, antisense to OR13C3 |
| ENSG00000170500 | LONRF2 | -0,710030157 | 0,61130736 | 0,1184 | 0,43614969 | LON peptidase N-terminal domain and ring finger 2 |
| ENSG00000160678 | S100A1 | -0,710185111 | 0,611241706 | 0,1838 | 0,50744379 | S100 calcium binding protein A1 |
| ENSG00000164556 | FAM183BP | -0,710296668 | 0,611194443 | 0,31427 | 0,60102501 | family with sequence similarity 183 member B, pseudogene |
| ENSG00000265678 | AC129510,1 | -0,710786058 | 0,610987149 | 0,12826 | 0,45171208 | novel transcript, antisense CCDC57 |
| ENSG00000140451 | PIF1 | -0,710811116 | 0,610976537 | 0,05264 | 0,30807312 | PIF1 5'-to-3' DNA helicase |
| ENSG00000004777 | ARHGAP33 | -0,710851114 | 0,610959598 | 0,01188 | 0,14209957 | Rho GTPase activating protein 33 |
| ENSG00000214530 | STARD10 | -0,711159663 | 0,610828947 | 0,16868 | 0,49492999 | StAR related lipid transfer domain containing 10 |
| ENSG00000141741 | MIEN1 | -0,711402943 | 0,610725952 | 0,07845 | 0,36604454 | migration and invasion enhancer 1 |
| ENSG00000284138 | ATP6V0CP4 | -0,711448311 | 0,610706747 | 0,0718 | 0,35363591 | ATPase H+ transporting V0 subunit c pseudogene 4 |
| ENSG00000284882 | AL359762,1 | -0,711490946 | 0,610688699 | 0,25845 | 0,5624548 | novel transcript |
| ENSG00000214708 | AC116407,1 | -0,711663181 | 0,610615797 | 0,22944 | 0,54294653 | novel transcript, antisense to RHOT1 |
| ENSG00000181704 | YIPF6 | -0,711719527 | 0,61059195 | 0,14344 | 0,47083258 | Yip1 domain family member 6 |
| ENSG00000187244 | BCAM | -0,712243956 | 0,610370036 | 0,15533 | 0,48334025 | basal cell adhesion molecule (Lutheran blood group) |
| ENSG00000223374 | AC005104,1 | -0,712521282 | 0,610252717 | 0,36101 | 0,63311211 | novel transcript |
| ENSG00000204909 | SPINK9 | -0,712522717 | 0,61025211 | 0,14226 | 0,46928068 | serine peptidase inhibitor Kazal type 9 |
| ENSG00000232640 | AL354892,2 | -0,712535277 | 0,610246797 | 0,16277 | 0,48945635 | novel transcript |
| ENSG00000281625 | AL354892,5 | -0,712535277 | 0,610246797 | 0,16277 | 0,48945635 | novel transcript |
| ENSG00000105427 | CNFN | -0,712640559 | 0,610202265 | 0,15731 | 0,48443207 | cornifelin |
| ENSG00000259200 | AC068722,1 | -0,712650292 | 0,610198149 | 0,32834 | 0,60973412 | novel transcript |
| ENSG00000156218 | ADAMTSL3 | -0,71288703 | 0,610098027 | 0,05131 | 0,30462001 | ADAMTS like 3 |
| ENSG00000267710 | EDDM13 | -0,7131098 | 0,610003828 | 0,15497 | 0,48292767 | epididymal protein 13 |
| ENSG00000277476 | AC005332,5 | -0,713249763 | 0,609944651 | 0,07849 | 0,36604454 | novel transcript |
| ENSG00000174136 | RGMB | -0,713276263 | 0,609933447 | 0,01465 | 0,15853855 | repulsive guidance molecule BMP co-receptor b |
| ENSG00000135747 | ZNF670-ZNF69 | -0,713401509 | 0,609880499 | 0,10116 | 0,40785299 | ZNF670-ZNF695 readthrough (NMD candidate) |

| | | | | | | |
|---|---|---|---|---|---|---|
| ENSG00000077454 | LRCH4 | -0,713518113 | 0,609831208 | 0,1041 | 0,41245634 | leucine rich repeats and calponin homology domain containing 4 |
| ENSG00000166823 | MESP1 | -0,713911514 | 0,609664939 | 0,23289 | 0,54547822 | mesoderm posterior bHLH transcription factor 1 |
| ENSG00000166347 | CYB5A | -0,713967019 | 0,609641484 | 0,09328 | 0,39486101 | cytochrome b5 type A |
| ENSG00000268047 | AC018766,1 | -0,714145798 | 0,609565942 | 0,36731 | 0,6373003 | novel transcript, antisense to PTOV1 |
| ENSG00000248360 | LINC00504 | -0,714234437 | 0,609528491 | 0,07241 | 0,35526408 | long intergenic non-protein coding RNA 504 |
| ENSG00000128294 | TPST2 | -0,714241899 | 0,609525339 | 0,15052 | 0,47863728 | tyrosylprotein sulfotransferase 2 |
| ENSG00000281693 | AC217775,1 | -0,714324109 | 0,609490607 | 0,11725 | 0,43455623 | novel transcript |
| ENSG00000177082 | WDR73 | -0,714377698 | 0,609467967 | 0,06876 | 0,34665178 | WD repeat domain 73 |
| ENSG00000180769 | WDFY3-AS2 | -0,714382256 | 0,609466042 | 0,10649 | 0,41625157 | WDFY3 antisense RNA 2 |
| ENSG00000286758 | AL445438,1 | -0,714729517 | 0,609319359 | 0,32403 | 0,60668177 | novel transcript |
| ENSG00000258498 | DIO3OS | -0,714925402 | 0,609236633 | 0,33205 | 0,61205241 | DIO3 opposite strand upstream RNA |
| ENSG00000154783 | FGD5 | -0,714990693 | 0,609209062 | 0,34237 | 0,62005694 | FYVE, RhoGEF and PH domain containing 5 |
| ENSG00000186193 | SAPCD2 | -0,715577432 | 0,608961349 | 0,03541 | 0,25492595 | suppressor APC domain containing 2 |
| ENSG00000149380 | P4HA3 | -0,715602349 | 0,608950832 | 0,26281 | 0,56582568 | prolyl 4-hydroxylase subunit alpha 3 |
| ENSG00000228186 | PRR3 | -0,715839171 | 0,608850879 | 0,15205 | 0,48055609 | proline rich 3 |
| ENSG00000144031 | ANKRD53 | -0,716134078 | 0,608726434 | 0,13691 | 0,4641716 | ankyrin repeat domain 53 |
| ENSG00000042832 | TG | -0,71659859 | 0,608530471 | 0,37105 | 0,64030792 | thyroglobulin |
| ENSG00000251056 | ANKRD20A17 | -0,716935882 | 0,608388218 | 0,56057 | 0,7708846 | ankyrin repeat domain 20 family member A17, pseudogene |
| ENSG00000286314 | AC010359,3 | -0,717216242 | 0,60827 | 0,12768 | 0,45102957 | novel transcript, sense intronic to ERBIN |
| ENSG00000286575 | AL356608,3 | -0,717525283 | 0,608139716 | 0,11732 | 0,43468107 | novel transcript, antisense to CNNM2and AS3MT |
| ENSG00000265205 | AC010761,3 | -0,718029096 | 0,607927381 | 0,06087 | 0,32750733 | novel transcript, antisense to SUPT6H |
| ENSG00000275963 | AC008115,4 | -0,718102137 | 0,607896603 | 0,10111 | 0,40780603 | novel transcript |
| ENSG00000106266 | SNX8 | -0,718105826 | 0,607895049 | 0,11853 | 0,43623197 | sorting nexin 8 |
| ENSG00000261992 | GATAD2B | -0,718230625 | 0,607842466 | 0,10496 | 0,41375163 | GATA zinc finger domain containing 2B |
| ENSG00000274447 | AC005790,1 | -0,718280983 | 0,607821249 | 0,25801 | 0,56210015 | novel transcript, antisense to PTPRS |
| ENSG00000253636 | AC022893,1 | -0,718333985 | 0,607798919 | 0,14264 | 0,47004319 | novel transcript |
| ENSG00000198203 | SULT1C2 | -0,718350284 | 0,607792053 | 0,32775 | 0,60960585 | sulfotransferase family 1C member 2 |
| ENSG00000250305 | TRMT9B | -0,718504845 | 0,607726941 | 0,09073 | 0,38981626 | tRNA methyltransferase 9B (putative) |
| ENSG00000205863 | C1QTNF9B | -0,718619959 | 0,607678452 | 0,51108 | 0,73835122 | C1q and TNF related 9B |
| ENSG00000274328 | AC007551,2 | -0,718756844 | 0,607620797 | 0,41444 | 0,67110928 | gem (nuclear organelle) associated protein 7 (GEMIN7) pseudogene |

| | | | | | | |
|---|---|---|---|---|---|---|
| ENSG00000160194 | NDUFV3 | -0,719102335 | 0,607475304 | 0,101 | 0,40770184 | NADH:ubiquinone oxidoreductase subunit V3 |
| ENSG00000163915 | IGF2BP2-AS1 | -0,719146034 | 0,607456904 | 0,08641 | 0,38089807 | IGF2BP2 antisense RNA 1 |
| ENSG00000173960 | UBXN2A | -0,719293102 | 0,607394983 | 0,08073 | 0,37077071 | UBX domain protein 2A |
| ENSG00000196366 | C9orf163 | -0,719499891 | 0,607307929 | 0,10836 | 0,41899991 | chromosome 9 open reading frame 163 |
| ENSG00000224717 | LEMD1-DT | -0,71988095 | 0,607147542 | 0,09367 | 0,39553462 | LEMD1 divergent transcript |
| ENSG00000166342 | NETO1 | -0,719959718 | 0,607114393 | 0,20451 | 0,52439728 | neuropilin and tolloid like 1 |
| ENSG00000250456 | LINC02260 | -0,719991605 | 0,607100975 | 0,26893 | 0,57085484 | long intergenic non-protein coding RNA 2260 |
| ENSG00000008323 | PLEKHG6 | -0,720072295 | 0,607067021 | 0,11835 | 0,43614969 | pleckstrin homology and RhoGEF domain containing G6 |
| ENSG00000164070 | HSPA4L | -0,720104169 | 0,607053609 | 0,08297 | 0,37497922 | heat shock protein family A (Hsp70) member 4 like |
| ENSG00000254431 | AC084083,1 | -0,720226925 | 0,607001958 | 0,18728 | 0,5105374 | novel transcript |
| ENSG00000092820 | EZR | -0,720251936 | 0,606991435 | 0,00311 | 0,06638089 | ezrin |
| ENSG00000196236 | XPNPEP3 | -0,720318722 | 0,606963336 | 0,09426 | 0,39648446 | X-prolyl aminopeptidase 3 |
| ENSG00000200169 | RNU5D-1 | -0,720552038 | 0,606865185 | 0,11566 | 0,43162108 | RNA, U5D small nuclear 1 |
| ENSG00000214331 | AC009053,1 | -0,720626755 | 0,606833756 | 0,10203 | 0,40934973 | pyruvate dehydrogenase phosphatase regulatory subunit (PDPR) pseudogene |
| ENSG00000121361 | KCNJ8 | -0,720654362 | 0,606822144 | 0,15476 | 0,48266733 | potassium inwardly rectifying channel subfamily J member 8 |
| ENSG00000287234 | AL592211,2 | -0,720660485 | 0,606819569 | 0,2121 | 0,5308403 | novel transcript |
| ENSG00000287546 | AL163534,1 | -0,720725772 | 0,606792108 | 0,41236 | 0,67001327 | novel transcript |
| ENSG00000171813 | PWWP2B | -0,720744909 | 0,60678406 | 0,08989 | 0,38777461 | PWWP domain containing 2B |
| ENSG00000287105 | AC090577,1 | -0,720861314 | 0,606735103 | 0,05006 | 0,30089233 | novel transcript, antisense to NKAIN3 |
| ENSG00000249485 | RBBP4P1 | -0,721107366 | 0,606631633 | 0,26195 | 0,56481243 | RBBP4 pseudogene 1 |
| ENSG00000180251 | SLC9A4 | -0,721198695 | 0,606593231 | 0,08485 | 0,37817642 | solute carrier family 9 member A4 |
| ENSG00000125375 | DMAC2L | -0,721360398 | 0,606525246 | 0,05993 | 0,32541248 | distal membrane arm assembly complex 2 like |
| ENSG00000151025 | GPR158 | -0,72165198 | 0,606402674 | 0,20694 | 0,52659911 | G protein-coupled receptor 158 |
| ENSG00000157168 | NRG1 | -0,721797048 | 0,606341701 | 0,02146 | 0,19472643 | neuregulin 1 |
| ENSG00000221990 | EXOC3-AS1 | -0,721937311 | 0,606282754 | 0,12679 | 0,44955467 | EXOC3 antisense RNA 1 |
| ENSG00000164897 | TMUB1 | -0,722074428 | 0,606225134 | 0,05281 | 0,30846816 | transmembrane and ubiquitin like domain containing 1 |
| ENSG00000275756 | AL354751,3 | -0,722201327 | 0,606171813 | 0,19044 | 0,51350457 | novel pseudogene |
| ENSG00000230327 | MTCO1P42 | -0,722754431 | 0,605939462 | 0,37306 | 0,64180957 | MT-CO1 pseudogene 42 |
| ENSG00000115252 | PDE1A | -0,722825855 | 0,605909464 | 0,34997 | 0,62563749 | phosphodiesterase 1A |
| ENSG00000169894 | MUC3A | -0,723047105 | 0,60581655 | 0,2911 | 0,58613153 | mucin 3A, cell surface associated |
| ENSG00000234520 | HRAT17 | -0,72311564 | 0,605787771 | 0,10147 | 0,40833508 | heart tissue-associated transcript 17 |

| | | | | | | |
|---|---|---|---|---|---|---|
| ENSG00000285492 | AL356417,3 | -0,723218391 | 0,605744628 | 0,30015 | 0,59160746 | uncharacterized LOC112267968 |
| ENSG00000003137 | CYP26B1 | -0,723495806 | 0,605628161 | 0,05081 | 0,30326674 | cytochrome P450 family 26 subfamily B member 1 |
| ENSG00000137842 | TMEM62 | -0,723605793 | 0,605581991 | 0,04606 | 0,28930497 | transmembrane protein 62 |
| ENSG00000146858 | ZC3HAV1L | -0,723784461 | 0,605506998 | 0,06556 | 0,33915908 | zinc finger CCCH-type containing, antiviral 1 like |
| ENSG00000089692 | LAG3 | -0,724027044 | 0,605405193 | 0,04133 | 0,27533587 | lymphocyte activating 3 |
| ENSG00000284299 | AL590132,1 | -0,724056567 | 0,605392805 | 0,19974 | 0,52077515 | novel protein |
| ENSG00000183323 | CCDC125 | -0,724256804 | 0,605308786 | 0,03712 | 0,26086671 | coiled-coil domain containing 125 |
| ENSG00000196705 | ZNF431 | -0,724316806 | 0,605283611 | 0,10502 | 0,41375455 | zinc finger protein 431 |
| ENSG00000260988 | AC090260,1 | -0,724853061 | 0,605058667 | 0,13698 | 0,46424215 | novel transcript |
| ENSG00000206493 | HLA-E | -0,724977244 | 0,605006587 | 0,22622 | 0,54121239 | major histocompatibility complex, class I, E |
| ENSG00000185002 | RFX6 | -0,725957438 | 0,604595674 | 0,39743 | 0,65990329 | regulatory factor X6 |
| ENSG00000262986 | SMIM10L1 | -0,725977479 | 0,604587276 | 0,69964 | 0,85489385 | small integral membrane protein 10 like 1 |
| ENSG00000205002 | AARD | -0,726112774 | 0,604530581 | 0,17951 | 0,50387897 | alanine and arginine rich domain containing protein |
| ENSG00000197753 | LHFPL5 | -0,72615253 | 0,604513922 | 0,11969 | 0,43822857 | LHFPL tetraspan subfamily member 5 |
| ENSG00000133250 | ZNF414 | -0,726229666 | 0,604481601 | 0,06903 | 0,34722131 | zinc finger protein 414 |
| ENSG00000286472 | AL035252,5 | -0,726340853 | 0,604435016 | 0,05483 | 0,31349896 | novel transcript |
| ENSG00000188779 | SKOR1 | -0,726341924 | 0,604434568 | 0,3265 | 0,60866572 | SKI family transcriptional corepressor 1 |
| ENSG00000170381 | SEMA3E | -0,726440547 | 0,60439325 | 0,36239 | 0,63379594 | semaphorin 3E |
| ENSG00000286965 | AC108066,2 | -0,726466304 | 0,60438246 | 0,40362 | 0,66399439 | novel transcript |
| ENSG00000275953 | AL359535,1 | -0,726521064 | 0,60435952 | 0,33263 | 0,61256072 | Yes-associated protein 1 (YAP1) pseudogene |
| ENSG00000259479 | SORD2P | -0,726525071 | 0,604357841 | 0,10335 | 0,41114168 | sorbitol dehydrogenase 2, pseudogene |
| ENSG00000175595 | ERCC4 | -0,726526274 | 0,604357337 | 0,10549 | 0,41477267 | ERCC excision repair 4, endonuclease catalytic subunit |
| ENSG00000260121 | AC138028,4 | -0,726610335 | 0,604322124 | 0,23229 | 0,54499308 | novel transcript, antisense to FAM38A |
| ENSG00000113448 | PDE4D | -0,726618763 | 0,604318594 | 0,10765 | 0,41827415 | phosphodiesterase 4D |
| ENSG00000253218 | KLF3P1 | -0,727009356 | 0,604155004 | 0,01982 | 0,18695096 | Kruppel like factor 3 pseudogene 1 |
| ENSG00000164916 | FOXK1 | -0,727085506 | 0,604123115 | 0,0849 | 0,3782577 | forkhead box K1 |
| ENSG00000135472 | FAIM2 | -0,727270158 | 0,604045798 | 0,05196 | 0,30634534 | Fas apoptotic inhibitory molecule 2 |
| ENSG00000278570 | NR2E3 | -0,727534308 | 0,60393521 | 0,17429 | 0,49875953 | nuclear receptor subfamily 2 group E member 3 |
| ENSG00000128683 | GAD1 | -0,727569074 | 0,603920657 | 0,14539 | 0,47300165 | glutamate decarboxylase 1 |
| ENSG00000255857 | PXN-AS1 | -0,727599336 | 0,603907989 | 0,07171 | 0,35351167 | PXN antisense RNA 1 |
| ENSG00000250771 | AC106865,1 | -0,727655038 | 0,603884673 | 0,03971 | 0,26947142 | toll-like receptor 2 pseudogene |
| ENSG00000258301 | VASH1-AS1 | -0,727907465 | 0,603779021 | 0,21619 | 0,53480193 | VASH1 antisense RNA 1 |
| ENSG00000164638 | SLC29A4 | -0,728002561 | 0,603739224 | 0,06623 | 0,3406271 | solute carrier family 29 member 4 |

| | | | | | | |
|---|---|---|---|---|---|---|
| ENSG00000258555 | SPECC1L-ADO | -0,728047015 | 0,603720621 | 0,51933 | 0,74394224 | SPECC1L-ADORA2A readthrough (NMD candidate) |
| ENSG00000270647 | TAF15 | -0,728110291 | 0,603694143 | 0,10121 | 0,40785299 | TATA-box binding protein associated factor 15 |
| ENSG00000286837 | AC087533,1 | -0,72821009 | 0,603652383 | 0,18658 | 0,509961 | novel transcript, antisense to THAP1 |
| ENSG00000275734 | AC010538,1 | -0,728469546 | 0,603543832 | 0,03879 | 0,26660248 | novel transcript, sense overlapping CDK10 |
| ENSG00000230438 | SERPINB9P1 | -0,728475228 | 0,603541455 | 0,19916 | 0,52017864 | serpin family B member 9 pseudogene 1 |
| ENSG00000223569 | USP17L15 | -0,728624958 | 0,603478819 | 0,46846 | 0,70810969 | ubiquitin specific peptidase 17 like family member 15 |
| ENSG00000168010 | ATG16L2 | -0,728729557 | 0,603435067 | 0,04272 | 0,27853965 | autophagy related 16 like 2 |
| ENSG00000244198 | ARHGEF35-AS | -0,728974458 | 0,603332641 | 0,17341 | 0,49804988 | ARHGEF35 antisense RNA 1 |
| ENSG00000274898 | AC001226,1 | -0,729106109 | 0,603277588 | 0,60433 | 0,79789086 | novel transcript |
| ENSG00000228336 | OR9H1P | -0,729130779 | 0,603267272 | 0,05994 | 0,32541248 | olfactory receptor family 9 subfamily H member 1 pseudogene |
| ENSG00000143434 | SEMA6C | -0,729136588 | 0,603264843 | 0,02666 | 0,22038242 | semaphorin 6C |
| ENSG00000285535 | AC021683,5 | -0,729258228 | 0,603213981 | 0,02761 | 0,22390566 | novel transcript |
| ENSG00000227782 | AC002553,1 | -0,729426218 | 0,603143746 | 0,07507 | 0,35949119 | novel transcript, antisense to NCOR1 |
| ENSG00000168763 | CNNM3 | -0,729496567 | 0,603114336 | 0,05916 | 0,32424653 | cyclin and CBS domain divalent metal cation transport mediator 3 |
| ENSG00000262473 | GART | -0,729511038 | 0,603108286 | 0,08619 | 0,38055991 | phosphoribosylglycinamide formyltransferase, phosphoribosylglycinamide synthetase, phosphoribosylaminoimidazole synthetase |
| ENSG00000160299 | PCNT | -0,729846084 | 0,602968239 | 0,02044 | 0,18959706 | pericentrin |
| ENSG00000110148 | CCKBR | -0,729907764 | 0,602942461 | 0,47621 | 0,71446062 | cholecystokinin B receptor |
| ENSG00000235299 | MRPL53P1 | -0,730046497 | 0,602884483 | 0,50997 | 0,73758064 | mitochondrial ribosomal protein L53 pseudogene 1 |
| ENSG00000213970 | AC006122,1 | -0,730065921 | 0,602876366 | 0,28858 | 0,58455903 | ribosomal protein S15 (RPS15) pseudogene |
| ENSG00000267372 | AC005330,1 | -0,730125683 | 0,602851393 | 0,13405 | 0,46018397 | novel transcript |
| ENSG00000233695 | GAS6-AS1 | -0,730350138 | 0,602757608 | 0,32514 | 0,60761222 | GAS6 antisense RNA 1 |
| ENSG00000160072 | ATAD3B | -0,730464647 | 0,602709769 | 0,06175 | 0,32900116 | ATPase family AAA domain containing 3B |
| ENSG00000231049 | OR52B5P | -0,730768508 | 0,602582839 | 0,15773 | 0,4847949 | olfactory receptor family 52 subfamily B member 5 pseudogene |
| ENSG00000205424 | PCBP3-AS1 | -0,730851325 | 0,602548249 | 0,18273 | 0,50616031 | PCBP3 antisense RNA 1 |
| ENSG00000276203 | ANKRD20A3P | -0,73115284 | 0,602422333 | 0,13836 | 0,46555669 | ankyrin repeat domain 20 family member A3, pseudogene |
| ENSG00000244356 | RN7SL398P | -0,731673981 | 0,602204761 | 0,41263 | 0,6702832 | RNA, 7SL, cytoplasmic 398, pseudogene |
| ENSG00000182676 | PPP1R27 | -0,731959568 | 0,602085564 | 0,32669 | 0,60877026 | protein phosphatase 1 regulatory subunit 27 |
| ENSG00000288186 | PPP1R27 | -0,731959568 | 0,602085564 | 0,32669 | 0,60877026 | protein phosphatase 1 regulatory subunit 27 |

| | | | | | | |
|---|---|---|---|---|---|---|
| ENSG00000270948 | MTDHP1 | -0,732066718 | 0,602040848 | 0,28052 | 0,57800643 | metadherin pseudogene 1 |
| ENSG00000206634 | SNORA22 | -0,732480376 | 0,601868252 | 0,17894 | 0,5033932 | small nucleolar RNA, H/ACA box 22 |
| ENSG00000232860 | SMG7-AS1 | -0,732607928 | 0,601815042 | 0,15389 | 0,48214138 | SMG7 antisense RNA 1 |
| ENSG00000172840 | PDP2 | -0,73294246 | 0,60167551 | 0,12593 | 0,448127 | pyruvate dehyrogenase phosphatase catalytic subunit 2 |
| ENSG00000130287 | NCAN | -0,733191293 | 0,601571743 | 0,03756 | 0,26227157 | neurocan |
| ENSG00000234509 | AP000253,1 | -0,733206089 | 0,601565573 | 0,10666 | 0,41648041 | novel transcript |
| ENSG00000101222 | SPEF1 | -0,733237542 | 0,601552458 | 0,22476 | 0,54036191 | sperm flagellar 1 |
| ENSG00000186453 | FAM228A | -0,733262403 | 0,601542092 | 0,10816 | 0,41884147 | family with sequence similarity 228 member A |
| ENSG00000275338 | RAB11FIP3 | -0,733404562 | 0,601482821 | 0,00598 | 0,09746931 | RAB11 family interacting protein 3 |
| ENSG00000130270 | ATP8B3 | -0,733484441 | 0,601449519 | 0,10238 | 0,40992908 | ATPase phospholipid transporting 8B3 |
| ENSG00000228573 | AL138962,1 | -0,733579344 | 0,601409956 | 0,47371 | 0,71294959 | novel transcript |
| ENSG00000177947 | ODF3 | -0,733671581 | 0,601371507 | 0,30918 | 0,59774312 | outer dense fiber of sperm tails 3 |
| ENSG00000227492 | AL138921,1 | -0,733726141 | 0,601348764 | 0,2809 | 0,57822547 | novel transcript |
| ENSG00000103034 | NDRG4 | -0,733884369 | 0,601282815 | 0,01499 | 0,16042172 | NDRG family member 4 |
| ENSG00000214491 | SEC14L6 | -0,734090043 | 0,601197101 | 0,15789 | 0,48493328 | SEC14 like lipid binding 6 |
| ENSG00000049769 | PPP1R3F | -0,734294568 | 0,601111877 | 0,07128 | 0,35265844 | protein phosphatase 1 regulatory subunit 3F |
| ENSG00000261267 | AC026470,2 | -0,734405717 | 0,601065568 | 0,50263 | 0,73209345 | novel transcript, antisense to N4BP1 |
| ENSG00000224066 | AL049795,1 | -0,734485976 | 0,601032131 | 0,345 | 0,62226948 | novel transcript |
| ENSG00000188626 | GOLGA8M | -0,734539854 | 0,601009686 | 0,44684 | 0,6933405 | golgin A8 family member M |
| ENSG00000135622 | SEMA4F | -0,734541935 | 0,601008819 | 0,00768 | 0,11151515 | ssemaphorin 4F |
| ENSG00000286861 | AL049779,4 | -0,734570454 | 0,600996938 | 0,1275 | 0,45074579 | novel transcript, antisense to ARG2 |
| ENSG00000236617 | AC127070,1 | -0,734592836 | 0,600987614 | 0,45305 | 0,69744022 | novel transcript |
| ENSG00000283317 | AL831711,1 | -0,734634502 | 0,600970258 | 0,24521 | 0,55295131 | novel transcript |
| ENSG00000267523 | AC008735,2 | -0,734770789 | 0,600913489 | 0,13323 | 0,45919282 | novel transcript, antisense to U2AF2A |
| ENSG00000149639 | SOGA1 | -0,73482348 | 0,600891542 | 0,04251 | 0,2782038 | suppressor of glucose, autophagy associated 1 |
| ENSG00000113327 | GABRG2 | -0,734857675 | 0,6008773 | 0,06639 | 0,34113137 | gamma-aminobutyric acid type A receptor subunit gamma2 |
| ENSG00000102934 | PLLP | -0,734939811 | 0,600843091 | 0,08941 | 0,38670267 | plasmolipin |
| ENSG00000256571 | AC079866,2 | -0,735446316 | 0,600632183 | 0,37293 | 0,64180957 | novel transcript |
| ENSG00000184154 | LRTOMT | -0,735955102 | 0,600420399 | 0,07012 | 0,34975886 | leucine rich transmembrane and O-methyltransferase domain containing |
| ENSG00000258038 | LINC02327 | -0,736031033 | 0,600388799 | 0,25049 | 0,55682585 | long intergenic non-protein coding RNA 2327 |
| ENSG00000121440 | PDZRN3 | -0,736066438 | 0,600374065 | 0,13442 | 0,46058584 | PDZ domain containing ring finger 3 |
| ENSG00000277594 | DDX52 | -0,736474435 | 0,600204302 | 0,31707 | 0,60268749 | DExD-box helicase 52 |

| | | | | | | |
|---|---|---|---|---|---|---|
| ENSG00000275811 | HTR1DP1 | -0,736940531 | 0,600010424 | 0,1578 | 0,48490111 | 5-hydroxytryptamine receptor 1D pseudogene 1 |
| ENSG00000110077 | MS4A6A | -0,7371052 | 0,599941942 | 0,2334 | 0,54563244 | membrane spanning 4-domains A6A |
| ENSG00000203734 | ECT2L | -0,737606429 | 0,599733543 | 0,26132 | 0,56432558 | epithelial cell transforming 2 like |
| ENSG00000129472 | RAB2B | -0,738058715 | 0,599545556 | 0,06251 | 0,33112961 | RAB2B, member RAS oncogene family |
| ENSG00000237283 | AL691515,1 | -0,738245435 | 0,599467965 | 0,44217 | 0,69003826 | novel transcript |
| ENSG00000284997 | TUBAP2 | -0,738349414 | 0,599424761 | 0,39022 | 0,65427905 | tubulin alpha pseudogene 2 |
| ENSG00000214391 | TUBAP2 | -0,738349414 | 0,599424761 | 0,39022 | 0,65427905 | tubulin alpha pseudogene 2 |
| ENSG00000225032 | AL162586,1 | -0,738482653 | 0,599369405 | 0,33175 | 0,6117373 | novel transcript |
| ENSG00000230696 | AC011753,2 | -0,73880194 | 0,599236771 | 0,41626 | 0,67307944 | novel transcript, sense intronic to SEPT10 |
| ENSG00000159147 | DONSON | -0,738869055 | 0,599208895 | 0,13711 | 0,4642578 | DNA replication fork stabilization factor DONSON |
| ENSG00000205212 | CCDC144NL | -0,739074242 | 0,599123679 | 0,07605 | 0,36217039 | CCDC144A N-terminal pseudogene |
| ENSG00000169696 | ASPSCR1 | -0,739131014 | 0,599100103 | 0,08709 | 0,3821708 | ASPSCR1 tether for SLC2A4, UBX domain containing |
| ENSG00000167371 | PRRT2 | -0,73917109 | 0,599083461 | 0,02115 | 0,19303449 | proline rich transmembrane protein 2 |
| ENSG00000230922 | AL079301,1 | -0,739480736 | 0,598954893 | 0,11054 | 0,42322615 | novel transcript |
| ENSG00000277452 | RN7SL473P | -0,739819347 | 0,59881433 | 0,3405 | 0,61897407 | RNA, 7SL, cytoplasmic 473, pseudogene |
| ENSG00000215297 | AL354941,1 | -0,739926322 | 0,598769931 | 0,24642 | 0,55411366 | ribosomal protein L7 (RPL7) pseudogene |
| ENSG00000126603 | GLIS2 | -0,740269296 | 0,598627601 | 0,09034 | 0,3889575 | GLIS family zinc finger 2 |
| ENSG00000167280 | ENGASE | -0,740321595 | 0,598605901 | 0,08781 | 0,38349699 | endo-beta-N-acetylglucosaminidase |
| ENSG00000164690 | SHH | -0,740726628 | 0,598437867 | 0,41763 | 0,67416605 | sonic hedgehog signaling molecule |
| ENSG00000236699 | ARHGEF38 | -0,740954787 | 0,598343233 | 0,3098 | 0,59807209 | Rho guanine nucleotide exchange factor 38 |
| ENSG00000285632 | AC084024,4 | -0,741118644 | 0,598275279 | 0,32076 | 0,60555414 | novel transcript |
| ENSG00000204054 | LINC00963 | -0,741289176 | 0,598204565 | 0,16598 | 0,49197859 | long intergenic non-protein coding RNA 963 |
| ENSG00000121897 | LIAS | -0,741292085 | 0,598203359 | 0,12794 | 0,45145929 | lipoic acid synthetase |
| ENSG00000153363 | LINC00467 | -0,741406455 | 0,598155937 | 0,11054 | 0,42322615 | long intergenic non-protein coding RNA 467 |
| ENSG00000163806 | SPDYA | -0,741851815 | 0,597971315 | 0,0968 | 0,40094776 | speedy/RINGO cell cycle regulator family member A |
| ENSG00000265962 | GACAT2 | -0,741923423 | 0,597941636 | 0,10894 | 0,42024426 | gastric cancer associated transcript 2 |
| ENSG00000026036 | RTEL1-TNFRSI | -0,742099513 | 0,597868658 | 0,07755 | 0,3645785 | RTEL1-TNFRSF6B readthrough (NMD candidate) |
| ENSG00000242087 | RPL36AP41 | -0,742131892 | 0,597855239 | 0,32825 | 0,60971379 | ribosomal protein L36a pseudogene 41 |
| ENSG00000276768 | AL121890,2 | -0,742256356 | 0,597803664 | 0,30614 | 0,59517372 | novel transcript |
| ENSG00000281801 | KRTAP5-AS1 | -0,742290362 | 0,597789573 | 0,21123 | 0,53012872 | KRTAP5-1/KRTAP5-2 antisense RNA 1 |
| ENSG00000228897 | AC012441,2 | -0,742375556 | 0,597754273 | 0,18098 | 0,50451145 | novel pseudogene |
| ENSG00000204666 | AC010624,1 | -0,742621431 | 0,597652408 | 0,50744 | 0,73566984 | novel transcript, sense overlapping ZNF473 |
| ENSG00000196878 | LAMB3 | -0,742701985 | 0,597619039 | 0,48363 | 0,71986841 | laminin subunit beta 3 |

| | | | | | | |
|---|---|---|---|---|---|---|
| ENSG00000120729 | MYOT | -0,742705767 | 0,597617472 | 0,37383 | 0,6423379 | myotilin |
| ENSG00000227544 | AC018647,1 | -0,743232437 | 0,597399346 | 0,34051 | 0,61897407 | novel transcript |
| ENSG00000149781 | FERMT3 | -0,743456137 | 0,597306722 | 0,09574 | 0,39858189 | fermitin family member 3 |
| ENSG00000276155 | MAPT | -0,743459561 | 0,597305305 | 0,03477 | 0,25220349 | microtubule associated protein tau |
| ENSG00000254705 | AP000722,1 | -0,7440704 | 0,597052458 | 0,45562 | 0,69957443 | novel transcript |
| ENSG00000262648 | PHBP15 | -0,744268853 | 0,596970335 | 0,63303 | 0,81528545 | prohibitin pseudogene 15 |
| ENSG00000200785 | SNORD8 | -0,74431572 | 0,596950942 | 0,27214 | 0,57306433 | small nucleolar RNA, C/D box 8 |
| ENSG00000105619 | TFPT | -0,744470512 | 0,596886897 | 0,10088 | 0,40770184 | TCF3 fusion partner |
| ENSG00000224774 | CSNK2B | -0,744764246 | 0,596765382 | 0,1856 | 0,50901954 | casein kinase 2 beta |
| ENSG00000225683 | PACRG-AS3 | -0,744811251 | 0,596745939 | 0,15462 | 0,48266733 | PACRG antisense RNA 3 |
| ENSG00000287595 | AC097369,4 | -0,744882198 | 0,596716594 | 0,49215 | 0,72542601 | novel transcript |
| ENSG00000228274 | AL021707,2 | -0,745016079 | 0,596661221 | 0,1192 | 0,43703483 | novel transcript, antisense to CBY1 |
| ENSG00000201998 | SNORA23 | -0,745208432 | 0,596581675 | 0,0564 | 0,31712073 | small nucleolar RNA, H/ACA box 23 |
| ENSG00000261529 | AC100774,1 | -0,745605199 | 0,596417627 | 0,02883 | 0,2296248 | novel transcript, sense intronic to UBE3A |
| ENSG00000214787 | MS4A4E | -0,745666546 | 0,596392266 | 0,05753 | 0,32010411 | membrane spanning 4-domains A4E |
| ENSG00000004799 | PDK4 | -0,745725001 | 0,596368102 | 0,31216 | 0,59925736 | pyruvate dehydrogenase kinase 4 |
| ENSG00000221968 | FADS3 | -0,746011091 | 0,596249853 | 0,04472 | 0,28565589 | fatty acid desaturase 3 |
| ENSG00000223553 | SMPD4P1 | -0,74601846 | 0,596246807 | 0,52497 | 0,74873866 | sphingomyelin phosphodiesterase 4 pseudogene 1 |
| ENSG00000131242 | RAB11FIP4 | -0,746091734 | 0,596216524 | 0,119 | 0,43696075 | RAB11 family interacting protein 4 |
| ENSG00000163541 | SUCLG1 | -0,746521347 | 0,596039006 | 0,09843 | 0,40393831 | succinate-CoA ligase GDP/ADP-forming subunit alpha |
| ENSG00000286647 | AC008459,1 | -0,746707814 | 0,595961974 | 0,23586 | 0,5471053 | novel transcript |
| ENSG00000206077 | ZDHHC11B | -0,746746217 | 0,59594611 | 0,01094 | 0,13517515 | zinc finger DHHC-type containing 11B |
| ENSG00000226419 | SLC16A1-AS1 | -0,746898611 | 0,595883163 | 0,12309 | 0,44322266 | SLC16A1 antisense RNA 1 |
| ENSG00000215304 | AC135983,1 | -0,747124259 | 0,59578997 | 0,28565 | 0,58214108 | novel transcript |
| ENSG00000078699 | CBFA2T2 | -0,747192481 | 0,595761797 | 0,06954 | 0,34816245 | CBFA2/RUNX1 partner transcriptional co-repressor 2 |
| ENSG00000270863 | DDX55P1 | -0,747275809 | 0,595727388 | 0,35141 | 0,62674766 | DEAD-box helicase 55 pseudogene 1 |
| ENSG00000151014 | NOCT | -0,747618529 | 0,595585886 | 0,13205 | 0,45711259 | nocturnin |
| ENSG00000270055 | AC127502,2 | -0,747620224 | 0,595585186 | 0,07175 | 0,35351167 | novel transcript |
| ENSG00000284189 | AC141424,2 | -0,747704508 | 0,595550393 | 0,34229 | 0,62000209 | novel transcript, antisense to C17orf97 |
| ENSG00000241525 | AC141424,1 | -0,747704508 | 0,595550393 | 0,34229 | 0,62000209 | novel transcript, antisense to C17orf97 |
| ENSG00000261599 | HERC2P8 | -0,747959096 | 0,595445307 | 0,1075 | 0,41794846 | hect domain and RLD 2 pseudogene 8 |
| ENSG00000240418 | AC020917,1 | -0,747990481 | 0,595432353 | 0,11354 | 0,42829885 | ribosomal protein S2 (RPS2) pseudogene |
| ENSG00000006128 | TAC1 | -0,748071003 | 0,595399121 | 0,26016 | 0,56382871 | tachykinin precursor 1 |

| | | | | | | |
|---|---|---|---|---|---|---|
| ENSG00000273399 | AL159169,3 | -0,748127365 | 0,595375861 | 0,26462 | 0,56738489 | novel transcript |
| ENSG00000167889 | MGAT5B | -0,748476934 | 0,595231617 | 0,00428 | 0,08037599 | alpha-1,6-mannosylglycoprotein 6-beta-N-acetylglucosaminyltransferase B |
| ENSG00000177732 | SOX12 | -0,74849788 | 0,595222975 | 0,10029 | 0,4067769 | SRY-box transcription factor 12 |
| ENSG00000234928 | LINC01659 | -0,748503922 | 0,595220482 | 0,32075 | 0,60555414 | long intergenic non-protein coding RNA 1659 |
| ENSG00000280986 | LINC01659 | -0,748503922 | 0,595220482 | 0,32075 | 0,60555414 | long intergenic non-protein coding RNA 1659 |
| ENSG00000106483 | SFRP4 | -0,748524749 | 0,59521189 | 0,3176 | 0,60294242 | secreted frizzled related protein 4 |
| ENSG00000282254 | AC004980,5 | -0,748525202 | 0,595211703 | 0,09849 | 0,40409952 | pseudogene similar to part of postmeiotic segregation increased 2-like 3 PMS2L3 |
| ENSG00000143028 | SYPL2 | -0,748568939 | 0,595193659 | 0,16361 | 0,48997962 | synaptophysin like 2 |
| ENSG00000166192 | SENP8 | -0,748868701 | 0,595070003 | 0,09809 | 0,40342986 | SUMO peptidase family member, NEDD8 specific |
| ENSG00000276781 | CHGA | -0,748891403 | 0,595060639 | 0,45602 | 0,69996535 | chromogranin A |
| ENSG00000215045 | GRID2IP | -0,749150635 | 0,594953724 | 0,34976 | 0,62563749 | Grid2 interacting protein |
| ENSG00000260367 | AC109460,1 | -0,749162686 | 0,594948755 | 0,26065 | 0,56400103 | novel transcript, antisense to SPNS1 |
| ENSG00000174928 | C3orf33 | -0,749377144 | 0,594860321 | 0,07357 | 0,35672172 | chromosome 3 open reading frame 33 |
| ENSG00000283041 | AC008038,1 | -0,749577014 | 0,594777916 | 0,14622 | 0,47428193 | eukaryotic translation elongation factor 1 gamma ( |
| ENSG00000124074 | ENKD1 | -0,74975484 | 0,594704608 | 0,03377 | 0,24838006 | enkurin domain containing 1 |
| ENSG00000100433 | KCNK10 | -0,749953205 | 0,594622844 | 0,0378 | 0,26327969 | potassium two pore domain channel subfamily K member 10 |
| ENSG00000231683 | AL033397,1 | -0,750032214 | 0,594590281 | 0,14452 | 0,47209141 | novel transcript |
| ENSG00000145217 | SLC26A1 | -0,750034185 | 0,594589468 | 0,26186 | 0,56477027 | solute carrier family 26 member 1 |
| ENSG00000227587 | ATP6V1G2 | -0,75023896 | 0,594505079 | 0,03609 | 0,25727995 | ATPase H+ transporting V1 subunit G2 |
| ENSG00000287288 | AL121762,2 | -0,750312443 | 0,594474799 | 0,29313 | 0,58745839 | novel transcript |
| ENSG00000274305 | LENG8 | -0,750317959 | 0,594472526 | 0,02803 | 0,22583197 | leukocyte receptor cluster member 8 |
| ENSG00000125966 | MMP24 | -0,75033734 | 0,59446454 | 0,03893 | 0,26698555 | matrix metallopeptidase 24 |
| ENSG00000259926 | AC007342,1 | -0,750438078 | 0,594423032 | 0,28032 | 0,57784691 | novel transcript |
| ENSG00000226688 | ENTPD1-AS1 | -0,750440602 | 0,594421992 | 0,10803 | 0,41857851 | ENTPD1 antisense RNA 1 |
| ENSG00000166006 | KCNC2 | -0,750763371 | 0,594289019 | 0,32504 | 0,60753927 | potassium voltage-gated channel subfamily C member 2 |
| ENSG00000286107 | AC007655,2 | -0,750773674 | 0,594284775 | 0,38393 | 0,65047768 | novel transcript |
| ENSG00000229809 | ZNF688 | -0,750821602 | 0,594265032 | 0,07329 | 0,35655334 | zinc finger protein 688 |
| ENSG00000233966 | UBE2SP1 | -0,750875632 | 0,594242777 | 0,12418 | 0,44504227 | ubiquitin conjugating enzyme E2 S pseudogene 1 |
| ENSG00000036565 | SLC18A1 | -0,751039753 | 0,59417518 | 0,24968 | 0,55657567 | solute carrier family 18 member A1 |
| ENSG00000249931 | GOLGA8K | -0,751460704 | 0,594001836 | 0,04913 | 0,29844668 | golgin A8 family member K |

| | | | | | | |
|---|---|---|---|---|---|---|
| ENSG00000275468 | AC004678,1 | -0,751511282 | 0,593981012 | 0,24664 | 0,55425129 | novel pseudogene |
| ENSG00000163618 | CADPS | -0,751548029 | 0,593965883 | 0,01905 | 0,18296265 | calcium dependent secretion activator |
| ENSG00000231829 | AL157834,1 | -0,751639689 | 0,593928147 | 0,08601 | 0,38030214 | novel transcript |
| ENSG00000166816 | LDHD | -0,751769119 | 0,593874866 | 0,17407 | 0,49860418 | lactate dehydrogenase D |
| ENSG00000167604 | NFKBID | -0,751871279 | 0,593832814 | 0,12665 | 0,44928017 | NFKB inhibitor delta |
| ENSG00000104714 | ERICH1 | -0,752112796 | 0,593733411 | 0,00928 | 0,12292799 | glutamate rich 1 |
| ENSG00000288312 | LINC02018 | -0,752217931 | 0,593690145 | 0,03247 | 0,24322152 | long intergenic non-protein coding RNA 2018 |
| ENSG00000271989 | AL139424,1 | -0,752243274 | 0,593679715 | 0,19038 | 0,51350457 | novel transcript, antisense to APITD1-CORT |
| ENSG00000100711 | ZFYVE21 | -0,75232718 | 0,593645189 | 0,09662 | 0,4005995 | zinc finger FYVE-type containing 21 |
| ENSG00000254352 | AC100854,1 | -0,752370066 | 0,593627542 | 0,30605 | 0,59512079 | chromosome 4 open reading frame 46 (C4orf46) pseudogene |
| ENSG00000277956 | MAPT | -0,752583086 | 0,593539897 | 0,05613 | 0,31646223 | microtubule associated protein tau |
| ENSG00000268061 | NAPA-AS1 | -0,752737066 | 0,593476551 | 0,02967 | 0,23170642 | NAPA antisense RNA 1 |
| ENSG00000159210 | SNF8 | -0,752737229 | 0,593476484 | 0,09553 | 0,39847312 | SNF8 subunit of ESCRT-II |
| ENSG00000280809 | LINC00836 | -0,75281345 | 0,59344513 | 0,05455 | 0,31264967 | long intergenic non-protein coding RNA 836 |
| ENSG00000198626 | RYR2 | -0,752895542 | 0,593411363 | 0,11363 | 0,42841156 | ryanodine receptor 2 |
| ENSG00000275017 | AL353748,2 | -0,753104298 | 0,593325503 | 0,13777 | 0,46481656 | developmental pluripotency associated 3 (DPPA3) pseudogene |
| ENSG00000274449 | AC008463,2 | -0,753127805 | 0,593315836 | 0,06965 | 0,34826564 | novel transcript |
| ENSG00000173611 | SCAI | -0,753210204 | 0,59328195 | 0,07864 | 0,3663904 | suppressor of cancer cell invasion |
| ENSG00000281654 | AC233280,28 | -0,753263316 | 0,593260109 | 0,25268 | 0,55796133 | novel transcript |
| ENSG00000280482 | AC233280,8 | -0,753263316 | 0,593260109 | 0,25268 | 0,55796133 | novel transcript |
| ENSG00000281564 | AC233280,25 | -0,753263316 | 0,593260109 | 0,25268 | 0,55796133 | novel transcript |
| ENSG00000280629 | AC233280,11 | -0,753263316 | 0,593260109 | 0,25268 | 0,55796133 | novel transcript |
| ENSG00000280631 | AC233280,12 | -0,753263316 | 0,593260109 | 0,25268 | 0,55796133 | novel transcript |
| ENSG00000280748 | AC233280,13 | -0,753263316 | 0,593260109 | 0,25268 | 0,55796133 | novel transcript |
| ENSG00000229178 | AC233280,1 | -0,753263316 | 0,593260109 | 0,25268 | 0,55796133 | novel transcript |
| ENSG00000281094 | AC233280,19 | -0,753263316 | 0,593260109 | 0,25268 | 0,55796133 | novel transcript |
| ENSG00000284495 | MIEF2 | -0,753273568 | 0,593255893 | 0,13878 | 0,46606421 | mitochondrial elongation factor 2 |
| ENSG00000251161 | AC020661,1 | -0,753280219 | 0,593253158 | 0,09308 | 0,39410499 | novel transcript |
| ENSG00000183628 | DGCR6 | -0,753302195 | 0,593244121 | 0,01345 | 0,15210212 | DiGeorge syndrome critical region gene 6 |
| ENSG00000251187 | AC010261,2 | -0,753302203 | 0,593244118 | 0,20788 | 0,52746553 | novel transcript |
| ENSG00000282944 | AC140481,3 | -0,753428574 | 0,593192156 | 0,15515 | 0,4829976 | novel transcript |

| | | | | | | |
|---|---|---|---|---|---|---|
| ENSG00000233323 | TNXB | -0,753618772 | 0,593113958 | 0,23375 | 0,54563244 | tenascin XB |
| ENSG00000068137 | PLEKHH3 | -0,753778347 | 0,593048357 | 0,13265 | 0,45811487 | pleckstrin homology, MyTH4 and FERM domain containing H3 |
| ENSG00000168887 | C2orf68 | -0,75414316 | 0,592898413 | 0,08354 | 0,37573154 | chromosome 2 open reading frame 68 |
| ENSG00000172113 | NME6 | -0,754299031 | 0,592834359 | 0,06644 | 0,34124715 | NME/NM23 nucleoside diphosphate kinase 6 |
| ENSG00000243601 | RPL12P24 | -0,75440252 | 0,592791834 | 0,30263 | 0,59309252 | ribosomal protein L12 pseudogene 24 |
| ENSG00000143365 | RORC | -0,754483851 | 0,592758417 | 0,30051 | 0,59186076 | RAR related orphan receptor C |
| ENSG00000258410 | AC087386,1 | -0,754499885 | 0,592751829 | 0,20029 | 0,52107152 | novel transcript |
| ENSG00000163013 | FBXO41 | -0,754601145 | 0,592710227 | 0,00953 | 0,12494325 | F-box protein 41 |
| ENSG00000273233 | AC097724,1 | -0,75460489 | 0,592708688 | 0,28575 | 0,58215967 | novel transcript |
| ENSG00000284672 | AC109583,4 | -0,754836036 | 0,592613733 | 0,08625 | 0,38072744 | novel transcript |
| ENSG00000259877 | AC009113,1 | -0,755434031 | 0,592368146 | 0,06549 | 0,33900891 | novel transcript, antisense to ZNF778 |
| ENSG00000132204 | LINC00470 | -0,755751847 | 0,592237666 | 0,15267 | 0,48061452 | long intergenic non-protein coding RNA 470 |
| ENSG00000260261 | AC124944,2 | -0,755993458 | 0,592138491 | 0,02335 | 0,20398797 | programmed cell death 6 (PDCD6) pseudogene |
| ENSG00000055118 | KCNH2 | -0,756401671 | 0,591970968 | 0,03305 | 0,2455282 | potassium voltage-gated channel subfamily H member 2 |
| ENSG00000132881 | CPLANE2 | -0,756405007 | 0,591969599 | 0,01692 | 0,17086866 | ciliogenesis and planar polarity effector 2 |
| ENSG00000135426 | TESPA1 | -0,756409656 | 0,591967692 | 0,29141 | 0,58649127 | thymocyte expressed, positive selection associated 1 |
| ENSG00000286731 | AP001360,2 | -0,756480242 | 0,591938729 | 0,50673 | 0,73517996 | novel transcript |
| ENSG00000248333 | CDK11B | -0,756701473 | 0,591847965 | 0,04151 | 0,27614259 | cyclin dependent kinase 11B |
| ENSG00000203877 | RIPPLY2 | -0,756793781 | 0,591810098 | 0,22927 | 0,54273374 | ripply transcriptional repressor 2 |
| ENSG00000118369 | USP35 | -0,756803683 | 0,591806036 | 0,06002 | 0,32549025 | ubiquitin specific peptidase 35 |
| ENSG00000271761 | AL021368,1 | -0,756852929 | 0,591785835 | 0,41974 | 0,67539312 | novel transcript |
| ENSG00000267707 | AC015961,2 | -0,756879542 | 0,591774919 | 0,171 | 0,49641416 | novel transcript, antisense to CELF4 |
| ENSG00000256427 | AC010175,1 | -0,757084797 | 0,591690732 | 0,39128 | 0,65504152 | novel transcript |
| ENSG00000204869 | IGFL4 | -0,757133946 | 0,591670575 | 0,05646 | 0,31712073 | IGF like family member 4 |
| ENSG00000227487 | NCAM1-AS1 | -0,757205759 | 0,591641124 | 0,07643 | 0,36251343 | NCAM1 antisense RNA1 |
| ENSG00000108785 | HSD17B1P1 | -0,75731543 | 0,591596151 | 0,22408 | 0,53995756 | hydroxysteroid 17-beta dehydrogenase 1 pseudogene 1 |
| ENSG00000118513 | MYB | -0,757551028 | 0,591499548 | 0,00319 | 0,06765368 | MYB proto-oncogene, transcription factor |
| ENSG00000207955 | AL359091,1 | -0,7576275 | 0,591468196 | 0,3109 | 0,59883009 | novel transcript |
| ENSG00000250654 | AC023794,3 | -0,75767003 | 0,59145076 | 0,31368 | 0,60035326 | novel pseudogene |
| ENSG00000229534 | HNRNPA1P53 | -0,757762305 | 0,591412932 | 0,22924 | 0,54273374 | heterogeneous nuclear ribonucleoprotein A1 pseudogene 53 |
| ENSG00000237737 | DCTN1-AS1 | -0,75784448 | 0,591379246 | 0,19097 | 0,51398916 | DCTN1 antisense RNA 1 |

| | | | | | | |
|---|---|---|---|---|---|---|
| ENSG00000197563 | PIGN | -0,758331149 | 0,591179788 | 0,07328 | 0,35655334 | phosphatidylinositol glycan anchor biosynthesis class N |
| ENSG00000262580 | AC087741,1 | -0,758416142 | 0,591144961 | 0,13967 | 0,4668467 | novel transcript, antisense to CARD14 |
| ENSG00000100033 | PRODH | -0,758437351 | 0,591136271 | 0,16993 | 0,49606609 | proline dehydrogenase 1 |
| ENSG00000253457 | SMIM18 | -0,758539666 | 0,591094349 | 0,22249 | 0,53862734 | small integral membrane protein 18 |
| ENSG00000106113 | CRHR2 | -0,758671026 | 0,591040531 | 0,18082 | 0,50437478 | corticotropin releasing hormone receptor 2 |
| ENSG00000248318 | AC104958,1 | -0,759288896 | 0,590787458 | 0,31332 | 0,60008086 | novel transcript |
| ENSG00000235914 | MACROD2-AS | -0,759310131 | 0,590778762 | 0,2718 | 0,57305058 | MACROD2 antisense RNA 1 |
| ENSG00000152229 | PSTPIP2 | -0,759414571 | 0,590735996 | 0,1162 | 0,43270708 | proline-serine-threonine phosphatase interacting protein 2 |
| ENSG00000275744 | SLC2A11 | -0,759659988 | 0,590635514 | 0,05242 | 0,30735033 | solute carrier family 2 member 11 |
| ENSG00000224812 | TMEM72-AS1 | -0,759861842 | 0,590552882 | 0,10015 | 0,4067769 | TMEM72 antisense RNA 1 |
| ENSG00000284830 | AL049557,1 | -0,759872846 | 0,590548377 | 0,06215 | 0,33018211 | novel transcript |
| ENSG00000287565 | AC108168,1 | -0,759977361 | 0,590505597 | 0,12176 | 0,44093327 | novel transcript, antisense to RP11-526A4,1 |
| ENSG00000185862 | EVI2B | -0,760120759 | 0,590446906 | 0,05289 | 0,30865611 | ecotropic viral integration site 2B |
| ENSG00000164929 | BAALC | -0,760186339 | 0,590420067 | 0,13379 | 0,45984774 | BAALC binder of MAP3K1 and KLF4 |
| ENSG00000242038 | EGFL8 | -0,760194209 | 0,590416846 | 0,32146 | 0,60577115 | EGF like domain multiple 8 |
| ENSG00000204580 | DDR1 | -0,760267255 | 0,590386953 | 0,03908 | 0,26760797 | discoidin domain receptor tyrosine kinase 1 |
| ENSG00000272844 | AC074044,1 | -0,760310824 | 0,590369124 | 0,46961 | 0,70911683 | novel transcript, antisense to GTPBP8 |
| ENSG00000137267 | TUBB2A | -0,760323551 | 0,590363916 | 0,08757 | 0,38314975 | tubulin beta 2A class IIa |
| ENSG00000205485 | AC004980,1 | -0,76060754 | 0,590247716 | 0,0928 | 0,39343521 | pseudogene similar to part of postmeiotic segregation increased 2-like 3 PMS2L3 |
| ENSG00000197976 | AKAP17A | -0,760730892 | 0,590197252 | 0,05725 | 0,31908628 | A-kinase anchoring protein 17A |
| ENSG00000185596 | WASH3P | -0,760814331 | 0,590163118 | 0,09222 | 0,39260493 | WASP family homolog 3, pseudogene |
| ENSG00000288207 | WASH3P | -0,760814331 | 0,590163118 | 0,09222 | 0,39260493 | WASP family homolog 3, pseudogene |
| ENSG00000128159 | TUBGCP6 | -0,760976097 | 0,590096949 | 0,02587 | 0,21707964 | tubulin gamma complex associated protein 6 |
| ENSG00000232977 | LINC00327 | -0,761529192 | 0,589870763 | 0,31613 | 0,60224605 | long intergenic non-protein coding RNA 327 |
| ENSG00000262514 | AC020978,6 | -0,761590086 | 0,589845866 | 0,1402 | 0,46710628 | novel transcript |
| ENSG00000182372 | CLN8 | -0,761840187 | 0,589743621 | 0,19924 | 0,52026761 | CLN8 transmembrane ER and ERGIC protein |
| ENSG00000163995 | ABLIM2 | -0,761850428 | 0,589739434 | 0,03885 | 0,26677873 | actin binding LIM protein family member 2 |
| ENSG00000270679 | AC138473,1 | -0,761963126 | 0,589693368 | 0,32845 | 0,60989517 | mitochondrial ribosomal protein S15 (MRPS15) pseudogene |
| ENSG00000247121 | AC009126,1 | -0,762066945 | 0,589650934 | 0,15905 | 0,48652925 | novel transcript, antisense to LNPEP and ERAP2 |
| ENSG00000239149 | SNORA59A | -0,762224779 | 0,589586429 | 0,07712 | 0,36384125 | small nucleolar RNA, H/ACA box 59A |

| | | | | | | |
|---|---|---|---|---|---|---|
| ENSG00000254229 | FAM90A12P | -0,762304551 | 0,589553829 | 0,18487 | 0,50844776 | family with sequence similarity 90 member A12, pseudogene |
| ENSG00000166979 | EVA1C | -0,762546042 | 0,589455153 | 0,08048 | 0,37018111 | eva-1 homolog C |
| ENSG00000242593 | AC006148,1 | -0,762729771 | 0,589380089 | 0,09098 | 0,39030416 | novel transcript |
| ENSG00000274570 | SPDYE10P | -0,762803466 | 0,589349984 | 0,08698 | 0,38205469 | speedy/RINGO cell cycle regulator family member E10, pseudogene |
| ENSG00000275331 | ABCC6 | -0,762852003 | 0,589330156 | 0,13584 | 0,46280747 | ATP binding cassette subfamily C member 6 |
| ENSG00000227227 | AC017101,1 | -0,763056814 | 0,589246498 | 0,38805 | 0,65289784 | novel transcript |
| ENSG00000171222 | SCAND1 | -0,763109169 | 0,589225116 | 0,08136 | 0,37218783 | SCAN domain containing 1 |
| ENSG00000285630 | AL590068,3 | -0,763205036 | 0,589185963 | 0,04122 | 0,27497038 | novel transcript |
| ENSG00000122696 | SLC25A51 | -0,763463132 | 0,589080568 | 0,06583 | 0,33965348 | solute carrier family 25 member 51 |
| ENSG00000225670 | CADM3-AS1 | -0,764456639 | 0,588675039 | 0,00712 | 0,10752796 | CADM3 antisense RNA 1 |
| ENSG00000169862 | CTNND2 | -0,7647783 | 0,588543804 | 0,00464 | 0,08393639 | catenin delta 2 |
| ENSG00000127586 | CHTF18 | -0,764909826 | 0,58849015 | 0,05912 | 0,32424653 | chromosome transmission fidelity factor 18 |
| ENSG00000005884 | ITGA3 | -0,764929947 | 0,588481943 | 0,07626 | 0,36242671 | integrin subunit alpha 3 |
| ENSG00000244050 | DEFB109F | -0,764977328 | 0,588462616 | 0,36661 | 0,63644059 | defensin beta 109F (pseudogene) |
| ENSG00000230091 | TMEM254-AS | -0,765047676 | 0,588433923 | 0,10199 | 0,40934973 | TMEM254 antisense RNA 1 |
| ENSG00000106236 | NPTX2 | -0,765393963 | 0,588292699 | 6,57E-05 | 0,00424641 | neuronal pentraxin 2 |
| ENSG00000286339 | Z99496,1 | -0,765461593 | 0,588265122 | 0,17721 | 0,50187296 | novel transcript, antisense to CEP85L |
| ENSG00000237636 | ANKRD26P3 | -0,765499918 | 0,588249495 | 0,485 | 0,72064814 | ankyrin repeat domain 26 pseudogene 3 |
| ENSG00000185088 | RPS27L | -0,765676701 | 0,588177417 | 0,03602 | 0,25706681 | ribosomal protein S27 like |
| ENSG00000274205 | MAPK15 | -0,765762131 | 0,588142589 | 0,24521 | 0,55295131 | mitogen-activated protein kinase 15 |
| ENSG00000286722 | AC009244,1 | -0,765836345 | 0,588112335 | 0,08268 | 0,37450725 | novel transcript, antisense to AHCYL2 |
| ENSG00000226620 | LINC00343 | -0,765851033 | 0,588106348 | 0,0741 | 0,35756831 | long intergenic non-protein coding RNA 343 |
| ENSG00000234232 | AC243772,3 | -0,765933643 | 0,588072673 | 0,28517 | 0,58171017 | phosphodiesterase 4D interacting protein (PDE4DIP) pseudogene |
| ENSG00000235439 | DDX39B | -0,766202507 | 0,587963089 | 0,13643 | 0,46362513 | DExD-box helicase 39B |
| ENSG00000143772 | ITPKB | -0,766344287 | 0,58790531 | 0,0255 | 0,21485835 | inositol-trisphosphate 3-kinase B |
| ENSG00000266651 | AC093484,4 | -0,766477686 | 0,587850952 | 0,36017 | 0,63261458 | novel transcript |
| ENSG00000177306 | OR7E125P | -0,766740587 | 0,587743838 | 0,40682 | 0,66618707 | olfactory receptor family 7 subfamily E member 125 pseudogene |
| ENSG00000167194 | C16orf92 | -0,766811779 | 0,587714835 | 0,39535 | 0,65820274 | chromosome 16 open reading frame 92 |
| ENSG00000229739 | PDC-AS1 | -0,766907623 | 0,587675793 | 0,09482 | 0,39731026 | PDC antisense RNA 1 |

| | | | | | | |
|---|---|---|---|---|---|---|
| ENSG00000176909 | MAMSTR | -0,766939417 | 0,587662841 | 0,03703 | 0,26086671 | MEF2 activating motif and SAP domain containing transcriptional regulator |
| ENSG00000265334 | AC130324,2 | -0,767140147 | 0,587581083 | 0,28397 | 0,58089793 | novel transcript, antisense ATAD5 |
| ENSG00000275084 | SNORD91B | -0,767453761 | 0,587453367 | 0,36322 | 0,63407828 | small nucleolar RNA, C/D box 91B |
| ENSG00000169957 | ZNF768 | -0,767875711 | 0,587281578 | 0,03721 | 0,26111083 | zinc finger protein 768 |
| ENSG00000257941 | AC011611,5 | -0,76793587 | 0,587257089 | 0,13593 | 0,46291909 | density-regulated protein (DENR) pseudogene |
| ENSG00000135502 | SLC26A10 | -0,768259485 | 0,587125375 | 0,15382 | 0,48214138 | solute carrier family 26 member 10 |
| ENSG00000239827 | SUGT1P3 | -0,768384281 | 0,58707459 | 0,05347 | 0,31052177 | SUGT1 pseudogene 3 |
| ENSG00000257557 | PPP1R12A-AS | -0,768784316 | 0,586911826 | 0,17568 | 0,49964116 | PPP1R12A antisense RNA 1 |
| ENSG00000274229 | SOCS7 | -0,769156749 | 0,586760334 | 0,03513 | 0,25353048 | suppressor of cytokine signaling 7 |
| ENSG00000104888 | SLC17A7 | -0,769178642 | 0,58675143 | 0,11921 | 0,43703483 | solute carrier family 17 member 7 |
| ENSG00000268049 | AC012313,2 | -0,769224179 | 0,58673291 | 0,12069 | 0,43948272 | novel transcript, antisense to ZNF497 |
| ENSG00000261466 | AC136944,4 | -0,769475376 | 0,586630759 | 0,24959 | 0,55657567 | novel transcript |
| ENSG00000251300 | AC104596,2 | -0,769514141 | 0,586614997 | 0,35244 | 0,62742225 | LysM, putative peptidoglycan-binding, domain containing 4 (LYSMD4) pseudogene |
| ENSG00000282814 | AC010724,9 | -0,769547845 | 0,586601292 | 0,45519 | 0,69925766 | novel transcript |
| ENSG00000262265 | AC002558,3 | -0,769773255 | 0,586509648 | 0,03461 | 0,25152438 | novel transcript, sense intronic to CDC27 |
| ENSG00000240889 | NDUFB2-AS1 | -0,769887283 | 0,586463293 | 0,17982 | 0,50407792 | NDUFB2 antisense RNA 1 |
| ENSG00000250020 | AC113430,1 | -0,769912038 | 0,58645323 | 0,32416 | 0,60675074 | Spi-C transcription factor (Spi-1/PU,1 related) (SPIC) pseudogene |
| ENSG00000233048 | LINC01722 | -0,770212956 | 0,58633092 | 0,15635 | 0,4841567 | long intergenic non-protein coding RNA 1722 |
| ENSG00000224195 | AC022400,1 | -0,770371196 | 0,586266613 | 0,29408 | 0,58837921 | novel transcript |
| ENSG00000273151 | AC073957,3 | -0,770433124 | 0,586241447 | 0,07802 | 0,36538634 | novel transcript, antisense to GET4 |
| ENSG00000232872 | CTAGE3P | -0,770632375 | 0,586160487 | 0,03045 | 0,23546694 | CTAGE family member 3, pseudogene |
| ENSG00000258749 | AL110504,1 | -0,77063497 | 0,586159433 | 0,59641 | 0,79393339 | novel transcript, antisense to CCDC85C |
| ENSG00000221887 | HMSD | -0,770915225 | 0,586045578 | 0,24743 | 0,5549105 | histocompatibility minor serpin domain containing |
| ENSG00000234694 | AL139289,1 | -0,771068705 | 0,585983235 | 0,38695 | 0,65213944 | novel transcript |
| ENSG00000123143 | PKN1 | -0,771151669 | 0,585949538 | 0,03725 | 0,26113308 | protein kinase N1 |
| ENSG00000196199 | MPHOSPH8 | -0,771227847 | 0,5859186 | 0,02584 | 0,21707964 | M-phase phosphoprotein 8 |
| ENSG00000285836 | AC092896,1 | -0,771343891 | 0,585871473 | 0,22881 | 0,54259385 | Novel trancript, antisense to CCDC191 |
| ENSG00000268006 | PTOV1-AS1 | -0,772008635 | 0,585601586 | 0,06778 | 0,34426294 | PTOV1 antisense RNA 1 |
| ENSG00000167136 | ENDOG | -0,772140586 | 0,585548028 | 0,08822 | 0,38444882 | endonuclease G |
| ENSG00000197457 | STMN3 | -0,772378107 | 0,585451633 | 0,02732 | 0,22320635 | stathmin 3 |

| | | | | | | |
|---|---|---|---|---|---|---|
| ENSG00000235145 | RPSAP16 | -0,77250008 | 0,585402138 | 0,13277 | 0,45830601 | ribosomal protein SA pseudogene 16 |
| ENSG00000233114 | AL358075,3 | -0,772582464 | 0,58536871 | 0,25937 | 0,56317998 | cytochrome c oxidase subunit VIIb (COX7B) pseudogene |
| ENSG00000227706 | AL713998,1 | -0,772592219 | 0,585364752 | 0,06927 | 0,34763785 | novel transcript |
| ENSG00000283235 | AC139493,2 | -0,772678176 | 0,585329877 | 0,37061 | 0,63997937 | novel transcript |
| ENSG00000165966 | PDZRN4 | -0,772887807 | 0,585244831 | 0,31102 | 0,59883009 | PDZ domain containing ring finger 4 |
| ENSG00000255021 | AC093496,1 | -0,7735653 | 0,584970064 | 0,18182 | 0,50517706 | novel transcript |
| ENSG00000105808 | RASA4 | -0,773599844 | 0,584956057 | 0,07342 | 0,35662549 | RAS p21 protein activator 4 |
| ENSG00000270981 | AC092944,2 | -0,773693364 | 0,584918139 | 0,37212 | 0,64114037 | poly(rC) binding protein 1, pseudogene |
| ENSG00000261570 | AC073878,1 | -0,773701887 | 0,584914684 | 0,2803 | 0,57784691 | novel transcript, antisense to AGK |
| ENSG00000127663 | KDM4B | -0,773806437 | 0,584872298 | 0,02334 | 0,20398797 | lysine demethylase 4B |
| ENSG00000225725 | FAM66E | -0,773838679 | 0,584859227 | 0,10462 | 0,41313952 | family with sequence similarity 66 member E |
| ENSG00000282873 | BCL7A | -0,773915057 | 0,584828264 | 0,36426 | 0,63495751 | BAF chromatin remodeling complex subunit BCL7A |
| ENSG00000175084 | DES | -0,773952772 | 0,584812976 | 0,28787 | 0,5840701 | desmin |
| ENSG00000132429 | POPDC3 | -0,774097966 | 0,584754123 | 0,28238 | 0,57972352 | popeye domain containing 3 |
| ENSG00000237582 | POU5F1 | -0,774708849 | 0,584506572 | 0,17112 | 0,49641416 | POU class 5 homeobox 1 |
| ENSG00000206454 | POU5F1 | -0,774708849 | 0,584506572 | 0,17112 | 0,49641416 | POU class 5 homeobox 1 |
| ENSG00000259221 | AC023905,1 | -0,774850038 | 0,584449372 | 0,13594 | 0,46291909 | novel transcript, antisense to SEMA6D |
| ENSG00000222515 | RN7SKP240 | -0,775360929 | 0,584242442 | 0,13355 | 0,45946836 | RN7SK pseudogene 240 |
| ENSG00000286369 | AP002813,1 | -0,775465481 | 0,584200104 | 0,06654 | 0,34124715 | novel transcript |
| ENSG00000276880 | IFI27L1 | -0,775630306 | 0,584133364 | 0,01988 | 0,18711351 | interferon alpha inducible protein 27 like 1 |
| ENSG00000184144 | CNTN2 | -0,776091061 | 0,583946838 | 0,02899 | 0,22977124 | contactin 2 |
| ENSG00000286195 | AC007666,2 | -0,776354818 | 0,583840089 | 0,05887 | 0,3241112 | novel transcript, antisense to CECR2 and SLC25A18 |
| ENSG00000228566 | AC013287,1 | -0,776393858 | 0,583824291 | 0,39617 | 0,65876705 | novel transcript |
| ENSG00000164100 | NDST3 | -0,77654286 | 0,583763996 | 0,00336 | 0,07000131 | N-deacetylase and N-sulfotransferase 3 |
| ENSG00000116117 | PARD3B | -0,776609265 | 0,583737127 | 0,0002 | 0,01066645 | par-3 family cell polarity regulator beta |
| ENSG00000206838 | SNORA5A | -0,776764239 | 0,583674426 | 0,34475 | 0,62213186 | small nucleolar RNA, H/ACA box 5A |
| ENSG00000248103 | AC008443,2 | -0,776836771 | 0,583645082 | 0,06509 | 0,33804854 | novel transcript |
| ENSG00000163016 | ALMS1P1 | -0,776929131 | 0,583607718 | 0,00844 | 0,11595565 | ALMS1 pseudogene 1 |
| ENSG00000240694 | PNMA2 | -0,777457241 | 0,583394123 | 0,00209 | 0,05143262 | PNMA family member 2 |
| ENSG00000197937 | ZNF347 | -0,777760906 | 0,583271341 | 0,07557 | 0,36093649 | zinc finger protein 347 |
| ENSG00000200091 | RN7SKP163 | -0,777898772 | 0,583215605 | 0,4381 | 0,68754292 | RN7SK pseudogene 163 |
| ENSG00000236609 | ZNF853 | -0,777902482 | 0,583214105 | 0,00372 | 0,07421558 | zinc finger protein 853 |
| ENSG00000149761 | NUDT22 | -0,778175106 | 0,583103907 | 0,05967 | 0,32521754 | nudix hydrolase 22 |

| | | | | | | |
|---|---|---|---|---|---|---|
| ENSG00000286739 | AC007387,1 | -0,778336046 | 0,583038862 | 0,03717 | 0,26108134 | novel transcript |
| ENSG00000107897 | ACBD5 | -0,778621381 | 0,582923561 | 0,0474 | 0,29245799 | acyl-CoA binding domain containing 5 |
| ENSG00000270066 | AL356488,2 | -0,779088615 | 0,582734804 | 0,73356 | 0,87368351 | small Cajal body-specific RNA 2 |
| ENSG00000287358 | AL096706,1 | -0,779096916 | 0,582731452 | 0,14642 | 0,47433542 | novel transcript |
| ENSG00000126070 | AGO3 | -0,779118561 | 0,582722709 | 0,06138 | 0,32865897 | argonaute RISC catalytic component 3 |
| ENSG00000253854 | AC010834,3 | -0,779347299 | 0,582630326 | 0,1206 | 0,43944057 | novel transcript |
| ENSG00000176435 | CLEC14A | -0,779384493 | 0,582615305 | 0,23311 | 0,54557342 | C-type lectin domain containing 14A |
| ENSG00000285671 | AL139405,1 | -0,779612506 | 0,582523232 | 0,07357 | 0,35672172 | novel transcript |
| ENSG00000228400 | AC079154,1 | -0,779794098 | 0,582449915 | 0,50938 | 0,73725396 | novel transcript |
| ENSG00000120498 | TEX11 | -0,779912219 | 0,582402229 | 0,55624 | 0,76838329 | testis expressed 11 |
| ENSG00000214525 | AC130709,1 | -0,779996281 | 0,582368294 | 0,31734 | 0,60276767 | chromosome 15 open reading frame 2 (C15orf2) pseudogene |
| ENSG00000225559 | AC083867,2 | -0,780048401 | 0,582347256 | 0,3048 | 0,59446033 | novel transcript |
| ENSG00000274756 | AC243732,1 | -0,78018695 | 0,582291333 | 0,19236 | 0,514872 | ribosomal protein S2 (RPS2) pseudogene |
| ENSG00000282707 | AC003042,2 | -0,78018695 | 0,582291333 | 0,19236 | 0,514872 | ribosomal protein S2 (RPS2) pseudogene |
| ENSG00000117280 | RAB29 | -0,780257176 | 0,582262989 | 0,15196 | 0,48039146 | RAB29, member RAS oncogene family |
| ENSG00000176244 | ACBD7 | -0,780397377 | 0,582206408 | 0,03679 | 0,25987876 | acyl-CoA binding domain containing 7 |
| ENSG00000257438 | AC011595,1 | -0,780473098 | 0,582175851 | 0,22894 | 0,54262488 | novel transcript, antisense to NUAK1 |
| ENSG00000152670 | DDX4 | -0,78053858 | 0,582149427 | 0,34645 | 0,62342969 | DEAD-box helicase 4 |
| ENSG00000174292 | TNK1 | -0,780604233 | 0,582122936 | 0,06181 | 0,32905359 | tyrosine kinase non receptor 1 |
| ENSG00000283781 | TNK1 | -0,780604233 | 0,582122936 | 0,06181 | 0,32905359 | tyrosine kinase non receptor 1 |
| ENSG00000281091 | AL117327,1 | -0,780652311 | 0,582103537 | 0,18044 | 0,50407792 | novel transcript |
| ENSG00000196581 | AJAP1 | -0,780740748 | 0,582067855 | 0,08751 | 0,38306995 | adherens junctions associated protein 1 |
| ENSG00000258938 | AL162311,3 | -0,780811698 | 0,582039231 | 0,04127 | 0,27520393 | novel transcript, antisense to BRMS1L |
| ENSG00000276727 | AC137834,2 | -0,780862606 | 0,582018693 | 0,20181 | 0,52216179 | novel transcript, antisense to SHMT2 |
| ENSG00000213755 | AC093270,1 | -0,780931428 | 0,581990929 | 0,32047 | 0,60537847 | ribosomal protein L29 (RPL29) pseudogene |
| ENSG00000167103 | PIP5KL1 | -0,781355597 | 0,581819842 | 0,06163 | 0,32867767 | phosphatidylinositol-4-phosphate 5-kinase like 1 |
| ENSG00000089639 | GMIP | -0,781417267 | 0,581794972 | 0,00144 | 0,04055806 | GEM interacting protein |
| ENSG00000185038 | MROH2A | -0,781442882 | 0,581784642 | 0,36485 | 0,63505392 | maestro heat like repeat family member 2A |
| ENSG00000196169 | KIF19 | -0,781669091 | 0,581693428 | 0,04363 | 0,28187766 | kinesin family member 19 |
| ENSG00000287833 | AL157997,1 | -0,781709369 | 0,581677188 | 0,02116 | 0,19303449 | novel transcript |
| ENSG00000255185 | PDXDC2P | -0,781744759 | 0,581662919 | 0,01438 | 0,15682082 | pyridoxal dependent decarboxylase domain containing 2, pseudogene |

| | | | | | | |
|---|---|---|---|---|---|---|
| ENSG00000254470 | AP5B1 | -0,781783288 | 0,581647385 | 0,10149 | 0,40833508 | adaptor related protein complex 5 subunit beta 1 |
| ENSG00000096171 | VARS1 | -0,781822491 | 0,58163158 | 0,03841 | 0,26521398 | valyl-tRNA synthetase 1 |
| ENSG00000166582 | CENPV | -0,781990639 | 0,581563794 | 0,04849 | 0,29588455 | centromere protein V |
| ENSG00000258526 | AL049828,1 | -0,782366984 | 0,581412106 | 0,06283 | 0,33170499 | novel transcript |
| ENSG00000143816 | WNT9A | -0,7823708 | 0,581410568 | 0,1237 | 0,44453761 | Wnt family member 9A |
| ENSG00000241170 | AP001992,1 | -0,782559317 | 0,5813346 | 0,04591 | 0,28908129 | ribosomal protein L31 (RPL31) pseudogene |
| ENSG00000196482 | ESRRG | -0,782691903 | 0,581281177 | 0,15932 | 0,4867988 | estrogen related receptor gamma |
| ENSG00000175787 | ZNF169 | -0,782827483 | 0,581226553 | 0,08875 | 0,38517778 | zinc finger protein 169 |
| ENSG00000260719 | AC009133,3 | -0,782851768 | 0,581216769 | 0,09521 | 0,3979112 | novel transcript, antisense to C16orf54 |
| ENSG00000162069 | BICDL2 | -0,783261484 | 0,58105173 | 0,41061 | 0,66856387 | BICD family like cargo adaptor 2 |
| ENSG00000170310 | STX8 | -0,783587684 | 0,580920367 | 0,08123 | 0,3719701 | syntaxin 8 |
| ENSG00000196353 | CPNE4 | -0,783647278 | 0,580896371 | 0,14088 | 0,46791293 | copine 4 |
| ENSG00000216723 | NUDT19P3 | -0,783690208 | 0,580879086 | 0,41843 | 0,67472778 | NUDT19 pseudogene 3 |
| ENSG00000288263 | GRK1 | -0,783899624 | 0,580794773 | 0,47784 | 0,71554067 | G protein-coupled receptor kinase 1 |
| ENSG00000185974 | GRK1 | -0,783899624 | 0,580794773 | 0,47784 | 0,71554067 | G protein-coupled receptor kinase 1 |
| ENSG00000261067 | AC109460,3 | -0,783974916 | 0,580764464 | 0,15856 | 0,48579688 | novel transcript |
| ENSG00000259030 | FPGT-TNNI3K | -0,7840534 | 0,58073287 | 0,01108 | 0,13609211 | FPGT-TNNI3K readthrough |
| ENSG00000124217 | MOCS3 | -0,784191288 | 0,580677368 | 0,11572 | 0,43162108 | molybdenum cofactor synthesis 3 |
| ENSG00000128394 | APOBEC3F | -0,784326245 | 0,580623051 | 0,15664 | 0,48425071 | apolipoprotein B mRNA editing enzyme catalytic subunit 3F |
| ENSG00000287896 | AC073548,2 | -0,784535475 | 0,580538851 | 0,20911 | 0,52816443 | novel transcript, antisense to KPTNand SLC8A2 |
| ENSG00000263163 | SLC27A3 | -0,78496973 | 0,580364134 | 0,08418 | 0,37695424 | solute carrier family 27 member 3 |
| ENSG00000152931 | PART1 | -0,785002571 | 0,580350923 | 0,17309 | 0,49783873 | prostate androgen-regulated transcript 1 |
| ENSG00000130517 | PGPEP1 | -0,785026766 | 0,58034119 | 0,07173 | 0,35351167 | pyroglutamyl-peptidase I |
| ENSG00000138622 | HCN4 | -0,785171027 | 0,580283162 | 0,04669 | 0,2905134 | hyperpolarization activated cyclic nucleotide gated potassium channel 4 |
| ENSG00000238230 | LINC00391 | -0,785258233 | 0,580248087 | 0,05177 | 0,30592031 | long intergenic non-protein coding RNA 391 |
| ENSG00000228509 | AC006460,1 | -0,785316319 | 0,580224726 | 0,08372 | 0,37587554 | novel transcript, antisense to NAB1 |
| ENSG00000244131 | RPSAP51 | -0,785917 | 0,579983193 | 0,20955 | 0,52856218 | ribosomal protein SA pseudogene 51 |
| ENSG00000091536 | MYO15A | -0,785917987 | 0,579982797 | 0,1213 | 0,44065523 | myosin XVA |
| ENSG00000280721 | LINC01943 | -0,786089743 | 0,579913752 | 0,07669 | 0,36291117 | long intergenic non-protein coding RNA 1943 |
| ENSG00000188365 | AC092171,1 | -0,786130188 | 0,579897495 | 0,08526 | 0,3785936 | novel transcript |
| ENSG00000250575 | AL732372,3 | -0,786149173 | 0,579889864 | 0,07793 | 0,36517053 | novel pseudogene |

| | | | | | | |
|---|---|---|---|---|---|---|
| ENSG00000224957 | LINC01266 | -0,786265317 | 0,579843182 | 0,2817 | 0,57904089 | long intergenic non-protein coding RNA 1266 |
| ENSG00000271716 | AC022001,2 | -0,786435174 | 0,579774918 | 0,15383 | 0,48214138 | novel transcript, antisense to VGLL4 |
| ENSG00000175894 | TSPEAR | -0,78695347 | 0,579566668 | 0,10962 | 0,42176178 | thrombospondin type laminin G domain and EAR repeats |
| ENSG00000207165 | SNORA70 | -0,78713216 | 0,579494888 | 0,0362 | 0,25752605 | small nucleolar RNA, H/ACA box 70 |
| ENSG00000130749 | ZC3H4 | -0,787461403 | 0,579362655 | 0,05198 | 0,30636099 | zinc finger CCCH-type containing 4 |
| ENSG00000266258 | LINC01909 | -0,787616751 | 0,579300273 | 0,19462 | 0,51562158 | long intergenic non-protein coding RNA 1909 |
| ENSG00000163644 | PPM1K | -0,787625219 | 0,579296872 | 0,06949 | 0,34813287 | protein phosphatase, Mg2+/Mn2+ dependent 1K |
| ENSG00000173200 | PARP15 | -0,787657604 | 0,579283869 | 0,0822 | 0,37359506 | poly(ADP-ribose) polymerase family member 15 |
| ENSG00000275493 | AL627230,1 | -0,787711299 | 0,579262309 | 0,2046 | 0,52443222 | |
| ENSG00000167670 | CHAF1A | -0,78798395 | 0,579152846 | 0,04772 | 0,29349263 | chromatin assembly factor 1 subunit A |
| ENSG00000280677 | FAM66B | -0,788057304 | 0,5791234 | 0,01215 | 0,14321473 | family with sequence similarity 66 member B |
| ENSG00000260669 | AL096870,2 | -0,788298561 | 0,579026563 | 0,47863 | 0,71634089 | novel transcript |
| ENSG00000233393 | AP000688,2 | -0,788566646 | 0,578918977 | 0,21513 | 0,5338459 | novel transcript |
| ENSG00000237975 | FLG-AS1 | -0,788831502 | 0,578812706 | 0,11332 | 0,42797362 | FLG antisense RNA 1 |
| ENSG00000244879 | GABPB1-AS1 | -0,789120066 | 0,578696945 | 0,04619 | 0,28930497 | GABPB1 antisense RNA 1 |
| ENSG00000206403 | ABHD16A | -0,789307093 | 0,57862193 | 0,40508 | 0,66481282 | abhydrolase domain containing 16A, phospholipase |
| ENSG00000228149 | RPL3P1 | -0,789443755 | 0,578567121 | 0,0974 | 0,40207852 | ribosomal protein L3 pseudogene 1 |
| ENSG00000128596 | CCDC136 | -0,789652232 | 0,578483521 | 0,00225 | 0,05400651 | coiled-coil domain containing 136 |
| ENSG00000177614 | PGBD5 | -0,789679359 | 0,578472644 | 0,02354 | 0,20483295 | piggyBac transposable element derived 5 |
| ENSG00000285596 | AC017116,2 | -0,790041454 | 0,578327474 | 0,03703 | 0,26086671 | novel transcript |
| ENSG00000145526 | CDH18 | -0,790308243 | 0,578220537 | 0,03091 | 0,23705504 | cadherin 18 |
| ENSG00000282189 | AC067930,6 | -0,790335328 | 0,578209682 | 0,14868 | 0,47609532 | novel transcript |
| ENSG00000234717 | TMEM212-AS | -0,790439601 | 0,578167893 | 0,26332 | 0,56608528 | TMEM212 antisense RNA 1 |
| ENSG00000278743 | AC087239,1 | -0,790640069 | 0,57808756 | 0,13485 | 0,46156017 | novel transcript |
| ENSG00000127415 | IDUA | -0,790792283 | 0,578026571 | 0,21658 | 0,53490925 | alpha-L-iduronidase |
| ENSG00000269901 | AC010531,5 | -0,790940535 | 0,577967176 | 0,37136 | 0,64049253 | novel transcript, antisense to FBXO31 |
| ENSG00000123600 | METTL8 | -0,791276148 | 0,577832739 | 0,08843 | 0,38472725 | methyltransferase like 8 |
| ENSG00000240553 | AL031428,1 | -0,791292807 | 0,577826067 | 0,12494 | 0,44652417 | novel transcript |
| ENSG00000268030 | AC005253,1 | -0,791375611 | 0,577792903 | 0,04043 | 0,27194236 | novel transcript, antisense to KXD1 |
| ENSG00000287405 | AC017028,2 | -0,791698103 | 0,577663761 | 0,01575 | 0,16461429 | novel transcript, antisense to HDAC4 |
| ENSG00000205704 | LINC00634 | -0,791861409 | 0,577598376 | 0,06369 | 0,33458308 | long intergenic non-protein coding RNA 634 |
| ENSG00000272153 | AL365330,1 | -0,792140095 | 0,577486812 | 0,11991 | 0,43853551 | novel transcript, antisense to LRRC47 |
| ENSG00000185352 | HS6ST3 | -0,792293642 | 0,577425353 | 0,05672 | 0,31752508 | heparan sulfate 6-O-sulfotransferase 3 |

| | | | | | | |
|---|---|---|---|---|---|---|
| ENSG00000075702 | WDR62 | -0,792762849 | 0,577237587 | 0,08682 | 0,3816108 | WD repeat domain 62 |
| ENSG00000237333 | MSH5 | -0,792861712 | 0,577198033 | 0,25343 | 0,55850509 | mutS homolog 5 |
| ENSG00000161513 | FDXR | -0,792961862 | 0,577157966 | 0,07296 | 0,35631792 | ferredoxin reductase |
| ENSG00000282002 | AC243807,1 | -0,793295717 | 0,577024421 | 0,04649 | 0,28999108 | novel transcript |
| ENSG00000276089 | AL137060,6 | -0,793598736 | 0,576903237 | 0,50538 | 0,73393676 | |
| ENSG00000173678 | SPDYE2B | -0,793833244 | 0,57680947 | 0,05642 | 0,31712073 | speedy/RINGO cell cycle regulator family member E2B |
| ENSG00000157927 | RADIL | -0,793842452 | 0,576805789 | 0,02681 | 0,22097601 | Rap associating with DIL domain |
| ENSG00000231170 | AC002451,1 | -0,793952986 | 0,576761598 | 0,25537 | 0,55992862 | novel transcript |
| ENSG00000281965 | PKD1P3 | -0,794112014 | 0,576698025 | 0,07281 | 0,35592834 | polycystin 1, transient receptor potential channel interacting pseudogene 3 |
| ENSG00000146232 | NFKBIE | -0,794272868 | 0,576633729 | 0,00724 | 0,10792958 | NFKB inhibitor epsilon |
| ENSG00000268655 | AC008687,4 | -0,794529439 | 0,576531189 | 0,42179 | 0,67643128 | novel protein |
| ENSG00000248202 | LINC02234 | -0,795574468 | 0,576113724 | 0,26751 | 0,56973003 | long intergenic non-protein coding RNA 2234 |
| ENSG00000248794 | AC026436,1 | -0,795593924 | 0,576105955 | 0,20778 | 0,52736343 | ribosomal protein S12 (RPS12) pseudogene |
| ENSG00000203362 | POLH-AS1 | -0,795685397 | 0,576069429 | 0,16544 | 0,49168875 | POLH antisense RNA 1 |
| ENSG00000144749 | LRIG1 | -0,795938734 | 0,57596828 | 0,03354 | 0,24724664 | leucine rich repeats and immunoglobulin like domains 1 |
| ENSG00000267298 | AC006116,5 | -0,796188121 | 0,575868726 | 0,14488 | 0,47244928 | novel transcript, sense intronic to ZSCAN5A |
| ENSG00000167178 | ISLR2 | -0,796387879 | 0,575788995 | 0,12407 | 0,44501797 | immunoglobulin superfamily containing leucine rich repeat 2 |
| ENSG00000165238 | WNK2 | -0,796400246 | 0,57578406 | 0,04382 | 0,28236923 | WNK lysine deficient protein kinase 2 |
| ENSG00000174456 | C12orf76 | -0,796559771 | 0,575720396 | 0,05938 | 0,32456197 | chromosome 12 open reading frame 76 |
| ENSG00000248968 | AC012640,1 | -0,797052203 | 0,57552392 | 0,30919 | 0,59774312 | novel transcript |
| ENSG00000188693 | CYP51A1-AS1 | -0,797107434 | 0,575501888 | 0,15578 | 0,48358141 | CYP51A1 antisense RNA 1 |
| ENSG00000161179 | YDJC | -0,797219594 | 0,575457148 | 0,083 | 0,37504719 | YdjC chitooligosaccharide deacetylase homolog |
| ENSG00000165644 | COMTD1 | -0,797481283 | 0,575352776 | 0,13657 | 0,46380939 | catechol-O-methyltransferase domain containing 1 |
| ENSG00000128165 | ADM2 | -0,797565229 | 0,575319299 | 0,19941 | 0,52039716 | adrenomedullin 2 |
| ENSG00000254254 | AC012349,1 | -0,798004212 | 0,575144268 | 0,03916 | 0,26783649 | novel transcript |
| ENSG00000129968 | ABHD17A | -0,798123742 | 0,575096618 | 0,06185 | 0,32919617 | abhydrolase domain containing 17A, depalmitoylase |
| ENSG00000286232 | AL365229,1 | -0,798223797 | 0,575056734 | 0,10278 | 0,41062092 | novel transcript |
| ENSG00000157335 | CLEC18C | -0,798353602 | 0,575004997 | 0,1263 | 0,44903997 | C-type lectin domain family 18 member C |
| ENSG00000267263 | AC111170,2 | -0,798839757 | 0,574811266 | 0,58026 | 0,78351044 | novel transcript, sense intronic to SEPT9 |
| ENSG00000162706 | CADM3 | -0,798850701 | 0,574806906 | 0,00605 | 0,09828178 | cell adhesion molecule 3 |
| ENSG00000288538 | AP002985,1 | -0,79892428 | 0,57477759 | 0,05876 | 0,32380736 | novel transcript, antisense to KCTD21 |

| | | | | | | |
|---|---|---|---|---|---|---|
| ENSG00000161149 | TUBA3FP | -0,799254249 | 0,574646144 | 0,10311 | 0,41078404 | tubulin alpha 3f pseudogene |
| ENSG00000223461 | AC004471,1 | -0,799289419 | 0,574632136 | 0,26844 | 0,57055808 | novel transcript |
| ENSG00000277449 | CEBPB-AS1 | -0,799412501 | 0,574583114 | 0,30229 | 0,59294633 | CEBPB antisense RNA 1 |
| ENSG00000261187 | AC079322,1 | -0,799450924 | 0,574567811 | 0,15289 | 0,48105429 | novel transcript |
| ENSG00000121289 | CEP89 | -0,799601412 | 0,574507881 | 0,05189 | 0,30606616 | centrosomal protein 89 |
| ENSG00000241534 | CFB | -0,799620247 | 0,57450038 | 0,18141 | 0,50490902 | complement factor B |
| ENSG00000092295 | TGM1 | -0,799782542 | 0,574435756 | 0,08772 | 0,3833014 | transglutaminase 1 |
| ENSG00000285348 | TGM1 | -0,799782542 | 0,574435756 | 0,08772 | 0,3833014 | transglutaminase 1 |
| ENSG00000170091 | NSG2 | -0,799935712 | 0,574374772 | 0,07482 | 0,35899472 | neuronal vesicle trafficking associated 2 |
| ENSG00000272918 | AC005070,3 | -0,800107494 | 0,574306385 | 0,09622 | 0,39969703 | novel transcript, antisense to MLL5 |
| ENSG00000131401 | NAPSB | -0,800159309 | 0,574285759 | 0,07165 | 0,35351167 | napsin B aspartic peptidase, pseudogene |
| ENSG00000185567 | AHNAK2 | -0,80017578 | 0,574279202 | 0,01604 | 0,16606232 | AHNAK nucleoprotein 2 |
| ENSG00000258053 | AC025575,2 | -0,800207791 | 0,57426646 | 0,29044 | 0,58563599 | novel transcript |
| ENSG00000129749 | CHRNA10 | -0,800541837 | 0,574133508 | 0,09399 | 0,39609058 | cholinergic receptor nicotinic alpha 10 subunit |
| ENSG00000235718 | MFRP | -0,800816512 | 0,574024209 | 0,50118 | 0,73120038 | membrane frizzled-related protein |
| ENSG00000286433 | AL020997,4 | -0,800877187 | 0,574000068 | 0,27147 | 0,57285347 | novel transcript, antisense to PPP1R8 |
| ENSG00000271259 | AC010201,1 | -0,80090342 | 0,573989631 | 0,22808 | 0,54220742 | novel transcript |
| ENSG00000197768 | STPG3 | -0,800909718 | 0,573987125 | 0,47038 | 0,70985356 | sperm-tail PG-rich repeat containing 3 |
| ENSG00000233058 | LINC00884 | -0,800958036 | 0,573967902 | 0,03847 | 0,26548082 | long intergenic non-protein coding RNA 884 |
| ENSG00000145214 | DGKQ | -0,80128645 | 0,573837259 | 0,09096 | 0,39027377 | diacylglycerol kinase theta |
| ENSG00000153982 | GDPD1 | -0,801355083 | 0,573809961 | 0,09182 | 0,39174496 | glycerophosphodiester phosphodiesterase domain containing 1 |
| ENSG00000266371 | AC079915,1 | -0,801620743 | 0,573704308 | 0,06461 | 0,33687219 | novel transcript, sense intronic NF1 |
| ENSG00000153002 | CPB1 | -0,801651522 | 0,573692069 | 0,44949 | 0,69516404 | carboxypeptidase B1 |
| ENSG00000184271 | POU6F1 | -0,802449537 | 0,573374823 | 0,00249 | 0,05782412 | POU class 6 homeobox 1 |
| ENSG00000230022 | FNTAP2 | -0,802524598 | 0,573344993 | 0,197 | 0,51824743 | farnesyltransferase, CAAX box, alpha pseudogene 2 |
| ENSG00000274893 | AL136317,2 | -0,802549261 | 0,573335191 | 0,14978 | 0,47782859 | novel transcript |
| ENSG00000134086 | VHL | -0,802627274 | 0,573304189 | 0,08916 | 0,38617197 | von Hippel-Lindau tumor suppressor |
| ENSG00000286318 | AC096576,7 | -0,802710119 | 0,573271269 | 0,37701 | 0,64554193 | novel transcript, sense intronic to KCNIP4 |
| ENSG00000210049 | MT-TF | -0,802718906 | 0,573267777 | 0,34259 | 0,62025394 | mitochondrially encoded tRNA-Phe (UUU/C) |
| ENSG00000259431 | THTPA | -0,802756059 | 0,573253014 | 0,04858 | 0,29611078 | thiamine triphosphatase |
| ENSG00000173681 | BCLAF3 | -0,80281835 | 0,573228263 | 0,03377 | 0,24838006 | BCLAF1 and THRAP3 family member 3 |
| ENSG00000241288 | LINC02614 | -0,802983371 | 0,573162699 | 0,00941 | 0,1242017 | long intergenic non-protein coding RNA 2614 |

| | | | | | | |
|---|---|---|---|---|---|---|
| ENSG00000100345 | MYH9 | -0,803155166 | 0,573094451 | 0,04425 | 0,28361109 | myosin heavy chain 9 |
| ENSG00000243053 | RPL31P58 | -0,803164209 | 0,573090859 | 0,38337 | 0,65009654 | ribosomal protein L31 pseudogene 58 |
| ENSG00000169750 | RAC3 | -0,803247087 | 0,573057938 | 0,04489 | 0,28635407 | Rac family small GTPase 3 |
| ENSG00000236859 | NIFK-AS1 | -0,803388298 | 0,57300185 | 0,07028 | 0,34990935 | NIFK antisense RNA 1 |
| ENSG00000229029 | CDCA4P1 | -0,803946332 | 0,572780256 | 0,21698 | 0,53505085 | cell division cycle associated 4 pseudogene 1 |
| ENSG00000133477 | FAM83F | -0,804062967 | 0,572733951 | 0,16507 | 0,49125639 | family with sequence similarity 83 member F |
| ENSG00000263818 | RDM1P5 | -0,804365988 | 0,572613668 | 0,0981 | 0,40342986 | RDM1 pseudogene 5 |
| ENSG00000213707 | HMGB1P10 | -0,804504921 | 0,572558527 | 0,35228 | 0,62740642 | high mobility group box 1 pseudogene 10 |
| ENSG00000239899 | RN7SL674P | -0,804521315 | 0,572552021 | 0,22357 | 0,53953208 | RNA, 7SL, cytoplasmic 674, pseudogene |
| ENSG00000275183 | LENG9 | -0,805054475 | 0,572340469 | 0,65734 | 0,82868311 | leukocyte receptor cluster member 9 |
| ENSG00000271428 | MPHOSPH6P1 | -0,805138945 | 0,572306959 | 0,22236 | 0,53859118 | MPHOSPH6 pseudogene 1 |
| ENSG00000204962 | PCDHA8 | -0,805407794 | 0,572200319 | 0,0248 | 0,21105933 | protocadherin alpha 8 |
| ENSG00000272970 | AC107294,2 | -0,805535404 | 0,572149708 | 0,08498 | 0,37843844 | novel transcript |
| ENSG00000134070 | IRAK2 | -0,805670573 | 0,572096105 | 0,08707 | 0,3821708 | interleukin 1 receptor associated kinase 2 |
| ENSG00000270124 | AC092127,2 | -0,805767072 | 0,57205784 | 0,30594 | 0,59512079 | novel transcript |
| ENSG00000128578 | STRIP2 | -0,805776919 | 0,572053935 | 0,09589 | 0,39884178 | striatin interacting protein 2 |
| ENSG00000239272 | RPL21P10 | -0,805777706 | 0,572053623 | 0,20279 | 0,52269757 | ribosomal protein L21 pseudogene 10 |
| ENSG00000225693 | LAGE3P1 | -0,805839873 | 0,572028973 | 0,19985 | 0,52086852 | L antigen family member 3 pseudogene 1 |
| ENSG00000234465 | PINLYP | -0,806089528 | 0,571929994 | 0,01057 | 0,1325438 | phospholipase A2 inhibitor and LY6/PLAUR domain containing |
| ENSG00000119946 | CNNM1 | -0,806312987 | 0,571841414 | 0,1068 | 0,41660052 | cyclin and CBS domain divalent metal cation transport mediator 1 |
| ENSG00000183186 | C2CD4C | -0,806525474 | 0,571757197 | 0,09563 | 0,39857837 | C2 calcium dependent domain containing 4C |
| ENSG00000125741 | OPA3 | -0,806734009 | 0,571674558 | 0,11098 | 0,42382951 | outer mitochondrial membrane lipid metabolism regulator OPA3 |
| ENSG00000225279 | AL121987,1 | -0,806869468 | 0,571620885 | 0,43315 | 0,68455289 | novel transcript |
| ENSG00000197210 | AP000550,1 | -0,806884615 | 0,571614883 | 0,06064 | 0,32698319 | novel transcript |
| ENSG00000074964 | ARHGEF10L | -0,807131661 | 0,571517009 | 0,03239 | 0,2428284 | Rho guanine nucleotide exchange factor 10 like |
| ENSG00000049192 | ADAMTS6 | -0,807244836 | 0,571472176 | 0,1917 | 0,51436129 | ADAM metallopeptidase with thrombospondin type 1 motif 6 |
| ENSG00000156395 | SORCS3 | -0,807954436 | 0,571191163 | 0,15079 | 0,47896194 | sortilin related VPS10 domain containing receptor 3 |
| ENSG00000116703 | PDC | -0,807980031 | 0,571181029 | 0,37517 | 0,64350562 | phosducin |
| ENSG00000259933 | AC009065,2 | -0,808002543 | 0,571172117 | 0,28335 | 0,58063542 | novel transcript, overlapping TSC2 |

| | | | | | | |
|---|---|---|---|---|---|---|
| ENSG00000227960 | AL138799,1 | -0,808321201 | 0,571045972 | 0,55535 | 0,76788267 | novel transcript |
| ENSG00000287045 | AC072028,1 | -0,80842273 | 0,571005786 | 0,09583 | 0,39879007 | novel transcript, antisense to PLS1 |
| ENSG00000254339 | AC064802,1 | -0,80883288 | 0,570843475 | 0,35791 | 0,63185282 | novel transcript |
| ENSG00000234042 | AL591438,2 | -0,809001552 | 0,570776739 | 0,38426 | 0,65052153 | novel pseudogene |
| ENSG00000231274 | SBK3 | -0,809115166 | 0,570731792 | 0,43738 | 0,68717176 | SH3 domain binding kinase family member 3 |
| ENSG00000260392 | AC090518,1 | -0,809128 | 0,570726715 | 0,17682 | 0,50114978 | novel transcript |
| ENSG00000242307 | RPS26P52 | -0,80941376 | 0,57061368 | 0,2733 | 0,57344244 | ribosomal protein S26 pseudogene 52 |
| ENSG00000080618 | CPB2 | -0,809451748 | 0,570598655 | 0,0593 | 0,32445511 | carboxypeptidase B2 |
| ENSG00000215041 | NEURL4 | -0,809536165 | 0,570565269 | 0,03182 | 0,24056445 | neuralized E3 ubiquitin protein ligase 4 |
| ENSG00000288301 | NEURL4 | -0,809536165 | 0,570565269 | 0,03182 | 0,24056445 | neuralized E3 ubiquitin protein ligase 4 |
| ENSG00000233968 | AL157895,1 | -0,809636233 | 0,570525695 | 0,05781 | 0,32108765 | novel transcript, antisense to C10orf112 |
| ENSG00000179240 | GVQW3 | -0,809814416 | 0,570455235 | 0,04067 | 0,27268567 | GVQW motif containing 3 |
| ENSG00000286681 | AF235103,3 | -0,809829772 | 0,570449163 | 0,05623 | 0,31679839 | novel transcript |
| ENSG00000248874 | C5orf17 | -0,809947511 | 0,57040261 | 0,20172 | 0,52214043 | chromosome 5 putative open reading frame 17 |
| ENSG00000287908 | AC025165,6 | -0,810064859 | 0,570356216 | 0,26948 | 0,57124604 | novel protein |
| ENSG00000272824 | AC245100,6 | -0,810340696 | 0,570247177 | 0,202 | 0,52230208 | novel transcript |
| ENSG00000117069 | ST6GALNAC5 | -0,810437934 | 0,570208743 | 0,15963 | 0,48693054 | ST6 N-acetylgalactosaminide alpha-2,6-sialyltransferase 5 |
| ENSG00000212170 | RNU1-77P | -0,810819144 | 0,570058094 | 0,27524 | 0,57491355 | RNA, U1 small nuclear 77, pseudogene |
| ENSG00000266908 | AL121757,2 | -0,811034459 | 0,569973022 | 0,34646 | 0,62342969 | novel transcript |
| ENSG00000253733 | LZTS1-AS1 | -0,811223897 | 0,569898185 | 0,22826 | 0,54221266 | LZTS1 antisense RNA 1 |
| ENSG00000256591 | AP003108,2 | -0,811281573 | 0,569875402 | 0,10737 | 0,41771747 | novel transcript |
| ENSG00000184524 | CEND1 | -0,811333472 | 0,569854902 | 0,08528 | 0,3785936 | cell cycle exit and neuronal differentiation 1 |
| ENSG00000167123 | CERCAM | -0,811448647 | 0,56980941 | 0,21542 | 0,5341454 | cerebral endothelial cell adhesion molecule |
| ENSG00000121413 | ZSCAN18 | -0,811456861 | 0,569806166 | 0,05921 | 0,32424653 | zinc finger and SCAN domain containing 18 |
| ENSG00000070444 | MNT | -0,811834555 | 0,569657012 | 0,10345 | 0,41135663 | MAX network transcriptional repressor |
| ENSG00000130957 | FBP2 | -0,812125878 | 0,569541993 | 0,23602 | 0,5471053 | fructose-bisphosphatase 2 |
| ENSG00000288569 | LINC01949 | -0,81219716 | 0,569513853 | 0,05137 | 0,30462001 | long intergenic non-protein coding RNA 1949 |
| ENSG00000142002 | DPP9 | -0,812311414 | 0,569468752 | 0,06911 | 0,34730156 | dipeptidyl peptidase 9 |
| ENSG00000131037 | EPS8L1 | -0,812337798 | 0,569458338 | 0,0717 | 0,35351167 | EPS8 like 1 |
| ENSG00000253203 | GUSBP3 | -0,812419375 | 0,569426139 | 0,07644 | 0,36251343 | GUSB pseudogene 3 |
| ENSG00000261373 | VPS9D1-AS1 | -0,81247908 | 0,569402574 | 0,22352 | 0,53953208 | VPS9D1 antisense RNA 1 |
| ENSG00000256288 | LINC02617 | -0,812787633 | 0,569280807 | 0,40588 | 0,66553796 | long intergenic non-protein coding RNA 2617 |
| ENSG00000171606 | ZNF274 | -0,812941142 | 0,569220237 | 0,06437 | 0,33619878 | zinc finger protein 274 |

| | | | | | | |
|---|---|---|---|---|---|---|
| ENSG00000248696 | AC011406,1 | -0,812945296 | 0,569218598 | 0,40916 | 0,66785537 | Transmembrane protein pseudogene |
| ENSG00000235545 | AC103923,1 | -0,81322984 | 0,569106341 | 0,08115 | 0,37189706 | novel transcript |
| ENSG00000161682 | FAM171A2 | -0,813293574 | 0,5690812 | 0,05756 | 0,32014909 | family with sequence similarity 171 member A2 |
| ENSG00000130827 | PLXNA3 | -0,813558054 | 0,568976884 | 0,02432 | 0,20842969 | plexin A3 |
| ENSG00000286710 | AC188617,1 | -0,813625702 | 0,568950205 | 0,43615 | 0,68658672 | novel transcript |
| ENSG00000225084 | AL450226,1 | -0,814006415 | 0,568800085 | 0,18962 | 0,51284737 | novel transcript, antisense to SMG6 |
| ENSG00000249412 | AC010285,1 | -0,814106382 | 0,568760673 | 0,04252 | 0,2782038 | novel transcript |
| ENSG00000120833 | SOCS2 | -0,814141151 | 0,568746966 | 0,10726 | 0,41745262 | suppressor of cytokine signaling 2 |
| ENSG00000173585 | CCR9 | -0,814452935 | 0,568624066 | 0,26274 | 0,56581069 | C-C motif chemokine receptor 9 |
| ENSG00000285680 | AL355481,1 | -0,814566373 | 0,568579357 | 0,14663 | 0,47449835 | novel transcript, antisense to SLITRK1 |
| ENSG00000163288 | GABRB1 | -0,814604611 | 0,568564287 | 0,42489 | 0,67857395 | gamma-aminobutyric acid type A receptor subunit beta1 |
| ENSG00000278907 | AC005609,2 | -0,81473196 | 0,568514102 | 0,65775 | 0,82881843 | novel transcript, antisense to PCDHA9 |
| ENSG00000130703 | OSBPL2 | -0,814835104 | 0,568473458 | 0,05777 | 0,32104013 | oxysterol binding protein like 2 |
| ENSG00000249502 | AC006160,1 | -0,814892422 | 0,568450873 | 0,15766 | 0,48476347 | novel transcript, antisense to LAP3 |
| ENSG00000196408 | NOXO1 | -0,81511793 | 0,568362025 | 0,30495 | 0,59446033 | NADPH oxidase organizer 1 |
| ENSG00000185105 | MYADML2 | -0,815256392 | 0,56830748 | 0,24892 | 0,55603752 | myeloid associated differentiation marker like 2 |
| ENSG00000230700 | CSNK2B | -0,815543044 | 0,568194573 | 0,22064 | 0,53767067 | casein kinase 2 beta |
| ENSG00000229368 | AC090587,1 | -0,815792174 | 0,568096463 | 0,19073 | 0,51380292 | novel transcript, sense overlapping STIM1 |
| ENSG00000229140 | CCDC26 | -0,815801027 | 0,568092977 | 0,02845 | 0,22807045 | CCDC26 long non-coding RNA |
| ENSG00000225539 | LINC01821 | -0,816135272 | 0,567961376 | 0,04867 | 0,29652102 | long intergenic non-protein coding RNA 1821 |
| ENSG00000175538 | KCNE3 | -0,816239135 | 0,567920489 | 0,09204 | 0,39230683 | potassium voltage-gated channel subfamily E regulatory subunit 3 |
| ENSG00000125931 | CITED1 | -0,816362074 | 0,567872095 | 0,05436 | 0,31205006 | Cbp/p300 interacting transactivator with Glu/Asp rich carboxy-terminal domain 1 |
| ENSG00000272157 | AC107373,2 | -0,816393627 | 0,567859676 | 0,32124 | 0,60573903 | novel transcript, antisense to NEFL |
| ENSG00000221955 | SLC12A8 | -0,817011808 | 0,567616405 | 0,0217 | 0,19569989 | solute carrier family 12 member 8 |
| ENSG00000214837 | LINC01347 | -0,817186124 | 0,567547826 | 0,13943 | 0,46666406 | long intergenic non-protein coding RNA 1347 |
| ENSG00000166762 | CATSPER2 | -0,817372064 | 0,567474683 | 0,02717 | 0,22241005 | cation channel sperm associated 2 |
| ENSG00000288505 | NANOS3 | -0,817571283 | 0,567396327 | 0,22633 | 0,54125836 | nanos C2HC-type zinc finger 3 |
| ENSG00000187556 | NANOS3 | -0,817571283 | 0,567396327 | 0,22633 | 0,54125836 | nanos C2HC-type zinc finger 3 |
| ENSG00000286044 | AP001977,1 | -0,817808735 | 0,567302947 | 0,043 | 0,27928585 | novel transcript |
| ENSG00000285130 | AL358113,1 | -0,818003751 | 0,567226267 | 0,02801 | 0,22579215 | novel protein |
| ENSG00000168907 | PLA2G4F | -0,81845913 | 0,567047254 | 0,22724 | 0,54168591 | phospholipase A2 group IVF |

| | | | | | | |
|---|---|---|---|---|---|---|
| ENSG00000166091 | CMTM5 | -0,818461173 | 0,567046451 | 0,24545 | 0,5532047 | CKLF like MARVEL transmembrane domain containing 5 |
| ENSG00000164794 | KCNV1 | -0,818476178 | 0,567040553 | 0,05283 | 0,30849176 | potassium voltage-gated channel modifier subfamily V member 1 |
| ENSG00000213722 | DDAH2 | -0,818633251 | 0,56697882 | 0,08193 | 0,37306719 | dimethylarginine dimethylaminohydrolase 2 |
| ENSG00000163873 | GRIK3 | -0,81866905 | 0,566964752 | 0,0041 | 0,07854299 | glutamate ionotropic receptor kainate type subunit 3 |
| ENSG00000169026 | SLC49A3 | -0,819123746 | 0,566786089 | 0,28411 | 0,58097626 | solute carrier family 49 member 3 |
| ENSG00000122432 | SPATA1 | -0,819187701 | 0,566760963 | 0,0292 | 0,2303054 | spermatogenesis associated 1 |
| ENSG00000244115 | DNAJC25-GN( | -0,819257387 | 0,566733588 | 0,03331 | 0,24664841 | DNAJC25-GNG10 readthrough |
| ENSG00000152784 | PRDM8 | -0,819489275 | 0,566642503 | 0,12586 | 0,448127 | PR/SET domain 8 |
| ENSG00000200913 | SNORD46 | -0,81950545 | 0,56663615 | 0,14181 | 0,46830665 | small nucleolar RNA, C/D box 46 |
| ENSG00000242085 | RPS20P33 | -0,819532358 | 0,566625581 | 0,24738 | 0,5549105 | ribosomal protein S20 pseudogene 33 |
| ENSG00000261279 | ULK4P1 | -0,819608151 | 0,566595814 | 0,10524 | 0,41429257 | ULK4 pseudogene 1 |
| ENSG00000119714 | GPR68 | -0,819678502 | 0,566568186 | 0,05422 | 0,31168784 | G protein-coupled receptor 68 |
| ENSG00000278901 | AC005609,1 | -0,819976064 | 0,566451341 | 0,17505 | 0,49919753 | novel transcript, antisense to PCDHA13 |
| ENSG00000166165 | CKB | -0,82026564 | 0,566337655 | 8,35E-05 | 0,00520428 | creatine kinase B |
| ENSG00000161920 | MED11 | -0,820317264 | 0,56631739 | 0,06564 | 0,33945082 | mediator complex subunit 11 |
| ENSG00000276757 | RN7SL192P | -0,820336168 | 0,566309969 | 0,21439 | 0,53334744 | RNA, 7SL, cytoplasmic 192, pseudogene |
| ENSG00000226053 | LINC01776 | -0,820383915 | 0,566291227 | 0,04225 | 0,27788591 | long intergenic non-protein coding RNA 1776 |
| ENSG00000166275 | BORCS7 | -0,820632621 | 0,566193613 | 0,03434 | 0,25054156 | BLOC-1 related complex subunit 7 |
| ENSG00000198468 | FLVCR1-DT | -0,820850044 | 0,56610829 | 0,09932 | 0,40577905 | FLVCR1 divergent transcript |
| ENSG00000229153 | EPHA1-AS1 | -0,820959112 | 0,566065494 | 0,32856 | 0,60992716 | EPHA1 antisense RNA 1 |
| ENSG00000225711 | AL512326,3 | -0,82103935 | 0,566034012 | 0,18075 | 0,50428279 | UDP-Gal:betaGlcNAc beta 1,3-galactosyltransferase, polypeptide 3 (B3GALT3) pseudogene |
| ENSG00000286812 | AP001120,5 | -0,821094004 | 0,566012569 | 0,05053 | 0,30240543 | novel transcript, antisense to GNAL |
| ENSG00000282176 | FLJ13224 | -0,821166793 | 0,565984012 | 0,06697 | 0,34279254 | uncharacterized LOC79857 |
| ENSG00000177340 | FLJ13224 | -0,821166793 | 0,565984012 | 0,06697 | 0,34279254 | uncharacterized LOC79857 |
| ENSG00000267075 | AC015818,8 | -0,821409929 | 0,565888636 | 0,06701 | 0,34281679 | novel transcript, piRNA host |
| ENSG00000205089 | CCNI2 | -0,821491315 | 0,565856714 | 0,20831 | 0,52766862 | cyclin I family member 2 |
| ENSG00000156232 | WHAMM | -0,821679073 | 0,565783076 | 0,051 | 0,30380236 | WASP homolog associated with actin, golgi membranes and microtubules |
| ENSG00000234003 | MTATP6P27 | -0,821997574 | 0,565658183 | 0,17091 | 0,49641416 | MT-ATP6 pseudogene 27 |
| ENSG00000267575 | AC006504,5 | -0,822110163 | 0,56561404 | 0,08175 | 0,3729912 | novel transcript |
| ENSG00000156500 | FAM122C | -0,822156302 | 0,565595951 | 0,05555 | 0,31503413 | family with sequence similarity 122C |

| | | | | | | |
|---|---|---|---|---|---|---|
| ENSG00000179168 | GGN | -0,822217157 | 0,565572094 | 0,17636 | 0,50050439 | gametogenetin |
| ENSG00000175003 | SLC22A1 | -0,822521843 | 0,565452662 | 0,20573 | 0,52518691 | solute carrier family 22 member 1 |
| ENSG00000125652 | ALKBH7 | -0,823081077 | 0,565233517 | 0,06147 | 0,32865897 | alkB homolog 7 |
| ENSG00000267152 | AC093227,1 | -0,823193472 | 0,565189484 | 0,0225 | 0,19974241 | novel transcript |
| ENSG00000248383 | PCDHAC1 | -0,823269782 | 0,56515959 | 0,00135 | 0,03911553 | protocadherin alpha subfamily C, 1 |
| ENSG00000256278 | AC048382,1 | -0,823396103 | 0,565110107 | 0,06005 | 0,32549025 | novel transcript, antisense to ZSCAN2 |
| ENSG00000002079 | MYH16 | -0,823805802 | 0,564949649 | 0,19205 | 0,5145257 | myosin heavy chain 16 pseudogene |
| ENSG00000142409 | ZNF787 | -0,823917911 | 0,564905749 | 0,09592 | 0,39891259 | zinc finger protein 787 |
| ENSG00000276072 | AATF | -0,823984833 | 0,564879546 | 0,37308 | 0,64180957 | apoptosis antagonizing transcription factor |
| ENSG00000225174 | OSTM1-AS1 | -0,824121419 | 0,564826069 | 0,35196 | 0,62703165 | OSTM1 antisense RNA 1 |
| ENSG00000135519 | KCNH3 | -0,824149326 | 0,564815143 | 0,1145 | 0,42992088 | potassium voltage-gated channel subfamily H member 3 |
| ENSG00000248485 | PCP4L1 | -0,824151549 | 0,564814273 | 0,29915 | 0,59122248 | Purkinje cell protein 4 like 1 |
| ENSG00000203392 | AC105020,1 | -0,824484902 | 0,56468378 | 0,13146 | 0,45657505 | novel transcript, antisense to CSPG4 |
| ENSG00000177606 | JUN | -0,82454951 | 0,564658493 | 0,05153 | 0,30518413 | Jun proto-oncogene, AP-1 transcription factor subunit |
| ENSG00000255411 | LINC02548 | -0,824716565 | 0,564593113 | 0,02305 | 0,2024534 | long intergenic non-protein coding RNA 2548 |
| ENSG00000059804 | SLC2A3 | -0,824731198 | 0,564587386 | 0,03302 | 0,24543613 | solute carrier family 2 member 3 |
| ENSG00000224975 | INE1 | -0,824765726 | 0,564573874 | 0,05937 | 0,32456197 | inactivation escape 1 |
| ENSG00000235222 | MSH5 | -0,824801606 | 0,564559833 | 0,05636 | 0,31702213 | mutS homolog 5 |
| ENSG00000185155 | MIXL1 | -0,824802713 | 0,5645594 | 0,08422 | 0,37695424 | Mix paired-like homeobox |
| ENSG00000247077 | PGAM5 | -0,824896943 | 0,564522527 | 0,10282 | 0,41062092 | PGAM family member 5, mitochondrial serine/threonine protein phosphatase |
| ENSG00000100867 | DHRS2 | -0,82492855 | 0,564510159 | 0,23976 | 0,5492136 | dehydrogenase/reductase 2 |
| ENSG00000167641 | PPP1R14A | -0,824983448 | 0,564488679 | 0,29316 | 0,58745839 | protein phosphatase 1 regulatory inhibitor subunit 14A |
| ENSG00000188452 | CERKL | -0,825304577 | 0,564363043 | 0,07987 | 0,36890401 | ceramide kinase like |
| ENSG00000003400 | CASP10 | -0,82548843 | 0,564291127 | 0,15354 | 0,48191866 | caspase 10 |
| ENSG00000278220 | CLN8 | -0,825521713 | 0,564278109 | 0,27722 | 0,57574407 | CLN8 transmembrane ER and ERGIC protein |
| ENSG00000152049 | KCNE4 | -0,825596159 | 0,564248992 | 0,08823 | 0,38444882 | potassium voltage-gated channel subfamily E regulatory subunit 4 |
| ENSG00000207279 | SNORD116-24 | -0,825663413 | 0,564222689 | 0,34984 | 0,62563749 | small nucleolar RNA, C/D box 116-24 |
| ENSG00000283945 | LINC00032 | -0,825869228 | 0,564142203 | 0,0349 | 0,25262813 | long intergenic non-protein coding RNA 32 |
| ENSG00000270690 | AC105129,3 | -0,825976021 | 0,564100444 | 0,27658 | 0,575468 | peroxiredoxin 3 (PRDX3) pseudogene |
| ENSG00000132000 | PODNL1 | -0,825987122 | 0,564096104 | 0,33079 | 0,6112353 | podocan like 1 |
| ENSG00000288195 | PODNL1 | -0,825987122 | 0,564096104 | 0,33079 | 0,6112353 | podocan like 1 |

| | | | | | | |
|---|---|---|---|---|---|---|
| ENSG00000224389 | C4B | -0,826116903 | 0,564045362 | 0,00548 | 0,09208977 | complement C4B (Chido blood group) |
| ENSG00000277925 | Telomerase-v | -0,826231994 | 0,564000367 | 0,01904 | 0,18296265 | Vertebrate telomerase RNA |
| ENSG00000268093 | AC022154,1 | -0,826604584 | 0,563854727 | 0,16222 | 0,48939373 | novel transcript, antisense to SPHK2 |
| ENSG00000146904 | EPHA1 | -0,826849631 | 0,563758963 | 0,15096 | 0,47923194 | EPH receptor A1 |
| ENSG00000284816 | EPHA1 | -0,826849631 | 0,563758963 | 0,15096 | 0,47923194 | EPH receptor A1 |
| ENSG00000227773 | ASH1L-IT1 | -0,826939288 | 0,563723928 | 0,43707 | 0,68709205 | ASH1L intronic transcript 1 |
| ENSG00000274717 | AL049757,1 | -0,826939352 | 0,563723904 | 0,45585 | 0,69982618 | ADAM metallopeptidase with thrombospondin type 1 motif, 7 (ADAMTS7) pseudogene |
| ENSG00000260698 | AL591848,2 | -0,827037103 | 0,563685709 | 0,26885 | 0,57085484 | novel transcript |
| ENSG00000277715 | AC079174,2 | -0,827185913 | 0,563627569 | 0,08248 | 0,37409152 | novel transcript, antisense to CKAP4 |
| ENSG00000277199 | LINC01632 | -0,827249352 | 0,563602786 | 0,08511 | 0,37850509 | long intergenic non-protein coding RNA 1632 |
| ENSG00000115641 | FHL2 | -0,827411171 | 0,563539573 | 0,06199 | 0,32961785 | four and a half LIM domains 2 |
| ENSG00000276505 | AP000892,3 | -0,82800267 | 0,563308572 | 0,37524 | 0,64359162 | novel transcript, sense intronic to BACE1 |
| ENSG00000259385 | AC066612,2 | -0,828038671 | 0,563294515 | 0,28916 | 0,58483331 | novel transcript, antisense to CTXN2 |
| ENSG00000214595 | EML6 | -0,828102027 | 0,563269779 | 0,01799 | 0,17701257 | EMAP like 6 |
| ENSG00000224699 | LAMTOR5-AS1 | -0,828395832 | 0,56315508 | 0,09962 | 0,40615251 | LAMTOR5 antisense RNA 1 |
| ENSG00000267222 | AC107993,1 | -0,828566423 | 0,563088494 | 0,45702 | 0,70056368 | novel transcript |
| ENSG00000286072 | AC002072,1 | -0,828721665 | 0,563027906 | 0,03707 | 0,26086671 | novel transcript |
| ENSG00000287997 | AL354950,3 | -0,828796774 | 0,562998595 | 0,28695 | 0,58337791 | novel transcript, antisense to EBF3 |
| ENSG00000235169 | SMIM1 | -0,828964526 | 0,562933135 | 0,13551 | 0,46223021 | small integral membrane protein 1 (Vel blood group) |
| ENSG00000187792 | ZNF70 | -0,828974224 | 0,56292935 | 0,08246 | 0,374054 | zinc finger protein 70 |
| ENSG00000273814 | OCLN | -0,82932563 | 0,562792251 | 0,08042 | 0,37007763 | occludin |
| ENSG00000214176 | PLEKHM1P1 | -0,829373994 | 0,562773385 | 0,05505 | 0,31407633 | pleckstrin homology and RUN domain containing M1 pseudogene 1 |
| ENSG00000167470 | MIDN | -0,829413256 | 0,562758069 | 0,06498 | 0,33787601 | midnolin |
| ENSG00000262480 | SAMD11P1 | -0,829425807 | 0,562753174 | 0,20995 | 0,52885604 | sterile alpha motif domain containing 11 pseudogene 1 |
| ENSG00000287049 | AC124862,1 | -0,829476458 | 0,562733417 | 0,23947 | 0,5491566 | novel transcript, antisense to ANKMY1 |
| ENSG00000265845 | AC024267,4 | -0,829513474 | 0,562718978 | 0,13278 | 0,45830601 | novel transcript, antisense to PHF12 |
| ENSG00000227741 | AL121987,2 | -0,829516578 | 0,562717768 | 0,14799 | 0,4751727 | novel transcript |
| ENSG00000235834 | Z93022,1 | -0,829618053 | 0,562678189 | 0,05913 | 0,32424653 | novel transcript |
| ENSG00000283696 | AL592295,4 | -0,829795059 | 0,562609158 | 0,06179 | 0,32905359 | novel transcript |
| ENSG00000149418 | ST14 | -0,830089141 | 0,562494486 | 0,19009 | 0,51331106 | ST14 transmembrane serine protease matriptase |
| ENSG00000160813 | PPP1R35 | -0,830382359 | 0,562380174 | 0,02355 | 0,20483295 | protein phosphatase 1 regulatory subunit 35 |

| | | | | | | |
|---|---|---|---|---|---|---|
| ENSG00000282744 | AC246789,2 | -0,830449363 | 0,562354056 | 0,09777 | 0,40289531 | novel transcript |
| ENSG00000124479 | NDP | -0,83057901 | 0,562303522 | 0,15543 | 0,48334025 | norrin cystine knot growth factor NDP |
| ENSG00000259294 | AC005096,1 | -0,830933669 | 0,562165308 | 0,14867 | 0,47609532 | novel transcript, antisense to CUX1 |
| ENSG00000285781 | AL117340,1 | -0,831024081 | 0,562130079 | 0,12527 | 0,44701864 | novel pseudogene similar to nuclear pore associated protein 1 (NPAP1) |
| ENSG00000287424 | AC080079,2 | -0,831296927 | 0,562023778 | 0,07947 | 0,3681274 | novel transcript |
| ENSG00000198208 | RPS6KL1 | -0,831429465 | 0,561972148 | 0,00609 | 0,09847604 | ribosomal protein S6 kinase like 1 |
| ENSG00000204682 | MIR1915HG | -0,831536866 | 0,561930314 | 0,00636 | 0,10059731 | MIR1915 host gene |
| ENSG00000272669 | AL021707,6 | -0,83225355 | 0,561651234 | 0,06012 | 0,32570742 | novel transcript, antisense to SUN2 |
| ENSG00000099377 | HSD3B7 | -0,832287054 | 0,561638191 | 0,11455 | 0,42992744 | hydroxy-delta-5-steroid dehydrogenase, 3 beta- and steroid delta-isomerase 7 |
| ENSG00000236559 | BCL2L1-AS1 | -0,832368902 | 0,561606328 | 0,3533 | 0,62807595 | BCL2L1 antisense RNA 1 |
| ENSG00000277675 | AC211486,5 | -0,832674188 | 0,5614875 | 0,0709 | 0,35165018 | novel transcript |
| ENSG00000278899 | AL358852,1 | -0,832799782 | 0,561438622 | 0,1157 | 0,43162108 | novel transcript, antisense to LATS1 |
| ENSG00000264229 | RNU4ATAC | -0,832900208 | 0,561399542 | 0,08861 | 0,38491339 | RNA, U4atac small nuclear (U12-dependent splicing) |
| ENSG00000280739 | EIF1B-AS1 | -0,833055183 | 0,56133924 | 0,05977 | 0,32535617 | EIF1B antisense RNA 1 |
| ENSG00000149609 | C20orf144 | -0,833072438 | 0,561332526 | 0,12547 | 0,44729629 | chromosome 20 open reading frame 144 |
| ENSG00000236942 | AC096533,1 | -0,83327972 | 0,561251881 | 0,29847 | 0,59082457 | GABA(A) receptor-associated protein (GABARAP) pseudogene |
| ENSG00000233334 | FAM53B-AS1 | -0,833499336 | 0,56116645 | 0,17107 | 0,49641416 | FAM53B antisense RNA 1 |
| ENSG00000138002 | IFT172 | -0,833785624 | 0,561055104 | 0,05604 | 0,31627719 | intraflagellar transport 172 |
| ENSG00000288293 | CC2D1A | -0,833826902 | 0,561039051 | 0,0535 | 0,31052177 | coiled-coil and C2 domain containing 1A |
| ENSG00000233296 | TMEM18-DT | -0,833959162 | 0,56098762 | 0,09974 | 0,40626368 | TMEM18 divergent transcript |
| ENSG00000073150 | PANX2 | -0,83407327 | 0,560943251 | 0,06852 | 0,34595829 | pannexin 2 |
| ENSG00000224358 | TMCO1-AS1 | -0,834491158 | 0,560780793 | 0,0858 | 0,37991409 | TMCO1 antisense RNA 1 |
| ENSG00000090534 | THPO | -0,834532564 | 0,560764699 | 0,17763 | 0,50205238 | thrombopoietin |
| ENSG00000006015 | REX1BD | -0,834616941 | 0,560731903 | 0,08955 | 0,38704804 | required for excision 1-B domain containing |
| ENSG00000164188 | RANBP3L | -0,834746919 | 0,560681386 | 0,1599 | 0,48710442 | RAN binding protein 3 like |
| ENSG00000281244 | ADAMTS13 | -0,834928648 | 0,560610765 | 0,05153 | 0,30518413 | ADAM metallopeptidase with thrombospondin type 1 motif 13 |
| ENSG00000004139 | SARM1 | -0,835790106 | 0,560276114 | 0,00075 | 0,02677782 | sterile alpha and TIR motif containing 1 |
| ENSG00000285803 | AL442003,1 | -0,835864174 | 0,56024735 | 0,06193 | 0,32950971 | novel transcript, antisense to AGAP6 |
| ENSG00000215193 | PEX26 | -0,836090534 | 0,560159454 | 0,06649 | 0,34124715 | peroxisomal biogenesis factor 26 |

| | | | | | | |
|---|---|---|---|---|---|---|
| ENSG00000061656 | SPAG4 | -0,836200804 | 0,560116641 | 0,12951 | 0,45306297 | sperm associated antigen 4 |
| ENSG00000129474 | AJUBA | -0,836234544 | 0,560103542 | 0,11797 | 0,43562473 | ajuba LIM protein |
| ENSG00000271078 | AC139426,3 | -0,836310275 | 0,560074141 | 0,14526 | 0,47289051 | NEDD4 binding protein 2 (N4BP2) pseudogene |
| ENSG00000180221 | TPT1P10 | -0,836492342 | 0,560003464 | 0,38437 | 0,65052153 | tumor protein, translationally-controlled 1 pseudogene 10 |
| ENSG00000235778 | AC130360,1 | -0,836638039 | 0,559946913 | 0,32121 | 0,60573903 | ubiquitin-specific protease (USP) pseudogene |
| ENSG00000242628 | AC009228,1 | -0,836714759 | 0,559917137 | 0,10437 | 0,4128259 | novel transcript |
| ENSG00000284048 | AC073111,2 | -0,836881962 | 0,559852248 | 0,55873 | 0,77025265 | novel transcript |
| ENSG00000266208 | AC080112,1 | -0,836921989 | 0,559836715 | 0,21356 | 0,53275614 | novel transcript, antisense to GJD3 |
| ENSG00000275964 | AL355001,2 | -0,837194165 | 0,559731108 | 0,01766 | 0,17511677 | novel transcript |
| ENSG00000196689 | TRPV1 | -0,837269886 | 0,559701731 | 0,08528 | 0,3785936 | transient receptor potential cation channel subfamily V member 1 |
| ENSG00000238272 | AL139142,1 | -0,837483136 | 0,559619005 | 0,15234 | 0,48061452 | novel transcript |
| ENSG00000213999 | MEF2B | -0,837607727 | 0,559570679 | 0,10265 | 0,41043823 | myocyte enhancer factor 2B |
| ENSG00000188315 | C3orf62 | -0,837647814 | 0,559555131 | 0,01652 | 0,16837263 | chromosome 3 open reading frame 62 |
| ENSG00000286661 | AC092933,2 | -0,837777995 | 0,559504642 | 0,05083 | 0,30326674 | novel transcript |
| ENSG00000145945 | FAM50B | -0,837974893 | 0,559428286 | 0,00764 | 0,11109228 | family with sequence similarity 50 member B |
| ENSG00000235618 | FAM21EP | -0,837995749 | 0,559420199 | 0,16909 | 0,49515165 | family with sequence similarity 21 member E, pseudogene |
| ENSG00000276504 | TFPT | -0,838049365 | 0,559399409 | 0,08636 | 0,38089807 | TCF3 fusion partner |
| ENSG00000103723 | AP3B2 | -0,838140681 | 0,559364003 | 0,04775 | 0,29349263 | adaptor related protein complex 3 subunit beta 2 |
| ENSG00000213363 | RPS3P6 | -0,838387645 | 0,559268258 | 0,33374 | 0,61370377 | ribosomal protein S3 pseudogene 6 |
| ENSG00000253309 | SERPINE3 | -0,838630255 | 0,559174217 | 0,21151 | 0,53032236 | serpin family E member 3 |
| ENSG00000275765 | AC091982,3 | -0,838726481 | 0,559136921 | 0,14382 | 0,47134131 | novel transcript |
| ENSG00000256810 | AC078925,1 | -0,838826719 | 0,559098074 | 0,1378 | 0,46485529 | novel transcript |
| ENSG00000205143 | ARID3C | -0,838865507 | 0,559083043 | 0,33843 | 0,61774255 | AT-rich interaction domain 3C |
| ENSG00000278291 | AL161772,1 | -0,838963192 | 0,559045188 | 0,05373 | 0,31090219 | novel transcript, antisense to IL17D |
| ENSG00000125430 | HS3ST3B1 | -0,839294532 | 0,558916809 | 0,21939 | 0,53676938 | heparan sulfate-glucosamine 3-sulfotransferase 3B1 |
| ENSG00000236130 | PTCSC2 | -0,839547536 | 0,558818801 | 0,11234 | 0,42601117 | papillary thyroid carcinoma susceptibility candidate 2 |
| ENSG00000235848 | RMDN2-AS1 | -0,839631593 | 0,558786242 | 0,2807 | 0,57812368 | RMDN2 antisense RNA 1 |
| ENSG00000114646 | CSPG5 | -0,839669442 | 0,558771583 | 0,04162 | 0,27635056 | chondroitin sulfate proteoglycan 5 |
| ENSG00000112041 | TULP1 | -0,839843868 | 0,55870403 | 0,053 | 0,30902552 | TUB like protein 1 |
| ENSG00000181965 | NEUROG1 | -0,839891387 | 0,558685628 | 0,0551 | 0,31415946 | neurogenin 1 |
| ENSG00000266171 | AP001020,3 | -0,840153419 | 0,558584165 | 0,14686 | 0,47471707 | novel transcript, antisense to YES1 |
| ENSG00000249614 | LINC02503 | -0,840456486 | 0,558466836 | 0,06867 | 0,34640847 | long intergenic non-protein coding RNA 2503 |

| | | | | | | |
|---|---|---|---|---|---|---|
| ENSG00000163328 | GPR155 | -0,840589898 | 0,558415194 | 0,05072 | 0,30287257 | G protein-coupled receptor 155 |
| ENSG00000220161 | LINC02076 | -0,840616936 | 0,558404729 | 0,14007 | 0,4669503 | long intergenic non-protein coding RNA 2076 |
| ENSG00000267632 | AC067852,3 | -0,840637409 | 0,558396805 | 0,04111 | 0,27462898 | novel transcript |
| ENSG00000255087 | AP001993,1 | -0,84087039 | 0,558306636 | 0,28635 | 0,58267001 | novel transcript |
| ENSG00000183688 | RFLNB | -0,840950262 | 0,558275728 | 0,07863 | 0,3663904 | refilin B |
| ENSG00000075340 | ADD2 | -0,841065511 | 0,558231132 | 0,00708 | 0,10712237 | adducin 2 |
| ENSG00000248161 | AC098487,1 | -0,841543438 | 0,558046235 | 0,04417 | 0,28322445 | novel transcript |
| ENSG00000140479 | PCSK6 | -0,841628724 | 0,558013247 | 0,18543 | 0,50888225 | proprotein convertase subtilisin/kexin type 6 |
| ENSG00000171169 | NAIF1 | -0,841686778 | 0,557990793 | 0,05425 | 0,31175187 | nuclear apoptosis inducing factor 1 |
| ENSG00000260005 | AC027601,1 | -0,842515355 | 0,557670416 | 0,10551 | 0,41477267 | novel transcript, antisense to C17orf56 |
| ENSG00000101546 | RBFA | -0,842543005 | 0,557659729 | 0,0712 | 0,3523325 | ribosome binding factor A |
| ENSG00000168004 | PLAAT5 | -0,842603397 | 0,557636385 | 0,08371 | 0,37587554 | phospholipase A and acyltransferase 5 |
| ENSG00000185046 | ANKS1B | -0,842685593 | 0,557604615 | 0,00131 | 0,03866806 | ankyrin repeat and sterile alpha motif domain containing 1B |
| ENSG00000244671 | RN7SL280P | -0,842806725 | 0,557557799 | 0,27612 | 0,57533851 | RNA, 7SL, cytoplasmic 280, pseudogene |
| ENSG00000152475 | ZNF837 | -0,842840133 | 0,557544888 | 0,05933 | 0,32455363 | zinc finger protein 837 |
| ENSG00000273424 | AL008582,1 | -0,843110477 | 0,557440421 | 0,27267 | 0,57315967 | novel transcript |
| ENSG00000270174 | AL022097,1 | -0,843216937 | 0,557399287 | 0,29535 | 0,58909567 | novel transcript, antisense to FARS2 |
| ENSG00000285224 | AL662854,2 | -0,843520353 | 0,557282072 | 0,63003 | 0,8132682 | novel protein |
| ENSG00000261342 | AC006538,1 | -0,843575002 | 0,557260963 | 0,0851 | 0,37850509 | novel transcript |
| ENSG00000259079 | AC005476,1 | -0,844001642 | 0,557096191 | 0,15566 | 0,48345635 | PDZ domain-containing protein 11 (PDZD11) pseudogene |
| ENSG00000176058 | TPRN | -0,844049117 | 0,557077859 | 0,04241 | 0,27820114 | taperin |
| ENSG00000234789 | AL590369,1 | -0,844097028 | 0,557059359 | 0,34533 | 0,62249219 | novel transcript |
| ENSG00000287243 | THCAT155 | -0,844528759 | 0,556892682 | 0,0456 | 0,2883604 | thyroid cancer-associated transcript 155 |
| ENSG00000176349 | AC104129,1 | -0,844595814 | 0,556866799 | 0,1779 | 0,50233162 | novel transcript, antisense to MAD1L1 |
| ENSG00000272549 | LINC02538 | -0,844861649 | 0,556764199 | 0,16666 | 0,49255702 | long intergenic non-protein coding RNA 2538 |
| ENSG00000158106 | RHPN1 | -0,844922853 | 0,55674058 | 0,06551 | 0,33905495 | rhophilin Rho GTPase binding protein 1 |
| ENSG00000227254 | VDAC1P12 | -0,844973883 | 0,556720887 | 0,34047 | 0,61897407 | voltage dependent anion channel 1 pseudogene 12 |
| ENSG00000196459 | TRAPPC2 | -0,845257645 | 0,556611397 | 0,07936 | 0,36796591 | trafficking protein particle complex 2 |
| ENSG00000154553 | PDLIM3 | -0,845729965 | 0,5564292 | 0,18308 | 0,50652733 | PDZ and LIM domain 3 |
| ENSG00000186409 | CCDC30 | -0,845849901 | 0,556382944 | 0,08937 | 0,38668036 | coiled-coil domain containing 30 |
| ENSG00000236534 | H3P36 | -0,845987851 | 0,556329745 | 0,31143 | 0,59890747 | H3 histone pseudogene 36 |
| ENSG00000287454 | AC008149,2 | -0,846039276 | 0,556309915 | 0,22216 | 0,53859118 | novel transcript, antisense to ELK3 |

| | | | | | | |
|---|---|---|---|---|---|---|
| ENSG00000254985 | RSF1-IT2 | -0,846212019 | 0,556243309 | 0,16241 | 0,48945635 | RSF1 intronic transcript 2 |
| ENSG00000270006 | AC010531,6 | -0,846320923 | 0,556201321 | 0,14336 | 0,47083258 | novel transcript, antisense to FBXO31 and C16orf95 readthrough |
| ENSG00000226644 | AL121899,1 | -0,846540529 | 0,556116663 | 0,12868 | 0,45191041 | novel transcript |
| ENSG00000152465 | NMT2 | -0,846604084 | 0,556092165 | 0,03793 | 0,26358169 | N-myristoyltransferase 2 |
| ENSG00000269892 | AC125494,2 | -0,846644264 | 0,556076678 | 0,14112 | 0,4679289 | novel transcript |
| ENSG00000011478 | QPCTL | -0,846691962 | 0,556058293 | 0,11257 | 0,42638197 | glutaminyl-peptide cyclotransferase like |
| ENSG00000213529 | AL359073,1 | -0,847006365 | 0,555937126 | 0,33072 | 0,6112353 | ribosomal protein L32 (RPL32) pseudogene |
| ENSG00000261786 | AC006058,1 | -0,847226491 | 0,555852308 | 0,04255 | 0,2782038 | novel transcript |
| ENSG00000232081 | LARGE-IT1 | -0,847410418 | 0,555781448 | 0,34059 | 0,61904269 | LARGE intronic transcript 1 |
| ENSG00000198796 | ALPK2 | -0,847445526 | 0,555767923 | 0,01339 | 0,15158202 | alpha kinase 2 |
| ENSG00000188037 | CLCN1 | -0,847645274 | 0,555690979 | 0,34085 | 0,61918628 | chloride voltage-gated channel 1 |
| ENSG00000105409 | ATP1A3 | -0,847682628 | 0,555676592 | 0,01182 | 0,1415704 | ATPase Na+/K+ transporting subunit alpha 3 |
| ENSG00000227659 | CLYBL-AS2 | -0,847710346 | 0,555665916 | 0,34456 | 0,62200601 | CLYBL antisense RNA 2 |
| ENSG00000287671 | AC068418,3 | -0,847780973 | 0,555638714 | 0,17197 | 0,49710979 | novel transcript |
| ENSG00000270035 | AL359881,2 | -0,847951435 | 0,555573066 | 0,1918 | 0,51436129 | novel transcript |
| ENSG00000167483 | NIBAN3 | -0,848090971 | 0,555519334 | 0,04195 | 0,27747877 | niban apoptosis regulator 3 |
| ENSG00000285022 | MFHAS1 | -0,8487121 | 0,555280216 | 0,10803 | 0,41857851 | malignant fibrous histiocytoma amplified sequence 1 |
| ENSG00000204296 | TSBP1 | -0,848780962 | 0,555253712 | 0,48784 | 0,72280177 | testis expressed basic protein 1 |
| ENSG00000189164 | ZNF527 | -0,849014905 | 0,555163681 | 0,07411 | 0,35756831 | zinc finger protein 527 |
| ENSG00000229388 | LINC01715 | -0,849259413 | 0,5550696 | 0,32081 | 0,60558211 | long intergenic non-protein coding RNA 1715 |
| ENSG00000237280 | AL136982,3 | -0,849366908 | 0,555028244 | 0,21049 | 0,5290538 | novel transcript |
| ENSG00000176887 | SOX11 | -0,849875876 | 0,55483247 | 0,00361 | 0,07319422 | SRY-box transcription factor 11 |
| ENSG00000276744 | AC105137,2 | -0,850149496 | 0,554727251 | 0,09833 | 0,40379437 | novel transcript, sense intronic to SIN3A |
| ENSG00000198846 | TOX | -0,85036803 | 0,554643229 | 0,03891 | 0,26698555 | thymocyte selection associated high mobility group box |
| ENSG00000284906 | AC091057,5 | -0,850387958 | 0,554635568 | 0,11162 | 0,42472977 | Rho GTPase-activating protein 11B |
| ENSG00000185499 | MUC1 | -0,850993398 | 0,554402859 | 0,01839 | 0,17938469 | mucin 1, cell surface associated |
| ENSG00000286366 | AC005052,2 | -0,851059429 | 0,554377485 | 0,14329 | 0,47082074 | novel transcript, antisense to UPF3B |
| ENSG00000283503 | LINC01902 | -0,851449582 | 0,554227583 | 0,045 | 0,28683057 | long intergenic non-protein coding RNA 1902 |
| ENSG00000229089 | ANKRD20A8P | -0,851491674 | 0,554211413 | 0,05445 | 0,31226071 | ankyrin repeat domain 20 family member A8, pseudogene |
| ENSG00000237940 | LINC01238 | -0,851698733 | 0,554131877 | 0,08645 | 0,38089807 | long intergenic non-protein coding RNA 1238 |
| ENSG00000106077 | ABHD11 | -0,852230623 | 0,553927618 | 0,06682 | 0,34224579 | abhydrolase domain containing 11 |
| ENSG00000108231 | LGI1 | -0,852352709 | 0,553880745 | 0,1843 | 0,50808005 | leucine rich glioma inactivated 1 |

| | | | | | | |
|---|---|---|---|---|---|---|
| ENSG00000286616 | AL008730,1 | -0,852588527 | 0,553790217 | 0,02644 | 0,21901972 | novel transcript |
| ENSG00000168096 | ANKS3 | -0,852736277 | 0,553733505 | 0,04377 | 0,28230121 | ankyrin repeat and sterile alpha motif domain containing 3 |
| ENSG00000054356 | PTPRN | -0,85285265 | 0,553688841 | 0,06583 | 0,33965348 | protein tyrosine phosphatase receptor type N |
| ENSG00000147642 | SYBU | -0,853024806 | 0,553622773 | 0,02067 | 0,1906977 | syntabulin |
| ENSG00000100299 | ARSA | -0,853084561 | 0,553599843 | 0,09105 | 0,39033853 | arylsulfatase A |
| ENSG00000165795 | NDRG2 | -0,85344513 | 0,5534615 | 0,06735 | 0,34355559 | NDRG family member 2 |
| ENSG00000213721 | HMGN2P30 | -0,853956938 | 0,55326519 | 0,33143 | 0,61155735 | high mobility group nucleosomal binding domain 2 pseudogene 30 |
| ENSG00000284734 | AC099063,4 | -0,854128793 | 0,553199289 | 0,07647 | 0,36251887 | novel transcript, antisense to GBP4 |
| ENSG00000263089 | AC007114,2 | -0,854313413 | 0,553128501 | 0,0484 | 0,29551007 | novel transcript, antisense to AKAP1 |
| ENSG00000262172 | AC116025,2 | -0,854324921 | 0,553124089 | 0,29358 | 0,58800349 | novel transcript |
| ENSG00000230638 | AL445933,1 | -0,854511755 | 0,553052462 | 0,42151 | 0,67643128 | novel pseudogene |
| ENSG00000264270 | AC087749,1 | -0,854533254 | 0,553044221 | 0,10429 | 0,41269448 | novel transcript, antisense to SAP30BP |
| ENSG00000135439 | AGAP2 | -0,854993039 | 0,552867994 | 0,02797 | 0,22566447 | ArfGAP with GTPase domain, ankyrin repeat and PH domain 2 |
| ENSG00000196535 | MYO18A | -0,855132181 | 0,552814675 | 0,01613 | 0,1663632 | myosin XVIIIA |
| ENSG00000230683 | DDTP1 | -0,855184159 | 0,552794758 | 0,34863 | 0,62475089 | D-dopachrome tautomerase pseudogene 1 |
| ENSG00000107099 | DOCK8 | -0,855348452 | 0,55273181 | 0,20955 | 0,52856218 | dedicator of cytokinesis 8 |
| ENSG00000272765 | NDUFA6 | -0,855382474 | 0,552718775 | 0,03313 | 0,24586374 | NADH:ubiquinone oxidoreductase subunit A6 |
| ENSG00000272768 | AC004854,2 | -0,855429988 | 0,552700573 | 0,13655 | 0,46380939 | novel transcript, antisense to PURB |
| ENSG00000249240 | AC069368,1 | -0,855830227 | 0,552547261 | 0,26719 | 0,56958284 | novel protein |
| ENSG00000226589 | VARS1 | -0,856087136 | 0,552448875 | 0,0242 | 0,20804286 | valyl-tRNA synthetase 1 |
| ENSG00000238273 | AC108058,1 | -0,856099926 | 0,552443977 | 0,10128 | 0,40805419 | novel transcript, antisense to FHL2 |
| ENSG00000214617 | SLC6A10P | -0,85618801 | 0,552410248 | 0,14649 | 0,47433542 | solute carrier family 6 member 10, pseudogene |
| ENSG00000183208 | GDPGP1 | -0,856543368 | 0,552274198 | 0,0742 | 0,35761188 | GDP-D-glucose phosphorylase 1 |
| ENSG00000136931 | NR5A1 | -0,856605637 | 0,552250361 | 0,46651 | 0,7067229 | nuclear receptor subfamily 5 group A member 1 |
| ENSG00000244490 | RWDD4P1 | -0,856652653 | 0,552232365 | 0,10254 | 0,41028143 | RWD domain containing 4 pseudogene 1 |
| ENSG00000285658 | AC127035,1 | -0,856751658 | 0,552194469 | 0,11898 | 0,43696075 | novel transcript, antisense to AMBRA1 |
| ENSG00000271239 | AC007423,1 | -0,856857232 | 0,552154062 | 0,36019 | 0,63261458 | novel transcript |
| ENSG00000226334 | AL359182,1 | -0,857104105 | 0,552059586 | 0,16406 | 0,4903483 | novel transcript |
| ENSG00000175906 | ARL4D | -0,857125088 | 0,552051556 | 0,05984 | 0,32535617 | ADP ribosylation factor like GTPase 4D |
| ENSG00000179761 | PIPOX | -0,85714472 | 0,552044044 | 0,01159 | 0,14000409 | pipecolic acid and sarcosine oxidase |
| ENSG00000246792 | AC106038,1 | -0,857453551 | 0,551925883 | 0,10209 | 0,40934973 | novel transcript |

| | | | | | | |
|---|---|---|---|---|---|---|
| ENSG00000148814 | LRRC27 | -0,857600188 | 0,551869788 | 0,03078 | 0,23675795 | leucine rich repeat containing 27 |
| ENSG00000183307 | TMEM121B | -0,857939023 | 0,551740189 | 0,0021 | 0,05154257 | transmembrane protein 121B |
| ENSG00000285619 | AL449214,1 | -0,858147812 | 0,551660347 | 0,13777 | 0,46481656 | Novel transcript, antisense to EEFSEC |
| ENSG00000260084 | AC126773,3 | -0,858336116 | 0,551588347 | 0,2794 | 0,57742338 | novel transcript, antisense to ZFP90 |
| ENSG00000287892 | AC090907,3 | -0,858357167 | 0,551580299 | 0,46321 | 0,70415312 | novel transcript, antisense to LRRK1 |
| ENSG00000034053 | APBA2 | -0,858513591 | 0,551520497 | 0,06676 | 0,34201747 | amyloid beta precursor protein binding family A member 2 |
| ENSG00000276833 | TAF15 | -0,858698281 | 0,551449897 | 0,02853 | 0,22839158 | TATA-box binding protein associated factor 15 |
| ENSG00000237352 | LINC01358 | -0,858921767 | 0,55136448 | 0,04416 | 0,28322445 | long intergenic non-protein coding RNA 1358 |
| ENSG00000277491 | AC087392,5 | -0,859065961 | 0,551309375 | 0,18944 | 0,51258824 | novel transcript |
| ENSG00000223510 | CDRT15 | -0,859149622 | 0,551277405 | 0,3582 | 0,63185282 | CMT1A duplicated region transcript 15 |
| ENSG00000287431 | AC027601,5 | -0,859283509 | 0,551226247 | 0,07298 | 0,35631792 | novel transcript |
| ENSG00000259744 | AC009269,4 | -0,859297724 | 0,551220816 | 0,3745 | 0,64298972 | novel transcript, sense intronic to LARP6 |
| ENSG00000282083 | AC015849,7 | -0,859551527 | 0,551123852 | 0,23318 | 0,54558536 | novel transcript, sense overlapping TAF15 |
| ENSG00000229589 | ACVR2B-AS1 | -0,859675067 | 0,551076661 | 0,07941 | 0,3679672 | ACVR2B antisense RNA 1 |
| ENSG00000172824 | CES4A | -0,859776448 | 0,551037937 | 0,03186 | 0,24068538 | carboxylesterase 4A |
| ENSG00000225914 | TSBP1-AS1 | -0,859790397 | 0,551032609 | 0,04929 | 0,29862897 | TSBP1 and BTNL2 antisense RNA 1 |
| ENSG00000226711 | FAM66C | -0,860106797 | 0,550911775 | 0,00482 | 0,08564971 | family with sequence similarity 66 member C |
| ENSG00000141258 | SGSM2 | -0,860240857 | 0,550860584 | 0,00052 | 0,02096524 | small G protein signaling modulator 2 |
| ENSG00000286957 | AC079776,5 | -0,860371623 | 0,550810657 | 0,16887 | 0,49495623 | novel transcript |
| ENSG00000281398 | SNHG4 | -0,860434606 | 0,550786611 | 0,14252 | 0,46989963 | small nucleolar RNA host gene 4 |
| ENSG00000151079 | KCNA6 | -0,860749365 | 0,550666456 | 0,00533 | 0,09086355 | potassium voltage-gated channel subfamily A member 6 |
| ENSG00000266947 | AC022916,1 | -0,860774362 | 0,550656915 | 0,04205 | 0,27770943 | novel transcript, antisense SLFN5 |
| ENSG00000234159 | RBPMSLP | -0,860786265 | 0,550652372 | 0,24249 | 0,55137204 | RNA binding protein with multiple splicing-like pseudogene |
| ENSG00000178665 | ZNF713 | -0,860847882 | 0,550628854 | 0,06242 | 0,33099652 | zinc finger protein 713 |
| ENSG00000198563 | DDX39B | -0,860854095 | 0,550626483 | 0,06065 | 0,32698319 | DExD-box helicase 39B |
| ENSG00000197093 | GAL3ST4 | -0,860892422 | 0,550611855 | 0,04568 | 0,28842796 | galactose-3-O-sulfotransferase 4 |
| ENSG00000285230 | RALY-AS1 | -0,861105016 | 0,550530724 | 0,0709 | 0,35165018 | RALY antisense RNA 1 |
| ENSG00000206811 | SNORA10 | -0,861126896 | 0,550522374 | 0,28149 | 0,57891473 | small nucleolar RNA, H/ACA box 10 |
| ENSG00000222460 | RN7SKP271 | -0,861327385 | 0,550445874 | 0,18208 | 0,5053975 | RN7SK pseudogene 271 |
| ENSG00000239556 | AC004951,2 | -0,861419784 | 0,550410622 | 0,18787 | 0,51107563 | uroplakin-like protein (UPLP) pseudogene |
| ENSG00000223496 | EXOSC6 | -0,861465954 | 0,550393007 | 0,0597 | 0,32521812 | exosome component 6 |
| ENSG00000172020 | GAP43 | -0,861855127 | 0,550244557 | 0,05351 | 0,31052177 | growth associated protein 43 |

| | | | | | | |
|---|---|---|---|---|---|---|
| ENSG00000176593 | AC008969,1 | -0,862142175 | 0,550135087 | 0,01942 | 0,18462928 | novel transcript, antisense to ZNF606 |
| ENSG00000139549 | DHH | -0,862392894 | 0,55003949 | 0,27528 | 0,57491355 | desert hedgehog signaling molecule |
| ENSG00000074706 | IPCEF1 | -0,862414396 | 0,550031292 | 0,02738 | 0,22321357 | interaction protein for cytohesin exchange factors 1 |
| ENSG00000090581 | GNPTG | -0,862696726 | 0,549923664 | 0,09447 | 0,39679147 | N-acetylglucosamine-1-phosphate transferase subunit gamma |
| ENSG00000260644 | HERC2P5 | -0,862977067 | 0,549816814 | 0,06956 | 0,34818658 | hect domain and RLD 2 pseudogene 5 |
| ENSG00000273759 | AL117379,1 | -0,863576065 | 0,549588581 | 0,10313 | 0,41083338 | novel transcript |
| ENSG00000272447 | AL135925,1 | -0,863784266 | 0,549509274 | 0,07869 | 0,36645437 | novel transcript |
| ENSG00000267117 | AC010525,1 | -0,864140982 | 0,549373421 | 0,48232 | 0,71892541 | novel transcript, antisense to EPN1 |
| ENSG00000085563 | ABCB1 | -0,864429873 | 0,549263423 | 0,18072 | 0,50425112 | ATP binding cassette subfamily B member 1 |
| ENSG00000246526 | LINC02481 | -0,865133584 | 0,548995571 | 0,21429 | 0,53334744 | long intergenic non-protein coding RNA 2481 |
| ENSG00000248508 | SRP14-AS1 | -0,865188483 | 0,548974681 | 0,04374 | 0,28225644 | SRP14 antisense RNA1 (head to head) |
| ENSG00000249102 | AC034223,1 | -0,865320476 | 0,548924457 | 0,03483 | 0,2523169 | novel transcript |
| ENSG00000198723 | TEX45 | -0,865355135 | 0,54891127 | 0,22123 | 0,53805234 | testis expressed 45 |
| ENSG00000267952 | AC008878,1 | -0,865367976 | 0,548906384 | 0,49715 | 0,72865531 | novel transcript |
| ENSG00000236069 | AL136987,1 | -0,865550412 | 0,548836976 | 0,1456 | 0,47304491 | novel transcript |
| ENSG00000128604 | IRF5 | -0,865936241 | 0,548690217 | 0,01701 | 0,17122091 | interferon regulatory factor 5 |
| ENSG00000179818 | PCBP1-AS1 | -0,866015693 | 0,548660001 | 0,06472 | 0,33727693 | PCBP1 antisense RNA 1 |
| ENSG00000177181 | RIMKLA | -0,866204449 | 0,548588221 | 0,02597 | 0,21717729 | ribosomal modification protein rimK like family member A |
| ENSG00000174276 | ZNHIT2 | -0,866228026 | 0,548579256 | 0,09479 | 0,39724142 | zinc finger HIT-type containing 2 |
| ENSG00000249278 | AC112249,1 | -0,86671117 | 0,548395573 | 0,00924 | 0,12269349 | ribosomal protein L18 (RPL18) pseudogene |
| ENSG00000277311 | AC012354,7 | -0,866790143 | 0,548365555 | 0,35002 | 0,62564488 | |
| ENSG00000164237 | CMBL | -0,867379352 | 0,548141643 | 0,10331 | 0,41114168 | carboxymethylenebutenolidase homolog |
| ENSG00000288061 | AP000926,2 | -0,86763635 | 0,548044007 | 0,15846 | 0,485647 | novel transcript, antisense to HSPA8and CLMP |
| ENSG00000168350 | DEGS2 | -0,867846185 | 0,547964302 | 0,03913 | 0,26771146 | delta 4-desaturase, sphingolipid 2 |
| ENSG00000237892 | KLF7-IT1 | -0,868085326 | 0,547873479 | 0,0131 | 0,14972794 | KLF7 intronic transcript 1 |
| ENSG00000283010 | AL589826,2 | -0,868165837 | 0,547842905 | 0,31732 | 0,60276767 | novel transcript |
| ENSG00000159335 | PTMS | -0,868221017 | 0,547821952 | 0,06164 | 0,32867767 | parathymosin |
| ENSG00000202058 | RN7SKP80 | -0,868370607 | 0,547765152 | 0,14801 | 0,4751727 | RN7SK pseudogene 80 |
| ENSG00000234618 | RPSAP9 | -0,868386713 | 0,547759037 | 0,2924 | 0,58694246 | ribosomal protein SA pseudogene 9 |
| ENSG00000275850 | AC005912,2 | -0,868421424 | 0,547745859 | 0,2066 | 0,52622584 | novel transcript |
| ENSG00000285938 | AC072022,2 | -0,868771269 | 0,547613049 | 0,28354 | 0,58068945 | BCL6 antisense 1 |
| ENSG00000254536 | AL360181,3 | -0,869058169 | 0,54750416 | 0,04305 | 0,27943834 | novel protein |

| | | | | | | |
|---|---|---|---|---|---|---|
| ENSG00000253357 | AC008708,1 | -0,869178078 | 0,547458656 | 0,1581 | 0,48511883 | novel transcript |
| ENSG00000181396 | OGFOD3 | -0,86928227 | 0,54741912 | 0,0416 | 0,27632896 | 2-oxoglutarate and iron dependent oxygenase domain containing 3 |
| ENSG00000146005 | PSD2 | -0,869291872 | 0,547415477 | 0,04616 | 0,28930497 | pleckstrin and Sec7 domain containing 2 |
| ENSG00000168970 | JMJD7-PLA2G | -0,869514161 | 0,547331138 | 0,03157 | 0,23991209 | JMJD7-PLA2G4B readthrough |
| ENSG00000233998 | SETP5 | -0,869649571 | 0,547279768 | 0,17735 | 0,50187931 | SET pseudogene 5 |
| ENSG00000234946 | SDHCP3 | -0,86971076 | 0,547256557 | 0,27943 | 0,57742338 | succinate dehydrogenase complex subunit C pseudogene 3 |
| ENSG00000287245 | AP001981,2 | -0,869760101 | 0,547237841 | 0,30346 | 0,59370687 | novel transcript, antisense to RDX |
| ENSG00000136938 | ANP32B | -0,869971183 | 0,54715778 | 0,07356 | 0,35672172 | acidic nuclear phosphoprotein 32 family member B |
| ENSG00000261889 | AC108134,2 | -0,870131025 | 0,547097161 | 0,05422 | 0,31168784 | novel transcript |
| ENSG00000277586 | NEFL | -0,870257499 | 0,547049202 | 0,00202 | 0,0505329 | neurofilament light |
| ENSG00000235488 | JARID2-AS1 | -0,870905205 | 0,546803656 | 0,18661 | 0,50999177 | JARID2 antisense RNA 1 |
| ENSG00000279159 | AC003681,1 | -0,871103132 | 0,546728644 | 0,06801 | 0,34456193 | novel transcript, antisense to MTMR3 |
| ENSG00000270049 | AC009061,2 | -0,871468763 | 0,546590101 | 0,0177 | 0,17522134 | novel transcript, antisense to AGRP |
| ENSG00000254370 | AC025871,2 | -0,871954453 | 0,54640612 | 0,25327 | 0,55836184 | novel transcript |
| ENSG00000267365 | KCNJ2-AS1 | -0,871963665 | 0,546402631 | 0,03208 | 0,24164117 | KCNJ2 antisense RNA 1 |
| ENSG00000166946 | CCNDBP1 | -0,872089741 | 0,546354883 | 0,03856 | 0,26560956 | cyclin D1 binding protein 1 |
| ENSG00000131944 | FAAP24 | -0,872234041 | 0,546300239 | 0,05469 | 0,31313954 | FA core complex associated protein 24 |
| ENSG00000135218 | CD36 | -0,872750092 | 0,546104863 | 0,15944 | 0,48684894 | CD36 molecule |
| ENSG00000263326 | AC133552,4 | -0,873738736 | 0,545730758 | 0,17253 | 0,49744497 | thyroid hormone receptor interactor 11 (TRIP11) pseudogene |
| ENSG00000233246 | PHC2-AS1 | -0,873817972 | 0,545700786 | 0,38283 | 0,64953876 | PHC2 antisense RNA 1 |
| ENSG00000283883 | VPS53 | -0,874458562 | 0,545458536 | 0,08633 | 0,38087445 | VPS53 subunit of GARP complex |
| ENSG00000215492 | HNRNPA1P7 | -0,874497152 | 0,545443947 | 0,07644 | 0,36251343 | heterogeneous nuclear ribonucleoprotein A1 pseudogene 7 |
| ENSG00000260852 | FBXL19-AS1 | -0,874563303 | 0,545418937 | 0,05762 | 0,32039326 | FBXL19 antisense RNA 1 |
| ENSG00000132024 | CC2D1A | -0,874599242 | 0,545405351 | 0,02207 | 0,19759515 | coiled-coil and C2 domain containing 1A |
| ENSG00000257534 | AC023794,4 | -0,874606942 | 0,545402439 | 0,25995 | 0,56377449 | novel transcript |
| ENSG00000173272 | MZT2A | -0,874933719 | 0,545278917 | 0,03153 | 0,23977194 | mitotic spindle organizing protein 2A |
| ENSG00000258839 | MC1R | -0,875215017 | 0,545172609 | 0,06103 | 0,32783036 | melanocortin 1 receptor |
| ENSG00000286449 | AC016590,4 | -0,875634659 | 0,545014055 | 0,2084 | 0,52769715 | novel transcript |
| ENSG00000128536 | CDHR3 | -0,875839745 | 0,544936584 | 0,17802 | 0,50235771 | cadherin related family member 3 |
| ENSG00000259438 | MAPK6-DT | -0,876091589 | 0,544841466 | 0,43764 | 0,68737508 | MAPK6 divergent transcript |

| | | | | | | |
|---|---|---|---|---|---|---|
| ENSG00000263740 | RN7SL4P | -0,876222659 | 0,544791969 | 0,05465 | 0,31304183 | RNA, 7SL, cytoplasmic 4, pseudogene |
| ENSG00000231431 | FAR2P4 | -0,876497634 | 0,544688143 | 0,09349 | 0,39516351 | fatty acyl-CoA reductase 2 pseudogene 4 |
| ENSG00000164418 | GRIK2 | -0,87661408 | 0,54464418 | 0,01889 | 0,18219113 | glutamate ionotropic receptor kainate type subunit 2 |
| ENSG00000237101 | AC092809,4 | -0,87670932 | 0,544608227 | 0,36852 | 0,63821729 | novel transcript |
| ENSG00000244642 | RN7SL396P | -0,876722865 | 0,544603113 | 0,27003 | 0,57184721 | RNA, 7SL, cytoplasmic 396, pseudogene |
| ENSG00000267321 | SNHG30 | -0,876730753 | 0,544600136 | 0,06837 | 0,34559843 | small nucleolar RNA host gene 30 |
| ENSG00000261826 | AC121333,1 | -0,877069308 | 0,54447235 | 0,05582 | 0,31585374 | novel transcript |
| ENSG00000251613 | AC063979,1 | -0,877370654 | 0,544358634 | 0,26395 | 0,56649862 | novel transcript |
| ENSG00000259111 | AL079307,2 | -0,877450064 | 0,544328672 | 0,55913 | 0,77044204 | novel transcript |
| ENSG00000254231 | AC103760,1 | -0,877451655 | 0,544328072 | 0,14933 | 0,47695763 | novel transcript |
| ENSG00000173013 | CCDC96 | -0,877495941 | 0,544311363 | 0,08648 | 0,38091118 | coiled-coil domain containing 96 |
| ENSG00000215325 | ASS1P10 | -0,87757202 | 0,54428266 | 0,05446 | 0,31226071 | argininosuccinate synthetase 1 pseudogene 10 |
| ENSG00000280433 | FP565260,6 | -0,877658288 | 0,544250115 | 0,13969 | 0,4668467 | novel protein, similar to trafficking protein particle complex 10 TRAPPC10 |
| ENSG00000188277 | C15orf62 | -0,877745928 | 0,544217054 | 0,09725 | 0,40176121 | chromosome 15 open reading frame 62 |
| ENSG00000238098 | ABCA17P | -0,877912819 | 0,544154103 | 0,197 | 0,51824743 | ATP binding cassette subfamily A member 17, pseudogene |
| ENSG00000282508 | LINC01002 | -0,877955478 | 0,544138013 | 0,06687 | 0,34241464 | long intergenic non-protein coding RNA 1002 |
| ENSG00000236567 | TCF3P1 | -0,877990659 | 0,544124744 | 0,28177 | 0,57909527 | transcription factor 3 pseudogene 1 |
| ENSG00000229227 | AL356056,1 | -0,878107058 | 0,544080845 | 0,09767 | 0,40273014 | novel transcript |
| ENSG00000238231 | AL512288,2 | -0,878209685 | 0,544042143 | 0,27152 | 0,57285347 | interferon-induced transmembrane protein pseudogene |
| ENSG00000118046 | STK11 | -0,878258828 | 0,544023611 | 0,05628 | 0,31687263 | serine/threonine kinase 11 |
| ENSG00000182463 | TSHZ2 | -0,878398824 | 0,543970823 | 0,00049 | 0,02013867 | teashirt zinc finger homeobox 2 |
| ENSG00000287093 | AC003093,1 | -0,87891107 | 0,543777714 | 0,32935 | 0,61040605 | novel transcript |
| ENSG00000232882 | PHKA1P1 | -0,879618158 | 0,543511265 | 0,05221 | 0,30683813 | phosphorylase kinase regulatory subunit alpha 1 pseudogene |
| ENSG00000259356 | AC022710,1 | -0,879842992 | 0,543426569 | 0,44671 | 0,69321646 | novel transcript, antisense to ADAMTS17 |
| ENSG00000176125 | UFSP1 | -0,879953899 | 0,543384795 | 0,12635 | 0,44903997 | UFM1 specific peptidase 1 (inactive) |
| ENSG00000008300 | CELSR3 | -0,880150863 | 0,543310614 | 0,00053 | 0,02135634 | cadherin EGF LAG seven-pass G-type receptor 3 |
| ENSG00000278921 | EPB41L4A-DT | -0,880166093 | 0,543304879 | 0,19414 | 0,51562158 | EPB41L4A divergent transcript |
| ENSG00000164199 | ADGRV1 | -0,880186237 | 0,543297293 | 0,13939 | 0,46666406 | adhesion G protein-coupled receptor V1 |
| ENSG00000198774 | RASSF9 | -0,880233786 | 0,543279387 | 0,15783 | 0,48490111 | Ras association domain family member 9 |
| ENSG00000187994 | RINL | -0,880515958 | 0,543173139 | 0,2282 | 0,54221266 | Ras and Rab interactor like |
| ENSG00000288087 | AC112770,1 | -0,880565961 | 0,543154313 | 0,14433 | 0,47189017 | novel transcript |

| | | | | | | |
|---|---|---|---|---|---|---|
| ENSG00000236939 | BAALC-AS2 | -0,880656836 | 0,543120101 | 0,2326 | 0,54529383 | BAALC antisense RNA 2 |
| ENSG00000172995 | ARPP21 | -0,880813951 | 0,543060957 | 0,20104 | 0,52181085 | cAMP regulated phosphoprotein 21 |
| ENSG00000263968 | RN7SL381P | -0,880916911 | 0,543022201 | 0,2183 | 0,535736 | RNA, 7SL, cytoplasmic 381, pseudogene |
| ENSG00000257543 | AC063948,1 | -0,881290959 | 0,54288143 | 0,33413 | 0,61394154 | novel transcript |
| ENSG00000214940 | NPIPA8 | -0,88157207 | 0,542775659 | 0,33992 | 0,618752 | nuclear pore complex interacting protein family member A8 |
| ENSG00000254186 | AC113414,1 | -0,881830281 | 0,542678523 | 0,04924 | 0,29853148 | novel transcript |
| ENSG00000015133 | CCDC88C | -0,882186142 | 0,54254468 | 0,01807 | 0,17747695 | coiled-coil domain containing 88C |
| ENSG00000205176 | REXO1L1P | -0,882198954 | 0,542539862 | 0,05965 | 0,32519243 | REXO1 like 1, pseudogene |
| ENSG00000261872 | AC002558,2 | -0,882616027 | 0,54238304 | 0,13319 | 0,45912454 | novel transcript, antisense to CDC27 |
| ENSG00000266998 | AC111182,1 | -0,882679612 | 0,542359136 | 0,12522 | 0,44695975 | novel transcript, antisense to SEPT9 |
| ENSG00000261220 | AC103706,1 | -0,882866603 | 0,542288844 | 0,16221 | 0,48939373 | novel transcript |
| ENSG00000278613 | NAIP | -0,882987423 | 0,542243432 | 0,08846 | 0,38476379 | NLR family apoptosis inhibitory protein |
| ENSG00000186960 | LINC01551 | -0,883013296 | 0,542233707 | 0,19227 | 0,51477601 | long intergenic non-protein coding RNA 1551 |
| ENSG00000143921 | ABCG8 | -0,883137572 | 0,542187 | 0,14489 | 0,47244928 | ATP binding cassette subfamily G member 8 |
| ENSG00000272040 | AC010245,2 | -0,883282207 | 0,542132647 | 0,19174 | 0,51436129 | novel transcript |
| ENSG00000264304 | AC024267,3 | -0,883328791 | 0,542115142 | 0,03717 | 0,26108134 | novel transcript, antisense to FLOT2 |
| ENSG00000287482 | AC124312,5 | -0,883709551 | 0,541972084 | 0,53113 | 0,75245598 | novel transcript |
| ENSG00000166997 | CNPY4 | -0,883767762 | 0,541950217 | 0,07745 | 0,36441467 | canopy FGF signaling regulator 4 |
| ENSG00000257568 | AC079035,1 | -0,88391313 | 0,541895612 | 0,06652 | 0,34124715 | novel transcript, antisense to MON2 |
| ENSG00000185245 | GP1BA | -0,883973895 | 0,541872788 | 0,08521 | 0,37857645 | glycoprotein Ib platelet subunit alpha |
| ENSG00000286652 | AL021918,5 | -0,884217083 | 0,541781455 | 0,33459 | 0,61433133 | novel transcript |
| ENSG00000225889 | AC012368,1 | -0,884352152 | 0,541730734 | 0,01655 | 0,16855868 | novel transcript, antisense to PELI1 |
| ENSG00000251441 | RTEL1P1 | -0,88438742 | 0,541717491 | 0,19529 | 0,51611615 | regulator of telomere elongation helicase 1 pseudogene 1 |
| ENSG00000164049 | FBXW12 | -0,885012186 | 0,541482949 | 0,07618 | 0,36240925 | F-box and WD repeat domain containing 12 |
| ENSG00000273186 | AL359091,4 | -0,886072849 | 0,541085 | 0,3723 | 0,64126849 | novel transcript, sense intronic to CERCAM |
| ENSG00000175182 | FAM131A | -0,886294907 | 0,541001723 | 0,0132 | 0,15012165 | family with sequence similarity 131 member A |
| ENSG00000237499 | WAKMAR2 | -0,886490958 | 0,54092821 | 0,03661 | 0,25940941 | wound and keratinocyte migration associated lncRNA 2 |
| ENSG00000274220 | AC009163,7 | -0,886569344 | 0,54089882 | 0,06011 | 0,32568394 | novel transcript |
| ENSG00000276480 | MYH11 | -0,886790743 | 0,540815819 | 0,08078 | 0,37087458 | myosin heavy chain 11 |
| ENSG00000182584 | ACTL10 | -0,886955011 | 0,540754245 | 0,02412 | 0,20758258 | actin like 10 |
| ENSG00000199377 | RNU5F-1 | -0,886966271 | 0,540750024 | 0,07626 | 0,36242671 | RNA, U5F small nuclear 1 |
| ENSG00000267547 | AC060766,4 | -0,886993309 | 0,540739889 | 0,01283 | 0,14814468 | novel transcript |

| | | | | | | |
|---|---|---|---|---|---|---|
| ENSG00000225361 | PPP1R26-AS1 | -0,887014351 | 0,540732003 | 0,04525 | 0,2876168 | PPP1R26 antisense RNA 1 |
| ENSG00000100505 | TRIM9 | -0,88728638 | 0,540630054 | 0,01999 | 0,18755241 | tripartite motif containing 9 |
| ENSG00000275936 | AC004263,1 | -0,887288608 | 0,540629219 | 0,1803 | 0,50407792 | novel transcript, antisense to PXN |
| ENSG00000071073 | MGAT4A | -0,887301307 | 0,54062446 | 0,00563 | 0,09352688 | alpha-1,3-mannosyl-glycoprotein 4-beta-N-acetylglucosaminyltransferase A |
| ENSG00000286011 | AC062037,3 | -0,8876719 | 0,540485605 | 0,28962 | 0,58522301 | novel transcript |
| ENSG00000204438 | GPANK1 | -0,887702163 | 0,540474268 | 0,09689 | 0,40110418 | G-patch domain and ankyrin repeats 1 |
| ENSG00000232117 | LINC00384 | -0,887956569 | 0,540378968 | 0,28036 | 0,57784691 | long intergenic non-protein coding RNA 384 |
| ENSG00000260785 | CASC17 | -0,888016959 | 0,540356349 | 0,01779 | 0,17571755 | cancer susceptibility 17 |
| ENSG00000104936 | DMPK | -0,888065475 | 0,540338178 | 0,03346 | 0,24712306 | DM1 protein kinase |
| ENSG00000153714 | LURAP1L | -0,888140417 | 0,54031011 | 0,03165 | 0,24008374 | leucine rich adaptor protein 1 like |
| ENSG00000182902 | SLC25A18 | -0,888200499 | 0,540287609 | 0,05326 | 0,30991339 | solute carrier family 25 member 18 |
| ENSG00000287016 | AC010996,1 | -0,888370108 | 0,540224095 | 0,13505 | 0,46159318 | novel transcript |
| ENSG00000286118 | AC234031,1 | -0,888918386 | 0,540018828 | 0,04522 | 0,2876168 | novel transcript, antisense to FAM156A |
| ENSG00000219445 | AC019155,2 | -0,888934781 | 0,540012692 | 0,33393 | 0,61382024 | novel transcript |
| ENSG00000244694 | PTCHD4 | -0,889260592 | 0,539890752 | 0,08399 | 0,37651068 | patched domain containing 4 |
| ENSG00000111305 | GSG1 | -0,889270666 | 0,539886982 | 0,05233 | 0,30711015 | germ cell associated 1 |
| ENSG00000287427 | AL390236,1 | -0,889668039 | 0,539738297 | 0,51235 | 0,73924429 | novel transcript, antisense to PTPRE |
| ENSG00000260369 | AC120024,1 | -0,889680276 | 0,539733718 | 0,16499 | 0,49122708 | novel transcript |
| ENSG00000205913 | SRRM2-AS1 | -0,889999867 | 0,539614168 | 0,04583 | 0,288874 | SRRM2 antisense RNA 1 |
| ENSG00000288385 | MG910337,1 | -0,890355224 | 0,53948127 | 0,24629 | 0,55395309 | |
| ENSG00000227039 | ITGB2-AS1 | -0,890464128 | 0,539440547 | 0,21567 | 0,53440695 | ITGB2 antisense RNA 1 |
| ENSG00000119820 | YIPF4 | -0,89063978 | 0,539374873 | 0,05556 | 0,31503413 | Yip1 domain family member 4 |
| ENSG00000116544 | DLGAP3 | -0,89066749 | 0,539364513 | 0,01156 | 0,1398032 | DLG associated protein 3 |
| ENSG00000228808 | HMGB3P4 | -0,890742324 | 0,539336537 | 0,15811 | 0,48511883 | high mobility group box 3 pseudogene 4 |
| ENSG00000068078 | FGFR3 | -0,890798671 | 0,539315473 | 0,14744 | 0,47492437 | fibroblast growth factor receptor 3 |
| ENSG00000064205 | CCN5 | -0,890900677 | 0,539277341 | 0,13605 | 0,46310781 | cellular communication network factor 5 |
| ENSG00000263717 | AC079336,1 | -0,891072607 | 0,539213078 | 0,36785 | 0,63780197 | novel transcript, sense intronic MYO1D |
| ENSG00000168591 | TMUB2 | -0,891195607 | 0,539167108 | 0,066 | 0,34008244 | transmembrane and ubiquitin like domain containing 2 |
| ENSG00000213058 | AL365357,1 | -0,891404593 | 0,539089011 | 0,37902 | 0,64689879 | ribosomal protein S14 (RPS14) pseudogene |
| ENSG00000139725 | RHOF | -0,891551064 | 0,539034282 | 0,10598 | 0,4154675 | ras homolog family member F, filopodia associated |
| ENSG00000258303 | AC012464,2 | -0,891841493 | 0,53892578 | 0,44408 | 0,6916877 | novel transcript |

| | | | | | | |
|---|---|---|---|---|---|---|
| ENSG00000260103 | AC012435,2 | -0,891865991 | 0,538916629 | 0,45322 | 0,69750019 | coiled-coil-helix-coiled-coil-helix domain-containing protein 7 (CHCHD7) pseudogene |
| ENSG00000154889 | MPPE1 | -0,891997167 | 0,538867631 | 0,07326 | 0,35655334 | metallophosphoesterase 1 |
| ENSG00000258654 | AC026495,1 | -0,892143662 | 0,538812915 | 0,07314 | 0,35655334 | novel transcript |
| ENSG00000285554 | AC242988,2 | -0,892270699 | 0,538765472 | 0,16406 | 0,4903483 | Novel transcript |
| ENSG00000214081 | CYP4F30P | -0,892366377 | 0,538729743 | 0,3261 | 0,6085098 | cytochrome P450 family 4 subfamily F member 30, pseudogene |
| ENSG00000281591 | DBET | -0,892665854 | 0,538617924 | 0,45216 | 0,69691985 | D4Z4 binding element transcript |
| ENSG00000288543 | AL035409,2 | -0,892792568 | 0,538570619 | 0,1451 | 0,47275989 | novel transcript |
| ENSG00000233115 | FAM90A11P | -0,892886266 | 0,538535642 | 0,05898 | 0,3241112 | family with sequence similarity 90 member A11, pseudogene |
| ENSG00000259242 | AC002306,1 | -0,892949897 | 0,53851189 | 0,24714 | 0,55483158 | novel transcript, antisense to PBX4 |
| ENSG00000134072 | CAMK1 | -0,893115184 | 0,538450197 | 0,00796 | 0,11367829 | calcium/calmodulin dependent protein kinase I |
| ENSG00000258924 | AC002094,1 | -0,893242561 | 0,538402659 | 0,15785 | 0,48490111 | novel transcript readthrough between TMEM199 and SARM1 |
| ENSG00000168309 | FAM107A | -0,893392038 | 0,538346878 | 0,06492 | 0,33778346 | family with sequence similarity 107 member A |
| ENSG00000254923 | AC130366,1 | -0,893395768 | 0,538345486 | 0,43001 | 0,68268477 | deubiquitinating enzyme 3 pseudogene |
| ENSG00000100726 | TELO2 | -0,893424605 | 0,538334725 | 0,0541 | 0,31168784 | telomere maintenance 2 |
| ENSG00000246273 | SBF2-AS1 | -0,893551344 | 0,538287435 | 0,03133 | 0,2388853 | SBF2 antisense RNA 1 |
| ENSG00000260233 | ZNRD2-AS1 | -0,893784704 | 0,538200373 | 0,08645 | 0,38089807 | ZNRD2 antisense RNA 1 (head to head) |
| ENSG00000106123 | EPHB6 | -0,893833182 | 0,538182289 | 0,07713 | 0,36384125 | EPH receptor B6 |
| ENSG00000184659 | FOXD4L4 | -0,894134981 | 0,538069717 | 0,07683 | 0,36335899 | forkhead box D4 like 4 |
| ENSG00000090971 | NAT14 | -0,894154413 | 0,53806247 | 0,05018 | 0,30107555 | N-acetyltransferase 14 (putative) |
| ENSG00000106617 | PRKAG2 | -0,89428204 | 0,538014873 | 0,05982 | 0,32535617 | protein kinase AMP-activated non-catalytic subunit gamma 2 |
| ENSG00000184274 | LINC00315 | -0,894566362 | 0,537908853 | 0,1639 | 0,4902029 | long intergenic non-protein coding RNA 315 |
| ENSG00000134146 | DPH6 | -0,894593517 | 0,537898728 | 0,05667 | 0,31746133 | diphthamine biosynthesis 6 |
| ENSG00000283843 | AC113189,6 | -0,894726872 | 0,53784901 | 0,52557 | 0,74917643 | novel transcript |
| ENSG00000262880 | AC113189,2 | -0,894726872 | 0,53784901 | 0,52557 | 0,74917643 | novel transcript |
| ENSG00000276570 | AC010327,6 | -0,89490111 | 0,537784057 | 0,04551 | 0,28824362 | novel transcript |
| ENSG00000250620 | LINC02515 | -0,895093948 | 0,537712178 | 0,1001 | 0,4066747 | long intergenic non-protein coding RNA 2515 |
| ENSG00000269356 | AL356740,2 | -0,895813451 | 0,537444077 | 0,27074 | 0,57239492 | novel transcript |
| ENSG00000240137 | ERICH6-AS1 | -0,896059842 | 0,537352297 | 0,05585 | 0,31590799 | ERICH6 antisense RNA 1 |

| | | | | | | |
|---|---|---|---|---|---|---|
| ENSG00000104731 | KLHDC4 | -0,896105223 | 0,537335394 | 0,03263 | 0,24385689 | kelch domain containing 4 |
| ENSG00000100478 | AP4S1 | -0,896119036 | 0,53733025 | 0,06214 | 0,33018211 | adaptor related protein complex 4 subunit sigma 1 |
| ENSG00000258527 | ASB9P1 | -0,896213115 | 0,537295211 | 0,21703 | 0,53505085 | ankyrin repeat and SOCS box containing 9 pseudogene 1 |
| ENSG00000196979 | GPRACR | -0,896455204 | 0,537205059 | 0,27238 | 0,57315552 | GPR107 adjacent cis regulating lncRNA |
| ENSG00000285548 | AC068725,1 | -0,896470777 | 0,53719926 | 0,26682 | 0,56923396 | Novel transcript, antisense to NRXN1 |
| ENSG00000281708 | ERC2-IT1 | -0,896504894 | 0,537186557 | 0,11416 | 0,42951506 | ERC2 intronic transcript 1 |
| ENSG00000172201 | ID4 | -0,896774041 | 0,537086349 | 0,009 | 0,12038221 | inhibitor of DNA binding 4, HLH protein |
| ENSG00000109265 | CRACD | -0,896989426 | 0,537006172 | 0,01096 | 0,13522945 | capping protein inhibiting regulator of actin dynamics |
| ENSG00000259953 | AL138756,1 | -0,89713053 | 0,536953652 | 0,01143 | 0,13882136 | novel transcript, overlapping SUSD1 |
| ENSG00000171931 | FBXW10 | -0,897293684 | 0,536892931 | 0,17686 | 0,50122202 | F-box and WD repeat domain containing 10 |
| ENSG00000169598 | DFFB | -0,897931512 | 0,536655619 | 0,10885 | 0,42013578 | DNA fragmentation factor subunit beta |
| ENSG00000223680 | DDR1 | -0,898080375 | 0,536600248 | 0,35188 | 0,62699036 | discoidin domain receptor tyrosine kinase 1 |
| ENSG00000166819 | PLIN1 | -0,898185016 | 0,536561328 | 0,36358 | 0,63439911 | perilipin 1 |
| ENSG00000105523 | FAM83E | -0,898402323 | 0,536480514 | 0,28463 | 0,58147833 | family with sequence similarity 83 member E |
| ENSG00000278600 | AC015871,3 | -0,898460439 | 0,536458904 | 0,02858 | 0,22861157 | novel transcript, sense intronic to ST20 |
| ENSG00000183696 | UPP1 | -0,898500468 | 0,53644402 | 0,0014 | 0,04004246 | uridine phosphorylase 1 |
| ENSG00000204347 | BTBD17 | -0,898523519 | 0,536435448 | 0,01354 | 0,15265267 | BTB domain containing 17 |
| ENSG00000259440 | NPM1P43 | -0,898749877 | 0,536351289 | 0,05487 | 0,31349896 | nucleophosmin 1 pseudogene 43 |
| ENSG00000269533 | AC003002,3 | -0,89875424 | 0,536349667 | 0,19857 | 0,51980353 | novel transcript |
| ENSG00000212232 | SNORD17 | -0,899020967 | 0,536250515 | 0,01638 | 0,16750744 | small nucleolar RNA, C/D box 17 |
| ENSG00000067842 | ATP2B3 | -0,899312499 | 0,536142163 | 0,17101 | 0,49641416 | ATPase plasma membrane Ca2+ transporting 3 |
| ENSG00000167625 | ZNF526 | -0,899395383 | 0,536111362 | 0,09123 | 0,39063588 | zinc finger protein 526 |
| ENSG00000142920 | AZIN2 | -0,899525167 | 0,536063136 | 0,07806 | 0,36538634 | antizyme inhibitor 2 |
| ENSG00000158545 | ZC3H18 | -0,899884574 | 0,535929608 | 0,04967 | 0,30003295 | zinc finger CCCH-type containing 18 |
| ENSG00000232489 | MFAP1P1 | -0,900475201 | 0,535710248 | 0,18149 | 0,50490902 | microfibril associated protein 1 pseudogene 1 |
| ENSG00000270804 | AC010326,4 | -0,900619753 | 0,535656575 | 0,06457 | 0,33679627 | ubiquitin-conjugating enzyme E2C (UBE2C) pseudogene |
| ENSG00000267470 | ZNF571-AS1 | -0,901167821 | 0,535453122 | 0,05604 | 0,31627719 | ZNF571 antisense RNA 1 |
| ENSG00000225177 | AL590617,2 | -0,901221832 | 0,535433076 | 0,03282 | 0,24447674 | novel transcript |
| ENSG00000197168 | NEK5 | -0,901558449 | 0,53530816 | 0,03007 | 0,23379082 | NIMA related kinase 5 |
| ENSG00000104879 | CKM | -0,901810636 | 0,535214595 | 0,42174 | 0,67643128 | creatine kinase, M-type |
| ENSG00000232233 | LINC02043 | -0,901894703 | 0,535183409 | 0,06764 | 0,34407703 | long intergenic non-protein coding RNA 2043 |
| ENSG00000130713 | EXOSC2 | -0,902012826 | 0,535139591 | 0,07764 | 0,36468824 | exosome component 2 |
| ENSG00000167700 | MFSD3 | -0,902594944 | 0,53492371 | 0,08711 | 0,3821708 | major facilitator superfamily domain containing 3 |

| | | | | | | |
|---|---|---|---|---|---|---|
| ENSG00000273654 | AC020904,2 | -0,902822142 | 0,534839476 | 0,35189 | 0,62699036 | citrate synthase (CS) pseudogene |
| ENSG00000231701 | BMS1P13 | -0,902922968 | 0,534802099 | 0,22885 | 0,54261107 | BMS1 pseudogene 13 |
| ENSG00000168067 | MAP4K2 | -0,903061471 | 0,534750758 | 0,08377 | 0,37595881 | mitogen-activated protein kinase kinase kinase kinase 2 |
| ENSG00000285243 | SLC39A4 | -0,903189313 | 0,534703375 | 0,23411 | 0,54593875 | solute carrier family 39 member 4 |
| ENSG00000103707 | MTFMT | -0,903498756 | 0,534588699 | 0,08875 | 0,38517778 | mitochondrial methionyl-tRNA formyltransferase |
| ENSG00000268575 | AL031282,2 | -0,904164685 | 0,534341996 | 0,04427 | 0,28372689 | novel transcript, readthrough between SLC35E2 and CDK11A |
| ENSG00000271049 | AC020910,4 | -0,904451004 | 0,534235961 | 0,41253 | 0,67020951 | small nuclear ribonucleoprotein polypeptide E (SNRPE) pseudogene |
| ENSG00000102575 | ACP5 | -0,904565959 | 0,534193394 | 0,297 | 0,59008525 | acid phosphatase 5, tartrate resistant |
| ENSG00000101298 | SNPH | -0,904607813 | 0,534177897 | 0,00835 | 0,11551322 | syntaphilin |
| ENSG00000069424 | KCNAB2 | -0,904819559 | 0,534099501 | 0,07299 | 0,35631792 | potassium voltage-gated channel subfamily A regulatory beta subunit 2 |
| ENSG00000285067 | AC270277,1 | -0,905532227 | 0,533835729 | 0,1611 | 0,48841163 | novel transcript, antisense to MTMR9 |
| ENSG00000280273 | AF131216,4 | -0,905532227 | 0,533835729 | 0,1611 | 0,48841163 | novel transcript, antisense to MTMR9 |
| ENSG00000270195 | AC016773,1 | -0,906049557 | 0,533644338 | 0,01489 | 0,16019538 | novel transcript |
| ENSG00000169660 | HEXD | -0,90605355 | 0,533642861 | 0,02589 | 0,21707964 | hexosaminidase D |
| ENSG00000080947 | CROCCP3 | -0,906090066 | 0,533629354 | 0,04253 | 0,2782038 | CROCC pseudogene 3 |
| ENSG00000103326 | CAPN15 | -0,906433283 | 0,533502419 | 0,03482 | 0,25226698 | calpain 15 |
| ENSG00000272219 | AC005072,1 | -0,906805676 | 0,533364727 | 0,07321 | 0,35655334 | novel transcript, sense intronic to CUX1 |
| ENSG00000280660 | AL157396,1 | -0,907449088 | 0,533126911 | 0,14092 | 0,46791293 | novel transcript, sense intronic to LGI1 |
| ENSG00000236581 | STARD13-AS | -0,907537518 | 0,533094234 | 0,07814 | 0,36540845 | STARD13 antisense RNA |
| ENSG00000267882 | AL031666,2 | -0,907644851 | 0,533054574 | 0,14162 | 0,46809977 | novel transcript, antisense to ZMYND8 |
| ENSG00000254048 | AC105150,1 | -0,90794843 | 0,532942418 | 0,2426 | 0,55152734 | novel transcript |
| ENSG00000225882 | LINC01456 | -0,908303208 | 0,532811376 | 0,21155 | 0,53034984 | long intergenic non-protein coding RNA 1456 |
| ENSG00000274475 | RN7SL650P | -0,908415092 | 0,532770057 | 0,56088 | 0,77115738 | RNA, 7SL, cytoplasmic 650, pseudogene |
| ENSG00000272202 | AC097358,2 | -0,908521147 | 0,532730894 | 0,13381 | 0,45985337 | novel transcript, antisense to ARHGEF3 |
| ENSG00000107816 | LZTS2 | -0,908754734 | 0,532644646 | 0,02493 | 0,21172372 | leucine zipper tumor suppressor 2 |
| ENSG00000168792 | ABHD15 | -0,908870089 | 0,532602059 | 0,03287 | 0,2447396 | abhydrolase domain containing 15 |
| ENSG00000285572 | AL136441,1 | -0,909294084 | 0,532445555 | 0,02822 | 0,22680636 | novel transcript |
| ENSG00000205822 | TPTE2P6 | -0,909880784 | 0,53222907 | 0,10616 | 0,41557034 | transmembrane phosphoinositide 3-phosphatase and tensin homolog 2 pseudogene 6 |
| ENSG00000174586 | ZNF497 | -0,910037777 | 0,532171156 | 0,00493 | 0,08701218 | zinc finger protein 497 |

| | | | | | | |
|---|---|---|---|---|---|---|
| ENSG00000240823 | RN7SL23P | -0,91027458 | 0,532083813 | 0,11466 | 0,43014087 | RNA, 7SL, cytoplasmic 23, pseudogene |
| ENSG00000162591 | MEGF6 | -0,910308759 | 0,532071208 | 0,07799 | 0,36536048 | multiple EGF like domains 6 |
| ENSG00000236854 | LINC00486 | -0,91042189 | 0,532029486 | 0,11763 | 0,43515458 | long intergenic non-protein coding RNA 486 |
| ENSG00000253797 | UTP14C | -0,910452178 | 0,532018317 | 0,11128 | 0,42416781 | UTP14C small subunit processome component |
| ENSG00000204305 | AGER | -0,910935404 | 0,531840149 | 0,10295 | 0,41078404 | advanced glycosylation end-product specific receptor |
| ENSG00000107742 | SPOCK2 | -0,911134011 | 0,531766939 | 0,13313 | 0,45903637 | SPARC (osteonectin), cwcv and kazal like domains proteoglycan 2 |
| ENSG00000267708 | RDM1P3 | -0,911293202 | 0,531708265 | 0,27652 | 0,575468 | RDM1 pseudogene 3 |
| ENSG00000233587 | LINC01884 | -0,911302238 | 0,531704935 | 0,54857 | 0,76372599 | long intergenic non-protein coding RNA 1884 |
| ENSG00000145423 | SFRP2 | -0,911500561 | 0,531631848 | 0,00288 | 0,06380901 | secreted frizzled related protein 2 |
| ENSG00000257838 | OTOAP1 | -0,911557874 | 0,531610729 | 0,34542 | 0,62262102 | OTOA pseudogene 1 |
| ENSG00000137825 | ITPKA | -0,912120572 | 0,531403424 | 0,15596 | 0,48369266 | inositol-trisphosphate 3-kinase A |
| ENSG00000127364 | TAS2R4 | -0,912197877 | 0,53137495 | 0,08552 | 0,37903997 | taste 2 receptor member 4 |
| ENSG00000286833 | AC097532,3 | -0,912260911 | 0,531351734 | 0,0151 | 0,16134382 | novel transcript |
| ENSG00000228113 | AC003991,1 | -0,913094575 | 0,53104478 | 0,04068 | 0,27268567 | novel transcript, antisense to STEAP4 |
| ENSG00000163346 | PBXIP1 | -0,913139231 | 0,531028343 | 0,04025 | 0,27130163 | PBX homeobox interacting protein 1 |
| ENSG00000233186 | KLF4P1 | -0,913954139 | 0,530728475 | 0,16073 | 0,48807078 | Kruppel like factor 4 pseudogene 1 |
| ENSG00000226026 | AC092802,1 | -0,914280436 | 0,530608453 | 0,0348 | 0,25226698 | novel transcript |
| ENSG00000229870 | RPL21P89 | -0,914759832 | 0,530432165 | 0,32322 | 0,6060085 | ribosomal protein L21 pseudogene 89 |
| ENSG00000100429 | HDAC10 | -0,914903123 | 0,530379484 | 0,04262 | 0,2782038 | histone deacetylase 10 |
| ENSG00000278077 | AL591926,6 | -0,915011198 | 0,530339754 | 0,33069 | 0,6112353 | zinc finger protein 829 (ZNF829) pseudogene |
| ENSG00000065621 | GSTO2 | -0,91516775 | 0,530282208 | 0,01408 | 0,15573084 | glutathione S-transferase omega 2 |
| ENSG00000112280 | COL9A1 | -0,915247761 | 0,5302528 | 0,07491 | 0,35918824 | collagen type IX alpha 1 chain |
| ENSG00000249593 | AC011405,1 | -0,915510304 | 0,530156313 | 0,02214 | 0,19795874 | novel transcript |
| ENSG00000273706 | LHX1 | -0,915582411 | 0,530129816 | 0,15889 | 0,48627932 | LIM homeobox 1 |
| ENSG00000227929 | SRGAP3-AS3 | -0,91604478 | 0,529959942 | 0,02363 | 0,20525328 | SRGAP3 antisense RNA 3 |
| ENSG00000182175 | RGMA | -0,916181306 | 0,529909793 | 1,07E-05 | 0,00088543 | repulsive guidance molecule BMP co-receptor a |
| ENSG00000154065 | ANKRD29 | -0,916206064 | 0,529900699 | 0,08275 | 0,37461566 | ankyrin repeat domain 29 |
| ENSG00000081913 | PHLPP1 | -0,916211433 | 0,529898727 | 0,00134 | 0,03910482 | PH domain and leucine rich repeat protein phosphatase 1 |
| ENSG00000150281 | CTF1 | -0,916549955 | 0,529774403 | 0,09497 | 0,39751728 | cardiotrophin 1 |
| ENSG00000141542 | RAB40B | -0,916796949 | 0,529683712 | 0,05542 | 0,31484345 | RAB40B, member RAS oncogene family |
| ENSG00000110046 | ATG2A | -0,91681691 | 0,529676384 | 0,06294 | 0,33203667 | autophagy related 2A |
| ENSG00000011590 | ZBTB32 | -0,91682608 | 0,529673017 | 0,0321 | 0,24164117 | zinc finger and BTB domain containing 32 |

| | | | | | | |
|---|---|---|---|---|---|---|
| ENSG00000147488 | ST18 | -0,91761631 | 0,52938297 | 0,03849 | 0,26548082 | ST18 C2H2C-type zinc finger transcription factor |
| ENSG00000250170 | RASA2-IT1 | -0,91767239 | 0,529362392 | 0,28886 | 0,58464073 | RASA2 intronic transcript 1 |
| ENSG00000282457 | CSPG4P10 | -0,918072635 | 0,529215552 | 0,00034 | 0,01561634 | chondroitin sulfate proteoglycan 4 pseudogene 10 |
| ENSG00000152467 | ZSCAN1 | -0,918191253 | 0,529172042 | 0,00913 | 0,12155654 | zinc finger and SCAN domain containing 1 |
| ENSG00000170734 | POLH | -0,918214341 | 0,529163574 | 0,04779 | 0,29355314 | DNA polymerase eta |
| ENSG00000184599 | TAFA3 | -0,918375643 | 0,529104414 | 0,28963 | 0,58522301 | TAFA chemokine like family member 3 |
| ENSG00000188283 | ZNF383 | -0,918612726 | 0,529017471 | 0,07639 | 0,36251343 | zinc finger protein 383 |
| ENSG00000229832 | AL360091,1 | -0,918708283 | 0,528982433 | 0,13963 | 0,4668467 | novel transcript, sense intronic to PPP2R5A |
| ENSG00000242252 | BGLAP | -0,918920563 | 0,528904603 | 0,18507 | 0,50871188 | bone gamma-carboxyglutamate protein |
| ENSG00000232957 | HLA-DOA | -0,919075621 | 0,528847761 | 0,23709 | 0,54809872 | major histocompatibility complex, class II, DO alpha |
| ENSG00000086730 | LAT2 | -0,919273625 | 0,528775183 | 0,04622 | 0,28930497 | linker for activation of T cells family member 2 |
| ENSG00000135424 | ITGA7 | -0,919369802 | 0,528739934 | 0,02663 | 0,22021571 | integrin subunit alpha 7 |
| ENSG00000198089 | SFI1 | -0,919529353 | 0,528681463 | 0,0372 | 0,26111083 | SFI1 centrin binding protein |
| ENSG00000143032 | BARHL2 | -0,919862121 | 0,528559532 | 0,05598 | 0,31614695 | BarH like homeobox 2 |
| ENSG00000234828 | IQCM | -0,920126621 | 0,528462637 | 0,40339 | 0,66399303 | IQ motif containing M |
| ENSG00000246339 | EXTL3-AS1 | -0,920428743 | 0,52835198 | 0,0426 | 0,2782038 | EXTL3 antisense RNA 1 |
| ENSG00000287269 | Z82215,1 | -0,920498436 | 0,528326458 | 0,07829 | 0,36569619 | novel transcript, antisense to ANKRD54and MYH9 |
| ENSG00000224536 | AC096677,1 | -0,920547019 | 0,528308666 | 0,19417 | 0,51562158 | novel transcript |
| ENSG00000130226 | DPP6 | -0,920576872 | 0,528297734 | 0,01748 | 0,17422253 | dipeptidyl peptidase like 6 |
| ENSG00000287895 | AL136379,1 | -0,920695263 | 0,528254383 | 0,09523 | 0,3979203 | novel transcript |
| ENSG00000230257 | NFE4 | -0,920697523 | 0,528253555 | 0,42358 | 0,67732256 | nuclear factor, erythroid 4 |
| ENSG00000198816 | ZNF358 | -0,920741634 | 0,528237404 | 0,03748 | 0,26187746 | zinc finger protein 358 |
| ENSG00000269296 | AC005614,1 | -0,920993335 | 0,528145253 | 0,0521 | 0,30670474 | novel transcript, antisense to ZNF780A |
| ENSG00000280797 | AC005614,3 | -0,920993335 | 0,528145253 | 0,0521 | 0,30670474 | novel transcript, antisense to ZNF780A |
| ENSG00000275445 | AC092119,3 | -0,921229948 | 0,52805864 | 0,39229 | 0,65563839 | novel transcript, antisense to C16orf52 |
| ENSG00000081760 | AACS | -0,921469921 | 0,527970812 | 0,01446 | 0,15737041 | acetoacetyl-CoA synthetase |
| ENSG00000248394 | FOSL1P1 | -0,9215997 | 0,52792332 | 0,2966 | 0,58979902 | FOSL1 pseudogene 1 |
| ENSG00000206312 | NOTCH4 | -0,921647981 | 0,527905652 | 0,02253 | 0,19992837 | notch receptor 4 |
| ENSG00000147509 | RGS20 | -0,921682563 | 0,527892999 | 0,02151 | 0,19510325 | regulator of G protein signaling 20 |
| ENSG00000181035 | SLC25A42 | -0,921737918 | 0,527872744 | 0,00423 | 0,07987039 | solute carrier family 25 member 42 |
| ENSG00000284820 | AC068946,2 | -0,921782312 | 0,527856501 | 0,02542 | 0,21440233 | novel protein |
| ENSG00000258779 | LINC01568 | -0,92193946 | 0,527799006 | 0,30268 | 0,59314261 | long intergenic non-protein coding RNA 1568 |
| ENSG00000188176 | SMTNL2 | -0,922038447 | 0,527762794 | 0,15444 | 0,48266733 | smoothelin like 2 |

| | | | | | | |
|---|---|---|---|---|---|---|
| ENSG00000259202 | AC012568,1 | -0,922103732 | 0,527738912 | 0,2163 | 0,53480193 | novel transcript, antisense to SMAD3 |
| ENSG00000165804 | ZNF219 | -0,922292368 | 0,527669914 | 0,03791 | 0,2635078 | zinc finger protein 219 |
| ENSG00000253880 | AC009435,1 | -0,922368389 | 0,527642109 | 0,10633 | 0,41594715 | novel transcript |
| ENSG00000274021 | AC024909,1 | -0,922539313 | 0,527579601 | 0,03265 | 0,24385689 | novel transcript, antisense to DUSP6 |
| ENSG00000123179 | EBPL | -0,922672353 | 0,527530951 | 0,07643 | 0,36251343 | EBP like |
| ENSG00000272630 | AL731563,3 | -0,922898526 | 0,527448256 | 0,0633 | 0,3332688 | novel transcript |
| ENSG00000235786 | ZNRF3-IT1 | -0,923215495 | 0,527332385 | 0,19564 | 0,51643293 | ZNRF3 intronic transcript 1 |
| ENSG00000237886 | NALT1 | -0,923220627 | 0,527330509 | 0,08342 | 0,37550713 | NOTCH1 associated lncRNA in T cell acute lymphoblastic leukemia 1 |
| ENSG00000124444 | ZNF576 | -0,923245592 | 0,527321384 | 0,06032 | 0,32601555 | zinc finger protein 576 |
| ENSG00000257355 | AC008770,1 | -0,923355088 | 0,527281364 | 0,44138 | 0,68972731 | novel protein |
| ENSG00000247157 | LINC01252 | -0,923365536 | 0,527277545 | 0,1043 | 0,41269448 | long intergenic non-protein coding RNA 1252 |
| ENSG00000167861 | HID1 | -0,923631195 | 0,527180461 | 0,02263 | 0,20056068 | HID1 domain containing |
| ENSG00000228998 | AC091167,1 | -0,923872086 | 0,527092443 | 0,03056 | 0,23584563 | PERP, TP53 apoptosis effector (PERP) pseudogene |
| ENSG00000223815 | DIAPH3-AS2 | -0,92418518 | 0,526978066 | 0,27475 | 0,57463533 | DIAPH3 antisense RNA 2 |
| ENSG00000146267 | FAXC | -0,924296088 | 0,526937556 | 0,01948 | 0,18486352 | failed axon connections homolog, metaxin like GST domain containing |
| ENSG00000172421 | EFCAB3 | -0,924302718 | 0,526935134 | 0,03463 | 0,25158473 | EF-hand calcium binding domain 3 |
| ENSG00000284400 | TBC1D9B | -0,925018019 | 0,52667394 | 0,06985 | 0,34882573 | TBC1 domain family member 9B |
| ENSG00000148408 | CACNA1B | -0,925087532 | 0,526648564 | 0,00224 | 0,05390184 | calcium voltage-gated channel subunit alpha1 B |
| ENSG00000266473 | AC007448,4 | -0,925476634 | 0,526506543 | 0,2371 | 0,54809872 | novel transcript |
| ENSG00000251192 | ZNF674 | -0,926157406 | 0,526258157 | 0,05204 | 0,30661383 | zinc finger protein 674 |
| ENSG00000139200 | PIANP | -0,926497448 | 0,526134133 | 0,03015 | 0,23425595 | PILR alpha associated neural protein |
| ENSG00000230596 | GPAA1P2 | -0,926596974 | 0,526097838 | 0,30489 | 0,59446033 | glycosylphosphatidylinositol anchor attachment 1 pseudogene 2 |
| ENSG00000267868 | AL356740,1 | -0,926880078 | 0,52599461 | 0,05348 | 0,31052177 | novel transcript |
| ENSG00000175985 | PLEKHD1 | -0,926991741 | 0,5259539 | 0,08004 | 0,36930228 | pleckstrin homology and coiled-coil domain containing D1 |
| ENSG00000150337 | FCGR1A | -0,927167515 | 0,525889824 | 0,4512 | 0,69626785 | Fc fragment of IgG receptor Ia |
| ENSG00000173175 | ADCY5 | -0,927231387 | 0,525866542 | 0,01715 | 0,17206613 | adenylate cyclase 5 |
| ENSG00000258465 | AL139011,2 | -0,927240917 | 0,525863068 | 0,74999 | 0,882436 | novel protein |
| ENSG00000234892 | Z82214,1 | -0,92752985 | 0,525757762 | 0,44601 | 0,69284796 | novel transcript |
| ENSG00000204055 | AL158151,1 | -0,928027087 | 0,525576586 | 0,16191 | 0,48914394 | novel transcript, antisense to IER5L |
| ENSG00000155980 | KIF5A | -0,928138008 | 0,525536179 | 0,00437 | 0,08120816 | kinesin family member 5A |

| | | | | | | |
|---|---|---|---|---|---|---|
| ENSG00000283243 | FO681507,1 | -0,928207122 | 0,525511003 | 0,79088 | 0,90292452 | |
| ENSG00000257704 | INAFM1 | -0,928254361 | 0,525493797 | 0,04569 | 0,28842796 | InaF motif containing 1 |
| ENSG00000260357 | DNM1P34 | -0,928876821 | 0,525267118 | 0,22231 | 0,53859118 | dynamin 1 pseudogene 34 |
| ENSG00000271601 | LIX1L | -0,928976872 | 0,525230692 | 0,00094 | 0,03102731 | limb and CNS expressed 1 like |
| ENSG00000273137 | AL022328,2 | -0,929153225 | 0,525166493 | 0,19286 | 0,51493916 | novel transcript, antisense to SELO |
| ENSG00000254109 | RBPMS-AS1 | -0,929475496 | 0,525049193 | 0,23803 | 0,54864017 | RBPMS antisense RNA 1 |
| ENSG00000249409 | AC025741,1 | -0,929961931 | 0,524872192 | 0,17168 | 0,49704622 | novel transcript |
| ENSG00000258768 | AL356019,2 | -0,93003015 | 0,524847373 | 0,14459 | 0,47220257 | novel transcript |
| ENSG00000277258 | PCGF2 | -0,930135845 | 0,524808923 | 0,05099 | 0,30380236 | polycomb group ring finger 2 |
| ENSG00000104866 | PPP1R37 | -0,930304981 | 0,5247474 | 0,03063 | 0,23611505 | protein phosphatase 1 regulatory subunit 37 |
| ENSG00000125285 | SOX21 | -0,930368092 | 0,524724445 | 0,00467 | 0,08412546 | SRY-box transcription factor 21 |
| ENSG00000272195 | AL356512,1 | -0,930390405 | 0,52471633 | 0,04218 | 0,27783166 | novel transcript, antisense to EFCAB2 |
| ENSG00000233547 | AL158212,2 | -0,930416896 | 0,524706695 | 0,22042 | 0,53750545 | novel transcript |
| ENSG00000253893 | FAM85B | -0,930759735 | 0,52458202 | 0,09674 | 0,40089337 | family with sequence similarity 85 member B |
| ENSG00000235884 | LINC00941 | -0,930868834 | 0,524542352 | 0,69072 | 0,84906099 | long intergenic non-protein coding RNA 941 |
| ENSG00000104361 | NIPAL2 | -0,931119676 | 0,524451157 | 0,01753 | 0,17446742 | NIPA like domain containing 2 |
| ENSG00000287617 | AC092902,6 | -0,931448153 | 0,524331762 | 0,03469 | 0,25183223 | novel transcript |
| ENSG00000242853 | RN7SL749P | -0,931453054 | 0,524329981 | 0,35377 | 0,62859222 | RNA, 7SL, cytoplasmic 749, pseudogene |
| ENSG00000271387 | AL445228,2 | -0,93180981 | 0,524200338 | 0,01596 | 0,16547046 | novel transcript |
| ENSG00000171368 | TPPP | -0,931959143 | 0,524146081 | 0,00453 | 0,08297896 | tubulin polymerization promoting protein |
| ENSG00000251359 | WWC2-AS2 | -0,932151855 | 0,524076072 | 0,12063 | 0,43944057 | WWC2 antisense RNA 2 |
| ENSG00000123999 | INHA | -0,93232705 | 0,524012434 | 0,07262 | 0,35556123 | inhibin subunit alpha |
| ENSG00000245213 | AC105285,1 | -0,932432985 | 0,523973958 | 0,06414 | 0,33569301 | novel transcript, antisense to GALNT7 |
| ENSG00000214659 | KRT8P26 | -0,932808809 | 0,52383748 | 0,09944 | 0,40604811 | keratin 8 pseudogene 26 |
| ENSG00000245482 | AC046130,1 | -0,932995608 | 0,523769658 | 0,52654 | 0,7497927 | novel transcript, antisense to ALG10 |
| ENSG00000226726 | TMEM256P2 | -0,933285722 | 0,523664343 | 0,16765 | 0,4939175 | transmembrane protein 256 pseudogene 2 |
| ENSG00000225465 | RFPL1S | -0,933595588 | 0,523551881 | 0,05988 | 0,32540929 | RFPL1 antisense RNA 1 |
| ENSG00000157992 | KRTCAP3 | -0,933863865 | 0,523454533 | 0,05094 | 0,30366564 | keratinocyte associated protein 3 |
| ENSG00000288212 | AC090826,7 | -0,933869365 | 0,523452537 | 0,12188 | 0,44093327 | novel transcript, antisense to SEMA7A |
| ENSG00000286696 | AC090826,3 | -0,933869365 | 0,523452537 | 0,12188 | 0,44093327 | novel transcript, antisense to SEMA7A |
| ENSG00000234709 | UPF3AP3 | -0,934031564 | 0,52339369 | 0,08401 | 0,37651068 | UPF3A pseudogene 3 |
| ENSG00000158683 | PKD1L1 | -0,934061698 | 0,523382758 | 0,18372 | 0,50744379 | polycystin 1 like 1, transient receptor potential channel interacting |

| | | | | | | |
|---|---|---|---|---|---|---|
| ENSG00000108786 | HSD17B1 | -0,934202052 | 0,523331842 | 0,15279 | 0,48083991 | hydroxysteroid 17-beta dehydrogenase 1 |
| ENSG00000139351 | SYCP3 | -0,934306763 | 0,52329386 | 0,04847 | 0,29584645 | synaptonemal complex protein 3 |
| ENSG00000270282 | AL096855,2 | -0,934366388 | 0,523272234 | 0,27942 | 0,57742338 | ferritin, light polypeptide (FTL) pseudogene |
| ENSG00000106686 | SPATA6L | -0,934667325 | 0,523163094 | 0,06964 | 0,34826564 | spermatogenesis associated 6 like |
| ENSG00000269001 | AC092070,2 | -0,935598481 | 0,522825538 | 0,0674 | 0,34363554 | zinc finger protein 818, pseudogene |
| ENSG00000243049 | RN7SL33P | -0,936012385 | 0,522675563 | 0,34269 | 0,62029024 | RNA, 7SL, cytoplasmic 33, pseudogene |
| ENSG00000280511 | AL591030,1 | -0,93602805 | 0,522669888 | 0,03317 | 0,24602205 | novel transcript |
| ENSG00000163157 | TMOD4 | -0,9360849 | 0,522649292 | 0,07847 | 0,36604454 | tropomodulin 4 |
| ENSG00000186862 | PDZD7 | -0,936110159 | 0,522640142 | 0,17098 | 0,49641416 | PDZ domain containing 7 |
| ENSG00000241169 | FO393419,2 | -0,936457888 | 0,522514186 | 0,31743 | 0,60284673 | novel transcript |
| ENSG00000274925 | ZKSCAN2-DT | -0,936559361 | 0,522477436 | 0,02317 | 0,20327138 | ZKSCAN2 divergent transcript |
| ENSG00000286944 | AL137018,1 | -0,936849796 | 0,522372264 | 0,06384 | 0,33492932 | novel transcript, antisense to RORB |
| ENSG00000230510 | PPP5D1 | -0,937049797 | 0,522299853 | 0,02756 | 0,22376406 | PPP5 tetratricopeptide repeat domain containing 1 |
| ENSG00000279082 | LINC01727 | -0,937350934 | 0,522190844 | 0,05056 | 0,30252802 | long intergenic non-protein coding RNA 1727 |
| ENSG00000272563 | AC016745,2 | -0,937382726 | 0,522179336 | 0,16275 | 0,48945635 | novel transcript |
| ENSG00000268798 | AC027307,3 | -0,937676624 | 0,522072972 | 0,08837 | 0,38462876 | novel transcript |
| ENSG00000264573 | RN7SL15P | -0,937843941 | 0,522012428 | 0,12385 | 0,4446254 | RNA, 7SL, cytoplasmic 15, pseudogene |
| ENSG00000234225 | AC244394,3 | -0,937858359 | 0,522007211 | 0,38879 | 0,65335633 | novel transcript |
| ENSG00000136002 | ARHGEF4 | -0,938351243 | 0,521828902 | 0,00845 | 0,11605624 | Rho guanine nucleotide exchange factor 4 |
| ENSG00000237159 | CNTFR-AS1 | -0,938424204 | 0,521802513 | 0,09672 | 0,40087593 | CNTFR antisense RNA 1 |
| ENSG00000099624 | ATP5F1D | -0,938482098 | 0,521781574 | 0,06586 | 0,33967813 | ATP synthase F1 subunit delta |
| ENSG00000186912 | P2RY4 | -0,939028508 | 0,52158399 | 0,22854 | 0,5424199 | pyrimidinergic receptor P2Y4 |
| ENSG00000237667 | LINC01115 | -0,939588024 | 0,521381745 | 0,38598 | 0,65145202 | long intergenic non-protein coding RNA 1115 |
| ENSG00000267260 | AC020928,1 | -0,939751622 | 0,521322625 | 0,0678 | 0,34426294 | novel transcript |
| ENSG00000101883 | RHOXF1 | -0,939912936 | 0,521264337 | 0,33351 | 0,61346096 | Rhox homeobox family member 1 |
| ENSG00000277007 | AC096642,1 | -0,939992099 | 0,521235735 | 0,1259 | 0,448127 | novel transcript |
| ENSG00000285878 | AP002961,1 | -0,940073232 | 0,521206423 | 0,22498 | 0,54037642 | novel transcript, antisense to DYNC2H1 |
| ENSG00000287732 | AL731574,1 | -0,940168554 | 0,521171987 | 0,40755 | 0,66665981 | novel transcript |
| ENSG00000205593 | DENND6B | -0,940362782 | 0,521101827 | 0,01838 | 0,17938469 | DENN domain containing 6B |
| ENSG00000246366 | LACTB2-AS1 | -0,940395966 | 0,521089841 | 0,07317 | 0,35655334 | LACTB2 antisense RNA 1 |
| ENSG00000179044 | EXOC3L1 | -0,940719107 | 0,520973138 | 0,21631 | 0,53480193 | exocyst complex component 3 like 1 |
| ENSG00000197106 | SLC6A17 | -0,941021404 | 0,520863987 | 0,34637 | 0,62342969 | solute carrier family 6 member 17 |
| ENSG00000269855 | RNF225 | -0,941797313 | 0,520583931 | 0,33156 | 0,61165554 | ring finger protein 225 |

| | | | | | | |
|---|---|---|---|---|---|---|
| ENSG00000170175 | CHRNB1 | -0,941816476 | 0,520577017 | 0,34461 | 0,62200601 | cholinergic receptor nicotinic beta 1 subunit |
| ENSG00000230961 | MSH5 | -0,942045269 | 0,520494467 | 0,06252 | 0,33112961 | mutS homolog 5 |
| ENSG00000241956 | AC109466,1 | -0,942152206 | 0,520455887 | 0,04631 | 0,28930497 | novel transcript |
| ENSG00000212978 | AC016747,1 | -0,942368773 | 0,520377766 | 0,02451 | 0,20943421 | novel transcript |
| ENSG00000105767 | CADM4 | -0,942551721 | 0,520311781 | 0,00617 | 0,09882554 | cell adhesion molecule 4 |
| ENSG00000234114 | CCHCR1 | -0,942615427 | 0,520288806 | 0,02527 | 0,21376722 | coiled-coil alpha-helical rod protein 1 |
| ENSG00000179862 | CITED4 | -0,942728216 | 0,520248132 | 0,07559 | 0,36093649 | Cbp/p300 interacting transactivator with Glu/Asp rich carboxy-terminal domain 4 |
| ENSG00000265494 | AC005304,2 | -0,942823193 | 0,520213883 | 0,05942 | 0,32465136 | novel transcript |
| ENSG00000260361 | AC106028,3 | -0,943075733 | 0,520122829 | 0,17869 | 0,50323303 | novel transcript |
| ENSG00000227678 | AL355581,1 | -0,94344323 | 0,519990356 | 0,00581 | 0,09564911 | novel transcript, antisense to L3MBTL3 |
| ENSG00000121716 | PILRB | -0,943779361 | 0,519869218 | 0,06245 | 0,33103715 | paired immunoglobin like type 2 receptor beta |
| ENSG00000037757 | MRI1 | -0,943859218 | 0,519840443 | 0,0525 | 0,30766476 | methylthioribose-1-phosphate isomerase 1 |
| ENSG00000157353 | FCSK | -0,944250526 | 0,519699463 | 0,03345 | 0,24712306 | fucose kinase |
| ENSG00000203809 | LIN28B-AS1 | -0,944354526 | 0,519662001 | 0,13533 | 0,46184564 | LIN28B antisense RNA 1 |
| ENSG00000111339 | ART4 | -0,94445581 | 0,519625519 | 0,10037 | 0,4067769 | ADP-ribosyltransferase 4 (Dombrock blood group) |
| ENSG00000134201 | GSTM5 | -0,944467994 | 0,519621131 | 0,07467 | 0,35882064 | glutathione S-transferase mu 5 |
| ENSG00000276876 | MYT1 | -0,944539586 | 0,519595346 | 0,04728 | 0,29217708 | myelin transcription factor 1 |
| ENSG00000140368 | PSTPIP1 | -0,944619228 | 0,519566663 | 0,17102 | 0,49641416 | proline-serine-threonine phosphatase interacting protein 1 |
| ENSG00000233061 | TTLL7-IT1 | -0,944669337 | 0,519548617 | 0,00945 | 0,12433437 | TTLL7 intronic transcript 1 |
| ENSG00000188525 | AC010969,1 | -0,944774318 | 0,519510813 | 0,21787 | 0,53550245 | novel transcript |
| ENSG00000177943 | MAMDC4 | -0,944923199 | 0,519457204 | 0,02454 | 0,20956669 | MAM domain containing 4 |
| ENSG00000253384 | AC124242,2 | -0,945098313 | 0,519394156 | 0,17122 | 0,49650902 | family with sequence similarity 175, member A (FAM175A) pseudogene |
| ENSG00000188185 | LINC00265 | -0,945195634 | 0,51935912 | 0,03592 | 0,25664143 | long intergenic non-protein coding RNA 265 |
| ENSG00000170604 | IRF2BP1 | -0,945297354 | 0,519322503 | 0,08364 | 0,37578481 | interferon regulatory factor 2 binding protein 1 |
| ENSG00000124191 | TOX2 | -0,94542883 | 0,519275178 | 0,05365 | 0,31073157 | TOX high mobility group box family member 2 |
| ENSG00000239528 | RPS14P8 | -0,945731457 | 0,519166264 | 0,08042 | 0,37007763 | ribosomal protein S14 pseudogene 8 |
| ENSG00000285676 | AL158212,5 | -0,945956872 | 0,519085153 | 0,27314 | 0,57344244 | novel transcript, antisense to VTI1A |
| ENSG00000204963 | PCDHA7 | -0,946340175 | 0,518947257 | 0,00553 | 0,09240758 | protocadherin alpha 7 |
| ENSG00000286431 | BX842570,1 | -0,947134651 | 0,518661558 | 0,00023 | 0,01192921 | novel transcript |

| | | | | | | |
|---|---|---|---|---|---|---|
| ENSG00000233585 | AC231533,2 | -0,947824585 | 0,51841358 | 0,23197 | 0,54471359 | acetyl-Coenzyme A acyltransferase 2 (mitochondrial 3-oxoacyl-Coenzyme A thiolase) (ACAA2) pseudogene |
| ENSG00000253154 | AC100801,1 | -0,947858419 | 0,518401422 | 0,17287 | 0,49768904 | novel transcript |
| ENSG00000250159 | AC106791,1 | -0,948057489 | 0,518329896 | 0,03942 | 0,2687016 | novel transcript |
| ENSG00000250874 | AC010595,1 | -0,948182538 | 0,51828497 | 0,13717 | 0,4642578 | novel transcript |
| ENSG00000272079 | AC004233,2 | -0,948215201 | 0,518273236 | 0,27654 | 0,575468 | novel transcript |
| ENSG00000273321 | AC023983,2 | -0,948423575 | 0,518198385 | 0,43738 | 0,68717176 | novel transcript |
| ENSG00000249388 | AC023794,2 | -0,949232181 | 0,517908025 | 0,16349 | 0,48977932 | novel transcript |
| ENSG00000239344 | AC090686,1 | -0,949289005 | 0,517887626 | 0,03649 | 0,25875342 | ribosomal protein S6 (RPS6) pseudogene |
| ENSG00000268257 | AIRN | -0,949470631 | 0,517822432 | 0,05876 | 0,32380736 | antisense of IGF2R non-protein coding RNA |
| ENSG00000261386 | AC027682,4 | -0,949734765 | 0,517727636 | 0,03196 | 0,24116556 | novel transcript |
| ENSG00000184905 | TCEAL2 | -0,949799588 | 0,517704374 | 0,0038 | 0,07519574 | transcription elongation factor A like 2 |
| ENSG00000234876 | NOTCH4 | -0,950184788 | 0,517566165 | 0,03682 | 0,26003223 | notch receptor 4 |
| ENSG00000199879 | RNVU1-22 | -0,950471602 | 0,517463281 | 0,09165 | 0,39158946 | RNA, variant U1 small nuclear 22 |
| ENSG00000175482 | POLD4 | -0,950546669 | 0,517436357 | 0,08215 | 0,373432 | DNA polymerase delta 4, accessory subunit |
| ENSG00000229240 | LINC00710 | -0,950755312 | 0,51736153 | 0,06035 | 0,3261119 | long intergenic non-protein coding RNA 710 |
| ENSG00000139160 | ETFBKMT | -0,950934887 | 0,517297137 | 0,04232 | 0,27814302 | electron transfer flavoprotein subunit beta lysine methyltransferase |
| ENSG00000275568 | ARHGAP11A | -0,951000165 | 0,517273732 | 0,55672 | 0,76859106 | Rho GTPase activating protein 11A |
| ENSG00000163673 | DCLK3 | -0,951974208 | 0,516924609 | 0,03706 | 0,26086671 | doublecortin like kinase 3 |
| ENSG00000287056 | AL606970,5 | -0,952260945 | 0,51682188 | 0,26025 | 0,56382982 | novel transcript |
| ENSG00000234653 | LRRTM4-AS1 | -0,952367005 | 0,516783888 | 0,20248 | 0,52259159 | LRRTM4 antisense RNA 1 |
| ENSG00000286450 | AC021573,1 | -0,952383349 | 0,516778033 | 0,05131 | 0,30462001 | novel transcript |
| ENSG00000114654 | EFCC1 | -0,952533273 | 0,516724332 | 0,15397 | 0,48229227 | EF-hand and coiled-coil domain containing 1 |
| ENSG00000166153 | DEPDC4 | -0,952760494 | 0,516642956 | 0,04886 | 0,29724532 | DEP domain containing 4 |
| ENSG00000127585 | FBXL16 | -0,952868123 | 0,516604415 | 0,03012 | 0,23411718 | F-box and leucine rich repeat protein 16 |
| ENSG00000262434 | AC087392,4 | -0,952933076 | 0,516581157 | 0,25002 | 0,55657567 | novel transcript, antisense to RNMTL1 |
| ENSG00000223356 | AL590666,1 | -0,952957897 | 0,516572269 | 0,32335 | 0,60611814 | novel transcript |
| ENSG00000248916 | NUP210P3 | -0,953257232 | 0,5164651 | 0,3608 | 0,63310436 | nucleoporin 210 pseudogene 3 |
| ENSG00000163435 | ELF3 | -0,953832365 | 0,516259251 | 0,1432 | 0,47082074 | E74 like ETS transcription factor 3 |
| ENSG00000100285 | NEFH | -0,954381757 | 0,516062692 | 0,00113 | 0,03501256 | neurofilament heavy |
| ENSG00000259295 | CSPG4P12 | -0,95453489 | 0,516007918 | 0,05685 | 0,31784869 | chondroitin sulfate proteoglycan 4 pseudogene 12 |
| ENSG00000230102 | LINC02028 | -0,954837357 | 0,515899747 | 0,26738 | 0,56973003 | long intergenic non-protein coding RNA 2028 |

| | | | | | | |
|---|---|---|---|---|---|---|
| ENSG00000159958 | TNFRSF13C | -0,955046885 | 0,515824826 | 0,10763 | 0,41827415 | TNF receptor superfamily member 13C |
| ENSG00000275211 | CYP2D6 | -0,955119877 | 0,515798729 | 0,24019 | 0,54948131 | cytochrome P450 family 2 subfamily D member 6 |
| ENSG00000287135 | AC010884,1 | -0,955150509 | 0,515787777 | 0,36257 | 0,63379594 | novel transcript |
| ENSG00000238151 | MLLT10P1 | -0,955224316 | 0,515761391 | 0,20479 | 0,52445677 | MLLT10 pseudogene 1 |
| ENSG00000242866 | STRC | -0,955250383 | 0,515752072 | 0,1111 | 0,42398965 | stereocilin |
| ENSG00000276075 | AC027682,6 | -0,955464695 | 0,515675463 | 0,18221 | 0,50570721 | novel transcript, antisense to FAM65A |
| ENSG00000231871 | IPO9-AS1 | -0,955511949 | 0,515658573 | 0,04198 | 0,27753946 | IPO9 antisense RNA 1 |
| ENSG00000135144 | DTX1 | -0,955918322 | 0,515513344 | 0,01778 | 0,1756157 | deltex E3 ubiquitin ligase 1 |
| ENSG00000271858 | Z84492,1 | -0,956171976 | 0,515422715 | 0,16637 | 0,49232464 | novel transcript, antisense to CACNA2D2 |
| ENSG00000215861 | AC245297,1 | -0,956314629 | 0,515371753 | 0,19951 | 0,52047306 | phosphodiesterase 4D interacting protein (myomegalin) (PDE4DIP) pseudogene |
| ENSG00000235221 | LINC00383 | -0,956381017 | 0,515348038 | 0,05715 | 0,31889093 | long intergenic non-protein coding RNA 383 |
| ENSG00000258733 | LINC02328 | -0,956569862 | 0,515280584 | 0,25089 | 0,55707572 | long intergenic non-protein coding RNA 2328 |
| ENSG00000116983 | HPCAL4 | -0,956746382 | 0,515217541 | 0,02989 | 0,23283161 | hippocalcin like 4 |
| ENSG00000260742 | AC009962,1 | -0,957124446 | 0,515082544 | 0,07469 | 0,35882064 | novel transcript, antisense to SSFA2 |
| ENSG00000253837 | AC090197,1 | -0,957196244 | 0,515056911 | 0,08025 | 0,36993659 | novel transcript |
| ENSG00000229635 | AL137159,1 | -0,957281296 | 0,515026547 | 0,22908 | 0,54263903 | pseudogene similar to part of F-box and WD-40 domain protein 1B (FBXW1B) |
| ENSG00000136267 | DGKB | -0,957448461 | 0,514966875 | 0,16922 | 0,49530908 | diacylglycerol kinase beta |
| ENSG00000205644 | AC025186,1 | -0,958003173 | 0,51476891 | 0,37956 | 0,64734351 | ribosomal protein S2 (RPS2) pseudogene |
| ENSG00000261442 | AC023830,1 | -0,958441565 | 0,51461251 | 0,13734 | 0,46435502 | novel transcript, antisense to MGRN1 |
| ENSG00000224159 | HMGB1P9 | -0,958463976 | 0,514604516 | 0,18029 | 0,50407792 | high mobility group box 1 pseudogene 9 |
| ENSG00000250251 | PKD1P6 | -0,958715031 | 0,514514974 | 0,07838 | 0,36597179 | polycystin 1, transient receptor potential channel interacting pseudogene 6 |
| ENSG00000171722 | SPATA46 | -0,958896469 | 0,514450271 | 0,02376 | 0,20581316 | spermatogenesis associated 46 |
| ENSG00000163705 | FANCD2OS | -0,95909013 | 0,514381218 | 0,1417 | 0,46826355 | FANCD2 opposite strand |
| ENSG00000269945 | AC004817,4 | -0,959109461 | 0,514374325 | 0,05563 | 0,31515737 | novel transcript |
| ENSG00000249494 | AC008629,1 | -0,9593658 | 0,514282939 | 0,02027 | 0,18910367 | novel transcript |
| ENSG00000258123 | LINC02444 | -0,959396241 | 0,514272088 | 0,01822 | 0,17852111 | long intergenic non-protein coding RNA 2444 |
| ENSG00000286642 | AC091646,1 | -0,959452458 | 0,514252049 | 0,08324 | 0,37550713 | novel transcript |
| ENSG00000114923 | SLC4A3 | -0,959860039 | 0,514106786 | 0,02072 | 0,19091508 | solute carrier family 4 member 3 |
| ENSG00000267943 | AC010328,1 | -0,960056196 | 0,51403689 | 0,0532 | 0,30971589 | novel transcript |
| ENSG00000278558 | TMEM191B | -0,960081594 | 0,514027841 | 0,19874 | 0,51984196 | transmembrane protein 191B |

| | | | | | | |
|---|---|---|---|---|---|---|
| ENSG00000125898 | FAM110A | -0,960475205 | 0,513887617 | 0,02087 | 0,19166705 | family with sequence similarity 110 member A |
| ENSG00000250698 | AC131182,1 | -0,960503387 | 0,513877579 | 0,00748 | 0,11001711 | novel transcript |
| ENSG00000181625 | SLX1B | -0,961701486 | 0,513451002 | 0,0826 | 0,37429151 | SLX1 homolog B, structure-specific endonuclease subunit |
| ENSG00000135407 | AVIL | -0,961702967 | 0,513450475 | 0,06778 | 0,34426294 | advillin |
| ENSG00000269318 | AC007292,2 | -0,961791171 | 0,513419084 | 0,08693 | 0,38197222 | novel transcript, antisense to MPND |
| ENSG00000223522 | AC093690,1 | -0,962132739 | 0,513297543 | 0,07347 | 0,356639 | novel transcript |
| ENSG00000276141 | WHAMMP3 | -0,962299383 | 0,513238256 | 0,02415 | 0,20778596 | WAS protein homolog associated with actin, golgi membranes and microtubules pseudogene 3 |
| ENSG00000085117 | CD82 | -0,962623719 | 0,513122887 | 0,12084 | 0,43982438 | CD82 molecule |
| ENSG00000103199 | ZNF500 | -0,962891986 | 0,513027481 | 0,06156 | 0,32866608 | zinc finger protein 500 |
| ENSG00000264290 | AC104564,3 | -0,963007332 | 0,512986465 | 0,15309 | 0,48120112 | novel transcript |
| ENSG00000052749 | RRP12 | -0,963317691 | 0,512876122 | 0,06204 | 0,32975861 | ribosomal RNA processing 12 homolog |
| ENSG00000286215 | AL356534,1 | -0,963509083 | 0,512808086 | 0,01827 | 0,17879941 | vimentin 2, pseudogene |
| ENSG00000234814 | SVIL2P | -0,963955168 | 0,512649549 | 0,01907 | 0,18296265 | supervillin family member 2, pseudogene |
| ENSG00000118242 | MREG | -0,964385769 | 0,512496562 | 0,09007 | 0,38819633 | melanoregulin |
| ENSG00000133454 | MYO18B | -0,964545392 | 0,512439861 | 0,02228 | 0,19831519 | myosin XVIIIB |
| ENSG00000153976 | HS3ST3A1 | -0,964642015 | 0,512405542 | 0,03307 | 0,24561547 | heparan sulfate-glucosamine 3-sulfotransferase 3A1 |
| ENSG00000258659 | TRIM34 | -0,964717343 | 0,512378788 | 0,10755 | 0,41807472 | tripartite motif containing 34 |
| ENSG00000104218 | CSPP1 | -0,965319369 | 0,512165021 | 0,02607 | 0,21736588 | centrosome and spindle pole associated protein 1 |
| ENSG00000223935 | LGALSL-DT | -0,965538095 | 0,512087378 | 0,0562 | 0,31666398 | LGALSL divergent transcript |
| ENSG00000253731 | PCDHGA6 | -0,965560942 | 0,512079269 | 0,09821 | 0,40370642 | protocadherin gamma subfamily A, 6 |
| ENSG00000168824 | NSG1 | -0,966062667 | 0,511901214 | 0,01271 | 0,1475797 | neuronal vesicle trafficking associated 1 |
| ENSG00000183018 | SPNS2 | -0,966075447 | 0,511896679 | 0,05308 | 0,30937107 | sphingolipid transporter 2 |
| ENSG00000187010 | RHD | -0,966270487 | 0,51182748 | 0,17385 | 0,49851138 | Rh blood group D antigen |
| ENSG00000233593 | LINC02609 | -0,966345736 | 0,511800785 | 0,02978 | 0,23233601 | long intergenic non-protein coding RNA 2609 |
| ENSG00000105672 | ETV2 | -0,96666678 | 0,511686906 | 0,03289 | 0,24484696 | ETS variant transcription factor 2 |
| ENSG00000137413 | TAF8 | -0,966790914 | 0,511642881 | 0,06512 | 0,33804854 | TATA-box binding protein associated factor 8 |
| ENSG00000287859 | AC008507,4 | -0,966849946 | 0,511621946 | 0,1694 | 0,49554671 | novel transcript |
| ENSG00000279566 | ZNF43 | -0,966876589 | 0,511612497 | 0,04625 | 0,28930497 | zinc finger protein 43 |
| ENSG00000218891 | ZNF579 | -0,967151575 | 0,51151499 | 0,03538 | 0,25487994 | zinc finger protein 579 |
| ENSG00000137497 | NUMA1 | -0,967860002 | 0,511263875 | 0,01734 | 0,17339933 | nuclear mitotic apparatus protein 1 |
| ENSG00000018610 | CXorf56 | -0,967874792 | 0,511258634 | 0,07978 | 0,36862496 | chromosome X open reading frame 56 |

| | | | | | | |
|---|---|---|---|---|---|---|
| ENSG00000230863 | AL137803,1 | -0,96810013 | 0,511178786 | 0,36631 | 0,63627876 | histidine acid phosphatase domain containing 1 (HISPPD1) pseudogene |
| ENSG00000272940 | U62317,2 | -0,968219323 | 0,511136555 | 0,39268 | 0,65593262 | novel transcript |
| ENSG00000264920 | AC018521,5 | -0,96833344 | 0,511096126 | 0,05045 | 0,30220996 | novel transcript |
| ENSG00000138030 | KHK | -0,968414811 | 0,5110673 | 0,00178 | 0,04660264 | ketohexokinase |
| ENSG00000182040 | USH1G | -0,968641535 | 0,51098699 | 0,41377 | 0,67085666 | USH1 protein network component sans |
| ENSG00000286784 | AC116611,1 | -0,968809281 | 0,51092758 | 0,02763 | 0,22402593 | novel transcript |
| ENSG00000205022 | PABPN1L | -0,969196733 | 0,510790383 | 0,19655 | 0,51769283 | PABPN1 like, cytoplasmic |
| ENSG00000245748 | AC097382,2 | -0,970617919 | 0,510287455 | 0,28341 | 0,58063542 | novel transcript, antisense to TBC1D14 and TADA2B |
| ENSG00000254768 | AC104009,1 | -0,970619178 | 0,51028701 | 0,04057 | 0,27233662 | novel transcript |
| ENSG00000286877 | AC090510,4 | -0,971024132 | 0,510143797 | 0,20537 | 0,52503195 | novel transcript, antisense to TTBK2 |
| ENSG00000204262 | COL5A2 | -0,971040497 | 0,51013801 | 0,0357 | 0,25626339 | collagen type V alpha 2 chain |
| ENSG00000277504 | AC010536,3 | -0,971441166 | 0,509996352 | 0,22722 | 0,54168591 | novel transcript, antisense to JPH3 |
| ENSG00000149596 | JPH2 | -0,97151916 | 0,509968782 | 0,20646 | 0,52609885 | junctophilin 2 |
| ENSG00000228782 | MRPL45P2 | -0,97174032 | 0,509890612 | 0,04051 | 0,27208001 | mitochondrial ribosomal protein L45 pseudogene 2 |
| ENSG00000231225 | MICA | -0,971763269 | 0,509882501 | 0,11444 | 0,42986344 | MHC class I polypeptide-related sequence A |
| ENSG00000285437 | POLR2J3 | -0,971827651 | 0,509859747 | 0,03034 | 0,2351139 | RNA polymerase II subunit J3 |
| ENSG00000109758 | HGFAC | -0,972519211 | 0,509615403 | 0,07743 | 0,3644095 | HGF activator |
| ENSG00000214626 | POLR3DP1 | -0,972881995 | 0,50948727 | 0,07526 | 0,36012885 | RNA polymerase III subunit D pseudogene 1 |
| ENSG00000274272 | AC069281,2 | -0,973314478 | 0,509334562 | 0,11368 | 0,42849514 | novel transcript |
| ENSG00000255474 | GAU1 | -0,973441287 | 0,509289795 | 0,0562 | 0,31666398 | GALNT8 antisense upstream 1 |
| ENSG00000273785 | CYP2D8P | -0,973895593 | 0,509129444 | 0,15474 | 0,48266733 | ccytochrome P450 family 2 subfamily D member 8, pseudogene |
| ENSG00000231920 | NEBL-AS1 | -0,974066253 | 0,509069221 | 0,29745 | 0,59034513 | NEBL antisense RNA 1 |
| ENSG00000144488 | ESPNL | -0,974310511 | 0,50898304 | 0,0912 | 0,39063588 | espin like |
| ENSG00000260903 | XKR7 | -0,974417096 | 0,508945438 | 0,00776 | 0,11202957 | XK related 7 |
| ENSG00000160214 | RRP1 | -0,974701504 | 0,508845116 | 0,05173 | 0,30583008 | ribosomal RNA processing 1 |
| ENSG00000080854 | IGSF9B | -0,974856697 | 0,508790382 | 0,02142 | 0,19439259 | immunoglobulin superfamily member 9B |
| ENSG00000176204 | LRRTM4 | -0,974995188 | 0,508741543 | 0,00197 | 0,04968365 | leucine rich repeat transmembrane neuronal 4 |
| ENSG00000272092 | AC087623,2 | -0,975148946 | 0,508687326 | 0,04256 | 0,2782038 | novel transcript |
| ENSG00000259230 | LINC02323 | -0,975260166 | 0,508648112 | 0,29596 | 0,58942061 | long intergenic non-protein coding RNA 2323 |
| ENSG00000185019 | UBOX5 | -0,97531434 | 0,508629012 | 0,07033 | 0,34995627 | U-box domain containing 5 |
| ENSG00000136868 | SLC31A1 | -0,975798129 | 0,508458479 | 0,07819 | 0,36540845 | solute carrier family 31 member 1 |

| | | | | | | |
|---|---|---|---|---|---|---|
| ENSG00000278788 | SBNO2 | -0,975806239 | 0,50845562 | 0,33122 | 0,61131843 | strawberry notch homolog 2 |
| ENSG00000101194 | SLC17A9 | -0,976003493 | 0,508386106 | 0,21017 | 0,52904355 | solute carrier family 17 member 9 |
| ENSG00000264015 | AC124254,2 | -0,97629197 | 0,508284461 | 0,01549 | 0,16309786 | novel transcript |
| ENSG00000227827 | PKD1P2 | -0,97638677 | 0,508251062 | 0,02034 | 0,18946827 | polycystin 1, transient receptor potential channel interacting pseudogene 2 |
| ENSG00000251279 | SMIM15-AS1 | -0,97649016 | 0,50821464 | 0,31183 | 0,59925736 | SMIM15 antisense RNA 1 |
| ENSG00000160949 | TONSL | -0,97652272 | 0,50820317 | 0,02134 | 0,19390172 | tonsoku like, DNA repair protein |
| ENSG00000152128 | TMEM163 | -0,97681677 | 0,508099599 | 0,00841 | 0,11567258 | transmembrane protein 163 |
| ENSG00000284977 | AL160272,1 | -0,976970155 | 0,508045581 | 0,08767 | 0,38329529 | novel transcript |
| ENSG00000166676 | TVP23A | -0,977037103 | 0,508022006 | 0,00305 | 0,06583598 | trans-golgi network vesicle protein 23 homolog A |
| ENSG00000284657 | AL031432,5 | -0,977457722 | 0,507873913 | 0,1163 | 0,43279558 | novel transcript |
| ENSG00000141098 | GFOD2 | -0,977618462 | 0,507817331 | 0,0321 | 0,24164117 | glucose-fructose oxidoreductase domain containing 2 |
| ENSG00000188917 | TRMT2B | -0,97765062 | 0,507806012 | 0,02767 | 0,22412082 | tRNA methyltransferase 2 homolog B |
| ENSG00000185215 | TNFAIP2 | -0,977916146 | 0,50771256 | 0,00509 | 0,08906183 | TNF alpha induced protein 2 |
| ENSG00000253327 | RAD21-AS1 | -0,978289978 | 0,507581018 | 0,07703 | 0,36376818 | RAD21 antisense RNA 1 |
| ENSG00000262655 | SPON1 | -0,978341093 | 0,507563034 | 0,05129 | 0,30462001 | spondin 1 |
| ENSG00000265802 | RN7SL49P | -0,978488554 | 0,507511158 | 0,12854 | 0,45191041 | RNA, 7SL, cytoplasmic 49, pseudogene |
| ENSG00000229463 | LYST-AS1 | -0,978733947 | 0,507424841 | 0,39441 | 0,65729552 | LYST antisense RNA 1 |
| ENSG00000188771 | PLET1 | -0,979148461 | 0,507279069 | 0,22778 | 0,54190354 | placenta expressed transcript 1 |
| ENSG00000228570 | NUTM2E | -0,979175069 | 0,507269713 | 0,02617 | 0,21796119 | NUT family member 2E |
| ENSG00000168955 | TM4SF20 | -0,979185045 | 0,507266205 | 0,06126 | 0,32857513 | transmembrane 4 L six family member 20 |
| ENSG00000288105 | AC117461,1 | -0,979427415 | 0,507180993 | 0,05919 | 0,32424653 | novel transcript, antisense to ROBO1 |
| ENSG00000237112 | GABBR1 | -0,979586986 | 0,507124899 | 0,10497 | 0,41375163 | gamma-aminobutyric acid type B receptor subunit 1 |
| ENSG00000206466 | GABBR1 | -0,979586986 | 0,507124899 | 0,10497 | 0,41375163 | gamma-aminobutyric acid type B receptor subunit 1 |
| ENSG00000151952 | TMEM132D | -0,979592265 | 0,507123043 | 0,04063 | 0,27256659 | transmembrane protein 132D |
| ENSG00000281356 | GOLGA8IP | -0,979625174 | 0,507111475 | 0,1072 | 0,41730934 | golgin A8 family member I, pseudogene |
| ENSG00000273356 | LINC02019 | -0,979844463 | 0,507034401 | 0,13154 | 0,45658336 | long intergenic non-protein coding RNA 2019 |
| ENSG00000278704 | BX004987,1 | -0,980940497 | 0,506649346 | 0,04881 | 0,29706388 | |
| ENSG00000171680 | PLEKHG5 | -0,981097967 | 0,506594048 | 0,00097 | 0,03190323 | pleckstrin homology and RhoGEF domain containing G5 |
| ENSG00000073350 | LLGL2 | -0,981144783 | 0,50657761 | 0,09143 | 0,39105147 | LLGL scribble cell polarity complex component 2 |
| ENSG00000262334 | RPH3AL | -0,981434432 | 0,506475914 | 0,00044 | 0,01893744 | rabphilin 3A like (without C2 domains) |
| ENSG00000223500 | RNF14P4 | -0,981495188 | 0,506454586 | 0,08608 | 0,3803347 | RNF14 pseudogene 4 |
| ENSG00000197813 | AC011450,1 | -0,981777565 | 0,506355468 | 0,00381 | 0,0752235 | novel transcript |

| | | | | | | |
|---|---|---|---|---|---|---|
| ENSG00000260163 | AC012508,1 | -0,981844826 | 0,506331861 | 0,07422 | 0,35761188 | novel transcript |
| ENSG00000102174 | PHEX | -0,981998662 | 0,506277873 | 0,00689 | 0,10589898 | phosphate regulating endopeptidase homolog X-linked |
| ENSG00000235100 | AL157400,3 | -0,982088615 | 0,506246308 | 0,12908 | 0,45248057 | novel transcript |
| ENSG00000112473 | SLC39A7 | -0,982417393 | 0,506130951 | 0,03723 | 0,26113308 | solute carrier family 39 member 7 |
| ENSG00000213965 | NUDT19 | -0,982447916 | 0,506120243 | 0,01099 | 0,1353263 | nudix hydrolase 19 |
| ENSG00000229183 | PGA4 | -0,982778523 | 0,506004275 | 0,3289 | 0,60998638 | pepsinogen A4 |
| ENSG00000231189 | AC013448,1 | -0,98292973 | 0,505951243 | 0,0431 | 0,27965963 | novel transcript |
| ENSG00000170962 | PDGFD | -0,982936084 | 0,505949015 | 0,04008 | 0,27058857 | platelet derived growth factor D |
| ENSG00000236438 | FAM157A | -0,982963085 | 0,505939546 | 0,12447 | 0,44546792 | family with sequence similarity 157 member A |
| ENSG00000180979 | LRRC57 | -0,983021088 | 0,505919205 | 0,04826 | 0,29506787 | leucine rich repeat containing 57 |
| ENSG00000260838 | AC022893,2 | -0,98325692 | 0,505836511 | 0,00512 | 0,08916539 | novel transcript |
| ENSG00000275023 | MLLT6 | -0,983461103 | 0,505764926 | 0,05538 | 0,31482368 | MLLT6, PHD finger containing |
| ENSG00000267439 | AD000671,3 | -0,983695119 | 0,505682894 | 0,3104 | 0,59844173 | novel transcript |
| ENSG00000264546 | AC008026,3 | -0,983975598 | 0,505584592 | 0,0571 | 0,31865533 | novel transcript |
| ENSG00000273992 | NLRP2 | -0,984131366 | 0,505530007 | 0,04261 | 0,2782038 | NLR family pyrin domain containing 2 |
| ENSG00000254303 | AC037486,1 | -0,984246481 | 0,505489671 | 0,15459 | 0,48266733 | novel transcript |
| ENSG00000203618 | GP1BB | -0,984441601 | 0,50542131 | 0,16261 | 0,48945635 | glycoprotein Ib platelet subunit beta |
| ENSG00000268650 | AC005759,1 | -0,984704483 | 0,505329223 | 0,18679 | 0,51023628 | novel transcript, antisense to PDE4C |
| ENSG00000240328 | AC091805,1 | -0,98474403 | 0,505315371 | 0,23358 | 0,54563244 | ribosomal protein L34 (RPL34) pseudogene |
| ENSG00000151413 | NUBPL | -0,985315393 | 0,505115286 | 0,05236 | 0,30715323 | nucleotide binding protein like |
| ENSG00000171847 | FAM90A1 | -0,985533536 | 0,505038916 | 0,03087 | 0,2370319 | family with sequence similarity 90 member A1 |
| ENSG00000260417 | AC092127,1 | -0,985746979 | 0,504964202 | 0,06882 | 0,34677117 | novel transcript |
| ENSG00000128655 | PDE11A | -0,985748435 | 0,504963693 | 0,02636 | 0,21871612 | phosphodiesterase 11A |
| ENSG00000104852 | SNRNP70 | -0,985781782 | 0,504952021 | 0,03919 | 0,26793262 | small nuclear ribonucleoprotein U1 subunit 70 |
| ENSG00000223723 | TFDP1P2 | -0,985791186 | 0,504948729 | 0,25124 | 0,55719434 | TFDP1 pseudogene 2 |
| ENSG00000267322 | SNHG22 | -0,985834813 | 0,50493346 | 0,04558 | 0,28834791 | small nucleolar RNA host gene 22 |
| ENSG00000148356 | LRSAM1 | -0,985990687 | 0,504878908 | 0,03534 | 0,25465323 | leucine rich repeat and sterile alpha motif containing 1 |
| ENSG00000275120 | AC048382,5 | -0,986747065 | 0,504614279 | 0,15964 | 0,48693054 | novel transcript, antisense to ZSCAN2 |
| ENSG00000177380 | PPFIA3 | -0,986862371 | 0,50457395 | 0,03781 | 0,26327969 | PTPRF interacting protein alpha 3 |
| ENSG00000280479 | CACFD1 | -0,986881499 | 0,50456726 | 0,05009 | 0,30089233 | calcium channel flower domain containing 1 |
| ENSG00000100311 | PDGFB | -0,98699505 | 0,504527548 | 0,13752 | 0,46456422 | platelet derived growth factor subunit B |
| ENSG00000215039 | CD27-AS1 | -0,987290206 | 0,504424339 | 0,00145 | 0,04066188 | CD27 antisense RNA 1 |
| ENSG00000198393 | ZNF26 | -0,987494532 | 0,504352904 | 0,05744 | 0,31984771 | zinc finger protein 26 |

| | | | | | | |
|---|---|---|---|---|---|---|
| ENSG00000177335 | C8orf31 | -0,98761871 | 0,504309494 | 0,00149 | 0,04142724 | chromosome 8 open reading frame 31 |
| ENSG00000162510 | MATN1 | -0,987679653 | 0,504288191 | 0,29224 | 0,58692263 | matrilin 1 |
| ENSG00000261227 | AC140912,1 | -0,987776649 | 0,504254288 | 0,26908 | 0,57085484 | novel transcript |
| ENSG00000150967 | ABCB9 | -0,987945072 | 0,504195423 | 0,02538 | 0,21424167 | ATP binding cassette subfamily B member 9 |
| ENSG00000227963 | RBM15-AS1 | -0,98795774 | 0,504190996 | 0,21984 | 0,53722186 | RBM15 antisense RNA 1 |
| ENSG00000205037 | AC134312,1 | -0,988008536 | 0,504173244 | 0,15556 | 0,48345635 | novel transcript |
| ENSG00000228218 | ATF4P3 | -0,988029261 | 0,504166002 | 0,16052 | 0,48780354 | activating transcription factor 4 pseudogene 3 |
| ENSG00000164430 | CGAS | -0,988670654 | 0,50394191 | 0,07813 | 0,36540845 | cyclic GMP-AMP synthase |
| ENSG00000221890 | NPTXR | -0,988699994 | 0,503931661 | 0,00307 | 0,06617279 | neuronal pentraxin receptor |
| ENSG00000236031 | AL591721,1 | -0,988790178 | 0,503900161 | 0,0796 | 0,36831832 | novel transcript |
| ENSG00000281618 | IFITM10 | -0,9888721 | 0,503871548 | 0,33352 | 0,61346096 | interferon induced transmembrane protein 10 |
| ENSG00000188732 | FAM221A | -0,988903734 | 0,5038605 | 0,05746 | 0,31984771 | family with sequence similarity 221 member A |
| ENSG00000260589 | STAM-AS1 | -0,988919659 | 0,503854938 | 0,08336 | 0,37550713 | STAM antisense RNA 1 (head to head) |
| ENSG00000164007 | CLDN19 | -0,989113204 | 0,503787348 | 0,11348 | 0,42823527 | claudin 19 |
| ENSG00000273711 | AC005520,5 | -0,989147038 | 0,503775533 | 0,07149 | 0,35329688 | novel transcript, antisense to PTGR2 |
| ENSG00000133627 | ACTR3B | -0,989499594 | 0,503652439 | 0,05957 | 0,32495341 | actin related protein 3B |
| ENSG00000063015 | SEZ6 | -0,989526037 | 0,503643208 | 0,06857 | 0,34608817 | seizure related 6 homolog |
| ENSG00000260774 | AC021087,3 | -0,989550156 | 0,503634788 | 0,06513 | 0,33804854 | novel transcript |
| ENSG00000265942 | RN7SL577P | -0,989857632 | 0,503527462 | 0,22034 | 0,53750545 | RNA, 7SL, cytoplasmic 577, pseudogene |
| ENSG00000254855 | AP001107,8 | -0,989877346 | 0,503520581 | 0,39876 | 0,6612889 | novel transcript |
| ENSG00000226091 | LINC00937 | -0,990092598 | 0,503445461 | 0,02922 | 0,2303054 | long intergenic non-protein coding RNA 937 |
| ENSG00000224608 | HLA-B | -0,990750605 | 0,503215894 | 0,14698 | 0,47471707 | major histocompatibility complex, class I, B |
| ENSG00000188687 | SLC4A5 | -0,990870825 | 0,503173963 | 0,03764 | 0,26251324 | solute carrier family 4 member 5 |
| ENSG00000196507 | TCEAL3 | -0,991013931 | 0,503124054 | 0,00086 | 0,02908139 | transcription elongation factor A like 3 |
| ENSG00000282740 | AL021920,3 | -0,99113577 | 0,503081565 | 0,71966 | 0,86592885 | novel transcript |
| ENSG00000214309 | MBLAC1 | -0,991372312 | 0,502999088 | 0,09052 | 0,38920435 | metallo-beta-lactamase domain containing 1 |
| ENSG00000130921 | C12orf65 | -0,991439202 | 0,502975767 | 0,05894 | 0,3241112 | chromosome 12 open reading frame 65 |
| ENSG00000095485 | CWF19L1 | -0,991500355 | 0,502954447 | 0,05009 | 0,30089233 | CWF19 like cell cycle control factor 1 |
| ENSG00000253807 | LINC01170 | -0,991648433 | 0,502902827 | 0,25243 | 0,55796133 | long intergenic non-protein coding RNA 1170 |
| ENSG00000228784 | LINC00954 | -0,99182449 | 0,502841459 | 0,01156 | 0,1398032 | long intergenic non-protein coding RNA 954 |
| ENSG00000204464 | C1orf195 | -0,991956827 | 0,502795336 | 0,5753 | 0,7801789 | chromosome 1 open reading frame 195 |
| ENSG00000173614 | NMNAT1 | -0,992513551 | 0,502601349 | 0,04658 | 0,29026543 | nicotinamide nucleotide adenylyltransferase 1 |
| ENSG00000229921 | KIF25-AS1 | -0,992553958 | 0,502587272 | 0,07779 | 0,36490452 | KIF25 antisense RNA 1 |

| | | | | | | |
|---|---|---|---|---|---|---|
| ENSG00000272512 | AL645608,7 | -0,992565947 | 0,502583096 | 0,10268 | 0,41043823 | novel transcript |
| ENSG00000225619 | MYT1L-AS1 | -0,992670602 | 0,502546639 | 0,09772 | 0,40284021 | MYT1L antisense RNA 1 |
| ENSG00000239686 | AL158801,1 | -0,992823906 | 0,50249324 | 0,19468 | 0,51562158 | ribosomal protein L34 (RPL34) pseudogene |
| ENSG00000185955 | C7orf61 | -0,99285421 | 0,502482685 | 0,22524 | 0,54058261 | chromosome 7 open reading frame 61 |
| ENSG00000188674 | C2orf80 | -0,993207148 | 0,502359774 | 0,19064 | 0,51369186 | chromosome 2 open reading frame 80 |
| ENSG00000206356 | AC114324,1 | -0,993279773 | 0,502334486 | 0,17223 | 0,49710979 | putative POM121-like protein 1-like |
| ENSG00000277559 | AC018553,1 | -0,993499183 | 0,502258095 | 0,32005 | 0,60496053 | novel transcript |
| ENSG00000230549 | USP17L1 | -0,993727794 | 0,502178513 | 0,35872 | 0,63189522 | ubiquitin specific peptidase 17 like family member 1 |
| ENSG00000183828 | NUDT14 | -0,99383073 | 0,502142684 | 0,05487 | 0,31349896 | nudix hydrolase 14 |
| ENSG00000228589 | SPCS2P4 | -0,993834016 | 0,50214154 | 0,31166 | 0,59911644 | signal peptidase complex subunit 2 pseudogene 4 |
| ENSG00000240922 | LSAMP-AS1 | -0,994020986 | 0,502076468 | 0,14565 | 0,47307324 | LSAMP antisense RNA 1 |
| ENSG00000158220 | ESYT3 | -0,994071789 | 0,502058788 | 0,08006 | 0,36931124 | extended synaptotagmin 3 |
| ENSG00000230293 | MSH5 | -0,994385347 | 0,501949682 | 0,13918 | 0,46648276 | mutS homolog 5 |
| ENSG00000105852 | PON3 | -0,99447378 | 0,501918915 | 0,24881 | 0,55591283 | paraoxonase 3 |
| ENSG00000236276 | NDP-AS1 | -0,994816682 | 0,501799632 | 0,15444 | 0,48266733 | NDP antisense RNA 1 |
| ENSG00000248125 | AC008700,1 | -0,995256625 | 0,501646634 | 0,3098 | 0,59807209 | novel transcript |
| ENSG00000263624 | AC055811,1 | -0,995804064 | 0,501456317 | 0,01713 | 0,17198199 | novel transcript |
| ENSG00000227370 | AC254562,1 | -0,995811559 | 0,501453712 | 0,1492 | 0,47682725 | chromosome 6 open reading frame 108 (C6orf108) pseudogene |
| ENSG00000206567 | AC022007,1 | -0,996322823 | 0,501276038 | 0,04625 | 0,28930497 | novel transcript |
| ENSG00000225663 | MCRIP1 | -0,996335502 | 0,501271632 | 0,04213 | 0,27783166 | MAPK regulated corepressor interacting protein 1 |
| ENSG00000253335 | AC009884,1 | -0,99651038 | 0,501210874 | 0,1217 | 0,44093327 | novel transcript |
| ENSG00000270894 | AC015849,4 | -0,996940833 | 0,501061351 | 0,1823 | 0,50576743 | novel transcript, sense overlapping TAF15 |
| ENSG00000073605 | GSDMB | -0,997018207 | 0,501034479 | 0,03199 | 0,24129103 | gasdermin B |
| ENSG00000206279 | DAXX | -0,997051039 | 0,501023077 | 0,09109 | 0,39033853 | death domain associated protein |
| ENSG00000288213 | MCRIP1 | -0,997402849 | 0,500900915 | 0,04204 | 0,27770943 | MAPK regulated corepressor interacting protein 1 |
| ENSG00000276467 | Z93929,1 | -0,997985526 | 0,500698651 | 0,15451 | 0,48266733 | zinc finger protein 831 (ZNF831) pseudogene |
| ENSG00000234019 | AL591212,1 | -0,998222866 | 0,500616287 | 0,24246 | 0,55137204 | novel transcript |
| ENSG00000259512 | HNRNPA1P5 | -0,998404866 | 0,500553137 | 0,11138 | 0,42416781 | heterogeneous nuclear ribonucleoprotein A1 pseudogene 5 |
| ENSG00000254598 | CSNK2A3 | -0,998411331 | 0,500550894 | 0,20732 | 0,52677688 | casein kinase 2 alpha 3 |
| ENSG00000228352 | AL354989,1 | -0,998647543 | 0,500468946 | 0,21068 | 0,52921344 | novel transcript |
| ENSG00000196218 | RYR1 | -0,998770892 | 0,500426158 | 0,00213 | 0,05214547 | ryanodine receptor 1 |

| | | | | | | |
|---|---|---|---|---|---|---|
| ENSG00000203710 | CR1 | -0,999159479 | 0,500291387 | 0,22533 | 0,54070011 | complement C3b/C4b receptor 1 (Knops blood group) |
| ENSG00000215302 | AC127502,1 | -0,999232862 | 0,50026594 | 0,03759 | 0,26238402 | WAS protein homolog associated with actin, golgi membranes and microtubules (WHAMM) pseudogene |
| ENSG00000174038 | C9orf131 | -0,999437019 | 0,500195152 | 0,00325 | 0,06841857 | chromosome 9 open reading frame 131 |
| ENSG00000228043 | AC114763,1 | -0,999557927 | 0,500153234 | 0,16819 | 0,49432269 | novel transcript |
| ENSG00000174776 | WDR49 | -0,999653424 | 0,500120129 | 0,02684 | 0,22105403 | WD repeat domain 49 |
| ENSG00000275226 | LINC00547 | -0,999991007 | 0,500003117 | 0,10424 | 0,41262354 | long intergenic non-protein coding RNA 547 |
| ENSG00000263823 | AC009831,1 | -1,000189019 | 0,499934495 | 0,10626 | 0,4157803 | novel transcript, antisense to KIAA1012 |
| ENSG00000163156 | SCNM1 | -1,000498061 | 0,499827415 | 0,06229 | 0,33075396 | sodium channel modifier 1 |
| ENSG00000184809 | B3GALT5-AS1 | -1,000627432 | 0,499782596 | 0,01156 | 0,1398032 | B3GALT5 antisense RNA 1 |
| ENSG00000285394 | C3orf52 | -1,00104383 | 0,499638367 | 0,24312 | 0,55199251 | chromosome 3 open reading frame 52 |
| ENSG00000160908 | ZNF394 | -1,00121761 | 0,499578186 | 0,0566 | 0,31740666 | zinc finger protein 394 |
| ENSG00000282725 | CORO7 | -1,001236768 | 0,499571553 | 0,23216 | 0,54496653 | coronin 7 |
| ENSG00000273062 | AL449106,1 | -1,001260084 | 0,499563479 | 0,14189 | 0,4683384 | novel transcript, antisense to RALGPS2 |
| ENSG00000229203 | AC103564,1 | -1,001329891 | 0,499539307 | 0,29403 | 0,58837921 | novel transcript |
| ENSG00000249740 | OSMR-AS1 | -1,001416848 | 0,499509199 | 0,16171 | 0,48914394 | OSMR antisense RNA 1 (head to head) |
| ENSG00000161618 | ALDH16A1 | -1,001485802 | 0,499485326 | 0,06777 | 0,34426294 | aldehyde dehydrogenase 16 family member A1 |
| ENSG00000176438 | SYNE3 | -1,001952083 | 0,499323917 | 0,20715 | 0,52670822 | spectrin repeat containing nuclear envelope family member 3 |
| ENSG00000244265 | SIAH2-AS1 | -1,002315248 | 0,49919824 | 0,26278 | 0,56582568 | SIAH2 antisense RNA 1 |
| ENSG00000249700 | SRD5A3-AS1 | -1,002416385 | 0,499163246 | 0,09574 | 0,39858189 | SRD5A3 antisense RNA 1 |
| ENSG00000241713 | LY6G5B | -1,002421658 | 0,499161421 | 0,10868 | 0,41972109 | lymphocyte antigen 6 family member G5B |
| ENSG00000237359 | AL354977,2 | -1,002534592 | 0,499122349 | 0,09611 | 0,39936437 | novel transcript |
| ENSG00000276805 | AL133216,2 | -1,002598897 | 0,499100102 | 0,01616 | 0,16645939 | general transcription factor IIi (GTF2I) pseudogene |
| ENSG00000272382 | AC025171,5 | -1,002603154 | 0,499098629 | 0,19477 | 0,51580179 | novel transcript |
| ENSG00000272692 | AC010997,4 | -1,002765151 | 0,499042589 | 0,15345 | 0,48185226 | novel transcript |
| ENSG00000287737 | AC132872,5 | -1,002879758 | 0,499002947 | 0,05564 | 0,31517819 | novel transcript |
| ENSG00000244357 | RN7SL145P | -1,002956139 | 0,498976529 | 0,24081 | 0,55001001 | RNA, 7SL, cytoplasmic 145, pseudogene |
| ENSG00000273374 | AC069222,1 | -1,003114444 | 0,49892178 | 0,10336 | 0,41114168 | novel transcript |
| ENSG00000179057 | IGSF22 | -1,003340259 | 0,498843694 | 0,16649 | 0,49245911 | immunoglobulin superfamily member 22 |
| ENSG00000287351 | AC092120,3 | -1,003556216 | 0,498769027 | 0,2919 | 0,58674733 | novel transcript, antisense to ANKRD11 |
| ENSG00000245468 | LINC02447 | -1,003787157 | 0,498689193 | 0,16342 | 0,48977932 | long intergenic non-protein coding RNA 2447 |
| ENSG00000259232 | AC105129,1 | -1,003797219 | 0,498685714 | 0,19865 | 0,51980353 | target of EGR1, member 1 (nuclear) (TOE1) pseudogene |

| | | | | | | |
|---|---|---|---|---|---|---|
| ENSG00000287165 | AL078621,2 | -1,003847917 | 0,498668191 | 0,20949 | 0,52856218 | novel transcript |
| ENSG00000274939 | TBC1D3B | -1,004156409 | 0,498561572 | 0,05873 | 0,32380167 | TBC1 domain family member 3B |
| ENSG00000273608 | SRCIN1 | -1,004272594 | 0,498521422 | 0,00134 | 0,03910482 | SRC kinase signaling inhibitor 1 |
| ENSG00000183476 | SH2D7 | -1,004671032 | 0,498383762 | 0,01629 | 0,16730761 | SH2 domain containing 7 |
| ENSG00000280543 | ASAP1-IT2 | -1,004761582 | 0,498352482 | 0,06027 | 0,32601555 | ASAP1 intronic transcript 2 |
| ENSG00000276879 | IFI27L2 | -1,005042906 | 0,498255313 | 0,15647 | 0,48425071 | interferon alpha inducible protein 27 like 2 |
| ENSG00000287000 | AC024651,3 | -1,005064607 | 0,498247818 | 0,05813 | 0,32179302 | novel transcript |
| ENSG00000188026 | RILPL1 | -1,005150817 | 0,498218046 | 0,0384 | 0,26521398 | Rab interacting lysosomal protein like 1 |
| ENSG00000271335 | AL117336,2 | -1,00516164 | 0,498214308 | 0,09965 | 0,40615251 | novel transcript, antisense to CCNY |
| ENSG00000204661 | C5orf60 | -1,005224247 | 0,498192688 | 0,39835 | 0,66095882 | chromosome 5 open reading frame 60 |
| ENSG00000287168 | AL512625,3 | -1,005235333 | 0,49818886 | 0,0046 | 0,08360185 | novel transcript |
| ENSG00000172273 | HINFP | -1,005464202 | 0,498109834 | 0,0556 | 0,31505767 | histone H4 transcription factor |
| ENSG00000259883 | EHD4-AS1 | -1,005696728 | 0,498029557 | 0,38805 | 0,65289784 | EHD4 antisense RNA 1 |
| ENSG00000181029 | TRAPPC5 | -1,005866194 | 0,49797106 | 0,0686 | 0,34614163 | trafficking protein particle complex 5 |
| ENSG00000181790 | ADGRB1 | -1,006048469 | 0,497908148 | 0,0007 | 0,02564289 | adhesion G protein-coupled receptor B1 |
| ENSG00000235586 | AC011247,1 | -1,006072436 | 0,497899877 | 0,22354 | 0,53953208 | novel transcript |
| ENSG00000186723 | OR10H1 | -1,006260421 | 0,497835004 | 0,04399 | 0,28274932 | olfactory receptor family 10 subfamily H member 1 |
| ENSG00000119559 | C19orf25 | -1,006372409 | 0,497796362 | 0,03503 | 0,25324761 | chromosome 19 open reading frame 25 |
| ENSG00000130830 | MPP1 | -1,006461731 | 0,497765542 | 0,00695 | 0,10609569 | membrane palmitoylated protein 1 |
| ENSG00000182749 | PAQR7 | -1,006521621 | 0,497744879 | 0,04571 | 0,28843251 | progestin and adipoQ receptor family member 7 |
| ENSG00000234961 | AL133415,1 | -1,006584591 | 0,497723155 | 0,07918 | 0,36743194 | novel transcript |
| ENSG00000255757 | AC087894,1 | -1,006802812 | 0,497647875 | 0,34802 | 0,62441612 | fibroblast growth factor receptor 1 (FGFR1) |
| ENSG00000101204 | CHRNA4 | -1,007086786 | 0,49754993 | 0,0494 | 0,29915481 | cholinergic receptor nicotinic alpha 4 subunit |
| ENSG00000275019 | LILRB3 | -1,007148751 | 0,49752856 | 0,23345 | 0,54563244 | leukocyte immunoglobulin like receptor B3 |
| ENSG00000197296 | FITM2 | -1,007317309 | 0,497470434 | 0,06848 | 0,3458068 | fat storage inducing transmembrane protein 2 |
| ENSG00000223802 | CERS1 | -1,007361799 | 0,497455093 | 0,03335 | 0,24678492 | ceramide synthase 1 |
| ENSG00000262966 | AC005695,1 | -1,007457217 | 0,497422193 | 0,01922 | 0,18350969 | novel transcript, antisense to NTN1 |
| ENSG00000273619 | AL121832,2 | -1,007464121 | 0,497419813 | 0,00483 | 0,08564971 | novel transcript, antisense to RPS21 |
| ENSG00000288012 | AP003307,1 | -1,007854883 | 0,497285103 | 0,09953 | 0,40609117 | novel transcript, antisense to CUL5 |
| ENSG00000081842 | PCDHA6 | -1,008112421 | 0,497196339 | 4,40E-05 | 0,00301003 | protocadherin alpha 6 |
| ENSG00000286721 | AC010623,1 | -1,009074495 | 0,49686489 | 0,01946 | 0,18482021 | novel transcript |
| ENSG00000288557 | AC080097,1 | -1,009175607 | 0,496830068 | 0,22139 | 0,53805234 | novel transcript |
| ENSG00000232518 | AC074366,1 | -1,009288175 | 0,496791304 | 0,2664 | 0,56917012 | novel transcript |

| | | | | | | |
|---|---|---|---|---|---|---|
| ENSG00000167110 | GOLGA2 | -1,00991652 | 0,496574981 | 0,02979 | 0,23233601 | golgin A2 |
| ENSG00000271803 | AL121906,1 | -1,01032555 | 0,496434213 | 0,15798 | 0,48506908 | novel transcript, antisense to NECAB3 |
| ENSG00000287099 | AL121750,1 | -1,010326285 | 0,49643396 | 0,10984 | 0,42203441 | novel transcript, antisense to ADD1 |
| ENSG00000129757 | CDKN1C | -1,010355966 | 0,496423746 | 0,03507 | 0,2533278 | cyclin dependent kinase inhibitor 1C |
| ENSG00000176842 | IRX5 | -1,010380673 | 0,496415245 | 0,14181 | 0,46830665 | iroquois homeobox 5 |
| ENSG00000278249 | SCARNA2 | -1,010435406 | 0,496396412 | 0,04841 | 0,29551886 | small Cajal body-specific RNA 2 |
| ENSG00000270409 | AC090950,1 | -1,011128599 | 0,496157959 | 0,16647 | 0,49245911 | chromosome 9 open reading frame 64 (C9orf64) pseudogene |
| ENSG00000152147 | GEMIN6 | -1,011371144 | 0,496074552 | 0,0588 | 0,32384911 | gem nuclear organelle associated protein 6 |
| ENSG00000243883 | RN7SL419P | -1,011817577 | 0,495921069 | 0,46302 | 0,70413675 | RNA, 7SL, cytoplasmic 419, pseudogene |
| ENSG00000262959 | RPL23AP86 | -1,012195078 | 0,495791321 | 0,26222 | 0,56507751 | ribosomal protein L23a pseudogene 86 |
| ENSG00000113100 | CDH9 | -1,012531637 | 0,495675674 | 0,19271 | 0,51490739 | cadherin 9 |
| ENSG00000231857 | MORF4L1P5 | -1,012691576 | 0,495620726 | 0,11421 | 0,42953156 | mortality factor 4 like 1 pseudogene 5 |
| ENSG00000232569 | GABBR1 | -1,01286485 | 0,495561203 | 0,14372 | 0,47123322 | gamma-aminobutyric acid type B receptor subunit 1 |
| ENSG00000273559 | CWC25 | -1,012969824 | 0,495525146 | 0,05521 | 0,31442055 | CWC25 spliceosome associated protein homolog |
| ENSG00000177045 | SIX5 | -1,012991071 | 0,495517848 | 0,055 | 0,31389132 | SIX homeobox 5 |
| ENSG00000119614 | VSX2 | -1,013003538 | 0,495513567 | 0,36315 | 0,63407828 | visual system homeobox 2 |
| ENSG00000179348 | GATA2 | -1,013301413 | 0,495411268 | 0,37756 | 0,64604267 | GATA binding protein 2 |
| ENSG00000277531 | PNMA8C | -1,013338877 | 0,495398403 | 0,02069 | 0,19083528 | PNMA family member 8C |
| ENSG00000215440 | NPEPL1 | -1,013844386 | 0,49522485 | 0,00206 | 0,0511876 | aminopeptidase like 1 |
| ENSG00000187609 | EXD3 | -1,014026331 | 0,495162399 | 0,01718 | 0,17228282 | exonuclease 3'-5' domain containing 3 |
| ENSG00000203258 | AC013549,1 | -1,014692453 | 0,494933825 | 0,209 | 0,5280176 | novel transcript |
| ENSG00000215014 | AL645728,1 | -1,014772141 | 0,494906488 | 0,07843 | 0,36604454 | novel transcript |
| ENSG00000197162 | ZNF785 | -1,014931758 | 0,494851735 | 0,02016 | 0,18846202 | zinc finger protein 785 |
| ENSG00000144401 | METTL21A | -1,015007291 | 0,494825827 | 0,04479 | 0,28590745 | methyltransferase like 21A |
| ENSG00000223812 | AC073365,1 | -1,015020309 | 0,494821363 | 0,30856 | 0,59736238 | novel transcript |
| ENSG00000188060 | RAB42 | -1,015320387 | 0,494718451 | 0,04767 | 0,29344099 | RAB42, member RAS oncogene family |
| ENSG00000248554 | C5orf34-AS1 | -1,015607876 | 0,494619877 | 0,04357 | 0,28174537 | C5orf34 antisense RNA 1 |
| ENSG00000257947 | AC025030,2 | -1,015715732 | 0,494582901 | 0,26063 | 0,56400103 | novel transcript, antisense to TMEM117 |
| ENSG00000232005 | PBX2 | -1,015754824 | 0,4945695 | 0,07868 | 0,36645437 | PBX homeobox 2 |
| ENSG00000223720 | AL109659,1 | -1,016220427 | 0,494409912 | 0,17102 | 0,49641416 | novel transcript |
| ENSG00000221949 | LINC01465 | -1,016494324 | 0,494316057 | 0,09916 | 0,4054333 | long intergenic non-protein coding RNA 1465 |
| ENSG00000230725 | AL035252,2 | -1,016585394 | 0,494284854 | 0,19357 | 0,51547251 | novel transcript |

| | | | | | | |
|---|---|---|---|---|---|---|
| ENSG00000260577 | AC126773,4 | -1,0173282 | 0,494030425 | 0,01659 | 0,16886598 | novel transcript, antisense to CDH3 |
| ENSG00000287481 | AL161663,1 | -1,01754394 | 0,493956554 | 0,32603 | 0,6085098 | novel transcript, antisense to STXBP6 |
| ENSG00000261423 | TMEM202-AS | -1,017639789 | 0,493923738 | 0,04034 | 0,27164865 | TMEM202 antisense RNA 1 |
| ENSG00000234118 | RPL13AP6 | -1,017655135 | 0,493918484 | 0,07724 | 0,36386039 | ribosomal protein L13a pseudogene 6 |
| ENSG00000249690 | AC110813,1 | -1,01783035 | 0,493858501 | 0,06939 | 0,34799331 | novel transcript, antisense to LRBA |
| ENSG00000236271 | FLOT1 | -1,018487861 | 0,493633476 | 0,06137 | 0,32865897 | flotillin 1 |
| ENSG00000232300 | FAM215B | -1,019179947 | 0,493396728 | 0,01192 | 0,14219678 | family with sequence similarity 215 member B |
| ENSG00000214381 | LINC00488 | -1,01933167 | 0,493344842 | 0,03355 | 0,24724664 | long intergenic non-protein coding RNA 488 |
| ENSG00000228876 | AC010745,2 | -1,019487889 | 0,493291424 | 0,09265 | 0,39335125 | novel transcript |
| ENSG00000286141 | AC097480,2 | -1,019626092 | 0,493244172 | 0,0694 | 0,34799331 | novel transcript |
| ENSG00000166492 | FAM86GP | -1,019727334 | 0,493209559 | 0,13934 | 0,46666406 | family with sequence similarity 86 member G, pseudogene |
| ENSG00000275538 | RNVU1-19 | -1,019742193 | 0,493204479 | 0,47398 | 0,71294959 | RNA, variant U1 small nuclear 19 |
| ENSG00000155052 | CNTNAP5 | -1,020632341 | 0,492900264 | 0,00325 | 0,06855329 | contactin associated protein family member 5 |
| ENSG00000249255 | PGAM1P9 | -1,020716349 | 0,492871563 | 0,1382 | 0,46543358 | phosphoglycerate mutase 1 pseudogene 9 |
| ENSG00000283062 | PGAM1P9 | -1,020716349 | 0,492871563 | 0,1382 | 0,46543358 | phosphoglycerate mutase 1 pseudogene 9 |
| ENSG00000182218 | HHIPL1 | -1,020967454 | 0,492785785 | 0,00339 | 0,07021313 | HHIP like 1 |
| ENSG00000203711 | C6orf99 | -1,021097399 | 0,492741401 | 0,04633 | 0,28930497 | chromosome 6 open reading frame 99 |
| ENSG00000225206 | MIR137HG | -1,021676622 | 0,492543612 | 0,00701 | 0,10660586 | MIR137 host gene |
| ENSG00000214842 | RAD51AP2 | -1,021700512 | 0,492535456 | 0,16331 | 0,48972962 | RAD51 associated protein 2 |
| ENSG00000134317 | GRHL1 | -1,022074284 | 0,492407867 | 0,00614 | 0,09864398 | grainyhead like transcription factor 1 |
| ENSG00000265491 | RNF115 | -1,022546717 | 0,492246647 | 0,0484 | 0,29551007 | ring finger protein 115 |
| ENSG00000165626 | BEND7 | -1,022917567 | 0,492120129 | 0,00812 | 0,11476366 | BEN domain containing 7 |
| ENSG00000231703 | AL354993,1 | -1,023157239 | 0,492038381 | 0,05752 | 0,32010411 | novel transcript, antisense to TSHZ2 |
| ENSG00000180488 | MIGA1 | -1,023840457 | 0,491805421 | 0,04794 | 0,29420465 | mitoguardin 1 |
| ENSG00000128510 | CPA4 | -1,023988629 | 0,491754913 | 0,15522 | 0,48317092 | carboxypeptidase A4 |
| ENSG00000254885 | AP001547,1 | -1,024086729 | 0,491721476 | 0,33229 | 0,61221238 | novel transcript |
| ENSG00000251666 | ZNF346-IT1 | -1,024180927 | 0,491689371 | 0,12474 | 0,44597546 | ZNF346 intronic transcript 1 |
| ENSG00000226031 | FGF13-AS1 | -1,024449562 | 0,491597825 | 0,04183 | 0,27702753 | FGF13 antisense RNA 1 |
| ENSG00000156959 | LHFPL4 | -1,024725956 | 0,491503653 | 0,11045 | 0,42311032 | LHFPL tetraspan subfamily member 4 |
| ENSG00000185250 | PPIL6 | -1,024793318 | 0,491480704 | 0,04237 | 0,27814338 | peptidylprolyl isomerase like 6 |
| ENSG00000196361 | ELAVL3 | -1,024904832 | 0,491442717 | 0,00344 | 0,07086238 | ELAV like RNA binding protein 3 |
| ENSG00000206417 | H1-10-AS1 | -1,02499335 | 0,491412564 | 0,01866 | 0,18080939 | H1-10 antisense RNA 1 |
| ENSG00000261884 | AC040162,1 | -1,025063358 | 0,491388719 | 0,02355 | 0,20483295 | novel transcript |

| | | | | | | |
|---|---|---|---|---|---|---|
| ENSG00000288352 | C1RL-AS1 | -1,025277549 | 0,49131577 | 0,10098 | 0,40770184 | C1RL antisense RNA 1 |
| ENSG00000254397 | AC132192,1 | -1,025306754 | 0,491305824 | 0,22489 | 0,5403718 | novel transcript |
| ENSG00000204390 | HSPA1L | -1,025453917 | 0,49125571 | 0,47488 | 0,71364583 | heat shock protein family A (Hsp70) member 1 like |
| ENSG00000253190 | AC084082,1 | -1,025733012 | 0,491160684 | 0,05594 | 0,3160797 | novel transcript |
| ENSG00000254622 | NAV2-AS4 | -1,026009877 | 0,491066436 | 0,29659 | 0,58979902 | NAV2 antisense RNA 4 |
| ENSG00000225043 | RPL18AP2 | -1,026627149 | 0,490856373 | 0,34568 | 0,62283284 | ribosomal protein L18a pseudogene 2 |
| ENSG00000188739 | RBM34 | -1,026629701 | 0,490855505 | 0,04608 | 0,28930497 | RNA binding motif protein 34 |
| ENSG00000288048 | AC104088,3 | -1,026762093 | 0,490810462 | 0,16532 | 0,49148455 | novel transcript |
| ENSG00000125817 | CENPB | -1,027071907 | 0,490705074 | 0,02551 | 0,21486896 | centromere protein B |
| ENSG00000249743 | AC116345,1 | -1,027284067 | 0,490632917 | 0,13967 | 0,4668467 | novel transcript |
| ENSG00000215769 | ARHGAP27P1- | -1,027464919 | 0,490571416 | 0,00065 | 0,02447582 | ARHGAP27P1-BPTFP1-KPNA2P3 readthrough, transcribed pseudogene |
| ENSG00000135127 | BICDL1 | -1,027471861 | 0,490569056 | 0,02592 | 0,21707964 | BICD family like cargo adaptor 1 |
| ENSG00000253438 | PCAT1 | -1,027864973 | 0,490435402 | 0,06722 | 0,34325886 | prostate cancer associated transcript 1 |
| ENSG00000242759 | LINC00882 | -1,028072956 | 0,490364704 | 0,03241 | 0,2429407 | long intergenic non-protein coding RNA 882 |
| ENSG00000124302 | CHST8 | -1,028158701 | 0,490335561 | 0,00895 | 0,11994782 | carbohydrate sulfotransferase 8 |
| ENSG00000155085 | AK9 | -1,028229782 | 0,490311403 | 0,03438 | 0,25062659 | adenylate kinase 9 |
| ENSG00000229953 | AL590666,2 | -1,028676301 | 0,490159673 | 0,25494 | 0,55945767 | novel transcript |
| ENSG00000239213 | NCK1-DT | -1,028806274 | 0,490115516 | 0,04167 | 0,2764623 | NCK1 divergent transcript |
| ENSG00000179922 | ZNF784 | -1,028817779 | 0,490111608 | 0,04387 | 0,28244553 | zinc finger protein 784 |
| ENSG00000205325 | AC005863,1 | -1,028915384 | 0,49007845 | 0,11032 | 0,42293017 | novel transcript |
| ENSG00000166257 | SCN3B | -1,028968421 | 0,490060434 | 0,01999 | 0,18755241 | sodium voltage-gated channel beta subunit 3 |
| ENSG00000235176 | POLR1H | -1,029150513 | 0,489998585 | 0,52531 | 0,74899497 | RNA polymerase I subunit H |
| ENSG00000215866 | LINC01356 | -1,02920257 | 0,489980904 | 0,02524 | 0,21358358 | long intergenic non-protein coding RNA 1356 |
| ENSG00000270059 | AC121493,1 | -1,029277093 | 0,489955595 | 0,03238 | 0,2428284 | novel transcript, intronic to MAGI1 |
| ENSG00000278128 | AL139093,1 | -1,029328089 | 0,489938276 | 0,06072 | 0,32716307 | nuclear pore associated protein 1 (NPAP1) pseudogene |
| ENSG00000206355 | CCHCR1 | -1,029478525 | 0,489887191 | 0,01766 | 0,17511677 | coiled-coil alpha-helical rod protein 1 |
| ENSG00000205181 | LINC00654 | -1,029976487 | 0,48971813 | 0,01181 | 0,14154028 | long intergenic non-protein coding RNA 654 |
| ENSG00000233871 | DLG5-AS1 | -1,030004224 | 0,489708715 | 0,058 | 0,32135028 | DLG5 antisense RNA 1 |
| ENSG00000103253 | HAGHL | -1,030105174 | 0,48967445 | 0,07089 | 0,35165018 | hydroxyacylglutathione hydrolase like |
| ENSG00000287740 | AC115485,1 | -1,030350554 | 0,489591171 | 0,47615 | 0,71446062 | novel transcript |
| ENSG00000103811 | CTSH | -1,030488933 | 0,489544213 | 0,02858 | 0,22861157 | cathepsin H |
| ENSG00000286373 | AC022387,2 | -1,030492117 | 0,489543133 | 0,09383 | 0,39569427 | novel transcript |

| | | | | | | |
|---|---|---|---|---|---|---|
| ENSG00000286320 | AC108053,1 | -1,030519655 | 0,489533788 | 0,01249 | 0,14623734 | novel transcript, antisense to MAML3 |
| ENSG00000130787 | HIP1R | -1,030528851 | 0,489530668 | 0,00953 | 0,12494325 | huntingtin interacting protein 1 related |
| ENSG00000223442 | TH2LCRR | -1,030568886 | 0,489517083 | 0,19255 | 0,51490739 | T helper type 2 locus control region associated RNA |
| ENSG00000130818 | ZNF426 | -1,030584148 | 0,489511905 | 0,05219 | 0,30681232 | zinc finger protein 426 |
| ENSG00000271314 | AL161729,2 | -1,030712528 | 0,489468347 | 0,09771 | 0,40284021 | novel transcript, antisense to PTCH1 |
| ENSG00000118729 | CASQ2 | -1,03164677 | 0,489151486 | 0,08838 | 0,38462876 | calsequestrin 2 |
| ENSG00000257097 | CLIP1-AS1 | -1,031691558 | 0,489136301 | 0,04722 | 0,29217708 | CLIP1 antisense RNA 1 |
| ENSG00000253208 | PSME2P5 | -1,031726551 | 0,489124437 | 0,02227 | 0,19831519 | proteasome activator subunit 2 pseudogene 5 |
| ENSG00000230286 | AC013472,1 | -1,031881093 | 0,489072044 | 0,37656 | 0,64494482 | novel transcript |
| ENSG00000120262 | CCDC170 | -1,032344086 | 0,488915115 | 0,11838 | 0,43614969 | coiled-coil domain containing 170 |
| ENSG00000152785 | BMP3 | -1,032530286 | 0,488852018 | 0,09717 | 0,40152057 | bone morphogenetic protein 3 |
| ENSG00000008710 | PKD1 | -1,032681948 | 0,488800631 | 0,01905 | 0,18296265 | polycystin 1, transient receptor potential channel interacting |
| ENSG00000189045 | ANKDD1B | -1,03308078 | 0,488665521 | 0,1855 | 0,5089355 | ankyrin repeat and death domain containing 1B |
| ENSG00000260884 | AC009120,3 | -1,033912569 | 0,488383861 | 0,08953 | 0,38703218 | novel transcript |
| ENSG00000264660 | AC107926,1 | -1,034208897 | 0,488283558 | 0,02086 | 0,19166705 | novel transcript |
| ENSG00000227354 | RBM26-AS1 | -1,034261475 | 0,488265763 | 0,04257 | 0,2782038 | RBM26 antisense RNA 1 |
| ENSG00000210194 | MT-TE | -1,034672398 | 0,48812671 | 0,07153 | 0,35335586 | mitochondrially encoded tRNA-Glu (GAA/G) |
| ENSG00000278156 | TSC22D1-AS1 | -1,034795138 | 0,488085183 | 0,03726 | 0,26113308 | TSC22D1 antisense RNA 1 |
| ENSG00000162989 | KCNJ3 | -1,035115916 | 0,487976671 | 0,24398 | 0,55253387 | potassium inwardly rectifying channel subfamily J member 3 |
| ENSG00000178722 | C5orf64 | -1,035144204 | 0,487967103 | 0,03042 | 0,23538999 | chromosome 5 open reading frame 64 |
| ENSG00000154188 | ANGPT1 | -1,035184753 | 0,487953388 | 0,03243 | 0,24300657 | angiopoietin 1 |
| ENSG00000101440 | ASIP | -1,035271977 | 0,487923888 | 0,28238 | 0,57972352 | agouti signaling protein |
| ENSG00000261787 | TCF24 | -1,035361203 | 0,487893713 | 0,21104 | 0,52983259 | transcription factor 24 |
| ENSG00000251602 | AL928654,1 | -1,035481421 | 0,487853059 | 0,05484 | 0,31349896 | novel transcript, antisense to MTA1 |
| ENSG00000206129 | AC006305,1 | -1,035503341 | 0,487845646 | 0,4987 | 0,729862 | novel transcript |
| ENSG00000241213 | LINC02024 | -1,036153944 | 0,487625696 | 0,18345 | 0,50703123 | long intergenic non-protein coding RNA 2024 |
| ENSG00000114859 | CLCN2 | -1,036385696 | 0,48754737 | 0,06654 | 0,34124715 | chloride voltage-gated channel 2 |
| ENSG00000224431 | AC063976,2 | -1,036387585 | 0,487546732 | 0,27311 | 0,57344244 | cytoplasmic linker associated protein 2 (CLASP2) pseudogene |
| ENSG00000229006 | TRIM27 | -1,036726284 | 0,487432285 | 0,29247 | 0,58699586 | tripartite motif containing 27 |
| ENSG00000275532 | AC006449,2 | -1,036744576 | 0,487426105 | 0,32805 | 0,60971379 | novel transcript, antisense to MLLT6 |

| | | | | | | |
|---|---|---|---|---|---|---|
| ENSG00000244165 | P2RY11 | -1,036966965 | 0,487350975 | 0,03181 | 0,24056445 | purinergic receptor P2Y11 |
| ENSG00000285275 | FAM167A-AS1 | -1,037241704 | 0,487258175 | 0,44907 | 0,6947028 | FAM167A antisense RNA 1 |
| ENSG00000241462 | AC100832,1 | -1,037630202 | 0,487126981 | 0,21691 | 0,53505085 | ribosomal protein S12 (RPS12) pseudogene |
| ENSG00000218676 | BRD7P4 | -1,037683801 | 0,487108883 | 0,05183 | 0,30603373 | bromodomain containing 7 pseudogene 4 |
| ENSG00000235513 | L3MBTL2-AS1 | -1,037777049 | 0,4870774 | 0,00809 | 0,11436636 | L3MBTL2 antisense RNA 1 |
| ENSG00000234663 | LINC01934 | -1,038137576 | 0,486955696 | 0,0488 | 0,29705113 | long intergenic non-protein coding RNA 1934 |
| ENSG00000228386 | AL031668,1 | -1,038450211 | 0,486850183 | 0,21825 | 0,53571161 | novel transcript |
| ENSG00000131969 | ABHD12B | -1,038568082 | 0,486810408 | 0,05794 | 0,32123433 | abhydrolase domain containing 12B |
| ENSG00000149527 | PLCH2 | -1,038761818 | 0,48674504 | 0,16185 | 0,48914394 | phospholipase C eta 2 |
| ENSG00000272282 | LINC02084 | -1,039001879 | 0,486664053 | 0,2859 | 0,58229607 | long intergenic non-protein coding RNA 2084 |
| ENSG00000277938 | AL035252,3 | -1,039591313 | 0,48646526 | 0,05041 | 0,30203302 | novel transcript |
| ENSG00000244945 | RUFY1-AS1 | -1,039841175 | 0,486381016 | 0,07018 | 0,34975886 | RUFY1 antisense RNA 1 |
| ENSG00000283808 | RUFY1-AS1 | -1,039841175 | 0,486381016 | 0,07018 | 0,34975886 | RUFY1 antisense RNA 1 |
| ENSG00000090932 | DLL3 | -1,039895156 | 0,486362818 | 0,02605 | 0,21736459 | delta like canonical Notch ligand 3 |
| ENSG00000198515 | CNGA1 | -1,04002639 | 0,486318578 | 0,25814 | 0,56220487 | cyclic nucleotide gated channel subunit alpha 1 |
| ENSG00000163735 | CXCL5 | -1,040094974 | 0,486295459 | 0,141 | 0,4679289 | C-X-C motif chemokine ligand 5 |
| ENSG00000276412 | AL591926,2 | -1,040383012 | 0,486198379 | 0,12755 | 0,45080057 | novel transcript |
| ENSG00000138496 | PARP9 | -1,040860917 | 0,486037348 | 0,0334 | 0,24709568 | poly(ADP-ribose) polymerase family member 9 |
| ENSG00000102172 | SMS | -1,041008709 | 0,48598756 | 0,00859 | 0,11699775 | spermine synthase |
| ENSG00000225632 | AL161644,1 | -1,041019515 | 0,48598392 | 0,07657 | 0,36266919 | novel transcript |
| ENSG00000273780 | ARHGAP23 | -1,041156934 | 0,485937632 | 0,03506 | 0,2533278 | Rho GTPase activating protein 23 |
| ENSG00000243319 | FGF14-IT1 | -1,041582736 | 0,485794232 | 0,04234 | 0,27814338 | FGF14 intronic transcript 1 |
| ENSG00000171786 | NHLH1 | -1,041599143 | 0,485788707 | 0,02327 | 0,20387451 | nescient helix-loop-helix 1 |
| ENSG00000243307 | POM121L6P | -1,041607647 | 0,485785843 | 0,06325 | 0,33311692 | POM121 transmembrane nucleoporin like 6, pseudogene |
| ENSG00000258620 | AL135838,1 | -1,041740229 | 0,485741202 | 0,03097 | 0,23719337 | novel transcript, intronic to WDR25 |
| ENSG00000215029 | TCP11X2 | -1,04188103 | 0,485693798 | 0,28733 | 0,58372981 | t-complex 11 family, X-linked 2 |
| ENSG00000256443 | AP003559,1 | -1,041905601 | 0,485685526 | 0,22132 | 0,53805234 | novel transcript |
| ENSG00000282614 | DND1P1 | -1,042499482 | 0,485485637 | 0,05407 | 0,31168784 | DND microRNA-mediated repression inhibitor 1 pseudogene 1 |
| ENSG00000264070 | DND1P1 | -1,042499482 | 0,485485637 | 0,05407 | 0,31168784 | DND microRNA-mediated repression inhibitor 1 pseudogene 1 |
| ENSG00000286024 | AC026803,3 | -1,042915237 | 0,48534575 | 0,18587 | 0,50915451 | novel transcript, antisense to BCAT2 |
| ENSG00000286539 | AC073973,1 | -1,043458067 | 0,485163167 | 0,29734 | 0,59031059 | novel transcript |

| | | | | | | |
|---|---|---|---|---|---|---|
| ENSG00000116871 | MAP7D1 | -1,043459021 | 0,485162847 | 0,03065 | 0,23611505 | MAP7 domain containing 1 |
| ENSG00000265912 | AC037487,1 | -1,043459172 | 0,485162796 | 0,17806 | 0,50236177 | novel transcript |
| ENSG00000259520 | AC051619,4 | -1,043746127 | 0,485066305 | 0,0118 | 0,14150455 | novel transcript, antisense to SLC28A2 |
| ENSG00000203688 | LINC02487 | -1,043864868 | 0,485026384 | 0,06116 | 0,32813211 | long intergenic non-protein coding RNA 2487 |
| ENSG00000235990 | RPL23AP20 | -1,044141146 | 0,484933509 | 0,16405 | 0,4903483 | ribosomal protein L23a pseudogene 20 |
| ENSG00000140527 | WDR93 | -1,044198298 | 0,484914299 | 0,14908 | 0,47668027 | WD repeat domain 93 |
| ENSG00000281706 | LINC01012 | -1,044334908 | 0,484868385 | 0,04507 | 0,2872055 | long intergenic non-protein coding RNA 1012 |
| ENSG00000278068 | AL136317,3 | -1,044383205 | 0,484852153 | 0,32544 | 0,60794826 | calponin 2 (CNN2) pseudogene |
| ENSG00000287303 | AP006248,5 | -1,044466924 | 0,484824018 | 0,389 | 0,65355561 | novel transcript, antisense to PDGFRL |
| ENSG00000148225 | WDR31 | -1,044489631 | 0,484816387 | 0,04416 | 0,28322445 | WD repeat domain 31 |
| ENSG00000197635 | DPP4 | -1,044688201 | 0,484749662 | 0,03643 | 0,25848105 | dipeptidyl peptidase 4 |
| ENSG00000246790 | AP000977,1 | -1,04469227 | 0,484748295 | 0,37423 | 0,64274924 | novel transcript, antisense to SORL1 |
| ENSG00000233622 | CYP2T1P | -1,045107559 | 0,484608777 | 0,04175 | 0,27670157 | cytochrome P450 family 2 subfamily T member 1, pseudogene |
| ENSG00000144182 | LIPT1 | -1,045959389 | 0,484322728 | 0,06148 | 0,32865897 | lipoyltransferase 1 |
| ENSG00000146001 | PCDHB18P | -1,046068254 | 0,484286182 | 0,00018 | 0,00987535 | protocadherin beta 18 pseudogene |
| ENSG00000260853 | AC109460,2 | -1,046180669 | 0,484248448 | 0,05124 | 0,30453389 | novel transcript, antisense to NFATC2IP |
| ENSG00000286364 | AL512308,1 | -1,046185511 | 0,484246823 | 0,05141 | 0,30462001 | novel transcript |
| ENSG00000162873 | KLHDC8A | -1,046382669 | 0,484180651 | 0,00992 | 0,12766438 | kelch domain containing 8A |
| ENSG00000144476 | ACKR3 | -1,046524293 | 0,484133123 | 0,00175 | 0,0459021 | atypical chemokine receptor 3 |
| ENSG00000018607 | ZNF806 | -1,046561025 | 0,484120797 | 0,00837 | 0,11551322 | zinc finger protein 806 |
| ENSG00000224822 | THRB-IT1 | -1,046838246 | 0,484027779 | 0,01732 | 0,17339159 | THRB intronic transcript 1 |
| ENSG00000183943 | PRKX | -1,046943909 | 0,48399233 | 2,26E-05 | 0,0017313 | protein kinase X-linked |
| ENSG00000229657 | AL391822,1 | -1,046947124 | 0,483991252 | 0,18177 | 0,50512193 | ribosomal protein L13a (RPL13A) pseudogene |
| ENSG00000168917 | SLC35G2 | -1,047002611 | 0,483972638 | 0,04134 | 0,27536199 | solute carrier family 35 member G2 |
| ENSG00000267288 | AC138150,2 | -1,047455596 | 0,483820701 | 0,14017 | 0,46710628 | novel transcript, antisense to HEXIM2 |
| ENSG00000111261 | MANSC1 | -1,04758717 | 0,483776579 | 0,22474 | 0,54036191 | MANSC domain containing 1 |
| ENSG00000125122 | LRRC29 | -1,048385661 | 0,483508896 | 0,02158 | 0,19535533 | leucine rich repeat containing 29 |
| ENSG00000276720 | CU151838,3 | -1,048714193 | 0,483398803 | 0,18946 | 0,51258824 | LSM14 homolog A (SCD6, S, cerevisiae) pseudogene |
| ENSG00000270716 | BNIP3P15 | -1,049133173 | 0,483258438 | 0,26853 | 0,57061705 | BCL2 interacting protein 3 pseudogene 15 |
| ENSG00000249346 | LINC01016 | -1,049148876 | 0,483253178 | 0,05067 | 0,30278705 | long intergenic non-protein coding RNA 1016 |
| ENSG00000171130 | ATP6V0E2 | -1,049235553 | 0,483224145 | 0,01392 | 0,15471973 | ATPase H+ transporting V0 subunit e2 |
| ENSG00000275118 | MBOAT7 | -1,049496073 | 0,483136893 | 0,06969 | 0,34826564 | membrane bound O-acyltransferase domain containing 7 |

| | | | | | | |
|---|---|---|---|---|---|---|
| ENSG00000178796 | RIIAD1 | -1,049610222 | 0,483098667 | 0,02396 | 0,20685571 | regulatory subunit of type II PKA R-subunit domain containing 1 |
| ENSG00000164483 | SAMD3 | -1,050211796 | 0,482897267 | 0,01533 | 0,16232837 | sterile alpha motif domain containing 3 |
| ENSG00000260773 | AC055855,1 | -1,050713445 | 0,482729385 | 0,12937 | 0,45283544 | novel transcript, antisense to DIS3L |
| ENSG00000277101 | ARHGEF26 | -1,050744472 | 0,482719003 | 0,05328 | 0,30991926 | Rho guanine nucleotide exchange factor 26 |
| ENSG00000250039 | AC096719,1 | -1,050898885 | 0,48266734 | 0,20026 | 0,52107152 | novel transcript |
| ENSG00000250579 | CTD-2297D10 | -1,051330206 | 0,482523059 | 0,03044 | 0,23546694 | uncharacterized LOC101929176 |
| ENSG00000213471 | TTLL13P | -1,051587286 | 0,482437084 | 0,14681 | 0,47471707 | tubulin tyrosine ligase like 13, pseudogene |
| ENSG00000275624 | PRB3 | -1,051944466 | 0,482317658 | 0,17923 | 0,50368273 | proline rich protein BstNI subfamily 3 |
| ENSG00000174151 | CYB561D1 | -1,052162468 | 0,482244782 | 0,02354 | 0,20483295 | cytochrome b561 family member D1 |
| ENSG00000236053 | LINC01067 | -1,052418366 | 0,482159251 | 0,01349 | 0,15241195 | long intergenic non-protein coding RNA 1067 |
| ENSG00000188107 | EYS | -1,052466455 | 0,48214318 | 0,08341 | 0,37550713 | eyes shut homolog |
| ENSG00000082684 | SEMA5B | -1,052702238 | 0,482064388 | 0,04077 | 0,27302816 | semaphorin 5B |
| ENSG00000224748 | TAP1 | -1,053058647 | 0,481945312 | 0,39189 | 0,65534953 | transporter 1, ATP binding cassette subfamily B member |
| ENSG00000255026 | AC136475,3 | -1,053204067 | 0,481896735 | 0,22155 | 0,53805234 | novel transcript |
| ENSG00000240015 | AC084024,1 | -1,053205924 | 0,481896115 | 0,24748 | 0,55497592 | novel transcript |
| ENSG00000253106 | AC090198,1 | -1,053441204 | 0,481817532 | 0,04699 | 0,29158646 | novel transcript, antisense to TATDN1 |
| ENSG00000278301 | GRAMD4P3 | -1,053479608 | 0,481804707 | 0,01243 | 0,14572968 | GRAM domain containing 4 pseudogene 3 |
| ENSG00000226328 | NUP50-DT | -1,053521581 | 0,48179069 | 0,07053 | 0,35061139 | NUP50 divergent transcript |
| ENSG00000285796 | AL162458,1 | -1,053684404 | 0,481736317 | 0,04715 | 0,2920993 | novel transcript |
| ENSG00000100209 | HSCB | -1,054033614 | 0,481619726 | 0,04524 | 0,2876168 | HscB mitochondrial iron-sulfur cluster cochaperone |
| ENSG00000261291 | AC007906,1 | -1,054463184 | 0,481476342 | 0,33023 | 0,61114402 | novel transcript |
| ENSG00000255933 | AC117500,2 | -1,054464139 | 0,481476023 | 0,11796 | 0,43562473 | novel transcript |
| ENSG00000070729 | CNGB1 | -1,054596925 | 0,48143171 | 0,00541 | 0,09148081 | cyclic nucleotide gated channel subunit beta 1 |
| ENSG00000286363 | AC087427,1 | -1,054802451 | 0,48136313 | 0,0609 | 0,32750733 | novel transcript, sense intronic to CNTN4 |
| ENSG00000179988 | PSTK | -1,054949515 | 0,481314064 | 0,05005 | 0,30089233 | phosphoseryl-tRNA kinase |
| ENSG00000167685 | ZNF444 | -1,055298123 | 0,481197775 | 0,03124 | 0,23849623 | zinc finger protein 444 |
| ENSG00000148948 | LRRC4C | -1,055443261 | 0,481149368 | 0,00209 | 0,05144845 | leucine rich repeat containing 4C |
| ENSG00000251535 | AC079140,4 | -1,055472941 | 0,48113947 | 0,30254 | 0,59306846 | pseudogene similar to part of zinc finger protein 74 (ZNF74) |
| ENSG00000105642 | KCNN1 | -1,055731464 | 0,48105326 | 0,0322 | 0,24207267 | potassium calcium-activated channel subfamily N member 1 |
| ENSG00000157637 | SLC38A10 | -1,05590537 | 0,480995276 | 0,01846 | 0,179662 | solute carrier family 38 member 10 |

| | | | | | | |
|---|---|---|---|---|---|---|
| ENSG00000235518 | AC011196,1 | -1,056346237 | 0,480848313 | 0,17394 | 0,49860418 | novel transcript |
| ENSG00000149926 | TLCD3B | -1,056804963 | 0,480695445 | 0,04316 | 0,27996046 | TLC domain containing 3B |
| ENSG00000225138 | SLC9A3-AS1 | -1,056988159 | 0,480634409 | 0,01996 | 0,18744234 | SLC9A3 antisense RNA 1 |
| ENSG00000287503 | AC027243,3 | -1,05699695 | 0,48063148 | 0,09229 | 0,39278042 | novel transcript, antisense to ETFA |
| ENSG00000140691 | ARMC5 | -1,057167168 | 0,480574776 | 0,06149 | 0,32865897 | armadillo repeat containing 5 |
| ENSG00000108448 | TRIM16L | -1,057172446 | 0,480573018 | 0,06834 | 0,34555414 | tripartite motif containing 16 like |
| ENSG00000285877 | AC007448,5 | -1,05717926 | 0,480570748 | 0,07343 | 0,35662549 | novel transcript, antisense to HELZ |
| ENSG00000261916 | AC027796,1 | -1,057238243 | 0,480551101 | 0,38377 | 0,65041894 | novel transcript, sense intronic to readthrough SHPK-TRPV1 |
| ENSG00000100767 | PAPLN | -1,057594341 | 0,480432502 | 0,05603 | 0,31627719 | papilin, proteoglycan like sulfated glycoprotein |
| ENSG00000155265 | GOLGA7B | -1,05764466 | 0,480415745 | 0,00627 | 0,09973971 | golgin A7 family member B |
| ENSG00000280789 | PAGR1 | -1,057977809 | 0,48030482 | 0,06936 | 0,34794943 | PAXIP1 associated glutamate rich protein 1 |
| ENSG00000266378 | AC005224,2 | -1,058184025 | 0,480236171 | 0,17378 | 0,49846579 | novel transcript |
| ENSG00000261003 | AL008628,1 | -1,058963238 | 0,479976861 | 0,32174 | 0,60577115 | novel transcript, overlapping to FAM120B |
| ENSG00000146830 | GIGYF1 | -1,059014372 | 0,479959849 | 0,03278 | 0,24429193 | GRB10 interacting GYF protein 1 |
| ENSG00000285750 | AL157385,2 | -1,059202999 | 0,4798971 | 0,20095 | 0,52181085 | novel transcript |
| ENSG00000236483 | MTND2P40 | -1,060070046 | 0,479608773 | 0,01846 | 0,179662 | MT-ND2 pseudogene 40 |
| ENSG00000232962 | HLA-DOA | -1,060173962 | 0,479574228 | 0,35747 | 0,63133849 | major histocompatibility complex, class II, DO alpha |
| ENSG00000183196 | CHST6 | -1,060204259 | 0,479564157 | 0,02543 | 0,21440233 | carbohydrate sulfotransferase 6 |
| ENSG00000185347 | TEDC1 | -1,061476477 | 0,479141448 | 0,09018 | 0,38841081 | tubulin epsilon and delta complex 1 |
| ENSG00000164037 | SLC9B1 | -1,061628209 | 0,479091058 | 0,07694 | 0,36356213 | solute carrier family 9 member B1 |
| ENSG00000198658 | ABHD17AP1 | -1,061844557 | 0,479019218 | 0,03959 | 0,26924564 | ABHD17A pseudogene 1 |
| ENSG00000251402 | FAM90A25P | -1,061881008 | 0,479007115 | 0,04047 | 0,27194236 | family with sequence similarity 90 member A25, pseudogene |
| ENSG00000198883 | PNMA5 | -1,061967733 | 0,478978322 | 0,25534 | 0,55992862 | PNMA family member 5 |
| ENSG00000283997 | TMEM159 | -1,062549586 | 0,478785184 | 0,01391 | 0,15466987 | transmembrane protein 159 |
| ENSG00000213563 | C8orf82 | -1,062664493 | 0,478747051 | 0,02543 | 0,21440233 | chromosome 8 open reading frame 82 |
| ENSG00000175745 | NR2F1 | -1,062675335 | 0,478743454 | 0,00165 | 0,0439245 | nuclear receptor subfamily 2 group F member 1 |
| ENSG00000255409 | RSF1-IT1 | -1,062756455 | 0,478716536 | 0,25453 | 0,55944852 | RSF1 intronic transcript 1 |
| ENSG00000162728 | KCNJ9 | -1,063072853 | 0,47861156 | 0,04472 | 0,28565589 | potassium inwardly rectifying channel subfamily J member 9 |
| ENSG00000269949 | AC069307,1 | -1,063239428 | 0,478556302 | 0,18468 | 0,50835796 | novel transcript |
| ENSG00000133597 | ADCK2 | -1,063300802 | 0,478535944 | 0,00506 | 0,08898421 | aarF domain containing kinase 2 |

| | | | | | | |
|---|---|---|---|---|---|---|
| ENSG00000168878 | SFTPB | -1,063411663 | 0,478499173 | 0,06414 | 0,33569301 | surfactant protein B |
| ENSG00000270030 | AC136475,7 | -1,063700858 | 0,478403265 | 0,30296 | 0,593345 | novel transcript |
| ENSG00000213658 | LAT | -1,063742997 | 0,478389292 | 0,10115 | 0,40785299 | linker for activation of T cells |
| ENSG00000175164 | ABO | -1,06387698 | 0,478344866 | 0,17293 | 0,49768904 | ABO, alpha 1-3-N-acetylgalactosaminyltransferase and alpha 1-3-galactosyltransferase |
| ENSG00000264853 | AC011933,2 | -1,063990626 | 0,478307187 | 0,00883 | 0,11905769 | novel transcript, sense intronic to GRB2 |
| ENSG00000287437 | AL161662,1 | -1,064553679 | 0,47812055 | 0,01835 | 0,17922726 | novel transcript, antisense to NRDE2 |
| ENSG00000256001 | AC079949,1 | -1,064825281 | 0,478030548 | 0,06432 | 0,33606729 | novel transcript |
| ENSG00000103528 | SYT17 | -1,065347945 | 0,477857397 | 0,00609 | 0,09844621 | synaptotagmin 17 |
| ENSG00000165655 | ZNF503 | -1,065443364 | 0,477825792 | 0,00124 | 0,03721366 | zinc finger protein 503 |
| ENSG00000226137 | BAIAP2-DT | -1,0655935 | 0,47777607 | 0,01393 | 0,15479024 | BAIAP2 divergent transcript |
| ENSG00000225738 | NRG3-AS1 | -1,065863913 | 0,477686525 | 0,13441 | 0,46058584 | NRG3 antisense RNA 1 |
| ENSG00000074276 | CDHR2 | -1,065933928 | 0,477663344 | 0,01056 | 0,1325438 | cadherin related family member 2 |
| ENSG00000260213 | AC092718,2 | -1,066613078 | 0,477438536 | 0,06106 | 0,32783036 | novel transcript, antisense to CENPN |
| ENSG00000276907 | ARHGAP27 | -1,066658609 | 0,477423468 | 0,06699 | 0,34281679 | Rho GTPase activating protein 27 |
| ENSG00000288046 | AL031123,5 | -1,066664005 | 0,477421683 | 0,24615 | 0,55394307 | novel transcript |
| ENSG00000281853 | AL034349,2 | -1,066919232 | 0,47733723 | 0,11801 | 0,43562845 | novel transcript, antisense to PTPRK |
| ENSG00000227533 | SLC2A1-AS1 | -1,066966468 | 0,477321601 | 0,00278 | 0,06217304 | SLC2A1 antisense RNA 1 |
| ENSG00000281880 | PAUPAR | -1,067000565 | 0,47731032 | 0,09953 | 0,40609117 | PAX6 upstream antisense RNA |
| ENSG00000170044 | ZPLD1 | -1,067113173 | 0,477273066 | 0,41865 | 0,67477592 | zona pellucida like domain containing 1 |
| ENSG00000284700 | AL049637,1 | -1,067255176 | 0,47722609 | 0,1799 | 0,50407792 | novel transcript |
| ENSG00000258910 | LINC01956 | -1,06751105 | 0,477141458 | 0,08671 | 0,38147501 | long intergenic non-protein coding RNA 1956 |
| ENSG00000255468 | AP001107,9 | -1,067543652 | 0,477130676 | 0,07012 | 0,34975886 | novel transcript, antisense to SLC29A2 |
| ENSG00000134962 | KLB | -1,067545044 | 0,477130215 | 0,07497 | 0,3592137 | klotho beta |
| ENSG00000197261 | C6orf141 | -1,067871335 | 0,477022316 | 0,1632 | 0,48972962 | chromosome 6 open reading frame 141 |
| ENSG00000249106 | AC097537,1 | -1,067969138 | 0,476989979 | 0,02908 | 0,2300184 | novel transcript, antisense to GPM6A |
| ENSG00000101463 | SYNDIG1 | -1,068523386 | 0,476806766 | 0,04815 | 0,29485684 | synapse differentiation inducing 1 |
| ENSG00000124006 | OBSL1 | -1,068621928 | 0,4767742 | 0,02689 | 0,22133474 | obscurin like cytoskeletal adaptor 1 |
| ENSG00000266980 | AC087289,1 | -1,068877386 | 0,476689785 | 0,05916 | 0,32424653 | novel transcript, antisense to UNK |
| ENSG00000243926 | TIPARP-AS1 | -1,069064739 | 0,476627884 | 0,07646 | 0,36251887 | TIPARP antisense RNA 1 |
| ENSG00000196391 | ZNF774 | -1,069230825 | 0,476573017 | 0,02471 | 0,21047862 | zinc finger protein 774 |
| ENSG00000239494 | RN7SL333P | -1,069931255 | 0,476341696 | 0,2565 | 0,56092974 | RNA, 7SL, cytoplasmic 333, pseudogene |
| ENSG00000110446 | SLC15A3 | -1,069989311 | 0,476322528 | 0,02102 | 0,19257726 | solute carrier family 15 member 3 |

| | | | | | | |
|---|---|---|---|---|---|---|
| ENSG00000245080 | MIR3150BHG | -1,070180947 | 0,476259262 | 0,07607 | 0,36217039 | MIR3150B host gene |
| ENSG00000187957 | DNER | -1,070306707 | 0,476217748 | 0,03751 | 0,26207258 | delta/notch like EGF repeat containing |
| ENSG00000151715 | TMEM45B | -1,071194426 | 0,475924811 | 0,05645 | 0,31712073 | transmembrane protein 45B |
| ENSG00000283632 | EXOC3L2 | -1,071362101 | 0,475869501 | 0,19567 | 0,51643293 | exocyst complex component 3 like 2 |
| ENSG00000235241 | BX284668,4 | -1,071447297 | 0,4758414 | 0,58096 | 0,78401175 | cyclin-dependent kinase 5 (CDK5) pseudogene |
| ENSG00000167555 | ZNF528 | -1,071527128 | 0,47581507 | 0,03998 | 0,27031997 | zinc finger protein 528 |
| ENSG00000136383 | ALPK3 | -1,071573135 | 0,475799897 | 0,01911 | 0,18296265 | alpha kinase 3 |
| ENSG00000234078 | DDR1 | -1,071674419 | 0,475766495 | 0,00203 | 0,05073405 | discoidin domain receptor tyrosine kinase 1 |
| ENSG00000215522 | DDR1 | -1,071674419 | 0,475766495 | 0,00203 | 0,05073405 | discoidin domain receptor tyrosine kinase 1 |
| ENSG00000229372 | SZT2-AS1 | -1,071987538 | 0,475663247 | 0,29229 | 0,58692263 | SZT2 antisense RNA 1 |
| ENSG00000187260 | WDR86 | -1,072286626 | 0,475564646 | 0,0247 | 0,21047862 | WD repeat domain 86 |
| ENSG00000225793 | TMEM30A-DT | -1,072675322 | 0,475436535 | 0,07254 | 0,35556123 | TMEM30A divergent transcript |
| ENSG00000226375 | AL022310,1 | -1,072746351 | 0,475413129 | 0,14545 | 0,47300165 | novel transcript |
| ENSG00000287922 | AC021979,3 | -1,072806243 | 0,475393393 | 0,02168 | 0,19558343 | novel transcript |
| ENSG00000188064 | WNT7B | -1,072961833 | 0,475342126 | 0,00933 | 0,12336347 | Wnt family member 7B |
| ENSG00000225822 | UBXN7-AS1 | -1,072983985 | 0,475334827 | 0,07231 | 0,35498653 | UBXN7 antisense RNA 1 |
| ENSG00000272449 | AL139246,5 | -1,072984288 | 0,475334727 | 0,24306 | 0,55199251 | novel transcript |
| ENSG00000251637 | LINC02754 | -1,073334787 | 0,47521926 | 0,02064 | 0,19052981 | long intergenic non-protein coding RNA 2754 |
| ENSG00000180660 | MAB21L1 | -1,073731395 | 0,475088637 | 0,00149 | 0,04144939 | mab-21 like 1 |
| ENSG00000183735 | TBK1 | -1,073889319 | 0,475036634 | 0,05627 | 0,31687103 | TANK binding kinase 1 |
| ENSG00000286337 | AC025370,2 | -1,073984831 | 0,475005186 | 0,05266 | 0,30807312 | novel transcript, antisense to FZD6 |
| ENSG00000232891 | AL450344,3 | -1,074005657 | 0,474998329 | 0,32894 | 0,60998638 | novel transcript |
| ENSG00000279243 | LINC02736 | -1,074069184 | 0,474977413 | 0,11678 | 0,43384482 | long intergenic non-protein coding RNA 2736 |
| ENSG00000199631 | SNORD33 | -1,074167182 | 0,474945151 | 0,01092 | 0,13517515 | small nucleolar RNA, C/D box 33 |
| ENSG00000270001 | AL121894,2 | -1,074459518 | 0,474848922 | 0,12247 | 0,4421901 | novel transcript |
| ENSG00000177875 | CCDC184 | -1,074848094 | 0,474721043 | 0,00748 | 0,11001711 | coiled-coil domain containing 184 |
| ENSG00000181896 | ZNF101 | -1,074968825 | 0,474681318 | 0,06538 | 0,33860034 | zinc finger protein 101 |
| ENSG00000169583 | CLIC3 | -1,07533054 | 0,47456232 | 0,43707 | 0,68709205 | chloride intracellular channel 3 |
| ENSG00000223525 | RABGAP1L-IT1 | -1,075393433 | 0,474541632 | 0,02203 | 0,19731935 | RABGAP1L intronic transcript 1 |
| ENSG00000167355 | OR51B5 | -1,07558539 | 0,474478496 | 0,17618 | 0,50029374 | olfactory receptor family 51 subfamily B member 5 |
| ENSG00000143847 | PPFIA4 | -1,075632339 | 0,474463056 | 0,01419 | 0,15631436 | PTPRF interacting protein alpha 4 |
| ENSG00000204052 | LRRC73 | -1,075706245 | 0,474438751 | 0,04324 | 0,28025067 | leucine rich repeat containing 73 |
| ENSG00000118257 | NRP2 | -1,075859849 | 0,47438824 | 0,04663 | 0,29026543 | neuropilin 2 |

| | | | | | | |
|---|---|---|---|---|---|---|
| ENSG00000286769 | AC064869,1 | -1,076002987 | 0,474341176 | 0,49331 | 0,72631 | novel transcript |
| ENSG00000168135 | KCNJ4 | -1,076057743 | 0,474323173 | 0,13832 | 0,46547269 | potassium inwardly rectifying channel subfamily J member 4 |
| ENSG00000214548 | MEG3 | -1,0762417 | 0,474262696 | 0,09958 | 0,40610985 | maternally expressed 3 |
| ENSG00000272767 | JMJD1C-AS1 | -1,076504728 | 0,474176238 | 0,04632 | 0,28930497 | JMJD1C antisense RNA 1 |
| ENSG00000272717 | AC112236,2 | -1,077063045 | 0,473992769 | 0,23616 | 0,54725688 | novel transcript, antisense to SETD7 |
| ENSG00000260280 | SLX1B-SULT1/ | -1,077172049 | 0,473956957 | 0,04991 | 0,30075789 | SLX1B-SULT1A4 readthrough (NMD candidate) |
| ENSG00000276517 | AL133243,3 | -1,077257753 | 0,473928803 | 0,02885 | 0,22968658 | novel transcript, sense intronic to BIRC6 |
| ENSG00000137857 | DUOX1 | -1,077489811 | 0,473852577 | 0,04047 | 0,27194236 | dual oxidase 1 |
| ENSG00000282205 | AC115090,1 | -1,077910456 | 0,473714437 | 0,03228 | 0,2425006 | novel transcript, antisense to ARHGAP23 |
| ENSG00000276170 | AC244153,1 | -1,077910456 | 0,473714437 | 0,03228 | 0,2425006 | novel transcript, antisense to ARHGAP23 |
| ENSG00000231113 | AL035587,1 | -1,077996718 | 0,473686113 | 0,02027 | 0,18910367 | novel transcript |
| ENSG00000252233 | RN7SKP253 | -1,078200428 | 0,473619233 | 0,21235 | 0,53117701 | RN7SK pseudogene 253 |
| ENSG00000188070 | C11orf95 | -1,078733099 | 0,473444395 | 0,02592 | 0,21707964 | chromosome 11 open reading frame 95 |
| ENSG00000257108 | NHLRC4 | -1,079701575 | 0,473126681 | 0,08718 | 0,38232685 | NHL repeat containing 4 |
| ENSG00000188186 | LAMTOR4 | -1,080010572 | 0,473025357 | 0,03036 | 0,23516009 | late endosomal/lysosomal adaptor, MAPK and MTOR activator 4 |
| ENSG00000203321 | CARNMT1-AS | -1,080075649 | 0,47300402 | 0,17931 | 0,50368273 | CARNMT1 antisense RNA 1 |
| ENSG00000213430 | HSPD1P1 | -1,080174016 | 0,472971771 | 0,22978 | 0,54324957 | heat shock protein family D (Hsp60) member 1 pseudogene 1 |
| ENSG00000227959 | AL451042,2 | -1,080297257 | 0,472931369 | 0,1679 | 0,49407144 | novel transcript |
| ENSG00000236673 | AL117378,1 | -1,080503359 | 0,472863811 | 0,05649 | 0,31717392 | novel transcript |
| ENSG00000280383 | Z95331,1 | -1,080905389 | 0,472732059 | 0,03098 | 0,23719337 | novel transcript, sense overlapping ATXN10 |
| ENSG00000182070 | OR52A1 | -1,081338477 | 0,472590169 | 0,01554 | 0,16340375 | olfactory receptor family 52 subfamily A member 1 |
| ENSG00000249328 | AC036214,1 | -1,081428965 | 0,472560528 | 0,12171 | 0,44093327 | novel transcript |
| ENSG00000285846 | AL358613,3 | -1,081886765 | 0,472410598 | 0,14455 | 0,47211995 | novel transcript |
| ENSG00000199719 | RN7SKP74 | -1,082200687 | 0,472307815 | 0,27185 | 0,57305058 | RN7SK pseudogene 74 |
| ENSG00000181291 | TMEM132E | -1,082617556 | 0,472171361 | 0,03331 | 0,24664841 | transmembrane protein 132E |
| ENSG00000253882 | AC099548,2 | -1,082682668 | 0,472150051 | 0,37336 | 0,64204597 | family with sequence similarity 115, member C (FAM115C) pseudogene |
| ENSG00000232499 | AL391058,1 | -1,083713431 | 0,471812834 | 0,03879 | 0,26660248 | heterogeneous nuclear ribonucleoprotein A3 (hnRNPA3) pseudogene |
| ENSG00000188643 | S100A16 | -1,084008732 | 0,47171627 | 0,06262 | 0,33123868 | S100 calcium binding protein A16 |

| | | | | | | |
|---|---|---|---|---|---|---|
| ENSG00000151834 | GABRA2 | -1,085231326 | 0,47131669 | 0,06665 | 0,34158908 | gamma-aminobutyric acid type A receptor subunit alpha2 |
| ENSG00000183918 | SH2D1A | -1,085275278 | 0,471302331 | 0,35844 | 0,63185282 | SH2 domain containing 1A |
| ENSG00000179921 | GPBAR1 | -1,085397814 | 0,471262302 | 0,06957 | 0,34818658 | G protein-coupled bile acid receptor 1 |
| ENSG00000276684 | RDH13 | -1,085837659 | 0,471118647 | 0,12284 | 0,44269063 | retinol dehydrogenase 13 |
| ENSG00000137364 | TPMT | -1,085936254 | 0,471086452 | 0,0359 | 0,25662951 | thiopurine S-methyltransferase |
| ENSG00000274628 | AL669942,1 | -1,086581893 | 0,470875677 | 0,19784 | 0,51942408 | novel cytochrome P450 family pseudogene |
| ENSG00000284944 | MED18 | -1,087000584 | 0,470739042 | 0,29283 | 0,58729524 | mediator complex subunit 18 |
| ENSG00000122912 | SLC25A16 | -1,087178736 | 0,470680916 | 0,03167 | 0,24008374 | solute carrier family 25 member 16 |
| ENSG00000154274 | C4orf19 | -1,087389944 | 0,470612014 | 0,09878 | 0,40461924 | chromosome 4 open reading frame 19 |
| ENSG00000227523 | RPS20P15 | -1,087473701 | 0,470584693 | 0,17646 | 0,50068603 | ribosomal protein S20 pseudogene 15 |
| ENSG00000175592 | FOSL1 | -1,087503851 | 0,470574859 | 0,0345 | 0,25106871 | FOS like 1, AP-1 transcription factor subunit |
| ENSG00000287968 | AC093606,1 | -1,087544384 | 0,470561638 | 0,32781 | 0,60960585 | novel transcript |
| ENSG00000287613 | AC069549,2 | -1,087619878 | 0,470537015 | 0,0057 | 0,09435135 | novel transcript, antisense to PLXDC2 |
| ENSG00000180902 | D2HGDH | -1,087797717 | 0,470479016 | 0,05276 | 0,30840541 | D-2-hydroxyglutarate dehydrogenase |
| ENSG00000008382 | MPND | -1,088134286 | 0,47036927 | 0,021 | 0,19244681 | MPN domain containing |
| ENSG00000253245 | MTND1P36 | -1,088300673 | 0,470315025 | 0,09141 | 0,39102429 | MT-ND1 pseudogene 36 |
| ENSG00000273722 | TMC4 | -1,088396196 | 0,470283886 | 0,32814 | 0,60971379 | transmembrane channel like 4 |
| ENSG00000286950 | AC087588,2 | -1,088739244 | 0,470172073 | 0,24062 | 0,54983486 | novel transcript, antisense to YARS2 |
| ENSG00000159905 | ZNF221 | -1,089349726 | 0,469973161 | 0,02989 | 0,23283161 | zinc finger protein 221 |
| ENSG00000285272 | TMEM249 | -1,089569405 | 0,469901603 | 0,10617 | 0,41557034 | transmembrane protein 249 |
| ENSG00000261587 | TMEM249 | -1,089569405 | 0,469901603 | 0,10617 | 0,41557034 | transmembrane protein 249 |
| ENSG00000278112 | AC145423,2 | -1,089580396 | 0,469898023 | 0,12168 | 0,44093327 | novel transcript, sense intronic to RILPL1 |
| ENSG00000237232 | ZNF295-AS1 | -1,089726347 | 0,469850488 | 0,11803 | 0,43562845 | ZNF295 antisense RNA 1 |
| ENSG00000236404 | VLDLR-AS1 | -1,089806453 | 0,469824401 | 0,02644 | 0,21901972 | VLDLR antisense RNA 1 |
| ENSG00000245832 | MIR4300HG | -1,089823684 | 0,469818789 | 0,31817 | 0,60349082 | MIR4300 host gene |
| ENSG00000260288 | AC019294,2 | -1,089846015 | 0,469811517 | 0,0548 | 0,31344524 | novel transcript |
| ENSG00000125611 | CHCHD5 | -1,089931104 | 0,469783809 | 0,03805 | 0,26375851 | coiled-coil-helix-coiled-coil-helix domain containing 5 |
| ENSG00000268157 | AC010524,1 | -1,089949627 | 0,469777777 | 0,13184 | 0,45708171 | novel transcript, antisense to TEAD2 |
| ENSG00000262732 | AC010401,1 | -1,090013571 | 0,469756956 | 0,01495 | 0,16037017 | novel transcript, antisense to ERCC4 |
| ENSG00000287434 | AC109133,1 | -1,090057837 | 0,469742543 | 0,13729 | 0,4642578 | novel transcript, antisense to DEPDC1B |
| ENSG00000252326 | SNORD116-25 | -1,090309493 | 0,46966061 | 0,25847 | 0,5624548 | small nucleolar RNA, C/D box 116-25 |
| ENSG00000236078 | LINC01447 | -1,090429271 | 0,469621619 | 0,00188 | 0,0481607 | long intergenic non-protein coding RNA 1447 |
| ENSG00000063127 | SLC6A16 | -1,090659939 | 0,469546539 | 0,00084 | 0,02886269 | solute carrier family 6 member 16 |

| | | | | | | |
|---|---|---|---|---|---|---|
| ENSG00000261801 | LOXL1-AS1 | -1,090699054 | 0,469533808 | 0,06794 | 0,34439007 | LOXL1 antisense RNA 1 |
| ENSG00000255745 | AC023796,1 | -1,090797506 | 0,469501767 | 0,28047 | 0,57795109 | novel transcript |
| ENSG00000224945 | AL353150,1 | -1,091189733 | 0,469374141 | 0,01828 | 0,17879941 | novel transcript |
| ENSG00000135736 | CCDC102A | -1,09150267 | 0,469272339 | 0,04851 | 0,29590716 | coiled-coil domain containing 102A |
| ENSG00000167543 | TP53I13 | -1,092120692 | 0,469071355 | 0,02342 | 0,20416051 | tumor protein p53 inducible protein 13 |
| ENSG00000090376 | IRAK3 | -1,092182846 | 0,469051147 | 0,00248 | 0,05777852 | interleukin 1 receptor associated kinase 3 |
| ENSG00000178175 | ZNF366 | -1,09224456 | 0,469031083 | 0,1177 | 0,43523522 | zinc finger protein 366 |
| ENSG00000250067 | YJEFN3 | -1,092905736 | 0,468816179 | 0,01448 | 0,15740445 | YjeF N-terminal domain containing 3 |
| ENSG00000215252 | GOLGA8B | -1,092942337 | 0,468804285 | 0,02445 | 0,20899529 | golgin A8 family member B |
| ENSG00000267787 | AC027097,2 | -1,092956587 | 0,468799655 | 0,05791 | 0,32123433 | novel transcript, antisense to ATP8B1 |
| ENSG00000163126 | ANKRD23 | -1,094001667 | 0,468460182 | 0,03787 | 0,26341726 | ankyrin repeat domain 23 |
| ENSG00000285755 | AC132153,1 | -1,094037258 | 0,468448625 | 0,08087 | 0,37109629 | Novel transcript |
| ENSG00000241520 | AC098820,1 | -1,094065949 | 0,468439309 | 0,17317 | 0,49786688 | novel transcript |
| ENSG00000167733 | HSD11B1L | -1,094266902 | 0,468374065 | 0,02997 | 0,23327096 | hydroxysteroid 11-beta dehydrogenase 1 like |
| ENSG00000075826 | SEC31B | -1,094277616 | 0,468370587 | 2,51E-05 | 0,00186998 | SEC31 homolog B, COPII coat complex component |
| ENSG00000151327 | FAM177A1 | -1,094454952 | 0,468313018 | 0,03852 | 0,26548082 | family with sequence similarity 177 member A1 |
| ENSG00000287869 | AC073349,5 | -1,094492091 | 0,468300963 | 0,21767 | 0,53547632 | novel transcript |
| ENSG00000101417 | PXMP4 | -1,094711787 | 0,468229654 | 0,03618 | 0,25742335 | peroxisomal membrane protein 4 |
| ENSG00000052344 | PRSS8 | -1,09504937 | 0,468120104 | 0,11069 | 0,42337179 | serine protease 8 |
| ENSG00000068400 | GRIPAP1 | -1,095304728 | 0,468037254 | 0,00863 | 0,11739798 | GRIP1 associated protein 1 |
| ENSG00000264549 | SNORD95 | -1,095359488 | 0,468019489 | 0,06844 | 0,34571361 | small nucleolar RNA, C/D box 95 |
| ENSG00000280441 | FP236383,3 | -1,095806874 | 0,467874376 | 0,03184 | 0,24060936 | novel transcript |
| ENSG00000072080 | SPP2 | -1,096056635 | 0,467793385 | 0,01783 | 0,17596434 | secreted phosphoprotein 2 |
| ENSG00000250896 | RNPS1P1 | -1,09639203 | 0,467684646 | 0,2196 | 0,53699318 | RNA binding protein with serine rich domain 1 pseudogene 1 |
| ENSG00000253919 | THAP12P7 | -1,096516326 | 0,467644354 | 0,26348 | 0,56614179 | THAP domain containing 12 pseudogene 7 |
| ENSG00000153721 | CNKSR3 | -1,097841666 | 0,467214947 | 0,04673 | 0,29062603 | CNKSR family member 3 |
| ENSG00000101004 | NINL | -1,097904254 | 0,467194678 | 0,02042 | 0,18959706 | ninein like |
| ENSG00000265739 | AC104984,3 | -1,098002565 | 0,467162843 | 0,1273 | 0,45051698 | novel transcript, antisense to CCDC55 |
| ENSG00000002822 | MAD1L1 | -1,09801298 | 0,46715947 | 0,02795 | 0,22565587 | mitotic arrest deficient 1 like 1 |
| ENSG00000286975 | AP005136,4 | -1,098437963 | 0,467021877 | 0,17978 | 0,50407792 | novel transcript |
| ENSG00000268087 | AC008764,2 | -1,099129178 | 0,466798174 | 0,08038 | 0,37007763 | novel transcript |
| ENSG00000198838 | RYR3 | -1,099348553 | 0,466727198 | 0,00354 | 0,07209525 | ryanodine receptor 3 |

| | | | | | | |
|---|---|---|---|---|---|---|
| ENSG00000239605 | STPG4 | -1,099595893 | 0,466647188 | 0,15388 | 0,48214138 | sperm-tail PG-rich repeat containing 4 |
| ENSG00000230332 | RNF39 | -1,099954837 | 0,4665311 | 0,23836 | 0,54875544 | ring finger protein 39 |
| ENSG00000204618 | RNF39 | -1,099954837 | 0,4665311 | 0,23836 | 0,54875544 | ring finger protein 39 |
| ENSG00000209582 | SNORA48 | -1,100030092 | 0,466506765 | 0,00428 | 0,08037599 | small nucleolar RNA, H/ACA box 48 |
| ENSG00000283189 | AC104452,1 | -1,100187805 | 0,46645577 | 0,01079 | 0,13440486 | novel protein |
| ENSG00000274148 | LILRA6 | -1,100379543 | 0,466393781 | 0,44507 | 0,69209822 | leukocyte immunoglobulin like receptor A6 |
| ENSG00000162365 | CYP4A22 | -1,100417576 | 0,466381486 | 0,19192 | 0,5145257 | cytochrome P450 family 4 subfamily A member 22 |
| ENSG00000197882 | OR7E13P | -1,100670037 | 0,46629988 | 0,04692 | 0,29141554 | olfactory receptor family 7 subfamily E member 13 pseudogene |
| ENSG00000235138 | AL445931,1 | -1,100671136 | 0,466299525 | 0,05191 | 0,30611375 | novel transcript, antisense to BRD3 |
| ENSG00000253404 | AC034243,1 | -1,100944498 | 0,466211178 | 0,20922 | 0,52826608 | novel transcript |
| ENSG00000285928 | AC103591,4 | -1,100969887 | 0,466202974 | 0,17561 | 0,49954066 | novel transcript |
| ENSG00000215867 | KRT18P57 | -1,101055084 | 0,466175444 | 0,00904 | 0,12067214 | keratin 18 pseudogene 57 |
| ENSG00000245680 | ZNF585B | -1,10199484 | 0,465871881 | 0,03989 | 0,27016182 | zinc finger protein 585B |
| ENSG00000161914 | ZNF653 | -1,102107991 | 0,465835344 | 0,01096 | 0,13522945 | zinc finger protein 653 |
| ENSG00000215458 | AATBC | -1,102135952 | 0,465826315 | 0,04925 | 0,29853148 | apoptosis associated transcript in bladder cancer |
| ENSG00000288050 | AC010727,1 | -1,102283723 | 0,465778605 | 0,0314 | 0,23928335 | novel transcript, antisense to RFTN1 |
| ENSG00000225792 | AC004540,2 | -1,102419995 | 0,465734611 | 0,01615 | 0,16645939 | novel transcript |
| ENSG00000233611 | AC019068,1 | -1,102421945 | 0,465733982 | 0,12853 | 0,45191041 | novel transcript, antisense to ASB18 and GBX2 |
| ENSG00000281521 | AC116351,4 | -1,102944467 | 0,46556533 | 0,0911 | 0,39033853 | novel transcript |
| ENSG00000272347 | AC116351,2 | -1,102944467 | 0,46556533 | 0,0911 | 0,39033853 | novel transcript |
| ENSG00000283918 | MRNIP | -1,103172087 | 0,465491882 | 0,06291 | 0,33195998 | MRN complex interacting protein |
| ENSG00000266946 | MRPL37P1 | -1,103803685 | 0,465288139 | 0,01018 | 0,12975801 | mitochondrial ribosomal protein L37 pseudogene 1 |
| ENSG00000168502 | MTCL1 | -1,103930391 | 0,465247276 | 0,02186 | 0,19657209 | microtubule crosslinking factor 1 |
| ENSG00000263142 | LRRC37A17P | -1,104013718 | 0,465220405 | 0,02593 | 0,21707964 | leucine rich repeat containing 37 member A17, pseudogene |
| ENSG00000277270 | AL160412,1 | -1,104057975 | 0,465206134 | 0,04338 | 0,28091273 | novel transcript |
| ENSG00000283538 | AC005972,3 | -1,104141296 | 0,465179268 | 0,05659 | 0,31739242 | novel transcript |
| ENSG00000227885 | AL590652,1 | -1,104226557 | 0,465151777 | 0,06962 | 0,34826564 | novel transcript |
| ENSG00000214338 | SOGA3 | -1,104260779 | 0,465140743 | 0,00543 | 0,0916301 | SOGA family member 3 |
| ENSG00000265179 | AP000894,2 | -1,104267037 | 0,465138726 | 0,04156 | 0,27616337 | novel transcript, antisense to ADCYAP1 |
| ENSG00000121281 | ADCY7 | -1,104817187 | 0,464961386 | 0,00021 | 0,01067558 | adenylate cyclase 7 |
| ENSG00000284076 | GOLGA8T | -1,10481988 | 0,464960518 | 4,65E-05 | 0,0031583 | golgin A8 family member T |

| | | | | | | |
|---|---|---|---|---|---|---|
| ENSG00000278475 | AC009464,1 | -1,105093347 | 0,464872392 | 0,09565 | 0,39857837 | novel transcript, sense intronic to ARID2 |
| ENSG00000187690 | EZHIP | -1,105567786 | 0,464719541 | 0,14787 | 0,4751727 | EZH inhibitory protein |
| ENSG00000274734 | ARHGAP11B | -1,105727103 | 0,464668224 | 0,12325 | 0,44354688 | Rho GTPase activating protein 11B |
| ENSG00000125503 | PPP1R12C | -1,106601579 | 0,464386656 | 0,0267 | 0,22056538 | protein phosphatase 1 regulatory subunit 12C |
| ENSG00000181524 | RPL24P4 | -1,106728938 | 0,464345662 | 0,10651 | 0,41625157 | RPL24 pseudogene 4 |
| ENSG00000198729 | PPP1R14C | -1,106780598 | 0,464329035 | 8,91E-06 | 0,00075488 | protein phosphatase 1 regulatory inhibitor subunit 14C |
| ENSG00000246763 | RGMB-AS1 | -1,106796602 | 0,464323884 | 0,00265 | 0,059998 | RGMB antisense RNA 1 |
| ENSG00000196553 | CCDC196 | -1,107487693 | 0,464101513 | 0,21689 | 0,53505085 | coiled-coil domain containing 196 |
| ENSG00000232807 | AL137186,2 | -1,107729784 | 0,464023641 | 0,05568 | 0,31531663 | novel transcript, antisense to FBXO18 |
| ENSG00000270540 | AC093915,1 | -1,108120264 | 0,463898066 | 0,11924 | 0,43706649 | novel transcript, antisense to IQCA1 |
| ENSG00000086696 | HSD17B2 | -1,108128684 | 0,463895358 | 0,11805 | 0,43562845 | hydroxysteroid 17-beta dehydrogenase 2 |
| ENSG00000130254 | SAFB2 | -1,108190344 | 0,463875532 | 0,01174 | 0,14104005 | scaffold attachment factor B2 |
| ENSG00000199363 | SNORA63E | -1,10826731 | 0,463850786 | 0,06572 | 0,33945082 | small nucleolar RNA, H/ACA box 63E |
| ENSG00000078269 | SYNJ2 | -1,108300192 | 0,463840214 | 0,0045 | 0,08275926 | synaptojanin 2 |
| ENSG00000282522 | PKD1P1 | -1,10833755 | 0,463828203 | 0,01143 | 0,13882136 | polycystin 1, transient receptor potential channel interacting pseudogene 1 |
| ENSG00000170088 | TMEM192 | -1,108569212 | 0,463753729 | 0,02445 | 0,20899529 | transmembrane protein 192 |
| ENSG00000237351 | ITPK1P1 | -1,108693026 | 0,463713931 | 0,03134 | 0,2388853 | ITPK1 pseudogene 1 |
| ENSG00000119004 | CYP20A1 | -1,108759053 | 0,463692709 | 0,04247 | 0,2782038 | cytochrome P450 family 20 subfamily A member 1 |
| ENSG00000130643 | CALY | -1,109224471 | 0,463543144 | 0,07182 | 0,35363591 | calcyon neuron specific vesicular protein |
| ENSG00000277459 | AP001527,2 | -1,109563964 | 0,463434077 | 0,08708 | 0,3821708 | novel transcript |
| ENSG00000224821 | COL4A2-AS2 | -1,10963005 | 0,463412849 | 0,14071 | 0,46776292 | COL4A2 antisense 2 |
| ENSG00000235149 | AC096659,1 | -1,109886582 | 0,463330455 | 0,1823 | 0,50576743 | novel transcript |
| ENSG00000263069 | RNF213-AS1 | -1,110071572 | 0,463271048 | 0,01876 | 0,18150273 | RNF213 antisense RNA 1 |
| ENSG00000276115 | AC026356,2 | -1,110094875 | 0,463263565 | 0,00445 | 0,08228165 | novel transcript, sense intronic to BICD1 |
| ENSG00000253559 | OSGEPL1-AS1 | -1,110120192 | 0,463255435 | 0,03064 | 0,23611505 | OSGEPL1 antisense RNA 1 |
| ENSG00000283973 | AC099795,1 | -1,110230684 | 0,463219957 | 0,24774 | 0,5551624 | novel transcript |
| ENSG00000177192 | PUS1 | -1,110234561 | 0,463218712 | 0,03399 | 0,248958 | pseudouridine synthase 1 |
| ENSG00000271848 | AC073389,2 | -1,110493564 | 0,463135559 | 0,11262 | 0,42641188 | novel transcript, antisense to SYNPO2L and AGAP5 |
| ENSG00000282202 | AC015849,10 | -1,110889869 | 0,463008355 | 0,15137 | 0,47956749 | novel transcript, antisense TAF15 |
| ENSG00000270871 | AC015849,3 | -1,110889869 | 0,463008355 | 0,15137 | 0,47956749 | novel transcript, antisense TAF15 |
| ENSG00000178597 | PSAPL1 | -1,111145225 | 0,46292641 | 0,10007 | 0,40666505 | prosaposin like 1 |
| ENSG00000236114 | AL450326,2 | -1,111480648 | 0,462818793 | 0,09929 | 0,4057206 | zinc finger protein pseudogene |

| | | | | | | |
|---|---|---|---|---|---|---|
| ENSG00000156968 | MPV17L | -1,111875424 | 0,462692165 | 0,0286 | 0,22866679 | MPV17 mitochondrial inner membrane protein like |
| ENSG00000259286 | AC087639,1 | -1,112956123 | 0,4623457 | 0,08798 | 0,38393623 | cytokine induced apoptosis inhibitor 1 (CIAPIN1) |
| ENSG00000235908 | RHOA-IT1 | -1,113193412 | 0,462269662 | 0,10175 | 0,40892507 | RHOA intronic transcript 1 |
| ENSG00000237505 | PKN2-AS1 | -1,113677919 | 0,462114442 | 0,04775 | 0,29349263 | PKN2 antisense RNA 1 |
| ENSG00000261888 | AC144831,1 | -1,113687917 | 0,462111239 | 0,12011 | 0,43868155 | novel transcript |
| ENSG00000237575 | PYY2 | -1,114129 | 0,461969977 | 0,10054 | 0,40713342 | peptide YY 2 (pseudogene) |
| ENSG00000204569 | PPP1R10 | -1,114135963 | 0,461967747 | 0,07252 | 0,35556123 | protein phosphatase 1 regulatory subunit 10 |
| ENSG00000198863 | RUNDC1 | -1,114156538 | 0,461961159 | 0,03275 | 0,24410674 | RUN domain containing 1 |
| ENSG00000163093 | BBS5 | -1,114190536 | 0,461950273 | 0,02747 | 0,22348662 | Bardet-Biedl syndrome 5 |
| ENSG00000239614 | HMGN1P7 | -1,114327412 | 0,461906447 | 0,37486 | 0,64325417 | high mobility group nucleosome binding domain 1 pseudogene 7 |
| ENSG00000172638 | EFEMP2 | -1,114789567 | 0,461758503 | 0,00019 | 0,01011637 | EGF containing fibulin extracellular matrix protein 2 |
| ENSG00000256218 | AC007848,2 | -1,114847808 | 0,461739862 | 0,0422 | 0,27783166 | novel transcript |
| ENSG00000161939 | RNASEK-C17o | -1,114924598 | 0,461715286 | 0,07307 | 0,35640071 | RNASEK-C17orf49 readthrough |
| ENSG00000251495 | PPIAP11 | -1,115089291 | 0,461662581 | 0,21892 | 0,53639845 | peptidylprolyl isomerase A pseudogene 11 |
| ENSG00000116852 | KIF21B | -1,115279809 | 0,46160162 | 0,00379 | 0,0751769 | kinesin family member 21B |
| ENSG00000180662 | RPL21P8 | -1,115311662 | 0,461591428 | 0,10018 | 0,4067769 | ribosomal protein L21 pseudogene 8 |
| ENSG00000218793 | AL049697,3 | -1,115685601 | 0,461471801 | 0,28305 | 0,58033644 | S-adenosylmethionine decarboxylase 1 (AMD1) pseudogene |
| ENSG00000167807 | AC011511,1 | -1,116250903 | 0,461291015 | 0,05266 | 0,30807312 | novel transcript |
| ENSG00000250536 | ABHD17AP3 | -1,116693623 | 0,46114948 | 0,08732 | 0,38254243 | ABHD17A pseudogene 3 |
| ENSG00000249992 | TMEM158 | -1,116712981 | 0,461143293 | 1,40E-06 | 0,00015251 | transmembrane protein 158 |
| ENSG00000224765 | AL035443,1 | -1,116826203 | 0,461107104 | 0,04011 | 0,27065903 | novel transcript |
| ENSG00000256671 | LIMS4 | -1,117172127 | 0,460996555 | 0,1937 | 0,51547251 | LIM zinc finger domain containing 4 |
| ENSG00000108556 | CHRNE | -1,117906802 | 0,460761857 | 0,11342 | 0,42803467 | cholinergic receptor nicotinic epsilon subunit |
| ENSG00000110723 | EXPH5 | -1,118139594 | 0,460687515 | 0,04484 | 0,28614848 | exophilin 5 |
| ENSG00000272108 | AC244517,1 | -1,118236803 | 0,460656475 | 0,00459 | 0,08349554 | novel transcript |
| ENSG00000213420 | GPC2 | -1,118457984 | 0,460585857 | 0,00685 | 0,10547716 | glypican 2 |
| ENSG00000239445 | ST3GAL6-AS1 | -1,118648191 | 0,460525136 | 0,02937 | 0,23068546 | ST3GAL6 antisense RNA 1 |
| ENSG00000178404 | CEP295NL | -1,118775438 | 0,460484519 | 0,00729 | 0,10833556 | CEP295 N-terminal like |
| ENSG00000104154 | SLC30A4 | -1,118781456 | 0,460482599 | 0,01319 | 0,15012165 | solute carrier family 30 member 4 |
| ENSG00000214832 | UPF3AP2 | -1,118914915 | 0,460440003 | 0,2214 | 0,53805234 | UPF3A pseudogene 2 |
| ENSG00000165091 | TMC1 | -1,119202381 | 0,460348266 | 0,06512 | 0,33804854 | transmembrane channel like 1 |

| | | | | | | |
|---|---|---|---|---|---|---|
| ENSG00000203730 | TEDDM1 | -1,119629335 | 0,46021205 | 0,23729 | 0,54824779 | transmembrane epididymal protein 1 |
| ENSG00000151208 | DLG5 | -1,119738484 | 0,460177234 | 0,03696 | 0,2607683 | discs large MAGUK scaffold protein 5 |
| ENSG00000185507 | IRF7 | -1,120063259 | 0,460073652 | 0,0896 | 0,38705587 | interferon regulatory factor 7 |
| ENSG00000142279 | WTIP | -1,120440218 | 0,459953456 | 0,04373 | 0,28225644 | WT1 interacting protein |
| ENSG00000275807 | AC145285,6 | -1,120466123 | 0,459945197 | 0,07438 | 0,35809404 | novel transcript, antisense to ATXN2L |
| ENSG00000117620 | SLC35A3 | -1,120685508 | 0,45987526 | 0,03598 | 0,25691454 | solute carrier family 35 member A3 |
| ENSG00000213152 | RPL7AP60 | -1,120999477 | 0,45977519 | 0,03785 | 0,26339938 | ribosomal protein L7a pseudogene 60 |
| ENSG00000229369 | AL450263,2 | -1,121142955 | 0,459729467 | 0,18901 | 0,51243448 | density-regulated protein (DENR) pseudogene |
| ENSG00000283294 | AP005212,4 | -1,122142355 | 0,459411108 | 0,04422 | 0,28352348 | novel transcript |
| ENSG00000237187 | NR2F1-AS1 | -1,122223738 | 0,459385193 | 0,00162 | 0,0433162 | NR2F1 antisense RNA 1 |
| ENSG00000100784 | RPS6KA5 | -1,122296901 | 0,459361897 | 0,04279 | 0,27879697 | ribosomal protein S6 kinase A5 |
| ENSG00000232104 | RFX3-AS1 | -1,123359739 | 0,459023608 | 0,00234 | 0,05563463 | RFX3 antisense RNA 1 |
| ENSG00000259892 | AC027139,1 | -1,1233674 | 0,459021171 | 0,21655 | 0,53490925 | amyloid beta (A4) precursor protein-binding, family A, member 2 (APBA2) pseudogene |
| ENSG00000279249 | AC007614,1 | -1,123381297 | 0,459016749 | 0,00291 | 0,06410301 | novel transcript, antisense to CBLN1 |
| ENSG00000220793 | RPL21P119 | -1,123539546 | 0,458966403 | 0,55741 | 0,76914186 | ribosomal protein L21 pseudogene 119 |
| ENSG00000287721 | AC061975,8 | -1,123580417 | 0,458953401 | 0,01206 | 0,14284116 | novel transcript |
| ENSG00000145476 | CYP4V2 | -1,123934893 | 0,458840648 | 0,04117 | 0,27486753 | cytochrome P450 family 4 subfamily V member 2 |
| ENSG00000153898 | MCOLN2 | -1,124627805 | 0,458620324 | 0,29722 | 0,59031059 | mucolipin 2 |
| ENSG00000154358 | OBSCN | -1,125023481 | 0,458494559 | 0,03831 | 0,264906 | obscurin, cytoskeletal calmodulin and titin-interacting RhoGEF |
| ENSG00000272356 | AL080317,2 | -1,125129294 | 0,458460933 | 0,05218 | 0,30681232 | novel transcript, antisense to REV3L |
| ENSG00000259869 | AL022344,1 | -1,125600928 | 0,458311081 | 0,27885 | 0,57700804 | novel transcript |
| ENSG00000101188 | NTSR1 | -1,126029163 | 0,458175061 | 0,08117 | 0,37191664 | neurotensin receptor 1 |
| ENSG00000104518 | GSDMD | -1,126177896 | 0,458127828 | 0,13957 | 0,46677981 | gasdermin D |
| ENSG00000234996 | ACTG1P25 | -1,126735716 | 0,457950727 | 0,0014 | 0,04000051 | actin gamma 1 pseudogene 25 |
| ENSG00000113645 | WWC1 | -1,126925787 | 0,457890397 | 0,00681 | 0,10528058 | WW and C2 domain containing 1 |
| ENSG00000105298 | CACTIN | -1,126942103 | 0,457885218 | 0,00964 | 0,12566531 | cactin, spliceosome C complex subunit |
| ENSG00000271860 | AL589740,1 | -1,126983197 | 0,457872176 | 0,00697 | 0,1063368 | novel transcript |
| ENSG00000265799 | AC090844,3 | -1,127041793 | 0,45785358 | 0,11911 | 0,43696075 | novel transcript |
| ENSG00000254477 | AP000640,1 | -1,127649078 | 0,457660892 | 0,09259 | 0,39335125 | novel transcript |
| ENSG00000226445 | BX322234,1 | -1,128257325 | 0,457467981 | 0,15132 | 0,47956749 | novel transcript |
| ENSG00000275329 | AL138781,1 | -1,128483791 | 0,457396176 | 0,08751 | 0,38306995 | novel transcript |

| | | | | | | |
|---|---|---|---|---|---|---|
| ENSG00000138834 | MAPK8IP3 | -1,128623286 | 0,457351952 | 0,00233 | 0,05562927 | mitogen-activated protein kinase 8 interacting protein 3 |
| ENSG00000266086 | AC015813,2 | -1,129030213 | 0,45722297 | 0,08464 | 0,37776115 | novel transcript |
| ENSG00000239908 | RN7SL75P | -1,129110615 | 0,457197489 | 0,15669 | 0,48425071 | RNA, 7SL, cytoplasmic 75, pseudogene |
| ENSG00000285935 | AL159978,1 | -1,129307625 | 0,45713506 | 0,0695 | 0,34813287 | novel transcript, antisense to WASF3 |
| ENSG00000269243 | AC008894,2 | -1,129942829 | 0,456933832 | 0,00408 | 0,07837165 | novel transcript, antisense RAB8A |
| ENSG00000224086 | PPM1F-AS1 | -1,130111244 | 0,456880494 | 0,03269 | 0,24399744 | PPM1F antisense RNA 1 |
| ENSG00000225342 | LRRK2-DT | -1,130729162 | 0,456684851 | 0,13589 | 0,46291909 | LRRK2 divergent transcript |
| ENSG00000142046 | TMEM91 | -1,130984237 | 0,456604114 | 0,03855 | 0,26560956 | transmembrane protein 91 |
| ENSG00000155428 | TRIM74 | -1,131280841 | 0,45651025 | 0,29766 | 0,59036937 | tripartite motif containing 74 |
| ENSG00000239389 | PCDHA13 | -1,132045854 | 0,456268242 | 0,00034 | 0,01573727 | protocadherin alpha 13 |
| ENSG00000266401 | AP002478,1 | -1,132452209 | 0,456139746 | 0,06432 | 0,33606729 | novel transcript, antisense to DLGAP1 |
| ENSG00000169884 | WNT10B | -1,132666061 | 0,456072137 | 0,09139 | 0,39102429 | Wnt family member 10B |
| ENSG00000228434 | AC004951,1 | -1,132747455 | 0,456046407 | 0,0239 | 0,20646034 | novel transcript |
| ENSG00000168101 | NUDT16L1 | -1,132877629 | 0,45600526 | 0,02601 | 0,21725277 | nudix hydrolase 16 like 1 |
| ENSG00000277558 | AL109923,1 | -1,133108807 | 0,455932195 | 0,16224 | 0,48939373 | novel transcript, sense intronic to ITCH |
| ENSG00000167995 | BEST1 | -1,133323378 | 0,45586439 | 0,06524 | 0,3380905 | bestrophin 1 |
| ENSG00000230587 | LINC02580 | -1,133370766 | 0,455849416 | 0,2408 | 0,55001001 | long intergenic non-protein coding RNA 2580 |
| ENSG00000270711 | AL162431,3 | -1,134472235 | 0,455501517 | 0,15384 | 0,48214138 | cell division cycle associated 8 (CDCA8) pseudogene |
| ENSG00000134531 | EMP1 | -1,134572733 | 0,455469788 | 0,0078 | 0,11235913 | epithelial membrane protein 1 |
| ENSG00000263924 | AC022960,1 | -1,13467832 | 0,455436455 | 0,2048 | 0,52445677 | novel transcript, antisense to KIAA1012 |
| ENSG00000288332 | PIMREGP3 | -1,134700507 | 0,455429451 | 0,17458 | 0,49884018 | PIMREG pseudogene 3 |
| ENSG00000217653 | PIMREGP3 | -1,134700507 | 0,455429451 | 0,17458 | 0,49884018 | PIMREG pseudogene 3 |
| ENSG00000227233 | CICP17 | -1,135457764 | 0,455190463 | 0,1678 | 0,49395055 | capicua transcriptional repressor pseudogene 17 |
| ENSG00000250320 | EDIL3-DT | -1,135719548 | 0,455107874 | 0,02536 | 0,21419157 | EDIL3 divergent transcript |
| ENSG00000285669 | AC026979,4 | -1,135796486 | 0,455083604 | 0,03876 | 0,26656704 | novel transcript |
| ENSG00000184163 | C1QTNF12 | -1,13581246 | 0,455078565 | 0,19278 | 0,51493916 | C1q and TNF related 12 |
| ENSG00000283946 | CHRNB1 | -1,135999201 | 0,455019664 | 0,01964 | 0,18592829 | cholinergic receptor nicotinic beta 1 subunit |
| ENSG00000276234 | TADA2A | -1,136130882 | 0,454978134 | 0,03166 | 0,24008374 | transcriptional adaptor 2A |
| ENSG00000204311 | PJVK | -1,136156142 | 0,454970168 | 0,02652 | 0,21946238 | pejvakin |
| ENSG00000286430 | AC020558,6 | -1,13617712 | 0,454963553 | 0,32647 | 0,60866572 | novel transcript, antisense to PEMT |
| ENSG00000258429 | PDF | -1,136205429 | 0,454954625 | 0,00925 | 0,12270888 | peptide deformylase, mitochondrial |
| ENSG00000213062 | Z99572,1 | -1,136289316 | 0,454928172 | 0,02872 | 0,2291366 | novel transcript |
| ENSG00000203804 | ADAMTSL4-AS | -1,136552743 | 0,454845113 | 0,05375 | 0,31090219 | ADAMTSL4 antisense RNA 1 |

| | | | | | | |
|---|---|---|---|---|---|---|
| ENSG00000236120 | AC110995,1 | -1,13657932 | 0,454836734 | 0,06521 | 0,33804854 | novel transcript |
| ENSG00000271437 | AL356423,1 | -1,136767018 | 0,454777563 | 0,18548 | 0,50893545 | high mobility group box 1 (HMGB1) pseudogene |
| ENSG00000226758 | DISC1-IT1 | -1,137492676 | 0,454548872 | 0,02533 | 0,21419157 | DISC1 intronic transcript 1 |
| ENSG00000215237 | AL592293,1 | -1,137548526 | 0,454531276 | 0,13054 | 0,45480419 | chromosome 4 open reading frame 27 (C4orf27) pseudogene |
| ENSG00000241357 | AC092849,1 | -1,137637054 | 0,454503385 | 0,27749 | 0,57588377 | novel transcript |
| ENSG00000173621 | LRFN4 | -1,137845091 | 0,454437851 | 0,01402 | 0,1554463 | leucine rich repeat and fibronectin type III domain containing 4 |
| ENSG00000164175 | SLC45A2 | -1,13792686 | 0,454412095 | 0,1878 | 0,51107563 | solute carrier family 45 member 2 |
| ENSG00000248568 | KRT8P48 | -1,137940244 | 0,454407879 | 0,1845 | 0,50835796 | keratin 8 pseudogene 48 |
| ENSG00000229656 | ITGB1-DT | -1,138084251 | 0,454362524 | 0,05548 | 0,31503413 | ITGB1 divergent transcript |
| ENSG00000103490 | PYCARD | -1,138235935 | 0,454314755 | 0,19009 | 0,51331106 | PYD and CARD domain containing |
| ENSG00000149922 | TBX6 | -1,138301649 | 0,454294061 | 0,097 | 0,40141222 | T-box transcription factor 6 |
| ENSG00000247774 | PCED1B-AS1 | -1,138622408 | 0,454193068 | 0,16505 | 0,49125639 | PCED1B antisense RNA 1 |
| ENSG00000204965 | PCDHA5 | -1,138712564 | 0,454164686 | 0,00018 | 0,0095631 | protocadherin alpha 5 |
| ENSG00000230498 | AL035409,1 | -1,13929968 | 0,453979897 | 0,00532 | 0,09086355 | novel transcript |
| ENSG00000157306 | ZFHX2-AS1 | -1,139602893 | 0,453884494 | 0,01694 | 0,17086866 | ZFHX2 antisense RNA 1 |
| ENSG00000188573 | FBLL1 | -1,139641901 | 0,453872222 | 0,00383 | 0,07538386 | fibrillarin like 1 |
| ENSG00000224360 | DDR1-DT | -1,139743826 | 0,453840157 | 0,14131 | 0,46803321 | DDR1 divergent transcript |
| ENSG00000223382 | LINC01778 | -1,139938069 | 0,453779057 | 0,47154 | 0,71092088 | long intergenic non-protein coding RNA 1778 |
| ENSG00000261462 | AC004023,1 | -1,140257206 | 0,453678688 | 0,25482 | 0,55944852 | novel transcript |
| ENSG00000253975 | AC018861,1 | -1,140892714 | 0,453478886 | 0,00839 | 0,11562524 | ubiquitin-conjugating enzyme E2L 3 (UBE2L3) pseudogene |
| ENSG00000239467 | AC007405,3 | -1,140892757 | 0,453478873 | 0,05814 | 0,32181744 | glutamate rich 2 |
| ENSG00000278341 | AC138028,6 | -1,141138291 | 0,453401701 | 0,13654 | 0,46380939 | novel transcript, antisense to CTU2 |
| ENSG00000240401 | AC012358,2 | -1,141462327 | 0,453299877 | 0,05557 | 0,31503413 | novel transcript, antisense to CCDC88A |
| ENSG00000258028 | AL135878,1 | -1,141912418 | 0,453158478 | 0,24807 | 0,55542983 | novel transcript |
| ENSG00000255737 | AGAP2-AS1 | -1,142579135 | 0,452949107 | 0,09356 | 0,39523994 | AGAP2 antisense RNA 1 |
| ENSG00000235535 | TRDN-AS1 | -1,142939149 | 0,452836091 | 0,09638 | 0,40013295 | TRDN antisense RNA 1 |
| ENSG00000143458 | GABPB2 | -1,143513097 | 0,452655975 | 0,03472 | 0,25192618 | GA binding protein transcription factor subunit beta 2 |
| ENSG00000177058 | SLC38A9 | -1,144148346 | 0,452456705 | 0,02942 | 0,23078264 | solute carrier family 38 member 9 |
| ENSG00000188910 | GJB3 | -1,14464333 | 0,452301495 | 0,0353 | 0,25450251 | gap junction protein beta 3 |
| ENSG00000287524 | AC007736,1 | -1,144703669 | 0,452282578 | 0,08672 | 0,38148201 | novel transcript, antisense to PARD3B |
| ENSG00000248730 | EGFLAM-AS4 | -1,144716087 | 0,452278686 | 0,35423 | 0,62892485 | EGFLAM antisense RNA 4 |

| | | | | | | |
|---|---|---|---|---|---|---|
| ENSG00000257896 | AC093012,1 | -1,144919924 | 0,452214788 | 0,02046 | 0,18969631 | La ribonucleoprotein domain family, member 7 (LARP7) |
| ENSG00000103184 | SEC14L5 | -1,14520478 | 0,452125508 | 0,11828 | 0,43607616 | SEC14 like lipid binding 5 |
| ENSG00000249628 | LINC00942 | -1,145538727 | 0,452020865 | 0,1569 | 0,48425071 | long intergenic non-protein coding RNA 942 |
| ENSG00000232732 | AC097717,1 | -1,145591374 | 0,45200437 | 0,11817 | 0,43593502 | novel transcript |
| ENSG00000149243 | KLHL35 | -1,14579772 | 0,451939725 | 0,00623 | 0,09947371 | kelch like family member 35 |
| ENSG00000233215 | LINC01687 | -1,145879337 | 0,451914159 | 0,05549 | 0,31503413 | long intergenic non-protein coding RNA 1687 |
| ENSG00000251171 | AC096741,1 | -1,146272576 | 0,451790996 | 0,1849 | 0,50844776 | novel transcript |
| ENSG00000246174 | KCTD21-AS1 | -1,146999271 | 0,451563483 | 0,04975 | 0,30012887 | KCTD21 antisense RNA 1 |
| ENSG00000170955 | CAVIN3 | -1,147416602 | 0,451432878 | 0,0237 | 0,20555253 | caveolae associated protein 3 |
| ENSG00000272787 | AC253536,6 | -1,147683527 | 0,451349362 | 0,18193 | 0,50521722 | novel transcript |
| ENSG00000281400 | AC253536,6 | -1,147683527 | 0,451349362 | 0,18193 | 0,50521722 | novel transcript |
| ENSG00000055813 | CCDC85A | -1,147957792 | 0,451263566 | 0,02793 | 0,22553251 | coiled-coil domain containing 85A |
| ENSG00000234906 | APOC2 | -1,148329994 | 0,451147159 | 0,24649 | 0,5541223 | apolipoprotein C2 |
| ENSG00000107249 | GLIS3 | -1,148730323 | 0,451021989 | 0,0001 | 0,00604054 | GLIS family zinc finger 3 |
| ENSG00000260206 | AC105020,2 | -1,148757706 | 0,451013428 | 0,24169 | 0,55047794 | novel transcript, antisense to IMP3 |
| ENSG00000231460 | AL365204,1 | -1,149124476 | 0,450898784 | 0,00184 | 0,04744946 | novel transcript |
| ENSG00000259683 | AC243562,2 | -1,149164324 | 0,45088633 | 0,09205 | 0,39230683 | golgin A6 family-like 9 (GOLGA6L9) pseudogene |
| ENSG00000130294 | KIF1A | -1,149598697 | 0,450750596 | 0,00398 | 0,07725019 | kinesin family member 1A |
| ENSG00000271992 | AL354872,1 | -1,150278839 | 0,450538144 | 0,13513 | 0,46165567 | novel transcript |
| ENSG00000256458 | AC080075,1 | -1,150461502 | 0,450481104 | 0,04666 | 0,29040463 | novel transcript, antisense to abl interactor 2 (ABI2) |
| ENSG00000170074 | FAM153A | -1,150710026 | 0,450403509 | 0,09134 | 0,3908781 | family with sequence similarity 153 member A |
| ENSG00000121270 | ABCC11 | -1,151103945 | 0,450280546 | 0,11503 | 0,43048326 | ATP binding cassette subfamily C member 11 |
| ENSG00000272277 | AL031963,3 | -1,15187809 | 0,450038992 | 0,03635 | 0,25801846 | novel transcript, antisense to RIPK1 |
| ENSG00000251031 | AC132803,1 | -1,152208293 | 0,449936 | 0,03913 | 0,26771146 | novel transcript |
| ENSG00000157578 | LCA5L | -1,152410275 | 0,449873011 | 0,05607 | 0,31627719 | lebercilin LCA5 like |
| ENSG00000251566 | HMGB1P35 | -1,152750321 | 0,449766988 | 0,13723 | 0,4642578 | high mobility group box 1 pseudogene 35 |
| ENSG00000287618 | AC025576,3 | -1,152885673 | 0,449724793 | 0,05233 | 0,30711015 | novel transcript, antisense to SRGAP1 |
| ENSG00000227516 | AL160163,1 | -1,152896608 | 0,449721385 | 0,2472 | 0,55483158 | novel transcript |
| ENSG00000232545 | AC253536,3 | -1,153649467 | 0,449486762 | 0,09498 | 0,39751728 | novel transcript |
| ENSG00000206342 | STK19 | -1,153669215 | 0,44948061 | 0,18311 | 0,50652733 | serine/threonine kinase 19 |
| ENSG00000241596 | AC092468,1 | -1,154251916 | 0,449299102 | 0,08111 | 0,37180968 | novel transcript |
| ENSG00000267265 | AC011476,3 | -1,15487536 | 0,449104985 | 0,04605 | 0,28930497 | novel transcript, antisense to RDH13 and GP6 |
| ENSG00000112474 | HSD17B8 | -1,155626311 | 0,448871278 | 0,13827 | 0,46543358 | hydroxysteroid 17-beta dehydrogenase 8 |

| | | | | | | |
|---|---|---|---|---|---|---|
| ENSG00000232357 | HSD17B8 | -1,155626311 | 0,448871278 | 0,13827 | 0,46543358 | hydroxysteroid 17-beta dehydrogenase 8 |
| ENSG00000225312 | HSD17B8 | -1,155626311 | 0,448871278 | 0,13827 | 0,46543358 | hydroxysteroid 17-beta dehydrogenase 8 |
| ENSG00000228712 | HSD17B8 | -1,155626311 | 0,448871278 | 0,13827 | 0,46543358 | hydroxysteroid 17-beta dehydrogenase 8 |
| ENSG00000228357 | HSD17B8 | -1,155626311 | 0,448871278 | 0,13827 | 0,46543358 | hydroxysteroid 17-beta dehydrogenase 8 |
| ENSG00000206579 | XKR4 | -1,155798844 | 0,4488176 | 0,01033 | 0,13101765 | XK related 4 |
| ENSG00000265015 | AP005901,3 | -1,1558173 | 0,448811858 | 0,03331 | 0,24664841 | kinase suppressor of ras 1 pseudogene |
| ENSG00000255176 | AP000941,1 | -1,155858043 | 0,448799184 | 0,0454 | 0,28785673 | novel transcript |
| ENSG00000213703 | AL138847,1 | -1,156043346 | 0,448741543 | 0,04915 | 0,29848184 | ribosomal protein L13a (RPL13A) pseudogene |
| ENSG00000266433 | TBC1D3P5 | -1,15606495 | 0,448734823 | 0,14298 | 0,47060958 | TBC1 domain family member 3 pseudogene 5 |
| ENSG00000225975 | LINC01534 | -1,156478997 | 0,448606056 | 0,0376 | 0,26242137 | long intergenic non-protein coding RNA 1534 |
| ENSG00000220848 | RPS18P9 | -1,156791013 | 0,448509046 | 0,02922 | 0,2303054 | ribosomal protein S18 pseudogene 9 |
| ENSG00000114853 | ZBTB47 | -1,157055736 | 0,448426755 | 0,02177 | 0,19608606 | zinc finger and BTB domain containing 47 |
| ENSG00000175471 | MCTP1 | -1,157245676 | 0,448367721 | 0,00377 | 0,07484606 | multiple C2 and transmembrane domain containing 1 |
| ENSG00000271816 | BMS1P4 | -1,157629127 | 0,448248566 | 0,05157 | 0,3052902 | BMS1 pseudogene 4 |
| ENSG00000075643 | MOCOS | -1,158073171 | 0,448110622 | 0,05187 | 0,30606616 | molybdenum cofactor sulfurase |
| ENSG00000182050 | MGAT4C | -1,158522938 | 0,447970943 | 0,32121 | 0,60573903 | MGAT4 family member C |
| ENSG00000131650 | KREMEN2 | -1,158900313 | 0,44785378 | 0,07205 | 0,354195 | kringle containing transmembrane protein 2 |
| ENSG00000248643 | RBM14-RBM4 | -1,159210259 | 0,447757574 | 0,05429 | 0,3118853 | RBM14-RBM4 readthrough |
| ENSG00000100604 | CHGA | -1,159264573 | 0,447740717 | 0,03805 | 0,26375851 | chromogranin A |
| ENSG00000100055 | CYTH4 | -1,160117552 | 0,447476073 | 0,25333 | 0,55839162 | cytohesin 4 |
| ENSG00000198049 | AVPR1B | -1,160623393 | 0,447319206 | 0,05008 | 0,30089233 | arginine vasopressin receptor 1B |
| ENSG00000260751 | AC008870,2 | -1,160711781 | 0,447291801 | 0,05236 | 0,30715323 | novel transcript |
| ENSG00000104863 | LIN7B | -1,160887455 | 0,447237338 | 0,03088 | 0,2370319 | lin-7 homolog B, crumbs cell polarity complex component |
| ENSG00000284693 | LINC02606 | -1,161168501 | 0,447150223 | 0,03065 | 0,23611505 | long intergenic non-protein coding RNA 2606 |
| ENSG00000101210 | EEF1A2 | -1,161606669 | 0,447014437 | 0,01324 | 0,15041844 | eukaryotic translation elongation factor 1 alpha 2 |
| ENSG00000269947 | AC135178,6 | -1,161636509 | 0,447005191 | 0,04884 | 0,29716844 | novel transcript |
| ENSG00000135437 | RDH5 | -1,16207733 | 0,446868628 | 0,0582 | 0,32195911 | retinol dehydrogenase 5 |
| ENSG00000259367 | AC087286,2 | -1,162326231 | 0,446791539 | 0,00666 | 0,10355399 | novel transcript, antisense to AKAP13 |
| ENSG00000261465 | AC099518,4 | -1,162713172 | 0,446671722 | 0,33376 | 0,61371201 | novel transcript, antisense to TMC7 and COQ7 |
| ENSG00000242330 | RN7SL683P | -1,16302695 | 0,446574584 | 0,34978 | 0,62563749 | RNA, 7SL, cytoplasmic 683, pseudogene |
| ENSG00000213176 | RPL13P6 | -1,163429304 | 0,446450056 | 0,19353 | 0,51547251 | ribosomal protein L13 pseudogene 6 |
| ENSG00000287808 | AL355999,1 | -1,163838059 | 0,446323582 | 0,05033 | 0,30174609 | novel transcript, antisense to DHX9 |
| ENSG00000229065 | AL354893,2 | -1,164800009 | 0,446026085 | 0,05972 | 0,32521812 | novel transcript |

| | | | | | | |
|---|---|---|---|---|---|---|
| ENSG00000167654 | ATCAY | -1,164839173 | 0,446013977 | 0,01094 | 0,13517515 | ATCAY kinesin light chain interacting caytaxin |
| ENSG00000162623 | TYW3 | -1,16528619 | 0,445875802 | 0,10233 | 0,40985316 | tRNA-yW synthesizing protein 3 homolog |
| ENSG00000270118 | AC138894,2 | -1,165465042 | 0,44582053 | 0,09489 | 0,39743157 | novel transcript |
| ENSG00000254166 | CASC19 | -1,165491753 | 0,445812276 | 0,0943 | 0,39648446 | cancer susceptibility 19 |
| ENSG00000080166 | DCT | -1,165590388 | 0,445781797 | 0,00353 | 0,07209525 | dopachrome tautomerase |
| ENSG00000235635 | AC016644,1 | -1,165818331 | 0,44571137 | 0,09981 | 0,40626368 | novel transcript |
| ENSG00000233478 | AL031280,1 | -1,165887784 | 0,445689913 | 0,15832 | 0,4853983 | novel transcript |
| ENSG00000235027 | AC068580,3 | -1,166017872 | 0,445649727 | 0,27889 | 0,57700804 | novel transcript |
| ENSG00000280496 | SNORA17B | -1,166669313 | 0,445448542 | 0,07604 | 0,36217039 | small nucleolar RNA, H/ACA box 17B |
| ENSG00000189431 | RASSF10 | -1,166701216 | 0,445438692 | 0,01481 | 0,15966819 | Ras association domain family member 10 |
| ENSG00000180616 | SSTR2 | -1,166822439 | 0,445401265 | 0,0098 | 0,12683981 | somatostatin receptor 2 |
| ENSG00000236980 | C3orf84 | -1,167077793 | 0,445322437 | 0,10594 | 0,4154675 | chromosome 3 open reading frame 84 |
| ENSG00000228084 | AC118553,1 | -1,167081573 | 0,44532127 | 0,1667 | 0,49257009 | novel transcript |
| ENSG00000028137 | TNFRSF1B | -1,16764983 | 0,445145899 | 0,028 | 0,22579215 | TNF receptor superfamily member 1B |
| ENSG00000280613 | NDUFA6-DT | -1,167699126 | 0,445130689 | 0,04869 | 0,29656979 | NDUFA6 divergent transcript |
| ENSG00000257279 | AC127164,1 | -1,167705769 | 0,445128639 | 0,05627 | 0,31687103 | novel transcript |
| ENSG00000102878 | HSF4 | -1,167877488 | 0,44507566 | 0,1083 | 0,4189697 | heat shock transcription factor 4 |
| ENSG00000273765 | AC024257,4 | -1,168751198 | 0,4448062 | 0,25813 | 0,56220487 | novel transcript |
| ENSG00000237149 | ZNF503-AS2 | -1,169048826 | 0,444714446 | 0,03702 | 0,26086671 | ZNF503 antisense RNA 2 |
| ENSG00000169871 | TRIM56 | -1,169087509 | 0,444702522 | 0,01173 | 0,14097305 | tripartite motif containing 56 |
| ENSG00000233370 | ACTR1AP1 | -1,169351948 | 0,444621017 | 0,11225 | 0,42587863 | ACTR1A pseudogene 1 |
| ENSG00000260426 | AC008060,4 | -1,169357877 | 0,44461919 | 0,29091 | 0,58590649 | novel transcript, overlapping novel protein |
| ENSG00000102996 | MMP15 | -1,169555306 | 0,44455835 | 0,00049 | 0,02013867 | matrix metallopeptidase 15 |
| ENSG00000169436 | COL22A1 | -1,170800342 | 0,444174864 | 0,03171 | 0,24023654 | collagen type XXII alpha 1 chain |
| ENSG00000271424 | GLRX5P3 | -1,171708231 | 0,443895432 | 0,16289 | 0,48945635 | GLRX5 pseudogene 3 |
| ENSG00000223972 | DDX11L1 | -1,171711097 | 0,443894551 | 0,0587 | 0,32371785 | DEAD/H-box helicase 11 like 1 (pseudogene) |
| ENSG00000274105 | AC084824,3 | -1,171989419 | 0,443808924 | 0,07263 | 0,35556123 | novel transcript, sense intronic to YARS2 |
| ENSG00000065320 | NTN1 | -1,1726806 | 0,44359635 | 0,00076 | 0,02699099 | netrin 1 |
| ENSG00000255556 | AC087203,2 | -1,172998681 | 0,443498558 | 0,07136 | 0,35283325 | ubiquitin specific peptidase 17-like 2 (USP17L2) pseudogene |
| ENSG00000274296 | DECR2 | -1,173856208 | 0,443235024 | 0,05337 | 0,31020602 | 2,4-dienoyl-CoA reductase 2 |
| ENSG00000261123 | AC009065,5 | -1,173997634 | 0,443191576 | 0,06948 | 0,34813287 | novel transcript, intronic to PKD1 |
| ENSG00000262833 | AC016245,1 | -1,17449362 | 0,443039237 | 0,04923 | 0,29853148 | novel transcript, antisense to RPTOR |

| | | | | | | |
|---|---|---|---|---|---|---|
| ENSG00000237301 | AL121992,1 | -1,17469822 | 0,442976411 | 0,08163 | 0,37277532 | novel transcript |
| ENSG00000260805 | AC092803,1 | -1,174840292 | 0,44293279 | 0,02223 | 0,19814016 | novel transcript |
| ENSG00000148655 | LRMDA | -1,175101212 | 0,44285269 | 0,11291 | 0,42702832 | leucine rich melanocyte differentiation associated |
| ENSG00000244378 | RPS2P45 | -1,175103078 | 0,442852118 | 0,06584 | 0,33965348 | ribosomal protein S2 pseudogene 45 |
| ENSG00000287527 | AC005753,3 | -1,175686613 | 0,442673031 | 0,03106 | 0,23745328 | novel transcript |
| ENSG00000256453 | DND1 | -1,175817748 | 0,442632795 | 0,24675 | 0,55445285 | DND microRNA-mediated repression inhibitor 1 |
| ENSG00000188677 | PARVB | -1,176023576 | 0,44256965 | 0,01036 | 0,1311124 | parvin beta |
| ENSG00000011201 | ANOS1 | -1,176422283 | 0,442447357 | 0,03862 | 0,26585242 | anosmin 1 |
| ENSG00000140522 | RLBP1 | -1,176966965 | 0,442280345 | 0,12639 | 0,44903997 | retinaldehyde binding protein 1 |
| ENSG00000282003 | TUBGCP3 | -1,177016113 | 0,442265278 | 0,39202 | 0,65540679 | tubulin gamma complex associated protein 3 |
| ENSG00000287355 | AL451123,2 | -1,177116154 | 0,442234611 | 0,02472 | 0,21047862 | novel transcript, sense intronic to MOB3B |
| ENSG00000107954 | NEURL1 | -1,177704731 | 0,442054229 | 0,0259 | 0,21707964 | neuralized E3 ubiquitin protein ligase 1 |
| ENSG00000071655 | MBD3 | -1,177784517 | 0,442029783 | 0,01936 | 0,18420145 | methyl-CpG binding domain protein 3 |
| ENSG00000236804 | RPS3AP12 | -1,178822954 | 0,441711729 | 0,14136 | 0,46803321 | RPS3A pseudogene 12 |
| ENSG00000280118 | AC090983,1 | -1,179641206 | 0,441461275 | 0,15606 | 0,48386631 | novel transcript |
| ENSG00000110619 | CARS1 | -1,179875418 | 0,441389612 | 0,00907 | 0,12087977 | cysteinyl-tRNA synthetase 1 |
| ENSG00000233006 | MIR3936HG | -1,180173346 | 0,441298471 | 0,01634 | 0,16750744 | MIR3936 host gene |
| ENSG00000154146 | NRGN | -1,180247246 | 0,441275867 | 0,0105 | 0,13230162 | neurogranin |
| ENSG00000227855 | DPY19L2P3 | -1,180407086 | 0,441226979 | 0,04696 | 0,29152224 | DPY19L2 pseudogene 3 |
| ENSG00000182103 | FAM181B | -1,180851774 | 0,441090999 | 0,00834 | 0,11548316 | family with sequence similarity 181 member B |
| ENSG00000163576 | EFHB | -1,180890364 | 0,441079201 | 0,05789 | 0,32123433 | EF-hand domain family member B |
| ENSG00000104140 | RHOV | -1,18149345 | 0,440894856 | 0,17692 | 0,5012673 | ras homolog family member V |
| ENSG00000227063 | RPL41P1 | -1,182195409 | 0,440680386 | 0,30035 | 0,59175704 | ribosomal protein L41 pseudogene 1 |
| ENSG00000133246 | PRAM1 | -1,182349042 | 0,44063346 | 0,07083 | 0,35145965 | PML-RARA regulated adaptor molecule 1 |
| ENSG00000236256 | DIAPH2-AS1 | -1,182761842 | 0,440507399 | 0,21046 | 0,5290538 | DIAPH2 antisense RNA 1 |
| ENSG00000103269 | RHBDL1 | -1,182798959 | 0,440496066 | 0,20275 | 0,52269757 | rhomboid like 1 |
| ENSG00000259772 | AC012236,1 | -1,182806729 | 0,440493694 | 0,1545 | 0,48266733 | uncharacterized LOC283710 |
| ENSG00000285852 | AL353147,1 | -1,182890325 | 0,440468171 | 0,06155 | 0,32865897 | novel transcript |
| ENSG00000198221 | AFDN-DT | -1,182978765 | 0,44044117 | 0,02193 | 0,19698513 | AFDN divergent transcript |
| ENSG00000260488 | AC007216,1 | -1,183525437 | 0,440274308 | 0,2811 | 0,57848427 | novel transcript, antisense to RUNDC2A |
| ENSG00000276952 | AL121772,3 | -1,185201629 | 0,439763073 | 0,02738 | 0,22321357 | novel transcript, antisense to PYGB |
| ENSG00000229173 | SRMP3 | -1,185715802 | 0,43960637 | 0,18301 | 0,50652064 | SRM pseudogene 3 |
| ENSG00000159263 | SIM2 | -1,185866721 | 0,439560386 | 0,03208 | 0,24164117 | SIM bHLH transcription factor 2 |

| | | | | | | |
|---|---|---|---|---|---|---|
| ENSG00000270022 | Z93241,1 | -1,186053072 | 0,439503612 | 0,04215 | 0,27783166 | RNA, U12 small nuclear |
| ENSG00000271395 | LINC02336 | -1,187225957 | 0,439146449 | 0,14477 | 0,47232606 | long intergenic non-protein coding RNA 2336 |
| ENSG00000234848 | AC008758,2 | -1,18740798 | 0,439091046 | 0,29808 | 0,59043587 | novel transcript, antisense to ZNF709 |
| ENSG00000185031 | SLC2A3P2 | -1,187725454 | 0,438994432 | 0,02904 | 0,22988377 | solute carrier family 2 member 3 pseudogene 2 |
| ENSG00000224877 | NDUFAF8 | -1,188467781 | 0,438768609 | 0,02423 | 0,20804286 | NADH:ubiquinone oxidoreductase complex assembly factor 8 |
| ENSG00000283633 | AP000547,3 | -1,188594102 | 0,438730193 | 0,01591 | 0,16521061 | novel transcript |
| ENSG00000051128 | HOMER3 | -1,188668703 | 0,438707506 | 0,00075 | 0,02678818 | homer scaffold protein 3 |
| ENSG00000267253 | WHSC1L2P | -1,188920186 | 0,43863104 | 0,05709 | 0,31865533 | Wolf-Hirschhorn syndrome candidate 1-like 2, pseudogene |
| ENSG00000287416 | AC093746,1 | -1,189148004 | 0,438561781 | 0,043 | 0,27928585 | novel transcript |
| ENSG00000226891 | LINC01359 | -1,189254771 | 0,438529326 | 0,01943 | 0,18463825 | long intergenic non-protein coding RNA 1359 |
| ENSG00000217227 | AL079341,1 | -1,189262071 | 0,438527107 | 0,10349 | 0,41144828 | TRK-fused gene (TFG) pseudogene |
| ENSG00000081138 | CDH7 | -1,189346761 | 0,438501365 | 0,01929 | 0,18393463 | cadherin 7 |
| ENSG00000237807 | AC022034,1 | -1,18945126 | 0,438469604 | 0,0204 | 0,18959706 | novel transcript |
| ENSG00000259146 | AC005476,2 | -1,190949107 | 0,438014609 | 0,25518 | 0,55966322 | novel transcript, antisense to SIPA1L1 |
| ENSG00000164853 | UNCX | -1,191033197 | 0,437989079 | 0,01424 | 0,15639135 | UNC homeobox |
| ENSG00000155749 | FLACC1 | -1,191191576 | 0,437940999 | 0,08194 | 0,37306719 | flagellum associated containing coiled-coil domains 1 |
| ENSG00000234776 | C11orf94 | -1,191673621 | 0,437794695 | 0,06069 | 0,32713591 | chromosome 11 open reading frame 94 |
| ENSG00000281197 | TNFRSF14-AS1 | -1,191723552 | 0,437779544 | 0,20472 | 0,52443222 | TNFRSF14 antisense RNA 1 |
| ENSG00000183715 | OPCML | -1,192137707 | 0,437653888 | 0,00524 | 0,09023148 | opioid binding protein/cell adhesion molecule like |
| ENSG00000176619 | LMNB2 | -1,192352934 | 0,437588602 | 0,01395 | 0,15492928 | lamin B2 |
| ENSG00000227888 | FAM66A | -1,192615298 | 0,437509031 | 0,02931 | 0,23063117 | family with sequence similarity 66 member A |
| ENSG00000269439 | AC010618,3 | -1,193647793 | 0,43719603 | 0,053 | 0,30902552 | novel transcript |
| ENSG00000141434 | MEP1B | -1,193756182 | 0,437163185 | 0,07333 | 0,35655334 | meprin A subunit beta |
| ENSG00000282509 | ULK4P3 | -1,194063796 | 0,437069983 | 0,00595 | 0,09720154 | ULK4 pseudogene 3 |
| ENSG00000178081 | ULK4P3 | -1,194063796 | 0,437069983 | 0,00595 | 0,09720154 | ULK4 pseudogene 3 |
| ENSG00000259660 | DNM1P47 | -1,194465297 | 0,436948363 | 0,00741 | 0,10952546 | dynamin 1 pseudogene 47 |
| ENSG00000130653 | PNPLA7 | -1,194667082 | 0,436887253 | 0,02129 | 0,19359286 | patatin like phospholipase domain containing 7 |
| ENSG00000172667 | ZMAT3 | -1,194730931 | 0,436867918 | 0,03091 | 0,23705504 | zinc finger matrin-type 3 |
| ENSG00000275557 | AC242842,1 | -1,195291185 | 0,436698298 | 0,08332 | 0,37550713 | novel transcript |
| ENSG00000248898 | AC022126,1 | -1,195342152 | 0,436682871 | 0,13825 | 0,46543358 | novel transcript |
| ENSG00000148832 | PAOX | -1,195707729 | 0,43657223 | 0,05518 | 0,31442055 | polyamine oxidase |
| ENSG00000247416 | AP000802,1 | -1,195730224 | 0,436565423 | 0,00808 | 0,11424852 | novel transcript, antisense to NCAM1 |

| | | | | | | |
|---|---|---|---|---|---|---|
| ENSG00000181085 | MAPK15 | -1,195814282 | 0,436539988 | 0,02397 | 0,20687236 | mitogen-activated protein kinase 15 |
| ENSG00000262050 | AC005696,1 | -1,195887439 | 0,436517852 | 0,22726 | 0,54168591 | novel transcript, antisense to CLUH |
| ENSG00000167971 | CASKIN1 | -1,196069326 | 0,436462822 | 0,00969 | 0,12603664 | CASK interacting protein 1 |
| ENSG00000237758 | BANF1P3 | -1,196320644 | 0,436386796 | 0,10035 | 0,4067769 | BANF1 pseudogene 3 |
| ENSG00000196277 | GRM7 | -1,196504272 | 0,436331256 | 0,03695 | 0,26075082 | glutamate metabotropic receptor 7 |
| ENSG00000137877 | SPTBN5 | -1,196950508 | 0,436196316 | 0,05506 | 0,31407633 | spectrin beta, non-erythrocytic 5 |
| ENSG00000276166 | AC092118,2 | -1,19788568 | 0,43591366 | 0,11536 | 0,43106723 | novel transcript, antisense to KIFC3 |
| ENSG00000153531 | ADPRHL1 | -1,19793168 | 0,435899762 | 0,03527 | 0,2543788 | ADP-ribosylhydrolase like 1 |
| ENSG00000216490 | IFI30 | -1,198610971 | 0,435694567 | 0,20463 | 0,52443222 | IFI30 lysosomal thiol reductase |
| ENSG00000235257 | ITGA9-AS1 | -1,198829687 | 0,43562852 | 0,04209 | 0,27778454 | ITGA9 antisense RNA 1 |
| ENSG00000182545 | RNASE10 | -1,199244185 | 0,435503378 | 0,02671 | 0,22056859 | ribonuclease A family member 10 (inactive) |
| ENSG00000267731 | AC005332,2 | -1,199462896 | 0,435437361 | 0,14649 | 0,47433542 | novel transcript |
| ENSG00000278823 | BMERB1 | -1,199599375 | 0,435396171 | 0,13747 | 0,46452141 | bMERB domain containing 1 |
| ENSG00000276191 | CRHR1 | -1,199605596 | 0,435394293 | 0,29451 | 0,58857672 | corticotropin releasing hormone receptor 1 |
| ENSG00000188735 | TMEM120B | -1,200128736 | 0,435236442 | 0,04631 | 0,28930497 | transmembrane protein 120B |
| ENSG00000231980 | AC011899,1 | -1,200168074 | 0,435224575 | 0,0305 | 0,23556309 | novel transcript |
| ENSG00000182255 | KCNA4 | -1,200697716 | 0,435064825 | 0,17528 | 0,49934416 | potassium voltage-gated channel subfamily A member 4 |
| ENSG00000161010 | MRNIP | -1,200961261 | 0,434985356 | 0,01307 | 0,14962749 | MRN complex interacting protein |
| ENSG00000238121 | LINC00426 | -1,20118607 | 0,43491758 | 0,0844 | 0,37731853 | long intergenic non-protein coding RNA 426 |
| ENSG00000168702 | LRP1B | -1,201504984 | 0,43482145 | 0,05182 | 0,30603373 | LDL receptor related protein 1B |
| ENSG00000241185 | AC005410,1 | -1,201838652 | 0,434720896 | 0,26827 | 0,5704222 | ribosomal protein L21 pseudogene 122 |
| ENSG00000229466 | LINC02627 | -1,201845379 | 0,434718868 | 0,14731 | 0,47481692 | long intergenic non-protein coding RNA 2627 |
| ENSG00000105443 | CYTH2 | -1,202151794 | 0,434626548 | 0,01125 | 0,13764117 | cytohesin 2 |
| ENSG00000287653 | AL355597,1 | -1,202262665 | 0,434593149 | 0,06278 | 0,33160043 | novel transcript |
| ENSG00000282739 | SLC25A26 | -1,202403758 | 0,434550648 | 3,22E-05 | 0,00229769 | solute carrier family 25 member 26 |
| ENSG00000228302 | AL512770,1 | -1,202546039 | 0,434507794 | 0,04538 | 0,28776102 | novel transcript |
| ENSG00000274253 | AC138649,1 | -1,202641364 | 0,434479085 | 0,00104 | 0,0330699 | novel transcript, sense overlapping NIPA1 |
| ENSG00000160613 | PCSK7 | -1,202930861 | 0,43439191 | 0,02702 | 0,22193038 | proprotein convertase subtilisin/kexin type 7 |
| ENSG00000140465 | CYP1A1 | -1,20304865 | 0,434356445 | 0,16917 | 0,49530908 | cytochrome P450 family 1 subfamily A member 1 |
| ENSG00000236810 | ELOA-AS1 | -1,203351611 | 0,434265241 | 0,0392 | 0,26793262 | ELOA antisense RNA 1 |
| ENSG00000151773 | CCDC122 | -1,203693577 | 0,434162318 | 0,03264 | 0,24385689 | coiled-coil domain containing 122 |
| ENSG00000274780 | PHRF1 | -1,203973027 | 0,434078229 | 0,00859 | 0,11699775 | PHD and ring finger domains 1 |
| ENSG00000188558 | OR2G6 | -1,204462567 | 0,433930961 | 0,03449 | 0,25106871 | olfactory receptor family 2 subfamily G member 6 |

| | | | | | | |
|---|---|---|---|---|---|---|
| ENSG00000129167 | TPH1 | -1,204757866 | 0,433842151 | 0,01494 | 0,16037017 | tryptophan hydroxylase 1 |
| ENSG00000102924 | CBLN1 | -1,204803019 | 0,433828573 | 0,00103 | 0,03299702 | cerebellin 1 precursor |
| ENSG00000285662 | FAM245B | -1,204850345 | 0,433814342 | 0,28626 | 0,58267001 | family with sequence similarity 245 member B |
| ENSG00000163683 | SMIM14 | -1,204929641 | 0,433790499 | 0,0318 | 0,24056445 | small integral membrane protein 14 |
| ENSG00000230593 | PPIAP40 | -1,205274507 | 0,433686816 | 0,11846 | 0,43614969 | peptidylprolyl isomerase A pseudogene 40 |
| ENSG00000288503 | DDX51 | -1,205312517 | 0,43367539 | 0,0657 | 0,33945082 | DEAD-box helicase 51 |
| ENSG00000204366 | ZBTB12 | -1,205426497 | 0,433641129 | 0,00681 | 0,10528058 | zinc finger and BTB domain containing 12 |
| ENSG00000076344 | RGS11 | -1,205836258 | 0,433517982 | 0,00225 | 0,05400651 | regulator of G protein signaling 11 |
| ENSG00000226899 | AL731577,1 | -1,205866585 | 0,433508869 | 0,07949 | 0,36813093 | novel transcript |
| ENSG00000177283 | FZD8 | -1,206426126 | 0,433340768 | 0,05899 | 0,3241112 | frizzled class receptor 8 |
| ENSG00000287769 | AL353748,3 | -1,206503501 | 0,433317527 | 0,01538 | 0,16247082 | novel transcript, antisense to SPIN1 |
| ENSG00000135702 | CHST5 | -1,206780771 | 0,433234257 | 0,03765 | 0,26251324 | carbohydrate sulfotransferase 5 |
| ENSG00000236719 | OVAAL | -1,207084353 | 0,433143102 | 0,22297 | 0,53908993 | ovarian adenocarcinoma amplified long non-coding RNA |
| ENSG00000118514 | ALDH8A1 | -1,207184952 | 0,4331129 | 0,07497 | 0,3592137 | aldehyde dehydrogenase 8 family member A1 |
| ENSG00000234710 | AC060834,2 | -1,207320094 | 0,433072331 | 0,10929 | 0,42113687 | novel transcript |
| ENSG00000235292 | AL109838,1 | -1,207844669 | 0,432914891 | 0,04994 | 0,30082943 | novel transcript |
| ENSG00000176654 | NANOGP1 | -1,20786729 | 0,432908103 | 0,17217 | 0,49710979 | Nanog homeobox pseudogene 1 |
| ENSG00000185477 | GPRIN3 | -1,207876542 | 0,432905327 | 0,02943 | 0,23080322 | GPRIN family member 3 |
| ENSG00000262666 | FAM189B | -1,208674433 | 0,432665972 | 0,06083 | 0,32750733 | family with sequence similarity 189 member B |
| ENSG00000235319 | AC012360,1 | -1,208763127 | 0,432639374 | 0,20339 | 0,52321822 | novel transcript |
| ENSG00000230733 | AC092171,2 | -1,208998569 | 0,432568774 | 0,72965 | 0,8715818 | novel transcript |
| ENSG00000257759 | AL357153,2 | -1,209479124 | 0,432424712 | 0,32508 | 0,60753927 | novel transcript |
| ENSG00000251143 | AP002490,1 | -1,209552154 | 0,432402823 | 0,02442 | 0,2089849 | novel transcript |
| ENSG00000228222 | AC073050,1 | -1,209669173 | 0,432367751 | 0,09048 | 0,38914225 | novel transcript (hCG1660379) |
| ENSG00000215893 | RPL23AP17 | -1,209722802 | 0,432351679 | 0,15019 | 0,47830235 | ribosomal protein L23a pseudogene 17 |
| ENSG00000130772 | MED18 | -1,210600894 | 0,43208861 | 0,25116 | 0,55719428 | mediator complex subunit 18 |
| ENSG00000260763 | AC106799,3 | -1,21076694 | 0,432038882 | 0,31695 | 0,60268749 | novel transcript |
| ENSG00000260318 | COX6CP1 | -1,21082811 | 0,432020564 | 0,22144 | 0,53805234 | cytochrome c oxidase subunit 6C pseudogene 1 |
| ENSG00000262583 | TMEM231P1 | -1,210878125 | 0,432005587 | 0,18989 | 0,51315563 | TMEM231 pseudogene 1 |
| ENSG00000246225 | AC006299,1 | -1,211261473 | 0,431890811 | 0,23374 | 0,54563244 | novel transcript, antisense to SVIP |
| ENSG00000241313 | WWTR1-AS1 | -1,211565337 | 0,431799855 | 0,04399 | 0,28274932 | WWTR1 antisense RNA 1 |
| ENSG00000250490 | LINC02145 | -1,211793852 | 0,431731465 | 0,27293 | 0,57331473 | long intergenic non-protein coding RNA 2145 |
| ENSG00000139629 | GALNT6 | -1,212102459 | 0,431639124 | 0,1847 | 0,50835796 | polypeptide N-acetylgalactosaminyltransferase 6 |

| | | | | | | |
|---|---|---|---|---|---|---|
| ENSG00000237004 | ZNRF2P1 | -1,212534788 | 0,431509795 | 0,01419 | 0,15631436 | zinc and ring finger 2 pseudogene 1 |
| ENSG00000130529 | TRPM4 | -1,212561846 | 0,431501702 | 0,0396 | 0,26924564 | transient receptor potential cation channel subfamily M member 4 |
| ENSG00000094755 | GABRP | -1,212707364 | 0,43145818 | 0,07262 | 0,35556123 | gamma-aminobutyric acid type A receptor subunit pi |
| ENSG00000226717 | PTPRD-AS2 | -1,212797945 | 0,431431092 | 0,19305 | 0,51505487 | PTPRD antisense RNA 2 (head to head) |
| ENSG00000137691 | CFAP300 | -1,213055169 | 0,431354177 | 0,01595 | 0,16543992 | cilia and flagella associated protein 300 |
| ENSG00000132274 | TRIM22 | -1,213388821 | 0,431254429 | 0,01907 | 0,18296265 | tripartite motif containing 22 |
| ENSG00000147127 | RAB41 | -1,214030552 | 0,431062644 | 0,03729 | 0,26130525 | RAB41, member RAS oncogene family |
| ENSG00000249854 | CDH18-AS1 | -1,214060492 | 0,431053698 | 0,01168 | 0,14071678 | CDH18 antisense RNA 1 |
| ENSG00000230555 | AL450326,1 | -1,214203355 | 0,431011015 | 0,16462 | 0,49065086 | novel transcript |
| ENSG00000198106 | AC025279,1 | -1,214550417 | 0,430907341 | 0,12649 | 0,4491539 | novel transcript |
| ENSG00000173011 | TADA2B | -1,214596618 | 0,430893542 | 0,02337 | 0,2040793 | transcriptional adaptor 2B |
| ENSG00000239791 | AC002310,2 | -1,214779096 | 0,430839045 | 0,11547 | 0,43130012 | novel transcript |
| ENSG00000186314 | PRELID2 | -1,214908395 | 0,430800433 | 0,01111 | 0,13644916 | PRELI domain containing 2 |
| ENSG00000269959 | SPACA6P-AS | -1,215157202 | 0,430726144 | 0,11893 | 0,43696075 | SPACA6P antisense RNA |
| ENSG00000142330 | CAPN10 | -1,215259484 | 0,430695608 | 0,00844 | 0,11598 | calpain 10 |
| ENSG00000234948 | LINC01524 | -1,215302759 | 0,430682689 | 0,06795 | 0,34439007 | long intergenic non-protein coding RNA 1524 |
| ENSG00000222009 | BTBD19 | -1,215518804 | 0,430618198 | 0,00765 | 0,11114888 | BTB domain containing 19 |
| ENSG00000287682 | AC020651,2 | -1,215713433 | 0,430560109 | 0,06725 | 0,34325886 | novel transcript |
| ENSG00000251520 | AL162431,2 | -1,215763041 | 0,430545304 | 0,06482 | 0,33757437 | pseudogene similar to part of major histocompatibility complex, class I-related MR1 |
| ENSG00000284639 | AP006294,2 | -1,215945282 | 0,430490921 | 0,26651 | 0,56917012 | RNA binding protein, fox-1 homolog (C, elegans) 1 (RBFOX1) pseudogene |
| ENSG00000256637 | LINC01965 | -1,216122263 | 0,430438114 | 0,12873 | 0,45196697 | long intergenic non-protein coding RNA 1965 |
| ENSG00000266897 | AC005546,1 | -1,216185862 | 0,43041914 | 0,19227 | 0,51477601 | novel transcript |
| ENSG00000287383 | AC110809,1 | -1,216349132 | 0,430370432 | 0,07372 | 0,35703856 | novel transcript |
| ENSG00000283141 | LINC02666 | -1,21660815 | 0,430293171 | 0,09087 | 0,39016581 | long intergenic non-protein coding RNA 2666 |
| ENSG00000270103 | AL360012,1 | -1,216798492 | 0,430236404 | 0,03566 | 0,2562416 | RNA, U11 small nuclear |
| ENSG00000179841 | AKAP5 | -1,216909724 | 0,430203234 | 0,00848 | 0,11625431 | A-kinase anchoring protein 5 |
| ENSG00000213862 | AC044787,1 | -1,217080159 | 0,430152414 | 0,06901 | 0,34717336 | ribosomal protein L7a (RPL7A) pseudogene |
| ENSG00000196440 | ARMCX4 | -1,217300668 | 0,430086673 | 0,00048 | 0,01972028 | armadillo repeat containing X-linked 4 |
| ENSG00000136213 | CHST12 | -1,217677726 | 0,429974281 | 0,03848 | 0,26548082 | carbohydrate sulfotransferase 12 |

| | | | | | | |
|---|---|---|---|---|---|---|
| ENSG00000233040 | FAM204BP | -1,217861979 | 0,429919371 | 0,01706 | 0,17158055 | family with sequence similarity 204 member B, pseudogene |
| ENSG00000187391 | MAGI2 | -1,218089011 | 0,429851722 | 0,00132 | 0,03878375 | membrane associated guanylate kinase, WW and PDZ domain containing 2 |
| ENSG00000161798 | AQP5 | -1,218389658 | 0,429762153 | 0,01781 | 0,1758522 | aquaporin 5 |
| ENSG00000285530 | AL445928,2 | -1,218426974 | 0,429751037 | 0,07432 | 0,3579004 | novel transcript, antisense to DBT |
| ENSG00000106012 | IQCE | -1,219281089 | 0,429496688 | 0,00771 | 0,11160063 | IQ motif containing E |
| ENSG00000244383 | FAM3D-AS1 | -1,219436582 | 0,429450399 | 0,06959 | 0,34818658 | FAM3D antisense RNA 1 |
| ENSG00000276914 | CEP20 | -1,219437749 | 0,429450052 | 0,13062 | 0,45494178 | centrosomal protein 20 |
| ENSG00000263072 | ZNF213-AS1 | -1,219504988 | 0,429430037 | 0,02865 | 0,22885696 | ZNF213 antisense RNA 1 (head to head) |
| ENSG00000123700 | KCNJ2 | -1,219789749 | 0,429345284 | 0,02844 | 0,22805204 | potassium inwardly rectifying channel subfamily J member 2 |
| ENSG00000117407 | ARTN | -1,219940004 | 0,429300571 | 0,00652 | 0,10239961 | artemin |
| ENSG00000270096 | AC078795,2 | -1,220334745 | 0,429183124 | 0,52253 | 0,74670504 | novel transcript, antisense to LRRC34 |
| ENSG00000141194 | OR4D1 | -1,220374558 | 0,429171281 | 0,23676 | 0,54767193 | olfactory receptor family 4 subfamily D member 1 |
| ENSG00000228137 | AP001469,2 | -1,22163869 | 0,428795393 | 0,0367 | 0,2596357 | novel transcript |
| ENSG00000260179 | AL162741,1 | -1,222083416 | 0,428663233 | 0,01523 | 0,16205591 | novel transcript |
| ENSG00000227315 | NEU1 | -1,222318204 | 0,428593476 | 0,28761 | 0,58395529 | neuraminidase 1 |
| ENSG00000287374 | AC239860,4 | -1,222544243 | 0,42852633 | 0,13026 | 0,45467139 | novel transcript |
| ENSG00000226636 | AC006020,1 | -1,222726731 | 0,428472129 | 0,2684 | 0,57051739 | tetratricopeptide repeat domain 14 (TTC14) pseudogene |
| ENSG00000214432 | VPS33B-DT | -1,223017426 | 0,428385803 | 0,07097 | 0,35173843 | VPS33B divergent transcript |
| ENSG00000004776 | HSPB6 | -1,223293099 | 0,428303954 | 0,03391 | 0,24870776 | heat shock protein family B (small) member 6 |
| ENSG00000188747 | NOXA1 | -1,223395266 | 0,428273624 | 0,04822 | 0,29503004 | NADPH oxidase activator 1 |
| ENSG00000161103 | AC008132,1 | -1,223660061 | 0,428195025 | 0,15588 | 0,4836223 | POM121 transmembrane nucleoporin (POM121) pseudogene |
| ENSG00000222019 | URAHP | -1,223962266 | 0,428105339 | 0,01911 | 0,18296265 | urate (hydroxyiso-) hydrolase, pseudogene |
| ENSG00000235673 | AL929472,4 | -1,223966211 | 0,428104169 | 0,25129 | 0,55720663 | ribosomal protein S2 (RPS2) pseudogene |
| ENSG00000042493 | CAPG | -1,224606331 | 0,427914262 | 7,27E-05 | 0,00461938 | capping actin protein, gelsolin like |
| ENSG00000178772 | CPN2 | -1,225245705 | 0,427724661 | 0,21036 | 0,5290538 | carboxypeptidase N subunit 2 |
| ENSG00000230732 | AC016949,1 | -1,225543848 | 0,427636278 | 0,04325 | 0,28025067 | novel transcript |
| ENSG00000233953 | AC009970,1 | -1,225815264 | 0,427555834 | 0,32682 | 0,60888005 | novel transcript |
| ENSG00000254688 | AC013549,3 | -1,225833037 | 0,427550566 | 0,24099 | 0,55003907 | novel transcript |
| ENSG00000268230 | AC012313,3 | -1,225889776 | 0,427533752 | 0,01118 | 0,13687226 | novel transcript |

| | | | | | | |
|---|---|---|---|---|---|---|
| ENSG00000259205 | PRKXP1 | -1,226622484 | 0,427316673 | 0,07318 | 0,35655334 | PRKX pseudogene 1 |
| ENSG00000224924 | LINC00320 | -1,227601985 | 0,42702665 | 0,0072 | 0,10776454 | long intergenic non-protein coding RNA 320 |
| ENSG00000243433 | AC010973,1 | -1,228199435 | 0,426849846 | 0,12374 | 0,4446254 | novel transcript |
| ENSG00000116254 | CHD5 | -1,228442451 | 0,426777951 | 0,01833 | 0,17905232 | chromodomain helicase DNA binding protein 5 |
| ENSG00000264860 | AC097641,1 | -1,228660126 | 0,426713563 | 0,23137 | 0,54433844 | novel transcript |
| ENSG00000287594 | AL356123,2 | -1,22919525 | 0,426555316 | 0,0303 | 0,23492717 | novel transcript |
| ENSG00000262380 | AC026401,2 | -1,229240418 | 0,426541962 | 0,18813 | 0,51128963 | novel transcript, sense to NDE1 |
| ENSG00000282255 | AC243742,1 | -1,229240418 | 0,426541962 | 0,18813 | 0,51128963 | novel transcript, sense to NDE1 |
| ENSG00000007968 | E2F2 | -1,229675633 | 0,426413307 | 0,18141 | 0,50490902 | E2F transcription factor 2 |
| ENSG00000283244 | FRG1-DT | -1,229744619 | 0,426392918 | 0,09919 | 0,40547136 | FRG1 divergent transcript |
| ENSG00000215481 | BCRP3 | -1,230289853 | 0,426231803 | 0,07017 | 0,34975886 | BCR pseudogene 3 |
| ENSG00000229196 | AC087071,1 | -1,231486364 | 0,42587845 | 0,20637 | 0,52602996 | novel transcript |
| ENSG00000287422 | AL442071,2 | -1,23164899 | 0,425830446 | 0,05467 | 0,31311579 | novel transcript, antisense to MACF1 |
| ENSG00000124194 | GDAP1L1 | -1,23205601 | 0,425710326 | 0,00784 | 0,11257373 | ganglioside induced differentiation associated protein 1 like 1 |
| ENSG00000176531 | PHLDB3 | -1,232122687 | 0,425690652 | 0,02668 | 0,22048053 | pleckstrin homology like domain family B member 3 |
| ENSG00000225292 | AL158212,1 | -1,232232262 | 0,425658321 | 0,43991 | 0,6885781 | novel transcript |
| ENSG00000107872 | FBXL15 | -1,23227683 | 0,425645171 | 0,00624 | 0,09963421 | F-box and leucine rich repeat protein 15 |
| ENSG00000144644 | GADL1 | -1,23254855 | 0,425565012 | 0,04207 | 0,27770943 | glutamate decarboxylase like 1 |
| ENSG00000213731 | RAB5CP1 | -1,23315485 | 0,425386204 | 0,13197 | 0,45708171 | RAB5C, member RAS oncogene family pseudogene 1 |
| ENSG00000141744 | PNMT | -1,233261378 | 0,425354795 | 0,04857 | 0,29611078 | phenylethanolamine N-methyltransferase |
| ENSG00000175611 | LINC00476 | -1,233639882 | 0,425243214 | 0,03038 | 0,23517553 | long intergenic non-protein coding RNA 476 |
| ENSG00000167657 | DAPK3 | -1,234252715 | 0,425062616 | 0,01531 | 0,16232837 | death associated protein kinase 3 |
| ENSG00000256108 | AC135586,1 | -1,234368309 | 0,42502856 | 0,04304 | 0,27943834 | tumor protein D52-like 2 (TPD52L2) pseudogene |
| ENSG00000278662 | GOLGA6L10 | -1,23445644 | 0,425002596 | 0,00258 | 0,05903475 | golgin A6 family like 10 |
| ENSG00000261838 | AC092718,6 | -1,235088344 | 0,424816485 | 0,25749 | 0,56160845 | novel transcript, antisense to C16orf46 |
| ENSG00000165233 | CARD19 | -1,235300675 | 0,424753966 | 0,02048 | 0,18980371 | caspase recruitment domain family member 19 |
| ENSG00000236753 | MKLN1-AS | -1,235506993 | 0,424693227 | 0,0289 | 0,22977124 | MKLN1 antisense RNA |
| ENSG00000177225 | GATD1 | -1,236089899 | 0,424521669 | 0,02583 | 0,2170617 | glutamine amidotransferase like class 1 domain containing 1 |
| ENSG00000268154 | Metazoa_SRP | -1,236295711 | 0,424461112 | 0,41023 | 0,66821634 | Metazoan signal recognition particle RNA |
| ENSG00000250069 | AC011379,1 | -1,236328838 | 0,424451366 | 0,00729 | 0,10839211 | novel transcript |
| ENSG00000162461 | SLC25A34 | -1,236329082 | 0,424451294 | 0,03272 | 0,24407309 | solute carrier family 25 member 34 |

| | | | | | | |
|---|---|---|---|---|---|---|
| ENSG00000251046 | ZNF969P | -1,236969238 | 0,424262997 | 0,00894 | 0,11991469 | zinc finger protein 969, pseudogene |
| ENSG00000056972 | TRAF3IP2 | -1,237026962 | 0,424246022 | 0,03735 | 0,26158694 | TRAF3 interacting protein 2 |
| ENSG00000267244 | AC012615,6 | -1,237154958 | 0,424208385 | 0,03353 | 0,24724664 | novel transcript, antisense to REXO1 |
| ENSG00000149483 | TMEM138 | -1,237735075 | 0,424037842 | 0,02429 | 0,20831626 | transmembrane protein 138 |
| ENSG00000277744 | AC011462,4 | -1,237862901 | 0,424000273 | 0,07909 | 0,36725857 | novel transcript, sense intronic TMEM91 |
| ENSG00000281732 | Z84468,1 | -1,237925374 | 0,423981913 | 0,0569 | 0,31803579 | novel transcript |
| ENSG00000234771 | SLC25A25-AS1 | -1,238198062 | 0,423901782 | 0,02216 | 0,19795874 | SLC25A25 antisense RNA 1 |
| ENSG00000164708 | PGAM2 | -1,238845595 | 0,423711563 | 0,11763 | 0,43515458 | phosphoglycerate mutase 2 |
| ENSG00000251138 | LINC02882 | -1,239035394 | 0,423655824 | 0,08122 | 0,3719701 | long intergenic non-protein coding RNA 2882 |
| ENSG00000286864 | AC012076,1 | -1,239100849 | 0,423636603 | 0,00222 | 0,05370447 | novel transcript, antisense to LRRFIP1 |
| ENSG00000277022 | AL031663,3 | -1,239168996 | 0,423616593 | 0,15639 | 0,48424189 | novel transcript |
| ENSG00000287760 | AC099521,3 | -1,239447034 | 0,42353496 | 0,05822 | 0,32195911 | novel transcript |
| ENSG00000270154 | AC130352,1 | -1,239588595 | 0,423493404 | 0,03161 | 0,23999626 | novel transcript |
| ENSG00000286981 | AL731553,1 | -1,239717682 | 0,423455513 | 0,09896 | 0,40492885 | novel transcript |
| ENSG00000264278 | ZNF236-DT | -1,239717831 | 0,42345547 | 0,01695 | 0,17088104 | ZNF236 divergent transcript |
| ENSG00000235975 | Z83313,1 | -1,240274386 | 0,423292143 | 0,03047 | 0,23546694 | tropomyosin 2 (beta) (TPM2) pseudogene |
| ENSG00000144063 | MALL | -1,240276982 | 0,423291381 | 0,02866 | 0,22888274 | mal, T cell differentiation protein like |
| ENSG00000285190 | AC018754,1 | -1,242099048 | 0,422757118 | 0,15877 | 0,48624756 | novel transcript, antisense to CCNH |
| ENSG00000123843 | C4BPB | -1,242516651 | 0,422634765 | 0,10032 | 0,4067769 | complement component 4 binding protein beta |
| ENSG00000053501 | USE1 | -1,242764281 | 0,422562228 | 0,02385 | 0,20623598 | unconventional SNARE in the ER 1 |
| ENSG00000223914 | LINC02471 | -1,24291847 | 0,422517069 | 0,30116 | 0,59252799 | long intergenic non-protein coding RNA 2471 |
| ENSG00000273327 | OR6L2P | -1,242994623 | 0,422494767 | 0,13622 | 0,463307 | olfactory receptor family 6 subfamily L member 2 pseudogene |
| ENSG00000244308 | RN7SL762P | -1,243033646 | 0,422483339 | 0,34332 | 0,62082523 | RNA, 7SL, cytoplasmic 762, pseudogene |
| ENSG00000100156 | SLC16A8 | -1,243525609 | 0,422339296 | 0,04073 | 0,2728706 | solute carrier family 16 member 8 |
| ENSG00000167393 | PPP2R3B | -1,243905038 | 0,422228235 | 0,01536 | 0,16236395 | protein phosphatase 2 regulatory subunit B''beta |
| ENSG00000258858 | AL583722,3 | -1,244169335 | 0,422150891 | 0,24521 | 0,55295131 | novel transcript, antisense to INF2 |
| ENSG00000245322 | H2AZ1-DT | -1,244717557 | 0,421990505 | 0,0404 | 0,27186146 | H2AZ1 divergent transcript |
| ENSG00000281123 | FCGBP | -1,244785605 | 0,421970601 | 0,01112 | 0,13644918 | Fc fragment of IgG binding protein |
| ENSG00000250282 | AC002401,2 | -1,24482322 | 0,4219596 | 0,21895 | 0,53640494 | novel transcript, antisense to PDK2 |
| ENSG00000146722 | AC211486,1 | -1,244842191 | 0,421954051 | 0,01565 | 0,164013 | FK506 binding protein 6, 36kDa (FKBP6 ) pseudogene |
| ENSG00000287013 | AC091646,2 | -1,245061986 | 0,421889771 | 0,16703 | 0,49293221 | novel transcript |
| ENSG00000241362 | RPL36AP43 | -1,245072969 | 0,421886559 | 0,18426 | 0,50802406 | ribosomal protein L36a pseudogene 43 |

| | | | | | | |
|---|---|---|---|---|---|---|
| ENSG00000223511 | AL683807,1 | -1,245207247 | 0,421847294 | 0,25172 | 0,55773568 | novel transcript |
| ENSG00000170439 | METTL7B | -1,245374205 | 0,421798478 | 0,07815 | 0,36540845 | methyltransferase like 7B |
| ENSG00000273793 | DUSP8 | -1,245805287 | 0,421672462 | 0,0039 | 0,07640394 | dual specificity phosphatase 8 |
| ENSG00000178031 | ADAMTSL1 | -1,245945473 | 0,421631491 | 0,03128 | 0,23867571 | ADAMTS like 1 |
| ENSG00000231444 | DDR1-DT | -1,24633396 | 0,421517969 | 0,51021 | 0,7378597 | DDR1 divergent transcript |
| ENSG00000258317 | AC034102,6 | -1,246434621 | 0,42148856 | 0,03047 | 0,23546694 | novel transcript, antisense to ZC3H10 and ESYT1 |
| ENSG00000284523 | AC004834,1 | -1,24650461 | 0,421468113 | 0,05345 | 0,31052104 | novel transcript |
| ENSG00000287069 | AC061958,1 | -1,24680394 | 0,421380676 | 0,03303 | 0,24547511 | novel transcript, antisense to KCNH8 |
| ENSG00000281207 | SLFNL1-AS1 | -1,246949797 | 0,421338076 | 0,04769 | 0,2934443 | SLFNL1 antisense RNA 1 |
| ENSG00000230415 | LINC01786 | -1,247193261 | 0,421266979 | 0,09748 | 0,40230004 | long intergenic non-protein coding RNA 1786 |
| ENSG00000278012 | AL031658,2 | -1,247347685 | 0,421221889 | 0,16515 | 0,49131595 | novel transcript, sense intronic to XKR7 |
| ENSG00000269352 | PTOV1-AS2 | -1,247467073 | 0,421187033 | 0,00975 | 0,12637434 | PTOV1 antisense RNA 2 |
| ENSG00000160991 | ORAI2 | -1,248360323 | 0,420926334 | 0,01385 | 0,15453657 | ORAI calcium release-activated calcium modulator 2 |
| ENSG00000255247 | NECTIN1-AS1 | -1,24840262 | 0,420913994 | 0,09435 | 0,39653034 | NECTIN1 antisense RNA 1 |
| ENSG00000177469 | CAVIN1 | -1,248640169 | 0,420844693 | 0,02985 | 0,23263128 | caveolae associated protein 1 |
| ENSG00000261377 | PDCD6IPP2 | -1,24972549 | 0,420528216 | 0,00846 | 0,11612498 | PDCD6IP pseudogene 2 |
| ENSG00000158023 | CFAP251 | -1,250477899 | 0,420308955 | 0,01629 | 0,16730761 | cilia and flagella associated protein 251 |
| ENSG00000270722 | RNVU1-31 | -1,250761869 | 0,420226233 | 0,02853 | 0,22839158 | RNA, variant U1 small nuclear 31 |
| ENSG00000284292 | AC004922,1 | -1,251363875 | 0,420050918 | 0,02676 | 0,22083401 | novel protein, ARPC1A and ARPC1B readthrough |
| ENSG00000287935 | AC136297,1 | -1,251401138 | 0,420040069 | 0,06795 | 0,34439007 | novel transcript |
| ENSG00000270419 | CAHM | -1,251973323 | 0,41987351 | 0,01352 | 0,15249214 | colon adenocarcinoma hypermethylated |
| ENSG00000226403 | AL392089,1 | -1,252822305 | 0,4196265 | 0,07169 | 0,35351167 | novel transcript |
| ENSG00000268218 | AC137932,3 | -1,254108922 | 0,419252438 | 0,01498 | 0,16040868 | novel transcript, antisense to ANKRD11 |
| ENSG00000239218 | RPS20P22 | -1,254262307 | 0,419207866 | 0,0264 | 0,21889454 | ribosomal protein S20 pseudogene 22 |
| ENSG00000227205 | PFN1P9 | -1,254994755 | 0,418995091 | 0,11134 | 0,42416781 | profilin 1 pseudogene 9 |
| ENSG00000286405 | AL132799,1 | -1,255393125 | 0,41887941 | 0,01387 | 0,15454988 | novel transcript |
| ENSG00000259804 | AC027682,1 | -1,255545976 | 0,418835033 | 0,10144 | 0,40833508 | novel transcript, antisense to CTCF |
| ENSG00000286230 | AC010327,7 | -1,255654799 | 0,418803441 | 0,03958 | 0,26924564 | novel transcript |
| ENSG00000261732 | AL031708,1 | -1,255932751 | 0,418722761 | 0,03803 | 0,26375851 | novel protein |
| ENSG00000241484 | ARHGAP8 | -1,256091425 | 0,418676711 | 0,11909 | 0,43696075 | Rho GTPase activating protein 8 |
| ENSG00000287380 | AL589935,3 | -1,25623534 | 0,418634948 | 0,01407 | 0,15573084 | novel transcript |
| ENSG00000183077 | AFMID | -1,25651073 | 0,418555044 | 0,03481 | 0,25226698 | arylformamidase |

| | | | | | | |
|---|---|---|---|---|---|---|
| ENSG00000270832 | AC092120,2 | -1,25662894 | 0,418520751 | 0,08035 | 0,37007763 | cytidine monophosphate (UMP-CMP) kinase 1, cytosolic (CMPK1) pseudogene |
| ENSG00000215908 | CROCCP2 | -1,257504717 | 0,418266768 | 0,01318 | 0,15012165 | CROCC pseudogene 2 |
| ENSG00000182759 | MAFA | -1,257570485 | 0,418247701 | 0,30026 | 0,59165391 | MAF bZIP transcription factor A |
| ENSG00000286966 | AL137800,1 | -1,257675314 | 0,418217311 | 0,00049 | 0,02010872 | novel transcript |
| ENSG00000213424 | KRT222 | -1,257819377 | 0,418175552 | 0,16782 | 0,49395055 | keratin 222 |
| ENSG00000262133 | AC087392,2 | -1,258011335 | 0,418119915 | 0,07244 | 0,3553822 | novel transcript, antisense to NXN |
| ENSG00000168334 | XIRP1 | -1,258400601 | 0,418007114 | 0,13484 | 0,46156017 | xin actin binding repeat containing 1 |
| ENSG00000256299 | AC073912,2 | -1,258743078 | 0,417907896 | 0,1571 | 0,48443207 | novel transcript |
| ENSG00000282715 | ANKRD20A9P | -1,258872238 | 0,417870484 | 0,28979 | 0,58528676 | ankyrin repeat domain 20 family member A9, pseudogene |
| ENSG00000118432 | CNR1 | -1,258916816 | 0,417857572 | 0,001 | 0,0323881 | cannabinoid receptor 1 |
| ENSG00000279110 | AL022323,3 | -1,259801395 | 0,417601444 | 0,03442 | 0,25073606 | novel transcript |
| ENSG00000286319 | AL135978,1 | -1,259956478 | 0,417556556 | 0,0514 | 0,30462001 | novel transcript, antisense to MPP5 |
| ENSG00000260388 | LINC00562 | -1,260398798 | 0,417428556 | 0,04045 | 0,27194236 | long intergenic non-protein coding RNA 562 |
| ENSG00000105321 | CCDC9 | -1,260421287 | 0,417422049 | 0,02419 | 0,20804286 | coiled-coil domain containing 9 |
| ENSG00000235529 | AGAP1-IT1 | -1,260589438 | 0,4173734 | 0,04379 | 0,28234289 | AGAP1 intronic transcript 1 |
| ENSG00000182601 | HS3ST4 | -1,260678109 | 0,417347748 | 0,04154 | 0,27616337 | heparan sulfate-glucosamine 3-sulfotransferase 4 |
| ENSG00000240498 | CDKN2B-AS1 | -1,260754632 | 0,417325612 | 0,00145 | 0,04066188 | CDKN2B antisense RNA 1 |
| ENSG00000236428 | PPP1R18 | -1,261116913 | 0,417220828 | 0,67025 | 0,83650134 | protein phosphatase 1 regulatory subunit 18 |
| ENSG00000276825 | DRD4 | -1,26118064 | 0,417202399 | 0,1211 | 0,44016175 | dopamine receptor D4 |
| ENSG00000224215 | AL606469,1 | -1,261278806 | 0,417174013 | 0,08981 | 0,38760032 | novel transcript |
| ENSG00000285589 | AC010422,8 | -1,261394345 | 0,417140604 | 0,02918 | 0,2303054 | novel transcript |
| ENSG00000163131 | CTSS | -1,261729541 | 0,417043697 | 0,14609 | 0,47410134 | cathepsin S |
| ENSG00000164972 | C9orf24 | -1,261778735 | 0,417029477 | 0,08257 | 0,37428581 | chromosome 9 open reading frame 24 |
| ENSG00000265801 | AC069366,2 | -1,26200919 | 0,416962866 | 0,01211 | 0,14307474 | novel transcript |
| ENSG00000287841 | AC096585,1 | -1,262113909 | 0,416932601 | 0,16173 | 0,48914394 | novel transcript |
| ENSG00000181409 | AATK | -1,262135775 | 0,416926282 | 0,02351 | 0,20467686 | apoptosis associated tyrosine kinase |
| ENSG00000224825 | RORB-AS1 | -1,262199186 | 0,416907957 | 0,06093 | 0,32753264 | RORB antisense RNA 1 |
| ENSG00000160360 | GPSM1 | -1,262586046 | 0,416796178 | 0,00101 | 0,03265007 | G protein signaling modulator 1 |
| ENSG00000281216 | KIF26B | -1,262663088 | 0,416773921 | 0,32178 | 0,60577115 | kinesin family member 26B |
| ENSG00000257576 | HSPD1P4 | -1,262714528 | 0,416759061 | 0,0084 | 0,11566397 | heat shock protein family D (Hsp60) member 1 pseudogene 4 |
| ENSG00000270497 | BX322635,1 | -1,262726969 | 0,416755467 | 0,03464 | 0,25161513 | ribosomal protein S3A (RPS3A) pseudogene |

| | | | | | | |
|---|---|---|---|---|---|---|
| ENSG00000215023 | AC131097,1 | -1,263072822 | 0,416655572 | 0,15759 | 0,48468257 | novel transcript |
| ENSG00000281082 | AC131097,4 | -1,263072822 | 0,416655572 | 0,15759 | 0,48468257 | novel transcript |
| ENSG00000141433 | ADCYAP1 | -1,264226122 | 0,416322628 | 0,02461 | 0,21002931 | adenylate cyclase activating polypeptide 1 |
| ENSG00000286679 | AC073225,1 | -1,264432083 | 0,416263197 | 0,12816 | 0,45171208 | novel transcript |
| ENSG00000277144 | AC092757,3 | -1,264987663 | 0,416102926 | 0,07598 | 0,36203962 | novel transcript, antisense to CCNB2 |
| ENSG00000253485 | PCDHGA5 | -1,265207756 | 0,416039451 | 0,05571 | 0,31543336 | protocadherin gamma subfamily A, 5 |
| ENSG00000116771 | AGMAT | -1,265212125 | 0,416038191 | 0,06588 | 0,3397198 | agmatinase |
| ENSG00000267060 | PTGES3L | -1,265427654 | 0,415976043 | 0,02984 | 0,23258582 | prostaglandin E synthase 3 like |
| ENSG00000272954 | AP000553,2 | -1,265899527 | 0,415840008 | 0,05584 | 0,31590444 | novel transcript, sense intronic to PPIL2 |
| ENSG00000245711 | NADK2-AS1 | -1,266696945 | 0,415610225 | 0,0466 | 0,29026543 | NADK2 antisense RNA 1 |
| ENSG00000173401 | GLIPR1L1 | -1,267007455 | 0,415520784 | 0,11846 | 0,43614969 | GLIPR1 like 1 |
| ENSG00000274574 | U1 | -1,267069452 | 0,415502928 | 0,00089 | 0,02988696 | U1 spliceosomal RNA |
| ENSG00000277918 | RNVU1-28 | -1,267069452 | 0,415502928 | 0,00089 | 0,02988696 | RNA, variant U1 small nuclear 28 |
| ENSG00000228999 | AC096570,1 | -1,267960732 | 0,415246314 | 0,17212 | 0,49710979 | long intergenic non-protein coding RNA 1830 |
| ENSG00000123901 | GPR83 | -1,268503471 | 0,415090129 | 0,10689 | 0,41675015 | G protein-coupled receptor 83 |
| ENSG00000262102 | DSEL | -1,268573556 | 0,415069965 | 0,00752 | 0,11001711 | dermatan sulfate epimerase like |
| ENSG00000269161 | AC010618,2 | -1,268615082 | 0,415058018 | 0,23187 | 0,5446505 | novel transcript, antisense PGLS |
| ENSG00000170893 | TRH | -1,269096166 | 0,414919635 | 0,0315 | 0,23963655 | thyrotropin releasing hormone |
| ENSG00000205363 | INSYN1 | -1,269326368 | 0,414853433 | 0,02324 | 0,20373153 | inhibitory synaptic factor 1 |
| ENSG00000177685 | CRACR2B | -1,269812338 | 0,414713714 | 0,04833 | 0,29522569 | calcium release activated channel regulator 2B |
| ENSG00000232577 | BX679671,1 | -1,270111863 | 0,414627623 | 0,48051 | 0,71764212 | novel transcript |
| ENSG00000233627 | C4A-AS1 | -1,270111863 | 0,414627623 | 0,48051 | 0,71764212 | C4A antisense RNA 1 |
| ENSG00000229776 | C4B-AS1 | -1,270111863 | 0,414627623 | 0,48051 | 0,71764212 | C4B antisense RNA 1 |
| ENSG00000187980 | PLA2G2C | -1,270424317 | 0,414537833 | 0,04013 | 0,27065903 | phospholipase A2 group IIC |
| ENSG00000134363 | FST | -1,271693535 | 0,414173302 | 0,01354 | 0,15265267 | follistatin |
| ENSG00000138642 | HERC6 | -1,271924268 | 0,414107068 | 0,01433 | 0,15653473 | HECT and RLD domain containing E3 ubiquitin protein ligase family member 6 |
| ENSG00000180767 | CHST13 | -1,271972706 | 0,414093165 | 0,12019 | 0,43872715 | carbohydrate sulfotransferase 13 |
| ENSG00000126653 | NSRP1 | -1,272710744 | 0,413881382 | 0,01285 | 0,14824976 | nuclear speckle splicing regulatory protein 1 |
| ENSG00000276116 | FUT8-AS1 | -1,273429647 | 0,413675194 | 0,01074 | 0,1339808 | FUT8 antisense RNA 1 |
| ENSG00000079112 | CDH17 | -1,273690825 | 0,413600311 | 0,20685 | 0,52649135 | cadherin 17 |
| ENSG00000206597 | SNORA57 | -1,27405116 | 0,413497021 | 0,00388 | 0,07610805 | small nucleolar RNA, H/ACA box 57 |
| ENSG00000235725 | AC007389,2 | -1,274158143 | 0,413466359 | 0,00992 | 0,12768018 | novel transcript |

| | | | | | | |
|---|---|---|---|---|---|---|
| ENSG00000222018 | FAM243A | -1,274553565 | 0,413353049 | 0,19151 | 0,51435195 | family with sequence similarity 243 member A |
| ENSG00000181847 | TIGIT | -1,274561436 | 0,413350794 | 0,00338 | 0,07008746 | T cell immunoreceptor with Ig and ITIM domains |
| ENSG00000225169 | BRI3P1 | -1,274907134 | 0,413251759 | 0,29006 | 0,58531219 | brain protein I3 pseudogene 1 |
| ENSG00000285451 | ADCK5 | -1,275070108 | 0,413205079 | 0,00475 | 0,08501852 | aarF domain containing kinase 5 |
| ENSG00000173137 | ADCK5 | -1,275070108 | 0,413205079 | 0,00475 | 0,08501852 | aarF domain containing kinase 5 |
| ENSG00000092051 | JPH4 | -1,275148436 | 0,413182645 | 0,02128 | 0,19349591 | junctophilin 4 |
| ENSG00000232260 | BTF3L4P1 | -1,275422814 | 0,413104072 | 0,08697 | 0,38205469 | basic transcription factor 3 like 4 pseudogene 1 |
| ENSG00000165553 | NGB | -1,276988864 | 0,41265589 | 0,22894 | 0,54262488 | neuroglobin |
| ENSG00000175920 | DOK7 | -1,277078912 | 0,412630134 | 0,0613 | 0,32863053 | docking protein 7 |
| ENSG00000187621 | TCL6 | -1,277591115 | 0,412483663 | 0,03144 | 0,23928335 | T cell leukemia/lymphoma 6 |
| ENSG00000227495 | KIF1C-AS1 | -1,277924478 | 0,412388361 | 0,14898 | 0,47654655 | KIF1C antisense RNA 1 |
| ENSG00000186951 | PPARA | -1,278490258 | 0,412226667 | 0,03439 | 0,25062659 | peroxisome proliferator activated receptor alpha |
| ENSG00000213188 | YBX1P4 | -1,278737372 | 0,412156065 | 0,0766 | 0,36266919 | Y-box binding protein 1 pseudogene 4 |
| ENSG00000186654 | PRR5 | -1,278797389 | 0,412138919 | 0,03552 | 0,25546549 | proline rich 5 |
| ENSG00000284783 | AC091057,4 | -1,278910249 | 0,412106679 | 0,23427 | 0,54609345 | OTU deubiquitinase 7A (OTUD7A), pseudogene |
| ENSG00000130758 | MAP3K10 | -1,279136905 | 0,41204194 | 0,00755 | 0,11001798 | mitogen-activated protein kinase kinase kinase 10 |
| ENSG00000225299 | AL139240,1 | -1,279562961 | 0,411920274 | 0,12182 | 0,44093327 | novel transcript |
| ENSG00000236890 | AC016027,5 | -1,279584444 | 0,41191414 | 0,26483 | 0,56748133 | ribosomal protein L18a (RPL18A) pseudogene |
| ENSG00000214919 | AC104472,1 | -1,279715871 | 0,411876617 | 0,0316 | 0,23997947 | novel transcript, sense intronic to PLCH1 |
| ENSG00000197558 | SSPOP | -1,279869875 | 0,411832653 | 0,0241 | 0,20752957 | SCO-spondin, pseudogene |
| ENSG00000236003 | AC007731,3 | -1,280446424 | 0,411668103 | 0,17052 | 0,49641416 | novel transcript |
| ENSG00000239732 | TLR9 | -1,281576531 | 0,411345757 | 0,4777 | 0,71545897 | toll like receptor 9 |
| ENSG00000223839 | FAM95B1 | -1,281676609 | 0,411317224 | 0,04813 | 0,29478742 | family with sequence similarity 95 member B1 |
| ENSG00000214691 | LINC01913 | -1,282015947 | 0,411220489 | 0,2013 | 0,52181085 | long intergenic non-protein coding RNA 1913 |
| ENSG00000260563 | AC132872,1 | -1,282290664 | 0,411142192 | 0,07484 | 0,35906132 | novel transcript |
| ENSG00000142459 | EVI5L | -1,282851833 | 0,4109823 | 0,01171 | 0,14084308 | ecotropic viral integration site 5 like |
| ENSG00000185163 | DDX51 | -1,282959514 | 0,410951626 | 0,09717 | 0,40152057 | DEAD-box helicase 51 |
| ENSG00000131044 | TTLL9 | -1,282998199 | 0,410940607 | 0,04167 | 0,2764623 | tubulin tyrosine ligase like 9 |
| ENSG00000227640 | SOX21-AS1 | -1,283410016 | 0,410823321 | 5,11E-05 | 0,00341912 | SOX21 antisense divergent transcript 1 |
| ENSG00000269758 | AC092070,4 | -1,283436721 | 0,410815716 | 0,06065 | 0,32698319 | novel transcript |
| ENSG00000233523 | PHBP5 | -1,283648367 | 0,410755453 | 0,0121 | 0,14307474 | prohibitin pseudogene 5 |
| ENSG00000268678 | AC005261,2 | -1,283686125 | 0,410744703 | 0,05838 | 0,32253335 | novel transcript, antisense to ZNF304 |
| ENSG00000223956 | LINC01767 | -1,283812903 | 0,41070861 | 0,07377 | 0,3570536 | long intergenic non-protein coding RNA 1767 |

| | | | | | | |
|---|---|---|---|---|---|---|
| ENSG00000141526 | SLC16A3 | -1,284127955 | 0,41061893 | 0,00593 | 0,09720154 | solute carrier family 16 member 3 |
| ENSG00000196421 | C20orf204 | -1,284273877 | 0,4105774 | 0,05114 | 0,30402651 | chromosome 20 open reading frame 204 |
| ENSG00000268531 | AC060814,2 | -1,28441891 | 0,410536127 | 0,24448 | 0,55253387 | novel transcript |
| ENSG00000286803 | AL109824,1 | -1,284756335 | 0,41044012 | 0,07366 | 0,35701531 | novel transcript, antisense to NCOA6 |
| ENSG00000259887 | FIGNL2-DT | -1,285115111 | 0,410338063 | 0,16521 | 0,49131595 | FIGNL2 divergent transcript |
| ENSG00000254653 | AC024475,1 | -1,286377931 | 0,409979043 | 0,09273 | 0,39335125 | novel transcript |
| ENSG00000142583 | SLC2A5 | -1,286429658 | 0,409964343 | 0,05012 | 0,30091295 | solute carrier family 2 member 5 |
| ENSG00000226824 | AC006001,2 | -1,286466324 | 0,409953924 | 0,00695 | 0,10609569 | novel transcript |
| ENSG00000188549 | CCDC9B | -1,286997958 | 0,409802884 | 0,15543 | 0,48334025 | coiled-coil domain containing 9B |
| ENSG00000136542 | GALNT5 | -1,287120114 | 0,409768187 | 0,13176 | 0,4570124 | polypeptide N-acetylgalactosaminyltransferase 5 |
| ENSG00000126001 | CEP250 | -1,287153078 | 0,409758824 | 0,00272 | 0,06123483 | centrosomal protein 250 |
| ENSG00000247903 | AC024896,1 | -1,287483516 | 0,409664983 | 0,01857 | 0,18012622 | novel transcript, antisense to TM7SF3 |
| ENSG00000230408 | BZW1-AS1 | -1,287703356 | 0,409602562 | 0,06054 | 0,32676053 | BZW1 antisense RNA 1 |
| ENSG00000234183 | LINC01952 | -1,287753008 | 0,409588465 | 0,12547 | 0,44729629 | long intergenic non-protein coding RNA 1952 |
| ENSG00000111879 | FAM184A | -1,287825915 | 0,409567767 | 0,01034 | 0,13101765 | family with sequence similarity 184 member A |
| ENSG00000255438 | AL354813,1 | -1,287893456 | 0,409548593 | 0,09499 | 0,39751728 | novel transcript, antisense to SULF2 |
| ENSG00000231711 | LINC00899 | -1,287908365 | 0,409544361 | 0,10446 | 0,41292721 | long intergenic non-protein coding RNA 899 |
| ENSG00000183454 | GRIN2A | -1,288193157 | 0,409463524 | 0,00545 | 0,09188602 | glutamate ionotropic receptor NMDA type subunit 2A |
| ENSG00000276017 | AC007325,1 | -1,288248842 | 0,409447719 | 0,0143 | 0,15653473 | |
| ENSG00000188004 | SNHG28 | -1,288488642 | 0,409379668 | 0,08501 | 0,37843844 | small nucleolar RNA host gene 28 |
| ENSG00000204536 | CCHCR1 | -1,28867133 | 0,409327832 | 0,03731 | 0,26136592 | coiled-coil alpha-helical rod protein 1 |
| ENSG00000248468 | AC107027,1 | -1,288779176 | 0,409297234 | 0,1688 | 0,49492999 | novel transcript |
| ENSG00000260917 | AL158212,3 | -1,289017631 | 0,40922959 | 0,02332 | 0,20398797 | novel transcript, overlapping to VTI1A |
| ENSG00000201229 | SNORA63D | -1,289767149 | 0,409017039 | 0,00979 | 0,12674119 | small nucleolar RNA, H/ACA box 63D |
| ENSG00000214049 | UCA1 | -1,28977425 | 0,409015026 | 0,0602 | 0,32601555 | urothelial cancer associated 1 |
| ENSG00000276653 | RN7SL856P | -1,289908133 | 0,408977071 | 0,09791 | 0,40323803 | RNA, 7SL, cytoplasmic 856, pseudogene |
| ENSG00000277949 | RTP5 | -1,289940618 | 0,408967862 | 0,12636 | 0,44903997 | receptor transporter protein 5 (putative) |
| ENSG00000188011 | RTP5 | -1,289940618 | 0,408967862 | 0,12636 | 0,44903997 | receptor transporter protein 5 (putative) |
| ENSG00000130600 | H19 | -1,290586548 | 0,408784798 | 0,15483 | 0,48266951 | H19 imprinted maternally expressed transcript |
| ENSG00000288237 | H19 | -1,290586548 | 0,408784798 | 0,15483 | 0,48266951 | H19 imprinted maternally expressed transcript |
| ENSG00000246100 | LINC00900 | -1,291001857 | 0,408667138 | 0,0082 | 0,11480072 | long intergenic non-protein coding RNA 900 |
| ENSG00000280798 | LINC00294 | -1,291049443 | 0,408653659 | 0,0295 | 0,23093703 | long intergenic non-protein coding RNA 294 |
| ENSG00000261519 | AC010542,4 | -1,291178837 | 0,408617009 | 0,10302 | 0,41078404 | novel transcript |

| | | | | | | |
|---|---|---|---|---|---|---|
| ENSG00000277463 | AC080038,2 | -1,291495454 | 0,408527343 | 0,17775 | 0,50207094 | novel transcript |
| ENSG00000148488 | ST8SIA6 | -1,29175952 | 0,408452574 | 0,07571 | 0,36122612 | ST8 alpha-N-acetyl-neuraminide alpha-2,8-sialyltransferase 6 |
| ENSG00000182870 | GALNT9 | -1,291837165 | 0,408430592 | 0,1816 | 0,50490902 | polypeptide N-acetylgalactosaminyltransferase 9 |
| ENSG00000260422 | Z97205,2 | -1,291858986 | 0,408424414 | 0,02719 | 0,22244887 | novel transcript |
| ENSG00000266111 | AC068025,2 | -1,292256941 | 0,40831177 | 0,10514 | 0,41407221 | novel transcript |
| ENSG00000188227 | ZNF793 | -1,29231689 | 0,408294803 | 0,02205 | 0,1974595 | zinc finger protein 793 |
| ENSG00000253982 | AC100810,1 | -1,29256224 | 0,408225373 | 0,02374 | 0,20579436 | novel transcript |
| ENSG00000277501 | AC243571,2 | -1,292656607 | 0,408198672 | 0,10231 | 0,40985316 | novel transcript, antisense DDX52 |
| ENSG00000185115 | NSMCE3 | -1,292694891 | 0,40818784 | 0,03412 | 0,24936671 | NSE3 homolog, SMC5-SMC6 complex component |
| ENSG00000104881 | PPP1R13L | -1,292722906 | 0,408179913 | 0,06368 | 0,33458308 | protein phosphatase 1 regulatory subunit 13 like |
| ENSG00000281614 | INPP5D | -1,293101229 | 0,408072889 | 0,09273 | 0,39335125 | inositol polyphosphate-5-phosphatase D |
| ENSG00000278780 | ZNF430 | -1,293371447 | 0,407996464 | 0,01371 | 0,15356644 | zinc finger protein 430 |
| ENSG00000279954 | BX324167,2 | -1,293702173 | 0,407902945 | 0,00291 | 0,06410301 | novel transcript |
| ENSG00000260886 | TAT-AS1 | -1,293750274 | 0,407889345 | 0,11981 | 0,43834197 | TAT antisense RNA 1 |
| ENSG00000215548 | FRG1JP | -1,293919579 | 0,407841481 | 0,04062 | 0,27256162 | FSHD region gene 1 family member J, pseudogene |
| ENSG00000128011 | LRFN1 | -1,294075421 | 0,407797427 | 0,0206 | 0,19042717 | leucine rich repeat and fibronectin type III domain containing 1 |
| ENSG00000206147 | RPL23AP57 | -1,29417953 | 0,407768001 | 0,13751 | 0,46456422 | ribosomal protein L23a pseudogene 57 |
| ENSG00000206281 | TAPBP | -1,294221815 | 0,407756049 | 0,03816 | 0,26448926 | TAP binding protein |
| ENSG00000242241 | RN7SL306P | -1,294293284 | 0,40773585 | 0,33191 | 0,61191629 | RNA, 7SL, cytoplasmic 306, pseudogene |
| ENSG00000110042 | DTX4 | -1,294379924 | 0,407711365 | 3,75E-16 | 1,91E-13 | deltex E3 ubiquitin ligase 4 |
| ENSG00000239419 | RN7SL535P | -1,294611397 | 0,407645955 | 0,03345 | 0,24712306 | RNA, 7SL, cytoplasmic 535, pseudogene |
| ENSG00000188981 | MSANTD1 | -1,294913236 | 0,407560676 | 0,07792 | 0,36517053 | Myb/SANT DNA binding domain containing 1 |
| ENSG00000258757 | AL133453,1 | -1,295021684 | 0,407530041 | 0,37135 | 0,64049253 | novel transcript, antisense to ERO1L |
| ENSG00000272175 | AL050343,3 | -1,295266256 | 0,407460961 | 0,11059 | 0,42326288 | novel transcript, antisense to NRD1 |
| ENSG00000196275 | GTF2IRD2 | -1,295342329 | 0,407439476 | 0,01128 | 0,13773024 | GTF2I repeat domain containing 2 |
| ENSG00000133048 | CHI3L1 | -1,297877215 | 0,406724213 | 0,05214 | 0,30680362 | chitinase 3 like 1 |
| ENSG00000141579 | ZNF750 | -1,297986344 | 0,406693449 | 0,02591 | 0,21707964 | zinc finger protein 750 |
| ENSG00000164362 | TERT | -1,298149207 | 0,40664754 | 0,07697 | 0,36357031 | telomerase reverse transcriptase |
| ENSG00000267682 | AC016590,2 | -1,298431951 | 0,406567852 | 0,06005 | 0,32549025 | novel transcript, sense intronic to HKR1 |
| ENSG00000165409 | TSHR | -1,298594802 | 0,406521961 | 0,05341 | 0,31039748 | thyroid stimulating hormone receptor |
| ENSG00000225420 | AC104134,1 | -1,298887938 | 0,40643937 | 0,33837 | 0,61774255 | novel transcript |

| | | | | | | |
|---|---|---|---|---|---|---|
| ENSG00000136297 | MMD2 | -1,298926174 | 0,406428598 | 0,08481 | 0,37815672 | monocyte to macrophage differentiation associated 2 |
| ENSG00000253138 | LINC00967 | -1,298947888 | 0,406422481 | 0,03951 | 0,2690131 | long intergenic non-protein coding RNA 967 |
| ENSG00000285238 | AC006064,6 | -1,299746363 | 0,406197605 | 0,7099 | 0,86019076 | novel transcript |
| ENSG00000142544 | CTU1 | -1,299753003 | 0,406195735 | 0,03927 | 0,26812486 | cytosolic thiouridylase subunit 1 |
| ENSG00000260796 | AC145285,3 | -1,299821916 | 0,406176333 | 0,08422 | 0,37695424 | novel transcript, antisense to ATXN2L |
| ENSG00000072195 | SPEG | -1,29998416 | 0,406130657 | 0,00523 | 0,09023148 | striated muscle enriched protein kinase |
| ENSG00000284801 | AC270301,2 | -1,300195877 | 0,406071062 | 0,05955 | 0,32490861 | novel transcript |
| ENSG00000253772 | AC112191,1 | -1,300249408 | 0,406055995 | 0,05979 | 0,32535617 | acid phosphatase 1, soluble (ACP1) pseudogene |
| ENSG00000275650 | TTYH1 | -1,300740986 | 0,40591766 | 0,02178 | 0,19610255 | tweety family member 1 |
| ENSG00000206406 | CSNK2B | -1,30082018 | 0,405895379 | 0,06007 | 0,32556609 | casein kinase 2 beta |
| ENSG00000236383 | CCDC200 | -1,302299581 | 0,40547937 | 0,04236 | 0,27814338 | coiled-coil domain containing 200 |
| ENSG00000243871 | RN7SL487P | -1,302375028 | 0,405458165 | 0,39449 | 0,65734592 | RNA, 7SL, cytoplasmic 487, pseudogene |
| ENSG00000069482 | GAL | -1,302635313 | 0,405385021 | 0,16154 | 0,48906157 | galanin and GMAP prepropeptide |
| ENSG00000130783 | CCDC62 | -1,303174714 | 0,405233482 | 0,01848 | 0,179662 | coiled-coil domain containing 62 |
| ENSG00000237529 | AL137847,2 | -1,303235663 | 0,405216363 | 0,0371 | 0,26086671 | novel transcript |
| ENSG00000279718 | SNX18P12 | -1,303334763 | 0,405188529 | 0,00852 | 0,11640065 | sorting nexin 18 pseudogene 12 |
| ENSG00000286138 | AC010336,7 | -1,303467843 | 0,405151155 | 0,06312 | 0,33273662 | novel transcript |
| ENSG00000172543 | CTSW | -1,303475998 | 0,405148865 | 0,18424 | 0,50802406 | cathepsin W |
| ENSG00000089199 | CHGB | -1,303944239 | 0,405017391 | 0,00086 | 0,02908139 | chromogranin B |
| ENSG00000163406 | SLC15A2 | -1,304858693 | 0,404760751 | 0,03395 | 0,24882853 | solute carrier family 15 member 2 |
| ENSG00000214193 | SH3D21 | -1,304966481 | 0,404730512 | 0,00456 | 0,08326903 | SH3 domain containing 21 |
| ENSG00000248522 | SBF1P1 | -1,30517765 | 0,404671275 | 0,13621 | 0,463307 | SET binding factor 1 pseudogene 1 |
| ENSG00000116783 | TNNI3K | -1,30542524 | 0,404601833 | 0,29841 | 0,59076323 | TNNI3 interacting kinase |
| ENSG00000287809 | AC005632,6 | -1,305536165 | 0,404570725 | 0,01896 | 0,18263335 | novel transcript |
| ENSG00000286098 | AC008770,4 | -1,305889598 | 0,404471625 | 0,57787 | 0,78171012 | zinc finger protein 844 |
| ENSG00000226539 | MLXP1 | -1,305891799 | 0,404471008 | 0,01361 | 0,1529196 | MLX pseudogene 1 |
| ENSG00000279483 | AC090498,1 | -1,306917145 | 0,404183646 | 0,07318 | 0,35655334 | ribosomal protein L41 (RPL41) pseudogene |
| ENSG00000226045 | AC234644,1 | -1,307184087 | 0,404108867 | 0,23584 | 0,5471053 | glutamyl-tRNA(Gln) amidotransferase, subunit C homolog (bacterial)(GATC) pseudogene |
| ENSG00000146674 | IGFBP3 | -1,307665002 | 0,403974182 | 0,05289 | 0,30865611 | insulin like growth factor binding protein 3 |
| ENSG00000285560 | AC103739,3 | -1,307915284 | 0,403904106 | 0,07135 | 0,35283325 | novel transcript, antisense to CRTC3 |
| ENSG00000211643 | IGLV5-52 | -1,309515319 | 0,4034564 | 0,08348 | 0,37564365 | immunoglobulin lambda variable 5-52 |
| ENSG00000004846 | ABCB5 | -1,310597498 | 0,403153877 | 0,05175 | 0,30583008 | ATP binding cassette subfamily B member 5 |

| | | | | | | |
|---|---|---|---|---|---|---|
| ENSG00000265293 | ARGFXP2 | -1,311181231 | 0,402990789 | 0,03405 | 0,24909651 | arginine-fifty homeobox pseudogene 2 |
| ENSG00000167619 | TMEM145 | -1,311189824 | 0,402988389 | 0,02937 | 0,23068546 | transmembrane protein 145 |
| ENSG00000110448 | CD5 | -1,311612453 | 0,402870353 | 0,04235 | 0,27814338 | CD5 molecule |
| ENSG00000128849 | CGNL1 | -1,311815723 | 0,402813594 | 0,00564 | 0,09352688 | cingulin like 1 |
| ENSG00000274904 | AC093512,1 | -1,311888376 | 0,402793309 | 0,2234 | 0,53953208 | novel transcript, antisense to ALDOA |
| ENSG00000057593 | F7 | -1,312014001 | 0,402758237 | 0,33035 | 0,6112353 | coagulation factor VII |
| ENSG00000053702 | NRIP2 | -1,312607323 | 0,402592633 | 0,01558 | 0,16367661 | nuclear receptor interacting protein 2 |
| ENSG00000231107 | LINC01508 | -1,313181912 | 0,402432322 | 0,00171 | 0,04508122 | long intergenic non-protein coding RNA 1508 |
| ENSG00000283658 | RPL23AP82 | -1,313380011 | 0,402377067 | 0,0525 | 0,30766476 | ribosomal protein L23a pseudogene 82 |
| ENSG00000265933 | LINC00668 | -1,314240114 | 0,40213725 | 0,02855 | 0,2284511 | long intergenic non-protein coding RNA 668 |
| ENSG00000125046 | SSUH2 | -1,314516374 | 0,402060253 | 0,00048 | 0,01999623 | ssu-2 homolog |
| ENSG00000248323 | LUCAT1 | -1,31454624 | 0,40205193 | 0,00632 | 0,10025151 | lung cancer associated transcript 1 |
| ENSG00000267106 | ZNF561-AS1 | -1,314736749 | 0,401998842 | 0,01981 | 0,18688735 | ZNF561 antisense RNA 1 (head to head) |
| ENSG00000244159 | RPS27AP13 | -1,314777922 | 0,40198737 | 0,14993 | 0,47794655 | RPS27A pseudogene 13 |
| ENSG00000183747 | ACSM2A | -1,31486315 | 0,401963623 | 0,06616 | 0,34052983 | acyl-CoA synthetase medium chain family member 2A |
| ENSG00000236790 | LINC00299 | -1,31488887 | 0,401956457 | 0,1788 | 0,50328925 | long intergenic non-protein coding RNA 299 |
| ENSG00000105227 | PRX | -1,31532754 | 0,401834255 | 0,04712 | 0,29208796 | periaxin |
| ENSG00000238288 | MTND3P23 | -1,315760803 | 0,401713596 | 0,31556 | 0,60172912 | MT-ND3 pseudogene 23 |
| ENSG00000180096 | SEPTIN1 | -1,316185876 | 0,401595254 | 0,02294 | 0,20217525 | septin 1 |
| ENSG00000231742 | LINC01273 | -1,316456518 | 0,401519924 | 0,05849 | 0,32289062 | long intergenic non-protein coding RNA 1273 |
| ENSG00000111424 | VDR | -1,316872974 | 0,401404036 | 0,1432 | 0,47082074 | vitamin D receptor |
| ENSG00000156381 | ANKRD9 | -1,316920914 | 0,401390697 | 0,0135 | 0,15241195 | ankyrin repeat domain 9 |
| ENSG00000204961 | PCDHA9 | -1,318348101 | 0,400993818 | 0,00066 | 0,02469756 | protocadherin alpha 9 |
| ENSG00000204427 | ABHD16A | -1,318687978 | 0,400899361 | 0,0559 | 0,31600461 | abhydrolase domain containing 16A, phospholipase |
| ENSG00000250222 | AC008443,4 | -1,318866944 | 0,400849632 | 0,04406 | 0,28277847 | novel transcript |
| ENSG00000248736 | AC094105,2 | -1,318965836 | 0,400822156 | 0,02507 | 0,21269649 | novel transcript |
| ENSG00000258807 | AL359237,1 | -1,319000444 | 0,400812541 | 0,07214 | 0,35442193 | novel transcript |
| ENSG00000233948 | TRIM27 | -1,319392324 | 0,400703683 | 0,00466 | 0,08400694 | tripartite motif containing 27 |
| ENSG00000250453 | AC008825,1 | -1,319617721 | 0,400641085 | 0,22667 | 0,54144 | novel transcript |
| ENSG00000272031 | ANKRD34A | -1,319665172 | 0,400627908 | 0,00272 | 0,06121178 | ankyrin repeat domain 34A |
| ENSG00000269834 | ZNF528-AS1 | -1,319770387 | 0,400598691 | 0,02588 | 0,21707964 | ZNF528 antisense RNA 1 |
| ENSG00000141013 | GAS8 | -1,320334836 | 0,400441989 | 1,47E-07 | 2,06E-05 | growth arrest specific 8 |
| ENSG00000130748 | TMEM160 | -1,321084057 | 0,400234085 | 0,01507 | 0,16106953 | transmembrane protein 160 |

| | | | | | | |
|---|---|---|---|---|---|---|
| ENSG00000225222 | CHCHD4P5 | -1,321105596 | 0,40022811 | 0,28723 | 0,58364712 | coiled-coil-helix-coiled-coil-helix domain containing 4 pseudogene 5 |
| ENSG00000115138 | POMC | -1,321647572 | 0,400077785 | 0,10027 | 0,4067769 | proopiomelanocortin |
| ENSG00000147655 | RSPO2 | -1,321676267 | 0,400069828 | 0,05009 | 0,30089233 | R-spondin 2 |
| ENSG00000268836 | Z69706,1 | -1,321862993 | 0,40001805 | 0,01436 | 0,15682082 | novel transcript |
| ENSG00000230358 | SPDYE21 | -1,321939076 | 0,399996955 | 0,05138 | 0,30462001 | speedy/RINGO cell cycle regulator family member E21 |
| ENSG00000237986 | CELF2-AS2 | -1,322228803 | 0,399916635 | 0,00457 | 0,08330333 | CELF2 antisense RNA 2 |
| ENSG00000228333 | RXRB | -1,322458535 | 0,399852958 | 0,00549 | 0,09213036 | retinoid X receptor beta |
| ENSG00000187808 | SOWAHD | -1,322810717 | 0,39975536 | 0,17703 | 0,50147587 | sosondowah ankyrin repeat domain family member D |
| ENSG00000152292 | SH2D6 | -1,323206962 | 0,39964558 | 0,16523 | 0,49131595 | SH2 domain containing 6 |
| ENSG00000273066 | AL355987,4 | -1,323509163 | 0,399561875 | 0,02079 | 0,19127841 | novel transcript |
| ENSG00000237682 | PRKCIP1 | -1,323646728 | 0,399523777 | 0,05421 | 0,31168784 | PRKCI pseudogene 1 |
| ENSG00000130038 | CRACR2A | -1,323690921 | 0,399511539 | 0,00139 | 0,03972417 | calcium release activated channel regulator 2A |
| ENSG00000287142 | AL355053,1 | -1,324309168 | 0,399340371 | 0,20368 | 0,52347929 | novel transcript, antisense to PRKD1 |
| ENSG00000164434 | FABP7 | -1,324673277 | 0,399239598 | 5,92E-05 | 0,00390694 | fatty acid binding protein 7 |
| ENSG00000275297 | AL162731,1 | -1,324849541 | 0,399190823 | 0,1097 | 0,42183783 | novel transcript |
| ENSG00000183397 | C19orf71 | -1,325249956 | 0,399080044 | 0,0691 | 0,34728769 | chromosome 19 open reading frame 71 |
| ENSG00000114204 | SERPINI2 | -1,325644117 | 0,398971026 | 0,04633 | 0,28930497 | serpin family I member 2 |
| ENSG00000171385 | KCND3 | -1,325849697 | 0,398914178 | 0,0254 | 0,21440233 | potassium voltage-gated channel subfamily D member 3 |
| ENSG00000101695 | RNF125 | -1,325884313 | 0,398904606 | 0,07998 | 0,36913629 | ring finger protein 125 |
| ENSG00000223797 | ENTPD3-AS1 | -1,326315746 | 0,398785333 | 0,0193 | 0,18400081 | ENTPD3 antisense RNA 1 |
| ENSG00000286579 | AC023483,1 | -1,326624376 | 0,398700031 | 0,12849 | 0,45191041 | novel transcript, antisense to SETMAR |
| ENSG00000275620 | FLJ16779 | -1,326701876 | 0,398678614 | 0,02379 | 0,20595388 | uncharacterized LOC100192386 |
| ENSG00000237599 | TAP2 | -1,326741636 | 0,398667627 | 0,61974 | 0,80726991 | transporter 2, ATP binding cassette subfamily B member |
| ENSG00000277425 | AL121890,4 | -1,327110447 | 0,398565724 | 0,08336 | 0,37550713 | novel transcript |
| ENSG00000146828 | SLC12A9 | -1,327280546 | 0,398518735 | 0,01628 | 0,16730761 | solute carrier family 12 member 9 |
| ENSG00000205482 | SPDYE18 | -1,327382488 | 0,398490576 | 0,02114 | 0,19303449 | speedy/RINGO cell cycle regulator family member E18 |
| ENSG00000204930 | FAM221B | -1,327461064 | 0,398468873 | 0,03121 | 0,23830231 | family with sequence similarity 221 member B |
| ENSG00000285518 | AC004900,1 | -1,328066975 | 0,398301557 | 0,13214 | 0,45719971 | novel transcript, antisense to SIPA1L1 |
| ENSG00000147573 | TRIM55 | -1,328173828 | 0,398272058 | 0,16556 | 0,49182669 | tripartite motif containing 55 |
| ENSG00000287452 | AL391477,1 | -1,328296787 | 0,398238115 | 0,01799 | 0,17701257 | novel transcript |
| ENSG00000281263 | AC136352,8 | -1,328651808 | 0,398140128 | 0,12612 | 0,44869905 | novel transcript |
| ENSG00000162144 | CYB561A3 | -1,32910631 | 0,398014719 | 0,02626 | 0,21852357 | cytochrome b561 family member A3 |

| | | | | | | |
|---|---|---|---|---|---|---|
| ENSG00000253102 | AC004707,1 | -1,329829182 | 0,397815341 | 0,23445 | 0,5462109 | novel transcript, antisense to LRRC59 |
| ENSG00000254538 | AC027018,1 | -1,330186785 | 0,397716746 | 0,05135 | 0,30462001 | novel transcript |
| ENSG00000228695 | CES1P1 | -1,330400791 | 0,397657754 | 0,21722 | 0,53520949 | carboxylesterase 1 pseudogene 1 |
| ENSG00000259138 | AL049780,2 | -1,332130526 | 0,397181264 | 0,38798 | 0,65289784 | novel transcript, antisense to NEK9 and TMED10 |
| ENSG00000231290 | APCDD1L-DT | -1,332402739 | 0,39710633 | 0,09333 | 0,39489949 | APCDD1L divergent transcript |
| ENSG00000227053 | MUC12-AS1 | -1,333014539 | 0,396937965 | 0,03047 | 0,23546694 | MUC12 antisense RNA 1 |
| ENSG00000273965 | AC243654,1 | -1,333063621 | 0,396924461 | 0,34429 | 0,6217351 | novel transcript, antisense TADA2A |
| ENSG00000178209 | PLEC | -1,33321881 | 0,396881767 | 0,02164 | 0,19544769 | plectin |
| ENSG00000275832 | ARHGAP23 | -1,333926137 | 0,396687231 | 0,0033 | 0,06918097 | Rho GTPase activating protein 23 |
| ENSG00000285889 | AL355312,4 | -1,333938697 | 0,396683777 | 0,06906 | 0,34726337 | novel transcript |
| ENSG00000258048 | AC073569,2 | -1,334057068 | 0,396651231 | 0,17861 | 0,50323303 | novel transcript |
| ENSG00000129990 | SYT5 | -1,334159966 | 0,396622942 | 0,01924 | 0,18363148 | synaptotagmin 5 |
| ENSG00000287945 | AL603824,1 | -1,334351376 | 0,396570323 | 0,06854 | 0,34596462 | novel transcript, antisense to CCNY |
| ENSG00000069399 | BCL3 | -1,334521262 | 0,396523627 | 0,00012 | 0,0070607 | BCL3 transcription coactivator |
| ENSG00000203808 | BVES-AS1 | -1,334533811 | 0,396520178 | 0,00654 | 0,1024996 | BVES antisense RNA 1 |
| ENSG00000054690 | PLEKHH1 | -1,335039693 | 0,396381163 | 0,01204 | 0,14273718 | pleckstrin homology, MyTH4 and FERM domain containing H1 |
| ENSG00000235590 | GNAS-AS1 | -1,335864017 | 0,396154744 | 0,01241 | 0,1456938 | GNAS antisense RNA 1 |
| ENSG00000166924 | NYAP1 | -1,336022258 | 0,396111294 | 0,00368 | 0,07408435 | neuronal tyrosine phosphorylated phosphoinositide-3-kinase adaptor 1 |
| ENSG00000286698 | AC027612,3 | -1,336165302 | 0,396072022 | 0,042 | 0,27753946 | novel transcript |
| ENSG00000178233 | TMEM151B | -1,336467112 | 0,395989172 | 0,00042 | 0,01813396 | transmembrane protein 151B |
| ENSG00000188603 | CLN3 | -1,336741352 | 0,395913907 | 0,01661 | 0,16891639 | CLN3 lysosomal/endosomal transmembrane protein, battenin |
| ENSG00000285549 | AC091973,1 | -1,336845385 | 0,395885358 | 0,3178 | 0,6032144 | Novel transcript |
| ENSG00000157322 | CLEC18A | -1,337103749 | 0,395814468 | 0,0388 | 0,26660248 | C-type lectin domain family 18 member A |
| ENSG00000178645 | C10orf53 | -1,337485485 | 0,395709749 | 0,21414 | 0,53334744 | chromosome 10 open reading frame 53 |
| ENSG00000177453 | NIM1K | -1,337748763 | 0,395637543 | 0,00237 | 0,05606528 | NIM1 serine/threonine protein kinase |
| ENSG00000251292 | ERVH-1 | -1,337915278 | 0,395591881 | 0,07934 | 0,36794989 | endogenous retrovirus group H member 1 |
| ENSG00000287536 | AC012500,1 | -1,33796915 | 0,395577109 | 0,18168 | 0,50497494 | novel transcript, antisense to PDE1A |
| ENSG00000276417 | AC092111,2 | -1,338014873 | 0,395564573 | 0,00514 | 0,08938358 | novel transcript |
| ENSG00000176723 | ZNF843 | -1,338048783 | 0,395555275 | 0,04817 | 0,29486819 | zinc finger protein 843 |
| ENSG00000233942 | AC004012,1 | -1,338280407 | 0,395491774 | 0,06792 | 0,34439007 | novel transcript |

| | | | | | | |
|---|---|---|---|---|---|---|
| ENSG00000249476 | AC008467,1 | -1,338453831 | 0,395444235 | 0,03551 | 0,25544114 | novel transcript |
| ENSG00000237015 | AL031186,1 | -1,338963148 | 0,395304656 | 0,09019 | 0,38841129 | novel transcript |
| ENSG00000249487 | LINC01586 | -1,339004583 | 0,395293303 | 0,0928 | 0,39343521 | long intergenic non-protein coding RNA 1586 |
| ENSG00000105617 | LENG1 | -1,339044669 | 0,395282319 | 0,36496 | 0,63519916 | leukocyte receptor cluster member 1 |
| ENSG00000124243 | BCAS4 | -1,339176113 | 0,395246307 | 0,01465 | 0,15853968 | breast carcinoma amplified sequence 4 |
| ENSG00000188937 | NYX | -1,339340742 | 0,395201207 | 0,0271 | 0,22199012 | nyctalopin |
| ENSG00000174165 | ZDHHC24 | -1,339535027 | 0,39514799 | 0,02368 | 0,20551719 | zinc finger DHHC-type containing 24 |
| ENSG00000172578 | KLHL6 | -1,339541655 | 0,395146174 | 0,01137 | 0,13841038 | kelch like family member 6 |
| ENSG00000106327 | TFR2 | -1,339619522 | 0,395124847 | 0,04404 | 0,28274932 | transferrin receptor 2 |
| ENSG00000285863 | AC009554,2 | -1,340229241 | 0,394957893 | 0,08199 | 0,37306719 | novel transcript, antisense to VPS13C |
| ENSG00000287713 | AC006517,2 | -1,340424908 | 0,39490433 | 0,17182 | 0,49710979 | novel transcript, antisense to SLC2A14 |
| ENSG00000116791 | CRYZ | -1,340579553 | 0,394862002 | 0,01124 | 0,1375342 | crystallin zeta |
| ENSG00000286125 | AC006115,2 | -1,340982502 | 0,394751731 | 0,01096 | 0,13522945 | novel transcript (contains ZIM2-AS1 and PEG3-AS1) |
| ENSG00000215559 | ANKRD20A11 | -1,341400216 | 0,394637452 | 0,22465 | 0,54036191 | ankyrin repeat domain 20 family member A11, pseudogene |
| ENSG00000278440 | AC243791,1 | -1,341684156 | 0,394559791 | 0,03255 | 0,24353935 | novel pseudogene KIAA2013 |
| ENSG00000261819 | AC138932,2 | -1,341684156 | 0,394559791 | 0,03255 | 0,24353935 | novel pseudogene KIAA2013 |
| ENSG00000177427 | MIEF2 | -1,342187841 | 0,394422063 | 0,08648 | 0,38091118 | mitochondrial elongation factor 2 |
| ENSG00000227012 | LINC02527 | -1,342580196 | 0,394314811 | 0,27274 | 0,57322782 | long intergenic non-protein coding RNA 2527 |
| ENSG00000227321 | MTND4P15 | -1,342640213 | 0,394298407 | 0,09141 | 0,39102429 | MT-ND4 pseudogene 15 |
| ENSG00000280620 | SCAANT1 | -1,342737977 | 0,394271688 | 0,14904 | 0,47664156 | SCA7/ATXN7 antisense RNA 1 |
| ENSG00000265702 | AC005821,1 | -1,342755455 | 0,394266912 | 0,02002 | 0,18762986 | novel transcript, antisense to MARCH10 |
| ENSG00000258711 | AL358334,2 | -1,34413225 | 0,393890834 | 0,01495 | 0,16037017 | novel transcript |
| ENSG00000197114 | ZGPAT | -1,344206882 | 0,393870458 | 0,02038 | 0,18949014 | zinc finger CCCH-type and G-patch domain containing |
| ENSG00000285831 | AC027329,1 | -1,344385898 | 0,393821588 | 0,0279 | 0,22539775 | novel transcript |
| ENSG00000253336 | AC018992,1 | -1,34469634 | 0,393736854 | 0,18812 | 0,51128963 | hydroxysteroid (17-beta) dehydrogenase 4 (HSD17B4) pseudogene |
| ENSG00000172339 | ALG14 | -1,344813438 | 0,393704897 | 0,021 | 0,19244681 | ALG14 UDP-N-acetylglucosaminyltransferase subunit |
| ENSG00000285766 | AC092042,4 | -1,345264621 | 0,39358179 | 0,12884 | 0,45214857 | Novel trranscript, antisense to POMGNT2 |
| ENSG00000232973 | CYP1B1-AS1 | -1,345331491 | 0,393563548 | 0,00948 | 0,12461039 | CYP1B1 antisense RNA 1 |
| ENSG00000229259 | LRRC37A12P | -1,34592685 | 0,393401169 | 0,14047 | 0,4673794 | leucine rich repeat containing 37 member A12, pseudogene |
| ENSG00000264324 | AC006030,1 | -1,347129588 | 0,393073337 | 0,10202 | 0,40934973 | novel protein |

| | | | | | | |
|---|---|---|---|---|---|---|
| ENSG00000169258 | GPRIN1 | -1,34740496 | 0,392998317 | 0,00037 | 0,01667641 | G protein regulated inducer of neurite outgrowth 1 |
| ENSG00000248516 | AC105415,1 | -1,347438328 | 0,392989228 | 0,19269 | 0,51490739 | novel transcript |
| ENSG00000164181 | ELOVL7 | -1,347614386 | 0,392941272 | 0,03176 | 0,24047554 | ELOVL fatty acid elongase 7 |
| ENSG00000255328 | AC136475,5 | -1,347977622 | 0,392842352 | 0,19378 | 0,51547251 | novel transcript |
| ENSG00000235903 | CPB2-AS1 | -1,348164496 | 0,39279147 | 0,0096 | 0,12527194 | CPB2 antisense RNA 1 |
| ENSG00000155034 | FBXL18 | -1,348843279 | 0,392606706 | 0,02331 | 0,20398797 | F-box and leucine rich repeat protein 18 |
| ENSG00000265727 | RN7SL648P | -1,349515918 | 0,392423701 | 0,04829 | 0,29514999 | RNA, 7SL, cytoplasmic 648, pseudogene |
| ENSG00000253708 | AC026979,1 | -1,350268777 | 0,392218971 | 0,17811 | 0,50236177 | novel transcript |
| ENSG00000280758 | AL356481,3 | -1,350597805 | 0,39212953 | 0,07897 | 0,3669053 | novel transcript, antisense to SPTAN1 |
| ENSG00000263276 | AC020978,7 | -1,350964189 | 0,392029958 | 0,02707 | 0,22193038 | novel transcript |
| ENSG00000243005 | RN7SL16P | -1,35112601 | 0,391985988 | 0,039 | 0,26725619 | RNA, 7SL, cytoplasmic 16, pseudogene |
| ENSG00000254897 | AP003035,1 | -1,351339735 | 0,391927922 | 0,22687 | 0,54168591 | DEAH (Asp-Glu-Ala-His) box polypeptide 9 (DHX9) pseudogene |
| ENSG00000224220 | DTNB-AS1 | -1,351822357 | 0,391796834 | 0,26918 | 0,5708709 | DTNB antisense RNA 1 |
| ENSG00000266644 | AC103810,5 | -1,352499154 | 0,391613077 | 0,01853 | 0,18002406 | novel transcript, antisense to LRRC37A3 |
| ENSG00000233483 | AC008105,2 | -1,352670845 | 0,391566475 | 0,21589 | 0,53454039 | novel pseudogene, ortholog of 1700023F06Rik (M, musculus) |
| ENSG00000116032 | GRIN3B | -1,353095181 | 0,391451321 | 0,06844 | 0,34571361 | glutamate ionotropic receptor NMDA type subunit 3B |
| ENSG00000181513 | ACBD4 | -1,353487443 | 0,391344902 | 0,02217 | 0,19795874 | acyl-CoA binding domain containing 4 |
| ENSG00000182095 | TNRC18 | -1,353808245 | 0,391257891 | 0,02957 | 0,23126047 | trinucleotide repeat containing 18 |
| ENSG00000230912 | AL021707,4 | -1,353821082 | 0,39125441 | 0,05676 | 0,31760957 | novel transcript |
| ENSG00000271032 | AC020907,4 | -1,353971859 | 0,391213522 | 0,09867 | 0,40453901 | novel transcript |
| ENSG00000143995 | MEIS1 | -1,354577226 | 0,3910494 | 0,00011 | 0,00659917 | Meis homeobox 1 |
| ENSG00000225383 | SFTA1P | -1,355185547 | 0,390884546 | 0,05484 | 0,31349896 | surfactant associated 1, lncRNA |
| ENSG00000157881 | PANK4 | -1,355306586 | 0,390851753 | 0,02965 | 0,23170642 | pantothenate kinase 4 (inactive) |
| ENSG00000260596 | DUX4 | -1,355583825 | 0,390776651 | 0,11898 | 0,43696075 | double homeobox 4 |
| ENSG00000277352 | AP001065,4 | -1,35655092 | 0,390514786 | 0,17275 | 0,49760599 | novel transcript |
| ENSG00000262304 | AC027796,3 | -1,356625914 | 0,390494487 | 0,04377 | 0,28230121 | novel readthrough SHPK-TRPV1 protein |
| ENSG00000251023 | AC114980,1 | -1,357441524 | 0,390273788 | 0,01747 | 0,17422253 | novel transcript |
| ENSG00000171532 | NEUROD2 | -1,357477279 | 0,390264116 | 0,15543 | 0,48334025 | neuronal differentiation 2 |
| ENSG00000146063 | TRIM41 | -1,35750352 | 0,390257018 | 0,01664 | 0,16922242 | tripartite motif containing 41 |
| ENSG00000259519 | AC051619,3 | -1,358010386 | 0,390119932 | 0,28035 | 0,57784691 | novel transcript, antisense to SHF |
| ENSG00000287853 | AL031668,2 | -1,358028708 | 0,390114977 | 0,00305 | 0,06592744 | novel transcript |

| | | | | | | |
|---|---|---|---|---|---|---|
| ENSG00000263874 | LINC00672 | -1,358365455 | 0,390023929 | 0,00319 | 0,06768121 | long intergenic non-protein coding RNA 672 |
| ENSG00000233783 | AP001442,1 | -1,359083226 | 0,389829932 | 0,01008 | 0,12883758 | novel transcript |
| ENSG00000262298 | AC004584,1 | -1,35984155 | 0,38962508 | 0,17493 | 0,49911246 | novel transcript, antisense to COIL |
| ENSG00000273145 | BX537318,1 | -1,359886075 | 0,389613055 | 0,09884 | 0,40464633 | novel transcript |
| ENSG00000067057 | PFKP | -1,360260265 | 0,389512015 | 7,28E-05 | 0,00461938 | phosphofructokinase, platelet |
| ENSG00000286447 | AC109782,1 | -1,36099989 | 0,389312375 | 0,02745 | 0,22348662 | novel transcript |
| ENSG00000200198 | RN7SKP211 | -1,361379288 | 0,389210008 | 0,14975 | 0,47782859 | RN7SK pseudogene 211 |
| ENSG00000169609 | C15orf40 | -1,361877632 | 0,389075588 | 0,01693 | 0,17086866 | chromosome 15 open reading frame 40 |
| ENSG00000274933 | TBC1D3I | -1,362295062 | 0,388963029 | 0,35925 | 0,6321355 | TBC1 domain family member 3I |
| ENSG00000234619 | RPL7P11 | -1,36247857 | 0,388913557 | 0,15265 | 0,48061452 | ribosomal protein L7 pseudogene 11 |
| ENSG00000105963 | ADAP1 | -1,362871489 | 0,38880765 | 0,01959 | 0,18572138 | ArfGAP with dual PH domains 1 |
| ENSG00000260518 | BMS1P8 | -1,363196909 | 0,388719959 | 0,0326 | 0,24377222 | BMS1 pseudogene 8 |
| ENSG00000141028 | CDRT15P1 | -1,363502526 | 0,388637622 | 0,26335 | 0,56608528 | CDRT15 pseudogene 1 |
| ENSG00000163083 | INHBB | -1,363531108 | 0,388629923 | 0,03508 | 0,2533278 | inhibin subunit beta B |
| ENSG00000230001 | AL359095,1 | -1,363574822 | 0,388618148 | 0,13773 | 0,46481656 | novel transcript |
| ENSG00000227115 | LINC01630 | -1,363992854 | 0,388505559 | 0,04783 | 0,29372502 | long intergenic non-protein coding RNA 1630 |
| ENSG00000240393 | AC092691,2 | -1,36488379 | 0,388265711 | 0,10209 | 0,40934973 | developmentally regulated GTP binding protein 1 (DRG1) pseudogene |
| ENSG00000264727 | AC005725,1 | -1,365136977 | 0,388197578 | 0,09677 | 0,40094776 | novel transcript |
| ENSG00000183578 | TNFAIP8L3 | -1,365506258 | 0,388098226 | 0,04741 | 0,29245799 | TNF alpha induced protein 8 like 3 |
| ENSG00000273253 | AL022328,4 | -1,365680883 | 0,388051253 | 0,04604 | 0,28930497 | novel transcript, antisense TRABD |
| ENSG00000232611 | AL683813,1 | -1,365964541 | 0,387974963 | 0,00807 | 0,11420372 | novel transcript |
| ENSG00000106038 | EVX1 | -1,366196758 | 0,387912519 | 0,25789 | 0,56199743 | even-skipped homeobox 1 |
| ENSG00000264365 | AC023983,1 | -1,366688243 | 0,387780391 | 0,14189 | 0,4683384 | novel transcript, antisense to OSBPL1A |
| ENSG00000069696 | DRD4 | -1,367000189 | 0,387696552 | 0,34396 | 0,62139417 | dopamine receptor D4 |
| ENSG00000249279 | LINC02057 | -1,367778053 | 0,387487573 | 0,12996 | 0,45419678 | long intergenic non-protein coding RNA 2057 |
| ENSG00000176884 | GRIN1 | -1,367872478 | 0,387462212 | 0,05242 | 0,30735033 | glutamate ionotropic receptor NMDA type subunit 1 |
| ENSG00000135604 | STX11 | -1,368059464 | 0,387411997 | 0,13702 | 0,4642578 | syntaxin 11 |
| ENSG00000254860 | TMEM9B-AS1 | -1,368812952 | 0,387209713 | 0,01855 | 0,18007753 | TMEM9B antisense RNA 1 |
| ENSG00000266936 | AC011442,1 | -1,36912468 | 0,387126057 | 0,0472 | 0,29217708 | novel transcript, antisense to SMARCA4 |
| ENSG00000273297 | AC009275,1 | -1,369235648 | 0,387096281 | 0,03 | 0,23340081 | novel transcript |
| ENSG00000265791 | AC127024,4 | -1,369295839 | 0,387080131 | 0,02302 | 0,20237428 | novel transcript, sense intronic CRLF3 |
| ENSG00000152939 | MARVELD2 | -1,369413237 | 0,387048634 | 0,17396 | 0,49860418 | MARVEL domain containing 2 |

| | | | | | | |
|---|---|---|---|---|---|---|
| ENSG00000242396 | AC096536,2 | -1,369451779 | 0,387038294 | 0,33676 | 0,61675087 | novel transcript |
| ENSG00000131584 | ACAP3 | -1,370031886 | 0,386882698 | 0,0042 | 0,07947406 | ArfGAP with coiled-coil, ankyrin repeat and PH domains 3 |
| ENSG00000225051 | HMGB3P22 | -1,370269323 | 0,38681903 | 0,19412 | 0,51562158 | high mobility group box 3 pseudogene 22 |
| ENSG00000260989 | AL133297,2 | -1,370306707 | 0,386809007 | 0,07094 | 0,35169607 | novel transcript, antisense to IFT140 |
| ENSG00000246985 | SOCS2-AS1 | -1,370445216 | 0,386771872 | 0,01357 | 0,15281979 | SOCS2 antisense RNA 1 |
| ENSG00000196341 | OR8D1 | -1,370794925 | 0,38667813 | 0,2287 | 0,54252903 | olfactory receptor family 8 subfamily D member 1 |
| ENSG00000154118 | JPH3 | -1,370799465 | 0,386676913 | 0,00067 | 0,02497236 | junctophilin 3 |
| ENSG00000174521 | TTC9B | -1,371146318 | 0,386583959 | 0,01114 | 0,13658037 | tetratricopeptide repeat domain 9B |
| ENSG00000188368 | PRR19 | -1,371338329 | 0,386532512 | 0,0275 | 0,22355667 | proline rich 19 |
| ENSG00000272772 | AC104109,3 | -1,37175302 | 0,386421422 | 0,10394 | 0,41230119 | novel protein |
| ENSG00000277758 | FO681492,1 | -1,371866623 | 0,386390995 | 0,02277 | 0,20142881 | synaptotagmin-15 |
| ENSG00000276486 | AC022960,3 | -1,371896564 | 0,386382976 | 0,14471 | 0,47224655 | family with sequence similarity 76, member A (FAM76A) pseudogene |
| ENSG00000092054 | MYH7 | -1,371903183 | 0,386381203 | 0,00892 | 0,11991469 | myosin heavy chain 7 |
| ENSG00000275470 | LENG8-AS1 | -1,372249163 | 0,386288555 | 0,17814 | 0,50236177 | LENG8 antisense RNA 1 |
| ENSG00000226264 | HLA-DMB | -1,37259306 | 0,386196485 | 0,14619 | 0,47428193 | major histocompatibility complex, class II, DM beta |
| ENSG00000286553 | AL357075,2 | -1,373012805 | 0,38608414 | 0,02949 | 0,23093703 | novel transcript, antisense to CNKSR3 |
| ENSG00000235961 | PNMA6A | -1,373328201 | 0,385999745 | 0,00616 | 0,09876116 | PNMA family member 6A |
| ENSG00000099814 | CEP170B | -1,373613196 | 0,385923501 | 0,00057 | 0,02226183 | centrosomal protein 170B |
| ENSG00000238086 | PPP1R26P1 | -1,373760095 | 0,385884207 | 0,01289 | 0,14857323 | protein phosphatase 1 regulatory subunit 26 pseudogene 1 |
| ENSG00000226404 | LY6G5C | -1,374328186 | 0,385732287 | 0,0466 | 0,29026543 | lymphocyte antigen 6 family member G5C |
| ENSG00000260325 | HSPB9 | -1,374494563 | 0,385687806 | 0,0766 | 0,36266919 | heat shock protein family B (small) member 9 |
| ENSG00000139266 | MARCHF9 | -1,374615112 | 0,38565558 | 0,00652 | 0,10241021 | membrane associated ring-CH-type finger 9 |
| ENSG00000255284 | AP006621,3 | -1,374791241 | 0,3856085 | 0,03177 | 0,24047554 | novel transcript |
| ENSG00000104368 | PLAT | -1,374935686 | 0,385569894 | 0,02248 | 0,19968345 | plasminogen activator, tissue type |
| ENSG00000250999 | AC136604,2 | -1,374966692 | 0,385561608 | 0,04922 | 0,29853148 | novel transcript |
| ENSG00000177839 | PCDHB9 | -1,374967122 | 0,385561493 | 0,01052 | 0,13246349 | protocadherin beta 9 |
| ENSG00000093010 | COMT | -1,375281172 | 0,385477572 | 2,32E-05 | 0,00175182 | catechol-O-methyltransferase |
| ENSG00000163630 | SYNPR | -1,375561164 | 0,385402767 | 0,02895 | 0,22977124 | synaptoporin |
| ENSG00000239511 | POM121L7P | -1,375782741 | 0,38534358 | 0,05373 | 0,31090219 | POM121 transmembrane nucleoporin like 7 pseudogene |
| ENSG00000236867 | Z98885,1 | -1,376216478 | 0,385227746 | 0,02221 | 0,19806119 | suppressor of cytokine signaling 2 (SOCS2) pseudogene |
| ENSG00000277753 | MUC20 | -1,376254398 | 0,385217621 | 0,15675 | 0,48425071 | mucin 20, cell surface associated |
| ENSG00000186567 | CEACAM19 | -1,376312491 | 0,385202109 | 0,0145 | 0,15751168 | CEA cell adhesion molecule 19 |

| | | | | | | |
|---|---|---|---|---|---|---|
| ENSG00000168928 | CTRB2 | -1,376541971 | 0,385140843 | 0,12154 | 0,44093327 | chymotrypsinogen B2 |
| ENSG00000136449 | MYCBPAP | -1,376932664 | 0,385036558 | 0,07691 | 0,36356213 | MYCBP associated protein |
| ENSG00000218672 | AC008060,1 | -1,377358613 | 0,384922894 | 0,14496 | 0,47257223 | novel transcript |
| ENSG00000270332 | SMC2-AS1 | -1,377757332 | 0,384816527 | 0,01075 | 0,13405812 | SMC2 antisense RNA 1 (head to head) |
| ENSG00000152086 | TUBA3E | -1,378039083 | 0,384741382 | 0,15226 | 0,48061452 | tubulin alpha 3e |
| ENSG00000285881 | AC105206,3 | -1,378082424 | 0,384729824 | 0,11199 | 0,42546126 | Novel transcript, antisense to BMP1 |
| ENSG00000233956 | BTF3P6 | -1,378169903 | 0,384706496 | 0,03445 | 0,25090278 | basic transcription factor 3 pseudogene 6 |
| ENSG00000157766 | ACAN | -1,378209181 | 0,384696023 | 0,08122 | 0,3719701 | aggrecan |
| ENSG00000272372 | AC009102,2 | -1,378397617 | 0,384645779 | 0,04767 | 0,29344099 | novel transcript, antisense to GNAO1 |
| ENSG00000262075 | DKFZP434A06 | -1,378666178 | 0,384574183 | 0,01437 | 0,15682082 | uncharacterized LOC26102 |
| ENSG00000185666 | SYN3 | -1,379160803 | 0,384442355 | 0,00237 | 0,05606528 | synapsin III |
| ENSG00000167207 | NOD2 | -1,37927473 | 0,384411998 | 0,10189 | 0,40926385 | nucleotide binding oligomerization domain containing 2 |
| ENSG00000244952 | AC123768,1 | -1,379863653 | 0,384255109 | 0,07882 | 0,36667857 | novel transcript |
| ENSG00000228701 | TNKS2-AS1 | -1,379878523 | 0,384251148 | 0,01838 | 0,17938469 | TNKS2 antisense RNA 1 (head to head) |
| ENSG00000261654 | AL360270,2 | -1,380484654 | 0,384089744 | 0,02872 | 0,2291366 | novel transcript, sense overlapping C1orf103 |
| ENSG00000207751 | AP000553,1 | -1,380487424 | 0,384089006 | 0,04439 | 0,28421599 | novel transcript |
| ENSG00000223494 | VARS2 | -1,380779714 | 0,384011198 | 0,05085 | 0,30326674 | valyl-tRNA synthetase 2, mitochondrial |
| ENSG00000254420 | AP003086,1 | -1,381023313 | 0,383946363 | 0,1068 | 0,41660052 | novel transcript |
| ENSG00000143382 | ADAMTSL4 | -1,381229266 | 0,383891556 | 0,02312 | 0,20293306 | ADAMTS like 4 |
| ENSG00000251687 | AC104791,1 | -1,38140825 | 0,383843933 | 0,0142 | 0,15631436 | novel transcript |
| ENSG00000177699 | AC011944,1 | -1,381474966 | 0,383826183 | 0,22854 | 0,5424199 | novel transcript, antisense to RASGRF1 |
| ENSG00000261560 | AC007216,3 | -1,382235318 | 0,383623946 | 0,01998 | 0,18752351 | novel transcript, sense intronic to GSPT1 |
| ENSG00000175984 | DENND2C | -1,382250201 | 0,383619989 | 0,01297 | 0,14899372 | DENN domain containing 2C |
| ENSG00000184988 | TMEM106A | -1,382319235 | 0,383601632 | 0,04697 | 0,29152224 | transmembrane protein 106A |
| ENSG00000238358 | AC004969,1 | -1,382330328 | 0,383598683 | 0,03403 | 0,24909651 | novel transcript |
| ENSG00000228486 | C2orf92 | -1,382420156 | 0,383574799 | 0,02243 | 0,19928952 | chromosome 2 open reading frame 92 |
| ENSG00000090565 | RAB11FIP3 | -1,382563377 | 0,383536723 | 0,00785 | 0,11266092 | RAB11 family interacting protein 3 |
| ENSG00000278812 | C8orf31 | -1,383082856 | 0,383398645 | 0,00904 | 0,12067214 | chromosome 8 open reading frame 31 |
| ENSG00000273165 | AL121652,1 | -1,383118846 | 0,383389081 | 0,06022 | 0,32601555 | novel transcript |
| ENSG00000013364 | MVP | -1,383744402 | 0,383222879 | 0,03803 | 0,26375851 | major vault protein |
| ENSG00000102003 | SYP | -1,383769348 | 0,383216252 | 0,00569 | 0,09424131 | synaptophysin |
| ENSG00000254896 | OPCML-IT1 | -1,383857355 | 0,383192876 | 0,11018 | 0,42271592 | OPCML intronic transcript 1 |
| ENSG00000286307 | AC017028,1 | -1,384058065 | 0,383139569 | 0,01035 | 0,13104019 | novel transcript, antisense to HDAC4 |

| | | | | | | |
|---|---|---|---|---|---|---|
| ENSG00000185880 | TRIM69 | -1,385317402 | 0,38280527 | 0,00525 | 0,09023573 | tripartite motif containing 69 |
| ENSG00000213600 | U73169,1 | -1,385400975 | 0,382783096 | 0,14632 | 0,47433542 | novel transcript, sense intronic to GNAI2 |
| ENSG00000242268 | LINC02082 | -1,385881994 | 0,382655491 | 0,05113 | 0,30402651 | long intergenic non-protein coding RNA 2082 |
| ENSG00000158122 | PRXL2C | -1,386322966 | 0,382538547 | 0,02638 | 0,2188126 | peroxiredoxin like 2C |
| ENSG00000261408 | TEN1-CDK3 | -1,386607028 | 0,382463233 | 0,02632 | 0,21861252 | TEN1-CDK3 readthrough (NMD candidate) |
| ENSG00000105613 | MAST1 | -1,386667326 | 0,382447248 | 0,00818 | 0,11480072 | microtubule associated serine/threonine kinase 1 |
| ENSG00000234722 | LINC01287 | -1,386667416 | 0,382447225 | 0,0187 | 0,18107187 | long intergenic non-protein coding RNA 1287 |
| ENSG00000206596 | RNU1-27P | -1,386996033 | 0,382360121 | 0,00823 | 0,11480072 | RNA, U1 small nuclear 27, pseudogene |
| ENSG00000207389 | RNU1-4 | -1,386996033 | 0,382360121 | 0,00823 | 0,11480072 | RNA, U1 small nuclear 4 |
| ENSG00000206652 | RNU1-1 | -1,386996033 | 0,382360121 | 0,00823 | 0,11480072 | RNA, U1 small nuclear 1 |
| ENSG00000273768 | RNVU1-29 | -1,386996033 | 0,382360121 | 0,00823 | 0,11480072 | RNA, variant U1 small nuclear 29 |
| ENSG00000206737 | RNVU1-18 | -1,386996033 | 0,382360121 | 0,00823 | 0,11480072 | RNA, variant U1 small nuclear 18 |
| ENSG00000206588 | RNU1-28P | -1,386996033 | 0,382360121 | 0,00823 | 0,11480072 | RNA, U1 small nuclear 28, pseudogene |
| ENSG00000275405 | U1 | -1,386996033 | 0,382360121 | 0,00823 | 0,11480072 | U1 spliceosomal RNA |
| ENSG00000207005 | RNU1-2 | -1,386996033 | 0,382360121 | 0,00823 | 0,11480072 | RNA, U1 small nuclear 2 |
| ENSG00000207513 | RNU1-3 | -1,386996033 | 0,382360121 | 0,00823 | 0,11480072 | RNA, U1 small nuclear 3 |
| ENSG00000171450 | CDK5R2 | -1,387128078 | 0,382325126 | 0,01619 | 0,16662112 | cyclin dependent kinase 5 regulatory subunit 2 |
| ENSG00000230536 | AL360268,1 | -1,38727183 | 0,382287033 | 0,06463 | 0,33689237 | novel transcript |
| ENSG00000183571 | PGPEP1L | -1,38751852 | 0,38222167 | 0,28806 | 0,58415113 | pyroglutamyl-peptidase I like |
| ENSG00000253112 | AC102945,1 | -1,38761515 | 0,38219607 | 0,18385 | 0,50754421 | novel transcript |
| ENSG00000204851 | PNMA8B | -1,388158513 | 0,382052151 | 0,05009 | 0,30089233 | PNMA family member 8B |
| ENSG00000172350 | ABCG4 | -1,388295225 | 0,382015948 | 0,00826 | 0,11480072 | ATP binding cassette subfamily G member 4 |
| ENSG00000231115 | RING1 | -1,389899488 | 0,381591387 | 0,05366 | 0,31073157 | ring finger protein 1 |
| ENSG00000228520 | RING1 | -1,389899488 | 0,381591387 | 0,05366 | 0,31073157 | ring finger protein 1 |
| ENSG00000103740 | ACSBG1 | -1,390176524 | 0,381518118 | 0,00369 | 0,0741128 | acyl-CoA synthetase bubblegum family member 1 |
| ENSG00000285830 | AL109628,2 | -1,391582653 | 0,381146451 | 0,01608 | 0,16620294 | novel transcript, antisense to TRAPPC6B |
| ENSG00000284874 | AC000093,1 | -1,391817517 | 0,381084407 | 0,02412 | 0,20758258 | novel transcript |
| ENSG00000286699 | AC084198,2 | -1,392118002 | 0,381005043 | 0,02105 | 0,19263105 | novel transcript |
| ENSG00000254423 | AC087203,1 | -1,392882935 | 0,380803083 | 0,06062 | 0,32697886 | ubiquitin specific peptidase 17-like 2 ( USP17L2) pseudogene |
| ENSG00000236801 | RPL24P8 | -1,393204134 | 0,380718311 | 0,16892 | 0,4950359 | RPL24 pseudogene 8 |
| ENSG00000105464 | GRIN2D | -1,394258868 | 0,380440075 | 0,00968 | 0,12598277 | glutamate ionotropic receptor NMDA type subunit 2D |
| ENSG00000268309 | AC008764,3 | -1,395243104 | 0,38018062 | 0,29489 | 0,58868965 | novel transcript, antisense SLC35E1 |

| | | | | | | |
|---|---|---|---|---|---|---|
| ENSG00000199785 | SNORA52 | -1,395387863 | 0,380142475 | 0,01547 | 0,16309786 | small nucleolar RNA, H/ACA box 52 |
| ENSG00000117400 | MPL | -1,395686831 | 0,380063707 | 0,09531 | 0,39794903 | MPL proto-oncogene, thrombopoietin receptor |
| ENSG00000167378 | IRGQ | -1,395904252 | 0,380006433 | 0,00889 | 0,11969635 | immunity related GTPase Q |
| ENSG00000244040 | IL12A-AS1 | -1,396023521 | 0,379975019 | 0,00529 | 0,09064667 | IL12A antisense RNA 1 |
| ENSG00000259425 | AC011767,1 | -1,396550418 | 0,379836271 | 0,1541 | 0,48235237 | Golgin subfamily A member 6 pseudogene |
| ENSG00000260236 | AC099778,1 | -1,396666867 | 0,379805613 | 0,01317 | 0,15012165 | novel transcript, antisense to PTPN23 |
| ENSG00000225952 | AL512444,1 | -1,397108509 | 0,379689364 | 0,09479 | 0,39724142 | novel transcript |
| ENSG00000243708 | PLA2G4B | -1,397739271 | 0,379523396 | 0,01264 | 0,14722977 | phospholipase A2 group IVB |
| ENSG00000106665 | CLIP2 | -1,398266053 | 0,379384843 | 0,0082 | 0,11480072 | CAP-Gly domain containing linker protein 2 |
| ENSG00000105639 | JAK3 | -1,398670805 | 0,37927842 | 0,01963 | 0,18592829 | Janus kinase 3 |
| ENSG00000198157 | HMGN5 | -1,39944477 | 0,379075003 | 0,01382 | 0,15438008 | high mobility group nucleosome binding domain 5 |
| ENSG00000224886 | AL132656,1 | -1,399566683 | 0,379042971 | 0,30566 | 0,59512079 | novel pseudogene |
| ENSG00000197475 | ADAM3A | -1,399890231 | 0,378957974 | 0,18869 | 0,5120698 | ADAM metallopeptidase domain 3A (pseudogene) |
| ENSG00000167676 | PLIN4 | -1,399949327 | 0,378942451 | 0,03128 | 0,23867571 | perilipin 4 |
| ENSG00000181873 | IBA57 | -1,400353428 | 0,378836324 | 0,01719 | 0,17234227 | iron-sulfur cluster assembly factor IBA57 |
| ENSG00000149488 | TMC2 | -1,400642425 | 0,378760444 | 0,02 | 0,187583 | transmembrane channel like 2 |
| ENSG00000081041 | CXCL2 | -1,401311031 | 0,378584951 | 0,07624 | 0,36242671 | C-X-C motif chemokine ligand 2 |
| ENSG00000213213 | CCDC183 | -1,402570734 | 0,37825453 | 0,01038 | 0,13132723 | coiled-coil domain containing 183 |
| ENSG00000226626 | NOP56P2 | -1,403154718 | 0,378101449 | 0,01274 | 0,14784043 | NOP56 ribonucleoprotein pseudogene 2 |
| ENSG00000183914 | DNAH2 | -1,404426916 | 0,377768178 | 0,0137 | 0,15356644 | dynein axonemal heavy chain 2 |
| ENSG00000168143 | FAM83B | -1,404520019 | 0,3777438 | 0,04285 | 0,27902746 | family with sequence similarity 83 member B |
| ENSG00000260372 | AQP4-AS1 | -1,404615913 | 0,377718692 | 0,0501 | 0,30089233 | AQP4 antisense RNA 1 |
| ENSG00000286470 | AL034427,1 | -1,404932122 | 0,377635913 | 0,04275 | 0,2786689 | novel transcript, antisense to PAK7 |
| ENSG00000271907 | SNORA35B | -1,405220564 | 0,377560419 | 0,01527 | 0,16217309 | small nucleolar RNA, H/ACA box 35B |
| ENSG00000285279 | AL353622,5 | -1,405708033 | 0,377432868 | 0,65399 | 0,82680123 | novel transcript |
| ENSG00000205930 | C21orf62-AS1 | -1,406006859 | 0,377354698 | 0,01447 | 0,15740445 | C21orf62 antisense RNA 1 |
| ENSG00000225377 | NRSN2-AS1 | -1,407280092 | 0,377021815 | 0,01669 | 0,16951872 | NRSN2 antisense RNA 1 |
| ENSG00000095713 | CRTAC1 | -1,407363248 | 0,377000085 | 0,02509 | 0,21271259 | cartilage acidic protein 1 |
| ENSG00000254440 | PBOV1 | -1,407534794 | 0,376955259 | 0,04706 | 0,29191438 | prostate and breast cancer overexpressed 1 |
| ENSG00000112394 | SLC16A10 | -1,408145539 | 0,376795714 | 0,01558 | 0,16367661 | solute carrier family 16 member 10 |
| ENSG00000244128 | LINC01322 | -1,408250377 | 0,376768334 | 0,00704 | 0,10676605 | long intergenic non-protein coding RNA 1322 |
| ENSG00000175602 | CCDC85B | -1,408384021 | 0,376733434 | 0,02164 | 0,19544769 | coiled-coil domain containing 85B |
| ENSG00000125650 | PSPN | -1,408585002 | 0,376680955 | 0,01856 | 0,18009781 | persephin |

| | | | | | | |
|---|---|---|---|---|---|---|
| ENSG00000240132 | ETF1P2 | -1,408628194 | 0,376669678 | 0,01969 | 0,18614972 | eukaryotic translation termination factor 1 pseudogene 2 |
| ENSG00000134539 | KLRD1 | -1,408669672 | 0,376658849 | 0,01637 | 0,16750744 | killer cell lectin like receptor D1 |
| ENSG00000166813 | KIF7 | -1,408708349 | 0,376648751 | 0,0075 | 0,11001711 | kinesin family member 7 |
| ENSG00000265519 | AC015922,2 | -1,409104419 | 0,376545362 | 0,01431 | 0,15653473 | novel transcript |
| ENSG00000104722 | NEFM | -1,409121834 | 0,376540817 | 8,50E-05 | 0,00527733 | neurofilament medium |
| ENSG00000140323 | DISP2 | -1,409838682 | 0,376353767 | 0,01577 | 0,16469616 | dispatched RND transporter family member 2 |
| ENSG00000231346 | LINC01160 | -1,410360205 | 0,376217743 | 0,02593 | 0,21707964 | long intergenic non-protein coding RNA 1160 |
| ENSG00000237436 | CAMTA1-DT | -1,410413534 | 0,376203836 | 0,11497 | 0,43043054 | CAMTA1 divergent transcript |
| ENSG00000270112 | AC090241,2 | -1,410560313 | 0,376165564 | 0,02955 | 0,23122134 | novel transcript, antisense to ST8SIA5 |
| ENSG00000255326 | AP001922,5 | -1,411147753 | 0,376012427 | 0,00994 | 0,12777099 | novel transcript |
| ENSG00000267734 | AL121989,1 | -1,411279214 | 0,375978165 | 0,05541 | 0,31484345 | novel transcript, antisense to HS2ST1 |
| ENSG00000038295 | TLL1 | -1,411384867 | 0,375950632 | 0,00464 | 0,08393639 | tolloid like 1 |
| ENSG00000262903 | AC027796,4 | -1,411506003 | 0,375919067 | 0,05459 | 0,3128315 | novel transcript, antisense to CTNS |
| ENSG00000286000 | AC234775,3 | -1,411801888 | 0,375841977 | 0,07942 | 0,3679672 | novel transcript |
| ENSG00000210077 | MT-TV | -1,413003401 | 0,375529096 | 0,10613 | 0,41557034 | mitochondrially encoded tRNA-Val (GUN) |
| ENSG00000271826 | PLS3-AS1 | -1,413981117 | 0,375274686 | 0,05844 | 0,32267557 | PLS3 antisense RNA 1 |
| ENSG00000206492 | GNL1 | -1,414894661 | 0,37503713 | 0,18757 | 0,51082796 | G protein nucleolar 1 (putative) |
| ENSG00000159314 | ARHGAP27 | -1,415035677 | 0,375000474 | 0,03976 | 0,2695995 | Rho GTPase activating protein 27 |
| ENSG00000246465 | AC138904,1 | -1,416212553 | 0,374694692 | 0,02595 | 0,21717729 | novel transcript LOC100506705 |
| ENSG00000242173 | ARHGDIG | -1,416516403 | 0,374615785 | 0,05277 | 0,30840541 | Rho GDP dissociation inhibitor gamma |
| ENSG00000254461 | AP001107,4 | -1,416664302 | 0,374577383 | 0,01311 | 0,14974648 | novel transcript |
| ENSG00000183458 | PKD1P3 | -1,416784788 | 0,374546102 | 0,00211 | 0,05175899 | polycystin 1, transient receptor potential channel interacting pseudogene 3 |
| ENSG00000167674 | HDGFL2 | -1,417009046 | 0,374487885 | 0,00957 | 0,12500859 | HDGF like 2 |
| ENSG00000170075 | GPR37L1 | -1,417016258 | 0,374486013 | 0,01484 | 0,15989783 | G protein-coupled receptor 37 like 1 |
| ENSG00000277903 | U2 | -1,417156481 | 0,374449617 | 0,00755 | 0,11001711 | U2 spliceosomal RNA |
| ENSG00000274862 | U2 | -1,417156481 | 0,374449617 | 0,00755 | 0,11001711 | U2 spliceosomal RNA |
| ENSG00000274062 | U2 | -1,417156481 | 0,374449617 | 0,00755 | 0,11001711 | U2 spliceosomal RNA |
| ENSG00000273709 | U2 | -1,417156481 | 0,374449617 | 0,00755 | 0,11001711 | U2 spliceosomal RNA |
| ENSG00000278048 | U2 | -1,417156481 | 0,374449617 | 0,00755 | 0,11001711 | U2 spliceosomal RNA |
| ENSG00000275219 | U2 | -1,417156481 | 0,374449617 | 0,00755 | 0,11001711 | U2 spliceosomal RNA |
| ENSG00000275616 | U2 | -1,417156481 | 0,374449617 | 0,00755 | 0,11001711 | U2 spliceosomal RNA |
| ENSG00000278591 | U2 | -1,417156481 | 0,374449617 | 0,00755 | 0,11001711 | U2 spliceosomal RNA |

| | | | | | | |
|---|---|---|---|---|---|---|
| ENSG00000274432 | U2 | -1,417156481 | 0,374449617 | 0,00755 | 0,11001711 | U2 spliceosomal RNA |
| ENSG00000276596 | U2 | -1,417156481 | 0,374449617 | 0,00755 | 0,11001711 | U2 spliceosomal RNA |
| ENSG00000274585 | RNU2-1 | -1,417156481 | 0,374449617 | 0,00755 | 0,11001711 | RNA, U2 small nuclear 1 |
| ENSG00000213888 | LINC01521 | -1,417332466 | 0,374403943 | 0,0164 | 0,16758111 | long intergenic non-protein coding RNA 1521 |
| ENSG00000255000 | AC139103,1 | -1,417988519 | 0,374233725 | 0,14888 | 0,47633105 | zinc finger protein pseudogene |
| ENSG00000287312 | AC127526,4 | -1,418434818 | 0,374117973 | 0,00151 | 0,04166341 | novel transcript |
| ENSG00000284713 | SMIM38 | -1,418779307 | 0,374028651 | 0,05316 | 0,30971127 | small integral membrane protein 38 |
| ENSG00000248483 | POU5F2 | -1,418851167 | 0,374010022 | 0,00701 | 0,10660586 | POU domain class 5, transcription factor 2 |
| ENSG00000144771 | LRTM1 | -1,419162884 | 0,37392922 | 0,25179 | 0,55773568 | leucine rich repeats and transmembrane domains 1 |
| ENSG00000226308 | AL122058,1 | -1,419216127 | 0,37391542 | 0,08448 | 0,37732439 | novel transcript |
| ENSG00000197774 | EME2 | -1,419324464 | 0,373887342 | 0,00478 | 0,08525507 | essential meiotic structure-specific endonuclease subunit 2 |
| ENSG00000103534 | TMC5 | -1,419804617 | 0,373762927 | 0,0897 | 0,38734232 | transmembrane channel like 5 |
| ENSG00000240322 | RN7SL481P | -1,419837325 | 0,373754453 | 0,06721 | 0,34325886 | RNA, 7SL, cytoplasmic 481, pseudogene |
| ENSG00000259564 | AC009554,1 | -1,420424776 | 0,373602295 | 0,13858 | 0,46578766 | novel transcript, antisense to VPS13C |
| ENSG00000224354 | MTND2P5 | -1,420523919 | 0,373576622 | 0,03624 | 0,25760275 | MT-ND2 pseudogene 5 |
| ENSG00000234912 | SNHG20 | -1,420824619 | 0,373498766 | 0,01916 | 0,18320398 | small nucleolar RNA host gene 20 |
| ENSG00000218475 | ACTG1P18 | -1,42097866 | 0,373458888 | 0,00983 | 0,12683981 | actin gamma 1 pseudogene 18 |
| ENSG00000153823 | PID1 | -1,421293532 | 0,373377389 | 0,04713 | 0,29208796 | phosphotyrosine interaction domain containing 1 |
| ENSG00000239974 | EGFL8 | -1,421690722 | 0,373274608 | 0,13004 | 0,45439235 | EGF like domain multiple 8 |
| ENSG00000280582 | AP003392,6 | -1,423122621 | 0,372904311 | 0,04624 | 0,28930497 | novel transcript, antisense to CCDC84 |
| ENSG00000254428 | AP003392,1 | -1,423122621 | 0,372904311 | 0,04624 | 0,28930497 | novel transcript, antisense to CCDC84 |
| ENSG00000167703 | SLC43A2 | -1,423439449 | 0,372822427 | 0,00938 | 0,12392004 | solute carrier family 43 member 2 |
| ENSG00000269399 | AC008764,6 | -1,423456036 | 0,37281814 | 0,01601 | 0,16581583 | novel transcript |
| ENSG00000228126 | FALEC | -1,423461989 | 0,372816602 | 0,08784 | 0,38352337 | focally amplified long non-coding RNA in epithelial cancer |
| ENSG00000187172 | BAGE2 | -1,423631335 | 0,372772843 | 0,04382 | 0,28236923 | BAGE family member 2 |
| ENSG00000221946 | FXYD7 | -1,423884566 | 0,372707417 | 0,01675 | 0,16990852 | FXYD domain containing ion transport regulator 7 |
| ENSG00000232445 | EMSLR | -1,425292039 | 0,372343986 | 0,01055 | 0,1325438 | E2F1 mRNA stabilizing lncRNA |
| ENSG00000286159 | AL035106,1 | -1,425921351 | 0,372181603 | 0,00602 | 0,09782483 | novel transcript, antisense to PREX1 |
| ENSG00000185361 | TNFAIP8L1 | -1,426355069 | 0,372069731 | 0,01413 | 0,15603075 | TNF alpha induced protein 8 like 1 |
| ENSG00000262772 | LINC01977 | -1,426557407 | 0,372017551 | 0,07619 | 0,36240925 | long intergenic non-protein coding RNA 1977 |
| ENSG00000286214 | AUXG010000! | -1,426724746 | 0,371974404 | 0,00482 | 0,08564971 | COPG2 imprinted transcript 1 |
| ENSG00000196183 | RPS2P4 | -1,426782516 | 0,371959509 | 0,16895 | 0,4950359 | ribosomal protein S2 pseudogene 4 |

| | | | | | | |
|---|---|---|---|---|---|---|
| ENSG00000287676 | AC096642,2 | -1,42686293 | 0,371938777 | 0,08893 | 0,38573292 | novel transcript |
| ENSG00000068079 | IFI35 | -1,427042569 | 0,371892467 | 0,05419 | 0,31168784 | interferon induced protein 35 |
| ENSG00000250328 | MGC32805 | -1,427109175 | 0,371875298 | 0,0037 | 0,07414958 | uncharacterized LOC153163 |
| ENSG00000273492 | AP000229,1 | -1,427914463 | 0,371667782 | 0,01421 | 0,15631436 | novel transcript |
| ENSG00000218416 | AC110619,1 | -1,428676257 | 0,37147158 | 0,00793 | 0,11357366 | novel transcript, antisense to GPC1 |
| ENSG00000185958 | FAM186A | -1,428874922 | 0,37142043 | 0,05691 | 0,31803579 | family with sequence similarity 186 member A |
| ENSG00000125787 | GNRH2 | -1,429161194 | 0,371346737 | 0,06907 | 0,34726337 | gonadotropin releasing hormone 2 |
| ENSG00000171798 | KNDC1 | -1,429550084 | 0,371246651 | 0,00033 | 0,01523174 | kinase non-catalytic C-lobe domain containing 1 |
| ENSG00000188130 | MAPK12 | -1,429552663 | 0,371245987 | 0,00127 | 0,03791398 | mitogen-activated protein kinase 12 |
| ENSG00000200320 | SNORA63 | -1,430039097 | 0,371120835 | 0,00239 | 0,05632389 | small nucleolar RNA, H/ACA box 63 |
| ENSG00000134574 | DDB2 | -1,430097749 | 0,371105748 | 0,01847 | 0,179662 | damage specific DNA binding protein 2 |
| ENSG00000226390 | KCNQ2-AS1 | -1,430108888 | 0,371102883 | 0,26288 | 0,56582568 | KCNQ2 antisense RNA 1 |
| ENSG00000281651 | KCNQ2-AS1 | -1,430108888 | 0,371102883 | 0,26288 | 0,56582568 | KCNQ2 antisense RNA 1 |
| ENSG00000223343 | AC137630,1 | -1,4302302 | 0,371071679 | 0,03988 | 0,27016182 | novel transcript |
| ENSG00000259448 | LINC02352 | -1,430471219 | 0,371009692 | 0,00038 | 0,01701402 | long intergenic non-protein coding RNA 2352 |
| ENSG00000264829 | AC087749,2 | -1,431887625 | 0,370645622 | 0,02195 | 0,19707985 | novel transcript, antisense to SAP30BP |
| ENSG00000239920 | AC104389,5 | -1,433152294 | 0,370320856 | 0,01794 | 0,17668892 | novel transcript |
| ENSG00000282005 | LMNTD2-AS1 | -1,43317951 | 0,37031387 | 0,22798 | 0,54213739 | LMNTD2 antisense RNA 1 |
| ENSG00000224323 | DPRXP1 | -1,43331682 | 0,370278626 | 0,11139 | 0,42416781 | divergent-paired related homeobox pseudogene 1 |
| ENSG00000233791 | LINC01136 | -1,43390398 | 0,370127958 | 0,23642 | 0,54746216 | long intergenic non-protein coding RNA 1136 |
| ENSG00000179698 | WDR97 | -1,4340774 | 0,370083469 | 0,02466 | 0,21034548 | WD repeat domain 97 |
| ENSG00000142513 | ACP4 | -1,43473998 | 0,369913542 | 0,04048 | 0,27198811 | acid phosphatase 4 |
| ENSG00000167984 | NLRC3 | -1,435140656 | 0,369810821 | 0,00714 | 0,1076344 | NLR family CARD domain containing 3 |
| ENSG00000275897 | AC021491,4 | -1,43554811 | 0,369706391 | 0,12739 | 0,45051698 | novel transcript |
| ENSG00000245904 | AC025164,1 | -1,436719893 | 0,369406231 | 0,05011 | 0,30089233 | novel transcript, antisense to BTG1 |
| ENSG00000278605 | TSEN34 | -1,437033819 | 0,369325858 | 0,07629 | 0,36242929 | tRNA splicing endonuclease subunit 34 |
| ENSG00000196811 | CHRNG | -1,437039215 | 0,369324477 | 0,0528 | 0,30846816 | cholinergic receptor nicotinic gamma subunit |
| ENSG00000288597 | AC234782,2 | -1,4371589 | 0,369293839 | 0,0135 | 0,15241195 | novel transcript |
| ENSG00000253477 | AC012213,1 | -1,437253414 | 0,369269647 | 0,02217 | 0,19795874 | novel transcript |
| ENSG00000249170 | AC096713,1 | -1,437285805 | 0,369261356 | 0,14215 | 0,46897057 | solute carrier family 35, member A4 (SLC35A4) pseudogene |
| ENSG00000223804 | AC244669,1 | -1,437439859 | 0,369221928 | 0,02904 | 0,22988377 | Poly [ADP-ribose] polymerase 8 (PARP8) pseudogene |
| ENSG00000243449 | C4orf48 | -1,437585388 | 0,369184685 | 0,01841 | 0,17944701 | chromosome 4 open reading frame 48 |
| ENSG00000265948 | AC015878,2 | -1,437861795 | 0,36911396 | 0,12758 | 0,45080057 | novel transcript |

| | | | | | | |
|---|---|---|---|---|---|---|
| ENSG00000109991 | P2RX3 | -1,43796921 | 0,369086478 | 0,02757 | 0,22376406 | purinergic receptor P2X 3 |
| ENSG00000183049 | CAMK1D | -1,438450772 | 0,3689633 | 0,01099 | 0,1353263 | calcium/calmodulin dependent protein kinase ID |
| ENSG00000089250 | NOS1 | -1,43894282 | 0,368837483 | 0,02333 | 0,20398797 | nitric oxide synthase 1 |
| ENSG00000248441 | LINC01197 | -1,439170472 | 0,368779286 | 0,05002 | 0,30089233 | long intergenic non-protein coding RNA 1197 |
| ENSG00000224861 | YBX1P1 | -1,439192872 | 0,36877356 | 0,10215 | 0,40949179 | Y-box binding protein 1 pseudogene 1 |
| ENSG00000258611 | YBX2P2 | -1,439494395 | 0,368696495 | 0,00037 | 0,01667641 | YBX2 pseudogene 2 |
| ENSG00000241057 | AC004951,3 | -1,439947597 | 0,368580692 | 0,1654 | 0,49167676 | RAS p21 protein activator 4 (RASA4) pseudogene |
| ENSG00000099864 | PALM | -1,440079388 | 0,368547024 | 0,00201 | 0,050435 | paralemmin |
| ENSG00000266714 | MYO15B | -1,440331088 | 0,36848273 | 0,00145 | 0,040729 | myosin XVB |
| ENSG00000141577 | CEP131 | -1,441642117 | 0,368148029 | 0,00139 | 0,03972417 | centrosomal protein 131 |
| ENSG00000160161 | CILP2 | -1,441719564 | 0,368128267 | 0,00119 | 0,03622879 | cartilage intermediate layer protein 2 |
| ENSG00000107281 | NPDC1 | -1,441842895 | 0,368096798 | 0,00072 | 0,02626224 | neural proliferation, differentiation and control 1 |
| ENSG00000215788 | TNFRSF25 | -1,441954008 | 0,368068449 | 0,00135 | 0,03912393 | TNF receptor superfamily member 25 |
| ENSG00000188157 | AGRN | -1,442107868 | 0,368029198 | 0,00814 | 0,11479236 | agrin |
| ENSG00000235204 | AL162724,2 | -1,44226775 | 0,367988414 | 0,09959 | 0,40610985 | novel transcript |
| ENSG00000256827 | AC068790,1 | -1,442462909 | 0,367938638 | 0,04229 | 0,27809282 | spindle and kinetochore associated complex subunit 2 (SKA2) |
| ENSG00000260495 | AC009148,1 | -1,442967302 | 0,367810023 | 0,02648 | 0,21924516 | novel transcript, antisense to GAN |
| ENSG00000099625 | CBARP | -1,443250143 | 0,36773792 | 0,00548 | 0,09208977 | CACN subunit beta associated regulatory protein |
| ENSG00000148204 | CRB2 | -1,443438453 | 0,367689924 | 0,00373 | 0,0744014 | crumbs cell polarity complex component 2 |
| ENSG00000275555 | DLL1 | -1,443697147 | 0,367623998 | 0,55219 | 0,76565817 | delta like canonical Notch ligand 1 |
| ENSG00000166448 | TMEM130 | -1,443712757 | 0,367620021 | 0,01917 | 0,18320398 | transmembrane protein 130 |
| ENSG00000260060 | AC009088,1 | -1,444174673 | 0,367502336 | 0,01947 | 0,18482021 | novel transcript, antisense to PYCARD |
| ENSG00000284227 | DCHS2 | -1,444689543 | 0,367371205 | 9,57E-05 | 0,00580109 | dachsous cadherin-related 2 |
| ENSG00000197410 | DCHS2 | -1,444689543 | 0,367371205 | 9,57E-05 | 0,00580109 | dachsous cadherin-related 2 |
| ENSG00000184545 | DUSP8 | -1,444791664 | 0,367345202 | 0,18924 | 0,51249536 | dual specificity phosphatase 8 |
| ENSG00000130222 | GADD45G | -1,445036255 | 0,367282928 | 0,01318 | 0,15012165 | growth arrest and DNA damage inducible gamma |
| ENSG00000196668 | LINC00173 | -1,445358666 | 0,367200858 | 0,02053 | 0,19002626 | long intergenic non-protein coding RNA 173 |
| ENSG00000286954 | AL033539,2 | -1,445806772 | 0,367086822 | 0,27287 | 0,57325456 | novel transcript |
| ENSG00000136546 | SCN7A | -1,446187159 | 0,366990047 | 0,00086 | 0,02908139 | sodium voltage-gated channel alpha subunit 7 |
| ENSG00000235829 | TBCAP2 | -1,446832869 | 0,366825829 | 0,06104 | 0,32783036 | tubulin folding cofactor A pseudogene 2 |
| ENSG00000142156 | COL6A1 | -1,446938494 | 0,366798973 | 0,02093 | 0,19204439 | collagen type VI alpha 1 chain |
| ENSG00000274251 | AP003419,2 | -1,447158992 | 0,366742917 | 0,09735 | 0,40201398 | novel transcript |

| | | | | | | |
|---|---|---|---|---|---|---|
| ENSG00000141934 | PLPP2 | -1,447216585 | 0,366728277 | 0,09681 | 0,40094776 | phospholipid phosphatase 2 |
| ENSG00000230805 | AL132709,7 | -1,44736485 | 0,36669059 | 0,01093 | 0,13517515 | novel transcript |
| ENSG00000249655 | AC008434,1 | -1,448830925 | 0,366318146 | 0,04564 | 0,2883604 | novel transcript |
| ENSG00000273416 | AL732292,2 | -1,448888707 | 0,366303475 | 0,21135 | 0,53032236 | novel transcript |
| ENSG00000279182 | CR559946,2 | -1,449384743 | 0,366177552 | 0,0191 | 0,18296265 | novel transcript, antisense to DENND6B |
| ENSG00000273831 | LRRC56 | -1,450292178 | 0,365947304 | 0,01257 | 0,1467975 | leucine rich repeat containing 56 |
| ENSG00000266853 | ITM2BP1 | -1,451043252 | 0,36575684 | 0,03871 | 0,26631237 | integral membrane protein 2B pseudogene 1 |
| ENSG00000243753 | HLA-L | -1,451099541 | 0,365742569 | 0,02116 | 0,19303449 | major histocompatibility complex, class I, L (pseudogene) |
| ENSG00000226448 | RPL7AP51 | -1,451534189 | 0,365632397 | 0,08103 | 0,37153397 | ribosomal protein L7a pseudogene 51 |
| ENSG00000256972 | AP000462,3 | -1,451891995 | 0,365541727 | 0,09991 | 0,40647621 | novel transcript |
| ENSG00000276612 | FP565260,2 | -1,451986801 | 0,365517706 | 0,29441 | 0,58857672 | novel protein |
| ENSG00000285649 | AL357079,2 | -1,452090021 | 0,365491555 | 0,0702 | 0,34975886 | novel transcript, antisense to DPH2 and ATP6V0B |
| ENSG00000224799 | AL139397,1 | -1,453096951 | 0,365236549 | 0,03913 | 0,26771146 | novel transcript |
| ENSG00000262633 | AC005670,2 | -1,453999679 | 0,365008084 | 0,6084 | 0,80115227 | novel protein |
| ENSG00000244586 | WNT5A-AS1 | -1,454583268 | 0,364860463 | 0,08176 | 0,3729912 | WNT5A antisense RNA 1 |
| ENSG00000054282 | SDCCAG8 | -1,455414016 | 0,364650426 | 0,0069 | 0,10589898 | SHH signaling and ciliogenesis regulator SDCCAG8 |
| ENSG00000204991 | SPIRE2 | -1,455433977 | 0,364645381 | 0,00784 | 0,11257373 | spire type actin nucleation factor 2 |
| ENSG00000205307 | SAP25 | -1,455596637 | 0,36460427 | 0,04825 | 0,29503004 | Sin3A associated protein 25 |
| ENSG00000214132 | AC094103,1 | -1,4556029 | 0,364602687 | 0,05386 | 0,31116022 | nucleoporin 50kDa (NUP50) pseudogene |
| ENSG00000255537 | AP000708,1 | -1,455793526 | 0,364554515 | 0,03628 | 0,25775574 | novel transcript, antisense to FEZ1 |
| ENSG00000225194 | LINC00092 | -1,456231067 | 0,364443969 | 0,03923 | 0,26801846 | long intergenic non-protein coding RNA 92 |
| ENSG00000269155 | AL009178,2 | -1,456269382 | 0,364434291 | 0,0104 | 0,13142144 | novel transcript |
| ENSG00000161610 | HCRT | -1,457298237 | 0,364174488 | 0,14705 | 0,47471707 | hypocretin neuropeptide precursor |
| ENSG00000272589 | ZSWIM8-AS1 | -1,457799067 | 0,364048087 | 0,14925 | 0,47691371 | ZSWIM8 antisense RNA 1 |
| ENSG00000255092 | AC010768,2 | -1,458743366 | 0,363809882 | 0,20486 | 0,52451442 | novel transcript |
| ENSG00000234338 | AC073349,2 | -1,458819629 | 0,36379065 | 0,00288 | 0,06379203 | endogenous retroviral family W, env(C7), member 2 (ERVWE2) pseudogene |
| ENSG00000255434 | AP001922,6 | -1,459089988 | 0,363722483 | 0,00082 | 0,02842937 | novel transcript |
| ENSG00000156096 | UGT2B4 | -1,459154069 | 0,363706328 | 0,06767 | 0,34407703 | UDP glucuronosyltransferase family 2 member B4 |
| ENSG00000254314 | AC021915,2 | -1,459236451 | 0,36368556 | 0,20543 | 0,52503195 | novel transcript |
| ENSG00000240634 | AC145285,1 | -1,459275294 | 0,363675768 | 0,15534 | 0,48334025 | ribosomal protein S15a (RPS15A) pseudogene |
| ENSG00000221676 | RNU6ATAC | -1,459428918 | 0,363637044 | 0,00667 | 0,10365198 | RNA, U6atac small nuclear (U12-dependent splicing) |
| ENSG00000237824 | RPL23AP33 | -1,459497796 | 0,363619684 | 0,32828 | 0,60971379 | ribosomal protein L23a pseudogene 33 |

| | | | | | | |
|---|---|---|---|---|---|---|
| ENSG00000238259 | PHF5GP | -1,459950814 | 0,363505522 | 0,06994 | 0,34916163 | PHD finger protein 5G pseudogene |
| ENSG00000259544 | AC021739,3 | -1,460829691 | 0,363284145 | 0,03025 | 0,23475421 | novel transcript, antisense to AKAP13 |
| ENSG00000013588 | GPRC5A | -1,461346296 | 0,363154082 | 0,1139 | 0,42900168 | G protein-coupled receptor class C group 5 member A |
| ENSG00000286176 | AC116317,1 | -1,461523248 | 0,363109543 | 0,01967 | 0,18612199 | novel transcript |
| ENSG00000287562 | AL109615,4 | -1,461832747 | 0,363031654 | 0,00837 | 0,11551322 | novel transcript |
| ENSG00000147465 | STAR | -1,46217845 | 0,362944674 | 0,01064 | 0,13305929 | steroidogenic acute regulatory protein |
| ENSG00000259361 | LINC00927 | -1,462808628 | 0,362786172 | 0,24591 | 0,55372423 | long intergenic non-protein coding RNA 927 |
| ENSG00000143994 | ABHD1 | -1,462926056 | 0,362756644 | 0,01072 | 0,1339093 | abhydrolase domain containing 1 |
| ENSG00000230475 | ABHD16A | -1,46314128 | 0,362702532 | 0,15402 | 0,4823167 | abhydrolase domain containing 16A, phospholipase |
| ENSG00000287569 | AL078587,2 | -1,463353967 | 0,362649065 | 0,05894 | 0,3241112 | novel transcript |
| ENSG00000278925 | AC005753,1 | -1,463392233 | 0,362639446 | 0,07809 | 0,36538634 | novel transcript |
| ENSG00000255201 | AC087623,1 | -1,463894218 | 0,362513288 | 0,08242 | 0,37394502 | novel transcript |
| ENSG00000215156 | AC138409,1 | -1,464133686 | 0,36245312 | 0,01107 | 0,1360165 | POM121-like protein 1 pseudogene |
| ENSG00000236283 | AC019197,1 | -1,464438562 | 0,362376533 | 0,00626 | 0,09970758 | novel transcript, antisense to SLC38A11 and SCN3A |
| ENSG00000179832 | MROH1 | -1,465054904 | 0,362221754 | 0,00198 | 0,05000974 | maestro heat like repeat family member 1 |
| ENSG00000274460 | AC092119,2 | -1,465552696 | 0,362096793 | 0,02834 | 0,22751771 | novel transcript |
| ENSG00000286112 | AL441992,2 | -1,466794225 | 0,36178532 | 0,01839 | 0,17938469 | kynurenine aminotransferase 1 |
| ENSG00000167800 | TBX10 | -1,467384357 | 0,361637363 | 0,10061 | 0,40721448 | T-box transcription factor 10 |
| ENSG00000213901 | SLC23A3 | -1,467695221 | 0,361559448 | 0,0667 | 0,34179768 | solute carrier family 23 member 3 |
| ENSG00000228672 | PROB1 | -1,468087381 | 0,36146118 | 0,01034 | 0,13101765 | proline rich basic protein 1 |
| ENSG00000128342 | LIF | -1,46888506 | 0,36126138 | 0,02948 | 0,23092879 | LIF interleukin 6 family cytokine |
| ENSG00000221475 | SNORA11D | -1,470723038 | 0,36080143 | 0,06088 | 0,32750733 | small nucleolar RNA, H/ACA box 11D |
| ENSG00000221705 | SNORA11E | -1,470723038 | 0,36080143 | 0,06088 | 0,32750733 | small nucleolar RNA, H/ACA box 11E |
| ENSG00000288596 | C8orf44 | -1,470942306 | 0,360746598 | 0,01579 | 0,16470677 | chromosome 8 open reading frame 44 |
| ENSG00000157214 | STEAP2 | -1,471029396 | 0,360724821 | 0,00982 | 0,12683981 | STEAP2 metalloreductase |
| ENSG00000259254 | AC022405,1 | -1,47106699 | 0,360715422 | 0,2366 | 0,54760364 | novel transcript, sense intronic to CASC5 |
| ENSG00000246203 | AL353807,3 | -1,471645847 | 0,36057072 | 0,001 | 0,0323881 | novel pseudogene |
| ENSG00000100987 | VSX1 | -1,472366441 | 0,360390668 | 0,11601 | 0,4324056 | visual system homeobox 1 |
| ENSG00000287177 | AL583825,1 | -1,473582704 | 0,360086969 | 0,01463 | 0,15842492 | novel transcript |
| ENSG00000284920 | RPS3AP34 | -1,473613738 | 0,360079223 | 0,0282 | 0,22670839 | RPS3A pseudogene 34 |
| ENSG00000235530 | AC087294,1 | -1,473695645 | 0,360058781 | 0,01892 | 0,18236662 | novel transcript, antisense to TMEM11 |
| ENSG00000261204 | AC004449,1 | -1,473782154 | 0,360037191 | 0,21824 | 0,53571161 | novel transcript |
| ENSG00000248008 | NRAV | -1,474317301 | 0,359903665 | 0,01446 | 0,15737041 | negative regulator of antiviral response |

| | | | | | | |
|---|---|---|---|---|---|---|
| ENSG00000221819 | GAS8-AS1 | -1,475608146 | 0,359581787 | 0,04167 | 0,2764623 | GAS8 antisense RNA 1 |
| ENSG00000261371 | PECAM1 | -1,475640198 | 0,359573798 | 0,01291 | 0,14873257 | platelet and endothelial cell adhesion molecule 1 |
| ENSG00000245685 | FRG1-DT | -1,476115658 | 0,359455315 | 0,07712 | 0,36384125 | FRG1 divergent transcript |
| ENSG00000181418 | DDN | -1,476459936 | 0,359369547 | 0,04852 | 0,29590716 | dendrin |
| ENSG00000248727 | LINC01948 | -1,476571083 | 0,359341862 | 0,01696 | 0,17088104 | long intergenic non-protein coding RNA 1948 |
| ENSG00000163794 | UCN | -1,476910936 | 0,359257222 | 0,06395 | 0,33512602 | urocortin |
| ENSG00000186628 | FSD2 | -1,477362248 | 0,359144855 | 0,01227 | 0,14430263 | fibronectin type III and SPRY domain containing 2 |
| ENSG00000162722 | TRIM58 | -1,477594296 | 0,359087093 | 0,00017 | 0,00936722 | tripartite motif containing 58 |
| ENSG00000226174 | TEX22 | -1,478108428 | 0,358959148 | 0,02934 | 0,23068546 | testis expressed 22 |
| ENSG00000234597 | AC010096,1 | -1,478110875 | 0,35895854 | 0,21615 | 0,53480193 | novel transcript |
| ENSG00000127241 | MASP1 | -1,478150872 | 0,358948588 | 0,00215 | 0,05242478 | mannan binding lectin serine peptidase 1 |
| ENSG00000235669 | CHN2-AS1 | -1,478203933 | 0,358935387 | 0,00376 | 0,07477699 | CHN2 antisense RNA 1 |
| ENSG00000277039 | Metazoa_SRP | -1,478294632 | 0,358912822 | 0,08992 | 0,38777461 | Metazoan signal recognition particle RNA |
| ENSG00000233974 | AC138951,1 | -1,480113925 | 0,358460504 | 0,01425 | 0,15639135 | POM121 membrane glycoprotein (POM121) pseudogene |
| ENSG00000104833 | TUBB4A | -1,480463741 | 0,358373598 | 0,00076 | 0,02696713 | tubulin beta 4A class IVa |
| ENSG00000254984 | FTLP6 | -1,480532196 | 0,358356593 | 0,06154 | 0,32865897 | ferritin light chain pseudogene 6 |
| ENSG00000276761 | CWC25 | -1,481114537 | 0,358211973 | 4,41E-05 | 0,00301003 | CWC25 spliceosome associated protein homolog |
| ENSG00000260220 | CCDC187 | -1,481137258 | 0,358206331 | 0,12348 | 0,44415944 | coiled-coil domain containing 187 |
| ENSG00000236625 | C4B | -1,481292663 | 0,358167748 | 0,11604 | 0,4324276 | complement C4B (Chido blood group) |
| ENSG00000252835 | SCARNA21 | -1,482201408 | 0,357942211 | 0,12091 | 0,43996704 | small Cajal body-specific RNA 21 |
| ENSG00000283448 | GPR1 | -1,482657696 | 0,357829021 | 0,04295 | 0,27922774 | G protein-coupled receptor 1 |
| ENSG00000173083 | HPSE | -1,482940848 | 0,357758798 | 0,04678 | 0,29089023 | heparanase |
| ENSG00000261780 | LINC02582 | -1,483302352 | 0,357669164 | 0,0184 | 0,17940189 | long intergenic non-protein coding RNA 2582 |
| ENSG00000276337 | AC105429,1 | -1,483304206 | 0,357668704 | 0,0036 | 0,07305642 | novel transcript, sense intronic to ZCCHC14 |
| ENSG00000178531 | CTXN1 | -1,483597314 | 0,357596045 | 0,01645 | 0,16797415 | cortexin 1 |
| ENSG00000277848 | MEGF11 | -1,483633272 | 0,357587132 | 0,104 | 0,41241152 | multiple EGF like domains 11 |
| ENSG00000241179 | AC010301,1 | -1,483804725 | 0,357544639 | 0,15883 | 0,48624756 | ribosomal protein L34 (RPL34) pseudogene |
| ENSG00000166664 | CHRFAM7A | -1,483925002 | 0,357514831 | 0,00271 | 0,06114469 | CHRNA7 (exons 5-10) and FAM7A (exons A-E) fusion |
| ENSG00000284748 | AL513220,1 | -1,483935687 | 0,357512184 | 0,09565 | 0,39857837 | novel transcript |
| ENSG00000175352 | NRIP3 | -1,484166297 | 0,357455041 | 0,01933 | 0,18418735 | nuclear receptor interacting protein 3 |
| ENSG00000272734 | ADIRF-AS1 | -1,484588436 | 0,357350463 | 0,00493 | 0,08699137 | ADIRF antisense RNA 1 |
| ENSG00000287509 | AP001033,3 | -1,484769466 | 0,357305626 | 0,00996 | 0,12783227 | novel transcript, sense intronic to TWSG1 |
| ENSG00000103254 | ANTKMT | -1,485011664 | 0,357245647 | 0,01433 | 0,15653473 | adenine nucleotide translocase lysine methyltransferase |

| | | | | | | |
|---|---|---|---|---|---|---|
| ENSG00000267491 | AC100788,1 | -1,485022348 | 0,357243001 | 0,07265 | 0,35556123 | novel transcript, antisense to TIMP2 |
| ENSG00000165023 | DIRAS2 | -1,485305944 | 0,357172783 | 0,02407 | 0,20739525 | DIRAS family GTPase 2 |
| ENSG00000287074 | AC137561,1 | -1,485802268 | 0,357049928 | 0,02121 | 0,19328293 | novel transcript, sense intronic to AEBP2 |
| ENSG00000285493 | NRL | -1,486138657 | 0,356966685 | 0,15151 | 0,47966555 | neural retina leucine zipper |
| ENSG00000154975 | CA10 | -1,486414296 | 0,35689849 | 0,07324 | 0,35655334 | carbonic anhydrase 10 |
| ENSG00000196660 | SLC30A10 | -1,48715014 | 0,356716501 | 0,0207 | 0,19083528 | solute carrier family 30 member 10 |
| ENSG00000261763 | AC110716,2 | -1,487556994 | 0,356615918 | 0,00814 | 0,11479236 | novel transcript |
| ENSG00000247982 | LINC00926 | -1,487961075 | 0,356516048 | 0,00724 | 0,10792958 | long intergenic non-protein coding RNA 926 |
| ENSG00000224437 | PIGUP1 | -1,48802206 | 0,356500978 | 0,05844 | 0,32267557 | phosphatidylinositol glycan anchor biosynthesis class U pseudogene 1 |
| ENSG00000197816 | CCDC180 | -1,488124303 | 0,356475714 | 0,01017 | 0,12964731 | coiled-coil domain containing 180 |
| ENSG00000162641 | AKNAD1 | -1,489038841 | 0,356249812 | 0,03711 | 0,26086671 | AKNA domain containing 1 |
| ENSG00000229492 | FO681548,1 | -1,490010178 | 0,356010037 | 0,00434 | 0,08101303 | Telomeric repeat-binding factor 2-interacting protein 1 (TERF2IP) pseudogene |
| ENSG00000226985 | LINC01203 | -1,491033571 | 0,355757587 | 0,13344 | 0,4594073 | long intergenic non-protein coding RNA 1203 |
| ENSG00000223799 | IL10RB-DT | -1,491561318 | 0,355627472 | 0,02214 | 0,19795874 | IL10RB divergent transcript |
| ENSG00000185386 | MAPK11 | -1,49203913 | 0,35550971 | 6,40E-05 | 0,00417691 | mitogen-activated protein kinase 11 |
| ENSG00000288184 | AC118282,7 | -1,492092556 | 0,355496545 | 0,01364 | 0,15297981 | |
| ENSG00000288249 | AC270106,1 | -1,492092556 | 0,355496545 | 0,01364 | 0,15297981 | |
| ENSG00000233220 | LINC00167 | -1,492102259 | 0,355494154 | 0,04893 | 0,2975639 | long intergenic non-protein coding RNA 167 |
| ENSG00000163380 | LMOD3 | -1,492118393 | 0,355490179 | 0,02424 | 0,20804286 | leiomodin 3 |
| ENSG00000119283 | TRIM67 | -1,492141297 | 0,355484535 | 0,00081 | 0,02810225 | tripartite motif containing 67 |
| ENSG00000178623 | GPR35 | -1,492231521 | 0,355462304 | 0,01829 | 0,17885917 | G protein-coupled receptor 35 |
| ENSG00000136918 | WDR38 | -1,492354187 | 0,355432082 | 0,04404 | 0,28274932 | WD repeat domain 38 |
| ENSG00000226235 | LEMD1-AS1 | -1,493335135 | 0,355190491 | 0,0177 | 0,17522134 | LEMD1 antisense RNA 1 |
| ENSG00000282895 | RINL | -1,493730261 | 0,355093225 | 0,08362 | 0,37578481 | Ras and Rab interactor like |
| ENSG00000059122 | FLYWCH1 | -1,494426026 | 0,354922016 | 0,01268 | 0,14729065 | FLYWCH-type zinc finger 1 |
| ENSG00000224493 | AC093156,1 | -1,495241129 | 0,354721547 | 0,00879 | 0,11878675 | pseudogene similar to part of nucleostemin (NS) |
| ENSG00000179253 | AL162457,1 | -1,495887364 | 0,35456269 | 0,11319 | 0,42776624 | novel transcript, antisense to CDH4 |
| ENSG00000235499 | AC073046,1 | -1,496088734 | 0,354513204 | 0,01088 | 0,13498814 | novel transcript (LOC100132318) |
| ENSG00000076826 | CAMSAP3 | -1,496316476 | 0,354457245 | 0,00369 | 0,0741128 | calmodulin regulated spectrin associated protein family member 3 |
| ENSG00000255105 | PTMAP15 | -1,496975837 | 0,354295283 | 0,15479 | 0,48266733 | prothymosin alpha pseudogene 15 |

| | | | | | | |
|---|---|---|---|---|---|---|
| ENSG00000089558 | KCNH4 | -1,49707178 | 0,354271722 | 0,12364 | 0,44437757 | potassium voltage-gated channel subfamily H member 4 |
| ENSG00000180176 | TH | -1,497548149 | 0,354154763 | 0,00273 | 0,06130397 | tyrosine hydroxylase |
| ENSG00000275216 | AL161431,1 | -1,498278157 | 0,353975605 | 0,01016 | 0,12964731 | novel transcript |
| ENSG00000202354 | RNY3 | -1,499307581 | 0,353723119 | 0,00017 | 0,00913443 | RNA, Ro60-associated Y3 |
| ENSG00000180610 | ZBTB12BP | -1,499391117 | 0,353702638 | 0,01017 | 0,12964731 | zinc finger and BTB domain containing 12B, pseudogene |
| ENSG00000285350 | FO681525,1 | -1,500085038 | 0,353532551 | 0,10238 | 0,40992908 | chymotrypsinogen B (CTRB) pseudogene |
| ENSG00000228740 | GABRG3-AS1 | -1,500127353 | 0,353522182 | 0,031 | 0,23728154 | GABRG3 antisense RNA 1 |
| ENSG00000159648 | TEPP | -1,500599823 | 0,353406426 | 0,02401 | 0,20710234 | testis, prostate and placenta expressed |
| ENSG00000214553 | LRRC37A11P | -1,500739646 | 0,353372176 | 0,08359 | 0,37573324 | leucine rich repeat containing 37 member A11, pseudogene |
| ENSG00000157765 | SLC34A2 | -1,502397501 | 0,352966336 | 0,05984 | 0,32535617 | solute carrier family 34 member 2 |
| ENSG00000271894 | AC007744,1 | -1,502488628 | 0,352944042 | 0,23912 | 0,5491566 | novel transcript |
| ENSG00000049283 | EPN3 | -1,502608019 | 0,352914835 | 0,01399 | 0,15521058 | epsin 3 |
| ENSG00000267016 | AC111170,1 | -1,502654703 | 0,352903415 | 0,01063 | 0,13304235 | novel transcript, antisense to SEPT9 |
| ENSG00000116711 | PLA2G4A | -1,503765651 | 0,352631766 | 0,02566 | 0,21591782 | phospholipase A2 group IVA |
| ENSG00000140986 | RPL3L | -1,504466705 | 0,352460452 | 0,04537 | 0,28776102 | ribosomal protein L3 like |
| ENSG00000225146 | AC073957,1 | -1,504653068 | 0,352414926 | 0,0222 | 0,19806119 | novel transcript |
| ENSG00000253210 | AC040970,1 | -1,505092115 | 0,352307693 | 0,02678 | 0,22093541 | novel transcript |
| ENSG00000180929 | GPR62 | -1,505277092 | 0,352262525 | 0,02233 | 0,19848029 | G protein-coupled receptor 62 |
| ENSG00000287498 | AC007749,2 | -1,505436604 | 0,352223579 | 0,19158 | 0,51435195 | novel transcript |
| ENSG00000249604 | AC096564,1 | -1,50577899 | 0,352139998 | 0,02459 | 0,20986664 | novel transcript |
| ENSG00000120088 | CRHR1 | -1,506214618 | 0,352033684 | 0,20637 | 0,52602996 | corticotropin releasing hormone receptor 1 |
| ENSG00000132329 | RAMP1 | -1,506693958 | 0,351916739 | 0,1346 | 0,46084956 | receptor activity modifying protein 1 |
| ENSG00000256193 | LINC00507 | -1,50679717 | 0,351891563 | 0,01279 | 0,14802902 | long intergenic non-protein coding RNA 507 |
| ENSG00000163623 | NKX6-1 | -1,507218589 | 0,351788789 | 0,06266 | 0,33130676 | NK6 homeobox 1 |
| ENSG00000141469 | SLC14A1 | -1,507950115 | 0,351610458 | 0,08852 | 0,38479442 | solute carrier family 14 member 1 (Kidd blood group) |
| ENSG00000134765 | DSC1 | -1,508102881 | 0,351573228 | 0,15774 | 0,4847949 | desmocollin 1 |
| ENSG00000273270 | AC090114,2 | -1,508160414 | 0,351559208 | 0,01094 | 0,13517515 | novel transcript |
| ENSG00000261864 | AC130462,2 | -1,508201726 | 0,351549141 | 0,09715 | 0,40152057 | novel transcript |
| ENSG00000273274 | ZBTB8B | -1,508330351 | 0,351517799 | 0,00988 | 0,12735196 | zinc finger and BTB domain containing 8B |
| ENSG00000287592 | AC073348,2 | -1,509230512 | 0,351298541 | 0,02628 | 0,21852536 | novel transcript |
| ENSG00000235994 | HGC6,3 | -1,509455437 | 0,351243775 | 0,14806 | 0,47519813 | uncharacterized LOC100128124 |
| ENSG00000215241 | LINC02449 | -1,509463622 | 0,351241783 | 0,01377 | 0,15401791 | long intergenic non-protein coding RNA 2449 |

| | | | | | | |
|---|---|---|---|---|---|---|
| ENSG00000272927 | AC107464,2 | -1,509594813 | 0,351209844 | 0,01059 | 0,13272448 | novel transcript, antisense to PDE6B |
| ENSG00000163632 | C3orf49 | -1,510375816 | 0,351019768 | 0,05755 | 0,32014909 | chromosome 3 open reading frame 49 |
| ENSG00000236213 | AC006369,1 | -1,510445181 | 0,351002891 | 0,05706 | 0,31859189 | novel transcript |
| ENSG00000286443 | AC091178,2 | -1,510477084 | 0,350995129 | 0,01204 | 0,14273718 | novel transcript, antisense to PLXDC1 |
| ENSG00000225614 | ZNF469 | -1,511146807 | 0,35083223 | 2,82E-05 | 0,00204966 | zinc finger protein 469 |
| ENSG00000111331 | OAS3 | -1,511702107 | 0,350697219 | 0,00133 | 0,03886499 | 2'-5'-oligoadenylate synthetase 3 |
| ENSG00000236510 | AC011284,1 | -1,513970975 | 0,350146125 | 0,01497 | 0,16040868 | novel transcript |
| ENSG00000059915 | PSD | -1,514273171 | 0,350072788 | 1,89E-05 | 0,00146726 | pleckstrin and Sec7 domain containing |
| ENSG00000258128 | MKRN9P | -1,515027559 | 0,349889783 | 0,27473 | 0,57463533 | makorin ring finger protein 9, pseudogene |
| ENSG00000159904 | ZNF890P | -1,516119563 | 0,349625044 | 0,17386 | 0,49851138 | zinc finger protein 890, pseudogene |
| ENSG00000234753 | FOXP4-AS1 | -1,516834601 | 0,349451804 | 0,01678 | 0,17000427 | FOXP4 antisense RNA 1 |
| ENSG00000287860 | AC079804,3 | -1,517206842 | 0,349361651 | 0,02587 | 0,21707964 | novel transcript |
| ENSG00000261064 | LINC02256 | -1,517287396 | 0,349342144 | 0,01534 | 0,16232837 | long intergenic non-protein coding RNA 2256 |
| ENSG00000112599 | GUCA1B | -1,517801313 | 0,349217724 | 0,01533 | 0,16232837 | guanylate cyclase activator 1B |
| ENSG00000170542 | SERPINB9 | -1,51794433 | 0,349183107 | 0,02065 | 0,19061908 | serpin family B member 9 |
| ENSG00000254610 | AC004923,1 | -1,518154019 | 0,349132358 | 0,06413 | 0,33569301 | RNA binding protein, fox-1 homolog (C, elegans) 1 (RBFOX1) pseudogene |
| ENSG00000099889 | ARVCF | -1,518533483 | 0,34904054 | 0,00123 | 0,03698156 | ARVCF delta catenin family member |
| ENSG00000241723 | AC074276,1 | -1,518637957 | 0,349015265 | 0,04003 | 0,27049438 | ribosomal protein L15 (RPL15) pseudogene |
| ENSG00000267765 | AC100793,3 | -1,519672895 | 0,348764984 | 0,35401 | 0,62877376 | novel transcript, antisense to CNTNAP1 |
| ENSG00000285664 | AL139317,5 | -1,520056316 | 0,348672306 | 0,02927 | 0,23044452 | novel transcript, antisense to FERMT2 |
| ENSG00000288023 | AL157936,1 | -1,521016622 | 0,348440295 | 0,16206 | 0,48917419 | novel transcript |
| ENSG00000225764 | P3H2-AS1 | -1,521094647 | 0,348421451 | 0,00894 | 0,11991469 | P3H2 antisense RNA 1 |
| ENSG00000188162 | OTOG | -1,521426835 | 0,348341234 | 0,09836 | 0,40379437 | otogelin |
| ENSG00000146216 | TTBK1 | -1,521739982 | 0,348265633 | 0,00274 | 0,06145058 | tau tubulin kinase 1 |
| ENSG00000273901 | AC012313,9 | -1,521917985 | 0,348222666 | 0,05478 | 0,31338925 | novel transcript, antisense to SLC27A5 |
| ENSG00000231560 | CLEC12A-AS1 | -1,521956515 | 0,348213366 | 0,00612 | 0,098625 | CLEC12A antisense RNA 1 |
| ENSG00000205045 | SLFN12L | -1,522674142 | 0,3480402 | 0,06146 | 0,32865897 | schlafen family member 12 like |
| ENSG00000129951 | PLPPR3 | -1,522710766 | 0,348031365 | 0,00569 | 0,09426358 | phospholipid phosphatase related 3 |
| ENSG00000224142 | AMMECR1-IT: | -1,52414968 | 0,347684419 | 0,11637 | 0,43288152 | AMMECR1 intronic transcript 1 |
| ENSG00000286585 | AC104472,4 | -1,524350626 | 0,347635995 | 0,0478 | 0,29355314 | novel transcript |
| ENSG00000185615 | PDIA2 | -1,524434279 | 0,347615838 | 0,01231 | 0,14467583 | protein disulfide isomerase family A member 2 |

| | | | | | | |
|---|---|---|---|---|---|---|
| ENSG00000237118 | CYP2F2P | -1,524516728 | 0,347595973 | 0,07595 | 0,36201732 | cytochrome P450 family 2 subfamily F member 2, pseudogene |
| ENSG00000275695 | UBE2Q2P6 | -1,524760044 | 0,347537354 | 0,08478 | 0,37808881 | ubiquitin conjugating enzyme E2 Q2 pseudogene 6 |
| ENSG00000237179 | LINC01797 | -1,525185723 | 0,347434826 | 0,09383 | 0,39569427 | long intergenic non-protein coding RNA 1797 |
| ENSG00000144908 | ALDH1L1 | -1,525382792 | 0,34738737 | 0,1253 | 0,44702572 | aldehyde dehydrogenase 1 family member L1 |
| ENSG00000241868 | RN7SL434P | -1,525388255 | 0,347386054 | 0,08728 | 0,38244677 | RNA, 7SL, cytoplasmic 434, pseudogene |
| ENSG00000124092 | CTCFL | -1,525417275 | 0,347379067 | 0,01638 | 0,16750744 | CCCTC-binding factor like |
| ENSG00000265458 | AC132938,3 | -1,526401616 | 0,347142133 | 0,01182 | 0,1415704 | novel transcript, antisense NARF |
| ENSG00000162692 | VCAM1 | -1,526405357 | 0,347141233 | 0,0002 | 0,01056153 | vascular cell adhesion molecule 1 |
| ENSG00000159842 | ABR | -1,526840489 | 0,347036548 | 0,01167 | 0,14066917 | ABR activator of RhoGEF and GTPase |
| ENSG00000197320 | AC060834,1 | -1,526996995 | 0,346998902 | 0,07586 | 0,36177044 | C3 and PZP-like, alpha-2-macroglobulin domain containing 8 (CPAMD8) pseudogene |
| ENSG00000235641 | LINC00484 | -1,527321385 | 0,346920889 | 0,13545 | 0,46214288 | long intergenic non-protein coding RNA 484 |
| ENSG00000224310 | LINC01567 | -1,528053317 | 0,346744928 | 0,01938 | 0,1843508 | long intergenic non-protein coding RNA 1567 |
| ENSG00000286290 | AC011593,1 | -1,528355792 | 0,346672237 | 0,15886 | 0,48624756 | novel transcript |
| ENSG00000257017 | HP | -1,528371453 | 0,346668473 | 0,03569 | 0,25626339 | haptoglobin |
| ENSG00000004848 | ARX | -1,528697192 | 0,34659021 | 0,01493 | 0,16037017 | aristaless related homeobox |
| ENSG00000248587 | GDNF-AS1 | -1,528957959 | 0,346527569 | 0,06708 | 0,34303817 | GDNF antisense RNA 1 |
| ENSG00000288642 | CDR1 | -1,529373784 | 0,346427705 | 0,0012 | 0,03627662 | cerebellar degeneration related protein 1 |
| ENSG00000169032 | MAP2K1 | -1,529374724 | 0,346427479 | 1,04E-06 | 0,00011857 | mitogen-activated protein kinase kinase 1 |
| ENSG00000284434 | AC270107,9 | -1,530328328 | 0,34619857 | 0,02443 | 0,2089849 | |
| ENSG00000176956 | LY6H | -1,531521405 | 0,34591239 | 0,11674 | 0,43375876 | lymphocyte antigen 6 family member H |
| ENSG00000230542 | LINC00102 | -1,531606144 | 0,345892073 | 0,04527 | 0,2876168 | long intergenic non-protein coding RNA 102 |
| ENSG00000253206 | AC090155,1 | -1,532898451 | 0,345582376 | 0,06786 | 0,34437708 | novel transcript |
| ENSG00000196600 | SLC22A25 | -1,533106755 | 0,345532482 | 0,18783 | 0,51107563 | solute carrier family 22 member 25 |
| ENSG00000287855 | AL023882,1 | -1,533895526 | 0,345343619 | 0,01548 | 0,16309786 | novel transcript, antisense to LMF1 |
| ENSG00000226669 | AL354977,1 | -1,534940549 | 0,345093559 | 0,03355 | 0,24724664 | novel transcript |
| ENSG00000105251 | SHD | -1,535499059 | 0,344959989 | 0,00467 | 0,08412546 | Src homology 2 domain containing transforming protein D |
| ENSG00000274161 | DLGAP2 | -1,535883275 | 0,344868132 | 0,02296 | 0,20217525 | DLG associated protein 2 |
| ENSG00000178222 | RNF212 | -1,536563055 | 0,344705672 | 0,00997 | 0,12783227 | ring finger protein 212 |
| ENSG00000227436 | FCF1P1 | -1,536814836 | 0,344645519 | 0,04531 | 0,2876168 | FCF1 pseudogene 1 |
| ENSG00000144504 | ANKMY1 | -1,537500988 | 0,344481643 | 0,00932 | 0,12336347 | ankyrin repeat and MYND domain containing 1 |
| ENSG00000167968 | DNASE1L2 | -1,537737226 | 0,34442524 | 0,02603 | 0,21733018 | deoxyribonuclease 1 like 2 |

| | | | | | | |
|---|---|---|---|---|---|---|
| ENSG00000203327 | AC012358,1 | -1,537787131 | 0,344413326 | 0,04572 | 0,28843251 | novel transcript |
| ENSG00000270324 | AC005972,1 | -1,537852701 | 0,344397673 | 0,17416 | 0,49867554 | ribosomal protein L29 (RPL29) pseudogene |
| ENSG00000233586 | AC246785,2 | -1,537909304 | 0,344384161 | 0,04606 | 0,28930497 | novel pseudogene (FLJ39739) |
| ENSG00000162004 | CCDC78 | -1,538050188 | 0,344350532 | 0,01973 | 0,18648554 | coiled-coil domain containing 78 |
| ENSG00000204257 | HLA-DMA | -1,53960049 | 0,343980696 | 0,12899 | 0,45236742 | major histocompatibility complex, class II, DM alpha |
| ENSG00000274461 | AC023282,1 | -1,539644168 | 0,343970282 | 0,01668 | 0,16951872 | novel transcript |
| ENSG00000196872 | CRACDL | -1,540515923 | 0,3437625 | 0,00456 | 0,08326903 | CRACD like |
| ENSG00000184925 | LCN12 | -1,541519302 | 0,343523499 | 0,0863 | 0,3808494 | lipocalin 12 |
| ENSG00000144485 | HES6 | -1,541849496 | 0,343444885 | 0,00411 | 0,07859127 | hes family bHLH transcription factor 6 |
| ENSG00000235597 | LINC01102 | -1,542199693 | 0,343361528 | 0,00112 | 0,03490596 | long intergenic non-protein coding RNA 1102 |
| ENSG00000241790 | ENO1P4 | -1,542838872 | 0,343209437 | 0,13571 | 0,462686 | enolase 1 pseudogene 4 |
| ENSG00000237775 | DDR1-DT | -1,543409342 | 0,343073752 | 0,40415 | 0,66426144 | DDR1 divergent transcript |
| ENSG00000259587 | AC114546,2 | -1,543493455 | 0,343053751 | 0,20485 | 0,52451442 | novel transcript |
| ENSG00000248340 | PIMREGP4 | -1,543494658 | 0,343053465 | 0,17928 | 0,50368273 | PIMREG pseudogene 4 |
| ENSG00000286671 | AC025253,2 | -1,543656101 | 0,343015078 | 0,01146 | 0,13897679 | novel transcript |
| ENSG00000287726 | AC005740,5 | -1,544152114 | 0,342897166 | 0,05019 | 0,30108624 | novel transcript |
| ENSG00000178685 | PARP10 | -1,544321078 | 0,342857009 | 0,01402 | 0,15545576 | poly(ADP-ribose) polymerase family member 10 |
| ENSG00000181392 | SYNE4 | -1,544474357 | 0,342820584 | 0,03497 | 0,252989 | spectrin repeat containing nuclear envelope family member 4 |
| ENSG00000220749 | RPL21P28 | -1,54596081 | 0,342467548 | 0,15774 | 0,4847949 | ribosomal protein L21 pseudogene 28 |
| ENSG00000182600 | SNORC | -1,5464149 | 0,342359773 | 0,00594 | 0,09720154 | secondary ossification center associated regulator of chondrocyte maturation |
| ENSG00000128482 | RNF112 | -1,546752235 | 0,342279731 | 0,01601 | 0,16581583 | ring finger protein 112 |
| ENSG00000226380 | AC016831,1 | -1,547114642 | 0,34219376 | 0,00973 | 0,12625914 | novel transcript |
| ENSG00000130612 | CYP2G1P | -1,547496305 | 0,342103245 | 0,01534 | 0,16232837 | cytochrome P450 family 2 subfamily G member 1, pseudogene |
| ENSG00000260249 | AC007608,3 | -1,547537072 | 0,342093578 | 0,03088 | 0,2370319 | novel transcript, antisense to SNX20 |
| ENSG00000159173 | TNNI1 | -1,547699031 | 0,342055177 | 0,06578 | 0,33961566 | troponin I1, slow skeletal type |
| ENSG00000149635 | OCSTAMP | -1,547703278 | 0,34205417 | 0,03911 | 0,26771146 | osteoclast stimulatory transmembrane protein |
| ENSG00000131152 | AC010531,1 | -1,547787978 | 0,342034088 | 0,23042 | 0,54355692 | novel protein |
| ENSG00000229263 | AC004691,1 | -1,547880501 | 0,342012154 | 0,11206 | 0,42549415 | novel transcript |
| ENSG00000250790 | AC127070,2 | -1,548118357 | 0,341955771 | 0,02523 | 0,21358358 | novel transcript |
| ENSG00000268970 | AC022150,2 | -1,548579811 | 0,341846412 | 0,01135 | 0,13831964 | novel transcript, sense intronic to ZNF83 |

| | | | | | | |
|---|---|---|---|---|---|---|
| ENSG00000106336 | FBXO24 | -1,549071579 | 0,341729908 | 0,02299 | 0,20221357 | F-box protein 24 |
| ENSG00000205116 | TMEM88B | -1,549902146 | 0,341533229 | 0,35408 | 0,62877376 | transmembrane protein 88B |
| ENSG00000267072 | NAGPA-AS1 | -1,550172979 | 0,34146912 | 0,09303 | 0,39400451 | NAGPA antisense RNA 1 |
| ENSG00000250273 | PSMC1P5 | -1,550677755 | 0,341349666 | 0,65763 | 0,82881843 | proteasome 26S subunit, ATPase 1 pseudogene 5 |
| ENSG00000282217 | AC006449,8 | -1,551090412 | 0,341252043 | 0,04555 | 0,28824362 | novel transcript, antisense to SRCIN1 |
| ENSG00000274996 | AC006449,1 | -1,551090412 | 0,341252043 | 0,04555 | 0,28824362 | novel transcript, antisense to SRCIN1 |
| ENSG00000184470 | TXNRD2 | -1,551425888 | 0,341172699 | 0,00017 | 0,00906182 | thioredoxin reductase 2 |
| ENSG00000173404 | INSM1 | -1,55287695 | 0,340829721 | 0,0012 | 0,03627662 | INSM transcriptional repressor 1 |
| ENSG00000228010 | AC073343,2 | -1,552939471 | 0,340814951 | 0,01848 | 0,179662 | novel transcript, antisense to ZNF12 |
| ENSG00000263477 | AC104982,2 | -1,553980961 | 0,340569003 | 0,12941 | 0,45292332 | novel transcript, antisense to SSH2 |
| ENSG00000228224 | NACA4P | -1,554241948 | 0,340507399 | 0,0942 | 0,39648446 | NACA family member 4, pseudogene |
| ENSG00000205018 | AC092384,1 | -1,55468264 | 0,340403402 | 0,04631 | 0,28930497 | novel transcript, antisense to CBFA2T3 |
| ENSG00000134955 | SLC37A2 | -1,554978467 | 0,340333609 | 0,00634 | 0,10046895 | solute carrier family 37 member 2 |
| ENSG00000261024 | GS1-279B7,1 | -1,555097546 | 0,340305519 | 0,02116 | 0,19303449 | microtubule associated protein 1 light chain 3 beta pseudogene |
| ENSG00000145198 | VWA5B2 | -1,555100213 | 0,34030489 | 0,01262 | 0,14717218 | von Willebrand factor A domain containing 5B2 |
| ENSG00000181908 | LINC02724 | -1,555105123 | 0,340303732 | 0,15186 | 0,48031011 | long intergenic non-protein coding RNA 2724 |
| ENSG00000235396 | NOTCH4 | -1,556515968 | 0,339971103 | 0,00153 | 0,0420401 | notch receptor 4 |
| ENSG00000116661 | FBXO2 | -1,55694562 | 0,339869871 | 0,04065 | 0,27263443 | F-box protein 2 |
| ENSG00000135116 | HRK | -1,557193424 | 0,339811498 | 0,03742 | 0,26177644 | harakiri, BCL2 interacting protein |
| ENSG00000230985 | VARS2 | -1,557286378 | 0,339789605 | 0,03092 | 0,237117 | valyl-tRNA synthetase 2, mitochondrial |
| ENSG00000287862 | AC114971,1 | -1,557483384 | 0,339743208 | 0,0342 | 0,24972789 | novel transcript |
| ENSG00000234630 | AC245060,2 | -1,557709737 | 0,339689908 | 0,02698 | 0,22180109 | novel transcript |
| ENSG00000261663 | AC009065,8 | -1,558583501 | 0,339484238 | 0,02926 | 0,23044452 | novel transcript, antisense to DCI |
| ENSG00000204969 | PCDHA2 | -1,559202808 | 0,339338539 | 7,70E-06 | 0,00067131 | protocadherin alpha 2 |
| ENSG00000147804 | SLC39A4 | -1,55923824 | 0,339330205 | 0,07637 | 0,36251343 | solute carrier family 39 member 4 |
| ENSG00000232998 | VPS13A-AS1 | -1,559607436 | 0,339243379 | 0,13049 | 0,45477153 | VPS13A antisense RNA 1 |
| ENSG00000172005 | MAL | -1,560060985 | 0,339136746 | 0,05557 | 0,31503413 | mal, T cell differentiation protein |
| ENSG00000268758 | ADGRE4P | -1,560087546 | 0,339130502 | 0,13375 | 0,459746 | adhesion G protein-coupled receptor E4, pseudogene |
| ENSG00000263934 | SNORD3A | -1,560103917 | 0,339126654 | 0,00801 | 0,11389729 | small nucleolar RNA, C/D box 3A |
| ENSG00000204923 | FBXO48 | -1,56029993 | 0,339080581 | 0,00851 | 0,11637365 | F-box protein 48 |
| ENSG00000232536 | AL365436,2 | -1,560529707 | 0,33902658 | 0,09405 | 0,39609058 | novel transcript |
| ENSG00000107485 | GATA3 | -1,560716551 | 0,338982676 | 0,10117 | 0,40785299 | GATA binding protein 3 |

| | | | | | | |
|---|---|---|---|---|---|---|
| ENSG00000261610 | AP000265,1 | -1,561761769 | 0,338737175 | 0,03852 | 0,26548082 | novel transcript |
| ENSG00000238009 | AL627309,1 | -1,561867885 | 0,338712261 | 0,02958 | 0,23126047 | novel transcript |
| ENSG00000067840 | PDZD4 | -1,562355837 | 0,33859772 | 0,00471 | 0,08459549 | PDZ domain containing 4 |
| ENSG00000220739 | AL513475,2 | -1,562416963 | 0,338583374 | 0,03086 | 0,2370319 | 5'-nucleotidase domain containing 1 (NT5DC1) pseudogene |
| ENSG00000286391 | AL592078,2 | -1,562430465 | 0,338580205 | 0,01057 | 0,1325438 | novel transcript |
| ENSG00000235389 | AL583805,2 | -1,562676058 | 0,338522573 | 0,06458 | 0,33679627 | novel transcript |
| ENSG00000259666 | LINGO1-AS1 | -1,56329775 | 0,338376727 | 0,06624 | 0,3406271 | LINGO1 antisense RNA 1 |
| ENSG00000272933 | AL391121,1 | -1,563487266 | 0,33833228 | 0,01001 | 0,12822568 | novel transcript |
| ENSG00000278718 | GSDMD | -1,563589552 | 0,338308293 | 0,13183 | 0,45708171 | gasdermin D |
| ENSG00000182256 | GABRG3 | -1,563693559 | 0,338283905 | 0,00206 | 0,0511876 | gamma-aminobutyric acid type A receptor subunit gamma3 |
| ENSG00000155158 | TTC39B | -1,564042786 | 0,338202027 | 0,0107 | 0,13371024 | tetratricopeptide repeat domain 39B |
| ENSG00000144460 | NYAP2 | -1,564098273 | 0,33818902 | 0,00777 | 0,11218615 | neuronal tyrosine-phosphorylated phosphoinositide-3-kinase adaptor 2 |
| ENSG00000169962 | TAS1R3 | -1,564156672 | 0,338175331 | 0,01967 | 0,18612199 | taste 1 receptor member 3 |
| ENSG00000239756 | PHF1 | -1,564328888 | 0,338134965 | 0,01458 | 0,15795863 | PHD finger protein 1 |
| ENSG00000170967 | DDI1 | -1,565289517 | 0,33790989 | 0,00147 | 0,04107269 | DNA damage inducible 1 homolog 1 |
| ENSG00000285667 | AC012291,3 | -1,565331221 | 0,337900123 | 0,03589 | 0,25662951 | novel transcript |
| ENSG00000258674 | AC011448,1 | -1,565895075 | 0,337768086 | 0,03296 | 0,24512211 | novel protein |
| ENSG00000254843 | XIAPP2 | -1,566006136 | 0,337742085 | 0,02265 | 0,20073789 | X-linked inhibitor of apoptosis pseudogene 2 |
| ENSG00000197177 | ADGRA1 | -1,566068435 | 0,3377275 | 0,00293 | 0,06450905 | adhesion G protein-coupled receptor A1 |
| ENSG00000198598 | MMP17 | -1,566258817 | 0,337682936 | 0,0013 | 0,0384253 | matrix metallopeptidase 17 |
| ENSG00000281818 | MANSC1 | -1,566553707 | 0,33761392 | 0,00068 | 0,02516175 | MANSC domain containing 1 |
| ENSG00000259554 | AC084882,1 | -1,56724427 | 0,337452356 | 0,12191 | 0,44093327 | novel transcript, intronic to SEMA6D |
| ENSG00000269473 | AC012313,8 | -1,567438729 | 0,337406874 | 0,01267 | 0,14729065 | novel transcript |
| ENSG00000287055 | AL359813,1 | -1,567531737 | 0,337385123 | 0,00462 | 0,08366397 | novel transcript |
| ENSG00000273990 | RNVU1-26 | -1,567811642 | 0,337319671 | 0,00238 | 0,05617483 | RNA, variant U1 small nuclear 26 |
| ENSG00000275291 | RNVU1-26 | -1,567811642 | 0,337319671 | 0,00238 | 0,05617483 | RNA, variant U1 small nuclear 26 |
| ENSG00000225246 | RPS2P1 | -1,568152121 | 0,337240072 | 0,05703 | 0,31850415 | ribosomal protein S2 pseudogene 1 |
| ENSG00000187848 | P2RX2 | -1,568421169 | 0,337177186 | 0,15132 | 0,47956749 | purinergic receptor P2X 2 |
| ENSG00000260588 | AC027702,1 | -1,568688723 | 0,337114661 | 0,0416 | 0,27632896 | novel transcript |
| ENSG00000288349 | AC139272,1 | -1,569003328 | 0,337041155 | 0,00319 | 0,06765368 | POM121 transmembrane nucleoporin C (POM121C) pseudogene |
| ENSG00000144290 | SLC4A10 | -1,569258983 | 0,336981435 | 0,01131 | 0,13803349 | solute carrier family 4 member 10 |

| | | | | | | |
|---|---|---|---|---|---|---|
| ENSG00000259682 | AC091231,1 | -1,569279881 | 0,336976554 | 0,09956 | 0,40609759 | novel transcript, sense intronic to ZNF609 |
| ENSG00000231024 | AC092431,1 | -1,569511662 | 0,33692242 | 0,16206 | 0,48917419 | novel transcript |
| ENSG00000286167 | AC105402,1 | -1,569512025 | 0,336922335 | 0,01874 | 0,18139566 | novel transcript, antisense to KIF5C |
| ENSG00000217702 | AC073263,1 | -1,570265078 | 0,336746515 | 0,07493 | 0,3592137 | novel transcript, antisense to BOLA3 |
| ENSG00000197629 | MPEG1 | -1,570311277 | 0,336735732 | 8,39E-06 | 0,00071762 | macrophage expressed 1 |
| ENSG00000125388 | GRK4 | -1,57046663 | 0,336699473 | 0,00819 | 0,11480072 | G protein-coupled receptor kinase 4 |
| ENSG00000188760 | TMEM198 | -1,571338183 | 0,33649613 | 0,00065 | 0,0244047 | transmembrane protein 198 |
| ENSG00000253431 | SRPK2P | -1,571489038 | 0,336460946 | 0,02914 | 0,23027064 | SRSF protein kinase 2 pseudogene |
| ENSG00000103319 | EEF2K | -1,571520085 | 0,336453705 | 0,03896 | 0,26712012 | eukaryotic elongation factor 2 kinase |
| ENSG00000188290 | HES4 | -1,571592902 | 0,336436724 | 0,01917 | 0,18320398 | hes family bHLH transcription factor 4 |
| ENSG00000228933 | AC107419,1 | -1,571903517 | 0,336364296 | 0,0253 | 0,21399985 | novel transcript |
| ENSG00000260714 | AC133552,1 | -1,571965178 | 0,33634992 | 0,04224 | 0,27788591 | thyroid hormone receptor interactor 11 (TRIP11) pseudogene |
| ENSG00000197134 | ZNF257 | -1,572409396 | 0,336246371 | 0,13577 | 0,462686 | zinc finger protein 257 |
| ENSG00000259994 | AL353796,1 | -1,572491011 | 0,33622735 | 0,02167 | 0,19556064 | novel transcript, overlapping to MTPAP |
| ENSG00000285936 | AL392024,1 | -1,573155784 | 0,336072457 | 0,03081 | 0,23691425 | novel transcript, antisense to DPF3 |
| ENSG00000204071 | TCEAL6 | -1,573164641 | 0,336070394 | 0,1107 | 0,42337179 | transcription elongation factor A like 6 |
| ENSG00000105219 | CCNP | -1,575493904 | 0,335528238 | 0,11111 | 0,42398965 | cyclin P |
| ENSG00000088340 | FER1L4 | -1,575655462 | 0,335490667 | 0,02777 | 0,22462771 | fer-1 like family member 4 (pseudogene) |
| ENSG00000130589 | HELZ2 | -1,575826269 | 0,335450949 | 0,00456 | 0,08326903 | helicase with zinc finger 2 |
| ENSG00000163285 | GABRG1 | -1,575925965 | 0,335427769 | 0,10694 | 0,41675015 | gamma-aminobutyric acid type A receptor subunit gamma1 |
| ENSG00000197530 | MIB2 | -1,576736261 | 0,335239427 | 0,00475 | 0,08501852 | mindbomb E3 ubiquitin protein ligase 2 |
| ENSG00000286305 | AC004893,2 | -1,57692089 | 0,335196528 | 0,0152 | 0,16197635 | novel transcript, antisense to TRRAP |
| ENSG00000179826 | MRGPRX3 | -1,577926719 | 0,334962914 | 0,01317 | 0,15012165 | MAS related GPR family member X3 |
| ENSG00000269405 | NXF2 | -1,579186979 | 0,334670437 | 0,31652 | 0,60239064 | nuclear RNA export factor 2 |
| ENSG00000141965 | FEM1A | -1,579205433 | 0,334666156 | 0,0113 | 0,13796554 | fem-1 homolog A |
| ENSG00000254528 | AP000757,1 | -1,579977318 | 0,334487147 | 0,02978 | 0,23233601 | novel transcript, antisense to FXYD6 |
| ENSG00000288398 | AL109627,1 | -1,579984536 | 0,334485474 | 0,00866 | 0,11761249 | novel transcript |
| ENSG00000263081 | AC211486,2 | -1,580165316 | 0,334443563 | 0,07772 | 0,36480631 | novel transcript |
| ENSG00000259986 | AC103876,1 | -1,580884921 | 0,334276787 | 0,04765 | 0,29344099 | novel transcript, antisense to HDGFRP3 |
| ENSG00000241404 | EGFL8 | -1,580991453 | 0,334252104 | 0,01553 | 0,16340375 | EGF like domain multiple 8 |
| ENSG00000222594 | RN7SKP235 | -1,581005492 | 0,334248851 | 0,13438 | 0,46058584 | RN7SK pseudogene 235 |
| ENSG00000123496 | IL13RA2 | -1,58104292 | 0,33424018 | 0,03616 | 0,25739586 | interleukin 13 receptor subunit alpha 2 |

| | | | | | | |
|---|---|---|---|---|---|---|
| ENSG00000275736 | OSCAR | -1,58118162 | 0,334208048 | 0,10691 | 0,41675015 | osteoclast associated Ig-like receptor |
| ENSG00000224675 | AC009227,1 | -1,581332772 | 0,334173035 | 0,1011 | 0,40780603 | novel transcript |
| ENSG00000205356 | TECPR1 | -1,582235732 | 0,333963946 | 0,00151 | 0,04166341 | tectonin beta-propeller repeat containing 1 |
| ENSG00000286534 | AC021683,6 | -1,582453411 | 0,333913561 | 0,00721 | 0,10777959 | novel transcript |
| ENSG00000230701 | FBXW4P1 | -1,584133636 | 0,333524897 | 0,01025 | 0,130304 | F-box and WD repeat domain containing 4 pseudogene 1 |
| ENSG00000231881 | AL109615,2 | -1,58428133 | 0,333490754 | 0,00562 | 0,09352688 | novel transcript |
| ENSG00000122728 | TAF1L | -1,585052107 | 0,33331263 | 0,07808 | 0,36538634 | TATA-box binding protein associated factor 1 like |
| ENSG00000075043 | KCNQ2 | -1,585403448 | 0,333231468 | 0,00191 | 0,04876303 | potassium voltage-gated channel subfamily Q member 2 |
| ENSG00000186994 | KANK3 | -1,585452421 | 0,333220157 | 0,05417 | 0,31168784 | KN motif and ankyrin repeat domains 3 |
| ENSG00000268170 | AC073342,1 | -1,585769235 | 0,33314699 | 0,01881 | 0,18167964 | novel transcript, antisense to TAS2R40 |
| ENSG00000281371 | INE2 | -1,586010295 | 0,333091329 | 0,24591 | 0,55372423 | inactivation escape 2 |
| ENSG00000134121 | CHL1 | -1,586100907 | 0,333070409 | 0,00073 | 0,02636261 | cell adhesion molecule L1 like |
| ENSG00000186676 | EEF1GP1 | -1,586820993 | 0,332904207 | 0,04949 | 0,29935562 | eukaryotic translation elongation factor 1 gamma pseudogene 1 |
| ENSG00000234222 | LIX1L-AS1 | -1,587414423 | 0,3327673 | 0,01016 | 0,12964731 | LIX1L antisense RNA 1 |
| ENSG00000253007 | SNORA50B | -1,587609179 | 0,332722381 | 0,05104 | 0,30386013 | small nucleolar RNA, H/ACA box 50B |
| ENSG00000116663 | FBXO6 | -1,587790278 | 0,332680618 | 0,00681 | 0,10528058 | F-box protein 6 |
| ENSG00000269680 | AC008760,1 | -1,588065847 | 0,332617079 | 0,17736 | 0,50187931 | novel transcript, antisense to TRIP10 |
| ENSG00000286302 | AC084724,1 | -1,588373494 | 0,332546157 | 0,04319 | 0,28011691 | novel transcript, antisense to NEK1 |
| ENSG00000269993 | KC877982,1 | -1,588651733 | 0,332482029 | 0,02431 | 0,20841902 | novel transcript |
| ENSG00000241882 | AC124893,1 | -1,58883174 | 0,332440547 | 0,09462 | 0,39709067 | novel transcript |
| ENSG00000074855 | ANO8 | -1,590342515 | 0,332092601 | 0,00236 | 0,05588694 | anoctamin 8 |
| ENSG00000125337 | KIF25 | -1,590380297 | 0,332083904 | 0,01771 | 0,17522134 | kinesin family member 25 |
| ENSG00000230068 | CDC42-IT1 | -1,590586736 | 0,332036389 | 0,05108 | 0,30389297 | CDC42 intronic transcript 1 |
| ENSG00000263338 | AC027796,5 | -1,590607718 | 0,33203156 | 0,07313 | 0,35655334 | novel transcript, sense intronic to TRPV1 |
| ENSG00000203446 | SUGCT-AS1 | -1,591330982 | 0,331865145 | 0,03995 | 0,27022885 | SUGCT antisense RNA 1 |
| ENSG00000108819 | PPP1R9B | -1,591422144 | 0,331844175 | 0,00725 | 0,10792958 | protein phosphatase 1 regulatory subunit 9B |
| ENSG00000159674 | SPON2 | -1,591830956 | 0,331750155 | 0,01281 | 0,14802902 | spondin 2 |
| ENSG00000258323 | AC073575,1 | -1,592611698 | 0,331570671 | 0,01288 | 0,14855349 | novel transcript, antisense to NAA25 |
| ENSG00000253326 | AL606534,4 | -1,593280974 | 0,331416888 | 0,0305 | 0,23556309 | novel pseudogene |
| ENSG00000284859 | MIR124-1HG | -1,593453942 | 0,331377156 | 0,00034 | 0,01561634 | MIR124-1 host gene |
| ENSG00000259891 | AC107375,1 | -1,593511739 | 0,331363881 | 0,01408 | 0,15573084 | novel transcript, antisense to CHRAC1 |
| ENSG00000281384 | AC093802,4 | -1,593563044 | 0,331352097 | 0,0671 | 0,34304078 | novel transcript |

| | | | | | | |
|---|---|---|---|---|---|---|
| ENSG00000235770 | LINC00607 | -1,594317686 | 0,33117882 | 0,01825 | 0,17860719 | long intergenic non-protein coding RNA 607 |
| ENSG00000214954 | LRRC69 | -1,59508926 | 0,331001748 | 0,05595 | 0,3160797 | leucine rich repeat containing 69 |
| ENSG00000224819 | AC093843,1 | -1,596548701 | 0,330667073 | 0,08976 | 0,38749519 | novel transcript |
| ENSG00000260409 | AC012414,4 | -1,597064233 | 0,330548934 | 0,00072 | 0,02620587 | novel transcript |
| ENSG00000276185 | AC005045,2 | -1,597122577 | 0,330535567 | 0,4311 | 0,6834044 | |
| ENSG00000288607 | ELOA3B | -1,597409098 | 0,330469928 | 0,5584 | 0,7700003 | elongin A3 family member B |
| ENSG00000281518 | FOXO6 | -1,598283154 | 0,330269774 | 0,04598 | 0,28917353 | forkhead box O6 |
| ENSG00000148411 | NACC2 | -1,598824948 | 0,330145767 | 0,00044 | 0,01864693 | NACC family member 2 |
| ENSG00000124839 | RAB17 | -1,599042395 | 0,33009601 | 0,09093 | 0,39026582 | RAB17, member RAS oncogene family |
| ENSG00000220868 | MRPL35P1 | -1,599374083 | 0,330020127 | 0,07659 | 0,36266919 | mitochondrial ribosomal protein L35 pseudogene 1 |
| ENSG00000267440 | LINC02594 | -1,600898079 | 0,329671693 | 0,08395 | 0,37651068 | long intergenic non-protein coding RNA 2594 |
| ENSG00000271983 | AC023302,1 | -1,602370541 | 0,329335391 | 0,14649 | 0,47433542 | novel transcript |
| ENSG00000048462 | TNFRSF17 | -1,602933652 | 0,32920687 | 0,03882 | 0,26668798 | TNF receptor superfamily member 17 |
| ENSG00000166317 | SYNPO2L | -1,603131128 | 0,329161811 | 0,01693 | 0,17086866 | synaptopodin 2 like |
| ENSG00000182405 | PGBD4 | -1,604242848 | 0,328908262 | 0,01333 | 0,15119996 | piggyBac transposable element derived 4 |
| ENSG00000132297 | HHLA1 | -1,604893684 | 0,328759916 | 0,00273 | 0,06130397 | HERV-H LTR-associating 1 |
| ENSG00000273533 | FAM181A | -1,60497032 | 0,328742453 | 0,00025 | 0,01275216 | family with sequence similarity 181 member A |
| ENSG00000140067 | FAM181A | -1,60497032 | 0,328742453 | 0,00025 | 0,01275216 | family with sequence similarity 181 member A |
| ENSG00000225647 | AC005487,1 | -1,605291589 | 0,328669255 | 0,13092 | 0,45558936 | novel transcript |
| ENSG00000286622 | AC034111,2 | -1,60531909 | 0,32866299 | 0,06453 | 0,33670487 | novel transcript, antisense to MSRA |
| ENSG00000263264 | AC119396,1 | -1,605695757 | 0,328577192 | 0,05139 | 0,30462001 | novel protein |
| ENSG00000242540 | AC010729,2 | -1,606015742 | 0,328504322 | 0,006 | 0,09769652 | novel transcript |
| ENSG00000177679 | SRRM3 | -1,606173891 | 0,328468313 | 0,00601 | 0,09775044 | serine/arginine repetitive matrix 3 |
| ENSG00000262079 | AC007638,1 | -1,606489319 | 0,328396506 | 0,19435 | 0,51562158 | novel transcript |
| ENSG00000233379 | AL139002,1 | -1,607984593 | 0,328056317 | 0,00272 | 0,06121178 | novel transcript |
| ENSG00000219491 | TPT1P8 | -1,608713193 | 0,327890681 | 0,31627 | 0,60226398 | tumor protein, translationally-controlled 1 pseudogene 8 |
| ENSG00000180509 | KCNE1 | -1,609528822 | 0,327705361 | 0,0593 | 0,32445511 | potassium voltage-gated channel subfamily E regulatory subunit 1 |
| ENSG00000279175 | AL033543,1 | -1,609596618 | 0,327689961 | 0,01189 | 0,14211147 | novel transcript, antisense to PARVB |
| ENSG00000282810 | MIR210HG | -1,609676517 | 0,327671814 | 0,0005 | 0,02023861 | MIR210 host gene |
| ENSG00000223734 | LAPTM4A-DT | -1,610102755 | 0,327575019 | 0,17207 | 0,49710979 | LAPTM4A divergent transcript |
| ENSG00000173898 | SPTBN2 | -1,610533452 | 0,32747724 | 0,00337 | 0,07000131 | spectrin beta, non-erythrocytic 2 |
| ENSG00000283057 | AC010653,3 | -1,612294961 | 0,327077639 | 0,2052 | 0,52486901 | parathymosin (PTMS) pseudogene |

| | | | | | | |
|---|---|---|---|---|---|---|
| ENSG00000261821 | AC090826,1 | -1,612496407 | 0,327031972 | 0,44653 | 0,69318358 | novel transcript, antisense to CYP11A1 |
| ENSG00000260228 | AC009119,1 | -1,612507856 | 0,327029377 | 0,18916 | 0,51249536 | novel transcript, antisense to CDH13 |
| ENSG00000237753 | AC079922,2 | -1,612785647 | 0,326966414 | 0,01753 | 0,17446742 | novel transcript (FLJ42351) |
| ENSG00000198483 | ANKRD35 | -1,612807192 | 0,326961531 | 0,00122 | 0,03698156 | ankyrin repeat domain 35 |
| ENSG00000132975 | GPR12 | -1,614503137 | 0,3265774 | 0,37262 | 0,64159479 | G protein-coupled receptor 12 |
| ENSG00000225216 | AC007362,1 | -1,616037955 | 0,326230154 | 0,006 | 0,09769652 | novel transcript |
| ENSG00000174343 | CHRNA9 | -1,61614015 | 0,326207046 | 0,04053 | 0,27217268 | cholinergic receptor nicotinic alpha 9 subunit |
| ENSG00000269425 | AC104521,1 | -1,616709418 | 0,326078355 | 0,12138 | 0,44078088 | novel transcript, antisense to STAP2 |
| ENSG00000243024 | AC012158,1 | -1,617966897 | 0,325794263 | 0,07778 | 0,36490452 | ribosomal protein S11 pseudogene 6 |
| ENSG00000268530 | AC008403,2 | -1,617975243 | 0,325792378 | 0,03346 | 0,24712306 | novel transcript, antisense to KCNJ14 |
| ENSG00000288602 | C8orf44-SGK3 | -1,618241123 | 0,325732342 | 0,05419 | 0,31168784 | C8orf44-SGK3 readthrough |
| ENSG00000278624 | AZU1 | -1,618317577 | 0,325715081 | 0,00487 | 0,08609281 | azurocidin 1 |
| ENSG00000238164 | TNFRSF14-AS: | -1,619174039 | 0,325521776 | 0,10368 | 0,4118325 | TNFRSF14 antisense RNA 1 |
| ENSG00000237328 | RAI1-AS1 | -1,619791575 | 0,325382468 | 0,01302 | 0,14934778 | RAI1 antisense RNA 1 |
| ENSG00000237483 | RPS27AP8 | -1,620410772 | 0,325242846 | 0,099 | 0,40500202 | RPS27A pseudogene 8 |
| ENSG00000162367 | TAL1 | -1,620573391 | 0,325206187 | 0,09042 | 0,3889575 | TAL bHLH transcription factor 1, erythroid differentiation factor |
| ENSG00000251158 | AC131392,1 | -1,620776258 | 0,325160461 | 0,0012 | 0,03627662 | POM121 membrane glycoprotein-like (POM121L1) pseudogene |
| ENSG00000255192 | NANOGP8 | -1,621494648 | 0,324998587 | 0,02818 | 0,22660674 | Nanog homeobox retrogene P8 |
| ENSG00000231720 | AL353747,3 | -1,621632805 | 0,324967466 | 0,08053 | 0,37032254 | novel transcript |
| ENSG00000270441 | LAMB2P1 | -1,621675544 | 0,324957839 | 0,03831 | 0,264906 | laminin subunit beta 2 pseudogene 1 |
| ENSG00000176155 | CCDC57 | -1,622245711 | 0,324829438 | 0,00727 | 0,10822218 | coiled-coil domain containing 57 |
| ENSG00000277383 | AC010331,1 | -1,622904617 | 0,324681116 | 0,03486 | 0,25246122 | novel transcript, antisense to GLTSCR1 |
| ENSG00000275014 | RN7SL166P | -1,622986318 | 0,32466273 | 0,09761 | 0,40268703 | RNA, 7SL, cytoplasmic 166, pseudogene |
| ENSG00000276458 | LENG8 | -1,623829685 | 0,324472995 | 0,025 | 0,21219706 | leukocyte receptor cluster member 8 |
| ENSG00000277303 | EPOP | -1,624085879 | 0,32441538 | 0,2408 | 0,55001001 | elongin BC and polycomb repressive complex 2 associated protein |
| ENSG00000228883 | LY6G5C | -1,624117344 | 0,324408304 | 0,0133 | 0,15097482 | lymphocyte antigen 6 family member G5C |
| ENSG00000228634 | AL136115,1 | -1,624142411 | 0,324402668 | 0,08488 | 0,37825613 | novel transcript |
| ENSG00000231344 | AL020997,1 | -1,62586246 | 0,32401613 | 0,15124 | 0,47956749 | novel pseudogene |
| ENSG00000121446 | RGSL1 | -1,625918175 | 0,324003617 | 0,25699 | 0,56155289 | regulator of G protein signaling like 1 |
| ENSG00000204588 | LINC01123 | -1,626038079 | 0,32397669 | 0,06375 | 0,3347372 | long intergenic non-protein coding RNA 1123 |

| | | | | | | |
|---|---|---|---|---|---|---|
| ENSG00000108309 | RUNDC3A | -1,626236134 | 0,323932217 | 0,00382 | 0,07526433 | RUN domain containing 3A |
| ENSG00000236017 | ASMTL-AS1 | -1,626427752 | 0,323889196 | 0,00225 | 0,05400651 | ASMTL antisense RNA 1 |
| ENSG00000237390 | AL139130,1 | -1,626586356 | 0,32385359 | 0,04222 | 0,27788591 | novel transcript |
| ENSG00000287756 | AL031293,1 | -1,626802586 | 0,323805055 | 0,03926 | 0,26812486 | novel transcript, antisense to KAZN |
| ENSG00000282676 | AC026369,8 | -1,626926606 | 0,323777221 | 0,00471 | 0,08459549 | |
| ENSG00000272068 | AL365181,2 | -1,62715021 | 0,323727042 | 0,06654 | 0,34124715 | novel transcript |
| ENSG00000254793 | FDPSP4 | -1,627272331 | 0,323699641 | 0,0192 | 0,18338313 | farnesyl diphosphate synthase pseudogene 4 |
| ENSG00000188505 | NCCRP1 | -1,627681805 | 0,323607779 | 0,03027 | 0,23475421 | NCCRP1, F-box associated domain containing |
| ENSG00000267795 | SMIM22 | -1,629146919 | 0,32327931 | 0,02825 | 0,22688194 | small integral membrane protein 22 |
| ENSG00000004660 | CAMKK1 | -1,629541645 | 0,323190872 | 0,00575 | 0,09488235 | calcium/calmodulin dependent protein kinase kinase 1 |
| ENSG00000257636 | G2E3-AS1 | -1,629611457 | 0,323175233 | 0,19501 | 0,51588752 | G2E3 antisense RNA 1 |
| ENSG00000248564 | AC079140,1 | -1,629832631 | 0,323125692 | 0,00691 | 0,10597793 | novel pseudogene |
| ENSG00000175467 | SART1 | -1,630853798 | 0,322897058 | 0,00669 | 0,10388673 | spliceosome associated factor 1, recruiter of U4/U6,U5 tri-snRNP |
| ENSG00000254676 | AP000873,4 | -1,631323432 | 0,322791964 | 0,01741 | 0,17391887 | novel transcript, sense overlapping PCF11 |
| ENSG00000242193 | CRYZL2P | -1,631488541 | 0,322755024 | 0,00156 | 0,04237714 | crystallin zeta like 2, pseudogene |
| ENSG00000261766 | AC133550,3 | -1,631775537 | 0,322690825 | 0,04723 | 0,29217708 | novel transcript, antisense to SH2B1 |
| ENSG00000225077 | LINC00337 | -1,633531215 | 0,322298367 | 0,02358 | 0,20492956 | long intergenic non-protein coding RNA 337 |
| ENSG00000286625 | AC113370,1 | -1,633873999 | 0,322221798 | 0,07656 | 0,36266919 | novel transcript |
| ENSG00000142700 | DMRTA2 | -1,633974767 | 0,322199293 | 0,00049 | 0,01999623 | DMRT like family A2 |
| ENSG00000286938 | AC092849,2 | -1,634918477 | 0,321988602 | 0,08916 | 0,38617197 | novel transcript |
| ENSG00000130433 | CACNG6 | -1,634984542 | 0,321973857 | 0,00527 | 0,09044255 | calcium voltage-gated channel auxiliary subunit gamma 6 |
| ENSG00000197744 | PTMAP2 | -1,635057203 | 0,321957641 | 0,05216 | 0,30680362 | prothymosin alpha pseudogene 2 |
| ENSG00000288110 | AC010941,1 | -1,635331544 | 0,321896424 | 0,21767 | 0,53547632 | novel transcript, antisense to CSMD1 |
| ENSG00000282951 | AC008537,4 | -1,638573183 | 0,321173957 | 0,05049 | 0,30230041 | novel transcript, antisense to RAB4B |
| ENSG00000259135 | AL445363,2 | -1,639629026 | 0,32093899 | 0,12764 | 0,45095465 | novel transcript, antisense to SNX6 |
| ENSG00000230319 | TTLL1-AS1 | -1,639715035 | 0,320919857 | 0,01733 | 0,17339933 | TTLL1 antisense RNA 1 |
| ENSG00000229852 | AC019205,1 | -1,640267221 | 0,32079705 | 0,00657 | 0,10284323 | novel transcript, antisense to KHDC1 |
| ENSG00000223626 | LINC01044 | -1,64059074 | 0,32072512 | 0,04357 | 0,28174537 | long intergenic non-protein coding RNA 1044 |
| ENSG00000131095 | GFAP | -1,641025745 | 0,320628429 | 0,00819 | 0,11480072 | glial fibrillary acidic protein |
| ENSG00000178965 | ERICH3 | -1,641344757 | 0,320557539 | 0,01964 | 0,18592829 | glutamate rich 3 |
| ENSG00000254402 | LRRC24 | -1,641838698 | 0,320447807 | 0,01148 | 0,1391122 | leucine rich repeat containing 24 |
| ENSG00000286596 | AC098595,1 | -1,642484742 | 0,320304342 | 0,03939 | 0,26865324 | novel transcript, antisense to PCDH7 |

| | | | | | | |
|---|---|---|---|---|---|---|
| ENSG00000226179 | LINC00685 | -1,642741771 | 0,320247282 | 0,02963 | 0,23163638 | long intergenic non-protein coding RNA 685 |
| ENSG00000225579 | EDNRB-AS1 | -1,643495462 | 0,320080022 | 0,01132 | 0,13803349 | EDNRB antisense RNA 1 |
| ENSG00000265055 | AC145343,1 | -1,643637804 | 0,320048443 | 0,0705 | 0,35057444 | novel transcript |
| ENSG00000271040 | AL390955,2 | -1,644089727 | 0,319948204 | 0,11064 | 0,42326288 | novel transcript |
| ENSG00000254153 | AC103957,2 | -1,644989274 | 0,319748773 | 0,00712 | 0,10752796 | novel transcript |
| ENSG00000285450 | AC270259,1 | -1,644989274 | 0,319748773 | 0,00712 | 0,10752796 | novel transcript |
| ENSG00000256925 | ADGRA1-AS1 | -1,645925822 | 0,31954127 | 0,02296 | 0,20217525 | ADGRA1 antisense RNA 1 |
| ENSG00000112414 | ADGRG6 | -1,646153409 | 0,319490866 | 0,00353 | 0,07205784 | adhesion G protein-coupled receptor G6 |
| ENSG00000282464 | AC137894,1 | -1,646332975 | 0,319451103 | 0,01167 | 0,14066917 | novel transcript |
| ENSG00000229052 | AL449283,1 | -1,646650295 | 0,319380847 | 0,05669 | 0,31746133 | GAJ protein pseudogene |
| ENSG00000239713 | APOBEC3G | -1,646667557 | 0,319377026 | 0,00436 | 0,08120816 | apolipoprotein B mRNA editing enzyme catalytic subunit 3G |
| ENSG00000149115 | TNKS1BP1 | -1,646905419 | 0,319324374 | 0,00612 | 0,098625 | tankyrase 1 binding protein 1 |
| ENSG00000204410 | MSH5 | -1,647654521 | 0,319158611 | 0,00303 | 0,06571644 | mutS homolog 5 |
| ENSG00000261777 | AC012184,3 | -1,647916427 | 0,319100677 | 0,00956 | 0,12500859 | novel transcript, antisense to DDX19B and DDX19A |
| ENSG00000287370 | AL356317,1 | -1,648132554 | 0,319052877 | 0,04396 | 0,28274932 | novel transcript |
| ENSG00000183682 | BMP8A | -1,648185698 | 0,319041124 | 0,00717 | 0,10764638 | bone morphogenetic protein 8a |
| ENSG00000177738 | AC025171,1 | -1,648556736 | 0,318959082 | 0,00608 | 0,09844621 | novel transcript, sense overlapping to ANXA2R |
| ENSG00000267746 | AC104985,1 | -1,649197557 | 0,318817437 | 0,0199 | 0,18715502 | novel transcript, antisense to NOL4 |
| ENSG00000272892 | AL133551,1 | -1,650339773 | 0,318565122 | 0,01673 | 0,16973453 | novel transcript |
| ENSG00000263958 | AC091138,1 | -1,650434685 | 0,318544165 | 0,03718 | 0,26111083 | novel transcript |
| ENSG00000251869 | SCARNA23 | -1,651524521 | 0,318303622 | 0,00135 | 0,03911553 | small Cajal body-specific RNA 23 |
| ENSG00000132164 | SLC6A11 | -1,65153908 | 0,31830041 | 0,0029 | 0,06399503 | solute carrier family 6 member 11 |
| ENSG00000236305 | SLC12A9-AS1 | -1,652841886 | 0,318013103 | 0,14495 | 0,47257223 | SLC12A9 antisense RNA 1 |
| ENSG00000079385 | CEACAM1 | -1,652996845 | 0,317978947 | 0,00815 | 0,11479236 | CEA cell adhesion molecule 1 |
| ENSG00000287518 | AC100778,4 | -1,654685475 | 0,31760698 | 0,05161 | 0,30535791 | novel transcript, antisense to DYM |
| ENSG00000233954 | UQCRHL | -1,655367108 | 0,317456955 | 0,00456 | 0,08326903 | ubiquinol-cytochrome c reductase hinge protein like |
| ENSG00000282164 | PEG13 | -1,656294668 | 0,317252916 | 0,00159 | 0,04261325 | paternally expressed 13 |
| ENSG00000225506 | CYP4A22-AS1 | -1,656875979 | 0,31712511 | 0,01502 | 0,16064277 | CYP4A22 antisense RNA 1 |
| ENSG00000131233 | GJA9 | -1,657005643 | 0,31709661 | 0,02366 | 0,20542049 | gap junction protein alpha 9 |
| ENSG00000183638 | RP1L1 | -1,657361614 | 0,317018379 | 0,01437 | 0,15682082 | RP1 like 1 |
| ENSG00000123080 | CDKN2C | -1,657851199 | 0,316910815 | 0,00012 | 0,00693199 | cyclin dependent kinase inhibitor 2C |
| ENSG00000236989 | AC130710,1 | -1,658009597 | 0,316876023 | 0,16552 | 0,49182669 | novel transcript |

| | | | | | | |
|---|---|---|---|---|---|---|
| ENSG00000227422 | DDR1-DT | -1,658083954 | 0,316859691 | 0,14714 | 0,47471707 | DDR1 divergent transcript |
| ENSG00000225493 | LINC01107 | -1,658859067 | 0,316689498 | 0,00264 | 0,059998 | long intergenic non-protein coding RNA 1107 |
| ENSG00000176595 | KBTBD11 | -1,659249922 | 0,316603712 | 0,02008 | 0,18801095 | kelch repeat and BTB domain containing 11 |
| ENSG00000221817 | PPP3CB-AS1 | -1,659587099 | 0,316529727 | 0,00826 | 0,11480072 | PPP3CB antisense RNA 1 (head to head) |
| ENSG00000267909 | CCDC177 | -1,660655024 | 0,316295509 | 0,00353 | 0,07205784 | coiled-coil domain containing 177 |
| ENSG00000267740 | AC024592,3 | -1,660666115 | 0,316293077 | 0,01818 | 0,17825548 | novel protein |
| ENSG00000162545 | CAMK2N1 | -1,661120514 | 0,316193472 | 2,21E-06 | 0,00022187 | calcium/calmodulin dependent protein kinase II inhibitor 1 |
| ENSG00000267394 | AC004596,1 | -1,662177446 | 0,31596191 | 0,08129 | 0,37209365 | novel transcript, antisense to ATXN7L3 |
| ENSG00000237495 | LY6G5C | -1,662702982 | 0,315846834 | 0,01276 | 0,14789627 | lymphocyte antigen 6 family member G5C |
| ENSG00000181215 | C4orf50 | -1,664147548 | 0,315530736 | 0,00134 | 0,03905892 | chromosome 4 open reading frame 50 |
| ENSG00000277631 | PGM5P3-AS1 | -1,664564769 | 0,315439499 | 0,07163 | 0,35351167 | PGM5P3 antisense RNA 1 |
| ENSG00000228810 | PABPC1P11 | -1,66533408 | 0,315271337 | 0,02956 | 0,23123936 | poly(A) binding protein cytoplasmic 1 pseudogene 11 |
| ENSG00000232815 | DUX4L50 | -1,665631888 | 0,315206264 | 0,00381 | 0,0752235 | double homeobox 4 like 50 (pseudogene) |
| ENSG00000273423 | OR13I1P | -1,665685514 | 0,315194548 | 0,10722 | 0,41734546 | olfactory receptor family 13 subfamily I member 1 pseudogene |
| ENSG00000264940 | SNORD3C | -1,666942453 | 0,314920056 | 0,01098 | 0,1353263 | small nucleolar RNA, C/D box 3C |
| ENSG00000233327 | USP32P2 | -1,667227858 | 0,314857763 | 0,00958 | 0,12507086 | ubiquitin specific peptidase 32 pseudogene 2 |
| ENSG00000253967 | AC022730,4 | -1,667229259 | 0,314857457 | 0,00547 | 0,0920741 | novel transcript |
| ENSG00000231390 | SNX18P8 | -1,667315596 | 0,314838615 | 0,24685 | 0,55453071 | sorting nexin 18 pseudogene 8 |
| ENSG00000283934 | IGSF6 | -1,667570287 | 0,314783039 | 0,01012 | 0,12924041 | immunoglobulin superfamily member 6 |
| ENSG00000240589 | RN7SL258P | -1,667616438 | 0,314772969 | 0,04545 | 0,2880569 | RNA, 7SL, cytoplasmic 258, pseudogene |
| ENSG00000266290 | AC015813,3 | -1,669006815 | 0,314469757 | 0,02322 | 0,20362407 | novel transcript |
| ENSG00000006747 | SCIN | -1,669016744 | 0,314467593 | 0,01204 | 0,14273718 | scinderin |
| ENSG00000269066 | AC020908,3 | -1,669139051 | 0,314440935 | 0,03974 | 0,26952004 | novel transcript, antisense MYO9B |
| ENSG00000206612 | SNORA2A | -1,670890913 | 0,314059342 | 0,00848 | 0,11625431 | small nucleolar RNA, H/ACA box 2A |
| ENSG00000240338 | AC009078,2 | -1,671709811 | 0,313881127 | 0,16449 | 0,49065086 | chymotrypsinogen B (CTRB) pseudogene |
| ENSG00000267665 | AC021683,3 | -1,672191549 | 0,313776334 | 0,02288 | 0,20189774 | novel transcript |
| ENSG00000254001 | AC084706,1 | -1,672941516 | 0,313613264 | 0,08538 | 0,37877491 | novel transcript |
| ENSG00000235768 | BRD7P5 | -1,673146675 | 0,31356867 | 0,01282 | 0,14814468 | bromodomain containing 7 pseudogene 5 |
| ENSG00000158315 | RHBDL2 | -1,67420382 | 0,313338984 | 0,02451 | 0,20943077 | rhomboid like 2 |
| ENSG00000281449 | BCL2L14 | -1,674549754 | 0,31326386 | 0,00482 | 0,08564971 | BCL2 like 14 |
| ENSG00000207118 | SNORD14D | -1,675523585 | 0,313052476 | 0,08071 | 0,37077071 | small nucleolar RNA, C/D box 14D |
| ENSG00000248165 | AC110760,1 | -1,675807844 | 0,3129908 | 0,10084 | 0,40770184 | novel transcript |

| | | | | | | |
|---|---|---|---|---|---|---|
| ENSG00000237638 | LINC02245 | -1,676245747 | 0,312895812 | 0,01706 | 0,17158055 | long intergenic non-protein coding RNA 2245 |
| ENSG00000238290 | AL034417,2 | -1,679217941 | 0,312251858 | 0,09574 | 0,39858189 | novel transcript |
| ENSG00000143851 | PTPN7 | -1,679385386 | 0,312215618 | 0,00096 | 0,03152018 | protein tyrosine phosphatase non-receptor type 7 |
| ENSG00000240373 | SEC62-AS1 | -1,680064369 | 0,312068713 | 0,08938 | 0,38668036 | SEC62 antisense RNA 1 |
| ENSG00000167772 | ANGPTL4 | -1,680559316 | 0,31196167 | 0,03788 | 0,26341956 | angiopoietin like 4 |
| ENSG00000287192 | AL031005,2 | -1,681105384 | 0,311843613 | 0,05403 | 0,31168784 | novel transcript |
| ENSG00000285434 | CRYGD | -1,681521819 | 0,311753612 | 0,00551 | 0,0922625 | crystallin gamma D |
| ENSG00000118231 | CRYGD | -1,681521819 | 0,311753612 | 0,00551 | 0,0922625 | crystallin gamma D |
| ENSG00000222035 | KIAA2012-AS1 | -1,682467706 | 0,311549281 | 0,05671 | 0,31748986 | KIAA2012 antisense RNA 1 |
| ENSG00000250138 | AC139495,2 | -1,682860534 | 0,311464462 | 0,00717 | 0,10764638 | POM121 membrane glycoprotein-like 1 (POM121L1) pseudogene |
| ENSG00000214279 | SCART1 | -1,683428622 | 0,311341841 | 0,0045 | 0,08275926 | scavenger receptor family member expressed on T cells 1 |
| ENSG00000268964 | ERVV-2 | -1,683673507 | 0,311288998 | 0,20367 | 0,52347929 | endogenous retrovirus group V member 2, envelope |
| ENSG00000288634 | AC079741,2 | -1,684574037 | 0,311094752 | 0,47182 | 0,71114958 | novel protein |
| ENSG00000285907 | AL161910,1 | -1,684899318 | 0,311024618 | 0,01063 | 0,13304235 | novel transcript |
| ENSG00000248405 | PRR5-ARHGAI | -1,68506837 | 0,310988175 | 0,27852 | 0,57673072 | PRR5-ARHGAP8 readthrough |
| ENSG00000255236 | AP002992,1 | -1,686345528 | 0,310712992 | 0,0189 | 0,18221517 | novel transcript |
| ENSG00000237017 | AC245052,4 | -1,68666048 | 0,310645168 | 0,07872 | 0,36645437 | novel transcript |
| ENSG00000275192 | LINC02197 | -1,686694678 | 0,310637805 | 0,16567 | 0,49182669 | long intergenic non-protein coding RNA 2197 |
| ENSG00000286483 | AL121762,1 | -1,687487914 | 0,310467054 | 0,01029 | 0,1306844 | novel transcript |
| ENSG00000232342 | AC022540,1 | -1,687540229 | 0,310455796 | 0,04253 | 0,2782038 | novel transcript |
| ENSG00000237224 | AL157832,2 | -1,688102335 | 0,310334859 | 0,07569 | 0,36122552 | novel transcript |
| ENSG00000255234 | AP000446,1 | -1,688210403 | 0,310311614 | 0,00191 | 0,04877274 | novel transcript, antisense to CCDC90B |
| ENSG00000286186 | AC018445,5 | -1,688343395 | 0,31028301 | 0,0247 | 0,21047862 | novel transcript, antisense to NFATC1 |
| ENSG00000185274 | GALNT17 | -1,689045726 | 0,310131995 | 1,02E-08 | 1,85E-06 | polypeptide N-acetylgalactosaminyltransferase 17 |
| ENSG00000234186 | C16orf82 | -1,689937998 | 0,309940245 | 0,01082 | 0,13459222 | chromosome 16 open reading frame 82 |
| ENSG00000270816 | LINC00221 | -1,690547833 | 0,309809259 | 0,13468 | 0,46109507 | long intergenic non-protein coding RNA 221 |
| ENSG00000116096 | SPR | -1,690878141 | 0,309738336 | 0,00717 | 0,10764638 | sepiapterin reductase |
| ENSG00000252634 | RN7SKP219 | -1,691808438 | 0,309538671 | 0,11338 | 0,42803427 | RN7SK pseudogene 219 |
| ENSG00000149646 | CNBD2 | -1,692342777 | 0,309424046 | 0,00716 | 0,10764638 | cyclic nucleotide binding domain containing 2 |
| ENSG00000267480 | AP001542,3 | -1,692366591 | 0,309418939 | 0,04772 | 0,29349263 | novel transcript, antisense IMPA2 |
| ENSG00000101203 | COL20A1 | -1,692446248 | 0,309401855 | 0,01404 | 0,15553198 | collagen type XX alpha 1 chain |
| ENSG00000224194 | AC008278,1 | -1,69390197 | 0,309089817 | 0,06318 | 0,332907 | novel transcript |

| | | | | | | |
|---|---|---|---|---|---|---|
| ENSG00000268001 | CARD8-AS1 | -1,694443372 | 0,308973846 | 0,02609 | 0,21750162 | CARD8 antisense RNA 1 |
| ENSG00000286086 | AC099661,1 | -1,694456509 | 0,308971033 | 0,05723 | 0,31908628 | novel transcript |
| ENSG00000182310 | SPACA6 | -1,694499811 | 0,308961759 | 0,01516 | 0,1618805 | sperm acrosome associated 6 |
| ENSG00000267703 | AC020917,2 | -1,694541225 | 0,30895289 | 0,19224 | 0,51477601 | novel transcript, antisense EPS15L1 |
| ENSG00000278126 | AC139768,1 | -1,695900344 | 0,308661972 | 0,00292 | 0,0643343 | novel transcript |
| ENSG00000278274 | SNORA61 | -1,695906373 | 0,308660682 | 0,01202 | 0,14273718 | small nucleolar RNA, H/ACA box 61 |
| ENSG00000284483 | TMEM256-PL: | -1,695935381 | 0,308654476 | 0,0335 | 0,2471482 | TMEM256-PLSCR3 readthrough (NMD candidate) |
| ENSG00000262481 | TMEM256-PL: | -1,695935381 | 0,308654476 | 0,0335 | 0,2471482 | TMEM256-PLSCR3 readthrough (NMD candidate) |
| ENSG00000267122 | AC004490,1 | -1,696271426 | 0,30858259 | 0,0047 | 0,08443369 | novel transcript, antisense to DOT1L |
| ENSG00000101213 | PTK6 | -1,696888467 | 0,308450638 | 0,00901 | 0,12042329 | protein tyrosine kinase 6 |
| ENSG00000287576 | AC067930,9 | -1,697869832 | 0,308240891 | 0,00878 | 0,11877365 | novel transcript |
| ENSG00000245293 | CYP2U1-AS1 | -1,698880158 | 0,308025104 | 0,01422 | 0,15638223 | CYP2U1 and SGMS2 antisense RNA 1 |
| ENSG00000133256 | PDE6B | -1,698971777 | 0,308005544 | 0,00367 | 0,07405288 | phosphodiesterase 6B |
| ENSG00000078081 | LAMP3 | -1,699056226 | 0,307987515 | 0,06089 | 0,32750733 | lysosomal associated membrane protein 3 |
| ENSG00000286763 | AC012533,1 | -1,69965578 | 0,307859548 | 0,25029 | 0,55661423 | novel transcript |
| ENSG00000249574 | AC226118,1 | -1,700096549 | 0,307765506 | 0,08185 | 0,37301152 | novel transcript |
| ENSG00000116035 | VAX2 | -1,700228012 | 0,307737463 | 0,00041 | 0,01788764 | ventral anterior homeobox 2 |
| ENSG00000259523 | AC022613,2 | -1,702142743 | 0,307329307 | 0,04713 | 0,29208796 | novel transcript, antisense to TJP1 |
| ENSG00000254419 | AL139349,1 | -1,702528387 | 0,307247167 | 0,04942 | 0,29915481 | novel transcript |
| ENSG00000230882 | AC005077,4 | -1,702741595 | 0,307201764 | 0,15753 | 0,48464679 | hypothetical protein LOC285908 (LOC285908) pseudogene |
| ENSG00000223576 | AL355001,1 | -1,702989102 | 0,307149065 | 0,066 | 0,34008244 | novel transcript |
| ENSG00000255002 | LINC02324 | -1,703881946 | 0,306959038 | 0,00383 | 0,07534426 | long intergenic non-protein coding RNA 2324 |
| ENSG00000286288 | AL109809,4 | -1,70402936 | 0,306927675 | 0,00453 | 0,08299378 | novel transcript |
| ENSG00000115756 | HPCAL1 | -1,704690301 | 0,306787094 | 0,01045 | 0,13188659 | hippocalcin like 1 |
| ENSG00000258472 | AC005726,1 | -1,704922034 | 0,306737821 | 0,00514 | 0,08937569 | novel protein |
| ENSG00000276087 | AC008073,3 | -1,706033235 | 0,306501654 | 0,00798 | 0,11389277 | novel transcript |
| ENSG00000259416 | LINC02883 | -1,706336811 | 0,306437166 | 0,00613 | 0,098625 | long intergenic non-protein coding RNA 2883 |
| ENSG00000033122 | LRRC7 | -1,70666437 | 0,306367598 | 0,00155 | 0,04237695 | leucine rich repeat containing 7 |
| ENSG00000155511 | GRIA1 | -1,706740387 | 0,306351456 | 2,56E-06 | 0,00025343 | glutamate ionotropic receptor AMPA type subunit 1 |
| ENSG00000266844 | AC093330,1 | -1,707292103 | 0,306234323 | 0,04563 | 0,2883604 | novel transcript, antisense to MBP |
| ENSG00000287182 | AC131953,2 | -1,707467088 | 0,306197182 | 0,02698 | 0,22180109 | novel transcript, antisense to NSUN7 |
| ENSG00000230537 | AL158071,2 | -1,70811326 | 0,30606007 | 0,03889 | 0,26695487 | novel transcript |
| ENSG00000167208 | SNX20 | -1,708287493 | 0,306023109 | 0,0877 | 0,3833014 | sorting nexin 20 |

| | | | | | | |
|---|---|---|---|---|---|---|
| ENSG00000267457 | AC004223,2 | -1,708615389 | 0,305953564 | 0,09596 | 0,39904722 | novel transcript |
| ENSG00000276786 | BX640515,1 | -1,70923461 | 0,305822274 | 0,06257 | 0,33113206 | novel transcript |
| ENSG00000243373 | RN7SL173P | -1,709360506 | 0,305795587 | 0,23383 | 0,54565913 | RNA, 7SL, cytoplasmic 173, pseudogene |
| ENSG00000104205 | SGK3 | -1,710385862 | 0,305578329 | 0,00815 | 0,11479236 | serum/glucocorticoid regulated kinase family member 3 |
| ENSG00000245479 | LINC01585 | -1,710653621 | 0,30552162 | 0,03804 | 0,26375851 | long intergenic non-protein coding RNA 1585 |
| ENSG00000260448 | LCMT1-AS1 | -1,71074984 | 0,305501244 | 0,01908 | 0,18296265 | LCMT1 antisense RNA 1 |
| ENSG00000129932 | DOHH | -1,71173497 | 0,305292707 | 0,01094 | 0,13517515 | deoxyhypusine hydroxylase |
| ENSG00000234855 | SLIT1-AS1 | -1,711904224 | 0,305256893 | 0,00982 | 0,12683981 | SLIT1 antisense RNA 1 |
| ENSG00000178078 | STAP2 | -1,712122978 | 0,30521061 | 0,01817 | 0,17825548 | signal transducing adaptor family member 2 |
| ENSG00000225761 | AL596247,1 | -1,712150151 | 0,305204862 | 0,17138 | 0,49665377 | novel transcript |
| ENSG00000255804 | OR6J1 | -1,712251131 | 0,3051835 | 0,01211 | 0,14307474 | olfactory receptor family 6 subfamily J member 1 |
| ENSG00000184388 | PABPC1L2B | -1,712651734 | 0,305098769 | 0,23806 | 0,54864017 | poly(A) binding protein cytoplasmic 1 like 2B |
| ENSG00000255182 | AC084125,2 | -1,713386331 | 0,304943458 | 0,00055 | 0,02167479 | novel transcript, antisense to PPP1R16A |
| ENSG00000268231 | AC008743,1 | -1,713427321 | 0,304934794 | 0,04382 | 0,28236923 | novel transcript, sense intronic to LRRC4B |
| ENSG00000177548 | RABEP2 | -1,71350708 | 0,304917936 | 0,0052 | 0,08998673 | rabaptin, RAB GTPase binding effector protein 2 |
| ENSG00000239039 | SNORD13 | -1,713717011 | 0,304873569 | 0,00168 | 0,04447133 | small nucleolar RNA, C/D box 13 |
| ENSG00000254973 | AC105219,3 | -1,715044149 | 0,304593245 | 0,1259 | 0,448127 | novel transcript |
| ENSG00000267395 | DM1-AS | -1,715260335 | 0,304547605 | 0,01618 | 0,16661451 | DM1 locus antisense RNA |
| ENSG00000284664 | AL161756,3 | -1,715337654 | 0,304531284 | 0,03658 | 0,25935019 | novel transcript |
| ENSG00000202252 | SNORD14C | -1,71652476 | 0,304280807 | 0,05702 | 0,31850415 | small nucleolar RNA, C/D box 14C |
| ENSG00000248503 | AL356235,1 | -1,716774076 | 0,304228228 | 0,00969 | 0,12603664 | chromosome 4 open reading frame 46 (C4orf46) pseudogene |
| ENSG00000171462 | DLK2 | -1,7188775 | 0,303784991 | 0,02269 | 0,20099577 | delta like non-canonical Notch ligand 2 |
| ENSG00000285051 | SLC7A14-AS1 | -1,718909944 | 0,30377816 | 0,01166 | 0,14066395 | SLC7A14 antisense RNA 1 |
| ENSG00000108622 | ICAM2 | -1,719704891 | 0,30361082 | 0,00597 | 0,09743057 | intercellular adhesion molecule 2 |
| ENSG00000265393 | AC084125,4 | -1,720066326 | 0,303534766 | 0,05742 | 0,3198382 | novel transcript, antisense to RECQL4 |
| ENSG00000174307 | PHLDA3 | -1,720711815 | 0,303398989 | 0,00014 | 0,00808329 | pleckstrin homology like domain family A member 3 |
| ENSG00000227218 | AL157935,1 | -1,721112296 | 0,30331478 | 0,00531 | 0,09082808 | novel transcript |
| ENSG00000288480 | AC270129,4 | -1,721706154 | 0,303189952 | 0,00354 | 0,07209525 | |
| ENSG00000163840 | DTX3L | -1,721833176 | 0,303163259 | 0,00085 | 0,02896396 | deltex E3 ubiquitin ligase 3L |
| ENSG00000236871 | LINC00106 | -1,721864492 | 0,303156678 | 0,02335 | 0,20398797 | long intergenic non-protein coding RNA 106 |
| ENSG00000213088 | ACKR1 | -1,72295724 | 0,302927143 | 0,00051 | 0,02068096 | atypical chemokine receptor 1 (Duffy blood group) |
| ENSG00000264569 | DCXR-DT | -1,72303147 | 0,302911557 | 0,0661 | 0,34035709 | DCXR divergent transcript |

| | | | | | | |
|---|---|---|---|---|---|---|
| ENSG00000175772 | LINC01106 | -1,723406413 | 0,302832844 | 0,03632 | 0,25797989 | long intergenic non-protein coding RNA 1106 |
| ENSG00000165714 | BORCS5 | -1,723498989 | 0,302813412 | 0,06506 | 0,33804854 | BLOC-1 related complex subunit 5 |
| ENSG00000285137 | MGAT4C | -1,723654602 | 0,302780752 | 0,09809 | 0,40342986 | MGAT4 family member C |
| ENSG00000273218 | AC005776,2 | -1,72676999 | 0,302127625 | 0,01645 | 0,16797415 | novel transcript |
| ENSG00000246379 | AC040168,1 | -1,727271187 | 0,302022684 | 0,00055 | 0,02167479 | novel transcript |
| ENSG00000042062 | RIPOR3 | -1,728006862 | 0,301868712 | 0,00451 | 0,08280499 | RIPOR family member 3 |
| ENSG00000287047 | AL133395,1 | -1,729778806 | 0,301498179 | 0,13672 | 0,46406381 | novel transcript, antisense to SORCS1 |
| ENSG00000219607 | PPP1R3G | -1,730056915 | 0,301440065 | 0,01056 | 0,1325438 | protein phosphatase 1 regulatory subunit 3G |
| ENSG00000287076 | AC092805,1 | -1,730875687 | 0,301269037 | 0,04714 | 0,29209764 | novel transcript |
| ENSG00000282935 | DUX4L34 | -1,730976549 | 0,301247976 | 0,00524 | 0,09023148 | double homeobox 4 like 34 (pseudogene) |
| ENSG00000011332 | DPF1 | -1,732018886 | 0,301030405 | 6,30E-05 | 0,00412036 | double PHD fingers 1 |
| ENSG00000260646 | AL031705,1 | -1,732109718 | 0,301011452 | 0,11329 | 0,42794837 | novel transcript |
| ENSG00000246308 | AC116535,1 | -1,732489461 | 0,300932231 | 0,0051 | 0,08907797 | novel transcript, antisense to ZBED5 |
| ENSG00000268401 | AC007785,3 | -1,733657261 | 0,300688738 | 0,01198 | 0,14258103 | novel transcript |
| ENSG00000205794 | AC098591,2 | -1,734835287 | 0,300443312 | 0,10197 | 0,40933045 | acyl-CoA thioesterase 7 (ACOT7) pseudogene |
| ENSG00000227695 | DNMBP-AS1 | -1,735111773 | 0,300385739 | 0,03725 | 0,26113308 | DNMBP antisense RNA 1 |
| ENSG00000209702 | SNORD41 | -1,736178891 | 0,300163635 | 0,02732 | 0,22320635 | small nucleolar RNA, C/D box 41 |
| ENSG00000139344 | AMDHD1 | -1,736850019 | 0,300024034 | 0,00694 | 0,10609569 | amidohydrolase domain containing 1 |
| ENSG00000276027 | RNU12 | -1,737027248 | 0,29998718 | 0,00538 | 0,09142613 | RNA, U12 small nuclear |
| ENSG00000182327 | GLTPD2 | -1,738584071 | 0,299663636 | 0,12701 | 0,44992777 | glycolipid transfer protein domain containing 2 |
| ENSG00000180875 | GREM2 | -1,738689099 | 0,299641821 | 2,32E-07 | 3,09E-05 | gremlin 2, DAN family BMP antagonist |
| ENSG00000261068 | AL512274,1 | -1,738732835 | 0,299632738 | 0,0518 | 0,30602123 | novel transcript |
| ENSG00000287872 | AC026956,2 | -1,739017555 | 0,29957361 | 0,00461 | 0,08360185 | novel transcript |
| ENSG00000231439 | WASIR2 | -1,740646988 | 0,299235452 | 0,04626 | 0,28930497 | WASH and IL9R antisense RNA 2 |
| ENSG00000226287 | TMEM191A | -1,740704301 | 0,299223564 | 0,01992 | 0,18715502 | transmembrane protein 191A (pseudogene) |
| ENSG00000177989 | ODF3B | -1,740756192 | 0,299212802 | 0,01694 | 0,17086866 | outer dense fiber of sperm tails 3B |
| ENSG00000226706 | AL161452,1 | -1,741863172 | 0,298983304 | 0,12549 | 0,44730555 | novel transcript |
| ENSG00000198576 | ARC | -1,741984701 | 0,29895812 | 0,00048 | 0,01978176 | activity regulated cytoskeleton associated protein |
| ENSG00000260293 | AC106820,4 | -1,742160283 | 0,298921737 | 0,00073 | 0,02636261 | novel transcript, intronic to TBC1D24 |
| ENSG00000267557 | AC008474,1 | -1,742330611 | 0,298886448 | 0,23547 | 0,54683669 | novel transcript, antisense to ANKRD27 |
| ENSG00000255097 | AP006259,1 | -1,742346134 | 0,298883232 | 0,04346 | 0,28117721 | novel transcript |
| ENSG00000207392 | SNORA20 | -1,744254612 | 0,298488114 | 0,10027 | 0,4067769 | small nucleolar RNA, H/ACA box 20 |
| ENSG00000214031 | CLDN7P1 | -1,744456368 | 0,298446374 | 0,16907 | 0,49515165 | CLDN7 pseudogene 1 |

| | | | | | | |
|---|---|---|---|---|---|---|
| ENSG00000266775 | AC011840,4 | -1,744463632 | 0,298444872 | 0,07263 | 0,35556123 | novel transcript, antisense to TBC1D29 |
| ENSG00000232871 | SEC1P | -1,746521955 | 0,298019378 | 0,03616 | 0,25739586 | secretory blood group 1, pseudogene |
| ENSG00000286932 | AC013716,1 | -1,747338301 | 0,297850792 | 0,06223 | 0,33046875 | novel transcript, sense intronic to LRRTM4 |
| ENSG00000240720 | LRRD1 | -1,747531139 | 0,297810982 | 0,00434 | 0,08101303 | leucine rich repeats and death domain containing 1 |
| ENSG00000171094 | ALK | -1,747711688 | 0,297773714 | 0,00015 | 0,00826026 | ALK receptor tyrosine kinase |
| ENSG00000286677 | AP006545,3 | -1,748179253 | 0,297677224 | 0,01536 | 0,16236395 | novel transcript |
| ENSG00000261467 | AC099398,1 | -1,749459063 | 0,297413273 | 0,0106 | 0,13274123 | novel transcript |
| ENSG00000271350 | AC103792,1 | -1,749599707 | 0,29738428 | 0,1064 | 0,41609301 | arsA arsenite transporter, ATP-binding, homolog 1 (bacterial), pseudogene |
| ENSG00000276359 | Metazoa_SRP | -1,749711059 | 0,297361328 | 0,146 | 0,47387591 | Metazoan signal recognition particle RNA |
| ENSG00000221963 | APOL6 | -1,749821036 | 0,297338661 | 0,01145 | 0,13897679 | apolipoprotein L6 |
| ENSG00000261714 | AC105137,1 | -1,751378335 | 0,297017876 | 0,07159 | 0,35351167 | high mobility group nucleosomal binding domain 2 (HMGN2) pseudogene |
| ENSG00000228590 | MIR4432HG | -1,751667287 | 0,296958393 | 0,15719 | 0,48443207 | MIR4432 host gene |
| ENSG00000152969 | JAKMIP1 | -1,752235068 | 0,296841546 | 0,0013 | 0,0384253 | janus kinase and microtubule interacting protein 1 |
| ENSG00000232229 | LINC00865 | -1,75453574 | 0,296368549 | 0,0001 | 0,00605632 | long intergenic non-protein coding RNA 865 |
| ENSG00000260834 | AC009055,2 | -1,756232826 | 0,296020127 | 0,00517 | 0,08972908 | novel transcript |
| ENSG00000008735 | MAPK8IP2 | -1,756678096 | 0,295928778 | 0,0011 | 0,03448302 | mitogen-activated protein kinase 8 interacting protein 2 |
| ENSG00000178342 | KCNG2 | -1,756797737 | 0,295904238 | 0,05929 | 0,32445511 | potassium voltage-gated channel modifier subfamily G member 2 |
| ENSG00000257298 | AC008147,2 | -1,756908827 | 0,295881454 | 0,00944 | 0,12431453 | novel transcript, sense intronic to LIMA1 |
| ENSG00000185101 | ANO9 | -1,756937727 | 0,295875526 | 0,04853 | 0,29590716 | anoctamin 9 |
| ENSG00000214855 | APOC1P1 | -1,757675796 | 0,295724198 | 0,08424 | 0,37695424 | apolipoprotein C1 pseudogene 1 |
| ENSG00000105971 | CAV2 | -1,757906674 | 0,295676876 | 0,01427 | 0,15652222 | caveolin 2 |
| ENSG00000260081 | AF274858,1 | -1,7585763 | 0,29553967 | 0,00954 | 0,1249879 | Morf4 family associated protein 1 (MRFAP1) pseudogene |
| ENSG00000251372 | LINC00499 | -1,759750292 | 0,295299273 | 0,0304 | 0,23530529 | long intergenic non-protein coding RNA 499 |
| ENSG00000130997 | POLN | -1,760325544 | 0,29518155 | 0,00715 | 0,1076344 | DNA polymerase nu |
| ENSG00000178715 | AL450998,1 | -1,760628337 | 0,295119604 | 0,04271 | 0,27853965 | ribosomal protein S16 (RPS16) pseudogene |
| ENSG00000260899 | AC106886,2 | -1,763331484 | 0,294567162 | 0,09983 | 0,40626368 | novel transcript |
| ENSG00000164303 | ENPP6 | -1,763487713 | 0,294535266 | 0,06028 | 0,32601555 | ectonucleotide pyrophosphatase/phosphodiesterase 6 |
| ENSG00000254966 | AC103974,1 | -1,763843767 | 0,294462584 | 0,0221 | 0,19768196 | novel transcript |
| ENSG00000237595 | LINC01275 | -1,764107537 | 0,294408752 | 0,1675 | 0,49362211 | long intergenic non-protein coding RNA 1275 |
| ENSG00000243404 | RPL35AP32 | -1,764600143 | 0,294308243 | 0,06419 | 0,33569685 | ribosomal protein L35a pseudogene 32 |

| | | | | | | |
|---|---|---|---|---|---|---|
| ENSG00000260979 | AC022167,3 | -1,764727547 | 0,294282254 | 0,0951 | 0,39772599 | novel transcript |
| ENSG00000203335 | AC006019,1 | -1,765004656 | 0,294225735 | 0,05887 | 0,3241112 | novel transcript |
| ENSG00000011052 | NME1-NME2 | -1,765512815 | 0,294122118 | 0,02817 | 0,22660674 | NME1-NME2 readthrough |
| ENSG00000206140 | TMEM191C | -1,766877439 | 0,293844044 | 0,00152 | 0,04181037 | transmembrane protein 191C |
| ENSG00000273264 | AL360219,1 | -1,766999657 | 0,293819152 | 0,06702 | 0,34281679 | novel transcript |
| ENSG00000281367 | AC044810,2 | -1,767489425 | 0,293719423 | 0,23347 | 0,54563244 | novel transcript, sense overlapping OR5P2, OR5P3 and OR5P1P |
| ENSG00000286349 | AC108471,3 | -1,767724828 | 0,293671501 | 0,01195 | 0,14252619 | novel transcript, antisense to SMIM14 |
| ENSG00000126461 | SCAF1 | -1,767783104 | 0,293659639 | 0,00749 | 0,11001711 | SR-related CTD associated factor 1 |
| ENSG00000137204 | SLC22A7 | -1,768661808 | 0,293480833 | 0,01713 | 0,17198199 | solute carrier family 22 member 7 |
| ENSG00000088899 | LZTS3 | -1,769759892 | 0,29325754 | 0,00368 | 0,07408435 | leucine zipper tumor suppressor family member 3 |
| ENSG00000201098 | RNY1 | -1,769966153 | 0,293215616 | 0,0004 | 0,01763971 | RNA, Ro60-associated Y1 |
| ENSG00000169683 | LRRC45 | -1,771218611 | 0,292961175 | 0,00069 | 0,02549506 | leucine rich repeat containing 45 |
| ENSG00000248409 | AP000344,2 | -1,771532264 | 0,29289749 | 0,01931 | 0,18404447 | coiled-coil domain containing 188 (CCDC188) pseudogene |
| ENSG00000118194 | TNNT2 | -1,772084481 | 0,2927854 | 0,07386 | 0,35715669 | troponin T2, cardiac type |
| ENSG00000234787 | LINC00458 | -1,772247495 | 0,292752319 | 0,04767 | 0,29344099 | long intergenic non-protein coding RNA 458 |
| ENSG00000204248 | COL11A2 | -1,774180105 | 0,292360416 | 0,3811 | 0,64852799 | collagen type XI alpha 2 chain |
| ENSG00000285367 | AC087564,1 | -1,774420027 | 0,2923118 | 0,02346 | 0,20434232 | novel transcript |
| ENSG00000202392 | RN7SKP292 | -1,774929009 | 0,292208691 | 0,1557 | 0,48346026 | RN7SK pseudogene 292 |
| ENSG00000160460 | SPTBN4 | -1,775083247 | 0,292177452 | 0,01028 | 0,13057158 | spectrin beta, non-erythrocytic 4 |
| ENSG00000228903 | RASA4CP | -1,775257814 | 0,292142101 | 0,00732 | 0,10869257 | RAS p21 protein activator 4C, pseudogene |
| ENSG00000179046 | TRIML2 | -1,775683307 | 0,292055952 | 0,09266 | 0,39335125 | tripartite motif family like 2 |
| ENSG00000285892 | AC006043,1 | -1,776011325 | 0,291989557 | 0,04029 | 0,27141252 | novel transcript |
| ENSG00000242559 | RN7SL144P | -1,776384821 | 0,291913974 | 0,18521 | 0,50876814 | RNA, 7SL, cytoplasmic 144, pseudogene |
| ENSG00000225108 | ZBTB45P1 | -1,777388232 | 0,291711015 | 0,17773 | 0,50205238 | zinc finger and BTB domain containing 45 pseudogene 1 |
| ENSG00000285636 | AL450163,1 | -1,777816779 | 0,291624376 | 0,00736 | 0,10888451 | Novel transcript, antisense to NOS1AP |
| ENSG00000273155 | AC092587,1 | -1,777946331 | 0,29159819 | 0,23195 | 0,54469907 | novel LIPT1-MRPL30 readthrough |
| ENSG00000281998 | LHX1-DT | -1,778224769 | 0,291541917 | 0,02176 | 0,19602491 | LHX1 divergent transcript |
| ENSG00000269194 | AC006942,1 | -1,778351456 | 0,291516317 | 0,019 | 0,18289762 | novel transcript, antisense to FUZ |
| ENSG00000286025 | FAM246C | -1,778598966 | 0,291466309 | 0,03752 | 0,26208613 | family with sequence similarity 246 member C (gene/pseudogene) |
| ENSG00000222005 | LINC01118 | -1,778765075 | 0,291432752 | 0,06422 | 0,33576796 | long intergenic non-protein coding RNA 1118 |
| ENSG00000173826 | KCNH6 | -1,77964035 | 0,291255995 | 0,20852 | 0,52778377 | potassium voltage-gated channel subfamily H member 6 |

| | | | | | | |
|---|---|---|---|---|---|---|
| ENSG00000263603 | AC127024,2 | -1,780087489 | 0,291165739 | 0,01469 | 0,15868223 | novel transcript |
| ENSG00000286940 | MICB-DT | -1,780255108 | 0,291131912 | 0,03802 | 0,26375851 | MICB divergent transcript |
| ENSG00000212607 | SNORA3B | -1,780466609 | 0,291089235 | 0,06439 | 0,33624556 | small nucleolar RNA, H/ACA box 3B |
| ENSG00000288425 | AC011509,6 | -1,780777319 | 0,29102655 | 0,23444 | 0,5462109 | novel transcript |
| ENSG00000265206 | AC004687,1 | -1,780964291 | 0,290988836 | 0,02063 | 0,19052318 | novel transcript, MIR142 host |
| ENSG00000188996 | HUS1B | -1,781647931 | 0,29085098 | 0,01587 | 0,16502318 | HUS1 checkpoint clamp component B |
| ENSG00000180539 | C9orf139 | -1,781756499 | 0,290829093 | 0,00406 | 0,07819491 | chromosome 9 open reading frame 139 |
| ENSG00000180846 | CSNK1G2-AS1 | -1,781828089 | 0,290814662 | 0,05591 | 0,31600461 | CSNK1G2 antisense RNA 1 |
| ENSG00000201403 | SNORD14B | -1,781974067 | 0,290785237 | 0,09834 | 0,40379437 | small nucleolar RNA, C/D box 14B |
| ENSG00000285417 | BX571818,1 | -1,783238211 | 0,290530552 | 0,01297 | 0,14899372 | novel pseudogene |
| ENSG00000230828 | AL583843,1 | -1,783802323 | 0,290416973 | 0,05272 | 0,30831416 | novel pseudogene |
| ENSG00000175718 | RBMXL3 | -1,784562603 | 0,290263968 | 0,00287 | 0,06351531 | RBMX like 3 |
| ENSG00000258150 | AC133555,3 | -1,784593312 | 0,290257789 | 0,10497 | 0,41375163 | coiled-coil domain containing 101 (CCDC101) pseudogene |
| ENSG00000268744 | AC008758,4 | -1,784816022 | 0,290212985 | 0,0141 | 0,15585442 | novel transcript |
| ENSG00000167094 | TTC16 | -1,786521572 | 0,289870099 | 0,02421 | 0,20804286 | tetratricopeptide repeat domain 16 |
| ENSG00000246145 | RRS1-AS1 | -1,786705306 | 0,289833185 | 0,0095 | 0,12474512 | RRS1 antisense RNA 1 (head to head) |
| ENSG00000206633 | SNORA80B | -1,787787376 | 0,289615882 | 0,05061 | 0,30267763 | small nucleolar RNA, H/ACA box 80B |
| ENSG00000228323 | MYADM-AS1 | -1,78802597 | 0,289567989 | 0,22178 | 0,53818364 | MYADM antisense RNA 1 |
| ENSG00000058453 | CROCC | -1,789348615 | 0,289302638 | 0,0031 | 0,06637856 | ciliary rootlet coiled-coil, rootletin |
| ENSG00000215277 | RNF212B | -1,789362607 | 0,289299833 | 0,14637 | 0,47433542 | ring finger protein 212B |
| ENSG00000243175 | RPSAP36 | -1,78946999 | 0,2892783 | 0,02277 | 0,20142881 | ribosomal protein SA pseudogene 36 |
| ENSG00000274918 | AC116447,1 | -1,789603085 | 0,289251614 | 0,02338 | 0,2040793 | novel transcript |
| ENSG00000282905 | NFKBIB | -1,790008649 | 0,289170312 | 0,00283 | 0,06289122 | NFKB inhibitor beta |
| ENSG00000287665 | AC092428,1 | -1,790323776 | 0,289107156 | 0,01102 | 0,13549982 | novel transcript, sense intronic to FOXK1 |
| ENSG00000273154 | AL121845,3 | -1,790827522 | 0,289006226 | 0,01079 | 0,13436765 | novel protein, ZGPAT-LIME1 readthrough |
| ENSG00000157005 | SST | -1,790891537 | 0,288993403 | 0,04923 | 0,29853148 | somatostatin |
| ENSG00000225096 | AL445250,1 | -1,792125319 | 0,288746363 | 0,03504 | 0,25324761 | novel transcript |
| ENSG00000254648 | AP000911,1 | -1,792402632 | 0,288690866 | 0,05491 | 0,31350641 | novel transcript |
| ENSG00000214248 | AC010336,1 | -1,793359982 | 0,288499359 | 0,00353 | 0,07209525 | novel transcript, antisense to LRRC8E |
| ENSG00000232520 | GCSIR | -1,793711354 | 0,288429103 | 0,03831 | 0,264906 | GPR55 cis regulatory suppressor of immune response RNA |
| ENSG00000284923 | SCRT1 | -1,794200254 | 0,288331376 | 0,00011 | 0,00631075 | scratch family transcriptional repressor 1 |
| ENSG00000261678 | SCRT1 | -1,794200254 | 0,288331376 | 0,00011 | 0,00631075 | scratch family transcriptional repressor 1 |
| ENSG00000213216 | AC007066,1 | -1,795032724 | 0,28816505 | 0,00695 | 0,10609569 | ribosomal protein S25 (RPS25) pseudogene |

| | | | | | | |
|---|---|---|---|---|---|---|
| ENSG00000150656 | CNDP1 | -1,795955296 | 0,287980834 | 0,03233 | 0,24253636 | carnosine dipeptidase 1 |
| ENSG00000282075 | AC093614,1 | -1,796094019 | 0,287953144 | 0,18117 | 0,50470865 | novel transcript |
| ENSG00000267980 | AC007292,1 | -1,796735416 | 0,287825154 | 0,00946 | 0,12436645 | novel transcript, antisense to SH3GL1 |
| ENSG00000203896 | LIME1 | -1,798043241 | 0,287564354 | 0,00863 | 0,11739798 | Lck interacting transmembrane adaptor 1 |
| ENSG00000261807 | LINC02141 | -1,798105955 | 0,287551854 | 0,00856 | 0,1168076 | long intergenic non-protein coding RNA 2141 |
| ENSG00000198286 | CARD11 | -1,800107613 | 0,287153169 | 0,00077 | 0,02713269 | caspase recruitment domain family member 11 |
| ENSG00000230991 | AC092625,1 | -1,802090833 | 0,286758701 | 0,06167 | 0,32870444 | novel transcript |
| ENSG00000148826 | NKX6-2 | -1,802117422 | 0,286753416 | 0,00304 | 0,06583598 | NK6 homeobox 2 |
| ENSG00000229057 | RPS3AP54 | -1,804391437 | 0,286301784 | 0,04005 | 0,27054267 | RPS3A pseudogene 54 |
| ENSG00000224165 | DNAJC27-AS1 | -1,804582546 | 0,286263861 | 0,0033 | 0,06918097 | DNAJC27 antisense RNA 1 |
| ENSG00000231325 | LY6G5C | -1,804716895 | 0,286237204 | 0,01387 | 0,15454988 | lymphocyte antigen 6 family member G5C |
| ENSG00000227025 | CR388205,1 | -1,8049058 | 0,286199727 | 0,18117 | 0,50470865 | HLA complex group 4 pseudogene 3 |
| ENSG00000223798 | CR759960,1 | -1,8049058 | 0,286199727 | 0,18117 | 0,50470865 | HLA complex group 4 pseudogene 3 |
| ENSG00000214456 | PLIN5 | -1,805857933 | 0,286010906 | 0,00144 | 0,04062048 | perilipin 5 |
| ENSG00000278996 | FP671120,4 | -1,805927538 | 0,285997108 | 0,60487 | 0,79819795 | novel transcript |
| ENSG00000162594 | IL23R | -1,805942945 | 0,285994053 | 0,06147 | 0,32865897 | interleukin 23 receptor |
| ENSG00000288018 | AP003120,1 | -1,806151311 | 0,285952751 | 0,03187 | 0,24068538 | novel transcript |
| ENSG00000226312 | CFLAR-AS1 | -1,806576439 | 0,2858685 | 0,01428 | 0,15652222 | CFLAR antisense RNA 1 |
| ENSG00000278703 | AC100847,1 | -1,806689824 | 0,285846033 | 0,13338 | 0,4594073 | novel transcript |
| ENSG00000262223 | AC110285,1 | -1,807778474 | 0,285630417 | 0,0186 | 0,18029394 | novel transcript |
| ENSG00000236539 | HNRNPA1P54 | -1,807815928 | 0,285623002 | 0,01081 | 0,13450293 | heterogeneous nuclear ribonucleoprotein A1 pseudogene 54 |
| ENSG00000277363 | SRCIN1 | -1,808936238 | 0,28540129 | 0,00289 | 0,06380901 | SRC kinase signaling inhibitor 1 |
| ENSG00000182272 | B4GALNT4 | -1,809493493 | 0,285291072 | 0,00262 | 0,05970947 | beta-1,4-N-acetyl-galactosaminyltransferase 4 |
| ENSG00000261634 | AC026992,2 | -1,809708164 | 0,285248625 | 0,1573 | 0,48443207 | novel transcript, antisense to PAQR5 |
| ENSG00000178403 | NEUROG2 | -1,80978042 | 0,285234339 | 7,06E-06 | 0,00062293 | neurogenin 2 |
| ENSG00000288115 | AC008731,2 | -1,810471151 | 0,285097807 | 0,04345 | 0,28117721 | novel transcript, antisense to TNRC6A |
| ENSG00000260969 | WWOX-AS1 | -1,810872874 | 0,285018432 | 0,04211 | 0,27783166 | WWOX antisense RNA 1 |
| ENSG00000282772 | AL358790,1 | -1,811145946 | 0,284964489 | 0,1343 | 0,46057566 | novel transcript, antisense to C10orf32 |
| ENSG00000288169 | ACAP1 | -1,811503043 | 0,284893963 | 0,08669 | 0,38144301 | ArfGAP with coiled-coil, ankyrin repeat and PH domains 1 |
| ENSG00000101082 | SLA2 | -1,812134588 | 0,284769277 | 0,007 | 0,10655525 | Src like adaptor 2 |
| ENSG00000271781 | AC026740,1 | -1,812466407 | 0,284703788 | 0,01266 | 0,14729065 | novel transcript, antisense to TPPP |
| ENSG00000204352 | C9orf129 | -1,812531568 | 0,284690929 | 0,00312 | 0,06664861 | chromosome 9 open reading frame 129 |

| | | | | | | |
|---|---|---|---|---|---|---|
| ENSG00000152578 | GRIA4 | -1,812626938 | 0,28467211 | 0,00868 | 0,11781444 | glutamate ionotropic receptor AMPA type subunit 4 |
| ENSG00000215397 | SCRT2 | -1,812933363 | 0,284611653 | 0,00028 | 0,01373045 | scratch family transcriptional repressor 2 |
| ENSG00000287264 | AP003778,1 | -1,813127672 | 0,284573323 | 0,02107 | 0,19278417 | novel transcript |
| ENSG00000278635 | AC141557,2 | -1,813194204 | 0,284560199 | 0,06137 | 0,32865897 | WAS protein family homolog 1 (WASH1) pseudogene |
| ENSG00000248455 | LINC02217 | -1,813651899 | 0,284469937 | 0,11618 | 0,43268574 | long intergenic non-protein coding RNA 2217 |
| ENSG00000183486 | MX2 | -1,814017582 | 0,284397841 | 0,00297 | 0,06492836 | MX dynamin like GTPase 2 |
| ENSG00000261649 | GOLGA6L7 | -1,814583108 | 0,284286381 | 0,0054 | 0,09146798 | golgin A6 family like 7 |
| ENSG00000232434 | AJM1 | -1,814743252 | 0,284254826 | 0,00406 | 0,07819491 | apical junction component 1 homolog |
| ENSG00000207088 | SNORA7B | -1,815813847 | 0,284043965 | 0,01081 | 0,13450293 | small nucleolar RNA, H/ACA box 7B |
| ENSG00000232491 | SAPCD2P3 | -1,816336851 | 0,283941012 | 0,04634 | 0,28930497 | suppressor APC domain containing 2 pseudogene 3 |
| ENSG00000267672 | AC010632,2 | -1,816388118 | 0,283930922 | 0,01454 | 0,15764463 | novel transcript |
| ENSG00000215246 | AC116351,1 | -1,81657491 | 0,283894163 | 0,00491 | 0,0866911 | novel transcript |
| ENSG00000187144 | SPATA21 | -1,817097734 | 0,2837913 | 0,06821 | 0,3453052 | spermatogenesis associated 21 |
| ENSG00000273448 | AC006480,2 | -1,818103676 | 0,283593491 | 0,00635 | 0,10053005 | novel transcript |
| ENSG00000278001 | AL591926,4 | -1,818305786 | 0,283553764 | 0,08933 | 0,38666749 | family with sequence similarity 27, member D1 (FAM27D1) pseudogene |
| ENSG00000250234 | AC025754,1 | -1,818386935 | 0,283537815 | 0,15927 | 0,4867709 | novel transcript |
| ENSG00000237331 | XIAP-AS1 | -1,8185815 | 0,28349958 | 0,00273 | 0,06124544 | XIAP antisense RNA 1 |
| ENSG00000149654 | CDH22 | -1,819431563 | 0,283332585 | 0,03388 | 0,24867791 | cadherin 22 |
| ENSG00000229010 | AL161672,1 | -1,819457197 | 0,283327551 | 0,00541 | 0,09151675 | novel pseudogene |
| ENSG00000267197 | AC011461,1 | -1,819917073 | 0,283237251 | 0,01728 | 0,17300352 | novel transcript, sense intronic with KEAP1 |
| ENSG00000280485 | AL358154,1 | -1,819932502 | 0,283234222 | 0,0389 | 0,26698555 | novel transcript, antisense to LGI1 |
| ENSG00000183401 | CCDC159 | -1,820014978 | 0,283218031 | 0,00417 | 0,07917294 | coiled-coil domain containing 159 |
| ENSG00000239265 | CLRN1-AS1 | -1,82027499 | 0,283166992 | 0,05937 | 0,32456197 | CLRN1 antisense RNA 1 |
| ENSG00000087258 | GNAO1 | -1,821915494 | 0,282845183 | 3,65E-06 | 0,00034801 | G protein subunit alpha o1 |
| ENSG00000258702 | AL137786,1 | -1,822020206 | 0,282824655 | 0,00188 | 0,0481607 | novel transcript |
| ENSG00000205041 | AC118344,1 | -1,823466098 | 0,282541345 | 0,00173 | 0,04563537 | novel transcript, sense intronic to AKT2 |
| ENSG00000185818 | NAT8L | -1,824086855 | 0,282419801 | 0,00956 | 0,12500859 | N-acetyltransferase 8 like |
| ENSG00000121410 | A1BG | -1,825128486 | 0,282215966 | 0,00359 | 0,07293271 | alpha-1-B glycoprotein |
| ENSG00000253227 | AC090192,2 | -1,826166663 | 0,282012954 | 0,06572 | 0,33945082 | novel transcript |
| ENSG00000122188 | LAX1 | -1,828258175 | 0,281604409 | 0,03001 | 0,23346163 | lymphocyte transmembrane adaptor 1 |
| ENSG00000282984 | AL357556,3 | -1,828360044 | 0,281584525 | 0,01085 | 0,13471553 | Tubulin-specific chaperone E |
| ENSG00000287463 | AC110620,2 | -1,82896566 | 0,281466346 | 0,028 | 0,22579215 | novel transcript, antisense to TMEM163 |

| | | | | | | |
|---|---|---|---|---|---|---|
| ENSG00000285625 | AC117378,1 | -1,829466365 | 0,281368677 | 0,04775 | 0,29349263 | novel protein |
| ENSG00000257474 | AC027288,1 | -1,829760778 | 0,281311263 | 0,00925 | 0,12270888 | novel transcript |
| ENSG00000243155 | AL162431,1 | -1,83028594 | 0,281208881 | 0,01194 | 0,14242284 | novel transcript |
| ENSG00000269106 | AC012313,7 | -1,83081278 | 0,281106208 | 0,28636 | 0,58267001 | novel transcript, antisense to ZNF446 |
| ENSG00000138675 | FGF5 | -1,830822275 | 0,281104358 | 0,02938 | 0,23068546 | fibroblast growth factor 5 |
| ENSG00000245025 | AC107959,1 | -1,831423709 | 0,280987195 | 0,00573 | 0,09462598 | novel transcript, antisense to RHOBTB2 |
| ENSG00000204241 | LINC02731 | -1,831635379 | 0,280945972 | 0,02084 | 0,19154368 | long intergenic non-protein coding RNA 2731 |
| ENSG00000269026 | AC003006,1 | -1,83436357 | 0,280415194 | 0,05792 | 0,32123433 | novel protein |
| ENSG00000185742 | C11orf87 | -1,834612979 | 0,280366721 | 0,00039 | 0,01713567 | chromosome 11 open reading frame 87 |
| ENSG00000261200 | AC136944,2 | -1,835467783 | 0,280200652 | 0,01243 | 0,14572968 | novel transcript |
| ENSG00000234945 | GTF3C2-AS1 | -1,836015684 | 0,280094259 | 0,00997 | 0,12783227 | GTF3C2 antisense RNA 1 |
| ENSG00000157150 | TIMP4 | -1,836096398 | 0,280078589 | 0,00086 | 0,02908139 | TIMP metallopeptidase inhibitor 4 |
| ENSG00000260059 | AC092620,2 | -1,836731735 | 0,279955274 | 0,01075 | 0,13405812 | novel transcript |
| ENSG00000266538 | AC005838,2 | -1,836757189 | 0,279950335 | 0,03556 | 0,25566849 | novel transcript |
| ENSG00000226842 | AL158834,1 | -1,837505258 | 0,279805212 | 0,16555 | 0,49182669 | novel transcript, antisense to EPC1 |
| ENSG00000251632 | LINC02172 | -1,837610308 | 0,279784839 | 0,132 | 0,45708171 | long intergenic non-protein coding RNA 2172 |
| ENSG00000266256 | LINC00683 | -1,837921603 | 0,279724475 | 0,00452 | 0,08293474 | long intergenic non-protein coding RNA 683 |
| ENSG00000180712 | LINC02363 | -1,838638232 | 0,279585562 | 0,01758 | 0,1746915 | long intergenic non-protein coding RNA 2363 |
| ENSG00000184343 | SRPK3 | -1,838653584 | 0,279582587 | 0,03708 | 0,26086671 | SRSF protein kinase 3 |
| ENSG00000167476 | JSRP1 | -1,839064568 | 0,279502953 | 0,12229 | 0,44183405 | junctional sarcoplasmic reticulum protein 1 |
| ENSG00000235020 | AL390783,1 | -1,840523084 | 0,279220528 | 0,01584 | 0,16482188 | novel transcript |
| ENSG00000259588 | AC012050,1 | -1,841259281 | 0,27907808 | 0,03498 | 0,252989 | novel transcript, antisense to SEMA6D |
| ENSG00000168405 | CMAHP | -1,841871871 | 0,278959605 | 0,10333 | 0,41114168 | cytidine monophospho-N-acetylneuraminic acid hydroxylase, pseudogene |
| ENSG00000166126 | AMN | -1,842444986 | 0,278848809 | 0,03994 | 0,27022885 | amnion associated transmembrane protein |
| ENSG00000060491 | OGFR | -1,843158454 | 0,278710942 | 0,00306 | 0,06608227 | opioid growth factor receptor |
| ENSG00000187857 | OR6C75 | -1,844996986 | 0,278355986 | 0,04953 | 0,29942963 | olfactory receptor family 6 subfamily C member 75 |
| ENSG00000261207 | AL031717,1 | -1,846386658 | 0,27808799 | 0,04256 | 0,2782038 | novel transcript, antisense to MAPK8IP3 |
| ENSG00000270165 | AC010530,1 | -1,846661375 | 0,278035041 | 0,03295 | 0,24506614 | novel transcript |
| ENSG00000207145 | SNORA18 | -1,848022963 | 0,277772761 | 0,13726 | 0,4642578 | small nucleolar RNA, H/ACA box 18 |
| ENSG00000259756 | AC100839,1 | -1,84967466 | 0,27745493 | 0,01311 | 0,14974648 | novel transcript, antisense to TLN2 |
| ENSG00000255475 | AP001007,2 | -1,850143793 | 0,277364722 | 0,19572 | 0,51646992 | novel transcript |
| ENSG00000277439 | GP6 | -1,850444689 | 0,277306879 | 0,04307 | 0,27954725 | glycoprotein VI platelet |

| | | | | | | |
|---|---|---|---|---|---|---|
| ENSG00000166111 | SVOP | -1,850732136 | 0,277251633 | 0,00395 | 0,07699664 | SV2 related protein |
| ENSG00000205744 | DENND1C | -1,85214161 | 0,276980898 | 0,05822 | 0,32195911 | DENN domain containing 1C |
| ENSG00000036530 | CYP46A1 | -1,853705317 | 0,276680847 | 0,00123 | 0,03698156 | cytochrome P450 family 46 subfamily A member 1 |
| ENSG00000271996 | AC019080,4 | -1,855970408 | 0,276246788 | 0,03422 | 0,24983232 | novel transcript, antisense to NFE2L2 |
| ENSG00000053524 | MCF2L2 | -1,856181118 | 0,276206444 | 0,00317 | 0,0674288 | MCF,2 cell line derived transforming sequence-like 2 |
| ENSG00000232110 | AL353751,1 | -1,856536874 | 0,276138342 | 0,00735 | 0,10881218 | novel transcript, antisense to LIPA |
| ENSG00000128692 | EIF2S2P4 | -1,856604963 | 0,27612531 | 0,08375 | 0,37595092 | eukaryotic translation initiation factor 2 subunit 2 beta pseudogene 4 |
| ENSG00000154237 | LRRK1 | -1,857190949 | 0,276013177 | 0,00147 | 0,04107269 | leucine rich repeat kinase 1 |
| ENSG00000222043 | AC079305,1 | -1,858181686 | 0,275823697 | 0,04446 | 0,2844409 | novel transcript |
| ENSG00000253720 | AC022973,3 | -1,858340449 | 0,275793345 | 0,01534 | 0,16232837 | novel transcript |
| ENSG00000183145 | RIPPLY3 | -1,858847458 | 0,27569644 | 0,00016 | 0,00902479 | ripply transcriptional repressor 3 |
| ENSG00000227733 | AC239809,3 | -1,859934628 | 0,275488762 | 4,97E-05 | 0,00335356 | novel transcript |
| ENSG00000227467 | LINC01537 | -1,860047881 | 0,275467136 | 0,00304 | 0,06579402 | long intergenic non-protein coding RNA 1537 |
| ENSG00000263982 | AC009716,1 | -1,862044876 | 0,275086096 | 0,18014 | 0,50407792 | novel transcript, antisense to ZFN156 |
| ENSG00000236272 | SCUBE1-AS2 | -1,862229236 | 0,275050945 | 0,12518 | 0,44690314 | SCUBE1 antisense RNA 2 |
| ENSG00000286228 | SPDYE11 | -1,86232362 | 0,275032951 | 0,00699 | 0,10650925 | speedy/RINGO cell cycle regulator family member E11 |
| ENSG00000273520 | SPDYE8 | -1,86232362 | 0,275032951 | 0,00699 | 0,10650925 | speedy/RINGO cell cycle regulator family member E8 |
| ENSG00000285544 | AC115220,1 | -1,863844012 | 0,274743259 | 0,15199 | 0,48041278 | novel transcript |
| ENSG00000111704 | NANOG | -1,863929969 | 0,27472689 | 0,01677 | 0,17000427 | Nanog homeobox |
| ENSG00000203690 | TCP10L3 | -1,864155256 | 0,274683993 | 0,02583 | 0,21706114 | t-complex 10 like 3, pseudogene |
| ENSG00000258730 | ITPK1-AS1 | -1,864406594 | 0,274636143 | 0,01768 | 0,17511677 | ITPK1 antisense RNA 1 |
| ENSG00000254687 | AC009646,2 | -1,864968916 | 0,274529118 | 0,06946 | 0,34812526 | novel transcript |
| ENSG00000254208 | AC011773,1 | -1,865138292 | 0,27449689 | 0,00508 | 0,08898421 | novel transcript, antisense to E2F5 |
| ENSG00000268756 | AC104534,1 | -1,865198749 | 0,274485387 | 0,07296 | 0,35631792 | novel transcript, antisense to ECH1 |
| ENSG00000271670 | AC010998,2 | -1,865295581 | 0,274466965 | 0,08468 | 0,37787496 | novel transcript, antisense to WDR11 |
| ENSG00000271851 | AC087501,4 | -1,865340183 | 0,274458479 | 0,00518 | 0,08972908 | novel transcript, sense intronic to STX8 |
| ENSG00000279668 | AC024610,2 | -1,865608676 | 0,274407406 | 0,00399 | 0,07731709 | novel transcript |
| ENSG00000287759 | AL353151,2 | -1,868157907 | 0,273922958 | 0,00529 | 0,09064667 | novel transcript, antisense to AK3 |
| ENSG00000222328 | RNU2-2P | -1,868749256 | 0,273810703 | 0,00076 | 0,02699099 | RNA, U2 small nuclear 2, pseudogene |
| ENSG00000089169 | RPH3A | -1,868975321 | 0,273767801 | 0,00659 | 0,10293464 | rabphilin 3A |
| ENSG00000258169 | LINC00485 | -1,869346149 | 0,273697441 | 0,01968 | 0,18612199 | long intergenic non-protein coding RNA 485 |
| ENSG00000203900 | AL121827,1 | -1,869459792 | 0,273675882 | 0,04244 | 0,2782038 | novel transcript, antisense to CHRNA4 |

| | | | | | | |
|---|---|---|---|---|---|---|
| ENSG00000155269 | GPR78 | -1,869865966 | 0,273598843 | 0,09647 | 0,40027716 | G protein-coupled receptor 78 |
| ENSG00000261505 | AL031714,1 | -1,871101219 | 0,273364684 | 0,00209 | 0,05143262 | novel transcript, antisense to UBE2I |
| ENSG00000180815 | MAP3K15 | -1,87114817 | 0,273355788 | 0,00592 | 0,09719731 | mitogen-activated protein kinase kinase kinase 15 |
| ENSG00000288577 | AL117334,1 | -1,872361336 | 0,273126019 | 0,03936 | 0,26861041 | novel transcript |
| ENSG00000155066 | PROM2 | -1,872385743 | 0,273121399 | 0,00151 | 0,04166341 | prominin 2 |
| ENSG00000122548 | KIAA0087 | -1,873226784 | 0,272962225 | 0,00185 | 0,04753043 | KIAA0087 lncRNA |
| ENSG00000249661 | TNRC18P1 | -1,874265468 | 0,272765773 | 0,00719 | 0,10764854 | trinucleotide repeat containing 18 pseudogene 1 |
| ENSG00000259555 | AC067863,1 | -1,874467024 | 0,272727669 | 0,00582 | 0,09586861 | novel transcript, antisense to CHRNB4 |
| ENSG00000166105 | GLB1L3 | -1,874672711 | 0,272688788 | 0,01804 | 0,17733967 | galactosidase beta 1 like 3 |
| ENSG00000228779 | Z69666,1 | -1,875266443 | 0,272576588 | 0,02747 | 0,22348662 | novel transcript |
| ENSG00000234255 | AC012370,2 | -1,87541352 | 0,272548801 | 0,13974 | 0,4668467 | novel transcript |
| ENSG00000274833 | AC110285,5 | -1,876143765 | 0,272410881 | 0,03673 | 0,25969721 | novel transcript |
| ENSG00000259954 | IL21R-AS1 | -1,876249495 | 0,272390918 | 0,00926 | 0,12271407 | IL21R antisense RNA 1 |
| ENSG00000175287 | PHYHD1 | -1,877031623 | 0,272243286 | 0,0078 | 0,11233096 | phytanoyl-CoA dioxygenase domain containing 1 |
| ENSG00000172940 | SLC22A13 | -1,877739593 | 0,272109722 | 0,06354 | 0,3341648 | solute carrier family 22 member 13 |
| ENSG00000268583 | AC011466,1 | -1,879093668 | 0,271854447 | 0,004 | 0,07736298 | novel transcript, antisense to CARD8 |
| ENSG00000179362 | HMGN2P46 | -1,879766787 | 0,271727637 | 0,01519 | 0,16197635 | high mobility group nucleosomal binding domain 2 pseudogene 46 |
| ENSG00000064787 | BCAS1 | -1,880089547 | 0,271666853 | 0,00991 | 0,12765439 | brain enriched myelin associated protein 1 |
| ENSG00000259055 | AL591770,1 | -1,880803341 | 0,271532475 | 0,03162 | 0,23999626 | novel transcript, antisense to TRIM9 |
| ENSG00000248873 | SERBP1P6 | -1,881059605 | 0,271484247 | 0,01132 | 0,13803349 | SERPINE1 mRNA binding protein 1 pseudogene 6 |
| ENSG00000204603 | LINC01257 | -1,881503775 | 0,271400677 | 0,08035 | 0,37007763 | long intergenic non-protein coding RNA 1257 |
| ENSG00000281076 | ZDHHC8P1 | -1,881992689 | 0,271308718 | 0,00424 | 0,07987039 | ZDHHC8 pseudogene 1 |
| ENSG00000284802 | AL136295,8 | -1,882264379 | 0,271257629 | 0,01303 | 0,14938099 | novel transcript |
| ENSG00000253507 | AC104257,1 | -1,883064633 | 0,271107206 | 0,0302 | 0,23452348 | novel transcript |
| ENSG00000264404 | LINC02675 | -1,884949647 | 0,270753211 | 0,05975 | 0,32533111 | long intergenic non-protein coding RNA 2675 |
| ENSG00000223493 | SKIV2L | -1,885136851 | 0,27071808 | 0,13522 | 0,46165567 | Ski2 like RNA helicase |
| ENSG00000269982 | AC018809,2 | -1,886062271 | 0,270544483 | 0,00832 | 0,11538752 | novel transcript, antisense to TMEM111 |
| ENSG00000273328 | AC099329,2 | -1,886519241 | 0,270458803 | 0,01654 | 0,16850885 | novel transcript |
| ENSG00000169085 | VXN | -1,886676513 | 0,270429321 | 0,00193 | 0,0490642 | vexin |
| ENSG00000236824 | BCYRN1 | -1,887416488 | 0,27029065 | 0,00203 | 0,05073405 | brain cytoplasmic RNA 1 |
| ENSG00000172602 | RND1 | -1,88753715 | 0,270268045 | 0,00193 | 0,0490642 | Rho family GTPase 1 |
| ENSG00000159251 | ACTC1 | -1,888288357 | 0,270127354 | 0,00098 | 0,0320617 | actin alpha cardiac muscle 1 |

| | | | | | | |
|---|---|---|---|---|---|---|
| ENSG00000232057 | AC093390,1 | -1,888738496 | 0,270043084 | 0,05106 | 0,30386013 | novel transcript |
| ENSG00000267908 | ZSCAN5DP | -1,88911555 | 0,269972516 | 0,10681 | 0,41660052 | zinc finger and SCAN domain containing 5D pseudogene |
| ENSG00000242686 | PDE6B-AS1 | -1,889176166 | 0,269961173 | 0,00067 | 0,02497236 | PDE6B antisense RNA 1 |
| ENSG00000242808 | SOX2-OT | -1,890712202 | 0,269673899 | 4,55E-09 | 9,05E-07 | SOX2 overlapping transcript |
| ENSG00000274808 | TBC1D3B | -1,890907469 | 0,269637401 | 0,00765 | 0,11109228 | TBC1 domain family member 3B |
| ENSG00000204301 | NOTCH4 | -1,89124541 | 0,269574248 | 0,00053 | 0,02118019 | notch receptor 4 |
| ENSG00000258928 | AL358333,2 | -1,891299436 | 0,269564153 | 0,02028 | 0,18915051 | novel transcript, antisense to GNG2 |
| ENSG00000286067 | AC004263,2 | -1,891913966 | 0,269449354 | 0,00556 | 0,09273388 | novel transcript, antisense to PXN |
| ENSG00000175264 | CHST1 | -1,893507285 | 0,269151937 | 0,00188 | 0,04829042 | carbohydrate sulfotransferase 1 |
| ENSG00000232628 | AC092809,2 | -1,893703847 | 0,269115269 | 0,07649 | 0,36256417 | novel transcript |
| ENSG00000286593 | AP006261,1 | -1,894557318 | 0,268956113 | 0,01618 | 0,16660955 | novel transcript |
| ENSG00000283199 | C13orf46 | -1,895290008 | 0,268819555 | 0,01524 | 0,16205694 | chromosome 13 open reading frame 46 |
| ENSG00000283302 | C13orf46 | -1,895290008 | 0,268819555 | 0,01524 | 0,16205694 | chromosome 13 open reading frame 46 |
| ENSG00000166573 | GALR1 | -1,896372257 | 0,268617973 | 0,0789 | 0,36683128 | galanin receptor 1 |
| ENSG00000268696 | ZNF723 | -1,89693831 | 0,268512599 | 0,11214 | 0,4256732 | zinc finger protein 723 |
| ENSG00000154262 | ABCA6 | -1,898141189 | 0,268288814 | 0,0357 | 0,25626339 | ATP binding cassette subfamily A member 6 |
| ENSG00000270255 | AC009884,2 | -1,900199017 | 0,267906406 | 0,01451 | 0,15751168 | chromosome 8 open reading frame 46 (C8orf46) pseudogene |
| ENSG00000286764 | AC188617,2 | -1,901998099 | 0,267572527 | 0,14074 | 0,46776292 | novel transcript |
| ENSG00000244203 | FOXP1-AS1 | -1,902803212 | 0,267423247 | 0,06217 | 0,33019735 | FOXP1 antisense RNA 1 |
| ENSG00000145920 | CPLX2 | -1,902930854 | 0,267399588 | 2,85E-07 | 3,75E-05 | complexin 2 |
| ENSG00000198010 | DLGAP2 | -1,903609818 | 0,267273773 | 0,09562 | 0,39857837 | DLG associated protein 2 |
| ENSG00000232367 | TAP1 | -1,904433914 | 0,267121145 | 0,08284 | 0,37470115 | transporter 1, ATP binding cassette subfamily B member |
| ENSG00000153292 | ADGRF1 | -1,90586642 | 0,266856042 | 0,0158 | 0,16482188 | adhesion G protein-coupled receptor F1 |
| ENSG00000276161 | SNORA17B | -1,905970722 | 0,26683675 | 0,00291 | 0,06410301 | small nucleolar RNA, H/ACA box 17B |
| ENSG00000287611 | AC007922,4 | -1,90718452 | 0,266612343 | 0,02262 | 0,20053455 | novel transcript |
| ENSG00000261327 | AC134312,5 | -1,90750567 | 0,266553001 | 0,05631 | 0,31687263 | novel transcript |
| ENSG00000135899 | SP110 | -1,907892707 | 0,266481501 | 0,00064 | 0,02439336 | SP110 nuclear body protein |
| ENSG00000286607 | AL135818,3 | -1,908542527 | 0,2663615 | 0,04764 | 0,29344099 | novel transcript |
| ENSG00000250302 | LINC01618 | -1,909371433 | 0,266208505 | 0,0349 | 0,25262813 | long intergenic non-protein coding RNA 1618 |
| ENSG00000216863 | LY86-AS1 | -1,909623699 | 0,26616196 | 0,01399 | 0,15521058 | LY86 antisense RNA 1 |
| ENSG00000215196 | BASP1-AS1 | -1,909774245 | 0,266134187 | 0,00376 | 0,07477699 | BASP1 antisense RNA 1 |
| ENSG00000261124 | AC135050,4 | -1,910355133 | 0,266027052 | 0,0342 | 0,24972789 | novel transcript, antisense to ZNF668 |

| | | | | | | |
|---|---|---|---|---|---|---|
| ENSG00000224315 | RPL7P7 | -1,910755005 | 0,265953328 | 0,01138 | 0,13851314 | ribosomal protein L7 pseudogene 7 |
| ENSG00000204128 | C2orf72 | -1,911299426 | 0,265852986 | 7,62E-06 | 0,00066823 | chromosome 2 open reading frame 72 |
| ENSG00000198517 | MAFK | -1,916576119 | 0,264882398 | 0,00938 | 0,12392004 | MAF bZIP transcription factor K |
| ENSG00000272622 | AC010735,2 | -1,916988863 | 0,264806628 | 0,00713 | 0,10752818 | novel transcript |
| ENSG00000184916 | JAG2 | -1,918583656 | 0,264514065 | 0,00019 | 0,0099537 | jagged canonical Notch ligand 2 |
| ENSG00000286565 | AC006348,1 | -1,919142977 | 0,264411535 | 0,05374 | 0,31090219 | novel transcript |
| ENSG00000250846 | EPHA5-AS1 | -1,919689963 | 0,264311305 | 0,0726 | 0,35556123 | EPHA5 antisense RNA 1 |
| ENSG00000233469 | ST6GALNAC4I | -1,921340284 | 0,264009128 | 0,13884 | 0,4661725 | ST6 N-acetylgalactosaminide alpha-2,6-sialyltransferase 4 pseudogene 1 |
| ENSG00000232725 | U52111,1 | -1,922188738 | 0,263853909 | 0,1105 | 0,42322208 | novel transcript |
| ENSG00000230077 | MTAPP2 | -1,923119163 | 0,263683799 | 0,07418 | 0,35759867 | methylthioadenosine phosphorylase pseudogene 2 |
| ENSG00000264491 | AC005544,1 | -1,924386926 | 0,263452189 | 0,01115 | 0,13658995 | novel transcript |
| ENSG00000244675 | AC108676,1 | -1,926198197 | 0,263121639 | 0,01595 | 0,16543992 | novel transcript, sense overlapping to ATP13A3 |
| ENSG00000265735 | RN7SL5P | -1,92640156 | 0,263084552 | 0,05529 | 0,31451946 | RNA, 7SL, cytoplasmic 5, pseudogene |
| ENSG00000223387 | LINC02068 | -1,927325337 | 0,262916149 | 0,08521 | 0,37857645 | long intergenic non-protein coding RNA 2068 |
| ENSG00000283462 | AC008505,1 | -1,928812433 | 0,262645281 | 0,02938 | 0,23068546 | novel transcript |
| ENSG00000274811 | CLPTM1L | -1,928829011 | 0,262642263 | 0,01491 | 0,16036692 | CLPTM1 like |
| ENSG00000259707 | AC105339,4 | -1,929205236 | 0,26257378 | 0,01053 | 0,1325438 | serine/arginine-rich splicing factor 9 (SRSF9) pseudogene |
| ENSG00000167701 | GPT | -1,929561927 | 0,262508869 | 0,00923 | 0,12259866 | glutamic--pyruvic transaminase |
| ENSG00000275116 | CU457734,1 | -1,930333955 | 0,262368431 | 0,03026 | 0,23475421 | LSM14 homolog A (SCD6, S, cerevisiae) pseudogene |
| ENSG00000172508 | CARNS1 | -1,931357436 | 0,262182367 | 0,00585 | 0,09617663 | carnosine synthase 1 |
| ENSG00000280913 | CTSLP3 | -1,931751309 | 0,262110798 | 0,00607 | 0,09835545 | cathepsin L pseudogene 3 |
| ENSG00000260975 | AC007333,2 | -1,931781173 | 0,262105372 | 0,0018 | 0,04684971 | novel transcript |
| ENSG00000280080 | TBC1D22A-AS | -1,932491625 | 0,26197633 | 0,00817 | 0,11480072 | TBC1D22A antisense RNA 1 |
| ENSG00000235997 | LINC01936 | -1,933991806 | 0,261704057 | 0,0503 | 0,30167993 | long intergenic non-protein coding RNA 1936 |
| ENSG00000240057 | AC078785,1 | -1,93416011 | 0,261673528 | 0,00725 | 0,10792958 | uncharacterized LOC107986114 |
| ENSG00000243193 | PRKAR2B-AS1 | -1,934433592 | 0,261623929 | 0,01755 | 0,17464254 | PRKAR2B antisense RNA 1 |
| ENSG00000230184 | SMYD3-IT1 | -1,934605901 | 0,261592684 | 0,18035 | 0,50407792 | SMYD3 intronic transcript 1 |
| ENSG00000281019 | AC116351,3 | -1,93613889 | 0,261314867 | 0,12652 | 0,44918536 | novel transcript |
| ENSG00000233198 | RNF224 | -1,936471745 | 0,261254584 | 0,06339 | 0,33352879 | ring finger protein 224 |
| ENSG00000255931 | AP003108,1 | -1,937900521 | 0,260995978 | 0,01636 | 0,16750744 | novel transcript |
| ENSG00000274287 | SCRIB | -1,939574636 | 0,260693292 | 0,00104 | 0,0330699 | scribble planar cell polarity protein |

| | | | | | | |
|---|---|---|---|---|---|---|
| ENSG00000230189 | AC008267,2 | -1,941113206 | 0,260415422 | 0,07062 | 0,3507077 | RAB guanine nucleotide exchange factor 1 (RABGEF1) pseudogene |
| ENSG00000205971 | AC074389,1 | -1,941564936 | 0,260333895 | 0,00644 | 0,1015592 | lncRNA radiation induced regulator of PLK1 and RAD51 |
| ENSG00000273769 | LCA10 | -1,941851914 | 0,260282115 | 0,21169 | 0,53056574 | lung carcinoma-associated 10 |
| ENSG00000284627 | AC079781,4 | -1,942136996 | 0,260230687 | 0,06568 | 0,33945082 | RNA binding protein, fox-1 homolog 1 (RBFOX1) pseudogene |
| ENSG00000272909 | AL122035,2 | -1,942983396 | 0,26007806 | 0,01127 | 0,13773024 | novel transcript, antisense to MTHFD1 |
| ENSG00000277095 | AC242852,2 | -1,943228997 | 0,260033788 | 0,32195 | 0,60577115 | otopetrin 1 (OTOP1) pseudogene |
| ENSG00000163833 | FBXO40 | -1,943916244 | 0,259909948 | 0,08546 | 0,37891248 | F-box protein 40 |
| ENSG00000150873 | C2orf50 | -1,944351534 | 0,259831539 | 0,00257 | 0,05898231 | chromosome 2 open reading frame 50 |
| ENSG00000248605 | AC022140,1 | -1,94463174 | 0,259781079 | 0,0287 | 0,22913601 | novel transcript |
| ENSG00000249509 | NEUROG2-AS | -1,944899313 | 0,259732902 | 0,02893 | 0,22977124 | NEUROG2 and ZGRF1 antisense RNA 1 |
| ENSG00000198765 | SYCP1 | -1,945036653 | 0,259708178 | 0,02197 | 0,19708179 | synaptonemal complex protein 1 |
| ENSG00000261630 | AC007496,2 | -1,945867155 | 0,259558717 | 0,01327 | 0,15062654 | novel transcript, antisense to FTO |
| ENSG00000171017 | LRRC8E | -1,946480545 | 0,259448384 | 0,00931 | 0,12325666 | leucine rich repeat containing 8 VRAC subunit E |
| ENSG00000231322 | RPL13AP17 | -1,9466463 | 0,259418577 | 0,00362 | 0,07330561 | ribosomal protein L13a pseudogene 17 |
| ENSG00000206159 | GYG2P1 | -1,947105329 | 0,25933605 | 0,0359 | 0,25662951 | glycogenin 2 pseudogene 1 |
| ENSG00000005379 | TSPOAP1 | -1,947115205 | 0,259334274 | 3,51E-05 | 0,00247834 | TSPO associated protein 1 |
| ENSG00000265443 | AC127024,3 | -1,9475648 | 0,259253469 | 0,0143 | 0,15653473 | novel transcript |
| ENSG00000275239 | FAM242F | -1,948228761 | 0,259134182 | 0,05416 | 0,31168784 | family with sequence similarity 242 member F |
| ENSG00000274041 | AC012314,2 | -1,949168765 | 0,258965396 | 0,03381 | 0,24846467 | novel transcript |
| ENSG00000178297 | TMPRSS9 | -1,950071193 | 0,258803459 | 0,0132 | 0,15012165 | transmembrane serine protease 9 |
| ENSG00000237637 | FRY-AS1 | -1,950246075 | 0,258772089 | 0,01843 | 0,17952781 | FRY antisense RNA 1 |
| ENSG00000226903 | LINC00354 | -1,950758109 | 0,258680264 | 0,00058 | 0,02269054 | long intergenic non-protein coding RNA 354 |
| ENSG00000150275 | PCDH15 | -1,951391969 | 0,258566635 | 0,01276 | 0,14789627 | protocadherin related 15 |
| ENSG00000230507 | RPL7AP8 | -1,952202722 | 0,258421369 | 0,27245 | 0,57315552 | ribosomal protein L7a pseudogene 8 |
| ENSG00000267612 | AC008759,2 | -1,952605619 | 0,258349211 | 0,09967 | 0,40615251 | novel transcript |
| ENSG00000261364 | AC063923,2 | -1,954868745 | 0,257944261 | 0,02872 | 0,2291366 | novel transcript |
| ENSG00000240993 | RN7SL459P | -1,95493003 | 0,257933304 | 0,07258 | 0,35556123 | RNA, 7SL, cytoplasmic 459, pseudogene |
| ENSG00000268987 | AC011462,3 | -1,956361109 | 0,257677575 | 0,03953 | 0,26903805 | novel transcript, sense intronic to BCKDHA |
| ENSG00000251075 | HTT-AS | -1,956498514 | 0,257653034 | 0,16599 | 0,49197859 | HTT antisense RNA |
| ENSG00000229122 | AGBL5-IT1 | -1,957281711 | 0,2575132 | 0,02165 | 0,19545722 | AGBL5 intronic transcript 1 |
| ENSG00000266541 | AP001381,1 | -1,958460273 | 0,257302919 | 0,00337 | 0,07000131 | novel transcript, antisense to PPP4R1 |

| | | | | | | |
|---|---|---|---|---|---|---|
| ENSG00000270316 | BORCS7-ASM[illegible] | -1,958515338 | 0,257293098 | 0,06197 | 0,32961785 | BORCS7-ASMT readthrough (NMD candidate) |
| ENSG00000259132 | AL132780,3 | -1,958762189 | 0,257249078 | 0,46057 | 0,70241437 | novel protein |
| ENSG00000167011 | NAT16 | -1,959725265 | 0,257077408 | 0,00347 | 0,07121616 | N-acetyltransferase 16 (putative) |
| ENSG00000284075 | AC015853,4 | -1,959886011 | 0,257048766 | 0,1448 | 0,4723264 | novel transcript, antisense to VPS53 |
| ENSG00000249684 | AC106795,2 | -1,960393145 | 0,256958424 | 0,02346 | 0,20434232 | novel transcript, antisense to PROP1 |
| ENSG00000227220 | AL133346,1 | -1,960554999 | 0,256929598 | 0,01743 | 0,17407115 | novel transcript |
| ENSG00000116299 | ELAPOR1 | -1,960569497 | 0,256927016 | 0,00453 | 0,08299378 | endosome-lysosome associated apoptosis and autophagy regulator 1 |
| ENSG00000274827 | LINC01297 | -1,961053863 | 0,256840771 | 0,02224 | 0,19814016 | long intergenic non-protein coding RNA 1297 |
| ENSG00000226239 | AL031658,1 | -1,961461119 | 0,256768278 | 0,00425 | 0,0801408 | novel transcript |
| ENSG00000198650 | TAT | -1,961925765 | 0,256685594 | 0,00707 | 0,10712237 | tyrosine aminotransferase |
| ENSG00000236501 | AC074286,1 | -1,961956937 | 0,256680048 | 0,0252 | 0,21343549 | novel transcript |
| ENSG00000284633 | AL031590,1 | -1,962541934 | 0,256575988 | 0,0775 | 0,36445727 | novel transcript, antisense to PDGFB |
| ENSG00000241231 | AC068308,1 | -1,965002787 | 0,256138711 | 0,00019 | 0,0103189 | novel transcript |
| ENSG00000243658 | MTND5P16 | -1,965779522 | 0,256000845 | 0,08809 | 0,38424528 | MT-ND5 pseudogene 16 |
| ENSG00000259810 | AC002519,1 | -1,968035655 | 0,255600816 | 0,02747 | 0,22348662 | novel transcript |
| ENSG00000141837 | CACNA1A | -1,968513602 | 0,255516152 | 0,00381 | 0,0752235 | calcium voltage-gated channel subunit alpha1 A |
| ENSG00000278631 | SOWAHCP5 | -1,970753215 | 0,255119801 | 0,02672 | 0,22056859 | SOWAHC pseudogene 5 |
| ENSG00000223573 | TINCR | -1,971414605 | 0,255002871 | 0,01909 | 0,18296265 | TINCR ubiquitin domain containing |
| ENSG00000272834 | AL022238,2 | -1,973688102 | 0,254601337 | 0,0886 | 0,38491339 | novel transcript, antisense to MKL1 |
| ENSG00000229999 | SGSM3-AS1 | -1,974534146 | 0,254452074 | 0,01627 | 0,16730761 | SGSM3 antisense RNA 1 |
| ENSG00000204118 | NAP1L6P | -1,976401139 | 0,254123 | 0,04206 | 0,27770943 | nucleosome assembly protein 1 like 6, pseudogene |
| ENSG00000168918 | INPP5D | -1,976864488 | 0,254041397 | 0,04693 | 0,29141554 | inositol polyphosphate-5-phosphatase D |
| ENSG00000210107 | MT-TQ | -1,977047581 | 0,254009158 | 0,04976 | 0,30012887 | mitochondrially encoded tRNA-Gln (CAA/G) |
| ENSG00000274124 | AC074029,3 | -1,979195763 | 0,253631218 | 0,11608 | 0,43251309 | novel transcript, antisense to ASB8 |
| ENSG00000274367 | AC004233,3 | -1,979369729 | 0,253600636 | 0,10516 | 0,41407649 | novel transcript |
| ENSG00000247033 | AC099508,1 | -1,979602128 | 0,253559788 | 0,57192 | 0,77838663 | novel transcript, antisense to ZNRF1 |
| ENSG00000262777 | AC032044,1 | -1,979661115 | 0,253549421 | 0,01073 | 0,1339456 | novel transcript, antisense to CRK |
| ENSG00000121858 | TNFSF10 | -1,980048208 | 0,2534814 | 2,29E-05 | 0,00173845 | TNF superfamily member 10 |
| ENSG00000236467 | KCNMA1-AS1 | -1,983535155 | 0,252869483 | 0,05898 | 0,3241112 | KCNMA1 antisense RNA 1 |
| ENSG00000198844 | ARHGEF15 | -1,983860532 | 0,252812459 | 0,07929 | 0,36776284 | Rho guanine nucleotide exchange factor 15 |
| ENSG00000275126 | H4C13 | -1,985527983 | 0,25252043 | 0,03319 | 0,24606495 | H4 clustered histone 13 |
| ENSG00000102195 | GPR50 | -1,986041579 | 0,252430549 | 6,44E-05 | 0,00418528 | G protein-coupled receptor 50 |

| | | | | | | |
|---|---|---|---|---|---|---|
| ENSG00000251393 | AC005280,1 | -1,986593472 | 0,252334002 | 0,15255 | 0,48061452 | novel transcript, antisense to NUMB and HEATR4 |
| ENSG00000267336 | EIF4A2P1 | -1,988218536 | 0,252049931 | 0,03471 | 0,25191864 | eukaryotic translation initiation factor 4A2 pseudogene 1 |
| ENSG00000226913 | BSN-DT | -1,988265161 | 0,252041785 | 0,0075 | 0,11001711 | BSN divergent transcript |
| ENSG00000237898 | HCG15 | -1,988817965 | 0,251945228 | 0,10588 | 0,4154675 | HLA complex group 15 |
| ENSG00000241132 | LY6G5B | -1,988922981 | 0,251926889 | 0,23629 | 0,54725688 | lymphocyte antigen 6 family member G5B |
| ENSG00000284871 | LINC00971 | -1,989691153 | 0,251792785 | 0,09876 | 0,40461924 | long intergenic non-protein coding RNA 971 |
| ENSG00000285887 | AL009176,1 | -1,990129891 | 0,251716224 | 0,00695 | 0,10609569 | novel transcript |
| ENSG00000286802 | AC104459,1 | -1,993631512 | 0,251106014 | 0,01744 | 0,17408809 | novel transcript, antisense to GBP7and GBP2 |
| ENSG00000266970 | AC061992,2 | -1,995343866 | 0,25080815 | 0,01945 | 0,18477688 | novel transcript |
| ENSG00000259762 | AC021739,4 | -1,997541211 | 0,250426439 | 0,00507 | 0,08898421 | novel transcript, antisense to AKAP13 |
| ENSG00000140044 | JDP2 | -1,998444064 | 0,250269769 | 0,01208 | 0,14294961 | Jun dimerization protein 2 |
| ENSG00000269313 | MAGIX | -1,99856967 | 0,25024798 | 0,00833 | 0,11543199 | MAGI family member, X-linked |
| ENSG00000237401 | LINC01304 | -2,000740304 | 0,249871748 | 0,07184 | 0,35364999 | long intergenic non-protein coding RNA 1304 |
| ENSG00000235881 | AC114776,1 | -2,001804108 | 0,249687567 | 0,15004 | 0,47814899 | novel transcript |
| ENSG00000284642 | AL139424,2 | -2,002036565 | 0,249647339 | 0,00195 | 0,04935825 | novel transcript |
| ENSG00000245017 | LINC02453 | -2,00321366 | 0,249443735 | 0,0057 | 0,09429746 | long intergenic non-protein coding RNA 2453 |
| ENSG00000175262 | C1orf127 | -2,003792759 | 0,249343628 | 0,01266 | 0,14729065 | chromosome 1 open reading frame 127 |
| ENSG00000285515 | CR936875,2 | -2,006164987 | 0,248933969 | 0,48448 | 0,72028271 | novel protein |
| ENSG00000285213 | AL773541,2 | -2,006164987 | 0,248933969 | 0,48448 | 0,72028271 | novel protein |
| ENSG00000254568 | AP003501,1 | -2,006939806 | 0,248800311 | 0,03362 | 0,24771686 | novel transcript |
| ENSG00000257956 | NOP56P3 | -2,007264227 | 0,248744369 | 0,10343 | 0,41135663 | NOP56 ribonucleoprotein pseudogene 3 |
| ENSG00000276130 | SLC22A18 | -2,007590038 | 0,2486882 | 0,01145 | 0,13897679 | solute carrier family 22 member 18 |
| ENSG00000268366 | AC010271,1 | -2,009241273 | 0,248403727 | 0,04013 | 0,27065903 | novel transcript, antisense to SERTAD3 |
| ENSG00000235294 | RPL7AP25 | -2,009812848 | 0,248305333 | 0,00429 | 0,08042226 | ribosomal protein L7a pseudogene 25 |
| ENSG00000008438 | PGLYRP1 | -2,010477541 | 0,248190957 | 0,06381 | 0,334882 | peptidoglycan recognition protein 1 |
| ENSG00000259198 | AC020658,2 | -2,010899501 | 0,248118377 | 0,17462 | 0,49886732 | novel transcript, antisense to PLCB2 |
| ENSG00000258778 | EIF2S2P1 | -2,012196333 | 0,247895445 | 0,39443 | 0,65729552 | eukaryotic translation initiation factor 2 subunit 2 beta pseudogene 1 |
| ENSG00000170827 | CELP | -2,013175292 | 0,247727289 | 0,00349 | 0,07162617 | carboxyl ester lipase pseudogene |
| ENSG00000181634 | TNFSF15 | -2,0132443 | 0,24771544 | 0,06193 | 0,32950971 | TNF superfamily member 15 |
| ENSG00000258765 | AC107958,2 | -2,014263585 | 0,247540487 | 0,00449 | 0,08275517 | novel transcript |
| ENSG00000197046 | SIGLEC15 | -2,014477368 | 0,247503809 | 0,03076 | 0,2367095 | sialic acid binding Ig like lectin 15 |
| ENSG00000224382 | LINC00703 | -2,014635213 | 0,247476731 | 0,02386 | 0,20623598 | long intergenic non-protein coding RNA 703 |

| | | | | | | |
|---|---|---|---|---|---|---|
| ENSG00000252906 | SCARNA3 | -2,014825342 | 0,247444119 | 0,09478 | 0,39724142 | small Cajal body-specific RNA 3 |
| ENSG00000132026 | RTBDN | -2,015059534 | 0,247403955 | 0,05204 | 0,30661383 | retbindin |
| ENSG00000250910 | AC097467,3 | -2,015220754 | 0,247376309 | 0,0067 | 0,10401251 | novel transcript, antisense to MAP9 |
| ENSG00000233388 | AL022334,1 | -2,016994596 | 0,247072338 | 0,06547 | 0,33894038 | tRNA methyltransferase 11 homolog (S, cerevisiae) (TRMT11) pseudogene |
| ENSG00000277789 | TMC4 | -2,017145945 | 0,24704642 | 0,01498 | 0,16040868 | transmembrane channel like 4 |
| ENSG00000254363 | AC011379,2 | -2,019998387 | 0,246558452 | 0,00644 | 0,1015592 | novel transcript |
| ENSG00000241269 | AC093620,1 | -2,021577589 | 0,246288712 | 0,00351 | 0,07177624 | novel transcript |
| ENSG00000225349 | ANKRD49P2 | -2,022962938 | 0,246052326 | 0,05127 | 0,30462001 | ANKRD49 pseudogene 2 |
| ENSG00000269177 | L34079,2 | -2,024162447 | 0,245847835 | 0,02489 | 0,21153603 | novel transcript, antisense to XRCC1 |
| ENSG00000227619 | AL391056,1 | -2,025513058 | 0,245617786 | 0,0715 | 0,35329893 | novel transcript |
| ENSG00000250427 | LINC02148 | -2,026538771 | 0,245443222 | 0,07371 | 0,35703856 | long intergenic non-protein coding RNA 2148 |
| ENSG00000186487 | MYT1L | -2,029036042 | 0,245018733 | 0,00129 | 0,03841105 | myelin transcription factor 1 like |
| ENSG00000234492 | RPL34-DT | -2,029959085 | 0,244862019 | 0,02977 | 0,23233601 | RPL34 divergent transcript |
| ENSG00000164265 | SCGB3A2 | -2,030667126 | 0,244741875 | 0,00838 | 0,1155444 | secretoglobin family 3A member 2 |
| ENSG00000258891 | AC005480,1 | -2,034184838 | 0,24414585 | 0,0035 | 0,07173085 | novel transcript, antisense to ZNF410 |
| ENSG00000234409 | CCDC188 | -2,034308275 | 0,244124962 | 0,02472 | 0,21047862 | coiled-coil domain containing 188 |
| ENSG00000285188 | AC008397,2 | -2,034778054 | 0,244045482 | 0,00525 | 0,09023573 | novel protein |
| ENSG00000243642 | RN7SL526P | -2,036714813 | 0,24371808 | 0,16368 | 0,49000347 | RNA, 7SL, cytoplasmic 526, pseudogene |
| ENSG00000196639 | HRH1 | -2,037707142 | 0,243550501 | 0,05361 | 0,31071932 | histamine receptor H1 |
| ENSG00000254746 | AC103855,2 | -2,038359945 | 0,243440322 | 0,01308 | 0,14966794 | novel transcript |
| ENSG00000284722 | AP003175,1 | -2,039204891 | 0,243297788 | 0,00522 | 0,09023148 | novel transcript |
| ENSG00000282024 | AL009031,1 | -2,039321165 | 0,24327818 | 0,00331 | 0,06922569 | novel transcript, antisense to GMPR |
| ENSG00000263257 | AC040173,1 | -2,03974343 | 0,243206985 | 0,22936 | 0,54284965 | novel transcript |
| ENSG00000261659 | Z92544,1 | -2,041215934 | 0,24295888 | 0,00302 | 0,06562119 | novel transcript, antisense to FBXL16 |
| ENSG00000204252 | HLA-DOA | -2,043390683 | 0,242592914 | 0,10464 | 0,41313952 | major histocompatibility complex, class II, DO alpha |
| ENSG00000258679 | AC121758,2 | -2,04372561 | 0,242536602 | 0,00973 | 0,12625914 | novel YWHA pseudogene |
| ENSG00000161281 | COX7A1 | -2,043898937 | 0,242507465 | 0,02705 | 0,22193038 | cytochrome c oxidase subunit 7A1 |
| ENSG00000232396 | AC004882,1 | -2,044180888 | 0,242460076 | 0,07355 | 0,35672172 | novel transcript |
| ENSG00000137033 | IL33 | -2,045418145 | 0,242252231 | 0,13061 | 0,45494178 | interleukin 33 |
| ENSG00000239268 | AC092691,1 | -2,045436907 | 0,24224908 | 0,01166 | 0,14066395 | novel transcript |
| ENSG00000258804 | LINC01148 | -2,046180691 | 0,24212422 | 0,01562 | 0,16386137 | long intergenic non-protein coding RNA 1148 |
| ENSG00000236423 | LINC01134 | -2,0462091 | 0,242119453 | 0,01394 | 0,1548079 | long intergenic non-protein coding RNA 1134 |

| | | | | | | |
|---|---|---|---|---|---|---|
| ENSG00000266268 | AP005203,1 | -2,046308289 | 0,242102807 | 0,0175 | 0,17433927 | novel transcript |
| ENSG00000288150 | AC090826,4 | -2,046373813 | 0,242091811 | 0,15424 | 0,48258262 | novel transcript, antisense to CYP11A1 |
| ENSG00000287743 | AL031289,2 | -2,046969838 | 0,241991816 | 0,03455 | 0,25132008 | novel transcript, antisense to RIMS3 |
| ENSG00000255046 | AC069185,1 | -2,047231734 | 0,241947891 | 0,01539 | 0,16247082 | novel transcript |
| ENSG00000287241 | AL359511,3 | -2,047559056 | 0,241893003 | 0,02386 | 0,20623598 | novel transcript, antisense to PCSK2 |
| ENSG00000286404 | AC011228,2 | -2,04942027 | 0,241581139 | 0,08229 | 0,37375431 | novel transcript |
| ENSG00000242893 | RN7SL413P | -2,049987718 | 0,241486138 | 0,05403 | 0,31168784 | RNA, 7SL, cytoplasmic 413, pseudogene |
| ENSG00000261143 | ADAMTS7P3 | -2,054228793 | 0,240777286 | 0,12418 | 0,44504227 | ADAMTS7 pseudogene 3 |
| ENSG00000249125 | AC093821,1 | -2,054359216 | 0,24075552 | 0,00263 | 0,05983445 | novel transcript |
| ENSG00000236418 | HLA-DQA1 | -2,054372859 | 0,240753244 | 0,22356 | 0,53953208 | major histocompatibility complex, class II, DQ alpha 1 |
| ENSG00000200792 | SNORA80A | -2,055663033 | 0,240538039 | 0,01196 | 0,14252936 | small nucleolar RNA, H/ACA box 80A |
| ENSG00000197599 | CCDC154 | -2,056135775 | 0,240459233 | 0,02707 | 0,22193038 | coiled-coil domain containing 154 |
| ENSG00000282846 | AC008982,4 | -2,057162354 | 0,24028819 | 0,00404 | 0,07795523 | novel transcript, sense intronic to HNRNPL |
| ENSG00000221858 | OR2A12 | -2,057885165 | 0,240167832 | 0,28831 | 0,58436774 | olfactory receptor family 2 subfamily A member 12 |
| ENSG00000168468 | ATF6B | -2,059115525 | 0,239963099 | 0,01364 | 0,15297981 | activating transcription factor 6 beta |
| ENSG00000255197 | AC090559,1 | -2,059294939 | 0,239933259 | 0,02021 | 0,18878814 | novel transcript |
| ENSG00000279345 | Z98885,3 | -2,059586211 | 0,239884823 | 0,00331 | 0,06920094 | novel transcript, antisense to BRD1 |
| ENSG00000181626 | ANKRD62 | -2,060881374 | 0,239669566 | 0,01427 | 0,15652222 | ankyrin repeat domain 62 |
| ENSG00000182223 | ZAR1 | -2,062234043 | 0,239444957 | 0,07115 | 0,35212832 | zygote arrest 1 |
| ENSG00000254990 | AP001781,1 | -2,064220394 | 0,239115508 | 0,06983 | 0,34878306 | novel transcript |
| ENSG00000107984 | DKK1 | -2,064827569 | 0,239014895 | 0,00615 | 0,09865643 | dickkopf WNT signaling pathway inhibitor 1 |
| ENSG00000287070 | AL499604,1 | -2,065187889 | 0,238955207 | 0,02115 | 0,19303449 | novel transcript, sense intronic to TRMO |
| ENSG00000258168 | AC025569,1 | -2,06569477 | 0,238871267 | 0,00682 | 0,10528058 | novel transcript, antisense to PTPRB |
| ENSG00000214650 | AC073592,1 | -2,069480263 | 0,238245313 | 0,02769 | 0,2242103 | novel transcript |
| ENSG00000249493 | ANKRD20A18 | -2,071281695 | 0,237948012 | 0,09208 | 0,39237469 | ankyrin repeat domain 20 family member A18, pseudogene |
| ENSG00000186765 | FSCN2 | -2,071325953 | 0,237940712 | 0,00651 | 0,10236887 | fascin actin-bundling protein 2, retinal |
| ENSG00000254501 | AP003068,1 | -2,071490004 | 0,237913657 | 0,36802 | 0,63798276 | novel transcript |
| ENSG00000231393 | AL357874,2 | -2,072055508 | 0,237820419 | 0,00843 | 0,11587456 | novel transcript |
| ENSG00000207014 | SNORD116-3 | -2,072990904 | 0,237666274 | 0,01758 | 0,1746915 | small nucleolar RNA, C/D box 116-3 |
| ENSG00000206727 | SNORD116-9 | -2,072990904 | 0,237666274 | 0,01758 | 0,1746915 | small nucleolar RNA, C/D box 116-9 |
| ENSG00000285382 | FO393400,1 | -2,07343838 | 0,237592569 | 0,00858 | 0,11699775 | novel protein |
| ENSG00000170790 | OR10A2 | -2,073630621 | 0,237560912 | 0,00606 | 0,09831822 | olfactory receptor family 10 subfamily A member 2 |

| | | | | | | |
|---|---|---|---|---|---|---|
| ENSG00000275379 | H3C11 | -2,073723204 | 0,237545667 | 0,00076 | 0,02699099 | H3 clustered histone 11 |
| ENSG00000245526 | LINC00461 | -2,07447304 | 0,237422235 | 4,21E-13 | 1,48E-10 | long intergenic non-protein coding RNA 461 |
| ENSG00000278214 | LINC02139 | -2,076076087 | 0,237158571 | 0,05823 | 0,32195911 | long intergenic non-protein coding RNA 2139 |
| ENSG00000275873 | LMNTD2 | -2,076600455 | 0,237072388 | 0,12867 | 0,45191041 | lamin tail domain containing 2 |
| ENSG00000255585 | AL590627,1 | -2,07687864 | 0,237026679 | 0,00137 | 0,03950275 | forkhead box H1 (FOXH1) pseudogene |
| ENSG00000231003 | MPIG6B | -2,079071336 | 0,236666705 | 0,55189 | 0,76553885 | megakaryocyte and platelet inhibitory receptor G6b |
| ENSG00000272636 | DOC2B | -2,080778571 | 0,236386808 | 0,39353 | 0,65651919 | double C2 domain beta |
| ENSG00000259207 | ITGB3 | -2,081375504 | 0,23628902 | 6,06E-06 | 0,00054275 | integrin subunit beta 3 |
| ENSG00000271199 | AL353804,2 | -2,082552949 | 0,236096253 | 0,05591 | 0,31600461 | novel transcript, sense overlapping FTX |
| ENSG00000271788 | AC008875,1 | -2,08370245 | 0,235908213 | 0,07061 | 0,3507077 | novel transcript |
| ENSG00000287920 | AL022098,1 | -2,08576493 | 0,235571199 | 0,01587 | 0,16502318 | novel transcript, antisense to NEDD9 |
| ENSG00000285820 | AL157829,1 | -2,08617053 | 0,23550498 | 0,03659 | 0,25938022 | Novel transcript, antisense to ASTN2 |
| ENSG00000223396 | RPS10P7 | -2,086426282 | 0,235463234 | 0,0042 | 0,07947406 | ribosomal protein S10 pseudogene 7 |
| ENSG00000267714 | AC008738,4 | -2,087233529 | 0,23533152 | 0,08332 | 0,37550713 | novel transcript, antisense to SLC7A10 |
| ENSG00000120280 | CXorf21 | -2,088406964 | 0,235140188 | 0,19861 | 0,51980353 | chromosome X open reading frame 21 |
| ENSG00000235023 | AP001626,1 | -2,090048533 | 0,234872786 | 0,00408 | 0,07837165 | novel transcript |
| ENSG00000285663 | AC108457,1 | -2,090939475 | 0,234727784 | 0,00127 | 0,03791245 | novel transcript |
| ENSG00000279548 | AL022323,4 | -2,092216198 | 0,234520152 | 0,0112 | 0,13709547 | novel transcript |
| ENSG00000285694 | AL162718,2 | -2,094069885 | 0,234219016 | 0,05333 | 0,31012262 | novel transcript |
| ENSG00000240481 | RN7SL38P | -2,094330744 | 0,23417667 | 0,07166 | 0,35351167 | RNA, 7SL, cytoplasmic 38, pseudogene |
| ENSG00000188763 | FZD9 | -2,096416813 | 0,233838306 | 5,92E-05 | 0,00390694 | frizzled class receptor 9 |
| ENSG00000285763 | AL358777,3 | -2,096891811 | 0,233761329 | 0,45303 | 0,69744022 | novel transcript, antisense to C6orf52 |
| ENSG00000259150 | LINC00929 | -2,097330026 | 0,233690335 | 0,05426 | 0,31175692 | long intergenic non-protein coding RNA 929 |
| ENSG00000288318 | PALM3 | -2,098297985 | 0,233533596 | 0,0055 | 0,09215976 | paralemmin 3 |
| ENSG00000187867 | PALM3 | -2,098297985 | 0,233533596 | 0,0055 | 0,09215976 | paralemmin 3 |
| ENSG00000272693 | AC073107,1 | -2,098479727 | 0,233504179 | 0,0025 | 0,05798721 | nuclear protein 2, transcriptional regulator (NUPR2) pseudogene |
| ENSG00000217325 | PRELID1P1 | -2,098969707 | 0,233424888 | 0,02379 | 0,20595388 | PRELID1 pseudogene 1 |
| ENSG00000255267 | LINC02687 | -2,0998757 | 0,233278346 | 0,04403 | 0,28274932 | long intergenic non-protein coding RNA 2687 |
| ENSG00000235091 | BX324167,1 | -2,100038443 | 0,233252032 | 0,00675 | 0,10456208 | novel transcript |
| ENSG00000180525 | PRR26 | -2,100407632 | 0,23319235 | 0,00242 | 0,05674593 | proline rich 26 |
| ENSG00000197405 | C5AR1 | -2,10050971 | 0,233175851 | 0,03344 | 0,24712306 | complement C5a receptor 1 |
| ENSG00000180871 | CXCR2 | -2,103039671 | 0,232767304 | 0,07266 | 0,35556123 | C-X-C motif chemokine receptor 2 |

| | | | | | | |
|---|---|---|---|---|---|---|
| ENSG00000082556 | OPRK1 | -2,103480629 | 0,23269617 | 2,55E-05 | 0,00188567 | opioid receptor kappa 1 |
| ENSG00000188585 | CLEC20A | -2,104037188 | 0,232606418 | 0,10989 | 0,42217079 | C-type lectin domain containing 20A |
| ENSG00000266527 | AC005697,2 | -2,104086335 | 0,232598495 | 0,01638 | 0,16750744 | novel transcript, antisense to C17orf108 |
| ENSG00000277324 | AC093462,1 | -2,105291032 | 0,232404348 | 0,0288 | 0,2296248 | novel transcript, antisense to POLI |
| ENSG00000259635 | AC100830,1 | -2,106054848 | 0,232281338 | 0,00702 | 0,10661861 | novel transcript, antisense OAZ2 |
| ENSG00000174939 | ASPHD1 | -2,10627805 | 0,232245404 | 0,00235 | 0,055751 | aspartate beta-hydroxylase domain containing 1 |
| ENSG00000239572 | AC108749,1 | -2,106633166 | 0,232188244 | 0,04971 | 0,30006068 | novel transcript |
| ENSG00000269032 | AC016629,1 | -2,107383784 | 0,232067471 | 0,03392 | 0,24870776 | interleukin 9 receptor (IL9R) pseudogene |
| ENSG00000248538 | AC022784,1 | -2,107992658 | 0,23196955 | 0,02186 | 0,19657209 | novel transcript |
| ENSG00000179082 | C9orf106 | -2,108255988 | 0,231927213 | 0,02626 | 0,21852357 | chromosome 9 putative open reading frame 106 |
| ENSG00000250334 | LINC00989 | -2,108469018 | 0,231892969 | 0,01054 | 0,1325438 | long intergenic non-protein coding RNA 989 |
| ENSG00000072858 | SIDT1 | -2,10906663 | 0,231796931 | 0,00398 | 0,07725019 | SID1 transmembrane family member 1 |
| ENSG00000281228 | AL132640,3 | -2,109505446 | 0,231726437 | 0,05439 | 0,31216639 | |
| ENSG00000111218 | PRMT8 | -2,10976674 | 0,231684472 | 0,00135 | 0,03912393 | protein arginine methyltransferase 8 |
| ENSG00000162592 | CCDC27 | -2,110768562 | 0,231523644 | 0,05916 | 0,32424653 | coiled-coil domain containing 27 |
| ENSG00000273079 | GRIN2B | -2,112460202 | 0,231252329 | 3,48E-08 | 5,68E-06 | glutamate ionotropic receptor NMDA type subunit 2B |
| ENSG00000131686 | CA6 | -2,112736755 | 0,231208004 | 0,09533 | 0,39794903 | carbonic anhydrase 6 |
| ENSG00000153029 | MR1 | -2,112841505 | 0,231191217 | 0,00393 | 0,0767581 | major histocompatibility complex, class I-related |
| ENSG00000274979 | AC020656,2 | -2,113368241 | 0,231106823 | 0,00341 | 0,07040998 | novel transcript |
| ENSG00000283077 | AC104534,3 | -2,115885209 | 0,230703979 | 0,04741 | 0,29245799 | novel transcript, antisense to ECH1 |
| ENSG00000255403 | HSPD1P13 | -2,1170865 | 0,230511958 | 0,04617 | 0,28930497 | heat shock protein family D (Hsp60) member 1 pseudogene 13 |
| ENSG00000202444 | RNU5E-6P | -2,117158833 | 0,230500401 | 0,00983 | 0,12683981 | RNA, U5E small nuclear 6, pseudogene |
| ENSG00000182521 | TBPL2 | -2,117775549 | 0,230401889 | 0,01938 | 0,1843508 | TATA-box binding protein like 2 |
| ENSG00000167702 | KIFC2 | -2,117814791 | 0,230395622 | 0,00027 | 0,01345586 | kinesin family member C2 |
| ENSG00000187021 | PNLIPRP1 | -2,118452773 | 0,23029376 | 0,08631 | 0,3808494 | pancreatic lipase related protein 1 |
| ENSG00000241983 | RN7SL566P | -2,118751206 | 0,230246127 | 0,04288 | 0,27914651 | RNA, 7SL, cytoplasmic 566, pseudogene |
| ENSG00000104537 | ANXA13 | -2,119101904 | 0,230190164 | 0,00727 | 0,10821167 | annexin A13 |
| ENSG00000144648 | ACKR2 | -2,119748309 | 0,23008705 | 0,04531 | 0,2876168 | atypical chemokine receptor 2 |
| ENSG00000284218 | NPIPB5 | -2,120347637 | 0,229991486 | 0,06781 | 0,34426294 | nuclear pore complex interacting protein family member B5 |
| ENSG00000264107 | AC138207,2 | -2,121440939 | 0,22981726 | 0,02833 | 0,22750748 | novel transcript, antisense to NF1 |
| ENSG00000141946 | ZIM3 | -2,123546667 | 0,229482069 | 0,16003 | 0,48727386 | zinc finger imprinted 3 |

| | | | | | | |
|---|---|---|---|---|---|---|
| ENSG00000183840 | GPR39 | -2,12384655 | 0,229434373 | 0,00646 | 0,10183581 | G protein-coupled receptor 39 |
| ENSG00000264151 | AC090403,1 | -2,124530554 | 0,22932562 | 0,0656 | 0,3393301 | novel transcript |
| ENSG00000171180 | OR2M4 | -2,126409304 | 0,229027175 | 0,02197 | 0,19708179 | olfactory receptor family 2 subfamily M member 4 |
| ENSG00000205976 | AC091951,1 | -2,126453186 | 0,229020209 | 0,03341 | 0,24709568 | golgi autoantigen, golgin subfamily a, 6-like 2 (GOLGA6L2) pseudogene |
| ENSG00000286417 | AL133334,1 | -2,127036501 | 0,22892763 | 0,01824 | 0,17858832 | novel transcript, antisense to SUPT3H |
| ENSG00000236683 | HMGA1P1 | -2,127311083 | 0,228884063 | 0,10494 | 0,41375163 | high mobility group AT-hook 1 pseudogene 1 |
| ENSG00000144596 | GRIP2 | -2,129150385 | 0,228592443 | 6,81E-05 | 0,00437444 | glutamate receptor interacting protein 2 |
| ENSG00000249623 | PALLD-AS1 | -2,130118826 | 0,228439047 | 0,07111 | 0,35204529 | PALLD antisense RNA 1 |
| ENSG00000285794 | AL031281,1 | -2,130248335 | 0,228418541 | 0,16897 | 0,495041 | SCAN domain containing 1 (SCAND1) pseudogene |
| ENSG00000284647 | AL109936,8 | -2,130304411 | 0,228409663 | 0,01022 | 0,1300932 | novel transcript |
| ENSG00000235172 | LINC01366 | -2,131648327 | 0,228196991 | 0,02772 | 0,22434063 | long intergenic non-protein coding RNA 1366 |
| ENSG00000261270 | AC012181,2 | -2,132197305 | 0,228110173 | 0,02489 | 0,21153603 | novel transcript, sense intronic to HERPUD1 |
| ENSG00000163273 | NPPC | -2,132199004 | 0,228109905 | 0,02736 | 0,22320635 | natriuretic peptide C |
| ENSG00000227801 | COL11A2 | -2,133825383 | 0,227852897 | 0,15076 | 0,47895084 | collagen type XI alpha 2 chain |
| ENSG00000240877 | RN7SL521P | -2,134883287 | 0,227685877 | 0,07477 | 0,35895201 | RNA, 7SL, cytoplasmic 521, pseudogene |
| ENSG00000258450 | AL139099,2 | -2,134897769 | 0,227683592 | 0,02285 | 0,20180274 | novel transcript |
| ENSG00000259654 | AC115102,1 | -2,135006975 | 0,227666358 | 0,0222 | 0,19806119 | novel transcript, antisense to SEC11A |
| ENSG00000176020 | AMIGO3 | -2,13538962 | 0,227605982 | 0,54116 | 0,75868526 | adhesion molecule with Ig like domain 3 |
| ENSG00000266605 | LONRF2P1 | -2,135872974 | 0,227529739 | 0,09152 | 0,39125372 | LONRF2 pseudogene 1 |
| ENSG00000099338 | CATSPERG | -2,136862954 | 0,227373661 | 0,00132 | 0,03867645 | cation channel sperm associated auxiliary subunit gamma |
| ENSG00000286592 | LINC02885 | -2,141576712 | 0,226631969 | 0,00663 | 0,10335057 | long intergenic non-protein coding RNA 2885 |
| ENSG00000261439 | DKFZP434H16 | -2,141785462 | 0,226599179 | 4,99E-05 | 0,00335711 | uncharacterized LOC26077 |
| ENSG00000228262 | LINC01320 | -2,142044353 | 0,226558519 | 0,00408 | 0,07837165 | long intergenic non-protein coding RNA 1320 |
| ENSG00000267178 | PHF5CP | -2,143029564 | 0,226403856 | 0,0418 | 0,27691618 | PHD finger protein 5C pseudogene |
| ENSG00000132688 | NES | -2,143400663 | 0,226345627 | 0,00035 | 0,01589581 | nestin |
| ENSG00000274422 | AC245060,5 | -2,143442718 | 0,226339029 | 0,00477 | 0,08525507 | novel transcript |
| ENSG00000244056 | RN7SL417P | -2,143952697 | 0,226259034 | 0,02125 | 0,19328293 | RNA, 7SL, cytoplasmic 417, pseudogene |
| ENSG00000275143 | SCARNA16 | -2,146833743 | 0,225807648 | 0,00139 | 0,03972417 | small Cajal body-specific RNA 16 |
| ENSG00000181101 | ENTR1P2 | -2,148030554 | 0,225620403 | 0,01673 | 0,16973453 | ENTR1 pseudogene 2 |
| ENSG00000164520 | RAET1E | -2,148751462 | 0,22550769 | 0,0153 | 0,16232837 | retinoic acid early transcript 1E |
| ENSG00000159640 | ACE | -2,149206004 | 0,225436652 | 0,00155 | 0,04237695 | angiotensin I converting enzyme |
| ENSG00000267048 | AC092143,3 | -2,149212235 | 0,225435678 | 0,00953 | 0,12494325 | novel transcript (MC1R-TUBB3 readthrough) |

| | | | | | | |
|---|---|---|---|---|---|---|
| ENSG00000228526 | MIR34AHG | -2,149979951 | 0,225315747 | 0,00258 | 0,05908361 | MIR34A host gene |
| ENSG00000171596 | NMUR1 | -2,150765731 | 0,22519306 | 0,03408 | 0,24923154 | neuromedin U receptor 1 |
| ENSG00000233008 | LINC01725 | -2,152497155 | 0,22492296 | 0,01448 | 0,15740445 | long intergenic non-protein coding RNA 1725 |
| ENSG00000147889 | CDKN2A | -2,153457811 | 0,224773239 | 2,10E-06 | 0,00021229 | cyclin dependent kinase inhibitor 2A |
| ENSG00000286582 | AL161753,1 | -2,156141976 | 0,224355433 | 0,1032 | 0,41099363 | novel transcript |
| ENSG00000104237 | RP1 | -2,156433677 | 0,224310074 | 0,03691 | 0,26055725 | RP1 axonemal microtubule associated |
| ENSG00000233557 | NFU1P2 | -2,158519852 | 0,223985951 | 0,01413 | 0,15603075 | NFU1 iron-sulfur cluster scaffold pseudogene 2 |
| ENSG00000210151 | MT-TS1 | -2,160835577 | 0,223626711 | 0,00372 | 0,07421558 | mitochondrially encoded tRNA-Ser (UCN) 1 |
| ENSG00000288242 | AC106845,2 | -2,161834936 | 0,223471857 | 0,05837 | 0,32253335 | novel transcript, antisense to DLC1 |
| ENSG00000152213 | ARL11 | -2,162319067 | 0,223396879 | 0,02523 | 0,21358358 | ADP ribosylation factor like GTPase 11 |
| ENSG00000145936 | KCNMB1 | -2,162819751 | 0,223319363 | 0,00994 | 0,12777099 | potassium calcium-activated channel subfamily M regulatory beta subunit 1 |
| ENSG00000112038 | OPRM1 | -2,163686245 | 0,223185276 | 0,00257 | 0,05898231 | opioid receptor mu 1 |
| ENSG00000253683 | AC027309,2 | -2,164192412 | 0,223106985 | 0,25748 | 0,56160845 | coiled-coil domain containing 72 (CCDC72) pseudogene |
| ENSG00000164241 | C5orf63 | -2,165269881 | 0,222940421 | 2,42E-13 | 8,69E-11 | chromosome 5 open reading frame 63 |
| ENSG00000235814 | ZNRD1ASP | -2,16548248 | 0,22290757 | 0,03099 | 0,23722256 | zinc ribbon domain containing 1 antisense, pseudogene |
| ENSG00000142408 | CACNG8 | -2,165697969 | 0,222874278 | 0,00423 | 0,07987039 | calcium voltage-gated channel auxiliary subunit gamma 8 |
| ENSG00000254681 | PKD1P5 | -2,166298105 | 0,222781586 | 0,0137 | 0,15356644 | polycystin 1, transient receptor potential channel interacting pseudogene 5 |
| ENSG00000259687 | LINC01220 | -2,167465309 | 0,222601418 | 0,01135 | 0,13828548 | long intergenic non-protein coding RNA 1220 |
| ENSG00000255171 | LINC01499 | -2,169972336 | 0,222214931 | 0,03819 | 0,26456946 | long intergenic non-protein coding RNA 1499 |
| ENSG00000227454 | MTND4P30 | -2,170298676 | 0,222164672 | 0,06889 | 0,34681583 | MT-ND4 pseudogene 30 |
| ENSG00000089847 | ANKRD24 | -2,17030551 | 0,222163619 | 0,01432 | 0,15653473 | ankyrin repeat domain 24 |
| ENSG00000167992 | VWCE | -2,170821025 | 0,222084248 | 0,0027 | 0,06097224 | von Willebrand factor C and EGF domains |
| ENSG00000248329 | APELA | -2,171443478 | 0,22198845 | 0,00054 | 0,02139887 | apelin receptor early endogenous ligand |
| ENSG00000288541 | AL353732,2 | -2,171738466 | 0,221943065 | 0,04591 | 0,28908129 | novel transcript |
| ENSG00000207051 | SNORA27 | -2,172246357 | 0,221864945 | 0,03577 | 0,25659799 | small nucleolar RNA, H/ACA box 27 |
| ENSG00000173442 | EHBP1L1 | -2,17270441 | 0,221794515 | 0,00022 | 0,01148455 | EH domain binding protein 1 like 1 |
| ENSG00000283635 | AC012485,3 | -2,173600059 | 0,221656864 | 0,04351 | 0,28147729 | novel transcript, antisense to PER2 |
| ENSG00000285737 | LINC02680 | -2,17374433 | 0,221634699 | 0,00613 | 0,098625 | long intergenic non-protein coding RNA 2680 |
| ENSG00000251131 | AC025171,3 | -2,174022124 | 0,221592027 | 0,01279 | 0,14802902 | novel transcript |
| ENSG00000285587 | AC012470,1 | -2,17446262 | 0,221524379 | 0,00057 | 0,02219024 | novel transcript |
| ENSG00000104899 | AMH | -2,175067796 | 0,221431474 | 0,00016 | 0,0090156 | anti-Mullerian hormone |

| | | | | | | |
|---|---|---|---|---|---|---|
| ENSG00000075429 | CACNG5 | -2,175450012 | 0,221372817 | 0,00138 | 0,03955033 | calcium voltage-gated channel auxiliary subunit gamma 5 |
| ENSG00000285280 | AL390957,1 | -2,176495278 | 0,221212486 | 0,11093 | 0,42371066 | novel transcript |
| ENSG00000239377 | ZNF775-AS1 | -2,179256132 | 0,220789561 | 0,01091 | 0,13517515 | ZNF775 antisense RNA 1 |
| ENSG00000267254 | ZNF790-AS1 | -2,180676327 | 0,220572322 | 3,06E-05 | 0,00220458 | ZNF790 antisense RNA 1 |
| ENSG00000141668 | CBLN2 | -2,181154986 | 0,220499152 | 0,0038 | 0,07518238 | cerebellin 2 precursor |
| ENSG00000224126 | UBE2SP2 | -2,181771616 | 0,220404928 | 0,05946 | 0,32472914 | ubiquitin conjugating enzyme E2 S pseudogene 2 |
| ENSG00000264456 | AC138207,4 | -2,185007374 | 0,219911145 | 0,00548 | 0,09208977 | novel transcript, sense intronic RNF135 |
| ENSG00000183484 | GPR132 | -2,185055074 | 0,219903874 | 0,04692 | 0,29141554 | G protein-coupled receptor 132 |
| ENSG00000164509 | IL31RA | -2,185606702 | 0,219819808 | 0,02599 | 0,21717729 | interleukin 31 receptor A |
| ENSG00000226763 | SRRM5 | -2,185978094 | 0,219763227 | 0,00391 | 0,07646656 | serine/arginine repetitive matrix 5 |
| ENSG00000281803 | AC136352,11 | -2,186839447 | 0,219632058 | 0,01148 | 0,1391122 | novel transcript |
| ENSG00000205231 | TTLL10-AS1 | -2,18752995 | 0,219526963 | 0,01957 | 0,18559269 | TTLL10 antisense RNA 1 |
| ENSG00000251003 | ZFPM2-AS1 | -2,189732893 | 0,219192009 | 0,00173 | 0,04563673 | ZFPM2 antisense RNA 1 |
| ENSG00000282044 | AC083849,3 | -2,190562433 | 0,219066011 | 0,02216 | 0,19795874 | novel transcript |
| ENSG00000152936 | LMNTD1 | -2,19124454 | 0,218962461 | 0,06029 | 0,32601555 | lamin tail domain containing 1 |
| ENSG00000152377 | SPOCK1 | -2,191304341 | 0,218953385 | 1,02E-09 | 2,31E-07 | SPARC (osteonectin), cwcv and kazal like domains proteoglycan 1 |
| ENSG00000233295 | FAM90A20P | -2,191682232 | 0,218896041 | 0,00881 | 0,11893447 | family with sequence similarity 90 member A20, pseudogene |
| ENSG00000276788 | SNORD26 | -2,192952932 | 0,218703326 | 0,00517 | 0,08972695 | small nucleolar RNA, C/D box 26 |
| ENSG00000230067 | HSPD1P6 | -2,193055359 | 0,2186878 | 0,07099 | 0,35177751 | heat shock protein family D (Hsp60) member 1 pseudogene 6 |
| ENSG00000271218 | AL033384,2 | -2,193645211 | 0,218598406 | 0,05555 | 0,31503413 | novel transcript, antisense to LRRC1 |
| ENSG00000197766 | CFD | -2,194862735 | 0,218414004 | 0,00562 | 0,09352688 | complement factor D |
| ENSG00000185028 | LRRC14B | -2,195414886 | 0,218330428 | 0,02109 | 0,19289451 | leucine rich repeat containing 14B |
| ENSG00000253348 | AC008514,1 | -2,195552834 | 0,218309553 | 0,00804 | 0,11395828 | novel transcript |
| ENSG00000259952 | AC009133,2 | -2,196248 | 0,218204385 | 0,05453 | 0,31262876 | novel transcript, antisense to MAZ |
| ENSG00000179520 | SLC17A8 | -2,196886215 | 0,218107878 | 0,05189 | 0,30606616 | solute carrier family 17 member 8 |
| ENSG00000198739 | LRRTM3 | -2,198170746 | 0,217913768 | 0,00034 | 0,01565603 | leucine rich repeat transmembrane neuronal 3 |
| ENSG00000285812 | AL390719,3 | -2,199563611 | 0,217703482 | 0,00309 | 0,0662736 | novel transcript, antisense to C1orf159 |
| ENSG00000167332 | OR51E2 | -2,20024466 | 0,217600736 | 0,10386 | 0,41222634 | olfactory receptor family 51 subfamily E member 2 |
| ENSG00000169402 | RSPH10B2 | -2,200291324 | 0,217593698 | 0,00324 | 0,06839437 | radial spoke head 10 homolog B2 |
| ENSG00000253334 | AC132219,1 | -2,202802742 | 0,217215244 | 0,054 | 0,31168784 | novel transcript, antisense to SNX16 |

| | | | | | | |
|---|---|---|---|---|---|---|
| ENSG00000281284 | MAPT-IT1 | -2,202845224 | 0,217208848 | 0,0031 | 0,06636757 | MAPT intronic transcript 1 |
| ENSG00000262299 | AL513523,6 | -2,209515071 | 0,216206969 | 0,0138 | 0,15426432 | immunoglobulin superfamily, DCC subclass, member 3 (IGDCC3) pseudogene |
| ENSG00000236532 | LINC01695 | -2,209798319 | 0,216164524 | 0,00142 | 0,04024594 | long intergenic non-protein coding RNA 1695 |
| ENSG00000261617 | LINC02177 | -2,210021891 | 0,216131028 | 0,00421 | 0,07968892 | long intergenic non-protein coding RNA 2177 |
| ENSG00000260949 | AP006545,1 | -2,21199276 | 0,215835973 | 0,00718 | 0,10764638 | novel transcript, overlapping to EIF4EBP1 |
| ENSG00000262003 | AC087392,1 | -2,213992644 | 0,215536985 | 0,00199 | 0,05000974 | novel transcript, antisense to NXN |
| ENSG00000254488 | AC007876,1 | -2,21454001 | 0,215455225 | 0,0052 | 0,08998673 | novel transcript |
| ENSG00000254101 | LINC02055 | -2,216341012 | 0,215186427 | 0,00206 | 0,0511876 | long intergenic non-protein coding RNA 2055 |
| ENSG00000164398 | ACSL6 | -2,216381264 | 0,215180423 | 0,00046 | 0,01927018 | acyl-CoA synthetase long chain family member 6 |
| ENSG00000255384 | AP001267,2 | -2,216994055 | 0,215089044 | 0,00506 | 0,08898421 | novel transcript |
| ENSG00000101276 | SLC52A3 | -2,217557348 | 0,21500508 | 0,11225 | 0,42587863 | solute carrier family 52 member 3 |
| ENSG00000229349 | ACTG1P9 | -2,217833484 | 0,214963931 | 0,03851 | 0,26548082 | actin gamma 1 pseudogene 9 |
| ENSG00000285261 | AC270285,6 | -2,21798095 | 0,21494196 | 0,00836 | 0,11551322 | novel transcript |
| ENSG00000239608 | RUVBL1-AS1 | -2,219257064 | 0,21475192 | 0,01202 | 0,14273718 | RUVBL1 antisense RNA 1 |
| ENSG00000285744 | AC083837,1 | -2,220619079 | 0,214549274 | 0,01691 | 0,17086866 | novel transcript |
| ENSG00000225556 | C2CD4D | -2,222937276 | 0,214204801 | 0,01055 | 0,1325438 | C2 calcium dependent domain containing 4D |
| ENSG00000214695 | NPAP1P2 | -2,223259953 | 0,214156897 | 0,00441 | 0,081876 | nuclear pore associated protein 1 pseudogene 2 |
| ENSG00000255372 | AC107886,1 | -2,223970959 | 0,21405138 | 0,00141 | 0,04015868 | novel transcript |
| ENSG00000241568 | RN7SL338P | -2,224785462 | 0,213930567 | 0,0471 | 0,29208796 | RNA, 7SL, cytoplasmic 338, pseudogene |
| ENSG00000287990 | AL358334,4 | -2,22652905 | 0,213672174 | 0,0019 | 0,04859118 | novel transcript, antisense to TRIM9 |
| ENSG00000267416 | AC025048,4 | -2,226843278 | 0,21362564 | 0,02599 | 0,21717729 | novel transcript |
| ENSG00000228022 | HCG20 | -2,230323058 | 0,213110996 | 0,10267 | 0,41043823 | HLA complex group 20 |
| ENSG00000113263 | ITK | -2,230463295 | 0,213090282 | 0,02672 | 0,22056859 | IL2 inducible T cell kinase |
| ENSG00000198835 | GJC2 | -2,230677906 | 0,213058586 | 0,01089 | 0,13508104 | gap junction protein gamma 2 |
| ENSG00000284973 | AC270285,2 | -2,232331622 | 0,212814503 | 0,02895 | 0,22977124 | novel transcript, sense intronic to CTSB |
| ENSG00000271727 | AL357054,2 | -2,234902974 | 0,212435536 | 0,25375 | 0,55879966 | novel transcript |
| ENSG00000173976 | RAX2 | -2,236988237 | 0,212128705 | 0,02543 | 0,21440233 | retina and anterior neural fold homeobox 2 |
| ENSG00000228415 | PTMAP1 | -2,237988151 | 0,211981732 | 0,0285 | 0,22823749 | prothymosin alpha pseudogene 1 |
| ENSG00000259134 | LINC00924 | -2,238514808 | 0,211904362 | 0,00309 | 0,06627153 | long intergenic non-protein coding RNA 924 |
| ENSG00000260198 | AC009102,1 | -2,238644092 | 0,211885374 | 0,00364 | 0,07353585 | novel transcript, sense intronic to GNAO1 |
| ENSG00000261170 | AC009053,3 | -2,239385037 | 0,211776581 | 0,00232 | 0,05549873 | novel transcript, antisense to CLEC18B |
| ENSG00000225091 | SNORA71A | -2,240192714 | 0,211658053 | 0,00406 | 0,07819491 | small nucleolar RNA, H/ACA box 71A |

| | | | | | | |
|---|---|---|---|---|---|---|
| ENSG00000250007 | AC087457,1 | -2,240216082 | 0,211654625 | 0,00116 | 0,03569587 | novel transcript, antisense to ACTC1 |
| ENSG00000081800 | SLC13A1 | -2,241077688 | 0,211528258 | 0,02914 | 0,23027064 | solute carrier family 13 member 1 |
| ENSG00000102362 | SYTL4 | -2,241952123 | 0,211400087 | 1,70E-07 | 2,36E-05 | synaptotagmin like 4 |
| ENSG00000231240 | KLF2P1 | -2,244430554 | 0,211037231 | 0,0048 | 0,08549364 | Kruppel like factor 2 pseudogene 1 |
| ENSG00000142910 | TINAGL1 | -2,24493503 | 0,210963449 | 0,16359 | 0,48997962 | tubulointerstitial nephritis antigen like 1 |
| ENSG00000283268 | TEX54 | -2,247134848 | 0,210642018 | 0,01531 | 0,16232837 | testis expressed 54 |
| ENSG00000231512 | SEPTIN14P21 | -2,247213285 | 0,210630566 | 0,05666 | 0,31746133 | septin 14 pseudogene 21 |
| ENSG00000230322 | AL136320,1 | -2,248066495 | 0,210506036 | 0,00254 | 0,0585725 | novel transcript |
| ENSG00000262585 | LINC01979 | -2,248067385 | 0,210505906 | 0,01649 | 0,16825348 | long intergenic non-protein coding RNA 1979 |
| ENSG00000151650 | VENTX | -2,249924922 | 0,210235044 | 0,02899 | 0,22977124 | VENT homeobox |
| ENSG00000265555 | LINC01903 | -2,251599924 | 0,209991098 | 0,07209 | 0,35432191 | long intergenic non-protein coding RNA 1903 |
| ENSG00000180828 | BHLHE22 | -2,252287362 | 0,209891062 | 8,06E-06 | 0,00069151 | basic helix-loop-helix family member e22 |
| ENSG00000213316 | LTC4S | -2,252288088 | 0,209890956 | 0,043 | 0,27928585 | leukotriene C4 synthase |
| ENSG00000283887 | LTC4S | -2,252288088 | 0,209890956 | 0,043 | 0,27928585 | leukotriene C4 synthase |
| ENSG00000272702 | AC010913,1 | -2,253337033 | 0,209738406 | 0,00123 | 0,03698641 | novel transcript |
| ENSG00000237918 | SAPCD1 | -2,255315873 | 0,20945092 | 0,01534 | 0,16232837 | suppressor APC domain containing 1 |
| ENSG00000224982 | TMEM233 | -2,25612179 | 0,20933395 | 0,03456 | 0,25132008 | transmembrane protein 233 |
| ENSG00000233485 | FHAD1-AS1 | -2,257098514 | 0,209192276 | 0,03709 | 0,26086671 | FHAD1 antisense RNA 1 |
| ENSG00000271771 | AC139792,1 | -2,257921682 | 0,209072949 | 0,00769 | 0,11151515 | novel transcript |
| ENSG00000226051 | ZNF503-AS1 | -2,258554197 | 0,208981307 | 0,11908 | 0,43696075 | ZNF503 antisense RNA 1 |
| ENSG00000287425 | AP003717,4 | -2,260038194 | 0,208766453 | 0,0028 | 0,06237064 | novel transcript |
| ENSG00000039068 | CDH1 | -2,262132118 | 0,20846367 | 0,02727 | 0,22298278 | cadherin 1 |
| ENSG00000254732 | AP001931,1 | -2,263904319 | 0,208207751 | 0,00753 | 0,11001711 | novel protein, C11orf31-CTNND1 readthrough |
| ENSG00000287025 | AC079117,1 | -2,26455654 | 0,208113644 | 0,00371 | 0,07418692 | novel transcript, sense intronic to LRRTM4 |
| ENSG00000208772 | SNORD94 | -2,264715855 | 0,208090664 | 0,00118 | 0,03613087 | small nucleolar RNA, C/D box 94 |
| ENSG00000285438 | SOX7 | -2,264766357 | 0,20808338 | 0,21051 | 0,5290538 | SRY-box transcription factor 7 |
| ENSG00000202347 | RNU1-16P | -2,264862553 | 0,208069506 | 0,00327 | 0,06873635 | RNA, U1 small nuclear 16, pseudogene |
| ENSG00000121380 | BCL2L14 | -2,265712494 | 0,207946961 | 4,73E-05 | 0,00320624 | BCL2 like 14 |
| ENSG00000286899 | AL139286,2 | -2,267001652 | 0,207761227 | 0,00099 | 0,03217084 | novel transcript, antisense to AGO1 |
| ENSG00000186407 | CD300E | -2,267383017 | 0,207706315 | 0,19726 | 0,51854293 | CD300e molecule |
| ENSG00000261602 | AC092115,2 | -2,267585787 | 0,207677124 | 0,09304 | 0,39400451 | novel transcript |
| ENSG00000281593 | AC006978,2 | -2,268102639 | 0,207602736 | 0,0824 | 0,37394502 | novel protein, ZNRF2-LINC01176 readthrough |
| ENSG00000261028 | AC012322,1 | -2,268321164 | 0,207571293 | 0,00277 | 0,06186307 | novel transcript |

| | | | | | | |
|---|---|---|---|---|---|---|
| ENSG00000199347 | RNU5E-1 | -2,269358148 | 0,207422148 | 0,00095 | 0,03123078 | RNA, U5E small nuclear 1 |
| ENSG00000284716 | AL034417,3 | -2,270628386 | 0,207239601 | 0,00525 | 0,09023148 | novel transcript |
| ENSG00000132702 | HAPLN2 | -2,271198708 | 0,207157692 | 0,0089 | 0,11977436 | hyaluronan and proteoglycan link protein 2 |
| ENSG00000189350 | TOGARAM2 | -2,271230273 | 0,207153159 | 0,00666 | 0,10355399 | TOG array regulator of axonemal microtubules 2 |
| ENSG00000277578 | Metazoa_SRP | -2,272369773 | 0,206989606 | 0,02914 | 0,23027064 | Metazoan signal recognition particle RNA |
| ENSG00000259402 | AC090515,5 | -2,273740808 | 0,206792991 | 0,00478 | 0,08525507 | novel transcript, sense intronic to FAM63B |
| ENSG00000158517 | NCF1 | -2,274564638 | 0,206674938 | 0,03666 | 0,25954613 | neutrophil cytosolic factor 1 |
| ENSG00000260115 | AC012174,1 | -2,274854373 | 0,206633436 | 0,11061 | 0,42326288 | novel transcript, antisense to CDH8 |
| ENSG00000162571 | TTLL10 | -2,275195359 | 0,206584603 | 0,00196 | 0,04956833 | tubulin tyrosine ligase like 10 |
| ENSG00000287262 | AC021148,2 | -2,276978867 | 0,206329375 | 0,00417 | 0,07918868 | novel transcript |
| ENSG00000184774 | MGAT4EP | -2,277141192 | 0,206306161 | 0,00966 | 0,12585684 | MGAT4 family member E, pseudogene |
| ENSG00000264346 | SNORA77B | -2,278274885 | 0,206144106 | 0,00015 | 0,00842649 | small nucleolar RNA, H/ACA box 77B |
| ENSG00000284543 | LINC01226 | -2,281147679 | 0,205734026 | 0,00176 | 0,04603979 | long intergenic non-protein coding RNA 1226 |
| ENSG00000261058 | AC099508,2 | -2,282599032 | 0,205527161 | 0,10211 | 0,40938532 | novel transcript |
| ENSG00000259771 | AC092756,1 | -2,283101131 | 0,205455644 | 0,16344 | 0,48977932 | novel transcript, sense intronic to MYO1E |
| ENSG00000111971 | LY6G5C | -2,284471085 | 0,20526064 | 0,03989 | 0,27016182 | lymphocyte antigen 6 family member G5C |
| ENSG00000249616 | CCNB3P1 | -2,284839086 | 0,205208289 | 0,075 | 0,35927361 | cyclin B3 pseudogene 1 |
| ENSG00000268945 | AC010422,2 | -2,286505005 | 0,204971466 | 0,00764 | 0,11109228 | novel transcript |
| ENSG00000229582 | AL358074,1 | -2,287363463 | 0,204849537 | 0,00907 | 0,12087977 | novel transcript |
| ENSG00000162068 | NTN3 | -2,287490965 | 0,204831434 | 0,00085 | 0,02896138 | netrin 3 |
| ENSG00000169208 | OR10G3 | -2,29156484 | 0,204253848 | 0,00483 | 0,08564971 | olfactory receptor family 10 subfamily G member 3 |
| ENSG00000248641 | HMGA1P2 | -2,293147252 | 0,204029936 | 0,09465 | 0,39712927 | high mobility group AT-hook 1 pseudogene 2 |
| ENSG00000142235 | LMTK3 | -2,29316734 | 0,204027095 | 0,0006 | 0,0232207 | lemur tyrosine kinase 3 |
| ENSG00000196132 | MYT1 | -2,294385981 | 0,203854826 | 0,00013 | 0,00727013 | myelin transcription factor 1 |
| ENSG00000277545 | AL356752,1 | -2,296216933 | 0,203596274 | 0,00583 | 0,09586861 | novel transcript, sense intronic to ATP11A |
| ENSG00000247699 | FABP6-AS1 | -2,297233933 | 0,203452804 | 0,04443 | 0,28434634 | FABP6 antisense RNA 1 |
| ENSG00000232564 | MRTFA-AS1 | -2,29745133 | 0,203422148 | 0,00127 | 0,03791245 | MRTFA antisense RNA 1 |
| ENSG00000267385 | AC011498,4 | -2,297827457 | 0,203369121 | 0,02122 | 0,19328293 | novel protein |
| ENSG00000258010 | AC016705,1 | -2,299034144 | 0,203199091 | 0,00456 | 0,08326903 | novel transcript, antisense to ARNT2 |
| ENSG00000236516 | KLF2P4 | -2,300615269 | 0,202976517 | 0,00108 | 0,03404843 | Kruppel like factor 2 pseudogene 4 |
| ENSG00000215483 | LINC00598 | -2,301150012 | 0,202901296 | 0,00665 | 0,10355399 | long intergenic non-protein coding RNA 598 |
| ENSG00000224884 | AC034187,1 | -2,301918327 | 0,202793269 | 0,14565 | 0,47307324 | novel transcript |
| ENSG00000221953 | C1orf229 | -2,302675201 | 0,202686906 | 0,00042 | 0,01799896 | chromosome 1 open reading frame 229 |

| | | | | | | |
|---|---|---|---|---|---|---|
| ENSG00000129226 | CD68 | -2,304215305 | 0,20247065 | 0,01912 | 0,18296697 | CD68 molecule |
| ENSG00000177508 | IRX3 | -2,30468163 | 0,202405215 | 0,00285 | 0,06340012 | iroquois homeobox 3 |
| ENSG00000249896 | LINC02495 | -2,305472306 | 0,202294317 | 0,01568 | 0,16409525 | long intergenic non-protein coding RNA 2495 |
| ENSG00000271275 | AC007326,1 | -2,306788456 | 0,202109851 | 0,02289 | 0,20192132 | POM121 transmembrane nucleoporin (POM121) pseudogene |
| ENSG00000250075 | AC104806,2 | -2,310654579 | 0,201568963 | 0,01212 | 0,14307474 | novel transcript |
| ENSG00000227195 | MIR663AHG | -2,31102158 | 0,201517694 | 0,01823 | 0,17856146 | MIR663A host gene |
| ENSG00000086506 | HBQ1 | -2,311523954 | 0,201447533 | 0,01388 | 0,15454988 | hemoglobin subunit theta 1 |
| ENSG00000230749 | MEIS1-AS2 | -2,312283196 | 0,201341546 | 4,77E-19 | 3,43E-16 | MEIS1 antisense RNA 2 |
| ENSG00000232719 | AC007272,1 | -2,313118108 | 0,20122506 | 0,02944 | 0,23083495 | novel transcript |
| ENSG00000261392 | AC087190,2 | -2,31483407 | 0,200985863 | 0,06899 | 0,34717336 | novel transcript |
| ENSG00000256906 | LINC02419 | -2,314850663 | 0,200983551 | 0,08173 | 0,37295776 | long intergenic non-protein coding RNA 2419 |
| ENSG00000119535 | CSF3R | -2,315836448 | 0,200846267 | 0,02823 | 0,2268256 | colony stimulating factor 3 receptor |
| ENSG00000226111 | MRPS18B | -2,31710181 | 0,200670186 | 0,02898 | 0,22977124 | mitochondrial ribosomal protein S18B |
| ENSG00000269560 | AC010422,4 | -2,317933673 | 0,200554512 | 0,00531 | 0,09082808 | novel transcript, sense intronic to ZNF564 |
| ENSG00000206258 | TNXB | -2,322136105 | 0,199971166 | 0,03842 | 0,26521398 | tenascin XB |
| ENSG00000231608 | TNXB | -2,322136105 | 0,199971166 | 0,03842 | 0,26521398 | tenascin XB |
| ENSG00000260473 | AC068987,2 | -2,32251087 | 0,199919226 | 0,0016 | 0,04288108 | novel transcript |
| ENSG00000268041 | ERFL | -2,322622185 | 0,199903802 | 0,07379 | 0,3570536 | ETS repressor factor like |
| ENSG00000165246 | NLGN4Y | -2,325699642 | 0,199477835 | 5,03E-07 | 6,11E-05 | neuroligin 4 Y-linked |
| ENSG00000115266 | APC2 | -2,32588103 | 0,199452757 | 0,00016 | 0,00890575 | APC regulator of WNT signaling pathway 2 |
| ENSG00000206245 | TSBP1 | -2,325981687 | 0,199438841 | 0,22546 | 0,5407493 | testis expressed basic protein 1 |
| ENSG00000133962 | CATSPERB | -2,326173366 | 0,199412345 | 0,06728 | 0,34325886 | cation channel sperm associated auxiliary subunit beta |
| ENSG00000196826 | AC008758,1 | -2,326324157 | 0,199391504 | 0,30983 | 0,59807209 | novel zinc finger protein |
| ENSG00000259704 | AC124248,1 | -2,326663954 | 0,199344547 | 0,04531 | 0,2876168 | novel transcript, sense overlapping 3' UTR of BLM |
| ENSG00000267033 | AC020911,1 | -2,329161858 | 0,198999697 | 0,01386 | 0,15454988 | novel transcript, antisense FAM32A |
| ENSG00000281833 | AL354892,6 | -2,330694154 | 0,198788451 | 0,31615 | 0,60224605 | novel transcript |
| ENSG00000275679 | PRH2 | -2,331109548 | 0,198731222 | 0,10076 | 0,40755636 | proline rich protein HaeIII subfamily 2 |
| ENSG00000272803 | PRH2 | -2,331109548 | 0,198731222 | 0,10076 | 0,40755636 | proline rich protein HaeIII subfamily 2 |
| ENSG00000232190 | LINC02181 | -2,338409466 | 0,197728198 | 0,01697 | 0,17097909 | long intergenic non-protein coding RNA 2181 |
| ENSG00000247627 | MTND4P12 | -2,338480984 | 0,197718396 | 0,00308 | 0,06624362 | MT-ND4 pseudogene 12 |
| ENSG00000275009 | GOLGA6L1 | -2,339425317 | 0,19758902 | 0,31302 | 0,59987747 | golgin A6 family like 1 |
| ENSG00000205403 | CFI | -2,33960326 | 0,197564651 | 0,01167 | 0,14066917 | complement factor I |

| | | | | | | |
|---|---|---|---|---|---|---|
| ENSG00000276975 | HYDIN2 | -2,339732421 | 0,197546964 | 0,00385 | 0,07555024 | HYDIN axonemal central pair apparatus protein 2 (pseudogene) |
| ENSG00000285621 | AL138999,2 | -2,342221846 | 0,197206383 | 0,10994 | 0,42223228 | novel transcript, antisense to STARD13 |
| ENSG00000268153 | AC005176,1 | -2,345720579 | 0,19672871 | 0,0099 | 0,12753789 | lectin, galactoside-binding, soluble, 14 (LGALS14) pseudogene |
| ENSG00000153303 | FRMD1 | -2,345754324 | 0,196724109 | 0,0603 | 0,32601555 | FERM domain containing 1 |
| ENSG00000286942 | AC017104,5 | -2,346077426 | 0,196680056 | 0,01785 | 0,17612603 | novel transcript |
| ENSG00000186529 | CYP4F3 | -2,347247978 | 0,196520541 | 0,08644 | 0,38089807 | cytochrome P450 family 4 subfamily F member 3 |
| ENSG00000050730 | TNIP3 | -2,349127637 | 0,196264665 | 0,0071 | 0,10744093 | TNFAIP3 interacting protein 3 |
| ENSG00000259006 | AC092143,2 | -2,349792721 | 0,196174208 | 0,28987 | 0,58531219 | novel transcript, antisense to MC1R |
| ENSG00000128564 | VGF | -2,35203862 | 0,195869053 | 0,00037 | 0,01681292 | VGF nerve growth factor inducible |
| ENSG00000249505 | AC108673,1 | -2,352612032 | 0,195791219 | 0,08137 | 0,37218783 | jumonji domain containing 4 (JMJD4) pseudogene |
| ENSG00000273188 | AL022328,3 | -2,352913302 | 0,195750337 | 0,03455 | 0,25132008 | novel transcript, antisense to TRABD |
| ENSG00000271959 | AC100803,4 | -2,353220284 | 0,195708689 | 0,02882 | 0,2296248 | novel transcript, antisense to MROH5 |
| ENSG00000266385 | AC005899,5 | -2,355684229 | 0,195374728 | 0,02406 | 0,20739525 | novel transcript |
| ENSG00000256705 | AL137779,1 | -2,355760098 | 0,195364454 | 0,0085 | 0,11637365 | novel transcript, antisense to PPP2R5C |
| ENSG00000109684 | CLNK | -2,356076874 | 0,195321562 | 0,00329 | 0,06904163 | cytokine dependent hematopoietic cell linker |
| ENSG00000263657 | AC023389,1 | -2,356613881 | 0,195248872 | 0,00588 | 0,0966472 | novel transcript, antisense to SSH2 |
| ENSG00000275139 | AL133492,1 | -2,361306156 | 0,194614869 | 0,00095 | 0,03138491 | novel transcript, antisense to PCBP3 |
| ENSG00000284887 | AC270254,1 | -2,361386946 | 0,194603971 | 0,25832 | 0,56226755 | |
| ENSG00000167664 | TMIGD2 | -2,361445161 | 0,194596119 | 0,00686 | 0,10559142 | transmembrane and immunoglobulin domain containing 2 |
| ENSG00000283959 | AP002851,1 | -2,361821847 | 0,194545316 | 0,01242 | 0,14572968 | novel transcript |
| ENSG00000249799 | SNRPCP13 | -2,363339204 | 0,194340811 | 0,02894 | 0,22977124 | small nuclear ribonucleoprotein polypeptide C pseudogene 13 |
| ENSG00000266237 | AC121320,1 | -2,364529132 | 0,194180585 | 0,10873 | 0,41976565 | novel transcript, sense intronic to TAF4B |
| ENSG00000276529 | AP001505,1 | -2,367081753 | 0,193837317 | 0,00161 | 0,04303245 | novel transcript |
| ENSG00000159961 | OR3A3 | -2,36769826 | 0,193754502 | 0,02435 | 0,2085577 | olfactory receptor family 3 subfamily A member 3 |
| ENSG00000250934 | AC016924,1 | -2,368011296 | 0,193712466 | 0,00152 | 0,04181037 | novel transcript |
| ENSG00000230490 | AL139383,1 | -2,369651193 | 0,1934924 | 0,00046 | 0,01927018 | novel transcript |
| ENSG00000269694 | AC005197,1 | -2,37031522 | 0,193403362 | 0,02704 | 0,22193038 | novel transcript, antisense to CERS1 |
| ENSG00000175604 | AC007601,1 | -2,370831419 | 0,193334174 | 0,00943 | 0,1242362 | novel transcript, antisense to SNX29 |
| ENSG00000273529 | HES5 | -2,373552556 | 0,192969861 | 2,28E-05 | 0,00173571 | hes family bHLH transcription factor 5 |
| ENSG00000183347 | GBP6 | -2,373872163 | 0,192927116 | 0,01988 | 0,18711351 | guanylate binding protein family member 6 |

| | | | | | | |
|---|---|---|---|---|---|---|
| ENSG00000233538 | AC017104,2 | -2,375160327 | 0,192754931 | 0,01511 | 0,16134382 | novel transcript |
| ENSG00000251733 | SCARNA8 | -2,375588161 | 0,192697778 | 0,30639 | 0,5953441 | small Cajal body-specific RNA 8 |
| ENSG00000267234 | AC092067,1 | -2,375961354 | 0,192647938 | 0,02199 | 0,19708179 | novel transcript, antisense to GATAD2A |
| ENSG00000229111 | MED4-AS1 | -2,376074046 | 0,19263289 | 0,00294 | 0,06466643 | MED4 antisense RNA 1 |
| ENSG00000198685 | LINC01565 | -2,376202296 | 0,192615766 | 0,01373 | 0,15370548 | long intergenic non-protein coding RNA 1565 |
| ENSG00000203499 | IQANK1 | -2,376655182 | 0,192555311 | 0,03596 | 0,25683797 | IQ motif and ankyrin repeat containing 1 |
| ENSG00000176920 | FUT2 | -2,376679383 | 0,192552081 | 0,00116 | 0,03569587 | fucosyltransferase 2 |
| ENSG00000133475 | GGT2 | -2,378064323 | 0,192367326 | 0,03131 | 0,23887055 | gamma-glutamyltransferase 2 |
| ENSG00000146147 | MLIP | -2,378694409 | 0,192283329 | 0,0088 | 0,11883496 | muscular LMNA interacting protein |
| ENSG00000165118 | C9orf64 | -2,382515593 | 0,191774713 | 0,01715 | 0,17207348 | chromosome 9 open reading frame 64 |
| ENSG00000239221 | RN7SL442P | -2,383445381 | 0,191651158 | 0,11721 | 0,43451125 | RNA, 7SL, cytoplasmic 442, pseudogene |
| ENSG00000254277 | AC009446,1 | -2,384013171 | 0,191575746 | 0,22093 | 0,53800198 | novel transcript |
| ENSG00000179886 | TIGD5 | -2,385274802 | 0,191408287 | 0,4814 | 0,71822668 | tigger transposable element derived 5 |
| ENSG00000242798 | AC073842,2 | -2,385639265 | 0,191359938 | 0,00382 | 0,0752235 | novel transcript |
| ENSG00000287560 | AL731533,3 | -2,386473458 | 0,191249323 | 0,00416 | 0,07917294 | novel transcript |
| ENSG00000287118 | AC008056,3 | -2,38805146 | 0,191040251 | 0,01259 | 0,1469008 | novel transcript, antisense to NRXN3 |
| ENSG00000277740 | Metazoa_SRP | -2,388460727 | 0,190986064 | 0,0188 | 0,18167964 | Metazoan signal recognition particle RNA |
| ENSG00000287550 | AL450345,2 | -2,38967624 | 0,19082522 | 0,00132 | 0,03881794 | novel transcript |
| ENSG00000236447 | ATG10-IT1 | -2,390972272 | 0,190653871 | 0,01677 | 0,17000427 | ATG10 intronic transcript 1 |
| ENSG00000203867 | RBM20 | -2,393051077 | 0,190379352 | 0,00024 | 0,01209726 | RNA binding motif protein 20 |
| ENSG00000231840 | TMEM139-AS | -2,393702348 | 0,190293429 | 0,01222 | 0,14377653 | TMEM139 antisense RNA 1 |
| ENSG00000224414 | AC010886,1 | -2,393827253 | 0,190276955 | 0,00636 | 0,10063153 | ATPase, H+/K+ transporting, nongastric, alpha polypeptide (ATP12A) pseudogene |
| ENSG00000287516 | AL359764,2 | -2,394542613 | 0,190182629 | 0,00223 | 0,05388971 | novel transcript |
| ENSG00000285795 | FAM90A23P | -2,394719498 | 0,190159313 | 0,03898 | 0,26717464 | family with sequence similarity 90 member A23, pseudogene |
| ENSG00000258735 | LINC00637 | -2,39537572 | 0,190072837 | 0,04478 | 0,28590745 | long intergenic non-protein coding RNA 637 |
| ENSG00000174680 | GRIK1-AS1 | -2,396129591 | 0,189973542 | 0,00303 | 0,06571644 | GRIK1 antisense RNA 1 |
| ENSG00000261411 | AL161938,1 | -2,396545817 | 0,189918741 | 0,07417 | 0,35759867 | novel transcript |
| ENSG00000274594 | AL391669,1 | -2,396666877 | 0,189902805 | 0,01084 | 0,13471553 | transmembrane protein 80 (TMEM80) pseudogene |
| ENSG00000243806 | RPL7P18 | -2,398292813 | 0,189688903 | 0,00339 | 0,07021313 | ribosomal protein L7 pseudogene 18 |
| ENSG00000228484 | AL137025,1 | -2,398483458 | 0,189663838 | 0,00839 | 0,11559293 | novel transcript |
| ENSG00000259881 | AC092384,2 | -2,400317839 | 0,189422835 | 0,01993 | 0,18715502 | novel transcript, antisense to CBFA2T3 |

| | | | | | | |
|---|---|---|---|---|---|---|
| ENSG00000260008 | AP001527,1 | -2,40198965 | 0,189203457 | 0,00382 | 0,07526433 | novel transcript |
| ENSG00000273382 | AL356488,3 | -2,403846757 | 0,188960061 | 0,00262 | 0,05975157 | novel transcript, antisense to TMEM167B |
| ENSG00000254251 | AC103770,1 | -2,405294277 | 0,188770564 | 0,05175 | 0,30583008 | novel transcript |
| ENSG00000187486 | KCNJ11 | -2,406746822 | 0,188580601 | 0,001 | 0,03233482 | potassium inwardly rectifying channel subfamily J member 11 |
| ENSG00000006534 | ALDH3B1 | -2,407301143 | 0,188508157 | 0,00309 | 0,06627153 | aldehyde dehydrogenase 3 family member B1 |
| ENSG00000249481 | SPATS1 | -2,408490342 | 0,188352836 | 0,00336 | 0,07000131 | spermatogenesis associated serine rich 1 |
| ENSG00000182177 | ASB18 | -2,408944157 | 0,188293597 | 0,0002 | 0,01061649 | ankyrin repeat and SOCS box containing 18 |
| ENSG00000200534 | SNORA33 | -2,41234992 | 0,187849617 | 0,00634 | 0,10046895 | small nucleolar RNA, H/ACA box 33 |
| ENSG00000261501 | AC079341,1 | -2,414655637 | 0,187549635 | 0,00607 | 0,09835545 | novel transcript |
| ENSG00000267691 | SHC1P2 | -2,416136836 | 0,187357179 | 0,04576 | 0,28848581 | SHC adaptor protein 1 pseudogene 2 |
| ENSG00000261513 | AC009097,4 | -2,416959227 | 0,187250409 | 0,0206 | 0,19042717 | novel transcript, antisense to PHLPP1 |
| ENSG00000241943 | RN7SL188P | -2,417261884 | 0,18721113 | 0,07276 | 0,35575672 | RNA, 7SL, cytoplasmic 188, pseudogene |
| ENSG00000174992 | ZG16 | -2,417526416 | 0,187176807 | 0,03585 | 0,25662951 | zymogen granule protein 16 |
| ENSG00000251330 | AC114939,1 | -2,419922199 | 0,186866233 | 0,00391 | 0,07646656 | novel transcript, antisense to FBXO38 |
| ENSG00000265992 | ESRG | -2,420693853 | 0,186766311 | 0,00038 | 0,01701402 | embryonic stem cell related |
| ENSG00000188620 | HMX3 | -2,420779187 | 0,186755264 | 0,00548 | 0,09208977 | H6 family homeobox 3 |
| ENSG00000282602 | Z82244,2 | -2,422087722 | 0,186585952 | 0,2235 | 0,53953208 | novel transcript |
| ENSG00000188396 | TCTEX1D4 | -2,423746387 | 0,186371558 | 0,00037 | 0,01667641 | Tctex1 domain containing 4 |
| ENSG00000275564 | FCAR | -2,426406889 | 0,186028183 | 0,4886 | 0,72350189 | Fc fragment of IgA receptor |
| ENSG00000188305 | PEAK3 | -2,428710937 | 0,185731325 | 0,00345 | 0,0708904 | PEAK family member 3 |
| ENSG00000230873 | STMND1 | -2,428804859 | 0,185719234 | 0,01191 | 0,14219678 | stathmin domain containing 1 |
| ENSG00000227124 | ZNF717 | -2,43343712 | 0,185123876 | 1,84E-07 | 2,53E-05 | zinc finger protein 717 |
| ENSG00000226028 | PFN1P12 | -2,434973657 | 0,184926815 | 0,0288 | 0,2296248 | profilin 1 pseudogene 12 |
| ENSG00000272473 | AC006273,1 | -2,437171306 | 0,184645332 | 0,00398 | 0,07725019 | novel transcript |
| ENSG00000166509 | CLEC3A | -2,43808977 | 0,184527818 | 0,04334 | 0,2807119 | C-type lectin domain family 3 member A |
| ENSG00000205791 | LOH12CR2 | -2,439011112 | 0,184410012 | 0,48418 | 0,72005601 | loss of heterozygosity on chromosome 12, region 2 |
| ENSG00000259776 | AC093426,1 | -2,439182391 | 0,18438812 | 0,00827 | 0,11480072 | novel transcript |
| ENSG00000251381 | LINC00958 | -2,439772211 | 0,184312751 | 0,00794 | 0,11357366 | long intergenic non-protein coding RNA 958 |
| ENSG00000176485 | PLAAT3 | -2,439922003 | 0,184293615 | 1,53E-05 | 0,00122771 | phospholipase A and acyltransferase 3 |
| ENSG00000009694 | TENM1 | -2,441872155 | 0,184044666 | 1,15E-05 | 0,00093166 | teneurin transmembrane protein 1 |
| ENSG00000278732 | Metazoa_SRP | -2,44330513 | 0,183861952 | 0,04432 | 0,28386225 | Metazoan signal recognition particle RNA |
| ENSG00000242430 | RN7SL274P | -2,443342172 | 0,183857232 | 0,00922 | 0,12259866 | RNA, 7SL, cytoplasmic 274, pseudogene |

| | | | | | | |
|---|---|---|---|---|---|---|
| ENSG00000263466 | AC006441,1 | -2,443673023 | 0,183815073 | 0,00848 | 0,11625431 | novel transcript, antisense to LASP1 |
| ENSG00000161929 | SCIMP | -2,443952828 | 0,183779426 | 0,00151 | 0,04166341 | SLP adaptor and CSK interacting membrane protein |
| ENSG00000237742 | AL365259,1 | -2,444747546 | 0,183678218 | 0,0592 | 0,32424653 | novel transcript, antisense to HEY2 |
| ENSG00000110090 | CPT1A | -2,447237259 | 0,183361511 | 2,81E-05 | 0,00204966 | carnitine palmitoyltransferase 1A |
| ENSG00000105664 | COMP | -2,453482103 | 0,182569528 | 0,00531 | 0,09082808 | cartilage oligomeric matrix protein |
| ENSG00000203772 | SPRN | -2,455596202 | 0,18230219 | 0,00042 | 0,01806535 | shadow of prion protein |
| ENSG00000270424 | AC145285,5 | -2,455999596 | 0,182251223 | 0,01792 | 0,17654822 | PRKC, apoptosis, WT1, regulator (PAWR) pseudogene |
| ENSG00000271420 | AL109936,2 | -2,456614815 | 0,182173521 | 0,06263 | 0,33123868 | novel transcript |
| ENSG00000283012 | AL109936,5 | -2,456614815 | 0,182173521 | 0,06263 | 0,33123868 | novel transcript |
| ENSG00000200084 | SNORD68 | -2,458645156 | 0,181917324 | 0,02022 | 0,18881231 | small nucleolar RNA, C/D box 68 |
| ENSG00000227214 | HCG15 | -2,459361213 | 0,181827055 | 0,48089 | 0,71795527 | HLA complex group 15 |
| ENSG00000182257 | PRR34 | -2,45980201 | 0,181771509 | 0,01113 | 0,13651927 | PRR34 long non-coding RNA |
| ENSG00000287346 | AL132712,2 | -2,460040091 | 0,181741514 | 0,00487 | 0,08609281 | novel transcript |
| ENSG00000231424 | BX284613,2 | -2,461872042 | 0,181510883 | 0,0024 | 0,05652636 | novel transcript, antisense to FMO1 |
| ENSG00000226572 | SNORD57 | -2,463124917 | 0,181353323 | 0,05097 | 0,30375806 | small nucleolar RNA, C/D box 57 |
| ENSG00000267287 | AC068473,3 | -2,464266914 | 0,181209825 | 0,07455 | 0,35852613 | novel transcript |
| ENSG00000142538 | PTH2 | -2,464346403 | 0,181199841 | 0,08207 | 0,37329007 | parathyroid hormone 2 |
| ENSG00000007933 | FMO3 | -2,464550599 | 0,181174196 | 0,01777 | 0,17559058 | flavin containing dimethylaniline monoxygenase 3 |
| ENSG00000286662 | AC087894,3 | -2,464752421 | 0,181148853 | 0,01974 | 0,1865312 | novel transcript |
| ENSG00000136160 | EDNRB | -2,46597585 | 0,180995301 | 6,56E-18 | 4,09E-15 | endothelin receptor type B |
| ENSG00000214770 | AL161756,1 | -2,466559864 | 0,180922048 | 0,00193 | 0,04912509 | novel transcript |
| ENSG00000286279 | AC108721,2 | -2,467025209 | 0,1808637 | 0,02436 | 0,2086544 | novel transcript |
| ENSG00000286641 | AC108073,3 | -2,469415234 | 0,180564323 | 0,0626 | 0,33123314 | novel transcript |
| ENSG00000258001 | AC126614,1 | -2,469703657 | 0,180528228 | 0,09228 | 0,39278042 | novel transcript |
| ENSG00000233151 | CYP21A2 | -2,471797033 | 0,180266468 | 0,06498 | 0,33787601 | cytochrome P450 family 21 subfamily A member 2 |
| ENSG00000196832 | OR11G2 | -2,473526171 | 0,18005054 | 0,00318 | 0,06765231 | olfactory receptor family 11 subfamily G member 2 |
| ENSG00000080573 | COL5A3 | -2,476146613 | 0,179723801 | 0,00027 | 0,01344733 | collagen type V alpha 3 chain |
| ENSG00000204642 | HLA-F | -2,476928683 | 0,179626401 | 0,02371 | 0,20555253 | major histocompatibility complex, class I, F |
| ENSG00000288470 | LINC02429 | -2,477729475 | 0,179526724 | 0,00866 | 0,11761249 | long intergenic non-protein coding RNA 2429 |
| ENSG00000269635 | AC004257,1 | -2,478709096 | 0,179404863 | 0,1087 | 0,41972109 | novel transcript |
| ENSG00000277448 | LINC02338 | -2,47885184 | 0,179387113 | 0,00615 | 0,09867005 | long intergenic non-protein coding RNA 2338 |
| ENSG00000166897 | ELFN2 | -2,479052711 | 0,179362138 | 0,01761 | 0,17483597 | extracellular leucine rich repeat and fibronectin type III domain containing 2 |

| | | | | | | |
|---|---|---|---|---|---|---|
| ENSG00000248050 | AC079061,1 | -2,480374134 | 0,179197929 | 0,00627 | 0,09970758 | novel transcript, antisense to SYBU |
| ENSG00000227189 | AC092535,2 | -2,481189188 | 0,179096719 | 0,00104 | 0,0330699 | novel transcript |
| ENSG00000287829 | AC002542,6 | -2,482051602 | 0,178989691 | 0,00263 | 0,05982247 | novel transcript, antisense to ST7 |
| ENSG00000232884 | AF127936,1 | -2,483317804 | 0,178832667 | 0,02735 | 0,22320635 | novel transcript |
| ENSG00000241461 | RN7SL182P | -2,483407209 | 0,178821584 | 0,17076 | 0,49641416 | RNA, 7SL, cytoplasmic 182, pseudogene |
| ENSG00000249816 | LINC00964 | -2,484495812 | 0,178686703 | 0,00637 | 0,10063153 | long intergenic non-protein coding RNA 964 |
| ENSG00000259884 | NR4A1AS | -2,48619982 | 0,178475776 | 0,07196 | 0,3539609 | NR4A1 antisense RNA |
| ENSG00000271806 | AL590822,2 | -2,486596478 | 0,178426712 | 0,00594 | 0,09720154 | novel transcript, antisense PRKCZ |
| ENSG00000282737 | LINC01044 | -2,488240721 | 0,178223475 | 0,00742 | 0,10960607 | long intergenic non-protein coding RNA 1044 |
| ENSG00000280213 | UCKL1-AS1 | -2,490838209 | 0,177902882 | 0,00182 | 0,04718641 | UCKL1 antisense RNA 1 |
| ENSG00000229206 | AL162408,1 | -2,491730193 | 0,177792923 | 0,00086 | 0,02911193 | novel transcript |
| ENSG00000258199 | AC073896,4 | -2,493363331 | 0,177591775 | 0,05282 | 0,30848882 | novel transcript, sense overlapping to SMARCC2 |
| ENSG00000230705 | TAP1 | -2,4936375 | 0,177558029 | 0,01905 | 0,18296265 | transporter 1, ATP binding cassette subfamily B member |
| ENSG00000278154 | LAIR1 | -2,494205888 | 0,177488089 | 0,0068 | 0,10528058 | leukocyte associated immunoglobulin like receptor 1 |
| ENSG00000283518 | LINC02775 | -2,495425718 | 0,177338082 | 0,04963 | 0,29985549 | long intergenic non-protein coding RNA 2775 |
| ENSG00000240915 | AC120042,1 | -2,496099632 | 0,177255263 | 0,00865 | 0,11751405 | novel transcript |
| ENSG00000277741 | GOLGA6L17P | -2,496980926 | 0,177147017 | 0,01848 | 0,179662 | golgin A6 family like 17, pseudogene |
| ENSG00000227420 | MRPS18B | -2,501444846 | 0,176599744 | 0,02092 | 0,19202875 | mitochondrial ribosomal protein S18B |
| ENSG00000277184 | SNORA9 | -2,502096671 | 0,176519972 | 0,07345 | 0,35662549 | small nucleolar RNA, H/ACA box 9 |
| ENSG00000276568 | Metazoa_SRP | -2,502817012 | 0,176431857 | 0,06513 | 0,33804854 | Metazoan signal recognition particle RNA |
| ENSG00000249860 | MTRNR2L5 | -2,505260737 | 0,176133259 | 0,00613 | 0,098625 | MT-RNR2 like 5 |
| ENSG00000283953 | FCAR | -2,506756105 | 0,17595079 | 0,13097 | 0,45558936 | Fc fragment of IgA receptor |
| ENSG00000234766 | LINC01496 | -2,509679096 | 0,175594663 | 0,00214 | 0,05235162 | long intergenic non-protein coding RNA 1496 |
| ENSG00000269487 | AC008635,1 | -2,509918641 | 0,17556551 | 0,04429 | 0,28377004 | novel transcript, antisense to STRN4 |
| ENSG00000259527 | LINC00052 | -2,512571204 | 0,175243009 | 0,00093 | 0,03069588 | long intergenic non-protein coding RNA 52 |
| ENSG00000267011 | AC011498,1 | -2,514720467 | 0,174982134 | 0,01978 | 0,18665735 | novel transcript |
| ENSG00000276110 | AC242988,1 | -2,515380686 | 0,174902075 | 0,03504 | 0,25324761 | novel transcript, antisense to CA14 |
| ENSG00000286804 | AC025871,3 | -2,515655203 | 0,174868798 | 0,01303 | 0,14938099 | novel transcript, sense intronic to FBXO16 |
| ENSG00000260001 | TGFBR3L | -2,522698558 | 0,174017155 | 0,02233 | 0,19848029 | transforming growth factor beta receptor 3 like |
| ENSG00000287539 | AC022966,2 | -2,522700776 | 0,174016887 | 0,00137 | 0,03936155 | novel transcript |
| ENSG00000233901 | LINC01503 | -2,52405962 | 0,173853062 | 0,00137 | 0,03936155 | long intergenic non-protein coding RNA 1503 |
| ENSG00000232599 | AL008707,1 | -2,525090236 | 0,173728911 | 0,0002 | 0,01061649 | odz, odd Oz\/ten-m homolog 1(Drosophila) (ODZ1) pseudogene |

| | | | | | | |
|---|---|---|---|---|---|---|
| ENSG00000272789 | AC010976,2 | -2,525505486 | 0,173678914 | 0,11749 | 0,43477245 | novel transcript, antisense to MYO7B |
| ENSG00000260792 | LINC02280 | -2,527847352 | 0,173397217 | 0,0068 | 0,10528058 | long intergenic non-protein coding RNA 2280 |
| ENSG00000229848 | AC139149,1 | -2,530665217 | 0,173058869 | 0,04631 | 0,28930497 | novel transcript, antisense to ACTG1 |
| ENSG00000275853 | Metazoa_SRP | -2,533676397 | 0,172698039 | 0,04245 | 0,2782038 | Metazoan signal recognition particle RNA |
| ENSG00000230454 | U73166,1 | -2,534914396 | 0,172549907 | 0,00109 | 0,03441486 | novel transcript |
| ENSG00000232065 | LINC01063 | -2,536666266 | 0,172340506 | 0,03333 | 0,2467172 | long intergenic non-protein coding RNA 1063 |
| ENSG00000181126 | HLA-V | -2,536982909 | 0,172302685 | 0,00997 | 0,12783227 | major histocompatibility complex, class I, V (pseudogene) |
| ENSG00000231437 | LINC01750 | -2,537899189 | 0,172193288 | 0,06702 | 0,34281679 | long intergenic non-protein coding RNA 1750 |
| ENSG00000287181 | AC107067,2 | -2,538130365 | 0,172165698 | 0,14281 | 0,47034016 | novel transcript |
| ENSG00000287897 | AP000465,1 | -2,539315861 | 0,172024283 | 0,00322 | 0,06803074 | novel transcript, sense intronic to CADM1 |
| ENSG00000228189 | LINC02843 | -2,539460017 | 0,172007095 | 0,11699 | 0,4343753 | long intergenic non-protein coding RNA 2843 |
| ENSG00000066735 | KIF26A | -2,539970964 | 0,171946188 | 5,04E-05 | 0,00338051 | kinesin family member 26A |
| ENSG00000261226 | AC092384,3 | -2,542884372 | 0,171599307 | 0,03784 | 0,26339938 | novel transcript |
| ENSG00000254790 | AP000842,1 | -2,543084494 | 0,171575505 | 0,02749 | 0,2235566 | novel transcript |
| ENSG00000196684 | HSH2D | -2,544858203 | 0,171364693 | 0,02188 | 0,19668785 | hematopoietic SH2 domain containing |
| ENSG00000248510 | LINC02267 | -2,545041868 | 0,171342878 | 0,00123 | 0,03698156 | long intergenic non-protein coding RNA 2267 |
| ENSG00000229861 | MRPS18B | -2,545131471 | 0,171332237 | 0,00437 | 0,08120816 | mitochondrial ribosomal protein S18B |
| ENSG00000198133 | TMEM229B | -2,546398923 | 0,171181782 | 5,04E-07 | 6,11E-05 | transmembrane protein 229B |
| ENSG00000228919 | AC097381,1 | -2,547403287 | 0,171062652 | 0,00272 | 0,06123483 | uncharacterized LOC389199 |
| ENSG00000235979 | AC004448,2 | -2,548049701 | 0,170986022 | 0,01858 | 0,18016932 | novel transcript |
| ENSG00000234857 | HNRNPUL2-BS | -2,548643263 | 0,170915689 | 0,00069 | 0,02550975 | HNRNPUL2-BSCL2 readthrough (NMD candidate) |
| ENSG00000251408 | AC113615,1 | -2,549711278 | 0,170789208 | 0,00308 | 0,06617471 | novel transcript |
| ENSG00000239774 | AC125618,1 | -2,550051054 | 0,170748989 | 0,01213 | 0,14313104 | novel transcript |
| ENSG00000253923 | AP002981,1 | -2,550665865 | 0,17067624 | 0,00182 | 0,04718641 | heat shock 10kDa protein 1 (chaperonin 10) (HSPE1) pseudogene |
| ENSG00000234173 | AL353784,1 | -2,551048337 | 0,170630998 | 0,01279 | 0,14802902 | novel transcript, antisense to PCDH15 |
| ENSG00000224404 | Z68323,1 | -2,554348081 | 0,170241175 | 0,02292 | 0,20210363 | novel transcript |
| ENSG00000101292 | PROKR2 | -2,554576816 | 0,170214186 | 0,02598 | 0,21717729 | prokineticin receptor 2 |
| ENSG00000282450 | AC012676,7 | -2,555118291 | 0,170150313 | 0,00704 | 0,10676605 | novel transcript, sense intronic with DNAJA3 |
| ENSG00000120055 | C10orf95 | -2,555177008 | 0,170143388 | 0,15309 | 0,48120112 | chromosome 10 open reading frame 95 |
| ENSG00000121900 | TMEM54 | -2,555373498 | 0,170120217 | 0,00011 | 0,00628762 | transmembrane protein 54 |
| ENSG00000286708 | AP003783,1 | -2,555492516 | 0,170106183 | 0,00537 | 0,09125239 | novel transcript, antisense to SHANK2 |
| ENSG00000279028 | AC010223,1 | -2,55560367 | 0,170093077 | 0,01526 | 0,16213925 | novel transcript, antisense to PCDHAC2 |

| | | | | | | |
|---|---|---|---|---|---|---|
| ENSG00000121895 | TMEM156 | -2,556070036 | 0,170038102 | 0,00685 | 0,10547716 | transmembrane protein 156 |
| ENSG00000283991 | AC017000,1 | -2,557269919 | 0,169896741 | 0,02164 | 0,19544769 | novel transcript, antisense to SDK1 |
| ENSG00000230615 | AL139220,2 | -2,557400761 | 0,169881333 | 0,0013 | 0,03841105 | novel transcript |
| ENSG00000253829 | AC067817,2 | -2,558781626 | 0,16971881 | 0,00742 | 0,10959823 | novel transcript |
| ENSG00000203624 | MRPS18B | -2,561107484 | 0,169445416 | 0,00496 | 0,08746202 | mitochondrial ribosomal protein S18B |
| ENSG00000275371 | AC012645,4 | -2,561866918 | 0,169356244 | 0,00314 | 0,06683654 | novel transcript, antisense to GDPD3 |
| ENSG00000269388 | AC018755,3 | -2,561869205 | 0,169355975 | 0,0022 | 0,05331646 | putative ATP-binding domain-containing protein 3-like protein (ABP3L) pseudogene |
| ENSG00000233671 | CENPNP1 | -2,561943694 | 0,169347231 | 0,01414 | 0,15607057 | CENPN pseudogene 1 |
| ENSG00000283643 | GPR1-AS | -2,56520824 | 0,168964464 | 0,00508 | 0,08900354 | GPR1 antisense RNA |
| ENSG00000245164 | LINC00861 | -2,566652461 | 0,168795405 | 0,02648 | 0,21924054 | long intergenic non-protein coding RNA 861 |
| ENSG00000258171 | LINC02412 | -2,566978406 | 0,168757274 | 0,02886 | 0,22968658 | long intergenic non-protein coding RNA 2412 |
| ENSG00000264739 | AC093484,2 | -2,566995932 | 0,168755224 | 0,00246 | 0,05743859 | novel transcript |
| ENSG00000274110 | LAIR1 | -2,567386464 | 0,168709549 | 0,00088 | 0,02961572 | leukocyte associated immunoglobulin like receptor 1 |
| ENSG00000199477 | SNORA31 | -2,567913541 | 0,168647923 | 0,01578 | 0,16470677 | small nucleolar RNA, H/ACA box 31 |
| ENSG00000286037 | AC011479,4 | -2,568279665 | 0,16860513 | 0,46237 | 0,70356854 | novel transcript, antisense to SPINT2 |
| ENSG00000238194 | PHACTR3-AS1 | -2,568472315 | 0,168582617 | 0,00137 | 0,03936155 | PHACTR3 antisense RNA 1 |
| ENSG00000102109 | PCSK1N | -2,571707326 | 0,168205021 | 0,00103 | 0,03289327 | proprotein convertase subtilisin/kexin type 1 inhibitor |
| ENSG00000105383 | CD33 | -2,57172869 | 0,16820253 | 0,0483 | 0,29516044 | CD33 molecule |
| ENSG00000279467 | AP000350,8 | -2,575887669 | 0,167718336 | 0,1094 | 0,42138503 | novel transcript, antisense to SLC2A11 |
| ENSG00000255366 | AC120036,4 | -2,576703209 | 0,167623554 | 0,00509 | 0,08904306 | novel transcript |
| ENSG00000262943 | ALOX12P2 | -2,578037246 | 0,167468627 | 0,00089 | 0,02981453 | arachidonate 12-lipoxygenase pseudogene 2 |
| ENSG00000276799 | WNT9B | -2,579179714 | 0,167336061 | 0,00927 | 0,12287842 | Wnt family member 9B |
| ENSG00000229170 | CR847861,1 | -2,579245767 | 0,1673284 | 0,04296 | 0,27922774 | chromosome 6 open reading frame 214 (putative) |
| ENSG00000122133 | PAEP | -2,579589035 | 0,167288591 | 0,08542 | 0,37882499 | progestagen associated endometrial protein |
| ENSG00000285966 | AC007686,4 | -2,580096159 | 0,167229798 | 0,00219 | 0,05329003 | novel transcript |
| ENSG00000260963 | AC026462,3 | -2,580627454 | 0,167168224 | 0,05444 | 0,31226071 | novel transcript |
| ENSG00000223775 | MRPS18B | -2,583972911 | 0,166781028 | 0,01567 | 0,16407493 | mitochondrial ribosomal protein S18B |
| ENSG00000134595 | SOX3 | -2,585736854 | 0,166577234 | 7,43E-16 | 3,48E-13 | SRY-box transcription factor 3 |
| ENSG00000276798 | LILRA4 | -2,588969222 | 0,166204434 | 0,05136 | 0,30462001 | leukocyte immunoglobulin like receptor A4 |
| ENSG00000251423 | AC016576,1 | -2,591483011 | 0,165915087 | 0,01253 | 0,1464509 | novel transcript |
| ENSG00000277985 | SNORA67 | -2,59619455 | 0,165374127 | 0,00046 | 0,01935233 | small nucleolar RNA, H/ACA box 67 |
| ENSG00000234913 | AC016027,2 | -2,597965384 | 0,165171264 | 0,00412 | 0,07868813 | novel transcript |

| | | | | | | |
|---|---|---|---|---|---|---|
| ENSG00000285652 | AL591428,1 | -2,599204004 | 0,165029518 | 0,00282 | 0,06285707 | novel transcript |
| ENSG00000112333 | NR2E1 | -2,599414315 | 0,165005462 | 0,00041 | 0,01788764 | nuclear receptor subfamily 2 group E member 1 |
| ENSG00000223703 | AC027612,1 | -2,602004865 | 0,164709439 | 0,00336 | 0,07000131 | pseudogene similar to part of immunoglobulin superfamily, member 3 (IGSF3) |
| ENSG00000119121 | TRPM6 | -2,602572397 | 0,164644657 | 0,0058 | 0,09561123 | transient receptor potential cation channel subfamily M member 6 |
| ENSG00000264116 | AC116003,1 | -2,604333447 | 0,164443804 | 0,00237 | 0,05606528 | novel transcript, antisense to SMIM21 |
| ENSG00000253686 | LINC01484 | -2,605086675 | 0,16435797 | 0,00129 | 0,03840911 | long intergenic non-protein coding RNA 1484 |
| ENSG00000261559 | FSCN1P1 | -2,609833194 | 0,163818115 | 0,00115 | 0,03540711 | fascin actin-bundling protein 1 pseudogene 1 |
| ENSG00000267522 | LINC01864 | -2,610724184 | 0,163716974 | 0,04631 | 0,28930497 | long intergenic non-protein coding RNA 1864 |
| ENSG00000230707 | AL589987,1 | -2,61544768 | 0,163181829 | 1,74E-05 | 0,00137044 | novel pseudogene |
| ENSG00000264019 | AC018521,2 | -2,617199239 | 0,162983832 | 0,0842 | 0,37695424 | novel transcript, antisense to PNPO |
| ENSG00000254344 | LINC01288 | -2,621686021 | 0,16247774 | 0,06518 | 0,33804854 | long intergenic non-protein coding RNA 1288 |
| ENSG00000287263 | AC008875,3 | -2,623007423 | 0,16232899 | 0,00563 | 0,09352688 | novel transcript |
| ENSG00000047662 | FAM184B | -2,624820017 | 0,162125169 | 0,00295 | 0,06468135 | family with sequence similarity 184 member B |
| ENSG00000115194 | SLC30A3 | -2,62488081 | 0,162118337 | 0,00628 | 0,09975871 | solute carrier family 30 member 3 |
| ENSG00000228692 | AL445307,1 | -2,625647343 | 0,162032224 | 0,10389 | 0,41228466 | novel transcript |
| ENSG00000198569 | SLC34A3 | -2,628480889 | 0,161714294 | 0,00522 | 0,09021858 | solute carrier family 34 member 3 |
| ENSG00000278330 | AC018529,2 | -2,631778459 | 0,161345086 | 0,00238 | 0,05617285 | novel transcript, antisense to MBP |
| ENSG00000247151 | CSTF3-DT | -2,63232828 | 0,161283608 | 0,00371 | 0,07418692 | CSTF3 divergent transcript |
| ENSG00000224259 | LINC01133 | -2,636157727 | 0,160856069 | 0,02949 | 0,23093703 | long intergenic non-protein coding RNA 1133 |
| ENSG00000255394 | C8orf49 | -2,637815813 | 0,160671304 | 0,12056 | 0,43944057 | chromosome 8 open reading frame 49 |
| ENSG00000168594 | ADAM29 | -2,638133089 | 0,160635973 | 0,02284 | 0,20179409 | ADAM metallopeptidase domain 29 |
| ENSG00000146955 | RAB19 | -2,638418017 | 0,160604251 | 0,03382 | 0,24846649 | RAB19, member RAS oncogene family |
| ENSG00000141485 | SLC13A5 | -2,638710786 | 0,160571662 | 7,51E-05 | 0,0047151 | solute carrier family 13 member 5 |
| ENSG00000125823 | CSTL1 | -2,641859428 | 0,160221601 | 0,12418 | 0,44504227 | cystatin like 1 |
| ENSG00000104894 | CD37 | -2,643864066 | 0,159999126 | 0,00011 | 0,00636759 | CD37 molecule |
| ENSG00000286933 | AL022170,1 | -2,645074794 | 0,15986491 | 0,00794 | 0,11357366 | novel transcript |
| ENSG00000274023 | AL360169,2 | -2,649697219 | 0,159353519 | 0,00516 | 0,08963917 | novel transcript, sense overlapping TULP4 |
| ENSG00000180432 | CYP8B1 | -2,650808601 | 0,159230808 | 0,02803 | 0,22583197 | cytochrome P450 family 8 subfamily B member 1 |
| ENSG00000176840 | MIR7-3HG | -2,652167794 | 0,159080864 | 0,00212 | 0,0519443 | MIR7-3 host gene |
| ENSG00000248802 | AC078850,2 | -2,654032579 | 0,158875373 | 0,01441 | 0,15696842 | novel transcript |
| ENSG00000282147 | FAM20C | -2,654776075 | 0,158793518 | 0,01988 | 0,18711351 | FAM20C golgi associated secretory pathway kinase |

| | | | | | | |
|---|---|---|---|---|---|---|
| ENSG00000255428 | AP002008,3 | -2,655178368 | 0,158749245 | 0,10129 | 0,40805419 | novel transcript, sense intronic to POU2AF1 |
| ENSG00000287400 | AC093151,7 | -2,657201941 | 0,158526734 | 0,08923 | 0,38636433 | novel transcript, antisense to SCMH1 |
| ENSG00000273203 | AC006946,2 | -2,657531773 | 0,158490495 | 0,10225 | 0,40974995 | novel transcript |
| ENSG00000145063 | AC062028,1 | -2,658082119 | 0,158430047 | 0,00823 | 0,11480072 | novel transcript |
| ENSG00000219201 | AC138392,1 | -2,658152555 | 0,158422312 | 0,09506 | 0,39768318 | novel pseudogene |
| ENSG00000242550 | SERPINB10 | -2,659274729 | 0,158299134 | 0,10252 | 0,41028143 | serpin family B member 10 |
| ENSG00000254791 | FAR1-IT1 | -2,660486401 | 0,15816624 | 0,00447 | 0,08248666 | FAR1 intronic transcript 1 |
| ENSG00000207445 | SNORD15B | -2,661439706 | 0,158061761 | 0,0006 | 0,02332263 | small nucleolar RNA, C/D box 15B |
| ENSG00000070190 | DAPP1 | -2,661970972 | 0,158003566 | 0,00112 | 0,03490596 | dual adaptor of phosphotyrosine and 3-phosphoinositides 1 |
| ENSG00000285772 | AC000120,2 | -2,670693556 | 0,157051153 | 0,0561 | 0,31638988 | novel protein |
| ENSG00000187871 | GFRAL | -2,671105664 | 0,157006298 | 0,10158 | 0,40840817 | GDNF family receptor alpha like |
| ENSG00000163064 | EN1 | -2,671524334 | 0,156960741 | 9,94E-10 | 2,27E-07 | engrailed homeobox 1 |
| ENSG00000273587 | SNORA78 | -2,673273836 | 0,156770516 | 0,03062 | 0,23611505 | small nucleolar RNA, H/ACA box 78 |
| ENSG00000200795 | RNU4-1 | -2,673885676 | 0,156704045 | 0,00087 | 0,02923181 | RNA, U4 small nuclear 1 |
| ENSG00000285565 | AL671762,1 | -2,674716697 | 0,156613806 | 0,30989 | 0,5981175 | novel transcript |
| ENSG00000246211 | P4HA3-AS1 | -2,674835457 | 0,156600915 | 0,01032 | 0,13096785 | P4HA3 antisense RNA 1 |
| ENSG00000285468 | EPHA1-AS1 | -2,678952168 | 0,156154693 | 0,03258 | 0,24369403 | EPHA1 antisense RNA 1 |
| ENSG00000286488 | AC103858,3 | -2,679293539 | 0,156117748 | 0,00221 | 0,05354564 | novel transcript, antisense to STARD5 |
| ENSG00000187847 | OR7E25P | -2,682660676 | 0,155753806 | 0,00156 | 0,04242446 | olfactory receptor family 7 subfamily E member 25 pseudogene |
| ENSG00000168967 | PMCHL1 | -2,686443011 | 0,155345999 | 0,01787 | 0,17623976 | pro-melanin concentrating hormone like 1 (pseudogene) |
| ENSG00000199266 | SNORA60 | -2,686955964 | 0,155290775 | 0,08508 | 0,37850509 | small nucleolar RNA, H/ACA box 60 |
| ENSG00000275918 | KIR3DX1 | -2,687139304 | 0,155271042 | 0,06389 | 0,33507507 | killer cell immunoglobulin like receptor, three Ig domains X1 (pseudogene) |
| ENSG00000257120 | AL356756,1 | -2,68723499 | 0,155260744 | 0,00628 | 0,09975871 | novel transcript, antisense to PRKD1 |
| ENSG00000288096 | AC109322,2 | -2,692110179 | 0,154736969 | 0,00725 | 0,10792958 | novel transcript |
| ENSG00000167419 | LPO | -2,69250764 | 0,154694345 | 0,01297 | 0,14899372 | lactoperoxidase |
| ENSG00000286446 | AL353688,1 | -2,693629337 | 0,154574116 | 0,0372 | 0,26111083 | novel transcript |
| ENSG00000248334 | WHAMMP2 | -2,695093688 | 0,154417301 | 0,00149 | 0,04142724 | WAS protein homolog associated with actin, golgi membranes and microtubules pseudogene 2 |
| ENSG00000256612 | CYP2B7P | -2,696140494 | 0,154305298 | 0,0032 | 0,06769433 | cytochrome P450 family 2 subfamily B member 7, pseudogene |

| | | | | | | |
|---|---|---|---|---|---|---|
| ENSG00000263847 | AP005899,1 | -2,696799594 | 0,15423482 | 0,00486 | 0,08609281 | novel transcript, antisense to NDUFV2 |
| ENSG00000223606 | HLA-V | -2,697512014 | 0,154158675 | 0,06434 | 0,33607088 | major histocompatibility complex, class I, V (pseudogene) |
| ENSG00000257431 | AC089998,1 | -2,700477729 | 0,1538421 | 0,10742 | 0,41771747 | novel transcript |
| ENSG00000282339 | AC010507,2 | -2,701803156 | 0,153700828 | 0,00196 | 0,04953228 | novel pseudogene |
| ENSG00000236635 | LINC02540 | -2,701833332 | 0,153697613 | 0,00202 | 0,05060501 | long intergenic non-protein coding RNA 2540 |
| ENSG00000254187 | AC008708,2 | -2,70630898 | 0,153221539 | 0,00045 | 0,01894901 | novel transcript |
| ENSG00000266896 | AL354892,3 | -2,706534988 | 0,153197537 | 0,00184 | 0,04744946 | novel transcript |
| ENSG00000144891 | AGTR1 | -2,708785325 | 0,152958764 | 0,00077 | 0,02712315 | angiotensin II receptor type 1 |
| ENSG00000223757 | VWA7 | -2,709637772 | 0,152868412 | 0,03698 | 0,26078413 | von Willebrand factor A domain containing 7 |
| ENSG00000205923 | CEMP1 | -2,711743174 | 0,152645485 | 0,25065 | 0,55690541 | cementum protein 1 |
| ENSG00000255886 | AC020611,2 | -2,712528488 | 0,152562417 | 0,00297 | 0,06492836 | novel transcript |
| ENSG00000187955 | COL14A1 | -2,715564073 | 0,152241747 | 1,64E-06 | 0,00017274 | collagen type XIV alpha 1 chain |
| ENSG00000244346 | AC092953,1 | -2,717594866 | 0,152027596 | 0,011 | 0,13537507 | chromosome 9 open reading frame 85 (C9orf85) pseudogene |
| ENSG00000214894 | LINC00243 | -2,718261225 | 0,151957393 | 0,09799 | 0,40323803 | long intergenic non-protein coding RNA 243 |
| ENSG00000119973 | PRLHR | -2,719974491 | 0,151777044 | 0,00322 | 0,06803074 | prolactin releasing hormone receptor |
| ENSG00000237806 | FAUP2 | -2,720315501 | 0,151741173 | 0,01549 | 0,16309786 | FAU pseudogene 2 |
| ENSG00000122787 | AKR1D1 | -2,722751632 | 0,151485159 | 0,02015 | 0,18846172 | aldo-keto reductase family 1 member D1 |
| ENSG00000255319 | ENPP7P8 | -2,723027052 | 0,151456243 | 0,0013 | 0,0384253 | ectonucleotide pyrophosphatase/phosphodiesterase 7 pseudogene 8 |
| ENSG00000286976 | AC084335,1 | -2,725348979 | 0,151212679 | 0,00094 | 0,03094099 | novel transcript |
| ENSG00000157388 | CACNA1D | -2,728369688 | 0,150896402 | 0,0011 | 0,03462398 | calcium voltage-gated channel subunit alpha1 D |
| ENSG00000223533 | CCHCR1 | -2,729403442 | 0,150788317 | 0,02013 | 0,18835147 | coiled-coil alpha-helical rod protein 1 |
| ENSG00000224180 | CCHCR1 | -2,729403442 | 0,150788317 | 0,02013 | 0,18835147 | coiled-coil alpha-helical rod protein 1 |
| ENSG00000225399 | AC121247,1 | -2,729792498 | 0,150747659 | 0,00635 | 0,10056798 | novel transcript |
| ENSG00000239856 | RN7SL225P | -2,73591112 | 0,150109676 | 0,06003 | 0,32549025 | RNA, 7SL, cytoplasmic 225, pseudogene |
| ENSG00000267412 | AC092068,2 | -2,742189495 | 0,149457842 | 0,00398 | 0,07725019 | novel transcript, sense intronic to GNG7 |
| ENSG00000230155 | CEP250-AS1 | -2,742492763 | 0,149426428 | 0,01582 | 0,16482188 | CEP250 antisense RNA 1 |
| ENSG00000261131 | AC012186,2 | -2,746995259 | 0,148960811 | 0,00613 | 0,098625 | novel transcript |
| ENSG00000213203 | GIMAP1 | -2,748411961 | 0,148814606 | 0,00249 | 0,05782412 | GTPase, IMAP family member 1 |
| ENSG00000080572 | DNAAF6 | -2,749845933 | 0,148666765 | 0,01339 | 0,15158202 | dynein axonemal assembly factor 6 |
| ENSG00000124772 | CPNE5 | -2,750898443 | 0,148558345 | 8,49E-05 | 0,00527733 | copine 5 |
| ENSG00000241933 | DENND6A-DT | -2,751447451 | 0,148501823 | 0,01382 | 0,15438008 | DENND6A divergent transcript |

| | | | | | | |
|---|---|---|---|---|---|---|
| ENSG00000285559 | AC022903,2 | -2,755535668 | 0,148081604 | 0,03711 | 0,26086671 | novel transcript, antisense to CCT6B |
| ENSG00000278192 | AL118505,1 | -2,75607074 | 0,148026693 | 0,07663 | 0,36266919 | novel transcript |
| ENSG00000284800 | AC235565,2 | -2,757536067 | 0,147876421 | 1,98E-06 | 0,00020155 | novel transcript |
| ENSG00000244677 | RN7SL706P | -2,759867171 | 0,147637675 | 0,02938 | 0,23068546 | RNA, 7SL, cytoplasmic 706, pseudogene |
| ENSG00000254042 | SLIT3-AS2 | -2,76126523 | 0,147494674 | 0,0011 | 0,03448302 | SLIT3 antisense RNA 2 |
| ENSG00000254202 | AC015522,1 | -2,763603701 | 0,147255793 | 0,03233 | 0,24253636 | novel transcript |
| ENSG00000182983 | ZNF662 | -2,765402031 | 0,147072352 | 0,00202 | 0,0505329 | zinc finger protein 662 |
| ENSG00000285599 | AC234917,2 | -2,766329438 | 0,14697784 | 0,01881 | 0,18167964 | Novel transcript, antisense to NRBP2 |
| ENSG00000205903 | ZNF316 | -2,766961664 | 0,146913445 | 0,0003 | 0,01424712 | zinc finger protein 316 |
| ENSG00000227906 | SNAP25-AS1 | -2,767704154 | 0,146837855 | 0,00156 | 0,04239659 | SNAP25 antisense RNA 1 |
| ENSG00000173727 | AP000769,1 | -2,769091953 | 0,146696672 | 0,0004 | 0,01750406 | Finkel-Biskis-Reilly murine sarcoma virus (FBR-MuSV) ubiquitously expressed (FAU) pseudogene |
| ENSG00000202538 | RNU4-2 | -2,772572526 | 0,146343186 | 0,00051 | 0,02072378 | RNA, U4 small nuclear 2 |
| ENSG00000251244 | AC107208,1 | -2,773425317 | 0,146256706 | 0,03273 | 0,24407309 | novel transcript |
| ENSG00000286004 | AC124944,3 | -2,774243827 | 0,146173751 | 0,00179 | 0,04676155 | novel transcript, antisense to TNK2 |
| ENSG00000107147 | KCNT1 | -2,775874934 | 0,146008581 | 0,0071 | 0,10744093 | potassium sodium-activated channel subfamily T member 1 |
| ENSG00000261765 | AC009127,1 | -2,776912353 | 0,145903626 | 0,00332 | 0,06940016 | novel transcript |
| ENSG00000260934 | AC130456,5 | -2,777685997 | 0,145825407 | 0,10681 | 0,41660052 | novel transcript, antisense to GDE1 |
| ENSG00000124256 | ZBP1 | -2,779024281 | 0,145690198 | 0,03054 | 0,2357794 | Z-DNA binding protein 1 |
| ENSG00000284601 | AL590440,2 | -2,780148782 | 0,145576685 | 0,06648 | 0,34124715 | novel transcript |
| ENSG00000203565 | AL450313,1 | -2,781538397 | 0,145436531 | 0,01902 | 0,18296265 | novel transcript |
| ENSG00000232541 | COL11A2 | -2,781559353 | 0,145434419 | 0,01953 | 0,18522413 | collagen type XI alpha 2 chain |
| ENSG00000281305 | LINC00574 | -2,781737691 | 0,145416442 | 0,00694 | 0,10609569 | long intergenic non-protein coding RNA 574 |
| ENSG00000243801 | RN7SL461P | -2,782851674 | 0,145304202 | 0,03771 | 0,26272136 | RNA, 7SL, cytoplasmic 461, pseudogene |
| ENSG00000145451 | GLRA3 | -2,784193731 | 0,145169096 | 0,00071 | 0,02609438 | glycine receptor alpha 3 |
| ENSG00000167751 | KLK2 | -2,784697332 | 0,145118431 | 0,00615 | 0,09865643 | kallikrein related peptidase 2 |
| ENSG00000236497 | LINC01744 | -2,79000957 | 0,144585064 | 0,15236 | 0,48061452 | long intergenic non-protein coding RNA 1744 |
| ENSG00000070388 | FGF22 | -2,790066809 | 0,144579328 | 0,0043 | 0,08060327 | fibroblast growth factor 22 |
| ENSG00000231812 | SNRPCP9 | -2,791099586 | 0,144475865 | 0,02006 | 0,18786958 | small nuclear ribonucleoprotein polypeptide C pseudogene 9 |
| ENSG00000199436 | SNORD9 | -2,793484862 | 0,144237194 | 0,00947 | 0,12448953 | small nucleolar RNA, C/D box 9 |
| ENSG00000264589 | MAPT-AS1 | -2,796500113 | 0,143936051 | 0,00072 | 0,02620601 | MAPT antisense RNA 1 |

| | | | | | | |
|---|---|---|---|---|---|---|
| ENSG00000285486 | AC003043,2 | -2,798611619 | 0,143725542 | 0,01182 | 0,1415704 | novel transcript |
| ENSG00000262692 | AC116914,2 | -2,79905901 | 0,143680979 | 0,00876 | 0,11870876 | novel transcript, antisense to ITGAE |
| ENSG00000249506 | ZEB2P1 | -2,800031174 | 0,143584192 | 0,00347 | 0,07121616 | zinc finger E-box binding homeobox 2 pseudogene 1 |
| ENSG00000182376 | AC138028,1 | -2,801469437 | 0,14344112 | 0,00301 | 0,06548828 | novel transcript, antisense to FAM38A |
| ENSG00000198692 | EIF1AY | -2,802179615 | 0,143370527 | 0,0046 | 0,08355176 | eukaryotic translation initiation factor 1A Y-linked |
| ENSG00000275517 | PRB1 | -2,802663494 | 0,143322449 | 0,03609 | 0,25727995 | proline rich protein BstNI subfamily 1 |
| ENSG00000236649 | PPT2 | -2,804300006 | 0,143159964 | 0,03706 | 0,26086671 | palmitoyl-protein thioesterase 2 |
| ENSG00000227517 | LINC01483 | -2,8056051 | 0,143030517 | 0,00141 | 0,04017644 | long intergenic non-protein coding RNA 1483 |
| ENSG00000267261 | AC099811,2 | -2,806336637 | 0,14295801 | 0,06945 | 0,34812526 | novel protein |
| ENSG00000236466 | AL035670,1 | -2,807992146 | 0,142794058 | 0,00242 | 0,05669968 | novel transcript |
| ENSG00000250584 | LINC01511 | -2,811424789 | 0,142454708 | 0,03024 | 0,23475421 | long intergenic non-protein coding RNA 1511 |
| ENSG00000287714 | AC025164,2 | -2,816338845 | 0,14197031 | 0,02747 | 0,22348662 | novel transcript |
| ENSG00000287254 | AC011509,3 | -2,818231394 | 0,141784193 | 0,01456 | 0,15778631 | novel transcript |
| ENSG00000248840 | AL645949,1 | -2,818633226 | 0,141744708 | 0,00459 | 0,08349554 | X antigen family, member 5 (XAGE5) pseudogene |
| ENSG00000253642 | AF279873,3 | -2,819159484 | 0,141693012 | 0,00391 | 0,07646656 | novel transcript |
| ENSG00000274841 | Metazoa_SRP | -2,819444693 | 0,141665003 | 0,02424 | 0,20804286 | Metazoan signal recognition particle RNA |
| ENSG00000238917 | SNORD10 | -2,81987334 | 0,141622919 | 0,00025 | 0,0124595 | small nucleolar RNA, C/D box 10 |
| ENSG00000275672 | AC025580,3 | -2,825892137 | 0,141033311 | 0,00077 | 0,02721863 | GATM antisense RNA 1 |
| ENSG00000161649 | CD300LG | -2,827491067 | 0,140877091 | 0,00111 | 0,03484471 | CD300 molecule like family member g |
| ENSG00000230013 | CT70 | -2,827702397 | 0,140856457 | 0,0085 | 0,11637365 | cancer/testis associated transcript 70 |
| ENSG00000279298 | Z99916,3 | -2,828944967 | 0,140735192 | 0,00184 | 0,04744946 | novel transcript |
| ENSG00000230060 | MPIG6B | -2,829656803 | 0,140665769 | 0,05069 | 0,30285446 | megakaryocyte and platelet inhibitory receptor G6b |
| ENSG00000197182 | MIRLET7BHG | -2,830376884 | 0,140595577 | 0,00157 | 0,04253303 | MIRLET7B host gene |
| ENSG00000261615 | LINC01858 | -2,832575302 | 0,140381497 | 0,00693 | 0,10609569 | long intergenic non-protein coding RNA 1858 |
| ENSG00000263558 | RN7SL716P | -2,833037305 | 0,140336549 | 0,05332 | 0,31010674 | RNA, 7SL, cytoplasmic 716, pseudogene |
| ENSG00000260911 | AC135050,3 | -2,836353882 | 0,140014303 | 0,03134 | 0,2388853 | novel transcript |
| ENSG00000228862 | AC068389,1 | -2,836638509 | 0,139986683 | 0,00067 | 0,02497236 | novel transcript |
| ENSG00000249175 | AC008534,1 | -2,837150042 | 0,139937057 | 0,02281 | 0,20153127 | novel transcript |
| ENSG00000273293 | AC093458,1 | -2,838191574 | 0,139836068 | 0,03857 | 0,26560956 | novel transcript |
| ENSG00000286569 | AC245427,2 | -2,838578104 | 0,139798608 | 0,04639 | 0,28939918 | novel transcript |
| ENSG00000200983 | SNORA3A | -2,83912203 | 0,13974591 | 0,00159 | 0,04261325 | small nucleolar RNA, H/ACA box 3A |
| ENSG00000264809 | Metazoa_SRP | -2,840800169 | 0,139583453 | 0,16006 | 0,48727386 | Metazoan signal recognition particle RNA |
| ENSG00000274414 | AL121772,1 | -2,841486893 | 0,139517027 | 0,00271 | 0,06117642 | novel transcript |

| | | | | | | |
|---|---|---|---|---|---|---|
| ENSG00000230651 | RGPD4-AS1 | -2,845953303 | 0,139085767 | 0,00112 | 0,03490596 | RGPD4 antisense RNA 1 (head to head) |
| ENSG00000228429 | AC004910,1 | -2,846962806 | 0,138988478 | 0,04865 | 0,29648386 | ribosomal protein L21 (RPL21) pseudogene |
| ENSG00000251434 | AC104071,1 | -2,848261753 | 0,138863394 | 0,00892 | 0,11991469 | novel transcript |
| ENSG00000284726 | AL109936,9 | -2,848837693 | 0,13880797 | 0,09017 | 0,38840544 | novel transcript |
| ENSG00000288225 | AL591926,7 | -2,849012609 | 0,138791141 | 0,01492 | 0,16037017 | family with sequence similarity 27 member E3 (FAM27E3) pseudogene |
| ENSG00000243468 | INGX | -2,850049503 | 0,138691425 | 0,00296 | 0,06492836 | inhibitor of growth family, X-linked (pseudogene) |
| ENSG00000163749 | CCDC158 | -2,85102976 | 0,138597221 | 0,00238 | 0,05617483 | coiled-coil domain containing 158 |
| ENSG00000221933 | OR2A25 | -2,854638424 | 0,138250977 | 0,0066 | 0,10299608 | olfactory receptor family 2 subfamily A member 25 |
| ENSG00000125207 | PIWIL1 | -2,858566065 | 0,137875109 | 0,01669 | 0,16951872 | piwi like RNA-mediated gene silencing 1 |
| ENSG00000232642 | AC008073,2 | -2,861241794 | 0,137619633 | 0,00824 | 0,11480072 | novel transcript |
| ENSG00000269437 | NXF2B | -2,861684339 | 0,137577425 | 0,05684 | 0,31784869 | nuclear RNA export factor 2B |
| ENSG00000274514 | Metazoa_SRP | -2,862805899 | 0,137470513 | 0,00386 | 0,07576608 | Metazoan signal recognition particle RNA |
| ENSG00000178803 | ADORA2A-AS1 | -2,865390631 | 0,137224441 | 0,00053 | 0,02125773 | ADORA2A antisense RNA 1 |
| ENSG00000221716 | SNORA11 | -2,865450823 | 0,137218716 | 0,07029 | 0,34990935 | small nucleolar RNA, H/ACA box 11 |
| ENSG00000259045 | MTCO1P2 | -2,866450086 | 0,137123706 | 0,00414 | 0,07886388 | MT-CO1 pseudogene 2 |
| ENSG00000198113 | TOR4A | -2,867988838 | 0,136977531 | 0,00031 | 0,01478654 | torsin family 4 member A |
| ENSG00000287271 | AC011455,8 | -2,870771623 | 0,136713572 | 0,00067 | 0,02497236 | novel transcript, antisense to FBXO17and FBXO27 |
| ENSG00000196166 | C8orf86 | -2,871326549 | 0,136660996 | 0,00363 | 0,07351896 | chromosome 8 open reading frame 86 |
| ENSG00000188039 | NWD1 | -2,872960562 | 0,1365063 | 0,00252 | 0,05844508 | NACHT and WD repeat domain containing 1 |
| ENSG00000087237 | CETP | -2,873889154 | 0,136418466 | 0,11877 | 0,43690527 | cholesteryl ester transfer protein |
| ENSG00000258986 | TMEM179 | -2,874797247 | 0,136332625 | 7,55E-05 | 0,00472998 | transmembrane protein 179 |
| ENSG00000254081 | LINC01299 | -2,879394178 | 0,135898913 | 0,00859 | 0,11703366 | long intergenic non-protein coding RNA 1299 |
| ENSG00000100427 | MLC1 | -2,882915151 | 0,135567649 | 9,75E-06 | 0,00081406 | modulator of VRAC current 1 |
| ENSG00000259616 | AC104574,2 | -2,88341127 | 0,135521037 | 0,0132 | 0,15012165 | novel transcript |
| ENSG00000278702 | Metazoa_SRP | -2,883919007 | 0,135473351 | 0,0357 | 0,25626339 | Metazoan signal recognition particle RNA |
| ENSG00000212588 | SNORA26 | -2,884942298 | 0,135377295 | 0,00337 | 0,07000131 | small nucleolar RNA, H/ACA box 26 |
| ENSG00000253313 | C1orf210 | -2,885291502 | 0,135344531 | 0,01022 | 0,1300932 | chromosome 1 open reading frame 210 |
| ENSG00000136944 | LMX1B | -2,886483031 | 0,135232795 | 1,07E-12 | 3,52E-10 | LIM homeobox transcription factor 1 beta |
| ENSG00000267151 | MIR2117HG | -2,886621087 | 0,135219855 | 0,0142 | 0,15631436 | MIR2117 host gene |
| ENSG00000251544 | MTND5P12 | -2,887118425 | 0,135173249 | 0,03461 | 0,25152438 | MT-ND5 pseudogene 12 |
| ENSG00000163898 | LIPH | -2,888075192 | 0,135083635 | 0,00119 | 0,03622756 | lipase H |
| ENSG00000267892 | AC022144,1 | -2,888295114 | 0,135063044 | 0,00236 | 0,05588694 | novel transcript, antisense to CAPN12 |

| | | | | | | |
|---|---|---|---|---|---|---|
| ENSG00000166473 | PKD1L2 | -2,896459208 | 0,134300892 | 0,00125 | 0,03740512 | polycystin 1 like 2 (gene/pseudogene) |
| ENSG00000142224 | IL19 | -2,896744691 | 0,134274319 | 0,00155 | 0,04237695 | interleukin 19 |
| ENSG00000276690 | AC117415,1 | -2,898256992 | 0,13413364 | 0,00754 | 0,11001711 | |
| ENSG00000259563 | AC025430,1 | -2,901093494 | 0,133870177 | 0,0025 | 0,05798721 | novel transcript, antisense to CTDSPL2 |
| ENSG00000154438 | ASZ1 | -2,901149619 | 0,133864969 | 0,08716 | 0,38232685 | ankyrin repeat, SAM and basic leucine zipper domain containing 1 |
| ENSG00000287364 | AL353709,1 | -2,902983589 | 0,133694907 | 0,00201 | 0,05041136 | uncharacterized LOC105371672 |
| ENSG00000184571 | PIWIL3 | -2,90319888 | 0,133674957 | 0,00354 | 0,07210091 | piwi like RNA-mediated gene silencing 3 |
| ENSG00000283769 | CR392000,2 | -2,912929113 | 0,132776422 | 0,0013 | 0,03846063 | novel transcript, antisense to a novel protein |
| ENSG00000274660 | Metazoa_SRP | -2,914291552 | 0,132651091 | 0,08906 | 0,38599819 | Metazoan signal recognition particle RNA |
| ENSG00000091592 | NLRP1 | -2,915444078 | 0,132545162 | 0,00175 | 0,0459021 | NLR family pyrin domain containing 1 |
| ENSG00000187268 | FAM9C | -2,917482019 | 0,132358062 | 0,00082 | 0,02832201 | family with sequence similarity 9 member C |
| ENSG00000238622 | SNORD97 | -2,920035466 | 0,132124007 | 3,26E-05 | 0,00231818 | small nucleolar RNA, C/D box 97 |
| ENSG00000258240 | AC002351,1 | -2,920159785 | 0,132112622 | 0,01332 | 0,15109916 | novel transcript |
| ENSG00000104848 | KCNA7 | -2,92130788 | 0,132007529 | 0,00062 | 0,02370104 | potassium voltage-gated channel subfamily A member 7 |
| ENSG00000111863 | ADTRP | -2,921730284 | 0,131968884 | 0,0414 | 0,27565808 | androgen dependent TFPI regulating protein |
| ENSG00000095752 | IL11 | -2,924330229 | 0,131731272 | 0,00038 | 0,0169515 | interleukin 11 |
| ENSG00000159337 | PLA2G4D | -2,930634184 | 0,131156918 | 0,00427 | 0,08031113 | phospholipase A2 group IVD |
| ENSG00000257495 | KRT73-AS1 | -2,931343225 | 0,131092475 | 0,06367 | 0,33458308 | KRT73 antisense RNA 1 |
| ENSG00000147113 | DIPK2B | -2,932340428 | 0,131001894 | 0,00103 | 0,03303848 | divergent protein kinase domain 2B |
| ENSG00000239822 | RN7SL754P | -2,933473116 | 0,130899082 | 0,00561 | 0,09344307 | RNA, 7SL, cytoplasmic 754, pseudogene |
| ENSG00000281951 | AC093627,11 | -2,934162566 | 0,130836541 | 0,39334 | 0,6563386 | novel transcript similar to family member with sequence similarity 157 |
| ENSG00000047936 | ROS1 | -2,938754189 | 0,130420794 | 0,00434 | 0,08101303 | ROS proto-oncogene 1, receptor tyrosine kinase |
| ENSG00000171772 | SYCE1 | -2,938895794 | 0,130407993 | 0,0142 | 0,15631436 | synaptonemal complex central element protein 1 |
| ENSG00000272180 | AC011306,1 | -2,940902774 | 0,130226705 | 0,01906 | 0,18296265 | novel transcript |
| ENSG00000228221 | LINC00578 | -2,941700504 | 0,130154716 | 0,0101 | 0,12907505 | long intergenic non-protein coding RNA 578 |
| ENSG00000159248 | GJD2 | -2,941886475 | 0,13013794 | 4,30E-12 | 1,36E-09 | gap junction protein delta 2 |
| ENSG00000088726 | TMEM40 | -2,942486305 | 0,130083844 | 0,01178 | 0,14140074 | transmembrane protein 40 |
| ENSG00000112212 | TSPO2 | -2,945421239 | 0,129819478 | 0,0051 | 0,08908408 | translocator protein 2 |
| ENSG00000258420 | LONRF2P3 | -2,946448934 | 0,129727034 | 0,00474 | 0,08501852 | LONRF2 pseudogene 3 |
| ENSG00000159708 | LRRC36 | -2,948217197 | 0,12956813 | 0,00076 | 0,02693893 | leucine rich repeat containing 36 |

| | | | | | | |
|---|---|---|---|---|---|---|
| ENSG00000099769 | IGFALS | -2,948867899 | 0,129509704 | 0,01184 | 0,1417348 | insulin like growth factor binding protein acid labile subunit |
| ENSG00000286020 | AC011997,2 | -2,948996912 | 0,129498123 | 0,02623 | 0,2183952 | novel transcript, antisense to PLCL1 |
| ENSG00000204421 | LY6G6C | -2,949145328 | 0,129484801 | 0,09463 | 0,39709067 | lymphocyte antigen 6 family member G6C |
| ENSG00000278760 | CORO7-PAM1 | -2,951045955 | 0,129314329 | 0,20507 | 0,52465509 | CORO7-PAM16 readthrough |
| ENSG00000103426 | CORO7-PAM1 | -2,951045955 | 0,129314329 | 0,20507 | 0,52465509 | CORO7-PAM16 readthrough |
| ENSG00000237222 | LINC01968 | -2,95203673 | 0,129225552 | 0,00943 | 0,1242251 | long intergenic non-protein coding RNA 1968 |
| ENSG00000280434 | AL031595,3 | -2,952215159 | 0,129209571 | 0,00031 | 0,01477593 | novel transcript, sense overlapping PARVB |
| ENSG00000199568 | RNU5A-1 | -2,952362953 | 0,129196335 | 0,00019 | 0,0102859 | RNA, U5A small nuclear 1 |
| ENSG00000246740 | PLA2G4E-AS1 | -2,953190204 | 0,129122274 | 0,04401 | 0,28274932 | PLA2G4E antisense RNA 1 |
| ENSG00000251340 | MTCYBP35 | -2,954003702 | 0,129049486 | 0,00855 | 0,11673898 | MT-CYB pseudogene 35 |
| ENSG00000021461 | CYP3A43 | -2,958307088 | 0,12866512 | 0,0156 | 0,16379098 | cytochrome P450 family 3 subfamily A member 43 |
| ENSG00000270866 | AC067773,1 | -2,959203361 | 0,128585212 | 0,0573 | 0,31930406 | novel transcript |
| ENSG00000238755 | LINC02006 | -2,959768389 | 0,128534862 | 0,00134 | 0,03909453 | long intergenic non-protein coding RNA 2006 |
| ENSG00000018280 | SLC11A1 | -2,964519967 | 0,128112223 | 0,00155 | 0,04237695 | solute carrier family 11 member 1 |
| ENSG00000260072 | AC008938,1 | -2,964742521 | 0,128092462 | 0,00059 | 0,02271395 | novel transcript |
| ENSG00000232118 | BACH1-AS1 | -2,965498434 | 0,128025364 | 0,00185 | 0,04753043 | BACH1 antisense RNA 1 |
| ENSG00000244414 | CFHR1 | -2,966701602 | 0,127918639 | 0,00457 | 0,08333243 | complement factor H related 1 |
| ENSG00000259048 | AL392023,2 | -2,968960281 | 0,127718526 | 0,03063 | 0,23611505 | novel transcript, antisense to TTC6 |
| ENSG00000232124 | AP001057,1 | -2,969433875 | 0,127676607 | 0,02059 | 0,19042717 | novel transcript |
| ENSG00000232983 | AL160270,2 | -2,978529921 | 0,126874151 | 0,03617 | 0,25739586 | novel pseudogene |
| ENSG00000253238 | AC034114,2 | -2,978617257 | 0,126866471 | 0,00226 | 0,05426803 | novel transcript |
| ENSG00000161640 | SIGLEC11 | -2,979482903 | 0,126790372 | 0,06748 | 0,34374257 | sialic acid binding Ig like lectin 11 |
| ENSG00000286381 | AL078622,1 | -2,980866301 | 0,126668851 | 0,02659 | 0,21992823 | novel transcript |
| ENSG00000074966 | TXK | -2,98086863 | 0,126668646 | 0,001 | 0,0323881 | TXK tyrosine kinase |
| ENSG00000225331 | LINC01678 | -2,981034495 | 0,126654084 | 0,01018 | 0,12975801 | long intergenic non-protein coding RNA 1678 |
| ENSG00000239381 | AC125613,1 | -2,981831617 | 0,126584124 | 0,00209 | 0,05143262 | novel transcript |
| ENSG00000203805 | PLPP4 | -2,983769256 | 0,126414227 | 0,00783 | 0,11257373 | phospholipid phosphatase 4 |
| ENSG00000267510 | AC011451,1 | -2,984432594 | 0,126356116 | 0,00654 | 0,1024996 | novel transcript, antisense to ZNF699 |
| ENSG00000269444 | AC011491,2 | -2,985259073 | 0,126283751 | 0,02917 | 0,2303054 | novel transcript, sense overlapping GTF2F1 |
| ENSG00000234336 | JAZF1-AS1 | -2,988264594 | 0,126020942 | 0,00094 | 0,03102731 | JAZF1 antisense RNA 1 |
| ENSG00000179673 | RPRML | -2,989300661 | 0,125930473 | 0,00144 | 0,04062048 | reprimo like |
| ENSG00000203588 | IGBP1-AS1 | -2,989628413 | 0,125901867 | 0,00256 | 0,05892196 | IGBP1 antisense RNA 1 |

| | | | | | | |
|---|---|---|---|---|---|---|
| ENSG00000212402 | SNORA74B | -2,989986678 | 0,125870606 | 0,00035 | 0,01575357 | small nucleolar RNA, H/ACA box 74B |
| ENSG00000233813 | MRPS18B | -2,990332811 | 0,125840411 | 0,00148 | 0,04128156 | mitochondrial ribosomal protein S18B |
| ENSG00000236393 | AC091806,1 | -2,990392239 | 0,125835227 | 0,00055 | 0,02167479 | novel transcript |
| ENSG00000261357 | AC099518,2 | -2,990418144 | 0,125832968 | 0,01265 | 0,14729065 | novel transcript |
| ENSG00000287108 | AC023302,2 | -2,990444221 | 0,125830693 | 0,00523 | 0,09023148 | novel transcript |
| ENSG00000259755 | AC090907,2 | -2,990782333 | 0,125801207 | 0,00744 | 0,1097853 | novel transcript, antisense to LRRK1 |
| ENSG00000284735 | AL139424,3 | -2,993989464 | 0,12552186 | 0,0004 | 0,01764485 | novel transcript |
| ENSG00000238266 | LINC00707 | -2,996602374 | 0,125294729 | 0,00231 | 0,05526704 | long intergenic non-protein coding RNA 707 |
| ENSG00000235262 | KDM5C-IT1 | -2,999014974 | 0,125085375 | 0,00049 | 0,02010872 | KDM5C intronic transcript 1 |
| ENSG00000258875 | AL135818,1 | -3,000342947 | 0,124970289 | 0,00046 | 0,01926552 | novel transcript, antisense to GPR68 |
| ENSG00000271171 | LINC00904 | -3,004153317 | 0,12464066 | 0,10091 | 0,40770184 | long intergenic non-protein coding RNA 904 |
| ENSG00000282307 | SERPINF1 | -3,005671878 | 0,124509534 | 0,01213 | 0,14313104 | serpin family F member 1 |
| ENSG00000110675 | ELMOD1 | -3,007855178 | 0,12432125 | 0,00041 | 0,01779212 | ELMO domain containing 1 |
| ENSG00000285833 | AL590989,1 | -3,008541289 | 0,12426214 | 0,07017 | 0,34975886 | novel transcript, antisense to DISP3 |
| ENSG00000259793 | AC013726,1 | -3,01045485 | 0,124097431 | 0,00101 | 0,03264954 | novel transcript, antisense to DGKD |
| ENSG00000267474 | AC008569,2 | -3,010889206 | 0,124060074 | 0,0005 | 0,02017094 | novel transcript, sense intronic to PKN1 |
| ENSG00000274775 | Z82195,2 | -3,012317301 | 0,12393733 | 0,04567 | 0,28842796 | fibrous sheath interacting protein 2 (FSIP2) pseudogene |
| ENSG00000263120 | AC004584,3 | -3,01551312 | 0,123663091 | 0,00082 | 0,0284201 | novel transcript, sense intronic to SCPEP1 |
| ENSG00000263588 | AC091043,1 | -3,015559175 | 0,123659144 | 0,01582 | 0,16482188 | novel transcript |
| ENSG00000186452 | TMPRSS12 | -3,01570206 | 0,123646897 | 0,00301 | 0,06548828 | transmembrane serine protease 12 |
| ENSG00000278522 | POTEB3 | -3,01630851 | 0,123594932 | 0,00038 | 0,01684107 | POTE ankyrin domain family member B3 |
| ENSG00000228661 | STIM1-AS1 | -3,019046981 | 0,123360551 | 0,00243 | 0,05680468 | STIM1 antisense RNA 1 |
| ENSG00000205488 | CALML3-AS1 | -3,020897965 | 0,12320238 | 0,00229 | 0,05474871 | CALML3 antisense RNA 1 |
| ENSG00000241490 | AC093010,1 | -3,023302272 | 0,12299723 | 0,00422 | 0,07971576 | novel transcript, antisense to ZNF80 |
| ENSG00000165810 | BTNL9 | -3,023734176 | 0,122960413 | 0,00374 | 0,07458671 | butyrophilin like 9 |
| ENSG00000181693 | OR8H1 | -3,024357442 | 0,122907304 | 0,05049 | 0,30230041 | olfactory receptor family 8 subfamily H member 1 |
| ENSG00000288084 | AL590608,1 | -3,026782845 | 0,122700851 | 0,00092 | 0,03058422 | novel transcript |
| ENSG00000260580 | AC009035,1 | -3,027877724 | 0,122607767 | 0,00718 | 0,10764638 | novel transcript |
| ENSG00000170683 | OR10A3 | -3,028704557 | 0,122537518 | 0,03675 | 0,25976579 | olfactory receptor family 10 subfamily A member 3 |
| ENSG00000271412 | PRR13P3 | -3,031064518 | 0,122337235 | 0,02441 | 0,20896203 | proline rich 13 pseudogene 3 |
| ENSG00000230709 | AC104024,1 | -3,032821617 | 0,122188328 | 0,05733 | 0,3193968 | novel transcript |
| ENSG00000229498 | AC105053,1 | -3,035065501 | 0,121998431 | 0,0017 | 0,0449436 | novel transcript |
| ENSG00000204832 | ST8SIA6-AS1 | -3,035113901 | 0,121994338 | 0,00129 | 0,0382843 | ST8SIA6 antisense RNA 1 |

| | | | | | | |
|---|---|---|---|---|---|---|
| ENSG00000282936 | AC004706,3 | -3,036344416 | 0,12189033 | 0,00011 | 0,00628762 | novel protein |
| ENSG00000256195 | AP002518,1 | -3,037155768 | 0,1218218 | 0,04109 | 0,27457447 | novel transcript |
| ENSG00000241286 | RPL29P25 | -3,037557359 | 0,121787894 | 0,0125 | 0,14623734 | ribosomal protein L29 pseudogene 25 |
| ENSG00000204183 | GDF5-AS1 | -3,038292447 | 0,121725856 | 0,01854 | 0,18002406 | GDF5 antisense RNA 1 |
| ENSG00000258375 | AC079363,1 | -3,040028107 | 0,1215795 | 0,00047 | 0,01961109 | transmembrane protein 66 (TMEM66) pseudogene |
| ENSG00000234962 | LINC00700 | -3,040654245 | 0,121526745 | 0,00208 | 0,05143262 | long intergenic non-protein coding RNA 700 |
| ENSG00000127533 | F2RL3 | -3,042222182 | 0,12139474 | 0,00096 | 0,03162131 | F2R like thrombin or trypsin receptor 3 |
| ENSG00000224116 | INHBA-AS1 | -3,045336449 | 0,121132975 | 0,00571 | 0,09435135 | INHBA antisense RNA 1 |
| ENSG00000260316 | AL008727,1 | -3,048726801 | 0,120848645 | 0,00103 | 0,03289327 | novel transcript, intronic to LMF1 |
| ENSG00000236081 | ELFN1-AS1 | -3,050142653 | 0,120730103 | 0,03591 | 0,25662951 | ELFN1 antisense RNA 1 |
| ENSG00000273566 | Metazoa_SRP | -3,050774411 | 0,120677246 | 0,02734 | 0,22320635 | Metazoan signal recognition particle RNA |
| ENSG00000283422 | LINC02452 | -3,051142663 | 0,120646447 | 0,00029 | 0,01389771 | long intergenic non-protein coding RNA 2452 |
| ENSG00000284240 | LINC02801 | -3,052118376 | 0,12056488 | 0,06922 | 0,34753175 | long intergenic non-protein coding RNA 2801 |
| ENSG00000181092 | ADIPOQ | -3,052366221 | 0,12054417 | 0,01583 | 0,16482188 | adiponectin, C1Q and collagen domain containing |
| ENSG00000286490 | AC067942,4 | -3,054349757 | 0,120378549 | 0,02815 | 0,22653994 | novel transcript |
| ENSG00000286503 | AC008462,1 | -3,05565643 | 0,12026957 | 0,03294 | 0,24506146 | novel transcript |
| ENSG00000266990 | AC004528,1 | -3,05628116 | 0,120217501 | 0,00098 | 0,0320617 | novel transcript, antisense to GRIN3B |
| ENSG00000275708 | MIR3648-1 | -3,057665266 | 0,120102221 | 0,04801 | 0,29426648 | microRNA 3648-1 |
| ENSG00000264462 | MIR3648-2 | -3,057665266 | 0,120102221 | 0,04801 | 0,29426648 | microRNA 3648-2 |
| ENSG00000174348 | PODN | -3,058137409 | 0,120062922 | 0,00084 | 0,02874882 | podocan |
| ENSG00000204364 | C2 | -3,058670935 | 0,12001853 | 0,0037 | 0,07414958 | complement C2 |
| ENSG00000154252 | GAL3ST2 | -3,059018758 | 0,119989598 | 0,16076 | 0,48807078 | galactose-3-O-sulfotransferase 2 |
| ENSG00000167895 | TMC8 | -3,06005919 | 0,119903095 | 0,00524 | 0,09023148 | transmembrane channel like 8 |
| ENSG00000248473 | LINC01962 | -3,062523728 | 0,119698441 | 0,00028 | 0,01366998 | long intergenic non-protein coding RNA 1962 |
| ENSG00000284995 | AC275565,7 | -3,063948092 | 0,119580322 | 0,03316 | 0,24602205 | RNA binding protein, fox-1 homolog (C, elegans) 1 (RBFOX1) pseudogene |
| ENSG00000285355 | AC005342,5 | -3,064314285 | 0,119549973 | 0,01355 | 0,152676 | novel transcript |
| ENSG00000179626 | OR6C4 | -3,064888005 | 0,119502441 | 0,06491 | 0,33778346 | olfactory receptor family 6 subfamily C member 4 |
| ENSG00000269103 | Metazoa_SRP | -3,06701614 | 0,119326292 | 0,00897 | 0,1201491 | Metazoan signal recognition particle RNA |
| ENSG00000280688 | CEP72 | -3,067573652 | 0,119280189 | 0,0739 | 0,35721215 | centrosomal protein 72 |
| ENSG00000188817 | SNTN | -3,071271273 | 0,118974865 | 0,00354 | 0,07209525 | sentan, cilia apical structure protein |
| ENSG00000263494 | AC004702,1 | -3,07217159 | 0,118900642 | 0,01197 | 0,14255836 | novel transcript |
| ENSG00000281680 | OR10A3 | -3,073371836 | 0,118801764 | 0,2667 | 0,56923396 | olfactory receptor family 10 subfamily A member 3 |

| | | | | | | |
|---|---|---|---|---|---|---|
| ENSG00000229961 | AL357143,1 | -3,076569412 | 0,118538745 | 0,00952 | 0,12493107 | novel SLAM family member pseudogene |
| ENSG00000100342 | APOL1 | -3,077457664 | 0,118465784 | 0,00072 | 0,0263314 | apolipoprotein L1 |
| ENSG00000274914 | LILRA5 | -3,078606611 | 0,118371477 | 0,00062 | 0,02365212 | leukocyte immunoglobulin like receptor A5 |
| ENSG00000233701 | PRR23C | -3,07911225 | 0,118329997 | 0,00995 | 0,12783227 | proline rich 23C |
| ENSG00000234199 | LINC01191 | -3,083702582 | 0,117954096 | 0,00191 | 0,04876303 | long intergenic non-protein coding RNA 1191 |
| ENSG00000282790 | AC068400,1 | -3,08637125 | 0,117736108 | 0,02071 | 0,19086086 | novel transcript, antisense TADA2A |
| ENSG00000240723 | RN7SL382P | -3,087740213 | 0,117624442 | 0,01724 | 0,17278665 | RNA, 7SL, cytoplasmic 382, pseudogene |
| ENSG00000249330 | AC095060,1 | -3,088123774 | 0,117593174 | 0,00609 | 0,09844621 | novel transcript |
| ENSG00000232328 | AC084030,1 | -3,088266501 | 0,117581541 | 0,00067 | 0,02497236 | novel transcript |
| ENSG00000244953 | AC087521,1 | -3,090313345 | 0,117414839 | 0,03446 | 0,25090278 | novel transcript |
| ENSG00000203987 | AL772363,1 | -3,092303929 | 0,117252946 | 0,04737 | 0,29244592 | novel transcript, antisense to CACNA1B and EHMT1 |
| ENSG00000267898 | AC026803,2 | -3,092990558 | 0,117197154 | 0,00358 | 0,07275302 | novel transcript |
| ENSG00000126262 | FFAR2 | -3,097950575 | 0,11679492 | 0,01268 | 0,14729065 | free fatty acid receptor 2 |
| ENSG00000254057 | AC084346,1 | -3,098927993 | 0,116715818 | 0,00028 | 0,01366998 | novel transcript |
| ENSG00000206478 | IER3 | -3,100716796 | 0,116571192 | 0,1513 | 0,47956749 | immediate early response 3 |
| ENSG00000263809 | AC135178,3 | -3,100753982 | 0,116568187 | 0,03512 | 0,25351219 | novel protein |
| ENSG00000224826 | USP21P1 | -3,101992486 | 0,11646816 | 0,00362 | 0,07336882 | USP21 pseudogene 1 |
| ENSG00000163694 | RBM47 | -3,104230711 | 0,116287609 | 1,08E-05 | 0,00088915 | RNA binding motif protein 47 |
| ENSG00000230930 | COL11A2 | -3,105035818 | 0,116222732 | 0,03348 | 0,2471482 | collagen type XI alpha 2 chain |
| ENSG00000164855 | TMEM184A | -3,105127428 | 0,116215353 | 0,00143 | 0,04046504 | transmembrane protein 184A |
| ENSG00000250264 | AL669918,1 | -3,108888433 | 0,115912782 | 0,00755 | 0,11001711 | novel protein, TAP2-HLA-DOB readthrough |
| ENSG00000268388 | FENDRR | -3,110078047 | 0,115817242 | 0,06141 | 0,32865897 | FOXF1 adjacent non-coding developmental regulatory RNA |
| ENSG00000281234 | LOH12CR2 | -3,110124771 | 0,115813491 | 0,1485 | 0,47602584 | loss of heterozygosity on chromosome 12, region 2 |
| ENSG00000165084 | C8orf34 | -3,111050002 | 0,115739241 | 3,98E-08 | 6,31E-06 | chromosome 8 open reading frame 34 |
| ENSG00000255595 | AC007368,1 | -3,114095766 | 0,115495155 | 0,01445 | 0,15735196 | uncharacterized LOC101927531 |
| ENSG00000252316 | RNY4 | -3,116938397 | 0,115267812 | 0,0413 | 0,27529279 | RNA, Ro60-associated Y4 |
| ENSG00000196566 | AL138767,1 | -3,122178921 | 0,114849866 | 0,00059 | 0,02271395 | novel transcript |
| ENSG00000244681 | MTHFD2P1 | -3,127183567 | 0,114452147 | 0,0335 | 0,2471482 | methylenetetrahydrofolate dehydrogenase (NADP+ dependent) 2, methenyltetrahydrofolate cyclohydrolase pseudogene 1 |
| ENSG00000261080 | RUNX2-AS1 | -3,133002868 | 0,113991419 | 0,00132 | 0,03880861 | RUNX2 antisense RNA 1 |
| ENSG00000221845 | C7orf65 | -3,134257531 | 0,113892328 | 0,00129 | 0,03841105 | chromosome 7 open reading frame 65 |
| ENSG00000269755 | AC008758,6 | -3,137977321 | 0,11359905 | 0,00141 | 0,04017644 | novel transcript |

| | | | | | | |
|---|---|---|---|---|---|---|
| ENSG00000285653 | AC006238,2 | -3,138603254 | 0,113549775 | 0,0015 | 0,04149527 | novel transcript, antisense to RAB31 |
| ENSG00000224008 | LINC01441 | -3,139753048 | 0,113459314 | 0,00064 | 0,0242766 | long intergenic non-protein coding RNA 1441 |
| ENSG00000226397 | C12orf77 | -3,139825689 | 0,113453601 | 0,01542 | 0,16268839 | chromosome 12 open reading frame 77 |
| ENSG00000267374 | MIR924HG | -3,141970674 | 0,113285045 | 1,13E-10 | 2,96E-08 | MIR924 host gene |
| ENSG00000279440 | AL008638,3 | -3,146611874 | 0,112921189 | 0,00835 | 0,11551322 | novel transcript |
| ENSG00000177324 | BEND2 | -3,149244889 | 0,112715288 | 0,00819 | 0,11480072 | BEN domain containing 2 |
| ENSG00000274711 | Metazoa_SRP | -3,150832537 | 0,112591316 | 0,16179 | 0,48914394 | Metazoan signal recognition particle RNA |
| ENSG00000287040 | AL590560,5 | -3,150992757 | 0,112578813 | 0,00072 | 0,02615353 | novel transcript |
| ENSG00000265313 | LINC01476 | -3,151585966 | 0,112532532 | 0,00118 | 0,03613087 | long intergenic non-protein coding RNA 1476 |
| ENSG00000204173 | LRRC37A5P | -3,152311721 | 0,112475936 | 0,03103 | 0,23730035 | leucine rich repeat containing 37 member A5, pseudogene |
| ENSG00000287226 | AC084740,1 | -3,155077781 | 0,112260494 | 0,00028 | 0,01349096 | novel transcript |
| ENSG00000162399 | BSND | -3,156766677 | 0,112129153 | 0,02017 | 0,18846202 | barttin CLCNK type accessory subunit beta |
| ENSG00000274064 | AC040978,1 | -3,158076183 | 0,112027422 | 0,01316 | 0,15012165 | glyceraldehyde-3-phosphate dehydrogenase (GAPDH) pseudogene |
| ENSG00000155026 | RSPH10B | -3,159138444 | 0,111944966 | 0,00074 | 0,02672199 | radial spoke head 10 homolog B |
| ENSG00000258851 | AL139300,2 | -3,164673921 | 0,111516267 | 0,00888 | 0,11966089 | novel transcript, antisense to BAG5 |
| ENSG00000249429 | AC016598,1 | -3,171564436 | 0,11098492 | 0,00684 | 0,10540607 | novel transcript |
| ENSG00000182352 | C17orf77 | -3,172099437 | 0,11094377 | 0,00647 | 0,10183667 | chromosome 17 open reading frame 77 |
| ENSG00000258881 | AC007040,2 | -3,174415499 | 0,110765807 | 0,00453 | 0,08299378 | novel protein |
| ENSG00000226923 | MCCD1 | -3,178523792 | 0,110450833 | 0,17538 | 0,49944679 | mitochondrial coiled-coil domain 1 |
| ENSG00000198077 | CYP2A7 | -3,180125744 | 0,110328258 | 0,00256 | 0,05892196 | cytochrome P450 family 2 subfamily A member 7 |
| ENSG00000233564 | PRR3 | -3,180206676 | 0,110322069 | 0,04219 | 0,27783166 | proline rich 3 |
| ENSG00000287711 | LINC02136 | -3,182976109 | 0,110110495 | 0,00049 | 0,02017094 | long intergenic non-protein coding RNA 2136 |
| ENSG00000260289 | AC093515,1 | -3,183764407 | 0,110050347 | 0,00734 | 0,10881218 | novel transcript |
| ENSG00000235200 | LINC01702 | -3,190475269 | 0,109539623 | 0,02376 | 0,20581316 | long intergenic non-protein coding RNA 1702 |
| ENSG00000179058 | C9orf50 | -3,192522768 | 0,109384273 | 0,00265 | 0,059998 | chromosome 9 open reading frame 50 |
| ENSG00000229235 | RPL37P18 | -3,196387685 | 0,10909163 | 0,00721 | 0,10777959 | ribosomal protein L37 pseudogene 18 |
| ENSG00000231336 | AC017006,1 | -3,20165266 | 0,108694236 | 0,04973 | 0,30011105 | novel transcript |
| ENSG00000135678 | CPM | -3,202451896 | 0,108634037 | 0,00037 | 0,01681292 | carboxypeptidase M |
| ENSG00000197614 | MFAP5 | -3,204743418 | 0,108461624 | 0,00031 | 0,01477593 | microfibril associated protein 5 |
| ENSG00000197334 | Metazoa_SRP | -3,205402928 | 0,108412053 | 0,015 | 0,16047226 | Metazoan signal recognition particle RNA |
| ENSG00000235158 | AC087430,1 | -3,205531178 | 0,108402416 | 0,02224 | 0,19814016 | novel transcript |
| ENSG00000205755 | CRLF2 | -3,210587092 | 0,108023186 | 0,00274 | 0,06145058 | cytokine receptor like factor 2 |

| | | | | | | |
|---|---|---|---|---|---|---|
| ENSG00000249768 | HSPE1P10 | -3,212554873 | 0,107875947 | 0,00426 | 0,08023657 | heat shock protein family E (Hsp10) member 1 pseudogene 10 |
| ENSG00000237261 | Metazoa_SRP | -3,214420346 | 0,107736548 | 0,00546 | 0,09204617 | Metazoan signal recognition particle RNA |
| ENSG00000250723 | AC016687,2 | -3,215390013 | 0,10766416 | 0,00061 | 0,02352135 | novel transcript |
| ENSG00000282806 | AC068400,2 | -3,215540104 | 0,10765296 | 0,00026 | 0,01283952 | novel transcript, sense intronic to C17orf78 |
| ENSG00000254951 | AC044810,2 | -3,216257539 | 0,107599439 | 0,01474 | 0,1589934 | novel transcript, sense overlapping OR5P2, OR5P3 and OR5P1P |
| ENSG00000267239 | AP001198,1 | -3,216982308 | 0,107545398 | 0,01207 | 0,14294961 | novel transcript, antisense C18orf1 |
| ENSG00000246394 | ZNF84-DT | -3,218017192 | 0,10746828 | 0,00303 | 0,06571644 | ZNF84 divergent transcript |
| ENSG00000276202 | Metazoa_SRP | -3,218161592 | 0,107457524 | 0,10496 | 0,41375163 | Metazoan signal recognition particle RNA |
| ENSG00000124882 | EREG | -3,218773869 | 0,107411929 | 0,00055 | 0,02167479 | epiregulin |
| ENSG00000196564 | BLOC1S6P1 | -3,218971145 | 0,107397242 | 0,00222 | 0,05370447 | BLOC1S6 pseudogene 1 |
| ENSG00000228090 | MPIG6B | -3,220074266 | 0,107315155 | 0,04801 | 0,29426648 | megakaryocyte and platelet inhibitory receptor G6b |
| ENSG00000285987 | AL157886,1 | -3,220167188 | 0,107308243 | 0,00795 | 0,11358895 | Novel transcript, antisense to SLC28A3 |
| ENSG00000106526 | ACTR3C | -3,220910983 | 0,107252934 | 0,0009 | 0,03006448 | actin related protein 3C |
| ENSG00000261873 | SMIM36 | -3,222212319 | 0,107156233 | 0,04046 | 0,27194236 | small integral membrane protein 36 |
| ENSG00000179709 | NLRP8 | -3,225176966 | 0,10693626 | 0,01589 | 0,16510852 | NLR family pyrin domain containing 8 |
| ENSG00000105398 | SULT2A1 | -3,228828902 | 0,106665912 | 0,0131 | 0,14972794 | sulfotransferase family 2A member 1 |
| ENSG00000285108 | AC103718,1 | -3,229707314 | 0,106600986 | 0,00463 | 0,08390289 | novel transcript |
| ENSG00000287490 | AC015818,9 | -3,231462071 | 0,106471405 | 0,00715 | 0,1076344 | novel transcript |
| ENSG00000250423 | KIAA1210 | -3,232625368 | 0,106385588 | 0,00295 | 0,06471154 | KIAA1210 |
| ENSG00000158901 | WFDC8 | -3,235115638 | 0,106202112 | 0,0008 | 0,02785281 | WAP four-disulfide core domain 8 |
| ENSG00000165828 | PRAP1 | -3,238090531 | 0,105983345 | 0,00711 | 0,10746892 | proline rich acidic protein 1 |
| ENSG00000206028 | Z99774,1 | -3,238870569 | 0,105926057 | 0,00068 | 0,02517055 | novel transcript |
| ENSG00000228141 | AC105339,1 | -3,239178852 | 0,105903425 | 0,00061 | 0,02341213 | novel pseudogene |
| ENSG00000228060 | PABPC4-AS1 | -3,241613183 | 0,105724879 | 0,00902 | 0,12060025 | PABPC4 antisense RNA 1 |
| ENSG00000233250 | AC233728,1 | -3,243334565 | 0,105598807 | 0,03771 | 0,26272136 | novel transcript |
| ENSG00000167769 | ACER1 | -3,244477064 | 0,105515214 | 0,00037 | 0,01667641 | alkaline ceramidase 1 |
| ENSG00000261459 | AC002310,4 | -3,247627126 | 0,105285077 | 0,22112 | 0,53805234 | novel transcript |
| ENSG00000273946 | RN7SL733P | -3,251211762 | 0,105023802 | 0,09995 | 0,40649331 | RNA, 7SL, cytoplasmic 733, pseudogene |
| ENSG00000281000 | SNORD3D | -3,25218033 | 0,104953317 | 0,02037 | 0,18949014 | small nucleolar RNA, C/D box 3D |
| ENSG00000197253 | TPSB2 | -3,257068419 | 0,10459832 | 0,01869 | 0,18100822 | tryptase beta 2 |
| ENSG00000267054 | AC008738,1 | -3,258621386 | 0,104485787 | 0,0268 | 0,2209591 | novel transcript, sense intronic to LRP3 |

| | | | | | | |
|---|---|---|---|---|---|---|
| ENSG00000288645 | AC084756,2 | -3,262859145 | 0,104179322 | 0,00064 | 0,0242766 | novel protein |
| ENSG00000066294 | CD84 | -3,263885232 | 0,104105253 | 0,00033 | 0,01523174 | CD84 molecule |
| ENSG00000254486 | LINC02547 | -3,264560162 | 0,104056561 | 0,00046 | 0,01934795 | long intergenic non-protein coding RNA 2547 |
| ENSG00000270755 | AL136141,1 | -3,264794815 | 0,104039638 | 0,02723 | 0,22269021 | novel transcript |
| ENSG00000148483 | TMEM236 | -3,265136132 | 0,104015027 | 0,00027 | 0,01332054 | transmembrane protein 236 |
| ENSG00000117215 | PLA2G2D | -3,268594407 | 0,103765991 | 0,00466 | 0,08400694 | phospholipase A2 group IID |
| ENSG00000203395 | AC015969,1 | -3,27060185 | 0,103621706 | 0,00112 | 0,03490449 | novel transcript, antisense to PLEK |
| ENSG00000122584 | NXPH1 | -3,272468681 | 0,103487708 | 3,79E-06 | 0,00035877 | neurexophilin 1 |
| ENSG00000224697 | NEFMP1 | -3,275218659 | 0,103290634 | 0,00214 | 0,05235162 | neurofilament medium pseudogene 1 |
| ENSG00000288584 | Z97205,3 | -3,276470959 | 0,103201013 | 0,00104 | 0,03307359 | novel transcript |
| ENSG00000253754 | AC087620,1 | -3,284393583 | 0,102635834 | 0,04216 | 0,27783166 | novel transcript |
| ENSG00000207344 | SNORA22C | -3,286727846 | 0,102469904 | 0,00246 | 0,05743859 | small nucleolar RNA, H/ACA box 22C |
| ENSG00000231002 | TRIM26 | -3,287500719 | 0,102415024 | 0,06466 | 0,33701981 | tripartite motif containing 26 |
| ENSG00000248801 | C8orf34-AS1 | -3,29136152 | 0,102141318 | 0,00047 | 0,01972028 | C8orf34 antisense RNA 1 |
| ENSG00000243313 | RN7SL285P | -3,297310618 | 0,101720995 | 0,00981 | 0,12683981 | RNA, 7SL, cytoplasmic 285, pseudogene |
| ENSG00000259976 | AC093010,2 | -3,298020296 | 0,101670969 | 0,27024 | 0,57196184 | novel transcript |
| ENSG00000253532 | AC107909,1 | -3,298524142 | 0,101635468 | 0,01989 | 0,18711351 | novel transcript |
| ENSG00000266677 | AC087164,1 | -3,300187006 | 0,10151839 | 0,00036 | 0,01613306 | novel transcript |
| ENSG00000286333 | AC004828,2 | -3,304348124 | 0,101226005 | 0,00152 | 0,04183999 | novel transcript, antisense to RGS6 |
| ENSG00000127084 | FGD3 | -3,308474173 | 0,100936917 | 2,63E-11 | 7,68E-09 | FYVE, RhoGEF and PH domain containing 3 |
| ENSG00000197644 | AC106870,1 | -3,312996223 | 0,100621031 | 0,0923 | 0,39278042 | novel transcript |
| ENSG00000234123 | RHBDF1P1 | -3,313317013 | 0,10059866 | 0,00685 | 0,10547948 | RHBDF1 pseudogene 1 |
| ENSG00000130592 | LSP1 | -3,313634534 | 0,100576521 | 0,01113 | 0,13651927 | lymphocyte specific protein 1 |
| ENSG00000169435 | RASSF6 | -3,314597649 | 0,100509401 | 0,0281 | 0,22622448 | Ras association domain family member 6 |
| ENSG00000214559 | AC019077,1 | -3,316615219 | 0,100368939 | 0,00065 | 0,02439336 | novel transcript |
| ENSG00000278767 | SERPINA10 | -3,323503888 | 0,099890834 | 0,11572 | 0,43162108 | serpin family A member 10 |
| ENSG00000229672 | AL450322,2 | -3,324811882 | 0,099800311 | 0,00286 | 0,06351531 | novel transcript |
| ENSG00000285025 | AL022312,1 | -3,33180702 | 0,099317584 | 0,33024 | 0,61114402 | MIEF1 upstream open reading frame protein |
| ENSG00000200463 | SNORD118 | -3,33223774 | 0,099287937 | 0,00898 | 0,12025942 | small nucleolar RNA, C/D box 118 |
| ENSG00000105697 | HAMP | -3,332298671 | 0,099283744 | 0,00031 | 0,01478522 | hepcidin antimicrobial peptide |
| ENSG00000140090 | SLC24A4 | -3,336104111 | 0,099022205 | 0,00028 | 0,01366998 | solute carrier family 24 member 4 |
| ENSG00000184454 | NCMAP | -3,336918233 | 0,098966342 | 0,00075 | 0,02681144 | non-compact myelin associated protein |
| ENSG00000272958 | AP000322,1 | -3,338248315 | 0,098875143 | 0,02807 | 0,22609619 | novel transcript |

| | | | | | | |
|---|---|---|---|---|---|---|
| ENSG00000265740 | RN7SL339P | -3,338805531 | 0,098836961 | 0,00115 | 0,03540527 | RNA, 7SL, cytoplasmic 339, pseudogene |
| ENSG00000274591 | AC025031,3 | -3,342893387 | 0,098557305 | 0,00393 | 0,07669432 | novel transcript, sense intronic to SLC38A1 |
| ENSG00000172689 | MS4A10 | -3,343004844 | 0,098549691 | 0,00018 | 0,00982279 | membrane spanning 4-domains A10 |
| ENSG00000242641 | LINC00971 | -3,344802315 | 0,098426983 | 0,00894 | 0,11991469 | long intergenic non-protein coding RNA 971 |
| ENSG00000213252 | AP001282,1 | -3,347724564 | 0,098227816 | 0,00066 | 0,02486898 | pyridine nucleotide-disulphide oxidoreductase domain 1 (PYROXD1) pseudogene |
| ENSG00000263300 | AC015853,3 | -3,348551422 | 0,098171535 | 0,01291 | 0,14873257 | novel transcript, antisense to VPS53 |
| ENSG00000157330 | C1orf158 | -3,348682832 | 0,098162593 | 0,00147 | 0,04106985 | chromosome 1 open reading frame 158 |
| ENSG00000281176 | KRTAP5-AS1 | -3,349474913 | 0,098108714 | 0,03274 | 0,24408093 | KRTAP5-1/KRTAP5-2 antisense RNA 1 |
| ENSG00000119125 | GDA | -3,351064973 | 0,098000643 | 3,37E-28 | 6,76E-25 | guanine deaminase |
| ENSG00000249736 | LINC02242 | -3,35649119 | 0,097632739 | 0,00119 | 0,03626584 | long intergenic non-protein coding RNA 2242 |
| ENSG00000284290 | ANKS4B | -3,360116276 | 0,097387723 | 0,05631 | 0,31687263 | ankyrin repeat and sterile alpha motif domain containing 4B |
| ENSG00000180264 | ADGRD2 | -3,36046806 | 0,097363979 | 0,00054 | 0,02167371 | adhesion G protein-coupled receptor D2 |
| ENSG00000188676 | IDO2 | -3,360724305 | 0,097346687 | 0,00614 | 0,09864398 | indoleamine 2,3-dioxygenase 2 |
| ENSG00000259462 | CPEB1-AS1 | -3,363405715 | 0,097165926 | 9,12E-05 | 0,00555949 | CPEB1 antisense RNA 1 |
| ENSG00000273710 | Metazoa_SRP | -3,364773851 | 0,097073825 | 0,0255 | 0,21485835 | Metazoan signal recognition particle RNA |
| ENSG00000135722 | FBXL8 | -3,372804127 | 0,096534997 | 0,0011 | 0,03460106 | F-box and leucine rich repeat protein 8 |
| ENSG00000261829 | AC009407,1 | -3,37281178 | 0,096534485 | 0,00564 | 0,09352688 | novel transcript |
| ENSG00000182261 | NLRP10 | -3,374102126 | 0,096448183 | 0,00034 | 0,01565603 | NLR family pyrin domain containing 10 |
| ENSG00000249806 | AC139720,1 | -3,374640442 | 0,096412202 | 0,01022 | 0,1300932 | novel transcript |
| ENSG00000182866 | LCK | -3,375003351 | 0,096387953 | 0,00259 | 0,05932226 | LCK proto-oncogene, Src family tyrosine kinase |
| ENSG00000285289 | LINC00596 | -3,375612091 | 0,096347291 | 0,01576 | 0,16469415 | long intergenic non-protein coding RNA 596 |
| ENSG00000284809 | AL365394,1 | -3,3773391 | 0,096232025 | 0,00958 | 0,12507086 | novel transcript |
| ENSG00000204655 | MOG | -3,378354552 | 0,096164315 | 0,01351 | 0,15245066 | myelin oligodendrocyte glycoprotein |
| ENSG00000160654 | CD3G | -3,379342334 | 0,096098496 | 0,00071 | 0,02613504 | CD3g molecule |
| ENSG00000287899 | AC055714,1 | -3,386535022 | 0,095620581 | 0,00112 | 0,03490596 | novel transcript |
| ENSG00000204117 | AL162293,1 | -3,3896785 | 0,095412461 | 0,00041 | 0,01788369 | novel transcript |
| ENSG00000275484 | AP003419,3 | -3,390587526 | 0,095352361 | 0,00195 | 0,04943508 | novel transcript |
| ENSG00000175886 | RPL7AP66 | -3,394081665 | 0,095121702 | 0,00104 | 0,03313249 | ribosomal protein L7a pseudogene 66 |
| ENSG00000248923 | MTND5P11 | -3,396345693 | 0,094972544 | 0,00042 | 0,01799896 | MT-ND5 pseudogene 11 |
| ENSG00000280356 | AL021877,2 | -3,397128077 | 0,094921054 | 0,00166 | 0,04403004 | novel transcript |
| ENSG00000113946 | CLDN16 | -3,397643797 | 0,094887128 | 0,00194 | 0,04920335 | claudin 16 |

| | | | | | | |
|---|---|---|---|---|---|---|
| ENSG00000147571 | CRH | -3,39853428 | 0,094828578 | 3,51E-08 | 5,68E-06 | corticotropin releasing hormone |
| ENSG00000228031 | AC078842,1 | -3,399874574 | 0,094740522 | 0,01466 | 0,15857384 | novel transcript |
| ENSG00000128285 | MCHR1 | -3,402719026 | 0,094553913 | 6,39E-08 | 9,69E-06 | melanin concentrating hormone receptor 1 |
| ENSG00000196188 | CTSE | -3,408382225 | 0,094183476 | 0,00824 | 0,11480072 | cathepsin E |
| ENSG00000268434 | AC011530,1 | -3,411895096 | 0,093954424 | 0,06103 | 0,32783036 | novel protein |
| ENSG00000228427 | AL590764,1 | -3,41227893 | 0,09392943 | 0,01494 | 0,16037017 | novel transcript |
| ENSG00000147697 | GSDMC | -3,413568758 | 0,093845491 | 0,03109 | 0,23760845 | gasdermin C |
| ENSG00000244306 | DUXAP10 | -3,419915246 | 0,093433567 | 0,0066 | 0,10299608 | double homeobox A pseudogene 10 |
| ENSG00000204396 | VWA7 | -3,420240092 | 0,093412531 | 0,00038 | 0,01694126 | von Willebrand factor A domain containing 7 |
| ENSG00000288253 | AC010332,3 | -3,421551695 | 0,093327645 | 0,07387 | 0,35715669 | novel transcript |
| ENSG00000197446 | CYP2F1 | -3,422186722 | 0,093286574 | 0,03147 | 0,2394635 | cytochrome P450 family 2 subfamily F member 1 |
| ENSG00000172782 | FADS6 | -3,42264019 | 0,093257257 | 0,00621 | 0,09929843 | fatty acid desaturase 6 |
| ENSG00000244921 | MTCYBP18 | -3,423139988 | 0,093224955 | 0,00208 | 0,05143262 | MT-CYB pseudogene 18 |
| ENSG00000163564 | PYHIN1 | -3,423452654 | 0,093204754 | 0,00107 | 0,03382756 | pyrin and HIN domain family member 1 |
| ENSG00000249345 | LINC02405 | -3,423880136 | 0,09317714 | 0,00957 | 0,12500859 | long intergenic non-protein coding RNA 2405 |
| ENSG00000287796 | AC016925,3 | -3,427571184 | 0,092939057 | 0,00706 | 0,10707107 | novel transcript |
| ENSG00000160013 | PTGIR | -3,435983671 | 0,092398698 | 0,00318 | 0,06765231 | prostaglandin I2 receptor |
| ENSG00000250493 | AP004147,1 | -3,440005686 | 0,092141463 | 7,25E-05 | 0,00461938 | novel transcript, antisense to GRIK4 |
| ENSG00000253894 | AC011124,2 | -3,440519423 | 0,092108658 | 0,00874 | 0,11841612 | novel transcript |
| ENSG00000251249 | AC097375,3 | -3,44078567 | 0,092091661 | 0,00034 | 0,01552867 | novel transcript |
| ENSG00000267659 | LINC01482 | -3,453061096 | 0,091311407 | 0,00234 | 0,05563463 | long intergenic non-protein coding RNA 1482 |
| ENSG00000149656 | LINC00266-1 | -3,458398477 | 0,090974216 | 0,05943 | 0,32465136 | long intergenic non-protein coding RNA 266-1 |
| ENSG00000250387 | LINC02197 | -3,459244703 | 0,09092087 | 0,06153 | 0,32865897 | long intergenic non-protein coding RNA 2197 |
| ENSG00000285016 | AC017002,3 | -3,463134042 | 0,090676088 | 0,01738 | 0,17367715 | novel transcript |
| ENSG00000261205 | NIFKP4 | -3,46493413 | 0,090563019 | 0,00098 | 0,0320617 | nucleolar protein interacting with the FHA domain of MKI67 pseudogene 4 |
| ENSG00000187714 | SLC18A3 | -3,466997739 | 0,090433572 | 2,99E-05 | 0,00216456 | solute carrier family 18 member A3 |
| ENSG00000231827 | AL513523,2 | -3,467191405 | 0,090421433 | 0,00011 | 0,00651858 | immunoglobulin superfamily, DCC subclass, member 3 (IGDCC3) pseudogene |
| ENSG00000285751 | AC021723,2 | -3,467480441 | 0,09040332 | 0,00508 | 0,08898421 | novel transcript |
| ENSG00000102962 | CCL22 | -3,474840986 | 0,089943262 | 0,00216 | 0,05259865 | C-C motif chemokine ligand 22 |
| ENSG00000125084 | WNT1 | -3,477168391 | 0,089798279 | 1,74E-05 | 0,00137044 | Wnt family member 1 |
| ENSG00000200156 | RNU5B-1 | -3,479767197 | 0,089636666 | 6,81E-05 | 0,00437444 | RNA, U5B small nuclear 1 |

| | | | | | | |
|---|---|---|---|---|---|---|
| ENSG00000288636 | AL049569,3 | -3,482540191 | 0,089464542 | 0,00856 | 0,11684463 | novel protein |
| ENSG00000125533 | BHLHE23 | -3,490000764 | 0,08900309 | 0,00647 | 0,10183667 | basic helix-loop-helix family member e23 |
| ENSG00000236188 | PRKX-AS1 | -3,490091986 | 0,088997463 | 0,00012 | 0,00677463 | PRKX antisense RNA 1 |
| ENSG00000215124 | AMD1P4 | -3,4930215 | 0,088816929 | 0,00486 | 0,08609281 | adenosylmethionine decarboxylase 1 pseudogene 4 |
| ENSG00000287666 | AP000821,2 | -3,493752638 | 0,088771929 | 0,00336 | 0,07000131 | novel transcript |
| ENSG00000236740 | AL033384,1 | -3,494402989 | 0,088731921 | 0,00627 | 0,09972852 | novel transcript |
| ENSG00000224058 | AC006509,1 | -3,496583985 | 0,088597882 | 0,01042 | 0,13153018 | novel pseudogene |
| ENSG00000281644 | AC106795,9 | -3,496992554 | 0,088572794 | 0,00143 | 0,04048455 | novel transcript, antisense to PROP1 |
| ENSG00000227674 | LINC00355 | -3,497136362 | 0,088563966 | 0,00012 | 0,0067978 | long intergenic non-protein coding RNA 355 |
| ENSG00000259773 | AC012100,2 | -3,49845415 | 0,088483107 | 0,00444 | 0,08217356 | novel transcript, antisense to SPPL2A |
| ENSG00000174792 | ODAPH | -3,499376776 | 0,088426538 | 0,00031 | 0,01478522 | odontogenesis associated phosphoprotein |
| ENSG00000262152 | LINC00514 | -3,501317171 | 0,088307686 | 0,00256 | 0,05892196 | long intergenic non-protein coding RNA 514 |
| ENSG00000168477 | TNXB | -3,501677675 | 0,088285623 | 7,27E-07 | 8,57E-05 | tenascin XB |
| ENSG00000284896 | OR2A5 | -3,504775421 | 0,08809626 | 0,0016 | 0,04295976 | olfactory receptor family 2 subfamily A member 5 |
| ENSG00000241985 | WWTR1-IT1 | -3,507702321 | 0,087917714 | 0,00183 | 0,04735593 | WWTR1 intronic transcript 1 |
| ENSG00000229031 | MTCO1P25 | -3,513159437 | 0,087585786 | 0,00063 | 0,02419154 | MT-CO1 pseudogene 25 |
| ENSG00000232756 | AC004990,1 | -3,517045778 | 0,087350164 | 0,00801 | 0,11389729 | novel transcript |
| ENSG00000260447 | AC009065,3 | -3,518815801 | 0,087243061 | 0,00921 | 0,12242453 | novel transcript, antisense to PKD1 |
| ENSG00000277825 | AC020917,3 | -3,523073438 | 0,086985971 | 0,00086 | 0,02908139 | novel transcript |
| ENSG00000206249 | AL359555,1 | -3,528418494 | 0,086664293 | 0,00083 | 0,02852749 | novel transcript |
| ENSG00000275152 | CCL16 | -3,528703671 | 0,086647163 | 0,00057 | 0,02219024 | C-C motif chemokine ligand 16 |
| ENSG00000185522 | LMNTD2 | -3,531558562 | 0,08647587 | 0,00656 | 0,1027687 | lamin tail domain containing 2 |
| ENSG00000279966 | AC008626,2 | -3,533424356 | 0,086364106 | 0,02508 | 0,21269649 | zinc finger protein 99 (ZNF99) pseudogene |
| ENSG00000265272 | RN7SL693P | -3,53547812 | 0,086241249 | 0,00014 | 0,00771723 | RNA, 7SL, cytoplasmic 693, pseudogene |
| ENSG00000277510 | Metazoa_SRP | -3,536767039 | 0,086164234 | 0,02295 | 0,20217525 | Metazoan signal recognition particle RNA |
| ENSG00000258080 | ARHGAP42P4 | -3,541816442 | 0,085863189 | 0,00253 | 0,0585725 | ARHGAP42 pseudogene 4 |
| ENSG00000253214 | AC079209,1 | -3,542559554 | 0,085818973 | 0,00405 | 0,07813668 | novel transcript |
| ENSG00000287289 | AP000428,2 | -3,545675076 | 0,085633846 | 0,00664 | 0,10348339 | novel transcript |
| ENSG00000277677 | Metazoa_SRP | -3,548148497 | 0,085487157 | 0,00318 | 0,06765368 | Metazoan signal recognition particle RNA |
| ENSG00000213373 | LINC00671 | -3,550141089 | 0,085369167 | 0,01217 | 0,14328772 | long intergenic non-protein coding RNA 671 |
| ENSG00000265916 | AC015818,5 | -3,551705422 | 0,08527665 | 0,02031 | 0,18924656 | novel pseudogene |
| ENSG00000275898 | AC125603,4 | -3,561503732 | 0,084699442 | 0,00135 | 0,03912393 | novel transcript |
| ENSG00000265052 | RN7SL622P | -3,563626735 | 0,084574893 | 0,03195 | 0,24116556 | RNA, 7SL, cytoplasmic 622, pseudogene |

| | | | | | | |
|---|---|---|---|---|---|---|
| ENSG00000137270 | GCM1 | -3,588325543 | 0,083139303 | 0,00659 | 0,10297277 | glial cells missing transcription factor 1 |
| ENSG00000244242 | IFITM10 | -3,594678636 | 0,082773993 | 0,01215 | 0,14321473 | interferon induced transmembrane protein 10 |
| ENSG00000166035 | LIPC | -3,604210671 | 0,082228899 | 0,00576 | 0,0949578 | lipase C, hepatic type |
| ENSG00000187513 | GJA4 | -3,6083609 | 0,08199269 | 2,09E-17 | 1,22E-14 | gap junction protein alpha 4 |
| ENSG00000184302 | SIX6 | -3,610153586 | 0,081890869 | 2,23E-10 | 5,57E-08 | SIX homeobox 6 |
| ENSG00000070748 | CHAT | -3,618002513 | 0,081446555 | 2,65E-08 | 4,46E-06 | choline O-acetyltransferase |
| ENSG00000234707 | SEC61G-DT | -3,621721672 | 0,081236862 | 6,21E-05 | 0,00407918 | SEC61G divergent transcript |
| ENSG00000240625 | RN7SL403P | -3,621856907 | 0,081229248 | 0,01228 | 0,14435261 | RNA, 7SL, cytoplasmic 403, pseudogene |
| ENSG00000287393 | AL591468,1 | -3,631471176 | 0,080689727 | 0,05164 | 0,3054345 | novel transcript |
| ENSG00000283689 | AC018553,2 | -3,633078322 | 0,08059989 | 0,00161 | 0,04305274 | novel transcript, antisense to IRX3 |
| ENSG00000253671 | AC027117,1 | -3,636143619 | 0,080428821 | 0,00082 | 0,02837345 | novel transcript |
| ENSG00000152034 | MCHR2 | -3,636810998 | 0,080391624 | 1,88E-05 | 0,00146354 | melanin concentrating hormone receptor 2 |
| ENSG00000257373 | AC012038,1 | -3,645584669 | 0,07990421 | 0,00397 | 0,07725019 | novel transcript |
| ENSG00000280936 | AL353658,1 | -3,647138728 | 0,079818184 | 0,04927 | 0,29853148 | novel transcript, antisense to KCNQ2 |
| ENSG00000281914 | AL353658,2 | -3,647138728 | 0,079818184 | 0,04927 | 0,29853148 | novel transcript, antisense to KCNQ2 |
| ENSG00000162896 | PIGR | -3,666780042 | 0,078738878 | 0,00053 | 0,02115883 | polymeric immunoglobulin receptor |
| ENSG00000253872 | AC083836,1 | -3,67014365 | 0,078555514 | 0,00023 | 0,01154256 | novel transcript |
| ENSG00000073067 | CYP2W1 | -3,677576759 | 0,078151818 | 0,00031 | 0,01465633 | cytochrome P450 family 2 subfamily W member 1 |
| ENSG00000223806 | LINC00114 | -3,685923985 | 0,077700948 | 0,00746 | 0,11001616 | long intergenic non-protein coding RNA 114 |
| ENSG00000257275 | AL139020,1 | -3,686131467 | 0,077689774 | 0,00244 | 0,05718802 | novel transcript, antisense to TCL1A |
| ENSG00000131746 | TNS4 | -3,68645457 | 0,077672377 | 0,00468 | 0,08422651 | tensin 4 |
| ENSG00000250377 | AC008525,1 | -3,705851342 | 0,076635075 | 6,99E-06 | 0,00061886 | novel transcript |
| ENSG00000171049 | FPR2 | -3,707883501 | 0,076527204 | 0,01199 | 0,14262383 | formyl peptide receptor 2 |
| ENSG00000276096 | Metazoa_SRP | -3,710377222 | 0,07639504 | 0,00483 | 0,08564971 | Metazoan signal recognition particle RNA |
| ENSG00000106648 | GALNTL5 | -3,718641821 | 0,075958655 | 0,00452 | 0,08293474 | polypeptide N-acetylgalactosaminyltransferase like 5 |
| ENSG00000264963 | RN7SL440P | -3,720060595 | 0,075883993 | 0,01039 | 0,13135977 | RNA, 7SL, cytoplasmic 440, pseudogene |
| ENSG00000269793 | ZIM2-AS1 | -3,72993175 | 0,075366555 | 0,08064 | 0,37072333 | ZIM2 antisense RNA 1 |
| ENSG00000262999 | AC099489,3 | -3,747345564 | 0,074462323 | 1,26E-06 | 0,00014061 | novel transcript, antisense to a novel protein gene |
| ENSG00000249222 | ATP5MGL | -3,749763793 | 0,074337615 | 0,00018 | 0,00974504 | ATP synthase membrane subunit g like |
| ENSG00000266445 | NARF-AS1 | -3,750718417 | 0,074288442 | 3,58E-05 | 0,00252386 | NARF antisense RNA 1 |
| ENSG00000237900 | ZBTB12 | -3,753154324 | 0,074163116 | 0,0417 | 0,276508 | zinc finger and BTB domain containing 12 |
| ENSG00000234852 | ZBTB12 | -3,753154324 | 0,074163116 | 0,0417 | 0,276508 | zinc finger and BTB domain containing 12 |
| ENSG00000272027 | AC073218,1 | -3,759016587 | 0,073862372 | 0,02198 | 0,19708179 | novel transcript |

| | | | | | | |
|---|---|---|---|---|---|---|
| ENSG00000274642 | AC244669,2 | -3,763422293 | 0,073637155 | 0,01164 | 0,14056963 | phosphodiesterase 4D interacting protein (PDE4DIP) pseudogene |
| ENSG00000148053 | NTRK2 | -3,767038622 | 0,073452804 | 3,35E-16 | 1,77E-13 | neurotrophic receptor tyrosine kinase 2 |
| ENSG00000280989 | LINC00581 | -3,768818456 | 0,073362242 | 0,08823 | 0,38444882 | long intergenic non-protein coding RNA 581 |
| ENSG00000286559 | AC099845,1 | -3,770325231 | 0,073285661 | 1,68E-10 | 4,28E-08 | novel transcript |
| ENSG00000281557 | LRIG2-DT | -3,771567829 | 0,073222567 | 0,00829 | 0,1150861 | LRIG2 divergent transcript |
| ENSG00000255439 | AC135050,2 | -3,781585199 | 0,072715907 | 0,00175 | 0,04596317 | novel protein, VKORC1 and PRSS53 readthrough |
| ENSG00000236434 | AL162430,2 | -3,781653537 | 0,072712462 | 0,0031 | 0,06637133 | novel transcript |
| ENSG00000239998 | LILRA2 | -3,792281953 | 0,072178754 | 0,00201 | 0,0505329 | leukocyte immunoglobulin like receptor A2 |
| ENSG00000105371 | ICAM4 | -3,793952743 | 0,072095212 | 0,01391 | 0,15466987 | intercellular adhesion molecule 4 (Landsteiner-Wiener blood group) |
| ENSG00000225249 | LINC00378 | -3,796183022 | 0,071983845 | 0,00092 | 0,03058125 | long intergenic non-protein coding RNA 378 |
| ENSG00000286519 | AC009236,1 | -3,798961889 | 0,071845326 | 0,00254 | 0,05861174 | novel transcript |
| ENSG00000276565 | AP006285,3 | -3,801194066 | 0,071734251 | 0,00051 | 0,02067067 | |
| ENSG00000234938 | AC012668,2 | -3,801942358 | 0,071697054 | 0,04596 | 0,28908951 | novel transcript |
| ENSG00000274965 | TRPM1 | -3,802769436 | 0,071655962 | 0,01172 | 0,14091628 | transient receptor potential cation channel subfamily M member 1 |
| ENSG00000203489 | HMGB1P39 | -3,807393165 | 0,071426678 | 0,00389 | 0,07611868 | high mobility group box 1 pseudogene 39 |
| ENSG00000250685 | AC009123,1 | -3,819601345 | 0,070824811 | 0,00466 | 0,08400694 | novel transcript, antisense to ADAD2 |
| ENSG00000243977 | H3P13 | -3,819708655 | 0,070819543 | 0,01048 | 0,13209363 | H3 histone pseudogene 13 |
| ENSG00000277278 | HERC2 | -3,820958658 | 0,070758209 | 0,03388 | 0,24867791 | HECT and RLD domain containing E3 ubiquitin protein ligase 2 |
| ENSG00000090659 | CD209 | -3,824738622 | 0,07057306 | 0,15025 | 0,47830235 | CD209 molecule |
| ENSG00000282620 | MAPT-IT1 | -3,846761849 | 0,06950392 | 0,03206 | 0,24164117 | MAPT intronic transcript 1 |
| ENSG00000259084 | AL133467,3 | -3,849462768 | 0,069373921 | 0,00034 | 0,01562182 | novel transcript |
| ENSG00000235482 | RPL21P135 | -3,855711546 | 0,06907409 | 0,00341 | 0,07040998 | ribosomal protein L21 pseudogene 135 |
| ENSG00000198488 | B3GNT6 | -3,86089687 | 0,06882627 | 0,00012 | 0,00671805 | UDP-GlcNAc:betaGal beta-1,3-N-acetylglucosaminyltransferase 6 |
| ENSG00000075290 | WNT8B | -3,861978107 | 0,068774707 | 3,79E-30 | 9,67E-27 | Wnt family member 8B |
| ENSG00000278700 | Metazoa_SRP | -3,868020276 | 0,068487273 | 4,04E-05 | 0,00279716 | Metazoan signal recognition particle RNA |
| ENSG00000257475 | AC068888,2 | -3,875112018 | 0,068151441 | 0,00081 | 0,02810225 | novel transcript |
| ENSG00000207405 | SNORA64 | -3,877647026 | 0,068031795 | 0,01537 | 0,16238205 | small nucleolar RNA, H/ACA box 64 |
| ENSG00000285476 | AC139491,7 | -3,8791956 | 0,06795881 | 0,02738 | 0,22321357 | novel transcript |

| | | | | | | |
|---|---|---|---|---|---|---|
| ENSG00000125999 | BPIFB1 | -3,881367236 | 0,067856591 | 0,00063 | 0,02415118 | BPI fold containing family B member 1 |
| ENSG00000260019 | LINC01992 | -3,888432026 | 0,067525114 | 0,00626 | 0,09970758 | long intergenic non-protein coding RNA 1992 |
| ENSG00000248148 | AC114954,1 | -3,888799812 | 0,067507902 | 0,01592 | 0,16525378 | novel transcript |
| ENSG00000263882 | RN7SL774P | -3,894164356 | 0,067257345 | 0,01926 | 0,18372297 | RNA, 7SL, cytoplasmic 774, pseudogene |
| ENSG00000178343 | SHISA3 | -3,894330018 | 0,067249623 | 0,00135 | 0,03912393 | shisa family member 3 |
| ENSG00000278794 | Metazoa_SRP | -3,902214929 | 0,066883079 | 0,06358 | 0,33422972 | Metazoan signal recognition particle RNA |
| ENSG00000287286 | AC024560,5 | -3,903438394 | 0,066826383 | 0,00026 | 0,01300234 | novel transcript, antisense to RUBCN |
| ENSG00000244296 | RN7SL168P | -3,905602992 | 0,066726193 | 0,00197 | 0,04969574 | RNA, 7SL, cytoplasmic 168, pseudogene |
| ENSG00000204576 | PRR3 | -3,906948622 | 0,066663985 | 0,02607 | 0,21736588 | proline rich 3 |
| ENSG00000256286 | AC078878,2 | -3,910957104 | 0,066479019 | 0,02164 | 0,19544769 | novel transcript |
| ENSG00000284607 | AL121936,1 | -3,930688854 | 0,065575974 | 0,00139 | 0,03972417 | novel transcript |
| ENSG00000276680 | AL109755,2 | -3,931365903 | 0,065545207 | 0,01142 | 0,13882136 | |
| ENSG00000269752 | AC008761,2 | -3,932575724 | 0,065490265 | 0,00145 | 0,04066188 | novel transcript, antisense UNC13A |
| ENSG00000235813 | AL672032,1 | -3,935771141 | 0,065345371 | 0,00101 | 0,03262591 | novel transcript |
| ENSG00000260818 | UNGP1 | -3,939605308 | 0,065171937 | 0,00051 | 0,02059488 | uracil-DNA glycosylase pseudogene 1 |
| ENSG00000223766 | PRR3 | -3,944006234 | 0,064973434 | 0,04635 | 0,28932018 | proline rich 3 |
| ENSG00000265648 | RN7SL279P | -3,945655706 | 0,064899191 | 0,00692 | 0,10602899 | RNA, 7SL, cytoplasmic 279, pseudogene |
| ENSG00000285134 | ITFG2-AS1 | -3,947535062 | 0,064814703 | 0,00045 | 0,01901843 | ITFG2 antisense RNA 1 |
| ENSG00000203262 | AL137845,1 | -3,948906852 | 0,064753103 | 0,0152 | 0,16197635 | zinc finger protein 265 (ZNF265) pseudogene |
| ENSG00000224736 | HLA-V | -3,949773277 | 0,064714227 | 0,00151 | 0,04166341 | major histocompatibility complex, class I, V (pseudogene) |
| ENSG00000281166 | NLRP10 | -3,952738654 | 0,064581347 | 0,02818 | 0,22660674 | NLR family pyrin domain containing 10 |
| ENSG00000204653 | ASPDH | -3,962533676 | 0,064144364 | 0,00064 | 0,0242212 | aspartate dehydrogenase domain containing |
| ENSG00000230647 | AC022816,1 | -3,964435261 | 0,064059873 | 0,00044 | 0,01864693 | novel transcript |
| ENSG00000188660 | LINC00319 | -3,973716469 | 0,063649083 | 0,01471 | 0,15877226 | long intergenic non-protein coding RNA 319 |
| ENSG00000225807 | AC069281,1 | -3,974416097 | 0,063618224 | 0,00199 | 0,05000974 | novel transcript |
| ENSG00000171564 | FGB | -3,978564691 | 0,063435547 | 0,00558 | 0,09311556 | fibrinogen beta chain |
| ENSG00000275363 | AC100757,3 | -3,981207665 | 0,063319441 | 0,02692 | 0,22156895 | M-phase phosphoprotein 10 (U3 small nucleolar ribonucleoprotein) (MPHOSPH10) pseudogene |
| ENSG00000268746 | AC010519,1 | -3,982929687 | 0,063243907 | 0,08192 | 0,37306719 | novel transcript |
| ENSG00000214772 | AC010168,1 | -3,987499672 | 0,063043888 | 0,00041 | 0,01776434 | novel transcript |
| ENSG00000274848 | Metazoa_SRP | -3,988596455 | 0,062995979 | 0,05948 | 0,32472914 | Metazoan signal recognition particle RNA |
| ENSG00000286272 | AC093843,2 | -3,99569144 | 0,062686933 | 0,00175 | 0,0459021 | novel transcript |
| ENSG00000229649 | LINC02681 | -4,010156388 | 0,062061553 | 3,15E-05 | 0,00225882 | long intergenic non-protein coding RNA 2681 |

| | | | | | | |
|---|---|---|---|---|---|---|
| ENSG00000267484 | AC027319,1 | -4,01904711 | 0,06168027 | 0,14004 | 0,46694389 | novel transcript, antisense to PLIN3 |
| ENSG00000276918 | Metazoa_SRP | -4,029329477 | 0,061242226 | 0,029 | 0,22977124 | Metazoan signal recognition particle RNA |
| ENSG00000263776 | SNORA4 | -4,043942365 | 0,060625041 | 0,00908 | 0,12099501 | small nucleolar RNA, H/ACA box 4 |
| ENSG00000177553 | C1orf167-AS1 | -4,048330873 | 0,060440907 | 0,00862 | 0,11734633 | C1orf167 antisense RNA 1 |
| ENSG00000266955 | AP001269,1 | -4,056283182 | 0,060108666 | 0,00787 | 0,11289244 | novel transcript, antisense IMPA2 |
| ENSG00000260235 | AC105020,3 | -4,057325144 | 0,060065269 | 0,03404 | 0,24909651 | novel transcript, antisense to SNUPN |
| ENSG00000277385 | Metazoa_SRP | -4,061307306 | 0,059899704 | 0,05963 | 0,325177 | Metazoan signal recognition particle RNA |
| ENSG00000274505 | Metazoa_SRP | -4,062444921 | 0,05985249 | 0,00805 | 0,11403014 | Metazoan signal recognition particle RNA |
| ENSG00000175311 | ANKS4B | -4,07162821 | 0,059472717 | 0,07747 | 0,36441467 | ankyrin repeat and sterile alpha motif domain containing 4B |
| ENSG00000279902 | AC073539,11 | -4,08665239 | 0,058856584 | 0,01127 | 0,13773024 | novel transcript |
| ENSG00000278007 | Metazoa_SRP | -4,105061453 | 0,058110334 | 0,01423 | 0,15639135 | Metazoan signal recognition particle RNA |
| ENSG00000264769 | AC145207,8 | -4,108537724 | 0,057970481 | 0,00159 | 0,04261325 | novel transcript, antisense MAFG |
| ENSG00000287765 | AL136295,22 | -4,113536528 | 0,057769967 | 0,04238 | 0,27814338 | novel transcript, antisense to TM9SF1 |
| ENSG00000287341 | AL136295,21 | -4,113536528 | 0,057769967 | 0,04238 | 0,27814338 | novel transcript, antisense to TM9SF1 |
| ENSG00000286144 | Z97056,2 | -4,139589542 | 0,056736087 | 0,00062 | 0,02380082 | novel transcript |
| ENSG00000154764 | WNT7A | -4,151230227 | 0,056280142 | 9,40E-09 | 1,73E-06 | Wnt family member 7A |
| ENSG00000282972 | GNG4 | -4,157290152 | 0,056044237 | 0,0808 | 0,37090142 | G protein subunit gamma 4 |
| ENSG00000185985 | SLITRK2 | -4,160752937 | 0,05590988 | 2,32E-13 | 8,47E-11 | SLIT and NTRK like family member 2 |
| ENSG00000264592 | RN7SL131P | -4,161062373 | 0,05589789 | 0,00183 | 0,04735721 | RNA, 7SL, cytoplasmic 131, pseudogene |
| ENSG00000275642 | Metazoa_SRP | -4,163719046 | 0,05579505 | 0,01467 | 0,15857384 | Metazoan signal recognition particle RNA |
| ENSG00000230885 | MOG | -4,1684918 | 0,055610773 | 0,01644 | 0,16797415 | myelin oligodendrocyte glycoprotein |
| ENSG00000258529 | AP001781,2 | -4,176791681 | 0,055291761 | 0,11783 | 0,43532264 | novel protein |
| ENSG00000282113 | AC068570,1 | -4,202085453 | 0,054330817 | 0,00327 | 0,06882129 | long intergenic non-protein coding RNA 976 |
| ENSG00000273572 | CDHR5 | -4,207703904 | 0,054119642 | 0,00032 | 0,01519301 | cadherin related family member 5 |
| ENSG00000224842 | AL161908,1 | -4,213972266 | 0,053885007 | 4,04E-06 | 0,00037775 | novel transcript, antisense to LIM1B |
| ENSG00000225005 | AC211485,1 | -4,217485924 | 0,053753931 | 0,01362 | 0,15293084 | ribosomal protein L7a (RPL7A) pseudogene |
| ENSG00000232531 | KMT5AP2 | -4,218262152 | 0,053725017 | 0,00236 | 0,05588694 | KMT5A pseudogene 2 |
| ENSG00000258593 | AL583810,1 | -4,224267773 | 0,053501837 | 0,00112 | 0,03490596 | novel transcript, antisense to KIAA0284 |
| ENSG00000181518 | OR8D4 | -4,225005003 | 0,053474504 | 0,00613 | 0,098625 | olfactory receptor family 8 subfamily D member 4 |
| ENSG00000264676 | RN7SL204P | -4,232590382 | 0,053194084 | 0,04095 | 0,27389081 | RNA, 7SL, cytoplasmic 204, pseudogene |
| ENSG00000258402 | AC016526,1 | -4,248016169 | 0,052628345 | 0,00982 | 0,12683981 | novel transcript |
| ENSG00000276493 | Metazoa_SRP | -4,251001491 | 0,052519555 | 0,00258 | 0,05909497 | Metazoan signal recognition particle RNA |

| | | | | | | |
|---|---|---|---|---|---|---|
| ENSG00000221737 | MIR548I1 | -4,283079541 | 0,05136468 | 0,00433 | 0,08098807 | microRNA 548i-1 |
| ENSG00000197408 | CYP2B6 | -4,284973009 | 0,05129731 | 0,0009 | 0,03001915 | cytochrome P450 family 2 subfamily B member 6 |
| ENSG00000263410 | RN7SL460P | -4,2957412 | 0,050915855 | 0,00771 | 0,11160063 | RNA, 7SL, cytoplasmic 460, pseudogene |
| ENSG00000176253 | OR4K13 | -4,332688679 | 0,049628454 | 0,0054 | 0,09146181 | olfactory receptor family 4 subfamily K member 13 |
| ENSG00000198868 | MTND4LP30 | -4,337837354 | 0,049451656 | 0,01499 | 0,16042172 | MT-ND4L pseudogene 30 |
| ENSG00000168925 | CTRB1 | -4,349243882 | 0,049062213 | 0,0054 | 0,09148081 | chymotrypsinogen B1 |
| ENSG00000182583 | VCX | -4,372601778 | 0,048274269 | 0,00073 | 0,02636261 | variable charge X-linked |
| ENSG00000282735 | GARRE1 | -4,373255084 | 0,048252414 | 9,86E-05 | 0,00592508 | granule associated Rac and RHOG effector 1 |
| ENSG00000277628 | Metazoa_SRP | -4,381081538 | 0,047991358 | 0,09712 | 0,40152057 | Metazoan signal recognition particle RNA |
| ENSG00000227163 | AL139220,1 | -4,389490773 | 0,047712438 | 0,00613 | 0,098625 | pseudogene similar to part of ATP synthase, H+ transporting, mitochondrial F0 complex, subunit c (subunit 9), isoform 1 (ATP5G1) |
| ENSG00000234096 | MOG | -4,390180119 | 0,047689646 | 0,01469 | 0,15868223 | myelin oligodendrocyte glycoprotein |
| ENSG00000242668 | RN7SL317P | -4,402550215 | 0,047282489 | 0,0079 | 0,11323874 | RNA, 7SL, cytoplasmic 317, pseudogene |
| ENSG00000209480 | SNORD83B | -4,406208811 | 0,047162735 | 0,00265 | 0,059998 | small nucleolar RNA, C/D box 83B |
| ENSG00000257127 | CLLU1 | -4,408370929 | 0,047092107 | 0,00961 | 0,12527194 | chronic lymphocytic leukemia up-regulated 1 |
| ENSG00000267559 | AL158154,3 | -4,412412039 | 0,046960382 | 0,05355 | 0,31056376 | novel transcript |
| ENSG00000147614 | ATP6V0D2 | -4,433492882 | 0,04627918 | 0,01888 | 0,1821523 | ATPase H+ transporting V0 subunit d2 |
| ENSG00000257563 | AC126755,2 | -4,442510137 | 0,045990824 | 0,09376 | 0,39568176 | microRNA 3180-3 |
| ENSG00000287827 | AC002529,1 | -4,466199462 | 0,045241812 | 0,00039 | 0,01713567 | novel transcript |
| ENSG00000269807 | AC007292,3 | -4,468800713 | 0,045160313 | 0,00449 | 0,08275517 | novel transcript, antisense to MPND |
| ENSG00000276848 | KIR3DX1 | -4,472738194 | 0,045037227 | 0,08979 | 0,38755464 | killer cell immunoglobulin like receptor, three Ig domains X1 (pseudogene) |
| ENSG00000274589 | Metazoa_SRP | -4,514538228 | 0,04375106 | 0,00416 | 0,07917257 | Metazoan signal recognition particle RNA |
| ENSG00000263490 | RN7SL155P | -4,543513935 | 0,04288111 | 0,00801 | 0,11389729 | RNA, 7SL, cytoplasmic 155, pseudogene |
| ENSG00000283464 | AC244452,35 | -4,575140169 | 0,041951315 | 0,03533 | 0,25465323 | novel pseudogene identical to IGHVII-44-2D |
| ENSG00000264519 | RN7SL596P | -4,581812299 | 0,041757747 | 0,00526 | 0,09039746 | RNA, 7SL, cytoplasmic 596, pseudogene |
| ENSG00000238284 | LINC01448 | -4,582732967 | 0,041731108 | 0,00464 | 0,08393639 | long intergenic non-protein coding RNA 1448 |
| ENSG00000278724 | Metazoa_SRP | -4,587380344 | 0,041596895 | 0,00626 | 0,09970758 | Metazoan signal recognition particle RNA |
| ENSG00000260905 | AC009021,1 | -4,592103558 | 0,041460934 | 0,00478 | 0,08525507 | novel transcript |
| ENSG00000168875 | SOX14 | -4,60168266 | 0,041186557 | 0,00111 | 0,03464369 | SRY-box transcription factor 14 |
| ENSG00000272021 | AC008592,4 | -4,601697431 | 0,041186135 | 0,0022 | 0,05335928 | novel transcript |
| ENSG00000173917 | HOXB2 | -4,627941659 | 0,040443687 | 2,35E-11 | 6,95E-09 | homeobox B2 |

| | | | | | | |
|---|---|---|---|---|---|---|
| ENSG00000284069 | AC270227,1 | -4,666286983 | 0,039382896 | 0,02318 | 0,20333236 | |
| ENSG00000284731 | AC083906,5 | -4,698639093 | 0,038509572 | 1,70E-05 | 0,00134398 | RNA binding protein, fox-1 homolog (C, elegans) 1 (RBFOX1) pseudogene |
| ENSG00000248774 | AC097534,1 | -4,706356242 | 0,03830413 | 0,03197 | 0,24119441 | novel transcript |
| ENSG00000248475 | AC016642,1 | -4,732728563 | 0,037610295 | 0,00092 | 0,03058422 | novel transcript |
| ENSG00000206941 | SNORD15A | -4,735152654 | 0,037547153 | 0,00271 | 0,06121178 | small nucleolar RNA, C/D box 15A |
| ENSG00000282446 | AC004675,1 | -4,737385295 | 0,037489092 | 0,17375 | 0,49846579 | novel transcript, antisense to CCL14 |
| ENSG00000188324 | OR6C6 | -4,819012639 | 0,035426859 | 0,00539 | 0,09146181 | olfactory receptor family 6 subfamily C member 6 |
| ENSG00000249667 | LINC01259 | -4,858467655 | 0,034471129 | 0,00805 | 0,11401251 | long intergenic non-protein coding RNA 1259 |
| ENSG00000274017 | Metazoa_SRP | -4,872169248 | 0,034145298 | 0,02104 | 0,19259837 | Metazoan signal recognition particle RNA |
| ENSG00000234793 | AC114730,2 | -4,894549997 | 0,033619685 | 0,00068 | 0,02517055 | novel transcript |
| ENSG00000264125 | AC104984,1 | -4,926311275 | 0,032887627 | 0,02134 | 0,19390172 | novel transcript, antisense to SLC6A4 |
| ENSG00000109851 | DBX1 | -4,946469272 | 0,032431301 | 1,33E-18 | 9,09E-16 | developing brain homeobox 1 |
| ENSG00000230246 | SPATA31C1 | -5,006916867 | 0,031100533 | 0,00052 | 0,0210367 | SPATA31 subfamily C member 1 |
| ENSG00000234623 | MOG | -5,039508408 | 0,030405826 | 0,0002 | 0,01061649 | myelin oligodendrocyte glycoprotein |
| ENSG00000254287 | AC007991,4 | -5,049074654 | 0,030204877 | 0,00903 | 0,12060929 | novel transcript |
| ENSG00000257746 | AC138123,2 | -5,073735825 | 0,029692949 | 0,0128 | 0,14802902 | novel transcript |
| ENSG00000284061 | FCAR | -5,100853265 | 0,029140041 | 0,12113 | 0,44021688 | Fc fragment of IgA receptor |
| ENSG00000188897 | AC099489,1 | -5,119658343 | 0,028762675 | 2,98E-22 | 3,21E-19 | novel lipoprotein amino terminal region containing protein |
| ENSG00000281734 | AL139246,13 | -5,221718609 | 0,026798228 | 0,13622 | 0,463307 | novel transcript |
| ENSG00000266036 | AC016888,1 | -5,28727498 | 0,025607763 | 0,00089 | 0,02988696 | novel transcript, antisense to SLC9A3R1 |
| ENSG00000182854 | OR4F15 | -5,410536068 | 0,023510743 | 0,01557 | 0,16365974 | olfactory receptor family 4 subfamily F member 15 |
| ENSG00000226829 | AC093655,1 | -5,454702265 | 0,022801898 | 0,03744 | 0,26177644 | novel transcript |
| ENSG00000235295 | LINC01634 | -5,465043128 | 0,022639044 | 0,00085 | 0,02896138 | long intergenic non-protein coding RNA 1634 |
| ENSG00000274370 | AC130371,2 | -5,534934125 | 0,021568443 | 0,03377 | 0,24838006 | novel transcript |
| ENSG00000278050 | AP000944,3 | -5,534934846 | 0,021568433 | 0,03381 | 0,24846467 | |
| ENSG00000227380 | HCG15 | -5,617546508 | 0,020368076 | 1,71E-09 | 3,63E-07 | HLA complex group 15 |
| ENSG00000267954 | AP000349,1 | -5,643187922 | 0,020009266 | 0,06117 | 0,32813211 | |
| ENSG00000264092 | RN7SL474P | -5,647490555 | 0,01994968 | 0,02905 | 0,22988377 | RNA, 7SL, cytoplasmic 474, pseudogene |
| ENSG00000179073 | TAAR3P | -5,772327881 | 0,018296 | 0,00342 | 0,07051554 | trace amine associated receptor 3, pseudogene |
| ENSG00000276950 | GSTT4 | -5,815794318 | 0,017752988 | 0,00837 | 0,11551322 | glutathione S-transferase theta 4 |
| ENSG00000265777 | RN7SL624P | -5,818363321 | 0,017721404 | 0,00044 | 0,01872839 | RNA, 7SL, cytoplasmic 624, pseudogene |

| | | | | | | |
|---|---|---|---|---|---|---|
| ENSG00000288604 | AC015871,8 | -5,897917203 | 0,016770654 | 0,00466 | 0,08400694 | SLAIN motif family member 2 (SLAIN2) pseudogene |
| ENSG00000233035 | HCG15 | -5,945177059 | 0,016230181 | 0,01376 | 0,15390159 | HLA complex group 15 |
| ENSG00000278348 | Metazoa_SRP | -5,955770803 | 0,016111439 | 0,0006 | 0,02321526 | Metazoan signal recognition particle RNA |
| ENSG00000277086 | MTMR10 | -6,027507273 | 0,015329907 | 0,00366 | 0,07390189 | myotubularin related protein 10 |
| ENSG00000281429 | FAM20C | -6,176453598 | 0,01382618 | 0,04218 | 0,27783166 | FAM20C golgi associated secretory pathway kinase |
| ENSG00000286085 | AL109653,3 | -6,211751026 | 0,013492009 | 0,00826 | 0,11480072 | novel transcript, antisense to SLITRK2 |
| ENSG00000171540 | OTP | -6,212419468 | 0,013485759 | 2,93E-15 | 1,35E-12 | orthopedia homeobox |
| ENSG00000186431 | FCAR | -6,243812476 | 0,013195479 | 0,00517 | 0,08972695 | Fc fragment of IgA receptor |
| ENSG00000263218 | AC127496,6 | -6,256662282 | 0,013078471 | 0,03802 | 0,26375851 | novel transcript, antisense to CHMP6 |
| ENSG00000267396 | AC090236,1 | -6,256662282 | 0,013078471 | 0,03802 | 0,26375851 | novel transcript, antisense to NEDD4L |
| ENSG00000276101 | AC027601,3 | -6,295154977 | 0,012734137 | 0,03613 | 0,25739586 | novel transcript, antisense to SLC38A10 |
| ENSG00000260727 | SLC7A5P1 | -6,33264676 | 0,012407474 | 0,03437 | 0,25062479 | solute carrier family 7 member 5 pseudogene 1 |
| ENSG00000206834 | SNORA1 | -6,369189421 | 0,012097146 | 0,03271 | 0,24406009 | small nucleolar RNA, H/ACA box 1 |
| ENSG00000283389 | Metazoa_SRP | -6,462012194 | 0,011343328 | 0,00136 | 0,03912393 | Metazoan signal recognition particle RNA |
| ENSG00000263999 | RN7SL42P | -6,489433228 | 0,011129764 | 0,00057 | 0,02219024 | RNA, 7SL, cytoplasmic 42, pseudogene |
| ENSG00000267007 | AC012615,2 | -6,603211177 | 0,010285736 | 0,00158 | 0,04260064 | novel transcript, antisense to REXO1 |
| ENSG00000284927 | AC270301,3 | -6,659327569 | 0,009893334 | 0,00445 | 0,08217507 | |
| ENSG00000214076 | CPSF1P1 | -6,738718952 | 0,009363613 | 0,00208 | 0,05143262 | CPSF1 pseudogene 1 |
| ENSG00000260892 | AC105020,4 | -6,760190519 | 0,009225287 | 0,00017 | 0,00905887 | novel transcript, antisense to CSPG4 |
| ENSG00000231641 | TRIM26 | -6,963159171 | 0,00801457 | 0,05543 | 0,31484345 | tripartite motif containing 26 |
| ENSG00000278144 | AP000944,4 | -6,991812128 | 0,007856965 | 0,00012 | 0,0070607 | |
| ENSG00000054803 | CBLN4 | -6,999220112 | 0,007816724 | 1,74E-06 | 0,00017951 | cerebellin 4 precursor |
| ENSG00000225368 | HCG15 | -7,259453794 | 0,006526595 | 0,02604 | 0,21733018 | HLA complex group 15 |
| ENSG00000281670 | CABIN1 | -7,372509959 | 0,006034668 | 1,98E-07 | 2,68E-05 | calcineurin binding protein 1 |
| ENSG00000257647 | AC124312,2 | -7,384279067 | 0,005985639 | 0,04215 | 0,27783166 | novel transcript |
| ENSG00000276858 | FCAR | -7,485706587 | 0,005579275 | 0,03939 | 0,26865324 | Fc fragment of IgA receptor |
| ENSG00000267125 | AC012615,3 | -8,232844887 | 0,003324044 | 0,02342 | 0,20416051 | novel transcript, antisense to KLF16 |
| ENSG00000262136 | AC092115,3 | -8,388308154 | 0,002984473 | 0,02092 | 0,19202875 | novel transcript |
| ENSG00000236561 | MOG | -8,482608501 | 0,002795635 | 0,01952 | 0,18522413 | myelin oligodendrocyte glycoprotein |
| ENSG00000257411 | AC034102,2 | -10,16719151 | 0,000869702 | 0,00511 | 0,08910452 | novel protein |
| ENSG00000273889 | FAM83H | -11,84455765 | 0,000271915 | 0,00111 | 0,03464369 | family with sequence similarity 83 member H |

**KO-NSC vs IC1-NSC**

| gene_ID | Gene_name | log2FoldChange | Fold_change | P-value | P_adjusted | Description |
|---|---|---|---|---|---|---|
| ENSG00000230907 | FKBPL | 21,98412913 | 4148415,971 | 1,73E-07 | 8,76E-06 | FKBP prolyl isomerase like |
| ENSG00000196954 | CASP4 | 10,3081192 | 1267,806804 | 1,45E-11 | 1,39E-09 | caspase 4 |
| ENSG00000263244 | AC087190,3 | 10,21712739 | 1190,314744 | 0,00765467 | 0,09470346 | novel transcript, 3' overlapping chromosome 16 open |
| ENSG00000111335 | OAS2 | 9,605559856 | 779,0433805 | 1,52E-08 | 9,23E-07 | 2'-5'-oligoadenylate synthetase 2 |
| ENSG00000179300 | RTL3 | 9,495955224 | 722,0501454 | 1,77E-09 | 1,26E-07 | retrotransposon Gag like 3 |
| ENSG00000250658 | AC097652,1 | 9,459406765 | 703,987872 | 1,99E-09 | 1,40E-07 | novel transcript |
| ENSG00000169429 | CXCL8 | 9,309906356 | 634,6891441 | 3,86E-08 | 2,17E-06 | C-X-C motif chemokine ligand 8 |
| ENSG00000285343 | PPP1R3B | 9,273204895 | 618,7466052 | 6,63E-10 | 5,08E-08 | protein phosphatase 1 regulatory subunit 3B |
| ENSG00000158714 | SLAMF8 | 9,068264287 | 536,8087165 | 3,72E-08 | 2,10E-06 | SLAM family member 8 |
| ENSG00000069122 | ADGRF5 | 9,038633142 | 525,8957918 | 1,38E-15 | 2,16E-13 | adhesion G protein-coupled receptor F5 |
| ENSG00000238034 | AL109807,1 | 8,991099177 | 508,8508992 | 8,31E-09 | 5,17E-07 | novel transcript |
| ENSG00000288622 | PDCD6-AHRR | 8,950499144 | 494,7305376 | 4,58E-09 | 3,01E-07 | PDCD6-AHRR readthrough (NMD candidate) |
| ENSG00000123610 | TNFAIP6 | 8,898344438 | 477,1650075 | 2,18E-09 | 1,52E-07 | TNF alpha induced protein 6 |
| ENSG00000243509 | TNFRSF6B | 8,760164054 | 433,5829015 | 2,93E-07 | 1,39E-05 | TNF receptor superfamily member 6b |
| ENSG00000205791 | LOH12CR2 | 8,74178054 | 428,093031 | 0,03771047 | 0,287183177 | loss of heterozygosity on chromosome 12, region 2 |
| ENSG00000108821 | COL1A1 | 8,684918023 | 411,5483214 | 1,81E-14 | 2,43E-12 | collagen type I alpha 1 chain |
| ENSG00000251493 | FOXD1 | 8,628302202 | 395,7106824 | 9,46E-21 | 2,29E-18 | forkhead box D1 |
| ENSG00000244721 | HLA-L | 8,626286419 | 395,158168 | 0,00366463 | 0,054792423 | major histocompatibility complex, class I, L (pseudogene) |
| ENSG00000205809 | KLRC2 | 8,54444058 | 373,364404 | 4,93E-07 | 2,24E-05 | killer cell lectin like receptor C2 |
| ENSG00000277117 | FP565260,3 | 8,527040721 | 368,8884278 | 1,67E-06 | 6,84E-05 | ICOS ligand |
| ENSG00000274538 | PHLDA2 | 8,51009825 | 364,5816778 | 0,04308895 | 0,312995545 | pleckstrin homology like domain family A member 2 |
| ENSG00000196090 | PTPRT | 8,497557026 | 361,426136 | 3,00E-07 | 1,42E-05 | protein tyrosine phosphatase receptor type T |
| ENSG00000163359 | COL6A3 | 8,496141952 | 361,0718032 | 2,96E-34 | 1,81E-31 | collagen type VI alpha 3 chain |
| ENSG00000285876 | AC093912,1 | 8,430854766 | 345,0961973 | 2,88E-08 | 1,67E-06 | novel transcript |
| ENSG00000255641 | AC068775,1 | 8,418241911 | 342,0923212 | 7,85E-08 | 4,27E-06 | novel protein |
| ENSG00000278444 | CDC42EP5 | 8,383087018 | 333,8571246 | 0,00538191 | 0,072847486 | CDC42 effector protein 5 |
| ENSG00000205810 | KLRC3 | 8,380008974 | 333,1455872 | 1,18E-07 | 6,23E-06 | killer cell lectin like receptor C3 |
| ENSG00000276476 | LINC00540 | 8,373081611 | 331,5497625 | 5,01E-08 | 2,80E-06 | long intergenic non-protein coding RNA 540 |
| ENSG00000143631 | FLG | 8,364155606 | 329,504785 | 0,00048552 | 0,010153285 | filaggrin |
| ENSG00000285032 | MTMR9 | 8,349110421 | 326,0863907 | 1,46E-07 | 7,57E-06 | myotubularin related protein 9 |
| ENSG00000128422 | KRT17 | 8,331928673 | 322,2259052 | 0,0007485 | 0,014598319 | keratin 17 |

| | | | | | | |
|---|---|---|---|---|---|---|
| ENSG00000182742 | HOXB4 | 8,315739228 | 318,6302055 | 0,00930705 | 0,108816623 | homeobox B4 |
| ENSG00000278996 | FP671120,4 | 8,260084246 | 306,5724501 | 0,0496074 | 0,33996618 | novel transcript |
| ENSG00000274847 | MAFIP | 8,242196006 | 302,7946712 | 1,80E-07 | 9,04E-06 | MAFF interacting protein (pseudogene) |
| ENSG00000124610 | H1-1 | 8,239591447 | 302,2485163 | 1,44E-31 | 7,01E-29 | H1,1 linker histone, cluster member |
| ENSG00000196611 | MMP1 | 8,13943132 | 281,9765386 | 2,60E-07 | 1,25E-05 | matrix metallopeptidase 1 |
| ENSG00000276033 | KIR3DX1 | 8,04754968 | 264,5780707 | 0,01300955 | 0,138910664 | killer cell immunoglobulin like receptor, three Ig domains X1 (pseudogene) |
| ENSG00000133110 | POSTN | 8,043618839 | 263,858169 | 1,10E-21 | 2,82E-19 | periostin |
| ENSG00000276886 | GREM1 | 8,02924394 | 261,2421616 | 1,59E-37 | 1,24E-34 | gremlin 1, DAN family BMP antagonist |
| ENSG00000104783 | KCNN4 | 8,005972059 | 257,0619124 | 7,68E-06 | 0,000271163 | potassium calcium-activated channel subfamily N member 4 |
| ENSG00000254139 | AC104051,2 | 7,953904719 | 247,9498803 | 3,23E-06 | 0,000125897 | novel transcript |
| ENSG00000286868 | AC074290,1 | 7,923357251 | 242,7550085 | 4,98E-07 | 2,25E-05 | novel transcript |
| ENSG00000288422 | ARHGEF35 | 7,890385199 | 237,2698864 | 9,83E-06 | 0,0003359 | Rho guanine nucleotide exchange factor 35 |
| ENSG00000011465 | DCN | 7,868271665 | 233,6607601 | 4,20E-39 | 3,65E-36 | decorin |
| ENSG00000255819 | KLRC4-KLRK1 | 7,810601922 | 224,5047131 | 9,21E-07 | 3,93E-05 | KLRC4-KLRK1 readthrough |
| ENSG00000188015 | S100A3 | 7,7979707 | 222,5476875 | 1,10E-06 | 4,61E-05 | S100 calcium binding protein A3 |
| ENSG00000170323 | FABP4 | 7,796795764 | 222,3665177 | 2,11E-06 | 8,51E-05 | fatty acid binding protein 4 |
| ENSG00000251643 | AC093763,2 | 7,782223475 | 220,1317564 | 4,21E-07 | 1,93E-05 | COP9 constitutive photomorphogenic homolog subunit 3 (Arabidopsis) (COPS3) pseudogene |
| ENSG00000263947 | RN7SL274P | 7,623407034 | 197,1851415 | 0,01200167 | 0,130693206 | RNA, 7SL, cytoplasmic 274, pseudogene |
| ENSG00000284798 | AC270272,1 | 7,621867505 | 196,9748335 | 0,00011984 | 0,003074697 | |
| ENSG00000135925 | WNT10A | 7,606668074 | 194,9105097 | 4,78E-06 | 0,000177239 | Wnt family member 10A |
| ENSG00000145824 | CXCL14 | 7,602803797 | 194,3891381 | 1,22E-61 | 4,78E-58 | C-X-C motif chemokine ligand 14 |
| ENSG00000113430 | IRX4 | 7,602390085 | 194,3334024 | 7,77E-16 | 1,22E-13 | iroquois homeobox 4 |
| ENSG00000126778 | SIX1 | 7,599842564 | 193,9905498 | 2,05E-34 | 1,28E-31 | SIX homeobox 1 |
| ENSG00000256268 | LINC02454 | 7,558495849 | 188,5098174 | 1,43E-06 | 5,94E-05 | long intergenic non-protein coding RNA 2454 |
| ENSG00000222032 | AC112721,2 | 7,538717765 | 185,9431466 | 3,39E-06 | 0,00013132 | novel protein (LOC728009) |
| ENSG00000283321 | AC019117,3 | 7,536088764 | 185,6046139 | 1,24E-05 | 0,000410214 | novel protein |
| ENSG00000133321 | PLAAT4 | 7,513731964 | 182,7505533 | 1,76E-05 | 0,000564097 | phospholipase A and acyltransferase 4 |
| ENSG00000278910 | BANCR | 7,504064568 | 181,5300487 | 6,94E-06 | 0,00024728 | BRAF-activated non-protein coding RNA |
| ENSG00000288297 | ECSCR | 7,491921901 | 180,0085848 | 1,56E-06 | 6,41E-05 | endothelial cell surface expressed chemotaxis and apoptosis regulator |

| | | | | | | |
|---|---|---|---|---|---|---|
| ENSG00000249751 | ECSCR | 7,491921901 | 180,0085848 | 1,56E-06 | 6,41E-05 | endothelial cell surface expressed chemotaxis and apoptosis regulator |
| ENSG00000256514 | AP003419,1 | 7,487537999 | 179,4624248 | 0,00801673 | 0,097789662 | novel protein |
| ENSG00000174600 | CMKLR1 | 7,472376646 | 177,5863207 | 9,37E-36 | 6,37E-33 | chemerin chemokine-like receptor 1 |
| ENSG00000092607 | TBX15 | 7,47217019 | 177,5609091 | 4,67E-29 | 2,00E-26 | T-box transcription factor 15 |
| ENSG00000230838 | LINC01614 | 7,468557587 | 177,1168411 | 1,33E-23 | 3,92E-21 | long intergenic non-protein coding RNA 1614 |
| ENSG00000043591 | ADRB1 | 7,468518601 | 177,1120549 | 9,46E-06 | 0,000326049 | adrenoceptor beta 1 |
| ENSG00000162761 | LMX1A | 7,461383479 | 176,2382762 | 1,90E-06 | 7,72E-05 | LIM homeobox transcription factor 1 alpha |
| ENSG00000104415 | CCN4 | 7,436828044 | 173,2639911 | 4,72E-19 | 1,00E-16 | cellular communication network factor 4 |
| ENSG00000227191 | TRGC2 | 7,430350424 | 172,4877892 | 8,64E-06 | 0,00030006 | T cell receptor gamma constant 2 |
| ENSG00000224642 | HCG15 | 7,428343006 | 172,2479504 | 0,07751934 | 0,440002866 | HLA complex group 15 |
| ENSG00000108511 | HOXB6 | 7,413162315 | 170,4449819 | 9,61E-05 | 0,002526223 | homeobox B6 |
| ENSG00000198542 | ITGBL1 | 7,340143133 | 162,0329188 | 4,39E-05 | 0,001260194 | integrin subunit beta like 1 |
| ENSG00000041982 | TNC | 7,328982338 | 160,7842564 | 5,07E-40 | 4,66E-37 | tenascin C |
| ENSG00000225813 | AC009299,1 | 7,305339411 | 158,1707927 | 2,39E-06 | 9,55E-05 | novel pseudogene |
| ENSG00000164035 | EMCN | 7,277947544 | 155,1959979 | 9,57E-06 | 0,000328779 | endomucin |
| ENSG00000163081 | CCDC140 | 7,258190195 | 153,0851143 | 3,16E-06 | 0,000123204 | CCDC140 long non-coding RNA |
| ENSG00000120149 | MSX2 | 7,248476157 | 152,0578152 | 6,56E-36 | 4,66E-33 | msh homeobox 2 |
| ENSG00000137673 | MMP7 | 7,206638926 | 147,7115599 | 2,66E-05 | 0,000808498 | matrix metallopeptidase 7 |
| ENSG00000259327 | AC023906,3 | 7,187136991 | 145,7282727 | 0,08767448 | 0,471734492 | novel transcript |
| ENSG00000120708 | TGFBI | 7,173273343 | 144,3345968 | 6,18E-41 | 6,44E-38 | transforming growth factor beta induced |
| ENSG00000118526 | TCF21 | 7,113413146 | 138,4684174 | 0,00033403 | 0,007427499 | transcription factor 21 |
| ENSG00000282046 | GREM1 | 7,068597881 | 134,2332142 | 1,85E-48 | 3,21E-45 | gremlin 1, DAN family BMP antagonist |
| ENSG00000183542 | KLRC4 | 7,065443209 | 133,9400135 | 4,00E-05 | 0,001155422 | killer cell lectin like receptor C4 |
| ENSG00000084636 | COL16A1 | 7,0353389 | 131,1740835 | 4,12E-22 | 1,09E-19 | collagen type XVI alpha 1 chain |
| ENSG00000278415 | FCAR | 7,034359624 | 131,0850751 | 0,09464559 | 0,488606312 | Fc fragment of IgA receptor |
| ENSG00000120075 | HOXB5 | 7,007022506 | 128,6245755 | 7,47E-08 | 4,09E-06 | homeobox B5 |
| ENSG00000265484 | AC091691,2 | 7,00166794 | 128,14807 | 1,00E-05 | 0,000342318 | novel transcript |
| ENSG00000278774 | U2 | 6,990603691 | 127,1690417 | 0,09672221 | 0,494090395 | U2 spliceosomal RNA |
| ENSG00000268854 | AC020909,3 | 6,989777845 | 127,0962669 | 0,09676176 | 0,494090395 | novel transcript, antisense to EMC10 |
| ENSG00000122691 | TWIST1 | 6,986845763 | 126,8382233 | 1,03E-16 | 1,77E-14 | twist family bHLH transcription factor 1 |
| ENSG00000168497 | CAVIN2 | 6,98221094 | 126,4313947 | 7,42E-14 | 9,40E-12 | caveolae associated protein 2 |
| ENSG00000248215 | LINC02505 | 6,964219127 | 124,8644629 | 4,47E-06 | 0,000166959 | long intergenic non-protein coding RNA 2505 |

| | | | | | | |
|---|---|---|---|---|---|---|
| ENSG00000251588 | AC125336,1 | 6,904220689 | 119,7781288 | 2,96E-05 | 0,000888988 | F-box protein 38 (FBXO38) pseudogene |
| ENSG00000227214 | HCG15 | 6,85618809 | 115,8559309 | 0,10333091 | 0,512638569 | HLA complex group 15 |
| ENSG00000179772 | FOXS1 | 6,849852494 | 115,348265 | 5,53E-13 | 6,37E-11 | forkhead box S1 |
| ENSG00000070047 | PHRF1 | 6,82020938 | 113,0023825 | 2,24E-07 | 1,10E-05 | PHD and ring finger domains 1 |
| ENSG00000117228 | GBP1 | 6,819679403 | 112,9608785 | 3,52E-53 | 8,46E-50 | guanylate binding protein 1 |
| ENSG00000286099 | AL390800,1 | 6,804567422 | 111,7838084 | 1,73E-09 | 1,24E-07 | novel transcript |
| ENSG00000182492 | BGN | 6,789608362 | 110,6307294 | 1,71E-70 | 1,07E-66 | biglycan |
| ENSG00000204261 | PSMB8-AS1 | 6,770532703 | 109,1775716 | 0,00047576 | 0,00996922 | PSMB8 antisense RNA 1 (head to head) |
| ENSG00000244563 | RPS26P19 | 6,765272186 | 108,7802009 | 0,00143831 | 0,025390852 | ribosomal protein S26 pseudogene 19 |
| ENSG00000024422 | EHD2 | 6,755099488 | 108,0158704 | 3,32E-29 | 1,44E-26 | EH domain containing 2 |
| ENSG00000178726 | THBD | 6,73840374 | 106,77305 | 5,17E-19 | 1,09E-16 | thrombomodulin |
| ENSG00000251254 | GTF2F2P1 | 6,734222446 | 106,4640428 | 7,53E-05 | 0,00204364 | general transcription factor IIF subunit 2 pseudogene 1 |
| ENSG00000158055 | GRHL3 | 6,732926038 | 106,368417 | 1,73E-25 | 5,89E-23 | grainyhead like transcription factor 3 |
| ENSG00000167767 | KRT80 | 6,732720634 | 106,3532739 | 6,39E-05 | 0,001765819 | keratin 80 |
| ENSG00000233911 | POU5F1 | 6,727571184 | 105,974341 | 0,00078266 | 0,015169893 | POU class 5 homeobox 1 |
| ENSG00000229094 | POU5F1 | 6,727571184 | 105,974341 | 0,00078266 | 0,015169893 | POU class 5 homeobox 1 |
| ENSG00000160180 | TFF3 | 6,726212268 | 105,8745677 | 3,47E-05 | 0,00102767 | trefoil factor 3 |
| ENSG00000112837 | TBX18 | 6,722583786 | 105,60862 | 9,19E-32 | 4,71E-29 | T-box transcription factor 18 |
| ENSG00000260549 | MT1L | 6,721190488 | 105,5066766 | 7,09E-05 | 0,001939353 | metallothionein 1L, pseudogene |
| ENSG00000258952 | SALRNA1 | 6,713822172 | 104,9691932 | 2,72E-05 | 0,000823721 | senescence associated long non-coding RNA 1 |
| ENSG00000222022 | AC112721,1 | 6,696197005 | 103,6946035 | 0,00012245 | 0,003127976 | novel transcript |
| ENSG00000163453 | IGFBP7 | 6,681297269 | 102,6291869 | 3,58E-83 | 3,73E-79 | insulin like growth factor binding protein 7 |
| ENSG00000140093 | SERPINA10 | 6,681192488 | 102,6217333 | 0,00108792 | 0,020102107 | serpin family A member 10 |
| ENSG00000187688 | TRPV2 | 6,664762734 | 101,459682 | 4,16E-26 | 1,46E-23 | transient receptor potential cation channel subfamily V member 2 |
| ENSG00000259807 | AC009093,1 | 6,640081459 | 99,73869756 | 3,21E-19 | 6,96E-17 | novel transcript |
| ENSG00000261683 | LINC00838 | 6,617562169 | 98,19394557 | 0,0001796 | 0,00433922 | long intergenic non-protein coding RNA 838 |
| ENSG00000288550 | AC018809,3 | 6,616176581 | 98,0996838 | 0,00074435 | 0,014535566 | novel protein |
| ENSG00000172061 | LRRC15 | 6,614954669 | 98,016632 | 5,05E-10 | 4,02E-08 | leucine rich repeat containing 15 |
| ENSG00000256861 | AC048338,1 | 6,598731507 | 96,9206051 | 0,11699121 | 0,546483353 | novel protein |
| ENSG00000259721 | AC090877,2 | 6,573692289 | 95,25297778 | 0,00023993 | 0,005560475 | novel transcript |
| ENSG00000282555 | AC090877,4 | 6,573692289 | 95,25297778 | 0,00023993 | 0,005560475 | novel transcript |

| | | | | | | |
|---|---|---|---|---|---|---|
| ENSG00000270617 | URGCP-MRPS2 | 6,571921588 | 95,13614011 | 0,00011605 | 0,002986058 | URGCP-MRPS24 readthrough |
| ENSG00000282143 | AL137798,2 | 6,569253978 | 94,96039149 | 0,11864212 | 0,550493797 | novel transcript |
| ENSG00000224712 | NPIPA3 | 6,562177998 | 94,49578001 | 0,00013538 | 0,003405111 | nuclear pore complex interacting protein family member A3 |
| ENSG00000276076 | CRYAA2 | 6,5387095 | 92,9710407 | 0,00041754 | 0,008953288 | crystallin alpha A2 |
| ENSG00000111799 | COL12A1 | 6,528306432 | 92,30305107 | 5,60E-108 | 1,75E-103 | collagen type XII alpha 1 chain |
| ENSG00000139329 | LUM | 6,519450825 | 91,73820823 | 1,85E-30 | 8,78E-28 | lumican |
| ENSG00000102359 | SRPX2 | 6,491036312 | 89,94906078 | 1,49E-60 | 5,19E-57 | sushi repeat containing protein X-linked 2 |
| ENSG00000232326 | TAP2 | 6,481617807 | 89,36374903 | 6,13E-05 | 0,001699043 | transporter 2, ATP binding cassette subfamily B member |
| ENSG00000271334 | LINC02104 | 6,478540045 | 89,17330882 | 8,74E-05 | 0,002330615 | long intergenic non-protein coding RNA 2104 |
| ENSG00000124564 | SLC17A3 | 6,465439891 | 88,36725048 | 0,00020628 | 0,004898559 | solute carrier family 17 member 3 |
| ENSG00000138316 | ADAMTS14 | 6,461761568 | 88,14223461 | 1,56E-19 | 3,40E-17 | ADAM metallopeptidase with thrombospondin type 1 motif 14 |
| ENSG00000228435 | C6orf47 | 6,456603697 | 87,82767433 | 0,12512071 | 0,565448648 | chromosome 6 open reading frame 47 |
| ENSG00000168542 | COL3A1 | 6,43401683 | 86,46335003 | 1,46E-29 | 6,44E-27 | collagen type III alpha 1 chain |
| ENSG00000260142 | AC068135,2 | 6,41179621 | 85,14183092 | 0,00017298 | 0,00418901 | novel transcript |
| ENSG00000267477 | AC008481,3 | 6,397269824 | 84,28884589 | 0,12864268 | 0,573081343 | novel protein |
| ENSG00000164283 | ESM1 | 6,387079861 | 83,69559918 | 2,64E-08 | 1,54E-06 | endothelial cell specific molecule 1 |
| ENSG00000243089 | AC107029,1 | 6,38622061 | 83,64576597 | 1,86E-08 | 1,12E-06 | novel transcript |
| ENSG00000277726 | AL109811,3 | 6,353725849 | 81,78281683 | 0,00294733 | 0,045752397 | novel protein |
| ENSG00000253414 | AC124067,2 | 6,344436097 | 81,25789556 | 0,00034437 | 0,007630317 | novel transcript |
| ENSG00000248461 | LINC02119 | 6,343802495 | 81,22221662 | 0,00217367 | 0,035937417 | long intergenic non-protein coding RNA 2119 |
| ENSG00000234421 | SLC25A5P4 | 6,333272633 | 80,63155461 | 0,00016709 | 0,004065362 | solute carrier family 25 member 5 pseudogene 4 |
| ENSG00000164438 | TLX3 | 6,330422543 | 80,47242166 | 0,00021736 | 0,005121088 | T cell leukemia homeobox 3 |
| ENSG00000204866 | IGFL2 | 6,321388604 | 79,97008987 | 0,00020651 | 0,004898559 | IGF like family member 2 |
| ENSG00000197467 | COL13A1 | 6,31306729 | 79,51015848 | 2,48E-21 | 6,21E-19 | collagen type XIII alpha 1 chain |
| ENSG00000144837 | PLA1A | 6,302989884 | 78,95670521 | 0,00012136 | 0,003104784 | phospholipase A1 member A |
| ENSG00000223485 | LINC01615 | 6,300974402 | 78,8464777 | 0,00032284 | 0,007209567 | long intergenic non-protein coding RNA 1615 |
| ENSG00000111262 | KCNA1 | 6,300441983 | 78,81738517 | 5,62E-22 | 1,46E-19 | potassium voltage-gated channel subfamily A member 1 |
| ENSG00000187094 | CCK | 6,299514647 | 78,76673918 | 0,00060244 | 0,012230969 | cholecystokinin |
| ENSG00000164920 | OSR2 | 6,288737885 | 78,18055303 | 1,79E-07 | 9,04E-06 | odd-skipped related transciption factor 2 |

| | | | | | | |
|---|---|---|---|---|---|---|
| ENSG00000168530 | MYL1 | 6,28808716 | 78,14529782 | 0,00014476 | 0,003606237 | myosin light chain 1 |
| ENSG00000188624 | IGFL3 | 6,286381511 | 78,05296389 | 0,00042314 | 0,00903617 | IGF like family member 3 |
| ENSG00000166923 | GREM1 | 6,281255582 | 77,77613197 | 3,44E-50 | 7,69E-47 | gremlin 1, DAN family BMP antagonist |
| ENSG00000140853 | NLRC5 | 6,279108533 | 77,66046998 | 7,17E-18 | 1,38E-15 | NLR family CARD domain containing 5 |
| ENSG00000222047 | C10orf55 | 6,272048193 | 77,28133937 | 5,78E-05 | 0,001612255 | chromosome 10 open reading frame 55 (putative) |
| ENSG00000115648 | MLPH | 6,26978453 | 77,16017607 | 4,58E-26 | 1,59E-23 | melanophilin |
| ENSG00000215523 | AL773551,1 | 6,260272481 | 76,65311314 | 0,13707009 | 0,590839562 | chromosome 6 open reading frame 214 (putative) |
| ENSG00000122641 | INHBA | 6,24598079 | 75,89751738 | 1,40E-67 | 7,28E-64 | inhibin subunit beta A |
| ENSG00000086967 | MYBPC2 | 6,232939027 | 75,21450612 | 9,87E-05 | 0,002587644 | myosin binding protein C2 |
| ENSG00000257219 | LNCOG | 6,204722984 | 73,75776238 | 0,00110619 | 0,020319143 | lncRNA osteogenesis associated |
| ENSG00000269590 | AC010422,5 | 6,197714458 | 73,40032054 | 0,14105793 | 0,598214217 | novel protein |
| ENSG00000135114 | OASL | 6,194430086 | 73,23341086 | 0,00049821 | 0,01039787 | 2'-5'-oligoadenylate synthetase like |
| ENSG00000259847 | LINC02126 | 6,188635703 | 72,93986896 | 0,0004271 | 0,009114566 | long intergenic non-protein coding RNA 2126 |
| ENSG00000204397 | CARD16 | 6,167195514 | 71,86390937 | 0,00013345 | 0,003367396 | caspase recruitment domain family member 16 |
| ENSG00000271945 | AL355612,1 | 6,139691226 | 70,50683552 | 0,00029399 | 0,006641104 | novel transcript |
| ENSG00000109132 | PHOX2B | 6,131132468 | 70,08979346 | 0,03032537 | 0,249039213 | paired like homeobox 2B |
| ENSG00000198798 | MAGEB3 | 6,127762531 | 69,92626444 | 0,00080333 | 0,015484207 | MAGE family member B3 |
| ENSG00000157613 | CREB3L1 | 6,123561936 | 69,72296109 | 4,07E-23 | 1,16E-20 | cAMP responsive element binding protein 3 like 1 |
| ENSG00000248751 | AC004997,1 | 6,103406835 | 68,75567233 | 0,00332786 | 0,050727562 | novel protein |
| ENSG00000275018 | AL121581,1 | 6,088823584 | 68,0641675 | 0,14821236 | 0,612762642 | novel transcript |
| ENSG00000196136 | SERPINA3 | 6,085855411 | 67,92427758 | 2,91E-07 | 1,39E-05 | serpin family A member 3 |
| ENSG00000175899 | A2M | 6,085826798 | 67,92293043 | 3,51E-63 | 1,57E-59 | alpha-2-macroglobulin |
| ENSG00000232641 | MOG | 6,079150656 | 67,60933989 | 0,14886113 | 0,613515281 | myelin oligodendrocyte glycoprotein |
| ENSG00000234362 | LINC01914 | 6,077106426 | 67,51360853 | 0,00032956 | 0,007338482 | long intergenic non-protein coding RNA 1914 |
| ENSG00000137709 | POU2F3 | 6,070477968 | 67,20412896 | 0,00129708 | 0,023279015 | POU class 2 homeobox 3 |
| ENSG00000276849 | TRBC2 | 6,062681395 | 66,84192582 | 5,56E-08 | 3,09E-06 | T cell receptor beta constant 2 |
| ENSG00000188662 | H1-9P | 6,059476651 | 66,69361068 | 0,00155179 | 0,027018437 | H1,9 linker histone, pseudogene |
| ENSG00000221420 | SNORA81 | 6,050597064 | 66,28438144 | 0,15078898 | 0,617373838 | small nucleolar RNA, H/ACA box 81 |
| ENSG00000256256 | AC019209,1 | 6,038331177 | 65,72321626 | 0,0010848 | 0,020056204 | novel transcript |
| ENSG00000197849 | OR8G1 | 6,031954915 | 65,43338109 | 0,00031219 | 0,007001545 | olfactory receptor family 8 subfamily G member 1 |
| ENSG00000285330 | AC126283,2 | 6,020533643 | 64,91741476 | 0,00623066 | 0,081266379 | novel protein |
| ENSG00000137752 | CASP1 | 6,017651834 | 64,7878705 | 0,00109847 | 0,020225367 | caspase 1 |
| ENSG00000008196 | TFAP2B | 6,014218036 | 64,63385047 | 2,51E-16 | 4,15E-14 | transcription factor AP-2 beta |

| | | | | | | |
|---|---|---|---|---|---|---|
| ENSG00000106991 | ENG | 6,013210376 | 64,58872231 | 1,26E-44 | 1,58E-41 | endoglin |
| ENSG00000258864 | AC008575,1 | 5,999078787 | 63,95914673 | 0,15431565 | 0,623607455 | novel protein |
| ENSG00000215912 | TTC34 | 5,997570086 | 63,89229629 | 2,61E-07 | 1,25E-05 | tetratricopeptide repeat domain 34 |
| ENSG00000225883 | BMS1P11 | 5,977256694 | 62,99898562 | 0,00418585 | 0,060783228 | BMS1 pseudogene 11 |
| ENSG00000183631 | PRR32 | 5,970452764 | 62,70257406 | 0,00213536 | 0,035454081 | proline rich 32 |
| ENSG00000084674 | APOB | 5,966866301 | 62,54689236 | 1,84E-53 | 4,80E-50 | apolipoprotein B |
| ENSG00000254779 | EGLN1P1 | 5,964058577 | 62,42528412 | 0,00489247 | 0,067740586 | egl-9 family hypoxia inducible factor 1 pseudogene 1 |
| ENSG00000266524 | GDF10 | 5,961249165 | 62,30383939 | 1,02E-29 | 4,63E-27 | growth differentiation factor 10 |
| ENSG00000163017 | ACTG2 | 5,944671188 | 61,59200499 | 1,07E-40 | 1,08E-37 | actin gamma 2, smooth muscle |
| ENSG00000124253 | PCK1 | 5,941635984 | 61,46256132 | 7,33E-15 | 1,03E-12 | phosphoenolpyruvate carboxykinase 1 |
| ENSG00000229637 | PRAC2 | 5,92293593 | 60,67103131 | 0,00094641 | 0,017835188 | PRAC2 small nuclear protein |
| ENSG00000122861 | PLAU | 5,921099871 | 60,59386687 | 9,77E-75 | 7,63E-71 | plasminogen activator, urokinase |
| ENSG00000277297 | ATP5F1AP10 | 5,9114071 | 60,18813044 | 0,00061093 | 0,012362556 | ATP synthase F1 subunit alpha pseudogene 10 |
| ENSG00000260534 | AC100827,3 | 5,898681344 | 59,65955634 | 0,16136896 | 0,637805201 | novel transcript |
| ENSG00000181649 | PHLDA2 | 5,896783309 | 59,58111879 | 1,32E-32 | 7,53E-30 | pleckstrin homology like domain family A member 2 |
| ENSG00000250384 | UBE2CP3 | 5,892280286 | 59,39544069 | 0,00117157 | 0,021307699 | ubiquitin conjugating enzyme E2 C pseudogene 3 |
| ENSG00000124491 | F13A1 | 5,885567358 | 59,1197129 | 1,71E-33 | 1,03E-30 | coagulation factor XIII A chain |
| ENSG00000145681 | HAPLN1 | 5,884980764 | 59,09567994 | 1,56E-28 | 6,33E-26 | hyaluronan and proteoglycan link protein 1 |
| ENSG00000286174 | AL596330,1 | 5,881151352 | 58,93902752 | 0,00081899 | 0,015747241 | novel transcript, antisense to CNIH3 |
| ENSG00000170498 | KISS1 | 5,877648809 | 58,79611023 | 0,00268343 | 0,042285674 | KiSS-1 metastasis suppressor |
| ENSG00000284811 | AC270285,1 | 5,864795548 | 58,27461037 | 0,00191799 | 0,032509055 | |
| ENSG00000261730 | AL034346,1 | 5,860282466 | 58,0925988 | 0,00210147 | 0,035002798 | novel transcript, antisense to FOXF2 |
| ENSG00000227510 | LINC01442 | 5,839509485 | 57,26213217 | 0,00132792 | 0,023750631 | long intergenic non-protein coding RNA 1442 |
| ENSG00000204644 | ZFP57 | 5,839139745 | 57,24745867 | 0,00468927 | 0,06577186 | ZFP57 zinc finger protein |
| ENSG00000260727 | SLC7A5P1 | 5,833133537 | 57,00962224 | 0,1661044 | 0,645424816 | solute carrier family 7 member 5 pseudogene 1 |
| ENSG00000078098 | FAP | 5,829596396 | 56,87001959 | 1,74E-14 | 2,34E-12 | fibroblast activation protein alpha |
| ENSG00000180644 | PRF1 | 5,822854474 | 56,6048778 | 0,00064944 | 0,012957299 | perforin 1 |
| ENSG00000142173 | COL6A2 | 5,817529064 | 56,39631775 | 3,07E-15 | 4,49E-13 | collagen type VI alpha 2 chain |
| ENSG00000159339 | PADI4 | 5,787479983 | 55,23381924 | 0,00555332 | 0,074739092 | peptidyl arginine deiminase 4 |
| ENSG00000170373 | CST1 | 5,780961341 | 54,98481489 | 0,00563003 | 0,075317553 | cystatin SN |
| ENSG00000113600 | C9 | 5,778825779 | 54,90348337 | 0,00110678 | 0,020319143 | complement C9 |
| ENSG00000254509 | AP000866,3 | 5,764465178 | 54,35968452 | 0,1711773 | 0,654843372 | endothelial cell adhesion molecule (ESAM) pseudogene |

| | | | | | | |
|---|---|---|---|---|---|---|
| ENSG00000185640 | KRT79 | 5,763105929 | 54,30849314 | 0,0034061 | 0,051668318 | keratin 79 |
| ENSG00000137273 | FOXF2 | 5,755589966 | 54,02629959 | 1,98E-20 | 4,77E-18 | forkhead box F2 |
| ENSG00000120093 | HOXB3 | 5,75411263 | 53,97100435 | 3,13E-10 | 2,54E-08 | homeobox B3 |
| ENSG00000164308 | ERAP2 | 5,748449225 | 53,75955256 | 1,63E-47 | 2,68E-44 | endoplasmic reticulum aminopeptidase 2 |
| ENSG00000100985 | MMP9 | 5,746602349 | 53,69077591 | 3,24E-20 | 7,50E-18 | matrix metallopeptidase 9 |
| ENSG00000267337 | LINC01478 | 5,721745434 | 52,77363428 | 0,00170646 | 0,029329691 | long intergenic non-protein coding RNA 1478 |
| ENSG00000163823 | CCR1 | 5,71425624 | 52,50039017 | 0,00044731 | 0,00950052 | C-C motif chemokine receptor 1 |
| ENSG00000008441 | NFIX | 5,709134863 | 52,31435112 | 2,40E-12 | 2,53E-10 | nuclear factor I X |
| ENSG00000268812 | AC004264,1 | 5,706491203 | 52,21857573 | 0,00247279 | 0,039727303 | novel transcript, antisense to LIF |
| ENSG00000224721 | AC007182,1 | 5,696236823 | 51,84873277 | 0,00712186 | 0,089636796 | novel transcript, antisense to FLVCR2 |
| ENSG00000167244 | IGF2 | 5,684967092 | 51,44528993 | 1,47E-40 | 1,39E-37 | insulin like growth factor 2 |
| ENSG00000262160 | AC020978,5 | 5,678899289 | 51,22937157 | 0,06852415 | 0,412543637 | novel transcript, antisense to NFATC3 |
| ENSG00000177409 | SAMD9L | 5,654035184 | 50,35402452 | 1,04E-27 | 3,92E-25 | sterile alpha motif domain containing 9 like |
| ENSG00000213654 | GPSM3 | 5,63515238 | 49,69925705 | 1,64E-05 | 0,000528182 | G protein signaling modulator 3 |
| ENSG00000272989 | LINC02012 | 5,624497977 | 49,33357652 | 0,03571865 | 0,27700703 | long intergenic non-protein coding RNA 2012 |
| ENSG00000236544 | AC008060,3 | 5,620195747 | 49,18667907 | 0,00343999 | 0,052056145 | novel transcript |
| ENSG00000159184 | HOXB13 | 5,612897061 | 48,9384689 | 0,00288778 | 0,04496196 | homeobox B13 |
| ENSG00000249669 | CARMN | 5,606860956 | 48,73414256 | 9,75E-56 | 2,77E-52 | cardiac mesoderm enhancer-associated non-coding RNA |
| ENSG00000125618 | PAX8 | 5,604771142 | 48,66359987 | 7,43E-35 | 4,84E-32 | paired box 8 |
| ENSG00000273471 | AC112229,4 | 5,583078215 | 47,93734873 | 0,04021925 | 0,299542756 | novel transcript |
| ENSG00000270249 | AC093668,1 | 5,577809576 | 47,76260358 | 0,01069819 | 0,120442319 | novel protein |
| ENSG00000267650 | AC008742,1 | 5,574722035 | 47,66049514 | 0,00308782 | 0,047710255 | novel transcript, antisense to COL5A3 |
| ENSG00000267387 | AC020931,1 | 5,564942333 | 47,33850806 | 0,00434536 | 0,062431658 | novel transcript, antisense to C19orf66 |
| ENSG00000248663 | LINC00992 | 5,553453182 | 46,96301679 | 0,00360517 | 0,054084458 | long intergenic non-protein coding RNA 992 |
| ENSG00000244081 | AL591379,2 | 5,550883526 | 46,8794431 | 0,00256134 | 0,040855994 | pseudogene similar to part of tripartite motif-containing 43 (TRIM43) |
| ENSG00000204264 | PSMB8 | 5,539942729 | 46,52527359 | 7,78E-06 | 0,000273554 | proteasome 20S subunit beta 8 |
| ENSG00000259375 | AC087286,3 | 5,537484781 | 46,44607506 | 0,07916571 | 0,446113303 | novel transcript, antisense to AKAP13 |
| ENSG00000185860 | CCDC190 | 5,525494535 | 46,06166111 | 0,00211556 | 0,03518124 | coiled-coil domain containing 190 |
| ENSG00000287352 | AC011029,1 | 5,509117442 | 45,54173791 | 0,00066726 | 0,013253727 | novel transcript |
| ENSG00000233656 | AL445430,1 | 5,503562383 | 45,36671782 | 0,02076369 | 0,191774313 | novel transcript |
| ENSG00000233101 | HOXB-AS3 | 5,495144381 | 45,10277771 | 0,00910957 | 0,106907512 | HOXB cluster antisense RNA 3 |

| | | | | | | |
|---|---|---|---|---|---|---|
| ENSG00000278122 | AQP7P5 | 5,494576075 | 45,08501434 | 0,03697037 | 0,283711768 | aquaporin 7 pseudogene 5 |
| ENSG00000151617 | EDNRA | 5,490258826 | 44,95029957 | 2,45E-46 | 3,19E-43 | endothelin receptor type A |
| ENSG00000240821 | RPL9P25 | 5,486274055 | 44,82631669 | 0,00378447 | 0,056261305 | ribosomal protein L9 pseudogene 25 |
| ENSG00000130711 | PRDM12 | 5,480846735 | 44,65799996 | 2,69E-16 | 4,43E-14 | PR/SET domain 12 |
| ENSG00000137745 | MMP13 | 5,477227475 | 44,54610773 | 0,00069775 | 0,01375439 | matrix metallopeptidase 13 |
| ENSG00000184344 | GDF3 | 5,475578103 | 44,49520917 | 0,00467345 | 0,065714391 | growth differentiation factor 3 |
| ENSG00000160185 | UBASH3A | 5,463968775 | 44,13859417 | 0,00195577 | 0,033033633 | ubiquitin associated and SH3 domain containing A |
| ENSG00000249239 | AC095059,2 | 5,459078915 | 43,98924439 | 0,00523711 | 0,071374436 | Guanine nucleotide-binding protein subunit beta-2-like 1 (GNB2L1) pseudogene |
| ENSG00000166825 | ANPEP | 5,457649354 | 43,94567721 | 1,31E-17 | 2,43E-15 | alanyl aminopeptidase, membrane |
| ENSG00000212643 | ZRSR2P1 | 5,456411576 | 43,90798963 | 0,0438642 | 0,31589834 | ZRSR2 pseudogene 1 |
| ENSG00000206485 | PPP1R18 | 5,455301401 | 43,8742148 | 0,01954032 | 0,183897798 | protein phosphatase 1 regulatory subunit 18 |
| ENSG00000186086 | NBPF6 | 5,454066533 | 43,83667693 | 0,01042176 | 0,118366144 | NBPF member 6 |
| ENSG00000275882 | IKBKGP1 | 5,452684131 | 43,79469241 | 0,0860997 | 0,467330023 | inhibitor of nuclear factor kappa B kinase subunit gamma pseudogene 1 |
| ENSG00000230141 | HLA-DOA | 5,452684131 | 43,79469241 | 0,0860997 | 0,467330023 | major histocompatibility complex, class II, DO alpha |
| ENSG00000235159 | AL121672,2 | 5,434108627 | 43,23442591 | 0,00420655 | 0,060998878 | novel transcript |
| ENSG00000177469 | CAVIN1 | 5,426694158 | 43,01280041 | 5,17E-14 | 6,68E-12 | caveolae associated protein 1 |
| ENSG00000248844 | LINC02753 | 5,418618074 | 42,7726911 | 0,04330795 | 0,314002702 | long intergenic non-protein coding RNA 2753 |
| ENSG00000111432 | FZD10 | 5,418292087 | 42,76302742 | 8,54E-37 | 6,36E-34 | frizzled class receptor 10 |
| ENSG00000236562 | IFITM3P1 | 5,411079764 | 42,54977989 | 0,00421175 | 0,061045942 | IFITM3 pseudogene 1 |
| ENSG00000273637 | AL139260,2 | 5,408337166 | 42,46896859 | 0,08989416 | 0,478293227 | novel transcript |
| ENSG00000286435 | AC025048,7 | 5,407126816 | 42,43335416 | 0,04105682 | 0,303420267 | novel transcript |
| ENSG00000267013 | LINC01929 | 5,405959941 | 42,39904725 | 0,00348756 | 0,052674011 | long intergenic non-protein coding RNA 1929 |
| ENSG00000171658 | NMRAL2P | 5,40311552 | 42,31553553 | 0,0118267 | 0,129464313 | NmrA like redox sensor 2, pseudogene |
| ENSG00000173253 | DMRT2 | 5,395625124 | 42,09640488 | 8,60E-06 | 0,000299033 | doublesex and mab-3 related transcription factor 2 |
| ENSG00000274185 | LILRA4 | 5,378141282 | 41,58932262 | 0,0143419 | 0,14921306 | leukocyte immunoglobulin like receptor A4 |
| ENSG00000259042 | AC244502,3 | 5,377315775 | 41,56553209 | 0,01455675 | 0,150888551 | novel transcript |
| ENSG00000108823 | SGCA | 5,375036437 | 41,49991386 | 0,0046797 | 0,065714391 | sarcoglycan alpha |
| ENSG00000233890 | TCF19 | 5,374042888 | 41,47134371 | 0,02178454 | 0,198207677 | transcription factor 19 |
| ENSG00000231010 | AL121672,1 | 5,373477134 | 41,45508389 | 0,00619971 | 0,081031602 | novel transcript |
| ENSG00000255191 | AP003555,2 | 5,370785039 | 41,37780008 | 0,04793071 | 0,333223418 | novel transcript |
| ENSG00000276278 | AC048382,6 | 5,370417143 | 41,36724984 | 0,04254922 | 0,310155915 | novel transcript, antisense to ZSCAN2 |

| | | | | | | |
|---|---|---|---|---|---|---|
| ENSG00000120337 | TNFSF18 | 5,368839219 | 41,32202983 | 0,00264915 | 0,041893209 | TNF superfamily member 18 |
| ENSG00000250042 | LINC02514 | 5,366710985 | 41,26111735 | 0,00340913 | 0,051689117 | long intergenic non-protein coding RNA 2514 |
| ENSG00000267280 | TBX2-AS1 | 5,362804343 | 41,14953843 | 5,12E-13 | 5,97E-11 | TBX2 antisense RNA 1 |
| ENSG00000274699 | TAS2R30 | 5,36258264 | 41,14321536 | 0,09397851 | 0,487240447 | taste 2 receptor member 30 |
| ENSG00000107796 | ACTA2 | 5,362542986 | 41,14208451 | 2,83E-36 | 2,06E-33 | actin alpha 2, smooth muscle |
| ENSG00000249116 | IRX4-AS1 | 5,359471164 | 41,05457694 | 0,00346334 | 0,052384115 | IRX4 antisense RNA 1 |
| ENSG00000183242 | WT1-AS | 5,351301786 | 40,82275904 | 0,01601968 | 0,161013527 | WT1 antisense RNA |
| ENSG00000135318 | NT5E | 5,350558981 | 40,80174589 | 1,59E-56 | 4,98E-53 | 5'-nucleotidase ecto |
| ENSG00000140563 | MCTP2 | 5,345500021 | 40,65892091 | 1,33E-07 | 6,96E-06 | multiple C2 and transmembrane domain containing 2 |
| ENSG00000204464 | C1orf195 | 5,34330151 | 40,59700831 | 0,00744278 | 0,092742506 | chromosome 1 open reading frame 195 |
| ENSG00000153294 | ADGRF4 | 5,338647342 | 40,46625241 | 0,00338893 | 0,051432725 | adhesion G protein-coupled receptor F4 |
| ENSG00000235180 | LINC00601 | 5,337035999 | 40,42108097 | 0,00521782 | 0,071173632 | long intergenic non-protein coding RNA 601 |
| ENSG00000211772 | TRBC2 | 5,315327732 | 39,81741706 | 0,09834915 | 0,49790609 | T cell receptor beta constant 2 |
| ENSG00000237633 | AC104623,1 | 5,309999473 | 39,67063189 | 0,00434408 | 0,062431658 | novel transcript, antisense to FAM49A |
| ENSG00000133816 | MICAL2 | 5,308228352 | 39,62196024 | 4,42E-28 | 1,70E-25 | microtubule associated monooxygenase, calponin and LIM domain containing 2 |
| ENSG00000278312 | LENG9 | 5,301214466 | 39,42979941 | 0,017542 | 0,171492516 | leukocyte receptor cluster member 9 |
| ENSG00000287369 | AC111197,1 | 5,298215549 | 39,34792217 | 0,00097384 | 0,01825309 | novel transcript |
| ENSG00000168685 | IL7R | 5,289463811 | 39,10995065 | 8,56E-35 | 5,46E-32 | interleukin 7 receptor |
| ENSG00000124875 | CXCL6 | 5,28782538 | 39,06555973 | 5,17E-13 | 6,01E-11 | C-X-C motif chemokine ligand 6 |
| ENSG00000012779 | ALOX5 | 5,285893664 | 39,01328737 | 6,86E-05 | 0,001877376 | arachidonate 5-lipoxygenase |
| ENSG00000285347 | AL359555,4 | 5,283520169 | 38,94915621 | 0,00776467 | 0,095647955 | novel transcript |
| ENSG00000113722 | CDX1 | 5,280399753 | 38,86500381 | 0,00166746 | 0,028754282 | caudal type homeobox 1 |
| ENSG00000284649 | AC009093,8 | 5,277409238 | 38,78452528 | 0,00226427 | 0,037043535 | BTG3 associated nuclear protein (BANP) pseudogene |
| ENSG00000157111 | TMEM171 | 5,276634796 | 38,76371124 | 0,00892096 | 0,105360946 | transmembrane protein 171 |
| ENSG00000221836 | OR2A5 | 5,270660233 | 38,60351291 | 0,01591923 | 0,160496195 | olfactory receptor family 2 subfamily A member 5 |
| ENSG00000262456 | AC006435,1 | 5,266472043 | 38,49160818 | 0,1030336 | 0,511633171 | novel transcript, antisense to MNT |
| ENSG00000262810 | AC090617,4 | 5,258467547 | 38,27863678 | 0,00584046 | 0,077338447 | novel transcript, antisense to SMG6 |
| ENSG00000284245 | FCAR | 5,252978536 | 38,13327505 | 0,04579785 | 0,324471967 | Fc fragment of IgA receptor |
| ENSG00000120949 | TNFRSF8 | 5,252151617 | 38,1114242 | 9,02E-05 | 0,002394839 | TNF receptor superfamily member 8 |
| ENSG00000250135 | AL049795,2 | 5,251993792 | 38,1072552 | 0,10438099 | 0,515175486 | novel transcript |
| ENSG00000164692 | COL1A2 | 5,235986673 | 37,68678113 | 2,10E-17 | 3,84E-15 | collagen type I alpha 2 chain |

| | | | | | | |
|---|---|---|---|---|---|---|
| ENSG00000283265 | AL356234,3 | 5,224474926 | 37,38726225 | 0,01043956 | 0,118383134 | novel transcript |
| ENSG00000254612 | DNAJB6P1 | 5,222819748 | 37,34439311 | 0,01118811 | 0,124301678 | DNAJB6 pseudogene 1 |
| ENSG00000139330 | KERA | 5,220930532 | 37,29552246 | 0,01587805 | 0,160236076 | keratocan |
| ENSG00000275871 | AC131025,2 | 5,219931944 | 37,26971662 | 0,02890217 | 0,241409949 | novel transcript |
| ENSG00000264578 | AC009630,3 | 5,215901954 | 37,16575354 | 0,10803796 | 0,523928283 | novel transcript, antisense to AGPAT6 |
| ENSG00000213877 | CFL1P7 | 5,209851806 | 37,01022 | 0,00591514 | 0,07796419 | cofilin 1 pseudogene 7 |
| ENSG00000163739 | CXCL1 | 5,205358052 | 36,8951187 | 2,76E-22 | 7,43E-20 | C-X-C motif chemokine ligand 1 |
| ENSG00000127743 | IL17B | 5,200050519 | 36,75963455 | 0,0074737 | 0,093016598 | interleukin 17B |
| ENSG00000176692 | FOXC2 | 5,189819989 | 36,49988439 | 1,46E-15 | 2,27E-13 | forkhead box C2 |
| ENSG00000273119 | AP005229,2 | 5,188835232 | 36,47497874 | 0,00534957 | 0,072653693 | novel transcript |
| ENSG00000266951 | AC093072,1 | 5,186003486 | 36,40345526 | 0,01344195 | 0,141928098 | novel transcript |
| ENSG00000147596 | PRDM14 | 5,175044021 | 36,12796332 | 0,00500833 | 0,068843113 | PR/SET domain 14 |
| ENSG00000145708 | CRHBP | 5,173460877 | 36,08833998 | 1,63E-08 | 9,82E-07 | corticotropin releasing hormone binding protein |
| ENSG00000232821 | AC003986,2 | 5,172702137 | 36,06936543 | 0,00197912 | 0,033391916 | novel transcript |
| ENSG00000155011 | DKK2 | 5,168080045 | 35,95399167 | 1,37E-29 | 6,11E-27 | dickkopf WNT signaling pathway inhibitor 2 |
| ENSG00000288508 | ISLR | 5,150660862 | 35,52249132 | 9,30E-20 | 2,06E-17 | immunoglobulin superfamily containing leucine rich repeat |
| ENSG00000129009 | ISLR | 5,150660862 | 35,52249132 | 9,30E-20 | 2,06E-17 | immunoglobulin superfamily containing leucine rich repeat |
| ENSG00000205795 | CYS1 | 5,150581467 | 35,52053649 | 2,29E-15 | 3,40E-13 | cystin 1 |
| ENSG00000274741 | AC138827,3 | 5,142618562 | 35,32502221 | 0,00831842 | 0,100528485 | occludin (OCLN) pseudogene |
| ENSG00000214285 | NPS | 5,141516783 | 35,29805495 | 0,02896385 | 0,241642594 | neuropeptide S |
| ENSG00000287792 | AC140118,2 | 5,141218748 | 35,29076376 | 0,01213853 | 0,131911723 | novel transcript |
| ENSG00000251194 | AL133330,1 | 5,137268488 | 35,19426589 | 0,02313726 | 0,207030121 | novel transcript |
| ENSG00000165474 | GJB2 | 5,134916015 | 35,13692451 | 2,42E-07 | 1,18E-05 | gap junction protein beta 2 |
| ENSG00000260139 | CSPG4P13 | 5,131682612 | 35,05826298 | 0,01133149 | 0,125514797 | chondroitin sulfate proteoglycan 4 pseudogene 13 |
| ENSG00000000971 | CFH | 5,12427171 | 34,87863585 | 4,00E-17 | 7,14E-15 | complement factor H |
| ENSG00000145649 | GZMA | 5,118347475 | 34,73570503 | 6,80E-13 | 7,79E-11 | granzyme A |
| ENSG00000258967 | HMGN1P3 | 5,111966339 | 34,58240599 | 0,03141315 | 0,255356429 | high mobility group nucleosome binding domain 1 pseudogene 3 |
| ENSG00000206298 | PSMB8 | 5,109176809 | 34,51560361 | 2,90E-05 | 0,000872002 | proteasome 20S subunit beta 8 |
| ENSG00000226201 | PSMB8 | 5,109176809 | 34,51560361 | 2,90E-05 | 0,000872002 | proteasome 20S subunit beta 8 |
| ENSG00000236443 | PSMB8 | 5,109176809 | 34,51560361 | 2,90E-05 | 0,000872002 | proteasome 20S subunit beta 8 |

| | | | | | | |
|---|---|---|---|---|---|---|
| ENSG00000230669 | PSMB8 | 5,109176809 | 34,51560361 | 2,90E-05 | 0,000872002 | proteasome 20S subunit beta 8 |
| ENSG00000284661 | AC105233,5 | 5,109106992 | 34,51393332 | 0,1192844 | 0,552100834 | RNA binding protein, fox-1 homolog (C, elegans) 1 (RBFOX1) pseudogene |
| ENSG00000258144 | LINC02406 | 5,105230356 | 34,42131614 | 0,00810009 | 0,098656485 | long intergenic non-protein coding RNA 2406 |
| ENSG00000120669 | SOHLH2 | 5,103879728 | 34,38910653 | 0,00702867 | 0,088859506 | spermatogenesis and oogenesis specific basic helix-loop-helix 2 |
| ENSG00000287294 | AL035694,1 | 5,099057946 | 34,27436294 | 0,006859 | 0,087388747 | novel transcript |
| ENSG00000261327 | AC134312,5 | 5,098366658 | 34,25794383 | 1,11E-09 | 8,32E-08 | novel transcript |
| ENSG00000188674 | C2orf80 | 5,08102857 | 33,84870115 | 1,32E-11 | 1,27E-09 | chromosome 2 open reading frame 80 |
| ENSG00000172232 | AZU1 | 5,079281596 | 33,80773821 | 0,1531228 | 0,621632863 | azurocidin 1 |
| ENSG00000136514 | RTP4 | 5,075364774 | 33,71607691 | 0,00554521 | 0,074662109 | receptor transporter protein 4 |
| ENSG00000165496 | RPL10L | 5,075137511 | 33,71076615 | 0,01047768 | 0,118729367 | ribosomal protein L10 like |
| ENSG00000178776 | C5orf46 | 5,073053891 | 33,66211435 | 0,01948595 | 0,183714864 | chromosome 5 open reading frame 46 |
| ENSG00000233980 | FDPSP2 | 5,071316624 | 33,62160344 | 0,06147231 | 0,386127478 | farnesyl diphosphate synthase pseudogene 2 |
| ENSG00000162595 | DIRAS3 | 5,071245326 | 33,6199419 | 5,35E-09 | 3,48E-07 | DIRAS family GTPase 3 |
| ENSG00000177234 | LINC01561 | 5,067009422 | 33,52137498 | 0,0048052 | 0,066828153 | long intergenic non-protein coding RNA 1561 |
| ENSG00000266728 | AC015688,4 | 5,059267461 | 33,34197041 | 0,06078508 | 0,383761049 | novel protein |
| ENSG00000187583 | PLEKHN1 | 5,054422022 | 33,23017584 | 0,00746018 | 0,09288537 | pleckstrin homology domain containing N1 |
| ENSG00000106366 | SERPINE1 | 5,053937108 | 33,21900849 | 3,63E-16 | 5,91E-14 | serpin family E member 1 |
| ENSG00000178394 | HTR1A | 5,048649661 | 33,09748437 | 2,85E-10 | 2,32E-08 | 5-hydroxytryptamine receptor 1A |
| ENSG00000137809 | ITGA11 | 5,04856668 | 33,09558071 | 1,01E-29 | 4,63E-27 | integrin subunit alpha 11 |
| ENSG00000175294 | CATSPER1 | 5,044482176 | 33,00201427 | 0,0076087 | 0,094283946 | cation channel sperm associated 1 |
| ENSG00000278960 | AL022329,3 | 5,043304569 | 32,97508719 | 0,00730821 | 0,091393601 | novel transcript, antisense to MYO18B |
| ENSG00000196427 | NBPF4 | 5,036853805 | 32,82797388 | 0,01035604 | 0,118005006 | NBPF member 4 |
| ENSG00000249842 | AC010486,3 | 5,032536092 | 32,72987283 | 0,02566609 | 0,222895766 | novel transcript |
| ENSG00000253647 | KCNIP1-OT1 | 5,030309093 | 32,67938873 | 0,00873662 | 0,103816706 | KCNIP1 overlapping transcript 1 |
| ENSG00000233728 | AL929472,3 | 5,03021456 | 32,67724747 | 0,06288612 | 0,391611013 | novel transcript |
| ENSG00000286251 | AC097528,1 | 5,028181243 | 32,63122499 | 0,01042849 | 0,118366144 | novel transcript |
| ENSG00000113721 | PDGFRB | 5,02616834 | 32,58572841 | 1,39E-46 | 1,97E-43 | platelet derived growth factor receptor beta |
| ENSG00000174332 | GLIS1 | 5,025445239 | 32,56940005 | 0,00012413 | 0,003160346 | GLIS family zinc finger 1 |
| ENSG00000133106 | EPSTI1 | 5,011563694 | 32,25752162 | 1,28E-14 | 1,75E-12 | epithelial stromal interaction 1 |
| ENSG00000251733 | SCARNA8 | 5,008579563 | 32,19086777 | 0,02277817 | 0,204838933 | small Cajal body-specific RNA 8 |
| ENSG00000277688 | AC243585,2 | 5,003331007 | 32,07396945 | 0,13169425 | 0,579654922 | novel transcript, antisense DDX52 |

| | | | | | | |
|---|---|---|---|---|---|---|
| ENSG00000254501 | AP003068,1 | 5,001110816 | 32,02464818 | 0,07627198 | 0,43651514 | novel transcript |
| ENSG00000272218 | AC108865,2 | 5,000528948 | 32,0117346 | 0,0249448 | 0,219004288 | novel transcript |
| ENSG00000268357 | VN1R81P | 4,996534402 | 31,92322282 | 0,07700899 | 0,438408786 | vomeronasal 1 receptor 81 pseudogene |
| ENSG00000285773 | AL161893,1 | 4,994309832 | 31,87403662 | 0,0844511 | 0,462476653 | novel transcript |
| ENSG00000121797 | CCRL2 | 4,988376187 | 31,74321143 | 0,0078483 | 0,096374424 | C-C motif chemokine receptor like 2 |
| ENSG00000170577 | SIX2 | 4,987962648 | 31,73411375 | 1,11E-24 | 3,59E-22 | SIX homeobox 2 |
| ENSG00000283436 | LINC01958 | 4,986103317 | 31,69324149 | 0,03129526 | 0,254596667 | long intergenic non-protein coding RNA 1958 |
| ENSG00000217236 | SP9 | 4,982570489 | 31,615727 | 1,01E-22 | 2,76E-20 | Sp9 transcription factor |
| ENSG00000197309 | OR10D3 | 4,980610841 | 31,57281172 | 7,76E-05 | 0,002096235 | olfactory receptor family 10 subfamily D member 3 (putative) |
| ENSG00000288243 | STING1 | 4,980168265 | 31,56312761 | 3,74E-14 | 4,87E-12 | stimulator of interferon response cGAMP interactor 1 |
| ENSG00000187546 | AGMO | 4,959677784 | 31,11800755 | 4,20E-05 | 0,001210559 | alkylglycerol monooxygenase |
| ENSG00000287496 | AC093227,3 | 4,957205866 | 31,06473551 | 0,02184012 | 0,198606627 | novel transcript |
| ENSG00000288136 | AC245175,2 | 4,952476891 | 30,96307588 | 0,0703721 | 0,417716603 | |
| ENSG00000164741 | DLC1 | 4,949485845 | 30,89894864 | 2,41E-41 | 2,59E-38 | DLC1 Rho GTPase activating protein |
| ENSG00000285955 | BX842242,1 | 4,944385736 | 30,7899098 | 0,02874113 | 0,240839934 | novel transcript |
| ENSG00000184584 | STING1 | 4,936213863 | 30,61599919 | 6,93E-15 | 9,85E-13 | stimulator of interferon response cGAMP interactor 1 |
| ENSG00000235431 | AC010991,1 | 4,935673959 | 30,60454382 | 0,01349144 | 0,142214228 | novel transcript |
| ENSG00000226147 | TUBBP10 | 4,93300987 | 30,54808144 | 0,02763343 | 0,234127782 | tubulin beta class I pseudogene 10 |
| ENSG00000268193 | AC002985,1 | 4,932209723 | 30,53114358 | 0,07768311 | 0,440457885 | novel transcript |
| ENSG00000259004 | LINC02285 | 4,93078247 | 30,50095415 | 0,01217103 | 0,131987045 | long intergenic non-protein coding RNA 2285 |
| ENSG00000173918 | C1QTNF1 | 4,930015198 | 30,48473707 | 7,19E-18 | 1,38E-15 | C1q and TNF related 1 |
| ENSG00000240216 | CPHL1P | 4,928389706 | 30,45040912 | 0,0325586 | 0,261251213 | ceruloplasmin and hephaestin like 1, pseudogene |
| ENSG00000225369 | CAPZA1P2 | 4,928367898 | 30,44994883 | 0,03539107 | 0,275308916 | CAPZA1 pseudogene 2 |
| ENSG00000281580 | LINC01060 | 4,921939362 | 30,31456807 | 0,01600897 | 0,161013527 | long intergenic non-protein coding RNA 1060 |
| ENSG00000274248 | AJ011932,1 | 4,921886452 | 30,31345634 | 0,0185924 | 0,177868046 | novel transcript |
| ENSG00000213876 | RPL7AP64 | 4,919994237 | 30,27372375 | 0,03957098 | 0,29639362 | ribosomal protein L7a pseudogene 64 |
| ENSG00000231259 | ANAPC1P2 | 4,918017494 | 30,23227191 | 0,14150161 | 0,599428784 | ANAPC1 pseudogene 2 |
| ENSG00000000005 | TNMD | 4,915995325 | 30,1899262 | 0,00373084 | 0,055649253 | tenomodulin |
| ENSG00000204539 | CDSN | 4,905711864 | 29,97549897 | 0,02064956 | 0,190898872 | corneodesmosin |
| ENSG00000189223 | PAX8-AS1 | 4,904198758 | 29,94407701 | 2,94E-27 | 1,08E-24 | PAX8 antisense RNA 1 |

| | | | | | | |
|---|---|---|---|---|---|---|
| ENSG00000287856 | AL445524,2 | 4,903380974 | 29,92710817 | 0,02923111 | 0,242859791 | novel protein |
| ENSG00000259977 | AL121578,1 | 4,901892469 | 29,8962467 | 0,00928089 | 0,108564101 | novel transcript |
| ENSG00000113739 | STC2 | 4,901201047 | 29,88192215 | 9,33E-10 | 7,06E-08 | stanniocalcin 2 |
| ENSG00000251574 | AC099520,1 | 4,898754896 | 29,83129899 | 0,02896657 | 0,241642594 | novel transcript |
| ENSG00000213981 | AC007277,1 | 4,890481215 | 29,66070975 | 0,0362744 | 0,279606213 | novel transcript, antisense to MYO3B |
| ENSG00000258955 | LINC00519 | 4,889292303 | 29,63627669 | 0,00268791 | 0,042334936 | long intergenic non-protein coding RNA 519 |
| ENSG00000130948 | HSD17B3 | 4,887739916 | 29,60440423 | 0,04597399 | 0,325335161 | hydroxysteroid 17-beta dehydrogenase 3 |
| ENSG00000115596 | WNT6 | 4,885352334 | 29,55545108 | 0,00725006 | 0,090884485 | Wnt family member 6 |
| ENSG00000237412 | PRSS56 | 4,884972961 | 29,54768016 | 0,01015509 | 0,116466907 | serine protease 56 |
| ENSG00000204540 | PSORS1C1 | 4,884784606 | 29,54382274 | 0,00248058 | 0,039770682 | psoriasis susceptibility 1 candidate 1 |
| ENSG00000241280 | AC106712,1 | 4,882602831 | 29,49917763 | 0,030605 | 0,250349254 | novel transcript |
| ENSG00000288673 | AC270289,4 | 4,878111968 | 29,4074945 | 2,38E-21 | 6,00E-19 | DLC1 Rho GTPase activating protein |
| ENSG00000258352 | IFITM3P6 | 4,871811907 | 29,2793558 | 0,04381081 | 0,31589834 | IFITM3P pseudogene 6 |
| ENSG00000255030 | OR8B5P | 4,870635002 | 29,25548036 | 0,01024609 | 0,117166653 | olfactory receptor family 8 subfamily B member 5 pseudogene |
| ENSG00000283375 | AC087521,4 | 4,86838192 | 29,20982723 | 0,14781892 | 0,612372753 | novel transcript, antisense HSD17B12 |
| ENSG00000284513 | AC006063,2 | 4,866199142 | 29,16566657 | 0,01348033 | 0,142189308 | novel transcript |
| ENSG00000164256 | PRDM9 | 4,864743164 | 29,13624722 | 0,01034836 | 0,117990971 | PR/SET domain 9 |
| ENSG00000282278 | AC058822,1 | 4,860105918 | 29,04274511 | 0,06737184 | 0,408200237 | novel FIP1L1-PDGFRA fusion protein |
| ENSG00000258945 | AL049775,3 | 4,854273462 | 28,92556957 | 0,07807643 | 0,441886578 | novel transcript |
| ENSG00000230630 | DNM3OS | 4,85156554 | 28,87132752 | 1,02E-13 | 1,27E-11 | DNM3 opposite strand/antisense RNA |
| ENSG00000211689 | TRGC1 | 4,851032377 | 28,86065979 | 7,52E-08 | 4,10E-06 | T cell receptor gamma constant 1 |
| ENSG00000183153 | GJD3 | 4,842892082 | 28,69827431 | 0,00017983 | 0,004341416 | gap junction protein delta 3 |
| ENSG00000240118 | PSMB9 | 4,840945087 | 28,65957056 | 0,00395455 | 0,05820865 | proteasome 20S subunit beta 9 |
| ENSG00000169908 | TM4SF1 | 4,830198176 | 28,44687315 | 5,20E-10 | 4,11E-08 | transmembrane 4 L six family member 1 |
| ENSG00000232126 | HLA-B | 4,827173883 | 28,38730298 | 7,22E-08 | 3,96E-06 | major histocompatibility complex, class I, B |
| ENSG00000226762 | LINC02668 | 4,815873883 | 28,16582614 | 0,0043213 | 0,062229857 | long intergenic non-protein coding RNA 2668 |
| ENSG00000174640 | SLCO2A1 | 4,813983079 | 28,12893603 | 5,07E-06 | 0,000187163 | solute carrier organic anion transporter family member 2A1 |
| ENSG00000275571 | TTC34 | 4,803438691 | 27,92409638 | 4,79E-07 | 2,17E-05 | tetratricopeptide repeat domain 34 |
| ENSG00000203805 | PLPP4 | 4,801564764 | 27,88784913 | 2,81E-14 | 3,69E-12 | phospholipid phosphatase 4 |
| ENSG00000151790 | TDO2 | 4,791221361 | 27,6886222 | 5,40E-09 | 3,51E-07 | tryptophan 2,3-dioxygenase |
| ENSG00000128815 | WDFY4 | 4,788600341 | 27,63836449 | 0,00013489 | 0,003398104 | WDFY family member 4 |

| | | | | | | |
|---|---|---|---|---|---|---|
| ENSG00000176510 | OR10AC1 | 4,78791463 | 27,62523114 | 0,00430783 | 0,062064547 | olfactory receptor family 10 subfamily AC member 1 (gene/pseudogene) |
| ENSG00000285252 | OR10AC1 | 4,78791463 | 27,62523114 | 0,00430783 | 0,062064547 | olfactory receptor family 10 subfamily AC member 1 (gene/pseudogene) |
| ENSG00000117226 | GBP3 | 4,77979274 | 27,47014728 | 1,15E-28 | 4,79E-26 | guanylate binding protein 3 |
| ENSG00000164488 | DACT2 | 4,777937153 | 27,43483801 | 2,75E-08 | 1,60E-06 | dishevelled binding antagonist of beta catenin 2 |
| ENSG00000130600 | H19 | 4,775344257 | 27,3855748 | 1,32E-11 | 1,27E-09 | H19 imprinted maternally expressed transcript |
| ENSG00000288237 | H19 | 4,775344257 | 27,3855748 | 1,32E-11 | 1,27E-09 | H19 imprinted maternally expressed transcript |
| ENSG00000213949 | ITGA1 | 4,761269747 | 27,11970813 | 2,08E-24 | 6,58E-22 | integrin subunit alpha 1 |
| ENSG00000164935 | DCSTAMP | 4,760942472 | 27,11355672 | 0,00424602 | 0,061411368 | dendrocyte expressed seven transmembrane protein |
| ENSG00000124205 | EDN3 | 4,75657397 | 27,03158065 | 1,61E-12 | 1,73E-10 | endothelin 3 |
| ENSG00000134107 | BHLHE40 | 4,752312868 | 26,95185876 | 7,16E-36 | 4,98E-33 | basic helix-loop-helix family member e40 |
| ENSG00000205413 | SAMD9 | 4,751531051 | 26,93725713 | 1,65E-14 | 2,23E-12 | sterile alpha motif domain containing 9 |
| ENSG00000102871 | TRADD | 4,750965678 | 26,92670284 | 3,29E-07 | 1,55E-05 | TNFRSF1A associated via death domain |
| ENSG00000133800 | LYVE1 | 4,742506855 | 26,76928786 | 1,95E-10 | 1,62E-08 | lymphatic vessel endothelial hyaluronan receptor 1 |
| ENSG00000123095 | BHLHE41 | 4,742336518 | 26,76612744 | 2,04E-09 | 1,43E-07 | basic helix-loop-helix family member e41 |
| ENSG00000005102 | MEOX1 | 4,737137999 | 26,66985358 | 2,02E-13 | 2,47E-11 | mesenchyme homeobox 1 |
| ENSG00000026508 | CD44 | 4,731834275 | 26,57198822 | 4,85E-19 | 1,02E-16 | CD44 molecule (Indian blood group) |
| ENSG00000020181 | ADGRA2 | 4,731729236 | 26,57005365 | 1,82E-28 | 7,30E-26 | adhesion G protein-coupled receptor A2 |
| ENSG00000121068 | TBX2 | 4,713299043 | 26,23278444 | 8,71E-05 | 0,002326706 | T-box transcription factor 2 |
| ENSG00000117586 | TNFSF4 | 4,711199023 | 26,19462717 | 5,41E-26 | 1,86E-23 | TNF superfamily member 4 |
| ENSG00000168481 | LGI3 | 4,701106442 | 26,01201836 | 0,00665589 | 0,085492961 | leucine rich repeat LGI family member 3 |
| ENSG00000249899 | AC025180,1 | 4,689402446 | 25,80184713 | 1,71E-09 | 1,23E-07 | novel transcript |
| ENSG00000134668 | SPOCD1 | 4,68650389 | 25,75006002 | 5,46E-10 | 4,29E-08 | SPOC domain containing 1 |
| ENSG00000138135 | CH25H | 4,659325509 | 25,26950518 | 0,00734999 | 0,091842517 | cholesterol 25-hydroxylase |
| ENSG00000166741 | NNMT | 4,658663294 | 25,25790883 | 4,15E-22 | 1,09E-19 | nicotinamide N-methyltransferase |
| ENSG00000240801 | AC132217,1 | 4,647258124 | 25,05902058 | 5,51E-11 | 4,88E-09 | novel transcript, 3' overlapping ncRNA IGF2 |
| ENSG00000287460 | AL442636,1 | 4,6443799 | 25,00907686 | 1,13E-23 | 3,38E-21 | novel transcript |
| ENSG00000013364 | MVP | 4,64168915 | 24,96247625 | 1,65E-12 | 1,77E-10 | major vault protein |
| ENSG00000184012 | TMPRSS2 | 4,636422089 | 24,87150818 | 6,27E-05 | 0,001735019 | transmembrane serine protease 2 |
| ENSG00000105974 | CAV1 | 4,62365784 | 24,65242803 | 2,37E-47 | 3,71E-44 | caveolin 1 |
| ENSG00000260418 | AL023284,4 | 4,61158738 | 24,44703133 | 0,00497197 | 0,068507629 | novel transcript |

| | | | | | | |
|---|---|---|---|---|---|---|
| ENSG00000226268 | AC135977,1 | 4,606459219 | 24,36028689 | 0,27299598 | 0,785672647 | solute carrier family 25, member 33 (SLC25A33) pseudogene |
| ENSG00000236841 | AC007750,1 | 4,601556736 | 24,27764766 | 0,01184424 | 0,129565539 | novel transcript |
| ENSG00000287382 | AC110611,2 | 4,600690382 | 24,26307305 | 0,00901186 | 0,106199263 | novel transcript |
| ENSG00000171631 | P2RY6 | 4,595468821 | 24,17541619 | 0,00819542 | 0,099387781 | pyrimidinergic receptor P2Y6 |
| ENSG00000230955 | AL929472,2 | 4,591650085 | 24,11150978 | 0,27454691 | 0,786383635 | novel transcript |
| ENSG00000277400 | AC145212,1 | 4,588578118 | 24,06022315 | 1,94E-15 | 2,91E-13 | MaFF-interacting protein |
| ENSG00000092758 | COL9A3 | 4,571125145 | 23,77090863 | 4,12E-09 | 2,74E-07 | collagen type IX alpha 3 chain |
| ENSG00000089327 | FXYD5 | 4,562346634 | 23,62670655 | 1,90E-12 | 2,01E-10 | FXYD domain containing ion transport regulator 5 |
| ENSG00000250208 | FZD10-AS1 | 4,560360926 | 23,59420941 | 1,55E-21 | 3,95E-19 | FZD10 antisense divergent transcript |
| ENSG00000107562 | CXCL12 | 4,553526534 | 23,48270232 | 1,58E-49 | 3,08E-46 | C-X-C motif chemokine ligand 12 |
| ENSG00000122585 | NPY | 4,553513939 | 23,48249731 | 9,81E-14 | 1,22E-11 | neuropeptide Y |
| ENSG00000138080 | EMILIN1 | 4,541793792 | 23,29250328 | 1,69E-09 | 1,22E-07 | elastin microfibril interfacer 1 |
| ENSG00000122862 | SRGN | 4,54127051 | 23,28405637 | 2,52E-06 | 0,000100354 | serglycin |
| ENSG00000226806 | LCT-AS1 | 4,540830673 | 23,2769588 | 5,62E-05 | 0,001580224 | LCT antisense RNA 1 |
| ENSG00000169715 | MT1E | 4,536814671 | 23,21225328 | 2,92E-08 | 1,69E-06 | metallothionein 1E |
| ENSG00000105246 | EBI3 | 4,535766792 | 23,19539954 | 0,00699092 | 0,088512268 | Epstein-Barr virus induced 3 |
| ENSG00000260873 | SNTB2 | 4,535621986 | 23,19307149 | 0,0353492 | 0,275051622 | syntrophin beta 2 |
| ENSG00000123572 | NRK | 4,534686283 | 23,17803382 | 2,77E-47 | 4,13E-44 | Nik related kinase |
| ENSG00000174885 | NLRP6 | 4,534131198 | 23,16911765 | 0,01019791 | 0,116701084 | NLR family pyrin domain containing 6 |
| ENSG00000135111 | TBX3 | 4,533276691 | 23,15539866 | 8,07E-09 | 5,04E-07 | T-box transcription factor 3 |
| ENSG00000161381 | PLXDC1 | 4,531867653 | 23,13279451 | 4,42E-18 | 8,74E-16 | plexin domain containing 1 |
| ENSG00000106034 | CPED1 | 4,523161166 | 22,99361134 | 4,05E-15 | 5,85E-13 | cadherin like and PC-esterase domain containing 1 |
| ENSG00000164093 | PITX2 | 4,521346339 | 22,96470488 | 1,85E-07 | 9,23E-06 | paired like homeodomain 2 |
| ENSG00000205578 | POM121B | 4,519010137 | 22,9275475 | 3,29E-19 | 7,10E-17 | POM121 transmembrane nucleoporin B (pseudogene) |
| ENSG00000164251 | F2RL1 | 4,513566492 | 22,84119921 | 1,33E-23 | 3,92E-21 | F2R like trypsin receptor 1 |
| ENSG00000227686 | VARS1 | 4,511254005 | 22,80461652 | 0,03018051 | 0,248045051 | valyl-tRNA synthetase 1 |
| ENSG00000020633 | RUNX3 | 4,505638015 | 22,71601738 | 0,0001116 | 0,002883465 | RUNX family transcription factor 3 |
| ENSG00000111424 | VDR | 4,500273724 | 22,63171053 | 3,83E-09 | 2,58E-07 | vitamin D receptor |
| ENSG00000105880 | DLX5 | 4,488191538 | 22,44296744 | 1,47E-28 | 6,05E-26 | distal-less homeobox 5 |
| ENSG00000100234 | TIMP3 | 4,488077361 | 22,44119133 | 4,24E-50 | 8,84E-47 | TIMP metallopeptidase inhibitor 3 |
| ENSG00000172137 | CALB2 | 4,486402658 | 22,41515636 | 6,85E-08 | 3,77E-06 | calbindin 2 |

| | | | | | | |
|---|---|---|---|---|---|---|
| ENSG00000282830 | CALB2 | 4,486402658 | 22,41515636 | 6,85E-08 | 3,77E-06 | calbindin 2 |
| ENSG00000224594 | RPL29P19 | 4,464664182 | 22,07993749 | 0,01115011 | 0,123967688 | ribosomal protein L29 pseudogene 19 |
| ENSG00000183160 | TMEM119 | 4,458468735 | 21,98532166 | 6,10E-15 | 8,75E-13 | transmembrane protein 119 |
| ENSG00000135636 | DYSF | 4,456685863 | 21,95816913 | 2,79E-31 | 1,34E-28 | dysferlin |
| ENSG00000250829 | AC108865,1 | 4,448500748 | 21,83394243 | 0,01224391 | 0,13259217 | novel transcript |
| ENSG00000276403 | AC105105,4 | 4,446698426 | 21,80668288 | 0,01080494 | 0,121294714 | novel transcript |
| ENSG00000285109 | GATA4 | 4,444038977 | 21,76652169 | 0,01565718 | 0,158672941 | GATA binding protein 4 |
| ENSG00000089012 | SIRPG | 4,440796111 | 21,71765021 | 0,00842836 | 0,101386842 | signal regulatory protein gamma |
| ENSG00000239836 | PSMB9 | 4,436412441 | 21,65176067 | 0,00907851 | 0,106703228 | proteasome 20S subunit beta 9 |
| ENSG00000243067 | PSMB9 | 4,436412441 | 21,65176067 | 0,00907851 | 0,106703228 | proteasome 20S subunit beta 9 |
| ENSG00000243594 | PSMB9 | 4,436412441 | 21,65176067 | 0,00907851 | 0,106703228 | proteasome 20S subunit beta 9 |
| ENSG00000156265 | MAP3K7CL | 4,430988143 | 21,57050642 | 1,05E-09 | 7,90E-08 | MAP3K7 C-terminal like |
| ENSG00000183421 | RIPK4 | 4,406341451 | 21,20513042 | 0,00055773 | 0,011456632 | receptor interacting serine/threonine kinase 4 |
| ENSG00000257178 | AC103702,1 | 4,402935323 | 21,15512527 | 0,03327339 | 0,264841921 | novel transcript, sense intronic to HOXB3 |
| ENSG00000196878 | LAMB3 | 4,40194608 | 21,14062436 | 1,19E-05 | 0,000394357 | laminin subunit beta 3 |
| ENSG00000138061 | CYP1B1 | 4,401388325 | 21,13245284 | 1,54E-44 | 1,85E-41 | cytochrome P450 family 1 subfamily B member 1 |
| ENSG00000143369 | ECM1 | 4,399421607 | 21,10366417 | 2,27E-32 | 1,27E-29 | extracellular matrix protein 1 |
| ENSG00000254818 | AP004607,3 | 4,399196437 | 21,10037066 | 0,00936895 | 0,109376713 | Cdon homolog (mouse) (CDON) pseudogene |
| ENSG00000108242 | CYP2C18 | 4,392242766 | 20,99891332 | 0,00902768 | 0,106345695 | cytochrome P450 family 2 subfamily C member 18 |
| ENSG00000228582 | TAP2 | 4,360896902 | 20,54758452 | 0,01583946 | 0,159949901 | transporter 2, ATP binding cassette subfamily B member |
| ENSG00000187479 | C11orf96 | 4,359443814 | 20,52689933 | 1,35E-08 | 8,24E-07 | chromosome 11 open reading frame 96 |
| ENSG00000248572 | EGFLAM-AS2 | 4,357734801 | 20,50259762 | 0,01478002 | 0,15245219 | EGFLAM antisense RNA 2 |
| ENSG00000081051 | AFP | 4,356663238 | 20,48737495 | 6,61E-05 | 0,001819667 | alpha fetoprotein |
| ENSG00000253974 | NRG1-IT1 | 4,356463864 | 20,48454388 | 0,00244503 | 0,03938249 | NRG1 intronic transcript 1 |
| ENSG00000136630 | HLX | 4,355966228 | 20,47747926 | 2,00E-05 | 0,000629467 | H2,0 like homeobox |
| ENSG00000247993 | FOXD1-AS1 | 4,352339467 | 20,426066 | 0,01993919 | 0,186138783 | FOXD1 antisense RNA 1 |
| ENSG00000137198 | GMPR | 4,345825727 | 20,3340506 | 3,59E-06 | 0,000138095 | guanosine monophosphate reductase |
| ENSG00000087303 | NID2 | 4,337361452 | 20,21510023 | 3,19E-49 | 5,87E-46 | nidogen 2 |
| ENSG00000286717 | AP001407,1 | 4,334342412 | 20,17284156 | 0,01977152 | 0,184933326 | novel transcript, antisense to KCNJ6 |
| ENSG00000105329 | TGFB1 | 4,325314421 | 20,04699958 | 1,53E-18 | 3,09E-16 | transforming growth factor beta 1 |
| ENSG00000233532 | LINC00460 | 4,30661103 | 19,78878361 | 2,41E-07 | 1,17E-05 | long intergenic non-protein coding RNA 460 |
| ENSG00000182601 | HS3ST4 | 4,306020759 | 19,7806888 | 1,69E-09 | 1,21E-07 | heparan sulfate-glucosamine 3-sulfotransferase 4 |

| | | | | | | |
|---|---|---|---|---|---|---|
| ENSG00000240065 | PSMB9 | 4,298634507 | 19,67967523 | 0,0218796 | 0,198849969 | proteasome 20S subunit beta 9 |
| ENSG00000196196 | HRCT1 | 4,280543203 | 19,43443422 | 1,58E-07 | 8,15E-06 | histidine rich carboxyl terminus 1 |
| ENSG00000171522 | PTGER4 | 4,280439234 | 19,43303371 | 4,60E-18 | 9,05E-16 | prostaglandin E receptor 4 |
| ENSG00000234155 | LINC02535 | 4,276192266 | 19,37591137 | 0,01103311 | 0,123104637 | long intergenic non-protein coding RNA 2535 |
| ENSG00000164176 | EDIL3 | 4,274811206 | 19,35737212 | 4,28E-38 | 3,62E-35 | EGF like repeats and discoidin domains 3 |
| ENSG00000198959 | TGM2 | 4,274385645 | 19,35166299 | 4,35E-16 | 7,04E-14 | transglutaminase 2 |
| ENSG00000088926 | F11 | 4,27390096 | 19,34516273 | 0,01548634 | 0,157554533 | coagulation factor XI |
| ENSG00000111348 | ARHGDIB | 4,27211112 | 19,32117757 | 1,18E-40 | 1,15E-37 | Rho GDP dissociation inhibitor beta |
| ENSG00000099860 | GADD45B | 4,263238725 | 19,20271935 | 2,43E-18 | 4,84E-16 | growth arrest and DNA damage inducible beta |
| ENSG00000135903 | PAX3 | 4,261684959 | 19,18204939 | 1,17E-27 | 4,35E-25 | paired box 3 |
| ENSG00000254646 | OR8B10P | 4,261162708 | 19,1751068 | 0,01353684 | 0,142544884 | olfactory receptor family 8 subfamily B member 10 pseudogene |
| ENSG00000075891 | PAX2 | 4,24937055 | 19,01901401 | 0,0036221 | 0,054312354 | paired box 2 |
| ENSG00000245662 | LINC02211 | 4,244430376 | 18,95399919 | 0,01501024 | 0,154317722 | long intergenic non-protein coding RNA 2211 |
| ENSG00000206235 | TAP2 | 4,23828362 | 18,87341544 | 0,01665283 | 0,165123398 | transporter 2, ATP binding cassette subfamily B member |
| ENSG00000039068 | CDH1 | 4,228624955 | 18,74748228 | 2,73E-11 | 2,54E-09 | cadherin 1 |
| ENSG00000139567 | ACVRL1 | 4,225282437 | 18,70409732 | 1,34E-16 | 2,27E-14 | activin A receptor like type 1 |
| ENSG00000119681 | LTBP2 | 4,224761689 | 18,69734721 | 9,53E-17 | 1,66E-14 | latent transforming growth factor beta binding protein 2 |
| ENSG00000115594 | IL1R1 | 4,20977341 | 18,50410445 | 1,33E-31 | 6,58E-29 | interleukin 1 receptor type 1 |
| ENSG00000176678 | FOXL1 | 4,207997936 | 18,48134612 | 1,85E-10 | 1,55E-08 | forkhead box L1 |
| ENSG00000184254 | ALDH1A3 | 4,20720355 | 18,47117259 | 1,23E-07 | 6,47E-06 | aldehyde dehydrogenase 1 family member A3 |
| ENSG00000135480 | KRT7 | 4,203940854 | 18,42944671 | 0,00072849 | 0,014252604 | keratin 7 |
| ENSG00000206450 | HLA-B | 4,203763093 | 18,42717608 | 3,05E-10 | 2,48E-08 | major histocompatibility complex, class I, B |
| ENSG00000145908 | ZNF300 | 4,20260935 | 18,4124455 | 4,17E-19 | 8,94E-17 | zinc finger protein 300 |
| ENSG00000147883 | CDKN2B | 4,195658579 | 18,32394941 | 1,17E-37 | 9,41E-35 | cyclin dependent kinase inhibitor 2B |
| ENSG00000197057 | DTHD1 | 4,167633087 | 17,97142729 | 2,35E-06 | 9,40E-05 | death domain containing 1 |
| ENSG00000179855 | GIPC3 | 4,154554238 | 17,80924236 | 3,99E-05 | 0,001154307 | GIPC PDZ domain containing family member 3 |
| ENSG00000162645 | GBP2 | 4,150391582 | 17,75793083 | 2,22E-11 | 2,08E-09 | guanylate binding protein 2 |
| ENSG00000154589 | LY96 | 4,150146128 | 17,75490983 | 0,01756991 | 0,171664029 | lymphocyte antigen 96 |
| ENSG00000115414 | FN1 | 4,136060812 | 17,58240859 | 4,56E-32 | 2,46E-29 | fibronectin 1 |
| ENSG00000089127 | OAS1 | 4,134254334 | 17,56040647 | 4,06E-06 | 0,000153851 | 2'-5'-oligoadenylate synthetase 1 |

| | | | | | | |
|---|---|---|---|---|---|---|
| ENSG00000228714 | AL691420,1 | 4,130220414 | 17,5113744 | 0,02507545 | 0,219719433 | novel transcript, antisense to DEC1 |
| ENSG00000151615 | POU4F2 | 4,121465029 | 17,4054237 | 1,72E-07 | 8,74E-06 | POU class 4 homeobox 2 |
| ENSG00000204381 | LAYN | 4,121449722 | 17,40523903 | 1,19E-14 | 1,66E-12 | layilin |
| ENSG00000106333 | PCOLCE | 4,121078959 | 17,40076657 | 4,02E-22 | 1,07E-19 | procollagen C-endopeptidase enhancer |
| ENSG00000174807 | CD248 | 4,111090716 | 17,28071152 | 3,49E-24 | 1,07E-21 | CD248 molecule |
| ENSG00000171992 | SYNPO | 4,110556433 | 17,27431303 | 8,82E-33 | 5,11E-30 | synaptopodin |
| ENSG00000274488 | LY6H | 4,105216984 | 17,21049854 | 0,00063777 | 0,012799522 | lymphocyte antigen 6 family member H |
| ENSG00000164690 | SHH | 4,104322918 | 17,19983617 | 8,68E-07 | 3,73E-05 | sonic hedgehog signaling molecule |
| ENSG00000163132 | MSX1 | 4,102221029 | 17,17479565 | 8,15E-38 | 6,70E-35 | msh homeobox 1 |
| ENSG00000268941 | LINC01711 | 4,100889825 | 17,15895543 | 0,02646965 | 0,227584468 | long intergenic non-protein coding RNA 1711 |
| ENSG00000052850 | ALX4 | 4,098940818 | 17,13579022 | 2,13E-13 | 2,58E-11 | ALX homeobox 4 |
| ENSG00000221869 | CEBPD | 4,096521463 | 17,1070781 | 4,00E-11 | 3,60E-09 | CCAAT enhancer binding protein delta |
| ENSG00000278788 | SBNO2 | 4,095095422 | 17,09017687 | 0,01453902 | 0,150812147 | strawberry notch homolog 2 |
| ENSG00000151468 | CCDC3 | 4,095059085 | 17,08974642 | 2,64E-25 | 8,78E-23 | coiled-coil domain containing 3 |
| ENSG00000232287 | SLC6A1-AS1 | 4,082591135 | 16,94269105 | 0,0184062 | 0,176657259 | SLC6A1 antisense RNA 1 |
| ENSG00000100097 | LGALS1 | 4,075441267 | 16,85893231 | 3,50E-26 | 1,24E-23 | galectin 1 |
| ENSG00000238279 | BX470102,1 | 4,074678365 | 16,8500196 | 0,02097954 | 0,192913017 | novel transcript |
| ENSG00000223532 | HLA-B | 4,065942691 | 16,7482992 | 8,00E-09 | 5,02E-07 | major histocompatibility complex, class I, B |
| ENSG00000150556 | LYPD6B | 4,052401459 | 16,59183398 | 1,15E-06 | 4,80E-05 | LY6/PLAUR domain containing 6B |
| ENSG00000116745 | RPE65 | 4,042319473 | 16,47628947 | 9,42E-12 | 9,26E-10 | retinoid isomerohydrolase RPE65 |
| ENSG00000169245 | CXCL10 | 4,040638206 | 16,45709976 | 0,00658322 | 0,08476847 | C-X-C motif chemokine ligand 10 |
| ENSG00000260517 | AC009093,2 | 4,035081564 | 16,39383599 | 6,77E-06 | 0,000242456 | novel transcript |
| ENSG00000253819 | LINC01151 | 4,033947498 | 16,38095427 | 2,53E-05 | 0,000775337 | long intergenic non-protein coding RNA 1151 |
| ENSG00000248498 | ASNSP1 | 4,033230205 | 16,37281185 | 0,00050364 | 0,010455365 | asparagine synthetase pseudogene 1 |
| ENSG00000198336 | MYL4 | 4,02563328 | 16,28682271 | 3,23E-08 | 1,84E-06 | myosin light chain 4 |
| ENSG00000226409 | AL450332,1 | 4,025394902 | 16,28413184 | 0,01943432 | 0,18350785 | novel transcript |
| ENSG00000223481 | TAP2 | 4,024876481 | 16,27828131 | 0,00122169 | 0,022001737 | transporter 2, ATP binding cassette subfamily B member |
| ENSG00000225967 | TAP2 | 4,024876481 | 16,27828131 | 0,00122169 | 0,022001737 | transporter 2, ATP binding cassette subfamily B member |
| ENSG00000099260 | PALMD | 4,019275265 | 16,21520396 | 2,00E-08 | 1,19E-06 | palmdelphin |
| ENSG00000164379 | FOXQ1 | 4,01440048 | 16,16050616 | 0,02061629 | 0,190694601 | forkhead box Q1 |
| ENSG00000165078 | CPA6 | 4,009730098 | 16,10827495 | 0,0195155 | 0,183719497 | carboxypeptidase A6 |

| | | | | | | |
|---|---|---|---|---|---|---|
| ENSG00000159618 | ADGRG5 | 4,009159093 | 16,10190071 | 0,01969817 | 0,18466078 | adhesion G protein-coupled receptor G5 |
| ENSG00000261040 | WFDC21P | 4,001483047 | 16,01645597 | 0,01942548 | 0,183479837 | WAP four-disulfide core domain 21, pseudogene |
| ENSG00000134853 | PDGFRA | 3,996115894 | 15,95698182 | 1,84E-16 | 3,07E-14 | platelet derived growth factor receptor alpha |
| ENSG00000172349 | IL16 | 3,991304151 | 15,90385001 | 5,05E-12 | 5,16E-10 | interleukin 16 |
| ENSG00000186795 | KCNK18 | 3,986723223 | 15,85343128 | 0,03209146 | 0,258851206 | potassium two pore domain channel subfamily K member 18 |
| ENSG00000237556 | KCND3-AS1 | 3,981927429 | 15,80081895 | 0,02913878 | 0,24236233 | KCND3 antisense RNA 1 |
| ENSG00000115165 | CYTIP | 3,977408857 | 15,75140764 | 0,02763328 | 0,234127782 | cytohesin 1 interacting protein |
| ENSG00000129354 | AP1M2 | 3,974202472 | 15,71643906 | 0,00391082 | 0,057755276 | adaptor related protein complex 1 subunit mu 2 |
| ENSG00000197461 | PDGFA | 3,973505915 | 15,70885273 | 6,09E-13 | 7,01E-11 | platelet derived growth factor subunit A |
| ENSG00000185112 | FAM43A | 3,972199441 | 15,69463357 | 1,03E-12 | 1,15E-10 | family with sequence similarity 43 member A |
| ENSG00000102007 | PLP2 | 3,968752231 | 15,65717721 | 3,89E-17 | 6,99E-15 | proteolipid protein 2 |
| ENSG00000050555 | LAMC3 | 3,968021092 | 15,64924436 | 2,33E-24 | 7,27E-22 | laminin subunit gamma 3 |
| ENSG00000111341 | MGP | 3,967365416 | 15,64213371 | 3,35E-07 | 1,57E-05 | matrix Gla protein |
| ENSG00000165935 | SMCO2 | 3,964705635 | 15,61332213 | 0,04670818 | 0,328080099 | single-pass membrane protein with coiled-coil domains 2 |
| ENSG00000180801 | ARSJ | 3,961962028 | 15,58365813 | 1,24E-19 | 2,74E-17 | arylsulfatase family member J |
| ENSG00000147041 | SYTL5 | 3,953221618 | 15,48953165 | 2,88E-13 | 3,46E-11 | synaptotagmin like 5 |
| ENSG00000165810 | BTNL9 | 3,953065922 | 15,48786012 | 1,93E-10 | 1,61E-08 | butyrophilin like 9 |
| ENSG00000087494 | PTHLH | 3,95126203 | 15,46850678 | 1,24E-13 | 1,54E-11 | parathyroid hormone like hormone |
| ENSG00000104413 | ESRP1 | 3,945381516 | 15,40558451 | 0,00745437 | 0,092849989 | epithelial splicing regulatory protein 1 |
| ENSG00000102802 | MEDAG | 3,940286537 | 15,35127455 | 1,52E-05 | 0,000490666 | mesenteric estrogen dependent adipogenesis |
| ENSG00000172819 | RARG | 3,937508108 | 15,3217386 | 1,03E-16 | 1,77E-14 | retinoic acid receptor gamma |
| ENSG00000128510 | CPA4 | 3,929913341 | 15,24129244 | 4,01E-13 | 4,73E-11 | carboxypeptidase A4 |
| ENSG00000280090 | OR8B4 | 3,929620236 | 15,23819625 | 0,02819281 | 0,237451497 | olfactory receptor family 8 subfamily B member 4 |
| ENSG00000171840 | NINJ2 | 3,929233419 | 15,23411112 | 0,00146814 | 0,025800885 | ninjurin 2 |
| ENSG00000136542 | GALNT5 | 3,928702267 | 15,22850346 | 3,52E-12 | 3,63E-10 | polypeptide N-acetylgalactosaminyltransferase 5 |
| ENSG00000135312 | HTR1B | 3,921608039 | 15,15380349 | 1,05E-17 | 1,97E-15 | 5-hydroxytryptamine receptor 1B |
| ENSG00000234710 | AC060834,2 | 3,91378461 | 15,07185003 | 3,89E-12 | 4,00E-10 | novel transcript |
| ENSG00000170801 | HTRA3 | 3,909041776 | 15,02238296 | 2,23E-28 | 8,81E-26 | HtrA serine peptidase 3 |
| ENSG00000216316 | AL022722,1 | 3,908890111 | 15,02080379 | 0,03101235 | 0,253151429 | 60S Ribosomal Protein L13A (RPL13) pseudogene |
| ENSG00000131620 | ANO1 | 3,904100641 | 14,97102029 | 1,76E-09 | 1,26E-07 | anoctamin 1 |
| ENSG00000258017 | AC011603,2 | 3,902257886 | 14,95191001 | 0,01071277 | 0,120476276 | novel transcript, antisense to TUBA1B |

| | | | | | | |
|---|---|---|---|---|---|---|
| ENSG00000185585 | OLFML2A | 3,893272159 | 14,85907248 | 7,80E-19 | 1,61E-16 | olfactomedin like 2A |
| ENSG00000287134 | AC106845,1 | 3,883646299 | 14,76026071 | 3,25E-06 | 0,000126662 | novel transcript, antisense to DLC1 |
| ENSG00000286785 | LINC01622 | 3,872435606 | 14,64600822 | 0,02621053 | 0,226054077 | long intergenic non-protein coding RNA 1622 |
| ENSG00000230937 | MIR205HG | 3,86557434 | 14,57651913 | 0,00437268 | 0,062681157 | MIR205 host gene |
| ENSG00000189120 | SP6 | 3,859652089 | 14,51680528 | 0,00026762 | 0,006107179 | Sp6 transcription factor |
| ENSG00000262352 | LINC02564 | 3,858855678 | 14,50879378 | 0,00392172 | 0,057834323 | long intergenic non-protein coding RNA 2564 |
| ENSG00000253563 | NKX2-1-AS1 | 3,856718633 | 14,48731801 | 0,00369198 | 0,055148628 | NKX2-1 antisense RNA 1 |
| ENSG00000224608 | HLA-B | 3,853591651 | 14,45595135 | 2,56E-06 | 0,000101851 | major histocompatibility complex, class I, B |
| ENSG00000198771 | RCSD1 | 3,853452872 | 14,45456084 | 2,76E-21 | 6,85E-19 | RCSD domain containing 1 |
| ENSG00000096696 | DSP | 3,853125786 | 14,45128409 | 4,94E-18 | 9,66E-16 | desmoplakin |
| ENSG00000136574 | GATA4 | 3,84982719 | 14,41828024 | 0,03432065 | 0,270005245 | GATA binding protein 4 |
| ENSG00000128284 | APOL3 | 3,848157787 | 14,40160589 | 9,56E-05 | 0,002522632 | apolipoprotein L3 |
| ENSG00000204118 | NAP1L6P | 3,84717268 | 14,39177548 | 1,06E-11 | 1,03E-09 | nucleosome assembly protein 1 like 6, pseudogene |
| ENSG00000267601 | AC022966,1 | 3,833723891 | 14,25823887 | 0,01349179 | 0,142214228 | novel transcript, antisense to TIMP2 |
| ENSG00000181195 | PENK | 3,832870296 | 14,24980524 | 5,43E-07 | 2,43E-05 | proenkephalin |
| ENSG00000217416 | ISCA1P1 | 3,823518853 | 14,15773776 | 0,28103572 | 0,791915358 | iron-sulfur cluster assembly 1 pseudogene 1 |
| ENSG00000124813 | RUNX2 | 3,823459185 | 14,15715223 | 4,05E-32 | 2,22E-29 | RUNX family transcription factor 2 |
| ENSG00000125148 | MT2A | 3,821492642 | 14,1378677 | 3,10E-13 | 3,71E-11 | metallothionein 2A |
| ENSG00000168830 | HTR1E | 3,821403687 | 14,13699601 | 0,0030134 | 0,046615966 | 5-hydroxytryptamine receptor 1E |
| ENSG00000138207 | RBP4 | 3,820489957 | 14,12804519 | 0,03940919 | 0,295677671 | retinol binding protein 4 |
| ENSG00000226816 | AC005082,1 | 3,818037748 | 14,10405155 | 6,90E-07 | 3,03E-05 | novel transcript |
| ENSG00000137507 | LRRC32 | 3,817818015 | 14,10190357 | 1,81E-15 | 2,74E-13 | leucine rich repeat containing 32 |
| ENSG00000279116 | OR8D2 | 3,815668518 | 14,08090854 | 0,00271533 | 0,042702264 | olfactory receptor family 8 subfamily D member 2 |
| ENSG00000224417 | AL606970,1 | 3,814309183 | 14,06764748 | 0,00849977 | 0,101736905 | novel transcript |
| ENSG00000230479 | AP000695,1 | 3,807302474 | 13,99949105 | 0,07327002 | 0,427132959 | novel transcript |
| ENSG00000154493 | C10orf90 | 3,805225623 | 13,9793524 | 0,00525875 | 0,071638184 | chromosome 10 open reading frame 90 |
| ENSG00000197355 | UAP1L1 | 3,804105559 | 13,96850347 | 2,09E-06 | 8,43E-05 | UDP-N-acetylglucosamine pyrophosphorylase 1 like 1 |
| ENSG00000160307 | S100B | 3,802133842 | 13,94942589 | 5,10E-07 | 2,30E-05 | S100 calcium binding protein B |
| ENSG00000124749 | COL21A1 | 3,795323859 | 13,88373529 | 5,91E-06 | 0,000213775 | collagen type XXI alpha 1 chain |
| ENSG00000168582 | CRYGA | 3,793760904 | 13,86870239 | 0,04370367 | 0,315756651 | crystallin gamma A |
| ENSG00000253706 | AC011632,1 | 3,79262426 | 13,85778007 | 4,44E-07 | 2,03E-05 | novel transcript, antisense to PI15 |
| ENSG00000266446 | AL449423,1 | 3,791160709 | 13,84372909 | 0,01723829 | 0,169264441 | novel transcript, sense intronic CDKN2B-AS1 |

| | | | | | | |
|---|---|---|---|---|---|---|
| ENSG00000163815 | CLEC3B | 3,784294388 | 13,77799819 | 0,07701068 | 0,438408786 | C-type lectin domain family 3 member B |
| ENSG00000159167 | STC1 | 3,780063685 | 13,73765339 | 2,70E-37 | 2,06E-34 | stanniocalcin 1 |
| ENSG00000278608 | MAMDC2 | 3,774686213 | 13,68654326 | 0,00056126 | 0,011521434 | MAM domain containing 2 |
| ENSG00000113578 | FGF1 | 3,773729875 | 13,67747369 | 1,71E-15 | 2,64E-13 | fibroblast growth factor 1 |
| ENSG00000118322 | ATP10B | 3,772678074 | 13,66750572 | 1,72E-07 | 8,74E-06 | ATPase phospholipid transporting 10B (putative) |
| ENSG00000065717 | TLE2 | 3,771912424 | 13,66025419 | 1,72E-07 | 8,74E-06 | TLE family member 2, transcriptional corepressor |
| ENSG00000275210 | AC245884,11 | 3,767673884 | 13,62018019 | 0,02692022 | 0,229957863 | novel transcript |
| ENSG00000213809 | KLRK1 | 3,766156114 | 13,60585877 | 0,00024303 | 0,005624148 | killer cell lectin like receptor K1 |
| ENSG00000261257 | AP000821,1 | 3,760125838 | 13,54910675 | 0,12557028 | 0,566089307 | novel transcript |
| ENSG00000235931 | LINC01553 | 3,759104517 | 13,53951839 | 0,0097651 | 0,113030801 | long intergenic non-protein coding RNA 1553 |
| ENSG00000247193 | AC104078,1 | 3,754555811 | 13,49689656 | 0,00051912 | 0,010748217 | novel transcript, antisense to CENTD1 |
| ENSG00000019549 | SNAI2 | 3,752960719 | 13,48198218 | 7,63E-17 | 1,33E-14 | snail family transcriptional repressor 2 |
| ENSG00000263257 | AC040173,1 | 3,752548165 | 13,47812742 | 0,07238505 | 0,424120881 | novel transcript |
| ENSG00000187498 | COL4A1 | 3,74226987 | 13,38244548 | 3,87E-09 | 2,58E-07 | collagen type IV alpha 1 chain |
| ENSG00000286318 | AC096576,7 | 3,742134006 | 13,38118527 | 3,92E-06 | 0,000149517 | novel transcript, sense intronic to KCNIP4 |
| ENSG00000287263 | AC008875,3 | 3,739617098 | 13,35786096 | 1,30E-10 | 1,10E-08 | novel transcript |
| ENSG00000164318 | EGFLAM | 3,739367726 | 13,35555224 | 4,32E-23 | 1,22E-20 | EGF like, fibronectin type III and laminin G domains |
| ENSG00000238097 | LINC02037 | 3,732959958 | 13,29636474 | 0,00683577 | 0,087222253 | long intergenic non-protein coding RNA 2037 |
| ENSG00000269586 | CT45A10 | 3,732814984 | 13,29502867 | 0,00430117 | 0,062025765 | cancer/testis antigen family 45 member A10 |
| ENSG00000283307 | OR5AK4P | 3,725510547 | 13,22788541 | 0,00616647 | 0,080698433 | olfactory receptor family 5 subfamily AK member 4 pseudogene |
| ENSG00000175084 | DES | 3,722277056 | 13,19827114 | 3,16E-10 | 2,55E-08 | desmin |
| ENSG00000116016 | EPAS1 | 3,722176325 | 13,19734965 | 4,38E-17 | 7,78E-15 | endothelial PAS domain protein 1 |
| ENSG00000185885 | IFITM1 | 3,720856367 | 13,18528058 | 3,43E-13 | 4,09E-11 | interferon induced transmembrane protein 1 |
| ENSG00000143520 | FLG2 | 3,711544864 | 13,10045363 | 0,03198797 | 0,258149711 | filaggrin family member 2 |
| ENSG00000167105 | TMEM92 | 3,711168682 | 13,09703814 | 2,85E-06 | 0,00011207 | transmembrane protein 92 |
| ENSG00000279141 | LINC01451 | 3,711058292 | 13,09603604 | 0,00093725 | 0,017705294 | long intergenic non-protein coding RNA 1451 |
| ENSG00000235387 | SPAAR | 3,710999115 | 13,09549887 | 3,75E-05 | 0,001095964 | small regulatory polypeptide of amino acid response |
| ENSG00000218728 | KRT18P44 | 3,710303766 | 13,08918864 | 0,04464943 | 0,319246353 | keratin 18 pseudogene 44 |
| ENSG00000186340 | THBS2 | 3,704908489 | 13,04033014 | 6,19E-12 | 6,22E-10 | thrombospondin 2 |
| ENSG00000116996 | ZP4 | 3,702358801 | 13,0173042 | 0,0016651 | 0,028729433 | zona pellucida glycoprotein 4 |
| ENSG00000109047 | RCVRN | 3,696371891 | 12,96339679 | 0,00822898 | 0,099666531 | recoverin |
| ENSG00000229415 | SFTA3 | 3,694209263 | 12,94397897 | 1,85E-07 | 9,23E-06 | surfactant associated 3 |

| | | | | | | |
|---|---|---|---|---|---|---|
| ENSG00000132481 | TRIM47 | 3,69311593 | 12,93417321 | 6,80E-06 | 0,000243082 | tripartite motif containing 47 |
| ENSG00000275287 | Metazoa_SRP | 3,693048084 | 12,93356496 | 0,10942091 | 0,527649315 | Metazoan signal recognition particle RNA |
| ENSG00000196639 | HRH1 | 3,691349316 | 12,9183447 | 5,00E-07 | 2,26E-05 | histamine receptor H1 |
| ENSG00000255225 | OR8B6P | 3,689473001 | 12,90155451 | 0,04100785 | 0,303331616 | olfactory receptor family 8 subfamily B member 6 pseudogene |
| ENSG00000197442 | MAP3K5 | 3,687709281 | 12,88579177 | 2,19E-09 | 1,53E-07 | mitogen-activated protein kinase kinase kinase 5 |
| ENSG00000204362 | LINC02783 | 3,670651221 | 12,73433062 | 0,04125851 | 0,304151397 | long intergenic non-protein coding RNA 2783 |
| ENSG00000281226 | LINC02783 | 3,670651221 | 12,73433062 | 0,04125851 | 0,304151397 | long intergenic non-protein coding RNA 2783 |
| ENSG00000231631 | PSMB8 | 3,669187822 | 12,72142008 | 3,26E-05 | 0,000969404 | proteasome 20S subunit beta 8 |
| ENSG00000260862 | AC099506,1 | 3,663125467 | 12,66807551 | 0,02632376 | 0,22653539 | novel transcript |
| ENSG00000157227 | MMP14 | 3,660325848 | 12,64351634 | 2,06E-14 | 2,75E-12 | matrix metallopeptidase 14 |
| ENSG00000281942 | AC068473,6 | 3,655514327 | 12,60141929 | 0,0911205 | 0,481283407 | novel transcript |
| ENSG00000137491 | SLCO2B1 | 3,652220777 | 12,57268413 | 3,44E-08 | 1,96E-06 | solute carrier organic anion transporter family member 2B1 |
| ENSG00000204248 | COL11A2 | 3,651742813 | 12,5685195 | 0,0031784 | 0,048806308 | collagen type XI alpha 2 chain |
| ENSG00000282697 | AC073488,11 | 3,650348488 | 12,55637824 | 0,03954014 | 0,296276117 | chromodomain protein, Y chromosome-like (CDYL) pseudogene |
| ENSG00000230148 | HOXB-AS1 | 3,635611596 | 12,42876977 | 1,85E-05 | 0,000586923 | HOXB cluster antisense RNA 1 |
| ENSG00000186895 | FGF3 | 3,632879051 | 12,40525126 | 0,06626027 | 0,40452277 | fibroblast growth factor 3 |
| ENSG00000198796 | ALPK2 | 3,627343822 | 12,35774687 | 4,45E-28 | 1,70E-25 | alpha kinase 2 |
| ENSG00000226037 | LINC01040 | 3,627310563 | 12,35746198 | 0,05750801 | 0,370798119 | long intergenic non-protein coding RNA 1040 |
| ENSG00000168404 | MLKL | 3,625111677 | 12,33864169 | 3,77E-13 | 4,46E-11 | mixed lineage kinase domain like pseudokinase |
| ENSG00000168621 | GDNF | 3,624368687 | 12,3322889 | 1,42E-07 | 7,40E-06 | glial cell derived neurotrophic factor |
| ENSG00000256824 | AP000721,2 | 3,622292932 | 12,31455792 | 0,04130977 | 0,304452496 | novel transcript, antisense to MACROD1 |
| ENSG00000277734 | TRAC | 3,61792342 | 12,27731706 | 7,48E-08 | 4,09E-06 | T cell receptor alpha constant |
| ENSG00000085741 | WNT11 | 3,616444423 | 12,26473726 | 9,69E-06 | 0,000332501 | Wnt family member 11 |
| ENSG00000163191 | S100A11 | 3,615677966 | 12,25822314 | 1,46E-35 | 9,71E-33 | S100 calcium binding protein A11 |
| ENSG00000176971 | FIBIN | 3,610515597 | 12,21443814 | 4,74E-20 | 1,08E-17 | fin bud initiation factor homolog |
| ENSG00000101825 | MXRA5 | 3,609241913 | 12,20365938 | 8,95E-44 | 1,04E-40 | matrix remodeling associated 5 |
| ENSG00000163131 | CTSS | 3,608251144 | 12,19528141 | 5,14E-07 | 2,31E-05 | cathepsin S |
| ENSG00000219992 | AL391422,1 | 3,604813185 | 12,16625452 | 0,14759216 | 0,611779236 | FUN14 domain containing 1 (FUNDC1) pseudogene |
| ENSG00000197646 | PDCD1LG2 | 3,604599255 | 12,16445058 | 3,22E-15 | 4,68E-13 | programmed cell death 1 ligand 2 |
| ENSG00000102034 | ELF4 | 3,604457019 | 12,16325133 | 2,86E-25 | 9,41E-23 | E74 like ETS transcription factor 4 |

| | | | | | | |
|---|---|---|---|---|---|---|
| ENSG00000177359 | AC024940,1 | 3,601891693 | 12,14164248 | 2,62E-05 | 0,000800485 | ovostatin 2 |
| ENSG00000235978 | AC018816,1 | 3,597823505 | 12,10745306 | 0,05254238 | 0,351978793 | novel transcript, antisense to ITPR1 |
| ENSG00000260592 | AC130456,3 | 3,577785248 | 11,94044954 | 0,0340767 | 0,268965908 | novel transcript, antisense to TMC5 |
| ENSG00000227120 | AC009238,1 | 3,576982603 | 11,9338083 | 0,04810201 | 0,334042943 | pseudogene similar to part of Cdon homolog (mouse) (CDON) |
| ENSG00000235959 | AC009237,6 | 3,576982603 | 11,9338083 | 0,04810201 | 0,334042943 | pseudogene similar to part of Cdon homolog (mouse) (CDON) |
| ENSG00000163814 | CDCP1 | 3,574119031 | 11,91014466 | 4,07E-06 | 0,000153996 | CUB domain containing protein 1 |
| ENSG00000270442 | AC008725,1 | 3,57383892 | 11,90783243 | 0,0526138 | 0,352155381 | peptidylprolyl isomerase G (cyclophilin G) (PPIG) pseudogene |
| ENSG00000080007 | DDX43 | 3,573534757 | 11,90532217 | 0,04467585 | 0,319329187 | DEAD-box helicase 43 |
| ENSG00000235649 | MXRA5Y | 3,571032761 | 11,88469324 | 9,13E-13 | 1,03E-10 | matrix remodeling associated 5 Y-linked (pseudogene) |
| ENSG00000107485 | GATA3 | 3,569284209 | 11,87029768 | 1,87E-08 | 1,12E-06 | GATA binding protein 3 |
| ENSG00000170891 | CYTL1 | 3,568909704 | 11,8672167 | 3,06E-11 | 2,83E-09 | cytokine like 1 |
| ENSG00000234889 | KPNB1P1 | 3,560723214 | 11,80006755 | 0,08646768 | 0,4687188 | KPNB1 pseudogene 1 |
| ENSG00000236765 | DXO | 3,555208468 | 11,75504753 | 0,04610053 | 0,325793402 | decapping exoribonuclease |
| ENSG00000176438 | SYNE3 | 3,552250232 | 11,73096859 | 4,89E-08 | 2,73E-06 | spectrin repeat containing nuclear envelope family member 3 |
| ENSG00000102755 | FLT1 | 3,547381403 | 11,69144547 | 3,45E-13 | 4,10E-11 | fms related receptor tyrosine kinase 1 |
| ENSG00000231574 | LINC02015 | 3,54726487 | 11,69050113 | 0,05991739 | 0,380820743 | long intergenic non-protein coding RNA 2015 |
| ENSG00000196569 | LAMA2 | 3,537944953 | 11,61522305 | 9,20E-14 | 1,15E-11 | laminin subunit alpha 2 |
| ENSG00000286648 | AC068931,1 | 3,537682588 | 11,61311092 | 0,00273615 | 0,042987437 | novel transcript, antisense to NRG1 |
| ENSG00000159231 | CBR3 | 3,536966939 | 11,60735167 | 8,62E-07 | 3,71E-05 | carbonyl reductase 3 |
| ENSG00000231981 | RPL7L1P12 | 3,536556084 | 11,60404656 | 0,09742036 | 0,495829213 | ribosomal protein L7 like 1 pseudogene 12 |
| ENSG00000142156 | COL6A1 | 3,53440348 | 11,5867454 | 0,00177183 | 0,030303261 | collagen type VI alpha 1 chain |
| ENSG00000288539 | AC022133,2 | 3,534189066 | 11,58502351 | 9,60E-05 | 0,002526223 | novel transcript |
| ENSG00000235941 | HSPA1A | 3,529959176 | 11,5511067 | 5,71E-11 | 5,04E-09 | heat shock protein family A (Hsp70) member 1A |
| ENSG00000230294 | LINC02370 | 3,525980744 | 11,5192968 | 0,10347602 | 0,513096613 | long intergenic non-protein coding RNA 2370 |
| ENSG00000283079 | LINC01515 | 3,524727307 | 11,50929299 | 1,53E-23 | 4,48E-21 | long intergenic non-protein coding RNA 1515 |
| ENSG00000238005 | AL391832,2 | 3,52458854 | 11,50818601 | 0,00879346 | 0,104096465 | novel transcript |
| ENSG00000286963 | AC074096,1 | 3,523305871 | 11,49795888 | 0,07097963 | 0,419165813 | novel transcript, antisense to ALK |
| ENSG00000260482 | AC008870,1 | 3,522624949 | 11,49253337 | 0,00079066 | 0,015306011 | novel transcript |

| | | | | | | |
|---|---|---|---|---|---|---|
| ENSG00000273686 | B2M | 3,52044298 | 11,4751649 | 0,00847204 | 0,101599441 | beta-2-microglobulin |
| ENSG00000058335 | RASGRF1 | 3,519694759 | 11,4692151 | 1,47E-09 | 1,08E-07 | Ras protein specific guanine nucleotide releasing factor 1 |
| ENSG00000162576 | MXRA8 | 3,516591275 | 11,44456938 | 2,21E-14 | 2,94E-12 | matrix remodeling associated 8 |
| ENSG00000128340 | RAC2 | 3,515417322 | 11,43526046 | 8,17E-06 | 0,000284636 | Rac family small GTPase 2 |
| ENSG00000136155 | SCEL | 3,514533373 | 11,42825615 | 0,09706519 | 0,494806155 | sciellin |
| ENSG00000288234 | BX255925,4 | 3,513601819 | 11,42087926 | 0,06694624 | 0,40664606 | novel transcript, antisense to NRARP |
| ENSG00000138395 | CDK15 | 3,513037144 | 11,41640997 | 1,35E-05 | 0,000441514 | cyclin dependent kinase 15 |
| ENSG00000160712 | IL6R | 3,510329871 | 11,39500673 | 6,18E-11 | 5,44E-09 | interleukin 6 receptor |
| ENSG00000285550 | AC110014,1 | 3,503969927 | 11,3448838 | 0,00675524 | 0,086413965 | novel transcript |
| ENSG00000136352 | NKX2-1 | 3,503903109 | 11,34435838 | 4,03E-11 | 3,62E-09 | NK2 homeobox 1 |
| ENSG00000286546 | AL079338,1 | 3,502791309 | 11,33561932 | 1,28E-12 | 1,41E-10 | novel transcript, antisense to MACROD2 |
| ENSG00000236700 | LINC01010 | 3,500113425 | 11,31459802 | 0,09915324 | 0,499826992 | long intergenic non-protein coding RNA 1010 |
| ENSG00000215895 | HSPA5P1 | 3,497055754 | 11,29064307 | 0,05704795 | 0,36911008 | HSPA5 pseudogene 1 |
| ENSG00000159450 | TCHH | 3,496862321 | 11,28912935 | 0,00022483 | 0,005273175 | trichohyalin |
| ENSG00000078401 | EDN1 | 3,494938735 | 11,27408727 | 9,19E-13 | 1,03E-10 | endothelin 1 |
| ENSG00000179046 | TRIML2 | 3,488253416 | 11,22196502 | 1,21E-05 | 0,000399808 | tripartite motif family like 2 |
| ENSG00000101425 | BPI | 3,487919535 | 11,21936824 | 0,01023031 | 0,117029035 | bactericidal permeability increasing protein |
| ENSG00000287712 | AC005865,2 | 3,487704601 | 11,21769689 | 0,06071604 | 0,38355752 | novel transcript |
| ENSG00000121743 | GJA3 | 3,481205049 | 11,1672732 | 0,01189433 | 0,129840887 | gap junction protein alpha 3 |
| ENSG00000286010 | AC108925,1 | 3,481054225 | 11,16610579 | 0,05972161 | 0,38023321 | novel transcript |
| ENSG00000254605 | AP003555,1 | 3,478829406 | 11,14889951 | 0,12749675 | 0,570260369 | novel transcript |
| ENSG00000110675 | ELMOD1 | 3,477776488 | 11,14076571 | 2,04E-08 | 1,21E-06 | ELMO domain containing 1 |
| ENSG00000187140 | FOXD3 | 3,47604913 | 11,12743471 | 0,10138781 | 0,506329296 | forkhead box D3 |
| ENSG00000165606 | DRGX | 3,459740537 | 11,00235564 | 0,00080903 | 0,015574903 | dorsal root ganglia homeobox |
| ENSG00000075429 | CACNG5 | 3,457567131 | 10,98579317 | 4,22E-09 | 2,79E-07 | calcium voltage-gated channel auxiliary subunit gamma 5 |
| ENSG00000282501 | AC073488,9 | 3,454379805 | 10,96154921 | 0,10886982 | 0,526415221 | chromodomain protein, Y chromosome-like (CDYL) pseudogene |
| ENSG00000087116 | ADAMTS2 | 3,453541695 | 10,95518312 | 1,36E-10 | 1,15E-08 | ADAM metallopeptidase with thrombospondin type 1 motif 2 |
| ENSG00000251321 | PCAT4 | 3,449378348 | 10,92361409 | 0,11553749 | 0,543019233 | prostate cancer associated transcript 4 |
| ENSG00000256022 | LINC02411 | 3,445641839 | 10,89535908 | 0,02365605 | 0,210168399 | long intergenic non-protein coding RNA 2411 |

| | | | | | | |
|---|---|---|---|---|---|---|
| ENSG00000038945 | MSR1 | 3,441476033 | 10,86394393 | 0,00446949 | 0,063517074 | macrophage scavenger receptor 1 |
| ENSG00000138772 | ANXA3 | 3,440783261 | 10,85872838 | 1,46E-17 | 2,69E-15 | annexin A3 |
| ENSG00000149591 | TAGLN | 3,439043098 | 10,8456386 | 1,78E-15 | 2,72E-13 | transgelin |
| ENSG00000188459 | WASF4P | 3,4363747 | 10,82559713 | 0,00848358 | 0,101620975 | WASP family member 4, pseudogene |
| ENSG00000225691 | HLA-C | 3,435378776 | 10,81812656 | 3,44E-09 | 2,34E-07 | major histocompatibility complex, class I, C |
| ENSG00000112175 | BMP5 | 3,435322433 | 10,81770408 | 1,46E-11 | 1,40E-09 | bone morphogenetic protein 5 |
| ENSG00000286194 | AC138207,9 | 3,431562879 | 10,78955066 | 0,06012487 | 0,381318641 | Novel transcript |
| ENSG00000133980 | VRTN | 3,429968671 | 10,77763457 | 0,00634648 | 0,082228016 | vertebrae development associated |
| ENSG00000171956 | FOXB1 | 3,418755883 | 10,69419425 | 3,09E-12 | 3,21E-10 | forkhead box B1 |
| ENSG00000090104 | RGS1 | 3,418640113 | 10,69333613 | 4,48E-07 | 2,04E-05 | regulator of G protein signaling 1 |
| ENSG00000054598 | FOXC1 | 3,414620057 | 10,66358073 | 7,22E-20 | 1,64E-17 | forkhead box C1 |
| ENSG00000228299 | HLA-C | 3,41407855 | 10,65957896 | 3,24E-07 | 1,53E-05 | major histocompatibility complex, class I, C |
| ENSG00000286153 | AP000331,1 | 3,412416633 | 10,6473067 | 7,59E-05 | 0,0020588 | novel transcript, antisense to RUNX1 |
| ENSG00000257809 | AC034102,5 | 3,410912433 | 10,63621126 | 0,11117657 | 0,532530151 | novel transcript, antisense to MYL6B |
| ENSG00000116774 | OLFML3 | 3,409079098 | 10,62270366 | 9,03E-29 | 3,81E-26 | olfactomedin like 3 |
| ENSG00000286423 | AC004817,5 | 3,408363994 | 10,61743958 | 0,07492395 | 0,432580317 | novel transcript |
| ENSG00000251129 | LINC02506 | 3,407779385 | 10,61313805 | 4,30E-05 | 0,001236416 | long intergenic non-protein coding RNA 2506 |
| ENSG00000187608 | ISG15 | 3,404586326 | 10,5896744 | 5,36E-10 | 4,22E-08 | ISG15 ubiquitin like modifier |
| ENSG00000273633 | CDC42EP5 | 3,399342626 | 10,55125444 | 7,42E-05 | 0,002018603 | CDC42 effector protein 5 |
| ENSG00000223749 | MIR503HG | 3,393690494 | 10,50999803 | 3,04E-20 | 7,09E-18 | MIR503 host gene |
| ENSG00000283802 | ADAMTS2 | 3,393258992 | 10,50685502 | 3,83E-11 | 3,48E-09 | ADAM metallopeptidase with thrombospondin type 1 motif 2 |
| ENSG00000124942 | AHNAK | 3,392800923 | 10,50351953 | 7,11E-12 | 7,12E-10 | AHNAK nucleoprotein |
| ENSG00000282540 | WHAMMP2 | 3,386860258 | 10,46035753 | 9,38E-11 | 8,08E-09 | WAS protein homolog associated with actin, golgi membranes and microtubules pseudogene 2 |
| ENSG00000166289 | PLEKHF1 | 3,385927067 | 10,45359356 | 2,49E-12 | 2,62E-10 | pleckstrin homology and FYVE domain containing 1 |
| ENSG00000276600 | RAB7B | 3,382426002 | 10,42825601 | 1,28E-12 | 1,41E-10 | RAB7B, member RAS oncogene family |
| ENSG00000170965 | PLAC1 | 3,380899452 | 10,41722746 | 0,01484918 | 0,153014099 | placenta enriched 1 |
| ENSG00000250243 | AC110296,1 | 3,379913709 | 10,41011217 | 0,06114135 | 0,384884976 | novel transcript |
| ENSG00000285520 | AL353586,1 | 3,378426832 | 10,39938878 | 0,07976105 | 0,447773275 | novel transcript, antisense to CCDC3 |
| ENSG00000285283 | AL035078,4 | 3,374246698 | 10,3693007 | 0,27988786 | 0,790565637 | novel protein |
| ENSG00000006377 | DLX6 | 3,37029509 | 10,34093757 | 6,47E-16 | 1,03E-13 | distal-less homeobox 6 |
| ENSG00000235085 | AC091153,2 | 3,369616839 | 10,33607715 | 0,06504122 | 0,400285189 | novel transcript, antisense to PELP1 |

| | | | | | | |
|---|---|---|---|---|---|---|
| ENSG00000134545 | KLRC1 | 3,366936492 | 10,31689184 | 0,08564908 | 0,465935763 | killer cell lectin like receptor C1 |
| ENSG00000259727 | AC103740,2 | 3,361045388 | 10,27484972 | 1,74E-07 | 8,81E-06 | novel transcript |
| ENSG00000118640 | VAMP8 | 3,358090639 | 10,25382758 | 1,55E-07 | 8,00E-06 | vesicle associated membrane protein 8 |
| ENSG00000255974 | CYP2A6 | 3,353485026 | 10,22114579 | 0,08684989 | 0,469592252 | cytochrome P450 family 2 subfamily A member 6 |
| ENSG00000276950 | GSTT4 | 3,352995867 | 10,2176808 | 0,13597123 | 0,587841041 | glutathione S-transferase theta 4 |
| ENSG00000164393 | ADGRF2 | 3,349189964 | 10,19076156 | 0,00023204 | 0,005415695 | adhesion G protein-coupled receptor F2 |
| ENSG00000012504 | NR1H4 | 3,346628107 | 10,17268144 | 0,11562348 | 0,543151268 | nuclear receptor subfamily 1 group H member 4 |
| ENSG00000286486 | AL590483,4 | 3,343240918 | 10,14882582 | 0,08737578 | 0,47120083 | novel transcript |
| ENSG00000130303 | BST2 | 3,342389385 | 10,14283737 | 3,55E-06 | 0,000136713 | bone marrow stromal cell antigen 2 |
| ENSG00000128342 | LIF | 3,341237951 | 10,13474547 | 0,00011002 | 0,002847281 | LIF interleukin 6 family cytokine |
| ENSG00000186910 | SERPINA11 | 3,339730554 | 10,12416173 | 0,11634867 | 0,544539657 | serpin family A member 11 |
| ENSG00000257042 | AC008011,2 | 3,338465181 | 10,11528582 | 0,0500979 | 0,342277237 | novel transcript, antisense to PTHLH |
| ENSG00000286123 | AL049828,2 | 3,338393827 | 10,11478554 | 8,68E-06 | 0,000300511 | novel transcript |
| ENSG00000232935 | AL359094,2 | 3,336299548 | 10,10011313 | 0,21358636 | 0,721033693 | novel transcript, antisense to DOCK1 |
| ENSG00000287136 | AC073578,5 | 3,331421116 | 10,06601757 | 0,12678622 | 0,568863994 | novel transcript |
| ENSG00000222033 | LINC01124 | 3,33006442 | 10,05655603 | 0,07778474 | 0,440668732 | long intergenic non-protein coding RNA 1124 |
| ENSG00000242807 | RPL26P31 | 3,317292729 | 9,967921652 | 0,11263463 | 0,536050184 | ribosomal protein L26 pseudogene 31 |
| ENSG00000075426 | FOSL2 | 3,316403112 | 9,961776975 | 1,25E-14 | 1,73E-12 | FOS like 2, AP-1 transcription factor subunit |
| ENSG00000126353 | CCR7 | 3,31292628 | 9,937798429 | 0,00422165 | 0,061132803 | C-C motif chemokine receptor 7 |
| ENSG00000134817 | APLNR | 3,310541371 | 9,921383901 | 4,21E-09 | 2,79E-07 | apelin receptor |
| ENSG00000196867 | ZFP28 | 3,306105639 | 9,890926311 | 1,27E-14 | 1,75E-12 | ZFP28 zinc finger protein |
| ENSG00000125378 | BMP4 | 3,305651952 | 9,887816377 | 6,25E-17 | 1,10E-14 | bone morphogenetic protein 4 |
| ENSG00000264501 | RN7SL731P | 3,298566904 | 9,839376541 | 0,13579022 | 0,587728418 | RNA, 7SL, cytoplasmic 731, pseudogene |
| ENSG00000237834 | MOG | 3,297115027 | 9,829479525 | 0,05619244 | 0,366305368 | myelin oligodendrocyte glycoprotein |
| ENSG00000278212 | AC134878,2 | 3,297090471 | 9,829312217 | 1,01E-05 | 0,000343651 | MAFF interacting protein (pseudogene) |
| ENSG00000106546 | AHR | 3,296505333 | 9,825326386 | 2,59E-20 | 6,09E-18 | aryl hydrocarbon receptor |
| ENSG00000226890 | Z97652,1 | 3,29167203 | 9,792464753 | 0,09460141 | 0,488518255 | novel transcript, antisense to CRAMP1L |
| ENSG00000124097 | HMGB1P1 | 3,288097489 | 9,768232159 | 0,01687709 | 0,166554751 | high mobility group box 1 pseudogene 1 |
| ENSG00000169248 | CXCL11 | 3,275725445 | 9,684821407 | 0,03391745 | 0,267979594 | C-X-C motif chemokine ligand 11 |
| ENSG00000229132 | EIF4A1P10 | 3,274812488 | 9,67869466 | 0,00522657 | 0,071261833 | eukaryotic translation initiation factor 4A1 pseudogene 10 |
| ENSG00000189058 | APOD | 3,271135351 | 9,654057042 | 0,01217114 | 0,131987045 | apolipoprotein D |
| ENSG00000159216 | RUNX1 | 3,270945886 | 9,652789284 | 2,08E-13 | 2,53E-11 | RUNX family transcription factor 1 |

| | | | | | | |
|---|---|---|---|---|---|---|
| ENSG00000237870 | AC073130,1 | 3,270220317 | 9,647935866 | 0,00370392 | 0,055300539 | uncharacterized LOC102724434 |
| ENSG00000136826 | KLF4 | 3,265550852 | 9,616759582 | 5,94E-08 | 3,29E-06 | Kruppel like factor 4 |
| ENSG00000204291 | COL15A1 | 3,265549135 | 9,616748134 | 3,95E-12 | 4,05E-10 | collagen type XV alpha 1 chain |
| ENSG00000248905 | FMN1 | 3,263494291 | 9,603060665 | 2,86E-23 | 8,20E-21 | formin 1 |
| ENSG00000123689 | G0S2 | 3,262859955 | 9,598839239 | 0,00632978 | 0,082213892 | G0/G1 switch 2 |
| ENSG00000253522 | MIR3142HG | 3,261386617 | 9,589041519 | 0,01626776 | 0,162750485 | MIR3142 host gene |
| ENSG00000281758 | KCNIP4 | 3,261138283 | 9,587391082 | 0,04622905 | 0,326327599 | potassium voltage-gated channel interacting protein 4 |
| ENSG00000103335 | PIEZO1 | 3,258664642 | 9,57096665 | 1,07E-07 | 5,71E-06 | piezo type mechanosensitive ion channel component 1 |
| ENSG00000138119 | MYOF | 3,256023639 | 9,55346203 | 1,20E-22 | 3,28E-20 | myoferlin |
| ENSG00000287303 | AP006248,5 | 3,254596422 | 9,544015735 | 0,01060678 | 0,119758169 | novel transcript, antisense to PDGFRL |
| ENSG00000234247 | DNAJB6P6 | 3,251288577 | 9,52215806 | 0,03264669 | 0,261468653 | DNAJB6 pseudogene 6 |
| ENSG00000168398 | BDKRB2 | 3,249159433 | 9,50811553 | 0,00015282 | 0,003761959 | bradykinin receptor B2 |
| ENSG00000126458 | RRAS | 3,247449803 | 9,496854847 | 1,89E-15 | 2,85E-13 | RAS related |
| ENSG00000185338 | SOCS1 | 3,246017303 | 9,487429781 | 1,66E-12 | 1,78E-10 | suppressor of cytokine signaling 1 |
| ENSG00000231826 | LINC01819 | 3,241672617 | 9,458901298 | 0,00249213 | 0,039914879 | long intergenic non-protein coding RNA 1819 |
| ENSG00000241186 | TDGF1 | 3,239530258 | 9,44486554 | 0,00089804 | 0,017046879 | teratocarcinoma-derived growth factor 1 |
| ENSG00000225028 | AC096541,1 | 3,236294251 | 9,423704176 | 0,00443673 | 0,063222453 | novel transcript |
| ENSG00000092969 | TGFB2 | 3,233634141 | 9,406344304 | 2,27E-13 | 2,74E-11 | transforming growth factor beta 2 |
| ENSG00000253953 | PCDHGB4 | 3,232322169 | 9,39779416 | 8,75E-33 | 5,11E-30 | protocadherin gamma subfamily B, 4 |
| ENSG00000277152 | AC110048,2 | 3,229650039 | 9,380403874 | 4,36E-07 | 1,99E-05 | novel transcript |
| ENSG00000043143 | JADE2 | 3,226499429 | 9,359940955 | 1,67E-07 | 8,57E-06 | jade family PHD finger 2 |
| ENSG00000133454 | MYO18B | 3,225396529 | 9,352788277 | 9,10E-08 | 4,90E-06 | myosin XVIIIB |
| ENSG00000225437 | PACRG-AS2 | 3,221917509 | 9,330261451 | 0,05237763 | 0,351288432 | PACRG antisense RNA 2 |
| ENSG00000284969 | AL049629,2 | 3,218488921 | 9,308114262 | 0,35943069 | 0,849982153 | novel protein |
| ENSG00000043462 | LCP2 | 3,213464115 | 9,275751143 | 0,00215976 | 0,035783057 | lymphocyte cytosolic protein 2 |
| ENSG00000226674 | TEX41 | 3,211133772 | 9,260780387 | 6,03E-10 | 4,70E-08 | testis expressed 41 |
| ENSG00000164125 | GASK1B | 3,200683611 | 9,193942297 | 9,57E-15 | 1,34E-12 | golgi associated kinase 1B |
| ENSG00000090339 | ICAM1 | 3,197372203 | 9,172863702 | 7,59E-09 | 4,83E-07 | intercellular adhesion molecule 1 |
| ENSG00000099869 | IGF2-AS | 3,195179026 | 9,158929758 | 0,00650651 | 0,083953588 | IGF2 antisense RNA |
| ENSG00000148848 | ADAM12 | 3,18933256 | 9,121888655 | 9,08E-20 | 2,04E-17 | ADAM metallopeptidase domain 12 |

| | | | | | | |
|---|---|---|---|---|---|---|
| ENSG00000260255 | AC093524,1 | 3,188832365 | 9,11872656 | 0,00794696 | 0,097242152 | TNF receptor-associated factor 7 (TRAF7) pseudogene |
| ENSG00000215512 | AP005901,1 | 3,188490243 | 9,116564398 | 0,0045165 | 0,064016873 | ankyrin repeat domain 30A (ANKRD30A) pseudogene |
| ENSG00000169885 | CALML6 | 3,185223791 | 9,095946639 | 0,03150436 | 0,255765345 | calmodulin like 6 |
| ENSG00000229970 | AC007128,1 | 3,184913958 | 9,093993407 | 1,04E-07 | 5,57E-06 | novel transcript, antisense to ICA1 |
| ENSG00000091136 | LAMB1 | 3,181825657 | 9,074547194 | 5,44E-25 | 1,77E-22 | laminin subunit beta 1 |
| ENSG00000088320 | REM1 | 3,178303493 | 9,052419816 | 0,07518383 | 0,433200749 | RRAD and GEM like GTPase 1 |
| ENSG00000276832 | CDRT15P6 | 3,177732599 | 9,048838362 | 0,11226238 | 0,535189241 | CDRT15 pseudogene 6 |
| ENSG00000224318 | CHL1-AS2 | 3,174456913 | 9,028315987 | 0,00219875 | 0,036256173 | CHL1 antisense RNA 2 |
| ENSG00000137033 | IL33 | 3,173471507 | 9,022151471 | 0,09188296 | 0,481984693 | interleukin 33 |
| ENSG00000110852 | CLEC2B | 3,170118742 | 9,0012087 | 0,00229436 | 0,037437843 | C-type lectin domain family 2 member B |
| ENSG00000197956 | S100A6 | 3,169143984 | 8,995129076 | 1,17E-09 | 8,65E-08 | S100 calcium binding protein A6 |
| ENSG00000275719 | AC008622,2 | 3,167800829 | 8,986758471 | 0,05506215 | 0,362489606 | novel transcript, antisense to SLC1A5 |
| ENSG00000225614 | ZNF469 | 3,166886246 | 8,981063209 | 1,45E-05 | 0,000469043 | zinc finger protein 469 |
| ENSG00000261189 | AL031058,1 | 3,162457833 | 8,953537726 | 1,06E-06 | 4,49E-05 | novel transcript, antisense to DSP |
| ENSG00000139211 | AMIGO2 | 3,161157394 | 8,945470682 | 3,28E-12 | 3,39E-10 | adhesion molecule with Ig like domain 2 |
| ENSG00000273622 | CDC42EP5 | 3,159945722 | 8,937960829 | 3,24E-05 | 0,000966403 | CDC42 effector protein 5 |
| ENSG00000231179 | MICB | 3,158608855 | 8,929682341 | 0,00035086 | 0,007727191 | MHC class I polypeptide-related sequence B |
| ENSG00000204516 | MICB | 3,158608855 | 8,929682341 | 0,00035086 | 0,007727191 | MHC class I polypeptide-related sequence B |
| ENSG00000240890 | AC020633,1 | 3,158171981 | 8,926978682 | 0,02097451 | 0,192913017 | G2/M-phase specific E3 ubiquitin ligase (G2E3) pseudogene |
| ENSG00000260228 | AC009119,1 | 3,155427859 | 8,91001499 | 1,20E-08 | 7,37E-07 | novel transcript, antisense to CDH13 |
| ENSG00000234393 | AL592546,2 | 3,149288803 | 8,872181029 | 0,01930802 | 0,182701508 | novel transcript |
| ENSG00000271857 | AL096865,1 | 3,147671375 | 8,862239865 | 2,98E-05 | 0,000892973 | novel transcript, antisense to RUNX2 |
| ENSG00000237586 | AC090192,1 | 3,146168122 | 8,853010437 | 0,02738534 | 0,232946937 | SRY (sex determining region Y)-box 2 (SOX2) pseudogene |
| ENSG00000115107 | STEAP3 | 3,136142082 | 8,791699537 | 1,04E-05 | 0,00035215 | STEAP3 metalloreductase |
| ENSG00000258998 | LINC02302 | 3,134431766 | 8,781283146 | 0,00576375 | 0,07654964 | long intergenic non-protein coding RNA 2302 |
| ENSG00000267765 | AC100793,3 | 3,131639261 | 8,764302386 | 0,05406557 | 0,35836452 | novel transcript, antisense to CNTNAP1 |
| ENSG00000178860 | MSC | 3,130292204 | 8,756122897 | 0,00018147 | 0,004374244 | musculin |
| ENSG00000244242 | IFITM10 | 3,127246101 | 8,737654747 | 0,00082877 | 0,015915568 | interferon induced transmembrane protein 10 |
| ENSG00000231453 | LINC01305 | 3,123942014 | 8,717666501 | 0,00019038 | 0,004578374 | long intergenic non-protein coding RNA 1305 |

| | | | | | | |
|---|---|---|---|---|---|---|
| ENSG00000288614 | AL031178,1 | 3,12032908 | 8,695862209 | 0,1060396 | 0,519559347 | novel protein |
| ENSG00000007908 | SELE | 3,113433027 | 8,654395347 | 0,04038357 | 0,300178807 | selectin E |
| ENSG00000234745 | HLA-B | 3,113190992 | 8,652943558 | 0,01017029 | 0,11653537 | major histocompatibility complex, class I, B |
| ENSG00000250131 | AC078881,1 | 3,112978442 | 8,651668828 | 1,19E-16 | 2,03E-14 | novel transcript |
| ENSG00000223611 | SUPT20HL2 | 3,110254403 | 8,635348504 | 0,0108923 | 0,121925143 | SUPT20H like 2 |
| ENSG00000134531 | EMP1 | 3,101274324 | 8,581764576 | 7,25E-13 | 8,28E-11 | epithelial membrane protein 1 |
| ENSG00000285882 | AC093516,1 | 3,10082864 | 8,579113866 | 0,01221799 | 0,132357286 | novel transcript, sense intronic to HS3ST4 |
| ENSG00000128274 | A4GALT | 3,096860805 | 8,555551202 | 1,72E-05 | 0,000551234 | alpha 1,4-galactosyltransferase (P blood group) |
| ENSG00000196468 | FGF16 | 3,095473694 | 8,547329233 | 0,07052984 | 0,418185447 | fibroblast growth factor 16 |
| ENSG00000171621 | SPSB1 | 3,095365398 | 8,54668765 | 6,57E-16 | 1,04E-13 | splA/ryanodine receptor domain and SOCS box containing 1 |
| ENSG00000271167 | LINC01109 | 3,095086165 | 8,545033604 | 0,00880524 | 0,104157077 | long intergenic non-protein coding RNA 1109 |
| ENSG00000130720 | FIBCD1 | 3,092835329 | 8,53171237 | 0,00040321 | 0,008675774 | fibrinogen C domain containing 1 |
| ENSG00000101276 | SLC52A3 | 3,092183625 | 8,527859237 | 0,00928198 | 0,108564101 | solute carrier family 52 member 3 |
| ENSG00000138435 | CHRNA1 | 3,091527131 | 8,523979545 | 5,36E-06 | 0,000197085 | cholinergic receptor nicotinic alpha 1 subunit |
| ENSG00000228065 | LINC01515 | 3,090931973 | 8,520463853 | 2,55E-20 | 6,04E-18 | long intergenic non-protein coding RNA 1515 |
| ENSG00000006116 | CACNG3 | 3,088185007 | 8,504255886 | 0,03339079 | 0,265327213 | calcium voltage-gated channel auxiliary subunit gamma 3 |
| ENSG00000249413 | AC116049,1 | 3,083960261 | 8,479388702 | 1,07E-06 | 4,50E-05 | novel transcript |
| ENSG00000258829 | AC026888,1 | 3,081163848 | 8,462968798 | 0,03018013 | 0,248045051 | novel transcript, antisense to NRXN3 |
| ENSG00000158270 | COLEC12 | 3,080979153 | 8,461885431 | 1,79E-26 | 6,44E-24 | collectin subfamily member 12 |
| ENSG00000233854 | POU6F2-AS2 | 3,078784948 | 8,449025472 | 0,00031707 | 0,007100888 | POU6F2 antisense RNA 2 |
| ENSG00000261742 | LINC00922 | 3,075031369 | 8,427071518 | 0,05153235 | 0,347838733 | long intergenic non-protein coding RNA 922 |
| ENSG00000198768 | APCDD1L | 3,072722862 | 8,41359785 | 0,00029199 | 0,006600644 | APC down-regulated 1 like |
| ENSG00000163762 | TM4SF18 | 3,071443535 | 8,406140294 | 1,04E-11 | 1,02E-09 | transmembrane 4 L six family member 18 |
| ENSG00000111087 | GLI1 | 3,069886279 | 8,397071541 | 1,20E-07 | 6,34E-06 | GLI family zinc finger 1 |
| ENSG00000230705 | TAP1 | 3,060878867 | 8,344808065 | 0,00020537 | 0,004886423 | transporter 1, ATP binding cassette subfamily B member |
| ENSG00000285838 | AL646090,2 | 3,060106032 | 8,340339045 | 0,01468384 | 0,15171041 | novel transcript |
| ENSG00000145536 | ADAMTS16 | 3,053566178 | 8,302617172 | 1,58E-11 | 1,51E-09 | ADAM metallopeptidase with thrombospondin type 1 motif 16 |
| ENSG00000152192 | POU4F1 | 3,050429236 | 8,284583881 | 1,24E-06 | 5,20E-05 | POU class 4 homeobox 1 |
| ENSG00000172986 | GXYLT2 | 3,049817184 | 8,28106996 | 5,19E-17 | 9,16E-15 | glucoside xylosyltransferase 2 |

| | | | | | | |
|---|---|---|---|---|---|---|
| ENSG00000075035 | WSCD2 | 3,049589845 | 8,27976514 | 1,44E-05 | 0,000467085 | WSC domain containing 2 |
| ENSG00000173917 | HOXB2 | 3,048380777 | 8,272829087 | 1,22E-07 | 6,46E-06 | homeobox B2 |
| ENSG00000136235 | GPNMB | 3,047155401 | 8,26580541 | 1,55E-08 | 9,37E-07 | glycoprotein nmb |
| ENSG00000112561 | TFEB | 3,045202086 | 8,254621625 | 6,39E-10 | 4,92E-08 | transcription factor EB |
| ENSG00000134640 | MTNR1B | 3,044699724 | 8,251747775 | 0,03890808 | 0,293396543 | melatonin receptor 1B |
| ENSG00000113083 | LOX | 3,044540689 | 8,250838198 | 3,70E-28 | 1,45E-25 | lysyl oxidase |
| ENSG00000168394 | TAP1 | 3,043909582 | 8,247229656 | 4,54E-06 | 0,000169377 | transporter 1, ATP binding cassette subfamily B member |
| ENSG00000280916 | FOXCUT | 3,039180435 | 8,220239538 | 0,01048783 | 0,118761539 | FOXC1 upstream transcript |
| ENSG00000169933 | FRMPD4 | 3,038413329 | 8,215869859 | 0,01497683 | 0,154075559 | FERM and PDZ domain containing 4 |
| ENSG00000274330 | AL160191,3 | 3,038040187 | 8,213745161 | 0,00225563 | 0,036960162 | ADAM metallopeptidase domain 20 (ADAM20) pseudogene |
| ENSG00000218672 | AC008060,1 | 3,03646356 | 8,204773801 | 6,89E-06 | 0,000246065 | novel transcript |
| ENSG00000257605 | MYG1-AS1 | 3,035466134 | 8,199103284 | 0,06959704 | 0,416357046 | MYG1 antisense RNA 1 |
| ENSG00000258815 | LINC02820 | 3,034420536 | 8,193163107 | 0,00022707 | 0,005320387 | long intergenic non-protein coding RNA 2820 |
| ENSG00000185774 | KCNIP4 | 3,034131003 | 8,191518993 | 1,83E-05 | 0,000581728 | potassium voltage-gated channel interacting protein 4 |
| ENSG00000105976 | MET | 3,031637628 | 8,177374019 | 3,29E-16 | 5,39E-14 | MET proto-oncogene, receptor tyrosine kinase |
| ENSG00000150361 | KLHL1 | 3,030185289 | 8,169146122 | 0,00054087 | 0,011152047 | kelch like family member 1 |
| ENSG00000134240 | HMGCS2 | 3,029329235 | 8,164300221 | 0,03169558 | 0,256717776 | 3-hydroxy-3-methylglutaryl-CoA synthase 2 |
| ENSG00000160219 | GAB3 | 3,028943454 | 8,162117354 | 5,43E-06 | 0,000199188 | GRB2 associated binding protein 3 |
| ENSG00000164509 | IL31RA | 3,028227596 | 8,158068359 | 3,67E-08 | 2,08E-06 | interleukin 31 receptor A |
| ENSG00000256654 | AC005906,2 | 3,022134294 | 8,123684971 | 0,00889036 | 0,105084407 | novel transcript, sense overlapping KCNA1 |
| ENSG00000274529 | SEBOX | 3,021007022 | 8,117339883 | 0,098246 | 0,497580255 | SEBOX homeobox |
| ENSG00000108691 | CCL2 | 3,020510768 | 8,114548185 | 2,38E-09 | 1,65E-07 | C-C motif chemokine ligand 2 |
| ENSG00000167757 | KLK11 | 3,018879382 | 8,105377516 | 0,07923873 | 0,446444361 | kallikrein related peptidase 11 |
| ENSG00000224982 | TMEM233 | 3,018514462 | 8,103327573 | 2,34E-05 | 0,000723898 | transmembrane protein 233 |
| ENSG00000268818 | AL049747,1 | 3,017849251 | 8,09959208 | 0,05702764 | 0,369056527 | novel transcript |
| ENSG00000124019 | FAM124B | 3,015885246 | 8,088573244 | 0,03110761 | 0,253664123 | family with sequence similarity 124 member B |
| ENSG00000274092 | AC106739,1 | 3,014138642 | 8,078786707 | 0,01757602 | 0,171664029 | novel transcript, antisense to IL4R |
| ENSG00000259863 | SH3RF3-AS1 | 3,014124936 | 8,078709959 | 0,00086888 | 0,016533644 | SH3RF3 antisense RNA 1 |
| ENSG00000245870 | LINC00682 | 3,012486466 | 8,069540168 | 0,12146984 | 0,556776403 | long intergenic non-protein coding RNA 682 |
| ENSG00000267120 | AD000671,2 | 3,012024258 | 8,066955278 | 0,01363494 | 0,143481291 | novel transcript |

| | | | | | | |
|---|---|---|---|---|---|---|
| ENSG00000261127 | AC133548,2 | 3,011856189 | 8,066015563 | 0,01031143 | 0,11765573 | protein phosphatase 2, regulatory subunit B'', beta (PPP2R3B) pseudogene |
| ENSG00000253687 | AC008456,1 | 3,011432333 | 8,06364616 | 0,06835256 | 0,411986588 | novel transcript |
| ENSG00000274354 | AC105105,3 | 3,010120767 | 8,056318762 | 2,30E-05 | 0,000714612 | novel transcript, sense intronic to ALPK2 |
| ENSG00000185404 | SP140L | 3,007665488 | 8,042619617 | 1,75E-15 | 2,69E-13 | SP140 nuclear body protein like |
| ENSG00000243349 | AL358394,2 | 3,004449389 | 8,024710736 | 0,05501041 | 0,362301551 | novel transcript |
| ENSG00000156466 | GDF6 | 3,003652544 | 8,020279664 | 1,29E-09 | 9,52E-08 | growth differentiation factor 6 |
| ENSG00000288175 | FO681492,2 | 3,003257821 | 8,01808561 | 0,27205515 | 0,78507773 | synaptotagmin-15 |
| ENSG00000166033 | HTRA1 | 2,997773073 | 7,987660821 | 4,44E-09 | 2,92E-07 | HtrA serine peptidase 1 |
| ENSG00000287278 | AL353600,2 | 2,99711684 | 7,984028331 | 0,0119867 | 0,130575624 | novel transcript |
| ENSG00000164220 | F2RL2 | 2,996367638 | 7,979883243 | 6,42E-16 | 1,03E-13 | coagulation factor II thrombin receptor like 2 |
| ENSG00000268089 | GABRQ | 2,993877687 | 7,966122618 | 1,44E-06 | 5,98E-05 | gamma-aminobutyric acid type A receptor subunit theta |
| ENSG00000270504 | AL391422,4 | 2,992748414 | 7,959889558 | 1,79E-06 | 7,26E-05 | novel transcript, antisense to PXDC1 |
| ENSG00000250420 | AACSP1 | 2,989740198 | 7,943309397 | 3,11E-05 | 0,000928763 | acetoacetyl-CoA synthetase pseudogene 1 |
| ENSG00000240508 | PSMB9 | 2,98714401 | 7,929027945 | 0,04843738 | 0,335106518 | proteasome 20S subunit beta 9 |
| ENSG00000242711 | PSMB9 | 2,98714401 | 7,929027945 | 0,04843738 | 0,335106518 | proteasome 20S subunit beta 9 |
| ENSG00000243958 | PSMB9 | 2,98714401 | 7,929027945 | 0,04843738 | 0,335106518 | proteasome 20S subunit beta 9 |
| ENSG00000137345 | MOG | 2,982802004 | 7,905200227 | 0,01116316 | 0,124068672 | myelin oligodendrocyte glycoprotein |
| ENSG00000286537 | AL512655,1 | 2,97763089 | 7,876915996 | 0,01770056 | 0,172556988 | novel transcript |
| ENSG00000277044 | OPRL1 | 2,975395775 | 7,864722025 | 0,09807657 | 0,497366739 | opioid related nociceptin receptor 1 |
| ENSG00000177699 | AC011944,1 | 2,97449709 | 7,859824452 | 0,00296002 | 0,045915666 | novel transcript, antisense to RASGRF1 |
| ENSG00000120820 | GLT8D2 | 2,97266096 | 7,84982755 | 5,19E-14 | 6,68E-12 | glycosyltransferase 8 domain containing 2 |
| ENSG00000165912 | PACSIN3 | 2,970655535 | 7,83892344 | 6,69E-05 | 0,001835492 | protein kinase C and casein kinase substrate in neurons 3 |
| ENSG00000221946 | FXYD7 | 2,969783937 | 7,834189019 | 8,95E-06 | 0,00030941 | FXYD domain containing ion transport regulator 7 |
| ENSG00000013725 | CD6 | 2,965080712 | 7,808690934 | 0,00145601 | 0,025616606 | CD6 molecule |
| ENSG00000159674 | SPON2 | 2,961161155 | 7,787504841 | 7,57E-06 | 0,00026753 | spondin 2 |
| ENSG00000008517 | IL32 | 2,958119595 | 7,771104141 | 0,00025902 | 0,005936939 | interleukin 32 |
| ENSG00000286502 | AL645768,1 | 2,957345544 | 7,766935814 | 5,63E-05 | 0,001581867 | novel transcript, antisense to COL5A1 |
| ENSG00000285547 | AL133500,1 | 2,956515816 | 7,762470152 | 0,00436522 | 0,062622653 | novel protein |
| ENSG00000053438 | NNAT | 2,955358529 | 7,756245827 | 5,24E-14 | 6,71E-12 | neuronatin |
| ENSG00000135046 | ANXA1 | 2,953759882 | 7,747655906 | 4,97E-32 | 2,63E-29 | annexin A1 |

| | | | | | | |
|---|---|---|---|---|---|---|
| ENSG00000229056 | HECW2-AS1 | 2,95293469 | 7,743225677 | 1,03E-06 | 4,37E-05 | HECW2 antisense RNA 1 |
| ENSG00000255004 | SLC1A2-AS1 | 2,950860337 | 7,732100221 | 0,03735498 | 0,285332555 | SLC1A2 antisense RNA 1 |
| ENSG00000184058 | TBX1 | 2,948797819 | 7,721054084 | 0,00078321 | 0,015171084 | T-box transcription factor 1 |
| ENSG00000260995 | AL512634,1 | 2,941163226 | 7,680302984 | 0,03469942 | 0,271754193 | novel transcript |
| ENSG00000164129 | NPY5R | 2,939844103 | 7,673283739 | 5,77E-05 | 0,001611277 | neuropeptide Y receptor Y5 |
| ENSG00000223382 | LINC01778 | 2,939356334 | 7,670689872 | 0,02281725 | 0,204988034 | long intergenic non-protein coding RNA 1778 |
| ENSG00000228459 | LINC01546 | 2,936215397 | 7,654007937 | 6,95E-07 | 3,05E-05 | long intergenic non-protein coding RNA 1546 |
| ENSG00000109846 | CRYAB | 2,93403958 | 7,642473159 | 0,01025139 | 0,117184454 | crystallin alpha B |
| ENSG00000176256 | HMGB4 | 2,933924981 | 7,641866111 | 0,039959 | 0,298370746 | high mobility group box 4 |
| ENSG00000204577 | LILRB3 | 2,933474553 | 7,639480592 | 0,04431478 | 0,31783836 | leukocyte immunoglobulin like receptor B3 |
| ENSG00000250722 | SELENOP | 2,931658106 | 7,629868042 | 2,83E-07 | 1,35E-05 | selenoprotein P |
| ENSG00000171234 | UGT2B7 | 2,931656583 | 7,629859987 | 0,000867 | 0,016517977 | UDP glucuronosyltransferase family 2 member B7 |
| ENSG00000240389 | EGFL8 | 2,923690189 | 7,5878449 | 0,00823116 | 0,099666531 | EGF like domain multiple 8 |
| ENSG00000134259 | NGF | 2,921973713 | 7,578822473 | 0,00454101 | 0,064298056 | nerve growth factor |
| ENSG00000113580 | NR3C1 | 2,919834063 | 7,567590711 | 2,33E-15 | 3,45E-13 | nuclear receptor subfamily 3 group C member 1 |
| ENSG00000185837 | HDHD5-AS1 | 2,918331219 | 7,559711714 | 0,10364809 | 0,513379913 | HDHD5 antisense RNA 1 |
| ENSG00000227496 | AC099066,2 | 2,918050247 | 7,558239566 | 0,00122945 | 0,022128683 | novel transcript |
| ENSG00000110719 | TCIRG1 | 2,917679608 | 7,55629805 | 5,16E-07 | 2,32E-05 | T cell immune regulator 1, ATPase H+ transporting V0 subunit a3 |
| ENSG00000196549 | MME | 2,917062262 | 7,553065309 | 1,66E-16 | 2,79E-14 | membrane metalloendopeptidase |
| ENSG00000165694 | FRMD7 | 2,912329322 | 7,528327147 | 0,00011024 | 0,002850725 | FERM domain containing 7 |
| ENSG00000107317 | PTGDS | 2,907793845 | 7,504697108 | 0,05198654 | 0,349829381 | prostaglandin D2 synthase |
| ENSG00000253161 | LINC01605 | 2,907239007 | 7,501811472 | 0,10424808 | 0,514802081 | long intergenic non-protein coding RNA 1605 |
| ENSG00000183378 | OVCH2 | 2,906602627 | 7,498503113 | 0,01102531 | 0,123085791 | ovochymase 2 |
| ENSG00000231437 | LINC01750 | 2,906025 | 7,49550146 | 0,00079959 | 0,015421594 | long intergenic non-protein coding RNA 1750 |
| ENSG00000105971 | CAV2 | 2,903620523 | 7,48301944 | 2,05E-24 | 6,55E-22 | caveolin 2 |
| ENSG00000145623 | OSMR | 2,903583578 | 7,482827816 | 9,18E-13 | 1,03E-10 | oncostatin M receptor |
| ENSG00000227365 | AC008060,2 | 2,902099648 | 7,475135072 | 0,00621575 | 0,081207299 | novel transcript, antisense to CNPY1 |
| ENSG00000133135 | RNF128 | 2,898723648 | 7,457663218 | 0,00065102 | 0,012972276 | ring finger protein 128 |
| ENSG00000277641 | WNT3 | 2,898682111 | 7,457448509 | 0,03523473 | 0,27443414 | Wnt family member 3 |
| ENSG00000122859 | NEUROG3 | 2,895740277 | 7,442257341 | 0,00234129 | 0,037987956 | neurogenin 3 |
| ENSG00000113389 | NPR3 | 2,894762316 | 7,437214159 | 1,03E-05 | 0,000350403 | natriuretic peptide receptor 3 |
| ENSG00000250509 | AC034213,1 | 2,893903064 | 7,43278596 | 0,03917016 | 0,294869635 | novel transcript |

| | | | | | | |
|---|---|---|---|---|---|---|
| ENSG00000156510 | HKDC1 | 2,89351907 | 7,43080788 | 0,00575247 | 0,076464763 | hexokinase domain containing 1 |
| ENSG00000170953 | OR8B12 | 2,893042334 | 7,428352791 | 0,06110629 | 0,384884976 | olfactory receptor family 8 subfamily B member 12 |
| ENSG00000150048 | CLEC1A | 2,891713771 | 7,421515244 | 0,00120273 | 0,021773145 | C-type lectin domain family 1 member A |
| ENSG00000260426 | AC008060,4 | 2,889930485 | 7,412347328 | 9,71E-05 | 0,002550558 | novel transcript, overlapping novel protein |
| ENSG00000275070 | Metazoa_SRP | 2,889192997 | 7,408559198 | 0,11373479 | 0,53883988 | Metazoan signal recognition particle RNA |
| ENSG00000154229 | PRKCA | 2,888881412 | 7,406959311 | 9,82E-23 | 2,72E-20 | protein kinase C alpha |
| ENSG00000225953 | SATB2-AS1 | 2,885778195 | 7,391044165 | 0,00074544 | 0,014547671 | SATB2 antisense RNA 1 |
| ENSG00000237599 | TAP2 | 2,885745897 | 7,390878705 | 0,08350491 | 0,460196964 | transporter 2, ATP binding cassette subfamily B member |
| ENSG00000206297 | TAP1 | 2,885172835 | 7,387943512 | 3,80E-05 | 0,001108776 | transporter 1, ATP binding cassette subfamily B member |
| ENSG00000167779 | IGFBP6 | 2,881162039 | 7,367433018 | 4,24E-06 | 0,000159062 | insulin like growth factor binding protein 6 |
| ENSG00000009709 | PAX7 | 2,881038035 | 7,366799793 | 3,24E-08 | 1,85E-06 | paired box 7 |
| ENSG00000162947 | LINC01931 | 2,880491356 | 7,364008829 | 0,07214899 | 0,423602654 | long intergenic non-protein coding RNA 1931 |
| ENSG00000182853 | VMO1 | 2,879515623 | 7,359030037 | 0,05521317 | 0,362784685 | vitelline membrane outer layer 1 homolog |
| ENSG00000259132 | AL132780,3 | 2,878178603 | 7,352213202 | 0,46754188 | 0,907940653 | novel protein |
| ENSG00000287692 | AC053545,1 | 2,877729797 | 7,349926368 | 0,03181533 | 0,257287766 | novel transcript |
| ENSG00000197125 | OR8B8 | 2,876503408 | 7,343681084 | 0,00020069 | 0,004787687 | olfactory receptor family 8 subfamily B member 8 |
| ENSG00000259417 | CTXND1 | 2,876155849 | 7,341912133 | 0,00035146 | 0,007727191 | cortexin domain containing 1 |
| ENSG00000144655 | CSRNP1 | 2,872046193 | 7,321027752 | 6,38E-10 | 4,92E-08 | cysteine and serine rich nuclear protein 1 |
| ENSG00000157502 | PWWP3B | 2,872035847 | 7,32097525 | 2,25E-11 | 2,10E-09 | PWWP domain containing 3B |
| ENSG00000275300 | MCCC2 | 2,870109707 | 7,311207549 | 0,02725597 | 0,232168734 | methylcrotonoyl-CoA carboxylase 2 |
| ENSG00000284669 | AC092053,3 | 2,866746767 | 7,294184881 | 1,51E-05 | 0,000488879 | novel transcript, antisense CSRNP1 |
| ENSG00000116132 | PRRX1 | 2,863870473 | 7,279656988 | 5,50E-12 | 5,58E-10 | paired related homeobox 1 |
| ENSG00000126785 | RHOJ | 2,859014602 | 7,255196063 | 5,55E-08 | 3,09E-06 | ras homolog family member J |
| ENSG00000224286 | LINC01142 | 2,858901301 | 7,254626307 | 0,07981188 | 0,447817422 | long intergenic non-protein coding RNA 1142 |
| ENSG00000255026 | AC136475,3 | 2,858236212 | 7,251282659 | 0,00909267 | 0,106789387 | novel transcript |
| ENSG00000282435 | AC246817,4 | 2,854065002 | 7,230347575 | 0,01170863 | 0,128601321 | novel transcript |
| ENSG00000227816 | TAP1 | 2,85156689 | 7,217838632 | 0,00023857 | 0,005545523 | transporter 1, ATP binding cassette subfamily B member |
| ENSG00000181273 | OR5AK2 | 2,851366392 | 7,21683561 | 0,05095882 | 0,345820358 | olfactory receptor family 5 subfamily AK member 2 |
| ENSG00000262327 | AGK | 2,850398312 | 7,211994577 | 0,00065501 | 0,013035069 | acylglycerol kinase |
| ENSG00000237022 | HLA-C | 2,846388303 | 7,191976479 | 0,00056395 | 0,011553977 | major histocompatibility complex, class I, C |

| | | | | | | |
|---|---|---|---|---|---|---|
| ENSG00000226171 | DHX16 | 2,845381152 | 7,186957489 | 0,00825395 | 0,099854406 | DEAH-box helicase 16 |
| ENSG00000256637 | LINC01965 | 2,839487777 | 7,157658818 | 2,84E-10 | 2,32E-08 | long intergenic non-protein coding RNA 1965 |
| ENSG00000123146 | ADGRE5 | 2,837905356 | 7,149812238 | 1,43E-08 | 8,74E-07 | adhesion G protein-coupled receptor E5 |
| ENSG00000215089 | KRT18P11 | 2,833596757 | 7,12849123 | 0,00553145 | 0,074508967 | keratin 18 pseudogene 11 |
| ENSG00000177468 | OLIG3 | 2,833007832 | 7,125581887 | 3,48E-07 | 1,62E-05 | oligodendrocyte transcription factor 3 |
| ENSG00000196460 | RFX8 | 2,831308703 | 7,117194699 | 0,00101407 | 0,018939078 | RFX family member 8, lacking RFX DNA binding domain |
| ENSG00000163565 | IFI16 | 2,8309969 | 7,115656657 | 2,30E-08 | 1,35E-06 | interferon gamma inducible protein 16 |
| ENSG00000285621 | AL138999,2 | 2,830901183 | 7,115184577 | 0,05434823 | 0,359759285 | novel transcript, antisense to STARD13 |
| ENSG00000285561 | AP001804,1 | 2,827884882 | 7,100324113 | 0,00028738 | 0,006510718 | novel transcript |
| ENSG00000160789 | LMNA | 2,827318266 | 7,097536023 | 2,75E-05 | 0,000832441 | lamin A/C |
| ENSG00000283849 | AC092053,2 | 2,827284665 | 7,097370721 | 0,05449903 | 0,360376004 | novel transcript, antisense CSRNP1 |
| ENSG00000109625 | CPZ | 2,822346875 | 7,073120675 | 0,00013156 | 0,003327728 | carboxypeptidase Z |
| ENSG00000278109 | Metazoa_SRP | 2,821731334 | 7,070103494 | 0,20079704 | 0,700325598 | Metazoan signal recognition particle RNA |
| ENSG00000136830 | NIBAN2 | 2,821051666 | 7,066773486 | 2,15E-06 | 8,68E-05 | niban apoptosis regulator 2 |
| ENSG00000257702 | LBX2-AS1 | 2,813189685 | 7,028367771 | 0,00072509 | 0,014221686 | LBX2 antisense RNA 1 |
| ENSG00000153071 | DAB2 | 2,812196744 | 7,023532131 | 8,83E-23 | 2,46E-20 | DAB adaptor protein 2 |
| ENSG00000196754 | S100A2 | 2,810180115 | 7,013721351 | 0,01016196 | 0,116502947 | S100 calcium binding protein A2 |
| ENSG00000165424 | ZCCHC24 | 2,809548154 | 7,010649721 | 4,53E-20 | 1,04E-17 | zinc finger CCHC-type containing 24 |
| ENSG00000175592 | FOSL1 | 2,809427504 | 7,010063458 | 0,00012823 | 0,003254008 | FOS like 1, AP-1 transcription factor subunit |
| ENSG00000234702 | VDAC1P3 | 2,809032527 | 7,008144525 | 0,00016445 | 0,004010503 | voltage dependent anion channel 1 pseudogene 3 |
| ENSG00000117152 | RGS4 | 2,807959344 | 7,002933288 | 8,27E-22 | 2,14E-19 | regulator of G protein signaling 4 |
| ENSG00000161544 | CYGB | 2,807932264 | 7,002801841 | 6,81E-07 | 3,00E-05 | cytoglobin |
| ENSG00000266846 | AP001021,3 | 2,80581619 | 6,992538007 | 3,86E-05 | 0,001123841 | novel transcript, antisense to L3MBTL4 |
| ENSG00000160255 | ITGB2 | 2,80464367 | 6,986857277 | 0,10541676 | 0,518119329 | integrin subunit beta 2 |
| ENSG00000230535 | BASP1P1 | 2,800996091 | 6,969214654 | 1,98E-05 | 0,000624413 | BASP1 pseudogene 1 |
| ENSG00000234091 | AL157392,2 | 2,800516312 | 6,966897373 | 0,04613314 | 0,325797698 | novel transcript |
| ENSG00000181577 | C6orf223 | 2,800039057 | 6,96459305 | 0,00075263 | 0,014669638 | chromosome 6 open reading frame 223 |
| ENSG00000123104 | ITPR2 | 2,79864373 | 6,957860381 | 1,11E-18 | 2,26E-16 | inositol 1,4,5-trisphosphate receptor type 2 |
| ENSG00000284461 | AC027644,4 | 2,798268584 | 6,956051354 | 0,00335699 | 0,051022338 | novel transcript |
| ENSG00000162631 | NTNG1 | 2,796126571 | 6,945731159 | 4,54E-08 | 2,54E-06 | netrin G1 |
| ENSG00000130300 | PLVAP | 2,793916614 | 6,935099655 | 0,01473054 | 0,15204212 | plasmalemma vesicle associated protein |
| ENSG00000223501 | VPS52 | 2,789736942 | 6,915036865 | 6,15E-05 | 0,001702946 | VPS52 subunit of GARP complex |

| | | | | | | |
|---|---|---|---|---|---|---|
| ENSG00000179954 | SSC5D | 2,787975715 | 6,906600207 | 1,66E-06 | 6,79E-05 | scavenger receptor cysteine rich family member with 5 domains |
| ENSG00000131015 | ULBP2 | 2,784057212 | 6,88786665 | 0,00046152 | 0,009742752 | UL16 binding protein 2 |
| ENSG00000196990 | FAM163B | 2,776663304 | 6,852656178 | 0,01775361 | 0,172912401 | family with sequence similarity 163 member B |
| ENSG00000280776 | LINC01202 | 2,77653493 | 6,852046445 | 0,08701852 | 0,470194775 | long intergenic non-protein coding RNA 1202 |
| ENSG00000112769 | LAMA4 | 2,775828565 | 6,848692399 | 2,76E-12 | 2,89E-10 | laminin subunit alpha 4 |
| ENSG00000146910 | CNPY1 | 2,773921753 | 6,839646455 | 0,00049279 | 0,01029155 | canopy FGF signaling regulator 1 |
| ENSG00000087589 | CASS4 | 2,773045603 | 6,835493989 | 2,66E-10 | 2,18E-08 | Cas scaffold protein family member 4 |
| ENSG00000006327 | TNFRSF12A | 2,766466857 | 6,804394831 | 4,17E-07 | 1,92E-05 | TNF receptor superfamily member 12A |
| ENSG00000188487 | INSC | 2,766331259 | 6,803755321 | 0,02991867 | 0,2466187 | INSC spindle orientation adaptor protein |
| ENSG00000127528 | KLF2 | 2,766104911 | 6,802687946 | 0,0005969 | 0,012133681 | Kruppel like factor 2 |
| ENSG00000154553 | PDLIM3 | 2,765068147 | 6,797801085 | 8,00E-24 | 2,43E-21 | PDZ and LIM domain 3 |
| ENSG00000183801 | OLFML1 | 2,758813399 | 6,768393279 | 6,07E-09 | 3,90E-07 | olfactomedin like 1 |
| ENSG00000287156 | AL138479,2 | 2,75733115 | 6,76144289 | 0,00073974 | 0,014454445 | novel transcript, antisense to BMP4 |
| ENSG00000205693 | MANSC4 | 2,755231874 | 6,751611421 | 0,00572057 | 0,076202714 | MANSC domain containing 4 |
| ENSG00000263310 | SALL3 | 2,75418017 | 6,746691386 | 5,67E-05 | 0,001591603 | spalt like transcription factor 3 |
| ENSG00000287023 | AC018978,1 | 2,752963177 | 6,741002579 | 0,07697882 | 0,438408786 | novel transcript |
| ENSG00000238107 | LINC02806 | 2,752761009 | 6,740058013 | 0,06279471 | 0,391611013 | long intergenic non-protein coding RNA 2806 |
| ENSG00000250379 | AC020659,2 | 2,750275415 | 6,728455685 | 0,10236645 | 0,509453154 | novel transcript |
| ENSG00000113209 | PCDHB5 | 2,748373546 | 6,71959157 | 4,73E-21 | 1,16E-18 | protocadherin beta 5 |
| ENSG00000164761 | TNFRSF11B | 2,743344338 | 6,696207979 | 9,79E-11 | 8,36E-09 | TNF receptor superfamily member 11b |
| ENSG00000110848 | CD69 | 2,743255456 | 6,695795449 | 0,04382275 | 0,31589834 | CD69 molecule |
| ENSG00000119917 | IFIT3 | 2,742434141 | 6,691984672 | 8,66E-09 | 5,38E-07 | interferon induced protein with tetratricopeptide repeats 3 |
| ENSG00000184374 | COLEC10 | 2,741478386 | 6,687552841 | 0,00500953 | 0,068843113 | collectin subfamily member 10 |
| ENSG00000100345 | MYH9 | 2,741265223 | 6,686564807 | 0,00035785 | 0,00783835 | myosin heavy chain 9 |
| ENSG00000134954 | ETS1 | 2,738588435 | 6,674170005 | 6,24E-11 | 5,48E-09 | ETS proto-oncogene 1, transcription factor |
| ENSG00000168899 | VAMP5 | 2,737026905 | 6,666949991 | 5,90E-10 | 4,61E-08 | vesicle associated membrane protein 5 |
| ENSG00000108798 | ABI3 | 2,733954313 | 6,652766103 | 0,00012712 | 0,003228531 | ABI family member 3 |
| ENSG00000115085 | ZAP70 | 2,732116784 | 6,644298018 | 0,00995193 | 0,114641543 | zeta chain of T cell receptor associated protein kinase 70 |
| ENSG00000237613 | FAM138A | 2,727525861 | 6,623188236 | 0,24056942 | 0,750839796 | family with sequence similarity 138 member A |
| ENSG00000102271 | KLHL4 | 2,726919476 | 6,620405002 | 1,89E-08 | 1,13E-06 | kelch like family member 4 |

| | | | | | | |
|---|---|---|---|---|---|---|
| ENSG00000135926 | TMBIM1 | 2,722438945 | 6,59987612 | 3,04E-12 | 3,16E-10 | transmembrane BAX inhibitor motif containing 1 |
| ENSG00000259856 | RAB43P1 | 2,722254257 | 6,599031284 | 0,01443021 | 0,149982094 | RAB43 pseudogene 1 |
| ENSG00000173068 | BNC2 | 2,72075959 | 6,592198066 | 5,11E-14 | 6,62E-12 | basonuclin 2 |
| ENSG00000168453 | HR | 2,720307343 | 6,590131909 | 6,15E-06 | 0,000221441 | HR lysine demethylase and nuclear receptor corepressor |
| ENSG00000196154 | S100A4 | 2,719578539 | 6,586803623 | 6,20E-07 | 2,74E-05 | S100 calcium binding protein A4 |
| ENSG00000005243 | COPZ2 | 2,719463011 | 6,586276188 | 2,61E-10 | 2,15E-08 | COPI coat complex subunit zeta 2 |
| ENSG00000204359 | CFB | 2,717221434 | 6,576050756 | 0,06912645 | 0,414811757 | complement factor B |
| ENSG00000224618 | AC096921,1 | 2,714303519 | 6,562763839 | 0,0101797 | 0,11653537 | ectodermal-neural cortex (ENC1) pseudogene |
| ENSG00000125740 | FOSB | 2,713716203 | 6,560092708 | 2,93E-07 | 1,39E-05 | FosB proto-oncogene, AP-1 transcription factor subunit |
| ENSG00000273936 | TNFRSF14 | 2,712837601 | 6,556098823 | 0,00234327 | 0,037997875 | TNF receptor superfamily member 14 |
| ENSG00000139926 | FRMD6 | 2,708788987 | 6,537726331 | 9,53E-18 | 1,81E-15 | FERM domain containing 6 |
| ENSG00000147256 | ARHGAP36 | 2,704661967 | 6,519051025 | 0,0015402 | 0,026885986 | Rho GTPase activating protein 36 |
| ENSG00000012211 | PRICKLE3 | 2,704456498 | 6,518122645 | 3,03E-06 | 0,000118632 | prickle planar cell polarity protein 3 |
| ENSG00000242829 | RPS26P21 | 2,704103333 | 6,516527232 | 0,02303533 | 0,206327482 | ribosomal protein S26 pseudogene 21 |
| ENSG00000142552 | RCN3 | 2,70249969 | 6,509287743 | 7,10E-06 | 0,000252236 | reticulocalbin 3 |
| ENSG00000240583 | AQP1 | 2,702196053 | 6,50791791 | 4,19E-10 | 3,36E-08 | aquaporin 1 (Colton blood group) |
| ENSG00000143867 | OSR1 | 2,701745588 | 6,505886204 | 2,46E-07 | 1,19E-05 | odd-skipped related transcription factor 1 |
| ENSG00000027075 | PRKCH | 2,701498172 | 6,504770569 | 9,52E-07 | 4,06E-05 | protein kinase C eta |
| ENSG00000258548 | LINC00645 | 2,701146348 | 6,503184471 | 2,57E-06 | 0,000101958 | long intergenic non-protein coding RNA 645 |
| ENSG00000186377 | CYP4X1 | 2,698447796 | 6,491031675 | 1,16E-05 | 0,00038575 | cytochrome P450 family 4 subfamily X member 1 |
| ENSG00000157873 | TNFRSF14 | 2,696479974 | 6,482184007 | 0,00251864 | 0,040298316 | TNF receptor superfamily member 14 |
| ENSG00000074410 | CA12 | 2,696432237 | 6,481969521 | 0,00014115 | 0,003524626 | carbonic anhydrase 12 |
| ENSG00000112297 | CRYBG1 | 2,696263677 | 6,481212231 | 1,02E-06 | 4,32E-05 | crystallin beta-gamma domain containing 1 |
| ENSG00000288335 | TWIST2 | 2,69335818 | 6,468172616 | 0,00468097 | 0,065714391 | twist family bHLH transcription factor 2 |
| ENSG00000233608 | TWIST2 | 2,69335818 | 6,468172616 | 0,00468097 | 0,065714391 | twist family bHLH transcription factor 2 |
| ENSG00000183770 | FOXL2 | 2,693026262 | 6,466684664 | 0,00458904 | 0,064860708 | forkhead box L2 |
| ENSG00000284609 | OR8B3 | 2,692045114 | 6,462288298 | 0,01833508 | 0,176288964 | olfactory receptor family 8 subfamily B member 3 |
| ENSG00000171246 | NPTX1 | 2,691378584 | 6,459303388 | 2,61E-06 | 0,000103305 | neuronal pentraxin 1 |
| ENSG00000108771 | DHX58 | 2,689110643 | 6,449157232 | 0,0036316 | 0,054387125 | DExH-box helicase 58 |
| ENSG00000203706 | SERTAD4-AS1 | 2,687827711 | 6,443424797 | 3,82E-07 | 1,77E-05 | SERTAD4 antisense RNA 1 |
| ENSG00000280963 | SERTAD4-AS1 | 2,687827711 | 6,443424797 | 3,82E-07 | 1,77E-05 | SERTAD4 antisense RNA 1 |

| | | | | | | |
|---|---|---|---|---|---|---|
| ENSG00000288295 | MCTS1 | 2,685655915 | 6,433732331 | 1,01E-09 | 7,58E-08 | MCTS1 re-initiation and release factor |
| ENSG00000267257 | AC105105,1 | 2,683524882 | 6,424235952 | 2,01E-09 | 1,42E-07 | novel transcript, antisense to ALPK2 |
| ENSG00000188643 | S100A16 | 2,682868199 | 6,421312446 | 1,04E-05 | 0,00035215 | S100 calcium binding protein A16 |
| ENSG00000170017 | ALCAM | 2,682300966 | 6,418788238 | 6,34E-18 | 1,23E-15 | activated leukocyte cell adhesion molecule |
| ENSG00000286378 | AL713852,2 | 2,68133393 | 6,414487176 | 0,00845398 | 0,101559908 | novel transcript |
| ENSG00000248187 | AC078850,1 | 2,680300044 | 6,409891976 | 0,00657108 | 0,084646994 | novel transcript |
| ENSG00000183246 | RIMBP3C | 2,678855263 | 6,40347603 | 0,01766249 | 0,172400906 | RIMS binding protein 3C |
| ENSG00000165757 | JCAD | 2,678588896 | 6,402293855 | 1,68E-07 | 8,60E-06 | junctional cadherin 5 associated |
| ENSG00000163501 | IHH | 2,677562847 | 6,397742146 | 0,1203231 | 0,553528761 | Indian hedgehog signaling molecule |
| ENSG00000075651 | PLD1 | 2,675758276 | 6,389744636 | 1,81E-09 | 1,28E-07 | phospholipase D1 |
| ENSG00000157601 | MX1 | 2,675258762 | 6,387532657 | 0,00014062 | 0,003514255 | MX dynamin like GTPase 1 |
| ENSG00000277013 | AC008556,1 | 2,675187145 | 6,387215583 | 7,72E-09 | 4,89E-07 | novel transcript |
| ENSG00000133067 | LGR6 | 2,671484305 | 6,370843088 | 0,0056426 | 0,075356774 | leucine rich repeat containing G protein-coupled receptor 6 |
| ENSG00000152137 | HSPB8 | 2,670363345 | 6,365894927 | 1,51E-12 | 1,64E-10 | heat shock protein family B (small) member 8 |
| ENSG00000140519 | RHCG | 2,669912941 | 6,363907832 | 0,05993446 | 0,380851824 | Rh family C glycoprotein |
| ENSG00000103241 | FOXF1 | 2,669564983 | 6,36237313 | 0,01970652 | 0,184683692 | forkhead box F1 |
| ENSG00000105464 | GRIN2D | 2,669383062 | 6,361570897 | 3,25E-07 | 1,53E-05 | glutamate ionotropic receptor NMDA type subunit 2D |
| ENSG00000275095 | CCL16 | 2,66892575 | 6,359554695 | 0,12422602 | 0,563196389 | C-C motif chemokine ligand 16 |
| ENSG00000206452 | HLA-C | 2,668347705 | 6,357007125 | 0,00092857 | 0,01757313 | major histocompatibility complex, class I, C |
| ENSG00000170955 | CAVIN3 | 2,667043341 | 6,351262245 | 5,30E-06 | 0,000194812 | caveolae associated protein 3 |
| ENSG00000236013 | AL357146,1 | 2,664254794 | 6,338997918 | 0,0162542 | 0,162718963 | novel transcript |
| ENSG00000154143 | PANX3 | 2,664003025 | 6,337891775 | 0,07597937 | 0,435457192 | pannexin 3 |
| ENSG00000026559 | KCNG1 | 2,66398612 | 6,337817512 | 5,12E-09 | 3,35E-07 | potassium voltage-gated channel modifier subfamily G member 1 |
| ENSG00000177575 | CD163 | 2,663423325 | 6,335345614 | 0,02898624 | 0,241642594 | CD163 molecule |
| ENSG00000230825 | AC005532,1 | 2,663158359 | 6,334182167 | 5,55E-06 | 0,000202913 | novel transcript |
| ENSG00000123609 | NMI | 2,654167685 | 6,294831171 | 2,13E-11 | 2,01E-09 | N-myc and STAT interactor |
| ENSG00000125965 | GDF5 | 2,649582991 | 6,27485878 | 0,00023773 | 0,005530041 | growth differentiation factor 5 |
| ENSG00000263955 | RN7SL850P | 2,647267472 | 6,264795741 | 0,15480874 | 0,624266783 | RNA, 7SL, cytoplasmic 850, pseudogene |
| ENSG00000287593 | AL022724,3 | 2,647214643 | 6,264566337 | 0,03784754 | 0,287759137 | novel transcript, antisense to ADTRP |
| ENSG00000288013 | AP003486,2 | 2,646436469 | 6,261188209 | 0,00013653 | 0,0034229 | novel transcript |

| | | | | | | |
|---|---|---|---|---|---|---|
| ENSG00000172985 | SH3RF3 | 2,64443237 | 6,25249661 | 6,14E-10 | 4,77E-08 | SH3 domain containing ring finger 3 |
| ENSG00000146038 | DCDC2 | 2,638705761 | 6,22772725 | 2,43E-07 | 1,18E-05 | doublecortin domain containing 2 |
| ENSG00000213928 | IRF9 | 2,635812122 | 6,215248705 | 1,34E-07 | 6,99E-06 | interferon regulatory factor 9 |
| ENSG00000166073 | GPR176 | 2,630595204 | 6,192814379 | 3,73E-08 | 2,11E-06 | G protein-coupled receptor 176 |
| ENSG00000225108 | ZBTB45P1 | 2,628611725 | 6,184306082 | 0,09397448 | 0,487240447 | zinc finger and BTB domain containing 45 pseudogene 1 |
| ENSG00000131981 | LGALS3 | 2,628461919 | 6,183663951 | 1,37E-08 | 8,36E-07 | galectin 3 |
| ENSG00000113905 | HRG | 2,628391232 | 6,183360984 | 0,09706169 | 0,494806155 | histidine rich glycoprotein |
| ENSG00000151090 | THRB | 2,625834363 | 6,172412 | 0,00049984 | 0,010411093 | thyroid hormone receptor beta |
| ENSG00000163513 | TGFBR2 | 2,623763223 | 6,163557211 | 2,75E-15 | 4,06E-13 | transforming growth factor beta receptor 2 |
| ENSG00000261499 | AC233699,1 | 2,621435825 | 6,153621998 | 0,18641427 | 0,67957739 | aminopeptidase puromycin sensitive (NPEPPS) pseudogene |
| ENSG00000282411 | AC233699,6 | 2,621435825 | 6,153621998 | 0,18641427 | 0,67957739 | aminopeptidase puromycin sensitive (NPEPPS) pseudogene |
| ENSG00000148680 | HTR7 | 2,619675602 | 6,146118583 | 0,00062745 | 0,012632275 | 5-hydroxytryptamine receptor 7 |
| ENSG00000141052 | MYOCD | 2,618673897 | 6,141852633 | 1,68E-06 | 6,86E-05 | myocardin |
| ENSG00000135547 | HEY2 | 2,618337581 | 6,140421035 | 5,62E-10 | 4,40E-08 | hes related family bHLH transcription factor with YRPW motif 2 |
| ENSG00000235523 | AL135924,2 | 2,616861128 | 6,134140146 | 0,13479982 | 0,585737556 | novel transcript |
| ENSG00000168938 | PPIC | 2,615553146 | 6,128581306 | 1,58E-09 | 1,14E-07 | peptidylprolyl isomerase C |
| ENSG00000273213 | H3-2 | 2,615369886 | 6,127802865 | 9,25E-18 | 1,76E-15 | H3,2 histone (putative) |
| ENSG00000182871 | COL18A1 | 2,614779022 | 6,125293705 | 3,52E-05 | 0,001038938 | collagen type XVIII alpha 1 chain |
| ENSG00000206486 | DHX16 | 2,614387362 | 6,123631047 | 0,11980483 | 0,552771274 | DEAH-box helicase 16 |
| ENSG00000284680 | OR8B2 | 2,613637287 | 6,120448123 | 0,08531238 | 0,465128991 | olfactory receptor family 8 subfamily B member 2 |
| ENSG00000174171 | AC020659,1 | 2,61338436 | 6,119375205 | 0,00010325 | 0,002692293 | novel transcript, antisense to SPTBN5 |
| ENSG00000115363 | EVA1A | 2,608970529 | 6,100681989 | 7,97E-06 | 0,000278696 | eva-1 homolog A, regulator of programmed cell death |
| ENSG00000007237 | GAS7 | 2,605550518 | 6,086237019 | 2,07E-05 | 0,000649718 | growth arrest specific 7 |
| ENSG00000259354 | AC025580,2 | 2,602745375 | 6,074414574 | 0,07351806 | 0,428147811 | novel transcript, antisense to SLC30A4 |
| ENSG00000148908 | RGS10 | 2,601269685 | 6,068204413 | 3,82E-07 | 1,77E-05 | regulator of G protein signaling 10 |
| ENSG00000262635 | TDO2 | 2,599151621 | 6,059302035 | 0,01950609 | 0,183714864 | tryptophan 2,3-dioxygenase |
| ENSG00000280944 | AC241851,2 | 2,598861914 | 6,058085394 | 0,0058613 | 0,07756432 | novel transcript |
| ENSG00000221963 | APOL6 | 2,595842818 | 6,045421026 | 3,08E-11 | 2,84E-09 | apolipoprotein L6 |

| | | | | | | |
|---|---|---|---|---|---|---|
| ENSG00000196341 | OR8D1 | 2,595637851 | 6,044562202 | 8,39E-07 | 3,63E-05 | olfactory receptor family 8 subfamily D member 1 |
| ENSG00000151117 | TMEM86A | 2,594636937 | 6,040370056 | 2,49E-06 | 9,92E-05 | transmembrane protein 86A |
| ENSG00000184937 | WT1 | 2,5943581 | 6,039202716 | 0,13599264 | 0,587841041 | WT1 transcription factor |
| ENSG00000274995 | AC013564,1 | 2,593899645 | 6,037283903 | 0,00083948 | 0,016052254 | novel transcript, sense intronic to SMAD6 |
| ENSG00000234218 | MICB | 2,591875876 | 6,028820917 | 0,00150461 | 0,026323598 | MHC class I polypeptide-related sequence B |
| ENSG00000147036 | LANCL3 | 2,590858959 | 6,024572861 | 0,00025641 | 0,005894434 | LanC like 3 |
| ENSG00000106511 | MEOX2 | 2,590725444 | 6,024015343 | 0,00257711 | 0,041012301 | mesenchyme homeobox 2 |
| ENSG00000285048 | IRF9 | 2,589344345 | 6,018251281 | 2,06E-07 | 1,01E-05 | interferon regulatory factor 9 |
| ENSG00000213253 | RPL12P42 | 2,58695774 | 6,008303709 | 0,08385734 | 0,461082665 | ribosomal protein L12 pseudogene 42 |
| ENSG00000122786 | CALD1 | 2,584327336 | 5,997359005 | 6,29E-14 | 8,00E-12 | caldesmon 1 |
| ENSG00000206408 | GPANK1 | 2,583887077 | 5,995529106 | 0,00024635 | 0,005684003 | G-patch domain and ankyrin repeats 1 |
| ENSG00000255571 | MIR9-3HG | 2,583829722 | 5,995290755 | 6,04E-07 | 2,67E-05 | MIR9-3 host gene |
| ENSG00000017483 | SLC38A5 | 2,58249677 | 5,989754071 | 1,70E-07 | 8,68E-06 | solute carrier family 38 member 5 |
| ENSG00000142089 | IFITM3 | 2,578643032 | 5,973775545 | 2,35E-16 | 3,91E-14 | interferon induced transmembrane protein 3 |
| ENSG00000137558 | PI15 | 2,575933667 | 5,962567391 | 5,79E-06 | 0,000209999 | peptidase inhibitor 15 |
| ENSG00000183762 | KREMEN1 | 2,574924603 | 5,95839845 | 1,24E-09 | 9,17E-08 | kringle containing transmembrane protein 1 |
| ENSG00000237724 | HSPA1A | 2,574552177 | 5,956860513 | 1,34E-10 | 1,14E-08 | heat shock protein family A (Hsp70) member 1A |
| ENSG00000169071 | ROR2 | 2,574370987 | 5,95611243 | 4,41E-10 | 3,52E-08 | receptor tyrosine kinase like orphan receptor 2 |
| ENSG00000139055 | ERP27 | 2,573936136 | 5,954317432 | 0,09809539 | 0,497381508 | endoplasmic reticulum protein 27 |
| ENSG00000267801 | AC087289,5 | 2,570690732 | 5,940937998 | 0,11329638 | 0,537503475 | novel transcript, antisense to TRIM47 |
| ENSG00000273055 | AC005046,1 | 2,570218325 | 5,938992969 | 1,49E-06 | 6,14E-05 | novel transcript, antisense to LAMB1 |
| ENSG00000184557 | SOCS3 | 2,570070554 | 5,938384686 | 4,80E-09 | 3,14E-07 | suppressor of cytokine signaling 3 |
| ENSG00000141753 | IGFBP4 | 2,568420787 | 5,931597839 | 6,82E-11 | 5,94E-09 | insulin like growth factor binding protein 4 |
| ENSG00000125844 | RRBP1 | 2,568265068 | 5,930957639 | 0,00109026 | 0,020109687 | ribosome binding protein 1 |
| ENSG00000227025 | CR388205,1 | 2,568151375 | 5,930490265 | 0,04826887 | 0,334607086 | HLA complex group 4 pseudogene 3 |
| ENSG00000223798 | CR759960,1 | 2,568151375 | 5,930490265 | 0,04826887 | 0,334607086 | HLA complex group 4 pseudogene 3 |
| ENSG00000105357 | MYH14 | 2,566961695 | 5,925601858 | 0,0078228 | 0,096174653 | myosin heavy chain 14 |
| ENSG00000162618 | ADGRL4 | 2,566841885 | 5,925109782 | 1,88E-07 | 9,35E-06 | adhesion G protein-coupled receptor L4 |
| ENSG00000253669 | GASAL1 | 2,566554227 | 5,923928495 | 0,00017526 | 0,004240873 | growth arrest associated lncRNA 1 |
| ENSG00000235394 | ZNRD1ASP | 2,562680833 | 5,908045076 | 0,00264826 | 0,041893209 | zinc ribbon domain containing 1 antisense, pseudogene |
| ENSG00000173826 | KCNH6 | 2,562225455 | 5,906180531 | 0,15372638 | 0,622713345 | potassium voltage-gated channel subfamily H member 6 |

| | | | | | | |
|---|---|---|---|---|---|---|
| ENSG00000062038 | CDH3 | 2,56059911 | 5,899526265 | 6,07E-10 | 4,72E-08 | cadherin 3 |
| ENSG00000285114 | GSDMC | 2,559719451 | 5,895930224 | 0,11164586 | 0,534286883 | gasdermin C |
| ENSG00000169594 | BNC1 | 2,559272129 | 5,894102416 | 2,17E-17 | 3,95E-15 | basonuclin 1 |
| ENSG00000284297 | AC208162,3 | 2,556210832 | 5,881608806 | 0,04159013 | 0,305946805 | novel transcript |
| ENSG00000170961 | HAS2 | 2,555245682 | 5,877675381 | 3,60E-07 | 1,68E-05 | hyaluronan synthase 2 |
| ENSG00000073146 | MOV10L1 | 2,551832752 | 5,863787215 | 0,01738648 | 0,170451797 | Mov10 like RISC complex RNA helicase 1 |
| ENSG00000137767 | SQOR | 2,549419779 | 5,853987963 | 0,00013135 | 0,003325174 | sulfide quinone oxidoreductase |
| ENSG00000206435 | HLA-C | 2,54914702 | 5,8528813 | 0,00118553 | 0,021499153 | major histocompatibility complex, class I, C |
| ENSG00000135074 | ADAM19 | 2,54703939 | 5,844337082 | 5,63E-12 | 5,70E-10 | ADAM metallopeptidase domain 19 |
| ENSG00000285735 | LINC02717 | 2,546640291 | 5,842720558 | 0,09394028 | 0,487240447 | long intergenic non-protein coding RNA 2717 |
| ENSG00000247287 | AL359220,1 | 2,546200694 | 5,840940521 | 0,07047467 | 0,418147371 | novel transcript, antisense to PRKCH |
| ENSG00000181036 | FCRL6 | 2,545870011 | 5,839601861 | 0,04781148 | 0,332623171 | Fc receptor like 6 |
| ENSG00000100065 | CARD10 | 2,545222125 | 5,83698 | 0,00184613 | 0,031402206 | caspase recruitment domain family member 10 |
| ENSG00000282765 | LINC01896 | 2,544859424 | 5,835512737 | 0,10660164 | 0,521482353 | long intergenic non-protein coding RNA 1896 |
| ENSG00000263146 | LINC01896 | 2,544859424 | 5,835512737 | 0,10660164 | 0,521482353 | long intergenic non-protein coding RNA 1896 |
| ENSG00000236056 | GAPDHP14 | 2,544719015 | 5,834944829 | 0,04368979 | 0,315756651 | glyceraldehyde-3-phosphate dehydrogenase pseudogene 14 |
| ENSG00000115604 | IL18R1 | 2,544040799 | 5,832202446 | 1,87E-05 | 0,000594202 | interleukin 18 receptor 1 |
| ENSG00000171223 | JUNB | 2,542560838 | 5,82622266 | 6,41E-08 | 3,54E-06 | JunB proto-oncogene, AP-1 transcription factor subunit |
| ENSG00000178732 | GP5 | 2,540800872 | 5,819119493 | 0,01742379 | 0,170657076 | glycoprotein V platelet |
| ENSG00000228692 | AL445307,1 | 2,53873778 | 5,810803948 | 0,01053937 | 0,119169172 | novel transcript |
| ENSG00000255394 | C8orf49 | 2,536670973 | 5,80248534 | 0,17289854 | 0,657443437 | chromosome 8 open reading frame 49 |
| ENSG00000285413 | AP001056,2 | 2,536321102 | 5,801078339 | 0,00078221 | 0,015169893 | novel transcript |
| ENSG00000288648 | AL139190,1 | 2,533310083 | 5,788983653 | 0,10755958 | 0,52347056 | novel transcript |
| ENSG00000288488 | SAMD1 | 2,533026152 | 5,787844459 | 0,05653601 | 0,367014478 | sterile alpha motif domain containing 1 |
| ENSG00000233345 | MSH5 | 2,53007789 | 5,776028619 | 0,00228736 | 0,037401667 | mutS homolog 5 |
| ENSG00000101463 | SYNDIG1 | 2,528317125 | 5,768983452 | 4,64E-06 | 0,000173085 | synapse differentiation inducing 1 |
| ENSG00000233077 | LINC01271 | 2,527592264 | 5,766085639 | 0,1153055 | 0,542418176 | long intergenic non-protein coding RNA 1271 |
| ENSG00000249406 | AC015909,1 | 2,526656744 | 5,762347816 | 0,04339947 | 0,314374628 | novel transcript |
| ENSG00000251075 | HTT-AS | 2,525920918 | 5,759409561 | 0,04558966 | 0,323511284 | HTT antisense RNA |
| ENSG00000279983 | AC244517,10 | 2,525428214 | 5,757442963 | 0,0416014 | 0,305957674 | novel protein |
| ENSG00000233519 | AL590490,1 | 2,523057608 | 5,747990224 | 0,11957389 | 0,552362303 | novel transcript, antisense to FMN2 |

| | | | | | | |
|---|---|---|---|---|---|---|
| ENSG00000131196 | NFATC1 | 2,520565004 | 5,738067757 | 2,35E-07 | 1,15E-05 | nuclear factor of activated T cells 1 |
| ENSG00000234546 | LNCTAM34A | 2,517671941 | 5,726572633 | 0,02030384 | 0,188783285 | long non coding transcriptional activator of miR34a |
| ENSG00000147606 | SLC26A7 | 2,516825466 | 5,723213657 | 0,10822105 | 0,524537831 | solute carrier family 26 member 7 |
| ENSG00000106031 | HOXA13 | 2,515434282 | 5,717697448 | 0,12383713 | 0,562657182 | homeobox A13 |
| ENSG00000120341 | SEC16B | 2,515049425 | 5,716172384 | 8,97E-07 | 3,85E-05 | SEC16 homolog B, endoplasmic reticulum export factor |
| ENSG00000162670 | BRINP3 | 2,509917672 | 5,695875734 | 0,00035802 | 0,00783835 | BMP/retinoic acid inducible neural specific 3 |
| ENSG00000277738 | AC126175,2 | 2,508990681 | 5,692217075 | 0,01033573 | 0,117889914 | novel transcript |
| ENSG00000100558 | PLEK2 | 2,508149392 | 5,688898701 | 0,00060432 | 0,0122526 | pleckstrin 2 |
| ENSG00000158458 | NRG2 | 2,505796322 | 5,679627533 | 3,26E-06 | 0,000126713 | neuregulin 2 |
| ENSG00000133710 | SPINK5 | 2,505291891 | 5,677642028 | 1,68E-12 | 1,79E-10 | serine peptidase inhibitor Kazal type 5 |
| ENSG00000140945 | CDH13 | 2,505193299 | 5,677254038 | 1,33E-09 | 9,80E-08 | cadherin 13 |
| ENSG00000272114 | AL136131,3 | 2,502169557 | 5,665367552 | 0,07082454 | 0,418829978 | novel transcript, antisense to VEGFA |
| ENSG00000233987 | AC106706,1 | 2,501855289 | 5,664133577 | 0,12603271 | 0,566777174 | novel transcript |
| ENSG00000271447 | MMP28 | 2,50152294 | 5,662828898 | 0,00180098 | 0,030740647 | matrix metallopeptidase 28 |
| ENSG00000140937 | CDH11 | 2,501103616 | 5,66118322 | 1,03E-17 | 1,93E-15 | cadherin 11 |
| ENSG00000112562 | SMOC2 | 2,500662195 | 5,659451333 | 0,00024525 | 0,005662876 | SPARC related modular calcium binding 2 |
| ENSG00000169129 | AFAP1L2 | 2,500395324 | 5,658404539 | 1,58E-12 | 1,71E-10 | actin filament associated protein 1 like 2 |
| ENSG00000172987 | HPSE2 | 2,498961797 | 5,652784885 | 0,00259643 | 0,041268249 | heparanase 2 (inactive) |
| ENSG00000178031 | ADAMTSL1 | 2,498505447 | 5,650997094 | 4,74E-06 | 0,000176069 | ADAMTS like 1 |
| ENSG00000242147 | LASTR | 2,498209382 | 5,649837531 | 0,12860636 | 0,573001179 | lncRNA associated with SART3 regulation of splicing |
| ENSG00000100249 | C22orf31 | 2,497565703 | 5,647317341 | 0,00223361 | 0,036693513 | chromosome 22 open reading frame 31 |
| ENSG00000187013 | LINC02875 | 2,497563501 | 5,647308721 | 0,00215516 | 0,035725886 | long intergenic non-protein coding RNA 2875 |
| ENSG00000188549 | CCDC9B | 2,49750839 | 5,647092996 | 0,00223466 | 0,036693513 | coiled-coil domain containing 9B |
| ENSG00000278546 | AC003101,2 | 2,495565192 | 5,639491923 | 0,12611982 | 0,567013255 | novel transcript |
| ENSG00000204103 | MAFB | 2,49503727 | 5,637428655 | 2,77E-15 | 4,06E-13 | MAF bZIP transcription factor B |
| ENSG00000188660 | LINC00319 | 2,494952565 | 5,637097672 | 0,13886742 | 0,594163262 | long intergenic non-protein coding RNA 319 |
| ENSG00000287427 | AL390236,1 | 2,492076396 | 5,625870703 | 0,04667836 | 0,32799008 | novel transcript, antisense to PTPRE |
| ENSG00000167360 | OR51Q1 | 2,489606857 | 5,616248833 | 0,10718113 | 0,522599956 | olfactory receptor family 51 subfamily Q member 1 |
| ENSG00000130592 | LSP1 | 2,488766073 | 5,612976711 | 0,00115856 | 0,021107938 | lymphocyte specific protein 1 |
| ENSG00000233221 | AC133785,1 | 2,486968664 | 5,60598803 | 0,15841259 | 0,631907243 | novel transcript, antisense to ARHGEF4 |
| ENSG00000285920 | AC087721,2 | 2,486664647 | 5,604806812 | 0,0287723 | 0,240839934 | novel protein |
| ENSG00000073598 | FNDC8 | 2,486642556 | 5,604720993 | 0,1716769 | 0,655397237 | fibronectin type III domain containing 8 |

| | | | | | | |
|---|---|---|---|---|---|---|
| ENSG00000091010 | POU4F3 | 2,485274808 | 5,599409951 | 0,10544539 | 0,518177398 | POU class 4 homeobox 3 |
| ENSG00000112343 | TRIM38 | 2,478006112 | 5,571269525 | 1,71E-15 | 2,64E-13 | tripartite motif containing 38 |
| ENSG00000259987 | DUX4L46 | 2,477412165 | 5,568976347 | 0,11347347 | 0,538172711 | double homeobox 4 like 46 (pseudogene) |
| ENSG00000121297 | TSHZ3 | 2,476541559 | 5,565616715 | 1,82E-13 | 2,24E-11 | teashirt zinc finger homeobox 3 |
| ENSG00000233754 | AP001628,2 | 2,476228989 | 5,564411016 | 0,13204557 | 0,580641112 | novel transcript |
| ENSG00000109743 | BST1 | 2,475786756 | 5,562705607 | 3,93E-11 | 3,55E-09 | bone marrow stromal cell antigen 1 |
| ENSG00000286156 | AC026273,1 | 2,475266859 | 5,56070136 | 6,09E-15 | 8,75E-13 | BMS1, ribosome biogenesis factor (BMS1) pseudogene |
| ENSG00000259446 | AC055874,1 | 2,473952462 | 5,555637477 | 0,08509075 | 0,464352786 | novel transcript, antisense to RYR3 |
| ENSG00000259447 | AC013652,2 | 2,47288189 | 5,551516368 | 0,06443453 | 0,397871785 | novel transcript |
| ENSG00000269898 | AC018695,2 | 2,471146409 | 5,54484222 | 0,20037946 | 0,700040616 | novel transcript, antisense to C16orf74 |
| ENSG00000103522 | IL21R | 2,470291765 | 5,541558461 | 1,45E-08 | 8,79E-07 | interleukin 21 receptor |
| ENSG00000058085 | LAMC2 | 2,467480596 | 5,530770949 | 0,00015696 | 0,003842736 | laminin subunit gamma 2 |
| ENSG00000105639 | JAK3 | 2,464337004 | 5,518732677 | 1,07E-07 | 5,73E-06 | Janus kinase 3 |
| ENSG00000270866 | AC067773,1 | 2,463152487 | 5,514203411 | 0,00674105 | 0,086338414 | novel transcript |
| ENSG00000251393 | AC005280,1 | 2,462486876 | 5,511659927 | 0,02330896 | 0,208089998 | novel transcript, antisense to NUMB and HEATR4 |
| ENSG00000197747 | S100A10 | 2,461041948 | 5,506142503 | 1,91E-13 | 2,34E-11 | S100 calcium binding protein A10 |
| ENSG00000163840 | DTX3L | 2,458654521 | 5,497038261 | 1,34E-07 | 6,99E-06 | deltex E3 ubiquitin ligase 3L |
| ENSG00000234661 | CHL1-AS1 | 2,456255996 | 5,487906858 | 0,03430186 | 0,270005245 | CHL1 antisense RNA 1 |
| ENSG00000184371 | CSF1 | 2,451205052 | 5,468727031 | 3,37E-07 | 1,58E-05 | colony stimulating factor 1 |
| ENSG00000285746 | AL450447,1 | 2,450912944 | 5,467619867 | 0,13232535 | 0,580797367 | putative UPF0607 protein ENSP00000383144 |
| ENSG00000288199 | LSP1 | 2,450734497 | 5,46694362 | 0,05821357 | 0,373407693 | lymphocyte specific protein 1 |
| ENSG00000182327 | GLTPD2 | 2,447577612 | 5,45499401 | 0,03476477 | 0,271925393 | glycolipid transfer protein domain containing 2 |
| ENSG00000182168 | UNC5C | 2,446938063 | 5,452576337 | 2,34E-08 | 1,37E-06 | unc-5 netrin receptor C |
| ENSG00000254550 | OMP | 2,446931435 | 5,452551288 | 0,09309708 | 0,484790551 | olfactory marker protein |
| ENSG00000137965 | IFI44 | 2,446456566 | 5,450756853 | 0,00138698 | 0,024693961 | interferon induced protein 44 |
| ENSG00000166710 | B2M | 2,443581986 | 5,439906994 | 1,80E-07 | 9,04E-06 | beta-2-microglobulin |
| ENSG00000166897 | ELFN2 | 2,441001305 | 5,430184833 | 0,00643296 | 0,083132372 | extracellular leucine rich repeat and fibronectin type III domain containing 2 |
| ENSG00000245164 | LINC00861 | 2,440358803 | 5,427767044 | 0,00459893 | 0,06494169 | long intergenic non-protein coding RNA 861 |
| ENSG00000156427 | FGF18 | 2,434404701 | 5,405412436 | 8,05E-05 | 0,002171169 | fibroblast growth factor 18 |
| ENSG00000155897 | ADCY8 | 2,433761951 | 5,403004751 | 0,00111184 | 0,020387378 | adenylate cyclase 8 |
| ENSG00000248869 | LINC02511 | 2,432249231 | 5,397342466 | 0,05243535 | 0,351487715 | long intergenic non-protein coding RNA 2511 |

| | | | | | | |
|---|---|---|---|---|---|---|
| ENSG00000275088 | AC026469,1 | 2,428141423 | 5,38199639 | 0,1398204 | 0,596038328 | novel transcript |
| ENSG00000275410 | HNF1B | 2,426785477 | 5,376940386 | 0,03212104 | 0,258956228 | HNF1 homeobox B |
| ENSG00000228964 | HLA-B | 2,422737968 | 5,361876416 | 0,22657872 | 0,735240631 | major histocompatibility complex, class I, B |
| ENSG00000264705 | AC090337,1 | 2,422657618 | 5,361577801 | 0,07796676 | 0,441433051 | novel transcript |
| ENSG00000228605 | GPANK1 | 2,422263987 | 5,360115124 | 0,00109135 | 0,02011794 | G-patch domain and ankyrin repeats 1 |
| ENSG00000019186 | CYP24A1 | 2,421158019 | 5,356007643 | 0,03367353 | 0,266716135 | cytochrome P450 family 24 subfamily A member 1 |
| ENSG00000250992 | AC139491,3 | 2,420665435 | 5,354179235 | 0,24218243 | 0,752479103 | novel transcript |
| ENSG00000232143 | GNL1 | 2,416850461 | 5,340039676 | 0,01211887 | 0,131831682 | G protein nucleolar 1 (putative) |
| ENSG00000228581 | GNL1 | 2,416850461 | 5,340039676 | 0,01211887 | 0,131831682 | G protein nucleolar 1 (putative) |
| ENSG00000204590 | GNL1 | 2,416850461 | 5,340039676 | 0,01211887 | 0,131831682 | G protein nucleolar 1 (putative) |
| ENSG00000119138 | KLF9 | 2,41656234 | 5,338973321 | 1,41E-06 | 5,85E-05 | Kruppel like factor 9 |
| ENSG00000178718 | RPP25 | 2,411395635 | 5,319887122 | 2,65E-06 | 0,000104622 | ribonuclease P and MRP subunit p25 |
| ENSG00000136110 | CNMD | 2,409551486 | 5,313091231 | 0,00035144 | 0,007727191 | chondromodulin |
| ENSG00000130052 | STARD8 | 2,407959662 | 5,307232168 | 1,93E-11 | 1,83E-09 | StAR related lipid transfer domain containing 8 |
| ENSG00000255118 | AP003306,2 | 2,404581691 | 5,294820186 | 0,19442175 | 0,691513253 | novel transcript |
| ENSG00000234571 | H2BP2 | 2,404016108 | 5,292744855 | 1,30E-08 | 7,96E-07 | H2B histone pseudogene 2 |
| ENSG00000171889 | MIR31HG | 2,402527019 | 5,287284724 | 0,03229164 | 0,260130339 | MIR31 host gene |
| ENSG00000077238 | IL4R | 2,401745257 | 5,284420447 | 1,62E-10 | 1,37E-08 | interleukin 4 receptor |
| ENSG00000235865 | GSN-AS1 | 2,401564757 | 5,283759339 | 8,46E-07 | 3,66E-05 | GSN antisense RNA 1 |
| ENSG00000099998 | GGT5 | 2,399528302 | 5,276306242 | 6,38E-10 | 4,92E-08 | gamma-glutamyltransferase 5 |
| ENSG00000228913 | UBD | 2,399140411 | 5,274887813 | 0,11613212 | 0,544096306 | ubiquitin D |
| ENSG00000106483 | SFRP4 | 2,397971057 | 5,270614068 | 0,00195315 | 0,033007107 | secreted frizzled related protein 4 |
| ENSG00000278718 | GSDMD | 2,397723058 | 5,269708127 | 0,01448619 | 0,150413865 | gasdermin D |
| ENSG00000216613 | AL023284,2 | 2,397059357 | 5,267284396 | 0,13774447 | 0,592433314 | phosphopantothenoylcysteine synthetase (PPCS) pseudogene |
| ENSG00000113924 | HGD | 2,396773475 | 5,266240745 | 3,70E-05 | 0,001083221 | homogentisate 1,2-dioxygenase |
| ENSG00000204277 | LINC01993 | 2,392894618 | 5,252100848 | 0,00355457 | 0,053492006 | long intergenic non-protein coding RNA 1993 |
| ENSG00000257614 | AL133304,1 | 2,391567906 | 5,247273199 | 0,13718871 | 0,591024932 | novel transcript, antisense to BRMS1L |
| ENSG00000120217 | CD274 | 2,390657933 | 5,24396455 | 7,02E-07 | 3,07E-05 | CD274 molecule |
| ENSG00000011422 | PLAUR | 2,390577927 | 5,243673751 | 3,65E-06 | 0,000140185 | plasminogen activator, urokinase receptor |
| ENSG00000151702 | FLI1 | 2,388661219 | 5,236711835 | 7,10E-05 | 0,001939353 | Fli-1 proto-oncogene, ETS transcription factor |
| ENSG00000282304 | AC140479,4 | 2,388008855 | 5,234344413 | 0,08972999 | 0,47775341 | novel transcript |
| ENSG00000143127 | ITGA10 | 2,387835977 | 5,23371722 | 8,65E-06 | 0,000300077 | integrin subunit alpha 10 |

| | | | | | | |
|---|---|---|---|---|---|---|
| ENSG00000050767 | COL23A1 | 2,387259622 | 5,231626772 | 0,00419513 | 0,060861508 | collagen type XXIII alpha 1 chain |
| ENSG00000288604 | AC015871,8 | 2,386534639 | 5,228998439 | 0,18727199 | 0,68119505 | SLAIN motif family member 2 (SLAIN2) pseudogene |
| ENSG00000158710 | TAGLN2 | 2,384721916 | 5,222432412 | 6,25E-09 | 4,01E-07 | transgelin 2 |
| ENSG00000268790 | AC008764,4 | 2,384449601 | 5,221446747 | 0,00985862 | 0,113944487 | novel protein |
| ENSG00000247970 | AL160313,1 | 2,384142314 | 5,220334724 | 0,14159358 | 0,599428784 | novel transcript |
| ENSG00000142910 | TINAGL1 | 2,380547584 | 5,207343528 | 0,02650118 | 0,227744064 | tubulointerstitial nephritis antigen like 1 |
| ENSG00000107819 | SFXN3 | 2,379807054 | 5,204671302 | 9,55E-11 | 8,21E-09 | sideroflexin 3 |
| ENSG00000138964 | PARVG | 2,378455959 | 5,199799368 | 0,00482198 | 0,066972197 | parvin gamma |
| ENSG00000213279 | Z97192,2 | 2,377753999 | 5,197269963 | 0,08516722 | 0,464688991 | novel transcript |
| ENSG00000171385 | KCND3 | 2,377423115 | 5,196078099 | 0,00014527 | 0,003616109 | potassium voltage-gated channel subfamily D member 3 |
| ENSG00000235030 | IER3 | 2,376789632 | 5,193797018 | 8,24E-12 | 8,15E-10 | immediate early response 3 |
| ENSG00000230128 | IER3 | 2,376789632 | 5,193797018 | 8,24E-12 | 8,15E-10 | immediate early response 3 |
| ENSG00000237155 | IER3 | 2,376789632 | 5,193797018 | 8,24E-12 | 8,15E-10 | immediate early response 3 |
| ENSG00000227231 | IER3 | 2,376789632 | 5,193797018 | 8,24E-12 | 8,15E-10 | immediate early response 3 |
| ENSG00000259952 | AC009133,2 | 2,376692537 | 5,193447481 | 0,00144999 | 0,025525005 | novel transcript, antisense to MAZ |
| ENSG00000114854 | TNNC1 | 2,372907251 | 5,17983898 | 0,00060331 | 0,012240058 | troponin C1, slow skeletal and cardiac type |
| ENSG00000104611 | SH2D4A | 2,371174673 | 5,173622081 | 1,54E-12 | 1,67E-10 | SH2 domain containing 4A |
| ENSG00000264630 | PRKCA-AS1 | 2,370581815 | 5,171496483 | 0,03304177 | 0,263861565 | PRKCA antisense RNA 1 |
| ENSG00000181381 | DDX60L | 2,370313088 | 5,170533289 | 1,26E-07 | 6,64E-06 | DExD/H-box 60 like |
| ENSG00000152779 | SLC16A12 | 2,369385372 | 5,167209481 | 0,00158814 | 0,027553613 | solute carrier family 16 member 12 |
| ENSG00000143786 | CNIH3 | 2,369363001 | 5,167129357 | 2,24E-05 | 0,000698223 | cornichon family AMPA receptor auxiliary protein 3 |
| ENSG00000215808 | LINC01139 | 2,367809192 | 5,161567258 | 2,77E-11 | 2,58E-09 | long intergenic non-protein coding RNA 1139 |
| ENSG00000122121 | XPNPEP2 | 2,36719719 | 5,159378148 | 6,68E-06 | 0,000239701 | X-prolyl aminopeptidase 2 |
| ENSG00000231185 | SPRY4-AS1 | 2,366349501 | 5,156347526 | 0,00021076 | 0,004980422 | SPRY4 antisense RNA 1 |
| ENSG00000157542 | KCNJ6 | 2,364666752 | 5,150336706 | 3,92E-05 | 0,001136945 | potassium inwardly rectifying channel subfamily J member 6 |
| ENSG00000278555 | LILRB4 | 2,363774123 | 5,147151059 | 0,11924107 | 0,552100834 | leukocyte immunoglobulin like receptor B4 |
| ENSG00000268916 | CSAG3 | 2,361850317 | 5,140292007 | 0,07577754 | 0,434777568 | CSAG family member 3 |
| ENSG00000171860 | C3AR1 | 2,356787157 | 5,122283692 | 0,00078112 | 0,015169893 | complement C3a receptor 1 |
| ENSG00000259848 | AC097374,1 | 2,355882294 | 5,119071986 | 2,24E-05 | 0,000697097 | pseudogene similar to part of asparagine synthetase (ASNS) |
| ENSG00000234869 | AL021392,1 | 2,355188614 | 5,116611212 | 0,02310316 | 0,206784218 | novel transcript |

| | | | | | | |
|---|---|---|---|---|---|---|
| ENSG00000278493 | AC039056,2 | 2,353646621 | 5,111145358 | 0,01808149 | 0,174744891 | novel transcript, sense intronic to EHD4 |
| ENSG00000282518 | AC240565,4 | 2,35162222 | 5,103978385 | 0,23476705 | 0,744321775 | novel transcript |
| ENSG00000254122 | PCDHGB7 | 2,351555238 | 5,103741425 | 3,44E-17 | 6,21E-15 | protocadherin gamma subfamily B, 7 |
| ENSG00000174016 | TENT5D | 2,350351572 | 5,099485058 | 0,15383174 | 0,622898002 | terminal nucleotidyltransferase 5D |
| ENSG00000179715 | PCED1B | 2,350049 | 5,09841567 | 0,00030425 | 0,006838394 | PC-esterase domain containing 1B |
| ENSG00000261121 | LINC02473 | 2,349732464 | 5,097297168 | 0,22473535 | 0,733203418 | long intergenic non-protein coding RNA 2473 |
| ENSG00000140839 | CLEC18B | 2,348220008 | 5,091956194 | 8,86E-05 | 0,002360504 | C-type lectin domain family 18 member B |
| ENSG00000188707 | ZBED6CL | 2,347785301 | 5,090422138 | 0,01315552 | 0,139782152 | ZBED6 C-terminal like |
| ENSG00000111199 | TRPV4 | 2,347199063 | 5,088354068 | 7,30E-05 | 0,001990701 | transient receptor potential cation channel subfamily V member 4 |
| ENSG00000162344 | FGF19 | 2,345256183 | 5,081506184 | 2,00E-08 | 1,19E-06 | fibroblast growth factor 19 |
| ENSG00000283128 | AC009403,2 | 2,344182515 | 5,077725885 | 0,00179312 | 0,030633997 | novel transcript |
| ENSG00000107831 | FGF8 | 2,343572017 | 5,075577625 | 2,02E-05 | 0,00063514 | fibroblast growth factor 8 |
| ENSG00000170345 | FOS | 2,343059481 | 5,073774781 | 6,40E-15 | 9,13E-13 | Fos proto-oncogene, AP-1 transcription factor subunit |
| ENSG00000111863 | ADTRP | 2,342060142 | 5,07026145 | 0,06216742 | 0,389109545 | androgen dependent TFPI regulating protein |
| ENSG00000019991 | HGF | 2,340649569 | 5,065306502 | 0,00033041 | 0,007352208 | hepatocyte growth factor |
| ENSG00000166147 | FBN1 | 2,336383186 | 5,050349346 | 1,68E-10 | 1,41E-08 | fibrillin 1 |
| ENSG00000135525 | MAP7 | 2,335541025 | 5,047402107 | 3,12E-09 | 2,13E-07 | microtubule associated protein 7 |
| ENSG00000144407 | PTH2R | 2,335245522 | 5,046368367 | 1,52E-09 | 1,11E-07 | parathyroid hormone 2 receptor |
| ENSG00000244444 | EGFL8 | 2,33446802 | 5,043649495 | 0,28421171 | 0,795731744 | EGF like domain multiple 8 |
| ENSG00000117298 | ECE1 | 2,332984031 | 5,038464151 | 4,72E-11 | 4,21E-09 | endothelin converting enzyme 1 |
| ENSG00000104371 | DKK4 | 2,331650833 | 5,033810243 | 0,01529569 | 0,156275953 | dickkopf WNT signaling pathway inhibitor 4 |
| ENSG00000133466 | C1QTNF6 | 2,330392889 | 5,029422974 | 1,53E-06 | 6,31E-05 | C1q and TNF related 6 |
| ENSG00000105643 | ARRDC2 | 2,329431363 | 5,026072087 | 0,00095885 | 0,018036986 | arrestin domain containing 2 |
| ENSG00000165072 | MAMDC2 | 2,325083599 | 5,010948113 | 7,91E-10 | 6,05E-08 | MAM domain containing 2 |
| ENSG00000184232 | OAF | 2,324890298 | 5,01027676 | 4,69E-16 | 7,56E-14 | out at first homolog |
| ENSG00000243738 | RN7SL181P | 2,32400167 | 5,007191632 | 0,15814934 | 0,631383073 | RNA, 7SL, cytoplasmic 181, pseudogene |
| ENSG00000163394 | CCKAR | 2,32364813 | 5,005964743 | 0,1316189 | 0,579455365 | cholecystokinin A receptor |
| ENSG00000170006 | TMEM154 | 2,3234742 | 5,005361265 | 6,62E-09 | 4,23E-07 | transmembrane protein 154 |
| ENSG00000287742 | AC068282,2 | 2,322468746 | 5,001874106 | 0,18507651 | 0,677443261 | novel transcript |
| ENSG00000249584 | LINC02225 | 2,321746413 | 4,999370379 | 0,1659033 | 0,645166759 | long intergenic non-protein coding RNA 2225 |

| | | | | | | |
|---|---|---|---|---|---|---|
| ENSG00000138642 | HERC6 | 2,320887178 | 4,996393758 | 0,0001935 | 0,004642882 | HECT and RLD domain containing E3 ubiquitin protein ligase family member 6 |
| ENSG00000287610 | AC009403,3 | 2,318643376 | 4,988628981 | 0,08604608 | 0,467282363 | novel transcript, antisense to CNPY1 |
| ENSG00000163431 | LMOD1 | 2,318626416 | 4,988570336 | 0,00012836 | 0,003254817 | leiomodin 1 |
| ENSG00000147408 | CSGALNACT1 | 2,316923223 | 4,982684487 | 9,03E-05 | 0,002395376 | chondroitin sulfate N-acetylgalactosaminyltransferase 1 |
| ENSG00000226173 | TAP1 | 2,316591456 | 4,981538785 | 0,00062541 | 0,012613995 | transporter 1, ATP binding cassette subfamily B member |
| ENSG00000232367 | TAP1 | 2,316591456 | 4,981538785 | 0,00062541 | 0,012613995 | transporter 1, ATP binding cassette subfamily B member |
| ENSG00000197550 | AL359955,1 | 2,315247368 | 4,976899893 | 8,58E-05 | 0,002298581 | zinc finger pseudogene |
| ENSG00000167641 | PPP1R14A | 2,314794652 | 4,97533839 | 0,00035111 | 0,007727191 | protein phosphatase 1 regulatory inhibitor subunit 14A |
| ENSG00000285105 | OR2A1 | 2,31417376 | 4,973197616 | 0,1791367 | 0,667603998 | olfactory receptor family 2 subfamily A member 1 |
| ENSG00000244681 | MTHFD2P1 | 2,31234839 | 4,966909257 | 0,12686201 | 0,568970141 | methylenetetrahydrofolate dehydrogenase (NADP+ dependent) 2, methenyltetrahydrofolate cyclohydrolase pseudogene 1 |
| ENSG00000165685 | TMEM52B | 2,311509455 | 4,964021813 | 0,16541849 | 0,644441588 | transmembrane protein 52B |
| ENSG00000129455 | KLK8 | 2,311153926 | 4,962798659 | 0,16432863 | 0,643240037 | kallikrein related peptidase 8 |
| ENSG00000285897 | AC004687,2 | 2,310631463 | 4,961001738 | 0,01645209 | 0,163860559 | novel transcript |
| ENSG00000231528 | FAM225A | 2,308973317 | 4,955303142 | 3,94E-05 | 0,00114066 | family with sequence similarity 225 member A |
| ENSG00000162849 | KIF26B | 2,308920467 | 4,955121617 | 3,87E-11 | 3,50E-09 | kinesin family member 26B |
| ENSG00000139874 | SSTR1 | 2,308660012 | 4,954227132 | 0,23245029 | 0,741488216 | somatostatin receptor 1 |
| ENSG00000101665 | SMAD7 | 2,307545531 | 4,950401471 | 2,68E-08 | 1,56E-06 | SMAD family member 7 |
| ENSG00000225449 | RAB6C-AS1 | 2,307449001 | 4,950070254 | 0,00556279 | 0,074829592 | RAB6C antisense RNA 1 |
| ENSG00000130176 | CNN1 | 2,304716986 | 4,940705235 | 1,82E-07 | 9,11E-06 | calponin 1 |
| ENSG00000189108 | IL1RAPL2 | 2,304227674 | 4,939029802 | 0,00329718 | 0,050358149 | interleukin 1 receptor accessory protein like 2 |
| ENSG00000148180 | GSN | 2,303344981 | 4,936008848 | 1,56E-08 | 9,42E-07 | gelsolin |
| ENSG00000240241 | AC108752,1 | 2,301963827 | 4,931285658 | 0,16121494 | 0,6373908 | novel transcript |
| ENSG00000148677 | ANKRD1 | 2,300242631 | 4,925405935 | 4,60E-10 | 3,67E-08 | ankyrin repeat domain 1 |
| ENSG00000198753 | PLXNB3 | 2,299623376 | 4,923292232 | 0,00519914 | 0,070980695 | plexin B3 |
| ENSG00000136099 | PCDH8 | 2,297790275 | 4,917040625 | 2,62E-05 | 0,000798461 | protocadherin 8 |
| ENSG00000267561 | AC093155,3 | 2,29733914 | 4,915503294 | 0,04379569 | 0,31589834 | novel protein |

| | | | | | | |
|---|---|---|---|---|---|---|
| ENSG00000286596 | AC098595,1 | 2,296257383 | 4,911818946 | 0,00174953 | 0,030003937 | novel transcript, antisense to PCDH7 |
| ENSG00000286235 | AL035461,3 | 2,296021299 | 4,911015239 | 0,42741751 | 0,892552417 | novel protein |
| ENSG00000228271 | LINC02048 | 2,294967589 | 4,90742966 | 0,03324926 | 0,264841921 | long intergenic non-protein coding RNA 2048 |
| ENSG00000147889 | CDKN2A | 2,294266121 | 4,905044145 | 8,42E-05 | 0,002261837 | cyclin dependent kinase inhibitor 2A |
| ENSG00000275400 | AC006001,4 | 2,2939806 | 4,904073496 | 0,00111595 | 0,020450817 | translocase of outer mitochondrial membrane 20 homolog (yeast) (TOMM20) pseudogene |
| ENSG00000254976 | OR8B7P | 2,293565149 | 4,902661478 | 0,03667286 | 0,28205173 | olfactory receptor family 8 subfamily B member 7 pseudogene |
| ENSG00000141905 | NFIC | 2,291455702 | 4,895498254 | 1,57E-07 | 8,09E-06 | nuclear factor I C |
| ENSG00000250234 | AC025754,1 | 2,290764534 | 4,893153476 | 0,31933315 | 0,824211281 | novel transcript |
| ENSG00000213605 | AC111200,1 | 2,289745546 | 4,889698619 | 0,07584718 | 0,434938753 | ribosomal protein S26 (RPS26) pseudogene |
| ENSG00000158525 | CPA5 | 2,289348525 | 4,888353187 | 0,03467075 | 0,271597663 | carboxypeptidase A5 |
| ENSG00000103888 | CEMIP | 2,289065988 | 4,887395949 | 4,59E-05 | 0,001313079 | cell migration inducing hyaluronidase 1 |
| ENSG00000063015 | SEZ6 | 2,286903977 | 4,880077221 | 2,51E-05 | 0,000771219 | seizure related 6 homolog |
| ENSG00000214039 | LINC02418 | 2,28598078 | 4,876955404 | 0,00043386 | 0,009252535 | long intergenic non-protein coding RNA 2418 |
| ENSG00000287125 | AC018695,9 | 2,28277683 | 4,866136635 | 0,07333525 | 0,427433518 | novel transcript, antisense to C16orf74 |
| ENSG00000285304 | Z83844,2 | 2,281850255 | 4,86301235 | 0,01219397 | 0,132188684 | novel protein |
| ENSG00000127863 | TNFRSF19 | 2,281384428 | 4,861442401 | 9,76E-09 | 6,03E-07 | TNF receptor superfamily member 19 |
| ENSG00000258018 | LINC02457 | 2,28072049 | 4,859205646 | 0,07844409 | 0,44332537 | long intergenic non-protein coding RNA 2457 |
| ENSG00000226673 | LINC01108 | 2,280714593 | 4,859185785 | 0,02869224 | 0,240556266 | long intergenic non-protein coding RNA 1108 |
| ENSG00000108379 | WNT3 | 2,279944535 | 4,856592821 | 0,11596639 | 0,544078746 | Wnt family member 3 |
| ENSG00000205359 | SLCO6A1 | 2,279821869 | 4,856179903 | 0,01279356 | 0,137355059 | solute carrier organic anion transporter family member 6A1 |
| ENSG00000223486 | AC092198,1 | 2,279789812 | 4,856072 | 7,67E-09 | 4,86E-07 | novel transcript |
| ENSG00000256630 | AC090424,1 | 2,278888981 | 4,853040772 | 0,05968861 | 0,380139452 | ribosomal protein S6 (RPS6) pseudogene |
| ENSG00000086205 | FOLH1 | 2,276898051 | 4,846348159 | 1,63E-06 | 6,67E-05 | folate hydrolase 1 |
| ENSG00000125089 | SH3TC1 | 2,276627228 | 4,845438486 | 0,00138539 | 0,024679641 | SH3 domain and tetratricopeptide repeats 1 |
| ENSG00000112559 | MDFI | 2,275828416 | 4,842756337 | 2,53E-07 | 1,22E-05 | MyoD family inhibitor |
| ENSG00000237819 | CDK6-AS1 | 2,273106897 | 4,833629508 | 3,78E-09 | 2,56E-07 | CDK6 antisense RNA 1 |
| ENSG00000237604 | AP001056,1 | 2,272128717 | 4,830353311 | 0,0779603 | 0,441433051 | novel transcript |
| ENSG00000267476 | AC104365,1 | 2,27121231 | 4,82728602 | 0,16159237 | 0,63813307 | novel transcript, antisense to MALT1 |
| ENSG00000124225 | PMEPA1 | 2,270090783 | 4,823534827 | 2,51E-10 | 2,07E-08 | prostate transmembrane protein, androgen induced 1 |

| | | | | | | |
|---|---|---|---|---|---|---|
| ENSG00000259583 | AC015712,2 | 2,269293252 | 4,820869083 | 0,00471767 | 0,066001114 | novel transcript, antisense to LRRK1 and ALDH1A3 |
| ENSG00000237181 | PRKAR1B-AS1 | 2,268993531 | 4,819867649 | 0,02527321 | 0,22064831 | PRKAR1B antisense RNA 1 |
| ENSG00000156804 | FBXO32 | 2,266151134 | 4,810380897 | 0,00036415 | 0,007955846 | F-box protein 32 |
| ENSG00000137834 | SMAD6 | 2,266035282 | 4,809994624 | 3,95E-07 | 1,82E-05 | SMAD family member 6 |
| ENSG00000111110 | PPM1H | 2,264590752 | 4,805180922 | 8,44E-10 | 6,42E-08 | protein phosphatase, Mg2+/Mn2+ dependent 1H |
| ENSG00000071205 | ARHGAP10 | 2,264374084 | 4,804459321 | 1,00E-18 | 2,06E-16 | Rho GTPase activating protein 10 |
| ENSG00000126709 | IFI6 | 2,261870451 | 4,79612896 | 0,00015062 | 0,003725447 | interferon alpha inducible protein 6 |
| ENSG00000258454 | AC016526,2 | 2,258866747 | 4,786153765 | 0,09209337 | 0,482440897 | novel transcript, sense overlapping GPATCH2L |
| ENSG00000162511 | LAPTM5 | 2,25595892 | 4,776516738 | 0,02494431 | 0,219004288 | lysosomal protein transmembrane 5 |
| ENSG00000102554 | KLF5 | 2,253733464 | 4,769154312 | 3,23E-07 | 1,52E-05 | Kruppel like factor 5 |
| ENSG00000100302 | RASD2 | 2,251821302 | 4,762837416 | 0,01613732 | 0,161911791 | RASD family member 2 |
| ENSG00000102290 | PCDH11X | 2,251493413 | 4,761755065 | 1,20E-05 | 0,000397951 | protocadherin 11 X-linked |
| ENSG00000263745 | AP005230,1 | 2,25134937 | 4,761279659 | 0,02531691 | 0,220844836 | novel transcript |
| ENSG00000223855 | HRAT92 | 2,251133787 | 4,760568233 | 0,06082078 | 0,383831435 | heart tissue-associated transcript 92 |
| ENSG00000134058 | CDK7 | 2,248195251 | 4,750881594 | 0,00050135 | 0,010435552 | cyclin dependent kinase 7 |
| ENSG00000224413 | AP001476,1 | 2,24666357 | 4,745840352 | 0,0643598 | 0,397835213 | novel transcript |
| ENSG00000162771 | FAM71A | 2,244491053 | 4,738699092 | 0,15875055 | 0,632735484 | family with sequence similarity 71 member A |
| ENSG00000275013 | PRAME | 2,244016766 | 4,7371415 | 0,00059002 | 0,01200944 | PRAME nuclear receptor transcriptional regulator |
| ENSG00000162241 | SLC25A45 | 2,243120053 | 4,734198025 | 0,00148046 | 0,025973673 | solute carrier family 25 member 45 |
| ENSG00000135905 | DOCK10 | 2,243060763 | 4,734003468 | 2,72E-09 | 1,87E-07 | dedicator of cytokinesis 10 |
| ENSG00000011105 | TSPAN9 | 2,241737926 | 4,729664751 | 2,20E-06 | 8,84E-05 | tetraspanin 9 |
| ENSG00000157554 | ERG | 2,24040935 | 4,725311216 | 0,00517554 | 0,07071409 | ETS transcription factor ERG |
| ENSG00000157103 | SLC6A1 | 2,240113533 | 4,724342413 | 2,58E-07 | 1,24E-05 | solute carrier family 6 member 1 |
| ENSG00000148175 | STOM | 2,238173692 | 4,717994352 | 9,92E-10 | 7,49E-08 | stomatin |
| ENSG00000111885 | MAN1A1 | 2,236760984 | 4,713376687 | 2,60E-12 | 2,73E-10 | mannosidase alpha class 1A member 1 |
| ENSG00000198626 | RYR2 | 2,234784604 | 4,706924147 | 1,07E-05 | 0,000361644 | ryanodine receptor 2 |
| ENSG00000255666 | LINC02700 | 2,233935936 | 4,704156106 | 0,00653959 | 0,08431079 | long intergenic non-protein coding RNA 2700 |
| ENSG00000179256 | SMCO3 | 2,233462583 | 4,702612908 | 0,15264564 | 0,621071504 | single-pass membrane protein with coiled-coil domains 3 |
| ENSG00000277646 | AL353684,1 | 2,233027277 | 4,701194198 | 0,03455379 | 0,271157069 | pyruvate dehydrogenase kinase, isozyme 1 (PDK1) pseudogene |
| ENSG00000282844 | ACTN4 | 2,230626611 | 4,693377846 | 3,35E-09 | 2,28E-07 | actinin alpha 4 |
| ENSG00000287929 | AL354953,1 | 2,229312439 | 4,689104527 | 0,0968821 | 0,494437146 | novel transcript, antisense to NMNAT2 |

| | | | | | | |
|---|---|---|---|---|---|---|
| ENSG00000170624 | SGCD | 2,227733905 | 4,683976719 | 4,88E-06 | 0,000180667 | sarcoglycan delta |
| ENSG00000163251 | FZD5 | 2,226312177 | 4,67936309 | 6,47E-07 | 2,86E-05 | frizzled class receptor 5 |
| ENSG00000285224 | AL662854,2 | 2,225673709 | 4,677292683 | 0,18616401 | 0,67937802 | novel protein |
| ENSG00000244491 | AL021707,5 | 2,223046193 | 4,668781891 | 0,12233847 | 0,559390308 | novel transcript, antisense to SUN2 |
| ENSG00000255282 | WTAPP1 | 2,22044254 | 4,66036367 | 0,00748781 | 0,093118073 | Wilms tumor 1 associated protein pseudogene 1 |
| ENSG00000198682 | PAPSS2 | 2,218247929 | 4,653279768 | 3,85E-21 | 9,48E-19 | 3'-phosphoadenosine 5'-phosphosulfate synthase 2 |
| ENSG00000164932 | CTHRC1 | 2,217913298 | 4,652200573 | 3,48E-10 | 2,79E-08 | collagen triple helix repeat containing 1 |
| ENSG00000060718 | COL11A1 | 2,21523259 | 4,643564229 | 9,08E-07 | 3,88E-05 | collagen type XI alpha 1 chain |
| ENSG00000108679 | LGALS3BP | 2,214912738 | 4,642534843 | 2,71E-06 | 0,000107043 | galectin 3 binding protein |
| ENSG00000283060 | ID3 | 2,211357136 | 4,631107148 | 3,35E-06 | 0,000129821 | inhibitor of DNA binding 3, HLH protein |
| ENSG00000117318 | ID3 | 2,211357136 | 4,631107148 | 3,35E-06 | 0,000129821 | inhibitor of DNA binding 3, HLH protein |
| ENSG00000139629 | GALNT6 | 2,210855493 | 4,629497134 | 0,00268145 | 0,042275714 | polypeptide N-acetylgalactosaminyltransferase 6 |
| ENSG00000146648 | EGFR | 2,207958031 | 4,620208731 | 1,58E-14 | 2,15E-12 | epidermal growth factor receptor |
| ENSG00000172403 | SYNPO2 | 2,20665722 | 4,616044782 | 2,92E-09 | 2,00E-07 | synaptopodin 2 |
| ENSG00000164161 | HHIP | 2,20642404 | 4,615298759 | 8,03E-10 | 6,13E-08 | hedgehog interacting protein |
| ENSG00000107731 | UNC5B | 2,206394043 | 4,615202797 | 2,94E-07 | 1,39E-05 | unc-5 netrin receptor B |
| ENSG00000287097 | AL590550,1 | 2,206028154 | 4,61403246 | 0,09359672 | 0,486323399 | uncharacterized LOC107986639 |
| ENSG00000183840 | GPR39 | 2,205081318 | 4,611005282 | 0,00865548 | 0,103087543 | G protein-coupled receptor 39 |
| ENSG00000182870 | GALNT9 | 2,203580093 | 4,606209702 | 0,00504416 | 0,069197242 | polypeptide N-acetylgalactosaminyltransferase 9 |
| ENSG00000182107 | TMEM30B | 2,203257285 | 4,605179163 | 0,01105159 | 0,12326687 | transmembrane protein 30B |
| ENSG00000259048 | AL392023,2 | 2,200774547 | 4,597260914 | 0,04863837 | 0,335726918 | novel transcript, antisense to TTC6 |
| ENSG00000047457 | CP | 2,197582521 | 4,587100514 | 7,78E-06 | 0,000273554 | ceruloplasmin |
| ENSG00000134569 | LRP4 | 2,195356204 | 4,580027321 | 1,01E-09 | 7,58E-08 | LDL receptor related protein 4 |
| ENSG00000231245 | C1DP1 | 2,195049596 | 4,579054057 | 0,20451208 | 0,705837666 | C1D nuclear receptor corepressor pseudogene 1 |
| ENSG00000136153 | LMO7 | 2,194575807 | 4,577550515 | 8,73E-14 | 1,10E-11 | LIM domain 7 |
| ENSG00000182326 | C1S | 2,193427677 | 4,573909045 | 5,20E-13 | 6,03E-11 | complement C1s |
| ENSG00000233621 | LINC01137 | 2,191520011 | 4,567864992 | 0,00021791 | 0,005129961 | long intergenic non-protein coding RNA 1137 |
| ENSG00000229647 | MYOSLID | 2,190746613 | 4,565416914 | 0,00223975 | 0,036757726 | myocardin-induced smooth muscle lncRNA, inducer of differentiation |
| ENSG00000285090 | AC002074,1 | 2,187920255 | 4,556481642 | 0,01667253 | 0,165266312 | COL1A2 antisense RNA 1 |
| ENSG00000137877 | SPTBN5 | 2,187003186 | 4,553586173 | 0,01042977 | 0,118366144 | spectrin beta, non-erythrocytic 5 |
| ENSG00000183779 | ZNF703 | 2,186270659 | 4,55127468 | 8,04E-09 | 5,04E-07 | zinc finger protein 703 |
| ENSG00000281453 | TGFB2-OT1 | 2,185220692 | 4,547963552 | 0,56600684 | 0,941107156 | TGFB2 overlapping transcript 1 |

| | | | | | | |
|---|---|---|---|---|---|---|
| ENSG00000225380 | IGKV1OR9-2 | 2,18508253 | 4,547528029 | 0,10966379 | 0,528090728 | immunoglobulin kappa variable 1/OR9-2 (pseudogene) |
| ENSG00000066813 | ACSM2B | 2,181875054 | 4,537428955 | 0,05434454 | 0,359759285 | acyl-CoA synthetase medium chain family member 2B |
| ENSG00000134013 | LOXL2 | 2,181628151 | 4,536652483 | 1,21E-05 | 0,000399808 | lysyl oxidase like 2 |
| ENSG00000116329 | OPRD1 | 2,180944069 | 4,534501851 | 0,00131102 | 0,023502099 | opioid receptor delta 1 |
| ENSG00000125492 | BARHL1 | 2,18070538 | 4,533751697 | 0,01464964 | 0,151489059 | BarH like homeobox 1 |
| ENSG00000232709 | MARK2P9 | 2,177913261 | 4,524985787 | 0,11224207 | 0,535189241 | microtubule affinity regulating kinase 2 pseudogene 9 |
| ENSG00000155465 | SLC7A7 | 2,177248453 | 4,52290111 | 3,91E-09 | 2,60E-07 | solute carrier family 7 member 7 |
| ENSG00000139174 | PRICKLE1 | 2,176377885 | 4,520172673 | 2,37E-14 | 3,12E-12 | prickle planar cell polarity protein 1 |
| ENSG00000092068 | SLC7A8 | 2,176124523 | 4,519378921 | 1,55E-10 | 1,30E-08 | solute carrier family 7 member 8 |
| ENSG00000182240 | BACE2 | 2,172292588 | 4,507390951 | 9,75E-14 | 1,22E-11 | beta-secretase 2 |
| ENSG00000007350 | TKTL1 | 2,169653692 | 4,499153822 | 1,75E-08 | 1,05E-06 | transketolase like 1 |
| ENSG00000148123 | PLPPR1 | 2,16898248 | 4,497061083 | 2,68E-05 | 0,000813297 | phospholipid phosphatase related 1 |
| ENSG00000166670 | MMP10 | 2,168515015 | 4,495604172 | 0,03115199 | 0,253879337 | matrix metallopeptidase 10 |
| ENSG00000204767 | INSYN2B | 2,166742343 | 4,490083714 | 0,00040651 | 0,008734801 | inhibitory synaptic factor family member 2B |
| ENSG00000088387 | DOCK9 | 2,166460219 | 4,489205749 | 6,14E-14 | 7,84E-12 | dedicator of cytokinesis 9 |
| ENSG00000144810 | COL8A1 | 2,166064031 | 4,487973106 | 6,72E-09 | 4,29E-07 | collagen type VIII alpha 1 chain |
| ENSG00000142178 | SIK1 | 2,164064651 | 4,481757693 | 1,09E-12 | 1,21E-10 | salt inducible kinase 1 |
| ENSG00000206299 | TAP2 | 2,164022048 | 4,481625346 | 0,07083864 | 0,418829978 | transporter 2, ATP binding cassette subfamily B member |
| ENSG00000197046 | SIGLEC15 | 2,162405045 | 4,476605059 | 0,01304563 | 0,139032628 | sialic acid binding Ig like lectin 15 |
| ENSG00000206208 | TAPBP | 2,162401843 | 4,476595125 | 0,17500712 | 0,660401055 | TAP binding protein |
| ENSG00000284075 | AC015853,4 | 2,162360588 | 4,476467115 | 0,2490846 | 0,761104998 | novel transcript, antisense to VPS53 |
| ENSG00000232392 | GOT2P7 | 2,160167908 | 4,469668724 | 0,17072721 | 0,6542799 | GOT2 pseudogene 7 |
| ENSG00000260235 | AC105020,3 | 2,159750804 | 4,468376664 | 0,22110175 | 0,729015508 | novel transcript, antisense to SNUPN |
| ENSG00000198848 | CES1 | 2,156821376 | 4,459312721 | 0,04972368 | 0,340424948 | carboxylesterase 1 |
| ENSG00000176170 | SPHK1 | 2,156018791 | 4,456832651 | 3,80E-06 | 0,000145319 | sphingosine kinase 1 |
| ENSG00000167994 | RAB3IL1 | 2,15489765 | 4,453370524 | 0,00026617 | 0,006082929 | RAB3A interacting protein like 1 |
| ENSG00000100292 | HMOX1 | 2,154656177 | 4,452625197 | 7,42E-07 | 3,23E-05 | heme oxygenase 1 |
| ENSG00000228031 | AC078842,1 | 2,153784424 | 4,449935496 | 0,06786767 | 0,410171049 | novel transcript |
| ENSG00000233817 | AL162727,1 | 2,153132798 | 4,447926037 | 0,00777283 | 0,095710835 | novel transcript |

| | | | | | | |
|---|---|---|---|---|---|---|
| ENSG00000256073 | URB1-AS1 | 2,15296856 | 4,447419709 | 8,44E-05 | 0,002266193 | URB1 antisense RNA 1 (head to head) |
| ENSG00000204267 | TAP2 | 2,151294507 | 4,44226207 | 0,00021062 | 0,004980422 | transporter 2, ATP binding cassette subfamily B member |
| ENSG00000235531 | MSC-AS1 | 2,151157074 | 4,441838915 | 0,00043556 | 0,00927602 | MSC antisense RNA 1 |
| ENSG00000232650 | LINC01780 | 2,150233878 | 4,438997445 | 0,06057712 | 0,383066986 | long intergenic non-protein coding RNA 1780 |
| ENSG00000109158 | GABRA4 | 2,150100086 | 4,438585802 | 0,01589989 | 0,160352922 | gamma-aminobutyric acid type A receptor subunit alpha4 |
| ENSG00000138496 | PARP9 | 2,149333597 | 4,436228253 | 0,00018061 | 0,004356848 | poly(ADP-ribose) polymerase family member 9 |
| ENSG00000227737 | OR8B1P | 2,14879867 | 4,434583679 | 0,06560997 | 0,402080164 | olfactory receptor family 8 subfamily B member 1 pseudogene |
| ENSG00000162733 | DDR2 | 2,148463743 | 4,433554292 | 1,01E-18 | 2,06E-16 | discoidin domain receptor tyrosine kinase 2 |
| ENSG00000160223 | ICOSLG | 2,146563681 | 4,427719046 | 0,00439984 | 0,062897441 | inducible T cell costimulator ligand |
| ENSG00000134242 | PTPN22 | 2,145665582 | 4,424963583 | 0,07730359 | 0,439421744 | protein tyrosine phosphatase non-receptor type 22 |
| ENSG00000113356 | POLR3G | 2,144665391 | 4,421896909 | 1,02E-05 | 0,000347006 | RNA polymerase III subunit G |
| ENSG00000279413 | AC112497,2 | 2,144276225 | 4,420704268 | 0,00076614 | 0,014914536 | |
| ENSG00000130513 | GDF15 | 2,144177416 | 4,420401506 | 0,00210545 | 0,035050408 | growth differentiation factor 15 |
| ENSG00000069399 | BCL3 | 2,142454514 | 4,415125705 | 0,00079705 | 0,015391523 | BCL3 transcription coactivator |
| ENSG00000142227 | EMP3 | 2,140735003 | 4,409866565 | 8,32E-08 | 4,51E-06 | epithelial membrane protein 3 |
| ENSG00000224729 | PCOLCE-AS1 | 2,139841079 | 4,407134966 | 0,0263186 | 0,22653539 | PCOLCE antisense RNA 1 |
| ENSG00000125347 | IRF1 | 2,139795132 | 4,40699461 | 6,71E-07 | 2,95E-05 | interferon regulatory factor 1 |
| ENSG00000101311 | FERMT1 | 2,139537923 | 4,406208983 | 9,82E-05 | 0,002578061 | fermitin family member 1 |
| ENSG00000288431 | AC187652,1 | 2,137875535 | 4,401134724 | 0,34586658 | 0,841168721 | |
| ENSG00000164694 | FNDC1 | 2,137527488 | 4,400073088 | 1,00E-05 | 0,000342318 | fibronectin type III domain containing 1 |
| ENSG00000223680 | DDR1 | 2,134402206 | 4,390551612 | 0,22093698 | 0,728635155 | discoidin domain receptor tyrosine kinase 1 |
| ENSG00000256574 | OR13A1 | 2,133927799 | 4,389108089 | 0,06494955 | 0,399805715 | olfactory receptor family 13 subfamily A member 1 |
| ENSG00000286338 | AC099487,2 | 2,131890378 | 4,38291402 | 0,05108918 | 0,345863615 | novel transcript |
| ENSG00000197587 | DMBX1 | 2,128518721 | 4,372682873 | 0,04886839 | 0,336895538 | diencephalon/mesencephalon homeobox 1 |
| ENSG00000238289 | MICB | 2,127231161 | 4,36878213 | 9,57E-05 | 0,002522632 | MHC class I polypeptide-related sequence B |
| ENSG00000288257 | STRA6 | 2,127101548 | 4,368389652 | 2,52E-09 | 1,74E-07 | signaling receptor and transporter of retinol STRA6 |
| ENSG00000235849 | HS6ST2-AS1 | 2,126396738 | 4,366256054 | 0,11979462 | 0,552771274 | HS6ST2 antisense RNA 1 |
| ENSG00000010610 | CD4 | 2,125898035 | 4,364747011 | 9,25E-11 | 7,99E-09 | CD4 molecule |
| ENSG00000118785 | SPP1 | 2,125584682 | 4,363799094 | 0,000566 | 0,011588267 | secreted phosphoprotein 1 |
| ENSG00000203307 | FAM78B-AS1 | 2,123326835 | 4,356974998 | 0,14242758 | 0,600682024 | FAM78B antisense RNA 1 |

| | | | | | | |
|---|---|---|---|---|---|---|
| ENSG00000271474 | AC106881,1 | 2,120617085 | 4,348799168 | 1,14E-05 | 0,000382594 | novel transcript, antisense to UNC5C |
| ENSG00000160973 | FOXH1 | 2,120388681 | 4,34811073 | 0,56103009 | 0,940489231 | forkhead box H1 |
| ENSG00000280445 | AL049536,1 | 2,118164059 | 4,34141115 | 0,12292757 | 0,560806566 | novel transcript |
| ENSG00000147119 | CHST7 | 2,118049408 | 4,341066155 | 0,00042273 | 0,009033644 | carbohydrate sulfotransferase 7 |
| ENSG00000172497 | ACOT12 | 2,117043461 | 4,338040316 | 0,01290798 | 0,138203846 | acyl-CoA thioesterase 12 |
| ENSG00000250072 | SH3TC2-DT | 2,115724811 | 4,334077079 | 0,12636275 | 0,567767162 | SH3TC2 divergent transcript |
| ENSG00000115194 | SLC30A3 | 2,11530276 | 4,332809358 | 0,00193211 | 0,032686991 | solute carrier family 30 member 3 |
| ENSG00000182667 | NTM | 2,114892818 | 4,331578363 | 2,32E-14 | 3,07E-12 | neurotrimin |
| ENSG00000162692 | VCAM1 | 2,114112247 | 4,329235394 | 1,46E-12 | 1,60E-10 | vascular cell adhesion molecule 1 |
| ENSG00000285275 | FAM167A-AS1 | 2,114029052 | 4,328985752 | 0,2381181 | 0,747367171 | FAM167A antisense RNA 1 |
| ENSG00000104059 | FAM189A1 | 2,113881137 | 4,328541938 | 0,00046848 | 0,009854107 | family with sequence similarity 189 member A1 |
| ENSG00000254578 | AC084125,1 | 2,109905985 | 4,316631637 | 0,30906895 | 0,816845699 | novel transcript, antisense to CYHR1 |
| ENSG00000277794 | Metazoa_SRP | 2,109881816 | 4,316559324 | 0,14357233 | 0,602664517 | Metazoan signal recognition particle RNA |
| ENSG00000251165 | F11-AS1 | 2,109794551 | 4,316298235 | 0,02158342 | 0,196898814 | F11 antisense RNA 1 |
| ENSG00000261359 | PYCARD-AS1 | 2,109003688 | 4,313932755 | 0,27138639 | 0,784254811 | PYCARD antisense RNA 1 |
| ENSG00000287071 | CDRT15P4 | 2,105667089 | 4,303967216 | 0,12929638 | 0,574683255 | CDRT15 pseudogene 4 |
| ENSG00000170425 | ADORA2B | 2,100914263 | 4,289811523 | 2,65E-09 | 1,83E-07 | adenosine A2b receptor |
| ENSG00000109099 | PMP22 | 2,098748719 | 4,283377172 | 4,42E-11 | 3,95E-09 | peripheral myelin protein 22 |
| ENSG00000166250 | CLMP | 2,098116082 | 4,281499276 | 8,96E-11 | 7,76E-09 | CXADR like membrane protein |
| ENSG00000134871 | COL4A2 | 2,09673871 | 4,277413588 | 0,00185571 | 0,031513885 | collagen type IV alpha 2 chain |
| ENSG00000275993 | SIK1B | 2,096490801 | 4,276678632 | 5,97E-09 | 3,85E-07 | salt inducible kinase 1B (putative) |
| ENSG00000237903 | AC004000,1 | 2,096260794 | 4,275996861 | 0,07077747 | 0,418829978 | novel transcript |
| ENSG00000260555 | AC092681,2 | 2,093172982 | 4,266854696 | 0,12571773 | 0,566345689 | novel transcript |
| ENSG00000267279 | AC090409,1 | 2,093032278 | 4,266438578 | 3,54E-05 | 0,001045459 | novel transcript |
| ENSG00000011028 | MRC2 | 2,092276106 | 4,264202958 | 6,75E-06 | 0,000241836 | mannose receptor C type 2 |
| ENSG00000197182 | MIRLET7BHG | 2,090777191 | 4,259774885 | 0,00729888 | 0,091345734 | MIRLET7B host gene |
| ENSG00000278771 | RN7SL3 | 2,089713867 | 4,256636418 | 1,49E-09 | 1,09E-07 | RNA component of signal recognition particle 7SL3 |
| ENSG00000132000 | PODNL1 | 2,088050772 | 4,251732324 | 0,0027512 | 0,043179425 | podocan like 1 |
| ENSG00000288195 | PODNL1 | 2,088050772 | 4,251732324 | 0,0027512 | 0,043179425 | podocan like 1 |
| ENSG00000286239 | AC093884,1 | 2,087991001 | 4,251556178 | 0,25903592 | 0,771947284 | novel protein, WDFY1-AP1S3 readthrough |
| ENSG00000196071 | OR2L13 | 2,08633605 | 4,246681909 | 0,13224128 | 0,580754493 | olfactory receptor family 2 subfamily L member 13 |
| ENSG00000153064 | BANK1 | 2,086120321 | 4,246046942 | 0,00793015 | 0,097133046 | B cell scaffold protein with ankyrin repeats 1 |
| ENSG00000185614 | INKA1 | 2,083365145 | 4,237945825 | 0,00046436 | 0,009789532 | inka box actin regulator 1 |

| | | | | | | |
|---|---|---|---|---|---|---|
| ENSG00000276729 | Metazoa_SRP | 2,081361986 | 4,232065588 | 0,31652471 | 0,821646322 | Metazoan signal recognition particle RNA |
| ENSG00000173559 | NABP1 | 2,080895923 | 4,230698637 | 2,31E-18 | 4,63E-16 | nucleic acid binding protein 1 |
| ENSG00000136997 | MYC | 2,072684384 | 4,206686726 | 2,45E-07 | 1,19E-05 | MYC proto-oncogene, bHLH transcription factor |
| ENSG00000013297 | CLDN11 | 2,07263883 | 4,206553901 | 9,78E-11 | 8,36E-09 | claudin 11 |
| ENSG00000286911 | AC009533,3 | 2,070560267 | 4,20049767 | 0,11611509 | 0,544096306 | novel transcript |
| ENSG00000277322 | GOLGA6L6 | 2,066162494 | 4,187712774 | 0,06639312 | 0,40505568 | golgin A6 family like 6 |
| ENSG00000132329 | RAMP1 | 2,066125719 | 4,187606028 | 0,03593123 | 0,277989137 | receptor activity modifying protein 1 |
| ENSG00000161249 | DMKN | 2,065802564 | 4,186668135 | 0,00728354 | 0,091227554 | dermokine |
| ENSG00000057657 | PRDM1 | 2,0653467 | 4,185345437 | 0,00016444 | 0,004010503 | PR/SET domain 1 |
| ENSG00000277829 | AL031651,2 | 2,064988519 | 4,18430646 | 0,01314549 | 0,139782152 | novel transcript, sense intronic to KIAA1755 |
| ENSG00000132031 | MATN3 | 2,064975211 | 4,184267862 | 7,85E-09 | 4,95E-07 | matrilin 3 |
| ENSG00000139044 | B4GALNT3 | 2,064305911 | 4,182327133 | 0,00715444 | 0,089938198 | beta-1,4-N-acetyl-galactosaminyltransferase 3 |
| ENSG00000146955 | RAB19 | 2,063699225 | 4,18056874 | 0,06944516 | 0,415766681 | RAB19, member RAS oncogene family |
| ENSG00000072682 | P4HA2 | 2,061604529 | 4,174503239 | 1,33E-06 | 5,53E-05 | prolyl 4-hydroxylase subunit alpha 2 |
| ENSG00000207405 | SNORA64 | 2,061500678 | 4,174202753 | 0,24847125 | 0,760720365 | small nucleolar RNA, H/ACA box 64 |
| ENSG00000254951 | AC044810,2 | 2,060715175 | 4,171930646 | 0,05592128 | 0,365161047 | novel transcript, sense overlapping OR5P2, OR5P3 and OR5P1P |
| ENSG00000079308 | TNS1 | 2,059039288 | 4,167087194 | 8,99E-07 | 3,85E-05 | tensin 1 |
| ENSG00000204661 | C5orf60 | 2,05897874 | 4,166912309 | 0,1420249 | 0,600058555 | chromosome 5 open reading frame 60 |
| ENSG00000113263 | ITK | 2,055436791 | 4,156694706 | 0,01360377 | 0,143201487 | IL2 inducible T cell kinase |
| ENSG00000185811 | IKZF1 | 2,055169735 | 4,155925334 | 1,54E-06 | 6,33E-05 | IKAROS family zinc finger 1 |
| ENSG00000280639 | LINC02204 | 2,054342557 | 4,153543193 | 0,02671711 | 0,228844863 | long intergenic non-protein coding RNA 2204 |
| ENSG00000073282 | TP63 | 2,054120658 | 4,152904391 | 0,04225474 | 0,308508641 | tumor protein p63 |
| ENSG00000256566 | AL049650,1 | 2,050971256 | 4,143848493 | 0,09906258 | 0,499450473 | novel protein |
| ENSG00000180787 | ZFP3 | 2,0495604 | 4,139798077 | 1,21E-05 | 0,000399877 | ZFP3 zinc finger protein |
| ENSG00000132530 | XAF1 | 2,049259404 | 4,138934461 | 7,85E-09 | 4,95E-07 | XIAP associated factor 1 |
| ENSG00000186867 | QRFPR | 2,049254725 | 4,138921036 | 0,00024424 | 0,005643754 | pyroglutamylated RFamide peptide receptor |
| ENSG00000271811 | Z97200,1 | 2,049019702 | 4,138246839 | 1,11E-07 | 5,90E-06 | novel transcript |
| ENSG00000082126 | MPP4 | 2,048667531 | 4,13723679 | 0,00755753 | 0,093795931 | membrane palmitoylated protein 4 |
| ENSG00000168122 | ZNF355P | 2,047916831 | 4,135084558 | 0,17042007 | 0,653594444 | zinc finger protein 355, pseudogene |
| ENSG00000241157 | AC104763,1 | 2,04782993 | 4,134835488 | 0,14571685 | 0,608454479 | ribosomal protein L32 (RPL32) pseudogene |
| ENSG00000102265 | TIMP1 | 2,047422474 | 4,133667863 | 3,81E-06 | 0,000145567 | TIMP metallopeptidase inhibitor 1 |
| ENSG00000283005 | CDC27P3 | 2,047092991 | 4,132723922 | 0,14803313 | 0,612582848 | cell division cycle 27 pseudogene 3 |

| | | | | | | |
|---|---|---|---|---|---|---|
| ENSG00000148942 | SLC5A12 | 2,045411563 | 4,127910132 | 0,00027412 | 0,006232838 | solute carrier family 5 member 12 |
| ENSG00000134996 | OSTF1 | 2,045049141 | 4,126873285 | 4,35E-09 | 2,88E-07 | osteoclast stimulating factor 1 |
| ENSG00000168594 | ADAM29 | 2,044458131 | 4,125183027 | 0,08753268 | 0,471587419 | ADAM metallopeptidase domain 29 |
| ENSG00000137868 | STRA6 | 2,043927242 | 4,123665306 | 3,18E-10 | 2,56E-08 | signaling receptor and transporter of retinol STRA6 |
| ENSG00000228261 | ITPRIP-AS1 | 2,043406212 | 4,12217631 | 0,01106981 | 0,123337997 | ITPRIP antisense RNA 1 |
| ENSG00000111077 | TNS2 | 2,041906125 | 4,117892377 | 1,52E-09 | 1,11E-07 | tensin 2 |
| ENSG00000154133 | ROBO4 | 2,04074593 | 4,114582158 | 0,00445635 | 0,063437482 | roundabout guidance receptor 4 |
| ENSG00000101438 | SLC32A1 | 2,039286904 | 4,110423102 | 0,00117832 | 0,021380662 | solute carrier family 32 member 1 |
| ENSG00000140044 | JDP2 | 2,036593986 | 4,10275779 | 0,0001207 | 0,003093101 | Jun dimerization protein 2 |
| ENSG00000163092 | XIRP2 | 2,034948058 | 4,098079745 | 0,05340632 | 0,355796083 | xin actin binding repeat containing 2 |
| ENSG00000087245 | MMP2 | 2,034655162 | 4,097247836 | 3,84E-09 | 2,58E-07 | matrix metallopeptidase 2 |
| ENSG00000283096 | AL731555,1 | 2,034203924 | 4,095966524 | 0,16722909 | 0,647913628 | novel transcript |
| ENSG00000227507 | LTB | 2,032428353 | 4,090928586 | 0,09713551 | 0,495003188 | lymphotoxin beta |
| ENSG00000079689 | SCGN | 2,029948415 | 4,083902476 | 0,08689548 | 0,469646795 | secretagogin, EF-hand calcium binding protein |
| ENSG00000164023 | SGMS2 | 2,029387894 | 4,082316092 | 3,05E-09 | 2,09E-07 | sphingomyelin synthase 2 |
| ENSG00000270962 | AC239800,3 | 2,028954722 | 4,081090553 | 0,13065736 | 0,577136255 | NHP2 non-histone chromosome protein 2-like 1 (S, cerevisiae) (NHP2L1) pseudogene |
| ENSG00000230131 | LINC02641 | 2,028472611 | 4,079726987 | 0,14640623 | 0,609811385 | long intergenic non-protein coding RNA 2641 |
| ENSG00000132170 | PPARG | 2,028100632 | 4,078675222 | 0,00023435 | 0,00546372 | peroxisome proliferator activated receptor gamma |
| ENSG00000119922 | IFIT2 | 2,027063145 | 4,075743173 | 0,00010429 | 0,002717183 | interferon induced protein with tetratricopeptide repeats 2 |
| ENSG00000236649 | PPT2 | 2,026943422 | 4,075404959 | 0,25780725 | 0,7701554 | palmitoyl-protein thioesterase 2 |
| ENSG00000235863 | B3GALT4 | 2,026873408 | 4,075207184 | 0,02410459 | 0,213244441 | beta-1,3-galactosyltransferase 4 |
| ENSG00000174145 | NWD2 | 2,025677434 | 4,071830295 | 0,0015182 | 0,026531575 | NACHT and WD repeat domain containing 2 |
| ENSG00000232591 | LINC02642 | 2,025403298 | 4,071056651 | 0,23379145 | 0,743112643 | long intergenic non-protein coding RNA 2642 |
| ENSG00000147614 | ATP6V0D2 | 2,021316628 | 4,059541036 | 0,24245978 | 0,752612359 | ATPase H+ transporting V0 subunit d2 |
| ENSG00000230045 | FAM90A15P | 2,019505855 | 4,05444897 | 0,32150955 | 0,824933727 | family with sequence similarity 90 member A15, pseudogene |
| ENSG00000237172 | B3GNT9 | 2,019271712 | 4,053791006 | 0,00068966 | 0,013620644 | UDP-GlcNAc:betaGal beta-1,3-N-acetylglucosaminyltransferase 9 |
| ENSG00000106688 | SLC1A1 | 2,018989126 | 4,052997052 | 3,73E-06 | 0,000142871 | solute carrier family 1 member 1 |
| ENSG00000177108 | ZDHHC22 | 2,018906709 | 4,052765523 | 0,000478 | 0,010009603 | zinc finger DHHC-type palmitoyltransferase 22 |
| ENSG00000100385 | IL2RB | 2,017886634 | 4,049900978 | 2,26E-05 | 0,000701675 | interleukin 2 receptor subunit beta |

| | | | | | | |
|---|---|---|---|---|---|---|
| ENSG00000144642 | RBMS3 | 2,017370342 | 4,048451916 | 1,00E-07 | 5,39E-06 | RNA binding motif single stranded interacting protein 3 |
| ENSG00000152207 | CYSLTR2 | 2,017083007 | 4,047645682 | 0,01438192 | 0,149579633 | cysteinyl leukotriene receptor 2 |
| ENSG00000231852 | CYP21A2 | 2,01622805 | 4,045247714 | 0,09298266 | 0,4846632 | cytochrome P450 family 21 subfamily A member 2 |
| ENSG00000172216 | CEBPB | 2,015641546 | 4,043603519 | 3,46E-05 | 0,001025182 | CCAAT enhancer binding protein beta |
| ENSG00000177301 | KCNA2 | 2,013193444 | 4,036747767 | 0,00088605 | 0,01683986 | potassium voltage-gated channel subfamily A member 2 |
| ENSG00000104825 | NFKBIB | 2,012135912 | 4,033789813 | 2,10E-06 | 8,49E-05 | NFKB inhibitor beta |
| ENSG00000149575 | SCN2B | 2,01213133 | 4,033777002 | 0,00012361 | 0,003152114 | sodium voltage-gated channel beta subunit 2 |
| ENSG00000196498 | NCOR2 | 2,011423444 | 4,031798237 | 0,00020024 | 0,004782621 | nuclear receptor corepressor 2 |
| ENSG00000100300 | TSPO | 2,011032709 | 4,030706426 | 5,03E-06 | 0,000185784 | translocator protein |
| ENSG00000112936 | C7 | 2,009365304 | 4,026050599 | 0,01122442 | 0,12466084 | complement C7 |
| ENSG00000285417 | BX571818,1 | 2,008310808 | 4,023108949 | 2,06E-05 | 0,000645612 | novel pseudogene |
| ENSG00000253144 | AC104117,2 | 2,006776693 | 4,018833179 | 0,2732283 | 0,785672647 | pseudogene similat to part of cold shock domain containing E1, RNA-binding CSDE1 |
| ENSG00000260779 | AP001180,1 | 2,006055379 | 4,016824358 | 0,15075079 | 0,617373838 | novel transcript, antisense to PIEZO2 |
| ENSG00000182118 | FAM89A | 2,004390724 | 4,012192216 | 1,76E-08 | 1,06E-06 | family with sequence similarity 89 member A |
| ENSG00000160691 | SHC1 | 2,003573527 | 4,009920202 | 1,04E-11 | 1,02E-09 | SHC adaptor protein 1 |
| ENSG00000260115 | AC012174,1 | 2,002866332 | 4,007955059 | 0,14749491 | 0,611575703 | novel transcript, antisense to CDH8 |
| ENSG00000171873 | ADRA1D | 2,002787523 | 4,007736127 | 0,02261752 | 0,203720621 | adrenoceptor alpha 1D |
| ENSG00000147010 | SH3KBP1 | 2,002357911 | 4,006542861 | 9,85E-08 | 5,30E-06 | SH3 domain containing kinase binding protein 1 |
| ENSG00000251504 | LINC01099 | 2,002037014 | 4,005651792 | 0,0299654 | 0,246731166 | long intergenic non-protein coding RNA 1099 |
| ENSG00000095752 | IL11 | 2,001917636 | 4,005320351 | 0,00674445 | 0,086346599 | interleukin 11 |
| ENSG00000228526 | MIR34AHG | 2,001370924 | 4,003802815 | 0,01246105 | 0,134570746 | MIR34A host gene |
| ENSG00000086717 | PPEF1 | 2,000528562 | 4,001465754 | 7,56E-06 | 0,000267298 | protein phosphatase with EF-hand domain 1 |
| ENSG00000231933 | MYO18B-AS1 | 1,998282185 | 3,995240041 | 0,04053456 | 0,300657756 | MYO18B antisense RNA 1 |
| ENSG00000074416 | MGLL | 1,997682891 | 3,993580765 | 0,00013407 | 0,003380354 | monoglyceride lipase |
| ENSG00000166501 | PRKCB | 1,997651108 | 3,993492787 | 0,00109728 | 0,020215388 | protein kinase C beta |
| ENSG00000096433 | ITPR3 | 1,997091784 | 3,991944834 | 0,00049889 | 0,01040156 | inositol 1,4,5-trisphosphate receptor type 3 |
| ENSG00000186704 | DTX2P1 | 1,997054127 | 3,991840638 | 0,00184771 | 0,031412115 | DTX2 pseudogene 1 |
| ENSG00000139832 | RAB20 | 1,995745329 | 3,988220923 | 0,0038408 | 0,056936307 | RAB20, member RAS oncogene family |
| ENSG00000239926 | PRDX3P4 | 1,994069129 | 3,983589887 | 0,01245365 | 0,134570746 | peroxiredoxin 3 pseudogene 4 |
| ENSG00000226985 | LINC01203 | 1,991961912 | 3,977775658 | 0,03431993 | 0,270005245 | long intergenic non-protein coding RNA 1203 |

| | | | | | | |
|---|---|---|---|---|---|---|
| ENSG00000254275 | LINC00824 | 1,99091569 | 3,974892076 | 0,14827051 | 0,612922036 | long intergenic non-protein coding RNA 824 |
| ENSG00000134121 | CHL1 | 1,989920875 | 3,972152122 | 0,00563426 | 0,075326629 | cell adhesion molecule L1 like |
| ENSG00000179528 | LBX2 | 1,989616184 | 3,97131331 | 0,06999796 | 0,417274281 | ladybird homeobox 2 |
| ENSG00000288520 | AL357075,4 | 1,987011095 | 3,96414874 | 0,35047793 | 0,843911129 | CNK3/IPCEF1 fusion protein |
| ENSG00000169435 | RASSF6 | 1,986423164 | 3,962533587 | 0,15599965 | 0,626944439 | Ras association domain family member 6 |
| ENSG00000105889 | STEAP1B | 1,984919183 | 3,958404878 | 2,57E-06 | 0,000102098 | STEAP family member 1B |
| ENSG00000123342 | MMP19 | 1,982362258 | 3,951395509 | 6,47E-05 | 0,00178784 | matrix metallopeptidase 19 |
| ENSG00000182022 | CHST15 | 1,982262659 | 3,951122728 | 5,33E-10 | 4,21E-08 | carbohydrate sulfotransferase 15 |
| ENSG00000231764 | DLX6-AS1 | 1,982132792 | 3,950767076 | 1,55E-07 | 8,00E-06 | DLX6 antisense RNA 1 |
| ENSG00000106624 | AEBP1 | 1,981791852 | 3,949833535 | 1,25E-05 | 0,000411563 | AE binding protein 1 |
| ENSG00000259319 | AF111167,2 | 1,980355565 | 3,9459032 | 0,06065735 | 0,383412341 | novel transcript, antisense to JDP2 |
| ENSG00000111801 | BTN3A3 | 1,97870264 | 3,941384888 | 5,53E-06 | 0,000202663 | butyrophilin subfamily 3 member A3 |
| ENSG00000288490 | GALNT9 | 1,974365144 | 3,929552819 | 0,01109358 | 0,123533851 | polypeptide N-acetylgalactosaminyltransferase 9 |
| ENSG00000128438 | TBC1D27P | 1,971605323 | 3,922042918 | 0,16181644 | 0,63836332 | TBC1 domain family member 27, pseudogene |
| ENSG00000236808 | POLR1H | 1,970883547 | 3,920081222 | 0,08357337 | 0,460330843 | RNA polymerase I subunit H |
| ENSG00000233795 | POLR1H | 1,970883547 | 3,920081222 | 0,08357337 | 0,460330843 | RNA polymerase I subunit H |
| ENSG00000235176 | POLR1H | 1,970883547 | 3,920081222 | 0,08357337 | 0,460330843 | RNA polymerase I subunit H |
| ENSG00000179097 | HTR1F | 1,970266214 | 3,918404168 | 0,07426335 | 0,430483363 | 5-hydroxytryptamine receptor 1F |
| ENSG00000229848 | AC139149,1 | 1,969448215 | 3,916183087 | 0,08292129 | 0,457949361 | novel transcript, antisense to ACTG1 |
| ENSG00000255298 | OR8G5 | 1,96893985 | 3,914803379 | 0,18678247 | 0,680285072 | olfactory receptor family 8 subfamily G member 5 |
| ENSG00000223725 | AC009226,1 | 1,967919777 | 3,912036354 | 0,16279467 | 0,640444531 | novel transcript |
| ENSG00000233009 | NALCN-AS1 | 1,967154728 | 3,909962385 | 0,26429773 | 0,775650458 | NALCN antisense RNA 1 |
| ENSG00000268864 | AC011487,2 | 1,965915468 | 3,906605208 | 0,07407543 | 0,429744686 | solute carrier family 7 (cationic amino acid transporter, y+ system), member 2 (SLC7A2) pseudogene |
| ENSG00000068079 | IFI35 | 1,963567989 | 3,900253751 | 0,00145687 | 0,025617312 | interferon induced protein 35 |
| ENSG00000115183 | TANC1 | 1,96279191 | 3,898156224 | 1,14E-08 | 7,03E-07 | tetratricopeptide repeat, ankyrin repeat and coiled-coil containing 1 |
| ENSG00000267592 | AC004134,1 | 1,962287802 | 3,896794364 | 0,33596375 | 0,835660292 | transmembrane protein 160 (TMEM160) pseudogene |
| ENSG00000261456 | TUBB8 | 1,962189276 | 3,896528251 | 2,04E-05 | 0,000641527 | tubulin beta 8 class VIII |
| ENSG00000125207 | PIWIL1 | 1,960839252 | 3,89288372 | 0,1288377 | 0,573541504 | piwi like RNA-mediated gene silencing 1 |
| ENSG00000231711 | LINC00899 | 1,959719975 | 3,8898647 | 3,72E-05 | 0,001087785 | long intergenic non-protein coding RNA 899 |
| ENSG00000198691 | ABCA4 | 1,95921613 | 3,888506445 | 2,29E-05 | 0,000712506 | ATP binding cassette subfamily A member 4 |

| | | | | | | |
|---|---|---|---|---|---|---|
| ENSG00000270210 | AC104695,2 | 1,958740766 | 3,887225403 | 0,18176796 | 0,671963282 | novel transcript |
| ENSG00000167880 | EVPL | 1,955245333 | 3,877818641 | 0,39460545 | 0,871872016 | envoplakin |
| ENSG00000175556 | LONRF3 | 1,954681519 | 3,876303463 | 0,00137311 | 0,024474889 | LON peptidase N-terminal domain and ring finger 3 |
| ENSG00000280721 | LINC01943 | 1,954607095 | 3,876103502 | 4,96E-05 | 0,001405597 | long intergenic non-protein coding RNA 1943 |
| ENSG00000081189 | MEF2C | 1,954256292 | 3,87516111 | 3,53E-08 | 2,01E-06 | myocyte enhancer factor 2C |
| ENSG00000156234 | CXCL13 | 1,952841772 | 3,871363492 | 0,19316218 | 0,689970564 | C-X-C motif chemokine ligand 13 |
| ENSG00000226440 | LAMA4-AS1 | 1,952839364 | 3,871357028 | 0,00685939 | 0,087388747 | LAMA4 antisense RNA 1 |
| ENSG00000233049 | DHX16 | 1,952259524 | 3,869801389 | 0,01858788 | 0,177868046 | DEAH-box helicase 16 |
| ENSG00000231377 | DHX16 | 1,952259524 | 3,869801389 | 0,01858788 | 0,177868046 | DEAH-box helicase 16 |
| ENSG00000277930 | AC010722,1 | 1,95012537 | 3,864081089 | 0,14052114 | 0,597347645 | novel transcript |
| ENSG00000180353 | HCLS1 | 1,947395436 | 3,85677621 | 0,01839487 | 0,176657259 | hematopoietic cell-specific Lyn substrate 1 |
| ENSG00000243179 | AC110769,1 | 1,946930467 | 3,855533404 | 0,06751564 | 0,408754627 | novel transcript |
| ENSG00000238189 | ENOX1-AS2 | 1,946716699 | 3,85496216 | 0,25518013 | 0,766521096 | ENOX1 antisense RNA 2 |
| ENSG00000271584 | LINC02550 | 1,943461241 | 3,846273201 | 0,16391278 | 0,642745357 | long intergenic non-protein coding RNA 2550 |
| ENSG00000234685 | NUS1P2 | 1,942796449 | 3,844501251 | 7,83E-06 | 0,00027488 | NUS1 pseudogene 2 |
| ENSG00000089041 | P2RX7 | 1,942455101 | 3,843591732 | 0,00561796 | 0,075243723 | purinergic receptor P2X 7 |
| ENSG00000111145 | ELK3 | 1,941306744 | 3,840533526 | 3,84E-09 | 2,58E-07 | ETS transcription factor ELK3 |
| ENSG00000170374 | SP7 | 1,939741755 | 3,8363697 | 0,01718674 | 0,168970493 | Sp7 transcription factor |
| ENSG00000261150 | EPPK1 | 1,939064378 | 3,834568861 | 0,02472122 | 0,217530026 | epiplakin 1 |
| ENSG00000130707 | ASS1 | 1,937379021 | 3,830091933 | 3,98E-05 | 0,001151515 | argininosuccinate synthase 1 |
| ENSG00000165886 | UBTD1 | 1,937002335 | 3,82909203 | 9,87E-05 | 0,002587659 | ubiquitin domain containing 1 |
| ENSG00000273769 | LCA10 | 1,935618793 | 3,825421697 | 0,17965844 | 0,668298509 | lung carcinoma-associated 10 |
| ENSG00000136541 | ERMN | 1,934998194 | 3,823776483 | 0,10603162 | 0,519559347 | ermin |
| ENSG00000196431 | CRYBA4 | 1,9337109 | 3,820366109 | 0,06245696 | 0,390296699 | crystallin beta A4 |
| ENSG00000248429 | FAM198B-AS1 | 1,933664875 | 3,820244233 | 0,03529441 | 0,2748305 | FAM198B antisense RNA 1 |
| ENSG00000284760 | AC246795,1 | 1,933612242 | 3,820104864 | 0,26877265 | 0,780939404 | RNA binding protein, fox-1 homolog (C, elegans) 1 (RBFOX1) pseudogene |
| ENSG00000072163 | LIMS2 | 1,933499395 | 3,81980607 | 1,54E-05 | 0,000495069 | LIM zinc finger domain containing 2 |
| ENSG00000273866 | Metazoa_SRP | 1,932998715 | 3,818480655 | 0,10129724 | 0,506227112 | Metazoan signal recognition particle RNA |
| ENSG00000215533 | LINC00189 | 1,932997433 | 3,818477263 | 0,1314107 | 0,579140697 | long intergenic non-protein coding RNA 189 |
| ENSG00000117600 | PLPPR4 | 1,932189374 | 3,816339118 | 0,00442936 | 0,063178325 | phospholipid phosphatase related 4 |
| ENSG00000173546 | CSPG4 | 1,932083975 | 3,816060318 | 0,00033783 | 0,007496035 | chondroitin sulfate proteoglycan 4 |
| ENSG00000213383 | AC104297,1 | 1,930906757 | 3,812947737 | 0,00599965 | 0,078887556 | pyrophosphatase (inorganic) 1 (PPA1) pseudogene |

| | | | | | | |
|---|---|---|---|---|---|---|
| ENSG00000104368 | PLAT | 1,930581068 | 3,81208706 | 5,56E-06 | 0,000203179 | plasminogen activator, tissue type |
| ENSG00000145632 | PLK2 | 1,930523376 | 3,811934622 | 7,68E-16 | 1,21E-13 | polo like kinase 2 |
| ENSG00000146072 | TNFRSF21 | 1,930474269 | 3,811804873 | 4,06E-13 | 4,76E-11 | TNF receptor superfamily member 21 |
| ENSG00000224243 | SOX1-OT | 1,928381621 | 3,806279807 | 2,40E-05 | 0,000740338 | SOX1 overlapping transcript |
| ENSG00000162641 | AKNAD1 | 1,927949421 | 3,8051397 | 0,00205688 | 0,0343696 | AKNA domain containing 1 |
| ENSG00000259518 | LINC01583 | 1,927822861 | 3,804805909 | 0,03012669 | 0,247798167 | long intergenic non-protein coding RNA 1583 |
| ENSG00000142698 | C1orf94 | 1,926795185 | 3,802096595 | 0,06066853 | 0,383412341 | chromosome 1 open reading frame 94 |
| ENSG00000167601 | AXL | 1,925013646 | 3,797404401 | 3,45E-05 | 0,001022847 | AXL receptor tyrosine kinase |
| ENSG00000261090 | AC127459,2 | 1,924907143 | 3,797124079 | 0,19570914 | 0,69375328 | novel transcript |
| ENSG00000161638 | ITGA5 | 1,923838926 | 3,794313609 | 3,21E-09 | 2,19E-07 | integrin subunit alpha 5 |
| ENSG00000141448 | GATA6 | 1,923457588 | 3,793310817 | 0,00791225 | 0,097045395 | GATA binding protein 6 |
| ENSG00000183166 | CALN1 | 1,923261934 | 3,792796415 | 0,00415563 | 0,060456726 | calneuron 1 |
| ENSG00000180287 | PLD5 | 1,92200881 | 3,789503423 | 9,45E-05 | 0,002500835 | phospholipase D family member 5 |
| ENSG00000281369 | AC093826,1 | 1,92197223 | 3,789407341 | 0,17732995 | 0,663639407 | |
| ENSG00000026103 | FAS | 1,921354292 | 3,787784603 | 3,51E-06 | 0,000135172 | Fas cell surface death receptor |
| ENSG00000233845 | AC093732,1 | 1,919762773 | 3,783608384 | 0,20011616 | 0,699646707 | novel transcript |
| ENSG00000154856 | APCDD1 | 1,919409602 | 3,782682273 | 8,96E-06 | 0,000309595 | APC down-regulated 1 |
| ENSG00000214944 | ARHGEF28 | 1,918096694 | 3,779241453 | 0,00022855 | 0,005348342 | Rho guanine nucleotide exchange factor 28 |
| ENSG00000163874 | ZC3H12A | 1,917893409 | 3,778708969 | 0,00010587 | 0,002751365 | zinc finger CCCH-type containing 12A |
| ENSG00000101188 | NTSR1 | 1,917156463 | 3,776779254 | 0,00286212 | 0,044740732 | neurotensin receptor 1 |
| ENSG00000010671 | BTK | 1,916311237 | 3,774567215 | 0,12990477 | 0,576076964 | Bruton tyrosine kinase |
| ENSG00000254966 | AC103974,1 | 1,916151211 | 3,774148557 | 0,00986871 | 0,114018922 | novel transcript |
| ENSG00000180269 | GPR139 | 1,915424909 | 3,772249 | 0,17009543 | 0,653300183 | G protein-coupled receptor 139 |
| ENSG00000091592 | NLRP1 | 1,914347783 | 3,769433664 | 0,00160688 | 0,027832381 | NLR family pyrin domain containing 1 |
| ENSG00000223695 | FO393418,1 | 1,913993167 | 3,768507248 | 0,1165853 | 0,545320605 | novel transcript |
| ENSG00000203469 | AL354956,1 | 1,913557155 | 3,7673685 | 0,20893296 | 0,713420724 | novel transcript, antisense to DFFA |
| ENSG00000277371 | Metazoa_SRP | 1,912714011 | 3,765167406 | 0,18344105 | 0,674559627 | Metazoan signal recognition particle RNA |
| ENSG00000282397 | AC104021,1 | 1,911117342 | 3,7610027 | 0,12158101 | 0,557102244 | novel transcript |
| ENSG00000173530 | TNFRSF10D | 1,910687871 | 3,759883266 | 5,22E-09 | 3,40E-07 | TNF receptor superfamily member 10d |
| ENSG00000131634 | TMEM204 | 1,909588477 | 3,757019168 | 2,05E-07 | 1,01E-05 | transmembrane protein 204 |
| ENSG00000223973 | PAFAH1B1P2 | 1,909209046 | 3,756031197 | 0,03897497 | 0,293830089 | platelet activating factor acetylhydrolase 1b regulatory subunit 1 pseudogene 2 |
| ENSG00000259757 | AC091891,1 | 1,908979839 | 3,755434507 | 0,05290381 | 0,353491098 | novel transcript |

| | | | | | | |
|---|---|---|---|---|---|---|
| ENSG00000082497 | SERTAD4 | 1,908591263 | 3,754423154 | 2,52E-05 | 0,000773457 | SERTA domain containing 4 |
| ENSG00000281230 | SERTAD4 | 1,908591263 | 3,754423154 | 2,52E-05 | 0,000773457 | SERTA domain containing 4 |
| ENSG00000233418 | DHX16 | 1,907677642 | 3,752046329 | 0,02886155 | 0,241135096 | DEAH-box helicase 16 |
| ENSG00000233561 | DHX16 | 1,907677642 | 3,752046329 | 0,02886155 | 0,241135096 | DEAH-box helicase 16 |
| ENSG00000183484 | GPR132 | 1,907643599 | 3,751957795 | 0,08177346 | 0,454743073 | G protein-coupled receptor 132 |
| ENSG00000131724 | IL13RA1 | 1,905182544 | 3,745562885 | 2,45E-07 | 1,19E-05 | interleukin 13 receptor subunit alpha 1 |
| ENSG00000275491 | LINC01730 | 1,904906881 | 3,74484727 | 0,18832212 | 0,682711355 | long intergenic non-protein coding RNA 1730 |
| ENSG00000267809 | NDUFV2P1 | 1,904563978 | 3,743957293 | 0,02814352 | 0,237109673 | NADH:ubiquinone oxidoreductase core subunit V2 pseudogene 1 |
| ENSG00000168743 | NPNT | 1,904033531 | 3,742580976 | 0,04062114 | 0,301085663 | nephronectin |
| ENSG00000162542 | TMCO4 | 1,903893092 | 3,742216671 | 0,00101596 | 0,018962991 | transmembrane and coiled-coil domains 4 |
| ENSG00000177363 | LRRN4CL | 1,903779482 | 3,74192199 | 0,0227987 | 0,204880322 | LRRN4 C-terminal like |
| ENSG00000238258 | AL121748,1 | 1,902898498 | 3,739637676 | 0,05146415 | 0,347585883 | novel transcript, antisense to NRP1 |
| ENSG00000288432 | AC270091,1 | 1,900889527 | 3,734433807 | 0,11768851 | 0,548330959 | |
| ENSG00000064012 | CASP8 | 1,900662552 | 3,733846327 | 1,52E-08 | 9,20E-07 | caspase 8 |
| ENSG00000164850 | GPER1 | 1,900209718 | 3,732674527 | 2,11E-09 | 1,48E-07 | G protein-coupled estrogen receptor 1 |
| ENSG00000282291 | SLC39A11 | 1,899583239 | 3,731053995 | 0,15145267 | 0,61895636 | solute carrier family 39 member 11 |
| ENSG00000196581 | AJAP1 | 1,899530965 | 3,73091881 | 0,00065787 | 0,013083699 | adherens junctions associated protein 1 |
| ENSG00000173868 | PHOSPHO1 | 1,897946809 | 3,72682431 | 0,19096856 | 0,68665222 | phosphoethanolamine/phosphocholine phosphatase 1 |
| ENSG00000103966 | EHD4 | 1,897290158 | 3,725128411 | 1,55E-07 | 8,00E-06 | EH domain containing 4 |
| ENSG00000249616 | CCNB3P1 | 1,896371897 | 3,722758158 | 0,12938263 | 0,57482144 | cyclin B3 pseudogene 1 |
| ENSG00000232504 | ST3GAL5-AS1 | 1,89504142 | 3,719326552 | 0,14209719 | 0,600058555 | ST3GAL5 antisense RNA 1 (head to head) |
| ENSG00000076706 | MCAM | 1,894302955 | 3,717423245 | 4,78E-06 | 0,000177239 | melanoma cell adhesion molecule |
| ENSG00000237512 | UNC5B-AS1 | 1,894219024 | 3,717206984 | 0,08051702 | 0,450041893 | UNC5B antisense RNA 1 |
| ENSG00000107807 | TLX1 | 1,894139313 | 3,717001611 | 0,12485949 | 0,564757949 | T cell leukemia homeobox 1 |
| ENSG00000104419 | NDRG1 | 1,893732883 | 3,715954618 | 2,07E-08 | 1,22E-06 | N-myc downstream regulated 1 |
| ENSG00000154127 | UBASH3B | 1,893195661 | 3,714571153 | 3,54E-14 | 4,63E-12 | ubiquitin associated and SH3 domain containing B |
| ENSG00000158869 | FCER1G | 1,892106464 | 3,711767806 | 0,16632839 | 0,646054255 | Fc fragment of IgE receptor Ig |
| ENSG00000154678 | PDE1C | 1,890391036 | 3,707356975 | 7,91E-09 | 4,97E-07 | phosphodiesterase 1C |
| ENSG00000111252 | SH2B3 | 1,890098032 | 3,706604106 | 1,35E-10 | 1,15E-08 | SH2B adaptor protein 3 |
| ENSG00000099139 | PCSK5 | 1,889734437 | 3,705670066 | 1,46E-06 | 6,03E-05 | proprotein convertase subtilisin/kexin type 5 |
| ENSG00000230581 | ACTG1P14 | 1,888948208 | 3,703651129 | 0,11288114 | 0,53680967 | actin gamma 1 pseudogene 14 |

| | | | | | | |
|---|---|---|---|---|---|---|
| ENSG00000137880 | GCHFR | 1,888222567 | 3,701788749 | 0,00728618 | 0,091227554 | GTP cyclohydrolase I feedback regulator |
| ENSG00000104327 | CALB1 | 1,888090206 | 3,701449141 | 4,80E-05 | 0,001365636 | calbindin 1 |
| ENSG00000235749 | AL390860,1 | 1,887866081 | 3,700874161 | 0,12331845 | 0,561682809 | novel transcript, antisense to OR13G1, OR6F1 and OR14A2 |
| ENSG00000281048 | AC114936,3 | 1,887710411 | 3,700474848 | 0,62640504 | 0,952793934 | novel transcript, antisense to PROP1 |
| ENSG00000270661 | Z99289,2 | 1,88653733 | 3,697467149 | 0,01616782 | 0,162061834 | novel transcript, sense intronic LAMA4 |
| ENSG00000168427 | KLHL30 | 1,884948287 | 3,693396853 | 0,07125802 | 0,420579013 | kelch like family member 30 |
| ENSG00000261175 | LINC02188 | 1,88453266 | 3,692332972 | 0,00058932 | 0,012002899 | long intergenic non-protein coding RNA 2188 |
| ENSG00000120833 | SOCS2 | 1,883889817 | 3,690688091 | 0,00021249 | 0,005013731 | suppressor of cytokine signaling 2 |
| ENSG00000171867 | PRNP | 1,882623015 | 3,687448792 | 2,77E-05 | 0,000838216 | prion protein |
| ENSG00000286263 | AL391832,4 | 1,882235251 | 3,686457823 | 0,13210402 | 0,580641112 | novel transcript |
| ENSG00000136010 | ALDH1L2 | 1,881968999 | 3,685777543 | 5,14E-11 | 4,56E-09 | aldehyde dehydrogenase 1 family member L2 |
| ENSG00000139278 | GLIPR1 | 1,881750269 | 3,685218777 | 3,74E-08 | 2,11E-06 | GLI pathogenesis related 1 |
| ENSG00000166145 | SPINT1 | 1,881573374 | 3,684766943 | 0,00761896 | 0,094373667 | serine peptidase inhibitor, Kunitz type 1 |
| ENSG00000264843 | AP001180,4 | 1,88092909 | 3,683121755 | 0,29233679 | 0,802771257 | novel transcript, antisense to PIEZO2 |
| ENSG00000204136 | GGTA1 | 1,880863306 | 3,682953816 | 0,02037706 | 0,189154353 | glycoprotein alpha-galactosyltransferase 1 (inactive) |
| ENSG00000076662 | ICAM3 | 1,880654563 | 3,682420968 | 0,00025269 | 0,005821706 | intercellular adhesion molecule 3 |
| ENSG00000031081 | ARHGAP31 | 1,880619823 | 3,682332298 | 8,50E-07 | 3,67E-05 | Rho GTPase activating protein 31 |
| ENSG00000213886 | UBD | 1,880037024 | 3,680845063 | 0,08899216 | 0,476005294 | ubiquitin D |
| ENSG00000010278 | CD9 | 1,878631253 | 3,677260172 | 5,24E-07 | 2,35E-05 | CD9 molecule |
| ENSG00000256377 | AC009509,1 | 1,878261437 | 3,676317674 | 0,19108375 | 0,686751133 | novel transcript |
| ENSG00000184716 | SERINC4 | 1,878244811 | 3,676275309 | 0,15724111 | 0,629552485 | serine incorporator 4 |
| ENSG00000234779 | BNC2-AS1 | 1,877528658 | 3,674450859 | 0,08274322 | 0,457565724 | BNC2 antisense RNA 1 |
| ENSG00000105851 | PIK3CG | 1,877180556 | 3,673564375 | 0,1688513 | 0,650760243 | phosphatidylinositol-4,5-bisphosphate 3-kinase catalytic subunit gamma |
| ENSG00000101096 | NFATC2 | 1,876375597 | 3,671515261 | 5,62E-05 | 0,001580755 | nuclear factor of activated T cells 2 |
| ENSG00000127666 | TICAM1 | 1,875968014 | 3,670478149 | 0,00208111 | 0,034737841 | toll like receptor adaptor molecule 1 |
| ENSG00000123685 | BATF3 | 1,874628119 | 3,667070798 | 0,03663265 | 0,281811808 | basic leucine zipper ATF-like transcription factor 3 |
| ENSG00000171227 | TMEM37 | 1,874284459 | 3,666197379 | 0,00039187 | 0,008490231 | transmembrane protein 37 |
| ENSG00000137959 | IFI44L | 1,873704413 | 3,664723655 | 0,00897708 | 0,105909272 | interferon induced protein 44 like |
| ENSG00000213694 | S1PR3 | 1,873662364 | 3,664616843 | 2,04E-09 | 1,43E-07 | sphingosine-1-phosphate receptor 3 |
| ENSG00000140416 | TPM1 | 1,873649551 | 3,664584298 | 5,19E-12 | 5,29E-10 | tropomyosin 1 |
| ENSG00000162520 | SYNC | 1,873414142 | 3,663986382 | 1,19E-05 | 0,00039411 | syncoilin, intermediate filament protein |

| | | | | | | |
|---|---|---|---|---|---|---|
| ENSG00000229983 | AL606468,1 | 1,872226981 | 3,660972613 | 0,34722605 | 0,841896177 | novel transcript |
| ENSG00000185686 | PRAME | 1,869530469 | 3,654136355 | 0,05100799 | 0,345820358 | PRAME nuclear receptor transcriptional regulator |
| ENSG00000132274 | TRIM22 | 1,867941193 | 3,650113166 | 3,93E-05 | 0,001139841 | tripartite motif containing 22 |
| ENSG00000248529 | AC026746,1 | 1,867296917 | 3,648483469 | 0,05020979 | 0,342742134 | novel transcript |
| ENSG00000235538 | AL078602,1 | 1,866952154 | 3,64761169 | 0,15413611 | 0,623607455 | novel transcript |
| ENSG00000105559 | PLEKHA4 | 1,866584687 | 3,646682728 | 0,00310892 | 0,047951302 | pleckstrin homology domain containing A4 |
| ENSG00000139269 | INHBE | 1,863805047 | 3,639663431 | 0,02668807 | 0,228721411 | inhibin subunit beta E |
| ENSG00000229931 | ATXN1-AS1 | 1,862523416 | 3,636431538 | 0,07438309 | 0,430889904 | ATXN1 antisense RNA 1 |
| ENSG00000242242 | NECTIN3-AS1 | 1,862312911 | 3,635900984 | 6,99E-15 | 9,89E-13 | NECTIN3 antisense RNA 1 |
| ENSG00000170545 | SMAGP | 1,857418872 | 3,623587852 | 0,00058594 | 0,011949606 | small cell adhesion glycoprotein |
| ENSG00000272668 | AL590560,3 | 1,855903541 | 3,619783823 | 0,00245375 | 0,039461957 | uncharacterized LOC107985216 |
| ENSG00000159247 | TUBBP5 | 1,855757102 | 3,61941642 | 0,00294417 | 0,045726071 | tubulin beta pseudogene 5 |
| ENSG00000277168 | Metazoa_SRP | 1,854714589 | 3,616801921 | 0,17771303 | 0,664232629 | Metazoan signal recognition particle RNA |
| ENSG00000274334 | VASN | 1,854462334 | 3,616169578 | 0,02093965 | 0,192886605 | vasorin |
| ENSG00000055732 | MCOLN3 | 1,853676815 | 3,614201183 | 0,06968978 | 0,416593001 | mucolipin 3 |
| ENSG00000283213 | AC130466,1 | 1,853384837 | 3,613469799 | 0,10563563 | 0,518704622 | novel transcript |
| ENSG00000168994 | PXDC1 | 1,853164861 | 3,612918876 | 0,00016828 | 0,004087798 | PX domain containing 1 |
| ENSG00000176134 | AL445665,1 | 1,853033033 | 3,612588757 | 1,11E-05 | 0,000372547 | methylenetetrahydrofolate dehydrogenase (NADP+ dependent) 1-like (MTHFD1L) pseudogene |
| ENSG00000231233 | CFAP58-DT | 1,852933357 | 3,61233917 | 0,13829834 | 0,593030588 | CFAP58 divergent transcript |
| ENSG00000197582 | GPX1P1 | 1,852704477 | 3,611766128 | 0,30496679 | 0,813769074 | glutathione peroxidase pseudogene 1 |
| ENSG00000258932 | AL390334,1 | 1,850888041 | 3,607221567 | 0,00067904 | 0,013444997 | ribosomal protein S27a (RPS27A) pseudogene |
| ENSG00000260428 | SCX | 1,850818033 | 3,607046528 | 0,02084311 | 0,192183647 | scleraxis bHLH transcription factor |
| ENSG00000196739 | COL27A1 | 1,849422352 | 3,603558714 | 0,00260598 | 0,041399037 | collagen type XXVII alpha 1 chain |
| ENSG00000183908 | LRRC55 | 1,849225506 | 3,603067066 | 5,74E-05 | 0,001605565 | leucine rich repeat containing 55 |
| ENSG00000230254 | HLA-E | 1,848993713 | 3,60248822 | 0,00013918 | 0,003481094 | major histocompatibility complex, class I, E |
| ENSG00000229252 | HLA-E | 1,848993713 | 3,60248822 | 0,00013918 | 0,003481094 | major histocompatibility complex, class I, E |
| ENSG00000259883 | EHD4-AS1 | 1,848391383 | 3,600984484 | 0,04257996 | 0,310307679 | EHD4 antisense RNA 1 |
| ENSG00000121075 | TBX4 | 1,847991721 | 3,599987061 | 0,12760788 | 0,570422178 | T-box transcription factor 4 |
| ENSG00000136244 | IL6 | 1,846657086 | 3,596658258 | 0,22895059 | 0,737173617 | interleukin 6 |
| ENSG00000198108 | CHSY3 | 1,845608381 | 3,594044773 | 2,19E-10 | 1,82E-08 | chondroitin sulfate synthase 3 |
| ENSG00000263904 | AC015563,1 | 1,844782127 | 3,591986996 | 0,11173199 | 0,534453626 | novel transcript, sense intronic to FAM59A |
| ENSG00000255433 | LINC02735 | 1,843570878 | 3,588972522 | 0,17322204 | 0,658193239 | long intergenic non-protein coding RNA 2735 |

| | | | | | | |
|---|---|---|---|---|---|---|
| ENSG00000237988 | OR2I1P | 1,84289396 | 3,587288959 | 0,06444364 | 0,397871785 | olfactory receptor family 2 subfamily I member 1 pseudogene |
| ENSG00000197594 | ENPP1 | 1,842734079 | 3,586891433 | 7,24E-07 | 3,16E-05 | ectonucleotide pyrophosphatase/phosphodiesterase 1 |
| ENSG00000187773 | DIPK1C | 1,842487543 | 3,586278537 | 0,04808805 | 0,334042943 | divergent protein kinase domain 1C |
| ENSG00000113140 | SPARC | 1,841768495 | 3,58449156 | 1,41E-09 | 1,04E-07 | secreted protein acidic and cysteine rich |
| ENSG00000148841 | ITPRIP | 1,84058754 | 3,581558584 | 5,62E-07 | 2,51E-05 | inositol 1,4,5-trisphosphate receptor interacting protein |
| ENSG00000264188 | AC106037,2 | 1,840178512 | 3,580543295 | 0,25454404 | 0,765935007 | novel transcript |
| ENSG00000135549 | PKIB | 1,839331216 | 3,578441057 | 9,55E-05 | 0,0025221 | cAMP-dependent protein kinase inhibitor beta |
| ENSG00000272430 | LINC02637 | 1,837964423 | 3,575052487 | 0,03824042 | 0,289829969 | long intergenic non-protein coding RNA 2637 |
| ENSG00000224505 | AC138150,1 | 1,837888216 | 3,574863648 | 0,43492418 | 0,896272827 | novel transcript, antisense to HEXIM1 and HEXIM2 |
| ENSG00000156049 | GNA14 | 1,836562448 | 3,571580027 | 0,01079207 | 0,121193719 | G protein subunit alpha 14 |
| ENSG00000007312 | CD79B | 1,836394369 | 3,571163951 | 0,18792562 | 0,682064851 | CD79b molecule |
| ENSG00000287283 | AL117353,1 | 1,836015562 | 3,570226397 | 0,10993408 | 0,528752437 | novel transcript, antisense to LRRC7 |
| ENSG00000141738 | GRB7 | 1,834240454 | 3,565836253 | 0,0602392 | 0,381629797 | growth factor receptor bound protein 7 |
| ENSG00000272459 | AC139795,3 | 1,831932336 | 3,560135954 | 0,05449435 | 0,360376004 | novel transcript |
| ENSG00000274840 | AC132807,2 | 1,830880108 | 3,557540319 | 0,05120394 | 0,346269002 | novel transcript |
| ENSG00000042493 | CAPG | 1,830613711 | 3,556883474 | 3,29E-05 | 0,000978652 | capping actin protein, gelsolin like |
| ENSG00000137962 | ARHGAP29 | 1,829451131 | 3,554018351 | 3,82E-07 | 1,77E-05 | Rho GTPase activating protein 29 |
| ENSG00000269113 | TRABD2B | 1,828905023 | 3,552673291 | 3,88E-05 | 0,001127746 | TraB domain containing 2B |
| ENSG00000126010 | GRPR | 1,828647825 | 3,552039993 | 0,00051646 | 0,010702297 | gastrin releasing peptide receptor |
| ENSG00000198838 | RYR3 | 1,82831089 | 3,551210526 | 0,00875244 | 0,103883368 | ryanodine receptor 3 |
| ENSG00000260103 | AC012435,2 | 1,827278368 | 3,548669869 | 0,07619274 | 0,436280172 | coiled-coil-helix-coiled-coil-helix domain-containing protein 7 (CHCHD7) pseudogene |
| ENSG00000139289 | PHLDA1 | 1,827035232 | 3,548071867 | 1,12E-09 | 8,34E-08 | pleckstrin homology like domain family A member 1 |
| ENSG00000168615 | ADAM9 | 1,824560333 | 3,541990476 | 3,81E-09 | 2,57E-07 | ADAM metallopeptidase domain 9 |
| ENSG00000282907 | Z98883,1 | 1,824443648 | 3,541704013 | 0,05260953 | 0,352155381 | novel transcript |
| ENSG00000130429 | ARPC1B | 1,823523695 | 3,539446319 | 8,96E-08 | 4,84E-06 | actin related protein 2/3 complex subunit 1B |
| ENSG00000134762 | DSC3 | 1,82295632 | 3,538054619 | 3,49E-11 | 3,19E-09 | desmocollin 3 |
| ENSG00000248636 | AC002070,1 | 1,822945304 | 3,538027604 | 0,16799636 | 0,649386526 | novel transcript |
| ENSG00000016402 | IL20RA | 1,822162994 | 3,536109608 | 0,26555787 | 0,776824277 | interleukin 20 receptor subunit alpha |
| ENSG00000077009 | NMRK2 | 1,821696109 | 3,534965437 | 0,08181413 | 0,454749316 | nicotinamide riboside kinase 2 |

| | | | | | | |
|---|---|---|---|---|---|---|
| ENSG00000128591 | FLNC | 1,821572333 | 3,534662167 | 0,00408208 | 0,059608622 | filamin C |
| ENSG00000261083 | LINC02516 | 1,821438513 | 3,534334318 | 0,21556797 | 0,722999018 | long intergenic non-protein coding RNA 2516 |
| ENSG00000233521 | LINC01638 | 1,821416958 | 3,534281513 | 0,04230422 | 0,308729941 | long intergenic non-protein coding RNA 1638 |
| ENSG00000233858 | LINC02599 | 1,821226879 | 3,533815892 | 0,02210442 | 0,200252843 | long intergenic non-protein coding RNA 2599 |
| ENSG00000140092 | FBLN5 | 1,819754119 | 3,530210274 | 8,18E-09 | 5,11E-07 | fibulin 5 |
| ENSG00000224057 | EGFR-AS1 | 1,819466365 | 3,529506224 | 0,00501489 | 0,068886379 | EGFR antisense RNA 1 |
| ENSG00000232762 | AL355483,2 | 1,81879641 | 3,527867581 | 0,13617325 | 0,588190164 | novel transcript |
| ENSG00000160932 | LY6E | 1,818334116 | 3,526737299 | 3,90E-07 | 1,80E-05 | lymphocyte antigen 6 family member E |
| ENSG00000254542 | NAV2-AS3 | 1,816483866 | 3,522217174 | 0,30870776 | 0,816730513 | NAV2 antisense RNA 3 |
| ENSG00000137801 | THBS1 | 1,815926902 | 3,520857655 | 1,29E-05 | 0,000424456 | thrombospondin 1 |
| ENSG00000238072 | CDC26P1 | 1,815715425 | 3,52034159 | 0,07771673 | 0,440494637 | cell division cycle 26 pseudogene 1 |
| ENSG00000281955 | AC073488,3 | 1,814958866 | 3,518495984 | 0,29223021 | 0,802771257 | chromodomain protein, Y chromosome-like (CDYL) pseudogene |
| ENSG00000229470 | GNL1 | 1,813946927 | 3,516028896 | 0,02364447 | 0,210168399 | G protein nucleolar 1 (putative) |
| ENSG00000261757 | AC005592,1 | 1,812598832 | 3,512744954 | 0,1893478 | 0,684603876 | novel transcript |
| ENSG00000177374 | HIC1 | 1,812123245 | 3,511587163 | 0,00362468 | 0,054324998 | HIC ZBTB transcriptional repressor 1 |
| ENSG00000101443 | WFDC2 | 1,810819377 | 3,508414919 | 1,27E-07 | 6,64E-06 | WAP four-disulfide core domain 2 |
| ENSG00000183876 | ARSI | 1,810160005 | 3,506811793 | 0,0011767 | 0,02136371 | arylsulfatase family member I |
| ENSG00000151917 | BEND6 | 1,809727177 | 3,505759859 | 2,96E-08 | 1,71E-06 | BEN domain containing 6 |
| ENSG00000138347 | MYPN | 1,809710027 | 3,505718187 | 0,10673276 | 0,521552542 | myopalladin |
| ENSG00000269374 | AC011497,2 | 1,808534797 | 3,502863565 | 0,00083012 | 0,015923883 | zinc finger, AN1-type domain 5 (ZFAND5) pseudogene |
| ENSG00000145040 | UCN2 | 1,806117077 | 3,496998257 | 0,11417419 | 0,540021451 | urocortin 2 |
| ENSG00000153233 | PTPRR | 1,805174566 | 3,494714419 | 0,01615141 | 0,162001207 | protein tyrosine phosphatase receptor type R |
| ENSG00000128016 | ZFP36 | 1,803971602 | 3,49180163 | 5,07E-08 | 2,83E-06 | ZFP36 ring finger protein |
| ENSG00000272444 | AL118558,4 | 1,803517033 | 3,490701597 | 0,21371617 | 0,721156449 | novel transcript |
| ENSG00000132205 | EMILIN2 | 1,803029483 | 3,489522134 | 4,17E-06 | 0,000157241 | elastin microfibril interfacer 2 |
| ENSG00000249306 | LINC01411 | 1,802184876 | 3,487479838 | 0,09819471 | 0,497481654 | long intergenic non-protein coding RNA 1411 |
| ENSG00000233343 | ATP6V1G1P4 | 1,801298897 | 3,485338784 | 0,08300945 | 0,458031678 | ATPase H+ transporting V1 subunit G1 pseudogene 4 |
| ENSG00000130203 | APOE | 1,800131268 | 3,482519108 | 1,94E-05 | 0,000612879 | apolipoprotein E |
| ENSG00000236138 | DUX4L26 | 1,799931154 | 3,482036084 | 0,22073171 | 0,728572782 | double homeobox 4 like 26 (pseudogene) |
| ENSG00000258045 | Metazoa_SRP | 1,799339309 | 3,480607923 | 0,37451764 | 0,861771195 | Metazoan signal recognition particle RNA |
| ENSG00000172164 | SNTB1 | 1,798066412 | 3,477538319 | 1,98E-07 | 9,83E-06 | syntrophin beta 1 |

| | | | | | | |
|---|---|---|---|---|---|---|
| ENSG00000251363 | LINC02315 | 1,797841562 | 3,476996372 | 2,87E-07 | 1,37E-05 | long intergenic non-protein coding RNA 2315 |
| ENSG00000179057 | IGSF22 | 1,797731359 | 3,476730786 | 0,03066489 | 0,250707942 | immunoglobulin superfamily member 22 |
| ENSG00000235721 | AC013268,3 | 1,797641118 | 3,476513321 | 0,20318838 | 0,703503738 | pseudogene similar to part of zinc finger protein 532 (ZNF532) |
| ENSG00000176753 | C15orf56 | 1,797037225 | 3,475058403 | 0,22288603 | 0,731402952 | chromosome 15 open reading frame 56 |
| ENSG00000245067 | IGFBP7-AS1 | 1,796994896 | 3,474956446 | 0,01178314 | 0,129168284 | IGFBP7 antisense RNA 1 |
| ENSG00000145451 | GLRA3 | 1,796746222 | 3,474357527 | 0,00598302 | 0,078759219 | glycine receptor alpha 3 |
| ENSG00000170381 | SEMA3E | 1,79671952 | 3,474293224 | 3,52E-05 | 0,001039995 | semaphorin 3E |
| ENSG00000286458 | AC083870,1 | 1,794755175 | 3,469565915 | 0,00027706 | 0,006290404 | novel transcript, antisense to AGBL3and CALD1 |
| ENSG00000101335 | MYL9 | 1,793163832 | 3,465740973 | 7,20E-09 | 4,59E-07 | myosin light chain 9 |
| ENSG00000230596 | GPAA1P2 | 1,792910065 | 3,46513141 | 0,01800137 | 0,174132087 | glycosylphosphatidylinositol anchor attachment 1 pseudogene 2 |
| ENSG00000115232 | ITGA4 | 1,791830147 | 3,462538582 | 2,30E-08 | 1,35E-06 | integrin subunit alpha 4 |
| ENSG00000132334 | PTPRE | 1,790719013 | 3,459872834 | 7,93E-06 | 0,000277579 | protein tyrosine phosphatase receptor type E |
| ENSG00000102760 | RGCC | 1,788870542 | 3,455442668 | 0,00216698 | 0,035883693 | regulator of cell cycle |
| ENSG00000233124 | LINC00456 | 1,788663135 | 3,454945936 | 0,04771184 | 0,332218942 | long intergenic non-protein coding RNA 456 |
| ENSG00000263961 | RHEX | 1,788406154 | 3,454330576 | 0,00233701 | 0,037975216 | regulator of hemoglobinization and erythroid cell expansion |
| ENSG00000287951 | AC026402,2 | 1,786975909 | 3,450907752 | 0,12662595 | 0,568389605 | novel transcript |
| ENSG00000228643 | AC079779,2 | 1,78537037 | 3,447069459 | 0,01683868 | 0,16634097 | novel transcript |
| ENSG00000139192 | TAPBPL | 1,785338702 | 3,446993795 | 0,00586967 | 0,077626585 | TAP binding protein like |
| ENSG00000230690 | AC013402,1 | 1,785141821 | 3,446523425 | 0,17398916 | 0,659185304 | novel transcript |
| ENSG00000275564 | FCAR | 1,784922128 | 3,44599863 | 0,39781465 | 0,874263835 | Fc fragment of IgA receptor |
| ENSG00000149150 | SLC43A1 | 1,784585309 | 3,445194203 | 0,00012373 | 0,003152571 | solute carrier family 43 member 1 |
| ENSG00000279851 | CR382287,1 | 1,782311568 | 3,439768726 | 0,04734283 | 0,330680606 | novel transcript |
| ENSG00000132975 | GPR12 | 1,781418912 | 3,437641056 | 0,17740654 | 0,663776507 | G protein-coupled receptor 12 |
| ENSG00000285479 | CACNA1C | 1,779873611 | 3,433960897 | 3,96E-06 | 0,000150731 | calcium voltage-gated channel subunit alpha1 C |
| ENSG00000151067 | CACNA1C | 1,779873611 | 3,433960897 | 3,96E-06 | 0,000150731 | calcium voltage-gated channel subunit alpha1 C |
| ENSG00000251563 | IARS2P1 | 1,779099537 | 3,432118907 | 0,30573113 | 0,814657623 | IARS2 pseudogene 1 |
| ENSG00000266076 | AC004805,1 | 1,778953936 | 3,431772546 | 0,07187796 | 0,422642955 | novel transcript |
| ENSG00000181019 | NQO1 | 1,778922972 | 3,431698893 | 2,15E-11 | 2,02E-09 | NAD(P)H quinone dehydrogenase 1 |
| ENSG00000140564 | FURIN | 1,778735183 | 3,431252234 | 3,70E-05 | 0,001083221 | furin, paired basic amino acid cleaving enzyme |
| ENSG00000171056 | SOX7 | 1,776996372 | 3,427119202 | 0,02755048 | 0,233868632 | SRY-box transcription factor 7 |

| | | | | | | |
|---|---|---|---|---|---|---|
| ENSG00000277971 | AC007731,4 | 1,776782792 | 3,42661188 | 0,17847789 | 0,665863105 | novel protein |
| ENSG00000222041 | CYTOR | 1,77672452 | 3,426473478 | 2,85E-11 | 2,65E-09 | cytoskeleton regulator RNA |
| ENSG00000168081 | PNOC | 1,776257799 | 3,42536517 | 0,14700138 | 0,610744318 | prepronociceptin |
| ENSG00000214286 | PDCL3P3 | 1,775964085 | 3,424667882 | 0,0324528 | 0,26082374 | PDCL3 pseudogene 3 |
| ENSG00000141338 | ABCA8 | 1,775350699 | 3,423212137 | 0,00574306 | 0,076404352 | ATP binding cassette subfamily A member 8 |
| ENSG00000149596 | JPH2 | 1,774982127 | 3,422337705 | 0,00411605 | 0,06004867 | junctophilin 2 |
| ENSG00000285778 | AL591463,1 | 1,774725678 | 3,421729415 | 0,01243123 | 0,134387939 | novel transcript |
| ENSG00000204446 | LINC02872 | 1,774432229 | 3,421033495 | 0,01321955 | 0,140068402 | long intergenic non-protein coding RNA 2872 |
| ENSG00000235220 | HLA-F | 1,774089995 | 3,420222058 | 0,32034679 | 0,824366234 | major histocompatibility complex, class I, F |
| ENSG00000175040 | CHST2 | 1,772503706 | 3,41646348 | 0,00068771 | 0,013590653 | carbohydrate sulfotransferase 2 |
| ENSG00000256463 | SALL3 | 1,772053644 | 3,415397849 | 0,00029906 | 0,00673626 | spalt like transcription factor 3 |
| ENSG00000277015 | SALL3 | 1,772053644 | 3,415397849 | 0,00029906 | 0,00673626 | spalt like transcription factor 3 |
| ENSG00000224888 | AC138028,2 | 1,771757974 | 3,414697962 | 0,07392152 | 0,429530795 | novel transcript, antisense to FAM38A |
| ENSG00000225492 | GBP1P1 | 1,771070222 | 3,413070516 | 0,16977529 | 0,652551597 | guanylate binding protein 1 pseudogene 1 |
| ENSG00000174307 | PHLDA3 | 1,770751337 | 3,412316195 | 0,00385839 | 0,057115888 | pleckstrin homology like domain family A member 3 |
| ENSG00000185652 | NTF3 | 1,769221086 | 3,408698707 | 0,00065282 | 0,012999839 | neurotrophin 3 |
| ENSG00000167716 | WDR81 | 1,768145887 | 3,406159248 | 0,00316835 | 0,048704482 | WD repeat domain 81 |
| ENSG00000254885 | AP001547,1 | 1,765814701 | 3,400659834 | 0,13478017 | 0,58573356 | novel transcript |
| ENSG00000254211 | LINC01485 | 1,76563562 | 3,400237736 | 0,16587901 | 0,645166759 | long intergenic non-protein coding RNA 1485 |
| ENSG00000273580 | AC087072,1 | 1,763685173 | 3,395643902 | 0,22737773 | 0,735964811 | zinc finger protein 383 (ZNF383) pseudogene |
| ENSG00000072952 | IRAG1 | 1,763284007 | 3,394699816 | 1,43E-09 | 1,05E-07 | inositol 1,4,5-triphosphate receptor associated 1 |
| ENSG00000145431 | PDGFC | 1,762545206 | 3,392961843 | 5,13E-07 | 2,31E-05 | platelet derived growth factor C |
| ENSG00000101000 | PROCR | 1,761392822 | 3,390252724 | 0,00288214 | 0,04491362 | protein C receptor |
| ENSG00000150722 | PPP1R1C | 1,761033176 | 3,389407681 | 0,00983688 | 0,113777361 | protein phosphatase 1 regulatory inhibitor subunit 1C |
| ENSG00000272958 | AP000322,1 | 1,758777982 | 3,384113562 | 0,38885826 | 0,870057277 | novel transcript |
| ENSG00000251694 | USP17L9P | 1,75847459 | 3,383401972 | 0,53463141 | 0,931649095 | ubiquitin specific peptidase 17 like family member 9, pseudogene |
| ENSG00000287542 | AC098582,1 | 1,756523562 | 3,378829524 | 0,36787847 | 0,857401877 | HECT and RLD domain containing E3 ubiquitin protein ligase 3 |
| ENSG00000155269 | GPR78 | 1,756488462 | 3,37874732 | 0,25146754 | 0,763892102 | G protein-coupled receptor 78 |
| ENSG00000270124 | AC092127,2 | 1,756243117 | 3,378172779 | 0,06568833 | 0,402209142 | novel transcript |
| ENSG00000164430 | CGAS | 1,755183248 | 3,375691929 | 1,26E-07 | 6,64E-06 | cyclic GMP-AMP synthase |

| | | | | | | |
|---|---|---|---|---|---|---|
| ENSG00000259697 | AC126323,2 | 1,754126454 | 3,373220095 | 0,11927928 | 0,552100834 | novel transcript |
| ENSG00000205572 | SERF1B | 1,753452944 | 3,371645703 | 0,00994802 | 0,114638731 | small EDRK-rich factor 1B |
| ENSG00000178033 | CALHM5 | 1,752589538 | 3,369628487 | 5,09E-10 | 4,04E-08 | calcium homeostasis modulator family member 5 |
| ENSG00000182718 | ANXA2 | 1,752413751 | 3,369217935 | 1,97E-15 | 2,94E-13 | annexin A2 |
| ENSG00000227063 | RPL41P1 | 1,751956907 | 3,368151206 | 0,25964325 | 0,772176335 | ribosomal protein L41 pseudogene 1 |
| ENSG00000282032 | AC243899,2 | 1,751618946 | 3,367362287 | 0,22470701 | 0,733203418 | C17orf58 pseudogene |
| ENSG00000261575 | AC005829,1 | 1,751618946 | 3,367362287 | 0,22470701 | 0,733203418 | C17orf58 pseudogene |
| ENSG00000177103 | DSCAML1 | 1,751116534 | 3,366189823 | 0,01186061 | 0,129653833 | DS cell adhesion molecule like 1 |
| ENSG00000285072 | AC270272,2 | 1,749630399 | 3,36272406 | 0,00984557 | 0,113835769 | |
| ENSG00000183199 | HSP90AB3P | 1,748314286 | 3,359657781 | 0,2369591 | 0,746456145 | heat shock protein 90 alpha family class B member 3, pseudogene |
| ENSG00000206457 | CCHCR1 | 1,747163791 | 3,35697965 | 0,10271396 | 0,510289111 | coiled-coil alpha-helical rod protein 1 |
| ENSG00000233841 | HLA-C | 1,746821232 | 3,356182649 | 0,00053738 | 0,011089529 | major histocompatibility complex, class I, C |
| ENSG00000250993 | AC020551,1 | 1,745939206 | 3,354131395 | 0,00839034 | 0,101084971 | novel transcript |
| ENSG00000154342 | WNT3A | 1,745389555 | 3,352853749 | 0,32060877 | 0,824366234 | Wnt family member 3A |
| ENSG00000256802 | AC022613,1 | 1,744896265 | 3,35170753 | 0,00427289 | 0,061690033 | novel transcript, antisense to TJP1 |
| ENSG00000248629 | AC092608,1 | 1,744513981 | 3,350819515 | 0,30090696 | 0,810566078 | novel transcript |
| ENSG00000262280 | LINC01896 | 1,743522083 | 3,348516514 | 0,01742162 | 0,170657076 | long intergenic non-protein coding RNA 1896 |
| ENSG00000231535 | LINC00278 | 1,743480957 | 3,34842106 | 0,06798873 | 0,410664541 | long intergenic non-protein coding RNA 278 |
| ENSG00000188089 | PLA2G4E | 1,743381443 | 3,348190102 | 0,06134735 | 0,38567538 | phospholipase A2 group IVE |
| ENSG00000173213 | TUBB8B | 1,743365447 | 3,348152979 | 0,01646232 | 0,163910156 | tubulin beta 8B |
| ENSG00000079337 | RAPGEF3 | 1,742249604 | 3,345564373 | 0,00044727 | 0,00950052 | Rap guanine nucleotide exchange factor 3 |
| ENSG00000231163 | CSMD2-AS1 | 1,742151112 | 3,345335981 | 0,16285336 | 0,640548464 | CSMD2 antisense RNA 1 |
| ENSG00000261371 | PECAM1 | 1,741669316 | 3,344218975 | 0,05804283 | 0,372771355 | platelet and endothelial cell adhesion molecule 1 |
| ENSG00000108684 | ASIC2 | 1,740597783 | 3,341736044 | 0,06222764 | 0,389221888 | acid sensing ion channel subunit 2 |
| ENSG00000149633 | KIAA1755 | 1,740483664 | 3,341471719 | 4,74E-06 | 0,000176069 | KIAA1755 |
| ENSG00000254154 | CRYZL2P-SEC16 | 1,740133691 | 3,340661234 | 4,24E-06 | 0,000159062 | CRYZL2P-SEC16B readthrough |
| ENSG00000254554 | AC080023,1 | 1,739892899 | 3,340103711 | 0,31138132 | 0,817751312 | novel transcript |
| ENSG00000269403 | AC008750,7 | 1,739870237 | 3,340051245 | 0,30344462 | 0,812669425 | novel protein |
| ENSG00000165970 | SLC6A5 | 1,738279097 | 3,336369551 | 0,07732579 | 0,439467994 | solute carrier family 6 member 5 |
| ENSG00000126561 | STAT5A | 1,737648669 | 3,334911946 | 0,0041605 | 0,060499544 | signal transducer and activator of transcription 5A |
| ENSG00000267263 | AC111170,2 | 1,736965383 | 3,333332846 | 0,18974714 | 0,684955795 | novel transcript, sense intronic to SEPT9 |
| ENSG00000262445 | AC099684,2 | 1,735963725 | 3,331019328 | 0,19169504 | 0,687672451 | novel transcript, antisense to RTN4RL1 |

| | | | | | | |
|---|---|---|---|---|---|---|
| ENSG00000147481 | SNTG1 | 1,734754995 | 3,328229676 | 0,00438421 | 0,062760045 | syntrophin gamma 1 |
| ENSG00000149260 | CAPN5 | 1,734655261 | 3,327999603 | 3,79E-05 | 0,001106769 | calpain 5 |
| ENSG00000104213 | PDGFRL | 1,734222105 | 3,32700055 | 0,00121756 | 0,021965248 | platelet derived growth factor receptor like |
| ENSG00000236444 | UBE2L5 | 1,73409596 | 3,326709661 | 0,24159795 | 0,751948055 | ubiquitin conjugating enzyme E2 L5 |
| ENSG00000270533 | CR382285,1 | 1,734047914 | 3,326598874 | 0,21703641 | 0,724629037 | ribosomal protein L7a (RPL7A) pseudogene |
| ENSG00000198889 | DCAF12L1 | 1,732802801 | 3,323729102 | 0,00045673 | 0,009661271 | DDB1 and CUL4 associated factor 12 like 1 |
| ENSG00000129667 | RHBDF2 | 1,732685533 | 3,323458947 | 0,00574548 | 0,076404352 | rhomboid 5 homolog 2 |
| ENSG00000240350 | SOCAR | 1,732330681 | 3,322641594 | 0,00184085 | 0,031329576 | serous ovarian cancer associated RNA |
| ENSG00000007129 | CEACAM21 | 1,731394614 | 3,320486457 | 1,68E-05 | 0,000538545 | CEA cell adhesion molecule 21 |
| ENSG00000123096 | SSPN | 1,730919148 | 3,319392311 | 0,00038179 | 0,008306423 | sarcospan |
| ENSG00000249740 | OSMR-AS1 | 1,730602933 | 3,318664834 | 0,03548463 | 0,27578235 | OSMR antisense RNA 1 (head to head) |
| ENSG00000258081 | AL110292,1 | 1,730488158 | 3,318400824 | 0,07220347 | 0,42371424 | novel transcript |
| ENSG00000151882 | CCL28 | 1,729142031 | 3,315305989 | 0,00416786 | 0,060578269 | C-C motif chemokine ligand 28 |
| ENSG00000148926 | ADM | 1,729043622 | 3,315079853 | 1,94E-07 | 9,63E-06 | adrenomedullin |
| ENSG00000197324 | LRP10 | 1,728793074 | 3,314504183 | 9,28E-05 | 0,002459412 | LDL receptor related protein 10 |
| ENSG00000276592 | PHBP6 | 1,728212889 | 3,313171512 | 0,27787112 | 0,789162936 | prohibitin pseudogene 6 |
| ENSG00000081320 | STK17B | 1,726936615 | 3,310241824 | 7,85E-07 | 3,41E-05 | serine/threonine kinase 17b |
| ENSG00000099250 | NRP1 | 1,726912953 | 3,310187534 | 5,72E-07 | 2,54E-05 | neuropilin 1 |
| ENSG00000123989 | CHPF | 1,726350662 | 3,308897638 | 0,00038647 | 0,008402299 | chondroitin polymerizing factor |
| ENSG00000225383 | SFTA1P | 1,726053352 | 3,308215812 | 0,00188867 | 0,032038803 | surfactant associated 1, lncRNA |
| ENSG00000065054 | SLC9A3R2 | 1,725544991 | 3,307050306 | 0,01176854 | 0,129098828 | SLC9A3 regulator 2 |
| ENSG00000198680 | TUSC1 | 1,725532218 | 3,307021026 | 1,07E-06 | 4,52E-05 | tumor suppressor candidate 1 |
| ENSG00000281019 | AC116351,3 | 1,725072213 | 3,305966745 | 0,14459044 | 0,605069656 | novel transcript |
| ENSG00000164292 | RHOBTB3 | 1,724372782 | 3,304364372 | 1,86E-05 | 0,000589837 | Rho related BTB domain containing 3 |
| ENSG00000010818 | HIVEP2 | 1,724344312 | 3,304299167 | 1,77E-09 | 1,26E-07 | HIVEP zinc finger 2 |
| ENSG00000278693 | ASB2 | 1,723902297 | 3,303286945 | 0,03325854 | 0,264841921 | ankyrin repeat and SOCS box containing 2 |
| ENSG00000156113 | KCNMA1 | 1,723753159 | 3,302945488 | 0,00022321 | 0,005242952 | potassium calcium-activated channel subfamily M alpha 1 |
| ENSG00000250387 | LINC02197 | 1,723458844 | 3,302271742 | 0,32031845 | 0,824366234 | long intergenic non-protein coding RNA 2197 |
| ENSG00000247317 | LY6E-DT | 1,723239964 | 3,301770773 | 0,18650629 | 0,679754349 | LY6E divergent transcript |
| ENSG00000214190 | RNF152P1 | 1,721423827 | 3,297616953 | 0,18471235 | 0,67694849 | ring finger protein 152 pseudogene 1 |
| ENSG00000176928 | GCNT4 | 1,720075619 | 3,294536748 | 0,00040275 | 0,008671976 | glucosaminyl (N-acetyl) transferase 4 |
| ENSG00000250584 | LINC01511 | 1,719178767 | 3,292489334 | 0,1581813 | 0,631383073 | long intergenic non-protein coding RNA 1511 |

| | | | | | | |
|---|---|---|---|---|---|---|
| ENSG00000122870 | BICC1 | 1,719149206 | 3,292421872 | 1,16E-05 | 0,00038575 | BicC family RNA binding protein 1 |
| ENSG00000095383 | TBC1D2 | 1,718254368 | 3,290380366 | 0,00113186 | 0,020718079 | TBC1 domain family member 2 |
| ENSG00000261522 | AL845331,2 | 1,718252753 | 3,290376682 | 0,00966228 | 0,112089569 | methylenetetrahydrofolate dehydrogenase (NADP+ dependent) 1-like (MTHFD1L) pseudogene |
| ENSG00000140470 | ADAMTS17 | 1,717101481 | 3,287752005 | 0,00310076 | 0,047849048 | ADAM metallopeptidase with thrombospondin type 1 motif 17 |
| ENSG00000170214 | ADRA1B | 1,717064604 | 3,287667967 | 0,04494174 | 0,320350768 | adrenoceptor alpha 1B |
| ENSG00000173482 | PTPRM | 1,716241242 | 3,285792195 | 1,73E-12 | 1,84E-10 | protein tyrosine phosphatase receptor type M |
| ENSG00000170356 | OR2A20P | 1,716046819 | 3,285349419 | 0,03850336 | 0,291183243 | olfactory receptor family 2 subfamily A member 20 pseudogene |
| ENSG00000282365 | LINC02091 | 1,713513661 | 3,279585896 | 0,21458758 | 0,722164151 | long intergenic non-protein coding RNA 2091 |
| ENSG00000175505 | CLCF1 | 1,713334444 | 3,279178519 | 0,0005439 | 0,011201867 | cardiotrophin like cytokine factor 1 |
| ENSG00000253586 | AC067817,1 | 1,713098802 | 3,278642958 | 0,12197075 | 0,55839706 | novel transcript |
| ENSG00000182771 | GRID1 | 1,712841448 | 3,278058152 | 0,00052656 | 0,010880544 | glutamate ionotropic receptor delta type subunit 1 |
| ENSG00000260807 | CEROX1 | 1,712424407 | 3,277110698 | 0,00063568 | 0,012780573 | cytoplasmic endogenous regulator of oxidative phosphorylation 1 |
| ENSG00000234487 | HLA-F | 1,712377326 | 3,277003754 | 0,39929829 | 0,875433675 | major histocompatibility complex, class I, F |
| ENSG00000140859 | KIFC3 | 1,711704907 | 3,275476747 | 0,00316485 | 0,04869389 | kinesin family member C3 |
| ENSG00000263494 | AC004702,1 | 1,710779658 | 3,273376748 | 0,22749707 | 0,736053199 | novel transcript |
| ENSG00000169347 | GP2 | 1,710426254 | 3,272574996 | 0,32857118 | 0,830659639 | glycoprotein 2 |
| ENSG00000163803 | PLB1 | 1,709866767 | 3,271306115 | 0,00031844 | 0,007121478 | phospholipase B1 |
| ENSG00000101955 | SRPX | 1,709547135 | 3,270581432 | 9,04E-09 | 5,61E-07 | sushi repeat containing protein X-linked |
| ENSG00000256100 | AP000721,1 | 1,708959201 | 3,26924886 | 0,24749026 | 0,759699102 | novel transcript |
| ENSG00000158815 | FGF17 | 1,70801057 | 3,267099902 | 0,00163304 | 0,028222909 | fibroblast growth factor 17 |
| ENSG00000235443 | POLR1H | 1,70791652 | 3,266886925 | 0,09643298 | 0,493240029 | RNA polymerase I subunit H |
| ENSG00000236949 | POLR1H | 1,70791652 | 3,266886925 | 0,09643298 | 0,493240029 | RNA polymerase I subunit H |
| ENSG00000266010 | GATA6-AS1 | 1,707910194 | 3,266872598 | 0,2185266 | 0,726304032 | GATA6 antisense RNA 1 (head to head) |
| ENSG00000234292 | AC123595,1 | 1,70772512 | 3,266453539 | 0,27661991 | 0,788353146 | novel transcript |
| ENSG00000077713 | SLC25A43 | 1,707355894 | 3,265617671 | 1,30E-12 | 1,43E-10 | solute carrier family 25 member 43 |
| ENSG00000225169 | BRI3P1 | 1,705582798 | 3,261606637 | 0,19337783 | 0,690313354 | brain protein I3 pseudogene 1 |
| ENSG00000185905 | C16orf54 | 1,70558053 | 3,26160151 | 0,01006147 | 0,11560521 | chromosome 16 open reading frame 54 |
| ENSG00000189431 | RASSF10 | 1,705000108 | 3,260289573 | 0,00115504 | 0,021056164 | Ras association domain family member 10 |
| ENSG00000230930 | COL11A2 | 1,704561536 | 3,259298613 | 0,17651296 | 0,662485145 | collagen type XI alpha 2 chain |

| | | | | | | |
|---|---|---|---|---|---|---|
| ENSG00000275234 | AC010503,4 | 1,703418439 | 3,25671718 | 0,30112121 | 0,810679941 | novel transcript, antisense to DENND1C |
| ENSG00000275009 | GOLGA6L1 | 1,702818196 | 3,255362484 | 0,52627242 | 0,929261495 | golgin A6 family like 1 |
| ENSG00000268388 | FENDRR | 1,699427631 | 3,247720843 | 0,16685874 | 0,647265026 | FOXF1 adjacent non-coding developmental regulatory RNA |
| ENSG00000133116 | KL | 1,699415767 | 3,247694134 | 0,071907 | 0,422734176 | klotho |
| ENSG00000159200 | RCAN1 | 1,698710687 | 3,246107296 | 3,33E-11 | 3,06E-09 | regulator of calcineurin 1 |
| ENSG00000278619 | MRM1 | 1,697789064 | 3,244034279 | 0,01814287 | 0,175067497 | mitochondrial rRNA methyltransferase 1 |
| ENSG00000227911 | LINC02344 | 1,6967203 | 3,241631952 | 0,30731264 | 0,816138329 | long intergenic non-protein coding RNA 2344 |
| ENSG00000282003 | TUBGCP3 | 1,694837673 | 3,237404584 | 0,33471645 | 0,834757602 | tubulin gamma complex associated protein 3 |
| ENSG00000228002 | DHX9P1 | 1,694022135 | 3,235575036 | 0,19725985 | 0,695943345 | DEAH-box helicase 9 pseudogene 1 |
| ENSG00000177606 | JUN | 1,693454303 | 3,234301792 | 0,00039006 | 0,008462812 | Jun proto-oncogene, AP-1 transcription factor subunit |
| ENSG00000141655 | TNFRSF11A | 1,692814801 | 3,232868444 | 0,00357788 | 0,05375238 | TNF receptor superfamily member 11a |
| ENSG00000225014 | KCTD9P1 | 1,691498823 | 3,229920875 | 0,15365919 | 0,622683206 | potassium channel tetramerization domain containing 9 pseudogene 1 |
| ENSG00000223701 | RAET1E-AS1 | 1,690631136 | 3,227978873 | 0,32805353 | 0,830076239 | RAET1E antisense RNA 1 |
| ENSG00000172965 | MIR4435-2HG | 1,68923003 | 3,224845468 | 6,81E-11 | 5,94E-09 | MIR4435-2 host gene |
| ENSG00000233904 | HLA-E | 1,688073969 | 3,22226237 | 6,73E-05 | 0,001844692 | major histocompatibility complex, class I, E |
| ENSG00000275565 | ALOX5 | 1,68802343 | 3,222149492 | 0,00994717 | 0,114638731 | arachidonate 5-lipoxygenase |
| ENSG00000164778 | EN2 | 1,687815461 | 3,221685043 | 0,17165942 | 0,655397237 | engrailed homeobox 2 |
| ENSG00000215146 | BX322639,1 | 1,686849795 | 3,219529335 | 0,00933924 | 0,10911135 | zinc finger protein pseudogene |
| ENSG00000267546 | AC015802,4 | 1,686787083 | 3,219389389 | 0,1135102 | 0,538265283 | novel transcript, antisense to PRCD |
| ENSG00000232310 | AL078590,2 | 1,686702654 | 3,219200989 | 0,05250005 | 0,351770604 | novel transcript |
| ENSG00000225932 | CTAGE4 | 1,68558025 | 3,216697454 | 0,06283978 | 0,391611013 | CTAGE family member 4 |
| ENSG00000206493 | HLA-E | 1,685529427 | 3,21658414 | 6,65E-05 | 0,001829897 | major histocompatibility complex, class I, E |
| ENSG00000230613 | HM13-AS1 | 1,685075097 | 3,215571339 | 0,19808606 | 0,696782178 | HM13 antisense RNA 1 |
| ENSG00000168675 | LDLRAD4 | 1,682979004 | 3,210902824 | 1,15E-07 | 6,13E-06 | low density lipoprotein receptor class A domain containing 4 |
| ENSG00000266835 | GAPLINC | 1,682760696 | 3,21041699 | 0,00426794 | 0,061660248 | gastric adenocarcinoma associated, positive CD44 regulator, long intergenic non-coding RNA |
| ENSG00000163492 | CCDC141 | 1,68161957 | 3,207878656 | 0,00020645 | 0,004898559 | coiled-coil domain containing 141 |
| ENSG00000172640 | OR10AD1 | 1,681438373 | 3,207475783 | 0,01582267 | 0,159883663 | olfactory receptor family 10 subfamily AD member 1 |
| ENSG00000266049 | AP001011,1 | 1,68068862 | 3,205809326 | 0,12189359 | 0,558242525 | novel transcript, antisense to SMCHD1 |

| | | | | | | |
|---|---|---|---|---|---|---|
| ENSG00000163637 | PRICKLE2 | 1,679656242 | 3,203516102 | 2,95E-08 | 1,71E-06 | prickle planar cell polarity protein 2 |
| ENSG00000268186 | ZNF114-AS1 | 1,67698562 | 3,197591449 | 0,14000867 | 0,596271774 | ZNF114 antisense RNA 1 |
| ENSG00000283511 | AC137936,2 | 1,67690998 | 3,197423806 | 0,23073198 | 0,739234644 | novel pseudogene |
| ENSG00000220256 | AC093802,1 | 1,67658961 | 3,196713852 | 0,0865209 | 0,468880561 | novel transcript |
| ENSG00000237530 | AL136140,1 | 1,675476702 | 3,194248829 | 0,18614848 | 0,67937802 | novel transcript |
| ENSG00000271099 | AL357315,2 | 1,675054911 | 3,193315084 | 0,20090374 | 0,700463311 | poly(A) binding protein, cytoplasmic 4 (inducible form) (PABPC4) pseudogene |
| ENSG00000181518 | OR8D4 | 1,674230704 | 3,191491276 | 0,1710259 | 0,654709629 | olfactory receptor family 8 subfamily D member 4 |
| ENSG00000176009 | ASCL3 | 1,674204308 | 3,191432885 | 0,26066993 | 0,772692192 | achaete-scute family bHLH transcription factor 3 |
| ENSG00000160685 | ZBTB7B | 1,674107395 | 3,191218507 | 0,00135029 | 0,02409561 | zinc finger and BTB domain containing 7B |
| ENSG00000142102 | PGGHG | 1,673912512 | 3,190787456 | 0,0043267 | 0,062250352 | protein-glucosylgalactosylhydroxylysine glucosidase |
| ENSG00000287787 | AC092275,1 | 1,673800292 | 3,190539272 | 0,00829482 | 0,10028199 | novel transcript, sense intronic to GSE1 |
| ENSG00000178882 | RFLNA | 1,673740296 | 3,190406593 | 0,00013518 | 0,003402789 | refilin A |
| ENSG00000226747 | FSIP2-AS2 | 1,671302169 | 3,18501942 | 0,06650091 | 0,405279626 | FSIP2 antisense RNA 2 |
| ENSG00000228153 | AL161793,1 | 1,670976975 | 3,184301575 | 0,31734025 | 0,822252078 | novel transcript |
| ENSG00000262528 | AL022341,2 | 1,669767849 | 3,181633922 | 0,27005531 | 0,782267151 | novel transcript, antisense to WDR90 |
| ENSG00000110660 | SLC35F2 | 1,669088261 | 3,180135552 | 5,66E-06 | 0,000205536 | solute carrier family 35 member F2 |
| ENSG00000221882 | OR3A2 | 1,66901555 | 3,179975279 | 0,08020215 | 0,44912052 | olfactory receptor family 3 subfamily A member 2 |
| ENSG00000166148 | AVPR1A | 1,668413737 | 3,178649046 | 0,00279816 | 0,043850441 | arginine vasopressin receptor 1A |
| ENSG00000251381 | LINC00958 | 1,668076792 | 3,17790675 | 0,00014835 | 0,003680878 | long intergenic non-protein coding RNA 958 |
| ENSG00000185274 | GALNT17 | 1,667847884 | 3,177402561 | 0,00821908 | 0,099597438 | polypeptide N-acetylgalactosaminyltransferase 17 |
| ENSG00000227000 | HSPD1P14 | 1,667656022 | 3,17698003 | 0,27755578 | 0,789067728 | heat shock protein family D (Hsp60) member 1 pseudogene 14 |
| ENSG00000080561 | MID2 | 1,666317402 | 3,174033604 | 3,60E-06 | 0,000138308 | midline 2 |
| ENSG00000198842 | STYXL2 | 1,66581951 | 3,172938393 | 0,02283754 | 0,205052511 | serine/threonine/tyrosine interacting like 2 |
| ENSG00000072110 | ACTN1 | 1,665490411 | 3,172214685 | 4,22E-06 | 0,000158782 | actinin alpha 1 |
| ENSG00000070540 | WIPI1 | 1,664164714 | 3,169301064 | 3,80E-11 | 3,46E-09 | WD repeat domain, phosphoinositide interacting 1 |
| ENSG00000168461 | RAB31 | 1,663984983 | 3,168906258 | 0,00010981 | 0,002846593 | RAB31, member RAS oncogene family |
| ENSG00000249565 | SERBP1P5 | 1,663396795 | 3,167614556 | 0,085872 | 0,466580174 | SERPINE1 mRNA binding protein 1 pseudogene 5 |
| ENSG00000124788 | ATXN1 | 1,663174156 | 3,167125762 | 2,01E-07 | 9,95E-06 | ataxin 1 |
| ENSG00000164465 | DCBLD1 | 1,662499919 | 3,165645966 | 4,27E-07 | 1,96E-05 | discoidin, CUB and LCCL domain containing 1 |
| ENSG00000240476 | LINC00973 | 1,661993398 | 3,164534722 | 0,05166648 | 0,34830577 | long intergenic non-protein coding RNA 973 |
| ENSG00000179348 | GATA2 | 1,66089052 | 3,162116497 | 0,05678616 | 0,367995736 | GATA binding protein 2 |

| | | | | | | |
|---|---|---|---|---|---|---|
| ENSG00000143507 | DUSP10 | 1,660369025 | 3,160973686 | 2,49E-05 | 0,000765057 | dual specificity phosphatase 10 |
| ENSG00000230555 | AL450326,1 | 1,660334149 | 3,160897271 | 0,02694037 | 0,230063809 | novel transcript |
| ENSG00000132470 | ITGB4 | 1,659199503 | 3,158412278 | 0,11432254 | 0,540477816 | integrin subunit beta 4 |
| ENSG00000207523 | SNORA66 | 1,658682586 | 3,157280822 | 0,3791483 | 0,864650895 | small nucleolar RNA, H/ACA box 66 |
| ENSG00000169242 | EFNA1 | 1,65778485 | 3,155316774 | 0,00070677 | 0,013905838 | ephrin A1 |
| ENSG00000227676 | LINC01068 | 1,657521089 | 3,154739955 | 0,18329904 | 0,674320815 | long intergenic non-protein coding RNA 1068 |
| ENSG00000228495 | LINC01013 | 1,657139383 | 3,153905389 | 0,00027307 | 0,006213418 | long intergenic non-protein coding RNA 1013 |
| ENSG00000080503 | SMARCA2 | 1,656673757 | 3,15288764 | 2,20E-07 | 1,08E-05 | SWI/SNF related, matrix associated, actin dependent regulator of chromatin, subfamily a, member 2 |
| ENSG00000267107 | PCAT19 | 1,656291717 | 3,152052833 | 0,01403002 | 0,146599474 | prostate cancer associated transcript 19 |
| ENSG00000256029 | AL590560,2 | 1,655987037 | 3,151387229 | 0,12165928 | 0,557379212 | novel protein |
| ENSG00000240666 | MME-AS1 | 1,655153166 | 3,149566267 | 0,33535456 | 0,835232738 | MME antisense RNA 1 |
| ENSG00000288560 | AL360015,1 | 1,654693405 | 3,148562717 | 0,32878771 | 0,830659639 | novel transcript |
| ENSG00000237525 | AC012668,3 | 1,653845136 | 3,146711985 | 0,30049223 | 0,810227614 | novel transcript |
| ENSG00000120658 | ENOX1 | 1,652892955 | 3,144635835 | 4,02E-07 | 1,85E-05 | ecto-NOX disulfide-thiol exchanger 1 |
| ENSG00000235619 | RPL36AP33 | 1,651849906 | 3,142363127 | 0,17473064 | 0,660057513 | ribosomal protein L36a pseudogene 33 |
| ENSG00000277639 | AC007906,2 | 1,651814024 | 3,142284972 | 4,23E-06 | 0,000159024 | novel protein |
| ENSG00000002586 | CD99 | 1,651597932 | 3,141814346 | 5,93E-06 | 0,000214004 | CD99 molecule (Xg blood group) |
| ENSG00000237424 | FOXD2-AS1 | 1,650581241 | 3,139601037 | 0,08774856 | 0,471937196 | FOXD2 adjacent opposite strand RNA 1 |
| ENSG00000200674 | RN7SKP160 | 1,647815748 | 3,133588519 | 0,33168505 | 0,832548086 | RN7SK pseudogene 160 |
| ENSG00000267519 | AC020916,1 | 1,647793913 | 3,133541095 | 0,00883141 | 0,104427089 | novel transcript |
| ENSG00000168484 | SFTPC | 1,646665919 | 3,131092042 | 0,27970914 | 0,790457076 | surfactant protein C |
| ENSG00000261634 | AC026992,2 | 1,646642066 | 3,131040274 | 0,37586683 | 0,862406526 | novel transcript, antisense to PAQR5 |
| ENSG00000166888 | STAT6 | 1,646037944 | 3,12972944 | 5,63E-09 | 3,65E-07 | signal transducer and activator of transcription 6 |
| ENSG00000171291 | ZNF439 | 1,645617567 | 3,128817622 | 3,15E-08 | 1,80E-06 | zinc finger protein 439 |
| ENSG00000125864 | BFSP1 | 1,64538312 | 3,128309212 | 0,00157925 | 0,027414558 | beaded filament structural protein 1 |
| ENSG00000224886 | AL132656,1 | 1,645179013 | 3,127866662 | 0,16878815 | 0,650757516 | novel pseudogene |
| ENSG00000264352 | RN7SL602P | 1,644579024 | 3,126566113 | 0,10394775 | 0,514131574 | RNA, 7SL, cytoplasmic 602, pseudogene |
| ENSG00000121858 | TNFSF10 | 1,643153938 | 3,123479233 | 0,02365459 | 0,210168399 | TNF superfamily member 10 |
| ENSG00000281680 | OR10A3 | 1,641750864 | 3,120443011 | 0,53586732 | 0,932139691 | olfactory receptor family 10 subfamily A member 3 |
| ENSG00000004399 | PLXND1 | 1,641158363 | 3,119161739 | 2,44E-06 | 9,72E-05 | plexin D1 |
| ENSG00000083444 | PLOD1 | 1,641150877 | 3,119145554 | 2,05E-06 | 8,31E-05 | procollagen-lysine,2-oxoglutarate 5-dioxygenase 1 |

| | | | | | | |
|---|---|---|---|---|---|---|
| ENSG00000275532 | AC006449,2 | 1,640935056 | 3,118678978 | 0,07216812 | 0,423632011 | novel transcript, antisense to MLLT6 |
| ENSG00000283979 | RFLNB | 1,640856 | 3,118508087 | 0,45553554 | 0,903545728 | refilin B |
| ENSG00000272533 | SNORA28 | 1,640689336 | 3,11814785 | 0,31898012 | 0,823977058 | small nucleolar RNA, H/ACA box 28 |
| ENSG00000135862 | LAMC1 | 1,640536327 | 3,117817163 | 7,24E-11 | 6,29E-09 | laminin subunit gamma 1 |
| ENSG00000082293 | COL19A1 | 1,639387181 | 3,115334726 | 0,0056876 | 0,075828159 | collagen type XIX alpha 1 chain |
| ENSG00000254035 | AC104117,3 | 1,639142578 | 3,114806578 | 0,22706659 | 0,735496246 | novel transcript |
| ENSG00000229060 | AL451124,1 | 1,638358841 | 3,113114934 | 0,08572419 | 0,466101036 | novel pseudogene |
| ENSG00000171502 | COL24A1 | 1,638348142 | 3,113091847 | 0,00084247 | 0,0160997 | collagen type XXIV alpha 1 chain |
| ENSG00000020577 | SAMD4A | 1,638065021 | 3,112480979 | 3,09E-07 | 1,46E-05 | sterile alpha motif domain containing 4A |
| ENSG00000282998 | AC009975,2 | 1,637998117 | 3,112336645 | 0,22058004 | 0,728572782 | novel transcript |
| ENSG00000112493 | TAPBP | 1,637365872 | 3,110972995 | 0,02307035 | 0,206549685 | TAP binding protein |
| ENSG00000248690 | HAS2-AS1 | 1,634710198 | 3,105251668 | 0,00130356 | 0,023381797 | HAS2 antisense RNA 1 |
| ENSG00000248854 | HNRNPH1P3 | 1,633889741 | 3,103486221 | 0,19854159 | 0,697657684 | heterogeneous nuclear ribonucleoprotein H1 pseudogene 3 |
| ENSG00000197712 | FAM114A1 | 1,633376599 | 3,102382561 | 1,76E-11 | 1,67E-09 | family with sequence similarity 114 member A1 |
| ENSG00000255400 | AC124276,2 | 1,632516109 | 3,100532708 | 0,30552861 | 0,814326218 | novel transcript |
| ENSG00000100122 | CRYBB1 | 1,631227131 | 3,09776377 | 0,24631691 | 0,758198269 | crystallin beta B1 |
| ENSG00000231367 | LINC02613 | 1,630623556 | 3,09646804 | 0,2112462 | 0,717012384 | long intergenic non-protein coding RNA 2613 |
| ENSG00000152952 | PLOD2 | 1,629541844 | 3,094147224 | 4,73E-05 | 0,001346654 | procollagen-lysine,2-oxoglutarate 5-dioxygenase 2 |
| ENSG00000263558 | RN7SL716P | 1,629163923 | 3,093336802 | 0,40970478 | 0,882177108 | RNA, 7SL, cytoplasmic 716, pseudogene |
| ENSG00000050165 | DKK3 | 1,628685315 | 3,092310772 | 2,99E-06 | 0,000117106 | dickkopf WNT signaling pathway inhibitor 3 |
| ENSG00000107821 | KAZALD1 | 1,628221437 | 3,091316643 | 5,12E-07 | 2,31E-05 | Kazal type serine peptidase inhibitor domain 1 |
| ENSG00000277782 | AC068870,2 | 1,627765601 | 3,090340061 | 0,03279176 | 0,262266996 | novel transcript |
| ENSG00000186470 | BTN3A2 | 1,626602905 | 3,0878505 | 6,56E-05 | 0,001809175 | butyrophilin subfamily 3 member A2 |
| ENSG00000171004 | HS6ST2 | 1,625799615 | 3,086131668 | 1,34E-06 | 5,55E-05 | heparan sulfate 6-O-sulfotransferase 2 |
| ENSG00000287175 | AL513303,1 | 1,625368823 | 3,085210281 | 0,08571385 | 0,466101036 | novel transcript |
| ENSG00000232023 | LINC01807 | 1,625169558 | 3,084784181 | 0,0889787 | 0,476005294 | long intergenic non-protein coding RNA 1807 |
| ENSG00000119630 | PGF | 1,625152075 | 3,084746798 | 0,00315964 | 0,048637582 | placental growth factor |
| ENSG00000244509 | APOBEC3C | 1,623744499 | 3,081738611 | 4,92E-05 | 0,001397948 | apolipoprotein B mRNA editing enzyme catalytic subunit 3C |
| ENSG00000003436 | TFPI | 1,622846251 | 3,07982046 | 1,01E-05 | 0,000343568 | tissue factor pathway inhibitor |
| ENSG00000268643 | AC006486,1 | 1,621452364 | 3,076846271 | 0,30706733 | 0,815892645 | novel protein |
| ENSG00000274721 | AC091825,1 | 1,620699539 | 3,075241134 | 0,16490059 | 0,644028999 | |

| | | | | | | |
|---|---|---|---|---|---|---|
| ENSG00000101986 | ABCD1 | 1,620404003 | 3,074611236 | 6,50E-06 | 0,000233788 | ATP binding cassette subfamily D member 1 |
| ENSG00000177663 | IL17RA | 1,620116382 | 3,073998331 | 3,01E-05 | 0,000899691 | interleukin 17 receptor A |
| ENSG00000232725 | U52111,1 | 1,620090267 | 3,073942689 | 0,15662994 | 0,627831388 | novel transcript |
| ENSG00000258402 | AC016526,1 | 1,620008332 | 3,073768115 | 0,34239836 | 0,840048641 | novel transcript |
| ENSG00000266970 | AC061992,2 | 1,619134687 | 3,071907312 | 0,0223625 | 0,202005507 | novel transcript |
| ENSG00000154783 | FGD5 | 1,617321707 | 3,068049388 | 0,00067385 | 0,013376072 | FYVE, RhoGEF and PH domain containing 5 |
| ENSG00000285748 | AC090337,2 | 1,617263408 | 3,067925412 | 0,1184558 | 0,550038851 | novel transcript |
| ENSG00000120907 | ADRA1A | 1,616844893 | 3,067035559 | 0,00220968 | 0,036397943 | adrenoceptor alpha 1A |
| ENSG00000227992 | AC108463,1 | 1,615587291 | 3,064363179 | 0,21794875 | 0,72555786 | pseudogene similar to part of ubiquitin carboxyl-terminal hydrolase L5 UCHL5 |
| ENSG00000203485 | INF2 | 1,615564782 | 3,064315369 | 0,002893 | 0,045020719 | inverted formin 2 |
| ENSG00000284116 | AL772307,1 | 1,615555219 | 3,064295057 | 0,00991861 | 0,11451092 | novel transcript |
| ENSG00000139155 | SLCO1C1 | 1,615315487 | 3,063785906 | 0,02759808 | 0,234018535 | solute carrier organic anion transporter family member 1C1 |
| ENSG00000233490 | GPSM3 | 1,613971511 | 3,060933094 | 3,59E-05 | 0,001056361 | G protein signaling modulator 3 |
| ENSG00000237582 | POU5F1 | 1,613898046 | 3,06077723 | 0,00711245 | 0,089581081 | POU class 5 homeobox 1 |
| ENSG00000206454 | POU5F1 | 1,613898046 | 3,06077723 | 0,00711245 | 0,089581081 | POU class 5 homeobox 1 |
| ENSG00000278787 | UBR7 | 1,613670365 | 3,060294227 | 0,01826084 | 0,1758801 | ubiquitin protein ligase E3 component n-recognin 7 |
| ENSG00000248730 | EGFLAM-AS4 | 1,611489196 | 3,055670953 | 0,23916134 | 0,748762296 | EGFLAM antisense RNA 4 |
| ENSG00000288595 | AC236668,1 | 1,609378757 | 3,051204247 | 0,15250904 | 0,620962584 | novel transcript |
| ENSG00000205946 | USP17L6P | 1,609222682 | 3,050874177 | 0,01076479 | 0,120930771 | ubiquitin specific peptidase 17 like family member 6, pseudogene |
| ENSG00000136068 | FLNB | 1,608999775 | 3,05040283 | 2,43E-05 | 0,000748493 | filamin B |
| ENSG00000267543 | AC015802,3 | 1,608988212 | 3,050378382 | 0,11548136 | 0,543019233 | novel transcript, sense intronic to PRCD |
| ENSG00000267228 | AC012254,2 | 1,608858394 | 3,050103913 | 0,02282626 | 0,205010087 | novel transcript |
| ENSG00000285280 | AL390957,1 | 1,608133535 | 3,048571822 | 0,2154226 | 0,72270101 | novel transcript |
| ENSG00000137076 | TLN1 | 1,607452838 | 3,047133774 | 0,00011268 | 0,002909149 | talin 1 |
| ENSG00000287079 | AL109806,2 | 1,606387907 | 3,044885351 | 0,14301121 | 0,601880115 | novel transcript |
| ENSG00000255219 | AP000755,2 | 1,606327236 | 3,044757303 | 0,09185854 | 0,481984693 | novel transcript |
| ENSG00000008516 | MMP25 | 1,60584298 | 3,04373547 | 0,00090537 | 0,017175593 | matrix metallopeptidase 25 |
| ENSG00000182759 | MAFA | 1,605718479 | 3,043472815 | 0,13984579 | 0,596065254 | MAF bZIP transcription factor A |
| ENSG00000283980 | GNG14 | 1,605484049 | 3,042978306 | 0,20296829 | 0,703345263 | G protein subunit gamma 14 |
| ENSG00000270002 | AC022028,2 | 1,605012108 | 3,041983037 | 0,25559283 | 0,767141836 | novel transcript, antisense to GRID1 |

| | | | | | | |
|---|---|---|---|---|---|---|
| ENSG00000286020 | AC011997,2 | 1,604323495 | 3,040531414 | 0,27367284 | 0,785719894 | novel transcript, antisense to PLCL1 |
| ENSG00000164344 | KLKB1 | 1,602585065 | 3,036869817 | 0,03670989 | 0,282267076 | kallikrein B1 |
| ENSG00000237238 | BMS1P10 | 1,601745853 | 3,03510379 | 0,00015275 | 0,003761959 | BMS1 pseudogene 10 |
| ENSG00000284337 | AC013271,1 | 1,601144828 | 3,033839635 | 0,26366716 | 0,774820034 | novel transcript |
| ENSG00000253688 | AC026904,2 | 1,600334855 | 3,032136822 | 0,07198333 | 0,422984135 | novel transcript |
| ENSG00000240750 | RN7SL559P | 1,599796302 | 3,031005147 | 0,27985422 | 0,790565637 | RNA, 7SL, cytoplasmic 559, pseudogene |
| ENSG00000285112 | SLC52A2 | 1,597230261 | 3,025618859 | 0,01633576 | 0,163324813 | solute carrier family 52 member 2 |
| ENSG00000257869 | AL139023,1 | 1,596607927 | 3,024313982 | 0,19355876 | 0,690424879 | novel transcript |
| ENSG00000160801 | PTH1R | 1,595702884 | 3,022417341 | 0,00098468 | 0,018445186 | parathyroid hormone 1 receptor |
| ENSG00000227117 | HORMAD2-AS1 | 1,595314601 | 3,021604004 | 0,0563294 | 0,366815744 | HORMAD2 and MTMR3 antisense RNA 1 |
| ENSG00000254721 | AP000879,1 | 1,593546423 | 3,017902973 | 0,68749052 | 0,965965731 | novel transcript |
| ENSG00000253882 | AC099548,2 | 1,59308703 | 3,016942145 | 0,05471148 | 0,361245992 | family with sequence similarity 115, member C (FAM115C) pseudogene |
| ENSG00000197320 | AC060834,1 | 1,592799199 | 3,016340296 | 0,03944992 | 0,295841289 | C3 and PZP-like, alpha-2-macroglobulin domain containing 8 (CPAMD8) pseudogene |
| ENSG00000161896 | IP6K3 | 1,59224378 | 3,015179267 | 0,11450777 | 0,540617781 | inositol hexakisphosphate kinase 3 |
| ENSG00000103316 | CRYM | 1,591030984 | 3,012645634 | 0,00509007 | 0,069582866 | crystallin mu |
| ENSG00000233117 | LINC00702 | 1,590810402 | 3,012185048 | 5,64E-06 | 0,000205034 | long intergenic non-protein coding RNA 702 |
| ENSG00000250754 | LINC02436 | 1,590319618 | 3,01116052 | 0,12416216 | 0,563164667 | long intergenic non-protein coding RNA 2436 |
| ENSG00000230276 | AL162388,1 | 1,589511604 | 3,009474524 | 0,22072352 | 0,728572782 | NHP2 non-histone chromosome protein 2-like 1 (S, cerevisiae) (NHP2L1) pseudogene |
| ENSG00000197019 | SERTAD1 | 1,588987017 | 3,00838043 | 0,00012977 | 0,003287837 | SERTA domain containing 1 |
| ENSG00000232852 | CICP4 | 1,588489155 | 3,007342443 | 0,34220438 | 0,840048641 | capicua transcriptional repressor pseudogene 4 |
| ENSG00000272896 | AL355987,3 | 1,588177276 | 3,006692391 | 0,11130425 | 0,532815219 | novel protein |
| ENSG00000175482 | POLD4 | 1,586955669 | 3,004147542 | 0,00079811 | 0,015402478 | DNA polymerase delta 4, accessory subunit |
| ENSG00000220008 | LINGO3 | 1,586012586 | 3,002184386 | 0,10382176 | 0,513912548 | leucine rich repeat and Ig domain containing 3 |
| ENSG00000278565 | CEACAM21 | 1,585362264 | 3,0008314 | 0,00035026 | 0,007727191 | CEA cell adhesion molecule 21 |
| ENSG00000284369 | COG5 | 1,583974691 | 2,99794661 | 0,02824997 | 0,237804851 | component of oligomeric golgi complex 5 |
| ENSG00000176381 | PRR18 | 1,583140919 | 2,996214519 | 0,12086581 | 0,555007274 | proline rich 18 |
| ENSG00000286198 | AL606753,2 | 1,583038178 | 2,996001151 | 0,16248897 | 0,639724866 | novel transcript |
| ENSG00000166106 | ADAMTS15 | 1,581930314 | 2,993701366 | 0,00300909 | 0,046603377 | ADAM metallopeptidase with thrombospondin type 1 motif 15 |
| ENSG00000166035 | LIPC | 1,581271306 | 2,992334187 | 0,13407635 | 0,584238558 | lipase C, hepatic type |

| | | | | | | |
|---|---|---|---|---|---|---|
| ENSG00000249628 | LINC00942 | 1,580940663 | 2,99164847 | 0,03453763 | 0,271098283 | long intergenic non-protein coding RNA 942 |
| ENSG00000152580 | IGSF10 | 1,57957678 | 2,988821587 | 0,00924538 | 0,108257566 | immunoglobulin superfamily member 10 |
| ENSG00000160963 | COL26A1 | 1,579382364 | 2,988418843 | 7,07E-06 | 0,000251409 | collagen type XXVI alpha 1 chain |
| ENSG00000286519 | AC009236,1 | 1,579352194 | 2,988356349 | 0,19356769 | 0,690424879 | novel transcript |
| ENSG00000197446 | CYP2F1 | 1,578545896 | 2,986686674 | 0,35363883 | 0,845346751 | cytochrome P450 family 2 subfamily F member 1 |
| ENSG00000151388 | ADAMTS12 | 1,577723404 | 2,984984426 | 1,06E-09 | 7,90E-08 | ADAM metallopeptidase with thrombospondin type 1 motif 12 |
| ENSG00000163520 | FBLN2 | 1,577661084 | 2,984855485 | 0,00020736 | 0,004914998 | fibulin 2 |
| ENSG00000144355 | DLX1 | 1,576819826 | 2,983115477 | 3,46E-07 | 1,61E-05 | distal-less homeobox 1 |
| ENSG00000152463 | OLAH | 1,575871123 | 2,981154452 | 0,13962342 | 0,595782615 | oleoyl-ACP hydrolase |
| ENSG00000280707 | HPAT5 | 1,575227525 | 2,979824833 | 0,02518728 | 0,220092673 | human pluripotency-associated transcript 5 |
| ENSG00000224149 | AC099786,2 | 1,573975876 | 2,977240726 | 0,20415166 | 0,70518148 | novel transcript, antisense to LHX8 |
| ENSG00000246985 | SOCS2-AS1 | 1,573947827 | 2,977182843 | 0,00043707 | 0,009301961 | SOCS2 antisense RNA 1 |
| ENSG00000257527 | AC126755,1 | 1,573738876 | 2,976751677 | 0,21588263 | 0,722999018 | microRNA 3179-3 |
| ENSG00000286645 | AC013267,1 | 1,572659283 | 2,974524957 | 0,31474664 | 0,81991112 | novel transcript |
| ENSG00000235934 | AC007405,2 | 1,572459455 | 2,974112984 | 0,02292482 | 0,205482064 | novel transcript |
| ENSG00000237781 | ADAMTSL4-AS2 | 1,570787264 | 2,970667761 | 0,26508836 | 0,776585686 | ADAMTSL4 antisense RNA 2 |
| ENSG00000101605 | MYOM1 | 1,570428201 | 2,969928503 | 9,92E-05 | 0,002596844 | myomesin 1 |
| ENSG00000173535 | TNFRSF10C | 1,569910976 | 2,968863937 | 0,04396499 | 0,316175438 | TNF receptor superfamily member 10c |
| ENSG00000170561 | IRX2 | 1,569794546 | 2,968624349 | 1,14E-05 | 0,000380349 | iroquois homeobox 2 |
| ENSG00000144668 | ITGA9 | 1,568949126 | 2,966885243 | 7,18E-07 | 3,14E-05 | integrin subunit alpha 9 |
| ENSG00000177098 | SCN4B | 1,568866612 | 2,96671556 | 0,03510043 | 0,273637274 | sodium voltage-gated channel beta subunit 4 |
| ENSG00000254894 | NAV2-AS1 | 1,567379629 | 2,963659348 | 0,25344799 | 0,765172364 | NAV2 antisense RNA 1 |
| ENSG00000185527 | PDE6G | 1,567251331 | 2,963395803 | 0,0539932 | 0,358091488 | phosphodiesterase 6G |
| ENSG00000278727 | AC000403,1 | 1,566984031 | 2,9628468 | 0,02941464 | 0,243736921 | novel transcript |
| ENSG00000183036 | PCP4 | 1,56661159 | 2,962082021 | 0,01315281 | 0,139782152 | Purkinje cell protein 4 |
| ENSG00000095596 | CYP26A1 | 1,56645658 | 2,961763778 | 0,19785712 | 0,696383642 | cytochrome P450 family 26 subfamily A member 1 |
| ENSG00000180139 | ACTA2-AS1 | 1,565253582 | 2,959295128 | 0,00067768 | 0,013427209 | ACTA2 antisense RNA 1 |
| ENSG00000259062 | ACTN1-AS1 | 1,564982911 | 2,958739972 | 0,20169825 | 0,701332924 | ACTN1 antisense RNA 1 |
| ENSG00000140682 | TGFB1I1 | 1,564971643 | 2,958716863 | 6,04E-05 | 0,001676594 | transforming growth factor beta 1 induced transcript 1 |
| ENSG00000227188 | MGAT3-AS1 | 1,564533266 | 2,957817964 | 0,21240301 | 0,719216058 | MGAT3 antisense RNA 1 |
| ENSG00000187553 | CYP26C1 | 1,561927489 | 2,952480413 | 0,06025576 | 0,381629797 | cytochrome P450 family 26 subfamily C member 1 |

| | | | | | | |
|---|---|---|---|---|---|---|
| ENSG00000253633 | AP002852,1 | 1,560963858 | 2,950508997 | 0,06634913 | 0,404907139 | novel transcript |
| ENSG00000171094 | ALK | 1,560955178 | 2,950491246 | 0,00162885 | 0,028166124 | ALK receptor tyrosine kinase |
| ENSG00000196109 | ZNF676 | 1,560832374 | 2,950240107 | 0,13097545 | 0,577911147 | zinc finger protein 676 |
| ENSG00000187536 | TPM3P7 | 1,56022225 | 2,948992697 | 0,12956813 | 0,575350249 | tropomyosin 3 pseudogene 7 |
| ENSG00000287818 | AL512283,3 | 1,559888254 | 2,94831006 | 0,09494461 | 0,489386708 | novel transcript, antisense to PKIB |
| ENSG00000196139 | AKR1C3 | 1,559187361 | 2,946878054 | 0,0399438 | 0,298328495 | aldo-keto reductase family 1 member C3 |
| ENSG00000259753 | AC068234,1 | 1,55890408 | 2,946299475 | 0,30804575 | 0,816424917 | novel protein |
| ENSG00000230456 | DDR1 | 1,558856655 | 2,946202625 | 0,18265449 | 0,673171032 | discoidin domain receptor tyrosine kinase 1 |
| ENSG00000263647 | BPTFP1 | 1,557870826 | 2,944190099 | 0,05398952 | 0,358091488 | bromodomain PHD finger transcription factor pseudogene 1 |
| ENSG00000237471 | AC073115,2 | 1,557774653 | 2,94399384 | 0,02899164 | 0,241642594 | novel transcript |
| ENSG00000274859 | AC131238,1 | 1,557491592 | 2,943416278 | 0,49858218 | 0,919683359 | novel transcript, antisense to VSIG10 |
| ENSG00000273573 | Metazoa_SRP | 1,557156204 | 2,94273209 | 0,50256535 | 0,920607749 | Metazoan signal recognition particle RNA |
| ENSG00000244953 | AC087521,1 | 1,557136709 | 2,942692327 | 0,33085739 | 0,832114797 | novel transcript |
| ENSG00000121904 | CSMD2 | 1,556766621 | 2,941937548 | 0,00013917 | 0,003481094 | CUB and Sushi multiple domains 2 |
| ENSG00000282800 | MIR22HG | 1,55674915 | 2,941901922 | 9,49E-06 | 0,000326416 | MIR22 host gene |
| ENSG00000186594 | MIR22HG | 1,55674915 | 2,941901922 | 9,49E-06 | 0,000326416 | MIR22 host gene |
| ENSG00000105227 | PRX | 1,554636462 | 2,937596943 | 0,04627975 | 0,326538078 | periaxin |
| ENSG00000203262 | AL137845,1 | 1,551496491 | 2,931210327 | 0,50611217 | 0,921780819 | zinc finger protein 265 (ZNF265) pseudogene |
| ENSG00000166592 | RRAD | 1,551447185 | 2,931110149 | 0,2378264 | 0,747090777 | RRAD, Ras related glycolysis inhibitor and calcium channel regulator |
| ENSG00000180061 | TMEM150B | 1,551286011 | 2,930782712 | 0,31396216 | 0,819877431 | transmembrane protein 150B |
| ENSG00000129226 | CD68 | 1,550597192 | 2,929383735 | 0,027297 | 0,232347819 | CD68 molecule |
| ENSG00000075223 | SEMA3C | 1,550236409 | 2,928651259 | 1,05E-05 | 0,000354451 | semaphorin 3C |
| ENSG00000278202 | AC243919,2 | 1,550024945 | 2,928222023 | 0,16713732 | 0,647757288 | novel transcript |
| ENSG00000165092 | ALDH1A1 | 1,549993616 | 2,928158435 | 0,21092804 | 0,716554846 | aldehyde dehydrogenase 1 family member A1 |
| ENSG00000257355 | AC008770,1 | 1,549537616 | 2,927233062 | 0,32018288 | 0,824366234 | novel protein |
| ENSG00000174791 | RIN1 | 1,549517825 | 2,927192908 | 0,00123975 | 0,022288326 | Ras and Rab interactor 1 |
| ENSG00000262533 | AC090617,3 | 1,549492906 | 2,927142347 | 0,43401169 | 0,896272827 | novel transcript, antisense to DPH1 and OVCA2 |
| ENSG00000235385 | LINC02154 | 1,549339628 | 2,926831373 | 0,07729862 | 0,439421744 | long intergenic non-protein coding RNA 2154 |
| ENSG00000166482 | MFAP4 | 1,54885295 | 2,925844202 | 0,00415262 | 0,060441045 | microfibril associated protein 4 |
| ENSG00000179673 | RPRML | 1,548346356 | 2,924816991 | 0,02994357 | 0,246681348 | reprimo like |
| ENSG00000248698 | LINC01085 | 1,54769863 | 2,923504131 | 0,18091968 | 0,670651294 | long intergenic non-protein coding RNA 1085 |

| | | | | | | |
|---|---|---|---|---|---|---|
| ENSG00000135898 | GPR55 | 1,547616668 | 2,923338047 | 0,08404646 | 0,461230231 | G protein-coupled receptor 55 |
| ENSG00000263528 | IKBKE | 1,54749921 | 2,92310005 | 1,89E-05 | 0,00059868 | inhibitor of nuclear factor kappa B kinase subunit epsilon |
| ENSG00000205927 | OLIG2 | 1,547271651 | 2,92263902 | 0,0023984 | 0,038751223 | oligodendrocyte transcription factor 2 |
| ENSG00000159733 | ZFYVE28 | 1,546915128 | 2,921916859 | 0,07225567 | 0,423827618 | zinc finger FYVE-type containing 28 |
| ENSG00000004776 | HSPB6 | 1,546718714 | 2,921519085 | 0,00685861 | 0,087388747 | heat shock protein family B (small) member 6 |
| ENSG00000223405 | DDR1-DT | 1,546616316 | 2,921311733 | 0,06516664 | 0,400662695 | DDR1 divergent transcript |
| ENSG00000174080 | CTSF | 1,546581519 | 2,921241272 | 1,13E-05 | 0,000378035 | cathepsin F |
| ENSG00000223953 | C1QTNF5 | 1,545548152 | 2,919149609 | 0,00467611 | 0,065714391 | C1q and TNF related 5 |
| ENSG00000286037 | AC011479,4 | 1,544907551 | 2,917853705 | 0,35292472 | 0,845031687 | novel transcript, antisense to SPINT2 |
| ENSG00000165899 | OTOGL | 1,54487169 | 2,917781177 | 0,00476206 | 0,066375821 | otogelin like |
| ENSG00000285338 | AC091564,7 | 1,544311919 | 2,916649286 | 0,25346521 | 0,765172364 | novel transcript |
| ENSG00000228358 | LINC02263 | 1,543512837 | 2,915034255 | 0,17982323 | 0,668568618 | long intergenic non-protein coding RNA 2263 |
| ENSG00000249341 | AC124017,1 | 1,542938435 | 2,91387388 | 0,40467181 | 0,878176155 | novel transcript |
| ENSG00000239593 | AL513122,2 | 1,541451762 | 2,910872728 | 0,1524918 | 0,620962584 | novel transcript |
| ENSG00000179082 | C9orf106 | 1,540760424 | 2,909478175 | 0,05105 | 0,345820358 | chromosome 9 putative open reading frame 106 |
| ENSG00000236683 | HMGA1P1 | 1,54019961 | 2,908347402 | 0,15986623 | 0,634614414 | high mobility group AT-hook 1 pseudogene 1 |
| ENSG00000205542 | TMSB4X | 1,539822027 | 2,907586328 | 8,16E-08 | 4,43E-06 | thymosin beta 4 X-linked |
| ENSG00000285599 | AC234917,2 | 1,539761161 | 2,907463663 | 0,14022731 | 0,596958956 | Novel transcript, antisense to NRBP2 |
| ENSG00000257225 | AC079601,1 | 1,539492005 | 2,906921282 | 0,02913591 | 0,24236233 | novel transcript, antisense to PRICKLE1 |
| ENSG00000043039 | BARX2 | 1,538873549 | 2,905675407 | 0,11750158 | 0,547966805 | BARX homeobox 2 |
| ENSG00000185972 | CCIN | 1,53832698 | 2,904574794 | 0,10438961 | 0,515175486 | calicin |
| ENSG00000058866 | DGKG | 1,538039786 | 2,903996644 | 0,01619638 | 0,16229599 | diacylglycerol kinase gamma |
| ENSG00000274135 | Metazoa_SRP | 1,53756898 | 2,903049114 | 0,23320086 | 0,742367548 | Metazoan signal recognition particle RNA |
| ENSG00000197734 | C14orf178 | 1,537472513 | 2,902855005 | 0,09227564 | 0,482910203 | chromosome 14 open reading frame 178 |
| ENSG00000131435 | PDLIM4 | 1,537195145 | 2,902296965 | 0,00557187 | 0,074892012 | PDZ and LIM domain 4 |
| ENSG00000183386 | FHL3 | 1,536592439 | 2,901084745 | 1,26E-06 | 5,26E-05 | four and a half LIM domains 3 |
| ENSG00000255921 | AC026310,1 | 1,536302926 | 2,900502627 | 0,13509636 | 0,586038109 | novel transcript, antisense to BCAT1 |
| ENSG00000187800 | PEAR1 | 1,53608668 | 2,900067902 | 0,0001051 | 0,002735807 | platelet endothelial aggregation receptor 1 |
| ENSG00000179776 | CDH5 | 1,534499612 | 2,896879375 | 0,00892387 | 0,105360946 | cadherin 5 |
| ENSG00000225213 | AC073367,1 | 1,533868583 | 2,895612568 | 0,14540548 | 0,607504591 | novel transcript, sense intronic to RSU1 |
| ENSG00000101892 | ATP1B4 | 1,533587272 | 2,895048007 | 0,08068781 | 0,450468504 | ATPase Na+/K+ transporting family member beta 4 |
| ENSG00000264028 | RN7SL744P | 1,533302786 | 2,894477187 | 0,13115614 | 0,578443778 | RNA, 7SL, cytoplasmic 744, pseudogene |

| | | | | | | |
|---|---|---|---|---|---|---|
| ENSG00000225778 | PROSER2-AS1 | 1,533203477 | 2,89427795 | 0,08247068 | 0,456832613 | PROSER2 antisense RNA 1 |
| ENSG00000260121 | AC138028,4 | 1,531044312 | 2,889949558 | 0,01445481 | 0,150137902 | novel transcript, antisense to FAM38A |
| ENSG00000179023 | KLHDC7A | 1,530813569 | 2,889487379 | 0,00330557 | 0,050445936 | kelch domain containing 7A |
| ENSG00000275935 | LENG1 | 1,530738924 | 2,889337882 | 0,06994606 | 0,417175017 | leukocyte receptor cluster member 1 |
| ENSG00000277457 | LENG1 | 1,530738924 | 2,889337882 | 0,06994606 | 0,417175017 | leukocyte receptor cluster member 1 |
| ENSG00000276628 | LENG1 | 1,530738924 | 2,889337882 | 0,06994606 | 0,417175017 | leukocyte receptor cluster member 1 |
| ENSG00000278257 | LENG1 | 1,530738924 | 2,889337882 | 0,06994606 | 0,417175017 | leukocyte receptor cluster member 1 |
| ENSG00000278426 | LENG1 | 1,530738924 | 2,889337882 | 0,06994606 | 0,417175017 | leukocyte receptor cluster member 1 |
| ENSG00000276774 | LENG1 | 1,530738924 | 2,889337882 | 0,06994606 | 0,417175017 | leukocyte receptor cluster member 1 |
| ENSG00000120498 | TEX11 | 1,530177727 | 2,888214171 | 0,22976557 | 0,73819656 | testis expressed 11 |
| ENSG00000270190 | AC068491,2 | 1,530086455 | 2,888031453 | 0,15621633 | 0,627034861 | novel transcript |
| ENSG00000022355 | GABRA1 | 1,529524266 | 2,886906265 | 0,29568132 | 0,805426883 | gamma-aminobutyric acid type A receptor subunit alpha1 |
| ENSG00000183087 | GAS6 | 1,528488917 | 2,884835223 | 1,44E-05 | 0,00046767 | growth arrest specific 6 |
| ENSG00000150636 | CCDC102B | 1,528466322 | 2,884790043 | 0,00010158 | 0,002653127 | coiled-coil domain containing 102B |
| ENSG00000179630 | LACC1 | 1,527903122 | 2,883664096 | 1,12E-05 | 0,000377999 | laccase domain containing 1 |
| ENSG00000182890 | GLUD2 | 1,527277296 | 2,882413464 | 5,79E-05 | 0,001612647 | glutamate dehydrogenase 2 |
| ENSG00000288118 | GLUD2 | 1,527277296 | 2,882413464 | 5,79E-05 | 0,001612647 | glutamate dehydrogenase 2 |
| ENSG00000151572 | ANO4 | 1,524796374 | 2,877460999 | 2,78E-05 | 0,000841352 | anoctamin 4 |
| ENSG00000232527 | LINC02802 | 1,524298339 | 2,876467837 | 4,77E-05 | 0,001359394 | long intergenic non-protein coding RNA 2802 |
| ENSG00000196975 | ANXA4 | 1,523719641 | 2,875314251 | 1,08E-06 | 4,54E-05 | annexin A4 |
| ENSG00000147234 | FRMPD3 | 1,523388332 | 2,874654023 | 7,74E-06 | 0,000272842 | FERM and PDZ domain containing 3 |
| ENSG00000065308 | TRAM2 | 1,523340642 | 2,874559 | 6,15E-08 | 3,40E-06 | translocation associated membrane protein 2 |
| ENSG00000157445 | CACNA2D3 | 1,5232768 | 2,874431797 | 0,0001707 | 0,004140237 | calcium voltage-gated channel auxiliary subunit alpha2delta 3 |
| ENSG00000184221 | OLIG1 | 1,523120511 | 2,874120423 | 0,00022719 | 0,005320387 | oligodendrocyte transcription factor 1 |
| ENSG00000163430 | FSTL1 | 1,522552242 | 2,872988548 | 2,68E-10 | 2,19E-08 | follistatin like 1 |
| ENSG00000179627 | ZBTB42 | 1,522333059 | 2,872552099 | 0,00672124 | 0,08612003 | zinc finger and BTB domain containing 42 |
| ENSG00000237462 | TRIM27 | 1,52215206 | 2,872191734 | 0,08371427 | 0,460944538 | tripartite motif containing 27 |
| ENSG00000128606 | LRRC17 | 1,521794184 | 2,871479345 | 8,29E-05 | 0,002231533 | leucine rich repeat containing 17 |
| ENSG00000162878 | PKDCC | 1,521436986 | 2,870768481 | 1,77E-06 | 7,21E-05 | protein kinase domain containing, cytoplasmic |
| ENSG00000278701 | Metazoa_SRP | 1,519445051 | 2,866807535 | 0,15508214 | 0,624724654 | Metazoan signal recognition particle RNA |
| ENSG00000168610 | STAT3 | 1,519057258 | 2,866037047 | 3,06E-08 | 1,75E-06 | signal transducer and activator of transcription 3 |

| | | | | | | |
|---|---|---|---|---|---|---|
| ENSG00000232977 | LINC00327 | 1,518979215 | 2,865882012 | 0,00039055 | 0,008467475 | long intergenic non-protein coding RNA 327 |
| ENSG00000133115 | STOML3 | 1,518592166 | 2,865113251 | 0,20640563 | 0,709439943 | stomatin like 3 |
| ENSG00000152804 | HHEX | 1,51781993 | 2,863580041 | 0,17257421 | 0,657085135 | hematopoietically expressed homeobox |
| ENSG00000186094 | AGBL4 | 1,517685481 | 2,863313189 | 0,00064237 | 0,012858692 | ATP/GTP binding protein like 4 |
| ENSG00000287446 | AC007731,5 | 1,516340027 | 2,860644115 | 0,22081772 | 0,728635155 | novel transcript |
| ENSG00000143387 | CTSK | 1,516312423 | 2,860589381 | 0,00010991 | 0,002846917 | cathepsin K |
| ENSG00000130147 | SH3BP4 | 1,515710335 | 2,859395804 | 0,00013651 | 0,0034229 | SH3 domain binding protein 4 |
| ENSG00000227934 | AL732414,1 | 1,515638837 | 2,859254099 | 0,18162741 | 0,67167359 | ring finger protein 4 (RNF4) pseudogene |
| ENSG00000273796 | BX322562,1 | 1,515480105 | 2,858939529 | 0,19131965 | 0,687045418 | novel transcript |
| ENSG00000253853 | AC246817,1 | 1,514421611 | 2,856842717 | 0,00244317 | 0,039372808 | novel transcript |
| ENSG00000253304 | TMEM200B | 1,512946235 | 2,853922652 | 0,0001178 | 0,003026113 | transmembrane protein 200B |
| ENSG00000138166 | DUSP5 | 1,512101163 | 2,85225143 | 4,59E-05 | 0,001313079 | dual specificity phosphatase 5 |
| ENSG00000284933 | ARHGEF5 | 1,512056847 | 2,852163817 | 0,00033621 | 0,007465363 | Rho guanine nucleotide exchange factor 5 |
| ENSG00000071967 | CYBRD1 | 1,511882178 | 2,851818523 | 0,00107054 | 0,019839552 | cytochrome b reductase 1 |
| ENSG00000229915 | AC016999,1 | 1,5116615 | 2,851382335 | 0,17602298 | 0,661519708 | novel transcript |
| ENSG00000283057 | AC010653,3 | 1,510437229 | 2,84896368 | 0,30536048 | 0,814326218 | parathymosin (PTMS) pseudogene |
| ENSG00000276539 | PLPPR1 | 1,510394358 | 2,84887902 | 0,02126159 | 0,194704806 | phospholipid phosphatase related 1 |
| ENSG00000274654 | AC022467,1 | 1,51024972 | 2,848593419 | 0,1297683 | 0,575798478 | novel transcript, sense intronic to FBN1 |
| ENSG00000178695 | KCTD12 | 1,509422932 | 2,846961399 | 2,07E-05 | 0,000649071 | potassium channel tetramerization domain containing 12 |
| ENSG00000224431 | AC063976,2 | 1,509294726 | 2,846708413 | 0,095733 | 0,491344168 | cytoplasmic linker associated protein 2 (CLASP2) pseudogene |
| ENSG00000267503 | AP005131,3 | 1,509105447 | 2,846334954 | 0,09245632 | 0,483289495 | novel transcript, antisense C18orf1 |
| ENSG00000170485 | NPAS2 | 1,508494023 | 2,845128914 | 0,00316866 | 0,048704482 | neuronal PAS domain protein 2 |
| ENSG00000288240 | AC020916,4 | 1,508051819 | 2,844256979 | 0,37602982 | 0,862512424 | |
| ENSG00000284702 | AL136115,3 | 1,506634166 | 2,841463466 | 0,19943359 | 0,698904803 | novel transcript |
| ENSG00000273262 | AL121928,1 | 1,505830056 | 2,83988017 | 0,37625466 | 0,86257156 | novel transcript, antisense to ACTR1A |
| ENSG00000186493 | C5orf38 | 1,505744113 | 2,839711 | 0,00071068 | 0,01397393 | chromosome 5 open reading frame 38 |
| ENSG00000241544 | LINC02029 | 1,505273464 | 2,838784756 | 0,19204397 | 0,68846033 | long intergenic non-protein coding RNA 2029 |
| ENSG00000267529 | AP005131,4 | 1,504850942 | 2,837953482 | 0,00536656 | 0,072776773 | novel transcript, antisense C18orf1 |
| ENSG00000228528 | AC068057,1 | 1,503327306 | 2,834957891 | 0,0876886 | 0,471734492 | novel transcript |
| ENSG00000155324 | GRAMD2B | 1,502926054 | 2,834169522 | 8,22E-05 | 0,002212308 | GRAM domain containing 2B |
| ENSG00000253496 | AC011586,2 | 1,502661719 | 2,833650286 | 0,19873169 | 0,697657684 | novel transcript |

| | | | | | | |
|---|---|---|---|---|---|---|
| ENSG00000197334 | Metazoa_SRP | 1,502539657 | 2,833410548 | 0,18632389 | 0,679406361 | Metazoan signal recognition particle RNA |
| ENSG00000103196 | CRISPLD2 | 1,502511473 | 2,833355197 | 0,00288177 | 0,04491362 | cysteine rich secretory protein LCCL domain containing 2 |
| ENSG00000287741 | AC009434,1 | 1,502434748 | 2,833204519 | 0,33566192 | 0,835387231 | novel transcript |
| ENSG00000231247 | PPP1R18 | 1,501526433 | 2,831421305 | 0,00079584 | 0,015377759 | protein phosphatase 1 regulatory subunit 18 |
| ENSG00000285243 | SLC39A4 | 1,501385081 | 2,831143902 | 0,05385331 | 0,357953158 | solute carrier family 39 member 4 |
| ENSG00000278709 | NKILA | 1,500567851 | 2,829540624 | 4,70E-05 | 0,001339483 | NF-kappaB interacting lncRNA |
| ENSG00000134363 | FST | 1,499158439 | 2,826777712 | 5,45E-05 | 0,00153701 | follistatin |
| ENSG00000285938 | AC072022,2 | 1,498980176 | 2,826428451 | 0,06445818 | 0,397871785 | BCL6 antisense 1 |
| ENSG00000186897 | C1QL4 | 1,498957463 | 2,826383952 | 0,0072743 | 0,091151848 | complement C1q like 4 |
| ENSG00000237410 | NRXN2-AS1 | 1,49786622 | 2,824246906 | 0,155299 | 0,625190083 | NRXN2 antisense RNA 1 |
| ENSG00000152104 | PTPN14 | 1,497681911 | 2,823886122 | 2,19E-08 | 1,29E-06 | protein tyrosine phosphatase non-receptor type 14 |
| ENSG00000123612 | ACVR1C | 1,497252316 | 2,823045371 | 0,024806 | 0,218153277 | activin A receptor type 1C |
| ENSG00000118777 | ABCG2 | 1,496429153 | 2,821435078 | 0,04505023 | 0,320904617 | ATP binding cassette subfamily G member 2 (Junior blood group) |
| ENSG00000079215 | SLC1A3 | 1,495551418 | 2,81971904 | 0,00292278 | 0,045416444 | solute carrier family 1 member 3 |
| ENSG00000165061 | ZMAT4 | 1,494901733 | 2,818449529 | 9,53E-05 | 0,002518276 | zinc finger matrin-type 4 |
| ENSG00000273047 | AL121845,2 | 1,494348467 | 2,817368875 | 0,20339593 | 0,703503738 | novel transcript, LIME1-SLC2A4RG readthrough |
| ENSG00000115267 | IFIH1 | 1,493567957 | 2,815845069 | 0,01884151 | 0,17950267 | interferon induced with helicase C domain 1 |
| ENSG00000066322 | ELOVL1 | 1,492868536 | 2,814480273 | 1,12E-07 | 5,95E-06 | ELOVL fatty acid elongase 1 |
| ENSG00000186335 | SLC36A2 | 1,492818672 | 2,814382997 | 0,1449332 | 0,606017324 | solute carrier family 36 member 2 |
| ENSG00000007944 | MYLIP | 1,492335909 | 2,813441389 | 5,45E-05 | 0,00153701 | myosin regulatory light chain interacting protein |
| ENSG00000213509 | PPIAP16 | 1,492313964 | 2,813398594 | 0,27822216 | 0,789609447 | peptidylprolyl isomerase A pseudogene 16 |
| ENSG00000134222 | PSRC1 | 1,49209505 | 2,812971723 | 4,05E-06 | 0,000153851 | proline and serine rich coiled-coil 1 |
| ENSG00000100196 | KDELR3 | 1,491813724 | 2,812423246 | 4,33E-08 | 2,43E-06 | KDEL endoplasmic reticulum protein retention receptor 3 |
| ENSG00000005513 | SOX8 | 1,491784048 | 2,812365395 | 0,00841971 | 0,101321672 | SRY-box transcription factor 8 |
| ENSG00000260910 | LINC00565 | 1,491522011 | 2,81185463 | 0,0039236 | 0,057834685 | long intergenic non-protein coding RNA 565 |
| ENSG00000284848 | LINC02359 | 1,491041516 | 2,810918287 | 0,07643514 | 0,436971724 | long intergenic non-protein coding RNA 2359 |
| ENSG00000224885 | EIPR1-IT1 | 1,490776391 | 2,810401772 | 0,11292852 | 0,53680967 | EIPR1 intronic transcript 1 |
| ENSG00000234474 | MIR3663HG | 1,490661992 | 2,810178928 | 0,02494292 | 0,219004288 | MIR3663 host gene |
| ENSG00000234964 | FABP5P7 | 1,489683389 | 2,808273386 | 0,02368931 | 0,210380919 | fatty acid binding protein 5 pseudogene 7 |
| ENSG00000086696 | HSD17B2 | 1,488114761 | 2,805221638 | 0,00374477 | 0,055793453 | hydroxysteroid 17-beta dehydrogenase 2 |

| | | | | | | |
|---|---|---|---|---|---|---|
| ENSG00000174132 | FAM174A | 1,487950619 | 2,804902493 | 3,15E-05 | 0,00093902 | family with sequence similarity 174 member A |
| ENSG00000145936 | KCNMB1 | 1,487421404 | 2,803873776 | 0,00521082 | 0,071109189 | potassium calcium-activated channel subfamily M regulatory beta subunit 1 |
| ENSG00000147869 | CER1 | 1,487259266 | 2,803558679 | 0,06601526 | 0,403499746 | cerberus 1, DAN family BMP antagonist |
| ENSG00000267117 | AC010525,1 | 1,487161689 | 2,803369066 | 0,35421926 | 0,84556089 | novel transcript, antisense to EPN1 |
| ENSG00000173110 | HSPA6 | 1,486373381 | 2,801837686 | 0,27112587 | 0,783770603 | heat shock protein family A (Hsp70) member 6 |
| ENSG00000282113 | AC068570,1 | 1,486138085 | 2,801380758 | 0,36287861 | 0,853138593 | long intergenic non-protein coding RNA 976 |
| ENSG00000251191 | LINC00589 | 1,485907711 | 2,800933461 | 0,22192387 | 0,729392589 | long intergenic non-protein coding RNA 589 |
| ENSG00000172780 | RAB43 | 1,485885844 | 2,800891006 | 0,09552006 | 0,490763275 | RAB43, member RAS oncogene family |
| ENSG00000172543 | CTSW | 1,485644519 | 2,800422531 | 0,07135314 | 0,420755215 | cathepsin W |
| ENSG00000224958 | PGM5-AS1 | 1,485410743 | 2,799968783 | 0,27585263 | 0,787819199 | PGM5 antisense RNA 1 |
| ENSG00000254405 | MSH5-SAPCD1 | 1,4848462 | 2,798873339 | 0,02178775 | 0,198207677 | MSH5-SAPCD1 readthrough (NMD candidate) |
| ENSG00000286573 | AL136225,1 | 1,484469699 | 2,79814301 | 0,26140066 | 0,77325059 | novel transcript |
| ENSG00000287095 | AF228727,1 | 1,483490059 | 2,796243619 | 0,33801931 | 0,837255242 | novel transcript, antisense to TMEM71 |
| ENSG00000134548 | SPX | 1,483461286 | 2,796187852 | 0,03954581 | 0,296276117 | spexin hormone |
| ENSG00000197256 | KANK2 | 1,483045617 | 2,79538233 | 0,00011988 | 0,003074697 | KN motif and ankyrin repeat domains 2 |
| ENSG00000138639 | ARHGAP24 | 1,482827977 | 2,79496066 | 8,10E-06 | 0,00028273 | Rho GTPase activating protein 24 |
| ENSG00000269994 | AL513318,1 | 1,482504157 | 2,794333388 | 5,81E-07 | 2,58E-05 | novel transcript |
| ENSG00000174130 | TLR6 | 1,481951162 | 2,793262507 | 5,78E-05 | 0,001612255 | toll like receptor 6 |
| ENSG00000249237 | AL359273,2 | 1,481878068 | 2,793120989 | 0,34523411 | 0,841168721 | novel pseudogene |
| ENSG00000165269 | AQP7 | 1,48133056 | 2,79206119 | 0,03835627 | 0,290314236 | aquaporin 7 |
| ENSG00000224227 | OR2L1P | 1,481019966 | 2,791460161 | 0,29246649 | 0,802851193 | olfactory receptor family 2 subfamily L member 1 pseudogene |
| ENSG00000237654 | AP003025,1 | 1,480835638 | 2,791103528 | 0,16518172 | 0,644240891 | novel transcript |
| ENSG00000283099 | CCER2 | 1,480826686 | 2,791086209 | 0,23869039 | 0,747998807 | coiled-coil glutamate rich protein 2 |
| ENSG00000262484 | CCER2 | 1,480826686 | 2,791086209 | 0,23869039 | 0,747998807 | coiled-coil glutamate rich protein 2 |
| ENSG00000216866 | RPS2P55 | 1,480695817 | 2,790833036 | 0,15314151 | 0,621632863 | ribosomal protein S2 pseudogene 55 |
| ENSG00000145244 | CORIN | 1,479511025 | 2,788542048 | 0,02588308 | 0,224198135 | corin, serine peptidase |
| ENSG00000035403 | VCL | 1,478430214 | 2,786453762 | 3,34E-11 | 3,06E-09 | vinculin |
| ENSG00000050628 | PTGER3 | 1,478326814 | 2,786254061 | 0,04902077 | 0,337648216 | prostaglandin E receptor 3 |
| ENSG00000062524 | LTK | 1,477247155 | 2,784169713 | 0,24277413 | 0,752892335 | leukocyte receptor tyrosine kinase |
| ENSG00000184731 | FAM110C | 1,476448336 | 2,782628546 | 0,00012129 | 0,003104784 | family with sequence similarity 110 member C |
| ENSG00000137948 | BRDT | 1,47625014 | 2,782246298 | 0,21652678 | 0,723777756 | bromodomain testis associated |

| | | | | | | |
|---|---|---|---|---|---|---|
| ENSG00000262788 | UTP4 | 1,475741045 | 2,781264678 | 0,00159757 | 0,027701853 | UTP4 small subunit processome component |
| ENSG00000140538 | NTRK3 | 1,474731749 | 2,779319611 | 7,41E-05 | 0,002018357 | neurotrophic receptor tyrosine kinase 3 |
| ENSG00000236404 | VLDLR-AS1 | 1,474590542 | 2,779047593 | 0,0214957 | 0,196273823 | VLDLR antisense RNA 1 |
| ENSG00000258385 | AL391516,1 | 1,474536289 | 2,778943088 | 0,07135505 | 0,420755215 | WD repeat-containing protein 75 (WDR75) pseudogene |
| ENSG00000267147 | LINC01842 | 1,473801689 | 2,77752845 | 0,07051898 | 0,418185447 | long intergenic non-protein coding RNA 1842 |
| ENSG00000256448 | AP000763,3 | 1,473748651 | 2,777426341 | 0,25534505 | 0,766910046 | novel transcript |
| ENSG00000159212 | CLIC6 | 1,473549178 | 2,77704235 | 0,05212107 | 0,350357588 | chloride intracellular channel 6 |
| ENSG00000237900 | ZBTB12 | 1,473536169 | 2,777017308 | 0,14255326 | 0,600806836 | zinc finger and BTB domain containing 12 |
| ENSG00000234852 | ZBTB12 | 1,473536169 | 2,777017308 | 0,14255326 | 0,600806836 | zinc finger and BTB domain containing 12 |
| ENSG00000102048 | ASB9 | 1,472541138 | 2,775102651 | 0,00215347 | 0,035716726 | ankyrin repeat and SOCS box containing 9 |
| ENSG00000076716 | GPC4 | 1,472413886 | 2,774857886 | 2,91E-09 | 2,00E-07 | glypican 4 |
| ENSG00000140807 | NKD1 | 1,470865221 | 2,771880805 | 1,35E-05 | 0,000441514 | NKD inhibitor of WNT signaling pathway 1 |
| ENSG00000272269 | AL138724,1 | 1,469916802 | 2,770059186 | 0,02934433 | 0,243477915 | novel transcript, antisense to NUP153 |
| ENSG00000260604 | AL590004,3 | 1,469815578 | 2,769864838 | 0,01705158 | 0,167958614 | novel transcript |
| ENSG00000198832 | SELENOM | 1,469243941 | 2,768767556 | 0,00163699 | 0,028275582 | selenoprotein M |
| ENSG00000254847 | AC009806,1 | 1,469237577 | 2,768755342 | 0,28174185 | 0,792816227 | novel transcript |
| ENSG00000141682 | PMAIP1 | 1,468124434 | 2,766619871 | 8,18E-05 | 0,002205954 | phorbol-12-myristate-13-acetate-induced protein 1 |
| ENSG00000173262 | SLC2A14 | 1,46770993 | 2,765825102 | 0,00060247 | 0,012230969 | solute carrier family 2 member 14 |
| ENSG00000162624 | LHX8 | 1,467028303 | 2,764518647 | 0,12604918 | 0,566777174 | LIM homeobox 8 |
| ENSG00000101017 | CD40 | 1,46697849 | 2,764423196 | 0,0087872 | 0,104076013 | CD40 molecule |
| ENSG00000285922 | AC104453,1 | 1,466712658 | 2,763913868 | 0,01675696 | 0,165787872 | novel transcript |
| ENSG00000235750 | KIAA0040 | 1,466608875 | 2,763715049 | 0,00840588 | 0,101233181 | KIAA0040 |
| ENSG00000161677 | JOSD2 | 1,465967807 | 2,762487252 | 0,00863208 | 0,102939044 | Josephin domain containing 2 |
| ENSG00000248927 | AC114284,1 | 1,465810181 | 2,762185445 | 0,26739679 | 0,779594822 | novel transcript |
| ENSG00000162975 | KCNF1 | 1,465284699 | 2,761179539 | 0,00491585 | 0,067999136 | potassium voltage-gated channel modifier subfamily F member 1 |
| ENSG00000242960 | FTH1P23 | 1,464322883 | 2,75933933 | 0,22030323 | 0,728572782 | ferritin heavy chain 1 pseudogene 23 |
| ENSG00000133636 | NTS | 1,464276594 | 2,759250798 | 0,08523129 | 0,464892813 | neurotensin |
| ENSG00000167600 | CYP2S1 | 1,463658987 | 2,758069835 | 0,0091143 | 0,106922843 | cytochrome P450 family 2 subfamily S member 1 |
| ENSG00000107438 | PDLIM1 | 1,462990327 | 2,756791822 | 3,99E-05 | 0,001154307 | PDZ and LIM domain 1 |
| ENSG00000132854 | KANK4 | 1,462800666 | 2,75642943 | 0,00465854 | 0,065605674 | KN motif and ankyrin repeat domains 4 |
| ENSG00000111331 | OAS3 | 1,462167816 | 2,755220564 | 0,0084782 | 0,101620975 | 2'-5'-oligoadenylate synthetase 3 |

| | | | | | | |
|---|---|---|---|---|---|---|
| ENSG00000240338 | AC009078,2 | 1,462126748 | 2,755142136 | 0,34359206 | 0,840901506 | chymotrypsinogen B (CTRB) pseudogene |
| ENSG00000268434 | AC011530,1 | 1,46190772 | 2,754723885 | 0,41039406 | 0,882614012 | novel protein |
| ENSG00000266863 | RN7SL123P | 1,459013011 | 2,749202185 | 0,33078677 | 0,832114797 | RNA, 7SL, cytoplasmic 123, pseudogene |
| ENSG00000280689 | BORCS5 | 1,458517265 | 2,748257653 | 0,05134861 | 0,346880488 | BLOC-1 related complex subunit 5 |
| ENSG00000258659 | TRIM34 | 1,458184266 | 2,74762338 | 0,00157809 | 0,027409682 | tripartite motif containing 34 |
| ENSG00000256340 | ABCC6P1 | 1,45801771 | 2,747306191 | 0,05202691 | 0,349872693 | ATP binding cassette subfamily C member 6 pseudogene 1 |
| ENSG00000258128 | MKRN9P | 1,457420516 | 2,746169197 | 0,30944539 | 0,816965091 | makorin ring finger protein 9, pseudogene |
| ENSG00000230082 | PRRT3-AS1 | 1,456973652 | 2,745318724 | 0,16823794 | 0,649838265 | PRRT3 antisense RNA 1 |
| ENSG00000285107 | CACNA1C-AS4 | 1,456938914 | 2,745252621 | 0,26141267 | 0,77325059 | CACNA1C antisense RNA 4 |
| ENSG00000256025 | CACNA1C-AS4 | 1,456938914 | 2,745252621 | 0,26141267 | 0,77325059 | CACNA1C antisense RNA 4 |
| ENSG00000274049 | INO80B-WBP1 | 1,456563908 | 2,744539129 | 0,44468393 | 0,900310737 | INO80B-WBP1 readthrough (NMD candidate) |
| ENSG00000172197 | MBOAT1 | 1,455403154 | 2,742331834 | 0,00045998 | 0,009721199 | membrane bound O-acyltransferase domain containing 1 |
| ENSG00000287944 | AC027584,1 | 1,455218617 | 2,741981082 | 0,12024436 | 0,553427919 | novel transcript |
| ENSG00000254339 | AC064802,1 | 1,454316147 | 2,740266386 | 0,08393476 | 0,461230231 | novel transcript |
| ENSG00000185269 | NOTUM | 1,452814605 | 2,737415829 | 0,27575269 | 0,787605704 | notum, palmitoleoyl-protein carboxylesterase |
| ENSG00000281166 | NLRP10 | 1,452695825 | 2,737190461 | 0,20675816 | 0,710261203 | NLR family pyrin domain containing 10 |
| ENSG00000261248 | AC009120,4 | 1,452357384 | 2,73654842 | 0,29406706 | 0,803927308 | novel transcript |
| ENSG00000062716 | VMP1 | 1,450876363 | 2,733740617 | 8,99E-07 | 3,85E-05 | vacuole membrane protein 1 |
| ENSG00000005020 | SKAP2 | 1,450068547 | 2,732210326 | 0,00045274 | 0,00958328 | src kinase associated phosphoprotein 2 |
| ENSG00000288455 | SEMA7A | 1,449261242 | 2,73068186 | 0,00486015 | 0,067352677 | semaphorin 7A (John Milton Hagen blood group) |
| ENSG00000138623 | SEMA7A | 1,449261242 | 2,73068186 | 0,00486015 | 0,067352677 | semaphorin 7A (John Milton Hagen blood group) |
| ENSG00000279440 | AL008638,3 | 1,448612089 | 2,729453443 | 0,14058185 | 0,597491964 | novel transcript |
| ENSG00000147526 | TACC1 | 1,447823744 | 2,727962371 | 7,38E-05 | 0,002012778 | transforming acidic coiled-coil containing protein 1 |
| ENSG00000275431 | AC244100,2 | 1,447425226 | 2,727208925 | 0,34635913 | 0,841168721 | novel transcript, antisense to CCL14 |
| ENSG00000228267 | C4B | 1,447225567 | 2,726831525 | 0,04604673 | 0,325702465 | complement C4B (Chido blood group) |
| ENSG00000063438 | AHRR | 1,447110922 | 2,726614843 | 0,06287305 | 0,391611013 | aryl-hydrocarbon receptor repressor |
| ENSG00000149403 | GRIK4 | 1,446960078 | 2,726329771 | 3,60E-05 | 0,001056416 | glutamate ionotropic receptor kainate type subunit 4 |
| ENSG00000240463 | RPS19P3 | 1,446889312 | 2,726196044 | 0,1423289 | 0,600436282 | ribosomal protein S19 pseudogene 3 |
| ENSG00000261594 | TPBGL | 1,446708891 | 2,725855132 | 0,03707434 | 0,284203181 | trophoblast glycoprotein like |
| ENSG00000271943 | AC098614,2 | 1,4466205 | 2,72568813 | 0,01927863 | 0,182533983 | tropomyosin 4 (TPM4) pseudogene |

| | | | | | | |
|---|---|---|---|---|---|---|
| ENSG00000273584 | AC096708,3 | 1,44639742 | 2,725266697 | 0,10134603 | 0,50630911 | novel transcript, sense intronic to CCDC102B |
| ENSG00000129038 | LOXL1 | 1,445543311 | 2,723653754 | 3,57E-05 | 0,001053388 | lysyl oxidase like 1 |
| ENSG00000173641 | HSPB7 | 1,444410996 | 2,721516902 | 0,19073935 | 0,685985838 | heat shock protein family B (small) member 7 |
| ENSG00000176076 | KCNE5 | 1,444090826 | 2,720912997 | 0,00155395 | 0,027035474 | potassium voltage-gated channel subfamily E regulatory subunit 5 |
| ENSG00000035862 | TIMP2 | 1,443641025 | 2,720064806 | 7,90E-06 | 0,000276902 | TIMP metallopeptidase inhibitor 2 |
| ENSG00000151726 | ACSL1 | 1,443552226 | 2,71989739 | 3,00E-05 | 0,000899604 | acyl-CoA synthetase long chain family member 1 |
| ENSG00000248587 | GDNF-AS1 | 1,443271083 | 2,719367407 | 0,02849794 | 0,23950529 | GDNF antisense RNA 1 |
| ENSG00000203531 | AC104772,1 | 1,443138113 | 2,719116779 | 0,31632549 | 0,821646322 | C-type lectin domain family 1, member A (CLEC1A) pseudogene |
| ENSG00000242559 | RN7SL144P | 1,443001043 | 2,71885845 | 0,30371084 | 0,812795736 | RNA, 7SL, cytoplasmic 144, pseudogene |
| ENSG00000226928 | RPS14P4 | 1,442152384 | 2,717259563 | 0,17281681 | 0,657292556 | ribosomal protein S14 pseudogene 4 |
| ENSG00000282837 | AC254788,1 | 1,441811102 | 2,716616847 | 0,5004913 | 0,920487088 | |
| ENSG00000100599 | RIN3 | 1,440951319 | 2,714998346 | 0,00015433 | 0,003790306 | Ras and Rab interactor 3 |
| ENSG00000232301 | LNCPRESS1 | 1,439296304 | 2,71188557 | 0,38130325 | 0,865900094 | lncRNA p53 regulated and ESC associated 1 |
| ENSG00000067082 | KLF6 | 1,439130477 | 2,711573877 | 1,72E-07 | 8,74E-06 | Kruppel like factor 6 |
| ENSG00000088002 | SULT2B1 | 1,438754387 | 2,7108671 | 0,25722158 | 0,769474266 | sulfotransferase family 2B member 1 |
| ENSG00000004948 | CALCR | 1,438633813 | 2,710640547 | 0,03910606 | 0,29449254 | calcitonin receptor |
| ENSG00000276988 | Metazoa_SRP | 1,438127998 | 2,709690351 | 0,18182154 | 0,672010829 | Metazoan signal recognition particle RNA |
| ENSG00000177692 | DNAJC28 | 1,437865525 | 2,709197416 | 0,05238162 | 0,351288432 | DnaJ heat shock protein family (Hsp40) member C28 |
| ENSG00000250748 | AC025419,1 | 1,43744397 | 2,708405905 | 0,10295071 | 0,51138402 | novel transcript |
| ENSG00000168487 | BMP1 | 1,437149777 | 2,707853665 | 1,54E-05 | 0,000495624 | bone morphogenetic protein 1 |
| ENSG00000255337 | AP001830,1 | 1,43609934 | 2,705882775 | 0,31214054 | 0,818547047 | novel transcript |
| ENSG00000107338 | SHB | 1,4359924 | 2,705682208 | 4,29E-07 | 1,96E-05 | SH2 domain containing adaptor protein B |
| ENSG00000231875 | AL359885,1 | 1,435913669 | 2,705534558 | 0,10903855 | 0,526890476 | novel pseudogene |
| ENSG00000229618 | AC011287,1 | 1,435879849 | 2,705471134 | 0,000348 | 0,007688978 | novel transcript |
| ENSG00000188223 | AD000671,1 | 1,435660192 | 2,705059246 | 0,47680073 | 0,91143003 | novel protein |
| ENSG00000243414 | TICAM2 | 1,435404275 | 2,704579442 | 0,07229457 | 0,423976244 | toll like receptor adaptor molecule 2 |
| ENSG00000234255 | AC012370,2 | 1,435308978 | 2,704400797 | 0,30010142 | 0,809942221 | novel transcript |
| ENSG00000258419 | AC022469,1 | 1,434747741 | 2,703348937 | 0,25173154 | 0,763986755 | novel transcript, sense intronic to NRXN3 |
| ENSG00000171587 | DSCAM | 1,434658332 | 2,703181406 | 0,00378343 | 0,056261305 | DS cell adhesion molecule |
| ENSG00000243339 | RN7SL738P | 1,434184453 | 2,702293644 | 0,34119214 | 0,839923707 | RNA, 7SL, cytoplasmic 738, pseudogene |
| ENSG00000237234 | Z99289,1 | 1,433327511 | 2,700688993 | 0,13756036 | 0,592217616 | novel transcript |

| | | | | | | |
|---|---|---|---|---|---|---|
| ENSG00000049249 | TNFRSF9 | 1,433022155 | 2,700117434 | 0,01062078 | 0,119872893 | TNF receptor superfamily member 9 |
| ENSG00000287661 | AL391844,1 | 1,431450148 | 2,697176902 | 0,34328103 | 0,840790189 | novel transcript, antisense to AGBL4 |
| ENSG00000178752 | ERFE | 1,431326189 | 2,696945166 | 0,00434929 | 0,062431658 | erythroferrone |
| ENSG00000287543 | AC013486,1 | 1,431298411 | 2,696893239 | 0,01035714 | 0,118005006 | novel transcript, antisense to PML |
| ENSG00000287906 | AC127526,5 | 1,430753823 | 2,695875409 | 0,25811371 | 0,770594635 | novel transcript |
| ENSG00000235200 | LINC01702 | 1,43019151 | 2,694824853 | 0,26950081 | 0,781602339 | long intergenic non-protein coding RNA 1702 |
| ENSG00000160183 | TMPRSS3 | 1,430129665 | 2,694709335 | 0,17563044 | 0,661199837 | transmembrane serine protease 3 |
| ENSG00000254880 | PKNOX2-AS1 | 1,42983611 | 2,694161079 | 0,22592824 | 0,734959876 | PKNOX2 antisense RNA 1 (head to head) |
| ENSG00000204851 | PNMA8B | 1,429634772 | 2,693785117 | 0,0669842 | 0,40671858 | PNMA family member 8B |
| ENSG00000216490 | IFI30 | 1,429466069 | 2,693470136 | 0,0896278 | 0,47775341 | IFI30 lysosomal thiol reductase |
| ENSG00000184481 | FOXO4 | 1,429420863 | 2,693385739 | 7,76E-07 | 3,37E-05 | forkhead box O4 |
| ENSG00000268964 | ERVV-2 | 1,429291154 | 2,693143595 | 0,14726151 | 0,611181688 | endogenous retrovirus group V member 2, envelope |
| ENSG00000260896 | ARLNC1 | 1,42823973 | 2,691181569 | 0,15901006 | 0,633144409 | androgen receptor regulated long noncoding RNA 1 |
| ENSG00000154330 | PGM5 | 1,427634249 | 2,69005235 | 0,00322401 | 0,049409506 | phosphoglucomutase 5 |
| ENSG00000273079 | GRIN2B | 1,426750648 | 2,68840529 | 0,00451138 | 0,063994488 | glutamate ionotropic receptor NMDA type subunit 2B |
| ENSG00000236824 | BCYRN1 | 1,426612326 | 2,688147545 | 0,03993927 | 0,298328495 | brain cytoplasmic RNA 1 |
| ENSG00000248340 | PIMREGP4 | 1,425915823 | 2,686850077 | 0,17013938 | 0,653308467 | PIMREG pseudogene 4 |
| ENSG00000072786 | STK10 | 1,425648261 | 2,68635182 | 0,00141577 | 0,02510636 | serine/threonine kinase 10 |
| ENSG00000285763 | AL358777,3 | 1,425474714 | 2,686028689 | 0,60463072 | 0,950878607 | novel transcript, antisense to C6orf52 |
| ENSG00000287469 | AC044798,3 | 1,424987803 | 2,685122304 | 0,19358574 | 0,690424879 | novel transcript |
| ENSG00000158955 | WNT9B | 1,424564053 | 2,684333743 | 0,00262014 | 0,041560616 | Wnt family member 9B |
| ENSG00000171570 | RAB4B-EGLN2 | 1,424519415 | 2,684250689 | 0,12424913 | 0,563219486 | RAB4B-EGLN2 readthrough (NMD candidate) |
| ENSG00000230578 | AC234781,5 | 1,424211633 | 2,683678097 | 0,1293374 | 0,57474603 | trimethyllysine hydroxylase, epsilon (TMLHE) pseudogene |
| ENSG00000226191 | CLK3P2 | 1,423522702 | 2,682396865 | 0,06828828 | 0,411780707 | CDC like kinase 3 pseudogene 2 |
| ENSG00000250189 | AC097504,1 | 1,422128441 | 2,679805773 | 0,26366781 | 0,774820034 | F-box and leucine-rich repeat protein 5 (FBXL5) pseudogene |
| ENSG00000228539 | AL157413,1 | 1,42208241 | 2,679720272 | 0,24448113 | 0,756029492 | novel transcript, antisense to SYNDIG1 |
| ENSG00000283199 | C13orf46 | 1,420469255 | 2,676725608 | 0,08385399 | 0,461082665 | chromosome 13 open reading frame 46 |
| ENSG00000283302 | C13orf46 | 1,420469255 | 2,676725608 | 0,08385399 | 0,461082665 | chromosome 13 open reading frame 46 |
| ENSG00000146122 | DAAM2 | 1,419330929 | 2,67461443 | 0,00016581 | 0,004037283 | dishevelled associated activator of morphogenesis 2 |
| ENSG00000288557 | AC080097,1 | 1,418585496 | 2,673232828 | 0,06593147 | 0,403224066 | novel transcript |

| | | | | | | |
|---|---|---|---|---|---|---|
| ENSG00000177800 | TMEM78 | 1,418364729 | 2,67282379 | 0,30346311 | 0,812669425 | transmembrane protein 78 |
| ENSG00000171812 | COL8A2 | 1,418001134 | 2,672150256 | 0,02680471 | 0,229343848 | collagen type VIII alpha 2 chain |
| ENSG00000228151 | AC004941,1 | 1,417799983 | 2,671777712 | 0,22207424 | 0,729759198 | uncharacterized LOC101928733 |
| ENSG00000155816 | FMN2 | 1,417495554 | 2,67121399 | 0,00011588 | 0,002984358 | formin 2 |
| ENSG00000286829 | AC093798,1 | 1,416458665 | 2,669294833 | 0,20310167 | 0,703503738 | novel transcript |
| ENSG00000164128 | NPY1R | 1,416238415 | 2,668887355 | 0,0063318 | 0,082213892 | neuropeptide Y receptor Y1 |
| ENSG00000272701 | MESTIT1 | 1,415585929 | 2,667680573 | 0,00029031 | 0,006567422 | MEST intronic transcript 1, antisense RNA |
| ENSG00000256977 | LIMS3 | 1,415155118 | 2,666884081 | 0,00149904 | 0,02624073 | LIM zinc finger domain containing 3 |
| ENSG00000232691 | ZNRD1ASP | 1,414143813 | 2,665015294 | 0,23446832 | 0,744090329 | zinc ribbon domain containing 1 antisense, pseudogene |
| ENSG00000187678 | SPRY4 | 1,413804901 | 2,664389313 | 1,39E-05 | 0,00045479 | sprouty RTK signaling antagonist 4 |
| ENSG00000250786 | SNHG18 | 1,413724278 | 2,664240423 | 0,00030238 | 0,006801198 | small nucleolar RNA host gene 18 |
| ENSG00000285106 | AC016831,6 | 1,41268592 | 2,662323567 | 0,00945913 | 0,110100586 | novel transcript |
| ENSG00000280004 | AC005086,4 | 1,412058487 | 2,661165964 | 0,20240522 | 0,702433811 | |
| ENSG00000239437 | RN7SL752P | 1,411945745 | 2,66095801 | 0,30840668 | 0,816625411 | RNA, 7SL, cytoplasmic 752, pseudogene |
| ENSG00000254715 | OR7E154P | 1,411311714 | 2,659788838 | 0,26360929 | 0,774793719 | olfactory receptor family 7 subfamily E member 154 pseudogene |
| ENSG00000158747 | NBL1 | 1,409494898 | 2,656441418 | 0,01303518 | 0,139032628 | NBL1, DAN family BMP antagonist |
| ENSG00000115844 | DLX2 | 1,408901094 | 2,655348269 | 3,26E-05 | 0,000969404 | distal-less homeobox 2 |
| ENSG00000187871 | GFRAL | 1,408436666 | 2,654493606 | 0,33227167 | 0,83293962 | GDNF family receptor alpha like |
| ENSG00000151136 | BTBD11 | 1,407620368 | 2,652992078 | 2,02E-05 | 0,000636848 | BTB domain containing 11 |
| ENSG00000182621 | PLCB1 | 1,407473558 | 2,652722121 | 2,36E-05 | 0,000729843 | phospholipase C beta 1 |
| ENSG00000124762 | CDKN1A | 1,405895408 | 2,649821921 | 0,00031731 | 0,007101185 | cyclin dependent kinase inhibitor 1A |
| ENSG00000130517 | PGPEP1 | 1,405386841 | 2,648887993 | 0,00229109 | 0,037418119 | pyroglutamyl-peptidase I |
| ENSG00000276002 | Metazoa_SRP | 1,405373962 | 2,648864347 | 0,20389469 | 0,704681782 | Metazoan signal recognition particle RNA |
| ENSG00000237918 | SAPCD1 | 1,405040475 | 2,648252117 | 0,12231489 | 0,559390308 | suppressor APC domain containing 1 |
| ENSG00000267366 | AP005131,2 | 1,404065872 | 2,646463711 | 0,05520806 | 0,362784685 | novel transcript, antisense C18orf1 |
| ENSG00000158516 | CPA2 | 1,403936149 | 2,646225761 | 0,07025207 | 0,417549035 | carboxypeptidase A2 |
| ENSG00000206505 | HLA-A | 1,403801256 | 2,645978348 | 0,25887093 | 0,771850203 | major histocompatibility complex, class I, A |
| ENSG00000275881 | Metazoa_SRP | 1,402729648 | 2,644013691 | 0,30842864 | 0,816625411 | Metazoan signal recognition particle RNA |
| ENSG00000148411 | NACC2 | 1,402522734 | 2,643634509 | 0,01840432 | 0,176657259 | NACC family member 2 |
| ENSG00000278250 | Metazoa_SRP | 1,401174351 | 2,641164849 | 0,45810312 | 0,904140862 | Metazoan signal recognition particle RNA |
| ENSG00000185222 | TCEAL9 | 1,400863329 | 2,640595518 | 2,66E-05 | 0,000807554 | transcription elongation factor A like 9 |

| | | | | | | |
|---|---|---|---|---|---|---|
| ENSG00000236540 | AC006547,1 | 1,400513884 | 2,639956 | 0,02017011 | 0,187832988 | novel transcript, antisense to C22orf25 |
| ENSG00000185340 | GAS2L1 | 1,400475618 | 2,639885978 | 0,00403016 | 0,059154371 | growth arrest specific 2 like 1 |
| ENSG00000249035 | CLMAT3 | 1,400101824 | 2,639202088 | 0,00401148 | 0,05893556 | colorectal liver metastasis associated transcript 3 |
| ENSG00000173391 | OLR1 | 1,399680748 | 2,638431901 | 0,37101441 | 0,858959913 | oxidized low density lipoprotein receptor 1 |
| ENSG00000153165 | RGPD3 | 1,399611396 | 2,638305073 | 0,13800534 | 0,592546669 | RANBP2 like and GRIP domain containing 3 |
| ENSG00000235997 | LINC01936 | 1,399446178 | 2,63800295 | 0,05042637 | 0,343470607 | long intergenic non-protein coding RNA 1936 |
| ENSG00000276259 | AC009118,3 | 1,399419485 | 2,637954141 | 0,11731746 | 0,547271437 | novel transcript, antisense to CNOT1 |
| ENSG00000151062 | CACNA2D4 | 1,398967425 | 2,637127684 | 0,00945131 | 0,1100828 | calcium voltage-gated channel auxiliary subunit alpha2delta 4 |
| ENSG00000284953 | CACNA2D4 | 1,398967425 | 2,637127684 | 0,00945131 | 0,1100828 | calcium voltage-gated channel auxiliary subunit alpha2delta 4 |
| ENSG00000204262 | COL5A2 | 1,39853904 | 2,636344748 | 0,00892095 | 0,105360946 | collagen type V alpha 2 chain |
| ENSG00000280651 | AC156455,2 | 1,398356927 | 2,636011979 | 0,00142161 | 0,025181433 | novel transcript |
| ENSG00000256546 | AC156455,1 | 1,398356927 | 2,636011979 | 0,00142161 | 0,025181433 | novel transcript |
| ENSG00000276396 | Metazoa_SRP | 1,398055893 | 2,635462004 | 0,3830167 | 0,867682628 | Metazoan signal recognition particle RNA |
| ENSG00000152208 | GRID2 | 1,397987968 | 2,635337926 | 0,00650639 | 0,083953588 | glutamate ionotropic receptor delta type subunit 2 |
| ENSG00000232006 | AC005537,1 | 1,39791967 | 2,635213169 | 0,00500498 | 0,068841073 | novel transcript |
| ENSG00000276130 | SLC22A18 | 1,397533551 | 2,634507983 | 0,01109924 | 0,123533851 | solute carrier family 22 member 18 |
| ENSG00000269110 | AC010636,2 | 1,39612109 | 2,631929948 | 0,16580078 | 0,645114847 | novel transcript |
| ENSG00000266578 | AP001025,1 | 1,395671866 | 2,631110549 | 0,15281243 | 0,62133458 | novel transcript |
| ENSG00000150594 | ADRA2A | 1,395528865 | 2,630849764 | 0,26866708 | 0,780823861 | adrenoceptor alpha 2A |
| ENSG00000154102 | C16orf74 | 1,395264811 | 2,630368289 | 0,19738403 | 0,695991044 | chromosome 16 open reading frame 74 |
| ENSG00000059378 | PARP12 | 1,394948084 | 2,629790884 | 0,00218306 | 0,036035485 | poly(ADP-ribose) polymerase family member 12 |
| ENSG00000162976 | SLC66A3 | 1,39491993 | 2,629739565 | 0,00142745 | 0,025249098 | solute carrier family 66 member 3 |
| ENSG00000090776 | EFNB1 | 1,394123061 | 2,628287436 | 8,18E-06 | 0,000284928 | ephrin B1 |
| ENSG00000250421 | AC106798,1 | 1,393491475 | 2,627137072 | 0,16434882 | 0,643240037 | novel transcript |
| ENSG00000229386 | OR8B9P | 1,392672203 | 2,625645606 | 0,01840933 | 0,176657259 | olfactory receptor family 8 subfamily B member 9 pseudogene |
| ENSG00000287684 | AL392172,1 | 1,392042286 | 2,624499434 | 0,17247021 | 0,656926048 | novel transcript, antisense to FAM177B |
| ENSG00000224700 | AP000844,1 | 1,391819282 | 2,624093784 | 0,13152269 | 0,579389234 | novel transcript |
| ENSG00000273564 | FAM189A1 | 1,391714061 | 2,623902406 | 7,68E-05 | 0,002079679 | family with sequence similarity 189 member A1 |
| ENSG00000117707 | PROX1 | 1,39126313 | 2,623082403 | 1,09E-06 | 4,59E-05 | prospero homeobox 1 |
| ENSG00000237339 | LINC01502 | 1,391194681 | 2,622957952 | 0,20275487 | 0,703075461 | long intergenic non-protein coding RNA 1502 |

| | | | | | | |
|---|---|---|---|---|---|---|
| ENSG00000213779 | AC107956,2 | 1,391017142 | 2,622635191 | 0,31054936 | 0,817751312 | ribosomal protein L29 (RPL29) pseudogene |
| ENSG00000166387 | PPFIBP2 | 1,390111272 | 2,620988952 | 4,45E-05 | 0,001275493 | PPFIA binding protein 2 |
| ENSG00000251199 | AC015631,1 | 1,389809226 | 2,620440271 | 0,17916525 | 0,667630798 | novel transcript |
| ENSG00000224940 | PRRT4 | 1,389642563 | 2,620137571 | 0,1977116 | 0,696048902 | proline rich transmembrane protein 4 |
| ENSG00000235688 | SNTG2-AS1 | 1,388821148 | 2,618646191 | 0,34645362 | 0,841168721 | SNTG2 antisense RNA 1 |
| ENSG00000225190 | PLEKHM1 | 1,388148289 | 2,617425164 | 0,01188499 | 0,12978427 | pleckstrin homology and RUN domain containing M1 |
| ENSG00000242641 | LINC00971 | 1,387955669 | 2,617075724 | 0,29146498 | 0,802058262 | long intergenic non-protein coding RNA 971 |
| ENSG00000273238 | TMEM271 | 1,387480892 | 2,616214612 | 0,03414732 | 0,26913287 | transmembrane protein 271 |
| ENSG00000165023 | DIRAS2 | 1,387007318 | 2,615355963 | 1,80E-05 | 0,000573309 | DIRAS family GTPase 2 |
| ENSG00000240233 | RN7SL587P | 1,385865765 | 2,613287344 | 0,33633208 | 0,835660292 | RNA, 7SL, cytoplasmic 587, pseudogene |
| ENSG00000075673 | ATP12A | 1,385698683 | 2,61298471 | 0,2674122 | 0,779594822 | ATPase H+/K+ transporting non-gastric alpha2 subunit |
| ENSG00000162745 | OLFML2B | 1,384953485 | 2,611635369 | 2,83E-06 | 0,000111132 | olfactomedin like 2B |
| ENSG00000143226 | FCGR2A | 1,384431051 | 2,610689805 | 0,09074828 | 0,480677819 | Fc fragment of IgG receptor IIa |
| ENSG00000256040 | PAPPA-AS1 | 1,384336306 | 2,61051836 | 0,62829442 | 0,952793934 | PAPPA antisense RNA 1 |
| ENSG00000280928 | AC013726,3 | 1,384063504 | 2,61002478 | 0,61852138 | 0,952793934 | novel transcript, antisense to DGKD |
| ENSG00000240654 | C1QTNF9 | 1,381894821 | 2,606104295 | 0,33162496 | 0,832548086 | C1q and TNF related 9 |
| ENSG00000124159 | MATN4 | 1,380669833 | 2,603892398 | 0,39461295 | 0,871872016 | matrilin 4 |
| ENSG00000196205 | EEF1A1P5 | 1,38043782 | 2,603473677 | 0,12338411 | 0,561691036 | eukaryotic translation elongation factor 1 alpha 1 pseudogene 5 |
| ENSG00000023445 | BIRC3 | 1,379617325 | 2,601993439 | 0,05646399 | 0,366903759 | baculoviral IAP repeat containing 3 |
| ENSG00000230943 | LINC02541 | 1,379565297 | 2,601899606 | 0,08059371 | 0,450103921 | long intergenic non-protein coding RNA 2541 |
| ENSG00000120659 | TNFSF11 | 1,379378206 | 2,60156221 | 0,30895325 | 0,816838423 | TNF superfamily member 11 |
| ENSG00000131982 | UBE2L2 | 1,378693009 | 2,600326911 | 0,19859666 | 0,697657684 | ubiquitin conjugating enzyme E2 L2 (pseudogene) |
| ENSG00000259106 | AC008056,2 | 1,378672835 | 2,600290549 | 0,25385005 | 0,76531122 | novel transcript, antisense to NRXN3 |
| ENSG00000225826 | LINC00626 | 1,378378759 | 2,599760565 | 0,18294872 | 0,673778851 | long intergenic non-protein coding RNA 626 |
| ENSG00000102287 | GABRE | 1,378344278 | 2,59969843 | 1,66E-05 | 0,000533988 | gamma-aminobutyric acid type A receptor subunit epsilon |
| ENSG00000277170 | AC012676,3 | 1,378056629 | 2,599180147 | 0,2845454 | 0,795919345 | novel transcript, sense intronic with DNAJA3 |
| ENSG00000183785 | TUBA8 | 1,378043892 | 2,599157199 | 0,13972867 | 0,595809743 | tubulin alpha 8 |
| ENSG00000005022 | SLC25A5 | 1,378033028 | 2,599137628 | 6,87E-06 | 0,000245289 | solute carrier family 25 member 5 |
| ENSG00000115170 | ACVR1 | 1,377886964 | 2,598874494 | 2,34E-08 | 1,37E-06 | activin A receptor type 1 |
| ENSG00000254233 | LINC02365 | 1,377768012 | 2,598660221 | 0,287882 | 0,799107075 | long intergenic non-protein coding RNA 2365 |

| | | | | | | |
|---|---|---|---|---|---|---|
| ENSG00000196188 | CTSE | 1,377558445 | 2,598282767 | 0,21666868 | 0,723942475 | cathepsin E |
| ENSG00000176183 | AC233964,1 | 1,377313914 | 2,597842404 | 0,28083903 | 0,791510309 | actin, beta (ACTB) pseudogene |
| ENSG00000100346 | CACNA1I | 1,376811091 | 2,596937136 | 0,04068622 | 0,301441804 | calcium voltage-gated channel subunit alpha1 I |
| ENSG00000114279 | FGF12 | 1,376463626 | 2,596311754 | 0,00046019 | 0,009721199 | fibroblast growth factor 12 |
| ENSG00000288242 | AC106845,2 | 1,376331561 | 2,596074097 | 0,21978687 | 0,727931767 | novel transcript, antisense to DLC1 |
| ENSG00000288169 | ACAP1 | 1,375884289 | 2,595269373 | 0,02016784 | 0,187832988 | ArfGAP with coiled-coil, ankyrin repeat and PH domains 1 |
| ENSG00000236206 | AL356441,1 | 1,375158406 | 2,593963908 | 0,28998305 | 0,800682868 | novel transcript |
| ENSG00000087253 | LPCAT2 | 1,37466417 | 2,593075423 | 1,95E-07 | 9,66E-06 | lysophosphatidylcholine acyltransferase 2 |
| ENSG00000008513 | ST3GAL1 | 1,374379355 | 2,592563552 | 1,12E-06 | 4,71E-05 | ST3 beta-galactoside alpha-2,3-sialyltransferase 1 |
| ENSG00000184313 | MROH7 | 1,372822544 | 2,589767429 | 0,06908393 | 0,414636184 | maestro heat like repeat family member 7 |
| ENSG00000274565 | AC080038,1 | 1,372743447 | 2,589625447 | 0,00137178 | 0,024465044 | novel transcript |
| ENSG00000225649 | AC064875,1 | 1,372450235 | 2,589099186 | 0,02681816 | 0,229377638 | novel transcript |
| ENSG00000273574 | LENG9 | 1,371745607 | 2,587834952 | 0,03157135 | 0,256176123 | leukocyte receptor cluster member 9 |
| ENSG00000065833 | ME1 | 1,371153528 | 2,586773128 | 8,48E-10 | 6,44E-08 | malic enzyme 1 |
| ENSG00000242198 | AC008897,1 | 1,369969876 | 2,584651693 | 0,24628551 | 0,758198269 | ribosomal protein S26 (RPS26) pseudogene |
| ENSG00000280466 | SCARNA4 | 1,369731358 | 2,584224412 | 0,2901008 | 0,800682868 | small Cajal body-specific RNA 4 |
| ENSG00000254337 | AC083967,1 | 1,368928973 | 2,582787541 | 0,09618145 | 0,492702847 | novel transcript |
| ENSG00000276916 | AL442125,2 | 1,367817085 | 2,580797749 | 0,18400129 | 0,675665532 | novel transcript, sense intronic |
| ENSG00000282307 | SERPINF1 | 1,367793919 | 2,580756308 | 0,17160523 | 0,655397237 | serpin family F member 1 |
| ENSG00000264423 | RN7SL718P | 1,367778846 | 2,580729345 | 0,32787052 | 0,829869166 | RNA, 7SL, cytoplasmic 718, pseudogene |
| ENSG00000214955 | AP000317,1 | 1,367488443 | 2,580209916 | 0,29071439 | 0,801555946 | novel transcript |
| ENSG00000264963 | RN7SL440P | 1,367393397 | 2,580039937 | 0,37233079 | 0,860223901 | RNA, 7SL, cytoplasmic 440, pseudogene |
| ENSG00000260305 | NTRK3-AS1 | 1,36721177 | 2,579715144 | 0,22796346 | 0,736341996 | NTRK3 antisense RNA 1 |
| ENSG00000277626 | WNT3 | 1,367051204 | 2,579428049 | 0,00023717 | 0,005521168 | Wnt family member 3 |
| ENSG00000277702 | AC239859,5 | 1,366714853 | 2,578826748 | 0,28325927 | 0,794558711 | lymphocyte-specific protein 1 (LSP1) pseudogene |
| ENSG00000197405 | C5AR1 | 1,366605775 | 2,578631778 | 0,15987646 | 0,634614414 | complement C5a receptor 1 |
| ENSG00000236714 | LINC01844 | 1,365549938 | 2,576745296 | 0,28857693 | 0,79998466 | long intergenic non-protein coding RNA 1844 |
| ENSG00000270106 | TSNAX-DISC1 | 1,365475825 | 2,57661293 | 0,02627866 | 0,226459876 | TSNAX-DISC1 readthrough (NMD candidate) |
| ENSG00000196502 | SULT1A1 | 1,364957104 | 2,575686674 | 3,42E-06 | 0,000132004 | sulfotransferase family 1A member 1 |
| ENSG00000280780 | JAKMIP2-AS1 | 1,364092122 | 2,574142859 | 7,49E-05 | 0,002034664 | JAKMIP2 antisense RNA 1 |
| ENSG00000167123 | CERCAM | 1,363171046 | 2,572499945 | 0,00233849 | 0,037979447 | cerebral endothelial cell adhesion molecule |
| ENSG00000220884 | MESTP1 | 1,362747668 | 2,571745121 | 0,13424958 | 0,584425029 | mesoderm specific transcript pseudogene 1 |

| | | | | | | |
|---|---|---|---|---|---|---|
| ENSG00000127124 | HIVEP3 | 1,362692631 | 2,571647014 | 0,00094436 | 0,017807303 | HIVEP zinc finger 3 |
| ENSG00000162888 | C1orf147 | 1,362442556 | 2,571201286 | 0,04435051 | 0,318021642 | chromosome 1 open reading frame 147 |
| ENSG00000215878 | MARCKSL1P2 | 1,362257281 | 2,570871106 | 0,23252539 | 0,74165209 | MARCKS like 1 pseudogene 2 |
| ENSG00000019169 | MARCO | 1,361535919 | 2,569585966 | 0,38778506 | 0,869820053 | macrophage receptor with collagenous structure |
| ENSG00000229896 | AL157373,2 | 1,360320172 | 2,567421511 | 0,18866943 | 0,682944308 | novel transcript |
| ENSG00000132386 | SERPINF1 | 1,360243354 | 2,567284809 | 0,00202399 | 0,033892951 | serpin family F member 1 |
| ENSG00000249465 | RBMXP4 | 1,359864093 | 2,566610001 | 0,32751636 | 0,829485803 | RBMX pseudogene 4 |
| ENSG00000213222 | TOMM40P4 | 1,359788648 | 2,566475784 | 0,01663529 | 0,165046811 | TOMM40 pseudogene 4 |
| ENSG00000281848 | TOMM40P4 | 1,359788648 | 2,566475784 | 0,01663529 | 0,165046811 | TOMM40 pseudogene 4 |
| ENSG00000165379 | LRFN5 | 1,35952619 | 2,566008927 | 0,0002747 | 0,006241398 | leucine rich repeat and fibronectin type III domain containing 5 |
| ENSG00000159840 | ZYX | 1,358873468 | 2,564848246 | 0,00036201 | 0,007914699 | zyxin |
| ENSG00000285443 | ZYX | 1,358873468 | 2,564848246 | 0,00036201 | 0,007914699 | zyxin |
| ENSG00000189152 | GRAPL | 1,358601148 | 2,564364155 | 0,20704575 | 0,710823213 | GRB2 related adaptor protein like |
| ENSG00000166750 | SLFN5 | 1,3585208 | 2,564221342 | 7,28E-05 | 0,001986917 | schlafen family member 5 |
| ENSG00000287367 | AC016583,2 | 1,358157733 | 2,563576115 | 0,11565244 | 0,543151268 | novel transcript |
| ENSG00000073008 | PVR | 1,357947706 | 2,563202936 | 5,21E-05 | 0,001472052 | PVR cell adhesion molecule |
| ENSG00000269608 | OSTCP3 | 1,35774795 | 2,562848059 | 0,18764867 | 0,681692777 | oligosaccharyltransferase complex subunit pseudogene 3 |
| ENSG00000286249 | AC016743,1 | 1,357579865 | 2,562549486 | 0,31298963 | 0,818922744 | novel transcript |
| ENSG00000235790 | PEF1-AS1 | 1,357085196 | 2,561670993 | 0,15452193 | 0,623626246 | PEF1 and COL16A1 antisense RNA 1 |
| ENSG00000269680 | AC008760,1 | 1,357010691 | 2,561538705 | 0,29008015 | 0,800682868 | novel transcript, antisense to TRIP10 |
| ENSG00000139508 | SLC46A3 | 1,356774581 | 2,561119521 | 0,00155114 | 0,027018437 | solute carrier family 46 member 3 |
| ENSG00000119699 | TGFB3 | 1,356702083 | 2,560990823 | 0,00327175 | 0,050075384 | transforming growth factor beta 3 |
| ENSG00000196917 | HCAR1 | 1,356130549 | 2,559976469 | 0,050257 | 0,342972789 | hydroxycarboxylic acid receptor 1 |
| ENSG00000261783 | AC009054,2 | 1,356045719 | 2,559825948 | 0,10524715 | 0,517691472 | novel transcript, sense intronic to CFDP1 |
| ENSG00000111057 | KRT18 | 1,356002636 | 2,559749505 | 0,00241086 | 0,038932326 | keratin 18 |
| ENSG00000176697 | BDNF | 1,354983491 | 2,557941892 | 0,00014902 | 0,003691955 | brain derived neurotrophic factor |
| ENSG00000101958 | GLRA2 | 1,35392515 | 2,556066109 | 0,02875646 | 0,240839934 | glycine receptor alpha 2 |
| ENSG00000213588 | ZBTB9 | 1,35364026 | 2,55556141 | 0,06643534 | 0,405191993 | zinc finger and BTB domain containing 9 |
| ENSG00000079112 | CDH17 | 1,353480128 | 2,555277773 | 0,16840042 | 0,650305178 | cadherin 17 |
| ENSG00000154864 | PIEZO2 | 1,353439752 | 2,555206259 | 1,39E-05 | 0,000454751 | piezo type mechanosensitive ion channel component 2 |

| | | | | | | |
|---|---|---|---|---|---|---|
| ENSG00000182179 | UBA7 | 1,353438481 | 2,555204009 | 0,00836101 | 0,100849048 | ubiquitin like modifier activating enzyme 7 |
| ENSG00000227694 | RPL23AP74 | 1,353181483 | 2,554748871 | 0,16654487 | 0,646493505 | ribosomal protein L23a pseudogene 74 |
| ENSG00000067182 | TNFRSF1A | 1,352522025 | 2,553581358 | 1,15E-06 | 4,80E-05 | TNF receptor superfamily member 1A |
| ENSG00000117643 | MAN1C1 | 1,351845801 | 2,552384718 | 0,00334846 | 0,050923041 | mannosidase alpha class 1C member 1 |
| ENSG00000167014 | TERB2 | 1,351212653 | 2,551264813 | 0,3329169 | 0,833301711 | telomere repeat binding bouquet formation protein 2 |
| ENSG00000178401 | DNAJC22 | 1,351204095 | 2,551249678 | 0,0106545 | 0,120123473 | DnaJ heat shock protein family (Hsp40) member C22 |
| ENSG00000228090 | MPIG6B | 1,351020229 | 2,550924552 | 0,0421101 | 0,307744302 | megakaryocyte and platelet inhibitory receptor G6b |
| ENSG00000196924 | FLNA | 1,351004584 | 2,550896889 | 0,00795662 | 0,097274568 | filamin A |
| ENSG00000282772 | AL358790,1 | 1,350866029 | 2,550651915 | 0,18714509 | 0,680971158 | novel transcript, antisense to C10orf32 |
| ENSG00000219201 | AC138392,1 | 1,350192697 | 2,549461757 | 0,40453381 | 0,878176155 | novel pseudogene |
| ENSG00000286163 | AC002377,1 | 1,350055967 | 2,549220145 | 0,1078158 | 0,523928283 | novel transcript |
| ENSG00000104953 | TLE6 | 1,348457974 | 2,54639808 | 0,03348869 | 0,265801029 | TLE family member 6, subcortical maternal complex member |
| ENSG00000123384 | LRP1 | 1,348227023 | 2,545990477 | 0,00217246 | 0,035936381 | LDL receptor related protein 1 |
| ENSG00000130813 | SHFL | 1,348086093 | 2,545741783 | 0,02767953 | 0,234454869 | shiftless antiviral inhibitor of ribosomal frameshifting |
| ENSG00000196141 | SPATS2L | 1,34776255 | 2,545170933 | 5,83E-06 | 0,000211342 | spermatogenesis associated serine rich 2 like |
| ENSG00000204054 | LINC00963 | 1,347626695 | 2,544931271 | 4,87E-05 | 0,001385017 | long intergenic non-protein coding RNA 963 |
| ENSG00000203286 | Metazoa_SRP | 1,34757694 | 2,544843503 | 0,3065302 | 0,815375074 | Metazoan signal recognition particle RNA |
| ENSG00000240592 | EGFL8 | 1,346366913 | 2,54270997 | 0,30014536 | 0,809990902 | EGF like domain multiple 8 |
| ENSG00000116717 | GADD45A | 1,346137927 | 2,542306421 | 0,00011507 | 0,002968346 | growth arrest and DNA damage inducible alpha |
| ENSG00000275038 | AC091980,2 | 1,346049678 | 2,542150913 | 0,2057706 | 0,708191543 | novel transcript, antisense to TLX3 |
| ENSG00000261717 | AC009163,5 | 1,345862461 | 2,541821043 | 0,36653135 | 0,856103393 | novel TMEM170A-CFDP1 readthrough protein |
| ENSG00000234243 | GPSM3 | 1,345717027 | 2,541564822 | 0,0913914 | 0,481302729 | G protein signaling modulator 3 |
| ENSG00000236697 | GPSM3 | 1,345717027 | 2,541564822 | 0,0913914 | 0,481302729 | G protein signaling modulator 3 |
| ENSG00000237052 | GPSM3 | 1,345717027 | 2,541564822 | 0,0913914 | 0,481302729 | G protein signaling modulator 3 |
| ENSG00000206314 | GPSM3 | 1,345717027 | 2,541564822 | 0,0913914 | 0,481302729 | G protein signaling modulator 3 |
| ENSG00000234508 | GPSM3 | 1,345717027 | 2,541564822 | 0,0913914 | 0,481302729 | G protein signaling modulator 3 |
| ENSG00000188004 | SNHG28 | 1,345435242 | 2,541068456 | 0,06212188 | 0,389024096 | small nucleolar RNA host gene 28 |
| ENSG00000115221 | ITGB6 | 1,344936307 | 2,540189816 | 0,30996571 | 0,817411297 | integrin subunit beta 6 |
| ENSG00000287988 | FO082814,1 | 1,344915358 | 2,540152932 | 0,28152729 | 0,792608024 | methylenetetrahydrofolate dehydrogenase (NADP+ dependent) 1 like (MTHFD1L) pseudogene |

| | | | | | | |
|---|---|---|---|---|---|---|
| ENSG00000234828 | IQCM | 1,344246642 | 2,538975796 | 0,21728883 | 0,725037995 | IQ motif containing M |
| ENSG00000278823 | BMERB1 | 1,344215654 | 2,538921261 | 0,0359787 | 0,278282126 | bMERB domain containing 1 |
| ENSG00000143318 | CASQ1 | 1,344210425 | 2,53891206 | 0,01577172 | 0,159575098 | calsequestrin 1 |
| ENSG00000228590 | MIR4432HG | 1,344162353 | 2,538827462 | 0,30604364 | 0,815073553 | MIR4432 host gene |
| ENSG00000184160 | ADRA2C | 1,3436523 | 2,537930039 | 0,14548726 | 0,60768385 | adrenoceptor alpha 2C |
| ENSG00000250846 | EPHA5-AS1 | 1,34363973 | 2,537907926 | 0,12566901 | 0,566207813 | EPHA5 antisense RNA 1 |
| ENSG00000287289 | AP000428,2 | 1,343169766 | 2,537081326 | 0,31191726 | 0,818226927 | novel transcript |
| ENSG00000158859 | ADAMTS4 | 1,342415382 | 2,535755036 | 0,00814616 | 0,098943848 | ADAM metallopeptidase with thrombospondin type 1 motif 4 |
| ENSG00000184867 | ARMCX2 | 1,342119949 | 2,53523582 | 2,80E-07 | 1,34E-05 | armadillo repeat containing X-linked 2 |
| ENSG00000143819 | EPHX1 | 1,34148273 | 2,534116288 | 0,00213799 | 0,035478809 | epoxide hydrolase 1 |
| ENSG00000197386 | HTT | 1,340807206 | 2,532929997 | 1,62E-09 | 1,17E-07 | huntingtin |
| ENSG00000154975 | CA10 | 1,340773838 | 2,532871415 | 0,07396413 | 0,42965711 | carbonic anhydrase 10 |
| ENSG00000073756 | PTGS2 | 1,340708805 | 2,532757241 | 0,03125369 | 0,254390894 | prostaglandin-endoperoxide synthase 2 |
| ENSG00000229205 | LINC00200 | 1,340347414 | 2,532122873 | 0,08639265 | 0,468476705 | long intergenic non-protein coding RNA 200 |
| ENSG00000264464 | LINC02837 | 1,340292744 | 2,532026922 | 0,14017637 | 0,59682337 | long intergenic non-protein coding RNA 2837 |
| ENSG00000226384 | GTF2H4 | 1,339619213 | 2,530845106 | 0,5374318 | 0,932883277 | general transcription factor IIH subunit 4 |
| ENSG00000239945 | AL627309,3 | 1,339424479 | 2,530503516 | 0,5504512 | 0,936204243 | novel transcript |
| ENSG00000187037 | GPR141 | 1,339328686 | 2,530335499 | 0,3448017 | 0,841168721 | G protein-coupled receptor 141 |
| ENSG00000242715 | CCDC169 | 1,339036511 | 2,529823108 | 0,00903637 | 0,106407889 | coiled-coil domain containing 169 |
| ENSG00000235335 | B3GALT1-AS1 | 1,338024542 | 2,528049202 | 0,23939148 | 0,749122637 | B3GALT1 antisense RNA 1 |
| ENSG00000214535 | RPS15AP1 | 1,337606378 | 2,527316554 | 0,28908451 | 0,800171579 | ribosomal protein S15a pseudogene 1 |
| ENSG00000184545 | DUSP8 | 1,337598369 | 2,527302525 | 0,02121653 | 0,194445863 | dual specificity phosphatase 8 |
| ENSG00000172380 | GNG12 | 1,337403233 | 2,526960709 | 8,99E-05 | 0,002387263 | G protein subunit gamma 12 |
| ENSG00000213234 | ST13P10 | 1,337243855 | 2,526681565 | 0,26275015 | 0,774160848 | ST13, Hsp70 interacting protein pseudogene 10 |
| ENSG00000281556 | AC243960,18 | 1,337072291 | 2,526381114 | 0,00743445 | 0,092677055 | novel transcript, sense intronic to CEACAM21 |
| ENSG00000156697 | UTP14A | 1,334964577 | 2,522692877 | 2,43E-08 | 1,42E-06 | UTP14A small subunit processome component |
| ENSG00000109667 | SLC2A9 | 1,334485897 | 2,521855997 | 0,16667682 | 0,646915725 | solute carrier family 2 member 9 |
| ENSG00000228221 | LINC00578 | 1,334477582 | 2,521841461 | 0,14321593 | 0,601976719 | long intergenic non-protein coding RNA 578 |
| ENSG00000241409 | AC064852,1 | 1,334467363 | 2,521823599 | 0,21991919 | 0,728200468 | novel transcript |
| ENSG00000145861 | C1QTNF2 | 1,334166798 | 2,521298268 | 0,00180352 | 0,030761259 | C1q and TNF related 2 |
| ENSG00000134243 | SORT1 | 1,334137751 | 2,521247504 | 1,11E-05 | 0,000372346 | sortilin 1 |
| ENSG00000283632 | EXOC3L2 | 1,333315956 | 2,519811749 | 0,0864755 | 0,4687188 | exocyst complex component 3 like 2 |

| | | | | | | |
|---|---|---|---|---|---|---|
| ENSG00000253227 | AC090192,2 | 1,332688454 | 2,518715992 | 0,14856199 | 0,613326406 | novel transcript |
| ENSG00000223731 | SUPT20HL1 | 1,331867317 | 2,517282825 | 0,32928059 | 0,831217482 | SUPT20H like 1 |
| ENSG00000234857 | HNRNPUL2-BS( | 1,331730601 | 2,517044288 | 0,08941863 | 0,477143535 | HNRNPUL2-BSCL2 readthrough (NMD candidate) |
| ENSG00000235831 | BHLHE40-AS1 | 1,330979965 | 2,515735007 | 0,04463893 | 0,319246353 | BHLHE40 antisense RNA 1 |
| ENSG00000181284 | TMEM102 | 1,330845773 | 2,515501017 | 0,08006562 | 0,448625211 | transmembrane protein 102 |
| ENSG00000284228 | TMEM102 | 1,330845773 | 2,515501017 | 0,08006562 | 0,448625211 | transmembrane protein 102 |
| ENSG00000132561 | MATN2 | 1,330722809 | 2,515286626 | 4,31E-05 | 0,001237371 | matrilin 2 |
| ENSG00000226040 | AC005740,1 | 1,329457262 | 2,513081157 | 0,35506819 | 0,846553187 | glycine cleavage system protein H (aminomethyl carrier) (GCSH) pseudogene |
| ENSG00000149380 | P4HA3 | 1,329444074 | 2,513058184 | 0,0419527 | 0,307024639 | prolyl 4-hydroxylase subunit alpha 3 |
| ENSG00000205403 | CFI | 1,328784779 | 2,511910008 | 0,02963261 | 0,244958706 | complement factor I |
| ENSG00000288299 | LDHA | 1,328235497 | 2,510953822 | 1,77E-07 | 8,95E-06 | lactate dehydrogenase A |
| ENSG00000234942 | GRID1-AS1 | 1,324892391 | 2,505142014 | 0,22464104 | 0,733203418 | GRID1 antisense RNA 1 |
| ENSG00000172554 | SNTG2 | 1,324691095 | 2,504792502 | 0,06116007 | 0,384884976 | syntrophin gamma 2 |
| ENSG00000102390 | PBDC1 | 1,324362519 | 2,504222096 | 3,21E-08 | 1,84E-06 | polysaccharide biosynthesis domain containing 1 |
| ENSG00000269752 | AC008761,2 | 1,32406698 | 2,503709154 | 0,29678574 | 0,806633854 | novel transcript, antisense UNC13A |
| ENSG00000018408 | WWTR1 | 1,323775833 | 2,503203938 | 2,13E-05 | 0,000665935 | WW domain containing transcription regulator 1 |
| ENSG00000147588 | PMP2 | 1,323752779 | 2,503163937 | 0,26315435 | 0,774637968 | peripheral myelin protein 2 |
| ENSG00000134333 | LDHA | 1,322990899 | 2,501842377 | 1,99E-07 | 9,83E-06 | lactate dehydrogenase A |
| ENSG00000261655 | AC100803,3 | 1,322770513 | 2,501460225 | 0,13433043 | 0,584428964 | novel transcript, overlapping to GPR20 |
| ENSG00000282573 | AC100803,3 | 1,322770513 | 2,501460225 | 0,13433043 | 0,584428964 | novel transcript, overlapping to GPR20 |
| ENSG00000112715 | VEGFA | 1,322240279 | 2,500541032 | 0,00204774 | 0,034235542 | vascular endothelial growth factor A |
| ENSG00000253102 | AC004707,1 | 1,322051983 | 2,500214691 | 0,26391796 | 0,775013078 | novel transcript, antisense to LRRC59 |
| ENSG00000285909 | AP002762,2 | 1,321234596 | 2,498798546 | 0,1414127 | 0,599297054 | novel transcript, antisense to CRTAM |
| ENSG00000186174 | BCL9L | 1,32051494 | 2,497552389 | 0,00255474 | 0,040813249 | BCL9 like |
| ENSG00000161011 | SQSTM1 | 1,32019438 | 2,496997506 | 0,0040455 | 0,059268345 | sequestosome 1 |
| ENSG00000231721 | LINC-PINT | 1,320149622 | 2,49692004 | 0,0012442 | 0,022355635 | long intergenic non-protein coding RNA, p53 induced transcript |
| ENSG00000225096 | AL445250,1 | 1,318464649 | 2,494005504 | 0,00683795 | 0,087222253 | novel transcript |
| ENSG00000215869 | AC092506,1 | 1,317949297 | 2,493114767 | 0,44653283 | 0,901644586 | THAP domain containing, apoptosis associated protein 3 (THAP3) pseudogene |
| ENSG00000116285 | ERRFI1 | 1,317551261 | 2,492427018 | 1,68E-06 | 6,85E-05 | ERBB receptor feedback inhibitor 1 |
| ENSG00000249119 | MTND6P4 | 1,316443516 | 2,490513992 | 0,43532743 | 0,896325752 | MT-ND6 pseudogene 4 |

| | | | | | | |
|---|---|---|---|---|---|---|
| ENSG00000125510 | OPRL1 | 1,31629551 | 2,490258503 | 0,00035409 | 0,007773988 | opioid related nociceptin receptor 1 |
| ENSG00000226149 | AL356124,1 | 1,316100412 | 2,489921764 | 0,00334883 | 0,050923041 | novel transcript |
| ENSG00000227619 | AL391056,1 | 1,314807559 | 2,487691452 | 0,19298097 | 0,689849199 | novel transcript |
| ENSG00000254307 | AP003692,1 | 1,314742416 | 2,487579126 | 0,41579616 | 0,885986046 | novel transcript |
| ENSG00000180964 | TCEAL8 | 1,314561301 | 2,487266856 | 0,00056364 | 0,011553977 | transcription elongation factor A like 8 |
| ENSG00000067066 | SP100 | 1,314348729 | 2,4869004 | 2,31E-06 | 9,27E-05 | SP100 nuclear antigen |
| ENSG00000286401 | AC069547,2 | 1,314058118 | 2,486399498 | 0,28310683 | 0,794386288 | novel transcript, antisense to SGMS1 |
| ENSG00000250986 | LINC02600 | 1,313949037 | 2,486211512 | 0,22897329 | 0,737173617 | long intergenic non-protein coding RNA 2600 |
| ENSG00000271454 | AL353729,2 | 1,31318406 | 2,484893567 | 0,09183243 | 0,481984693 | CAS1 domain containing 1 (CASD1) pseudogene |
| ENSG00000126391 | FRMD8 | 1,312813744 | 2,484255818 | 0,00788475 | 0,096783964 | FERM domain containing 8 |
| ENSG00000240634 | AC145285,1 | 1,312712152 | 2,484080888 | 0,17052927 | 0,653761763 | ribosomal protein S15a (RPS15A) pseudogene |
| ENSG00000108819 | PPP1R9B | 1,312216074 | 2,483226872 | 0,08189196 | 0,454835736 | protein phosphatase 1 regulatory subunit 9B |
| ENSG00000185022 | MAFF | 1,31176028 | 2,482442463 | 0,01181991 | 0,129435298 | MAF bZIP transcription factor F |
| ENSG00000165462 | PHOX2A | 1,31014686 | 2,479667806 | 0,13427526 | 0,584428964 | paired like homeobox 2A |
| ENSG00000102038 | SMARCA1 | 1,309975493 | 2,479373282 | 3,06E-06 | 0,00011966 | SWI/SNF related, matrix associated, actin dependent regulator of chromatin, subfamily a, member 1 |
| ENSG00000135697 | BCO1 | 1,30993417 | 2,479302266 | 0,25852616 | 0,771444876 | beta-carotene oxygenase 1 |
| ENSG00000235483 | GYG2-AS1 | 1,309758876 | 2,479001038 | 0,39241028 | 0,870909282 | GYG2 antisense RNA 1 |
| ENSG00000267691 | SHC1P2 | 1,309530667 | 2,478608935 | 0,34175486 | 0,840048641 | SHC adaptor protein 1 pseudogene 2 |
| ENSG00000205730 | ITPRIPL2 | 1,308822596 | 2,477392739 | 2,88E-05 | 0,000868564 | ITPRIP like 2 |
| ENSG00000109854 | HTATIP2 | 1,307475223 | 2,475080114 | 0,00019133 | 0,004594306 | HIV-1 Tat interactive protein 2 |
| ENSG00000248356 | AC093864,1 | 1,307098932 | 2,474434634 | 0,30263258 | 0,812334076 | novel transcript |
| ENSG00000167157 | PRRX2 | 1,307098497 | 2,474433889 | 0,08871517 | 0,475336951 | paired related homeobox 2 |
| ENSG00000258461 | AC012651,1 | 1,305490507 | 2,471677485 | 0,06917354 | 0,415014699 | novel transcript |
| ENSG00000268027 | AC243960,1 | 1,305106573 | 2,471019803 | 0,02332943 | 0,208213349 | novel transcript, sense intronic to CEACAM21 |
| ENSG00000118515 | SGK1 | 1,304656758 | 2,47024949 | 8,66E-06 | 0,000300077 | serum/glucocorticoid regulated kinase 1 |
| ENSG00000165376 | CLDN2 | 1,304385621 | 2,46978528 | 0,03258942 | 0,261251213 | claudin 2 |
| ENSG00000142733 | MAP3K6 | 1,304261028 | 2,469571996 | 0,00348349 | 0,052637854 | mitogen-activated protein kinase kinase kinase 6 |
| ENSG00000250290 | NCAPGP1 | 1,304194051 | 2,469457347 | 0,38826521 | 0,869820053 | non-SMC condensin I complex subunit G pseudogene 1 |
| ENSG00000165806 | CASP7 | 1,304135357 | 2,469356883 | 2,44E-05 | 0,000751131 | caspase 7 |
| ENSG00000180432 | CYP8B1 | 1,303988332 | 2,469105244 | 0,31628127 | 0,821646322 | cytochrome P450 family 8 subfamily B member 1 |

| | | | | | | |
|---|---|---|---|---|---|---|
| ENSG00000261453 | LINC01735 | 1,303795345 | 2,468774978 | 0,32523571 | 0,827324367 | long intergenic non-protein coding RNA 1735 |
| ENSG00000254560 | BBOX1-AS1 | 1,302079111 | 2,465839862 | 0,00121983 | 0,021993581 | BBOX1 antisense RNA 1 |
| ENSG00000287614 | AL158198,2 | 1,301828475 | 2,465411516 | 0,3554386 | 0,846840543 | novel transcript |
| ENSG00000135842 | NIBAN1 | 1,301700827 | 2,465193389 | 0,01559058 | 0,158151823 | niban apoptosis regulator 1 |
| ENSG00000173706 | HEG1 | 1,301665469 | 2,465132972 | 5,95E-09 | 3,84E-07 | heart development protein with EGF like domains 1 |
| ENSG00000170509 | HSD17B13 | 1,299598902 | 2,461604356 | 0,06110888 | 0,384884976 | hydroxysteroid 17-beta dehydrogenase 13 |
| ENSG00000232334 | AL683842,1 | 1,299307845 | 2,461107789 | 0,20330676 | 0,703503738 | novel pseudogene |
| ENSG00000149489 | ROM1 | 1,298910047 | 2,460429276 | 0,01941903 | 0,183474302 | retinal outer segment membrane protein 1 |
| ENSG00000185684 | EP400P1 | 1,298742323 | 2,460143249 | 0,01245916 | 0,134570746 | EP400 pseudogene 1 |
| ENSG00000213301 | HMGB3P11 | 1,297642404 | 2,458268337 | 0,38374988 | 0,868000076 | high mobility group box 3 pseudogene 11 |
| ENSG00000253500 | AF121898,1 | 1,297332716 | 2,457740703 | 0,42109504 | 0,888848798 | novel transcript, antisense to CNBD1 |
| ENSG00000287091 | AC009961,3 | 1,297264419 | 2,457624356 | 0,13787676 | 0,592546669 | novel transcript, sense intronic to CD302and LY75-CD302 |
| ENSG00000149090 | PAMR1 | 1,296686215 | 2,456639586 | 0,01860102 | 0,177872097 | peptidase domain containing associated with muscle regeneration 1 |
| ENSG00000246582 | AC100861,1 | 1,296582534 | 2,456463043 | 0,1090225 | 0,526890476 | novel transcript, antisense to TNFRSF10A |
| ENSG00000196119 | OR8A1 | 1,296443112 | 2,456225661 | 0,00154348 | 0,026913159 | olfactory receptor family 8 subfamily A member 1 |
| ENSG00000226994 | AC012593,1 | 1,296185217 | 2,455786628 | 0,26448249 | 0,775800337 | novel transcript |
| ENSG00000182508 | LHFPL1 | 1,296176914 | 2,455772494 | 0,02508358 | 0,219729073 | LHFPL tetraspan subfamily member 1 |
| ENSG00000143603 | KCNN3 | 1,295907397 | 2,455313762 | 0,00019105 | 0,004591078 | potassium calcium-activated channel subfamily N member 3 |
| ENSG00000145391 | SETD7 | 1,295486542 | 2,454597617 | 9,53E-09 | 5,90E-07 | SET domain containing 7, histone lysine methyltransferase |
| ENSG00000266934 | AC005746,1 | 1,295158193 | 2,454039028 | 0,13737914 | 0,591519288 | novel transcript, sense intronic to BCAS3 |
| ENSG00000214110 | LDHAP4 | 1,295086184 | 2,453916543 | 0,00783984 | 0,096308297 | lactate dehydrogenase A pseudogene 4 |
| ENSG00000260027 | HOXB7 | 1,294732042 | 2,453314247 | 0,40225093 | 0,87718264 | homeobox B7 |
| ENSG00000282057 | AC092807,3 | 1,294381363 | 2,452717988 | 0,00010959 | 0,002843437 | novel transcript |
| ENSG00000256969 | AC084375,1 | 1,294121825 | 2,452276788 | 0,34858792 | 0,842757646 | novel transcript |
| ENSG00000150540 | HNMT | 1,293182346 | 2,450680393 | 0,00612313 | 0,080231988 | histamine N-methyltransferase |
| ENSG00000272479 | AC010857,1 | 1,292746842 | 2,44994072 | 0,22641602 | 0,735064413 | novel transcript |
| ENSG00000070404 | FSTL3 | 1,292560416 | 2,449624159 | 0,015668 | 0,158731178 | follistatin like 3 |
| ENSG00000223694 | ADH5P3 | 1,292539199 | 2,449588134 | 0,02285973 | 0,205192794 | ADH5 pseudogene 3 |
| ENSG00000168454 | TXNDC2 | 1,291545213 | 2,447901 | 0,11105141 | 0,532146382 | thioredoxin domain containing 2 |

| | | | | | | |
|---|---|---|---|---|---|---|
| ENSG00000176014 | TUBB6 | 1,291247199 | 2,447395395 | 1,42E-05 | 0,000460802 | tubulin beta 6 class V |
| ENSG00000278384 | AL354822,1 | 1,291217929 | 2,447345742 | 0,00334475 | 0,050910539 | |
| ENSG00000286231 | AL445423,3 | 1,291090414 | 2,44712944 | 0,03496178 | 0,272988286 | novel protein |
| ENSG00000163873 | GRIK3 | 1,291031016 | 2,447028689 | 0,00517735 | 0,07071409 | glutamate ionotropic receptor kainate type subunit 3 |
| ENSG00000186312 | CA5BP1 | 1,290911375 | 2,446825768 | 1,43E-05 | 0,000465762 | carbonic anhydrase 5B pseudogene 1 |
| ENSG00000243742 | RPLP0P2 | 1,290393861 | 2,445948218 | 0,07086609 | 0,418838091 | ribosomal protein lateral stalk subunit P0 pseudogene 2 |
| ENSG00000288023 | AL157936,1 | 1,289106367 | 2,443766371 | 0,19883725 | 0,697657684 | novel transcript |
| ENSG00000166444 | DENND2B | 1,288217039 | 2,442260411 | 0,00011644 | 0,002993721 | DENN domain containing 2B |
| ENSG00000100351 | GRAP2 | 1,288158718 | 2,442161685 | 0,39968807 | 0,875433675 | GRB2 related adaptor protein 2 |
| ENSG00000203618 | GP1BB | 1,288025525 | 2,441936229 | 0,0314875 | 0,255694831 | glycoprotein Ib platelet subunit beta |
| ENSG00000065357 | DGKA | 1,287200736 | 2,440540573 | 0,00058228 | 0,011882718 | diacylglycerol kinase alpha |
| ENSG00000258175 | LINC02300 | 1,287075351 | 2,440328473 | 0,50043651 | 0,920485547 | long intergenic non-protein coding RNA 2300 |
| ENSG00000081803 | CADPS2 | 1,286216827 | 2,438876707 | 8,50E-05 | 0,002279955 | calcium dependent secretion activator 2 |
| ENSG00000236141 | AC013727,2 | 1,285912284 | 2,438361931 | 0,22114259 | 0,72907327 | novel transcript |
| ENSG00000224232 | KCTD5P1 | 1,28564712 | 2,437913807 | 0,2702082 | 0,782492524 | KCTD5 pseudogene 1 |
| ENSG00000101638 | ST8SIA5 | 1,283411198 | 2,434138399 | 0,05489926 | 0,362021772 | ST8 alpha-N-acetyl-neuraminide alpha-2,8-sialyltransferase 5 |
| ENSG00000198879 | SFMBT2 | 1,282743993 | 2,43301294 | 0,00567794 | 0,075796418 | Scm like with four mbt domains 2 |
| ENSG00000228960 | OR2A9P | 1,282503647 | 2,432607646 | 0,00019662 | 0,004706919 | olfactory receptor family 2 subfamily A member 9 pseudogene |
| ENSG00000260412 | AL353746,1 | 1,28234194 | 2,432334997 | 0,03183273 | 0,25735202 | novel transcript, overlapping LINGO2 |
| ENSG00000007384 | RHBDF1 | 1,281839166 | 2,431487485 | 0,00060811 | 0,012313456 | rhomboid 5 homolog 1 |
| ENSG00000182866 | LCK | 1,281720458 | 2,431287426 | 0,2132867 | 0,720652258 | LCK proto-oncogene, Src family tyrosine kinase |
| ENSG00000254670 | RASSF10-DT | 1,279859495 | 2,428153278 | 0,23564644 | 0,744839106 | RASSF10 divergent transcript |
| ENSG00000114812 | VIPR1 | 1,279820624 | 2,428087856 | 0,1528305 | 0,62133458 | vasoactive intestinal peptide receptor 1 |
| ENSG00000135362 | PRR5L | 1,279450075 | 2,427464294 | 2,36E-05 | 0,00072957 | proline rich 5 like |
| ENSG00000135916 | ITM2C | 1,279213218 | 2,427065793 | 5,56E-07 | 2,48E-05 | integral membrane protein 2C |
| ENSG00000200394 | SNORA38B | 1,278315584 | 2,425556161 | 0,15620622 | 0,627034861 | small nucleolar RNA, H/ACA box 38B |
| ENSG00000177548 | RABEP2 | 1,278003199 | 2,425031015 | 0,0722412 | 0,423822299 | rabaptin, RAB GTPase binding effector protein 2 |
| ENSG00000231579 | RPL7P21 | 1,277408843 | 2,424032167 | 0,39538351 | 0,871996943 | ribosomal protein L7 pseudogene 21 |
| ENSG00000127507 | ADGRE2 | 1,277292416 | 2,423836553 | 0,031611 | 0,256419452 | adhesion G protein-coupled receptor E2 |

| | | | | | | |
|---|---|---|---|---|---|---|
| ENSG00000134247 | PTGFRN | 1,277279689 | 2,42381517 | 1,14E-05 | 0,000381175 | prostaglandin F2 receptor inhibitor |
| ENSG00000124915 | MYRF-AS1 | 1,277018757 | 2,423376829 | 0,22165681 | 0,729254784 | MYRF antisense RNA 1 |
| ENSG00000117525 | F3 | 1,276577 | 2,422634897 | 0,00017809 | 0,004306186 | coagulation factor III, tissue factor |
| ENSG00000251467 | AC020893,1 | 1,276278524 | 2,422133735 | 0,15962986 | 0,634138231 | dullard homolog (Xenopus laevis) (DULLARD) pseudogene |
| ENSG00000230737 | AC106870,2 | 1,275927111 | 2,421543822 | 0,07861887 | 0,443831782 | novel transcript |
| ENSG00000206341 | HLA-H | 1,275090264 | 2,420139593 | 0,2842457 | 0,795731744 | major histocompatibility complex, class I, H (pseudogene) |
| ENSG00000111321 | LTBR | 1,274946497 | 2,419898434 | 0,03065051 | 0,250655898 | lymphotoxin beta receptor |
| ENSG00000256632 | AC135586,2 | 1,274632195 | 2,419371298 | 0,36415472 | 0,85421167 | novel transcript |
| ENSG00000264540 | RN7SL405P | 1,274502981 | 2,419154619 | 0,31889221 | 0,823818062 | RNA, 7SL, cytoplasmic 405, pseudogene |
| ENSG00000002745 | WNT16 | 1,274331926 | 2,418867805 | 0,32770502 | 0,829743891 | Wnt family member 16 |
| ENSG00000280118 | AC090983,1 | 1,274284619 | 2,41878849 | 0,01331797 | 0,140952253 | novel transcript |
| ENSG00000106211 | HSPB1 | 1,274229953 | 2,41869684 | 0,00091993 | 0,01742009 | heat shock protein family B (small) member 1 |
| ENSG00000243978 | RTL9 | 1,273602702 | 2,417645475 | 0,00072849 | 0,014252604 | retrotransposon Gag like 9 |
| ENSG00000287644 | AC080112,4 | 1,272655894 | 2,416059349 | 0,30916895 | 0,816845699 | novel transcript, antisense to RARA |
| ENSG00000174099 | MSRB3 | 1,272628127 | 2,416012849 | 2,33E-07 | 1,14E-05 | methionine sulfoxide reductase B3 |
| ENSG00000188517 | COL25A1 | 1,272346388 | 2,41554108 | 0,02045279 | 0,189631051 | collagen type XXV alpha 1 chain |
| ENSG00000160886 | LY6K | 1,272053457 | 2,415050668 | 0,15396645 | 0,623201341 | lymphocyte antigen 6 family member K |
| ENSG00000119686 | FLVCR2 | 1,271873754 | 2,414749866 | 0,00063638 | 0,012786531 | FLVCR heme transporter 2 |
| ENSG00000237797 | MACORIS | 1,271870261 | 2,414744019 | 0,22718799 | 0,735737019 | macrophage enriched lincRNA repressor of IFN-gamma signaling |
| ENSG00000185304 | RGPD2 | 1,27163275 | 2,414346513 | 0,22606511 | 0,734959876 | RANBP2 like and GRIP domain containing 2 |
| ENSG00000280109 | PLAC4 | 1,270863926 | 2,41306023 | 0,00093351 | 0,017645276 | placenta enriched 4 |
| ENSG00000197971 | MBP | 1,270608407 | 2,412632886 | 0,00026463 | 0,0060522 | myelin basic protein |
| ENSG00000134508 | CABLES1 | 1,269722164 | 2,411151268 | 5,64E-06 | 0,000205034 | Cdk5 and Abl enzyme substrate 1 |
| ENSG00000269987 | AC004542,2 | 1,269476116 | 2,410740087 | 0,4650685 | 0,907804479 | novel transcript |
| ENSG00000264982 | AC015563,2 | 1,26898078 | 2,409912524 | 0,03055141 | 0,250041727 | novel transcript, antisense to FAM59A |
| ENSG00000267226 | AC104971,1 | 1,268459507 | 2,409041934 | 0,04241308 | 0,309296116 | novel transcript, antisense to MALT1 |
| ENSG00000109943 | CRTAM | 1,26830963 | 2,408791681 | 0,20211211 | 0,701895011 | cytotoxic and regulatory T cell molecule |
| ENSG00000183963 | SMTN | 1,268157699 | 2,408538021 | 0,01423224 | 0,148389909 | smoothelin |
| ENSG00000138448 | ITGAV | 1,268107801 | 2,40845472 | 0,00015398 | 0,003784716 | integrin subunit alpha V |
| ENSG00000182253 | SYNM | 1,268068289 | 2,408388759 | 0,02796939 | 0,236352093 | synemin |

| | | | | | | |
|---|---|---|---|---|---|---|
| ENSG00000236166 | AL021408,1 | 1,267991793 | 2,408261063 | 0,10789427 | 0,523928283 | novel transcript |
| ENSG00000169994 | MYO7B | 1,26733644 | 2,407167343 | 0,35369995 | 0,845346751 | myosin VIIB |
| ENSG00000205562 | AL049775,1 | 1,266277169 | 2,405400577 | 0,0053633 | 0,072776773 | novel transcript |
| ENSG00000084453 | SLCO1A2 | 1,265340726 | 2,403839755 | 0,00583208 | 0,077292963 | solute carrier organic anion transporter family member 1A2 |
| ENSG00000167117 | ANKRD40CL | 1,26505303 | 2,403360438 | 0,08389254 | 0,461113993 | ANKRD40 C-terminal like |
| ENSG00000269304 | FKBP1AP1 | 1,264895866 | 2,403098637 | 0,29558701 | 0,805426883 | FKBP prolyl isomerase 1A pseudogene 1 |
| ENSG00000288510 | AC148477,19 | 1,264842765 | 2,403010188 | 0,09150602 | 0,481302729 | novel transcript |
| ENSG00000256542 | AC148477,2 | 1,264842765 | 2,403010188 | 0,09150602 | 0,481302729 | novel transcript |
| ENSG00000186197 | EDARADD | 1,264607811 | 2,402618871 | 0,02163322 | 0,197183925 | EDAR associated death domain |
| ENSG00000117598 | PLPPR5 | 1,264034488 | 2,401664266 | 0,05919799 | 0,377938736 | phospholipid phosphatase related 5 |
| ENSG00000075275 | CELSR1 | 1,263920438 | 2,401474414 | 0,01219989 | 0,132207041 | cadherin EGF LAG seven-pass G-type receptor 1 |
| ENSG00000239713 | APOBEC3G | 1,263820737 | 2,40130846 | 0,00052262 | 0,010813506 | apolipoprotein B mRNA editing enzyme catalytic subunit 3G |
| ENSG00000272717 | AC112236,2 | 1,263058205 | 2,400039592 | 0,16729062 | 0,647940282 | novel transcript, antisense to SETD7 |
| ENSG00000286669 | AC008532,1 | 1,262289317 | 2,398760824 | 0,1085619 | 0,525236658 | novel transcript |
| ENSG00000170549 | IRX1 | 1,261165023 | 2,396892196 | 0,00133032 | 0,02378002 | iroquois homeobox 1 |
| ENSG00000274737 | AC004466,2 | 1,260891597 | 2,39643797 | 0,09134928 | 0,481302729 | novel transcript, sense intronic to HDAC7 |
| ENSG00000288156 | AC104530,1 | 1,260678526 | 2,396084066 | 0,28298126 | 0,794176498 | novel transcript |
| ENSG00000203463 | SLC44A4 | 1,26059827 | 2,395950778 | 0,29281528 | 0,803295982 | solute carrier family 44 member 4 |
| ENSG00000165633 | VSTM4 | 1,260437693 | 2,395684116 | 0,00108364 | 0,020056204 | V-set and transmembrane domain containing 4 |
| ENSG00000261592 | AC010531,3 | 1,260336059 | 2,395515352 | 0,2888334 | 0,800014136 | novel transcript, antisense to MAP1LC3B |
| ENSG00000227294 | AC016994,1 | 1,260212325 | 2,395309906 | 0,263896 | 0,775013078 | novel transcript |
| ENSG00000167173 | C15orf39 | 1,259990282 | 2,394941277 | 0,00012542 | 0,003187922 | chromosome 15 open reading frame 39 |
| ENSG00000110628 | SLC22A18 | 1,259373797 | 2,393918102 | 0,05084411 | 0,345412912 | solute carrier family 22 member 18 |
| ENSG00000198517 | MAFK | 1,258825574 | 2,393008587 | 0,08558939 | 0,465692069 | MAF bZIP transcription factor K |
| ENSG00000204334 | ERICH2 | 1,25857418 | 2,392591634 | 0,18369074 | 0,674921526 | glutamate rich 2 |
| ENSG00000285407 | AL033530,1 | 1,25813056 | 2,39185604 | 0,00392957 | 0,057895474 | novel transcript |
| ENSG00000010379 | SLC6A13 | 1,257229195 | 2,390362127 | 0,1449855 | 0,606154944 | solute carrier family 6 member 13 |
| ENSG00000188419 | CHM | 1,257133828 | 2,39020412 | 0,00286922 | 0,044824652 | CHM Rab escort protein |
| ENSG00000183878 | UTY | 1,256772783 | 2,389606029 | 0,21775417 | 0,72540192 | ubiquitously transcribed tetratricopeptide repeat containing, Y-linked |
| ENSG00000198467 | TPM2 | 1,256671721 | 2,389438642 | 0,01979774 | 0,185094647 | tropomyosin 2 |

| | | | | | | |
|---|---|---|---|---|---|---|
| ENSG00000237773 | AC073332,1 | 1,256250784 | 2,388741573 | 0,01076107 | 0,120930771 | novel transcript |
| ENSG00000234557 | PPIAP90 | 1,255550682 | 2,387582662 | 0,25540897 | 0,766913772 | peptidylprolyl isomerase A pseudogene 90 |
| ENSG00000277288 | LINC02881 | 1,254794913 | 2,386332231 | 0,39059539 | 0,870057277 | long intergenic non-protein coding RNA 2881 |
| ENSG00000226085 | UQCRFS1P1 | 1,254155278 | 2,385274459 | 0,05831743 | 0,373902954 | ubiquinol-cytochrome c reductase, Rieske iron-sulfur polypeptide 1 pseudogene 1 |
| ENSG00000227060 | LINC00629 | 1,252821002 | 2,383069459 | 0,11313075 | 0,537199228 | long intergenic non-protein coding RNA 629 |
| ENSG00000167306 | MYO5B | 1,252373675 | 2,382330671 | 0,00243211 | 0,039235021 | myosin VB |
| ENSG00000104313 | EYA1 | 1,251765212 | 2,381326123 | 0,00824087 | 0,099745442 | EYA transcriptional coactivator and phosphatase 1 |
| ENSG00000243914 | RPL5P14 | 1,251747801 | 2,381297385 | 0,25805458 | 0,770501859 | ribosomal protein L5 pseudogene 14 |
| ENSG00000274657 | FAM230I | 1,251425099 | 2,380764797 | 0,32001702 | 0,824366234 | family with sequence similarity 230 member I |
| ENSG00000178248 | FAM230I | 1,251425099 | 2,380764797 | 0,32001702 | 0,824366234 | family with sequence similarity 230 member I |
| ENSG00000264910 | RN7SL525P | 1,251199629 | 2,38039275 | 0,20196346 | 0,701576182 | RNA, 7SL, cytoplasmic 525, pseudogene |
| ENSG00000248050 | AC079061,1 | 1,250267743 | 2,37885567 | 0,08854367 | 0,474721701 | novel transcript, antisense to SYBU |
| ENSG00000111732 | AICDA | 1,250223483 | 2,378782691 | 0,21829216 | 0,725947652 | activation induced cytidine deaminase |
| ENSG00000128283 | CDC42EP1 | 1,250187895 | 2,378724013 | 0,00590324 | 0,077905832 | CDC42 effector protein 1 |
| ENSG00000138944 | SHISAL1 | 1,250106079 | 2,378589118 | 4,73E-06 | 0,000175945 | shisa like 1 |
| ENSG00000282572 | AC215522,2 | 1,249548675 | 2,377670296 | 0,20018906 | 0,69971898 | novel transcript similar to family member with sequence similarity 157 |
| ENSG00000065361 | ERBB3 | 1,24871801 | 2,376301691 | 0,00480304 | 0,066828153 | erb-b2 receptor tyrosine kinase 3 |
| ENSG00000157514 | TSC22D3 | 1,248077349 | 2,375246676 | 4,40E-09 | 2,91E-07 | TSC22 domain family member 3 |
| ENSG00000139985 | ADAM21 | 1,248061504 | 2,375220589 | 0,07754243 | 0,440002866 | ADAM metallopeptidase domain 21 |
| ENSG00000248641 | HMGA1P2 | 1,246926525 | 2,373352719 | 0,27416066 | 0,7862019 | high mobility group AT-hook 1 pseudogene 2 |
| ENSG00000188649 | CC2D2B | 1,246775467 | 2,373104231 | 0,16390227 | 0,642745357 | coiled-coil and C2 domain containing 2B |
| ENSG00000160396 | HIPK4 | 1,246745968 | 2,373055707 | 0,14840894 | 0,613326406 | homeodomain interacting protein kinase 4 |
| ENSG00000132359 | RAP1GAP2 | 1,246652064 | 2,372901252 | 0,00010275 | 0,002681498 | RAP1 GTPase activating protein 2 |
| ENSG00000265399 | AP005329,2 | 1,246352248 | 2,372408175 | 0,01124424 | 0,124829844 | novel transcript, antisense to MYOM1 |
| ENSG00000185561 | TLCD2 | 1,245761363 | 2,371436706 | 0,0476106 | 0,331735648 | TLC domain containing 2 |
| ENSG00000235700 | CYCSP52 | 1,244908145 | 2,370034639 | 0,23899241 | 0,748458245 | CYCS pseudogene 52 |
| ENSG00000181191 | PJA1 | 1,244656442 | 2,369621182 | 8,89E-08 | 4,81E-06 | praja ring finger ubiquitin ligase 1 |
| ENSG00000067057 | PFKP | 1,244372167 | 2,369154308 | 0,02414771 | 0,213444732 | phosphofructokinase, platelet |
| ENSG00000278817 | AC007325,4 | 1,243702547 | 2,368054932 | 0,03603677 | 0,27830728 | protein DGCR6 |
| ENSG00000238241 | CCR12P | 1,243668012 | 2,367998246 | 0,26233958 | 0,774068771 | C-C motif chemokine receptor 12, pseudogene |
| ENSG00000070882 | OSBPL3 | 1,243381643 | 2,367528255 | 0,00170469 | 0,029315377 | oxysterol binding protein like 3 |

| | | | | | | |
|---|---|---|---|---|---|---|
| ENSG00000268751 | SCGB1B2P | 1,243145004 | 2,367139951 | 0,28005811 | 0,790779983 | secretoglobin family 1B member 2, pseudogene |
| ENSG00000261465 | AC099518,4 | 1,242210069 | 2,365606429 | 0,33303154 | 0,833301711 | novel transcript, antisense to TMC7 and COQ7 |
| ENSG00000162746 | FCRLB | 1,242055967 | 2,365353759 | 0,02006142 | 0,18705645 | Fc receptor like B |
| ENSG00000103496 | STX4 | 1,241037414 | 2,363684391 | 0,01577721 | 0,15957902 | syntaxin 4 |
| ENSG00000250900 | TRIM7-AS1 | 1,240924111 | 2,363498765 | 0,25507452 | 0,76642479 | TRIM7 antisense RNA 1 |
| ENSG00000277900 | Metazoa_SRP | 1,240812027 | 2,36331515 | 0,22402711 | 0,732710917 | Metazoan signal recognition particle RNA |
| ENSG00000188783 | PRELP | 1,239954253 | 2,361910427 | 0,03729349 | 0,285054324 | proline and arginine rich end leucine rich repeat protein |
| ENSG00000110811 | P3H3 | 1,239773731 | 2,361614904 | 0,00573366 | 0,076312149 | prolyl 3-hydroxylase 3 |
| ENSG00000213352 | GAPDHP44 | 1,23956468 | 2,361272723 | 0,2590223 | 0,771947284 | glyceraldehyde 3 phosphate dehydrogenase pseudogene 44 |
| ENSG00000226380 | AC016831,1 | 1,237777111 | 2,358348804 | 0,00266031 | 0,042048519 | novel transcript |
| ENSG00000250320 | EDIL3-DT | 1,23718064 | 2,357373964 | 0,0004739 | 0,009937001 | EDIL3 divergent transcript |
| ENSG00000274874 | AC068790,8 | 1,2367873 | 2,356731332 | 0,41440976 | 0,885579089 | novel protein |
| ENSG00000249532 | MIR302CHG | 1,236304057 | 2,355942057 | 0,14367757 | 0,602944354 | miR-302/367 cluster host gene |
| ENSG00000061337 | LZTS1 | 1,235450037 | 2,354547843 | 0,0137622 | 0,144528528 | leucine zipper tumor suppressor 1 |
| ENSG00000288584 | Z97205,3 | 1,234899053 | 2,353648781 | 0,1733252 | 0,658330849 | novel transcript |
| ENSG00000030582 | GRN | 1,234854107 | 2,353575457 | 0,00249207 | 0,039914879 | granulin precursor |
| ENSG00000134070 | IRAK2 | 1,234541545 | 2,353065607 | 0,02035959 | 0,189112948 | interleukin 1 receptor associated kinase 2 |
| ENSG00000188959 | C9orf152 | 1,234047975 | 2,35226072 | 0,17082911 | 0,654404445 | chromosome 9 open reading frame 152 |
| ENSG00000273445 | AC133644,1 | 1,233995518 | 2,352175194 | 0,02811474 | 0,237067462 | novel transcript |
| ENSG00000229867 | STEAP3-AS1 | 1,233150298 | 2,350797547 | 0,24475272 | 0,756644825 | STEAP3 antisense RNA 1 |
| ENSG00000288658 | AC010980,1 | 1,232883644 | 2,350363089 | 0,06445489 | 0,397871785 | novel protein, ortholog of Gm2102 (M, musculus) |
| ENSG00000162302 | RPS6KA4 | 1,232291527 | 2,34939864 | 0,01536808 | 0,156675544 | ribosomal protein S6 kinase A4 |
| ENSG00000203999 | LINC01270 | 1,232207748 | 2,349262211 | 0,01891184 | 0,180041477 | long intergenic non-protein coding RNA 1270 |
| ENSG00000259275 | AC087477,2 | 1,232078256 | 2,349051358 | 0,0786911 | 0,444135046 | novel transcript |
| ENSG00000110375 | UPK2 | 1,231045232 | 2,347369952 | 0,20802223 | 0,712494639 | uroplakin 2 |
| ENSG00000257718 | CPNE8-AS1 | 1,23087972 | 2,347100666 | 0,15431828 | 0,623607455 | CPNE8 antisense RNA 1 |
| ENSG00000250995 | TMEM30BP1 | 1,230173184 | 2,345951494 | 0,16859173 | 0,650506523 | TMEM30B pseudogene 1 |
| ENSG00000177324 | BEND2 | 1,229826446 | 2,345387734 | 0,32555413 | 0,827465638 | BEN domain containing 2 |
| ENSG00000184906 | AMYH0202086 | 1,229383204 | 2,344667267 | 0,04739008 | 0,330875511 | methylenetetrahydrofolate dehydrogenase (NADP+ dependent) 1 like pseudogene |
| ENSG00000145248 | SLC10A4 | 1,228614636 | 2,343418523 | 0,00247636 | 0,039747347 | solute carrier family 10 member 4 |

| | | | | | | |
|---|---|---|---|---|---|---|
| ENSG00000154252 | GAL3ST2 | 1,227964283 | 2,342362371 | 0,41266976 | 0,884576962 | galactose-3-O-sulfotransferase 2 |
| ENSG00000243370 | RN7SL775P | 1,227901607 | 2,342260614 | 0,21350379 | 0,720983667 | RNA, 7SL, cytoplasmic 775, pseudogene |
| ENSG00000166311 | SMPD1 | 1,227878335 | 2,34222283 | 0,000582 | 0,011882718 | sphingomyelin phosphodiesterase 1 |
| ENSG00000226690 | AC013470,2 | 1,227730386 | 2,341982646 | 0,26032613 | 0,772692192 | novel protein |
| ENSG00000118523 | CCN2 | 1,227338392 | 2,341346393 | 0,00357095 | 0,05367413 | cellular communication network factor 2 |
| ENSG00000224079 | AC091729,1 | 1,227303625 | 2,341289972 | 0,38323199 | 0,867740551 | novel transcript |
| ENSG00000179388 | EGR3 | 1,226738098 | 2,340372382 | 0,03138308 | 0,255178302 | early growth response 3 |
| ENSG00000220925 | IGBP1-AS2 | 1,224544657 | 2,336816836 | 0,35109659 | 0,844035671 | IGBP1 antisense RNA 2 |
| ENSG00000155366 | RHOC | 1,224007586 | 2,335947073 | 1,33E-05 | 0,000436144 | ras homolog family member C |
| ENSG00000238203 | VWA7 | 1,222498941 | 2,333505618 | 0,40498452 | 0,878176155 | von Willebrand factor A domain containing 7 |
| ENSG00000103257 | SLC7A5 | 1,22244312 | 2,333415332 | 0,01063399 | 0,119935439 | solute carrier family 7 member 5 |
| ENSG00000244682 | FCGR2C | 1,222286559 | 2,333162125 | 0,43702703 | 0,896948268 | Fc fragment of IgG receptor IIc (gene/pseudogene) |
| ENSG00000261916 | AC027796,1 | 1,221867316 | 2,332484212 | 0,31846968 | 0,823270714 | novel transcript, sense intronic to readthrough SHPK-TRPV1 |
| ENSG00000240882 | AC063952,2 | 1,221807402 | 2,332387348 | 0,36542618 | 0,855323027 | prostate tumor overexpressed 1 (PTOV1) pseudogene |
| ENSG00000066248 | NGEF | 1,221270854 | 2,331520079 | 0,01503086 | 0,154428164 | neuronal guanine nucleotide exchange factor |
| ENSG00000257038 | AP002761,3 | 1,220994222 | 2,331073061 | 0,14489946 | 0,605968311 | novel transcript |
| ENSG00000135063 | FAM189A2 | 1,220468546 | 2,33022384 | 0,02070994 | 0,1913909 | family with sequence similarity 189 member A2 |
| ENSG00000152953 | STK32B | 1,220234747 | 2,329846242 | 0,00148759 | 0,02608417 | serine/threonine kinase 32B |
| ENSG00000288602 | C8orf44-SGK3 | 1,219870709 | 2,32925842 | 0,17355808 | 0,658349911 | C8orf44-SGK3 readthrough |
| ENSG00000172794 | RAB37 | 1,219637256 | 2,328881537 | 0,11129466 | 0,532815219 | RAB37, member RAS oncogene family |
| ENSG00000203266 | AL157414,1 | 1,219328896 | 2,328383817 | 0,30716624 | 0,815953743 | novel transcript |
| ENSG00000137975 | CLCA2 | 1,218827075 | 2,327574063 | 0,42776416 | 0,892705337 | chloride channel accessory 2 |
| ENSG00000153246 | PLA2R1 | 1,217594801 | 2,325586821 | 2,19E-05 | 0,000683867 | phospholipase A2 receptor 1 |
| ENSG00000251632 | LINC02172 | 1,216432561 | 2,323714074 | 0,45141555 | 0,902062594 | long intergenic non-protein coding RNA 2172 |
| ENSG00000203585 | LINC02408 | 1,216308725 | 2,323514623 | 0,32284189 | 0,824955371 | long intergenic non-protein coding RNA 2408 |
| ENSG00000287544 | AC097518,2 | 1,216182974 | 2,323312105 | 0,2563111 | 0,768515424 | novel transcript |
| ENSG00000128918 | ALDH1A2 | 1,215432772 | 2,322104296 | 0,04640325 | 0,327114174 | aldehyde dehydrogenase 1 family member A2 |
| ENSG00000261105 | LMO7-AS1 | 1,214932261 | 2,321298834 | 0,04157916 | 0,305946805 | LMO7 antisense RNA 1 |
| ENSG00000232618 | AL355304,1 | 1,214099117 | 2,319958691 | 0,02587072 | 0,224174655 | novel transcript |
| ENSG00000186056 | MATN1-AS1 | 1,213420289 | 2,318867343 | 0,04664371 | 0,32799008 | MATN1 antisense RNA 1 |
| ENSG00000181234 | TMEM132C | 1,213056234 | 2,318282266 | 8,90E-05 | 0,002366751 | transmembrane protein 132C |

| | | | | | | |
|---|---|---|---|---|---|---|
| ENSG00000134538 | SLCO1B1 | 1,212881466 | 2,318001445 | 0,35378487 | 0,845346751 | solute carrier organic anion transporter family member 1B1 |
| ENSG00000005884 | ITGA3 | 1,212806013 | 2,317880217 | 0,01709252 | 0,168308818 | integrin subunit alpha 3 |
| ENSG00000214336 | FOXI3 | 1,212696044 | 2,317703544 | 0,15220743 | 0,620418906 | forkhead box I3 |
| ENSG00000147164 | SNX12 | 1,212479578 | 2,317355816 | 1,16E-07 | 6,17E-06 | sorting nexin 12 |
| ENSG00000171243 | SOSTDC1 | 1,212464408 | 2,317331449 | 0,01340085 | 0,141637695 | sclerostin domain containing 1 |
| ENSG00000094841 | UPRT | 1,21177874 | 2,316230355 | 0,00180726 | 0,030808162 | uracil phosphoribosyltransferase homolog |
| ENSG00000100344 | PNPLA3 | 1,211585413 | 2,315919991 | 7,27E-06 | 0,00025768 | patatin like phospholipase domain containing 3 |
| ENSG00000263360 | RN7SL134P | 1,211136609 | 2,31519965 | 0,43084647 | 0,89530663 | RNA, 7SL, cytoplasmic 134, pseudogene |
| ENSG00000259223 | AC009654,1 | 1,210256037 | 2,313786962 | 0,07474253 | 0,432166624 | novel transcript |
| ENSG00000248746 | ACTN3 | 1,209982628 | 2,313348512 | 0,19727083 | 0,695943345 | actinin alpha 3 |
| ENSG00000127824 | TUBA4A | 1,208790916 | 2,311438401 | 0,06334926 | 0,393826088 | tubulin alpha 4a |
| ENSG00000187634 | SAMD11 | 1,208059243 | 2,310266436 | 0,00149725 | 0,026224089 | sterile alpha motif domain containing 11 |
| ENSG00000249609 | AC080188,1 | 1,206668736 | 2,308040814 | 0,10702498 | 0,522327361 | novel transcript |
| ENSG00000225603 | AC244394,1 | 1,206371279 | 2,307564987 | 0,34670087 | 0,8414568 | insulin-like growth factor 2 mRNA binding protein 2 (IGF2BP2) pseudogene |
| ENSG00000254467 | AP002765,1 | 1,205817961 | 2,306680135 | 0,21816984 | 0,725855256 | novel transcript |
| ENSG00000178996 | SNX18 | 1,205686557 | 2,306470047 | 9,98E-07 | 4,24E-05 | sorting nexin 18 |
| ENSG00000139178 | C1RL | 1,205667166 | 2,306439047 | 0,00224867 | 0,036865517 | complement C1r subcomponent like |
| ENSG00000286294 | AC107398,5 | 1,205563897 | 2,306273956 | 0,33266319 | 0,83323041 | novel transcript |
| ENSG00000281833 | AL354892,6 | 1,205493486 | 2,3061614 | 0,26200932 | 0,773902615 | novel transcript |
| ENSG00000231107 | LINC01508 | 1,205256246 | 2,305782201 | 0,0056238 | 0,075266531 | long intergenic non-protein coding RNA 1508 |
| ENSG00000139117 | CPNE8 | 1,205000836 | 2,305374029 | 0,00439675 | 0,062897441 | copine 8 |
| ENSG00000232719 | AC007272,1 | 1,204910287 | 2,305229341 | 0,2931442 | 0,803365244 | novel transcript |
| ENSG00000113070 | HBEGF | 1,204574371 | 2,304692654 | 0,00479599 | 0,066789281 | heparin binding EGF like growth factor |
| ENSG00000162783 | IER5 | 1,204362208 | 2,304353752 | 0,00407803 | 0,059577367 | immediate early response 5 |
| ENSG00000090989 | EXOC1 | 1,203684987 | 2,303272309 | 0,00096416 | 0,018126011 | exocyst complex component 1 |
| ENSG00000171561 | OR2AT4 | 1,203617136 | 2,303163987 | 0,05621344 | 0,366342158 | olfactory receptor family 2 subfamily AT member 4 |
| ENSG00000166681 | BEX3 | 1,203584741 | 2,303112271 | 1,77E-09 | 1,26E-07 | brain expressed X-linked 3 |
| ENSG00000224599 | BMS1P12 | 1,203107036 | 2,302349792 | 0,00669722 | 0,085882668 | BMS1 pseudogene 12 |
| ENSG00000172572 | PDE3A | 1,203084228 | 2,302313394 | 0,00013319 | 0,003363566 | phosphodiesterase 3A |
| ENSG00000168505 | GBX2 | 1,203039743 | 2,302242403 | 0,00798152 | 0,097436235 | gastrulation brain homeobox 2 |
| ENSG00000234257 | SOD2P1 | 1,202485051 | 2,3013574 | 0,21359077 | 0,721033693 | superoxide dismutase 2 pseudogene 1 |

| | | | | | | |
|---|---|---|---|---|---|---|
| ENSG00000287502 | AC092418,2 | 1,201722605 | 2,300141483 | 0,41278174 | 0,884576962 | novel transcript |
| ENSG00000282413 | AL133461,1 | 1,200793799 | 2,298661131 | 0,00755335 | 0,093783925 | VMA21 vacuolar H+-ATPase homolog (S, cerevisiae) (VMA21) pseudogene |
| ENSG00000134470 | IL15RA | 1,200709953 | 2,298527541 | 0,4686738 | 0,907940653 | interleukin 15 receptor subunit alpha |
| ENSG00000058668 | ATP2B4 | 1,200537387 | 2,298252623 | 4,93E-05 | 0,001398352 | ATPase plasma membrane Ca2+ transporting 4 |
| ENSG00000171772 | SYCE1 | 1,200290868 | 2,297859944 | 0,3268146 | 0,828612831 | synaptonemal complex central element protein 1 |
| ENSG00000078596 | ITM2A | 1,199661005 | 2,296856946 | 0,00849424 | 0,101709666 | integral membrane protein 2A |
| ENSG00000132692 | BCAN | 1,199531429 | 2,296650662 | 0,07994801 | 0,448292044 | brevican |
| ENSG00000196923 | PDLIM7 | 1,199255702 | 2,296211771 | 0,00104457 | 0,019427394 | PDZ and LIM domain 7 |
| ENSG00000259171 | AL163636,1 | 1,198504812 | 2,295016956 | 0,75239882 | 0,975901808 | novel protein, ANG-RNASE4 readthrough |
| ENSG00000150961 | SEC24D | 1,19812025 | 2,294405282 | 6,15E-06 | 0,000221441 | SEC24 homolog D, COPII coat complex component |
| ENSG00000165816 | VWA2 | 1,197912556 | 2,294074996 | 0,00775129 | 0,095596237 | von Willebrand factor A domain containing 2 |
| ENSG00000184163 | C1QTNF12 | 1,19758055 | 2,293547124 | 0,32997758 | 0,831651927 | C1q and TNF related 12 |
| ENSG00000260571 | BNIP3P5 | 1,197424963 | 2,293299791 | 0,36819288 | 0,857548638 | BCL2 interacting protein 3 pseudogene 5 |
| ENSG00000274799 | Metazoa_SRP | 1,196900403 | 2,292466105 | 0,29076401 | 0,801555946 | Metazoan signal recognition particle RNA |
| ENSG00000261101 | AC234775,2 | 1,196204641 | 2,291360793 | 0,01635129 | 0,163324813 | novel transcript, overlapping to ARMCX3 |
| ENSG00000264186 | SNRPCP4 | 1,195711296 | 2,290577372 | 0,30625031 | 0,815375074 | small nuclear ribonucleoprotein polypeptide C pseudogene 4 |
| ENSG00000240163 | AC087385,1 | 1,195619639 | 2,290431852 | 0,14271498 | 0,601245287 | ribosomal protein L36a (RPL36A) pseudogene |
| ENSG00000105388 | CEACAM5 | 1,195245669 | 2,289838213 | 0,47169384 | 0,908896785 | CEA cell adhesion molecule 5 |
| ENSG00000276048 | AC012354,4 | 1,195215122 | 2,289789729 | 0,37474439 | 0,861771195 | |
| ENSG00000254321 | AC016813,1 | 1,195143303 | 2,289675743 | 0,41796221 | 0,886992289 | novel transcript |
| ENSG00000125997 | BPIFB9P | 1,194856257 | 2,289220223 | 0,2822326 | 0,793283115 | BPI fold containing family B member 9, pseudogene |
| ENSG00000225969 | ABHD11-AS1 | 1,194454844 | 2,288583363 | 0,31520445 | 0,820597202 | ABHD11 antisense RNA 1 (tail to tail) |
| ENSG00000183066 | WBP2NL | 1,192585408 | 2,285619751 | 0,46082364 | 0,906029574 | WBP2 N-terminal like |
| ENSG00000205794 | AC098591,2 | 1,19244612 | 2,285399091 | 0,25239665 | 0,76432863 | acyl-CoA thioesterase 7 (ACOT7) pseudogene |
| ENSG00000091073 | DTX2 | 1,19200864 | 2,284706176 | 0,00039462 | 0,008543835 | deltex E3 ubiquitin ligase 2 |
| ENSG00000154027 | AK5 | 1,191344767 | 2,283655085 | 0,00099169 | 0,018554184 | adenylate kinase 5 |
| ENSG00000227751 | RCC2-AS1 | 1,190581991 | 2,282447997 | 0,11770244 | 0,548330959 | RCC2 antisense RNA 1 |
| ENSG00000281391 | RCC2-AS1 | 1,190581991 | 2,282447997 | 0,11770244 | 0,548330959 | RCC2 antisense RNA 1 |
| ENSG00000150907 | FOXO1 | 1,190276735 | 2,281965112 | 0,0014154 | 0,02510636 | forkhead box O1 |
| ENSG00000129911 | KLF16 | 1,189257384 | 2,280353336 | 0,01439224 | 0,149637139 | Kruppel like factor 16 |
| ENSG00000142871 | CCN1 | 1,18798784 | 2,278347551 | 0,00046869 | 0,009854107 | cellular communication network factor 1 |

| | | | | | | |
|---|---|---|---|---|---|---|
| ENSG00000258303 | AC012464,2 | 1,187454162 | 2,277504906 | 0,45579195 | 0,903545728 | novel transcript |
| ENSG00000275074 | NUDT18 | 1,187275218 | 2,277222434 | 0,04608615 | 0,325793402 | nudix hydrolase 18 |
| ENSG00000285676 | AL158212,5 | 1,187120661 | 2,276978488 | 0,1199224 | 0,552987303 | novel transcript, antisense to VTI1A |
| ENSG00000288007 | AC092803,4 | 1,18695411 | 2,276715638 | 0,43109488 | 0,89530663 | novel transcript |
| ENSG00000077044 | DGKD | 1,186865128 | 2,27657522 | 0,14278229 | 0,601366788 | diacylglycerol kinase delta |
| ENSG00000240875 | LINC00886 | 1,186281134 | 2,275653862 | 0,01070812 | 0,120467271 | long intergenic non-protein coding RNA 886 |
| ENSG00000269755 | AC008758,6 | 1,186156812 | 2,275457771 | 0,19573311 | 0,69375328 | novel transcript |
| ENSG00000086289 | EPDR1 | 1,186110646 | 2,275384958 | 1,31E-06 | 5,48E-05 | ependymin related 1 |
| ENSG00000177459 | ERICH5 | 1,186027814 | 2,275254321 | 0,22660602 | 0,735240631 | glutamate rich 5 |
| ENSG00000165240 | ATP7A | 1,185219193 | 2,273979413 | 0,00161195 | 0,027904786 | ATPase copper transporting alpha |
| ENSG00000287064 | AL606500,1 | 1,185098136 | 2,273788611 | 0,00154114 | 0,026887445 | novel transcript |
| ENSG00000278488 | NAPRT | 1,18500541 | 2,273642473 | 0,14234937 | 0,600436282 | nicotinate phosphoribosyltransferase |
| ENSG00000169499 | PLEKHA2 | 1,184347438 | 2,272605765 | 0,01087265 | 0,121820431 | pleckstrin homology domain containing A2 |
| ENSG00000259129 | LINC00648 | 1,184322281 | 2,272566137 | 1,50E-05 | 0,000486546 | long intergenic non-protein coding RNA 648 |
| ENSG00000229780 | UBE2Q1-AS1 | 1,183796694 | 2,271738372 | 0,46537564 | 0,90786072 | UBE2Q1 antisense RNA 1 |
| ENSG00000270069 | MIR222HG | 1,183484822 | 2,271247335 | 0,00047389 | 0,009937001 | miR222/221 cluster host gene |
| ENSG00000128294 | TPST2 | 1,183328703 | 2,271001569 | 0,0034131 | 0,051724216 | tyrosylprotein sulfotransferase 2 |
| ENSG00000263879 | SNORA38 | 1,182327117 | 2,269425482 | 0,39088439 | 0,870057277 | small nucleolar RNA, H/ACA box 38 |
| ENSG00000200816 | SNORA38 | 1,182327117 | 2,269425482 | 0,39088439 | 0,870057277 | small nucleolar RNA, H/ACA box 38 |
| ENSG00000266847 | SNORA38 | 1,182327117 | 2,269425482 | 0,39088439 | 0,870057277 | small nucleolar RNA, H/ACA box 38 |
| ENSG00000264153 | SNORA38 | 1,182327117 | 2,269425482 | 0,39088439 | 0,870057277 | small nucleolar RNA, H/ACA box 38 |
| ENSG00000263994 | SNORA38 | 1,182327117 | 2,269425482 | 0,39088439 | 0,870057277 | small nucleolar RNA, H/ACA box 38 |
| ENSG00000116525 | TRIM62 | 1,182128678 | 2,269113349 | 3,75E-05 | 0,001095964 | tripartite motif containing 62 |
| ENSG00000152592 | DMP1 | 1,181207095 | 2,26766432 | 0,34413512 | 0,840938551 | dentin matrix acidic phosphoprotein 1 |
| ENSG00000203987 | AL772363,1 | 1,180993577 | 2,267328732 | 0,40578283 | 0,87874173 | novel transcript, antisense to CACNA1B and EHMT1 |
| ENSG00000123562 | MORF4L2 | 1,18030921 | 2,26625344 | 0,00035484 | 0,007785167 | mortality factor 4 like 2 |
| ENSG00000226625 | RBM17P1 | 1,180301337 | 2,266241072 | 0,13272973 | 0,581510979 | RNA binding motif protein 17 pseudogene 1 |
| ENSG00000223766 | PRR3 | 1,180281026 | 2,266209169 | 0,51683429 | 0,925606197 | proline rich 3 |
| ENSG00000166292 | TMEM100 | 1,180120984 | 2,265957785 | 0,08111757 | 0,452221784 | transmembrane protein 100 |
| ENSG00000145741 | BTF3 | 1,179669412 | 2,265248638 | 6,98E-06 | 0,000248656 | basic transcription factor 3 |
| ENSG00000266441 | AP005205,2 | 1,179410709 | 2,264842471 | 0,13959719 | 0,595782615 | novel transcript, antisense to ARHGAP28, |
| ENSG00000288637 | AC013477,1 | 1,179031497 | 2,264247236 | 0,1354504 | 0,586770311 | novel protein |
| ENSG00000287923 | AL138701,2 | 1,178935604 | 2,264096743 | 0,22284071 | 0,731402952 | novel transcript |

| | | | | | | |
|---|---|---|---|---|---|---|
| ENSG00000142408 | CACNG8 | 1,178026315 | 2,262670197 | 0,11874177 | 0,550651408 | calcium voltage-gated channel auxiliary subunit gamma 8 |
| ENSG00000206281 | TAPBP | 1,177726176 | 2,262199519 | 0,01563553 | 0,158504956 | TAP binding protein |
| ENSG00000188133 | TMEM215 | 1,17728875 | 2,261513722 | 0,26913655 | 0,781347689 | transmembrane protein 215 |
| ENSG00000206412 | GNL1 | 1,176323427 | 2,260001025 | 0,10691727 | 0,521964616 | G protein nucleolar 1 (putative) |
| ENSG00000266891 | AP000902,1 | 1,176141925 | 2,259716718 | 0,46169585 | 0,906174847 | ribosomal protein L27a (RPL27A) pseudogene |
| ENSG00000089356 | FXYD3 | 1,175531133 | 2,258760226 | 0,4752433 | 0,911082311 | FXYD domain containing ion transport regulator 3 |
| ENSG00000169851 | PCDH7 | 1,175237635 | 2,258300758 | 2,19E-05 | 0,000683873 | protocadherin 7 |
| ENSG00000227129 | NEU1 | 1,17510662 | 2,258095684 | 0,00139205 | 0,024770103 | neuraminidase 1 |
| ENSG00000131171 | SH3BGRL | 1,175020811 | 2,257961381 | 0,01819506 | 0,175462769 | SH3 domain binding glutamate rich protein like |
| ENSG00000248714 | AC091180,2 | 1,174925941 | 2,257812905 | 0,04439459 | 0,318259511 | novel transcript |
| ENSG00000073712 | FERMT2 | 1,174454626 | 2,257075419 | 2,64E-05 | 0,00080382 | fermitin family member 2 |
| ENSG00000153563 | CD8A | 1,17418489 | 2,25665346 | 0,15130588 | 0,618627212 | CD8a molecule |
| ENSG00000245651 | AC083805,1 | 1,173972124 | 2,256320677 | 0,05128983 | 0,34665805 | novel transcript |
| ENSG00000173077 | DELEC1 | 1,17354331 | 2,255650128 | 0,30893895 | 0,816838423 | deleted in esophageal cancer 1 |
| ENSG00000170927 | PKHD1 | 1,173191367 | 2,255099933 | 0,02763335 | 0,234127782 | PKHD1 ciliary IPT domain containing fibrocystin/polyductin |
| ENSG00000174236 | REP15 | 1,172931702 | 2,254694083 | 0,01539066 | 0,156734063 | RAB15 effector protein |
| ENSG00000267072 | NAGPA-AS1 | 1,172800816 | 2,254489539 | 0,2308655 | 0,739268552 | NAGPA antisense RNA 1 |
| ENSG00000101811 | CSTF2 | 1,172787941 | 2,25446942 | 2,98E-08 | 1,72E-06 | cleavage stimulation factor subunit 2 |
| ENSG00000148484 | RSU1 | 1,172511444 | 2,254037385 | 3,26E-06 | 0,000126713 | Ras suppressor protein 1 |
| ENSG00000241111 | PRICKLE2-AS1 | 1,172127869 | 2,253438174 | 0,01966091 | 0,184422056 | PRICKLE2 antisense RNA 1 |
| ENSG00000177807 | KCNJ10 | 1,171946433 | 2,253154796 | 0,04313294 | 0,313242318 | potassium inwardly rectifying channel subfamily J member 10 |
| ENSG00000168056 | LTBP3 | 1,171937652 | 2,253141083 | 0,01551133 | 0,157757432 | latent transforming growth factor beta binding protein 3 |
| ENSG00000254198 | AC113191,1 | 1,171934773 | 2,253136586 | 0,16502645 | 0,644037807 | Family with sequence similarity 9, member C (FAM9C) pseudogene |
| ENSG00000089289 | IGBP1 | 1,17166172 | 2,252710185 | 8,32E-05 | 0,00223542 | immunoglobulin binding protein 1 |
| ENSG00000230143 | FLOT1 | 1,171419197 | 2,252331527 | 0,0022631 | 0,037043535 | flotillin 1 |
| ENSG00000272874 | AL034548,1 | 1,171241259 | 2,252053747 | 0,2694125 | 0,781544346 | novel transcript |
| ENSG00000274768 | CISD3 | 1,171052781 | 2,251759551 | 0,03737594 | 0,28542293 | CDGSH iron sulfur domain 3 |
| ENSG00000149257 | SERPINH1 | 1,170422971 | 2,250776758 | 4,24E-05 | 0,001221696 | serpin family H member 1 |

| | | | | | | |
|---|---|---|---|---|---|---|
| ENSG00000198892 | SHISA4 | 1,170263194 | 2,2505275 | 0,00443075 | 0,063178325 | shisa family member 4 |
| ENSG00000153495 | TEX29 | 1,170186802 | 2,250408336 | 0,34978204 | 0,843534837 | testis expressed 29 |
| ENSG00000225406 | ELF2P4 | 1,170184551 | 2,250404826 | 0,09577952 | 0,491457584 | ELF2 pseudogene 4 |
| ENSG00000105810 | CDK6 | 1,169599163 | 2,249491886 | 0,00014415 | 0,00359387 | cyclin dependent kinase 6 |
| ENSG00000167657 | DAPK3 | 1,169011844 | 2,248576308 | 0,03309551 | 0,264223189 | death associated protein kinase 3 |
| ENSG00000248634 | AC008588,1 | 1,168810207 | 2,248262059 | 0,318449 | 0,823270714 | novel transcript |
| ENSG00000140968 | IRF8 | 1,167981874 | 2,246971575 | 0,21650384 | 0,723777756 | interferon regulatory factor 8 |
| ENSG00000282978 | AC110994,2 | 1,167968322 | 2,246950468 | 0,13393845 | 0,584034405 | basic transcription factor 3 (BTF3) pseudogene |
| ENSG00000280641 | CDH4 | 1,167468621 | 2,246172335 | 0,01919594 | 0,182079045 | cadherin 4 |
| ENSG00000203411 | Metazoa_SRP | 1,167335767 | 2,245965501 | 0,32530713 | 0,827332801 | Metazoan signal recognition particle RNA |
| ENSG00000088386 | SLC15A1 | 1,167191489 | 2,245740902 | 0,17198367 | 0,655879181 | solute carrier family 15 member 1 |
| ENSG00000269902 | AC234772,1 | 1,167061155 | 2,24553803 | 0,03375708 | 0,267117518 | novel transcript |
| ENSG00000254453 | NAV2-AS2 | 1,166257161 | 2,24428697 | 0,36885552 | 0,857836723 | NAV2 antisense RNA 2 |
| ENSG00000125872 | LRRN4 | 1,166162402 | 2,244139566 | 0,15253943 | 0,620962584 | leucine rich repeat neuronal 4 |
| ENSG00000174343 | CHRNA9 | 1,165325436 | 2,242838027 | 0,09009575 | 0,478714084 | cholinergic receptor nicotinic alpha 9 subunit |
| ENSG00000082781 | ITGB5 | 1,165324863 | 2,242837137 | 0,00062578 | 0,012613995 | integrin subunit beta 5 |
| ENSG00000147100 | SLC16A2 | 1,165288926 | 2,242781268 | 1,08E-05 | 0,000364352 | solute carrier family 16 member 2 |
| ENSG00000253194 | AL137009,1 | 1,165272585 | 2,242755867 | 0,06617278 | 0,404225451 | novel transcript, antisense to FAM184A |
| ENSG00000286656 | AC008875,2 | 1,165165297 | 2,242589086 | 0,3664451 | 0,856103393 | novel transcript |
| ENSG00000050327 | ARHGEF5 | 1,164960253 | 2,24227038 | 0,08884653 | 0,475558385 | Rho guanine nucleotide exchange factor 5 |
| ENSG00000171864 | PRND | 1,16457609 | 2,241673383 | 0,14334026 | 0,602337365 | prion like protein doppel |
| ENSG00000047365 | ARAP2 | 1,164272209 | 2,24120126 | 0,01594931 | 0,160695804 | ArfGAP with RhoGAP domain, ankyrin repeat and PH domain 2 |
| ENSG00000149798 | CDC42EP2 | 1,162979493 | 2,239193948 | 0,09981686 | 0,501267282 | CDC42 effector protein 2 |
| ENSG00000281766 | RYBP | 1,162940506 | 2,239133438 | 0,00176019 | 0,030137218 | RING1 and YY1 binding protein |
| ENSG00000285379 | RIPK3 | 1,162776308 | 2,238878609 | 0,3448057 | 0,841168721 | receptor interacting serine/threonine kinase 3 |
| ENSG00000129465 | RIPK3 | 1,162776308 | 2,238878609 | 0,3448057 | 0,841168721 | receptor interacting serine/threonine kinase 3 |
| ENSG00000213553 | RPLP0P6 | 1,162507181 | 2,238460997 | 0,3883456 | 0,869820053 | ribosomal protein lateral stalk subunit P0 pseudogene 6 |
| ENSG00000258935 | AL096869,1 | 1,161452069 | 2,236824502 | 0,43304101 | 0,895777109 | novel transcript, antisense & intronic to TTC7B |
| ENSG00000135899 | SP110 | 1,161345849 | 2,23665982 | 0,00355541 | 0,053492006 | SP110 nuclear body protein |
| ENSG00000286060 | AL365256,1 | 1,161056792 | 2,236211729 | 0,18952504 | 0,684878649 | novel transcript, antisense to IGSF1 |
| ENSG00000174482 | LINGO2 | 1,160898765 | 2,235966796 | 0,03743889 | 0,28576405 | leucine rich repeat and Ig domain containing 2 |

| | | | | | | |
|---|---|---|---|---|---|---|
| ENSG00000140332 | TLE3 | 1,160678885 | 2,235626041 | 1,39E-05 | 0,000454062 | TLE family member 3, transcriptional corepressor |
| ENSG00000197043 | ANXA6 | 1,160156078 | 2,234816037 | 5,21E-06 | 0,00019179 | annexin A6 |
| ENSG00000256739 | AP002892,2 | 1,159788803 | 2,234247179 | 0,35018635 | 0,843789477 | novel transcript |
| ENSG00000149654 | CDH22 | 1,158773451 | 2,232675296 | 0,14739546 | 0,611530401 | cadherin 22 |
| ENSG00000124313 | IQSEC2 | 1,158544355 | 2,232320781 | 4,23E-06 | 0,000159062 | IQ motif and Sec7 domain ArfGEF 2 |
| ENSG00000226308 | AL122058,1 | 1,158513767 | 2,232273452 | 0,08683204 | 0,469592252 | novel transcript |
| ENSG00000102032 | RENBP | 1,157569068 | 2,230812202 | 0,05562867 | 0,36392023 | renin binding protein |
| ENSG00000274442 | PRPF8 | 1,157470885 | 2,23066039 | 0,04706751 | 0,32967633 | pre-mRNA processing factor 8 |
| ENSG00000122012 | SV2C | 1,157447962 | 2,230624947 | 1,76E-05 | 0,000562581 | synaptic vesicle glycoprotein 2C |
| ENSG00000177096 | PHETA2 | 1,157444558 | 2,230619684 | 0,00668966 | 0,085820922 | PH domain containing endocytic trafficking adaptor 2 |
| ENSG00000125845 | BMP2 | 1,157436324 | 2,230606953 | 0,01014595 | 0,11641031 | bone morphogenetic protein 2 |
| ENSG00000071282 | LMCD1 | 1,156914722 | 2,22980063 | 0,00077935 | 0,015152741 | LIM and cysteine rich domains 1 |
| ENSG00000227536 | SOCS5P4 | 1,15681858 | 2,22965204 | 0,13101988 | 0,577988644 | suppressor of cytokine signaling 5 pseudogene 4 |
| ENSG00000116729 | WLS | 1,156551436 | 2,229239212 | 1,03E-06 | 4,36E-05 | Wnt ligand secretion mediator |
| ENSG00000235128 | AC013474,1 | 1,156298037 | 2,228847698 | 0,06537653 | 0,401164726 | novel pseudogene |
| ENSG00000122126 | OCRL | 1,155439187 | 2,227521238 | 0,0001502 | 0,0037181 | OCRL inositol polyphosphate-5-phosphatase |
| ENSG00000135723 | FHOD1 | 1,155382181 | 2,227433222 | 0,00870893 | 0,103566409 | formin homology 2 domain containing 1 |
| ENSG00000196358 | NTNG2 | 1,155278524 | 2,227273188 | 0,00203228 | 0,033995352 | netrin G2 |
| ENSG00000111907 | TPD52L1 | 1,155271359 | 2,227262127 | 0,0021188 | 0,035216523 | TPD52 like 1 |
| ENSG00000273186 | AL359091,4 | 1,154902508 | 2,22669276 | 0,22660476 | 0,735240631 | novel transcript, sense intronic to CERCAM |
| ENSG00000287963 | AC010476,2 | 1,154325108 | 2,225801763 | 0,06325701 | 0,393511843 | novel transcript |
| ENSG00000134716 | CYP2J2 | 1,154130161 | 2,225501018 | 0,08215969 | 0,455998678 | cytochrome P450 family 2 subfamily J member 2 |
| ENSG00000263873 | THY1-AS1 | 1,15388068 | 2,225116202 | 0,04423842 | 0,3175092 | THY1 antisense RNA 1 |
| ENSG00000233559 | LINC00513 | 1,153694842 | 2,224829597 | 0,07554661 | 0,434139326 | long intergenic non-protein coding RNA 513 |
| ENSG00000155254 | MARVELD1 | 1,153648149 | 2,224757591 | 0,0001364 | 0,0034229 | MARVEL domain containing 1 |
| ENSG00000213699 | SLC35F6 | 1,153605975 | 2,224692555 | 0,00037194 | 0,008097746 | solute carrier family 35 member F6 |
| ENSG00000275549 | STPG3-AS1 | 1,15345576 | 2,224460931 | 0,35092936 | 0,844035671 | STPG3 antisense RNA 1 |
| ENSG00000147485 | PXDNL | 1,152768515 | 2,223401534 | 0,01170687 | 0,128601321 | peroxidasin like |
| ENSG00000105939 | ZC3HAV1 | 1,152442951 | 2,22289985 | 0,00155211 | 0,027018437 | zinc finger CCCH-type containing, antiviral 1 |
| ENSG00000117385 | P3H1 | 1,152134871 | 2,222425212 | 0,00014591 | 0,003626251 | prolyl 3-hydroxylase 1 |
| ENSG00000042832 | TG | 1,152014737 | 2,222240158 | 0,17040446 | 0,653594444 | thyroglobulin |
| ENSG00000237457 | LINC01351 | 1,151248261 | 2,221059838 | 0,08470513 | 0,463056695 | long intergenic non-protein coding RNA 1351 |
| ENSG00000184743 | ATL3 | 1,151087885 | 2,220812949 | 3,60E-05 | 0,001057694 | atlastin GTPase 3 |

| | | | | | | |
|---|---|---|---|---|---|---|
| ENSG00000238222 | MKRN4P | 1,151025787 | 2,220717361 | 0,05513623 | 0,362604555 | makorin ring finger protein 4, pseudogene |
| ENSG00000185745 | IFIT1 | 1,150918933 | 2,220552889 | 0,02663396 | 0,228445591 | interferon induced protein with tetratricopeptide repeats 1 |
| ENSG00000205111 | CDKL4 | 1,15068577 | 2,22019404 | 0,09679364 | 0,494147005 | cyclin dependent kinase like 4 |
| ENSG00000274383 | AC103691,1 | 1,150184928 | 2,219423418 | 0,05684235 | 0,368162243 | novel transcript, antisense to ANKDD1A |
| ENSG00000248747 | AC107220,1 | 1,149796662 | 2,218826193 | 0,51949893 | 0,926800715 | novel transcript |
| ENSG00000221926 | TRIM16 | 1,149704315 | 2,218684171 | 0,00121026 | 0,021896773 | tripartite motif containing 16 |
| ENSG00000245146 | MALINC1 | 1,149430838 | 2,218263637 | 0,06678738 | 0,406075591 | mitosis associated long intergenic non-coding RNA 1 |
| ENSG00000149218 | ENDOD1 | 1,149304545 | 2,21806946 | 3,88E-05 | 0,001127746 | endonuclease domain containing 1 |
| ENSG00000288542 | AL133318,1 | 1,149185447 | 2,21788636 | 0,50616363 | 0,921780819 | novel transcript |
| ENSG00000181143 | MUC16 | 1,148952818 | 2,217528764 | 0,35906764 | 0,849814998 | mucin 16, cell surface associated |
| ENSG00000274811 | CLPTM1L | 1,14862238 | 2,217020915 | 0,11989937 | 0,552962686 | CLPTM1 like |
| ENSG00000204173 | LRRC37A5P | 1,148515702 | 2,216856986 | 0,40040963 | 0,876148288 | leucine rich repeat containing 37 member A5, pseudogene |
| ENSG00000261582 | AL121753,1 | 1,148407192 | 2,216690255 | 0,53950291 | 0,933350549 | novel transcript |
| ENSG00000254648 | AP000911,1 | 1,148225059 | 2,216410427 | 0,09084352 | 0,480889225 | novel transcript |
| ENSG00000185903 | OR11N1P | 1,147934923 | 2,215964735 | 0,18448162 | 0,676713995 | olfactory receptor family 11 subfamily N member 1 pseudogene |
| ENSG00000182257 | PRR34 | 1,147862884 | 2,215854087 | 0,1306438 | 0,577136255 | PRR34 long non-coding RNA |
| ENSG00000262094 | AC139099,1 | 1,147756145 | 2,215690152 | 0,29987531 | 0,809891287 | novel transcript |
| ENSG00000146070 | PLA2G7 | 1,147600607 | 2,215451289 | 0,09021778 | 0,478874121 | phospholipase A2 group VII |
| ENSG00000140478 | GOLGA6D | 1,147551844 | 2,215376409 | 0,56670167 | 0,941311563 | golgin A6 family member D |
| ENSG00000243905 | RN7SL679P | 1,147328009 | 2,215032718 | 0,25343396 | 0,765172364 | RNA, 7SL, cytoplasmic 679, pseudogene |
| ENSG00000281092 | AL627313,4 | 1,147328009 | 2,215032718 | 0,25343396 | 0,765172364 | |
| ENSG00000272369 | AC008035,1 | 1,146406811 | 2,213618814 | 0,43542114 | 0,89634451 | novel transcript |
| ENSG00000288499 | FAM20C | 1,146156298 | 2,21323447 | 0,01261769 | 0,135653188 | FAM20C golgi associated secretory pathway kinase |
| ENSG00000177706 | FAM20C | 1,146156298 | 2,21323447 | 0,01261769 | 0,135653188 | FAM20C golgi associated secretory pathway kinase |
| ENSG00000273733 | AC011472,4 | 1,145851545 | 2,212766998 | 0,38968654 | 0,870057277 | novel transcript, antisense to DOCK6 |
| ENSG00000237188 | AC242426,2 | 1,145527404 | 2,212269895 | 0,04704407 | 0,329624762 | novel transcript, antisense to FMO5 |
| ENSG00000234343 | NEU1 | 1,145389498 | 2,212058436 | 0,00374185 | 0,055786943 | neuraminidase 1 |
| ENSG00000204498 | NFKBIL1 | 1,144487036 | 2,210675141 | 0,52957553 | 0,930619378 | NFKB inhibitor like 1 |
| ENSG00000196758 | AC079612,1 | 1,14404672 | 2,210000537 | 0,25070114 | 0,762963147 | uncharacterized LOC401040 |
| ENSG00000137648 | TMPRSS4 | 1,14284405 | 2,208158987 | 0,21250504 | 0,719332768 | transmembrane serine protease 4 |

| | | | | | | |
|---|---|---|---|---|---|---|
| ENSG00000182095 | TNRC18 | 1,1422903 | 2,207311591 | 0,08923144 | 0,476498052 | trinucleotide repeat containing 18 |
| ENSG00000064489 | BORCS8-MEF2I | 1,141581363 | 2,206227191 | 0,06617169 | 0,404225451 | BORCS8-MEF2B readthrough |
| ENSG00000175489 | LRRC25 | 1,141561548 | 2,206196889 | 0,32020441 | 0,824366234 | leucine rich repeat containing 25 |
| ENSG00000251230 | MIR3945HG | 1,140803954 | 2,205038666 | 0,22757173 | 0,736053199 | MIR3945 host gene |
| ENSG00000143248 | RGS5 | 1,14006101 | 2,20390343 | 0,00918333 | 0,107611518 | regulator of G protein signaling 5 |
| ENSG00000277721 | Metazoa_SRP | 1,139797724 | 2,203501264 | 0,41735169 | 0,886750438 | Metazoan signal recognition particle RNA |
| ENSG00000233347 | ERP29P1 | 1,139350479 | 2,20281827 | 0,33244052 | 0,83293962 | endoplasmic reticulum lumenal protein 29 pseudogene 1 |
| ENSG00000284237 | LINC02767 | 1,138292995 | 2,201204212 | 0,2018907 | 0,701480895 | long intergenic non-protein coding RNA 2767 |
| ENSG00000248231 | KRT8P4 | 1,138093556 | 2,200899939 | 0,04701756 | 0,329512866 | keratin 8 pseudogene 4 |
| ENSG00000266821 | AC018521,7 | 1,13779729 | 2,200448017 | 0,17935242 | 0,66800978 | novel transcript |
| ENSG00000151789 | ZNF385D | 1,136937788 | 2,199137465 | 0,01884365 | 0,17950267 | zinc finger protein 385D |
| ENSG00000284730 | TMDD1 | 1,136723981 | 2,198811578 | 0,29456089 | 0,804714394 | transmembrane and death domain 1 |
| ENSG00000242732 | RTL5 | 1,136365037 | 2,198264579 | 0,00020651 | 0,004898559 | retrotransposon Gag like 5 |
| ENSG00000014914 | MTMR11 | 1,136197569 | 2,198009419 | 0,00083428 | 0,015972491 | myotubularin related protein 11 |
| ENSG00000235172 | LINC01366 | 1,135751961 | 2,197330621 | 0,24890791 | 0,760912969 | long intergenic non-protein coding RNA 1366 |
| ENSG00000261693 | AC134682,1 | 1,135401851 | 2,196797443 | 0,21527481 | 0,722575157 | novel transcript, antisense to TSNARE1 |
| ENSG00000267346 | EIF5AP3 | 1,135019214 | 2,196214876 | 0,31966941 | 0,824366234 | eukaryotic translation initiation factor 5A pseudogene 3 |
| ENSG00000287259 | AL359732,1 | 1,134034614 | 2,194716531 | 0,44327519 | 0,899789356 | novel transcript |
| ENSG00000145901 | TNIP1 | 1,134015097 | 2,194686842 | 0,00676527 | 0,086506916 | TNFAIP3 interacting protein 1 |
| ENSG00000270640 | AC104695,3 | 1,132731113 | 2,192734462 | 0,37958579 | 0,865157837 | novel transcript, sense intronic to FOSL2 |
| ENSG00000174125 | TLR1 | 1,132594862 | 2,192527385 | 0,11607846 | 0,544096306 | toll like receptor 1 |
| ENSG00000237742 | AL365259,1 | 1,132357584 | 2,192166813 | 0,37614228 | 0,862539384 | novel transcript, antisense to HEY2 |
| ENSG00000132669 | RIN2 | 1,132353431 | 2,192160503 | 0,00064385 | 0,012870387 | Ras and Rab interactor 2 |
| ENSG00000162399 | BSND | 1,13230188 | 2,192082172 | 0,44471896 | 0,90032335 | barttin CLCNK type accessory subunit beta |
| ENSG00000251279 | SMIM15-AS1 | 1,132172647 | 2,19188582 | 0,36355468 | 0,853893276 | SMIM15 antisense RNA 1 |
| ENSG00000236006 | AL662848,4 | 1,132108161 | 2,191787849 | 0,40333697 | 0,877589651 | chromosome 6 open reading frame 214 (putative) |
| ENSG00000140511 | HAPLN3 | 1,131829639 | 2,19136475 | 0,00406315 | 0,059443318 | hyaluronan and proteoglycan link protein 3 |
| ENSG00000144908 | ALDH1L1 | 1,131766131 | 2,191268288 | 0,31246676 | 0,818574551 | aldehyde dehydrogenase 1 family member L1 |
| ENSG00000163702 | IL17RC | 1,131682696 | 2,191141565 | 0,01974504 | 0,184776117 | interleukin 17 receptor C |
| ENSG00000074660 | SCARF1 | 1,131666348 | 2,191116734 | 0,0288063 | 0,240931045 | scavenger receptor class F member 1 |
| ENSG00000153179 | RASSF3 | 1,131643793 | 2,191082479 | 0,00045096 | 0,009558531 | Ras association domain family member 3 |

| | | | | | | |
|---|---|---|---|---|---|---|
| ENSG00000260951 | AC005100,1 | 1,130520955 | 2,189377841 | 0,55372573 | 0,937338325 | novel transcript |
| ENSG00000126790 | L3HYPDH | 1,130215942 | 2,188915014 | 3,88E-05 | 0,001127746 | trans-L-3-hydroxyproline dehydratase |
| ENSG00000141934 | PLPP2 | 1,129555655 | 2,187913429 | 0,05326608 | 0,355152621 | phospholipid phosphatase 2 |
| ENSG00000261653 | AC092337,1 | 1,12937745 | 2,187643189 | 0,21870767 | 0,72663937 | novel transcript |
| ENSG00000110799 | VWF | 1,129068789 | 2,187175198 | 0,01087905 | 0,121820431 | von Willebrand factor |
| ENSG00000171388 | APLN | 1,128617474 | 2,186491098 | 0,00470999 | 0,065944067 | apelin |
| ENSG00000108370 | RGS9 | 1,128096931 | 2,185702325 | 2,93E-05 | 0,000880956 | regulator of G protein signaling 9 |
| ENSG00000184986 | TMEM121 | 1,127987412 | 2,185536409 | 0,02992019 | 0,2466187 | transmembrane protein 121 |
| ENSG00000225824 | HLA-DQB1 | 1,127728627 | 2,185144411 | 0,13061645 | 0,577136255 | major histocompatibility complex, class II, DQ beta 1 |
| ENSG00000272670 | DOC2B | 1,127672691 | 2,18505969 | 0,64688727 | 0,956899963 | double C2 domain beta |
| ENSG00000147255 | IGSF1 | 1,127498929 | 2,184796532 | 0,03920877 | 0,294869635 | immunoglobulin superfamily member 1 |
| ENSG00000171476 | HOPX | 1,126674627 | 2,183548577 | 0,15954184 | 0,633929525 | HOP homeobox |
| ENSG00000198835 | GJC2 | 1,126603717 | 2,183441256 | 0,22809394 | 0,736341996 | gap junction protein gamma 2 |
| ENSG00000238250 | ST6GAL2-IT1 | 1,126400955 | 2,183134409 | 0,31129139 | 0,817751312 | ST6GAL2 intronic transcript 1 |
| ENSG00000162885 | B3GALNT2 | 1,125551393 | 2,181849201 | 0,00798999 | 0,097501565 | beta-1,3-N-acetylgalactosaminyltransferase 2 |
| ENSG00000248571 | AC106882,1 | 1,125399623 | 2,181619685 | 0,3836941 | 0,868000076 | novel transcript |
| ENSG00000115850 | LCT | 1,123334056 | 2,178498403 | 0,16229467 | 0,639443053 | lactase |
| ENSG00000281721 | LINC01080 | 1,123086529 | 2,178124665 | 0,46532835 | 0,90786072 | long intergenic non-protein coding RNA 1080 |
| ENSG00000276504 | TFPT | 1,122915679 | 2,177866737 | 0,0350846 | 0,273605654 | TCF3 fusion partner |
| ENSG00000105619 | TFPT | 1,122915679 | 2,177866737 | 0,0350846 | 0,273605654 | TCF3 fusion partner |
| ENSG00000132357 | CARD6 | 1,122503863 | 2,177245156 | 0,04600104 | 0,325452954 | caspase recruitment domain family member 6 |
| ENSG00000265485 | LINC01915 | 1,122481121 | 2,177210835 | 0,41701944 | 0,886616509 | long intergenic non-protein coding RNA 1915 |
| ENSG00000233639 | PANTR1 | 1,12117888 | 2,175246474 | 0,00437629 | 0,06270406 | POU3F3 adjacent non-coding transcript 1 |
| ENSG00000235776 | AC000089,1 | 1,120910262 | 2,174841498 | 0,39908136 | 0,875433675 | ribosomal protein L7a (RPL7A) pseudogene |
| ENSG00000111704 | NANOG | 1,120756515 | 2,17460974 | 0,02836559 | 0,238585384 | Nanog homeobox |
| ENSG00000237115 | AL139805,1 | 1,120547253 | 2,174294337 | 0,214438 | 0,721889709 | T-cell activation leucine repeat-rich protein (TA-LRRP) pseudogene |
| ENSG00000106852 | LHX6 | 1,120208096 | 2,173783252 | 0,46091792 | 0,906029574 | LIM homeobox 6 |
| ENSG00000275696 | Metazoa_SRP | 1,119699496 | 2,173017053 | 0,40459157 | 0,878176155 | Metazoan signal recognition particle RNA |
| ENSG00000258930 | AC005230,1 | 1,11954111 | 2,172778501 | 0,33299302 | 0,833301711 | tetratricopeptide repeat domain 7B (TTC7B) pseudogene |
| ENSG00000025039 | RRAGD | 1,119438635 | 2,172624174 | 0,0036725 | 0,054883819 | Ras related GTP binding D |
| ENSG00000249395 | CASC9 | 1,119242849 | 2,17232935 | 0,0003555 | 0,007794133 | cancer susceptibility 9 |

| | | | | | | |
|---|---|---|---|---|---|---|
| ENSG00000230065 | AC010974,1 | 1,119197897 | 2,172261666 | 0,28456968 | 0,795919345 | novel transcript |
| ENSG00000116741 | RGS2 | 1,118150926 | 2,170685816 | 0,00110682 | 0,020319143 | regulator of G protein signaling 2 |
| ENSG00000102057 | KCND1 | 1,117958864 | 2,170396858 | 0,04828438 | 0,334640427 | potassium voltage-gated channel subfamily D member 1 |
| ENSG00000106538 | RARRES2 | 1,117855791 | 2,1702418 | 0,00560846 | 0,075200114 | retinoic acid receptor responder 2 |
| ENSG00000232888 | RPS11P5 | 1,117788999 | 2,170141326 | 0,16532805 | 0,644399285 | ribosomal protein S11 pseudogene 5 |
| ENSG00000167925 | GHDC | 1,117689028 | 2,169990953 | 0,00697653 | 0,088412767 | GH3 domain containing |
| ENSG00000101966 | XIAP | 1,11756194 | 2,169799806 | 3,88E-07 | 1,80E-05 | X-linked inhibitor of apoptosis |
| ENSG00000268460 | AC006262,1 | 1,11710906 | 2,169118785 | 0,25781878 | 0,7701554 | novel transcript |
| ENSG00000119042 | SATB2 | 1,116957382 | 2,168890746 | 0,00025765 | 0,005918594 | SATB homeobox 2 |
| ENSG00000244716 | BX679664,3 | 1,11688559 | 2,168782819 | 0,23299686 | 0,742171568 | ribosomal protein L17 (RPL17) pseudogene |
| ENSG00000196656 | AC004057,1 | 1,116483873 | 2,168179008 | 0,01475145 | 0,152207703 | ribosomal protein S26 (RPS26) pseudogene |
| ENSG00000267535 | LINC00868 | 1,11573491 | 2,167053709 | 0,56420303 | 0,940865625 | long intergenic non-protein coding RNA 868 |
| ENSG00000178573 | MAF | 1,115690317 | 2,166986728 | 0,00875552 | 0,103883368 | MAF bZIP transcription factor |
| ENSG00000143344 | RGL1 | 1,1154539 | 2,166631649 | 0,00030468 | 0,006842963 | ral guanine nucleotide dissociation stimulator like 1 |
| ENSG00000286389 | AL731702,1 | 1,115450439 | 2,166626451 | 0,1108505 | 0,531538341 | novel transcript |
| ENSG00000253831 | ETV3L | 1,115131808 | 2,166147987 | 0,33745629 | 0,836578328 | ETS variant transcription factor 3 like |
| ENSG00000264769 | AC145207,8 | 1,114971398 | 2,16590715 | 0,51266819 | 0,923541244 | novel transcript, antisense MAFG |
| ENSG00000258177 | AC008149,1 | 1,114786867 | 2,165630134 | 0,39574427 | 0,872227623 | novel transcript, antisense to ELK3 |
| ENSG00000089159 | PXN | 1,114723468 | 2,165534967 | 0,0024137 | 0,038958175 | paxillin |
| ENSG00000253317 | AC078906,1 | 1,114512195 | 2,165217863 | 0,25685303 | 0,769107677 | novel transcript |
| ENSG00000274779 | NOMO1 | 1,113973467 | 2,164409482 | 3,41E-06 | 0,000131589 | NODAL modulator 1 |
| ENSG00000130055 | GDPD2 | 1,113746038 | 2,164068309 | 5,73E-05 | 0,001603546 | glycerophosphodiester phosphodiesterase domain containing 2 |
| ENSG00000273375 | AC055764,2 | 1,113564007 | 2,163795275 | 0,2085202 | 0,713245584 | novel transcript |
| ENSG00000287717 | AC133637,2 | 1,113443994 | 2,163615285 | 0,30249431 | 0,812236791 | novel transcript |
| ENSG00000125730 | C3 | 1,113414221 | 2,163570635 | 0,16891425 | 0,650892223 | complement C3 |
| ENSG00000060491 | OGFR | 1,112425623 | 2,162088568 | 0,10669524 | 0,521491876 | opioid growth factor receptor |
| ENSG00000248468 | AC107027,1 | 1,11148762 | 2,16068329 | 0,19386819 | 0,690644393 | novel transcript |
| ENSG00000188483 | IER5L | 1,110881045 | 2,159775032 | 0,00034628 | 0,007656431 | immediate early response 5 like |
| ENSG00000277559 | AC018553,1 | 1,110678829 | 2,159472327 | 0,22466368 | 0,733203418 | novel transcript |
| ENSG00000255970 | LINC02421 | 1,110656697 | 2,159439199 | 0,18689916 | 0,680474679 | long intergenic non-protein coding RNA 2421 |
| ENSG00000244586 | WNT5A-AS1 | 1,110037947 | 2,158513247 | 0,07405055 | 0,4296801 | WNT5A antisense RNA 1 |

| | | | | | | |
|---|---|---|---|---|---|---|
| ENSG00000170923 | OR7G2 | 1,109205478 | 2,157268093 | 0,35311366 | 0,845063255 | olfactory receptor family 7 subfamily G member 2 |
| ENSG00000065485 | PDIA5 | 1,108959573 | 2,156900423 | 0,00089643 | 0,017026728 | protein disulfide isomerase family A member 5 |
| ENSG00000261600 | AC233266,2 | 1,108780118 | 2,156632145 | 0,47203904 | 0,909171312 | novel transcript |
| ENSG00000214976 | VDAC2P1 | 1,108234395 | 2,155816518 | 0,3461836 | 0,841168721 | VDAC2 pseudogene 1 |
| ENSG00000185432 | METTL7A | 1,107855994 | 2,155251148 | 0,02876974 | 0,240839934 | methyltransferase like 7A |
| ENSG00000198910 | L1CAM | 1,10779643 | 2,155162167 | 0,00099921 | 0,018683827 | L1 cell adhesion molecule |
| ENSG00000168468 | ATF6B | 1,107051658 | 2,154049881 | 0,13896505 | 0,594174432 | activating transcription factor 6 beta |
| ENSG00000102962 | CCL22 | 1,10701463 | 2,153994596 | 0,18550762 | 0,678110103 | C-C motif chemokine ligand 22 |
| ENSG00000156535 | CD109 | 1,106646389 | 2,153444869 | 0,0063395 | 0,082213892 | CD109 molecule |
| ENSG00000127152 | BCL11B | 1,106368479 | 2,153030086 | 0,0217162 | 0,197882657 | BAF chromatin remodeling complex subunit BCL11B |
| ENSG00000145012 | LPP | 1,105839317 | 2,152240527 | 3,69E-06 | 0,000141416 | LIM domain containing preferred translocation partner in lipoma |
| ENSG00000091428 | RAPGEF4 | 1,105754398 | 2,152113847 | 0,07123126 | 0,420501137 | Rap guanine nucleotide exchange factor 4 |
| ENSG00000280249 | AL691477,1 | 1,105614198 | 2,151904715 | 0,48988075 | 0,916063942 | |
| ENSG00000260577 | AC126773,4 | 1,105079851 | 2,151107838 | 0,03558621 | 0,276263058 | novel transcript, antisense to CDH3 |
| ENSG00000285774 | AL133444,1 | 1,104621795 | 2,150424969 | 0,13208897 | 0,580641112 | novel transcript, antisense to SAMD4A |
| ENSG00000063660 | GPC1 | 1,104360859 | 2,150036063 | 0,01460892 | 0,151236194 | glypican 1 |
| ENSG00000104689 | TNFRSF10A | 1,104300009 | 2,149945382 | 0,03358073 | 0,266328762 | TNF receptor superfamily member 10a |
| ENSG00000230397 | SPTLC1P1 | 1,103823979 | 2,149236105 | 0,13464481 | 0,585389409 | serine palmitoyltransferase long chain base subunit 1 pseudogene 1 |
| ENSG00000131482 | G6PC | 1,103608633 | 2,148915319 | 0,50242734 | 0,920585659 | glucose-6-phosphatase catalytic subunit |
| ENSG00000116871 | MAP7D1 | 1,103454704 | 2,148686052 | 0,06008075 | 0,381239005 | MAP7 domain containing 1 |
| ENSG00000137573 | SULF1 | 1,103277805 | 2,148422603 | 0,0007067 | 0,013905838 | sulfatase 1 |
| ENSG00000129910 | CDH15 | 1,103238601 | 2,148364223 | 0,20140224 | 0,701094807 | cadherin 15 |
| ENSG00000283148 | AC137936,1 | 1,102008068 | 2,146532577 | 0,48209233 | 0,912981134 | novel transcript |
| ENSG00000251201 | TMED7-TICAM2 | 1,101660462 | 2,146015449 | 0,00015102 | 0,003729453 | TMED7-TICAM2 readthrough |
| ENSG00000204209 | DAXX | 1,101335898 | 2,145532713 | 0,33349996 | 0,833923279 | death domain associated protein |
| ENSG00000269952 | AL117336,1 | 1,100311651 | 2,144010023 | 0,37745776 | 0,862854083 | novel transcript, sense intronic to CREM |
| ENSG00000288498 | AC020916,5 | 1,099970804 | 2,143503547 | 0,38261261 | 0,867188687 | novel transcript |
| ENSG00000233953 | AC009970,1 | 1,098924638 | 2,141949754 | 0,4498874 | 0,902062594 | novel transcript |
| ENSG00000261915 | AC026954,2 | 1,098912957 | 2,141932413 | 0,17902875 | 0,667360786 | novel protein |
| ENSG00000288456 | AC026954,6 | 1,098912957 | 2,141932413 | 0,17902875 | 0,667360786 | novel protein |
| ENSG00000251128 | WWC2-AS1 | 1,098676243 | 2,141580997 | 0,43853025 | 0,897220559 | WWC2 antisense RNA 1 |

| | | | | | | |
|---|---|---|---|---|---|---|
| ENSG00000162267 | ITIH3 | 1,098596094 | 2,141462026 | 0,23619147 | 0,745253314 | inter-alpha-trypsin inhibitor heavy chain 3 |
| ENSG00000145757 | SPATA9 | 1,098343194 | 2,141086666 | 0,20049822 | 0,700064356 | spermatogenesis associated 9 |
| ENSG00000284099 | SQSTM1 | 1,098027592 | 2,140618336 | 0,03717725 | 0,284809996 | sequestosome 1 |
| ENSG00000235659 | AL391987,4 | 1,097881371 | 2,14040139 | 0,23406655 | 0,743586269 | novel transcript |
| ENSG00000179242 | CDH4 | 1,097162256 | 2,139334768 | 0,01272633 | 0,136680168 | cadherin 4 |
| ENSG00000284618 | AL391294,1 | 1,097160986 | 2,139332885 | 0,1971139 | 0,695927373 | novel transcript |
| ENSG00000162458 | FBLIM1 | 1,096604136 | 2,138507306 | 0,01571635 | 0,159066362 | filamin binding LIM protein 1 |
| ENSG00000263238 | CTSO | 1,096599312 | 2,138500155 | 0,00536794 | 0,072776773 | cathepsin O |
| ENSG00000258376 | AC004846,1 | 1,096226433 | 2,13794751 | 0,19766194 | 0,696048902 | novel transcript, antisense to PAPLN |
| ENSG00000079931 | MOXD1 | 1,096164439 | 2,137855643 | 0,00313285 | 0,048248943 | monooxygenase DBH like 1 |
| ENSG00000272555 | AC009974,1 | 1,096062736 | 2,137704939 | 0,38800498 | 0,869820053 | novel transcript |
| ENSG00000226264 | HLA-DMB | 1,096004629 | 2,137618842 | 0,12338249 | 0,561691036 | major histocompatibility complex, class II, DM beta |
| ENSG00000196843 | ARID5A | 1,095008228 | 2,136142998 | 0,01537894 | 0,156675544 | AT-rich interaction domain 5A |
| ENSG00000255320 | AP000759,1 | 1,094975661 | 2,136094778 | 0,26345107 | 0,774637968 | novel transcript |
| ENSG00000166006 | KCNC2 | 1,094635308 | 2,135590902 | 0,21900382 | 0,727236912 | potassium voltage-gated channel subfamily C member 2 |
| ENSG00000026025 | VIM | 1,094574982 | 2,135501605 | 0,00025349 | 0,00583579 | vimentin |
| ENSG00000276021 | WDR81 | 1,09434455 | 2,135160543 | 0,02987784 | 0,246399596 | WD repeat domain 81 |
| ENSG00000081923 | ATP8B1 | 1,094126165 | 2,134837361 | 0,00054112 | 0,011152047 | ATPase phospholipid transporting 8B1 |
| ENSG00000228401 | AL590226,1 | 1,094083301 | 2,134773935 | 0,3149121 | 0,820109271 | novel transcript, antisense to EGFL7 (HSPC324) |
| ENSG00000071127 | WDR1 | 1,093865577 | 2,134451789 | 4,48E-05 | 0,001282567 | WD repeat domain 1 |
| ENSG00000267645 | AC105052,3 | 1,093823049 | 2,13438887 | 0,53833793 | 0,933011647 | novel protein, POLR2J2-UPK3BL readthrough |
| ENSG00000237225 | OR13K1P | 1,093643251 | 2,134122885 | 0,03918366 | 0,294869635 | olfactory receptor family 13 subfamily K member 1 pseudogene |
| ENSG00000153823 | PID1 | 1,09352261 | 2,133944434 | 0,09430202 | 0,487879903 | phosphotyrosine interaction domain containing 1 |
| ENSG00000227950 | AL590128,1 | 1,093186803 | 2,133447786 | 0,4217297 | 0,889720796 | ribosomal protein L37 (RPL37) pseudogene |
| ENSG00000287633 | AL161640,3 | 1,092161328 | 2,13193186 | 0,21303198 | 0,72028467 | novel transcript |
| ENSG00000163254 | CRYGC | 1,091877264 | 2,131512128 | 0,40334647 | 0,877589651 | crystallin gamma C |
| ENSG00000285011 | CRYGC | 1,091877264 | 2,131512128 | 0,40334647 | 0,877589651 | crystallin gamma C |
| ENSG00000248975 | AL133372,2 | 1,091463409 | 2,130900765 | 0,3694063 | 0,857991124 | novel transcript |
| ENSG00000167460 | TPM4 | 1,091462623 | 2,130899604 | 1,60E-07 | 8,22E-06 | tropomyosin 4 |
| ENSG00000287647 | AC093575,1 | 1,090680995 | 2,129745431 | 0,32265812 | 0,824955371 | novel transcript, antisense to AK5 |
| ENSG00000104856 | RELB | 1,090557228 | 2,12956273 | 0,02339556 | 0,208565361 | RELB proto-oncogene, NF-kB subunit |

| | | | | | | |
|---|---|---|---|---|---|---|
| ENSG00000112773 | TENT5A | 1,089996014 | 2,128734484 | 0,03399147 | 0,268360931 | terminal nucleotidyltransferase 5A |
| ENSG00000173193 | PARP14 | 1,089746087 | 2,128365742 | 0,00434862 | 0,062431658 | poly(ADP-ribose) polymerase family member 14 |
| ENSG00000242971 | RN7SL233P | 1,089457034 | 2,127939353 | 0,35067288 | 0,844035671 | RNA, 7SL, cytoplasmic 233, pseudogene |
| ENSG00000114626 | ABTB1 | 1,089094712 | 2,127405003 | 0,00783439 | 0,096279218 | ankyrin repeat and BTB domain containing 1 |
| ENSG00000232021 | LEF1-AS1 | 1,088888682 | 2,127101213 | 0,04521674 | 0,32172423 | LEF1 antisense RNA 1 |
| ENSG00000205786 | LINC01531 | 1,088447888 | 2,126451408 | 0,4378813 | 0,897156497 | long intergenic non-protein coding RNA 1531 |
| ENSG00000135406 | PRPH | 1,08755337 | 2,125133347 | 0,0904573 | 0,479775816 | peripherin |
| ENSG00000163032 | VSNL1 | 1,087063906 | 2,124412474 | 0,00813184 | 0,098882383 | visinin like 1 |
| ENSG00000184185 | KCNJ12 | 1,086359088 | 2,123374862 | 0,05681897 | 0,368087068 | potassium inwardly rectifying channel subfamily J member 12 |
| ENSG00000276500 | BMS1P14 | 1,085845342 | 2,122618859 | 0,08182761 | 0,454749316 | BMS1 pseudogene 14 |
| ENSG00000228818 | AL359918,1 | 1,08577205 | 2,122511029 | 0,02426314 | 0,214222733 | proteasome 26S non-ATPase subunit 2 (PSMD2) pseudogene |
| ENSG00000144230 | GPR17 | 1,085568106 | 2,122211005 | 0,14103942 | 0,598214217 | G protein-coupled receptor 17 |
| ENSG00000271983 | AC023302,1 | 1,085171263 | 2,121627327 | 0,27308654 | 0,785672647 | novel transcript |
| ENSG00000090659 | CD209 | 1,085055866 | 2,12145763 | 0,53959037 | 0,933350549 | CD209 molecule |
| ENSG00000255565 | AC073651,1 | 1,084363431 | 2,120439662 | 0,10705335 | 0,522384265 | novel transcript |
| ENSG00000269210 | AC019171,1 | 1,084305504 | 2,120354524 | 0,18589082 | 0,678777213 | novel transcript, antisense to ARHGEF33 |
| ENSG00000205220 | PSMB10 | 1,084133085 | 2,120101132 | 0,00645598 | 0,083370406 | proteasome 20S subunit beta 10 |
| ENSG00000229246 | LINC00377 | 1,083851525 | 2,119687409 | 0,25221062 | 0,76432863 | long intergenic non-protein coding RNA 377 |
| ENSG00000166535 | A2ML1 | 1,083467763 | 2,119123638 | 0,22314615 | 0,731704543 | alpha-2-macroglobulin like 1 |
| ENSG00000273319 | AC058791,1 | 1,083436151 | 2,119077205 | 0,01066032 | 0,120145713 | novel transcript |
| ENSG00000272549 | LINC02538 | 1,083418458 | 2,119051217 | 0,02139502 | 0,195525884 | long intergenic non-protein coding RNA 2538 |
| ENSG00000287108 | AC023302,2 | 1,083223771 | 2,118765277 | 0,24870744 | 0,760720365 | novel transcript |
| ENSG00000042980 | ADAM28 | 1,083040932 | 2,118496774 | 0,12238756 | 0,559485996 | ADAM metallopeptidase domain 28 |
| ENSG00000110090 | CPT1A | 1,082288747 | 2,117392531 | 0,00086797 | 0,016526335 | carnitine palmitoyltransferase 1A |
| ENSG00000183239 | AL109615,1 | 1,082226887 | 2,117301743 | 0,44949266 | 0,902062594 | ribosomal protein L29 (RPL29) pseudogene |
| ENSG00000254441 | EIF2S2P6 | 1,081775085 | 2,116638782 | 0,43298392 | 0,895777109 | eukaryotic translation initiation factor 2 subunit 2 beta pseudogene 6 |
| ENSG00000287693 | AC091691,3 | 1,081620041 | 2,116411322 | 0,09124366 | 0,481302729 | novel transcript |
| ENSG00000182631 | RXFP3 | 1,080850161 | 2,115282221 | 0,32069094 | 0,824366234 | relaxin family peptide receptor 3 |
| ENSG00000105472 | CLEC11A | 1,080790359 | 2,115194541 | 0,00587727 | 0,077671698 | C-type lectin domain containing 11A |
| ENSG00000169618 | PROKR1 | 1,080716139 | 2,115085727 | 0,09622071 | 0,492709242 | prokineticin receptor 1 |

| | | | | | | |
|---|---|---|---|---|---|---|
| ENSG00000171428 | NAT1 | 1,080508216 | 2,114780921 | 0,04692178 | 0,329207362 | N-acetyltransferase 1 |
| ENSG00000288384 | AC148477,16 | 1,0802563 | 2,11441168 | 0,38505853 | 0,868764521 | novel transcript |
| ENSG00000277011 | AC148477,4 | 1,0802563 | 2,11441168 | 0,38505853 | 0,868764521 | novel transcript |
| ENSG00000261468 | AC096921,2 | 1,080255389 | 2,114410345 | 0,09307018 | 0,484790551 | novel transcript |
| ENSG00000114779 | ABHD14B | 1,079933027 | 2,113937945 | 0,00012246 | 0,003127976 | abhydrolase domain containing 14B |
| ENSG00000251348 | HSPD1P11 | 1,079891629 | 2,113877286 | 0,0068194 | 0,087129511 | heat shock protein family D (Hsp60) member 1 pseudogene 11 |
| ENSG00000171444 | MCC | 1,079772291 | 2,113702437 | 0,00222966 | 0,036688402 | MCC regulator of WNT signaling pathway |
| ENSG00000164687 | FABP5 | 1,07859864 | 2,111983612 | 0,00413757 | 0,060306354 | fatty acid binding protein 5 |
| ENSG00000254835 | RNF185-AS1 | 1,078287442 | 2,111528093 | 0,23130197 | 0,739268552 | RNF185 antisense RNA 1 |
| ENSG00000128052 | KDR | 1,078265334 | 2,111495736 | 0,03849763 | 0,291183243 | kinase insert domain receptor |
| ENSG00000227477 | STK4-AS1 | 1,078136265 | 2,111306842 | 0,21581997 | 0,722999018 | STK4 antisense RNA 1 (head to head) |
| ENSG00000185070 | FLRT2 | 1,078101185 | 2,111255505 | 0,01718405 | 0,168970493 | fibronectin leucine rich transmembrane protein 2 |
| ENSG00000101928 | MOSPD1 | 1,078002442 | 2,111111009 | 0,04058102 | 0,300930985 | motile sperm domain containing 1 |
| ENSG00000131759 | RARA | 1,077991762 | 2,11109538 | 0,00223196 | 0,036693513 | retinoic acid receptor alpha |
| ENSG00000078808 | SDF4 | 1,077915683 | 2,110984057 | 0,01214956 | 0,131911723 | stromal cell derived factor 4 |
| ENSG00000135631 | RAB11FIP5 | 1,077889562 | 2,110945836 | 0,02097638 | 0,192913017 | RAB11 family interacting protein 5 |
| ENSG00000232952 | AL512844,1 | 1,07730747 | 2,110094294 | 0,2971381 | 0,806635068 | matrin 3 (MATR3) pseudogene |
| ENSG00000101608 | MYL12A | 1,076493838 | 2,108904606 | 0,00121392 | 0,021937506 | myosin light chain 12A |
| ENSG00000198417 | MT1F | 1,076160512 | 2,108417412 | 0,02234417 | 0,201956662 | metallothionein 1F |
| ENSG00000240793 | UBA52P8 | 1,076133149 | 2,108377424 | 0,26503244 | 0,776509969 | ubiquitin A-52 residue ribosomal protein fusion product 1 pseudogene 8 |
| ENSG00000189171 | S100A13 | 1,075687205 | 2,107725815 | 0,00553088 | 0,074508967 | S100 calcium binding protein A13 |
| ENSG00000282876 | Z98752,4 | 1,075527188 | 2,107492048 | 0,3144879 | 0,819877431 | novel transcript |
| ENSG00000204131 | NHSL2 | 1,075147922 | 2,106938089 | 0,00142785 | 0,025249098 | NHS like 2 |
| ENSG00000233476 | EEF1A1P6 | 1,075059918 | 2,10680957 | 0,01191733 | 0,130001219 | eukaryotic translation elongation factor 1 alpha 1 pseudogene 6 |
| ENSG00000258943 | AL049871,1 | 1,074835261 | 2,106481523 | 0,2786363 | 0,789775666 | novel transcript |
| ENSG00000132294 | EFR3A | 1,07458364 | 2,106114163 | 0,00354752 | 0,053424696 | EFR3 homolog A |
| ENSG00000272636 | DOC2B | 1,074527663 | 2,106032446 | 0,61922781 | 0,952793934 | double C2 domain beta |
| ENSG00000091106 | NLRC4 | 1,07444633 | 2,105913721 | 0,09400372 | 0,487240447 | NLR family CARD domain containing 4 |
| ENSG00000276858 | FCAR | 1,073319126 | 2,104268974 | 0,6094949 | 0,951793902 | Fc fragment of IgA receptor |
| ENSG00000263669 | RN7SL470P | 1,073075729 | 2,103913992 | 0,38797849 | 0,869820053 | RNA, 7SL, cytoplasmic 470, pseudogene |

| | | | | | | |
|---|---|---|---|---|---|---|
| ENSG00000224481 | AC245100,1 | 1,071955332 | 2,102280728 | 0,27253852 | 0,785492261 | novel transcript |
| ENSG00000110841 | PPFIBP1 | 1,071775952 | 2,102019353 | 2,09E-05 | 0,00065555 | PPFIA binding protein 1 |
| ENSG00000120251 | GRIA2 | 1,071604462 | 2,101769505 | 0,08831918 | 0,473946233 | glutamate ionotropic receptor AMPA type subunit 2 |
| ENSG00000186577 | SMIM29 | 1,071150283 | 2,101107946 | 0,00722115 | 0,090558361 | small integral membrane protein 29 |
| ENSG00000064218 | DMRT3 | 1,070766219 | 2,100548678 | 0,25805799 | 0,770501859 | doublesex and mab-3 related transcription factor 3 |
| ENSG00000107957 | SH3PXD2A | 1,070755423 | 2,100532959 | 0,00322763 | 0,049440738 | SH3 and PX domains 2A |
| ENSG00000225879 | AL513210,1 | 1,070569603 | 2,100262426 | 0,45936551 | 0,904862549 | novel transcript |
| ENSG00000261478 | AC013391,3 | 1,070078567 | 2,099547703 | 0,42727288 | 0,89251416 | novel transcript |
| ENSG00000140297 | GCNT3 | 1,069849538 | 2,099214424 | 0,19551077 | 0,693689284 | glucosaminyl (N-acetyl) transferase 3, mucin type |
| ENSG00000286549 | AP001631,2 | 1,069806694 | 2,099152084 | 0,33874066 | 0,837449623 | novel transcript, sense intronic to MX2 |
| ENSG00000285517 | AC010198,2 | 1,069641885 | 2,098912297 | 0,00349615 | 0,052778242 | long intergenic non-protein coding RNA 941 |
| ENSG00000259300 | TUBBP8 | 1,069106348 | 2,098133312 | 0,47697526 | 0,91143003 | tubulin beta class I pseudogene 8 |
| ENSG00000108669 | CYTH1 | 1,068918188 | 2,097859688 | 0,00072125 | 0,014155072 | cytohesin 1 |
| ENSG00000157782 | CABP1 | 1,068717291 | 2,097567578 | 0,06209462 | 0,389024096 | calcium binding protein 1 |
| ENSG00000104951 | IL4I1 | 1,068209485 | 2,096829397 | 0,28264901 | 0,793528967 | interleukin 4 induced 1 |
| ENSG00000223685 | LINC00571 | 1,068185121 | 2,096793987 | 0,09877256 | 0,498792641 | long intergenic non-protein coding RNA 571 |
| ENSG00000132429 | POPDC3 | 1,068142546 | 2,096732109 | 0,05481855 | 0,361697451 | popeye domain containing 3 |
| ENSG00000142185 | TRPM2 | 1,068053413 | 2,096602572 | 0,30414734 | 0,813056239 | transient receptor potential cation channel subfamily M member 2 |
| ENSG00000240755 | ERLEC1P1 | 1,06795123 | 2,09645408 | 0,55676985 | 0,938264532 | endoplasmic reticulum lectin 1 pseudogene 1 |
| ENSG00000176654 | NANOGP1 | 1,067826042 | 2,096272171 | 0,06595229 | 0,403272526 | Nanog homeobox pseudogene 1 |
| ENSG00000249992 | TMEM158 | 1,067728667 | 2,096130687 | 0,00450619 | 0,063949825 | transmembrane protein 158 |
| ENSG00000288084 | AL590608,1 | 1,067432282 | 2,095700105 | 0,12501678 | 0,565142382 | novel transcript |
| ENSG00000106714 | CNTNAP3 | 1,067248861 | 2,09543368 | 0,00028257 | 0,006410885 | contactin associated protein family member 3 |
| ENSG00000127415 | IDUA | 1,067143147 | 2,095280142 | 0,08550918 | 0,465660842 | alpha-L-iduronidase |
| ENSG00000171914 | TLN2 | 1,066998457 | 2,095070013 | 0,00010579 | 0,002751365 | talin 2 |
| ENSG00000150093 | ITGB1 | 1,06601103 | 2,09363657 | 6,65E-05 | 0,001829897 | integrin subunit beta 1 |
| ENSG00000223947 | AC016738,1 | 1,065928267 | 2,093516469 | 0,34441232 | 0,841161388 | novel transcript |
| ENSG00000273173 | SNURF | 1,065527877 | 2,092935537 | 0,08534427 | 0,465128991 | SNRPN upstream reading frame |
| ENSG00000175538 | KCNE3 | 1,064507144 | 2,09145527 | 0,01425956 | 0,148554089 | potassium voltage-gated channel subfamily E regulatory subunit 3 |
| ENSG00000242361 | HLA-DMA | 1,064326662 | 2,091193644 | 0,03177076 | 0,257126831 | major histocompatibility complex, class II, DM alpha |
| ENSG00000213659 | RSU1P3 | 1,064226298 | 2,091048172 | 0,10970853 | 0,528090728 | Ras suppressor protein 1 pseudogene 3 |

| | | | | | | |
|---|---|---|---|---|---|---|
| ENSG00000266473 | AC007448,4 | 1,063778359 | 2,090399028 | 0,19371868 | 0,690594523 | novel transcript |
| ENSG00000186815 | TPCN1 | 1,063459805 | 2,089937509 | 0,00157056 | 0,027309155 | two pore segment channel 1 |
| ENSG00000129932 | DOHH | 1,063200357 | 2,089561697 | 0,16350078 | 0,642091247 | deoxyhypusine hydroxylase |
| ENSG00000179820 | MYADM | 1,063009296 | 2,089284988 | 0,00667132 | 0,085630448 | myeloid associated differentiation marker |
| ENSG00000183337 | BCOR | 1,063002035 | 2,089274473 | 6,60E-06 | 0,000237157 | BCL6 corepressor |
| ENSG00000286984 | AC090578,3 | 1,06280258 | 2,088985646 | 0,19115787 | 0,686867719 | novel transcript |
| ENSG00000101160 | CTSZ | 1,062275595 | 2,088222725 | 0,00021225 | 0,005012017 | cathepsin Z |
| ENSG00000286882 | AC116428,1 | 1,062132722 | 2,088015935 | 0,34293556 | 0,840419133 | novel transcript, antisense to FER |
| ENSG00000149115 | TNKS1BP1 | 1,061960878 | 2,087767239 | 0,15321751 | 0,62169919 | tankyrase 1 binding protein 1 |
| ENSG00000239393 | AC146507,1 | 1,060697652 | 2,085939988 | 0,48558985 | 0,913941424 | ribosomal protein S20 (RPS20) pseudogene |
| ENSG00000243709 | LEFTY1 | 1,060393945 | 2,085500914 | 0,33630737 | 0,835660292 | left-right determination factor 1 |
| ENSG00000152518 | ZFP36L2 | 1,059991787 | 2,084919653 | 0,00023143 | 0,005407583 | ZFP36 ring finger protein like 2 |
| ENSG00000149177 | PTPRJ | 1,05898905 | 2,083471045 | 1,27E-05 | 0,000418742 | protein tyrosine phosphatase receptor type J |
| ENSG00000285132 | AC270285,3 | 1,058788626 | 2,083181622 | 0,00536586 | 0,072776773 | cathepsin B |
| ENSG00000236890 | AC016027,5 | 1,0584877 | 2,082747145 | 0,46312829 | 0,906879806 | ribosomal protein L18a (RPL18A) pseudogene |
| ENSG00000267136 | AP005131,1 | 1,058366787 | 2,082572596 | 0,01536794 | 0,156675544 | novel transcript, antisense C18orf1 |
| ENSG00000267571 | AC104532,2 | 1,058269739 | 2,082432509 | 0,24197102 | 0,752285407 | novel transcript, antisense to CAPS |
| ENSG00000285881 | AC105206,3 | 1,057676655 | 2,081576608 | 0,27458831 | 0,786383635 | Novel transcript, antisense to BMP1 |
| ENSG00000113916 | BCL6 | 1,057651914 | 2,081540911 | 0,01700269 | 0,167582635 | BCL6 transcription repressor |
| ENSG00000232913 | PLCE1-AS2 | 1,057432781 | 2,081224768 | 0,24723096 | 0,759350486 | PLCE1 antisense RNA 2 |
| ENSG00000179219 | LINC00311 | 1,057310129 | 2,081047838 | 0,1927537 | 0,689581368 | long intergenic non-protein coding RNA 311 |
| ENSG00000223932 | GPANK1 | 1,057211219 | 2,080905167 | 0,01949611 | 0,183714864 | G-patch domain and ankyrin repeats 1 |
| ENSG00000119599 | DCAF4 | 1,056683888 | 2,080144699 | 0,00121321 | 0,021937365 | DDB1 and CUL4 associated factor 4 |
| ENSG00000283959 | AP002851,1 | 1,056659918 | 2,080110138 | 0,14361953 | 0,602781713 | novel transcript |
| ENSG00000236345 | SCAT8 | 1,056176437 | 2,079413161 | 0,38050964 | 0,865900094 | S-phase cancer associated transcript 8 |
| ENSG00000136237 | RAPGEF5 | 1,05591028 | 2,079029573 | 0,0103631 | 0,118029873 | Rap guanine nucleotide exchange factor 5 |
| ENSG00000256056 | AC087257,2 | 1,055898026 | 2,079011914 | 0,16388043 | 0,642745357 | chromosome 12 open reading frame 71 (C12orf71) pseudogene |
| ENSG00000244502 | HDAC11-AS1 | 1,05586792 | 2,078968531 | 0,50224643 | 0,920520138 | HDAC11 antisense RNA 1 |
| ENSG00000242861 | AL591895,1 | 1,054917317 | 2,077599132 | 0,51359967 | 0,92410971 | novel transcript |
| ENSG00000165617 | DACT1 | 1,054856856 | 2,077512066 | 0,00432412 | 0,062241853 | dishevelled binding antagonist of beta catenin 1 |
| ENSG00000276266 | FP325318,1 | 1,054751801 | 2,07736079 | 0,02821919 | 0,237609688 | novel transcript |
| ENSG00000278716 | AC133540,1 | 1,054664423 | 2,077234977 | 0,56813867 | 0,94163744 | novel transcript |

| | | | | | | |
|---|---|---|---|---|---|---|
| ENSG00000120693 | SMAD9 | 1,054171822 | 2,076525837 | 0,02106888 | 0,193450047 | SMAD family member 9 |
| ENSG00000154319 | FAM167A | 1,053573321 | 2,075664569 | 0,00126324 | 0,022684575 | family with sequence similarity 167 member A |
| ENSG00000273076 | AL021707,7 | 1,053430312 | 2,075458826 | 0,54134741 | 0,933429141 | novel transcript, antisense to SUN2 |
| ENSG00000128335 | APOL2 | 1,053332792 | 2,07531854 | 0,00775838 | 0,095645933 | apolipoprotein L2 |
| ENSG00000281828 | HERC2P7 | 1,053012719 | 2,074858165 | 0,61818476 | 0,952793934 | hect domain and RLD 2 pseudogene 7 |
| ENSG00000288291 | LINC02361 | 1,052687065 | 2,074389868 | 0,34456167 | 0,841168721 | long intergenic non-protein coding RNA 2361 |
| ENSG00000256576 | LINC02361 | 1,052687065 | 2,074389868 | 0,34456167 | 0,841168721 | long intergenic non-protein coding RNA 2361 |
| ENSG00000283809 | AC007326,4 | 1,052270061 | 2,073790362 | 0,50838457 | 0,922427526 | novel protein |
| ENSG00000163932 | PRKCD | 1,052031587 | 2,073447599 | 0,00655264 | 0,0844443 | protein kinase C delta |
| ENSG00000104154 | SLC30A4 | 1,051291361 | 2,072384015 | 0,00139577 | 0,024808062 | solute carrier family 30 member 4 |
| ENSG00000109113 | RAB34 | 1,050997551 | 2,07196201 | 0,00106224 | 0,019709205 | RAB34, member RAS oncogene family |
| ENSG00000275441 | AC020765,2 | 1,050357556 | 2,071043069 | 0,37382182 | 0,861319894 | novel transcript, antisense to CCDC101 |
| ENSG00000233478 | AL031280,1 | 1,04973896 | 2,070155242 | 0,30156347 | 0,811179119 | novel transcript |
| ENSG00000276053 | LAIR1 | 1,049355918 | 2,06960568 | 0,20617995 | 0,709131999 | leukocyte associated immunoglobulin like receptor 1 |
| ENSG00000231995 | AL590399,4 | 1,049046056 | 2,069161217 | 0,26886201 | 0,781126457 | pseudogene similar to part of ribosomal protein L7a (RPL7A) |
| ENSG00000167705 | RILP | 1,048574112 | 2,06848445 | 0,02178804 | 0,198207677 | Rab interacting lysosomal protein |
| ENSG00000108556 | CHRNE | 1,048232848 | 2,067995216 | 0,10186535 | 0,50776757 | cholinergic receptor nicotinic epsilon subunit |
| ENSG00000249679 | AC106897,1 | 1,047962871 | 2,06760826 | 0,08464324 | 0,462904176 | novel transcript, antisense to CCDC110 |
| ENSG00000180347 | ITPRID1 | 1,047892075 | 2,0675068 | 0,41860626 | 0,887094957 | ITPR interacting domain containing 1 |
| ENSG00000204196 | RPL12P16 | 1,047205027 | 2,066522436 | 0,46782909 | 0,907940653 | ribosomal protein L12 pseudogene 16 |
| ENSG00000261801 | LOXL1-AS1 | 1,04713665 | 2,066424495 | 0,0004499 | 0,009549075 | LOXL1 antisense RNA 1 |
| ENSG00000144591 | GMPPA | 1,047065359 | 2,066322385 | 0,00021939 | 0,005157114 | GDP-mannose pyrophosphorylase A |
| ENSG00000227267 | AC072039,1 | 1,046797489 | 2,065938759 | 0,46065351 | 0,905949965 | downstream neighbor of SON (DONSON) pseudogene |
| ENSG00000078549 | ADCYAP1R1 | 1,046338806 | 2,06528203 | 0,06648819 | 0,405279626 | ADCYAP receptor type I |
| ENSG00000128394 | APOBEC3F | 1,046225949 | 2,065120476 | 0,01008407 | 0,115779813 | apolipoprotein B mRNA editing enzyme catalytic subunit 3F |
| ENSG00000247675 | LRP4-AS1 | 1,046148264 | 2,065009279 | 0,00391301 | 0,057760366 | LRP4 antisense RNA 1 |
| ENSG00000237257 | MTND2P8 | 1,045896722 | 2,064649265 | 0,28439733 | 0,795731744 | MT-ND2 pseudogene 8 |
| ENSG00000198034 | RPS4X | 1,045889259 | 2,064638583 | 0,00256718 | 0,040907452 | ribosomal protein S4 X-linked |
| ENSG00000163817 | SLC6A20 | 1,045744935 | 2,064432053 | 0,00878665 | 0,104076013 | solute carrier family 6 member 20 |

| | | | | | | |
|---|---|---|---|---|---|---|
| ENSG00000139304 | PTPRQ | 1,045670133 | 2,064325016 | 0,30320233 | 0,812669425 | protein tyrosine phosphatase receptor type Q |
| ENSG00000226572 | SNORD57 | 1,045332726 | 2,063842284 | 0,39253595 | 0,870909282 | small nucleolar RNA, C/D box 57 |
| ENSG00000243094 | AC079203,1 | 1,045105307 | 2,063516976 | 0,40819962 | 0,881221125 | ribosomal protein L32 (RPL32) pseudogene |
| ENSG00000259498 | TPM1-AS | 1,044915073 | 2,063244898 | 0,00473802 | 0,066158733 | TPM1 antisense RNA |
| ENSG00000223493 | SKIV2L | 1,044549545 | 2,062722211 | 0,29929283 | 0,809131439 | Ski2 like RNA helicase |
| ENSG00000101935 | AMMECR1 | 1,044497493 | 2,062647789 | 0,01306887 | 0,139163869 | AMMECR nuclear protein 1 |
| ENSG00000160888 | IER2 | 1,044468538 | 2,062606392 | 0,01257978 | 0,135431952 | immediate early response 2 |
| ENSG00000159197 | KCNE2 | 1,04409553 | 2,062073176 | 0,47704734 | 0,91143003 | potassium voltage-gated channel subfamily E regulatory subunit 2 |
| ENSG00000082397 | EPB41L3 | 1,043764804 | 2,061600517 | 0,01959003 | 0,184067141 | erythrocyte membrane protein band 4,1 like 3 |
| ENSG00000274965 | TRPM1 | 1,043496467 | 2,061217101 | 0,4967734 | 0,918457927 | transient receptor potential cation channel subfamily M member 1 |
| ENSG00000231977 | AL096828,1 | 1,043230346 | 2,060836921 | 0,08570816 | 0,466101036 | novel transcript |
| ENSG00000241211 | IQCJ-SCHIP1-AS | 1,042771141 | 2,060181069 | 0,35598692 | 0,847322051 | IQCJ-SCHIP1 readthrough antisense RNA 1 |
| ENSG00000126803 | HSPA2 | 1,04253274 | 2,059840659 | 0,00460525 | 0,065001531 | heat shock protein family A (Hsp70) member 2 |
| ENSG00000118985 | ELL2 | 1,042320604 | 2,059537799 | 0,00071997 | 0,014138946 | elongation factor for RNA polymerase II 2 |
| ENSG00000211829 | TRDC | 1,042249025 | 2,059435617 | 0,14303071 | 0,601880115 | T cell receptor delta constant |
| ENSG00000223396 | RPS10P7 | 1,042034429 | 2,059129306 | 0,09196007 | 0,482073274 | ribosomal protein S10 pseudogene 7 |
| ENSG00000064666 | CNN2 | 1,041228984 | 2,057980032 | 3,58E-05 | 0,001054604 | calponin 2 |
| ENSG00000175602 | CCDC85B | 1,041072618 | 2,05775699 | 0,09870831 | 0,498721099 | coiled-coil domain containing 85B |
| ENSG00000156515 | HK1 | 1,040942587 | 2,057571532 | 0,00086474 | 0,016494994 | hexokinase 1 |
| ENSG00000278071 | AL161669,3 | 1,040364195 | 2,056746795 | 0,27141879 | 0,784254811 | novel transcript |
| ENSG00000250644 | AC068580,4 | 1,040151229 | 2,056443206 | 0,01508382 | 0,154803784 | novel protein |
| ENSG00000280515 | SALRNA2 | 1,039767499 | 2,055896304 | 0,46172666 | 0,906178422 | senescence associated long non-coding RNA 2 |
| ENSG00000105894 | PTN | 1,039146737 | 2,055011884 | 0,0007933 | 0,015347569 | pleiotrophin |
| ENSG00000288541 | AL353732,2 | 1,038913621 | 2,054679855 | 0,30268143 | 0,812334076 | novel transcript |
| ENSG00000166126 | AMN | 1,038668296 | 2,054330494 | 0,19307229 | 0,689931655 | amnion associated transmembrane protein |
| ENSG00000133195 | SLC39A11 | 1,038519636 | 2,054118819 | 0,00098844 | 0,018504493 | solute carrier family 39 member 11 |
| ENSG00000087077 | TRIP6 | 1,03824262 | 2,053724439 | 0,01017556 | 0,11653537 | thyroid hormone receptor interactor 6 |
| ENSG00000123240 | OPTN | 1,038186137 | 2,053644036 | 0,0001648 | 0,004015881 | optineurin |
| ENSG00000092531 | SNAP23 | 1,038008067 | 2,053390573 | 0,00106153 | 0,019707591 | synaptosome associated protein 23 |
| ENSG00000065320 | NTN1 | 1,037561762 | 2,052755444 | 0,08289278 | 0,457872768 | netrin 1 |
| ENSG00000175793 | SFN | 1,037543331 | 2,05272922 | 0,33465645 | 0,834757602 | stratifin |

| | | | | | | |
|---|---|---|---|---|---|---|
| ENSG00000163564 | PYHIN1 | 1,037418516 | 2,052551636 | 0,29169149 | 0,802285739 | pyrin and HIN domain family member 1 |
| ENSG00000256995 | AC084816,1 | 1,03721284 | 2,052259037 | 0,37622012 | 0,86257156 | novel transcript |
| ENSG00000275909 | Metazoa_SRP | 1,036877609 | 2,05178222 | 0,52759802 | 0,929546857 | Metazoan signal recognition particle RNA |
| ENSG00000122359 | ANXA11 | 1,03672817 | 2,051569701 | 0,00147078 | 0,02583285 | annexin A11 |
| ENSG00000170421 | KRT8 | 1,035929774 | 2,050434665 | 0,03165127 | 0,256491777 | keratin 8 |
| ENSG00000115828 | QPCT | 1,035479038 | 2,049794154 | 0,03204747 | 0,258563162 | glutaminyl-peptide cyclotransferase |
| ENSG00000120690 | ELF1 | 1,03509201 | 2,049244335 | 0,00499498 | 0,06873368 | E74 like ETS transcription factor 1 |
| ENSG00000228097 | MTATP6P11 | 1,034447126 | 2,048328529 | 0,4164845 | 0,886262687 | MT-ATP6 pseudogene 11 |
| ENSG00000119227 | PIGZ | 1,034234242 | 2,048026299 | 0,05156672 | 0,347838733 | phosphatidylinositol glycan anchor biosynthesis class Z |
| ENSG00000181634 | TNFSF15 | 1,033848235 | 2,047478404 | 0,27784052 | 0,789162936 | TNF superfamily member 15 |
| ENSG00000282851 | BISPR | 1,033766719 | 2,04736272 | 0,09381555 | 0,486996636 | BST2 interferon stimulated positive regulator |
| ENSG00000265413 | AP001094,2 | 1,033631881 | 2,047171376 | 0,10942441 | 0,527649315 | novel transcript, sense intronic to PTPRM |
| ENSG00000251273 | LINC02228 | 1,033015505 | 2,046296932 | 0,52333056 | 0,927995032 | long intergenic non-protein coding RNA 2228 |
| ENSG00000276096 | Metazoa_SRP | 1,032841242 | 2,046049775 | 0,3994629 | 0,875433675 | Metazoan signal recognition particle RNA |
| ENSG00000140464 | PML | 1,032742833 | 2,045910214 | 0,00682841 | 0,08719714 | PML nuclear body scaffold |
| ENSG00000099715 | PCDH11Y | 1,032626654 | 2,045745466 | 0,03927998 | 0,294920606 | protocadherin 11 Y-linked |
| ENSG00000205054 | LINC01121 | 1,032283635 | 2,045259121 | 0,31102052 | 0,817751312 | long intergenic non-protein coding RNA 1121 |
| ENSG00000223957 | NEU1 | 1,032257264 | 2,045221737 | 0,00104552 | 0,01943341 | neuraminidase 1 |
| ENSG00000287454 | AC008149,2 | 1,032193166 | 2,04513087 | 0,07166252 | 0,421847873 | novel transcript, antisense to ELK3 |
| ENSG00000106397 | PLOD3 | 1,031980773 | 2,044829809 | 0,00637294 | 0,082502575 | procollagen-lysine,2-oxoglutarate 5-dioxygenase 3 |
| ENSG00000186439 | TRDN | 1,031955911 | 2,044794571 | 0,34485796 | 0,841168721 | triadin |
| ENSG00000147852 | VLDLR | 1,03054351 | 2,042793694 | 0,03262619 | 0,261468653 | very low density lipoprotein receptor |
| ENSG00000171517 | LPAR3 | 1,030539772 | 2,0427884 | 0,49740739 | 0,918759905 | lysophosphatidic acid receptor 3 |
| ENSG00000261312 | AC002550,1 | 1,030509093 | 2,04274496 | 0,37550888 | 0,862024808 | novel transcript, antisense to C16orf88 |
| ENSG00000187098 | MITF | 1,030490908 | 2,042719212 | 0,01600637 | 0,161013527 | melanocyte inducing transcription factor |
| ENSG00000129116 | PALLD | 1,030070531 | 2,042124085 | 1,77E-06 | 7,21E-05 | palladin, cytoskeletal associated protein |
| ENSG00000140575 | IQGAP1 | 1,030069925 | 2,042123228 | 5,11E-06 | 0,000188529 | IQ motif containing GTPase activating protein 1 |
| ENSG00000151491 | EPS8 | 1,029881897 | 2,041857093 | 0,00113896 | 0,020823628 | epidermal growth factor receptor pathway substrate 8 |
| ENSG00000242539 | AC007620,2 | 1,029480421 | 2,04128896 | 0,0908816 | 0,480927969 | novel transcript |
| ENSG00000125868 | DSTN | 1,029305706 | 2,041041767 | 0,00526134 | 0,071642267 | destrin, actin depolymerizing factor |
| ENSG00000269378 | ITGB1P1 | 1,029199624 | 2,040891695 | 0,00014755 | 0,003663932 | integrin subunit beta 1 pseudogene 1 |

| | | | | | | |
|---|---|---|---|---|---|---|
| ENSG00000263096 | AC007638,2 | 1,028853098 | 2,040401545 | 0,41574228 | 0,885986046 | novel transcript, antisense to HLF |
| ENSG00000204706 | MAMDC2-AS1 | 1,028503678 | 2,03990742 | 0,01316489 | 0,139782152 | MAMDC2 antisense RNA 1 |
| ENSG00000143641 | GALNT2 | 1,028481213 | 2,039875656 | 0,00281856 | 0,044125856 | polypeptide N-acetylgalactosaminyltransferase 2 |
| ENSG00000281371 | INE2 | 1,027769657 | 2,038869811 | 0,48607193 | 0,91402339 | inactivation escape 2 |
| ENSG00000231953 | AL031432,1 | 1,027443966 | 2,038409585 | 0,41412028 | 0,885579089 | novel transcript |
| ENSG00000125354 | SEPTIN6 | 1,027180643 | 2,038037564 | 0,00015694 | 0,003842736 | septin 6 |
| ENSG00000164877 | MICALL2 | 1,026938393 | 2,037695377 | 0,04995839 | 0,341697436 | MICAL like 2 |
| ENSG00000285815 | GET1-SH3BGR | 1,026834039 | 2,037547989 | 0,36704707 | 0,856406773 | GET1-SH3BGR readthrough |
| ENSG00000283448 | GPR1 | 1,026783912 | 2,037477196 | 0,03831251 | 0,290235613 | G protein-coupled receptor 1 |
| ENSG00000276725 | CEP170 | 1,026314458 | 2,036814307 | 0,13765784 | 0,592433314 | centrosomal protein 170 |
| ENSG00000228037 | AL139246,3 | 1,025904631 | 2,036235791 | 0,36912067 | 0,857942508 | novel transcript |
| ENSG00000248029 | AC034238,2 | 1,025899313 | 2,036228284 | 0,65696927 | 0,957596503 | novel transcript |
| ENSG00000141526 | SLC16A3 | 1,025673342 | 2,035909373 | 0,04962724 | 0,339981227 | solute carrier family 16 member 3 |
| ENSG00000227315 | NEU1 | 1,025073501 | 2,035063061 | 0,0251623 | 0,219987188 | neuraminidase 1 |
| ENSG00000261411 | AL161938,1 | 1,024883872 | 2,03479559 | 0,47956469 | 0,912721304 | novel transcript |
| ENSG00000140848 | CPNE2 | 1,024869679 | 2,034775571 | 0,01047376 | 0,118727922 | copine 2 |
| ENSG00000153822 | KCNJ16 | 1,024814898 | 2,03469831 | 0,23671223 | 0,746175744 | potassium inwardly rectifying channel subfamily J member 16 |
| ENSG00000084112 | SSH1 | 1,02465915 | 2,034478663 | 1,59E-05 | 0,000512165 | slingshot protein phosphatase 1 |
| ENSG00000205683 | DPF3 | 1,024649046 | 2,034464415 | 0,02550071 | 0,22195269 | double PHD fingers 3 |
| ENSG00000183911 | RPL21P132 | 1,023830604 | 2,033310589 | 0,44197593 | 0,899273096 | ribosomal protein L21 pseudogene 132 |
| ENSG00000258256 | AC011773,3 | 1,023821219 | 2,033297362 | 0,26729323 | 0,779594822 | novel transcript, antisense to CA13 |
| ENSG00000125931 | CITED1 | 1,023341869 | 2,03262189 | 0,01506446 | 0,154671635 | Cbp/p300 interacting transactivator with Glu/Asp rich carboxy-terminal domain 1 |
| ENSG00000119121 | TRPM6 | 1,023209323 | 2,032435153 | 0,18926552 | 0,684385512 | transient receptor potential cation channel subfamily M member 6 |
| ENSG00000172716 | SLFN11 | 1,02275855 | 2,031800214 | 0,00200869 | 0,033752576 | schlafen family member 11 |
| ENSG00000204231 | RXRB | 1,022148797 | 2,030941659 | 0,01459344 | 0,151125944 | retinoid X receptor beta |
| ENSG00000134339 | SAA2 | 1,02200892 | 2,030744757 | 0,40301788 | 0,877589651 | serum amyloid A2 |
| ENSG00000184494 | NEU1 | 1,021906532 | 2,030600641 | 0,00116762 | 0,021248274 | neuraminidase 1 |
| ENSG00000243038 | AL117692,1 | 1,021703543 | 2,030314953 | 0,21950776 | 0,727838473 | ribosomal protein S15a (RPS15A) pseudogene |
| ENSG00000264438 | RN7SL560P | 1,021601086 | 2,030170769 | 0,45885035 | 0,90450802 | RNA, 7SL, cytoplasmic 560, pseudogene |
| ENSG00000266964 | FXYD1 | 1,021349626 | 2,029816945 | 0,07631182 | 0,436562264 | FXYD domain containing ion transport regulator 1 |

| | | | | | | |
|---|---|---|---|---|---|---|
| ENSG00000236813 | BTF3P8 | 1,021249171 | 2,029675613 | 0,48098567 | 0,912907448 | basic transcription factor 3 pseudogene 8 |
| ENSG00000238290 | AL034417,2 | 1,021118802 | 2,02949221 | 0,31379576 | 0,819877431 | novel transcript |
| ENSG00000237928 | NFIA-AS2 | 1,020756685 | 2,02898287 | 0,01365617 | 0,143656275 | NFIA antisense RNA 2 |
| ENSG00000156709 | AIFM1 | 1,020637464 | 2,028815206 | 5,78E-07 | 2,57E-05 | apoptosis inducing factor mitochondria associated 1 |
| ENSG00000167588 | GPD1 | 1,020483714 | 2,028599004 | 0,31444271 | 0,819877431 | glycerol-3-phosphate dehydrogenase 1 |
| ENSG00000286781 | AC099332,1 | 1,020248509 | 2,028268305 | 0,13370579 | 0,583882708 | novel transcript |
| ENSG00000240184 | PCDHGC3 | 1,020133866 | 2,028107136 | 0,00069408 | 0,013690664 | protocadherin gamma subfamily C, 3 |
| ENSG00000230102 | LINC02028 | 1,020062265 | 2,028006484 | 0,15441033 | 0,623607455 | long intergenic non-protein coding RNA 2028 |
| ENSG00000196184 | OR10J1 | 1,019785121 | 2,027616938 | 0,42519127 | 0,891853867 | olfactory receptor family 10 subfamily J member 1 |
| ENSG00000041353 | RAB27B | 1,019637387 | 2,027409318 | 0,03372764 | 0,266952134 | RAB27B, member RAS oncogene family |
| ENSG00000142347 | MYO1F | 1,019601705 | 2,027359175 | 0,12753176 | 0,570260369 | myosin IF |
| ENSG00000233471 | KRT18P62 | 1,01943069 | 2,027118869 | 0,2597203 | 0,772220217 | keratin 18 pseudogene 62 |
| ENSG00000259415 | AC012291,2 | 1,019290627 | 2,026922077 | 0,45561131 | 0,903545728 | novel transcript |
| ENSG00000264472 | AC096708,2 | 1,019057089 | 2,026593993 | 0,27073963 | 0,783073289 | novel transcript, antisense to CCDC102B |
| ENSG00000276386 | CNTNAP3P2 | 1,019044325 | 2,026576064 | 0,00256537 | 0,040899377 | CNTNAP3 pseudogene 2 |
| ENSG00000258565 | BLZF2P | 1,018975469 | 2,026479343 | 0,28013952 | 0,790779983 | basic leucine zipper nuclear factor 2, pseudogene |
| ENSG00000236445 | LINC00608 | 1,018727164 | 2,026130592 | 0,33396016 | 0,834406649 | long intergenic non-protein coding RNA 608 |
| ENSG00000276823 | Metazoa_SRP | 1,017787417 | 2,024811234 | 0,34773838 | 0,842293776 | Metazoan signal recognition particle RNA |
| ENSG00000262966 | AC005695,1 | 1,017628339 | 2,024587982 | 0,06255821 | 0,390552401 | novel transcript, antisense to NTN1 |
| ENSG00000255513 | AC005363,1 | 1,017254332 | 2,024063192 | 0,459229 | 0,904855066 | ribosomal protein L26 (RPL26) pseudogene |
| ENSG00000230389 | HCP5 | 1,017097073 | 2,023842573 | 0,56031248 | 0,940070839 | HLA complex P5 |
| ENSG00000287252 | AL009179,1 | 1,017038507 | 2,023760417 | 0,14179693 | 0,599761485 | novel transcript |
| ENSG00000276438 | PPP2R3B | 1,01649695 | 2,023000883 | 0,41715469 | 0,886651836 | protein phosphatase 2 regulatory subunit B''beta |
| ENSG00000150893 | FREM2 | 1,016050003 | 2,022374255 | 0,02485794 | 0,218487143 | FRAS1 related extracellular matrix 2 |
| ENSG00000269883 | AC007375,3 | 1,015803956 | 2,022029375 | 0,23113874 | 0,739268552 | novel transcript, antisense to ZDHHC22 |
| ENSG00000123575 | FAM199X | 1,015670601 | 2,021842479 | 0,01969203 | 0,184658561 | family with sequence similarity 199, X-linked |
| ENSG00000257956 | NOP56P3 | 1,01520146 | 2,021185114 | 0,39652249 | 0,873142625 | NOP56 ribonucleoprotein pseudogene 3 |
| ENSG00000233485 | FHAD1-AS1 | 1,015155692 | 2,021120996 | 0,37179039 | 0,859864971 | FHAD1 antisense RNA 1 |
| ENSG00000145934 | TENM2 | 1,014583335 | 2,020319321 | 0,01367464 | 0,143802252 | teneurin transmembrane protein 2 |
| ENSG00000259639 | AC021351,1 | 1,01448674 | 2,020184055 | 0,41368317 | 0,885121175 | novel transcript |
| ENSG00000287910 | AC104024,4 | 1,014328424 | 2,01996238 | 0,39111288 | 0,870057277 | novel transcript |
| ENSG00000231105 | ECE1-AS1 | 1,014271224 | 2,019882293 | 0,18124487 | 0,670981593 | ECE1 antisense RNA 1 |
| ENSG00000287477 | AL356495,1 | 1,014021777 | 2,019533079 | 0,01444405 | 0,150076058 | novel transcript, antisense to TRPC4 |

| | | | | | | |
|---|---|---|---|---|---|---|
| ENSG00000177752 | YIPF7 | 1,013886611 | 2,019343878 | 0,37098267 | 0,858959913 | Yip1 domain family member 7 |
| ENSG00000198312 | BMS1P9 | 1,013111837 | 2,018259716 | 0,02046792 | 0,189715124 | BMS1 pseudogene 9 |
| ENSG00000151623 | NR3C2 | 1,012719675 | 2,017711175 | 0,04722434 | 0,33044357 | nuclear receptor subfamily 3 group C member 2 |
| ENSG00000175600 | SUGCT | 1,012531913 | 2,017448594 | 0,00455572 | 0,064447972 | succinyl-CoA:glutarate-CoA transferase |
| ENSG00000119535 | CSF3R | 1,012341998 | 2,017183037 | 0,38288279 | 0,867549469 | colony stimulating factor 3 receptor |
| ENSG00000229981 | LINC01435 | 1,012074662 | 2,016809281 | 0,28054759 | 0,791456745 | long intergenic non-protein coding RNA 1435 |
| ENSG00000229172 | AC073065,1 | 1,011843527 | 2,016486193 | 0,32086062 | 0,824366234 | ATP-binding cassette, sub-family G (WHITE), member 2 (ABCG2) pseudogene |
| ENSG00000204603 | LINC01257 | 1,011794534 | 2,016417715 | 0,39090319 | 0,870057277 | long intergenic non-protein coding RNA 1257 |
| ENSG00000137225 | CAPN11 | 1,011740244 | 2,016341836 | 0,27019932 | 0,782492524 | calpain 11 |
| ENSG00000076043 | REXO2 | 1,011650409 | 2,016216286 | 5,71E-05 | 0,00160022 | RNA exonuclease 2 |
| ENSG00000168918 | INPP5D | 1,011494214 | 2,01599801 | 0,34626276 | 0,841168721 | inositol polyphosphate-5-phosphatase D |
| ENSG00000068366 | ACSL4 | 1,01130676 | 2,015736081 | 0,0417669 | 0,306597901 | acyl-CoA synthetase long chain family member 4 |
| ENSG00000182021 | AL591379,1 | 1,011282272 | 2,015701867 | 0,01654045 | 0,164530876 | methylenetetrahydrofolate dehydrogenase (NADP+ dependent) 1 like pseudogene |
| ENSG00000174652 | ZNF266 | 1,010842172 | 2,015087062 | 0,00036924 | 0,008044632 | zinc finger protein 266 |
| ENSG00000175772 | LINC01106 | 1,010683475 | 2,014865413 | 0,17720374 | 0,663325859 | long intergenic non-protein coding RNA 1106 |
| ENSG00000141469 | SLC14A1 | 1,010551442 | 2,014681025 | 0,25558173 | 0,767141836 | solute carrier family 14 member 1 (Kidd blood group) |
| ENSG00000168961 | LGALS9 | 1,010398426 | 2,014467355 | 0,09216203 | 0,482558005 | galectin 9 |
| ENSG00000106560 | GIMAP2 | 1,010375783 | 2,014435737 | 0,23359058 | 0,742832118 | GTPase, IMAP family member 2 |
| ENSG00000186710 | CFAP73 | 1,010072377 | 2,014012137 | 0,39527197 | 0,871981986 | cilia and flagella associated protein 73 |
| ENSG00000287666 | AP000821,2 | 1,009877498 | 2,013740102 | 0,35099112 | 0,844035671 | novel transcript |
| ENSG00000265170 | RN7SL667P | 1,009855341 | 2,013709176 | 0,51397615 | 0,924227101 | RNA, 7SL, cytoplasmic 667, pseudogene |
| ENSG00000101194 | SLC17A9 | 1,009693753 | 2,013483643 | 0,21766623 | 0,72540192 | solute carrier family 17 member 9 |
| ENSG00000226850 | ATP6V1G2 | 1,009636302 | 2,013403465 | 0,48481274 | 0,913690171 | ATPase H+ transporting V1 subunit G2 |
| ENSG00000234920 | ATP6V1G2 | 1,009636302 | 2,013403465 | 0,48481274 | 0,913690171 | ATPase H+ transporting V1 subunit G2 |
| ENSG00000206445 | ATP6V1G2 | 1,009636302 | 2,013403465 | 0,48481274 | 0,913690171 | ATPase H+ transporting V1 subunit G2 |
| ENSG00000230900 | ATP6V1G2 | 1,009636302 | 2,013403465 | 0,48481274 | 0,913690171 | ATPase H+ transporting V1 subunit G2 |
| ENSG00000234668 | ATP6V1G2 | 1,009636302 | 2,013403465 | 0,48481274 | 0,913690171 | ATPase H+ transporting V1 subunit G2 |
| ENSG00000125462 | MIR9-1HG | 1,009387773 | 2,013056652 | 0,11549516 | 0,543019233 | MIR9-1 host gene |
| ENSG00000021762 | OSBPL5 | 1,00933129 | 2,012977839 | 0,02339219 | 0,208565361 | oxysterol binding protein like 5 |
| ENSG00000232280 | FLOT1 | 1,008467778 | 2,011773351 | 0,00058211 | 0,011882718 | flotillin 1 |

| | | | | | | |
|---|---|---|---|---|---|---|
| ENSG00000258001 | AC126614,1 | 1,008361927 | 2,011625752 | 0,51541926 | 0,924901769 | novel transcript |
| ENSG00000286367 | AC000072,1 | 1,008267229 | 2,011493713 | 0,39403848 | 0,871635019 | novel transcript, antisense to CLTCL1 |
| ENSG00000232599 | AL008707,1 | 1,008174338 | 2,011364203 | 0,10826056 | 0,524537831 | odz, odd Oz\/ten-m homolog 1(Drosophila) (ODZ1) pseudogene |
| ENSG00000128692 | EIF2S2P4 | 1,008103619 | 2,011265611 | 0,28928925 | 0,800260154 | eukaryotic translation initiation factor 2 subunit 2 beta pseudogene 4 |
| ENSG00000100342 | APOL1 | 1,007670605 | 2,010662035 | 0,16743584 | 0,648316613 | apolipoprotein L1 |
| ENSG00000287797 | AC093577,1 | 1,007542305 | 2,010483234 | 0,35717817 | 0,848693232 | novel transcript |
| ENSG00000161791 | FMNL3 | 1,007029063 | 2,009768127 | 5,09E-05 | 0,001440283 | formin like 3 |
| ENSG00000226772 | AL021393,1 | 1,006696214 | 2,0093045 | 0,24184135 | 0,752128472 | novel transcript |
| ENSG00000206480 | FLOT1 | 1,006430455 | 2,0089344 | 0,00772688 | 0,095370355 | flotillin 1 |
| ENSG00000206379 | FLOT1 | 1,006430455 | 2,0089344 | 0,00772688 | 0,095370355 | flotillin 1 |
| ENSG00000274346 | AC016999,2 | 1,006296575 | 2,008747983 | 0,35969363 | 0,850125614 | novel zinc finger protein pseudogene |
| ENSG00000260573 | AC007493,1 | 1,005778372 | 2,008026589 | 0,26241556 | 0,774068771 | novel transcript |
| ENSG00000182511 | FES | 1,005761413 | 2,008002984 | 0,05492715 | 0,362021772 | FES proto-oncogene, tyrosine kinase |
| ENSG00000242020 | RN7SL68P | 1,005547277 | 2,007704963 | 0,34521886 | 0,841168721 | RNA, 7SL, cytoplasmic 68, pseudogene |
| ENSG00000123243 | ITIH5 | 1,00554213 | 2,0076978 | 0,07186014 | 0,422642955 | inter-alpha-trypsin inhibitor heavy chain 5 |
| ENSG00000253894 | AC011124,2 | 1,005392874 | 2,007490101 | 0,45263994 | 0,902285042 | novel transcript |
| ENSG00000265073 | AC010761,2 | 1,005248015 | 2,007288542 | 0,3160889 | 0,821394376 | novel transcript, antisense to NEK8 |
| ENSG00000258479 | LINC00640 | 1,00498949 | 2,006928877 | 0,5644365 | 0,940865625 | long intergenic non-protein coding RNA 640 |
| ENSG00000006534 | ALDH3B1 | 1,004979354 | 2,006914776 | 0,11453864 | 0,540681869 | aldehyde dehydrogenase 3 family member B1 |
| ENSG00000262712 | AC012676,1 | 1,004754885 | 2,006602545 | 0,20348293 | 0,703726811 | novel transcript |
| ENSG00000205959 | AC105345,1 | 1,004547181 | 2,006313676 | 0,14740574 | 0,611530401 | novel transcript |
| ENSG00000246350 | AL049543,1 | 1,003761452 | 2,005221283 | 0,47213704 | 0,909171312 | novel transcript |
| ENSG00000224821 | COL4A2-AS2 | 1,003753572 | 2,005210331 | 0,15765171 | 0,630235931 | COL4A2 antisense 2 |
| ENSG00000184988 | TMEM106A | 1,003246132 | 2,004505161 | 0,02934445 | 0,243477915 | transmembrane protein 106A |
| ENSG00000260641 | AC114811,2 | 1,003085558 | 2,004282069 | 0,0215315 | 0,196543348 | novel transcript, antisense TSPAN5 |
| ENSG00000283239 | AC019257,8 | 1,00224125 | 2,003109447 | 0,08665671 | 0,468969234 | novel transcript |
| ENSG00000225472 | AL136366,1 | 1,00222036 | 2,003080442 | 0,14599653 | 0,608997405 | novel transcript |
| ENSG00000120051 | CFAP58 | 1,001466046 | 2,002033405 | 0,04523513 | 0,32178181 | cilia and flagella associated protein 58 |
| ENSG00000284898 | AL136419,1 | 1,001117685 | 2,001550041 | 0,21137083 | 0,717057897 | novel transcript |
| ENSG00000285467 | AL136419,1 | 1,001117685 | 2,001550041 | 0,21137083 | 0,717057897 | novel transcript |
| ENSG00000088256 | GNA11 | 1,000752229 | 2,001043083 | 0,01670554 | 0,165411048 | G protein subunit alpha 11 |

| | | | | | | |
|---|---|---|---|---|---|---|
| ENSG00000182916 | TCEAL7 | 1,000696642 | 2,000965984 | 0,03510616 | 0,273637274 | transcription elongation factor A like 7 |
| ENSG00000260941 | LINC00622 | 1,000479006 | 2,000664154 | 0,10549415 | 0,518254119 | long intergenic non-protein coding RNA 622 |
| ENSG00000104814 | MAP4K1 | 1,000468209 | 2,000649181 | 0,09828775 | 0,49771109 | mitogen-activated protein kinase kinase kinase kinase 1 |
| ENSG00000250682 | LINC00491 | 1,000364201 | 2,000504954 | 0,00045207 | 0,009575492 | long intergenic non-protein coding RNA 491 |
| ENSG00000149499 | EML3 | 1,000009819 | 2,000013612 | 0,02606911 | 0,22520713 | EMAP like 3 |
| ENSG00000120738 | EGR1 | 0,999946053 | 1,999925215 | 0,00224737 | 0,036863452 | early growth response 1 |
| ENSG00000237844 | AC016766,1 | 0,999838919 | 1,999776706 | 0,49465216 | 0,91758518 | novel transcript |
| ENSG00000287257 | AC111182,2 | 0,999364638 | 1,999119395 | 0,29612638 | 0,805962851 | novel transcript, antisense to SEPT9 |
| ENSG00000285602 | AC097625,1 | 0,998972352 | 1,998575885 | 0,42271302 | 0,890403187 | novel protein |
| ENSG00000100336 | APOL4 | 0,998906618 | 1,998484825 | 0,01632261 | 0,163246983 | apolipoprotein L4 |
| ENSG00000168032 | ENTPD3 | 0,998774303 | 1,998301544 | 0,14743427 | 0,611567589 | ectonucleoside triphosphate diphosphohydrolase 3 |
| ENSG00000175220 | ARHGAP1 | 0,997531834 | 1,996581321 | 0,00328888 | 0,050255868 | Rho GTPase activating protein 1 |
| ENSG00000138678 | GPAT3 | 0,996895317 | 1,995700623 | 0,57150759 | 0,942436428 | glycerol-3-phosphate acyltransferase 3 |
| ENSG00000183715 | OPCML | 0,996271926 | 1,994838464 | 0,06714372 | 0,40729169 | opioid binding protein/cell adhesion molecule like |
| ENSG00000146376 | ARHGAP18 | 0,996227747 | 1,994777377 | 0,00201843 | 0,033817896 | Rho GTPase activating protein 18 |
| ENSG00000090615 | GOLGA3 | 0,995986889 | 1,994444378 | 0,05203774 | 0,349872693 | golgin A3 |
| ENSG00000276110 | AC242988,1 | 0,995667978 | 1,994003549 | 0,351494 | 0,84405766 | novel transcript, antisense to CA14 |
| ENSG00000124782 | RREB1 | 0,995626595 | 1,993946353 | 0,00024401 | 0,005642477 | ras responsive element binding protein 1 |
| ENSG00000120280 | CXorf21 | 0,99542902 | 1,993673303 | 0,54048721 | 0,933385624 | chromosome X open reading frame 21 |
| ENSG00000288586 | AL357874,3 | 0,995133002 | 1,993264276 | 0,12764649 | 0,570513207 | novel transcript |
| ENSG00000260167 | AC093249,2 | 0,995006608 | 1,993089655 | 0,27305182 | 0,785672647 | novel transcript, antisense to ZNF689 |
| ENSG00000015568 | RGPD5 | 0,993898598 | 1,991559522 | 0,04659438 | 0,327821686 | RANBP2 like and GRIP domain containing 5 |
| ENSG00000277594 | DDX52 | 0,993369873 | 1,99082978 | 0,0205039 | 0,189935954 | DExD-box helicase 52 |
| ENSG00000285695 | AC016383,2 | 0,993291539 | 1,990721687 | 0,29653033 | 0,806142623 | novel transcript |
| ENSG00000228380 | MYL6BP1 | 0,992992696 | 1,990309367 | 0,4330468 | 0,895777109 | myosin light chain 6B pseudogene 1 |
| ENSG00000274441 | AC008957,2 | 0,992783869 | 1,990021296 | 0,08883917 | 0,475558385 | novel transcript, antisense SLC1A3 |
| ENSG00000073060 | SCARB1 | 0,99257868 | 1,989738282 | 0,00046561 | 0,009809192 | scavenger receptor class B member 1 |
| ENSG00000176046 | NUPR1 | 0,992440412 | 1,989547595 | 0,2277488 | 0,736053199 | nuclear protein 1, transcriptional regulator |
| ENSG00000287490 | AC015818,9 | 0,991929729 | 1,988843462 | 0,39697934 | 0,873609704 | novel transcript |
| ENSG00000102181 | CD99L2 | 0,991265053 | 1,987927376 | 9,75E-06 | 0,000334115 | CD99 molecule like 2 |
| ENSG00000236342 | ABCF1 | 0,990870294 | 1,987383502 | 0,17564112 | 0,661199837 | ATP binding cassette subfamily F member 1 |

| | | | | | | |
|---|---|---|---|---|---|---|
| ENSG00000177519 | RPRM | 0,990718057 | 1,987173799 | 0,00025409 | 0,00584535 | reprimo, TP53 dependent G2 arrest mediator homolog |
| ENSG00000267633 | AC008686,1 | 0,990617719 | 1,987035598 | 0,60553705 | 0,950927937 | novel transcript, antisense to MRI1 |
| ENSG00000175352 | NRIP3 | 0,990384626 | 1,986714583 | 0,00739688 | 0,092339375 | nuclear receptor interacting protein 3 |
| ENSG00000256232 | LINC02387 | 0,990071253 | 1,986283089 | 0,32684918 | 0,82862575 | long intergenic non-protein coding RNA 2387 |
| ENSG00000235174 | RPL39P3 | 0,990070825 | 1,986282499 | 0,64196365 | 0,956548736 | ribosomal protein L39 pseudogene 3 |
| ENSG00000189149 | CRYM-AS1 | 0,989895302 | 1,986040856 | 0,24261779 | 0,752798972 | CRYM antisense RNA 1 |
| ENSG00000172927 | MYEOV | 0,98965857 | 1,985714993 | 0,33054159 | 0,832114797 | myeloma overexpressed |
| ENSG00000268366 | AC010271,1 | 0,989558709 | 1,98557755 | 0,4435907 | 0,900079149 | novel transcript, antisense to SERTAD3 |
| ENSG00000205414 | AC007608,1 | 0,989525636 | 1,985532032 | 0,07385497 | 0,429465432 | novel transcript, antisense to NKD1 |
| ENSG00000157637 | SLC38A10 | 0,989370245 | 1,985318185 | 0,11629909 | 0,544389084 | solute carrier family 38 member 10 |
| ENSG00000128564 | VGF | 0,988800403 | 1,984534171 | 0,15598694 | 0,626944439 | VGF nerve growth factor inducible |
| ENSG00000234409 | CCDC188 | 0,988645863 | 1,9843216 | 0,21734287 | 0,725037995 | coiled-coil domain containing 188 |
| ENSG00000131459 | GFPT2 | 0,988484769 | 1,98410004 | 0,00200294 | 0,033702909 | glutamine-fructose-6-phosphate transaminase 2 |
| ENSG00000187808 | SOWAHD | 0,988242935 | 1,983767479 | 0,29238601 | 0,802771257 | sosondowah ankyrin repeat domain family member D |
| ENSG00000060138 | YBX3 | 0,988109962 | 1,983584644 | 0,00386993 | 0,057259514 | Y-box binding protein 3 |
| ENSG00000183055 | FAM133CP | 0,987977339 | 1,983402306 | 0,31726513 | 0,822252078 | family with sequence similarity 133 member C, pseudogene |
| ENSG00000265474 | AC010761,4 | 0,987782172 | 1,983134012 | 0,05304594 | 0,354145183 | novel transcript, antisense to TRAF4 |
| ENSG00000242893 | RN7SL413P | 0,987718142 | 1,983045998 | 0,36834361 | 0,857548638 | RNA, 7SL, cytoplasmic 413, pseudogene |
| ENSG00000006638 | TBXA2R | 0,987611405 | 1,982899288 | 0,07541761 | 0,433690715 | thromboxane A2 receptor |
| ENSG00000133392 | MYH11 | 0,98753654 | 1,982796393 | 0,19953012 | 0,698904803 | myosin heavy chain 11 |
| ENSG00000228430 | AL162726,3 | 0,987443362 | 1,982668337 | 0,49087316 | 0,916995341 | serine palmitoyltransferase long chain base subunit 1 (SPTLC1) pseudogene |
| ENSG00000287636 | AC005229,5 | 0,987104941 | 1,982203306 | 0,34379694 | 0,840901506 | novel transcript, antisense to C7orf33 |
| ENSG00000182310 | SPACA6 | 0,987095946 | 1,982190948 | 0,08958254 | 0,477709449 | sperm acrosome associated 6 |
| ENSG00000235568 | NFAM1 | 0,986768586 | 1,981741222 | 0,18755866 | 0,681556081 | NFAT activating protein with ITAM motif 1 |
| ENSG00000163710 | PCOLCE2 | 0,98638251 | 1,981210964 | 0,03746098 | 0,285862844 | procollagen C-endopeptidase enhancer 2 |
| ENSG00000282873 | BCL7A | 0,986209033 | 1,980972747 | 0,49465885 | 0,91758518 | BAF chromatin remodeling complex subunit BCL7A |
| ENSG00000274772 | Metazoa_SRP | 0,985896266 | 1,980543332 | 0,35650517 | 0,847845555 | Metazoan signal recognition particle RNA |
| ENSG00000225652 | KCNMA1-AS3 | 0,985715509 | 1,980295203 | 0,48640069 | 0,914091794 | KCNMA1 antisense RNA 3 |

| | | | | | | |
|---|---|---|---|---|---|---|
| ENSG00000183148 | ANKRD20A2P | 0,985168158 | 1,979544031 | 0,13589331 | 0,587841041 | ankyrin repeat domain 20 family member A2, pseudogene |
| ENSG00000183971 | NPW | 0,98479416 | 1,979030929 | 0,5162689 | 0,925176589 | neuropeptide W |
| ENSG00000182752 | PAPPA | 0,98420679 | 1,978225363 | 0,04615713 | 0,325893531 | pappalysin 1 |
| ENSG00000170004 | CHD3 | 0,98410236 | 1,978082174 | 0,04277767 | 0,311526734 | chromodomain helicase DNA binding protein 3 |
| ENSG00000238186 | AL603839,2 | 0,98375909 | 1,977611571 | 0,39143874 | 0,870284501 | novel transcript |
| ENSG00000254319 | AC246817,2 | 0,983592593 | 1,977383355 | 0,00772067 | 0,09536899 | novel transcript |
| ENSG00000092841 | MYL6 | 0,98354091 | 1,977312518 | 0,00056724 | 0,011606083 | myosin light chain 6 |
| ENSG00000100678 | SLC8A3 | 0,983353099 | 1,977055127 | 0,03240966 | 0,260687565 | solute carrier family 8 member A3 |
| ENSG00000274162 | SNX18P1Y | 0,983264778 | 1,976934097 | 0,36650732 | 0,856103393 | sorting nexin 18 pseudogene 1, Y-linked |
| ENSG00000272255 | AC113361,1 | 0,982697059 | 1,976156301 | 0,24248542 | 0,752612359 | novel transcript |
| ENSG00000227064 | Z82205,1 | 0,982658989 | 1,976104155 | 0,45478145 | 0,903424009 | N-acetyltransferase 5 (ARD1 homolog, S, cerevisiae) (NAT5) pseudogene |
| ENSG00000163346 | PBXIP1 | 0,982411395 | 1,975765046 | 0,06286862 | 0,391611013 | PBX homeobox interacting protein 1 |
| ENSG00000187823 | RTL4 | 0,982333991 | 1,975659045 | 0,27655924 | 0,788324941 | retrotransposon Gag like 4 |
| ENSG00000233208 | LINC00642 | 0,982263347 | 1,975562306 | 0,44536424 | 0,901068127 | long intergenic non-protein coding RNA 642 |
| ENSG00000185989 | RASA3 | 0,982142556 | 1,975396907 | 0,00117651 | 0,02136371 | RAS p21 protein activator 3 |
| ENSG00000280477 | RASA3 | 0,982142556 | 1,975396907 | 0,00117651 | 0,02136371 | RAS p21 protein activator 3 |
| ENSG00000262454 | MIR193BHG | 0,981911694 | 1,975080827 | 0,10253142 | 0,509949455 | MIR193B host gene |
| ENSG00000102575 | ACP5 | 0,981056402 | 1,97391026 | 0,24701759 | 0,759068012 | acid phosphatase 5, tartrate resistant |
| ENSG00000050030 | NEXMIF | 0,980745893 | 1,973485464 | 0,00941212 | 0,109821064 | neurite extension and migration factor |
| ENSG00000267360 | AC012309,1 | 0,980372038 | 1,972974128 | 0,28067005 | 0,791510309 | novel protein |
| ENSG00000104324 | CPQ | 0,980263981 | 1,972826359 | 0,00309683 | 0,047812073 | carboxypeptidase Q |
| ENSG00000254718 | AL157756,1 | 0,980213702 | 1,972757606 | 0,35852844 | 0,849814998 | novel transcript, antisense to PPM1A |
| ENSG00000248605 | AC022140,1 | 0,980177054 | 1,972707494 | 0,18785824 | 0,681978628 | novel transcript |
| ENSG00000137265 | IRF4 | 0,980008298 | 1,972476754 | 0,1947247 | 0,692275751 | interferon regulatory factor 4 |
| ENSG00000157933 | SKI | 0,978966582 | 1,971053017 | 0,00508405 | 0,06953097 | SKI proto-oncogene |
| ENSG00000002834 | LASP1 | 0,978903356 | 1,970966638 | 0,00020076 | 0,004787687 | LIM and SH3 protein 1 |
| ENSG00000090554 | FLT3LG | 0,978721836 | 1,970718665 | 0,14101173 | 0,598214217 | fms related receptor tyrosine kinase 3 ligand |
| ENSG00000172548 | NIPAL4 | 0,978501699 | 1,970417982 | 0,00910572 | 0,106902479 | NIPA like domain containing 4 |
| ENSG00000276547 | PCDHGB5 | 0,978274939 | 1,9701083 | 0,00609953 | 0,080000234 | protocadherin gamma subfamily B, 5 |
| ENSG00000260078 | AC007342,3 | 0,978168123 | 1,96996244 | 0,35531816 | 0,846840543 | u3 small nucleolar ribonucleoprotein protein MPP10-like |

| | | | | | | |
|---|---|---|---|---|---|---|
| ENSG00000108797 | CNTNAP1 | 0,978103644 | 1,969874398 | 0,01145504 | 0,126436253 | contactin associated protein 1 |
| ENSG00000264083 | AC005899,1 | 0,978036386 | 1,969782564 | 0,36088826 | 0,851667053 | novel transcript |
| ENSG00000286710 | AC188617,1 | 0,977685725 | 1,969303849 | 0,44623147 | 0,901348735 | novel transcript |
| ENSG00000285245 | AL162417,1 | 0,976158407 | 1,967220136 | 0,71983939 | 0,972014805 | novel protein |
| ENSG00000261771 | DNAAF4-CCPG1 | 0,975704228 | 1,966600927 | 0,80504732 | 0,98109454 | DNAAF4-CCPG1 readthrough (NMD candidate) |
| ENSG00000203684 | IBA57-DT | 0,975591666 | 1,966447494 | 0,47473075 | 0,91069468 | IBA57 divergent transcript |
| ENSG00000160683 | CXCR5 | 0,975476361 | 1,966290336 | 0,18346087 | 0,674559627 | C-X-C motif chemokine receptor 5 |
| ENSG00000136048 | DRAM1 | 0,975263432 | 1,96600015 | 0,01179515 | 0,129254616 | DNA damage regulated autophagy modulator 1 |
| ENSG00000274001 | AL512506,1 | 0,975155078 | 1,965852499 | 0,51725015 | 0,925746338 | novel transcript, sense intronic to CCDC122 |
| ENSG00000185164 | NOMO2 | 0,974884738 | 1,965484162 | 0,00015521 | 0,003808848 | NODAL modulator 2 |
| ENSG00000146112 | PPP1R18 | 0,974180902 | 1,964525511 | 0,03481858 | 0,272210084 | protein phosphatase 1 regulatory subunit 18 |
| ENSG00000223511 | AL683807,1 | 0,974005322 | 1,964286437 | 0,31377904 | 0,819877431 | novel transcript |
| ENSG00000158290 | CUL4B | 0,974000099 | 1,964279325 | 0,02115363 | 0,194057264 | cullin 4B |
| ENSG00000157600 | TMEM164 | 0,973923305 | 1,96417477 | 4,15E-06 | 0,000156723 | transmembrane protein 164 |
| ENSG00000256943 | AC148477,3 | 0,973921535 | 1,964172361 | 0,42426049 | 0,890949676 | novel transcript |
| ENSG00000288209 | AC148477,12 | 0,973921535 | 1,964172361 | 0,42426049 | 0,890949676 | novel transcript |
| ENSG00000272865 | AL591686,2 | 0,973886982 | 1,964125319 | 0,43350514 | 0,896073042 | novel transcript, antisense to PLD5 |
| ENSG00000233355 | CHRM3-AS2 | 0,973740202 | 1,963925499 | 0,1340811 | 0,584238558 | CHRM3 antisense RNA 2 |
| ENSG00000264592 | RN7SL131P | 0,973588517 | 1,963719022 | 0,61264239 | 0,952413269 | RNA, 7SL, cytoplasmic 131, pseudogene |
| ENSG00000286329 | AC108690,1 | 0,973017687 | 1,962942193 | 0,2563934 | 0,768541051 | novel transcript |
| ENSG00000125675 | GRIA3 | 0,97296701 | 1,962873242 | 0,00904598 | 0,106453868 | glutamate ionotropic receptor AMPA type subunit 3 |
| ENSG00000172935 | MRGPRF | 0,972724642 | 1,962543514 | 0,16990132 | 0,652875483 | MAS related GPR family member F |
| ENSG00000253490 | LINC02099 | 0,972716016 | 1,962531779 | 0,24542114 | 0,757479368 | long intergenic non-protein coding RNA 2099 |
| ENSG00000285159 | AL627422,2 | 0,972673079 | 1,962473373 | 0,41394504 | 0,885456174 | novel transcript |
| ENSG00000229870 | RPL21P89 | 0,972254159 | 1,961903606 | 0,31230372 | 0,818547047 | ribosomal protein L21 pseudogene 89 |
| ENSG00000223571 | DHRSX-IT1 | 0,972045475 | 1,961619839 | 0,38231871 | 0,866930927 | DHRSX intronic transcript 1 |
| ENSG00000168792 | ABHD15 | 0,972029454 | 1,961598056 | 0,02438879 | 0,215149831 | abhydrolase domain containing 15 |
| ENSG00000197702 | PARVA | 0,972010705 | 1,961572563 | 5,54E-06 | 0,000202663 | parvin alpha |
| ENSG00000165704 | HPRT1 | 0,971317825 | 1,960630709 | 0,03048075 | 0,249663376 | hypoxanthine phosphoribosyltransferase 1 |
| ENSG00000256116 | AP001453,1 | 0,970791827 | 1,959916005 | 0,49595304 | 0,918105073 | novel transcript |
| ENSG00000124493 | GRM4 | 0,970565762 | 1,959608918 | 0,14631307 | 0,609656943 | glutamate metabotropic receptor 4 |
| ENSG00000102362 | SYTL4 | 0,970475162 | 1,959485859 | 0,00610715 | 0,080056171 | synaptotagmin like 4 |
| ENSG00000106266 | SNX8 | 0,97012041 | 1,95900409 | 0,07027709 | 0,417549035 | sorting nexin 8 |

| | | | | | | |
|---|---|---|---|---|---|---|
| ENSG00000109458 | GAB1 | 0,969909829 | 1,958718168 | 0,01527981 | 0,156164763 | GRB2 associated binding protein 1 |
| ENSG00000224281 | SLC25A5-AS1 | 0,969877781 | 1,958674657 | 0,00398104 | 0,058515839 | SLC25A5 antisense RNA 1 |
| ENSG00000255381 | AP001258,1 | 0,969812917 | 1,958586597 | 0,47025107 | 0,908536 | protein kinase, cAMP-dependent, regulatory, type II, beta (PRKAR2B) pseudogene |
| ENSG00000229544 | NKX1-2 | 0,969805448 | 1,958576457 | 0,35055465 | 0,843970352 | NK1 homeobox 2 |
| ENSG00000187942 | LDLRAD2 | 0,969803005 | 1,958573141 | 0,09420077 | 0,487613759 | low density lipoprotein receptor class A domain containing 2 |
| ENSG00000243049 | RN7SL33P | 0,969393947 | 1,958017891 | 0,28618097 | 0,797420923 | RNA, 7SL, cytoplasmic 33, pseudogene |
| ENSG00000266210 | RN7SL371P | 0,969269104 | 1,957848462 | 0,32495453 | 0,826983176 | RNA, 7SL, cytoplasmic 371, pseudogene |
| ENSG00000183134 | PTGDR2 | 0,968755429 | 1,957151489 | 0,11007265 | 0,529025546 | prostaglandin D2 receptor 2 |
| ENSG00000102309 | PIN4 | 0,968468724 | 1,956762585 | 0,00897358 | 0,10590787 | peptidylprolyl cis/trans isomerase, NIMA-interacting 4 |
| ENSG00000278268 | MOAP1 | 0,968291052 | 1,956521619 | 0,17985887 | 0,668621595 | modulator of apoptosis 1 |
| ENSG00000223305 | RN7SKP30 | 0,968243877 | 1,956457643 | 0,29286506 | 0,803304991 | RN7SK pseudogene 30 |
| ENSG00000228133 | AC099684,1 | 0,96804529 | 1,956188355 | 0,49125839 | 0,916995341 | novel transcript, antisense to RTN4RL1 |
| ENSG00000228502 | EEF1A1P11 | 0,967922956 | 1,956022487 | 0,42247096 | 0,890275827 | eukaryotic translation elongation factor 1 alpha 1 pseudogene 11 |
| ENSG00000103313 | MEFV | 0,967860745 | 1,955938142 | 0,23341427 | 0,74254697 | MEFV innate immuity regulator, pyrin |
| ENSG00000106153 | CHCHD2 | 0,967707995 | 1,955731062 | 7,81E-06 | 0,000274334 | coiled-coil-helix-coiled-coil-helix domain containing 2 |
| ENSG00000284161 | EEF2K | 0,967201868 | 1,955045072 | 0,01171085 | 0,128601321 | eukaryotic elongation factor 2 kinase |
| ENSG00000214650 | AC073592,1 | 0,967188692 | 1,955027217 | 0,23452714 | 0,74410295 | novel transcript |
| ENSG00000204392 | LSM2 | 0,967181789 | 1,955017863 | 0,00034837 | 0,007691734 | LSM2 homolog, U6 small nuclear RNA and mRNA degradation associated |
| ENSG00000223883 | LINC01707 | 0,966229616 | 1,953727984 | 0,3191367 | 0,82414099 | long intergenic non-protein coding RNA 1707 |
| ENSG00000010704 | HFE | 0,96613712 | 1,953602727 | 0,00764079 | 0,094569166 | homeostatic iron regulator |
| ENSG00000254087 | LYN | 0,966134992 | 1,953599847 | 0,0134517 | 0,141982678 | LYN proto-oncogene, Src family tyrosine kinase |
| ENSG00000273253 | AL022328,4 | 0,966123009 | 1,95358362 | 0,15434085 | 0,623607455 | novel transcript, antisense TRABD |
| ENSG00000272211 | AC114760,2 | 0,965344037 | 1,952529082 | 0,04579794 | 0,324471967 | novel transcript, antisense to STK17B |
| ENSG00000273474 | AL157392,4 | 0,965098243 | 1,952196456 | 0,28765128 | 0,799000743 | novel transcript, antisense to FRMD4A |
| ENSG00000177628 | GBA | 0,965026185 | 1,952098952 | 0,00063824 | 0,012799522 | glucosylceramidase beta |
| ENSG00000025708 | TYMP | 0,964902669 | 1,95193183 | 0,1872183 | 0,681158265 | thymidine phosphorylase |
| ENSG00000198759 | EGFL6 | 0,964753815 | 1,951730445 | 0,25350391 | 0,765172364 | EGF like domain multiple 6 |

| | | | | | | |
|---|---|---|---|---|---|---|
| ENSG00000077721 | UBE2A | 0,964453526 | 1,951324246 | 0,0114596 | 0,126436253 | ubiquitin conjugating enzyme E2 A |
| ENSG00000287670 | AC108479,1 | 0,964414902 | 1,951272005 | 0,06024022 | 0,381629797 | novel transcript |
| ENSG00000235426 | AL133481,1 | 0,964231352 | 1,951023766 | 0,40086564 | 0,876599365 | novel transcript |
| ENSG00000235927 | NEXN-AS1 | 0,964161097 | 1,95092876 | 0,19567993 | 0,69375328 | NEXN antisense RNA 1 |
| ENSG00000236708 | SDK1-AS1 | 0,964143022 | 1,950904317 | 0,17609973 | 0,661639583 | SDK1 antisense RNA 1 |
| ENSG00000206289 | RXRB | 0,96412054 | 1,950873916 | 0,05278461 | 0,352845398 | retinoid X receptor beta |
| ENSG00000124920 | MYRF | 0,963890817 | 1,950563299 | 0,00412095 | 0,060092113 | myelin regulatory factor |
| ENSG00000224563 | LPP-AS1 | 0,963795147 | 1,950433954 | 0,32870524 | 0,830659639 | LPP antisense RNA 1 |
| ENSG00000240563 | L1TD1 | 0,963675781 | 1,950272586 | 0,07317915 | 0,426957349 | LINE1 type transposase domain containing 1 |
| ENSG00000138829 | FBN2 | 0,963547474 | 1,950099145 | 1,13E-05 | 0,000378035 | fibrillin 2 |
| ENSG00000120913 | PDLIM2 | 0,963498464 | 1,950032899 | 0,01950151 | 0,183714864 | PDZ and LIM domain 2 |
| ENSG00000120725 | SIL1 | 0,963430928 | 1,949941616 | 0,01287274 | 0,137968237 | SIL1 nucleotide exchange factor |
| ENSG00000234961 | AL133415,1 | 0,963358422 | 1,949843619 | 0,01382819 | 0,14497809 | novel transcript |
| ENSG00000101888 | NXT2 | 0,963317263 | 1,949787992 | 0,13978083 | 0,595950877 | nuclear transport factor 2 like export factor 2 |
| ENSG00000203546 | AL139353,1 | 0,963174436 | 1,949594973 | 0,16031334 | 0,635722509 | novel protein |
| ENSG00000135535 | CD164 | 0,962688999 | 1,948939085 | 0,03466509 | 0,271597663 | CD164 molecule |
| ENSG00000233087 | RAB6D | 0,962655589 | 1,948893951 | 0,25288671 | 0,76514565 | RAB6D, member RAS oncogene family |
| ENSG00000177144 | NUDT4B | 0,962221712 | 1,948307928 | 0,07616431 | 0,43619729 | nudix hydrolase 4B |
| ENSG00000147130 | ZMYM3 | 0,9621577 | 1,948221484 | 3,03E-06 | 0,000118663 | zinc finger MYM-type containing 3 |
| ENSG00000227301 | AL356489,1 | 0,96203347 | 1,948053731 | 0,42429795 | 0,890949676 | novel transcript |
| ENSG00000128165 | ADM2 | 0,960662667 | 1,946203631 | 0,16526703 | 0,644333269 | adrenomedullin 2 |
| ENSG00000090013 | BLVRB | 0,96063747 | 1,94616964 | 0,02412524 | 0,213366798 | biliverdin reductase B |
| ENSG00000225043 | RPL18AP2 | 0,960608878 | 1,94613107 | 0,4241751 | 0,890949676 | ribosomal protein L18a pseudogene 2 |
| ENSG00000248810 | LINC02432 | 0,960547713 | 1,946048563 | 0,06028046 | 0,381654162 | long intergenic non-protein coding RNA 2432 |
| ENSG00000254028 | AC083843,1 | 0,960397904 | 1,945846497 | 0,46875373 | 0,907940653 | novel transcript |
| ENSG00000267095 | AC025048,1 | 0,959633041 | 1,944815155 | 0,50918214 | 0,922427526 | novel transcript |
| ENSG00000271204 | AC016831,4 | 0,959592718 | 1,944760799 | 0,03355161 | 0,266165357 | novel transcript |
| ENSG00000277778 | PGM5P2 | 0,959412594 | 1,944518006 | 0,03185489 | 0,257407954 | phosphoglucomutase 5 pseudogene 2 |
| ENSG00000269032 | AC016629,1 | 0,958812732 | 1,943709658 | 0,3558769 | 0,847256893 | interleukin 9 receptor (IL9R) pseudogene |
| ENSG00000278243 | GFUS | 0,958588833 | 1,943408027 | 0,03729045 | 0,285054324 | GDP-L-fucose synthase |
| ENSG00000214652 | ZNF727 | 0,958496073 | 1,943283077 | 0,40347868 | 0,877691113 | zinc finger protein 727 |
| ENSG00000287689 | AC009704,3 | 0,95834656 | 1,943081696 | 0,64565987 | 0,956899963 | novel transcript |
| ENSG00000198049 | AVPR1B | 0,958074331 | 1,942715081 | 0,0949846 | 0,489386708 | arginine vasopressin receptor 1B |

| | | | | | | |
|---|---|---|---|---|---|---|
| ENSG00000272662 | AC073352,1 | 0,958056957 | 1,942691686 | 0,3809882 | 0,865900094 | novel transcript |
| ENSG00000155657 | TTN | 0,957912035 | 1,942496548 | 0,00404486 | 0,059268345 | titin |
| ENSG00000229539 | PANK2-AS1 | 0,95774621 | 1,942273288 | 0,10969576 | 0,528090728 | PANK2 antisense RNA 1 |
| ENSG00000204314 | PRRT1 | 0,957668736 | 1,94216899 | 0,38794623 | 0,869820053 | proline rich transmembrane protein 1 |
| ENSG00000258050 | AL139316,1 | 0,957608566 | 1,94208799 | 0,32435782 | 0,82642149 | novel transcript |
| ENSG00000264448 | MIR378D2HG | 0,956798187 | 1,940997402 | 0,37203507 | 0,859937929 | MIR378D2 host gene |
| ENSG00000285125 | AL136295,13 | 0,955940659 | 1,93984403 | 0,14692547 | 0,610674677 | novel protein |
| ENSG00000259522 | AL136295,4 | 0,955940659 | 1,93984403 | 0,14692547 | 0,610674677 | novel protein |
| ENSG00000280700 | AC136687,1 | 0,955799126 | 1,939653734 | 0,35491828 | 0,846374661 | M-phase phosphoprotein 10 (U3 small nucleolar ribonucleoprotein) (MPHOSPH10) pseudogene |
| ENSG00000125912 | NCLN | 0,955501834 | 1,939254077 | 0,02738974 | 0,232946937 | nicalin |
| ENSG00000102385 | DRP2 | 0,955345137 | 1,939043457 | 2,61E-05 | 0,000798461 | dystrophin related protein 2 |
| ENSG00000131016 | AKAP12 | 0,955160896 | 1,938795845 | 0,03271589 | 0,261798703 | A-kinase anchoring protein 12 |
| ENSG00000239557 | PPP2R5CP | 0,954734902 | 1,938223449 | 0,24640645 | 0,758198269 | protein phosphatase 2 regulatory subunit B'gamma pseudogene |
| ENSG00000155090 | KLF10 | 0,954726239 | 1,938211811 | 0,00084492 | 0,016132236 | Kruppel like factor 10 |
| ENSG00000105355 | PLIN3 | 0,954122302 | 1,937400612 | 0,0062813 | 0,081714521 | perilipin 3 |
| ENSG00000165807 | PPP1R36 | 0,953969312 | 1,937195171 | 0,28009291 | 0,790779983 | protein phosphatase 1 regulatory subunit 36 |
| ENSG00000169184 | MN1 | 0,953738116 | 1,936884756 | 0,00957789 | 0,111275759 | MN1 proto-oncogene, transcriptional regulator |
| ENSG00000124067 | SLC12A4 | 0,952810377 | 1,935639624 | 0,03311389 | 0,264235013 | solute carrier family 12 member 4 |
| ENSG00000267177 | AP002505,1 | 0,952809253 | 1,935638115 | 0,39086272 | 0,870057277 | novel transcript, antisense C18orf1 |
| ENSG00000146373 | RNF217 | 0,952636581 | 1,935406458 | 0,00560872 | 0,075200114 | ring finger protein 217 |
| ENSG00000235597 | LINC01102 | 0,952598553 | 1,935355443 | 0,01083747 | 0,121616202 | long intergenic non-protein coding RNA 1102 |
| ENSG00000214514 | KRT42P | 0,951279764 | 1,933587114 | 0,26134506 | 0,77325059 | keratin 42 pseudogene |
| ENSG00000115884 | SDC1 | 0,950879812 | 1,93305115 | 0,00440324 | 0,062917285 | syndecan 1 |
| ENSG00000164300 | SERINC5 | 0,950776933 | 1,932913307 | 0,00239827 | 0,038751223 | serine incorporator 5 |
| ENSG00000111319 | SCNN1A | 0,950756484 | 1,93288591 | 0,33070684 | 0,832114797 | sodium channel epithelial 1 subunit alpha |
| ENSG00000068697 | LAPTM4A | 0,950753639 | 1,932882098 | 0,02004239 | 0,186934787 | lysosomal protein transmembrane 4 alpha |
| ENSG00000235837 | AC073333,1 | 0,95064791 | 1,932740451 | 0,40597805 | 0,878981828 | novel transcript |
| ENSG00000033100 | CHPF2 | 0,949736985 | 1,931520494 | 0,00588663 | 0,077752195 | chondroitin polymerizing factor 2 |
| ENSG00000183479 | TREX2 | 0,949036294 | 1,930582618 | 0,4830429 | 0,91363215 | three prime repair exonuclease 2 |
| ENSG00000131236 | CAP1 | 0,948961248 | 1,930482194 | 0,00021845 | 0,005139013 | cyclase associated actin cytoskeleton regulatory protein 1 |

| | | | | | | |
|---|---|---|---|---|---|---|
| ENSG00000164733 | CTSB | 0,94872596 | 1,93016738 | 0,00442814 | 0,063178325 | cathepsin B |
| ENSG00000185758 | CLDN24 | 0,948561622 | 1,929947525 | 0,18243688 | 0,6727656 | claudin 24 |
| ENSG00000144802 | NFKBIZ | 0,94814756 | 1,929393698 | 0,00083901 | 0,016052254 | NFKB inhibitor zeta |
| ENSG00000286602 | AC021660,4 | 0,947353774 | 1,928332418 | 0,42205128 | 0,889775006 | novel transcript |
| ENSG00000207304 | SNORA8 | 0,946707033 | 1,927468166 | 0,67436557 | 0,963103112 | small nucleolar RNA, H/ACA box 8 |
| ENSG00000137871 | ZNF280D | 0,94653475 | 1,927238006 | 0,06332868 | 0,393776438 | zinc finger protein 280D |
| ENSG00000232756 | AC004990,1 | 0,946188358 | 1,926775332 | 0,47406522 | 0,910491353 | novel transcript |
| ENSG00000250337 | PURPL | 0,94599541 | 1,926517659 | 0,3871941 | 0,869441674 | p53 upregulated regulator of p53 levels |
| ENSG00000171155 | C1GALT1C1 | 0,945318967 | 1,925614575 | 0,01909193 | 0,181315309 | C1GALT1 specific chaperone 1 |
| ENSG00000288368 | C1GALT1C1 | 0,945318967 | 1,925614575 | 0,01909193 | 0,181315309 | C1GALT1 specific chaperone 1 |
| ENSG00000204694 | OR11A1 | 0,945285068 | 1,925569329 | 0,51003056 | 0,922427526 | olfactory receptor family 11 subfamily A member 1 |
| ENSG00000133121 | STARD13 | 0,945184663 | 1,925435324 | 0,0280541 | 0,23672966 | StAR related lipid transfer domain containing 13 |
| ENSG00000288046 | AL031123,5 | 0,944071919 | 1,923950817 | 0,26838507 | 0,780709773 | novel transcript |
| ENSG00000243695 | AL353729,1 | 0,943824272 | 1,923620587 | 0,16482008 | 0,643964432 | TGF beta-inducible nuclear protein 1 (TINP1) pseudogene |
| ENSG00000235779 | AC079779,3 | 0,94324888 | 1,922853541 | 0,18207511 | 0,672311616 | novel transcript |
| ENSG00000099331 | MYO9B | 0,943039622 | 1,922574658 | 0,02780444 | 0,235385294 | myosin IXB |
| ENSG00000185347 | TEDC1 | 0,942777425 | 1,922225278 | 0,12306244 | 0,561176198 | tubulin epsilon and delta complex 1 |
| ENSG00000172346 | CSDC2 | 0,942504335 | 1,921861452 | 0,04328616 | 0,313927122 | cold shock domain containing C2 |
| ENSG00000269911 | FAM226B | 0,942443759 | 1,921780758 | 0,00835159 | 0,100812358 | family with sequence similarity 226 member B |
| ENSG00000250669 | AC112206,4 | 0,942378644 | 1,921694022 | 0,3665769 | 0,856103393 | mitochondrial ribosomal protein L39 (MRPL39) pseudogene |
| ENSG00000102024 | PLS3 | 0,942360912 | 1,921670402 | 0,00165691 | 0,028603891 | plastin 3 |
| ENSG00000243359 | RN7SL815P | 0,942158925 | 1,921401374 | 0,3884212 | 0,869820053 | RNA, 7SL, cytoplasmic 815, pseudogene |
| ENSG00000275620 | FLJ16779 | 0,942032307 | 1,92123275 | 0,05040021 | 0,343470607 | uncharacterized LOC100192386 |
| ENSG00000129566 | TEP1 | 0,941800989 | 1,92092473 | 0,00471829 | 0,066001114 | telomerase associated protein 1 |
| ENSG00000258677 | AC022826,2 | 0,941699256 | 1,920789279 | 0,14158671 | 0,599428784 | novel protein, UBE2W-STAU2 readthrough |
| ENSG00000121060 | TRIM25 | 0,94152979 | 1,920563667 | 0,0254867 | 0,221892605 | tripartite motif containing 25 |
| ENSG00000116604 | MEF2D | 0,941492933 | 1,920514603 | 0,00149169 | 0,026141383 | myocyte enhancer factor 2D |
| ENSG00000257921 | AC025165,3 | 0,94089178 | 1,919714514 | 0,71318481 | 0,970812834 | novel protein |
| ENSG00000135373 | EHF | 0,940890929 | 1,919713382 | 0,54289513 | 0,933663748 | ETS homologous factor |
| ENSG00000255745 | AC023796,1 | 0,940854434 | 1,919664821 | 0,39456642 | 0,871872016 | novel transcript |
| ENSG00000185624 | P4HB | 0,940849181 | 1,919657831 | 0,00107031 | 0,019839552 | prolyl 4-hydroxylase subunit beta |

| | | | | | | |
|---|---|---|---|---|---|---|
| ENSG00000243313 | RN7SL285P | 0,940772066 | 1,919555225 | 0,50248975 | 0,920588312 | RNA, 7SL, cytoplasmic 285, pseudogene |
| ENSG00000164111 | ANXA5 | 0,94066804 | 1,919416818 | 0,00045093 | 0,009558531 | annexin A5 |
| ENSG00000235308 | AL445991,1 | 0,940014972 | 1,918548149 | 0,43255334 | 0,89560871 | novel pseudogene |
| ENSG00000256950 | AC069503,2 | 0,939584425 | 1,917975677 | 0,20501494 | 0,706634751 | novel transcript |
| ENSG00000207392 | SNORA20 | 0,939361826 | 1,917679768 | 0,47782 | 0,911721977 | small nucleolar RNA, H/ACA box 20 |
| ENSG00000130402 | ACTN4 | 0,93932361 | 1,917628971 | 0,01067467 | 0,120264106 | actinin alpha 4 |
| ENSG00000189401 | OTUD6A | 0,939269704 | 1,917557321 | 0,33705822 | 0,836223219 | OTU deubiquitinase 6A |
| ENSG00000166573 | GALR1 | 0,93866118 | 1,916748672 | 0,2085663 | 0,713259317 | galanin receptor 1 |
| ENSG00000254910 | AC136475,2 | 0,938285388 | 1,916249464 | 0,19452002 | 0,691630774 | novel transcript |
| ENSG00000082175 | PGR | 0,938014132 | 1,915889203 | 0,10369792 | 0,513464006 | progesterone receptor |
| ENSG00000125945 | ZNF436 | 0,937950159 | 1,91580425 | 0,01198594 | 0,130575624 | zinc finger protein 436 |
| ENSG00000283009 | ZNF436 | 0,937950159 | 1,91580425 | 0,01198594 | 0,130575624 | zinc finger protein 436 |
| ENSG00000231194 | FARP1-AS1 | 0,937919091 | 1,915762994 | 0,48316705 | 0,91363215 | FARP1 antisense RNA 1 |
| ENSG00000256671 | LIMS4 | 0,937894709 | 1,915730617 | 0,22434944 | 0,73307493 | LIM zinc finger domain containing 4 |
| ENSG00000169515 | CCDC8 | 0,937752223 | 1,915541422 | 0,0844978 | 0,462651338 | coiled-coil domain containing 8 |
| ENSG00000188460 | ACTBP11 | 0,937568256 | 1,915297175 | 0,11942734 | 0,552333761 | ACTB pseudogene 11 |
| ENSG00000277773 | SMN2 | 0,937568009 | 1,915296847 | 0,21961214 | 0,727931767 | survival of motor neuron 2, centromeric |
| ENSG00000233564 | PRR3 | 0,937557871 | 1,915283388 | 0,55086586 | 0,936366211 | proline rich 3 |
| ENSG00000236514 | AL135791,1 | 0,937542976 | 1,915263614 | 0,18232183 | 0,672658692 | novel transcript |
| ENSG00000254349 | MIR2052HG | 0,937416545 | 1,915095776 | 0,04030208 | 0,299940355 | MIR2052 host gene |
| ENSG00000153976 | HS3ST3A1 | 0,937275026 | 1,914907927 | 0,04991474 | 0,341473625 | heparan sulfate-glucosamine 3-sulfotransferase 3A1 |
| ENSG00000140832 | MARVELD3 | 0,937195855 | 1,914802845 | 0,0417953 | 0,306600121 | MARVEL domain containing 3 |
| ENSG00000198223 | CSF2RA | 0,937172512 | 1,914771863 | 0,23868602 | 0,747998807 | colony stimulating factor 2 receptor subunit alpha |
| ENSG00000254887 | AC010247,1 | 0,936836593 | 1,914326078 | 0,4569739 | 0,903711377 | novel transcript, antisense to POU2F2 |
| ENSG00000111261 | MANSC1 | 0,936823622 | 1,914308867 | 0,08794189 | 0,472326951 | MANSC domain containing 1 |
| ENSG00000248475 | AC016642,1 | 0,936508396 | 1,91389064 | 0,43859558 | 0,897246228 | novel transcript |
| ENSG00000130066 | SAT1 | 0,936242008 | 1,91353728 | 0,00563552 | 0,075326629 | spermidine/spermine N1-acetyltransferase 1 |
| ENSG00000203472 | AC009163,1 | 0,936109353 | 1,913361339 | 0,32391662 | 0,825946442 | novel transcript, antisense to CHST6 |
| ENSG00000161217 | PCYT1A | 0,936032236 | 1,913259066 | 0,0002606 | 0,005968773 | phosphate cytidylyltransferase 1, choline, alpha |
| ENSG00000112782 | CLIC5 | 0,935579714 | 1,912659039 | 0,25902587 | 0,771947284 | chloride intracellular channel 5 |
| ENSG00000234370 | GTF2H4 | 0,935548822 | 1,912618085 | 0,24418791 | 0,755571155 | general transcription factor IIH subunit 4 |
| ENSG00000106536 | POU6F2 | 0,935378122 | 1,912391796 | 0,00341626 | 0,051747052 | POU class 6 homeobox 2 |
| ENSG00000287260 | AL357075,3 | 0,93526034 | 1,912235674 | 0,1971194 | 0,695927373 | novel transcript |

| | | | | | | |
|---|---|---|---|---|---|---|
| ENSG00000131187 | F12 | 0,934658453 | 1,911438063 | 0,21696525 | 0,724546211 | coagulation factor XII |
| ENSG00000182220 | ATP6AP2 | 0,934656836 | 1,911435921 | 0,01313348 | 0,13975671 | ATPase H+ transporting accessory protein 2 |
| ENSG00000264019 | AC018521,2 | 0,934598185 | 1,911358216 | 0,56457261 | 0,940865625 | novel transcript, antisense to PNPO |
| ENSG00000238188 | AL049733,1 | 0,934227965 | 1,910867791 | 0,39473638 | 0,871888817 | novel pseudogene |
| ENSG00000114315 | HES1 | 0,934074783 | 1,910664911 | 0,00209877 | 0,034976441 | hes family bHLH transcription factor 1 |
| ENSG00000164742 | ADCY1 | 0,9340232 | 1,910596597 | 0,03682803 | 0,282977152 | adenylate cyclase 1 |
| ENSG00000178685 | PARP10 | 0,933081675 | 1,909350119 | 0,16504752 | 0,644039657 | poly(ADP-ribose) polymerase family member 10 |
| ENSG00000166823 | MESP1 | 0,932730978 | 1,908886042 | 0,09276132 | 0,484139956 | mesoderm posterior bHLH transcription factor 1 |
| ENSG00000123427 | EEF1AKMT3 | 0,932221611 | 1,908212198 | 0,01095004 | 0,122483758 | EEF1A lysine methyltransferase 3 |
| ENSG00000188937 | NYX | 0,932131028 | 1,908092389 | 0,22297216 | 0,731402952 | nyctalopin |
| ENSG00000131174 | COX7B | 0,931980306 | 1,907893057 | 0,03773442 | 0,287183177 | cytochrome c oxidase subunit 7B |
| ENSG00000288099 | AL359259,1 | 0,93181229 | 1,907670877 | 0,57105262 | 0,942436428 | novel transcript |
| ENSG00000269220 | LINC00528 | 0,93147951 | 1,907230894 | 0,39166174 | 0,870347042 | long intergenic non-protein coding RNA 528 |
| ENSG00000136869 | TLR4 | 0,931458722 | 1,907203413 | 0,04242137 | 0,309296116 | toll like receptor 4 |
| ENSG00000205922 | ONECUT3 | 0,931235907 | 1,90690888 | 0,21033271 | 0,715586415 | one cut homeobox 3 |
| ENSG00000267439 | AD000671,3 | 0,931185605 | 1,906842392 | 0,3705975 | 0,858545694 | novel transcript |
| ENSG00000143341 | HMCN1 | 0,931172013 | 1,906824428 | 0,00561969 | 0,075243723 | hemicentin 1 |
| ENSG00000002822 | MAD1L1 | 0,930685409 | 1,906181388 | 0,1260443 | 0,566777174 | mitotic arrest deficient 1 like 1 |
| ENSG00000076928 | ARHGEF1 | 0,930525526 | 1,905970151 | 0,0512935 | 0,34665805 | Rho guanine nucleotide exchange factor 1 |
| ENSG00000274215 | AC106028,4 | 0,930464065 | 1,905888956 | 0,43637117 | 0,896850871 | novel transcript |
| ENSG00000073803 | MAP3K13 | 0,93041885 | 1,905829225 | 0,00491767 | 0,067999136 | mitogen-activated protein kinase kinase kinase 13 |
| ENSG00000285471 | AC007846,2 | 0,93029086 | 1,905660155 | 0,59883109 | 0,948854865 | novel protein |
| ENSG00000183671 | GPR1 | 0,930101874 | 1,905410539 | 0,06262316 | 0,390866548 | G protein-coupled receptor 1 |
| ENSG00000258080 | ARHGAP42P4 | 0,930014766 | 1,905295497 | 0,44694762 | 0,901798674 | ARHGAP42 pseudogene 4 |
| ENSG00000236751 | LINC01186 | 0,929434426 | 1,904529224 | 0,38627338 | 0,869312875 | long intergenic non-protein coding RNA 1186 |
| ENSG00000234288 | OR1AA1P | 0,929417828 | 1,904507313 | 0,47538233 | 0,91115935 | olfactory receptor family 1 subfamily AA member 1 pseudogene |
| ENSG00000264558 | AC015674,1 | 0,929326699 | 1,904387018 | 0,08046699 | 0,449985752 | novel transcript, antisense to KPNB1 |
| ENSG00000258870 | EIF4EBP1P1 | 0,92893096 | 1,903864705 | 0,477954 | 0,911872483 | eukaryotic translation initiation factor 4E binding protein 1 pseudogene 1 |
| ENSG00000233198 | RNF224 | 0,92885001 | 1,903757883 | 0,5558229 | 0,9379924 | ring finger protein 224 |
| ENSG00000260475 | AL353719,1 | 0,928273413 | 1,902997166 | 0,2636092 | 0,774793719 | novel transcript |
| ENSG00000141959 | PFKL | 0,92819056 | 1,902887881 | 0,03241084 | 0,260687565 | phosphofructokinase, liver type |

| | | | | | | |
|---|---|---|---|---|---|---|
| ENSG00000136603 | SKIL | 0,928138383 | 1,902819061 | 0,0411382 | 0,30369413 | SKI like proto-oncogene |
| ENSG00000145794 | MEGF10 | 0,928018417 | 1,902660841 | 0,00662719 | 0,085219165 | multiple EGF like domains 10 |
| ENSG00000131370 | SH3BP5 | 0,928000914 | 1,902637758 | 0,03475546 | 0,271925393 | SH3 domain binding protein 5 |
| ENSG00000279483 | AC090498,1 | 0,927978695 | 1,902608456 | 0,21971812 | 0,727931767 | ribosomal protein L41 (RPL41) pseudogene |
| ENSG00000050405 | LIMA1 | 0,927976662 | 1,902605775 | 0,00102272 | 0,01905494 | LIM domain and actin binding 1 |
| ENSG00000267373 | AC008894,1 | 0,92788746 | 1,90248814 | 0,47605285 | 0,911413793 | novel transcript |
| ENSG00000085185 | BCORL1 | 0,927875049 | 1,902471774 | 1,16E-05 | 0,000386157 | BCL6 corepressor like 1 |
| ENSG00000260626 | AC136944,1 | 0,927684756 | 1,902220852 | 0,25535867 | 0,766910046 | novel transcript |
| ENSG00000285942 | AC018362,3 | 0,927452208 | 1,901914258 | 0,81484145 | 0,981521413 | novel protein |
| ENSG00000241907 | RPS20P4 | 0,927244701 | 1,901640719 | 0,34446176 | 0,841168721 | ribosomal protein S20 pseudogene 4 |
| ENSG00000152939 | MARVELD2 | 0,927237018 | 1,901630593 | 0,07779077 | 0,440668732 | MARVEL domain containing 2 |
| ENSG00000231225 | MICA | 0,927118531 | 1,90147442 | 0,03857423 | 0,29137102 | MHC class I polypeptide-related sequence A |
| ENSG00000285744 | AC083837,1 | 0,927013275 | 1,901335698 | 0,29107187 | 0,801689099 | novel transcript |
| ENSG00000274653 | AC106782,5 | 0,92697112 | 1,901280143 | 0,25120561 | 0,7635322 | novel transcript, antisense to TBC1D10B |
| ENSG00000168672 | LRATD2 | 0,926889382 | 1,901172425 | 0,00039658 | 0,00857444 | LRAT domain containing 2 |
| ENSG00000240893 | LINC02042 | 0,926714722 | 1,900942274 | 0,41465923 | 0,885579089 | long intergenic non-protein coding RNA 2042 |
| ENSG00000151135 | TMEM263 | 0,926283109 | 1,900373652 | 0,01320949 | 0,140068402 | transmembrane protein 263 |
| ENSG00000061455 | PRDM6 | 0,926138284 | 1,900182892 | 0,15502523 | 0,624656368 | PR/SET domain 6 |
| ENSG00000246130 | AC107959,2 | 0,926056348 | 1,900074976 | 0,41456338 | 0,885579089 | novel transcript |
| ENSG00000198569 | SLC34A3 | 0,925919515 | 1,899894772 | 0,36349066 | 0,85380707 | solute carrier family 34 member 3 |
| ENSG00000287754 | AL109837,3 | 0,925697692 | 1,899602674 | 0,53211242 | 0,931291085 | novel transcript |
| ENSG00000170275 | CRTAP | 0,925413923 | 1,89922907 | 0,00028874 | 0,006536611 | cartilage associated protein |
| ENSG00000225066 | MDM4P1 | 0,925412854 | 1,899227663 | 0,34858443 | 0,842757646 | MDM4 pseudogene 1 |
| ENSG00000189129 | PLAC9 | 0,924264696 | 1,897716779 | 0,09624623 | 0,492720165 | placenta associated 9 |
| ENSG00000262152 | LINC00514 | 0,923517787 | 1,896734552 | 0,47000362 | 0,908378389 | long intergenic non-protein coding RNA 514 |
| ENSG00000270919 | AC108451,2 | 0,923397403 | 1,896576287 | 0,15747494 | 0,629899768 | novel transcript, antisense to ARNT2 |
| ENSG00000224985 | AL590714,1 | 0,923232568 | 1,896359606 | 0,08280196 | 0,457613669 | novel transcript |
| ENSG00000165702 | GFI1B | 0,923165815 | 1,896271864 | 0,41833788 | 0,887094957 | growth factor independent 1B transcriptional repressor |
| ENSG00000174705 | SH3PXD2B | 0,922941134 | 1,895976568 | 0,00236452 | 0,038282935 | SH3 and PX domains 2B |
| ENSG00000197114 | ZGPAT | 0,922846177 | 1,89585178 | 0,13284455 | 0,581850928 | zinc finger CCCH-type and G-patch domain containing |
| ENSG00000236039 | AC019117,1 | 0,922660607 | 1,895607937 | 0,21589896 | 0,722999018 | novel transcript |

| | | | | | | |
|---|---|---|---|---|---|---|
| ENSG00000284952 | AC104472,3 | 0,922564634 | 1,895481839 | 0,1302576 | 0,576741782 | novel transcript |
| ENSG00000185551 | NR2F2 | 0,922442155 | 1,895320927 | 0,00015824 | 0,003867985 | nuclear receptor subfamily 2 group F member 2 |
| ENSG00000259234 | ANKRD34C-AS1 | 0,922406862 | 1,895274562 | 0,3033924 | 0,812669425 | ANKRD34C antisense RNA 1 |
| ENSG00000287952 | AC105328,1 | 0,921788463 | 1,894462343 | 0,46130934 | 0,906165604 | novel transcript, antisense to COX6C |
| ENSG00000106128 | GHRHR | 0,92170698 | 1,894355347 | 0,35643832 | 0,847792901 | growth hormone releasing hormone receptor |
| ENSG00000283563 | AC098650,1 | 0,920613108 | 1,892919564 | 0,06159315 | 0,38671794 | novel protein |
| ENSG00000115415 | STAT1 | 0,920411987 | 1,892655697 | 0,01383583 | 0,145009495 | signal transducer and activator of transcription 1 |
| ENSG00000226791 | LINC02611 | 0,919799969 | 1,891852968 | 0,2878817 | 0,799107075 | long intergenic non-protein coding RNA 2611 |
| ENSG00000115306 | SPTBN1 | 0,91958726 | 1,891574056 | 0,00301195 | 0,046615966 | spectrin beta, non-erythrocytic 1 |
| ENSG00000108039 | XPNPEP1 | 0,919182282 | 1,891043148 | 0,00046641 | 0,009819355 | X-prolyl aminopeptidase 1 |
| ENSG00000225249 | LINC00378 | 0,919174469 | 1,891032908 | 0,40822433 | 0,881221125 | long intergenic non-protein coding RNA 378 |
| ENSG00000286353 | AC115282,1 | 0,919097041 | 1,89093142 | 0,46435662 | 0,907500217 | novel transcript, antisense to CACNA2D3 |
| ENSG00000241169 | FO393419,2 | 0,918343717 | 1,889944301 | 0,39931152 | 0,875433675 | novel transcript |
| ENSG00000258240 | AC002351,1 | 0,918293955 | 1,889879113 | 0,5406973 | 0,933385624 | novel transcript |
| ENSG00000167034 | NKX3-1 | 0,918055889 | 1,889567282 | 0,17144047 | 0,65533907 | NK3 homeobox 1 |
| ENSG00000231851 | MRPS9-AS1 | 0,917635524 | 1,889016789 | 0,42410923 | 0,890949676 | MRPS9 antisense RNA 1 |
| ENSG00000261888 | AC144831,1 | 0,91751404 | 1,888857729 | 0,15028553 | 0,615957904 | novel transcript |
| ENSG00000125753 | VASP | 0,917427061 | 1,888743855 | 0,00505862 | 0,069304383 | vasodilator stimulated phosphoprotein |
| ENSG00000287192 | AL031005,2 | 0,91721532 | 1,888466668 | 0,19310819 | 0,689970564 | novel transcript |
| ENSG00000235233 | MICA | 0,917128353 | 1,888352832 | 0,00451706 | 0,064016873 | MHC class I polypeptide-related sequence A |
| ENSG00000269053 | AC010319,3 | 0,916884707 | 1,88803395 | 0,3968688 | 0,873609704 | novel transcript, antisense FAM125A |
| ENSG00000171700 | RGS19 | 0,916725814 | 1,88782602 | 0,00288231 | 0,04491362 | regulator of G protein signaling 19 |
| ENSG00000286410 | AC089985,1 | 0,916645638 | 1,887721109 | 0,4327656 | 0,89560871 | novel transcript, antisense to SLC41A2 |
| ENSG00000269959 | SPACA6P-AS | 0,916519753 | 1,887556401 | 0,10927523 | 0,527544915 | SPACA6P antisense RNA |
| ENSG00000239697 | TNFSF12 | 0,916426421 | 1,887434294 | 0,07211543 | 0,423561288 | TNF superfamily member 12 |
| ENSG00000147224 | PRPS1 | 0,916245021 | 1,887196987 | 0,00025853 | 0,005931065 | phosphoribosyl pyrophosphate synthetase 1 |
| ENSG00000237631 | AL161454,1 | 0,916127573 | 1,887043359 | 0,35076124 | 0,844035671 | parkinson disease 7 (PARK7) pseudogene |
| ENSG00000260537 | AC012184,2 | 0,915981246 | 1,886851975 | 0,5084182 | 0,922427526 | novel protein, DDX19B and DDX19A readthrough |
| ENSG00000147140 | NONO | 0,915967889 | 1,886834504 | 9,30E-06 | 0,000320787 | non-POU domain containing octamer binding |
| ENSG00000112655 | PTK7 | 0,915741512 | 1,886538461 | 0,00247756 | 0,039747347 | protein tyrosine kinase 7 (inactive) |
| ENSG00000249252 | C1QTNF7-AS1 | 0,915693144 | 1,886475213 | 0,47295909 | 0,909532026 | C1QTNF7 antisense RNA 1 |
| ENSG00000198597 | ZNF536 | 0,91515009 | 1,885765246 | 0,00766461 | 0,094788909 | zinc finger protein 536 |
| ENSG00000142694 | EVA1B | 0,915069638 | 1,885660089 | 0,05797757 | 0,372428727 | eva-1 homolog B |

| | | | | | | |
|---|---|---|---|---|---|---|
| ENSG00000287493 | AC069503,5 | 0,915023585 | 1,885599897 | 0,54983378 | 0,936121729 | novel transcript |
| ENSG00000107165 | TYRP1 | 0,914913482 | 1,885455998 | 0,41276636 | 0,884576962 | tyrosinase related protein 1 |
| ENSG00000173210 | ABLIM3 | 0,914810089 | 1,885320878 | 0,0311746 | 0,253879337 | actin binding LIM protein family member 3 |
| ENSG00000281015 | LINC01596 | 0,914498512 | 1,884913752 | 0,49531915 | 0,917777383 | long intergenic non-protein coding RNA 1596 |
| ENSG00000187398 | LUZP2 | 0,913503397 | 1,88361406 | 0,02375126 | 0,210728318 | leucine zipper protein 2 |
| ENSG00000081041 | CXCL2 | 0,913481974 | 1,88358609 | 0,1935683 | 0,690424879 | C-X-C motif chemokine ligand 2 |
| ENSG00000275031 | METRNL | 0,913434505 | 1,883524115 | 0,09199236 | 0,482073274 | meteorin like, glial cell differentiation regulator |
| ENSG00000277527 | LINC01796 | 0,913319887 | 1,883374481 | 0,54748593 | 0,935404974 | long intergenic non-protein coding RNA 1796 |
| ENSG00000249349 | AC114928,1 | 0,912778488 | 1,882667841 | 0,45944119 | 0,904873969 | novel transcript |
| ENSG00000245330 | AP005717,1 | 0,912747023 | 1,882626781 | 0,29366528 | 0,803461221 | novel transcript |
| ENSG00000228222 | AC073050,1 | 0,912422614 | 1,882203495 | 0,22674032 | 0,735383523 | novel transcript (hCG1660379) |
| ENSG00000115641 | FHL2 | 0,912405959 | 1,882181766 | 0,02207842 | 0,200191372 | four and a half LIM domains 2 |
| ENSG00000240280 | TCAM1P | 0,911369875 | 1,880830547 | 0,23620506 | 0,745253314 | testicular cell adhesion molecule 1, pseudogene |
| ENSG00000234906 | APOC2 | 0,911102476 | 1,880481973 | 0,43412284 | 0,896272827 | apolipoprotein C2 |
| ENSG00000263278 | AC233701,1 | 0,910619755 | 1,879852874 | 0,49569393 | 0,918046024 | novel transcript, sense intronic to RBFOX3 |
| ENSG00000170340 | B3GNT2 | 0,909957344 | 1,878989942 | 0,02502078 | 0,219551856 | UDP-GlcNAc:betaGal beta-1,3-N-acetylglucosaminyltransferase 2 |
| ENSG00000197471 | SPN | 0,909732353 | 1,878696932 | 0,18246559 | 0,672792109 | sialophorin |
| ENSG00000282895 | RINL | 0,909719931 | 1,878680756 | 0,15207561 | 0,620332737 | Ras and Rab interactor like |
| ENSG00000214575 | CPEB1 | 0,909505261 | 1,878401233 | 0,04863782 | 0,335726918 | cytoplasmic polyadenylation element binding protein 1 |
| ENSG00000271303 | SRXN1 | 0,909471326 | 1,87835705 | 0,00044049 | 0,009368268 | sulfiredoxin 1 |
| ENSG00000250390 | AP001790,1 | 0,909419079 | 1,878289028 | 0,30237094 | 0,812035949 | novel transcript |
| ENSG00000137831 | UACA | 0,90935472 | 1,878205238 | 0,00020533 | 0,004886423 | uveal autoantigen with coiled-coil domains and ankyrin repeats |
| ENSG00000159176 | CSRP1 | 0,90924324 | 1,878060111 | 0,00039751 | 0,008588647 | cysteine and glycine rich protein 1 |
| ENSG00000176463 | SLCO3A1 | 0,909217496 | 1,878026599 | 0,00084521 | 0,016132236 | solute carrier organic anion transporter family member 3A1 |
| ENSG00000267048 | AC092143,3 | 0,908583732 | 1,877201778 | 0,24275405 | 0,752892335 | novel transcript (MC1R-TUBB3 readthrough) |
| ENSG00000085117 | CD82 | 0,908436197 | 1,877009819 | 0,03446347 | 0,270819603 | CD82 molecule |
| ENSG00000244672 | LY6G5B | 0,908245364 | 1,876761553 | 0,35581506 | 0,8472355 | lymphocyte antigen 6 family member G5B |
| ENSG00000145242 | EPHA5 | 0,908179991 | 1,876676514 | 0,10465353 | 0,515975114 | EPH receptor A5 |
| ENSG00000179218 | CALR | 0,908136734 | 1,876620245 | 0,00068227 | 0,013491822 | calreticulin |

| | | | | | | |
|---|---|---|---|---|---|---|
| ENSG00000245667 | AC006064,1 | 0,907570719 | 1,875884132 | 0,35893416 | 0,849814998 | novel transcript |
| ENSG00000261052 | SULT1A3 | 0,907455557 | 1,875734397 | 0,00534423 | 0,072612755 | sulfotransferase family 1A member 3 |
| ENSG00000102409 | BEX4 | 0,907216678 | 1,875423841 | 0,02576488 | 0,223443317 | brain expressed X-linked 4 |
| ENSG00000227516 | AL160163,1 | 0,906995322 | 1,875136113 | 0,48186827 | 0,912981134 | novel transcript |
| ENSG00000169239 | CA5B | 0,906748165 | 1,874814899 | 0,00402926 | 0,059154371 | carbonic anhydrase 5B |
| ENSG00000273010 | AL360270,3 | 0,905989692 | 1,873829505 | 0,07737736 | 0,439601283 | novel transcript, antisense to LRIF1 |
| ENSG00000102010 | BMX | 0,90576722 | 1,873540572 | 0,54991068 | 0,93612918 | BMX non-receptor tyrosine kinase |
| ENSG00000283958 | MIR1248 | 0,905258669 | 1,872880264 | 0,70409328 | 0,969077997 | microRNA 1248 |
| ENSG00000269694 | AC005197,1 | 0,905048379 | 1,87260729 | 0,45319626 | 0,902540855 | novel transcript, antisense to CERS1 |
| ENSG00000279998 | FP475955,3 | 0,904790874 | 1,872273079 | 0,43588124 | 0,896422069 | novel transcript |
| ENSG00000196839 | ADA | 0,904670052 | 1,872116288 | 0,02735821 | 0,232805386 | adenosine deaminase |
| ENSG00000124570 | SERPINB6 | 0,904463512 | 1,87184829 | 0,00779764 | 0,095940774 | serpin family B member 6 |
| ENSG00000285460 | BCAR1 | 0,904364104 | 1,871719316 | 0,03960581 | 0,296512476 | BCAR1 scaffold protein, Cas family member |
| ENSG00000197872 | CYRIA | 0,904209783 | 1,871519115 | 0,01921894 | 0,182079045 | CYFIP related Rac1 interactor A |
| ENSG00000129103 | SUMF2 | 0,904149539 | 1,871440966 | 0,0049494 | 0,068317084 | sulfatase modifying factor 2 |
| ENSG00000246363 | LINC02458 | 0,903995603 | 1,871241293 | 0,12685121 | 0,568970141 | long intergenic non-protein coding RNA 2458 |
| ENSG00000105281 | SLC1A5 | 0,903957826 | 1,871192294 | 0,00226042 | 0,037019304 | solute carrier family 1 member 5 |
| ENSG00000188000 | OR7D2 | 0,90364384 | 1,870785095 | 0,28018681 | 0,790779983 | olfactory receptor family 7 subfamily D member 2 |
| ENSG00000163807 | KIAA1143 | 0,903384264 | 1,870448526 | 0,14378681 | 0,603159913 | KIAA1143 |
| ENSG00000115963 | RND3 | 0,903323284 | 1,870369468 | 0,01958297 | 0,184067141 | Rho family GTPase 3 |
| ENSG00000157557 | ETS2 | 0,903173298 | 1,87017503 | 0,00032402 | 0,00722599 | ETS proto-oncogene 2, transcription factor |
| ENSG00000214354 | PAFAH1B1P1 | 0,903077206 | 1,870050468 | 0,37057225 | 0,858545694 | platelet activating factor acetylhydrolase 1b regulatory subunit 1 pseudogene 1 |
| ENSG00000226355 | AL353803,1 | 0,902687916 | 1,869545932 | 0,23398362 | 0,743537585 | novel transcript |
| ENSG00000167800 | TBX10 | 0,902628036 | 1,869468336 | 0,21579415 | 0,722999018 | T-box transcription factor 10 |
| ENSG00000285022 | MFHAS1 | 0,902172057 | 1,868877565 | 0,22469924 | 0,733203418 | malignant fibrous histiocytoma amplified sequence 1 |
| ENSG00000108604 | SMARCD2 | 0,902127062 | 1,868819279 | 0,00107167 | 0,019848716 | SWI/SNF related, matrix associated, actin dependent regulator of chromatin, subfamily d, member 2 |
| ENSG00000114853 | ZBTB47 | 0,902122066 | 1,868812806 | 0,09457206 | 0,488468697 | zinc finger and BTB domain containing 47 |
| ENSG00000113248 | PCDHB15 | 0,902064596 | 1,868738364 | 0,00132595 | 0,023728925 | protocadherin beta 15 |
| ENSG00000158486 | DNAH3 | 0,90192163 | 1,868553188 | 0,20371436 | 0,704215578 | dynein axonemal heavy chain 3 |

| | | | | | | |
|---|---|---|---|---|---|---|
| ENSG00000264113 | RN7SL784P | 0,901915635 | 1,868545422 | 0,36875412 | 0,857685565 | RNA, 7SL, cytoplasmic 784, pseudogene |
| ENSG00000176720 | BOK | 0,901355101 | 1,867819574 | 0,02093099 | 0,192865038 | BCL2 family apoptosis regulator BOK |
| ENSG00000278139 | P3R3URF-PIK3F | 0,900929176 | 1,86726822 | 0,81971548 | 0,98249617 | P3R3URF-PIK3R3 readthrough |
| ENSG00000116017 | ARID3A | 0,900794617 | 1,867094071 | 0,02682333 | 0,229377638 | AT-rich interaction domain 3A |
| ENSG00000172243 | CLEC7A | 0,900608421 | 1,866853116 | 0,18818893 | 0,682491978 | C-type lectin domain containing 7A |
| ENSG00000213977 | TAX1BP3 | 0,900361051 | 1,866533046 | 0,00255799 | 0,040844283 | Tax1 binding protein 3 |
| ENSG00000185201 | IFITM2 | 0,900352755 | 1,866522312 | 7,71E-05 | 0,002085919 | interferon induced transmembrane protein 2 |
| ENSG00000151364 | KCTD14 | 0,900223207 | 1,866354715 | 0,06932521 | 0,415287103 | potassium channel tetramerization domain containing 14 |
| ENSG00000115935 | WIPF1 | 0,899977265 | 1,866036577 | 0,00121688 | 0,021965248 | WAS/WASL interacting protein family member 1 |
| ENSG00000180422 | LINC00304 | 0,899973965 | 1,866032308 | 0,40511717 | 0,878176155 | long intergenic non-protein coding RNA 304 |
| ENSG00000100276 | RASL10A | 0,899604493 | 1,865554481 | 0,14581 | 0,608479758 | RAS like family 10 member A |
| ENSG00000197927 | NBEAP2 | 0,899465084 | 1,86537422 | 0,5331787 | 0,931291085 | neurobeachin pseudogene 2 |
| ENSG00000280025 | AL008638,6 | 0,899267228 | 1,865118414 | 0,46450122 | 0,907547847 | novel transcript |
| ENSG00000119777 | TMEM214 | 0,898827239 | 1,864549681 | 0,00052491 | 0,010853767 | transmembrane protein 214 |
| ENSG00000119408 | NEK6 | 0,898772702 | 1,864479199 | 0,00457397 | 0,064676905 | NIMA related kinase 6 |
| ENSG00000213398 | LCAT | 0,89765156 | 1,863030845 | 0,06218612 | 0,389148682 | lecithin-cholesterol acyltransferase |
| ENSG00000255363 | LINC02757 | 0,897068703 | 1,862278321 | 0,50261814 | 0,920607749 | long intergenic non-protein coding RNA 2757 |
| ENSG00000165841 | CYP2C19 | 0,897036807 | 1,86223715 | 0,41541114 | 0,885986046 | cytochrome P450 family 2 subfamily C member 19 |
| ENSG00000134030 | CTIF | 0,896935273 | 1,862106094 | 0,00081881 | 0,015747241 | cap binding complex dependent translation initiation factor |
| ENSG00000101972 | STAG2 | 0,896757708 | 1,861876923 | 0,02511462 | 0,219776434 | stromal antigen 2 |
| ENSG00000204434 | POTEKP | 0,896589524 | 1,861659885 | 0,26915404 | 0,781347689 | POTE ankyrin domain family member K, pseudogene |
| ENSG00000182158 | CREB3L2 | 0,896491349 | 1,861533203 | 1,93E-05 | 0,00061048 | cAMP responsive element binding protein 3 like 2 |
| ENSG00000136436 | CALCOCO2 | 0,896386738 | 1,861398226 | 0,00496272 | 0,068419936 | calcium binding and coiled-coil domain 2 |
| ENSG00000229802 | SLC39A7 | 0,896106443 | 1,861036618 | 0,017976 | 0,174048239 | solute carrier family 39 member 7 |
| ENSG00000206288 | SLC39A7 | 0,896106443 | 1,861036618 | 0,017976 | 0,174048239 | solute carrier family 39 member 7 |
| ENSG00000227402 | SLC39A7 | 0,896106443 | 1,861036618 | 0,017976 | 0,174048239 | solute carrier family 39 member 7 |
| ENSG00000224399 | SLC39A7 | 0,896106443 | 1,861036618 | 0,017976 | 0,174048239 | solute carrier family 39 member 7 |
| ENSG00000226614 | SLC39A7 | 0,896106443 | 1,861036618 | 0,017976 | 0,174048239 | solute carrier family 39 member 7 |
| ENSG00000188706 | ZDHHC9 | 0,895911157 | 1,860784722 | 3,39E-05 | 0,001007311 | zinc finger DHHC-type palmitoyltransferase 9 |
| ENSG00000204869 | IGFL4 | 0,895857049 | 1,860714936 | 0,16262857 | 0,639889157 | IGF like family member 4 |
| ENSG00000187848 | P2RX2 | 0,895642891 | 1,860438746 | 0,35684403 | 0,848328773 | purinergic receptor P2X 2 |

| | | | | | | |
|---|---|---|---|---|---|---|
| ENSG00000286281 | AL353152,2 | 0,895167466 | 1,859825758 | 0,27212741 | 0,785213785 | novel transcript |
| ENSG00000178951 | ZBTB7A | 0,895118133 | 1,859762162 | 0,04179663 | 0,306600121 | zinc finger and BTB domain containing 7A |
| ENSG00000101938 | CHRDL1 | 0,895079085 | 1,859711828 | 0,0552729 | 0,362959872 | chordin like 1 |
| ENSG00000271631 | AL139041,1 | 0,894607148 | 1,859103574 | 0,4675384 | 0,907940653 | novel transcript, antisense to SLC46A2 |
| ENSG00000105397 | TYK2 | 0,894603119 | 1,859098382 | 0,01377712 | 0,144636673 | tyrosine kinase 2 |
| ENSG00000270137 | AF230666,2 | 0,894589741 | 1,859081144 | 0,36058085 | 0,851271174 | novel transcript, intronic to PHF20L1 |
| ENSG00000276261 | AC009509,4 | 0,894321223 | 1,858735159 | 0,51086035 | 0,922693987 | novel transcript, antisense to a lncRNA |
| ENSG00000242479 | AC109992,1 | 0,894297269 | 1,858704298 | 0,13430661 | 0,584428964 | eukaryotic translation initiation factor 2-alpha kinase 1 (EIF2AK1) pseudogene |
| ENSG00000243811 | APOBEC3D | 0,894192255 | 1,858569006 | 0,16497341 | 0,644037807 | apolipoprotein B mRNA editing enzyme catalytic subunit 3D |
| ENSG00000154511 | DIPK1A | 0,894152589 | 1,858517908 | 0,01043861 | 0,118383134 | divergent protein kinase domain 1A |
| ENSG00000213363 | RPS3P6 | 0,893859334 | 1,858140167 | 0,24938877 | 0,761381055 | ribosomal protein S3 pseudogene 6 |
| ENSG00000231945 | VARS1 | 0,893752399 | 1,858002443 | 0,00696324 | 0,088341044 | valyl-tRNA synthetase 1 |
| ENSG00000165949 | IFI27 | 0,893711984 | 1,857950395 | 0,10733761 | 0,522893717 | interferon alpha inducible protein 27 |
| ENSG00000145819 | ARHGAP26 | 0,893612563 | 1,857822361 | 0,00976299 | 0,113030801 | Rho GTPase activating protein 26 |
| ENSG00000206811 | SNORA10 | 0,893605699 | 1,857813523 | 0,49443209 | 0,91758518 | small nucleolar RNA, H/ACA box 10 |
| ENSG00000111846 | GCNT2 | 0,893288118 | 1,857404606 | 0,1196056 | 0,552362303 | glucosaminyl (N-acetyl) transferase 2 (I blood group) |
| ENSG00000259704 | AC124248,1 | 0,89283598 | 1,856822591 | 0,43150469 | 0,89530663 | novel transcript, sense overlapping 3' UTR of BLM |
| ENSG00000144063 | MALL | 0,892622216 | 1,856547486 | 0,0263243 | 0,22653539 | mal, T cell differentiation protein like |
| ENSG00000241155 | ARHGAP31-AS1 | 0,892226268 | 1,856038026 | 0,23514885 | 0,744654651 | ARHGAP31 antisense RNA 1 |
| ENSG00000226067 | LINC00623 | 0,892196639 | 1,855999908 | 0,00587805 | 0,077671698 | long intergenic non-protein coding RNA 623 |
| ENSG00000181789 | COPG1 | 0,891991879 | 1,855736507 | 0,00014237 | 0,003552374 | COPI coat complex subunit gamma 1 |
| ENSG00000287932 | AC000065,2 | 0,891968558 | 1,855706509 | 0,0376342 | 0,286834587 | novel transcript, sense intronic to CDK6 |
| ENSG00000257275 | AL139020,1 | 0,891829498 | 1,855527648 | 0,45963676 | 0,905088088 | novel transcript, antisense to TCL1A |
| ENSG00000182704 | TSKU | 0,891718424 | 1,855384796 | 0,0023298 | 0,037897375 | tsukushi, small leucine rich proteoglycan |
| ENSG00000105499 | PLA2G4C | 0,89099699 | 1,854457224 | 0,11287236 | 0,53680967 | phospholipase A2 group IVC |
| ENSG00000261751 | AC007603,1 | 0,89091134 | 1,854347131 | 0,45345329 | 0,902754541 | novel transcript |
| ENSG00000264169 | RN7SL665P | 0,89087734 | 1,85430343 | 0,45378142 | 0,902912642 | RNA, 7SL, cytoplasmic 665, pseudogene |
| ENSG00000287842 | AL161714,1 | 0,890726112 | 1,854109067 | 0,31250562 | 0,818574551 | novel transcript, sense intronic to ENOX1 |
| ENSG00000261239 | ANKRD26P1 | 0,890685483 | 1,854056853 | 0,49148476 | 0,916995341 | ankyrin repeat domain 26 pseudogene 1 |
| ENSG00000005893 | LAMP2 | 0,890627062 | 1,853981775 | 0,03234853 | 0,260320302 | lysosomal associated membrane protein 2 |
| ENSG00000249302 | FTH1P24 | 0,890601841 | 1,853949364 | 0,31501869 | 0,820318541 | ferritin heavy chain 1 pseudogene 24 |

| | | | | | | |
|---|---|---|---|---|---|---|
| ENSG00000117151 | CTBS | 0,890535252 | 1,853863796 | 0,05545148 | 0,363656395 | chitobiase |
| ENSG00000156413 | FUT6 | 0,890329512 | 1,853599438 | 0,30433382 | 0,813156808 | fucosyltransferase 6 |
| ENSG00000151929 | BAG3 | 0,890308198 | 1,853572054 | 0,00144082 | 0,025420869 | BAG cochaperone 3 |
| ENSG00000267423 | AC005616,1 | 0,889501212 | 1,85253553 | 0,25956618 | 0,772176335 | novel transcript |
| ENSG00000266442 | MSH5-SAPCD1 | 0,889488566 | 1,852519291 | 0,40729021 | 0,880314995 | MSH5-SAPCD1 readthrough (NMD candidate) |
| ENSG00000206279 | DAXX | 0,888937112 | 1,851811321 | 0,01384717 | 0,145079743 | death domain associated protein |
| ENSG00000138669 | PRKG2 | 0,888922007 | 1,851791933 | 0,15011602 | 0,615828266 | protein kinase cGMP-dependent 2 |
| ENSG00000230863 | AL137803,1 | 0,88882925 | 1,851672877 | 0,40747966 | 0,880449269 | histidine acid phosphatase domain containing 1 (HISPPD1) pseudogene |
| ENSG00000100867 | DHRS2 | 0,888815366 | 1,851655057 | 0,22028927 | 0,728572782 | dehydrogenase/reductase 2 |
| ENSG00000186567 | CEACAM19 | 0,888241379 | 1,850918509 | 0,10404992 | 0,514473624 | CEA cell adhesion molecule 19 |
| ENSG00000261821 | AC090826,1 | 0,888038008 | 1,850657611 | 0,579801 | 0,943828882 | novel transcript, antisense to CYP11A1 |
| ENSG00000206941 | SNORD15A | 0,887977789 | 1,850580365 | 0,6310366 | 0,953647802 | small nucleolar RNA, C/D box 15A |
| ENSG00000204217 | BMPR2 | 0,887815739 | 1,850372511 | 0,00689592 | 0,087746857 | bone morphogenetic protein receptor type 2 |
| ENSG00000185129 | PURA | 0,88757418 | 1,850062718 | 0,0043379 | 0,062382733 | purine rich element binding protein A |
| ENSG00000120332 | TNN | 0,88738401 | 1,849818868 | 0,13362312 | 0,583882708 | tenascin N |
| ENSG00000286090 | AC034110,1 | 0,887084312 | 1,849434635 | 0,48533696 | 0,913885618 | novel transcript |
| ENSG00000164099 | PRSS12 | 0,886923845 | 1,849228939 | 0,0773524 | 0,439539362 | serine protease 12 |
| ENSG00000170775 | GPR37 | 0,886795699 | 1,849064691 | 0,00443162 | 0,063178325 | G protein-coupled receptor 37 |
| ENSG00000229520 | LINC00404 | 0,886757258 | 1,849015422 | 0,13335731 | 0,583524541 | long intergenic non-protein coding RNA 404 |
| ENSG00000279914 | AL008638,5 | 0,886500764 | 1,848686718 | 0,30238221 | 0,812035949 | novel transcript |
| ENSG00000069702 | TGFBR3 | 0,88647981 | 1,848659868 | 0,01554103 | 0,157881532 | transforming growth factor beta receptor 3 |
| ENSG00000021645 | NRXN3 | 0,885986797 | 1,848028233 | 0,0035428 | 0,053389744 | neurexin 3 |
| ENSG00000153303 | FRMD1 | 0,885788186 | 1,847773838 | 0,46782451 | 0,907940653 | FERM domain containing 1 |
| ENSG00000129151 | BBOX1 | 0,885775948 | 1,847758163 | 0,16876617 | 0,650757516 | gamma-butyrobetaine hydroxylase 1 |
| ENSG00000185499 | MUC1 | 0,885503491 | 1,847409243 | 0,07155506 | 0,421457662 | mucin 1, cell surface associated |
| ENSG00000215841 | AP002761,1 | 0,885218211 | 1,84704397 | 0,36537016 | 0,855323027 | novel transcript |
| ENSG00000232412 | AL121601,1 | 0,884978484 | 1,84673708 | 0,10385426 | 0,513912548 | novel transcript |
| ENSG00000076641 | PAG1 | 0,884207739 | 1,845750743 | 0,01086224 | 0,121796931 | phosphoprotein membrane anchor with glycosphingolipid microdomains 1 |
| ENSG00000197165 | SULT1A2 | 0,88407676 | 1,845583179 | 0,13867924 | 0,593837907 | sulfotransferase family 1A member 2 |
| ENSG00000106384 | MOGAT3 | 0,884046379 | 1,845544314 | 0,38836438 | 0,869820053 | monoacylglycerol O-acyltransferase 3 |
| ENSG00000172318 | B3GALT1 | 0,88404089 | 1,845537292 | 0,01670225 | 0,165411048 | beta-1,3-galactosyltransferase 1 |

| | | | | | | |
|---|---|---|---|---|---|---|
| ENSG00000157399 | ARSL | 0,884033617 | 1,845527988 | 0,00025857 | 0,005931065 | arylsulfatase L |
| ENSG00000225217 | HSPA7 | 0,883918418 | 1,845380629 | 0,54976747 | 0,936108121 | heat shock protein family A (Hsp70) member 7 (pseudogene) |
| ENSG00000150551 | LYPD1 | 0,883911138 | 1,845371317 | 0,0336078 | 0,266408273 | LY6/PLAUR domain containing 1 |
| ENSG00000272695 | GAS6-DT | 0,883334375 | 1,844633719 | 0,21944968 | 0,727838473 | GAS6 divergent transcript |
| ENSG00000145246 | ATP10D | 0,883093754 | 1,844326086 | 0,00119539 | 0,02166534 | ATPase phospholipid transporting 10D (putative) |
| ENSG00000250620 | LINC02515 | 0,882717553 | 1,843845217 | 0,15862665 | 0,632402923 | long intergenic non-protein coding RNA 2515 |
| ENSG00000258215 | AC078886,1 | 0,88261146 | 1,843709629 | 0,35259646 | 0,844912682 | tripartite motif containing 61 (TRIM61) pseudogene |
| ENSG00000176349 | AC104129,1 | 0,882376919 | 1,843409919 | 0,32274673 | 0,824955371 | novel transcript, antisense to MAD1L1 |
| ENSG00000167778 | SPRYD3 | 0,882079132 | 1,84302946 | 0,05405698 | 0,35836452 | SPRY domain containing 3 |
| ENSG00000288216 | SLC5A11 | 0,881992802 | 1,842919178 | 0,46775843 | 0,907940653 | solute carrier family 5 member 11 |
| ENSG00000158865 | SLC5A11 | 0,881992802 | 1,842919178 | 0,46775843 | 0,907940653 | solute carrier family 5 member 11 |
| ENSG00000245059 | AC092718,1 | 0,881480925 | 1,842265414 | 0,23058968 | 0,739234644 | novel transcript |
| ENSG00000282735 | GARRE1 | 0,88132027 | 1,842060275 | 0,36179132 | 0,851979035 | granule associated Rac and RHOG effector 1 |
| ENSG00000267264 | AC006504,3 | 0,880752054 | 1,841334909 | 0,37588232 | 0,862406526 | vomeronasal receptor pseudogene |
| ENSG00000253532 | AC107909,1 | 0,880725344 | 1,841300819 | 0,60282709 | 0,950352286 | novel transcript |
| ENSG00000214031 | CLDN7P1 | 0,880676547 | 1,841238542 | 0,50659021 | 0,921824694 | CLDN7 pseudogene 1 |
| ENSG00000282041 | Z82244,1 | 0,880519696 | 1,841038372 | 0,33412314 | 0,834518413 | novel transcript |
| ENSG00000253767 | PCDHGA8 | 0,880192312 | 1,840620641 | 0,0270444 | 0,230637202 | protocadherin gamma subfamily A, 8 |
| ENSG00000175467 | SART1 | 0,879993023 | 1,840366401 | 0,2171619 | 0,724864986 | spliceosome associated factor 1, recruiter of U4/U6,U5 tri-snRNP |
| ENSG00000169756 | LIMS1 | 0,879726237 | 1,840026108 | 6,55E-05 | 0,001806572 | LIM zinc finger domain containing 1 |
| ENSG00000232312 | GPANK1 | 0,879510051 | 1,839750404 | 0,04926907 | 0,338538066 | G-patch domain and ankyrin repeats 1 |
| ENSG00000236383 | CCDC200 | 0,879313925 | 1,839500317 | 0,18686032 | 0,680474679 | coiled-coil domain containing 200 |
| ENSG00000205622 | AP001042,1 | 0,879149016 | 1,839290062 | 0,41033963 | 0,882614012 | novel transcript |
| ENSG00000049089 | COL9A2 | 0,87913619 | 1,839273711 | 0,09795734 | 0,497096776 | collagen type IX alpha 2 chain |
| ENSG00000034152 | MAP2K3 | 0,879135862 | 1,839273293 | 0,02060622 | 0,1906578 | mitogen-activated protein kinase kinase 3 |
| ENSG00000227308 | LINC02832 | 0,878947236 | 1,839032831 | 0,25453644 | 0,765935007 | long intergenic non-protein coding RNA 2832 |
| ENSG00000108846 | ABCC3 | 0,878809652 | 1,838857459 | 0,2361547 | 0,745253314 | ATP binding cassette subfamily C member 3 |
| ENSG00000197976 | AKAP17A | 0,878695315 | 1,838711731 | 0,11189139 | 0,534735697 | A-kinase anchoring protein 17A |
| ENSG00000128965 | CHAC1 | 0,878231058 | 1,838120132 | 0,11477035 | 0,541181798 | ChaC glutathione specific gamma-glutamylcyclotransferase 1 |
| ENSG00000124208 | PEDS1-UBE2V1 | 0,878045375 | 1,83788357 | 0,46743022 | 0,907940653 | PEDS1-UBE2V1 readthrough |

| | | | | | | |
|---|---|---|---|---|---|---|
| ENSG00000137449 | CPEB2 | 0,877960831 | 1,837775871 | 0,02189818 | 0,198960988 | cytoplasmic polyadenylation element binding protein 2 |
| ENSG00000104964 | TLE5 | 0,877796374 | 1,83756639 | 0,02454881 | 0,216317369 | TLE family member 5, transcriptional modulator |
| ENSG00000198825 | INPP5F | 0,877377083 | 1,837032415 | 0,02619667 | 0,225996859 | inositol polyphosphate-5-phosphatase F |
| ENSG00000162989 | KCNJ3 | 0,877350322 | 1,836998339 | 0,27840879 | 0,789733543 | potassium inwardly rectifying channel subfamily J member 3 |
| ENSG00000136717 | BIN1 | 0,877136364 | 1,836725924 | 0,02699673 | 0,230419233 | bridging integrator 1 |
| ENSG00000049540 | ELN | 0,876638576 | 1,836092289 | 0,05020934 | 0,342742134 | elastin |
| ENSG00000255276 | RRM1-AS1 | 0,876428283 | 1,835824672 | 0,37941894 | 0,86496672 | RRM1 antisense RNA 1 |
| ENSG00000161835 | TAMALIN | 0,876297359 | 1,83565808 | 0,07552197 | 0,434109026 | trafficking regulator and scaffold protein tamalin |
| ENSG00000118971 | CCND2 | 0,875994435 | 1,835272685 | 0,00296225 | 0,045915666 | cyclin D2 |
| ENSG00000250635 | CXXC5-AS1 | 0,875434394 | 1,834560387 | 0,30838883 | 0,816625411 | CXXC5 antisense RNA 1 |
| ENSG00000049860 | HEXB | 0,875299711 | 1,83438913 | 0,01138828 | 0,126077616 | hexosaminidase subunit beta |
| ENSG00000273293 | AC093458,1 | 0,875165774 | 1,834218836 | 0,5931839 | 0,947416795 | novel transcript |
| ENSG00000285527 | AC136285,3 | 0,875048849 | 1,834070186 | 0,288342 | 0,799673937 | novel transcript |
| ENSG00000267325 | LINC01415 | 0,87490693 | 1,833889777 | 0,14984738 | 0,61544661 | long intergenic non-protein coding RNA 1415 |
| ENSG00000227893 | LINC00352 | 0,874846931 | 1,83381351 | 0,38850409 | 0,869820053 | long intergenic non-protein coding RNA 352 |
| ENSG00000213997 | PGAM1P7 | 0,874753471 | 1,833694716 | 0,32700525 | 0,828735859 | phosphoglycerate mutase 1 pseudogene 7 |
| ENSG00000114251 | WNT5A | 0,874746446 | 1,833685787 | 0,04104386 | 0,303420267 | Wnt family member 5A |
| ENSG00000233038 | PTPRN2-AS1 | 0,874734472 | 1,833670568 | 0,59184333 | 0,947416795 | PTPRN2 antisense RNA 1 |
| ENSG00000255200 | PGAM1P8 | 0,874653992 | 1,83356828 | 0,25225263 | 0,76432863 | phosphoglycerate mutase 1 pseudogene 8 |
| ENSG00000144057 | ST6GAL2 | 0,874624765 | 1,833531135 | 0,01455736 | 0,150888551 | ST6 beta-galactoside alpha-2,6-sialyltransferase 2 |
| ENSG00000124201 | ZNFX1 | 0,874493739 | 1,833364621 | 0,00627667 | 0,081714521 | zinc finger NFX1-type containing 1 |
| ENSG00000200488 | RN7SKP203 | 0,874463402 | 1,833326069 | 0,43582433 | 0,896422069 | RN7SK pseudogene 203 |
| ENSG00000286662 | AC087894,3 | 0,874323342 | 1,833148096 | 0,40321018 | 0,877589651 | novel transcript |
| ENSG00000236739 | CLIC4P1 | 0,8742431 | 1,833046139 | 0,37275362 | 0,860564851 | chloride intracellular channel 4 pseudogene 1 |
| ENSG00000285508 | AL034430,1 | 0,874229106 | 1,833028359 | 0,10665318 | 0,521491876 | novel protein |
| ENSG00000278087 | NOMO3 | 0,872660963 | 1,831037024 | 0,00072825 | 0,014252604 | NODAL modulator 3 |
| ENSG00000258743 | LINC02301 | 0,872616686 | 1,83098083 | 0,00908472 | 0,106736016 | long intergenic non-protein coding RNA 2301 |
| ENSG00000287309 | AL583859,3 | 0,872598628 | 1,830957912 | 0,25435829 | 0,765826923 | novel transcript |
| ENSG00000161091 | MFSD12 | 0,872273497 | 1,830545326 | 0,03316466 | 0,264437603 | major facilitator superfamily domain containing 12 |
| ENSG00000101882 | NKAP | 0,87199922 | 1,830197347 | 0,00113367 | 0,020739016 | NFKB activating protein |
| ENSG00000226445 | BX322234,1 | 0,871981681 | 1,830175097 | 0,1880031 | 0,68218766 | novel transcript |

| | | | | | | |
|---|---|---|---|---|---|---|
| ENSG00000180611 | MB21D2 | 0,871946768 | 1,830130808 | 0,00273621 | 0,042987437 | Mab-21 domain containing 2 |
| ENSG00000237732 | CT75 | 0,871596116 | 1,829686042 | 0,21121578 | 0,716986998 | cancer/testis associated transcript 75 |
| ENSG00000126461 | SCAF1 | 0,871535257 | 1,82960886 | 0,15493252 | 0,624591393 | SR-related CTD associated factor 1 |
| ENSG00000215915 | ATAD3C | 0,871419245 | 1,829461741 | 0,21808807 | 0,72573766 | ATPase family AAA domain containing 3C |
| ENSG00000189221 | MAOA | 0,871342203 | 1,829364048 | 0,02328742 | 0,207957138 | monoamine oxidase A |
| ENSG00000142669 | SH3BGRL3 | 0,871052388 | 1,828996593 | 0,00047095 | 0,009894953 | SH3 domain binding glutamate rich protein like 3 |
| ENSG00000259341 | AC015660,1 | 0,87094338 | 1,828858403 | 0,29747505 | 0,806684007 | novel transcript |
| ENSG00000154134 | ROBO3 | 0,870824065 | 1,828707157 | 0,01658703 | 0,164889286 | roundabout guidance receptor 3 |
| ENSG00000257986 | LINC02306 | 0,870714389 | 1,828568141 | 0,1605499 | 0,636125619 | long intergenic non-protein coding RNA 2306 |
| ENSG00000247774 | PCED1B-AS1 | 0,870655791 | 1,828493871 | 0,23500749 | 0,744654651 | PCED1B antisense RNA 1 |
| ENSG00000166333 | ILK | 0,870501723 | 1,828298614 | 5,98E-05 | 0,001661705 | integrin linked kinase |
| ENSG00000206432 | TMEM200C | 0,870456401 | 1,828241179 | 0,01181666 | 0,129435298 | transmembrane protein 200C |
| ENSG00000014216 | CAPN1 | 0,870063209 | 1,827742978 | 0,01581909 | 0,159883663 | calpain 1 |
| ENSG00000139083 | ETV6 | 0,869400074 | 1,826903048 | 0,00076975 | 0,014975415 | ETS variant transcription factor 6 |
| ENSG00000227210 | AC079145,1 | 0,869204539 | 1,826655457 | 0,08655872 | 0,468880561 | novel transcript |
| ENSG00000215397 | SCRT2 | 0,869185108 | 1,826630855 | 0,00838388 | 0,101046165 | scratch family transcriptional repressor 2 |
| ENSG00000253671 | AC027117,1 | 0,869165329 | 1,826605813 | 0,4099526 | 0,882453744 | novel transcript |
| ENSG00000124260 | MAGEA10 | 0,868581298 | 1,825866516 | 0,47618026 | 0,911413793 | MAGE family member A10 |
| ENSG00000198157 | HMGN5 | 0,868088147 | 1,825242494 | 0,07458734 | 0,431593284 | high mobility group nucleosome binding domain 5 |
| ENSG00000243260 | RN7SL558P | 0,867795075 | 1,824871748 | 0,46208221 | 0,906277266 | RNA, 7SL, cytoplasmic 558, pseudogene |
| ENSG00000267082 | AC011472,1 | 0,867413498 | 1,824389153 | 0,59511681 | 0,947639622 | novel transcript, antisense to DOCK6 |
| ENSG00000114654 | EFCC1 | 0,867362668 | 1,824324877 | 0,11434766 | 0,540514852 | EF-hand and coiled-coil domain containing 1 |
| ENSG00000106648 | GALNTL5 | 0,867324011 | 1,824275995 | 0,56397312 | 0,940865625 | polypeptide N-acetylgalactosaminyltransferase like 5 |
| ENSG00000270547 | LINC01235 | 0,867020161 | 1,823891819 | 0,244795 | 0,756700694 | long intergenic non-protein coding RNA 1235 |
| ENSG00000204642 | HLA-F | 0,866983236 | 1,823845137 | 0,25859056 | 0,771444876 | major histocompatibility complex, class I, F |
| ENSG00000270872 | SRGAP2D | 0,866662544 | 1,823439767 | 0,00058791 | 0,011981944 | SLIT-ROBO Rho GTPase activating protein 2D (pseudogene) |
| ENSG00000168418 | KCNG4 | 0,866644945 | 1,823417523 | 0,38600948 | 0,869112622 | potassium voltage-gated channel modifier subfamily G member 4 |
| ENSG00000224536 | AC096677,1 | 0,866552399 | 1,823300558 | 0,23137611 | 0,739268552 | novel transcript |
| ENSG00000185950 | IRS2 | 0,86653876 | 1,823283321 | 0,00832265 | 0,100540653 | insulin receptor substrate 2 |
| ENSG00000287056 | AL606970,5 | 0,866446329 | 1,82316651 | 0,2843779 | 0,795731744 | novel transcript |

| | | | | | | |
|---|---|---|---|---|---|---|
| ENSG00000166025 | AMOTL1 | 0,866308591 | 1,822992456 | 0,00012491 | 0,003177629 | angiomotin like 1 |
| ENSG00000234551 | LINC01309 | 0,866267283 | 1,822940259 | 0,36975689 | 0,858104601 | long intergenic non-protein coding RNA 1309 |
| ENSG00000273387 | AC005005,3 | 0,866116434 | 1,822749661 | 0,16011176 | 0,635191158 | novel transcript, antisense to SMTN |
| ENSG00000154928 | EPHB1 | 0,86600131 | 1,822604216 | 0,02987443 | 0,246399596 | EPH receptor B1 |
| ENSG00000268173 | AC007192,1 | 0,865444129 | 1,821900447 | 0,46540058 | 0,90786072 | novel protein, readthrough between PIK3R2 and IFI30 |
| ENSG00000233701 | PRR23C | 0,865438554 | 1,821893407 | 0,52818238 | 0,92998848 | proline rich 23C |
| ENSG00000171408 | PDE7B | 0,865362853 | 1,821797811 | 0,2298717 | 0,738309937 | phosphodiesterase 7B |
| ENSG00000150938 | CRIM1 | 0,865238579 | 1,821640888 | 0,0047454 | 0,066232244 | cysteine rich transmembrane BMP regulator 1 |
| ENSG00000175806 | MSRA | 0,864942622 | 1,821267232 | 0,18576688 | 0,678777213 | methionine sulfoxide reductase A |
| ENSG00000078124 | ACER3 | 0,864356092 | 1,820526943 | 0,01931595 | 0,182721243 | alkaline ceramidase 3 |
| ENSG00000267563 | AC011471,2 | 0,864266562 | 1,820413969 | 0,51091834 | 0,922715081 | novel transcript, antisense to MLLT1 |
| ENSG00000165675 | ENOX2 | 0,864072071 | 1,820168574 | 0,00334099 | 0,050879833 | ecto-NOX disulfide-thiol exchanger 2 |
| ENSG00000235986 | GNL1 | 0,864056325 | 1,820148709 | 0,04124679 | 0,304151397 | G protein nucleolar 1 (putative) |
| ENSG00000287430 | AL136090,2 | 0,863973463 | 1,82004417 | 0,51086978 | 0,922693987 | novel transcript |
| ENSG00000282992 | LGALS4 | 0,863716527 | 1,819720059 | 0,43894366 | 0,897402199 | galectin 4 |
| ENSG00000125733 | TRIP10 | 0,863386282 | 1,819303557 | 0,01105791 | 0,123293364 | thyroid hormone receptor interactor 10 |
| ENSG00000238018 | AC093110,1 | 0,862950744 | 1,818754408 | 0,09314262 | 0,484931061 | novel transcript, antisense to SPTBN1 |
| ENSG00000253288 | AC046195,1 | 0,862737551 | 1,818485663 | 0,53867183 | 0,933279924 | novel transcript |
| ENSG00000166426 | CRABP1 | 0,862717073 | 1,818459851 | 0,00191876 | 0,032509055 | cellular retinoic acid binding protein 1 |
| ENSG00000077942 | FBLN1 | 0,8624304 | 1,818098547 | 2,62E-05 | 0,000798461 | fibulin 1 |
| ENSG00000277800 | TBX22 | 0,862199 | 1,817806957 | 0,4858177 | 0,913997663 | T-box transcription factor 22 |
| ENSG00000129514 | FOXA1 | 0,861927539 | 1,817464947 | 0,39060621 | 0,870057277 | forkhead box A1 |
| ENSG00000095637 | SORBS1 | 0,861834752 | 1,81734806 | 0,00096626 | 0,018143718 | sorbin and SH3 domain containing 1 |
| ENSG00000250770 | AC005865,1 | 0,861614943 | 1,81707119 | 0,225491 | 0,734461165 | tetraspanin 11 (TSPAN11) pseudogene |
| ENSG00000267033 | AC020911,1 | 0,86148035 | 1,816901679 | 0,25109928 | 0,76347766 | novel transcript, antisense FAM32A |
| ENSG00000288607 | ELOA3B | 0,861439616 | 1,816850379 | 0,58953649 | 0,946941469 | elongin A3 family member B |
| ENSG00000185736 | ADARB2 | 0,861386786 | 1,816783849 | 0,10852281 | 0,52521009 | adenosine deaminase RNA specific B2 (inactive) |
| ENSG00000287038 | AL162388,2 | 0,861372759 | 1,816766185 | 0,0369681 | 0,283711768 | novel transcript |
| ENSG00000282454 | AC023315,1 | 0,861320636 | 1,816700548 | 0,55120607 | 0,936366211 | novel transcript, antisense to LHX1 |
| ENSG00000117280 | RAB29 | 0,861308817 | 1,816685666 | 0,06570876 | 0,402255481 | RAB29, member RAS oncogene family |
| ENSG00000162704 | ARPC5 | 0,861072867 | 1,816388574 | 0,00161378 | 0,027921043 | actin related protein 2/3 complex subunit 5 |
| ENSG00000060069 | CTDP1 | 0,861000474 | 1,816297432 | 0,11100867 | 0,532051958 | CTD phosphatase subunit 1 |

| | | | | | | |
|---|---|---|---|---|---|---|
| ENSG00000275568 | ARHGAP11A | 0,860512832 | 1,815683614 | 0,2593031 | 0,772081162 | Rho GTPase activating protein 11A |
| ENSG00000198142 | SOWAHC | 0,860478649 | 1,815640593 | 0,00079461 | 0,015363444 | sosondowah ankyrin repeat domain family member C |
| ENSG00000185896 | LAMP1 | 0,86036409 | 1,815496426 | 0,0006064 | 0,012286697 | lysosomal associated membrane protein 1 |
| ENSG00000274364 | AL110115,1 | 0,860250989 | 1,815354105 | 0,3073696 | 0,816143546 | novel transcript, sense intronic to HM13 |
| ENSG00000232633 | AC079465,1 | 0,859984294 | 1,815018551 | 0,12310961 | 0,561282565 | novel transcript |
| ENSG00000163378 | EOGT | 0,859862239 | 1,814865003 | 0,0235747 | 0,209803448 | EGF domain specific O-linked N-acetylglucosamine transferase |
| ENSG00000286234 | AC079790,2 | 0,859861116 | 1,814863591 | 0,4348509 | 0,896272827 | novel transcript |
| ENSG00000173040 | EVC2 | 0,859781877 | 1,814763913 | 0,02983592 | 0,246313735 | EvC ciliary complex subunit 2 |
| ENSG00000273189 | AC010619,2 | 0,859164117 | 1,813987 | 0,56000891 | 0,940033215 | novel transcript |
| ENSG00000227067 | DPPA3P1 | 0,859091581 | 1,813895799 | 0,4268375 | 0,892261855 | developmental pluripotency associated 3 pseudogene 1 |
| ENSG00000161133 | USP41 | 0,858662484 | 1,813356376 | 0,47201597 | 0,909171312 | ubiquitin specific peptidase 41 |
| ENSG00000048140 | TSPAN17 | 0,858053478 | 1,812591065 | 0,00317507 | 0,048779097 | tetraspanin 17 |
| ENSG00000166862 | CACNG2 | 0,858035909 | 1,812568991 | 0,30500723 | 0,813769074 | calcium voltage-gated channel auxiliary subunit gamma 2 |
| ENSG00000088726 | TMEM40 | 0,857594366 | 1,812014332 | 0,49715433 | 0,918759905 | transmembrane protein 40 |
| ENSG00000225032 | AL162586,1 | 0,857537075 | 1,811942376 | 0,28830913 | 0,799653716 | novel transcript |
| ENSG00000159387 | IRX6 | 0,857368803 | 1,811731049 | 0,1356394 | 0,587344894 | iroquois homeobox 6 |
| ENSG00000235527 | HIPK1-AS1 | 0,857312923 | 1,811660875 | 0,17966508 | 0,668298509 | HIPK1 antisense RNA 1 |
| ENSG00000287774 | AC025614,2 | 0,857180949 | 1,811495158 | 0,3161412 | 0,821394376 | novel transcript |
| ENSG00000261227 | AC140912,1 | 0,857139692 | 1,811443354 | 0,47070029 | 0,908869097 | novel transcript |
| ENSG00000172667 | ZMAT3 | 0,857123415 | 1,811422917 | 0,00346697 | 0,052413557 | zinc finger matrin-type 3 |
| ENSG00000286011 | AC062037,3 | 0,856753117 | 1,810958037 | 0,28235622 | 0,793419467 | novel transcript |
| ENSG00000120500 | ARR3 | 0,855791456 | 1,809751304 | 0,51736538 | 0,9257552 | arrestin 3 |
| ENSG00000248996 | AC145098,1 | 0,855753501 | 1,809703694 | 0,55926334 | 0,939727587 | novel transcript |
| ENSG00000050820 | BCAR1 | 0,855475368 | 1,80935484 | 0,04392268 | 0,316175438 | BCAR1 scaffold protein, Cas family member |
| ENSG00000276879 | IFI27L2 | 0,855331692 | 1,809174657 | 0,13893945 | 0,594174432 | interferon alpha inducible protein 27 like 2 |
| ENSG00000187024 | PTRH1 | 0,855249268 | 1,809071298 | 0,11395364 | 0,53948382 | peptidyl-tRNA hydrolase 1 homolog |
| ENSG00000285864 | AP000593,4 | 0,855114222 | 1,808901965 | 0,44062252 | 0,8985469 | novel transcript |
| ENSG00000113073 | SLC4A9 | 0,855009858 | 1,808771114 | 0,4851507 | 0,913690171 | solute carrier family 4 member 9 |
| ENSG00000225216 | AC007362,1 | 0,854899158 | 1,808632331 | 0,16603409 | 0,645311996 | novel transcript |

| | | | | | | |
|---|---|---|---|---|---|---|
| ENSG00000224819 | AC093843,1 | 0,854760206 | 1,808458141 | 0,34368138 | 0,840901506 | novel transcript |
| ENSG00000163947 | ARHGEF3 | 0,854477862 | 1,80810425 | 0,00943503 | 0,109999648 | Rho guanine nucleotide exchange factor 3 |
| ENSG00000248712 | CCDC153 | 0,854000907 | 1,807506589 | 0,14225281 | 0,600366679 | coiled-coil domain containing 153 |
| ENSG00000278843 | MMP28 | 0,853777707 | 1,807226971 | 0,53622921 | 0,932274935 | matrix metallopeptidase 28 |
| ENSG00000154188 | ANGPT1 | 0,853776961 | 1,807226036 | 0,1015611 | 0,506654908 | angiopoietin 1 |
| ENSG00000186111 | PIP5K1C | 0,853541212 | 1,806930745 | 0,01883403 | 0,17950267 | phosphatidylinositol-4-phosphate 5-kinase type 1 gamma |
| ENSG00000221955 | SLC12A8 | 0,853417511 | 1,80677582 | 0,02937759 | 0,243559001 | solute carrier family 12 member 8 |
| ENSG00000182687 | GALR2 | 0,853250047 | 1,806566106 | 0,52909722 | 0,930207681 | galanin receptor 2 |
| ENSG00000234203 | AC004771,1 | 0,852930658 | 1,806166206 | 0,46308366 | 0,906849208 | novel transcript, antisense to CAMTA2 |
| ENSG00000164116 | GUCY1A1 | 0,852678892 | 1,805851039 | 0,09969073 | 0,500760119 | guanylate cyclase 1 soluble subunit alpha 1 |
| ENSG00000064601 | CTSA | 0,85267822 | 1,805850198 | 0,00591489 | 0,07796419 | cathepsin A |
| ENSG00000174667 | OR7D4 | 0,851645653 | 1,804558175 | 0,48335445 | 0,913690171 | olfactory receptor family 7 subfamily D member 4 |
| ENSG00000189292 | ALKAL2 | 0,851589138 | 1,804487485 | 0,00998029 | 0,11484133 | ALK and LTK ligand 2 |
| ENSG00000263535 | AK4P1 | 0,851429139 | 1,804287374 | 0,25435479 | 0,765826923 | adenylate kinase 4 pseudogene 1 |
| ENSG00000282997 | AL162456,1 | 0,851268937 | 1,804087031 | 0,27080951 | 0,783073289 | novel transcript |
| ENSG00000168040 | FADD | 0,851065499 | 1,80383265 | 0,00748096 | 0,093069966 | Fas associated via death domain |
| ENSG00000152784 | PRDM8 | 0,850885044 | 1,803607037 | 0,02706499 | 0,230749905 | PR/SET domain 8 |
| ENSG00000130309 | COLGALT1 | 0,850846991 | 1,803559465 | 0,00561163 | 0,075200114 | collagen beta(1-O)galactosyltransferase 1 |
| ENSG00000186352 | ANKRD37 | 0,850845318 | 1,803557373 | 0,1084881 | 0,525152765 | ankyrin repeat domain 37 |
| ENSG00000188501 | LCTL | 0,850371778 | 1,802965484 | 0,00553018 | 0,074508967 | lactase like |
| ENSG00000203645 | LINC00501 | 0,850262106 | 1,80282843 | 0,40233798 | 0,87718264 | long intergenic non-protein coding RNA 501 |
| ENSG00000183397 | C19orf71 | 0,850005675 | 1,802508015 | 0,35130913 | 0,84405766 | chromosome 19 open reading frame 71 |
| ENSG00000007866 | TEAD3 | 0,849889939 | 1,80236342 | 0,00061534 | 0,012435622 | TEA domain transcription factor 3 |
| ENSG00000086062 | B4GALT1 | 0,849765316 | 1,802207735 | 0,00396984 | 0,058378646 | beta-1,4-galactosyltransferase 1 |
| ENSG00000288096 | AC109322,2 | 0,849690336 | 1,802114073 | 0,42647629 | 0,892138574 | novel transcript |
| ENSG00000288055 | AC107373,3 | 0,849173575 | 1,801468686 | 0,1046609 | 0,515975114 | novel transcript |
| ENSG00000287219 | AL021707,9 | 0,849164056 | 1,801456801 | 0,52051241 | 0,927042262 | novel transcript |
| ENSG00000222014 | RAB6C | 0,848789249 | 1,800988849 | 0,29849326 | 0,807973435 | RAB6C, member RAS oncogene family |
| ENSG00000228308 | LINC01209 | 0,848685411 | 1,800859228 | 0,29773557 | 0,806960585 | long intergenic non-protein coding RNA 1209 |
| ENSG00000243385 | AC094085,1 | 0,848435668 | 1,800547511 | 0,54177152 | 0,933491609 | ribosomal protein L13 (RPL13) pseudogene |
| ENSG00000271046 | AL512631,1 | 0,848288282 | 1,800363575 | 0,38333662 | 0,867760194 | novel transcript |
| ENSG00000250423 | KIAA1210 | 0,847667691 | 1,799589296 | 0,40315109 | 0,877589651 | KIAA1210 |

| | | | | | | |
|---|---|---|---|---|---|---|
| ENSG00000121440 | PDZRN3 | 0,847505862 | 1,799387445 | 0,02936389 | 0,243557543 | PDZ domain containing ring finger 3 |
| ENSG00000197769 | MAP1LC3C | 0,847497203 | 1,799376645 | 0,24764465 | 0,759874608 | microtubule associated protein 1 light chain 3 gamma |
| ENSG00000253519 | AC106801,1 | 0,847246425 | 1,799063894 | 0,07505663 | 0,432866745 | novel transcript |
| ENSG00000275586 | Metazoa_SRP | 0,847037154 | 1,798802948 | 0,35231379 | 0,844912682 | Metazoan signal recognition particle RNA |
| ENSG00000177112 | IRAG1-AS1 | 0,846911837 | 1,798646706 | 0,4707022 | 0,908869097 | IRAG1 antisense RNA 1 |
| ENSG00000265158 | LRRC37A7P | 0,846908826 | 1,798642952 | 0,32195204 | 0,824955371 | leucine rich repeat containing 37 member A7, pseudogene |
| ENSG00000256980 | KHDC1L | 0,846701236 | 1,798384163 | 0,40851046 | 0,88153752 | KH domain containing 1 like |
| ENSG00000249484 | LINC01470 | 0,846319827 | 1,797908782 | 0,3597885 | 0,85022129 | long intergenic non-protein coding RNA 1470 |
| ENSG00000188050 | RNF133 | 0,846232008 | 1,797799345 | 0,41118156 | 0,883335414 | ring finger protein 133 |
| ENSG00000111961 | SASH1 | 0,845996443 | 1,797505821 | 0,00702652 | 0,088859506 | SAM and SH3 domain containing 1 |
| ENSG00000077943 | ITGA8 | 0,845886541 | 1,797368896 | 0,11367881 | 0,538699894 | integrin subunit alpha 8 |
| ENSG00000281857 | SAG | 0,845739437 | 1,797185637 | 0,59916741 | 0,948894897 | S-antigen visual arrestin |
| ENSG00000287886 | AC005914,1 | 0,845696342 | 1,797131954 | 0,37775854 | 0,863195661 | novel transcript |
| ENSG00000147133 | TAF1 | 0,845325729 | 1,796670348 | 8,71E-05 | 0,002326706 | TATA-box binding protein associated factor 1 |
| ENSG00000178878 | APOLD1 | 0,844709893 | 1,795903577 | 0,01092174 | 0,122210917 | apolipoprotein L domain containing 1 |
| ENSG00000183486 | MX2 | 0,844673737 | 1,795858569 | 0,15807496 | 0,631249902 | MX dynamin like GTPase 2 |
| ENSG00000230230 | TRIM26 | 0,844179534 | 1,795243494 | 0,07522921 | 0,433332501 | tripartite motif containing 26 |
| ENSG00000257524 | AL157935,2 | 0,844013539 | 1,795036947 | 0,24181951 | 0,752128472 | novel protein |
| ENSG00000236081 | ELFN1-AS1 | 0,843892709 | 1,794886614 | 0,60728838 | 0,950993594 | ELFN1 antisense RNA 1 |
| ENSG00000271590 | AC108463,3 | 0,843761679 | 1,794723604 | 0,0704848 | 0,418147371 | novel transcript |
| ENSG00000259459 | LINC02568 | 0,843119878 | 1,793925376 | 0,34039183 | 0,839010589 | long intergenic non-protein coding RNA 2568 |
| ENSG00000227695 | DNMBP-AS1 | 0,843095986 | 1,793895667 | 0,41964442 | 0,888090642 | DNMBP antisense RNA 1 |
| ENSG00000265666 | RARA-AS1 | 0,842878909 | 1,793625767 | 0,13786351 | 0,592546669 | RARA antisense RNA 1 |
| ENSG00000226652 | PSMD10P2 | 0,842706667 | 1,79341164 | 0,48309538 | 0,91363215 | proteasome 26S subunit, non-ATPase, 10 pseudogene 2 |
| ENSG00000200972 | RNU5A-8P | 0,842521726 | 1,793181756 | 0,49298298 | 0,917090312 | RNA, U5A small nuclear 8, pseudogene |
| ENSG00000281499 | AL034351,2 | 0,842521726 | 1,793181756 | 0,49298298 | 0,917090312 | |
| ENSG00000058262 | SEC61A1 | 0,841985748 | 1,792515691 | 8,59E-05 | 0,00230006 | SEC61 translocon subunit alpha 1 |
| ENSG00000157322 | CLEC18A | 0,841819524 | 1,792309174 | 0,18666779 | 0,680025828 | C-type lectin domain family 18 member A |
| ENSG00000118855 | MFSD1 | 0,841642299 | 1,792089015 | 0,00662913 | 0,085219165 | major facilitator superfamily domain containing 1 |
| ENSG00000266613 | COP1P1 | 0,841370917 | 1,791751941 | 0,47099745 | 0,908869685 | COP1 pseudogene 1 |

| | | | | | | |
|---|---|---|---|---|---|---|
| ENSG00000119508 | NR4A3 | 0,841365418 | 1,791745111 | 0,03116504 | 0,253879337 | nuclear receptor subfamily 4 group A member 3 |
| ENSG00000230215 | LINC01840 | 0,840655552 | 1,790863715 | 0,25102752 | 0,763388331 | long intergenic non-protein coding RNA 1840 |
| ENSG00000266767 | AP000897,2 | 0,84060039 | 1,790795242 | 0,39952557 | 0,875433675 | novel transcript, antisense to PTPRM |
| ENSG00000228300 | FAM174C | 0,840567789 | 1,790754775 | 0,05598723 | 0,36527227 | family with sequence similarity 174 member C |
| ENSG00000260860 | AC126773,5 | 0,840554966 | 1,790738858 | 0,54304983 | 0,933827063 | calponin 2 (CNN2) pseudogene |
| ENSG00000201512 | SNORA71C | 0,839906817 | 1,789934527 | 0,2999371 | 0,809916717 | small nucleolar RNA, H/ACA box 71C |
| ENSG00000152936 | LMNTD1 | 0,839733314 | 1,789719276 | 0,50498485 | 0,921400408 | lamin tail domain containing 1 |
| ENSG00000260492 | CISTR | 0,839618445 | 1,789576783 | 0,3450099 | 0,841168721 | chondrogenesis-associated transcript |
| ENSG00000169247 | SH3TC2 | 0,839575328 | 1,7895233 | 0,00943638 | 0,109999648 | SH3 domain and tetratricopeptide repeats 2 |
| ENSG00000132677 | RHBG | 0,838921957 | 1,78871304 | 0,42655994 | 0,892152052 | Rh family B glycoprotein |
| ENSG00000136026 | CKAP4 | 0,838800444 | 1,78856239 | 0,00790653 | 0,097013276 | cytoskeleton associated protein 4 |
| ENSG00000203601 | LINC00970 | 0,838606819 | 1,788322361 | 0,30113606 | 0,810679941 | long intergenic non-protein coding RNA 970 |
| ENSG00000106236 | NPTX2 | 0,838313795 | 1,787959174 | 0,0612804 | 0,38540944 | neuronal pentraxin 2 |
| ENSG00000132109 | TRIM21 | 0,838069087 | 1,787655929 | 0,00628332 | 0,081714521 | tripartite motif containing 21 |
| ENSG00000105321 | CCDC9 | 0,837962166 | 1,787523448 | 0,20836273 | 0,713031122 | coiled-coil domain containing 9 |
| ENSG00000187510 | PLEKHG7 | 0,83783011 | 1,787359835 | 0,41426146 | 0,885579089 | pleckstrin homology and RhoGEF domain containing G7 |
| ENSG00000219951 | AL391416,1 | 0,837816379 | 1,787342823 | 0,49586334 | 0,918105073 | glyceraldehyde-3-phosphate dehydrogenase (GAPDH) pseudogene |
| ENSG00000233086 | LNCARSR | 0,837756406 | 1,787268525 | 0,37810277 | 0,863761554 | lncRNA regulator of Akt signaling associated with HCC and RCC |
| ENSG00000038427 | VCAN | 0,837413983 | 1,786844369 | 0,02041723 | 0,189467657 | versican |
| ENSG00000286679 | AC073225,1 | 0,837120707 | 1,786481169 | 0,27916587 | 0,790013761 | novel transcript |
| ENSG00000237222 | LINC01968 | 0,837093682 | 1,786447705 | 0,54257497 | 0,933663748 | long intergenic non-protein coding RNA 1968 |
| ENSG00000238099 | LINC01625 | 0,836540011 | 1,785762241 | 0,07971658 | 0,447697198 | long intergenic non-protein coding RNA 1625 |
| ENSG00000268001 | CARD8-AS1 | 0,836326739 | 1,785498273 | 0,09351062 | 0,485956782 | CARD8 antisense RNA 1 |
| ENSG00000107201 | DDX58 | 0,83606914 | 1,785179495 | 0,02303893 | 0,206327482 | DExD/H-box helicase 58 |
| ENSG00000224017 | LINC01449 | 0,835930415 | 1,785007846 | 0,49661798 | 0,918455764 | long intergenic non-protein coding RNA 1449 |
| ENSG00000143878 | RHOB | 0,835827867 | 1,78488097 | 0,0125191 | 0,135057725 | ras homolog family member B |
| ENSG00000206680 | SNORD21 | 0,835361523 | 1,784304109 | 0,43064009 | 0,895267028 | small nucleolar RNA, C/D box 21 |
| ENSG00000181126 | HLA-V | 0,835293865 | 1,784220433 | 0,35858344 | 0,849814998 | major histocompatibility complex, class I, V (pseudogene) |
| ENSG00000095380 | NANS | 0,835022737 | 1,783885153 | 0,00683041 | 0,08719714 | N-acetylneuraminate synthase |

| | | | | | | |
|---|---|---|---|---|---|---|
| ENSG00000257704 | INAFM1 | 0,834969795 | 1,783819692 | 0,12771295 | 0,570728673 | InaF motif containing 1 |
| ENSG00000104805 | NUCB1 | 0,834964264 | 1,783812853 | 0,03075172 | 0,251352083 | nucleobindin 1 |
| ENSG00000185532 | PRKG1 | 0,834962682 | 1,783810897 | 0,06188844 | 0,388245132 | protein kinase cGMP-dependent 1 |
| ENSG00000204421 | LY6G6C | 0,834745278 | 1,783542109 | 0,58320709 | 0,944174844 | lymphocyte antigen 6 family member G6C |
| ENSG00000141522 | ARHGDIA | 0,834654193 | 1,783429509 | 0,0194556 | 0,183597917 | Rho GDP dissociation inhibitor alpha |
| ENSG00000274542 | AC243734,1 | 0,834586456 | 1,783345776 | 0,83055127 | 0,984156028 | cholinergic receptor nicotinic alpha 7 subunit |
| ENSG00000004939 | SLC4A1 | 0,834390152 | 1,783103135 | 0,36427403 | 0,854302928 | solute carrier family 4 member 1 (Diego blood group) |
| ENSG00000147065 | MSN | 0,834377307 | 1,78308726 | 0,0010188 | 0,019004674 | moesin |
| ENSG00000227877 | MRLN | 0,834355756 | 1,783060625 | 0,41251563 | 0,884523593 | myoregulin |
| ENSG00000250120 | PCDHA10 | 0,834336023 | 1,783036236 | 6,74E-05 | 0,001847711 | protocadherin alpha 10 |
| ENSG00000271278 | ELOCP33 | 0,834016453 | 1,782641321 | 0,38231378 | 0,866930927 | elongin C pseudogene 33 |
| ENSG00000156504 | FAM122B | 0,833875616 | 1,782467307 | 0,01652567 | 0,164488529 | family with sequence similarity 122B |
| ENSG00000287572 | AC270284,1 | 0,833814009 | 1,782391193 | 0,33923571 | 0,838276153 | novel transcript |
| ENSG00000285972 | CERNA2 | 0,833695449 | 1,782244723 | 0,12544388 | 0,565907826 | competing endogenous lncRNA 2 for microRNA let-7b |
| ENSG00000226017 | PRICKLE2-AS3 | 0,83360272 | 1,782130173 | 0,2300179 | 0,738551869 | PRICKLE2 antisense RNA 3 |
| ENSG00000253837 | AC090197,1 | 0,833593789 | 1,78211914 | 0,12875502 | 0,573336703 | novel transcript |
| ENSG00000102401 | ARMCX3 | 0,833476662 | 1,781974462 | 0,0044696 | 0,063517074 | armadillo repeat containing X-linked 3 |
| ENSG00000234380 | LINC01426 | 0,833473729 | 1,78197084 | 0,11483496 | 0,541181798 | long intergenic non-protein coding RNA 1426 |
| ENSG00000204386 | NEU1 | 0,833226514 | 1,781665515 | 0,31584274 | 0,821233148 | neuraminidase 1 |
| ENSG00000250075 | AC104806,2 | 0,832717414 | 1,781036909 | 0,26147949 | 0,773332221 | novel transcript |
| ENSG00000215475 | SIAH3 | 0,83247748 | 1,78074073 | 0,08720627 | 0,470802426 | siah E3 ubiquitin protein ligase family member 3 |
| ENSG00000130766 | SESN2 | 0,832426624 | 1,780677959 | 0,02243871 | 0,202518437 | sestrin 2 |
| ENSG00000285069 | SESN2 | 0,832426624 | 1,780677959 | 0,02243871 | 0,202518437 | sestrin 2 |
| ENSG00000235643 | LINC01647 | 0,832266402 | 1,780480212 | 0,23468137 | 0,744134035 | long intergenic non-protein coding RNA 1647 |
| ENSG00000085733 | CTTN | 0,831395629 | 1,779405885 | 0,01602719 | 0,161013527 | cortactin |
| ENSG00000288401 | CTTN | 0,831395629 | 1,779405885 | 0,01602719 | 0,161013527 | cortactin |
| ENSG00000285633 | AL132633,1 | 0,831321089 | 1,779313951 | 0,09984741 | 0,501305496 | novel transcript |
| ENSG00000197063 | MAFG | 0,831247842 | 1,779223615 | 0,01683947 | 0,16634097 | MAF bZIP transcription factor G |
| ENSG00000125968 | ID1 | 0,831205109 | 1,779170916 | 0,04193682 | 0,307024639 | inhibitor of DNA binding 1, HLH protein |
| ENSG00000285966 | AC007686,4 | 0,830969924 | 1,778880902 | 0,28306363 | 0,794336345 | novel transcript |
| ENSG00000120549 | KIAA1217 | 0,830691532 | 1,778537671 | 0,00419067 | 0,060825029 | KIAA1217 |

| | | | | | | |
|---|---|---|---|---|---|---|
| ENSG00000254615 | AC027031,2 | 0,830610003 | 1,778437165 | 0,00529306 | 0,072011359 | novel transcript |
| ENSG00000126005 | MMP24OS | 0,830450534 | 1,778240595 | 0,02604315 | 0,225107295 | MMP24 opposite strand |
| ENSG00000185924 | RTN4RL1 | 0,830352294 | 1,778119511 | 0,15185597 | 0,619956253 | reticulon 4 receptor like 1 |
| ENSG00000255175 | LINC02759 | 0,830350182 | 1,778116908 | 0,38590328 | 0,869112622 | long intergenic non-protein coding RNA 2759 |
| ENSG00000159871 | LYPD5 | 0,830292304 | 1,778045575 | 0,42632252 | 0,892138574 | LY6/PLAUR domain containing 5 |
| ENSG00000250892 | AC108475,1 | 0,830281705 | 1,778032513 | 0,43460975 | 0,896272827 | pseudogene similar to part of WEE1 homolog (WEE1) |
| ENSG00000224468 | LAMC1-AS1 | 0,83024931 | 1,777992588 | 0,40707563 | 0,880078305 | LAMC1 antisense RNA 1 |
| ENSG00000250111 | AC107982,1 | 0,830000719 | 1,777686249 | 0,50222467 | 0,920520138 | coiled-coil domain containing 144B (pseudogene) (CCDC144B) pseudogene |
| ENSG00000263155 | MYZAP | 0,829993254 | 1,77767705 | 0,46151809 | 0,906165604 | myocardial zonula adherens protein |
| ENSG00000163466 | ARPC2 | 0,82996204 | 1,777638589 | 0,0005519 | 0,011351768 | actin related protein 2/3 complex subunit 2 |
| ENSG00000224655 | LINC01823 | 0,829386425 | 1,776929478 | 0,56976981 | 0,94221458 | long intergenic non-protein coding RNA 1823 |
| ENSG00000266074 | BAHCC1 | 0,82905217 | 1,776517832 | 0,07165413 | 0,421847873 | BAH domain and coiled-coil containing 1 |
| ENSG00000230427 | ALG14-AS1 | 0,828994596 | 1,776446937 | 0,38862911 | 0,869977116 | ALG14 antisense RNA 1 |
| ENSG00000183535 | COL18A1-AS1 | 0,828834168 | 1,776249407 | 0,380964 | 0,865900094 | COL18A1 antisense RNA 1 |
| ENSG00000075420 | FNDC3B | 0,828827903 | 1,776241693 | 0,0019883 | 0,033510694 | fibronectin type III domain containing 3B |
| ENSG00000112576 | CCND3 | 0,82882152 | 1,776233835 | 0,0010262 | 0,019108459 | cyclin D3 |
| ENSG00000276298 | AC234782,1 | 0,828692002 | 1,776074381 | 0,28136519 | 0,792415204 | HYDIN, axonemal central pair apparatus protein (HYDIN) pseudogene |
| ENSG00000166401 | SERPINB8 | 0,828577408 | 1,775933312 | 0,01111454 | 0,123616096 | serpin family B member 8 |
| ENSG00000166833 | NAV2 | 0,828498897 | 1,775836669 | 0,02352536 | 0,209423932 | neuron navigator 2 |
| ENSG00000241985 | WWTR1-IT1 | 0,828464377 | 1,775794178 | 0,41563624 | 0,885986046 | WWTR1 intronic transcript 1 |
| ENSG00000161509 | GRIN2C | 0,828211163 | 1,775482527 | 0,03145415 | 0,255623198 | glutamate ionotropic receptor NMDA type subunit 2C |
| ENSG00000273143 | AL355512,1 | 0,827933367 | 1,775140686 | 0,41252682 | 0,884523593 | novel transcript |
| ENSG00000257512 | AC124947,2 | 0,827607321 | 1,774739552 | 0,32029465 | 0,824366234 | anaphase promoting complex subunit 10 (ANAPC10) pseudogene |
| ENSG00000268894 | PLCE1-AS1 | 0,827600302 | 1,774730918 | 0,10025649 | 0,502713526 | PLCE1 antisense RNA 1 |
| ENSG00000189114 | BLOC1S3 | 0,827042409 | 1,774044759 | 0,0195542 | 0,183917723 | biogenesis of lysosomal organelles complex 1 subunit 3 |
| ENSG00000270426 | AC099343,2 | 0,82696564 | 1,773950361 | 0,26013403 | 0,772692192 | novel transcript, sense intronic to IRF2 |
| ENSG00000185015 | CA13 | 0,826881882 | 1,773847374 | 0,06189958 | 0,388245132 | carbonic anhydrase 13 |

| | | | | | | |
|---|---|---|---|---|---|---|
| ENSG00000225968 | ELFN1 | 0,826827529 | 1,773780546 | 0,03232824 | 0,260224035 | extracellular leucine rich repeat and fibronectin type III domain containing 1 |
| ENSG00000262298 | AC004584,1 | 0,826762283 | 1,773700328 | 0,42775453 | 0,892705337 | novel transcript, antisense to COIL |
| ENSG00000088826 | SMOX | 0,826403813 | 1,773259667 | 0,04677124 | 0,328449224 | spermine oxidase |
| ENSG00000287387 | AC010883,3 | 0,825774847 | 1,772486755 | 0,05735079 | 0,370182715 | novel transcript, antisense to ZFP36L2 |
| ENSG00000180509 | KCNE1 | 0,825562702 | 1,772226135 | 0,39440674 | 0,871799523 | potassium voltage-gated channel subfamily E regulatory subunit 1 |
| ENSG00000266498 | AC055811,3 | 0,825357894 | 1,771974563 | 0,3357058 | 0,83542993 | novel transcript |
| ENSG00000272834 | AL022238,2 | 0,825080633 | 1,771634053 | 0,54812479 | 0,935404974 | novel transcript, antisense to MKL1 |
| ENSG00000274580 | FCAR | 0,824926958 | 1,77144535 | 0,37399617 | 0,861369706 | Fc fragment of IgA receptor |
| ENSG00000225807 | AC069281,1 | 0,824091388 | 1,770419673 | 0,55315889 | 0,936992468 | novel transcript |
| ENSG00000287830 | AC007218,3 | 0,824089762 | 1,770417677 | 0,43469911 | 0,896272827 | novel transcript, antisense to GRIN2A |
| ENSG00000237548 | TTLL11-IT1 | 0,823928448 | 1,770219731 | 0,50460663 | 0,921400408 | TTLL11 intronic transcript 1 |
| ENSG00000034510 | TMSB10 | 0,823923119 | 1,770213192 | 0,00546971 | 0,073868213 | thymosin beta 10 |
| ENSG00000108381 | ASPA | 0,823761117 | 1,770014424 | 0,41601227 | 0,885986046 | aspartoacylase |
| ENSG00000160703 | NLRX1 | 0,823726789 | 1,769972308 | 0,02547669 | 0,221867198 | NLR family member X1 |
| ENSG00000182347 | PDSS1P1 | 0,823580136 | 1,769792396 | 0,25041048 | 0,762598222 | decaprenyl diphosphate synthase subunit 1 pseudogene 1 |
| ENSG00000260494 | AC002310,3 | 0,82356869 | 1,769778355 | 0,3386126 | 0,837371637 | ribosomal protein L7 (RPL7) pseudogene |
| ENSG00000065621 | GSTO2 | 0,823372477 | 1,769537674 | 0,03637043 | 0,280277294 | glutathione S-transferase omega 2 |
| ENSG00000150990 | DHX37 | 0,823024031 | 1,769110339 | 0,05391509 | 0,357953158 | DEAH-box helicase 37 |
| ENSG00000276230 | RNH1 | 0,823001096 | 1,769082215 | 0,05466409 | 0,361009299 | ribonuclease/angiogenin inhibitor 1 |
| ENSG00000165568 | AKR1E2 | 0,822576612 | 1,768561774 | 0,01369703 | 0,143989223 | aldo-keto reductase family 1 member E2 |
| ENSG00000275799 | AP001059,1 | 0,822383375 | 1,768324906 | 0,53583626 | 0,932139691 | novel transcript |
| ENSG00000257411 | AC034102,2 | 0,822168686 | 1,768061779 | 0,83542148 | 0,985034063 | novel protein |
| ENSG00000042304 | C2orf83 | 0,82206903 | 1,767939653 | 0,55549133 | 0,937793851 | chromosome 2 open reading frame 83 |
| ENSG00000104361 | NIPAL2 | 0,821909303 | 1,767743927 | 0,01374075 | 0,144400323 | NIPA like domain containing 2 |
| ENSG00000184634 | MED12 | 0,821676117 | 1,767458226 | 0,00062749 | 0,012632275 | mediator complex subunit 12 |
| ENSG00000154734 | ADAMTS1 | 0,821637964 | 1,767411484 | 0,01431935 | 0,149077648 | ADAM metallopeptidase with thrombospondin type 1 motif 1 |
| ENSG00000236222 | WDR46 | 0,821525871 | 1,767274168 | 0,13516116 | 0,586080761 | WD repeat domain 46 |
| ENSG00000285082 | AL160272,2 | 0,821422685 | 1,767147771 | 0,31444226 | 0,819877431 | novel protein |
| ENSG00000165801 | ARHGEF40 | 0,821356738 | 1,767066996 | 0,0700321 | 0,417362474 | Rho guanine nucleotide exchange factor 40 |

| | | | | | | |
|---|---|---|---|---|---|---|
| ENSG00000166510 | CCDC68 | 0,821265148 | 1,766954815 | 0,15114286 | 0,618336884 | coiled-coil domain containing 68 |
| ENSG00000235082 | SUMO1P3 | 0,821111265 | 1,766766356 | 0,51969422 | 0,926822772 | SUMO1 pseudogene 3 |
| ENSG00000285301 | BOP1 | 0,821050113 | 1,766691469 | 0,13162598 | 0,579455365 | BOP1 ribosomal biogenesis factor |
| ENSG00000261236 | BOP1 | 0,821050113 | 1,766691469 | 0,13162598 | 0,579455365 | BOP1 ribosomal biogenesis factor |
| ENSG00000101199 | ARFGAP1 | 0,820963014 | 1,766584813 | 0,04655117 | 0,327787303 | ADP ribosylation factor GTPase activating protein 1 |
| ENSG00000281005 | LINC00921 | 0,820843604 | 1,766438601 | 0,2455831 | 0,757563881 | long intergenic non-protein coding RNA 921 |
| ENSG00000283516 | LITAFD | 0,820820754 | 1,766410624 | 0,38292334 | 0,867578459 | LITAF domain containing |
| ENSG00000213170 | SAR1AP2 | 0,820607618 | 1,766149683 | 0,40089104 | 0,876599365 | secretion associated Ras related GTPase 1A pseudogene 2 |
| ENSG00000054219 | LY75 | 0,820477544 | 1,765990453 | 0,18540894 | 0,677967843 | lymphocyte antigen 75 |
| ENSG00000171766 | GATM | 0,820294839 | 1,76576682 | 0,14854475 | 0,613326406 | glycine amidinotransferase |
| ENSG00000228055 | FAM245A | 0,820177394 | 1,76562308 | 0,5391595 | 0,933350549 | family with sequence similarity 245 member A |
| ENSG00000238061 | AL356273,2 | 0,820158013 | 1,765599361 | 0,21690562 | 0,724544002 | proteasome (prosome, macropain) 26S subunit, non-ATPase, 7 (Mov34 homolog) (PSMD7) pseudogene |
| ENSG00000255774 | LINC02747 | 0,819905068 | 1,76528983 | 0,52223786 | 0,927617924 | long intergenic non-protein coding RNA 2747 |
| ENSG00000231290 | APCDD1L-DT | 0,819904043 | 1,765288575 | 0,34870499 | 0,842757646 | APCDD1L divergent transcript |
| ENSG00000184500 | PROS1 | 0,819891954 | 1,765273783 | 0,09337094 | 0,485553749 | protein S |
| ENSG00000188039 | NWD1 | 0,819879432 | 1,765258461 | 0,34697138 | 0,841625668 | NACHT and WD repeat domain containing 1 |
| ENSG00000275880 | AL139385,1 | 0,819429569 | 1,764708102 | 0,14153717 | 0,599428784 | novel transcript, antisense to CARKD |
| ENSG00000142634 | EFHD2 | 0,819340219 | 1,764598812 | 0,04492507 | 0,320304999 | EF-hand domain family member D2 |
| ENSG00000224320 | HLA-A | 0,819224638 | 1,764457448 | 0,02852195 | 0,239578272 | major histocompatibility complex, class I, A |
| ENSG00000102580 | DNAJC3 | 0,819154498 | 1,764371667 | 0,0099438 | 0,114638731 | DnaJ heat shock protein family (Hsp40) member C3 |
| ENSG00000265388 | RN7SL391P | 0,819081483 | 1,764282374 | 0,35902706 | 0,849814998 | RNA, 7SL, cytoplasmic 391, pseudogene |
| ENSG00000233901 | LINC01503 | 0,818983238 | 1,764162233 | 0,15641974 | 0,62744507 | long intergenic non-protein coding RNA 1503 |
| ENSG00000117425 | PTCH2 | 0,818555455 | 1,763639207 | 0,03031613 | 0,249039213 | patched 2 |
| ENSG00000232406 | AL121895,1 | 0,818471971 | 1,763537154 | 0,2727625 | 0,78561201 | novel transcript, antisense to EPB41L1 |
| ENSG00000229215 | HLA-A | 0,818332889 | 1,76336715 | 0,0286194 | 0,240203215 | major histocompatibility complex, class I, A |
| ENSG00000282053 | AC019257,1 | 0,818311842 | 1,763341426 | 0,45917445 | 0,904804618 | novel transcript |
| ENSG00000168264 | IRF2BP2 | 0,817809491 | 1,762727531 | 0,00038941 | 0,008455021 | interferon regulatory factor 2 binding protein 2 |
| ENSG00000236548 | RNF217-AS1 | 0,81691311 | 1,761632647 | 0,0803152 | 0,449444373 | RNF217 antisense RNA 1 (head to head) |
| ENSG00000214264 | KCTD9P4 | 0,816817309 | 1,761515671 | 0,52648798 | 0,929261495 | potassium channel tetramerization domain containing 9 pseudogene 4 |
| ENSG00000258096 | AC025031,2 | 0,816797184 | 1,761491099 | 0,46941269 | 0,908150886 | novel transcript, antisense to SLC38A2 |

| | | | | | | |
|---|---|---|---|---|---|---|
| ENSG00000286947 | AC022467,2 | 0,816604471 | 1,761255816 | 0,29998942 | 0,809919611 | novel transcript, antisense to FBN1 |
| ENSG00000117472 | TSPAN1 | 0,816515229 | 1,761146873 | 0,28021721 | 0,790779983 | tetraspanin 1 |
| ENSG00000177239 | MAN1B1 | 0,816481416 | 1,761105597 | 0,01662051 | 0,165046811 | mannosidase alpha class 1B member 1 |
| ENSG00000234789 | AL590369,1 | 0,816286957 | 1,760868235 | 0,55510513 | 0,937793851 | novel transcript |
| ENSG00000255346 | NOX5 | 0,815973528 | 1,760485724 | 0,24740574 | 0,759663439 | NADPH oxidase 5 |
| ENSG00000272755 | AC245297,2 | 0,815253762 | 1,759607629 | 0,36687345 | 0,856313634 | novel transcript |
| ENSG00000119147 | ECRG4 | 0,814127688 | 1,758234731 | 0,13581516 | 0,587728418 | ECRG4 augurin precursor |
| ENSG00000237552 | LINC02567 | 0,814079296 | 1,758175755 | 0,46957621 | 0,908293218 | long intergenic non-protein coding RNA 2567 |
| ENSG00000230638 | AL445933,1 | 0,813912168 | 1,757972093 | 0,45993976 | 0,905341196 | novel pseudogene |
| ENSG00000288577 | AL117334,1 | 0,813845179 | 1,757890466 | 0,15603425 | 0,626944439 | novel transcript |
| ENSG00000128567 | PODXL | 0,813498206 | 1,757467738 | 0,01834664 | 0,176288964 | podocalyxin like |
| ENSG00000197879 | MYO1C | 0,813477856 | 1,757442948 | 0,02675736 | 0,229082684 | myosin IC |
| ENSG00000164054 | SHISA5 | 0,813461885 | 1,757423493 | 0,00217057 | 0,035924072 | shisa family member 5 |
| ENSG00000154310 | TNIK | 0,813242303 | 1,757156029 | 0,02337257 | 0,208479303 | TRAF2 and NCK interacting kinase |
| ENSG00000275214 | IFI27 | 0,813184581 | 1,757085727 | 0,13221555 | 0,580737262 | interferon alpha inducible protein 27 |
| ENSG00000267059 | AC005943,1 | 0,813029351 | 1,75689668 | 0,17367411 | 0,658552229 | novel transcript, readthrough between UQCR11 and MBD3 |
| ENSG00000257529 | RPL36A-HNRNI | 0,812911665 | 1,756753369 | 0,2014025 | 0,701094807 | RPL36A-HNRNPH2 readthrough |
| ENSG00000271503 | CCL5 | 0,812688077 | 1,756481129 | 0,31730797 | 0,822252078 | C-C motif chemokine ligand 5 |
| ENSG00000260051 | AL031600,1 | 0,812566314 | 1,756332889 | 0,23827958 | 0,747710567 | novel transcript, antisense to CLCN7 |
| ENSG00000179284 | DAND5 | 0,812483737 | 1,756232363 | 0,45044169 | 0,902062594 | DAN domain BMP antagonist family member 5 |
| ENSG00000261997 | AC007336,1 | 0,81233098 | 1,756046418 | 0,149749 | 0,615291456 | novel transcript |
| ENSG00000259414 | AC242376,1 | 0,812242613 | 1,755938861 | 0,25662655 | 0,769018737 | hect domain and RLD 2 pseudogene 7 |
| ENSG00000173218 | VANGL1 | 0,811915213 | 1,755540419 | 0,00108859 | 0,020102644 | VANGL planar cell polarity protein 1 |
| ENSG00000258646 | AL049780,1 | 0,811889686 | 1,755509357 | 0,51262881 | 0,923540891 | novel transcript, antisense to EIF2B2 |
| ENSG00000253507 | AC104257,1 | 0,811166744 | 1,754629883 | 0,46064503 | 0,905949965 | novel transcript |
| ENSG00000285395 | XYLT1 | 0,811047824 | 1,754485256 | 0,19648981 | 0,694444647 | xylosyltransferase 1 |
| ENSG00000176845 | METRNL | 0,810979294 | 1,754401918 | 0,01069259 | 0,120422602 | meteorin like, glial cell differentiation regulator |
| ENSG00000111652 | COPS7A | 0,810975106 | 1,754396825 | 0,00258223 | 0,041063507 | COP9 signalosome subunit 7A |
| ENSG00000177105 | RHOG | 0,810690334 | 1,75405056 | 0,00841643 | 0,101321206 | ras homolog family member G |
| ENSG00000149577 | SIDT2 | 0,810567723 | 1,753901494 | 0,00115156 | 0,021017096 | SID1 transmembrane family member 2 |
| ENSG00000267530 | LINC01836 | 0,810435509 | 1,753740768 | 0,2951 | 0,805254732 | long intergenic non-protein coding RNA 1836 |
| ENSG00000148426 | PROSER2 | 0,810276335 | 1,753547286 | 0,16058423 | 0,636125619 | proline and serine rich 2 |

| | | | | | | |
|---|---|---|---|---|---|---|
| ENSG00000196114 | IFITM3P3 | 0,810252991 | 1,753518912 | 0,42915476 | 0,894012535 | IFITM3 pseudogene 3 |
| ENSG00000248099 | INSL3 | 0,81022717 | 1,753487528 | 0,58826867 | 0,946176848 | insulin like 3 |
| ENSG00000103490 | PYCARD | 0,810109015 | 1,753343926 | 0,30278444 | 0,812344697 | PYD and CARD domain containing |
| ENSG00000075624 | ACTB | 0,809924065 | 1,753119167 | 0,00091898 | 0,017412738 | actin beta |
| ENSG00000141194 | OR4D1 | 0,809682073 | 1,75282513 | 0,43687095 | 0,896866043 | olfactory receptor family 4 subfamily D member 1 |
| ENSG00000277278 | HERC2 | 0,809524847 | 1,752634116 | 0,48870713 | 0,91557151 | HECT and RLD domain containing E3 ubiquitin protein ligase 2 |
| ENSG00000255874 | PRECSIT | 0,809272461 | 1,752327536 | 0,04124197 | 0,304151397 | p53 regulated carcinoma associated Stat3 activating long intergenic non-protein coding transcript |
| ENSG00000224864 | AC011447,1 | 0,809243532 | 1,752292399 | 0,72914579 | 0,974009551 | ribosomal protein S16 (RPS16) pseudogene |
| ENSG00000260029 | AC007608,2 | 0,80831555 | 1,751165637 | 0,08914463 | 0,47641326 | novel transcript, antisense to NKD1 |
| ENSG00000129538 | RNASE1 | 0,808110697 | 1,750917001 | 0,50115127 | 0,920520138 | ribonuclease A family member 1, pancreatic |
| ENSG00000268996 | MAN1B1-DT | 0,808100888 | 1,750905097 | 0,24785565 | 0,760164991 | MAN1B1 divergent transcript |
| ENSG00000256453 | DND1 | 0,808020927 | 1,750808056 | 0,33451026 | 0,834713776 | DND microRNA-mediated repression inhibitor 1 |
| ENSG00000149428 | HYOU1 | 0,80801584 | 1,750801882 | 0,06565184 | 0,402080164 | hypoxia up-regulated 1 |
| ENSG00000280682 | HYOU1 | 0,80801584 | 1,750801882 | 0,06565184 | 0,402080164 | hypoxia up-regulated 1 |
| ENSG00000269980 | AC068768,1 | 0,807733092 | 1,750458783 | 0,35337169 | 0,84514549 | novel transcript, antisense to CDK2AP1 |
| ENSG00000175093 | SPSB4 | 0,807391701 | 1,750044613 | 0,00026839 | 0,006120202 | splA/ryanodine receptor domain and SOCS box containing 4 |
| ENSG00000168993 | CPLX1 | 0,807363418 | 1,750010306 | 0,05735559 | 0,370182715 | complexin 1 |
| ENSG00000276626 | 7SK | 0,80692271 | 1,749475802 | 0,4554506 | 0,903545728 | 7SK RNA |
| ENSG00000117791 | MTARC2 | 0,806903577 | 1,749452601 | 0,0575006 | 0,370798119 | mitochondrial amidoxime reducing component 2 |
| ENSG00000287001 | AC010624,5 | 0,806740862 | 1,749255299 | 0,0039492 | 0,058157244 | novel transcript |
| ENSG00000173517 | PEAK1 | 0,806458542 | 1,748913022 | 0,00303664 | 0,046952225 | pseudopodium enriched atypical kinase 1 |
| ENSG00000186654 | PRR5 | 0,80639953 | 1,748841486 | 0,08669218 | 0,469007996 | proline rich 5 |
| ENSG00000285588 | AL160254,1 | 0,80616968 | 1,748562883 | 0,36800563 | 0,857401877 | novel transcript, antisense to PCDH9 |
| ENSG00000271225 | BNIP3P4 | 0,806161985 | 1,748553557 | 0,5341874 | 0,93151582 | BCL2 interacting protein 3 pseudogene 4 |
| ENSG00000267690 | LDLRAD4-AS1 | 0,805894011 | 1,7482288 | 0,33717272 | 0,836323083 | LDLRAD4 antisense RNA 1 |
| ENSG00000137193 | PIM1 | 0,805861958 | 1,74818996 | 0,00064251 | 0,012858692 | Pim-1 proto-oncogene, serine/threonine kinase |
| ENSG00000287413 | AL449363,2 | 0,805694068 | 1,747986531 | 0,50385377 | 0,921212789 | novel transcript |
| ENSG00000206403 | ABHD16A | 0,805363917 | 1,747586562 | 0,38599267 | 0,869112622 | abhydrolase domain containing 16A, phospholipase |
| ENSG00000116690 | PRG4 | 0,804905904 | 1,747031842 | 0,39532602 | 0,871981986 | proteoglycan 4 |
| ENSG00000230183 | CNOT6LP1 | 0,804778403 | 1,746877452 | 0,49275093 | 0,917090312 | CNOT6L pseudogene 1 |

| | | | | | | |
|---|---|---|---|---|---|---|
| ENSG00000226103 | C6orf47 | 0,804719633 | 1,746806292 | 0,14424838 | 0,60428531 | chromosome 6 open reading frame 47 |
| ENSG00000203623 | C6orf47 | 0,804719633 | 1,746806292 | 0,14424838 | 0,60428531 | chromosome 6 open reading frame 47 |
| ENSG00000226531 | C6orf47 | 0,804719633 | 1,746806292 | 0,14424838 | 0,60428531 | chromosome 6 open reading frame 47 |
| ENSG00000234271 | Z98752,2 | 0,804437751 | 1,746465025 | 0,5483917 | 0,935602633 | transmembrane protein 14C (TMEM14C) pseudogene |
| ENSG00000197245 | FAM110D | 0,804271822 | 1,746264169 | 0,12119252 | 0,555892438 | family with sequence similarity 110 member D |
| ENSG00000287360 | AC006296,3 | 0,803581254 | 1,745428493 | 0,35878455 | 0,849814998 | novel transcript |
| ENSG00000148700 | ADD3 | 0,80323054 | 1,745004237 | 0,0538767 | 0,357953158 | adducin 3 |
| ENSG00000145734 | BDP1 | 0,803162848 | 1,744922363 | 0,23938502 | 0,749122637 | B double prime 1, subunit of RNA polymerase III transcription initiation factor IIIB |
| ENSG00000239468 | RN7SL569P | 0,803019752 | 1,744749299 | 0,539857 | 0,933385624 | RNA, 7SL, cytoplasmic 569, pseudogene |
| ENSG00000253404 | AC034243,1 | 0,802686302 | 1,744346082 | 0,31676185 | 0,821646322 | novel transcript |
| ENSG00000234096 | MOG | 0,802534365 | 1,744162386 | 0,66075671 | 0,959226315 | myelin oligodendrocyte glycoprotein |
| ENSG00000182372 | CLN8 | 0,802280657 | 1,74385569 | 0,1097279 | 0,528090728 | CLN8 transmembrane ER and ERGIC protein |
| ENSG00000158104 | HPD | 0,802176537 | 1,743729839 | 0,60197564 | 0,949939978 | 4-hydroxyphenylpyruvate dioxygenase |
| ENSG00000256500 | AL139300,1 | 0,802064933 | 1,743594953 | 0,01891749 | 0,180041477 | novel protein |
| ENSG00000130477 | UNC13A | 0,801951258 | 1,743457575 | 0,04332692 | 0,314067408 | unc-13 homolog A |
| ENSG00000228140 | AL031283,1 | 0,801922407 | 1,743422709 | 0,29879408 | 0,808507705 | novel transcript |
| ENSG00000285867 | BX470102,2 | 0,801652733 | 1,743096852 | 0,33927139 | 0,838276153 | Novel transcript |
| ENSG00000152684 | PELO | 0,801475368 | 1,74288257 | 0,0001703 | 0,00413376 | pelota mRNA surveillance and ribosome rescue factor |
| ENSG00000265753 | RN7SL444P | 0,801449727 | 1,742851593 | 0,26805623 | 0,780680587 | RNA, 7SL, cytoplasmic 444, pseudogene |
| ENSG00000170608 | FOXA3 | 0,801367447 | 1,742752198 | 0,5893667 | 0,946844281 | forkhead box A3 |
| ENSG00000237713 | AC011287,2 | 0,801293144 | 1,742662443 | 0,60457056 | 0,950864972 | novel transcript |
| ENSG00000173281 | PPP1R3B | 0,801259402 | 1,742621686 | 0,00388437 | 0,057418855 | protein phosphatase 1 regulatory subunit 3B |
| ENSG00000281980 | AC246817,3 | 0,801137823 | 1,742474838 | 0,09495045 | 0,489386708 | novel transcript |
| ENSG00000112511 | PHF1 | 0,800924537 | 1,742217252 | 0,20243615 | 0,702433811 | PHD finger protein 1 |
| ENSG00000273084 | AC092171,5 | 0,800897803 | 1,742184967 | 0,0458335 | 0,324561392 | novel transcript |
| ENSG00000205356 | TECPR1 | 0,800791732 | 1,742056882 | 0,14928564 | 0,614476095 | tectonin beta-propeller repeat containing 1 |
| ENSG00000249353 | NPM1P27 | 0,800760464 | 1,742019126 | 0,31090847 | 0,817751312 | nucleophosmin 1 pseudogene 27 |
| ENSG00000224116 | INHBA-AS1 | 0,800743183 | 1,74199826 | 0,44659389 | 0,901644586 | INHBA antisense RNA 1 |
| ENSG00000090382 | LYZ | 0,800662182 | 1,741900457 | 0,28626268 | 0,797444206 | lysozyme |
| ENSG00000123395 | ATG101 | 0,800534077 | 1,741745792 | 0,05982413 | 0,380382656 | autophagy related 101 |

| | | | | | | |
|---|---|---|---|---|---|---|
| ENSG00000147862 | NFIB | 0,800238258 | 1,741388689 | 0,05908982 | 0,377556505 | nuclear factor I B |
| ENSG00000167674 | HDGFL2 | 0,800180782 | 1,741319315 | 0,20679494 | 0,710309478 | HDGF like 2 |
| ENSG00000263940 | RN7SL275P | 0,800084247 | 1,741202802 | 0,51080655 | 0,922693987 | RNA, 7SL, cytoplasmic 275, pseudogene |
| ENSG00000175197 | DDIT3 | 0,800079399 | 1,741196951 | 0,00752679 | 0,093528464 | DNA damage inducible transcript 3 |
| ENSG00000136478 | TEX2 | 0,799724823 | 1,740769064 | 0,00200693 | 0,033751863 | testis expressed 2 |
| ENSG00000146094 | DOK3 | 0,799701633 | 1,740741083 | 0,16844731 | 0,650405917 | docking protein 3 |
| ENSG00000276840 | PMS2P10 | 0,799642619 | 1,740669878 | 0,18480472 | 0,677025387 | PMS1 homolog 2, mismatch repair system component pseudogene 10 |
| ENSG00000156642 | NPTN | 0,799638937 | 1,740665436 | 0,00439924 | 0,062897441 | neuroplastin |
| ENSG00000248673 | LINC01331 | 0,799625041 | 1,74064867 | 0,48515881 | 0,913690171 | long intergenic non-protein coding RNA 1331 |
| ENSG00000164889 | SLC4A2 | 0,799434129 | 1,740418345 | 0,08219341 | 0,456024762 | solute carrier family 4 member 2 |
| ENSG00000254294 | IMPDH1P6 | 0,799420349 | 1,740401722 | 0,59822026 | 0,948608142 | inosine monophosphate dehydrogenase 1 pseudogene 6 |
| ENSG00000255650 | FAM222A-AS1 | 0,799158731 | 1,740086145 | 0,49139403 | 0,916995341 | FAM222A antisense RNA 1 |
| ENSG00000121064 | SCPEP1 | 0,799117636 | 1,740036581 | 0,00449453 | 0,063842344 | serine carboxypeptidase 1 |
| ENSG00000198909 | MAP3K3 | 0,799081297 | 1,739992753 | 0,00173353 | 0,029745949 | mitogen-activated protein kinase kinase kinase 3 |
| ENSG00000171608 | PIK3CD | 0,798790008 | 1,739641473 | 0,12869538 | 0,573234424 | phosphatidylinositol-4,5-bisphosphate 3-kinase catalytic subunit delta |
| ENSG00000275363 | AC100757,3 | 0,798730383 | 1,739569576 | 0,61133826 | 0,952062377 | M-phase phosphoprotein 10 (U3 small nucleolar ribonucleoprotein) (MPHOSPH10) pseudogene |
| ENSG00000121966 | CXCR4 | 0,798121607 | 1,738835683 | 0,03428401 | 0,270005245 | C-X-C motif chemokine receptor 4 |
| ENSG00000213270 | RPL6P25 | 0,797605558 | 1,738213816 | 0,56900631 | 0,942177697 | ribosomal protein L6 pseudogene 25 |
| ENSG00000267391 | MIR122HG | 0,797221902 | 1,737751635 | 0,26579956 | 0,77713706 | MIR122 host gene |
| ENSG00000225349 | ANKRD49P2 | 0,796562774 | 1,736957885 | 0,54380713 | 0,934259495 | ANKRD49 pseudogene 2 |
| ENSG00000013293 | SLC7A14 | 0,796461083 | 1,736835456 | 0,10259936 | 0,510071005 | solute carrier family 7 member 14 |
| ENSG00000264596 | AP000897,1 | 0,796351757 | 1,736703846 | 0,16640484 | 0,646270935 | novel transcript, sense intronic to PTPRM |
| ENSG00000128595 | CALU | 0,796201758 | 1,736523288 | 0,01265903 | 0,136004094 | calumenin |
| ENSG00000126903 | SLC10A3 | 0,796093337 | 1,736392789 | 0,02239657 | 0,202254907 | solute carrier family 10 member 3 |
| ENSG00000249736 | LINC02242 | 0,79593632 | 1,736203818 | 0,45868348 | 0,904420115 | long intergenic non-protein coding RNA 2242 |
| ENSG00000240808 | AC126389,1 | 0,795470185 | 1,735642941 | 0,34906969 | 0,843141089 | ribosomal protein L34 (RPL34) pseudogene |
| ENSG00000235310 | GXYLT1P6 | 0,795013709 | 1,735093862 | 0,59738199 | 0,948282297 | GXYLT1 pseudogene 6 |
| ENSG00000250358 | LINC02200 | 0,79475434 | 1,734781953 | 0,2295022 | 0,737904201 | long intergenic non-protein coding RNA 2200 |
| ENSG00000270959 | LPP-AS2 | 0,794443616 | 1,73440836 | 0,00718017 | 0,090189178 | LPP antisense RNA 2 |

| | | | | | | |
|---|---|---|---|---|---|---|
| ENSG00000189127 | ANKRD34B | 0,79434612 | 1,734291154 | 0,33553677 | 0,835320086 | ankyrin repeat domain 34B |
| ENSG00000278936 | AC244517,4 | 0,794264054 | 1,734192504 | 0,64885451 | 0,956899963 | novel transcript, antisense to PCDHB16, PCDHB9, PCDHB10 and PCDHB11 |
| ENSG00000288640 | AC005192,1 | 0,794119137 | 1,734018315 | 0,14202987 | 0,600058555 | novel protein |
| ENSG00000268189 | AC005785,1 | 0,794052982 | 1,733938803 | 0,38589681 | 0,869112622 | novel transcript, antisense to AKAP8 and AKAP8L |
| ENSG00000159289 | GOLGA6A | 0,794042856 | 1,733926633 | 0,36089064 | 0,851667053 | golgin A6 family member A |
| ENSG00000253341 | PCBP2P2 | 0,794033636 | 1,733915553 | 0,60088156 | 0,94964552 | poly(rC) binding protein 2 pseudogene 2 |
| ENSG00000151012 | SLC7A11 | 0,793979612 | 1,733850624 | 0,00446463 | 0,063504172 | solute carrier family 7 member 11 |
| ENSG00000184009 | ACTG1 | 0,793716909 | 1,733534933 | 0,00024185 | 0,005600879 | actin gamma 1 |
| ENSG00000185339 | TCN2 | 0,793054003 | 1,732738571 | 0,00404273 | 0,059268345 | transcobalamin 2 |
| ENSG00000254650 | MTCYBP41 | 0,792828868 | 1,732468196 | 0,60786042 | 0,951015776 | MT-CYB pseudogene 41 |
| ENSG00000285980 | AP003025,2 | 0,792670938 | 1,732278555 | 0,45653633 | 0,903545728 | novel transcript |
| ENSG00000056972 | TRAF3IP2 | 0,79255935 | 1,732144574 | 0,05042615 | 0,343470607 | TRAF3 interacting protein 2 |
| ENSG00000251018 | HMMR-AS1 | 0,792384372 | 1,731934503 | 0,5166801 | 0,925489094 | HMMR antisense RNA 1 |
| ENSG00000287362 | AC103563,9 | 0,792383739 | 1,731933743 | 0,44362537 | 0,900090367 | novel transcript |
| ENSG00000130511 | SSBP4 | 0,792330573 | 1,731869918 | 0,00974458 | 0,112876842 | single stranded DNA binding protein 4 |
| ENSG00000282114 | LINC00824 | 0,791946167 | 1,731408524 | 0,61370881 | 0,952413269 | long intergenic non-protein coding RNA 824 |
| ENSG00000256043 | CTSO | 0,791852109 | 1,731295646 | 0,67643576 | 0,963580269 | cathepsin O |
| ENSG00000253591 | KCNIP1-AS1 | 0,791683191 | 1,73109295 | 0,43141559 | 0,89530663 | KCNIP1 antisense RNA 1 |
| ENSG00000156030 | MIDEAS | 0,791640904 | 1,73104221 | 0,00863996 | 0,102981152 | mitotic deacetylase associated SANT domain protein |
| ENSG00000237914 | SIRPG-AS1 | 0,791500652 | 1,730873934 | 0,40092649 | 0,876604367 | SIRPG antisense RNA 1 |
| ENSG00000166407 | LMO1 | 0,791468668 | 1,730835562 | 0,0226428 | 0,203830812 | LIM domain only 1 |
| ENSG00000263477 | AC104982,2 | 0,790687416 | 1,729898529 | 0,40455853 | 0,878176155 | novel transcript, antisense to SSH2 |
| ENSG00000143545 | RAB13 | 0,789897159 | 1,728951212 | 0,01342907 | 0,14188797 | RAB13, member RAS oncogene family |
| ENSG00000167419 | LPO | 0,78985324 | 1,728898579 | 0,47942264 | 0,912714933 | lactoperoxidase |
| ENSG00000259744 | AC009269,4 | 0,789770661 | 1,72879962 | 0,4528234 | 0,902361309 | novel transcript, sense intronic to LARP6 |
| ENSG00000177721 | ANXA2R | 0,789280572 | 1,728212441 | 0,15985398 | 0,634614414 | annexin A2 receptor |
| ENSG00000110876 | SELPLG | 0,789151019 | 1,728057256 | 0,19535383 | 0,693577833 | selectin P ligand |
| ENSG00000148734 | NPFFR1 | 0,788931527 | 1,727794368 | 0,26529238 | 0,77668189 | neuropeptide FF receptor 1 |
| ENSG00000230985 | VARS2 | 0,788656539 | 1,727465069 | 0,28426271 | 0,795731744 | valyl-tRNA synthetase 2, mitochondrial |
| ENSG00000259660 | DNM1P47 | 0,788502738 | 1,72728092 | 0,28724058 | 0,798816004 | dynamin 1 pseudogene 47 |
| ENSG00000253733 | LZTS1-AS1 | 0,788388429 | 1,727144068 | 0,29218705 | 0,802771257 | LZTS1 antisense RNA 1 |

| | | | | | | |
|---|---|---|---|---|---|---|
| ENSG00000183514 | TDGF1P2 | 0,787894088 | 1,726552362 | 0,55551515 | 0,937793851 | teratocarcinoma-derived growth factor 1 pseudogene 2 |
| ENSG00000082805 | ERC1 | 0,787570924 | 1,726165657 | 0,00244873 | 0,039401463 | ELKS/RAB6-interacting/CAST family member 1 |
| ENSG00000254109 | RBPMS-AS1 | 0,78708282 | 1,725581745 | 0,35949197 | 0,849982153 | RBPMS antisense RNA 1 |
| ENSG00000164588 | HCN1 | 0,786982916 | 1,725462256 | 0,11137873 | 0,53309009 | hyperpolarization activated cyclic nucleotide gated potassium channel 1 |
| ENSG00000140939 | NOL3 | 0,786857394 | 1,725312138 | 0,21254874 | 0,719402819 | nucleolar protein 3 |
| ENSG00000162104 | ADCY9 | 0,786738963 | 1,725170514 | 0,03385189 | 0,267664545 | adenylate cyclase 9 |
| ENSG00000205500 | MAPRE3-AS1 | 0,786600835 | 1,725005349 | 0,33840535 | 0,837263082 | MAPRE3 antisense RNA 1 |
| ENSG00000101144 | BMP7 | 0,786259728 | 1,724597541 | 0,00015202 | 0,003748322 | bone morphogenetic protein 7 |
| ENSG00000147044 | CASK | 0,786218819 | 1,724548639 | 0,01932816 | 0,182760901 | calcium/calmodulin dependent serine protein kinase |
| ENSG00000228013 | IL6R-AS1 | 0,786179365 | 1,724501477 | 0,58043316 | 0,943832501 | IL6R antisense RNA 1 |
| ENSG00000164136 | IL15 | 0,786076468 | 1,724378485 | 0,02436598 | 0,215070018 | interleukin 15 |
| ENSG00000288293 | CC2D1A | 0,785927014 | 1,72419986 | 0,05478092 | 0,36155177 | coiled-coil and C2 domain containing 1A |
| ENSG00000254489 | MPPED2-AS1 | 0,785847006 | 1,724104244 | 0,30777205 | 0,816424917 | MPPED2 antisense RNA 1 |
| ENSG00000173821 | RNF213 | 0,785600792 | 1,723810029 | 0,00568554 | 0,075828159 | ring finger protein 213 |
| ENSG00000187492 | CDHR4 | 0,785305092 | 1,723456746 | 0,31145475 | 0,817751312 | cadherin related family member 4 |
| ENSG00000104970 | KIR3DX1 | 0,784906941 | 1,722981177 | 0,29414115 | 0,804046662 | killer cell immunoglobulin like receptor, three Ig domains X1 (pseudogene) |
| ENSG00000171729 | TMEM51 | 0,784690543 | 1,722722757 | 0,02248576 | 0,202767423 | transmembrane protein 51 |
| ENSG00000130294 | KIF1A | 0,784640934 | 1,72266352 | 0,09526738 | 0,489967108 | kinesin family member 1A |
| ENSG00000286650 | AL139234,1 | 0,784605194 | 1,722620845 | 0,26555693 | 0,776824277 | novel transcript |
| ENSG00000258111 | AC079316,1 | 0,784491922 | 1,7224856 | 0,43767994 | 0,897021798 | ATP synthase, H+ transporting, mitochondrial Fo complex, subunit C1 (subunit 9) (ATP5G1) pseudogene |
| ENSG00000121281 | ADCY7 | 0,784306765 | 1,722264548 | 0,06484619 | 0,399478103 | adenylate cyclase 7 |
| ENSG00000182287 | AP1S2 | 0,784217474 | 1,722157958 | 0,13582196 | 0,587728418 | adaptor related protein complex 1 subunit sigma 2 |
| ENSG00000167202 | TBC1D2B | 0,784165536 | 1,72209596 | 0,00120053 | 0,021745953 | TBC1 domain family member 2B |
| ENSG00000198715 | GLMP | 0,784057602 | 1,721967127 | 0,0086736 | 0,103264064 | glycosylated lysosomal membrane protein |
| ENSG00000125398 | SOX9 | 0,783357267 | 1,721131427 | 0,00810356 | 0,098656485 | SRY-box transcription factor 9 |
| ENSG00000275978 | Metazoa_SRP | 0,782873402 | 1,720554274 | 0,40423439 | 0,878176155 | Metazoan signal recognition particle RNA |
| ENSG00000282733 | NSFP1 | 0,78269198 | 1,720337925 | 0,01255361 | 0,135289937 | N-ethylmaleimide-sensitive factor pseudogene 1 |

| | | | | | | |
|---|---|---|---|---|---|---|
| ENSG00000133401 | PDZD2 | 0,782562829 | 1,720183925 | 0,00396814 | 0,058378646 | PDZ domain containing 2 |
| ENSG00000141756 | FKBP10 | 0,781991292 | 1,719502594 | 0,01717675 | 0,168970493 | FKBP prolyl isomerase 10 |
| ENSG00000255478 | AP000944,1 | 0,781939763 | 1,719441178 | 0,60413255 | 0,950672921 | novel transcript |
| ENSG00000260170 | AC090527,2 | 0,781815776 | 1,719293414 | 0,72201326 | 0,972516173 | novel protein |
| ENSG00000274297 | AC009269,5 | 0,781712217 | 1,719170005 | 0,44375786 | 0,900184646 | novel transcript, sense intronic to UACA |
| ENSG00000266527 | AC005697,2 | 0,781649865 | 1,719095705 | 0,25367947 | 0,765248462 | novel transcript, antisense to C17orf108 |
| ENSG00000233821 | ENOX1-AS1 | 0,78152577 | 1,718947842 | 0,16397572 | 0,642745357 | ENOX1 antisense RNA 1 |
| ENSG00000286019 | NOTCH2NLB | 0,781485957 | 1,718900406 | 0,12604417 | 0,566777174 | notch 2 N-terminal like B |
| ENSG00000167914 | GSDMA | 0,781459978 | 1,718869453 | 0,37850466 | 0,864031013 | gasdermin A |
| ENSG00000135722 | FBXL8 | 0,781440579 | 1,718846342 | 0,41266133 | 0,884576962 | F-box and leucine rich repeat protein 8 |
| ENSG00000258410 | AC087386,1 | 0,781177079 | 1,718532433 | 0,19742686 | 0,695991044 | novel transcript |
| ENSG00000274598 | AC087893,2 | 0,780927995 | 1,71823575 | 0,20862528 | 0,713284351 | novel transcript, sense intronic to SSH1 |
| ENSG00000188305 | PEAK3 | 0,780565153 | 1,717803664 | 0,41629642 | 0,886060654 | PEAK family member 3 |
| ENSG00000185761 | ADAMTSL5 | 0,780125884 | 1,71728071 | 0,55475006 | 0,937793851 | ADAMTS like 5 |
| ENSG00000230547 | HMGB1P11 | 0,780048723 | 1,717188865 | 0,56566244 | 0,940866081 | high mobility group box 1 pseudogene 11 |
| ENSG00000253313 | C1orf210 | 0,779808863 | 1,716903392 | 0,49531348 | 0,917777383 | chromosome 1 open reading frame 210 |
| ENSG00000079385 | CEACAM1 | 0,779746051 | 1,716828643 | 0,08934241 | 0,476981049 | CEA cell adhesion molecule 1 |
| ENSG00000243015 | RN7SL737P | 0,77964872 | 1,716712822 | 0,55509956 | 0,937793851 | RNA, 7SL, cytoplasmic 737, pseudogene |
| ENSG00000229373 | LINC00452 | 0,779283031 | 1,71627773 | 0,43800165 | 0,897177404 | long intergenic non-protein coding RNA 452 |
| ENSG00000266308 | RN7SL510P | 0,779154353 | 1,716124658 | 0,52000607 | 0,926822772 | RNA, 7SL, cytoplasmic 510, pseudogene |
| ENSG00000257949 | TEN1 | 0,779125274 | 1,716090069 | 0,02359621 | 0,20993506 | TEN1 subunit of CST complex |
| ENSG00000274206 | Metazoa_SRP | 0,779026652 | 1,715972761 | 0,39559628 | 0,872227623 | Metazoan signal recognition particle RNA |
| ENSG00000230795 | HLA-K | 0,778677155 | 1,715557113 | 0,56030896 | 0,940070839 | major histocompatibility complex, class I, K (pseudogene) |
| ENSG00000198113 | TOR4A | 0,778676128 | 1,715555891 | 0,38974743 | 0,870057277 | torsin family 4 member A |
| ENSG00000197454 | OR2L5 | 0,778585978 | 1,715448693 | 0,44350583 | 0,900020258 | olfactory receptor family 2 subfamily L member 5 |
| ENSG00000171695 | LKAAEAR1 | 0,778509691 | 1,715357987 | 0,59087305 | 0,947416795 | LKAAEAR motif containing 1 |
| ENSG00000100055 | CYTH4 | 0,778342237 | 1,715158897 | 0,45646816 | 0,903545728 | cytohesin 4 |
| ENSG00000233538 | AC017104,2 | 0,778299216 | 1,715107751 | 0,32962143 | 0,831406561 | novel transcript |
| ENSG00000204745 | AC083899,1 | 0,778281567 | 1,71508677 | 0,10978592 | 0,528215911 | anaphase promoting complex subunit 1 (ANAPC1) pseudogene |
| ENSG00000079432 | CIC | 0,778196094 | 1,714985161 | 0,09859612 | 0,498520872 | capicua transcriptional repressor |
| ENSG00000115525 | ST3GAL5 | 0,777979879 | 1,714728159 | 0,00100463 | 0,018773878 | ST3 beta-galactoside alpha-2,3-sialyltransferase 5 |

| | | | | | | |
|---|---|---|---|---|---|---|
| ENSG00000181211 | HECW1-IT1 | 0,777926896 | 1,714665186 | 0,2147026 | 0,722164151 | HECW1 intronic transcript 1 |
| ENSG00000224875 | EML4-AS1 | 0,777664335 | 1,714353156 | 0,38471292 | 0,868536368 | EML4 antisense RNA 1 |
| ENSG00000100253 | MIOX | 0,777599381 | 1,714275973 | 0,22603756 | 0,734959876 | myo-inositol oxygenase |
| ENSG00000277941 | Z99774,4 | 0,777108917 | 1,713693281 | 0,58257284 | 0,944174844 | |
| ENSG00000249240 | AC069368,1 | 0,776935674 | 1,713487508 | 0,13411052 | 0,584263748 | novel protein |
| ENSG00000124818 | OPN5 | 0,776763887 | 1,713283488 | 0,63804683 | 0,956018518 | opsin 5 |
| ENSG00000229456 | RLIMP1 | 0,776561435 | 1,713043083 | 0,24148656 | 0,751948055 | ring finger protein, LIM domain interacting pseudogene 1 |
| ENSG00000164241 | C5orf63 | 0,776428404 | 1,71288513 | 0,14845904 | 0,613326406 | chromosome 5 open reading frame 63 |
| ENSG00000186416 | NKRF | 0,776249659 | 1,712672923 | 0,01386539 | 0,145173348 | NFKB repressing factor |
| ENSG00000126368 | NR1D1 | 0,775992085 | 1,712367175 | 0,11423643 | 0,540234136 | nuclear receptor subfamily 1 group D member 1 |
| ENSG00000231129 | ABCF1 | 0,775774669 | 1,712109138 | 0,24556422 | 0,757563881 | ATP binding cassette subfamily F member 1 |
| ENSG00000106665 | CLIP2 | 0,775744306 | 1,712073105 | 0,21066649 | 0,716131719 | CAP-Gly domain containing linker protein 2 |
| ENSG00000186765 | FSCN2 | 0,775083585 | 1,711289194 | 0,38147384 | 0,865900094 | fascin actin-bundling protein 2, retinal |
| ENSG00000125657 | TNFSF9 | 0,775060103 | 1,711261341 | 0,23151935 | 0,73957278 | TNF superfamily member 9 |
| ENSG00000184840 | TMED9 | 0,775039412 | 1,711236798 | 0,03327466 | 0,264841921 | transmembrane p24 trafficking protein 9 |
| ENSG00000205363 | INSYN1 | 0,77497526 | 1,711160708 | 0,14281172 | 0,601409695 | inhibitory synaptic factor 1 |
| ENSG00000105656 | ELL | 0,774768032 | 1,710914935 | 0,02037717 | 0,189154353 | elongation factor for RNA polymerase II |
| ENSG00000067221 | STOML1 | 0,774610241 | 1,710727818 | 0,04826128 | 0,334607086 | stomatin like 1 |
| ENSG00000204183 | GDF5-AS1 | 0,774386563 | 1,710462604 | 0,51026307 | 0,922499069 | GDF5 antisense RNA 1 |
| ENSG00000197157 | SND1 | 0,774354733 | 1,710424867 | 0,00036923 | 0,008044632 | staphylococcal nuclease and tudor domain containing 1 |
| ENSG00000270933 | AC010719,1 | 0,773775082 | 1,709737784 | 0,34597427 | 0,841168721 | novel transcript |
| ENSG00000161714 | PLCD3 | 0,773523086 | 1,70943917 | 0,11786996 | 0,548937448 | phospholipase C delta 3 |
| ENSG00000188277 | C15orf62 | 0,773203702 | 1,709060776 | 0,15363853 | 0,622683206 | chromosome 15 open reading frame 62 |
| ENSG00000100647 | SUSD6 | 0,773005342 | 1,708825809 | 0,01423601 | 0,148389909 | sushi domain containing 6 |
| ENSG00000145214 | DGKQ | 0,772945829 | 1,70875532 | 0,09656825 | 0,493721941 | diacylglycerol kinase theta |
| ENSG00000288642 | CDR1 | 0,772801858 | 1,708584806 | 0,16056628 | 0,636125619 | cerebellar degeneration related protein 1 |
| ENSG00000136114 | THSD1 | 0,77257833 | 1,708320102 | 0,14152774 | 0,599428784 | thrombospondin type 1 domain containing 1 |
| ENSG00000275325 | PDCD6IPP1 | 0,772456693 | 1,708176076 | 0,60875202 | 0,951601152 | PDCD6IP pseudogene 1 |
| ENSG00000132382 | MYBBP1A | 0,772328786 | 1,708024638 | 0,13135517 | 0,578977575 | MYB binding protein 1a |
| ENSG00000141505 | ASGR1 | 0,771783858 | 1,707379613 | 0,05925778 | 0,378088799 | asialoglycoprotein receptor 1 |
| ENSG00000072840 | EVC | 0,771649867 | 1,707221047 | 0,07581456 | 0,434831495 | EvC ciliary complex subunit 1 |

| | | | | | | |
|---|---|---|---|---|---|---|
| ENSG00000227741 | AL121987,2 | 0,77100527 | 1,70645843 | 0,14234401 | 0,600436282 | novel transcript |
| ENSG00000286444 | AL035661,2 | 0,770988311 | 1,70643837 | 0,31476748 | 0,81991112 | novel transcript, antisense to APMAP |
| ENSG00000197380 | DACT3 | 0,770692466 | 1,706088477 | 0,00950634 | 0,110608911 | dishevelled binding antagonist of beta catenin 3 |
| ENSG00000172738 | TMEM217 | 0,770678777 | 1,706072289 | 0,29951939 | 0,809349546 | transmembrane protein 217 |
| ENSG00000242767 | ZBTB20-AS4 | 0,770620362 | 1,706003211 | 0,61208348 | 0,952285547 | ZBTB20 antisense RNA 4 |
| ENSG00000236064 | AL109946,1 | 0,770591019 | 1,705968513 | 0,32375717 | 0,825794844 | novel transcript, antisense to MID2 |
| ENSG00000141985 | SH3GL1 | 0,770510285 | 1,705873048 | 0,04360136 | 0,315398641 | SH3 domain containing GRB2 like 1, endophilin A2 |
| ENSG00000109762 | SNX25 | 0,770364123 | 1,705700232 | 0,00445157 | 0,063404996 | sorting nexin 25 |
| ENSG00000269335 | IKBKG | 0,770321446 | 1,705649776 | 0,10556325 | 0,518512104 | inhibitor of nuclear factor kappa B kinase regulatory subunit gamma |
| ENSG00000285108 | AC103718,1 | 0,770303261 | 1,705628276 | 0,53290608 | 0,931291085 | novel transcript |
| ENSG00000243446 | RN7SL284P | 0,769900295 | 1,705151936 | 0,59131102 | 0,947416795 | RNA, 7SL, cytoplasmic 284, pseudogene |
| ENSG00000276516 | RN7SL284P | 0,769900295 | 1,705151936 | 0,59131102 | 0,947416795 | RNA, 7SL, cytoplasmic 284, pseudogene |
| ENSG00000145335 | SNCA | 0,769595361 | 1,704791566 | 0,01017716 | 0,11653537 | synuclein alpha |
| ENSG00000183336 | BOLA2 | 0,769371303 | 1,704526823 | 0,22092224 | 0,728635155 | bolA family member 2 |
| ENSG00000146192 | FGD2 | 0,76925377 | 1,704387966 | 0,42423806 | 0,890949676 | FYVE, RhoGEF and PH domain containing 2 |
| ENSG00000026751 | SLAMF7 | 0,76912672 | 1,704237876 | 0,54125143 | 0,933422598 | SLAM family member 7 |
| ENSG00000227006 | AL136988,2 | 0,769111157 | 1,704219492 | 0,29691834 | 0,806635068 | novel transcript |
| ENSG00000064300 | NGFR | 0,769016478 | 1,704107653 | 0,04611685 | 0,325797698 | nerve growth factor receptor |
| ENSG00000177485 | ZBTB33 | 0,768796333 | 1,703847639 | 0,13891264 | 0,594174432 | zinc finger and BTB domain containing 33 |
| ENSG00000013016 | EHD3 | 0,768683873 | 1,703714826 | 0,01450181 | 0,150526119 | EH domain containing 3 |
| ENSG00000252835 | SCARNA21 | 0,768674135 | 1,703703327 | 0,46022254 | 0,905443183 | small Cajal body-specific RNA 21 |
| ENSG00000184408 | KCND2 | 0,768321754 | 1,703287246 | 0,17783045 | 0,664478432 | potassium voltage-gated channel subfamily D member 2 |
| ENSG00000122254 | HS3ST2 | 0,768200639 | 1,703144259 | 0,35931148 | 0,849982153 | heparan sulfate-glucosamine 3-sulfotransferase 2 |
| ENSG00000258628 | RHPN2P1 | 0,76813693 | 1,70306905 | 0,56675303 | 0,941311563 | rhophilin Rho GTPase binding protein 2 pseudogene 1 |
| ENSG00000108375 | RNF43 | 0,767969906 | 1,702871893 | 0,06359328 | 0,394965288 | ring finger protein 43 |
| ENSG00000214803 | AC090921,1 | 0,767919267 | 1,702812124 | 0,50145789 | 0,920520138 | novel transcript |
| ENSG00000206376 | EHMT2 | 0,76785999 | 1,70274216 | 0,17622407 | 0,661798106 | euchromatic histone lysine methyltransferase 2 |
| ENSG00000262097 | LINC02185 | 0,7678221 | 1,702697442 | 0,3419302 | 0,840048641 | long intergenic non-protein coding RNA 2185 |
| ENSG00000158716 | DUSP23 | 0,767760235 | 1,702624428 | 0,01977457 | 0,184933326 | dual specificity phosphatase 23 |
| ENSG00000135506 | OS9 | 0,767304635 | 1,702086828 | 0,02684937 | 0,229474736 | OS9 endoplasmic reticulum lectin |

| | | | | | | |
|---|---|---|---|---|---|---|
| ENSG00000236936 | AL031005,1 | 0,767247217 | 1,702019088 | 0,49244011 | 0,917008021 | novel transcript |
| ENSG00000130589 | HELZ2 | 0,767110802 | 1,70185816 | 0,18699068 | 0,680567707 | helicase with zinc finger 2 |
| ENSG00000227630 | LINC01132 | 0,766810385 | 1,701503814 | 0,23334837 | 0,742507431 | long intergenic non-protein coding RNA 1132 |
| ENSG00000278330 | AC018529,2 | 0,766691822 | 1,701363987 | 0,3692374 | 0,857942508 | novel transcript, antisense to MBP |
| ENSG00000260190 | AL807752,5 | 0,766597644 | 1,701252927 | 0,2810932 | 0,792005925 | novel transcript, overlapping FBXW5 |
| ENSG00000196083 | IL1RAP | 0,766547609 | 1,701193926 | 0,05887026 | 0,37676968 | interleukin 1 receptor accessory protein |
| ENSG00000281644 | AC106795,9 | 0,766443196 | 1,701070809 | 0,44808069 | 0,902062594 | novel transcript, antisense to PROP1 |
| ENSG00000260580 | AC009035,1 | 0,766064283 | 1,700624094 | 0,54885156 | 0,936029622 | novel transcript |
| ENSG00000232569 | GABBR1 | 0,766037781 | 1,700592854 | 0,37876371 | 0,864112469 | gamma-aminobutyric acid type B receptor subunit 1 |
| ENSG00000272822 | AC073610,2 | 0,765879876 | 1,700406732 | 0,19632336 | 0,694046542 | novel protein |
| ENSG00000171067 | C11orf24 | 0,765838948 | 1,700358494 | 0,00042085 | 0,009000416 | chromosome 11 open reading frame 24 |
| ENSG00000224714 | LINC02671 | 0,76578538 | 1,70029536 | 0,53999509 | 0,933385624 | long intergenic non-protein coding RNA 2671 |
| ENSG00000089060 | SLC8B1 | 0,765465892 | 1,699918867 | 0,07005629 | 0,41742708 | solute carrier family 8 member B1 |
| ENSG00000162434 | JAK1 | 0,765005715 | 1,699376731 | 0,00175618 | 0,030101606 | Janus kinase 1 |
| ENSG00000249188 | ENPP7P1 | 0,764489998 | 1,698769366 | 0,54565878 | 0,935146964 | ectonucleotide pyrophosphatase/phosphodiesterase 7 pseudogene 1 |
| ENSG00000246223 | LINC01550 | 0,764068479 | 1,698273101 | 0,36176081 | 0,851979035 | long intergenic non-protein coding RNA 1550 |
| ENSG00000159128 | IFNGR2 | 0,763941981 | 1,698124201 | 0,12049792 | 0,553925447 | interferon gamma receptor 2 |
| ENSG00000260608 | AC024270,3 | 0,763853575 | 1,698020145 | 0,5755313 | 0,942700009 | novel transcript, antisense to BTBD1 |
| ENSG00000182534 | MXRA7 | 0,763811639 | 1,697970789 | 0,0026231 | 0,0415655 | matrix remodeling associated 7 |
| ENSG00000128228 | SDF2L1 | 0,763809796 | 1,697968619 | 0,04624969 | 0,326399584 | stromal cell derived factor 2 like 1 |
| ENSG00000108518 | PFN1 | 0,763779944 | 1,697933486 | 0,00168063 | 0,028933421 | profilin 1 |
| ENSG00000225916 | AC007879,2 | 0,762916856 | 1,696918006 | 0,52553573 | 0,92884555 | novel transcript |
| ENSG00000116254 | CHD5 | 0,762893382 | 1,696890395 | 0,23439727 | 0,744054861 | chromodomain helicase DNA binding protein 5 |
| ENSG00000197448 | GSTK1 | 0,762579814 | 1,696521618 | 0,00296223 | 0,045915666 | glutathione S-transferase kappa 1 |
| ENSG00000146904 | EPHA1 | 0,762468345 | 1,696390543 | 0,28261168 | 0,793528967 | EPH receptor A1 |
| ENSG00000284816 | EPHA1 | 0,762468345 | 1,696390543 | 0,28261168 | 0,793528967 | EPH receptor A1 |
| ENSG00000205929 | C21orf62 | 0,762348907 | 1,696250107 | 0,07687869 | 0,438361367 | chromosome 21 open reading frame 62 |
| ENSG00000250155 | AC008957,1 | 0,761819663 | 1,695627962 | 0,25950389 | 0,772164235 | novel transcript |
| ENSG00000038382 | TRIO | 0,761786062 | 1,695588471 | 0,00232039 | 0,037803447 | trio Rho guanine nucleotide exchange factor |
| ENSG00000114270 | COL7A1 | 0,761422629 | 1,695161385 | 0,28622881 | 0,797420923 | collagen type VII alpha 1 chain |
| ENSG00000125726 | CD70 | 0,761410058 | 1,695146614 | 0,56778081 | 0,94163744 | CD70 molecule |

| | | | | | | |
|---|---|---|---|---|---|---|
| ENSG00000140678 | ITGAX | 0,760846003 | 1,694483986 | 0,42636059 | 0,892138574 | integrin subunit alpha X |
| ENSG00000141384 | TAF4B | 0,760715571 | 1,694330797 | 0,09087656 | 0,480927969 | TATA-box binding protein associated factor 4b |
| ENSG00000113732 | ATP6V0E1 | 0,760161619 | 1,69368035 | 0,00488206 | 0,067626397 | ATPase H+ transporting V0 subunit e1 |
| ENSG00000106080 | FKBP14 | 0,760109453 | 1,69361911 | 0,00059163 | 0,012034278 | FKBP prolyl isomerase 14 |
| ENSG00000224891 | ARL14EPP1 | 0,760095833 | 1,693603121 | 0,24123858 | 0,751659063 | ARL14EP pseudogene 1 |
| ENSG00000225366 | TDGF1P3 | 0,760080103 | 1,693584655 | 0,28228233 | 0,793283115 | teratocarcinoma-derived growth factor 1 pseudogene 3 |
| ENSG00000203688 | LINC02487 | 0,759969568 | 1,693454903 | 0,09738839 | 0,495806966 | long intergenic non-protein coding RNA 2487 |
| ENSG00000175662 | TOM1L2 | 0,759418253 | 1,692807885 | 0,02537853 | 0,221166029 | target of myb1 like 2 membrane trafficking protein |
| ENSG00000254510 | AP001107,5 | 0,75897149 | 1,69228375 | 0,0492503 | 0,338502855 | novel transcript |
| ENSG00000213176 | RPL13P6 | 0,758844742 | 1,692135081 | 0,36116591 | 0,851821025 | ribosomal protein L13 pseudogene 6 |
| ENSG00000198547 | C20orf203 | 0,758781376 | 1,692060761 | 0,59324231 | 0,947416795 | chromosome 20 open reading frame 203 |
| ENSG00000184785 | SMIM10 | 0,758711721 | 1,691979069 | 0,07127135 | 0,420579013 | small integral membrane protein 10 |
| ENSG00000276184 | UPK3B | 0,758299294 | 1,691495447 | 0,39621993 | 0,872845252 | uroplakin 3B |
| ENSG00000269646 | AC010487,1 | 0,758216951 | 1,691398906 | 0,40267095 | 0,877341338 | novel transcript |
| ENSG00000248909 | HMGB1P21 | 0,757923491 | 1,691054893 | 0,25433079 | 0,765826923 | high mobility group box 1 pseudogene 21 |
| ENSG00000139645 | ANKRD52 | 0,7575576 | 1,690626067 | 0,01463362 | 0,151391619 | ankyrin repeat domain 52 |
| ENSG00000229743 | LINC01159 | 0,757168676 | 1,690170367 | 0,00936762 | 0,109376713 | long intergenic non-protein coding RNA 1159 |
| ENSG00000197785 | ATAD3A | 0,757114681 | 1,690107112 | 0,12003499 | 0,553128623 | ATPase family AAA domain containing 3A |
| ENSG00000227714 | MTND6P18 | 0,757054511 | 1,690036624 | 0,59318096 | 0,947416795 | MT-ND6 pseudogene 18 |
| ENSG00000198844 | ARHGEF15 | 0,756719542 | 1,689644273 | 0,59291393 | 0,947416795 | Rho guanine nucleotide exchange factor 15 |
| ENSG00000257086 | AP001453,2 | 0,756696999 | 1,689617871 | 0,26916308 | 0,781347689 | novel transcript |
| ENSG00000080572 | DNAAF6 | 0,756571715 | 1,68947115 | 0,53072951 | 0,931265667 | dynein axonemal assembly factor 6 |
| ENSG00000285518 | AC004900,1 | 0,756321175 | 1,689177781 | 0,45145121 | 0,902062594 | novel transcript, antisense to SIPA1L1 |
| ENSG00000250942 | ENPP7P11 | 0,756127122 | 1,688950589 | 0,58107805 | 0,943919685 | ectonucleotide pyrophosphatase/phosphodiesterase 7 pseudogene 11 |
| ENSG00000283378 | CNTNAP3C | 0,756043665 | 1,688852889 | 0,18236005 | 0,672670499 | contactin associated protein family member 3C |
| ENSG00000006062 | MAP3K14 | 0,755964186 | 1,688759852 | 0,03452041 | 0,271098283 | mitogen-activated protein kinase kinase kinase 14 |
| ENSG00000042753 | AP2S1 | 0,755496776 | 1,68821281 | 0,01407035 | 0,146925708 | adaptor related protein complex 2 subunit sigma 1 |
| ENSG00000263809 | AC135178,3 | 0,755319914 | 1,688005862 | 0,57683622 | 0,943261034 | novel protein |
| ENSG00000271208 | Z99289,3 | 0,754961322 | 1,687586348 | 0,54268154 | 0,933663748 | novel transcript, sense intronic LAMA4 |
| ENSG00000285732 | AC121334,3 | 0,754881239 | 1,687492674 | 0,51363403 | 0,92410971 | novel transcript, antisense to ABCD2 |

| | | | | | | |
|---|---|---|---|---|---|---|
| ENSG00000153902 | LGI4 | 0,754754332 | 1,687344239 | 0,12774937 | 0,570756819 | leucine rich repeat LGI family member 4 |
| ENSG00000286241 | AC080089,2 | 0,754572365 | 1,687131429 | 0,3325048 | 0,83303391 | novel transcript |
| ENSG00000158816 | VWA5B1 | 0,754504035 | 1,687051523 | 0,36649409 | 0,856103393 | von Willebrand factor A domain containing 5B1 |
| ENSG00000183597 | TANGO2 | 0,7542518 | 1,686756591 | 0,07994112 | 0,448292044 | transport and golgi organization 2 homolog |
| ENSG00000158813 | EDA | 0,754243792 | 1,68674723 | 0,04415918 | 0,317145854 | ectodysplasin A |
| ENSG00000115556 | PLCD4 | 0,754169923 | 1,686660866 | 0,12890969 | 0,573624329 | phospholipase C delta 4 |
| ENSG00000078269 | SYNJ2 | 0,753835966 | 1,686270481 | 0,08994687 | 0,478492263 | synaptojanin 2 |
| ENSG00000102387 | TAF7L | 0,753625572 | 1,686024583 | 0,17826169 | 0,665463944 | TATA-box binding protein associated factor 7 like |
| ENSG00000188868 | ZNF563 | 0,753592565 | 1,68598601 | 0,07413995 | 0,429959479 | zinc finger protein 563 |
| ENSG00000266208 | AC080112,1 | 0,753446048 | 1,685814794 | 0,49938782 | 0,920135598 | novel transcript, antisense to GJD3 |
| ENSG00000245748 | AC097382,2 | 0,753268599 | 1,685607454 | 0,48008614 | 0,912847708 | novel transcript, antisense to TBC1D14 and TADA2B |
| ENSG00000217643 | PTGES3P2 | 0,753048163 | 1,685349922 | 0,37144981 | 0,859476158 | prostaglandin E synthase 3 pseudogene 2 |
| ENSG00000161652 | IZUMO2 | 0,753028987 | 1,685327521 | 0,44494771 | 0,900436606 | IZUMO family member 2 |
| ENSG00000158008 | EXTL1 | 0,752651723 | 1,684886866 | 0,06523876 | 0,400948417 | exostosin like glycosyltransferase 1 |
| ENSG00000065154 | OAT | 0,752644851 | 1,684878841 | 0,03422057 | 0,269624964 | ornithine aminotransferase |
| ENSG00000124006 | OBSL1 | 0,752612862 | 1,684841482 | 0,13966514 | 0,595782615 | obscurin like cytoskeletal adaptor 1 |
| ENSG00000234162 | AC009505,1 | 0,752240184 | 1,684406309 | 0,55035746 | 0,936146662 | novel transcript, antisense to NCK2 |
| ENSG00000286373 | AC022387,2 | 0,752078993 | 1,684218122 | 0,14855062 | 0,613326406 | novel transcript |
| ENSG00000203437 | AC022080,1 | 0,751708424 | 1,683785571 | 0,39780778 | 0,874263835 | TAF10 RNA polymerase II, TATA box binding protein (TBP)-associated factor, 30kDa (TAF10) pseudogene |
| ENSG00000239856 | RN7SL225P | 0,751697822 | 1,683773197 | 0,61681316 | 0,952793934 | RNA, 7SL, cytoplasmic 225, pseudogene |
| ENSG00000096968 | JAK2 | 0,751651285 | 1,683718885 | 0,06289275 | 0,391611013 | Janus kinase 2 |
| ENSG00000122642 | FKBP9 | 0,751584212 | 1,683640609 | 0,00671456 | 0,086069735 | FKBP prolyl isomerase 9 |
| ENSG00000204839 | MROH6 | 0,75141109 | 1,683438585 | 0,23880712 | 0,748055366 | maestro heat like repeat family member 6 |
| ENSG00000239329 | HLA-DMB | 0,751386075 | 1,683409396 | 0,32000539 | 0,824366234 | major histocompatibility complex, class II, DM beta |
| ENSG00000171303 | KCNK3 | 0,751230266 | 1,683227601 | 0,11829263 | 0,549606291 | potassium two pore domain channel subfamily K member 3 |
| ENSG00000276045 | ORAI1 | 0,751033922 | 1,682998537 | 0,04732528 | 0,330631884 | ORAI calcium release-activated calcium modulator 1 |
| ENSG00000286272 | AC093843,2 | 0,750596592 | 1,682488439 | 0,57285821 | 0,942475308 | novel transcript |
| ENSG00000051108 | HERPUD1 | 0,750526771 | 1,682407015 | 0,00086619 | 0,016512546 | homocysteine inducible ER protein with ubiquitin like domain 1 |
| ENSG00000123130 | ACOT9 | 0,750482618 | 1,682355527 | 0,00032404 | 0,00722599 | acyl-CoA thioesterase 9 |
| ENSG00000053108 | FSTL4 | 0,749855212 | 1,681624056 | 0,07386245 | 0,429465432 | follistatin like 4 |

| | | | | | | |
|---|---|---|---|---|---|---|
| ENSG00000099290 | WASHC2A | 0,749581066 | 1,681304537 | 0,0024474 | 0,039400443 | WASH complex subunit 2A |
| ENSG00000254477 | AP000640,1 | 0,749140596 | 1,680791296 | 0,26082633 | 0,772952333 | novel transcript |
| ENSG00000196526 | AFAP1 | 0,748693652 | 1,68027067 | 0,01955392 | 0,183917723 | actin filament associated protein 1 |
| ENSG00000215720 | MFFP1 | 0,748631085 | 1,680197802 | 0,61808401 | 0,952793934 | MFF pseudogene 1 |
| ENSG00000234787 | LINC00458 | 0,748613126 | 1,680176887 | 0,50583237 | 0,921780819 | long intergenic non-protein coding RNA 458 |
| ENSG00000244021 | AC093591,1 | 0,748404912 | 1,679934416 | 0,60755424 | 0,950993594 | ribosomal protein L21 (RPL21) pseudogene |
| ENSG00000122176 | FMOD | 0,748362439 | 1,67988496 | 0,06033483 | 0,381843747 | fibromodulin |
| ENSG00000163377 | TAFA4 | 0,748078601 | 1,679554489 | 0,33035539 | 0,831982503 | TAFA chemokine like family member 4 |
| ENSG00000167110 | GOLGA2 | 0,748058962 | 1,679531626 | 0,14894529 | 0,613619689 | golgin A2 |
| ENSG00000119457 | SLC46A2 | 0,747682752 | 1,679093713 | 0,55672726 | 0,93825245 | solute carrier family 46 member 2 |
| ENSG00000231672 | DIRC3 | 0,747638552 | 1,679042272 | 0,12858788 | 0,573000492 | disrupted in renal carcinoma 3 |
| ENSG00000284391 | AL139398,1 | 0,747638058 | 1,679041696 | 0,16877399 | 0,650757516 | uncharacterized LOC105373244 |
| ENSG00000271959 | AC100803,4 | 0,747624952 | 1,679026443 | 0,49488982 | 0,917622746 | novel transcript, antisense to MROH5 |
| ENSG00000259151 | CAP2P1 | 0,74755816 | 1,678948712 | 0,31657947 | 0,821646322 | cyclase associated actin cytoskeleton regulatory protein 2 pseudogene 1 |
| ENSG00000272381 | LINC02664 | 0,747082462 | 1,678395206 | 0,30063992 | 0,81031109 | long intergenic non-protein coding RNA 2664 |
| ENSG00000224271 | AL117329,1 | 0,747037497 | 1,678342895 | 0,33102491 | 0,832186595 | novel transcript |
| ENSG00000103742 | IGDCC4 | 0,747004173 | 1,678304129 | 0,02786133 | 0,235803123 | immunoglobulin superfamily DCC subclass member 4 |
| ENSG00000277250 | Metazoa_SRP | 0,74698242 | 1,678278824 | 0,52106749 | 0,927046149 | Metazoan signal recognition particle RNA |
| ENSG00000183638 | RP1L1 | 0,746857718 | 1,678133765 | 0,35887403 | 0,849814998 | RP1 like 1 |
| ENSG00000112699 | GMDS | 0,746669468 | 1,677914808 | 0,00740354 | 0,092364048 | GDP-mannose 4,6-dehydratase |
| ENSG00000198756 | COLGALT2 | 0,746516873 | 1,677737343 | 0,0742716 | 0,430483363 | collagen beta(1-O)galactosyltransferase 2 |
| ENSG00000137501 | SYTL2 | 0,74645028 | 1,677659902 | 0,04868917 | 0,33595633 | synaptotagmin like 2 |
| ENSG00000179750 | APOBEC3B | 0,746351851 | 1,677545447 | 0,07039634 | 0,417781142 | apolipoprotein B mRNA editing enzyme catalytic subunit 3B |
| ENSG00000130772 | MED18 | 0,746216031 | 1,677387525 | 0,74829031 | 0,975901808 | mediator complex subunit 18 |
| ENSG00000226964 | RHEBP2 | 0,746086884 | 1,677237375 | 0,14771226 | 0,612152172 | RHEB pseudogene 2 |
| ENSG00000184925 | LCN12 | 0,745991679 | 1,677126696 | 0,46779773 | 0,907940653 | lipocalin 12 |
| ENSG00000105137 | SYDE1 | 0,745984328 | 1,67711815 | 0,05197333 | 0,349829381 | synapse defective Rho GTPase homolog 1 |
| ENSG00000224691 | AL133553,1 | 0,745826631 | 1,676934839 | 0,57077861 | 0,942436428 | novel transcript |
| ENSG00000229118 | AC068491,1 | 0,745619522 | 1,676694121 | 0,40506758 | 0,878176155 | novel transcript |
| ENSG00000135404 | CD63 | 0,745515131 | 1,676572802 | 0,00793284 | 0,097133046 | CD63 molecule |

| | | | | | | |
|---|---|---|---|---|---|---|
| ENSG00000244036 | AC073320,1 | 0,745304123 | 1,676327606 | 0,3575674 | 0,848693232 | novel transcript, antisense to ZC3HC1 |
| ENSG00000286742 | AC000065,1 | 0,745159184 | 1,676159203 | 0,0299372 | 0,246681348 | novel transcript, antisense to CDK6 |
| ENSG00000284418 | AL391117,1 | 0,744879621 | 1,675834431 | 0,6499778 | 0,956899963 | novel transcript |
| ENSG00000273423 | OR13I1P | 0,7447929 | 1,675733699 | 0,52282074 | 0,92790906 | olfactory receptor family 13 subfamily I member 1 pseudogene |
| ENSG00000203650 | LINC01285 | 0,744524307 | 1,67542175 | 0,06468045 | 0,398771343 | long intergenic non-protein coding RNA 1285 |
| ENSG00000261177 | AC135012,1 | 0,74429589 | 1,675156506 | 0,39015061 | 0,870057277 | novel transcript |
| ENSG00000149925 | ALDOA | 0,744286441 | 1,675145536 | 0,00898246 | 0,105928449 | aldolase, fructose-bisphosphate A |
| ENSG00000275454 | AC105020,6 | 0,743974538 | 1,674783417 | 0,19600333 | 0,694033976 | novel transcript, antisense to IMP3 |
| ENSG00000177951 | BET1L | 0,743950927 | 1,674756007 | 0,01760735 | 0,171916321 | Bet1 golgi vesicular membrane trafficking protein like |
| ENSG00000076770 | MBNL3 | 0,743644692 | 1,674400552 | 0,18877852 | 0,683164698 | muscleblind like splicing regulator 3 |
| ENSG00000145075 | CCDC39 | 0,743518467 | 1,67425406 | 0,65771086 | 0,957864535 | coiled-coil domain containing 39 |
| ENSG00000259330 | INAFM2 | 0,743353914 | 1,674063107 | 0,02105824 | 0,193441015 | InaF motif containing 2 |
| ENSG00000148248 | SURF4 | 0,743349003 | 1,674057408 | 0,00067772 | 0,013427209 | surfeit 4 |
| ENSG00000280951 | SURF4 | 0,743349003 | 1,674057408 | 0,00067772 | 0,013427209 | surfeit 4 |
| ENSG00000286797 | AC009315,1 | 0,742908507 | 1,673546349 | 0,08480979 | 0,463466776 | novel transcript |
| ENSG00000273805 | AC083805,3 | 0,742900681 | 1,673537271 | 0,48156531 | 0,912981134 | novel transcript, antisense to Linc |
| ENSG00000203867 | RBM20 | 0,742307861 | 1,672849736 | 0,07060919 | 0,418488309 | RNA binding motif protein 20 |
| ENSG00000274414 | AL121772,1 | 0,742254123 | 1,672787426 | 0,40378186 | 0,877691457 | novel transcript |
| ENSG00000110080 | ST3GAL4 | 0,742011552 | 1,672506192 | 0,00269445 | 0,04239523 | ST3 beta-galactoside alpha-2,3-sialyltransferase 4 |
| ENSG00000087086 | FTL | 0,741915253 | 1,672394557 | 0,00875121 | 0,103883368 | ferritin light chain |
| ENSG00000182132 | KCNIP1 | 0,741903722 | 1,672381191 | 0,13805045 | 0,592614201 | potassium voltage-gated channel interacting protein 1 |
| ENSG00000131686 | CA6 | 0,741114198 | 1,67146622 | 0,72815507 | 0,973888243 | carbonic anhydrase 6 |
| ENSG00000259218 | LINC00928 | 0,740846866 | 1,671156524 | 0,54434814 | 0,934381824 | long intergenic non-protein coding RNA 928 |
| ENSG00000228750 | LINC01672 | 0,740744902 | 1,671038418 | 0,51618137 | 0,925176589 | long intergenic non-protein coding RNA 1672 |
| ENSG00000188112 | C6orf132 | 0,740714238 | 1,671002901 | 0,20633753 | 0,709417043 | chromosome 6 open reading frame 132 |
| ENSG00000236251 | HSPA1L | 0,740704762 | 1,670991926 | 0,77199218 | 0,978033855 | heat shock protein family A (Hsp70) member 1 like |
| ENSG00000206383 | HSPA1L | 0,740704762 | 1,670991926 | 0,77199218 | 0,978033855 | heat shock protein family A (Hsp70) member 1 like |
| ENSG00000184164 | CRELD2 | 0,740690855 | 1,670975817 | 0,05584833 | 0,36489907 | cysteine rich with EGF like domains 2 |
| ENSG00000168306 | ACOX2 | 0,740690854 | 1,670975817 | 0,14749125 | 0,611575703 | acyl-CoA oxidase 2 |
| ENSG00000282142 | AC245187,2 | 0,740666463 | 1,670947566 | 0,3434762 | 0,840901506 | novel transcript |

| | | | | | | |
|---|---|---|---|---|---|---|
| ENSG00000239677 | PDZRN3-AS1 | 0,740634788 | 1,670910881 | 0,17755722 | 0,664013033 | PDZRN3 antisense RNA 1 |
| ENSG00000272582 | AL031587,3 | 0,740502434 | 1,670757597 | 0,44922489 | 0,902062594 | novel transcript, antisense to C22orf23 |
| ENSG00000228604 | AL121759,1 | 0,74042738 | 1,670670681 | 0,66706282 | 0,9611116 | novel transcript |
| ENSG00000018610 | CXorf56 | 0,740339981 | 1,670569474 | 0,0036323 | 0,054387125 | chromosome X open reading frame 56 |
| ENSG00000169682 | SPNS1 | 0,740128493 | 1,670324599 | 0,07232765 | 0,424090732 | sphingolipid transporter 1 (putative) |
| ENSG00000185825 | BCAP31 | 0,740002786 | 1,670179064 | 0,01532533 | 0,156527661 | B cell receptor associated protein 31 |
| ENSG00000168282 | MGAT2 | 0,739885957 | 1,670043819 | 0,00232955 | 0,037897375 | alpha-1,6-mannosyl-glycoprotein 2-beta-N-acetylglucosaminyltransferase |
| ENSG00000110104 | CCDC86 | 0,739804488 | 1,669949514 | 0,03622589 | 0,279341761 | coiled-coil domain containing 86 |
| ENSG00000258896 | SCOCP1 | 0,739614958 | 1,669730144 | 0,54085025 | 0,933385624 | short coiled-coil protein pseudogene 1 |
| ENSG00000229122 | AGBL5-IT1 | 0,739391348 | 1,669471364 | 0,36026746 | 0,850774364 | AGBL5 intronic transcript 1 |
| ENSG00000270066 | AL356488,2 | 0,738678753 | 1,668646961 | 0,51850451 | 0,926739362 | small Cajal body-specific RNA 2 |
| ENSG00000108784 | NAGLU | 0,73831511 | 1,668226418 | 0,04932712 | 0,338713596 | N-acetyl-alpha-glucosaminidase |
| ENSG00000165288 | BRWD3 | 0,738075638 | 1,667949533 | 0,06428817 | 0,397764749 | bromodomain and WD repeat domain containing 3 |
| ENSG00000179918 | SEPHS2 | 0,738026567 | 1,667892801 | 0,00288324 | 0,04491362 | selenophosphate synthetase 2 |
| ENSG00000271121 | NUDT4P2 | 0,737948768 | 1,66780286 | 0,04755045 | 0,33161214 | nudix hydrolase 4 pseudogene 2 |
| ENSG00000229859 | PGA3 | 0,737864018 | 1,66770489 | 0,71396027 | 0,970867497 | pepsinogen A3 |
| ENSG00000111817 | DSE | 0,737710268 | 1,66752717 | 0,021541 | 0,196572635 | dermatan sulfate epimerase |
| ENSG00000166794 | PPIB | 0,737460362 | 1,667238343 | 0,00110115 | 0,020262677 | peptidylprolyl isomerase B |
| ENSG00000282987 | AC099753,1 | 0,737453924 | 1,667230903 | 0,65427303 | 0,957444211 | novel transcript |
| ENSG00000070159 | PTPN3 | 0,737067866 | 1,66678482 | 0,03270447 | 0,261798703 | protein tyrosine phosphatase non-receptor type 3 |
| ENSG00000287067 | AC106791,3 | 0,736730496 | 1,666395093 | 0,30912944 | 0,816845699 | zinc finger BED-type containing 9 (ZBED9) pseudogene |
| ENSG00000142494 | SLC47A1 | 0,736608832 | 1,66625457 | 0,39752239 | 0,874236071 | solute carrier family 47 member 1 |
| ENSG00000133107 | TRPC4 | 0,736581762 | 1,666223305 | 0,08298633 | 0,458031678 | transient receptor potential cation channel subfamily C member 4 |
| ENSG00000047662 | FAM184B | 0,736508197 | 1,666138345 | 0,42342544 | 0,890929178 | family with sequence similarity 184 member B |
| ENSG00000287654 | AP002982,2 | 0,736406162 | 1,66602051 | 0,54804249 | 0,935404974 | novel transcript |
| ENSG00000213228 | RPL12P38 | 0,736229335 | 1,665816323 | 0,4996237 | 0,920190594 | ribosomal protein L12 pseudogene 38 |
| ENSG00000283108 | AC011451,3 | 0,736067212 | 1,665629138 | 0,5561764 | 0,938046927 | novel transcript |
| ENSG00000271699 | SNX29P2 | 0,735815914 | 1,665339033 | 0,49148034 | 0,916995341 | sorting nexin 29 pseudogene 2 |
| ENSG00000163141 | BNIPL | 0,735760831 | 1,665275451 | 0,19199042 | 0,688347294 | BCL2 interacting protein like |
| ENSG00000231646 | FSIP2-AS1 | 0,735569652 | 1,665054791 | 0,11398911 | 0,53948382 | FSIP2 antisense RNA 1 |

| | | | | | | |
|---|---|---|---|---|---|---|
| ENSG00000173221 | GLRX | 0,735420296 | 1,664882423 | 0,02335187 | 0,208354147 | glutaredoxin |
| ENSG00000197381 | ADARB1 | 0,735341683 | 1,664791706 | 0,02562889 | 0,22266762 | adenosine deaminase RNA specific B1 |
| ENSG00000239705 | AL354710,2 | 0,735238075 | 1,664672152 | 0,51126033 | 0,922893865 | novel transcript |
| ENSG00000168754 | FAM178B | 0,735179989 | 1,664605131 | 0,41467422 | 0,885579089 | family with sequence similarity 178 member B |
| ENSG00000166340 | TPP1 | 0,735113756 | 1,664528712 | 0,0150278 | 0,154428164 | tripeptidyl peptidase 1 |
| ENSG00000187554 | TLR5 | 0,734999962 | 1,664397425 | 0,18821794 | 0,682491978 | toll like receptor 5 |
| ENSG00000269823 | AL513523,9 | 0,734837492 | 1,664209999 | 0,17353783 | 0,658349911 | |
| ENSG00000286606 | AC097468,3 | 0,734611018 | 1,663948772 | 0,45451653 | 0,903383444 | novel transcript |
| ENSG00000204673 | AKT1S1 | 0,73452567 | 1,663850338 | 0,01518911 | 0,155491887 | AKT1 substrate 1 |
| ENSG00000183853 | KIRREL1 | 0,733809311 | 1,663024371 | 0,00031399 | 0,007036967 | kirre like nephrin family adhesion molecule 1 |
| ENSG00000239300 | AC080162,1 | 0,733757254 | 1,662964364 | 0,5443587 | 0,934381824 | novel transcript |
| ENSG00000286205 | AC139887,5 | 0,733494577 | 1,662661611 | 0,36925775 | 0,857942508 | novel transcript |
| ENSG00000236801 | RPL24P8 | 0,733484261 | 1,662649721 | 0,45742196 | 0,903831777 | RPL24 pseudogene 8 |
| ENSG00000286000 | AC234775,3 | 0,733260368 | 1,662391713 | 0,33074542 | 0,832114797 | novel transcript |
| ENSG00000259764 | PCSK6-AS1 | 0,733214377 | 1,66233872 | 0,34435119 | 0,84111511 | PCSK6 antisense RNA 1 |
| ENSG00000125820 | NKX2-2 | 0,732955955 | 1,662040981 | 0,50735318 | 0,921988479 | NK2 homeobox 2 |
| ENSG00000202406 | RN7SKP187 | 0,732855118 | 1,661924817 | 0,28951588 | 0,800444324 | RN7SK pseudogene 187 |
| ENSG00000145063 | AC062028,1 | 0,732718115 | 1,661767002 | 0,38151467 | 0,865900094 | novel transcript |
| ENSG00000160318 | CLDND2 | 0,73265553 | 1,661694915 | 0,29399135 | 0,803927308 | claudin domain containing 2 |
| ENSG00000285974 | AC026624,1 | 0,732589144 | 1,661618454 | 0,40530375 | 0,878251762 | novel transcript, antisense to EFL1 |
| ENSG00000107719 | PALD1 | 0,731625217 | 1,660508626 | 0,04638024 | 0,327048609 | phosphatase domain containing paladin 1 |
| ENSG00000189409 | MMP23B | 0,730765825 | 1,65951978 | 0,26487886 | 0,776408453 | matrix metallopeptidase 23B |
| ENSG00000258232 | AC125611,3 | 0,730553052 | 1,659275047 | 0,03323269 | 0,264777507 | novel transcript, antisense to TUBA1C |
| ENSG00000226493 | RPL26P27 | 0,729975586 | 1,658611024 | 0,58131492 | 0,943919685 | ribosomal protein L26 pseudogene 27 |
| ENSG00000197106 | SLC6A17 | 0,729975246 | 1,658610633 | 0,47238967 | 0,909229736 | solute carrier family 6 member 17 |
| ENSG00000205755 | CRLF2 | 0,72979549 | 1,658403987 | 0,50733763 | 0,921988479 | cytokine receptor like factor 2 |
| ENSG00000158863 | FAM160B2 | 0,729512978 | 1,658079266 | 0,02027823 | 0,188628172 | family with sequence similarity 160 member B2 |
| ENSG00000198354 | DCAF12L2 | 0,729500929 | 1,658065418 | 0,09982376 | 0,501267282 | DDB1 and CUL4 associated factor 12 like 2 |
| ENSG00000235482 | RPL21P135 | 0,729491331 | 1,658054388 | 0,50757887 | 0,922140988 | ribosomal protein L21 pseudogene 135 |
| ENSG00000102078 | SLC25A14 | 0,729311989 | 1,657848287 | 0,0769571 | 0,438408786 | solute carrier family 25 member 14 |
| ENSG00000239704 | CDRT4 | 0,728873932 | 1,657344978 | 0,06884548 | 0,413761082 | CMT1A duplicated region transcript 4 |
| ENSG00000287149 | AC092640,1 | 0,728865161 | 1,657334902 | 0,45265916 | 0,902285042 | novel transcript, antisense to CCDC141and TTN |

| | | | | | | |
|---|---|---|---|---|---|---|
| ENSG00000256254 | AP000619,3 | 0,728617613 | 1,657050549 | 0,65042363 | 0,956899963 | spastic ataxia of Charlevoix-Saguenay (SACS) pseudogene |
| ENSG00000143028 | SYPL2 | 0,728368212 | 1,656764117 | 0,20183123 | 0,701461582 | synaptophysin like 2 |
| ENSG00000286100 | AL008725,1 | 0,72822478 | 1,65659941 | 0,40949798 | 0,882177108 | novel transcript |
| ENSG00000075240 | GRAMD4 | 0,728224651 | 1,656599262 | 0,02592429 | 0,224390102 | GRAM domain containing 4 |
| ENSG00000248538 | AC022784,1 | 0,728008493 | 1,656351073 | 0,2700546 | 0,782267151 | novel transcript |
| ENSG00000283703 | VSIG10L2 | 0,727922269 | 1,656252083 | 0,28691374 | 0,798400609 | V-set and immunoglobulin domain containing 10 like 2 |
| ENSG00000164342 | TLR3 | 0,727921172 | 1,656250824 | 0,10953946 | 0,527842443 | toll like receptor 3 |
| ENSG00000227573 | AC004986,1 | 0,727565924 | 1,65584304 | 0,58749263 | 0,946068138 | v-rel reticuloendotheliosis viral oncogene homolog A (avian) (RELA) pseudogene |
| ENSG00000235112 | HSPE1P27 | 0,727493345 | 1,655759741 | 0,56242514 | 0,940865625 | heat shock protein family E (Hsp10) member 1 pseudogene 27 |
| ENSG00000278603 | AC245033,3 | 0,727476443 | 1,655740342 | 0,26959871 | 0,781791089 | dynamin 1 (DNM1) pseudogene |
| ENSG00000237426 | ZIK1P1 | 0,727377819 | 1,655627157 | 0,52795195 | 0,929699769 | zinc finger protein interacting with K protein 1 pseudogene 1 |
| ENSG00000056291 | NPFFR2 | 0,726911633 | 1,655092253 | 0,40957603 | 0,882177108 | neuropeptide FF receptor 2 |
| ENSG00000243175 | RPSAP36 | 0,726761323 | 1,654919822 | 0,17644666 | 0,662315819 | ribosomal protein SA pseudogene 36 |
| ENSG00000228634 | AL136115,1 | 0,726710208 | 1,654861189 | 0,47979273 | 0,912847708 | novel transcript |
| ENSG00000285278 | TFAP2A-AS2 | 0,726399002 | 1,654504255 | 0,1080401 | 0,523928283 | TFAP2A antisense RNA 2 |
| ENSG00000282419 | TEX13D | 0,726209095 | 1,654286481 | 0,31072943 | 0,817751312 | TEX13 family member D |
| ENSG00000226650 | KIF4B | 0,726172445 | 1,654244457 | 0,15657285 | 0,627737077 | kinesin family member 4B |
| ENSG00000272716 | AL121658,1 | 0,725979183 | 1,65402287 | 0,35800691 | 0,849032846 | novel transcript |
| ENSG00000103152 | MPG | 0,725761845 | 1,653773716 | 0,06518272 | 0,400682776 | N-methylpurine DNA glycosylase |
| ENSG00000143502 | SUSD4 | 0,725753693 | 1,65376437 | 0,09048899 | 0,479824907 | sushi domain containing 4 |
| ENSG00000139354 | GAS2L3 | 0,725708934 | 1,653713064 | 0,058986 | 0,377278711 | growth arrest specific 2 like 3 |
| ENSG00000276786 | BX640515,1 | 0,725621265 | 1,653612575 | 0,50182665 | 0,920520138 | novel transcript |
| ENSG00000167323 | STIM1 | 0,725549869 | 1,653530743 | 0,00640353 | 0,082829913 | stromal interaction molecule 1 |
| ENSG00000236723 | AL606760,2 | 0,72546063 | 1,653428466 | 0,28930491 | 0,800260154 | novel transcript, antisense to CPT2 |
| ENSG00000272933 | AL391121,1 | 0,725112031 | 1,653028996 | 0,19267763 | 0,689545952 | novel transcript |
| ENSG00000278590 | RN7SL113P | 0,72510523 | 1,653021203 | 0,48320457 | 0,91363215 | RNA, 7SL, cytoplasmic 113, pseudogene |
| ENSG00000181092 | ADIPOQ | 0,725069221 | 1,652979945 | 0,58996752 | 0,947252221 | adiponectin, C1Q and collagen domain containing |
| ENSG00000136859 | ANGPTL2 | 0,724973179 | 1,652869907 | 0,04155737 | 0,305921747 | angiopoietin like 2 |

| | | | | | | |
|---|---|---|---|---|---|---|
| ENSG00000168502 | MTCL1 | 0,724816027 | 1,652689871 | 0,18300402 | 0,673903158 | microtubule crosslinking factor 1 |
| ENSG00000008086 | CDKL5 | 0,724698866 | 1,652555662 | 0,00825626 | 0,099854406 | cyclin dependent kinase like 5 |
| ENSG00000285928 | AC103591,4 | 0,724172542 | 1,651952886 | 0,30763093 | 0,816380056 | novel transcript |
| ENSG00000141524 | TMC6 | 0,724014353 | 1,651771763 | 0,04325933 | 0,313869373 | transmembrane channel like 6 |
| ENSG00000123992 | DNPEP | 0,723881342 | 1,651619483 | 0,01379226 | 0,144680458 | aspartyl aminopeptidase |
| ENSG00000122863 | CHST3 | 0,723780904 | 1,651504504 | 0,02900706 | 0,241642594 | carbohydrate sulfotransferase 3 |
| ENSG00000253015 | RN7SKP64 | 0,723720932 | 1,651435853 | 0,33426182 | 0,83453565 | RN7SK pseudogene 64 |
| ENSG00000288553 | AC106870,3 | 0,723695022 | 1,651406195 | 0,57552386 | 0,942700009 | novel transcript |
| ENSG00000181788 | SIAH2 | 0,723270051 | 1,650919816 | 0,01854173 | 0,177678855 | siah E3 ubiquitin protein ligase 2 |
| ENSG00000103145 | HCFC1R1 | 0,723174289 | 1,650810236 | 0,02528465 | 0,22068654 | host cell factor C1 regulator 1 |
| ENSG00000121895 | TMEM156 | 0,723018666 | 1,650632173 | 0,4342871 | 0,896272827 | transmembrane protein 156 |
| ENSG00000162946 | DISC1 | 0,722970908 | 1,650577532 | 0,00792436 | 0,097117653 | DISC1 scaffold protein |
| ENSG00000136205 | TNS3 | 0,722924389 | 1,650524312 | 0,01660596 | 0,164996341 | tensin 3 |
| ENSG00000112033 | PPARD | 0,722895928 | 1,650491751 | 0,01070777 | 0,120467271 | peroxisome proliferator activated receptor delta |
| ENSG00000227057 | WDR46 | 0,72252249 | 1,65006458 | 0,36433518 | 0,854320801 | WD repeat domain 46 |
| ENSG00000154654 | NCAM2 | 0,721644546 | 1,649060748 | 0,08031748 | 0,449444373 | neural cell adhesion molecule 2 |
| ENSG00000255933 | AC117500,2 | 0,721509045 | 1,648905873 | 0,34937094 | 0,843141089 | novel transcript |
| ENSG00000011052 | NME1-NME2 | 0,721449125 | 1,648837389 | 0,2652951 | 0,77668189 | NME1-NME2 readthrough |
| ENSG00000240441 | AC007622,1 | 0,72133315 | 1,648704848 | 0,58263178 | 0,944174844 | ribosomal protein L17 (RPL17) pseudogene |
| ENSG00000279727 | LINC02033 | 0,721183661 | 1,648534022 | 0,59195463 | 0,947416795 | long intergenic non-protein coding RNA 2033 |
| ENSG00000112902 | SEMA5A | 0,721110887 | 1,648450867 | 0,01263056 | 0,1357449 | semaphorin 5A |
| ENSG00000206491 | PRR3 | 0,720945221 | 1,648261584 | 0,03756647 | 0,286458107 | proline rich 3 |
| ENSG00000287269 | Z82215,1 | 0,720845908 | 1,648148124 | 0,28262456 | 0,793528967 | novel transcript, antisense to ANKRD54and MYH9 |
| ENSG00000283654 | LMLN2 | 0,720801586 | 1,648097491 | 0,44814303 | 0,902062594 | leishmanolysin like peptidase 2 |
| ENSG00000165124 | SVEP1 | 0,72079964 | 1,648095269 | 0,05068858 | 0,344431146 | sushi, von Willebrand factor type A, EGF and pentraxin domain containing 1 |
| ENSG00000276836 | ARHGAP27 | 0,720268735 | 1,647488889 | 0,16522273 | 0,644240891 | Rho GTPase activating protein 27 |
| ENSG00000150991 | UBC | 0,720260456 | 1,647479435 | 0,00080785 | 0,015561719 | ubiquitin C |
| ENSG00000225077 | LINC00337 | 0,720017572 | 1,647202097 | 0,14500776 | 0,606166963 | long intergenic non-protein coding RNA 337 |
| ENSG00000145888 | GLRA1 | 0,719745994 | 1,646892052 | 0,32703374 | 0,828735859 | glycine receptor alpha 1 |
| ENSG00000235554 | SRSF6P2 | 0,719393113 | 1,646489273 | 0,4133269 | 0,884678298 | SRSF6 pseudogene 2 |
| ENSG00000273873 | BDP1 | 0,719391238 | 1,646487133 | 0,28056919 | 0,791456745 | B double prime 1, subunit of RNA polymerase III transcription initiation factor IIIB |

| | | | | | | |
|---|---|---|---|---|---|---|
| ENSG00000203804 | ADAMTSL4-AS1 | 0,719201716 | 1,646270854 | 0,17317712 | 0,658102544 | ADAMTSL4 antisense RNA 1 |
| ENSG00000277459 | AP001527,2 | 0,719141545 | 1,646202194 | 0,2471737 | 0,759323829 | novel transcript |
| ENSG00000244486 | SCARF2 | 0,719128338 | 1,646187124 | 0,1648295 | 0,643964432 | scavenger receptor class F member 2 |
| ENSG00000157350 | ST3GAL2 | 0,719050185 | 1,64609795 | 0,0398938 | 0,298118347 | ST3 beta-galactoside alpha-2,3-sialyltransferase 2 |
| ENSG00000168003 | SLC3A2 | 0,718934145 | 1,645965555 | 0,0358129 | 0,277297887 | solute carrier family 3 member 2 |
| ENSG00000133142 | TCEAL4 | 0,718929088 | 1,645959786 | 0,00875269 | 0,103883368 | transcription elongation factor A like 4 |
| ENSG00000214706 | IFRD2 | 0,71856471 | 1,645544122 | 0,07673103 | 0,437679087 | interferon related developmental regulator 2 |
| ENSG00000176472 | ZNF575 | 0,718561826 | 1,645540833 | 0,12407221 | 0,563164667 | zinc finger protein 575 |
| ENSG00000235912 | AL031729,1 | 0,718398899 | 1,645355008 | 0,51254378 | 0,923529987 | novel pseudogene |
| ENSG00000071859 | FAM50A | 0,718324098 | 1,645269701 | 0,04730636 | 0,330573569 | family with sequence similarity 50 member A |
| ENSG00000197081 | IGF2R | 0,718321986 | 1,645267293 | 0,01040401 | 0,118280376 | insulin like growth factor 2 receptor |
| ENSG00000244356 | RN7SL398P | 0,718015333 | 1,644917619 | 0,44149646 | 0,898921877 | RNA, 7SL, cytoplasmic 398, pseudogene |
| ENSG00000103381 | CPPED1 | 0,718008824 | 1,644910198 | 0,00105015 | 0,019507958 | calcineurin like phosphoesterase domain containing 1 |
| ENSG00000188042 | ARL4C | 0,717882329 | 1,64476598 | 0,00480431 | 0,066828153 | ADP ribosylation factor like GTPase 4C |
| ENSG00000185634 | SHC4 | 0,717534894 | 1,644369929 | 0,0128498 | 0,137769634 | SHC adaptor protein 4 |
| ENSG00000258285 | TESC-AS1 | 0,717243097 | 1,644037375 | 0,50852459 | 0,922427526 | TESC antisense RNA 1 (head to head) |
| ENSG00000269226 | TMSB15B | 0,717014135 | 1,64377648 | 0,05374447 | 0,357655859 | thymosin beta 15B |
| ENSG00000110455 | ACCS | 0,716970506 | 1,64372677 | 0,02779144 | 0,235339037 | 1-aminocyclopropane-1-carboxylate synthase homolog (inactive) |
| ENSG00000246211 | P4HA3-AS1 | 0,716572289 | 1,643273127 | 0,53224427 | 0,931291085 | P4HA3 antisense RNA 1 |
| ENSG00000110880 | CORO1C | 0,716423494 | 1,643103654 | 0,00144287 | 0,025442636 | coronin 1C |
| ENSG00000167553 | TUBA1C | 0,716404833 | 1,6430824 | 0,01256289 | 0,135303249 | tubulin alpha 1c |
| ENSG00000163735 | CXCL5 | 0,716018662 | 1,64264265 | 0,18630041 | 0,679406361 | C-X-C motif chemokine ligand 5 |
| ENSG00000153814 | JAZF1 | 0,715966479 | 1,642583236 | 0,07404343 | 0,4296801 | JAZF zinc finger 1 |
| ENSG00000285751 | AC021723,2 | 0,715669031 | 1,64224461 | 0,5889538 | 0,946604802 | novel transcript |
| ENSG00000258839 | MC1R | 0,715518497 | 1,642073265 | 0,18584474 | 0,678777213 | melanocortin 1 receptor |
| ENSG00000284740 | AL645728,2 | 0,715496383 | 1,642048094 | 0,64135718 | 0,956465487 | ATPase family, AAA domain containing 3C (ATAD3C) pseudogene |
| ENSG00000287512 | AC019131,3 | 0,715385392 | 1,641921771 | 0,61305184 | 0,952413269 | novel transcript |
| ENSG00000285053 | AL357556,3 | 0,715313673 | 1,64184015 | 0,33041265 | 0,832059703 | Tubulin-specific chaperone E |
| ENSG00000227455 | AL450345,1 | 0,71528624 | 1,641808931 | 0,52710625 | 0,92927181 | novel transcript |
| ENSG00000138675 | FGF5 | 0,714963179 | 1,641441324 | 0,34723638 | 0,841896177 | fibroblast growth factor 5 |

| | | | | | | |
|---|---|---|---|---|---|---|
| ENSG00000172725 | CORO1B | 0,714898575 | 1,641367821 | 0,02196916 | 0,199432031 | coronin 1B |
| ENSG00000142627 | EPHA2 | 0,714882479 | 1,641349509 | 0,05101376 | 0,345820358 | EPH receptor A2 |
| ENSG00000229672 | AL450322,2 | 0,714799945 | 1,641255613 | 0,54975154 | 0,936108121 | novel transcript |
| ENSG00000281548 | LINC00895 | 0,714741426 | 1,641189041 | 0,36490931 | 0,855057583 | long intergenic non-protein coding RNA 895 |
| ENSG00000272167 | AL606537,1 | 0,714728886 | 1,641174776 | 0,46281223 | 0,906849208 | PROX1 antisense RNA 1 |
| ENSG00000266998 | AC111182,1 | 0,714330872 | 1,640722067 | 0,31426943 | 0,819877431 | novel transcript, antisense to SEPT9 |
| ENSG00000151914 | DST | 0,713894978 | 1,640226416 | 0,01461697 | 0,151269455 | dystonin |
| ENSG00000164294 | GPX8 | 0,713546117 | 1,639829838 | 0,04653194 | 0,327767757 | glutathione peroxidase 8 (putative) |
| ENSG00000114850 | SSR3 | 0,71350987 | 1,639788638 | 0,00282285 | 0,044170984 | signal sequence receptor subunit 3 |
| ENSG00000283018 | MRPS12 | 0,713500432 | 1,639777911 | 0,08418518 | 0,461585627 | mitochondrial ribosomal protein S12 |
| ENSG00000166452 | AKIP1 | 0,713440424 | 1,639709707 | 0,00250838 | 0,040154681 | A-kinase interacting protein 1 |
| ENSG00000074181 | NOTCH3 | 0,712833534 | 1,639020086 | 0,08343494 | 0,45997353 | notch receptor 3 |
| ENSG00000181885 | CLDN7 | 0,71245246 | 1,638587211 | 0,48976804 | 0,916063942 | claudin 7 |
| ENSG00000288292 | CLDN7 | 0,71245246 | 1,638587211 | 0,48976804 | 0,916063942 | claudin 7 |
| ENSG00000286042 | LCAL1 | 0,712402981 | 1,638531014 | 0,39411115 | 0,871635019 | lung cancer associated lncRNA 1 |
| ENSG00000131773 | KHDRBS3 | 0,712402707 | 1,638530704 | 0,01193632 | 0,130162906 | KH RNA binding domain containing, signal transduction associated 3 |
| ENSG00000107104 | KANK1 | 0,712377796 | 1,638502411 | 0,01319927 | 0,140027857 | KN motif and ankyrin repeat domains 1 |
| ENSG00000124134 | KCNS1 | 0,712215417 | 1,638318004 | 0,10424553 | 0,514802081 | potassium voltage-gated channel modifier subfamily S member 1 |
| ENSG00000278220 | CLN8 | 0,71207966 | 1,638163846 | 0,2748371 | 0,786383635 | CLN8 transmembrane ER and ERGIC protein |
| ENSG00000142192 | APP | 0,711905902 | 1,637966558 | 0,00405348 | 0,05935735 | amyloid beta precursor protein |
| ENSG00000172375 | C2CD2L | 0,711770891 | 1,63781328 | 0,0228739 | 0,205211335 | C2CD2 like |
| ENSG00000235802 | HCFC1-AS1 | 0,711730122 | 1,637766998 | 0,35340428 | 0,84514549 | HCFC1 antisense RNA 1 |
| ENSG00000268658 | LINC00664 | 0,711651206 | 1,637677414 | 0,01793871 | 0,174004579 | long intergenic non-protein coding RNA 664 |
| ENSG00000287207 | AC096861,2 | 0,711564098 | 1,637578537 | 0,58821479 | 0,946176848 | novel transcript |
| ENSG00000154096 | THY1 | 0,711123943 | 1,637079 | 0,11404196 | 0,53948382 | Thy-1 cell surface antigen |
| ENSG00000186810 | CXCR3 | 0,711069357 | 1,63701706 | 0,72552885 | 0,97326334 | C-X-C motif chemokine receptor 3 |
| ENSG00000203620 | AL354919,1 | 0,711046451 | 1,636991069 | 0,2756149 | 0,787284069 | novel transcript |
| ENSG00000247402 | AC099487,1 | 0,710997554 | 1,636935587 | 0,62521446 | 0,952793934 | novel transcript |
| ENSG00000232215 | OR2L6P | 0,710966189 | 1,636900001 | 0,53001767 | 0,931030033 | olfactory receptor family 2 subfamily L member 6 pseudogene |
| ENSG00000254245 | PCDHGA3 | 0,710801844 | 1,636713543 | 0,22338656 | 0,731984265 | protocadherin gamma subfamily A, 3 |

| | | | | | | |
|---|---|---|---|---|---|---|
| ENSG00000088367 | EPB41L1 | 0,710794614 | 1,636705341 | 0,0043666 | 0,062622653 | erythrocyte membrane protein band 4,1 like 1 |
| ENSG00000089847 | ANKRD24 | 0,710749782 | 1,636654481 | 0,50706916 | 0,921951873 | ankyrin repeat domain 24 |
| ENSG00000230870 | FBXW11P1 | 0,710425617 | 1,636286776 | 0,50306853 | 0,920888747 | F-box and WD repeat domain containing 11 pseudogene 1 |
| ENSG00000186480 | INSIG1 | 0,709585846 | 1,635334594 | 0,03265013 | 0,261468653 | insulin induced gene 1 |
| ENSG00000231651 | DLG3-AS1 | 0,709585751 | 1,635334487 | 0,31780193 | 0,822700957 | DLG3 antisense RNA 1 |
| ENSG00000146147 | MLIP | 0,709512513 | 1,635251472 | 0,3562981 | 0,847740014 | muscular LMNA interacting protein |
| ENSG00000125503 | PPP1R12C | 0,709372113 | 1,63509234 | 0,17858352 | 0,666018763 | protein phosphatase 1 regulatory subunit 12C |
| ENSG00000068323 | TFE3 | 0,709155789 | 1,634847186 | 0,01670657 | 0,165411048 | transcription factor binding to IGHM enhancer 3 |
| ENSG00000247596 | TWF2 | 0,708938776 | 1,634601287 | 0,07727169 | 0,439400181 | twinfilin actin binding protein 2 |
| ENSG00000274829 | PARN | 0,708877219 | 1,634531544 | 0,23447985 | 0,744090329 | poly(A)-specific ribonuclease |
| ENSG00000166016 | ABTB2 | 0,708662123 | 1,634287864 | 0,10266271 | 0,510196647 | ankyrin repeat and BTB domain containing 2 |
| ENSG00000213866 | YBX1P10 | 0,708598665 | 1,634215981 | 0,6286972 | 0,952793934 | Y-box binding protein 1 pseudogene 10 |
| ENSG00000224185 | SNX18P9 | 0,708310584 | 1,633889689 | 0,26696229 | 0,779386557 | sorting nexin 18 pseudogene 9 |
| ENSG00000164930 | FZD6 | 0,708183085 | 1,633745299 | 0,04053282 | 0,300657756 | frizzled class receptor 6 |
| ENSG00000258325 | ITFG2-AS1 | 0,707388034 | 1,63284521 | 0,3734445 | 0,861319894 | ITFG2 antisense RNA 1 |
| ENSG00000099377 | HSD3B7 | 0,70729506 | 1,632739986 | 0,13801576 | 0,592546669 | hydroxy-delta-5-steroid dehydrogenase, 3 beta- and steroid delta-isomerase 7 |
| ENSG00000285446 | Z84488,1 | 0,706996934 | 1,632402624 | 0,79052553 | 0,979462318 | novel protein |
| ENSG00000260914 | AC026464,4 | 0,706933325 | 1,632330651 | 0,00472406 | 0,066050442 | novel protein |
| ENSG00000182798 | MAGEB17 | 0,706688083 | 1,632053197 | 0,48177819 | 0,912981134 | MAGE family member B17 |
| ENSG00000173705 | SUSD5 | 0,706570472 | 1,631920155 | 0,07538422 | 0,433690715 | sushi domain containing 5 |
| ENSG00000269696 | AC005498,3 | 0,706513077 | 1,631855233 | 0,19627194 | 0,694046542 | novel transcript, antisense to ZFP28 |
| ENSG00000234199 | LINC01191 | 0,706217699 | 1,63152116 | 0,43751589 | 0,897021798 | long intergenic non-protein coding RNA 1191 |
| ENSG00000275557 | AC242842,1 | 0,705901612 | 1,631163742 | 0,42925532 | 0,894029049 | novel transcript |
| ENSG00000108829 | LRRC59 | 0,705820189 | 1,631071684 | 0,00237 | 0,038351788 | leucine rich repeat containing 59 |
| ENSG00000229455 | RPS10P18 | 0,70568693 | 1,630921033 | 0,62900165 | 0,952793934 | ribosomal protein S10 pseudogene 18 |
| ENSG00000253302 | STAU2-AS1 | 0,705686879 | 1,630920975 | 0,37841971 | 0,864011808 | STAU2 antisense RNA 1 |
| ENSG00000231256 | CFAP97D1 | 0,705668147 | 1,630899799 | 0,41383988 | 0,885395886 | CFAP97 domain containing 1 |
| ENSG00000287796 | AC016925,3 | 0,705590987 | 1,630812576 | 0,71761613 | 0,971697135 | novel transcript |
| ENSG00000124191 | TOX2 | 0,705307133 | 1,63049174 | 0,06993425 | 0,417175017 | TOX high mobility group box family member 2 |
| ENSG00000285241 | PCK2 | 0,705208139 | 1,630379864 | 0,09076852 | 0,480677819 | phosphoenolpyruvate carboxykinase 2, mitochondrial |

| | | | | | | |
|---|---|---|---|---|---|---|
| ENSG00000100889 | PCK2 | 0,705208139 | 1,630379864 | 0,09076852 | 0,480677819 | phosphoenolpyruvate carboxykinase 2, mitochondrial |
| ENSG00000261829 | AC009407,1 | 0,704897291 | 1,630028615 | 0,58314437 | 0,944174844 | novel transcript |
| ENSG00000242612 | DECR2 | 0,704644437 | 1,629742953 | 0,14726304 | 0,611181688 | 2,4-dienoyl-CoA reductase 2 |
| ENSG00000129250 | KIF1C | 0,704591309 | 1,629682938 | 0,12799008 | 0,571345532 | kinesin family member 1C |
| ENSG00000237207 | RBM17P3 | 0,704465205 | 1,629540495 | 0,30841105 | 0,816625411 | RNA binding motif protein 17 pseudogene 3 |
| ENSG00000274843 | MUC20 | 0,70421715 | 1,629260339 | 0,58755277 | 0,946068138 | mucin 20, cell surface associated |
| ENSG00000147459 | DOCK5 | 0,70414739 | 1,629181559 | 0,00642786 | 0,083110242 | dedicator of cytokinesis 5 |
| ENSG00000230399 | RBBP8P1 | 0,70409132 | 1,629118243 | 0,33228748 | 0,83293962 | RBBP8 pseudogene 1 |
| ENSG00000248890 | HHIP-AS1 | 0,703329711 | 1,628258447 | 0,2064926 | 0,709660852 | HHIP antisense RNA 1 |
| ENSG00000135604 | STX11 | 0,703313942 | 1,62824065 | 0,4336026 | 0,896108974 | syntaxin 11 |
| ENSG00000267413 | LINC01901 | 0,703296123 | 1,628220539 | 0,23761099 | 0,746907875 | long intergenic non-protein coding RNA 1901 |
| ENSG00000117400 | MPL | 0,702728386 | 1,627579919 | 0,3797081 | 0,865238161 | MPL proto-oncogene, thrombopoietin receptor |
| ENSG00000108176 | DNAJC12 | 0,702587194 | 1,627420641 | 0,05018349 | 0,342712245 | DnaJ heat shock protein family (Hsp40) member C12 |
| ENSG00000188976 | NOC2L | 0,702422064 | 1,627234379 | 0,10420072 | 0,514802081 | NOC2 like nucleolar associated transcriptional repressor |
| ENSG00000260963 | AC026462,3 | 0,702417958 | 1,627229747 | 0,70908329 | 0,970282889 | novel transcript |
| ENSG00000144118 | RALB | 0,702101403 | 1,626872741 | 0,01411573 | 0,147301149 | RAS like proto-oncogene B |
| ENSG00000198488 | B3GNT6 | 0,702029299 | 1,626791433 | 0,47701559 | 0,91143003 | UDP-GlcNAc:betaGal beta-1,3-N-acetylglucosaminyltransferase 6 |
| ENSG00000236467 | KCNMA1-AS1 | 0,701975969 | 1,626731299 | 0,37838934 | 0,864005583 | KCNMA1 antisense RNA 1 |
| ENSG00000253829 | AC067817,2 | 0,701865459 | 1,626606698 | 0,4256874 | 0,892133432 | novel transcript |
| ENSG00000226386 | PARD3-AS1 | 0,70174856 | 1,626474902 | 0,52029483 | 0,926865961 | PARD3 antisense RNA 1 |
| ENSG00000015532 | XYLT2 | 0,701730819 | 1,6264549 | 0,08007409 | 0,448625211 | xylosyltransferase 2 |
| ENSG00000200354 | SNORA71D | 0,701674962 | 1,62639193 | 0,3474708 | 0,842200946 | small nucleolar RNA, H/ACA box 71D |
| ENSG00000172175 | MALT1 | 0,701360479 | 1,626037443 | 0,04110012 | 0,303484684 | MALT1 paracaspase |
| ENSG00000180900 | SCRIB | 0,701264903 | 1,625929725 | 0,32168095 | 0,824933727 | scribble planar cell polarity protein |
| ENSG00000249021 | AC008549,2 | 0,701256175 | 1,625919887 | 0,60753539 | 0,950993594 | novel transcript |
| ENSG00000236699 | ARHGEF38 | 0,700790084 | 1,625394688 | 0,25763859 | 0,769984975 | Rho guanine nucleotide exchange factor 38 |
| ENSG00000136720 | HS6ST1 | 0,70070733 | 1,625301456 | 0,05351558 | 0,356208443 | heparan sulfate 6-O-sulfotransferase 1 |
| ENSG00000158195 | WASF2 | 0,700571049 | 1,625147933 | 0,00236294 | 0,038277257 | WASP family member 2 |
| ENSG00000240497 | AC092919,1 | 0,70035336 | 1,624902732 | 0,3510533 | 0,844035671 | novel transcript |
| ENSG00000146054 | TRIM7 | 0,700326946 | 1,624872983 | 0,33862876 | 0,837371637 | tripartite motif containing 7 |

| | | | | | | |
|---|---|---|---|---|---|---|
| ENSG00000157978 | LDLRAP1 | 0,700069534 | 1,624583091 | 0,04920513 | 0,338321763 | low density lipoprotein receptor adaptor protein 1 |
| ENSG00000266651 | AC093484,4 | 0,699953024 | 1,624451897 | 0,49722469 | 0,918759905 | novel transcript |
| ENSG00000228649 | SNHG26 | 0,699947734 | 1,624445942 | 0,08400736 | 0,461230231 | small nucleolar RNA host gene 26 |
| ENSG00000248771 | SMIM31 | 0,699900292 | 1,624392524 | 0,48055305 | 0,912847708 | small integral membrane protein 31 |
| ENSG00000233562 | RPS3AP8 | 0,699887128 | 1,624377702 | 0,57939845 | 0,943828882 | RPS3A pseudogene 8 |
| ENSG00000173867 | AC013489,1 | 0,699823615 | 1,624306192 | 0,39258294 | 0,870909282 | novel transcript |
| ENSG00000287151 | C2orf27A | 0,69978589 | 1,624263718 | 0,0183257 | 0,176287633 | chromosome 2 open reading frame 27A |
| ENSG00000242125 | SNHG3 | 0,699749004 | 1,624222191 | 0,0011418 | 0,020863304 | small nucleolar RNA host gene 3 |
| ENSG00000274630 | AC125257,2 | 0,699343896 | 1,623766174 | 0,58582804 | 0,945246954 | novel transcript, antisense to ACLY |
| ENSG00000178826 | TMEM139 | 0,699270793 | 1,623683898 | 0,1610539 | 0,637192669 | transmembrane protein 139 |
| ENSG00000106733 | NMRK1 | 0,699222451 | 1,623629492 | 0,07543533 | 0,433690715 | nicotinamide riboside kinase 1 |
| ENSG00000186471 | AKAP14 | 0,699198942 | 1,623603035 | 0,6184172 | 0,952793934 | A-kinase anchoring protein 14 |
| ENSG00000213025 | COX20P1 | 0,698854583 | 1,623215541 | 0,52652041 | 0,929261495 | COX20 pseudogene 1 |
| ENSG00000276375 | AL157712,1 | 0,698854174 | 1,623215081 | 0,50655298 | 0,921824694 | zinc finger protein pseudogene |
| ENSG00000258466 | AL049779,1 | 0,698761536 | 1,623110854 | 0,49529435 | 0,917777383 | novel protein |
| ENSG00000213638 | ADAT3 | 0,698731965 | 1,623077586 | 0,1961092 | 0,694033976 | adenosine deaminase tRNA specific 3 |
| ENSG00000270030 | AC136475,7 | 0,698278669 | 1,622567694 | 0,55703046 | 0,938292968 | novel transcript |
| ENSG00000284889 | PSME2 | 0,698010246 | 1,622265832 | 0,01643978 | 0,163842262 | proteasome activator subunit 2 |
| ENSG00000100911 | PSME2 | 0,698010246 | 1,622265832 | 0,01643978 | 0,163842262 | proteasome activator subunit 2 |
| ENSG00000227593 | PHBP16 | 0,697692954 | 1,621909087 | 0,60504741 | 0,950927937 | prohibitin pseudogene 16 |
| ENSG00000266278 | LINC01910 | 0,697646423 | 1,621856776 | 0,45155304 | 0,902062594 | long intergenic non-protein coding RNA 1910 |
| ENSG00000100938 | GMPR2 | 0,697335944 | 1,621507778 | 0,01292012 | 0,138286373 | guanosine monophosphate reductase 2 |
| ENSG00000128298 | BAIAP2L2 | 0,69680753 | 1,62091398 | 0,37173134 | 0,859817753 | BAR/IMD domain containing adaptor protein 2 like 2 |
| ENSG00000173166 | RAPH1 | 0,696774157 | 1,620876484 | 0,00473789 | 0,066158733 | Ras association (RalGDS/AF-6) and pleckstrin homology domains 1 |
| ENSG00000234177 | LINC01114 | 0,696484296 | 1,620550856 | 0,10005133 | 0,50208745 | long intergenic non-protein coding RNA 1114 |
| ENSG00000235027 | AC068580,3 | 0,69641181 | 1,620469436 | 0,53569382 | 0,932139691 | novel transcript |
| ENSG00000221866 | PLXNA4 | 0,696191672 | 1,620222191 | 0,07476904 | 0,432166624 | plexin A4 |
| ENSG00000273587 | SNORA78 | 0,696190752 | 1,620221158 | 0,68217691 | 0,964699524 | small nucleolar RNA, H/ACA box 78 |
| ENSG00000235204 | AL162724,2 | 0,696182157 | 1,620211505 | 0,53388811 | 0,931381467 | novel transcript |
| ENSG00000256925 | ADGRA1-AS1 | 0,69615837 | 1,620184791 | 0,13962738 | 0,595782615 | ADGRA1 antisense RNA 1 |
| ENSG00000226053 | LINC01776 | 0,696078892 | 1,620095538 | 0,05343675 | 0,355858082 | long intergenic non-protein coding RNA 1776 |

| | | | | | | |
|---|---|---|---|---|---|---|
| ENSG00000219102 | HNRNPA3P12 | 0,696019171 | 1,620028475 | 0,38559062 | 0,869008435 | heterogeneous nuclear ribonucleoprotein A3 pseudogene 12 |
| ENSG00000183010 | PYCR1 | 0,695995509 | 1,620001904 | 0,04006521 | 0,298878256 | pyrroline-5-carboxylate reductase 1 |
| ENSG00000273729 | AC007686,3 | 0,695943213 | 1,619943183 | 0,06839033 | 0,412055338 | novel transcript, antisense to IRF2BPL |
| ENSG00000197191 | CYSRT1 | 0,695928341 | 1,619926483 | 0,23468403 | 0,744134035 | cysteine rich tail 1 |
| ENSG00000214093 | AL096701,1 | 0,695868308 | 1,619859077 | 0,56676205 | 0,941311563 | ribosomal protein S18 (RPS18) pseudogene |
| ENSG00000196365 | LONP1 | 0,695667946 | 1,619634125 | 0,15913673 | 0,633144664 | lon peptidase 1, mitochondrial |
| ENSG00000225138 | SLC9A3-AS1 | 0,695661772 | 1,619627194 | 0,16240414 | 0,639629762 | SLC9A3 antisense RNA 1 |
| ENSG00000183888 | SRARP | 0,695424633 | 1,619360995 | 0,56963346 | 0,94221458 | steroid receptor associated and regulated protein |
| ENSG00000287287 | AL133476,1 | 0,695255024 | 1,619170627 | 0,3936814 | 0,871635019 | novel transcript, antisense to RUSC2 |
| ENSG00000260979 | AC022167,3 | 0,695246778 | 1,619161373 | 0,54135525 | 0,933429141 | novel transcript |
| ENSG00000268949 | MRPS17P1 | 0,69511682 | 1,619015525 | 0,43064456 | 0,895267028 | mitochondrial ribosomal protein S17 pseudogene 1 |
| ENSG00000134352 | IL6ST | 0,695063906 | 1,618956145 | 0,07900641 | 0,445376198 | interleukin 6 signal transducer |
| ENSG00000262246 | CORO7 | 0,695049641 | 1,618940138 | 0,19126413 | 0,686939473 | coronin 7 |
| ENSG00000223804 | AC244669,1 | 0,694713898 | 1,618563423 | 0,30353175 | 0,812685668 | Poly [ADP-ribose] polymerase 8 (PARP8) pseudogene |
| ENSG00000121769 | FABP3 | 0,694638209 | 1,618478509 | 0,11758126 | 0,54817483 | fatty acid binding protein 3 |
| ENSG00000002746 | HECW1 | 0,694614161 | 1,618451532 | 0,07137349 | 0,420784584 | HECT, C2 and WW domain containing E3 ubiquitin protein ligase 1 |
| ENSG00000253282 | AP001330,1 | 0,693984189 | 1,617744967 | 0,45594936 | 0,903545728 | novel transcript |
| ENSG00000272410 | AC022384,1 | 0,692725786 | 1,61633449 | 0,02279743 | 0,204880322 | novel protein |
| ENSG00000266903 | AC243964,3 | 0,692612284 | 1,616207333 | 0,24145099 | 0,751948055 | novel transcript, antisense to CEACAM16, CEACAM19, PVR & IGSF23 |
| ENSG00000241568 | RN7SL338P | 0,692331861 | 1,615893214 | 0,58300261 | 0,944174844 | RNA, 7SL, cytoplasmic 338, pseudogene |
| ENSG00000276194 | HNF1B | 0,692301178 | 1,615858848 | 0,45589796 | 0,903545728 | HNF1 homeobox B |
| ENSG00000131446 | MGAT1 | 0,692204861 | 1,615750974 | 0,04688794 | 0,329047041 | alpha-1,3-mannosyl-glycoprotein 2-beta-N-acetylglucosaminyltransferase |
| ENSG00000227144 | VWA7 | 0,691965171 | 1,615482554 | 0,60327567 | 0,950352286 | von Willebrand factor A domain containing 7 |
| ENSG00000287200 | AC022506,2 | 0,69183759 | 1,615339699 | 0,34602594 | 0,841168721 | novel transcript, antisense to INHBEand GLI1 |
| ENSG00000274589 | Metazoa_SRP | 0,691738288 | 1,615228518 | 0,66822679 | 0,961289023 | Metazoan signal recognition particle RNA |
| ENSG00000105519 | CAPS | 0,691720604 | 1,615208719 | 0,1089245 | 0,5265021 | calcyphosine |
| ENSG00000251992 | SCARNA17 | 0,691452968 | 1,614909108 | 0,74958264 | 0,975901808 | small Cajal body-specific RNA 17 |
| ENSG00000287582 | AC099796,2 | 0,691448223 | 1,614903797 | 0,62830887 | 0,952793934 | novel transcript |

| | | | | | | |
|---|---|---|---|---|---|---|
| ENSG00000123064 | DDX54 | 0,691224537 | 1,61465343 | 0,20484378 | 0,706459522 | DEAD-box helicase 54 |
| ENSG00000172752 | COL6A5 | 0,691144806 | 1,614564198 | 0,51385858 | 0,924227101 | collagen type VI alpha 5 chain |
| ENSG00000287724 | AC096536,3 | 0,691033759 | 1,614439927 | 0,40369597 | 0,877691113 | novel transcript, antisense to DHCR24 |
| ENSG00000227528 | DIAPH3-AS1 | 0,691012723 | 1,614416386 | 0,48745252 | 0,91491358 | DIAPH3 antisense RNA 1 |
| ENSG00000271780 | AL118558,3 | 0,690375609 | 1,613703595 | 0,28603046 | 0,797420923 | novel transcript |
| ENSG00000114737 | CISH | 0,690303929 | 1,61362342 | 0,01004191 | 0,115425382 | cytokine inducible SH2 containing protein |
| ENSG00000242735 | RPSAP26 | 0,690301594 | 1,613620809 | 0,42326183 | 0,890801612 | ribosomal protein SA pseudogene 26 |
| ENSG00000163347 | CLDN1 | 0,689986021 | 1,613267887 | 0,18393077 | 0,675485923 | claudin 1 |
| ENSG00000237753 | AC079922,2 | 0,689940047 | 1,613216478 | 0,06230629 | 0,389548927 | novel transcript (FLJ42351) |
| ENSG00000125817 | CENPB | 0,689707893 | 1,612956905 | 0,19350889 | 0,690424879 | centromere protein B |
| ENSG00000088298 | EDEM2 | 0,689682418 | 1,612928424 | 0,01318679 | 0,139942956 | ER degradation enhancing alpha-mannosidase like protein 2 |
| ENSG00000259589 | AC073167,1 | 0,689529094 | 1,612757017 | 0,48034109 | 0,912847708 | novel transcript, antisense to HERC1 |
| ENSG00000061273 | HDAC7 | 0,689467248 | 1,612687882 | 0,04158085 | 0,305946805 | histone deacetylase 7 |
| ENSG00000203506 | RBMS3-AS2 | 0,689253758 | 1,612449254 | 0,218961 | 0,727171948 | RBMS3 antisense RNA 2 |
| ENSG00000238184 | CD81-AS1 | 0,689099981 | 1,612277393 | 0,28806716 | 0,799375215 | CD81 antisense RNA 1 |
| ENSG00000241388 | HNF1A-AS1 | 0,688994689 | 1,612159728 | 0,37373757 | 0,861319894 | HNF1A antisense RNA 1 |
| ENSG00000173153 | ESRRA | 0,688935587 | 1,612093685 | 0,16745242 | 0,648316613 | estrogen related receptor alpha |
| ENSG00000273913 | Metazoa_SRP | 0,688439267 | 1,611539184 | 0,54281298 | 0,933663748 | Metazoan signal recognition particle RNA |
| ENSG00000267239 | AP001198,1 | 0,688431521 | 1,611530531 | 0,73326938 | 0,974353474 | novel transcript, antisense C18orf1 |
| ENSG00000100417 | PMM1 | 0,688326472 | 1,611413193 | 0,06893969 | 0,41408848 | phosphomannomutase 1 |
| ENSG00000072864 | NDE1 | 0,688202563 | 1,611274799 | 0,01055838 | 0,119335457 | nudE neurodevelopment protein 1 |
| ENSG00000276839 | Metazoa_SRP | 0,688027701 | 1,611079516 | 0,47947896 | 0,912714933 | Metazoan signal recognition particle RNA |
| ENSG00000181467 | RAP2B | 0,68802506 | 1,611076566 | 0,00063867 | 0,012799522 | RAP2B, member of RAS oncogene family |
| ENSG00000072609 | CHFR | 0,68795601 | 1,61099946 | 0,03034655 | 0,249117099 | checkpoint with forkhead and ring finger domains |
| ENSG00000140650 | PMM2 | 0,687665526 | 1,610675121 | 0,00235236 | 0,038125556 | phosphomannomutase 2 |
| ENSG00000226704 | HSPA1L | 0,687664737 | 1,610674239 | 0,22193935 | 0,729392589 | heat shock protein family A (Hsp70) member 1 like |
| ENSG00000274351 | AL137000,1 | 0,687659174 | 1,610668028 | 0,59506154 | 0,947636708 | protein-kinase, interferon-inducible double stranded RNA dependent inhibitor, repressor of (P58 repressor) PRKRIR |
| ENSG00000179796 | LRRC3B | 0,687621561 | 1,610626037 | 0,22680256 | 0,735383523 | leucine rich repeat containing 3B |
| ENSG00000272525 | AC099522,2 | 0,687162522 | 1,610113647 | 0,1733846 | 0,658330849 | novel transcript |
| ENSG00000230798 | FOXD3-AS1 | 0,687101225 | 1,610045238 | 0,52167731 | 0,927472429 | FOXD3 antisense RNA 1 |

| | | | | | | |
|---|---|---|---|---|---|---|
| ENSG00000129625 | REEP5 | 0,687042381 | 1,609979569 | 0,06859453 | 0,412873795 | receptor accessory protein 5 |
| ENSG00000197614 | MFAP5 | 0,686735188 | 1,609636793 | 0,31018134 | 0,817687618 | microfibril associated protein 5 |
| ENSG00000251535 | AC079140,4 | 0,686669845 | 1,609563891 | 0,65900344 | 0,95828296 | pseudogene similar to part of zinc finger protein 74 (ZNF74) |
| ENSG00000260144 | AC100827,2 | 0,686660234 | 1,609553169 | 0,73738542 | 0,974353474 | golgin A6 family, member A (GOLGA6A) pseudogene |
| ENSG00000226153 | CHP1P3 | 0,685993493 | 1,608809486 | 0,56381313 | 0,940865625 | CHP1 pseudogene 3 |
| ENSG00000287836 | AC016168,4 | 0,685768422 | 1,608558519 | 0,42279229 | 0,8904128 | novel transcript |
| ENSG00000212952 | BX088651,1 | 0,685668119 | 1,608446689 | 0,39381239 | 0,871635019 | novel pseudogene |
| ENSG00000226028 | PFN1P12 | 0,685429797 | 1,608181008 | 0,54113773 | 0,933385624 | profilin 1 pseudogene 12 |
| ENSG00000175115 | PACS1 | 0,685188422 | 1,607911968 | 0,03035876 | 0,249117099 | phosphofurin acidic cluster sorting protein 1 |
| ENSG00000171671 | SHANK2-AS3 | 0,684905071 | 1,607596199 | 0,45543266 | 0,903545728 | SHANK2 antisense RNA 3 |
| ENSG00000258458 | AL160314,2 | 0,684832494 | 1,607515328 | 0,34674028 | 0,8414568 | novel transcript, antisense to OXA1L |
| ENSG00000166349 | RAG1 | 0,684687773 | 1,607354082 | 0,10422301 | 0,514802081 | recombination activating 1 |
| ENSG00000156869 | FRRS1 | 0,684245209 | 1,606861082 | 0,00590644 | 0,077915148 | ferric chelate reductase 1 |
| ENSG00000288580 | AC023296,1 | 0,683990252 | 1,606577138 | 0,45080307 | 0,902062594 | novel transcript |
| ENSG00000255139 | AP000442,1 | 0,683754017 | 1,60631409 | 0,37542925 | 0,862022464 | novel transcript |
| ENSG00000202252 | SNORD14C | 0,683737455 | 1,606295649 | 0,37924579 | 0,86476118 | small nucleolar RNA, C/D box 14C |
| ENSG00000196187 | TMEM63A | 0,68355108 | 1,606088153 | 0,11746748 | 0,547889487 | transmembrane protein 63A |
| ENSG00000106789 | CORO2A | 0,683169439 | 1,605663345 | 0,03602575 | 0,278307135 | coronin 2A |
| ENSG00000087884 | AAMDC | 0,683099009 | 1,605584961 | 0,02518845 | 0,220092673 | adipogenesis associated Mth938 domain containing |
| ENSG00000228830 | AL160408,2 | 0,682764822 | 1,605213085 | 0,05730325 | 0,370150588 | novel transcript |
| ENSG00000259094 | AC013457,1 | 0,682709916 | 1,605151996 | 0,55058264 | 0,936249395 | novel transcript, antisense to MARCO |
| ENSG00000164749 | HNF4G | 0,682503133 | 1,604921944 | 0,24611667 | 0,758198269 | hepatocyte nuclear factor 4 gamma |
| ENSG00000159110 | IFNAR2 | 0,682069026 | 1,604439095 | 0,00610032 | 0,080000234 | interferon alpha and beta receptor subunit 2 |
| ENSG00000238266 | LINC00707 | 0,681929693 | 1,604284149 | 0,383618 | 0,868000076 | long intergenic non-protein coding RNA 707 |
| ENSG00000122705 | CLTA | 0,681927485 | 1,604281694 | 0,00709707 | 0,089504999 | clathrin light chain A |
| ENSG00000171132 | PRKCE | 0,681904897 | 1,604256576 | 0,01677785 | 0,165942007 | protein kinase C epsilon |
| ENSG00000104903 | LYL1 | 0,681890248 | 1,604240286 | 0,22900144 | 0,737173617 | LYL1 basic helix-loop-helix family member |
| ENSG00000129521 | EGLN3 | 0,681577608 | 1,603892676 | 0,15294138 | 0,621569237 | egl-9 family hypoxia inducible factor 3 |
| ENSG00000166562 | SEC11C | 0,68135831 | 1,603648894 | 0,13795075 | 0,592546669 | SEC11 homolog C, signal peptidase complex subunit |
| ENSG00000100934 | SEC23A | 0,681258822 | 1,603538311 | 0,04754947 | 0,33161214 | Sec23 homolog A, COPII coat complex component |
| ENSG00000248931 | MTCYBP40 | 0,681155996 | 1,603424025 | 0,51174446 | 0,923100549 | MT-CYB pseudogene 40 |
| ENSG00000187193 | MT1X | 0,681037125 | 1,603291916 | 0,05954157 | 0,379434925 | metallothionein 1X |

| | | | | | | |
|---|---|---|---|---|---|---|
| ENSG00000230666 | CEACAM22P | 0,680921432 | 1,603163349 | 0,38904651 | 0,870057277 | CEA cell adhesion molecule 22, pseudogene |
| ENSG00000271646 | IRF2-DT | 0,680907268 | 1,603147611 | 0,16395737 | 0,642745357 | IRF2 divergent transcript |
| ENSG00000251533 | LINC00605 | 0,680858449 | 1,603093363 | 0,70994914 | 0,970482408 | long intergenic non-protein coding RNA 605 |
| ENSG00000049245 | VAMP3 | 0,680707804 | 1,602925978 | 0,06066533 | 0,383412341 | vesicle associated membrane protein 3 |
| ENSG00000173442 | EHBP1L1 | 0,680599173 | 1,602805286 | 0,37851103 | 0,864031013 | EH domain binding protein 1 like 1 |
| ENSG00000091137 | SLC26A4 | 0,680508241 | 1,602704266 | 0,47515437 | 0,911082311 | solute carrier family 26 member 4 |
| ENSG00000253722 | AC010834,1 | 0,680461382 | 1,602652211 | 0,80537978 | 0,981213877 | novel transcript |
| ENSG00000261437 | AC108860,2 | 0,680194928 | 1,602356241 | 0,5305623 | 0,931265667 | novel transcript, antisense to ESRP1 |
| ENSG00000263847 | AP005899,1 | 0,680040801 | 1,602185066 | 0,43571085 | 0,896422069 | novel transcript, antisense to NDUFV2 |
| ENSG00000197496 | SLC2A10 | 0,67991684 | 1,602047407 | 0,01297958 | 0,138732823 | solute carrier family 2 member 10 |
| ENSG00000184838 | PRR16 | 0,679892683 | 1,602020582 | 0,13509982 | 0,586038109 | proline rich 16 |
| ENSG00000277270 | AL160412,1 | 0,679829102 | 1,601949981 | 0,04300116 | 0,31257571 | novel transcript |
| ENSG00000229962 | AP000221,1 | 0,679790046 | 1,601906615 | 0,49358211 | 0,917289924 | novel transcript |
| ENSG00000238141 | BRWD1-AS1 | 0,67958312 | 1,601676869 | 0,43546433 | 0,89634451 | BRWD1 antisense RNA 1 |
| ENSG00000275355 | Metazoa_SRP | 0,679497476 | 1,60158179 | 0,47612418 | 0,911413793 | Metazoan signal recognition particle RNA |
| ENSG00000105650 | PDE4C | 0,679399763 | 1,60147332 | 0,39106443 | 0,870057277 | phosphodiesterase 4C |
| ENSG00000279066 | HEXD-IT1 | 0,679395806 | 1,601468927 | 0,46247781 | 0,906571346 | HEXD intronic transcript 1 |
| ENSG00000228415 | PTMAP1 | 0,679214312 | 1,601267471 | 0,5376183 | 0,932883277 | prothymosin alpha pseudogene 1 |
| ENSG00000197694 | SPTAN1 | 0,679112301 | 1,601154252 | 0,0550069 | 0,362301551 | spectrin alpha, non-erythrocytic 1 |
| ENSG00000276054 | AC243654,3 | 0,679040763 | 1,601074859 | 0,58059422 | 0,943832501 | novel transcript, sense intronic to C17orf78 |
| ENSG00000090861 | AARS1 | 0,678991747 | 1,601020463 | 0,02560156 | 0,222582632 | alanyl-tRNA synthetase 1 |
| ENSG00000269404 | SPIB | 0,67860668 | 1,600593194 | 0,35019635 | 0,843789477 | Spi-B transcription factor |
| ENSG00000100100 | PIK3IP1 | 0,67854288 | 1,600522414 | 0,02514885 | 0,219931089 | phosphoinositide-3-kinase interacting protein 1 |
| ENSG00000115993 | TRAK2 | 0,678496569 | 1,600471037 | 0,01027345 | 0,117393679 | trafficking kinesin protein 2 |
| ENSG00000228589 | SPCS2P4 | 0,678214572 | 1,60015823 | 0,11910282 | 0,551730733 | signal peptidase complex subunit 2 pseudogene 4 |
| ENSG00000133488 | SEC14L4 | 0,678147505 | 1,600083846 | 0,32034434 | 0,824366234 | SEC14 like lipid binding 4 |
| ENSG00000157064 | NMNAT2 | 0,677427387 | 1,599285366 | 0,06132407 | 0,385606542 | nicotinamide nucleotide adenylyltransferase 2 |
| ENSG00000130733 | YIPF2 | 0,677108031 | 1,598931387 | 0,04957214 | 0,339816563 | Yip1 domain family member 2 |
| ENSG00000177675 | CD163L1 | 0,676311989 | 1,598049381 | 0,2932572 | 0,80339933 | CD163 molecule like 1 |
| ENSG00000273125 | LINC01990 | 0,676052277 | 1,597761728 | 0,48431095 | 0,913690171 | long intergenic non-protein coding RNA 1990 |
| ENSG00000227201 | CNN2P1 | 0,675920323 | 1,597615598 | 0,26036913 | 0,772692192 | calponin 2 pseudogene 1 |
| ENSG00000229950 | TFAP2A-AS1 | 0,675771522 | 1,597450826 | 0,48166012 | 0,912981134 | TFAP2A antisense RNA 1 |
| ENSG00000115091 | ACTR3 | 0,675566437 | 1,597223759 | 0,0590379 | 0,377379048 | actin related protein 3 |

| | | | | | | |
|---|---|---|---|---|---|---|
| ENSG00000223494 | VARS2 | 0,675557051 | 1,597213366 | 0,31656338 | 0,821646322 | valyl-tRNA synthetase 2, mitochondrial |
| ENSG00000210151 | MT-TS1 | 0,675471223 | 1,597118349 | 0,47111168 | 0,908869685 | mitochondrially encoded tRNA-Ser (UCN) 1 |
| ENSG00000166272 | WBP1L | 0,675419399 | 1,597060979 | 0,00376769 | 0,056091926 | WW domain binding protein 1 like |
| ENSG00000286334 | AL161430,1 | 0,675361249 | 1,596996609 | 0,54578979 | 0,935146964 | novel transcript, antisense to MBNL2 |
| ENSG00000103174 | NAGPA | 0,675224118 | 1,596844818 | 0,07403458 | 0,4296801 | N-acetylglucosamine-1-phosphodiester alpha-N-acetylglucosaminidase |
| ENSG00000219404 | AL359694,1 | 0,674891932 | 1,596477181 | 0,52711286 | 0,92927181 | neugrin, neurite outgrowth associated (NGRN) pseudogene |
| ENSG00000258130 | AC106782,1 | 0,674866809 | 1,59644938 | 0,36683136 | 0,856313634 | coiled-coil domain containing 101 (CCDC101) pseudogene |
| ENSG00000003400 | CASP10 | 0,674808951 | 1,596385357 | 0,15607421 | 0,626979676 | caspase 10 |
| ENSG00000277440 | AC012676,4 | 0,674799072 | 1,596374426 | 0,50125494 | 0,920520138 | novel transcript, sense intronic with DNAJA3 |
| ENSG00000282590 | AC012676,8 | 0,674799072 | 1,596374426 | 0,50125494 | 0,920520138 | novel transcript, sense intronic with DNAJA3 |
| ENSG00000101346 | POFUT1 | 0,674711463 | 1,596277487 | 0,00311154 | 0,047967995 | protein O-fucosyltransferase 1 |
| ENSG00000070388 | FGF22 | 0,674585425 | 1,596138038 | 0,42127201 | 0,889067654 | fibroblast growth factor 22 |
| ENSG00000253842 | AP003472,2 | 0,674333744 | 1,595859613 | 0,4261791 | 0,892133432 | novel transcript |
| ENSG00000274233 | CCL5 | 0,674285123 | 1,595805831 | 0,4052449 | 0,878185962 | C-C motif chemokine ligand 5 |
| ENSG00000130164 | LDLR | 0,674023226 | 1,595516166 | 0,00898549 | 0,105928449 | low density lipoprotein receptor |
| ENSG00000196338 | NLGN3 | 0,673846534 | 1,595320769 | 0,00696805 | 0,088341044 | neuroligin 3 |
| ENSG00000154529 | CNTNAP3B | 0,6734644 | 1,594898264 | 0,02485235 | 0,218487143 | contactin associated protein family member 3B |
| ENSG00000107521 | HPS1 | 0,673324144 | 1,594743219 | 0,04396488 | 0,316175438 | HPS1 biogenesis of lysosomal organelles complex 3 subunit 1 |
| ENSG00000087510 | TFAP2C | 0,67314382 | 1,594543903 | 0,31831599 | 0,823093059 | transcription factor AP-2 gamma |
| ENSG00000084234 | APLP2 | 0,673102115 | 1,594497809 | 0,00391907 | 0,05782249 | amyloid beta precursor like protein 2 |
| ENSG00000259513 | CYCSP38 | 0,672824544 | 1,594191061 | 0,61983714 | 0,952793934 | CYCS pseudogene 38 |
| ENSG00000255303 | OR5BA1P | 0,672639153 | 1,593986215 | 0,30489358 | 0,813743624 | olfactory receptor family 5 subfamily BA member 1 pseudogene |
| ENSG00000233673 | ANAPC1P1 | 0,672555612 | 1,593893916 | 0,5801113 | 0,943828882 | anaphase promoting complex subunit 1 pseudogene 1 |
| ENSG00000262700 | AC133552,3 | 0,672489405 | 1,593820772 | 0,606149 | 0,950927937 | thyroid hormone receptor interactor 11 (TRIP11) pseudogene |
| ENSG00000287274 | AL627232,1 | 0,67220774 | 1,593509632 | 0,60168208 | 0,949939978 | novel transcript, antisense to STARD13 |
| ENSG00000137313 | TRIM26 | 0,672107386 | 1,593398791 | 0,17041007 | 0,653594444 | tripartite motif containing 26 |

| | | | | | | |
|---|---|---|---|---|---|---|
| ENSG00000286313 | AL360178,1 | 0,672105916 | 1,593397168 | 0,58151216 | 0,944092866 | novel transcript, antisense to PDE7B |
| ENSG00000134597 | RBMX2 | 0,671933137 | 1,593206353 | 0,01280402 | 0,137391732 | RNA binding motif protein X-linked 2 |
| ENSG00000287120 | AL138773,1 | 0,67170758 | 1,592957283 | 0,39563671 | 0,872227623 | novel transcript |
| ENSG00000075618 | FSCN1 | 0,670937502 | 1,592107226 | 0,04234575 | 0,308960928 | fascin actin-bundling protein 1 |
| ENSG00000134153 | EMC7 | 0,670855497 | 1,592016731 | 0,01791008 | 0,173950874 | ER membrane protein complex subunit 7 |
| ENSG00000267327 | AC009271,1 | 0,670719786 | 1,59186698 | 0,09385491 | 0,487017413 | novel transcript |
| ENSG00000184162 | NR2C2AP | 0,670653168 | 1,591793475 | 0,02404416 | 0,212830338 | nuclear receptor 2C2 associated protein |
| ENSG00000206028 | Z99774,1 | 0,670428865 | 1,59154601 | 0,37545291 | 0,862022464 | novel transcript |
| ENSG00000034677 | RNF19A | 0,67031532 | 1,591420756 | 0,04068847 | 0,301441804 | ring finger protein 19A, RBR E3 ubiquitin protein ligase |
| ENSG00000100299 | ARSA | 0,670019901 | 1,591094916 | 0,0724278 | 0,424244694 | arylsulfatase A |
| ENSG00000155660 | PDIA4 | 0,669933779 | 1,590999937 | 0,00264585 | 0,041883474 | protein disulfide isomerase family A member 4 |
| ENSG00000076356 | PLXNA2 | 0,669624368 | 1,590658757 | 0,15897122 | 0,633144409 | plexin A2 |
| ENSG00000173805 | HAP1 | 0,669468443 | 1,590486849 | 0,34913694 | 0,843141089 | huntingtin associated protein 1 |
| ENSG00000203721 | LINC00862 | 0,669209749 | 1,590201679 | 0,51906384 | 0,926800715 | long intergenic non-protein coding RNA 862 |
| ENSG00000163659 | TIPARP | 0,66919713 | 1,590187771 | 0,03732981 | 0,285209938 | TCDD inducible poly(ADP-ribose) polymerase |
| ENSG00000164120 | HPGD | 0,669171608 | 1,590159639 | 0,37052766 | 0,858545694 | 15-hydroxyprostaglandin dehydrogenase |
| ENSG00000271437 | AL356423,1 | 0,66910561 | 1,590086897 | 0,50671725 | 0,921824694 | high mobility group box 1 (HMGB1) pseudogene |
| ENSG00000286427 | AC009486,2 | 0,669081888 | 1,590060752 | 0,20556254 | 0,708021064 | novel transcript |
| ENSG00000165192 | ASB11 | 0,668947364 | 1,589912494 | 0,46758948 | 0,907940653 | ankyrin repeat and SOCS box containing 11 |
| ENSG00000131408 | NR1H2 | 0,668888163 | 1,589847252 | 0,06021497 | 0,381629797 | nuclear receptor subfamily 1 group H member 2 |
| ENSG00000074842 | MYDGF | 0,668417686 | 1,589328873 | 0,03215802 | 0,259187498 | myeloid derived growth factor |
| ENSG00000140030 | GPR65 | 0,668295456 | 1,589194225 | 0,31346362 | 0,819801802 | G protein-coupled receptor 65 |
| ENSG00000278224 | PRICKLE4 | 0,667884109 | 1,588741172 | 0,41559133 | 0,885986046 | prickle planar cell polarity protein 4 |
| ENSG00000170788 | DYDC1 | 0,66787704 | 1,588733388 | 0,49282801 | 0,917090312 | DPY30 domain containing 1 |
| ENSG00000232533 | AC093673,1 | 0,667775731 | 1,588621828 | 0,02973679 | 0,245625065 | novel transcript |
| ENSG00000106392 | C1GALT1 | 0,667612384 | 1,588441969 | 0,18013129 | 0,669236428 | core 1 synthase, glycoprotein-N-acetylgalactosamine 3-beta-galactosyltransferase 1 |
| ENSG00000162909 | CAPN2 | 0,667266547 | 1,588061239 | 0,0085413 | 0,102155722 | calpain 2 |
| ENSG00000204531 | POU5F1 | 0,667198335 | 1,587986156 | 0,32067121 | 0,824366234 | POU class 5 homeobox 1 |
| ENSG00000112077 | RHAG | 0,667174858 | 1,587960314 | 0,48926981 | 0,915936198 | Rh associated glycoprotein |
| ENSG00000248318 | AC104958,1 | 0,667025269 | 1,587795672 | 0,47843495 | 0,912212078 | novel transcript |
| ENSG00000088756 | ARHGAP28 | 0,666584669 | 1,587310832 | 0,14454572 | 0,604963494 | Rho GTPase activating protein 28 |

| | | | | | | |
|---|---|---|---|---|---|---|
| ENSG00000083857 | FAT1 | 0,666502491 | 1,587220419 | 0,00699288 | 0,088512268 | FAT atypical cadherin 1 |
| ENSG00000235631 | RNF148 | 0,666463296 | 1,587177298 | 0,43944836 | 0,897557559 | ring finger protein 148 |
| ENSG00000180448 | ARHGAP45 | 0,666392655 | 1,587099585 | 0,34936158 | 0,843141089 | Rho GTPase activating protein 45 |
| ENSG00000276408 | AC025287,2 | 0,665780562 | 1,586426368 | 0,44273548 | 0,899437949 | novel transcript |
| ENSG00000228330 | AL355355,1 | 0,665384248 | 1,58599063 | 0,46960247 | 0,908293218 | pseudogene similar to part of COX10 homolog, cytochrome c oxidase assembly protein, heme A: farnesyltransferase (yeast) (COX10) |
| ENSG00000237441 | RGL2 | 0,665209486 | 1,585798522 | 0,04479076 | 0,319566021 | ral guanine nucleotide dissociation stimulator like 2 |
| ENSG00000170382 | LRRN2 | 0,665184152 | 1,585770675 | 0,08624744 | 0,46788826 | leucine rich repeat neuronal 2 |
| ENSG00000183741 | CBX6 | 0,665141676 | 1,585723988 | 0,06381817 | 0,395590561 | chromobox 6 |
| ENSG00000204950 | LRRC10B | 0,664916208 | 1,585476186 | 0,027406 | 0,232982843 | leucine rich repeat containing 10B |
| ENSG00000029993 | HMGB3 | 0,664488973 | 1,585006737 | 0,01672175 | 0,165491831 | high mobility group box 3 |
| ENSG00000282933 | RHOXF1P3 | 0,66438808 | 1,584895896 | 0,22061078 | 0,728572782 | Rhox homeobox family member 1 pseudogene 3 |
| ENSG00000167578 | RAB4B | 0,664214803 | 1,584705551 | 0,18403234 | 0,675673608 | RAB4B, member RAS oncogene family |
| ENSG00000197768 | STPG3 | 0,664052914 | 1,584527737 | 0,68320943 | 0,96498124 | sperm-tail PG-rich repeat containing 3 |
| ENSG00000288369 | RBP5 | 0,663985414 | 1,584453602 | 0,43450599 | 0,896272827 | retinol binding protein 5 |
| ENSG00000139194 | RBP5 | 0,663985414 | 1,584453602 | 0,43450599 | 0,896272827 | retinol binding protein 5 |
| ENSG00000240661 | AC063952,1 | 0,663836211 | 1,584289747 | 0,0636841 | 0,395357349 | novel V-set domain-containing pseudogene |
| ENSG00000105404 | RABAC1 | 0,663699485 | 1,584139609 | 0,07325681 | 0,427132959 | Rab acceptor 1 |
| ENSG00000198918 | RPL39 | 0,663500245 | 1,58392085 | 0,0994076 | 0,50040065 | ribosomal protein L39 |
| ENSG00000109944 | JHY | 0,663389603 | 1,583799382 | 0,07022672 | 0,417549035 | junctional cadherin complex regulator |
| ENSG00000227474 | RPL6P24 | 0,66318655 | 1,583576486 | 0,42111145 | 0,888848798 | ribosomal protein L6 pseudogene 24 |
| ENSG00000099282 | TSPAN15 | 0,663172244 | 1,583560783 | 0,11519501 | 0,542308656 | tetraspanin 15 |
| ENSG00000003096 | KLHL13 | 0,662987446 | 1,583357954 | 0,1139659 | 0,53948382 | kelch like family member 13 |
| ENSG00000130997 | POLN | 0,66282471 | 1,583179361 | 0,10760976 | 0,52347056 | DNA polymerase nu |
| ENSG00000008405 | CRY1 | 0,662663659 | 1,583002637 | 0,02199571 | 0,199557087 | cryptochrome circadian regulator 1 |
| ENSG00000270490 | AC110048,1 | 0,662460627 | 1,582779876 | 0,48044247 | 0,912847708 | leucine zipper-EF-hand containing transmembrane protein 1 (LETM1) pseudogene |
| ENSG00000224184 | MIR3681HG | 0,662376513 | 1,582687596 | 0,21370487 | 0,721156449 | MIR3681 host gene |
| ENSG00000110013 | SIAE | 0,662329906 | 1,582636468 | 0,03247102 | 0,260903141 | sialic acid acetylesterase |
| ENSG00000223923 | TNS1-AS1 | 0,662163216 | 1,582453619 | 0,4176091 | 0,886857227 | TNS1 antisense RNA 1 |
| ENSG00000167941 | SOST | 0,662147658 | 1,582436554 | 0,51889166 | 0,926800715 | sclerostin |
| ENSG00000165915 | SLC39A13 | 0,662080594 | 1,582362996 | 0,07489317 | 0,432482457 | solute carrier family 39 member 13 |

| | | | | | | |
|---|---|---|---|---|---|---|
| ENSG00000130508 | PXDN | 0,662034726 | 1,582312689 | 0,01152299 | 0,127029148 | peroxidasin |
| ENSG00000239774 | AC125618,1 | 0,661946273 | 1,582215678 | 0,65229668 | 0,957257009 | novel transcript |
| ENSG00000261789 | AC023830,2 | 0,661937187 | 1,582205714 | 0,50208233 | 0,920520138 | novel transcript, antisense to C16orf96 |
| ENSG00000163116 | STPG2 | 0,661932639 | 1,582200726 | 0,38033558 | 0,865730737 | sperm tail PG-rich repeat containing 2 |
| ENSG00000259993 | AC135731,1 | 0,661787511 | 1,582041573 | 0,31248662 | 0,818574551 | hyperpolarization activated cyclic nucleotide-gated potassium channel 2 (HCN2) pseudogene |
| ENSG00000067225 | PKM | 0,661774787 | 1,58202762 | 0,03117257 | 0,253879337 | pyruvate kinase M1/2 |
| ENSG00000090020 | SLC9A1 | 0,661555664 | 1,581787353 | 0,02953103 | 0,244377517 | solute carrier family 9 member A1 |
| ENSG00000244161 | FLNB-AS1 | 0,661513252 | 1,581740852 | 0,09848848 | 0,498280077 | FLNB antisense RNA 1 |
| ENSG00000206284 | WDR46 | 0,661224963 | 1,581424809 | 0,24027942 | 0,750234283 | WD repeat domain 46 |
| ENSG00000282283 | PLPPR3 | 0,661141582 | 1,581333413 | 0,57907933 | 0,943828882 | phospholipid phosphatase related 3 |
| ENSG00000285694 | AL162718,2 | 0,660907769 | 1,581077152 | 0,59737532 | 0,948282297 | novel transcript |
| ENSG00000285781 | AL117340,1 | 0,660870358 | 1,581036153 | 0,32208163 | 0,824955371 | novel pseudogene similar to nuclear pore associated protein 1 (NPAP1) |
| ENSG00000205903 | ZNF316 | 0,660633088 | 1,580776153 | 0,37333565 | 0,861257555 | zinc finger protein 316 |
| ENSG00000181072 | CHRM2 | 0,660383739 | 1,580502962 | 0,24539632 | 0,757479368 | cholinergic receptor muscarinic 2 |
| ENSG00000254017 | IGHEP2 | 0,660378812 | 1,580497565 | 0,45728607 | 0,903831777 | immunoglobulin heavy constant epsilon P2 (pseudogene) |
| ENSG00000171206 | TRIM8 | 0,660196244 | 1,580297571 | 0,05107 | 0,345820358 | tripartite motif containing 8 |
| ENSG00000198153 | ZNF849P | 0,660057325 | 1,580145409 | 0,52532296 | 0,928752911 | zinc finger protein 849, pseudogene |
| ENSG00000233557 | NFU1P2 | 0,660045104 | 1,580132024 | 0,39036098 | 0,870057277 | NFU1 iron-sulfur cluster scaffold pseudogene 2 |
| ENSG00000188917 | TRMT2B | 0,659469772 | 1,579502009 | 0,01006721 | 0,115628657 | tRNA methyltransferase 2 homolog B |
| ENSG00000235569 | MSH5 | 0,6592796 | 1,579293817 | 0,16883225 | 0,650760243 | mutS homolog 5 |
| ENSG00000231298 | MANCR | 0,658915059 | 1,578894811 | 0,39037377 | 0,870057277 | mitotically associated long non coding RNA |
| ENSG00000164088 | PPM1M | 0,658792972 | 1,578761203 | 0,05932973 | 0,378470661 | protein phosphatase, Mg2+/Mn2+ dependent 1M |
| ENSG00000166927 | MS4A7 | 0,658751034 | 1,578715311 | 0,47219217 | 0,909171312 | membrane spanning 4-domains A7 |
| ENSG00000230417 | LINC00595 | 0,658514375 | 1,57845636 | 0,46850365 | 0,907940653 | long intergenic non-protein coding RNA 595 |
| ENSG00000115325 | DOK1 | 0,658422739 | 1,578356104 | 0,14909018 | 0,614037073 | docking protein 1 |
| ENSG00000180209 | MYLPF | 0,658216265 | 1,578130231 | 0,55115952 | 0,936366211 | myosin light chain, phosphorylatable, fast skeletal muscle |
| ENSG00000259250 | AC018904,1 | 0,658196429 | 1,578108533 | 0,42979636 | 0,894607 | novel transcript, antisense to ADAM10 |
| ENSG00000248235 | AC037459,1 | 0,658155029 | 1,578063248 | 0,21580745 | 0,722999018 | novel protein |
| ENSG00000285005 | PLEKHM3 | 0,658137713 | 1,578044308 | 0,68686519 | 0,965769168 | pleckstrin homology domain containing M3 |

| | | | | | | |
|---|---|---|---|---|---|---|
| ENSG00000286130 | AC006040,1 | 0,657978516 | 1,577870185 | 0,36924926 | 0,857942508 | novel transcript |
| ENSG00000286264 | AP001453,3 | 0,657552298 | 1,577404101 | 0,2495944 | 0,761448016 | Uncharacterized protein LOC114841035 |
| ENSG00000108786 | HSD17B1 | 0,657299865 | 1,577128121 | 0,24971818 | 0,761653149 | hydroxysteroid 17-beta dehydrogenase 1 |
| ENSG00000236829 | Z97634,1 | 0,657244347 | 1,57706743 | 0,50781978 | 0,922300318 | ribosomal protein L23a (RPL23A) pseudogene |
| ENSG00000136240 | KDELR2 | 0,657032912 | 1,57683632 | 0,02975068 | 0,245674909 | KDEL endoplasmic reticulum protein retention receptor 2 |
| ENSG00000277863 | AL162574,1 | 0,656847783 | 1,57663399 | 0,69476413 | 0,967417311 | novel transcript |
| ENSG00000112335 | SNX3 | 0,656747648 | 1,576524563 | 0,06159977 | 0,38671794 | sorting nexin 3 |
| ENSG00000135077 | HAVCR2 | 0,656508441 | 1,576263188 | 0,36894456 | 0,857916156 | hepatitis A virus cellular receptor 2 |
| ENSG00000142798 | HSPG2 | 0,65648233 | 1,57623466 | 0,14250541 | 0,600806836 | heparan sulfate proteoglycan 2 |
| ENSG00000202260 | RN7SKP69 | 0,656341293 | 1,576080576 | 0,64844268 | 0,956899963 | RN7SK pseudogene 69 |
| ENSG00000234883 | MIR155HG | 0,656318697 | 1,576055891 | 0,33824944 | 0,837263082 | MIR155 host gene |
| ENSG00000260385 | AL356961,1 | 0,656306988 | 1,5760431 | 0,36969011 | 0,858104601 | novel transcript |
| ENSG00000287652 | AC000082,1 | 0,65623536 | 1,575964852 | 0,73202668 | 0,974353474 | novel transcript, antisense to DGCR8and CDC45 |
| ENSG00000277363 | SRCIN1 | 0,656072956 | 1,575787456 | 0,16975603 | 0,652551597 | SRC kinase signaling inhibitor 1 |
| ENSG00000158428 | CATIP | 0,655939708 | 1,575641922 | 0,42567436 | 0,892133432 | ciliogenesis associated TTC17 interacting protein |
| ENSG00000196235 | SUPT5H | 0,655359177 | 1,575008023 | 0,19012235 | 0,685541073 | SPT5 homolog, DSIF elongation factor subunit |
| ENSG00000278191 | CARS1 | 0,65494706 | 1,574558173 | 0,01167352 | 0,128326667 | cysteinyl-tRNA synthetase 1 |
| ENSG00000249150 | AC010267,1 | 0,654861294 | 1,574464571 | 0,39868643 | 0,875433675 | novel transcript |
| ENSG00000160695 | VPS11 | 0,654859899 | 1,574463049 | 0,01299023 | 0,138779332 | VPS11 core subunit of CORVET and HOPS complexes |
| ENSG00000182580 | EPHB3 | 0,65376263 | 1,573266016 | 0,0686054 | 0,412873795 | EPH receptor B3 |
| ENSG00000260916 | CCPG1 | 0,653645103 | 1,573137858 | 0,03600758 | 0,278307135 | cell cycle progression 1 |
| ENSG00000229491 | AC136489,1 | 0,653316926 | 1,572780049 | 0,62137025 | 0,952793934 | novel transcript |
| ENSG00000170604 | IRF2BP1 | 0,65331363 | 1,572776455 | 0,21386505 | 0,721188427 | interferon regulatory factor 2 binding protein 1 |
| ENSG00000236353 | PBX2 | 0,653059715 | 1,572499671 | 0,37221205 | 0,860223901 | PBX homeobox 2 |
| ENSG00000239627 | RPL12P20 | 0,653009118 | 1,572444522 | 0,585715 | 0,945246954 | ribosomal protein L12 pseudogene 20 |
| ENSG00000213303 | AC008481,1 | 0,652671171 | 1,572076225 | 0,54232885 | 0,933491609 | ribosomal protein S12 (RPS12) pseudogene |
| ENSG00000278154 | LAIR1 | 0,652250811 | 1,571618233 | 0,42642363 | 0,892138574 | leukocyte associated immunoglobulin like receptor 1 |
| ENSG00000118680 | MYL12B | 0,652230175 | 1,571595753 | 0,03767356 | 0,287047407 | myosin light chain 12B |
| ENSG00000284744 | AL591163,1 | 0,652163246 | 1,571522847 | 0,51016624 | 0,922427526 | novel transcript |
| ENSG00000181826 | RELL1 | 0,652052214 | 1,571401905 | 0,07402075 | 0,4296801 | RELT like 1 |
| ENSG00000023839 | ABCC2 | 0,651783261 | 1,571108984 | 0,10145939 | 0,506329296 | ATP binding cassette subfamily C member 2 |

| | | | | | | |
|---|---|---|---|---|---|---|
| ENSG00000103042 | SLC38A7 | 0,651752309 | 1,571075278 | 0,01828114 | 0,176021407 | solute carrier family 38 member 7 |
| ENSG00000111058 | ACSS3 | 0,651723843 | 1,571044279 | 0,05839173 | 0,374166639 | acyl-CoA synthetase short chain family member 3 |
| ENSG00000277047 | AL390964,1 | 0,651713594 | 1,571033119 | 0,35791901 | 0,849032846 | novel transcript |
| ENSG00000235245 | AL360181,2 | 0,65165237 | 1,57096645 | 0,38538391 | 0,868895792 | novel transcript |
| ENSG00000225989 | ABCF1 | 0,651377639 | 1,570667321 | 0,14454192 | 0,604963494 | ATP binding cassette subfamily F member 1 |
| ENSG00000287265 | AL365272,1 | 0,651358534 | 1,570646521 | 0,34794981 | 0,842451708 | novel transcript, antisense to DUSP22 |
| ENSG00000227344 | HAUS6P1 | 0,650993855 | 1,570249549 | 0,59013291 | 0,947262972 | HAUS augmin like complex subunit 6 pseudogene 1 |
| ENSG00000095139 | ARCN1 | 0,650948197 | 1,570199854 | 0,02701937 | 0,23048668 | archain 1 |
| ENSG00000254066 | LINC01938 | 0,65066602 | 1,569892769 | 0,56033201 | 0,940070839 | long intergenic non-protein coding RNA 1938 |
| ENSG00000255301 | AP002893,1 | 0,650318771 | 1,56951495 | 0,53844032 | 0,933082867 | novel transcript |
| ENSG00000214140 | PRCD | 0,650317693 | 1,569513777 | 0,36345298 | 0,85380707 | photoreceptor disc component |
| ENSG00000235026 | DPP10-AS1 | 0,65025457 | 1,569445107 | 0,37002607 | 0,85830893 | DPP10 antisense RNA 1 |
| ENSG00000256091 | MTRF1LP1 | 0,650186239 | 1,569370774 | 0,69510672 | 0,967519493 | mitochondrial translational release factor 1 like pseudogene 1 |
| ENSG00000263620 | AC129492,3 | 0,649742309 | 1,56888794 | 0,52070569 | 0,927046149 | novel protein |
| ENSG00000163749 | CCDC158 | 0,649686249 | 1,568826977 | 0,41565839 | 0,885986046 | coiled-coil domain containing 158 |
| ENSG00000138821 | SLC39A8 | 0,649499079 | 1,568623456 | 0,11725109 | 0,547082438 | solute carrier family 39 member 8 |
| ENSG00000188779 | SKOR1 | 0,649145698 | 1,568239277 | 0,41143141 | 0,883655718 | SKI family transcriptional corepressor 1 |
| ENSG00000153820 | SPHKAP | 0,648875669 | 1,567945778 | 0,71499257 | 0,971041579 | SPHK1 interactor, AKAP domain containing |
| ENSG00000184451 | CCR10 | 0,648434996 | 1,567466919 | 0,27255054 | 0,785492261 | C-C motif chemokine receptor 10 |
| ENSG00000260390 | AL360014,1 | 0,648178128 | 1,567187861 | 0,46400055 | 0,907337579 | novel transcript |
| ENSG00000268931 | AC136469,1 | 0,647994078 | 1,566987942 | 0,46506106 | 0,907804479 | novel transcript, antisense to MARCH2 |
| ENSG00000182700 | IGIP | 0,647887395 | 1,566872071 | 0,17772217 | 0,664232629 | IgA inducing protein |
| ENSG00000159388 | BTG2 | 0,647872561 | 1,566855961 | 0,0491381 | 0,33801251 | BTG anti-proliferation factor 2 |
| ENSG00000234667 | ACTBP13 | 0,647801723 | 1,566779029 | 0,46887682 | 0,907940653 | ACTB pseudogene 13 |
| ENSG00000196544 | BORCS6 | 0,647676683 | 1,56664324 | 0,11618657 | 0,544242846 | BLOC-1 related complex subunit 6 |
| ENSG00000126247 | CAPNS1 | 0,647491868 | 1,56644256 | 0,00459702 | 0,06494169 | calpain small subunit 1 |
| ENSG00000144847 | IGSF11 | 0,647465078 | 1,566413472 | 0,17345904 | 0,658349911 | immunoglobulin superfamily member 11 |
| ENSG00000224596 | ZMIZ1-AS1 | 0,647459762 | 1,5664077 | 0,37635568 | 0,862638114 | ZMIZ1 antisense RNA 1 |
| ENSG00000233614 | DDX11L10 | 0,647348359 | 1,566286749 | 0,34161488 | 0,840048641 | DEAD/H-box helicase 11 like 10 (pseudogene) |
| ENSG00000240521 | AC092979,1 | 0,647299857 | 1,566234092 | 0,47300659 | 0,909532026 | novel transcript |
| ENSG00000144583 | MARCHF4 | 0,646991218 | 1,565899061 | 0,19153528 | 0,687341458 | membrane associated ring-CH-type finger 4 |
| ENSG00000163820 | FYCO1 | 0,646695287 | 1,565577891 | 0,12062024 | 0,554057131 | FYVE and coiled-coil domain autophagy adaptor 1 |

| | | | | | | |
|---|---|---|---|---|---|---|
| ENSG00000187840 | EIF4EBP1 | 0,646582044 | 1,565455007 | 0,06002059 | 0,381011893 | eukaryotic translation initiation factor 4E binding protein 1 |
| ENSG00000253932 | AC019270,1 | 0,646192836 | 1,565032738 | 0,41618446 | 0,885986046 | novel transcript |
| ENSG00000233912 | AC026202,2 | 0,645851916 | 1,564662952 | 0,61195956 | 0,952187506 | novel transcript, antisense to ARL8B |
| ENSG00000219747 | AL133260,1 | 0,645559348 | 1,564345682 | 0,6096616 | 0,951793902 | 60S ribosomal protein L32 (RPL32) pseudogene |
| ENSG00000159720 | ATP6V0D1 | 0,645358563 | 1,564127982 | 0,0033411 | 0,050879833 | ATPase H+ transporting V0 subunit d1 |
| ENSG00000141639 | MAPK4 | 0,645336021 | 1,564103542 | 0,06042645 | 0,382191519 | mitogen-activated protein kinase 4 |
| ENSG00000124429 | POF1B | 0,645325609 | 1,564092254 | 0,40918122 | 0,882068652 | POF1B actin binding protein |
| ENSG00000000003 | TSPAN6 | 0,645296915 | 1,564061146 | 0,22576616 | 0,734785882 | tetraspanin 6 |
| ENSG00000229582 | AL358074,1 | 0,644969918 | 1,56370668 | 0,24762813 | 0,759874608 | novel transcript |
| ENSG00000267750 | RUNDC3A-AS1 | 0,644962921 | 1,563699097 | 0,40181284 | 0,877131444 | RUNDC3A antisense RNA 1 |
| ENSG00000236320 | SLFN14 | 0,644855945 | 1,563583153 | 0,34174013 | 0,840048641 | schlafen family member 14 |
| ENSG00000005108 | THSD7A | 0,644855 | 1,563582129 | 0,10756618 | 0,52347056 | thrombospondin type 1 domain containing 7A |
| ENSG00000286572 | AL592146,2 | 0,64434858 | 1,56303337 | 0,51665945 | 0,925489094 | novel transcript, antisense to SOX13 |
| ENSG00000169991 | IFFO2 | 0,644278216 | 1,562957139 | 0,13259709 | 0,581174167 | intermediate filament family orphan 2 |
| ENSG00000167114 | SLC27A4 | 0,644242293 | 1,562918222 | 0,07980213 | 0,447817422 | solute carrier family 27 member 4 |
| ENSG00000267272 | LINC01140 | 0,644084317 | 1,562747091 | 0,22097591 | 0,728677442 | long intergenic non-protein coding RNA 1140 |
| ENSG00000105875 | WDR91 | 0,643850235 | 1,562493551 | 0,01736195 | 0,170318153 | WD repeat domain 91 |
| ENSG00000144746 | ARL6IP5 | 0,643839076 | 1,562481466 | 0,08880207 | 0,475558385 | ADP ribosylation factor like GTPase 6 interacting protein 5 |
| ENSG00000119673 | ACOT2 | 0,64379555 | 1,562434326 | 0,02860838 | 0,240203215 | acyl-CoA thioesterase 2 |
| ENSG00000082512 | TRAF5 | 0,643760248 | 1,562396094 | 0,01225974 | 0,13267178 | TNF receptor associated factor 5 |
| ENSG00000226784 | PGAM4 | 0,643491317 | 1,562104877 | 0,35336406 | 0,84514549 | phosphoglycerate mutase family member 4 |
| ENSG00000226237 | GAS1RR | 0,643190898 | 1,561779627 | 0,18862304 | 0,682944308 | GAS1 adjacent regulatory RNA |
| ENSG00000260949 | AP006545,1 | 0,643078776 | 1,561658254 | 0,4276414 | 0,892656446 | novel transcript, overlapping to EIF4EBP1 |
| ENSG00000165029 | ABCA1 | 0,642864659 | 1,561426499 | 0,02793825 | 0,236266271 | ATP binding cassette subfamily A member 1 |
| ENSG00000100399 | CHADL | 0,642835903 | 1,561395377 | 0,33127632 | 0,832357374 | chondroadherin like |
| ENSG00000237898 | HCG15 | 0,642548609 | 1,561084476 | 0,81275996 | 0,981246822 | HLA complex group 15 |
| ENSG00000157191 | NECAP2 | 0,642419257 | 1,560944515 | 0,00845655 | 0,101559908 | NECAP endocytosis associated 2 |
| ENSG00000272382 | AC025171,5 | 0,642242555 | 1,560753342 | 0,4768343 | 0,91143003 | novel transcript |
| ENSG00000277586 | NEFL | 0,64198013 | 1,560469468 | 0,12645531 | 0,567868238 | neurofilament light |
| ENSG00000104870 | FCGRT | 0,641750956 | 1,560221605 | 0,14387282 | 0,603329615 | Fc fragment of IgG receptor and transporter |
| ENSG00000276742 | AL731566,3 | 0,641636876 | 1,560098237 | 0,54353017 | 0,934259495 | novel transcript, antisense to NSMCE4A |

| | | | | | | |
|---|---|---|---|---|---|---|
| ENSG00000251664 | PCDHA12 | 0,641614375 | 1,560073905 | 0,02079347 | 0,191935899 | protocadherin alpha 12 |
| ENSG00000172037 | LAMB2 | 0,641573686 | 1,560029906 | 0,08052502 | 0,450041893 | laminin subunit beta 2 |
| ENSG00000224114 | AL591846,1 | 0,64152502 | 1,559977283 | 0,40684556 | 0,880078305 | ribosomal protein S14 (RPS14) pseudogene |
| ENSG00000081087 | OSTM1 | 0,640777816 | 1,559169544 | 0,07887987 | 0,444823281 | osteoclastogenesis associated transmembrane protein 1 |
| ENSG00000271075 | AL139317,4 | 0,640699357 | 1,559084754 | 0,32769934 | 0,829743891 | golgi membrane protein 1 (GOLM1) pseudogene |
| ENSG00000234724 | HDAC1P1 | 0,640617676 | 1,558996485 | 0,48603727 | 0,914018932 | histone deacetylase 1 pseudogene 1 |
| ENSG00000272247 | AC080013,4 | 0,640558885 | 1,558932956 | 0,60529447 | 0,950927937 | novel transcript, antisense to MFSD1 |
| ENSG00000267179 | AC008770,2 | 0,640430007 | 1,558793702 | 0,4515492 | 0,902062594 | novel protein |
| ENSG00000105383 | CD33 | 0,640159469 | 1,558501419 | 0,62013684 | 0,952793934 | CD33 molecule |
| ENSG00000255008 | LINC02739 | 0,640085656 | 1,558421683 | 0,40822323 | 0,881221125 | long intergenic non-protein coding RNA 2739 |
| ENSG00000178852 | EFCAB13 | 0,640045929 | 1,558378771 | 0,15639309 | 0,627418665 | EF-hand calcium binding domain 13 |
| ENSG00000214309 | MBLAC1 | 0,639819953 | 1,558134694 | 0,21247824 | 0,719319956 | metallo-beta-lactamase domain containing 1 |
| ENSG00000086991 | NOX4 | 0,639360737 | 1,557638811 | 0,06001392 | 0,381011893 | NADPH oxidase 4 |
| ENSG00000123159 | GIPC1 | 0,639341174 | 1,55761769 | 0,05713294 | 0,369432136 | GIPC PDZ domain containing family member 1 |
| ENSG00000106355 | LSM5 | 0,639331791 | 1,55760756 | 0,1810207 | 0,670752303 | LSM5 homolog, U6 small nuclear RNA and mRNA degradation associated |
| ENSG00000152661 | GJA1 | 0,63917292 | 1,557436044 | 0,04008319 | 0,298941023 | gap junction protein alpha 1 |
| ENSG00000133169 | BEX1 | 0,639143869 | 1,557404683 | 0,01510208 | 0,154803784 | brain expressed X-linked 1 |
| ENSG00000214513 | NOTO | 0,639030645 | 1,557282461 | 0,51152965 | 0,922893865 | notochord homeobox |
| ENSG00000255192 | NANOGP8 | 0,638874792 | 1,557114238 | 0,26453918 | 0,775800337 | Nanog homeobox retrogene P8 |
| ENSG00000273509 | CNTNAP3P1 | 0,638824997 | 1,557060495 | 0,63326128 | 0,954594048 | CNTNAP3 pseudogene 1 |
| ENSG00000236830 | CBR3-AS1 | 0,638731053 | 1,556959106 | 0,39113354 | 0,870057277 | CBR3 antisense RNA 1 |
| ENSG00000129221 | AIPL1 | 0,638662147 | 1,556884746 | 0,46691033 | 0,907940653 | aryl hydrocarbon receptor interacting protein like 1 |
| ENSG00000213265 | TSGA13 | 0,638467591 | 1,556674804 | 0,40524529 | 0,878185962 | testis specific 13 |
| ENSG00000276914 | CEP20 | 0,638443833 | 1,55664917 | 0,25260768 | 0,764671439 | centrosomal protein 20 |
| ENSG00000100605 | ITPK1 | 0,637694521 | 1,555840881 | 0,05345156 | 0,355858082 | inositol-tetrakisphosphate 1-kinase |
| ENSG00000140320 | BAHD1 | 0,637694509 | 1,555840869 | 0,08173623 | 0,454697768 | bromo adjacent homology domain containing 1 |
| ENSG00000214063 | TSPAN4 | 0,637473151 | 1,555602169 | 0,0783006 | 0,442692864 | tetraspanin 4 |
| ENSG00000214013 | GANC | 0,637427292 | 1,555552721 | 0,03166037 | 0,256499062 | glucosidase alpha, neutral C |
| ENSG00000264235 | AP005329,1 | 0,636921278 | 1,555007219 | 0,28319332 | 0,794486366 | novel transcript, antisense to MYL12A |
| ENSG00000138193 | PLCE1 | 0,636917221 | 1,555002846 | 0,01555384 | 0,157881532 | phospholipase C epsilon 1 |
| ENSG00000173486 | FKBP2 | 0,636835681 | 1,554914961 | 0,04724123 | 0,330487755 | FKBP prolyl isomerase 2 |

| | | | | | | |
|---|---|---|---|---|---|---|
| ENSG00000120324 | PCDHB10 | 0,636803103 | 1,55487985 | 0,0279557 | 0,236345822 | protocadherin beta 10 |
| ENSG00000174370 | C11orf45 | 0,636788898 | 1,55486454 | 0,36006888 | 0,850433889 | chromosome 11 open reading frame 45 |
| ENSG00000171302 | CANT1 | 0,636724689 | 1,55479534 | 0,03339504 | 0,265327213 | calcium activated nucleotidase 1 |
| ENSG00000131100 | ATP6V1E1 | 0,636647064 | 1,554711686 | 0,02851498 | 0,239578272 | ATPase H+ transporting V1 subunit E1 |
| ENSG00000216867 | RPL22P12 | 0,636349677 | 1,554391241 | 0,4644053 | 0,907500217 | ribosomal protein L22 pseudogene 12 |
| ENSG00000245522 | LINC02709 | 0,636236699 | 1,554269522 | 0,30799881 | 0,816424917 | long intergenic non-protein coding RNA 2709 |
| ENSG00000151474 | FRMD4A | 0,636061432 | 1,554080712 | 0,00384466 | 0,056966512 | FERM domain containing 4A |
| ENSG00000203364 | AL390760,1 | 0,636032744 | 1,554049809 | 0,4358434 | 0,896422069 | novel transcript |
| ENSG00000009765 | IYD | 0,636022904 | 1,554039209 | 0,50392185 | 0,921268512 | iodotyrosine deiodinase |
| ENSG00000173698 | ADGRG2 | 0,635884527 | 1,55389016 | 0,10541166 | 0,518119329 | adhesion G protein-coupled receptor G2 |
| ENSG00000167207 | NOD2 | 0,63586984 | 1,553874341 | 0,47128127 | 0,908869685 | nucleotide binding oligomerization domain containing 2 |
| ENSG00000145949 | MYLK4 | 0,635847579 | 1,553850364 | 0,21584361 | 0,722999018 | myosin light chain kinase family member 4 |
| ENSG00000248516 | AC105415,1 | 0,635753847 | 1,553749413 | 0,57407088 | 0,942475308 | novel transcript |
| ENSG00000230155 | CEP250-AS1 | 0,635674964 | 1,553664461 | 0,59183544 | 0,947416795 | CEP250 antisense RNA 1 |
| ENSG00000240015 | AC084024,1 | 0,635638939 | 1,553625665 | 0,50221026 | 0,920520138 | novel transcript |
| ENSG00000260588 | AC027702,1 | 0,635443004 | 1,553414679 | 0,28175027 | 0,792816227 | novel transcript |
| ENSG00000248787 | AC092903,2 | 0,635312031 | 1,553273661 | 0,35678883 | 0,848268504 | novel transcript |
| ENSG00000102683 | SGCG | 0,635189063 | 1,553141273 | 0,32076411 | 0,824366234 | sarcoglycan gamma |
| ENSG00000185298 | CCDC137 | 0,634973533 | 1,552909261 | 0,1480513 | 0,612582848 | coiled-coil domain containing 137 |
| ENSG00000023191 | RNH1 | 0,634816589 | 1,552740336 | 0,0993973 | 0,50040065 | ribonuclease/angiogenin inhibitor 1 |
| ENSG00000090581 | GNPTG | 0,634769955 | 1,552690146 | 0,11934767 | 0,552128662 | N-acetylglucosamine-1-phosphate transferase subunit gamma |
| ENSG00000099992 | TBC1D10A | 0,634650645 | 1,552561744 | 0,12963351 | 0,575441236 | TBC1 domain family member 10A |
| ENSG00000236352 | AC005220,1 | 0,634594245 | 1,55250105 | 0,46198959 | 0,906277266 | novel transcript |
| ENSG00000175591 | P2RY2 | 0,634517612 | 1,552418588 | 0,41897393 | 0,887452634 | purinergic receptor P2Y2 |
| ENSG00000137393 | RNF144B | 0,634405783 | 1,552298258 | 0,27561019 | 0,787284069 | ring finger protein 144B |
| ENSG00000148120 | AOPEP | 0,634265382 | 1,552147198 | 0,02701673 | 0,23048668 | aminopeptidase O (putative) |
| ENSG00000259175 | AC108451,1 | 0,634215419 | 1,552093445 | 0,35461091 | 0,846034781 | novel transcript, antisense to ARNT2 |
| ENSG00000166189 | HPS6 | 0,634099681 | 1,551968936 | 0,18320419 | 0,674243186 | HPS6 biogenesis of lysosomal organelles complex 2 subunit 3 |
| ENSG00000286333 | AC004828,2 | 0,633820288 | 1,55166841 | 0,56196404 | 0,940865625 | novel transcript, antisense to RGS6 |
| ENSG00000230105 | AL354793,1 | 0,633722285 | 1,551563008 | 0,29102001 | 0,801689099 | novel transcript |

| | | | | | | |
|---|---|---|---|---|---|---|
| ENSG00000230521 | AL645929,1 | 0,633493064 | 1,551316509 | 0,6414488 | 0,956465487 | HLA complex group 4 pseudogene 7 |
| ENSG00000197208 | SLC22A4 | 0,633145193 | 1,550942492 | 0,31753234 | 0,822351863 | solute carrier family 22 member 4 |
| ENSG00000260289 | AC093515,1 | 0,633105915 | 1,550900267 | 0,64646792 | 0,956899963 | novel transcript |
| ENSG00000002549 | LAP3 | 0,633046421 | 1,550836313 | 0,00198107 | 0,033406759 | leucine aminopeptidase 3 |
| ENSG00000241945 | PWP2 | 0,632837109 | 1,550611328 | 0,04966937 | 0,340167192 | PWP2 small subunit processome component |
| ENSG00000101977 | MCF2 | 0,632817859 | 1,550590637 | 0,24281645 | 0,752892335 | MCF,2 cell line derived transforming sequence |
| ENSG00000274475 | RN7SL650P | 0,632657138 | 1,550417907 | 0,62216226 | 0,952793934 | RNA, 7SL, cytoplasmic 650, pseudogene |
| ENSG00000278469 | Metazoa_SRP | 0,632545665 | 1,550298114 | 0,6636754 | 0,959920886 | Metazoan signal recognition particle RNA |
| ENSG00000153885 | KCTD15 | 0,632117931 | 1,549838546 | 0,01028624 | 0,11746904 | potassium channel tetramerization domain containing 15 |
| ENSG00000128683 | GAD1 | 0,632057007 | 1,549773099 | 0,15117877 | 0,618402864 | glutamate decarboxylase 1 |
| ENSG00000115677 | HDLBP | 0,631984964 | 1,549695711 | 0,00505351 | 0,06927661 | high density lipoprotein binding protein |
| ENSG00000236609 | ZNF853 | 0,631979782 | 1,549690144 | 0,15250399 | 0,620962584 | zinc finger protein 853 |
| ENSG00000129194 | SOX15 | 0,631777994 | 1,549473406 | 0,19838293 | 0,697413278 | SRY-box transcription factor 15 |
| ENSG00000119669 | IRF2BPL | 0,631687734 | 1,549376469 | 0,0754036 | 0,433690715 | interferon regulatory factor 2 binding protein like |
| ENSG00000126821 | SGPP1 | 0,631556589 | 1,549235633 | 0,09239306 | 0,48312034 | sphingosine-1-phosphate phosphatase 1 |
| ENSG00000285281 | SGPP1 | 0,631556589 | 1,549235633 | 0,09239306 | 0,48312034 | sphingosine-1-phosphate phosphatase 1 |
| ENSG00000139364 | TMEM132B | 0,631484003 | 1,549157688 | 0,19121186 | 0,686939473 | transmembrane protein 132B |
| ENSG00000274963 | RN7SL600P | 0,631441801 | 1,549112372 | 0,46713976 | 0,907940653 | RNA, 7SL, cytoplasmic 600, pseudogene |
| ENSG00000116260 | QSOX1 | 0,631361928 | 1,54902661 | 0,17643493 | 0,662315819 | quiescin sulfhydryl oxidase 1 |
| ENSG00000220161 | LINC02076 | 0,631295543 | 1,548955334 | 0,38345648 | 0,867918747 | long intergenic non-protein coding RNA 2076 |
| ENSG00000230885 | MOG | 0,631287113 | 1,548946283 | 0,58245146 | 0,944174844 | myelin oligodendrocyte glycoprotein |
| ENSG00000287494 | AC069368,2 | 0,631226035 | 1,548880708 | 0,58405831 | 0,944526397 | novel transcript |
| ENSG00000285854 | AC010197,2 | 0,63120306 | 1,548856042 | 0,3421713 | 0,840048641 | novel transcript |
| ENSG00000231898 | MYO3B-AS1 | 0,631058971 | 1,548701358 | 0,53984582 | 0,933385624 | MYO3B antisense RNA 1 |
| ENSG00000102100 | SLC35A2 | 0,630909749 | 1,54854118 | 0,01059552 | 0,119674239 | solute carrier family 35 member A2 |
| ENSG00000116962 | NID1 | 0,630881903 | 1,548511292 | 0,00941404 | 0,109821064 | nidogen 1 |
| ENSG00000105854 | PON2 | 0,630860868 | 1,548488714 | 0,14721211 | 0,611181688 | paraoxonase 2 |
| ENSG00000110917 | MLEC | 0,630328291 | 1,547917188 | 0,0153799 | 0,156675544 | malectin |
| ENSG00000142002 | DPP9 | 0,630133052 | 1,547707724 | 0,09343861 | 0,485744051 | dipeptidyl peptidase 9 |
| ENSG00000139514 | SLC7A1 | 0,629999144 | 1,547564076 | 0,01419396 | 0,148068051 | solute carrier family 7 member 1 |
| ENSG00000277424 | HMOX2 | 0,629998252 | 1,547563118 | 0,05773006 | 0,371372961 | heme oxygenase 2 |
| ENSG00000273987 | AC121761,2 | 0,629930319 | 1,54749025 | 0,45309235 | 0,902489761 | novel transcript, sense intronic to CAPS2 |

| | | | | | | |
|---|---|---|---|---|---|---|
| ENSG00000225655 | BX255923,1 | 0,629844945 | 1,547398677 | 0,67599766 | 0,963560994 | novel transcript |
| ENSG00000013288 | MAN2B2 | 0,62969228 | 1,547234941 | 0,04205328 | 0,307400909 | mannosidase alpha class 2B member 2 |
| ENSG00000059122 | FLYWCH1 | 0,629685461 | 1,547227628 | 0,24582378 | 0,757752583 | FLYWCH-type zinc finger 1 |
| ENSG00000142188 | TMEM50B | 0,629657609 | 1,547197758 | 0,06255338 | 0,390552401 | transmembrane protein 50B |
| ENSG00000165895 | ARHGAP42 | 0,629486334 | 1,547014087 | 0,09622833 | 0,492709242 | Rho GTPase activating protein 42 |
| ENSG00000255348 | AP001775,2 | 0,629374866 | 1,546894564 | 0,53720555 | 0,932883277 | novel transcript |
| ENSG00000115520 | COQ10B | 0,629283555 | 1,54679666 | 0,05670603 | 0,367607368 | coenzyme Q10B |
| ENSG00000103175 | WFDC1 | 0,628865657 | 1,546348673 | 0,31229768 | 0,818547047 | WAP four-disulfide core domain 1 |
| ENSG00000234353 | AP000346,2 | 0,628679666 | 1,546149332 | 0,45877617 | 0,904420115 | immunoglobulin lambda-like polypeptide 1 (IGLL1) pseudogene |
| ENSG00000136147 | PHF11 | 0,628634182 | 1,546100587 | 0,0611566 | 0,384884976 | PHD finger protein 11 |
| ENSG00000282147 | FAM20C | 0,62819877 | 1,545634037 | 0,60861331 | 0,951557131 | FAM20C golgi associated secretory pathway kinase |
| ENSG00000155886 | SLC24A2 | 0,628090694 | 1,545518254 | 0,32062092 | 0,824366234 | solute carrier family 24 member 2 |
| ENSG00000231007 | CDC20P1 | 0,628085722 | 1,545512928 | 0,15703072 | 0,629249971 | cell division cycle 20 pseudogene 1 |
| ENSG00000138071 | ACTR2 | 0,627897784 | 1,545311609 | 0,08062634 | 0,450205741 | actin related protein 2 |
| ENSG00000091436 | MAP3K20 | 0,627470931 | 1,544854463 | 0,01495634 | 0,153915406 | mitogen-activated protein kinase kinase kinase 20 |
| ENSG00000174917 | MICOS13 | 0,627453288 | 1,54483557 | 0,03414487 | 0,26913287 | mitochondrial contact site and cristae organizing system subunit 13 |
| ENSG00000227359 | AC017074,1 | 0,627451017 | 1,544833138 | 0,61802352 | 0,952793934 | novel transcript |
| ENSG00000246331 | AC010198,1 | 0,627233617 | 1,544600365 | 0,61767079 | 0,952793934 | novel transcript |
| ENSG00000137460 | FHDC1 | 0,62722123 | 1,544587103 | 0,13277386 | 0,581622796 | FH2 domain containing 1 |
| ENSG00000186635 | ARAP1 | 0,627209032 | 1,544574043 | 0,17753552 | 0,664011316 | ArfGAP with RhoGAP domain, ankyrin repeat and PH domain 1 |
| ENSG00000241570 | PAQR9-AS1 | 0,626839045 | 1,544177979 | 0,3431181 | 0,840759015 | PAQR9 antisense RNA 1 |
| ENSG00000105419 | MEIS3 | 0,62683219 | 1,544170642 | 0,06165366 | 0,386978526 | Meis homeobox 3 |
| ENSG00000078902 | TOLLIP | 0,626697469 | 1,544026452 | 0,08534385 | 0,465128991 | toll interacting protein |
| ENSG00000232453 | LINC02777 | 0,626573878 | 1,543894186 | 0,26800144 | 0,780680587 | long intergenic non-protein coding RNA 2777 |
| ENSG00000156675 | RAB11FIP1 | 0,626395854 | 1,543703686 | 0,06701563 | 0,406751428 | RAB11 family interacting protein 1 |
| ENSG00000147206 | NXF3 | 0,626356339 | 1,543661405 | 0,62157637 | 0,952793934 | nuclear RNA export factor 3 |
| ENSG00000275779 | NTAN1 | 0,62616056 | 1,543451939 | 0,06925882 | 0,415287103 | N-terminal asparagine amidase |
| ENSG00000260596 | DUX4 | 0,626159875 | 1,543451207 | 0,50414694 | 0,921400408 | double homeobox 4 |
| ENSG00000074966 | TXK | 0,62614541 | 1,543435731 | 0,43563164 | 0,896422069 | TXK tyrosine kinase |
| ENSG00000256030 | CBX3P4 | 0,626086895 | 1,543373131 | 0,66855508 | 0,961351573 | chromobox 3 pseudogene 4 |

| | | | | | | |
|---|---|---|---|---|---|---|
| ENSG00000196970 | NXF4 | 0,625741582 | 1,543003765 | 0,62186345 | 0,952793934 | nuclear RNA export factor 4 pseudogene |
| ENSG00000264548 | AC132872,2 | 0,625613349 | 1,542866622 | 0,44259609 | 0,899437949 | novel transcript, antisense CCDC57 |
| ENSG00000136960 | ENPP2 | 0,625434501 | 1,542675368 | 0,18774409 | 0,681801755 | ectonucleotide pyrophosphatase/phosphodiesterase 2 |
| ENSG00000227857 | AL358075,2 | 0,625243072 | 1,542470686 | 0,69899468 | 0,968522643 | novel transcript |
| ENSG00000173083 | HPSE | 0,625124734 | 1,542344169 | 0,22834313 | 0,736649232 | heparanase |
| ENSG00000204850 | AC011484,1 | 0,625114212 | 1,542332921 | 0,60726656 | 0,950993594 | novel transcript, antisense to PPP5D1 |
| ENSG00000157020 | SEC13 | 0,625030991 | 1,542243955 | 0,00239648 | 0,038751223 | SEC13 homolog, nuclear pore and COPII coat complex component |
| ENSG00000133519 | ZDHHC8P1 | 0,624926121 | 1,542131853 | 0,24332296 | 0,753715365 | ZDHHC8 pseudogene 1 |
| ENSG00000140545 | MFGE8 | 0,624901193 | 1,542105207 | 0,04853047 | 0,335601999 | milk fat globule EGF and factor V/VIII domain containing |
| ENSG00000069482 | GAL | 0,624890464 | 1,542093738 | 0,40054346 | 0,876257137 | galanin and GMAP prepropeptide |
| ENSG00000073417 | PDE8A | 0,624885753 | 1,542088703 | 0,02223621 | 0,20109717 | phosphodiesterase 8A |
| ENSG00000182544 | MFSD5 | 0,624720044 | 1,541911587 | 0,02572625 | 0,223230867 | major facilitator superfamily domain containing 5 |
| ENSG00000165169 | DYNLT3 | 0,624705151 | 1,54189567 | 0,23126612 | 0,739268552 | dynein light chain Tctex-type 3 |
| ENSG00000109956 | B3GAT1 | 0,624642587 | 1,541828806 | 0,05547202 | 0,363656395 | beta-1,3-glucuronyltransferase 1 |
| ENSG00000119812 | FAM98A | 0,624424443 | 1,54159569 | 0,02684196 | 0,229474164 | family with sequence similarity 98 member A |
| ENSG00000086288 | NME8 | 0,624339875 | 1,541505327 | 0,46200698 | 0,906277266 | NME/NM23 family member 8 |
| ENSG00000178764 | ZHX2 | 0,62414724 | 1,541299512 | 0,03409601 | 0,269045724 | zinc fingers and homeoboxes 2 |
| ENSG00000114302 | PRKAR2A | 0,624141764 | 1,541293662 | 0,00181079 | 0,030851518 | protein kinase cAMP-dependent type II regulatory subunit alpha |
| ENSG00000213901 | SLC23A3 | 0,624024285 | 1,541168159 | 0,37305287 | 0,861064958 | solute carrier family 23 member 3 |
| ENSG00000262823 | AC127521,1 | 0,62374405 | 1,540868825 | 0,43049402 | 0,895238505 | novel transcript, antisense to SPNS3 |
| ENSG00000288478 | NIPA1 | 0,623610108 | 1,540725776 | 0,35042869 | 0,843857545 | NIPA magnesium transporter 1 |
| ENSG00000196209 | SIRPB2 | 0,623426264 | 1,540529452 | 0,46922717 | 0,907940653 | signal regulatory protein beta 2 |
| ENSG00000264313 | RN7SL644P | 0,623415028 | 1,540517454 | 0,5956784 | 0,947991313 | RNA, 7SL, cytoplasmic 644, pseudogene |
| ENSG00000104915 | STX10 | 0,623106496 | 1,540188037 | 0,04716715 | 0,330117289 | syntaxin 10 |
| ENSG00000174851 | YIF1A | 0,623069633 | 1,540148684 | 0,00756329 | 0,093795931 | Yip1 interacting factor homolog A, membrane trafficking protein |
| ENSG00000170190 | SLC16A5 | 0,622731491 | 1,539787742 | 0,15705197 | 0,629254479 | solute carrier family 16 member 5 |
| ENSG00000287616 | AL357522,1 | 0,622486225 | 1,539525992 | 0,59471903 | 0,947469769 | novel transcript |
| ENSG00000197815 | AC122129,1 | 0,622280005 | 1,539305947 | 0,31241969 | 0,818574551 | novel transcript, antisense to TOM1L2 |

| | | | | | | |
|---|---|---|---|---|---|---|
| ENSG00000070669 | ASNS | 0,622215496 | 1,53923712 | 0,06513751 | 0,400662695 | asparagine synthetase (glutamine-hydrolyzing) |
| ENSG00000013619 | MAMLD1 | 0,622153163 | 1,539170617 | 0,09101306 | 0,481283407 | mastermind like domain containing 1 |
| ENSG00000082641 | NFE2L1 | 0,621966139 | 1,538971099 | 0,10687226 | 0,521837326 | nuclear factor, erythroid 2 like 1 |
| ENSG00000204241 | LINC02731 | 0,621884076 | 1,538883563 | 0,21416652 | 0,721518909 | long intergenic non-protein coding RNA 2731 |
| ENSG00000144455 | SUMF1 | 0,621850114 | 1,538847336 | 0,01829853 | 0,176080387 | sulfatase modifying factor 1 |
| ENSG00000274780 | PHRF1 | 0,621834601 | 1,53883079 | 0,35302453 | 0,845031687 | PHD and ring finger domains 1 |
| ENSG00000226847 | PRPS1P1 | 0,621824013 | 1,538819495 | 0,67515341 | 0,963390425 | phosphoribosyl pyrophosphate synthetase 1 pseudogene 1 |
| ENSG00000287313 | AC016995,1 | 0,621671316 | 1,538656634 | 0,38103661 | 0,865900094 | novel transcript |
| ENSG00000271860 | AL589740,1 | 0,621563829 | 1,538542001 | 0,1583414 | 0,631749049 | novel transcript |
| ENSG00000266004 | SNORA38 | 0,62092295 | 1,537858697 | 0,58649934 | 0,945395317 | small nucleolar RNA, H/ACA box 38 |
| ENSG00000265754 | SNORA38 | 0,62092295 | 1,537858697 | 0,58649934 | 0,945395317 | small nucleolar RNA, H/ACA box 38 |
| ENSG00000157087 | ATP2B2 | 0,620852867 | 1,537783993 | 0,31079329 | 0,817751312 | ATPase plasma membrane Ca2+ transporting 2 |
| ENSG00000277101 | ARHGEF26 | 0,620748679 | 1,537672941 | 0,47144101 | 0,908896785 | Rho guanine nucleotide exchange factor 26 |
| ENSG00000239577 | RN7SL388P | 0,620710553 | 1,537632306 | 0,44568616 | 0,901173082 | RNA, 7SL, cytoplasmic 388, pseudogene |
| ENSG00000072201 | LNX1 | 0,620637848 | 1,537554819 | 0,11319608 | 0,537264645 | ligand of numb-protein X 1 |
| ENSG00000272109 | AC008906,2 | 0,620565979 | 1,537478226 | 0,52859019 | 0,930033144 | novel transcript, antisense to ERAP1 |
| ENSG00000180537 | RNF182 | 0,620457681 | 1,537362817 | 0,07608903 | 0,435925861 | ring finger protein 182 |
| ENSG00000267339 | LINC00906 | 0,620370852 | 1,537270294 | 0,38541554 | 0,868895792 | long intergenic non-protein coding RNA 906 |
| ENSG00000274041 | AC012314,2 | 0,620259477 | 1,537151622 | 0,62759876 | 0,952793934 | novel transcript |
| ENSG00000243570 | CFB | 0,620177057 | 1,537063808 | 0,43225267 | 0,89558934 | complement factor B |
| ENSG00000242335 | CFB | 0,620177057 | 1,537063808 | 0,43225267 | 0,89558934 | complement factor B |
| ENSG00000274422 | AC245060,5 | 0,620011424 | 1,536887351 | 0,3905298 | 0,870057277 | novel transcript |
| ENSG00000267532 | MIR497HG | 0,619943102 | 1,53681457 | 0,34385768 | 0,840901506 | mir-497-195 cluster host gene |
| ENSG00000242802 | AP5Z1 | 0,619940433 | 1,536811727 | 0,09374771 | 0,486865208 | adaptor related protein complex 5 subunit zeta 1 |
| ENSG00000160471 | COX6B2 | 0,619738375 | 1,536596502 | 0,43482828 | 0,896272827 | cytochrome c oxidase subunit 6B2 |
| ENSG00000172995 | ARPP21 | 0,619718555 | 1,536575393 | 0,32402974 | 0,82610013 | cAMP regulated phosphoprotein 21 |
| ENSG00000231321 | RXRB | 0,619478175 | 1,536319391 | 0,28014274 | 0,790779983 | retinoid X receptor beta |
| ENSG00000246394 | ZNF84-DT | 0,619419526 | 1,536256938 | 0,6287674 | 0,952793934 | ZNF84 divergent transcript |
| ENSG00000282846 | AC008982,4 | 0,619261029 | 1,536088171 | 0,47756676 | 0,911578505 | novel transcript, sense intronic to HNRNPL |
| ENSG00000287033 | AL590101,1 | 0,618838833 | 1,53563871 | 0,61060178 | 0,951841373 | novel transcript |
| ENSG00000244229 | RPL26P35 | 0,618654607 | 1,535442628 | 0,46408426 | 0,90737035 | ribosomal protein L26 pseudogene 35 |
| ENSG00000152689 | RASGRP3 | 0,618573654 | 1,535356473 | 0,35926455 | 0,849982153 | RAS guanyl releasing protein 3 |

| | | | | | | |
|---|---|---|---|---|---|---|
| ENSG00000206503 | HLA-A | 0,618153004 | 1,534908871 | 0,08415048 | 0,461477024 | major histocompatibility complex, class I, A |
| ENSG00000179165 | PXT1 | 0,61806107 | 1,534811064 | 0,5629557 | 0,940865625 | peroxisomal testis enriched protein 1 |
| ENSG00000117984 | CTSD | 0,618059861 | 1,534809778 | 0,04293281 | 0,312296718 | cathepsin D |
| ENSG00000145990 | GFOD1 | 0,618005618 | 1,534752073 | 0,09436796 | 0,4879788 | glucose-fructose oxidoreductase domain containing 1 |
| ENSG00000101347 | SAMHD1 | 0,617811692 | 1,534545786 | 0,06579049 | 0,402522813 | SAM and HD domain containing deoxynucleoside triphosphate triphosphohydrolase 1 |
| ENSG00000103264 | FBXO31 | 0,617734277 | 1,534463445 | 0,08911278 | 0,47641326 | F-box protein 31 |
| ENSG00000110092 | CCND1 | 0,617064816 | 1,533751565 | 0,00256127 | 0,040855994 | cyclin D1 |
| ENSG00000280395 | AL034546,1 | 0,616984402 | 1,533666078 | 0,37435032 | 0,861515531 | novel transcript, sense intronic to MOV10L1 |
| ENSG00000126945 | HNRNPH2 | 0,616955969 | 1,533635852 | 0,02504315 | 0,219619974 | heterogeneous nuclear ribonucleoprotein H2 |
| ENSG00000146966 | DENND2A | 0,616636946 | 1,533296757 | 0,099564 | 0,500686631 | DENN domain containing 2A |
| ENSG00000174460 | ZCCHC12 | 0,616544573 | 1,533198586 | 0,09784486 | 0,496997855 | zinc finger CCHC-type containing 12 |
| ENSG00000246225 | AC006299,1 | 0,616397544 | 1,533042342 | 0,56313775 | 0,940865625 | novel transcript, antisense to SVIP |
| ENSG00000272273 | IER3-AS1 | 0,616338358 | 1,532979451 | 0,56516568 | 0,940865625 | IER3 antisense RNA 1 |
| ENSG00000177963 | RIC8A | 0,615694524 | 1,532295478 | 0,04191858 | 0,306990518 | RIC8 guanine nucleotide exchange factor A |
| ENSG00000243224 | AC006252,1 | 0,615684431 | 1,532284758 | 0,31388112 | 0,819877431 | novel transcript, antisense to TWF2 |
| ENSG00000239650 | GUSBP4 | 0,615584901 | 1,532179051 | 0,52356532 | 0,928038684 | GUSB pseudogene 4 |
| ENSG00000254325 | AC018607,1 | 0,615512621 | 1,53210229 | 0,62004435 | 0,952793934 | novel transcript |
| ENSG00000227038 | GTF2IP7 | 0,615426354 | 1,532010679 | 0,3464318 | 0,841168721 | general transcription factor IIi pseudogene 7 |
| ENSG00000042445 | RETSAT | 0,61536389 | 1,53194435 | 0,0379778 | 0,288398822 | retinol saturase |
| ENSG00000255165 | AC134775,1 | 0,614932723 | 1,531486578 | 0,42019046 | 0,888457169 | novel transcript |
| ENSG00000110711 | AIP | 0,614854394 | 1,53140343 | 0,01584551 | 0,159959274 | aryl hydrocarbon receptor interacting protein |
| ENSG00000248008 | NRAV | 0,614433959 | 1,530957209 | 0,08959891 | 0,477709449 | negative regulator of antiviral response |
| ENSG00000100979 | PLTP | 0,614382476 | 1,530902577 | 0,04397169 | 0,316175438 | phospholipid transfer protein |
| ENSG00000230873 | STMND1 | 0,614300635 | 1,530815734 | 0,5143125 | 0,924319732 | stathmin domain containing 1 |
| ENSG00000140829 | DHX38 | 0,614173733 | 1,530681087 | 0,23874765 | 0,747998807 | DEAH-box helicase 38 |
| ENSG00000168884 | TNIP2 | 0,613971806 | 1,53046686 | 0,11868332 | 0,550565054 | TNFAIP3 interacting protein 2 |
| ENSG00000085063 | CD59 | 0,613858038 | 1,530346176 | 0,03910994 | 0,29449254 | CD59 molecule (CD59 blood group) |
| ENSG00000258610 | AF111169,1 | 0,613689066 | 1,530166948 | 0,59765614 | 0,9482846 | novel transcript |
| ENSG00000187239 | FNBP1 | 0,613505961 | 1,529972754 | 0,05557693 | 0,363892597 | formin binding protein 1 |
| ENSG00000135677 | GNS | 0,613427926 | 1,52989 | 0,03920098 | 0,294869635 | glucosamine (N-acetyl)-6-sulfatase |
| ENSG00000257880 | AC078789,1 | 0,612962362 | 1,529396377 | 0,59675396 | 0,948282297 | novel transcript, sense intronic to FAM19A2 |

| | | | | | | |
|---|---|---|---|---|---|---|
| ENSG00000137819 | PAQR5 | 0,61291222 | 1,529343223 | 0,0873426 | 0,471131638 | progestin and adipoQ receptor family member 5 |
| ENSG00000255409 | RSF1-IT1 | 0,612739727 | 1,529160381 | 0,53210042 | 0,931291085 | RSF1 intronic transcript 1 |
| ENSG00000259746 | HSPE1P3 | 0,612601212 | 1,529013571 | 0,5283577 | 0,930033144 | heat shock protein family E (Hsp10) member 1 pseudogene 3 |
| ENSG00000264041 | RN7SL670P | 0,612541102 | 1,528949867 | 0,40259415 | 0,877321369 | RNA, 7SL, cytoplasmic 670, pseudogene |
| ENSG00000068831 | RASGRP2 | 0,612441902 | 1,528844739 | 0,33424136 | 0,83453565 | RAS guanyl releasing protein 2 |
| ENSG00000260792 | LINC02280 | 0,612341579 | 1,528738429 | 0,43852419 | 0,897220559 | long intergenic non-protein coding RNA 2280 |
| ENSG00000268375 | AC010325,2 | 0,61211917 | 1,528502773 | 0,74181817 | 0,975176043 | novel transcript, antisense to ACPT |
| ENSG00000226828 | LINC02774 | 0,611875201 | 1,528244316 | 0,54218194 | 0,933491609 | long intergenic non-protein coding RNA 2774 |
| ENSG00000168925 | CTRB1 | 0,611779355 | 1,52814279 | 0,66447019 | 0,96046247 | chymotrypsinogen B1 |
| ENSG00000131323 | TRAF3 | 0,61177344 | 1,528136524 | 0,03043041 | 0,249541473 | TNF receptor associated factor 3 |
| ENSG00000274287 | SCRIB | 0,611714386 | 1,528073974 | 0,3594777 | 0,849982153 | scribble planar cell polarity protein |
| ENSG00000127022 | CANX | 0,611608023 | 1,52796132 | 0,04861058 | 0,335726918 | calnexin |
| ENSG00000254788 | CKLF-CMTM1 | 0,611393876 | 1,527734534 | 0,20817025 | 0,71274406 | CKLF-CMTM1 readthrough |
| ENSG00000273264 | AL360219,1 | 0,611223408 | 1,527554029 | 0,45145356 | 0,902062594 | novel transcript |
| ENSG00000134115 | CNTN6 | 0,611172419 | 1,527500041 | 0,08536688 | 0,465128991 | contactin 6 |
| ENSG00000172661 | WASHC2C | 0,610934577 | 1,527248239 | 0,0071269 | 0,089664163 | WASH complex subunit 2C |
| ENSG00000260273 | AL359711,2 | 0,610778123 | 1,527082624 | 0,52910322 | 0,930207681 | novel transcript, antisense to CD164 |
| ENSG00000235499 | AC073046,1 | 0,610394746 | 1,526676877 | 0,27650458 | 0,788265979 | novel transcript (LOC100132318) |
| ENSG00000262412 | AL160291,1 | 0,610293448 | 1,526569686 | 0,56258131 | 0,940865625 | novel transcript |
| ENSG00000150867 | PIP4K2A | 0,610119144 | 1,526385259 | 0,02220312 | 0,200856008 | phosphatidylinositol-5-phosphate 4-kinase type 2 alpha |
| ENSG00000249923 | AC007663,2 | 0,609927683 | 1,526182705 | 0,44928977 | 0,902062594 | novel transcript |
| ENSG00000126088 | UROD | 0,609918902 | 1,526173416 | 0,01426793 | 0,14859184 | uroporphyrinogen decarboxylase |
| ENSG00000285641 | AL358472,6 | 0,609852651 | 1,526103333 | 0,17945085 | 0,668296783 | novel protein |
| ENSG00000215277 | RNF212B | 0,609765777 | 1,526011439 | 0,61009768 | 0,951801946 | ring finger protein 212B |
| ENSG00000196411 | EPHB4 | 0,609758229 | 1,526003456 | 0,06286097 | 0,391611013 | EPH receptor B4 |
| ENSG00000069812 | HES2 | 0,609599338 | 1,525835399 | 0,47643293 | 0,91143003 | hes family bHLH transcription factor 2 |
| ENSG00000267436 | AC005786,3 | 0,609539028 | 1,525771615 | 0,16414256 | 0,642888389 | novel transcript, antisense to MFSD12 |
| ENSG00000141696 | P3H4 | 0,609222608 | 1,52543701 | 0,09410614 | 0,487592685 | prolyl 3-hydroxylase family member 4 (inactive) |
| ENSG00000160214 | RRP1 | 0,609025734 | 1,525228859 | 0,25506248 | 0,76642479 | ribosomal RNA processing 1 |
| ENSG00000234902 | AC007879,3 | 0,608766561 | 1,524954884 | 0,58783975 | 0,946129322 | novel transcript |
| ENSG00000262079 | AC007638,1 | 0,608454517 | 1,524625084 | 0,63747808 | 0,956017934 | novel transcript |

| | | | | | | |
|---|---|---|---|---|---|---|
| ENSG00000019144 | PHLDB1 | 0,608404574 | 1,524572305 | 0,18497454 | 0,677431213 | pleckstrin homology like domain family B member 1 |
| ENSG00000125637 | PSD4 | 0,608390383 | 1,524557309 | 0,16256268 | 0,639794628 | pleckstrin and Sec7 domain containing 4 |
| ENSG00000135956 | TMEM127 | 0,608360352 | 1,524525574 | 0,02217633 | 0,200729778 | transmembrane protein 127 |
| ENSG00000286940 | MICB-DT | 0,608316022 | 1,524478731 | 0,42946457 | 0,894343583 | MICB divergent transcript |
| ENSG00000143314 | MRPL24 | 0,608103712 | 1,524254401 | 0,03836932 | 0,290314236 | mitochondrial ribosomal protein L24 |
| ENSG00000090539 | CHRD | 0,607760097 | 1,523891405 | 0,19047807 | 0,685949202 | chordin |
| ENSG00000278840 | AC009878,1 | 0,607678033 | 1,523804725 | 0,64276245 | 0,956633898 | novel transcript, sense intronic to GABRB3 |
| ENSG00000203876 | ADD3-AS1 | 0,607667133 | 1,523793212 | 0,22562684 | 0,734461165 | ADD3 antisense RNA 1 |
| ENSG00000268598 | VN1R80P | 0,60745786 | 1,523572191 | 0,58516251 | 0,945013723 | vomeronasal 1 receptor 80 pseudogene |
| ENSG00000255737 | AGAP2-AS1 | 0,607116126 | 1,523211341 | 0,40355938 | 0,877691113 | AGAP2 antisense RNA 1 |
| ENSG00000170581 | STAT2 | 0,60685879 | 1,522939668 | 0,03346252 | 0,265660811 | signal transducer and activator of transcription 2 |
| ENSG00000260507 | AC133919,1 | 0,606695417 | 1,522767218 | 0,53963548 | 0,933350549 | novel transcript |
| ENSG00000228728 | PPIAP91 | 0,606584739 | 1,522650401 | 0,43149272 | 0,89530663 | peptidylprolyl isomerase A pseudogene 91 |
| ENSG00000235183 | SRP14P3 | 0,606518869 | 1,522580883 | 0,42184621 | 0,889720796 | signal recognition particle 14 pseudogene 3 |
| ENSG00000134830 | C5AR2 | 0,606494894 | 1,522555581 | 0,41806374 | 0,887087337 | complement component 5a receptor 2 |
| ENSG00000166130 | IKBIP | 0,606446883 | 1,522504912 | 0,02901488 | 0,241642594 | IKBKB interacting protein |
| ENSG00000154803 | FLCN | 0,606252858 | 1,522300167 | 0,09419702 | 0,487613759 | folliculin |
| ENSG00000142405 | NLRP12 | 0,606056584 | 1,522093078 | 0,43585282 | 0,896422069 | NLR family pyrin domain containing 12 |
| ENSG00000009694 | TENM1 | 0,605738253 | 1,521757265 | 0,27994079 | 0,790611456 | teneurin transmembrane protein 1 |
| ENSG00000230149 | AL021707,3 | 0,605703222 | 1,521720314 | 0,51976474 | 0,926822772 | novel transcript, antisense to SUN2 |
| ENSG00000110108 | TMEM109 | 0,605616764 | 1,521629124 | 0,02861573 | 0,240203215 | transmembrane protein 109 |
| ENSG00000255015 | AP000755,1 | 0,605592829 | 1,521603879 | 0,6236635 | 0,952793934 | novel transcript |
| ENSG00000249896 | LINC02495 | 0,605378087 | 1,521377409 | 0,40978286 | 0,882209844 | long intergenic non-protein coding RNA 2495 |
| ENSG00000267714 | AC008738,4 | 0,605350018 | 1,521347809 | 0,57890088 | 0,943828882 | novel transcript, antisense to SLC7A10 |
| ENSG00000288185 | IL27RA | 0,605035253 | 1,52101592 | 0,19324173 | 0,689985107 | interleukin 27 receptor subunit alpha |
| ENSG00000104998 | IL27RA | 0,605035253 | 1,52101592 | 0,19324173 | 0,689985107 | interleukin 27 receptor subunit alpha |
| ENSG00000253112 | AC102945,1 | 0,604825789 | 1,5207951 | 0,63988725 | 0,956018518 | novel transcript |
| ENSG00000275857 | AC009133,4 | 0,604820342 | 1,520789359 | 0,87126869 | 0,989217334 | novel transcript, antisense to KIF22 |
| ENSG00000229344 | MTCO2P12 | 0,604818719 | 1,520787647 | 0,61344836 | 0,952413269 | MT-CO2 pseudogene 12 |
| ENSG00000178722 | C5orf64 | 0,604716343 | 1,520679734 | 0,05034987 | 0,343323549 | chromosome 5 open reading frame 64 |
| ENSG00000255038 | AP006287,2 | 0,60465234 | 1,520612273 | 0,60026097 | 0,949340299 | novel transcript, antisense to SF3B2 |
| ENSG00000255277 | ABCC6P2 | 0,604493908 | 1,520445293 | 0,44929841 | 0,902062594 | ATP binding cassette subfamily C member 6 pseudogene 2 |

| | | | | | | |
|---|---|---|---|---|---|---|
| ENSG00000223551 | TMSB4XP4 | 0,604467743 | 1,520417718 | 0,23993318 | 0,749799162 | TMSB4X pseudogene 4 |
| ENSG00000118194 | TNNT2 | 0,604359485 | 1,520303633 | 0,66459374 | 0,96046247 | troponin T2, cardiac type |
| ENSG00000173080 | RXFP4 | 0,604253405 | 1,52019185 | 0,33645233 | 0,835713011 | relaxin family peptide/INSL5 receptor 4 |
| ENSG00000066027 | PPP2R5A | 0,604211498 | 1,520147693 | 0,02758672 | 0,233985684 | protein phosphatase 2 regulatory subunit B'alpha |
| ENSG00000239264 | TXNDC5 | 0,604003632 | 1,519928683 | 0,01905623 | 0,181086311 | thioredoxin domain containing 5 |
| ENSG00000273294 | C1QTNF3-AMA | 0,603985017 | 1,519909072 | 0,35542583 | 0,846840543 | C1QTNF3-AMACR readthrough (NMD candidate) |
| ENSG00000235814 | ZNRD1ASP | 0,603633927 | 1,519539236 | 0,42905808 | 0,893964132 | zinc ribbon domain containing 1 antisense, pseudogene |
| ENSG00000091879 | ANGPT2 | 0,603438145 | 1,519333039 | 0,02520486 | 0,220174572 | angiopoietin 2 |
| ENSG00000183114 | FAM43B | 0,603426676 | 1,519320961 | 0,30817011 | 0,816424917 | family with sequence similarity 43 member B |
| ENSG00000006283 | CACNA1G | 0,603410589 | 1,519304021 | 0,11519665 | 0,542308656 | calcium voltage-gated channel subunit alpha1 G |
| ENSG00000179058 | C9orf50 | 0,603236887 | 1,519121106 | 0,53244594 | 0,931291085 | chromosome 9 open reading frame 50 |
| ENSG00000018625 | ATP1A2 | 0,603158316 | 1,519038375 | 0,08346126 | 0,460037525 | ATPase Na+/K+ transporting subunit alpha 2 |
| ENSG00000259203 | AC016044,1 | 0,602512979 | 1,51835904 | 0,31114844 | 0,817751312 | novel transcript |
| ENSG00000104635 | SLC39A14 | 0,602287334 | 1,51812158 | 0,01798769 | 0,174107492 | solute carrier family 39 member 14 |
| ENSG00000274776 | AC090241,3 | 0,602272809 | 1,518106295 | 0,26976594 | 0,781791089 | novel transcript |
| ENSG00000171054 | OR13H1 | 0,601921729 | 1,517736909 | 0,2134019 | 0,720944157 | olfactory receptor family 13 subfamily H member 1 |
| ENSG00000262434 | AC087392,4 | 0,601803264 | 1,517612286 | 0,51724895 | 0,925746338 | novel transcript, antisense to RNMTL1 |
| ENSG00000137275 | RIPK1 | 0,601567141 | 1,517363923 | 0,01187234 | 0,129691422 | receptor interacting serine/threonine kinase 1 |
| ENSG00000258337 | AC130895,1 | 0,601554809 | 1,517350952 | 0,50502551 | 0,921400408 | novel transcript, sense intronic to MED13L |
| ENSG00000215861 | AC245297,1 | 0,601462147 | 1,517253498 | 0,44452215 | 0,900210555 | phosphodiesterase 4D interacting protein (myomegalin) (PDE4DIP) pseudogene |
| ENSG00000105366 | SIGLEC8 | 0,601368083 | 1,517154576 | 0,54979585 | 0,936108121 | sialic acid binding Ig like lectin 8 |
| ENSG00000171604 | CXXC5 | 0,601338952 | 1,517123942 | 0,00308871 | 0,047710255 | CXXC finger protein 5 |
| ENSG00000205971 | AC074389,1 | 0,601310141 | 1,517093645 | 0,56450672 | 0,940865625 | lncRNA radiation induced regulator of PLK1 and RAD51 |
| ENSG00000136250 | AOAH | 0,601272468 | 1,51705403 | 0,38116704 | 0,865900094 | acyloxyacyl hydrolase |
| ENSG00000235806 | AF241728,1 | 0,601209103 | 1,516987401 | 0,69875547 | 0,968522643 | novel transcript |
| ENSG00000275438 | AL031008,1 | 0,600945664 | 1,516710421 | 0,43774688 | 0,897021798 | novel transcript |
| ENSG00000262185 | LINC02861 | 0,600901936 | 1,51666445 | 0,44704619 | 0,901822945 | long intergenic non-protein coding RNA 2861 |
| ENSG00000261873 | SMIM36 | 0,600584346 | 1,516330614 | 0,72138919 | 0,972142678 | small integral membrane protein 36 |
| ENSG00000275678 | AL133320,2 | 0,600341751 | 1,516075658 | 0,53581819 | 0,932139691 | novel transcript, sense intronic to C1orf141 |
| ENSG00000184979 | USP18 | 0,599992315 | 1,515708492 | 0,12856148 | 0,572964502 | ubiquitin specific peptidase 18 |

| | | | | | | |
|---|---|---|---|---|---|---|
| ENSG00000139410 | SDSL | 0,5997539 | 1,515458032 | 0,20335716 | 0,703503738 | serine dehydratase like |
| ENSG00000137486 | ARRB1 | 0,599723826 | 1,515426442 | 0,02074256 | 0,191635713 | arrestin beta 1 |
| ENSG00000229097 | CALM2P2 | 0,599578942 | 1,515274261 | 0,26665696 | 0,778772823 | calmodulin 2 pseudogene 2 |
| ENSG00000233448 | PMS2P9 | 0,599442017 | 1,515130454 | 0,62445511 | 0,952793934 | PMS1 homolog 2, mismatch repair system component pseudogene 9 |
| ENSG00000168159 | RNF187 | 0,59879077 | 1,514446664 | 0,04198322 | 0,307032357 | ring finger protein 187 |
| ENSG00000285426 | RNF187 | 0,59879077 | 1,514446664 | 0,04198322 | 0,307032357 | ring finger protein 187 |
| ENSG00000166557 | TMED3 | 0,598703993 | 1,514355574 | 0,00695537 | 0,088323639 | transmembrane p24 trafficking protein 3 |
| ENSG00000117602 | RCAN3 | 0,598682561 | 1,514333077 | 0,02375951 | 0,210728318 | RCAN family member 3 |
| ENSG00000258010 | AC016705,1 | 0,598576942 | 1,514222218 | 0,43821568 | 0,897177404 | novel transcript, antisense to ARNT2 |
| ENSG00000232994 | RPL7P14 | 0,598545401 | 1,514189113 | 0,58231784 | 0,944174844 | ribosomal protein L7 pseudogene 14 |
| ENSG00000071909 | MYO3B | 0,598528206 | 1,514171066 | 0,23191196 | 0,740375324 | myosin IIIB |
| ENSG00000105609 | LILRB5 | 0,598500265 | 1,514141742 | 0,51430265 | 0,924319732 | leukocyte immunoglobulin like receptor B5 |
| ENSG00000277258 | PCGF2 | 0,59835029 | 1,513984348 | 0,02661801 | 0,228371442 | polycomb group ring finger 2 |
| ENSG00000185668 | POU3F1 | 0,598343628 | 1,513977356 | 0,11893527 | 0,551281265 | POU class 3 homeobox 1 |
| ENSG00000162881 | OXER1 | 0,598167604 | 1,513792647 | 0,44882807 | 0,902062594 | oxoeicosanoid receptor 1 |
| ENSG00000005844 | ITGAL | 0,598073553 | 1,513693964 | 0,46828067 | 0,907940653 | integrin subunit alpha L |
| ENSG00000265043 | AC068418,2 | 0,597832406 | 1,513440971 | 0,40862594 | 0,88153752 | novel transcript |
| ENSG00000251045 | SLC25A48-AS1 | 0,597668307 | 1,513268834 | 0,59752947 | 0,948282297 | SLC25A48 antisense RNA 1 |
| ENSG00000106992 | AK1 | 0,597636789 | 1,513235775 | 0,03115893 | 0,253879337 | adenylate kinase 1 |
| ENSG00000145217 | SLC26A1 | 0,597537318 | 1,513131444 | 0,53693486 | 0,932883277 | solute carrier family 26 member 1 |
| ENSG00000247345 | AC092343,1 | 0,597535248 | 1,513129272 | 0,65684892 | 0,957476526 | novel transcript |
| ENSG00000240373 | SEC62-AS1 | 0,597498817 | 1,513091063 | 0,42055016 | 0,888563911 | SEC62 antisense RNA 1 |
| ENSG00000044574 | HSPA5 | 0,597347927 | 1,512932819 | 0,00405958 | 0,059418825 | heat shock protein family A (Hsp70) member 5 |
| ENSG00000249464 | LINC01091 | 0,597072212 | 1,512643709 | 0,13096997 | 0,577911147 | long intergenic non-protein coding RNA 1091 |
| ENSG00000213145 | CRIP1 | 0,596591765 | 1,512140051 | 0,24111095 | 0,751551821 | cysteine rich protein 1 |
| ENSG00000006210 | CX3CL1 | 0,596337 | 1,511873046 | 0,15804512 | 0,631211378 | C-X3-C motif chemokine ligand 1 |
| ENSG00000260136 | AC008915,2 | 0,596163767 | 1,511691517 | 0,28683452 | 0,798325866 | novel transcript |
| ENSG00000286341 | AP002762,3 | 0,596038047 | 1,51155979 | 0,43207253 | 0,89558934 | novel transcript, antisense to UBASH3B |
| ENSG00000153283 | CD96 | 0,595838323 | 1,511350548 | 0,38371086 | 0,868000076 | CD96 molecule |
| ENSG00000137310 | TCF19 | 0,595734709 | 1,511242007 | 0,62352797 | 0,952793934 | transcription factor 19 |
| ENSG00000285250 | MSRA | 0,59548167 | 1,510976969 | 0,10968366 | 0,528090728 | methionine sulfoxide reductase A |
| ENSG00000259322 | AC090607,1 | 0,595277958 | 1,51076363 | 0,49751399 | 0,918759905 | novel transcript, antisense to WDR61 |

| | | | | | | |
|---|---|---|---|---|---|---|
| ENSG00000229153 | EPHA1-AS1 | 0,595213142 | 1,510695758 | 0,64160038 | 0,956465487 | EPHA1 antisense RNA 1 |
| ENSG00000138594 | TMOD3 | 0,595144185 | 1,510623552 | 0,01640823 | 0,163632161 | tropomodulin 3 |
| ENSG00000183308 | AC005037,1 | 0,595070914 | 1,510546833 | 0,43266663 | 0,89560871 | novel transcript |
| ENSG00000167508 | MVD | 0,59484203 | 1,510307203 | 0,11621469 | 0,544242846 | mevalonate diphosphate decarboxylase |
| ENSG00000142684 | ZNF593 | 0,594764842 | 1,5102264 | 0,17350476 | 0,658349911 | zinc finger protein 593 |
| ENSG00000277463 | AC080038,2 | 0,594696826 | 1,510155202 | 0,47043769 | 0,908757071 | novel transcript |
| ENSG00000167987 | VPS37C | 0,594603365 | 1,510057373 | 0,0576797 | 0,371278162 | VPS37C subunit of ESCRT-I |
| ENSG00000128536 | CDHR3 | 0,594428939 | 1,509874815 | 0,20735735 | 0,711147468 | cadherin related family member 3 |
| ENSG00000151116 | UEVLD | 0,59422913 | 1,509665716 | 0,12374501 | 0,562483871 | UEV and lactate/malate dehyrogenase domains |
| ENSG00000104341 | LAPTM4B | 0,594131919 | 1,509563996 | 0,07077746 | 0,418829978 | lysosomal protein transmembrane 4 beta |
| ENSG00000264879 | RN7SL690P | 0,594055924 | 1,509484481 | 0,72142452 | 0,972142678 | RNA, 7SL, cytoplasmic 690, pseudogene |
| ENSG00000167565 | SERTAD3 | 0,594048723 | 1,509476946 | 0,03680047 | 0,282894027 | SERTA domain containing 3 |
| ENSG00000287594 | AL356123,2 | 0,594010293 | 1,509436737 | 0,1716463 | 0,655397237 | novel transcript |
| ENSG00000110446 | SLC15A3 | 0,593802261 | 1,509219097 | 0,3221247 | 0,824955371 | solute carrier family 15 member 3 |
| ENSG00000224081 | SLC44A3-AS1 | 0,593738723 | 1,509152631 | 0,07741465 | 0,439733188 | SLC44A3 antisense RNA 1 |
| ENSG00000157330 | C1orf158 | 0,593716777 | 1,509129674 | 0,64821357 | 0,956899963 | chromosome 1 open reading frame 158 |
| ENSG00000203952 | CCDC160 | 0,593670871 | 1,509081655 | 0,2615715 | 0,773396846 | coiled-coil domain containing 160 |
| ENSG00000227036 | LINC00511 | 0,593542037 | 1,508946899 | 0,2166358 | 0,723909983 | long intergenic non-protein coding RNA 511 |
| ENSG00000138604 | GLCE | 0,593517819 | 1,508921569 | 0,17156835 | 0,655397237 | glucuronic acid epimerase |
| ENSG00000276870 | PLA2G10 | 0,593358276 | 1,508754711 | 0,50676294 | 0,921824694 | phospholipase A2 group X |
| ENSG00000254081 | LINC01299 | 0,593246504 | 1,508637826 | 0,57515439 | 0,942667371 | long intergenic non-protein coding RNA 1299 |
| ENSG00000132613 | MTSS2 | 0,593109041 | 1,508494086 | 0,13528173 | 0,586275832 | MTSS I-BAR domain containing 2 |
| ENSG00000205045 | SLFN12L | 0,593100645 | 1,508485308 | 0,38397255 | 0,868113415 | schlafen family member 12 like |
| ENSG00000169291 | SHE | 0,593072876 | 1,508456272 | 0,39072745 | 0,870057277 | Src homology 2 domain containing E |
| ENSG00000165495 | PKNOX2 | 0,59302752 | 1,50840885 | 0,01110404 | 0,123543331 | PBX/knotted 1 homeobox 2 |
| ENSG00000144136 | SLC20A1 | 0,593015928 | 1,50839673 | 0,00437996 | 0,062727859 | solute carrier family 20 member 1 |
| ENSG00000197665 | AC007952,1 | 0,593007945 | 1,508388383 | 0,58022036 | 0,943828882 | novel transcript, antisense to GRAPL |
| ENSG00000131203 | IDO1 | 0,592870739 | 1,508244937 | 0,40218708 | 0,87718264 | indoleamine 2,3-dioxygenase 1 |
| ENSG00000255197 | AC090559,1 | 0,592793703 | 1,508164402 | 0,48441429 | 0,913690171 | novel transcript |
| ENSG00000203335 | AC006019,1 | 0,592555581 | 1,507915495 | 0,49179809 | 0,916995341 | novel transcript |
| ENSG00000226377 | AC084809,1 | 0,592321622 | 1,507670979 | 0,41181462 | 0,883966506 | novel transcript |
| ENSG00000144285 | SCN1A | 0,591860958 | 1,507189644 | 0,24437939 | 0,755789604 | sodium voltage-gated channel alpha subunit 1 |
| ENSG00000287389 | AC007106,2 | 0,59165801 | 1,50697764 | 0,53386731 | 0,931381467 | novel transcript |

| | | | | | | |
|---|---|---|---|---|---|---|
| ENSG00000254990 | AP001781,1 | 0,591610399 | 1,506927907 | 0,66940571 | 0,961867165 | novel transcript |
| ENSG00000236498 | AC107081,1 | 0,591527415 | 1,506841231 | 0,2910109 | 0,801689099 | novel transcript |
| ENSG00000125741 | OPA3 | 0,591406578 | 1,506715027 | 0,13145954 | 0,579274292 | outer mitochondrial membrane lipid metabolism regulator OPA3 |
| ENSG00000206190 | ATP10A | 0,591241385 | 1,506542513 | 0,16449135 | 0,643636727 | ATPase phospholipid transporting 10A (putative) |
| ENSG00000229791 | AL356259,1 | 0,591165501 | 1,506463273 | 0,5858844 | 0,945246954 | novel transcript |
| ENSG00000119714 | GPR68 | 0,591133799 | 1,50643017 | 0,40237259 | 0,87718264 | G protein-coupled receptor 68 |
| ENSG00000188157 | AGRN | 0,591053398 | 1,50634622 | 0,29820816 | 0,807621278 | agrin |
| ENSG00000145022 | TCTA | 0,590797318 | 1,506078865 | 0,02611859 | 0,225509966 | T cell leukemia translocation altered |
| ENSG00000169223 | LMAN2 | 0,590795114 | 1,506076564 | 0,06344371 | 0,394334795 | lectin, mannose binding 2 |
| ENSG00000276359 | Metazoa_SRP | 0,59074743 | 1,506026786 | 0,665499 | 0,960946962 | Metazoan signal recognition particle RNA |
| ENSG00000245928 | SDAD1-AS1 | 0,590669267 | 1,505945194 | 0,48579703 | 0,913997663 | SDAD1 antisense RNA 1 |
| ENSG00000151414 | NEK7 | 0,590589317 | 1,505861741 | 0,15042434 | 0,616446007 | NIMA related kinase 7 |
| ENSG00000112893 | MAN2A1 | 0,590570104 | 1,505841687 | 0,06931804 | 0,415287103 | mannosidase alpha class 2A member 1 |
| ENSG00000149782 | PLCB3 | 0,590564 | 1,505835316 | 0,12687264 | 0,568970141 | phospholipase C beta 3 |
| ENSG00000252699 | SNORA21B | 0,590326618 | 1,505587565 | 0,42613173 | 0,892133432 | small nucleolar RNA, H/ACA box 21B |
| ENSG00000087460 | GNAS | 0,590279824 | 1,505538733 | 0,00837089 | 0,100928472 | GNAS complex locus |
| ENSG00000203573 | Metazoa_SRP | 0,590252935 | 1,505510672 | 0,63969108 | 0,956018518 | Metazoan signal recognition particle RNA |
| ENSG00000272414 | FAM47E-STBD1 | 0,590105834 | 1,505357174 | 0,15284942 | 0,62133458 | FAM47E-STBD1 readthrough |
| ENSG00000114450 | GNB4 | 0,589652128 | 1,504883836 | 0,09515001 | 0,489674048 | G protein subunit beta 4 |
| ENSG00000214212 | C19orf38 | 0,589565893 | 1,504793887 | 0,54092316 | 0,933385624 | chromosome 19 open reading frame 38 |
| ENSG00000151718 | WWC2 | 0,589513226 | 1,504738953 | 0,00756332 | 0,093795931 | WW and C2 domain containing 2 |
| ENSG00000262815 | AC087501,2 | 0,589469193 | 1,504693028 | 0,5123322 | 0,923445033 | novel transcript, antisense to STX8 |
| ENSG00000152291 | TGOLN2 | 0,589376105 | 1,504595943 | 0,00957461 | 0,111275759 | trans-golgi network protein 2 |
| ENSG00000010327 | STAB1 | 0,589207316 | 1,504419922 | 0,54425557 | 0,934381824 | stabilin 1 |
| ENSG00000285392 | AC270276,1 | 0,588846924 | 1,504044158 | 0,27497219 | 0,786383635 | novel transcript |
| ENSG00000162889 | MAPKAPK2 | 0,588677167 | 1,503867192 | 0,01322102 | 0,140068402 | MAPK activated protein kinase 2 |
| ENSG00000229750 | AC096649,1 | 0,58848442 | 1,503666286 | 0,68230806 | 0,964749521 | novel transcript |
| ENSG00000150281 | CTF1 | 0,588286407 | 1,503459918 | 0,30443688 | 0,813181541 | cardiotrophin 1 |
| ENSG00000221838 | AP4M1 | 0,588221934 | 1,503392732 | 0,03381706 | 0,267456737 | adaptor related protein complex 4 subunit mu 1 |
| ENSG00000179104 | TMTC2 | 0,588141816 | 1,503309245 | 0,10502796 | 0,517182893 | transmembrane O-mannosyltransferase targeting cadherins 2 |
| ENSG00000013563 | DNASE1L1 | 0,587839766 | 1,502994538 | 0,06467333 | 0,398771343 | deoxyribonuclease 1 like 1 |

| | | | | | | |
|---|---|---|---|---|---|---|
| ENSG00000175283 | DOLK | 0,587801864 | 1,502955052 | 0,04414328 | 0,317117513 | dolichol kinase |
| ENSG00000032444 | PNPLA6 | 0,587676037 | 1,502823975 | 0,15874832 | 0,632735484 | patatin like phospholipase domain containing 6 |
| ENSG00000196564 | BLOC1S6P1 | 0,587623303 | 1,502769045 | 0,62028568 | 0,952793934 | BLOC1S6 pseudogene 1 |
| ENSG00000172350 | ABCG4 | 0,587608474 | 1,502753598 | 0,18758903 | 0,681556081 | ATP binding cassette subfamily G member 4 |
| ENSG00000163364 | LINC01116 | 0,587563656 | 1,502706915 | 0,53153085 | 0,931291085 | long intergenic non-protein coding RNA 1116 |
| ENSG00000187630 | DHRS4L2 | 0,587187381 | 1,502315039 | 0,14784383 | 0,612372753 | dehydrogenase/reductase 4 like 2 |
| ENSG00000100219 | XBP1 | 0,587176573 | 1,502303784 | 0,00481164 | 0,066887986 | X-box binding protein 1 |
| ENSG00000250565 | ATP6V1E2 | 0,587057891 | 1,502180204 | 0,07166257 | 0,421847873 | ATPase H+ transporting V1 subunit E2 |
| ENSG00000103226 | NOMO3 | 0,587044573 | 1,502166337 | 0,17762922 | 0,664043981 | NODAL modulator 3 |
| ENSG00000122224 | LY9 | 0,586967868 | 1,502086472 | 0,51297532 | 0,923725914 | lymphocyte antigen 9 |
| ENSG00000089820 | ARHGAP4 | 0,586837558 | 1,501950804 | 0,2161073 | 0,72322574 | Rho GTPase activating protein 4 |
| ENSG00000257246 | PHBP19 | 0,586610993 | 1,501714951 | 0,35701701 | 0,848610955 | prohibitin pseudogene 19 |
| ENSG00000271806 | AL590822,2 | 0,586553791 | 1,50165541 | 0,50852699 | 0,922427526 | novel transcript, antisense PRKCZ |
| ENSG00000286379 | AL591845,1 | 0,586095397 | 1,501178358 | 0,33387165 | 0,83425218 | novel transcript |
| ENSG00000269903 | AC025165,4 | 0,586050352 | 1,501131488 | 0,47997655 | 0,912847708 | novel transcript |
| ENSG00000171310 | CHST11 | 0,585996583 | 1,501075543 | 0,04384166 | 0,31589834 | carbohydrate sulfotransferase 11 |
| ENSG00000278920 | AC005005,4 | 0,585980862 | 1,501059186 | 0,45091719 | 0,902062594 | novel transcript, sense overlapping SMTN |
| ENSG00000278016 | TIGD5 | 0,585960058 | 1,50103754 | 0,18758697 | 0,681556081 | tigger transposable element derived 5 |
| ENSG00000236490 | TAPBP | 0,585767277 | 1,500836976 | 0,70565542 | 0,969485452 | TAP binding protein |
| ENSG00000250602 | AC093535,1 | 0,585541448 | 1,500602064 | 0,34802402 | 0,842479505 | novel transcript, antisense to GRAMD3 |
| ENSG00000141367 | CLTC | 0,58533663 | 1,500389041 | 0,06823275 | 0,411581814 | clathrin heavy chain |
| ENSG00000213648 | SULT1A4 | 0,58525499 | 1,500304138 | 0,4709413 | 0,908869685 | sulfotransferase family 1A member 4 |
| ENSG00000257900 | AL162632,1 | 0,585219877 | 1,500267623 | 0,43725043 | 0,897021798 | novel transcript |
| ENSG00000173421 | IHO1 | 0,585062299 | 1,500103766 | 0,47110581 | 0,908869685 | interactor of HORMAD1 1 |
| ENSG00000114353 | GNAI2 | 0,585045923 | 1,500086738 | 0,02127346 | 0,194756461 | G protein subunit alpha i2 |
| ENSG00000169641 | LUZP1 | 0,585010656 | 1,500050069 | 0,0038826 | 0,057418855 | leucine zipper protein 1 |
| ENSG00000141540 | TTYH2 | 0,584998391 | 1,500037317 | 0,03747447 | 0,285896037 | tweety family member 2 |
| ENSG00000234042 | AL591438,2 | 0,584918597 | 1,499954353 | 0,58113302 | 0,943919685 | novel pseudogene |
| ENSG00000167208 | SNX20 | 0,584913581 | 1,499949138 | 0,58415453 | 0,94460573 | sorting nexin 20 |
| ENSG00000026950 | BTN3A1 | 0,584767568 | 1,499797338 | 0,09711818 | 0,494995584 | butyrophilin subfamily 3 member A1 |
| ENSG00000132874 | SLC14A2 | 0,58471397 | 1,49974162 | 0,39006769 | 0,870057277 | solute carrier family 14 member 2 |
| ENSG00000275342 | PRAG1 | 0,584415952 | 1,499431849 | 0,1631771 | 0,641545388 | PEAK1 related, kinase-activating pseudokinase 1 |
| ENSG00000164081 | TEX264 | 0,5844007 | 1,499415998 | 0,05318526 | 0,354689414 | testis expressed 264, ER-phagy receptor |

| | | | | | | |
|---|---|---|---|---|---|---|
| ENSG00000275086 | TFPT | 0,584354927 | 1,499368426 | 0,23125131 | 0,739268552 | TCF3 fusion partner |
| ENSG00000278161 | TFPT | 0,584354927 | 1,499368426 | 0,23125131 | 0,739268552 | TCF3 fusion partner |
| ENSG00000276022 | TFPT | 0,584354927 | 1,499368426 | 0,23125131 | 0,739268552 | TCF3 fusion partner |
| ENSG00000276323 | TFPT | 0,584354927 | 1,499368426 | 0,23125131 | 0,739268552 | TCF3 fusion partner |
| ENSG00000276296 | TFPT | 0,584354927 | 1,499368426 | 0,23125131 | 0,739268552 | TCF3 fusion partner |
| ENSG00000274073 | TFPT | 0,584354927 | 1,499368426 | 0,23125131 | 0,739268552 | TCF3 fusion partner |
| ENSG00000276263 | TFPT | 0,584354927 | 1,499368426 | 0,23125131 | 0,739268552 | TCF3 fusion partner |
| ENSG00000273833 | TFPT | 0,584354927 | 1,499368426 | 0,23125131 | 0,739268552 | TCF3 fusion partner |
| ENSG00000176635 | HORMAD2 | 0,584247251 | 1,499256524 | 0,73122921 | 0,974353474 | HORMA domain containing 2 |
| ENSG00000136732 | GYPC | 0,584074 | 1,499076491 | 0,01821636 | 0,175614018 | glycophorin C (Gerbich blood group) |
| ENSG00000185453 | ZSWIM9 | 0,584070418 | 1,499072769 | 0,46830593 | 0,907940653 | zinc finger SWIM-type containing 9 |
| ENSG00000177551 | NHLH2 | 0,583921555 | 1,498918098 | 0,31023695 | 0,817687618 | nescient helix-loop-helix 2 |
| ENSG00000261117 | AC009486,1 | 0,58383687 | 1,498830114 | 0,32692263 | 0,82867756 | novel transcript |
| ENSG00000285684 | AL137781,1 | 0,583834845 | 1,498828011 | 0,4494168 | 0,902062594 | novel transcript |
| ENSG00000253712 | LINC02886 | 0,583737154 | 1,498726522 | 0,40833437 | 0,881397793 | long intergenic non-protein coding RNA 2886 |
| ENSG00000226416 | MRPL23-AS1 | 0,583682921 | 1,498670184 | 0,68972612 | 0,966868992 | MRPL23 antisense RNA 1 |
| ENSG00000288320 | MRPL23-AS1 | 0,583682921 | 1,498670184 | 0,68972612 | 0,966868992 | MRPL23 antisense RNA 1 |
| ENSG00000228742 | LINC02577 | 0,583667952 | 1,498654634 | 0,35216809 | 0,844912682 | long intergenic non-protein coding RNA 2577 |
| ENSG00000186603 | HPDL | 0,583453332 | 1,498431707 | 0,12194538 | 0,558362693 | 4-hydroxyphenylpyruvate dioxygenase like |
| ENSG00000137266 | SLC22A23 | 0,583329058 | 1,498302636 | 0,03465075 | 0,271577099 | solute carrier family 22 member 23 |
| ENSG00000168374 | ARF4 | 0,583013603 | 1,497975058 | 0,11387832 | 0,539356413 | ADP ribosylation factor 4 |
| ENSG00000108384 | RAD51C | 0,582972392 | 1,497932269 | 0,10094725 | 0,504961114 | RAD51 paralog C |
| ENSG00000154655 | L3MBTL4 | 0,582934857 | 1,497893297 | 0,29120611 | 0,801926163 | L3MBTL histone methyl-lysine binding protein 4 |
| ENSG00000130287 | NCAN | 0,582855199 | 1,497810594 | 0,04550944 | 0,323249559 | neurocan |
| ENSG00000078795 | PKD2L2 | 0,582445765 | 1,497385578 | 0,39472135 | 0,871888817 | polycystin 2 like 2, transient receptor potential cation channel |
| ENSG00000261512 | AC092368,3 | 0,582384588 | 1,497322084 | 0,3122217 | 0,818547047 | novel transcript |
| ENSG00000105429 | MEGF8 | 0,582304571 | 1,497239039 | 0,22777141 | 0,736053199 | multiple EGF like domains 8 |
| ENSG00000236882 | LINC01554 | 0,582295075 | 1,497229184 | 0,35388978 | 0,845346751 | long intergenic non-protein coding RNA 1554 |
| ENSG00000100284 | TOM1 | 0,582160178 | 1,497089195 | 0,12782879 | 0,570756819 | target of myb1 membrane trafficking protein |
| ENSG00000283741 | SMCR8 | 0,582024589 | 1,4969485 | 0,76875635 | 0,977381742 | SMCR8-C9orf72 complex subunit |
| ENSG00000244038 | DDOST | 0,581839643 | 1,496756611 | 0,00770814 | 0,095251919 | dolichyl-diphosphooligosaccharide--protein glycosyltransferase non-catalytic subunit |

| | | | | | | |
|---|---|---|---|---|---|---|
| ENSG00000189238 | LINC00943 | 0,581823476 | 1,496739839 | 0,47718501 | 0,91143003 | long intergenic non-protein coding RNA 943 |
| ENSG00000131873 | CHSY1 | 0,581487102 | 1,496390904 | 0,01600136 | 0,161013527 | chondroitin sulfate synthase 1 |
| ENSG00000214210 | AC084017,1 | 0,581469014 | 1,496372143 | 0,48118263 | 0,912911021 | neugrin, neurite outgrowth associated (NGRN) pseudogene |
| ENSG00000282825 | CTIF | 0,581451902 | 1,496354395 | 0,06463028 | 0,398619244 | cap binding complex dependent translation initiation factor |
| ENSG00000164051 | CCDC51 | 0,581313967 | 1,496211336 | 0,07489119 | 0,432482457 | coiled-coil domain containing 51 |
| ENSG00000283149 | AC068631,2 | 0,581174173 | 1,496066364 | 0,40151481 | 0,876898854 | novel protein |
| ENSG00000245685 | FRG1-DT | 0,58106017 | 1,495948149 | 0,47638293 | 0,91143003 | FRG1 divergent transcript |
| ENSG00000140279 | DUOX2 | 0,580963529 | 1,495847943 | 0,398336 | 0,875040521 | dual oxidase 2 |
| ENSG00000265099 | AC090774,1 | 0,580953833 | 1,49583789 | 0,58920789 | 0,946705487 | novel pseudogene |
| ENSG00000134594 | RAB33A | 0,580875823 | 1,495757009 | 0,08237658 | 0,45655405 | RAB33A, member RAS oncogene family |
| ENSG00000272754 | AL133245,1 | 0,580845555 | 1,495725628 | 0,49977952 | 0,920310866 | novel transcript |
| ENSG00000182111 | ZNF716 | 0,580827439 | 1,495706846 | 0,58448145 | 0,944792307 | zinc finger protein 716 |
| ENSG00000157510 | AFAP1L1 | 0,580817677 | 1,495696725 | 0,19116031 | 0,686867719 | actin filament associated protein 1 like 1 |
| ENSG00000258884 | LINC02321 | 0,580743661 | 1,495619992 | 0,49196934 | 0,916995341 | long intergenic non-protein coding RNA 2321 |
| ENSG00000267221 | C17orf113 | 0,580248627 | 1,495106886 | 0,41333469 | 0,884678298 | chromosome 17 open reading frame 113 |
| ENSG00000231252 | AC099792,1 | 0,579877565 | 1,494722393 | 0,2160121 | 0,723046035 | novel transcript |
| ENSG00000261796 | ISY1-RAB43 | 0,579824092 | 1,494666992 | 0,06662855 | 0,40559155 | ISY1-RAB43 readthrough |
| ENSG00000026036 | RTEL1-TNFRSF6 | 0,57981161 | 1,494654061 | 0,16327224 | 0,641596686 | RTEL1-TNFRSF6B readthrough (NMD candidate) |
| ENSG00000256288 | LINC02617 | 0,579553595 | 1,494386778 | 0,48879858 | 0,91557151 | long intergenic non-protein coding RNA 2617 |
| ENSG00000187796 | CARD9 | 0,57948336 | 1,494314028 | 0,20801847 | 0,712494639 | caspase recruitment domain family member 9 |
| ENSG00000113407 | TARS1 | 0,579366553 | 1,494193046 | 0,01882942 | 0,17950267 | threonyl-tRNA synthetase 1 |
| ENSG00000071575 | TRIB2 | 0,578730074 | 1,493533993 | 0,0495747 | 0,339816563 | tribbles pseudokinase 2 |
| ENSG00000176840 | MIR7-3HG | 0,578673813 | 1,493475751 | 0,40725541 | 0,880314995 | MIR7-3 host gene |
| ENSG00000116521 | SCAMP3 | 0,578633943 | 1,493434478 | 0,01339504 | 0,141624131 | secretory carrier membrane protein 3 |
| ENSG00000204219 | TCEA3 | 0,578534058 | 1,493331084 | 0,31232241 | 0,818547047 | transcription elongation factor A3 |
| ENSG00000282941 | TCEA3 | 0,578534058 | 1,493331084 | 0,31232241 | 0,818547047 | transcription elongation factor A3 |
| ENSG00000248079 | DPH6-DT | 0,57850959 | 1,493305757 | 0,57217863 | 0,942475308 | DPH6 divergent transcript |
| ENSG00000259207 | ITGB3 | 0,578493673 | 1,493289282 | 0,13592855 | 0,587841041 | integrin subunit beta 3 |
| ENSG00000249129 | SUDS3P1 | 0,578324948 | 1,49311465 | 0,69379623 | 0,967340633 | SDS3 homolog, SIN3A corepressor complex component pseudogene 1 |

| | | | | | | |
|---|---|---|---|---|---|---|
| ENSG00000274681 | SUDS3P1 | 0,578324948 | 1,49311465 | 0,69379623 | 0,967340633 | SDS3 homolog, SIN3A corepressor complex component pseudogene 1 |
| ENSG00000169490 | TM2D2 | 0,578177759 | 1,492962325 | 0,03178807 | 0,257200368 | TM2 domain containing 2 |
| ENSG00000205352 | PRR13 | 0,578135223 | 1,492918307 | 0,02915542 | 0,24236233 | proline rich 13 |
| ENSG00000200693 | U3 | 0,578116165 | 1,492898586 | 0,51932833 | 0,926800715 | Small nucleolar RNA U3 |
| ENSG00000235070 | AC062015,1 | 0,57810012 | 1,492881983 | 0,51549467 | 0,924901769 | novel transcript |
| ENSG00000113319 | RASGRF2 | 0,578061449 | 1,492841966 | 0,24019423 | 0,750118118 | Ras protein specific guanine nucleotide releasing factor 2 |
| ENSG00000230013 | CT70 | 0,577782267 | 1,492553109 | 0,65557704 | 0,957444211 | cancer/testis associated transcript 70 |
| ENSG00000119523 | ALG2 | 0,57777816 | 1,492548859 | 0,01683088 | 0,16634097 | ALG2 alpha-1,3/1,6-mannosyltransferase |
| ENSG00000126432 | PRDX5 | 0,577706398 | 1,492474619 | 0,05531351 | 0,363130125 | peroxiredoxin 5 |
| ENSG00000162105 | SHANK2 | 0,57721081 | 1,491962019 | 0,05272495 | 0,35267286 | SH3 and multiple ankyrin repeat domains 2 |
| ENSG00000274145 | RILP | 0,57711185 | 1,491859684 | 0,62689635 | 0,952793934 | Rab interacting lysosomal protein |
| ENSG00000163888 | CAMK2N2 | 0,577049578 | 1,491795291 | 0,06122134 | 0,385192985 | calcium/calmodulin dependent protein kinase II inhibitor 2 |
| ENSG00000136156 | ITM2B | 0,577034178 | 1,491779367 | 0,20376388 | 0,70430889 | integral membrane protein 2B |
| ENSG00000281000 | SNORD3D | 0,577022881 | 1,491767686 | 0,41608349 | 0,885986046 | small nucleolar RNA, C/D box 3D |
| ENSG00000213967 | ZNF726 | 0,576955931 | 1,491698461 | 0,00863034 | 0,102939044 | zinc finger protein 726 |
| ENSG00000230916 | MTCO1P53 | 0,576953956 | 1,491696418 | 0,42262784 | 0,890403187 | MT-CO1 pseudogene 53 |
| ENSG00000177697 | CD151 | 0,576871971 | 1,491611651 | 0,17226539 | 0,656553102 | CD151 molecule (Raph blood group) |
| ENSG00000272682 | AC004471,2 | 0,576696562 | 1,491430305 | 0,61484908 | 0,952793934 | novel transcript |
| ENSG00000198752 | CDC42BPB | 0,576554234 | 1,491283177 | 0,07701485 | 0,438408786 | CDC42 binding protein kinase beta |
| ENSG00000213614 | HEXA | 0,57651276 | 1,491240307 | 0,03288348 | 0,262798865 | hexosaminidase subunit alpha |
| ENSG00000101307 | SIRPB1 | 0,576327755 | 1,491049089 | 0,4757788 | 0,911331229 | signal regulatory protein beta 1 |
| ENSG00000234619 | RPL7P11 | 0,576109628 | 1,490823668 | 0,43506575 | 0,896272827 | ribosomal protein L7 pseudogene 11 |
| ENSG00000220891 | LL22NC03-63E[illegible] | 0,575943288 | 1,490651788 | 0,68188845 | 0,964654539 | uncharacterized LOC648691 |
| ENSG00000197780 | TAF13 | 0,575881875 | 1,490588336 | 0,07153581 | 0,421423698 | TATA-box binding protein associated factor 13 |
| ENSG00000102158 | MAGT1 | 0,575753575 | 1,490455782 | 0,00505437 | 0,06927661 | magnesium transporter 1 |
| ENSG00000283384 | AL138694,1 | 0,575648488 | 1,49034722 | 0,43590283 | 0,896422069 | novel transcript |
| ENSG00000232748 | AC135050,1 | 0,57559796 | 1,490295024 | 0,22412886 | 0,732890339 | novel transcript, antisense to ZNF668 |
| ENSG00000173013 | CCDC96 | 0,575324014 | 1,490012066 | 0,23912636 | 0,748727732 | coiled-coil domain containing 96 |
| ENSG00000126254 | RBM42 | 0,575160751 | 1,489843458 | 0,13990575 | 0,596239558 | RNA binding motif protein 42 |
| ENSG00000215208 | KRT18P60 | 0,57514903 | 1,489831355 | 0,54673285 | 0,935404974 | keratin 18 pseudogene 60 |

| | | | | | | |
|---|---|---|---|---|---|---|
| ENSG00000155876 | RRAGA | 0,575126619 | 1,489808211 | 0,0145165 | 0,15062851 | Ras related GTP binding A |
| ENSG00000138413 | IDH1 | 0,57508493 | 1,489765161 | 0,16477399 | 0,643964432 | isocitrate dehydrogenase (NADP(+)) 1 |
| ENSG00000257594 | GALNT4 | 0,574909271 | 1,489583782 | 0,04876629 | 0,336414216 | polypeptide N-acetylgalactosaminyltransferase 4 |
| ENSG00000176054 | RPL23P2 | 0,574787699 | 1,489458264 | 0,68284224 | 0,964914784 | ribosomal protein L23 pseudogene 2 |
| ENSG00000267283 | AC005306,1 | 0,574771499 | 1,48944154 | 0,7381679 | 0,974353474 | novel transcript, antisense to BTBD2 |
| ENSG00000104142 | VPS18 | 0,574585815 | 1,489249851 | 0,11553451 | 0,543019233 | VPS18 core subunit of CORVET and HOPS complexes |
| ENSG00000259687 | LINC01220 | 0,574472977 | 1,489133377 | 0,36035762 | 0,850858747 | long intergenic non-protein coding RNA 1220 |
| ENSG00000173039 | RELA | 0,574274757 | 1,488928791 | 0,01750961 | 0,171283038 | RELA proto-oncogene, NF-kB subunit |
| ENSG00000259219 | AC084855,2 | 0,574211547 | 1,488863556 | 0,50095362 | 0,920520138 | novel transcript, antisense to ADAMTS17 |
| ENSG00000239389 | PCDHA13 | 0,57418451 | 1,488835654 | 0,07962496 | 0,447576616 | protocadherin alpha 13 |
| ENSG00000229498 | AC105053,1 | 0,573840832 | 1,488481027 | 0,53537606 | 0,932037868 | novel transcript |
| ENSG00000168917 | SLC35G2 | 0,573742378 | 1,488379452 | 0,01965328 | 0,184405835 | solute carrier family 35 member G2 |
| ENSG00000179041 | RRS1 | 0,573685487 | 1,48832076 | 0,20439537 | 0,70555556 | ribosome biogenesis regulator 1 homolog |
| ENSG00000171443 | ZNF524 | 0,573665284 | 1,488299919 | 0,2617553 | 0,773497425 | zinc finger protein 524 |
| ENSG00000218682 | AC064847,1 | 0,573648365 | 1,488282465 | 0,59642686 | 0,948282297 | enolase 2 (gamma, neuronal) (ENO2) pseudogene |
| ENSG00000159202 | UBE2Z | 0,573602917 | 1,488235581 | 0,01386233 | 0,145173348 | ubiquitin conjugating enzyme E2 Z |
| ENSG00000105369 | CD79A | 0,573590161 | 1,488222423 | 0,67836553 | 0,9640473 | CD79a molecule |
| ENSG00000132467 | UTP3 | 0,573514452 | 1,488144327 | 0,04581049 | 0,324471967 | UTP3 small subunit processome component |
| ENSG00000107281 | NPDC1 | 0,573262104 | 1,487884051 | 0,19316929 | 0,689970564 | neural proliferation, differentiation and control 1 |
| ENSG00000269927 | AC004817,3 | 0,572650294 | 1,487253212 | 0,56057658 | 0,940170934 | novel transcript |
| ENSG00000143761 | ARF1 | 0,572457077 | 1,487054041 | 0,00691203 | 0,087880377 | ADP ribosylation factor 1 |
| ENSG00000079263 | SP140 | 0,572348615 | 1,486942248 | 0,48006122 | 0,912847708 | SP140 nuclear body protein |
| ENSG00000235641 | LINC00484 | 0,572094382 | 1,486680241 | 0,59880447 | 0,948854865 | long intergenic non-protein coding RNA 484 |
| ENSG00000196700 | ZNF512B | 0,571985888 | 1,486568443 | 0,26564266 | 0,776863208 | zinc finger protein 512B |
| ENSG00000106400 | ZNHIT1 | 0,571815539 | 1,486392924 | 0,07078847 | 0,418829978 | zinc finger HIT-type containing 1 |
| ENSG00000179562 | GCC1 | 0,571704498 | 1,486278525 | 0,07235601 | 0,424120881 | GRIP and coiled-coil domain containing 1 |
| ENSG00000287176 | AC099517,1 | 0,571621715 | 1,486193243 | 0,54662843 | 0,935401821 | novel transcript |
| ENSG00000163959 | SLC51A | 0,571602374 | 1,486173319 | 0,1300262 | 0,576206833 | solute carrier family 51 subunit alpha |
| ENSG00000287519 | AC004921,2 | 0,571567089 | 1,486136971 | 0,53165997 | 0,931291085 | novel transcript |
| ENSG00000126231 | PROZ | 0,571498496 | 1,486066314 | 0,59207305 | 0,947416795 | protein Z, vitamin K dependent plasma glycoprotein |
| ENSG00000047644 | WWC3 | 0,571437558 | 1,486003546 | 0,01757572 | 0,171664029 | WWC family member 3 |
| ENSG00000132688 | NES | 0,571382401 | 1,485946734 | 0,41794592 | 0,886992289 | nestin |
| ENSG00000006756 | ARSD | 0,571352774 | 1,485916218 | 0,05958969 | 0,379664157 | arylsulfatase D |

| | | | | | | |
|---|---|---|---|---|---|---|
| ENSG00000167703 | SLC43A2 | 0,571141774 | 1,485698914 | 0,20927127 | 0,713678247 | solute carrier family 43 member 2 |
| ENSG00000281180 | AP000350,11 | 0,571111662 | 1,485667904 | 0,75380969 | 0,975901808 | novel transcript, antisense to SLC2A11 |
| ENSG00000262102 | DSEL | 0,570997158 | 1,485549994 | 0,27401209 | 0,7862019 | dermatan sulfate epimerase like |
| ENSG00000105929 | ATP6V0A4 | 0,570761715 | 1,485307577 | 0,70006517 | 0,968522643 | ATPase H+ transporting V0 subunit a4 |
| ENSG00000267121 | AC008105,3 | 0,570608195 | 1,485149531 | 0,35956366 | 0,849982153 | novel transcript, antisense to FMNL1 |
| ENSG00000140691 | ARMC5 | 0,570527014 | 1,485065963 | 0,22398934 | 0,732710917 | armadillo repeat containing 5 |
| ENSG00000288585 | AC010184,1 | 0,570490709 | 1,485028593 | 0,35513209 | 0,846584484 | novel transcript |
| ENSG00000269313 | MAGIX | 0,570306418 | 1,484838906 | 0,3105393 | 0,817751312 | MAGI family member, X-linked |
| ENSG00000084652 | TXLNA | 0,570249694 | 1,484780527 | 0,03682939 | 0,282977152 | taxilin alpha |
| ENSG00000040633 | PHF23 | 0,570177571 | 1,484706301 | 0,0208096 | 0,192028119 | PHD finger protein 23 |
| ENSG00000175544 | CABP4 | 0,570084415 | 1,484610436 | 0,44010328 | 0,897906863 | calcium binding protein 4 |
| ENSG00000122218 | COPA | 0,570066844 | 1,484592354 | 0,00379094 | 0,056324176 | COPI coat complex subunit alpha |
| ENSG00000142409 | ZNF787 | 0,569986092 | 1,484509259 | 0,22001149 | 0,728200468 | zinc finger protein 787 |
| ENSG00000103266 | STUB1 | 0,569851388 | 1,484370658 | 0,04746695 | 0,331177592 | STIP1 homology and U-box containing protein 1 |
| ENSG00000068903 | SIRT2 | 0,56977338 | 1,484290398 | 0,11291101 | 0,53680967 | sirtuin 2 |
| ENSG00000158806 | NPM2 | 0,569754894 | 1,48427138 | 0,42042132 | 0,888563911 | nucleophosmin/nucleoplasmin 2 |
| ENSG00000267262 | AC011444,2 | 0,569477778 | 1,483986305 | 0,63499698 | 0,955158181 | novel transcript, antisense to RFX2 |
| ENSG00000259515 | AL117209,1 | 0,569380813 | 1,483886568 | 0,32645389 | 0,828450693 | novel transcript, antisense to CDC42BPB |
| ENSG00000237686 | AL109615,3 | 0,569164269 | 1,483663858 | 0,46538942 | 0,90786072 | novel transcript, antisense to C6orf223 |
| ENSG00000287170 | AC010327,8 | 0,569102229 | 1,483600058 | 0,42796341 | 0,89282532 | novel transcript, antisense to PPP1R12C |
| ENSG00000275019 | LILRB3 | 0,569029854 | 1,483525633 | 0,57903191 | 0,943828882 | leukocyte immunoglobulin like receptor B3 |
| ENSG00000144857 | BOC | 0,568555507 | 1,483037941 | 0,01042304 | 0,118366144 | BOC cell adhesion associated, oncogene regulated |
| ENSG00000158552 | ZFAND2B | 0,568410833 | 1,482889228 | 0,04037433 | 0,300178807 | zinc finger AN1-type containing 2B |
| ENSG00000282826 | FRG1CP | 0,568139438 | 1,482610299 | 0,03780592 | 0,287579371 | FSHD region gene 1 family member C, pseudogene |
| ENSG00000142686 | C1orf216 | 0,568091597 | 1,482561135 | 0,09607342 | 0,492399878 | chromosome 1 open reading frame 216 |
| ENSG00000116685 | KIAA2013 | 0,568041893 | 1,482510059 | 0,06538128 | 0,401164726 | KIAA2013 |
| ENSG00000182827 | ACBD3 | 0,568023477 | 1,482491135 | 0,01810259 | 0,17489473 | acyl-CoA binding domain containing 3 |
| ENSG00000184178 | SCFD2 | 0,567933444 | 1,482398621 | 0,01039936 | 0,118280376 | sec1 family domain containing 2 |
| ENSG00000107077 | KDM4C | 0,567725272 | 1,482184735 | 0,04325944 | 0,313869373 | lysine demethylase 4C |
| ENSG00000267432 | DNAH17-AS1 | 0,567705409 | 1,482164328 | 0,52209617 | 0,927599011 | DNAH17 antisense RNA 1 |
| ENSG00000114767 | RRP9 | 0,567574494 | 1,482029838 | 0,17437478 | 0,65976679 | ribosomal RNA processing 9, U3 small nucleolar RNA binding protein |
| ENSG00000138073 | PREB | 0,567552346 | 1,482007086 | 0,0400449 | 0,298798028 | prolactin regulatory element binding |

| | | | | | | |
|---|---|---|---|---|---|---|
| ENSG00000243646 | IL10RB | 0,567356179 | 1,481805588 | 0,03722264 | 0,284946761 | interleukin 10 receptor subunit beta |
| ENSG00000286659 | Z99716,2 | 0,567246187 | 1,481692618 | 0,53387084 | 0,931381467 | novel transcript, antisense to POLDIP3 |
| ENSG00000267281 | ATF7-NPFF | 0,567169971 | 1,481614344 | 0,11059028 | 0,530942044 | ATF7-NPFF readthrough |
| ENSG00000172000 | ZNF556 | 0,567116383 | 1,481559312 | 0,46134784 | 0,906165604 | zinc finger protein 556 |
| ENSG00000226437 | TRIM39 | 0,566990794 | 1,481430344 | 0,66086647 | 0,959296562 | tripartite motif containing 39 |
| ENSG00000087088 | BAX | 0,566911486 | 1,48134891 | 0,06007507 | 0,381239005 | BCL2 associated X, apoptosis regulator |
| ENSG00000257763 | OR5BK1P | 0,566721664 | 1,481154014 | 0,52771191 | 0,929546857 | olfactory receptor family 5 subfamily BK member 1 pseudogene |
| ENSG00000172940 | SLC22A13 | 0,56653705 | 1,480964491 | 0,5805516 | 0,943832501 | solute carrier family 22 member 13 |
| ENSG00000112658 | SRF | 0,566330879 | 1,480752867 | 0,09319392 | 0,485117333 | serum response factor |
| ENSG00000163792 | TCF23 | 0,566244691 | 1,480664408 | 0,41306366 | 0,884663685 | transcription factor 23 |
| ENSG00000277797 | AL359693,1 | 0,565978874 | 1,48039162 | 0,76783445 | 0,977309816 | novel transcript |
| ENSG00000171219 | CDC42BPG | 0,565814164 | 1,480222616 | 0,45345629 | 0,902754541 | CDC42 binding protein kinase gamma |
| ENSG00000228334 | AC024028,1 | 0,565611982 | 1,48001519 | 0,4856579 | 0,913997663 | novel transcript |
| ENSG00000272689 | AC004832,4 | 0,56539627 | 1,479793913 | 0,53599992 | 0,932139691 | novel transcript, sense intonic to |
| ENSG00000274516 | FAM74A6 | 0,565342401 | 1,47973866 | 0,488734 | 0,91557151 | family with sequence similarity 74 member A6 |
| ENSG00000140553 | UNC45A | 0,565217327 | 1,479610381 | 0,14250305 | 0,600806836 | unc-45 myosin chaperone A |
| ENSG00000142224 | IL19 | 0,565020153 | 1,479408175 | 0,4600542 | 0,905443183 | interleukin 19 |
| ENSG00000234290 | AC116366,1 | 0,565002016 | 1,479389576 | 0,31869082 | 0,823501926 | novel transcript |
| ENSG00000287059 | AC090004,2 | 0,564880786 | 1,479265268 | 0,62722184 | 0,952793934 | novel transcript |
| ENSG00000100242 | SUN2 | 0,564851625 | 1,479235368 | 0,1747436 | 0,660057513 | Sad1 and UNC84 domain containing 2 |
| ENSG00000155926 | SLA | 0,564833428 | 1,47921671 | 0,50409919 | 0,921400408 | Src like adaptor |
| ENSG00000149451 | ADAM33 | 0,564612934 | 1,478990652 | 0,17477195 | 0,660057513 | ADAM metallopeptidase domain 33 |
| ENSG00000285540 | AC022023,2 | 0,564341947 | 1,478712874 | 0,44948299 | 0,902062594 | novel transcript |
| ENSG00000124479 | NDP | 0,564300068 | 1,47866995 | 0,30120452 | 0,810679941 | norrin cystine knot growth factor NDP |
| ENSG00000228484 | AL137025,1 | 0,564157296 | 1,478523625 | 0,53707396 | 0,932883277 | novel transcript |
| ENSG00000216819 | TUBB2BP1 | 0,564114684 | 1,478479955 | 0,54012003 | 0,933385624 | tubulin beta 2B class IIb pseudogene 1 |
| ENSG00000197099 | AC068631,1 | 0,564095162 | 1,478459949 | 0,48721022 | 0,914678448 | novel transcript, antisense to KNG1, HRG, FETUB, AHSG and DNAJB11 |
| ENSG00000164951 | PDP1 | 0,564003932 | 1,47836646 | 0,05724577 | 0,369932153 | pyruvate dehyrogenase phosphatase catalytic subunit 1 |
| ENSG00000064932 | SBNO2 | 0,563975622 | 1,478337451 | 0,15772073 | 0,63039898 | strawberry notch homolog 2 |
| ENSG00000236252 | AL512605,1 | 0,563898506 | 1,478258432 | 0,56948541 | 0,94221458 | uncharacterized LOC101928381 |

| | | | | | | |
|---|---|---|---|---|---|---|
| ENSG00000282416 | AC092299,1 | 0,563757848 | 1,478114314 | 0,83533646 | 0,984987524 | novel pseudogene |
| ENSG00000228444 | AL137244,1 | 0,563577229 | 1,477929271 | 0,60118131 | 0,949882873 | novel transcript, sense intronic to LMO7 |
| ENSG00000259426 | AC027237,3 | 0,563527708 | 1,477878542 | 0,16902265 | 0,651019362 | novel transcript, antisense to PAQR5 |
| ENSG00000256812 | CAPNS2 | 0,563384595 | 1,477731946 | 0,58718871 | 0,945891795 | calpain small subunit 2 |
| ENSG00000254027 | AC009902,2 | 0,56314668 | 1,477488273 | 0,36700417 | 0,856397852 | novel transcript |
| ENSG00000169327 | OR5AU1 | 0,563019 | 1,47735752 | 0,46815659 | 0,907940653 | olfactory receptor family 5 subfamily AU member 1 |
| ENSG00000274958 | ITPK1 | 0,562813265 | 1,477146857 | 0,05313602 | 0,354512283 | inositol-tetrakisphosphate 1-kinase |
| ENSG00000130054 | FAM155B | 0,562754139 | 1,477086321 | 0,17137766 | 0,655246551 | family with sequence similarity 155 member B |
| ENSG00000254519 | AC044839,1 | 0,562692148 | 1,477022853 | 0,50994986 | 0,922427526 | novel transcript |
| ENSG00000004700 | RECQL | 0,562450553 | 1,476775531 | 0,08308705 | 0,458378944 | RecQ like helicase |
| ENSG00000274860 | Metazoa_SRP | 0,562266348 | 1,476586985 | 0,46906099 | 0,907940653 | Metazoan signal recognition particle RNA |
| ENSG00000233586 | AC246785,2 | 0,562251946 | 1,476572246 | 0,44629217 | 0,901348735 | novel pseudogene (FLJ39739) |
| ENSG00000087074 | PPP1R15A | 0,562233696 | 1,476553567 | 0,20705466 | 0,710823213 | protein phosphatase 1 regulatory subunit 15A |
| ENSG00000169241 | SLC50A1 | 0,562203313 | 1,476522471 | 0,04175805 | 0,306597901 | solute carrier family 50 member 1 |
| ENSG00000267273 | AC008567,2 | 0,562075983 | 1,476392161 | 0,76064276 | 0,97624106 | novel transcript |
| ENSG00000137936 | BCAR3 | 0,562046466 | 1,476361955 | 0,16643124 | 0,646293152 | BCAR3 adaptor protein, NSP family member |
| ENSG00000143337 | TOR1AIP1 | 0,56164741 | 1,475953643 | 0,03773505 | 0,287183177 | torsin 1A interacting protein 1 |
| ENSG00000228594 | FNDC10 | 0,561571891 | 1,475876385 | 0,07376427 | 0,429054144 | fibronectin type III domain containing 10 |
| ENSG00000271155 | AL161729,1 | 0,561546461 | 1,475850371 | 0,15437114 | 0,623607455 | novel transcript, antisense to PTCH1 |
| ENSG00000162444 | RBP7 | 0,561544263 | 1,475848122 | 0,66652909 | 0,9611116 | retinol binding protein 7 |
| ENSG00000277449 | CEBPB-AS1 | 0,561518854 | 1,475822129 | 0,40872728 | 0,88153752 | CEBPB antisense RNA 1 |
| ENSG00000124783 | SSR1 | 0,561493076 | 1,47579576 | 0,09371847 | 0,486794191 | signal sequence receptor subunit 1 |
| ENSG00000100600 | LGMN | 0,561243999 | 1,47554099 | 0,02267601 | 0,204071065 | legumain |
| ENSG00000047849 | MAP4 | 0,561134522 | 1,475429025 | 0,00863313 | 0,102939044 | microtubule associated protein 4 |
| ENSG00000249378 | LINC01060 | 0,560991061 | 1,475282317 | 0,42247058 | 0,890275827 | long intergenic non-protein coding RNA 1060 |
| ENSG00000186334 | SLC36A3 | 0,560977318 | 1,475268262 | 0,5608014 | 0,940368987 | solute carrier family 36 member 3 |
| ENSG00000274272 | AC069281,2 | 0,560928799 | 1,475218649 | 0,48070165 | 0,912907448 | novel transcript |
| ENSG00000169439 | SDC2 | 0,560925923 | 1,475215709 | 0,04972879 | 0,340424948 | syndecan 2 |
| ENSG00000261103 | AC009063,3 | 0,560870318 | 1,475158851 | 0,4515842 | 0,902062594 | novel transcript, antisense to CDH13 |
| ENSG00000230162 | CT45A11P | 0,560834576 | 1,475122305 | 0,56748431 | 0,941510797 | cancer/testis antigen family 45 member A11, pseudogene |
| ENSG00000104853 | CLPTM1 | 0,560699314 | 1,47498401 | 0,02048315 | 0,189800033 | CLPTM1 regulator of GABA type A receptor forward trafficking |

| | | | | | | |
|---|---|---|---|---|---|---|
| ENSG00000241553 | ARPC4 | 0,560623808 | 1,474906816 | 0,03183974 | 0,25735202 | actin related protein 2/3 complex subunit 4 |
| ENSG00000139722 | VPS37B | 0,560622796 | 1,474905781 | 0,1306441 | 0,577136255 | VPS37B subunit of ESCRT-I |
| ENSG00000227712 | AL359915,1 | 0,560596129 | 1,474878518 | 0,54278178 | 0,933663748 | novel transcript |
| ENSG00000156463 | SH3RF2 | 0,560562298 | 1,474843933 | 0,27888701 | 0,789775666 | SH3 domain containing ring finger 2 |
| ENSG00000161904 | LEMD2 | 0,560240225 | 1,47451472 | 0,08536067 | 0,465128991 | LEM domain nuclear envelope protein 2 |
| ENSG00000111671 | SPSB2 | 0,560232589 | 1,474506916 | 0,10887713 | 0,526415221 | splA/ryanodine receptor domain and SOCS box containing 2 |
| ENSG00000284721 | AL662907,3 | 0,56002609 | 1,474295879 | 0,44422027 | 0,900210555 | novel transcript |
| ENSG00000166140 | ZFYVE19 | 0,560014667 | 1,474284205 | 0,08404585 | 0,461230231 | zinc finger FYVE-type containing 19 |
| ENSG00000248333 | CDK11B | 0,559996626 | 1,47426577 | 0,23855454 | 0,747998807 | cyclin dependent kinase 11B |
| ENSG00000285296 | AL096870,5 | 0,559875343 | 1,474141838 | 0,78053181 | 0,978597905 | novel transcript |
| ENSG00000105063 | PPP6R1 | 0,559714383 | 1,473977378 | 0,12851997 | 0,572897424 | protein phosphatase 6 regulatory subunit 1 |
| ENSG00000131398 | KCNC3 | 0,559703532 | 1,473966292 | 0,07573363 | 0,434766111 | potassium voltage-gated channel subfamily C member 3 |
| ENSG00000169105 | CHST14 | 0,559634728 | 1,473895998 | 0,0417491 | 0,306597901 | carbohydrate sulfotransferase 14 |
| ENSG00000281309 | SURF6 | 0,559539467 | 1,47379868 | 0,2326835 | 0,741684508 | surfeit 6 |
| ENSG00000148296 | SURF6 | 0,559539467 | 1,47379868 | 0,2326835 | 0,741684508 | surfeit 6 |
| ENSG00000170525 | PFKFB3 | 0,559524864 | 1,473783763 | 0,07757471 | 0,440002866 | 6-phosphofructo-2-kinase/fructose-2,6-biphosphatase 3 |
| ENSG00000157368 | IL34 | 0,559398679 | 1,473654865 | 0,50317468 | 0,920888747 | interleukin 34 |
| ENSG00000144567 | RETREG2 | 0,559304469 | 1,473558635 | 0,07965979 | 0,447606687 | reticulophagy regulator family member 2 |
| ENSG00000164303 | ENPP6 | 0,559274903 | 1,473528438 | 0,56127916 | 0,940567693 | ectonucleotide pyrophosphatase/phosphodiesterase 6 |
| ENSG00000265369 | PCAT18 | 0,558957291 | 1,473204073 | 0,51735925 | 0,9257552 | prostate cancer associated transcript 18 |
| ENSG00000230201 | ATP6V0CP1 | 0,558917802 | 1,47316375 | 0,68156849 | 0,964654539 | ATPase H+ transporting V0 subunit c pseudogene 1 |
| ENSG00000238120 | LINC01589 | 0,55870729 | 1,472948808 | 0,44736532 | 0,902062594 | long intergenic non-protein coding RNA 1589 |
| ENSG00000185988 | PLK5 | 0,558637216 | 1,472877266 | 0,59129084 | 0,947416795 | polo like kinase 5 (inactive) |
| ENSG00000104880 | ARHGEF18 | 0,558618642 | 1,472858303 | 0,20633772 | 0,709417043 | Rho/Rac guanine nucleotide exchange factor 18 |
| ENSG00000270978 | AC091982,2 | 0,558391951 | 1,472626891 | 0,73039308 | 0,974321178 | glutamate-ammonia ligase (GLUL) pseudogene |
| ENSG00000204117 | AL162293,1 | 0,558305956 | 1,472539115 | 0,60273407 | 0,950352286 | novel transcript |
| ENSG00000259005 | AC005479,1 | 0,558221159 | 1,472452565 | 0,68121327 | 0,964654539 | novel transcript |
| ENSG00000197142 | ACSL5 | 0,558101246 | 1,472330184 | 0,23053379 | 0,739234644 | acyl-CoA synthetase long chain family member 5 |
| ENSG00000177508 | IRX3 | 0,557823754 | 1,472047019 | 0,35272555 | 0,845031687 | iroquois homeobox 3 |

| | | | | | | |
|---|---|---|---|---|---|---|
| ENSG00000168785 | TSPAN5 | 0,55768244 | 1,471902838 | 0,06012989 | 0,381318641 | tetraspanin 5 |
| ENSG00000224361 | AC011239,1 | 0,557558289 | 1,471776178 | 0,58348213 | 0,944174844 | novel transcript, antisense to KLHL29 |
| ENSG00000117877 | POLR1G | 0,557472057 | 1,471688211 | 0,27886584 | 0,789775666 | RNA polymerase I subunit G |
| ENSG00000224255 | PDCL3P6 | 0,557467353 | 1,471683413 | 0,52094014 | 0,927046149 | PDCL3 pseudogene 6 |
| ENSG00000118689 | FOXO3 | 0,55741288 | 1,471627846 | 0,01554722 | 0,157881532 | forkhead box O3 |
| ENSG00000286196 | AL162586,2 | 0,557200779 | 1,471411507 | 0,44225293 | 0,899331131 | Novel transcript, antisense to SH2D3C |
| ENSG00000174576 | NPAS4 | 0,557170636 | 1,471380764 | 0,23462351 | 0,744134035 | neuronal PAS domain protein 4 |
| ENSG00000288352 | C1RL-AS1 | 0,557132591 | 1,471341963 | 0,31187717 | 0,818208978 | C1RL antisense RNA 1 |
| ENSG00000147099 | HDAC8 | 0,55711563 | 1,471324666 | 0,06766795 | 0,409359696 | histone deacetylase 8 |
| ENSG00000167996 | FTH1 | 0,55696203 | 1,471168026 | 0,03516647 | 0,273970725 | ferritin heavy chain 1 |
| ENSG00000274734 | ARHGAP11B | 0,556820034 | 1,471023235 | 0,52582891 | 0,929056124 | Rho GTPase activating protein 11B |
| ENSG00000283154 | IQCJ-SCHIP1 | 0,556666486 | 1,47086668 | 0,01916171 | 0,181867586 | IQCJ-SCHIP1 readthrough |
| ENSG00000186866 | POFUT2 | 0,556435113 | 1,470630808 | 0,04071064 | 0,30153461 | protein O-fucosyltransferase 2 |
| ENSG00000157766 | ACAN | 0,55632288 | 1,470516406 | 0,47529915 | 0,911082311 | aggrecan |
| ENSG00000261033 | AC005730,2 | 0,556289561 | 1,470482445 | 0,41867672 | 0,887094957 | novel transcript, antisense to SPECC1 |
| ENSG00000155506 | LARP1 | 0,556205436 | 1,470396702 | 0,05239438 | 0,351288432 | La ribonucleoprotein 1, translational regulator |
| ENSG00000232118 | BACH1-AS1 | 0,556202382 | 1,47039359 | 0,54808255 | 0,935404974 | BACH1 antisense RNA 1 |
| ENSG00000240499 | AC004594,1 | 0,556146678 | 1,470336817 | 0,44800185 | 0,902062594 | novel transcript, antisense to CADPS2 |
| ENSG00000285152 | RNF31 | 0,555798308 | 1,469981815 | 0,13082232 | 0,577524567 | ring finger protein 31 |
| ENSG00000270580 | PKD1P6-NPIPP1 | 0,555760401 | 1,469943192 | 0,29754816 | 0,806812288 | PKD1P6-NPIPP1 readthrough |
| ENSG00000235411 | DRG1P1 | 0,555721368 | 1,469903422 | 0,58048439 | 0,943832501 | DRG1 pseudogene 1 |
| ENSG00000182902 | SLC25A18 | 0,555357146 | 1,469532378 | 0,18530854 | 0,677679985 | solute carrier family 25 member 18 |
| ENSG00000100243 | CYB5R3 | 0,555312295 | 1,469486694 | 0,04935586 | 0,338781394 | cytochrome b5 reductase 3 |
| ENSG00000259999 | AC009054,1 | 0,555285666 | 1,46945957 | 0,5841891 | 0,94461278 | novel transcript, sense intronic to CFDP1 |
| ENSG00000267015 | AC023090,1 | 0,555284791 | 1,469458679 | 0,71253804 | 0,970812834 | novel transcript, antisense to ATP9B |
| ENSG00000186148 | AC013268,1 | 0,555212419 | 1,469384966 | 0,43919529 | 0,897434993 | two-pore calcium channel 3 (TPCN3) pseudogene |
| ENSG00000197599 | CCDC154 | 0,555071917 | 1,469241872 | 0,61344891 | 0,952413269 | coiled-coil domain containing 154 |
| ENSG00000161921 | CXCL16 | 0,554887421 | 1,469053993 | 0,47281851 | 0,909509054 | C-X-C motif chemokine ligand 16 |
| ENSG00000277592 | Metazoa_SRP | 0,554720604 | 1,468884137 | 0,77825263 | 0,978587255 | Metazoan signal recognition particle RNA |
| ENSG00000280011 | AL031595,2 | 0,554712949 | 1,468876344 | 0,46419145 | 0,90737035 | novel transcript, antisense to PARVG |
| ENSG00000274667 | AC090517,2 | 0,554689251 | 1,468852216 | 0,48545781 | 0,913907266 | novel transcript, sense intronic to |
| ENSG00000286266 | AC104370,1 | 0,554673369 | 1,468836046 | 0,55713611 | 0,938334502 | novel transcript |
| ENSG00000177406 | NINJ2-AS1 | 0,554672706 | 1,468835371 | 0,17495051 | 0,66036665 | NINJ2 antisense RNA 1 |

| | | | | | | |
|---|---|---|---|---|---|---|
| ENSG00000288657 | AL139280,3 | 0,554653599 | 1,468815919 | 0,62843781 | 0,952793934 | novel transcript |
| ENSG00000286874 | AC072054,1 | 0,554403037 | 1,468560842 | 0,65014563 | 0,956899963 | novel transcript |
| ENSG00000175311 | ANKS4B | 0,554252647 | 1,468407764 | 0,88804607 | 0,990717729 | ankyrin repeat and sterile alpha motif domain containing 4B |
| ENSG00000245156 | AP001107,1 | 0,554219667 | 1,468374196 | 0,38151006 | 0,865900094 | novel transcript |
| ENSG00000164713 | BRI3 | 0,55415315 | 1,468306496 | 0,09875177 | 0,498768217 | brain protein I3 |
| ENSG00000198001 | IRAK4 | 0,554053441 | 1,468205021 | 0,02473825 | 0,217618609 | interleukin 1 receptor associated kinase 4 |
| ENSG00000287096 | AC005165,3 | 0,553958985 | 1,468108898 | 0,46672914 | 0,907940653 | novel transcript |
| ENSG00000148357 | HMCN2 | 0,553855451 | 1,468003544 | 0,50594707 | 0,921780819 | hemicentin 2 |
| ENSG00000159579 | RSPRY1 | 0,553822979 | 1,467970503 | 0,08268243 | 0,45739309 | ring finger and SPRY domain containing 1 |
| ENSG00000185215 | TNFAIP2 | 0,553780044 | 1,467926816 | 0,10793686 | 0,523928283 | TNF alpha induced protein 2 |
| ENSG00000112763 | BTN2A1 | 0,553656388 | 1,467801003 | 0,01086526 | 0,121796931 | butyrophilin subfamily 2 member A1 |
| ENSG00000002933 | TMEM176A | 0,553563882 | 1,46770689 | 0,42606078 | 0,892133432 | transmembrane protein 176A |
| ENSG00000253230 | MIR124-1HG | 0,55356141 | 1,467704375 | 0,18894972 | 0,683322632 | MIR124-1 host gene |
| ENSG00000163577 | EIF5A2 | 0,553465281 | 1,467606583 | 0,13010649 | 0,576317557 | eukaryotic translation initiation factor 5A2 |
| ENSG00000197385 | ZNF860 | 0,553450893 | 1,467591946 | 0,30343794 | 0,812669425 | zinc finger protein 860 |
| ENSG00000267452 | LINC02073 | 0,553291856 | 1,467430174 | 0,29085684 | 0,80167048 | long intergenic non-protein coding RNA 2073 |
| ENSG00000124155 | PIGT | 0,55326028 | 1,467398057 | 0,02096649 | 0,192913017 | phosphatidylinositol glycan anchor biosynthesis class T |
| ENSG00000262223 | AC110285,1 | 0,553227164 | 1,467364375 | 0,47825995 | 0,912116096 | novel transcript |
| ENSG00000284652 | BX323043,1 | 0,55305981 | 1,467194168 | 0,48168784 | 0,912981134 | novel transcript |
| ENSG00000115461 | IGFBP5 | 0,55285598 | 1,466986892 | 0,09120127 | 0,481302729 | insulin like growth factor binding protein 5 |
| ENSG00000286933 | AL022170,1 | 0,552838422 | 1,466969039 | 0,5755164 | 0,942700009 | novel transcript |
| ENSG00000119771 | KLHL29 | 0,55256308 | 1,466689091 | 0,07995383 | 0,448292044 | kelch like family member 29 |
| ENSG00000286615 | AC011416,4 | 0,552515764 | 1,466640988 | 0,2787701 | 0,789775666 | novel transcript |
| ENSG00000100504 | PYGL | 0,5523993 | 1,466522596 | 0,09890647 | 0,499216097 | glycogen phosphorylase L |
| ENSG00000228838 | AL355483,1 | 0,552379142 | 1,466502105 | 0,47999062 | 0,912847708 | novel transcript |
| ENSG00000113100 | CDH9 | 0,552373167 | 1,466496032 | 0,41827877 | 0,887094957 | cadherin 9 |
| ENSG00000213368 | ST13P11 | 0,552275801 | 1,466397063 | 0,41434202 | 0,885579089 | ST13, Hsp70 interacting protein pseudogene 11 |
| ENSG00000140743 | CDR2 | 0,552257626 | 1,466378589 | 0,01401515 | 0,146544982 | cerebellar degeneration related protein 2 |
| ENSG00000262795 | IFNGR2 | 0,552234209 | 1,466354788 | 0,22694917 | 0,735420703 | interferon gamma receptor 2 |
| ENSG00000101695 | RNF125 | 0,552183215 | 1,466302959 | 0,30488326 | 0,813743624 | ring finger protein 125 |
| ENSG00000262172 | AC116025,2 | 0,551914071 | 1,466029436 | 0,49877786 | 0,919683359 | novel transcript |

| | | | | | | |
|---|---|---|---|---|---|---|
| ENSG00000230528 | NOS2P3 | 0,551854536 | 1,465968939 | 0,69140621 | 0,967201016 | nitric oxide synthase 2 pseudogene 3 |
| ENSG00000262691 | AC040160,1 | 0,551843139 | 1,465957359 | 0,47352117 | 0,91012947 | novel transcript |
| ENSG00000143473 | KCNH1 | 0,551782128 | 1,465895365 | 0,15177371 | 0,619863279 | potassium voltage-gated channel subfamily H member 1 |
| ENSG00000168476 | REEP4 | 0,551536897 | 1,465646212 | 0,0591468 | 0,377766175 | receptor accessory protein 4 |
| ENSG00000287961 | AC020911,3 | 0,551425326 | 1,465532869 | 0,54776698 | 0,935404974 | novel transcript, antisense to AP1M1 |
| ENSG00000153113 | CAST | 0,551419852 | 1,465527309 | 0,05539982 | 0,36341169 | calpastatin |
| ENSG00000269889 | AC078802,1 | 0,55092247 | 1,465022142 | 0,57755947 | 0,94337651 | novel transcript, antisense to ACTRT3 |
| ENSG00000184384 | MAML2 | 0,550656915 | 1,464752503 | 0,12449947 | 0,56384187 | mastermind like transcriptional coactivator 2 |
| ENSG00000221716 | SNORA11 | 0,550598243 | 1,464692935 | 0,75440353 | 0,975901808 | small nucleolar RNA, H/ACA box 11 |
| ENSG00000124103 | FAM209A | 0,550543806 | 1,464637669 | 0,62792225 | 0,952793934 | family with sequence similarity 209 member A |
| ENSG00000162066 | AMDHD2 | 0,550408123 | 1,464499929 | 0,25715383 | 0,769358275 | amidohydrolase domain containing 2 |
| ENSG00000269779 | AC010329,2 | 0,550361296 | 1,464452395 | 0,58380861 | 0,944276183 | zinc finger protein 85 pseudogene |
| ENSG00000130368 | MAS1 | 0,550327268 | 1,464417854 | 0,48592319 | 0,914018575 | MAS1 proto-oncogene, G protein-coupled receptor |
| ENSG00000150995 | ITPR1 | 0,55014311 | 1,464230935 | 0,03197507 | 0,258112191 | inositol 1,4,5-trisphosphate receptor type 1 |
| ENSG00000272416 | AC025175,1 | 0,550113559 | 1,464200944 | 0,65487861 | 0,957444211 | novel transcript |
| ENSG00000273090 | AC007378,1 | 0,550042255 | 1,464128578 | 0,59946494 | 0,949137642 | novel transcript |
| ENSG00000206342 | STK19 | 0,549783814 | 1,463866321 | 0,45072812 | 0,902062594 | serine/threonine kinase 19 |
| ENSG00000114316 | USP4 | 0,549780129 | 1,463862582 | 0,01306592 | 0,139163869 | ubiquitin specific peptidase 4 |
| ENSG00000144867 | SRPRB | 0,549776344 | 1,463858741 | 0,0082092 | 0,099516288 | SRP receptor subunit beta |
| ENSG00000158545 | ZC3H18 | 0,549760326 | 1,463842488 | 0,30309672 | 0,812669425 | zinc finger CCCH-type containing 18 |
| ENSG00000285292 | ABCF2-H2BE1 | 0,549737246 | 1,463819071 | 0,01332639 | 0,140993633 | ABCF2-H2BE1 readthrough |
| ENSG00000166579 | NDEL1 | 0,549659054 | 1,463739736 | 0,00933334 | 0,10908314 | nudE neurodevelopment protein 1 like 1 |
| ENSG00000229975 | LIPT1P1 | 0,549510608 | 1,463589132 | 0,53569816 | 0,932139691 | lipoyltransferase 1 pseudogene 1 |
| ENSG00000180264 | ADGRD2 | 0,54931558 | 1,463391292 | 0,60946392 | 0,951793902 | adhesion G protein-coupled receptor D2 |
| ENSG00000213358 | AC092933,1 | 0,549245503 | 1,463320212 | 0,52311547 | 0,92790906 | N-acetyltransferase 13 (GCN5-related) (NAT13) pseudogene |
| ENSG00000144645 | OSBPL10 | 0,548856264 | 1,462925462 | 0,164646 | 0,643811841 | oxysterol binding protein like 10 |
| ENSG00000173545 | ZNF622 | 0,548846716 | 1,46291578 | 0,06717149 | 0,407302071 | zinc finger protein 622 |
| ENSG00000287535 | AC009630,4 | 0,548629498 | 1,462695534 | 0,48473247 | 0,913690171 | novel transcript |
| ENSG00000241369 | LINC01192 | 0,548618163 | 1,462684042 | 0,58241704 | 0,944174844 | long intergenic non-protein coding RNA 1192 |
| ENSG00000103512 | NOMO1 | 0,548559019 | 1,46262408 | 0,62513006 | 0,952793934 | NODAL modulator 1 |
| ENSG00000140749 | IGSF6 | 0,548436064 | 1,462499432 | 0,17509345 | 0,660487662 | immunoglobulin superfamily member 6 |

| | | | | | | |
|---|---|---|---|---|---|---|
| ENSG00000261780 | LINC02582 | 0,548052859 | 1,462111018 | 0,39736216 | 0,874053739 | long intergenic non-protein coding RNA 2582 |
| ENSG00000165271 | NOL6 | 0,547925336 | 1,461981784 | 0,11106241 | 0,532146382 | nucleolar protein 6 |
| ENSG00000257303 | AC073896,2 | 0,547874992 | 1,461930768 | 0,08005554 | 0,448625211 | novel transcript, antisense to CNPY2 and CS |
| ENSG00000069974 | RAB27A | 0,547782896 | 1,461837448 | 0,02113558 | 0,193948538 | RAB27A, member RAS oncogene family |
| ENSG00000012171 | SEMA3B | 0,547627637 | 1,461680136 | 0,12345145 | 0,561884706 | semaphorin 3B |
| ENSG00000272472 | AL512283,1 | 0,547413657 | 1,461463357 | 0,45699287 | 0,903711377 | novel transcript |
| ENSG00000172081 | MOB3A | 0,546877326 | 1,46092015 | 0,08730105 | 0,470988771 | MOB kinase activator 3A |
| ENSG00000121351 | IAPP | 0,546821346 | 1,460863463 | 0,46592234 | 0,907940653 | islet amyloid polypeptide |
| ENSG00000100897 | DCAF11 | 0,546795695 | 1,460837489 | 0,27631221 | 0,788263115 | DDB1 and CUL4 associated factor 11 |
| ENSG00000240764 | PCDHGC5 | 0,546792555 | 1,46083431 | 0,12655958 | 0,568173268 | protocadherin gamma subfamily C, 5 |
| ENSG00000072071 | ADGRL1 | 0,546625807 | 1,460665476 | 0,04726757 | 0,330524125 | adhesion G protein-coupled receptor L1 |
| ENSG00000288324 | ADGRL1 | 0,546625807 | 1,460665476 | 0,04726757 | 0,330524125 | adhesion G protein-coupled receptor L1 |
| ENSG00000136802 | LRRC8A | 0,546399201 | 1,460436065 | 0,10042942 | 0,503419184 | leucine rich repeat containing 8 VRAC subunit A |
| ENSG00000235967 | COX5BP3 | 0,546343633 | 1,460379815 | 0,63340216 | 0,954714356 | cytochrome c oxidase subunit 5B pseudogene 3 |
| ENSG00000138109 | CYP2C9 | 0,546334474 | 1,460370543 | 0,54617408 | 0,935341056 | cytochrome P450 family 2 subfamily C member 9 |
| ENSG00000179967 | PPP1R14BP3 | 0,546265709 | 1,460300937 | 0,62196961 | 0,952793934 | protein phosphatase 1 regulatory inhibitor subunit 14B pseudogene 3 |
| ENSG00000198074 | AKR1B10 | 0,546249575 | 1,460284607 | 0,72051643 | 0,972142678 | aldo-keto reductase family 1 member B10 |
| ENSG00000105088 | OLFM2 | 0,546226144 | 1,46026089 | 0,1387154 | 0,593837907 | olfactomedin 2 |
| ENSG00000258048 | AC073569,2 | 0,546104592 | 1,460137864 | 0,5944584 | 0,947469769 | novel transcript |
| ENSG00000225408 | AL136980,1 | 0,545994688 | 1,460026635 | 0,61616104 | 0,952793934 | novel transcript |
| ENSG00000241735 | FABP5P3 | 0,545896996 | 1,459927772 | 0,43232582 | 0,89558934 | fatty acid binding protein 5 pseudogene 3 |
| ENSG00000178607 | ERN1 | 0,545753881 | 1,459782956 | 0,0401238 | 0,299101235 | endoplasmic reticulum to nucleus signaling 1 |
| ENSG00000267414 | SETBP1-DT | 0,545723986 | 1,459752707 | 0,20044603 | 0,700064356 | SETBP1 divergent transcript |
| ENSG00000125826 | RBCK1 | 0,545720825 | 1,459749508 | 0,10537961 | 0,518119329 | RANBP2-type and C3HC4-type zinc finger containing 1 |
| ENSG00000151014 | NOCT | 0,545509764 | 1,459535967 | 0,09068362 | 0,480530948 | nocturnin |
| ENSG00000237025 | AL121601,2 | 0,545460134 | 1,459485759 | 0,22476168 | 0,733203418 | par-6 partitioning defective 6 homolog beta (C, elegans) (PARD6B) pseudogene |
| ENSG00000240498 | CDKN2B-AS1 | 0,54544938 | 1,45947488 | 0,09012778 | 0,478721526 | CDKN2B antisense RNA 1 |
| ENSG00000120314 | WDR55 | 0,545274366 | 1,459297841 | 0,04530087 | 0,322102876 | WD repeat domain 55 |
| ENSG00000276533 | AC018926,2 | 0,545057576 | 1,459078572 | 0,608806 | 0,951637955 | novel transcript, sense intronic to RAB27A |
| ENSG00000243250 | RPS6P16 | 0,545022745 | 1,459043347 | 0,59431336 | 0,947469769 | ribosomal protein S6 pseudogene 16 |

| | | | | | | |
|---|---|---|---|---|---|---|
| ENSG00000125877 | ITPA | 0,544950521 | 1,458970306 | 0,04125426 | 0,304151397 | inosine triphosphatase |
| ENSG00000183617 | MRPL54 | 0,544918087 | 1,458937506 | 0,06968474 | 0,416593001 | mitochondrial ribosomal protein L54 |
| ENSG00000103066 | PLA2G15 | 0,544877328 | 1,45889629 | 0,109719 | 0,528090728 | phospholipase A2 group XV |
| ENSG00000118257 | NRP2 | 0,544625212 | 1,458641365 | 0,2740754 | 0,7862019 | neuropilin 2 |
| ENSG00000226849 | AL109811,1 | 0,544608813 | 1,458624784 | 0,40004313 | 0,87583672 | novel transcript |
| ENSG00000249577 | AC010424,1 | 0,5445705 | 1,458586049 | 0,60774405 | 0,950993594 | ARP3 actin-related protein 3 homolog (yeast) pseudogene |
| ENSG00000244357 | RN7SL145P | 0,544428189 | 1,458442178 | 0,5686782 | 0,941793403 | RNA, 7SL, cytoplasmic 145, pseudogene |
| ENSG00000217555 | CKLF | 0,544368084 | 1,458381417 | 0,04198176 | 0,307032357 | chemokine like factor |
| ENSG00000109111 | SUPT6H | 0,544236869 | 1,458248782 | 0,1338786 | 0,584034405 | SPT6 homolog, histone chaperone and transcription elongation factor |
| ENSG00000204569 | PPP1R10 | 0,544091735 | 1,45810209 | 0,50374186 | 0,921200643 | protein phosphatase 1 regulatory subunit 10 |
| ENSG00000253276 | CCDC71L | 0,544058156 | 1,458068153 | 0,01956489 | 0,183962882 | coiled-coil domain containing 71 like |
| ENSG00000286366 | AC005052,2 | 0,543957862 | 1,457966794 | 0,29047092 | 0,801030501 | novel transcript, antisense to UPF3B |
| ENSG00000263466 | AC006441,1 | 0,54369676 | 1,457702952 | 0,5258769 | 0,929056124 | novel transcript, antisense to LASP1 |
| ENSG00000117036 | ETV3 | 0,543590112 | 1,457595199 | 0,01973735 | 0,184776117 | ETS variant transcription factor 3 |
| ENSG00000197180 | CH17-340M24, | 0,543543424 | 1,45754803 | 0,12234874 | 0,559390308 | uncharacterized protein BC009467 |
| ENSG00000143382 | ADAMTSL4 | 0,543503595 | 1,45750779 | 0,33214972 | 0,83293962 | ADAMTS like 4 |
| ENSG00000183691 | NOG | 0,543210442 | 1,457211658 | 0,26450914 | 0,775800337 | noggin |
| ENSG00000030110 | BAK1 | 0,543063239 | 1,457062982 | 0,03398401 | 0,268360931 | BCL2 antagonist/killer 1 |
| ENSG00000286181 | AC232271,3 | 0,543049143 | 1,457048745 | 0,40929098 | 0,882177108 | FOXP3 regulating long intergenic non-coding RNA |
| ENSG00000198873 | GRK5 | 0,542910651 | 1,456908882 | 0,1000803 | 0,502152202 | G protein-coupled receptor kinase 5 |
| ENSG00000258354 | AC138932,1 | 0,542881481 | 1,456879424 | 0,83863617 | 0,985672131 | microRNA 3180-1 |
| ENSG00000243568 | AC020779,1 | 0,542866757 | 1,456864556 | 0,46090803 | 0,906029574 | ribosomal protein L21 (RPL21) pseudogene |
| ENSG00000118508 | RAB32 | 0,542710943 | 1,45670722 | 0,06403793 | 0,396451853 | RAB32, member RAS oncogene family |
| ENSG00000130522 | JUND | 0,542588936 | 1,456584033 | 0,11606789 | 0,544096306 | JunD proto-oncogene, AP-1 transcription factor subunit |
| ENSG00000075785 | RAB7A | 0,542165162 | 1,456156243 | 0,04595792 | 0,325324163 | RAB7A, member RAS oncogene family |
| ENSG00000259038 | AL121820,2 | 0,54211801 | 1,456108651 | 0,7259697 | 0,973344786 | novel transcript, antisense to RAD51L1 |
| ENSG00000215199 | YWHAZP6 | 0,542097162 | 1,45608761 | 0,20719931 | 0,711073462 | tyrosine 3-monooxygenase/tryptophan 5-monooxygenase activation protein zeta pseudogene 6 |
| ENSG00000010626 | LRRC23 | 0,542006361 | 1,455995968 | 0,06664533 | 0,40559155 | leucine rich repeat containing 23 |

| | | | | | | |
|---|---|---|---|---|---|---|
| ENSG00000205885 | C1RL-AS1 | 0,541922355 | 1,45591119 | 0,31993573 | 0,824366234 | C1RL antisense RNA 1 |
| ENSG00000183214 | MICA | 0,541660744 | 1,455647207 | 0,54829725 | 0,935543592 | MHC class I polypeptide-related sequence A |
| ENSG00000228956 | SATB1-AS1 | 0,541489201 | 1,455474134 | 0,38727792 | 0,869513953 | SATB1 antisense RNA 1 |
| ENSG00000248954 | AC027801,2 | 0,54148116 | 1,455466022 | 0,73849337 | 0,974353474 | novel transcript |
| ENSG00000134910 | STT3A | 0,541357937 | 1,455341714 | 0,08136066 | 0,452914752 | STT3 oligosaccharyltransferase complex catalytic subunit A |
| ENSG00000145506 | NKD2 | 0,541333532 | 1,455317094 | 0,55074412 | 0,9363105 | NKD inhibitor of WNT signaling pathway 2 |
| ENSG00000232186 | TERF1P7 | 0,541267642 | 1,45525063 | 0,42579195 | 0,892133432 | TERF1 pseudogene 7 |
| ENSG00000162882 | HAAO | 0,54117875 | 1,455160967 | 0,21089355 | 0,716515521 | 3-hydroxyanthranilate 3,4-dioxygenase |
| ENSG00000122884 | P4HA1 | 0,540826664 | 1,454805882 | 0,13116005 | 0,578443778 | prolyl 4-hydroxylase subunit alpha 1 |
| ENSG00000283530 | MGAT4C | 0,540807417 | 1,454786474 | 0,47486389 | 0,91069468 | MGAT4 family member C |
| ENSG00000135116 | HRK | 0,540784919 | 1,454763788 | 0,23235548 | 0,741261398 | harakiri, BCL2 interacting protein |
| ENSG00000197859 | ADAMTSL2 | 0,540615165 | 1,454592623 | 0,27253638 | 0,785492261 | ADAMTS like 2 |
| ENSG00000182154 | MRPL41 | 0,540420394 | 1,45439626 | 0,15609766 | 0,626979676 | mitochondrial ribosomal protein L41 |
| ENSG00000284879 | AC133644,2 | 0,540411347 | 1,454387139 | 0,0656544 | 0,402080164 | novel transcript |
| ENSG00000130810 | PPAN | 0,540006973 | 1,453979544 | 0,24101817 | 0,75149019 | peter pan homolog |
| ENSG00000236426 | AL583824,1 | 0,539783579 | 1,453754421 | 0,57234455 | 0,942475308 | novel transcript |
| ENSG00000150457 | LATS2 | 0,539702328 | 1,45367255 | 0,01485518 | 0,153025461 | large tumor suppressor kinase 2 |
| ENSG00000139549 | DHH | 0,539661658 | 1,453631571 | 0,51449491 | 0,924434989 | desert hedgehog signaling molecule |
| ENSG00000100029 | PES1 | 0,539585821 | 1,45355516 | 0,18118856 | 0,670965888 | pescadillo ribosomal biogenesis factor 1 |
| ENSG00000123728 | RAP2C | 0,539571012 | 1,453540241 | 0,13757921 | 0,592217616 | RAP2C, member of RAS oncogene family |
| ENSG00000136560 | TANK | 0,539513912 | 1,453482712 | 0,19052717 | 0,685949202 | TRAF family member associated NFKB activator |
| ENSG00000184454 | NCMAP | 0,539471851 | 1,453440338 | 0,61485751 | 0,952793934 | non-compact myelin associated protein |
| ENSG00000284693 | LINC02606 | 0,53926738 | 1,453234358 | 0,18351878 | 0,674606849 | long intergenic non-protein coding RNA 2606 |
| ENSG00000143375 | CGN | 0,539078135 | 1,453043742 | 0,35857392 | 0,849814998 | cingulin |
| ENSG00000260585 | AL359851,1 | 0,538984918 | 1,452949861 | 0,48452233 | 0,913690171 | novel transcript |
| ENSG00000196183 | RPS2P4 | 0,538981137 | 1,452946052 | 0,68602092 | 0,965740648 | ribosomal protein S2 pseudogene 4 |
| ENSG00000068028 | RASSF1 | 0,538930841 | 1,4528954 | 0,08658214 | 0,468880561 | Ras association domain family member 1 |
| ENSG00000235681 | AC245884,3 | 0,538808247 | 1,452771945 | 0,65621306 | 0,957444211 | LSM6 homolog, U6 small nuclear RNA associated (S, cerevisiae) (LSM6) pseudogene |
| ENSG00000282485 | LINC00235 | 0,538493071 | 1,452454601 | 0,22626627 | 0,734959876 | long intergenic non-protein coding RNA 235 |
| ENSG00000277142 | LINC00235 | 0,538493071 | 1,452454601 | 0,22626627 | 0,734959876 | long intergenic non-protein coding RNA 235 |
| ENSG00000257497 | AC121761,1 | 0,538350358 | 1,452310931 | 0,24297566 | 0,753092752 | novel transcript, antisense to GLIPR1 |

| | | | | | | |
|---|---|---|---|---|---|---|
| ENSG00000283183 | AC116565,1 | 0,538266363 | 1,452226378 | 0,32081589 | 0,824366234 | novel transcript |
| ENSG00000261270 | AC012181,2 | 0,538196875 | 1,452156433 | 0,56452786 | 0,940865625 | novel transcript, sense intronic to HERPUD1 |
| ENSG00000267676 | THA1P | 0,538075165 | 1,45203393 | 0,70657454 | 0,969503793 | threonine aldolase 1, pseudogene |
| ENSG00000260460 | AL365181,1 | 0,537792438 | 1,4517494 | 0,51230078 | 0,923445033 | novel transcript |
| ENSG00000105401 | CDC37 | 0,537737353 | 1,451693971 | 0,11962785 | 0,552362303 | cell division cycle 37, HSP90 cochaperone |
| ENSG00000142544 | CTU1 | 0,537654905 | 1,451611011 | 0,29390372 | 0,803761875 | cytosolic thiouridylase subunit 1 |
| ENSG00000272328 | AC007128,2 | 0,537423099 | 1,45137779 | 0,1640864 | 0,642888389 | novel transcript, antisense to ICA1 |
| ENSG00000143753 | DEGS1 | 0,537390529 | 1,451345025 | 0,09419108 | 0,487613759 | delta 4-desaturase, sphingolipid 1 |
| ENSG00000183783 | KCTD8 | 0,537358393 | 1,451312696 | 0,31264096 | 0,818683344 | potassium channel tetramerization domain containing 8 |
| ENSG00000198857 | HSD3BP5 | 0,53730456 | 1,451258543 | 0,35141015 | 0,84405766 | hydroxy-delta-5-steroid dehydrogenase, 3 beta, pseudogene 5 |
| ENSG00000285424 | AL162718,1 | 0,537252174 | 1,451205847 | 0,62972852 | 0,952812183 | novel transcript |
| ENSG00000262772 | LINC01977 | 0,537204943 | 1,451158339 | 0,38098206 | 0,865900094 | long intergenic non-protein coding RNA 1977 |
| ENSG00000214967 | NPIPA7 | 0,537188169 | 1,451141466 | 0,22354947 | 0,731984265 | nuclear pore complex interacting protein family member A7 |
| ENSG00000280441 | FP236383,3 | 0,537119735 | 1,451072634 | 0,14960879 | 0,614957829 | novel transcript |
| ENSG00000213676 | ATF6B | 0,537089004 | 1,451041724 | 0,3653258 | 0,855323027 | activating transcription factor 6 beta |
| ENSG00000179253 | AL162457,1 | 0,536969335 | 1,450921368 | 0,59260981 | 0,947416795 | novel transcript, antisense to CDH4 |
| ENSG00000287383 | AC110809,1 | 0,536790478 | 1,450741502 | 0,19424631 | 0,691291421 | novel transcript |
| ENSG00000100034 | PPM1F | 0,536679324 | 1,450629733 | 0,18973911 | 0,684955795 | protein phosphatase, Mg2+/Mn2+ dependent 1F |
| ENSG00000091490 | SEL1L3 | 0,53667837 | 1,450628773 | 0,21407156 | 0,72150508 | SEL1L family member 3 |
| ENSG00000160408 | ST6GALNAC6 | 0,536562159 | 1,450511928 | 0,07484573 | 0,432448167 | ST6 N-acetylgalactosaminide alpha-2,6-sialyltransferase 6 |
| ENSG00000269621 | AL589765,7 | 0,536457775 | 1,450406982 | 0,56907016 | 0,942177697 | novel transcript, antisense to MRPL9 |
| ENSG00000229951 | FLJ31356 | 0,536414153 | 1,450363128 | 0,48927007 | 0,915936198 | uncharacterized protein FLJ31356 |
| ENSG00000269533 | AC003002,3 | 0,535791925 | 1,449737727 | 0,4883085 | 0,915394579 | novel transcript |
| ENSG00000110651 | CD81 | 0,535447381 | 1,449391543 | 0,092744 | 0,484139956 | CD81 molecule |
| ENSG00000188997 | KCTD21 | 0,535442432 | 1,449386571 | 0,05632656 | 0,366815744 | potassium channel tetramerization domain containing 21 |
| ENSG00000109066 | TMEM104 | 0,535295478 | 1,449238942 | 0,04820771 | 0,334553986 | transmembrane protein 104 |
| ENSG00000052749 | RRP12 | 0,535189383 | 1,44913237 | 0,27068417 | 0,783073289 | ribosomal RNA processing 12 homolog |
| ENSG00000184697 | CLDN6 | 0,535062098 | 1,449004523 | 0,32272853 | 0,824955371 | claudin 6 |

| | | | | | | |
|---|---|---|---|---|---|---|
| ENSG00000110046 | ATG2A | 0,534837014 | 1,448778472 | 0,21972621 | 0,727931767 | autophagy related 2A |
| ENSG00000228981 | AC097658,1 | 0,534764454 | 1,448705608 | 0,36706278 | 0,856406773 | ribosomal protein S2 (RPS2) pseudogene |
| ENSG00000228022 | HCG20 | 0,534659481 | 1,448600201 | 0,6389387 | 0,956018518 | HLA complex group 20 |
| ENSG00000254486 | LINC02547 | 0,534628754 | 1,448569349 | 0,50631433 | 0,921780819 | long intergenic non-protein coding RNA 2547 |
| ENSG00000137628 | DDX60 | 0,534392403 | 1,448332055 | 0,2100819 | 0,71523009 | DExD/H-box helicase 60 |
| ENSG00000223431 | MTND6P21 | 0,53435038 | 1,448289868 | 0,64249058 | 0,956633898 | MT-ND6 pseudogene 21 |
| ENSG00000157168 | NRG1 | 0,534127001 | 1,44806564 | 0,19554387 | 0,693689284 | neuregulin 1 |
| ENSG00000273888 | FRMD6-AS1 | 0,53398706 | 1,447925185 | 0,23346291 | 0,74254697 | FRMD6 antisense RNA 1 |
| ENSG00000287336 | AL137157,1 | 0,533964275 | 1,447902318 | 0,52255324 | 0,927822837 | novel transcript, antisense to DNM3 |
| ENSG00000167004 | PDIA3 | 0,533675585 | 1,447612615 | 0,02915164 | 0,24236233 | protein disulfide isomerase family A member 3 |
| ENSG00000177946 | CENPBD1 | 0,533645984 | 1,447582914 | 0,06190513 | 0,388245132 | CENPB DNA-binding domain containing 1 |
| ENSG00000180891 | CUEDC1 | 0,533614129 | 1,44755095 | 0,14382349 | 0,603232838 | CUE domain containing 1 |
| ENSG00000196557 | CACNA1H | 0,533593201 | 1,447529953 | 0,18755183 | 0,681556081 | calcium voltage-gated channel subunit alpha1 H |
| ENSG00000164659 | ELAPOR2 | 0,533517026 | 1,447453525 | 0,2157821 | 0,722999018 | endosome-lysosome associated apoptosis and autophagy regulator family member 2 |
| ENSG00000141582 | CBX4 | 0,533444789 | 1,447381051 | 0,23773286 | 0,746907875 | chromobox 4 |
| ENSG00000181104 | F2R | 0,533394526 | 1,447330625 | 0,05546265 | 0,363656395 | coagulation factor II thrombin receptor |
| ENSG00000181804 | SLC9A9 | 0,533182717 | 1,447118152 | 0,30073137 | 0,810383166 | solute carrier family 9 member A9 |
| ENSG00000206113 | CFAP99 | 0,533087761 | 1,447022907 | 0,64541298 | 0,956899963 | cilia and flagella associated protein 99 |
| ENSG00000168297 | PXK | 0,533071839 | 1,447006937 | 0,09108578 | 0,481283407 | PX domain containing serine/threonine kinase like |
| ENSG00000118960 | HS1BP3 | 0,53304231 | 1,44697732 | 0,09129701 | 0,481302729 | HCLS1 binding protein 3 |
| ENSG00000062096 | ARSF | 0,533040109 | 1,446975113 | 0,6549234 | 0,957444211 | arylsulfatase F |
| ENSG00000099219 | ERMP1 | 0,533021571 | 1,44695652 | 0,18725686 | 0,68119505 | endoplasmic reticulum metallopeptidase 1 |
| ENSG00000146592 | CREB5 | 0,532998348 | 1,446933229 | 0,09289168 | 0,484503945 | cAMP responsive element binding protein 5 |
| ENSG00000280693 | SH3PXD2A-AS1 | 0,532994109 | 1,446928977 | 0,44584902 | 0,901173082 | SH3PXD2A antisense RNA 1 |
| ENSG00000151876 | FBXO4 | 0,532695054 | 1,446629076 | 0,16753294 | 0,648316613 | F-box protein 4 |
| ENSG00000143549 | TPM3 | 0,532606352 | 1,446540135 | 0,00737696 | 0,09214276 | tropomyosin 3 |
| ENSG00000127903 | ZNF835 | 0,532585435 | 1,446519162 | 0,63931604 | 0,956018518 | zinc finger protein 835 |
| ENSG00000236857 | RAP1BP1 | 0,532556016 | 1,446489665 | 0,71135012 | 0,970635643 | RAP1B, member of RAS oncogene family pseudogene 1 |
| ENSG00000236276 | NDP-AS1 | 0,532438225 | 1,44637157 | 0,46796666 | 0,907940653 | NDP antisense RNA 1 |
| ENSG00000276718 | AC005840,4 | 0,532289507 | 1,44622248 | 0,31711812 | 0,822224319 | novel transcript, antisense to VAMP1 |
| ENSG00000286878 | AC100805,1 | 0,532270613 | 1,44620354 | 0,45212791 | 0,902062594 | novel transcript |

| | | | | | | |
|---|---|---|---|---|---|---|
| ENSG00000030304 | MUSK | 0,53225653 | 1,446189424 | 0,40759326 | 0,880588468 | muscle associated receptor tyrosine kinase |
| ENSG00000145675 | PIK3R1 | 0,532115674 | 1,446048233 | 0,10580774 | 0,519142056 | phosphoinositide-3-kinase regulatory subunit 1 |
| ENSG00000152642 | GPD1L | 0,531878523 | 1,44581055 | 0,11669549 | 0,545591104 | glycerol-3-phosphate dehydrogenase 1 like |
| ENSG00000239763 | AC009120,1 | 0,531860069 | 1,445792056 | 0,33840332 | 0,837263082 | ribosomal protein L21 (RPL21) pseudogene |
| ENSG00000107745 | MICU1 | 0,531713896 | 1,445645577 | 0,06829206 | 0,411780707 | mitochondrial calcium uptake 1 |
| ENSG00000256940 | PPP1R14B-AS1 | 0,531603795 | 1,445535255 | 0,25773905 | 0,7701554 | PPP1R14B antisense RNA 1 |
| ENSG00000126562 | WNK4 | 0,531503232 | 1,445434497 | 0,46485581 | 0,907804479 | WNK lysine deficient protein kinase 4 |
| ENSG00000144589 | STK11IP | 0,531357634 | 1,445288631 | 0,14440289 | 0,604689513 | serine/threonine kinase 11 interacting protein |
| ENSG00000165233 | CARD19 | 0,531303959 | 1,44523486 | 0,21796441 | 0,72555786 | caspase recruitment domain family member 19 |
| ENSG00000134684 | YARS1 | 0,531299467 | 1,44523036 | 0,03379949 | 0,267385469 | tyrosyl-tRNA synthetase 1 |
| ENSG00000283100 | SIRT2 | 0,531007292 | 1,444937702 | 0,08406472 | 0,461249431 | sirtuin 2 |
| ENSG00000130202 | NECTIN2 | 0,530975407 | 1,444905767 | 0,08117591 | 0,452385684 | nectin cell adhesion molecule 2 |
| ENSG00000251450 | RASGRF2-AS1 | 0,530839342 | 1,4447695 | 0,58571338 | 0,945246954 | RASGRF2 antisense RNA 1 |
| ENSG00000259225 | LINC02345 | 0,530764032 | 1,444694084 | 0,68312798 | 0,964965151 | long intergenic non-protein coding RNA 2345 |
| ENSG00000198925 | ATG9A | 0,530555088 | 1,444484865 | 0,07702723 | 0,438408786 | autophagy related 9A |
| ENSG00000281468 | AC006504,7 | 0,530529231 | 1,444458977 | 0,63351105 | 0,95478642 | novel transcript, sense intronic to novel transcript |
| ENSG00000245812 | LINC02202 | 0,530477948 | 1,444407632 | 0,46408846 | 0,90737035 | long intergenic non-protein coding RNA 2202 |
| ENSG00000286670 | AL450467,1 | 0,530458529 | 1,444388189 | 0,45497732 | 0,903424009 | novel transcript, antisense to VLDLR |
| ENSG00000163297 | ANTXR2 | 0,530451523 | 1,444381176 | 0,07798502 | 0,441449172 | ANTXR cell adhesion molecule 2 |
| ENSG00000260367 | AC109460,1 | 0,530423319 | 1,444352939 | 0,41924123 | 0,88768414 | novel transcript, antisense to SPNS1 |
| ENSG00000223549 | MTND5P28 | 0,530380412 | 1,444309983 | 0,57537285 | 0,942700009 | MT-ND5 pseudogene 28 |
| ENSG00000114200 | BCHE | 0,530307505 | 1,444236996 | 0,29259364 | 0,803059213 | butyrylcholinesterase |
| ENSG00000232804 | HSPA1B | 0,5298783 | 1,443807397 | 0,39589251 | 0,872308364 | heat shock protein family A (Hsp70) member 1B |
| ENSG00000158473 | CD1D | 0,529803073 | 1,443732114 | 0,64542439 | 0,956899963 | CD1d molecule |
| ENSG00000274352 | ZNF707 | 0,529648486 | 1,443577424 | 0,40123368 | 0,876659568 | zinc finger protein 707 |
| ENSG00000249337 | SNX18P25 | 0,529575738 | 1,443504633 | 0,64466612 | 0,956633898 | sorting nexin 18 pseudogene 25 |
| ENSG00000196535 | MYO18A | 0,529354554 | 1,443283342 | 0,31414283 | 0,819877431 | myosin XVIIIA |
| ENSG00000148396 | SEC16A | 0,529277776 | 1,443206535 | 0,03699817 | 0,283855382 | SEC16 homolog A, endoplasmic reticulum export factor |
| ENSG00000066455 | GOLGA5 | 0,52924599 | 1,443174738 | 0,0976734 | 0,496691781 | golgin A5 |
| ENSG00000237667 | LINC01115 | 0,529230685 | 1,443159428 | 0,64398156 | 0,956633898 | long intergenic non-protein coding RNA 1115 |
| ENSG00000215375 | MYL5 | 0,529202861 | 1,443131595 | 0,09570511 | 0,491317678 | myosin light chain 5 |
| ENSG00000225689 | AL008633,1 | 0,52907094 | 1,44299964 | 0,16426872 | 0,643087554 | novel transcript |

| | | | | | | |
|---|---|---|---|---|---|---|
| ENSG00000130940 | CASZ1 | 0,52905801 | 1,442986707 | 0,0597755 | 0,380382656 | castor zinc finger 1 |
| ENSG00000188163 | FAM166A | 0,529048184 | 1,44297688 | 0,5020438 | 0,920520138 | family with sequence similarity 166 member A |
| ENSG00000197321 | SVIL | 0,528996474 | 1,44292516 | 0,10781703 | 0,523928283 | supervillin |
| ENSG00000132361 | CLUH | 0,528992067 | 1,442920753 | 0,19678647 | 0,695257351 | clustered mitochondria homolog |
| ENSG00000253394 | LINC00534 | 0,528875386 | 1,442804058 | 0,43867613 | 0,89732861 | long intergenic non-protein coding RNA 534 |
| ENSG00000249476 | AC008467,1 | 0,528608891 | 1,442537568 | 0,18616365 | 0,67937802 | novel transcript |
| ENSG00000105393 | BABAM1 | 0,52854071 | 1,442469396 | 0,04295694 | 0,312399557 | BRISC and BRCA1 A complex member 1 |
| ENSG00000286771 | AC097658,3 | 0,527809475 | 1,44173846 | 0,34253218 | 0,840113448 | novel transcript, antisense to GAB1 |
| ENSG00000168491 | CCDC110 | 0,527612218 | 1,441541347 | 0,4518404 | 0,902062594 | coiled-coil domain containing 110 |
| ENSG00000168209 | DDIT4 | 0,527599846 | 1,441528986 | 0,09693985 | 0,494570395 | DNA damage inducible transcript 4 |
| ENSG00000124574 | ABCC10 | 0,527552924 | 1,441482102 | 0,29418788 | 0,804046662 | ATP binding cassette subfamily C member 10 |
| ENSG00000242372 | EIF6 | 0,527515447 | 1,441444657 | 0,07018233 | 0,417549035 | eukaryotic translation initiation factor 6 |
| ENSG00000242038 | EGFL8 | 0,527507534 | 1,441436751 | 0,34618004 | 0,841168721 | EGF like domain multiple 8 |
| ENSG00000249779 | AC106800,1 | 0,527389694 | 1,441319019 | 0,70292349 | 0,968742626 | spermine synthase (SMS) pseudogene |
| ENSG00000064042 | LIMCH1 | 0,527004626 | 1,44093437 | 0,20912359 | 0,71355079 | LIM and calponin homology domains 1 |
| ENSG00000125841 | NRSN2 | 0,526980512 | 1,440910285 | 0,05895857 | 0,377180408 | neurensin 2 |
| ENSG00000198947 | DMD | 0,526909781 | 1,440839643 | 0,1354081 | 0,586668333 | dystrophin |
| ENSG00000130244 | FAM98C | 0,526801375 | 1,44073138 | 0,32279522 | 0,824955371 | family with sequence similarity 98 member C |
| ENSG00000227500 | SCAMP4 | 0,526687558 | 1,440617723 | 0,08017762 | 0,449063597 | secretory carrier membrane protein 4 |
| ENSG00000269243 | AC008894,2 | 0,52659287 | 1,440523175 | 0,3847618 | 0,868536368 | novel transcript, antisense RAB8A |
| ENSG00000123094 | RASSF8 | 0,526339783 | 1,44027049 | 0,17500609 | 0,660401055 | Ras association domain family member 8 |
| ENSG00000261207 | AL031717,1 | 0,526278295 | 1,440209107 | 0,57478301 | 0,942533938 | novel transcript, antisense to MAPK8IP3 |
| ENSG00000148834 | GSTO1 | 0,526267003 | 1,440197835 | 0,04938772 | 0,338828378 | glutathione S-transferase omega 1 |
| ENSG00000154262 | ABCA6 | 0,52624305 | 1,440173924 | 0,53856947 | 0,933154278 | ATP binding cassette subfamily A member 6 |
| ENSG00000105298 | CACTIN | 0,526033689 | 1,439964943 | 0,38561137 | 0,869008435 | cactin, spliceosome C complex subunit |
| ENSG00000141858 | SAMD1 | 0,525785983 | 1,439717727 | 0,03136947 | 0,255133979 | sterile alpha motif domain containing 1 |
| ENSG00000275118 | MBOAT7 | 0,525721067 | 1,439652947 | 0,29176933 | 0,802346409 | membrane bound O-acyltransferase domain containing 7 |
| ENSG00000114867 | EIF4G1 | 0,525585966 | 1,439518137 | 0,09574171 | 0,491344168 | eukaryotic translation initiation factor 4 gamma 1 |
| ENSG00000204364 | C2 | 0,525483915 | 1,439416315 | 0,60999182 | 0,951801946 | complement C2 |
| ENSG00000255284 | AP006621,3 | 0,525443882 | 1,439376372 | 0,33956358 | 0,838481794 | novel transcript |
| ENSG00000267317 | AC027307,2 | 0,525421521 | 1,439354064 | 0,14625781 | 0,609635114 | novel transcript, antisense to APC2 |
| ENSG00000109062 | SLC9A3R1 | 0,525403167 | 1,439335752 | 0,08136646 | 0,452914752 | SLC9A3 regulator 1 |

| | | | | | | |
|---|---|---|---|---|---|---|
| ENSG00000140367 | UBE2Q2 | 0,525295993 | 1,439228831 | 0,20077037 | 0,700325598 | ubiquitin conjugating enzyme E2 Q2 |
| ENSG00000224910 | LINC01766 | 0,525264788 | 1,439197702 | 0,73536691 | 0,974353474 | long intergenic non-protein coding RNA 1766 |
| ENSG00000287087 | AC025175,2 | 0,525244053 | 1,439177017 | 0,49950351 | 0,920186067 | novel transcript, antisense to ARL15 |
| ENSG00000147649 | MTDH | 0,525042936 | 1,438976405 | 0,0244493 | 0,215622841 | metadherin |
| ENSG00000285665 | TOP1P1 | 0,524935747 | 1,438869496 | 0,66160964 | 0,959582738 | DNA topoisomerase I pseudogene 1 |
| ENSG00000141252 | VPS53 | 0,524834056 | 1,438768078 | 0,20969696 | 0,714468807 | VPS53 subunit of GARP complex |
| ENSG00000246898 | LINC00920 | 0,524803308 | 1,438737414 | 0,43507886 | 0,896272827 | long intergenic non-protein coding RNA 920 |
| ENSG00000256167 | ATF4P4 | 0,524744237 | 1,438678507 | 0,66063674 | 0,959226315 | activating transcription factor 4 pseudogene 4 |
| ENSG00000183287 | CCBE1 | 0,524688333 | 1,43862276 | 0,31784664 | 0,822748587 | collagen and calcium binding EGF domains 1 |
| ENSG00000165458 | INPPL1 | 0,524535684 | 1,438470549 | 0,05333345 | 0,355526001 | inositol polyphosphate phosphatase like 1 |
| ENSG00000126524 | SBDS | 0,524423313 | 1,438358512 | 0,08175708 | 0,454732877 | SBDS ribosome maturation factor |
| ENSG00000254535 | PABPC4L | 0,524394402 | 1,438329688 | 0,11525936 | 0,542364351 | poly(A) binding protein cytoplasmic 4 like |
| ENSG00000110047 | EHD1 | 0,524124247 | 1,438060375 | 0,10786942 | 0,523928283 | EH domain containing 1 |
| ENSG00000141002 | TCF25 | 0,523858611 | 1,437795617 | 0,0855699 | 0,465670107 | transcription factor 25 |
| ENSG00000257386 | AC025257,1 | 0,52368983 | 1,437627419 | 0,37236167 | 0,860231676 | novel transcript, antisense to ATXN7L3B |
| ENSG00000250182 | EEF1A1P13 | 0,523586712 | 1,437524667 | 0,53363401 | 0,931381467 | eukaryotic translation elongation factor 1 alpha 1 pseudogene 13 |
| ENSG00000204576 | PRR3 | 0,523252588 | 1,437191779 | 0,79484025 | 0,979776981 | proline rich 3 |
| ENSG00000277558 | AL109923,1 | 0,523137812 | 1,437077445 | 0,57401475 | 0,942475308 | novel transcript, sense intronic to ITCH |
| ENSG00000163645 | ERICH6 | 0,5230069 | 1,43694705 | 0,64668148 | 0,956899963 | glutamate rich 6 |
| ENSG00000116903 | EXOC8 | 0,522750068 | 1,436691263 | 0,03548722 | 0,27578235 | exocyst complex component 8 |
| ENSG00000120910 | PPP3CC | 0,522605037 | 1,436546843 | 0,06706136 | 0,406871024 | protein phosphatase 3 catalytic subunit gamma |
| ENSG00000126062 | TMEM115 | 0,522532285 | 1,436474403 | 0,12643314 | 0,567868238 | transmembrane protein 115 |
| ENSG00000167797 | CDK2AP2 | 0,522526618 | 1,436468761 | 0,08210117 | 0,45575481 | cyclin dependent kinase 2 associated protein 2 |
| ENSG00000144579 | CTDSP1 | 0,522503554 | 1,436445796 | 0,09567036 | 0,491219916 | CTD small phosphatase 1 |
| ENSG00000236390 | AC092800,1 | 0,522433975 | 1,43637652 | 0,52411848 | 0,928404671 | novel transcript |
| ENSG00000266677 | AC087164,1 | 0,522411108 | 1,436353754 | 0,50595356 | 0,921780819 | novel transcript |
| ENSG00000113790 | EHHADH | 0,522399768 | 1,436342463 | 0,21571855 | 0,722999018 | enoyl-CoA hydratase and 3-hydroxyacyl CoA dehydrogenase |
| ENSG00000215301 | DDX3X | 0,52234909 | 1,436292009 | 0,1541831 | 0,623607455 | DEAD-box helicase 3 X-linked |
| ENSG00000173991 | TCAP | 0,522300382 | 1,436243518 | 0,51206456 | 0,923307981 | titin-cap |
| ENSG00000260658 | AC138305,1 | 0,52225498 | 1,43619832 | 0,60738828 | 0,950993594 | novel transcript |
| ENSG00000119321 | FKBP15 | 0,522221572 | 1,436165063 | 0,0246244 | 0,216800136 | FKBP prolyl isomerase 15 |

| | | | | | | |
|---|---|---|---|---|---|---|
| ENSG00000263234 | AC010401,2 | 0,52207626 | 1,436020416 | 0,49044524 | 0,916897686 | novel transcript |
| ENSG00000170266 | GLB1 | 0,522034832 | 1,435979181 | 0,10506468 | 0,517282232 | galactosidase beta 1 |
| ENSG00000229751 | AC044781,1 | 0,521973015 | 1,435917652 | 0,57672249 | 0,943261034 | novel transcript |
| ENSG00000137824 | RMDN3 | 0,521602285 | 1,435548711 | 0,11835796 | 0,549825725 | regulator of microtubule dynamics 3 |
| ENSG00000254802 | AC022182,2 | 0,521566515 | 1,435513119 | 0,53332116 | 0,931381467 | novel transcript |
| ENSG00000267505 | AC005180,2 | 0,521072216 | 1,435021365 | 0,53214536 | 0,931291085 | novel transcript |
| ENSG00000286503 | AC008462,1 | 0,520959707 | 1,434909458 | 0,74227797 | 0,975176043 | novel transcript |
| ENSG00000100109 | TFIP11 | 0,520867323 | 1,434817576 | 0,02939953 | 0,243676239 | tuftelin interacting protein 11 |
| ENSG00000285577 | AC019127,1 | 0,520860425 | 1,434810715 | 0,45826606 | 0,904140862 | novel transcript, antisense to LTBP1 |
| ENSG00000157654 | PALM2AKAP2 | 0,520700734 | 1,434651906 | 0,01686503 | 0,166488266 | PALM2 and AKAP2 fusion |
| ENSG00000140497 | SCAMP2 | 0,520412672 | 1,434365479 | 0,04551389 | 0,323249559 | secretory carrier membrane protein 2 |
| ENSG00000204469 | PRRC2A | 0,520391969 | 1,434344896 | 0,27343177 | 0,785672647 | proline rich coiled-coil 2A |
| ENSG00000222009 | BTBD19 | 0,520296263 | 1,434249747 | 0,22612063 | 0,734959876 | BTB domain containing 19 |
| ENSG00000225636 | HNRNPA3P15 | 0,520163149 | 1,434117418 | 0,69247097 | 0,967236299 | heterogeneous nuclear ribonucleoprotein A3 pseudogene 15 |
| ENSG00000285692 | AC023509,6 | 0,52008766 | 1,43404238 | 0,62087517 | 0,952793934 | Novel transcript, antisense to ATF7 |
| ENSG00000277086 | MTMR10 | 0,520076633 | 1,434031419 | 0,89545023 | 0,991303328 | myotubularin related protein 10 |
| ENSG00000112038 | OPRM1 | 0,519956967 | 1,433912476 | 0,31461328 | 0,819877431 | opioid receptor mu 1 |
| ENSG00000181097 | LINC02878 | 0,51991719 | 1,433872942 | 0,62722209 | 0,952793934 | long intergenic non-protein coding RNA 2878 |
| ENSG00000286635 | AC006112,1 | 0,51990305 | 1,433858888 | 0,62230376 | 0,952793934 | novel transcript, sense intronic to RBFOX1 |
| ENSG00000274400 | AC015967,1 | 0,51981624 | 1,433772613 | 0,77260979 | 0,978310326 | novel transcript |
| ENSG00000131069 | ACSS2 | 0,519661093 | 1,433618434 | 0,04700952 | 0,329512866 | acyl-CoA synthetase short chain family member 2 |
| ENSG00000095739 | BAMBI | 0,519504146 | 1,433462483 | 0,2733811 | 0,785672647 | BMP and activin membrane bound inhibitor |
| ENSG00000182983 | ZNF662 | 0,519447102 | 1,433405805 | 0,55225024 | 0,936619966 | zinc finger protein 662 |
| ENSG00000174327 | SLC16A13 | 0,519270948 | 1,433230796 | 0,11923739 | 0,552100834 | solute carrier family 16 member 13 |
| ENSG00000163827 | LRRC2 | 0,519256521 | 1,433216463 | 0,4993375 | 0,920135598 | leucine rich repeat containing 2 |
| ENSG00000161013 | MGAT4B | 0,519157904 | 1,433118497 | 0,13395066 | 0,584034405 | alpha-1,3-mannosyl-glycoprotein 4-beta-N-acetylglucosaminyltransferase B |
| ENSG00000284501 | MGAT4B | 0,519157904 | 1,433118497 | 0,13395066 | 0,584034405 | alpha-1,3-mannosyl-glycoprotein 4-beta-N-acetylglucosaminyltransferase B |
| ENSG00000153956 | CACNA2D1 | 0,519022951 | 1,432984447 | 0,19913437 | 0,698344029 | calcium voltage-gated channel auxiliary subunit alpha2delta 1 |
| ENSG00000125351 | UPF3B | 0,518928693 | 1,432890826 | 0,05853154 | 0,374908869 | UPF3B regulator of nonsense mediated mRNA decay |

| | | | | | | |
|---|---|---|---|---|---|---|
| ENSG00000251359 | WWC2-AS2 | 0,518875728 | 1,432838222 | 0,19807548 | 0,696782178 | WWC2 antisense RNA 2 |
| ENSG00000189377 | CXCL17 | 0,518778764 | 1,432741924 | 0,66007475 | 0,958859629 | C-X-C motif chemokine ligand 17 |
| ENSG00000166780 | BMERB1 | 0,518748092 | 1,432711464 | 0,07168877 | 0,421847873 | bMERB domain containing 1 |
| ENSG00000105647 | PIK3R2 | 0,518664479 | 1,432628432 | 0,08741571 | 0,47120083 | phosphoinositide-3-kinase regulatory subunit 2 |
| ENSG00000065989 | PDE4A | 0,518660444 | 1,432624425 | 0,16394159 | 0,642745357 | phosphodiesterase 4A |
| ENSG00000177556 | ATOX1 | 0,518293913 | 1,432260498 | 0,051024 | 0,345820358 | antioxidant 1 copper chaperone |
| ENSG00000111981 | ULBP1 | 0,518127116 | 1,432094918 | 0,22144553 | 0,729254784 | UL16 binding protein 1 |
| ENSG00000163902 | RPN1 | 0,518084759 | 1,432052873 | 0,0186207 | 0,177975438 | ribophorin I |
| ENSG00000272855 | AC104458,1 | 0,51787516 | 1,431844834 | 0,69971574 | 0,968522643 | novel transcript |
| ENSG00000171451 | DSEL | 0,517840697 | 1,431810631 | 0,24369944 | 0,754507765 | dermatan sulfate epimerase like |
| ENSG00000269940 | AL049840,4 | 0,517808537 | 1,431778714 | 0,15554156 | 0,625769067 | novel transcript, sense intronic to KLC1 |
| ENSG00000198055 | GRK6 | 0,517746243 | 1,431716893 | 0,14080929 | 0,597646146 | G protein-coupled receptor kinase 6 |
| ENSG00000271156 | AC090578,2 | 0,517733029 | 1,43170378 | 0,70042628 | 0,968522643 | novel transcript |
| ENSG00000184860 | SDR42E1 | 0,517728442 | 1,431699228 | 0,51083709 | 0,922693987 | short chain dehydrogenase/reductase family 42E, member 1 |
| ENSG00000257520 | SFTA3 | 0,517549811 | 1,431521969 | 0,82195629 | 0,982952211 | surfactant associated 3 |
| ENSG00000206249 | AL359555,1 | 0,517337779 | 1,431311594 | 0,76459484 | 0,976640468 | novel transcript |
| ENSG00000189060 | H1-0 | 0,517319271 | 1,431293233 | 0,0517251 | 0,348595265 | H1,0 linker histone |
| ENSG00000186591 | UBE2H | 0,517288264 | 1,431262471 | 0,04022136 | 0,299542756 | ubiquitin conjugating enzyme E2 H |
| ENSG00000283635 | AC012485,3 | 0,517154847 | 1,431130117 | 0,68233564 | 0,964749521 | novel transcript, antisense to PER2 |
| ENSG00000112210 | RAB23 | 0,517138485 | 1,431113887 | 0,14155572 | 0,599428784 | RAB23, member RAS oncogene family |
| ENSG00000186907 | RTN4RL2 | 0,517114207 | 1,431089803 | 0,27316221 | 0,785672647 | reticulon 4 receptor like 2 |
| ENSG00000167799 | NUDT8 | 0,517100017 | 1,431075728 | 0,23154136 | 0,73957278 | nudix hydrolase 8 |
| ENSG00000184216 | IRAK1 | 0,517054092 | 1,431030173 | 0,19710966 | 0,695927373 | interleukin 1 receptor associated kinase 1 |
| ENSG00000288516 | PRKACA | 0,516951953 | 1,430928864 | 0,02018912 | 0,187910875 | protein kinase cAMP-activated catalytic subunit alpha |
| ENSG00000110906 | KCTD10 | 0,516919231 | 1,430896409 | 0,0353272 | 0,274948876 | potassium channel tetramerization domain containing 10 |
| ENSG00000126822 | PLEKHG3 | 0,516908036 | 1,430885305 | 0,19491616 | 0,692641391 | pleckstrin homology and RhoGEF domain containing G3 |
| ENSG00000250251 | PKD1P6 | 0,516905622 | 1,430882912 | 0,29207716 | 0,802771257 | polycystin 1, transient receptor potential channel interacting pseudogene 6 |
| ENSG00000174282 | ZBTB4 | 0,516809706 | 1,430787784 | 0,13015814 | 0,576383034 | zinc finger and BTB domain containing 4 |

| | | | | | | |
|---|---|---|---|---|---|---|
| ENSG00000283868 | ZBTB4 | 0,516809706 | 1,430787784 | 0,13015814 | 0,576383034 | zinc finger and BTB domain containing 4 |
| ENSG00000172757 | CFL1 | 0,51674259 | 1,430721224 | 0,0498246 | 0,340931531 | cofilin 1 |
| ENSG00000287401 | AC034139,1 | 0,516462903 | 1,430443885 | 0,65263961 | 0,957275891 | novel transcript, antisense to STBD1and FAM47E |
| ENSG00000198792 | TMEM184B | 0,5164442 | 1,430425341 | 0,07023733 | 0,417549035 | transmembrane protein 184B |
| ENSG00000285533 | RELA-DT | 0,516347577 | 1,430329543 | 0,22965043 | 0,738054186 | RELA divergent transcript |
| ENSG00000284968 | AC093827,4 | 0,516323114 | 1,43030529 | 0,3998404 | 0,875699486 | Novel transcript, antisense to AFF1 |
| ENSG00000175764 | TTLL11 | 0,516320353 | 1,430302552 | 0,14679162 | 0,610674677 | tubulin tyrosine ligase like 11 |
| ENSG00000141503 | MINK1 | 0,516291128 | 1,430273579 | 0,17966106 | 0,668298509 | misshapen like kinase 1 |
| ENSG00000241839 | PLEKHO2 | 0,516139483 | 1,430123247 | 0,11473986 | 0,541181798 | pleckstrin homology domain containing O2 |
| ENSG00000200087 | SNORA73B | 0,516013112 | 1,429997983 | 0,09310013 | 0,484790551 | small nucleolar RNA, H/ACA box 73B |
| ENSG00000275793 | RIMBP3 | 0,515792534 | 1,429779362 | 0,26242299 | 0,774068771 | RIMS binding protein 3 |
| ENSG00000235158 | AC087430,1 | 0,515739107 | 1,429726415 | 0,8307594 | 0,984156028 | novel transcript |
| ENSG00000244479 | OR2A1-AS1 | 0,515562166 | 1,429551076 | 0,41633152 | 0,886060654 | OR2A1 antisense RNA 1 |
| ENSG00000116678 | LEPR | 0,515517559 | 1,429506875 | 0,21568452 | 0,722999018 | leptin receptor |
| ENSG00000250107 | CACNA1G-AS1 | 0,515293964 | 1,429285342 | 0,60935727 | 0,951793902 | CACNA1G antisense RNA 1 |
| ENSG00000166925 | TSC22D4 | 0,515281157 | 1,429272654 | 0,17390212 | 0,659015241 | TSC22 domain family member 4 |
| ENSG00000178950 | GAK | 0,515222493 | 1,429214537 | 0,12038251 | 0,553609804 | cyclin G associated kinase |
| ENSG00000275700 | AATF | 0,515124571 | 1,429117533 | 0,26266983 | 0,774111884 | apoptosis antagonizing transcription factor |
| ENSG00000285851 | AL359762,3 | 0,515087945 | 1,429081252 | 0,27876401 | 0,789775666 | novel transcript |
| ENSG00000277161 | PIGW | 0,515066942 | 1,429060448 | 0,08922284 | 0,476498052 | phosphatidylinositol glycan anchor biosynthesis class W |
| ENSG00000244219 | TMEM225B | 0,514731017 | 1,428727736 | 0,54406274 | 0,934259495 | transmembrane protein 225B |
| ENSG00000129451 | KLK10 | 0,514706707 | 1,428703662 | 0,50184339 | 0,920520138 | kallikrein related peptidase 10 |
| ENSG00000254473 | AL354920,1 | 0,514705223 | 1,428702192 | 0,19751983 | 0,696048902 | novel transcript, antisense to UBQLN1 |
| ENSG00000181830 | SLC35C1 | 0,514483002 | 1,428482143 | 0,07975941 | 0,447773275 | solute carrier family 35 member C1 |
| ENSG00000101255 | TRIB3 | 0,514288197 | 1,42828927 | 0,17476563 | 0,660057513 | tribbles pseudokinase 3 |
| ENSG00000235092 | ID2-AS1 | 0,514192683 | 1,428194713 | 0,21561117 | 0,722999018 | ID2 antisense RNA 1 |
| ENSG00000180066 | LINC02870 | 0,514110933 | 1,428113787 | 0,67130634 | 0,962026636 | long intergenic non-protein coding RNA 2870 |
| ENSG00000259362 | AC046168,1 | 0,514095734 | 1,428098742 | 0,4669815 | 0,907940653 | novel transcript |
| ENSG00000123595 | RAB9A | 0,514065093 | 1,428068411 | 0,18490713 | 0,677321217 | RAB9A, member RAS oncogene family |
| ENSG00000242737 | AC012170,1 | 0,514047626 | 1,428051122 | 0,58122954 | 0,943919685 | ribosomal protein S20 (RPS20) pseudogene |
| ENSG00000172890 | NADSYN1 | 0,513649592 | 1,427657182 | 0,09467359 | 0,488670145 | NAD synthetase 1 |
| ENSG00000285662 | FAM245B | 0,513509808 | 1,427518862 | 0,70110955 | 0,968677171 | family with sequence similarity 245 member B |

| | | | | | | |
|---|---|---|---|---|---|---|
| ENSG00000247151 | CSTF3-DT | 0,513429613 | 1,427439512 | 0,515391 | 0,924901769 | CSTF3 divergent transcript |
| ENSG00000135686 | KLHL36 | 0,513386244 | 1,427396603 | 0,02631022 | 0,22653539 | kelch like family member 36 |
| ENSG00000239215 | RPL27P12 | 0,513330871 | 1,427341818 | 0,59767647 | 0,9482846 | ribosomal protein L27 pseudogene 12 |
| ENSG00000258676 | AC091544,2 | 0,513207032 | 1,427219302 | 0,60778541 | 0,950993594 | novel transcript |
| ENSG00000227512 | AL592301,1 | 0,513115196 | 1,427128454 | 0,53848396 | 0,933082867 | novel transcript |
| ENSG00000233922 | LINC01694 | 0,51302672 | 1,427040935 | 0,4093221 | 0,882177108 | long intergenic non-protein coding RNA 1694 |
| ENSG00000090273 | NUDC | 0,512897715 | 1,426913336 | 0,20730497 | 0,711123838 | nuclear distribution C, dynein complex regulator |
| ENSG00000046653 | GPM6B | 0,512720236 | 1,42673781 | 0,2774072 | 0,789067728 | glycoprotein M6B |
| ENSG00000180015 | AC093909,1 | 0,512325033 | 1,426347031 | 0,53210577 | 0,931291085 | mitochondrial translational release factor 1-like (MTRF1L) pseudogene |
| ENSG00000267140 | AC007998,2 | 0,512269683 | 1,426292309 | 0,61299658 | 0,952413269 | novel protein |
| ENSG00000156531 | PHF6 | 0,512152169 | 1,426176136 | 0,29778023 | 0,806960585 | PHD finger protein 6 |
| ENSG00000160741 | CRTC2 | 0,512091621 | 1,426116283 | 0,07639191 | 0,436940481 | CREB regulated transcription coactivator 2 |
| ENSG00000151470 | C4orf33 | 0,512009174 | 1,426034785 | 0,19662487 | 0,694800312 | chromosome 4 open reading frame 33 |
| ENSG00000204065 | TCEAL5 | 0,511821535 | 1,425849325 | 0,16946268 | 0,651991274 | transcription elongation factor A like 5 |
| ENSG00000262292 | AC007952,5 | 0,511586609 | 1,425617162 | 0,45526027 | 0,903545728 | novel transcript |
| ENSG00000125266 | EFNB2 | 0,511477966 | 1,425509808 | 0,08835526 | 0,474058486 | ephrin B2 |
| ENSG00000101856 | PGRMC1 | 0,511437968 | 1,425470288 | 0,23810178 | 0,747367171 | progesterone receptor membrane component 1 |
| ENSG00000132507 | EIF5A | 0,511349505 | 1,425382883 | 0,0392449 | 0,294869635 | eukaryotic translation initiation factor 5A |
| ENSG00000288145 | EIF5A | 0,511349505 | 1,425382883 | 0,0392449 | 0,294869635 | eukaryotic translation initiation factor 5A |
| ENSG00000074800 | ENO1 | 0,511346638 | 1,425380051 | 0,03738866 | 0,285450288 | enolase 1 |
| ENSG00000205710 | C17orf107 | 0,51123911 | 1,425273818 | 0,46013767 | 0,905443183 | chromosome 17 open reading frame 107 |
| ENSG00000105523 | FAM83E | 0,511188488 | 1,425223808 | 0,61268136 | 0,952413269 | family with sequence similarity 83 member E |
| ENSG00000133063 | CHIT1 | 0,511097342 | 1,425133768 | 0,62130688 | 0,952793934 | chitinase 1 |
| ENSG00000277203 | F8A1 | 0,5108405 | 1,424880075 | 0,0602644 | 0,381629797 | coagulation factor VIII associated 1 |
| ENSG00000068137 | PLEKHH3 | 0,510807599 | 1,424847581 | 0,23306985 | 0,742177197 | pleckstrin homology, MyTH4 and FERM domain containing H3 |
| ENSG00000100258 | LMF2 | 0,510783553 | 1,424823832 | 0,12312167 | 0,561282565 | lipase maturation factor 2 |
| ENSG00000185345 | PRKN | 0,510603935 | 1,42464645 | 0,1488044 | 0,613515281 | parkin RBR E3 ubiquitin protein ligase |
| ENSG00000048828 | FAM120A | 0,510546067 | 1,424589308 | 0,01568684 | 0,158870544 | family with sequence similarity 120A |
| ENSG00000116096 | SPR | 0,510390157 | 1,424435363 | 0,26513187 | 0,776598285 | sepiapterin reductase |
| ENSG00000126001 | CEP250 | 0,510324132 | 1,424370175 | 0,43423006 | 0,896272827 | centrosomal protein 250 |
| ENSG00000169504 | CLIC4 | 0,510161556 | 1,424209673 | 0,26896049 | 0,781267382 | chloride intracellular channel 4 |

| | | | | | | |
|---|---|---|---|---|---|---|
| ENSG00000259663 | AC010478,1 | 0,51006774 | 1,424117061 | 0,42650346 | 0,892138574 | novel transcript, antisense to ISL1 |
| ENSG00000108175 | ZMIZ1 | 0,509994777 | 1,42404504 | 0,03718697 | 0,28481462 | zinc finger MIZ-type containing 1 |
| ENSG00000177156 | TALDO1 | 0,509951418 | 1,424002243 | 0,05340811 | 0,355796083 | transaldolase 1 |
| ENSG00000148516 | ZEB1 | 0,509919448 | 1,423970687 | 0,11066802 | 0,531152136 | zinc finger E-box binding homeobox 1 |
| ENSG00000126107 | HECTD3 | 0,509801129 | 1,423853909 | 0,09638297 | 0,493177918 | HECT domain E3 ubiquitin protein ligase 3 |
| ENSG00000152583 | SPARCL1 | 0,509639278 | 1,42369418 | 0,47598472 | 0,911390632 | SPARC like 1 |
| ENSG00000282489 | AC016482,1 | 0,509349961 | 1,423408702 | 0,70068135 | 0,968522643 | ribosomal protein L18a (RPL18A) pseudogene |
| ENSG00000276810 | AC244093,2 | 0,509349961 | 1,423408702 | 0,70068135 | 0,968522643 | ribosomal protein L18a (RPL18A) pseudogene |
| ENSG00000279389 | GARRE1 | 0,509254842 | 1,423314857 | 0,59557362 | 0,947923966 | granule associated Rac and RHOG effector 1 |
| ENSG00000148400 | NOTCH1 | 0,509173058 | 1,423234175 | 0,21068896 | 0,716131719 | notch receptor 1 |
| ENSG00000287190 | AC239799,2 | 0,509145396 | 1,423206886 | 0,38714805 | 0,869441674 | novel transcript |
| ENSG00000117758 | STX12 | 0,509136393 | 1,423198005 | 0,15385947 | 0,622929595 | syntaxin 12 |
| ENSG00000178187 | ZNF454 | 0,509020297 | 1,423083483 | 0,27186856 | 0,784843944 | zinc finger protein 454 |
| ENSG00000196131 | VN1R2 | 0,508992739 | 1,423056299 | 0,48634618 | 0,914091794 | vomeronasal 1 receptor 2 |
| ENSG00000200084 | SNORD68 | 0,508838619 | 1,422904285 | 0,65645046 | 0,957444211 | small nucleolar RNA, C/D box 68 |
| ENSG00000284648 | AC097493,3 | 0,508620102 | 1,422688782 | 0,64269199 | 0,956633898 | RNA binding protein, fox-1 homolog (C, elegans) 1 (RBFOX1) pseudogene |
| ENSG00000180539 | C9orf139 | 0,508544436 | 1,422614167 | 0,33934311 | 0,838276153 | chromosome 9 open reading frame 139 |
| ENSG00000112238 | PRDM13 | 0,50811148 | 1,422187302 | 0,6268248 | 0,952793934 | PR/SET domain 13 |
| ENSG00000231206 | HNRNPA1P25 | 0,508001867 | 1,422079251 | 0,48793365 | 0,915157336 | heterogeneous nuclear ribonucleoprotein A1 pseudogene 25 |
| ENSG00000198356 | GET3 | 0,507932687 | 1,422011061 | 0,05557459 | 0,363892597 | guided entry of tail-anchored proteins factor 3, ATPase |
| ENSG00000231532 | LINC01249 | 0,50786413 | 1,421943489 | 0,55148698 | 0,936366211 | long intergenic non-protein coding RNA 1249 |
| ENSG00000268051 | AC008395,1 | 0,507806403 | 1,421886593 | 0,53768318 | 0,932883277 | novel transcript, sense intronic to SIPA1L3 |
| ENSG00000189067 | LITAF | 0,507686075 | 1,421768005 | 0,2006281 | 0,700152419 | lipopolysaccharide induced TNF factor |
| ENSG00000174177 | CTU2 | 0,507649337 | 1,421731801 | 0,21227372 | 0,719057938 | cytosolic thiouridylase subunit 2 |
| ENSG00000134757 | DSG3 | 0,507477228 | 1,421562203 | 0,49432807 | 0,91758518 | desmoglein 3 |
| ENSG00000156194 | PPEF2 | 0,507401765 | 1,421487847 | 0,68658493 | 0,965769168 | protein phosphatase with EF-hand domain 2 |
| ENSG00000149761 | NUDT22 | 0,50694671 | 1,421039552 | 0,19474986 | 0,692283541 | nudix hydrolase 22 |
| ENSG00000180758 | GPR157 | 0,506907158 | 1,421000594 | 0,33621706 | 0,835660292 | G protein-coupled receptor 157 |
| ENSG00000277635 | Metazoa_SRP | 0,506849955 | 1,420944252 | 0,64848993 | 0,956899963 | Metazoan signal recognition particle RNA |
| ENSG00000148335 | NTMT1 | 0,506741265 | 1,420837206 | 0,08299869 | 0,458031678 | N-terminal Xaa-Pro-Lys N-methyltransferase 1 |

| | | | | | | |
|---|---|---|---|---|---|---|
| ENSG00000256589 | ENPP7P5 | 0,50656732 | 1,420665907 | 0,63689795 | 0,955881979 | ectonucleotide pyrophosphatase/phosphodiesterase 7 pseudogene 5 |
| ENSG00000272772 | AC104109,3 | 0,506372902 | 1,42047447 | 0,45749787 | 0,903831777 | novel protein |
| ENSG00000126500 | FLRT1 | 0,506197055 | 1,420301342 | 0,22769133 | 0,736053199 | fibronectin leucine rich transmembrane protein 1 |
| ENSG00000274895 | AC011700,1 | 0,50597237 | 1,420080162 | 0,50134028 | 0,920520138 | novel transcript, antisense to PROX1 |
| ENSG00000131725 | WDR44 | 0,505889989 | 1,419999074 | 0,15004076 | 0,615757437 | WD repeat domain 44 |
| ENSG00000182472 | CAPN12 | 0,505604765 | 1,419718365 | 0,49830328 | 0,919439944 | calpain 12 |
| ENSG00000198960 | ARMCX6 | 0,505559806 | 1,419674123 | 0,03329122 | 0,264906253 | armadillo repeat containing X-linked 6 |
| ENSG00000115756 | HPCAL1 | 0,505519479 | 1,41963444 | 0,23644298 | 0,745627731 | hippocalcin like 1 |
| ENSG00000121690 | DEPDC7 | 0,505482311 | 1,419597867 | 0,22881392 | 0,737173617 | DEP domain containing 7 |
| ENSG00000284607 | AL121936,1 | 0,505469045 | 1,419584813 | 0,64366659 | 0,956633898 | novel transcript |
| ENSG00000198774 | RASSF9 | 0,505363388 | 1,419480852 | 0,12805592 | 0,571345532 | Ras association domain family member 9 |
| ENSG00000236297 | AC048351,1 | 0,505290699 | 1,419409334 | 0,70714823 | 0,969565008 | eukaryotic translation elongation factor 1 alpha 1 (EEF1A1) pseudogene |
| ENSG00000127954 | STEAP4 | 0,50492725 | 1,419051797 | 0,42580331 | 0,892133432 | STEAP4 metalloreductase |
| ENSG00000285884 | AL022345,4 | 0,504922889 | 1,419047507 | 0,66286978 | 0,959897678 | novel transcript |
| ENSG00000237611 | Metazoa_SRP | 0,504829596 | 1,418955747 | 0,50679129 | 0,921824694 | Metazoan signal recognition particle RNA |
| ENSG00000214353 | VAC14-AS1 | 0,504742863 | 1,418870443 | 0,18261302 | 0,673171032 | VAC14 antisense RNA 1 |
| ENSG00000138279 | ANXA7 | 0,504729759 | 1,418857555 | 0,03502162 | 0,273319002 | annexin A7 |
| ENSG00000260360 | AL353708,1 | 0,504357689 | 1,41849168 | 0,50771983 | 0,922172348 | novel transcript |
| ENSG00000161203 | AP2M1 | 0,504338744 | 1,418473054 | 0,03163002 | 0,256419452 | adaptor related protein complex 2 subunit mu 1 |
| ENSG00000255136 | AP001972,3 | 0,50416803 | 1,418305215 | 0,65581628 | 0,957444211 | novel transcript |
| ENSG00000259767 | AC022558,1 | 0,504086214 | 1,418224785 | 0,58836264 | 0,946199432 | novel transcript, sense intronic to BTBD1 |
| ENSG00000266397 | AP000919,2 | 0,503734462 | 1,417879041 | 0,58798765 | 0,946139044 | novel transcript, antisense to LPIN2 |
| ENSG00000184571 | PIWIL3 | 0,503692894 | 1,417838189 | 0,61982083 | 0,952793934 | piwi like RNA-mediated gene silencing 3 |
| ENSG00000277150 | F8A3 | 0,503578491 | 1,417725762 | 0,10632078 | 0,520515619 | coagulation factor VIII associated 3 |
| ENSG00000259869 | AL022344,1 | 0,503530512 | 1,417678613 | 0,71633142 | 0,971438758 | novel transcript |
| ENSG00000160216 | AGPAT3 | 0,503465165 | 1,417614401 | 0,10761742 | 0,52347056 | 1-acylglycerol-3-phosphate O-acyltransferase 3 |
| ENSG00000197226 | TBC1D9B | 0,503443932 | 1,417593538 | 0,09796911 | 0,497096776 | TBC1 domain family member 9B |
| ENSG00000262446 | GBA | 0,503238156 | 1,417391357 | 0,43685958 | 0,896866043 | glucosylceramidase beta |
| ENSG00000171989 | LDHAL6B | 0,503204791 | 1,417358577 | 0,48960865 | 0,916063942 | lactate dehydrogenase A like 6B |

| | | | | | | |
|---|---|---|---|---|---|---|
| ENSG00000168924 | LETM1 | 0,503128436 | 1,417283564 | 0,20003167 | 0,699646707 | leucine zipper and EF-hand containing transmembrane protein 1 |
| ENSG00000182376 | AC138028,1 | 0,503117868 | 1,417273183 | 0,66414357 | 0,960088568 | novel transcript, antisense to FAM38A |
| ENSG00000155393 | HEATR3 | 0,50282119 | 1,416981763 | 0,01583758 | 0,159949901 | HEAT repeat containing 3 |
| ENSG00000143257 | NR1I3 | 0,502776344 | 1,416937717 | 0,41953767 | 0,887991693 | nuclear receptor subfamily 1 group I member 3 |
| ENSG00000042286 | AIFM2 | 0,502601111 | 1,416765623 | 0,13494935 | 0,586038109 | apoptosis inducing factor mitochondria associated 2 |
| ENSG00000213468 | FIRRE | 0,502472276 | 1,416639109 | 0,26980039 | 0,781794063 | firre intergenic repeating RNA element |
| ENSG00000186364 | NUDT17 | 0,50229366 | 1,416463729 | 0,1431572 | 0,601880115 | nudix hydrolase 17 |
| ENSG00000277277 | FAM243B | 0,502214502 | 1,416386013 | 0,44866837 | 0,902062594 | family with sequence similarity 243 member B |
| ENSG00000169371 | SNUPN | 0,502045203 | 1,416219811 | 0,02377316 | 0,210789592 | snurportin 1 |
| ENSG00000105509 | HAS1 | 0,501852908 | 1,416031058 | 0,63269402 | 0,954291097 | hyaluronan synthase 1 |
| ENSG00000204655 | MOG | 0,501668284 | 1,415849857 | 0,77790212 | 0,978587255 | myelin oligodendrocyte glycoprotein |
| ENSG00000162438 | CTRC | 0,501528554 | 1,415712734 | 0,54110028 | 0,933385624 | chymotrypsin C |
| ENSG00000169221 | TBC1D10B | 0,501435344 | 1,41562127 | 0,22950935 | 0,737904201 | TBC1 domain family member 10B |
| ENSG00000170270 | GON7 | 0,501422623 | 1,415608788 | 0,1524965 | 0,620962584 | GON7 subunit of KEOPS complex |
| ENSG00000177238 | TRIM72 | 0,501036075 | 1,415229548 | 0,51058151 | 0,922693987 | tripartite motif containing 72 |
| ENSG00000277111 | PLEKHM1 | 0,501017561 | 1,415211387 | 0,17303178 | 0,657758111 | pleckstrin homology and RUN domain containing M1 |
| ENSG00000234127 | TRIM26 | 0,500884488 | 1,415080855 | 0,20508831 | 0,706634751 | tripartite motif containing 26 |
| ENSG00000285816 | AP000944,5 | 0,500786947 | 1,414985184 | 0,29039518 | 0,800962934 | novel protein, POLA2-CDC42EP2 readthrough |
| ENSG00000159335 | PTMS | 0,50074245 | 1,414941542 | 0,25497559 | 0,76642479 | parathymosin |
| ENSG00000234352 | AC009264,1 | 0,500642068 | 1,414843095 | 0,3418042 | 0,840048641 | novel transcript, antisense to CHRM2 |
| ENSG00000099624 | ATP5F1D | 0,500604019 | 1,414805781 | 0,30065278 | 0,81031109 | ATP synthase F1 subunit delta |
| ENSG00000126581 | BECN1 | 0,500506154 | 1,414709811 | 0,02972458 | 0,245589145 | beclin 1 |
| ENSG00000090432 | MUL1 | 0,50048522 | 1,414689283 | 0,05663019 | 0,367473274 | mitochondrial E3 ubiquitin protein ligase 1 |
| ENSG00000117868 | ESYT2 | 0,500479544 | 1,414683718 | 0,01989632 | 0,185849586 | extended synaptotagmin 2 |
| ENSG00000172742 | OR4D9 | 0,500476876 | 1,414681101 | 0,55260699 | 0,936881074 | olfactory receptor family 4 subfamily D member 9 |
| ENSG00000149091 | DGKZ | 0,500381999 | 1,41458807 | 0,15664643 | 0,627831388 | diacylglycerol kinase zeta |
| ENSG00000136840 | ST6GALNAC4 | 0,50035469 | 1,414561293 | 0,26461629 | 0,775800337 | ST6 N-acetylgalactosaminide alpha-2,6-sialyltransferase 4 |
| ENSG00000158856 | DMTN | 0,5003234 | 1,414530613 | 0,26260833 | 0,774111884 | dematin actin binding protein |
| ENSG00000197746 | PSAP | 0,500291889 | 1,414499718 | 0,07202281 | 0,423096806 | prosaposin |
| ENSG00000204740 | MALRD1 | 0,500197379 | 1,414407058 | 0,124183 | 0,563164667 | MAM and LDL receptor class A domain containing 1 |
| ENSG00000160785 | SLC25A44 | 0,500075386 | 1,414287462 | 0,0333939 | 0,265327213 | solute carrier family 25 member 44 |

| | | | | | | |
|---|---|---|---|---|---|---|
| ENSG00000152292 | SH2D6 | 0,49988827 | 1,414104042 | 0,59003294 | 0,947252221 | SH2 domain containing 6 |
| ENSG00000116954 | RRAGC | 0,499847261 | 1,414063847 | 0,19766217 | 0,696048902 | Ras related GTP binding C |
| ENSG00000276720 | CU151838,3 | 0,499785945 | 1,414003748 | 0,54040742 | 0,933385624 | LSM14 homolog A (SCD6, S, cerevisiae) pseudogene |
| ENSG00000168679 | SLC16A4 | 0,499720944 | 1,413940042 | 0,10072593 | 0,504258684 | solute carrier family 16 member 4 |
| ENSG00000185436 | IFNLR1 | 0,499690324 | 1,413910033 | 0,42025805 | 0,888457169 | interferon lambda receptor 1 |
| ENSG00000178449 | COX14 | 0,499598712 | 1,413820252 | 0,142922 | 0,601785046 | cytochrome c oxidase assembly factor COX14 |
| ENSG00000120937 | NPPB | 0,499439483 | 1,413664218 | 0,62263467 | 0,952793934 | natriuretic peptide B |
| ENSG00000227370 | AC254562,1 | 0,499398037 | 1,413623606 | 0,54009265 | 0,933385624 | chromosome 6 open reading frame 108 (C6orf108) pseudogene |
| ENSG00000268087 | AC008764,2 | 0,499266114 | 1,413494347 | 0,39714018 | 0,873764299 | novel transcript |
| ENSG00000250490 | LINC02145 | 0,499206392 | 1,413435836 | 0,72016437 | 0,972142678 | long intergenic non-protein coding RNA 2145 |
| ENSG00000158109 | TPRG1L | 0,49918356 | 1,413413467 | 0,04325051 | 0,313869373 | tumor protein p63 regulated 1 like |
| ENSG00000129219 | PLD2 | 0,499082261 | 1,413314228 | 0,07934544 | 0,446804019 | phospholipase D2 |
| ENSG00000146909 | NOM1 | 0,498941997 | 1,413176827 | 0,19383573 | 0,690637119 | nucleolar protein with MIF4G domain 1 |
| ENSG00000271784 | AL031055,1 | 0,498817747 | 1,413055124 | 0,59683546 | 0,948282297 | novel transcript, antisense to TP53RK |
| ENSG00000105669 | COPE | 0,498757811 | 1,412996421 | 0,08255506 | 0,45713804 | COPI coat complex subunit epsilon |
| ENSG00000124145 | SDC4 | 0,498579811 | 1,412822095 | 0,1371231 | 0,590986594 | syndecan 4 |
| ENSG00000239467 | AC007405,3 | 0,49856077 | 1,412803449 | 0,28598466 | 0,797420923 | glutamate rich 2 |
| ENSG00000185669 | SNAI3 | 0,498321038 | 1,412568704 | 0,27334545 | 0,785672647 | snail family transcriptional repressor 3 |
| ENSG00000181625 | SLX1B | 0,498272932 | 1,412521603 | 0,20887846 | 0,713420724 | SLX1 homolog B, structure-specific endonuclease subunit |
| ENSG00000235097 | LINC00330 | 0,498184166 | 1,412434696 | 0,69411434 | 0,967359949 | long intergenic non-protein coding RNA 330 |
| ENSG00000265882 | RN7SL73P | 0,498156487 | 1,412407598 | 0,52660331 | 0,929261495 | RNA, 7SL, cytoplasmic 73, pseudogene |
| ENSG00000189306 | RRP7A | 0,498125162 | 1,412376931 | 0,20071887 | 0,700287312 | ribosomal RNA processing 7 homolog A |
| ENSG00000131386 | GALNT15 | 0,4980767 | 1,412329488 | 0,50211086 | 0,920520138 | polypeptide N-acetylgalactosaminyltransferase 15 |
| ENSG00000265727 | RN7SL648P | 0,498023475 | 1,412277384 | 0,52296407 | 0,92790906 | RNA, 7SL, cytoplasmic 648, pseudogene |
| ENSG00000140961 | OSGIN1 | 0,49795222 | 1,412207633 | 0,34011163 | 0,838849007 | oxidative stress induced growth inhibitor 1 |
| ENSG00000249330 | AC095060,1 | 0,4979078 | 1,412164153 | 0,69626304 | 0,96810276 | novel transcript |
| ENSG00000150760 | DOCK1 | 0,4977956 | 1,412054331 | 0,04102941 | 0,303392509 | dedicator of cytokinesis 1 |
| ENSG00000108312 | UBTF | 0,497602046 | 1,411864901 | 0,15665662 | 0,627831388 | upstream binding transcription factor |
| ENSG00000237232 | ZNF295-AS1 | 0,497579094 | 1,41184244 | 0,34459203 | 0,841168721 | ZNF295 antisense RNA 1 |
| ENSG00000228875 | CSNK2B | 0,497562167 | 1,411825874 | 0,2342029 | 0,743739908 | casein kinase 2 beta |
| ENSG00000230424 | EMC1-AS1 | 0,497517972 | 1,411782626 | 0,43847917 | 0,897220559 | EMC1 antisense RNA 1 |

| | | | | | | |
|---|---|---|---|---|---|---|
| ENSG00000145685 | LHFPL2 | 0,497410515 | 1,411677476 | 0,05650081 | 0,366903759 | LHFPL tetraspan subfamily member 2 |
| ENSG00000170122 | FOXD4 | 0,497339902 | 1,411608382 | 0,475757 | 0,911331229 | forkhead box D4 |
| ENSG00000284792 | PTEN | 0,497262242 | 1,411532397 | 0,09110607 | 0,481283407 | phosphatase and tensin homolog |
| ENSG00000115419 | GLS | 0,497172405 | 1,411444504 | 0,1600731 | 0,635172652 | glutaminase |
| ENSG00000263551 | AP000829,1 | 0,497109964 | 1,411383417 | 0,2770738 | 0,788874154 | novel transcript |
| ENSG00000185960 | SHOX | 0,497088663 | 1,411362578 | 0,69152076 | 0,967201016 | short stature homeobox |
| ENSG00000008710 | PKD1 | 0,497015423 | 1,41129093 | 0,31676401 | 0,821646322 | polycystin 1, transient receptor potential channel interacting |
| ENSG00000138835 | RGS3 | 0,496969792 | 1,411246294 | 0,0581805 | 0,373272114 | regulator of G protein signaling 3 |
| ENSG00000284981 | UPK3BL2 | 0,496944962 | 1,411222005 | 0,15262894 | 0,621071504 | uroplakin 3B like 2 |
| ENSG00000126705 | AHDC1 | 0,496866685 | 1,411145438 | 0,20767579 | 0,712005258 | AT-hook DNA binding motif containing 1 |
| ENSG00000166173 | LARP6 | 0,496761527 | 1,411042583 | 0,08456139 | 0,462844615 | La ribonucleoprotein 6, translational regulator |
| ENSG00000173905 | GOLIM4 | 0,496747056 | 1,411028429 | 0,09497713 | 0,489386708 | golgi integral membrane protein 4 |
| ENSG00000143515 | ATP8B2 | 0,496621508 | 1,410905643 | 0,03368076 | 0,266716135 | ATPase phospholipid transporting 8B2 |
| ENSG00000233010 | RPEP4 | 0,496587139 | 1,410872031 | 0,50888197 | 0,922427526 | ribulose-5-phosphate-3-epimerase pseudogene 4 |
| ENSG00000077150 | NFKB2 | 0,496573416 | 1,410858611 | 0,20469749 | 0,706286529 | nuclear factor kappa B subunit 2 |
| ENSG00000272477 | AC144521,1 | 0,496277911 | 1,410569656 | 0,5422987 | 0,933491609 | novel transcript, sense intronic to SATB1 |
| ENSG00000163110 | PDLIM5 | 0,496110038 | 1,410405531 | 0,05558442 | 0,363892597 | PDZ and LIM domain 5 |
| ENSG00000006282 | SPATA20 | 0,495974279 | 1,410272818 | 0,09303397 | 0,48476901 | spermatogenesis associated 20 |
| ENSG00000229433 | LINC02069 | 0,495965929 | 1,410264655 | 0,61084339 | 0,951873283 | long intergenic non-protein coding RNA 2069 |
| ENSG00000070831 | CDC42 | 0,495944712 | 1,410243915 | 0,11570539 | 0,543236711 | cell division cycle 42 |
| ENSG00000127530 | OR7C1 | 0,495871347 | 1,410172202 | 0,50111843 | 0,920520138 | olfactory receptor family 7 subfamily C member 1 |
| ENSG00000177738 | AC025171,1 | 0,495812348 | 1,410114534 | 0,26802306 | 0,780680587 | novel transcript, sense overlapping to ANXA2R |
| ENSG00000285766 | AC092042,4 | 0,49580845 | 1,410110725 | 0,60447513 | 0,950813598 | Novel trranscript, antisense to POMGNT2 |
| ENSG00000237574 | LINC01856 | 0,495753512 | 1,410057028 | 0,66216456 | 0,959874876 | long intergenic non-protein coding RNA 1856 |
| ENSG00000259776 | AC093426,1 | 0,495478906 | 1,409788661 | 0,55175239 | 0,936366211 | novel transcript |
| ENSG00000269242 | AC010422,3 | 0,495437376 | 1,409748078 | 0,47933212 | 0,912714933 | novel transcript |
| ENSG00000277383 | AC010331,1 | 0,495261151 | 1,409575888 | 0,47748847 | 0,911574108 | novel transcript, antisense to GLTSCR1 |
| ENSG00000166473 | PKD1L2 | 0,495207772 | 1,409523736 | 0,54648712 | 0,935401821 | polycystin 1 like 2 (gene/pseudogene) |
| ENSG00000111913 | RIPOR2 | 0,495157053 | 1,409474184 | 0,23872669 | 0,747998807 | RHO family interacting cell polarization regulator 2 |
| ENSG00000165006 | UBAP1 | 0,495067655 | 1,409386846 | 0,0408016 | 0,301993643 | ubiquitin associated protein 1 |
| ENSG00000135047 | CTSL | 0,494943114 | 1,409265186 | 0,03759132 | 0,28657768 | cathepsin L |
| ENSG00000157483 | MYO1E | 0,494819012 | 1,409143965 | 0,06097898 | 0,38428679 | myosin IE |

| | | | | | | |
|---|---|---|---|---|---|---|
| ENSG00000224941 | TCF19 | 0,494642821 | 1,408971882 | 0,27727345 | 0,789067728 | transcription factor 19 |
| ENSG00000224472 | TCF19 | 0,494642821 | 1,408971882 | 0,27727345 | 0,789067728 | transcription factor 19 |
| ENSG00000283228 | AC068547,1 | 0,494578072 | 1,408908648 | 0,79643011 | 0,980015022 | novel transcript |
| ENSG00000111684 | LPCAT3 | 0,49457719 | 1,408907787 | 0,08233601 | 0,456410098 | lysophosphatidylcholine acyltransferase 3 |
| ENSG00000248837 | AC097512,1 | 0,494513286 | 1,40884538 | 0,71869786 | 0,971869926 | novel transcript |
| ENSG00000141551 | CSNK1D | 0,4944982 | 1,408830649 | 0,08969788 | 0,47775341 | casein kinase 1 delta |
| ENSG00000198746 | GPATCH3 | 0,494176944 | 1,408516968 | 0,25381773 | 0,76531122 | G-patch domain containing 3 |
| ENSG00000130561 | SAG | 0,494128228 | 1,408469407 | 0,53569139 | 0,932139691 | S-antigen visual arrestin |
| ENSG00000154222 | CC2D1B | 0,494123949 | 1,40846523 | 0,20137381 | 0,701094807 | coiled-coil and C2 domain containing 1B |
| ENSG00000169083 | AR | 0,494027033 | 1,408370616 | 0,03561157 | 0,276336597 | androgen receptor |
| ENSG00000280384 | FP325332,1 | 0,493970872 | 1,408315793 | 0,63266427 | 0,954291097 | novel transcript, sense intronic to PPARA |
| ENSG00000087076 | HSD17B14 | 0,493915354 | 1,408261599 | 0,16096418 | 0,637001317 | hydroxysteroid 17-beta dehydrogenase 14 |
| ENSG00000161204 | ABCF3 | 0,493872307 | 1,40821958 | 0,11236159 | 0,535454522 | ATP binding cassette subfamily F member 3 |
| ENSG00000264812 | AC132938,2 | 0,493493131 | 1,407849513 | 0,49662534 | 0,918455764 | novel transcript, sense intronic C17orf101 |
| ENSG00000276561 | IRF7 | 0,493484497 | 1,407841088 | 0,2847965 | 0,796197593 | interferon regulatory factor 7 |
| ENSG00000280165 | PCDH20 | 0,493377692 | 1,407736867 | 0,22551395 | 0,734461165 | protocadherin 20 |
| ENSG00000226696 | LENG8-AS1 | 0,493243333 | 1,40760577 | 0,34695399 | 0,841625668 | LENG8 antisense RNA 1 |
| ENSG00000106263 | EIF3B | 0,49313691 | 1,40750194 | 0,12597111 | 0,566777174 | eukaryotic translation initiation factor 3 subunit B |
| ENSG00000263120 | AC004584,3 | 0,492979686 | 1,407348559 | 0,49889101 | 0,919683359 | novel transcript, sense intronic to SCPEP1 |
| ENSG00000188770 | OPTC | 0,492925503 | 1,407295704 | 0,70909137 | 0,970282889 | opticin |
| ENSG00000100726 | TELO2 | 0,492904638 | 1,407275352 | 0,30860277 | 0,816730513 | telomere maintenance 2 |
| ENSG00000103485 | QPRT | 0,492852413 | 1,407224409 | 0,06529975 | 0,401164726 | quinolinate phosphoribosyltransferase |
| ENSG00000262619 | LINC00621 | 0,492689993 | 1,407065992 | 0,09562776 | 0,49111852 | long intergenic non-protein coding RNA 621 |
| ENSG00000266627 | RN7SL202P | 0,492578261 | 1,406957024 | 0,64731109 | 0,956899963 | RNA, 7SL, cytoplasmic 202, pseudogene |
| ENSG00000171596 | NMUR1 | 0,492521006 | 1,406901188 | 0,651833 | 0,957160627 | neuromedin U receptor 1 |
| ENSG00000255690 | TRIL | 0,492509242 | 1,406889716 | 0,13067193 | 0,577136255 | TLR4 interactor with leucine rich repeats |
| ENSG00000118046 | STK11 | 0,492488745 | 1,406869727 | 0,22167199 | 0,729254784 | serine/threonine kinase 11 |
| ENSG00000258716 | AL139193,2 | 0,492254449 | 1,406641268 | 0,49161616 | 0,916995341 | novel transcript, antisense to TTC7B |
| ENSG00000131061 | ZNF341 | 0,492246886 | 1,406633894 | 0,25950372 | 0,772164235 | zinc finger protein 341 |
| ENSG00000124357 | NAGK | 0,492223556 | 1,406611147 | 0,12544113 | 0,565907826 | N-acetylglucosamine kinase |
| ENSG00000144031 | ANKRD53 | 0,492050787 | 1,40644271 | 0,2555134 | 0,767141836 | ankyrin repeat domain 53 |
| ENSG00000285904 | AC006452,1 | 0,492013243 | 1,406406109 | 0,48088216 | 0,912907448 | novel transcript |
| ENSG00000109501 | WFS1 | 0,492011251 | 1,406404168 | 0,18789526 | 0,682033821 | wolframin ER transmembrane glycoprotein |

| | | | | | | |
|---|---|---|---|---|---|---|
| ENSG00000110436 | SLC1A2 | 0,491923056 | 1,406318194 | 0,14178924 | 0,599761485 | solute carrier family 1 member 2 |
| ENSG00000248115 | AC023154,1 | 0,491743698 | 1,40614337 | 0,41244921 | 0,884523593 | novel transcript |
| ENSG00000101224 | CDC25B | 0,491726516 | 1,406126623 | 0,07510088 | 0,43291799 | cell division cycle 25B |
| ENSG00000160193 | WDR4 | 0,491647786 | 1,40604989 | 0,3679023 | 0,857401877 | WD repeat domain 4 |
| ENSG00000283608 | AL031056,2 | 0,49157566 | 1,405979598 | 0,67708909 | 0,963580269 | novel transcript |
| ENSG00000263080 | AC009121,3 | 0,49144921 | 1,405856372 | 0,48754122 | 0,914936533 | novel transcript, antisense to RMI2 |
| ENSG00000161405 | IKZF3 | 0,491334623 | 1,405744715 | 0,43970202 | 0,897665148 | IKAROS family zinc finger 3 |
| ENSG00000178773 | CPNE7 | 0,49128727 | 1,405698576 | 0,64458839 | 0,956633898 | copine 7 |
| ENSG00000260848 | AC092114,1 | 0,491192254 | 1,405605999 | 0,4893292 | 0,915936198 | novel transcript |
| ENSG00000212663 | AC003035,1 | 0,491110987 | 1,405526824 | 0,49066464 | 0,916995341 | novel transcript |
| ENSG00000213463 | SYNJ2BP | 0,491098996 | 1,405515142 | 0,15125588 | 0,618627212 | synaptojanin 2 binding protein |
| ENSG00000053501 | USE1 | 0,491054973 | 1,405472254 | 0,18207207 | 0,672311616 | unconventional SNARE in the ER 1 |
| ENSG00000268750 | AC010522,1 | 0,490926149 | 1,405346759 | 0,46550734 | 0,907899037 | novel protein |
| ENSG00000232662 | LDHBP1 | 0,490853889 | 1,405276372 | 0,553844 | 0,937388968 | lactate dehydrogenase B pseudogene 1 |
| ENSG00000274079 | Metazoa_SRP | 0,490846703 | 1,405269372 | 0,76621136 | 0,977110125 | Metazoan signal recognition particle RNA |
| ENSG00000285449 | H2BU1 | 0,490791172 | 1,405215282 | 0,72612668 | 0,973344786 | H2B,U histone 1 |
| ENSG00000130175 | PRKCSH | 0,490775854 | 1,405200363 | 0,16521236 | 0,644240891 | protein kinase C substrate 80K-H |
| ENSG00000179115 | FARSA | 0,490621147 | 1,405049684 | 0,20133877 | 0,701094807 | phenylalanyl-tRNA synthetase subunit alpha |
| ENSG00000233231 | HNRNPA1P49 | 0,490611395 | 1,405040187 | 0,35386522 | 0,845346751 | heterogeneous nuclear ribonucleoprotein A1 pseudogene 49 |
| ENSG00000239254 | AC009220,2 | 0,490487381 | 1,404919415 | 0,45103866 | 0,902062594 | Male-specific lethal-3 homolog 1 (Msl3l1) pseudogene |
| ENSG00000074755 | ZZEF1 | 0,490463865 | 1,404896515 | 0,05537937 | 0,363353754 | zinc finger ZZ-type and EF-hand domain containing 1 |
| ENSG00000165929 | TC2N | 0,490385765 | 1,404820463 | 0,70449977 | 0,969132783 | tandem C2 domains, nuclear |
| ENSG00000168016 | TRANK1 | 0,489963519 | 1,404409363 | 0,05643978 | 0,366903759 | tetratricopeptide repeat and ankyrin repeat containing 1 |
| ENSG00000244693 | CTAGE8 | 0,489943728 | 1,404390097 | 0,72331138 | 0,972913663 | CTAGE family member 8 |
| ENSG00000169026 | SLC49A3 | 0,489655733 | 1,404109777 | 0,62837631 | 0,952793934 | solute carrier family 49 member 3 |
| ENSG00000138434 | ITPRID2 | 0,489577499 | 1,404033637 | 0,051671 | 0,34830577 | ITPR interacting domain containing 2 |
| ENSG00000135617 | PRADC1 | 0,489556487 | 1,404013188 | 0,09123236 | 0,481302729 | protease associated domain containing 1 |
| ENSG00000115295 | CLIP4 | 0,489542122 | 1,403999208 | 0,12447537 | 0,563836263 | CAP-Gly domain containing linker protein family member 4 |
| ENSG00000108828 | VAT1 | 0,489524716 | 1,40398227 | 0,04350967 | 0,315026964 | vesicle amine transport 1 |

| | | | | | | |
|---|---|---|---|---|---|---|
| ENSG00000246100 | LINC00900 | 0,489468668 | 1,403927727 | 0,14260015 | 0,600923438 | long intergenic non-protein coding RNA 900 |
| ENSG00000114698 | PLSCR4 | 0,489444297 | 1,40390401 | 0,2603544 | 0,772692192 | phospholipid scramblase 4 |
| ENSG00000106178 | CCL24 | 0,48940542 | 1,403866179 | 0,5770041 | 0,943288205 | C-C motif chemokine ligand 24 |
| ENSG00000226312 | CFLAR-AS1 | 0,489346231 | 1,403808584 | 0,34609276 | 0,841168721 | CFLAR antisense RNA 1 |
| ENSG00000179119 | SPTY2D1 | 0,489325895 | 1,403788797 | 0,1125281 | 0,535800356 | SPT2 chromatin protein domain containing 1 |
| ENSG00000160256 | FAM207A | 0,489200765 | 1,403667046 | 0,34607921 | 0,841168721 | family with sequence similarity 207 member A |
| ENSG00000128791 | TWSG1 | 0,488829705 | 1,403306071 | 0,1239533 | 0,562842145 | twisted gastrulation BMP signaling modulator 1 |
| ENSG00000198561 | CTNND1 | 0,488639645 | 1,403121212 | 0,04864475 | 0,335726918 | catenin delta 1 |
| ENSG00000142208 | AKT1 | 0,488598205 | 1,403080909 | 0,0809954 | 0,451779326 | AKT serine/threonine kinase 1 |
| ENSG00000286688 | AP002748,6 | 0,488589634 | 1,403072574 | 0,31292496 | 0,818890608 | novel transcript, antisense to BBS1 |
| ENSG00000164574 | GALNT10 | 0,488565509 | 1,403049111 | 0,09696017 | 0,494593368 | polypeptide N-acetylgalactosaminyltransferase 10 |
| ENSG00000274114 | ALOX15P1 | 0,48821276 | 1,402706098 | 0,50857572 | 0,922427526 | arachidonate 15-lipoxygenase pseudogene 1 |
| ENSG00000258233 | ARHGAP42P5 | 0,487625413 | 1,402135147 | 0,76283281 | 0,97647393 | ARHGAP42 pseudogene 5 |
| ENSG00000138674 | SEC31A | 0,487615389 | 1,402125405 | 0,07196487 | 0,422984135 | SEC31 homolog A, COPII coat complex component |
| ENSG00000144815 | NXPE3 | 0,48760616 | 1,402116436 | 0,04155096 | 0,305921747 | neurexophilin and PC-esterase domain family member 3 |
| ENSG00000203593 | AC005342,1 | 0,487520067 | 1,402032767 | 0,70303141 | 0,968742626 | novel transcript |
| ENSG00000175463 | TBC1D10C | 0,487428946 | 1,401944217 | 0,2990342 | 0,808877417 | TBC1 domain family member 10C |
| ENSG00000235770 | LINC00607 | 0,487295625 | 1,401814668 | 0,29623258 | 0,805971225 | long intergenic non-protein coding RNA 607 |
| ENSG00000156219 | ART3 | 0,487227589 | 1,401748561 | 0,52112601 | 0,927046149 | ADP-ribosyltransferase 3 |
| ENSG00000261578 | AP003119,3 | 0,487202242 | 1,401723934 | 0,25327359 | 0,765172364 | novel transcript, overlapping to TSKU |
| ENSG00000233330 | AL078581,2 | 0,487180403 | 1,401702715 | 0,53708938 | 0,932883277 | novel transcript |
| ENSG00000103249 | CLCN7 | 0,487067364 | 1,401592892 | 0,18334671 | 0,674370766 | chloride voltage-gated channel 7 |
| ENSG00000108774 | RAB5C | 0,487024607 | 1,401551354 | 0,03341942 | 0,265453414 | RAB5C, member RAS oncogene family |
| ENSG00000274303 | 7SK | 0,486968212 | 1,401496569 | 0,68821966 | 0,966426338 | 7SK RNA |
| ENSG00000179431 | FJX1 | 0,486944926 | 1,401473948 | 0,14865287 | 0,613346323 | four-jointed box kinase 1 |
| ENSG00000095370 | SH2D3C | 0,48693303 | 1,401462391 | 0,20847859 | 0,713245584 | SH2 domain containing 3C |
| ENSG00000124587 | PEX6 | 0,486861086 | 1,401392505 | 0,25200759 | 0,764267906 | peroxisomal biogenesis factor 6 |
| ENSG00000133065 | SLC41A1 | 0,486708691 | 1,401244482 | 0,15647886 | 0,627521192 | solute carrier family 41 member 1 |
| ENSG00000159479 | MED8 | 0,486481469 | 1,401023805 | 0,02530567 | 0,220808367 | mediator complex subunit 8 |
| ENSG00000198914 | POU3F3 | 0,486465023 | 1,401007834 | 0,25200669 | 0,764267906 | POU class 3 homeobox 3 |
| ENSG00000175416 | CLTB | 0,486419538 | 1,400963664 | 0,08655752 | 0,468880561 | clathrin light chain B |
| ENSG00000186951 | PPARA | 0,486395614 | 1,400940432 | 0,17856751 | 0,666018763 | peroxisome proliferator activated receptor alpha |

| | | | | | | |
|---|---|---|---|---|---|---|
| ENSG00000146834 | MEPCE | 0,486356818 | 1,400902759 | 0,18529457 | 0,677679985 | methylphosphate capping enzyme |
| ENSG00000179761 | PIPOX | 0,486215595 | 1,400765634 | 0,15729059 | 0,629565097 | pipecolic acid and sarcosine oxidase |
| ENSG00000224830 | OR2X1P | 0,485901028 | 1,400460243 | 0,69496283 | 0,967509655 | olfactory receptor family 2 subfamily X member 1 pseudogene |
| ENSG00000227383 | RPL35AP2 | 0,48588257 | 1,400442325 | 0,62775455 | 0,952793934 | ribosomal protein L35a pseudogene 2 |
| ENSG00000272715 | AL157904,1 | 0,485714994 | 1,400279666 | 0,74530807 | 0,9756851 | novel transcript |
| ENSG00000281484 | BTG3 | 0,48569531 | 1,400260562 | 0,3199177 | 0,824366234 | BTG anti-proliferation factor 3 |
| ENSG00000125952 | MAX | 0,485626657 | 1,40019393 | 0,02647533 | 0,227584468 | MYC associated factor X |
| ENSG00000184432 | COPB2 | 0,485563432 | 1,400132569 | 0,10994817 | 0,528752437 | COPI coat complex subunit beta 2 |
| ENSG00000162004 | CCDC78 | 0,485531025 | 1,400101118 | 0,53311362 | 0,931291085 | coiled-coil domain containing 78 |
| ENSG00000288235 | FAM106C | 0,485497419 | 1,400068505 | 0,49866716 | 0,919683359 | family with sequence similarity 106 member C |
| ENSG00000269397 | AC011503,2 | 0,485464931 | 1,400036977 | 0,42386837 | 0,890949676 | novel transcript, sense intronic to ZNF726 |
| ENSG00000100207 | TCF20 | 0,485285231 | 1,399862601 | 0,21064781 | 0,716131719 | transcription factor 20 |
| ENSG00000147251 | DOCK11 | 0,485150772 | 1,39973214 | 0,28393116 | 0,79541433 | dedicator of cytokinesis 11 |
| ENSG00000203995 | ZYG11A | 0,485141064 | 1,399722721 | 0,32604341 | 0,828085941 | zyg-11 family member A, cell cycle regulator |
| ENSG00000229273 | BX664615,1 | 0,485134653 | 1,399716501 | 0,50955754 | 0,922427526 | novel pseudogene |
| ENSG00000196576 | PLXNB2 | 0,485124579 | 1,399706728 | 0,16551972 | 0,644635638 | plexin B2 |
| ENSG00000100139 | MICALL1 | 0,485064586 | 1,399648523 | 0,18820824 | 0,682491978 | MICAL like 1 |
| ENSG00000132026 | RTBDN | 0,484970261 | 1,399557016 | 0,56559073 | 0,940865625 | retbindin |
| ENSG00000248610 | HSPA8P4 | 0,484904567 | 1,399493287 | 0,44699027 | 0,901815148 | heat shock protein family A (Hsp70) member 8 pseudogene 4 |
| ENSG00000197622 | CDC42SE1 | 0,484846133 | 1,399436605 | 0,03905007 | 0,294183453 | CDC42 small effector 1 |
| ENSG00000131165 | CHMP1A | 0,484817153 | 1,399408493 | 0,11208253 | 0,535066156 | charged multivesicular body protein 1A |
| ENSG00000071553 | ATP6AP1 | 0,48477553 | 1,39936812 | 0,04226403 | 0,308508641 | ATPase H+ transporting accessory protein 1 |
| ENSG00000163584 | RPL22L1 | 0,484754494 | 1,399347716 | 0,25902411 | 0,771947284 | ribosomal protein L22 like 1 |
| ENSG00000148110 | MFSD14B | 0,484663586 | 1,399259542 | 0,14886537 | 0,613515281 | major facilitator superfamily domain containing 14B |
| ENSG00000205790 | DPP9-AS1 | 0,484654031 | 1,399250275 | 0,42096119 | 0,888848798 | DPP9 antisense RNA 1 |
| ENSG00000286702 | AC011455,7 | 0,484647126 | 1,399243578 | 0,47296172 | 0,909532026 | novel transcript, antisense to FBXO17and FBXO27 |
| ENSG00000232536 | AL365436,2 | 0,484547648 | 1,399147099 | 0,71973811 | 0,972003986 | novel transcript |
| ENSG00000151689 | INPP1 | 0,484488622 | 1,399089856 | 0,11947195 | 0,552362303 | inositol polyphosphate-1-phosphatase |
| ENSG00000285564 | AL031121,2 | 0,484383215 | 1,398987638 | 0,44626419 | 0,901348735 | novel transcript |
| ENSG00000007376 | RPUSD1 | 0,484031215 | 1,398646344 | 0,20885573 | 0,713420724 | RNA pseudouridine synthase domain containing 1 |
| ENSG00000214655 | ZSWIM8 | 0,484021402 | 1,398636831 | 0,12339093 | 0,561691036 | zinc finger SWIM-type containing 8 |

| | | | | | | |
|---|---|---|---|---|---|---|
| ENSG00000100906 | NFKBIA | 0,483857213 | 1,398477666 | 0,11655156 | 0,545320605 | NFKB inhibitor alpha |
| ENSG00000275374 | AC113385,3 | 0,483753893 | 1,398377515 | 0,62213368 | 0,952793934 | glucuronidase, beta (GUSB) pseudogene |
| ENSG00000002726 | AOC1 | 0,483593946 | 1,398222491 | 0,68173454 | 0,964654539 | amine oxidase copper containing 1 |
| ENSG00000275873 | LMNTD2 | 0,483553219 | 1,39818302 | 0,73820965 | 0,974353474 | lamin tail domain containing 2 |
| ENSG00000113262 | GRM6 | 0,483548975 | 1,398178907 | 0,45042855 | 0,902062594 | glutamate metabotropic receptor 6 |
| ENSG00000259967 | AC087761,1 | 0,483495104 | 1,398126698 | 0,73546508 | 0,974353474 | novel transcript |
| ENSG00000135253 | KCP | 0,483256076 | 1,397895074 | 0,22885684 | 0,737173617 | kielin cysteine rich BMP regulator |
| ENSG00000120254 | MTHFD1L | 0,483177162 | 1,397818612 | 0,04462973 | 0,319246353 | methylenetetrahydrofolate dehydrogenase (NADP+ dependent) 1 like |
| ENSG00000160226 | CFAP410 | 0,48313939 | 1,397782016 | 0,25910091 | 0,771989931 | cilia and flagella associated protein 410 |
| ENSG00000133789 | SWAP70 | 0,48300144 | 1,397648367 | 0,01843006 | 0,176747665 | switching B cell complex subunit SWAP70 |
| ENSG00000286240 | AL034422,1 | 0,482907667 | 1,397557524 | 0,686579 | 0,965769168 | novel transcript |
| ENSG00000260063 | AL512408,1 | 0,482851192 | 1,397502818 | 0,26258192 | 0,774111884 | novel transcript, antisense to ARID1A |
| ENSG00000278516 | LENG1 | 0,482779525 | 1,397433398 | 0,34672638 | 0,8414568 | leukocyte receptor cluster member 1 |
| ENSG00000124228 | DDX27 | 0,482727275 | 1,397382788 | 0,14957485 | 0,614957829 | DEAD-box helicase 27 |
| ENSG00000285372 | TNKS | 0,482676344 | 1,397333457 | 0,16920738 | 0,651287551 | tankyrase |
| ENSG00000181817 | LSM10 | 0,482518836 | 1,39718091 | 0,0546498 | 0,361009299 | LSM10, U7 small nuclear RNA associated |
| ENSG00000272734 | ADIRF-AS1 | 0,482365031 | 1,397031966 | 0,30644323 | 0,815375074 | ADIRF antisense RNA 1 |
| ENSG00000038532 | CLEC16A | 0,48232999 | 1,396998034 | 0,04498559 | 0,320517213 | C-type lectin domain containing 16A |
| ENSG00000120889 | TNFRSF10B | 0,482258155 | 1,396928476 | 0,14314351 | 0,601880115 | TNF receptor superfamily member 10b |
| ENSG00000094661 | OR1I1 | 0,482205804 | 1,396877787 | 0,56741788 | 0,941450549 | olfactory receptor family 1 subfamily I member 1 |
| ENSG00000008128 | CDK11A | 0,482172378 | 1,396845422 | 0,32307829 | 0,825150495 | cyclin dependent kinase 11A |
| ENSG00000138758 | SEPTIN11 | 0,482153247 | 1,396826899 | 0,1479969 | 0,612582848 | septin 11 |
| ENSG00000280537 | AC068946,1 | 0,482111284 | 1,396786271 | 0,62614267 | 0,952793934 | novel protein, SLC23A3-NHEJ1 readthrough |
| ENSG00000101474 | APMAP | 0,481855261 | 1,396538417 | 0,02750174 | 0,233645239 | adipocyte plasma membrane associated protein |
| ENSG00000171241 | SHCBP1 | 0,481737912 | 1,396424827 | 0,02513492 | 0,219870742 | SHC binding and spindle associated 1 |
| ENSG00000280789 | PAGR1 | 0,481706355 | 1,396394283 | 0,22132737 | 0,729254784 | PAXIP1 associated glutamate rich protein 1 |
| ENSG00000134046 | MBD2 | 0,481665901 | 1,396355128 | 0,05751816 | 0,370798119 | methyl-CpG binding domain protein 2 |
| ENSG00000105671 | DDX49 | 0,481431178 | 1,396127962 | 0,09599296 | 0,492148866 | DEAD-box helicase 49 |
| ENSG00000280711 | JADRR | 0,481294024 | 1,395995241 | 0,5258926 | 0,929056124 | JADE1 adjacent regulatory RNA |
| ENSG00000286116 | AL157394,3 | 0,481236804 | 1,395939874 | 0,4070416 | 0,880078305 | FAS antisense RNA 1 |
| ENSG00000186575 | NF2 | 0,481180474 | 1,395885372 | 0,13491693 | 0,586038109 | neurofibromin 2 |
| ENSG00000129562 | DAD1 | 0,481139001 | 1,395845245 | 0,11035582 | 0,530142011 | defender against cell death 1 |

| | | | | | | |
|---|---|---|---|---|---|---|
| ENSG00000049768 | FOXP3 | 0,480146017 | 1,394884837 | 0,54340184 | 0,934226837 | forkhead box P3 |
| ENSG00000152822 | GRM1 | 0,48008682 | 1,394827603 | 0,35102919 | 0,844035671 | glutamate metabotropic receptor 1 |
| ENSG00000109063 | MYH3 | 0,480038945 | 1,394781318 | 0,48384623 | 0,913690171 | myosin heavy chain 3 |
| ENSG00000126934 | MAP2K2 | 0,480038596 | 1,39478098 | 0,14979274 | 0,615346044 | mitogen-activated protein kinase kinase 2 |
| ENSG00000126456 | IRF3 | 0,479987411 | 1,394731495 | 0,20125028 | 0,701094807 | interferon regulatory factor 3 |
| ENSG00000281089 | ATG16L1 | 0,47989953 | 1,394646539 | 0,05121379 | 0,346269002 | autophagy related 16 like 1 |
| ENSG00000085978 | ATG16L1 | 0,47989953 | 1,394646539 | 0,05121379 | 0,346269002 | autophagy related 16 like 1 |
| ENSG00000269886 | AC022382,1 | 0,479880035 | 1,394627693 | 0,62898344 | 0,952793934 | novel transcript, antisense to TTLL3 |
| ENSG00000262420 | AC007613,1 | 0,479808666 | 1,394558704 | 0,4577301 | 0,90388627 | novel transcript, antisense to TXNDC11 |
| ENSG00000156096 | UGT2B4 | 0,479808506 | 1,394558549 | 0,5517817 | 0,936366211 | UDP glucuronosyltransferase family 2 member B4 |
| ENSG00000280571 | AC006059,2 | 0,479509494 | 1,394269544 | 0,2495149 | 0,761442412 | novel protein |
| ENSG00000128849 | CGNL1 | 0,479474742 | 1,394235959 | 0,36873733 | 0,857685565 | cingulin like 1 |
| ENSG00000089169 | RPH3A | 0,479470587 | 1,394231944 | 0,60525814 | 0,950927937 | rabphilin 3A |
| ENSG00000110801 | PSMD9 | 0,479287813 | 1,39405532 | 0,09528531 | 0,489967108 | proteasome 26S subunit, non-ATPase 9 |
| ENSG00000167183 | PRR15L | 0,479227652 | 1,393997189 | 0,69909432 | 0,968522643 | proline rich 15 like |
| ENSG00000137672 | TRPC6 | 0,479052524 | 1,393827982 | 0,62158165 | 0,952793934 | transient receptor potential cation channel subfamily C member 6 |
| ENSG00000166598 | HSP90B1 | 0,478990528 | 1,393768088 | 0,03258153 | 0,261251213 | heat shock protein 90 beta family member 1 |
| ENSG00000238082 | NUDCP2 | 0,47891287 | 1,393693065 | 0,55749124 | 0,938578675 | nuclear distribution C pseudogene 2 |
| ENSG00000277801 | AL138478,1 | 0,478825143 | 1,39360832 | 0,50045283 | 0,920485547 | novel transcript |
| ENSG00000284194 | SCO2 | 0,478796215 | 1,393580378 | 0,44278236 | 0,899437949 | synthesis of cytochrome C oxidase 2 |
| ENSG00000150403 | TMCO3 | 0,478712892 | 1,393499894 | 0,05305834 | 0,354145183 | transmembrane and coiled-coil domains 3 |
| ENSG00000155849 | ELMO1 | 0,478661336 | 1,393450096 | 0,12518912 | 0,565471028 | engulfment and cell motility 1 |
| ENSG00000231370 | PRRC2A | 0,478547187 | 1,393339848 | 0,38883852 | 0,870057277 | proline rich coiled-coil 2A |
| ENSG00000171872 | KLF17 | 0,478455951 | 1,393251736 | 0,67719119 | 0,963580269 | Kruppel like factor 17 |
| ENSG00000003989 | SLC7A2 | 0,478141575 | 1,392948167 | 0,23156595 | 0,73957278 | solute carrier family 7 member 2 |
| ENSG00000008130 | NADK | 0,478116059 | 1,392923531 | 0,11364422 | 0,538655668 | NAD kinase |
| ENSG00000196566 | AL138767,1 | 0,478034216 | 1,392844514 | 0,55374746 | 0,937338325 | novel transcript |
| ENSG00000286622 | AC034111,2 | 0,477788942 | 1,392607736 | 0,58115253 | 0,943919685 | novel transcript, antisense to MSRA |
| ENSG00000259342 | AC025580,1 | 0,477734402 | 1,39255509 | 0,50632491 | 0,921780819 | novel transcript, antisense to C15orf48 |
| ENSG00000163219 | ARHGAP25 | 0,477575444 | 1,392401665 | 0,23201773 | 0,740561742 | Rho GTPase activating protein 25 |
| ENSG00000106089 | STX1A | 0,477535632 | 1,392363241 | 0,27509128 | 0,786506974 | syntaxin 1A |
| ENSG00000181908 | LINC02724 | 0,47748856 | 1,392317812 | 0,57107945 | 0,942436428 | long intergenic non-protein coding RNA 2724 |

| | | | | | | |
|---|---|---|---|---|---|---|
| ENSG00000227959 | AL451042,2 | 0,477418739 | 1,39225043 | 0,65602766 | 0,957444211 | novel transcript |
| ENSG00000156599 | ZDHHC5 | 0,477409601 | 1,392241613 | 0,03387948 | 0,267747212 | zinc finger DHHC-type palmitoyltransferase 5 |
| ENSG00000243468 | INGX | 0,47718378 | 1,392023706 | 0,54952783 | 0,936083284 | inhibitor of growth family, X-linked (pseudogene) |
| ENSG00000240970 | RPL23AP64 | 0,477082642 | 1,391926123 | 0,49129287 | 0,916995341 | ribosomal protein L23a pseudogene 64 |
| ENSG00000281913 | RPL23AP64 | 0,477082642 | 1,391926123 | 0,49129287 | 0,916995341 | ribosomal protein L23a pseudogene 64 |
| ENSG00000162729 | IGSF8 | 0,477081699 | 1,391925214 | 0,27436905 | 0,786239603 | immunoglobulin superfamily member 8 |
| ENSG00000214305 | Metazoa_SRP | 0,476995687 | 1,391842231 | 0,64710047 | 0,956899963 | Metazoan signal recognition particle RNA |
| ENSG00000157916 | RER1 | 0,476879746 | 1,391730381 | 0,02257682 | 0,203412591 | retention in endoplasmic reticulum sorting receptor 1 |
| ENSG00000213590 | AL807752,1 | 0,476822738 | 1,391675388 | 0,44252424 | 0,899432962 | ribosomal protein L30 (RPL30) pseudogene |
| ENSG00000165102 | HGSNAT | 0,476771792 | 1,391626245 | 0,04290245 | 0,312148546 | heparan-alpha-glucosaminide N-acetyltransferase |
| ENSG00000107581 | EIF3A | 0,47670658 | 1,391563343 | 0,27219001 | 0,785277096 | eukaryotic translation initiation factor 3 subunit A |
| ENSG00000157593 | SLC35B2 | 0,476580481 | 1,391441718 | 0,09064746 | 0,480420766 | solute carrier family 35 member B2 |
| ENSG00000236069 | AL136987,1 | 0,476488308 | 1,391352822 | 0,45530844 | 0,903545728 | novel transcript |
| ENSG00000288407 | CCDC33 | 0,476389107 | 1,391257155 | 0,27265137 | 0,785565607 | coiled-coil domain containing 33 |
| ENSG00000140481 | CCDC33 | 0,476389107 | 1,391257155 | 0,27265137 | 0,785565607 | coiled-coil domain containing 33 |
| ENSG00000277173 | AC004241,4 | 0,47633701 | 1,391206916 | 0,73048949 | 0,974321178 | novel transcript, sense intronic to SLC48A1 |
| ENSG00000102466 | FGF14 | 0,476089929 | 1,390968673 | 0,41914646 | 0,887577549 | fibroblast growth factor 14 |
| ENSG00000147231 | RADX | 0,476084625 | 1,39096356 | 0,33985189 | 0,838611198 | RPA1 related single stranded DNA binding protein, X-linked |
| ENSG00000113716 | HMGXB3 | 0,475940605 | 1,390824711 | 0,08380454 | 0,461082665 | HMG-box containing 3 |
| ENSG00000236271 | FLOT1 | 0,4758376 | 1,390725413 | 0,39701824 | 0,873618956 | flotillin 1 |
| ENSG00000272502 | AC104958,2 | 0,475833642 | 1,390721597 | 0,74222801 | 0,975176043 | novel transcript |
| ENSG00000185920 | PTCH1 | 0,47574172 | 1,390632989 | 0,06256034 | 0,390552401 | patched 1 |
| ENSG00000135835 | KIAA1614 | 0,475470488 | 1,39037157 | 0,27197391 | 0,784915764 | KIAA1614 |
| ENSG00000249664 | AC027338,1 | 0,475328859 | 1,390235085 | 0,64185035 | 0,95651945 | novel transcript |
| ENSG00000276890 | Metazoa_SRP | 0,475323761 | 1,390230172 | 0,70380732 | 0,968976619 | Metazoan signal recognition particle RNA |
| ENSG00000284916 | PSME1 | 0,475206433 | 1,390117116 | 0,20850103 | 0,713245584 | proteasome activator subunit 1 |
| ENSG00000092010 | PSME1 | 0,475206433 | 1,390117116 | 0,20850103 | 0,713245584 | proteasome activator subunit 1 |
| ENSG00000090975 | PITPNM2 | 0,475153719 | 1,390066323 | 0,22889856 | 0,737173617 | phosphatidylinositol transfer protein membrane associated 2 |
| ENSG00000258465 | AL139011,2 | 0,474883979 | 1,389806448 | 0,87233229 | 0,989217334 | novel protein |
| ENSG00000084070 | SMAP2 | 0,474671935 | 1,389602192 | 0,03997908 | 0,298449351 | small ArfGAP2 |

| | | | | | | |
|---|---|---|---|---|---|---|
| ENSG00000228501 | RPL15P18 | 0,474628402 | 1,389560262 | 0,64067441 | 0,956348702 | ribosomal protein L15 pseudogene 18 |
| ENSG00000124098 | FAM210B | 0,47456529 | 1,389499476 | 0,22351882 | 0,731984265 | family with sequence similarity 210 member B |
| ENSG00000228691 | NEU1 | 0,474552401 | 1,389487062 | 0,24330032 | 0,753715365 | neuraminidase 1 |
| ENSG00000258904 | AL157871,5 | 0,474459331 | 1,389397427 | 0,57143075 | 0,942436428 | novel transcript, intronic to YY1 |
| ENSG00000228485 | GRK5-IT1 | 0,474318636 | 1,389261937 | 0,68681333 | 0,965769168 | GRK5 intronic transcript 1 |
| ENSG00000110002 | VWA5A | 0,474181089 | 1,389129491 | 0,11246725 | 0,535800356 | von Willebrand factor A domain containing 5A |
| ENSG00000255872 | AL138752,2 | 0,474170441 | 1,389119237 | 0,4829158 | 0,91363215 | novel transcript |
| ENSG00000147804 | SLC39A4 | 0,474022027 | 1,388976342 | 0,50822194 | 0,922427526 | solute carrier family 39 member 4 |
| ENSG00000167807 | AC011511,1 | 0,473958832 | 1,388915502 | 0,29339961 | 0,803461221 | novel transcript |
| ENSG00000100557 | CCDC198 | 0,473939703 | 1,388897086 | 0,26535116 | 0,776700553 | coiled-coil domain containing 198 |
| ENSG00000173812 | EIF1 | 0,473854234 | 1,388814806 | 0,08082665 | 0,451082539 | eukaryotic translation initiation factor 1 |
| ENSG00000103260 | METRN | 0,473842729 | 1,388803731 | 0,21425582 | 0,721587186 | meteorin, glial cell differentiation regulator |
| ENSG00000236187 | GJA6P | 0,473744112 | 1,388708802 | 0,61917103 | 0,952793934 | gap junction protein alpha 6, pseudogene |
| ENSG00000185043 | CIB1 | 0,473729202 | 1,38869445 | 0,09116429 | 0,481283407 | calcium and integrin binding 1 |
| ENSG00000249550 | LINC01234 | 0,473714487 | 1,388680285 | 0,2571582 | 0,769358275 | long intergenic non-protein coding RNA 1234 |
| ENSG00000288503 | DDX51 | 0,473552423 | 1,388524298 | 0,44391313 | 0,900210555 | DEAD-box helicase 51 |
| ENSG00000101298 | SNPH | 0,473532651 | 1,388505268 | 0,29571991 | 0,805426883 | syntaphilin |
| ENSG00000143036 | SLC44A3 | 0,473521604 | 1,388494636 | 0,19902611 | 0,698148483 | solute carrier family 44 member 3 |
| ENSG00000277481 | PKD1L3 | 0,473416586 | 1,388393568 | 0,67435781 | 0,963103112 | polycystin 1 like 3, transient receptor potential channel interacting |
| ENSG00000267484 | AC027319,1 | 0,473379632 | 1,388358004 | 0,78773362 | 0,979462318 | novel transcript, antisense to PLIN3 |
| ENSG00000123136 | DDX39A | 0,473379173 | 1,388357563 | 0,05460164 | 0,360901847 | DExD-box helicase 39A |
| ENSG00000258216 | AC009522,1 | 0,473347984 | 1,388327549 | 0,48975414 | 0,916063942 | novel transcript |
| ENSG00000175497 | DPP10 | 0,473220333 | 1,388204714 | 0,37079938 | 0,858780045 | dipeptidyl peptidase like 10 |
| ENSG00000270755 | AL136141,1 | 0,473202798 | 1,388187841 | 0,75653777 | 0,975901808 | novel transcript |
| ENSG00000269103 | Metazoa_SRP | 0,47313204 | 1,388119759 | 0,71223257 | 0,970812834 | Metazoan signal recognition particle RNA |
| ENSG00000285653 | AC006238,2 | 0,473108588 | 1,388097194 | 0,67633734 | 0,963580269 | novel transcript, antisense to RAB31 |
| ENSG00000273382 | AL356488,3 | 0,473103293 | 1,388092099 | 0,37458443 | 0,861771195 | novel transcript, antisense to TMEM167B |
| ENSG00000172638 | EFEMP2 | 0,473036281 | 1,388027625 | 0,18586998 | 0,678777213 | EGF containing fibulin extracellular matrix protein 2 |
| ENSG00000248740 | LINC02428 | 0,473013409 | 1,388005619 | 0,23270627 | 0,741684508 | long intergenic non-protein coding RNA 2428 |
| ENSG00000011304 | PTBP1 | 0,472960699 | 1,387954909 | 0,03048123 | 0,249663376 | polypyrimidine tract binding protein 1 |
| ENSG00000212747 | RTL8B | 0,472948859 | 1,387943518 | 0,02698449 | 0,230377648 | retrotransposon Gag like 8B |

| | | | | | | |
|---|---|---|---|---|---|---|
| ENSG00000212989 | AC124293,1 | 0,472724973 | 1,387728145 | 0,69479583 | 0,967417311 | nucleolar pre-rRNA processing protein, homolog (S, cerevisiae) (ESF1) pseudogene |
| ENSG00000126262 | FFAR2 | 0,472603327 | 1,387611139 | 0,75925642 | 0,976120207 | free fatty acid receptor 2 |
| ENSG00000255655 | AC007570,1 | 0,472436358 | 1,387450555 | 0,59707247 | 0,948282297 | novel transcript, antisense to MYO1H |
| ENSG00000115738 | ID2 | 0,472396254 | 1,387411987 | 0,45621536 | 0,903545728 | inhibitor of DNA binding 2 |
| ENSG00000282367 | PTDSS2 | 0,472372854 | 1,387389483 | 0,29853087 | 0,808005284 | phosphatidylserine synthase 2 |
| ENSG00000288212 | AC090826,7 | 0,472006583 | 1,387037298 | 0,58040397 | 0,943832501 | novel transcript, antisense to SEMA7A |
| ENSG00000286696 | AC090826,3 | 0,472006583 | 1,387037298 | 0,58040397 | 0,943832501 | novel transcript, antisense to SEMA7A |
| ENSG00000267123 | SCAT1 | 0,47193834 | 1,386971689 | 0,65271579 | 0,95732962 | S-phase cancer associated transcript 1 |
| ENSG00000260316 | AL008727,1 | 0,471761037 | 1,386801245 | 0,63146554 | 0,953823688 | novel transcript, intronic to LMF1 |
| ENSG00000135924 | DNAJB2 | 0,471666316 | 1,386710197 | 0,18039638 | 0,66974377 | DnaJ heat shock protein family (Hsp40) member B2 |
| ENSG00000118503 | TNFAIP3 | 0,471602768 | 1,386649117 | 0,28431863 | 0,795731744 | TNF alpha induced protein 3 |
| ENSG00000090863 | GLG1 | 0,471240467 | 1,386300934 | 0,03001824 | 0,247101211 | golgi glycoprotein 1 |
| ENSG00000285661 | AC127520,1 | 0,471232341 | 1,386293125 | 0,65047084 | 0,956899963 | novel transcript |
| ENSG00000261002 | AC036103,1 | 0,471163974 | 1,386227432 | 0,6511387 | 0,957089277 | novel transcript, antisense to TMEM87A |
| ENSG00000179846 | NKPD1 | 0,471104153 | 1,386169954 | 0,43639456 | 0,896850871 | NTPase KAP family P-loop domain containing 1 |
| ENSG00000285219 | HULC | 0,470915143 | 1,385988362 | 0,35329434 | 0,84514549 | hepatocellular carcinoma up-regulated long non-coding RNA |
| ENSG00000241563 | CORT | 0,470767088 | 1,385846133 | 0,50498564 | 0,921400408 | cortistatin |
| ENSG00000142459 | EVI5L | 0,470721008 | 1,38580187 | 0,39698609 | 0,873609704 | ecotropic viral integration site 5 like |
| ENSG00000133424 | LARGE1 | 0,470711291 | 1,385792536 | 0,07612794 | 0,436068882 | LARGE xylosyl- and glucuronyltransferase 1 |
| ENSG00000103415 | HMOX2 | 0,470599327 | 1,385684992 | 0,08394775 | 0,461230231 | heme oxygenase 2 |
| ENSG00000110274 | CEP164 | 0,470481479 | 1,385571806 | 0,33412503 | 0,834518413 | centrosomal protein 164 |
| ENSG00000197536 | IRF1-AS1 | 0,47046159 | 1,385552705 | 0,25075255 | 0,762971073 | IRF1 antisense RNA 1 |
| ENSG00000263823 | AC009831,1 | 0,470388268 | 1,385482288 | 0,44542161 | 0,901068127 | novel transcript, antisense to KIAA1012 |
| ENSG00000142675 | CNKSR1 | 0,47034142 | 1,385437299 | 0,41884709 | 0,887244078 | connector enhancer of kinase suppressor of Ras 1 |
| ENSG00000203588 | IGBP1-AS1 | 0,470297422 | 1,385395048 | 0,61626351 | 0,952793934 | IGBP1 antisense RNA 1 |
| ENSG00000188822 | CNR2 | 0,470269205 | 1,385367952 | 0,4791463 | 0,912714933 | cannabinoid receptor 2 |
| ENSG00000275971 | AC018659,4 | 0,470121012 | 1,385225654 | 0,89761235 | 0,991708791 | |
| ENSG00000101577 | LPIN2 | 0,470105267 | 1,385210537 | 0,09770314 | 0,496762239 | lipin 2 |
| ENSG00000275397 | DHRS11 | 0,470027041 | 1,38513543 | 0,3907675 | 0,870057277 | dehydrogenase/reductase 11 |
| ENSG00000092098 | RNF31 | 0,469971134 | 1,385081755 | 0,18783744 | 0,681978628 | ring finger protein 31 |
| ENSG00000228725 | MTND2P12 | 0,469650192 | 1,384773663 | 0,71770981 | 0,971697135 | MT-ND2 pseudogene 12 |

| | | | | | | |
|---|---|---|---|---|---|---|
| ENSG00000048162 | NOP16 | 0,469625363 | 1,384749832 | 0,09934463 | 0,50040065 | NOP16 nucleolar protein |
| ENSG00000285262 | MCPH1 | 0,469576484 | 1,384702917 | 0,35737339 | 0,848693232 | microcephalin 1 |
| ENSG00000259235 | AC066612,1 | 0,469436144 | 1,384568224 | 0,69232134 | 0,967236299 | novel transcript |
| ENSG00000257438 | AC011595,1 | 0,469213705 | 1,384354764 | 0,51644769 | 0,925337876 | novel transcript, antisense to NUAK1 |
| ENSG00000137845 | ADAM10 | 0,469155275 | 1,384298698 | 0,23075886 | 0,739234644 | ADAM metallopeptidase domain 10 |
| ENSG00000162430 | SELENON | 0,469118251 | 1,384263173 | 0,07831265 | 0,442692864 | selenoprotein N |
| ENSG00000240481 | RN7SL38P | 0,469090833 | 1,384236865 | 0,70367271 | 0,968975668 | RNA, 7SL, cytoplasmic 38, pseudogene |
| ENSG00000227964 | LINC01429 | 0,468984074 | 1,384134436 | 0,66616485 | 0,961020116 | long intergenic non-protein coding RNA 1429 |
| ENSG00000102393 | GLA | 0,468979207 | 1,384129767 | 0,165392 | 0,644418684 | galactosidase alpha |
| ENSG00000242247 | ARFGAP3 | 0,468951817 | 1,384103489 | 0,16859821 | 0,650506523 | ADP ribosylation factor GTPase activating protein 3 |
| ENSG00000123066 | MED13L | 0,46888891 | 1,384043138 | 0,06223517 | 0,389221888 | mediator complex subunit 13L |
| ENSG00000185385 | OR7A17 | 0,468817351 | 1,38397449 | 0,50669638 | 0,921824694 | olfactory receptor family 7 subfamily A member 17 |
| ENSG00000237261 | Metazoa_SRP | 0,468660033 | 1,383823583 | 0,70592798 | 0,969503793 | Metazoan signal recognition particle RNA |
| ENSG00000271010 | AC011475,1 | 0,468600521 | 1,383766501 | 0,63924211 | 0,956018518 | LSM2 homolog, U6 small nuclear RNA associated (S, cerevisiae) (LSM2) pseudogene |
| ENSG00000129347 | KRI1 | 0,468528786 | 1,383697698 | 0,34992832 | 0,843677595 | KRI1 homolog |
| ENSG00000201428 | RN7SKP71 | 0,468507585 | 1,383677365 | 0,45764532 | 0,903831777 | RN7SK pseudogene 71 |
| ENSG00000120509 | PDZD11 | 0,468430176 | 1,383603124 | 0,10262112 | 0,510071005 | PDZ domain containing 11 |
| ENSG00000278518 | AL161645,1 | 0,468405147 | 1,38357912 | 0,67983309 | 0,96448882 | novel transcript, antisense to CYP2E1 |
| ENSG00000286782 | AL355377,3 | 0,46811434 | 1,383300257 | 0,52070164 | 0,927046149 | novel transcript |
| ENSG00000077549 | CAPZB | 0,467910727 | 1,38310504 | 0,03438391 | 0,27043482 | capping actin protein of muscle Z-line subunit beta |
| ENSG00000166770 | ZNF667-AS1 | 0,467753273 | 1,382954098 | 0,56481976 | 0,940865625 | ZNF667 antisense RNA 1 (head to head) |
| ENSG00000270111 | CELF2-DT | 0,467692986 | 1,382896309 | 0,53379811 | 0,931381467 | CELF2 divergent transript |
| ENSG00000225556 | C2CD4D | 0,467474552 | 1,382686945 | 0,6870577 | 0,965769168 | C2 calcium dependent domain containing 4D |
| ENSG00000010295 | IFFO1 | 0,467309201 | 1,382528481 | 0,14269677 | 0,601245287 | intermediate filament family orphan 1 |
| ENSG00000287185 | AC041005,1 | 0,467195625 | 1,382419646 | 0,63466243 | 0,955149453 | novel transcript |
| ENSG00000008282 | SYPL1 | 0,467162269 | 1,382387683 | 0,34232847 | 0,840048641 | synaptophysin like 1 |
| ENSG00000230317 | LINC01284 | 0,467048804 | 1,382278966 | 0,54168997 | 0,933491609 | long intergenic non-protein coding RNA 1284 |
| ENSG00000255498 | AC068385,1 | 0,466998143 | 1,382230428 | 0,67829716 | 0,9640473 | novel transcript |
| ENSG00000162836 | ACP6 | 0,466894754 | 1,382131375 | 0,20832772 | 0,712989352 | acid phosphatase 6, lysophosphatidic |
| ENSG00000227322 | RXRB | 0,466882182 | 1,38211933 | 0,05287233 | 0,353356233 | retinoid X receptor beta |
| ENSG00000130706 | ADRM1 | 0,466701715 | 1,381946452 | 0,13499971 | 0,586038109 | adhesion regulating molecule 1 |
| ENSG00000130208 | APOC1 | 0,466628881 | 1,381876687 | 0,15496165 | 0,624591393 | apolipoprotein C1 |

| | | | | | | |
|---|---|---|---|---|---|---|
| ENSG00000261615 | LINC01858 | 0,466611329 | 1,381859875 | 0,62785788 | 0,952793934 | long intergenic non-protein coding RNA 1858 |
| ENSG00000243232 | PCDHAC2 | 0,466405097 | 1,381662353 | 0,16347569 | 0,642091247 | protocadherin alpha subfamily C, 2 |
| ENSG00000116406 | EDEM3 | 0,466163081 | 1,381430595 | 0,16458828 | 0,643811841 | ER degradation enhancing alpha-mannosidase like protein 3 |
| ENSG00000225831 | RPS18P1 | 0,466044277 | 1,381316841 | 0,73754753 | 0,974353474 | ribosomal protein S18 pseudogene 1 |
| ENSG00000135824 | RGS8 | 0,466023184 | 1,381296645 | 0,21922581 | 0,727587652 | regulator of G protein signaling 8 |
| ENSG00000184378 | ACTRT3 | 0,466003213 | 1,381277524 | 0,23039987 | 0,739095163 | actin related protein T3 |
| ENSG00000204520 | MICA | 0,465997648 | 1,381272196 | 0,40122291 | 0,876659568 | MHC class I polypeptide-related sequence A |
| ENSG00000188761 | BCL2L15 | 0,465920241 | 1,381198087 | 0,48289636 | 0,91363215 | BCL2 like 15 |
| ENSG00000263968 | RN7SL381P | 0,46566716 | 1,380955815 | 0,6130145 | 0,952413269 | RNA, 7SL, cytoplasmic 381, pseudogene |
| ENSG00000053254 | FOXN3 | 0,465640772 | 1,380930557 | 0,07628962 | 0,43651514 | forkhead box N3 |
| ENSG00000110171 | TRIM3 | 0,465512367 | 1,380807654 | 0,09002766 | 0,478514897 | tripartite motif containing 3 |
| ENSG00000079313 | REXO1 | 0,465502761 | 1,380798461 | 0,23503529 | 0,744654651 | RNA exonuclease 1 homolog |
| ENSG00000188313 | PLSCR1 | 0,465445276 | 1,380743443 | 0,22614805 | 0,734959876 | phospholipid scramblase 1 |
| ENSG00000065000 | AP3D1 | 0,465389352 | 1,380689921 | 0,19725493 | 0,695943345 | adaptor related protein complex 3 subunit delta 1 |
| ENSG00000159363 | ATP13A2 | 0,465383074 | 1,380683913 | 0,22913813 | 0,737173617 | ATPase cation transporting 13A2 |
| ENSG00000085265 | FCN1 | 0,465359674 | 1,380661519 | 0,72056214 | 0,972142678 | ficolin 1 |
| ENSG00000240849 | PEDS1 | 0,465272191 | 1,380577801 | 0,0614265 | 0,386017694 | plasmanylethanolamine desaturase 1 |
| ENSG00000144711 | IQSEC1 | 0,465053347 | 1,380368395 | 0,04928883 | 0,338599406 | IQ motif and Sec7 domain ArfGEF 1 |
| ENSG00000284686 | AC119674,2 | 0,465033921 | 1,380349808 | 0,59703927 | 0,948282297 | novel transcript |
| ENSG00000213970 | AC006122,1 | 0,464971732 | 1,380290307 | 0,543916 | 0,934259495 | ribosomal protein S15 (RPS15) pseudogene |
| ENSG00000286879 | AC093639,3 | 0,464936851 | 1,380256936 | 0,49206292 | 0,916995341 | novel transcript |
| ENSG00000139438 | FAM222A | 0,464768274 | 1,380095664 | 0,1698369 | 0,652708149 | family with sequence similarity 222 member A |
| ENSG00000286232 | AL365229,1 | 0,464724128 | 1,380053435 | 0,46913244 | 0,907940653 | novel transcript |
| ENSG00000254093 | PINX1 | 0,464294307 | 1,379642337 | 0,04698955 | 0,329464272 | PIN2 (TERF1) interacting telomerase inhibitor 1 |
| ENSG00000226390 | KCNQ2-AS1 | 0,464187078 | 1,379539799 | 0,71412599 | 0,970867497 | KCNQ2 antisense RNA 1 |
| ENSG00000281651 | KCNQ2-AS1 | 0,464187078 | 1,379539799 | 0,71412599 | 0,970867497 | KCNQ2 antisense RNA 1 |
| ENSG00000242931 | RPL7P49 | 0,464030515 | 1,379390098 | 0,47657039 | 0,91143003 | ribosomal protein L7 pseudogene 49 |
| ENSG00000277126 | LINC02878 | 0,463952987 | 1,379315974 | 0,66144249 | 0,959582738 | long intergenic non-protein coding RNA 2878 |
| ENSG00000189337 | KAZN | 0,463873837 | 1,379240302 | 0,22805988 | 0,736341996 | kazrin, periplakin interacting protein |
| ENSG00000079156 | OSBPL6 | 0,463851648 | 1,37921909 | 0,17599732 | 0,661519708 | oxysterol binding protein like 6 |
| ENSG00000224531 | SMIM13 | 0,46381848 | 1,379187382 | 0,22482753 | 0,733203418 | small integral membrane protein 13 |
| ENSG00000116704 | SLC35D1 | 0,463772681 | 1,3791436 | 0,19374375 | 0,690594523 | solute carrier family 35 member D1 |

| | | | | | | |
|---|---|---|---|---|---|---|
| ENSG00000256879 | AC129102,1 | 0,463712795 | 1,379086352 | 0,78886829 | 0,979462318 | novel transcript |
| ENSG00000076201 | PTPN23 | 0,463470404 | 1,378854668 | 0,12190128 | 0,558242525 | protein tyrosine phosphatase non-receptor type 23 |
| ENSG00000255966 | AC006064,3 | 0,463457258 | 1,378842104 | 0,65713894 | 0,957709856 | novel transcript |
| ENSG00000104976 | SNAPC2 | 0,463439693 | 1,378825317 | 0,20361516 | 0,704097757 | small nuclear RNA activating complex polypeptide 2 |
| ENSG00000068971 | PPP2R5B | 0,463356895 | 1,378746186 | 0,06755064 | 0,408854824 | protein phosphatase 2 regulatory subunit B'beta |
| ENSG00000251675 | AC010260,1 | 0,463336817 | 1,378726998 | 0,76921269 | 0,977508554 | novel transcript |
| ENSG00000133742 | CA1 | 0,46333494 | 1,378725204 | 0,70016114 | 0,968522643 | carbonic anhydrase 1 |
| ENSG00000274756 | AC243732,1 | 0,463159957 | 1,37855799 | 0,58331235 | 0,944174844 | ribosomal protein S2 (RPS2) pseudogene |
| ENSG00000282707 | AC003042,2 | 0,463159957 | 1,37855799 | 0,58331235 | 0,944174844 | ribosomal protein S2 (RPS2) pseudogene |
| ENSG00000219200 | RNASEK | 0,463158868 | 1,37855695 | 0,06717091 | 0,407302071 | ribonuclease K |
| ENSG00000258791 | LINC00520 | 0,462859539 | 1,378270958 | 0,61477304 | 0,952793934 | long intergenic non-protein coding RNA 520 |
| ENSG00000163399 | ATP1A1 | 0,462813699 | 1,378227165 | 0,03814829 | 0,289219826 | ATPase Na+/K+ transporting subunit alpha 1 |
| ENSG00000198692 | EIF1AY | 0,462811415 | 1,378224984 | 0,5962517 | 0,948282297 | eukaryotic translation initiation factor 1A Y-linked |
| ENSG00000171813 | PWWP2B | 0,462764675 | 1,378180333 | 0,31429361 | 0,819877431 | PWWP domain containing 2B |
| ENSG00000160606 | TLCD1 | 0,462719566 | 1,378137242 | 0,13648628 | 0,589216672 | TLC domain containing 1 |
| ENSG00000242282 | AC108488,1 | 0,462462347 | 1,377891554 | 0,30188802 | 0,811368958 | novel transcript |
| ENSG00000100625 | SIX4 | 0,462441812 | 1,377871943 | 0,15423096 | 0,623607455 | SIX homeobox 4 |
| ENSG00000203414 | BTBD7P1 | 0,46240928 | 1,377840872 | 0,61524736 | 0,952793934 | BTB domain containing 7 pseudogene 1 |
| ENSG00000088876 | ZNF343 | 0,462301908 | 1,377738331 | 0,03350902 | 0,26589493 | zinc finger protein 343 |
| ENSG00000005981 | ASB4 | 0,462291997 | 1,377728867 | 0,49345172 | 0,917289924 | ankyrin repeat and SOCS box containing 4 |
| ENSG00000148803 | FUOM | 0,462279498 | 1,37771693 | 0,2583813 | 0,771153141 | fucose mutarotase |
| ENSG00000012124 | CD22 | 0,462011938 | 1,377461445 | 0,67909866 | 0,964151912 | CD22 molecule |
| ENSG00000170915 | PAQR8 | 0,461776353 | 1,377236531 | 0,25389058 | 0,76531122 | progestin and adipoQ receptor family member 8 |
| ENSG00000109089 | CDR2L | 0,461737926 | 1,377199847 | 0,30030175 | 0,810130728 | cerebellar degeneration related protein 2 like |
| ENSG00000047249 | ATP6V1H | 0,461645745 | 1,377111854 | 0,04450089 | 0,318734455 | ATPase H+ transporting V1 subunit H |
| ENSG00000112149 | CD83 | 0,4615069 | 1,376979327 | 0,14388516 | 0,603329615 | CD83 molecule |
| ENSG00000149930 | TAOK2 | 0,46149311 | 1,376966165 | 0,31679007 | 0,821646322 | TAO kinase 2 |
| ENSG00000127463 | EMC1 | 0,461417358 | 1,376893866 | 0,05642364 | 0,366903759 | ER membrane protein complex subunit 1 |
| ENSG00000273999 | RBM17P2 | 0,461412913 | 1,376889624 | 0,28828062 | 0,799645584 | RNA binding motif protein 17 pseudogene 2 |
| ENSG00000272081 | AC008972,2 | 0,461389834 | 1,376867598 | 0,40336097 | 0,877589651 | novel transcript |
| ENSG00000061938 | TNK2 | 0,461370655 | 1,376849294 | 0,48482016 | 0,913690171 | tyrosine kinase non receptor 2 |
| ENSG00000173852 | DPY19L1 | 0,461221481 | 1,376706936 | 0,23791394 | 0,747090777 | dpy-19 like C-mannosyltransferase 1 |

| | | | | | | |
|---|---|---|---|---|---|---|
| ENSG00000181392 | SYNE4 | 0,461188445 | 1,376675412 | 0,39459305 | 0,871872016 | spectrin repeat containing nuclear envelope family member 4 |
| ENSG00000167085 | PHB | 0,461042397 | 1,376536054 | 0,08276402 | 0,457565724 | prohibitin |
| ENSG00000168495 | POLR3D | 0,460891506 | 1,37639209 | 0,06388183 | 0,395798941 | RNA polymerase III subunit D |
| ENSG00000137070 | IL11RA | 0,460799818 | 1,376304619 | 0,09662142 | 0,493760245 | interleukin 11 receptor subunit alpha |
| ENSG00000168575 | SLC20A2 | 0,46074807 | 1,376255253 | 0,03257008 | 0,261251213 | solute carrier family 20 member 2 |
| ENSG00000089682 | RBM41 | 0,46073311 | 1,376240982 | 0,19950577 | 0,698904803 | RNA binding motif protein 41 |
| ENSG00000105738 | SIPA1L3 | 0,46063355 | 1,376146011 | 0,15989182 | 0,634614414 | signal induced proliferation associated 1 like 3 |
| ENSG00000066230 | SLC9A3 | 0,460539219 | 1,376056034 | 0,15514101 | 0,624800779 | solute carrier family 9 member A3 |
| ENSG00000165272 | AQP3 | 0,460460015 | 1,37598049 | 0,37472677 | 0,861771195 | aquaporin 3 (Gill blood group) |
| ENSG00000197177 | ADGRA1 | 0,460320349 | 1,375847289 | 0,26350674 | 0,774637968 | adhesion G protein-coupled receptor A1 |
| ENSG00000258388 | PPT2-EGFL8 | 0,460314549 | 1,375841758 | 0,34203025 | 0,840048641 | PPT2-EGFL8 readthrough (NMD candidate) |
| ENSG00000158427 | TMSB15B | 0,460261025 | 1,375790716 | 0,25149357 | 0,763892102 | thymosin beta 15B |
| ENSG00000168874 | ATOH8 | 0,460259845 | 1,37578959 | 0,26224125 | 0,774068771 | atonal bHLH transcription factor 8 |
| ENSG00000172354 | GNB2 | 0,460254451 | 1,375784447 | 0,13633874 | 0,588661013 | G protein subunit beta 2 |
| ENSG00000172425 | TTC36 | 0,460235082 | 1,375765976 | 0,39876885 | 0,875433675 | tetratricopeptide repeat domain 36 |
| ENSG00000249471 | ZNF324B | 0,460202202 | 1,375734622 | 0,22752394 | 0,736053199 | zinc finger protein 324B |
| ENSG00000267544 | HIKESHIP2 | 0,460196617 | 1,375729296 | 0,73670956 | 0,974353474 | HIKESHI pseudogene 2 |
| ENSG00000254044 | AC034154,1 | 0,460024884 | 1,375565544 | 0,19927056 | 0,698588788 | novel transcript, sense overlapping to STPG2 |
| ENSG00000137502 | RAB30 | 0,459920569 | 1,375466086 | 0,26935682 | 0,781544346 | RAB30, member RAS oncogene family |
| ENSG00000260464 | AL049796,1 | 0,45975637 | 1,375309548 | 0,41094451 | 0,882983885 | novel transcript |
| ENSG00000130956 | HABP4 | 0,459626477 | 1,375185728 | 0,10045988 | 0,503435287 | hyaluronan binding protein 4 |
| ENSG00000287054 | AC113410,5 | 0,459543285 | 1,37510643 | 0,69201859 | 0,967225449 | novel transcript, antisense to ZCCHC10 |
| ENSG00000187079 | TEAD1 | 0,459449933 | 1,375017455 | 0,13595847 | 0,587841041 | TEA domain transcription factor 1 |
| ENSG00000101294 | HM13 | 0,459378824 | 1,374949684 | 0,02864566 | 0,240299088 | histocompatibility minor 13 |
| ENSG00000118094 | TREH | 0,459374697 | 1,37494575 | 0,41675868 | 0,886368152 | trehalase |
| ENSG00000276429 | PLCH2 | 0,459043893 | 1,374630517 | 0,56223751 | 0,940865625 | phospholipase C eta 2 |
| ENSG00000100221 | JOSD1 | 0,458941405 | 1,374532868 | 0,03946415 | 0,295877055 | Josephin domain containing 1 |
| ENSG00000174564 | IL20RB | 0,458606587 | 1,374213906 | 0,28027869 | 0,790779983 | interleukin 20 receptor subunit beta |
| ENSG00000106105 | GARS1 | 0,458546735 | 1,374156896 | 0,03724979 | 0,284946761 | glycyl-tRNA synthetase 1 |
| ENSG00000259308 | FABP5P8 | 0,458506922 | 1,374118975 | 0,69148511 | 0,967201016 | fatty acid binding protein 5 pseudogene 8 |
| ENSG00000105698 | USF2 | 0,458436561 | 1,374051961 | 0,0987143 | 0,498721099 | upstream transcription factor 2, c-fos interacting |
| ENSG00000232284 | GNG12-AS1 | 0,458302203 | 1,373924001 | 0,24634611 | 0,758198269 | GNG12, DIRAS3 and WLS antisense RNA 1 |

| | | | | | | |
|---|---|---|---|---|---|---|
| ENSG00000236212 | AC009262,1 | 0,458164024 | 1,373792415 | 0,52652714 | 0,929261495 | novel transcript |
| ENSG00000167995 | BEST1 | 0,458112125 | 1,373742996 | 0,14809357 | 0,612595581 | bestrophin 1 |
| ENSG00000089505 | CMTM1 | 0,45808775 | 1,373719786 | 0,25033465 | 0,762441549 | CKLF like MARVEL transmembrane domain containing 1 |
| ENSG00000250493 | AP004147,1 | 0,458016032 | 1,373651498 | 0,53159218 | 0,931291085 | novel transcript, antisense to GRIK4 |
| ENSG00000232706 | NUTM2HP | 0,457665045 | 1,373317349 | 0,65213192 | 0,957241719 | NUT family member 2H, pseudogene |
| ENSG00000283658 | RPL23AP82 | 0,457608696 | 1,373263711 | 0,40657036 | 0,879715952 | ribosomal protein L23a pseudogene 82 |
| ENSG00000102030 | NAA10 | 0,457563943 | 1,373221113 | 0,09248757 | 0,483372031 | N-alpha-acetyltransferase 10, NatA catalytic subunit |
| ENSG00000124508 | BTN2A2 | 0,457336035 | 1,373004197 | 0,09290562 | 0,484503945 | butyrophilin subfamily 2 member A2 |
| ENSG00000143774 | GUK1 | 0,457229053 | 1,372902386 | 0,16511284 | 0,644200984 | guanylate kinase 1 |
| ENSG00000178980 | SELENOW | 0,456959297 | 1,372645704 | 0,06952678 | 0,416095977 | selenoprotein W |
| ENSG00000236908 | LINC02827 | 0,456857898 | 1,372549231 | 0,63253562 | 0,954288117 | long intergenic non-protein coding RNA 2827 |
| ENSG00000226916 | WDR46 | 0,456842792 | 1,37253486 | 0,10668423 | 0,521491876 | WD repeat domain 46 |
| ENSG00000204221 | WDR46 | 0,456842792 | 1,37253486 | 0,10668423 | 0,521491876 | WD repeat domain 46 |
| ENSG00000127838 | PNKD | 0,456562228 | 1,372267967 | 0,08419981 | 0,461585627 | PNKD metallo-beta-lactamase domain containing |
| ENSG00000229530 | AL136097,1 | 0,456534854 | 1,372241929 | 0,71109938 | 0,970482408 | novel pseudogene |
| ENSG00000116044 | NFE2L2 | 0,456488403 | 1,372197747 | 0,14925245 | 0,614462554 | nuclear factor, erythroid 2 like 2 |
| ENSG00000137161 | CNPY3 | 0,456462369 | 1,372172985 | 0,15299975 | 0,621569237 | canopy FGF signaling regulator 3 |
| ENSG00000280055 | TMEM75 | 0,456450611 | 1,372161802 | 0,23538513 | 0,744654651 | transmembrane protein 75 |
| ENSG00000156140 | ADAMTS3 | 0,456450589 | 1,372161781 | 0,32362061 | 0,825794844 | ADAM metallopeptidase with thrombospondin type 1 motif 3 |
| ENSG00000228862 | AC068389,1 | 0,456294396 | 1,372013233 | 0,5339118 | 0,931381467 | novel transcript |
| ENSG00000111676 | ATN1 | 0,456272085 | 1,371992015 | 0,31398978 | 0,819877431 | atrophin 1 |
| ENSG00000278376 | AP004609,3 | 0,456213589 | 1,371936387 | 0,33785076 | 0,83719797 | novel transcript |
| ENSG00000160710 | ADAR | 0,456199407 | 1,3719229 | 0,06580393 | 0,402522813 | adenosine deaminase RNA specific |
| ENSG00000130702 | LAMA5 | 0,456102839 | 1,371831072 | 0,36853965 | 0,857548638 | laminin subunit alpha 5 |
| ENSG00000158352 | SHROOM4 | 0,455997793 | 1,37173119 | 0,08985444 | 0,478293227 | shroom family member 4 |
| ENSG00000170390 | DCLK2 | 0,455671051 | 1,371420555 | 0,25103619 | 0,763388331 | doublecortin like kinase 2 |
| ENSG00000258651 | SEC23A-AS1 | 0,455581612 | 1,371335537 | 0,74859567 | 0,975901808 | SEC23A antisense RNA 1 |
| ENSG00000175348 | TMEM9B | 0,455468719 | 1,371228233 | 0,17495322 | 0,66036665 | TMEM9 domain family member B |
| ENSG00000260209 | AP000842,3 | 0,455450664 | 1,371211072 | 0,75505198 | 0,975901808 | novel transcript |
| ENSG00000158792 | SPATA2L | 0,4553284 | 1,371094871 | 0,30829629 | 0,816620804 | spermatogenesis associated 2 like |
| ENSG00000280802 | ITPK1-AS1 | 0,455248879 | 1,371019299 | 0,61512948 | 0,952793934 | ITPK1 antisense RNA 1 |

| | | | | | | |
|---|---|---|---|---|---|---|
| ENSG00000258730 | ITPK1-AS1 | 0,455248879 | 1,371019299 | 0,61512948 | 0,952793934 | ITPK1 antisense RNA 1 |
| ENSG00000090674 | MCOLN1 | 0,455241577 | 1,37101236 | 0,09413469 | 0,487613759 | mucolipin 1 |
| ENSG00000183049 | CAMK1D | 0,455111693 | 1,370888935 | 0,17115138 | 0,654843372 | calcium/calmodulin dependent protein kinase ID |
| ENSG00000165476 | REEP3 | 0,455090127 | 1,370868443 | 0,29514502 | 0,805254732 | receptor accessory protein 3 |
| ENSG00000286506 | FP325317,2 | 0,454783367 | 1,370576986 | 0,65965218 | 0,958691226 | methylenetetrahydrofolate dehydrogenase (NADP+ dependent) 1 like (MTHFD1L) pseudogene |
| ENSG00000117305 | HMGCL | 0,454780106 | 1,370573888 | 0,08636987 | 0,468471143 | 3-hydroxy-3-methylglutaryl-CoA lyase |
| ENSG00000239636 | TFAP2E-AS1 | 0,454769776 | 1,370564075 | 0,71480896 | 0,971041579 | TFAP2E antisense RNA 1 |
| ENSG00000209480 | SNORD83B | 0,454646921 | 1,370447367 | 0,79930388 | 0,980161849 | small nucleolar RNA, C/D box 83B |
| ENSG00000141741 | MIEN1 | 0,454449657 | 1,370259994 | 0,05513091 | 0,362604555 | migration and invasion enhancer 1 |
| ENSG00000198951 | NAGA | 0,454351981 | 1,370167225 | 0,06198078 | 0,388629081 | alpha-N-acetylgalactosaminidase |
| ENSG00000203280 | KIAA1671-AS1 | 0,454071865 | 1,369901217 | 0,61328375 | 0,952413269 | KIAA1671 antisense RNA 1 |
| ENSG00000285818 | AL591893,1 | 0,45402248 | 1,369854325 | 0,55636907 | 0,938061341 | putative UPF0607 protein ENSP00000383783 |
| ENSG00000267586 | LINC00907 | 0,453989369 | 1,369822886 | 0,59321236 | 0,947416795 | long intergenic non-protein coding RNA 907 |
| ENSG00000103549 | RNF40 | 0,453913556 | 1,369750905 | 0,32602324 | 0,828085941 | ring finger protein 40 |
| ENSG00000111405 | ENDOU | 0,453872135 | 1,369711578 | 0,65227403 | 0,957257009 | endonuclease, poly(U) specific |
| ENSG00000162413 | KLHL21 | 0,453675529 | 1,369524931 | 0,17714574 | 0,663320644 | kelch like family member 21 |
| ENSG00000160094 | ZNF362 | 0,45366137 | 1,36951149 | 0,05649858 | 0,366903759 | zinc finger protein 362 |
| ENSG00000286039 | AC093849,2 | 0,453630499 | 1,369482185 | 0,6408189 | 0,956410077 | novel transcript |
| ENSG00000263567 | AC007923,1 | 0,453546477 | 1,36940243 | 0,69893999 | 0,968522643 | novel transcript |
| ENSG00000231393 | AL357874,2 | 0,453522336 | 1,369379515 | 0,61524552 | 0,952793934 | novel transcript |
| ENSG00000165507 | DEPP1 | 0,453392944 | 1,369256704 | 0,23032381 | 0,738927002 | DEPP1 autophagy regulator |
| ENSG00000160799 | CCDC12 | 0,453259959 | 1,369130495 | 0,23345852 | 0,74254697 | coiled-coil domain containing 12 |
| ENSG00000188677 | PARVB | 0,453166378 | 1,369041688 | 0,25066648 | 0,762931919 | parvin beta |
| ENSG00000105438 | KDELR1 | 0,453121332 | 1,368998943 | 0,05269564 | 0,352552191 | KDEL endoplasmic reticulum protein retention receptor 1 |
| ENSG00000283526 | PRRT1B | 0,453087111 | 1,36896647 | 0,7245989 | 0,97326334 | proline rich transmembrane protein 1B |
| ENSG00000159899 | NPR2 | 0,453052207 | 1,36893335 | 0,05131668 | 0,346739721 | natriuretic peptide receptor 2 |
| ENSG00000170889 | RPS9 | 0,45294896 | 1,368835386 | 0,2262529 | 0,734959876 | ribosomal protein S9 |
| ENSG00000123983 | ACSL3 | 0,452905766 | 1,368794403 | 0,19619754 | 0,694046542 | acyl-CoA synthetase long chain family member 3 |
| ENSG00000145050 | MANF | 0,452733959 | 1,368631407 | 0,07467895 | 0,432043455 | mesencephalic astrocyte derived neurotrophic factor |
| ENSG00000107959 | PITRM1 | 0,452684145 | 1,368584151 | 0,05394426 | 0,357994781 | pitrilysin metallopeptidase 1 |

| | | | | | | |
|---|---|---|---|---|---|---|
| ENSG00000165996 | HACD1 | 0,452456037 | 1,368367778 | 0,08917235 | 0,476479989 | 3-hydroxyacyl-CoA dehydratase 1 |
| ENSG00000129474 | AJUBA | 0,452437903 | 1,368350578 | 0,18978039 | 0,684955795 | ajuba LIM protein |
| ENSG00000280634 | THRIL | 0,452415113 | 1,368328962 | 0,39324582 | 0,871294303 | TNF and HNRNPL related immunoregulatory long non-coding RNA |
| ENSG00000142186 | SCYL1 | 0,452051649 | 1,367984278 | 0,12878597 | 0,573392824 | SCY1 like pseudokinase 1 |
| ENSG00000274600 | RIMBP3B | 0,452035953 | 1,367969394 | 0,57004053 | 0,942251611 | RIMS binding protein 3B |
| ENSG00000166734 | GOLM2 | 0,451995491 | 1,367931028 | 0,17024099 | 0,653377651 | golgi membrane protein 2 |
| ENSG00000123901 | GPR83 | 0,451959745 | 1,367897136 | 0,55967585 | 0,939811058 | G protein-coupled receptor 83 |
| ENSG00000200959 | SNORA74A | 0,451954949 | 1,367892588 | 0,38714072 | 0,869441674 | small nucleolar RNA, H/ACA box 74A |
| ENSG00000250979 | AC022905,1 | 0,451776267 | 1,367723181 | 0,34050978 | 0,83910285 | transient receptor potential cation channel, subfamily A, member 1 (TRPA1) pseudogene |
| ENSG00000172765 | TMCC1 | 0,451668173 | 1,367620708 | 0,08196432 | 0,455075899 | transmembrane and coiled-coil domain family 1 |
| ENSG00000125999 | BPIFB1 | 0,451499373 | 1,367460701 | 0,61769125 | 0,952793934 | BPI fold containing family B member 1 |
| ENSG00000253642 | AF279873,3 | 0,451401573 | 1,367368005 | 0,59734152 | 0,948282297 | novel transcript |
| ENSG00000287475 | AL844892,2 | 0,451319634 | 1,367290346 | 0,61617378 | 0,952793934 | novel transcript |
| ENSG00000230647 | AC022816,1 | 0,451293626 | 1,367265698 | 0,75090585 | 0,975901808 | novel transcript |
| ENSG00000286928 | AC012325,1 | 0,451118112 | 1,367099371 | 0,52756461 | 0,92954125 | novel transcript |
| ENSG00000155367 | PPM1J | 0,450866031 | 1,366860519 | 0,30119378 | 0,810679941 | protein phosphatase, Mg2+/Mn2+ dependent 1J |
| ENSG00000172900 | ACTE1P | 0,450861585 | 1,366856307 | 0,5010684 | 0,920520138 | actin epsilon 1, pseudogene |
| ENSG00000006712 | PAF1 | 0,450828541 | 1,366825 | 0,24696101 | 0,759015651 | PAF1 homolog, Paf1/RNA polymerase II complex component |
| ENSG00000285707 | AL008720,1 | 0,450821854 | 1,366818665 | 0,64468206 | 0,956633898 | novel transcript |
| ENSG00000177885 | GRB2 | 0,450614634 | 1,366622357 | 0,04187651 | 0,306904544 | growth factor receptor bound protein 2 |
| ENSG00000148704 | VAX1 | 0,450609463 | 1,366617459 | 0,0486035 | 0,335726918 | ventral anterior homeobox 1 |
| ENSG00000109686 | SH3D19 | 0,450588879 | 1,366597961 | 0,11312546 | 0,537199228 | SH3 domain containing 19 |
| ENSG00000143772 | ITPKB | 0,450522833 | 1,3665354 | 0,29387098 | 0,803761875 | inositol-trisphosphate 3-kinase B |
| ENSG00000236567 | TCF3P1 | 0,450505594 | 1,366519071 | 0,57773088 | 0,94337651 | transcription factor 3 pseudogene 1 |
| ENSG00000123643 | SLC36A1 | 0,450425229 | 1,366442951 | 0,0325742 | 0,261251213 | solute carrier family 36 member 1 |
| ENSG00000055483 | USP36 | 0,450283917 | 1,366309115 | 0,13355349 | 0,583811016 | ubiquitin specific peptidase 36 |
| ENSG00000100994 | PYGB | 0,450168149 | 1,36619948 | 0,16956012 | 0,652045207 | glycogen phosphorylase B |
| ENSG00000155714 | PDZD9 | 0,450067992 | 1,366104638 | 0,54573249 | 0,935146964 | PDZ domain containing 9 |
| ENSG00000174715 | PPIAP72 | 0,449998426 | 1,366038767 | 0,64891083 | 0,956899963 | peptidylprolyl isomerase A pseudogene 72 |
| ENSG00000175727 | MLXIP | 0,449961707 | 1,366003999 | 0,12034456 | 0,553546035 | MLX interacting protein |

| | | | | | | |
|---|---|---|---|---|---|---|
| ENSG00000104375 | STK3 | 0,449844094 | 1,365892642 | 0,188156 | 0,682491978 | serine/threonine kinase 3 |
| ENSG00000253372 | AC016405,1 | 0,449818433 | 1,365868348 | 0,61258252 | 0,952413269 | novel transcript |
| ENSG00000185359 | HGS | 0,449552224 | 1,365616338 | 0,15588182 | 0,626815323 | hepatocyte growth factor-regulated tyrosine kinase substrate |
| ENSG00000281358 | RASSF1-AS1 | 0,449498266 | 1,365565264 | 0,6802168 | 0,964580199 | RASSF1 antisense RNA 1 |
| ENSG00000252316 | RNY4 | 0,449475382 | 1,365543604 | 0,8193709 | 0,98249617 | RNA, Ro60-associated Y4 |
| ENSG00000162944 | RFTN2 | 0,449243828 | 1,36532445 | 0,26560307 | 0,776856049 | raftlin family member 2 |
| ENSG00000253807 | LINC01170 | 0,44922832 | 1,365309774 | 0,73292422 | 0,974353474 | long intergenic non-protein coding RNA 1170 |
| ENSG00000125977 | EIF2S2 | 0,449131806 | 1,36521844 | 0,07016237 | 0,417549035 | eukaryotic translation initiation factor 2 subunit beta |
| ENSG00000188981 | MSANTD1 | 0,449070823 | 1,365160733 | 0,60762142 | 0,950993594 | Myb/SANT DNA binding domain containing 1 |
| ENSG00000213467 | HMGB1P37 | 0,449028674 | 1,36512085 | 0,51705246 | 0,925746338 | high mobility group box 1 pseudogene 37 |
| ENSG00000144218 | AFF3 | 0,448923874 | 1,365021689 | 0,11806727 | 0,549146283 | AF4/FMR2 family member 3 |
| ENSG00000132963 | POMP | 0,448728883 | 1,364837208 | 0,17115834 | 0,654843372 | proteasome maturation protein |
| ENSG00000167608 | TMC4 | 0,448531199 | 1,364650205 | 0,76403036 | 0,976640468 | transmembrane channel like 4 |
| ENSG00000131378 | RFTN1 | 0,448380195 | 1,364507378 | 0,08436575 | 0,462252136 | raftlin, lipid raft linker 1 |
| ENSG00000262380 | AC026401,2 | 0,448366633 | 1,364494551 | 0,70224289 | 0,968742626 | novel transcript, sense to NDE1 |
| ENSG00000282255 | AC243742,1 | 0,448366633 | 1,364494551 | 0,70224289 | 0,968742626 | novel transcript, sense to NDE1 |
| ENSG00000273171 | AC002094,3 | 0,448360038 | 1,364488313 | 0,78061323 | 0,978597905 | novel protein, readthrough between VTN and SEBOX |
| ENSG00000262903 | AC027796,4 | 0,448207928 | 1,364344456 | 0,60317314 | 0,950352286 | novel transcript, antisense to CTNS |
| ENSG00000259782 | AC008915,1 | 0,448100222 | 1,364242604 | 0,56357575 | 0,940865625 | novel transcript |
| ENSG00000259539 | AC051619,5 | 0,448086107 | 1,364229256 | 0,50236277 | 0,920585659 | novel transcript |
| ENSG00000148488 | ST8SIA6 | 0,448000999 | 1,364148779 | 0,5460006 | 0,935201756 | ST8 alpha-N-acetyl-neuraminide alpha-2,8-sialyltransferase 6 |
| ENSG00000182199 | SHMT2 | 0,447978578 | 1,364127579 | 0,11197987 | 0,534821079 | serine hydroxymethyltransferase 2 |
| ENSG00000257767 | AC002996,1 | 0,447946783 | 1,364097516 | 0,36536976 | 0,855323027 | novel protein |
| ENSG00000227373 | RABGAP1L-DT | 0,447928834 | 1,364080545 | 0,49810925 | 0,919274055 | RABGAP1L divergent transcript |
| ENSG00000169692 | AGPAT2 | 0,447925769 | 1,364077647 | 0,20872114 | 0,713398665 | 1-acylglycerol-3-phosphate O-acyltransferase 2 |
| ENSG00000187068 | C3orf70 | 0,447890154 | 1,364043973 | 0,14345418 | 0,602438935 | chromosome 3 open reading frame 70 |
| ENSG00000134294 | SLC38A2 | 0,447763073 | 1,363923826 | 0,29996244 | 0,809916717 | solute carrier family 38 member 2 |
| ENSG00000095951 | HIVEP1 | 0,447722311 | 1,363885289 | 0,11449041 | 0,540617781 | HIVEP zinc finger 1 |
| ENSG00000105379 | ETFB | 0,447708243 | 1,36387199 | 0,21481981 | 0,722164151 | electron transfer flavoprotein subunit beta |
| ENSG00000206353 | SKIV2L | 0,447540134 | 1,363713076 | 0,22142578 | 0,729254784 | Ski2 like RNA helicase |
| ENSG00000228896 | SKIV2L | 0,447540134 | 1,363713076 | 0,22142578 | 0,729254784 | Ski2 like RNA helicase |

| | | | | | | |
|---|---|---|---|---|---|---|
| ENSG00000232616 | SKIV2L | 0,447540134 | 1,363713076 | 0,22142578 | 0,729254784 | Ski2 like RNA helicase |
| ENSG00000111229 | ARPC3 | 0,447501411 | 1,363676473 | 0,22736527 | 0,735964811 | actin related protein 2/3 complex subunit 3 |
| ENSG00000176619 | LMNB2 | 0,447379677 | 1,363561411 | 0,42639002 | 0,892138574 | lamin B2 |
| ENSG00000100439 | ABHD4 | 0,447287023 | 1,363473842 | 0,19570292 | 0,69375328 | abhydrolase domain containing 4, N-acyl phospholipase B |
| ENSG00000099991 | CABIN1 | 0,447217581 | 1,363408215 | 0,10742656 | 0,523062468 | calcineurin binding protein 1 |
| ENSG00000207088 | SNORA7B | 0,447115012 | 1,363311287 | 0,41152075 | 0,883760466 | small nucleolar RNA, H/ACA box 7B |
| ENSG00000126767 | ELK1 | 0,447100257 | 1,363297343 | 0,14210701 | 0,600058555 | ETS transcription factor ELK1 |
| ENSG00000237938 | AL450998,2 | 0,446707876 | 1,362926607 | 0,53678985 | 0,932810731 | novel transcript, antisense to SLC25A34 and PLEKHM2 |
| ENSG00000272895 | AC009303,3 | 0,446565094 | 1,362791726 | 0,51910128 | 0,926800715 | novel transcript, antisense to CCDC93 |
| ENSG00000169359 | SLC33A1 | 0,446559042 | 1,362786009 | 0,07295408 | 0,426006068 | solute carrier family 33 member 1 |
| ENSG00000254873 | AP001267,1 | 0,446525806 | 1,362754615 | 0,63715789 | 0,955985674 | novel transcript, antisense to ATP5L and UBE4A |
| ENSG00000236535 | RC3H1-IT1 | 0,446374789 | 1,362611973 | 0,54561767 | 0,935146964 | RC3H1 intronic transcript 1 |
| ENSG00000282493 | AC100810,2 | 0,446297837 | 1,362539295 | 0,59692943 | 0,948282297 | DNA fragmentation factor, 45kDa, alpha polypeptide (DFFA) pseudogene |
| ENSG00000282021 | AC100810,3 | 0,446297837 | 1,362539295 | 0,59692943 | 0,948282297 | novel transcript |
| ENSG00000141569 | TRIM65 | 0,446295413 | 1,362537006 | 0,21028675 | 0,715586415 | tripartite motif containing 65 |
| ENSG00000102230 | PCYT1B | 0,446289744 | 1,362531651 | 0,24786374 | 0,760164991 | phosphate cytidylyltransferase 1, choline, beta |
| ENSG00000288051 | AC008060,5 | 0,446285292 | 1,362527447 | 0,66590806 | 0,961020116 | novel transcript |
| ENSG00000154328 | NEIL2 | 0,446275549 | 1,362518245 | 0,13894891 | 0,594174432 | nei like DNA glycosylase 2 |
| ENSG00000206357 | NELFE | 0,446169762 | 1,362418342 | 0,29573382 | 0,805426883 | negative elongation factor complex member E |
| ENSG00000231044 | NELFE | 0,446169762 | 1,362418342 | 0,29573382 | 0,805426883 | negative elongation factor complex member E |
| ENSG00000229363 | NELFE | 0,446169762 | 1,362418342 | 0,29573382 | 0,805426883 | negative elongation factor complex member E |
| ENSG00000233801 | NELFE | 0,446169762 | 1,362418342 | 0,29573382 | 0,805426883 | negative elongation factor complex member E |
| ENSG00000206268 | NELFE | 0,446169762 | 1,362418342 | 0,29573382 | 0,805426883 | negative elongation factor complex member E |
| ENSG00000264943 | SH3GL1P2 | 0,446053981 | 1,362309007 | 0,48467519 | 0,913690171 | SH3 domain containing GRB2 like 1, endophilin A2 pseudogene 2 |
| ENSG00000100024 | UPB1 | 0,445979802 | 1,362238963 | 0,57101678 | 0,942436428 | beta-ureidopropionase 1 |
| ENSG00000131871 | SELENOS | 0,445912769 | 1,362175669 | 0,04899015 | 0,337586055 | selenoprotein S |
| ENSG00000145349 | CAMK2D | 0,445791797 | 1,362061454 | 0,14490209 | 0,605968311 | calcium/calmodulin dependent protein kinase II delta |
| ENSG00000149131 | SERPING1 | 0,44567461 | 1,361950821 | 0,10012762 | 0,502228437 | serpin family G member 1 |

| | | | | | | |
|---|---|---|---|---|---|---|
| ENSG00000256343 | AC095350,1 | 0,445534706 | 1,361818754 | 0,77287788 | 0,978310326 | novel transcript |
| ENSG00000064763 | FAR2 | 0,445367067 | 1,361660521 | 0,25159279 | 0,763892102 | fatty acyl-CoA reductase 2 |
| ENSG00000198948 | MFAP3L | 0,445361091 | 1,361654881 | 0,18598461 | 0,679040373 | microfibril associated protein 3 like |
| ENSG00000142303 | ADAMTS10 | 0,445351802 | 1,361646114 | 0,19231257 | 0,688712519 | ADAM metallopeptidase with thrombospondin type 1 motif 10 |
| ENSG00000174243 | DDX23 | 0,445151397 | 1,361456981 | 0,21978894 | 0,727931767 | DEAD-box helicase 23 |
| ENSG00000242610 | OR5BH1P | 0,445094583 | 1,361403367 | 0,63154437 | 0,953845566 | olfactory receptor family 5 subfamily BH member 1 pseudogene |
| ENSG00000254202 | AC015522,1 | 0,44495797 | 1,361274458 | 0,78146571 | 0,978597905 | novel transcript |
| ENSG00000231728 | TMSB15B-AS1 | 0,444896362 | 1,361216328 | 0,1682245 | 0,649838265 | TMSB15B antisense RNA 1 |
| ENSG00000257557 | PPP1R12A-AS1 | 0,444810208 | 1,361135042 | 0,18514254 | 0,677548457 | PPP1R12A antisense RNA 1 |
| ENSG00000270077 | AP003117,1 | 0,444802522 | 1,361127791 | 0,72058391 | 0,972142678 | novel transcript, sense intronic to PGCP |
| ENSG00000206129 | AC006305,1 | 0,444794897 | 1,361120597 | 0,70029849 | 0,968522643 | novel transcript |
| ENSG00000237170 | RPS7P15 | 0,444634794 | 1,360969555 | 0,66364068 | 0,959920886 | ribosomal protein S7 pseudogene 15 |
| ENSG00000233862 | AC016907,2 | 0,444620878 | 1,360956427 | 0,5409018 | 0,933385624 | novel transcript |
| ENSG00000250899 | AC125807,2 | 0,44449793 | 1,36084045 | 0,3931328 | 0,871240371 | novel transcript |
| ENSG00000103472 | RRN3P2 | 0,444396927 | 1,360745181 | 0,57337926 | 0,942475308 | RRN3 pseudogene 2 |
| ENSG00000224957 | LINC01266 | 0,44426864 | 1,360624187 | 0,5780446 | 0,943425812 | long intergenic non-protein coding RNA 1266 |
| ENSG00000144843 | ADPRH | 0,44423687 | 1,360594225 | 0,13294309 | 0,582037931 | ADP-ribosylarginine hydrolase |
| ENSG00000243883 | RN7SL419P | 0,444234707 | 1,360592184 | 0,71921002 | 0,971869926 | RNA, 7SL, cytoplasmic 419, pseudogene |
| ENSG00000006555 | TTC22 | 0,444226891 | 1,360584813 | 0,70214727 | 0,968742626 | tetratricopeptide repeat domain 22 |
| ENSG00000102144 | PGK1 | 0,444175298 | 1,360536157 | 0,03314419 | 0,264359088 | phosphoglycerate kinase 1 |
| ENSG00000154889 | MPPE1 | 0,444121941 | 1,36048584 | 0,1131293 | 0,537199228 | metallophosphoesterase 1 |
| ENSG00000213523 | SRA1 | 0,443974573 | 1,360346877 | 0,03583646 | 0,277393225 | steroid receptor RNA activator 1 |
| ENSG00000168036 | CTNNB1 | 0,443927911 | 1,360302879 | 0,095029 | 0,489534774 | catenin beta 1 |
| ENSG00000226007 | BX005266,2 | 0,443890799 | 1,360267887 | 0,50940329 | 0,922427526 | novel transcript |
| ENSG00000229300 | SNHG32 | 0,44378904 | 1,360171945 | 0,23031377 | 0,738927002 | small nucleolar RNA host gene 32 |
| ENSG00000144040 | SFXN5 | 0,443659395 | 1,360049721 | 0,23131851 | 0,739268552 | sideroflexin 5 |
| ENSG00000281300 | NXN | 0,443607261 | 1,360000574 | 0,68518406 | 0,965415049 | nucleoredoxin |
| ENSG00000280563 | NXN | 0,443607261 | 1,360000574 | 0,68518406 | 0,965415049 | nucleoredoxin |
| ENSG00000114745 | GORASP1 | 0,443484834 | 1,35988517 | 0,08876588 | 0,475527164 | golgi reassembly stacking protein 1 |
| ENSG00000070756 | PABPC1 | 0,443453178 | 1,359855331 | 0,10773376 | 0,523905449 | poly(A) binding protein cytoplasmic 1 |
| ENSG00000177725 | AC105206,1 | 0,443363413 | 1,359770723 | 0,31763871 | 0,822414621 | novel transcript, sense overlapping BMP1 |

| | | | | | | |
|---|---|---|---|---|---|---|
| ENSG00000235032 | BMP7-AS1 | 0,443239399 | 1,359653842 | 0,71316072 | 0,970812834 | BMP7 antisense RNA 1 |
| ENSG00000254180 | AC004083,1 | 0,443220135 | 1,359635688 | 0,61027908 | 0,951841373 | novel transcript |
| ENSG00000127249 | ATP13A4 | 0,443069827 | 1,35949404 | 0,41234441 | 0,884523593 | ATPase 13A4 |
| ENSG00000067560 | RHOA | 0,443067526 | 1,359491872 | 0,06514447 | 0,400662695 | ras homolog family member A |
| ENSG00000143537 | ADAM15 | 0,443014724 | 1,359442116 | 0,21588447 | 0,722999018 | ADAM metallopeptidase domain 15 |
| ENSG00000110048 | OSBP | 0,442949633 | 1,359380783 | 0,04695586 | 0,329301904 | oxysterol binding protein |
| ENSG00000011132 | APBA3 | 0,442749716 | 1,359192424 | 0,21912648 | 0,727412416 | amyloid beta precursor protein binding family A member 3 |
| ENSG00000286538 | AC008798,2 | 0,442632813 | 1,359082291 | 0,56736683 | 0,941425553 | novel transcript |
| ENSG00000253872 | AC083836,1 | 0,442450709 | 1,358910753 | 0,61363529 | 0,952413269 | novel transcript |
| ENSG00000287085 | AC012066,1 | 0,442390798 | 1,358854322 | 0,37976276 | 0,865245798 | novel transcript, antisense to FMNL2 |
| ENSG00000164902 | PHAX | 0,442355053 | 1,358820655 | 0,04205147 | 0,307400909 | phosphorylated adaptor for RNA export |
| ENSG00000186222 | BLOC1S4 | 0,442334596 | 1,358801387 | 0,06009578 | 0,38125698 | biogenesis of lysosomal organelles complex 1 subunit 4 |
| ENSG00000285532 | LINC02788 | 0,442327043 | 1,358794273 | 0,65532575 | 0,957444211 | long intergenic non-protein coding RNA 2788 |
| ENSG00000137872 | SEMA6D | 0,442218132 | 1,3586917 | 0,31755857 | 0,822351863 | semaphorin 6D |
| ENSG00000166166 | TRMT61A | 0,441981901 | 1,358469242 | 0,3402007 | 0,838936323 | tRNA methyltransferase 61A |
| ENSG00000230836 | LINC01293 | 0,441972609 | 1,358460493 | 0,63988554 | 0,956018518 | long intergenic non-protein coding RNA 1293 |
| ENSG00000143816 | WNT9A | 0,441919446 | 1,358410435 | 0,44397941 | 0,900210555 | Wnt family member 9A |
| ENSG00000248599 | FLJ42969 | 0,441611744 | 1,35812074 | 0,80263718 | 0,98082499 | uncharacterized LOC441374 |
| ENSG00000090520 | DNAJB11 | 0,441191115 | 1,357724827 | 0,09901603 | 0,499376869 | DnaJ heat shock protein family (Hsp40) member B11 |
| ENSG00000248559 | AC109454,2 | 0,440926372 | 1,3574757 | 0,48239921 | 0,913158695 | novel transcript |
| ENSG00000130749 | ZC3H4 | 0,440917789 | 1,357467623 | 0,39413716 | 0,871635019 | zinc finger CCCH-type containing 4 |
| ENSG00000264026 | LINC02003 | 0,440864425 | 1,357417413 | 0,51408056 | 0,924227101 | long intergenic non-protein coding RNA 2003 |
| ENSG00000250731 | TPM3P6 | 0,44074977 | 1,35730954 | 0,61695749 | 0,952793934 | tropomyosin 3 pseudogene 6 |
| ENSG00000243437 | RN7SL370P | 0,440654561 | 1,357219968 | 0,78979238 | 0,979462318 | RNA, 7SL, cytoplasmic 370, pseudogene |
| ENSG00000206560 | ANKRD28 | 0,440596558 | 1,357165403 | 0,17403676 | 0,659285746 | ankyrin repeat domain 28 |
| ENSG00000204580 | DDR1 | 0,440510883 | 1,357084809 | 0,24549558 | 0,757479368 | discoidin domain receptor tyrosine kinase 1 |
| ENSG00000244207 | C4A | 0,440462823 | 1,357039602 | 0,73503123 | 0,974353474 | complement C4A (Rodgers blood group) |
| ENSG00000231114 | AC078842,2 | 0,440425318 | 1,357004325 | 0,13734066 | 0,591516506 | novel transcript |
| ENSG00000269944 | AP001267,4 | 0,44032317 | 1,356908247 | 0,68992343 | 0,96699391 | novel transcript |
| ENSG00000148339 | SLC25A25 | 0,44029842 | 1,356884969 | 0,24621776 | 0,758198269 | solute carrier family 25 member 25 |
| ENSG00000273145 | BX537318,1 | 0,440239844 | 1,356829878 | 0,52328687 | 0,927995032 | novel transcript |

| | | | | | | |
|---|---|---|---|---|---|---|
| ENSG00000276463 | TAF9 | 0,440129551 | 1,356726153 | 0,27178605 | 0,784843944 | TATA-box binding protein associated factor 9 |
| ENSG00000237333 | MSH5 | 0,440102427 | 1,356700646 | 0,63876023 | 0,956018518 | mutS homolog 5 |
| ENSG00000143933 | CALM2 | 0,439988335 | 1,356593359 | 0,28786734 | 0,799107075 | calmodulin 2 |
| ENSG00000138185 | ENTPD1 | 0,439773153 | 1,356391034 | 0,24591099 | 0,757752922 | ectonucleoside triphosphate diphosphohydrolase 1 |
| ENSG00000252326 | SNORD116-25 | 0,439749526 | 1,35636882 | 0,75980585 | 0,97614835 | small nucleolar RNA, C/D box 116-25 |
| ENSG00000103502 | CDIPT | 0,439311917 | 1,355957459 | 0,12147428 | 0,556776403 | CDP-diacylglycerol--inositol 3-phosphatidyltransferase |
| ENSG00000171150 | SOCS5 | 0,439298706 | 1,355945042 | 0,27862102 | 0,789775666 | suppressor of cytokine signaling 5 |
| ENSG00000166960 | CCDC178 | 0,438965433 | 1,355631846 | 0,46499778 | 0,907804479 | coiled-coil domain containing 178 |
| ENSG00000131944 | FAAP24 | 0,438937003 | 1,355605132 | 0,16850187 | 0,650506523 | FA core complex associated protein 24 |
| ENSG00000261490 | AC005674,1 | 0,43892088 | 1,355589982 | 0,15277058 | 0,62133458 | novel transcript, overlapping WDR1 |
| ENSG00000198075 | SULT1C4 | 0,438756785 | 1,355435803 | 0,22402161 | 0,732710917 | sulfotransferase family 1C member 4 |
| ENSG00000175582 | RAB6A | 0,438713119 | 1,355394779 | 0,28762376 | 0,799000743 | RAB6A, member RAS oncogene family |
| ENSG00000198106 | AC025279,1 | 0,438660648 | 1,355345484 | 0,47111503 | 0,908869685 | novel transcript |
| ENSG00000107798 | LIPA | 0,438537216 | 1,35522953 | 0,26626056 | 0,77776044 | lipase A, lysosomal acid type |
| ENSG00000198363 | ASPH | 0,438423642 | 1,355122846 | 0,11253748 | 0,535800356 | aspartate beta-hydroxylase |
| ENSG00000169744 | LDB2 | 0,43840281 | 1,355103279 | 0,3738087 | 0,861319894 | LIM domain binding 2 |
| ENSG00000229194 | HLA-V | 0,438368539 | 1,355071088 | 0,62823311 | 0,952793934 | major histocompatibility complex, class I, V (pseudogene) |
| ENSG00000164880 | INTS1 | 0,438037053 | 1,354759771 | 0,33680192 | 0,836162806 | integrator complex subunit 1 |
| ENSG00000151322 | NPAS3 | 0,438028793 | 1,354752015 | 0,20952447 | 0,71402241 | neuronal PAS domain protein 3 |
| ENSG00000100095 | SEZ6L | 0,437991896 | 1,354717368 | 0,28375232 | 0,7950558 | seizure related 6 homolog like |
| ENSG00000111737 | RAB35 | 0,437882182 | 1,354614348 | 0,08890192 | 0,475766813 | RAB35, member RAS oncogene family |
| ENSG00000176087 | SLC35A4 | 0,437814083 | 1,354550408 | 0,06926398 | 0,415287103 | solute carrier family 35 member A4 |
| ENSG00000187650 | VMAC | 0,437686307 | 1,354430445 | 0,22621017 | 0,734959876 | vimentin type intermediate filament associated coiled-coil protein |
| ENSG00000136141 | LRCH1 | 0,437648018 | 1,354394498 | 0,11876265 | 0,550651408 | leucine rich repeats and calponin homology domain containing 1 |
| ENSG00000160285 | LSS | 0,43764254 | 1,354389355 | 0,14328027 | 0,602166199 | lanosterol synthase |
| ENSG00000158014 | SLC30A2 | 0,437384704 | 1,354147323 | 0,59669985 | 0,948282297 | solute carrier family 30 member 2 |
| ENSG00000266946 | MRPL37P1 | 0,437362956 | 1,35412691 | 0,48649884 | 0,914166353 | mitochondrial ribosomal protein L37 pseudogene 1 |
| ENSG00000060656 | PTPRU | 0,437231799 | 1,35400381 | 0,21831729 | 0,725947652 | protein tyrosine phosphatase receptor type U |
| ENSG00000268400 | AC008763,2 | 0,437200219 | 1,353974172 | 0,77799533 | 0,978587255 | novel transcript |

| | | | | | | |
|---|---|---|---|---|---|---|
| ENSG00000184281 | TSSC4 | 0,437196925 | 1,353971081 | 0,31736263 | 0,822252078 | tumor suppressing subtransferable candidate 4 |
| ENSG00000243678 | NME2 | 0,437169509 | 1,353945351 | 0,09804213 | 0,497317904 | NME/NM23 nucleoside diphosphate kinase 2 |
| ENSG00000286883 | AC005041,5 | 0,437142754 | 1,353920242 | 0,42459956 | 0,891157406 | novel transcript, antisense to CCDC142and MRPL53and MOGS |
| ENSG00000215227 | NUTF2P5 | 0,437104098 | 1,353883965 | 0,72346524 | 0,972913663 | nuclear transport factor 2 pseudogene 5 |
| ENSG00000229656 | ITGB1-DT | 0,437074704 | 1,353856381 | 0,44842085 | 0,902062594 | ITGB1 divergent transcript |
| ENSG00000172830 | SSH3 | 0,437059374 | 1,353841995 | 0,37652055 | 0,862818448 | slingshot protein phosphatase 3 |
| ENSG00000254561 | AP003393,1 | 0,436935885 | 1,353726117 | 0,55600936 | 0,938031417 | novel transcript, antisense to PVRL1 |
| ENSG00000013561 | RNF14 | 0,436797762 | 1,353596517 | 0,0809106 | 0,451470483 | ring finger protein 14 |
| ENSG00000170037 | CNTROB | 0,436730996 | 1,353533877 | 0,3250374 | 0,826983176 | centrobin, centriole duplication and spindle assembly protein |
| ENSG00000280026 | AC005618,3 | 0,436672648 | 1,353479135 | 0,56615209 | 0,941123407 | novel transcript, antisense to PCDHGB1 |
| ENSG00000228208 | LINC02869 | 0,436654827 | 1,353462416 | 0,67941825 | 0,964375386 | long intergenic non-protein coding RNA 2869 |
| ENSG00000255104 | AC005324,3 | 0,436524155 | 1,353339833 | 0,50011195 | 0,92041337 | novel transcript |
| ENSG00000259775 | AL138976,2 | 0,436388466 | 1,353212553 | 0,46358982 | 0,907101775 | novel transcript, antisense to EIF5 |
| ENSG00000198853 | RUSC2 | 0,436186769 | 1,35302338 | 0,15395951 | 0,623201341 | RUN and SH3 domain containing 2 |
| ENSG00000108479 | GALK1 | 0,43612557 | 1,352965986 | 0,2253227 | 0,734259846 | galactokinase 1 |
| ENSG00000136938 | ANP32B | 0,436003902 | 1,35285189 | 0,29236892 | 0,802771257 | acidic nuclear phosphoprotein 32 family member B |
| ENSG00000125037 | EMC3 | 0,435876224 | 1,352732169 | 0,09872649 | 0,498721099 | ER membrane protein complex subunit 3 |
| ENSG00000063241 | ISOC2 | 0,435740481 | 1,352604896 | 0,22561735 | 0,734461165 | isochorismatase domain containing 2 |
| ENSG00000143870 | PDIA6 | 0,435693817 | 1,352561146 | 0,07743387 | 0,439762503 | protein disulfide isomerase family A member 6 |
| ENSG00000071655 | MBD3 | 0,435628821 | 1,352500213 | 0,37185983 | 0,859898346 | methyl-CpG binding domain protein 3 |
| ENSG00000103326 | CAPN15 | 0,435576601 | 1,352451258 | 0,42797516 | 0,89282532 | calpain 15 |
| ENSG00000233348 | BAG6 | 0,435505528 | 1,352384632 | 0,37873644 | 0,864112469 | BAG cochaperone 6 |
| ENSG00000173511 | VEGFB | 0,435463965 | 1,352345673 | 0,30384487 | 0,812957289 | vascular endothelial growth factor B |
| ENSG00000107833 | NPM3 | 0,435371614 | 1,352259108 | 0,10716886 | 0,522599956 | nucleophosmin/nucleoplasmin 3 |
| ENSG00000115216 | NRBP1 | 0,435354463 | 1,352243032 | 0,05855115 | 0,37495765 | nuclear receptor binding protein 1 |
| ENSG00000068438 | FTSJ1 | 0,435303338 | 1,352195113 | 0,04371145 | 0,315756651 | FtsJ RNA 2'-O-methyltransferase 1 |
| ENSG00000251354 | AC024451,1 | 0,435261972 | 1,352156342 | 0,6762701 | 0,963580269 | ribosomal protein S4, X-linked (RPS4X) pseudogene |
| ENSG00000196664 | TLR7 | 0,435225516 | 1,352122174 | 0,5501476 | 0,93612918 | toll like receptor 7 |
| ENSG00000127419 | TMEM175 | 0,435113623 | 1,35201731 | 0,1210582 | 0,555610668 | transmembrane protein 175 |
| ENSG00000243849 | CFAP44-AS1 | 0,435055879 | 1,351963196 | 0,7029704 | 0,968742626 | CFAP44 antisense RNA 1 |
| ENSG00000196428 | TSC22D2 | 0,435041652 | 1,351949864 | 0,03727377 | 0,285054324 | TSC22 domain family member 2 |

| | | | | | | |
|---|---|---|---|---|---|---|
| ENSG00000227908 | FLJ31104 | 0,434896412 | 1,351813767 | 0,52012492 | 0,926827337 | uncharacterized LOC441072 |
| ENSG00000017797 | RALBP1 | 0,434772003 | 1,3516972 | 0,03232753 | 0,260224035 | ralA binding protein 1 |
| ENSG00000108509 | CAMTA2 | 0,434646948 | 1,351580038 | 0,22970512 | 0,738154077 | calmodulin binding transcription activator 2 |
| ENSG00000258056 | AC009779,2 | 0,434630616 | 1,351564738 | 0,15724716 | 0,629552485 | novel transcript, antisense to CD63 |
| ENSG00000070193 | FGF10 | 0,434602004 | 1,351537933 | 0,49759467 | 0,918759905 | fibroblast growth factor 10 |
| ENSG00000071889 | FAM3A | 0,434561743 | 1,351500216 | 0,16712986 | 0,647757288 | FAM3 metabolism regulating signaling molecule A |
| ENSG00000259439 | LINC01833 | 0,434422478 | 1,351369761 | 0,28432568 | 0,795731744 | long intergenic non-protein coding RNA 1833 |
| ENSG00000277873 | AC131159,2 | 0,434405746 | 1,351354088 | 0,44201288 | 0,899273096 | novel transcript, antisense to RFC5 |
| ENSG00000117118 | SDHB | 0,434382853 | 1,351332645 | 0,10181489 | 0,507596965 | succinate dehydrogenase complex iron sulfur subunit B |
| ENSG00000022267 | FHL1 | 0,434378868 | 1,351328912 | 0,24115813 | 0,751551821 | four and a half LIM domains 1 |
| ENSG00000137216 | TMEM63B | 0,434356226 | 1,351307704 | 0,07696158 | 0,438408786 | transmembrane protein 63B |
| ENSG00000260924 | LINC01311 | 0,434286473 | 1,351242371 | 0,29359199 | 0,803461221 | long intergenic non-protein coding RNA 1311 |
| ENSG00000136261 | BZW2 | 0,434282751 | 1,351238885 | 0,10784567 | 0,523928283 | basic leucine zipper and W2 domains 2 |
| ENSG00000123131 | PRDX4 | 0,434171283 | 1,351134487 | 0,07214957 | 0,423602654 | peroxiredoxin 4 |
| ENSG00000186834 | HEXIM1 | 0,434070227 | 1,351039848 | 0,19018584 | 0,685594362 | HEXIM P-TEFb complex subunit 1 |
| ENSG00000233806 | LINC01237 | 0,433939055 | 1,350917015 | 0,61966741 | 0,952793934 | long intergenic non-protein coding RNA 1237 |
| ENSG00000185627 | PSMD13 | 0,433831499 | 1,350816305 | 0,07764116 | 0,440299906 | proteasome 26S subunit, non-ATPase 13 |
| ENSG00000147454 | SLC25A37 | 0,433807733 | 1,350794053 | 0,05039357 | 0,343470607 | solute carrier family 25 member 37 |
| ENSG00000158882 | TOMM40L | 0,433787505 | 1,350775113 | 0,05526184 | 0,362959872 | translocase of outer mitochondrial membrane 40 like |
| ENSG00000253183 | AC005531,1 | 0,433708161 | 1,350700827 | 0,53385944 | 0,931381467 | suppression of tumorigenicity 13 (colon carcinoma) (Hsp70 interacting protein) (ST13) pseudogene |
| ENSG00000226203 | LINC01733 | 0,433707875 | 1,350700559 | 0,72206344 | 0,972531855 | long intergenic non-protein coding RNA 1733 |
| ENSG00000156860 | FBRS | 0,433672906 | 1,350667821 | 0,22483129 | 0,733203418 | fibrosin |
| ENSG00000270615 | SGO1P1 | 0,433671747 | 1,350666735 | 0,77649339 | 0,978587255 | shugoshin 1 pseudogene 1 |
| ENSG00000087085 | ACHE | 0,433603463 | 1,350602809 | 0,30998044 | 0,817411297 | acetylcholinesterase (Cartwright blood group) |
| ENSG00000136932 | TRMO | 0,433434756 | 1,35044488 | 0,15955704 | 0,633929525 | tRNA methyltransferase O |
| ENSG00000234766 | LINC01496 | 0,433417337 | 1,350428575 | 0,52501528 | 0,928660697 | long intergenic non-protein coding RNA 1496 |
| ENSG00000113594 | LIFR | 0,433391609 | 1,350404493 | 0,33506593 | 0,834858938 | LIF receptor subunit alpha |
| ENSG00000278869 | BX539320,1 | 0,433246392 | 1,350268572 | 0,76144896 | 0,976285211 | novel transcript |
| ENSG00000099864 | PALM | 0,433217584 | 1,350241611 | 0,32753334 | 0,829485803 | paralemmin |
| ENSG00000039560 | RAI14 | 0,433054789 | 1,350089256 | 0,07026084 | 0,417549035 | retinoic acid induced 14 |

| | | | | | | |
|---|---|---|---|---|---|---|
| ENSG00000173418 | NAA20 | 0,433011498 | 1,350048745 | 0,15453021 | 0,623626246 | N-alpha-acetyltransferase 20, NatB catalytic subunit |
| ENSG00000240682 | ISY1 | 0,432971955 | 1,350011742 | 0,09456502 | 0,488468697 | ISY1 splicing factor homolog |
| ENSG00000232891 | AL450344,3 | 0,432931397 | 1,34997379 | 0,58796233 | 0,946139044 | novel transcript |
| ENSG00000250264 | AL669918,1 | 0,43272032 | 1,349776293 | 0,81185628 | 0,981213877 | novel protein, TAP2-HLA-DOB readthrough |
| ENSG00000286496 | AL359764,1 | 0,432672243 | 1,349731313 | 0,57595396 | 0,942955084 | novel transcript |
| ENSG00000234754 | LINC02817 | 0,432651913 | 1,349712293 | 0,44353289 | 0,900020258 | long intergenic non-protein coding RNA 2817 |
| ENSG00000100266 | PACSIN2 | 0,432507277 | 1,349576986 | 0,13071129 | 0,57719744 | protein kinase C and casein kinase substrate in neurons 2 |
| ENSG00000249631 | AC005699,1 | 0,432224078 | 1,349312092 | 0,76388232 | 0,976640468 | novel transcript |
| ENSG00000170348 | TMED10 | 0,432169759 | 1,34926129 | 0,23557913 | 0,744706368 | transmembrane p24 trafficking protein 10 |
| ENSG00000257452 | AC004551,1 | 0,432163585 | 1,349255516 | 0,71075449 | 0,970482408 | novel transcript, antisense to OAS1, OAS2 and OAS3 |
| ENSG00000141293 | SKAP1 | 0,432131977 | 1,349225956 | 0,73561965 | 0,974353474 | src kinase associated phosphoprotein 1 |
| ENSG00000235513 | L3MBTL2-AS1 | 0,431984381 | 1,349087928 | 0,17052085 | 0,653761763 | L3MBTL2 antisense RNA 1 |
| ENSG00000236635 | LINC02540 | 0,431951796 | 1,349057458 | 0,57999912 | 0,943828882 | long intergenic non-protein coding RNA 2540 |
| ENSG00000172432 | GTPBP2 | 0,431901668 | 1,349010585 | 0,09171991 | 0,481764036 | GTP binding protein 2 |
| ENSG00000113273 | ARSB | 0,431762859 | 1,348880796 | 0,04037451 | 0,300178807 | arylsulfatase B |
| ENSG00000284290 | ANKS4B | 0,431687903 | 1,348810716 | 0,87597053 | 0,990109814 | ankyrin repeat and sterile alpha motif domain containing 4B |
| ENSG00000287169 | AC107050,1 | 0,431524182 | 1,348657657 | 0,3045079 | 0,813200217 | novel transcript, antisense to FAT1 |
| ENSG00000166224 | SGPL1 | 0,431422668 | 1,348562764 | 0,03922673 | 0,294869635 | sphingosine-1-phosphate lyase 1 |
| ENSG00000261644 | AC007728,2 | 0,431071017 | 1,348234097 | 0,68370825 | 0,965206123 | novel transcript, antisense to CYLD and NOD2 |
| ENSG00000287374 | AC239860,4 | 0,430966102 | 1,348136055 | 0,60570466 | 0,950927937 | novel transcript |
| ENSG00000169188 | APEX2 | 0,43096163 | 1,348131876 | 0,09143081 | 0,481302729 | apurinic/apyrimidinic endodeoxyribonuclease 2 |
| ENSG00000226789 | WBP11P2 | 0,430915623 | 1,348088886 | 0,58071972 | 0,943832501 | WBP11 pseudogene 2 |
| ENSG00000070495 | JMJD6 | 0,430742856 | 1,347927458 | 0,10576347 | 0,51904616 | jumonji domain containing 6, arginine demethylase and lysine hydroxylase |
| ENSG00000288213 | MCRIP1 | 0,430652887 | 1,347843402 | 0,31459003 | 0,819877431 | MAPK regulated corepressor interacting protein 1 |
| ENSG00000225663 | MCRIP1 | 0,430652887 | 1,347843402 | 0,31459003 | 0,819877431 | MAPK regulated corepressor interacting protein 1 |
| ENSG00000131188 | PRR7 | 0,430649745 | 1,347840466 | 0,38083263 | 0,865900094 | proline rich 7, synaptic |
| ENSG00000175213 | ZNF408 | 0,43064473 | 1,34783578 | 0,19126817 | 0,686939473 | zinc finger protein 408 |
| ENSG00000197905 | TEAD4 | 0,430577497 | 1,347772969 | 0,18631668 | 0,679406361 | TEA domain transcription factor 4 |
| ENSG00000163576 | EFHB | 0,430411004 | 1,34761744 | 0,35472153 | 0,84617839 | EF-hand domain family member B |

| | | | | | | |
|---|---|---|---|---|---|---|
| ENSG00000227013 | OR7E96P | 0,430364164 | 1,347573687 | 0,72655715 | 0,973344786 | olfactory receptor family 7 subfamily E member 96 pseudogene |
| ENSG00000126012 | KDM5C | 0,430356202 | 1,347566251 | 0,21752344 | 0,725248255 | lysine demethylase 5C |
| ENSG00000275265 | AC127002,2 | 0,430330982 | 1,347542694 | 0,74004354 | 0,974724313 | novel transcript |
| ENSG00000280474 | AL356481,2 | 0,430318252 | 1,347530803 | 0,5357539 | 0,932139691 | novel transcript, sense intronic to SPTAN1 |
| ENSG00000070214 | SLC44A1 | 0,430150693 | 1,347374306 | 0,30371638 | 0,812795736 | solute carrier family 44 member 1 |
| ENSG00000149922 | TBX6 | 0,430131162 | 1,347356066 | 0,47825286 | 0,912116096 | T-box transcription factor 6 |
| ENSG00000254505 | CHMP4A | 0,430044222 | 1,347274873 | 0,11725696 | 0,547082438 | charged multivesicular body protein 4A |
| ENSG00000155252 | PI4K2A | 0,429939525 | 1,347177104 | 0,09611841 | 0,49254976 | phosphatidylinositol 4-kinase type 2 alpha |
| ENSG00000160211 | G6PD | 0,429876296 | 1,347118063 | 0,13423146 | 0,584425029 | glucose-6-phosphate dehydrogenase |
| ENSG00000126522 | ASL | 0,429782769 | 1,347030735 | 0,26704161 | 0,779386557 | argininosuccinate lyase |
| ENSG00000241484 | ARHGAP8 | 0,429770575 | 1,34701935 | 0,5596962 | 0,939811058 | Rho GTPase activating protein 8 |
| ENSG00000278341 | AC138028,6 | 0,429747839 | 1,346998122 | 0,56917943 | 0,94219857 | novel transcript, antisense to CTU2 |
| ENSG00000287372 | AL627316,1 | 0,429386683 | 1,346660964 | 0,63227419 | 0,954164226 | novel transcript |
| ENSG00000181481 | RNF135 | 0,429376329 | 1,3466513 | 0,17240805 | 0,656917487 | ring finger protein 135 |
| ENSG00000109016 | DHRS7B | 0,429316513 | 1,346595466 | 0,08183277 | 0,454749316 | dehydrogenase/reductase 7B |
| ENSG00000182195 | LDOC1 | 0,429291582 | 1,346572197 | 0,08050467 | 0,450041893 | LDOC1 regulator of NFKB signaling |
| ENSG00000158315 | RHBDL2 | 0,429202761 | 1,346489296 | 0,44858586 | 0,902062594 | rhomboid like 2 |
| ENSG00000119280 | C1orf198 | 0,429067806 | 1,346363346 | 0,04454079 | 0,318908336 | chromosome 1 open reading frame 198 |
| ENSG00000244405 | ETV5 | 0,429019683 | 1,346318437 | 0,18198873 | 0,672311616 | ETS variant transcription factor 5 |
| ENSG00000197150 | ABCB8 | 0,428932087 | 1,346236696 | 0,19007235 | 0,685541073 | ATP binding cassette subfamily B member 8 |
| ENSG00000237766 | GGTA2P | 0,428788783 | 1,34610298 | 0,60994127 | 0,951801946 | glycoprotein alpha-galactosyltransferase 2, pseudogene |
| ENSG00000176871 | WSB2 | 0,428720989 | 1,346039726 | 0,08849547 | 0,474566472 | WD repeat and SOCS box containing 2 |
| ENSG00000260072 | AC008938,1 | 0,428621521 | 1,345946925 | 0,55027901 | 0,936146662 | novel transcript |
| ENSG00000228628 | ATF6B | 0,428552116 | 1,345882176 | 0,56937709 | 0,94221458 | activating transcription factor 6 beta |
| ENSG00000160401 | CFAP157 | 0,428510154 | 1,34584303 | 0,39360038 | 0,8715972 | cilia and flagella associated protein 157 |
| ENSG00000118412 | CASP8AP2 | 0,428493117 | 1,345827137 | 0,56764795 | 0,941632387 | caspase 8 associated protein 2 |
| ENSG00000227935 | AL390243,1 | 0,428473182 | 1,345808541 | 0,58951293 | 0,946941469 | novel transcript |
| ENSG00000196968 | FUT11 | 0,428377978 | 1,345719733 | 0,05201284 | 0,349872693 | fucosyltransferase 11 |
| ENSG00000128272 | ATF4 | 0,428325974 | 1,345671226 | 0,03516629 | 0,273970725 | activating transcription factor 4 |
| ENSG00000262766 | AC135050,5 | 0,428040263 | 1,345404755 | 0,43826863 | 0,897177404 | novel transcript |
| ENSG00000265313 | LINC01476 | 0,427966629 | 1,345336089 | 0,59197841 | 0,947416795 | long intergenic non-protein coding RNA 1476 |

| | | | | | | |
|---|---|---|---|---|---|---|
| ENSG00000227200 | AL162724,1 | 0,427923099 | 1,345295498 | 0,68516645 | 0,965415049 | novel transcript |
| ENSG00000203356 | LINC01562 | 0,42790773 | 1,345281166 | 0,63120777 | 0,953705762 | long intergenic non-protein coding RNA 1562 |
| ENSG00000281951 | AC093627,11 | 0,427892711 | 1,345267161 | 0,84511264 | 0,986639902 | novel transcript similar to family member with sequence similarity 157 |
| ENSG00000162390 | ACOT11 | 0,427870122 | 1,345246098 | 0,28955236 | 0,800474391 | acyl-CoA thioesterase 11 |
| ENSG00000182934 | SRPRA | 0,427870041 | 1,345246022 | 0,03866258 | 0,291967859 | SRP receptor subunit alpha |
| ENSG00000117595 | IRF6 | 0,427786981 | 1,345168575 | 0,54998664 | 0,93612918 | interferon regulatory factor 6 |
| ENSG00000270000 | AC005479,2 | 0,427784659 | 1,34516641 | 0,75230567 | 0,975901808 | novel transcript |
| ENSG00000184916 | JAG2 | 0,427722463 | 1,345108419 | 0,49461533 | 0,91758518 | jagged canonical Notch ligand 2 |
| ENSG00000182446 | NPLOC4 | 0,427720616 | 1,345106698 | 0,09505563 | 0,489591294 | NPL4 homolog, ubiquitin recognition factor |
| ENSG00000103534 | TMC5 | 0,42754483 | 1,344942812 | 0,51245516 | 0,92347658 | transmembrane channel like 5 |
| ENSG00000287916 | AC091607,2 | 0,427499119 | 1,344900199 | 0,52313604 | 0,92790906 | novel transcript, antisense to SSR3 |
| ENSG00000273616 | NDUFA6-DT | 0,427445921 | 1,344850608 | 0,3669329 | 0,856359365 | NDUFA6 divergent transcript |
| ENSG00000229052 | AL449283,1 | 0,427439405 | 1,344844535 | 0,56838099 | 0,94163744 | GAJ protein pseudogene |
| ENSG00000136280 | CCM2 | 0,427215701 | 1,344636019 | 0,15796743 | 0,630981681 | CCM2 scaffold protein |
| ENSG00000205464 | ATP6AP1L | 0,427122922 | 1,344549549 | 0,1917802 | 0,687672451 | ATPase H+ transporting accessory protein 1 like |
| ENSG00000167625 | ZNF526 | 0,427113396 | 1,344540671 | 0,3154685 | 0,82082095 | zinc finger protein 526 |
| ENSG00000116663 | FBXO6 | 0,426998464 | 1,344433563 | 0,39944666 | 0,875433675 | F-box protein 6 |
| ENSG00000228697 | AL023755,1 | 0,426877071 | 1,344320442 | 0,67963798 | 0,964459885 | novel transcript |
| ENSG00000181016 | LSMEM1 | 0,426846183 | 1,344291662 | 0,56463753 | 0,940865625 | leucine rich single-pass membrane protein 1 |
| ENSG00000163406 | SLC15A2 | 0,426701488 | 1,344156842 | 0,23977727 | 0,749639658 | solute carrier family 15 member 2 |
| ENSG00000092529 | CAPN3 | 0,426673195 | 1,344130482 | 0,39240097 | 0,870909282 | calpain 3 |
| ENSG00000284874 | AC000093,1 | 0,426542771 | 1,344008974 | 0,48767093 | 0,914936533 | novel transcript |
| ENSG00000039523 | RIPOR1 | 0,426402658 | 1,343878451 | 0,21971103 | 0,727931767 | RHO family interacting cell polarization regulator 1 |
| ENSG00000235319 | AC012360,1 | 0,42638803 | 1,343864826 | 0,64960847 | 0,956899963 | novel transcript |
| ENSG00000106636 | YKT6 | 0,426348916 | 1,343828391 | 0,04124593 | 0,304151397 | YKT6 v-SNARE homolog |
| ENSG00000111729 | CLEC4A | 0,426318859 | 1,343800394 | 0,62906743 | 0,952793934 | C-type lectin domain family 4 member A |
| ENSG00000117519 | CNN3 | 0,42626977 | 1,343754671 | 0,2479127 | 0,760164991 | calponin 3 |
| ENSG00000135709 | KIAA0513 | 0,426206528 | 1,343695768 | 0,23063306 | 0,739234644 | KIAA0513 |
| ENSG00000163344 | PMVK | 0,426141245 | 1,343634966 | 0,35342349 | 0,84514549 | phosphomevalonate kinase |
| ENSG00000236871 | LINC00106 | 0,426074645 | 1,34357294 | 0,66966383 | 0,962016728 | long intergenic non-protein coding RNA 106 |
| ENSG00000104872 | PIH1D1 | 0,426045312 | 1,343545623 | 0,1149185 | 0,541330721 | PIH1 domain containing 1 |
| ENSG00000274636 | GLIS2 | 0,425950826 | 1,343457634 | 0,18955993 | 0,684882433 | GLIS family zinc finger 2 |

| | | | | | | |
|---|---|---|---|---|---|---|
| ENSG00000272434 | AC137630,3 | 0,42590285 | 1,343412958 | 0,6966142 | 0,968300554 | novel transcript, antisense to IMPDH2 and QRICH1 |
| ENSG00000241132 | LY6G5B | 0,425881579 | 1,343393151 | 0,44452975 | 0,900210555 | lymphocyte antigen 6 family member G5B |
| ENSG00000178537 | SLC25A20 | 0,425794224 | 1,343311811 | 0,10046484 | 0,503435287 | solute carrier family 25 member 20 |
| ENSG00000103855 | CD276 | 0,425779737 | 1,343298322 | 0,15016776 | 0,615923738 | CD276 molecule |
| ENSG00000272768 | AC004854,2 | 0,425731869 | 1,343253753 | 0,57598558 | 0,942955084 | novel transcript, antisense to PURB |
| ENSG00000170175 | CHRNB1 | 0,425678958 | 1,34320449 | 0,6531423 | 0,95732962 | cholinergic receptor nicotinic beta 1 subunit |
| ENSG00000173786 | CNP | 0,42566483 | 1,343191337 | 0,17102781 | 0,654709629 | 2',3'-cyclic nucleotide 3' phosphodiesterase |
| ENSG00000188986 | NELFB | 0,425610577 | 1,343140826 | 0,32810408 | 0,830076239 | negative elongation factor complex member B |
| ENSG00000082014 | SMARCD3 | 0,425570329 | 1,343103356 | 0,06089801 | 0,383945238 | SWI/SNF related, matrix associated, actin dependent regulator of chromatin, subfamily d, member 3 |
| ENSG00000153292 | ADGRF1 | 0,425545196 | 1,343079958 | 0,43712129 | 0,897021798 | adhesion G protein-coupled receptor F1 |
| ENSG00000253626 | EIF5AL1 | 0,425537656 | 1,343072939 | 0,67669373 | 0,963580269 | eukaryotic translation initiation factor 5A like 1 |
| ENSG00000214273 | AGGF1P1 | 0,425494371 | 1,343032644 | 0,46870896 | 0,907940653 | angiogenic factor with G-patch and FHA domains 1 pseudogene 1 |
| ENSG00000284859 | MIR124-1HG | 0,425468497 | 1,343008557 | 0,18078368 | 0,670312865 | MIR124-1 host gene |
| ENSG00000117682 | DHDDS | 0,425432962 | 1,342975478 | 0,07647637 | 0,436971724 | dehydrodolichyl diphosphate synthase subunit |
| ENSG00000285170 | AC009152,2 | 0,425267031 | 1,342821025 | 0,78709228 | 0,979424965 | novel transcript, sense intronic to XYLT1 |
| ENSG00000275927 | AC009152,1 | 0,425267031 | 1,342821025 | 0,78709228 | 0,979424965 | novel transcript, sense intronic to XYLT1 |
| ENSG00000124541 | RRP36 | 0,425263478 | 1,342817718 | 0,05814152 | 0,373098594 | ribosomal RNA processing 36 |
| ENSG00000172269 | DPAGT1 | 0,425249799 | 1,342804986 | 0,09333975 | 0,485472307 | dolichyl-phosphate N-acetylglucosaminephosphotransferase 1 |
| ENSG00000106367 | AP1S1 | 0,425242276 | 1,342797984 | 0,18157507 | 0,671647294 | adaptor related protein complex 1 subunit sigma 1 |
| ENSG00000125434 | SLC25A35 | 0,425229793 | 1,342786365 | 0,24142333 | 0,751948055 | solute carrier family 25 member 35 |
| ENSG00000241061 | RPL5P1 | 0,425019966 | 1,342591083 | 0,50970259 | 0,922427526 | ribosomal protein L5 pseudogene 1 |
| ENSG00000276191 | CRHR1 | 0,425014859 | 1,342586331 | 0,75777276 | 0,976120207 | corticotropin releasing hormone receptor 1 |
| ENSG00000179071 | CCDC89 | 0,424829366 | 1,342413721 | 0,54433476 | 0,934381824 | coiled-coil domain containing 89 |
| ENSG00000226479 | TMEM185B | 0,424760076 | 1,342349248 | 0,06326096 | 0,393511843 | transmembrane protein 185B |
| ENSG00000238227 | TMEM250 | 0,424741699 | 1,34233215 | 0,25793885 | 0,770351899 | transmembrane protein 250 |
| ENSG00000087269 | NOP14 | 0,424652954 | 1,342249581 | 0,22677285 | 0,735383523 | NOP14 nucleolar protein |
| ENSG00000162512 | SDC3 | 0,424443234 | 1,342054476 | 0,1963548 | 0,694046542 | syndecan 3 |
| ENSG00000287391 | AC103871,1 | 0,424401062 | 1,342015247 | 0,55872139 | 0,939635125 | novel transcript |
| ENSG00000272341 | AL137003,1 | 0,42430469 | 1,341925603 | 0,55547593 | 0,937793851 | novel transcript |

| | | | | | | |
|---|---|---|---|---|---|---|
| ENSG00000107036 | RIC1 | 0,424061456 | 1,341699378 | 0,18194448 | 0,672297868 | RIC1 homolog, RAB6A GEF complex partner 1 |
| ENSG00000170734 | POLH | 0,424027732 | 1,341668015 | 0,1271998 | 0,569656846 | DNA polymerase eta |
| ENSG00000038295 | TLL1 | 0,423846147 | 1,341499156 | 0,3288498 | 0,830667565 | tolloid like 1 |
| ENSG00000228124 | AL096678,1 | 0,423767051 | 1,34142561 | 0,61445071 | 0,952793934 | novel transcript |
| ENSG00000086015 | MAST2 | 0,423735555 | 1,341396325 | 0,10776123 | 0,523928283 | microtubule associated serine/threonine kinase 2 |
| ENSG00000242337 | INHCAP | 0,423719793 | 1,341381671 | 0,68575922 | 0,965740648 | inhibitor of carbonic anhydrase pseudogene |
| ENSG00000276488 | AC008735,4 | 0,423618728 | 1,341287705 | 0,58180352 | 0,944145586 | novel transcript, antisense to ZNF865 |
| ENSG00000054282 | SDCCAG8 | 0,423550029 | 1,341223837 | 0,41833616 | 0,887094957 | SHH signaling and ciliogenesis regulator SDCCAG8 |
| ENSG00000224063 | AC007319,1 | 0,423458005 | 1,341138288 | 0,51009951 | 0,922427526 | novel transcript, antisense to TFPI and CALCRL |
| ENSG00000173976 | RAX2 | 0,423444495 | 1,34112573 | 0,729628 | 0,974009551 | retina and anterior neural fold homeobox 2 |
| ENSG00000255237 | AC138230,1 | 0,42342872 | 1,341111065 | 0,60020031 | 0,949340299 | novel transcript |
| ENSG00000070182 | SPTB | 0,423390382 | 1,341075427 | 0,50982242 | 0,922427526 | spectrin beta, erythrocytic |
| ENSG00000273820 | USP27X | 0,423348635 | 1,341036621 | 0,15581479 | 0,626626384 | ubiquitin specific peptidase 27 X-linked |
| ENSG00000253480 | AC122713,1 | 0,42324345 | 1,340938851 | 0,72143978 | 0,972142678 | chromosome 4 open reading frame 27 (C4orf27) pseudogene |
| ENSG00000164949 | GEM | 0,423216921 | 1,340914194 | 0,28810073 | 0,799375215 | GTP binding protein overexpressed in skeletal muscle |
| ENSG00000169220 | RGS14 | 0,423106242 | 1,340811327 | 0,26277764 | 0,774168869 | regulator of G protein signaling 14 |
| ENSG00000103876 | FAH | 0,423091367 | 1,340797503 | 0,14067032 | 0,597543965 | fumarylacetoacetate hydrolase |
| ENSG00000226247 | SUPT4H1P1 | 0,423072516 | 1,340779983 | 0,7349362 | 0,974353474 | SPT4 homolog, DSIF elongation factor subunit pseudogene |
| ENSG00000168646 | AXIN2 | 0,422917885 | 1,340636283 | 0,06453333 | 0,39817842 | axin 2 |
| ENSG00000165861 | ZFYVE1 | 0,422903516 | 1,340622931 | 0,07978438 | 0,447817422 | zinc finger FYVE-type containing 1 |
| ENSG00000135045 | C9orf40 | 0,422886628 | 1,340607238 | 0,20545325 | 0,707800596 | chromosome 9 open reading frame 40 |
| ENSG00000135736 | CCDC102A | 0,422857383 | 1,340580063 | 0,4975803 | 0,918759905 | coiled-coil domain containing 102A |
| ENSG00000182985 | CADM1 | 0,422856096 | 1,340578866 | 0,24092392 | 0,751467261 | cell adhesion molecule 1 |
| ENSG00000056277 | ZNF280C | 0,422791589 | 1,340518927 | 0,32459928 | 0,826676737 | zinc finger protein 280C |
| ENSG00000268282 | AC010328,2 | 0,422639334 | 1,340377463 | 0,77224718 | 0,97820249 | actin related protein 2/3 complex, subunit 1A, 41kDa pseudogene |
| ENSG00000260996 | BX255925,1 | 0,422559285 | 1,340303092 | 0,66086372 | 0,959296562 | novel transcript, overlapping NRARP |
| ENSG00000139734 | DIAPH3 | 0,422552575 | 1,340296859 | 0,07392789 | 0,429530795 | diaphanous related formin 3 |
| ENSG00000270409 | AC090950,1 | 0,422490838 | 1,340239506 | 0,48488311 | 0,913690171 | chromosome 9 open reading frame 64 (C9orf64) pseudogene |

| | | | | | | |
|---|---|---|---|---|---|---|
| ENSG00000174151 | CYB561D1 | 0,422487905 | 1,34023678 | 0,19832937 | 0,69732 | cytochrome b561 family member D1 |
| ENSG00000116863 | ADPRS | 0,422418063 | 1,3401719 | 0,18312943 | 0,674047379 | ADP-ribosylserine hydrolase |
| ENSG00000221974 | GTF2H4 | 0,422406386 | 1,340161053 | 0,39098661 | 0,870057277 | general transcription factor IIH subunit 4 |
| ENSG00000233149 | GTF2H4 | 0,422406386 | 1,340161053 | 0,39098661 | 0,870057277 | general transcription factor IIH subunit 4 |
| ENSG00000260750 | AC092720,1 | 0,422301013 | 1,340063173 | 0,69685772 | 0,968300554 | novel transcript |
| ENSG00000229721 | AC104115,1 | 0,422264139 | 1,340028922 | 0,6077321 | 0,950993594 | ribosomal protein S8 (RPS8) pseudogene |
| ENSG00000117597 | UTP25 | 0,422095492 | 1,339872286 | 0,05569685 | 0,364179132 | UTP25 small subunit processor component |
| ENSG00000171189 | GRIK1 | 0,421835385 | 1,339630739 | 0,45653895 | 0,903545728 | glutamate ionotropic receptor kainate type subunit 1 |
| ENSG00000168310 | IRF2 | 0,421745617 | 1,339547386 | 0,18031532 | 0,66974377 | interferon regulatory factor 2 |
| ENSG00000242525 | OR7E100P | 0,421562017 | 1,339376923 | 0,70348452 | 0,968940012 | olfactory receptor family 7 subfamily E member 100 pseudogene |
| ENSG00000174775 | HRAS | 0,421561866 | 1,339376783 | 0,24855507 | 0,760720365 | HRas proto-oncogene, GTPase |
| ENSG00000143393 | PI4KB | 0,421511647 | 1,339330162 | 0,08261601 | 0,457152163 | phosphatidylinositol 4-kinase beta |
| ENSG00000255663 | AP002373,1 | 0,42140483 | 1,339231001 | 0,87733552 | 0,990109814 | novel transcript, RBM7-REXO2 readthrough |
| ENSG00000270689 | BUD13P1 | 0,421392534 | 1,339219587 | 0,74482167 | 0,975649186 | BUD13 homolog pseudogene 1 |
| ENSG00000148356 | LRSAM1 | 0,421390853 | 1,339218027 | 0,27630546 | 0,788263115 | leucine rich repeat and sterile alpha motif containing 1 |
| ENSG00000105325 | FZR1 | 0,421352316 | 1,339182254 | 0,25558075 | 0,767141836 | fizzy and cell division cycle 20 related 1 |
| ENSG00000178021 | TSPYL6 | 0,421324651 | 1,339156575 | 0,62953749 | 0,952812183 | TSPY like 6 |
| ENSG00000277144 | AC092757,3 | 0,421242549 | 1,339080367 | 0,56138028 | 0,940567693 | novel transcript, antisense to CCNB2 |
| ENSG00000185088 | RPS27L | 0,421217711 | 1,339057313 | 0,19911053 | 0,69833874 | ribosomal protein S27 like |
| ENSG00000140931 | CMTM3 | 0,421196271 | 1,339037414 | 0,09795916 | 0,497096776 | CKLF like MARVEL transmembrane domain containing 3 |
| ENSG00000261404 | AC138627,1 | 0,421145197 | 1,33899001 | 0,54116013 | 0,933385624 | novel transcript |
| ENSG00000185507 | IRF7 | 0,421105771 | 1,338953418 | 0,62474781 | 0,952793934 | interferon regulatory factor 7 |
| ENSG00000166411 | IDH3A | 0,421046306 | 1,338898231 | 0,16406735 | 0,642888389 | isocitrate dehydrogenase (NAD(+)) 3 catalytic subunit alpha |
| ENSG00000273230 | AC102953,2 | 0,421036992 | 1,338889587 | 0,49440947 | 0,91758518 | novel transcript |
| ENSG00000125651 | GTF2F1 | 0,421008761 | 1,338863387 | 0,31272562 | 0,818683344 | general transcription factor IIF subunit 1 |
| ENSG00000282977 | PCBP2-OT1 | 0,420975148 | 1,338832194 | 0,34451704 | 0,841168721 | PCBP2 overlapping transcript 1 |
| ENSG00000234553 | AC022431,1 | 0,420829447 | 1,338696989 | 0,68123309 | 0,964654539 | novel transcript |
| ENSG00000099953 | MMP11 | 0,420775289 | 1,338646736 | 0,3324186 | 0,83293962 | matrix metallopeptidase 11 |

| | | | | | | |
|---|---|---|---|---|---|---|
| ENSG00000047936 | ROS1 | 0,420690382 | 1,338567955 | 0,66709961 | 0,9611116 | ROS proto-oncogene 1, receptor tyrosine kinase |
| ENSG00000104866 | PPP1R37 | 0,420558021 | 1,338445153 | 0,34177089 | 0,840048641 | protein phosphatase 1 regulatory subunit 37 |
| ENSG00000261670 | AC012213,2 | 0,420389715 | 1,338289018 | 0,61091087 | 0,951873283 | novel transcript |
| ENSG00000155850 | SLC26A2 | 0,420221966 | 1,338133418 | 0,19235649 | 0,688790885 | solute carrier family 26 member 2 |
| ENSG00000224424 | PRKAR2A-AS1 | 0,419911676 | 1,337845648 | 0,51608882 | 0,925176589 | PRKAR2A antisense RNA 1 |
| ENSG00000057019 | DCBLD2 | 0,419849854 | 1,337788319 | 0,11657032 | 0,545320605 | discoidin, CUB and LCCL domain containing 2 |
| ENSG00000204348 | DXO | 0,419843331 | 1,337782271 | 0,12943804 | 0,574985916 | decapping exoribonuclease |
| ENSG00000288505 | NANOS3 | 0,419796836 | 1,337739158 | 0,52170324 | 0,927472429 | nanos C2HC-type zinc finger 3 |
| ENSG00000187556 | NANOS3 | 0,419796836 | 1,337739158 | 0,52170324 | 0,927472429 | nanos C2HC-type zinc finger 3 |
| ENSG00000132432 | SEC61G | 0,419776576 | 1,337720372 | 0,32995006 | 0,831651927 | SEC61 translocon subunit gamma |
| ENSG00000189045 | ANKDD1B | 0,419632745 | 1,337587013 | 0,57237109 | 0,942475308 | ankyrin repeat and death domain containing 1B |
| ENSG00000119632 | IFI27L2 | 0,419631161 | 1,337585545 | 0,31593157 | 0,821327495 | interferon alpha inducible protein 27 like 2 |
| ENSG00000087152 | ATXN7L3 | 0,419527214 | 1,337489174 | 0,10812475 | 0,524257484 | ataxin 7 like 3 |
| ENSG00000278229 | RPS17 | 0,419445551 | 1,337413468 | 0,12730542 | 0,569803406 | ribosomal protein S17 |
| ENSG00000055332 | EIF2AK2 | 0,419336225 | 1,337312124 | 0,19972435 | 0,698923433 | eukaryotic translation initiation factor 2 alpha kinase 2 |
| ENSG00000130158 | DOCK6 | 0,419324909 | 1,337301635 | 0,27190678 | 0,784843944 | dedicator of cytokinesis 6 |
| ENSG00000068912 | ERLEC1 | 0,419248843 | 1,337231128 | 0,26444574 | 0,775793521 | endoplasmic reticulum lectin 1 |
| ENSG00000113719 | ERGIC1 | 0,419117307 | 1,337109213 | 0,10456994 | 0,515821166 | endoplasmic reticulum-golgi intermediate compartment 1 |
| ENSG00000263020 | AL662899,2 | 0,419092252 | 1,337085992 | 0,68269449 | 0,964820564 | novel protein |
| ENSG00000123836 | PFKFB2 | 0,419051392 | 1,337048124 | 0,14369881 | 0,602952565 | 6-phosphofructo-2-kinase/fructose-2,6-biphosphatase 2 |
| ENSG00000228980 | LINC01205 | 0,418950508 | 1,33695463 | 0,64921729 | 0,956899963 | long intergenic non-protein coding RNA 1205 |
| ENSG00000284807 | DHRS4 | 0,41888415 | 1,336893137 | 0,19275101 | 0,689581368 | dehydrogenase/reductase 4 |
| ENSG00000157326 | DHRS4 | 0,41888415 | 1,336893137 | 0,19275101 | 0,689581368 | dehydrogenase/reductase 4 |
| ENSG00000164402 | SEPTIN8 | 0,418825663 | 1,336838941 | 0,0385574 | 0,291314266 | septin 8 |
| ENSG00000147383 | NSDHL | 0,418594137 | 1,33662442 | 0,10391413 | 0,514127437 | NAD(P) dependent steroid dehydrogenase-like |
| ENSG00000240668 | KRT8P36 | 0,418581457 | 1,336612672 | 0,61840034 | 0,952793934 | keratin 8 pseudogene 36 |
| ENSG00000258926 | AL355916,2 | 0,418573847 | 1,336605621 | 0,71369447 | 0,970812834 | novel transcript, antisense to PRKCH |
| ENSG00000157107 | FCHO2 | 0,418555491 | 1,336588616 | 0,34586748 | 0,841168721 | FCH and mu domain containing endocytic adaptor 2 |
| ENSG00000270479 | BNIP3P37 | 0,418497084 | 1,336534505 | 0,42836179 | 0,893058343 | BCL2 interacting protein 3 pseudogene 37 |
| ENSG00000237720 | AC011995,2 | 0,41827588 | 1,336329594 | 0,75934829 | 0,976120207 | novel transcript |

| | | | | | | |
|---|---|---|---|---|---|---|
| ENSG00000119383 | PTPA | 0,417637855 | 1,33573874 | 0,16255186 | 0,639794628 | protein phosphatase 2 phosphatase activator |
| ENSG00000274023 | AL360169,2 | 0,417617485 | 1,33571988 | 0,61895283 | 0,952793934 | novel transcript, sense overlapping TULP4 |
| ENSG00000234913 | AC016027,2 | 0,417563171 | 1,335669594 | 0,62889671 | 0,952793934 | novel transcript |
| ENSG00000115234 | SNX17 | 0,417554311 | 1,335661392 | 0,09644244 | 0,493240029 | sorting nexin 17 |
| ENSG00000234729 | AGER | 0,417422354 | 1,33553923 | 0,40105981 | 0,876659568 | advanced glycosylation end-product specific receptor |
| ENSG00000276931 | AC009041,3 | 0,417413611 | 1,335531136 | 0,55724661 | 0,938419505 | novel transcript, sense intronic to LMF1 |
| ENSG00000175573 | C11orf68 | 0,417319337 | 1,335443868 | 0,21571126 | 0,722999018 | chromosome 11 open reading frame 68 |
| ENSG00000119715 | ESRRB | 0,417170763 | 1,335306346 | 0,5649554 | 0,940865625 | estrogen related receptor beta |
| ENSG00000146223 | RPL7L1 | 0,417059039 | 1,335202943 | 0,04653791 | 0,327767757 | ribosomal protein L7 like 1 |
| ENSG00000134375 | TIMM17A | 0,416973073 | 1,335123384 | 0,1305398 | 0,577136255 | translocase of inner mitochondrial membrane 17A |
| ENSG00000105701 | FKBP8 | 0,416970442 | 1,335120949 | 0,27245749 | 0,785492261 | FKBP prolyl isomerase 8 |
| ENSG00000100949 | RABGGTA | 0,416911865 | 1,335066741 | 0,24432535 | 0,755755471 | Rab geranylgeranyltransferase subunit alpha |
| ENSG00000285193 | RABGGTA | 0,416911865 | 1,335066741 | 0,24432535 | 0,755755471 | Rab geranylgeranyltransferase subunit alpha |
| ENSG00000143845 | ETNK2 | 0,416899936 | 1,335055702 | 0,31966184 | 0,824366234 | ethanolamine kinase 2 |
| ENSG00000269480 | AC020913,2 | 0,416896389 | 1,33505242 | 0,61075148 | 0,951873283 | novel transcript, antisense MYO9B |
| ENSG00000102934 | PLLP | 0,416748306 | 1,334915392 | 0,19445616 | 0,691556971 | plasmolipin |
| ENSG00000160200 | CBS | 0,415892062 | 1,334123351 | 0,30336313 | 0,812669425 | cystathionine beta-synthase |
| ENSG00000276063 | AL391262,1 | 0,415786959 | 1,334026161 | 0,72162156 | 0,972195829 | novel transcript |
| ENSG00000259299 | AC061965,2 | 0,415758617 | 1,333999954 | 0,59308224 | 0,947416795 | DDB1 and CUL4 associated factor 6 (DCAF6) pseudogene |
| ENSG00000228146 | CASP16P | 0,415739981 | 1,333982723 | 0,62081795 | 0,952793934 | caspase 16, pseudogene |
| ENSG00000115486 | GGCX | 0,415712984 | 1,33395776 | 0,04841515 | 0,335106518 | gamma-glutamyl carboxylase |
| ENSG00000272788 | AP000864,1 | 0,415644151 | 1,333894117 | 0,38582662 | 0,869112622 | novel transcript, sense intronic to SOGA2 |
| ENSG00000126895 | AVPR2 | 0,415526965 | 1,333785772 | 0,80635156 | 0,981213877 | arginine vasopressin receptor 2 |
| ENSG00000234814 | SVIL2P | 0,415418189 | 1,333685212 | 0,47689684 | 0,91143003 | supervillin family member 2, pseudogene |
| ENSG00000170043 | TRAPPC1 | 0,415301651 | 1,333577483 | 0,06301311 | 0,39228231 | trafficking protein particle complex 1 |
| ENSG00000176022 | B3GALT6 | 0,415212374 | 1,333494961 | 0,21373579 | 0,721156449 | beta-1,3-galactosyltransferase 6 |
| ENSG00000134516 | DOCK2 | 0,415171137 | 1,333456846 | 0,35985024 | 0,850238665 | dedicator of cytokinesis 2 |
| ENSG00000187944 | C2orf66 | 0,415137229 | 1,333425507 | 0,53410093 | 0,93151582 | chromosome 2 open reading frame 66 |
| ENSG00000286979 | AC005234,2 | 0,415037547 | 1,333333377 | 0,72726211 | 0,973344786 | novel transcript, antisense to ADD2 |
| ENSG00000100304 | TTLL12 | 0,414924719 | 1,333229106 | 0,18064219 | 0,670229021 | tubulin tyrosine ligase like 12 |
| ENSG00000271360 | AL512631,2 | 0,414724601 | 1,333044185 | 0,68612709 | 0,965740648 | novel transcript, antisense to USP6NL |

| | | | | | | |
|---|---|---|---|---|---|---|
| ENSG00000079805 | DNM2 | 0,41470384 | 1,333025002 | 0,0956349 | 0,49111852 | dynamin 2 |
| ENSG00000101844 | ATG4A | 0,414548724 | 1,332881685 | 0,20239138 | 0,702433811 | autophagy related 4A cysteine peptidase |
| ENSG00000169926 | KLF13 | 0,414519662 | 1,332854836 | 0,124858 | 0,564757949 | Kruppel like factor 13 |
| ENSG00000077800 | FKBP6 | 0,414426215 | 1,332768507 | 0,50747663 | 0,922098508 | FKBP prolyl isomerase 6 |
| ENSG00000139998 | RAB15 | 0,414391636 | 1,332736563 | 0,0741312 | 0,429959479 | RAB15, member RAS oncogene family |
| ENSG00000169764 | UGP2 | 0,414355001 | 1,33270272 | 0,26201679 | 0,773902615 | UDP-glucose pyrophosphorylase 2 |
| ENSG00000196092 | PAX5 | 0,414062288 | 1,332432351 | 0,38546101 | 0,868895792 | paired box 5 |
| ENSG00000163898 | LIPH | 0,413941388 | 1,332320696 | 0,5501004 | 0,93612918 | lipase H |
| ENSG00000186862 | PDZD7 | 0,413856532 | 1,332242334 | 0,49725695 | 0,918759905 | PDZ domain containing 7 |
| ENSG00000177822 | TENM3-AS1 | 0,413825745 | 1,332213905 | 0,15936177 | 0,633636845 | TENM3 antisense RNA 1 |
| ENSG00000170379 | TCAF2 | 0,413811532 | 1,33220078 | 0,26702224 | 0,779386557 | TRPM8 channel associated factor 2 |
| ENSG00000179304 | FAM156B | 0,413743314 | 1,332137789 | 0,28606929 | 0,797420923 | family with sequence similarity 156 member B |
| ENSG00000084093 | REST | 0,413717157 | 1,332113636 | 0,27853577 | 0,789775666 | RE1 silencing transcription factor |
| ENSG00000044115 | CTNNA1 | 0,41362556 | 1,332029063 | 0,05062521 | 0,344150161 | catenin alpha 1 |
| ENSG00000273797 | AL133445,2 | 0,413486905 | 1,331901049 | 0,59414081 | 0,947469769 | novel transcript, sense intronic to KIAA0247 |
| ENSG00000127418 | FGFRL1 | 0,413477146 | 1,331892041 | 0,29301287 | 0,803365244 | fibroblast growth factor receptor like 1 |
| ENSG00000120656 | TAF12 | 0,413453308 | 1,331870033 | 0,18121916 | 0,670965888 | TATA-box binding protein associated factor 12 |
| ENSG00000128045 | RASL11B | 0,413378126 | 1,331800628 | 0,17121074 | 0,654848602 | RAS like family 11 member B |
| ENSG00000261114 | AC012181,1 | 0,413365807 | 1,331789256 | 0,58320614 | 0,944174844 | novel transcript, sense intronic to HERPUD1 |
| ENSG00000105726 | ATP13A1 | 0,413110325 | 1,331553435 | 0,17847565 | 0,665863105 | ATPase 13A1 |
| ENSG00000282101 | AC019257,5 | 0,413033039 | 1,331482105 | 0,78982357 | 0,979462318 | novel transcript, antisense to ARHGEF10 |
| ENSG00000267009 | AC007780,1 | 0,413016182 | 1,331466547 | 0,62382145 | 0,952793934 | novel transcript |
| ENSG00000215305 | VPS16 | 0,412999947 | 1,331451565 | 0,08538322 | 0,465136959 | VPS16 core subunit of CORVET and HOPS complexes |
| ENSG00000251076 | AC104126,1 | 0,412992448 | 1,331444644 | 0,70586972 | 0,969503793 | novel transcript |
| ENSG00000121388 | AL139327,1 | 0,412990341 | 1,331442699 | 0,77923161 | 0,978587255 | CDC42 binding protein kinase beta (DMPK-like) (CDC42BPB) pseudogene |
| ENSG00000109991 | P2RX3 | 0,412879445 | 1,331340359 | 0,32859078 | 0,830659639 | purinergic receptor P2X 3 |
| ENSG00000262664 | OVCA2 | 0,4128395 | 1,331303497 | 0,14067053 | 0,597543965 | OVCA2 serine hydrolase domain containing |
| ENSG00000141971 | MVB12A | 0,412836776 | 1,331300984 | 0,20063541 | 0,700152419 | multivesicular body subunit 12A |
| ENSG00000288675 | AP006621,6 | 0,412756132 | 1,331226568 | 0,47040818 | 0,908757071 | Proapoptotic nucleolar protein 1 |
| ENSG00000259699 | HMGB1P8 | 0,412738049 | 1,331209883 | 0,29841806 | 0,807839821 | high mobility group box 1 pseudogene 8 |
| ENSG00000265489 | AC005358,1 | 0,41269841 | 1,331173308 | 0,55680292 | 0,938264532 | novel transcript, antisense to MYOCD |
| ENSG00000164619 | BMPER | 0,412677693 | 1,331154192 | 0,75640198 | 0,975901808 | BMP binding endothelial regulator |

| | | | | | | |
|---|---|---|---|---|---|---|
| ENSG00000224051 | CPTP | 0,412631158 | 1,331111255 | 0,24894896 | 0,76093812 | ceramide-1-phosphate transfer protein |
| ENSG00000244217 | RPS4XP10 | 0,412566958 | 1,331052022 | 0,6191448 | 0,952793934 | ribosomal protein S4X pseudogene 10 |
| ENSG00000198841 | KTI12 | 0,412549948 | 1,331036328 | 0,16726031 | 0,647913628 | KTI12 chromatin associated homolog |
| ENSG00000248968 | AC012640,1 | 0,412501693 | 1,33099181 | 0,65390837 | 0,957444211 | novel transcript |
| ENSG00000135828 | RNASEL | 0,412341459 | 1,33084399 | 0,08723343 | 0,470867736 | ribonuclease L |
| ENSG00000265692 | LINC01970 | 0,412285726 | 1,330792579 | 0,60102857 | 0,94978555 | long intergenic non-protein coding RNA 1970 |
| ENSG00000275340 | FGD5P1 | 0,412272336 | 1,330780227 | 0,66091858 | 0,959312917 | FYVE, RhoGEF and PH domain containing 5 pseudogene 1 |
| ENSG00000233220 | LINC00167 | 0,412115894 | 1,330635929 | 0,57819703 | 0,943425812 | long intergenic non-protein coding RNA 167 |
| ENSG00000231401 | AC023481,1 | 0,412018303 | 1,330545922 | 0,51062327 | 0,922693987 | novel transcript |
| ENSG00000277200 | AC005696,4 | 0,411967242 | 1,33049883 | 0,55072792 | 0,9363105 | novel transcript |
| ENSG00000214999 | AC129492,1 | 0,411909287 | 1,330445384 | 0,67732372 | 0,963580269 | novel transcript, antisense to ALOX12B |
| ENSG00000275832 | ARHGAP23 | 0,411796151 | 1,330341054 | 0,44482875 | 0,900384201 | Rho GTPase activating protein 23 |
| ENSG00000285493 | NRL | 0,41176577 | 1,33031304 | 0,6629931 | 0,959920886 | neural retina leucine zipper |
| ENSG00000177427 | MIEF2 | 0,411764981 | 1,330312312 | 0,18865746 | 0,682944308 | mitochondrial elongation factor 2 |
| ENSG00000260025 | CRIM1-DT | 0,411721638 | 1,330272345 | 0,5032723 | 0,920888747 | CRIM1 divergent transcript |
| ENSG00000183117 | CSMD1 | 0,411615977 | 1,330174922 | 0,43151794 | 0,89530663 | CUB and Sushi multiple domains 1 |
| ENSG00000162614 | NEXN | 0,411587943 | 1,330149075 | 0,25297556 | 0,76515438 | nexilin F-actin binding protein |
| ENSG00000187066 | TMEM262 | 0,411268507 | 1,329854591 | 0,40494747 | 0,878176155 | transmembrane protein 262 |
| ENSG00000182329 | KIAA2012 | 0,411267919 | 1,329854049 | 0,57162788 | 0,942436428 | KIAA2012 |
| ENSG00000227007 | LINC01247 | 0,411191443 | 1,329783556 | 0,52972898 | 0,930679791 | long intergenic non-protein coding RNA 1247 |
| ENSG00000196979 | GPRACR | 0,411121693 | 1,329719267 | 0,61503909 | 0,952793934 | GPR107 adjacent cis regulating lncRNA |
| ENSG00000267222 | AC107993,1 | 0,41109094 | 1,329690923 | 0,76528974 | 0,976830764 | novel transcript |
| ENSG00000162734 | PEA15 | 0,411084006 | 1,329684531 | 0,11236941 | 0,535454522 | proliferation and apoptosis adaptor protein 15 |
| ENSG00000181135 | ZNF707 | 0,411036075 | 1,329640355 | 0,44424635 | 0,900210555 | zinc finger protein 707 |
| ENSG00000088970 | KIZ | 0,411017633 | 1,329623359 | 0,23388656 | 0,743339368 | kizuna centrosomal protein |
| ENSG00000261399 | AL031710,1 | 0,411010425 | 1,329616716 | 0,69285648 | 0,967271667 | novel transcript, antisense to MAPK8IP3 |
| ENSG00000175206 | NPPA | 0,411008435 | 1,329614882 | 0,64222277 | 0,956633898 | natriuretic peptide A |
| ENSG00000140406 | TLNRD1 | 0,410647568 | 1,329282342 | 0,26534208 | 0,776700553 | talin rod domain containing 1 |
| ENSG00000226065 | ZBTB45P2 | 0,410610208 | 1,329247919 | 0,84173048 | 0,98571999 | zinc finger and BTB domain containing 45 pseudogene 2 |
| ENSG00000099330 | OCEL1 | 0,410606502 | 1,329244504 | 0,20363523 | 0,704097757 | occludin/ELL domain containing 1 |
| ENSG00000134287 | ARF3 | 0,41060111 | 1,329239537 | 0,05692355 | 0,368611841 | ADP ribosylation factor 3 |

| | | | | | | |
|---|---|---|---|---|---|---|
| ENSG00000183688 | RFLNB | 0,4104987 | 1,329145183 | 0,8410314 | 0,985672131 | refilin B |
| ENSG00000169814 | BTD | 0,410468691 | 1,329117537 | 0,07854267 | 0,443561759 | biotinidase |
| ENSG00000196961 | AP2A1 | 0,41046522 | 1,329114339 | 0,10915287 | 0,527070407 | adaptor related protein complex 2 subunit alpha 1 |
| ENSG00000217195 | AL513475,1 | 0,410432431 | 1,329084132 | 0,8071545 | 0,981213877 | 5'-nucleotidase domain containing 1 (NT5DC1) pseudogene |
| ENSG00000132915 | PDE6A | 0,410423962 | 1,32907633 | 0,53452412 | 0,93162317 | phosphodiesterase 6A |
| ENSG00000103978 | TMEM87A | 0,410360501 | 1,329017868 | 0,09271956 | 0,484139956 | transmembrane protein 87A |
| ENSG00000206487 | C6orf136 | 0,410261639 | 1,328926798 | 0,35258605 | 0,844912682 | chromosome 6 open reading frame 136 |
| ENSG00000242759 | LINC00882 | 0,410248125 | 1,328914351 | 0,2095431 | 0,71402241 | long intergenic non-protein coding RNA 882 |
| ENSG00000258768 | AL356019,2 | 0,41022832 | 1,328896108 | 0,27933274 | 0,790200177 | novel transcript |
| ENSG00000286110 | AC090983,2 | 0,410169571 | 1,328841994 | 0,28618285 | 0,797420923 | novel transcript |
| ENSG00000182749 | PAQR7 | 0,410134616 | 1,328809798 | 0,35779912 | 0,848920986 | progestin and adipoQ receptor family member 7 |
| ENSG00000172250 | SERHL | 0,410030918 | 1,328714289 | 0,73521587 | 0,974353474 | serine hydrolase like (pseudogene) |
| ENSG00000153443 | UBALD1 | 0,409922972 | 1,328614875 | 0,27952734 | 0,790302992 | UBA like domain containing 1 |
| ENSG00000221970 | OR2A1 | 0,409902317 | 1,328595854 | 0,50426976 | 0,921400408 | olfactory receptor family 2 subfamily A member 1 |
| ENSG00000262304 | AC027796,3 | 0,409700219 | 1,328409752 | 0,51654339 | 0,92540328 | novel readthrough SHPK-TRPV1 protein |
| ENSG00000286916 | ABBA0104507 | 0,409694154 | 1,328404168 | 0,75767434 | 0,976120207 | novel tripartite motif-containing protein family pseudogene |
| ENSG00000273559 | CWC25 | 0,409534705 | 1,328257358 | 0,2844008 | 0,795731744 | CWC25 spliceosome associated protein homolog |
| ENSG00000167281 | RBFOX3 | 0,409358525 | 1,328095163 | 0,44669297 | 0,901650934 | RNA binding fox-1 homolog 3 |
| ENSG00000167118 | URM1 | 0,4093495 | 1,328086855 | 0,12303849 | 0,561148863 | ubiquitin related modifier 1 |
| ENSG00000035681 | NSMAF | 0,40932323 | 1,328062672 | 0,22439124 | 0,733134896 | neutral sphingomyelinase activation associated factor |
| ENSG00000271375 | AC134879,2 | 0,409319964 | 1,328059665 | 0,72937681 | 0,974009551 | immunoglobulin kappa variable 1D-39 (IGKV1D-39) pseudogene |
| ENSG00000177106 | EPS8L2 | 0,40912535 | 1,327880528 | 0,54718243 | 0,935404974 | EPS8 like 2 |
| ENSG00000103245 | CIAO3 | 0,409019364 | 1,32778298 | 0,22544067 | 0,734457344 | cytosolic iron-sulfur assembly component 3 |
| ENSG00000223575 | RBMX2P3 | 0,408874675 | 1,327649823 | 0,64514814 | 0,956899963 | RBMX2 pseudogene 3 |
| ENSG00000089597 | GANAB | 0,408753979 | 1,327538756 | 0,06329231 | 0,393628542 | glucosidase II alpha subunit |
| ENSG00000234788 | HSPA8P3 | 0,408733748 | 1,32752014 | 0,58437082 | 0,944741266 | heat shock protein family A (Hsp70) member 8 pseudogene 3 |
| ENSG00000156509 | FBXO43 | 0,408669758 | 1,32746126 | 0,207941 | 0,712494639 | F-box protein 43 |
| ENSG00000161618 | ALDH16A1 | 0,408651034 | 1,327444031 | 0,4372612 | 0,897021798 | aldehyde dehydrogenase 16 family member A1 |

| | | | | | | |
|---|---|---|---|---|---|---|
| ENSG00000173950 | XXYLT1 | 0,408586572 | 1,32738472 | 0,08027788 | 0,449383651 | xyloside xylosyltransferase 1 |
| ENSG00000279457 | WASH9P | 0,408317422 | 1,327137105 | 0,34632259 | 0,841168721 | WAS protein family homolog 9, pseudogene |
| ENSG00000198026 | ZNF335 | 0,40829236 | 1,327114051 | 0,10858161 | 0,525250741 | zinc finger protein 335 |
| ENSG00000265451 | AC012447,1 | 0,408286641 | 1,327108791 | 0,62840214 | 0,952793934 | novel transcript, antisense to CLASP1 |
| ENSG00000102316 | MAGED2 | 0,408082195 | 1,326920737 | 0,05596124 | 0,365178946 | MAGE family member D2 |
| ENSG00000236449 | AC010894,2 | 0,408074198 | 1,326913382 | 0,33541284 | 0,835232738 | novel transcript, antisense to WIPF1 and CHRNA1 |
| ENSG00000152767 | FARP1 | 0,407900816 | 1,326753925 | 0,03813167 | 0,289216019 | FERM, ARH/RhoGEF and pleckstrin domain protein 1 |
| ENSG00000177380 | PPFIA3 | 0,407877106 | 1,32673212 | 0,41716593 | 0,886651836 | PTPRF interacting protein alpha 3 |
| ENSG00000009335 | UBE3C | 0,407624815 | 1,326500128 | 0,07023807 | 0,417549035 | ubiquitin protein ligase E3C |
| ENSG00000105722 | ERF | 0,407562951 | 1,326443248 | 0,23424622 | 0,743801925 | ETS2 repressor factor |
| ENSG00000274266 | SNORA73A | 0,40752641 | 1,326409652 | 0,12000401 | 0,553128623 | small nucleolar RNA, H/ACA box 73A |
| ENSG00000172689 | MS4A10 | 0,407391153 | 1,326285303 | 0,61811164 | 0,952793934 | membrane spanning 4-domains A10 |
| ENSG00000124575 | H1-3 | 0,407341032 | 1,326239226 | 0,20419139 | 0,705240783 | H1,3 linker histone, cluster member |
| ENSG00000037897 | METTL1 | 0,407316166 | 1,326216368 | 0,0899851 | 0,478514897 | methyltransferase like 1 |
| ENSG00000141933 | TPGS1 | 0,407297516 | 1,326199224 | 0,44483541 | 0,900384201 | tubulin polyglutamylase complex subunit 1 |
| ENSG00000232169 | ABCF1 | 0,407247799 | 1,326153522 | 0,26837371 | 0,780709773 | ATP binding cassette subfamily F member 1 |
| ENSG00000264491 | AC005544,1 | 0,407241358 | 1,326147602 | 0,7052413 | 0,969427597 | novel transcript |
| ENSG00000203387 | SNF8P1 | 0,407061593 | 1,325982369 | 0,605031 | 0,950927937 | SNF8 pseudogene 1 |
| ENSG00000011275 | RNF216 | 0,407030484 | 1,325953777 | 0,04838752 | 0,335106518 | ring finger protein 216 |
| ENSG00000271519 | AL590065,1 | 0,406986945 | 1,325913762 | 0,5450567 | 0,934923038 | CDC42 binding protein kinase beta (DMPK-like) (CDC42BPB) pseudogene |
| ENSG00000105676 | ARMC6 | 0,406965461 | 1,325894018 | 0,18064539 | 0,670229021 | armadillo repeat containing 6 |
| ENSG00000282503 | AC243686,1 | 0,406944399 | 1,32587466 | 0,42802155 | 0,89282532 | novel transcript |
| ENSG00000259690 | AC107980,1 | 0,406944399 | 1,32587466 | 0,42802155 | 0,89282532 | novel transcript |
| ENSG00000131149 | GSE1 | 0,406909233 | 1,325842342 | 0,10086665 | 0,504639949 | Gse1 coiled-coil protein |
| ENSG00000267834 | AL592211,1 | 0,406888027 | 1,325822855 | 0,54666832 | 0,935404974 | novel transcript |
| ENSG00000009950 | MLXIPL | 0,406836536 | 1,325775535 | 0,58755209 | 0,946068138 | MLX interacting protein like |
| ENSG00000229770 | AL022337,1 | 0,406616552 | 1,325573395 | 0,72376204 | 0,973025002 | novel transcript |
| ENSG00000283847 | HBP1 | 0,406615235 | 1,325572185 | 0,22093978 | 0,728635155 | HMG-box transcription factor 1 |
| ENSG00000178093 | TSSK6 | 0,406586369 | 1,325545662 | 0,46839804 | 0,907940653 | testis specific serine kinase 6 |
| ENSG00000165175 | MID1IP1 | 0,406569584 | 1,32553024 | 0,07336856 | 0,427547926 | MID1 interacting protein 1 |
| ENSG00000214717 | ZBED1 | 0,406569399 | 1,325530071 | 0,19530791 | 0,693577833 | zinc finger BED-type containing 1 |
| ENSG00000036054 | TBC1D23 | 0,406501031 | 1,325467256 | 0,34530012 | 0,841168721 | TBC1 domain family member 23 |

| | | | | | | |
|---|---|---|---|---|---|---|
| ENSG00000196453 | ZNF777 | 0,406434454 | 1,325406091 | 0,21172441 | 0,718113208 | zinc finger protein 777 |
| ENSG00000106785 | TRIM14 | 0,406112744 | 1,325110568 | 0,35587825 | 0,847256893 | tripartite motif containing 14 |
| ENSG00000088836 | SLC4A11 | 0,406108232 | 1,325106423 | 0,58441561 | 0,944741266 | solute carrier family 4 member 11 |
| ENSG00000023909 | GCLM | 0,406082839 | 1,325083101 | 0,12778103 | 0,570756819 | glutamate-cysteine ligase modifier subunit |
| ENSG00000260615 | RPL23AP97 | 0,405736426 | 1,324764966 | 0,67450592 | 0,963174978 | ribosomal protein L23a pseudogene 97 |
| ENSG00000226125 | LINC01907 | 0,405608309 | 1,324647327 | 0,68146068 | 0,964654539 | long intergenic non-protein coding RNA 1907 |
| ENSG00000049656 | CLPTM1L | 0,405597536 | 1,324637436 | 0,11122736 | 0,532691822 | CLPTM1 like |
| ENSG00000134215 | VAV3 | 0,405541343 | 1,324585842 | 0,22340297 | 0,731984265 | vav guanine nucleotide exchange factor 3 |
| ENSG00000242265 | PEG10 | 0,405534299 | 1,324579375 | 0,10207504 | 0,508625897 | paternally expressed 10 |
| ENSG00000076067 | RBMS2 | 0,405524197 | 1,3245701 | 0,05650722 | 0,366903759 | RNA binding motif single stranded interacting protein 2 |
| ENSG00000145016 | RUBCN | 0,405490712 | 1,324539357 | 0,12677699 | 0,568863994 | rubicon autophagy regulator |
| ENSG00000285316 | AC275401,5 | 0,405456904 | 1,324508318 | 0,58011648 | 0,943828882 | |
| ENSG00000227337 | RPL23AP43 | 0,405352329 | 1,324412313 | 0,67851807 | 0,9640473 | ribosomal protein L23a pseudogene 43 |
| ENSG00000155034 | FBXL18 | 0,405302225 | 1,324366319 | 0,33800975 | 0,837255242 | F-box and leucine rich repeat protein 18 |
| ENSG00000116815 | CD58 | 0,405080766 | 1,324163038 | 0,47321429 | 0,909724186 | CD58 molecule |
| ENSG00000008256 | CYTH3 | 0,405068185 | 1,324151491 | 0,07026841 | 0,417549035 | cytohesin 3 |
| ENSG00000185518 | SV2B | 0,405048212 | 1,324133159 | 0,5330852 | 0,931291085 | synaptic vesicle glycoprotein 2B |
| ENSG00000244187 | TMEM141 | 0,405012652 | 1,324100522 | 0,21575077 | 0,722999018 | transmembrane protein 141 |
| ENSG00000234432 | AC092171,3 | 0,404957852 | 1,324050228 | 0,24722353 | 0,759350486 | uncharacterized LOC100129484 |
| ENSG00000135740 | SLC9A5 | 0,404857458 | 1,323958093 | 0,40545782 | 0,878367308 | solute carrier family 9 member A5 |
| ENSG00000203857 | HSD3B1 | 0,404681113 | 1,323796272 | 0,73622828 | 0,974353474 | hydroxy-delta-5-steroid dehydrogenase, 3 beta- and steroid delta-isomerase 1 |
| ENSG00000055070 | SZRD1 | 0,404543479 | 1,323669987 | 0,06848586 | 0,412469927 | SUZ RNA binding domain containing 1 |
| ENSG00000099337 | KCNK6 | 0,404540768 | 1,323667499 | 0,54531486 | 0,935085411 | potassium two pore domain channel subfamily K member 6 |
| ENSG00000189050 | RNFT1 | 0,404528829 | 1,323656546 | 0,23324703 | 0,742438913 | ring finger protein, transmembrane 1 |
| ENSG00000073921 | PICALM | 0,404427355 | 1,323563448 | 0,30336257 | 0,812669425 | phosphatidylinositol binding clathrin assembly protein |
| ENSG00000239263 | RBM43P1 | 0,404131569 | 1,323292114 | 0,75353339 | 0,975901808 | RNA binding motif protein 43 pseudogene 1 |
| ENSG00000257740 | AC073896,3 | 0,404085017 | 1,323249416 | 0,45670917 | 0,90366076 | novel transcript, antisense to CNPY2 |
| ENSG00000197858 | GPAA1 | 0,404068604 | 1,323234362 | 0,19218287 | 0,688538797 | glycosylphosphatidylinositol anchor attachment 1 |
| ENSG00000269482 | Z69720,1 | 0,403984515 | 1,323157238 | 0,62541879 | 0,952793934 | novel transcript, antisense to NPRL3 |

| | | | | | | |
|---|---|---|---|---|---|---|
| ENSG00000253463 | HMGB1P19 | 0,403936812 | 1,323113488 | 0,66676493 | 0,9611116 | high mobility group box 1 pseudogene 19 |
| ENSG00000101150 | TPD52L2 | 0,403801151 | 1,322989078 | 0,09604827 | 0,492351692 | TPD52 like 2 |
| ENSG00000285737 | LINC02680 | 0,4037037 | 1,322899715 | 0,48414083 | 0,913690171 | long intergenic non-protein coding RNA 2680 |
| ENSG00000250614 | AC005522,1 | 0,403564071 | 1,322771687 | 0,80226385 | 0,980710009 | novel transcript, antisense to DTX2 |
| ENSG00000187715 | KBTBD12 | 0,403530679 | 1,322741071 | 0,54017322 | 0,933385624 | kelch repeat and BTB domain containing 12 |
| ENSG00000074621 | SLC24A1 | 0,403507378 | 1,322719707 | 0,18848312 | 0,68281995 | solute carrier family 24 member 1 |
| ENSG00000179271 | GADD45GIP1 | 0,40350576 | 1,322718224 | 0,25955395 | 0,772176335 | GADD45G interacting protein 1 |
| ENSG00000139112 | GABARAPL1 | 0,403451943 | 1,322668884 | 0,18425769 | 0,676130552 | GABA type A receptor associated protein like 1 |
| ENSG00000260908 | AC009093,3 | 0,40337126 | 1,322594916 | 0,56322005 | 0,940865625 | phospholipase A2, group X (PLA2G10) pseudogene |
| ENSG00000089472 | HEPH | 0,40329362 | 1,32252374 | 0,20146044 | 0,701094807 | hephaestin |
| ENSG00000101842 | VSIG1 | 0,403188475 | 1,322427357 | 0,55811322 | 0,939078529 | V-set and immunoglobulin domain containing 1 |
| ENSG00000224208 | ZCRB1P1 | 0,403145317 | 1,322387798 | 0,61415104 | 0,952705073 | ZCRB1 pseudogene 1 |
| ENSG00000254846 | AL355075,1 | 0,40311023 | 1,322355637 | 0,75243837 | 0,975901808 | novel transcript |
| ENSG00000272476 | AL024507,2 | 0,403087452 | 1,322334759 | 0,31665084 | 0,821646322 | novel transcript, antisense to SEC63 |
| ENSG00000241015 | TPM3P9 | 0,4029619 | 1,322219687 | 0,15441208 | 0,623607455 | tropomyosin 3 pseudogene 9 |
| ENSG00000233654 | AC108047,1 | 0,402899768 | 1,322162744 | 0,39276434 | 0,871063652 | novel transcript |
| ENSG00000286599 | AC110611,1 | 0,402832273 | 1,322100889 | 0,42543181 | 0,892125569 | novel transcript, antisense to EXOC1 |
| ENSG00000182196 | ARL6IP4 | 0,402775022 | 1,322048425 | 0,24640973 | 0,758198269 | ADP ribosylation factor like GTPase 6 interacting protein 4 |
| ENSG00000226404 | LY6G5C | 0,402644943 | 1,321929229 | 0,57105862 | 0,942436428 | lymphocyte antigen 6 family member G5C |
| ENSG00000234383 | CTBP2P8 | 0,40263067 | 1,321916151 | 0,49804904 | 0,919274055 | CTBP2 pseudogene 8 |
| ENSG00000185955 | C7orf61 | 0,402599397 | 1,321887497 | 0,61529278 | 0,952793934 | chromosome 7 open reading frame 61 |
| ENSG00000156959 | LHFPL4 | 0,402517689 | 1,321812633 | 0,44264611 | 0,899437949 | LHFPL tetraspan subfamily member 4 |
| ENSG00000232192 | KIF26B-AS1 | 0,402511698 | 1,321807144 | 0,77174471 | 0,978033855 | KIF26B antisense RNA 1 |
| ENSG00000115685 | PPP1R7 | 0,402453369 | 1,321753703 | 0,050656 | 0,344284595 | protein phosphatase 1 regulatory subunit 7 |
| ENSG00000214954 | LRRC69 | 0,402420183 | 1,321723299 | 0,58874612 | 0,946498624 | leucine rich repeat containing 69 |
| ENSG00000171657 | GPR82 | 0,402367873 | 1,321675377 | 0,22025015 | 0,728572782 | G protein-coupled receptor 82 |
| ENSG00000060971 | ACAA1 | 0,402339184 | 1,321649095 | 0,16460537 | 0,643811841 | acetyl-CoA acyltransferase 1 |
| ENSG00000167785 | ZNF558 | 0,402320052 | 1,321631568 | 0,57957703 | 0,943828882 | zinc finger protein 558 |
| ENSG00000198369 | SPRED2 | 0,402254317 | 1,32157135 | 0,12052572 | 0,553971792 | sprouty related EVH1 domain containing 2 |
| ENSG00000224093 | BCAR3-AS1 | 0,402201242 | 1,321522733 | 0,52448233 | 0,928616707 | BCAR3 antisense RNA 1 |
| ENSG00000184271 | POU6F1 | 0,402147479 | 1,321473486 | 0,30809593 | 0,816424917 | POU class 6 homeobox 1 |
| ENSG00000273311 | DGCR11 | 0,402086404 | 1,321417543 | 0,23136711 | 0,739268552 | DiGeorge syndrome critical region gene 11 |

| | | | | | | |
|---|---|---|---|---|---|---|
| ENSG00000122965 | RBM19 | 0,401918715 | 1,32126396 | 0,26554109 | 0,776824277 | RNA binding motif protein 19 |
| ENSG00000278310 | GOLGA8F | 0,401917597 | 1,321262936 | 0,77889405 | 0,978587255 | golgin A8 family member F |
| ENSG00000213221 | DNLZ | 0,401842128 | 1,321193822 | 0,35289167 | 0,845031687 | DNL-type zinc finger |
| ENSG00000248431 | AC021134,1 | 0,401841923 | 1,321193634 | 0,48662657 | 0,914296452 | novel transcript |
| ENSG00000173156 | RHOD | 0,401798234 | 1,321153625 | 0,37775294 | 0,863195661 | ras homolog family member D |
| ENSG00000172379 | ARNT2 | 0,401765706 | 1,321123837 | 0,19380255 | 0,690637119 | aryl hydrocarbon receptor nuclear translocator 2 |
| ENSG00000137203 | TFAP2A | 0,401612449 | 1,320983502 | 0,31438292 | 0,819877431 | transcription factor AP-2 alpha |
| ENSG00000134109 | EDEM1 | 0,401496889 | 1,320877695 | 0,09046438 | 0,479775816 | ER degradation enhancing alpha-mannosidase like protein 1 |
| ENSG00000117395 | EBNA1BP2 | 0,401436305 | 1,320822228 | 0,09404827 | 0,487373632 | EBNA1 binding protein 2 |
| ENSG00000206394 | CLIC1 | 0,401393878 | 1,320783386 | 0,21844763 | 0,726161274 | chloride intracellular channel 1 |
| ENSG00000122420 | PTGFR | 0,40134997 | 1,320743189 | 0,50988348 | 0,922427526 | prostaglandin F receptor |
| ENSG00000109610 | SOD3 | 0,401327439 | 1,320722563 | 0,67607963 | 0,963560994 | superoxide dismutase 3 |
| ENSG00000226054 | MEMO1P1 | 0,401279515 | 1,320678691 | 0,79891698 | 0,980161849 | mediator of cell motility 1 pseudogene 1 |
| ENSG00000079459 | FDFT1 | 0,401039298 | 1,320458809 | 0,20478203 | 0,706459522 | farnesyl-diphosphate farnesyltransferase 1 |
| ENSG00000234350 | AC007405,1 | 0,400961386 | 1,3203875 | 0,47758006 | 0,911578505 | novel transcript (FLJ13453) |
| ENSG00000169826 | CSGALNACT2 | 0,400870692 | 1,320304498 | 0,21651538 | 0,723777756 | chondroitin sulfate N-acetylgalactosaminyltransferase 2 |
| ENSG00000197181 | PIWIL2 | 0,400834534 | 1,320271408 | 0,61519567 | 0,952793934 | piwi like RNA-mediated gene silencing 2 |
| ENSG00000147166 | ITGB1BP2 | 0,400824088 | 1,320261848 | 0,26436342 | 0,775697617 | integrin subunit beta 1 binding protein 2 |
| ENSG00000276536 | HRAS | 0,400695201 | 1,320143904 | 0,46095694 | 0,906029574 | HRas proto-oncogene, GTPase |
| ENSG00000224008 | LINC01441 | 0,400673883 | 1,320124397 | 0,57552358 | 0,942700009 | long intergenic non-protein coding RNA 1441 |
| ENSG00000197982 | C1orf122 | 0,400666846 | 1,320117958 | 0,22974309 | 0,73819656 | chromosome 1 open reading frame 122 |
| ENSG00000197361 | FBXL22 | 0,400648309 | 1,320100996 | 0,44400117 | 0,900210555 | F-box and leucine rich repeat protein 22 |
| ENSG00000272431 | AC104117,5 | 0,400629741 | 1,320084006 | 0,7355179 | 0,974353474 | novel transcript |
| ENSG00000275618 | BFAR | 0,400626369 | 1,32008092 | 0,21950717 | 0,727838473 | bifunctional apoptosis regulator |
| ENSG00000177337 | DLGAP1-AS1 | 0,400385599 | 1,319860632 | 0,08479269 | 0,46345429 | DLGAP1 antisense RNA 1 |
| ENSG00000204758 | AC008429,1 | 0,400282585 | 1,319766392 | 0,53084171 | 0,931265667 | novel transcript, antisense to RPL26L1 |
| ENSG00000258592 | AL391152,1 | 0,400182038 | 1,319674416 | 0,2226381 | 0,731227818 | novel transcript |
| ENSG00000130764 | LRRC47 | 0,400149635 | 1,319644776 | 0,26317339 | 0,774637968 | leucine rich repeat containing 47 |
| ENSG00000175931 | UBE2O | 0,399907663 | 1,319423461 | 0,22878062 | 0,737173617 | ubiquitin conjugating enzyme E2 O |
| ENSG00000166889 | PATL1 | 0,39988024 | 1,319398381 | 0,14689818 | 0,610674677 | PAT1 homolog 1, processing body mRNA decay factor |

| | | | | | | |
|---|---|---|---|---|---|---|
| ENSG00000240549 | THOC7-AS1 | 0,399823612 | 1,319346594 | 0,61863387 | 0,952793934 | THOC7 antisense RNA 1 |
| ENSG00000273659 | AC108125,1 | 0,399804126 | 1,319328774 | 0,65115247 | 0,957089277 | novel transcript |
| ENSG00000167930 | FAM234A | 0,399790723 | 1,319316517 | 0,19029098 | 0,685793354 | family with sequence similarity 234 member A |
| ENSG00000167332 | OR51E2 | 0,399711221 | 1,319243816 | 0,75845905 | 0,976120207 | olfactory receptor family 51 subfamily E member 2 |
| ENSG00000174326 | SLC16A11 | 0,399532209 | 1,319080133 | 0,61477622 | 0,952793934 | solute carrier family 16 member 11 |
| ENSG00000239791 | AC002310,2 | 0,399488109 | 1,319039811 | 0,62168771 | 0,952793934 | novel transcript |
| ENSG00000106638 | TBL2 | 0,39937646 | 1,318937736 | 0,06405555 | 0,396482442 | transducin beta like 2 |
| ENSG00000149541 | B3GAT3 | 0,399234436 | 1,318807902 | 0,29568379 | 0,805426883 | beta-1,3-glucuronyltransferase 3 |
| ENSG00000236149 | ABCF1 | 0,399072642 | 1,318660009 | 0,30170426 | 0,811326505 | ATP binding cassette subfamily F member 1 |
| ENSG00000168077 | SCARA3 | 0,398929707 | 1,31852937 | 0,30037856 | 0,810130728 | scavenger receptor class A member 3 |
| ENSG00000122728 | TAF1L | 0,39890669 | 1,318508334 | 0,63659868 | 0,95557086 | TATA-box binding protein associated factor 1 like |
| ENSG00000141965 | FEM1A | 0,398871814 | 1,31847646 | 0,41444408 | 0,885579089 | fem-1 homolog A |
| ENSG00000267205 | AC005954,2 | 0,398779816 | 1,318392386 | 0,63207213 | 0,953935145 | novel transcript, antisense to APBA3 |
| ENSG00000249816 | LINC00964 | 0,39876084 | 1,318375045 | 0,62832517 | 0,952793934 | long intergenic non-protein coding RNA 964 |
| ENSG00000286402 | AC090629,1 | 0,39864528 | 1,318269447 | 0,61965038 | 0,952793934 | novel transcript |
| ENSG00000236423 | LINC01134 | 0,398576605 | 1,318206697 | 0,55902927 | 0,939635125 | long intergenic non-protein coding RNA 1134 |
| ENSG00000103319 | EEF2K | 0,398497875 | 1,318134763 | 0,57950513 | 0,943828882 | eukaryotic elongation factor 2 kinase |
| ENSG00000273154 | AL121845,3 | 0,398474806 | 1,318113686 | 0,60871663 | 0,951601152 | novel protein, ZGPAT-LIME1 readthrough |
| ENSG00000130340 | SNX9 | 0,39841518 | 1,31805921 | 0,10339535 | 0,512859293 | sorting nexin 9 |
| ENSG00000107902 | LHPP | 0,39818657 | 1,317850366 | 0,1858511 | 0,678777213 | phospholysine phosphohistidine inorganic pyrophosphate phosphatase |
| ENSG00000162076 | FLYWCH2 | 0,398003729 | 1,317683358 | 0,19281946 | 0,689737701 | FLYWCH family member 2 |
| ENSG00000286675 | AC068700,2 | 0,397996772 | 1,317677003 | 0,50500304 | 0,921400408 | novel transcript |
| ENSG00000205702 | CYP2D7 | 0,397991644 | 1,31767232 | 0,76401063 | 0,976640468 | cytochrome P450 family 2 subfamily D member 7 (gene/pseudogene) |
| ENSG00000262938 | C21orf62 | 0,397975641 | 1,317657704 | 0,68532104 | 0,965434008 | chromosome 21 open reading frame 62 |
| ENSG00000163931 | TKT | 0,397972771 | 1,317655083 | 0,08661774 | 0,468880561 | transketolase |
| ENSG00000227560 | RPS15AP30 | 0,397967695 | 1,317650447 | 0,64111567 | 0,956465487 | ribosomal protein S15a pseudogene 30 |
| ENSG00000226651 | CLIC1 | 0,397823375 | 1,317518641 | 0,22125899 | 0,729227317 | chloride intracellular channel 1 |
| ENSG00000260070 | AC006960,3 | 0,397790277 | 1,317488416 | 0,56451073 | 0,940865625 | novel transcript, antisense to KIAA0895 |
| ENSG00000119899 | SLC17A5 | 0,397784434 | 1,31748308 | 0,16108903 | 0,637192669 | solute carrier family 17 member 5 |
| ENSG00000131094 | C1QL1 | 0,397759699 | 1,317460492 | 0,46855291 | 0,907940653 | complement C1q like 1 |
| ENSG00000130876 | SLC7A10 | 0,397570825 | 1,317288025 | 0,45752199 | 0,903831777 | solute carrier family 7 member 10 |

| | | | | | | |
|---|---|---|---|---|---|---|
| ENSG00000260886 | TAT-AS1 | 0,397338987 | 1,317076356 | 0,62334605 | 0,952793934 | TAT antisense RNA 1 |
| ENSG00000169946 | ZFPM2 | 0,397172675 | 1,316924534 | 0,20030026 | 0,699920362 | zinc finger protein, FOG family member 2 |
| ENSG00000235726 | AC010148,1 | 0,397132134 | 1,316887528 | 0,33931427 | 0,838276153 | novel transcript |
| ENSG00000112186 | CAP2 | 0,397118135 | 1,316874749 | 0,23304451 | 0,742172102 | cyclase associated actin cytoskeleton regulatory protein 2 |
| ENSG00000012822 | CALCOCO1 | 0,397072525 | 1,316833118 | 0,10139821 | 0,506329296 | calcium binding and coiled-coil domain 1 |
| ENSG00000124198 | ARFGEF2 | 0,397031533 | 1,316795703 | 0,09943355 | 0,50040065 | ADP ribosylation factor guanine nucleotide exchange factor 2 |
| ENSG00000266933 | AC005775,1 | 0,396734316 | 1,316524451 | 0,53599526 | 0,932139691 | novel transcript, antisense to MADCAM1 and TPGS1 |
| ENSG00000213453 | FTH1P3 | 0,396701621 | 1,316494616 | 0,59071246 | 0,947416795 | ferritin heavy chain 1 pseudogene 3 |
| ENSG00000233164 | C6orf136 | 0,396635297 | 1,316434094 | 0,48090858 | 0,912907448 | chromosome 6 open reading frame 136 |
| ENSG00000263818 | RDM1P5 | 0,396622463 | 1,316422384 | 0,54212541 | 0,933491609 | RDM1 pseudogene 5 |
| ENSG00000237440 | ZNF737 | 0,396539512 | 1,316346696 | 0,32954635 | 0,831379626 | zinc finger protein 737 |
| ENSG00000160321 | ZNF208 | 0,396436248 | 1,316252478 | 0,74028663 | 0,974833668 | zinc finger protein 208 |
| ENSG00000198894 | CIPC | 0,396430615 | 1,316247339 | 0,12957527 | 0,575350249 | CLOCK interacting pacemaker |
| ENSG00000196998 | WDR45 | 0,396351454 | 1,316175118 | 0,15591918 | 0,626884945 | WD repeat domain 45 |
| ENSG00000198673 | TAFA2 | 0,396329672 | 1,316155247 | 0,22059423 | 0,728572782 | TAFA chemokine like family member 2 |
| ENSG00000154736 | ADAMTS5 | 0,396314686 | 1,316141576 | 0,30654647 | 0,815375074 | ADAM metallopeptidase with thrombospondin type 1 motif 5 |
| ENSG00000253660 | AC008464,1 | 0,396255685 | 1,316087751 | 0,69890491 | 0,968522643 | novel transcript |
| ENSG00000105245 | NUMBL | 0,39621539 | 1,316050993 | 0,30497614 | 0,813769074 | NUMB like endocytic adaptor protein |
| ENSG00000184702 | SEPTIN5 | 0,396194891 | 1,316032294 | 0,22663108 | 0,735240631 | septin 5 |
| ENSG00000241253 | CFB | 0,396192366 | 1,31602999 | 0,65030208 | 0,956899963 | complement factor B |
| ENSG00000007080 | CCDC124 | 0,396078559 | 1,315926179 | 0,22217908 | 0,730026977 | coiled-coil domain containing 124 |
| ENSG00000274943 | AC079684,1 | 0,395994001 | 1,315849053 | 0,68828616 | 0,966426338 | novel transcript, sense intronic to PPHLN1 |
| ENSG00000174744 | BRMS1 | 0,39597925 | 1,3158356 | 0,20638124 | 0,709434078 | BRMS1 transcriptional repressor and anoikis regulator |
| ENSG00000129925 | PGAP6 | 0,39593422 | 1,315794529 | 0,21853704 | 0,726304032 | post-glycosylphosphatidylinositol attachment to proteins 6 |
| ENSG00000215244 | LINC02649 | 0,395933707 | 1,315794062 | 0,59876628 | 0,948854865 | long intergenic non-protein coding RNA 2649 |
| ENSG00000281560 | LSINCT5 | 0,395865452 | 1,315731812 | 0,45388292 | 0,902912642 | long stress-induced non-coding transcript 5 |
| ENSG00000183020 | AP2A2 | 0,395808717 | 1,315680071 | 0,11807072 | 0,549146283 | adaptor related protein complex 2 subunit alpha 2 |
| ENSG00000240823 | RN7SL23P | 0,395737973 | 1,315615557 | 0,66143746 | 0,959582738 | RNA, 7SL, cytoplasmic 23, pseudogene |

| | | | | | | |
|---|---|---|---|---|---|---|
| ENSG00000158683 | PKD1L1 | 0,395686479 | 1,3155686 | 0,496231 | 0,918259281 | polycystin 1 like 1, transient receptor potential channel interacting |
| ENSG00000140577 | CRTC3 | 0,395684334 | 1,315566644 | 0,08855479 | 0,474721701 | CREB regulated transcription coactivator 3 |
| ENSG00000121578 | B4GALT4 | 0,395657066 | 1,315541779 | 0,13568428 | 0,587376532 | beta-1,4-galactosyltransferase 4 |
| ENSG00000116649 | SRM | 0,395643595 | 1,315529495 | 0,23849383 | 0,747998807 | spermidine synthase |
| ENSG00000251003 | ZFPM2-AS1 | 0,395523298 | 1,315419806 | 0,35752691 | 0,848693232 | ZFPM2 antisense RNA 1 |
| ENSG00000286358 | AC096647,1 | 0,39551619 | 1,315413325 | 0,07658661 | 0,43728459 | novel transcript |
| ENSG00000204361 | NXPE2 | 0,395349596 | 1,315261438 | 0,73349117 | 0,974353474 | neurexophilin and PC-esterase domain family member 2 |
| ENSG00000106244 | PDAP1 | 0,39534563 | 1,315257822 | 0,14464119 | 0,605120065 | PDGFA associated protein 1 |
| ENSG00000116337 | AMPD2 | 0,395225445 | 1,315148258 | 0,16213734 | 0,639225966 | adenosine monophosphate deaminase 2 |
| ENSG00000285784 | AL353595,1 | 0,39518857 | 1,315114643 | 0,6944961 | 0,967417311 | Novel transcript |
| ENSG00000175215 | CTDSP2 | 0,395084168 | 1,315019477 | 0,12999363 | 0,576206833 | CTD small phosphatase 2 |
| ENSG00000280893 | AC009133,6 | 0,395047234 | 1,314985812 | 0,28732751 | 0,798896538 | novel transcript |
| ENSG00000198053 | SIRPA | 0,394964825 | 1,314910701 | 0,13119055 | 0,578496667 | signal regulatory protein alpha |
| ENSG00000140521 | POLG | 0,394944283 | 1,314891978 | 0,23623826 | 0,745282842 | DNA polymerase gamma, catalytic subunit |
| ENSG00000046647 | GEMIN8 | 0,394749555 | 1,314714512 | 0,13317098 | 0,582952249 | gem nuclear organelle associated protein 8 |
| ENSG00000254901 | BORCS8 | 0,394730306 | 1,314696971 | 0,19019239 | 0,685594362 | BLOC-1 related complex subunit 8 |
| ENSG00000224570 | AC097063,1 | 0,394705186 | 1,31467408 | 0,67208548 | 0,962290553 | UDP-N-acetyl-alpha-D-galactosamine:polypeptide N-acetylgalactosaminyltransferase 11 (GalNAc-T11) (GALNT11) pseudogene |
| ENSG00000157833 | GAREM2 | 0,394615678 | 1,314592517 | 0,18351656 | 0,674606849 | GRB2 associated regulator of MAPK1 subtype 2 |
| ENSG00000101337 | TM9SF4 | 0,394599058 | 1,314577373 | 0,10132925 | 0,506306167 | transmembrane 9 superfamily member 4 |
| ENSG00000233012 | HDAC1P2 | 0,39459027 | 1,314569366 | 0,34345388 | 0,840901506 | histone deacetylase 1 pseudogene 2 |
| ENSG00000164916 | FOXK1 | 0,394566984 | 1,314548147 | 0,29346271 | 0,803461221 | forkhead box K1 |
| ENSG00000203710 | CR1 | 0,394538571 | 1,314522258 | 0,54265328 | 0,933663748 | complement C3b/C4b receptor 1 (Knops blood group) |
| ENSG00000286067 | AC004263,2 | 0,39451289 | 1,314498859 | 0,49213154 | 0,916995341 | novel transcript, antisense to PXN |
| ENSG00000087274 | ADD1 | 0,394409751 | 1,314404888 | 0,07874935 | 0,4442685 | adducin 1 |
| ENSG00000185291 | IL3RA | 0,394323037 | 1,314325888 | 0,39298529 | 0,871121102 | interleukin 3 receptor subunit alpha |
| ENSG00000006025 | OSBPL7 | 0,394256867 | 1,314265606 | 0,22157243 | 0,729254784 | oxysterol binding protein like 7 |
| ENSG00000267394 | AC004596,1 | 0,394252109 | 1,314261272 | 0,69989453 | 0,968522643 | novel transcript, antisense to ATXN7L3 |
| ENSG00000221823 | PPP3R1 | 0,394231728 | 1,314242706 | 0,27065802 | 0,783073289 | protein phosphatase 3 regulatory subunit B, alpha |

| | | | | | | |
|---|---|---|---|---|---|---|
| ENSG00000088832 | FKBP1A | 0,394207715 | 1,314220831 | 0,12523418 | 0,565471028 | FKBP prolyl isomerase 1A |
| ENSG00000105223 | PLD3 | 0,394164623 | 1,314181577 | 0,13655699 | 0,589440545 | phospholipase D family member 3 |
| ENSG00000187905 | LRRC74B | 0,394127626 | 1,314147876 | 0,63146887 | 0,953823688 | leucine rich repeat containing 74B |
| ENSG00000147570 | DNAJC5B | 0,393927268 | 1,313965383 | 0,72866288 | 0,973955059 | DnaJ heat shock protein family (Hsp40) member C5 beta |
| ENSG00000182057 | OGFRP1 | 0,393850505 | 1,313895472 | 0,239611 | 0,749431667 | opioid growth factor receptor pseudogene 1 |
| ENSG00000197136 | PCNX3 | 0,393733033 | 1,313788491 | 0,28950566 | 0,800444324 | pecanex 3 |
| ENSG00000170854 | RIOX2 | 0,393716957 | 1,313773852 | 0,20158614 | 0,701122367 | ribosomal oxygenase 2 |
| ENSG00000177370 | TIMM22 | 0,393688423 | 1,313747868 | 0,09676674 | 0,494090395 | translocase of inner mitochondrial membrane 22 |
| ENSG00000248522 | SBF1P1 | 0,393682305 | 1,313742297 | 0,62364855 | 0,952793934 | SET binding factor 1 pseudogene 1 |
| ENSG00000267011 | AC011498,1 | 0,393638576 | 1,313702478 | 0,72167791 | 0,972229848 | novel transcript |
| ENSG00000169193 | CCDC126 | 0,393469549 | 1,313548572 | 0,40484402 | 0,878176155 | coiled-coil domain containing 126 |
| ENSG00000166900 | STX3 | 0,393346743 | 1,313436764 | 0,24031955 | 0,750284634 | syntaxin 3 |
| ENSG00000090054 | SPTLC1 | 0,393299253 | 1,31339353 | 0,28089969 | 0,791603381 | serine palmitoyltransferase long chain base subunit 1 |
| ENSG00000162236 | STX5 | 0,393269576 | 1,313366513 | 0,12592748 | 0,566777174 | syntaxin 5 |
| ENSG00000172936 | MYD88 | 0,39320089 | 1,313303986 | 0,11919036 | 0,552054434 | MYD88 innate immune signal transduction adaptor |
| ENSG00000151239 | TWF1 | 0,393140891 | 1,313249369 | 0,43463262 | 0,896272827 | twinfilin actin binding protein 1 |
| ENSG00000269858 | EGLN2 | 0,392886436 | 1,313017765 | 0,36401104 | 0,854207909 | egl-9 family hypoxia inducible factor 2 |
| ENSG00000198865 | CCDC152 | 0,392824139 | 1,312961068 | 0,48891128 | 0,915617728 | coiled-coil domain containing 152 |
| ENSG00000288031 | AC093838,2 | 0,39278197 | 1,312922693 | 0,39219643 | 0,870909282 | novel pseudogene |
| ENSG00000118705 | RPN2 | 0,392712085 | 1,312859096 | 0,10124322 | 0,506118967 | ribophorin II |
| ENSG00000136827 | TOR1A | 0,392589269 | 1,312747337 | 0,12928786 | 0,574683255 | torsin family 1 member A |
| ENSG00000072310 | SREBF1 | 0,392450668 | 1,312621226 | 0,22292925 | 0,731402952 | sterol regulatory element binding transcription factor 1 |
| ENSG00000107331 | ABCA2 | 0,392432557 | 1,312604748 | 0,29925198 | 0,809131439 | ATP binding cassette subfamily A member 2 |
| ENSG00000065911 | MTHFD2 | 0,392418678 | 1,312592121 | 0,16571277 | 0,645029358 | methylenetetrahydrofolate dehydrogenase (NADP+ dependent) 2, methenyltetrahydrofolate cyclohydrolase |
| ENSG00000198042 | MAK16 | 0,392311037 | 1,31249419 | 0,17545838 | 0,661199837 | MAK16 homolog |
| ENSG00000086598 | TMED2 | 0,392250772 | 1,312439366 | 0,2774793 | 0,789067728 | transmembrane p24 trafficking protein 2 |
| ENSG00000286805 | AL138716,1 | 0,392216587 | 1,312408267 | 0,74281014 | 0,975201495 | novel transcript, antisense to PARK2 |
| ENSG00000261200 | AC136944,2 | 0,392180834 | 1,312375744 | 0,50046102 | 0,920485547 | novel transcript |

| | | | | | | |
|---|---|---|---|---|---|---|
| ENSG00000104946 | TBC1D17 | 0,392136143 | 1,31233509 | 0,23829926 | 0,747710567 | TBC1 domain family member 17 |
| ENSG00000170419 | VSTM2A | 0,392130446 | 1,312329907 | 0,48545609 | 0,913907266 | V-set and transmembrane domain containing 2A |
| ENSG00000236719 | OVAAL | 0,392043252 | 1,312250595 | 0,7699596 | 0,977742359 | ovarian adenocarcinoma amplified long non-coding RNA |
| ENSG00000090238 | YPEL3 | 0,391881785 | 1,312103736 | 0,17471995 | 0,660057513 | yippee like 3 |
| ENSG00000140941 | MAP1LC3B | 0,391858537 | 1,312082592 | 0,33413835 | 0,834518413 | microtubule associated protein 1 light chain 3 beta |
| ENSG00000204816 | FGF7P5 | 0,391829116 | 1,312055835 | 0,81792233 | 0,982323352 | fibroblast growth factor 7 pseudogene 5 |
| ENSG00000181029 | TRAPPC5 | 0,391655113 | 1,311897598 | 0,213807 | 0,721188427 | trafficking protein particle complex 5 |
| ENSG00000132793 | LPIN3 | 0,391649823 | 1,311892788 | 0,34833287 | 0,842757646 | lipin 3 |
| ENSG00000177707 | NECTIN3 | 0,391628541 | 1,311873435 | 0,35221099 | 0,844912682 | nectin cell adhesion molecule 3 |
| ENSG00000104129 | DNAJC17 | 0,391602108 | 1,311849399 | 0,20334211 | 0,703503738 | DnaJ heat shock protein family (Hsp40) member C17 |
| ENSG00000068120 | COASY | 0,391600593 | 1,311848022 | 0,15780944 | 0,630592312 | Coenzyme A synthase |
| ENSG00000161671 | EMC10 | 0,391549735 | 1,311801778 | 0,21805092 | 0,72573766 | ER membrane protein complex subunit 10 |
| ENSG00000196353 | CPNE4 | 0,391546341 | 1,311798691 | 0,48535558 | 0,913885618 | copine 4 |
| ENSG00000214176 | PLEKHM1P1 | 0,391489819 | 1,311747298 | 0,30683351 | 0,815610044 | pleckstrin homology and RUN domain containing M1 pseudogene 1 |
| ENSG00000255508 | AP002990,1 | 0,391466715 | 1,311726292 | 0,28330303 | 0,794558711 | novel protein |
| ENSG00000100991 | TRPC4AP | 0,39141221 | 1,311676736 | 0,07526451 | 0,433425987 | transient receptor potential cation channel subfamily C member 4 associated protein |
| ENSG00000282379 | DTX2 | 0,391374328 | 1,311642295 | 0,57647699 | 0,943216284 | deltex E3 ubiquitin ligase 2 |
| ENSG00000115956 | PLEK | 0,391320094 | 1,311592988 | 0,72579404 | 0,973344786 | pleckstrin |
| ENSG00000171940 | ZNF217 | 0,391110193 | 1,311402175 | 0,19553591 | 0,693689284 | zinc finger protein 217 |
| ENSG00000277791 | PSMB3 | 0,391088106 | 1,311382098 | 0,05981531 | 0,380382656 | proteasome 20S subunit beta 3 |
| ENSG00000196166 | C8orf86 | 0,391064949 | 1,31136105 | 0,65009604 | 0,956899963 | chromosome 8 open reading frame 86 |
| ENSG00000287358 | AL096706,1 | 0,391021388 | 1,311321454 | 0,44958703 | 0,902062594 | novel transcript |
| ENSG00000131018 | SYNE1 | 0,390988057 | 1,311291159 | 0,1636344 | 0,642293268 | spectrin repeat containing nuclear envelope protein 1 |
| ENSG00000196421 | C20orf204 | 0,390879391 | 1,311192394 | 0,56665253 | 0,941311563 | chromosome 20 open reading frame 204 |
| ENSG00000105556 | MIER2 | 0,390852412 | 1,311167875 | 0,27419663 | 0,7862019 | MIER family member 2 |
| ENSG00000287041 | AC079354,2 | 0,390778246 | 1,311100472 | 0,58241117 | 0,944174844 | novel transcript, antisense to KIAA2012 |
| ENSG00000154175 | ABI3BP | 0,390754392 | 1,311078794 | 0,40229022 | 0,87718264 | ABI family member 3 binding protein |
| ENSG00000067829 | IDH3G | 0,390689762 | 1,311020061 | 0,13220047 | 0,580737262 | isocitrate dehydrogenase (NAD(+)) 3 non-catalytic subunit gamma |

| | | | | | | |
|---|---|---|---|---|---|---|
| ENSG00000055211 | GINM1 | 0,390673659 | 1,311005428 | 0,12059625 | 0,55405161 | glycoprotein integral membrane 1 |
| ENSG00000167522 | ANKRD11 | 0,390593602 | 1,310932681 | 0,37618789 | 0,862569716 | ankyrin repeat domain 11 |
| ENSG00000166509 | CLEC3A | 0,390451034 | 1,31080314 | 0,81210343 | 0,981213877 | C-type lectin domain family 3 member A |
| ENSG00000204588 | LINC01123 | 0,390059178 | 1,310447156 | 0,6746684 | 0,963215238 | long intergenic non-protein coding RNA 1123 |
| ENSG00000240405 | SAMMSON | 0,390043533 | 1,310432946 | 0,63383506 | 0,954929949 | survival associated mitochondrial melanoma specific oncogenic non-coding RNA |
| ENSG00000163249 | CCNYL1 | 0,389937317 | 1,310336471 | 0,16368657 | 0,642417397 | cyclin Y like 1 |
| ENSG00000207014 | SNORD116-3 | 0,389909746 | 1,310311429 | 0,71328515 | 0,970812834 | small nucleolar RNA, C/D box 116-3 |
| ENSG00000206727 | SNORD116-9 | 0,389909746 | 1,310311429 | 0,71328515 | 0,970812834 | small nucleolar RNA, C/D box 116-9 |
| ENSG00000182197 | EXT1 | 0,389758023 | 1,310173636 | 0,10947509 | 0,527694919 | exostosin glycosyltransferase 1 |
| ENSG00000167900 | TK1 | 0,389607579 | 1,310037018 | 0,17929839 | 0,66796771 | thymidine kinase 1 |
| ENSG00000234674 | TCF19 | 0,389590262 | 1,310021293 | 0,3068451 | 0,815610044 | transcription factor 19 |
| ENSG00000206455 | TCF19 | 0,389590262 | 1,310021293 | 0,3068451 | 0,815610044 | transcription factor 19 |
| ENSG00000166986 | MARS1 | 0,389519893 | 1,309957397 | 0,07099165 | 0,419165813 | methionyl-tRNA synthetase 1 |
| ENSG00000165280 | VCP | 0,389515704 | 1,309953594 | 0,07558003 | 0,434203245 | valosin containing protein |
| ENSG00000227544 | AC018647,1 | 0,389501524 | 1,309940718 | 0,6633283 | 0,959920886 | novel transcript |
| ENSG00000237693 | IRGM | 0,389495203 | 1,30993498 | 0,71221745 | 0,970812834 | immunity related GTPase M |
| ENSG00000250001 | AC027335,1 | 0,389340845 | 1,309794833 | 0,64819743 | 0,956899963 | novel transcript |
| ENSG00000186350 | RXRA | 0,38929034 | 1,309748981 | 0,21240716 | 0,719216058 | retinoid X receptor alpha |
| ENSG00000263843 | AC022211,2 | 0,389239374 | 1,309702713 | 0,35086536 | 0,844035671 | novel transcript, antisense to SLC25A19 |
| ENSG00000129680 | MAP7D3 | 0,389195941 | 1,309663284 | 0,12500012 | 0,565142382 | MAP7 domain containing 3 |
| ENSG00000284207 | HMGB3P22 | 0,389179264 | 1,309648145 | 0,57738412 | 0,94337651 | high mobility group box 3 pseudogene 22 |
| ENSG00000136279 | DBNL | 0,389028061 | 1,309510893 | 0,32424977 | 0,826368833 | drebrin like |
| ENSG00000131779 | PEX11B | 0,389004811 | 1,30948979 | 0,09152541 | 0,481302729 | peroxisomal biogenesis factor 11 beta |
| ENSG00000244414 | CFHR1 | 0,388991847 | 1,309478023 | 0,72712568 | 0,973344786 | complement factor H related 1 |
| ENSG00000102858 | MGRN1 | 0,388868189 | 1,309365788 | 0,22048956 | 0,728572782 | mahogunin ring finger 1 |
| ENSG00000275371 | AC012645,4 | 0,388862702 | 1,309360808 | 0,52225981 | 0,927617924 | novel transcript, antisense to GDPD3 |
| ENSG00000137710 | RDX | 0,388846812 | 1,309346387 | 0,17582951 | 0,661199837 | radixin |
| ENSG00000095203 | EPB41L4B | 0,388768933 | 1,309275708 | 0,22564039 | 0,734461165 | erythrocyte membrane protein band 4,1 like 4B |
| ENSG00000206195 | DUXAP8 | 0,388761053 | 1,309268557 | 0,44741434 | 0,902062594 | double homeobox A pseudogene 8 |
| ENSG00000240342 | RPS2P5 | 0,388509361 | 1,309040162 | 0,26393181 | 0,775013078 | ribosomal protein S2 pseudogene 5 |
| ENSG00000248540 | AC010931,1 | 0,388450089 | 1,308986382 | 0,18859684 | 0,682944308 | novel transcript |
| ENSG00000255256 | AL136146,2 | 0,388420626 | 1,308959651 | 0,7389 | 0,974448853 | novel transcript |

| | | | | | | |
|---|---|---|---|---|---|---|
| ENSG00000230537 | AL158071,2 | 0,388406244 | 1,308946602 | 0,58856578 | 0,946306024 | novel transcript |
| ENSG00000218018 | RBM38-AS1 | 0,388385201 | 1,308927509 | 0,58071346 | 0,943832501 | RBM38 antisense RNA 1 |
| ENSG00000171314 | PGAM1 | 0,388210985 | 1,308769456 | 0,09892028 | 0,499216097 | phosphoglycerate mutase 1 |
| ENSG00000261553 | AL137782,1 | 0,388193057 | 1,308753193 | 0,30828955 | 0,816620804 | novel transcript |
| ENSG00000150347 | ARID5B | 0,388167789 | 1,308730271 | 0,27841745 | 0,789733543 | AT-rich interaction domain 5B |
| ENSG00000277423 | AC069234,5 | 0,388152558 | 1,308716455 | 0,66342909 | 0,959920886 | novel transcript |
| ENSG00000254854 | AP003390,1 | 0,387947185 | 1,308530167 | 0,76035411 | 0,97624106 | novel transcript |
| ENSG00000228150 | AL357140,2 | 0,387902065 | 1,308489244 | 0,71210043 | 0,970812834 | novel transcript |
| ENSG00000267085 | AL512605,2 | 0,387817394 | 1,308412452 | 0,78244857 | 0,978628303 | NADH dehydrogenase pseudogene |
| ENSG00000278270 | RPS9 | 0,387799297 | 1,308396039 | 0,3140367 | 0,819877431 | ribosomal protein S9 |
| ENSG00000188676 | IDO2 | 0,387692038 | 1,308298768 | 0,80677521 | 0,981213877 | indoleamine 2,3-dioxygenase 2 |
| ENSG00000264569 | DCXR-DT | 0,387489002 | 1,308114659 | 0,71150631 | 0,970739528 | DCXR divergent transcript |
| ENSG00000288225 | AL591926,7 | 0,387387111 | 1,308022277 | 0,75573556 | 0,975901808 | family with sequence similarity 27 member E3 (FAM27E3) pseudogene |
| ENSG00000107130 | NCS1 | 0,387260054 | 1,307907085 | 0,1396446 | 0,595782615 | neuronal calcium sensor 1 |
| ENSG00000109736 | MFSD10 | 0,387162962 | 1,307819067 | 0,33609438 | 0,835660292 | major facilitator superfamily domain containing 10 |
| ENSG00000164265 | SCGB3A2 | 0,387139146 | 1,307797478 | 0,67488108 | 0,963268551 | secretoglobin family 3A member 2 |
| ENSG00000005187 | ACSM3 | 0,387069728 | 1,307734552 | 0,46900943 | 0,907940653 | acyl-CoA synthetase medium chain family member 3 |
| ENSG00000148840 | PPRC1 | 0,386936621 | 1,307613903 | 0,22188989 | 0,729383401 | PPARG related coactivator 1 |
| ENSG00000162585 | FAAP20 | 0,386857175 | 1,307541897 | 0,33076547 | 0,832114797 | FA core complex associated protein 20 |
| ENSG00000179889 | PDXDC1 | 0,386833937 | 1,307520836 | 0,092786 | 0,484139956 | pyridoxal dependent decarboxylase domain containing 1 |
| ENSG00000260088 | DDX59-AS1 | 0,386833935 | 1,307520835 | 0,60071245 | 0,949489681 | DDX59 antisense RNA 1 |
| ENSG00000283297 | TEX52 | 0,386677572 | 1,307379129 | 0,79642339 | 0,980015022 | testis expressed 52 |
| ENSG00000165948 | IFI27L1 | 0,3866434 | 1,307348163 | 0,27686936 | 0,788777434 | interferon alpha inducible protein 27 like 1 |
| ENSG00000182950 | ODF3L1 | 0,386600181 | 1,307308999 | 0,66589888 | 0,961020116 | outer dense fiber of sperm tails 3 like 1 |
| ENSG00000237588 | AL590666,3 | 0,386584328 | 1,307294634 | 0,57294573 | 0,942475308 | novel transcript |
| ENSG00000284323 | AC015853,6 | 0,386552509 | 1,307265802 | 0,74939909 | 0,975901808 | novel transcript, antisense to VPS53 |
| ENSG00000262905 | AC015853,1 | 0,386552509 | 1,307265802 | 0,74939909 | 0,975901808 | novel transcript, antisense to VPS53 |
| ENSG00000109445 | ZNF330 | 0,386517478 | 1,30723406 | 0,10758057 | 0,52347056 | zinc finger protein 330 |
| ENSG00000164007 | CLDN19 | 0,386456595 | 1,307178894 | 0,5803739 | 0,943832501 | claudin 19 |
| ENSG00000169507 | SLC38A11 | 0,386388351 | 1,307117062 | 0,74840978 | 0,975901808 | solute carrier family 38 member 11 |
| ENSG00000269743 | SLC25A53 | 0,386263744 | 1,30700417 | 0,23277279 | 0,741684508 | solute carrier family 25 member 53 |

| | | | | | | |
|---|---|---|---|---|---|---|
| ENSG00000172009 | THOP1 | 0,386243774 | 1,306986079 | 0,38536184 | 0,868895792 | thimet oligopeptidase 1 |
| ENSG00000203441 | LINC00449 | 0,386216635 | 1,306961492 | 0,56348518 | 0,940865625 | long intergenic non-protein coding RNA 449 |
| ENSG00000197705 | KLHL14 | 0,386192992 | 1,306940074 | 0,50948107 | 0,922427526 | kelch like family member 14 |
| ENSG00000160218 | TRAPPC10 | 0,386145825 | 1,306897346 | 0,05592683 | 0,365161047 | trafficking protein particle complex 10 |
| ENSG00000244203 | FOXP1-AS1 | 0,386131345 | 1,306884229 | 0,74942487 | 0,975901808 | FOXP1 antisense RNA 1 |
| ENSG00000223768 | LINC00205 | 0,386088634 | 1,306845539 | 0,18033517 | 0,66974377 | long intergenic non-protein coding RNA 205 |
| ENSG00000254614 | AP003068,2 | 0,386055178 | 1,306815234 | 0,44453201 | 0,900210555 | novel transcript, antisense to CAPN1 |
| ENSG00000225921 | NOL7 | 0,385989392 | 1,306755645 | 0,07496414 | 0,432645898 | nucleolar protein 7 |
| ENSG00000169249 | ZRSR2 | 0,385905448 | 1,306679613 | 0,21750559 | 0,725248255 | zinc finger CCCH-type, RNA binding motif and serine/arginine rich 2 |
| ENSG00000134590 | RTL8C | 0,385825634 | 1,306607326 | 0,11725945 | 0,547082438 | retrotransposon Gag like 8C |
| ENSG00000060982 | BCAT1 | 0,385766495 | 1,306553767 | 0,29089282 | 0,801689099 | branched chain amino acid transaminase 1 |
| ENSG00000079819 | EPB41L2 | 0,385655887 | 1,3064536 | 0,0993687 | 0,50040065 | erythrocyte membrane protein band 4,1 like 2 |
| ENSG00000187051 | RPS19BP1 | 0,38563596 | 1,306435555 | 0,25724653 | 0,769475277 | ribosomal protein S19 binding protein 1 |
| ENSG00000145817 | YIPF5 | 0,385619458 | 1,306420611 | 0,34294485 | 0,840419133 | Yip1 domain family member 5 |
| ENSG00000241666 | AL031733,2 | 0,385583609 | 1,306388149 | 0,71442839 | 0,971029555 | novel transcript |
| ENSG00000188735 | TMEM120B | 0,385514228 | 1,306325325 | 0,38443283 | 0,868418217 | transmembrane protein 120B |
| ENSG00000143575 | HAX1 | 0,385384754 | 1,306208095 | 0,09787829 | 0,497051225 | HCLS1 associated protein X-1 |
| ENSG00000241278 | ENPP7P4 | 0,385353205 | 1,306179531 | 0,5148366 | 0,924720046 | ectonucleotide pyrophosphatase/phosphodiesterase 7 pseudogene 4 |
| ENSG00000186479 | RGS7BP | 0,385285872 | 1,306118571 | 0,33551811 | 0,835320086 | regulator of G protein signaling 7 binding protein |
| ENSG00000287463 | AC110620,2 | 0,385282071 | 1,306115129 | 0,57475975 | 0,942533938 | novel transcript, antisense to TMEM163 |
| ENSG00000179583 | CIITA | 0,385176661 | 1,306019702 | 0,35215625 | 0,844912682 | class II major histocompatibility complex transactivator |
| ENSG00000149503 | INCENP | 0,385116787 | 1,305965502 | 0,4070753 | 0,880078305 | inner centromere protein |
| ENSG00000131653 | TRAF7 | 0,385038649 | 1,30589477 | 0,19384406 | 0,690637119 | TNF receptor associated factor 7 |
| ENSG00000176248 | ANAPC2 | 0,385003869 | 1,305863289 | 0,3949585 | 0,871888817 | anaphase promoting complex subunit 2 |
| ENSG00000204351 | SKIV2L | 0,384804715 | 1,305683036 | 0,18472829 | 0,67694849 | Ski2 like RNA helicase |
| ENSG00000262959 | RPL23AP86 | 0,384803632 | 1,305682056 | 0,71313442 | 0,970812834 | ribosomal protein L23a pseudogene 86 |
| ENSG00000070601 | FRMPD1 | 0,384673145 | 1,305563967 | 0,57200135 | 0,942475308 | FERM and PDZ domain containing 1 |
| ENSG00000279080 | AL022322,2 | 0,384620378 | 1,305516216 | 0,56090411 | 0,940374625 | novel transcript, antisense to PLA2G6 |
| ENSG00000178297 | TMPRSS9 | 0,384497428 | 1,305404961 | 0,72754221 | 0,973502227 | transmembrane serine protease 9 |

| | | | | | | |
|---|---|---|---|---|---|---|
| ENSG00000026297 | RNASET2 | 0,384357173 | 1,30527806 | 0,37706834 | 0,862818448 | ribonuclease T2 |
| ENSG00000122912 | SLC25A16 | 0,384266673 | 1,305196183 | 0,13381864 | 0,583990205 | solute carrier family 25 member 16 |
| ENSG00000258679 | AC121758,2 | 0,384233423 | 1,305166102 | 0,51758131 | 0,925883285 | novel YWHA pseudogene |
| ENSG00000072121 | ZFYVE26 | 0,384045577 | 1,304996174 | 0,13044804 | 0,577136255 | zinc finger FYVE-type containing 26 |
| ENSG00000130304 | SLC27A1 | 0,383988593 | 1,30494463 | 0,40364973 | 0,877691113 | solute carrier family 27 member 1 |
| ENSG00000132681 | ATP1A4 | 0,383783775 | 1,304759382 | 0,54805019 | 0,935404974 | ATPase Na+/K+ transporting subunit alpha 4 |
| ENSG00000254879 | PTPRJ-AS1 | 0,38372481 | 1,304706055 | 0,72908852 | 0,974009551 | PTPRJ antisense RNA 1 |
| ENSG00000231841 | FAM192BP | 0,383687763 | 1,304672552 | 0,69588982 | 0,967806915 | family with sequence similarity 192 member B, pseudogene |
| ENSG00000286025 | FAM246C | 0,383643682 | 1,304632688 | 0,62871891 | 0,952793934 | family with sequence similarity 246 member C (gene/pseudogene) |
| ENSG00000224641 | HCG20 | 0,383423579 | 1,304433664 | 0,70192961 | 0,968742626 | HLA complex group 20 |
| ENSG00000243986 | ENO1P3 | 0,383398176 | 1,304410696 | 0,54050417 | 0,933385624 | enolase 1 pseudogene 3 |
| ENSG00000255150 | EID3 | 0,383376862 | 1,304391425 | 0,23399649 | 0,743537585 | EP300 interacting inhibitor of differentiation 3 |
| ENSG00000173264 | GPR137 | 0,38334699 | 1,304364416 | 0,3237307 | 0,825794844 | G protein-coupled receptor 137 |
| ENSG00000244218 | RN7SL81P | 0,383209702 | 1,304240299 | 0,65218896 | 0,957257009 | RNA, 7SL, cytoplasmic 81, pseudogene |
| ENSG00000205213 | LGR4 | 0,383148063 | 1,304184576 | 0,29014717 | 0,800682868 | leucine rich repeat containing G protein-coupled receptor 4 |
| ENSG00000099364 | FBXL19 | 0,383145156 | 1,304181948 | 0,53350037 | 0,931381467 | F-box and leucine rich repeat protein 19 |
| ENSG00000225194 | LINC00092 | 0,383136788 | 1,304174384 | 0,55116466 | 0,936366211 | long intergenic non-protein coding RNA 92 |
| ENSG00000135838 | NPL | 0,383094336 | 1,304136008 | 0,4346287 | 0,896272827 | N-acetylneuraminate pyruvate lyase |
| ENSG00000157881 | PANK4 | 0,383066676 | 1,304111005 | 0,54117591 | 0,933385624 | pantothenate kinase 4 (inactive) |
| ENSG00000139433 | GLTP | 0,382994413 | 1,304045686 | 0,09482176 | 0,489192643 | glycolipid transfer protein |
| ENSG00000170876 | TMEM43 | 0,382861579 | 1,303925623 | 0,07088879 | 0,418874878 | transmembrane protein 43 |
| ENSG00000230876 | LINC00486 | 0,382833285 | 1,30390005 | 0,59444662 | 0,947469769 | long intergenic non-protein coding RNA 486 |
| ENSG00000271643 | AC112220,2 | 0,382641526 | 1,303726751 | 0,37419553 | 0,861369706 | novel transcript |
| ENSG00000153250 | RBMS1 | 0,382640822 | 1,303726115 | 0,32275613 | 0,824955371 | RNA binding motif single stranded interacting protein 1 |
| ENSG00000251513 | LIX1-AS1 | 0,382634099 | 1,30372004 | 0,62702922 | 0,952793934 | LIX1 and RIOK2 antisense RNA 1 |
| ENSG00000223720 | AL109659,1 | 0,382592111 | 1,303682097 | 0,69672478 | 0,968300554 | novel transcript |
| ENSG00000159348 | CYB5R1 | 0,382580916 | 1,303671981 | 0,11718904 | 0,547080491 | cytochrome b5 reductase 1 |
| ENSG00000266187 | RN7SL480P | 0,38251929 | 1,303616294 | 0,76442467 | 0,976640468 | RNA, 7SL, cytoplasmic 480, pseudogene |
| ENSG00000165527 | ARF6 | 0,382475605 | 1,303576821 | 0,11660494 | 0,545330857 | ADP ribosylation factor 6 |

| | | | | | | |
|---|---|---|---|---|---|---|
| ENSG00000183576 | SETD3 | 0,382375913 | 1,303486746 | 0,10738807 | 0,523028605 | SET domain containing 3, actin histidine methyltransferase |
| ENSG00000285095 | AC025887,2 | 0,38216253 | 1,303293966 | 0,27219959 | 0,785277096 | novel transcript |
| ENSG00000197989 | SNHG12 | 0,382145517 | 1,303278597 | 0,14581405 | 0,608479758 | small nucleolar RNA host gene 12 |
| ENSG00000152380 | FAM151B | 0,382144967 | 1,303278101 | 0,39713916 | 0,873764299 | family with sequence similarity 151 member B |
| ENSG00000258609 | LINC-ROR | 0,382109272 | 1,303245856 | 0,6418285 | 0,95651945 | long intergenic non-protein coding RNA, regulator of reprogramming |
| ENSG00000132824 | SERINC3 | 0,38205329 | 1,303195286 | 0,21262796 | 0,719437301 | serine incorporator 3 |
| ENSG00000275756 | AL354751,3 | 0,381997812 | 1,303145173 | 0,59205711 | 0,947416795 | novel pseudogene |
| ENSG00000136295 | TTYH3 | 0,381885522 | 1,303043748 | 0,25127195 | 0,763579382 | tweety family member 3 |
| ENSG00000207181 | SNORA14B | 0,381824556 | 1,302988685 | 0,80669978 | 0,981213877 | small nucleolar RNA, H/ACA box 14B |
| ENSG00000125533 | BHLHE23 | 0,381824298 | 1,302988452 | 0,76125314 | 0,97624106 | basic helix-loop-helix family member e23 |
| ENSG00000244165 | P2RY11 | 0,38175584 | 1,302926625 | 0,58497536 | 0,944857905 | purinergic receptor P2Y11 |
| ENSG00000160877 | NACC1 | 0,381715677 | 1,302890354 | 0,24078983 | 0,751152776 | nucleus accumbens associated 1 |
| ENSG00000097007 | ABL1 | 0,38170627 | 1,302881858 | 0,19522269 | 0,693415365 | ABL proto-oncogene 1, non-receptor tyrosine kinase |
| ENSG00000109320 | NFKB1 | 0,381617728 | 1,302801899 | 0,12725546 | 0,569742888 | nuclear factor kappa B subunit 1 |
| ENSG00000148158 | SNX30 | 0,381516091 | 1,302710121 | 0,16349429 | 0,642091247 | sorting nexin family member 30 |
| ENSG00000156958 | GALK2 | 0,381464713 | 1,302663729 | 0,16231945 | 0,639460073 | galactokinase 2 |
| ENSG00000281538 | AC254562,3 | 0,381384304 | 1,302591126 | 0,72806188 | 0,973888243 | novel transcript |
| ENSG00000282969 | AC235563,1 | 0,381384304 | 1,302591126 | 0,72806188 | 0,973888243 | novel transcript |
| ENSG00000282874 | AC254847,1 | 0,381384304 | 1,302591126 | 0,72806188 | 0,973888243 | novel transcript |
| ENSG00000164896 | FASTK | 0,381363259 | 1,302572125 | 0,28212722 | 0,793203731 | Fas activated serine/threonine kinase |
| ENSG00000281165 | SLC2A6 | 0,38135837 | 1,302567712 | 0,32008465 | 0,824366234 | solute carrier family 2 member 6 |
| ENSG00000125352 | RNF113A | 0,381281129 | 1,302497974 | 0,14168889 | 0,599581043 | ring finger protein 113A |
| ENSG00000243896 | OR2A7 | 0,38125069 | 1,302470494 | 0,5197956 | 0,926822772 | olfactory receptor family 2 subfamily A member 7 |
| ENSG00000136750 | GAD2 | 0,381235586 | 1,302456858 | 0,3958687 | 0,872308364 | glutamate decarboxylase 2 |
| ENSG00000167962 | ZNF598 | 0,381228214 | 1,302450203 | 0,27853175 | 0,789775666 | zinc finger protein 598, E3 ubiquitin ligase |
| ENSG00000163352 | LENEP | 0,380986422 | 1,302231934 | 0,6896051 | 0,966868992 | lens epithelial protein |
| ENSG00000284944 | MED18 | 0,380808303 | 1,302071167 | 0,67064895 | 0,962026636 | mediator complex subunit 18 |
| ENSG00000089351 | GRAMD1A | 0,380736177 | 1,302006072 | 0,25234956 | 0,76432863 | GRAM domain containing 1A |
| ENSG00000217128 | FNIP1 | 0,380415472 | 1,301716674 | 0,28128597 | 0,792337715 | folliculin interacting protein 1 |
| ENSG00000276461 | TCF20 | 0,380209752 | 1,30153107 | 0,36201609 | 0,852161624 | transcription factor 20 |
| ENSG00000262024 | TCF20 | 0,380209752 | 1,30153107 | 0,36201609 | 0,852161624 | transcription factor 20 |

| | | | | | | |
|---|---|---|---|---|---|---|
| ENSG00000261701 | HPR | 0,380207114 | 1,30152869 | 0,61821129 | 0,952793934 | haptoglobin-related protein |
| ENSG00000174276 | ZNHIT2 | 0,380052725 | 1,301389415 | 0,41828953 | 0,887094957 | zinc finger HIT-type containing 2 |
| ENSG00000165312 | OTUD1 | 0,379983457 | 1,301326933 | 0,27092598 | 0,783337629 | OTU deubiquitinase 1 |
| ENSG00000281692 | PACRG-AS1 | 0,379869821 | 1,301224437 | 0,69024737 | 0,967144432 | PACRG antisense RNA 1 |
| ENSG00000169410 | PTPN9 | 0,379710628 | 1,301080862 | 0,0974686 | 0,495973057 | protein tyrosine phosphatase non-receptor type 9 |
| ENSG00000281998 | LHX1-DT | 0,379532574 | 1,300920295 | 0,58611325 | 0,945246954 | LHX1 divergent transcript |
| ENSG00000157014 | TATDN2 | 0,3795025 | 1,300893177 | 0,17022597 | 0,653377651 | TatD DNase domain containing 2 |
| ENSG00000186395 | KRT10 | 0,379466444 | 1,300860665 | 0,44618435 | 0,901348735 | keratin 10 |
| ENSG00000225272 | AL451074,2 | 0,379360825 | 1,300765434 | 0,72544229 | 0,97326334 | ribosomal protein L21 (RPL21) pseudogene |
| ENSG00000125810 | CD93 | 0,37932466 | 1,300732826 | 0,49119924 | 0,916995341 | CD93 molecule |
| ENSG00000248936 | AC027607,1 | 0,379271849 | 1,300685214 | 0,67468298 | 0,963215238 | putatitive novel transcript |
| ENSG00000241420 | RN7SL505P | 0,379248622 | 1,300664273 | 0,77480167 | 0,978553937 | RNA, 7SL, cytoplasmic 505, pseudogene |
| ENSG00000226033 | STK19 | 0,379244802 | 1,300660829 | 0,40546945 | 0,878367308 | serine/threonine kinase 19 |
| ENSG00000236250 | STK19 | 0,379244802 | 1,300660829 | 0,40546945 | 0,878367308 | serine/threonine kinase 19 |
| ENSG00000228566 | AC013287,1 | 0,37908943 | 1,300520761 | 0,65257873 | 0,957275891 | novel transcript |
| ENSG00000165502 | RPL36AL | 0,378987114 | 1,300428531 | 0,13463646 | 0,585389409 | ribosomal protein L36a like |
| ENSG00000103126 | AXIN1 | 0,378933224 | 1,300379956 | 0,3702957 | 0,858545694 | axin 1 |
| ENSG00000134438 | RAX | 0,37892231 | 1,300370119 | 0,60733116 | 0,950993594 | retina and anterior neural fold homeobox |
| ENSG00000213856 | VDAC1P2 | 0,378815902 | 1,300274211 | 0,63941786 | 0,956018518 | voltage dependent anion channel 1 pseudogene 2 |
| ENSG00000109689 | STIM2 | 0,378755036 | 1,300219356 | 0,2562664 | 0,768515424 | stromal interaction molecule 2 |
| ENSG00000138311 | ZNF365 | 0,378619163 | 1,300096907 | 0,40773004 | 0,880762246 | zinc finger protein 365 |
| ENSG00000119943 | PYROXD2 | 0,378162654 | 1,299685585 | 0,15893489 | 0,633144409 | pyridine nucleotide-disulphide oxidoreductase domain 2 |
| ENSG00000144821 | MYH15 | 0,378142137 | 1,299667102 | 0,43076548 | 0,89530663 | myosin heavy chain 15 |
| ENSG00000228386 | AL031668,1 | 0,378114276 | 1,299642003 | 0,69391987 | 0,967340633 | novel transcript |
| ENSG00000166394 | CYB5R2 | 0,378094405 | 1,299624103 | 0,13768435 | 0,592433314 | cytochrome b5 reductase 2 |
| ENSG00000179262 | RAD23A | 0,378089314 | 1,299619517 | 0,09931001 | 0,50040065 | RAD23 homolog A, nucleotide excision repair protein |
| ENSG00000100280 | AP1B1 | 0,377636326 | 1,299211516 | 0,14792986 | 0,612582848 | adaptor related protein complex 1 subunit beta 1 |
| ENSG00000257298 | AC008147,2 | 0,377596321 | 1,299175491 | 0,4038514 | 0,877749224 | novel transcript, sense intronic to LIMA1 |
| ENSG00000287299 | AC012459,1 | 0,377584485 | 1,299164832 | 0,20163051 | 0,701198711 | novel transcript, antisense to SPATS2L |
| ENSG00000103005 | USB1 | 0,377488825 | 1,299078692 | 0,17574456 | 0,661199837 | U6 snRNA biogenesis phosphodiesterase 1 |
| ENSG00000054116 | TRAPPC3 | 0,377465218 | 1,299057436 | 0,12059207 | 0,55405161 | trafficking protein particle complex 3 |

| | | | | | | |
|---|---|---|---|---|---|---|
| ENSG00000129083 | COPB1 | 0,377232397 | 1,298847812 | 0,22463918 | 0,733203418 | COPI coat complex subunit beta 1 |
| ENSG00000243388 | RPL3P3 | 0,377192967 | 1,298812313 | 0,77142839 | 0,978033855 | ribosomal protein L3 pseudogene 3 |
| ENSG00000128655 | PDE11A | 0,377191528 | 1,298811017 | 0,39337138 | 0,87137239 | phosphodiesterase 11A |
| ENSG00000168405 | CMAHP | 0,377177581 | 1,298798462 | 0,72328048 | 0,972913663 | cytidine monophospho-N-acetylneuraminic acid hydroxylase, pseudogene |
| ENSG00000007923 | DNAJC11 | 0,377144211 | 1,29876842 | 0,17920275 | 0,667690954 | DnaJ heat shock protein family (Hsp40) member C11 |
| ENSG00000286990 | AC016026,2 | 0,37709515 | 1,298724255 | 0,65164486 | 0,957089277 | novel transcript, antisense to MICAL3 |
| ENSG00000166359 | WDR88 | 0,376934998 | 1,298580093 | 0,54766625 | 0,935404974 | WD repeat domain 88 |
| ENSG00000184441 | AP001062,1 | 0,376919126 | 1,298565807 | 0,3181998 | 0,823060685 | novel transcript, antisense to C21orf2 |
| ENSG00000146701 | MDH2 | 0,376799391 | 1,298458037 | 0,1732873 | 0,658330849 | malate dehydrogenase 2 |
| ENSG00000132356 | PRKAA1 | 0,376745688 | 1,298409704 | 0,25890147 | 0,771850203 | protein kinase AMP-activated catalytic subunit alpha 1 |
| ENSG00000156026 | MCU | 0,376690151 | 1,298359723 | 0,26706771 | 0,779389991 | mitochondrial calcium uniporter |
| ENSG00000259763 | LINC02157 | 0,376678961 | 1,298349652 | 0,78574082 | 0,979320406 | long intergenic non-protein coding RNA 2157 |
| ENSG00000187164 | SHTN1 | 0,376671758 | 1,29834317 | 0,24227799 | 0,752491461 | shootin 1 |
| ENSG00000173327 | MAP3K11 | 0,376614065 | 1,29829125 | 0,36158266 | 0,851893059 | mitogen-activated protein kinase kinase kinase 11 |
| ENSG00000109790 | KLHL5 | 0,376562675 | 1,298245006 | 0,26051926 | 0,772692192 | kelch like family member 5 |
| ENSG00000099949 | LZTR1 | 0,376559507 | 1,298242155 | 0,12702126 | 0,569312214 | leucine zipper like transcription regulator 1 |
| ENSG00000112685 | EXOC2 | 0,376343487 | 1,298047779 | 0,22357569 | 0,731984265 | exocyst complex component 2 |
| ENSG00000063587 | ZNF275 | 0,376303777 | 1,29801205 | 0,07771777 | 0,440494637 | zinc finger protein 275 |
| ENSG00000173662 | TAS1R1 | 0,376055269 | 1,297788484 | 0,7007415 | 0,968522643 | taste 1 receptor member 1 |
| ENSG00000280759 | AP2A2 | 0,375936979 | 1,297682079 | 0,17286663 | 0,657402061 | adaptor related protein complex 2 subunit alpha 2 |
| ENSG00000277997 | AC079779,4 | 0,375826023 | 1,29758228 | 0,68043679 | 0,964654539 | novel transcript |
| ENSG00000137727 | ARHGAP20 | 0,375692406 | 1,297462109 | 0,57368729 | 0,942475308 | Rho GTPase activating protein 20 |
| ENSG00000227039 | ITGB2-AS1 | 0,375647458 | 1,297421686 | 0,70940594 | 0,970411233 | ITGB2 antisense RNA 1 |
| ENSG00000134061 | CD180 | 0,375623463 | 1,297400107 | 0,57456564 | 0,942475308 | CD180 molecule |
| ENSG00000171148 | TADA3 | 0,375547569 | 1,297331858 | 0,23438905 | 0,744054861 | transcriptional adaptor 3 |
| ENSG00000287549 | AC092268,1 | 0,375491228 | 1,297281195 | 0,66731699 | 0,9611116 | novel transcript |
| ENSG00000231638 | LUARIS | 0,375327863 | 1,297134305 | 0,42315682 | 0,890714198 | lncRNA upregulator of antiviral response interferon signaling |
| ENSG00000166913 | YWHAB | 0,37526498 | 1,297077767 | 0,22829006 | 0,736631691 | tyrosine 3-monooxygenase/tryptophan 5-monooxygenase activation protein beta |
| ENSG00000248866 | USP46-DT | 0,375221026 | 1,297038251 | 0,39347491 | 0,871526705 | USP46 divergent transcript |

| | | | | | | |
|---|---|---|---|---|---|---|
| ENSG00000214992 | AKAP17BP | 0,375209637 | 1,297028011 | 0,51046931 | 0,922663618 | A-kinase anchoring protein 17B, pseudogene |
| ENSG00000016082 | ISL1 | 0,374790172 | 1,296650954 | 0,47091631 | 0,908869685 | ISL LIM homeobox 1 |
| ENSG00000253214 | AC079209,1 | 0,374689804 | 1,29656075 | 0,77860599 | 0,978587255 | novel transcript |
| ENSG00000139793 | MBNL2 | 0,374020147 | 1,295959063 | 0,40605601 | 0,87908976 | muscleblind like splicing regulator 2 |
| ENSG00000108557 | RAI1 | 0,37401972 | 1,295958679 | 0,31366196 | 0,819846908 | retinoic acid induced 1 |
| ENSG00000244730 | AC139451,1 | 0,373952173 | 1,295898005 | 0,64673009 | 0,956899963 | PHD finger protein 5A (PHF5A) pseudogene |
| ENSG00000087842 | PIR | 0,373933565 | 1,29588129 | 0,17658593 | 0,662599918 | pirin |
| ENSG00000263606 | CHORDC1P4 | 0,37379618 | 1,295757891 | 0,48079271 | 0,912907448 | CHORDC1 pseudogene 4 |
| ENSG00000148343 | MIGA2 | 0,373763161 | 1,295728236 | 0,30583888 | 0,81473644 | mitoguardin 2 |
| ENSG00000255176 | AP000941,1 | 0,373734155 | 1,295702185 | 0,553262 | 0,936992468 | novel transcript |
| ENSG00000119729 | RHOQ | 0,373634485 | 1,295612673 | 0,32853671 | 0,830659639 | ras homolog family member Q |
| ENSG00000236155 | ZPLD2P | 0,373595243 | 1,295577432 | 0,66067457 | 0,959226315 | zona pellucida like domain containing 2, pseudogene |
| ENSG00000100227 | POLDIP3 | 0,373544603 | 1,295531957 | 0,1071619 | 0,522599956 | DNA polymerase delta interacting protein 3 |
| ENSG00000206283 | PFDN6 | 0,373526638 | 1,295515824 | 0,27602105 | 0,788074712 | prefoldin subunit 6 |
| ENSG00000224955 | KCTD10P1 | 0,373459154 | 1,295455227 | 0,67589126 | 0,963549282 | potassium channel tetramerization domain containing 10 pseudogene 1 |
| ENSG00000087263 | OGFOD1 | 0,373456393 | 1,295452747 | 0,12115804 | 0,55581584 | 2-oxoglutarate and iron dependent oxygenase domain containing 1 |
| ENSG00000257771 | LINC02395 | 0,373456346 | 1,295452705 | 0,62259071 | 0,952793934 | long intergenic non-protein coding RNA 2395 |
| ENSG00000277595 | AC007546,1 | 0,373383178 | 1,295387007 | 0,67533472 | 0,963390425 | novel transcript, sense intronic to ANKRD13A |
| ENSG00000235893 | BX284632,1 | 0,373380632 | 1,295384721 | 0,65068444 | 0,956899963 | novel pseudogene |
| ENSG00000178199 | ZC3H12D | 0,373369528 | 1,29537475 | 0,61306672 | 0,952413269 | zinc finger CCCH-type containing 12D |
| ENSG00000136238 | RAC1 | 0,373318369 | 1,295328816 | 0,22508321 | 0,733632361 | Rac family small GTPase 1 |
| ENSG00000109917 | ZPR1 | 0,373277756 | 1,295292352 | 0,0781407 | 0,442170262 | ZPR1 zinc finger |
| ENSG00000287381 | AC012433,2 | 0,373065213 | 1,295101539 | 0,69953879 | 0,968522643 | novel transcript, antisense to MAPK4 |
| ENSG00000225760 | LINC00431 | 0,373054919 | 1,295092298 | 0,75619908 | 0,975901808 | long intergenic non-protein coding RNA 431 |
| ENSG00000269891 | ARHGAP19-SLI | 0,372952171 | 1,295000066 | 0,39773193 | 0,874263835 | ARHGAP19-SLIT1 readthrough (NMD candidate) |
| ENSG00000107175 | CREB3 | 0,372828149 | 1,294888745 | 0,13266696 | 0,581398927 | cAMP responsive element binding protein 3 |
| ENSG00000283026 | TCF20 | 0,37281304 | 1,294875184 | 0,38100808 | 0,865900094 | transcription factor 20 |
| ENSG00000181274 | FRAT2 | 0,372791867 | 1,29485618 | 0,27770318 | 0,789067728 | FRAT regulator of WNT signaling pathway 2 |
| ENSG00000256533 | AP002373,2 | 0,372514858 | 1,294607581 | 0,602708 | 0,950352286 | CCR4 carbon catabolite repression 4-like (S, cerevisiae) (CCRN4L) pseudogene |

| | | | | | | |
|---|---|---|---|---|---|---|
| ENSG00000257252 | AC124947,1 | 0,372481061 | 1,294577255 | 0,73773353 | 0,974353474 | novel transcript, antisense to NUDT4 |
| ENSG00000135269 | TES | 0,372426278 | 1,294528097 | 0,23453142 | 0,74410295 | testin LIM domain protein |
| ENSG00000183401 | CCDC159 | 0,372294179 | 1,29440957 | 0,45766101 | 0,903831777 | coiled-coil domain containing 159 |
| ENSG00000263322 | AC110597,2 | 0,372284678 | 1,294401045 | 0,42107563 | 0,888848798 | novel transcript |
| ENSG00000121577 | POPDC2 | 0,372245624 | 1,294366006 | 0,52514521 | 0,928752911 | popeye domain containing 2 |
| ENSG00000197530 | MIB2 | 0,372196949 | 1,294322336 | 0,55689141 | 0,938264532 | mindbomb E3 ubiquitin protein ligase 2 |
| ENSG00000141012 | GALNS | 0,372170494 | 1,294298603 | 0,15720109 | 0,629552485 | galactosamine (N-acetyl)-6-sulfatase |
| ENSG00000138315 | OIT3 | 0,372141698 | 1,294272769 | 0,76631172 | 0,977154522 | oncoprotein induced transcript 3 |
| ENSG00000134780 | DAGLA | 0,372072614 | 1,294210793 | 0,28294796 | 0,794166737 | diacylglycerol lipase alpha |
| ENSG00000133985 | TTC9 | 0,372055129 | 1,294195108 | 0,17374395 | 0,658735233 | tetratricopeptide repeat domain 9 |
| ENSG00000122787 | AKR1D1 | 0,37195715 | 1,294107217 | 0,71754675 | 0,971697135 | aldo-keto reductase family 1 member D1 |
| ENSG00000232713 | RPS12P3 | 0,371936696 | 1,29408887 | 0,71856474 | 0,971869926 | ribosomal protein S12 pseudogene 3 |
| ENSG00000101084 | RAB5IF | 0,371828188 | 1,293991542 | 0,11837503 | 0,549825725 | RAB5 interacting factor |
| ENSG00000261226 | AC092384,3 | 0,371715753 | 1,293890701 | 0,69521332 | 0,967597791 | novel transcript |
| ENSG00000286806 | AC080005,1 | 0,371656416 | 1,293837485 | 0,60467047 | 0,950878607 | novel transcript |
| ENSG00000168894 | RNF181 | 0,371633917 | 1,293817308 | 0,09553433 | 0,490763275 | ring finger protein 181 |
| ENSG00000233892 | PAIP1P1 | 0,371604616 | 1,29379103 | 0,70102244 | 0,968677171 | poly(A) binding protein interacting protein 1 pseudogene 1 |
| ENSG00000164591 | MYOZ3 | 0,37151922 | 1,293714451 | 0,29702895 | 0,806635068 | myozenin 3 |
| ENSG00000115318 | LOXL3 | 0,371401773 | 1,293609136 | 0,25515496 | 0,766521096 | lysyl oxidase like 3 |
| ENSG00000086619 | ERO1B | 0,371241418 | 1,29346536 | 0,38013282 | 0,86564734 | endoplasmic reticulum oxidoreductase 1 beta |
| ENSG00000281416 | AC006359,4 | 0,371053846 | 1,293297201 | 0,6139239 | 0,952494509 | novel transcript |
| ENSG00000275746 | KLF13 | 0,370908781 | 1,293167164 | 0,09160381 | 0,481464668 | Kruppel like factor 13 |
| ENSG00000160072 | ATAD3B | 0,370897048 | 1,293156648 | 0,41611048 | 0,885986046 | ATPase family AAA domain containing 3B |
| ENSG00000185332 | TMEM105 | 0,37086631 | 1,293129096 | 0,69361853 | 0,967340633 | TMEM105 long non-coding RNA |
| ENSG00000007520 | TSR3 | 0,370595374 | 1,292886272 | 0,39155814 | 0,870333649 | TSR3 ribosome maturation factor |
| ENSG00000139436 | GIT2 | 0,370512184 | 1,292811722 | 0,22402545 | 0,732710917 | GIT ArfGAP 2 |
| ENSG00000274726 | ARHGEF10 | 0,370444048 | 1,292750666 | 0,13965672 | 0,595782615 | Rho guanine nucleotide exchange factor 10 |
| ENSG00000163481 | RNF25 | 0,370438417 | 1,29274562 | 0,11657818 | 0,545320605 | ring finger protein 25 |
| ENSG00000196116 | TDRD7 | 0,370332969 | 1,292651136 | 0,36617341 | 0,856056641 | tudor domain containing 7 |
| ENSG00000253923 | AP002981,1 | 0,370326497 | 1,292645336 | 0,67915145 | 0,964171781 | heat shock 10kDa protein 1 (chaperonin 10) (HSPE1) pseudogene |
| ENSG00000134256 | CD101 | 0,370235101 | 1,29256345 | 0,38071488 | 0,865900094 | CD101 molecule |

| | | | | | | |
|---|---|---|---|---|---|---|
| ENSG00000123178 | SPRYD7 | 0,369943308 | 1,292302047 | 0,12976519 | 0,575798478 | SPRY domain containing 7 |
| ENSG00000031698 | SARS1 | 0,369907937 | 1,292270364 | 0,11606943 | 0,544096306 | seryl-tRNA synthetase 1 |
| ENSG00000106484 | MEST | 0,369785788 | 1,292160956 | 0,3068181 | 0,815610044 | mesoderm specific transcript |
| ENSG00000116786 | PLEKHM2 | 0,369703179 | 1,292086969 | 0,2080666 | 0,712550568 | pleckstrin homology and RUN domain containing M2 |
| ENSG00000183621 | ZNF438 | 0,36967229 | 1,292059305 | 0,31535166 | 0,82082095 | zinc finger protein 438 |
| ENSG00000286001 | AC021087,5 | 0,369558461 | 1,291957365 | 0,75917515 | 0,976120207 | novel protein |
| ENSG00000282033 | AC074387,1 | 0,369499097 | 1,291904204 | 0,67255514 | 0,962290553 | uncharacterized LOC105373553 |
| ENSG00000104695 | PPP2CB | 0,369451671 | 1,291861736 | 0,20910292 | 0,71355079 | protein phosphatase 2 catalytic subunit beta |
| ENSG00000162517 | PEF1 | 0,36928863 | 1,29171575 | 0,11201469 | 0,534905665 | penta-EF-hand domain containing 1 |
| ENSG00000158125 | XDH | 0,369240457 | 1,291672618 | 0,42942015 | 0,89431066 | xanthine dehydrogenase |
| ENSG00000048392 | RRM2B | 0,369126623 | 1,291570705 | 0,36307364 | 0,853318246 | ribonucleotide reductase regulatory TP53 inducible subunit M2B |
| ENSG00000138698 | RAP1GDS1 | 0,369073388 | 1,291523047 | 0,33419537 | 0,83453565 | Rap1 GTPase-GDP dissociation stimulator 1 |
| ENSG00000078814 | MYH7B | 0,368979024 | 1,291438573 | 0,5695878 | 0,94221458 | myosin heavy chain 7B |
| ENSG00000187601 | MAGEH1 | 0,368969976 | 1,291430474 | 0,20251995 | 0,702572531 | MAGE family member H1 |
| ENSG00000151348 | EXT2 | 0,368936572 | 1,291400573 | 0,12433255 | 0,563515931 | exostosin glycosyltransferase 2 |
| ENSG00000264573 | RN7SL15P | 0,368920768 | 1,291386426 | 0,56681743 | 0,941353523 | RNA, 7SL, cytoplasmic 15, pseudogene |
| ENSG00000169562 | GJB1 | 0,368823977 | 1,291299789 | 0,5695131 | 0,94221458 | gap junction protein beta 1 |
| ENSG00000140688 | RUSF1 | 0,368617932 | 1,29111538 | 0,25447173 | 0,765935007 | RUS family member 1 |
| ENSG00000103647 | CORO2B | 0,36855061 | 1,291055133 | 0,21544183 | 0,72270101 | coronin 2B |
| ENSG00000133059 | DSTYK | 0,368546795 | 1,291051718 | 0,10141617 | 0,506329296 | dual serine/threonine and tyrosine protein kinase |
| ENSG00000250723 | AC016687,2 | 0,368447627 | 1,290962977 | 0,63950813 | 0,956018518 | novel transcript |
| ENSG00000214553 | LRRC37A11P | 0,368409813 | 1,290929141 | 0,64132406 | 0,956465487 | leucine rich repeat containing 37 member A11, pseudogene |
| ENSG00000186529 | CYP4F3 | 0,36834077 | 1,290867362 | 0,78551756 | 0,979320406 | cytochrome P450 family 4 subfamily F member 3 |
| ENSG00000283681 | TCF20 | 0,368274575 | 1,290808135 | 0,38695606 | 0,869441674 | transcription factor 20 |
| ENSG00000282892 | TCF20 | 0,368274575 | 1,290808135 | 0,38695606 | 0,869441674 | transcription factor 20 |
| ENSG00000280467 | TCF20 | 0,368274575 | 1,290808135 | 0,38695606 | 0,869441674 | transcription factor 20 |
| ENSG00000281897 | TCF20 | 0,368274575 | 1,290808135 | 0,38695606 | 0,869441674 | transcription factor 20 |
| ENSG00000068489 | PRR11 | 0,368256537 | 1,290791996 | 0,10577159 | 0,51904616 | proline rich 11 |
| ENSG00000197774 | EME2 | 0,368191452 | 1,290733765 | 0,4478883 | 0,902062594 | essential meiotic structure-specific endonuclease subunit 2 |
| ENSG00000225742 | LINC02036 | 0,367957657 | 1,290524613 | 0,53540939 | 0,932037868 | long intergenic non-protein coding RNA 2036 |

| | | | | | | |
|---|---|---|---|---|---|---|
| ENSG00000287291 | AC068286,2 | 0,367922429 | 1,290493101 | 0,84188221 | 0,98571999 | novel transcript |
| ENSG00000273403 | AC107294,3 | 0,367867948 | 1,290444369 | 0,75724834 | 0,975901808 | novel transcript |
| ENSG00000124496 | TRERF1 | 0,367782147 | 1,290367625 | 0,14445569 | 0,604748614 | transcriptional regulating factor 1 |
| ENSG00000189375 | TBC1D28 | 0,367744461 | 1,290333918 | 0,77192746 | 0,978033855 | TBC1 domain family member 28 |
| ENSG00000223639 | CLIC1 | 0,367740203 | 1,29033011 | 0,24849365 | 0,760720365 | chloride intracellular channel 1 |
| ENSG00000226248 | CLIC1 | 0,367740203 | 1,29033011 | 0,24849365 | 0,760720365 | chloride intracellular channel 1 |
| ENSG00000226417 | CLIC1 | 0,367740203 | 1,29033011 | 0,24849365 | 0,760720365 | chloride intracellular channel 1 |
| ENSG00000213719 | CLIC1 | 0,367740203 | 1,29033011 | 0,24849365 | 0,760720365 | chloride intracellular channel 1 |
| ENSG00000156232 | WHAMM | 0,36760064 | 1,290205293 | 0,22890345 | 0,737173617 | WASP homolog associated with actin, golgi membranes and microtubules |
| ENSG00000182263 | FIGN | 0,367581826 | 1,290188467 | 0,41940391 | 0,887822044 | fidgetin, microtubule severing factor |
| ENSG00000229625 | AC016877,1 | 0,367514855 | 1,290128577 | 0,55885037 | 0,939635125 | novel transcript |
| ENSG00000183570 | PCBP3 | 0,367371112 | 1,290000041 | 0,36131009 | 0,851821025 | poly(rC) binding protein 3 |
| ENSG00000206418 | RAB12 | 0,367099064 | 1,28975681 | 0,24537195 | 0,757479368 | RAB12, member RAS oncogene family |
| ENSG00000175137 | SH3BP5L | 0,367045929 | 1,289709309 | 0,28807409 | 0,799375215 | SH3 binding domain protein 5 like |
| ENSG00000224379 | TCF19 | 0,367032512 | 1,289697315 | 0,20001303 | 0,699646707 | transcription factor 19 |
| ENSG00000206492 | GNL1 | 0,366959993 | 1,289632488 | 0,18038695 | 0,66974377 | G protein nucleolar 1 (putative) |
| ENSG00000269439 | AC010618,3 | 0,366814547 | 1,289502479 | 0,50300529 | 0,920831325 | novel transcript |
| ENSG00000277781 | MROH6 | 0,366705359 | 1,28940489 | 0,53930097 | 0,933350549 | maestro heat like repeat family member 6 |
| ENSG00000212961 | HNRNPA1P40 | 0,366547853 | 1,289264127 | 0,60156022 | 0,949939978 | heterogeneous nuclear ribonucleoprotein A1 pseudogene 40 |
| ENSG00000172531 | PPP1CA | 0,36650231 | 1,289223428 | 0,10495199 | 0,516971651 | protein phosphatase 1 catalytic subunit alpha |
| ENSG00000172159 | FRMD3 | 0,366410398 | 1,289141296 | 0,40882328 | 0,881578006 | FERM domain containing 3 |
| ENSG00000146858 | ZC3HAV1L | 0,366362612 | 1,289098596 | 0,17270689 | 0,657107871 | zinc finger CCCH-type containing, antiviral 1 like |
| ENSG00000134533 | RERG | 0,366348108 | 1,289085636 | 0,57980811 | 0,943828882 | RAS like estrogen regulated growth inhibitor |
| ENSG00000122390 | NAA60 | 0,366304219 | 1,289046421 | 0,10934182 | 0,527621964 | N-alpha-acetyltransferase 60, NatF catalytic subunit |
| ENSG00000258675 | LINC02308 | 0,3662842 | 1,289028534 | 0,80168077 | 0,980623173 | long intergenic non-protein coding RNA 2308 |
| ENSG00000229243 | LINC01981 | 0,366217762 | 1,288969175 | 0,49597609 | 0,918105073 | long intergenic non-protein coding RNA 1981 |
| ENSG00000181350 | LRRC75A | 0,36620215 | 1,288955226 | 0,28061659 | 0,791510309 | leucine rich repeat containing 75A |
| ENSG00000100325 | ASCC2 | 0,366078803 | 1,288845028 | 0,10020653 | 0,502543647 | activating signal cointegrator 1 complex subunit 2 |
| ENSG00000142082 | SIRT3 | 0,36606842 | 1,288835752 | 0,17354919 | 0,658349911 | sirtuin 3 |
| ENSG00000103227 | LMF1 | 0,366014459 | 1,288787547 | 0,27918994 | 0,790013761 | lipase maturation factor 1 |
| ENSG00000135929 | CYP27A1 | 0,365981282 | 1,28875791 | 0,30961696 | 0,816986489 | cytochrome P450 family 27 subfamily A member 1 |

| | | | | | | |
|---|---|---|---|---|---|---|
| ENSG00000101280 | ANGPT4 | 0,365861952 | 1,288651317 | 0,57335718 | 0,942475308 | angiopoietin 4 |
| ENSG00000241490 | AC093010,1 | 0,365851778 | 1,288642229 | 0,72520428 | 0,97326334 | novel transcript, antisense to ZNF80 |
| ENSG00000147457 | CHMP7 | 0,36576172 | 1,28856179 | 0,11265896 | 0,536050184 | charged multivesicular body protein 7 |
| ENSG00000136933 | RABEPK | 0,365753413 | 1,288554371 | 0,1494604 | 0,614794108 | Rab9 effector protein with kelch motifs |
| ENSG00000168071 | CCDC88B | 0,365697024 | 1,288504007 | 0,53799531 | 0,932883277 | coiled-coil domain containing 88B |
| ENSG00000203799 | CCDC162P | 0,365636898 | 1,288450309 | 0,45028776 | 0,902062594 | coiled-coil domain containing 162, pseudogene |
| ENSG00000264860 | AC097641,1 | 0,365534499 | 1,288358861 | 0,7742695 | 0,978503783 | novel transcript |
| ENSG00000222043 | AC079305,1 | 0,365532547 | 1,288357117 | 0,71833926 | 0,971789526 | novel transcript |
| ENSG00000126351 | THRA | 0,36550381 | 1,288331455 | 0,24260262 | 0,752798972 | thyroid hormone receptor alpha |
| ENSG00000267462 | AC090377,1 | 0,365246522 | 1,288101716 | 0,8124579 | 0,981213877 | novel transcript |
| ENSG00000258384 | RCCD1-AS1 | 0,365157311 | 1,288022067 | 0,54645888 | 0,935401821 | RCCD1 and UNC45A antisense RNA 1 |
| ENSG00000230685 | CLIC1 | 0,365119805 | 1,287988583 | 0,21994657 | 0,728200468 | chloride intracellular channel 1 |
| ENSG00000228395 | AL356481,1 | 0,364905952 | 1,287797677 | 0,28641518 | 0,797584768 | novel transcript |
| ENSG00000148498 | PARD3 | 0,364837036 | 1,287736161 | 0,15107742 | 0,618311888 | par-3 family cell polarity regulator |
| ENSG00000257950 | P2RX5-TAX1BP | 0,364803872 | 1,287706559 | 0,444619 | 0,900295889 | P2RX5-TAX1BP3 readthrough (NMD candidate) |
| ENSG00000182208 | MOB2 | 0,364768507 | 1,287674995 | 0,24488374 | 0,756825336 | MOB kinase activator 2 |
| ENSG00000287851 | AP001885,2 | 0,364703158 | 1,287616669 | 0,75167183 | 0,975901808 | novel transcript |
| ENSG00000259917 | HNRNPLP2 | 0,364573158 | 1,287500648 | 0,43872897 | 0,897377994 | heterogeneous nuclear ribonucleoprotein L pseudogene 2 |
| ENSG00000253925 | AC011369,1 | 0,364446134 | 1,287387293 | 0,62207337 | 0,952793934 | novel transcript |
| ENSG00000108439 | PNPO | 0,364360832 | 1,287311176 | 0,46260392 | 0,906730926 | pyridoxamine 5'-phosphate oxidase |
| ENSG00000162419 | GMEB1 | 0,364358343 | 1,287308955 | 0,09968274 | 0,500760119 | glucocorticoid modulatory element binding protein 1 |
| ENSG00000284430 | AC020912,1 | 0,364316101 | 1,287271264 | 0,66095837 | 0,959312917 | novel transcript |
| ENSG00000161036 | LRWD1 | 0,364313749 | 1,287269165 | 0,39642114 | 0,873099971 | leucine rich repeats and WD repeat domain containing 1 |
| ENSG00000124299 | PEPD | 0,364184023 | 1,287153419 | 0,17338149 | 0,658330849 | peptidase D |
| ENSG00000163798 | SLC4A1AP | 0,364141298 | 1,287115301 | 0,09890627 | 0,499216097 | solute carrier family 4 member 1 adaptor protein |
| ENSG00000106803 | SEC61B | 0,36411085 | 1,287088138 | 0,36710073 | 0,85643141 | SEC61 translocon subunit beta |
| ENSG00000138814 | PPP3CA | 0,363938564 | 1,286934443 | 0,28364943 | 0,794981235 | protein phosphatase 3 catalytic subunit alpha |
| ENSG00000230523 | LINC01756 | 0,363862569 | 1,286866655 | 0,74102262 | 0,975146552 | long intergenic non-protein coding RNA 1756 |
| ENSG00000232818 | RPS2P32 | 0,363733451 | 1,286751488 | 0,37382714 | 0,861319894 | ribosomal protein S2 pseudogene 32 |
| ENSG00000187994 | RINL | 0,363708798 | 1,2867295 | 0,43034674 | 0,895167022 | Ras and Rab interactor like |

| | | | | | | |
|---|---|---|---|---|---|---|
| ENSG00000279927 | Z95114,4 | 0,363636705 | 1,286665202 | 0,69890137 | 0,968522643 | novel transcript |
| ENSG00000225080 | PFN1P4 | 0,363620985 | 1,286651183 | 0,542883 | 0,933663748 | profilin 1 pseudogene 4 |
| ENSG00000196378 | ZNF34 | 0,363587714 | 1,286621511 | 0,42344931 | 0,890929178 | zinc finger protein 34 |
| ENSG00000130768 | SMPDL3B | 0,363537068 | 1,286576345 | 0,77872165 | 0,978587255 | sphingomyelin phosphodiesterase acid like 3B |
| ENSG00000068394 | GPKOW | 0,363536531 | 1,286575866 | 0,30186689 | 0,811368958 | G-patch domain and KOW motifs |
| ENSG00000131788 | PIAS3 | 0,363368039 | 1,286425615 | 0,12087527 | 0,555007274 | protein inhibitor of activated STAT 3 |
| ENSG00000235189 | AL034405,1 | 0,363238999 | 1,286310558 | 0,54661097 | 0,935401821 | novel transcript |
| ENSG00000164961 | WASHC5 | 0,363192336 | 1,286268954 | 0,33622414 | 0,835660292 | WASH complex subunit 5 |
| ENSG00000137497 | NUMA1 | 0,363143971 | 1,286225834 | 0,51345607 | 0,924108601 | nuclear mitotic apparatus protein 1 |
| ENSG00000162813 | BPNT1 | 0,363101456 | 1,28618793 | 0,13689039 | 0,590472005 | 3'(2'), 5'-bisphosphate nucleotidase 1 |
| ENSG00000235065 | RPL24P2 | 0,362950082 | 1,286052985 | 0,62567504 | 0,952793934 | RPL24 pseudogene 2 |
| ENSG00000250571 | GLI4 | 0,362819067 | 1,2859362 | 0,35949362 | 0,849982153 | GLI family zinc finger 4 |
| ENSG00000259065 | AC005520,2 | 0,36277764 | 1,285899275 | 0,56945228 | 0,94221458 | novel transcript, antisense to C14orf43 |
| ENSG00000266990 | AC004528,1 | 0,362754374 | 1,285878537 | 0,62573398 | 0,952793934 | novel transcript, antisense to GRIN3B |
| ENSG00000103187 | COTL1 | 0,362691121 | 1,285822161 | 0,17351252 | 0,658349911 | coactosin like F-actin binding protein 1 |
| ENSG00000106526 | ACTR3C | 0,36256701 | 1,28571155 | 0,66288182 | 0,959897678 | actin related protein 3C |
| ENSG00000224066 | AL049795,1 | 0,362564078 | 1,285708937 | 0,66786283 | 0,961113267 | novel transcript |
| ENSG00000130962 | PRRG1 | 0,362362233 | 1,285529068 | 0,4455564 | 0,901085146 | proline rich and Gla domain 1 |
| ENSG00000100142 | POLR2F | 0,362180773 | 1,285367387 | 0,25401639 | 0,76531122 | RNA polymerase II, I and III subunit F |
| ENSG00000103489 | XYLT1 | 0,361812831 | 1,285039612 | 0,2032881 | 0,703503738 | xylosyltransferase 1 |
| ENSG00000179010 | MRFAP1 | 0,361811112 | 1,28503808 | 0,08939487 | 0,47709818 | Morf4 family associated protein 1 |
| ENSG00000114480 | GBE1 | 0,361693001 | 1,28493288 | 0,28470291 | 0,796099998 | 1,4-alpha-glucan branching enzyme 1 |
| ENSG00000134285 | FKBP11 | 0,36167925 | 1,284920633 | 0,29429885 | 0,804089131 | FKBP prolyl isomerase 11 |
| ENSG00000105248 | YJU2 | 0,361642627 | 1,284888015 | 0,2280796 | 0,736341996 | YJU2 splicing factor homolog |
| ENSG00000283247 | CCDC201 | 0,361636834 | 1,284882857 | 0,76233219 | 0,976315588 | coiled-coil domain containing 201 |
| ENSG00000160613 | PCSK7 | 0,36156344 | 1,284817493 | 0,34137062 | 0,840048641 | proprotein convertase subtilisin/kexin type 7 |
| ENSG00000104907 | TRMT1 | 0,361468567 | 1,284733004 | 0,25712316 | 0,769358275 | tRNA methyltransferase 1 |
| ENSG00000146674 | IGFBP3 | 0,361324596 | 1,284604803 | 0,17932709 | 0,667995017 | insulin like growth factor binding protein 3 |
| ENSG00000132821 | VSTM2L | 0,36120292 | 1,284496465 | 0,28964122 | 0,800507706 | V-set and transmembrane domain containing 2 like |
| ENSG00000242798 | AC073842,2 | 0,360988415 | 1,284305495 | 0,60964756 | 0,951793902 | novel transcript |
| ENSG00000177694 | NAALADL2 | 0,360810253 | 1,284146904 | 0,38901505 | 0,870057277 | N-acetylated alpha-linked acidic dipeptidase like 2 |
| ENSG00000004142 | POLDIP2 | 0,360722011 | 1,284068362 | 0,09763025 | 0,496553115 | DNA polymerase delta interacting protein 2 |
| ENSG00000104904 | OAZ1 | 0,360711936 | 1,284059394 | 0,14848051 | 0,613326406 | ornithine decarboxylase antizyme 1 |

| | | | | | | |
|---|---|---|---|---|---|---|
| ENSG00000260037 | AC104938,1 | 0,36062327 | 1,28398048 | 0,67851119 | 0,9640473 | novel transcript, antisense to MYO9A |
| ENSG00000113758 | DBN1 | 0,360584604 | 1,283946069 | 0,28618816 | 0,797420923 | drebrin 1 |
| ENSG00000141564 | RPTOR | 0,360565042 | 1,283928659 | 0,25944288 | 0,772164235 | regulatory associated protein of MTOR complex 1 |
| ENSG00000266869 | AC005993,1 | 0,36049968 | 1,283870491 | 0,73674774 | 0,974353474 | novel transcript |
| ENSG00000108582 | CPD | 0,360339068 | 1,283727569 | 0,30759264 | 0,816380056 | carboxypeptidase D |
| ENSG00000235078 | AC231981,1 | 0,360298375 | 1,28369136 | 0,62482773 | 0,952793934 | novel transcript, antisense to ADI1 |
| ENSG00000230982 | DSTNP1 | 0,36028202 | 1,283676808 | 0,40797645 | 0,880990181 | DSTN pseudogene 1 |
| ENSG00000149932 | TMEM219 | 0,360272058 | 1,283667944 | 0,21948781 | 0,727838473 | transmembrane protein 219 |
| ENSG00000254986 | DPP3 | 0,360228163 | 1,283628888 | 0,27304776 | 0,785672647 | dipeptidyl peptidase 3 |
| ENSG00000099917 | MED15 | 0,360206869 | 1,283609942 | 0,24152757 | 0,751948055 | mediator complex subunit 15 |
| ENSG00000250033 | SLC7A11-AS1 | 0,36004899 | 1,28346948 | 0,51809937 | 0,926332996 | SLC7A11 antisense RNA 1 |
| ENSG00000278247 | GOLGA6L1 | 0,359927164 | 1,283361104 | 0,89987555 | 0,991845106 | golgin A6 family like 1 |
| ENSG00000285465 | TM9SF1 | 0,359750204 | 1,283203697 | 0,139987 | 0,596260721 | transmembrane 9 superfamily member 1 |
| ENSG00000100926 | TM9SF1 | 0,359750204 | 1,283203697 | 0,139987 | 0,596260721 | transmembrane 9 superfamily member 1 |
| ENSG00000228436 | AL139260,1 | 0,359512141 | 1,28299197 | 0,45653454 | 0,903545728 | novel transcript |
| ENSG00000122042 | UBL3 | 0,359384697 | 1,282878639 | 0,28705238 | 0,798434649 | ubiquitin like 3 |
| ENSG00000196821 | ILRUN | 0,35936289 | 1,282859247 | 0,09497955 | 0,489386708 | inflammation and lipid regulator with UBA-like and NBR1-like domains |
| ENSG00000101134 | DOK5 | 0,359343652 | 1,282842141 | 0,4967024 | 0,918457927 | docking protein 5 |
| ENSG00000018280 | SLC11A1 | 0,359182721 | 1,282699049 | 0,66358471 | 0,959920886 | solute carrier family 11 member 1 |
| ENSG00000106948 | AKNA | 0,359018572 | 1,282553113 | 0,32295849 | 0,825051003 | AT-hook transcription factor |
| ENSG00000143924 | EML4 | 0,358975003 | 1,282514381 | 0,30842471 | 0,816625411 | EMAP like 4 |
| ENSG00000166317 | SYNPO2L | 0,358832142 | 1,282387388 | 0,56737271 | 0,941425553 | synaptopodin 2 like |
| ENSG00000145107 | TM4SF19 | 0,358802303 | 1,282360865 | 0,58548401 | 0,945213113 | transmembrane 4 L six family member 19 |
| ENSG00000187566 | NHLRC1 | 0,358793526 | 1,282353063 | 0,31541872 | 0,82082095 | NHL repeat containing E3 ubiquitin protein ligase 1 |
| ENSG00000288307 | CTDNEP1 | 0,358734572 | 1,282300662 | 0,15303661 | 0,621569237 | CTD nuclear envelope phosphatase 1 |
| ENSG00000175826 | CTDNEP1 | 0,358734572 | 1,282300662 | 0,15303661 | 0,621569237 | CTD nuclear envelope phosphatase 1 |
| ENSG00000171603 | CLSTN1 | 0,358719968 | 1,282287682 | 0,16722851 | 0,647913628 | calsyntenin 1 |
| ENSG00000100376 | FAM118A | 0,358692011 | 1,282262834 | 0,15459018 | 0,623707245 | family with sequence similarity 118 member A |
| ENSG00000071462 | BUD23 | 0,358639174 | 1,282215873 | 0,20409092 | 0,705049569 | BUD23 rRNA methyltransferase and ribosome maturation factor |
| ENSG00000179241 | LDLRAD3 | 0,358549705 | 1,282136359 | 0,25230338 | 0,76432863 | low density lipoprotein receptor class A domain containing 3 |

| | | | | | | |
|---|---|---|---|---|---|---|
| ENSG00000223687 | ZNF311 | 0,358541081 | 1,282128695 | 0,47589955 | 0,911390632 | zinc finger protein 311 |
| ENSG00000071626 | DAZAP1 | 0,358364996 | 1,281972217 | 0,12888115 | 0,573624329 | DAZ associated protein 1 |
| ENSG00000277566 | AC089999,2 | 0,358296145 | 1,281911038 | 0,79552246 | 0,979879951 | novel transcript, sense intronic to TPCN1 |
| ENSG00000225339 | AL354740,1 | 0,358271867 | 1,281889465 | 0,36967224 | 0,858104601 | novel transcript, antisense to C6orf1 |
| ENSG00000227205 | PFN1P9 | 0,358088495 | 1,281726543 | 0,70582148 | 0,969503793 | profilin 1 pseudogene 9 |
| ENSG00000272405 | AL365181,3 | 0,357785378 | 1,281457275 | 0,50953531 | 0,922427526 | novel transcript, antisense to BCAN |
| ENSG00000182551 | ADI1 | 0,357748848 | 1,281424827 | 0,1460555 | 0,609080854 | acireductone dioxygenase 1 |
| ENSG00000103423 | DNAJA3 | 0,35774265 | 1,281419322 | 0,17594412 | 0,661519708 | DnaJ heat shock protein family (Hsp40) member A3 |
| ENSG00000015676 | NUDCD3 | 0,357685726 | 1,281368763 | 0,17614062 | 0,661643674 | NudC domain containing 3 |
| ENSG00000137200 | CMTR1 | 0,357609901 | 1,281301419 | 0,12952273 | 0,575280402 | cap methyltransferase 1 |
| ENSG00000236782 | AL391650,1 | 0,357553978 | 1,281251753 | 0,44960365 | 0,902062594 | novel protein |
| ENSG00000100395 | L3MBTL2 | 0,357492208 | 1,281196896 | 0,22495576 | 0,733369856 | L3MBTL histone methyl-lysine binding protein 2 |
| ENSG00000166816 | LDHD | 0,357460684 | 1,281168901 | 0,52095644 | 0,927046149 | lactate dehydrogenase D |
| ENSG00000197586 | ENTPD6 | 0,357415636 | 1,281128897 | 0,24877191 | 0,760720365 | ectonucleoside triphosphate diphosphohydrolase 6 |
| ENSG00000230475 | ABHD16A | 0,357329381 | 1,281052305 | 0,7183112 | 0,971789526 | abhydrolase domain containing 16A, phospholipase |
| ENSG00000225727 | AL139184,1 | 0,357275171 | 1,281004169 | 0,7573784 | 0,975917168 | filamin A interacting protein 1-like (FILIP1L) pseudogene |
| ENSG00000134086 | VHL | 0,357261501 | 1,280992031 | 0,16704257 | 0,647621408 | von Hippel-Lindau tumor suppressor |
| ENSG00000001617 | SEMA3F | 0,357013184 | 1,280771565 | 0,37113422 | 0,859119168 | semaphorin 3F |
| ENSG00000182183 | SHISAL2A | 0,356956563 | 1,2807213 | 0,69158709 | 0,967201016 | shisa like 2A |
| ENSG00000065618 | COL17A1 | 0,356940657 | 1,28070718 | 0,58459762 | 0,944857905 | collagen type XVII alpha 1 chain |
| ENSG00000226686 | LINC01535 | 0,356934 | 1,280701271 | 0,60774102 | 0,950993594 | long intergenic non-protein coding RNA 1535 |
| ENSG00000158122 | PRXL2C | 0,356874805 | 1,280648724 | 0,21827896 | 0,725947652 | peroxiredoxin like 2C |
| ENSG00000167881 | SRP68 | 0,356842649 | 1,28062018 | 0,09811719 | 0,497396091 | signal recognition particle 68 |
| ENSG00000258983 | AL162171,2 | 0,356812027 | 1,280592998 | 0,61990848 | 0,952793934 | novel transcript, antisense to PTPN21 |
| ENSG00000137411 | VARS2 | 0,356739418 | 1,28052855 | 0,55124824 | 0,936366211 | valyl-tRNA synthetase 2, mitochondrial |
| ENSG00000162298 | SYVN1 | 0,356657272 | 1,280455639 | 0,18505762 | 0,677443261 | synoviolin 1 |
| ENSG00000133612 | AGAP3 | 0,356588055 | 1,280394207 | 0,27403011 | 0,7862019 | ArfGAP with GTPase domain, ankyrin repeat and PH domain 3 |
| ENSG00000115307 | AUP1 | 0,356531567 | 1,280344075 | 0,17691113 | 0,66278604 | AUP1 lipid droplet regulating VLDL assembly factor |
| ENSG00000124659 | TBCC | 0,356530827 | 1,280343418 | 0,13342267 | 0,583647193 | tubulin folding cofactor C |
| ENSG00000119487 | MAPKAP1 | 0,356519114 | 1,280333024 | 0,10802914 | 0,523928283 | MAPK associated protein 1 |
| ENSG00000167491 | GATAD2A | 0,356473236 | 1,280292309 | 0,0889324 | 0,47584845 | GATA zinc finger domain containing 2A |

| | | | | | | |
|---|---|---|---|---|---|---|
| ENSG00000090316 | MAEA | 0,356294304 | 1,280133529 | 0,21661308 | 0,723909983 | macrophage erythroblast attacher, E3 ubiquitin ligase |
| ENSG00000254858 | MPV17L2 | 0,356270253 | 1,280112189 | 0,2012206 | 0,701094807 | MPV17 mitochondrial inner membrane protein like 2 |
| ENSG00000274372 | LINC02804 | 0,356255456 | 1,280099059 | 0,71416876 | 0,970867497 | long intergenic non-protein coding RNA 2804 |
| ENSG00000269964 | MEI4 | 0,356241783 | 1,280086927 | 0,55822791 | 0,939161457 | meiotic double-stranded break formation protein 4 |
| ENSG00000181693 | OR8H1 | 0,356162642 | 1,280016708 | 0,82423268 | 0,983374097 | olfactory receptor family 8 subfamily H member 1 |
| ENSG00000146063 | TRIM41 | 0,356100135 | 1,279961251 | 0,49839121 | 0,919491485 | tripartite motif containing 41 |
| ENSG00000063245 | EPN1 | 0,35604571 | 1,279912965 | 0,33428953 | 0,83453565 | epsin 1 |
| ENSG00000143970 | ASXL2 | 0,355853848 | 1,279742763 | 0,1950361 | 0,69283135 | ASXL transcriptional regulator 2 |
| ENSG00000173992 | CCS | 0,355822585 | 1,279715032 | 0,28331396 | 0,794558711 | copper chaperone for superoxide dismutase |
| ENSG00000185963 | BICD2 | 0,35581751 | 1,27971053 | 0,28076655 | 0,791510309 | BICD cargo adaptor 2 |
| ENSG00000159337 | PLA2G4D | 0,35578333 | 1,279680212 | 0,74504073 | 0,975666349 | phospholipase A2 group IVD |
| ENSG00000284753 | EEF1AKMT4 | 0,355728314 | 1,279631413 | 0,40750067 | 0,880449269 | EEF1A lysine methyltransferase 4 |
| ENSG00000261205 | NIFKP4 | 0,355650145 | 1,279562081 | 0,73626085 | 0,974353474 | nucleolar protein interacting with the FHA domain of MKI67 pseudogene 4 |
| ENSG00000281221 | PSMC4 | 0,355637125 | 1,279550534 | 0,08968091 | 0,47775341 | proteasome 26S subunit, ATPase 4 |
| ENSG00000013275 | PSMC4 | 0,355637125 | 1,279550534 | 0,08968091 | 0,47775341 | proteasome 26S subunit, ATPase 4 |
| ENSG00000229334 | AC046143,1 | 0,355589969 | 1,279508711 | 0,4632745 | 0,906995673 | novel transcript |
| ENSG00000180879 | SSR4 | 0,355538949 | 1,279463462 | 0,16248225 | 0,639724866 | signal sequence receptor subunit 4 |
| ENSG00000071537 | SEL1L | 0,35541823 | 1,279356406 | 0,32649928 | 0,828450693 | SEL1L adaptor subunit of ERAD E3 ubiquitin ligase |
| ENSG00000003056 | M6PR | 0,355132488 | 1,279103041 | 0,24124163 | 0,751659063 | mannose-6-phosphate receptor, cation dependent |
| ENSG00000227082 | LINC02798 | 0,355103178 | 1,279077054 | 0,36789769 | 0,857401877 | long intergenic non-protein coding RNA 2798 |
| ENSG00000151176 | PLBD2 | 0,355074113 | 1,279051286 | 0,16573442 | 0,645029358 | phospholipase B domain containing 2 |
| ENSG00000272420 | AL513477,2 | 0,354956392 | 1,278946922 | 0,5475956 | 0,935404974 | novel transcript, sense intronic to MORN1 |
| ENSG00000168807 | SNTB2 | 0,354895573 | 1,278893007 | 0,16513008 | 0,644200984 | syntrophin beta 2 |
| ENSG00000285549 | AC091973,1 | 0,354808087 | 1,278815457 | 0,84939066 | 0,987036295 | Novel transcript |
| ENSG00000169902 | TPST1 | 0,354715853 | 1,278733703 | 0,21407306 | 0,72150508 | tyrosylprotein sulfotransferase 1 |
| ENSG00000099821 | POLRMT | 0,354569436 | 1,278603933 | 0,48581896 | 0,913997663 | RNA polymerase mitochondrial |
| ENSG00000258053 | AC025575,2 | 0,354545212 | 1,278582464 | 0,65073727 | 0,956899963 | novel transcript |
| ENSG00000130038 | CRACR2A | 0,35451922 | 1,278559429 | 0,44805388 | 0,902062594 | calcium release activated channel regulator 2A |
| ENSG00000283010 | AL589826,2 | 0,354501585 | 1,278543801 | 0,64904637 | 0,956899963 | novel transcript |
| ENSG00000275576 | AL049539,1 | 0,354456045 | 1,278503442 | 0,69720533 | 0,968522643 | novel transcript, sense intronic to TM9SF4 |
| ENSG00000272909 | AL122035,2 | 0,354390658 | 1,278445498 | 0,45671519 | 0,90366076 | novel transcript, antisense to MTHFD1 |

| | | | | | | |
|---|---|---|---|---|---|---|
| ENSG00000185633 | NDUFA4L2 | 0,354285821 | 1,2783526 | 0,35898112 | 0,849814998 | NDUFA4 mitochondrial complex associated like 2 |
| ENSG00000260422 | Z97205,2 | 0,354268517 | 1,278337268 | 0,54388283 | 0,934259495 | novel transcript |
| ENSG00000249786 | EAF1-AS1 | 0,354213282 | 1,278288326 | 0,59094852 | 0,947416795 | EAF1 antisense RNA 1 |
| ENSG00000265905 | RN7SL96P | 0,354191565 | 1,278269084 | 0,73084048 | 0,974353474 | RNA, 7SL, cytoplasmic 96, pseudogene |
| ENSG00000115896 | PLCL1 | 0,354177376 | 1,278256512 | 0,32466203 | 0,826700906 | phospholipase C like 1 (inactive) |
| ENSG00000108592 | FTSJ3 | 0,35400258 | 1,278101649 | 0,39963433 | 0,875433675 | FtsJ RNA 2'-O-methyltransferase 3 |
| ENSG00000111412 | SPRING1 | 0,353753741 | 1,277881219 | 0,20151453 | 0,701094807 | SREBF pathway regulator in golgi 1 |
| ENSG00000174225 | ARL13A | 0,353747992 | 1,277876126 | 0,76726959 | 0,977201891 | ADP ribosylation factor like GTPase 13A |
| ENSG00000274148 | LILRA6 | 0,353548274 | 1,277699237 | 0,81026903 | 0,981213877 | leukocyte immunoglobulin like receptor A6 |
| ENSG00000286201 | AC011751,1 | 0,353414517 | 1,277580782 | 0,63298575 | 0,954408281 | novel transcript |
| ENSG00000234947 | STK19 | 0,353398614 | 1,2775667 | 0,22427146 | 0,73307493 | serine/threonine kinase 19 |
| ENSG00000142207 | URB1 | 0,35336524 | 1,277537146 | 0,14679525 | 0,610674677 | URB1 ribosome biogenesis homolog |
| ENSG00000196933 | RPS26P11 | 0,353344277 | 1,277518583 | 0,71092201 | 0,970482408 | ribosomal protein S26 pseudogene 11 |
| ENSG00000277825 | AC020917,3 | 0,353080254 | 1,27728481 | 0,70463428 | 0,969219816 | novel transcript |
| ENSG00000231760 | AL355312,2 | 0,352998103 | 1,27721208 | 0,63546147 | 0,955314044 | novel transcript |
| ENSG00000108946 | PRKAR1A | 0,352979541 | 1,277195647 | 0,22326389 | 0,73194107 | protein kinase cAMP-dependent type I regulatory subunit alpha |
| ENSG00000267827 | AC011468,2 | 0,352947059 | 1,277166892 | 0,67894019 | 0,964151912 | novel transcript |
| ENSG00000163602 | RYBP | 0,352905876 | 1,277130434 | 0,45940642 | 0,904862549 | RING1 and YY1 binding protein |
| ENSG00000288310 | AL133216,7 | 0,352859259 | 1,277089168 | 0,82441836 | 0,983374097 | general transcription factor IIi (GTF2I) pseudogene |
| ENSG00000286388 | AC026748,7 | 0,352785063 | 1,27702349 | 0,21049753 | 0,716025966 | novel transcript |
| ENSG00000175756 | AURKAIP1 | 0,352662818 | 1,276915288 | 0,29291444 | 0,803304991 | aurora kinase A interacting protein 1 |
| ENSG00000130119 | GNL3L | 0,352598981 | 1,276858788 | 0,13885524 | 0,594163262 | G protein nucleolar 3 like |
| ENSG00000187997 | C17orf99 | 0,352493144 | 1,27676512 | 0,68357116 | 0,965143332 | chromosome 17 open reading frame 99 |
| ENSG00000130037 | KCNA5 | 0,352459419 | 1,276735274 | 0,65984822 | 0,958837121 | potassium voltage-gated channel subfamily A member 5 |
| ENSG00000131966 | ACTR10 | 0,352387491 | 1,276671622 | 0,34624209 | 0,841168721 | actin related protein 10 |
| ENSG00000138246 | DNAJC13 | 0,352360785 | 1,276647989 | 0,27040799 | 0,782859427 | DnaJ heat shock protein family (Hsp40) member C13 |
| ENSG00000167671 | UBXN6 | 0,352282334 | 1,27657857 | 0,25464409 | 0,766088613 | UBX domain protein 6 |
| ENSG00000106404 | CLDN15 | 0,352199451 | 1,276505232 | 0,35522154 | 0,846725094 | claudin 15 |
| ENSG00000127603 | MACF1 | 0,35201928 | 1,276345826 | 0,10394785 | 0,514131574 | microtubule actin crosslinking factor 1 |
| ENSG00000288191 | MISP3 | 0,351923596 | 1,276261178 | 0,4771614 | 0,91143003 | MISP family member 3 |
| ENSG00000141854 | MISP3 | 0,351923596 | 1,276261178 | 0,4771614 | 0,91143003 | MISP family member 3 |

| | | | | | | |
|---|---|---|---|---|---|---|
| ENSG00000160654 | CD3G | 0,351921545 | 1,276259362 | 0,73122118 | 0,974353474 | CD3g molecule |
| ENSG00000280582 | AP003392,6 | 0,351900699 | 1,276240922 | 0,56227094 | 0,940865625 | novel transcript, antisense to CCDC84 |
| ENSG00000254428 | AP003392,1 | 0,351900699 | 1,276240922 | 0,56227094 | 0,940865625 | novel transcript, antisense to CCDC84 |
| ENSG00000215483 | LINC00598 | 0,351861969 | 1,276206661 | 0,60564183 | 0,950927937 | long intergenic non-protein coding RNA 598 |
| ENSG00000104983 | CCDC61 | 0,351858184 | 1,276203313 | 0,44619221 | 0,901348735 | coiled-coil domain containing 61 |
| ENSG00000206260 | PRR23A | 0,351660827 | 1,276028744 | 0,83365102 | 0,984905093 | proline rich 23A |
| ENSG00000101421 | CHMP4B | 0,351525442 | 1,275909004 | 0,11449205 | 0,540617781 | charged multivesicular body protein 4B |
| ENSG00000233008 | LINC01725 | 0,351478452 | 1,275867447 | 0,70206044 | 0,968742626 | long intergenic non-protein coding RNA 1725 |
| ENSG00000236466 | AL035670,1 | 0,351333605 | 1,275739356 | 0,69567515 | 0,967723599 | novel transcript |
| ENSG00000267057 | LINC01905 | 0,351174994 | 1,275599108 | 0,77871295 | 0,978587255 | long intergenic non-protein coding RNA 1905 |
| ENSG00000078487 | ZCWPW1 | 0,35110621 | 1,275538293 | 0,29026803 | 0,800682868 | zinc finger CW-type and PWWP domain containing 1 |
| ENSG00000204568 | MRPS18B | 0,351104261 | 1,275536569 | 0,14052878 | 0,597347645 | mitochondrial ribosomal protein S18B |
| ENSG00000283118 | AC009951,5 | 0,351043799 | 1,275483114 | 0,76663825 | 0,977201891 | novel transcript, antisense to ZEB2 |
| ENSG00000091986 | CCDC80 | 0,351040498 | 1,275480195 | 0,28419233 | 0,795731744 | coiled-coil domain containing 80 |
| ENSG00000226533 | BTBD9-AS1 | 0,351024779 | 1,275466298 | 0,72306351 | 0,972838903 | BTBD9 antisense RNA 1 |
| ENSG00000099341 | PSMD8 | 0,351014441 | 1,275457158 | 0,11953545 | 0,552362303 | proteasome 26S subunit, non-ATPase 8 |
| ENSG00000178381 | ZFAND2A | 0,350911085 | 1,275365787 | 0,14293935 | 0,601785046 | zinc finger AN1-type containing 2A |
| ENSG00000173714 | WFIKKN2 | 0,350716782 | 1,275194031 | 0,59724329 | 0,948282297 | WAP, follistatin/kazal, immunoglobulin, kunitz and netrin domain containing 2 |
| ENSG00000184530 | C6orf58 | 0,350713444 | 1,275191081 | 0,71399488 | 0,970867497 | chromosome 6 open reading frame 58 |
| ENSG00000103043 | VAC14 | 0,35069208 | 1,275172198 | 0,23302949 | 0,742172102 | VAC14 component of PIKFYVE complex |
| ENSG00000089154 | GCN1 | 0,350568665 | 1,275063118 | 0,29493757 | 0,805040011 | GCN1 activator of EIF2AK4 |
| ENSG00000224259 | LINC01133 | 0,350543866 | 1,275041201 | 0,79263476 | 0,979471509 | long intergenic non-protein coding RNA 1133 |
| ENSG00000253154 | AC100801,1 | 0,350517832 | 1,275018192 | 0,56621371 | 0,941123407 | novel transcript |
| ENSG00000172732 | MUS81 | 0,350508793 | 1,275010203 | 0,15854742 | 0,632328936 | MUS81 structure-specific endonuclease subunit |
| ENSG00000141456 | PELP1 | 0,350418973 | 1,274930826 | 0,42319181 | 0,890714198 | proline, glutamate and leucine rich protein 1 |
| ENSG00000159069 | FBXW5 | 0,350329714 | 1,274851949 | 0,24041533 | 0,750433792 | F-box and WD repeat domain containing 5 |
| ENSG00000249481 | SPATS1 | 0,350144191 | 1,27468802 | 0,63947907 | 0,956018518 | spermatogenesis associated serine rich 1 |
| ENSG00000229688 | CRPPA-AS1 | 0,350091563 | 1,274641522 | 0,69980479 | 0,968522643 | CRPPA antisense RNA 1 |
| ENSG00000145740 | SLC30A5 | 0,350000699 | 1,274561245 | 0,2446442 | 0,75645892 | solute carrier family 30 member 5 |
| ENSG00000231840 | TMEM139-AS1 | 0,349886474 | 1,274460335 | 0,74757872 | 0,975901808 | TMEM139 antisense RNA 1 |
| ENSG00000115053 | NCL | 0,349835063 | 1,274414921 | 0,44882783 | 0,902062594 | nucleolin |
| ENSG00000256235 | SMIM3 | 0,349808412 | 1,274391379 | 0,33648783 | 0,835713011 | small integral membrane protein 3 |

| | | | | | | |
|---|---|---|---|---|---|---|
| ENSG00000108576 | SLC6A4 | 0,349624326 | 1,274228779 | 0,59248992 | 0,947416795 | solute carrier family 6 member 4 |
| ENSG00000135424 | ITGA7 | 0,349450181 | 1,274074978 | 0,4531854 | 0,902540855 | integrin subunit alpha 7 |
| ENSG00000167723 | TRPV3 | 0,349407879 | 1,27403762 | 0,30194312 | 0,81141049 | transient receptor potential cation channel subfamily V member 3 |
| ENSG00000136810 | TXN | 0,349397734 | 1,274028662 | 0,36562035 | 0,855531368 | thioredoxin |
| ENSG00000173809 | TDRD12 | 0,349346386 | 1,273983317 | 0,59636614 | 0,948282297 | tudor domain containing 12 |
| ENSG00000197670 | AL157838,1 | 0,349316556 | 1,273956976 | 0,2951821 | 0,805285623 | novel transcript, antisense to ZNF217 |
| ENSG00000180902 | D2HGDH | 0,349277141 | 1,273922172 | 0,51206618 | 0,923307981 | D-2-hydroxyglutarate dehydrogenase |
| ENSG00000273064 | AC017083,1 | 0,349058066 | 1,27372874 | 0,62056553 | 0,952793934 | novel transcript |
| ENSG00000128739 | SNRPN | 0,349045058 | 1,273717255 | 0,11972354 | 0,552559297 | small nuclear ribonucleoprotein polypeptide N |
| ENSG00000283020 | DUX4L37 | 0,348991978 | 1,273670393 | 0,60676118 | 0,950927937 | double homeobox 4 like 37 (pseudogene) |
| ENSG00000188827 | SLX4 | 0,348947231 | 1,273630889 | 0,25563812 | 0,767141836 | SLX4 structure-specific endonuclease subunit |
| ENSG00000260669 | AL096870,2 | 0,348824284 | 1,273522355 | 0,87589806 | 0,990109814 | novel transcript |
| ENSG00000284612 | AL591543,1 | 0,348811954 | 1,273511471 | 0,7597215 | 0,976120207 | novel transcript |
| ENSG00000233735 | AL359918,2 | 0,348770797 | 1,273475141 | 0,72676341 | 0,973344786 | novel transcript |
| ENSG00000140675 | SLC5A2 | 0,348720442 | 1,273430693 | 0,5398688 | 0,933385624 | solute carrier family 5 member 2 |
| ENSG00000226060 | TRIM26 | 0,348631077 | 1,273351815 | 0,56087067 | 0,940368987 | tripartite motif containing 26 |
| ENSG00000228881 | TRIM26 | 0,348631077 | 1,273351815 | 0,56087067 | 0,940368987 | tripartite motif containing 26 |
| ENSG00000162129 | CLPB | 0,348457335 | 1,273198476 | 0,16826672 | 0,649869138 | caseinolytic mitochondrial matrix peptidase chaperone subunit B |
| ENSG00000119723 | COQ6 | 0,348365647 | 1,273117563 | 0,33297612 | 0,833301711 | coenzyme Q6, monooxygenase |
| ENSG00000258012 | AC026111,1 | 0,348362739 | 1,273114996 | 0,57620912 | 0,943123502 | methionine adenosyltransferase II, alpha (MAT2A) pseudogene |
| ENSG00000176842 | IRX5 | 0,348349837 | 1,273103611 | 0,48804435 | 0,915188998 | iroquois homeobox 5 |
| ENSG00000115561 | CHMP3 | 0,348301295 | 1,273060776 | 0,1715531 | 0,655397237 | charged multivesicular body protein 3 |
| ENSG00000274505 | Metazoa_SRP | 0,348246014 | 1,273011996 | 0,84410845 | 0,98632753 | Metazoan signal recognition particle RNA |
| ENSG00000143862 | ARL8A | 0,348179299 | 1,272953129 | 0,13200478 | 0,580641112 | ADP ribosylation factor like GTPase 8A |
| ENSG00000235917 | MTCO2P11 | 0,347953733 | 1,272754117 | 0,75501957 | 0,975901808 | MT-CO2 pseudogene 11 |
| ENSG00000022277 | RTF2 | 0,347923795 | 1,272727706 | 0,11611396 | 0,544096306 | replication termination factor 2 |
| ENSG00000070010 | UFD1 | 0,3478761 | 1,272685631 | 0,14433063 | 0,604548856 | ubiquitin recognition factor in ER associated degradation 1 |
| ENSG00000125875 | TBC1D20 | 0,347754017 | 1,272577939 | 0,13050183 | 0,577136255 | TBC1 domain family member 20 |
| ENSG00000163479 | SSR2 | 0,34773466 | 1,272560865 | 0,09174844 | 0,481764036 | signal sequence receptor subunit 2 |

| | | | | | | |
|---|---|---|---|---|---|---|
| ENSG00000213018 | PABPN1P1 | 0,347681078 | 1,272513602 | 0,65997845 | 0,958859629 | PABPN1 pseudogene 1 |
| ENSG00000106268 | NUDT1 | 0,347636399 | 1,272474195 | 0,25960335 | 0,772176335 | nudix hydrolase 1 |
| ENSG00000276345 | AC004556,3 | 0,347559643 | 1,272406497 | 0,2881095 | 0,799375215 | 39S ribosomal protein L23, mitochondrial |
| ENSG00000253898 | LINC01419 | 0,347548063 | 1,272396284 | 0,48465298 | 0,913690171 | long intergenic non-protein coding RNA 1419 |
| ENSG00000109079 | TNFAIP1 | 0,347438126 | 1,272299327 | 0,15018505 | 0,615923738 | TNF alpha induced protein 1 |
| ENSG00000273398 | AC017083,3 | 0,347383316 | 1,272250992 | 0,58115185 | 0,943919685 | novel protein |
| ENSG00000197930 | ERO1A | 0,347336517 | 1,272209723 | 0,24746042 | 0,759682112 | endoplasmic reticulum oxidoreductase 1 alpha |
| ENSG00000267646 | AC008543,5 | 0,347095191 | 1,271996932 | 0,68486255 | 0,965415049 | novel transcript, antisense to ZNF491 |
| ENSG00000114638 | UPK1B | 0,347000028 | 1,271913031 | 0,6405972 | 0,956348702 | uroplakin 1B |
| ENSG00000107404 | DVL1 | 0,346893575 | 1,271819184 | 0,39253052 | 0,870909282 | dishevelled segment polarity protein 1 |
| ENSG00000072364 | AFF4 | 0,34686489 | 1,271793896 | 0,30657371 | 0,815375074 | AF4/FMR2 family member 4 |
| ENSG00000242156 | AC000041,1 | 0,346838107 | 1,271770286 | 0,69741214 | 0,968522643 | adaptor-related protein complex 1, beta 1 subunit (AP1B1) pseudogene |
| ENSG00000249141 | AL159163,1 | 0,346685207 | 1,271635508 | 0,70363597 | 0,968967751 | novel protein |
| ENSG00000285697 | AC002059,3 | 0,3466801 | 1,271631007 | 0,69924438 | 0,968522643 | chromosome 22 open reading frame 42 (C22orf42) pseudogene |
| ENSG00000250280 | AC026124,1 | 0,346646029 | 1,271600976 | 0,29025504 | 0,800682868 | novel transcript |
| ENSG00000161533 | ACOX1 | 0,346625633 | 1,271582999 | 0,14275459 | 0,601331098 | acyl-CoA oxidase 1 |
| ENSG00000188641 | DPYD | 0,346616412 | 1,271574871 | 0,27870197 | 0,789775666 | dihydropyrimidine dehydrogenase |
| ENSG00000196972 | SMIM10L2B | 0,346600287 | 1,271560659 | 0,26837673 | 0,780709773 | small integral membrane protein 10 like 2B |
| ENSG00000283638 | AC002407,1 | 0,346557439 | 1,271522895 | 0,21010341 | 0,71523009 | novel transcript |
| ENSG00000224055 | GAPDHP55 | 0,346507051 | 1,271478486 | 0,71859529 | 0,971869926 | glyceraldehyde 3 phosphate dehydrogenase pseudogene 55 |
| ENSG00000184227 | ACOT1 | 0,346493646 | 1,271466672 | 0,35038886 | 0,843857545 | acyl-CoA thioesterase 1 |
| ENSG00000005810 | MYCBP2 | 0,34634865 | 1,271338891 | 0,19094703 | 0,68665222 | MYC binding protein 2 |
| ENSG00000185753 | CXorf38 | 0,34634318 | 1,271334071 | 0,14633015 | 0,609656943 | chromosome X open reading frame 38 |
| ENSG00000239998 | LILRA2 | 0,346310309 | 1,271305105 | 0,77802319 | 0,978587255 | leukocyte immunoglobulin like receptor A2 |
| ENSG00000181638 | ZFP41 | 0,346291969 | 1,271288944 | 0,32212065 | 0,824955371 | ZFP41 zinc finger protein |
| ENSG00000228592 | D21S2088E | 0,346196291 | 1,271204635 | 0,67456161 | 0,963183892 | D21S2088E |
| ENSG00000288198 | PPFIA1 | 0,346108596 | 1,271127368 | 0,26717812 | 0,779493925 | PTPRF interacting protein alpha 1 |
| ENSG00000287621 | AC021851,2 | 0,346046251 | 1,271072438 | 0,65607263 | 0,957444211 | novel transcript |
| ENSG00000253361 | AC069120,1 | 0,345999304 | 1,271031076 | 0,64994401 | 0,956899963 | novel transcript |
| ENSG00000267303 | AC011511,4 | 0,345717553 | 1,270782874 | 0,86189129 | 0,989074806 | novel transcript |

| | | | | | | |
|---|---|---|---|---|---|---|
| ENSG00000100075 | SLC25A1 | 0,345707018 | 1,270773595 | 0,14677715 | 0,610674677 | solute carrier family 25 member 1 |
| ENSG00000141527 | CARD14 | 0,3456943 | 1,270762392 | 0,53380959 | 0,931381467 | caspase recruitment domain family member 14 |
| ENSG00000150782 | IL18 | 0,345682308 | 1,27075183 | 0,67867412 | 0,9640473 | interleukin 18 |
| ENSG00000259073 | FOXN3-AS2 | 0,345494276 | 1,270586218 | 0,73169429 | 0,974353474 | FOXN3 antisense RNA 2 |
| ENSG00000107862 | GBF1 | 0,345338539 | 1,270449068 | 0,09994497 | 0,501634237 | golgi brefeldin A resistant guanine nucleotide exchange factor 1 |
| ENSG00000166603 | MC4R | 0,345324705 | 1,270436886 | 0,77005032 | 0,977778135 | melanocortin 4 receptor |
| ENSG00000065970 | FOXJ2 | 0,345173419 | 1,27030367 | 0,13500871 | 0,586038109 | forkhead box J2 |
| ENSG00000234962 | LINC00700 | 0,34517109 | 1,27030162 | 0,72150482 | 0,972142678 | long intergenic non-protein coding RNA 700 |
| ENSG00000103671 | TRIP4 | 0,345151345 | 1,270284234 | 0,13497006 | 0,586038109 | thyroid hormone receptor interactor 4 |
| ENSG00000286867 | AC008798,3 | 0,345116418 | 1,270253482 | 0,72833019 | 0,973955059 | novel transcript |
| ENSG00000182585 | EPGN | 0,34510894 | 1,270246897 | 0,63400336 | 0,954929949 | epithelial mitogen |
| ENSG00000124593 | AL365205,1 | 0,345104709 | 1,270243172 | 0,32980875 | 0,831610673 | novel protein |
| ENSG00000138172 | CALHM2 | 0,345094891 | 1,270234527 | 0,31168064 | 0,817962189 | calcium homeostasis modulator family member 2 |
| ENSG00000174607 | UGT8 | 0,344877688 | 1,270043303 | 0,42518302 | 0,891853867 | UDP glycosyltransferase 8 |
| ENSG00000117983 | MUC5B | 0,344849948 | 1,270018883 | 0,79754131 | 0,980161849 | mucin 5B, oligomeric mucus/gel-forming |
| ENSG00000220517 | ASS1P1 | 0,344849907 | 1,270018847 | 0,69457069 | 0,967417311 | argininosuccinate synthetase 1 pseudogene 1 |
| ENSG00000268069 | AC004466,1 | 0,344820733 | 1,269993166 | 0,4704527 | 0,908757071 | novel transcript, antisense to HDAC7 |
| ENSG00000134970 | TMED7 | 0,344764638 | 1,269943787 | 0,50212182 | 0,920520138 | transmembrane p24 trafficking protein 7 |
| ENSG00000236780 | LINC01829 | 0,344701835 | 1,269888504 | 0,72352363 | 0,972913663 | long intergenic non-protein coding RNA 1829 |
| ENSG00000095261 | PSMD5 | 0,344621864 | 1,269818114 | 0,2812796 | 0,792337715 | proteasome 26S subunit, non-ATPase 5 |
| ENSG00000257221 | AC007569,1 | 0,344584289 | 1,269785042 | 0,48383539 | 0,913690171 | novel transcript, antisense to SELPLG |
| ENSG00000130182 | ZSCAN10 | 0,344579422 | 1,269780759 | 0,67156747 | 0,962098952 | zinc finger and SCAN domain containing 10 |
| ENSG00000157335 | CLEC18C | 0,344532921 | 1,269739832 | 0,54689239 | 0,935404974 | C-type lectin domain family 18 member C |
| ENSG00000106682 | EIF4H | 0,344468928 | 1,269683512 | 0,10546912 | 0,51821258 | eukaryotic translation initiation factor 4H |
| ENSG00000159461 | AMFR | 0,344382078 | 1,269607079 | 0,13425475 | 0,584425029 | autocrine motility factor receptor |
| ENSG00000189229 | AC069277,1 | 0,344322133 | 1,269554328 | 0,47713502 | 0,91143003 | novel transcript |
| ENSG00000119403 | PHF19 | 0,344228183 | 1,269471655 | 0,25177037 | 0,763986755 | PHD finger protein 19 |
| ENSG00000102189 | EEA1 | 0,344193645 | 1,269441265 | 0,32916306 | 0,831116147 | early endosome antigen 1 |
| ENSG00000255458 | AC108471,2 | 0,344022705 | 1,269290862 | 0,47802272 | 0,911886158 | novel transcript |
| ENSG00000100056 | ESS2 | 0,344003349 | 1,269273832 | 0,30638426 | 0,815375074 | ess-2 splicing factor homolog |
| ENSG00000105447 | GRWD1 | 0,343780248 | 1,269077564 | 0,38417332 | 0,868125769 | glutamate rich WD repeat containing 1 |
| ENSG00000160209 | PDXK | 0,343555471 | 1,268879853 | 0,26804358 | 0,780680587 | pyridoxal kinase |

| | | | | | | |
|---|---|---|---|---|---|---|
| ENSG00000177689 | MAGEB10 | 0,343553519 | 1,268878136 | 0,84183809 | 0,98571999 | MAGE family member B10 |
| ENSG00000166716 | ZNF592 | 0,343487672 | 1,268820225 | 0,1260122 | 0,566777174 | zinc finger protein 592 |
| ENSG00000107815 | TWNK | 0,343345809 | 1,268695465 | 0,49140391 | 0,916995341 | twinkle mtDNA helicase |
| ENSG00000213625 | LEPROT | 0,34329474 | 1,268650556 | 0,29430629 | 0,804089131 | leptin receptor overlapping transcript |
| ENSG00000265933 | LINC00668 | 0,343290196 | 1,26864656 | 0,50386191 | 0,921212789 | long intergenic non-protein coding RNA 668 |
| ENSG00000070785 | EIF2B3 | 0,343204207 | 1,268570947 | 0,14308363 | 0,601880115 | eukaryotic translation initiation factor 2B subunit gamma |
| ENSG00000155368 | DBI | 0,34319418 | 1,26856213 | 0,32037855 | 0,824366234 | diazepam binding inhibitor, acyl-CoA binding protein |
| ENSG00000049239 | H6PD | 0,343160011 | 1,268532086 | 0,2742176 | 0,7862019 | hexose-6-phosphate dehydrogenase/glucose 1-dehydrogenase |
| ENSG00000272573 | MUSTN1 | 0,343081295 | 1,268462875 | 0,84490393 | 0,986557476 | musculoskeletal, embryonic nuclear protein 1 |
| ENSG00000182223 | ZAR1 | 0,343054454 | 1,268439275 | 0,74497577 | 0,975666349 | zygote arrest 1 |
| ENSG00000136718 | IMP4 | 0,342992887 | 1,268385146 | 0,20019787 | 0,69971898 | IMP U3 small nucleolar ribonucleoprotein 4 |
| ENSG00000238280 | AC067751,1 | 0,342947778 | 1,268345487 | 0,85148553 | 0,987259878 | novel transcript |
| ENSG00000110422 | HIPK3 | 0,342946127 | 1,268344036 | 0,32068495 | 0,824366234 | homeodomain interacting protein kinase 3 |
| ENSG00000223944 | AL353604,1 | 0,342885086 | 1,268290373 | 0,64953425 | 0,956899963 | novel transcript |
| ENSG00000176182 | MYPOP | 0,342837593 | 1,268248622 | 0,33125914 | 0,832357374 | Myb related transcription factor, partner of profilin |
| ENSG00000255067 | AC013549,4 | 0,342744578 | 1,268166857 | 0,7354021 | 0,974353474 | novel transcript |
| ENSG00000259562 | AC090607,2 | 0,342612703 | 1,268050941 | 0,51189614 | 0,923184606 | zinc finger CCHC-type and RNA binding motif 1 (ZCRB1) pseudogene |
| ENSG00000224973 | LARGE-AS1 | 0,342472141 | 1,267927401 | 0,56771608 | 0,94163744 | LARGE antisense RNA 1 |
| ENSG00000104848 | KCNA7 | 0,342453161 | 1,26791072 | 0,66669756 | 0,9611116 | potassium voltage-gated channel subfamily A member 7 |
| ENSG00000112697 | TMEM30A | 0,342417357 | 1,267879254 | 0,42370695 | 0,890949676 | transmembrane protein 30A |
| ENSG00000262227 | AC004771,2 | 0,342389349 | 1,26785464 | 0,66366659 | 0,959920886 | novel transcript, antisense to INCA1 |
| ENSG00000136463 | TACO1 | 0,342366542 | 1,267834596 | 0,15710416 | 0,629382925 | translational activator of cytochrome c oxidase I |
| ENSG00000207496 | SNORA7A | 0,342325416 | 1,267798456 | 0,69534915 | 0,967602563 | small nucleolar RNA, H/ACA box 7A |
| ENSG00000110057 | UNC93B1 | 0,342245297 | 1,267728051 | 0,50907513 | 0,922427526 | unc-93 homolog B1, TLR signaling regulator |
| ENSG00000160233 | LRRC3 | 0,342188236 | 1,267677912 | 0,30875569 | 0,816730513 | leucine rich repeat containing 3 |
| ENSG00000259762 | AC021739,4 | 0,342170272 | 1,267662127 | 0,72481823 | 0,97326334 | novel transcript, antisense to AKAP13 |
| ENSG00000283777 | CANX | 0,342116846 | 1,267615184 | 0,20656131 | 0,709779284 | calnexin |
| ENSG00000188229 | TUBB4B | 0,342097914 | 1,267598549 | 0,12989862 | 0,576076964 | tubulin beta 4B class IVb |

| | | | | | | |
|---|---|---|---|---|---|---|
| ENSG00000261739 | GOLGA8S | 0,342058647 | 1,267564048 | 0,82172957 | 0,982952211 | golgin A8 family member S |
| ENSG00000259230 | LINC02323 | 0,341883095 | 1,267409816 | 0,75281091 | 0,975901808 | long intergenic non-protein coding RNA 2323 |
| ENSG00000198171 | DDRGK1 | 0,341823409 | 1,267357384 | 0,4476869 | 0,902062594 | DDRGK domain containing 1 |
| ENSG00000119392 | GLE1 | 0,341811576 | 1,267346989 | 0,12529335 | 0,56557482 | GLE1 RNA export mediator |
| ENSG00000127399 | LRRC61 | 0,341697667 | 1,267246929 | 0,82344337 | 0,983374097 | leucine rich repeat containing 61 |
| ENSG00000115762 | PLEKHB2 | 0,341552124 | 1,267119092 | 0,12473805 | 0,564453675 | pleckstrin homology domain containing B2 |
| ENSG00000261994 | WEE2-AS1 | 0,341451405 | 1,267030633 | 0,65748236 | 0,957853146 | WEE2 antisense RNA 1 |
| ENSG00000154957 | ZNF18 | 0,341393986 | 1,266980207 | 0,13506425 | 0,586038109 | zinc finger protein 18 |
| ENSG00000197296 | FITM2 | 0,341380373 | 1,266968252 | 0,22327668 | 0,73194107 | fat storage inducing transmembrane protein 2 |
| ENSG00000259895 | AC106820,2 | 0,341235037 | 1,266840625 | 0,66715947 | 0,9611116 | novel transcript, antisense to C16orf59 |
| ENSG00000279263 | OR2L8 | 0,341163013 | 1,266777381 | 0,79001842 | 0,979462318 | olfactory receptor family 2 subfamily L member 8 |
| ENSG00000040531 | CTNS | 0,341151174 | 1,266766986 | 0,16344942 | 0,642091247 | cystinosin, lysosomal cystine transporter |
| ENSG00000128487 | SPECC1 | 0,341148721 | 1,266764832 | 0,1407264 | 0,597618856 | sperm antigen with calponin homology and coiled-coil domains 1 |
| ENSG00000163156 | SCNM1 | 0,34112915 | 1,266747648 | 0,30253638 | 0,812236791 | sodium channel modifier 1 |
| ENSG00000178605 | GTPBP6 | 0,341119631 | 1,26673929 | 0,35872022 | 0,849814998 | GTP binding protein 6 (putative) |
| ENSG00000119004 | CYP20A1 | 0,341101415 | 1,266723295 | 0,27241328 | 0,785492261 | cytochrome P450 family 20 subfamily A member 1 |
| ENSG00000126603 | GLIS2 | 0,341062377 | 1,26668902 | 0,43201711 | 0,89558934 | GLIS family zinc finger 2 |
| ENSG00000180438 | TPRXL | 0,341003537 | 1,266637359 | 0,78711783 | 0,979424965 | tetrapeptide repeat homeobox like (pseudogene) |
| ENSG00000158042 | MRPL17 | 0,340806969 | 1,266464791 | 0,11932599 | 0,552110068 | mitochondrial ribosomal protein L17 |
| ENSG00000107937 | GTPBP4 | 0,340711399 | 1,266380898 | 0,15911642 | 0,633144409 | GTP binding protein 4 |
| ENSG00000160953 | PWWP3A | 0,340697357 | 1,266368572 | 0,21506757 | 0,722298056 | PWWP domain containing 3A, DNA repair factor |
| ENSG00000173890 | GPR160 | 0,340632976 | 1,266312061 | 0,56666093 | 0,941311563 | G protein-coupled receptor 160 |
| ENSG00000181585 | TMIE | 0,340602121 | 1,266284979 | 0,70437188 | 0,969132783 | transmembrane inner ear |
| ENSG00000114395 | CYB561D2 | 0,340486058 | 1,266183111 | 0,23526107 | 0,744654651 | cytochrome b561 family member D2 |
| ENSG00000241785 | RN7SL390P | 0,340356708 | 1,266069593 | 0,65920417 | 0,958394501 | RNA, 7SL, cytoplasmic 390, pseudogene |
| ENSG00000053372 | MRTO4 | 0,340340223 | 1,266055126 | 0,18078141 | 0,670312865 | MRT4 homolog, ribosome maturation factor |
| ENSG00000127586 | CHTF18 | 0,340296282 | 1,266016565 | 0,51123836 | 0,922893865 | chromosome transmission fidelity factor 18 |
| ENSG00000233314 | PPP1R11 | 0,340270646 | 1,265994069 | 0,24550724 | 0,757479368 | protein phosphatase 1 regulatory inhibitor subunit 11 |
| ENSG00000234058 | PPP1R11 | 0,340270646 | 1,265994069 | 0,24550724 | 0,757479368 | protein phosphatase 1 regulatory inhibitor subunit 11 |

| | | | | | | |
|---|---|---|---|---|---|---|
| ENSG00000227720 | PPP1R11 | 0,340270646 | 1,265994069 | 0,24550724 | 0,757479368 | protein phosphatase 1 regulatory inhibitor subunit 11 |
| ENSG00000237403 | PPP1R11 | 0,340270646 | 1,265994069 | 0,24550724 | 0,757479368 | protein phosphatase 1 regulatory inhibitor subunit 11 |
| ENSG00000270518 | AC008626,1 | 0,340261681 | 1,265986202 | 0,77903361 | 0,978587255 | zinc finger protein 99 (ZNF99) pseudogene |
| ENSG00000275145 | FRG1 | 0,340136673 | 1,26587651 | 0,2512944 | 0,763579382 | FSHD region gene 1 |
| ENSG00000273592 | MBOAT7 | 0,340116176 | 1,265858525 | 0,36957326 | 0,858037898 | membrane bound O-acyltransferase domain containing 7 |
| ENSG00000170776 | AKAP13 | 0,340053603 | 1,265803624 | 0,15157037 | 0,61913273 | A-kinase anchoring protein 13 |
| ENSG00000236754 | AC007666,1 | 0,339944654 | 1,265708037 | 0,70715787 | 0,969565008 | novel transcript |
| ENSG00000264151 | AC090403,1 | 0,339882469 | 1,265653482 | 0,80079566 | 0,980505307 | novel transcript |
| ENSG00000242288 | BMS1P4-AGAP. | 0,339810704 | 1,265590525 | 0,56288403 | 0,940865625 | BMS1P4-AGAP5 readthough |
| ENSG00000167792 | NDUFV1 | 0,339745182 | 1,265533048 | 0,27300154 | 0,785672647 | NADH:ubiquinone oxidoreductase core subunit V1 |
| ENSG00000229173 | SRMP3 | 0,33972602 | 1,265516239 | 0,72243532 | 0,972594421 | SRM pseudogene 3 |
| ENSG00000225987 | PBX2 | 0,339656152 | 1,265454953 | 0,48375553 | 0,913690171 | PBX homeobox 2 |
| ENSG00000134452 | FBH1 | 0,339551648 | 1,265363291 | 0,10803163 | 0,523928283 | F-box DNA helicase 1 |
| ENSG00000181031 | RPH3AL | 0,339395615 | 1,265226444 | 0,37879162 | 0,864112469 | rabphilin 3A like (without C2 domains) |
| ENSG00000005812 | FBXL3 | 0,339350686 | 1,265187043 | 0,4237367 | 0,890949676 | F-box and leucine rich repeat protein 3 |
| ENSG00000183955 | KMT5A | 0,339266242 | 1,265112991 | 0,16056451 | 0,636125619 | lysine methyltransferase 5A |
| ENSG00000118369 | USP35 | 0,339043531 | 1,264917709 | 0,36376897 | 0,854139916 | ubiquitin specific peptidase 35 |
| ENSG00000198663 | C6orf89 | 0,338870352 | 1,264765879 | 0,19875701 | 0,697657684 | chromosome 6 open reading frame 89 |
| ENSG00000177398 | UMODL1 | 0,338840099 | 1,264739357 | 0,72707873 | 0,973344786 | uromodulin like 1 |
| ENSG00000260280 | SLX1B-SULT1A4 | 0,338807192 | 1,264710509 | 0,47241325 | 0,909229736 | SLX1B-SULT1A4 readthrough (NMD candidate) |
| ENSG00000162493 | PDPN | 0,338753596 | 1,264663526 | 0,37117809 | 0,859147985 | podoplanin |
| ENSG00000136811 | ODF2 | 0,338637506 | 1,264561766 | 0,20118675 | 0,701094807 | outer dense fiber of sperm tails 2 |
| ENSG00000101182 | PSMA7 | 0,338614555 | 1,264541649 | 0,10827124 | 0,524537831 | proteasome 20S subunit alpha 7 |
| ENSG00000188738 | FSIP2 | 0,338559285 | 1,264493205 | 0,30427466 | 0,813156808 | fibrous sheath interacting protein 2 |
| ENSG00000169895 | SYAP1 | 0,338515332 | 1,264454682 | 0,10566651 | 0,518774794 | synapse associated protein 1 |
| ENSG00000265740 | RN7SL339P | 0,338337741 | 1,264299042 | 0,80588573 | 0,981213877 | RNA, 7SL, cytoplasmic 339, pseudogene |
| ENSG00000286185 | AC242842,3 | 0,338217426 | 1,264193609 | 0,24942974 | 0,761381055 | novel protein, identical to neuroblastoma breakpoint family, member 19 NBPF19 |
| ENSG00000232396 | AC004882,1 | 0,338042411 | 1,264040257 | 0,78701998 | 0,979424965 | novel transcript |
| ENSG00000115539 | PDCL3 | 0,338030434 | 1,264029763 | 0,13231511 | 0,580797367 | phosducin like 3 |

| | | | | | | |
|---|---|---|---|---|---|---|
| ENSG00000104331 | BPNT2 | 0,338028224 | 1,264027826 | 0,32232428 | 0,824955371 | 3'(2'), 5'-bisphosphate nucleotidase 2 |
| ENSG00000230805 | AL132709,7 | 0,338022207 | 1,264022555 | 0,4601853 | 0,905443183 | novel transcript |
| ENSG00000143164 | DCAF6 | 0,337861159 | 1,26388146 | 0,26594739 | 0,777251371 | DDB1 and CUL4 associated factor 6 |
| ENSG00000132819 | RBM38 | 0,337844222 | 1,263866622 | 0,17051792 | 0,653761763 | RNA binding motif protein 38 |
| ENSG00000111859 | NEDD9 | 0,337739811 | 1,263775157 | 0,24859158 | 0,760720365 | neural precursor cell expressed, developmentally down-regulated 9 |
| ENSG00000129467 | ADCY4 | 0,337689481 | 1,26373107 | 0,47344034 | 0,910083046 | adenylate cyclase 4 |
| ENSG00000198399 | ITSN2 | 0,337687069 | 1,263728957 | 0,21820837 | 0,725906202 | intersectin 2 |
| ENSG00000266412 | NCOA4 | 0,337675106 | 1,263718478 | 0,31002853 | 0,817469158 | nuclear receptor coactivator 4 |
| ENSG00000261051 | AC107021,2 | 0,337587799 | 1,263642004 | 0,76233668 | 0,976315588 | novel transcript, overlapping to PLOD2 |
| ENSG00000286271 | AC008945,2 | 0,337575672 | 1,263631382 | 0,5284223 | 0,930033144 | novel transcript, antisense to SEPP1 |
| ENSG00000254681 | PKD1P5 | 0,337548076 | 1,263607212 | 0,55361355 | 0,937334216 | polycystin 1, transient receptor potential channel interacting pseudogene 5 |
| ENSG00000134490 | TMEM241 | 0,337535027 | 1,263595782 | 0,15740407 | 0,629849164 | transmembrane protein 241 |
| ENSG00000254855 | AP001107,8 | 0,337469151 | 1,263538086 | 0,78394953 | 0,979279135 | novel transcript |
| ENSG00000260830 | AL135744,1 | 0,33734166 | 1,263426432 | 0,70484963 | 0,969335769 | novel transcript, antisense to CHD8 and SUPT16H |
| ENSG00000284260 | RUFY1 | 0,337253515 | 1,263349243 | 0,22053971 | 0,728572782 | RUN and FYVE domain containing 1 |
| ENSG00000100241 | SBF1 | 0,337207503 | 1,26330895 | 0,44466055 | 0,900310737 | SET binding factor 1 |
| ENSG00000166813 | KIF7 | 0,337023323 | 1,263147682 | 0,57494222 | 0,942533938 | kinesin family member 7 |
| ENSG00000175967 | LINC02880 | 0,336957836 | 1,263090346 | 0,61265846 | 0,952413269 | long intergenic non-protein coding RNA 2880 |
| ENSG00000260032 | NORAD | 0,336833883 | 1,262981829 | 0,31690991 | 0,821888971 | non-coding RNA activated by DNA damage |
| ENSG00000173171 | MTX1 | 0,336825946 | 1,262974881 | 0,23767489 | 0,746907875 | metaxin 1 |
| ENSG00000146535 | GNA12 | 0,336796854 | 1,262949413 | 0,15570298 | 0,626262282 | G protein subunit alpha 12 |
| ENSG00000269982 | AC018809,2 | 0,336587577 | 1,262766223 | 0,50831836 | 0,922427526 | novel transcript, antisense to TMEM111 |
| ENSG00000174032 | SLC25A30 | 0,336546819 | 1,262730549 | 0,15455423 | 0,623642664 | solute carrier family 25 member 30 |
| ENSG00000115946 | PNO1 | 0,336369735 | 1,262575565 | 0,22485919 | 0,733203418 | partner of NOB1 homolog |
| ENSG00000105392 | CRX | 0,336301311 | 1,262515684 | 0,64092131 | 0,956410077 | cone-rod homeobox |
| ENSG00000167693 | NXN | 0,3362834 | 1,26250001 | 0,24798979 | 0,760187537 | nucleoredoxin |
| ENSG00000221916 | C19orf73 | 0,336120008 | 1,262357034 | 0,51445829 | 0,924434989 | chromosome 19 open reading frame 73 |
| ENSG00000124126 | PREX1 | 0,336111756 | 1,262349814 | 0,33459167 | 0,834757602 | phosphatidylinositol-3,4,5-trisphosphate dependent Rac exchange factor 1 |
| ENSG00000251485 | NRBF2P6 | 0,336066989 | 1,262310644 | 0,42892871 | 0,893763475 | NRBF2 pseudogene 6 |
| ENSG00000177830 | CHID1 | 0,336020481 | 1,262269951 | 0,28335174 | 0,79457429 | chitinase domain containing 1 |

| | | | | | | |
|---|---|---|---|---|---|---|
| ENSG00000112308 | C6orf62 | 0,336018289 | 1,262268034 | 0,32373264 | 0,825794844 | chromosome 6 open reading frame 62 |
| ENSG00000274605 | PCCA-DT | 0,335898699 | 1,262163404 | 0,19011895 | 0,685541073 | PCCA divergent transcript |
| ENSG00000165238 | WNK2 | 0,335818351 | 1,262093112 | 0,44248638 | 0,899414493 | WNK lysine deficient protein kinase 2 |
| ENSG00000253347 | AC040934,1 | 0,335677289 | 1,261969716 | 0,77970311 | 0,978587255 | novel transcript |
| ENSG00000108448 | TRIM16L | 0,335674173 | 1,26196699 | 0,37004689 | 0,85830893 | tripartite motif containing 16 like |
| ENSG00000176029 | C11orf16 | 0,335645211 | 1,261941656 | 0,55611774 | 0,938046927 | chromosome 11 open reading frame 16 |
| ENSG00000104863 | LIN7B | 0,335469223 | 1,261787726 | 0,50979728 | 0,922427526 | lin-7 homolog B, crumbs cell polarity complex component |
| ENSG00000146828 | SLC12A9 | 0,335429987 | 1,261753411 | 0,44975539 | 0,902062594 | solute carrier family 12 member 9 |
| ENSG00000288620 | AC092032,2 | 0,335344214 | 1,261678398 | 0,66169164 | 0,959603271 | ring finger protein 216 pseudogene 1 |
| ENSG00000109971 | HSPA8 | 0,335288944 | 1,261630064 | 0,1917373 | 0,687672451 | heat shock protein family A (Hsp70) member 8 |
| ENSG00000197021 | EOLA2 | 0,335259817 | 1,261604592 | 0,20937362 | 0,713678247 | endothelium and lymphocyte associated ASCH domain 2 |
| ENSG00000170075 | GPR37L1 | 0,335103439 | 1,261467851 | 0,56855534 | 0,9417884 | G protein-coupled receptor 37 like 1 |
| ENSG00000236682 | MAP3K2-DT | 0,33508987 | 1,261455986 | 0,50721508 | 0,921951873 | MAP3K2 divergent transcript |
| ENSG00000288275 | DNAJC18 | 0,335035483 | 1,261408433 | 0,37308302 | 0,861070989 | DnaJ heat shock protein family (Hsp40) member C18 |
| ENSG00000235173 | HGH1 | 0,334987214 | 1,26136623 | 0,42485894 | 0,891582081 | HGH1 homolog |
| ENSG00000230772 | VN1R108P | 0,334981444 | 1,261361185 | 0,69539206 | 0,967602563 | vomeronasal 1 receptor 108 pseudogene |
| ENSG00000010165 | EEF1AKNMT | 0,334753248 | 1,261161687 | 0,2965506 | 0,806142623 | eEF1A lysine and N-terminal methyltransferase |
| ENSG00000184208 | C22orf46 | 0,334687042 | 1,261103813 | 0,5846137 | 0,944857905 | chromosome 22 open reading frame 46 |
| ENSG00000103507 | BCKDK | 0,334672597 | 1,261091186 | 0,21504985 | 0,722298056 | branched chain keto acid dehydrogenase kinase |
| ENSG00000089737 | DDX24 | 0,334646667 | 1,26106852 | 0,34702301 | 0,841678758 | DEAD-box helicase 24 |
| ENSG00000214359 | RPL18P10 | 0,33464334 | 1,261065612 | 0,77020367 | 0,977853702 | ribosomal protein L18 pseudogene 10 |
| ENSG00000272320 | AL445309,1 | 0,334616213 | 1,261041901 | 0,25992156 | 0,77245129 | novel transcript, antisense to SLC22A23 |
| ENSG00000179403 | VWA1 | 0,334589512 | 1,261018562 | 0,38833427 | 0,869820053 | von Willebrand factor A domain containing 1 |
| ENSG00000129255 | MPDU1 | 0,334565235 | 1,260997343 | 0,26328088 | 0,774637968 | mannose-P-dolichol utilization defect 1 |
| ENSG00000158887 | MPZ | 0,334551611 | 1,260985435 | 0,4328341 | 0,89560871 | myelin protein zero |
| ENSG00000116771 | AGMAT | 0,334516021 | 1,260954327 | 0,54276628 | 0,933663748 | agmatinase |
| ENSG00000173846 | PLK3 | 0,334508373 | 1,260947643 | 0,30643996 | 0,815375074 | polo like kinase 3 |
| ENSG00000153904 | DDAH1 | 0,334502987 | 1,260942935 | 0,32384806 | 0,82583898 | dimethylarginine dimethylaminohydrolase 1 |
| ENSG00000160323 | ADAMTS13 | 0,334271024 | 1,260740211 | 0,53491226 | 0,931774961 | ADAM metallopeptidase with thrombospondin type 1 motif 13 |
| ENSG00000278513 | AC091046,2 | 0,334238979 | 1,260712208 | 0,67233253 | 0,962290553 | zinc finger protein 551 (ZNF551) pseudogene |

| | | | | | | |
|---|---|---|---|---|---|---|
| ENSG00000145703 | IQGAP2 | 0,334194677 | 1,260673495 | 0,36536141 | 0,855323027 | IQ motif containing GTPase activating protein 2 |
| ENSG00000153815 | CMIP | 0,334152935 | 1,26063702 | 0,13854744 | 0,593533053 | c-Maf inducing protein |
| ENSG00000244198 | ARHGEF35-AS1 | 0,334068708 | 1,260563424 | 0,24190578 | 0,752203493 | ARHGEF35 antisense RNA 1 |
| ENSG00000280613 | NDUFA6-DT | 0,33393264 | 1,260444539 | 0,67468816 | 0,963215238 | NDUFA6 divergent transcript |
| ENSG00000224401 | RPL7P57 | 0,333916253 | 1,260430223 | 0,62669968 | 0,952793934 | ribosomal protein L7 pseudogene 57 |
| ENSG00000173801 | JUP | 0,33381273 | 1,260339782 | 0,26823806 | 0,780692104 | junction plakoglobin |
| ENSG00000186318 | BACE1 | 0,333776902 | 1,260308483 | 0,11801792 | 0,549146283 | beta-secretase 1 |
| ENSG00000276262 | NSF | 0,333699094 | 1,260240513 | 0,27366767 | 0,785719894 | N-ethylmaleimide sensitive factor, vesicle fusing ATPase |
| ENSG00000073969 | NSF | 0,333699094 | 1,260240513 | 0,27366767 | 0,785719894 | N-ethylmaleimide sensitive factor, vesicle fusing ATPase |
| ENSG00000100239 | PPP6R2 | 0,333620331 | 1,260171713 | 0,32965605 | 0,831426808 | protein phosphatase 6 regulatory subunit 2 |
| ENSG00000185361 | TNFAIP8L1 | 0,333609144 | 1,260161941 | 0,42953224 | 0,894365375 | TNF alpha induced protein 8 like 1 |
| ENSG00000011478 | QPCTL | 0,333558919 | 1,260118072 | 0,39018674 | 0,870057277 | glutaminyl-peptide cyclotransferase like |
| ENSG00000247595 | SPTY2D1OS | 0,333546997 | 1,260107659 | 0,44997488 | 0,902062594 | SPTY2D1 opposite strand |
| ENSG00000183207 | RUVBL2 | 0,333427185 | 1,260003014 | 0,23613285 | 0,745253314 | RuvB like AAA ATPase 2 |
| ENSG00000277222 | BTBD7 | 0,333194742 | 1,259800022 | 0,39907574 | 0,875433675 | BTB domain containing 7 |
| ENSG00000143384 | MCL1 | 0,33315456 | 1,259764935 | 0,11266587 | 0,536050184 | MCL1 apoptosis regulator, BCL2 family member |
| ENSG00000172803 | SNX32 | 0,333152265 | 1,259762931 | 0,26822258 | 0,780692104 | sorting nexin 32 |
| ENSG00000119866 | BCL11A | 0,333089156 | 1,259707825 | 0,30211882 | 0,811603618 | BAF chromatin remodeling complex subunit BCL11A |
| ENSG00000116584 | ARHGEF2 | 0,333053454 | 1,259676652 | 0,27079028 | 0,783073289 | Rho/Rac guanine nucleotide exchange factor 2 |
| ENSG00000229619 | MBNL1-AS1 | 0,333004796 | 1,259634167 | 0,29233821 | 0,802771257 | MBNL1 antisense RNA 1 |
| ENSG00000072858 | SIDT1 | 0,332941901 | 1,259579255 | 0,52390676 | 0,928328101 | SID1 transmembrane family member 1 |
| ENSG00000179833 | SERTAD2 | 0,332903782 | 1,259545974 | 0,31856951 | 0,823392614 | SERTA domain containing 2 |
| ENSG00000231940 | RPS7P3 | 0,332785344 | 1,259442576 | 0,71407405 | 0,970867497 | ribosomal protein S7 pseudogene 3 |
| ENSG00000064102 | INTS13 | 0,332746945 | 1,259409055 | 0,33603446 | 0,835660292 | integrator complex subunit 13 |
| ENSG00000276934 | AC009704,2 | 0,332545815 | 1,259233489 | 0,71287082 | 0,970812834 | novel transcript, sense intronic to CNDP2 |
| ENSG00000228474 | OST4 | 0,332499448 | 1,25919302 | 0,33538444 | 0,835232738 | oligosaccharyltransferase complex subunit 4, non-catalytic |
| ENSG00000171532 | NEUROD2 | 0,332497939 | 1,259191703 | 0,75735169 | 0,975917168 | neuronal differentiation 2 |
| ENSG00000158050 | DUSP2 | 0,332448669 | 1,2591487 | 0,63905947 | 0,956018518 | dual specificity phosphatase 2 |
| ENSG00000183111 | ARHGEF37 | 0,332415375 | 1,259119642 | 0,32869715 | 0,830659639 | Rho guanine nucleotide exchange factor 37 |
| ENSG00000228262 | LINC01320 | 0,332413936 | 1,259118386 | 0,61473515 | 0,952793934 | long intergenic non-protein coding RNA 1320 |

| | | | | | | |
|---|---|---|---|---|---|---|
| ENSG00000167658 | EEF2 | 0,332387407 | 1,259095233 | 0,20662865 | 0,709894291 | eukaryotic translation elongation factor 2 |
| ENSG00000197291 | RAMP2-AS1 | 0,332342578 | 1,25905611 | 0,60500697 | 0,950927937 | RAMP2 antisense RNA 1 |
| ENSG00000265656 | AC106037,3 | 0,332296288 | 1,259015712 | 0,73311155 | 0,974353474 | novel transcript, antisense to ABHD3 |
| ENSG00000251138 | LINC02882 | 0,332287655 | 1,259008178 | 0,49759498 | 0,918759905 | long intergenic non-protein coding RNA 2882 |
| ENSG00000236729 | HCG17 | 0,332211389 | 1,258941625 | 0,77534064 | 0,978587255 | HLA complex group 17 |
| ENSG00000197265 | GTF2E2 | 0,332143273 | 1,258882186 | 0,16572138 | 0,645029358 | general transcription factor IIE subunit 2 |
| ENSG00000119801 | YPEL5 | 0,332099029 | 1,25884358 | 0,33988499 | 0,838611198 | yippee like 5 |
| ENSG00000170906 | NDUFA3 | 0,332024671 | 1,258778699 | 0,37540442 | 0,862022464 | NADH:ubiquinone oxidoreductase subunit A3 |
| ENSG00000170832 | USP32 | 0,332023623 | 1,258777785 | 0,31044864 | 0,817751312 | ubiquitin specific peptidase 32 |
| ENSG00000196914 | ARHGEF12 | 0,332006084 | 1,258762482 | 0,20573174 | 0,708191543 | Rho guanine nucleotide exchange factor 12 |
| ENSG00000277140 | MARF1 | 0,331893112 | 1,258663917 | 0,37683988 | 0,862818448 | meiosis regulator and mRNA stability factor 1 |
| ENSG00000270647 | TAF15 | 0,331862888 | 1,258637548 | 0,34187644 | 0,840048641 | TATA-box binding protein associated factor 15 |
| ENSG00000286209 | AC005252,4 | 0,331859248 | 1,258634373 | 0,66055145 | 0,959226315 | novel transcript |
| ENSG00000254866 | DEFB109D | 0,331828511 | 1,258607557 | 0,72584981 | 0,973344786 | defensin beta 109D (pseudogene) |
| ENSG00000116141 | MARK1 | 0,331771877 | 1,258558151 | 0,36986781 | 0,858104601 | microtubule affinity regulating kinase 1 |
| ENSG00000243989 | ACY1 | 0,331591949 | 1,258401198 | 0,32276703 | 0,824955371 | aminoacylase 1 |
| ENSG00000121931 | LRIF1 | 0,331502213 | 1,258322927 | 0,46708897 | 0,907940653 | ligand dependent nuclear receptor interacting factor 1 |
| ENSG00000153767 | GTF2E1 | 0,33133285 | 1,258175217 | 0,10479997 | 0,51630419 | general transcription factor IIE subunit 1 |
| ENSG00000179933 | C14orf119 | 0,331295341 | 1,258142506 | 0,36931543 | 0,857991124 | chromosome 14 open reading frame 119 |
| ENSG00000226107 | AC004383,1 | 0,331055 | 1,257932928 | 0,81977729 | 0,98249617 | ribosomal protein L36a (RPL36A) pseudogene |
| ENSG00000141644 | MBD1 | 0,330958683 | 1,257848948 | 0,19875765 | 0,697657684 | methyl-CpG binding domain protein 1 |
| ENSG00000286228 | SPDYE11 | 0,330946991 | 1,257838754 | 0,78672601 | 0,979320406 | speedy/RINGO cell cycle regulator family member E11 |
| ENSG00000273520 | SPDYE8 | 0,330946991 | 1,257838754 | 0,78672601 | 0,979320406 | speedy/RINGO cell cycle regulator family member E8 |
| ENSG00000101152 | DNAJC5 | 0,330811699 | 1,257720803 | 0,20044068 | 0,700064356 | DnaJ heat shock protein family (Hsp40) member C5 |
| ENSG00000161956 | SENP3 | 0,330674191 | 1,257600931 | 0,20010807 | 0,699646707 | SUMO specific peptidase 3 |
| ENSG00000119682 | AREL1 | 0,330558538 | 1,25750012 | 0,16448993 | 0,643636727 | apoptosis resistant E3 ubiquitin protein ligase 1 |
| ENSG00000248724 | NPHP3-AS1 | 0,330497114 | 1,257446582 | 0,64453743 | 0,956633898 | NPHP3 antisense RNA 1 |
| ENSG00000243156 | MICAL3 | 0,330476152 | 1,257428312 | 0,41057918 | 0,882829958 | microtubule associated monooxygenase, calponin and LIM domain containing 3 |
| ENSG00000198189 | HSD17B11 | 0,330428958 | 1,257387179 | 0,43892298 | 0,897402199 | hydroxysteroid 17-beta dehydrogenase 11 |

| | | | | | | |
|---|---|---|---|---|---|---|
| ENSG00000186973 | FAM183A | 0,330352897 | 1,25732089 | 0,59240608 | 0,947416795 | family with sequence similarity 183 member A |
| ENSG00000167772 | ANGPTL4 | 0,330311763 | 1,257285041 | 0,78017404 | 0,978587255 | angiopoietin like 4 |
| ENSG00000272079 | AC004233,2 | 0,330259053 | 1,257239106 | 0,72473115 | 0,97326334 | novel transcript |
| ENSG00000214211 | CTAGE14P | 0,330178351 | 1,25716878 | 0,49154014 | 0,916995341 | CTAGE family member 14, pseudogene |
| ENSG00000203395 | AC015969,1 | 0,330138527 | 1,257134078 | 0,80848183 | 0,981213877 | novel transcript, antisense to PLEK |
| ENSG00000198331 | HYLS1 | 0,330138023 | 1,257133639 | 0,44933337 | 0,902062594 | HYLS1 centriolar and ciliogenesis associated |
| ENSG00000139505 | MTMR6 | 0,330111373 | 1,257110417 | 0,41306759 | 0,884663685 | myotubularin related protein 6 |
| ENSG00000089558 | KCNH4 | 0,330082572 | 1,257085321 | 0,65573615 | 0,957444211 | potassium voltage-gated channel subfamily H member 4 |
| ENSG00000170445 | HARS1 | 0,329887553 | 1,256915403 | 0,20399869 | 0,704808813 | histidyl-tRNA synthetase 1 |
| ENSG00000253356 | AC084024,3 | 0,329883477 | 1,256911852 | 0,79978331 | 0,980161849 | novel transcript |
| ENSG00000132581 | SDF2 | 0,329823248 | 1,256859381 | 0,13820536 | 0,592953528 | stromal cell derived factor 2 |
| ENSG00000271947 | AC017076,1 | 0,3298104 | 1,256848187 | 0,76070539 | 0,97624106 | novel transcript |
| ENSG00000267151 | MIR2117HG | 0,329723157 | 1,256772185 | 0,79296784 | 0,979565591 | MIR2117 host gene |
| ENSG00000235257 | ITGA9-AS1 | 0,329717703 | 1,256767435 | 0,38261164 | 0,867188687 | ITGA9 antisense RNA 1 |
| ENSG00000153214 | TMEM87B | 0,329708177 | 1,256759136 | 0,33628482 | 0,835660292 | transmembrane protein 87B |
| ENSG00000162341 | TPCN2 | 0,329520024 | 1,256595244 | 0,36077273 | 0,851517344 | two pore segment channel 2 |
| ENSG00000259182 | AC019254,1 | 0,329443039 | 1,25652819 | 0,77567704 | 0,978587255 | novel transcript |
| ENSG00000169047 | IRS1 | 0,32942799 | 1,256515084 | 0,2435642 | 0,754313147 | insulin receptor substrate 1 |
| ENSG00000169230 | PRELID1 | 0,329384487 | 1,256477196 | 0,17597084 | 0,661519708 | PRELI domain containing 1 |
| ENSG00000182318 | ZSCAN22 | 0,32933997 | 1,256438425 | 0,43229989 | 0,89558934 | zinc finger and SCAN domain containing 22 |
| ENSG00000254531 | FLJ20021 | 0,329225356 | 1,256338612 | 0,49273678 | 0,917090312 | uncharacterized LOC90024 |
| ENSG00000166548 | TK2 | 0,329055262 | 1,256190498 | 0,21000032 | 0,715112736 | thymidine kinase 2 |
| ENSG00000197977 | ELOVL2 | 0,329024395 | 1,256163622 | 0,48573944 | 0,913997663 | ELOVL fatty acid elongase 2 |
| ENSG00000143751 | SDE2 | 0,329009549 | 1,256150696 | 0,13451381 | 0,584982573 | SDE2 telomere maintenance homolog |
| ENSG00000238009 | AL627309,1 | 0,328988492 | 1,256132361 | 0,43893346 | 0,897402199 | novel transcript |
| ENSG00000167566 | NCKAP5L | 0,328873471 | 1,256032218 | 0,31734096 | 0,822252078 | NCK associated protein 5 like |
| ENSG00000277753 | MUC20 | 0,328728558 | 1,255906061 | 0,73619453 | 0,974353474 | mucin 20, cell surface associated |
| ENSG00000160282 | FTCD | 0,328697828 | 1,25587931 | 0,83440969 | 0,984905093 | formimidoyltransferase cyclodeaminase |
| ENSG00000162368 | CMPK1 | 0,328338019 | 1,255566132 | 0,41551927 | 0,885986046 | cytidine/uridine monophosphate kinase 1 |
| ENSG00000067208 | EVI5 | 0,328308419 | 1,255540372 | 0,23578981 | 0,744966003 | ecotropic viral integration site 5 |
| ENSG00000084623 | EIF3I | 0,328201024 | 1,255446912 | 0,13057875 | 0,577136255 | eukaryotic translation initiation factor 3 subunit I |
| ENSG00000241790 | ENO1P4 | 0,328141398 | 1,255395026 | 0,63914893 | 0,956018518 | enolase 1 pseudogene 4 |

| | | | | | | |
|---|---|---|---|---|---|---|
| ENSG00000142546 | NOSIP | 0,32805745 | 1,255321979 | 0,33149289 | 0,832548086 | nitric oxide synthase interacting protein |
| ENSG00000108861 | DUSP3 | 0,328011236 | 1,255281768 | 0,12670387 | 0,568657756 | dual specificity phosphatase 3 |
| ENSG00000182247 | UBE2E2 | 0,327976945 | 1,255251932 | 0,35729651 | 0,848693232 | ubiquitin conjugating enzyme E2 E2 |
| ENSG00000181626 | ANKRD62 | 0,327932059 | 1,255212878 | 0,72693072 | 0,973344786 | ankyrin repeat domain 62 |
| ENSG00000164304 | CAGE1 | 0,327890079 | 1,255176354 | 0,73565262 | 0,974353474 | cancer antigen 1 |
| ENSG00000173011 | TADA2B | 0,327873718 | 1,25516212 | 0,34551866 | 0,841168721 | transcriptional adaptor 2B |
| ENSG00000234367 | PFN1P3 | 0,327733699 | 1,255040307 | 0,75514513 | 0,975901808 | profilin 1 pseudogene 3 |
| ENSG00000224167 | LINC01357 | 0,327648878 | 1,254966521 | 0,62313706 | 0,952793934 | long intergenic non-protein coding RNA 1357 |
| ENSG00000270757 | HSPE1-MOB4 | 0,327483696 | 1,254822841 | 0,37029594 | 0,858545694 | HSPE1-MOB4 readthrough |
| ENSG00000280538 | AC208876,1 | 0,327377418 | 1,254730407 | 0,82032916 | 0,982609114 | novel transcript |
| ENSG00000101417 | PXMP4 | 0,327360926 | 1,254716064 | 0,2860171 | 0,797420923 | peroxisomal membrane protein 4 |
| ENSG00000177757 | FAM87B | 0,327114464 | 1,254501733 | 0,5528116 | 0,936894267 | family with sequence similarity 87 member B |
| ENSG00000183571 | PGPEP1L | 0,327067453 | 1,254460856 | 0,85202726 | 0,98732978 | pyroglutamyl-peptidase I like |
| ENSG00000119185 | ITGB1BP1 | 0,327055655 | 1,254450597 | 0,21685094 | 0,724474005 | integrin subunit beta 1 binding protein 1 |
| ENSG00000161243 | FBXO27 | 0,327019288 | 1,254418975 | 0,61951472 | 0,952793934 | F-box protein 27 |
| ENSG00000141968 | VAV1 | 0,326896061 | 1,254311834 | 0,74344725 | 0,975310206 | vav guanine nucleotide exchange factor 1 |
| ENSG00000274378 | AC005702,3 | 0,326862185 | 1,254282382 | 0,73335152 | 0,974353474 | double C2-like domains, beta (DOC2B) pseudogene |
| ENSG00000204843 | DCTN1 | 0,326852033 | 1,254273556 | 0,39569834 | 0,872227623 | dynactin subunit 1 |
| ENSG00000178927 | CYBC1 | 0,326801318 | 1,254229465 | 0,34612962 | 0,841168721 | cytochrome b-245 chaperone 1 |
| ENSG00000162639 | HENMT1 | 0,326762435 | 1,254195662 | 0,33465794 | 0,834757602 | HEN methyltransferase 1 |
| ENSG00000159210 | SNF8 | 0,326751272 | 1,254185957 | 0,21912198 | 0,727412416 | SNF8 subunit of ESCRT-II |
| ENSG00000244731 | C4A | 0,326746334 | 1,254181665 | 0,5288504 | 0,930077007 | complement C4A (Rodgers blood group) |
| ENSG00000100865 | CINP | 0,326726313 | 1,25416426 | 0,1322055 | 0,580737262 | cyclin dependent kinase 2 interacting protein |
| ENSG00000172500 | FIBP | 0,326517026 | 1,253982336 | 0,15907681 | 0,633144409 | FGF1 intracellular binding protein |
| ENSG00000196449 | YRDC | 0,326450031 | 1,253924106 | 0,29230174 | 0,802771257 | yrdC N6-threonylcarbamoyltransferase domain containing |
| ENSG00000160058 | BSDC1 | 0,326358349 | 1,253844422 | 0,18888491 | 0,683246297 | BSD domain containing 1 |
| ENSG00000264271 | RN7SL488P | 0,326267486 | 1,253765456 | 0,76482496 | 0,976722333 | RNA, 7SL, cytoplasmic 488, pseudogene |
| ENSG00000238273 | AC108058,1 | 0,326118828 | 1,253636272 | 0,56492561 | 0,940865625 | novel transcript, antisense to FHL2 |
| ENSG00000145832 | SLC25A48 | 0,32610359 | 1,253623031 | 0,72708651 | 0,973344786 | solute carrier family 25 member 48 |
| ENSG00000266306 | AC061975,5 | 0,326082271 | 1,253604506 | 0,75415597 | 0,975901808 | leucine rich repeat containing 37, member A3 (LRRC37A3) pseudogene |
| ENSG00000257475 | AC068888,2 | 0,325936208 | 1,253477594 | 0,75200559 | 0,975901808 | novel transcript |

| | | | | | | |
|---|---|---|---|---|---|---|
| ENSG00000105618 | PRPF31 | 0,325894446 | 1,25344131 | 0,32652475 | 0,828450693 | pre-mRNA processing factor 31 |
| ENSG00000100401 | RANGAP1 | 0,325847657 | 1,253400659 | 0,44477767 | 0,900383899 | Ran GTPase activating protein 1 |
| ENSG00000276657 | PYCR3 | 0,325792701 | 1,253352915 | 0,57067713 | 0,942407031 | pyrroline-5-carboxylate reductase 3 |
| ENSG00000260450 | LINC02134 | 0,325762089 | 1,253326321 | 0,81235185 | 0,981213877 | long intergenic non-protein coding RNA 2134 |
| ENSG00000184056 | VPS33B | 0,325719791 | 1,253289575 | 0,37332629 | 0,861257555 | VPS33B late endosome and lysosome associated |
| ENSG00000076604 | TRAF4 | 0,325716321 | 1,253286561 | 0,27275737 | 0,78561201 | TNF receptor associated factor 4 |
| ENSG00000218891 | ZNF579 | 0,325574808 | 1,253163633 | 0,47620088 | 0,911413793 | zinc finger protein 579 |
| ENSG00000214832 | UPF3AP2 | 0,325556187 | 1,253147458 | 0,76448583 | 0,976640468 | UPF3A pseudogene 2 |
| ENSG00000095587 | TLL2 | 0,325551021 | 1,253142971 | 0,38470792 | 0,868536368 | tolloid like 2 |
| ENSG00000272333 | KMT2B | 0,32551184 | 1,253108938 | 0,44925473 | 0,902062594 | lysine methyltransferase 2B |
| ENSG00000213700 | RPL17P50 | 0,325499037 | 1,253097818 | 0,66789608 | 0,961113267 | ribosomal protein L17 pseudogene 50 |
| ENSG00000124535 | WRNIP1 | 0,3254513 | 1,253056355 | 0,22734293 | 0,735964811 | WRN helicase interacting protein 1 |
| ENSG00000229110 | AC006355,1 | 0,325432723 | 1,25304022 | 0,79543149 | 0,979879951 | heterogeneous nuclear ribonucleoprotein A1-like 3 (HNRPA1L3) pseudogene |
| ENSG00000125388 | GRK4 | 0,325309589 | 1,252933278 | 0,30353539 | 0,812685668 | G protein-coupled receptor kinase 4 |
| ENSG00000179295 | PTPN11 | 0,325185576 | 1,252825581 | 0,19298836 | 0,689849199 | protein tyrosine phosphatase non-receptor type 11 |
| ENSG00000179950 | PUF60 | 0,325106371 | 1,252756802 | 0,52493101 | 0,928616707 | poly(U) binding splicing factor 60 |
| ENSG00000096080 | MRPS18A | 0,324959334 | 1,252629129 | 0,13442421 | 0,584751412 | mitochondrial ribosomal protein S18A |
| ENSG00000224470 | ATXN1L | 0,324685327 | 1,252391244 | 0,13407469 | 0,584238558 | ataxin 1 like |
| ENSG00000163472 | TMEM79 | 0,324588423 | 1,252307125 | 0,3502167 | 0,843789477 | transmembrane protein 79 |
| ENSG00000160588 | MPZL3 | 0,324569274 | 1,252290503 | 0,38969965 | 0,870057277 | myelin protein zero like 3 |
| ENSG00000226310 | AL022157,1 | 0,324524844 | 1,252251938 | 0,5984886 | 0,948793037 | novel transcript |
| ENSG00000241852 | C8orf58 | 0,324503471 | 1,252233386 | 0,28080972 | 0,791510309 | chromosome 8 open reading frame 58 |
| ENSG00000134278 | SPIRE1 | 0,324452489 | 1,252189136 | 0,27412149 | 0,7862019 | spire type actin nucleation factor 1 |
| ENSG00000280418 | AC007326,2 | 0,324432309 | 1,25217162 | 0,78628284 | 0,979320406 | novel transcript |
| ENSG00000215529 | EFCAB8 | 0,32441936 | 1,252160381 | 0,80783814 | 0,981213877 | EF-hand calcium binding domain 8 |
| ENSG00000230604 | TSEN15P2 | 0,324386774 | 1,2521321 | 0,77887728 | 0,978587255 | tRNA splicing endonuclease subunit 15 pseudogene 2 |
| ENSG00000226257 | STK19 | 0,324366878 | 1,252114832 | 0,43534757 | 0,896325752 | serine/threonine kinase 19 |
| ENSG00000159352 | PSMD4 | 0,324266132 | 1,252027397 | 0,17814954 | 0,665432182 | proteasome 26S subunit, non-ATPase 4 |
| ENSG00000267508 | ZNF285 | 0,324157804 | 1,25193339 | 0,38448505 | 0,868434303 | zinc finger protein 285 |
| ENSG00000170027 | YWHAG | 0,324028275 | 1,251820992 | 0,12376739 | 0,562503814 | tyrosine 3-monooxygenase/tryptophan 5-monooxygenase activation protein gamma |

| | | | | | | |
|---|---|---|---|---|---|---|
| ENSG00000284708 | AL031731,1 | 0,323903569 | 1,251712791 | 0,77154407 | 0,978033855 | novel transcript |
| ENSG00000169598 | DFFB | 0,323844925 | 1,251661911 | 0,31736572 | 0,822252078 | DNA fragmentation factor subunit beta |
| ENSG00000277369 | AC010654,1 | 0,323834529 | 1,251652891 | 0,45654245 | 0,903545728 | novel transcript |
| ENSG00000228421 | AC005013,1 | 0,323739166 | 1,251570159 | 0,81777462 | 0,982323352 | novel transcript |
| ENSG00000244045 | TMEM199 | 0,323686321 | 1,251524316 | 0,18415446 | 0,675910428 | transmembrane protein 199 |
| ENSG00000136848 | DAB2IP | 0,323616425 | 1,251463683 | 0,32990012 | 0,831639839 | DAB2 interacting protein |
| ENSG00000050393 | MCUR1 | 0,323553415 | 1,251409026 | 0,23489496 | 0,744576287 | mitochondrial calcium uniporter regulator 1 |
| ENSG00000090924 | PLEKHG2 | 0,323506269 | 1,251368132 | 0,23934903 | 0,749122637 | pleckstrin homology and RhoGEF domain containing G2 |
| ENSG00000025800 | KPNA6 | 0,323404505 | 1,251279867 | 0,12758026 | 0,570380291 | karyopherin subunit alpha 6 |
| ENSG00000265334 | AC130324,2 | 0,323374518 | 1,251253859 | 0,64274501 | 0,956633898 | novel transcript, antisense ATAD5 |
| ENSG00000221968 | FADS3 | 0,323339584 | 1,25122356 | 0,57218465 | 0,942475308 | fatty acid desaturase 3 |
| ENSG00000244306 | DUXAP10 | 0,323305934 | 1,251194377 | 0,81355243 | 0,981436305 | double homeobox A pseudogene 10 |
| ENSG00000184304 | PRKD1 | 0,323291732 | 1,25118206 | 0,32376217 | 0,825794844 | protein kinase D1 |
| ENSG00000162627 | SNX7 | 0,32321491 | 1,251115438 | 0,42617636 | 0,892133432 | sorting nexin 7 |
| ENSG00000239617 | AC073610,1 | 0,323172865 | 1,251078976 | 0,73182444 | 0,974353474 | ribosomal protein L32 (RPL32) pseudogene |
| ENSG00000261466 | AC136944,4 | 0,323142094 | 1,251052293 | 0,66510211 | 0,960681381 | novel transcript |
| ENSG00000179546 | HTR1D | 0,323115474 | 1,251029209 | 0,57764042 | 0,94337651 | 5-hydroxytryptamine receptor 1D |
| ENSG00000163945 | UVSSA | 0,323085496 | 1,251003214 | 0,33020325 | 0,831800372 | UV stimulated scaffold protein A |
| ENSG00000225479 | PLCB1-IT1 | 0,322838935 | 1,250789432 | 0,79949432 | 0,980161849 | PLCB1 intronic transcript 1 |
| ENSG00000254112 | AP003467,2 | 0,322704149 | 1,25067258 | 0,82693219 | 0,983548538 | ATP synthase, H+ transporting, mitochondrial F0 complex, subunit F6 (ATP5J) pseudogene |
| ENSG00000174306 | ZHX3 | 0,322663714 | 1,250637528 | 0,19715419 | 0,695927373 | zinc fingers and homeoboxes 3 |
| ENSG00000115310 | RTN4 | 0,322653369 | 1,25062856 | 0,17306613 | 0,657760693 | reticulon 4 |
| ENSG00000287637 | AP001437,2 | 0,322637821 | 1,250615082 | 0,83413002 | 0,984905093 | novel transcript, antisense to DYRK1A |
| ENSG00000101306 | MYLK2 | 0,322619756 | 1,250599422 | 0,58160456 | 0,944139372 | myosin light chain kinase 2 |
| ENSG00000132694 | ARHGEF11 | 0,322586804 | 1,250570858 | 0,16256808 | 0,639794628 | Rho guanine nucleotide exchange factor 11 |
| ENSG00000277954 | AC092376,2 | 0,322565638 | 1,250552511 | 0,48037039 | 0,912847708 | novel transcript, antisense to WWOX |
| ENSG00000066294 | CD84 | 0,322535847 | 1,250526688 | 0,70300135 | 0,968742626 | CD84 molecule |
| ENSG00000136754 | ABI1 | 0,322525386 | 1,25051762 | 0,37999202 | 0,865511891 | abl interactor 1 |
| ENSG00000259571 | BLID | 0,3224553 | 1,250456871 | 0,54835982 | 0,935599286 | BH3-like motif containing, cell death inducer |
| ENSG00000015475 | BID | 0,322442574 | 1,250445842 | 0,21761976 | 0,72540192 | BH3 interacting domain death agonist |
| ENSG00000124333 | VAMP7 | 0,322428099 | 1,250433296 | 0,50446917 | 0,921400408 | vesicle associated membrane protein 7 |

| | | | | | | |
|---|---|---|---|---|---|---|
| ENSG00000070614 | NDST1 | 0,322415097 | 1,250422026 | 0,21313448 | 0,720449383 | N-deacetylase and N-sulfotransferase 1 |
| ENSG00000266127 | ZNF415P1 | 0,322390931 | 1,250401081 | 0,73875444 | 0,974367986 | zinc finger protein 415 pseudogene 1 |
| ENSG00000131771 | PPP1R1B | 0,322321777 | 1,250341146 | 0,67267034 | 0,962355986 | protein phosphatase 1 regulatory inhibitor subunit 1B |
| ENSG00000130724 | CHMP2A | 0,322284745 | 1,250309052 | 0,17670575 | 0,662651108 | charged multivesicular body protein 2A |
| ENSG00000102878 | HSF4 | 0,322279963 | 1,250304908 | 0,76608912 | 0,977102081 | heat shock transcription factor 4 |
| ENSG00000012232 | EXTL3 | 0,322146209 | 1,250188995 | 0,29008287 | 0,800682868 | exostosin like glycosyltransferase 3 |
| ENSG00000112378 | PERP | 0,322130412 | 1,250175307 | 0,36591142 | 0,855700097 | p53 apoptosis effector related to PMP22 |
| ENSG00000174792 | ODAPH | 0,322092031 | 1,250142048 | 0,74168827 | 0,975176043 | odontogenesis associated phosphoprotein |
| ENSG00000070444 | MNT | 0,322071364 | 1,250124139 | 0,38996562 | 0,870057277 | MAX network transcriptional repressor |
| ENSG00000213772 | EIF1P7 | 0,322025308 | 1,250084232 | 0,82806299 | 0,983595244 | eukaryotic translation initiation factor 1 pseudogene 7 |
| ENSG00000105479 | CCDC114 | 0,321981963 | 1,250046674 | 0,62900357 | 0,952793934 | coiled-coil domain containing 114 |
| ENSG00000064989 | CALCRL | 0,321927597 | 1,249999568 | 0,53118754 | 0,931265667 | calcitonin receptor like receptor |
| ENSG00000089248 | ERP29 | 0,321922028 | 1,249994744 | 0,14818253 | 0,612760682 | endoplasmic reticulum protein 29 |
| ENSG00000183813 | CCR4 | 0,321902619 | 1,249977927 | 0,65572937 | 0,957444211 | C-C motif chemokine receptor 4 |
| ENSG00000205625 | AL591499,1 | 0,321839151 | 1,249922938 | 0,80573586 | 0,981213877 | stathmin-like 2 (STMN2) pseudogene |
| ENSG00000100353 | EIF3D | 0,321726916 | 1,249825704 | 0,15777829 | 0,630548432 | eukaryotic translation initiation factor 3 subunit D |
| ENSG00000087008 | ACOX3 | 0,321508227 | 1,249636265 | 0,29726073 | 0,806635068 | acyl-CoA oxidase 3, pristanoyl |
| ENSG00000160213 | CSTB | 0,321439347 | 1,249576604 | 0,21110948 | 0,716859662 | cystatin B |
| ENSG00000087365 | SF3B2 | 0,321433443 | 1,24957149 | 0,25858102 | 0,771444876 | splicing factor 3b subunit 2 |
| ENSG00000111452 | ADGRD1 | 0,321386973 | 1,249531242 | 0,39283691 | 0,871101009 | adhesion G protein-coupled receptor D1 |
| ENSG00000272970 | AC107294,2 | 0,321315503 | 1,249469342 | 0,36534942 | 0,855323027 | novel transcript |
| ENSG00000177173 | NAP1L4P1 | 0,32124202 | 1,249405703 | 0,48713287 | 0,914593485 | nucleosome assembly protein 1 like 4 pseudogene 1 |
| ENSG00000005007 | UPF1 | 0,320890522 | 1,249101335 | 0,36511181 | 0,855173479 | UPF1 RNA helicase and ATPase |
| ENSG00000100612 | DHRS7 | 0,320881248 | 1,249093305 | 0,1818453 | 0,672010829 | dehydrogenase/reductase 7 |
| ENSG00000133398 | MED10 | 0,320870631 | 1,249084113 | 0,14225679 | 0,600366679 | mediator complex subunit 10 |
| ENSG00000162398 | LEXM | 0,320853447 | 1,249069235 | 0,68699662 | 0,965769168 | lymphocyte expansion molecule |
| ENSG00000266086 | AC015813,2 | 0,32077969 | 1,249005379 | 0,4551196 | 0,903545728 | novel transcript |
| ENSG00000101350 | KIF3B | 0,320617637 | 1,248865091 | 0,12006694 | 0,553128623 | kinesin family member 3B |
| ENSG00000223427 | RBM7P1 | 0,320397241 | 1,24867432 | 0,70003168 | 0,968522643 | RBM7 pseudogene 1 |
| ENSG00000136270 | TBRG4 | 0,320126311 | 1,248439848 | 0,26910536 | 0,781347689 | transforming growth factor beta regulator 4 |
| ENSG00000103047 | TANGO6 | 0,32009834 | 1,248415643 | 0,1813393 | 0,671070342 | transport and golgi organization 6 homolog |

| | | | | | | |
|---|---|---|---|---|---|---|
| ENSG00000236017 | ASMTL-AS1 | 0,320093276 | 1,248411261 | 0,67733088 | 0,963580269 | ASMTL antisense RNA 1 |
| ENSG00000165916 | PSMC3 | 0,320047788 | 1,248371899 | 0,15496915 | 0,624591393 | proteasome 26S subunit, ATPase 3 |
| ENSG00000173848 | NET1 | 0,320043037 | 1,248367789 | 0,28727575 | 0,798842742 | neuroepithelial cell transforming 1 |
| ENSG00000144560 | VGLL4 | 0,320026426 | 1,248353414 | 0,2340463 | 0,743586269 | vestigial like family member 4 |
| ENSG00000234592 | AL359845,1 | 0,319843802 | 1,248195401 | 0,70887834 | 0,970282889 | tight junction protein 4 (peripheral) (TJP4) pseudogene |
| ENSG00000160445 | ZER1 | 0,319798542 | 1,248156244 | 0,21830305 | 0,725947652 | zyg-11 related cell cycle regulator |
| ENSG00000168843 | FSTL5 | 0,319798139 | 1,248155896 | 0,56895685 | 0,942177697 | follistatin like 5 |
| ENSG00000047648 | ARHGAP6 | 0,319690027 | 1,248062365 | 0,24466935 | 0,756461896 | Rho GTPase activating protein 6 |
| ENSG00000139631 | CSAD | 0,319609718 | 1,247992893 | 0,43377394 | 0,896272827 | cysteine sulfinic acid decarboxylase |
| ENSG00000141994 | DUS3L | 0,319583396 | 1,247970123 | 0,36420652 | 0,854269087 | dihydrouridine synthase 3 like |
| ENSG00000251423 | AC016576,1 | 0,319519177 | 1,247914574 | 0,75268507 | 0,975901808 | novel transcript |
| ENSG00000146826 | MAP11 | 0,31948233 | 1,247882701 | 0,35255 | 0,844912682 | microtubule associated protein 11 |
| ENSG00000257122 | RRN3P3 | 0,319473212 | 1,247874815 | 0,32230267 | 0,824955371 | RRN3 pseudogene 3 |
| ENSG00000138760 | SCARB2 | 0,319402458 | 1,247813617 | 0,38473997 | 0,868536368 | scavenger receptor class B member 2 |
| ENSG00000148795 | CYP17A1 | 0,319399826 | 1,24781134 | 0,77801399 | 0,978587255 | cytochrome P450 family 17 subfamily A member 1 |
| ENSG00000125775 | SDCBP2 | 0,319304159 | 1,247728599 | 0,55323534 | 0,936992468 | syndecan binding protein 2 |
| ENSG00000109787 | KLF3 | 0,319239999 | 1,247673111 | 0,39900636 | 0,875433675 | Kruppel like factor 3 |
| ENSG00000010438 | PRSS3 | 0,319184502 | 1,247625117 | 0,82793385 | 0,983595244 | serine protease 3 |
| ENSG00000177311 | ZBTB38 | 0,319167035 | 1,247610012 | 0,30761818 | 0,816380056 | zinc finger and BTB domain containing 38 |
| ENSG00000259921 | AC022819,1 | 0,31901192 | 1,247475879 | 0,57298679 | 0,942475308 | novel transcript, sense intronic to TTC23 |
| ENSG00000107147 | KCNT1 | 0,318966289 | 1,247436423 | 0,80439615 | 0,980875302 | potassium sodium-activated channel subfamily T member 1 |
| ENSG00000177731 | FLII | 0,318769696 | 1,247266449 | 0,34232779 | 0,840048641 | FLII actin remodeling protein |
| ENSG00000284571 | FLII | 0,318769696 | 1,247266449 | 0,34232779 | 0,840048641 | FLII actin remodeling protein |
| ENSG00000173208 | ABCD2 | 0,318767152 | 1,24726425 | 0,55020218 | 0,936137195 | ATP binding cassette subfamily D member 2 |
| ENSG00000286905 | AC108488,2 | 0,318688226 | 1,247196017 | 0,55467381 | 0,937793851 | novel protein |
| ENSG00000118898 | PPL | 0,318673876 | 1,247183611 | 0,57216486 | 0,942475308 | periplakin |
| ENSG00000108788 | MLX | 0,318649792 | 1,247162791 | 0,1431737 | 0,601880115 | MAX dimerization protein MLX |
| ENSG00000285531 | Z83840,1 | 0,318636304 | 1,247151132 | 0,61148379 | 0,952155617 | LLP homolog, long-term synaptic facilitation ( LLPH) pseudogene |
| ENSG00000144749 | LRIG1 | 0,318551718 | 1,247078013 | 0,37426767 | 0,86138873 | leucine rich repeats and immunoglobulin like domains 1 |

| | | | | | | |
|---|---|---|---|---|---|---|
| ENSG00000163734 | CXCL3 | 0,318540731 | 1,247068516 | 0,62962891 | 0,952812183 | C-X-C motif chemokine ligand 3 |
| ENSG00000196388 | INCA1 | 0,318525187 | 1,24705508 | 0,59068631 | 0,947416795 | inhibitor of CDK, cyclin A1 interacting protein 1 |
| ENSG00000148634 | HERC4 | 0,31843824 | 1,246979925 | 0,23073564 | 0,739234644 | HECT and RLD domain containing E3 ubiquitin protein ligase 4 |
| ENSG00000213658 | LAT | 0,318437411 | 1,246979209 | 0,55450523 | 0,937793851 | linker for activation of T cells |
| ENSG00000069667 | RORA | 0,318415855 | 1,246960578 | 0,44564226 | 0,901173082 | RAR related orphan receptor A |
| ENSG00000040487 | SLC66A1 | 0,318284502 | 1,24684705 | 0,33158235 | 0,832548086 | solute carrier family 66 member 1 |
| ENSG00000223505 | RPL35AP5 | 0,318281528 | 1,24684448 | 0,79839703 | 0,980161849 | ribosomal protein L35a pseudogene 5 |
| ENSG00000196440 | ARMCX4 | 0,318234996 | 1,246804266 | 0,43510617 | 0,896272827 | armadillo repeat containing X-linked 4 |
| ENSG00000142687 | KIAA0319L | 0,318106022 | 1,246692809 | 0,12121665 | 0,555921553 | KIAA0319 like |
| ENSG00000272854 | AC004839,2 | 0,317981736 | 1,246585413 | 0,79252523 | 0,979471509 | novel transcript, antisense to BCAP29 |
| ENSG00000284582 | AC004839,4 | 0,317981736 | 1,246585413 | 0,79252523 | 0,979471509 | novel transcript, antisense to BCAP29 |
| ENSG00000218996 | ARL4AP5 | 0,317941701 | 1,24655082 | 0,67701238 | 0,963580269 | ADP ribosylation factor like GTPase 4A pseudogene 5 |
| ENSG00000276291 | FRG1HP | 0,317884021 | 1,246500983 | 0,21099706 | 0,716555746 | FSHD region gene 1 family member H, pseudogene |
| ENSG00000284400 | TBC1D9B | 0,317747247 | 1,246382815 | 0,52488256 | 0,928616707 | TBC1 domain family member 9B |
| ENSG00000276493 | Metazoa_SRP | 0,317734528 | 1,246371827 | 0,87148322 | 0,989217334 | Metazoan signal recognition particle RNA |
| ENSG00000134440 | NARS1 | 0,317681943 | 1,246326398 | 0,20864533 | 0,713284351 | asparaginyl-tRNA synthetase 1 |
| ENSG00000240694 | PNMA2 | 0,317655176 | 1,246303275 | 0,19635497 | 0,694046542 | PNMA family member 2 |
| ENSG00000143167 | GPA33 | 0,31764833 | 1,246297361 | 0,62556968 | 0,952793934 | glycoprotein A33 |
| ENSG00000228352 | AL354989,1 | 0,317615415 | 1,246268927 | 0,76268093 | 0,97639149 | novel transcript |
| ENSG00000172519 | OR10H5 | 0,317458719 | 1,246133573 | 0,63763069 | 0,956017934 | olfactory receptor family 10 subfamily H member 5 |
| ENSG00000065802 | ASB1 | 0,317380204 | 1,246065757 | 0,31661349 | 0,821646322 | ankyrin repeat and SOCS box containing 1 |
| ENSG00000132471 | WBP2 | 0,317308333 | 1,246003684 | 0,23007807 | 0,738593358 | WW domain binding protein 2 |
| ENSG00000068308 | OTUD5 | 0,317173007 | 1,245886812 | 0,2147237 | 0,722164151 | OTU deubiquitinase 5 |
| ENSG00000064961 | HMG20B | 0,317022986 | 1,245757263 | 0,30652492 | 0,815375074 | high mobility group 20B |
| ENSG00000166682 | TMPRSS5 | 0,316968084 | 1,245709857 | 0,46349366 | 0,907101775 | transmembrane serine protease 5 |
| ENSG00000196810 | CTBP1-DT | 0,316914793 | 1,245663843 | 0,15499984 | 0,624634583 | CTBP1 divergent transcript |
| ENSG00000242028 | HYPK | 0,316909069 | 1,245658901 | 0,25228601 | 0,76432863 | huntingtin interacting protein K |
| ENSG00000138646 | HERC5 | 0,316851545 | 1,245609235 | 0,50319869 | 0,920888747 | HECT and RLD domain containing E3 ubiquitin protein ligase 5 |
| ENSG00000170468 | RIOX1 | 0,316658201 | 1,245442315 | 0,16914452 | 0,651208888 | ribosomal oxygenase 1 |
| ENSG00000122545 | SEPTIN7 | 0,316572736 | 1,245368536 | 0,3460878 | 0,841168721 | septin 7 |
| ENSG00000127311 | HELB | 0,316535322 | 1,245336241 | 0,19869289 | 0,697657684 | DNA helicase B |

| | | | | | | |
|---|---|---|---|---|---|---|
| ENSG00000135100 | HNF1A | 0,316512464 | 1,24531651 | 0,73611166 | 0,974353474 | HNF1 homeobox A |
| ENSG00000213411 | RBM22P2 | 0,316458927 | 1,245270298 | 0,65286088 | 0,95732962 | RNA binding motif protein 22 pseudogene 2 |
| ENSG00000179222 | MAGED1 | 0,316089717 | 1,244951653 | 0,1407709 | 0,597646146 | MAGE family member D1 |
| ENSG00000226766 | FABP7P1 | 0,315732334 | 1,244643292 | 0,6808137 | 0,964654539 | fatty acid binding protein 7 pseudogene 1 |
| ENSG00000132254 | ARFIP2 | 0,315693285 | 1,244609605 | 0,1795217 | 0,668298509 | ADP ribosylation factor interacting protein 2 |
| ENSG00000105576 | TNPO2 | 0,315555109 | 1,244490406 | 0,23723408 | 0,746624292 | transportin 2 |
| ENSG00000226741 | LINC02554 | 0,315528218 | 1,244467211 | 0,62254707 | 0,952793934 | long intergenic non-protein coding RNA 2554 |
| ENSG00000144848 | ATG3 | 0,315475566 | 1,244421793 | 0,3481752 | 0,84258453 | autophagy related 3 |
| ENSG00000224153 | LINC02054 | 0,31544821 | 1,244398197 | 0,76902753 | 0,977432179 | long intergenic non-protein coding RNA 2054 |
| ENSG00000111181 | SLC6A12 | 0,315257383 | 1,24423361 | 0,8106348 | 0,981213877 | solute carrier family 6 member 12 |
| ENSG00000177192 | PUS1 | 0,315207131 | 1,244190271 | 0,58441965 | 0,944741266 | pseudouridine synthase 1 |
| ENSG00000277631 | PGM5P3-AS1 | 0,31517313 | 1,24416095 | 0,72282775 | 0,972707638 | PGM5P3 antisense RNA 1 |
| ENSG00000184911 | DMRTC1B | 0,315164121 | 1,24415318 | 0,54923398 | 0,936040289 | DMRT like family C1B |
| ENSG00000141376 | BCAS3 | 0,315118951 | 1,244114227 | 0,25704836 | 0,769358275 | BCAS3 microtubule associated cell migration factor |
| ENSG00000265939 | UBE2CP2 | 0,315077567 | 1,244078539 | 0,82386656 | 0,983374097 | ubiquitin conjugating enzyme E2 C pseudogene 2 |
| ENSG00000184640 | SEPTIN9 | 0,315057897 | 1,244061578 | 0,35393594 | 0,845346751 | septin 9 |
| ENSG00000264772 | AC016876,2 | 0,314959948 | 1,243977118 | 0,19219799 | 0,688538797 | novel transcript |
| ENSG00000260743 | AC007823,1 | 0,314829974 | 1,243865051 | 0,58672853 | 0,945395317 | novel transcript |
| ENSG00000267612 | AC008759,2 | 0,314818851 | 1,243855461 | 0,84446571 | 0,986340494 | novel transcript |
| ENSG00000163166 | IWS1 | 0,314808798 | 1,243846794 | 0,22717754 | 0,735737019 | interacts with SUPT6H, CTD assembly factor 1 |
| ENSG00000072422 | RHOBTB1 | 0,314798072 | 1,243837546 | 0,20321469 | 0,703503738 | Rho related BTB domain containing 1 |
| ENSG00000246016 | LINC01513 | 0,314775381 | 1,243817983 | 0,77694818 | 0,978587255 | long intergenic non-protein coding RNA 1513 |
| ENSG00000070423 | RNF126 | 0,314689267 | 1,243743743 | 0,30562971 | 0,814456808 | ring finger protein 126 |
| ENSG00000159322 | ADPGK | 0,314674983 | 1,243731428 | 0,14723997 | 0,611181688 | ADP dependent glucokinase |
| ENSG00000171703 | TCEA2 | 0,314590347 | 1,243658467 | 0,40232809 | 0,87718264 | transcription elongation factor A2 |
| ENSG00000111912 | NCOA7 | 0,314571112 | 1,243641885 | 0,33834719 | 0,837263082 | nuclear receptor coactivator 7 |
| ENSG00000136197 | C7orf25 | 0,314523117 | 1,243600513 | 0,42716278 | 0,892343796 | chromosome 7 open reading frame 25 |
| ENSG00000127580 | WDR24 | 0,314383367 | 1,243480055 | 0,3519932 | 0,844762053 | WD repeat domain 24 |
| ENSG00000175820 | CCDC168 | 0,314355193 | 1,243455772 | 0,61810665 | 0,952793934 | coiled-coil domain containing 168 |
| ENSG00000132437 | DDC | 0,314336065 | 1,243439285 | 0,57664394 | 0,943261034 | dopa decarboxylase |
| ENSG00000110237 | ARHGEF17 | 0,314258697 | 1,243372605 | 0,30587618 | 0,814766371 | Rho guanine nucleotide exchange factor 17 |
| ENSG00000143570 | SLC39A1 | 0,314251176 | 1,243366123 | 0,18404671 | 0,675673608 | solute carrier family 39 member 1 |
| ENSG00000162526 | TSSK3 | 0,314203567 | 1,243325092 | 0,60929408 | 0,951793902 | testis specific serine kinase 3 |

| | | | | | | |
|---|---|---|---|---|---|---|
| ENSG00000196155 | PLEKHG4 | 0,314176651 | 1,243301896 | 0,61693949 | 0,952793934 | pleckstrin homology and RhoGEF domain containing G4 |
| ENSG00000078804 | TP53INP2 | 0,31416147 | 1,243288813 | 0,17578314 | 0,661199837 | tumor protein p53 inducible nuclear protein 2 |
| ENSG00000126804 | ZBTB1 | 0,31416034 | 1,243287839 | 0,43277867 | 0,89560871 | zinc finger and BTB domain containing 1 |
| ENSG00000079950 | STX7 | 0,314137383 | 1,243268055 | 0,52308289 | 0,92790906 | syntaxin 7 |
| ENSG00000227825 | SLC9A7P1 | 0,314105122 | 1,243240254 | 0,32474446 | 0,826793081 | solute carrier family 9 member 7 pseudogene 1 |
| ENSG00000259845 | HERC2P10 | 0,314104846 | 1,243240017 | 0,51495389 | 0,924720046 | hect domain and RLD 2 pseudogene 10 |
| ENSG00000238142 | BX284668,5 | 0,314050745 | 1,243193396 | 0,3414938 | 0,840048641 | novel transcript |
| ENSG00000138867 | GUCD1 | 0,313686726 | 1,242879754 | 0,1695935 | 0,652093372 | guanylyl cyclase domain containing 1 |
| ENSG00000115275 | MOGS | 0,313660008 | 1,242856737 | 0,28796654 | 0,799270774 | mannosyl-oligosaccharide glucosidase |
| ENSG00000285715 | AC063960,1 | 0,313419554 | 1,242649607 | 0,55048498 | 0,936210753 | novel transcript, antisense to TACC2 |
| ENSG00000232490 | OSBPL10-AS1 | 0,313292432 | 1,242540117 | 0,81800914 | 0,982323352 | OSBPL10 antisense RNA 1 |
| ENSG00000132016 | C19orf57 | 0,313218666 | 1,242476587 | 0,37338424 | 0,861257555 | chromosome 19 open reading frame 57 |
| ENSG00000288152 | C19orf57 | 0,313218666 | 1,242476587 | 0,37338424 | 0,861257555 | chromosome 19 open reading frame 57 |
| ENSG00000230158 | MTND1P28 | 0,31315338 | 1,242420362 | 0,77727165 | 0,978587255 | MT-ND1 pseudogene 28 |
| ENSG00000251449 | MTND1P19 | 0,313128916 | 1,242399295 | 0,833472 | 0,984905093 | MT-ND1 pseudogene 19 |
| ENSG00000140157 | NIPA2 | 0,312997575 | 1,242286194 | 0,30007851 | 0,809942221 | NIPA magnesium transporter 2 |
| ENSG00000179526 | SHARPIN | 0,312980196 | 1,242271229 | 0,36815831 | 0,857548638 | SHANK associated RH domain interactor |
| ENSG00000106565 | TMEM176B | 0,312812151 | 1,242126538 | 0,62925789 | 0,952793934 | transmembrane protein 176B |
| ENSG00000078237 | TIGAR | 0,312672114 | 1,242005975 | 0,24942659 | 0,761381055 | TP53 induced glycolysis regulatory phosphatase |
| ENSG00000187135 | VSTM2B | 0,312530536 | 1,241884098 | 0,59397238 | 0,947469769 | V-set and transmembrane domain containing 2B |
| ENSG00000278032 | LY6E | 0,31251502 | 1,241870741 | 0,58001169 | 0,943828882 | lymphocyte antigen 6 family member E |
| ENSG00000168005 | SPINDOC | 0,312463524 | 1,241826414 | 0,30056684 | 0,81031109 | spindlin interactor and repressor of chromatin binding |
| ENSG00000204370 | SDHD | 0,312378173 | 1,241752949 | 0,42362385 | 0,890949676 | succinate dehydrogenase complex subunit D |
| ENSG00000174915 | PTDSS2 | 0,312351029 | 1,241729586 | 0,28349371 | 0,794687304 | phosphatidylserine synthase 2 |
| ENSG00000124786 | SLC35B3 | 0,312220385 | 1,241617145 | 0,45197845 | 0,902062594 | solute carrier family 35 member B3 |
| ENSG00000127511 | SIN3B | 0,312056331 | 1,241475965 | 0,3531579 | 0,845063255 | SIN3 transcription regulator family member B |
| ENSG00000171853 | TRAPPC12 | 0,311955215 | 1,241388955 | 0,34564858 | 0,841168721 | trafficking protein particle complex 12 |
| ENSG00000177993 | ZNRF3-AS1 | 0,311934177 | 1,241370852 | 0,63400166 | 0,954929949 | ZNRF3 antisense RNA 1 |
| ENSG00000180104 | EXOC3 | 0,311830226 | 1,241281411 | 0,27141782 | 0,784254811 | exocyst complex component 3 |
| ENSG00000214078 | CPNE1 | 0,311790193 | 1,241246968 | 0,19605249 | 0,694033976 | copine 1 |
| ENSG00000101181 | MTG2 | 0,311789001 | 1,241245942 | 0,35286364 | 0,845031687 | mitochondrial ribosome associated GTPase 2 |

| | | | | | | |
|---|---|---|---|---|---|---|
| ENSG00000100209 | HSCB | 0,311745756 | 1,241208736 | 0,25120438 | 0,7635322 | HscB mitochondrial iron-sulfur cluster cochaperone |
| ENSG00000169957 | ZNF768 | 0,311700269 | 1,241169602 | 0,50342457 | 0,921005254 | zinc finger protein 768 |
| ENSG00000104892 | KLC3 | 0,311523475 | 1,241017513 | 0,67377523 | 0,962923244 | kinesin light chain 3 |
| ENSG00000167741 | GGT6 | 0,311486346 | 1,240985575 | 0,65528318 | 0,957444211 | gamma-glutamyltransferase 6 |
| ENSG00000154978 | VOPP1 | 0,311432559 | 1,240939309 | 0,12637832 | 0,567767162 | VOPP1 WW domain binding protein |
| ENSG00000132313 | MRPL35 | 0,311330868 | 1,240851842 | 0,46678431 | 0,907940653 | mitochondrial ribosomal protein L35 |
| ENSG00000173163 | COMMD1 | 0,311241505 | 1,240774984 | 0,21406229 | 0,72150508 | copper metabolism domain containing 1 |
| ENSG00000242512 | LINC01206 | 0,31119166 | 1,240732116 | 0,59867123 | 0,948854865 | long intergenic non-protein coding RNA 1206 |
| ENSG00000275278 | AC012150,2 | 0,311155592 | 1,240701098 | 0,80281032 | 0,980866933 | novel transcript, antisense to FAR2 |
| ENSG00000163171 | CDC42EP3 | 0,311049172 | 1,240609581 | 0,25043891 | 0,762610495 | CDC42 effector protein 3 |
| ENSG00000266538 | AC005838,2 | 0,310831189 | 1,240422146 | 0,64405751 | 0,956633898 | novel transcript |
| ENSG00000258034 | AC012157,2 | 0,310826555 | 1,240418162 | 0,72691477 | 0,973344786 | novel transcript, antisense to MED13L |
| ENSG00000257446 | ZNF878 | 0,310748436 | 1,240350997 | 0,49743855 | 0,918759905 | zinc finger protein 878 |
| ENSG00000258524 | NT5CP1 | 0,310700241 | 1,240309563 | 0,75270901 | 0,975901808 | 5',3'-nucleotidase, cytosolic pseudogene 1 |
| ENSG00000116667 | C1orf21 | 0,310670092 | 1,240283644 | 0,40698362 | 0,880078305 | chromosome 1 open reading frame 21 |
| ENSG00000112078 | KCTD20 | 0,310648609 | 1,240265175 | 0,31067043 | 0,817751312 | potassium channel tetramerization domain containing 20 |
| ENSG00000125746 | EML2 | 0,310557688 | 1,240187014 | 0,43916492 | 0,897434993 | EMAP like 2 |
| ENSG00000176358 | TAC4 | 0,310543273 | 1,240174622 | 0,74336291 | 0,975310206 | tachykinin precursor 4 |
| ENSG00000271723 | MROH7-TTC4 | 0,310520389 | 1,24015495 | 0,74207648 | 0,975176043 | MROH7-TTC4 readthrough (NMD candidate) |
| ENSG00000125656 | CLPP | 0,310448938 | 1,240093532 | 0,23376348 | 0,743099279 | caseinolytic mitochondrial matrix peptidase proteolytic subunit |
| ENSG00000172775 | PSME3IP1 | 0,310410547 | 1,240060533 | 0,22182853 | 0,729383401 | proteasome activator subunit 3 interacting protein 1 |
| ENSG00000213881 | NPM1P6 | 0,310356642 | 1,240014201 | 0,77116947 | 0,978033855 | nucleophosmin 1 pseudogene 6 |
| ENSG00000226471 | Z93930,2 | 0,310273034 | 1,239942341 | 0,5777383 | 0,94337651 | novel transcript |
| ENSG00000166707 | ZCCHC18 | 0,310211142 | 1,239889147 | 0,33505581 | 0,834858938 | zinc finger CCHC-type containing 18 |
| ENSG00000131831 | RAI2 | 0,31020121 | 1,239880612 | 0,58487105 | 0,944857905 | retinoic acid induced 2 |
| ENSG00000132326 | PER2 | 0,310158384 | 1,239843806 | 0,36122835 | 0,851821025 | period circadian regulator 2 |
| ENSG00000136448 | NMT1 | 0,310141348 | 1,239829166 | 0,14026328 | 0,597018607 | N-myristoyltransferase 1 |
| ENSG00000214145 | LINC00887 | 0,310131518 | 1,239820719 | 0,6479653 | 0,956899963 | long intergenic non-protein coding RNA 887 |
| ENSG00000021355 | SERPINB1 | 0,309853681 | 1,239581974 | 0,3595824 | 0,849982153 | serpin family B member 1 |
| ENSG00000188916 | INSYN2A | 0,309851619 | 1,239580203 | 0,38415749 | 0,868125769 | inhibitory synaptic factor 2A |
| ENSG00000158286 | RNF207 | 0,309829795 | 1,239561452 | 0,51086328 | 0,922693987 | ring finger protein 207 |

| | | | | | | |
|---|---|---|---|---|---|---|
| ENSG00000248919 | ATP5MF-PTCD1 | 0,309799404 | 1,23953534 | 0,37093087 | 0,858959913 | ATP5MF-PTCD1 readthrough |
| ENSG00000189343 | RPS2P46 | 0,309778243 | 1,239517159 | 0,74242091 | 0,975176043 | ribosomal protein S2 pseudogene 46 |
| ENSG00000174992 | ZG16 | 0,309586046 | 1,23935204 | 0,76549722 | 0,97683946 | zymogen granule protein 16 |
| ENSG00000278961 | SMIM34B | 0,309561811 | 1,239331222 | 0,79067878 | 0,979466731 | small integral membrane protein 34B |
| ENSG00000234851 | RPL23AP42 | 0,309467908 | 1,239250558 | 0,22429343 | 0,73307493 | ribosomal protein L23a pseudogene 42 |
| ENSG00000133316 | WDR74 | 0,309386621 | 1,239180736 | 0,28361106 | 0,794944971 | WD repeat domain 74 |
| ENSG00000166997 | CNPY4 | 0,309299649 | 1,239106035 | 0,33920158 | 0,838276153 | canopy FGF signaling regulator 4 |
| ENSG00000100554 | ATP6V1D | 0,309249298 | 1,239062789 | 0,28261163 | 0,793528967 | ATPase H+ transporting V1 subunit D |
| ENSG00000260630 | SNAI3-AS1 | 0,309244418 | 1,239058598 | 0,40873746 | 0,88153752 | SNAI3 antisense RNA 1 |
| ENSG00000183291 | SELENOF | 0,309197724 | 1,239018496 | 0,42197409 | 0,889720796 | selenoprotein F |
| ENSG00000173540 | GMPPB | 0,30919744 | 1,239018252 | 0,41614127 | 0,885986046 | GDP-mannose pyrophosphorylase B |
| ENSG00000224831 | TMEM183B | 0,309191213 | 1,239012904 | 0,72398202 | 0,973153353 | transmembrane protein 183B |
| ENSG00000138162 | TACC2 | 0,309102235 | 1,238936491 | 0,23265965 | 0,741684508 | transforming acidic coiled-coil containing protein 2 |
| ENSG00000240244 | GAPDHP33 | 0,309038444 | 1,238881711 | 0,76199565 | 0,976315588 | glyceraldehyde 3 phosphate dehydrogenase pseudogene 33 |
| ENSG00000011426 | ANLN | 0,309007761 | 1,238855362 | 0,34351969 | 0,840901506 | anillin actin binding protein |
| ENSG00000153989 | NUS1 | 0,308931116 | 1,238789548 | 0,39180394 | 0,87047742 | NUS1 dehydrodolichyl diphosphate synthase subunit |
| ENSG00000197965 | MPZL1 | 0,308898824 | 1,23876182 | 0,3331167 | 0,833301711 | myelin protein zero like 1 |
| ENSG00000284713 | SMIM38 | 0,308889778 | 1,238754053 | 0,62781623 | 0,952793934 | small integral membrane protein 38 |
| ENSG00000111696 | NT5DC3 | 0,30885428 | 1,238723574 | 0,34710636 | 0,841757143 | 5'-nucleotidase domain containing 3 |
| ENSG00000155729 | KCTD18 | 0,308451404 | 1,238377705 | 0,17003416 | 0,653256728 | potassium channel tetramerization domain containing 18 |
| ENSG00000270557 | AC013731,1 | 0,308353499 | 1,238293669 | 0,58286639 | 0,944174844 | novel transcript |
| ENSG00000119929 | CUTC | 0,308349182 | 1,238289964 | 0,31505336 | 0,820340495 | cutC copper transporter |
| ENSG00000175895 | PLEKHF2 | 0,308158389 | 1,238126214 | 0,46068994 | 0,905949965 | pleckstrin homology and FYVE domain containing 2 |
| ENSG00000175029 | CTBP2 | 0,308101759 | 1,238077614 | 0,21891305 | 0,72708995 | C-terminal binding protein 2 |
| ENSG00000137509 | PRCP | 0,308065537 | 1,23804653 | 0,44312239 | 0,899712842 | prolylcarboxypeptidase |
| ENSG00000248268 | AC010275,1 | 0,307900357 | 1,237904789 | 0,60197706 | 0,949939978 | novel transcript |
| ENSG00000234640 | AL390763,1 | 0,307841433 | 1,23785423 | 0,87276126 | 0,989262536 | novel transcript |
| ENSG00000137409 | MTCH1 | 0,307812166 | 1,237829119 | 0,14911422 | 0,614055206 | mitochondrial carrier 1 |
| ENSG00000160439 | RDH13 | 0,307787482 | 1,237807941 | 0,45155855 | 0,902062594 | retinol dehydrogenase 13 |
| ENSG00000121270 | ABCC11 | 0,307711901 | 1,237743095 | 0,71420744 | 0,970867497 | ATP binding cassette subfamily C member 11 |

| | | | | | | |
|---|---|---|---|---|---|---|
| ENSG00000142327 | RNPEPL1 | 0,307643428 | 1,237684351 | 0,48022118 | 0,912847708 | arginyl aminopeptidase like 1 |
| ENSG00000004961 | HCCS | 0,307527535 | 1,23758493 | 0,33164794 | 0,832548086 | holocytochrome c synthase |
| ENSG00000162510 | MATN1 | 0,307253672 | 1,237350025 | 0,71473051 | 0,971041579 | matrilin 1 |
| ENSG00000105402 | NAPA | 0,3071682 | 1,237276721 | 0,19006334 | 0,685541073 | NSF attachment protein alpha |
| ENSG00000164197 | RNF180 | 0,307167034 | 1,237275721 | 0,4783665 | 0,912208437 | ring finger protein 180 |
| ENSG00000175550 | DRAP1 | 0,307109516 | 1,237226394 | 0,27502106 | 0,786439192 | DR1 associated protein 1 |
| ENSG00000285051 | SLC7A14-AS1 | 0,307058733 | 1,237182845 | 0,46381166 | 0,907251937 | SLC7A14 antisense RNA 1 |
| ENSG00000148824 | MTG1 | 0,307014402 | 1,237144829 | 0,34747793 | 0,842200946 | mitochondrial ribosome associated GTPase 1 |
| ENSG00000213973 | ZNF99 | 0,306941255 | 1,237082105 | 0,78401613 | 0,97928555 | zinc finger protein 99 |
| ENSG00000259523 | AC022613,2 | 0,306889136 | 1,237037415 | 0,62708736 | 0,952793934 | novel transcript, antisense to TJP1 |
| ENSG00000198917 | SPOUT1 | 0,30683312 | 1,236989385 | 0,49190806 | 0,916995341 | SPOUT domain containing methyltransferase 1 |
| ENSG00000226627 | SHANK2-AS1 | 0,306713068 | 1,236886455 | 0,64879322 | 0,956899963 | SHANK2 antisense RNA 1 |
| ENSG00000142765 | SYTL1 | 0,306648102 | 1,236830757 | 0,63387283 | 0,954929949 | synaptotagmin like 1 |
| ENSG00000182810 | DDX28 | 0,306582878 | 1,236774842 | 0,20935687 | 0,713678247 | DEAD-box helicase 28 |
| ENSG00000158828 | PINK1 | 0,306571923 | 1,236765451 | 0,21941908 | 0,727838473 | PTEN induced kinase 1 |
| ENSG00000226453 | LINC02542 | 0,30650909 | 1,236711588 | 0,75112366 | 0,975901808 | long intergenic non-protein coding RNA 2542 |
| ENSG00000287606 | LINC02542 | 0,30650909 | 1,236711588 | 0,75112366 | 0,975901808 | long intergenic non-protein coding RNA 2542 |
| ENSG00000174437 | ATP2A2 | 0,306433335 | 1,23664665 | 0,1285281 | 0,572897424 | ATPase sarcoplasmic/endoplasmic reticulum Ca2+ transporting 2 |
| ENSG00000281376 | ABALON | 0,306424063 | 1,236638703 | 0,57999361 | 0,943828882 | apoptotic BCL2L1-antisense long non-coding RNA |
| ENSG00000082458 | DLG3 | 0,306423133 | 1,236637905 | 0,21647305 | 0,723777756 | discs large MAGUK scaffold protein 3 |
| ENSG00000077348 | EXOSC5 | 0,306202236 | 1,236448573 | 0,38934426 | 0,870057277 | exosome component 5 |
| ENSG00000128891 | CCDC32 | 0,30612197 | 1,236379784 | 0,23988416 | 0,749799162 | coiled-coil domain containing 32 |
| ENSG00000248668 | OXCT1-AS1 | 0,306051472 | 1,236319369 | 0,546003 | 0,935201756 | OXCT1 antisense RNA 1 |
| ENSG00000276814 | AC004801,6 | 0,306031018 | 1,236301841 | 0,6264665 | 0,952793934 | novel transcript |
| ENSG00000107816 | LZTS2 | 0,305867055 | 1,236161343 | 0,52743519 | 0,929454507 | leucine zipper tumor suppressor 2 |
| ENSG00000255851 | Metazoa_SRP | 0,305832618 | 1,236131836 | 0,73947425 | 0,974715504 | Metazoan signal recognition particle RNA |
| ENSG00000101812 | H2BW2 | 0,305810052 | 1,236112501 | 0,50090867 | 0,920520138 | H2B,W histone 2 |
| ENSG00000206159 | GYG2P1 | 0,305794808 | 1,23609944 | 0,7545709 | 0,975901808 | glycogenin 2 pseudogene 1 |
| ENSG00000112599 | GUCA1B | 0,305771824 | 1,236079748 | 0,56159868 | 0,940681545 | guanylate cyclase activator 1B |
| ENSG00000159228 | CBR1 | 0,305764655 | 1,236073605 | 0,24622497 | 0,758198269 | carbonyl reductase 1 |
| ENSG00000243649 | CFB | 0,305744561 | 1,236056389 | 0,57303107 | 0,942475308 | complement factor B |
| ENSG00000288095 | AC005618,4 | 0,305674374 | 1,235996257 | 0,6574183 | 0,957853146 | novel transcript |

| | | | | | | |
|---|---|---|---|---|---|---|
| ENSG00000111328 | CDK2AP1 | 0,305366118 | 1,235732194 | 0,35591707 | 0,847284806 | cyclin dependent kinase 2 associated protein 1 |
| ENSG00000175287 | PHYHD1 | 0,305356001 | 1,235723528 | 0,48409918 | 0,913690171 | phytanoyl-CoA dioxygenase domain containing 1 |
| ENSG00000159346 | ADIPOR1 | 0,30522376 | 1,235610264 | 0,18039478 | 0,66974377 | adiponectin receptor 1 |
| ENSG00000054611 | TBC1D22A | 0,305195635 | 1,235586176 | 0,28784865 | 0,799107075 | TBC1 domain family member 22A |
| ENSG00000183258 | DDX41 | 0,305190747 | 1,23558199 | 0,21425461 | 0,721587186 | DEAD-box helicase 41 |
| ENSG00000235142 | LINC02532 | 0,305187377 | 1,235579104 | 0,79463862 | 0,979776981 | long intergenic non-protein coding RNA 2532 |
| ENSG00000064787 | BCAS1 | 0,305146918 | 1,235544453 | 0,51013411 | 0,922427526 | brain enriched myelin associated protein 1 |
| ENSG00000285351 | TCAF2P1 | 0,30514686 | 1,235544404 | 0,80084359 | 0,980505307 | TRPM8 channel associated factor 2 pseudogene 1 |
| ENSG00000037474 | NSUN2 | 0,305116928 | 1,23551877 | 0,16581829 | 0,645114847 | NOP2/Sun RNA methyltransferase 2 |
| ENSG00000288663 | AC073611,1 | 0,304890218 | 1,235324631 | 0,55855094 | 0,939452149 | novel protein, readthrough PFDN5-C12orf10 |
| ENSG00000167447 | SMG8 | 0,304868934 | 1,235306407 | 0,23617984 | 0,745253314 | SMG8 nonsense mediated mRNA decay factor |
| ENSG00000258656 | AL160236,1 | 0,3048603 | 1,235299014 | 0,83872309 | 0,985672131 | high mobility group nucleosomal binding domain 2 (HMGN2) pseudogene |
| ENSG00000154845 | PPP4R1 | 0,304851588 | 1,235291554 | 0,32167338 | 0,824933727 | protein phosphatase 4 regulatory subunit 1 |
| ENSG00000244921 | MTCYBP18 | 0,304764865 | 1,235217301 | 0,76316238 | 0,976607938 | MT-CYB pseudogene 18 |
| ENSG00000204257 | HLA-DMA | 0,304561635 | 1,23504331 | 0,79225684 | 0,979471509 | major histocompatibility complex, class II, DM alpha |
| ENSG00000141385 | AFG3L2 | 0,30449447 | 1,234985814 | 0,13122705 | 0,578575994 | AFG3 like matrix AAA peptidase subunit 2 |
| ENSG00000174442 | ZWILCH | 0,304349359 | 1,234861601 | 0,47361344 | 0,910190245 | zwilch kinetochore protein |
| ENSG00000268655 | AC008687,4 | 0,304272929 | 1,234796184 | 0,7902669 | 0,979462318 | novel protein |
| ENSG00000139173 | TMEM117 | 0,304181603 | 1,234718021 | 0,45657035 | 0,903545728 | transmembrane protein 117 |
| ENSG00000165511 | ZNF22-AS1 | 0,304165013 | 1,234703823 | 0,41914333 | 0,887577549 | ZNF22 antisense RNA 1 |
| ENSG00000260913 | LINC01254 | 0,304055095 | 1,234609754 | 0,77686924 | 0,978587255 | long intergenic non-protein coding RNA 1254 |
| ENSG00000115145 | STAM2 | 0,304035353 | 1,23459286 | 0,46191189 | 0,906277266 | signal transducing adaptor molecule 2 |
| ENSG00000260602 | HMGN2P40 | 0,303855286 | 1,234438776 | 0,81019198 | 0,981213877 | high mobility group nucleosomal binding domain 2 pseudogene 40 |
| ENSG00000265496 | AC100778,3 | 0,303798181 | 1,234389916 | 0,7814879 | 0,978597905 | microRNA 1539 |
| ENSG00000156273 | BACH1 | 0,303675465 | 1,234284922 | 0,346815 | 0,841572812 | BTB domain and CNC homolog 1 |
| ENSG00000074590 | NUAK1 | 0,303652553 | 1,234265321 | 0,28511901 | 0,796602794 | NUAK family kinase 1 |
| ENSG00000100036 | SLC35E4 | 0,303607636 | 1,234226893 | 0,44650973 | 0,901644586 | solute carrier family 35 member E4 |
| ENSG00000266718 | AC079336,5 | 0,303540537 | 1,234169492 | 0,64316548 | 0,956633898 | novel transcript, antisense MYO1D |
| ENSG00000137996 | RTCA | 0,303433587 | 1,234078003 | 0,28958859 | 0,800503785 | RNA 3'-terminal phosphate cyclase |
| ENSG00000182050 | MGAT4C | 0,3034275 | 1,234072797 | 0,78109055 | 0,978597905 | MGAT4 family member C |
| ENSG00000187713 | TMEM203 | 0,303426777 | 1,234072178 | 0,21774733 | 0,72540192 | transmembrane protein 203 |

| | | | | | | |
|---|---|---|---|---|---|---|
| ENSG00000113441 | LNPEP | 0,303391424 | 1,234041938 | 0,43498228 | 0,896272827 | leucyl and cystinyl aminopeptidase |
| ENSG00000169710 | FASN | 0,303383331 | 1,234035015 | 0,48141127 | 0,912981134 | fatty acid synthase |
| ENSG00000105287 | PRKD2 | 0,303380158 | 1,234032301 | 0,3071832 | 0,815953743 | protein kinase D2 |
| ENSG00000120370 | GORAB | 0,303355061 | 1,234010834 | 0,36123024 | 0,851821025 | golgin, RAB6 interacting |
| ENSG00000231787 | H2BP9 | 0,303107075 | 1,233798738 | 0,67752333 | 0,9635902 | H2B histone pseudogene 9 |
| ENSG00000287086 | AC034193,1 | 0,303100001 | 1,233792688 | 0,71610282 | 0,971360663 | novel transcript, antisense to VHL |
| ENSG00000261732 | AL031708,1 | 0,303064098 | 1,233761984 | 0,50978616 | 0,922427526 | novel protein |
| ENSG00000288656 | AC020765,6 | 0,302993987 | 1,233702028 | 0,86553904 | 0,989074806 | Sulfotransferase 1A1 |
| ENSG00000230701 | FBXW4P1 | 0,302990239 | 1,233698823 | 0,70068057 | 0,968522643 | F-box and WD repeat domain containing 4 pseudogene 1 |
| ENSG00000141867 | BRD4 | 0,302977222 | 1,233687691 | 0,27025462 | 0,78255445 | bromodomain containing 4 |
| ENSG00000185189 | NRBP2 | 0,302942779 | 1,233658239 | 0,35133897 | 0,84405766 | nuclear receptor binding protein 2 |
| ENSG00000198355 | PIM3 | 0,302925467 | 1,233643436 | 0,4423015 | 0,899331131 | Pim-3 proto-oncogene, serine/threonine kinase |
| ENSG00000109794 | FAM149A | 0,302894648 | 1,233617082 | 0,24725739 | 0,759357074 | family with sequence similarity 149 member A |
| ENSG00000142655 | PEX14 | 0,302889598 | 1,233612764 | 0,32850475 | 0,830659639 | peroxisomal biogenesis factor 14 |
| ENSG00000287665 | AC092428,1 | 0,302803439 | 1,233539094 | 0,68621829 | 0,965740648 | novel transcript, sense intronic to FOXK1 |
| ENSG00000165732 | DDX21 | 0,302716488 | 1,233464751 | 0,24268959 | 0,752892335 | DExD-box helicase 21 |
| ENSG00000270711 | AL162431,3 | 0,302701654 | 1,233452069 | 0,73941367 | 0,974715504 | cell division cycle associated 8 (CDCA8) pseudogene |
| ENSG00000272787 | AC253536,6 | 0,302652474 | 1,233410022 | 0,74513078 | 0,975666349 | novel transcript |
| ENSG00000281400 | AC253536,6 | 0,302652474 | 1,233410022 | 0,74513078 | 0,975666349 | novel transcript |
| ENSG00000176915 | ANKLE2 | 0,302594095 | 1,233360112 | 0,25355437 | 0,765172364 | ankyrin repeat and LEM domain containing 2 |
| ENSG00000237927 | AL078604,2 | 0,302557674 | 1,233328977 | 0,80140751 | 0,980557464 | novel transcript |
| ENSG00000058272 | PPP1R12A | 0,302525663 | 1,233301611 | 0,30042388 | 0,810183053 | protein phosphatase 1 regulatory subunit 12A |
| ENSG00000253649 | PRSS51 | 0,302468196 | 1,233252486 | 0,62368081 | 0,952793934 | serine protease 51 |
| ENSG00000143149 | ALDH9A1 | 0,302388902 | 1,233184706 | 0,40068374 | 0,876441365 | aldehyde dehydrogenase 9 family member A1 |
| ENSG00000130811 | EIF3G | 0,30229415 | 1,233103716 | 0,20013244 | 0,699646707 | eukaryotic translation initiation factor 3 subunit G |
| ENSG00000215077 | BRD2 | 0,302259617 | 1,233074201 | 0,18419154 | 0,675967184 | bromodomain containing 2 |
| ENSG00000134250 | NOTCH2 | 0,302201373 | 1,233024421 | 0,14626641 | 0,609635114 | notch receptor 2 |
| ENSG00000066382 | MPPED2 | 0,302016696 | 1,232866593 | 0,43658836 | 0,896866043 | metallophosphoesterase domain containing 2 |
| ENSG00000131471 | AOC3 | 0,301984728 | 1,232839274 | 0,64812954 | 0,956899963 | amine oxidase copper containing 3 |
| ENSG00000115457 | IGFBP2 | 0,301961307 | 1,232819261 | 0,40439086 | 0,878176155 | insulin like growth factor binding protein 2 |
| ENSG00000248593 | DSTNP2 | 0,301852898 | 1,232726626 | 0,34236236 | 0,840048641 | DSTN pseudogene 2 |
| ENSG00000068078 | FGFR3 | 0,301511997 | 1,232435374 | 0,23826845 | 0,747710567 | fibroblast growth factor receptor 3 |

| | | | | | | |
|---|---|---|---|---|---|---|
| ENSG00000204959 | ARHGEF34P | 0,301407657 | 1,232346243 | 0,52585174 | 0,929056124 | Rho guanine nucleotide exchange factor 34, pseudogene |
| ENSG00000203740 | METTL11B | 0,301372023 | 1,232315806 | 0,73099276 | 0,974353474 | methyltransferase like 11B |
| ENSG00000236779 | AL512306,1 | 0,301359583 | 1,23230518 | 0,64592093 | 0,956899963 | PEF protein with a long N-terminal hydrophobic domain (peflin) (PEF) pseudogene |
| ENSG00000184428 | TOP1MT | 0,301267428 | 1,232226467 | 0,48170322 | 0,912981134 | DNA topoisomerase I mitochondrial |
| ENSG00000116209 | TMEM59 | 0,301196752 | 1,232166102 | 0,35079682 | 0,844035671 | transmembrane protein 59 |
| ENSG00000114019 | AMOTL2 | 0,301145775 | 1,232122565 | 0,33256258 | 0,833045148 | angiomotin like 2 |
| ENSG00000272485 | AL392183,1 | 0,301127162 | 1,232106669 | 0,81880494 | 0,98249617 | novel transcript |
| ENSG00000104365 | IKBKB | 0,301068484 | 1,232056557 | 0,2780719 | 0,789397971 | inhibitor of nuclear factor kappa B kinase subunit beta |
| ENSG00000168291 | PDHB | 0,301052615 | 1,232043005 | 0,22772304 | 0,736053199 | pyruvate dehydrogenase E1 subunit beta |
| ENSG00000201793 | RN7SKP9 | 0,301028027 | 1,232022007 | 0,61676769 | 0,952793934 | RN7SK pseudogene 9 |
| ENSG00000149357 | LAMTOR1 | 0,301007541 | 1,232004513 | 0,26060936 | 0,772692192 | late endosomal/lysosomal adaptor, MAPK and MTOR activator 1 |
| ENSG00000131748 | STARD3 | 0,300984349 | 1,231984708 | 0,31548275 | 0,82082095 | StAR related lipid transfer domain containing 3 |
| ENSG00000258414 | AL121790,1 | 0,300980177 | 1,231981146 | 0,81992495 | 0,982564818 | novel transcript |
| ENSG00000188626 | GOLGA8M | 0,300979737 | 1,23198077 | 0,73325237 | 0,974353474 | golgin A8 family member M |
| ENSG00000265790 | RNASEH1P1 | 0,300889833 | 1,231903999 | 0,5095765 | 0,922427526 | ribonuclease H1 pseudogene 1 |
| ENSG00000185085 | INTS5 | 0,300791008 | 1,231819616 | 0,29824922 | 0,807662492 | integrator complex subunit 5 |
| ENSG00000102452 | NALCN | 0,300776126 | 1,23180691 | 0,41023859 | 0,882614012 | sodium leak channel, non-selective |
| ENSG00000120327 | PCDHB14 | 0,300722901 | 1,231761466 | 0,35148077 | 0,84405766 | protocadherin beta 14 |
| ENSG00000189420 | ZFP92 | 0,300655046 | 1,231703533 | 0,7921456 | 0,979471509 | ZFP92 zinc finger protein |
| ENSG00000223764 | LINC02593 | 0,30065399 | 1,231702632 | 0,46825347 | 0,907940653 | long intergenic non-protein coding RNA 2593 |
| ENSG00000008323 | PLEKHG6 | 0,300443522 | 1,231522957 | 0,62891817 | 0,952793934 | pleckstrin homology and RhoGEF domain containing G6 |
| ENSG00000161057 | PSMC2 | 0,300306876 | 1,231406318 | 0,27259214 | 0,785539753 | proteasome 26S subunit, ATPase 2 |
| ENSG00000285084 | SLC35G5 | 0,300294311 | 1,231395593 | 0,71785926 | 0,971762632 | solute carrier family 35 member G5 |
| ENSG00000081985 | IL12RB2 | 0,300176126 | 1,231294722 | 0,48573974 | 0,913997663 | interleukin 12 receptor subunit beta 2 |
| ENSG00000079150 | FKBP7 | 0,300151225 | 1,23127347 | 0,49572841 | 0,918046024 | FKBP prolyl isomerase 7 |
| ENSG00000125122 | LRRC29 | 0,300121486 | 1,23124809 | 0,62752322 | 0,952793934 | leucine rich repeat containing 29 |
| ENSG00000254565 | MTND2P26 | 0,30009925 | 1,231229113 | 0,76478758 | 0,976722333 | MT-ND2 pseudogene 26 |
| ENSG00000163380 | LMOD3 | 0,300086524 | 1,231218252 | 0,47964901 | 0,912816326 | leiomodin 3 |

| | | | | | | |
|---|---|---|---|---|---|---|
| ENSG00000141542 | RAB40B | 0,300081068 | 1,231213596 | 0,2360579 | 0,745253314 | RAB40B, member RAS oncogene family |
| ENSG00000167895 | TMC8 | 0,30002488 | 1,231165645 | 0,78858529 | 0,979462318 | transmembrane channel like 8 |
| ENSG00000126777 | KTN1 | 0,299961189 | 1,231111294 | 0,22084039 | 0,728635155 | kinectin 1 |
| ENSG00000063761 | ADCK1 | 0,299956452 | 1,231107252 | 0,32797856 | 0,830008227 | aarF domain containing kinase 1 |
| ENSG00000160194 | NDUFV3 | 0,299940284 | 1,231093455 | 0,25254122 | 0,76458964 | NADH:ubiquinone oxidoreductase subunit V3 |
| ENSG00000282854 | ASAP3 | 0,299892731 | 1,231052877 | 0,23790976 | 0,747090777 | ArfGAP with SH3 domain, ankyrin repeat and PH domain 3 |
| ENSG00000088280 | ASAP3 | 0,299892731 | 1,231052877 | 0,23790976 | 0,747090777 | ArfGAP with SH3 domain, ankyrin repeat and PH domain 3 |
| ENSG00000067167 | TRAM1 | 0,299868129 | 1,231031885 | 0,47911914 | 0,912714933 | translocation associated membrane protein 1 |
| ENSG00000091128 | LAMB4 | 0,299866929 | 1,230030861 | 0,70027676 | 0,968522643 | laminin subunit beta 4 |
| ENSG00000101365 | IDH3B | 0,299816086 | 1,230987477 | 0,16414977 | 0,642888389 | isocitrate dehydrogenase (NAD(+)) 3 non-catalytic subunit beta |
| ENSG00000110442 | COMMD9 | 0,299801161 | 1,230974743 | 0,24227276 | 0,752491461 | COMM domain containing 9 |
| ENSG00000178404 | CEP295NL | 0,299791269 | 1,230966303 | 0,70218199 | 0,968742626 | CEP295 N-terminal like |
| ENSG00000135124 | P2RX4 | 0,299713826 | 1,230900227 | 0,47021815 | 0,908528529 | purinergic receptor P2X 4 |
| ENSG00000177788 | AL162595,1 | 0,299617847 | 1,230818341 | 0,31362885 | 0,819846908 | novel transcript |
| ENSG00000171815 | PCDHB1 | 0,299611221 | 1,230812688 | 0,62893839 | 0,952793934 | protocadherin beta 1 |
| ENSG00000203363 | GPR160P1 | 0,299547942 | 1,230758703 | 0,77776912 | 0,978587255 | GPR160 pseudogene 1 |
| ENSG00000075461 | CACNG4 | 0,299403672 | 1,230635634 | 0,23504751 | 0,744654651 | calcium voltage-gated channel auxiliary subunit gamma 4 |
| ENSG00000273381 | AL158071,4 | 0,299386264 | 1,230620784 | 0,60394556 | 0,950551453 | novel transcript |
| ENSG00000188002 | AC026412,1 | 0,299339471 | 1,23058087 | 0,3833268 | 0,867760194 | programmed cell death 6 (PDCD6) pseudogene |
| ENSG00000103653 | CSK | 0,299290356 | 1,230538977 | 0,25303996 | 0,76515438 | C-terminal Src kinase |
| ENSG00000266967 | AARSD1 | 0,299198419 | 1,230460563 | 0,33049688 | 0,832114797 | alanyl-tRNA synthetase domain containing 1 |
| ENSG00000125991 | ERGIC3 | 0,299152969 | 1,230421799 | 0,19602625 | 0,694033976 | ERGIC and golgi 3 |
| ENSG00000287262 | AC021148,2 | 0,299089945 | 1,23036805 | 0,59819496 | 0,948608142 | novel transcript |
| ENSG00000160310 | PRMT2 | 0,299077574 | 1,230357499 | 0,21002964 | 0,715134683 | protein arginine methyltransferase 2 |
| ENSG00000265992 | ESRG | 0,298992148 | 1,230284649 | 0,61064107 | 0,951841373 | embryonic stem cell related |
| ENSG00000169871 | TRIM56 | 0,298937058 | 1,23023767 | 0,5417061 | 0,933491609 | tripartite motif containing 56 |
| ENSG00000138834 | MAPK8IP3 | 0,298891809 | 1,230199086 | 0,53202199 | 0,931291085 | mitogen-activated protein kinase 8 interacting protein 3 |
| ENSG00000229349 | ACTG1P9 | 0,29885133 | 1,23016457 | 0,80901861 | 0,981213877 | actin gamma 1 pseudogene 9 |

| | | | | | | |
|---|---|---|---|---|---|---|
| ENSG00000187054 | TMPRSS11A | 0,298782392 | 1,230105788 | 0,74589593 | 0,975707496 | transmembrane serine protease 11A |
| ENSG00000185052 | SLC24A3 | 0,298767079 | 1,230092732 | 0,52397642 | 0,928352631 | solute carrier family 24 member 3 |
| ENSG00000274569 | AC024884,1 | 0,298707611 | 1,230042029 | 0,74961673 | 0,975901808 | apolipoprotein F (APOF) pseudogene |
| ENSG00000286433 | AL020997,4 | 0,29869247 | 1,230029119 | 0,67231681 | 0,962290553 | novel transcript, antisense to PPP1R8 |
| ENSG00000100403 | ZC3H7B | 0,298630964 | 1,229976681 | 0,37043356 | 0,858545694 | zinc finger CCCH-type containing 7B |
| ENSG00000233321 | LINC02669 | 0,298629998 | 1,229975858 | 0,54432849 | 0,934381824 | long intergenic non-protein coding RNA 2669 |
| ENSG00000162804 | SNED1 | 0,298604828 | 1,229954399 | 0,48214402 | 0,912981134 | sushi, nidogen and EGF like domains 1 |
| ENSG00000154478 | GPR26 | 0,298512818 | 1,229875959 | 0,80198747 | 0,980710009 | G protein-coupled receptor 26 |
| ENSG00000122068 | FYTTD1 | 0,298329636 | 1,229719809 | 0,4348879 | 0,896272827 | forty-two-three domain containing 1 |
| ENSG00000260493 | AC011773,4 | 0,298320398 | 1,229711935 | 0,58760079 | 0,946068138 | novel transcript, antisense to E2F5 |
| ENSG00000152217 | SETBP1 | 0,298304839 | 1,229698673 | 0,22135396 | 0,729254784 | SET binding protein 1 |
| ENSG00000140443 | IGF1R | 0,298283213 | 1,22968024 | 0,171317 | 0,655094738 | insulin like growth factor 1 receptor |
| ENSG00000227906 | SNAP25-AS1 | 0,298238436 | 1,229642075 | 0,66913802 | 0,961703264 | SNAP25 antisense RNA 1 |
| ENSG00000179168 | GGN | 0,29823827 | 1,229641934 | 0,65324847 | 0,957395476 | gametogenetin |
| ENSG00000106025 | TSPAN12 | 0,298156857 | 1,229572545 | 0,3991881 | 0,875433675 | tetraspanin 12 |
| ENSG00000287795 | AC026341,2 | 0,297996582 | 1,229435955 | 0,71929694 | 0,971869926 | novel transcript |
| ENSG00000070759 | TESK2 | 0,297867802 | 1,229326216 | 0,39496269 | 0,871888817 | testis associated actin remodelling kinase 2 |
| ENSG00000153066 | TXNDC11 | 0,297727961 | 1,229207062 | 0,20562148 | 0,708146065 | thioredoxin domain containing 11 |
| ENSG00000274373 | AC148476,1 | 0,297708899 | 1,229190821 | 0,60987262 | 0,951801946 | novel transcript |
| ENSG00000124882 | EREG | 0,297706172 | 1,229188498 | 0,76995587 | 0,977742359 | epiregulin |
| ENSG00000277491 | AC087392,5 | 0,297661629 | 1,229150547 | 0,70649532 | 0,969503793 | novel transcript |
| ENSG00000196767 | POU3F4 | 0,29764661 | 1,229137751 | 0,37934147 | 0,864853196 | POU class 3 homeobox 4 |
| ENSG00000267673 | FDX2 | 0,297600319 | 1,229098313 | 0,40168359 | 0,87705851 | ferredoxin 2 |
| ENSG00000135521 | LTV1 | 0,297595675 | 1,229094357 | 0,26847141 | 0,780709773 | LTV1 ribosome biogenesis factor |
| ENSG00000264449 | AP001021,1 | 0,297529774 | 1,229038215 | 0,60218874 | 0,950130148 | novel transcript, antisense to TMEM200C |
| ENSG00000215124 | AMD1P4 | 0,297388571 | 1,228917929 | 0,81204674 | 0,981213877 | adenosylmethionine decarboxylase 1 pseudogene 4 |
| ENSG00000108239 | TBC1D12 | 0,297290852 | 1,228834692 | 0,24924534 | 0,761274564 | TBC1 domain family member 12 |
| ENSG00000177150 | FAM210A | 0,297265886 | 1,228813427 | 0,43449314 | 0,896272827 | family with sequence similarity 210 member A |
| ENSG00000088888 | MAVS | 0,297202588 | 1,228759515 | 0,24114904 | 0,751551821 | mitochondrial antiviral signaling protein |
| ENSG00000260930 | LINC01416 | 0,297158942 | 1,228722342 | 0,65368088 | 0,957444211 | long intergenic non-protein coding RNA 1416 |
| ENSG00000273102 | AP000569,1 | 0,297111098 | 1,228681594 | 0,8440172 | 0,986273696 | novel transcript |
| ENSG00000147202 | DIAPH2 | 0,29706633 | 1,228643468 | 0,38974481 | 0,870057277 | diaphanous related formin 2 |
| ENSG00000131148 | EMC8 | 0,296806241 | 1,228421988 | 0,18055549 | 0,670229021 | ER membrane protein complex subunit 8 |

| | | | | | | |
|---|---|---|---|---|---|---|
| ENSG00000076321 | KLHL20 | 0,296766435 | 1,228388094 | 0,28251364 | 0,793528967 | kelch like family member 20 |
| ENSG00000278311 | GGNBP2 | 0,296695567 | 1,228327755 | 0,23512538 | 0,744654651 | gametogenetin binding protein 2 |
| ENSG00000164077 | MON1A | 0,296650152 | 1,228289089 | 0,32714234 | 0,82883128 | MON1 homolog A, secretory trafficking associated |
| ENSG00000023892 | DEF6 | 0,296568918 | 1,22821993 | 0,54408556 | 0,934259495 | DEF6 guanine nucleotide exchange factor |
| ENSG00000139641 | ESYT1 | 0,296551634 | 1,228205215 | 0,20572087 | 0,708191543 | extended synaptotagmin 1 |
| ENSG00000197548 | ATG7 | 0,296452719 | 1,228121009 | 0,24873561 | 0,760720365 | autophagy related 7 |
| ENSG00000233968 | AL157895,1 | 0,296439022 | 1,228109349 | 0,5767474 | 0,943261034 | novel transcript, antisense to C10orf112 |
| ENSG00000136040 | PLXNC1 | 0,29628666 | 1,227979657 | 0,36923399 | 0,857942508 | plexin C1 |
| ENSG00000027001 | MIPEP | 0,2962625 | 1,227959092 | 0,29353882 | 0,803461221 | mitochondrial intermediate peptidase |
| ENSG00000235754 | APOM | 0,296257967 | 1,227955234 | 0,49023069 | 0,916554718 | apolipoprotein M |
| ENSG00000175874 | CREG2 | 0,296222316 | 1,227924889 | 0,68825268 | 0,966426338 | cellular repressor of E1A stimulated genes 2 |
| ENSG00000236268 | LINC01361 | 0,296162399 | 1,227873894 | 0,56500229 | 0,940865625 | long intergenic non-protein coding RNA 1361 |
| ENSG00000283008 | CYP2D7 | 0,296160499 | 1,227872277 | 0,5124846 | 0,92347658 | cytochrome P450 family 2 subfamily D member 7 (gene/pseudogene) |
| ENSG00000272031 | ANKRD34A | 0,29611475 | 1,22783334 | 0,42622151 | 0,892133432 | ankyrin repeat domain 34A |
| ENSG00000130529 | TRPM4 | 0,2960974 | 1,227818574 | 0,49802286 | 0,919274055 | transient receptor potential cation channel subfamily M member 4 |
| ENSG00000140264 | SERF2 | 0,296079733 | 1,227803539 | 0,18026862 | 0,669667075 | small EDRK-rich factor 2 |
| ENSG00000120896 | SORBS3 | 0,296052559 | 1,227780413 | 0,31756186 | 0,822351863 | sorbin and SH3 domain containing 3 |
| ENSG00000256663 | AC112777,1 | 0,295931705 | 1,227677566 | 0,55476896 | 0,937793851 | ubiquitin-like with PHD and ring finger domains 1 (UHRF1) pseudogene |
| ENSG00000167112 | TRUB2 | 0,295909663 | 1,22765881 | 0,27478961 | 0,786383635 | TruB pseudouridine synthase family member 2 |
| ENSG00000258317 | AC034102,6 | 0,295907229 | 1,227656739 | 0,62469238 | 0,952793934 | novel transcript, antisense to ZC3H10 and ESYT1 |
| ENSG00000164970 | FAM219A | 0,295888359 | 1,227640681 | 0,2582566 | 0,770947602 | family with sequence similarity 219 member A |
| ENSG00000284726 | AL109936,9 | 0,295841275 | 1,227600616 | 0,69251447 | 0,967243233 | novel transcript |
| ENSG00000271659 | AL161729,4 | 0,295708503 | 1,227487644 | 0,7000718 | 0,968522643 | novel transcript, antisense to PTCH1 |
| ENSG00000206337 | HCP5 | 0,295647296 | 1,227435569 | 0,63490797 | 0,955158181 | HLA complex P5 |
| ENSG00000254991 | AC124276,1 | 0,29562503 | 1,227416625 | 0,78576158 | 0,979320406 | novel transcript |
| ENSG00000169251 | NMD3 | 0,295611924 | 1,227405475 | 0,35070058 | 0,844035671 | NMD3 ribosome export adaptor |
| ENSG00000136868 | SLC31A1 | 0,295569426 | 1,22736932 | 0,24842675 | 0,760720365 | solute carrier family 31 member 1 |
| ENSG00000107929 | LARP4B | 0,29556431 | 1,227364967 | 0,15757531 | 0,630220619 | La ribonucleoprotein 4B |
| ENSG00000227692 | MED28P3 | 0,295446766 | 1,227264971 | 0,79495188 | 0,979776981 | mediator complex subunit 28 pseudogene 3 |
| ENSG00000179157 | RPS2P28 | 0,295397351 | 1,227222936 | 0,72099752 | 0,972142678 | ribosomal protein S2 pseudogene 28 |

| | | | | | | |
|---|---|---|---|---|---|---|
| ENSG00000167543 | TP53I13 | 0,295303822 | 1,227143378 | 0,5210981 | 0,927046149 | tumor protein p53 inducible protein 13 |
| ENSG00000103544 | VPS35L | 0,295095062 | 1,226965822 | 0,16362926 | 0,642293268 | VPS35 endosomal protein sorting factor like |
| ENSG00000214760 | RPL21P1 | 0,295087881 | 1,226959715 | 0,78085134 | 0,978597905 | ribosomal protein L21 pseudogene 1 |
| ENSG00000176584 | DMBT1L1 | 0,295070546 | 1,226944972 | 0,76687541 | 0,977201891 | deleted in malignant brain tumors 1 like 1 (pseudogene) |
| ENSG00000286104 | AC016629,3 | 0,295020062 | 1,226902038 | 0,26213319 | 0,773902615 | Novel transcript |
| ENSG00000267284 | AC022031,2 | 0,295008894 | 1,226892541 | 0,38830421 | 0,869820053 | novel transcript |
| ENSG00000131462 | TUBG1 | 0,294877975 | 1,22678121 | 0,1799993 | 0,668905046 | tubulin gamma 1 |
| ENSG00000236529 | AC093582,1 | 0,294738538 | 1,226662647 | 0,77737839 | 0,978587255 | novel transcript |
| ENSG00000230454 | U73166,1 | 0,294521956 | 1,22647851 | 0,63785301 | 0,956018518 | novel transcript |
| ENSG00000273761 | DDX24 | 0,294444857 | 1,226412968 | 0,1849805 | 0,677431213 | DEAD-box helicase 24 |
| ENSG00000122756 | CNTFR | 0,294434348 | 1,226404035 | 0,36599217 | 0,85572078 | ciliary neurotrophic factor receptor |
| ENSG00000107738 | VSIR | 0,294408672 | 1,226382208 | 0,75151099 | 0,975901808 | V-set immunoregulatory receptor |
| ENSG00000074319 | TSG101 | 0,294393284 | 1,226369127 | 0,3984326 | 0,875129737 | tumor susceptibility 101 |
| ENSG00000285367 | AC087564,1 | 0,294345586 | 1,226328582 | 0,66511155 | 0,960681381 | novel transcript |
| ENSG00000230457 | PA2G4P4 | 0,294334516 | 1,226319172 | 0,78244832 | 0,978628303 | proliferation-associated 2G4 pseudogene 4 |
| ENSG00000230490 | AL139383,1 | 0,294258016 | 1,226254147 | 0,67104984 | 0,962026636 | novel transcript |
| ENSG00000278239 | AC243967,3 | 0,294245492 | 1,226243503 | 0,78432867 | 0,979320406 | novel transcript |
| ENSG00000264071 | RN7SL531P | 0,294226415 | 1,226227288 | 0,72439709 | 0,973250998 | RNA, 7SL, cytoplasmic 531, pseudogene |
| ENSG00000235993 | AC007559,1 | 0,294199213 | 1,226204167 | 0,78663754 | 0,979320406 | thioredoxin domain containing 17 (TXNDC17) pseudogene |
| ENSG00000113282 | CLINT1 | 0,294168584 | 1,226178135 | 0,37539556 | 0,862022464 | clathrin interactor 1 |
| ENSG00000235090 | RPL7L1P3 | 0,294055516 | 1,22608204 | 0,682682 | 0,964820564 | ribosomal protein L7 like 1 pseudogene 3 |
| ENSG00000187838 | PLSCR3 | 0,293924275 | 1,225970509 | 0,39024241 | 0,870057277 | phospholipid scramblase 3 |
| ENSG00000284009 | PLSCR3 | 0,293924275 | 1,225970509 | 0,39024241 | 0,870057277 | phospholipid scramblase 3 |
| ENSG00000141873 | SLC39A3 | 0,29388225 | 1,225934798 | 0,41434335 | 0,885579089 | solute carrier family 39 member 3 |
| ENSG00000248544 | AC008676,1 | 0,293720342 | 1,225797224 | 0,6154606 | 0,952793934 | novel transcript, antisense to CYFIP2 |
| ENSG00000228223 | HCG11 | 0,293714443 | 1,225792211 | 0,27070551 | 0,783073289 | HLA complex group 11 |
| ENSG00000177666 | PNPLA2 | 0,293662741 | 1,225748283 | 0,4322311 | 0,89558934 | patatin like phospholipase domain containing 2 |
| ENSG00000171943 | SRGAP2C | 0,293622053 | 1,225713714 | 0,31820415 | 0,823060685 | SLIT-ROBO Rho GTPase activating protein 2C |
| ENSG00000157379 | DHRS1 | 0,293612339 | 1,225705461 | 0,36387829 | 0,854140004 | dehydrogenase/reductase 1 |
| ENSG00000236393 | AC091806,1 | 0,293559967 | 1,225660967 | 0,68296183 | 0,964931888 | novel transcript |
| ENSG00000172382 | PRSS27 | 0,293480142 | 1,225593152 | 0,54347516 | 0,934259495 | serine protease 27 |

| | | | | | | |
|---|---|---|---|---|---|---|
| ENSG00000059691 | GATB | 0,293333791 | 1,225468831 | 0,21856237 | 0,726311 | glutamyl-tRNA amidotransferase subunit B |
| ENSG00000125843 | AP5S1 | 0,293219716 | 1,225371937 | 0,32420252 | 0,826338461 | adaptor related protein complex 5 subunit sigma 1 |
| ENSG00000269604 | AC005523,2 | 0,293169668 | 1,225329428 | 0,24313292 | 0,753350591 | novel transcript, antisense to FEM1A |
| ENSG00000287356 | AL590822,3 | 0,293083078 | 1,225255887 | 0,65302394 | 0,95732962 | novel transcript, antisense to MORN1 |
| ENSG00000127526 | SLC35E1 | 0,293072549 | 1,225246945 | 0,16896128 | 0,650934541 | solute carrier family 35 member E1 |
| ENSG00000169217 | CD2BP2 | 0,293020114 | 1,225202413 | 0,38144187 | 0,865900094 | CD2 cytoplasmic tail binding protein 2 |
| ENSG00000113645 | WWC1 | 0,293012466 | 1,225195919 | 0,4413436 | 0,898885877 | WW and C2 domain containing 1 |
| ENSG00000286084 | AL096794,1 | 0,292921819 | 1,22511894 | 0,6207428 | 0,952793934 | novel transcript |
| ENSG00000224738 | AC099850,1 | 0,292905425 | 1,225105018 | 0,61980954 | 0,952793934 | novel transcript |
| ENSG00000277452 | RN7SL473P | 0,2928672 | 1,225072559 | 0,79206363 | 0,979471509 | RNA, 7SL, cytoplasmic 473, pseudogene |
| ENSG00000285018 | AC245140,3 | 0,292837924 | 1,225047699 | 0,78474176 | 0,979320406 | novel transcript, antisense to FLNA |
| ENSG00000248192 | ATG10-AS1 | 0,292808644 | 1,225022836 | 0,78593608 | 0,979320406 | ATG10 antisense RNA 1 |
| ENSG00000104067 | TJP1 | 0,292707549 | 1,224936997 | 0,20704303 | 0,710823213 | tight junction protein 1 |
| ENSG00000176108 | CHMP6 | 0,292704164 | 1,224934123 | 0,36508484 | 0,855173479 | charged multivesicular body protein 6 |
| ENSG00000027697 | IFNGR1 | 0,292698268 | 1,224929117 | 0,46513055 | 0,907804479 | interferon gamma receptor 1 |
| ENSG00000285877 | AC007448,5 | 0,292641908 | 1,224881266 | 0,61999585 | 0,952793934 | novel transcript, antisense to HELZ |
| ENSG00000139324 | TMTC3 | 0,292627311 | 1,224868873 | 0,50323167 | 0,920888747 | transmembrane O-mannosyltransferase targeting cadherins 3 |
| ENSG00000279217 | Z95114,1 | 0,292607548 | 1,224852094 | 0,6515791 | 0,957089277 | novel transcript |
| ENSG00000177294 | FBXO39 | 0,292521713 | 1,224779222 | 0,73870871 | 0,974353474 | F-box protein 39 |
| ENSG00000169604 | ANTXR1 | 0,292487 | 1,224749753 | 0,37021227 | 0,858545694 | ANTXR cell adhesion molecule 1 |
| ENSG00000275365 | MMP11 | 0,292457201 | 1,224724455 | 0,64246192 | 0,956633898 | matrix metallopeptidase 11 |
| ENSG00000100412 | ACO2 | 0,292380016 | 1,224658933 | 0,21925584 | 0,727610082 | aconitase 2 |
| ENSG00000162654 | GBP4 | 0,292333624 | 1,224619554 | 0,61729431 | 0,952793934 | guanylate binding protein 4 |
| ENSG00000285091 | AL109840,2 | 0,292271468 | 1,224566794 | 0,77502161 | 0,978587255 | novel transcript |
| ENSG00000100522 | GNPNAT1 | 0,292248168 | 1,224547017 | 0,42000163 | 0,888425645 | glucosamine-phosphate N-acetyltransferase 1 |
| ENSG00000002919 | SNX11 | 0,292230334 | 1,22453188 | 0,21813055 | 0,725801782 | sorting nexin 11 |
| ENSG00000198373 | WWP2 | 0,292201971 | 1,224507806 | 0,26767887 | 0,780226749 | WW domain containing E3 ubiquitin protein ligase 2 |
| ENSG00000128203 | ASPHD2 | 0,292190384 | 1,224497971 | 0,30429156 | 0,813156808 | aspartate beta-hydroxylase domain containing 2 |
| ENSG00000225885 | AC023590,1 | 0,292181747 | 1,224490641 | 0,6933964 | 0,967340633 | novel transcript, antisense to SAMD12 |
| ENSG00000282130 | NSMCE3 | 0,29205533 | 1,224383349 | 0,26040725 | 0,772692192 | NSE3 homolog, SMC5-SMC6 complex component |
| ENSG00000250151 | ARPC4-TTLL3 | 0,292021054 | 1,22435426 | 0,33270166 | 0,833260009 | ARPC4-TTLL3 readthrough |
| ENSG00000166133 | RPUSD2 | 0,291964344 | 1,224306133 | 0,22875461 | 0,737173617 | RNA pseudouridine synthase domain containing 2 |

| | | | | | | |
|---|---|---|---|---|---|---|
| ENSG00000228639 | ROCR | 0,291874861 | 1,224230199 | 0,81710384 | 0,981979397 | regulator of chondrogenesis RNA |
| ENSG00000140943 | MBTPS1 | 0,291799705 | 1,224166425 | 0,20312717 | 0,703503738 | membrane bound transcription factor peptidase, site 1 |
| ENSG00000277802 | AF252830,2 | 0,291749055 | 1,224123448 | 0,86536306 | 0,989074806 | novel ubiquitin specific peptidase pseudogene |
| ENSG00000203896 | LIME1 | 0,291739321 | 1,224115189 | 0,69502295 | 0,967509655 | Lck interacting transmembrane adaptor 1 |
| ENSG00000281886 | TGM4 | 0,291667881 | 1,224054574 | 0,72263384 | 0,972594421 | transglutaminase 4 |
| ENSG00000163810 | TGM4 | 0,291667881 | 1,224054574 | 0,72263384 | 0,972594421 | transglutaminase 4 |
| ENSG00000181027 | FKRP | 0,291664164 | 1,22405142 | 0,43968148 | 0,897665148 | fukutin related protein |
| ENSG00000104388 | RAB2A | 0,291597312 | 1,223994701 | 0,5166554 | 0,925489094 | RAB2A, member RAS oncogene family |
| ENSG00000188026 | RILPL1 | 0,29153036 | 1,223937899 | 0,46739762 | 0,907940653 | Rab interacting lysosomal protein like 1 |
| ENSG00000275488 | AC023509,5 | 0,291469383 | 1,22388617 | 0,69527853 | 0,967599788 | zinc finger protein 337 (ZNF337) pseudogene |
| ENSG00000184887 | BTBD6 | 0,291433327 | 1,223855582 | 0,28970784 | 0,800522458 | BTB domain containing 6 |
| ENSG00000105371 | ICAM4 | 0,291397751 | 1,223825404 | 0,87806052 | 0,990109814 | intercellular adhesion molecule 4 (Landsteiner-Wiener blood group) |
| ENSG00000261221 | ZNF865 | 0,291332634 | 1,223770166 | 0,50228861 | 0,920543477 | zinc finger protein 865 |
| ENSG00000106686 | SPATA6L | 0,291303635 | 1,223745568 | 0,48045197 | 0,912847708 | spermatogenesis associated 6 like |
| ENSG00000164845 | FAM86FP | 0,291275472 | 1,223721679 | 0,77561168 | 0,978587255 | family with sequence similarity 86 member F, pseudogene |
| ENSG00000141068 | KSR1 | 0,29127251 | 1,223719167 | 0,34763818 | 0,842276143 | kinase suppressor of ras 1 |
| ENSG00000173598 | NUDT4 | 0,291267784 | 1,223715158 | 0,48804985 | 0,915188998 | nudix hydrolase 4 |
| ENSG00000270605 | AL353622,1 | 0,291224791 | 1,223678691 | 0,86955189 | 0,989074806 | novel transcript |
| ENSG00000241889 | AC079944,2 | 0,291192611 | 1,223651397 | 0,6560044 | 0,957444211 | dullard homolog (Xenopus laevis) (DULLARD) pseudogene |
| ENSG00000287236 | Z73420,1 | 0,291035305 | 1,223517982 | 0,72352089 | 0,972913663 | novel transcript, antisense to ANKRD54 |
| ENSG00000188215 | DCUN1D3 | 0,290917221 | 1,223417842 | 0,20907611 | 0,71355079 | defective in cullin neddylation 1 domain containing 3 |
| ENSG00000130803 | ZNF317 | 0,290890884 | 1,223395508 | 0,17639163 | 0,662268274 | zinc finger protein 317 |
| ENSG00000122376 | SHLD2 | 0,290678825 | 1,223215696 | 0,49368161 | 0,917289924 | shieldin complex subunit 2 |
| ENSG00000080189 | SLC35C2 | 0,290548244 | 1,223104986 | 0,2680593 | 0,780680587 | solute carrier family 35 member C2 |
| ENSG00000236810 | ELOA-AS1 | 0,290535875 | 1,2230945 | 0,54159779 | 0,933491609 | ELOA antisense RNA 1 |
| ENSG00000101333 | PLCB4 | 0,290505691 | 1,22306891 | 0,38016671 | 0,865661467 | phospholipase C beta 4 |
| ENSG00000113615 | SEC24A | 0,290459885 | 1,223030078 | 0,4312691 | 0,89530663 | SEC24 homolog A, COPII coat complex component |
| ENSG00000224846 | NQO2-AS1 | 0,290290659 | 1,222886627 | 0,7815658 | 0,978597905 | NQO2 antisense RNA 1 |

| | | | | | | |
|---|---|---|---|---|---|---|
| ENSG00000064651 | SLC12A2 | 0,290249849 | 1,222852036 | 0,37056222 | 0,858545694 | solute carrier family 12 member 2 |
| ENSG00000225156 | AC012354,1 | 0,290191701 | 1,222802749 | 0,53504907 | 0,931805607 | novel transcript |
| ENSG00000133193 | FAM104A | 0,290173655 | 1,222787454 | 0,39252967 | 0,870909282 | family with sequence similarity 104 member A |
| ENSG00000284602 | AL031432,4 | 0,290171642 | 1,222785747 | 0,53926228 | 0,933350549 | novel transcript |
| ENSG00000136045 | PWP1 | 0,290123006 | 1,222744526 | 0,3338051 | 0,834152569 | PWP1 homolog, endonuclein |
| ENSG00000115155 | OTOF | 0,290110961 | 1,222734318 | 0,74581642 | 0,975707496 | otoferlin |
| ENSG00000237356 | AL365295,1 | 0,290067458 | 1,222697447 | 0,50838368 | 0,922427526 | novel transcript |
| ENSG00000129968 | ABHD17A | 0,290059208 | 1,222690455 | 0,49158353 | 0,916995341 | abhydrolase domain containing 17A, depalmitoylase |
| ENSG00000084073 | ZMPSTE24 | 0,290017108 | 1,222654776 | 0,45455264 | 0,903383444 | zinc metallopeptidase STE24 |
| ENSG00000250501 | AC093766,1 | 0,289992975 | 1,222634325 | 0,76875625 | 0,977381742 | novel transcript |
| ENSG00000278112 | AC145423,2 | 0,289979609 | 1,222622997 | 0,74975221 | 0,975901808 | novel transcript, sense intronic to RILPL1 |
| ENSG00000198961 | PJA2 | 0,289951354 | 1,222599053 | 0,41701548 | 0,886616509 | praja ring finger ubiquitin ligase 2 |
| ENSG00000187815 | ZFP69 | 0,289856429 | 1,222518612 | 0,38182342 | 0,866412204 | ZFP69 zinc finger protein |
| ENSG00000167548 | KMT2D | 0,289730592 | 1,222411984 | 0,44857714 | 0,902062594 | lysine methyltransferase 2D |
| ENSG00000108883 | EFTUD2 | 0,289705346 | 1,222390593 | 0,17301234 | 0,657758111 | elongation factor Tu GTP binding domain containing 2 |
| ENSG00000113312 | TTC1 | 0,289684521 | 1,222372948 | 0,29474289 | 0,804859966 | tetratricopeptide repeat domain 1 |
| ENSG00000278720 | AL669942,2 | 0,289582625 | 1,222286616 | 0,84144485 | 0,985672131 | zinc finger protein 114 (ZNF114) pseudogene |
| ENSG00000215399 | HMGB3P7 | 0,289545304 | 1,222254997 | 0,82541232 | 0,983383773 | high mobility group box 3 pseudogene 7 |
| ENSG00000153786 | ZDHHC7 | 0,289388937 | 1,22212253 | 0,18693438 | 0,680521261 | zinc finger DHHC-type palmitoyltransferase 7 |
| ENSG00000240210 | AL122013,1 | 0,289202411 | 1,221964532 | 0,82457892 | 0,983374097 | ribosomal protein L7a (RPL7A) pseudogene |
| ENSG00000255159 | AC026894,1 | 0,289174943 | 1,221941267 | 0,66847642 | 0,961351573 | novel transcript |
| ENSG00000163463 | KRTCAP2 | 0,289036641 | 1,221824133 | 0,21312368 | 0,720449383 | keratinocyte associated protein 2 |
| ENSG00000167193 | CRK | 0,289008264 | 1,2218001 | 0,32849743 | 0,830659639 | CRK proto-oncogene, adaptor protein |
| ENSG00000260563 | AC132872,1 | 0,288961531 | 1,221760524 | 0,65657565 | 0,957444211 | novel transcript |
| ENSG00000108021 | TASOR2 | 0,288868793 | 1,22168199 | 0,29577489 | 0,805426883 | transcription activation suppressor family member 2 |
| ENSG00000175324 | LSM1 | 0,288746247 | 1,221578221 | 0,35679154 | 0,848268504 | LSM1 homolog, mRNA degradation associated |
| ENSG00000236003 | AC007731,3 | 0,288738977 | 1,221572066 | 0,78171078 | 0,978597905 | novel transcript |
| ENSG00000232626 | AC099336,1 | 0,288642827 | 1,221490655 | 0,69270815 | 0,967271667 | FK506 binding protein 8, 38kDa (FKBP8) pseudogene |
| ENSG00000182858 | ALG12 | 0,288566785 | 1,221426275 | 0,22292006 | 0,731402952 | ALG12 alpha-1,6-mannosyltransferase |
| ENSG00000254003 | SRI-AS1 | 0,28843842 | 1,221317602 | 0,6652495 | 0,960743128 | SRI antisense RNA 1 |
| ENSG00000206490 | ABCF1 | 0,288401042 | 1,22128596 | 0,43448611 | 0,896272827 | ATP binding cassette subfamily F member 1 |
| ENSG00000162736 | NCSTN | 0,288374261 | 1,221263289 | 0,27664708 | 0,788353146 | nicastrin |

| | | | | | | |
|---|---|---|---|---|---|---|
| ENSG00000100644 | HIF1A | 0,288246747 | 1,221155352 | 0,39429301 | 0,871794663 | hypoxia inducible factor 1 subunit alpha |
| ENSG00000271275 | AC007326,1 | 0,288015605 | 1,220959719 | 0,73258613 | 0,974353474 | POM121 transmembrane nucleoporin (POM121) pseudogene |
| ENSG00000247077 | PGAM5 | 0,287883059 | 1,22084755 | 0,39024926 | 0,870057277 | PGAM family member 5, mitochondrial serine/threonine protein phosphatase |
| ENSG00000172590 | MRPL52 | 0,287839441 | 1,22081064 | 0,23757372 | 0,746907875 | mitochondrial ribosomal protein L52 |
| ENSG00000287115 | AC009951,6 | 0,287807915 | 1,220783963 | 0,7176937 | 0,971697135 | novel transcript |
| ENSG00000184414 | IRS3P | 0,287790807 | 1,220769486 | 0,68109782 | 0,964654539 | insulin receptor substrate 3, pseudogene |
| ENSG00000262826 | INTS3 | 0,287752498 | 1,220737071 | 0,23761078 | 0,746907875 | integrator complex subunit 3 |
| ENSG00000168591 | TMUB2 | 0,287706175 | 1,220697875 | 0,34750693 | 0,842205934 | transmembrane and ubiquitin like domain containing 2 |
| ENSG00000110921 | MVK | 0,287655604 | 1,220655087 | 0,22592414 | 0,734959876 | mevalonate kinase |
| ENSG00000100591 | AHSA1 | 0,287567815 | 1,220580811 | 0,15887017 | 0,63313153 | activator of HSP90 ATPase activity 1 |
| ENSG00000244005 | NFS1 | 0,287499968 | 1,220523411 | 0,21191691 | 0,718553023 | NFS1 cysteine desulfurase |
| ENSG00000147955 | SIGMAR1 | 0,287483359 | 1,22050936 | 0,26551961 | 0,776824277 | sigma non-opioid intracellular receptor 1 |
| ENSG00000232926 | AC000078,1 | 0,287475247 | 1,220502498 | 0,78658196 | 0,979320406 | ribosomal protein L8 (RPL8) pseudogene |
| ENSG00000106635 | BCL7B | 0,287241729 | 1,22030496 | 0,24236437 | 0,752610231 | BAF chromatin remodeling complex subunit BCL7B |
| ENSG00000101246 | ARFRP1 | 0,287229703 | 1,220294788 | 0,53458673 | 0,93162317 | ADP ribosylation factor related protein 1 |
| ENSG00000101542 | CDH20 | 0,287223318 | 1,220289387 | 0,37354022 | 0,861319894 | cadherin 20 |
| ENSG00000173456 | RNF26 | 0,287203921 | 1,22027298 | 0,30176177 | 0,811332971 | ring finger protein 26 |
| ENSG00000137760 | ALKBH8 | 0,287099965 | 1,220185055 | 0,31341851 | 0,819801802 | alkB homolog 8, tRNA methyltransferase |
| ENSG00000188542 | DUSP28 | 0,287055875 | 1,220147765 | 0,31126124 | 0,817751312 | dual specificity phosphatase 28 |
| ENSG00000165355 | FBXO33 | 0,287021824 | 1,220118967 | 0,46152151 | 0,906165604 | F-box protein 33 |
| ENSG00000177119 | ANO6 | 0,286922652 | 1,220035098 | 0,36937486 | 0,857991124 | anoctamin 6 |
| ENSG00000274180 | NATD1 | 0,286865381 | 1,219986667 | 0,2967798 | 0,806633854 | N-acetyltransferase domain containing 1 |
| ENSG00000010017 | RANBP9 | 0,286806972 | 1,219937276 | 0,40413713 | 0,878036356 | RAN binding protein 9 |
| ENSG00000271797 | AC008494,3 | 0,286763424 | 1,219900452 | 0,5614915 | 0,940653209 | novel transcript, antisense to PGGT1B |
| ENSG00000140995 | DEF8 | 0,286660604 | 1,219813514 | 0,33823383 | 0,837263082 | differentially expressed in FDCP 8 homolog |
| ENSG00000197555 | SIPA1L1 | 0,286538873 | 1,219710593 | 0,36920104 | 0,857942508 | signal induced proliferation associated 1 like 1 |
| ENSG00000168936 | TMEM129 | 0,286533349 | 1,219705923 | 0,44584448 | 0,901173082 | transmembrane protein 129, E3 ubiquitin ligase |
| ENSG00000135392 | DNAJC14 | 0,286514476 | 1,219689967 | 0,30070215 | 0,810374278 | DnaJ heat shock protein family (Hsp40) member C14 |
| ENSG00000111711 | GOLT1B | 0,286491244 | 1,219670327 | 0,60059689 | 0,949439306 | golgi transport 1B |
| ENSG00000106052 | TAX1BP1 | 0,286457226 | 1,219641568 | 0,35743809 | 0,848693232 | Tax1 binding protein 1 |

| | | | | | | |
|---|---|---|---|---|---|---|
| ENSG00000111358 | GTF2H3 | 0,286307565 | 1,219515052 | 0,18356037 | 0,674680375 | general transcription factor IIH subunit 3 |
| ENSG00000126756 | UXT | 0,286295374 | 1,219504747 | 0,33493508 | 0,834858938 | ubiquitously expressed prefoldin like chaperone |
| ENSG00000089693 | MLF2 | 0,286247931 | 1,219464644 | 0,25414456 | 0,765569035 | myeloid leukemia factor 2 |
| ENSG00000237004 | ZNRF2P1 | 0,286198147 | 1,219422564 | 0,61452491 | 0,952793934 | zinc and ring finger 2 pseudogene 1 |
| ENSG00000157693 | TMEM268 | 0,286096105 | 1,219336317 | 0,20337993 | 0,703503738 | transmembrane protein 268 |
| ENSG00000152778 | IFIT5 | 0,286017252 | 1,219269674 | 0,50006679 | 0,92041337 | interferon induced protein with tetratricopeptide repeats 5 |
| ENSG00000184698 | OR51M1 | 0,285985148 | 1,219242543 | 0,76339766 | 0,976640468 | olfactory receptor family 51 subfamily M member 1 |
| ENSG00000170265 | ZNF282 | 0,285906118 | 1,219175754 | 0,47383344 | 0,910337894 | zinc finger protein 282 |
| ENSG00000284967 | FDFT1 | 0,285879467 | 1,219153233 | 0,34239958 | 0,840048641 | farnesyl-diphosphate farnesyltransferase 1 |
| ENSG00000103254 | ANTKMT | 0,285839684 | 1,219119615 | 0,54334989 | 0,934188893 | adenine nucleotide translocase lysine methyltransferase |
| ENSG00000166886 | NAB2 | 0,285815106 | 1,219098846 | 0,42797735 | 0,89282532 | NGFI-A binding protein 2 |
| ENSG00000216639 | AL133406,1 | 0,285753334 | 1,219046649 | 0,75872717 | 0,976120207 | ribosomal protein L7 (RPL7) pseudogene |
| ENSG00000283189 | AC104452,1 | 0,285720167 | 1,219018623 | 0,49500855 | 0,917691373 | novel protein |
| ENSG00000079999 | KEAP1 | 0,28570116 | 1,219002564 | 0,32170999 | 0,824933727 | kelch like ECH associated protein 1 |
| ENSG00000285248 | IPO4 | 0,28562463 | 1,218937902 | 0,47484674 | 0,91069468 | importin 4 |
| ENSG00000196497 | IPO4 | 0,28562463 | 1,218937902 | 0,47484674 | 0,91069468 | importin 4 |
| ENSG00000286691 | AL590009,1 | 0,285613694 | 1,218928662 | 0,82669438 | 0,983548538 | novel transcript |
| ENSG00000148600 | CDHR1 | 0,285490148 | 1,218824282 | 0,44145289 | 0,898921877 | cadherin related family member 1 |
| ENSG00000073169 | SELENOO | 0,285406055 | 1,21875324 | 0,5112925 | 0,922893865 | selenoprotein O |
| ENSG00000104529 | EEF1D | 0,285322519 | 1,218682673 | 0,42916262 | 0,894012535 | eukaryotic translation elongation factor 1 delta |
| ENSG00000180881 | CAPS2 | 0,285317341 | 1,218678299 | 0,44617407 | 0,901348735 | calcyphosine 2 |
| ENSG00000220842 | RPL21P16 | 0,285284115 | 1,218650233 | 0,89980017 | 0,991845106 | ribosomal protein L21 pseudogene 16 |
| ENSG00000189350 | TOGARAM2 | 0,285161327 | 1,218546518 | 0,66363613 | 0,959920886 | TOG array regulator of axonemal microtubules 2 |
| ENSG00000247228 | AC009060,1 | 0,285010582 | 1,218419201 | 0,56448753 | 0,940865625 | novel transcript, antisense to PDPR & CLEC18C |
| ENSG00000238198 | LRIG2-DT | 0,284956617 | 1,218373625 | 0,66067019 | 0,959226315 | LRIG2 divergent transcript |
| ENSG00000164776 | PHKG1 | 0,284715589 | 1,218170091 | 0,56174727 | 0,940728834 | phosphorylase kinase catalytic subunit gamma 1 |
| ENSG00000164933 | SLC25A32 | 0,284712016 | 1,218167075 | 0,42102261 | 0,888848798 | solute carrier family 25 member 32 |
| ENSG00000198087 | CD2AP | 0,284703258 | 1,21815968 | 0,32673698 | 0,82854287 | CD2 associated protein |
| ENSG00000285840 | AL353648,1 | 0,2844499 | 1,217945772 | 0,73326583 | 0,974353474 | novel transcript, antisense to HMGB1 |
| ENSG00000173599 | PC | 0,28443307 | 1,217931564 | 0,28882413 | 0,800014136 | pyruvate carboxylase |
| ENSG00000169627 | BOLA2B | 0,284426322 | 1,217925867 | 0,31547576 | 0,82082095 | bolA family member 2B |

| | | | | | | |
|---|---|---|---|---|---|---|
| ENSG00000075568 | TMEM131 | 0,284389283 | 1,217894599 | 0,35808655 | 0,849151846 | transmembrane protein 131 |
| ENSG00000267338 | AP000365,1 | 0,284382985 | 1,217889283 | 0,76897646 | 0,977406998 | novel transcript |
| ENSG00000224141 | MIR548XHG | 0,284291313 | 1,217811898 | 0,69437429 | 0,967417311 | MIR548X host gene |
| ENSG00000186162 | CIDECP1 | 0,284273134 | 1,217796553 | 0,35960574 | 0,849982153 | cell death inducing DFFA like effector c pseudogene 1 |
| ENSG00000169100 | SLC25A6 | 0,284262726 | 1,217787767 | 0,20602212 | 0,708745123 | solute carrier family 25 member 6 |
| ENSG00000227780 | ZBTB22 | 0,284232839 | 1,21776254 | 0,34622355 | 0,841168721 | zinc finger and BTB domain containing 22 |
| ENSG00000229253 | ZBTB22 | 0,284232839 | 1,21776254 | 0,34622355 | 0,841168721 | zinc finger and BTB domain containing 22 |
| ENSG00000160714 | UBE2Q1 | 0,284198176 | 1,217733282 | 0,19267338 | 0,689545952 | ubiquitin conjugating enzyme E2 Q1 |
| ENSG00000228760 | BAG6 | 0,284127424 | 1,217673563 | 0,38559037 | 0,869008435 | BAG cochaperone 6 |
| ENSG00000152601 | MBNL1 | 0,284123977 | 1,217670654 | 0,41448359 | 0,885579089 | muscleblind like splicing regulator 1 |
| ENSG00000127334 | DYRK2 | 0,284039123 | 1,217599037 | 0,43451761 | 0,896272827 | dual specificity tyrosine phosphorylation regulated kinase 2 |
| ENSG00000129675 | ARHGEF6 | 0,284010286 | 1,2175747 | 0,39778206 | 0,874263835 | Rac/Cdc42 guanine nucleotide exchange factor 6 |
| ENSG00000288146 | NOC4L | 0,283991671 | 1,217558989 | 0,39237196 | 0,870909282 | nucleolar complex associated 4 homolog |
| ENSG00000184967 | NOC4L | 0,283991671 | 1,217558989 | 0,39237196 | 0,870909282 | nucleolar complex associated 4 homolog |
| ENSG00000119616 | FCF1 | 0,283956192 | 1,217529047 | 0,30580446 | 0,814714157 | FCF1 rRNA-processing protein |
| ENSG00000282928 | MAP4K1 | 0,283908412 | 1,217488725 | 0,687128 | 0,965769168 | mitogen-activated protein kinase kinase kinase kinase 1 |
| ENSG00000268635 | AP003680,1 | 0,28385689 | 1,217445247 | 0,6085391 | 0,951511094 | novel transcript, antisense to PAK1 |
| ENSG00000138303 | ASCC1 | 0,283812508 | 1,217407795 | 0,35023849 | 0,843789477 | activating signal cointegrator 1 complex subunit 1 |
| ENSG00000240057 | AC078785,1 | 0,283784253 | 1,217383952 | 0,65299835 | 0,95732962 | uncharacterized LOC107986114 |
| ENSG00000171462 | DLK2 | 0,283686521 | 1,217301486 | 0,66992916 | 0,962026636 | delta like non-canonical Notch ligand 2 |
| ENSG00000115806 | GORASP2 | 0,283660881 | 1,217279852 | 0,36166171 | 0,851893059 | golgi reassembly stacking protein 2 |
| ENSG00000087266 | SH3BP2 | 0,283645946 | 1,217267251 | 0,36636634 | 0,856103393 | SH3 domain binding protein 2 |
| ENSG00000236184 | TCEA1P4 | 0,283630072 | 1,217253857 | 0,60544058 | 0,950927937 | transcription elongation factor A1 pseudogene 4 |
| ENSG00000258315 | C17orf49 | 0,28349171 | 1,217137122 | 0,22149766 | 0,729254784 | chromosome 17 open reading frame 49 |
| ENSG00000138031 | ADCY3 | 0,28347937 | 1,217126711 | 0,34942925 | 0,843190352 | adenylate cyclase 3 |
| ENSG00000228056 | CFL1P3 | 0,283461191 | 1,217111374 | 0,81212256 | 0,981213877 | cofilin 1 pseudogene 3 |
| ENSG00000104881 | PPP1R13L | 0,283441814 | 1,217095027 | 0,56496748 | 0,940865625 | protein phosphatase 1 regulatory subunit 13 like |
| ENSG00000147124 | ZNF41 | 0,283380256 | 1,217043097 | 0,36679883 | 0,856302171 | zinc finger protein 41 |
| ENSG00000287271 | AC011455,8 | 0,283214607 | 1,216903364 | 0,7025641 | 0,968742626 | novel transcript, antisense to FBXO17and FBXO27 |
| ENSG00000249096 | LINC02362 | 0,283086359 | 1,216795193 | 0,6834962 | 0,965081074 | long intergenic non-protein coding RNA 2362 |

| | | | | | | |
|---|---|---|---|---|---|---|
| ENSG00000131143 | COX4I1 | 0,283069973 | 1,216781373 | 0,1823707 | 0,672670499 | cytochrome c oxidase subunit 4I1 |
| ENSG00000044090 | CUL7 | 0,283036209 | 1,216752897 | 0,38948832 | 0,870057277 | cullin 7 |
| ENSG00000143256 | PFDN2 | 0,283023568 | 1,216742235 | 0,26739844 | 0,779594822 | prefoldin subunit 2 |
| ENSG00000142784 | WDTC1 | 0,282960941 | 1,216689418 | 0,22043388 | 0,728572782 | WD and tetratricopeptide repeats 1 |
| ENSG00000128512 | DOCK4 | 0,282940635 | 1,216672293 | 0,34157885 | 0,840048641 | dedicator of cytokinesis 4 |
| ENSG00000264619 | MSH5-SAPCD1 | 0,282836788 | 1,216584718 | 0,41319689 | 0,884663685 | MSH5-SAPCD1 readthrough (NMD candidate) |
| ENSG00000154608 | CEP170P1 | 0,282830889 | 1,216579745 | 0,71739058 | 0,971697135 | centrosomal protein 170 pseudogene 1 |
| ENSG00000285971 | AC023095,1 | 0,282809406 | 1,216561628 | 0,60022647 | 0,949340299 | novel transcript |
| ENSG00000143437 | ARNT | 0,282805969 | 1,21655873 | 0,32265926 | 0,824955371 | aryl hydrocarbon receptor nuclear translocator |
| ENSG00000214253 | FIS1 | 0,282742647 | 1,216505334 | 0,22679222 | 0,735383523 | fission, mitochondrial 1 |
| ENSG00000182117 | NOP10 | 0,282716613 | 1,216483382 | 0,39493852 | 0,871888817 | NOP10 ribonucleoprotein |
| ENSG00000079739 | PGM1 | 0,282584499 | 1,216371989 | 0,24583366 | 0,757752583 | phosphoglucomutase 1 |
| ENSG00000119772 | DNMT3A | 0,282206866 | 1,216053639 | 0,26729535 | 0,779594822 | DNA methyltransferase 3 alpha |
| ENSG00000280303 | ERICD | 0,282167254 | 1,21602025 | 0,53888403 | 0,933348247 | E2F1-regulated inhibitor of cell death |
| ENSG00000125780 | TGM3 | 0,282138229 | 1,215995786 | 0,82638346 | 0,983471137 | transglutaminase 3 |
| ENSG00000185019 | UBOX5 | 0,282132214 | 1,215990716 | 0,39578946 | 0,872265727 | U-box domain containing 5 |
| ENSG00000171862 | PTEN | 0,28208202 | 1,21594841 | 0,63483054 | 0,955158181 | phosphatase and tensin homolog |
| ENSG00000285943 | AC112128,1 | 0,282071423 | 1,215939479 | 0,32897741 | 0,830787537 | novel protein |
| ENSG00000132676 | DAP3 | 0,281939379 | 1,215828194 | 0,25030358 | 0,762441549 | death associated protein 3 |
| ENSG00000171109 | MFN1 | 0,281891899 | 1,215788181 | 0,39002309 | 0,870057277 | mitofusin 1 |
| ENSG00000282790 | AC068400,1 | 0,281854908 | 1,215757008 | 0,88020011 | 0,990268682 | novel transcript, antisense TADA2A |
| ENSG00000286835 | AL133477,1 | 0,281838348 | 1,215743053 | 0,57234746 | 0,942475308 | novel transcript, sense intronic to ZNF367 |
| ENSG00000135624 | CCT7 | 0,281668781 | 1,215600169 | 0,20993113 | 0,71495498 | chaperonin containing TCP1 subunit 7 |
| ENSG00000106615 | RHEB | 0,281611601 | 1,215551991 | 0,32043625 | 0,824366234 | Ras homolog, mTORC1 binding |
| ENSG00000268307 | LINC02560 | 0,281536161 | 1,21548843 | 0,8164852 | 0,981900869 | long intergenic non-protein coding RNA 2560 |
| ENSG00000272508 | AL136982,6 | 0,281391546 | 1,215366596 | 0,7135046 | 0,970812834 | novel transcript |
| ENSG00000108854 | SMURF2 | 0,28138952 | 1,21536489 | 0,25681536 | 0,769107677 | SMAD specific E3 ubiquitin protein ligase 2 |
| ENSG00000275672 | AC025580,3 | 0,281302849 | 1,215291878 | 0,66667027 | 0,9611116 | GATM antisense RNA 1 |
| ENSG00000259192 | AC109631,1 | 0,281283212 | 1,215275336 | 0,69459542 | 0,967417311 | CTD (carboxy-terminal domain, RNA polymerase II, polypeptide A) small phosphatase 2 (CTDSP2) pseudogene |
| ENSG00000284254 | HNRNPH1 | 0,281233123 | 1,215233144 | 0,36407928 | 0,854207909 | heterogeneous nuclear ribonucleoprotein H1 |
| ENSG00000121680 | PEX16 | 0,281224241 | 1,215225662 | 0,39031786 | 0,870057277 | peroxisomal biogenesis factor 16 |

| | | | | | | |
|---|---|---|---|---|---|---|
| ENSG00000263990 | AC004253,1 | 0,281202069 | 1,215206986 | 0,72329265 | 0,972913663 | novel transcript |
| ENSG00000231084 | RPL22P24 | 0,28101268 | 1,215047471 | 0,68456199 | 0,965415049 | ribosomal protein L22 pseudogene 24 |
| ENSG00000272638 | AC006027,1 | 0,281007176 | 1,215042836 | 0,60658331 | 0,950927937 | novel transcript, antisense to NOD1 |
| ENSG00000235410 | AL157896,1 | 0,280947639 | 1,214992694 | 0,77883132 | 0,978587255 | novel transcript |
| ENSG00000127837 | AAMP | 0,280944089 | 1,214989704 | 0,42582626 | 0,892133432 | angio associated migratory cell protein |
| ENSG00000287707 | AC115284,4 | 0,280915407 | 1,21496555 | 0,67635282 | 0,963580269 | novel transcript |
| ENSG00000001629 | ANKIB1 | 0,280907646 | 1,214959014 | 0,33721497 | 0,836323083 | ankyrin repeat and IBR domain containing 1 |
| ENSG00000269068 | AC009955,4 | 0,280878579 | 1,214934536 | 0,64243851 | 0,956633898 | novel transcript, antisense to OBSL1 |
| ENSG00000175183 | CSRP2 | 0,28087504 | 1,214931555 | 0,28773522 | 0,799107075 | cysteine and glycine rich protein 2 |
| ENSG00000259658 | AC027559,1 | 0,280833884 | 1,214896897 | 0,76468565 | 0,976640468 | ubiquitin-conjugating enzyme E2Q family member 2 (UBE2Q2) pseudogene |
| ENSG00000084754 | HADHA | 0,28081487 | 1,214880886 | 0,31050113 | 0,817751312 | hydroxyacyl-CoA dehydrogenase trifunctional multienzyme complex subunit alpha |
| ENSG00000144043 | TEX261 | 0,280788552 | 1,214858724 | 0,19072976 | 0,685985838 | testis expressed 261 |
| ENSG00000284738 | AL358472,5 | 0,280774991 | 1,214847304 | 0,62821202 | 0,952793934 | novel transcript, antisense to DENND4B |
| ENSG00000270800 | RPS10-NUDT3 | 0,280591662 | 1,214692939 | 0,35699339 | 0,848610955 | RPS10-NUDT3 readthrough |
| ENSG00000235920 | THRAP3P3 | 0,280519277 | 1,214631995 | 0,55322494 | 0,936992468 | THRAP3 pseudogene 3 |
| ENSG00000143110 | C1orf162 | 0,28037774 | 1,214512838 | 0,74434537 | 0,975641819 | chromosome 1 open reading frame 162 |
| ENSG00000267322 | SNHG22 | 0,280376717 | 1,214511977 | 0,48410968 | 0,913690171 | small nucleolar RNA host gene 22 |
| ENSG00000267049 | AC002398,1 | 0,280343354 | 1,214483891 | 0,70615105 | 0,969503793 | novel transcript |
| ENSG00000203499 | IQANK1 | 0,280290329 | 1,214439254 | 0,75702494 | 0,975901808 | IQ motif and ankyrin repeat containing 1 |
| ENSG00000241399 | CD302 | 0,280256162 | 1,214410493 | 0,49476455 | 0,91758518 | CD302 molecule |
| ENSG00000287118 | AC008056,3 | 0,280247706 | 1,214403376 | 0,76532468 | 0,976830764 | novel transcript, antisense to NRXN3 |
| ENSG00000101161 | PRPF6 | 0,280167402 | 1,214335781 | 0,45808045 | 0,904140862 | pre-mRNA processing factor 6 |
| ENSG00000068654 | POLR1A | 0,280137141 | 1,21431031 | 0,29732959 | 0,806635068 | RNA polymerase I subunit A |
| ENSG00000257315 | ZBED6 | 0,2801094 | 1,214286961 | 0,43148078 | 0,89530663 | zinc finger BED-type containing 6 |
| ENSG00000270909 | CR788268,1 | 0,280104239 | 1,214282616 | 0,51700392 | 0,925746338 | CAS1 domain containing 1 (CASD1) pseudogene |
| ENSG00000260628 | AC142381,3 | 0,280079969 | 1,214262189 | 0,83650174 | 0,985423106 | Rho GTPase activating protein 23 (ARHGAP23) pseudogene |
| ENSG00000113583 | C5orf15 | 0,279851591 | 1,214069987 | 0,53256126 | 0,931291085 | chromosome 5 open reading frame 15 |
| ENSG00000104852 | SNRNP70 | 0,279830735 | 1,214052437 | 0,53823533 | 0,932900071 | small nuclear ribonucleoprotein U1 subunit 70 |
| ENSG00000127720 | METTL25 | 0,279822534 | 1,214045535 | 0,43853721 | 0,897220559 | methyltransferase like 25 |
| ENSG00000091640 | SPAG7 | 0,279812693 | 1,214037254 | 0,29715114 | 0,806635068 | sperm associated antigen 7 |

| | | | | | | |
|---|---|---|---|---|---|---|
| ENSG00000120451 | SNX19 | 0,279775679 | 1,214006107 | 0,15637705 | 0,627418665 | sorting nexin 19 |
| ENSG00000160201 | U2AF1 | 0,27973743 | 1,213973921 | 0,46845455 | 0,907940653 | U2 small nuclear RNA auxiliary factor 1 |
| ENSG00000118900 | UBN1 | 0,27959675 | 1,21385555 | 0,24754042 | 0,759778489 | ubinuclein 1 |
| ENSG00000164983 | TMEM65 | 0,279550948 | 1,213817014 | 0,43098485 | 0,89530663 | transmembrane protein 65 |
| ENSG00000253618 | GRPEL2-AS1 | 0,279401437 | 1,213691228 | 0,81228655 | 0,981213877 | GRPEL2 antisense RNA 1 |
| ENSG00000166963 | MAP1A | 0,279389248 | 1,213680974 | 0,42619368 | 0,892133432 | microtubule associated protein 1A |
| ENSG00000132256 | TRIM5 | 0,279373492 | 1,21366772 | 0,2033116 | 0,703503738 | tripartite motif containing 5 |
| ENSG00000174903 | RAB1B | 0,279311061 | 1,213615201 | 0,30370177 | 0,812795736 | RAB1B, member RAS oncogene family |
| ENSG00000112079 | STK38 | 0,279252755 | 1,213566154 | 0,33474143 | 0,834757602 | serine/threonine kinase 38 |
| ENSG00000203435 | E2F3P2 | 0,279183001 | 1,21350748 | 0,67274717 | 0,962371283 | E2F transcription factor 3 pseudogene 2 |
| ENSG00000261043 | AC019294,3 | 0,27914012 | 1,213471412 | 0,82626474 | 0,983471137 | novel transcript |
| ENSG00000257851 | HNRNPA3P10 | 0,279114661 | 1,213449998 | 0,68403608 | 0,965403332 | heterogeneous nuclear ribonucleoprotein A3 pseudogene 10 |
| ENSG00000277177 | LILRA6 | 0,279098859 | 1,213436707 | 0,73725494 | 0,974353474 | leukocyte immunoglobulin like receptor A6 |
| ENSG00000137825 | ITPKA | 0,27909584 | 1,213434168 | 0,66869492 | 0,961375797 | inositol-trisphosphate 3-kinase A |
| ENSG00000124217 | MOCS3 | 0,27903963 | 1,213386891 | 0,22451314 | 0,733203418 | molybdenum cofactor synthesis 3 |
| ENSG00000139370 | SLC15A4 | 0,278984897 | 1,213340858 | 0,2485265 | 0,760720365 | solute carrier family 15 member 4 |
| ENSG00000132603 | NIP7 | 0,278875006 | 1,213248441 | 0,2399482 | 0,749799162 | nucleolar pre-rRNA processing protein NIP7 |
| ENSG00000158805 | ZNF276 | 0,278651942 | 1,213060867 | 0,44275513 | 0,899437949 | zinc finger protein 276 |
| ENSG00000168517 | HEXIM2 | 0,278647691 | 1,213057293 | 0,4018521 | 0,877155908 | HEXIM P-TEFb complex subunit 2 |
| ENSG00000105700 | KXD1 | 0,278616169 | 1,213030788 | 0,25210769 | 0,764267906 | KxDL motif containing 1 |
| ENSG00000226328 | NUP50-DT | 0,278566199 | 1,212988774 | 0,37694069 | 0,862818448 | NUP50 divergent transcript |
| ENSG00000266648 | SETP3 | 0,278406114 | 1,212854185 | 0,69393653 | 0,967340633 | SET pseudogene 3 |
| ENSG00000142166 | IFNAR1 | 0,278316974 | 1,212779249 | 0,33017265 | 0,831800372 | interferon alpha and beta receptor subunit 1 |
| ENSG00000231509 | AL162253,1 | 0,278268712 | 1,212738679 | 0,86011739 | 0,989074806 | novel transcript |
| ENSG00000072062 | PRKACA | 0,278219278 | 1,212697125 | 0,27590978 | 0,787910415 | protein kinase cAMP-activated catalytic subunit alpha |
| ENSG00000277022 | AL031663,3 | 0,278218174 | 1,212696198 | 0,75307517 | 0,975901808 | novel transcript |
| ENSG00000091947 | TMEM101 | 0,278180132 | 1,212664221 | 0,37224205 | 0,860223901 | transmembrane protein 101 |
| ENSG00000184575 | XPOT | 0,278173029 | 1,21265825 | 0,47976576 | 0,912847708 | exportin for tRNA |
| ENSG00000047579 | DTNBP1 | 0,278127985 | 1,212620389 | 0,21034542 | 0,715586415 | dystrobrevin binding protein 1 |
| ENSG00000072195 | SPEG | 0,278127239 | 1,212619762 | 0,58097935 | 0,943914064 | striated muscle enriched protein kinase |
| ENSG00000099203 | TMED1 | 0,278117869 | 1,212611886 | 0,40741828 | 0,880392941 | transmembrane p24 trafficking protein 1 |

| | | | | | | |
|---|---|---|---|---|---|---|
| ENSG00000224994 | TRIM39 | 0,278074017 | 1,212575028 | 0,33369134 | 0,834001599 | tripartite motif containing 39 |
| ENSG00000229929 | TRIM39 | 0,278074017 | 1,212575028 | 0,33369134 | 0,834001599 | tripartite motif containing 39 |
| ENSG00000232839 | TRIM39 | 0,278074017 | 1,212575028 | 0,33369134 | 0,834001599 | tripartite motif containing 39 |
| ENSG00000204599 | TRIM39 | 0,278074017 | 1,212575028 | 0,33369134 | 0,834001599 | tripartite motif containing 39 |
| ENSG00000230308 | TRIM39 | 0,278074017 | 1,212575028 | 0,33369134 | 0,834001599 | tripartite motif containing 39 |
| ENSG00000248840 | AL645949,1 | 0,278055298 | 1,212559295 | 0,79198912 | 0,979471509 | X antigen family, member 5 (XAGE5) pseudogene |
| ENSG00000159792 | PSKH1 | 0,278005696 | 1,212517606 | 0,3461996 | 0,841168721 | protein serine kinase H1 |
| ENSG00000234636 | MED14OS | 0,27800567 | 1,212517584 | 0,53076547 | 0,931265667 | MED14 opposite strand |
| ENSG00000131480 | AOC2 | 0,277941276 | 1,212463465 | 0,48505156 | 0,913690171 | amine oxidase copper containing 2 |
| ENSG00000077454 | LRCH4 | 0,27792833 | 1,212452585 | 0,43315877 | 0,895845815 | leucine rich repeats and calponin homology domain containing 4 |
| ENSG00000196371 | FUT4 | 0,277928246 | 1,212452515 | 0,31969572 | 0,824366234 | fucosyltransferase 4 |
| ENSG00000180917 | CMTR2 | 0,277694025 | 1,212255689 | 0,49230228 | 0,916995341 | cap methyltransferase 2 |
| ENSG00000152082 | MZT2B | 0,277653513 | 1,212221649 | 0,42264151 | 0,890403187 | mitotic spindle organizing protein 2B |
| ENSG00000163644 | PPM1K | 0,277647557 | 1,212216645 | 0,23691968 | 0,746456145 | protein phosphatase, Mg2+/Mn2+ dependent 1K |
| ENSG00000091527 | CDV3 | 0,277628948 | 1,212201009 | 0,4241853 | 0,890949676 | CDV3 homolog |
| ENSG00000224675 | AC009227,1 | 0,277502257 | 1,212094563 | 0,77362629 | 0,978415098 | novel transcript |
| ENSG00000149534 | MS4A2 | 0,277371636 | 1,211984826 | 0,74683161 | 0,975796253 | membrane spanning 4-domains A2 |
| ENSG00000173548 | SNX33 | 0,277264122 | 1,211894509 | 0,32908616 | 0,830995037 | sorting nexin 33 |
| ENSG00000169040 | PMCHL2 | 0,277238745 | 1,211873191 | 0,88857141 | 0,990717729 | pro-melanin concentrating hormone like 2 (pseudogene) |
| ENSG00000175634 | RPS6KB2 | 0,277101024 | 1,21175751 | 0,28589305 | 0,797420923 | ribosomal protein S6 kinase B2 |
| ENSG00000188878 | FBF1 | 0,277067409 | 1,211729276 | 0,57324515 | 0,942475308 | Fas binding factor 1 |
| ENSG00000167670 | CHAF1A | 0,277021862 | 1,211691022 | 0,54871504 | 0,935949972 | chromatin assembly factor 1 subunit A |
| ENSG00000255153 | TOLLIP-AS1 | 0,276963443 | 1,211641957 | 0,58329114 | 0,944174844 | TOLLIP antisense RNA 1 (head to head) |
| ENSG00000120805 | ARL1 | 0,276961922 | 1,211640681 | 0,48775455 | 0,914986092 | ADP ribosylation factor like GTPase 1 |
| ENSG00000023228 | NDUFS1 | 0,276935394 | 1,211618401 | 0,50759643 | 0,922140988 | NADH:ubiquinone oxidoreductase core subunit S1 |
| ENSG00000205853 | RFPL3S | 0,27691345 | 1,211599972 | 0,66851138 | 0,961351573 | RFPL3 antisense |
| ENSG00000183722 | LHFPL6 | 0,276903928 | 1,211591975 | 0,32262706 | 0,824955371 | LHFPL tetraspan subfamily member 6 |
| ENSG00000182400 | TRAPPC6B | 0,276881295 | 1,211572968 | 0,59900462 | 0,948894897 | trafficking protein particle complex 6B |
| ENSG00000167977 | KCTD5 | 0,276817722 | 1,211519581 | 0,26245374 | 0,774068771 | potassium channel tetramerization domain containing 5 |
| ENSG00000234156 | AL359636,2 | 0,276756108 | 1,211467841 | 0,79483862 | 0,979776981 | novel transcript |

| | | | | | | |
|---|---|---|---|---|---|---|
| ENSG00000176853 | FAM91A1 | 0,276725133 | 1,21144183 | 0,46952996 | 0,908265361 | family with sequence similarity 91 member A1 |
| ENSG00000148218 | ALAD | 0,276716995 | 1,211434997 | 0,32616659 | 0,828184588 | aminolevulinate dehydratase |
| ENSG00000033170 | FUT8 | 0,276673337 | 1,211398338 | 0,40213204 | 0,87718264 | fucosyltransferase 8 |
| ENSG00000197892 | KIF13B | 0,27665631 | 1,21138404 | 0,32460967 | 0,826676737 | kinesin family member 13B |
| ENSG00000011114 | BTBD7 | 0,27664428 | 1,21137394 | 0,4824647 | 0,913223171 | BTB domain containing 7 |
| ENSG00000204560 | DHX16 | 0,276638108 | 1,211368757 | 0,3988191 | 0,875433675 | DEAH-box helicase 16 |
| ENSG00000196588 | MRTFA | 0,276637947 | 1,211368622 | 0,44945357 | 0,902062594 | myocardin related transcription factor A |
| ENSG00000121871 | SLITRK3 | 0,276603473 | 1,211339676 | 0,63359018 | 0,954859653 | SLIT and NTRK like family member 3 |
| ENSG00000270019 | AC110769,2 | 0,276495773 | 1,21124925 | 0,49477517 | 0,91758518 | novel transcript |
| ENSG00000269506 | AC110792,2 | 0,276362907 | 1,211137705 | 0,6724458 | 0,962290553 | novel transcript, antisense to CHIC2 |
| ENSG00000116977 | LGALS8 | 0,276357517 | 1,21113318 | 0,45644498 | 0,903545728 | galectin 8 |
| ENSG00000224774 | CSNK2B | 0,27633816 | 1,21111693 | 0,67923268 | 0,964199528 | casein kinase 2 beta |
| ENSG00000206406 | CSNK2B | 0,27633816 | 1,21111693 | 0,67923268 | 0,964199528 | casein kinase 2 beta |
| ENSG00000110076 | NRXN2 | 0,276240779 | 1,211035183 | 0,25497607 | 0,76642479 | neurexin 2 |
| ENSG00000243916 | AC092183,1 | 0,27618244 | 1,210986213 | 0,835875 | 0,985102385 | ribosomal protein S2 (RPS2) pseudogene |
| ENSG00000272779 | AC245060,4 | 0,276156119 | 1,210964119 | 0,26135066 | 0,77325059 | BMS1, ribosome biogenesis factor (BSM1) pseudogene |
| ENSG00000254936 | AF131215,4 | 0,276137031 | 1,210948098 | 0,67098719 | 0,962026636 | novel transcript |
| ENSG00000075945 | KIFAP3 | 0,276126225 | 1,210939027 | 0,42417084 | 0,890949676 | kinesin associated protein 3 |
| ENSG00000148331 | ASB6 | 0,275982522 | 1,210818415 | 0,33303691 | 0,833301711 | ankyrin repeat and SOCS box containing 6 |
| ENSG00000174945 | AMZ1 | 0,275685363 | 1,210569042 | 0,61161185 | 0,952155617 | archaelysin family metallopeptidase 1 |
| ENSG00000281856 | AC106017,2 | 0,275494467 | 1,210408872 | 0,76805135 | 0,977377956 | novel transcript |
| ENSG00000185245 | GP1BA | 0,275421195 | 1,210347399 | 0,63198367 | 0,953935145 | glycoprotein Ib platelet subunit alpha |
| ENSG00000174165 | ZDHHC24 | 0,275411142 | 1,210338965 | 0,4587114 | 0,904420115 | zinc finger DHHC-type containing 24 |
| ENSG00000287829 | AC002542,6 | 0,275343445 | 1,210282172 | 0,7416639 | 0,975176043 | novel transcript, antisense to ST7 |
| ENSG00000231530 | AL157932,1 | 0,27528022 | 1,210229134 | 0,77599247 | 0,978587255 | novel transcript |
| ENSG00000175376 | EIF1AD | 0,275211993 | 1,210171901 | 0,17698799 | 0,662915139 | eukaryotic translation initiation factor 1A domain containing |
| ENSG00000155287 | SLC25A28 | 0,275197047 | 1,210159365 | 0,25922926 | 0,771989931 | solute carrier family 25 member 28 |
| ENSG00000266680 | AL135905,1 | 0,275192393 | 1,210155461 | 0,68203168 | 0,964699524 | novel transcript, antisense to PTP4A1 |
| ENSG00000189241 | TSPYL1 | 0,275186505 | 1,210150522 | 0,19408532 | 0,691024186 | TSPY like 1 |
| ENSG00000120686 | UFM1 | 0,275161621 | 1,210129649 | 0,55602004 | 0,938031417 | ubiquitin fold modifier 1 |
| ENSG00000070371 | CLTCL1 | 0,275032934 | 1,210021712 | 0,27701307 | 0,788874154 | clathrin heavy chain like 1 |

| | | | | | | |
|---|---|---|---|---|---|---|
| ENSG00000095321 | CRAT | 0,274998563 | 1,209992884 | 0,35736856 | 0,848693232 | carnitine O-acetyltransferase |
| ENSG00000146555 | SDK1 | 0,274932247 | 1,209937266 | 0,37399183 | 0,861369706 | sidekick cell adhesion molecule 1 |
| ENSG00000166341 | DCHS1 | 0,2749139 | 1,209921879 | 0,38032273 | 0,865730737 | dachsous cadherin-related 1 |
| ENSG00000169093 | ASMTL | 0,274801091 | 1,209827275 | 0,41116393 | 0,883335414 | acetylserotonin O-methyltransferase like |
| ENSG00000182809 | CRIP2 | 0,274691629 | 1,209735485 | 0,39926048 | 0,875433675 | cysteine rich protein 2 |
| ENSG00000140105 | WARS1 | 0,274685407 | 1,209730268 | 0,26594958 | 0,777251371 | tryptophanyl-tRNA synthetase 1 |
| ENSG00000270300 | PHACTR2P1 | 0,27449014 | 1,209566543 | 0,45801379 | 0,904140862 | phosphatase and actin regulator 2 pseudogene 1 |
| ENSG00000187097 | ENTPD5 | 0,274306722 | 1,209412774 | 0,31955179 | 0,824366234 | ectonucleoside triphosphate diphosphohydrolase 5 (inactive) |
| ENSG00000078061 | ARAF | 0,274289419 | 1,209398269 | 0,33846219 | 0,837263082 | A-Raf proto-oncogene, serine/threonine kinase |
| ENSG00000183735 | TBK1 | 0,274177295 | 1,20930428 | 0,21352315 | 0,720983667 | TANK binding kinase 1 |
| ENSG00000130544 | ZNF557 | 0,274170245 | 1,20929837 | 0,23515207 | 0,744654651 | zinc finger protein 557 |
| ENSG00000178772 | CPN2 | 0,274130562 | 1,209265108 | 0,82218052 | 0,982952211 | carboxypeptidase N subunit 2 |
| ENSG00000068400 | GRIPAP1 | 0,274124648 | 1,209260151 | 0,55248642 | 0,936760964 | GRIP1 associated protein 1 |
| ENSG00000158186 | MRAS | 0,274114589 | 1,209251719 | 0,23416644 | 0,743699674 | muscle RAS oncogene homolog |
| ENSG00000130311 | DDA1 | 0,274044866 | 1,20919328 | 0,19811023 | 0,69678458 | DET1 and DDB1 associated 1 |
| ENSG00000231566 | LINC02595 | 0,273893007 | 1,209066006 | 0,57766327 | 0,94337651 | long intergenic non-protein coding RNA 2595 |
| ENSG00000224897 | POT1-AS1 | 0,273848122 | 1,209028391 | 0,45695995 | 0,903711377 | POT1 antisense RNA 1 |
| ENSG00000112110 | MRPL18 | 0,273838163 | 1,209020044 | 0,34695259 | 0,841625668 | mitochondrial ribosomal protein L18 |
| ENSG00000163541 | SUCLG1 | 0,273754777 | 1,208950167 | 0,2208773 | 0,728635155 | succinate-CoA ligase GDP/ADP-forming subunit alpha |
| ENSG00000188599 | NPIPP1 | 0,273750014 | 1,208946175 | 0,38276978 | 0,867356268 | nuclear pore complex interacting protein pseudogene 1 |
| ENSG00000140992 | PDPK1 | 0,273684569 | 1,208891334 | 0,19068546 | 0,685985838 | 3-phosphoinositide dependent protein kinase 1 |
| ENSG00000178342 | KCNG2 | 0,273574505 | 1,208799111 | 0,78662876 | 0,979320406 | potassium voltage-gated channel modifier subfamily G member 2 |
| ENSG00000117242 | PINK1-AS | 0,273447006 | 1,208692288 | 0,20212259 | 0,701895011 | PINK1 antisense RNA |
| ENSG00000171421 | MRPL36 | 0,273340099 | 1,208602724 | 0,4085192 | 0,88153752 | mitochondrial ribosomal protein L36 |
| ENSG00000188167 | TMPPE | 0,27327914 | 1,208551658 | 0,32872173 | 0,830659639 | transmembrane protein with metallophosphoesterase domain |
| ENSG00000213754 | KPNA4P1 | 0,273226011 | 1,208507152 | 0,68559791 | 0,965606498 | KPNA4 pseudogene 1 |
| ENSG00000135144 | DTX1 | 0,273142609 | 1,20843729 | 0,37916977 | 0,864650895 | deltex E3 ubiquitin ligase 1 |
| ENSG00000131507 | NDFIP1 | 0,273076308 | 1,208381756 | 0,44391321 | 0,900210555 | Nedd4 family interacting protein 1 |

| | | | | | | |
|---|---|---|---|---|---|---|
| ENSG00000148337 | CIZ1 | 0,273062975 | 1,208370589 | 0,26016896 | 0,772692192 | CDKN1A interacting zinc finger protein 1 |
| ENSG00000143322 | ABL2 | 0,272990021 | 1,208309486 | 0,18478477 | 0,677025387 | ABL proto-oncogene 2, non-receptor tyrosine kinase |
| ENSG00000061918 | GUCY1B1 | 0,272961092 | 1,208285256 | 0,54624212 | 0,93535572 | guanylate cyclase 1 soluble subunit beta 1 |
| ENSG00000130985 | UBA1 | 0,272928805 | 1,208258216 | 0,23460844 | 0,744134035 | ubiquitin like modifier activating enzyme 1 |
| ENSG00000231154 | MORF4L2-AS1 | 0,27275237 | 1,208110461 | 0,61923631 | 0,952793934 | MORF4L2 antisense RNA 1 |
| ENSG00000003402 | CFLAR | 0,272694134 | 1,208061695 | 0,24921488 | 0,761255882 | CASP8 and FADD like apoptosis regulator |
| ENSG00000135241 | PNPLA8 | 0,272609566 | 1,207990883 | 0,540456 | 0,933385624 | patatin like phospholipase domain containing 8 |
| ENSG00000144791 | LIMD1 | 0,272546803 | 1,207938331 | 0,41611793 | 0,885986046 | LIM domains containing 1 |
| ENSG00000170412 | GPRC5C | 0,272530275 | 1,207924493 | 0,46209589 | 0,906277266 | G protein-coupled receptor class C group 5 member C |
| ENSG00000230415 | LINC01786 | 0,272490425 | 1,207891128 | 0,79729748 | 0,980161849 | long intergenic non-protein coding RNA 1786 |
| ENSG00000147145 | LPAR4 | 0,272430512 | 1,207840968 | 0,61161561 | 0,952155617 | lysophosphatidic acid receptor 4 |
| ENSG00000262580 | AC087741,1 | 0,272393201 | 1,20780973 | 0,44274002 | 0,899437949 | novel transcript, antisense to CARD14 |
| ENSG00000240370 | RPL13P5 | 0,272345824 | 1,207770067 | 0,46490134 | 0,907804479 | ribosomal protein L13 pseudogene 5 |
| ENSG00000197893 | NRAP | 0,272280793 | 1,207715627 | 0,65285046 | 0,95732962 | nebulin related anchoring protein |
| ENSG00000232721 | AC239800,2 | 0,272241453 | 1,207682695 | 0,58026603 | 0,943828882 | novel transcript |
| ENSG00000255419 | MSH5-SAPCD1 | 0,272169994 | 1,207622878 | 0,43422416 | 0,896272827 | MSH5-SAPCD1 readthrough (NMD candidate) |
| ENSG00000115616 | SLC9A2 | 0,272136961 | 1,207595228 | 0,70239562 | 0,968742626 | solute carrier family 9 member A2 |
| ENSG00000088812 | ATRN | 0,272067766 | 1,207537311 | 0,27662028 | 0,788353146 | attractin |
| ENSG00000151458 | ANKRD50 | 0,271996766 | 1,207477885 | 0,44701348 | 0,901815148 | ankyrin repeat domain 50 |
| ENSG00000250282 | AC002401,2 | 0,271903502 | 1,207399829 | 0,83383105 | 0,984905093 | novel transcript, antisense to PDK2 |
| ENSG00000152127 | MGAT5 | 0,271705819 | 1,207234398 | 0,26419015 | 0,775595866 | alpha-1,6-mannosylglycoprotein 6-beta-N-acetylglucosaminyltransferase |
| ENSG00000242498 | ARPIN | 0,271691184 | 1,207222152 | 0,24969779 | 0,761653149 | actin related protein 2/3 complex inhibitor |
| ENSG00000106976 | DNM1 | 0,271668525 | 1,207203192 | 0,38360078 | 0,868000076 | dynamin 1 |
| ENSG00000258811 | AL583810,2 | 0,271654806 | 1,207191712 | 0,83493418 | 0,984905093 | novel transcript |
| ENSG00000116171 | SCP2 | 0,27154221 | 1,207097499 | 0,36509541 | 0,855173479 | sterol carrier protein 2 |
| ENSG00000023330 | ALAS1 | 0,271541252 | 1,207096698 | 0,2079227 | 0,712494639 | 5'-aminolevulinate synthase 1 |
| ENSG00000102125 | TAZ | 0,271531689 | 1,207088696 | 0,46299095 | 0,906849208 | tafazzin |
| ENSG00000167083 | GNGT2 | 0,271467953 | 1,20703537 | 0,77639982 | 0,978587255 | G protein subunit gamma transducin 2 |
| ENSG00000263368 | AC069366,1 | 0,271413442 | 1,206989765 | 0,78438678 | 0,979320406 | chromosome 9 open reading frame 30 (C9orf30) pseudogene |
| ENSG00000259751 | TUBAP11 | 0,271386308 | 1,206967064 | 0,68017524 | 0,964580199 | tubulin alpha pseudogene 11 |

| | | | | | | |
|---|---|---|---|---|---|---|
| ENSG00000182405 | PGBD4 | 0,271349716 | 1,206936451 | 0,49599865 | 0,918105073 | piggyBac transposable element derived 4 |
| ENSG00000203362 | POLH-AS1 | 0,271234553 | 1,206840112 | 0,63073179 | 0,953375088 | POLH antisense RNA 1 |
| ENSG00000258843 | AL133485,2 | 0,271179945 | 1,206794432 | 0,74705146 | 0,975796253 | novel transcript, antisense to NIN |
| ENSG00000224715 | Z82186,1 | 0,271077001 | 1,206708324 | 0,72759648 | 0,973533177 | novel transcript |
| ENSG00000248202 | LINC02234 | 0,270896984 | 1,206557762 | 0,71765077 | 0,971697135 | long intergenic non-protein coding RNA 2234 |
| ENSG00000251370 | SEMA5A-AS1 | 0,270876716 | 1,206540812 | 0,58225108 | 0,944174844 | SEMA5A antisense RNA 1 |
| ENSG00000198019 | FCGR1B | 0,270864257 | 1,206530392 | 0,76679421 | 0,977201891 | Fc fragment of IgG receptor Ib |
| ENSG00000155760 | FZD7 | 0,270776233 | 1,20645678 | 0,24977324 | 0,761653149 | frizzled class receptor 7 |
| ENSG00000266433 | TBC1D3P5 | 0,270758027 | 1,206441555 | 0,77963319 | 0,978587255 | TBC1 domain family member 3 pseudogene 5 |
| ENSG00000284705 | AL596257,1 | 0,270723879 | 1,206412999 | 0,80923239 | 0,981213877 | novel transcript |
| ENSG00000223502 | AL731537,1 | 0,270514228 | 1,206237697 | 0,84981127 | 0,987036295 | novel transcript |
| ENSG00000136950 | ARPC5L | 0,270507475 | 1,206232051 | 0,22606646 | 0,734959876 | actin related protein 2/3 complex subunit 5 like |
| ENSG00000264668 | AC138696,1 | 0,270473899 | 1,206203979 | 0,81528433 | 0,981521413 | novel protein |
| ENSG00000177369 | FLJ40194 | 0,270453583 | 1,206186993 | 0,74905355 | 0,975901808 | uncharacterized FLJ40194 |
| ENSG00000271826 | PLS3-AS1 | 0,270450561 | 1,206184467 | 0,64366019 | 0,956633898 | PLS3 antisense RNA 1 |
| ENSG00000101945 | SUV39H1 | 0,270449198 | 1,206183327 | 0,30874369 | 0,816730513 | suppressor of variegation 3-9 homolog 1 |
| ENSG00000260919 | AC100835,1 | 0,270416517 | 1,206156004 | 0,840992 | 0,985672131 | novel transcript, antisense to CLK3 |
| ENSG00000010404 | IDS | 0,270413626 | 1,206153587 | 0,24896482 | 0,76093812 | iduronate 2-sulfatase |
| ENSG00000175414 | ARL10 | 0,270356568 | 1,206105885 | 0,22904841 | 0,737173617 | ADP ribosylation factor like GTPase 10 |
| ENSG00000077147 | TM9SF3 | 0,270300848 | 1,206059303 | 0,49673087 | 0,918457927 | transmembrane 9 superfamily member 3 |
| ENSG00000267504 | AC090236,2 | 0,27013803 | 1,205923199 | 0,65279981 | 0,95732962 | novel transcript, sense intronic to NEDD4L |
| ENSG00000257294 | AC080011,1 | 0,270117552 | 1,205906082 | 0,7276899 | 0,973616511 | novel transcript |
| ENSG00000110777 | POU2AF1 | 0,270001127 | 1,20580877 | 0,69319533 | 0,967340633 | POU class 2 homeobox associating factor 1 |
| ENSG00000260834 | AC009055,2 | 0,269813054 | 1,205651588 | 0,65677934 | 0,957476526 | novel transcript |
| ENSG00000198937 | CCDC167 | 0,269807445 | 1,205646901 | 0,31474601 | 0,81991112 | coiled-coil domain containing 167 |
| ENSG00000100711 | ZFYVE21 | 0,269736164 | 1,205587333 | 0,34231214 | 0,840048641 | zinc finger FYVE-type containing 21 |
| ENSG00000103111 | MON1B | 0,269691572 | 1,20555007 | 0,22955261 | 0,73796738 | MON1 homolog B, secretory trafficking associated |
| ENSG00000034053 | APBA2 | 0,269587923 | 1,205463462 | 0,52449745 | 0,928616707 | amyloid beta precursor protein binding family A member 2 |
| ENSG00000214900 | LINC01588 | 0,269482272 | 1,205375187 | 0,61591889 | 0,952793934 | long intergenic non-protein coding RNA 1588 |
| ENSG00000125356 | NDUFA1 | 0,269345372 | 1,205260812 | 0,24630949 | 0,758198269 | NADH:ubiquinone oxidoreductase subunit A1 |
| ENSG00000286708 | AP003783,1 | 0,269305018 | 1,2052271 | 0,70725713 | 0,969565008 | novel transcript, antisense to SHANK2 |
| ENSG00000270761 | AL355353,1 | 0,269268398 | 1,205196508 | 0,82319619 | 0,983243765 | novel transcript |

| | | | | | | |
|---|---|---|---|---|---|---|
| ENSG00000259224 | SLC35G6 | 0,269160797 | 1,205106624 | 0,80129667 | 0,980557464 | solute carrier family 35 member G6 |
| ENSG00000284515 | SLC35G6 | 0,269160797 | 1,205106624 | 0,80129667 | 0,980557464 | solute carrier family 35 member G6 |
| ENSG00000185947 | ZNF267 | 0,269070591 | 1,205031276 | 0,4883436 | 0,915394579 | zinc finger protein 267 |
| ENSG00000272620 | AFAP1-AS1 | 0,269031985 | 1,20499903 | 0,41805312 | 0,887087337 | AFAP1 antisense RNA 1 |
| ENSG00000101782 | RIOK3 | 0,269010974 | 1,204981481 | 0,34417997 | 0,840938551 | RIO kinase 3 |
| ENSG00000103342 | GSPT1 | 0,268977209 | 1,204953279 | 0,35569434 | 0,847012548 | G1 to S phase transition 1 |
| ENSG00000130204 | TOMM40 | 0,268774189 | 1,204783728 | 0,36800063 | 0,857401877 | translocase of outer mitochondrial membrane 40 |
| ENSG00000130748 | TMEM160 | 0,2687134 | 1,204732963 | 0,58933703 | 0,946844281 | transmembrane protein 160 |
| ENSG00000248923 | MTND5P11 | 0,268658491 | 1,204687113 | 0,80370251 | 0,980875302 | MT-ND5 pseudogene 11 |
| ENSG00000099889 | ARVCF | 0,268643567 | 1,204674651 | 0,59957215 | 0,949259316 | ARVCF delta catenin family member |
| ENSG00000132704 | FCRL2 | 0,268440977 | 1,204505496 | 0,82771541 | 0,983595244 | Fc receptor like 2 |
| ENSG00000214770 | AL161756,1 | 0,268279188 | 1,204370426 | 0,64916726 | 0,956899963 | novel transcript |
| ENSG00000022840 | RNF10 | 0,26813421 | 1,204249404 | 0,25233935 | 0,76432863 | ring finger protein 10 |
| ENSG00000227440 | ATP5MC1P4 | 0,268112183 | 1,204231018 | 0,79183172 | 0,979471509 | ATP synthase membrane subunit c locus 1 pseudogene 4 |
| ENSG00000159173 | TNNI1 | 0,268109811 | 1,204229038 | 0,6956053 | 0,967669485 | troponin I1, slow skeletal type |
| ENSG00000125971 | DYNLRB1 | 0,26809621 | 1,204217685 | 0,25047947 | 0,762632703 | dynein light chain roadblock-type 1 |
| ENSG00000154099 | DNAAF1 | 0,267992642 | 1,20413124 | 0,6879922 | 0,966192982 | dynein axonemal assembly factor 1 |
| ENSG00000112531 | QKI | 0,267978443 | 1,204119389 | 0,51316995 | 0,92377973 | QKI, KH domain containing RNA binding |
| ENSG00000174238 | PITPNA | 0,267976656 | 1,204117897 | 0,22065197 | 0,728572782 | phosphatidylinositol transfer protein alpha |
| ENSG00000158417 | EIF5B | 0,267973449 | 1,204115221 | 0,44222854 | 0,899331131 | eukaryotic translation initiation factor 5B |
| ENSG00000110888 | CAPRIN2 | 0,267921344 | 1,204071734 | 0,29634629 | 0,806070152 | caprin family member 2 |
| ENSG00000226491 | FTOP1 | 0,267884311 | 1,204040826 | 0,75884455 | 0,976120207 | FTO pseudogene 1 |
| ENSG00000260772 | AC012321,1 | 0,267849052 | 1,2040114 | 0,81807421 | 0,982323352 | novel transcript |
| ENSG00000133422 | MORC2 | 0,267602031 | 1,203805264 | 0,37420227 | 0,861369706 | MORC family CW-type zinc finger 2 |
| ENSG00000111906 | HDDC2 | 0,267422791 | 1,203655713 | 0,39195423 | 0,870625724 | HD domain containing 2 |
| ENSG00000279738 | AL022311,1 | 0,267410417 | 1,20364539 | 0,63510267 | 0,955159215 | novel transcript, sense overlapping to EIF3L |
| ENSG00000233622 | CYP2T1P | 0,267236603 | 1,203500384 | 0,49730501 | 0,918759905 | cytochrome P450 family 2 subfamily T member 1, pseudogene |
| ENSG00000197744 | PTMAP2 | 0,267160148 | 1,203436608 | 0,81163219 | 0,981213877 | prothymosin alpha pseudogene 2 |
| ENSG00000285973 | AC063960,2 | 0,266918692 | 1,203235212 | 0,56825224 | 0,94163744 | novel transcript, antisense to TACC2 |
| ENSG00000140987 | ZSCAN32 | 0,266912322 | 1,203229899 | 0,25752283 | 0,769908546 | zinc finger and SCAN domain containing 32 |
| ENSG00000183340 | JRKL | 0,266907419 | 1,20322581 | 0,34798559 | 0,842451708 | JRK like |

| | | | | | | |
|---|---|---|---|---|---|---|
| ENSG00000224020 | MIR181A2HG | 0,266789526 | 1,203127489 | 0,48808327 | 0,915188998 | MIR181A2 host gene |
| ENSG00000136819 | C9orf78 | 0,266742766 | 1,203088495 | 0,2376545 | 0,746907875 | chromosome 9 open reading frame 78 |
| ENSG00000164244 | PRRC1 | 0,266668611 | 1,203026657 | 0,42065378 | 0,888569577 | proline rich coiled-coil 1 |
| ENSG00000155229 | MMS19 | 0,266622065 | 1,202987844 | 0,25394533 | 0,76531122 | MMS19 homolog, cytosolic iron-sulfur assembly component |
| ENSG00000166685 | COG1 | 0,266436726 | 1,20283331 | 0,24096275 | 0,751467261 | component of oligomeric golgi complex 1 |
| ENSG00000197102 | DYNC1H1 | 0,266372041 | 1,20277938 | 0,26136115 | 0,77325059 | dynein cytoplasmic 1 heavy chain 1 |
| ENSG00000100997 | ABHD12 | 0,266242977 | 1,202671784 | 0,28826516 | 0,799645584 | abhydrolase domain containing 12, lysophospholipase |
| ENSG00000142961 | MOB3C | 0,266129741 | 1,202577391 | 0,48475929 | 0,913690171 | MOB kinase activator 3C |
| ENSG00000163812 | ZDHHC3 | 0,266102291 | 1,202554511 | 0,23310028 | 0,742198503 | zinc finger DHHC-type palmitoyltransferase 3 |
| ENSG00000274166 | GOLGA6L10 | 0,266040149 | 1,202502713 | 0,67787913 | 0,963853615 | golgin A6 family like 10 |
| ENSG00000223760 | MED15P9 | 0,265825198 | 1,202323562 | 0,72366947 | 0,972971932 | mediator complex subunit 15 pseudogene 9 |
| ENSG00000099904 | ZDHHC8 | 0,26573333 | 1,202247003 | 0,46729616 | 0,907940653 | zinc finger DHHC-type palmitoyltransferase 8 |
| ENSG00000056558 | TRAF1 | 0,265672196 | 1,202196059 | 0,58293986 | 0,944174844 | TNF receptor associated factor 1 |
| ENSG00000102699 | PARP4 | 0,265599338 | 1,202135349 | 0,4014991 | 0,876898854 | poly(ADP-ribose) polymerase family member 4 |
| ENSG00000179832 | MROH1 | 0,265568268 | 1,202109459 | 0,55478206 | 0,937793851 | maestro heat like repeat family member 1 |
| ENSG00000229081 | LINC01165 | 0,265315851 | 1,201899154 | 0,77776711 | 0,978587255 | long intergenic non-protein coding RNA 1165 |
| ENSG00000106123 | EPHB6 | 0,265229473 | 1,201827196 | 0,59814323 | 0,948582224 | EPH receptor B6 |
| ENSG00000029534 | ANK1 | 0,265212552 | 1,201813099 | 0,54940325 | 0,936051084 | ankyrin 1 |
| ENSG00000115977 | AAK1 | 0,265206649 | 1,201808182 | 0,23496421 | 0,744644825 | AP2 associated kinase 1 |
| ENSG00000236035 | NSA2P1 | 0,265204786 | 1,20180663 | 0,75718385 | 0,975901808 | NSA2 pseudogene 1 |
| ENSG00000169180 | XPO6 | 0,265136734 | 1,201749942 | 0,24338908 | 0,753845477 | exportin 6 |
| ENSG00000259619 | AC010867,1 | 0,265091061 | 1,201711897 | 0,76039581 | 0,97624106 | intraflagellar transport 80 homolog (Chlamydomonas) (IFT80) pseudogene |
| ENSG00000117143 | UAP1 | 0,265059946 | 1,20168598 | 0,33775561 | 0,837066292 | UDP-N-acetylglucosamine pyrophosphorylase 1 |
| ENSG00000160818 | GPATCH4 | 0,265050522 | 1,201678131 | 0,49584889 | 0,918105073 | G-patch domain containing 4 |
| ENSG00000086544 | ITPKC | 0,26504114 | 1,201670316 | 0,49200542 | 0,916995341 | inositol-trisphosphate 3-kinase C |
| ENSG00000233143 | DIRC3-AS1 | 0,264949148 | 1,201593695 | 0,59237345 | 0,947416795 | DIRC3 antisense RNA 1 |
| ENSG00000243477 | NAA80 | 0,264792287 | 1,201463056 | 0,40327514 | 0,877589651 | N-alpha-acetyltransferase 80, NatH catalytic subunit |
| ENSG00000225386 | AC099754,1 | 0,264753469 | 1,201430729 | 0,71024545 | 0,970482408 | novel transcript |
| ENSG00000186716 | BCR | 0,264648236 | 1,201343097 | 0,44975614 | 0,902062594 | BCR activator of RhoGEF and GTPase |
| ENSG00000184014 | DENND5A | 0,264645343 | 1,201340688 | 0,21348467 | 0,720983667 | DENN domain containing 5A |

| | | | | | | |
|---|---|---|---|---|---|---|
| ENSG00000277578 | Metazoa_SRP | 0,264613614 | 1,201314268 | 0,81005046 | 0,981213877 | Metazoan signal recognition particle RNA |
| ENSG00000135407 | AVIL | 0,264471065 | 1,201195575 | 0,46763622 | 0,907940653 | advillin |
| ENSG00000203685 | STUM | 0,264425273 | 1,201157448 | 0,57283098 | 0,942475308 | stum, mechanosensory transduction mediator homolog |
| ENSG00000109519 | GRPEL1 | 0,264380829 | 1,201120446 | 0,26611584 | 0,77755567 | GrpE like 1, mitochondrial |
| ENSG00000085644 | ZNF213 | 0,26432588 | 1,201074699 | 0,5048586 | 0,921400408 | zinc finger protein 213 |
| ENSG00000116688 | MFN2 | 0,264275602 | 1,201032842 | 0,31740963 | 0,822252078 | mitofusin 2 |
| ENSG00000164100 | NDST3 | 0,264203052 | 1,200972447 | 0,47004825 | 0,908378389 | N-deacetylase and N-sulfotransferase 3 |
| ENSG00000183255 | PTTG1IP | 0,264137861 | 1,200918179 | 0,21228585 | 0,719057938 | PTTG1 interacting protein |
| ENSG00000127481 | UBR4 | 0,264092084 | 1,200880074 | 0,20913082 | 0,71355079 | ubiquitin protein ligase E3 component n-recognin 4 |
| ENSG00000117601 | SERPINC1 | 0,264022599 | 1,200822237 | 0,75848709 | 0,976120207 | serpin family C member 1 |
| ENSG00000248121 | SMURF2P1 | 0,264017003 | 1,20081758 | 0,75425342 | 0,975901808 | SMAD specific E3 ubiquitin protein ligase 2 pseudogene 1 |
| ENSG00000071242 | RPS6KA2 | 0,26391132 | 1,200729618 | 0,43360851 | 0,896108974 | ribosomal protein S6 kinase A2 |
| ENSG00000108963 | DPH1 | 0,263863139 | 1,200689519 | 0,33179047 | 0,832645466 | diphthamide biosynthesis 1 |
| ENSG00000071539 | TRIP13 | 0,263806211 | 1,200642142 | 0,37837114 | 0,864005583 | thyroid hormone receptor interactor 13 |
| ENSG00000175866 | BAIAP2 | 0,263723874 | 1,200573621 | 0,45303269 | 0,902428411 | BAR/IMD domain containing adaptor protein 2 |
| ENSG00000237125 | HAND2-AS1 | 0,263685348 | 1,20054156 | 0,71334108 | 0,970812834 | HAND2 antisense RNA 1 |
| ENSG00000253536 | AC138832,1 | 0,263633177 | 1,200498147 | 0,79398053 | 0,979776981 | sterol carrier protein (SCP2) pseudogene |
| ENSG00000274506 | AC138832,3 | 0,263633177 | 1,200498147 | 0,79398053 | 0,979776981 | sterol carrier protein (SCP2) pseudogene |
| ENSG00000162069 | BICDL2 | 0,26353707 | 1,200418177 | 0,81726325 | 0,982048938 | BICD family like cargo adaptor 2 |
| ENSG00000243155 | AL162431,1 | 0,263293678 | 1,200215676 | 0,6661417 | 0,961020116 | novel transcript |
| ENSG00000100593 | ISM2 | 0,263035463 | 1,200000879 | 0,83688048 | 0,985437134 | isthmin 2 |
| ENSG00000262171 | AC130651,1 | 0,262807956 | 1,199811659 | 0,80489201 | 0,98109454 | novel transcript, antisense to C16orf63 |
| ENSG00000282412 | AC243629,3 | 0,262807956 | 1,199811659 | 0,80489201 | 0,98109454 | novel transcript, antisense to C16orf63 |
| ENSG00000168546 | GFRA2 | 0,262798194 | 1,199803541 | 0,52851795 | 0,930033144 | GDNF family receptor alpha 2 |
| ENSG00000105135 | ILVBL | 0,262707999 | 1,199728533 | 0,45374791 | 0,902912642 | ilvB acetolactate synthase like |
| ENSG00000282951 | AC008537,4 | 0,262624172 | 1,199658825 | 0,75008663 | 0,975901808 | novel transcript, antisense to RAB4B |
| ENSG00000213445 | SIPA1 | 0,262462309 | 1,199524238 | 0,56493317 | 0,940865625 | signal-induced proliferation-associated 1 |
| ENSG00000167397 | VKORC1 | 0,262382607 | 1,199457971 | 0,26331419 | 0,774637968 | vitamin K epoxide reductase complex subunit 1 |
| ENSG00000275322 | AC103746,1 | 0,262307962 | 1,199395913 | 0,70531299 | 0,969441015 | novel transcript |
| ENSG00000229154 | KCNQ5-AS1 | 0,262184175 | 1,199293006 | 0,77016441 | 0,977843577 | KCNQ5 antisense RNA 1 |
| ENSG00000125144 | MT1G | 0,262080659 | 1,199206958 | 0,89396348 | 0,991135494 | metallothionein 1G |

| | | | | | | |
|---|---|---|---|---|---|---|
| ENSG00000226288 | OR52I2 | 0,26207937 | 1,199205886 | 0,72665727 | 0,973344786 | olfactory receptor family 52 subfamily I member 2 |
| ENSG00000182612 | TSPAN10 | 0,261970321 | 1,199115245 | 0,68502792 | 0,965415049 | tetraspanin 10 |
| ENSG00000144741 | SLC25A26 | 0,261776961 | 1,198954542 | 0,63873111 | 0,956018518 | solute carrier family 25 member 26 |
| ENSG00000144644 | GADL1 | 0,261709548 | 1,19889852 | 0,56235401 | 0,940865625 | glutamate decarboxylase like 1 |
| ENSG00000156103 | MMP16 | 0,261639457 | 1,198840275 | 0,54486623 | 0,93489368 | matrix metallopeptidase 16 |
| ENSG00000259728 | LINC00933 | 0,26154948 | 1,198765509 | 0,75372875 | 0,975901808 | long intergenic non-protein coding RNA 933 |
| ENSG00000267457 | AC004223,2 | 0,261454664 | 1,198686727 | 0,82709495 | 0,983567657 | novel transcript |
| ENSG00000137802 | MAPKBP1 | 0,261363126 | 1,198610673 | 0,37770506 | 0,863195661 | mitogen-activated protein kinase binding protein 1 |
| ENSG00000142867 | BCL10 | 0,261348452 | 1,198598482 | 0,45566231 | 0,903545728 | BCL10 immune signaling adaptor |
| ENSG00000182872 | RBM10 | 0,261343654 | 1,198594496 | 0,41250712 | 0,884523593 | RNA binding motif protein 10 |
| ENSG00000103710 | RASL12 | 0,261285979 | 1,19854658 | 0,57796949 | 0,943425812 | RAS like family 12 |
| ENSG00000072506 | HSD17B10 | 0,261231888 | 1,198501644 | 0,28863316 | 0,79998466 | hydroxysteroid 17-beta dehydrogenase 10 |
| ENSG00000068024 | HDAC4 | 0,261084251 | 1,198379003 | 0,38674223 | 0,869441674 | histone deacetylase 4 |
| ENSG00000286699 | AC084198,2 | 0,261059788 | 1,198358683 | 0,47078961 | 0,908869685 | novel transcript |
| ENSG00000260005 | AC027601,1 | 0,261009051 | 1,198316539 | 0,64429689 | 0,956633898 | novel transcript, antisense to C17orf56 |
| ENSG00000169299 | PGM2 | 0,26099459 | 1,198304528 | 0,40207111 | 0,87718264 | phosphoglucomutase 2 |
| ENSG00000178096 | BOLA1 | 0,260990228 | 1,198300905 | 0,4128461 | 0,884576962 | bolA family member 1 |
| ENSG00000221990 | EXOC3-AS1 | 0,260771528 | 1,198119267 | 0,45684742 | 0,903699469 | EXOC3 antisense RNA 1 |
| ENSG00000119725 | ZNF410 | 0,260679265 | 1,198042647 | 0,28671691 | 0,79806958 | zinc finger protein 410 |
| ENSG00000134001 | EIF2S1 | 0,26053377 | 1,197921831 | 0,38205354 | 0,866612629 | eukaryotic translation initiation factor 2 subunit alpha |
| ENSG00000285212 | AL009178,3 | 0,260473634 | 1,197871899 | 0,7592401 | 0,976120207 | novel transcript |
| ENSG00000204524 | ZNF805 | 0,260473143 | 1,197871491 | 0,21456537 | 0,722164151 | zinc finger protein 805 |
| ENSG00000225234 | TRAPPC12-AS1 | 0,260462613 | 1,197862749 | 0,54582607 | 0,935154874 | TRAPPC12 antisense RNA 1 |
| ENSG00000177426 | TGIF1 | 0,260414281 | 1,19782262 | 0,33588546 | 0,835660292 | TGFB induced factor homeobox 1 |
| ENSG00000176986 | SEC24C | 0,260392974 | 1,197804929 | 0,21361975 | 0,721033693 | SEC24 homolog C, COPII coat complex component |
| ENSG00000135940 | COX5B | 0,260228028 | 1,19766799 | 0,28648263 | 0,797701552 | cytochrome c oxidase subunit 5B |
| ENSG00000118729 | CASQ2 | 0,26018241 | 1,19763012 | 0,75319669 | 0,975901808 | calsequestrin 2 |
| ENSG00000144566 | RAB5A | 0,260171232 | 1,197620841 | 0,57526469 | 0,942667371 | RAB5A, member RAS oncogene family |
| ENSG00000177025 | C19orf18 | 0,260144653 | 1,197598777 | 0,7089464 | 0,970282889 | chromosome 19 open reading frame 18 |
| ENSG00000144747 | TMF1 | 0,260112865 | 1,19757239 | 0,3951595 | 0,871981986 | TATA element modulatory factor 1 |
| ENSG00000234264 | DEPDC1-AS1 | 0,26006562 | 1,197533173 | 0,78648158 | 0,979320406 | DEPDC1 antisense RNA 1 |
| ENSG00000158526 | TSR2 | 0,260052919 | 1,19752263 | 0,42975214 | 0,894607 | TSR2 ribosome maturation factor |

| | | | | | | |
|---|---|---|---|---|---|---|
| ENSG00000149972 | CNTN5 | 0,260018963 | 1,197494445 | 0,69576567 | 0,96776072 | contactin 5 |
| ENSG00000267898 | AC026803,2 | 0,25998796 | 1,197468711 | 0,78548889 | 0,979320406 | novel transcript |
| ENSG00000185475 | TMEM179B | 0,259826071 | 1,197334347 | 0,32617618 | 0,828184588 | transmembrane protein 179B |
| ENSG00000127870 | RNF6 | 0,259708053 | 1,197236405 | 0,44629204 | 0,901348735 | ring finger protein 6 |
| ENSG00000105219 | CCNP | 0,259703116 | 1,197232308 | 0,80640284 | 0,981213877 | cyclin P |
| ENSG00000168701 | TMEM208 | 0,259640827 | 1,197180618 | 0,27839976 | 0,789733543 | transmembrane protein 208 |
| ENSG00000151148 | UBE3B | 0,259590722 | 1,19713904 | 0,20181443 | 0,701461582 | ubiquitin protein ligase E3B |
| ENSG00000167695 | TLCD3A | 0,259584403 | 1,197133797 | 0,26815528 | 0,780692104 | TLC domain containing 3A |
| ENSG00000102471 | NDFIP2 | 0,259513297 | 1,197074795 | 0,54516424 | 0,934923038 | Nedd4 family interacting protein 2 |
| ENSG00000277401 | TJP1 | 0,259480187 | 1,197047323 | 0,56518428 | 0,940865625 | tight junction protein 1 |
| ENSG00000230225 | MTND5P14 | 0,259464445 | 1,197034261 | 0,84188125 | 0,98571999 | MT-ND5 pseudogene 14 |
| ENSG00000272068 | AL365181,2 | 0,259408011 | 1,196987438 | 0,75288822 | 0,975901808 | novel transcript |
| ENSG00000100319 | ZMAT5 | 0,259365385 | 1,196952071 | 0,41284996 | 0,884576962 | zinc finger matrin-type 5 |
| ENSG00000134108 | ARL8B | 0,259356299 | 1,196944534 | 0,52634889 | 0,929261495 | ADP ribosylation factor like GTPase 8B |
| ENSG00000136643 | RPS6KC1 | 0,259308157 | 1,196904593 | 0,42623455 | 0,892133432 | ribosomal protein S6 kinase C1 |
| ENSG00000260439 | LMF1-AS1 | 0,259300222 | 1,196898009 | 0,68006303 | 0,964580199 | LMF1 antisense RNA 1 |
| ENSG00000109680 | TBC1D19 | 0,259275955 | 1,196877877 | 0,44048216 | 0,898377869 | TBC1 domain family member 19 |
| ENSG00000103274 | NUBP1 | 0,259155115 | 1,196777631 | 0,24655792 | 0,758400921 | nucleotide binding protein 1 |
| ENSG00000171617 | ENC1 | 0,259002005 | 1,196650626 | 0,31601558 | 0,821394376 | ectodermal-neural cortex 1 |
| ENSG00000261253 | AC137932,2 | 0,258993552 | 1,196643615 | 0,6249561 | 0,952793934 | novel transcript, antisense to ANKRD11 |
| ENSG00000274671 | MARVELD2 | 0,258968497 | 1,196622833 | 0,69089079 | 0,967179044 | MARVEL domain containing 2 |
| ENSG00000156983 | BRPF1 | 0,25895034 | 1,196607773 | 0,2485756 | 0,760720365 | bromodomain and PHD finger containing 1 |
| ENSG00000174740 | PABPC5 | 0,258857486 | 1,196530761 | 0,44689328 | 0,901798674 | poly(A) binding protein cytoplasmic 5 |
| ENSG00000178307 | TMEM11 | 0,258856166 | 1,196529666 | 0,24203609 | 0,752344083 | transmembrane protein 11 |
| ENSG00000185670 | ZBTB3 | 0,258765023 | 1,196454076 | 0,3647012 | 0,854852166 | zinc finger and BTB domain containing 3 |
| ENSG00000268257 | AIRN | 0,258756305 | 1,196446847 | 0,60536599 | 0,950927937 | antisense of IGF2R non-protein coding RNA |
| ENSG00000267784 | AC010680,1 | 0,258741808 | 1,196434824 | 0,65543358 | 0,957444211 | novel transcript, antisense to titin |
| ENSG00000144736 | SHQ1 | 0,258592956 | 1,196311386 | 0,30325474 | 0,812669425 | SHQ1, H/ACA ribonucleoprotein assembly factor |
| ENSG00000255031 | AP002807,1 | 0,258591786 | 1,196310416 | 0,76435468 | 0,976640468 | novel transcript, antisense to CHKA |
| ENSG00000158079 | PTPDC1 | 0,258447414 | 1,196190706 | 0,44180408 | 0,899138311 | protein tyrosine phosphatase domain containing 1 |
| ENSG00000161920 | MED11 | 0,258425635 | 1,196172649 | 0,33844541 | 0,837263082 | mediator complex subunit 11 |
| ENSG00000259370 | AC103740,1 | 0,258403114 | 1,196153976 | 0,37797901 | 0,863636321 | uncharacterized LOC105370854 |
| ENSG00000161681 | SHANK1 | 0,258399439 | 1,19615093 | 0,53939738 | 0,933350549 | SH3 and multiple ankyrin repeat domains 1 |

| | | | | | | |
|---|---|---|---|---|---|---|
| ENSG00000188266 | HYKK | 0,258383929 | 1,19613807 | 0,63385809 | 0,954929949 | hydroxylysine kinase |
| ENSG00000262228 | AC087392,3 | 0,258380979 | 1,196135624 | 0,61281101 | 0,952413269 | novel transcript |
| ENSG00000135966 | TGFBRAP1 | 0,258335066 | 1,196097558 | 0,30414938 | 0,813056239 | transforming growth factor beta receptor associated protein 1 |
| ENSG00000268240 | AC123912,2 | 0,258273653 | 1,196046643 | 0,77633818 | 0,978587255 | novel transcript |
| ENSG00000108344 | PSMD3 | 0,258248898 | 1,196026121 | 0,24637419 | 0,758198269 | proteasome 26S subunit, non-ATPase 3 |
| ENSG00000188554 | NBR1 | 0,258248573 | 1,196025852 | 0,35393284 | 0,845346751 | NBR1 autophagy cargo receptor |
| ENSG00000171540 | OTP | 0,258240565 | 1,196019213 | 0,67095262 | 0,962026636 | orthopedia homeobox |
| ENSG00000286408 | AC113410,4 | 0,258144691 | 1,195939735 | 0,61168763 | 0,952161039 | novel transcript |
| ENSG00000283045 | AC103703,1 | 0,25811674 | 1,195916564 | 0,65244258 | 0,957275891 | novel transcript, sense intronic to GPATCH8 |
| ENSG00000113734 | BNIP1 | 0,258112277 | 1,195912864 | 0,33961409 | 0,838481794 | BCL2 interacting protein 1 |
| ENSG00000284814 | ADCY4 | 0,258047498 | 1,195859167 | 0,56050822 | 0,940170934 | adenylate cyclase 4 |
| ENSG00000162144 | CYB561A3 | 0,258013519 | 1,195831003 | 0,48352429 | 0,913690171 | cytochrome b561 family member A3 |
| ENSG00000112679 | DUSP22 | 0,257961259 | 1,195787686 | 0,32011201 | 0,824366234 | dual specificity phosphatase 22 |
| ENSG00000172270 | BSG | 0,257955331 | 1,195782772 | 0,32255521 | 0,824955371 | basigin (Ok blood group) |
| ENSG00000256262 | USP30-AS1 | 0,257922531 | 1,195755586 | 0,76873353 | 0,977381742 | USP30 antisense RNA 1 |
| ENSG00000171953 | ATPAF2 | 0,257913131 | 1,195747795 | 0,26945581 | 0,781544346 | ATP synthase mitochondrial F1 complex assembly factor 2 |
| ENSG00000234492 | RPL34-DT | 0,257871012 | 1,195712886 | 0,78259012 | 0,978647973 | RPL34 divergent transcript |
| ENSG00000099381 | SETD1A | 0,257792761 | 1,195648033 | 0,43886253 | 0,897402199 | SET domain containing 1A, histone lysine methyltransferase |
| ENSG00000109452 | INPP4B | 0,257682137 | 1,195556356 | 0,59759275 | 0,9482846 | inositol polyphosphate-4-phosphatase type II B |
| ENSG00000131467 | PSME3 | 0,257666371 | 1,195543291 | 0,24230881 | 0,752512424 | proteasome activator subunit 3 |
| ENSG00000175166 | PSMD2 | 0,257460404 | 1,19537262 | 0,32427559 | 0,826368833 | proteasome 26S subunit, non-ATPase 2 |
| ENSG00000188396 | TCTEX1D4 | 0,257435344 | 1,195351857 | 0,68544332 | 0,965475755 | Tctex1 domain containing 4 |
| ENSG00000155846 | PPARGC1B | 0,257291042 | 1,195232301 | 0,42211946 | 0,889775006 | PPARG coactivator 1 beta |
| ENSG00000151612 | ZNF827 | 0,257258435 | 1,195205287 | 0,21737511 | 0,725063004 | zinc finger protein 827 |
| ENSG00000182087 | TMEM259 | 0,257250278 | 1,195198529 | 0,42816076 | 0,892936968 | transmembrane protein 259 |
| ENSG00000154370 | TRIM11 | 0,25723549 | 1,195186278 | 0,5004328 | 0,920485547 | tripartite motif containing 11 |
| ENSG00000279058 | AGAP14P | 0,257159969 | 1,195123716 | 0,68259945 | 0,964817084 | ArfGAP with GTPase domain, ankyrin repeat and PH domain 14, pseudogene |
| ENSG00000178997 | EXD1 | 0,257140446 | 1,195107543 | 0,70697044 | 0,969565008 | exonuclease 3'-5' domain containing 1 |
| ENSG00000141485 | SLC13A5 | 0,257065243 | 1,195045247 | 0,64961513 | 0,956899963 | solute carrier family 13 member 5 |

| | | | | | | |
|---|---|---|---|---|---|---|
| ENSG00000261996 | AC004706,1 | 0,257033717 | 1,195019133 | 0,7286393 | 0,973955059 | novel transcript |
| ENSG00000276302 | AL021997,2 | 0,257026049 | 1,195012782 | 0,71727634 | 0,971697135 | novel protein |
| ENSG00000168090 | COPS6 | 0,257007258 | 1,194997217 | 0,24590253 | 0,757752922 | COP9 signalosome subunit 6 |
| ENSG00000230615 | AL139220,2 | 0,256997845 | 1,19498942 | 0,66423956 | 0,960180581 | novel transcript |
| ENSG00000132840 | BHMT2 | 0,256997313 | 1,194988979 | 0,68925548 | 0,966707784 | betaine--homocysteine S-methyltransferase 2 |
| ENSG00000276497 | APBA1 | 0,256968827 | 1,194965385 | 0,68948407 | 0,966868992 | amyloid beta precursor protein binding family A member 1 |
| ENSG00000124222 | STX16 | 0,256815614 | 1,194838487 | 0,22653035 | 0,735240631 | syntaxin 16 |
| ENSG00000185798 | WDR53 | 0,256815585 | 1,194838463 | 0,35595165 | 0,8473026 | WD repeat domain 53 |
| ENSG00000213706 | RHOG2P | 0,256811957 | 1,194835458 | 0,82888192 | 0,983795764 | RHOG family member 2, pseudogene |
| ENSG00000169122 | FAM110B | 0,256753774 | 1,194787272 | 0,29146729 | 0,802058262 | family with sequence similarity 110 member B |
| ENSG00000108100 | CCNY | 0,256727899 | 1,194765844 | 0,30811378 | 0,816424917 | cyclin Y |
| ENSG00000151240 | DIP2C | 0,256722791 | 1,194761613 | 0,26331125 | 0,774637968 | disco interacting protein 2 homolog C |
| ENSG00000151304 | SRFBP1 | 0,256686496 | 1,194731557 | 0,39294384 | 0,871121102 | serum response factor binding protein 1 |
| ENSG00000215041 | NEURL4 | 0,256618163 | 1,194674969 | 0,57376422 | 0,942475308 | neuralized E3 ubiquitin protein ligase 4 |
| ENSG00000288301 | NEURL4 | 0,256618163 | 1,194674969 | 0,57376422 | 0,942475308 | neuralized E3 ubiquitin protein ligase 4 |
| ENSG00000261231 | LINC02133 | 0,256374732 | 1,194473404 | 0,72279969 | 0,972707638 | long intergenic non-protein coding RNA 2133 |
| ENSG00000275116 | CU457734,1 | 0,256351594 | 1,194454247 | 0,79213415 | 0,979471509 | LSM14 homolog A (SCD6, S, cerevisiae) pseudogene |
| ENSG00000228141 | AC105339,1 | 0,256235132 | 1,194357829 | 0,74788128 | 0,975901808 | novel pseudogene |
| ENSG00000122490 | SLC66A2 | 0,255943357 | 1,194116303 | 0,49467473 | 0,91758518 | solute carrier family 66 member 2 |
| ENSG00000180846 | CSNK1G2-AS1 | 0,255930212 | 1,194105423 | 0,82966133 | 0,984081562 | CSNK1G2 antisense RNA 1 |
| ENSG00000232518 | AC074366,1 | 0,255784376 | 1,193984722 | 0,78998613 | 0,979462318 | novel transcript |
| ENSG00000108588 | CCDC47 | 0,255683694 | 1,1939014 | 0,25868358 | 0,771485596 | coiled-coil domain containing 47 |
| ENSG00000219642 | BMPR1AP1 | 0,255574163 | 1,193810761 | 0,8071581 | 0,981213877 | bone morphogenetic protein receptor type 1A pseudogene 1 |
| ENSG00000231628 | AL133406,2 | 0,25555861 | 1,193797891 | 0,61502014 | 0,952793934 | novel transcript |
| ENSG00000131504 | DIAPH1 | 0,255370443 | 1,193642197 | 0,20287568 | 0,703260357 | diaphanous related formin 1 |
| ENSG00000006715 | VPS41 | 0,255294676 | 1,193579511 | 0,36222754 | 0,85231292 | VPS41 subunit of HOPS complex |
| ENSG00000261420 | AL022069,1 | 0,255152625 | 1,193461995 | 0,69098834 | 0,967179456 | novel transcript, antisense to BRP44L |
| ENSG00000205611 | LINC01597 | 0,25512165 | 1,193436371 | 0,52306111 | 0,92790906 | long intergenic non-protein coding RNA 1597 |
| ENSG00000285043 | AC093512,2 | 0,255065556 | 1,193389969 | 0,35480426 | 0,846311161 | uncharaterized LOC112694756 |
| ENSG00000166855 | CLPX | 0,254999181 | 1,193335065 | 0,2662266 | 0,777733913 | caseinolytic mitochondrial matrix peptidase chaperone subunit X |

| | | | | | | |
|---|---|---|---|---|---|---|
| ENSG00000172315 | TP53RK | 0,254991747 | 1,193328917 | 0,41364776 | 0,885121175 | TP53 regulating kinase |
| ENSG00000187531 | SIRT7 | 0,254991649 | 1,193328836 | 0,48223349 | 0,91306612 | sirtuin 7 |
| ENSG00000105771 | SMG9 | 0,254988154 | 1,193325944 | 0,3164473 | 0,821646322 | SMG9 nonsense mediated mRNA decay factor |
| ENSG00000272100 | AL513218,1 | 0,254946701 | 1,193291657 | 0,75484061 | 0,975901808 | novel transcript, antisense to CC2D1B |
| ENSG00000256972 | AP000462,3 | 0,254786 | 1,193158745 | 0,79718114 | 0,980161849 | novel transcript |
| ENSG00000113971 | NPHP3 | 0,254695452 | 1,193083861 | 0,37573872 | 0,862362023 | nephrocystin 3 |
| ENSG00000143319 | ISG20L2 | 0,254634106 | 1,19303313 | 0,28493763 | 0,796433333 | interferon stimulated exonuclease gene 20 like 2 |
| ENSG00000277656 | GSTT1 | 0,25458251 | 1,192990463 | 0,32092184 | 0,824428958 | glutathione S-transferase theta 1 |
| ENSG00000119718 | EIF2B2 | 0,254564062 | 1,192975208 | 0,23319968 | 0,742367548 | eukaryotic translation initiation factor 2B subunit beta |
| ENSG00000078369 | GNB1 | 0,254545407 | 1,192959782 | 0,23215705 | 0,740855165 | G protein subunit beta 1 |
| ENSG00000173626 | TRAPPC3L | 0,254533449 | 1,192949895 | 0,76676644 | 0,977201891 | trafficking protein particle complex 3 like |
| ENSG00000113621 | TXNDC15 | 0,254445386 | 1,192877078 | 0,27640432 | 0,788263115 | thioredoxin domain containing 15 |
| ENSG00000008438 | PGLYRP1 | 0,254341386 | 1,19279109 | 0,81532183 | 0,981521413 | peptidoglycan recognition protein 1 |
| ENSG00000204822 | MRPL53 | 0,254299453 | 1,192756421 | 0,26289581 | 0,774444028 | mitochondrial ribosomal protein L53 |
| ENSG00000236756 | DNAJC9-AS1 | 0,254257518 | 1,192721752 | 0,70904482 | 0,970282889 | DNAJC9 antisense RNA 1 |
| ENSG00000214203 | RPS4XP1 | 0,254209269 | 1,192681864 | 0,81507049 | 0,981521413 | ribosomal protein S4X pseudogene 1 |
| ENSG00000104812 | GYS1 | 0,254082106 | 1,192576742 | 0,33730985 | 0,836427495 | glycogen synthase 1 |
| ENSG00000266962 | AC067852,2 | 0,253697155 | 1,192258572 | 0,39902397 | 0,875433675 | novel transcript, antisense to HSD17B1 |
| ENSG00000274523 | RCC1L | 0,253682927 | 1,192246814 | 0,39698006 | 0,873609704 | RCC1 like |
| ENSG00000052802 | MSMO1 | 0,253608233 | 1,192185088 | 0,56477811 | 0,940865625 | methylsterol monooxygenase 1 |
| ENSG00000144659 | SLC25A38 | 0,253596146 | 1,1921751 | 0,28018861 | 0,790779983 | solute carrier family 25 member 38 |
| ENSG00000110756 | HPS5 | 0,253482495 | 1,192081188 | 0,28850332 | 0,799943767 | HPS5 biogenesis of lysosomal organelles complex 2 subunit 2 |
| ENSG00000185127 | C6orf120 | 0,25342722 | 1,192035515 | 0,46156937 | 0,906165604 | chromosome 6 open reading frame 120 |
| ENSG00000228079 | AC012368,2 | 0,253420048 | 1,192029589 | 0,86422443 | 0,989074806 | novel transcript |
| ENSG00000275084 | SNORD91B | 0,25329411 | 1,191925538 | 0,78266846 | 0,978678077 | small nucleolar RNA, C/D box 91B |
| ENSG00000185122 | HSF1 | 0,253207693 | 1,191854144 | 0,3460843 | 0,841168721 | heat shock transcription factor 1 |
| ENSG00000284774 | HSF1 | 0,253207693 | 1,191854144 | 0,3460843 | 0,841168721 | heat shock transcription factor 1 |
| ENSG00000221890 | NPTXR | 0,253071953 | 1,19174201 | 0,49603127 | 0,918105073 | neuronal pentraxin receptor |
| ENSG00000167136 | ENDOG | 0,253050378 | 1,191724188 | 0,57484855 | 0,942533938 | endonuclease G |
| ENSG00000174840 | PDE12 | 0,252996074 | 1,191679331 | 0,2687079 | 0,780823861 | phosphodiesterase 12 |
| ENSG00000177352 | CCDC71 | 0,252985959 | 1,191670977 | 0,33652408 | 0,835736667 | coiled-coil domain containing 71 |

| | | | | | | |
|---|---|---|---|---|---|---|
| ENSG00000204323 | SMIM5 | 0,252980621 | 1,191666567 | 0,64072671 | 0,956348702 | small integral membrane protein 5 |
| ENSG00000282228 | PAM16 | 0,252672279 | 1,191411904 | 0,70726076 | 0,969565008 | presequence translocase associated motor 16 |
| ENSG00000136051 | WASHC4 | 0,252666144 | 1,191406838 | 0,50143248 | 0,920520138 | WASH complex subunit 4 |
| ENSG00000281690 | ADAMTS12 | 0,252588192 | 1,191342465 | 0,55172615 | 0,936366211 | ADAM metallopeptidase with thrombospondin type 1 motif 12 |
| ENSG00000229638 | RPL4P4 | 0,252548677 | 1,191309835 | 0,81912209 | 0,98249617 | ribosomal protein L4 pseudogene 4 |
| ENSG00000143374 | TARS2 | 0,252523336 | 1,19128891 | 0,36843003 | 0,857548638 | threonyl-tRNA synthetase 2, mitochondrial |
| ENSG00000153406 | NMRAL1 | 0,252475124 | 1,1912491 | 0,47171697 | 0,908896785 | NmrA like redox sensor 1 |
| ENSG00000106608 | URGCP | 0,252427803 | 1,191210027 | 0,46568876 | 0,907940653 | upregulator of cell proliferation |
| ENSG00000163638 | ADAMTS9 | 0,252425287 | 1,191207949 | 0,45327864 | 0,902630791 | ADAM metallopeptidase with thrombospondin type 1 motif 9 |
| ENSG00000199691 | RN7SKP173 | 0,252405795 | 1,191191856 | 0,84020461 | 0,985672131 | RN7SK pseudogene 173 |
| ENSG00000112146 | FBXO9 | 0,252378314 | 1,191169165 | 0,42179729 | 0,889720796 | F-box protein 9 |
| ENSG00000268756 | AC104534,1 | 0,252140993 | 1,190973236 | 0,83574359 | 0,985034063 | novel transcript, antisense to ECH1 |
| ENSG00000283077 | AC104534,3 | 0,252140993 | 1,190973236 | 0,83574359 | 0,985034063 | novel transcript, antisense to ECH1 |
| ENSG00000011260 | UTP18 | 0,252010472 | 1,190865494 | 0,2557129 | 0,767236865 | UTP18 small subunit processome component |
| ENSG00000139180 | NDUFA9 | 0,251996664 | 1,190854096 | 0,3333912 | 0,833718 | NADH:ubiquinone oxidoreductase subunit A9 |
| ENSG00000143614 | GATAD2B | 0,251820335 | 1,190708556 | 0,26812992 | 0,780692104 | GATA zinc finger domain containing 2B |
| ENSG00000115361 | ACADL | 0,251800341 | 1,190692055 | 0,62199837 | 0,952793934 | acyl-CoA dehydrogenase long chain |
| ENSG00000138600 | SPPL2A | 0,251793372 | 1,190686302 | 0,37061581 | 0,858545694 | signal peptide peptidase like 2A |
| ENSG00000263797 | AP005210,1 | 0,251789116 | 1,19068279 | 0,77093356 | 0,978033855 | novel transcript, antisense to ARHGAP28 |
| ENSG00000248896 | AC105001,1 | 0,251659881 | 1,190576135 | 0,56009722 | 0,940070839 | novel transcript, antisense to SOX7 and PINX1 |
| ENSG00000165689 | ENTR1 | 0,251574068 | 1,19050532 | 0,30102993 | 0,810679941 | endosome associated trafficking regulator 1 |
| ENSG00000204227 | RING1 | 0,251539149 | 1,190476506 | 0,50428008 | 0,921400408 | ring finger protein 1 |
| ENSG00000261080 | RUNX2-AS1 | 0,251512038 | 1,190454134 | 0,76728254 | 0,977201891 | RUNX2 antisense RNA 1 |
| ENSG00000277324 | AC093462,1 | 0,251455817 | 1,190407744 | 0,78901428 | 0,979462318 | novel transcript, antisense to POLI |
| ENSG00000135678 | CPM | 0,25142103 | 1,190379041 | 0,69582565 | 0,96776072 | carboxypeptidase M |
| ENSG00000108958 | AC130689,1 | 0,251419304 | 1,190377617 | 0,80087974 | 0,980505307 | succinate dehydrogenase complex, subunit C, integral membrane protein, 15kDa (SDHC) pseudogene |
| ENSG00000285302 | CHMP4A | 0,251405405 | 1,190366148 | 0,54896439 | 0,936040289 | charged multivesicular body protein 4A |
| ENSG00000108528 | SLC25A11 | 0,251393477 | 1,190356307 | 0,2728179 | 0,785672647 | solute carrier family 25 member 11 |
| ENSG00000183579 | ZNRF3 | 0,25136192 | 1,19033027 | 0,26391864 | 0,775013078 | zinc and ring finger 3 |

| | | | | | | |
|---|---|---|---|---|---|---|
| ENSG00000271848 | AC073389,2 | 0,251325798 | 1,190300466 | 0,7121624 | 0,970812834 | novel transcript, antisense to SYNPO2L and AGAP5 |
| ENSG00000115084 | SLC35F5 | 0,25125837 | 1,190244837 | 0,28548583 | 0,796984464 | solute carrier family 35 member F5 |
| ENSG00000150051 | MKX | 0,251231376 | 1,190222566 | 0,76165229 | 0,976285211 | mohawk homeobox |
| ENSG00000174123 | TLR10 | 0,251085179 | 1,190101959 | 0,71645318 | 0,971438758 | toll like receptor 10 |
| ENSG00000173465 | ZNRD2 | 0,250884248 | 1,18993622 | 0,45940645 | 0,904862549 | zinc ribbon domain containing 2 |
| ENSG00000148483 | TMEM236 | 0,250877356 | 1,189930536 | 0,74664141 | 0,975796253 | transmembrane protein 236 |
| ENSG00000072778 | ACADVL | 0,250841181 | 1,189900699 | 0,38245784 | 0,867026464 | acyl-CoA dehydrogenase very long chain |
| ENSG00000152402 | GUCY1A2 | 0,250764634 | 1,189837566 | 0,58624908 | 0,945256674 | guanylate cyclase 1 soluble subunit alpha 2 |
| ENSG00000148300 | REXO4 | 0,250761391 | 1,189834892 | 0,43665839 | 0,896866043 | REX4 homolog, 3'-5' exonuclease |
| ENSG00000280706 | REXO4 | 0,250761391 | 1,189834892 | 0,43665839 | 0,896866043 | REX4 homolog, 3'-5' exonuclease |
| ENSG00000105229 | PIAS4 | 0,250760634 | 1,189834268 | 0,34514093 | 0,841168721 | protein inhibitor of activated STAT 4 |
| ENSG00000150783 | TEX12 | 0,250737871 | 1,189815494 | 0,87248684 | 0,989217334 | testis expressed 12 |
| ENSG00000134463 | ECHDC3 | 0,250721289 | 1,189801819 | 0,51156889 | 0,922893865 | enoyl-CoA hydratase domain containing 3 |
| ENSG00000123415 | SMUG1 | 0,250581152 | 1,189686252 | 0,34118204 | 0,839923707 | single-strand-selective monofunctional uracil-DNA glycosylase 1 |
| ENSG00000214185 | XPOTP1 | 0,250500461 | 1,189619714 | 0,78557807 | 0,979320406 | exportin for tRNA pseudogene 1 |
| ENSG00000271741 | AC114490,2 | 0,250381814 | 1,189521884 | 0,42107988 | 0,888848798 | novel transcript, ZMYM6-ZMYM6NB readthrough |
| ENSG00000124216 | SNAI1 | 0,250355603 | 1,189500273 | 0,65017945 | 0,956899963 | snail family transcriptional repressor 1 |
| ENSG00000234737 | KRT18P15 | 0,250213437 | 1,189383063 | 0,83554546 | 0,985034063 | keratin 18 pseudogene 15 |
| ENSG00000282936 | AC004706,3 | 0,250086571 | 1,189278477 | 0,77525002 | 0,978587255 | novel protein |
| ENSG00000270248 | AC011468,4 | 0,250016652 | 1,189220842 | 0,81503728 | 0,981521413 | transmembrane protein 50B (TMEM50B) pseudogene |
| ENSG00000120053 | GOT1 | 0,249999192 | 1,189206449 | 0,22793946 | 0,736341996 | glutamic-oxaloacetic transaminase 1 |
| ENSG00000047621 | C12orf4 | 0,249982741 | 1,189192888 | 0,43646322 | 0,896850871 | chromosome 12 open reading frame 4 |
| ENSG00000133265 | HSPBP1 | 0,249975252 | 1,189186715 | 0,49704496 | 0,918742673 | HSPA (Hsp70) binding protein 1 |
| ENSG00000171222 | SCAND1 | 0,249961404 | 1,189175301 | 0,51527819 | 0,924832497 | SCAN domain containing 1 |
| ENSG00000133112 | TPT1 | 0,24991438 | 1,189136541 | 0,57063595 | 0,9423888 | tumor protein, translationally-controlled 1 |
| ENSG00000112320 | SOBP | 0,249907474 | 1,189130849 | 0,35146479 | 0,84405766 | sine oculis binding protein homolog |
| ENSG00000263531 | AC130324,1 | 0,249853741 | 1,189086561 | 0,75410649 | 0,975901808 | novel transcript, antisense ATAD5 |
| ENSG00000165171 | METTL27 | 0,249851077 | 1,189084365 | 0,70539036 | 0,969485452 | methyltransferase like 27 |
| ENSG00000151327 | FAM177A1 | 0,249779208 | 1,189025131 | 0,32271173 | 0,824955371 | family with sequence similarity 177 member A1 |
| ENSG00000102524 | TNFSF13B | 0,249645867 | 1,188915241 | 0,75473212 | 0,975901808 | TNF superfamily member 13b |
| ENSG00000244265 | SIAH2-AS1 | 0,249569143 | 1,188852014 | 0,82225288 | 0,982952211 | SIAH2 antisense RNA 1 |

| | | | | | | |
|---|---|---|---|---|---|---|
| ENSG00000213585 | VDAC1 | 0,249529233 | 1,188819127 | 0,46686941 | 0,907940653 | voltage dependent anion channel 1 |
| ENSG00000257636 | G2E3-AS1 | 0,249469651 | 1,188770031 | 0,86072093 | 0,989074806 | G2E3 antisense RNA 1 |
| ENSG00000092931 | MFSD11 | 0,2494623 | 1,188763974 | 0,27847577 | 0,789775666 | major facilitator superfamily domain containing 11 |
| ENSG00000270367 | AC092053,1 | 0,249414621 | 1,188724687 | 0,71491471 | 0,971041579 | Adenosine deaminase domain-containing protein 1, pseudogene |
| ENSG00000287039 | AC006478,2 | 0,24938352 | 1,188699062 | 0,79979522 | 0,980161849 | novel transcript |
| ENSG00000211448 | DIO2 | 0,249318723 | 1,188645674 | 0,63306466 | 0,954408281 | iodothyronine deiodinase 2 |
| ENSG00000275023 | MLLT6 | 0,249297503 | 1,188628191 | 0,64299449 | 0,956633898 | MLLT6, PHD finger containing |
| ENSG00000126947 | ARMCX1 | 0,249294693 | 1,188625876 | 0,34459189 | 0,841168721 | armadillo repeat containing X-linked 1 |
| ENSG00000164086 | DUSP7 | 0,249201383 | 1,188549001 | 0,32933185 | 0,83127975 | dual specificity phosphatase 7 |
| ENSG00000137547 | MRPL15 | 0,249199477 | 1,188547431 | 0,33097763 | 0,832186595 | mitochondrial ribosomal protein L15 |
| ENSG00000110344 | UBE4A | 0,248831552 | 1,188244359 | 0,46291397 | 0,906849208 | ubiquitination factor E4A |
| ENSG00000196642 | RABL6 | 0,248656588 | 1,188100262 | 0,57354167 | 0,942475308 | RAB, member RAS oncogene family like 6 |
| ENSG00000138785 | INTS12 | 0,248651733 | 1,188096264 | 0,42050321 | 0,888563911 | integrator complex subunit 12 |
| ENSG00000226578 | AL132657,1 | 0,248626004 | 1,188075075 | 0,75869231 | 0,976120207 | novel transcript |
| ENSG00000232320 | MXRA7P1 | 0,248613603 | 1,188064863 | 0,89438102 | 0,991135494 | MXRA7 pseudogene 1 |
| ENSG00000267858 | MZF1-AS1 | 0,248581061 | 1,188038065 | 0,56794002 | 0,94163744 | MZF1 antisense RNA 1 |
| ENSG00000180185 | FAHD1 | 0,248436928 | 1,187919379 | 0,32451803 | 0,82667087 | fumarylacetoacetate hydrolase domain containing 1 |
| ENSG00000006607 | FARP2 | 0,248373388 | 1,187867061 | 0,31255943 | 0,818619247 | FERM, ARH/RhoGEF and pleckstrin domain protein 2 |
| ENSG00000027644 | INSRR | 0,248354629 | 1,187851616 | 0,77367382 | 0,978415098 | insulin receptor related receptor |
| ENSG00000234678 | ELF3-AS1 | 0,248309105 | 1,187814134 | 0,64463682 | 0,956633898 | ELF3 antisense RNA 1 |
| ENSG00000108830 | RND2 | 0,248227674 | 1,187747091 | 0,28847549 | 0,799943767 | Rho family GTPase 2 |
| ENSG00000102910 | LONP2 | 0,248151048 | 1,187684008 | 0,29984831 | 0,809888342 | lon peptidase 2, peroxisomal |
| ENSG00000288325 | GPS2 | 0,248120162 | 1,187658582 | 0,31390973 | 0,819877431 | G protein pathway suppressor 2 |
| ENSG00000132522 | GPS2 | 0,248120162 | 1,187658582 | 0,31390973 | 0,819877431 | G protein pathway suppressor 2 |
| ENSG00000242073 | AC211429,1 | 0,248028882 | 1,18758344 | 0,58377248 | 0,944276183 | postmeiotic segregation increased 2-like 3 (PMS2L3) pseudogene |
| ENSG00000175604 | AC007601,1 | 0,248014855 | 1,187571894 | 0,75911029 | 0,976120207 | novel transcript, antisense to SNX29 |
| ENSG00000168802 | CHTF8 | 0,24800598 | 1,187564588 | 0,36164418 | 0,851893059 | chromosome transmission fidelity factor 8 |
| ENSG00000203392 | AC105020,1 | 0,247884763 | 1,187464812 | 0,69288555 | 0,967271667 | novel transcript, antisense to CSPG4 |
| ENSG00000101361 | NOP56 | 0,247846574 | 1,18743338 | 0,21497653 | 0,722298056 | NOP56 ribonucleoprotein |
| ENSG00000183292 | PRSS40A | 0,247799401 | 1,187394554 | 0,75686678 | 0,975901808 | serine protease 40A (pseudogene) |
| ENSG00000065427 | KARS1 | 0,247793404 | 1,187389618 | 0,2728628 | 0,785672647 | lysyl-tRNA synthetase 1 |

| | | | | | | |
|---|---|---|---|---|---|---|
| ENSG00000057252 | SOAT1 | 0,247741573 | 1,18734696 | 0,44215578 | 0,899331131 | sterol O-acyltransferase 1 |
| ENSG00000256195 | AP002518,1 | 0,247731637 | 1,187338783 | 0,83995796 | 0,985672131 | novel transcript |
| ENSG00000151465 | CDC123 | 0,2477193 | 1,18732863 | 0,43764863 | 0,897021798 | cell division cycle 123 |
| ENSG00000280433 | FP565260,6 | 0,247717315 | 1,187326996 | 0,44177273 | 0,899133035 | novel protein, similar to trafficking protein particle complex 10 TRAPPC10 |
| ENSG00000168288 | MMADHC | 0,247572183 | 1,187207559 | 0,48152817 | 0,912981134 | metabolism of cobalamin associated D |
| ENSG00000106100 | NOD1 | 0,247507742 | 1,187154531 | 0,3265542 | 0,828450693 | nucleotide binding oligomerization domain containing 1 |
| ENSG00000187912 | CLEC17A | 0,247501196 | 1,187149144 | 0,76953593 | 0,977683747 | C-type lectin domain containing 17A |
| ENSG00000132376 | INPP5K | 0,247478006 | 1,187130062 | 0,31863821 | 0,823434042 | inositol polyphosphate-5-phosphatase K |
| ENSG00000283175 | AC007920,2 | 0,247191669 | 1,186894472 | 0,72561212 | 0,97326334 | novel transcript, antisense to MASP1 |
| ENSG00000034693 | PEX3 | 0,247106905 | 1,18682474 | 0,55982652 | 0,939880787 | peroxisomal biogenesis factor 3 |
| ENSG00000279375 | AC244517,7 | 0,247106399 | 1,186824323 | 0,74775554 | 0,975901808 | novel transcript, antisense to PCDHB10 |
| ENSG00000250615 | AC008581,1 | 0,24709615 | 1,186815892 | 0,74898601 | 0,975901808 | novel transcript |
| ENSG00000257800 | FNBP1P1 | 0,247036193 | 1,186766569 | 0,41031958 | 0,882614012 | formin binding protein 1 pseudogene 1 |
| ENSG00000214160 | ALG3 | 0,247027643 | 1,186759537 | 0,36947733 | 0,857991124 | ALG3 alpha-1,3- mannosyltransferase |
| ENSG00000130307 | USHBP1 | 0,246997802 | 1,18673499 | 0,74250581 | 0,975176043 | USH1 protein network component harmonin binding protein 1 |
| ENSG00000253915 | MAPRE1P1 | 0,246951846 | 1,186697187 | 0,69114496 | 0,967179456 | MAPRE1 pseudogene 1 |
| ENSG00000286544 | AL359502,2 | 0,246864252 | 1,186625139 | 0,87498389 | 0,990078774 | novel transcript |
| ENSG00000142279 | WTIP | 0,246746645 | 1,18652841 | 0,59454213 | 0,947469769 | WT1 interacting protein |
| ENSG00000280486 | AC005329,3 | 0,246536155 | 1,186355308 | 0,66375287 | 0,959920886 | novel transcript, antisense to NDUFS7 |
| ENSG00000085998 | POMGNT1 | 0,246466211 | 1,186297793 | 0,37589604 | 0,862406526 | protein O-linked mannose N-acetylglucosaminyltransferase 1 (beta 1,2-) |
| ENSG00000143418 | CERS2 | 0,246369598 | 1,186218353 | 0,30891342 | 0,816838423 | ceramide synthase 2 |
| ENSG00000110514 | MADD | 0,246318412 | 1,186176267 | 0,28315789 | 0,794458239 | MAP kinase activating death domain |
| ENSG00000100416 | TRMU | 0,246291989 | 1,186154543 | 0,34046232 | 0,839052015 | tRNA 5-methylaminomethyl-2-thiouridylate methyltransferase |
| ENSG00000228824 | MIR4500HG | 0,246266269 | 1,186133396 | 0,51720437 | 0,925746338 | MIR4500 host gene |
| ENSG00000260569 | AC090398,1 | 0,246163803 | 1,186049156 | 0,81747771 | 0,98215965 | novel transcript |
| ENSG00000133131 | MORC4 | 0,246020239 | 1,185931136 | 0,32232767 | 0,824955371 | MORC family CW-type zinc finger 4 |
| ENSG00000231346 | LINC01160 | 0,245944957 | 1,185869254 | 0,72662528 | 0,973344786 | long intergenic non-protein coding RNA 1160 |
| ENSG00000275807 | AC145285,6 | 0,245857794 | 1,18579761 | 0,57798216 | 0,943425812 | novel transcript, antisense to ATXN2L |

| | | | | | | |
|---|---|---|---|---|---|---|
| ENSG00000274746 | ZNF100 | 0,245829014 | 1,185773955 | 0,58730829 | 0,945938196 | zinc finger protein 100 |
| ENSG00000236559 | BCL2L1-AS1 | 0,245822981 | 1,185768997 | 0,79180123 | 0,979471509 | BCL2L1 antisense RNA 1 |
| ENSG00000096155 | BAG6 | 0,245806004 | 1,185755043 | 0,36256801 | 0,85266484 | BAG cochaperone 6 |
| ENSG00000269918 | AF131215,6 | 0,245691824 | 1,185661202 | 0,56499815 | 0,940865625 | novel transcript |
| ENSG00000124257 | NEURL2 | 0,245680847 | 1,185652181 | 0,74315222 | 0,975305642 | neuralized E3 ubiquitin protein ligase 2 |
| ENSG00000161202 | DVL3 | 0,245639564 | 1,185618254 | 0,23728319 | 0,746695699 | dishevelled segment polarity protein 3 |
| ENSG00000260236 | AC099778,1 | 0,245612636 | 1,185596124 | 0,66697506 | 0,9611116 | novel transcript, antisense to PTPN23 |
| ENSG00000251815 | SNORD116-26 | 0,245602881 | 1,185588107 | 0,86794687 | 0,989074806 | small nucleolar RNA, C/D box 116-26 |
| ENSG00000135317 | SNX14 | 0,245574356 | 1,185564666 | 0,53559213 | 0,932139691 | sorting nexin 14 |
| ENSG00000107282 | APBA1 | 0,245548627 | 1,185543523 | 0,48976781 | 0,916063942 | amyloid beta precursor protein binding family A member 1 |
| ENSG00000101940 | WDR13 | 0,24542402 | 1,185441131 | 0,41066031 | 0,882835102 | WD repeat domain 13 |
| ENSG00000027847 | B4GALT7 | 0,245417823 | 1,185436039 | 0,43799068 | 0,897177404 | beta-1,4-galactosyltransferase 7 |
| ENSG00000254416 | LINC02732 | 0,245381872 | 1,185406499 | 0,60122794 | 0,949908544 | long intergenic non-protein coding RNA 2732 |
| ENSG00000160838 | LRRC71 | 0,245371781 | 1,185398208 | 0,75709579 | 0,975901808 | leucine rich repeat containing 71 |
| ENSG00000141429 | GALNT1 | 0,245310366 | 1,185347747 | 0,50988768 | 0,922427526 | polypeptide N-acetylgalactosaminyltransferase 1 |
| ENSG00000167302 | TEPSIN | 0,245265987 | 1,185311285 | 0,49427109 | 0,91758518 | TEPSIN adaptor related protein complex 4 accessory protein |
| ENSG00000124549 | BTN2A3P | 0,24525125 | 1,185299177 | 0,33004605 | 0,831739454 | butyrophilin subfamily 2 member A3, pseudogene |
| ENSG00000114439 | BBX | 0,245247054 | 1,185295729 | 0,45036739 | 0,902062594 | BBX high mobility group box domain containing |
| ENSG00000261423 | TMEM202-AS1 | 0,245243677 | 1,185292955 | 0,61372681 | 0,952413269 | TMEM202 antisense RNA 1 |
| ENSG00000260033 | AC023813,2 | 0,245212252 | 1,185267137 | 0,80219488 | 0,980710009 | ribosomal L24 domain containing 1 (RSL24D1) pseudogene |
| ENSG00000174527 | MYO1H | 0,245210737 | 1,185265892 | 0,55204531 | 0,936552095 | myosin IH |
| ENSG00000276802 | HERC2 | 0,245155986 | 1,185220912 | 0,51769044 | 0,925915783 | HECT and RLD domain containing E3 ubiquitin protein ligase 2 |
| ENSG00000121644 | DESI2 | 0,245075043 | 1,185154416 | 0,57333349 | 0,942475308 | desumoylating isopeptidase 2 |
| ENSG00000133983 | COX16 | 0,245043302 | 1,185128341 | 0,27829426 | 0,789652735 | cytochrome c oxidase assembly factor COX16 |
| ENSG00000107263 | RAPGEF1 | 0,244984219 | 1,185079808 | 0,30178645 | 0,811332971 | Rap guanine nucleotide exchange factor 1 |
| ENSG00000135083 | CCNJL | 0,244921764 | 1,185028506 | 0,25436114 | 0,765826923 | cyclin J like |
| ENSG00000119396 | RAB14 | 0,244917403 | 1,185024924 | 0,49876135 | 0,919683359 | RAB14, member RAS oncogene family |
| ENSG00000257108 | NHLRC4 | 0,244871332 | 1,184987082 | 0,66606642 | 0,961020116 | NHL repeat containing 4 |
| ENSG00000122694 | GLIPR2 | 0,244869989 | 1,184985979 | 0,42662577 | 0,892175781 | GLI pathogenesis related 2 |

| | | | | | | |
|---|---|---|---|---|---|---|
| ENSG00000138801 | PAPSS1 | 0,244768911 | 1,184902959 | 0,38938208 | 0,870057277 | 3'-phosphoadenosine 5'-phosphosulfate synthase 1 |
| ENSG00000250496 | ABT1P1 | 0,244727853 | 1,184869239 | 0,75574213 | 0,975901808 | activator of basal transcription 1 pseudogene 1 |
| ENSG00000287711 | LINC02136 | 0,244689536 | 1,18483777 | 0,77313215 | 0,978363975 | long intergenic non-protein coding RNA 2136 |
| ENSG00000100983 | GSS | 0,244456102 | 1,184646073 | 0,29427336 | 0,804089131 | glutathione synthetase |
| ENSG00000130165 | ELOF1 | 0,244360349 | 1,18456745 | 0,41546096 | 0,885986046 | elongation factor 1 homolog |
| ENSG00000260927 | AC009107,2 | 0,244085294 | 1,18434163 | 0,31353538 | 0,819801802 | uncharacterized LOC101927556 |
| ENSG00000181396 | OGFOD3 | 0,244060835 | 1,184321551 | 0,56589975 | 0,940979133 | 2-oxoglutarate and iron dependent oxygenase domain containing 3 |
| ENSG00000186367 | MINAR2 | 0,24405647 | 1,184317968 | 0,76825435 | 0,977381742 | membrane integral NOTCH2 associated receptor 2 |
| ENSG00000257550 | AC023509,2 | 0,243971979 | 1,184248611 | 0,71547196 | 0,971152196 | novel transcript, antisense to ATF7 |
| ENSG00000091732 | ZC3HC1 | 0,243771802 | 1,184084305 | 0,29304785 | 0,803365244 | zinc finger C3HC-type containing 1 |
| ENSG00000163623 | NKX6-1 | 0,243769881 | 1,184082728 | 0,69242124 | 0,967236299 | NK6 homeobox 1 |
| ENSG00000235613 | NSRP1P1 | 0,243724381 | 1,184045385 | 0,61660851 | 0,952793934 | nuclear speckle splicing regulatory protein 1 pseudogene 1 |
| ENSG00000226137 | BAIAP2-DT | 0,24366967 | 1,184000484 | 0,52046083 | 0,927003219 | BAIAP2 divergent transcript |
| ENSG00000171307 | ZDHHC16 | 0,243652015 | 1,183985994 | 0,24859694 | 0,760720365 | zinc finger DHHC-type palmitoyltransferase 16 |
| ENSG00000125304 | TM9SF2 | 0,243574075 | 1,183922033 | 0,55157235 | 0,936366211 | transmembrane 9 superfamily member 2 |
| ENSG00000100033 | PRODH | 0,243550623 | 1,183902787 | 0,63567501 | 0,955389018 | proline dehydrogenase 1 |
| ENSG00000107140 | TESK1 | 0,243494019 | 1,183856338 | 0,31121732 | 0,817751312 | testis associated actin remodelling kinase 1 |
| ENSG00000181610 | MRPS23 | 0,243456295 | 1,183825382 | 0,33685915 | 0,836162806 | mitochondrial ribosomal protein S23 |
| ENSG00000125650 | PSPN | 0,243404182 | 1,183782621 | 0,5469828 | 0,935404974 | persephin |
| ENSG00000159147 | DONSON | 0,243393019 | 1,183773462 | 0,27148727 | 0,784380175 | DNA replication fork stabilization factor DONSON |
| ENSG00000158615 | PPP1R15B | 0,243360535 | 1,183746807 | 0,37413968 | 0,861369706 | protein phosphatase 1 regulatory subunit 15B |
| ENSG00000166342 | NETO1 | 0,24335797 | 1,183744703 | 0,64768655 | 0,956899963 | neuropilin and tolloid like 1 |
| ENSG00000198856 | OSTC | 0,243350882 | 1,183738888 | 0,57103915 | 0,942436428 | oligosaccharyltransferase complex non-catalytic subunit |
| ENSG00000206355 | CCHCR1 | 0,243290724 | 1,183689528 | 0,67703696 | 0,963580269 | coiled-coil alpha-helical rod protein 1 |
| ENSG00000287004 | AC007686,5 | 0,24328615 | 1,183685776 | 0,81609813 | 0,981708551 | novel transcript |
| ENSG00000114923 | SLC4A3 | 0,243147186 | 1,183571766 | 0,55428567 | 0,937726571 | solute carrier family 4 member 3 |
| ENSG00000288552 | AL162573,1 | 0,243097947 | 1,183531371 | 0,82079358 | 0,982664102 | novel transcript |
| ENSG00000135919 | SERPINE2 | 0,243005197 | 1,183455285 | 0,33463776 | 0,834757602 | serpin family E member 2 |
| ENSG00000242258 | LINC00996 | 0,242936617 | 1,18339903 | 0,81606469 | 0,981708551 | long intergenic non-protein coding RNA 996 |
| ENSG00000267244 | AC012615,6 | 0,242847072 | 1,183325581 | 0,65519055 | 0,957444211 | novel transcript, antisense to REXO1 |

| | | | | | | |
|---|---|---|---|---|---|---|
| ENSG00000122729 | ACO1 | 0,242816921 | 1,18330085 | 0,3565601 | 0,847911681 | aconitase 1 |
| ENSG00000108094 | CUL2 | 0,242751063 | 1,183246835 | 0,55644143 | 0,938061341 | cullin 2 |
| ENSG00000129245 | FXR2 | 0,242661183 | 1,183173121 | 0,42763426 | 0,892656446 | FMR1 autosomal homolog 2 |
| ENSG00000206440 | NFKBIL1 | 0,242502943 | 1,183043353 | 0,65038525 | 0,956899963 | NFKB inhibitor like 1 |
| ENSG00000236346 | NFKBIL1 | 0,242502943 | 1,183043353 | 0,65038525 | 0,956899963 | NFKB inhibitor like 1 |
| ENSG00000234530 | NFKBIL1 | 0,242502943 | 1,183043353 | 0,65038525 | 0,956899963 | NFKB inhibitor like 1 |
| ENSG00000227565 | NFKBIL1 | 0,242502943 | 1,183043353 | 0,65038525 | 0,956899963 | NFKB inhibitor like 1 |
| ENSG00000236196 | NFKBIL1 | 0,242502943 | 1,183043353 | 0,65038525 | 0,956899963 | NFKB inhibitor like 1 |
| ENSG00000213197 | NUDCP1 | 0,242490075 | 1,183032801 | 0,72930765 | 0,974009551 | nuclear distribution C pseudogene 1 |
| ENSG00000139637 | MYG1 | 0,24247004 | 1,183016372 | 0,33214248 | 0,83293962 | MYG1 exonuclease |
| ENSG00000196600 | SLC22A25 | 0,242346356 | 1,182914955 | 0,84288202 | 0,986208256 | solute carrier family 22 member 25 |
| ENSG00000259488 | AC023355,1 | 0,242325478 | 1,182897836 | 0,59172954 | 0,947416795 | novel transcript, antisense to DUT |
| ENSG00000115514 | TXNDC9 | 0,242303517 | 1,182879831 | 0,55180435 | 0,936366211 | thioredoxin domain containing 9 |
| ENSG00000123154 | WDR83 | 0,242273738 | 1,182855414 | 0,61741682 | 0,952793934 | WD repeat domain 83 |
| ENSG00000216906 | AL355312,1 | 0,242099783 | 1,182712799 | 0,77696259 | 0,978587255 | multimerin 2 (MMRN2) pseudogene |
| ENSG00000177045 | SIX5 | 0,241983853 | 1,182617764 | 0,65567208 | 0,957444211 | SIX homeobox 5 |
| ENSG00000223459 | TCAF1P1 | 0,241854069 | 1,182511381 | 0,55616473 | 0,938046927 | TRPM8 channel associated factor 1 pseudogene 1 |
| ENSG00000154217 | PITPNC1 | 0,241718811 | 1,182400522 | 0,35866477 | 0,849814998 | phosphatidylinositol transfer protein cytoplasmic 1 |
| ENSG00000134899 | ERCC5 | 0,241700631 | 1,182385622 | 0,28813201 | 0,799375215 | ERCC excision repair 5, endonuclease |
| ENSG00000168758 | SEMA4C | 0,241589559 | 1,182294595 | 0,4776713 | 0,911605141 | semaphorin 4C |
| ENSG00000151208 | DLG5 | 0,241541367 | 1,182255101 | 0,73048774 | 0,974321178 | discs large MAGUK scaffold protein 5 |
| ENSG00000261028 | AC012322,1 | 0,241494314 | 1,182216543 | 0,82856077 | 0,983721455 | novel transcript |
| ENSG00000272417 | AC034229,4 | 0,241460444 | 1,182188789 | 0,81316491 | 0,981246822 | novel transcript |
| ENSG00000183798 | EMILIN3 | 0,24140788 | 1,182145717 | 0,49547716 | 0,917815326 | elastin microfibril interfacer 3 |
| ENSG00000114054 | PCCB | 0,24135141 | 1,182099446 | 0,3005775 | 0,81031109 | propionyl-CoA carboxylase subunit beta |
| ENSG00000131116 | ZNF428 | 0,241183965 | 1,181962255 | 0,47713672 | 0,91143003 | zinc finger protein 428 |
| ENSG00000233051 | MICA | 0,241179374 | 1,181958494 | 0,43496852 | 0,896272827 | MHC class I polypeptide-related sequence A |
| ENSG00000167461 | RAB8A | 0,241161547 | 1,181943889 | 0,40356449 | 0,877691113 | RAB8A, member RAS oncogene family |
| ENSG00000134824 | FADS2 | 0,241091245 | 1,181886294 | 0,29906242 | 0,808883765 | fatty acid desaturase 2 |
| ENSG00000104763 | ASAH1 | 0,240950105 | 1,181770675 | 0,53727245 | 0,932883277 | N-acylsphingosine amidohydrolase 1 |
| ENSG00000223528 | AL359094,1 | 0,240687123 | 1,181555276 | 0,64437345 | 0,956633898 | novel transcript |
| ENSG00000155097 | ATP6V1C1 | 0,240648915 | 1,181523984 | 0,53296718 | 0,931291085 | ATPase H+ transporting V1 subunit C1 |
| ENSG00000240356 | RPL23AP7 | 0,240644223 | 1,181520141 | 0,40456213 | 0,878176155 | ribosomal protein L23a pseudogene 7 |

| | | | | | | |
|---|---|---|---|---|---|---|
| ENSG00000105202 | FBL | 0,240607766 | 1,181490285 | 0,45223049 | 0,902062594 | fibrillarin |
| ENSG00000154832 | CXXC1 | 0,240560354 | 1,181451458 | 0,46974666 | 0,908378389 | CXXC finger protein 1 |
| ENSG00000287215 | AC231759,2 | 0,24046288 | 1,181371637 | 0,68406505 | 0,965403332 | novel transcript |
| ENSG00000250726 | AC107222,1 | 0,240458762 | 1,181368265 | 0,78871844 | 0,979462318 | low density lipoprotein receptor-related protein 8, apolipoprotein e receptor (LRP8) pseudogene |
| ENSG00000130725 | UBE2M | 0,240451207 | 1,181362078 | 0,29650212 | 0,806142623 | ubiquitin conjugating enzyme E2 M |
| ENSG00000249743 | AC116345,1 | 0,240285714 | 1,18122657 | 0,80872116 | 0,981213877 | novel transcript |
| ENSG00000184343 | SRPK3 | 0,240260489 | 1,181205917 | 0,79332349 | 0,979753729 | SRSF protein kinase 3 |
| ENSG00000119471 | HSDL2 | 0,240250555 | 1,181197784 | 0,54757951 | 0,935404974 | hydroxysteroid dehydrogenase like 2 |
| ENSG00000065413 | ANKRD44 | 0,240090183 | 1,181066488 | 0,4107626 | 0,882835102 | ankyrin repeat domain 44 |
| ENSG00000163950 | SLBP | 0,240035246 | 1,181021514 | 0,27913921 | 0,790013761 | stem-loop binding protein |
| ENSG00000215795 | AL390728,2 | 0,240019538 | 1,181008655 | 0,67206942 | 0,962290553 | peroxisomal biogenesis factor 5 (PEX5) pseudogene |
| ENSG00000140299 | BNIP2 | 0,239986184 | 1,180981351 | 0,55599731 | 0,938031417 | BCL2 interacting protein 2 |
| ENSG00000204677 | FAM153CP | 0,239945603 | 1,180948133 | 0,7192737 | 0,971869926 | family with sequence similarity 153 member C, pseudogene |
| ENSG00000121073 | SLC35B1 | 0,239882283 | 1,180896302 | 0,36130826 | 0,851821025 | solute carrier family 35 member B1 |
| ENSG00000185596 | WASH3P | 0,23987456 | 1,180889981 | 0,57952256 | 0,943828882 | WASP family homolog 3, pseudogene |
| ENSG00000288207 | WASH3P | 0,23987456 | 1,180889981 | 0,57952256 | 0,943828882 | WASP family homolog 3, pseudogene |
| ENSG00000114993 | RTKN | 0,239819179 | 1,18084465 | 0,41395303 | 0,885456174 | rhotekin |
| ENSG00000139405 | RITA1 | 0,239815101 | 1,180841312 | 0,34686533 | 0,841625668 | RBPJ interacting and tubulin associated 1 |
| ENSG00000161526 | SAP30BP | 0,23975342 | 1,180790828 | 0,3266826 | 0,828503246 | SAP30 binding protein |
| ENSG00000272321 | AP003355,2 | 0,239594627 | 1,180660869 | 0,75611096 | 0,975901808 | novel transcript, antisense to STK3 |
| ENSG00000165219 | GAPVD1 | 0,239529759 | 1,180607784 | 0,4059206 | 0,878918323 | GTPase activating protein and VPS9 domains 1 |
| ENSG00000110025 | SNX15 | 0,239309228 | 1,18042733 | 0,4728202 | 0,909509054 | sorting nexin 15 |
| ENSG00000271869 | AC026979,2 | 0,239295884 | 1,180416412 | 0,67025164 | 0,962026636 | novel transcript |
| ENSG00000144369 | FAM171B | 0,239285598 | 1,180407996 | 0,62189466 | 0,952793934 | family with sequence similarity 171 member B |
| ENSG00000174059 | CD34 | 0,239257199 | 1,180384759 | 0,70509195 | 0,969413904 | CD34 molecule |
| ENSG00000269983 | AC146944,2 | 0,239191604 | 1,180331093 | 0,83401554 | 0,984905093 | novel transcript |
| ENSG00000125734 | GPR108 | 0,239135114 | 1,180284877 | 0,36199229 | 0,852161624 | G protein-coupled receptor 108 |
| ENSG00000262814 | MRPL12 | 0,239118838 | 1,180271561 | 0,4745118 | 0,91069468 | mitochondrial ribosomal protein L12 |
| ENSG00000231903 | AC079354,1 | 0,239106102 | 1,180261141 | 0,84939516 | 0,987036295 | novel transcript |
| ENSG00000260086 | AC007611,1 | 0,239072807 | 1,180233904 | 0,72556487 | 0,97326334 | novel transcript |
| ENSG00000260340 | LINC02176 | 0,239071387 | 1,180232742 | 0,7867391 | 0,979320406 | long intergenic non-protein coding RNA 2176 |

| | | | | | | |
|---|---|---|---|---|---|---|
| ENSG00000144045 | DQX1 | 0,239014569 | 1,180186261 | 0,79652219 | 0,980064923 | DEAQ-box RNA dependent ATPase 1 |
| ENSG00000049759 | NEDD4L | 0,239007991 | 1,18018088 | 0,46782183 | 0,907940653 | NEDD4 like E3 ubiquitin protein ligase |
| ENSG00000123143 | PKN1 | 0,238900929 | 1,180093303 | 0,59002283 | 0,947252221 | protein kinase N1 |
| ENSG00000144840 | RABL3 | 0,238889252 | 1,180083751 | 0,51502057 | 0,924720046 | RAB, member of RAS oncogene family like 3 |
| ENSG00000272945 | AL356299,2 | 0,238651794 | 1,179889533 | 0,7975024 | 0,980161849 | novel transcript |
| ENSG00000074695 | LMAN1 | 0,238629177 | 1,179871036 | 0,47925852 | 0,912714933 | lectin, mannose binding 1 |
| ENSG00000249258 | AC079193,2 | 0,238628735 | 1,179870674 | 0,72231788 | 0,972594421 | novel transcript |
| ENSG00000090372 | STRN4 | 0,238609391 | 1,179854855 | 0,41659038 | 0,886367338 | striatin 4 |
| ENSG00000189077 | TMEM120A | 0,238548494 | 1,179805054 | 0,44005429 | 0,897906863 | transmembrane protein 120A |
| ENSG00000186001 | LRCH3 | 0,238489439 | 1,179756761 | 0,27533283 | 0,78698169 | leucine rich repeats and calponin homology domain containing 3 |
| ENSG00000287100 | AL078600,1 | 0,238118418 | 1,179453399 | 0,83548786 | 0,985034063 | novel transcript |
| ENSG00000169718 | DUS1L | 0,238060603 | 1,179406135 | 0,52943 | 0,930491829 | dihydrouridine synthase 1 like |
| ENSG00000259692 | AC104041,1 | 0,238047939 | 1,179395782 | 0,37257675 | 0,860420124 | novel transcript |
| ENSG00000140400 | MAN2C1 | 0,238030165 | 1,179381252 | 0,52868825 | 0,930033144 | mannosidase alpha class 2C member 1 |
| ENSG00000198805 | PNP | 0,238011941 | 1,179366354 | 0,44870945 | 0,902062594 | purine nucleoside phosphorylase |
| ENSG00000249500 | LINC01179 | 0,237989307 | 1,179347851 | 0,79753438 | 0,980161849 | long intergenic non-protein coding RNA 1179 |
| ENSG00000249859 | PVT1 | 0,2379157 | 1,179287682 | 0,48681644 | 0,914444641 | Pvt1 oncogene |
| ENSG00000179965 | ZNF771 | 0,237874471 | 1,179253981 | 0,50970411 | 0,922427526 | zinc finger protein 771 |
| ENSG00000141750 | STAC2 | 0,23776057 | 1,179160882 | 0,76536841 | 0,976830764 | SH3 and cysteine rich domain 2 |
| ENSG00000185186 | LINC00313 | 0,237681184 | 1,179096 | 0,88579613 | 0,990717729 | long intergenic non-protein coding RNA 313 |
| ENSG00000168496 | FEN1 | 0,237589753 | 1,179021276 | 0,27423016 | 0,7862019 | flap structure-specific endonuclease 1 |
| ENSG00000284703 | AL805961,1 | 0,237580266 | 1,179013523 | 0,71936432 | 0,971869926 | novel transcript |
| ENSG00000124370 | MCEE | 0,23757504 | 1,179009253 | 0,53253747 | 0,931291085 | methylmalonyl-CoA epimerase |
| ENSG00000074603 | DPP8 | 0,2375738 | 1,179008239 | 0,5222521 | 0,927617924 | dipeptidyl peptidase 8 |
| ENSG00000173261 | PLAC8L1 | 0,237543636 | 1,178983589 | 0,63125978 | 0,953738253 | PLAC8 like 1 |
| ENSG00000232164 | LINC01873 | 0,237536715 | 1,178977932 | 0,42683816 | 0,892261855 | long intergenic non-protein coding RNA 1873 |
| ENSG00000167315 | ACAA2 | 0,237531796 | 1,178973913 | 0,37378311 | 0,861319894 | acetyl-CoA acyltransferase 2 |
| ENSG00000183309 | ZNF623 | 0,237521543 | 1,178965534 | 0,44426343 | 0,900210555 | zinc finger protein 623 |
| ENSG00000132740 | IGHMBP2 | 0,237484193 | 1,178935012 | 0,53774051 | 0,932883277 | immunoglobulin mu DNA binding protein 2 |
| ENSG00000155275 | TRMT44 | 0,237468884 | 1,178922502 | 0,43820722 | 0,897177404 | tRNA methyltransferase 44 homolog |
| ENSG00000225553 | PHF1 | 0,237428321 | 1,178889355 | 0,35463427 | 0,846034781 | PHD finger protein 1 |
| ENSG00000213671 | OLA1P2 | 0,237347548 | 1,178823354 | 0,77911396 | 0,978587255 | OLA1 pseudogene 2 |

| | | | | | | |
|---|---|---|---|---|---|---|
| ENSG00000028310 | BRD9 | 0,237191479 | 1,178695838 | 0,50201994 | 0,920520138 | bromodomain containing 9 |
| ENSG00000265298 | AC132812,1 | 0,237160393 | 1,17867044 | 0,70503456 | 0,969413904 | small nuclear ribonucleoprotein 200kDa (U5) (SNRNP200) pseudogene |
| ENSG00000171100 | MTM1 | 0,237138609 | 1,178652643 | 0,53890682 | 0,933348247 | myotubularin 1 |
| ENSG00000182473 | EXOC7 | 0,237043247 | 1,178574737 | 0,25415305 | 0,765569035 | exocyst complex component 7 |
| ENSG00000150527 | MIA2 | 0,237036454 | 1,178569187 | 0,41366015 | 0,885121175 | MIA SH3 domain ER export factor 2 |
| ENSG00000243725 | TTC4 | 0,23702299 | 1,178558188 | 0,28226028 | 0,793283115 | tetratricopeptide repeat domain 4 |
| ENSG00000184203 | PPP1R2 | 0,237017323 | 1,178553559 | 0,54074474 | 0,933385624 | protein phosphatase 1 regulatory inhibitor subunit 2 |
| ENSG00000223887 | PRR3 | 0,236917191 | 1,178471763 | 0,51237849 | 0,923445033 | proline rich 3 |
| ENSG00000213015 | ZNF580 | 0,236747029 | 1,178332773 | 0,58658924 | 0,945395317 | zinc finger protein 580 |
| ENSG00000172977 | KAT5 | 0,236636055 | 1,178242138 | 0,32108867 | 0,824586518 | lysine acetyltransferase 5 |
| ENSG00000223510 | CDRT15 | 0,236635446 | 1,178241641 | 0,78270965 | 0,978678077 | CMT1A duplicated region transcript 15 |
| ENSG00000106290 | TAF6 | 0,236595507 | 1,178209024 | 0,463758 | 0,907251937 | TATA-box binding protein associated factor 6 |
| ENSG00000275797 | AC008746,4 | 0,236577091 | 1,178193983 | 0,72501374 | 0,97326334 | novel transcript |
| ENSG00000267838 | AC245884,8 | 0,236577091 | 1,178193983 | 0,72501374 | 0,97326334 | novel transcript |
| ENSG00000144233 | AMMECR1L | 0,236565318 | 1,178184369 | 0,28128698 | 0,792337715 | AMMECR1 like |
| ENSG00000185024 | BRF1 | 0,236549324 | 1,178171307 | 0,61098927 | 0,951873283 | BRF1 RNA polymerase III transcription initiation factor subunit |
| ENSG00000143367 | TUFT1 | 0,236516778 | 1,178144729 | 0,49455766 | 0,91758518 | tuftelin 1 |
| ENSG00000116661 | FBXO2 | 0,236467964 | 1,178104867 | 0,74353419 | 0,975310206 | F-box protein 2 |
| ENSG00000139865 | TTC6 | 0,236395621 | 1,178045793 | 0,76883686 | 0,977381742 | tetratricopeptide repeat domain 6 |
| ENSG00000187446 | CHP1 | 0,236393321 | 1,178043915 | 0,34548499 | 0,841168721 | calcineurin like EF-hand protein 1 |
| ENSG00000146007 | ZMAT2 | 0,236352556 | 1,178010629 | 0,31660786 | 0,821646322 | zinc finger matrin-type 2 |
| ENSG00000123349 | PFDN5 | 0,236331865 | 1,177993734 | 0,40695543 | 0,880078305 | prefoldin subunit 5 |
| ENSG00000273350 | AC004832,5 | 0,236306511 | 1,177973032 | 0,73978622 | 0,974715504 | novel transcript, sense intronic to SEC14L2 |
| ENSG00000176142 | TMEM39A | 0,236274973 | 1,177947281 | 0,46216852 | 0,906306011 | transmembrane protein 39A |
| ENSG00000287624 | AL590666,5 | 0,236237767 | 1,177916903 | 0,68093678 | 0,964654539 | novel transcript, antisense to PRCC |
| ENSG00000107551 | RASSF4 | 0,236178546 | 1,177868552 | 0,30650506 | 0,815375074 | Ras association domain family member 4 |
| ENSG00000101310 | SEC23B | 0,236108972 | 1,177811751 | 0,30203564 | 0,811449876 | SEC23 homolog B, COPII coat complex component |
| ENSG00000285625 | AC117378,1 | 0,236096893 | 1,17780189 | 0,78702441 | 0,979424965 | novel protein |
| ENSG00000160781 | PAQR6 | 0,236065297 | 1,177776095 | 0,55457161 | 0,937793851 | progestin and adipoQ receptor family member 6 |
| ENSG00000229917 | RPL7P46 | 0,235790423 | 1,177551717 | 0,80816942 | 0,981213877 | ribosomal protein L7 pseudogene 46 |
| ENSG00000165475 | CRYL1 | 0,235498927 | 1,177313817 | 0,62972471 | 0,952812183 | crystallin lambda 1 |

| | | | | | | |
|---|---|---|---|---|---|---|
| ENSG00000120675 | DNAJC15 | 0,235488271 | 1,177305121 | 0,60195223 | 0,949939978 | DnaJ heat shock protein family (Hsp40) member C15 |
| ENSG00000164405 | UQCRQ | 0,235457875 | 1,177280317 | 0,30576314 | 0,814673493 | ubiquinol-cytochrome c reductase complex III subunit VII |
| ENSG00000101460 | MAP1LC3A | 0,2354233 | 1,177252103 | 0,38845709 | 0,869820053 | microtubule associated protein 1 light chain 3 alpha |
| ENSG00000161265 | U2AF1L4 | 0,235420626 | 1,177249921 | 0,40288535 | 0,877551313 | U2 small nuclear RNA auxiliary factor 1 like 4 |
| ENSG00000267560 | AC027514,2 | 0,235390166 | 1,177225065 | 0,75231358 | 0,975901808 | novel transcript, antisense to KIAA1468 |
| ENSG00000173120 | KDM2A | 0,235267891 | 1,177125295 | 0,33929406 | 0,838276153 | lysine demethylase 2A |
| ENSG00000130558 | OLFM1 | 0,235222262 | 1,177088066 | 0,40521889 | 0,878185962 | olfactomedin 1 |
| ENSG00000157680 | DGKI | 0,235158412 | 1,177035972 | 0,50092411 | 0,920520138 | diacylglycerol kinase iota |
| ENSG00000118564 | FBXL5 | 0,235080716 | 1,176972585 | 0,50885989 | 0,922427526 | F-box and leucine rich repeat protein 5 |
| ENSG00000272735 | AC007881,3 | 0,235070932 | 1,176964603 | 0,80164773 | 0,980623173 | novel transcript |
| ENSG00000108091 | CCDC6 | 0,23499776 | 1,17690491 | 0,50590053 | 0,921780819 | coiled-coil domain containing 6 |
| ENSG00000162300 | ZFPL1 | 0,234920772 | 1,176842107 | 0,28543832 | 0,796948543 | zinc finger protein like 1 |
| ENSG00000170310 | STX8 | 0,2348641 | 1,176795879 | 0,30884625 | 0,816831843 | syntaxin 8 |
| ENSG00000228392 | AL162391,1 | 0,234861254 | 1,176793558 | 0,73401376 | 0,974353474 | novel transcript |
| ENSG00000151812 | SLC35F4 | 0,234834513 | 1,176771746 | 0,77143941 | 0,978033855 | solute carrier family 35 member F4 |
| ENSG00000162819 | BROX | 0,234821228 | 1,17676091 | 0,61738145 | 0,952793934 | BRO1 domain and CAAX motif containing |
| ENSG00000070413 | DGCR2 | 0,234803045 | 1,176746079 | 0,45392326 | 0,902912642 | DiGeorge syndrome critical region gene 2 |
| ENSG00000244257 | PKD1P1 | 0,234784292 | 1,176730782 | 0,76459812 | 0,976640468 | polycystin 1, transient receptor potential channel interacting pseudogene 1 |
| ENSG00000165637 | VDAC2 | 0,23478167 | 1,176728644 | 0,43258119 | 0,89560871 | voltage dependent anion channel 2 |
| ENSG00000174446 | SNAPC5 | 0,234777503 | 1,176725245 | 0,44318738 | 0,899727948 | small nuclear RNA activating complex polypeptide 5 |
| ENSG00000107566 | ERLIN1 | 0,234698495 | 1,176660804 | 0,44766683 | 0,902062594 | ER lipid raft associated 1 |
| ENSG00000235125 | NFKBIL1 | 0,234578242 | 1,17656273 | 0,62923389 | 0,952793934 | NFKB inhibitor like 1 |
| ENSG00000267308 | LINC01764 | 0,234540914 | 1,176532288 | 0,75534716 | 0,975901808 | long intergenic non-protein coding RNA 1764 |
| ENSG00000197345 | MRPL21 | 0,234539932 | 1,176531488 | 0,37563695 | 0,862191735 | mitochondrial ribosomal protein L21 |
| ENSG00000149182 | ARFGAP2 | 0,234346992 | 1,176374154 | 0,37627146 | 0,86257156 | ADP ribosylation factor GTPase activating protein 2 |
| ENSG00000260249 | AC007608,3 | 0,234334579 | 1,176364032 | 0,64118654 | 0,956465487 | novel transcript, antisense to SNX20 |
| ENSG00000067445 | TRO | 0,234256056 | 1,176300007 | 0,30937404 | 0,816845699 | trophinin |
| ENSG00000136213 | CHST12 | 0,234078551 | 1,176155287 | 0,57917629 | 0,943828882 | carbohydrate sulfotransferase 12 |
| ENSG00000125485 | DDX31 | 0,234054572 | 1,176135738 | 0,33172504 | 0,832548086 | DEAD-box helicase 31 |
| ENSG00000100764 | PSMC1 | 0,233973719 | 1,176069826 | 0,32605354 | 0,828085941 | proteasome 26S subunit, ATPase 1 |
| ENSG00000033122 | LRRC7 | 0,233971406 | 1,176067941 | 0,63902997 | 0,956018518 | leucine rich repeat containing 7 |

| | | | | | | |
|---|---|---|---|---|---|---|
| ENSG00000278068 | AL136317,3 | 0,233937421 | 1,176040237 | 0,8021632 | 0,980710009 | calponin 2 (CNN2) pseudogene |
| ENSG00000260740 | AC026471,3 | 0,23392455 | 1,176029745 | 0,9154115 | 0,993361299 | novel transcript, antisense to SLC5A2 |
| ENSG00000225387 | AL591896,1 | 0,233872124 | 1,17598701 | 0,85872283 | 0,989074806 | novel transcript |
| ENSG00000100023 | PPIL2 | 0,233808712 | 1,175935322 | 0,30775709 | 0,816424917 | peptidylprolyl isomerase like 2 |
| ENSG00000180581 | SRP9P1 | 0,233775495 | 1,175908247 | 0,60179224 | 0,949939978 | signal recognition particle 9 pseudogene 1 |
| ENSG00000167107 | ACSF2 | 0,233662943 | 1,175816512 | 0,39353465 | 0,8715972 | acyl-CoA synthetase family member 2 |
| ENSG00000141699 | RETREG3 | 0,233636822 | 1,175795223 | 0,27913539 | 0,790013761 | reticulophagy regulator family member 3 |
| ENSG00000133250 | ZNF414 | 0,233600208 | 1,175765384 | 0,51563079 | 0,924926785 | zinc finger protein 414 |
| ENSG00000198818 | SFT2D1 | 0,233575325 | 1,175745104 | 0,50627174 | 0,921780819 | SFT2 domain containing 1 |
| ENSG00000173614 | NMNAT1 | 0,233558191 | 1,175731141 | 0,43692598 | 0,896866043 | nicotinamide nucleotide adenylyltransferase 1 |
| ENSG00000224029 | AL450344,1 | 0,233461693 | 1,175652502 | 0,80566644 | 0,981213877 | novel transcript |
| ENSG00000116809 | ZBTB17 | 0,233255979 | 1,175484877 | 0,57949959 | 0,943828882 | zinc finger and BTB domain containing 17 |
| ENSG00000277118 | Metazoa_SRP | 0,233230363 | 1,175464006 | 0,78863601 | 0,979462318 | Metazoan signal recognition particle RNA |
| ENSG00000134575 | ACP2 | 0,233108095 | 1,17536439 | 0,35265627 | 0,844991243 | acid phosphatase 2, lysosomal |
| ENSG00000225937 | PCA3 | 0,233104285 | 1,175361286 | 0,69964343 | 0,968522643 | prostate cancer associated 3 |
| ENSG00000203402 | AL591503,1 | 0,233077677 | 1,175339609 | 0,74457411 | 0,975649186 | novel transcript |
| ENSG00000154122 | ANKH | 0,232993133 | 1,175270734 | 0,38604679 | 0,869112622 | ANKH inorganic pyrophosphate transport regulator |
| ENSG00000160447 | PKN3 | 0,232969721 | 1,175251662 | 0,59811727 | 0,948582224 | protein kinase N3 |
| ENSG00000232628 | AC092809,2 | 0,232800663 | 1,175113952 | 0,83808928 | 0,985672131 | novel transcript |
| ENSG00000115761 | NOL10 | 0,232784539 | 1,175100819 | 0,40735059 | 0,880314995 | nucleolar protein 10 |
| ENSG00000160908 | ZNF394 | 0,232741613 | 1,175065855 | 0,4904851 | 0,916917372 | zinc finger protein 394 |
| ENSG00000142444 | TIMM29 | 0,232705618 | 1,175036537 | 0,46341647 | 0,907101775 | translocase of inner mitochondrial membrane 29 |
| ENSG00000136731 | UGGT1 | 0,232665932 | 1,175004215 | 0,25947467 | 0,772164235 | UDP-glucose glycoprotein glucosyltransferase 1 |
| ENSG00000258818 | RNASE4 | 0,232453973 | 1,174831597 | 0,67739275 | 0,963580269 | ribonuclease A family member 4 |
| ENSG00000273850 | PPP2R3B | 0,232297018 | 1,174703791 | 0,54357584 | 0,934259495 | protein phosphatase 2 regulatory subunit B''beta |
| ENSG00000186103 | ARGFX | 0,232258964 | 1,174672807 | 0,75348492 | 0,975901808 | arginine-fifty homeobox |
| ENSG00000149929 | HIRIP3 | 0,232257979 | 1,174672004 | 0,5946951 | 0,947469769 | HIRA interacting protein 3 |
| ENSG00000165389 | SPTSSA | 0,232196425 | 1,174621887 | 0,63762437 | 0,956017934 | serine palmitoyltransferase small subunit A |
| ENSG00000167701 | GPT | 0,231973725 | 1,174440581 | 0,78183425 | 0,978597905 | glutamic--pyruvic transaminase |
| ENSG00000120992 | LYPLA1 | 0,231925938 | 1,174401681 | 0,63886346 | 0,956018518 | lysophospholipase 1 |
| ENSG00000169972 | PUSL1 | 0,23192526 | 1,174401129 | 0,53378096 | 0,931381467 | pseudouridine synthase like 1 |
| ENSG00000274587 | LILRB3 | 0,231915497 | 1,174393182 | 0,86570295 | 0,989074806 | leukocyte immunoglobulin like receptor B3 |
| ENSG00000143543 | JTB | 0,231867852 | 1,174354398 | 0,30295923 | 0,81253474 | jumping translocation breakpoint |

| | | | | | | |
|---|---|---|---|---|---|---|
| ENSG00000228302 | AL512770,1 | 0,23176239 | 1,174268555 | 0,56394321 | 0,940865625 | novel transcript |
| ENSG00000196932 | TMEM26 | 0,231761498 | 1,174267829 | 0,77773358 | 0,978587255 | transmembrane protein 26 |
| ENSG00000198954 | KIFBP | 0,231725393 | 1,174238442 | 0,37647151 | 0,862818448 | kinesin family binding protein |
| ENSG00000267642 | AC091198,1 | 0,231707102 | 1,174223555 | 0,82977365 | 0,984144281 | novel transcript |
| ENSG00000253623 | AC009879,1 | 0,231669304 | 1,17419279 | 0,84384658 | 0,986273696 | TAF9 RNA polymerase II, TATA box binding protein (TBP)-associated factor, 32kDa (TAF9) pseudogene |
| ENSG00000288052 | AL356952,1 | 0,231656003 | 1,174181965 | 0,79903888 | 0,980161849 | novel transcript |
| ENSG00000260267 | AC026471,1 | 0,231638093 | 1,174167388 | 0,49129771 | 0,916995341 | novel transcript, antisense to ARMC5 |
| ENSG00000170653 | ATF7 | 0,23158445 | 1,174123731 | 0,34205703 | 0,840048641 | activating transcription factor 7 |
| ENSG00000270387 | AC115621,1 | 0,23156877 | 1,17411097 | 0,85660614 | 0,988299298 | ret finger protein-like 4A (RFPL4A) pseudogene |
| ENSG00000254783 | AP003084,1 | 0,231362916 | 1,173943452 | 0,70149673 | 0,968742626 | immunoglobin superfamily, member 21 (IGSF21) pseudogene |
| ENSG00000104312 | RIPK2 | 0,231303985 | 1,1738955 | 0,50476053 | 0,921400408 | receptor interacting serine/threonine kinase 2 |
| ENSG00000177189 | RPS6KA3 | 0,231236773 | 1,173840812 | 0,45457372 | 0,903383444 | ribosomal protein S6 kinase A3 |
| ENSG00000009830 | POMT2 | 0,231153375 | 1,173772958 | 0,32414085 | 0,826248637 | protein O-mannosyltransferase 2 |
| ENSG00000108433 | GOSR2 | 0,231147099 | 1,173767851 | 0,33243298 | 0,83293962 | golgi SNAP receptor complex member 2 |
| ENSG00000213839 | TMX2P1 | 0,231120323 | 1,173746066 | 0,45892445 | 0,904597057 | thioredoxin related transmembrane protein 2 pseudogene 1 |
| ENSG00000204220 | PFDN6 | 0,231106648 | 1,173734941 | 0,46163317 | 0,906165604 | prefoldin subunit 6 |
| ENSG00000266775 | AC011840,4 | 0,230988436 | 1,173638771 | 0,84874259 | 0,987036295 | novel transcript, antisense to TBC1D29 |
| ENSG00000273270 | AC090114,2 | 0,230896376 | 1,173563882 | 0,51690286 | 0,92567597 | novel transcript |
| ENSG00000236942 | AC096533,1 | 0,230870904 | 1,173543162 | 0,80480575 | 0,98109454 | GABA(A) receptor-associated protein (GABARAP) pseudogene |
| ENSG00000075391 | RASAL2 | 0,230861282 | 1,173535335 | 0,44009423 | 0,897906863 | RAS protein activator like 2 |
| ENSG00000135823 | STX6 | 0,23085704 | 1,173531884 | 0,37077247 | 0,858780045 | syntaxin 6 |
| ENSG00000105058 | FAM32A | 0,230794324 | 1,173480871 | 0,34402195 | 0,840938551 | family with sequence similarity 32 member A |
| ENSG00000161267 | BDH1 | 0,230736814 | 1,173434093 | 0,61561975 | 0,952793934 | 3-hydroxybutyrate dehydrogenase 1 |
| ENSG00000224945 | AL353150,1 | 0,230517265 | 1,173255533 | 0,64010936 | 0,956157624 | novel transcript |
| ENSG00000196976 | LAGE3 | 0,230506431 | 1,173246723 | 0,56129451 | 0,940567693 | L antigen family member 3 |
| ENSG00000154930 | ACSS1 | 0,230404375 | 1,173163731 | 0,69922155 | 0,968522643 | acyl-CoA synthetase short chain family member 1 |
| ENSG00000144677 | CTDSPL | 0,230367305 | 1,173133586 | 0,46351469 | 0,907101775 | CTD small phosphatase like |
| ENSG00000234769 | WASH4P | 0,230320646 | 1,173095647 | 0,66041071 | 0,959163386 | WASP family homolog 4, pseudogene |
| ENSG00000184985 | SORCS2 | 0,230297212 | 1,173076592 | 0,53106452 | 0,931265667 | sortilin related VPS10 domain containing receptor 2 |

| | | | | | | |
|---|---|---|---|---|---|---|
| ENSG00000278638 | AC244093,5 | 0,230263309 | 1,173049025 | 0,8131753 | 0,981246822 | novel transcript, antisense AATF |
| ENSG00000139291 | TMEM19 | 0,230253706 | 1,173041217 | 0,39644726 | 0,873099971 | transmembrane protein 19 |
| ENSG00000151502 | VPS26B | 0,230245424 | 1,173034483 | 0,30725811 | 0,816083392 | VPS26, retromer complex component B |
| ENSG00000253978 | CTB-178M22,2 | 0,230203398 | 1,173000313 | 0,72015107 | 0,972142678 | uncharacterized LOC101927862 |
| ENSG00000112208 | BAG2 | 0,23010458 | 1,17291997 | 0,42349343 | 0,890929178 | BAG cochaperone 2 |
| ENSG00000160410 | SHKBP1 | 0,23004951 | 1,172875199 | 0,41783893 | 0,886857227 | SH3KBP1 binding protein 1 |
| ENSG00000288258 | BX470111,1 | 0,230041598 | 1,172868767 | 0,79458156 | 0,979776981 | novel protein |
| ENSG00000157895 | C12orf43 | 0,230030251 | 1,172859542 | 0,41874832 | 0,887094957 | chromosome 12 open reading frame 43 |
| ENSG00000105778 | AVL9 | 0,229978362 | 1,172817359 | 0,43547137 | 0,89634451 | AVL9 cell migration associated |
| ENSG00000170634 | ACYP2 | 0,229847677 | 1,172711125 | 0,36857795 | 0,857573944 | acylphosphatase 2 |
| ENSG00000119977 | TCTN3 | 0,229800076 | 1,172672433 | 0,38851217 | 0,869820053 | tectonic family member 3 |
| ENSG00000230516 | AL589642,2 | 0,229756548 | 1,172637052 | 0,83206077 | 0,984653014 | novel transcript |
| ENSG00000250303 | LINC02762 | 0,229750012 | 1,17263174 | 0,57936646 | 0,943828882 | long intergenic non-protein coding RNA 2762 |
| ENSG00000215845 | TSTD1 | 0,229696491 | 1,172588239 | 0,77998747 | 0,978587255 | thiosulfate sulfurtransferase like domain containing 1 |
| ENSG00000172663 | TMEM134 | 0,229678581 | 1,172573681 | 0,48694128 | 0,914557983 | transmembrane protein 134 |
| ENSG00000285339 | VPS28 | 0,229552171 | 1,172470944 | 0,33508503 | 0,834858938 | VPS28 subunit of ESCRT-I |
| ENSG00000124116 | WFDC3 | 0,229515102 | 1,172440819 | 0,79731887 | 0,980161849 | WAP four-disulfide core domain 3 |
| ENSG00000111727 | HCFC2 | 0,229500663 | 1,172429085 | 0,59360308 | 0,947469769 | host cell factor C2 |
| ENSG00000127663 | KDM4B | 0,22948 | 1,172412293 | 0,50182865 | 0,920520138 | lysine demethylase 4B |
| ENSG00000225489 | AL354707,1 | 0,229453408 | 1,172390683 | 0,7144509 | 0,971029555 | novel transcript |
| ENSG00000116514 | RNF19B | 0,229371373 | 1,17232402 | 0,30144615 | 0,811013605 | ring finger protein 19B |
| ENSG00000222404 | RN7SKP281 | 0,22914952 | 1,172143758 | 0,72820306 | 0,973888243 | RN7SK pseudogene 281 |
| ENSG00000280071 | GATD3B | 0,229138127 | 1,172134501 | 0,40963021 | 0,882177108 | glutamine amidotransferase like class 1 domain containing 3B |
| ENSG00000166002 | SMCO4 | 0,229095258 | 1,172099672 | 0,41095026 | 0,882983885 | single-pass membrane protein with coiled-coil domains 4 |
| ENSG00000196663 | TECPR2 | 0,229071028 | 1,172079987 | 0,34389361 | 0,840901506 | tectonin beta-propeller repeat containing 2 |
| ENSG00000267348 | GEMIN7-AS1 | 0,229057032 | 1,172068616 | 0,5578628 | 0,938984031 | GEMIN7 antisense RNA 1 |
| ENSG00000057663 | ATG5 | 0,228998636 | 1,172021175 | 0,53789858 | 0,932883277 | autophagy related 5 |
| ENSG00000066827 | ZFAT | 0,22899299 | 1,172016589 | 0,34636689 | 0,841168721 | zinc finger and AT-hook domain containing |
| ENSG00000149635 | OCSTAMP | 0,228965006 | 1,171993855 | 0,6752232 | 0,963390425 | osteoclast stimulatory transmembrane protein |
| ENSG00000266289 | AC012213,3 | 0,228898725 | 1,171940013 | 0,77593811 | 0,978587255 | novel transcript |

| | | | | | | |
|---|---|---|---|---|---|---|
| ENSG00000158669 | GPAT4 | 0,228873304 | 1,171919362 | 0,35285553 | 0,845031687 | glycerol-3-phosphate acyltransferase 4 |
| ENSG00000245750 | DRAIC | 0,228853707 | 1,171903443 | 0,65248751 | 0,957275891 | downregulated RNA in cancer, inhibitor of cell invasion and migration |
| ENSG00000139974 | SLC38A6 | 0,228845324 | 1,171896634 | 0,49572284 | 0,918046024 | solute carrier family 38 member 6 |
| ENSG00000175595 | ERCC4 | 0,228800003 | 1,17185982 | 0,28342352 | 0,79458592 | ERCC excision repair 4, endonuclease catalytic subunit |
| ENSG00000101849 | TBL1X | 0,228745023 | 1,171815163 | 0,33064833 | 0,832114797 | transducin beta like 1 X-linked |
| ENSG00000170759 | KIF5B | 0,228714401 | 1,17179029 | 0,50571545 | 0,921780819 | kinesin family member 5B |
| ENSG00000185900 | POMK | 0,228646055 | 1,17173478 | 0,29927734 | 0,809131439 | protein O-mannose kinase |
| ENSG00000177981 | ASB8 | 0,228614332 | 1,171709015 | 0,33948613 | 0,838364485 | ankyrin repeat and SOCS box containing 8 |
| ENSG00000110321 | EIF4G2 | 0,228585729 | 1,171685785 | 0,44016896 | 0,897914806 | eukaryotic translation initiation factor 4 gamma 2 |
| ENSG00000261238 | AC009166,1 | 0,22851966 | 1,171632128 | 0,81779737 | 0,982323352 | novel transcript, antisense to SALL1 |
| ENSG00000176058 | TPRN | 0,228394592 | 1,171530563 | 0,60564567 | 0,950927937 | taperin |
| ENSG00000198837 | DENND4B | 0,228326193 | 1,171475021 | 0,39814087 | 0,874759093 | DENN domain containing 4B |
| ENSG00000276141 | WHAMMP3 | 0,228316382 | 1,171467055 | 0,35536272 | 0,846840543 | WAS protein homolog associated with actin, golgi membranes and microtubules pseudogene 3 |
| ENSG00000235910 | APOA1-AS | 0,228309191 | 1,171461216 | 0,68237428 | 0,964760518 | APOA1 antisense RNA |
| ENSG00000106683 | LIMK1 | 0,228278605 | 1,171436381 | 0,467083 | 0,907940653 | LIM domain kinase 1 |
| ENSG00000233942 | AC004012,1 | 0,228265053 | 1,171425377 | 0,71298209 | 0,970812834 | novel transcript |
| ENSG00000152147 | GEMIN6 | 0,228237946 | 1,171403367 | 0,33916142 | 0,838276153 | gem nuclear organelle associated protein 6 |
| ENSG00000177700 | POLR2L | 0,228235346 | 1,171401256 | 0,37241486 | 0,86029098 | RNA polymerase II, I and III subunit L |
| ENSG00000228816 | AK3P5 | 0,22809193 | 1,171284815 | 0,6005139 | 0,949439306 | adenylate kinase 3 pseudogene 5 |
| ENSG00000260778 | AC009065,4 | 0,228023605 | 1,171229345 | 0,76700853 | 0,977201891 | novel transcript |
| ENSG00000125787 | GNRH2 | 0,22774774 | 1,171005409 | 0,77484202 | 0,978553937 | gonadotropin releasing hormone 2 |
| ENSG00000161270 | NPHS1 | 0,227737914 | 1,170997433 | 0,8149921 | 0,981521413 | NPHS1 adhesion molecule, nephrin |
| ENSG00000138768 | USO1 | 0,227688258 | 1,17095713 | 0,52492425 | 0,928616707 | USO1 vesicle transport factor |
| ENSG00000251602 | AL928654,1 | 0,22753658 | 1,170834027 | 0,54543298 | 0,935108602 | novel transcript, antisense to MTA1 |
| ENSG00000230333 | AC004160,1 | 0,227513777 | 1,170815522 | 0,68547437 | 0,965476003 | novel transcript, antisense to THSD7A |
| ENSG00000269955 | FMC1-LUC7L2 | 0,227399948 | 1,170723148 | 0,38764915 | 0,869820053 | FMC1-LUC7L2 readthrough |
| ENSG00000163956 | LRPAP1 | 0,227330109 | 1,170666476 | 0,53253936 | 0,931291085 | LDL receptor related protein associated protein 1 |
| ENSG00000165406 | MARCHF8 | 0,227186424 | 1,170549889 | 0,47455582 | 0,91069468 | membrane associated ring-CH-type finger 8 |
| ENSG00000287586 | Z98886,1 | 0,227158619 | 1,170527329 | 0,7624655 | 0,976315588 | novel transcript, antisense to AJAP1 |
| ENSG00000148358 | GPR107 | 0,227127517 | 1,170502095 | 0,32681758 | 0,828612831 | G protein-coupled receptor 107 |

| | | | | | | |
|---|---|---|---|---|---|---|
| ENSG00000115526 | CHST10 | 0,22707033 | 1,170455699 | 0,33327897 | 0,833504026 | carbohydrate sulfotransferase 10 |
| ENSG00000287254 | AC011509,3 | 0,227064095 | 1,17045064 | 0,85765865 | 0,988893234 | novel transcript |
| ENSG00000227080 | AC012441,1 | 0,22702655 | 1,17042018 | 0,68450652 | 0,965415049 | novel pseudogene |
| ENSG00000090565 | RAB11FIP3 | 0,226944008 | 1,170353218 | 0,62449416 | 0,952793934 | RAB11 family interacting protein 3 |
| ENSG00000129559 | NEDD8 | 0,226937307 | 1,170347783 | 0,43219246 | 0,89558934 | NEDD8 ubiquitin like modifier |
| ENSG00000167862 | MRPL58 | 0,226876434 | 1,170298402 | 0,34218589 | 0,840048641 | mitochondrial ribosomal protein L58 |
| ENSG00000176055 | MBLAC2 | 0,226855093 | 1,17028109 | 0,62222327 | 0,952793934 | metallo-beta-lactamase domain containing 2 |
| ENSG00000214866 | DCDC2C | 0,226843823 | 1,170271948 | 0,8164178 | 0,981862057 | doublecortin domain containing 2C |
| ENSG00000129244 | ATP1B2 | 0,226801274 | 1,170237435 | 0,62516157 | 0,952793934 | ATPase Na+/K+ transporting subunit beta 2 |
| ENSG00000275183 | LENG9 | 0,226795457 | 1,170232716 | 0,95259298 | 0,996448973 | leukocyte receptor cluster member 9 |
| ENSG00000284722 | AP003175,1 | 0,226767459 | 1,170210006 | 0,71271985 | 0,970812834 | novel transcript |
| ENSG00000287302 | AL133477,2 | 0,226686834 | 1,17014461 | 0,84987487 | 0,987036295 | novel transcript |
| ENSG00000070814 | TCOF1 | 0,226681849 | 1,170140567 | 0,6190941 | 0,952793934 | treacle ribosome biogenesis factor 1 |
| ENSG00000213859 | KCTD11 | 0,226515224 | 1,170005429 | 0,48300969 | 0,91363215 | potassium channel tetramerization domain containing 11 |
| ENSG00000288399 | KCTD11 | 0,226515224 | 1,170005429 | 0,48300969 | 0,91363215 | potassium channel tetramerization domain containing 11 |
| ENSG00000126953 | TIMM8A | 0,226483069 | 1,169979352 | 0,44833012 | 0,902062594 | translocase of inner mitochondrial membrane 8A |
| ENSG00000275198 | AL512791,2 | 0,226479787 | 1,16997669 | 0,58964459 | 0,947017802 | novel transcript |
| ENSG00000212402 | SNORA74B | 0,226470214 | 1,169968927 | 0,81203197 | 0,981213877 | small nucleolar RNA, H/ACA box 74B |
| ENSG00000169084 | DHRSX | 0,226450216 | 1,169952709 | 0,35645591 | 0,847792901 | dehydrogenase/reductase X-linked |
| ENSG00000251550 | AC091133,5 | 0,22644312 | 1,169946955 | 0,67017537 | 0,962026636 | COX assembly mitochondrial protein homolog (S, cerevisiae) (CMC1) pseudogene |
| ENSG00000274751 | AC120498,9 | 0,226415279 | 1,169924378 | 0,73131567 | 0,974353474 | novel transcript, antisense to UBE2I |
| ENSG00000288318 | PALM3 | 0,22633293 | 1,1698576 | 0,80628973 | 0,981213877 | paralemmin 3 |
| ENSG00000187867 | PALM3 | 0,22633293 | 1,1698576 | 0,80628973 | 0,981213877 | paralemmin 3 |
| ENSG00000150753 | CCT5 | 0,226291631 | 1,169824112 | 0,3727357 | 0,860564851 | chaperonin containing TCP1 subunit 5 |
| ENSG00000140932 | CMTM2 | 0,226261299 | 1,169799517 | 0,76238359 | 0,976315588 | CKLF like MARVEL transmembrane domain containing 2 |
| ENSG00000160991 | ORAI2 | 0,226243486 | 1,169785074 | 0,58050578 | 0,943832501 | ORAI calcium release-activated calcium modulator 2 |
| ENSG00000198221 | AFDN-DT | 0,226235935 | 1,169778951 | 0,40680407 | 0,880078305 | AFDN divergent transcript |
| ENSG00000139219 | COL2A1 | 0,226121127 | 1,169685865 | 0,65154719 | 0,957089277 | collagen type II alpha 1 chain |
| ENSG00000281388 | SCARNA6 | 0,226098388 | 1,169667429 | 0,51002272 | 0,922427526 | small Cajal body-specific RNA 6 |

| | | | | | | |
|---|---|---|---|---|---|---|
| ENSG00000251791 | SCARNA6 | 0,226098388 | 1,169667429 | 0,51002272 | 0,922427526 | small Cajal body-specific RNA 6 |
| ENSG00000279571 | AL162426,1 | 0,226051574 | 1,169629476 | 0,69680947 | 0,968300554 | novel transcript |
| ENSG00000261487 | AC135048,1 | 0,226050815 | 1,16962886 | 0,67720706 | 0,963580269 | novel transcript |
| ENSG00000124831 | LRRFIP1 | 0,22597101 | 1,169564163 | 0,27765031 | 0,789067728 | LRR binding FLII interacting protein 1 |
| ENSG00000275294 | LINC02340 | 0,225928716 | 1,169529876 | 0,83696032 | 0,985452772 | long intergenic non-protein coding RNA 2340 |
| ENSG00000276935 | MBOAT7 | 0,225863887 | 1,169477323 | 0,4558173 | 0,903545728 | membrane bound O-acyltransferase domain containing 7 |
| ENSG00000246705 | H2AJ | 0,225822173 | 1,169443509 | 0,40362898 | 0,877691113 | H2A,J histone |
| ENSG00000162998 | FRZB | 0,2258021 | 1,169427238 | 0,67845477 | 0,9640473 | frizzled related protein |
| ENSG00000214128 | TMEM213 | 0,225794972 | 1,16942146 | 0,75401737 | 0,975901808 | transmembrane protein 213 |
| ENSG00000233426 | EIF3FP3 | 0,225741581 | 1,169378183 | 0,46803109 | 0,907940653 | eukaryotic translation initiation factor 3 subunit F pseudogene 3 |
| ENSG00000143294 | PRCC | 0,225704737 | 1,16934832 | 0,33596444 | 0,835660292 | proline rich mitotic checkpoint control factor |
| ENSG00000272969 | AC024243,1 | 0,225657577 | 1,169310096 | 0,75922186 | 0,976120207 | novel transcript, antisense to clock |
| ENSG00000273312 | AL121749,1 | 0,225583615 | 1,169250151 | 0,79833137 | 0,980161849 | novel transcript |
| ENSG00000037757 | MRI1 | 0,22551704 | 1,169196196 | 0,54104761 | 0,933385624 | methylthioribose-1-phosphate isomerase 1 |
| ENSG00000153574 | RPIA | 0,225512415 | 1,169192448 | 0,33470243 | 0,834757602 | ribose 5-phosphate isomerase A |
| ENSG00000196369 | SRGAP2B | 0,225480606 | 1,169166669 | 0,42309874 | 0,890698195 | SLIT-ROBO Rho GTPase activating protein 2B |
| ENSG00000183283 | DAZAP2 | 0,225475377 | 1,169162432 | 0,55002926 | 0,93612918 | DAZ associated protein 2 |
| ENSG00000131473 | ACLY | 0,225443121 | 1,169136291 | 0,32268949 | 0,824955371 | ATP citrate lyase |
| ENSG00000136371 | MTHFS | 0,225434206 | 1,169129067 | 0,50558501 | 0,921780819 | methenyltetrahydrofolate synthetase |
| ENSG00000162194 | LBHD1 | 0,225347572 | 1,169058862 | 0,44826007 | 0,902062594 | LBH domain containing 1 |
| ENSG00000272030 | AL162258,2 | 0,225341567 | 1,169053996 | 0,68880865 | 0,966556275 | novel transcript, antisense to S100A1 and CHTOP |
| ENSG00000171865 | RNASEH1 | 0,225227802 | 1,168961813 | 0,33236809 | 0,83293962 | ribonuclease H1 |
| ENSG00000075886 | TUBA3D | 0,225221762 | 1,16895692 | 0,71231196 | 0,970812834 | tubulin alpha 3d |
| ENSG00000122882 | ECD | 0,225012685 | 1,168787526 | 0,48627357 | 0,914091794 | ecdysoneless cell cycle regulator |
| ENSG00000101670 | LIPG | 0,225005377 | 1,168781605 | 0,31269573 | 0,818683344 | lipase G, endothelial type |
| ENSG00000004866 | ST7 | 0,224982919 | 1,16876341 | 0,4975252 | 0,918759905 | suppression of tumorigenicity 7 |
| ENSG00000071051 | NCK2 | 0,224982403 | 1,168762992 | 0,34592938 | 0,841168721 | NCK adaptor protein 2 |
| ENSG00000143811 | PYCR2 | 0,224936555 | 1,168725851 | 0,38547801 | 0,868895792 | pyrroline-5-carboxylate reductase 2 |
| ENSG00000115839 | RAB3GAP1 | 0,224749767 | 1,168574543 | 0,37491722 | 0,861771195 | RAB3 GTPase activating protein catalytic subunit 1 |
| ENSG00000197498 | RPF2 | 0,224744178 | 1,168570016 | 0,48165907 | 0,912981134 | ribosome production factor 2 homolog |
| ENSG00000172889 | EGFL7 | 0,224734432 | 1,168562122 | 0,6288181 | 0,952793934 | EGF like domain multiple 7 |

| | | | | | | |
|---|---|---|---|---|---|---|
| ENSG00000281794 | MUC20-OT1 | 0,22472361 | 1,168553357 | 0,53032332 | 0,931172326 | MUC20 overlapping transcript |
| ENSG00000173406 | DAB1 | 0,224705442 | 1,168538641 | 0,50035077 | 0,920485547 | DAB adaptor protein 1 |
| ENSG00000011007 | ELOA | 0,224646629 | 1,168491006 | 0,28073452 | 0,791510309 | elongin A |
| ENSG00000197696 | NMB | 0,224633934 | 1,168480723 | 0,60668309 | 0,950927937 | neuromedin B |
| ENSG00000127445 | PIN1 | 0,22463287 | 1,168479861 | 0,39921179 | 0,875433675 | peptidylprolyl cis/trans isomerase, NIMA-interacting 1 |
| ENSG00000286106 | NOTCH2NLR | 0,224621518 | 1,168470667 | 0,59455962 | 0,947469769 | notch 2 N-terminal like R (pseudogene) |
| ENSG00000114125 | RNF7 | 0,224592286 | 1,168446992 | 0,43455467 | 0,896272827 | ring finger protein 7 |
| ENSG00000047932 | GOPC | 0,224568519 | 1,168427743 | 0,56578844 | 0,940866081 | golgi associated PDZ and coiled-coil motif containing |
| ENSG00000271874 | AC025754,2 | 0,224507385 | 1,168378232 | 0,83422141 | 0,984905093 | novel transcript |
| ENSG00000180440 | SERTM1 | 0,224505488 | 1,168376696 | 0,8041675 | 0,980875302 | serine rich and transmembrane domain containing 1 |
| ENSG00000183751 | TBL3 | 0,224387057 | 1,168280787 | 0,55799012 | 0,939078529 | transducin beta like 3 |
| ENSG00000131381 | RBSN | 0,224306926 | 1,1682159 | 0,34076857 | 0,839409751 | rabenosyn, RAB effector |
| ENSG00000025434 | NR1H3 | 0,224281549 | 1,168195351 | 0,4466117 | 0,901644586 | nuclear receptor subfamily 1 group H member 3 |
| ENSG00000232059 | AL451007,1 | 0,224226107 | 1,168150459 | 0,87247103 | 0,989217334 | ribosomal protein L37 (RPL37) pseudogene |
| ENSG00000138079 | SLC3A1 | 0,224132567 | 1,168074723 | 0,55285782 | 0,936894267 | solute carrier family 3 member 1 |
| ENSG00000286591 | AC006197,2 | 0,22411305 | 1,16805892 | 0,68421607 | 0,965415049 | novel transcript |
| ENSG00000283761 | AC118553,2 | 0,224075777 | 1,168028743 | 0,47624711 | 0,911416789 | novel protein |
| ENSG00000111667 | USP5 | 0,224066931 | 1,168021581 | 0,42648752 | 0,892138574 | ubiquitin specific peptidase 5 |
| ENSG00000143153 | ATP1B1 | 0,224064154 | 1,168019333 | 0,54675439 | 0,935404974 | ATPase Na+/K+ transporting subunit beta 1 |
| ENSG00000174028 | FAM3C2P | 0,223965912 | 1,167939798 | 0,43930454 | 0,897434993 | family with sequence similarity 3 member C2, pseudogene |
| ENSG00000071246 | VASH1 | 0,223951955 | 1,167928499 | 0,58758435 | 0,946068138 | vasohibin 1 |
| ENSG00000232341 | RPL4P2 | 0,223868525 | 1,167860961 | 0,79973039 | 0,980161849 | ribosomal protein L4 pseudogene 2 |
| ENSG00000205138 | SDHAF1 | 0,22363532 | 1,167672196 | 0,49983619 | 0,920310866 | succinate dehydrogenase complex assembly factor 1 |
| ENSG00000125447 | GGA3 | 0,223621025 | 1,167660627 | 0,30966255 | 0,816986489 | golgi associated, gamma adaptin ear containing, ARF binding protein 3 |
| ENSG00000227232 | WASH7P | 0,223609284 | 1,167651124 | 0,43801672 | 0,897177404 | WASP family homolog 7, pseudogene |
| ENSG00000176788 | BASP1 | 0,223582027 | 1,167629064 | 0,35912765 | 0,849846046 | brain abundant membrane attached signal protein 1 |
| ENSG00000284376 | AC092017,3 | 0,223519705 | 1,167578626 | 0,86143095 | 0,989074806 | novel transcript |
| ENSG00000177051 | FBXO46 | 0,223476474 | 1,167543639 | 0,43957523 | 0,89764124 | F-box protein 46 |
| ENSG00000213780 | GTF2H4 | 0,223444824 | 1,167518025 | 0,65244363 | 0,957275891 | general transcription factor IIH subunit 4 |
| ENSG00000142039 | CCDC97 | 0,223303714 | 1,167403836 | 0,43980288 | 0,897753805 | coiled-coil domain containing 97 |

| | | | | | | |
|---|---|---|---|---|---|---|
| ENSG00000134851 | TMEM165 | 0,223289385 | 1,167392241 | 0,41777022 | 0,886857227 | transmembrane protein 165 |
| ENSG00000204410 | MSH5 | 0,22317948 | 1,167303312 | 0,56909172 | 0,942177697 | mutS homolog 5 |
| ENSG00000160294 | MCM3AP | 0,223133987 | 1,167266504 | 0,36130884 | 0,851821025 | minichromosome maintenance complex component 3 associated protein |
| ENSG00000238171 | AC068196,1 | 0,223106118 | 1,167243955 | 0,88249589 | 0,990717729 | novel transcript |
| ENSG00000159658 | EFCAB14 | 0,223078586 | 1,16722168 | 0,33288923 | 0,833301711 | EF-hand calcium binding domain 14 |
| ENSG00000274646 | RPS9 | 0,223010882 | 1,167166905 | 0,69308703 | 0,967333542 | ribosomal protein S9 |
| ENSG00000112619 | PRPH2 | 0,222945374 | 1,16711391 | 0,68251257 | 0,964781531 | peripherin 2 |
| ENSG00000145293 | ENOPH1 | 0,222937558 | 1,167107587 | 0,46491633 | 0,907804479 | enolase-phosphatase 1 |
| ENSG00000187003 | ACTL7A | 0,222925782 | 1,16709806 | 0,86426217 | 0,989074806 | actin like 7A |
| ENSG00000138629 | UBL7 | 0,22292453 | 1,167097047 | 0,34910983 | 0,843141089 | ubiquitin like 7 |
| ENSG00000288408 | UBL7 | 0,22292453 | 1,167097047 | 0,34910983 | 0,843141089 | ubiquitin like 7 |
| ENSG00000265490 | AP001178,2 | 0,222924188 | 1,167096771 | 0,89553517 | 0,991303328 | novel transcript, antisense to ENOSF1 |
| ENSG00000170248 | PDCD6IP | 0,222882221 | 1,167062821 | 0,52120898 | 0,927046149 | programmed cell death 6 interacting protein |
| ENSG00000249790 | AC092490,1 | 0,22285267 | 1,167038916 | 0,50315641 | 0,920888747 | novel transcript |
| ENSG00000229183 | PGA4 | 0,222789 | 1,166987412 | 0,84903038 | 0,987036295 | pepsinogen A4 |
| ENSG00000129473 | BCL2L2 | 0,222585742 | 1,16682301 | 0,59211953 | 0,947416795 | BCL2 like 2 |
| ENSG00000178896 | EXOSC4 | 0,222584738 | 1,166822198 | 0,59249088 | 0,947416795 | exosome component 4 |
| ENSG00000138069 | RAB1A | 0,22248726 | 1,166743363 | 0,61056205 | 0,951841373 | RAB1A, member RAS oncogene family |
| ENSG00000205861 | PCOTH | 0,222404848 | 1,166676716 | 0,69129977 | 0,967201016 | Pro-X-Gly collagen triple helix like repeat containing |
| ENSG00000162599 | NFIA | 0,222191287 | 1,166504026 | 0,60775719 | 0,950993594 | nuclear factor I A |
| ENSG00000063046 | EIF4B | 0,222183638 | 1,166497841 | 0,34806044 | 0,842502453 | eukaryotic translation initiation factor 4B |
| ENSG00000285850 | AC009152,3 | 0,222179449 | 1,166494455 | 0,80451295 | 0,980875302 | novel transcript |
| ENSG00000285903 | AC009152,4 | 0,222179449 | 1,166494455 | 0,80451295 | 0,980875302 | novel transcript |
| ENSG00000091542 | ALKBH5 | 0,221989413 | 1,16634081 | 0,33786229 | 0,83719797 | alkB homolog 5, RNA demethylase |
| ENSG00000011021 | CLCN6 | 0,221981454 | 1,166334377 | 0,2914913 | 0,802058262 | chloride voltage-gated channel 6 |
| ENSG00000230651 | RGPD4-AS1 | 0,221858144 | 1,166234692 | 0,75648804 | 0,975901808 | RGPD4 antisense RNA 1 (head to head) |
| ENSG00000143183 | TMCO1 | 0,22184499 | 1,166224059 | 0,6098828 | 0,951801946 | transmembrane and coiled-coil domains 1 |
| ENSG00000237335 | PFDN6 | 0,221601608 | 1,166027333 | 0,53793793 | 0,932883277 | prefoldin subunit 6 |
| ENSG00000224782 | PFDN6 | 0,221601608 | 1,166027333 | 0,53793793 | 0,932883277 | prefoldin subunit 6 |
| ENSG00000124209 | RAB22A | 0,221600657 | 1,166026565 | 0,59717654 | 0,948282297 | RAB22A, member RAS oncogene family |
| ENSG00000124356 | STAMBP | 0,221586075 | 1,166014779 | 0,43382758 | 0,896272827 | STAM binding protein |
| ENSG00000264522 | OTUD7B | 0,221564984 | 1,165997733 | 0,49887519 | 0,919683359 | OTU deubiquitinase 7B |

| | | | | | | |
|---|---|---|---|---|---|---|
| ENSG00000107164 | FUBP3 | 0,22152384 | 1,165964481 | 0,48388247 | 0,913690171 | far upstream element binding protein 3 |
| ENSG00000132842 | AP3B1 | 0,221497042 | 1,165942823 | 0,47954815 | 0,912721304 | adaptor related protein complex 3 subunit beta 1 |
| ENSG00000117500 | TMED5 | 0,221492129 | 1,165938853 | 0,64443227 | 0,956633898 | transmembrane p24 trafficking protein 5 |
| ENSG00000121053 | EPX | 0,221412164 | 1,16587423 | 0,81905472 | 0,98249617 | eosinophil peroxidase |
| ENSG00000250624 | MTCYBP43 | 0,221377372 | 1,165846114 | 0,71601701 | 0,971360663 | MT-CYB pseudogene 43 |
| ENSG00000185883 | ATP6V0C | 0,221326632 | 1,165805111 | 0,3284039 | 0,830613989 | ATPase H+ transporting V0 subunit c |
| ENSG00000172465 | TCEAL1 | 0,221291862 | 1,165777015 | 0,55227329 | 0,936619966 | transcription elongation factor A like 1 |
| ENSG00000041802 | LSG1 | 0,221263222 | 1,165753872 | 0,29473599 | 0,804859966 | large 60S subunit nuclear export GTPase 1 |
| ENSG00000213189 | BTF3L4P2 | 0,221229837 | 1,165726896 | 0,43065859 | 0,895267028 | basic transcription factor 3 like 4 pseudogene 2 |
| ENSG00000251249 | AC097375,3 | 0,221174409 | 1,16568211 | 0,80737509 | 0,981213877 | novel transcript |
| ENSG00000111275 | ALDH2 | 0,221168517 | 1,16567735 | 0,33318209 | 0,833395069 | aldehyde dehydrogenase 2 family member |
| ENSG00000136935 | GOLGA1 | 0,221106692 | 1,165627397 | 0,33091852 | 0,832153798 | golgin A1 |
| ENSG00000124151 | NCOA3 | 0,221025697 | 1,165561959 | 0,51920439 | 0,926800715 | nuclear receptor coactivator 3 |
| ENSG00000243701 | DUBR | 0,220984255 | 1,165528478 | 0,50496922 | 0,921400408 | DPPA2 upstream binding RNA |
| ENSG00000159377 | PSMB4 | 0,220871167 | 1,165437119 | 0,3872955 | 0,869513953 | proteasome 20S subunit beta 4 |
| ENSG00000278494 | RRN3 | 0,220783733 | 1,165366491 | 0,7443361 | 0,975641819 | RRN3 homolog, RNA polymerase I transcription factor |
| ENSG00000273618 | AL135999,2 | 0,220699421 | 1,165298388 | 0,80446361 | 0,980875302 | TEC |
| ENSG00000103852 | TTC23 | 0,220676562 | 1,165279925 | 0,42555609 | 0,892133432 | tetratricopeptide repeat domain 23 |
| ENSG00000234405 | Z69733,1 | 0,220672902 | 1,165276968 | 0,66328469 | 0,959920886 | novel transcript |
| ENSG00000288445 | HPS5 | 0,220660904 | 1,165267278 | 0,38248771 | 0,867031314 | HPS5 biogenesis of lysosomal organelles complex 2 subunit 2 |
| ENSG00000257501 | PAFAH1B2P2 | 0,220555648 | 1,165182265 | 0,78784868 | 0,979462318 | PAFAH1B2 pseudogene 2 |
| ENSG00000258366 | RTEL1 | 0,22046392 | 1,165108184 | 0,54644058 | 0,935401821 | regulator of telomere elongation helicase 1 |
| ENSG00000225748 | PRRC2A | 0,220458497 | 1,165103804 | 0,45556347 | 0,903545728 | proline rich coiled-coil 2A |
| ENSG00000105398 | SULT2A1 | 0,220431173 | 1,165081738 | 0,88797493 | 0,990717729 | sulfotransferase family 2A member 1 |
| ENSG00000164715 | LMTK2 | 0,220420582 | 1,165073186 | 0,27883253 | 0,789775666 | lemur tyrosine kinase 2 |
| ENSG00000272345 | AL031775,1 | 0,22032031 | 1,164992211 | 0,6923622 | 0,967236299 | novel transcript, antisense to ACOT13 |
| ENSG00000103253 | HAGHL | 0,220300768 | 1,164976432 | 0,67132343 | 0,962026636 | hydroxyacylglutathione hydrolase like |
| ENSG00000159788 | RGS12 | 0,220262995 | 1,16494593 | 0,51361992 | 0,92410971 | regulator of G protein signaling 12 |
| ENSG00000188186 | LAMTOR4 | 0,220249823 | 1,164935294 | 0,54938197 | 0,936051084 | late endosomal/lysosomal adaptor, MAPK and MTOR activator 4 |
| ENSG00000286193 | AC025449,2 | 0,220229831 | 1,164919151 | 0,72978795 | 0,974106992 | novel transcript, antisense to CPLANE1 |

| | | | | | | |
|---|---|---|---|---|---|---|
| ENSG00000179588 | ZFPM1 | 0,220054724 | 1,164777768 | 0,59019896 | 0,947275534 | zinc finger protein, FOG family member 1 |
| ENSG00000279175 | AL033543,1 | 0,219841271 | 1,164605446 | 0,70651925 | 0,969503793 | novel transcript, antisense to PARVB |
| ENSG00000130313 | PGLS | 0,21982908 | 1,164595606 | 0,50178684 | 0,920520138 | 6-phosphogluconolactonase |
| ENSG00000135632 | SMYD5 | 0,21968889 | 1,164482444 | 0,36183223 | 0,851979035 | SMYD family member 5 |
| ENSG00000105127 | AKAP8 | 0,219678797 | 1,164474298 | 0,42533831 | 0,892049171 | A-kinase anchoring protein 8 |
| ENSG00000166477 | LEO1 | 0,219566229 | 1,164383442 | 0,38792543 | 0,869820053 | LEO1 homolog, Paf1/RNA polymerase II complex component |
| ENSG00000092330 | TINF2 | 0,219464412 | 1,16430127 | 0,34066303 | 0,839282032 | TERF1 interacting nuclear factor 2 |
| ENSG00000131043 | AAR2 | 0,219331324 | 1,164193868 | 0,33506856 | 0,834858938 | AAR2 splicing factor |
| ENSG00000075413 | MARK3 | 0,219236933 | 1,164117701 | 0,3478496 | 0,842383417 | microtubule affinity regulating kinase 3 |
| ENSG00000116288 | PARK7 | 0,219196042 | 1,164084707 | 0,28687241 | 0,79836024 | Parkinsonism associated deglycase |
| ENSG00000267197 | AC011461,1 | 0,219152238 | 1,164049362 | 0,81781308 | 0,982323352 | novel transcript, sense intronic with KEAP1 |
| ENSG00000235447 | TRAPPC13P1 | 0,219052227 | 1,16396867 | 0,73028085 | 0,974321178 | trafficking protein particle complex 13 pseudogene 1 |
| ENSG00000286849 | AC009152,6 | 0,218863784 | 1,163816645 | 0,81448458 | 0,981521413 | novel transcript, antisense to XYLT1 |
| ENSG00000125450 | NUP85 | 0,218851482 | 1,16380672 | 0,32634258 | 0,828416252 | nucleoporin 85 |
| ENSG00000232940 | HCG25 | 0,218830214 | 1,163789564 | 0,69630843 | 0,96810276 | HLA complex group 25 |
| ENSG00000234473 | ZNF101P2 | 0,218769278 | 1,16374041 | 0,75298002 | 0,975901808 | zinc finger protein 101 pseudogene 2 |
| ENSG00000144959 | NCEH1 | 0,218709856 | 1,163692478 | 0,43256167 | 0,89560871 | neutral cholesterol ester hydrolase 1 |
| ENSG00000100714 | MTHFD1 | 0,218675815 | 1,16366502 | 0,35722444 | 0,848693232 | methylenetetrahydrofolate dehydrogenase, cyclohydrolase and formyltetrahydrofolate synthetase 1 |
| ENSG00000264727 | AC005725,1 | 0,218671415 | 1,163661472 | 0,75557286 | 0,975901808 | novel transcript |
| ENSG00000130150 | MOSPD2 | 0,218611152 | 1,163612865 | 0,59423779 | 0,947469769 | motile sperm domain containing 2 |
| ENSG00000225092 | AL355336,1 | 0,21860408 | 1,163607161 | 0,67256304 | 0,962290553 | novel transcript |
| ENSG00000064726 | BTBD1 | 0,21860097 | 1,163604652 | 0,59429042 | 0,947469769 | BTB domain containing 1 |
| ENSG00000272153 | AL365330,1 | 0,218555964 | 1,163568353 | 0,71812225 | 0,97178302 | novel transcript, antisense to LRRC47 |
| ENSG00000076513 | ANKRD13A | 0,218500571 | 1,163523679 | 0,36945439 | 0,857991124 | ankyrin repeat domain 13A |
| ENSG00000179409 | GEMIN4 | 0,218414432 | 1,16345421 | 0,4039021 | 0,877769731 | gem nuclear organelle associated protein 4 |
| ENSG00000267568 | AC016168,2 | 0,218339129 | 1,163393484 | 0,83951369 | 0,985672131 | novel transcript |
| ENSG00000251372 | LINC00499 | 0,218319476 | 1,163377636 | 0,74198049 | 0,975176043 | long intergenic non-protein coding RNA 499 |
| ENSG00000143106 | PSMA5 | 0,218309506 | 1,163369596 | 0,4707185 | 0,908869097 | proteasome 20S subunit alpha 5 |
| ENSG00000272540 | AL662797,1 | 0,218297503 | 1,163359917 | 0,51417592 | 0,924286789 | novel transcript, antisense to TUBB |
| ENSG00000165983 | PTER | 0,218270798 | 1,163338383 | 0,58652377 | 0,945395317 | phosphotriesterase related |

| | | | | | | |
|---|---|---|---|---|---|---|
| ENSG00000080608 | PUM3 | 0,218257114 | 1,163327348 | 0,43505111 | 0,896272827 | pumilio RNA binding family member 3 |
| ENSG00000006015 | REX1BD | 0,218225963 | 1,16330223 | 0,60345094 | 0,950352286 | required for excision 1-B domain containing |
| ENSG00000164305 | CASP3 | 0,218140399 | 1,163233239 | 0,59989207 | 0,949340299 | caspase 3 |
| ENSG00000230021 | AL669831,3 | 0,217923124 | 1,163058065 | 0,7303236 | 0,974321178 | septin 14 (SEPT14) pseudogene |
| ENSG00000112183 | RBM24 | 0,217881548 | 1,163024547 | 0,46858407 | 0,907940653 | RNA binding motif protein 24 |
| ENSG00000020426 | MNAT1 | 0,217868471 | 1,163014006 | 0,47112701 | 0,908869685 | MNAT1 component of CDK activating kinase |
| ENSG00000286370 | AC064856,1 | 0,217548921 | 1,162756433 | 0,71489978 | 0,971041579 | novel transcript |
| ENSG00000267249 | AP005482,2 | 0,217505255 | 1,16272124 | 0,73524054 | 0,974353474 | novel transcript, sense intronic CEP76 |
| ENSG00000090487 | SPG21 | 0,217465956 | 1,162689568 | 0,46429498 | 0,907500217 | SPG21 abhydrolase domain containing, maspardin |
| ENSG00000254653 | AC024475,1 | 0,217417486 | 1,162650506 | 0,75968234 | 0,976120207 | novel transcript |
| ENSG00000138792 | ENPEP | 0,217302506 | 1,162557848 | 0,61383002 | 0,952443383 | glutamyl aminopeptidase |
| ENSG00000231441 | DST-AS1 | 0,217301653 | 1,162557161 | 0,82143966 | 0,982950156 | DST antisense RNA 1 |
| ENSG00000267169 | AC022098,1 | 0,217272453 | 1,162533632 | 0,57635367 | 0,943158459 | novel transcript, antisense to LPHN1 |
| ENSG00000288348 | AC022098,6 | 0,217272453 | 1,162533632 | 0,57635367 | 0,943158459 | novel transcript, antisense to LPHN1 |
| ENSG00000236156 | CHCHD4P3 | 0,21721346 | 1,162486095 | 0,79572046 | 0,979879951 | coiled-coil-helix-coiled-coil-helix domain containing 4 pseudogene 3 |
| ENSG00000101384 | JAG1 | 0,217167241 | 1,162448854 | 0,29627968 | 0,805997082 | jagged canonical Notch ligand 1 |
| ENSG00000148572 | NRBF2 | 0,217118139 | 1,162409291 | 0,572478 | 0,942475308 | nuclear receptor binding factor 2 |
| ENSG00000234114 | CCHCR1 | 0,217106116 | 1,162399603 | 0,70825388 | 0,97011892 | coiled-coil alpha-helical rod protein 1 |
| ENSG00000104957 | CCDC130 | 0,217045183 | 1,16235051 | 0,44109475 | 0,898864957 | coiled-coil domain containing 130 |
| ENSG00000138035 | PNPT1 | 0,216833914 | 1,162180308 | 0,48042215 | 0,912847708 | polyribonucleotide nucleotidyltransferase 1 |
| ENSG00000188603 | CLN3 | 0,216812987 | 1,162163449 | 0,54755647 | 0,935404974 | CLN3 lysosomal/endosomal transmembrane protein, battenin |
| ENSG00000176723 | ZNF843 | 0,216777389 | 1,162134774 | 0,62562658 | 0,952793934 | zinc finger protein 843 |
| ENSG00000196396 | PTPN1 | 0,216766637 | 1,162126113 | 0,38604492 | 0,869112622 | protein tyrosine phosphatase non-receptor type 1 |
| ENSG00000080200 | CRYBG3 | 0,216741211 | 1,162105631 | 0,54410742 | 0,934259495 | crystallin beta-gamma domain containing 3 |
| ENSG00000117408 | IPO13 | 0,216667434 | 1,162046205 | 0,35038153 | 0,843857545 | importin 13 |
| ENSG00000067596 | DHX8 | 0,216648733 | 1,162031142 | 0,29098333 | 0,801689099 | DEAH-box helicase 8 |
| ENSG00000119401 | TRIM32 | 0,216639657 | 1,162023832 | 0,37364918 | 0,861319894 | tripartite motif containing 32 |
| ENSG00000274929 | AL157813,1 | 0,216622706 | 1,162010179 | 0,75653166 | 0,975901808 | novel transcript, sense intronic to RCBTB2 |
| ENSG00000168872 | DDX19A | 0,21646541 | 1,161883493 | 0,31122083 | 0,817751312 | DEAD-box helicase 19A |
| ENSG00000204442 | FAM155A | 0,216271202 | 1,161727096 | 0,47299589 | 0,909532026 | family with sequence similarity 155 member A |
| ENSG00000284126 | SMURF1 | 0,21625374 | 1,161713035 | 0,33201183 | 0,832927044 | SMAD specific E3 ubiquitin protein ligase 1 |

| | | | | | | |
|---|---|---|---|---|---|---|
| ENSG00000165733 | BMS1 | 0,21607385 | 1,16156819 | 0,4702181 | 0,908528529 | BMS1 ribosome biogenesis factor |
| ENSG00000114988 | LMAN2L | 0,215930043 | 1,161452412 | 0,32317342 | 0,825262912 | lectin, mannose binding 2 like |
| ENSG00000176101 | SSNA1 | 0,215913579 | 1,161439157 | 0,49074214 | 0,916995341 | SS nuclear autoantigen 1 |
| ENSG00000013306 | SLC25A39 | 0,215860148 | 1,161396143 | 0,46680274 | 0,907940653 | solute carrier family 25 member 39 |
| ENSG00000136634 | IL10 | 0,215841288 | 1,161380961 | 0,74152064 | 0,975176043 | interleukin 10 |
| ENSG00000258169 | LINC00485 | 0,215837831 | 1,161378178 | 0,67975542 | 0,964459885 | long intergenic non-protein coding RNA 485 |
| ENSG00000075073 | TACR2 | 0,215449453 | 1,161065573 | 0,75961561 | 0,976120207 | tachykinin receptor 2 |
| ENSG00000175147 | TMEM51-AS1 | 0,215413842 | 1,161036914 | 0,56325112 | 0,940865625 | TMEM51 antisense RNA 1 |
| ENSG00000268584 | AC073389,1 | 0,215294076 | 1,160940534 | 0,64976425 | 0,956899963 | novel transcript, antisense to USP54 |
| ENSG00000077235 | GTF3C1 | 0,215239628 | 1,16089672 | 0,43915023 | 0,897434993 | general transcription factor IIIC subunit 1 |
| ENSG00000180071 | ANKRD18A | 0,215131416 | 1,160809648 | 0,66943606 | 0,961867165 | ankyrin repeat domain 18A |
| ENSG00000170892 | TSEN34 | 0,215080265 | 1,160768493 | 0,61349161 | 0,952413269 | tRNA splicing endonuclease subunit 34 |
| ENSG00000078747 | ITCH | 0,215063761 | 1,160755214 | 0,54929573 | 0,936040289 | itchy E3 ubiquitin protein ligase |
| ENSG00000261659 | Z92544,1 | 0,215057977 | 1,16075056 | 0,69453433 | 0,967417311 | novel transcript, antisense to FBXL16 |
| ENSG00000137054 | POLR1E | 0,215034226 | 1,160731451 | 0,37521119 | 0,862022464 | RNA polymerase I subunit E |
| ENSG00000138587 | MNS1 | 0,214995846 | 1,160700572 | 0,51992996 | 0,926822772 | meiosis specific nuclear structural 1 |
| ENSG00000134574 | DDB2 | 0,214968127 | 1,160678272 | 0,54645239 | 0,935401821 | damage specific DNA binding protein 2 |
| ENSG00000106327 | TFR2 | 0,214966583 | 1,16067703 | 0,73469952 | 0,974353474 | transferrin receptor 2 |
| ENSG00000131080 | EDA2R | 0,214896939 | 1,160621001 | 0,60008467 | 0,949340299 | ectodysplasin A2 receptor |
| ENSG00000143486 | EIF2D | 0,214851392 | 1,16058436 | 0,37200942 | 0,859937929 | eukaryotic translation initiation factor 2D |
| ENSG00000111325 | OGFOD2 | 0,214841193 | 1,160576155 | 0,46189569 | 0,906277266 | 2-oxoglutarate and iron dependent oxygenase domain containing 2 |
| ENSG00000131351 | HAUS8 | 0,214786419 | 1,160532093 | 0,33731058 | 0,836427495 | HAUS augmin like complex subunit 8 |
| ENSG00000280434 | AL031595,3 | 0,214636587 | 1,160411572 | 0,73713902 | 0,974353474 | novel transcript, sense overlapping PARVB |
| ENSG00000152348 | ATG10 | 0,214625302 | 1,160402495 | 0,50890046 | 0,922427526 | autophagy related 10 |
| ENSG00000126856 | PRDM7 | 0,214566515 | 1,160355211 | 0,68464416 | 0,965415049 | PR/SET domain 7 |
| ENSG00000233325 | MIPEPP3 | 0,214519767 | 1,160317612 | 0,76267355 | 0,97639149 | mitochondrial intermediate peptidase pseudogene 3 |
| ENSG00000131475 | VPS25 | 0,214434561 | 1,160249086 | 0,35251189 | 0,844912682 | vacuolar protein sorting 25 homolog |
| ENSG00000136856 | SLC2A8 | 0,214419471 | 1,16023695 | 0,56813676 | 0,94163744 | solute carrier family 2 member 8 |
| ENSG00000007968 | E2F2 | 0,214341954 | 1,160174612 | 0,8811208 | 0,990662809 | E2F transcription factor 2 |
| ENSG00000139437 | TCHP | 0,214336082 | 1,16016989 | 0,60582306 | 0,950927937 | trichoplein keratin filament binding |
| ENSG00000147789 | ZNF7 | 0,214184337 | 1,160047868 | 0,3076598 | 0,816387366 | zinc finger protein 7 |

| | | | | | | |
|---|---|---|---|---|---|---|
| ENSG00000233367 | AL133480,1 | 0,214151825 | 1,160021725 | 0,79154342 | 0,979471509 | novel transcript |
| ENSG00000288079 | AC026329,1 | 0,214113699 | 1,15999107 | 0,51190917 | 0,923184606 | novel transcript, antisense to BOC |
| ENSG00000229404 | LINC00858 | 0,214052634 | 1,159941972 | 0,58578956 | 0,945246954 | long intergenic non-protein coding RNA 858 |
| ENSG00000104131 | EIF3J | 0,214033786 | 1,159926818 | 0,4838067 | 0,913690171 | eukaryotic translation initiation factor 3 subunit J |
| ENSG00000204929 | AC007389,1 | 0,214018579 | 1,159914592 | 0,78573725 | 0,979320406 | novel transcript |
| ENSG00000245694 | CRNDE | 0,213902913 | 1,159821601 | 0,67336244 | 0,962685359 | colorectal neoplasia differentially expressed |
| ENSG00000037241 | RPL26L1 | 0,213888666 | 1,159810147 | 0,55033676 | 0,936146662 | ribosomal protein L26 like 1 |
| ENSG00000144895 | EIF2A | 0,213844133 | 1,159774347 | 0,6001819 | 0,949340299 | eukaryotic translation initiation factor 2A |
| ENSG00000232632 | GABBR1 | 0,213784434 | 1,159726356 | 0,70527222 | 0,969427597 | gamma-aminobutyric acid type B receptor subunit 1 |
| ENSG00000169020 | ATP5ME | 0,213723007 | 1,159676979 | 0,32001711 | 0,824366234 | ATP synthase membrane subunit e |
| ENSG00000273018 | FAM106A | 0,213654594 | 1,159621987 | 0,60417478 | 0,950672921 | family with sequence similarity 106 member A |
| ENSG00000148719 | DNAJB12 | 0,213608928 | 1,159585282 | 0,32066716 | 0,824366234 | DnaJ heat shock protein family (Hsp40) member B12 |
| ENSG00000165879 | FRAT1 | 0,213526843 | 1,159519307 | 0,61761644 | 0,952793934 | FRAT regulator of WNT signaling pathway 1 |
| ENSG00000115657 | ABCB6 | 0,213426523 | 1,159438681 | 0,38900814 | 0,870057277 | ATP binding cassette subfamily B member 6 (Langereis blood group) |
| ENSG00000165688 | PMPCA | 0,213377842 | 1,159399559 | 0,42695476 | 0,892273007 | peptidase, mitochondrial processing subunit alpha |
| ENSG00000075336 | TIMM21 | 0,213376283 | 1,159398306 | 0,5488468 | 0,936029622 | translocase of inner mitochondrial membrane 21 |
| ENSG00000116039 | ATP6V1B1 | 0,213373516 | 1,159396082 | 0,8118454 | 0,981213877 | ATPase H+ transporting V1 subunit B1 |
| ENSG00000279494 | AL117328,2 | 0,213288133 | 1,159327468 | 0,82101383 | 0,982753437 | novel transcript |
| ENSG00000254470 | AP5B1 | 0,213228927 | 1,159279892 | 0,52084487 | 0,927046149 | adaptor related protein complex 5 subunit beta 1 |
| ENSG00000213639 | PPP1CB | 0,213181886 | 1,159242093 | 0,54512718 | 0,934923038 | protein phosphatase 1 catalytic subunit beta |
| ENSG00000263465 | SRSF8 | 0,213111584 | 1,159185604 | 0,47081143 | 0,908869685 | serine and arginine rich splicing factor 8 |
| ENSG00000287548 | AC021269,3 | 0,213060213 | 1,159144329 | 0,77095285 | 0,978033855 | novel transcript |
| ENSG00000104980 | TIMM44 | 0,213048477 | 1,1591349 | 0,46305883 | 0,906849208 | translocase of inner mitochondrial membrane 44 |
| ENSG00000272447 | AL135925,1 | 0,213013962 | 1,159107169 | 0,46718854 | 0,907940653 | novel transcript |
| ENSG00000076650 | GPATCH1 | 0,21300221 | 1,159097728 | 0,33998842 | 0,838677456 | G-patch domain containing 1 |
| ENSG00000187550 | SBK2 | 0,212994926 | 1,159091875 | 0,79841315 | 0,980161849 | SH3 domain binding kinase family member 2 |
| ENSG00000099385 | BCL7C | 0,212977878 | 1,159078179 | 0,52929719 | 0,93044404 | BAF chromatin remodeling complex subunit BCL7C |
| ENSG00000183644 | HOATZ | 0,212957876 | 1,159062109 | 0,80165754 | 0,980623173 | HOATZ cilia and flagella associated protein |
| ENSG00000231389 | HLA-DPA1 | 0,212939131 | 1,15904705 | 0,84773722 | 0,987035044 | major histocompatibility complex, class II, DP alpha 1 |
| ENSG00000229685 | HLA-DPA1 | 0,212939131 | 1,15904705 | 0,84773722 | 0,987035044 | major histocompatibility complex, class II, DP alpha 1 |

| | | | | | | |
|---|---|---|---|---|---|---|
| ENSG00000170185 | USP38 | 0,212929624 | 1,159039412 | 0,49737114 | 0,918759905 | ubiquitin specific peptidase 38 |
| ENSG00000268157 | AC010524,1 | 0,212739511 | 1,158886687 | 0,78921133 | 0,979462318 | novel transcript, antisense to TEAD2 |
| ENSG00000174943 | KCTD13 | 0,212602286 | 1,158776463 | 0,45910541 | 0,90476838 | potassium channel tetramerization domain containing 13 |
| ENSG00000073734 | ABCB11 | 0,212541162 | 1,158727369 | 0,73627326 | 0,974353474 | ATP binding cassette subfamily B member 11 |
| ENSG00000251417 | AC145285,2 | 0,212510441 | 1,158702695 | 0,75170614 | 0,975901808 | novel transcript |
| ENSG00000108599 | AKAP10 | 0,212502251 | 1,158696117 | 0,45931789 | 0,904862549 | A-kinase anchoring protein 10 |
| ENSG00000155875 | SAXO1 | 0,212421229 | 1,158631047 | 0,7163524 | 0,971438758 | stabilizer of axonemal microtubules 1 |
| ENSG00000281183 | NPTN-IT1 | 0,212367967 | 1,158588273 | 0,40998682 | 0,882457617 | NPTN intronic transcript 1 |
| ENSG00000025770 | NCAPH2 | 0,212289304 | 1,158525102 | 0,50004353 | 0,92041337 | non-SMC condensin II complex subunit H2 |
| ENSG00000280627 | SURF1 | 0,212201662 | 1,158454725 | 0,41482367 | 0,885579089 | SURF1 cytochrome c oxidase assembly factor |
| ENSG00000148290 | SURF1 | 0,212201662 | 1,158454725 | 0,41482367 | 0,885579089 | SURF1 cytochrome c oxidase assembly factor |
| ENSG00000006744 | ELAC2 | 0,212149888 | 1,158413153 | 0,32074331 | 0,824366234 | elaC ribonuclease Z 2 |
| ENSG00000155026 | RSPH10B | 0,212119568 | 1,158388807 | 0,77629711 | 0,978587255 | radial spoke head 10 homolog B |
| ENSG00000168488 | ATXN2L | 0,212097505 | 1,158371092 | 0,36921342 | 0,857942508 | ataxin 2 like |
| ENSG00000174912 | METTL15P1 | 0,212028795 | 1,158315925 | 0,68130196 | 0,964654539 | methyltransferase like 15 pseudogene 1 |
| ENSG00000126464 | PRR12 | 0,211965609 | 1,158265195 | 0,61738662 | 0,952793934 | proline rich 12 |
| ENSG00000155158 | TTC39B | 0,211899171 | 1,158211857 | 0,50287207 | 0,920749191 | tetratricopeptide repeat domain 39B |
| ENSG00000203772 | SPRN | 0,211861101 | 1,158181294 | 0,78965065 | 0,979462318 | shadow of prion protein |
| ENSG00000128829 | EIF2AK4 | 0,211837991 | 1,158162742 | 0,38950277 | 0,870057277 | eukaryotic translation initiation factor 2 alpha kinase 4 |
| ENSG00000113240 | CLK4 | 0,211808472 | 1,158139045 | 0,53746686 | 0,932883277 | CDC like kinase 4 |
| ENSG00000152894 | PTPRK | 0,211721409 | 1,158069156 | 0,48055523 | 0,912847708 | protein tyrosine phosphatase receptor type K |
| ENSG00000127990 | SGCE | 0,211708642 | 1,158058908 | 0,59739482 | 0,948282297 | sarcoglycan epsilon |
| ENSG00000063978 | RNF4 | 0,211670173 | 1,158028029 | 0,34378095 | 0,840901506 | ring finger protein 4 |
| ENSG00000281903 | LINC02246 | 0,211663363 | 1,158022563 | 0,80327803 | 0,980866933 | long intergenic non-protein coding RNA 2246 |
| ENSG00000174013 | FBXO45 | 0,211623588 | 1,157990636 | 0,36623333 | 0,856103393 | F-box protein 45 |
| ENSG00000236515 | ZBTB9 | 0,211573882 | 1,15795074 | 0,40260949 | 0,877321369 | zinc finger and BTB domain containing 9 |
| ENSG00000175701 | MTLN | 0,21152247 | 1,157909477 | 0,56397082 | 0,940865625 | mitoregulin |
| ENSG00000142751 | GPN2 | 0,211504128 | 1,157894755 | 0,4343563 | 0,896272827 | GPN-loop GTPase 2 |
| ENSG00000237872 | POU5F1P4 | 0,211433361 | 1,15783796 | 0,71261153 | 0,970812834 | POU class 5 homeobox 1 pseudogene 4 |
| ENSG00000145362 | ANK2 | 0,21138 | 1,157795135 | 0,46849623 | 0,907940653 | ankyrin 2 |
| ENSG00000277147 | LINC00869 | 0,211286083 | 1,157719768 | 0,44083701 | 0,898675567 | long intergenic non-protein coding RNA 869 |

| | | | | | | |
|---|---|---|---|---|---|---|
| ENSG00000285563 | AC106872,12 | 0,211254309 | 1,15769427 | 0,76515694 | 0,976830764 | novel transcript |
| ENSG00000119541 | VPS4B | 0,211196655 | 1,157648006 | 0,47660756 | 0,91143003 | vacuolar protein sorting 4 homolog B |
| ENSG00000011638 | TMEM159 | 0,211152026 | 1,157612195 | 0,71080453 | 0,970482408 | transmembrane protein 159 |
| ENSG00000255296 | AP002490,2 | 0,211093172 | 1,157564972 | 0,84489117 | 0,986557476 | ribosomal protein L17 (RPL17) pseudogene |
| ENSG00000211455 | STK38L | 0,211001054 | 1,157491063 | 0,56534133 | 0,940865625 | serine/threonine kinase 38 like |
| ENSG00000180035 | ZNF48 | 0,210930009 | 1,157434064 | 0,48234165 | 0,913113923 | zinc finger protein 48 |
| ENSG00000285226 | PAK1IP1 | 0,210760122 | 1,157297776 | 0,54211483 | 0,933491609 | PAK1 interacting protein 1 |
| ENSG00000111845 | PAK1IP1 | 0,210760122 | 1,157297776 | 0,54211483 | 0,933491609 | PAK1 interacting protein 1 |
| ENSG00000164068 | RNF123 | 0,210753865 | 1,157292757 | 0,45823471 | 0,904140862 | ring finger protein 123 |
| ENSG00000273209 | AC069148,1 | 0,210674861 | 1,157229384 | 0,6396791 | 0,956018518 | novel transcript |
| ENSG00000204237 | OXLD1 | 0,210624745 | 1,157189185 | 0,49226302 | 0,916995341 | oxidoreductase like domain containing 1 |
| ENSG00000260576 | EIF5A2P1 | 0,210557066 | 1,1571349 | 0,86370215 | 0,989074806 | eukaryotic translation initiation factor 5A2 pseudogene 1 |
| ENSG00000100359 | SGSM3 | 0,21054105 | 1,157122055 | 0,51507223 | 0,924720046 | small G protein signaling modulator 3 |
| ENSG00000235618 | FAM21EP | 0,210536562 | 1,157118456 | 0,69792983 | 0,968522643 | family with sequence similarity 21 member E, pseudogene |
| ENSG00000173402 | DAG1 | 0,210517591 | 1,15710324 | 0,37721431 | 0,862818448 | dystroglycan 1 |
| ENSG00000198203 | SULT1C2 | 0,210489873 | 1,157081009 | 0,67855617 | 0,9640473 | sulfotransferase family 1C member 2 |
| ENSG00000280994 | LINC01347 | 0,210484952 | 1,157077062 | 0,60920102 | 0,951793902 | long intergenic non-protein coding RNA 1347 |
| ENSG00000214837 | LINC01347 | 0,210484952 | 1,157077062 | 0,60920102 | 0,951793902 | long intergenic non-protein coding RNA 1347 |
| ENSG00000277050 | AL122125,1 | 0,210346508 | 1,156966032 | 0,81503138 | 0,981521413 | novel transcript, antisense to FRMD6 |
| ENSG00000050748 | MAPK9 | 0,210231067 | 1,156873458 | 0,5071964 | 0,921951873 | mitogen-activated protein kinase 9 |
| ENSG00000138495 | COX17 | 0,210155221 | 1,15681264 | 0,4021473 | 0,87718264 | cytochrome c oxidase copper chaperone COX17 |
| ENSG00000244398 | AC116533,1 | 0,210113957 | 1,156779553 | 0,54544014 | 0,935108602 | ribosomal protein L36a (RPL36A) pseudogene |
| ENSG00000166483 | WEE1 | 0,210084446 | 1,156755891 | 0,50354218 | 0,921005254 | WEE1 G2 checkpoint kinase |
| ENSG00000149609 | C20orf144 | 0,210076256 | 1,156749324 | 0,70214945 | 0,968742626 | chromosome 20 open reading frame 144 |
| ENSG00000142599 | RERE | 0,210068308 | 1,156742952 | 0,35953556 | 0,849982153 | arginine-glutamic acid dipeptide repeats |
| ENSG00000235650 | RPS18 | 0,209975059 | 1,156668187 | 0,56981125 | 0,94221458 | ribosomal protein S18 |
| ENSG00000120885 | CLU | 0,209952859 | 1,156650389 | 0,51397655 | 0,924227101 | clusterin |
| ENSG00000099797 | TECR | 0,209833246 | 1,156554496 | 0,37747106 | 0,862854083 | trans-2,3-enoyl-CoA reductase |
| ENSG00000138942 | RNF185 | 0,209823742 | 1,156546877 | 0,47411717 | 0,910491353 | ring finger protein 185 |
| ENSG00000130741 | EIF2S3 | 0,20978533 | 1,156516084 | 0,55899894 | 0,939635125 | eukaryotic translation initiation factor 2 subunit gamma |

| | | | | | | |
|---|---|---|---|---|---|---|
| ENSG00000230219 | CIBAR1P2 | 0,209628651 | 1,156390491 | 0,82613664 | 0,983471137 | CIBAR1 pseudogene 2 |
| ENSG00000169231 | THBS3 | 0,209620571 | 1,156384015 | 0,38810384 | 0,869820053 | thrombospondin 3 |
| ENSG00000113643 | RARS1 | 0,209558229 | 1,156334046 | 0,5503995 | 0,936167238 | arginyl-tRNA synthetase 1 |
| ENSG00000165828 | PRAP1 | 0,20955235 | 1,156329334 | 0,85523935 | 0,988016959 | proline rich acidic protein 1 |
| ENSG00000255310 | AF131215,5 | 0,20954928 | 1,156326873 | 0,50079978 | 0,920520138 | novel transcript |
| ENSG00000146676 | PURB | 0,20936762 | 1,156181281 | 0,50959578 | 0,922427526 | purine rich element binding protein B |
| ENSG00000111670 | GNPTAB | 0,209367111 | 1,156180873 | 0,50947813 | 0,922427526 | N-acetylglucosamine-1-phosphate transferase subunits alpha and beta |
| ENSG00000261069 | AC124312,3 | 0,209320442 | 1,156143473 | 0,39770721 | 0,874263835 | novel transcript, SNORD host |
| ENSG00000007047 | MARK4 | 0,209305154 | 1,156131222 | 0,39963527 | 0,875433675 | microtubule affinity regulating kinase 4 |
| ENSG00000081760 | AACS | 0,209202085 | 1,156048628 | 0,46158469 | 0,906165604 | acetoacetyl-CoA synthetase |
| ENSG00000201403 | SNORD14B | 0,209188905 | 1,156038067 | 0,86483517 | 0,989074806 | small nucleolar RNA, C/D box 14B |
| ENSG00000232075 | MRPL35P2 | 0,209163783 | 1,156017937 | 0,8587249 | 0,989074806 | mitochondrial ribosomal protein L35 pseudogene 2 |
| ENSG00000249364 | AC112206,2 | 0,209096142 | 1,155963738 | 0,73935866 | 0,974715504 | novel transcript |
| ENSG00000120705 | ETF1 | 0,209060085 | 1,155934848 | 0,54565842 | 0,935146964 | eukaryotic translation termination factor 1 |
| ENSG00000161960 | EIF4A1 | 0,209054395 | 1,155930288 | 0,3287762 | 0,830659639 | eukaryotic translation initiation factor 4A1 |
| ENSG00000259079 | AC005476,1 | 0,208928244 | 1,155829217 | 0,7825424 | 0,978647973 | PDZ domain-containing protein 11 (PDZD11) pseudogene |
| ENSG00000114648 | KLHL18 | 0,208916689 | 1,15581996 | 0,3385883 | 0,837371637 | kelch like family member 18 |
| ENSG00000137166 | FOXP4 | 0,208775345 | 1,155706727 | 0,53601297 | 0,932139691 | forkhead box P4 |
| ENSG00000214026 | MRPL23 | 0,20874865 | 1,155685342 | 0,40493488 | 0,878176155 | mitochondrial ribosomal protein L23 |
| ENSG00000288141 | MRPL23 | 0,20874865 | 1,155685342 | 0,40493488 | 0,878176155 | mitochondrial ribosomal protein L23 |
| ENSG00000213593 | TMX2 | 0,20873818 | 1,155676955 | 0,46783201 | 0,907940653 | thioredoxin related transmembrane protein 2 |
| ENSG00000164610 | RP9 | 0,208713239 | 1,155656977 | 0,44546503 | 0,901068127 | RP9 pre-mRNA splicing factor |
| ENSG00000095066 | HOOK2 | 0,208709178 | 1,155653723 | 0,72025515 | 0,972142678 | hook microtubule tethering protein 2 |
| ENSG00000267796 | LIN37 | 0,208698532 | 1,155645196 | 0,45559094 | 0,903545728 | lin-37 DREAM MuvB core complex component |
| ENSG00000138085 | ATRAID | 0,20864063 | 1,155598816 | 0,43239229 | 0,895608365 | all-trans retinoic acid induced differentiation factor |
| ENSG00000265566 | RN7SL605P | 0,208581713 | 1,155551624 | 0,77065747 | 0,977966587 | RNA, 7SL, cytoplasmic 605, pseudogene |
| ENSG00000198406 | BZW1P2 | 0,208557261 | 1,155532038 | 0,58004984 | 0,943828882 | basic leucine zipper and W2 domains 1 pseudogene 2 |
| ENSG00000181852 | RNF41 | 0,208540324 | 1,155518473 | 0,35096398 | 0,844035671 | ring finger protein 41 |
| ENSG00000111669 | TPI1 | 0,208338053 | 1,155356477 | 0,37134787 | 0,859476158 | triosephosphate isomerase 1 |
| ENSG00000083937 | CHMP2B | 0,208276987 | 1,155307574 | 0,62024603 | 0,952793934 | charged multivesicular body protein 2B |

| | | | | | | |
|---|---|---|---|---|---|---|
| ENSG00000265354 | TIMM23 | 0,208222544 | 1,155263977 | 0,52278785 | 0,92790906 | translocase of inner mitochondrial membrane 23 |
| ENSG00000288109 | AC079331,3 | 0,208207776 | 1,155252151 | 0,83547176 | 0,985034063 | novel transcript, antisense to PITPNC1 |
| ENSG00000204619 | PPP1R11 | 0,208183868 | 1,155233007 | 0,36154886 | 0,851893059 | protein phosphatase 1 regulatory inhibitor subunit 11 |
| ENSG00000103199 | ZNF500 | 0,208078923 | 1,155148975 | 0,61316032 | 0,952413269 | zinc finger protein 500 |
| ENSG00000173085 | COQ2 | 0,208056485 | 1,15513101 | 0,58837843 | 0,946199432 | coenzyme Q2, polyprenyltransferase |
| ENSG00000136003 | ISCU | 0,208040117 | 1,155117905 | 0,45776104 | 0,90388627 | iron-sulfur cluster assembly enzyme |
| ENSG00000273973 | AC025162,2 | 0,208014733 | 1,155097581 | 0,84088059 | 0,985672131 | novel transcript, sense intronic to DGKA |
| ENSG00000062598 | ELMO2 | 0,207973797 | 1,155064806 | 0,36495293 | 0,855057583 | engulfment and cell motility 2 |
| ENSG00000259781 | HMGB1P6 | 0,207919421 | 1,155021271 | 0,49131549 | 0,916995341 | high mobility group box 1 pseudogene 6 |
| ENSG00000172613 | RAD9A | 0,207908041 | 1,155012161 | 0,66997529 | 0,962026636 | RAD9 checkpoint clamp component A |
| ENSG00000106799 | TGFBR1 | 0,207890963 | 1,154998488 | 0,60650976 | 0,950927937 | transforming growth factor beta receptor 1 |
| ENSG00000198863 | RUNDC1 | 0,207827474 | 1,154947661 | 0,53596552 | 0,932139691 | RUN domain containing 1 |
| ENSG00000122707 | RECK | 0,207792228 | 1,154919445 | 0,58581091 | 0,945246954 | reversion inducing cysteine rich protein with kazal motifs |
| ENSG00000108424 | KPNB1 | 0,207748726 | 1,154884621 | 0,35240673 | 0,844912682 | karyopherin subunit beta 1 |
| ENSG00000148660 | CAMK2G | 0,207732081 | 1,154871297 | 0,43782688 | 0,89711755 | calcium/calmodulin dependent protein kinase II gamma |
| ENSG00000131828 | PDHA1 | 0,207724738 | 1,154865419 | 0,31744663 | 0,822257775 | pyruvate dehydrogenase E1 subunit alpha 1 |
| ENSG00000132128 | LRRC41 | 0,207723244 | 1,154864222 | 0,3867549 | 0,869441674 | leucine rich repeat containing 41 |
| ENSG00000260612 | AC009097,3 | 0,207650468 | 1,154805967 | 0,81889228 | 0,98249617 | novel transcript |
| ENSG00000187605 | TET3 | 0,207606918 | 1,154771109 | 0,41633287 | 0,886060654 | tet methylcytosine dioxygenase 3 |
| ENSG00000087111 | PIGS | 0,207488666 | 1,154676461 | 0,32527735 | 0,827324367 | phosphatidylinositol glycan anchor biosynthesis class S |
| ENSG00000225715 | PSMA1P1 | 0,207481119 | 1,15467042 | 0,80967794 | 0,981213877 | PSMA1 pseudogene 1 |
| ENSG00000234032 | VARS2 | 0,207466275 | 1,15465854 | 0,55302998 | 0,936971186 | valyl-tRNA synthetase 2, mitochondrial |
| ENSG00000005700 | IBTK | 0,207438234 | 1,154636097 | 0,58678979 | 0,945395317 | inhibitor of Bruton tyrosine kinase |
| ENSG00000092621 | PHGDH | 0,207415738 | 1,154618093 | 0,54031903 | 0,933385624 | phosphoglycerate dehydrogenase |
| ENSG00000229385 | CTNNA2-AS1 | 0,207398286 | 1,154604126 | 0,76639442 | 0,977167878 | CTNNA2 antisense RNA 1 |
| ENSG00000221914 | PPP2R2A | 0,207350718 | 1,154566057 | 0,61004097 | 0,951801946 | protein phosphatase 2 regulatory subunit Balpha |
| ENSG00000182108 | DEXI | 0,207322353 | 1,154543357 | 0,48309164 | 0,91363215 | Dexi homolog |
| ENSG00000133961 | NUMB | 0,207316553 | 1,154538716 | 0,44127122 | 0,898885877 | NUMB endocytic adaptor protein |
| ENSG00000183154 | AC138356,1 | 0,207299024 | 1,154524688 | 0,79178581 | 0,979471509 | novel transcript, antisense to ERLIN2 |

| | | | | | | |
|---|---|---|---|---|---|---|
| ENSG00000104884 | ERCC2 | 0,207175475 | 1,154425822 | 0,46790532 | 0,907940653 | ERCC excision repair 2, TFIIH core complex helicase subunit |
| ENSG00000245937 | LINC01184 | 0,207091346 | 1,154358505 | 0,43896549 | 0,897402199 | long intergenic non-protein coding RNA 1184 |
| ENSG00000130479 | MAP1S | 0,207039504 | 1,154317024 | 0,59286867 | 0,947416795 | microtubule associated protein 1S |
| ENSG00000213085 | CFAP45 | 0,206937155 | 1,154235137 | 0,78819934 | 0,979462318 | cilia and flagella associated protein 45 |
| ENSG00000153093 | ACOXL | 0,206868808 | 1,154180457 | 0,58619929 | 0,945256674 | acyl-CoA oxidase like |
| ENSG00000205155 | PSENEN | 0,206773809 | 1,154104458 | 0,34860193 | 0,842757646 | presenilin enhancer, gamma-secretase subunit |
| ENSG00000286226 | AC093849,3 | 0,206703637 | 1,154048325 | 0,87240376 | 0,989217334 | novel transcript |
| ENSG00000204681 | GABBR1 | 0,206664859 | 1,154017306 | 0,6597744 | 0,958824284 | gamma-aminobutyric acid type B receptor subunit 1 |
| ENSG00000091317 | CMTM6 | 0,206566303 | 1,153938473 | 0,62275931 | 0,952793934 | CKLF like MARVEL transmembrane domain containing 6 |
| ENSG00000287964 | AC066580,1 | 0,206562808 | 1,153935678 | 0,75997324 | 0,97624106 | novel transcript |
| ENSG00000274340 | AC032011,1 | 0,206548675 | 1,153924373 | 0,791592 | 0,979471509 | novel transcript |
| ENSG00000214960 | CRPPA | 0,206491155 | 1,153878367 | 0,50326673 | 0,920888747 | CDP-L-ribitol pyrophosphorylase A |
| ENSG00000267939 | AC008946,1 | 0,206457366 | 1,153851343 | 0,74700845 | 0,975796253 | novel transcript |
| ENSG00000008952 | SEC62 | 0,20643167 | 1,153830792 | 0,56053089 | 0,940170934 | SEC62 homolog, preprotein translocation factor |
| ENSG00000176974 | SHMT1 | 0,206339639 | 1,153757191 | 0,56859972 | 0,941793403 | serine hydroxymethyltransferase 1 |
| ENSG00000273594 | EEF1D | 0,206337532 | 1,153755506 | 0,45292629 | 0,902428411 | eukaryotic translation elongation factor 1 delta |
| ENSG00000184110 | EIF3C | 0,206269959 | 1,153701467 | 0,43189605 | 0,895589185 | eukaryotic translation initiation factor 3 subunit C |
| ENSG00000136867 | SLC31A2 | 0,206240361 | 1,153677798 | 0,76334313 | 0,976640468 | solute carrier family 31 member 2 |
| ENSG00000111647 | UHRF1BP1L | 0,206208325 | 1,15365218 | 0,6057843 | 0,950927937 | UHRF1 binding protein 1 like |
| ENSG00000143858 | SYT2 | 0,206187533 | 1,153635554 | 0,56320814 | 0,940865625 | synaptotagmin 2 |
| ENSG00000286154 | AC009974,2 | 0,206140511 | 1,153597954 | 0,79446736 | 0,979776981 | novel transcript, antisense to CYP27A1 |
| ENSG00000176783 | RUFY1 | 0,206110314 | 1,153573809 | 0,41813393 | 0,887094957 | RUN and FYVE domain containing 1 |
| ENSG00000029364 | SLC39A9 | 0,206049474 | 1,153525162 | 0,5243454 | 0,928579043 | solute carrier family 39 member 9 |
| ENSG00000162896 | PIGR | 0,205966907 | 1,153459146 | 0,85272894 | 0,98761642 | polymeric immunoglobulin receptor |
| ENSG00000072958 | AP1M1 | 0,20592484 | 1,153425514 | 0,39890835 | 0,875433675 | adaptor related protein complex 1 subunit mu 1 |
| ENSG00000248476 | BACH1-IT1 | 0,205864228 | 1,153377055 | 0,67845145 | 0,9640473 | BACH1 intronic transcript 1 |
| ENSG00000176393 | RNPEP | 0,205745425 | 1,153282081 | 0,42695766 | 0,892273007 | arginyl aminopeptidase |
| ENSG00000086065 | CHMP5 | 0,205736508 | 1,153274953 | 0,54705948 | 0,935404974 | charged multivesicular body protein 5 |
| ENSG00000105568 | PPP2R1A | 0,205628944 | 1,153188971 | 0,44935263 | 0,902062594 | protein phosphatase 2 scaffold subunit Aalpha |
| ENSG00000171777 | RASGRP4 | 0,205555594 | 1,153130342 | 0,76164334 | 0,976285211 | RAS guanyl releasing protein 4 |
| ENSG00000100413 | POLR3H | 0,205553385 | 1,153128576 | 0,53090346 | 0,931265667 | RNA polymerase III subunit H |

| | | | | | | |
|---|---|---|---|---|---|---|
| ENSG00000287655 | AC023283,1 | 0,205392193 | 1,152999744 | 0,81568686 | 0,981590958 | novel transcript, antisense to SHTN1 |
| ENSG00000156639 | ZFAND3 | 0,205378525 | 1,152988821 | 0,41829999 | 0,887094957 | zinc finger AN1-type containing 3 |
| ENSG00000122145 | TBX22 | 0,205344576 | 1,15296169 | 0,87223105 | 0,989217334 | T-box transcription factor 22 |
| ENSG00000103051 | COG4 | 0,205328626 | 1,152948943 | 0,31945875 | 0,824366234 | component of oligomeric golgi complex 4 |
| ENSG00000237945 | LINC00649 | 0,20528863 | 1,15291698 | 0,44167526 | 0,899078209 | long intergenic non-protein coding RNA 649 |
| ENSG00000165671 | NSD1 | 0,205081036 | 1,152751095 | 0,35561699 | 0,846892875 | nuclear receptor binding SET domain protein 1 |
| ENSG00000139624 | CERS5 | 0,205023038 | 1,152704754 | 0,32690261 | 0,82867756 | ceramide synthase 5 |
| ENSG00000183935 | HTR7P1 | 0,205010855 | 1,15269502 | 0,47595823 | 0,911390632 | 5-hydroxytryptamine receptor 7 pseudogene 1 |
| ENSG00000272509 | AC087752,4 | 0,204958735 | 1,152653377 | 0,78479969 | 0,979320406 | novel transcript, antisense to CCNE2 |
| ENSG00000186297 | GABRA5 | 0,204814882 | 1,15253845 | 0,69782375 | 0,968522643 | gamma-aminobutyric acid type A receptor subunit alpha5 |
| ENSG00000087087 | SRRT | 0,204798143 | 1,152525079 | 0,59477173 | 0,947469769 | serrate, RNA effector molecule |
| ENSG00000148672 | GLUD1 | 0,204718705 | 1,152461619 | 0,44237121 | 0,89933631 | glutamate dehydrogenase 1 |
| ENSG00000275045 | GTF2H2 | 0,204705612 | 1,15245116 | 0,62865872 | 0,952793934 | general transcription factor IIH subunit 2 |
| ENSG00000130640 | TUBGCP2 | 0,204660146 | 1,152414842 | 0,36291885 | 0,853169027 | tubulin gamma complex associated protein 2 |
| ENSG00000273599 | AL731571,1 | 0,20460524 | 1,152370984 | 0,65665042 | 0,957444211 | novel transcript, antisense to CTBP2 |
| ENSG00000164237 | CMBL | 0,20450361 | 1,152289808 | 0,42345433 | 0,890929178 | carboxymethylenebutenolidase homolog |
| ENSG00000257979 | SNRPGP18 | 0,20423345 | 1,152074051 | 0,90240542 | 0,992230188 | small nuclear ribonucleoprotein polypeptide G pseudogene 18 |
| ENSG00000104969 | SGTA | 0,20415047 | 1,152007788 | 0,43450863 | 0,896272827 | small glutamine rich tetratricopeptide repeat containing alpha |
| ENSG00000154240 | CEP112 | 0,204126189 | 1,1519884 | 0,3992898 | 0,875433675 | centrosomal protein 112 |
| ENSG00000196634 | LUADT1 | 0,204104148 | 1,1519708 | 0,77547024 | 0,978587255 | lung adenocarcinoma associated transcript 1 |
| ENSG00000187268 | FAM9C | 0,204098454 | 1,151966253 | 0,76028268 | 0,97624106 | family with sequence similarity 9 member C |
| ENSG00000225973 | PIGBOS1 | 0,204002184 | 1,151889386 | 0,47913039 | 0,912714933 | PIGB opposite strand 1 |
| ENSG00000203325 | AL445248,1 | 0,203996506 | 1,151884853 | 0,75706819 | 0,975901808 | novel transcript |
| ENSG00000107960 | STN1 | 0,203845851 | 1,151764572 | 0,45000103 | 0,902062594 | STN1 subunit of CST complex |
| ENSG00000254162 | AC009812,3 | 0,203836489 | 1,151757098 | 0,70767558 | 0,969831641 | novel transcript |
| ENSG00000235844 | HLA-DPA1 | 0,203829184 | 1,151751266 | 0,86346259 | 0,989074806 | major histocompatibility complex, class II, DP alpha 1 |
| ENSG00000168384 | HLA-DPA1 | 0,203829184 | 1,151751266 | 0,86346259 | 0,989074806 | major histocompatibility complex, class II, DP alpha 1 |
| ENSG00000272523 | LINC01023 | 0,203797824 | 1,151726231 | 0,77976528 | 0,978587255 | long intergenic non-protein coding RNA 1023 |

| | | | | | | |
|---|---|---|---|---|---|---|
| ENSG00000240219 | AL512306,2 | 0,203685024 | 1,151636185 | 0,68513828 | 0,965415049 | novel transcript |
| ENSG00000214216 | IQCJ | 0,203681533 | 1,151633397 | 0,87510196 | 0,990078774 | IQ motif containing J |
| ENSG00000170234 | PWWP2A | 0,203630112 | 1,151592351 | 0,5028505 | 0,920749191 | PWWP domain containing 2A |
| ENSG00000111641 | NOP2 | 0,203626717 | 1,151589642 | 0,3689365 | 0,857916156 | NOP2 nucleolar protein |
| ENSG00000068354 | TBC1D25 | 0,203605897 | 1,151573023 | 0,50078356 | 0,920520138 | TBC1 domain family member 25 |
| ENSG00000287530 | AL445985,2 | 0,203603009 | 1,151570718 | 0,77761753 | 0,978587255 | novel transcript |
| ENSG00000074964 | ARHGEF10L | 0,203594991 | 1,151564318 | 0,59745992 | 0,948282297 | Rho guanine nucleotide exchange factor 10 like |
| ENSG00000271816 | BMS1P4 | 0,203516878 | 1,151501969 | 0,61617364 | 0,952793934 | BMS1 pseudogene 4 |
| ENSG00000160633 | SAFB | 0,203392963 | 1,151403069 | 0,65160693 | 0,957089277 | scaffold attachment factor B |
| ENSG00000287226 | AC084740,1 | 0,203361723 | 1,151378137 | 0,79488486 | 0,979776981 | novel transcript |
| ENSG00000275035 | CCDC92 | 0,203318983 | 1,151344027 | 0,40047317 | 0,876197594 | coiled-coil domain containing 92 |
| ENSG00000181904 | C5orf24 | 0,203293928 | 1,151324033 | 0,63026035 | 0,953103521 | chromosome 5 open reading frame 24 |
| ENSG00000136100 | VPS36 | 0,203180024 | 1,151233137 | 0,48092993 | 0,912907448 | vacuolar protein sorting 36 homolog |
| ENSG00000186063 | AIDA | 0,20314109 | 1,151202069 | 0,57094158 | 0,942436428 | axin interactor, dorsalization associated |
| ENSG00000169696 | ASPSCR1 | 0,203005835 | 1,151094147 | 0,63847043 | 0,956018518 | ASPSCR1 tether for SLC2A4, UBX domain containing |
| ENSG00000189051 | RNF222 | 0,202948221 | 1,151048179 | 0,75773589 | 0,976120207 | ring finger protein 222 |
| ENSG00000256028 | AC026362,1 | 0,202939243 | 1,151041016 | 0,42690039 | 0,892272623 | novel transcript |
| ENSG00000239382 | ALKBH6 | 0,202907911 | 1,151016018 | 0,51619822 | 0,925176589 | alkB homolog 6 |
| ENSG00000101558 | VAPA | 0,202880522 | 1,150994167 | 0,62126772 | 0,952793934 | VAMP associated protein A |
| ENSG00000254687 | AC009646,2 | 0,20275693 | 1,150895569 | 0,8201409 | 0,982609114 | novel transcript |
| ENSG00000132612 | VPS4A | 0,202734122 | 1,150877373 | 0,35151272 | 0,84405766 | vacuolar protein sorting 4 homolog A |
| ENSG00000134802 | SLC43A3 | 0,202701429 | 1,150851294 | 0,34610952 | 0,841168721 | solute carrier family 43 member 3 |
| ENSG00000286449 | AC016590,4 | 0,202618898 | 1,15078546 | 0,75306812 | 0,975901808 | novel transcript |
| ENSG00000259735 | AC092868,2 | 0,202564589 | 1,150742141 | 0,89133814 | 0,990820775 | novel transcript, sense intronic to MYO1E |
| ENSG00000054179 | ENTPD2 | 0,202552718 | 1,150732672 | 0,92040363 | 0,993787755 | ectonucleoside triphosphate diphosphohydrolase 2 |
| ENSG00000230280 | HNRNPA1P59 | 0,202451214 | 1,150651712 | 0,67120325 | 0,962026636 | heterogeneous nuclear ribonucleoprotein A1 pseudogene 59 |
| ENSG00000169855 | ROBO1 | 0,202288919 | 1,150522277 | 0,54574283 | 0,935146964 | roundabout guidance receptor 1 |
| ENSG00000285667 | AC012291,3 | 0,202248054 | 1,150489689 | 0,70806433 | 0,970109258 | novel transcript |
| ENSG00000178184 | PARD6G | 0,20216123 | 1,150420453 | 0,34494627 | 0,841168721 | par-6 family cell polarity regulator gamma |
| ENSG00000210135 | MT-TN | 0,202139048 | 1,150402764 | 0,73601832 | 0,974353474 | mitochondrially encoded tRNA-Asn (AAU/C) |
| ENSG00000163781 | TOPBP1 | 0,202049723 | 1,15033154 | 0,50573453 | 0,921780819 | DNA topoisomerase II binding protein 1 |
| ENSG00000233178 | AL161457,2 | 0,201999421 | 1,150291432 | 0,59413097 | 0,947469769 | novel transcript, antisense to PGM5 |

| | | | | | | |
|---|---|---|---|---|---|---|
| ENSG00000112941 | TENT4A | 0,201910037 | 1,150220166 | 0,38118873 | 0,865900094 | terminal nucleotidyltransferase 4A |
| ENSG00000112249 | ASCC3 | 0,201808671 | 1,150139352 | 0,56059109 | 0,940170934 | activating signal cointegrator 1 complex subunit 3 |
| ENSG00000136930 | PSMB7 | 0,201799863 | 1,150132331 | 0,40080896 | 0,876592597 | proteasome 20S subunit beta 7 |
| ENSG00000258790 | AL121594,1 | 0,201736271 | 1,150081635 | 0,68657122 | 0,965769168 | novel transcript |
| ENSG00000237390 | AL139130,1 | 0,201683685 | 1,150039716 | 0,84186252 | 0,98571999 | novel transcript |
| ENSG00000236753 | MKLN1-AS | 0,201599958 | 1,149972976 | 0,58540787 | 0,945213113 | MKLN1 antisense RNA |
| ENSG00000197497 | ZNF665 | 0,201526647 | 1,149914541 | 0,7107249 | 0,970482408 | zinc finger protein 665 |
| ENSG00000160087 | UBE2J2 | 0,201495606 | 1,1498898 | 0,40118862 | 0,876659568 | ubiquitin conjugating enzyme E2 J2 |
| ENSG00000226617 | RPL21P110 | 0,201367365 | 1,149787591 | 0,85165506 | 0,987289464 | ribosomal protein L21 pseudogene 110 |
| ENSG00000188191 | PRKAR1B | 0,201321324 | 1,149750898 | 0,56630589 | 0,941153917 | protein kinase cAMP-dependent type I regulatory subunit beta |
| ENSG00000108018 | SORCS1 | 0,201307534 | 1,149739908 | 0,62207336 | 0,952793934 | sortilin related VPS10 domain containing receptor 1 |
| ENSG00000067798 | NAV3 | 0,201270119 | 1,149710091 | 0,44861322 | 0,902062594 | neuron navigator 3 |
| ENSG00000169228 | RAB24 | 0,201192284 | 1,149648064 | 0,49805608 | 0,919274055 | RAB24, member RAS oncogene family |
| ENSG00000234160 | AL513165,1 | 0,201191234 | 1,149647227 | 0,76415895 | 0,976640468 | novel transcript |
| ENSG00000087502 | ERGIC2 | 0,201182787 | 1,149640496 | 0,64126722 | 0,956465487 | ERGIC and golgi 2 |
| ENSG00000241685 | ARPC1A | 0,201021616 | 1,149512071 | 0,41016049 | 0,882597401 | actin related protein 2/3 complex subunit 1A |
| ENSG00000177732 | SOX12 | 0,20095209 | 1,149456676 | 0,56327962 | 0,940865625 | SRY-box transcription factor 12 |
| ENSG00000092929 | UNC13D | 0,200901151 | 1,149416091 | 0,77730086 | 0,978587255 | unc-13 homolog D |
| ENSG00000224877 | NDUFAF8 | 0,20084273 | 1,149369547 | 0,52072216 | 0,927046149 | NADH:ubiquinone oxidoreductase complex assembly factor 8 |
| ENSG00000125505 | MBOAT7 | 0,20079208 | 1,149329196 | 0,64133937 | 0,956465487 | membrane bound O-acyltransferase domain containing 7 |
| ENSG00000161643 | SIGLEC16 | 0,200734049 | 1,149282966 | 0,76341232 | 0,976640468 | sialic acid binding Ig like lectin 16 |
| ENSG00000136152 | COG3 | 0,20063422 | 1,149203442 | 0,53736393 | 0,932883277 | component of oligomeric golgi complex 3 |
| ENSG00000256269 | HMBS | 0,200605416 | 1,149180499 | 0,44070625 | 0,898600418 | hydroxymethylbilane synthase |
| ENSG00000109171 | SLAIN2 | 0,200486101 | 1,149085462 | 0,56415837 | 0,940865625 | SLAIN motif family member 2 |
| ENSG00000272899 | ATP6V1FNB | 0,200348345 | 1,148975746 | 0,63509881 | 0,955159215 | ATP6V1F neighbor |
| ENSG00000258136 | AC007622,2 | 0,200337287 | 1,14896694 | 0,57981447 | 0,943828882 | novel transcript, antisense to PRDM4 |
| ENSG00000142632 | ARHGEF19 | 0,200322758 | 1,148955369 | 0,64685829 | 0,956899963 | Rho guanine nucleotide exchange factor 19 |
| ENSG00000160293 | VAV2 | 0,200316016 | 1,14895 | 0,5376181 | 0,932883277 | vav guanine nucleotide exchange factor 2 |
| ENSG00000197024 | ZNF398 | 0,200289655 | 1,148929006 | 0,44119347 | 0,898885877 | zinc finger protein 398 |
| ENSG00000275464 | FP565260,1 | 0,200175527 | 1,148838121 | 0,60635807 | 0,950927937 | periodic tryptophan protein 2 homolog |

| | | | | | | |
|---|---|---|---|---|---|---|
| ENSG00000107758 | PPP3CB | 0,200157609 | 1,148823853 | 0,56229681 | 0,940865625 | protein phosphatase 3 catalytic subunit beta |
| ENSG00000163719 | MTMR14 | 0,200145276 | 1,148814032 | 0,38266822 | 0,867251844 | myotubularin related protein 14 |
| ENSG00000263826 | AC112907,3 | 0,200111502 | 1,148787139 | 0,51774756 | 0,925915783 | novel transcript, antisense to eukaryotic translation initiation factor 4A, isoform 2 EIF4A2 |
| ENSG00000256746 | MADD-AS1 | 0,20006335 | 1,148748797 | 0,81924537 | 0,98249617 | MADD antisense RNA 1 |
| ENSG00000287193 | AC132938,6 | 0,200029773 | 1,148722061 | 0,7650115 | 0,976739398 | novel transcript |
| ENSG00000114423 | CBLB | 0,200019371 | 1,148713779 | 0,52745592 | 0,929454507 | Cbl proto-oncogene B |
| ENSG00000239857 | GET4 | 0,199951511 | 1,148659748 | 0,53702753 | 0,932883277 | guided entry of tail-anchored proteins factor 4 |
| ENSG00000160299 | PCNT | 0,199824018 | 1,148558244 | 0,69012331 | 0,967053457 | pericentrin |
| ENSG00000089195 | TRMT6 | 0,199549709 | 1,148339881 | 0,50564693 | 0,921780819 | tRNA methyltransferase 6 |
| ENSG00000132002 | DNAJB1 | 0,199490988 | 1,148293142 | 0,34646013 | 0,841168721 | DnaJ heat shock protein family (Hsp40) member B1 |
| ENSG00000169592 | INO80E | 0,199448021 | 1,148258944 | 0,50843418 | 0,922427526 | INO80 complex subunit E |
| ENSG00000107854 | TNKS2 | 0,199358715 | 1,148187866 | 0,59835518 | 0,948677746 | tankyrase 2 |
| ENSG00000213923 | CSNK1E | 0,199338328 | 1,148171642 | 0,38768297 | 0,869820053 | casein kinase 1 epsilon |
| ENSG00000168172 | HOOK3 | 0,199333445 | 1,148167755 | 0,4534044 | 0,902754541 | hook microtubule tethering protein 3 |
| ENSG00000163126 | ANKRD23 | 0,199295213 | 1,148137329 | 0,70817621 | 0,97011892 | ankyrin repeat domain 23 |
| ENSG00000250427 | LINC02148 | 0,19920386 | 1,148064629 | 0,8670983 | 0,989074806 | long intergenic non-protein coding RNA 2148 |
| ENSG00000105254 | TBCB | 0,199195627 | 1,148058078 | 0,37710181 | 0,862818448 | tubulin folding cofactor B |
| ENSG00000233215 | LINC01687 | 0,199125603 | 1,148002356 | 0,72337154 | 0,972913663 | long intergenic non-protein coding RNA 1687 |
| ENSG00000136816 | TOR1B | 0,199045578 | 1,147938679 | 0,39988123 | 0,875727577 | torsin family 1 member B |
| ENSG00000155438 | NIFK | 0,198930197 | 1,147846876 | 0,61643577 | 0,952793934 | nucleolar protein interacting with the FHA domain of MKI67 |
| ENSG00000114999 | TTL | 0,198855479 | 1,147787429 | 0,46779706 | 0,907940653 | tubulin tyrosine ligase |
| ENSG00000237949 | LINC00844 | 0,198742807 | 1,147697792 | 0,79670892 | 0,980146131 | long intergenic non-protein coding RNA 844 |
| ENSG00000227001 | NBPF2P | 0,198705081 | 1,147667781 | 0,874343 | 0,98990147 | NBPF member 2, pseudogene |
| ENSG00000285317 | DNAJC8 | 0,198699966 | 1,147663712 | 0,46390003 | 0,907292403 | DnaJ heat shock protein family (Hsp40) member C8 |
| ENSG00000126698 | DNAJC8 | 0,198699966 | 1,147663712 | 0,46390003 | 0,907292403 | DnaJ heat shock protein family (Hsp40) member C8 |
| ENSG00000065135 | GNAI3 | 0,198680448 | 1,147648186 | 0,57172674 | 0,942436428 | G protein subunit alpha i3 |
| ENSG00000254397 | AC132192,1 | 0,198651181 | 1,147624904 | 0,82439314 | 0,983374097 | novel transcript |
| ENSG00000276116 | FUT8-AS1 | 0,198536751 | 1,147533882 | 0,6245757 | 0,952793934 | FUT8 antisense RNA 1 |
| ENSG00000165813 | CCDC186 | 0,198426155 | 1,147445916 | 0,58605102 | 0,945246954 | coiled-coil domain containing 186 |
| ENSG00000132024 | CC2D1A | 0,198414793 | 1,147436879 | 0,6627921 | 0,959897678 | coiled-coil and C2 domain containing 1A |
| ENSG00000111266 | DUSP16 | 0,198328204 | 1,147368013 | 0,49159817 | 0,916995341 | dual specificity phosphatase 16 |

| | | | | | | |
|---|---|---|---|---|---|---|
| ENSG00000280962 | DUSP16 | 0,198328204 | 1,147368013 | 0,49159817 | 0,916995341 | dual specificity phosphatase 16 |
| ENSG00000218416 | AC110619,1 | 0,198289696 | 1,147337388 | 0,78137984 | 0,978597905 | novel transcript, antisense to GPC1 |
| ENSG00000264954 | PRR29-AS1 | 0,198275352 | 1,147325981 | 0,83103858 | 0,984300284 | PRR29 antisense RNA 1 |
| ENSG00000126267 | COX6B1 | 0,198122957 | 1,147204793 | 0,34336579 | 0,840873784 | cytochrome c oxidase subunit 6B1 |
| ENSG00000230992 | FAM201B | 0,198070692 | 1,147163234 | 0,77039247 | 0,977891334 | family with sequence similarity 201 member B |
| ENSG00000258044 | AC073569,1 | 0,198008142 | 1,147113498 | 0,78020943 | 0,978587255 | novel transcript, antisense to PPP1R12A |
| ENSG00000107021 | TBC1D13 | 0,197996559 | 1,147104288 | 0,42617784 | 0,892133432 | TBC1 domain family member 13 |
| ENSG00000116489 | CAPZA1 | 0,197882793 | 1,147013835 | 0,67038567 | 0,962026636 | capping actin protein of muscle Z-line subunit alpha 1 |
| ENSG00000087191 | PSMC5 | 0,197867138 | 1,147001389 | 0,3655333 | 0,855455735 | proteasome 26S subunit, ATPase 5 |
| ENSG00000014164 | ZC3H3 | 0,197821902 | 1,146965425 | 0,45004985 | 0,902062594 | zinc finger CCCH-type containing 3 |
| ENSG00000102743 | SLC25A15 | 0,197812636 | 1,146958058 | 0,38450096 | 0,868434303 | solute carrier family 25 member 15 |
| ENSG00000286305 | AC004893,2 | 0,19776987 | 1,146924059 | 0,66357633 | 0,959920886 | novel transcript, antisense to TRRAP |
| ENSG00000145390 | USP53 | 0,197713516 | 1,146879259 | 0,61172936 | 0,952161039 | ubiquitin specific peptidase 53 |
| ENSG00000260676 | LINC01541 | 0,197683156 | 1,146855125 | 0,86382249 | 0,989074806 | long intergenic non-protein coding RNA 1541 |
| ENSG00000185504 | FAAP100 | 0,197672178 | 1,146846398 | 0,60603206 | 0,950927937 | FA core complex associated protein 100 |
| ENSG00000143891 | GALM | 0,197591767 | 1,146782478 | 0,52339229 | 0,927995032 | galactose mutarotase |
| ENSG00000281691 | RBM5-AS1 | 0,197544238 | 1,146744699 | 0,5686212 | 0,941793403 | RBM5 antisense RNA 1 |
| ENSG00000185800 | DMWD | 0,19747521 | 1,146689833 | 0,5658013 | 0,940866081 | DM1 locus, WD repeat containing |
| ENSG00000267321 | SNHG30 | 0,197449898 | 1,146669714 | 0,45263064 | 0,902285042 | small nucleolar RNA host gene 30 |
| ENSG00000155256 | ZFYVE27 | 0,197365059 | 1,146602285 | 0,48744892 | 0,91491358 | zinc finger FYVE-type containing 27 |
| ENSG00000273607 | CHCHD10 | 0,197344922 | 1,146586281 | 0,65999899 | 0,958859629 | coiled-coil-helix-coiled-coil-helix domain containing 10 |
| ENSG00000147475 | ERLIN2 | 0,197282797 | 1,146536908 | 0,45610531 | 0,903545728 | ER lipid raft associated 2 |
| ENSG00000123352 | SPATS2 | 0,197267186 | 1,146524501 | 0,45070642 | 0,902062594 | spermatogenesis associated serine rich 2 |
| ENSG00000113742 | CPEB4 | 0,197258397 | 1,146517517 | 0,54904506 | 0,936040289 | cytoplasmic polyadenylation element binding protein 4 |
| ENSG00000206476 | VARS2 | 0,197252659 | 1,146512956 | 0,57266335 | 0,942475308 | valyl-tRNA synthetase 2, mitochondrial |
| ENSG00000221994 | ZNF630 | 0,197234723 | 1,146498703 | 0,56572994 | 0,940866081 | zinc finger protein 630 |
| ENSG00000287912 | AP003464,1 | 0,197205628 | 1,146475582 | 0,87035248 | 0,989074806 | novel transcript |
| ENSG00000221883 | ARIH2OS | 0,197180504 | 1,146455617 | 0,68229754 | 0,964749521 | ARIH2 opposite strand lncRNA |
| ENSG00000166228 | PCBD1 | 0,197159048 | 1,146438566 | 0,52868037 | 0,930033144 | pterin-4 alpha-carbinolamine dehydratase 1 |
| ENSG00000255537 | AP000708,1 | 0,197094911 | 1,146387601 | 0,77958839 | 0,978587255 | novel transcript, antisense to FEZ1 |

| | | | | | | |
|---|---|---|---|---|---|---|
| ENSG00000163930 | BAP1 | 0,197040456 | 1,146344331 | 0,47559181 | 0,911264385 | BRCA1 associated protein 1 |
| ENSG00000228340 | MIR646HG | 0,197012317 | 1,146321973 | 0,77678192 | 0,978587255 | MIR646 host gene |
| ENSG00000176018 | LYSMD3 | 0,196974601 | 1,146292005 | 0,73940972 | 0,974715504 | LysM domain containing 3 |
| ENSG00000243697 | AC009108,1 | 0,196930683 | 1,14625711 | 0,8388098 | 0,985672131 | ribosomal protein L7a (RPL7A) pseudogene |
| ENSG00000185267 | CDNF | 0,196915833 | 1,146245312 | 0,79249984 | 0,979471509 | cerebral dopamine neurotrophic factor |
| ENSG00000253710 | ALG11 | 0,196872533 | 1,14621091 | 0,42159413 | 0,889627345 | ALG11 alpha-1,2-mannosyltransferase |
| ENSG00000264672 | SEPTIN4-AS1 | 0,196864204 | 1,146204292 | 0,77496674 | 0,978575875 | SEPTIN4 antisense RNA 1 |
| ENSG00000172409 | CLP1 | 0,196812589 | 1,146163285 | 0,4536396 | 0,902908855 | cleavage factor polyribonucleotide kinase subunit 1 |
| ENSG00000285682 | AC005343,5 | 0,196663302 | 1,146044689 | 0,76633994 | 0,977154522 | novel transcript, antisense to ADIPOR2 |
| ENSG00000002330 | BAD | 0,19655506 | 1,145958708 | 0,55442681 | 0,937793851 | BCL2 associated agonist of cell death |
| ENSG00000233401 | PRKAR1AP1 | 0,196504918 | 1,14591888 | 0,85791097 | 0,988965329 | protein kinase cAMP-dependent type I regulatory subunit alpha pseudogene 1 |
| ENSG00000276570 | AC010327,6 | 0,196459707 | 1,14588297 | 0,70924522 | 0,970282889 | novel transcript |
| ENSG00000105085 | MED26 | 0,196446176 | 1,145872222 | 0,47670429 | 0,91143003 | mediator complex subunit 26 |
| ENSG00000213995 | NAXD | 0,196422181 | 1,145853164 | 0,37729293 | 0,862818448 | NAD(P)HX dehydratase |
| ENSG00000174516 | PELI3 | 0,196325265 | 1,145776192 | 0,50156707 | 0,920520138 | pellino E3 ubiquitin protein ligase family member 3 |
| ENSG00000164253 | WDR41 | 0,196298419 | 1,145754871 | 0,47891522 | 0,912714933 | WD repeat domain 41 |
| ENSG00000170873 | MTSS1 | 0,196245137 | 1,145712556 | 0,5428176 | 0,933663748 | MTSS I-BAR domain containing 1 |
| ENSG00000132589 | FLOT2 | 0,196233824 | 1,145703572 | 0,55140287 | 0,936366211 | flotillin 2 |
| ENSG00000213626 | LBH | 0,196223328 | 1,145695237 | 0,60675764 | 0,950927937 | LBH regulator of WNT signaling pathway |
| ENSG00000115902 | SLC1A4 | 0,196150694 | 1,145637557 | 0,38695581 | 0,869441674 | solute carrier family 1 member 4 |
| ENSG00000141580 | WDR45B | 0,196132761 | 1,145623317 | 0,39926735 | 0,875433675 | WD repeat domain 45B |
| ENSG00000103035 | PSMD7 | 0,196111368 | 1,145606329 | 0,47725052 | 0,91143003 | proteasome 26S subunit, non-ATPase 7 |
| ENSG00000108515 | ENO3 | 0,196058493 | 1,145564343 | 0,51726929 | 0,925746338 | enolase 3 |
| ENSG00000135378 | PRRG4 | 0,19598275 | 1,145504201 | 0,81422094 | 0,981521413 | proline rich and Gla domain 4 |
| ENSG00000101997 | CCDC22 | 0,195977604 | 1,145500116 | 0,52122624 | 0,927046149 | coiled-coil domain containing 22 |
| ENSG00000138107 | ACTR1A | 0,195974319 | 1,145497507 | 0,36142633 | 0,851893059 | actin related protein 1A |
| ENSG00000107223 | EDF1 | 0,195956168 | 1,145483095 | 0,37524803 | 0,862022464 | endothelial differentiation related factor 1 |
| ENSG00000250506 | CDK3 | 0,195900669 | 1,14543903 | 0,5480552 | 0,935404974 | cyclin dependent kinase 3 |
| ENSG00000170502 | NUDT9 | 0,195884032 | 1,145425822 | 0,43768123 | 0,897021798 | nudix hydrolase 9 |
| ENSG00000261736 | AC002551,1 | 0,195860907 | 1,145407462 | 0,89722365 | 0,991644439 | novel transcript |
| ENSG00000061936 | SFSWAP | 0,195807517 | 1,145365074 | 0,37536552 | 0,862022464 | splicing factor SWAP |
| ENSG00000105705 | SUGP1 | 0,195754914 | 1,145323313 | 0,45234462 | 0,90209238 | SURP and G-patch domain containing 1 |

| | | | | | | |
|---|---|---|---|---|---|---|
| ENSG00000138744 | NAAA | 0,195730487 | 1,145303921 | 0,73019777 | 0,974303405 | N-acylethanolamine acid amidase |
| ENSG00000280548 | FBL | 0,195707685 | 1,14528582 | 0,53307516 | 0,931291085 | fibrillarin |
| ENSG00000137693 | YAP1 | 0,195677821 | 1,145262112 | 0,58398992 | 0,944486102 | Yes1 associated transcriptional regulator |
| ENSG00000196943 | NOP9 | 0,195649585 | 1,145239698 | 0,62399311 | 0,952793934 | NOP9 nucleolar protein |
| ENSG00000285326 | NOP9 | 0,195649585 | 1,145239698 | 0,62399311 | 0,952793934 | NOP9 nucleolar protein |
| ENSG00000074071 | MRPS34 | 0,195568918 | 1,145175664 | 0,4954461 | 0,917815326 | mitochondrial ribosomal protein S34 |
| ENSG00000061676 | NCKAP1 | 0,195370359 | 1,145018064 | 0,61567278 | 0,952793934 | NCK associated protein 1 |
| ENSG00000149792 | MRPL49 | 0,195269536 | 1,144938047 | 0,35765154 | 0,848828488 | mitochondrial ribosomal protein L49 |
| ENSG00000113141 | IK | 0,195143596 | 1,144838104 | 0,45217819 | 0,902062594 | IK cytokine |
| ENSG00000054356 | PTPRN | 0,195075701 | 1,144784228 | 0,59262134 | 0,947416795 | protein tyrosine phosphatase receptor type N |
| ENSG00000286070 | AP000356,5 | 0,195053469 | 1,144766587 | 0,39969103 | 0,875433675 | novel protein |
| ENSG00000112855 | HARS2 | 0,195018533 | 1,144738866 | 0,34015051 | 0,838878717 | histidyl-tRNA synthetase 2, mitochondrial |
| ENSG00000184898 | RBM43 | 0,194964063 | 1,144695646 | 0,655165 | 0,957444211 | RNA binding motif protein 43 |
| ENSG00000152818 | UTRN | 0,194929234 | 1,144668012 | 0,38625332 | 0,869312875 | utrophin |
| ENSG00000116005 | PCYOX1 | 0,194887035 | 1,144634531 | 0,60363663 | 0,950352286 | prenylcysteine oxidase 1 |
| ENSG00000126067 | PSMB2 | 0,194772315 | 1,144543515 | 0,35754796 | 0,848693232 | proteasome 20S subunit beta 2 |
| ENSG00000214194 | SMIM30 | 0,194761781 | 1,144535158 | 0,67561858 | 0,963449137 | small integral membrane protein 30 |
| ENSG00000249915 | PDCD6 | 0,194722878 | 1,144504296 | 0,42553307 | 0,892133432 | programmed cell death 6 |
| ENSG00000149923 | PPP4C | 0,194692864 | 1,144480485 | 0,46261723 | 0,906730926 | protein phosphatase 4 catalytic subunit |
| ENSG00000118308 | IRAG2 | 0,194691031 | 1,144479032 | 0,82562599 | 0,983383773 | inositol 1,4,5-triphosphate receptor associated 2 |
| ENSG00000123144 | TRIR | 0,194570844 | 1,144383692 | 0,4737117 | 0,910271829 | telomerase RNA component interacting RNase |
| ENSG00000276219 | ZNF676 | 0,19447793 | 1,144309993 | 0,91747672 | 0,993361299 | zinc finger protein 676 |
| ENSG00000139718 | SETD1B | 0,194468515 | 1,144302525 | 0,60314414 | 0,950352286 | SET domain containing 1B, histone lysine methyltransferase |
| ENSG00000114861 | FOXP1 | 0,194421787 | 1,144265463 | 0,48633414 | 0,914091794 | forkhead box P1 |
| ENSG00000135930 | EIF4E2 | 0,194387933 | 1,144238612 | 0,37816394 | 0,863806148 | eukaryotic translation initiation factor 4E family member 2 |
| ENSG00000175334 | BANF1 | 0,194319672 | 1,144184473 | 0,35180335 | 0,844406182 | BAF nuclear assembly factor 1 |
| ENSG00000128563 | PRKRIP1 | 0,194309691 | 1,144176557 | 0,57446226 | 0,942475308 | PRKR interacting protein 1 |
| ENSG00000127928 | GNGT1 | 0,194234751 | 1,144117126 | 0,85320766 | 0,987861659 | G protein subunit gamma transducin 1 |
| ENSG00000235758 | AGPAT1 | 0,194112925 | 1,144020516 | 0,45092379 | 0,902062594 | 1-acylglycerol-3-phosphate O-acyltransferase 1 |
| ENSG00000206324 | AGPAT1 | 0,194112925 | 1,144020516 | 0,45092379 | 0,902062594 | 1-acylglycerol-3-phosphate O-acyltransferase 1 |
| ENSG00000226467 | AGPAT1 | 0,194112925 | 1,144020516 | 0,45092379 | 0,902062594 | 1-acylglycerol-3-phosphate O-acyltransferase 1 |

| | | | | | | |
|---|---|---|---|---|---|---|
| ENSG00000180957 | PITPNB | 0,194109572 | 1,144017858 | 0,58899383 | 0,946604802 | phosphatidylinositol transfer protein beta |
| ENSG00000118454 | ANKRD13C | 0,194109173 | 1,144017541 | 0,51794561 | 0,926164002 | ankyrin repeat domain 13C |
| ENSG00000283000 | LINC02635 | 0,194034874 | 1,143958626 | 0,82024886 | 0,982609114 | long intergenic non-protein coding RNA 2635 |
| ENSG00000175318 | GRAMD2A | 0,193844616 | 1,143807774 | 0,7923246 | 0,979471509 | GRAM domain containing 2A |
| ENSG00000176125 | UFSP1 | 0,193842155 | 1,143805823 | 0,76647827 | 0,977174732 | UFM1 specific peptidase 1 (inactive) |
| ENSG00000251095 | AC093866,1 | 0,193827841 | 1,143794474 | 0,83441398 | 0,984905093 | novel transcript |
| ENSG00000167967 | E4F1 | 0,193801604 | 1,143773673 | 0,63810645 | 0,956018518 | E4F transcription factor 1 |
| ENSG00000286136 | AC107071,1 | 0,193790921 | 1,143765204 | 0,71073832 | 0,970482408 | novel transcript |
| ENSG00000250510 | GPR162 | 0,193693195 | 1,143687729 | 0,50072197 | 0,920520138 | G protein-coupled receptor 162 |
| ENSG00000166086 | JAM3 | 0,193664839 | 1,143665251 | 0,49799741 | 0,919274055 | junctional adhesion molecule 3 |
| ENSG00000234215 | CT66 | 0,193639517 | 1,143645178 | 0,79495448 | 0,979776981 | cancer/testis associated transcript 66 |
| ENSG00000242255 | RPL39P34 | 0,193589212 | 1,1436053 | 0,8998367 | 0,991845106 | ribosomal protein L39 pseudogene 34 |
| ENSG00000130560 | UBAC1 | 0,193510222 | 1,143542688 | 0,36663154 | 0,856103393 | UBA domain containing 1 |
| ENSG00000085377 | PREP | 0,193486741 | 1,143524076 | 0,49146709 | 0,916995341 | prolyl endopeptidase |
| ENSG00000089902 | RCOR1 | 0,193443995 | 1,143490195 | 0,49057907 | 0,916995341 | REST corepressor 1 |
| ENSG00000102908 | NFAT5 | 0,193430941 | 1,143479848 | 0,59327042 | 0,947416795 | nuclear factor of activated T cells 5 |
| ENSG00000184857 | TMEM186 | 0,193391988 | 1,143448975 | 0,48874395 | 0,91557151 | transmembrane protein 186 |
| ENSG00000138131 | LOXL4 | 0,193301777 | 1,143377478 | 0,59750777 | 0,948282297 | lysyl oxidase like 4 |
| ENSG00000136636 | KCTD3 | 0,193277475 | 1,143358217 | 0,54410814 | 0,934259495 | potassium channel tetramerization domain containing 3 |
| ENSG00000073578 | SDHA | 0,193225606 | 1,143317112 | 0,41862198 | 0,887094957 | succinate dehydrogenase complex flavoprotein subunit A |
| ENSG00000010539 | ZNF200 | 0,193182179 | 1,143282696 | 0,45821291 | 0,904140862 | zinc finger protein 200 |
| ENSG00000179134 | SAMD4B | 0,193171541 | 1,143274266 | 0,42760426 | 0,892656446 | sterile alpha motif domain containing 4B |
| ENSG00000033050 | ABCF2 | 0,19315815 | 1,143263654 | 0,44710858 | 0,901890609 | ATP binding cassette subfamily F member 2 |
| ENSG00000276358 | PLEKHM1 | 0,193130226 | 1,143241526 | 0,75014517 | 0,975901808 | pleckstrin homology and RUN domain containing M1 |
| ENSG00000230189 | AC008267,2 | 0,193124791 | 1,143237219 | 0,91425089 | 0,993361299 | RAB guanine nucleotide exchange factor 1 (RABGEF1) pseudogene |
| ENSG00000261408 | TEN1-CDK3 | 0,193124338 | 1,14323686 | 0,76097648 | 0,97624106 | TEN1-CDK3 readthrough (NMD candidate) |
| ENSG00000116670 | MAD2L2 | 0,193076577 | 1,143199014 | 0,40258867 | 0,877321369 | mitotic arrest deficient 2 like 2 |
| ENSG00000110497 | AMBRA1 | 0,192924782 | 1,143078737 | 0,38402685 | 0,868125769 | autophagy and beclin 1 regulator 1 |
| ENSG00000259345 | AC013652,1 | 0,192877354 | 1,14304116 | 0,60277702 | 0,950352286 | novel transcript |
| ENSG00000215817 | ZC3H11B | 0,192812993 | 1,142990168 | 0,79277876 | 0,979471509 | zinc finger CCCH-type containing 11B |

| | | | | | | |
|---|---|---|---|---|---|---|
| ENSG00000254050 | AC018861,2 | 0,192747436 | 1,142938231 | 0,86788188 | 0,989074806 | novel transcript |
| ENSG00000106336 | FBXO24 | 0,192740205 | 1,142932502 | 0,73519173 | 0,974353474 | F-box protein 24 |
| ENSG00000229502 | AL391863,1 | 0,192711451 | 1,142909722 | 0,81115643 | 0,981213877 | novel transcript |
| ENSG00000287845 | AC093765,5 | 0,192656968 | 1,142866562 | 0,84010301 | 0,985672131 | novel transcript |
| ENSG00000132670 | PTPRA | 0,192597151 | 1,142819178 | 0,52379914 | 0,928295231 | protein tyrosine phosphatase receptor type A |
| ENSG00000115020 | PIKFYVE | 0,192561921 | 1,142791271 | 0,57639117 | 0,943158459 | phosphoinositide kinase, FYVE-type zinc finger containing |
| ENSG00000162923 | WDR26 | 0,192471534 | 1,142719675 | 0,55658148 | 0,938062556 | WD repeat domain 26 |
| ENSG00000272092 | AC087623,2 | 0,192260693 | 1,142552686 | 0,65884934 | 0,958240464 | novel transcript |
| ENSG00000099817 | POLR2E | 0,192197184 | 1,14250239 | 0,41137307 | 0,883655718 | RNA polymerase II, I and III subunit E |
| ENSG00000110063 | DCPS | 0,192108878 | 1,142432462 | 0,47933076 | 0,912714933 | decapping enzyme, scavenger |
| ENSG00000239305 | RNF103 | 0,192075724 | 1,142406208 | 0,53117783 | 0,931265667 | ring finger protein 103 |
| ENSG00000230408 | BZW1-AS1 | 0,192007128 | 1,142351891 | 0,78566459 | 0,979320406 | BZW1 antisense RNA 1 |
| ENSG00000261519 | AC010542,4 | 0,19198633 | 1,142335423 | 0,82753792 | 0,983595244 | novel transcript |
| ENSG00000106609 | TMEM248 | 0,191934069 | 1,142294043 | 0,47565646 | 0,911264385 | transmembrane protein 248 |
| ENSG00000169136 | ATF5 | 0,191918466 | 1,142281689 | 0,57059859 | 0,9423888 | activating transcription factor 5 |
| ENSG00000171425 | ZNF581 | 0,191875657 | 1,142247795 | 0,55705143 | 0,938292968 | zinc finger protein 581 |
| ENSG00000137955 | RABGGTB | 0,191837729 | 1,142217765 | 0,61374968 | 0,952413269 | Rab geranylgeranyltransferase subunit beta |
| ENSG00000141101 | NOB1 | 0,191746201 | 1,142145303 | 0,45231641 | 0,90209238 | NIN1 (RPN12) binding protein 1 homolog |
| ENSG00000167264 | DUS2 | 0,191716187 | 1,142121542 | 0,48335762 | 0,913690171 | dihydrouridine synthase 2 |
| ENSG00000181873 | IBA57 | 0,19169968 | 1,142108474 | 0,70310377 | 0,968742626 | iron-sulfur cluster assembly factor IBA57 |
| ENSG00000162604 | TM2D1 | 0,191689724 | 1,142100592 | 0,44791118 | 0,902062594 | TM2 domain containing 1 |
| ENSG00000169181 | GSG1L | 0,191596482 | 1,14202678 | 0,86450838 | 0,989074806 | GSG1 like |
| ENSG00000165282 | PIGO | 0,191525685 | 1,141970739 | 0,50942205 | 0,922427526 | phosphatidylinositol glycan anchor biosynthesis class O |
| ENSG00000160961 | ZNF333 | 0,191514938 | 1,141962232 | 0,41045864 | 0,88263147 | zinc finger protein 333 |
| ENSG00000275484 | AP003419,3 | 0,191506825 | 1,141955811 | 0,87059964 | 0,989074806 | novel transcript |
| ENSG00000183431 | SF3A3 | 0,191442761 | 1,141905103 | 0,41864062 | 0,887094957 | splicing factor 3a subunit 3 |
| ENSG00000233719 | GOT2P3 | 0,191403858 | 1,141874311 | 0,90373736 | 0,992462326 | GOT2 pseudogene 3 |
| ENSG00000135655 | USP15 | 0,191373325 | 1,141850145 | 0,62067922 | 0,952793934 | ubiquitin specific peptidase 15 |
| ENSG00000231023 | LINC00326 | 0,191331215 | 1,141816816 | 0,79995299 | 0,980161849 | long intergenic non-protein coding RNA 326 |
| ENSG00000131584 | ACAP3 | 0,191311895 | 1,141801526 | 0,68051207 | 0,964654539 | ArfGAP with coiled-coil, ankyrin repeat and PH domains 3 |

| | | | | | | |
|---|---|---|---|---|---|---|
| ENSG00000283196 | LSP1P5 | 0,191288468 | 1,141782985 | 0,42688205 | 0,892272623 | LSP1 pseudogene 5 |
| ENSG00000110717 | NDUFS8 | 0,191243987 | 1,141747782 | 0,4475641 | 0,902062594 | NADH:ubiquinone oxidoreductase core subunit S8 |
| ENSG00000198064 | NPIPB13 | 0,191236139 | 1,141741571 | 0,73816456 | 0,974353474 | nuclear pore complex interacting protein family, member B13 |
| ENSG00000102178 | UBL4A | 0,191192575 | 1,141707095 | 0,52874176 | 0,930033144 | ubiquitin like 4A |
| ENSG00000198855 | FICD | 0,191181647 | 1,141698448 | 0,42699062 | 0,892282255 | FIC domain protein adenylyltransferase |
| ENSG00000264940 | SNORD3C | 0,191121705 | 1,141651012 | 0,78317072 | 0,978919905 | small nucleolar RNA, C/D box 3C |
| ENSG00000164332 | UBLCP1 | 0,191069255 | 1,141609508 | 0,67546601 | 0,963400586 | ubiquitin like domain containing CTD phosphatase 1 |
| ENSG00000280496 | SNORA17B | 0,191064403 | 1,141605668 | 0,75802315 | 0,976120207 | small nucleolar RNA, H/ACA box 17B |
| ENSG00000106351 | AGFG2 | 0,191049359 | 1,141593764 | 0,56318338 | 0,940865625 | ArfGAP with FG repeats 2 |
| ENSG00000175224 | ATG13 | 0,190991788 | 1,141548209 | 0,42347064 | 0,890929178 | autophagy related 13 |
| ENSG00000272836 | AL022328,1 | 0,190952451 | 1,141517084 | 0,83525335 | 0,984926668 | novel transcript, antisense to SELO |
| ENSG00000121964 | GTDC1 | 0,190950654 | 1,141515662 | 0,3929382 | 0,871121102 | glycosyltransferase like domain containing 1 |
| ENSG00000223374 | AC005104,1 | 0,190867447 | 1,141449827 | 0,78185622 | 0,978597905 | novel transcript |
| ENSG00000112651 | MRPL2 | 0,190824472 | 1,141415827 | 0,37971132 | 0,865238161 | mitochondrial ribosomal protein L2 |
| ENSG00000254999 | BRK1 | 0,19081373 | 1,141407328 | 0,59612478 | 0,948282297 | BRICK1 subunit of SCAR/WAVE actin nucleating complex |
| ENSG00000260448 | LCMT1-AS1 | 0,19081177 | 1,141405777 | 0,72153437 | 0,972142678 | LCMT1 antisense RNA 1 |
| ENSG00000202538 | RNU4-2 | 0,190776752 | 1,141378072 | 0,77614412 | 0,978587255 | RNA, U4 small nuclear 2 |
| ENSG00000103222 | ABCC1 | 0,190694208 | 1,14131277 | 0,55111525 | 0,936366211 | ATP binding cassette subfamily C member 1 |
| ENSG00000176105 | YES1 | 0,190644288 | 1,14127328 | 0,62273382 | 0,952793934 | YES proto-oncogene 1, Src family tyrosine kinase |
| ENSG00000264164 | AC090616,2 | 0,190397079 | 1,141077736 | 0,86700863 | 0,989074806 | bromodomain PHD finger transcription factor (BPTF) pseudogene |
| ENSG00000028277 | POU2F2 | 0,190379643 | 1,141063946 | 0,60043384 | 0,949439306 | POU class 2 homeobox 2 |
| ENSG00000133019 | CHRM3 | 0,190369724 | 1,141056101 | 0,59218573 | 0,947416795 | cholinergic receptor muscarinic 3 |
| ENSG00000105171 | POP4 | 0,190303189 | 1,141003478 | 0,53122415 | 0,931265667 | POP4 homolog, ribonuclease P/MRP subunit |
| ENSG00000106617 | PRKAG2 | 0,190220324 | 1,140937943 | 0,41460523 | 0,885579089 | protein kinase AMP-activated non-catalytic subunit gamma 2 |
| ENSG00000054118 | THRAP3 | 0,190132781 | 1,140868713 | 0,36448076 | 0,854463603 | thyroid hormone receptor associated protein 3 |
| ENSG00000274081 | PUF60 | 0,189956692 | 1,140729472 | 0,37174247 | 0,859817753 | poly(U) binding splicing factor 60 |
| ENSG00000102900 | NUP93 | 0,189908783 | 1,140691591 | 0,39060849 | 0,870057277 | nucleoporin 93 |
| ENSG00000230882 | AC005077,4 | 0,189897742 | 1,140682862 | 0,8711474 | 0,989217334 | hypothetical protein LOC285908 (LOC285908) pseudogene |

| | | | | | | |
|---|---|---|---|---|---|---|
| ENSG00000197483 | ZNF628 | 0,189882074 | 1,140670473 | 0,60655488 | 0,950927937 | zinc finger protein 628 |
| ENSG00000234028 | EIF2AK3-DT | 0,189841351 | 1,140638276 | 0,71517467 | 0,971041579 | EIF2AK3 divergent transcript |
| ENSG00000274008 | Metazoa_SRP | 0,189793313 | 1,140600296 | 0,89826476 | 0,9918188 | Metazoan signal recognition particle RNA |
| ENSG00000260371 | AC026464,3 | 0,189746687 | 1,140563435 | 0,71906836 | 0,971869926 | novel protein |
| ENSG00000176946 | THAP4 | 0,189686097 | 1,140515535 | 0,48000947 | 0,912847708 | THAP domain containing 4 |
| ENSG00000118873 | RAB3GAP2 | 0,189652105 | 1,140488662 | 0,57026774 | 0,9423888 | RAB3 GTPase activating non-catalytic protein subunit 2 |
| ENSG00000168818 | STX18 | 0,189628311 | 1,140469853 | 0,4434674 | 0,900014899 | syntaxin 18 |
| ENSG00000100982 | PCIF1 | 0,189585157 | 1,14043574 | 0,42566855 | 0,892133432 | PDX1 C-terminal inhibiting factor 1 |
| ENSG00000119125 | GDA | 0,189493881 | 1,140363589 | 0,66143896 | 0,959582738 | guanine deaminase |
| ENSG00000103067 | ESRP2 | 0,189470427 | 1,14034505 | 0,70857957 | 0,970181083 | epithelial splicing regulatory protein 2 |
| ENSG00000153046 | CDYL | 0,189459138 | 1,140336127 | 0,49254836 | 0,917016235 | chromodomain Y like |
| ENSG00000233461 | AL445524,1 | 0,189399572 | 1,140289046 | 0,66161368 | 0,959582738 | novel transcript |
| ENSG00000283267 | FAM237B | 0,189267416 | 1,140184596 | 0,87374791 | 0,989633545 | family with sequence similarity 237 member B |
| ENSG00000227053 | MUC12-AS1 | 0,189262467 | 1,140180685 | 0,78025639 | 0,978587255 | MUC12 antisense RNA 1 |
| ENSG00000136758 | YME1L1 | 0,189256227 | 1,140175753 | 0,44210405 | 0,899331131 | YME1 like 1 ATPase |
| ENSG00000103064 | SLC7A6 | 0,189218173 | 1,140145679 | 0,55710303 | 0,938329338 | solute carrier family 7 member 6 |
| ENSG00000077463 | SIRT6 | 0,189206045 | 1,140136095 | 0,58877725 | 0,946499989 | sirtuin 6 |
| ENSG00000134042 | MRO | 0,189162243 | 1,140101479 | 0,77561614 | 0,978587255 | maestro |
| ENSG00000121289 | CEP89 | 0,189039407 | 1,140004411 | 0,48147124 | 0,912981134 | centrosomal protein 89 |
| ENSG00000112365 | ZBTB24 | 0,189004182 | 1,139976577 | 0,56474623 | 0,940865625 | zinc finger and BTB domain containing 24 |
| ENSG00000187783 | TMEM72 | 0,188996617 | 1,1399706 | 0,73838583 | 0,974353474 | transmembrane protein 72 |
| ENSG00000276371 | SINHCAF | 0,188852363 | 1,139856621 | 0,53307554 | 0,931291085 | SIN3-HDAC complex associated factor |
| ENSG00000242114 | MTFP1 | 0,188679865 | 1,13972034 | 0,56524438 | 0,940865625 | mitochondrial fission process 1 |
| ENSG00000023902 | PLEKHO1 | 0,188672374 | 1,139714422 | 0,50851057 | 0,922427526 | pleckstrin homology domain containing O1 |
| ENSG00000285799 | AL645929,2 | 0,188670226 | 1,139712725 | 0,70778713 | 0,969856974 | MHC class I polypeptide-related sequence F pseudogene |
| ENSG00000241973 | PI4KA | 0,188666916 | 1,13971011 | 0,4313018 | 0,89530663 | phosphatidylinositol 4-kinase alpha |
| ENSG00000125995 | ROMO1 | 0,188650194 | 1,1396969 | 0,57987038 | 0,943828882 | reactive oxygen species modulator 1 |
| ENSG00000259682 | AC091231,1 | 0,188590937 | 1,139650089 | 0,8860584 | 0,990717729 | novel transcript, sense intronic to ZNF609 |
| ENSG00000000419 | DPM1 | 0,188541634 | 1,139611144 | 0,65564757 | 0,957444211 | dolichyl-phosphate mannosyltransferase subunit 1, catalytic |
| ENSG00000153560 | UBP1 | 0,188465621 | 1,139551101 | 0,55333034 | 0,936992468 | upstream binding protein 1 |

| | | | | | | |
|---|---|---|---|---|---|---|
| ENSG00000115841 | RMDN2 | 0,188287857 | 1,139410698 | 0,62513468 | 0,952793934 | regulator of microtubule dynamics 2 |
| ENSG00000067177 | PHKA1 | 0,188237421 | 1,139370865 | 0,61881484 | 0,952793934 | phosphorylase kinase regulatory subunit alpha 1 |
| ENSG00000197562 | RAB40C | 0,188204664 | 1,139344996 | 0,59614941 | 0,948282297 | RAB40C, member RAS oncogene family |
| ENSG00000143776 | CDC42BPA | 0,188192808 | 1,139335632 | 0,46214654 | 0,906306011 | CDC42 binding protein kinase alpha |
| ENSG00000166747 | AP1G1 | 0,188072545 | 1,139240662 | 0,5412621 | 0,933422598 | adaptor related protein complex 1 subunit gamma 1 |
| ENSG00000189283 | FHIT | 0,188044332 | 1,139218384 | 0,57522975 | 0,942667371 | fragile histidine triad diadenosine triphosphatase |
| ENSG00000282885 | AL627171,2 | 0,188017583 | 1,139197261 | 0,52105295 | 0,927046149 | novel transcript |
| ENSG00000274626 | RPS9 | 0,187973437 | 1,139162403 | 0,52009272 | 0,926822772 | ribosomal protein S9 |
| ENSG00000277359 | RPS9 | 0,187973437 | 1,139162403 | 0,52009272 | 0,926822772 | ribosomal protein S9 |
| ENSG00000274005 | RPS9 | 0,187973437 | 1,139162403 | 0,52009272 | 0,926822772 | ribosomal protein S9 |
| ENSG00000278081 | RPS9 | 0,187973437 | 1,139162403 | 0,52009272 | 0,926822772 | ribosomal protein S9 |
| ENSG00000275323 | RPS9 | 0,187973437 | 1,139162403 | 0,52009272 | 0,926822772 | ribosomal protein S9 |
| ENSG00000277079 | RPS9 | 0,187973437 | 1,139162403 | 0,52009272 | 0,926822772 | ribosomal protein S9 |
| ENSG00000013374 | NUB1 | 0,187947697 | 1,139142079 | 0,46100675 | 0,906029574 | negative regulator of ubiquitin like proteins 1 |
| ENSG00000144724 | PTPRG | 0,187938511 | 1,139134825 | 0,57804767 | 0,943425812 | protein tyrosine phosphatase receptor type G |
| ENSG00000175221 | MED16 | 0,187834341 | 1,139052577 | 0,6145682 | 0,952793934 | mediator complex subunit 16 |
| ENSG00000171126 | KCNG3 | 0,187818153 | 1,139039796 | 0,85470243 | 0,987861659 | potassium voltage-gated channel modifier subfamily G member 3 |
| ENSG00000177225 | GATD1 | 0,187810563 | 1,139033804 | 0,62753635 | 0,952793934 | glutamine amidotransferase like class 1 domain containing 1 |
| ENSG00000034533 | ASTE1 | 0,187799529 | 1,139025092 | 0,59539469 | 0,947781041 | asteroid homolog 1 |
| ENSG00000146700 | SSC4D | 0,187720729 | 1,13896288 | 0,67851472 | 0,9640473 | scavenger receptor cysteine rich family member with 4 domains |
| ENSG00000179091 | CYC1 | 0,187703101 | 1,138948964 | 0,48882804 | 0,915571519 | cytochrome c1 |
| ENSG00000136940 | PDCL | 0,187565653 | 1,138840459 | 0,50971616 | 0,922427526 | phosducin like |
| ENSG00000183347 | GBP6 | 0,187500282 | 1,138788857 | 0,87803739 | 0,990109814 | guanylate binding protein family member 6 |
| ENSG00000255248 | MIR100HG | 0,18745242 | 1,138751078 | 0,62313708 | 0,952793934 | mir-100-let-7a-2-mir-125b-1 cluster host gene |
| ENSG00000107185 | RGP1 | 0,187397573 | 1,138707787 | 0,58141764 | 0,944037449 | RGP1 homolog, RAB6A GEF complex partner 1 |
| ENSG00000097033 | SH3GLB1 | 0,187392106 | 1,138703472 | 0,60339311 | 0,950352286 | SH3 domain containing GRB2 like, endophilin B1 |
| ENSG00000110768 | GTF2H1 | 0,187379055 | 1,138693171 | 0,57039181 | 0,9423888 | general transcription factor IIH subunit 1 |
| ENSG00000214595 | EML6 | 0,187336375 | 1,138659485 | 0,66256415 | 0,959897678 | EMAP like 6 |
| ENSG00000103479 | RBL2 | 0,187301642 | 1,138632072 | 0,60527305 | 0,950927937 | RB transcriptional corepressor like 2 |
| ENSG00000286161 | AC104066,4 | 0,187295408 | 1,138627152 | 0,81490637 | 0,981521413 | novel transcript |

| | | | | | | |
|---|---|---|---|---|---|---|
| ENSG00000164855 | TMEM184A | 0,18714225 | 1,138506281 | 0,86874799 | 0,989074806 | transmembrane protein 184A |
| ENSG00000270722 | RNVU1-31 | 0,187008765 | 1,138400945 | 0,75403968 | 0,975901808 | RNA, variant U1 small nuclear 31 |
| ENSG00000272791 | AC073389,3 | 0,186841136 | 1,13826868 | 0,80830241 | 0,981213877 | novel transcript |
| ENSG00000283068 | IBTK | 0,186813574 | 1,138246935 | 0,62459915 | 0,952793934 | inhibitor of Bruton tyrosine kinase |
| ENSG00000153060 | TEKT5 | 0,186794701 | 1,138232045 | 0,82245022 | 0,98299282 | tektin 5 |
| ENSG00000273855 | AC020658,5 | 0,1865988 | 1,138077497 | 0,87539475 | 0,990078774 | novel transcript, sense intronic to PLCB2 |
| ENSG00000063322 | MED29 | 0,186597921 | 1,138076803 | 0,38655659 | 0,869441674 | mediator complex subunit 29 |
| ENSG00000167985 | SDHAF2 | 0,186573175 | 1,138057282 | 0,50491089 | 0,921400408 | succinate dehydrogenase complex assembly factor 2 |
| ENSG00000278702 | Metazoa_SRP | 0,186470037 | 1,137975925 | 0,91214165 | 0,993361299 | Metazoan signal recognition particle RNA |
| ENSG00000100731 | PCNX1 | 0,186451315 | 1,137961158 | 0,50982481 | 0,922427526 | pecanex 1 |
| ENSG00000131845 | ZNF304 | 0,186440842 | 1,137952897 | 0,50538778 | 0,921762096 | zinc finger protein 304 |
| ENSG00000285589 | AC010422,8 | 0,186437593 | 1,137950334 | 0,59292834 | 0,947416795 | novel transcript |
| ENSG00000272463 | AL357054,4 | 0,186425694 | 1,137940949 | 0,78942276 | 0,979462318 | novel transcript |
| ENSG00000120063 | GNA13 | 0,186425312 | 1,137940648 | 0,67184595 | 0,962161102 | G protein subunit alpha 13 |
| ENSG00000185838 | GNB1L | 0,186414386 | 1,13793203 | 0,66979455 | 0,962026636 | G protein subunit beta 1 like |
| ENSG00000108262 | GIT1 | 0,186233591 | 1,137789436 | 0,50114521 | 0,920520138 | GIT ArfGAP 1 |
| ENSG00000278440 | AC243791,1 | 0,18623145 | 1,137787747 | 0,81462086 | 0,981521413 | novel pseudogene KIAA2013 |
| ENSG00000261819 | AC138932,2 | 0,18623145 | 1,137787747 | 0,81462086 | 0,981521413 | novel pseudogene KIAA2013 |
| ENSG00000233246 | PHC2-AS1 | 0,186204916 | 1,137766821 | 0,88573478 | 0,990717729 | PHC2 antisense RNA 1 |
| ENSG00000137364 | TPMT | 0,186193675 | 1,137757957 | 0,43070207 | 0,895267028 | thiopurine S-methyltransferase |
| ENSG00000095713 | CRTAC1 | 0,186170407 | 1,137739606 | 0,77462978 | 0,978553937 | cartilage acidic protein 1 |
| ENSG00000137947 | GTF2B | 0,186063092 | 1,137654978 | 0,66006804 | 0,958859629 | general transcription factor IIB |
| ENSG00000224743 | TEX26-AS1 | 0,186003786 | 1,137608213 | 0,8022942 | 0,980710009 | TEX26 antisense RNA 1 |
| ENSG00000181751 | MACIR | 0,185961621 | 1,137574966 | 0,6817711 | 0,964654539 | macrophage immunometabolism regulator |
| ENSG00000225164 | PRRC2A | 0,185887094 | 1,137516202 | 0,50750221 | 0,922098508 | proline rich coiled-coil 2A |
| ENSG00000204946 | ZNF783 | 0,18587964 | 1,137510325 | 0,64919719 | 0,956899963 | zinc finger family member 783 |
| ENSG00000228645 | PHKG1P2 | 0,185867231 | 1,137500541 | 0,87861759 | 0,990109814 | phosphorylase kinase catalytic subunit gamma 1 pseudogene 2 |
| ENSG00000284802 | AL136295,8 | 0,185784424 | 1,137435253 | 0,71306454 | 0,970812834 | novel transcript |
| ENSG00000155463 | OXA1L | 0,185731525 | 1,137393548 | 0,52687945 | 0,92927181 | OXA1L mitochondrial inner membrane protein |
| ENSG00000124557 | BTN1A1 | 0,185716569 | 1,137381756 | 0,86064713 | 0,989074806 | butyrophilin subfamily 1 member A1 |
| ENSG00000223839 | FAM95B1 | 0,185688098 | 1,137359311 | 0,77169339 | 0,978033855 | family with sequence similarity 95 member B1 |
| ENSG00000100461 | RBM23 | 0,185600656 | 1,137290378 | 0,39309891 | 0,871240371 | RNA binding motif protein 23 |

| | | | | | | |
|---|---|---|---|---|---|---|
| ENSG00000066032 | CTNNA2 | 0,185560775 | 1,137258939 | 0,63814136 | 0,956018518 | catenin alpha 2 |
| ENSG00000197170 | PSMD12 | 0,185533559 | 1,137237485 | 0,54116255 | 0,933385624 | proteasome 26S subunit, non-ATPase 12 |
| ENSG00000287581 | AC016549,1 | 0,185507871 | 1,137217237 | 0,72423741 | 0,973178901 | novel transcript, antisense to TRIO |
| ENSG00000224870 | MRPL20-AS1 | 0,185506469 | 1,137216131 | 0,5268345 | 0,92927181 | MRPL20 antisense RNA 1 |
| ENSG00000183706 | OR4N4 | 0,185345101 | 1,137088939 | 0,8926169 | 0,991048031 | olfactory receptor family 4 subfamily N member 4 |
| ENSG00000278621 | AC037198,2 | 0,185316203 | 1,137066162 | 0,78542127 | 0,979320406 | novel transcript, antisense to THBS1 |
| ENSG00000104549 | SQLE | 0,185308032 | 1,137059723 | 0,5118169 | 0,923177951 | squalene epoxidase |
| ENSG00000235655 | H3P6 | 0,185292419 | 1,137047417 | 0,80715052 | 0,981213877 | H3 histone pseudogene 6 |
| ENSG00000168385 | SEPTIN2 | 0,185272858 | 1,137032001 | 0,53422718 | 0,93151582 | septin 2 |
| ENSG00000147439 | BIN3 | 0,185159523 | 1,136942682 | 0,45640339 | 0,903545728 | bridging integrator 3 |
| ENSG00000091039 | OSBPL8 | 0,185049922 | 1,136856312 | 0,63439273 | 0,955097925 | oxysterol binding protein like 8 |
| ENSG00000196110 | ZNF699 | 0,184976495 | 1,136798452 | 0,52492403 | 0,928616707 | zinc finger protein 699 |
| ENSG00000048544 | MRPS10 | 0,184896191 | 1,136735177 | 0,63442038 | 0,955097925 | mitochondrial ribosomal protein S10 |
| ENSG00000204316 | MRPL38 | 0,18489558 | 1,136734696 | 0,55841897 | 0,939334338 | mitochondrial ribosomal protein L38 |
| ENSG00000248898 | AC022126,1 | 0,184879814 | 1,136722273 | 0,8204509 | 0,982609114 | novel transcript |
| ENSG00000101439 | CST3 | 0,184808565 | 1,136666136 | 0,40255076 | 0,877321369 | cystatin C |
| ENSG00000173320 | STOX2 | 0,184781768 | 1,136645024 | 0,58773787 | 0,946124093 | storkhead box 2 |
| ENSG00000123360 | PDE1B | 0,184752459 | 1,136621932 | 0,58291708 | 0,944174844 | phosphodiesterase 1B |
| ENSG00000273372 | SFTPD-AS1 | 0,1847392 | 1,136611486 | 0,77789762 | 0,978587255 | SFTPD antisense RNA 1 |
| ENSG00000228536 | LYPLAL1-AS1 | 0,184513276 | 1,136433508 | 0,78403196 | 0,97928555 | LYPLAL1 antisense RNA 1 |
| ENSG00000231877 | AL359551,1 | 0,184457662 | 1,136389701 | 0,80618081 | 0,981213877 | novel transcript |
| ENSG00000079332 | SAR1A | 0,18442135 | 1,136361099 | 0,57258524 | 0,942475308 | secretion associated Ras related GTPase 1A |
| ENSG00000236375 | POU5F1P5 | 0,184415133 | 1,136356202 | 0,86131609 | 0,989074806 | POU class 5 homeobox 1 pseudogene 5 |
| ENSG00000146433 | TMEM181 | 0,184395723 | 1,136340914 | 0,53937746 | 0,933350549 | transmembrane protein 181 |
| ENSG00000122873 | CISD1 | 0,18430195 | 1,136267056 | 0,59708057 | 0,948282297 | CDGSH iron sulfur domain 1 |
| ENSG00000278183 | ABCC1 | 0,184265789 | 1,136238576 | 0,43720722 | 0,897021798 | ATP binding cassette subfamily C member 1 |
| ENSG00000165152 | PGAP4 | 0,18425892 | 1,136233166 | 0,51912004 | 0,926800715 | post-GPI attachment to proteins GalNAc transferase 4 |
| ENSG00000106305 | AIMP2 | 0,184253016 | 1,136228516 | 0,45285469 | 0,902361309 | aminoacyl tRNA synthetase complex interacting multifunctional protein 2 |
| ENSG00000127452 | FBXL12 | 0,184224336 | 1,136205929 | 0,44429631 | 0,900210555 | F-box and leucine rich repeat protein 12 |
| ENSG00000151445 | VIPAS39 | 0,184212449 | 1,136196567 | 0,47581452 | 0,91134381 | VPS33B interacting protein, apical-basolateral polarity regulator, spe-39 homolog |

| | | | | | | |
|---|---|---|---|---|---|---|
| ENSG00000239732 | TLR9 | 0,184196764 | 1,136184215 | 0,90990336 | 0,992951496 | toll like receptor 9 |
| ENSG00000285246 | NEDD8 | 0,184087683 | 1,136098312 | 0,46285124 | 0,906849208 | NEDD8 ubiquitin like modifier |
| ENSG00000143368 | SF3B4 | 0,184078917 | 1,136091409 | 0,46440415 | 0,907500217 | splicing factor 3b subunit 4 |
| ENSG00000140983 | RHOT2 | 0,184052103 | 1,136070294 | 0,63874056 | 0,956018518 | ras homolog family member T2 |
| ENSG00000269693 | AC010422,6 | 0,184007154 | 1,136034898 | 0,73766656 | 0,974353474 | novel transcript |
| ENSG00000281385 | AP2A2 | 0,183986844 | 1,136018905 | 0,41315857 | 0,884663685 | adaptor related protein complex 2 subunit alpha 2 |
| ENSG00000117450 | PRDX1 | 0,183959705 | 1,135997536 | 0,53993675 | 0,933385624 | peroxiredoxin 1 |
| ENSG00000203605 | AC096543,1 | 0,183771554 | 1,135849393 | 0,82192648 | 0,982952211 | novel transcript |
| ENSG00000119689 | DLST | 0,183769976 | 1,13584815 | 0,37053044 | 0,858545694 | dihydrolipoamide S-succinyltransferase |
| ENSG00000150394 | CDH8 | 0,183761001 | 1,135841084 | 0,48012628 | 0,912847708 | cadherin 8 |
| ENSG00000103197 | TSC2 | 0,183718299 | 1,135807465 | 0,54846372 | 0,935674429 | TSC complex subunit 2 |
| ENSG00000244675 | AC108676,1 | 0,183623901 | 1,135733149 | 0,73486179 | 0,974353474 | novel transcript, sense overlapping to ATP13A3 |
| ENSG00000116237 | ICMT | 0,183599059 | 1,135713594 | 0,43676412 | 0,896866043 | isoprenylcysteine carboxyl methyltransferase |
| ENSG00000160360 | GPSM1 | 0,183592524 | 1,135708449 | 0,7105733 | 0,970482408 | G protein signaling modulator 1 |
| ENSG00000108651 | UTP6 | 0,18349112 | 1,135628625 | 0,38082587 | 0,865900094 | UTP6 small subunit processome component |
| ENSG00000182272 | B4GALNT4 | 0,183468641 | 1,135610931 | 0,7307732 | 0,974336839 | beta-1,4-N-acetyl-galactosaminyltransferase 4 |
| ENSG00000224295 | OLFM5P | 0,183352891 | 1,135519823 | 0,82489213 | 0,983374097 | olfactomedin family member 5, pseudogene |
| ENSG00000179943 | FIZ1 | 0,183254251 | 1,135442188 | 0,56462232 | 0,940865625 | FLT3 interacting zinc finger 1 |
| ENSG00000135218 | CD36 | 0,183229454 | 1,135422672 | 0,73556415 | 0,974353474 | CD36 molecule |
| ENSG00000136698 | CFC1 | 0,183204246 | 1,135402833 | 0,88845764 | 0,990717729 | cripto, FRL-1, cryptic family 1 |
| ENSG00000225783 | MIAT | 0,183176137 | 1,135380711 | 0,54287361 | 0,933663748 | myocardial infarction associated transcript |
| ENSG00000100601 | ALKBH1 | 0,183133814 | 1,135347404 | 0,6066725 | 0,950927937 | alkB homolog 1, histone H2A dioxygenase |
| ENSG00000236502 | SIX3-AS1 | 0,183091934 | 1,135314446 | 0,7466394 | 0,975796253 | SIX3 antisense RNA 1 |
| ENSG00000236434 | AL162430,2 | 0,183015937 | 1,135254643 | 0,91470475 | 0,993361299 | novel transcript |
| ENSG00000235897 | TM4SF19-AS1 | 0,182982768 | 1,135228543 | 0,81748204 | 0,98215965 | TM4SF19 antisense RNA 1 |
| ENSG00000134504 | KCTD1 | 0,182925808 | 1,135183723 | 0,43836919 | 0,897220559 | potassium channel tetramerization domain containing 1 |
| ENSG00000198301 | SDAD1 | 0,182912853 | 1,135173529 | 0,54518238 | 0,934923038 | SDA1 domain containing 1 |
| ENSG00000180535 | BHLHA15 | 0,182774383 | 1,13506458 | 0,77978042 | 0,978587255 | basic helix-loop-helix family member a15 |
| ENSG00000147650 | LRP12 | 0,18276929 | 1,135060573 | 0,5409999 | 0,933385624 | LDL receptor related protein 12 |
| ENSG00000274825 | AL023803,2 | 0,182678788 | 1,134989372 | 0,80800805 | 0,981213877 | novel transcript |
| ENSG00000127952 | STYXL1 | 0,182673351 | 1,134985094 | 0,45613658 | 0,903545728 | serine/threonine/tyrosine interacting like 1 |
| ENSG00000241520 | AC098820,1 | 0,182650832 | 1,134967379 | 0,79051469 | 0,979462318 | novel transcript |

| | | | | | | |
|---|---|---|---|---|---|---|
| ENSG00000185262 | UBALD2 | 0,182643874 | 1,134961905 | 0,57434364 | 0,942475308 | UBA like domain containing 2 |
| ENSG00000275160 | AL354718,1 | 0,182497153 | 1,134846486 | 0,85334346 | 0,987861659 | ribosomal protein L7a (RPL7A) pseudogene |
| ENSG00000242193 | CRYZL2P | 0,182487332 | 1,13483876 | 0,57549353 | 0,942700009 | crystallin zeta like 2, pseudogene |
| ENSG00000173212 | MAB21L3 | 0,182364541 | 1,134742176 | 0,86829926 | 0,989074806 | mab-21 like 3 |
| ENSG00000089053 | ANAPC5 | 0,18225733 | 1,134657853 | 0,44724573 | 0,902050878 | anaphase promoting complex subunit 5 |
| ENSG00000107651 | SEC23IP | 0,182235398 | 1,134640604 | 0,54505243 | 0,934923038 | SEC23 interacting protein |
| ENSG00000160563 | MED27 | 0,182131616 | 1,134558985 | 0,38383426 | 0,868069622 | mediator complex subunit 27 |
| ENSG00000286460 | AC006538,4 | 0,182098577 | 1,134533003 | 0,91632718 | 0,993361299 | novel transcript |
| ENSG00000188385 | JAKMIP3 | 0,182065929 | 1,134507329 | 0,71001746 | 0,970482408 | Janus kinase and microtubule interacting protein 3 |
| ENSG00000258603 | AC005225,2 | 0,181852676 | 1,134339643 | 0,79500328 | 0,979776981 | novel transcript, antisense to ACOT6 |
| ENSG00000162910 | MRPL55 | 0,181806465 | 1,13430331 | 0,5472105 | 0,935404974 | mitochondrial ribosomal protein L55 |
| ENSG00000249363 | AC011411,1 | 0,181577796 | 1,134123536 | 0,84152302 | 0,98570213 | transient receptor potential cation channel family (TRPC7) pseudogene |
| ENSG00000144730 | IL17RD | 0,181571334 | 1,134118456 | 0,61271995 | 0,952413269 | interleukin 17 receptor D |
| ENSG00000218739 | CEBPZOS | 0,181500227 | 1,134062559 | 0,47525333 | 0,911082311 | CEBPZ opposite strand |
| ENSG00000177602 | HASPIN | 0,181448819 | 1,134022149 | 0,46347566 | 0,907101775 | histone H3 associated protein kinase |
| ENSG00000175938 | ORAI3 | 0,181381571 | 1,133969291 | 0,59117377 | 0,947416795 | ORAI calcium release-activated calcium modulator 3 |
| ENSG00000170684 | ZNF296 | 0,181252227 | 1,13386763 | 0,74946717 | 0,975901808 | zinc finger protein 296 |
| ENSG00000136271 | DDX56 | 0,181201312 | 1,133827615 | 0,42736406 | 0,892552417 | DEAD-box helicase 56 |
| ENSG00000249894 | AC024581,1 | 0,181149673 | 1,133787032 | 0,66720029 | 0,9611116 | novel transcript |
| ENSG00000165644 | COMTD1 | 0,18101929 | 1,133684571 | 0,72983246 | 0,974106992 | catechol-O-methyltransferase domain containing 1 |
| ENSG00000099899 | TRMT2A | 0,180987875 | 1,133659885 | 0,59039557 | 0,947347931 | tRNA methyltransferase 2 homolog A |
| ENSG00000144744 | UBA3 | 0,18091987 | 1,133606449 | 0,62413737 | 0,952793934 | ubiquitin like modifier activating enzyme 3 |
| ENSG00000004059 | ARF5 | 0,180917117 | 1,133604285 | 0,47713024 | 0,91143003 | ADP ribosylation factor 5 |
| ENSG00000198874 | TYW1 | 0,180889323 | 1,133582447 | 0,4271267 | 0,892343796 | tRNA-yW synthesizing protein 1 homolog |
| ENSG00000148308 | GTF3C5 | 0,180877922 | 1,133573488 | 0,51028975 | 0,922499069 | general transcription factor IIIC subunit 5 |
| ENSG00000140505 | CYP1A2 | 0,180753734 | 1,133475913 | 0,8659382 | 0,989074806 | cytochrome P450 family 1 subfamily A member 2 |
| ENSG00000234899 | SOX9-AS1 | 0,180717278 | 1,133447272 | 0,56619272 | 0,941123407 | SOX9 antisense RNA 1 |
| ENSG00000200795 | RNU4-1 | 0,180713355 | 1,13344419 | 0,78594066 | 0,979320406 | RNA, U4 small nuclear 1 |
| ENSG00000282019 | GPI | 0,180699402 | 1,133433228 | 0,48203286 | 0,912981134 | glucose-6-phosphate isomerase |
| ENSG00000014641 | MDH1 | 0,180639267 | 1,133385984 | 0,5416876 | 0,933491609 | malate dehydrogenase 1 |
| ENSG00000114670 | NEK11 | 0,180579806 | 1,133339272 | 0,48466346 | 0,913690171 | NIMA related kinase 11 |
| ENSG00000274659 | LINC02371 | 0,180517084 | 1,133290001 | 0,78271708 | 0,978678077 | long intergenic non-protein coding RNA 2371 |

| | | | | | | |
|---|---|---|---|---|---|---|
| ENSG00000127191 | TRAF2 | 0,180498683 | 1,133275547 | 0,64955533 | 0,956899963 | TNF receptor associated factor 2 |
| ENSG00000245468 | LINC02447 | 0,180383429 | 1,133185015 | 0,76930569 | 0,97758701 | long intergenic non-protein coding RNA 2447 |
| ENSG00000150967 | ABCB9 | 0,180373013 | 1,133176834 | 0,63347725 | 0,954781501 | ATP binding cassette subfamily B member 9 |
| ENSG00000173898 | SPTBN2 | 0,180330937 | 1,133143785 | 0,7530069 | 0,975901808 | spectrin beta, non-erythrocytic 2 |
| ENSG00000278970 | HEIH | 0,180309083 | 1,133126621 | 0,49137171 | 0,916995341 | hepatocellular carcinoma up-regulated EZH2-associated long non-coding RNA |
| ENSG00000101191 | DIDO1 | 0,180027936 | 1,132905822 | 0,50847678 | 0,922427526 | death inducer-obliterator 1 |
| ENSG00000106299 | WASL | 0,179999659 | 1,132883618 | 0,65828522 | 0,957906876 | WASP like actin nucleation promoting factor |
| ENSG00000254503 | AC010319,1 | 0,179924189 | 1,132824356 | 0,81394369 | 0,981521413 | high-mobility group box 3 (HMGB3) pseudogene |
| ENSG00000123268 | ATF1 | 0,179921118 | 1,132821945 | 0,66850667 | 0,961351573 | activating transcription factor 1 |
| ENSG00000153827 | TRIP12 | 0,17991747 | 1,13281908 | 0,59691367 | 0,948282297 | thyroid hormone receptor interactor 12 |
| ENSG00000198169 | ZNF251 | 0,179748042 | 1,132686051 | 0,49234531 | 0,916995341 | zinc finger protein 251 |
| ENSG00000134815 | DHX34 | 0,179742098 | 1,132681385 | 0,54484446 | 0,93489368 | DExH-box helicase 34 |
| ENSG00000183092 | BEGAIN | 0,179725019 | 1,132667975 | 0,67000652 | 0,962026636 | brain enriched guanylate kinase associated |
| ENSG00000162222 | TTC9C | 0,179664953 | 1,132620819 | 0,49340719 | 0,917289924 | tetratricopeptide repeat domain 9C |
| ENSG00000129128 | SPCS3 | 0,179651167 | 1,132609995 | 0,67549024 | 0,963400586 | signal peptidase complex subunit 3 |
| ENSG00000120158 | RCL1 | 0,179624772 | 1,132589274 | 0,46774831 | 0,907940653 | RNA terminal phosphate cyclase like 1 |
| ENSG00000072182 | ASIC4 | 0,179579535 | 1,132553761 | 0,69979093 | 0,968522643 | acid sensing ion channel subunit family member 4 |
| ENSG00000259519 | AC051619,3 | 0,179524873 | 1,132510851 | 0,87708623 | 0,990109814 | novel transcript, antisense to SHF |
| ENSG00000198382 | UVRAG | 0,179515261 | 1,132503306 | 0,53480911 | 0,931751025 | UV radiation resistance associated |
| ENSG00000267419 | ZNF56 | 0,179476237 | 1,132472672 | 0,73598999 | 0,974353474 | zinc finger protein 56 |
| ENSG00000146410 | MTFR2 | 0,179463132 | 1,132462385 | 0,63945155 | 0,956018518 | mitochondrial fission regulator 2 |
| ENSG00000233016 | SNHG7 | 0,179461613 | 1,132461193 | 0,51917098 | 0,926800715 | small nucleolar RNA host gene 7 |
| ENSG00000270607 | AC009549,1 | 0,179422976 | 1,132430865 | 0,75941864 | 0,976120207 | novel transcript |
| ENSG00000122696 | SLC25A51 | 0,179353703 | 1,132376491 | 0,43153069 | 0,89530663 | solute carrier family 25 member 51 |
| ENSG00000100030 | MAPK1 | 0,17923608 | 1,132284172 | 0,57724283 | 0,94337651 | mitogen-activated protein kinase 1 |
| ENSG00000215012 | RTL10 | 0,179172203 | 1,13223404 | 0,5302572 | 0,931157231 | retrotransposon Gag like 10 |
| ENSG00000272921 | AC005832,4 | 0,179139019 | 1,132207997 | 0,66708002 | 0,9611116 | novel protein |
| ENSG00000232830 | AC110994,1 | 0,179118882 | 1,132192194 | 0,86529478 | 0,989074806 | family with sequence similarity 136, member A (FAM136A) pseudogene |
| ENSG00000283504 | AC006974,1 | 0,179116482 | 1,13219031 | 0,89062545 | 0,990717729 | novel transcript |
| ENSG00000119574 | ZBTB45 | 0,179082878 | 1,13216394 | 0,62086845 | 0,952793934 | zinc finger and BTB domain containing 45 |
| ENSG00000141627 | DYM | 0,17908154 | 1,132162889 | 0,60824855 | 0,951332922 | dymeclin |

| | | | | | | |
|---|---|---|---|---|---|---|
| ENSG00000198901 | PRC1 | 0,179072062 | 1,132155451 | 0,41700526 | 0,886616509 | protein regulator of cytokinesis 1 |
| ENSG00000259797 | AC020978,1 | 0,178863599 | 1,131991872 | 0,85006892 | 0,987036295 | novel transcript, intronic to PRMT7 |
| ENSG00000160392 | C19orf47 | 0,178853346 | 1,131983827 | 0,60305062 | 0,950352286 | chromosome 19 open reading frame 47 |
| ENSG00000197217 | ENTPD4 | 0,178834998 | 1,131969431 | 0,44763886 | 0,902062594 | ectonucleoside triphosphate diphosphohydrolase 4 |
| ENSG00000115649 | CNPPD1 | 0,178779867 | 1,131926175 | 0,51324162 | 0,923829003 | cyclin Pas1/PHO80 domain containing 1 |
| ENSG00000183682 | BMP8A | 0,178489629 | 1,131698479 | 0,75531219 | 0,975901808 | bone morphogenetic protein 8a |
| ENSG00000235217 | TSPY26P | 0,178488728 | 1,131697773 | 0,46803878 | 0,907940653 | testis specific protein Y-linked 26, pseudogene |
| ENSG00000236032 | OR5H14 | 0,178451018 | 1,131668192 | 0,74190622 | 0,975176043 | olfactory receptor family 5 subfamily H member 14 |
| ENSG00000214062 | RPL7P17 | 0,178440101 | 1,131659629 | 0,84272293 | 0,986183927 | ribosomal protein L7 pseudogene 17 |
| ENSG00000130254 | SAFB2 | 0,178410666 | 1,131636539 | 0,71886799 | 0,971869926 | scaffold attachment factor B2 |
| ENSG00000287735 | AL355075,6 | 0,178402976 | 1,131630508 | 0,65746825 | 0,957853146 | novel transcript, antisense to CCNB1IP1 |
| ENSG00000166887 | VPS39 | 0,178401922 | 1,131629681 | 0,41777151 | 0,886857227 | VPS39 subunit of HOPS complex |
| ENSG00000172057 | ORMDL3 | 0,178376734 | 1,131609924 | 0,53966082 | 0,933350549 | ORMDL sphingolipid biosynthesis regulator 3 |
| ENSG00000013375 | PGM3 | 0,1781662 | 1,131444799 | 0,62736476 | 0,952793934 | phosphoglucomutase 3 |
| ENSG00000100883 | SRP54 | 0,177926027 | 1,131256457 | 0,64340047 | 0,956633898 | signal recognition particle 54 |
| ENSG00000136699 | SMPD4 | 0,177700059 | 1,131079283 | 0,44904065 | 0,902062594 | sphingomyelin phosphodiesterase 4 |
| ENSG00000204356 | NELFE | 0,177573919 | 1,130980393 | 0,46435038 | 0,907500217 | negative elongation factor complex member E |
| ENSG00000167077 | MEI1 | 0,177561098 | 1,130970343 | 0,78894974 | 0,979462318 | meiotic double-stranded break formation protein 1 |
| ENSG00000282164 | PEG13 | 0,177514014 | 1,130933433 | 0,64123943 | 0,956465487 | paternally expressed 13 |
| ENSG00000010932 | FMO1 | 0,177477874 | 1,130905103 | 0,87805226 | 0,990109814 | flavin containing dimethylaniline monoxygenase 1 |
| ENSG00000261924 | AC127496,1 | 0,177388127 | 1,130834754 | 0,75430206 | 0,975901808 | novel transcript, antisense to RPTOR |
| ENSG00000234176 | HSPA8P1 | 0,177359075 | 1,130811982 | 0,86948007 | 0,989074806 | heat shock protein family A (Hsp70) member 8 pseudogene 1 |
| ENSG00000173621 | LRFN4 | 0,177325473 | 1,130785644 | 0,7086148 | 0,970181083 | leucine rich repeat and fibronectin type III domain containing 4 |
| ENSG00000183569 | SERHL2 | 0,177310909 | 1,130774229 | 0,67074066 | 0,962026636 | serine hydrolase like 2 |
| ENSG00000228793 | AL033523,1 | 0,177268558 | 1,130741035 | 0,63408676 | 0,954940644 | novel transcript |
| ENSG00000177169 | ULK1 | 0,177224401 | 1,130706427 | 0,56155786 | 0,940663581 | unc-51 like autophagy activating kinase 1 |
| ENSG00000152359 | POC5 | 0,177080174 | 1,130593395 | 0,63889014 | 0,956018518 | POC5 centriolar protein |
| ENSG00000197948 | FCHSD1 | 0,177029833 | 1,130553945 | 0,63231025 | 0,954172582 | FCH and double SH3 domains 1 |
| ENSG00000284647 | AL109936,8 | 0,177005753 | 1,130535075 | 0,9258437 | 0,993915537 | novel transcript |
| ENSG00000173692 | PSMD1 | 0,176955655 | 1,130495818 | 0,57010732 | 0,942262392 | proteasome 26S subunit, non-ATPase 1 |
| ENSG00000260490 | MYL12BP1 | 0,176913275 | 1,130462609 | 0,87828852 | 0,990109814 | myosin light chain 12B pseudogene 1 |

| | | | | | | |
|---|---|---|---|---|---|---|
| ENSG00000257964 | AC020612,3 | 0,176826671 | 1,13039475 | 0,8247299 | 0,983374097 | novel transcript, antisense to PRPF40B |
| ENSG00000137124 | ALDH1B1 | 0,17681421 | 1,130384987 | 0,42294678 | 0,890541302 | aldehyde dehydrogenase 1 family member B1 |
| ENSG00000143793 | C1orf35 | 0,17677695 | 1,130355793 | 0,61589385 | 0,952793934 | chromosome 1 open reading frame 35 |
| ENSG00000225177 | AL590617,2 | 0,176770279 | 1,130350566 | 0,53033695 | 0,931172326 | novel transcript |
| ENSG00000147912 | FBXO10 | 0,176707787 | 1,130301605 | 0,44538341 | 0,901068127 | F-box protein 10 |
| ENSG00000219438 | TAFA5 | 0,176694923 | 1,130291527 | 0,5315469 | 0,931291085 | TAFA chemokine like family member 5 |
| ENSG00000149781 | FERMT3 | 0,17668679 | 1,130285155 | 0,71790923 | 0,971762632 | fermitin family member 3 |
| ENSG00000232024 | LSM12P1 | 0,176683315 | 1,130282432 | 0,73978395 | 0,974715504 | LSM12 pseudogene 1 |
| ENSG00000280870 | MIR325HG | 0,176642833 | 1,130250717 | 0,75332604 | 0,975901808 | MIR325 host gene |
| ENSG00000091513 | TF | 0,176608705 | 1,130223981 | 0,76172977 | 0,976285211 | transferrin |
| ENSG00000278211 | TRIM69 | 0,17657089 | 1,130194356 | 0,79374717 | 0,979776981 | tripartite motif containing 69 |
| ENSG00000166546 | BEAN1 | 0,176445967 | 1,130096497 | 0,75080235 | 0,975901808 | brain expressed associated with NEDD4 1 |
| ENSG00000163374 | YY1AP1 | 0,176278105 | 1,129965015 | 0,4297353 | 0,894607 | YY1 associated protein 1 |
| ENSG00000187621 | TCL6 | 0,176200727 | 1,129904411 | 0,73406712 | 0,974353474 | T cell leukemia/lymphoma 6 |
| ENSG00000198870 | STKLD1 | 0,17618441 | 1,129891632 | 0,73769947 | 0,974353474 | serine/threonine kinase like domain containing 1 |
| ENSG00000271989 | AL139424,1 | 0,176110293 | 1,129833586 | 0,80313253 | 0,980866933 | novel transcript, antisense to APITD1-CORT |
| ENSG00000221995 | TIAF1 | 0,175986516 | 1,129736656 | 0,60048274 | 0,949439306 | TGFB1-induced anti-apoptotic factor 1 |
| ENSG00000119559 | C19orf25 | 0,175974944 | 1,129727594 | 0,67587194 | 0,963549282 | chromosome 19 open reading frame 25 |
| ENSG00000103932 | RPAP1 | 0,175942857 | 1,129702468 | 0,57845338 | 0,943425812 | RNA polymerase II associated protein 1 |
| ENSG00000248713 | C4orf54 | 0,175897737 | 1,129667137 | 0,82392986 | 0,983374097 | chromosome 4 open reading frame 54 |
| ENSG00000158796 | DEDD | 0,17588407 | 1,129656435 | 0,4855088 | 0,913907266 | death effector domain containing |
| ENSG00000271746 | AL031848,2 | 0,175880864 | 1,129653925 | 0,79906235 | 0,980161849 | novel transcript, antisense to ACOT7 |
| ENSG00000172932 | ANKRD13D | 0,175872417 | 1,129647311 | 0,57106199 | 0,942436428 | ankyrin repeat domain 13D |
| ENSG00000134697 | GNL2 | 0,175859002 | 1,129636807 | 0,42076441 | 0,888709349 | G protein nucleolar 2 |
| ENSG00000275740 | AC091959,1 | 0,175855088 | 1,129633743 | 0,594429 | 0,947469769 | novel readthrough transcript |
| ENSG00000168028 | RPSA | 0,175830715 | 1,129614659 | 0,4844032 | 0,913690171 | ribosomal protein SA |
| ENSG00000063854 | HAGH | 0,175828637 | 1,129613031 | 0,51157084 | 0,922893865 | hydroxyacylglutathione hydrolase |
| ENSG00000181588 | MEX3D | 0,175810697 | 1,129598985 | 0,60356974 | 0,950352286 | mex-3 RNA binding family member D |
| ENSG00000284868 | DHRS1 | 0,175807956 | 1,129596838 | 0,51993461 | 0,926822772 | dehydrogenase/reductase 1 |
| ENSG00000011376 | LARS2 | 0,175769528 | 1,129566751 | 0,398077 | 0,874717411 | leucyl-tRNA synthetase 2, mitochondrial |
| ENSG00000104714 | ERICH1 | 0,175438592 | 1,129307672 | 0,70520426 | 0,96941942 | glutamate rich 1 |
| ENSG00000198276 | UCKL1 | 0,175384543 | 1,129265365 | 0,51310681 | 0,923745908 | uridine-cytidine kinase 1 like 1 |
| ENSG00000003393 | ALS2 | 0,175329954 | 1,129222636 | 0,52731131 | 0,929454507 | alsin Rho guanine nucleotide exchange factor ALS2 |

| | | | | | | |
|---|---|---|---|---|---|---|
| ENSG00000136628 | EPRS1 | 0,175282408 | 1,129185421 | 0,53017111 | 0,931131717 | glutamyl-prolyl-tRNA synthetase 1 |
| ENSG00000188152 | NUTM2G | 0,175274216 | 1,12917901 | 0,71359014 | 0,970812834 | NUT family member 2G |
| ENSG00000236047 | RPL13AP12 | 0,175188061 | 1,129111579 | 0,72126135 | 0,972142678 | ribosomal protein L13a pseudogene 12 |
| ENSG00000090061 | CCNK | 0,17515225 | 1,129083553 | 0,53573398 | 0,932139691 | cyclin K |
| ENSG00000122140 | MRPS2 | 0,175054237 | 1,129006848 | 0,554113 | 0,937637409 | mitochondrial ribosomal protein S2 |
| ENSG00000064195 | DLX3 | 0,175033956 | 1,128990977 | 0,88631119 | 0,990717729 | distal-less homeobox 3 |
| ENSG00000181404 | WASHC1 | 0,175032999 | 1,128990228 | 0,56361555 | 0,940865625 | WASH complex subunit 1 |
| ENSG00000269894 | AC018809,1 | 0,175031496 | 1,128989052 | 0,83191887 | 0,984596888 | novel transcript, sense intronic to CRELD1 |
| ENSG00000078967 | UBE2D4 | 0,174971454 | 1,128942067 | 0,45100701 | 0,902062594 | ubiquitin conjugating enzyme E2 D4 (putative) |
| ENSG00000138592 | USP8 | 0,17489782 | 1,128884448 | 0,41286294 | 0,884576962 | ubiquitin specific peptidase 8 |
| ENSG00000084676 | NCOA1 | 0,174864139 | 1,128858094 | 0,5574893 | 0,938578675 | nuclear receptor coactivator 1 |
| ENSG00000137478 | FCHSD2 | 0,174851058 | 1,128847858 | 0,62865017 | 0,952793934 | FCH and double SH3 domains 2 |
| ENSG00000164010 | ERMAP | 0,174772568 | 1,128786445 | 0,46757588 | 0,907940653 | erythroblast membrane associated protein (Scianna blood group) |
| ENSG00000286004 | AC124944,3 | 0,174763672 | 1,128779484 | 0,84690881 | 0,986939308 | novel transcript, antisense to TNK2 |
| ENSG00000286657 | AC008453,2 | 0,174749089 | 1,128768074 | 0,80627184 | 0,981213877 | novel transcript, antisense to SYNPO |
| ENSG00000186132 | C2orf76 | 0,174732429 | 1,128755039 | 0,62722418 | 0,952793934 | chromosome 2 open reading frame 76 |
| ENSG00000235946 | H2BW3P | 0,174672337 | 1,128708025 | 0,68021124 | 0,964580199 | H2B,W histone 3, pseudogene |
| ENSG00000117362 | APH1A | 0,174647316 | 1,128688449 | 0,49181002 | 0,916995341 | aph-1 homolog A, gamma-secretase subunit |
| ENSG00000260793 | AC003102,1 | 0,174629063 | 1,12867417 | 0,70141547 | 0,968742626 | novel transcript, antisense to UBTF |
| ENSG00000124422 | USP22 | 0,174598528 | 1,128650281 | 0,39519192 | 0,871981986 | ubiquitin specific peptidase 22 |
| ENSG00000215717 | TMEM167B | 0,174590938 | 1,128644343 | 0,65776831 | 0,957864535 | transmembrane protein 167B |
| ENSG00000091262 | ABCC6 | 0,174465104 | 1,128545906 | 0,79699035 | 0,980161849 | ATP binding cassette subfamily C member 6 |
| ENSG00000277287 | AL109976,1 | 0,174432613 | 1,12852049 | 0,85450543 | 0,987861659 | novel transcript |
| ENSG00000126106 | TMEM53 | 0,174398516 | 1,128493818 | 0,49252147 | 0,917016235 | transmembrane protein 53 |
| ENSG00000281296 | LL22NC03-63E9 | 0,174386809 | 1,128484661 | 0,93119695 | 0,994433362 | uncharacterized LOC648691 |
| ENSG00000120800 | UTP20 | 0,174326369 | 1,128437386 | 0,57008043 | 0,942262392 | UTP20 small subunit processome component |
| ENSG00000234206 | AL121957,1 | 0,174196436 | 1,12833576 | 0,80292265 | 0,980866933 | novel transcript |
| ENSG00000096996 | IL12RB1 | 0,174084863 | 1,128248502 | 0,84180716 | 0,98571999 | interleukin 12 receptor subunit beta 1 |
| ENSG00000177674 | AGTRAP | 0,174010697 | 1,128190503 | 0,51626301 | 0,925176589 | angiotensin II receptor associated protein |
| ENSG00000177535 | OR2B11 | 0,174004838 | 1,12818592 | 0,78919471 | 0,979462318 | olfactory receptor family 2 subfamily B member 11 |
| ENSG00000149823 | VPS51 | 0,173772557 | 1,128004292 | 0,62143905 | 0,952793934 | VPS51 subunit of GARP complex |
| ENSG00000286713 | AC113348,3 | 0,173741358 | 1,127979898 | 0,83027947 | 0,984156028 | novel transcript |

| | | | | | | |
|---|---|---|---|---|---|---|
| ENSG00000104859 | CLASRP | 0,173729707 | 1,127970789 | 0,62737328 | 0,952793934 | CLK4 associating serine/arginine rich protein |
| ENSG00000223573 | TINCR | 0,17369841 | 1,12794632 | 0,81245182 | 0,981213877 | TINCR ubiquitin domain containing |
| ENSG00000172809 | RPL38 | 0,173584335 | 1,127857135 | 0,43756148 | 0,897021798 | ribosomal protein L38 |
| ENSG00000223508 | RPL23AP53 | 0,173584092 | 1,127856946 | 0,59488271 | 0,947469769 | ribosomal protein L23a pseudogene 53 |
| ENSG00000204149 | AGAP6 | 0,173351149 | 1,127674853 | 0,52876597 | 0,930033144 | ArfGAP with GTPase domain, ankyrin repeat and PH domain 6 |
| ENSG00000187778 | MCRS1 | 0,173326861 | 1,127655868 | 0,45382137 | 0,902912642 | microspherule protein 1 |
| ENSG00000128731 | HERC2 | 0,173255156 | 1,127599823 | 0,58512422 | 0,945000709 | HECT and RLD domain containing E3 ubiquitin protein ligase 2 |
| ENSG00000137207 | YIPF3 | 0,173233761 | 1,1275831 | 0,47934123 | 0,912714933 | Yip1 domain family member 3 |
| ENSG00000205293 | LINC01602 | 0,173130094 | 1,12750208 | 0,86788022 | 0,989074806 | long intergenic non-protein coding RNA 1602 |
| ENSG00000168904 | LRRC28 | 0,173087396 | 1,12746871 | 0,43439144 | 0,896272827 | leucine rich repeat containing 28 |
| ENSG00000163530 | DPPA2 | 0,173004249 | 1,127403732 | 0,81121428 | 0,981213877 | developmental pluripotency associated 2 |
| ENSG00000283192 | AC007383,6 | 0,172945587 | 1,127357892 | 0,70045575 | 0,968522643 | novel transcript |
| ENSG00000227946 | AC007383,1 | 0,172945587 | 1,127357892 | 0,70045575 | 0,968522643 | novel transcript |
| ENSG00000274317 | LINC02334 | 0,1729231 | 1,12734032 | 0,63735547 | 0,956017912 | long intergenic non-protein coding RNA 2334 |
| ENSG00000270115 | AL513327,2 | 0,172867883 | 1,127297174 | 0,72956809 | 0,974009551 | novel transcript |
| ENSG00000132591 | ERAL1 | 0,172844122 | 1,127278607 | 0,4251978 | 0,891853867 | Era like 12S mitochondrial rRNA chaperone 1 |
| ENSG00000274684 | NMRAL1 | 0,172797904 | 1,127242495 | 0,67119879 | 0,962026636 | NmrA like redox sensor 1 |
| ENSG00000067900 | ROCK1 | 0,172753303 | 1,127207646 | 0,48629638 | 0,914091794 | Rho associated coiled-coil containing protein kinase 1 |
| ENSG00000198598 | MMP17 | 0,172711976 | 1,127175357 | 0,78004715 | 0,978587255 | matrix metallopeptidase 17 |
| ENSG00000284869 | EEFSEC | 0,172711331 | 1,127174853 | 0,51527706 | 0,924832497 | eukaryotic elongation factor, selenocysteine-tRNA specific |
| ENSG00000130703 | OSBPL2 | 0,172694757 | 1,127161904 | 0,5317296 | 0,931291085 | oxysterol binding protein like 2 |
| ENSG00000135297 | MTO1 | 0,172599201 | 1,127087249 | 0,47160934 | 0,908896785 | mitochondrial tRNA translation optimization 1 |
| ENSG00000262333 | HNRNPA1P16 | 0,172515501 | 1,127021862 | 0,61248338 | 0,952413269 | heterogeneous nuclear ribonucleoprotein A1 pseudogene 16 |
| ENSG00000165724 | ZMYND19 | 0,172489956 | 1,127001907 | 0,45680404 | 0,903699469 | zinc finger MYND-type containing 19 |
| ENSG00000241749 | RPSAP52 | 0,172466818 | 1,126983832 | 0,57853528 | 0,943425812 | ribosomal protein SA pseudogene 52 |
| ENSG00000186073 | CDIN1 | 0,172422413 | 1,126949145 | 0,64478715 | 0,956699983 | CDAN1 interacting nuclease 1 |
| ENSG00000196655 | TRAPPC4 | 0,172352253 | 1,126894341 | 0,48841949 | 0,915409567 | trafficking protein particle complex 4 |
| ENSG00000228049 | POLR2J2 | 0,172308065 | 1,126859826 | 0,66073313 | 0,959226315 | RNA polymerase II subunit J2 |

| | | | | | | |
|---|---|---|---|---|---|---|
| ENSG00000067955 | CBFB | 0,172219525 | 1,126790672 | 0,63144329 | 0,953823688 | core-binding factor subunit beta |
| ENSG00000166479 | TMX3 | 0,172136384 | 1,126725737 | 0,71491649 | 0,971041579 | thioredoxin related transmembrane protein 3 |
| ENSG00000243319 | FGF14-IT1 | 0,172135464 | 1,126725019 | 0,76655338 | 0,977201891 | FGF14 intronic transcript 1 |
| ENSG00000171236 | LRG1 | 0,172113546 | 1,126707901 | 0,82411118 | 0,983374097 | leucine rich alpha-2-glycoprotein 1 |
| ENSG00000282534 | IMPDH1P5 | 0,172043263 | 1,126653014 | 0,76462305 | 0,976640468 | inosine monophosphate dehydrogenase 1 pseudogene 5 |
| ENSG00000213513 | IMPDH1P5 | 0,172043263 | 1,126653014 | 0,76462305 | 0,976640468 | inosine monophosphate dehydrogenase 1 pseudogene 5 |
| ENSG00000284153 | RPS4XP17 | 0,171919538 | 1,126556397 | 0,87647489 | 0,990109814 | ribosomal protein S4X pseudogene 17 |
| ENSG00000244097 | RPS4XP17 | 0,171919538 | 1,126556397 | 0,87647489 | 0,990109814 | ribosomal protein S4X pseudogene 17 |
| ENSG00000105821 | DNAJC2 | 0,171898156 | 1,1265397 | 0,47221883 | 0,909171312 | DnaJ heat shock protein family (Hsp40) member C2 |
| ENSG00000168439 | STIP1 | 0,171855172 | 1,126506136 | 0,44135473 | 0,898885877 | stress induced phosphoprotein 1 |
| ENSG00000282189 | AC067930,6 | 0,171841035 | 1,126495097 | 0,74268578 | 0,975201495 | novel transcript |
| ENSG00000101189 | MRGBP | 0,171829725 | 1,126486266 | 0,43680052 | 0,896866043 | MRG domain binding protein |
| ENSG00000124469 | CEACAM8 | 0,17180488 | 1,126466867 | 0,83878907 | 0,985672131 | CEA cell adhesion molecule 8 |
| ENSG00000251026 | LINC02163 | 0,171745695 | 1,126420656 | 0,71107572 | 0,970482408 | long intergenic non-protein coding RNA 2163 |
| ENSG00000105053 | VRK3 | 0,171673056 | 1,126363943 | 0,48453732 | 0,913690171 | VRK serine/threonine kinase 3 |
| ENSG00000136813 | ECPAS | 0,171621012 | 1,12632331 | 0,48489633 | 0,913690171 | Ecm29 proteasome adaptor and scaffold |
| ENSG00000287077 | AL157893,2 | 0,171582836 | 1,126293507 | 0,78848716 | 0,979462318 | novel transcript |
| ENSG00000154710 | RABGEF1 | 0,171568566 | 1,126282367 | 0,50347596 | 0,921005254 | RAB guanine nucleotide exchange factor 1 |
| ENSG00000017260 | ATP2C1 | 0,171551496 | 1,12626904 | 0,5527167 | 0,936894267 | ATPase secretory pathway Ca2+ transporting 1 |
| ENSG00000285768 | LINC02849 | 0,171486433 | 1,126218249 | 0,87589634 | 0,990109814 | long intergenic non-protein coding RNA 2849 |
| ENSG00000155363 | MOV10 | 0,171437862 | 1,126180334 | 0,53639864 | 0,932338187 | Mov10 RISC complex RNA helicase |
| ENSG00000170802 | FOXN2 | 0,171376164 | 1,126132172 | 0,7230222 | 0,972825184 | forkhead box N2 |
| ENSG00000038358 | EDC4 | 0,171318981 | 1,126087537 | 0,51767814 | 0,925915783 | enhancer of mRNA decapping 4 |
| ENSG00000219712 | AL357054,1 | 0,171316308 | 1,126085451 | 0,77651926 | 0,978587255 | novel serine/threonine kinase pseudogene |
| ENSG00000136807 | CDK9 | 0,171220068 | 1,126010334 | 0,51487567 | 0,924720046 | cyclin dependent kinase 9 |
| ENSG00000005238 | FAM214B | 0,171183803 | 1,12598203 | 0,50024127 | 0,920485547 | family with sequence similarity 214 member B |
| ENSG00000187824 | TMEM220 | 0,171125663 | 1,125936654 | 0,53775292 | 0,932883277 | transmembrane protein 220 |
| ENSG00000130305 | NSUN5 | 0,171052877 | 1,125879851 | 0,5859269 | 0,945246954 | NOP2/Sun RNA methyltransferase 5 |
| ENSG00000236554 | ASNSP3 | 0,171027005 | 1,12585966 | 0,8629825 | 0,989074806 | asparagine synthetase pseudogene 3 |
| ENSG00000186501 | TMEM222 | 0,170804323 | 1,125685896 | 0,51279405 | 0,923640376 | transmembrane protein 222 |
| ENSG00000237836 | PHKA2-AS1 | 0,170756341 | 1,125648458 | 0,89540089 | 0,991303328 | PHKA2 antisense RNA 1 |

| | | | | | | |
|---|---|---|---|---|---|---|
| ENSG00000271868 | AC114810,1 | 0,170711441 | 1,125613425 | 0,88087321 | 0,990566588 | novel transcript |
| ENSG00000282522 | PKD1P1 | 0,170582801 | 1,125513063 | 0,74235191 | 0,975176043 | polycystin 1, transient receptor potential channel interacting pseudogene 1 |
| ENSG00000140718 | FTO | 0,170573879 | 1,125506102 | 0,46717595 | 0,907940653 | FTO alpha-ketoglutarate dependent dioxygenase |
| ENSG00000171824 | EXOSC10 | 0,170534152 | 1,12547511 | 0,4818394 | 0,912981134 | exosome component 10 |
| ENSG00000258007 | AC063947,1 | 0,170484624 | 1,125436473 | 0,8659122 | 0,989074806 | novel transcript, antisense to ANO4 |
| ENSG00000114520 | SNX4 | 0,170399831 | 1,125370328 | 0,65914786 | 0,958379996 | sorting nexin 4 |
| ENSG00000165410 | CFL2 | 0,170399105 | 1,125369762 | 0,73599382 | 0,974353474 | cofilin 2 |
| ENSG00000173020 | GRK2 | 0,170281303 | 1,125277875 | 0,58091679 | 0,943914064 | G protein-coupled receptor kinase 2 |
| ENSG00000110435 | PDHX | 0,170267654 | 1,125267229 | 0,61540385 | 0,952793934 | pyruvate dehydrogenase complex component X |
| ENSG00000128383 | APOBEC3A | 0,170229562 | 1,125237519 | 0,82113618 | 0,982810827 | apolipoprotein B mRNA editing enzyme catalytic subunit 3A |
| ENSG00000112276 | BVES | 0,17020334 | 1,125217067 | 0,52118022 | 0,927046149 | blood vessel epicardial substance |
| ENSG00000185813 | PCYT2 | 0,170179238 | 1,125198269 | 0,58369187 | 0,944213336 | phosphate cytidylyltransferase 2, ethanolamine |
| ENSG00000226800 | CACTIN-AS1 | 0,170071262 | 1,125114058 | 0,88884421 | 0,990717729 | CACTIN antisense RNA 1 |
| ENSG00000143222 | UFC1 | 0,170009591 | 1,125065964 | 0,55544456 | 0,937793851 | ubiquitin-fold modifier conjugating enzyme 1 |
| ENSG00000278818 | Metazoa_SRP | 0,1699154 | 1,124992513 | 0,91077374 | 0,993222655 | Metazoan signal recognition particle RNA |
| ENSG00000237399 | PITRM1-AS1 | 0,169912338 | 1,124990125 | 0,68168908 | 0,964654539 | PITRM1 antisense RNA 1 |
| ENSG00000164209 | SLC25A46 | 0,169894033 | 1,124975851 | 0,7101806 | 0,970482408 | solute carrier family 25 member 46 |
| ENSG00000283782 | AC116366,2 | 0,169871335 | 1,124958152 | 0,93208858 | 0,994479239 | novel protein |
| ENSG00000048471 | SNX29 | 0,1698337 | 1,124928807 | 0,45172579 | 0,902062594 | sorting nexin 29 |
| ENSG00000248510 | LINC02267 | 0,169815709 | 1,124914778 | 0,81127409 | 0,981213877 | long intergenic non-protein coding RNA 2267 |
| ENSG00000084733 | RAB10 | 0,16975481 | 1,124867294 | 0,56463613 | 0,940865625 | RAB10, member RAS oncogene family |
| ENSG00000275338 | RAB11FIP3 | 0,169683112 | 1,124811393 | 0,74329621 | 0,975310206 | RAB11 family interacting protein 3 |
| ENSG00000113194 | FAF2 | 0,169679214 | 1,124808354 | 0,55384625 | 0,937388968 | Fas associated factor family member 2 |
| ENSG00000280852 | AC025048,6 | 0,169564474 | 1,124718899 | 0,67834257 | 0,9640473 | adaptor-related protein complex 1, sigma 2 subunit (AP1S2) pseudogene |
| ENSG00000272368 | AC074032,1 | 0,169546095 | 1,124704571 | 0,78781417 | 0,979462318 | novel transcript, antisense to CERS5 |
| ENSG00000159692 | CTBP1 | 0,169517131 | 1,124681992 | 0,53812234 | 0,932900071 | C-terminal binding protein 1 |
| ENSG00000154174 | TOMM70 | 0,169424041 | 1,124609423 | 0,66726505 | 0,9611116 | translocase of outer mitochondrial membrane 70 |
| ENSG00000126749 | EMG1 | 0,169393456 | 1,124585582 | 0,42616122 | 0,892133432 | EMG1 N1-specific pseudouridine methyltransferase |
| ENSG00000277605 | Metazoa_SRP | 0,169386599 | 1,124580237 | 0,88620563 | 0,990717729 | Metazoan signal recognition particle RNA |
| ENSG00000284832 | POLR2A | 0,169269931 | 1,124489298 | 0,47778714 | 0,911721977 | RNA polymerase II subunit A |

| | | | | | | |
|---|---|---|---|---|---|---|
| ENSG00000004975 | DVL2 | 0,169266518 | 1,124486638 | 0,42715977 | 0,892343796 | dishevelled segment polarity protein 2 |
| ENSG00000163755 | HPS3 | 0,169263919 | 1,124484612 | 0,6340631 | 0,954940644 | HPS3 biogenesis of lysosomal organelles complex 2 subunit 1 |
| ENSG00000117481 | NSUN4 | 0,169161624 | 1,124404883 | 0,4657122 | 0,907940653 | NOP2/Sun RNA methyltransferase 4 |
| ENSG00000164520 | RAET1E | 0,16915677 | 1,1244011 | 0,85062492 | 0,987147801 | retinoic acid early transcript 1E |
| ENSG00000136908 | DPM2 | 0,169066268 | 1,124330567 | 0,49628413 | 0,918259281 | dolichyl-phosphate mannosyltransferase subunit 2, regulatory |
| ENSG00000241684 | ADAMTS9-AS2 | 0,169061176 | 1,124326598 | 0,7100997 | 0,970482408 | ADAMTS9 antisense RNA 2 |
| ENSG00000236107 | SCN1A-AS1 | 0,169030243 | 1,124302492 | 0,73056733 | 0,974321178 | SCN1A and SCN9A antisense RNA 1 |
| ENSG00000128185 | DGCR6L | 0,168957096 | 1,12424549 | 0,59090644 | 0,947416795 | DiGeorge syndrome critical region gene 6 like |
| ENSG00000226721 | EEF1DP2 | 0,168956764 | 1,124245231 | 0,83152116 | 0,984456953 | eukaryotic translation elongation factor 1 delta pseudogene 2 |
| ENSG00000011243 | AKAP8L | 0,168937036 | 1,124229858 | 0,57632508 | 0,943158459 | A-kinase anchoring protein 8 like |
| ENSG00000241680 | RPL31P49 | 0,168682663 | 1,124031653 | 0,89537102 | 0,991303328 | ribosomal protein L31 pseudogene 49 |
| ENSG00000113838 | TBCCD1 | 0,168667111 | 1,124019536 | 0,49206587 | 0,916995341 | TBCC domain containing 1 |
| ENSG00000223675 | AC093117,1 | 0,168617487 | 1,123980874 | 0,87859056 | 0,990109814 | novel transcript |
| ENSG00000274429 | DLG5 | 0,168603691 | 1,123970126 | 0,59813121 | 0,948582224 | discs large MAGUK scaffold protein 5 |
| ENSG00000213212 | NCLP1 | 0,168523418 | 1,123907589 | 0,77429612 | 0,978503783 | nucleolin pseudogene 1 |
| ENSG00000277959 | AL162274,2 | 0,168493431 | 1,123884229 | 0,82331298 | 0,983308136 | novel transcript, antisense to BNIP3 |
| ENSG00000249125 | AC093821,1 | 0,168481522 | 1,123874951 | 0,73192035 | 0,974353474 | novel transcript |
| ENSG00000281618 | IFITM10 | 0,168416363 | 1,123824193 | 0,93107874 | 0,994433362 | interferon induced transmembrane protein 10 |
| ENSG00000108107 | RPL28 | 0,168269657 | 1,123709918 | 0,4848043 | 0,913690171 | ribosomal protein L28 |
| ENSG00000171311 | EXOSC1 | 0,168223406 | 1,123673894 | 0,46836697 | 0,907940653 | exosome component 1 |
| ENSG00000125459 | MSTO1 | 0,168176451 | 1,123637323 | 0,43932944 | 0,897434993 | misato mitochondrial distribution and morphology regulator 1 |
| ENSG00000156486 | KCNS2 | 0,168139482 | 1,12360853 | 0,73589139 | 0,974353474 | potassium voltage-gated channel modifier subfamily S member 2 |
| ENSG00000224746 | AC015987,1 | 0,16812659 | 1,123598489 | 0,82337831 | 0,983348593 | novel transcript, antisense to MTPN |
| ENSG00000280213 | UCKL1-AS1 | 0,168095978 | 1,123574648 | 0,79154919 | 0,979471509 | UCKL1 antisense RNA 1 |
| ENSG00000116473 | RAP1A | 0,168066594 | 1,123551765 | 0,69049938 | 0,967179044 | RAP1A, member of RAS oncogene family |
| ENSG00000196832 | OR11G2 | 0,168047233 | 1,123536686 | 0,78942107 | 0,979462318 | olfactory receptor family 11 subfamily G member 2 |
| ENSG00000119231 | SENP5 | 0,168019074 | 1,123514757 | 0,55639023 | 0,938061341 | SUMO specific peptidase 5 |
| ENSG00000177879 | AP3S1 | 0,168016755 | 1,123512951 | 0,73340094 | 0,974353474 | adaptor related protein complex 3 subunit sigma 1 |

| | | | | | | |
|---|---|---|---|---|---|---|
| ENSG00000001461 | NIPAL3 | 0,167996017 | 1,123496801 | 0,59964227 | 0,94932226 | NIPA like domain containing 3 |
| ENSG00000076924 | XAB2 | 0,167907721 | 1,123428043 | 0,52642566 | 0,929261495 | XPA binding protein 2 |
| ENSG00000111361 | EIF2B1 | 0,167872666 | 1,123400746 | 0,56816847 | 0,94163744 | eukaryotic translation initiation factor 2B subunit alpha |
| ENSG00000206538 | VGLL3 | 0,167860601 | 1,123391351 | 0,69556833 | 0,967669485 | vestigial like family member 3 |
| ENSG00000186684 | CYP27C1 | 0,167810626 | 1,123352437 | 0,59798858 | 0,948577552 | cytochrome P450 family 27 subfamily C member 1 |
| ENSG00000225737 | SKIV2L | 0,167761648 | 1,123314302 | 0,70621018 | 0,969503793 | Ski2 like RNA helicase |
| ENSG00000105849 | POLR1F | 0,167579618 | 1,123172578 | 0,69679155 | 0,968300554 | RNA polymerase I subunit F |
| ENSG00000151923 | TIAL1 | 0,167536623 | 1,123139106 | 0,62750228 | 0,952793934 | TIA1 cytotoxic granule associated RNA binding protein like 1 |
| ENSG00000107554 | DNMBP | 0,16745568 | 1,123076093 | 0,46323471 | 0,906974564 | dynamin binding protein |
| ENSG00000283771 | TMEM256 | 0,167425745 | 1,123052791 | 0,58334964 | 0,944174844 | transmembrane protein 256 |
| ENSG00000205544 | TMEM256 | 0,167425745 | 1,123052791 | 0,58334964 | 0,944174844 | transmembrane protein 256 |
| ENSG00000262771 | SSBP1 | 0,167387059 | 1,123022676 | 0,59084301 | 0,947416795 | single stranded DNA binding protein 1 |
| ENSG00000165322 | ARHGAP12 | 0,167334907 | 1,12298208 | 0,68998582 | 0,967038 | Rho GTPase activating protein 12 |
| ENSG00000099940 | SNAP29 | 0,167225006 | 1,122896538 | 0,57307356 | 0,942475308 | synaptosome associated protein 29 |
| ENSG00000198898 | CAPZA2 | 0,167182798 | 1,122863686 | 0,65744702 | 0,957853146 | capping actin protein of muscle Z-line subunit alpha 2 |
| ENSG00000139725 | RHOF | 0,167144694 | 1,12283403 | 0,71547302 | 0,971152196 | ras homolog family member F, filopodia associated |
| ENSG00000258555 | SPECC1L-ADOR | 0,167111831 | 1,122808453 | 0,78615892 | 0,979320406 | SPECC1L-ADORA2A readthrough (NMD candidate) |
| ENSG00000198176 | TFDP1 | 0,167060421 | 1,122768444 | 0,45757351 | 0,903831777 | transcription factor Dp-1 |
| ENSG00000232541 | COL11A2 | 0,166945519 | 1,122679025 | 0,89381797 | 0,991135494 | collagen type XI alpha 2 chain |
| ENSG00000274717 | AL049757,1 | 0,166854883 | 1,122608496 | 0,86368435 | 0,989074806 | ADAM metallopeptidase with thrombospondin type 1 motif, 7 (ADAMTS7) pseudogene |
| ENSG00000122783 | CYREN | 0,166844853 | 1,122600691 | 0,53037945 | 0,931194643 | cell cycle regulator of NHEJ |
| ENSG00000198933 | TBKBP1 | 0,166777777 | 1,122548499 | 0,69069231 | 0,967179044 | TBK1 binding protein 1 |
| ENSG00000237100 | C6orf136 | 0,166750653 | 1,122527394 | 0,75366558 | 0,975901808 | chromosome 6 open reading frame 136 |
| ENSG00000233641 | C6orf136 | 0,166750653 | 1,122527394 | 0,75366558 | 0,975901808 | chromosome 6 open reading frame 136 |
| ENSG00000224120 | C6orf136 | 0,166750653 | 1,122527394 | 0,75366558 | 0,975901808 | chromosome 6 open reading frame 136 |
| ENSG00000237012 | C6orf136 | 0,166750653 | 1,122527394 | 0,75366558 | 0,975901808 | chromosome 6 open reading frame 136 |
| ENSG00000116198 | CEP104 | 0,166724415 | 1,122506979 | 0,45421913 | 0,90318049 | centrosomal protein 104 |
| ENSG00000134324 | LPIN1 | 0,166671486 | 1,122465798 | 0,55329314 | 0,936992468 | lipin 1 |
| ENSG00000084092 | NOA1 | 0,166628549 | 1,122432392 | 0,59217907 | 0,947416795 | nitric oxide associated 1 |

| | | | | | | |
|---|---|---|---|---|---|---|
| ENSG00000225697 | SLC26A6 | 0,166600333 | 1,12241044 | 0,57722835 | 0,94337651 | solute carrier family 26 member 6 |
| ENSG00000230068 | CDC42-IT1 | 0,16652327 | 1,122350487 | 0,79226273 | 0,979471509 | CDC42 intronic transcript 1 |
| ENSG00000078618 | NRDC | 0,166449625 | 1,122293196 | 0,46716943 | 0,907940653 | nardilysin convertase |
| ENSG00000144320 | LNPK | 0,166406408 | 1,122259577 | 0,66600134 | 0,961020116 | lunapark, ER junction formation factor |
| ENSG00000147003 | CLTRN | 0,166309196 | 1,122183959 | 0,70279731 | 0,968742626 | collectrin, amino acid transport regulator |
| ENSG00000093009 | CDC45 | 0,166247587 | 1,122136038 | 0,43964748 | 0,897665148 | cell division cycle 45 |
| ENSG00000004766 | VPS50 | 0,166247093 | 1,122135654 | 0,61719039 | 0,952793934 | VPS50 subunit of EARP/GARPII complex |
| ENSG00000276421 | AC012314,11 | 0,166156398 | 1,122065113 | 0,86904484 | 0,989074806 | pre-mRNA processing factor 31 |
| ENSG00000274894 | AC012314,6 | 0,166156398 | 1,122065113 | 0,86904484 | 0,989074806 | pre-mRNA processing factor 31 |
| ENSG00000274651 | AC012314,5 | 0,166156398 | 1,122065113 | 0,86904484 | 0,989074806 | pre-mRNA processing factor 31 |
| ENSG00000275117 | AC012314,7 | 0,166156398 | 1,122065113 | 0,86904484 | 0,989074806 | pre-mRNA processing factor 31 |
| ENSG00000277953 | AC012314,14 | 0,166156398 | 1,122065113 | 0,86904484 | 0,989074806 | pre-mRNA processing factor 31 |
| ENSG00000274144 | AC012314,3 | 0,166156398 | 1,122065113 | 0,86904484 | 0,989074806 | pre-mRNA processing factor 31 |
| ENSG00000277707 | AC012314,13 | 0,166156398 | 1,122065113 | 0,86904484 | 0,989074806 | pre-mRNA processing factor 31 |
| ENSG00000277154 | AC012314,12 | 0,166156398 | 1,122065113 | 0,86904484 | 0,989074806 | pre-mRNA processing factor 31 |
| ENSG00000257727 | CNPY2 | 0,166146825 | 1,122057668 | 0,44168827 | 0,899078209 | canopy FGF signaling regulator 2 |
| ENSG00000133026 | MYH10 | 0,166117326 | 1,122034726 | 0,6494976 | 0,956899963 | myosin heavy chain 10 |
| ENSG00000258858 | AL583722,3 | 0,166074811 | 1,12200166 | 0,87924885 | 0,990148595 | novel transcript, antisense to INF2 |
| ENSG00000056998 | GYG2 | 0,166064977 | 1,121994012 | 0,45114706 | 0,902062594 | glycogenin 2 |
| ENSG00000125520 | SLC2A4RG | 0,166063033 | 1,1219925 | 0,64278051 | 0,956633898 | SLC2A4 regulator |
| ENSG00000184007 | PTP4A2 | 0,166038515 | 1,121973433 | 0,65786917 | 0,957864535 | protein tyrosine phosphatase 4A2 |
| ENSG00000185324 | CDK10 | 0,165978483 | 1,121926747 | 0,62651003 | 0,952793934 | cyclin dependent kinase 10 |
| ENSG00000162636 | FAM102B | 0,165956823 | 1,121909904 | 0,67280416 | 0,962371283 | family with sequence similarity 102 member B |
| ENSG00000136111 | TBC1D4 | 0,165943716 | 1,121899711 | 0,5974818 | 0,948282297 | TBC1 domain family member 4 |
| ENSG00000125779 | PANK2 | 0,165925449 | 1,121885506 | 0,46503701 | 0,907804479 | pantothenate kinase 2 |
| ENSG00000108561 | C1QBP | 0,165811304 | 1,121796746 | 0,46500459 | 0,907804479 | complement C1q binding protein |
| ENSG00000275016 | AC015574,1 | 0,165790258 | 1,121780382 | 0,81962085 | 0,98249617 | novel transcript |
| ENSG00000229190 | AC069542,1 | 0,165789125 | 1,121779501 | 0,86992008 | 0,989074806 | novel transcript |
| ENSG00000062485 | CS | 0,165689553 | 1,121702081 | 0,52025192 | 0,926865961 | citrate synthase |
| ENSG00000228107 | AP000692,1 | 0,165671765 | 1,12168825 | 0,81452612 | 0,981521413 | novel transcript, sense overlapping MORC3 |
| ENSG00000100347 | SAMM50 | 0,165631026 | 1,121656576 | 0,46460744 | 0,907616025 | SAMM50 sorting and assembly machinery component |
| ENSG00000213930 | GALT | 0,165607631 | 1,121638388 | 0,49516039 | 0,917777383 | galactose-1-phosphate uridylyltransferase |

| | | | | | | |
|---|---|---|---|---|---|---|
| ENSG00000114573 | ATP6V1A | 0,165583629 | 1,121619727 | 0,65007901 | 0,956899963 | ATPase H+ transporting V1 subunit A |
| ENSG00000095059 | DHPS | 0,165576147 | 1,12161391 | 0,47626075 | 0,911416789 | deoxyhypusine synthase |
| ENSG00000171097 | KYAT1 | 0,165564954 | 1,121605208 | 0,56488268 | 0,940865625 | kynurenine aminotransferase 1 |
| ENSG00000257913 | DDN-AS1 | 0,165537063 | 1,121583525 | 0,66626913 | 0,961070325 | DDN and PRKAG1 antisense RNA 1 |
| ENSG00000170515 | PA2G4 | 0,165519114 | 1,121569572 | 0,47616677 | 0,911413793 | proliferation-associated 2G4 |
| ENSG00000116157 | GPX7 | 0,165515966 | 1,121567124 | 0,68336893 | 0,96503213 | glutathione peroxidase 7 |
| ENSG00000287155 | AC022215,2 | 0,165453273 | 1,121518387 | 0,89968137 | 0,991845106 | novel transcript, antisense to PXYLP1 |
| ENSG00000103056 | SMPD3 | 0,165142916 | 1,121277148 | 0,65659062 | 0,957444211 | sphingomyelin phosphodiesterase 3 |
| ENSG00000101216 | GMEB2 | 0,16513101 | 1,121267895 | 0,5827897 | 0,944174844 | glucocorticoid modulatory element binding protein 2 |
| ENSG00000013810 | TACC3 | 0,165128206 | 1,121265715 | 0,57748501 | 0,94337651 | transforming acidic coiled-coil containing protein 3 |
| ENSG00000254827 | SLC22A18AS | 0,165063657 | 1,121215549 | 0,77304412 | 0,978360981 | solute carrier family 22 member 18 antisense |
| ENSG00000111249 | CUX2 | 0,164974059 | 1,121145919 | 0,55149441 | 0,936366211 | cut like homeobox 2 |
| ENSG00000287040 | AL590560,5 | 0,164920776 | 1,121104512 | 0,86882873 | 0,989074806 | novel transcript |
| ENSG00000166484 | MAPK7 | 0,164851446 | 1,121050638 | 0,54059931 | 0,933385624 | mitogen-activated protein kinase 7 |
| ENSG00000274104 | AC020910,5 | 0,164846403 | 1,121046719 | 0,84834334 | 0,987036295 | novel transcript |
| ENSG00000067334 | DNTTIP2 | 0,164818387 | 1,121024949 | 0,56906182 | 0,942177697 | deoxynucleotidyltransferase terminal interacting protein 2 |
| ENSG00000260081 | AF274858,1 | 0,164816785 | 1,121023705 | 0,74162625 | 0,975176043 | Morf4 family associated protein 1 (MRFAP1) pseudogene |
| ENSG00000146109 | ABT1 | 0,16474842 | 1,120970584 | 0,52406811 | 0,928403516 | activator of basal transcription 1 |
| ENSG00000119421 | NDUFA8 | 0,164705034 | 1,120936873 | 0,60908142 | 0,951793902 | NADH:ubiquinone oxidoreductase subunit A8 |
| ENSG00000068383 | INPP5A | 0,164701637 | 1,120934234 | 0,49983409 | 0,920310866 | inositol polyphosphate-5-phosphatase A |
| ENSG00000227195 | MIR663AHG | 0,164700437 | 1,120933302 | 0,85964814 | 0,989074806 | MIR663A host gene |
| ENSG00000128590 | DNAJB9 | 0,164675223 | 1,120913712 | 0,72100287 | 0,972142678 | DnaJ heat shock protein family (Hsp40) member B9 |
| ENSG00000285863 | AC009554,2 | 0,164553379 | 1,120819048 | 0,76335745 | 0,976640468 | novel transcript, antisense to VPS13C |
| ENSG00000163444 | TMEM183A | 0,164520296 | 1,120793346 | 0,5016022 | 0,920520138 | transmembrane protein 183A |
| ENSG00000247626 | MARS2 | 0,164424412 | 1,120718859 | 0,61133218 | 0,952062377 | methionyl-tRNA synthetase 2, mitochondrial |
| ENSG00000258140 | AC127894,1 | 0,164312445 | 1,120631884 | 0,91819157 | 0,993361299 | novel transcript |
| ENSG00000229058 | AGER | 0,164303738 | 1,120625121 | 0,78946296 | 0,979462318 | advanced glycosylation end-product specific receptor |
| ENSG00000237405 | AGER | 0,164303738 | 1,120625121 | 0,78946296 | 0,979462318 | advanced glycosylation end-product specific receptor |

| | | | | | | |
|---|---|---|---|---|---|---|
| ENSG00000206320 | AGER | 0,164303738 | 1,120625121 | 0,78946296 | 0,979462318 | advanced glycosylation end-product specific receptor |
| ENSG00000284915 | TINF2 | 0,164296429 | 1,120619443 | 0,46588222 | 0,907940653 | TERF1 interacting nuclear factor 2 |
| ENSG00000234937 | AL139128,1 | 0,164249063 | 1,120582652 | 0,82587363 | 0,983383773 | proteasome (prosome, macropain) 26S subunit, non-ATPase, 7 (Mov34 homolog) (PSMD7) pseudogene |
| ENSG00000149634 | SPATA25 | 0,164213491 | 1,120555022 | 0,80715956 | 0,981213877 | spermatogenesis associated 25 |
| ENSG00000112474 | HSD17B8 | 0,164153024 | 1,120508058 | 0,86814451 | 0,989074806 | hydroxysteroid 17-beta dehydrogenase 8 |
| ENSG00000232357 | HSD17B8 | 0,164153024 | 1,120508058 | 0,86814451 | 0,989074806 | hydroxysteroid 17-beta dehydrogenase 8 |
| ENSG00000225312 | HSD17B8 | 0,164153024 | 1,120508058 | 0,86814451 | 0,989074806 | hydroxysteroid 17-beta dehydrogenase 8 |
| ENSG00000228712 | HSD17B8 | 0,164153024 | 1,120508058 | 0,86814451 | 0,989074806 | hydroxysteroid 17-beta dehydrogenase 8 |
| ENSG00000228357 | HSD17B8 | 0,164153024 | 1,120508058 | 0,86814451 | 0,989074806 | hydroxysteroid 17-beta dehydrogenase 8 |
| ENSG00000249699 | LINC02261 | 0,164094337 | 1,120462478 | 0,82693002 | 0,983548538 | long intergenic non-protein coding RNA 2261 |
| ENSG00000111540 | RAB5B | 0,164079911 | 1,120451275 | 0,53536797 | 0,932037868 | RAB5B, member RAS oncogene family |
| ENSG00000008083 | JARID2 | 0,164072389 | 1,120445432 | 0,47460451 | 0,91069468 | jumonji and AT-rich interaction domain containing 2 |
| ENSG00000273841 | TAF9 | 0,164062308 | 1,120437603 | 0,80883924 | 0,981213877 | TATA-box binding protein associated factor 9 |
| ENSG00000162039 | MEIOB | 0,164038241 | 1,120418913 | 0,85610938 | 0,988185425 | meiosis specific with OB-fold |
| ENSG00000140280 | LYSMD2 | 0,163985861 | 1,120378234 | 0,79344273 | 0,979776981 | LysM domain containing 2 |
| ENSG00000197779 | ZNF81 | 0,16398297 | 1,120375989 | 0,62247356 | 0,952793934 | zinc finger protein 81 |
| ENSG00000166171 | DPCD | 0,163832707 | 1,120259303 | 0,47074326 | 0,908869097 | deleted in primary ciliary dyskinesia homolog (mouse) |
| ENSG00000242338 | BMS1P4 | 0,163797251 | 1,120231772 | 0,73871227 | 0,974353474 | BMS1 pseudogene 4 |
| ENSG00000287141 | AC093675,2 | 0,163762156 | 1,120204521 | 0,82955653 | 0,984081562 | novel transcript, antisense to TMEM87B |
| ENSG00000134049 | IER3IP1 | 0,163651886 | 1,120118903 | 0,77433963 | 0,978503783 | immediate early response 3 interacting protein 1 |
| ENSG00000189280 | GJB5 | 0,163556924 | 1,120045177 | 0,87037384 | 0,989074806 | gap junction protein beta 5 |
| ENSG00000162377 | COA7 | 0,16354772 | 1,120038031 | 0,47434963 | 0,910676852 | cytochrome c oxidase assembly factor 7 (putative) |
| ENSG00000097021 | ACOT7 | 0,163534303 | 1,120027615 | 0,51923045 | 0,926800715 | acyl-CoA thioesterase 7 |
| ENSG00000274507 | AL121772,2 | 0,163514118 | 1,120011944 | 0,88162015 | 0,990717729 | novel transcript, antisense to PYGB |
| ENSG00000167291 | TBC1D16 | 0,163491924 | 1,119994714 | 0,54536672 | 0,935085411 | TBC1 domain family member 16 |
| ENSG00000145982 | FARS2 | 0,163490985 | 1,119993986 | 0,47522653 | 0,911082311 | phenylalanyl-tRNA synthetase 2, mitochondrial |
| ENSG00000099795 | NDUFB7 | 0,163443646 | 1,119957236 | 0,64315399 | 0,956633898 | NADH:ubiquinone oxidoreductase subunit B7 |
| ENSG00000280007 | AC008079,1 | 0,163370286 | 1,119900288 | 0,7853443 | 0,979320406 | novel transcript, antisense to TUBA8 |
| ENSG00000106246 | PTCD1 | 0,16332794 | 1,119867418 | 0,60776597 | 0,950993594 | pentatricopeptide repeat domain 1 |
| ENSG00000100804 | PSMB5 | 0,163270138 | 1,119822551 | 0,4811741 | 0,912911021 | proteasome 20S subunit beta 5 |

| | | | | | | |
|---|---|---|---|---|---|---|
| ENSG00000238132 | CASC4P1 | 0,163256971 | 1,11981233 | 0,86704238 | 0,989074806 | cancer susceptibility 4 pseudogene 1 |
| ENSG00000224848 | AL589843,1 | 0,163216148 | 1,119780644 | 0,78279987 | 0,978687331 | novel transcript |
| ENSG00000120697 | ALG5 | 0,163190203 | 1,119760506 | 0,66966124 | 0,962016728 | ALG5 dolichyl-phosphate beta-glucosyltransferase |
| ENSG00000160075 | SSU72 | 0,163008242 | 1,119619285 | 0,45480451 | 0,903424009 | SSU72 homolog, RNA polymerase II CTD phosphatase |
| ENSG00000177138 | FAM9B | 0,162912019 | 1,119544613 | 0,91318322 | 0,993361299 | family with sequence similarity 9 member B |
| ENSG00000099910 | KLHL22 | 0,162824459 | 1,119476667 | 0,56826689 | 0,94163744 | kelch like family member 22 |
| ENSG00000018699 | TTC27 | 0,162741176 | 1,119412045 | 0,51772756 | 0,925915783 | tetratricopeptide repeat domain 27 |
| ENSG00000179151 | EDC3 | 0,162734724 | 1,119407039 | 0,44772374 | 0,902062594 | enhancer of mRNA decapping 3 |
| ENSG00000080845 | DLGAP4 | 0,162595846 | 1,119299286 | 0,45186102 | 0,902062594 | DLG associated protein 4 |
| ENSG00000162191 | UBXN1 | 0,162577102 | 1,119284744 | 0,54595109 | 0,935201756 | UBX domain protein 1 |
| ENSG00000154359 | LONRF1 | 0,16251622 | 1,119237511 | 0,47911638 | 0,912714933 | LON peptidase N-terminal domain and ring finger 1 |
| ENSG00000137463 | MGARP | 0,162405604 | 1,119151699 | 0,86478861 | 0,989074806 | mitochondria localized glutamic acid rich protein |
| ENSG00000136710 | CCDC115 | 0,162401571 | 1,119148571 | 0,509274 | 0,922427526 | coiled-coil domain containing 115 |
| ENSG00000075711 | DLG1 | 0,162385112 | 1,119135802 | 0,56622023 | 0,941123407 | discs large MAGUK scaffold protein 1 |
| ENSG00000178464 | RPL10P16 | 0,162371598 | 1,11912532 | 0,91078905 | 0,993222655 | ribosomal protein L10 pseudogene 16 |
| ENSG00000130958 | SLC35D2 | 0,162282452 | 1,119056169 | 0,53432508 | 0,931584152 | solute carrier family 35 member D2 |
| ENSG00000100348 | TXN2 | 0,162265372 | 1,119042921 | 0,48760968 | 0,914936533 | thioredoxin 2 |
| ENSG00000153029 | MR1 | 0,162248544 | 1,119029868 | 0,7817661 | 0,978597905 | major histocompatibility complex, class I-related |
| ENSG00000006194 | ZNF263 | 0,162203973 | 1,118995297 | 0,49566566 | 0,918046024 | zinc finger protein 263 |
| ENSG00000278622 | TSEN34 | 0,162203426 | 1,118994873 | 0,59188139 | 0,947416795 | tRNA splicing endonuclease subunit 34 |
| ENSG00000115289 | PCGF1 | 0,162161389 | 1,118962269 | 0,55151372 | 0,936366211 | polycomb group ring finger 1 |
| ENSG00000138029 | HADHB | 0,162059667 | 1,118883375 | 0,62470173 | 0,952793934 | hydroxyacyl-CoA dehydrogenase trifunctional multienzyme complex subunit beta |
| ENSG00000244687 | UBE2V1 | 0,16203094 | 1,118861096 | 0,59664308 | 0,948282297 | ubiquitin conjugating enzyme E2 V1 |
| ENSG00000180370 | PAK2 | 0,161953717 | 1,118801209 | 0,61942877 | 0,952793934 | p21 (RAC1) activated kinase 2 |
| ENSG00000163382 | NAXE | 0,161928998 | 1,118782039 | 0,45845478 | 0,90420131 | NAD(P)HX epimerase |
| ENSG00000252464 | RN7SKP70 | 0,161892612 | 1,118753823 | 0,8453992 | 0,986639902 | RN7SK pseudogene 70 |
| ENSG00000145029 | NICN1 | 0,161867783 | 1,118734569 | 0,57450969 | 0,942475308 | nicolin 1 |
| ENSG00000128908 | INO80 | 0,161866739 | 1,11873376 | 0,43764299 | 0,897021798 | INO80 complex ATPase subunit |
| ENSG00000143179 | UCK2 | 0,161847327 | 1,118718707 | 0,50028345 | 0,920485547 | uridine-cytidine kinase 2 |
| ENSG00000114331 | ACAP2 | 0,161847306 | 1,118718691 | 0,67306767 | 0,962528022 | ArfGAP with coiled-coil, ankyrin repeat and PH domains 2 |
| ENSG00000286071 | AL136171,2 | 0,161733704 | 1,118630602 | 0,6318128 | 0,953915351 | disrupted in schizophrenia 2 |

| | | | | | | |
|---|---|---|---|---|---|---|
| ENSG00000103044 | HAS3 | 0,16169192 | 1,118598204 | 0,60071987 | 0,949489681 | hyaluronan synthase 3 |
| ENSG00000025293 | PHF20 | 0,161585951 | 1,118516045 | 0,43404819 | 0,896272827 | PHD finger protein 20 |
| ENSG00000198858 | R3HDM4 | 0,161578064 | 1,11850993 | 0,56523476 | 0,940865625 | R3H domain containing 4 |
| ENSG00000155749 | FLACC1 | 0,16156274 | 1,118498049 | 0,792372 | 0,979471509 | flagellum associated containing coiled-coil domains 1 |
| ENSG00000241674 | HLA-DMB | 0,161559846 | 1,118495805 | 0,69910948 | 0,968522643 | major histocompatibility complex, class II, DM beta |
| ENSG00000288114 | GTF2H1 | 0,161528547 | 1,11847154 | 0,68117324 | 0,964654539 | general transcription factor IIH subunit 1 |
| ENSG00000105221 | AKT2 | 0,161524224 | 1,118468188 | 0,54969773 | 0,936108121 | AKT serine/threonine kinase 2 |
| ENSG00000139644 | TMBIM6 | 0,161504515 | 1,118452909 | 0,55902833 | 0,939635125 | transmembrane BAX inhibitor motif containing 6 |
| ENSG00000286033 | CR381670,2 | 0,161362531 | 1,118342841 | 0,73152551 | 0,974353474 | novel transcript |
| ENSG00000244296 | RN7SL168P | 0,161354978 | 1,118336987 | 0,90022879 | 0,991845106 | RNA, 7SL, cytoplasmic 168, pseudogene |
| ENSG00000204859 | ZBTB48 | 0,161318065 | 1,118308372 | 0,65654004 | 0,957444211 | zinc finger and BTB domain containing 48 |
| ENSG00000099999 | RNF215 | 0,161295254 | 1,118290691 | 0,6430568 | 0,956633898 | ring finger protein 215 |
| ENSG00000100083 | GGA1 | 0,161288445 | 1,118285413 | 0,56114293 | 0,940567693 | golgi associated, gamma adaptin ear containing, ARF binding protein 1 |
| ENSG00000154914 | USP43 | 0,161164463 | 1,118189314 | 0,65224831 | 0,957257009 | ubiquitin specific peptidase 43 |
| ENSG00000155858 | LSM11 | 0,161153318 | 1,118180676 | 0,52005378 | 0,926822772 | LSM11, U7 small nuclear RNA associated |
| ENSG00000285486 | AC003043,2 | 0,161139887 | 1,118170266 | 0,89365886 | 0,991135494 | novel transcript |
| ENSG00000094914 | AAAS | 0,161117351 | 1,118152799 | 0,46846166 | 0,907940653 | aladin WD repeat nucleoporin |
| ENSG00000287553 | AC090617,10 | 0,161058167 | 1,118106931 | 0,83100968 | 0,984300284 | novel transcript, antisense to DPH1 |
| ENSG00000225695 | HNRNPA1P35 | 0,160958967 | 1,118030051 | 0,80350573 | 0,980875302 | heterogeneous nuclear ribonucleoprotein A1 pseudogene 35 |
| ENSG00000274384 | TMC4 | 0,160909817 | 1,117991963 | 0,81389876 | 0,981521413 | transmembrane channel like 4 |
| ENSG00000104228 | TRIM35 | 0,160866405 | 1,117958322 | 0,54258618 | 0,933663748 | tripartite motif containing 35 |
| ENSG00000156671 | SAMD8 | 0,16076982 | 1,11788348 | 0,65890834 | 0,958240464 | sterile alpha motif domain containing 8 |
| ENSG00000186300 | ZNF555 | 0,160721083 | 1,117845716 | 0,51752911 | 0,925842873 | zinc finger protein 555 |
| ENSG00000197566 | ZNF624 | 0,160690102 | 1,117821712 | 0,70021262 | 0,968522643 | zinc finger protein 624 |
| ENSG00000162869 | PPP1R21 | 0,160571745 | 1,117730011 | 0,59230499 | 0,947416795 | protein phosphatase 1 regulatory subunit 21 |
| ENSG00000163376 | KBTBD8 | 0,160457268 | 1,117641323 | 0,78502959 | 0,979320406 | kelch repeat and BTB domain containing 8 |
| ENSG00000235944 | ZNF815P | 0,160451046 | 1,117636503 | 0,69826494 | 0,968522643 | zinc finger protein 815, pseudogene |
| ENSG00000203930 | LINC00632 | 0,16040616 | 1,117601731 | 0,50189483 | 0,920520138 | long intergenic non-protein coding RNA 632 |
| ENSG00000058600 | POLR3E | 0,160381912 | 1,117582947 | 0,47805199 | 0,911886357 | RNA polymerase III subunit E |
| ENSG00000065613 | SLK | 0,160360061 | 1,11756602 | 0,57171685 | 0,942436428 | STE20 like kinase |

| | | | | | | |
|---|---|---|---|---|---|---|
| ENSG00000227794 | RPS18 | 0,160357473 | 1,117564015 | 0,69852223 | 0,968522643 | ribosomal protein S18 |
| ENSG00000239884 | RN7SL608P | 0,16034304 | 1,117552835 | 0,87562942 | 0,990078774 | RNA, 7SL, cytoplasmic 608, pseudogene |
| ENSG00000130396 | AFDN | 0,160188826 | 1,117433383 | 0,50319283 | 0,920888747 | afadin, adherens junction formation factor |
| ENSG00000262319 | AC007952,6 | 0,160178403 | 1,11742531 | 0,83112857 | 0,984369574 | novel transcript, antisense to GRAPL |
| ENSG00000122122 | SASH3 | 0,16014936 | 1,117402815 | 0,87159143 | 0,989217334 | SAM and SH3 domain containing 3 |
| ENSG00000270813 | NANOGNBP3 | 0,160065925 | 1,117338195 | 0,86656953 | 0,989074806 | NANOGNB pseudogene 3 |
| ENSG00000249534 | LINC01258 | 0,159975199 | 1,117267931 | 0,86197233 | 0,989074806 | long intergenic non-protein coding RNA 1258 |
| ENSG00000231133 | HAR1B | 0,159974033 | 1,117267028 | 0,91427883 | 0,993361299 | highly accelerated region 1B |
| ENSG00000274471 | HERC2P7 | 0,159948511 | 1,117247263 | 0,62318156 | 0,952793934 | hect domain and RLD 2 pseudogene 7 |
| ENSG00000225558 | UBE2D3P2 | 0,159906574 | 1,117214787 | 0,88988545 | 0,990717729 | ubiquitin conjugating enzyme E2 D3 pseudogene 2 |
| ENSG00000105373 | NOP53 | 0,159901763 | 1,117211061 | 0,63848288 | 0,956018518 | NOP53 ribosome biogenesis factor |
| ENSG00000172482 | AGXT | 0,159899323 | 1,117209172 | 0,87169826 | 0,989217334 | alanine--glyoxylate and serine--pyruvate aminotransferase |
| ENSG00000277292 | PIP4K2B | 0,159887185 | 1,117199772 | 0,4564312 | 0,903545728 | phosphatidylinositol-5-phosphate 4-kinase type 2 beta |
| ENSG00000276293 | PIP4K2B | 0,159887185 | 1,117199772 | 0,4564312 | 0,903545728 | phosphatidylinositol-5-phosphate 4-kinase type 2 beta |
| ENSG00000163468 | CCT3 | 0,159862315 | 1,117180513 | 0,53947624 | 0,933350549 | chaperonin containing TCP1 subunit 3 |
| ENSG00000272040 | AC010245,2 | 0,15975767 | 1,117099483 | 0,82680223 | 0,983548538 | novel transcript |
| ENSG00000274514 | Metazoa_SRP | 0,15972112 | 1,117071182 | 0,88862596 | 0,990717729 | Metazoan signal recognition particle RNA |
| ENSG00000167972 | ABCA3 | 0,159521888 | 1,116916928 | 0,55512153 | 0,937793851 | ATP binding cassette subfamily A member 3 |
| ENSG00000255910 | AC024901,1 | 0,159452195 | 1,116862974 | 0,87454221 | 0,989995204 | novel transcript |
| ENSG00000179520 | SLC17A8 | 0,159400239 | 1,116822753 | 0,88266327 | 0,990717729 | solute carrier family 17 member 8 |
| ENSG00000083750 | RRAGB | 0,159339165 | 1,116775475 | 0,61853805 | 0,952793934 | Ras related GTP binding B |
| ENSG00000184205 | TSPYL2 | 0,159319318 | 1,116760112 | 0,57131414 | 0,942436428 | TSPY like 2 |
| ENSG00000220848 | RPS18P9 | 0,159266214 | 1,116719006 | 0,67162943 | 0,962143623 | ribosomal protein S18 pseudogene 9 |
| ENSG00000141560 | FN3KRP | 0,15924222 | 1,116700433 | 0,50271771 | 0,92068224 | fructosamine 3 kinase related protein |
| ENSG00000112640 | PPP2R5D | 0,159188224 | 1,116658639 | 0,47904929 | 0,912714933 | protein phosphatase 2 regulatory subunit B'delta |
| ENSG00000250874 | AC010595,1 | 0,159151465 | 1,116630188 | 0,8032085 | 0,980866933 | novel transcript |
| ENSG00000100104 | SRRD | 0,159073504 | 1,116569849 | 0,53264749 | 0,931291085 | SRR1 domain containing |
| ENSG00000072042 | RDH11 | 0,159067425 | 1,116565144 | 0,65923342 | 0,958394501 | retinol dehydrogenase 11 |
| ENSG00000215838 | AL451074,1 | 0,159049454 | 1,116551236 | 0,88455144 | 0,990717729 | carbonic anhydrase XIV (CA14) pseudogene |
| ENSG00000142330 | CAPN10 | 0,158988886 | 1,11650436 | 0,75120154 | 0,975901808 | calpain 10 |

| | | | | | | |
|---|---|---|---|---|---|---|
| ENSG00000174353 | STAG3L3 | 0,158940798 | 1,116467146 | 0,63939703 | 0,956018518 | stromal antigen 3-like 3 (pseudogene) |
| ENSG00000237493 | AC034102,1 | 0,158891503 | 1,116428998 | 0,79959252 | 0,980161849 | RAB13 member RAS oncogene family pseudogene |
| ENSG00000198590 | C3orf35 | 0,158804076 | 1,116361345 | 0,77467411 | 0,978553937 | chromosome 3 open reading frame 35 |
| ENSG00000235120 | BSN-AS1 | 0,158750619 | 1,116319981 | 0,88664829 | 0,990717729 | BSN antisense RNA 1 |
| ENSG00000143842 | SOX13 | 0,158743689 | 1,116314619 | 0,69852981 | 0,968522643 | SRY-box transcription factor 13 |
| ENSG00000137312 | FLOT1 | 0,158696719 | 1,116278275 | 0,56702075 | 0,941425553 | flotillin 1 |
| ENSG00000276663 | AC009090,3 | 0,158657102 | 1,116247622 | 0,80404303 | 0,980875302 | novel transcript, antisense to ARL2BP |
| ENSG00000108298 | RPL19 | 0,158555814 | 1,116169256 | 0,51841899 | 0,926651255 | ribosomal protein L19 |
| ENSG00000169018 | FEM1B | 0,15850405 | 1,116129208 | 0,67863006 | 0,9640473 | fem-1 homolog B |
| ENSG00000273562 | NAP1L4 | 0,15849953 | 1,116125711 | 0,45912712 | 0,90476838 | nucleosome assembly protein 1 like 4 |
| ENSG00000164296 | TIGD6 | 0,15846588 | 1,116099679 | 0,6063588 | 0,950927937 | tigger transposable element derived 6 |
| ENSG00000112144 | CILK1 | 0,158463356 | 1,116097726 | 0,66633884 | 0,961082184 | ciliogenesis associated kinase 1 |
| ENSG00000173275 | ZNF449 | 0,158392024 | 1,116042544 | 0,68365549 | 0,965175217 | zinc finger protein 449 |
| ENSG00000118217 | ATF6 | 0,158371407 | 1,116026595 | 0,60352283 | 0,950352286 | activating transcription factor 6 |
| ENSG00000198162 | MAN1A2 | 0,158319085 | 1,115986121 | 0,67142768 | 0,962085713 | mannosidase alpha class 1A member 2 |
| ENSG00000157353 | FCSK | 0,158247947 | 1,115931094 | 0,68541652 | 0,965475755 | fucose kinase |
| ENSG00000266885 | AC132825,4 | 0,158209454 | 1,11590132 | 0,74975126 | 0,975901808 | chromosome 17 open reading frame 51 (C17orf51) pseudogene |
| ENSG00000129646 | QRICH2 | 0,158132446 | 1,115841757 | 0,62914765 | 0,952793934 | glutamine rich 2 |
| ENSG00000135932 | CAB39 | 0,158125195 | 1,115836149 | 0,62872715 | 0,952793934 | calcium binding protein 39 |
| ENSG00000104447 | TRPS1 | 0,158030844 | 1,115763176 | 0,64816873 | 0,956899963 | transcriptional repressor GATA binding 1 |
| ENSG00000104412 | EMC2 | 0,157930532 | 1,115685599 | 0,66463254 | 0,96046247 | ER membrane protein complex subunit 2 |
| ENSG00000117480 | FAAH | 0,157914753 | 1,115673397 | 0,6859324 | 0,965740648 | fatty acid amide hydrolase |
| ENSG00000165660 | ABRAXAS2 | 0,157901 | 1,115662761 | 0,65160082 | 0,957089277 | abraxas 2, BRISC complex subunit |
| ENSG00000197724 | PHF2 | 0,157895815 | 1,115658751 | 0,47402228 | 0,910491353 | PHD finger protein 2 |
| ENSG00000286409 | AL391839,3 | 0,15788231 | 1,115648308 | 0,89887376 | 0,991832796 | novel transcript |
| ENSG00000163161 | ERCC3 | 0,157833583 | 1,115610628 | 0,46290819 | 0,906849208 | ERCC excision repair 3, TFIIH core complex helicase subunit |
| ENSG00000287723 | AP005901,6 | 0,157791759 | 1,115578286 | 0,89375053 | 0,991135494 | novel transcript |
| ENSG00000279467 | AP000350,8 | 0,157650985 | 1,115469437 | 0,93503923 | 0,994479239 | novel transcript, antisense to SLC2A11 |
| ENSG00000282986 | EIF3K | 0,157612938 | 1,11544002 | 0,46917951 | 0,907940653 | eukaryotic translation initiation factor 3 subunit K |
| ENSG00000236397 | DDX11L2 | 0,157588498 | 1,115421124 | 0,65051781 | 0,956899963 | DEAD/H-box helicase 11 like 2 (pseudogene) |
| ENSG00000261342 | AC006538,1 | 0,157557746 | 1,115397348 | 0,67863755 | 0,9640473 | novel transcript |

| | | | | | | |
|---|---|---|---|---|---|---|
| ENSG00000105323 | HNRNPUL1 | 0,15755638 | 1,115396292 | 0,50616262 | 0,921780819 | heterogeneous nuclear ribonucleoprotein U like 1 |
| ENSG00000151287 | TEX30 | 0,157538908 | 1,115382784 | 0,72501332 | 0,97326334 | testis expressed 30 |
| ENSG00000140694 | PARN | 0,157535856 | 1,115380424 | 0,50665692 | 0,921824694 | poly(A)-specific ribonuclease |
| ENSG00000254051 | AC011853,1 | 0,157501654 | 1,115353982 | 0,84964942 | 0,987036295 | NADH dehydrogenase [ubiquinone] flavoprotein 2, mitochondrial (NDUFV2) pseudogene |
| ENSG00000147113 | DIPK2B | 0,157499636 | 1,115352422 | 0,84956175 | 0,987036295 | divergent protein kinase domain 2B |
| ENSG00000178188 | SH2B1 | 0,157430877 | 1,115299265 | 0,64281608 | 0,956633898 | SH2B adaptor protein 1 |
| ENSG00000198468 | FLVCR1-DT | 0,157396353 | 1,115272576 | 0,67703269 | 0,963580269 | FLVCR1 divergent transcript |
| ENSG00000279463 | AL159972,1 | 0,157381856 | 1,11526137 | 0,86881867 | 0,989074806 | novel transcript, sense intronic to KLF12 |
| ENSG00000260394 | LINC02867 | 0,157377886 | 1,1152583 | 0,90720775 | 0,992590661 | long intergenic non-protein coding RNA 2867 |
| ENSG00000245105 | A2M-AS1 | 0,157206997 | 1,115126205 | 0,70288303 | 0,968742626 | A2M antisense RNA 1 |
| ENSG00000137818 | RPLP1 | 0,157188125 | 1,115111618 | 0,55396515 | 0,937524781 | ribosomal protein lateral stalk subunit P1 |
| ENSG00000068724 | TTC7A | 0,157178259 | 1,115103992 | 0,6439041 | 0,956633898 | tetratricopeptide repeat domain 7A |
| ENSG00000071054 | MAP4K4 | 0,15716826 | 1,115096264 | 0,49890704 | 0,919683359 | mitogen-activated protein kinase kinase kinase kinase 4 |
| ENSG00000119446 | RBM18 | 0,157082096 | 1,115029667 | 0,70576871 | 0,969503793 | RNA binding motif protein 18 |
| ENSG00000259954 | IL21R-AS1 | 0,157031969 | 1,114990926 | 0,84322558 | 0,986218419 | IL21R antisense RNA 1 |
| ENSG00000279712 | FP325330,3 | 0,15690599 | 1,114893567 | 0,80269898 | 0,980832528 | novel transcript |
| ENSG00000095319 | NUP188 | 0,156905735 | 1,11489337 | 0,43566446 | 0,896422069 | nucleoporin 188 |
| ENSG00000218189 | POM121L14P | 0,156840342 | 1,114842836 | 0,83860568 | 0,985672131 | POM121 transmembrane nucleoporin like 14, pseudogene |
| ENSG00000249028 | AC004672,1 | 0,156651931 | 1,114697251 | 0,82071622 | 0,982609114 | novel transcript |
| ENSG00000163632 | C3orf49 | 0,156580252 | 1,11464187 | 0,85045095 | 0,987075267 | chromosome 3 open reading frame 49 |
| ENSG00000115866 | DARS1 | 0,156571543 | 1,114635141 | 0,67723573 | 0,963580269 | aspartyl-tRNA synthetase 1 |
| ENSG00000266554 | LINC01443 | 0,156482241 | 1,114566148 | 0,88494087 | 0,990717729 | long intergenic non-protein coding RNA 1443 |
| ENSG00000179364 | PACS2 | 0,156384129 | 1,114490353 | 0,61060241 | 0,951841373 | phosphofurin acidic cluster sorting protein 2 |
| ENSG00000234345 | ELF2P1 | 0,156365058 | 1,114475621 | 0,71482634 | 0,971041579 | ELF2 pseudogene 1 |
| ENSG00000204814 | AL935212,1 | 0,156336529 | 1,114453582 | 0,90328029 | 0,992311172 | novel transcript |
| ENSG00000150316 | CWC15 | 0,156322907 | 1,11444306 | 0,593387 | 0,947416795 | CWC15 spliceosome associated protein homolog |
| ENSG00000276077 | CU633904,1 | 0,156162859 | 1,114319434 | 0,64717259 | 0,956899963 | uncharacterized LOC102724951 |
| ENSG00000161542 | PRPSAP1 | 0,156162772 | 1,114319367 | 0,53306355 | 0,931291085 | phosphoribosyl pyrophosphate synthetase associated protein 1 |
| ENSG00000144868 | TMEM108 | 0,156109767 | 1,114278427 | 0,56571443 | 0,940866081 | transmembrane protein 108 |

| | | | | | | |
|---|---|---|---|---|---|---|
| ENSG00000173727 | AP000769,1 | 0,156079222 | 1,114254836 | 0,76551998 | 0,97683946 | Finkel-Biskis-Reilly murine sarcoma virus (FBR-MuSV) ubiquitously expressed (FAU) pseudogene |
| ENSG00000128159 | TUBGCP6 | 0,156054627 | 1,11423584 | 0,67033177 | 0,962026636 | tubulin gamma complex associated protein 6 |
| ENSG00000270264 | NDUFB8P2 | 0,156034816 | 1,114220539 | 0,87833712 | 0,990109814 | NADH:ubiquinone oxidoreductase subunit B8 pseudogene 2 |
| ENSG00000106009 | BRAT1 | 0,15601814 | 1,114207661 | 0,70516924 | 0,969413904 | BRCA1 associated ATM activator 1 |
| ENSG00000285036 | DHRS4L1 | 0,15595225 | 1,114156774 | 0,86533603 | 0,989074806 | dehydrogenase/reductase 4 like 1 |
| ENSG00000225766 | DHRS4L1 | 0,15595225 | 1,114156774 | 0,86533603 | 0,989074806 | dehydrogenase/reductase 4 like 1 |
| ENSG00000153094 | BCL2L11 | 0,155909011 | 1,114123383 | 0,67041528 | 0,962026636 | BCL2 like 11 |
| ENSG00000236255 | AC009404,1 | 0,155884526 | 1,114104474 | 0,63859043 | 0,956018518 | novel transcript |
| ENSG00000178105 | DDX10 | 0,155862133 | 1,114087181 | 0,51007918 | 0,922427526 | DEAD-box helicase 10 |
| ENSG00000105887 | MTPN | 0,15584909 | 1,114077109 | 0,66147856 | 0,959582738 | myotrophin |
| ENSG00000288089 | AC011303,1 | 0,155835817 | 1,11406686 | 0,88597765 | 0,990717729 | novel transcript |
| ENSG00000149806 | FAU | 0,1558318 | 1,114063758 | 0,54511346 | 0,934923038 | FAU ubiquitin like and ribosomal protein S30 fusion |
| ENSG00000277443 | MARCKS | 0,155804993 | 1,114043057 | 0,43385434 | 0,896272827 | myristoylated alanine rich protein kinase C substrate |
| ENSG00000198677 | TTC37 | 0,155789326 | 1,114030959 | 0,6948361 | 0,967417311 | tetratricopeptide repeat domain 37 |
| ENSG00000165684 | SNAPC4 | 0,155711381 | 1,113970772 | 0,72562037 | 0,97326334 | small nuclear RNA activating complex polypeptide 4 |
| ENSG00000168386 | FILIP1L | 0,15570892 | 1,113968872 | 0,46123235 | 0,906165604 | filamin A interacting protein 1 like |
| ENSG00000228522 | AL845321,1 | 0,155692079 | 1,113955868 | 0,83563828 | 0,985034063 | methylenetetrahydrofolate dehydrogenase (NADP+ dependent) 1-like (MTHFD1L) pseudogene |
| ENSG00000109738 | GLRB | 0,155655287 | 1,113927461 | 0,73192315 | 0,974353474 | glycine receptor beta |
| ENSG00000156110 | ADK | 0,155556163 | 1,113850928 | 0,59325241 | 0,947416795 | adenosine kinase |
| ENSG00000183520 | UTP11 | 0,155509629 | 1,113815001 | 0,60647245 | 0,950927937 | UTP11 small subunit processome component |
| ENSG00000129460 | NGDN | 0,155500457 | 1,11380792 | 0,51005361 | 0,922427526 | neuroguidin |
| ENSG00000264456 | AC138207,4 | 0,155463234 | 1,113779184 | 0,76238171 | 0,976315588 | novel transcript, sense intronic RNF135 |
| ENSG00000171456 | ASXL1 | 0,155396021 | 1,113727295 | 0,45709732 | 0,90378766 | ASXL transcriptional regulator 1 |
| ENSG00000284747 | AL034417,4 | 0,155382233 | 1,113716651 | 0,76866658 | 0,977381742 | novel transcript, antisense to ERRFI1 |
| ENSG00000075702 | WDR62 | 0,155345788 | 1,113688517 | 0,66988296 | 0,962026636 | WD repeat domain 62 |
| ENSG00000285184 | AC244033,2 | 0,155314704 | 1,113664522 | 0,68194459 | 0,964654539 | novel transcript |
| ENSG00000117475 | BLZF1 | 0,155261383 | 1,113623363 | 0,67735863 | 0,963580269 | basic leucine zipper nuclear factor 1 |
| ENSG00000260282 | EIF4EBP2P2 | 0,155127198 | 1,113519789 | 0,91989311 | 0,993758187 | eukaryotic translation initiation factor 4E binding protein 2 pseudogene 2 |
| ENSG00000083720 | OXCT1 | 0,155118502 | 1,113513078 | 0,62431255 | 0,952793934 | 3-oxoacid CoA-transferase 1 |

| | | | | | | |
|---|---|---|---|---|---|---|
| ENSG00000172296 | SPTLC3 | 0,15497898 | 1,113405396 | 0,68345628 | 0,965068292 | serine palmitoyltransferase long chain base subunit 3 |
| ENSG00000164976 | MYORG | 0,154965943 | 1,113395334 | 0,47322302 | 0,909724186 | myogenesis regulating glycosidase (putative) |
| ENSG00000226763 | SRRM5 | 0,154905698 | 1,113348842 | 0,79940553 | 0,980161849 | serine/arginine repetitive matrix 5 |
| ENSG00000171475 | WIPF2 | 0,154853959 | 1,113308915 | 0,48139289 | 0,912981134 | WAS/WASL interacting protein family member 2 |
| ENSG00000156299 | TIAM1 | 0,154818602 | 1,11328163 | 0,5722565 | 0,942475308 | TIAM Rac1 associated GEF 1 |
| ENSG00000237499 | WAKMAR2 | 0,15478231 | 1,113253626 | 0,7461754 | 0,975758222 | wound and keratinocyte migration associated lncRNA 2 |
| ENSG00000196230 | TUBB | 0,154741983 | 1,113222508 | 0,45367097 | 0,902908855 | tubulin beta class I |
| ENSG00000125458 | NT5C | 0,154734312 | 1,113216588 | 0,55057951 | 0,936249395 | 5', 3'-nucleotidase, cytosolic |
| ENSG00000112039 | FANCE | 0,154695006 | 1,113186259 | 0,55554063 | 0,937793851 | FA complementation group E |
| ENSG00000146066 | HIGD2A | 0,154620928 | 1,113129102 | 0,58220303 | 0,944174844 | HIG1 hypoxia inducible domain family member 2A |
| ENSG00000288398 | AL109627,1 | 0,154614051 | 1,113123796 | 0,74530027 | 0,9756851 | novel transcript |
| ENSG00000239569 | KMT2E-AS1 | 0,154613071 | 1,11312304 | 0,71892513 | 0,971869926 | KMT2E antisense RNA 1 |
| ENSG00000118600 | RXYLT1 | 0,15460875 | 1,113119706 | 0,64568667 | 0,956899963 | ribitol xylosyltransferase 1 |
| ENSG00000281540 | RCC2 | 0,154605055 | 1,113116855 | 0,46783636 | 0,907940653 | regulator of chromosome condensation 2 |
| ENSG00000179051 | RCC2 | 0,154605055 | 1,113116855 | 0,46783636 | 0,907940653 | regulator of chromosome condensation 2 |
| ENSG00000092203 | TOX4 | 0,154575026 | 1,113093686 | 0,55943219 | 0,939727587 | TOX high mobility group box family member 4 |
| ENSG00000133313 | CNDP2 | 0,154564842 | 1,113085829 | 0,45268841 | 0,902285042 | carnosine dipeptidase 2 |
| ENSG00000133773 | CCDC59 | 0,154512663 | 1,113045572 | 0,70662862 | 0,969503793 | coiled-coil domain containing 59 |
| ENSG00000183665 | TRMT12 | 0,154490719 | 1,113028642 | 0,48984609 | 0,916063942 | tRNA methyltransferase 12 homolog |
| ENSG00000203635 | AC144450,1 | 0,154485838 | 1,113024877 | 0,83350636 | 0,984905093 | novel transcript |
| ENSG00000135272 | MDFIC | 0,154439419 | 1,112989065 | 0,65193208 | 0,957216199 | MyoD family inhibitor domain containing |
| ENSG00000176410 | DNAJC30 | 0,15438912 | 1,112950262 | 0,5195171 | 0,926800715 | DnaJ heat shock protein family (Hsp40) member C30 |
| ENSG00000215193 | PEX26 | 0,154344909 | 1,112916157 | 0,57167366 | 0,942436428 | peroxisomal biogenesis factor 26 |
| ENSG00000187664 | HAPLN4 | 0,154196267 | 1,112801498 | 0,79035541 | 0,979462318 | hyaluronan and proteoglycan link protein 4 |
| ENSG00000215190 | LINC00680 | 0,154064908 | 1,112700181 | 0,53608872 | 0,932162269 | long intergenic non-protein coding RNA 680 |
| ENSG00000285070 | ADIPOR2 | 0,154062707 | 1,112698483 | 0,61491082 | 0,952793934 | adiponectin receptor 2 |
| ENSG00000006831 | ADIPOR2 | 0,154062707 | 1,112698483 | 0,61491082 | 0,952793934 | adiponectin receptor 2 |
| ENSG00000288332 | PIMREGP3 | 0,154047639 | 1,112686862 | 0,87228733 | 0,989217334 | PIMREG pseudogene 3 |
| ENSG00000217653 | PIMREGP3 | 0,154047639 | 1,112686862 | 0,87228733 | 0,989217334 | PIMREG pseudogene 3 |
| ENSG00000213443 | NPM1P7 | 0,154015743 | 1,112662262 | 0,79961991 | 0,980161849 | nucleophosmin 1 pseudogene 7 |
| ENSG00000232489 | MFAP1P1 | 0,154007066 | 1,11265557 | 0,8776256 | 0,990109814 | microfibril associated protein 1 pseudogene 1 |

| | | | | | | |
|---|---|---|---|---|---|---|
| ENSG00000171722 | SPATA46 | 0,153988559 | 1,112641297 | 0,77276411 | 0,978310326 | spermatogenesis associated 46 |
| ENSG00000285664 | AL139317,5 | 0,153946855 | 1,112609135 | 0,67189506 | 0,962171454 | novel transcript, antisense to FERMT2 |
| ENSG00000072518 | MARK2 | 0,153820196 | 1,112511459 | 0,53739107 | 0,932883277 | microtubule affinity regulating kinase 2 |
| ENSG00000186522 | SEPTIN10 | 0,153815046 | 1,112507487 | 0,6900682 | 0,967053457 | septin 10 |
| ENSG00000285580 | CR769776,3 | 0,153776934 | 1,112478099 | 0,93471262 | 0,994479239 | Novel transcript |
| ENSG00000287913 | AL132986,1 | 0,153767418 | 1,112470761 | 0,86509892 | 0,989074806 | novel transcript |
| ENSG00000008838 | MED24 | 0,153712843 | 1,112428678 | 0,53892025 | 0,933348247 | mediator complex subunit 24 |
| ENSG00000263165 | AC087292,1 | 0,153677615 | 1,112401515 | 0,77699424 | 0,978587255 | novel transcript, antisense to ANKFY1 |
| ENSG00000137522 | RNF121 | 0,153663224 | 1,112390419 | 0,46851552 | 0,907940653 | ring finger protein 121 |
| ENSG00000063244 | U2AF2 | 0,153607212 | 1,112347232 | 0,61422685 | 0,952775398 | U2 small nuclear RNA auxiliary factor 2 |
| ENSG00000234136 | AC055764,1 | 0,153550505 | 1,11230351 | 0,88278109 | 0,990717729 | novel transcript |
| ENSG00000280495 | TRAPPC4 | 0,1535467 | 1,112300577 | 0,55205298 | 0,936552095 | trafficking protein particle complex 4 |
| ENSG00000218226 | TATDN2P2 | 0,153484458 | 1,11225259 | 0,71638061 | 0,971438758 | TatD DNase domain containing 2 pseudogene 2 |
| ENSG00000241219 | AC078785,2 | 0,153483314 | 1,112251708 | 0,8348786 | 0,984905093 | novel transcript |
| ENSG00000135766 | EGLN1 | 0,153438523 | 1,112217177 | 0,59140289 | 0,947416795 | egl-9 family hypoxia inducible factor 1 |
| ENSG00000047056 | WDR37 | 0,153310813 | 1,112118726 | 0,47123737 | 0,908869685 | WD repeat domain 37 |
| ENSG00000158411 | MITD1 | 0,153299911 | 1,112110322 | 0,68129235 | 0,964654539 | microtubule interacting and trafficking domain containing 1 |
| ENSG00000174109 | C16orf91 | 0,15317197 | 1,112011702 | 0,65540136 | 0,957444211 | chromosome 16 open reading frame 91 |
| ENSG00000278901 | AC005609,1 | 0,153162383 | 1,112004312 | 0,81970321 | 0,98249617 | novel transcript, antisense to PCDHA13 |
| ENSG00000117862 | TXNDC12 | 0,153149161 | 1,111994121 | 0,58998935 | 0,947252221 | thioredoxin domain containing 12 |
| ENSG00000182968 | SOX1 | 0,1531012 | 1,111957154 | 0,47021348 | 0,908528529 | SRY-box transcription factor 1 |
| ENSG00000260738 | LINC00554 | 0,15307644 | 1,111938071 | 0,87223048 | 0,989217334 | long intergenic non-protein coding RNA 554 |
| ENSG00000172071 | EIF2AK3 | 0,153074318 | 1,111936436 | 0,63470624 | 0,955149453 | eukaryotic translation initiation factor 2 alpha kinase 3 |
| ENSG00000284976 | BX255925,3 | 0,153062349 | 1,111927211 | 0,73790601 | 0,974353474 | novel RING finger protein |
| ENSG00000007202 | KIAA0100 | 0,153050735 | 1,111918259 | 0,44919868 | 0,902062594 | KIAA0100 |
| ENSG00000261126 | RBFADN | 0,153049109 | 1,111917006 | 0,80332535 | 0,980866933 | RBFA downstream neighbor |
| ENSG00000239775 | AC017116,1 | 0,153040778 | 1,111910586 | 0,91721593 | 0,993361299 | novel transcript, sense overlapping DBNL |
| ENSG00000177646 | ACAD9 | 0,15301432 | 1,111890194 | 0,51155546 | 0,922893865 | acyl-CoA dehydrogenase family member 9 |
| ENSG00000182712 | CMC4 | 0,152971079 | 1,111856868 | 0,71226103 | 0,970812834 | C-X9-C motif containing 4 |
| ENSG00000151650 | VENTX | 0,152897906 | 1,111800477 | 0,87614794 | 0,990109814 | VENT homeobox |
| ENSG00000015520 | NPC1L1 | 0,152897567 | 1,111800216 | 0,92305908 | 0,993825008 | NPC1 like intracellular cholesterol transporter 1 |

| | | | | | | |
|---|---|---|---|---|---|---|
| ENSG00000125454 | SLC25A19 | 0,152810863 | 1,1117334 | 0,64406132 | 0,956633898 | solute carrier family 25 member 19 |
| ENSG00000248206 | AC074194,1 | 0,152719917 | 1,11166332 | 0,7079478 | 0,969992117 | novel transcript |
| ENSG00000197223 | C1D | 0,152695556 | 1,111644549 | 0,75135396 | 0,975901808 | C1D nuclear receptor corepressor |
| ENSG00000280195 | AC245140,2 | 0,15255956 | 1,111539764 | 0,64767718 | 0,956899963 | novel transcript, antisense to RPL10 |
| ENSG00000239605 | STPG4 | 0,152536195 | 1,111521762 | 0,80036254 | 0,980237962 | sperm-tail PG-rich repeat containing 4 |
| ENSG00000145247 | OCIAD2 | 0,152524997 | 1,111513135 | 0,61448624 | 0,952793934 | OCIA domain containing 2 |
| ENSG00000250329 | KDELC1P1 | 0,152476842 | 1,111476035 | 0,81124664 | 0,981213877 | KDEL motif containing 1 pseudogene 1 |
| ENSG00000251577 | AC105460,2 | 0,152393671 | 1,111411961 | 0,91657082 | 0,993361299 | novel transcript, antisense to TACR3 |
| ENSG00000111886 | GABRR2 | 0,152389291 | 1,111408586 | 0,78075362 | 0,978597905 | gamma-aminobutyric acid type A receptor subunit rho2 |
| ENSG00000196305 | IARS1 | 0,152380997 | 1,111402197 | 0,58206789 | 0,944174844 | isoleucyl-tRNA synthetase 1 |
| ENSG00000100220 | RTCB | 0,15231227 | 1,111349253 | 0,48021007 | 0,912847708 | RNA 2',3'-cyclic phosphate and 5'-OH ligase |
| ENSG00000212807 | OR2A42 | 0,152213811 | 1,11127341 | 0,71457452 | 0,971041579 | olfactory receptor family 2 subfamily A member 42 |
| ENSG00000266094 | RASSF5 | 0,152152881 | 1,111226478 | 0,76562791 | 0,976862445 | Ras association domain family member 5 |
| ENSG00000067365 | METTL22 | 0,152062034 | 1,111156506 | 0,60903648 | 0,951793902 | methyltransferase like 22 |
| ENSG00000008018 | PSMB1 | 0,152043206 | 1,111142004 | 0,5725699 | 0,942475308 | proteasome 20S subunit beta 1 |
| ENSG00000258485 | SRMP2 | 0,151945938 | 1,111067093 | 0,77979421 | 0,978587255 | SRM pseudogene 2 |
| ENSG00000101082 | SLA2 | 0,151945313 | 1,111066612 | 0,79256827 | 0,979471509 | Src like adaptor 2 |
| ENSG00000284842 | FAM91A2P | 0,151904295 | 1,111035022 | 0,87031842 | 0,989074806 | family with sequence similarity 91 member A2, pseudogene |
| ENSG00000139842 | CUL4A | 0,151875811 | 1,111013088 | 0,50577412 | 0,921780819 | cullin 4A |
| ENSG00000144426 | NBEAL1 | 0,151839801 | 1,110985356 | 0,65073123 | 0,956899963 | neurobeachin like 1 |
| ENSG00000100099 | HPS4 | 0,151736787 | 1,11090603 | 0,46756653 | 0,907940653 | HPS4 biogenesis of lysosomal organelles complex 3 subunit 2 |
| ENSG00000121039 | RDH10 | 0,151724379 | 1,110896476 | 0,56016131 | 0,940070839 | retinol dehydrogenase 10 |
| ENSG00000169021 | UQCRFS1 | 0,151658516 | 1,110845762 | 0,55100239 | 0,936366211 | ubiquinol-cytochrome c reductase, Rieske iron-sulfur polypeptide 1 |
| ENSG00000183747 | ACSM2A | 0,151617568 | 1,110814233 | 0,81190811 | 0,981213877 | acyl-CoA synthetase medium chain family member 2A |
| ENSG00000116691 | MIIP | 0,151615311 | 1,110812495 | 0,69998808 | 0,968522643 | migration and invasion inhibitory protein |
| ENSG00000267696 | ERVK-28 | 0,151557576 | 1,110768043 | 0,80820721 | 0,981213877 | endogenous retrovirus group K member 28 |
| ENSG00000182324 | KCNJ14 | 0,151399298 | 1,110646188 | 0,73920078 | 0,974715504 | potassium inwardly rectifying channel subfamily J member 14 |

| | | | | | | |
|---|---|---|---|---|---|---|
| ENSG00000271755 | AL031118,1 | 0,151341743 | 1,11060188 | 0,75394223 | 0,975901808 | novel transcript |
| ENSG00000157538 | VPS26C | 0,151247327 | 1,1105292 | 0,57003495 | 0,942251611 | VPS26 endosomal protein sorting factor C |
| ENSG00000148384 | INPP5E | 0,151203894 | 1,110495767 | 0,63645993 | 0,955547532 | inositol polyphosphate-5-phosphatase E |
| ENSG00000268015 | AC010320,2 | 0,151159273 | 1,110461422 | 0,90578503 | 0,992590661 | novel transcript, antisense to PPP2R1A |
| ENSG00000176994 | SMCR8 | 0,151106369 | 1,110420702 | 0,93072797 | 0,994405002 | SMCR8-C9orf72 complex subunit |
| ENSG00000270839 | AC091393,1 | 0,151088315 | 1,110406805 | 0,8617489 | 0,989074806 | nuclear transcription factor Y, beta (NFYB) pseudogene |
| ENSG00000164924 | YWHAZ | 0,151015829 | 1,110351016 | 0,70071782 | 0,968522643 | tyrosine 3-monooxygenase/tryptophan 5-monooxygenase activation protein zeta |
| ENSG00000102119 | EMD | 0,150966103 | 1,110312746 | 0,50379633 | 0,921200643 | emerin |
| ENSG00000183654 | MARCHF11 | 0,150920466 | 1,110277624 | 0,86701825 | 0,989074806 | membrane associated ring-CH-type finger 11 |
| ENSG00000079277 | MKNK1 | 0,150877132 | 1,110244275 | 0,52594086 | 0,929088883 | MAPK interacting serine/threonine kinase 1 |
| ENSG00000163961 | RNF168 | 0,150792442 | 1,110179102 | 0,53433503 | 0,931584152 | ring finger protein 168 |
| ENSG00000267734 | AL121989,1 | 0,150775509 | 1,110166072 | 0,80135179 | 0,980557464 | novel transcript, antisense to HS2ST1 |
| ENSG00000184599 | TAFA3 | 0,150709565 | 1,110115329 | 0,8661764 | 0,989074806 | TAFA chemokine like family member 3 |
| ENSG00000169905 | TOR1AIP2 | 0,150674443 | 1,110088304 | 0,56551421 | 0,940865625 | torsin 1A interacting protein 2 |
| ENSG00000259049 | AL139317,3 | 0,150664272 | 1,110080478 | 0,93536306 | 0,994479239 | novel transcript, antisense to GNPNAT1 |
| ENSG00000236895 | GTF2H4 | 0,150636482 | 1,110059095 | 0,74583378 | 0,975707496 | general transcription factor IIH subunit 4 |
| ENSG00000275066 | SYNRG | 0,150607244 | 1,110036599 | 0,65562716 | 0,957444211 | synergin gamma |
| ENSG00000157954 | WIPI2 | 0,150570657 | 1,110008448 | 0,51277686 | 0,923640376 | WD repeat domain, phosphoinositide interacting 2 |
| ENSG00000187792 | ZNF70 | 0,150471797 | 1,109932388 | 0,60387837 | 0,950551453 | zinc finger protein 70 |
| ENSG00000131373 | HACL1 | 0,15029684 | 1,109797794 | 0,60854223 | 0,951511094 | 2-hydroxyacyl-CoA lyase 1 |
| ENSG00000239463 | HLA-DMA | 0,150285559 | 1,109789116 | 0,77919803 | 0,978587255 | major histocompatibility complex, class II, DM alpha |
| ENSG00000243215 | HLA-DMA | 0,150285559 | 1,109789116 | 0,77919803 | 0,978587255 | major histocompatibility complex, class II, DM alpha |
| ENSG00000243189 | HLA-DMA | 0,150285559 | 1,109789116 | 0,77919803 | 0,978587255 | major histocompatibility complex, class II, DM alpha |
| ENSG00000242685 | HLA-DMA | 0,150285559 | 1,109789116 | 0,77919803 | 0,978587255 | major histocompatibility complex, class II, DM alpha |
| ENSG00000243719 | HLA-DMA | 0,150285559 | 1,109789116 | 0,77919803 | 0,978587255 | major histocompatibility complex, class II, DM alpha |
| ENSG00000196683 | TOMM7 | 0,150237831 | 1,109752402 | 0,50951895 | 0,922427526 | translocase of outer mitochondrial membrane 7 |
| ENSG00000155111 | CDK19 | 0,150223133 | 1,109741096 | 0,69297207 | 0,967271667 | cyclin dependent kinase 19 |
| ENSG00000230069 | LRRC37A15P | 0,150208565 | 1,10972989 | 0,88516616 | 0,990717729 | leucine rich repeat containing 37 member A15, pseudogene |
| ENSG00000171490 | RSL1D1 | 0,150157699 | 1,109690764 | 0,51484045 | 0,924720046 | ribosomal L1 domain containing 1 |
| ENSG00000203722 | RAET1G | 0,150120163 | 1,109661893 | 0,88961258 | 0,990717729 | retinoic acid early transcript 1G |

| | | | | | | |
|---|---|---|---|---|---|---|
| ENSG00000225087 | AL583808,1 | 0,150095236 | 1,10964272 | 0,7556864 | 0,975901808 | novel transcript |
| ENSG00000101457 | DNTTIP1 | 0,150091436 | 1,109639797 | 0,47223749 | 0,909171312 | deoxynucleotidyltransferase terminal interacting protein 1 |
| ENSG00000221949 | LINC01465 | 0,150074071 | 1,109626441 | 0,75990656 | 0,976197503 | long intergenic non-protein coding RNA 1465 |
| ENSG00000132881 | CPLANE2 | 0,150054401 | 1,109611312 | 0,75426783 | 0,975901808 | ciliogenesis and planar polarity effector 2 |
| ENSG00000164106 | SCRG1 | 0,150042804 | 1,109602393 | 0,7113408 | 0,970635643 | stimulator of chondrogenesis 1 |
| ENSG00000166441 | RPL27A | 0,15003577 | 1,109596983 | 0,55984109 | 0,939880787 | ribosomal protein L27a |
| ENSG00000104671 | DCTN6 | 0,149995751 | 1,109566204 | 0,75232644 | 0,975901808 | dynactin subunit 6 |
| ENSG00000185651 | UBE2L3 | 0,14997345 | 1,109549052 | 0,62750888 | 0,952793934 | ubiquitin conjugating enzyme E2 L3 |
| ENSG00000163162 | RNF149 | 0,149950829 | 1,109531655 | 0,53287974 | 0,931291085 | ring finger protein 149 |
| ENSG00000102317 | RBM3 | 0,149813804 | 1,109426279 | 0,56130742 | 0,940567693 | RNA binding motif protein 3 |
| ENSG00000230896 | AL604028,1 | 0,149803944 | 1,109418697 | 0,79886776 | 0,980161849 | novel transcript |
| ENSG00000198912 | C1orf174 | 0,149803785 | 1,109418574 | 0,50441071 | 0,921400408 | chromosome 1 open reading frame 174 |
| ENSG00000237854 | LINC00674 | 0,149702768 | 1,109340896 | 0,7130417 | 0,970812834 | long intergenic non-protein coding RNA 674 |
| ENSG00000286764 | AC188617,2 | 0,149683736 | 1,109326262 | 0,9158924 | 0,993361299 | novel transcript |
| ENSG00000223799 | IL10RB-DT | 0,149612537 | 1,109271516 | 0,64954175 | 0,956899963 | IL10RB divergent transcript |
| ENSG00000274541 | AL121955,1 | 0,149596995 | 1,109259566 | 0,92640057 | 0,993915537 | siah E3 ubiquitin protein ligase family member 3 (SIAH3) pseudogene |
| ENSG00000197016 | ZNF470 | 0,149540086 | 1,109215811 | 0,63568329 | 0,955389018 | zinc finger protein 470 |
| ENSG00000114209 | PDCD10 | 0,149532432 | 1,109209926 | 0,74347279 | 0,975310206 | programmed cell death 10 |
| ENSG00000064545 | TMEM161A | 0,149406555 | 1,10911315 | 0,70467695 | 0,969219816 | transmembrane protein 161A |
| ENSG00000240731 | AL139287,1 | 0,149388926 | 1,109099598 | 0,80022468 | 0,980161849 | novel transcript |
| ENSG00000164484 | TMEM200A | 0,149357184 | 1,109075195 | 0,59689137 | 0,948282297 | transmembrane protein 200A |
| ENSG00000277581 | AL023803,3 | 0,149333321 | 1,109056851 | 0,86111296 | 0,989074806 | novel transcript, sense intronic to DHX35 |
| ENSG00000127325 | BEST3 | 0,14933198 | 1,10905582 | 0,89843037 | 0,991832796 | bestrophin 3 |
| ENSG00000083838 | ZNF446 | 0,149318361 | 1,109045351 | 0,60932468 | 0,951793902 | zinc finger protein 446 |
| ENSG00000117410 | ATP6V0B | 0,14923106 | 1,108978242 | 0,53457986 | 0,93162317 | ATPase H+ transporting V0 subunit b |
| ENSG00000165630 | PRPF18 | 0,149179515 | 1,108938621 | 0,64262518 | 0,956633898 | pre-mRNA processing factor 18 |
| ENSG00000219529 | AP000580,1 | 0,149108041 | 1,108883683 | 0,79242353 | 0,979471509 | ribosomal protein S20 (RPS20) pseudogene |
| ENSG00000214087 | ARL16 | 0,149089868 | 1,108869715 | 0,49128132 | 0,916995341 | ADP ribosylation factor like GTPase 16 |
| ENSG00000166391 | MOGAT2 | 0,14907812 | 1,108860685 | 0,9252156 | 0,993915537 | monoacylglycerol O-acyltransferase 2 |
| ENSG00000229719 | MIR194-2HG | 0,149044482 | 1,108834832 | 0,85953719 | 0,989074806 | MIR194-2 host gene |
| ENSG00000133703 | KRAS | 0,148921147 | 1,108740042 | 0,76051047 | 0,97624106 | KRAS proto-oncogene, GTPase |

| | | | | | | |
|---|---|---|---|---|---|---|
| ENSG00000168286 | THAP11 | 0,148919123 | 1,108738486 | 0,52824436 | 0,92998848 | THAP domain containing 11 |
| ENSG00000283924 | TNPO1P2 | 0,148886279 | 1,108713246 | 0,90138935 | 0,992041918 | transportin 1 pseudogene 2 |
| ENSG00000230662 | TNPO1P2 | 0,148886279 | 1,108713246 | 0,90138935 | 0,992041918 | transportin 1 pseudogene 2 |
| ENSG00000184319 | RPL23AP82 | 0,148866278 | 1,108697875 | 0,65427108 | 0,957444211 | ribosomal protein L23a pseudogene 82 |
| ENSG00000154144 | TBRG1 | 0,148863688 | 1,108695884 | 0,57936054 | 0,943828882 | transforming growth factor beta regulator 1 |
| ENSG00000256417 | AC006206,2 | 0,148840834 | 1,108678322 | 0,76768528 | 0,977309816 | novel transcript |
| ENSG00000109083 | IFT20 | 0,148790555 | 1,108639684 | 0,64646888 | 0,956899963 | intraflagellar transport 20 |
| ENSG00000198265 | HELZ | 0,148763162 | 1,108618633 | 0,66812301 | 0,961289023 | helicase with zinc finger |
| ENSG00000204564 | C6orf136 | 0,148713235 | 1,108580269 | 0,77381246 | 0,978415098 | chromosome 6 open reading frame 136 |
| ENSG00000233554 | B4GALT1-AS1 | 0,148670583 | 1,108547495 | 0,86770575 | 0,989074806 | B4GALT1 antisense RNA 1 |
| ENSG00000166908 | PIP4K2C | 0,148612121 | 1,108502575 | 0,5090909 | 0,922427526 | phosphatidylinositol-5-phosphate 4-kinase type 2 gamma |
| ENSG00000184226 | PCDH9 | 0,148310553 | 1,108270887 | 0,72674892 | 0,973344786 | protocadherin 9 |
| ENSG00000225210 | DUXAP9 | 0,148310467 | 1,108270822 | 0,90726101 | 0,992590661 | double homeobox A pseudogene 9 |
| ENSG00000166188 | ZNF319 | 0,148220853 | 1,108201983 | 0,56212484 | 0,940865625 | zinc finger protein 319 |
| ENSG00000102225 | CDK16 | 0,148183151 | 1,108173022 | 0,48685994 | 0,914460148 | cyclin dependent kinase 16 |
| ENSG00000164307 | ERAP1 | 0,148119786 | 1,108124351 | 0,62960245 | 0,952812183 | endoplasmic reticulum aminopeptidase 1 |
| ENSG00000171861 | MRM3 | 0,148101277 | 1,108110135 | 0,60007536 | 0,949340299 | mitochondrial rRNA methyltransferase 3 |
| ENSG00000012061 | ERCC1 | 0,148057625 | 1,108076606 | 0,57760919 | 0,94337651 | ERCC excision repair 1, endonuclease non-catalytic subunit |
| ENSG00000172273 | HINFP | 0,148025415 | 1,108051868 | 0,59383458 | 0,947469769 | histone H4 transcription factor |
| ENSG00000247049 | AC008443,1 | 0,147971085 | 1,108010141 | 0,92434451 | 0,993915537 | novel transcript |
| ENSG00000196850 | PPTC7 | 0,147936001 | 1,107983196 | 0,69923446 | 0,968522643 | protein phosphatase targeting COQ7 |
| ENSG00000226087 | AC073283,2 | 0,147916354 | 1,107968107 | 0,83646554 | 0,985423106 | novel transcript |
| ENSG00000241014 | AC114490,1 | 0,147873385 | 1,107935108 | 0,87526992 | 0,990078774 | novel platelet-activating factor receptor family pseudogene |
| ENSG00000120253 | NUP43 | 0,147840639 | 1,10790996 | 0,56673216 | 0,941311563 | nucleoporin 43 |
| ENSG00000039650 | PNKP | 0,147769797 | 1,107855559 | 0,65590917 | 0,957444211 | polynucleotide kinase 3'-phosphatase |
| ENSG00000173267 | SNCG | 0,147706908 | 1,107807267 | 0,70849512 | 0,970181083 | synuclein gamma |
| ENSG00000138795 | LEF1 | 0,147660628 | 1,107771731 | 0,72876266 | 0,974009551 | lymphoid enhancer binding factor 1 |
| ENSG00000214274 | ANG | 0,14756741 | 1,107700156 | 0,71250946 | 0,970812834 | angiogenin |
| ENSG00000284238 | TOP3A | 0,147564042 | 1,10769757 | 0,70822623 | 0,97011892 | DNA topoisomerase III alpha |
| ENSG00000212283 | SNORD89 | 0,147555206 | 1,107690786 | 0,79277077 | 0,979471509 | small nucleolar RNA, C/D box 89 |

| | | | | | | |
|---|---|---|---|---|---|---|
| ENSG00000230993 | RPL12P15 | 0,147539148 | 1,107678456 | 0,86151845 | 0,989074806 | ribosomal protein L12 pseudogene 15 |
| ENSG00000229320 | KRT8P12 | 0,147531358 | 1,107672475 | 0,61907962 | 0,952793934 | keratin 8 pseudogene 12 |
| ENSG00000236673 | AL117378,1 | 0,147528208 | 1,107670057 | 0,79843521 | 0,980161849 | novel transcript |
| ENSG00000107863 | ARHGAP21 | 0,147503753 | 1,107651281 | 0,65773763 | 0,957864535 | Rho GTPase activating protein 21 |
| ENSG00000167333 | TRIM68 | 0,147494475 | 1,107644157 | 0,51401483 | 0,924227101 | tripartite motif containing 68 |
| ENSG00000239789 | MRPS17 | 0,147483967 | 1,10763609 | 0,51739127 | 0,9257552 | mitochondrial ribosomal protein S17 |
| ENSG00000282034 | AC106886,5 | 0,147418149 | 1,107585559 | 0,65306533 | 0,95732962 | novel transcript |
| ENSG00000157800 | SLC37A3 | 0,147411111 | 1,107580156 | 0,57746521 | 0,94337651 | solute carrier family 37 member 3 |
| ENSG00000205302 | SNX2 | 0,14735682 | 1,107538477 | 0,70446442 | 0,969132783 | sorting nexin 2 |
| ENSG00000136444 | RSAD1 | 0,147351816 | 1,107534635 | 0,54702042 | 0,935404974 | radical S-adenosyl methionine domain containing 1 |
| ENSG00000164022 | AIMP1 | 0,147336983 | 1,107523248 | 0,59592982 | 0,948198368 | aminoacyl tRNA synthetase complex interacting multifunctional protein 1 |
| ENSG00000103148 | NPRL3 | 0,147318512 | 1,107509069 | 0,60058637 | 0,949439306 | NPR3 like, GATOR1 complex subunit |
| ENSG00000239280 | AC108693,1 | 0,147312327 | 1,107504321 | 0,79067658 | 0,979466731 | pseudogene similar to part of dihydrolipoamide branched chain transacylase E2 DBT |
| ENSG00000272720 | AL022322,1 | 0,147189577 | 1,107410094 | 0,8085704 | 0,981213877 | novel transcript, antisense to BAIAP2L2 |
| ENSG00000136943 | CTSV | 0,147145788 | 1,107376482 | 0,72019797 | 0,972142678 | cathepsin V |
| ENSG00000038002 | AGA | 0,147138135 | 1,107370608 | 0,63209188 | 0,953935145 | aspartylglucosaminidase |
| ENSG00000135845 | PIGC | 0,147088116 | 1,107332216 | 0,66394027 | 0,959985685 | phosphatidylinositol glycan anchor biosynthesis class C |
| ENSG00000259155 | STK16P1 | 0,14706997 | 1,107318288 | 0,8273883 | 0,983595244 | serine/threonine kinase 16 pseudogene 1 |
| ENSG00000274474 | YWHAE | 0,14704576 | 1,107299705 | 0,50562623 | 0,921780819 | tyrosine 3-monooxygenase/tryptophan 5-monooxygenase activation protein epsilon |
| ENSG00000126775 | ATG14 | 0,147034235 | 1,10729086 | 0,52652586 | 0,929261495 | autophagy related 14 |
| ENSG00000100603 | SNW1 | 0,147020165 | 1,107280062 | 0,4812487 | 0,912970838 | SNW domain containing 1 |
| ENSG00000276467 | Z93929,1 | 0,147016929 | 1,107277577 | 0,86566695 | 0,989074806 | zinc finger protein 831 (ZNF831) pseudogene |
| ENSG00000123358 | NR4A1 | 0,146956983 | 1,10723157 | 0,6829893 | 0,964931888 | nuclear receptor subfamily 4 group A member 1 |
| ENSG00000095485 | CWF19L1 | 0,146924542 | 1,107206672 | 0,5185687 | 0,926748138 | CWF19 like cell cycle control factor 1 |
| ENSG00000224195 | AC022400,1 | 0,146866816 | 1,107162371 | 0,86191042 | 0,989074806 | novel transcript |
| ENSG00000125962 | ARMCX5 | 0,146855364 | 1,107153582 | 0,60765503 | 0,950993594 | armadillo repeat containing X-linked 5 |
| ENSG00000276690 | AC117415,1 | 0,146825103 | 1,10713036 | 0,89677893 | 0,991644439 | |
| ENSG00000259088 | AL137779,2 | 0,146817734 | 1,107124705 | 0,8430383 | 0,986218419 | novel transcript, antisense to PPP2R5C |

| | | | | | | |
|---|---|---|---|---|---|---|
| ENSG00000169372 | CRADD | 0,146813131 | 1,107121173 | 0,60329012 | 0,950352286 | CASP2 and RIPK1 domain containing adaptor with death domain |
| ENSG00000092096 | SLC22A17 | 0,146808429 | 1,107117564 | 0,61783243 | 0,952793934 | solute carrier family 22 member 17 |
| ENSG00000225518 | LINC01703 | 0,146715969 | 1,107046613 | 0,84158321 | 0,98571999 | long intergenic non-protein coding RNA 1703 |
| ENSG00000143252 | SDHC | 0,14671448 | 1,10704547 | 0,57906668 | 0,943828882 | succinate dehydrogenase complex subunit C |
| ENSG00000272589 | ZSWIM8-AS1 | 0,146652189 | 1,106997673 | 0,82568379 | 0,983383773 | ZSWIM8 antisense RNA 1 |
| ENSG00000158864 | NDUFS2 | 0,146570654 | 1,106935112 | 0,51129125 | 0,922893865 | NADH:ubiquinone oxidoreductase core subunit S2 |
| ENSG00000070778 | PTPN21 | 0,146479913 | 1,106865491 | 0,50167586 | 0,920520138 | protein tyrosine phosphatase non-receptor type 21 |
| ENSG00000181523 | SGSH | 0,146477057 | 1,1068633 | 0,6636052 | 0,959920886 | N-sulfoglucosamine sulfohydrolase |
| ENSG00000116898 | MRPS15 | 0,146467255 | 1,106855779 | 0,55624182 | 0,938046927 | mitochondrial ribosomal protein S15 |
| ENSG00000023318 | ERP44 | 0,146455317 | 1,106846621 | 0,72477788 | 0,97326334 | endoplasmic reticulum protein 44 |
| ENSG00000249504 | PCDHA14 | 0,146306312 | 1,106732309 | 0,95770335 | 0,996608711 | protocadherin alpha 14 pseudogene |
| ENSG00000170852 | KBTBD2 | 0,146241528 | 1,106682613 | 0,6496292 | 0,956899963 | kelch repeat and BTB domain containing 2 |
| ENSG00000019485 | PRDM11 | 0,146212432 | 1,106660293 | 0,50004873 | 0,92041337 | PR/SET domain 11 |
| ENSG00000100916 | BRMS1L | 0,146200122 | 1,10665085 | 0,66126197 | 0,959582738 | BRMS1 like transcriptional repressor |
| ENSG00000147573 | TRIM55 | 0,146143949 | 1,106607763 | 0,86271295 | 0,989074806 | tripartite motif containing 55 |
| ENSG00000174405 | LIG4 | 0,146126597 | 1,106594453 | 0,72888086 | 0,974009551 | DNA ligase 4 |
| ENSG00000196507 | TCEAL3 | 0,146096408 | 1,106571298 | 0,62775873 | 0,952793934 | transcription elongation factor A like 3 |
| ENSG00000171346 | KRT15 | 0,146082563 | 1,106560678 | 0,80921254 | 0,981213877 | keratin 15 |
| ENSG00000269318 | AC007292,2 | 0,146031776 | 1,106521724 | 0,83918994 | 0,985672131 | novel transcript, antisense to MPND |
| ENSG00000178982 | EIF3K | 0,145914858 | 1,106432054 | 0,50161282 | 0,920520138 | eukaryotic translation initiation factor 3 subunit K |
| ENSG00000121900 | TMEM54 | 0,145897703 | 1,106418898 | 0,81569003 | 0,981590958 | transmembrane protein 54 |
| ENSG00000105518 | TMEM205 | 0,145760736 | 1,106313861 | 0,60447707 | 0,950813598 | transmembrane protein 205 |
| ENSG00000285253 | AC090517,4 | 0,14571038 | 1,106275247 | 0,74303469 | 0,975240824 | zinc finger protein 280D |
| ENSG00000162616 | DNAJB4 | 0,145678207 | 1,106250576 | 0,61028061 | 0,951841373 | DnaJ heat shock protein family (Hsp40) member B4 |
| ENSG00000159403 | C1R | 0,14564911 | 1,106228265 | 0,7095598 | 0,970482408 | complement C1r |
| ENSG00000134901 | POGLUT2 | 0,145610179 | 1,106198414 | 0,6686686 | 0,961375797 | protein O-glucosyltransferase 2 |
| ENSG00000175061 | SNHG29 | 0,14560268 | 1,106192665 | 0,55975577 | 0,939860604 | small nucleolar RNA host gene 29 |
| ENSG00000284896 | OR2A5 | 0,14552368 | 1,106132093 | 0,9328214 | 0,994479239 | olfactory receptor family 2 subfamily A member 5 |
| ENSG00000196417 | ZNF765 | 0,145351822 | 1,106000334 | 0,59668667 | 0,948282297 | zinc finger protein 765 |
| ENSG00000285397 | AC270273,5 | 0,14534435 | 1,105994606 | 0,89149784 | 0,990820775 | family with sequence similarity 86 member B1 |
| ENSG00000163683 | SMIM14 | 0,145327741 | 1,105981873 | 0,53264517 | 0,931291085 | small integral membrane protein 14 |
| ENSG00000259315 | ACTG1P17 | 0,145229529 | 1,105906587 | 0,81900302 | 0,98249617 | actin gamma 1 pseudogene 17 |

| | | | | | | |
|---|---|---|---|---|---|---|
| ENSG00000225364 | ATP6V0E1P1 | 0,145174237 | 1,105864203 | 0,88029294 | 0,990315537 | ATPase H+ transporting V0 subunit e1 pseudogene 1 |
| ENSG00000124459 | ZNF45 | 0,145149794 | 1,105845467 | 0,5670734 | 0,941425553 | zinc finger protein 45 |
| ENSG00000095397 | WHRN | 0,145056323 | 1,105773822 | 0,62387775 | 0,952793934 | whirlin |
| ENSG00000286950 | AC087588,2 | 0,145045479 | 1,10576551 | 0,88207665 | 0,990717729 | novel transcript, antisense to YARS2 |
| ENSG00000275493 | AL627230,1 | 0,145027078 | 1,105751407 | 0,81851273 | 0,9824772 | |
| ENSG00000185803 | SLC52A2 | 0,145019527 | 1,105745619 | 0,70216004 | 0,968742626 | solute carrier family 52 member 2 |
| ENSG00000136457 | CHAD | 0,145006477 | 1,105735617 | 0,90257387 | 0,992230188 | chondroadherin |
| ENSG00000264714 | KIAA0895LP1 | 0,144970868 | 1,105708326 | 0,87848913 | 0,990109814 | KIAA00895L pseudogene 1 |
| ENSG00000167378 | IRGQ | 0,144963657 | 1,105702799 | 0,72721739 | 0,973344786 | immunity related GTPase Q |
| ENSG00000244501 | RN7SL623P | 0,144888227 | 1,10564499 | 0,89546615 | 0,991303328 | RNA, 7SL, cytoplasmic 623, pseudogene |
| ENSG00000188566 | NDOR1 | 0,144855059 | 1,105619571 | 0,6869652 | 0,965769168 | NADPH dependent diflavin oxidoreductase 1 |
| ENSG00000236308 | AL138921,2 | 0,144793165 | 1,105572139 | 0,85095428 | 0,987259878 | novel transcript, antisense to CHUK |
| ENSG00000205209 | SCGB2B2 | 0,144758208 | 1,105545351 | 0,87068501 | 0,989102723 | secretoglobin family 2B member 2 |
| ENSG00000160014 | CALM3 | 0,144734895 | 1,105527486 | 0,48588284 | 0,913997663 | calmodulin 3 |
| ENSG00000124440 | HIF3A | 0,144691465 | 1,105494207 | 0,65535619 | 0,957444211 | hypoxia inducible factor 3 subunit alpha |
| ENSG00000185442 | FAM174B | 0,144597814 | 1,105422447 | 0,67228338 | 0,962290553 | family with sequence similarity 174 member B |
| ENSG00000165025 | SYK | 0,144517664 | 1,105361037 | 0,69706174 | 0,968488939 | spleen associated tyrosine kinase |
| ENSG00000129515 | SNX6 | 0,144473976 | 1,105327564 | 0,7165021 | 0,971438758 | sorting nexin 6 |
| ENSG00000171503 | ETFDH | 0,144389532 | 1,105262869 | 0,64234482 | 0,956633898 | electron transfer flavoprotein dehydrogenase |
| ENSG00000115317 | HTRA2 | 0,144331974 | 1,105218774 | 0,53107315 | 0,931265667 | HtrA serine peptidase 2 |
| ENSG00000095777 | MYO3A | 0,144328971 | 1,105216474 | 0,78138197 | 0,978597905 | myosin IIIA |
| ENSG00000229622 | MTND5P2 | 0,14426385 | 1,105166587 | 0,86014462 | 0,989074806 | MT-ND5 pseudogene 2 |
| ENSG00000240915 | AC120042,1 | 0,144222755 | 1,105135106 | 0,87110172 | 0,989217334 | novel transcript |
| ENSG00000074582 | BCS1L | 0,144162414 | 1,105088885 | 0,57074812 | 0,942436428 | BCS1 homolog, ubiquinol-cytochrome c reductase complex chaperone |
| ENSG00000213885 | RPL13AP7 | 0,144051572 | 1,105003984 | 0,90592274 | 0,992590661 | ribosomal protein L13a pseudogene 7 |
| ENSG00000118242 | MREG | 0,144042046 | 1,104996689 | 0,59274532 | 0,947416795 | melanoregulin |
| ENSG00000156508 | EEF1A1 | 0,144028976 | 1,104986678 | 0,58964096 | 0,947017802 | eukaryotic translation elongation factor 1 alpha 1 |
| ENSG00000244060 | RPS2P41 | 0,143816835 | 1,104824207 | 0,8696126 | 0,989074806 | ribosomal protein S2 pseudogene 41 |
| ENSG00000145868 | FBXO38 | 0,143795056 | 1,104807529 | 0,63905431 | 0,956018518 | F-box protein 38 |
| ENSG00000107736 | CDH23 | 0,143640784 | 1,104689394 | 0,76504588 | 0,976739398 | cadherin related 23 |
| ENSG00000162441 | LZIC | 0,143601239 | 1,104659115 | 0,54056804 | 0,933385624 | leucine zipper and CTNNBIP1 domain containing |
| ENSG00000125686 | MED1 | 0,143448886 | 1,104542466 | 0,64042089 | 0,956251063 | mediator complex subunit 1 |

| | | | | | | |
|---|---|---|---|---|---|---|
| ENSG00000285880 | AC144573,1 | 0,143446876 | 1,104540926 | 0,94186413 | 0,995373376 | ADRB3-GOT1L1 readthrough |
| ENSG00000284770 | TBCE | 0,143401981 | 1,104506555 | 0,66388097 | 0,959985685 | tubulin folding cofactor E |
| ENSG00000117632 | STMN1 | 0,14333255 | 1,104453401 | 0,58625022 | 0,945256674 | stathmin 1 |
| ENSG00000223597 | LINC02142 | 0,143292433 | 1,10442269 | 0,87481749 | 0,990078774 | long intergenic non-protein coding RNA 2142 |
| ENSG00000184574 | LPAR5 | 0,143290758 | 1,104421408 | 0,88305156 | 0,990717729 | lysophosphatidic acid receptor 5 |
| ENSG00000240861 | AC009153,1 | 0,143255452 | 1,10439438 | 0,90729359 | 0,992590661 | ribosomal protein S10 (RPS10) pseudogene |
| ENSG00000169740 | ZNF32 | 0,143211044 | 1,104360386 | 0,63108306 | 0,953655598 | zinc finger protein 32 |
| ENSG00000267943 | AC010328,1 | 0,143187791 | 1,104342586 | 0,75387805 | 0,975901808 | novel transcript |
| ENSG00000198887 | SMC5 | 0,143133379 | 1,104300936 | 0,64977618 | 0,956899963 | structural maintenance of chromosomes 5 |
| ENSG00000236051 | MYCBP2-AS1 | 0,143112756 | 1,104285151 | 0,86777037 | 0,989074806 | MYCBP2 antisense RNA 1 |
| ENSG00000152492 | CCDC50 | 0,142967706 | 1,104174131 | 0,70038275 | 0,968522643 | coiled-coil domain containing 50 |
| ENSG00000198246 | SLC29A3 | 0,142880565 | 1,104107439 | 0,68212992 | 0,964699524 | solute carrier family 29 member 3 |
| ENSG00000231500 | RPS18 | 0,142860295 | 1,104091926 | 0,69500373 | 0,967509655 | ribosomal protein S18 |
| ENSG00000117640 | MTFR1L | 0,142851845 | 1,104085459 | 0,53278404 | 0,931291085 | mitochondrial fission regulator 1 like |
| ENSG00000010810 | FYN | 0,142824918 | 1,104064852 | 0,59699257 | 0,948282297 | FYN proto-oncogene, Src family tyrosine kinase |
| ENSG00000158850 | B4GALT3 | 0,142803285 | 1,104048298 | 0,49586964 | 0,918105073 | beta-1,4-galactosyltransferase 3 |
| ENSG00000277496 | AL357033,4 | 0,142688497 | 1,103960457 | 0,897653 | 0,991708791 | novel transcript, antisense to SLCO4A1 |
| ENSG00000259617 | AC020661,3 | 0,142674161 | 1,103949487 | 0,90944638 | 0,992864002 | novel transcript, antisense to INO80 |
| ENSG00000185880 | TRIM69 | 0,142574311 | 1,103873085 | 0,66723987 | 0,9611116 | tripartite motif containing 69 |
| ENSG00000090686 | USP48 | 0,142522105 | 1,10383314 | 0,61951009 | 0,952793934 | ubiquitin specific peptidase 48 |
| ENSG00000229010 | AL161672,1 | 0,142477937 | 1,103799347 | 0,85917443 | 0,989074806 | novel pseudogene |
| ENSG00000089006 | SNX5 | 0,142284146 | 1,103651088 | 0,64137195 | 0,956465487 | sorting nexin 5 |
| ENSG00000259933 | AC009065,2 | 0,142256643 | 1,103630049 | 0,88226276 | 0,990717729 | novel transcript, overlapping TSC2 |
| ENSG00000197798 | FAM118B | 0,142224613 | 1,103605547 | 0,69171873 | 0,967204896 | family with sequence similarity 118 member B |
| ENSG00000020129 | NCDN | 0,142194269 | 1,103582335 | 0,69015151 | 0,967053457 | neurochondrin |
| ENSG00000255224 | AC109322,1 | 0,142070911 | 1,103487977 | 0,72421211 | 0,973178901 | TEC |
| ENSG00000253908 | AC104115,2 | 0,142064274 | 1,103482901 | 0,83127127 | 0,98438942 | zinc finger protein pseudogene |
| ENSG00000241135 | LINC00881 | 0,142025254 | 1,103453056 | 0,90760607 | 0,992590661 | long intergenic non-protein coding RNA 881 |
| ENSG00000087157 | PGS1 | 0,142017449 | 1,103447086 | 0,61638373 | 0,952793934 | phosphatidylglycerophosphate synthase 1 |
| ENSG00000147416 | ATP6V1B2 | 0,14196788 | 1,103409173 | 0,62965084 | 0,952812183 | ATPase H+ transporting V1 subunit B2 |
| ENSG00000164609 | SLU7 | 0,141876137 | 1,103339009 | 0,66864454 | 0,961375797 | SLU7 homolog, splicing factor |
| ENSG00000041880 | PARP3 | 0,141780175 | 1,103265622 | 0,6492446 | 0,956899963 | poly(ADP-ribose) polymerase family member 3 |

| | | | | | | |
|---|---|---|---|---|---|---|
| ENSG00000144647 | POMGNT2 | 0,141777317 | 1,103263436 | 0,5820566 | 0,944174844 | protein O-linked mannose N-acetylglucosaminyltransferase 2 (beta 1,4-) |
| ENSG00000172534 | HCFC1 | 0,141606399 | 1,103132738 | 0,64027899 | 0,956251063 | host cell factor C1 |
| ENSG00000274265 | AC245297,3 | 0,141604847 | 1,103131552 | 0,64461423 | 0,956633898 | novel transcript |
| ENSG00000224280 | AC005014,1 | 0,14156957 | 1,103104578 | 0,88727245 | 0,990717729 | ataxin 2-like (ATXN2L) pseudogene |
| ENSG00000232097 | AC079781,3 | 0,141552131 | 1,103091245 | 0,89796573 | 0,991711719 | novel pseudogene |
| ENSG00000125818 | PSMF1 | 0,141528209 | 1,103072954 | 0,51584935 | 0,925114101 | proteasome inhibitor subunit 1 |
| ENSG00000101004 | NINL | 0,141410706 | 1,102983115 | 0,76943324 | 0,977683747 | ninein like |
| ENSG00000175984 | DENND2C | 0,141285993 | 1,102887773 | 0,71735199 | 0,971697135 | DENN domain containing 2C |
| ENSG00000070610 | GBA2 | 0,141247698 | 1,102858498 | 0,5270858 | 0,92927181 | glucosylceramidase beta 2 |
| ENSG00000261812 | TUBB8P7 | 0,141232983 | 1,102847249 | 0,8908981 | 0,990717729 | tubulin beta 8 class VIII pseudogene 7 |
| ENSG00000086504 | MRPL28 | 0,141078456 | 1,10272913 | 0,59142571 | 0,947416795 | mitochondrial ribosomal protein L28 |
| ENSG00000131791 | PRKAB2 | 0,14104731 | 1,102705323 | 0,69463587 | 0,967417311 | protein kinase AMP-activated non-catalytic subunit beta 2 |
| ENSG00000120278 | PLEKHG1 | 0,140924461 | 1,102611429 | 0,61640639 | 0,952793934 | pleckstrin homology and RhoGEF domain containing G1 |
| ENSG00000099800 | TIMM13 | 0,140651277 | 1,102402662 | 0,65189912 | 0,957212753 | translocase of inner mitochondrial membrane 13 |
| ENSG00000188107 | EYS | 0,140621674 | 1,102380042 | 0,75663959 | 0,975901808 | eyes shut homolog |
| ENSG00000131943 | C19orf12 | 0,140502691 | 1,102289129 | 0,53234902 | 0,931291085 | chromosome 19 open reading frame 12 |
| ENSG00000005194 | CIAPIN1 | 0,140499577 | 1,10228675 | 0,49361016 | 0,917289924 | cytokine induced apoptosis inhibitor 1 |
| ENSG00000233452 | STXBP5-AS1 | 0,140451867 | 1,102250298 | 0,81033035 | 0,981213877 | STXBP5 antisense RNA 1 |
| ENSG00000174227 | PIGG | 0,140384681 | 1,102198968 | 0,52343254 | 0,928013771 | phosphatidylinositol glycan anchor biosynthesis class G |
| ENSG00000261857 | MIA | 0,140371098 | 1,10218859 | 0,87880811 | 0,990109814 | MIA SH3 domain containing |
| ENSG00000125249 | RAP2A | 0,140315684 | 1,102146256 | 0,75517678 | 0,975901808 | RAP2A, member of RAS oncogene family |
| ENSG00000111639 | MRPL51 | 0,140312672 | 1,102143955 | 0,55471641 | 0,937793851 | mitochondrial ribosomal protein L51 |
| ENSG00000197728 | RPS26 | 0,140288123 | 1,102125201 | 0,57796106 | 0,943425812 | ribosomal protein S26 |
| ENSG00000086666 | ZFAND6 | 0,140238988 | 1,102087666 | 0,74222708 | 0,975176043 | zinc finger AN1-type containing 6 |
| ENSG00000104979 | C19orf53 | 0,140197532 | 1,102055998 | 0,58259843 | 0,944174844 | chromosome 19 open reading frame 53 |
| ENSG00000129636 | ITFG1 | 0,14019352 | 1,102052933 | 0,7608159 | 0,97624106 | integrin alpha FG-GAP repeat containing 1 |
| ENSG00000282355 | CARS1-AS1 | 0,140175014 | 1,102038797 | 0,78663957 | 0,979320406 | CARS1 antisense RNA 1 |
| ENSG00000088038 | CNOT3 | 0,140112805 | 1,101991278 | 0,57311227 | 0,942475308 | CCR4-NOT transcription complex subunit 3 |
| ENSG00000118507 | AKAP7 | 0,140058104 | 1,101949496 | 0,73511776 | 0,974353474 | A-kinase anchoring protein 7 |

| | | | | | | |
|---|---|---|---|---|---|---|
| ENSG00000167645 | YIF1B | 0,139994988 | 1,101901288 | 0,61060652 | 0,951841373 | Yip1 interacting factor homolog B, membrane trafficking protein |
| ENSG00000183760 | ACP7 | 0,139992603 | 1,101899466 | 0,91065713 | 0,993222655 | acid phosphatase 7, tartrate resistant (putative) |
| ENSG00000224501 | HSPA1B | 0,139949201 | 1,101866317 | 0,77059234 | 0,977966587 | heat shock protein family A (Hsp70) member 1B |
| ENSG00000224950 | AL390066,1 | 0,139927359 | 1,101849635 | 0,83856281 | 0,985672131 | novel transcript |
| ENSG00000119655 | NPC2 | 0,139899904 | 1,101828667 | 0,66401262 | 0,959985685 | NPC intracellular cholesterol transporter 2 |
| ENSG00000136193 | SCRN1 | 0,139898145 | 1,101827324 | 0,64007091 | 0,956157624 | secernin 1 |
| ENSG00000114529 | C3orf52 | 0,139884165 | 1,101816647 | 0,82479439 | 0,983374097 | chromosome 3 open reading frame 52 |
| ENSG00000283765 | AC131160,1 | 0,139873516 | 1,101808514 | 0,78640211 | 0,979320406 | presenilin associated rhomboid like |
| ENSG00000016391 | CHDH | 0,139850388 | 1,101790851 | 0,75543079 | 0,975901808 | choline dehydrogenase |
| ENSG00000128928 | IVD | 0,13979385 | 1,101747674 | 0,63888538 | 0,956018518 | isovaleryl-CoA dehydrogenase |
| ENSG00000104731 | KLHDC4 | 0,139752984 | 1,101716466 | 0,77190528 | 0,978033855 | kelch domain containing 4 |
| ENSG00000233315 | LINC02569 | 0,139743468 | 1,101709198 | 0,89708464 | 0,991644439 | long intergenic non-protein coding RNA 2569 |
| ENSG00000232523 | LINC02569 | 0,139743468 | 1,101709198 | 0,89708464 | 0,991644439 | long intergenic non-protein coding RNA 2569 |
| ENSG00000225382 | LINC02569 | 0,139743468 | 1,101709198 | 0,89708464 | 0,991644439 | long intergenic non-protein coding RNA 2569 |
| ENSG00000236934 | LINC02569 | 0,139743468 | 1,101709198 | 0,89708464 | 0,991644439 | long intergenic non-protein coding RNA 2569 |
| ENSG00000223827 | LINC02569 | 0,139743468 | 1,101709198 | 0,89708464 | 0,991644439 | long intergenic non-protein coding RNA 2569 |
| ENSG00000152464 | RPP38 | 0,139625054 | 1,101618776 | 0,65264635 | 0,957275891 | ribonuclease P/MRP subunit p38 |
| ENSG00000122678 | POLM | 0,139602441 | 1,101601509 | 0,62939447 | 0,952812183 | DNA polymerase mu |
| ENSG00000122034 | GTF3A | 0,139530851 | 1,101546846 | 0,63604911 | 0,955399772 | general transcription factor IIIA |
| ENSG00000188690 | UROS | 0,139523811 | 1,101541471 | 0,57373352 | 0,942475308 | uroporphyrinogen III synthase |
| ENSG00000204310 | AGPAT1 | 0,139518159 | 1,101537155 | 0,59437893 | 0,947469769 | 1-acylglycerol-3-phosphate O-acyltransferase 1 |
| ENSG00000254635 | WAC-AS1 | 0,139491014 | 1,10151643 | 0,61763168 | 0,952793934 | WAC antisense RNA 1 (head to head) |
| ENSG00000273899 | NOL12 | 0,139466356 | 1,101497604 | 0,59198723 | 0,947416795 | nucleolar protein 12 |
| ENSG00000144339 | TMEFF2 | 0,139351143 | 1,101409642 | 0,59424139 | 0,947469769 | transmembrane protein with EGF like and two follistatin like domains 2 |
| ENSG00000178809 | TRIM73 | 0,139277385 | 1,101353333 | 0,82228278 | 0,982952211 | tripartite motif containing 73 |
| ENSG00000139352 | ASCL1 | 0,139254421 | 1,101335803 | 0,70385936 | 0,968976619 | achaete-scute family bHLH transcription factor 1 |
| ENSG00000183048 | SLC25A10 | 0,139249271 | 1,101331872 | 0,67390755 | 0,963037987 | solute carrier family 25 member 10 |
| ENSG00000276168 | RN7SL1 | 0,139156226 | 1,101260844 | 0,58970924 | 0,947051012 | RNA component of signal recognition particle 7SL1 |
| ENSG00000198952 | SMG5 | 0,139154579 | 1,101259587 | 0,56626703 | 0,94113936 | SMG5 nonsense mediated mRNA decay factor |
| ENSG00000116221 | MRPL37 | 0,139123918 | 1,101236183 | 0,55524009 | 0,937793851 | mitochondrial ribosomal protein L37 |
| ENSG00000176454 | LPCAT4 | 0,139110916 | 1,101226259 | 0,60319735 | 0,950352286 | lysophosphatidylcholine acyltransferase 4 |

| | | | | | | |
|---|---|---|---|---|---|---|
| ENSG00000104936 | DMPK | 0,139109559 | 1,101225223 | 0,75026646 | 0,975901808 | DM1 protein kinase |
| ENSG00000264107 | AC138207,2 | 0,139092395 | 1,101212121 | 0,8545038 | 0,987861659 | novel transcript, antisense to NF1 |
| ENSG00000286904 | AC093675,1 | 0,139038701 | 1,101171137 | 0,8078845 | 0,981213877 | novel transcript, antisense to TMEM87B |
| ENSG00000113384 | GOLPH3 | 0,139013389 | 1,101151817 | 0,63266544 | 0,954291097 | golgi phosphoprotein 3 |
| ENSG00000110448 | CD5 | 0,138877965 | 1,101048458 | 0,8318963 | 0,984596888 | CD5 molecule |
| ENSG00000267493 | CIRBP-AS1 | 0,138861084 | 1,101035575 | 0,73244287 | 0,974353474 | CIRBP antisense RNA 1 |
| ENSG00000236230 | AL356108,1 | 0,138845879 | 1,101023971 | 0,86179094 | 0,989074806 | novel transcript |
| ENSG00000109586 | GALNT7 | 0,138796034 | 1,100985932 | 0,72402317 | 0,973156872 | polypeptide N-acetylgalactosaminyltransferase 7 |
| ENSG00000165632 | TAF3 | 0,138775459 | 1,10097023 | 0,62091577 | 0,952793934 | TATA-box binding protein associated factor 3 |
| ENSG00000067113 | PLPP1 | 0,138765712 | 1,100962792 | 0,68736824 | 0,965965731 | phospholipid phosphatase 1 |
| ENSG00000253982 | AC100810,1 | 0,138719701 | 1,10092768 | 0,73497731 | 0,974353474 | novel transcript |
| ENSG00000131931 | THAP1 | 0,138700583 | 1,100913091 | 0,75588994 | 0,975901808 | THAP domain containing 1 |
| ENSG00000227403 | LINC01806 | 0,138699959 | 1,100912615 | 0,87056475 | 0,989074806 | long intergenic non-protein coding RNA 1806 |
| ENSG00000116353 | MECR | 0,13867996 | 1,100897354 | 0,58274216 | 0,944174844 | mitochondrial trans-2-enoyl-CoA reductase |
| ENSG00000133030 | MPRIP | 0,138578806 | 1,100820168 | 0,66704929 | 0,9611116 | myosin phosphatase Rho interacting protein |
| ENSG00000115163 | CENPA | 0,13852279 | 1,100777426 | 0,67413938 | 0,96304746 | centromere protein A |
| ENSG00000179270 | PCARE | 0,138436294 | 1,100711431 | 0,91079135 | 0,993222655 | photoreceptor cilium actin regulator |
| ENSG00000223461 | AC004471,1 | 0,138344348 | 1,100641283 | 0,88390325 | 0,990717729 | novel transcript |
| ENSG00000272950 | AC093799,1 | 0,13831735 | 1,100620686 | 0,90862741 | 0,992603769 | novel transcript, antisense to BAIAP2L1 |
| ENSG00000069998 | HDHD5 | 0,138250167 | 1,100569434 | 0,64190017 | 0,956529637 | haloacid dehalogenase like hydrolase domain containing 5 |
| ENSG00000164897 | TMUB1 | 0,138238499 | 1,100560533 | 0,72523701 | 0,97326334 | transmembrane and ubiquitin like domain containing 1 |
| ENSG00000262001 | DLGAP1-AS2 | 0,138176167 | 1,100512985 | 0,60134943 | 0,949939978 | DLGAP1 antisense RNA 2 |
| ENSG00000170855 | TRIAP1 | 0,138096917 | 1,100452533 | 0,68620162 | 0,965740648 | TP53 regulated inhibitor of apoptosis 1 |
| ENSG00000174021 | GNG5 | 0,138022458 | 1,100395739 | 0,71054868 | 0,970482408 | G protein subunit gamma 5 |
| ENSG00000118620 | ZNF430 | 0,138016562 | 1,100391242 | 0,68878243 | 0,966556275 | zinc finger protein 430 |
| ENSG00000083799 | CYLD | 0,138010761 | 1,100386817 | 0,6283009 | 0,952793934 | CYLD lysine 63 deubiquitinase |
| ENSG00000283753 | RPL21P121 | 0,137924392 | 1,100320943 | 0,90025552 | 0,991845106 | ribosomal protein L21 pseudogene 121 |
| ENSG00000237382 | RPL21P121 | 0,137924392 | 1,100320943 | 0,90025552 | 0,991845106 | ribosomal protein L21 pseudogene 121 |
| ENSG00000131469 | RPL27 | 0,137923748 | 1,100320451 | 0,69610261 | 0,968059789 | ribosomal protein L27 |
| ENSG00000258435 | AC048337,1 | 0,137892155 | 1,100296356 | 0,8969145 | 0,991644439 | novel transcript |
| ENSG00000204335 | SP5 | 0,137852364 | 1,10026601 | 0,78256796 | 0,978647973 | Sp5 transcription factor |

| | | | | | | |
|---|---|---|---|---|---|---|
| ENSG00000140526 | ABHD2 | 0,137780381 | 1,100211113 | 0,59211983 | 0,947416795 | abhydrolase domain containing 2, acylglycerol lipase |
| ENSG00000224455 | VPS52 | 0,137767761 | 1,10020149 | 0,88241739 | 0,990717729 | VPS52 subunit of GARP complex |
| ENSG00000236014 | VPS52 | 0,137767761 | 1,10020149 | 0,88241739 | 0,990717729 | VPS52 subunit of GARP complex |
| ENSG00000075415 | SLC25A3 | 0,137743748 | 1,100183177 | 0,57699072 | 0,943288205 | solute carrier family 25 member 3 |
| ENSG00000253540 | FAM86HP | 0,137642171 | 1,100105718 | 0,78444675 | 0,979320406 | family with sequence similarity 86 member H, pseudogene |
| ENSG00000180423 | HARBI1 | 0,137611669 | 1,10008246 | 0,61098802 | 0,951873283 | harbinger transposase derived 1 |
| ENSG00000287738 | AL359764,3 | 0,137575026 | 1,100054519 | 0,92808049 | 0,994108531 | novel transcript |
| ENSG00000015171 | ZMYND11 | 0,137509551 | 1,100004596 | 0,68713209 | 0,965769168 | zinc finger MYND-type containing 11 |
| ENSG00000162396 | PARS2 | 0,13749468 | 1,099993257 | 0,60305521 | 0,950352286 | prolyl-tRNA synthetase 2, mitochondrial |
| ENSG00000100216 | TOMM22 | 0,137492183 | 1,099991353 | 0,52486129 | 0,928616707 | translocase of outer mitochondrial membrane 22 |
| ENSG00000198911 | SREBF2 | 0,137435126 | 1,099947851 | 0,53519979 | 0,931905243 | sterol regulatory element binding transcription factor 2 |
| ENSG00000228000 | RPL7AP65 | 0,137381032 | 1,099906609 | 0,84984598 | 0,987036295 | ribosomal protein L7a pseudogene 65 |
| ENSG00000284212 | RPL7AP65 | 0,137381032 | 1,099906609 | 0,84984598 | 0,987036295 | ribosomal protein L7a pseudogene 65 |
| ENSG00000198833 | UBE2J1 | 0,13735961 | 1,099890277 | 0,71919437 | 0,971869926 | ubiquitin conjugating enzyme E2 J1 |
| ENSG00000268670 | AC016586,1 | 0,137352417 | 1,099884793 | 0,87883303 | 0,990109814 | novel transcript, sense intronic to ZBTB7A |
| ENSG00000227176 | NSA2P5 | 0,137337443 | 1,099873377 | 0,86741614 | 0,989074806 | NSA2 pseudogene 5 |
| ENSG00000274270 | AL137060,3 | 0,137320064 | 1,099860128 | 0,86902523 | 0,989074806 | novel transcript, sense intronic to DLEU1 |
| ENSG00000137806 | NDUFAF1 | 0,137280041 | 1,099829616 | 0,56524314 | 0,940865625 | NADH:ubiquinone oxidoreductase complex assembly factor 1 |
| ENSG00000225564 | LINC01492 | 0,137268724 | 1,099820988 | 0,85210249 | 0,98732978 | long intergenic non-protein coding RNA 1492 |
| ENSG00000182552 | RWDD4 | 0,137215953 | 1,09978076 | 0,75614695 | 0,975901808 | RWD domain containing 4 |
| ENSG00000152253 | SPC25 | 0,137210538 | 1,099776632 | 0,75531351 | 0,975901808 | SPC25 component of NDC80 kinetochore complex |
| ENSG00000168397 | ATG4B | 0,137185376 | 1,099757451 | 0,60333763 | 0,950352286 | autophagy related 4B cysteine peptidase |
| ENSG00000119950 | MXI1 | 0,137174163 | 1,099748904 | 0,75781042 | 0,976120207 | MAX interactor 1, dimerization protein |
| ENSG00000126246 | IGFLR1 | 0,137164464 | 1,09974151 | 0,6961693 | 0,96810276 | IGF like family receptor 1 |
| ENSG00000134248 | LAMTOR5 | 0,137157246 | 1,099736008 | 0,68597481 | 0,965740648 | late endosomal/lysosomal adaptor, MAPK and MTOR activator 5 |
| ENSG00000169813 | HNRNPF | 0,137133669 | 1,099718036 | 0,6103362 | 0,951841373 | heterogeneous nuclear ribonucleoprotein F |
| ENSG00000223606 | HLA-V | 0,137095796 | 1,099689167 | 0,89848151 | 0,991832796 | major histocompatibility complex, class I, V (pseudogene) |
| ENSG00000171564 | FGB | 0,137080293 | 1,09967735 | 0,93381513 | 0,994479239 | fibrinogen beta chain |

| | | | | | | |
|---|---|---|---|---|---|---|
| ENSG00000136143 | SUCLA2 | 0,136924302 | 1,099558454 | 0,71517433 | 0,971041579 | succinate-CoA ligase ADP-forming subunit beta |
| ENSG00000287844 | AC018953,2 | 0,136884094 | 1,09952781 | 0,89662229 | 0,991644439 | novel transcript |
| ENSG00000143569 | UBAP2L | 0,136848068 | 1,099500354 | 0,51909093 | 0,926800715 | ubiquitin associated protein 2 like |
| ENSG00000159640 | ACE | 0,136833045 | 1,099488905 | 0,82654894 | 0,983471137 | angiotensin I converting enzyme |
| ENSG00000285582 | AC016042,1 | 0,136828819 | 1,099485684 | 0,79020766 | 0,979462318 | novel transcript, antisense to ATRNL1 |
| ENSG00000232176 | AL161909,1 | 0,136753647 | 1,099428396 | 0,90317258 | 0,992311172 | ribosomal protein S6 (RPS6) pseudogene |
| ENSG00000122687 | MRM2 | 0,136745956 | 1,099422535 | 0,51991509 | 0,926822772 | mitochondrial rRNA methyltransferase 2 |
| ENSG00000159208 | CIART | 0,136606542 | 1,099316298 | 0,70121093 | 0,96870039 | circadian associated repressor of transcription |
| ENSG00000169857 | AVEN | 0,136605701 | 1,099315657 | 0,55892192 | 0,939635125 | apoptosis and caspase activation inhibitor |
| ENSG00000167721 | TSR1 | 0,136562994 | 1,099283116 | 0,53798852 | 0,932883277 | TSR1 ribosome maturation factor |
| ENSG00000205571 | SMN2 | 0,136535828 | 1,099262416 | 0,69079595 | 0,967179044 | survival of motor neuron 2, centromeric |
| ENSG00000128311 | TST | 0,136484234 | 1,099223105 | 0,67590837 | 0,963549282 | thiosulfate sulfurtransferase |
| ENSG00000077585 | GPR137B | 0,136481347 | 1,099220905 | 0,7104165 | 0,970482408 | G protein-coupled receptor 137B |
| ENSG00000257379 | AC023509,1 | 0,136464831 | 1,099208322 | 0,77105658 | 0,978033855 | novel transcript |
| ENSG00000275125 | PRXL2B | 0,136464071 | 1,099207742 | 0,753038 | 0,975901808 | peroxiredoxin like 2B |
| ENSG00000233186 | KLF4P1 | 0,136456498 | 1,099201972 | 0,86839921 | 0,989074806 | Kruppel like factor 4 pseudogene 1 |
| ENSG00000229308 | AC010737,1 | 0,136347043 | 1,099118581 | 0,92067282 | 0,993787755 | novel transcript |
| ENSG00000149639 | SOGA1 | 0,136325609 | 1,099102252 | 0,7286622 | 0,973955059 | suppressor of glucose, autophagy associated 1 |
| ENSG00000110011 | DNAJC4 | 0,136294699 | 1,099078703 | 0,65183227 | 0,957160627 | DnaJ heat shock protein family (Hsp40) member C4 |
| ENSG00000141452 | RMC1 | 0,136282763 | 1,09906961 | 0,64869337 | 0,956899963 | regulator of MON1-CCZ1 |
| ENSG00000154545 | MAGED4 | 0,13623251 | 1,099031327 | 0,56143512 | 0,94060917 | MAGE family member D4 |
| ENSG00000187010 | RHD | 0,136176577 | 1,098988719 | 0,77548932 | 0,978587255 | Rh blood group D antigen |
| ENSG00000085872 | CHERP | 0,136171747 | 1,09898504 | 0,57034499 | 0,9423888 | calcium homeostasis endoplasmic reticulum protein |
| ENSG00000126746 | ZNF384 | 0,136168356 | 1,098982456 | 0,5557563 | 0,937930638 | zinc finger protein 384 |
| ENSG00000132792 | CTNNBL1 | 0,136093666 | 1,098925562 | 0,62573288 | 0,952793934 | catenin beta like 1 |
| ENSG00000156711 | MAPK13 | 0,136077196 | 1,098913017 | 0,81020929 | 0,981213877 | mitogen-activated protein kinase 13 |
| ENSG00000282988 | AL031777,2 | 0,136029758 | 1,098876884 | 0,67244678 | 0,962290553 | novel protein |
| ENSG00000232196 | MTRNR2L4 | 0,136021939 | 1,098870928 | 0,89971657 | 0,991845106 | MT-RNR2 like 4 |
| ENSG00000239920 | AC104389,5 | 0,135983933 | 1,09884198 | 0,73534327 | 0,974353474 | novel transcript |
| ENSG00000114023 | FAM162A | 0,135975939 | 1,098835891 | 0,72812508 | 0,973888243 | family with sequence similarity 162 member A |
| ENSG00000258289 | CHURC1 | 0,135951954 | 1,098817623 | 0,74827248 | 0,975901808 | churchill domain containing 1 |
| ENSG00000166105 | GLB1L3 | 0,135822149 | 1,098718763 | 0,83271531 | 0,984905093 | galactosidase beta 1 like 3 |
| ENSG00000167978 | SRRM2 | 0,135753107 | 1,098666183 | 0,53475812 | 0,931714104 | serine/arginine repetitive matrix 2 |

| | | | | | | |
|---|---|---|---|---|---|---|
| ENSG00000136273 | HUS1 | 0,135741507 | 1,098657349 | 0,68461344 | 0,965415049 | HUS1 checkpoint clamp component |
| ENSG00000286629 | AC005803,1 | 0,135682671 | 1,098612545 | 0,88726286 | 0,990717729 | novel transcript, antisense to ERN1 |
| ENSG00000232712 | KIZ-AS1 | 0,135587588 | 1,098540141 | 0,8252269 | 0,983383773 | KIZ antisense RNA 1 |
| ENSG00000165782 | PIP4P1 | 0,135587094 | 1,098539765 | 0,64471201 | 0,956633898 | phosphatidylinositol-4,5-bisphosphate 4-phosphatase 1 |
| ENSG00000168765 | GSTM4 | 0,135584121 | 1,098537501 | 0,61609186 | 0,952793934 | glutathione S-transferase mu 4 |
| ENSG00000105963 | ADAP1 | 0,135504096 | 1,098476569 | 0,78375572 | 0,979226738 | ArfGAP with dual PH domains 1 |
| ENSG00000228894 | HCG18 | 0,135440929 | 1,098428474 | 0,76204781 | 0,976315588 | HLA complex group 18 |
| ENSG00000224619 | HCG18 | 0,135440929 | 1,098428474 | 0,76204781 | 0,976315588 | HLA complex group 18 |
| ENSG00000105723 | GSK3A | 0,135398453 | 1,098396134 | 0,51729748 | 0,925746338 | glycogen synthase kinase 3 alpha |
| ENSG00000171931 | FBXW10 | 0,135398433 | 1,098396119 | 0,83920004 | 0,985672131 | F-box and WD repeat domain containing 10 |
| ENSG00000135249 | RINT1 | 0,135314556 | 1,098332261 | 0,69176114 | 0,967204896 | RAD50 interactor 1 |
| ENSG00000136059 | VILL | 0,135282467 | 1,098307832 | 0,74545431 | 0,975707496 | villin like |
| ENSG00000179344 | HLA-DQB1 | 0,135249011 | 1,098282362 | 0,87540983 | 0,990078774 | major histocompatibility complex, class II, DQ beta 1 |
| ENSG00000172086 | KRCC1 | 0,135243564 | 1,098278216 | 0,76827062 | 0,977381742 | lysine rich coiled-coil 1 |
| ENSG00000200997 | RNVU1-34 | 0,135212309 | 1,098254423 | 0,89983138 | 0,991845106 | RNA, variant U1 small nuclear 34 |
| ENSG00000196743 | GM2A | 0,135195863 | 1,098241903 | 0,59346191 | 0,947454078 | GM2 ganglioside activator |
| ENSG00000233927 | RPS28 | 0,135031205 | 1,098116565 | 0,59881975 | 0,948854865 | ribosomal protein S28 |
| ENSG00000213465 | ARL2 | 0,134982479 | 1,098079478 | 0,64271287 | 0,956633898 | ADP ribosylation factor like GTPase 2 |
| ENSG00000180447 | GAS1 | 0,134962548 | 1,098064308 | 0,60851414 | 0,951511094 | growth arrest specific 1 |
| ENSG00000110200 | ANAPC15 | 0,134960746 | 1,098062937 | 0,58258567 | 0,944174844 | anaphase promoting complex subunit 15 |
| ENSG00000266338 | NBPF15 | 0,134831611 | 1,097964653 | 0,57157962 | 0,942436428 | NBPF member 15 |
| ENSG00000133872 | SARAF | 0,134722985 | 1,097881987 | 0,67412635 | 0,96304746 | store-operated calcium entry associated regulatory factor |
| ENSG00000229924 | FAM90A26 | 0,134721107 | 1,097880558 | 0,90737291 | 0,992590661 | family with sequence similarity 90 member A26 |
| ENSG00000102218 | RP2 | 0,134584193 | 1,097776372 | 0,74363185 | 0,975310206 | RP2 activator of ARL3 GTPase |
| ENSG00000118007 | STAG1 | 0,134499904 | 1,097712237 | 0,66312248 | 0,959920886 | stromal antigen 1 |
| ENSG00000152133 | GPATCH11 | 0,134485895 | 1,097701578 | 0,68791442 | 0,966127143 | G-patch domain containing 11 |
| ENSG00000139668 | WDFY2 | 0,134359178 | 1,097605167 | 0,58056724 | 0,943832501 | WD repeat and FYVE domain containing 2 |
| ENSG00000161888 | SPC24 | 0,134339109 | 1,097589898 | 0,62874488 | 0,952793934 | SPC24 component of NDC80 kinetochore complex |
| ENSG00000173926 | MARCHF3 | 0,134303789 | 1,097563027 | 0,73249178 | 0,974353474 | membrane associated ring-CH-type finger 3 |
| ENSG00000116580 | GON4L | 0,134244566 | 1,097517973 | 0,56361166 | 0,940865625 | gon-4 like |

| | | | | | | |
|---|---|---|---|---|---|---|
| ENSG00000159921 | GNE | 0,134229755 | 1,097506706 | 0,50768573 | 0,922163985 | glucosamine (UDP-N-acetyl)-2-epimerase/N-acetylmannosamine kinase |
| ENSG00000267064 | UXT-AS1 | 0,134199328 | 1,09748356 | 0,77493246 | 0,97857211 | UXT antisense RNA 1 |
| ENSG00000186787 | SPIN2B | 0,134197454 | 1,097482134 | 0,70160623 | 0,968742626 | spindlin family member 2B |
| ENSG00000258964 | AL137129,1 | 0,134172882 | 1,097463442 | 0,97208599 | 0,998214946 | novel transcript |
| ENSG00000165731 | RET | 0,134156133 | 1,097450701 | 0,74192918 | 0,975176043 | ret proto-oncogene |
| ENSG00000276524 | AC010999,2 | 0,134106215 | 1,097412729 | 0,89028024 | 0,990717729 | novel transcript |
| ENSG00000164151 | ICE1 | 0,134050223 | 1,097370139 | 0,64858411 | 0,956899963 | interactor of little elongation complex ELL subunit 1 |
| ENSG00000274695 | AC108704,1 | 0,134028766 | 1,097353817 | 0,86750009 | 0,989074806 | novel transcript |
| ENSG00000253190 | AC084082,1 | 0,133978496 | 1,097315581 | 0,79833845 | 0,980161849 | novel transcript |
| ENSG00000253438 | PCAT1 | 0,133929327 | 1,097278184 | 0,79884296 | 0,980161849 | prostate cancer associated transcript 1 |
| ENSG00000168135 | KCNJ4 | 0,133863895 | 1,097228419 | 0,86861421 | 0,989074806 | potassium inwardly rectifying channel subfamily J member 4 |
| ENSG00000174799 | CEP135 | 0,133858664 | 1,097224441 | 0,66288184 | 0,959897678 | centrosomal protein 135 |
| ENSG00000258749 | AL110504,1 | 0,133849876 | 1,097217757 | 0,93469423 | 0,994479239 | novel transcript, antisense to CCDC85C |
| ENSG00000264066 | AC024267,1 | 0,133834119 | 1,097205774 | 0,92055017 | 0,993787755 | novel transcript |
| ENSG00000168826 | ZBTB49 | 0,133820507 | 1,097195422 | 0,65147437 | 0,957089277 | zinc finger and BTB domain containing 49 |
| ENSG00000175691 | ZNF77 | 0,133738573 | 1,097133111 | 0,62247296 | 0,952793934 | zinc finger protein 77 |
| ENSG00000123338 | NCKAP1L | 0,133713836 | 1,097114299 | 0,79396865 | 0,979776981 | NCK associated protein 1 like |
| ENSG00000108587 | GOSR1 | 0,133712347 | 1,097113167 | 0,50531606 | 0,921715363 | golgi SNAP receptor complex member 1 |
| ENSG00000169727 | GPS1 | 0,13369217 | 1,097097823 | 0,63659896 | 0,95557086 | G protein pathway suppressor 1 |
| ENSG00000164743 | C8orf48 | 0,133685265 | 1,097092572 | 0,74678582 | 0,975796253 | chromosome 8 open reading frame 48 |
| ENSG00000278550 | SLC43A2 | 0,13368094 | 1,097089283 | 0,68500835 | 0,965415049 | solute carrier family 43 member 2 |
| ENSG00000286733 | LINC02348 | 0,13366846 | 1,097079793 | 0,86696041 | 0,989074806 | long intergenic non-protein coding RNA 2348 |
| ENSG00000119986 | AVPI1 | 0,133623531 | 1,097045628 | 0,78307657 | 0,978917309 | arginine vasopressin induced 1 |
| ENSG00000113712 | CSNK1A1 | 0,133612232 | 1,097037036 | 0,68438714 | 0,965415049 | casein kinase 1 alpha 1 |
| ENSG00000112062 | MAPK14 | 0,133599718 | 1,09702752 | 0,63117442 | 0,953701469 | mitogen-activated protein kinase 14 |
| ENSG00000113068 | PFDN1 | 0,133581071 | 1,097013341 | 0,61800743 | 0,952793934 | prefoldin subunit 1 |
| ENSG00000204435 | CSNK2B | 0,133563547 | 1,097000016 | 0,55092734 | 0,936366211 | casein kinase 2 beta |
| ENSG00000228288 | PCAT6 | 0,133541417 | 1,096983189 | 0,72330054 | 0,972913663 | prostate cancer associated transcript 6 |
| ENSG00000148344 | PTGES | 0,133537951 | 1,096980553 | 0,81076767 | 0,981213877 | prostaglandin E synthase |
| ENSG00000121350 | PYROXD1 | 0,133532668 | 1,096976536 | 0,77886169 | 0,978587255 | pyridine nucleotide-disulphide oxidoreductase domain 1 |

| | | | | | | |
|---|---|---|---|---|---|---|
| ENSG00000110400 | NECTIN1 | 0,133460391 | 1,096921581 | 0,59919022 | 0,948894897 | nectin cell adhesion molecule 1 |
| ENSG00000105711 | SCN1B | 0,133402809 | 1,096877801 | 0,81556046 | 0,981590958 | sodium voltage-gated channel beta subunit 1 |
| ENSG00000287710 | AC067852,5 | 0,133306239 | 1,096804381 | 0,79936425 | 0,980161849 | novel transcript, antisense to PSMC3IP |
| ENSG00000204922 | UQCC3 | 0,133274992 | 1,096780626 | 0,74185475 | 0,975176043 | ubiquinol-cytochrome c reductase complex assembly factor 3 |
| ENSG00000109861 | CTSC | 0,133179787 | 1,09670825 | 0,72821376 | 0,973888243 | cathepsin C |
| ENSG00000067248 | DHX29 | 0,133132748 | 1,096672493 | 0,71902572 | 0,971869926 | DExH-box helicase 29 |
| ENSG00000133835 | HSD17B4 | 0,133104539 | 1,09665105 | 0,67290056 | 0,962421114 | hydroxysteroid 17-beta dehydrogenase 4 |
| ENSG00000142507 | PSMB6 | 0,133048718 | 1,096608619 | 0,56015651 | 0,940070839 | proteasome 20S subunit beta 6 |
| ENSG00000266466 | AC233702,7 | 0,133011628 | 1,096580427 | 0,88984007 | 0,990717729 | novel transcript, antisense to C17orf51 |
| ENSG00000108671 | PSMD11 | 0,132874789 | 1,096476422 | 0,59534959 | 0,947757499 | proteasome 26S subunit, non-ATPase 11 |
| ENSG00000163636 | PSMD6 | 0,132808674 | 1,096426174 | 0,70828859 | 0,97011892 | proteasome 26S subunit, non-ATPase 6 |
| ENSG00000135596 | MICAL1 | 0,132732512 | 1,096368294 | 0,6628039 | 0,959897678 | microtubule associated monooxygenase, calponin and LIM domain containing 1 |
| ENSG00000175073 | VCPIP1 | 0,132711256 | 1,09635214 | 0,68285375 | 0,964914784 | valosin containing protein interacting protein 1 |
| ENSG00000188739 | RBM34 | 0,132661427 | 1,096314274 | 0,58848491 | 0,946292796 | RNA binding motif protein 34 |
| ENSG00000261761 | LINC02616 | 0,132613876 | 1,096278141 | 0,85175374 | 0,987289464 | long intergenic non-protein coding RNA 2616 |
| ENSG00000107897 | ACBD5 | 0,132605449 | 1,096271737 | 0,52865664 | 0,930033144 | acyl-CoA binding domain containing 5 |
| ENSG00000099246 | RAB18 | 0,132560833 | 1,096237835 | 0,77396476 | 0,978415098 | RAB18, member RAS oncogene family |
| ENSG00000132275 | RRP8 | 0,132543667 | 1,096224791 | 0,55289468 | 0,936894267 | ribosomal RNA processing 8 |
| ENSG00000275055 | AC011468,5 | 0,132527346 | 1,09621239 | 0,76236722 | 0,976315588 | novel transcript, antisense to ZNF432 |
| ENSG00000006576 | PHTF2 | 0,132511511 | 1,096200358 | 0,75472957 | 0,975901808 | putative homeodomain transcription factor 2 |
| ENSG00000174574 | AKIRIN1 | 0,132362806 | 1,096087374 | 0,7251346 | 0,97326334 | akirin 1 |
| ENSG00000165914 | TTC7B | 0,132362161 | 1,096086884 | 0,51937089 | 0,926800715 | tetratricopeptide repeat domain 7B |
| ENSG00000064490 | RFXANK | 0,132352579 | 1,096079604 | 0,67477187 | 0,963268551 | regulatory factor X associated ankyrin containing protein |
| ENSG00000109118 | PHF12 | 0,132347057 | 1,096075408 | 0,58341296 | 0,944174844 | PHD finger protein 12 |
| ENSG00000265735 | RN7SL5P | 0,13233965 | 1,096069781 | 0,88802172 | 0,990717729 | RNA, 7SL, cytoplasmic 5, pseudogene |
| ENSG00000227666 | CYCSP24 | 0,132302585 | 1,096041622 | 0,88006788 | 0,990268593 | CYCS pseudogene 24 |
| ENSG00000213888 | LINC01521 | 0,132285293 | 1,096028485 | 0,76174774 | 0,976285211 | long intergenic non-protein coding RNA 1521 |
| ENSG00000135148 | TRAFD1 | 0,132283807 | 1,096027356 | 0,56272149 | 0,940865625 | TRAF-type zinc finger domain containing 1 |
| ENSG00000236337 | FMR1-IT1 | 0,132281927 | 1,096025927 | 0,88220006 | 0,990717729 | FMR1 intronic transcript 1 |

| | | | | | | |
|---|---|---|---|---|---|---|
| ENSG00000173457 | PPP1R14B | 0,132195678 | 1,095960406 | 0,6407241 | 0,956348702 | protein phosphatase 1 regulatory inhibitor subunit 14B |
| ENSG00000060237 | WNK1 | 0,132164113 | 1,095936427 | 0,63545867 | 0,955314044 | WNK lysine deficient protein kinase 1 |
| ENSG00000155428 | TRIM74 | 0,13210339 | 1,0958903 | 0,93841396 | 0,994549499 | tripartite motif containing 74 |
| ENSG00000286069 | AC076968,2 | 0,132087597 | 1,095878303 | 0,7658039 | 0,976949161 | novel transcript |
| ENSG00000177452 | AL161742,1 | 0,131968934 | 1,09578817 | 0,88491504 | 0,990717729 | ribosomal protein L19 (RPL19) pseudogene |
| ENSG00000119711 | ALDH6A1 | 0,131858817 | 1,095704535 | 0,64042697 | 0,956251063 | aldehyde dehydrogenase 6 family member A1 |
| ENSG00000116731 | PRDM2 | 0,13184107 | 1,095691056 | 0,65736124 | 0,957853146 | PR/SET domain 2 |
| ENSG00000261737 | CLCA4-AS1 | 0,131819383 | 1,095674586 | 0,85107683 | 0,987259878 | CLCA4 antisense RNA 1 |
| ENSG00000148481 | MINDY3 | 0,131646176 | 1,095543049 | 0,73721081 | 0,974353474 | MINDY lysine 48 deubiquitinase 3 |
| ENSG00000211460 | TSN | 0,131582877 | 1,095494983 | 0,748338 | 0,975901808 | translin |
| ENSG00000243312 | AC093827,1 | 0,131563756 | 1,095480463 | 0,76068172 | 0,97624106 | ribosomal protein L6 (RPL6) pseudogene |
| ENSG00000233994 | GDI2P2 | 0,131499512 | 1,095431682 | 0,89916995 | 0,991832796 | GDP dissociation inhibitor 2 pseudogene 2 |
| ENSG00000178202 | POGLUT3 | 0,131485999 | 1,095421422 | 0,63499188 | 0,955158181 | protein O-glucosyltransferase 3 |
| ENSG00000230549 | USP17L1 | 0,131235906 | 1,095231546 | 0,88680908 | 0,990717729 | ubiquitin specific peptidase 17 like family member 1 |
| ENSG00000184005 | ST6GALNAC3 | 0,131217192 | 1,095217339 | 0,8213788 | 0,982950156 | ST6 N-acetylgalactosaminide alpha-2,6-sialyltransferase 3 |
| ENSG00000158169 | FANCC | 0,1311508 | 1,095166939 | 0,55954762 | 0,939727587 | FA complementation group C |
| ENSG00000175104 | TRAF6 | 0,131109474 | 1,095135568 | 0,68915704 | 0,966685038 | TNF receptor associated factor 6 |
| ENSG00000196937 | FAM3C | 0,131089106 | 1,095120107 | 0,77638055 | 0,978587255 | FAM3 metabolism regulating signaling molecule C |
| ENSG00000134809 | TIMM10 | 0,131067715 | 1,09510387 | 0,55945554 | 0,939727587 | translocase of inner mitochondrial membrane 10 |
| ENSG00000101557 | USP14 | 0,130969214 | 1,095029104 | 0,72493257 | 0,97326334 | ubiquitin specific peptidase 14 |
| ENSG00000168528 | SERINC2 | 0,130958746 | 1,095021158 | 0,56719902 | 0,941425553 | serine incorporator 2 |
| ENSG00000148843 | PDCD11 | 0,130958684 | 1,095021111 | 0,65734637 | 0,957853146 | programmed cell death 11 |
| ENSG00000134313 | KIDINS220 | 0,130895986 | 1,094973523 | 0,7152614 | 0,971041579 | kinase D interacting substrate 220 |
| ENSG00000281566 | AL157778,1 | 0,130750289 | 1,094862948 | 0,75133144 | 0,975901808 | novel transcript |
| ENSG00000134909 | ARHGAP32 | 0,130748074 | 1,094861267 | 0,67974773 | 0,964459885 | Rho GTPase activating protein 32 |
| ENSG00000177479 | ARIH2 | 0,130738179 | 1,094853758 | 0,57595955 | 0,942955084 | ariadne RBR E3 ubiquitin protein ligase 2 |
| ENSG00000256646 | AC010132,3 | 0,130715968 | 1,094836903 | 0,71729249 | 0,971697135 | UPF0415 protein C7orf25 |
| ENSG00000185437 | SH3BGR | 0,130560036 | 1,094718574 | 0,74281274 | 0,975201495 | SH3 domain binding glutamate rich protein |
| ENSG00000230700 | CSNK2B | 0,130534683 | 1,094699337 | 0,74699209 | 0,975796253 | casein kinase 2 beta |
| ENSG00000110900 | TSPAN11 | 0,13053147 | 1,094696899 | 0,66716415 | 0,9611116 | tetraspanin 11 |
| ENSG00000239917 | RPS10P16 | 0,13052776 | 1,094694084 | 0,923242 | 0,993825008 | ribosomal protein S10 pseudogene 16 |

| | | | | | | |
|---|---|---|---|---|---|---|
| ENSG00000226919 | AL365184,1 | 0,130527118 | 1,094693597 | 0,86632848 | 0,989074806 | novel transcript, antisense to RGS7 |
| ENSG00000081177 | EXD2 | 0,130469977 | 1,09465024 | 0,60600687 | 0,950927937 | exonuclease 3'-5' domain containing 2 |
| ENSG00000100225 | FBXO7 | 0,130458341 | 1,094641411 | 0,65876668 | 0,958206085 | F-box protein 7 |
| ENSG00000164828 | SUN1 | 0,130454824 | 1,094638743 | 0,5527262 | 0,936894267 | Sad1 and UNC84 domain containing 1 |
| ENSG00000277159 | AL139384,2 | 0,130404529 | 1,094600582 | 0,74549687 | 0,975707496 | novel transcript |
| ENSG00000105492 | SIGLEC6 | 0,130368822 | 1,094573491 | 0,86649322 | 0,989074806 | sialic acid binding Ig like lectin 6 |
| ENSG00000055955 | ITIH4 | 0,130346546 | 1,094556591 | 0,7993718 | 0,980161849 | inter-alpha-trypsin inhibitor heavy chain 4 |
| ENSG00000277965 | Metazoa_SRP | 0,13033686 | 1,094549242 | 0,90795771 | 0,992590661 | Metazoan signal recognition particle RNA |
| ENSG00000108590 | MED31 | 0,130293076 | 1,094516024 | 0,7320029 | 0,974353474 | mediator complex subunit 31 |
| ENSG00000161103 | AC008132,1 | 0,130287747 | 1,094511981 | 0,86606873 | 0,989074806 | POM121 transmembrane nucleoporin (POM121) pseudogene |
| ENSG00000131409 | LRRC4B | 0,130270574 | 1,094498952 | 0,69154246 | 0,967201016 | leucine rich repeat containing 4B |
| ENSG00000259240 | MIR4713HG | 0,130248055 | 1,094481869 | 0,93029136 | 0,994336072 | MIR4713 host gene |
| ENSG00000284735 | AL139424,3 | 0,13022641 | 1,094465448 | 0,81772659 | 0,982323352 | novel transcript |
| ENSG00000205323 | SARNP | 0,130218613 | 1,094459533 | 0,54018555 | 0,933385624 | SAP domain containing ribonucleoprotein |
| ENSG00000239419 | RN7SL535P | 0,130182525 | 1,094432157 | 0,82623346 | 0,983471137 | RNA, 7SL, cytoplasmic 535, pseudogene |
| ENSG00000277763 | AL138995,1 | 0,130164868 | 1,094418762 | 0,89589148 | 0,99150948 | novel transcript |
| ENSG00000065183 | WDR3 | 0,130159924 | 1,094415011 | 0,59455525 | 0,947469769 | WD repeat domain 3 |
| ENSG00000104938 | CLEC4M | 0,130090196 | 1,094362117 | 0,90198964 | 0,992182254 | C-type lectin domain family 4 member M |
| ENSG00000166169 | POLL | 0,130070217 | 1,094346963 | 0,63475518 | 0,955149453 | DNA polymerase lambda |
| ENSG00000169062 | UPF3A | 0,130043708 | 1,094326855 | 0,60020838 | 0,949340299 | UPF3A regulator of nonsense mediated mRNA decay |
| ENSG00000064607 | SUGP2 | 0,130022036 | 1,094310416 | 0,54029704 | 0,933385624 | SURP and G-patch domain containing 2 |
| ENSG00000121022 | COPS5 | 0,129961955 | 1,094264844 | 0,63587323 | 0,955399772 | COP9 signalosome subunit 5 |
| ENSG00000259644 | AC022613,3 | 0,1299039 | 1,094220811 | 0,7321962 | 0,974353474 | novel transcript |
| ENSG00000140365 | COMMD4 | 0,129887093 | 1,094208064 | 0,61160185 | 0,952155617 | COMM domain containing 4 |
| ENSG00000276131 | AC009118,2 | 0,129870605 | 1,094195559 | 0,87766376 | 0,990109814 | novel transcript, antisense to GINS3 |
| ENSG00000168259 | DNAJC7 | 0,129801322 | 1,094143013 | 0,53686682 | 0,932883277 | DnaJ heat shock protein family (Hsp40) member C7 |
| ENSG00000186141 | POLR3C | 0,129780824 | 1,094127467 | 0,63589801 | 0,955399772 | RNA polymerase III subunit C |
| ENSG00000115286 | NDUFS7 | 0,1297324 | 1,094090744 | 0,65563473 | 0,957444211 | NADH:ubiquinone oxidoreductase core subunit S7 |
| ENSG00000176407 | KCMF1 | 0,129717664 | 1,094079569 | 0,67154719 | 0,962098952 | potassium channel modulatory factor 1 |
| ENSG00000133275 | CSNK1G2 | 0,129631586 | 1,094014292 | 0,60677415 | 0,950927937 | casein kinase 1 gamma 2 |
| ENSG00000156471 | PTDSS1 | 0,129610316 | 1,093998163 | 0,62142295 | 0,952793934 | phosphatidylserine synthase 1 |
| ENSG00000161970 | RPL26 | 0,129598016 | 1,093988836 | 0,73957158 | 0,974715504 | ribosomal protein L26 |

| | | | | | | |
|---|---|---|---|---|---|---|
| ENSG00000164944 | VIRMA | 0,129556862 | 1,09395763 | 0,71677777 | 0,971438758 | vir like m6A methyltransferase associated |
| ENSG00000169738 | DCXR | 0,129528636 | 1,093936227 | 0,70701096 | 0,969565008 | dicarbonyl and L-xylulose reductase |
| ENSG00000172301 | COPRS | 0,129422991 | 1,093856124 | 0,59579385 | 0,948096995 | coordinator of PRMT5 and differentiation stimulator |
| ENSG00000214331 | AC009053,1 | 0,12940751 | 1,093844386 | 0,68744134 | 0,965965731 | pyruvate dehydrogenase phosphatase regulatory subunit (PDPR) pseudogene |
| ENSG00000148334 | PTGES2 | 0,129226051 | 1,093706813 | 0,69631931 | 0,96810276 | prostaglandin E synthase 2 |
| ENSG00000174197 | MGA | 0,129206954 | 1,093692336 | 0,67604433 | 0,963560994 | MAX dimerization protein MGA |
| ENSG00000106772 | PRUNE2 | 0,129194598 | 1,093682969 | 0,68173353 | 0,964654539 | prune homolog 2 with BCH domain |
| ENSG00000246695 | RASSF8-AS1 | 0,129151818 | 1,093650539 | 0,6109423 | 0,951873283 | RASSF8 antisense RNA 1 |
| ENSG00000005175 | RPAP3 | 0,129061346 | 1,093581958 | 0,763983 | 0,976640468 | RNA polymerase II associated protein 3 |
| ENSG00000163877 | SNIP1 | 0,129057248 | 1,093578851 | 0,60655585 | 0,950927937 | Smad nuclear interacting protein 1 |
| ENSG00000132517 | SLC52A1 | 0,129036128 | 1,093562842 | 0,84756623 | 0,987035044 | solute carrier family 52 member 1 |
| ENSG00000136158 | SPRY2 | 0,129003658 | 1,09353823 | 0,62988156 | 0,952812183 | sprouty RTK signaling antagonist 2 |
| ENSG00000270917 | AL161911,1 | 0,128977466 | 1,093518377 | 0,92845122 | 0,994108531 | chromatin assembly factor 1, subunit A (p150) (CHAF1A) pseudogene |
| ENSG00000153936 | HS2ST1 | 0,128969407 | 1,093512269 | 0,76466554 | 0,976640468 | heparan sulfate 2-O-sulfotransferase 1 |
| ENSG00000119760 | SUPT7L | 0,12894936 | 1,093497074 | 0,64965349 | 0,956899963 | SPT7 like, STAGA complex subunit gamma |
| ENSG00000197852 | INKA2 | 0,128943305 | 1,093492485 | 0,7113683 | 0,970635643 | inka box actin regulator 2 |
| ENSG00000100138 | SNU13 | 0,128936638 | 1,093487431 | 0,6336874 | 0,954914102 | small nuclear ribonucleoprotein 13 |
| ENSG00000274616 | CNOT3 | 0,128902406 | 1,093461485 | 0,82483096 | 0,983374097 | CCR4-NOT transcription complex subunit 3 |
| ENSG00000229717 | AC110615,1 | 0,12887101 | 1,09343769 | 0,8992893 | 0,991832796 | novel transcript |
| ENSG00000273780 | ARHGAP23 | 0,12885298 | 1,093424025 | 0,83058501 | 0,984156028 | Rho GTPase activating protein 23 |
| ENSG00000130816 | DNMT1 | 0,128821122 | 1,09339988 | 0,58941541 | 0,946844281 | DNA methyltransferase 1 |
| ENSG00000155100 | OTUD6B | 0,128779868 | 1,093368614 | 0,71170377 | 0,97080996 | OTU deubiquitinase 6B |
| ENSG00000214248 | AC010336,1 | 0,128774866 | 1,093364823 | 0,80796924 | 0,981213877 | novel transcript, antisense to LRRC8E |
| ENSG00000247121 | AC009126,1 | 0,128754236 | 1,093349189 | 0,74040054 | 0,974833668 | novel transcript, antisense to LNPEP and ERAP2 |
| ENSG00000282105 | RPL41P2 | 0,128739959 | 1,093338369 | 0,87969967 | 0,990261566 | ribosomal protein L41 pseudogene 2 |
| ENSG00000198964 | SGMS1 | 0,128707216 | 1,093313555 | 0,67739617 | 0,963580269 | sphingomyelin synthase 1 |
| ENSG00000170296 | GABARAP | 0,128705937 | 1,093312586 | 0,62653493 | 0,952793934 | GABA type A receptor-associated protein |
| ENSG00000262435 | WASH8P | 0,128675608 | 1,093289602 | 0,67090326 | 0,962026636 | WAS protein family homolog 8, pseudogene |
| ENSG00000226210 | WASH8P | 0,128675608 | 1,093289602 | 0,67090326 | 0,962026636 | WAS protein family homolog 8, pseudogene |
| ENSG00000275236 | AC009120,5 | 0,128571658 | 1,093210831 | 0,84124916 | 0,985672131 | novel transcript |
| ENSG00000106245 | BUD31 | 0,128500898 | 1,093157213 | 0,57084411 | 0,942436428 | BUD31 homolog |

| | | | | | | |
|---|---|---|---|---|---|---|
| ENSG00000127314 | RAP1B | 0,128386262 | 1,093070354 | 0,75035363 | 0,975901808 | RAP1B, member of RAS oncogene family |
| ENSG00000257923 | CUX1 | 0,128381467 | 1,093066722 | 0,53849841 | 0,933082867 | cut like homeobox 1 |
| ENSG00000287249 | AC005400,1 | 0,128375775 | 1,093062409 | 0,85118541 | 0,987259878 | novel transcript, antisense to DPY19L1 |
| ENSG00000162851 | TFB2M | 0,128371694 | 1,093059317 | 0,6420235 | 0,956548736 | transcription factor B2, mitochondrial |
| ENSG00000286457 | AC009560,3 | 0,128316891 | 1,093017796 | 0,81767943 | 0,982321308 | novel transcript, antisense to RORA |
| ENSG00000155893 | PXYLP1 | 0,128290547 | 1,092997838 | 0,71037143 | 0,970482408 | 2-phosphoxylose phosphatase 1 |
| ENSG00000232653 | GOLGA8N | 0,128281882 | 1,092991273 | 0,7714222 | 0,978033855 | golgin A8 family member N |
| ENSG00000089818 | NECAP1 | 0,128275497 | 1,092986436 | 0,62623965 | 0,952793934 | NECAP endocytosis associated 1 |
| ENSG00000128524 | ATP6V1F | 0,128246604 | 1,092964546 | 0,59635681 | 0,948282297 | ATPase H+ transporting V1 subunit F |
| ENSG00000136631 | VPS45 | 0,128234361 | 1,092955272 | 0,71852766 | 0,971869926 | vacuolar protein sorting 45 homolog |
| ENSG00000197037 | ZSCAN25 | 0,128171726 | 1,092907822 | 0,62767637 | 0,952793934 | zinc finger and SCAN domain containing 25 |
| ENSG00000251131 | AC025171,3 | 0,128111365 | 1,092862097 | 0,8651162 | 0,989074806 | novel transcript |
| ENSG00000169019 | COMMD8 | 0,128077699 | 1,092836594 | 0,79949767 | 0,980161849 | COMM domain containing 8 |
| ENSG00000255046 | AC069185,1 | 0,127981946 | 1,092764064 | 0,86931991 | 0,989074806 | novel transcript |
| ENSG00000174137 | FAM53A | 0,127942941 | 1,09273452 | 0,8365586 | 0,985423106 | family with sequence similarity 53 member A |
| ENSG00000224398 | CSNK2B | 0,127934333 | 1,092728 | 0,62221563 | 0,952793934 | casein kinase 2 beta |
| ENSG00000088179 | PTPN4 | 0,127910584 | 1,092710013 | 0,79961337 | 0,980161849 | protein tyrosine phosphatase non-receptor type 4 |
| ENSG00000079691 | CARMIL1 | 0,1278779 | 1,092685258 | 0,6697538 | 0,962026636 | capping protein regulator and myosin 1 linker 1 |
| ENSG00000023734 | STRAP | 0,127821647 | 1,092642653 | 0,71449186 | 0,971041579 | serine/threonine kinase receptor associated protein |
| ENSG00000254413 | CHKB-CPT1B | 0,12775706 | 1,092593739 | 0,67037456 | 0,962026636 | CHKB-CPT1B readthrough (NMD candidate) |
| ENSG00000180398 | MCFD2 | 0,127732805 | 1,09257537 | 0,69451581 | 0,967417311 | multiple coagulation factor deficiency 2, ER cargo receptor complex subunit |
| ENSG00000199347 | RNU5E-1 | 0,127666189 | 1,092524921 | 0,85466654 | 0,987861659 | RNA, U5E small nuclear 1 |
| ENSG00000261934 | PCDHGA9 | 0,127623422 | 1,092492535 | 0,73738192 | 0,974353474 | protocadherin gamma subfamily A, 9 |
| ENSG00000221983 | UBA52 | 0,127615465 | 1,09248651 | 0,63413264 | 0,954940644 | ubiquitin A-52 residue ribosomal protein fusion product 1 |
| ENSG00000103495 | MAZ | 0,127590546 | 1,092467639 | 0,65169257 | 0,957089277 | MYC associated zinc finger protein |
| ENSG00000262666 | FAM189B | 0,12757575 | 1,092456436 | 0,69389466 | 0,967340633 | family with sequence similarity 189 member B |
| ENSG00000227671 | AL390728,4 | 0,127555782 | 1,092441315 | 0,68220771 | 0,964699524 | zinc finger protein pseudogene |
| ENSG00000058799 | YIPF1 | 0,12754515 | 1,092433265 | 0,64909645 | 0,956899963 | Yip1 domain family member 1 |
| ENSG00000258568 | RHOQP1 | 0,127532837 | 1,092423941 | 0,77517394 | 0,978587255 | ras homolog family member Q pseudogene 1 |
| ENSG00000203436 | Metazoa_SRP | 0,12739739 | 1,092321384 | 0,91087984 | 0,993222655 | Metazoan signal recognition particle RNA |
| ENSG00000050344 | NFE2L3 | 0,127364415 | 1,092296418 | 0,71351607 | 0,970812834 | nuclear factor, erythroid 2 like 3 |

| | | | | | | |
|---|---|---|---|---|---|---|
| ENSG00000143067 | ZNF697 | 0,127343526 | 1,092280602 | 0,62280625 | 0,952793934 | zinc finger protein 697 |
| ENSG00000270184 | AC018695,4 | 0,127324861 | 1,092266471 | 0,84107595 | 0,985672131 | novel transcript, antisense to COX4NB |
| ENSG00000242485 | MRPL20 | 0,127269615 | 1,092224645 | 0,53011744 | 0,93110065 | mitochondrial ribosomal protein L20 |
| | | | | | | RNA binding protein, fox-1 homolog (C, elegans) 1 (RBFOX1) |
| ENSG00000284727 | AC116562,3 | 0,127234859 | 1,092198333 | 0,71521496 | 0,971041579 | pseudogene |
| ENSG00000196199 | MPHOSPH8 | 0,127197585 | 1,092170114 | 0,67148886 | 0,962085713 | M-phase phosphoprotein 8 |
| ENSG00000123843 | C4BPB | 0,127142964 | 1,092128765 | 0,87544466 | 0,990078774 | complement component 4 binding protein beta |
| ENSG00000159882 | ZNF230 | 0,127060142 | 1,09206607 | 0,69077171 | 0,967179044 | zinc finger protein 230 |
| ENSG00000168653 | NDUFS5 | 0,12704599 | 1,092055357 | 0,63871476 | 0,956018518 | NADH:ubiquinone oxidoreductase subunit S5 |
| ENSG00000100462 | PRMT5 | 0,12701662 | 1,092033126 | 0,56580166 | 0,940866081 | protein arginine methyltransferase 5 |
| ENSG00000104356 | POP1 | 0,126938808 | 1,091974229 | 0,63606834 | 0,955399772 | POP1 homolog, ribonuclease P/MRP subunit |
| ENSG00000154316 | TDH | 0,126924675 | 1,091963532 | 0,91976933 | 0,993699236 | L-threonine dehydrogenase (pseudogene) |
| ENSG00000284856 | TDH | 0,126924675 | 1,091963532 | 0,91976933 | 0,993699236 | L-threonine dehydrogenase (pseudogene) |
| ENSG00000277809 | AC012354,8 | 0,126855811 | 1,09191141 | 0,77828161 | 0,978587255 | |
| ENSG00000224103 | HLA-DPA1 | 0,126762609 | 1,091840872 | 0,9369526 | 0,994479239 | major histocompatibility complex, class II, DP alpha 1 |
| ENSG00000165304 | MELK | 0,126657451 | 1,09176129 | 0,76038953 | 0,97624106 | maternal embryonic leucine zipper kinase |
| ENSG00000105993 | DNAJB6 | 0,126577936 | 1,091701119 | 0,58666675 | 0,945395317 | DnaJ heat shock protein family (Hsp40) member B6 |
| ENSG00000124120 | TTPAL | 0,126560638 | 1,09168803 | 0,55921676 | 0,939727587 | alpha tocopherol transfer protein like |
| ENSG00000151553 | FAM160B1 | 0,126518226 | 1,091655937 | 0,75371848 | 0,975901808 | family with sequence similarity 160 member B1 |
| ENSG00000182149 | IST1 | 0,126489304 | 1,091634053 | 0,57743088 | 0,94337651 | IST1 factor associated with ESCRT-III |
| ENSG00000163312 | HELQ | 0,126487816 | 1,091632926 | 0,67497199 | 0,963268551 | helicase, POLQ like |
| ENSG00000188710 | QRFP | 0,126460713 | 1,091612419 | 0,80208445 | 0,980710009 | pyroglutamylated RFamide peptide |
| ENSG00000169554 | ZEB2 | 0,126334649 | 1,091517037 | 0,68386954 | 0,965385216 | zinc finger E-box binding homeobox 2 |
| ENSG00000058091 | CDK14 | 0,126330693 | 1,091514044 | 0,712663 | 0,970812834 | cyclin dependent kinase 14 |
| ENSG00000227212 | PFN1P6 | 0,126320819 | 1,091506574 | 0,86847254 | 0,989074806 | profilin 1 pseudogene 6 |
| ENSG00000226419 | SLC16A1-AS1 | 0,126320271 | 1,091506159 | 0,69299413 | 0,967271667 | SLC16A1 antisense RNA 1 |
| ENSG00000115464 | USP34 | 0,12622492 | 1,091434021 | 0,75489792 | 0,975901808 | ubiquitin specific peptidase 34 |
| ENSG00000115661 | STK16 | 0,126222353 | 1,091432079 | 0,61830598 | 0,952793934 | serine/threonine kinase 16 |
| ENSG00000173401 | GLIPR1L1 | 0,126179879 | 1,091399947 | 0,90081286 | 0,991865079 | GLIPR1 like 1 |
| ENSG00000180694 | TMEM64 | 0,126133656 | 1,09136498 | 0,76287026 | 0,97647393 | transmembrane protein 64 |
| ENSG00000277196 | AC007325,2 | 0,126109793 | 1,091346929 | 0,85046208 | 0,987075267 | proline dehydrogenase 1, mitochondrial |
| ENSG00000103429 | BFAR | 0,126069312 | 1,091316306 | 0,70054137 | 0,968522643 | bifunctional apoptosis regulator |

| | | | | | | |
|---|---|---|---|---|---|---|
| ENSG00000148154 | UGCG | 0,126061856 | 1,091310666 | 0,77452344 | 0,978553937 | UDP-glucose ceramide glucosyltransferase |
| ENSG00000108773 | KAT2A | 0,126057942 | 1,091307706 | 0,62689394 | 0,952793934 | lysine acetyltransferase 2A |
| ENSG00000151693 | ASAP2 | 0,126025693 | 1,091283311 | 0,64352615 | 0,956633898 | ArfGAP with SH3 domain, ankyrin repeat and PH domain 2 |
| ENSG00000277301 | AL034550,2 | 0,126012485 | 1,091273321 | 0,87198571 | 0,989217334 | novel transcript, antisense to NOL4L |
| ENSG00000181991 | MRPS11 | 0,125961419 | 1,091234695 | 0,58558831 | 0,945213113 | mitochondrial ribosomal protein S11 |
| ENSG00000218690 | H2AC10P | 0,125807625 | 1,091118373 | 0,85223139 | 0,987405951 | H2A clustered histone 10, pseudogene |
| ENSG00000160752 | FDPS | 0,125796763 | 1,091110158 | 0,56726544 | 0,941425553 | farnesyl diphosphate synthase |
| ENSG00000235683 | AC018442,1 | 0,125776381 | 1,091094743 | 0,88589203 | 0,990717729 | leucine zipper-EF-hand containing transmembrane protein 1 (LETM1) pseudogene |
| ENSG00000276234 | TADA2A | 0,125717316 | 1,091050074 | 0,64822826 | 0,956899963 | transcriptional adaptor 2A |
| ENSG00000149658 | YTHDF1 | 0,125602252 | 1,090963059 | 0,63036324 | 0,953166783 | YTH N6-methyladenosine RNA binding protein 1 |
| ENSG00000163093 | BBS5 | 0,125550015 | 1,090923559 | 0,77805757 | 0,978587255 | Bardet-Biedl syndrome 5 |
| ENSG00000285628 | AC126755,5 | 0,125503516 | 1,090888398 | 0,76381085 | 0,976640468 | novel transcript |
| ENSG00000178078 | STAP2 | 0,125499813 | 1,090885598 | 0,80491886 | 0,98109454 | signal transducing adaptor family member 2 |
| ENSG00000170619 | COMMD5 | 0,125370111 | 1,090787529 | 0,65936715 | 0,958425175 | COMM domain containing 5 |
| ENSG00000278759 | TBCD | 0,125315877 | 1,090746525 | 0,74596443 | 0,975707496 | tubulin folding cofactor D |
| ENSG00000137776 | SLTM | 0,12519588 | 1,090655805 | 0,66617285 | 0,961020116 | SAFB like transcription modulator |
| ENSG00000127080 | IPPK | 0,125186756 | 1,090648907 | 0,60210519 | 0,950046267 | inositol-pentakisphosphate 2-kinase |
| ENSG00000016864 | GLT8D1 | 0,125090917 | 1,090576458 | 0,701272 | 0,968741948 | glycosyltransferase 8 domain containing 1 |
| ENSG00000141664 | ZCCHC2 | 0,125060003 | 1,090553089 | 0,67014286 | 0,962026636 | zinc finger CCHC-type containing 2 |
| ENSG00000085871 | MGST2 | 0,12500718 | 1,09051316 | 0,78088564 | 0,978597905 | microsomal glutathione S-transferase 2 |
| ENSG00000183873 | SCN5A | 0,124995918 | 1,090504647 | 0,81815281 | 0,982323806 | sodium voltage-gated channel alpha subunit 5 |
| ENSG00000007168 | PAFAH1B1 | 0,124932972 | 1,090457068 | 0,6709456 | 0,962026636 | platelet activating factor acetylhydrolase 1b regulatory subunit 1 |
| ENSG00000286355 | AL355834,2 | 0,124928237 | 1,09045349 | 0,88810577 | 0,990717729 | novel transcript |
| ENSG00000207001 | SNORD116-2 | 0,124845901 | 1,090391258 | 0,89643928 | 0,991638165 | small nucleolar RNA, C/D box 116-2 |
| ENSG00000287009 | AC021037,1 | 0,124787236 | 1,09034692 | 0,75379099 | 0,975901808 | novel transcript, antisense to ZNF518A |
| ENSG00000173660 | UQCRH | 0,124751419 | 1,090319851 | 0,64937993 | 0,956899963 | ubiquinol-cytochrome c reductase hinge protein |
| ENSG00000181026 | AEN | 0,124733803 | 1,090306537 | 0,63709083 | 0,955945807 | apoptosis enhancing nuclease |
| ENSG00000257207 | AC112229,3 | 0,124623671 | 1,090223309 | 0,90086322 | 0,991865079 | novel transcript |
| ENSG00000167981 | ZNF597 | 0,124590689 | 1,090198386 | 0,74872621 | 0,975901808 | zinc finger protein 597 |
| ENSG00000094804 | CDC6 | 0,124571038 | 1,090183536 | 0,62824558 | 0,952793934 | cell division cycle 6 |

| | | | | | | |
|---|---|---|---|---|---|---|
| ENSG00000148341 | SH3GLB2 | 0,124537497 | 1,090158191 | 0,72224874 | 0,972594421 | SH3 domain containing GRB2 like, endophilin B2 |
| ENSG00000197620 | EOLA1 | 0,124380156 | 1,090039304 | 0,59890005 | 0,948894897 | endothelium and lymphocyte associated ASCH domain 1 |
| ENSG00000241218 | CSP2 | 0,124311274 | 1,089987261 | 0,91950262 | 0,993617097 | CS pseudogene 2 |
| ENSG00000285898 | AC007317,1 | 0,124245973 | 1,089937926 | 0,79374421 | 0,979776981 | Novel transcript |
| ENSG00000108061 | SHOC2 | 0,124135924 | 1,089854789 | 0,78259909 | 0,978647973 | SHOC2 leucine rich repeat scaffold protein |
| ENSG00000179362 | HMGN2P46 | 0,124124941 | 1,089846492 | 0,88961102 | 0,990717729 | high mobility group nucleosomal binding domain 2 pseudogene 46 |
| ENSG00000281091 | AL117327,1 | 0,124063981 | 1,089800442 | 0,73654576 | 0,974353474 | novel transcript |
| ENSG00000149100 | EIF3M | 0,124063237 | 1,08979988 | 0,6504082 | 0,956899963 | eukaryotic translation initiation factor 3 subunit M |
| ENSG00000160007 | ARHGAP35 | 0,124042091 | 1,089783906 | 0,5372929 | 0,932883277 | Rho GTPase activating protein 35 |
| ENSG00000225951 | ODF2-AS1 | 0,124038005 | 1,08978082 | 0,84632605 | 0,986939308 | ODF2 antisense RNA 1 |
| ENSG00000284858 | AC091185,2 | 0,123996531 | 1,089749492 | 0,93724036 | 0,994479239 | novel oocyte-secreted protein family member pseudogene |
| ENSG00000269026 | AC003006,1 | 0,12397987 | 1,089736907 | 0,88766613 | 0,990717729 | novel protein |
| ENSG00000144048 | DUSP11 | 0,123892619 | 1,089671004 | 0,72420787 | 0,973178901 | dual specificity phosphatase 11 |
| ENSG00000101166 | PRELID3B | 0,123783899 | 1,089588891 | 0,79959907 | 0,980161849 | PRELI domain containing 3B |
| ENSG00000100316 | RPL3 | 0,123780807 | 1,089586556 | 0,62604154 | 0,952793934 | ribosomal protein L3 |
| ENSG00000100124 | ANKRD54 | 0,123769984 | 1,089578382 | 0,60874963 | 0,951601152 | ankyrin repeat domain 54 |
| ENSG00000261338 | AC021016,1 | 0,123758483 | 1,089569696 | 0,72915196 | 0,974009551 | novel transcript, overlapping to ARPC2 and GPBAR1 |
| ENSG00000047578 | KATNIP | 0,123750199 | 1,089563439 | 0,60323615 | 0,950352286 | katanin interacting protein |
| ENSG00000110851 | PRDM4 | 0,123742285 | 1,089557463 | 0,61740351 | 0,952793934 | PR/SET domain 4 |
| ENSG00000105928 | GSDME | 0,123598813 | 1,089449114 | 0,67804873 | 0,963963232 | gasdermin E |
| ENSG00000141232 | TOB1 | 0,123586811 | 1,089440051 | 0,7256142 | 0,97326334 | transducer of ERBB2, 1 |
| ENSG00000102901 | CENPT | 0,123522402 | 1,089391414 | 0,61252465 | 0,952413269 | centromere protein T |
| ENSG00000213380 | COG8 | 0,12349242 | 1,089368775 | 0,57132507 | 0,942436428 | component of oligomeric golgi complex 8 |
| ENSG00000138433 | CIR1 | 0,123486248 | 1,089364114 | 0,70401249 | 0,969016754 | corepressor interacting with RBPJ, CIR1 |
| ENSG00000264047 | RN7SL455P | 0,123483998 | 1,089362416 | 0,87964326 | 0,990261566 | RNA, 7SL, cytoplasmic 455, pseudogene |
| ENSG00000230678 | BRD2 | 0,123471136 | 1,089352703 | 0,68762187 | 0,966063455 | bromodomain containing 2 |
| ENSG00000268030 | AC005253,1 | 0,123455292 | 1,08934074 | 0,66466589 | 0,96046247 | novel transcript, antisense to KXD1 |
| ENSG00000234539 | ATF6B | 0,123433519 | 1,0893243 | 0,68416112 | 0,965415049 | activating transcription factor 6 beta |
| ENSG00000199293 | SNORA21 | 0,123370097 | 1,089276414 | 0,87117826 | 0,989217334 | small nucleolar RNA, H/ACA box 21 |
| ENSG00000254166 | CASC19 | 0,123270321 | 1,089201082 | 0,8502138 | 0,987036295 | cancer susceptibility 19 |

| | | | | | | |
|---|---|---|---|---|---|---|
| ENSG00000136942 | RPL35 | 0,123240998 | 1,089178944 | 0,54801146 | 0,935404974 | ribosomal protein L35 |
| ENSG00000203950 | RTL8A | 0,123017803 | 1,089010454 | 0,55924341 | 0,939727587 | retrotransposon Gag like 8A |
| ENSG00000198231 | DDX42 | 0,123002385 | 1,088998816 | 0,57758922 | 0,94337651 | DEAD-box helicase 42 |
| ENSG00000206206 | DAXX | 0,123001276 | 1,088997978 | 0,62392037 | 0,952793934 | death domain associated protein |
| ENSG00000227046 | DAXX | 0,123001276 | 1,088997978 | 0,62392037 | 0,952793934 | death domain associated protein |
| ENSG00000065526 | SPEN | 0,12283705 | 1,088874022 | 0,54957624 | 0,936091063 | spen family transcriptional repressor |
| ENSG00000267255 | AC011498,3 | 0,122836246 | 1,088873415 | 0,8777526 | 0,990109814 | novel transcript, antisense to CHAF1A |
| ENSG00000100226 | GTPBP1 | 0,122798859 | 1,088845198 | 0,66355957 | 0,959920886 | GTP binding protein 1 |
| ENSG00000105197 | TIMM50 | 0,122562636 | 1,088666928 | 0,67033155 | 0,962026636 | translocase of inner mitochondrial membrane 50 |
| ENSG00000254772 | EEF1G | 0,122500192 | 1,088619808 | 0,59978718 | 0,949340299 | eukaryotic translation elongation factor 1 gamma |
| ENSG00000259793 | AC013726,1 | 0,122495593 | 1,088616338 | 0,91513303 | 0,993361299 | novel transcript, antisense to DGKD |
| ENSG00000183742 | MACC1 | 0,122387561 | 1,088534823 | 0,82803449 | 0,983595244 | MET transcriptional regulator MACC1 |
| ENSG00000138663 | COPS4 | 0,12237986 | 1,088529013 | 0,75878588 | 0,976120207 | COP9 signalosome subunit 4 |
| ENSG00000064201 | TSPAN32 | 0,122359929 | 1,088513974 | 0,91488072 | 0,993361299 | tetraspanin 32 |
| ENSG00000262179 | MYMX | 0,122346775 | 1,08850405 | 0,87257531 | 0,989217334 | myomixer, myoblast fusion factor |
| ENSG00000163681 | SLMAP | 0,122332209 | 1,08849306 | 0,6496166 | 0,956899963 | sarcolemma associated protein |
| ENSG00000102302 | FGD1 | 0,122302769 | 1,088470848 | 0,63518729 | 0,95524053 | FYVE, RhoGEF and PH domain containing 1 |
| ENSG00000085982 | USP40 | 0,122265405 | 1,088442658 | 0,55965745 | 0,939811058 | ubiquitin specific peptidase 40 |
| ENSG00000275092 | AL031710,2 | 0,122198584 | 1,088392246 | 0,85944362 | 0,989074806 | novel transcript, sense intronic to MAPK8IP3 |
| ENSG00000273494 | PANK4 | 0,122157224 | 1,088361044 | 0,61681304 | 0,952793934 | pantothenate kinase 4 (inactive) |
| ENSG00000265185 | SNORD3B-1 | 0,122120785 | 1,088333555 | 0,65769491 | 0,957864535 | small nucleolar RNA, C/D box 3B-1 |
| ENSG00000111897 | SERINC1 | 0,12211277 | 1,088327509 | 0,77286389 | 0,978310326 | serine incorporator 1 |
| ENSG00000125449 | ARMC7 | 0,122087299 | 1,088308294 | 0,64719523 | 0,956899963 | armadillo repeat containing 7 |
| ENSG00000229519 | AL591866,1 | 0,122077717 | 1,088301067 | 0,88299887 | 0,990717729 | ribosomal protein L34 (RPL34) pseudogene |
| ENSG00000147403 | RPL10 | 0,122033716 | 1,088267874 | 0,65418474 | 0,957444211 | ribosomal protein L10 |
| ENSG00000263677 | AC012588,1 | 0,122007766 | 1,088248299 | 0,89376873 | 0,991135494 | novel transcript |
| ENSG00000165699 | TSC1 | 0,121968278 | 1,088218513 | 0,59141346 | 0,947416795 | TSC complex subunit 1 |
| ENSG00000143740 | SNAP47 | 0,121966344 | 1,088217055 | 0,62338388 | 0,952793934 | synaptosome associated protein 47 |
| ENSG00000223496 | EXOSC6 | 0,121905803 | 1,08817139 | 0,64058735 | 0,956348702 | exosome component 6 |
| ENSG00000287920 | AL022098,1 | 0,12182916 | 1,088113583 | 0,87271119 | 0,989262536 | novel transcript, antisense to NEDD9 |
| ENSG00000037042 | TUBG2 | 0,121802666 | 1,0880936 | 0,629261 | 0,952793934 | tubulin gamma 2 |
| ENSG00000253250 | C8orf88 | 0,121792374 | 1,088085838 | 0,79900186 | 0,980161849 | chromosome 8 open reading frame 88 |
| ENSG00000127920 | GNG11 | 0,121792041 | 1,088085587 | 0,73055006 | 0,974321178 | G protein subunit gamma 11 |

| | | | | | | |
|---|---|---|---|---|---|---|
| ENSG00000152700 | SAR1B | 0,121763695 | 1,088064209 | 0,6154527 | 0,952793934 | secretion associated Ras related GTPase 1B |
| ENSG00000165487 | MICU2 | 0,121587707 | 1,087931489 | 0,74918705 | 0,975901808 | mitochondrial calcium uptake 2 |
| ENSG00000125834 | STK35 | 0,121560584 | 1,087911036 | 0,56550369 | 0,940865625 | serine/threonine kinase 35 |
| ENSG00000275003 | DDT | 0,12152086 | 1,08788108 | 0,69299065 | 0,967271667 | D-dopachrome tautomerase |
| ENSG00000258701 | LINC00638 | 0,121497341 | 1,087863346 | 0,80257724 | 0,980798762 | long intergenic non-protein coding RNA 638 |
| ENSG00000111215 | PRR4 | 0,12146352 | 1,087837843 | 0,86746134 | 0,989074806 | proline rich 4 |
| ENSG00000196204 | RNF216P1 | 0,121282957 | 1,087701702 | 0,56135764 | 0,940567693 | ring finger protein 216 pseudogene 1 |
| ENSG00000188021 | UBQLN2 | 0,121274232 | 1,087695124 | 0,72057971 | 0,972142678 | ubiquilin 2 |
| ENSG00000234728 | SNHG32 | 0,121113896 | 1,087574248 | 0,69097084 | 0,967179456 | small nucleolar RNA host gene 32 |
| ENSG00000206380 | SNHG32 | 0,121113896 | 1,087574248 | 0,69097084 | 0,967179456 | small nucleolar RNA host gene 32 |
| ENSG00000142583 | SLC2A5 | 0,121088938 | 1,087555433 | 0,85207284 | 0,98732978 | solute carrier family 2 member 5 |
| ENSG00000103269 | RHBDL1 | 0,121010205 | 1,087496083 | 0,8990812 | 0,991832796 | rhomboid like 1 |
| ENSG00000186854 | TRABD2A | 0,120948501 | 1,087449572 | 0,78681317 | 0,979373608 | TraB domain containing 2A |
| ENSG00000163785 | RYK | 0,120923622 | 1,08743082 | 0,75206913 | 0,975901808 | receptor like tyrosine kinase |
| ENSG00000055147 | FAM114A2 | 0,120885435 | 1,087402036 | 0,70414534 | 0,969077997 | family with sequence similarity 114 member A2 |
| ENSG00000109065 | NAT9 | 0,120797145 | 1,087335491 | 0,72048635 | 0,972142678 | N-acetyltransferase 9 (putative) |
| ENSG00000156564 | LRFN2 | 0,120751559 | 1,087301135 | 0,90491309 | 0,992590661 | leucine rich repeat and fibronectin type III domain containing 2 |
| ENSG00000259420 | AC046168,2 | 0,120694041 | 1,087257787 | 0,89396219 | 0,991135494 | novel transcript |
| ENSG00000119636 | BBOF1 | 0,12057887 | 1,087170994 | 0,73859133 | 0,974353474 | basal body orientation factor 1 |
| ENSG00000165055 | METTL2B | 0,120495752 | 1,087108361 | 0,60543873 | 0,950927937 | methyltransferase like 2B |
| ENSG00000133794 | ARNTL | 0,120449926 | 1,08707383 | 0,72155101 | 0,972142678 | aryl hydrocarbon receptor nuclear translocator like |
| ENSG00000163848 | ZNF148 | 0,12037656 | 1,08701855 | 0,76071894 | 0,97624106 | zinc finger protein 148 |
| ENSG00000130584 | ZBTB46 | 0,120347312 | 1,086996513 | 0,69185634 | 0,967204896 | zinc finger and BTB domain containing 46 |
| ENSG00000135953 | MFSD9 | 0,120331939 | 1,08698493 | 0,6581054 | 0,957906876 | major facilitator superfamily domain containing 9 |
| ENSG00000257612 | MIR4307HG | 0,120303753 | 1,086963694 | 0,92508034 | 0,993915537 | MIR4307 host gene |
| ENSG00000278058 | AC009159,3 | 0,120296572 | 1,086958283 | 0,85391254 | 0,987861659 | novel transcript |
| ENSG00000088881 | EBF4 | 0,120285138 | 1,086949669 | 0,69916613 | 0,968522643 | EBF family member 4 |
| ENSG00000273284 | AP001033,2 | 0,120281864 | 1,086947202 | 0,87227565 | 0,989217334 | novel transcript, sense intronic to ANKRD12 |
| ENSG00000155008 | APOOL | 0,120196397 | 1,086882812 | 0,66468034 | 0,96046247 | apolipoprotein O like |
| ENSG00000259332 | ST20-MTHFS | 0,120151492 | 1,086848982 | 0,78867056 | 0,979462318 | ST20-MTHFS readthrough |
| ENSG00000267648 | AC060766,5 | 0,120130532 | 1,086833192 | 0,81487772 | 0,981521413 | DTW domain containing 2 (DTWD2) pseudogene |
| ENSG00000133884 | DPF2 | 0,12011698 | 1,086822983 | 0,60543538 | 0,950927937 | double PHD fingers 2 |

| | | | | | | |
|---|---|---|---|---|---|---|
| ENSG00000237328 | RAI1-AS1 | 0,1200195 | 1,086749551 | 0,88468042 | 0,990717729 | RAI1 antisense RNA 1 |
| ENSG00000261560 | AC007216,3 | 0,119998553 | 1,086733772 | 0,82930316 | 0,984034229 | novel transcript, sense intronic to GSPT1 |
| ENSG00000168256 | NKIRAS2 | 0,119929765 | 1,086681958 | 0,57839638 | 0,943425812 | NFKB inhibitor interacting Ras like 2 |
| ENSG00000274914 | LILRA5 | 0,119905842 | 1,086663939 | 0,87889649 | 0,990109814 | leukocyte immunoglobulin like receptor A5 |
| ENSG00000232057 | AC093390,1 | 0,119896163 | 1,086656649 | 0,925868 | 0,993915537 | novel transcript |
| ENSG00000148362 | PAXX | 0,119852798 | 1,086623986 | 0,71900143 | 0,971869926 | PAXX non-homologous end joining factor |
| ENSG00000118197 | DDX59 | 0,119837961 | 1,08661281 | 0,74982025 | 0,975901808 | DEAD-box helicase 59 |
| ENSG00000144524 | COPS7B | 0,119836089 | 1,086611401 | 0,57446267 | 0,942475308 | COP9 signalosome subunit 7B |
| ENSG00000182670 | TTC3 | 0,119831025 | 1,086607587 | 0,7270287 | 0,973344786 | tetratricopeptide repeat domain 3 |
| ENSG00000115241 | PPM1G | 0,119785687 | 1,086573439 | 0,59903867 | 0,948894897 | protein phosphatase, Mg2+/Mn2+ dependent 1G |
| ENSG00000136888 | ATP6V1G1 | 0,119743636 | 1,086541769 | 0,66894283 | 0,961599545 | ATPase H+ transporting V1 subunit G1 |
| ENSG00000161654 | LSM12 | 0,119678077 | 1,086492396 | 0,65595611 | 0,957444211 | LSM12 homolog |
| ENSG00000261786 | AC006058,1 | 0,119658961 | 1,086478 | 0,77662084 | 0,978587255 | novel transcript |
| ENSG00000169564 | PCBP1 | 0,119585066 | 1,086422351 | 0,66400079 | 0,959985685 | poly(rC) binding protein 1 |
| ENSG00000058063 | ATP11B | 0,11957481 | 1,086414628 | 0,77884458 | 0,978587255 | ATPase phospholipid transporting 11B (putative) |
| ENSG00000272142 | LYRM4-AS1 | 0,1195576 | 1,086401669 | 0,69085381 | 0,967179044 | LYRM4 antisense RNA 1 |
| ENSG00000100372 | SLC25A17 | 0,119518928 | 1,086372547 | 0,63285511 | 0,954396141 | solute carrier family 25 member 17 |
| ENSG00000112578 | BYSL | 0,119471792 | 1,086337054 | 0,59901256 | 0,948894897 | bystin like |
| ENSG00000155329 | ZCCHC10 | 0,119458813 | 1,086327281 | 0,79901508 | 0,980161849 | zinc finger CCHC-type containing 10 |
| ENSG00000178234 | GALNT11 | 0,119392423 | 1,086277291 | 0,58344794 | 0,944174844 | polypeptide N-acetylgalactosaminyltransferase 11 |
| ENSG00000232938 | RPL23AP87 | 0,119289965 | 1,086200148 | 0,80124051 | 0,980557464 | ribosomal protein L23a pseudogene 87 |
| ENSG00000225526 | MKRN2OS | 0,119216984 | 1,086145203 | 0,86579027 | 0,989074806 | MKRN2 opposite strand |
| ENSG00000287070 | AL499604,1 | 0,119176754 | 1,086114915 | 0,87335142 | 0,989614687 | novel transcript, sense intronic to TRMO |
| ENSG00000141552 | ANAPC11 | 0,119146541 | 1,08609217 | 0,67397054 | 0,963037987 | anaphase promoting complex subunit 11 |
| ENSG00000008382 | MPND | 0,119074147 | 1,086037671 | 0,80332887 | 0,980866933 | MPN domain containing |
| ENSG00000066933 | MYO9A | 0,119060137 | 1,086027125 | 0,70169154 | 0,968742626 | myosin IXA |
| ENSG00000275498 | PDXDC1 | 0,119030273 | 1,086004644 | 0,71512565 | 0,971041579 | pyridoxal dependent decarboxylase domain containing 1 |
| ENSG00000286960 | AC009869,1 | 0,118953705 | 1,085947008 | 0,93709139 | 0,994479239 | novel transcript |
| ENSG00000179930 | ZNF648 | 0,118946116 | 1,085941297 | 0,88509325 | 0,990717729 | zinc finger protein 648 |
| ENSG00000127884 | ECHS1 | 0,118918015 | 1,085920144 | 0,57714719 | 0,94337651 | enoyl-CoA hydratase, short chain 1 |
| ENSG00000179912 | R3HDM2 | 0,118848208 | 1,085867602 | 0,65767931 | 0,957864535 | R3H domain containing 2 |
| ENSG00000035664 | DAPK2 | 0,118838812 | 1,08586053 | 0,79541391 | 0,979879951 | death associated protein kinase 2 |

| | | | | | | |
|---|---|---|---|---|---|---|
| ENSG00000117751 | PPP1R8 | 0,118815307 | 1,085842838 | 0,69842169 | 0,968522643 | protein phosphatase 1 regulatory subunit 8 |
| ENSG00000198624 | CCDC69 | 0,118750957 | 1,085794407 | 0,75450782 | 0,975901808 | coiled-coil domain containing 69 |
| ENSG00000159199 | ATP5MC1 | 0,118623679 | 1,085698619 | 0,62851546 | 0,952793934 | ATP synthase membrane subunit c locus 1 |
| ENSG00000286177 | AC011462,5 | 0,118509906 | 1,085613003 | 0,72634685 | 0,973344786 | novel transcript, antisense to CCDC97 and HNRNPUL1 |
| ENSG00000129187 | DCTD | 0,118449855 | 1,085567817 | 0,61431746 | 0,952793934 | dCMP deaminase |
| ENSG00000118263 | KLF7 | 0,118414245 | 1,085541021 | 0,65604004 | 0,957444211 | Kruppel like factor 7 |
| ENSG00000151233 | GXYLT1 | 0,118374197 | 1,085510888 | 0,7731836 | 0,978363975 | glucoside xylosyltransferase 1 |
| ENSG00000100604 | CHGA | 0,118309236 | 1,085462012 | 0,85403314 | 0,987861659 | chromogranin A |
| ENSG00000196652 | ZKSCAN5 | 0,118286539 | 1,085444934 | 0,66214927 | 0,959874876 | zinc finger with KRAB and SCAN domains 5 |
| ENSG00000187957 | DNER | 0,118270024 | 1,085432509 | 0,80435806 | 0,980875302 | delta/notch like EGF repeat containing |
| ENSG00000101057 | MYBL2 | 0,118267969 | 1,085430963 | 0,64651134 | 0,956899963 | MYB proto-oncogene like 2 |
| ENSG00000164818 | DNAAF5 | 0,118234723 | 1,08540595 | 0,63758276 | 0,956017934 | dynein axonemal assembly factor 5 |
| ENSG00000282012 | AP000357,2 | 0,118218463 | 1,085393717 | 0,88717988 | 0,990717729 | novel transcript |
| ENSG00000198242 | RPL23A | 0,118183239 | 1,085367218 | 0,72599735 | 0,973344786 | ribosomal protein L23a |
| ENSG00000278128 | AL139093,1 | 0,118180926 | 1,085365477 | 0,85036536 | 0,987036295 | nuclear pore associated protein 1 (NPAP1) pseudogene |
| ENSG00000156831 | NSMCE2 | 0,118179644 | 1,085364512 | 0,74035704 | 0,974833668 | NSE2 (MMS21) homolog, SMC5-SMC6 complex SUMO ligase |
| ENSG00000243279 | PRAF2 | 0,118064417 | 1,085277829 | 0,64144143 | 0,956465487 | PRA1 domain family member 2 |
| ENSG00000281603 | MUC20-OT1 | 0,118062633 | 1,085276487 | 0,69187312 | 0,967204896 | MUC20 overlapping transcript |
| ENSG00000176171 | BNIP3 | 0,11798016 | 1,085214448 | 0,63554173 | 0,955314044 | BCL2 interacting protein 3 |
| ENSG00000106012 | IQCE | 0,1179707 | 1,085207332 | 0,80544191 | 0,981213877 | IQ motif containing E |
| ENSG00000038210 | PI4K2B | 0,117903119 | 1,085156498 | 0,73601692 | 0,974353474 | phosphatidylinositol 4-kinase type 2 beta |
| ENSG00000174444 | RPL4 | 0,117844297 | 1,085112254 | 0,64118467 | 0,956465487 | ribosomal protein L4 |
| ENSG00000105220 | GPI | 0,117844263 | 1,085112229 | 0,63928831 | 0,956018518 | glucose-6-phosphate isomerase |
| ENSG00000196517 | SLC6A9 | 0,117800736 | 1,085079491 | 0,77945369 | 0,978587255 | solute carrier family 6 member 9 |
| ENSG00000237529 | AL137847,2 | 0,117776221 | 1,085061053 | 0,85376311 | 0,987861659 | novel transcript |
| ENSG00000285023 | CHORDC1 | 0,11777617 | 1,085061015 | 0,74055002 | 0,974885509 | cysteine and histidine rich domain containing 1 |
| ENSG00000243403 | AC090543,2 | 0,117774115 | 1,085059469 | 0,94315371 | 0,995402494 | ribosomal protein S26 (RPS26) pseudogene |
| ENSG00000156931 | VPS8 | 0,117764216 | 1,085052024 | 0,75726542 | 0,975901808 | VPS8 subunit of CORVET complex |
| ENSG00000239614 | HMGN1P7 | 0,117751604 | 1,085042538 | 0,93476931 | 0,994479239 | high mobility group nucleosome binding domain 1 pseudogene 7 |

| | | | | | | |
|---|---|---|---|---|---|---|
| ENSG00000090971 | NAT14 | 0,117715298 | 1,085015233 | 0,76113091 | 0,97624106 | N-acetyltransferase 14 (putative) |
| ENSG00000233695 | GAS6-AS1 | 0,117600325 | 1,084928769 | 0,90901327 | 0,992814265 | GAS6 antisense RNA 1 |
| ENSG00000145545 | SRD5A1 | 0,117504184 | 1,084856471 | 0,63597757 | 0,955399772 | steroid 5 alpha-reductase 1 |
| ENSG00000162959 | MEMO1 | 0,117499726 | 1,084853119 | 0,73061623 | 0,974321178 | mediator of cell motility 1 |
| ENSG00000177030 | DEAF1 | 0,117377468 | 1,084761189 | 0,66685426 | 0,9611116 | DEAF1 transcription factor |
| ENSG00000164124 | TMEM144 | 0,117373492 | 1,0847582 | 0,83382105 | 0,984905093 | transmembrane protein 144 |
| ENSG00000163795 | ZNF513 | 0,117293425 | 1,084698 | 0,74428951 | 0,975641819 | zinc finger protein 513 |
| ENSG00000127616 | SMARCA4 | 0,117279682 | 1,084687667 | 0,76240574 | 0,976315588 | SWI/SNF related, matrix associated, actin dependent regulator of chromatin, subfamily a, member 4 |
| ENSG00000167280 | ENGASE | 0,117255698 | 1,084669634 | 0,76303943 | 0,976576876 | endo-beta-N-acetylglucosaminidase |
| ENSG00000214223 | HNRNPA1P10 | 0,117244546 | 1,08466125 | 0,78931078 | 0,979462318 | heterogeneous nuclear ribonucleoprotein A1 pseudogene 10 |
| ENSG00000186767 | SPIN4 | 0,117185412 | 1,084616792 | 0,75933728 | 0,976120207 | spindlin family member 4 |
| ENSG00000128309 | MPST | 0,117185005 | 1,084616487 | 0,73830809 | 0,974353474 | mercaptopyruvate sulfurtransferase |
| ENSG00000178719 | GRINA | 0,117129232 | 1,084574558 | 0,62437196 | 0,952793934 | glutamate ionotropic receptor NMDA type subunit associated protein 1 |
| ENSG00000100890 | PRORP | 0,117107928 | 1,084558541 | 0,65322987 | 0,957395476 | protein only RNase P catalytic subunit |
| ENSG00000229227 | AL356056,1 | 0,117025963 | 1,084496925 | 0,75191464 | 0,975901808 | novel transcript |
| ENSG00000254593 | OR7E126P | 0,11699457 | 1,084473327 | 0,76413086 | 0,976640468 | olfactory receptor family 7 subfamily E member 126 pseudogene |
| ENSG00000139579 | NABP2 | 0,116981896 | 1,0844638 | 0,58358739 | 0,944174844 | nucleic acid binding protein 2 |
| ENSG00000143079 | CTTNBP2NL | 0,116963173 | 1,084449726 | 0,70625513 | 0,969503793 | CTTNBP2 N-terminal like |
| ENSG00000135637 | CCDC142 | 0,116960613 | 1,084447802 | 0,77549166 | 0,978587255 | coiled-coil domain containing 142 |
| ENSG00000118655 | DCLRE1B | 0,11693954 | 1,084431962 | 0,6596307 | 0,958691226 | DNA cross-link repair 1B |
| ENSG00000166902 | MRPL16 | 0,116873148 | 1,084382058 | 0,69079293 | 0,967179044 | mitochondrial ribosomal protein L16 |
| ENSG00000137154 | RPS6 | 0,116868182 | 1,084378326 | 0,62149531 | 0,952793934 | ribosomal protein S6 |
| ENSG00000141458 | NPC1 | 0,116800819 | 1,084327695 | 0,62942549 | 0,952812183 | NPC intracellular cholesterol transporter 1 |
| ENSG00000197451 | HNRNPAB | 0,116773536 | 1,084307188 | 0,60137401 | 0,949939978 | heterogeneous nuclear ribonucleoprotein A/B |
| ENSG00000134265 | NAPG | 0,116724002 | 1,08426996 | 0,74177027 | 0,975176043 | NSF attachment protein gamma |
| ENSG00000113845 | TIMMDC1 | 0,116699664 | 1,084251669 | 0,60416132 | 0,950672921 | translocase of inner mitochondrial membrane domain containing 1 |
| ENSG00000151657 | KIN | 0,116698175 | 1,08425055 | 0,66290371 | 0,959897678 | Kin17 DNA and RNA binding protein |

| | | | | | | |
|---|---|---|---|---|---|---|
| ENSG00000185100 | ADSS1 | 0,116696026 | 1,084248935 | 0,74945774 | 0,975901808 | adenylosuccinate synthase 1 |
| ENSG00000126882 | FAM78A | 0,116675242 | 1,084233315 | 0,74165584 | 0,975176043 | family with sequence similarity 78 member A |
| ENSG00000198121 | LPAR1 | 0,11666177 | 1,08422319 | 0,80821115 | 0,981213877 | lysophosphatidic acid receptor 1 |
| ENSG00000143882 | ATP6V1C2 | 0,116617724 | 1,084190089 | 0,7009282 | 0,968652222 | ATPase H+ transporting V1 subunit C2 |
| ENSG00000229563 | LINC01204 | 0,11660111 | 1,084177604 | 0,88750622 | 0,990717729 | long intergenic non-protein coding RNA 1204 |
| ENSG00000243056 | EIF4EBP3 | 0,116586021 | 1,084166264 | 0,78859981 | 0,979462318 | eukaryotic translation initiation factor 4E binding protein 3 |
| ENSG00000116455 | WDR77 | 0,116572087 | 1,084155793 | 0,5931004 | 0,947416795 | WD repeat domain 77 |
| ENSG00000144029 | MRPS5 | 0,116493434 | 1,084096689 | 0,60702709 | 0,950993594 | mitochondrial ribosomal protein S5 |
| ENSG00000023516 | AKAP11 | 0,116285721 | 1,083940617 | 0,76706305 | 0,977201891 | A-kinase anchoring protein 11 |
| ENSG00000128626 | MRPS12 | 0,116242168 | 1,083907894 | 0,71747065 | 0,971697135 | mitochondrial ribosomal protein S12 |
| ENSG00000179922 | ZNF784 | 0,116175615 | 1,083857894 | 0,83419136 | 0,984905093 | zinc finger protein 784 |
| ENSG00000086758 | HUWE1 | 0,116044448 | 1,083759356 | 0,57217825 | 0,942475308 | HECT, UBA and WWE domain containing E3 ubiquitin protein ligase 1 |
| ENSG00000132321 | IQCA1 | 0,116031377 | 1,083749537 | 0,78186147 | 0,978597905 | IQ motif containing with AAA domain 1 |
| ENSG00000162981 | LRATD1 | 0,116023811 | 1,083743854 | 0,61881941 | 0,952793934 | LRAT domain containing 1 |
| ENSG00000160049 | DFFA | 0,115954923 | 1,083692106 | 0,59892862 | 0,948894897 | DNA fragmentation factor subunit alpha |
| ENSG00000103363 | ELOB | 0,115944524 | 1,083684295 | 0,65768056 | 0,957864535 | elongin B |
| ENSG00000236617 | AC127070,1 | 0,115900649 | 1,083651339 | 0,90053456 | 0,991865079 | novel transcript |
| ENSG00000055208 | TAB2 | 0,115843912 | 1,083608723 | 0,76778299 | 0,977309816 | TGF-beta activated kinase 1 (MAP3K7) binding protein 2 |
| ENSG00000130726 | TRIM28 | 0,115834577 | 1,083601711 | 0,70662953 | 0,969503793 | tripartite motif containing 28 |
| ENSG00000198783 | ZNF830 | 0,115796008 | 1,083572743 | 0,63051671 | 0,953306622 | zinc finger protein 830 |
| ENSG00000112851 | ERBIN | 0,115729255 | 1,083522607 | 0,77958222 | 0,978587255 | erbb2 interacting protein |
| ENSG00000169118 | CSNK1G1 | 0,115574293 | 1,083406231 | 0,68147504 | 0,964654539 | casein kinase 1 gamma 1 |
| ENSG00000166199 | ALKBH3 | 0,115505302 | 1,083354423 | 0,63079506 | 0,953404722 | alkB homolog 3, alpha-ketoglutarate dependent dioxygenase |
| ENSG00000129472 | RAB2B | 0,115489221 | 1,083342347 | 0,65583203 | 0,957444211 | RAB2B, member RAS oncogene family |
| ENSG00000272622 | AC010735,2 | 0,11544425 | 1,083308578 | 0,85031483 | 0,987036295 | novel transcript |
| ENSG00000288283 | RFX1 | 0,115425972 | 1,083294853 | 0,67910662 | 0,964151912 | regulatory factor X1 |
| ENSG00000132005 | RFX1 | 0,115425972 | 1,083294853 | 0,67910662 | 0,964151912 | regulatory factor X1 |
| ENSG00000108839 | ALOX12 | 0,115421412 | 1,083291429 | 0,86975269 | 0,989074806 | arachidonate 12-lipoxygenase, 12S type |
| ENSG00000260818 | UNGP1 | 0,115339135 | 1,083229651 | 0,91631196 | 0,993361299 | uracil-DNA glycosylase pseudogene 1 |

| | | | | | | |
|---|---|---|---|---|---|---|
| ENSG00000085465 | OVGP1 | 0,115310585 | 1,083208214 | 0,82808375 | 0,983595244 | oviductal glycoprotein 1 |
| ENSG00000233665 | AC060234,2 | 0,115268953 | 1,083176957 | 0,79129264 | 0,979471509 | novel transcript |
| ENSG00000237754 | AC093655,2 | 0,115199724 | 1,083124981 | 0,90964155 | 0,992948341 | septin 13 (SEPT13) pseudogene |
| ENSG00000273233 | AC097724,1 | 0,115087828 | 1,083040977 | 0,9033327 | 0,992311172 | novel transcript |
| ENSG00000164463 | CREBRF | 0,11504347 | 1,083007677 | 0,73461718 | 0,974353474 | CREB3 regulatory factor |
| ENSG00000140395 | WDR61 | 0,115018583 | 1,082988995 | 0,62149967 | 0,952793934 | WD repeat domain 61 |
| ENSG00000267711 | AC060766,6 | 0,115011834 | 1,082983929 | 0,90063424 | 0,991865079 | novel transcript |
| ENSG00000254706 | AL512785,2 | 0,114998297 | 1,082973767 | 0,77897093 | 0,978587255 | novel protein |
| ENSG00000168556 | ING2 | 0,114780983 | 1,08281065 | 0,78431956 | 0,979320406 | inhibitor of growth family member 2 |
| ENSG00000242575 | TUBAP13 | 0,114710284 | 1,082757589 | 0,86729617 | 0,989074806 | tubulin alpha pseudogene 13 |
| ENSG00000225505 | AC104332,1 | 0,114698087 | 1,082748435 | 0,84614004 | 0,986939308 | novel pseudogene |
| ENSG00000275223 | AL121906,2 | 0,114615565 | 1,082686503 | 0,87244481 | 0,989217334 | novel transcript |
| ENSG00000112406 | HECA | 0,114514667 | 1,082610786 | 0,74919236 | 0,975901808 | hdc homolog, cell cycle regulator |
| ENSG00000259135 | AL445363,2 | 0,114486325 | 1,082589518 | 0,90890961 | 0,992756615 | novel transcript, antisense to SNX6 |
| ENSG00000169925 | BRD3 | 0,11441514 | 1,082536103 | 0,6015364 | 0,949939978 | bromodomain containing 3 |
| ENSG00000169891 | REPS2 | 0,114218206 | 1,082388342 | 0,72896003 | 0,974009551 | RALBP1 associated Eps domain containing 2 |
| ENSG00000138641 | HERC3 | 0,114189739 | 1,082366985 | 0,72982762 | 0,974106992 | HECT and RLD domain containing E3 ubiquitin protein ligase 3 |
| ENSG00000282092 | MED16 | 0,114152604 | 1,082339125 | 0,76430564 | 0,976640468 | mediator complex subunit 16 |
| ENSG00000262634 | SKA1 | 0,114140214 | 1,08232983 | 0,64262312 | 0,956633898 | spindle and kinetochore associated complex subunit 1 |
| ENSG00000255152 | MSH5-SAPCD1 | 0,114139242 | 1,082329101 | 0,88784295 | 0,990717729 | MSH5-SAPCD1 readthrough (NMD candidate) |
| ENSG00000286732 | AC091181,2 | 0,114017816 | 1,082238009 | 0,90398659 | 0,992484155 | novel transcript, antisense to MSI2 |
| ENSG00000213613 | RPL11P3 | 0,113964294 | 1,08219786 | 0,8561273 | 0,988185425 | ribosomal protein L11 pseudogene 3 |
| ENSG00000283120 | RPL11P3 | 0,113964294 | 1,08219786 | 0,8561273 | 0,988185425 | ribosomal protein L11 pseudogene 3 |
| ENSG00000116133 | DHCR24 | 0,113958974 | 1,08219387 | 0,59876658 | 0,948854865 | 24-dehydrocholesterol reductase |
| ENSG00000115652 | UXS1 | 0,113911349 | 1,082158146 | 0,68001495 | 0,964580199 | UDP-glucuronate decarboxylase 1 |
| ENSG00000229431 | MED8-AS1 | 0,113818473 | 1,082088482 | 0,91472603 | 0,993361299 | MED8 antisense RNA 1 |
| ENSG00000088833 | NSFL1C | 0,113768568 | 1,082051052 | 0,60713041 | 0,950993594 | NSFL1 cofactor |
| ENSG00000117505 | DR1 | 0,113737933 | 1,082028075 | 0,74699837 | 0,975796253 | down-regulator of transcription 1 |
| ENSG00000169403 | PTAFR | 0,113662421 | 1,081971443 | 0,86496685 | 0,989074806 | platelet activating factor receptor |
| ENSG00000270060 | AC090589,3 | 0,113651371 | 1,081963156 | 0,86045215 | 0,989074806 | novel transcript |

| | | | | | | |
|---|---|---|---|---|---|---|
| ENSG00000254527 | ENPP7P12 | 0,11358108 | 1,081910441 | 0,82467836 | 0,983374097 | ectonucleotide pyrophosphatase/phosphodiesterase 7 pseudogene 12 |
| ENSG00000233560 | KRT8P39 | 0,11353766 | 1,08187788 | 0,77399444 | 0,978415098 | keratin 8 pseudogene 39 |
| ENSG00000249673 | NOP14-AS1 | 0,113440098 | 1,081804721 | 0,65044083 | 0,956899963 | NOP14 antisense RNA 1 |
| ENSG00000269148 | AC092301,1 | 0,113395858 | 1,081771547 | 0,85753913 | 0,988893234 | novel transcript, antisense to SYMPK |
| ENSG00000285360 | LONRF1 | 0,113390229 | 1,081767327 | 0,7832996 | 0,978936626 | LON peptidase N-terminal domain and ring finger 1 |
| ENSG00000164024 | METAP1 | 0,113204273 | 1,081627902 | 0,75466861 | 0,975901808 | methionyl aminopeptidase 1 |
| ENSG00000255680 | AC091564,6 | 0,113078383 | 1,081533522 | 0,93447134 | 0,994479239 | novel transcript |
| ENSG00000163689 | CFAP20DC | 0,11305924 | 1,081519172 | 0,74402832 | 0,975509237 | CFAP20 domain containing |
| ENSG00000164168 | TMEM184C | 0,113023838 | 1,081492633 | 0,73456461 | 0,974353474 | transmembrane protein 184C |
| ENSG00000261402 | AL591222,1 | 0,113023325 | 1,081492249 | 0,90468622 | 0,992590661 | novel transcript |
| ENSG00000171161 | ZNF672 | 0,112958369 | 1,081443557 | 0,61524531 | 0,952793934 | zinc finger protein 672 |
| ENSG00000285236 | AC270291,5 | 0,112921713 | 1,08141608 | 0,88159023 | 0,990717729 | novel transcript |
| ENSG00000176340 | COX8A | 0,112880249 | 1,081385 | 0,6372184 | 0,955985674 | cytochrome c oxidase subunit 8A |
| ENSG00000167395 | ZNF646 | 0,112819916 | 1,081339777 | 0,78556397 | 0,979320406 | zinc finger protein 646 |
| ENSG00000006125 | AP2B1 | 0,112706843 | 1,081255029 | 0,68457442 | 0,965415049 | adaptor related protein complex 2 subunit beta 1 |
| ENSG00000024048 | UBR2 | 0,112636396 | 1,081202233 | 0,72522157 | 0,97326334 | ubiquitin protein ligase E3 component n-recognin 2 |
| ENSG00000273706 | LHX1 | 0,112616979 | 1,081187681 | 0,8276598 | 0,983595244 | LIM homeobox 1 |
| ENSG00000126460 | PRRG2 | 0,112586295 | 1,081164686 | 0,87168189 | 0,989217334 | proline rich and Gla domain 2 |
| ENSG00000238090 | AC006195,1 | 0,112557427 | 1,081143053 | 0,92841597 | 0,994108531 | RAB3A interacting protein (rabin3)-like 1 (RAB3IL1) pseudogene |
| ENSG00000104774 | MAN2B1 | 0,112539398 | 1,081129541 | 0,69381074 | 0,967340633 | mannosidase alpha class 2B member 1 |
| ENSG00000274129 | TSEN34 | 0,112479701 | 1,081084807 | 0,69155923 | 0,967201016 | tRNA splicing endonuclease subunit 34 |
| ENSG00000198324 | PHETA1 | 0,112429374 | 1,081047095 | 0,75068049 | 0,975901808 | PH domain containing endocytic trafficking adaptor 1 |
| ENSG00000115129 | TP53I3 | 0,112337914 | 1,080978564 | 0,64760219 | 0,956899963 | tumor protein p53 inducible protein 3 |
| ENSG00000170365 | SMAD1 | 0,112281397 | 1,080936217 | 0,68140248 | 0,964654539 | SMAD family member 1 |
| ENSG00000167863 | ATP5PD | 0,112279217 | 1,080934584 | 0,68265314 | 0,964820564 | ATP synthase peripheral stalk subunit d |
| ENSG00000143412 | ANXA9 | 0,112276135 | 1,080932275 | 0,82260879 | 0,98299282 | annexin A9 |
| ENSG00000109265 | CRACD | 0,112265 | 1,080923932 | 0,8210568 | 0,982753437 | capping protein inhibiting regulator of actin dynamics |
| ENSG00000166012 | TAF1D | 0,112243657 | 1,080907941 | 0,75587257 | 0,975901808 | TATA-box binding protein associated factor, RNA polymerase I subunit D |

| | | | | | | |
|---|---|---|---|---|---|---|
| ENSG00000115694 | STK25 | 0,112140951 | 1,080830994 | 0,65937715 | 0,958425175 | serine/threonine kinase 25 |
| ENSG00000109929 | SC5D | 0,112137148 | 1,080828145 | 0,8289855 | 0,983844005 | sterol-C5-desaturase |
| ENSG00000249249 | AC010226,1 | 0,112077118 | 1,080783173 | 0,7584719 | 0,976120207 | novel transcript |
| ENSG00000106348 | IMPDH1 | 0,112010149 | 1,080733005 | 0,77291391 | 0,978310326 | inosine monophosphate dehydrogenase 1 |
| ENSG00000248275 | TRIM52-AS1 | 0,111943593 | 1,080683148 | 0,65442453 | 0,957444211 | TRIM52 antisense RNA 1 (head to head) |
| ENSG00000281518 | FOXO6 | 0,111905144 | 1,080654347 | 0,88272732 | 0,990717729 | forkhead box O6 |
| ENSG00000092094 | OSGEP | 0,111863335 | 1,080623031 | 0,5938196 | 0,947469769 | O-sialoglycoprotein endopeptidase |
| ENSG00000085760 | MTIF2 | 0,111822016 | 1,080592082 | 0,70899064 | 0,970282889 | mitochondrial translational initiation factor 2 |
| ENSG00000273523 | AL139082,1 | 0,111792245 | 1,080569784 | 0,88726767 | 0,990717729 | novel transcript, antisense to NEK3 & NEK5 |
| ENSG00000083290 | ULK2 | 0,111788113 | 1,080566689 | 0,70784536 | 0,969894262 | unc-51 like autophagy activating kinase 2 |
| ENSG00000103994 | ZNF106 | 0,111778822 | 1,08055973 | 0,68980338 | 0,966868992 | zinc finger protein 106 |
| ENSG00000166689 | PLEKHA7 | 0,111729091 | 1,080522483 | 0,71982196 | 0,972014805 | pleckstrin homology domain containing A7 |
| ENSG00000235692 | PFDN6 | 0,111725896 | 1,08052009 | 0,74714932 | 0,975807355 | prefoldin subunit 6 |
| ENSG00000163870 | TPRA1 | 0,111552074 | 1,080389912 | 0,75924798 | 0,976120207 | transmembrane protein adipocyte associated 1 |
| ENSG00000140259 | MFAP1 | 0,111550257 | 1,080388551 | 0,7126025 | 0,970812834 | microfibril associated protein 1 |
| ENSG00000180329 | CCDC43 | 0,11154434 | 1,08038412 | 0,71416615 | 0,970867497 | coiled-coil domain containing 43 |
| ENSG00000227773 | ASH1L-IT1 | 0,11149578 | 1,080347756 | 0,91631373 | 0,993361299 | ASH1L intronic transcript 1 |
| ENSG00000132640 | BTBD3 | 0,11141368 | 1,080286278 | 0,77742622 | 0,978587255 | BTB domain containing 3 |
| ENSG00000276727 | AC137834,2 | 0,111390174 | 1,080268677 | 0,86743005 | 0,989074806 | novel transcript, antisense to SHMT2 |
| ENSG00000189136 | UBE2Q2P1 | 0,111368803 | 1,080252675 | 0,83066852 | 0,984156028 | ubiquitin conjugating enzyme E2 Q2 pseudogene 1 |
| ENSG00000160993 | ALKBH4 | 0,111361787 | 1,080247422 | 0,74646482 | 0,975796253 | alkB homolog 4, lysine demethylase |
| ENSG00000113749 | HRH2 | 0,111287977 | 1,080192156 | 0,74377022 | 0,975346349 | histamine receptor H2 |
| ENSG00000145860 | RNF145 | 0,111244629 | 1,0801597 | 0,69715574 | 0,968522643 | ring finger protein 145 |
| ENSG00000143420 | ENSA | 0,111235419 | 1,080152805 | 0,62070679 | 0,952793934 | endosulfine alpha |
| ENSG00000196459 | TRAPPC2 | 0,111216277 | 1,080138474 | 0,64342405 | 0,956633898 | trafficking protein particle complex 2 |
| ENSG00000261642 | AL713866,1 | 0,111073412 | 1,080031516 | 0,86254153 | 0,989074806 | novel transcript |
| ENSG00000235408 | SNORA71B | 0,110982067 | 1,079963136 | 0,86441037 | 0,989074806 | small nucleolar RNA, H/ACA box 71B |
| ENSG00000264247 | LINC00909 | 0,110979708 | 1,079961369 | 0,68333528 | 0,965028201 | long intergenic non-protein coding RNA 909 |
| ENSG00000282853 | ECH1 | 0,110960557 | 1,079947034 | 0,68686398 | 0,965769168 | enoyl-CoA hydratase 1 |
| ENSG00000104823 | ECH1 | 0,110960557 | 1,079947034 | 0,68686398 | 0,965769168 | enoyl-CoA hydratase 1 |
| ENSG00000124172 | ATP5F1E | 0,110929215 | 1,079923573 | 0,76169458 | 0,976285211 | ATP synthase F1 subunit epsilon |
| ENSG00000129084 | PSMA1 | 0,110857434 | 1,079869843 | 0,81659139 | 0,981919736 | proteasome 20S subunit alpha 1 |
| ENSG00000136878 | USP20 | 0,110788959 | 1,07981859 | 0,73555764 | 0,974353474 | ubiquitin specific peptidase 20 |

| | | | | | | |
|---|---|---|---|---|---|---|
| ENSG00000172428 | COPS9 | 0,110783828 | 1,079814749 | 0,62349043 | 0,952793934 | COP9 signalosome subunit 9 |
| ENSG00000238133 | MAP3K20-AS1 | 0,11070588 | 1,079756409 | 0,88549033 | 0,990717729 | MAP3K20 antisense RNA 1 |
| ENSG00000278107 | AC027575,2 | 0,110668629 | 1,07972853 | 0,86679355 | 0,989074806 | novel transcript |
| ENSG00000282339 | AC010507,2 | 0,110529586 | 1,079624473 | 0,85103608 | 0,987259878 | novel pseudogene |
| ENSG00000185485 | SDHAP1 | 0,110524754 | 1,079620857 | 0,67060092 | 0,962026636 | succinate dehydrogenase complex flavoprotein subunit A pseudogene 1 |
| ENSG00000172893 | DHCR7 | 0,110472219 | 1,079581545 | 0,65309376 | 0,95732962 | 7-dehydrocholesterol reductase |
| ENSG00000230176 | LINC01433 | 0,110407843 | 1,079533372 | 0,93874105 | 0,994705993 | long intergenic non-protein coding RNA 1433 |
| ENSG00000163485 | ADORA1 | 0,110352389 | 1,079491878 | 0,79730338 | 0,980161849 | adenosine A1 receptor |
| ENSG00000059573 | ALDH18A1 | 0,110311521 | 1,079461299 | 0,61739253 | 0,952793934 | aldehyde dehydrogenase 18 family member A1 |
| ENSG00000164885 | CDK5 | 0,110143418 | 1,079335528 | 0,66370697 | 0,959920886 | cyclin dependent kinase 5 |
| ENSG00000287804 | Z98745,2 | 0,110132403 | 1,079327287 | 0,95205978 | 0,996432861 | novel transcript |
| ENSG00000175203 | DCTN2 | 0,110100891 | 1,079303712 | 0,61495413 | 0,952793934 | dynactin subunit 2 |
| ENSG00000287977 | AC068630,1 | 0,110064584 | 1,07927655 | 0,91139938 | 0,993222655 | tigger transposable element derived 1 (TIGD1) pseudogene |
| ENSG00000219545 | UMAD1 | 0,109944058 | 1,079186389 | 0,80325846 | 0,980866933 | UBAP1-MVB12-associated (UMA) domain containing 1 |
| ENSG00000129691 | ASH2L | 0,109923342 | 1,079170893 | 0,6655338 | 0,960946962 | ASH2 like, histone lysine methyltransferase complex subunit |
| ENSG00000234073 | AC011816,1 | 0,109892284 | 1,079147661 | 0,93470291 | 0,994479239 | chromobox homolog 3 (CBX3) pseudogene |
| ENSG00000116539 | ASH1L | 0,109750625 | 1,079041704 | 0,68777605 | 0,966064784 | ASH1 like histone lysine methyltransferase |
| ENSG00000111962 | UST | 0,109747352 | 1,079039256 | 0,73466412 | 0,974353474 | uronyl 2-sulfotransferase |
| ENSG00000174989 | FBXW8 | 0,10971597 | 1,079015785 | 0,61839219 | 0,952793934 | F-box and WD repeat domain containing 8 |
| ENSG00000280081 | LINC01667 | 0,109600673 | 1,078929555 | 0,81859525 | 0,9824772 | long intergenic non-protein coding RNA 1667 |
| ENSG00000206286 | VPS52 | 0,109599875 | 1,078928959 | 0,79304819 | 0,97960721 | VPS52 subunit of GARP complex |
| ENSG00000260470 | AC023794,5 | 0,109521183 | 1,07887011 | 0,94263444 | 0,995390305 | novel transcript |
| ENSG00000160113 | NR2F6 | 0,109491022 | 1,078847555 | 0,72813368 | 0,973888243 | nuclear receptor subfamily 2 group F member 6 |
| ENSG00000064115 | TM7SF3 | 0,10939831 | 1,078778228 | 0,65996611 | 0,958859629 | transmembrane 7 superfamily member 3 |
| ENSG00000167182 | SP2 | 0,109386239 | 1,078769202 | 0,64659048 | 0,956899963 | Sp2 transcription factor |
| ENSG00000248100 | AC087257,1 | 0,109382513 | 1,078766416 | 0,79134598 | 0,979471509 | novel transcript |
| ENSG00000149201 | CCDC81 | 0,109355262 | 1,078746039 | 0,75251984 | 0,975901808 | coiled-coil domain containing 81 |
| ENSG00000168411 | RFWD3 | 0,109347363 | 1,078740132 | 0,65047064 | 0,956899963 | ring finger and WD repeat domain 3 |
| ENSG00000006016 | CRLF1 | 0,109309627 | 1,078711917 | 0,7719483 | 0,978033855 | cytokine receptor like factor 1 |

| | | | | | | |
|---|---|---|---|---|---|---|
| ENSG00000109220 | CHIC2 | 0,109219449 | 1,078644492 | 0,71032603 | 0,970482408 | cysteine rich hydrophobic domain 2 |
| ENSG00000147872 | PLIN2 | 0,109121261 | 1,078571084 | 0,68192593 | 0,964654539 | perilipin 2 |
| ENSG00000112530 | PACRG | 0,109108243 | 1,078561351 | 0,78290397 | 0,978755191 | parkin coregulated |
| ENSG00000287809 | AC005632,6 | 0,109045275 | 1,078514277 | 0,85945992 | 0,989074806 | novel transcript |
| ENSG00000077380 | DYNC1I2 | 0,108931504 | 1,078429229 | 0,7034818 | 0,968940012 | dynein cytoplasmic 1 intermediate chain 2 |
| ENSG00000138686 | BBS7 | 0,108915291 | 1,07841711 | 0,77318404 | 0,978363975 | Bardet-Biedl syndrome 7 |
| ENSG00000224411 | HSP90AA2P | 0,108849811 | 1,078368164 | 0,78932409 | 0,979462318 | heat shock protein 90 alpha family class A member 2, pseudogene |
| ENSG00000104537 | ANXA13 | 0,108746266 | 1,07829077 | 0,88757539 | 0,990717729 | annexin A13 |
| ENSG00000146574 | CCZ1B | 0,108702756 | 1,078258251 | 0,72265418 | 0,972594421 | CCZ1 homolog B, vacuolar protein trafficking and biogenesis associated |
| ENSG00000132478 | UNK | 0,108653466 | 1,078221413 | 0,64340175 | 0,956633898 | unk zinc finger |
| ENSG00000111237 | VPS29 | 0,108651942 | 1,078220274 | 0,80870219 | 0,981213877 | VPS29 retromer complex component |
| ENSG00000115944 | COX7A2L | 0,108624963 | 1,078200111 | 0,69756793 | 0,968522643 | cytochrome c oxidase subunit 7A2 like |
| ENSG00000265681 | RPL17 | 0,108585055 | 1,078170286 | 0,73177206 | 0,974353474 | ribosomal protein L17 |
| ENSG00000143373 | ZNF687 | 0,108584786 | 1,078170085 | 0,69800208 | 0,968522643 | zinc finger protein 687 |
| ENSG00000117308 | GALE | 0,108582271 | 1,078168205 | 0,69509158 | 0,967519493 | UDP-galactose-4-epimerase |
| ENSG00000272595 | OR10AH1P | 0,108577707 | 1,078164795 | 0,9185823 | 0,993481057 | olfactory receptor family 10 subfamily AH member 1 pseudogene |
| ENSG00000130827 | PLXNA3 | 0,108529489 | 1,078128761 | 0,76540933 | 0,976830764 | plexin A3 |
| ENSG00000068878 | PSME4 | 0,108395255 | 1,078028452 | 0,72453116 | 0,97326334 | proteasome activator subunit 4 |
| ENSG00000162063 | CCNF | 0,10838847 | 1,078023382 | 0,70902825 | 0,970282889 | cyclin F |
| ENSG00000110169 | HPX | 0,10833546 | 1,077983772 | 0,82708887 | 0,983567657 | hemopexin |
| ENSG00000254001 | AC084706,1 | 0,108230789 | 1,077905565 | 0,93737085 | 0,994479239 | novel transcript |
| ENSG00000147799 | ARHGAP39 | 0,108205444 | 1,077886629 | 0,76958754 | 0,977683747 | Rho GTPase activating protein 39 |
| ENSG00000205086 | C2orf91 | 0,108187755 | 1,077873412 | 0,82229184 | 0,982952211 | chromosome 2 putative open reading frame 91 |
| ENSG00000226318 | RPS3AP38 | 0,108167091 | 1,077857974 | 0,84769422 | 0,987035044 | RPS3A pseudogene 38 |
| ENSG00000137288 | UQCC2 | 0,108163858 | 1,077855558 | 0,60526888 | 0,950927937 | ubiquinol-cytochrome c reductase complex assembly factor 2 |
| ENSG00000285807 | AL353768,2 | 0,108151471 | 1,077846304 | 0,91693217 | 0,993361299 | Novel transcript, antisense to C9orf3 |
| ENSG00000162337 | LRP5 | 0,108106728 | 1,077812877 | 0,76713034 | 0,977201891 | LDL receptor related protein 5 |
| ENSG00000175470 | PPP2R2D | 0,107973761 | 1,077713544 | 0,60748797 | 0,950993594 | protein phosphatase 2 regulatory subunit Bdelta |
| ENSG00000167470 | MIDN | 0,107946397 | 1,077693103 | 0,74384606 | 0,975394113 | midnolin |

| | | | | | | |
|---|---|---|---|---|---|---|
| ENSG00000220305 | HNRNPH1P1 | 0,107872634 | 1,077638004 | 0,86072962 | 0,989074806 | heterogeneous nuclear ribonucleoprotein H1 pseudogene 1 |
| ENSG00000275176 | ACACA | 0,107826959 | 1,077603887 | 0,79415513 | 0,979776981 | acetyl-CoA carboxylase alpha |
| ENSG00000069345 | DNAJA2 | 0,107741328 | 1,077539927 | 0,75906477 | 0,976120207 | DnaJ heat shock protein family (Hsp40) member A2 |
| ENSG00000086102 | NFX1 | 0,107652501 | 1,077473585 | 0,62951001 | 0,952812183 | nuclear transcription factor, X-box binding 1 |
| ENSG00000165338 | HECTD2 | 0,107585349 | 1,077423433 | 0,83705217 | 0,985462684 | HECT domain E3 ubiquitin protein ligase 2 |
| ENSG00000203993 | ARRDC1-AS1 | 0,107487641 | 1,077350466 | 0,70207072 | 0,968742626 | ARRDC1 antisense RNA 1 |
| ENSG00000276043 | UHRF1 | 0,107479903 | 1,077344688 | 0,80245997 | 0,980770465 | ubiquitin like with PHD and ring finger domains 1 |
| ENSG00000122033 | MTIF3 | 0,107433934 | 1,077310361 | 0,78013612 | 0,978587255 | mitochondrial translational initiation factor 3 |
| ENSG00000169208 | OR10G3 | 0,10741852 | 1,077298851 | 0,87530248 | 0,990078774 | olfactory receptor family 10 subfamily G member 3 |
| ENSG00000067064 | IDI1 | 0,107403458 | 1,077287603 | 0,79910466 | 0,980161849 | isopentenyl-diphosphate delta isomerase 1 |
| ENSG00000261170 | AC009053,3 | 0,107401552 | 1,07728618 | 0,86609443 | 0,989074806 | novel transcript, antisense to CLEC18B |
| ENSG00000175309 | PHYKPL | 0,107330077 | 1,07723281 | 0,70706335 | 0,969565008 | 5-phosphohydroxy-L-lysine phospho-lyase |
| ENSG00000116918 | TSNAX | 0,107317664 | 1,077223542 | 0,77243389 | 0,978261893 | translin associated factor X |
| ENSG00000230438 | SERPINB9P1 | 0,107236204 | 1,077162719 | 0,82134578 | 0,982950156 | serpin family B member 9 pseudogene 1 |
| ENSG00000180096 | SEPTIN1 | 0,10721966 | 1,077150367 | 0,84103137 | 0,985672131 | septin 1 |
| ENSG00000164338 | UTP15 | 0,10717391 | 1,07711621 | 0,79181265 | 0,979471509 | UTP15 small subunit processome component |
| ENSG00000129493 | HEATR5A | 0,106950892 | 1,076949717 | 0,64033273 | 0,956251063 | HEAT repeat containing 5A |
| ENSG00000102898 | NUTF2 | 0,1069506 | 1,076949499 | 0,72844133 | 0,973955059 | nuclear transport factor 2 |
| ENSG00000196262 | PPIA | 0,106936396 | 1,076938896 | 0,69285416 | 0,967271667 | peptidylprolyl isomerase A |
| ENSG00000235872 | AC078777,1 | 0,1069121 | 1,07692076 | 0,82154933 | 0,982952211 | novel transcript |
| ENSG00000286467 | AC108052,1 | 0,106883053 | 1,076899077 | 0,88572856 | 0,990717729 | novel transcript |
| ENSG00000183060 | LYSMD4 | 0,106869353 | 1,076888851 | 0,64823272 | 0,956899963 | LysM domain containing 4 |
| ENSG00000005302 | MSL3 | 0,106851894 | 1,076875819 | 0,71066219 | 0,970482408 | MSL complex subunit 3 |
| ENSG00000169758 | TMEM266 | 0,106784592 | 1,076825584 | 0,83949874 | 0,985672131 | transmembrane protein 266 |
| ENSG00000120438 | TCP1 | 0,106752248 | 1,076801443 | 0,72660979 | 0,973344786 | t-complex 1 |
| ENSG00000092020 | PPP2R3C | 0,106663721 | 1,076735369 | 0,79724129 | 0,980161849 | protein phosphatase 2 regulatory subunit B''gamma |
| ENSG00000154723 | ATP5PF | 0,106585738 | 1,07667717 | 0,80800121 | 0,981213877 | ATP synthase peripheral stalk subunit F6 |
| ENSG00000116459 | ATP5PB | 0,106388174 | 1,076529739 | 0,79570575 | 0,979879951 | ATP synthase peripheral stalk-membrane subunit b |
| ENSG00000253967 | AC022730,4 | 0,10628819 | 1,076455134 | 0,88539214 | 0,990717729 | novel transcript |
| ENSG00000276163 | LAIR1 | 0,106261535 | 1,076435246 | 0,84276114 | 0,986191719 | leukocyte associated immunoglobulin like receptor 1 |
| ENSG00000114554 | PLXNA1 | 0,106220482 | 1,076404616 | 0,73813472 | 0,974353474 | plexin A1 |

| | | | | | | |
|---|---|---|---|---|---|---|
| ENSG00000105641 | SLC5A5 | 0,106215476 | 1,07640088 | 0,89406812 | 0,991135494 | solute carrier family 5 member 5 |
| ENSG00000188672 | RHCE | 0,106197812 | 1,076387702 | 0,87188196 | 0,989217334 | Rh blood group CcEe antigens |
| ENSG00000102780 | DGKH | 0,106180461 | 1,076374756 | 0,69812494 | 0,968522643 | diacylglycerol kinase eta |
| ENSG00000143155 | TIPRL | 0,106138333 | 1,076343325 | 0,78984238 | 0,979462318 | TOR signaling pathway regulator |
| ENSG00000175305 | CCNE2 | 0,106035062 | 1,076266281 | 0,80817663 | 0,981213877 | cyclin E2 |
| ENSG00000245864 | MEF2C-AS2 | 0,106019973 | 1,076255025 | 0,88386435 | 0,990717729 | MEF2C antisense RNA 2 |
| ENSG00000138442 | WDR12 | 0,106007222 | 1,076245512 | 0,65398808 | 0,957444211 | WD repeat domain 12 |
| ENSG00000100445 | SDR39U1 | 0,105845388 | 1,076124792 | 0,64542666 | 0,956899963 | short chain dehydrogenase/reductase family 39U member 1 |
| ENSG00000272483 | AC022001,3 | 0,10581233 | 1,076100133 | 0,86937579 | 0,989074806 | novel transcript, antisense to VGLL4 |
| ENSG00000242083 | RPL7AP31 | 0,105777023 | 1,076073798 | 0,93798285 | 0,994488324 | ribosomal protein L7a pseudogene 31 |
| ENSG00000151413 | NUBPL | 0,105715919 | 1,076028223 | 0,66319497 | 0,959920886 | nucleotide binding protein like |
| ENSG00000178057 | NDUFAF3 | 0,10554545 | 1,075901087 | 0,68018578 | 0,964580199 | NADH:ubiquinone oxidoreductase complex assembly factor 3 |
| ENSG00000247400 | DNAJC3-DT | 0,105525103 | 1,075885914 | 0,7623892 | 0,976315588 | DNAJC3 divergent transcript |
| ENSG00000183077 | AFMID | 0,105479801 | 1,07585213 | 0,79319995 | 0,979659177 | arylformamidase |
| ENSG00000286652 | AL021918,5 | 0,105394142 | 1,075788254 | 0,91804103 | 0,993361299 | novel transcript |
| ENSG00000144283 | PKP4 | 0,105349582 | 1,075755027 | 0,71351859 | 0,970812834 | plakophilin 4 |
| ENSG00000149636 | DSN1 | 0,105302591 | 1,075719988 | 0,64593086 | 0,956899963 | DSN1 component of MIS12 kinetochore complex |
| ENSG00000167664 | TMIGD2 | 0,105294126 | 1,075713676 | 0,90067431 | 0,991865079 | transmembrane and immunoglobulin domain containing 2 |
| ENSG00000139428 | MMAB | 0,105286008 | 1,075707624 | 0,60362735 | 0,950352286 | metabolism of cobalamin associated B |
| ENSG00000186283 | TOR3A | 0,10528505 | 1,075706909 | 0,6683196 | 0,961351573 | torsin family 3 member A |
| ENSG00000139899 | CBLN3 | 0,105274538 | 1,075699071 | 0,8095076 | 0,981213877 | cerebellin 3 precursor |
| ENSG00000167515 | TRAPPC2L | 0,105265952 | 1,07569267 | 0,62917629 | 0,952793934 | trafficking protein particle complex 2 like |
| ENSG00000156253 | RWDD2B | 0,10525711 | 1,075686077 | 0,7313643 | 0,974353474 | RWD domain containing 2B |
| ENSG00000213024 | NUP62 | 0,105255862 | 1,075685147 | 0,64598007 | 0,956899963 | nucleoporin 62 |
| ENSG00000127993 | RBM48 | 0,105214322 | 1,075654174 | 0,71264077 | 0,970812834 | RNA binding motif protein 48 |
| ENSG00000226826 | HLA-DPB1 | 0,10519997 | 1,075643474 | 0,81543919 | 0,981521413 | major histocompatibility complex, class II, DP beta 1 |
| ENSG00000237710 | HLA-DPB1 | 0,10519997 | 1,075643474 | 0,81543919 | 0,981521413 | major histocompatibility complex, class II, DP beta 1 |
| ENSG00000260230 | FRRS1L | 0,105192841 | 1,075638158 | 0,74592239 | 0,975707496 | ferric chelate reductase 1 like |
| ENSG00000265100 | AC005332,1 | 0,105175693 | 1,075625373 | 0,89468158 | 0,991223105 | novel transcript, antisense to AMZ2 |
| ENSG00000141179 | PCTP | 0,105174187 | 1,075624251 | 0,69236725 | 0,967236299 | phosphatidylcholine transfer protein |

| | | | | | | |
|---|---|---|---|---|---|---|
| ENSG00000188938 | FAM120AOS | 0,105158407 | 1,075612485 | 0,61873984 | 0,952793934 | family with sequence similarity 120A opposite strand |
| ENSG00000174485 | DENND4A | 0,105157294 | 1,075611656 | 0,79793477 | 0,980161849 | DENN domain containing 4A |
| ENSG00000166831 | RBPMS2 | 0,105149827 | 1,075606089 | 0,63946477 | 0,956018518 | RNA binding protein, mRNA processing factor 2 |
| ENSG00000231762 | AL590426,1 | 0,105121855 | 1,075585234 | 0,92595969 | 0,993915537 | novel pseudogene |
| ENSG00000244026 | FAM86DP | 0,105014888 | 1,075505489 | 0,65788847 | 0,957864535 | family with sequence similarity 86 member D, pseudogene |
| ENSG00000288323 | FAM86DP | 0,105014888 | 1,075505489 | 0,65788847 | 0,957864535 | family with sequence similarity 86 member D, pseudogene |
| ENSG00000261534 | AL596244,1 | 0,104993018 | 1,075489186 | 0,78998749 | 0,979462318 | novel transcript, overlapping TTLL11 |
| ENSG00000271361 | HTATSF1P2 | 0,104909097 | 1,075426627 | 0,85392506 | 0,987861659 | HIV-1 Tat specific factor 1 pseudogene 2 |
| ENSG00000273073 | AC073869,1 | 0,10489214 | 1,075413986 | 0,91669278 | 0,993361299 | novel transcript |
| ENSG00000242252 | BGLAP | 0,104848872 | 1,075381734 | 0,92278487 | 0,993797669 | bone gamma-carboxyglutamate protein |
| ENSG00000234928 | LINC01659 | 0,104839415 | 1,075374685 | 0,91499577 | 0,993361299 | long intergenic non-protein coding RNA 1659 |
| ENSG00000280986 | LINC01659 | 0,104839415 | 1,075374685 | 0,91499577 | 0,993361299 | long intergenic non-protein coding RNA 1659 |
| ENSG00000023171 | GRAMD1B | 0,104805244 | 1,075349214 | 0,68330377 | 0,965027283 | GRAM domain containing 1B |
| ENSG00000228166 | MTND1P11 | 0,104709386 | 1,075277767 | 0,88553586 | 0,990717729 | MT-ND1 pseudogene 11 |
| ENSG00000173230 | GOLGB1 | 0,104641086 | 1,075226862 | 0,67114081 | 0,962026636 | golgin B1 |
| ENSG00000101367 | MAPRE1 | 0,104634531 | 1,075221976 | 0,71895228 | 0,971869926 | microtubule associated protein RP/EB family member 1 |
| ENSG00000212719 | LINC02693 | 0,104467021 | 1,075097141 | 0,64509489 | 0,956899963 | long intergenic non-protein coding RNA 2693 |
| ENSG00000206530 | CFAP44 | 0,104466641 | 1,075096857 | 0,7706474 | 0,977966587 | cilia and flagella associated protein 44 |
| ENSG00000230409 | TCEA1P2 | 0,104421922 | 1,075063534 | 0,80888245 | 0,981213877 | transcription elongation factor A1 pseudogene 2 |
| ENSG00000137133 | HINT2 | 0,104374344 | 1,07502808 | 0,66981257 | 0,962026636 | histidine triad nucleotide binding protein 2 |
| ENSG00000054965 | FAM168A | 0,104290668 | 1,074965731 | 0,67734416 | 0,963580269 | family with sequence similarity 168 member A |
| ENSG00000255529 | POLR2M | 0,104083817 | 1,074811615 | 0,77859919 | 0,978587255 | RNA polymerase II subunit M |
| ENSG00000183475 | ASB7 | 0,104071693 | 1,074802583 | 0,73794481 | 0,974353474 | ankyrin repeat and SOCS box containing 7 |
| ENSG00000269091 | AC010624,3 | 0,104046396 | 1,074783736 | 0,83471289 | 0,984905093 | novel transcript, antisense to ZNF473 |
| ENSG00000139970 | RTN1 | 0,104041592 | 1,074780157 | 0,66154876 | 0,959582738 | reticulon 1 |
| ENSG00000088356 | PDRG1 | 0,103991071 | 1,074742521 | 0,66339671 | 0,959920886 | p53 and DNA damage regulated 1 |
| ENSG00000157870 | PRXL2B | 0,103931927 | 1,074698463 | 0,74075243 | 0,975059523 | peroxiredoxin like 2B |
| ENSG00000250685 | AC009123,1 | 0,103908775 | 1,074681216 | 0,95902528 | 0,996608711 | novel transcript, antisense to ADAD2 |
| ENSG00000104728 | ARHGEF10 | 0,103813972 | 1,074610598 | 0,71680274 | 0,971438758 | Rho guanine nucleotide exchange factor 10 |

| | | | | | | |
|---|---|---|---|---|---|---|
| ENSG00000287516 | AL359764,2 | 0,103587697 | 1,074442068 | 0,8673558 | 0,989074806 | novel transcript |
| ENSG00000266744 | AC005304,3 | 0,103558117 | 1,074420038 | 0,93145469 | 0,994433362 | novel transcript |
| ENSG00000165140 | FBP1 | 0,103534505 | 1,074402454 | 0,90739954 | 0,992590661 | fructose-bisphosphatase 1 |
| ENSG00000249019 | AC108471,1 | 0,103440841 | 1,074332703 | 0,90758258 | 0,992590661 | ribosomal protein S7 (RPS7) pseudogene |
| ENSG00000276856 | TUBGCP5 | 0,103424866 | 1,074320807 | 0,85543101 | 0,988075747 | tubulin gamma complex associated protein 5 |
| ENSG00000091157 | WDR7 | 0,103421101 | 1,074318003 | 0,75395956 | 0,975901808 | WD repeat domain 7 |
| ENSG00000110107 | PRPF19 | 0,103390517 | 1,074295228 | 0,61003842 | 0,951801946 | pre-mRNA processing factor 19 |
| ENSG00000090857 | PDPR | 0,103314279 | 1,07423846 | 0,68077555 | 0,964654539 | pyruvate dehydrogenase phosphatase regulatory subunit |
| ENSG00000274447 | AC005790,1 | 0,103201671 | 1,074154614 | 0,88610914 | 0,990717729 | novel transcript, antisense to PTPRS |
| ENSG00000229020 | AKR7A2P1 | 0,10318447 | 1,074141808 | 0,88879212 | 0,990717729 | AKR7A2 pseudogene 1 |
| ENSG00000280751 | AKR7A2P1 | 0,10318447 | 1,074141808 | 0,88879212 | 0,990717729 | AKR7A2 pseudogene 1 |
| ENSG00000069849 | ATP1B3 | 0,103152051 | 1,074117671 | 0,7774063 | 0,978587255 | ATPase Na+/K+ transporting subunit beta 3 |
| ENSG00000150401 | DCUN1D2 | 0,103129003 | 1,074100511 | 0,8092069 | 0,981213877 | defective in cullin neddylation 1 domain containing 2 |
| ENSG00000215252 | GOLGA8B | 0,103055686 | 1,074045927 | 0,77397542 | 0,978415098 | golgin A8 family member B |
| ENSG00000287686 | AC062029,1 | 0,10302699 | 1,074024564 | 0,87440048 | 0,98990147 | novel transcript |
| ENSG00000284633 | AL031590,1 | 0,10297026 | 1,073982332 | 0,92593442 | 0,993915537 | novel transcript, antisense to PDGFB |
| ENSG00000133619 | KRBA1 | 0,102904638 | 1,073933483 | 0,80397405 | 0,980875302 | KRAB-A domain containing 1 |
| ENSG00000229082 | OR5AW1P | 0,102890779 | 1,073923166 | 0,91709242 | 0,993361299 | olfactory receptor family 5 subfamily AW member 1 pseudogene |
| ENSG00000121361 | KCNJ8 | 0,102847239 | 1,073890756 | 0,77701336 | 0,978587255 | potassium inwardly rectifying channel subfamily J member 8 |
| ENSG00000169446 | MMGT1 | 0,102817647 | 1,073868729 | 0,81850255 | 0,9824772 | membrane magnesium transporter 1 |
| ENSG00000075142 | SRI | 0,10280435 | 1,073858832 | 0,79419117 | 0,979776981 | sorcin |
| ENSG00000261123 | AC009065,5 | 0,102733688 | 1,073806236 | 0,88016721 | 0,990268593 | novel transcript, intronic to PKD1 |
| ENSG00000271853 | AL162258,1 | 0,102719922 | 1,07379599 | 0,85902407 | 0,989074806 | novel transcript |
| ENSG00000183172 | SMDT1 | 0,102666745 | 1,073756411 | 0,75355539 | 0,975901808 | single-pass membrane protein with aspartate rich tail 1 |
| ENSG00000249042 | AC008771,1 | 0,102631828 | 1,073730423 | 0,81747889 | 0,98215965 | novel transcript |
| ENSG00000141568 | FOXK2 | 0,102627739 | 1,07372738 | 0,62102561 | 0,952793934 | forkhead box K2 |
| ENSG00000116701 | NCF2 | 0,102556821 | 1,073674601 | 0,84916109 | 0,987036295 | neutrophil cytosolic factor 2 |
| ENSG00000172046 | USP19 | 0,102544448 | 1,073665393 | 0,70604837 | 0,969503793 | ubiquitin specific peptidase 19 |

| | | | | | | |
|---|---|---|---|---|---|---|
| ENSG00000241343 | RPL36A | 0,102529949 | 1,073654603 | 0,68618328 | 0,965740648 | ribosomal protein L36a |
| ENSG00000148737 | TCF7L2 | 0,102428601 | 1,073579181 | 0,73144997 | 0,974353474 | transcription factor 7 like 2 |
| ENSG00000104081 | BMF | 0,102317371 | 1,073496413 | 0,78628065 | 0,979320406 | Bcl2 modifying factor |
| ENSG00000234665 | LERFS | 0,102248695 | 1,073445313 | 0,88009955 | 0,990268593 | lncRNA negative regulator of fibroblast-like synoviocyte migration, SYNCRIP interacting |
| ENSG00000232577 | BX679671,1 | 0,102161242 | 1,073380245 | 0,96143389 | 0,996608711 | novel transcript |
| ENSG00000233627 | C4A-AS1 | 0,102161242 | 1,073380245 | 0,96143389 | 0,996608711 | C4A antisense RNA 1 |
| ENSG00000229776 | C4B-AS1 | 0,102161242 | 1,073380245 | 0,96143389 | 0,996608711 | C4B antisense RNA 1 |
| ENSG00000234911 | TEX21P | 0,10213072 | 1,073357537 | 0,83447123 | 0,984905093 | testis expressed 21, pseudogene |
| ENSG00000183684 | ALYREF | 0,102111636 | 1,073343338 | 0,66342092 | 0,959920886 | Aly/REF export factor |
| ENSG00000215014 | AL645728,1 | 0,102003485 | 1,073262879 | 0,75936586 | 0,976120207 | novel transcript |
| ENSG00000159433 | STARD9 | 0,101996947 | 1,073258015 | 0,68872662 | 0,966556275 | StAR related lipid transfer domain containing 9 |
| ENSG00000115459 | ELMOD3 | 0,101865046 | 1,073159895 | 0,65450158 | 0,957444211 | ELMO domain containing 3 |
| ENSG00000161513 | FDXR | 0,101837105 | 1,073139111 | 0,84362235 | 0,986273696 | ferredoxin reductase |
| ENSG00000185818 | NAT8L | 0,101778154 | 1,073095262 | 0,84085914 | 0,985672131 | N-acetyltransferase 8 like |
| ENSG00000180488 | MIGA1 | 0,101716122 | 1,073049123 | 0,66263874 | 0,959897678 | mitoguardin 1 |
| ENSG00000284901 | RUVBL1 | 0,101674093 | 1,073017863 | 0,69197694 | 0,967225449 | RuvB like AAA ATPase 1 |
| ENSG00000169330 | MINAR1 | 0,101656485 | 1,073004767 | 0,717956 | 0,971780083 | membrane integral NOTCH2 associated receptor 1 |
| ENSG00000116273 | PHF13 | 0,101623036 | 1,072979889 | 0,69424127 | 0,967417311 | PHD finger protein 13 |
| ENSG00000185721 | DRG1 | 0,101618139 | 1,072976247 | 0,68453756 | 0,965415049 | developmentally regulated GTP binding protein 1 |
| ENSG00000234807 | LINC01135 | 0,101614902 | 1,07297384 | 0,90577224 | 0,992590661 | long intergenic non-protein coding RNA 1135 |
| ENSG00000106077 | ABHD11 | 0,101550227 | 1,07292574 | 0,72746574 | 0,973441576 | abhydrolase domain containing 11 |
| ENSG00000166337 | TAF10 | 0,101380687 | 1,072799661 | 0,62182274 | 0,952793934 | TATA-box binding protein associated factor 10 |
| ENSG00000037749 | MFAP3 | 0,101333906 | 1,072764875 | 0,77665448 | 0,978587255 | microfibril associated protein 3 |
| ENSG00000184922 | FMNL1 | 0,101312027 | 1,072748607 | 0,81257399 | 0,981224866 | formin like 1 |
| ENSG00000228575 | MDC1 | 0,101271425 | 1,072718416 | 0,7341462 | 0,974353474 | mediator of DNA damage checkpoint 1 |
| ENSG00000106591 | MRPL32 | 0,101230191 | 1,072687757 | 0,80874116 | 0,981213877 | mitochondrial ribosomal protein L32 |
| ENSG00000075856 | SART3 | 0,10116549 | 1,072639651 | 0,67286353 | 0,962412177 | spliceosome associated factor 3, U4/U6 recycling protein |
| ENSG00000148468 | FAM171A1 | 0,101137374 | 1,072618747 | 0,61541924 | 0,952793934 | family with sequence similarity 171 member A1 |
| ENSG00000253348 | AC008514,1 | 0,101133914 | 1,072616174 | 0,89884721 | 0,991832796 | novel transcript |
| ENSG00000108443 | RPS6KB1 | 0,1010422 | 1,072547989 | 0,78970618 | 0,979462318 | ribosomal protein S6 kinase B1 |
| ENSG00000163884 | KLF15 | 0,100970231 | 1,072494487 | 0,79125703 | 0,979471509 | Kruppel like factor 15 |

| | | | | | | |
|---|---|---|---|---|---|---|
| ENSG00000198642 | KLHL9 | 0,100955652 | 1,072483649 | 0,81080293 | 0,981213877 | kelch like family member 9 |
| ENSG00000145476 | CYP4V2 | 0,100805278 | 1,072371868 | 0,71241251 | 0,970812834 | cytochrome P450 family 4 subfamily V member 2 |
| ENSG00000167685 | ZNF444 | 0,100788505 | 1,072359401 | 0,81692506 | 0,981977498 | zinc finger protein 444 |
| ENSG00000284454 | NPIPB3 | 0,100741468 | 1,072324439 | 0,75667723 | 0,975901808 | nuclear pore complex interacting protein family member B3 |
| ENSG00000011566 | MAP4K3 | 0,100685779 | 1,072283047 | 0,79618505 | 0,980015022 | mitogen-activated protein kinase kinase kinase kinase 3 |
| ENSG00000273327 | OR6L2P | 0,100667731 | 1,072269632 | 0,92874031 | 0,994173587 | olfactory receptor family 6 subfamily L member 2 pseudogene |
| ENSG00000278213 | CNTNAP3P5 | 0,100664749 | 1,072267416 | 0,94756479 | 0,996107107 | CNTNAP3 pseudogene 5 |
| ENSG00000267719 | LINC02078 | 0,100658626 | 1,072262865 | 0,89288448 | 0,991120066 | long intergenic non-protein coding RNA 2078 |
| ENSG00000133935 | ERG28 | 0,100630252 | 1,072241777 | 0,72999514 | 0,974178338 | ergosterol biosynthesis 28 homolog |
| ENSG00000135801 | TAF5L | 0,100538885 | 1,072173874 | 0,7401158 | 0,974724313 | TATA-box binding protein associated factor 5 like |
| ENSG00000259358 | DEPDC1P1 | 0,100474812 | 1,072126257 | 0,87956699 | 0,990261566 | DEP domain containing 1 pseudogene 1 |
| ENSG00000253305 | PCDHGB6 | 0,100465225 | 1,072119132 | 0,7505625 | 0,975901808 | protocadherin gamma subfamily B, 6 |
| ENSG00000118946 | PCDH17 | 0,100391486 | 1,072064336 | 0,72201487 | 0,972516173 | protocadherin 17 |
| ENSG00000140199 | SLC12A6 | 0,100373272 | 1,072050801 | 0,66304317 | 0,959920886 | solute carrier family 12 member 6 |
| ENSG00000248367 | AC008610,1 | 0,100334034 | 1,072021644 | 0,92040554 | 0,993787755 | novel transcript |
| ENSG00000135334 | AKIRIN2 | 0,100331025 | 1,072019408 | 0,70730896 | 0,969565008 | akirin 2 |
| ENSG00000173915 | ATP5MD | 0,100315287 | 1,072007714 | 0,84320204 | 0,986218419 | ATP synthase membrane subunit DAPIT |
| ENSG00000135426 | TESPA1 | 0,100286423 | 1,071986267 | 0,92143906 | 0,993787755 | thymocyte expressed, positive selection associated 1 |
| ENSG00000102226 | USP11 | 0,100257799 | 1,071964998 | 0,62897873 | 0,952793934 | ubiquitin specific peptidase 11 |
| ENSG00000171169 | NAIF1 | 0,100225785 | 1,071941211 | 0,76610519 | 0,977102081 | nuclear apoptosis inducing factor 1 |
| ENSG00000010219 | DYRK4 | 0,100187562 | 1,071912811 | 0,76975376 | 0,977719248 | dual specificity tyrosine phosphorylation regulated kinase 4 |
| ENSG00000186912 | P2RY4 | 0,100076835 | 1,071830544 | 0,92555533 | 0,993915537 | pyrimidinergic receptor P2Y4 |
| ENSG00000147854 | UHRF2 | 0,100026082 | 1,071792839 | 0,75494507 | 0,975901808 | ubiquitin like with PHD and ring finger domains 2 |
| ENSG00000153975 | ZUP1 | 0,099968997 | 1,071750431 | 0,8265506 | 0,983471137 | zinc finger containing ubiquitin peptidase 1 |
| ENSG00000270673 | YTHDF3-AS1 | 0,099956857 | 1,071741412 | 0,83074112 | 0,984156028 | YTHDF3 antisense RNA 1 (head to head) |
| ENSG00000143061 | IGSF3 | 0,099953589 | 1,071738985 | 0,63963913 | 0,956018518 | immunoglobulin superfamily member 3 |
| ENSG00000156521 | TYSND1 | 0,099929301 | 1,071720942 | 0,74443392 | 0,975649186 | trypsin domain containing 1 |
| ENSG00000275552 | AC243965,2 | 0,099810833 | 1,07163294 | 0,90807239 | 0,992590661 | novel transcript, antisense ABHD4 |

| | | | | | | |
|---|---|---|---|---|---|---|
| ENSG00000108349 | CASC3 | 0,099718579 | 1,071564416 | 0,62845766 | 0,952793934 | CASC3 exon junction complex subunit |
| ENSG00000130723 | PRRC2B | 0,099652775 | 1,071515541 | 0,78197907 | 0,978597905 | proline rich coiled-coil 2B |
| ENSG00000218713 | AL512378,1 | 0,099647806 | 1,071511851 | 0,91489931 | 0,993361299 | regulator of chromosome condensation 1 (RCC1) pseudogene |
| ENSG00000177984 | LCN15 | 0,099641366 | 1,071507067 | 0,93180688 | 0,994433362 | lipocalin 15 |
| ENSG00000139131 | YARS2 | 0,099635202 | 1,07150249 | 0,74257984 | 0,975176043 | tyrosyl-tRNA synthetase 2 |
| ENSG00000154582 | ELOC | 0,099635065 | 1,071502388 | 0,80435003 | 0,980875302 | elongin C |
| ENSG00000151292 | CSNK1G3 | 0,099590805 | 1,071469516 | 0,8070061 | 0,981213877 | casein kinase 1 gamma 3 |
| ENSG00000134186 | PRPF38B | 0,099519736 | 1,071416735 | 0,72404689 | 0,973156872 | pre-mRNA processing factor 38B |
| ENSG00000103642 | LACTB | 0,099470538 | 1,071380199 | 0,71699657 | 0,971583263 | lactamase beta |
| ENSG00000066697 | MSANTD3 | 0,099455072 | 1,071368714 | 0,73059915 | 0,974321178 | Myb/SANT DNA binding domain containing 3 |
| ENSG00000108932 | SLC16A6 | 0,09943353 | 1,071352717 | 0,82464826 | 0,983374097 | solute carrier family 16 member 6 |
| ENSG00000284973 | AC270285,2 | 0,099424992 | 1,071346376 | 0,93031203 | 0,994336072 | novel transcript, sense intronic to CTSB |
| ENSG00000151651 | ADAM8 | 0,099359152 | 1,071297485 | 0,83982965 | 0,985672131 | ADAM metallopeptidase domain 8 |
| ENSG00000256966 | AL513165,2 | 0,099351584 | 1,071291865 | 0,87361885 | 0,989633545 | novel transcript |
| ENSG00000108469 | RECQL5 | 0,099351079 | 1,07129149 | 0,7551896 | 0,975901808 | RecQ like helicase 5 |
| ENSG00000235813 | AL672032,1 | 0,099346095 | 1,071287789 | 0,92256321 | 0,993797669 | novel transcript |
| ENSG00000081692 | JMJD4 | 0,099320981 | 1,07126914 | 0,78911528 | 0,979462318 | jumonji domain containing 4 |
| ENSG00000234742 | AC144530,1 | 0,099297057 | 1,071251376 | 0,88355704 | 0,990717729 | ribosomal protein L17 (RPL17) pseudogene |
| ENSG00000133895 | MEN1 | 0,099267037 | 1,071229085 | 0,71685803 | 0,971471587 | menin 1 |
| ENSG00000163904 | SENP2 | 0,099233706 | 1,071204336 | 0,70202528 | 0,968742626 | SUMO specific peptidase 2 |
| ENSG00000213178 | RPL22P1 | 0,099166811 | 1,071154668 | 0,7061467 | 0,969503793 | ribosomal protein L22 pseudogene 1 |
| ENSG00000168827 | GFM1 | 0,09914106 | 1,071135549 | 0,77200357 | 0,978033855 | G elongation factor mitochondrial 1 |
| ENSG00000225442 | MPRIP-AS1 | 0,099137057 | 1,071132577 | 0,84777546 | 0,987035044 | MPRIP antisense RNA 1 |
| ENSG00000248473 | LINC01962 | 0,099082449 | 1,071092034 | 0,88875791 | 0,990717729 | long intergenic non-protein coding RNA 1962 |
| ENSG00000226710 | SNHG32 | 0,099064269 | 1,071078536 | 0,79808676 | 0,980161849 | small nucleolar RNA host gene 32 |
| ENSG00000249193 | HSPD1P5 | 0,098991923 | 1,071024827 | 0,92576515 | 0,993915537 | heat shock protein family D (Hsp60) member 1 pseudogene 5 |
| ENSG00000171467 | ZNF318 | 0,098984184 | 1,071019081 | 0,68673128 | 0,965769168 | zinc finger protein 318 |
| ENSG00000164587 | RPS14 | 0,098964619 | 1,071004558 | 0,67902069 | 0,964151912 | ribosomal protein S14 |
| ENSG00000275476 | AC009318,4 | 0,098947709 | 1,070992004 | 0,91219893 | 0,993361299 | novel transcript, antisense to FAR2 |
| ENSG00000116984 | MTR | 0,098918365 | 1,070970221 | 0,69958733 | 0,968522643 | 5-methyltetrahydrofolate-homocysteine methyltransferase |

| | | | | | | |
|---|---|---|---|---|---|---|
| ENSG00000100003 | SEC14L2 | 0,098870777 | 1,070934895 | 0,74044212 | 0,974833668 | SEC14 like lipid binding 2 |
| ENSG00000285991 | AL355312,5 | 0,098776981 | 1,070865271 | 0,83849873 | 0,985672131 | novel transcript |
| ENSG00000206256 | PPT2 | 0,098716363 | 1,070820277 | 0,74687661 | 0,975796253 | palmitoyl-protein thioesterase 2 |
| ENSG00000134897 | BIVM | 0,098622372 | 1,070750516 | 0,78244859 | 0,978628303 | basic, immunoglobulin-like variable motif containing |
| ENSG00000288565 | AC068057,3 | 0,09860206 | 1,07073544 | 0,85033122 | 0,987036295 | novel transcript |
| ENSG00000186376 | ZNF75D | 0,098551513 | 1,070697926 | 0,6923148 | 0,967236299 | zinc finger protein 75D |
| ENSG00000083844 | ZNF264 | 0,098491958 | 1,070653729 | 0,67983752 | 0,96448882 | zinc finger protein 264 |
| ENSG00000279070 | AC073263,2 | 0,098469072 | 1,070636744 | 0,93347545 | 0,994479239 | novel transcript |
| ENSG00000139679 | LPAR6 | 0,098466853 | 1,070635098 | 0,83615119 | 0,985159643 | lysophosphatidic acid receptor 6 |
| ENSG00000108010 | GLRX3 | 0,098445709 | 1,070619406 | 0,73433466 | 0,974353474 | glutaredoxin 3 |
| ENSG00000105953 | OGDH | 0,098443064 | 1,070617443 | 0,692061 | 0,967225449 | oxoglutarate dehydrogenase |
| ENSG00000174365 | SNHG11 | 0,098405024 | 1,070589215 | 0,75100339 | 0,975901808 | small nucleolar RNA host gene 11 |
| ENSG00000009954 | BAZ1B | 0,098373448 | 1,070565783 | 0,6846696 | 0,965415049 | bromodomain adjacent to zinc finger domain 1B |
| ENSG00000234832 | NXT1-AS1 | 0,098361433 | 1,070556868 | 0,91765387 | 0,993361299 | NXT1 antisense RNA 1 |
| ENSG00000107290 | SETX | 0,098171738 | 1,070416113 | 0,77297793 | 0,978316811 | senataxin |
| ENSG00000132323 | ILKAP | 0,098161149 | 1,070408256 | 0,63796615 | 0,956018518 | ILK associated serine/threonine phosphatase |
| ENSG00000204778 | CBWD4P | 0,098140292 | 1,070392782 | 0,87977685 | 0,990261566 | COBW domain containing 4, pseudogene |
| ENSG00000063176 | SPHK2 | 0,098127681 | 1,070383425 | 0,79215817 | 0,979471509 | sphingosine kinase 2 |
| ENSG00000163811 | WDR43 | 0,09812694 | 1,070382875 | 0,77569997 | 0,978587255 | WD repeat domain 43 |
| ENSG00000165650 | PDZD8 | 0,098084754 | 1,070351577 | 0,76537761 | 0,976830764 | PDZ domain containing 8 |
| ENSG00000286443 | AC091178,2 | 0,097909173 | 1,070221319 | 0,86766004 | 0,989074806 | novel transcript, antisense to PLXDC1 |
| ENSG00000123416 | TUBA1B | 0,097749235 | 1,07010268 | 0,65401728 | 0,957444211 | tubulin alpha 1b |
| ENSG00000106070 | GRB10 | 0,097749128 | 1,070102601 | 0,73072699 | 0,974336839 | growth factor receptor bound protein 10 |
| ENSG00000008056 | SYN1 | 0,097745289 | 1,070099753 | 0,73407966 | 0,974353474 | synapsin I |
| ENSG00000005882 | PDK2 | 0,097721755 | 1,070082298 | 0,69271719 | 0,967271667 | pyruvate dehydrogenase kinase 2 |
| ENSG00000229117 | RPL41 | 0,097689935 | 1,070058696 | 0,70393403 | 0,968994082 | ribosomal protein L41 |
| ENSG00000158092 | NCK1 | 0,097681323 | 1,070052308 | 0,82063888 | 0,982609114 | NCK adaptor protein 1 |
| ENSG00000287737 | AC132872,5 | 0,097656703 | 1,070034048 | 0,79794995 | 0,980161849 | novel transcript |
| ENSG00000077782 | FGFR1 | 0,097533127 | 1,069942396 | 0,73857126 | 0,974353474 | fibroblast growth factor receptor 1 |
| ENSG00000276684 | RDH13 | 0,097509076 | 1,06992456 | 0,81315398 | 0,981246822 | retinol dehydrogenase 13 |
| ENSG00000268573 | AC011815,1 | 0,097450025 | 1,069880768 | 0,78935564 | 0,979462318 | novel transcript |
| ENSG00000172020 | GAP43 | 0,097434518 | 1,069869268 | 0,72823565 | 0,973888243 | growth associated protein 43 |

| | | | | | | |
|---|---|---|---|---|---|---|
| ENSG00000204842 | ATXN2 | 0,097386727 | 1,069833827 | 0,70656458 | 0,969503793 | ataxin 2 |
| ENSG00000278742 | LRRC37A | 0,097375168 | 1,069825256 | 0,74874677 | 0,975901808 | leucine rich repeat containing 37A |
| ENSG00000261158 | AC109597,2 | 0,097346835 | 1,069804246 | 0,90706597 | 0,992590661 | novel transcript, antisense to CPPED1 |
| ENSG00000025772 | TOMM34 | 0,097319076 | 1,069783662 | 0,67010895 | 0,962026636 | translocase of outer mitochondrial membrane 34 |
| ENSG00000271871 | AC005740,4 | 0,097308293 | 1,069775666 | 0,91815323 | 0,993361299 | novel transcript, antisense to PCDH12 |
| ENSG00000149547 | EI24 | 0,097296937 | 1,069767245 | 0,72816653 | 0,973888243 | EI24 autophagy associated transmembrane protein |
| ENSG00000177613 | CSTF2T | 0,097266891 | 1,069744966 | 0,69921605 | 0,968522643 | cleavage stimulation factor subunit 2 tau variant |
| ENSG00000280036 | AC020661,4 | 0,097158301 | 1,069664451 | 0,85508268 | 0,988016959 | novel transcript, sense intronic to INO80 |
| ENSG00000198836 | OPA1 | 0,097110384 | 1,069628924 | 0,78309627 | 0,978917309 | OPA1 mitochondrial dynamin like GTPase |
| ENSG00000136738 | STAM | 0,097093141 | 1,069616141 | 0,82440759 | 0,983374097 | signal transducing adaptor molecule |
| ENSG00000279145 | AC011912,1 | 0,097081911 | 1,069607815 | 0,82955236 | 0,984081562 | TEC |
| ENSG00000101190 | TCFL5 | 0,097053649 | 1,069586861 | 0,75450945 | 0,975901808 | transcription factor like 5 |
| ENSG00000272157 | AC107373,2 | 0,097036924 | 1,069574462 | 0,90708747 | 0,992590661 | novel transcript, antisense to NEFL |
| ENSG00000147059 | SPIN2A | 0,097023516 | 1,069564521 | 0,75078084 | 0,975901808 | spindlin family member 2A |
| ENSG00000187555 | USP7 | 0,097012323 | 1,069556224 | 0,76110446 | 0,97624106 | ubiquitin specific peptidase 7 |
| ENSG00000270118 | AC138894,2 | 0,097012019 | 1,069555998 | 0,87955165 | 0,990261566 | novel transcript |
| ENSG00000100767 | PAPLN | 0,096936734 | 1,069500186 | 0,85931623 | 0,989074806 | papilin, proteoglycan like sulfated glycoprotein |
| ENSG00000254905 | AP001318,1 | 0,096907801 | 1,069478738 | 0,89419323 | 0,991135494 | novel transcript, antisense to TIRAP |
| ENSG00000227258 | SMIM2-AS1 | 0,096777164 | 1,0693819 | 0,87841988 | 0,990109814 | SMIM2 antisense RNA 1 |
| ENSG00000248554 | C5orf34-AS1 | 0,096758877 | 1,069368346 | 0,87602727 | 0,990109814 | C5orf34 antisense RNA 1 |
| ENSG00000278615 | C11orf98 | 0,096712453 | 1,069333935 | 0,70244644 | 0,968742626 | chromosome 11 open reading frame 98 |
| ENSG00000169976 | SF3B5 | 0,096707962 | 1,069330606 | 0,66656869 | 0,9611116 | splicing factor 3b subunit 5 |
| ENSG00000271664 | AC004890,3 | 0,096695636 | 1,06932147 | 0,87997966 | 0,990268593 | SECIS binding protein 2, pseudogene |
| ENSG00000121316 | PLBD1 | 0,096687613 | 1,069315523 | 0,84194918 | 0,98572121 | phospholipase B domain containing 1 |
| ENSG00000268047 | AC018766,1 | 0,096665171 | 1,06929889 | 0,91364392 | 0,993361299 | novel transcript, antisense to PTOV1 |
| ENSG00000250021 | ARPIN-AP3S2 | 0,096516749 | 1,069188888 | 0,70771296 | 0,969840367 | ARPIN-AP3S2 readthrough |
| ENSG00000255468 | AP001107,9 | 0,096349011 | 1,069064584 | 0,8356825 | 0,985034063 | novel transcript, antisense to SLC29A2 |
| ENSG00000245025 | AC107959,1 | 0,096228786 | 1,068975499 | 0,84676216 | 0,986939308 | novel transcript, antisense to RHOBTB2 |
| ENSG00000158156 | XKR8 | 0,096220985 | 1,068969718 | 0,70817666 | 0,97011892 | XK related 8 |
| ENSG00000170471 | RALGAPB | 0,096203808 | 1,068956991 | 0,76148253 | 0,976285211 | Ral GTPase activating protein non-catalytic subunit beta |
| ENSG00000117222 | RBBP5 | 0,096138487 | 1,068908593 | 0,65303177 | 0,95732962 | RB binding protein 5, histone lysine methyltransferase complex subunit |

| | | | | | | |
|---|---|---|---|---|---|---|
| ENSG00000277476 | AC005332,5 | 0,096068413 | 1,068856675 | 0,85946644 | 0,989074806 | novel transcript |
| ENSG00000274386 | TMEM269 | 0,09606509 | 1,068854213 | 0,84688549 | 0,986939308 | transmembrane protein 269 |
| ENSG00000224614 | TNK2-AS1 | 0,095929822 | 1,068754002 | 0,89248635 | 0,991048031 | TNK2 antisense RNA 1 |
| ENSG00000198689 | SLC9A6 | 0,095927809 | 1,068752511 | 0,76403886 | 0,976640468 | solute carrier family 9 member A6 |
| ENSG00000115268 | RPS15 | 0,095920503 | 1,068747098 | 0,68584358 | 0,965740648 | ribosomal protein S15 |
| ENSG00000140374 | ETFA | 0,095894252 | 1,068727652 | 0,76915799 | 0,97747878 | electron transfer flavoprotein subunit alpha |
| ENSG00000047634 | SCML1 | 0,095885114 | 1,068720882 | 0,86225415 | 0,989074806 | Scm polycomb group protein like 1 |
| ENSG00000213203 | GIMAP1 | 0,09584366 | 1,068690175 | 0,94318919 | 0,995402494 | GTPase, IMAP family member 1 |
| ENSG00000286627 | AP005900,1 | 0,095835385 | 1,068684045 | 0,8474595 | 0,987035044 | novel transcript, antisense to PTPRM |
| ENSG00000183530 | PRR14L | 0,095774116 | 1,06863866 | 0,69310513 | 0,967333542 | proline rich 14 like |
| ENSG00000038219 | BOD1L1 | 0,095666285 | 1,06855879 | 0,73447535 | 0,974353474 | biorientation of chromosomes in cell division 1 like 1 |
| ENSG00000125445 | MRPS7 | 0,09565357 | 1,068549373 | 0,71379461 | 0,970812834 | mitochondrial ribosomal protein S7 |
| ENSG00000198000 | NOL8 | 0,0956343 | 1,0685351 | 0,75121051 | 0,975901808 | nucleolar protein 8 |
| ENSG00000225401 | TGIF2P1 | 0,095588978 | 1,068501533 | 0,95169518 | 0,996432861 | TGFB induced factor homeobox 2 pseudogene 1 |
| ENSG00000158006 | PAFAH2 | 0,095584531 | 1,06849824 | 0,68250865 | 0,964781531 | platelet activating factor acetylhydrolase 2 |
| ENSG00000234650 | PCCA-AS1 | 0,095554403 | 1,068475926 | 0,91509039 | 0,993361299 | PCCA antisense RNA 1 |
| ENSG00000129484 | PARP2 | 0,095495558 | 1,068432346 | 0,7833497 | 0,978960107 | poly(ADP-ribose) polymerase 2 |
| ENSG00000135052 | GOLM1 | 0,095418581 | 1,06837534 | 0,66306806 | 0,959920886 | golgi membrane protein 1 |
| ENSG00000178913 | TAF7 | 0,095412161 | 1,068370585 | 0,75791942 | 0,976120207 | TATA-box binding protein associated factor 7 |
| ENSG00000151881 | TMEM267 | 0,095263114 | 1,068260216 | 0,80942327 | 0,981213877 | transmembrane protein 267 |
| ENSG00000108826 | MRPL27 | 0,095261509 | 1,068259027 | 0,63937088 | 0,956018518 | mitochondrial ribosomal protein L27 |
| ENSG00000161980 | POLR3K | 0,095258133 | 1,068256528 | 0,75166535 | 0,975901808 | RNA polymerase III subunit K |
| ENSG00000174255 | ZNF80 | 0,09515438 | 1,068179706 | 0,90573353 | 0,992590661 | zinc finger protein 80 |
| ENSG00000244968 | LIFR-AS1 | 0,095090405 | 1,068132339 | 0,78100345 | 0,978597905 | LIFR antisense RNA 1 |
| ENSG00000189332 | SLC25A51P4 | 0,095005963 | 1,068069823 | 0,8123226 | 0,981213877 | SLC25A51 pseudogene 4 |
| ENSG00000163349 | HIPK1 | 0,094994964 | 1,06806168 | 0,76503071 | 0,976739398 | homeodomain interacting protein kinase 1 |
| ENSG00000164849 | GPR146 | 0,094984655 | 1,068054048 | 0,90042636 | 0,991865079 | G protein-coupled receptor 146 |
| ENSG00000174721 | FGFBP3 | 0,094939903 | 1,068020918 | 0,72955907 | 0,974009551 | fibroblast growth factor binding protein 3 |
| ENSG00000197912 | SPG7 | 0,094893024 | 1,067986214 | 0,77437758 | 0,978503783 | SPG7 matrix AAA peptidase subunit, paraplegin |
| ENSG00000255112 | CHMP1B | 0,094868649 | 1,06796817 | 0,66364378 | 0,959920886 | charged multivesicular body protein 1B |
| ENSG00000147324 | MFHAS1 | 0,094842599 | 1,067948887 | 0,75608698 | 0,975901808 | malignant fibrous histiocytoma amplified sequence 1 |

| | | | | | | |
|---|---|---|---|---|---|---|
| ENSG00000100815 | TRIP11 | 0,094793806 | 1,067912768 | 0,70316263 | 0,968742626 | thyroid hormone receptor interactor 11 |
| ENSG00000182521 | TBPL2 | 0,094731968 | 1,067866995 | 0,92505062 | 0,993915537 | TATA-box binding protein like 2 |
| ENSG00000133812 | SBF2 | 0,094729928 | 1,067865485 | 0,70244622 | 0,968742626 | SET binding factor 2 |
| ENSG00000278545 | MARCHF8 | 0,094688751 | 1,067835007 | 0,73780768 | 0,974353474 | membrane associated ring-CH-type finger 8 |
| ENSG00000143494 | VASH2 | 0,094588271 | 1,067760637 | 0,7470784 | 0,975796253 | vasohibin 2 |
| ENSG00000169967 | MAP3K2 | 0,094562557 | 1,067741606 | 0,81789769 | 0,982323352 | mitogen-activated protein kinase kinase kinase 2 |
| ENSG00000140382 | HMG20A | 0,094513763 | 1,067705495 | 0,72264258 | 0,972594421 | high mobility group 20A |
| ENSG00000159884 | CCDC107 | 0,094505387 | 1,067699296 | 0,72542712 | 0,97326334 | coiled-coil domain containing 107 |
| ENSG00000276726 | DNAJA3 | 0,094442417 | 1,067652695 | 0,67326258 | 0,962605657 | DnaJ heat shock protein family (Hsp40) member A3 |
| ENSG00000169436 | COL22A1 | 0,094439258 | 1,067650357 | 0,88679875 | 0,990717729 | collagen type XXII alpha 1 chain |
| ENSG00000274020 | LINC01138 | 0,094421155 | 1,06763696 | 0,70007964 | 0,968522643 | long intergenic non-protein coding RNA 1138 |
| ENSG00000171135 | JAGN1 | 0,094392228 | 1,067615553 | 0,73577311 | 0,974353474 | jagunal homolog 1 |
| ENSG00000167699 | GLOD4 | 0,094379614 | 1,067606219 | 0,66755802 | 0,9611116 | glyoxalase domain containing 4 |
| ENSG00000214447 | FAM187A | 0,094379257 | 1,067605954 | 0,81127077 | 0,981213877 | family with sequence similarity 187 member A |
| ENSG00000161800 | RACGAP1 | 0,094377731 | 1,067604825 | 0,75799153 | 0,976120207 | Rac GTPase activating protein 1 |
| ENSG00000136783 | NIPSNAP3A | 0,094355884 | 1,067588659 | 0,82698136 | 0,983556324 | nipsnap homolog 3A |
| ENSG00000065548 | ZC3H15 | 0,094343782 | 1,067579703 | 0,72530964 | 0,97326334 | zinc finger CCCH-type containing 15 |
| ENSG00000146233 | CYP39A1 | 0,094313107 | 1,067557004 | 0,84558739 | 0,986639902 | cytochrome P450 family 39 subfamily A member 1 |
| ENSG00000132801 | ZSWIM3 | 0,094309137 | 1,067554066 | 0,77951769 | 0,978587255 | zinc finger SWIM-type containing 3 |
| ENSG00000147669 | POLR2K | 0,094288329 | 1,067538669 | 0,83894199 | 0,985672131 | RNA polymerase II, I and III subunit K |
| ENSG00000041357 | PSMA4 | 0,094259813 | 1,067517569 | 0,80849543 | 0,981213877 | proteasome 20S subunit alpha 4 |
| ENSG00000158480 | SPATA2 | 0,094225913 | 1,067492485 | 0,66909396 | 0,961684145 | spermatogenesis associated 2 |
| ENSG00000136490 | LIMD2 | 0,094222281 | 1,067489797 | 0,74083161 | 0,97509197 | LIM domain containing 2 |
| ENSG00000160062 | ZBTB8A | 0,094219824 | 1,067487979 | 0,71009911 | 0,970482408 | zinc finger and BTB domain containing 8A |
| ENSG00000113361 | CDH6 | 0,094205205 | 1,067477163 | 0,80712167 | 0,981213877 | cadherin 6 |
| ENSG00000269842 | AC011453,1 | 0,094172006 | 1,067452598 | 0,9005087 | 0,991865079 | novel transcript, host to microRNA 519a-2 |
| ENSG00000236611 | LINC02556 | 0,094108995 | 1,067405977 | 0,93207519 | 0,994479239 | long intergenic non-protein coding RNA 2556 |
| ENSG00000198931 | APRT | 0,094102631 | 1,067401269 | 0,78829355 | 0,979462318 | adenine phosphoribosyltransferase |
| ENSG00000138271 | GPR87 | 0,093990755 | 1,067318498 | 0,94436348 | 0,995542144 | G protein-coupled receptor 87 |
| ENSG00000271425 | NBPF10 | 0,093927862 | 1,06727197 | 0,7346076 | 0,974353474 | NBPF member 10 |
| ENSG00000261971 | MMP25-AS1 | 0,093910198 | 1,067258904 | 0,83975267 | 0,985672131 | MMP25 antisense RNA 1 |
| ENSG00000283918 | MRNIP | 0,093898146 | 1,067249988 | 0,81638061 | 0,981855101 | MRN complex interacting protein |
| ENSG00000101158 | NELFCD | 0,093702045 | 1,06710493 | 0,69051351 | 0,967179044 | negative elongation factor complex member C/D |

| | | | | | | |
|---|---|---|---|---|---|---|
| ENSG00000184381 | PLA2G6 | 0,093684237 | 1,067091758 | 0,7372836 | 0,974353474 | phospholipase A2 group VI |
| ENSG00000259462 | CPEB1-AS1 | 0,09363056 | 1,067052056 | 0,90386322 | 0,992462326 | CPEB1 antisense RNA 1 |
| ENSG00000164258 | NDUFS4 | 0,093607094 | 1,067034701 | 0,77688949 | 0,978587255 | NADH:ubiquinone oxidoreductase subunit S4 |
| ENSG00000224740 | FLOT1 | 0,093543219 | 1,066987459 | 0,84460451 | 0,986428908 | flotillin 1 |
| ENSG00000128923 | MINDY2 | 0,09340006 | 1,066881587 | 0,77347057 | 0,978415098 | MINDY lysine 48 deubiquitinase 2 |
| ENSG00000178690 | DYNAP | 0,093377865 | 1,066865173 | 0,86114586 | 0,989074806 | dynactin associated protein |
| ENSG00000285752 | CDC42-AS1 | 0,093331853 | 1,066831148 | 0,81236368 | 0,981213877 | CDC42 antisense RNA 1 |
| ENSG00000267248 | AC025048,2 | 0,093327143 | 1,066827665 | 0,77466783 | 0,978553937 | novel transcript |
| ENSG00000259654 | AC115102,1 | 0,093207066 | 1,066738876 | 0,93495828 | 0,994479239 | novel transcript, antisense to SEC11A |
| ENSG00000096654 | ZNF184 | 0,093118161 | 1,066673141 | 0,81928992 | 0,98249617 | zinc finger protein 184 |
| ENSG00000227252 | AC012063,1 | 0,093098254 | 1,066658422 | 0,72860732 | 0,973955059 | novel transcript, antisense to COPS8 |
| ENSG00000105176 | URI1 | 0,093067673 | 1,066635813 | 0,74865256 | 0,975901808 | URI1 prefoldin like chaperone |
| ENSG00000275216 | AL161431,1 | 0,092993396 | 1,066580899 | 0,85305349 | 0,98780934 | novel transcript |
| ENSG00000099995 | SF3A1 | 0,09292513 | 1,066530431 | 0,66150875 | 0,959582738 | splicing factor 3a subunit 1 |
| ENSG00000077157 | PPP1R12B | 0,092908773 | 1,066518339 | 0,70640145 | 0,969503793 | protein phosphatase 1 regulatory subunit 12B |
| ENSG00000178015 | GPR150 | 0,092861103 | 1,066483099 | 0,93007509 | 0,994331958 | G protein-coupled receptor 150 |
| ENSG00000256603 | AP003170,4 | 0,092833824 | 1,066462934 | 0,9194122 | 0,993598006 | novel transcript |
| ENSG00000233420 | AC002069,2 | 0,092826237 | 1,066457325 | 0,93917063 | 0,994857716 | novel transcript |
| ENSG00000224587 | MDC1 | 0,092713043 | 1,066373654 | 0,75050372 | 0,975901808 | mediator of DNA damage checkpoint 1 |
| ENSG00000237095 | MDC1 | 0,092713043 | 1,066373654 | 0,75050372 | 0,975901808 | mediator of DNA damage checkpoint 1 |
| ENSG00000124104 | SNX21 | 0,092582073 | 1,066276852 | 0,79632383 | 0,980015022 | sorting nexin family member 21 |
| ENSG00000136379 | ABHD17C | 0,092565693 | 1,066264745 | 0,81684842 | 0,981977498 | abhydrolase domain containing 17C, depalmitoylase |
| ENSG00000234751 | AP002381,1 | 0,092540407 | 1,066246057 | 0,9174185 | 0,993361299 | ribosomal protein L15 (RPL15) pseudogene |
| ENSG00000140990 | NDUFB10 | 0,092446621 | 1,066176746 | 0,71432974 | 0,970991522 | NADH:ubiquinone oxidoreductase subunit B10 |
| ENSG00000257953 | AC083805,2 | 0,092445696 | 1,066176062 | 0,85348168 | 0,987861659 | novel transcript, antisense to CTDSP2 |
| ENSG00000151948 | GLT1D1 | 0,092418862 | 1,066156232 | 0,70659787 | 0,969503793 | glycosyltransferase 1 domain containing 1 |
| ENSG00000103168 | TAF1C | 0,092415217 | 1,066153538 | 0,78770646 | 0,979462318 | TATA-box binding protein associated factor, RNA polymerase I subunit C |
| ENSG00000164306 | PRIMPOL | 0,092385161 | 1,066131327 | 0,72907639 | 0,974009551 | primase and DNA directed polymerase |
| ENSG00000119537 | KDSR | 0,092277531 | 1,066051793 | 0,70680774 | 0,969565008 | 3-ketodihydrosphingosine reductase |
| ENSG00000161847 | RAVER1 | 0,092259417 | 1,066038408 | 0,76069889 | 0,97624106 | ribonucleoprotein, PTB binding 1 |
| ENSG00000235786 | ZNRF3-IT1 | 0,092246494 | 1,066028859 | 0,89409772 | 0,991135494 | ZNRF3 intronic transcript 1 |
| ENSG00000204956 | PCDHGA1 | 0,092185086 | 1,065983484 | 0,85271991 | 0,98761642 | protocadherin gamma subfamily A, 1 |

| | | | | | | |
|---|---|---|---|---|---|---|
| ENSG00000149600 | COMMD7 | 0,092150012 | 1,065957569 | 0,75220645 | 0,975901808 | COMM domain containing 7 |
| ENSG00000166949 | SMAD3 | 0,092137348 | 1,065948212 | 0,73984386 | 0,974715504 | SMAD family member 3 |
| ENSG00000146576 | C7orf26 | 0,092091175 | 1,065914098 | 0,74361017 | 0,975310206 | chromosome 7 open reading frame 26 |
| ENSG00000115350 | POLE4 | 0,092082995 | 1,065908053 | 0,79150412 | 0,979471509 | DNA polymerase epsilon 4, accessory subunit |
| ENSG00000236973 | GAPDHP51 | 0,092076474 | 1,065903236 | 0,92828222 | 0,994108531 | glyceraldehyde 3 phosphate dehydrogenase pseudogene 51 |
| ENSG00000282788 | AC243791,4 | 0,092071456 | 1,065899528 | 0,8225122 | 0,98299282 | novel transcript |
| ENSG00000213782 | DDX47 | 0,092050573 | 1,065884099 | 0,75070978 | 0,975901808 | DEAD-box helicase 47 |
| ENSG00000179335 | CLK3 | 0,092045875 | 1,065880628 | 0,71749247 | 0,971697135 | CDC like kinase 3 |
| ENSG00000277564 | RBFOX2 | 0,092035523 | 1,065872981 | 0,81332645 | 0,981282128 | RNA binding fox-1 homolog 2 |
| ENSG00000198932 | GPRASP1 | 0,092004798 | 1,065850281 | 0,80436596 | 0,980875302 | G protein-coupled receptor associated sorting protein 1 |
| ENSG00000083807 | SLC27A5 | 0,09200325 | 1,065849137 | 0,83753301 | 0,98566906 | solute carrier family 27 member 5 |
| ENSG00000169570 | DTWD2 | 0,09199021 | 1,065839504 | 0,72369144 | 0,972971932 | DTW domain containing 2 |
| ENSG00000163964 | PIGX | 0,091970947 | 1,065825272 | 0,75208547 | 0,975901808 | phosphatidylinositol glycan anchor biosynthesis class X |
| ENSG00000160602 | NEK8 | 0,091933224 | 1,065797404 | 0,80192361 | 0,980710009 | NIMA related kinase 8 |
| ENSG00000065268 | WDR18 | 0,091890403 | 1,06576577 | 0,75674729 | 0,975901808 | WD repeat domain 18 |
| ENSG00000287130 | AC068616,3 | 0,091789819 | 1,065691468 | 0,84111721 | 0,985672131 | novel transcript |
| ENSG00000215174 | NLRP2B | 0,091779145 | 1,065683583 | 0,9482841 | 0,99635783 | NLR family pyrin domain containing 2B |
| ENSG00000100393 | EP300 | 0,091742735 | 1,065656689 | 0,68073932 | 0,964654539 | E1A binding protein p300 |
| ENSG00000166200 | COPS2 | 0,09172655 | 1,065644734 | 0,83488166 | 0,984905093 | COP9 signalosome subunit 2 |
| ENSG00000267882 | AL031666,2 | 0,091680838 | 1,065610969 | 0,88274259 | 0,990717729 | novel transcript, antisense to ZMYND8 |
| ENSG00000167615 | LENG8 | 0,091665079 | 1,065599329 | 0,70582353 | 0,969503793 | leukocyte receptor cluster member 8 |
| ENSG00000104687 | GSR | 0,091655681 | 1,065592387 | 0,70598795 | 0,969503793 | glutathione-disulfide reductase |
| ENSG00000231726 | HMGN2P38 | 0,091624095 | 1,065569058 | 0,95541668 | 0,996608711 | high mobility group nucleosomal binding domain 2 pseudogene 38 |
| ENSG00000272617 | AC026464,6 | 0,091616428 | 1,065563395 | 0,80240557 | 0,980742323 | novel protein, COG8-PDF readthrough |
| ENSG00000139625 | MAP3K12 | 0,09160372 | 1,065554009 | 0,73377594 | 0,974353474 | mitogen-activated protein kinase kinase kinase 12 |
| ENSG00000134882 | UBAC2 | 0,091588987 | 1,065543127 | 0,70272647 | 0,968742626 | UBA domain containing 2 |
| ENSG00000280858 | RPL7A | 0,091579771 | 1,065536321 | 0,69110394 | 0,967179456 | ribosomal protein L7a |
| ENSG00000148303 | RPL7A | 0,091579771 | 1,065536321 | 0,69110394 | 0,967179456 | ribosomal protein L7a |
| ENSG00000157388 | CACNA1D | 0,091570912 | 1,065529778 | 0,88708551 | 0,990717729 | calcium voltage-gated channel subunit alpha1 D |

| | | | | | | |
|---|---|---|---|---|---|---|
| ENSG00000241506 | PSMC1P1 | 0,091527439 | 1,06549767 | 0,96360515 | 0,996733015 | proteasome 26S subunit, ATPase 1 pseudogene 1 |
| ENSG00000137691 | CFAP300 | 0,091517706 | 1,065490482 | 0,76608908 | 0,977102081 | cilia and flagella associated protein 300 |
| ENSG00000229023 | RAB1AP1 | 0,091473515 | 1,065457846 | 0,93587782 | 0,994479239 | RAB1A pseudogene 1 |
| ENSG00000260868 | LINC01960 | 0,091472237 | 1,065456902 | 0,94194478 | 0,995373376 | long intergenic non-protein coding RNA 1960 |
| ENSG00000228021 | AL158835,1 | 0,091427603 | 1,065423939 | 0,9111922 | 0,993222655 | novel transcript |
| ENSG00000135452 | TSPAN31 | 0,091410522 | 1,065411325 | 0,674902 | 0,963268551 | tetraspanin 31 |
| ENSG00000119013 | NDUFB3 | 0,091398686 | 1,065402585 | 0,79476656 | 0,979776981 | NADH:ubiquinone oxidoreductase subunit B3 |
| ENSG00000106853 | PTGR1 | 0,091353123 | 1,065368938 | 0,74059675 | 0,974905965 | prostaglandin reductase 1 |
| ENSG00000175193 | PARL | 0,091312917 | 1,065339248 | 0,76037862 | 0,97624106 | presenilin associated rhomboid like |
| ENSG00000266028 | SRGAP2 | 0,091298406 | 1,065328533 | 0,69384157 | 0,967340633 | SLIT-ROBO Rho GTPase activating protein 2 |
| ENSG00000267740 | AC024592,3 | 0,091289086 | 1,06532165 | 0,9036133 | 0,992431891 | novel protein |
| ENSG00000160959 | LRRC14 | 0,091274766 | 1,065311076 | 0,77251754 | 0,978288572 | leucine rich repeat containing 14 |
| ENSG00000286895 | AC092745,5 | 0,091219524 | 1,065270285 | 0,86099666 | 0,989074806 | novel transcript |
| ENSG00000240152 | LINC02271 | 0,091123482 | 1,065199371 | 0,92685459 | 0,993915537 | long intergenic non-protein coding RNA 2271 |
| ENSG00000145782 | ATG12 | 0,091070694 | 1,065160397 | 0,82180049 | 0,982952211 | autophagy related 12 |
| ENSG00000124226 | RNF114 | 0,090994507 | 1,065104148 | 0,73233367 | 0,974353474 | ring finger protein 114 |
| ENSG00000196357 | ZNF565 | 0,090991302 | 1,065101782 | 0,75842779 | 0,976120207 | zinc finger protein 565 |
| ENSG00000157617 | C2CD2 | 0,09098644 | 1,065098192 | 0,72065009 | 0,972142678 | C2 calcium dependent domain containing 2 |
| ENSG00000123700 | KCNJ2 | 0,090752243 | 1,064925306 | 0,82581809 | 0,983383773 | potassium inwardly rectifying channel subfamily J member 2 |
| ENSG00000069509 | FUNDC1 | 0,090730218 | 1,064909048 | 0,81711214 | 0,981979397 | FUN14 domain containing 1 |
| ENSG00000161649 | CD300LG | 0,090727631 | 1,064907139 | 0,90750016 | 0,992590661 | CD300 molecule like family member g |
| ENSG00000189143 | CLDN4 | 0,090722402 | 1,064903279 | 0,86839294 | 0,989074806 | claudin 4 |
| ENSG00000146556 | WASH2P | 0,090605202 | 1,064816773 | 0,73322664 | 0,974353474 | WASP family homolog 2, pseudogene |
| ENSG00000233266 | HMGB1P31 | 0,090588336 | 1,064804324 | 0,88873954 | 0,990717729 | high mobility group box 1 pseudogene 31 |
| ENSG00000243667 | WDR92 | 0,090579633 | 1,064797901 | 0,7070891 | 0,969565008 | WD repeat domain 92 |
| ENSG00000124702 | KLHDC3 | 0,090560035 | 1,064783437 | 0,71370839 | 0,970812834 | kelch domain containing 3 |
| ENSG00000074047 | GLI2 | 0,090510854 | 1,06474714 | 0,75800544 | 0,976120207 | GLI family zinc finger 2 |
| ENSG00000274950 | RPS9 | 0,090421646 | 1,064681304 | 0,77434697 | 0,978503783 | ribosomal protein S9 |
| ENSG00000249456 | AL731577,2 | 0,090354716 | 1,064631912 | 0,81509239 | 0,981521413 | novel transcript, sense overlapping ZRANB1 |
| ENSG00000263017 | AC036164,1 | 0,090350961 | 1,064629141 | 0,94983524 | 0,996432861 | chromosome 19 open reading frame 70 (C19orf70) pseudogene |
| ENSG00000287241 | AL359511,3 | 0,090334617 | 1,06461708 | 0,92856592 | 0,994108531 | novel transcript, antisense to PCSK2 |

| | | | | | | |
|---|---|---|---|---|---|---|
| ENSG00000186871 | ERCC6L | 0,090283284 | 1,0645792 | 0,8265625 | 0,983471137 | ERCC excision repair 6 like, spindle assembly checkpoint helicase |
| ENSG00000253893 | FAM85B | 0,090260466 | 1,064562362 | 0,88554847 | 0,990717729 | family with sequence similarity 85 member B |
| ENSG00000282806 | AC068400,2 | 0,090254483 | 1,064557947 | 0,92515325 | 0,993915537 | novel transcript, sense intronic to C17orf78 |
| ENSG00000048707 | VPS13D | 0,090236169 | 1,064544434 | 0,6811732 | 0,964654539 | vacuolar protein sorting 13 homolog D |
| ENSG00000166965 | RCCD1 | 0,090186233 | 1,064507587 | 0,74268675 | 0,975201495 | RCC1 domain containing 1 |
| ENSG00000203875 | SNHG5 | 0,090184291 | 1,064506154 | 0,75044656 | 0,975901808 | small nucleolar RNA host gene 5 |
| ENSG00000272625 | AP000919,3 | 0,090113899 | 1,064454217 | 0,90016467 | 0,991845106 | novel transcript, antisense to LPIN2 |
| ENSG00000181472 | ZBTB2 | 0,090082268 | 1,064430879 | 0,79563057 | 0,979879951 | zinc finger and BTB domain containing 2 |
| ENSG00000139719 | VPS33A | 0,090066702 | 1,064419394 | 0,70146759 | 0,968742626 | VPS33A core subunit of CORVET and HOPS complexes |
| ENSG00000196363 | WDR5 | 0,090029667 | 1,06439207 | 0,67241484 | 0,962290553 | WD repeat domain 5 |
| ENSG00000217624 | YWHAZP10 | 0,090012853 | 1,064379665 | 0,893998 | 0,991135494 | YWHAZ pseudogene 10 |
| ENSG00000174206 | C12orf66 | 0,08987861 | 1,064280629 | 0,75692997 | 0,975901808 | chromosome 12 open reading frame 66 |
| ENSG00000198612 | COPS8 | 0,089846103 | 1,064256649 | 0,8399764 | 0,985672131 | COP9 signalosome subunit 8 |
| ENSG00000272669 | AL021707,6 | 0,089829772 | 1,064244602 | 0,86418751 | 0,989074806 | novel transcript, antisense to SUN2 |
| ENSG00000163069 | SGCB | 0,089817232 | 1,064235351 | 0,82508706 | 0,983383773 | sarcoglycan beta |
| ENSG00000059804 | SLC2A3 | 0,089781461 | 1,064208964 | 0,80336866 | 0,980866933 | solute carrier family 2 member 3 |
| ENSG00000181929 | PRKAG1 | 0,089779115 | 1,064207233 | 0,77292354 | 0,978310326 | protein kinase AMP-activated non-catalytic subunit gamma 1 |
| ENSG00000112514 | CUTA | 0,089760986 | 1,064193861 | 0,71335075 | 0,970812834 | cutA divalent cation tolerance homolog |
| ENSG00000215784 | FAM72D | 0,089756241 | 1,06419036 | 0,85634611 | 0,988229014 | family with sequence similarity 72 member D |
| ENSG00000066777 | ARFGEF1 | 0,089712571 | 1,064158149 | 0,78446611 | 0,979320406 | ADP ribosylation factor guanine nucleotide exchange factor 1 |
| ENSG00000161016 | RPL8 | 0,089664809 | 1,064122919 | 0,70649368 | 0,969503793 | ribosomal protein L8 |
| ENSG00000101109 | STK4 | 0,089600462 | 1,064075458 | 0,74737697 | 0,975901808 | serine/threonine kinase 4 |
| ENSG00000127922 | SEM1 | 0,089594927 | 1,064071376 | 0,70068163 | 0,968522643 | SEM1 26S proteasome complex subunit |
| ENSG00000267023 | LRRC37A16P | 0,089461314 | 1,063972833 | 0,75075001 | 0,975901808 | leucine rich repeat containing 37 member A16, pseudogene |
| ENSG00000075218 | GTSE1 | 0,089431303 | 1,0639507 | 0,68021103 | 0,964580199 | G2 and S-phase expressed 1 |
| ENSG00000134262 | AP4B1 | 0,089404132 | 1,063930663 | 0,70001993 | 0,968522643 | adaptor related protein complex 4 subunit beta 1 |
| ENSG00000253676 | TAGLN2P1 | 0,089377376 | 1,063910931 | 0,93689668 | 0,994479239 | transgelin 2 pseudogene 1 |
| ENSG00000253716 | MINCR | 0,089353307 | 1,063893182 | 0,79196636 | 0,979471509 | MYC-induced long non-coding RNA |

| | | | | | | |
|---|---|---|---|---|---|---|
| ENSG00000138449 | SLC40A1 | 0,089343929 | 1,063886266 | 0,86760211 | 0,989074806 | solute carrier family 40 member 1 |
| ENSG00000161860 | SYCE2 | 0,089162247 | 1,063752297 | 0,83782465 | 0,98566906 | synaptonemal complex central element protein 2 |
| ENSG00000233822 | H2BC15 | 0,089154914 | 1,06374689 | 0,73859184 | 0,974353474 | H2B clustered histone 15 |
| ENSG00000169981 | ZNF35 | 0,089149837 | 1,063743146 | 0,71274469 | 0,970812834 | zinc finger protein 35 |
| ENSG00000281306 | ZNF35 | 0,089149837 | 1,063743146 | 0,71274469 | 0,970812834 | zinc finger protein 35 |
| ENSG00000236511 | LINC01231 | 0,08913855 | 1,063734824 | 0,91010188 | 0,992965454 | long intergenic non-protein coding RNA 1231 |
| ENSG00000284468 | SLC25A24 | 0,089120663 | 1,063721636 | 0,8276339 | 0,983595244 | solute carrier family 25 member 24 |
| ENSG00000085491 | SLC25A24 | 0,089120663 | 1,063721636 | 0,8276339 | 0,983595244 | solute carrier family 25 member 24 |
| ENSG00000148832 | PAOX | 0,0889888 | 1,063624416 | 0,91698481 | 0,993361299 | polyamine oxidase |
| ENSG00000114030 | KPNA1 | 0,088965608 | 1,063607318 | 0,80018305 | 0,980161849 | karyopherin subunit alpha 1 |
| ENSG00000277978 | AC010542,5 | 0,08884645 | 1,063519474 | 0,86196232 | 0,989074806 | novel transcript, antisense to CKLF |
| ENSG00000033178 | UBA6 | 0,088795269 | 1,063481745 | 0,83862914 | 0,985672131 | ubiquitin like modifier activating enzyme 6 |
| ENSG00000275014 | RN7SL166P | 0,088782432 | 1,063472282 | 0,93055818 | 0,994344567 | RNA, 7SL, cytoplasmic 166, pseudogene |
| ENSG00000131238 | PPT1 | 0,088724272 | 1,063429411 | 0,7814472 | 0,978597905 | palmitoyl-protein thioesterase 1 |
| ENSG00000231087 | FDPSP7 | 0,088722608 | 1,063428184 | 0,91462196 | 0,993361299 | farnesyl diphosphate synthase pseudogene 7 |
| ENSG00000214369 | CYB5AP2 | 0,08867546 | 1,063393431 | 0,91692772 | 0,993361299 | cytochrome b5 type A pseudogene 2 |
| ENSG00000284820 | AC068946,2 | 0,088530489 | 1,06328658 | 0,87900337 | 0,990109814 | novel protein |
| ENSG00000213965 | NUDT19 | 0,088511387 | 1,063272502 | 0,74363113 | 0,975310206 | nudix hydrolase 19 |
| ENSG00000160305 | DIP2A | 0,088476939 | 1,063247114 | 0,68963387 | 0,966868992 | disco interacting protein 2 homolog A |
| ENSG00000037637 | FBXO42 | 0,088472931 | 1,06324416 | 0,69114409 | 0,967179456 | F-box protein 42 |
| ENSG00000103248 | MTHFSD | 0,088400516 | 1,063190793 | 0,69558436 | 0,967669485 | methenyltetrahydrofolate synthetase domain containing |
| ENSG00000167676 | PLIN4 | 0,088352813 | 1,063155639 | 0,89994265 | 0,991845106 | perilipin 4 |
| ENSG00000164164 | OTUD4 | 0,088288172 | 1,063108005 | 0,75427498 | 0,975901808 | OTU deubiquitinase 4 |
| ENSG00000156500 | FAM122C | 0,088235972 | 1,063069539 | 0,82579365 | 0,983383773 | family with sequence similarity 122C |
| ENSG00000127955 | GNAI1 | 0,088233316 | 1,063067583 | 0,82355799 | 0,983374097 | G protein subunit alpha i1 |
| ENSG00000285314 | AC002470,2 | 0,088178116 | 1,063026908 | 0,97464311 | 0,99837368 | novel transcript |
| ENSG00000168612 | ZSWIM1 | 0,088090018 | 1,062961997 | 0,72264452 | 0,972594421 | zinc finger SWIM-type containing 1 |
| ENSG00000204604 | ZNF468 | 0,088027147 | 1,062915675 | 0,77767967 | 0,978587255 | zinc finger protein 468 |
| ENSG00000164002 | EXO5 | 0,087968447 | 1,062872428 | 0,69930692 | 0,968522643 | exonuclease 5 |
| ENSG00000099937 | SERPIND1 | 0,08793693 | 1,062849209 | 0,86305477 | 0,989074806 | serpin family D member 1 |
| ENSG00000134283 | PPHLN1 | 0,087856606 | 1,062790035 | 0,78917239 | 0,979462318 | periphilin 1 |

| | | | | | | |
|---|---|---|---|---|---|---|
| ENSG00000255836 | AC131206,1 | 0,087853803 | 1,06278797 | 0,87937144 | 0,990178196 | M-phase phosphoprotein 8 (MPHOSPH8) pseudogene |
| ENSG00000114354 | TFG | 0,087834755 | 1,062773938 | 0,82957384 | 0,984081562 | trafficking from ER to golgi regulator |
| ENSG00000153774 | CFDP1 | 0,087808575 | 1,062754653 | 0,70332727 | 0,968926756 | craniofacial development protein 1 |
| ENSG00000203327 | AC012358,1 | 0,087808127 | 1,062754322 | 0,87816609 | 0,990109814 | novel transcript |
| ENSG00000260774 | AC021087,3 | 0,087745495 | 1,062708186 | 0,81055055 | 0,981213877 | novel transcript |
| ENSG00000134330 | IAH1 | 0,087704436 | 1,062677942 | 0,76540611 | 0,976830764 | isoamyl acetate hydrolyzing esterase 1 (putative) |
| ENSG00000100811 | YY1 | 0,087703036 | 1,062676911 | 0,74289189 | 0,975201495 | YY1 transcription factor |
| ENSG00000259950 | AC034105,3 | 0,087649722 | 1,062637641 | 0,94290172 | 0,995402494 | protein tyrosine phosphatase, receptor type, f polypeptide (PTPRF), interacting protein (liprin), alpha 1 (PPFIA1) pseudogene |
| ENSG00000103707 | MTFMT | 0,087632002 | 1,062624589 | 0,7554532 | 0,975901808 | mitochondrial methionyl-tRNA formyltransferase |
| ENSG00000249700 | SRD5A3-AS1 | 0,087596579 | 1,062598498 | 0,85183172 | 0,98730877 | SRD5A3 antisense RNA 1 |
| ENSG00000227321 | MTND4P15 | 0,087590409 | 1,062593954 | 0,91480441 | 0,993361299 | MT-ND4 pseudogene 15 |
| ENSG00000100441 | KHNYN | 0,08749877 | 1,062526461 | 0,75686904 | 0,975901808 | KH and NYN domain containing |
| ENSG00000183155 | RABIF | 0,087475847 | 1,062509578 | 0,79499185 | 0,979776981 | RAB interacting factor |
| ENSG00000285821 | AC011899,3 | 0,087403867 | 1,062456568 | 0,93342506 | 0,994479239 | novel transcript, antisense to PTPRN2 |
| ENSG00000108960 | MMD | 0,0871754 | 1,06228833 | 0,86121954 | 0,989074806 | monocyte to macrophage differentiation associated |
| ENSG00000101193 | GID8 | 0,087153591 | 1,062272271 | 0,67632515 | 0,963580269 | GID complex subunit 8 homolog |
| ENSG00000154945 | ANKRD40 | 0,087007496 | 1,062164706 | 0,71453525 | 0,971041579 | ankyrin repeat domain 40 |
| ENSG00000120265 | PCMT1 | 0,087000034 | 1,062159212 | 0,82249298 | 0,98299282 | protein-L-isoaspartate (D-aspartate) O-methyltransferase |
| ENSG00000136144 | RCBTB1 | 0,086989403 | 1,062151384 | 0,81521036 | 0,981521413 | RCC1 and BTB domain containing protein 1 |
| ENSG00000249807 | AC034229,1 | 0,086966046 | 1,062134189 | 0,90248882 | 0,992230188 | novel transcript |
| ENSG00000099840 | IZUMO4 | 0,086956113 | 1,062126876 | 0,87019258 | 0,989074806 | IZUMO family member 4 |
| ENSG00000163933 | RFT1 | 0,08693461 | 1,062111045 | 0,69447333 | 0,967417311 | RFT1 homolog |
| ENSG00000178952 | TUFM | 0,086926465 | 1,062105049 | 0,70078432 | 0,968539008 | Tu translation elongation factor, mitochondrial |
| ENSG00000135090 | TAOK3 | 0,086813595 | 1,062021958 | 0,76975043 | 0,977719248 | TAO kinase 3 |
| ENSG00000125755 | SYMPK | 0,086806761 | 1,062016927 | 0,79403402 | 0,979776981 | symplekin |
| ENSG00000225329 | LHFPL3-AS2 | 0,086762129 | 1,061984072 | 0,90202178 | 0,992182704 | LHFPL3 antisense RNA 2 |
| ENSG00000214719 | AC005562,1 | 0,086723054 | 1,061955309 | 0,81603696 | 0,981708551 | novel transcript |
| ENSG00000104886 | PLEKHJ1 | 0,086660535 | 1,061909291 | 0,78827811 | 0,979462318 | pleckstrin homology domain containing J1 |
| ENSG00000103353 | UBFD1 | 0,086571279 | 1,061843595 | 0,7174567 | 0,971697135 | ubiquitin family domain containing 1 |

| | | | | | | |
|---|---|---|---|---|---|---|
| ENSG00000185236 | RAB11B | 0,086530515 | 1,061813593 | 0,69189923 | 0,967204896 | RAB11B, member RAS oncogene family |
| ENSG00000204272 | NBDY | 0,086529556 | 1,061812887 | 0,79658478 | 0,980103371 | negative regulator of P-body association |
| ENSG00000081870 | HSPB11 | 0,086521039 | 1,061806618 | 0,82715843 | 0,983579842 | heat shock protein family B (small) member 11 |
| ENSG00000229809 | ZNF688 | 0,086479555 | 1,061776087 | 0,82792483 | 0,983595244 | zinc finger protein 688 |
| ENSG00000116991 | SIPA1L2 | 0,086456566 | 1,061759168 | 0,73391047 | 0,974353474 | signal induced proliferation associated 1 like 2 |
| ENSG00000099977 | DDT | 0,086361615 | 1,061689291 | 0,78046739 | 0,978597905 | D-dopachrome tautomerase |
| ENSG00000143093 | STRIP1 | 0,086332507 | 1,06166787 | 0,68610173 | 0,965740648 | striatin interacting protein 1 |
| ENSG00000283668 | FRG1-DT | 0,08627027 | 1,061622071 | 0,87258227 | 0,989217334 | FRG1 divergent transcript |
| ENSG00000266708 | AP001793,1 | 0,086216207 | 1,061582289 | 0,90795589 | 0,992590661 | novel transcript, antisense to RAB12 |
| ENSG00000092445 | TYRO3 | 0,08619292 | 1,061565154 | 0,77323425 | 0,978363975 | TYRO3 protein tyrosine kinase |
| ENSG00000219891 | ZSCAN12P1 | 0,086186259 | 1,061560253 | 0,87241909 | 0,989217334 | zinc finger and SCAN domain containing 12 pseudogene 1 |
| ENSG00000282532 | AC016482,2 | 0,086178082 | 1,061554236 | 0,91176383 | 0,993357417 | ribosomal protein S2 (RPS2) pseudogene |
| ENSG00000274308 | AC244093,1 | 0,086178082 | 1,061554236 | 0,91176383 | 0,993357417 | ribosomal protein S2 (RPS2) pseudogene |
| ENSG00000118200 | CAMSAP2 | 0,086156689 | 1,061538495 | 0,82918533 | 0,983957145 | calmodulin regulated spectrin associated protein family member 2 |
| ENSG00000102935 | ZNF423 | 0,086055923 | 1,061464353 | 0,6891811 | 0,966685038 | zinc finger protein 423 |
| ENSG00000261889 | AC108134,2 | 0,085919431 | 1,061363934 | 0,82497236 | 0,983374097 | novel transcript |
| ENSG00000117010 | ZNF684 | 0,085905721 | 1,061353848 | 0,83256906 | 0,98489473 | zinc finger protein 684 |
| ENSG00000147394 | ZNF185 | 0,085884953 | 1,06133857 | 0,80396038 | 0,980875302 | zinc finger protein 185 with LIM domain |
| ENSG00000009790 | TRAF3IP3 | 0,0858598 | 1,061320065 | 0,92167215 | 0,993787755 | TRAF3 interacting protein 3 |
| ENSG00000156976 | EIF4A2 | 0,085795063 | 1,061272443 | 0,80017705 | 0,980161849 | eukaryotic translation initiation factor 4A2 |
| ENSG00000156097 | GPR61 | 0,085790216 | 1,061268877 | 0,88307145 | 0,990717729 | G protein-coupled receptor 61 |
| ENSG00000182776 | AC239585,1 | 0,085695417 | 1,061199144 | 0,93238372 | 0,994479239 | novel transcript |
| ENSG00000179028 | AC245177,1 | 0,085695417 | 1,061199144 | 0,93238372 | 0,994479239 | novel transcript |
| ENSG00000131711 | MAP1B | 0,085673127 | 1,061182749 | 0,69063051 | 0,967179044 | microtubule associated protein 1B |
| ENSG00000110944 | IL23A | 0,08561411 | 1,061139339 | 0,92929682 | 0,994293848 | interleukin 23 subunit alpha |
| ENSG00000165802 | NSMF | 0,085502361 | 1,061057148 | 0,76304439 | 0,976576876 | NMDA receptor synaptonuclear signaling and neuronal migration factor |
| ENSG00000113575 | PPP2CA | 0,085492771 | 1,061050094 | 0,8244171 | 0,983374097 | protein phosphatase 2 catalytic subunit alpha |
| ENSG00000237679 | VDAC1P11 | 0,085428697 | 1,061002971 | 0,91650071 | 0,993361299 | voltage dependent anion channel 1 pseudogene 11 |
| ENSG00000269897 | COMMD3-BMI | 0,085413992 | 1,060992157 | 0,87551573 | 0,990078774 | COMMD3-BMI1 readthrough |
| ENSG00000227674 | LINC00355 | 0,085407905 | 1,060987681 | 0,91042959 | 0,99320976 | long intergenic non-protein coding RNA 355 |

| | | | | | | |
|---|---|---|---|---|---|---|
| ENSG00000213514 | AL731556,1 | 0,085350686 | 1,060945601 | 0,91638133 | 0,993361299 | novel zinc finger pseudogene (KIAA1629) |
| ENSG00000126243 | LRFN3 | 0,085343889 | 1,060940603 | 0,78550054 | 0,979320406 | leucine rich repeat and fibronectin type III domain containing 3 |
| ENSG00000117020 | AKT3 | 0,085324206 | 1,060926129 | 0,77804684 | 0,978587255 | AKT serine/threonine kinase 3 |
| ENSG00000134183 | GNAT2 | 0,085317948 | 1,060921526 | 0,90175328 | 0,992136926 | G protein subunit alpha transducin 2 |
| ENSG00000281965 | PKD1P3 | 0,085309002 | 1,060914948 | 0,85533655 | 0,988074261 | polycystin 1, transient receptor potential channel interacting pseudogene 3 |
| ENSG00000150779 | TIMM8B | 0,085283141 | 1,060895931 | 0,71375792 | 0,970812834 | translocase of inner mitochondrial membrane 8 homolog B |
| ENSG00000219607 | PPP1R3G | 0,085281678 | 1,060894855 | 0,90323017 | 0,992311172 | protein phosphatase 1 regulatory subunit 3G |
| ENSG00000094755 | GABRP | 0,085209135 | 1,060841511 | 0,89793443 | 0,991711719 | gamma-aminobutyric acid type A receptor subunit pi |
| ENSG00000143622 | RIT1 | 0,085188549 | 1,060826374 | 0,80286361 | 0,980866933 | Ras like without CAAX 1 |
| ENSG00000151748 | SAV1 | 0,085172905 | 1,060814871 | 0,81862956 | 0,9824772 | salvador family WW domain containing protein 1 |
| ENSG00000278845 | MRPL45 | 0,085146176 | 1,060795218 | 0,79051059 | 0,979462318 | mitochondrial ribosomal protein L45 |
| ENSG00000122591 | FAM126A | 0,085120527 | 1,060776358 | 0,79431717 | 0,979776981 | family with sequence similarity 126 member A |
| ENSG00000137575 | SDCBP | 0,085081173 | 1,060747423 | 0,8439191 | 0,986273696 | syndecan binding protein |
| ENSG00000164091 | WDR82 | 0,085078891 | 1,060745745 | 0,7735016 | 0,978415098 | WD repeat domain 82 |
| ENSG00000174827 | PDZK1 | 0,08501115 | 1,060695939 | 0,87193467 | 0,989217334 | PDZ domain containing 1 |
| ENSG00000230312 | SLC25A53P1 | 0,084899878 | 1,060614133 | 0,88994849 | 0,990717729 | SLC25A53 pseudogene 1 |
| ENSG00000259920 | AC007938,1 | 0,084857787 | 1,06058319 | 0,92126648 | 0,993787755 | novel transcript |
| ENSG00000226179 | LINC00685 | 0,084747575 | 1,060502172 | 0,91606287 | 0,993361299 | long intergenic non-protein coding RNA 685 |
| ENSG00000128191 | DGCR8 | 0,084569561 | 1,060371324 | 0,72895799 | 0,974009551 | DGCR8 microprocessor complex subunit |
| ENSG00000107968 | MAP3K8 | 0,084548653 | 1,060355957 | 0,81457332 | 0,981521413 | mitogen-activated protein kinase kinase kinase 8 |
| ENSG00000151532 | VTI1A | 0,084489637 | 1,060312582 | 0,7358191 | 0,974353474 | vesicle transport through interaction with t-SNAREs 1A |
| ENSG00000226937 | CEP164P1 | 0,084452307 | 1,060285147 | 0,89601572 | 0,991528124 | centrosomal protein 164 pseudogene 1 |
| ENSG00000197958 | RPL12 | 0,084360511 | 1,060217685 | 0,79012406 | 0,979462318 | ribosomal protein L12 |
| ENSG00000076053 | RBM7 | 0,08418065 | 1,060085516 | 0,81500475 | 0,981521413 | RNA binding motif protein 7 |
| ENSG00000241878 | PISD | 0,084166014 | 1,060074761 | 0,73431054 | 0,974353474 | phosphatidylserine decarboxylase |
| ENSG00000101473 | ACOT8 | 0,084095296 | 1,0600228 | 0,74052216 | 0,974885509 | acyl-CoA thioesterase 8 |
| ENSG00000270804 | AC010326,4 | 0,08398446 | 1,059941366 | 0,82304214 | 0,983204348 | ubiquitin-conjugating enzyme E2C (UBE2C) pseudogene |
| ENSG00000078081 | LAMP3 | 0,083963836 | 1,059926214 | 0,93086097 | 0,994433362 | lysosomal associated membrane protein 3 |

| | | | | | | |
|---|---|---|---|---|---|---|
| ENSG00000183726 | TMEM50A | 0,083871688 | 1,059858516 | 0,83672027 | 0,985426146 | transmembrane protein 50A |
| ENSG00000167130 | DOLPP1 | 0,083832107 | 1,059829439 | 0,71236718 | 0,970812834 | dolichyldiphosphatase 1 |
| ENSG00000161277 | THAP8 | 0,083801632 | 1,059807052 | 0,73616178 | 0,974353474 | THAP domain containing 8 |
| ENSG00000114948 | ADAM23 | 0,083624414 | 1,059676875 | 0,80226345 | 0,980710009 | ADAM metallopeptidase domain 23 |
| ENSG00000092054 | MYH7 | 0,083581453 | 1,05964532 | 0,91229025 | 0,993361299 | myosin heavy chain 7 |
| ENSG00000176095 | IP6K1 | 0,083573832 | 1,059639722 | 0,6897364 | 0,966868992 | inositol hexakisphosphate kinase 1 |
| ENSG00000267106 | ZNF561-AS1 | 0,083534379 | 1,059610745 | 0,8179234 | 0,982323352 | ZNF561 antisense RNA 1 (head to head) |
| ENSG00000116750 | UCHL5 | 0,083463723 | 1,059558852 | 0,84117515 | 0,985672131 | ubiquitin C-terminal hydrolase L5 |
| ENSG00000168061 | SAC3D1 | 0,083300363 | 1,059438882 | 0,82206838 | 0,982952211 | SAC3 domain containing 1 |
| ENSG00000286159 | AL035106,1 | 0,083254486 | 1,059405194 | 0,86579503 | 0,989074806 | novel transcript, antisense to PREX1 |
| ENSG00000159459 | UBR1 | 0,083249666 | 1,059401654 | 0,80450488 | 0,980875302 | ubiquitin protein ligase E3 component n-recognin 1 |
| ENSG00000228069 | MTCO3P29 | 0,083188679 | 1,059356871 | 0,92605484 | 0,993915537 | MT-CO3 pseudogene 29 |
| ENSG00000061794 | MRPS35 | 0,08318024 | 1,059350674 | 0,8196632 | 0,98249617 | mitochondrial ribosomal protein S35 |
| ENSG00000181894 | ZNF329 | 0,083167381 | 1,059341232 | 0,73496301 | 0,974353474 | zinc finger protein 329 |
| ENSG00000141371 | C17orf64 | 0,083099601 | 1,059291463 | 0,93524204 | 0,994479239 | chromosome 17 open reading frame 64 |
| ENSG00000151151 | IPMK | 0,083088796 | 1,05928353 | 0,86320237 | 0,989074806 | inositol polyphosphate multikinase |
| ENSG00000183580 | FBXL7 | 0,083074178 | 1,059272797 | 0,76746779 | 0,977201891 | F-box and leucine rich repeat protein 7 |
| ENSG00000257918 | AC079385,3 | 0,083058271 | 1,059261118 | 0,84747011 | 0,987035044 | novel transcript, antisense to RFX4 |
| ENSG00000031823 | RANBP3 | 0,083042273 | 1,059249372 | 0,70114493 | 0,968677171 | RAN binding protein 3 |
| ENSG00000119979 | DENND10 | 0,083005613 | 1,059222456 | 0,80201816 | 0,980710009 | DENN domain containing 10 |
| ENSG00000226101 | LINC02097 | 0,082966126 | 1,059193465 | 0,89407485 | 0,991135494 | long intergenic non-protein coding RNA 2097 |
| ENSG00000012174 | MBTPS2 | 0,082945043 | 1,059177986 | 0,79353412 | 0,979776981 | membrane bound transcription factor peptidase, site 2 |
| ENSG00000258891 | AC005480,1 | 0,082942244 | 1,059175931 | 0,87631923 | 0,990109814 | novel transcript, antisense to ZNF410 |
| ENSG00000033011 | ALG1 | 0,082923246 | 1,059161984 | 0,79736691 | 0,980161849 | ALG1 chitobiosyldiphosphodolichol beta-mannosyltransferase |
| ENSG00000064995 | TAF11 | 0,082857496 | 1,059113714 | 0,76353202 | 0,976640468 | TATA-box binding protein associated factor 11 |
| ENSG00000237153 | AL162725,2 | 0,082790076 | 1,059064221 | 0,9221348 | 0,993797669 | novel transcript |
| ENSG00000196367 | TRRAP | 0,082718227 | 1,059011479 | 0,67842728 | 0,9640473 | transformation/transcription domain associated protein |
| ENSG00000122692 | SMU1 | 0,082680846 | 1,05898404 | 0,77294151 | 0,978310326 | SMU1 DNA replication regulator and spliceosomal factor |
| ENSG00000082929 | LINC01587 | 0,082651639 | 1,058962601 | 0,95995575 | 0,996608711 | long intergenic non-protein coding RNA 1587 |

| | | | | | | |
|---|---|---|---|---|---|---|
| ENSG00000121716 | PILRB | 0,082647477 | 1,058959546 | 0,84975206 | 0,987036295 | paired immunoglobin like type 2 receptor beta |
| ENSG00000230587 | LINC02580 | 0,08255005 | 1,058888036 | 0,94224192 | 0,995373376 | long intergenic non-protein coding RNA 2580 |
| ENSG00000196756 | SNHG17 | 0,082511609 | 1,058859822 | 0,79281152 | 0,979471509 | small nucleolar RNA host gene 17 |
| ENSG00000253704 | AC023632,2 | 0,08248621 | 1,05884118 | 0,88783508 | 0,990717729 | novel transcript |
| ENSG00000184719 | RNLS | 0,082477102 | 1,058834495 | 0,80837027 | 0,981213877 | renalase, FAD dependent amine oxidase |
| ENSG00000198843 | SELENOT | 0,082393067 | 1,058772822 | 0,85889532 | 0,989074806 | selenoprotein T |
| ENSG00000109436 | TBC1D9 | 0,082382968 | 1,05876541 | 0,7199408 | 0,972067771 | TBC1 domain family member 9 |
| ENSG00000076984 | MAP2K7 | 0,082360239 | 1,05874873 | 0,80902544 | 0,981213877 | mitogen-activated protein kinase kinase 7 |
| ENSG00000232828 | AC231533,1 | 0,082343273 | 1,058736279 | 0,95815406 | 0,996608711 | novel transcript |
| ENSG00000010310 | GIPR | 0,082334485 | 1,05872983 | 0,92010493 | 0,993787755 | gastric inhibitory polypeptide receptor |
| ENSG00000272657 | AP000317,2 | 0,082260992 | 1,058675898 | 0,94989886 | 0,996432861 | novel transcript |
| ENSG00000004478 | FKBP4 | 0,082138786 | 1,058586225 | 0,74234651 | 0,975176043 | FKBP prolyl isomerase 4 |
| ENSG00000278053 | DDX52 | 0,082129212 | 1,0585792 | 0,78317001 | 0,978919905 | DExD-box helicase 52 |
| ENSG00000103591 | AAGAB | 0,082121944 | 1,058573867 | 0,80885517 | 0,981213877 | alpha and gamma adaptin binding protein |
| ENSG00000123353 | ORMDL2 | 0,082038259 | 1,058512465 | 0,74581804 | 0,975707496 | ORMDL sphingolipid biosynthesis regulator 2 |
| ENSG00000133704 | IPO8 | 0,081797747 | 1,058336015 | 0,7958362 | 0,979879951 | importin 8 |
| ENSG00000105835 | NAMPT | 0,08176748 | 1,058313812 | 0,85911406 | 0,989074806 | nicotinamide phosphoribosyltransferase |
| ENSG00000237492 | OR2L9P | 0,081764233 | 1,05831143 | 0,95041965 | 0,996432861 | olfactory receptor family 2 subfamily L member 9 pseudogene |
| ENSG00000236095 | AL807757,2 | 0,081649228 | 1,05822707 | 0,94764758 | 0,996113938 | novel transcript |
| ENSG00000172456 | FGGY | 0,081611755 | 1,058199583 | 0,74909841 | 0,975901808 | FGGY carbohydrate kinase domain containing |
| ENSG00000253238 | AC034114,2 | 0,081582456 | 1,058178093 | 0,92534739 | 0,993915537 | novel transcript |
| ENSG00000275850 | AC005912,2 | 0,081534018 | 1,058142566 | 0,92862887 | 0,994132759 | novel transcript |
| ENSG00000216895 | AC009403,1 | 0,08149246 | 1,058112085 | 0,85817592 | 0,989067345 | novel transcript, antisense to RBM33 |
| ENSG00000110330 | BIRC2 | 0,081438804 | 1,058072733 | 0,84074612 | 0,985672131 | baculoviral IAP repeat containing 2 |
| ENSG00000110925 | CSRNP2 | 0,081423796 | 1,058061727 | 0,69410481 | 0,967359949 | cysteine and serine rich nuclear protein 2 |
| ENSG00000007541 | PIGQ | 0,081307151 | 1,057976183 | 0,82577441 | 0,983383773 | phosphatidylinositol glycan anchor biosynthesis class Q |
| ENSG00000242488 | AF107885,1 | 0,081278093 | 1,057954874 | 0,9186941 | 0,993485783 | ribosomal protein S2 (RPS2) pseudogene |
| ENSG00000257270 | AL928654,2 | 0,081276079 | 1,057953398 | 0,92393924 | 0,993915537 | novel transcript, antisense to MTA1 |
| ENSG00000102241 | HTATSF1 | 0,081222519 | 1,057914122 | 0,82548618 | 0,983383773 | HIV-1 Tat specific factor 1 |
| ENSG00000067842 | ATP2B3 | 0,081151306 | 1,057861904 | 0,90342837 | 0,992361703 | ATPase plasma membrane Ca2+ transporting 3 |
| ENSG00000130382 | MLLT1 | 0,081054566 | 1,057790971 | 0,74655579 | 0,975796253 | MLLT1 super elongation complex subunit |

| | | | | | | |
|---|---|---|---|---|---|---|
| ENSG00000285377 | EMC9 | 0,081042808 | 1,05778235 | 0,80016993 | 0,980161849 | ER membrane protein complex subunit 9 |
| ENSG00000100908 | EMC9 | 0,081042808 | 1,05778235 | 0,80016993 | 0,980161849 | ER membrane protein complex subunit 9 |
| ENSG00000259630 | FABP5P9 | 0,081022457 | 1,057767429 | 0,90958227 | 0,992918304 | fatty acid binding protein 5 pseudogene 9 |
| ENSG00000173064 | HECTD4 | 0,080975935 | 1,05773332 | 0,7000838 | 0,968522643 | HECT domain E3 ubiquitin protein ligase 4 |
| ENSG00000130177 | CDC16 | 0,080949516 | 1,05771395 | 0,8100427 | 0,981213877 | cell division cycle 16 |
| ENSG00000067836 | ROGDI | 0,080934231 | 1,057702745 | 0,79548865 | 0,979879951 | rogdi atypical leucine zipper |
| ENSG00000170191 | NANP | 0,080887991 | 1,057668845 | 0,85144388 | 0,987259878 | N-acetylneuraminic acid phosphatase |
| ENSG00000285672 | AL160396,2 | 0,080795692 | 1,05760118 | 0,88841916 | 0,990717729 | novel transcript |
| ENSG00000250397 | AP006623,1 | 0,080782407 | 1,057591441 | 0,91424865 | 0,993361299 | novel transcript |
| ENSG00000272457 | AC113194,1 | 0,08072663 | 1,057550554 | 0,90815984 | 0,992595473 | novel transcript, antisense to RGS20 |
| ENSG00000132600 | PRMT7 | 0,080696448 | 1,05752843 | 0,77818354 | 0,978587255 | protein arginine methyltransferase 7 |
| ENSG00000268129 | AC026304,1 | 0,080653928 | 1,057497262 | 0,87853987 | 0,990109814 | novel transcript |
| ENSG00000198466 | ZNF587 | 0,080649012 | 1,057493658 | 0,70298377 | 0,968742626 | zinc finger protein 587 |
| ENSG00000165526 | RPUSD4 | 0,080561863 | 1,05742978 | 0,72993723 | 0,974178338 | RNA pseudouridine synthase D4 |
| ENSG00000276464 | POMZP3 | 0,080535696 | 1,057410601 | 0,75417265 | 0,975901808 | POM121 and ZP3 fusion |
| ENSG00000134982 | APC | 0,080510771 | 1,057392333 | 0,82774385 | 0,983595244 | APC regulator of WNT signaling pathway |
| ENSG00000274080 | AC005089,1 | 0,080493065 | 1,057379356 | 0,9493226 | 0,996432861 | novel transcript |
| ENSG00000175110 | MRPS22 | 0,080490342 | 1,05737736 | 0,82711672 | 0,983567657 | mitochondrial ribosomal protein S22 |
| ENSG00000162695 | SLC30A7 | 0,080468964 | 1,057361692 | 0,83457298 | 0,984905093 | solute carrier family 30 member 7 |
| ENSG00000228423 | AL357552,2 | 0,080371271 | 1,057290094 | 0,92528322 | 0,993915537 | novel transcript |
| ENSG00000176715 | ACSF3 | 0,080348009 | 1,057273046 | 0,83751647 | 0,98566906 | acyl-CoA synthetase family member 3 |
| ENSG00000261291 | AC007906,1 | 0,080300592 | 1,057238298 | 0,94400858 | 0,995542144 | novel transcript |
| ENSG00000107874 | CUEDC2 | 0,080198225 | 1,057163284 | 0,77814644 | 0,978587255 | CUE domain containing 2 |
| ENSG00000278605 | TSEN34 | 0,080173998 | 1,057145531 | 0,94886315 | 0,996432861 | tRNA splicing endonuclease subunit 34 |
| ENSG00000228817 | BACH1-IT2 | 0,080168252 | 1,057141321 | 0,8763392 | 0,990109814 | BACH1 intronic transcript 2 |
| ENSG00000161912 | ADCY10P1 | 0,080118744 | 1,057105044 | 0,86177289 | 0,989074806 | ADCY10 pseudogene 1 |
| ENSG00000104325 | DECR1 | 0,080100055 | 1,05709135 | 0,81259253 | 0,981224866 | 2,4-dienoyl-CoA reductase 1 |
| ENSG00000230303 | AL353743,2 | 0,079974976 | 1,056999707 | 0,8965222 | 0,991644439 | novel transcript |
| ENSG00000175746 | C15orf54 | 0,079942724 | 1,056976077 | 0,939907 | 0,994949796 | chromosome 15 putative open reading frame 54 |
| ENSG00000164331 | ANKRA2 | 0,079928054 | 1,056965329 | 0,81290771 | 0,981246822 | ankyrin repeat family A member 2 |
| ENSG00000130699 | TAF4 | 0,07991733 | 1,056957472 | 0,7837823 | 0,979226738 | TATA-box binding protein associated factor 4 |
| ENSG00000265808 | SEC22B | 0,079913647 | 1,056954774 | 0,84319281 | 0,986218419 | SEC22 homolog B, vesicle trafficking protein |

| | | | | | | |
|---|---|---|---|---|---|---|
| ENSG00000204152 | TIMM23B | 0,079905614 | 1,056948889 | 0,75082462 | 0,975901808 | translocase of inner mitochondrial membrane 23 homolog B |
| ENSG00000154781 | CCDC174 | 0,079873652 | 1,056925474 | 0,72653017 | 0,973344786 | coiled-coil domain containing 174 |
| ENSG00000066135 | KDM4A | 0,079820043 | 1,0568862 | 0,70867533 | 0,970181083 | lysine demethylase 4A |
| ENSG00000080603 | SRCAP | 0,079763498 | 1,056844777 | 0,79442869 | 0,979776981 | Snf2 related CREBBP activator protein |
| ENSG00000168268 | NT5DC2 | 0,079646138 | 1,056758809 | 0,74482945 | 0,975649186 | 5'-nucleotidase domain containing 2 |
| ENSG00000273136 | NBPF26 | 0,079642055 | 1,056755818 | 0,73331951 | 0,974353474 | NBPF member 26 |
| ENSG00000287834 | AP005210,2 | 0,079635415 | 1,056750954 | 0,88877306 | 0,990717729 | novel transcript, antisense to ARHGAP28 |
| ENSG00000197601 | FAR1 | 0,079579526 | 1,056710017 | 0,86864691 | 0,989074806 | fatty acyl-CoA reductase 1 |
| ENSG00000165194 | PCDH19 | 0,079531465 | 1,056674815 | 0,82283036 | 0,983107285 | protocadherin 19 |
| ENSG00000274612 | AC011447,5 | 0,079530472 | 1,056674088 | 0,94849628 | 0,996431033 | BCL2/adenovirus E1B 19kDa interacting protein 3 (BNIP3) pseudogene |
| ENSG00000168890 | TMEM150A | 0,079529677 | 1,056673506 | 0,80266144 | 0,98082499 | transmembrane protein 150A |
| ENSG00000148606 | POLR3A | 0,079520745 | 1,056666964 | 0,71060428 | 0,970482408 | RNA polymerase III subunit A |
| ENSG00000185608 | MRPL40 | 0,079505465 | 1,056655772 | 0,77570324 | 0,978587255 | mitochondrial ribosomal protein L40 |
| ENSG00000112305 | SMAP1 | 0,079493091 | 1,056646709 | 0,80836456 | 0,981213877 | small ArfGAP 1 |
| ENSG00000060642 | PIGV | 0,079488652 | 1,056643458 | 0,75567417 | 0,975901808 | phosphatidylinositol glycan anchor biosynthesis class V |
| ENSG00000288045 | AL352979,4 | 0,079437069 | 1,056605679 | 0,84047386 | 0,985672131 | novel transcript, antisense to FERMT2 |
| ENSG00000284845 | AL136295,10 | 0,079429624 | 1,056600226 | 0,83594109 | 0,985106004 | novel transcript |
| ENSG00000259529 | AL136295,5 | 0,079429624 | 1,056600226 | 0,83594109 | 0,985106004 | novel transcript |
| ENSG00000275835 | TUBGCP5 | 0,079421364 | 1,056594177 | 0,79219955 | 0,979471509 | tubulin gamma complex associated protein 5 |
| ENSG00000103404 | USP31 | 0,079419241 | 1,056592622 | 0,77091545 | 0,978033855 | ubiquitin specific peptidase 31 |
| ENSG00000226167 | AP4B1-AS1 | 0,079415505 | 1,056589886 | 0,87176688 | 0,989217334 | AP4B1 antisense RNA 1 |
| ENSG00000151503 | NCAPD3 | 0,079415498 | 1,056589881 | 0,76574301 | 0,976911351 | non-SMC condensin II complex subunit D3 |
| ENSG00000155380 | SLC16A1 | 0,079265244 | 1,056479845 | 0,83905741 | 0,985672131 | solute carrier family 16 member 1 |
| ENSG00000134453 | RBM17 | 0,079212905 | 1,056441518 | 0,74230914 | 0,975176043 | RNA binding motif protein 17 |
| ENSG00000278126 | AC139768,1 | 0,079204882 | 1,056435643 | 0,83346862 | 0,984905093 | novel transcript |
| ENSG00000182600 | SNORC | 0,07917536 | 1,056414025 | 0,90379598 | 0,992462326 | secondary ossification center associated regulator of chondrocyte maturation |
| ENSG00000231889 | TRAF3IP2-AS1 | 0,079127118 | 1,0563787 | 0,76294872 | 0,976534376 | TRAF3IP2 antisense RNA 1 |
| ENSG00000198900 | TOP1 | 0,07903785 | 1,056313338 | 0,75893012 | 0,976120207 | DNA topoisomerase I |
| ENSG00000158435 | CNOT11 | 0,079015101 | 1,056296682 | 0,79058732 | 0,979462318 | CCR4-NOT transcription complex subunit 11 |

| | | | | | | |
|---|---|---|---|---|---|---|
| ENSG00000239900 | ADSL | 0,078978119 | 1,056269605 | 0,75017742 | 0,975901808 | adenylosuccinate lyase |
| ENSG00000104044 | OCA2 | 0,078950419 | 1,056249325 | 0,9001405 | 0,991845106 | OCA2 melanosomal transmembrane protein |
| ENSG00000232860 | SMG7-AS1 | 0,07891099 | 1,056220458 | 0,87086209 | 0,98919606 | SMG7 antisense RNA 1 |
| ENSG00000164087 | POC1A | 0,078895264 | 1,056208945 | 0,76641292 | 0,977167878 | POC1 centriolar protein A |
| ENSG00000283244 | FRG1-DT | 0,07888477 | 1,056201262 | 0,90044574 | 0,991865079 | FRG1 divergent transcript |
| ENSG00000274891 | TRAPPC12 | 0,078832788 | 1,056163207 | 0,8258757 | 0,983383773 | trafficking protein particle complex 12 |
| ENSG00000173113 | TRMT112 | 0,078820797 | 1,056154428 | 0,72487488 | 0,97326334 | tRNA methyltransferase subunit 11-2 |
| ENSG00000227268 | KLLN | 0,078802328 | 1,056140908 | 0,79103045 | 0,979471509 | killin, p53 regulated DNA replication inhibitor |
| ENSG00000270553 | AC011921,3 | 0,078743677 | 1,056097972 | 0,94241296 | 0,995373376 | splicing factor 3B, 14 kDa subunit (SF3B14) pseudogene |
| ENSG00000268592 | RAET1E-AS1 | 0,078715305 | 1,056077203 | 0,93287144 | 0,994479239 | RAET1E antisense RNA 1 |
| ENSG00000172795 | DCP2 | 0,078683865 | 1,056054189 | 0,83848123 | 0,985672131 | decapping mRNA 2 |
| ENSG00000107771 | CCSER2 | 0,078521565 | 1,055935392 | 0,83027001 | 0,984156028 | coiled-coil serine rich protein 2 |
| ENSG00000150712 | MTMR12 | 0,078499612 | 1,055919324 | 0,72097705 | 0,972142678 | myotubularin related protein 12 |
| ENSG00000158163 | DZIP1L | 0,078389187 | 1,055838506 | 0,8071827 | 0,981213877 | DAZ interacting zinc finger protein 1 like |
| ENSG00000227115 | LINC01630 | 0,078381358 | 1,055832776 | 0,87248613 | 0,989217334 | long intergenic non-protein coding RNA 1630 |
| ENSG00000237190 | CDKN2AIPNL | 0,078257217 | 1,055741929 | 0,77023728 | 0,977856668 | CDKN2A interacting protein N-terminal like |
| ENSG00000154124 | OTULIN | 0,078179533 | 1,055685082 | 0,7955242 | 0,979879951 | OTU deubiquitinase with linear linkage specificity |
| ENSG00000115112 | TFCP2L1 | 0,078146002 | 1,055660546 | 0,93979461 | 0,994943406 | transcription factor CP2 like 1 |
| ENSG00000143851 | PTPN7 | 0,078110322 | 1,055634438 | 0,88462683 | 0,990717729 | protein tyrosine phosphatase non-receptor type 7 |
| ENSG00000100350 | FOXRED2 | 0,078002837 | 1,055555794 | 0,75117235 | 0,975901808 | FAD dependent oxidoreductase domain containing 2 |
| ENSG00000286285 | AL138778,1 | 0,077968722 | 1,055530833 | 0,94418955 | 0,995542144 | novel transcript |
| ENSG00000057593 | F7 | 0,077932668 | 1,055504455 | 0,93781757 | 0,994479239 | coagulation factor VII |
| ENSG00000113108 | APBB3 | 0,077874884 | 1,05546218 | 0,80815877 | 0,981213877 | amyloid beta precursor protein binding family B member 3 |
| ENSG00000168724 | DNAJC21 | 0,077861552 | 1,055452427 | 0,77951455 | 0,978587255 | DnaJ heat shock protein family (Hsp40) member C21 |
| ENSG00000151164 | RAD9B | 0,077850754 | 1,055444527 | 0,85891485 | 0,989074806 | RAD9 checkpoint clamp component B |
| ENSG00000186432 | KPNA4 | 0,077844718 | 1,055440111 | 0,81530062 | 0,981521413 | karyopherin subunit alpha 4 |
| ENSG00000273943 | CNOT3 | 0,07774409 | 1,055366496 | 0,89040381 | 0,990717729 | CCR4-NOT transcription complex subunit 3 |
| ENSG00000274176 | CNOT3 | 0,07774409 | 1,055366496 | 0,89040381 | 0,990717729 | CCR4-NOT transcription complex subunit 3 |
| ENSG00000274941 | CNOT3 | 0,07774409 | 1,055366496 | 0,89040381 | 0,990717729 | CCR4-NOT transcription complex subunit 3 |
| ENSG00000276082 | CNOT3 | 0,07774409 | 1,055366496 | 0,89040381 | 0,990717729 | CCR4-NOT transcription complex subunit 3 |
| ENSG00000275979 | CNOT3 | 0,07774409 | 1,055366496 | 0,89040381 | 0,990717729 | CCR4-NOT transcription complex subunit 3 |

| | | | | | | |
|---|---|---|---|---|---|---|
| ENSG00000277615 | CNOT3 | 0,07774409 | 1,055366496 | 0,89040381 | 0,990717729 | CCR4-NOT transcription complex subunit 3 |
| ENSG00000271105 | SCML2P2 | 0,07774126 | 1,055364427 | 0,91730983 | 0,993361299 | SCML2 pseudogene 2 |
| ENSG00000204920 | ZNF155 | 0,077661325 | 1,055305953 | 0,83018327 | 0,984156028 | zinc finger protein 155 |
| ENSG00000141837 | CACNA1A | 0,077574992 | 1,055242805 | 0,86282921 | 0,989074806 | calcium voltage-gated channel subunit alpha1 A |
| ENSG00000198160 | MIER1 | 0,077570282 | 1,055239359 | 0,85077102 | 0,987147801 | MIER1 transcriptional regulator |
| ENSG00000268747 | AC022432,1 | 0,07750247 | 1,055189761 | 0,95043858 | 0,996432861 | COX16 cytochrome c oxidase assembly homolog (S, cerevisiae) (COX16) pseudogene |
| ENSG00000122299 | ZC3H7A | 0,077453225 | 1,055153743 | 0,79513029 | 0,97981137 | zinc finger CCCH-type containing 7A |
| ENSG00000133858 | ZFC3H1 | 0,077433856 | 1,055139578 | 0,79041242 | 0,979462318 | zinc finger C3H1-type containing |
| ENSG00000164188 | RANBP3L | 0,077317417 | 1,055054421 | 0,87493566 | 0,990078774 | RAN binding protein 3 like |
| ENSG00000259112 | NDUFC2-KCTD1 | 0,077285337 | 1,055030961 | 0,88677681 | 0,990717729 | NDUFC2-KCTD14 readthrough |
| ENSG00000110721 | CHKA | 0,077218114 | 1,054981802 | 0,80504648 | 0,98109454 | choline kinase alpha |
| ENSG00000253715 | AC083841,2 | 0,077156949 | 1,054937076 | 0,91343623 | 0,993361299 | novel transcript |
| ENSG00000287870 | AC093433,2 | 0,077067676 | 1,054871799 | 0,95942037 | 0,996608711 | novel transcript |
| ENSG00000237522 | NONOP2 | 0,076983261 | 1,054810079 | 0,88505542 | 0,990717729 | non-POU domain containing, octamer-binding pseudogene 2 |
| ENSG00000213246 | SUPT4H1 | 0,076981148 | 1,054808533 | 0,76854906 | 0,977381742 | SPT4 homolog, DSIF elongation factor subunit |
| ENSG00000258099 | ATXN2-AS | 0,076973236 | 1,054802749 | 0,93678731 | 0,994479239 | ATXN2 antisense RNA |
| ENSG00000206527 | HACD2 | 0,076946227 | 1,054783002 | 0,81046063 | 0,981213877 | 3-hydroxyacyl-CoA dehydratase 2 |
| ENSG00000182952 | HMGN4 | 0,076880725 | 1,054735113 | 0,82282981 | 0,983107285 | high mobility group nucleosomal binding domain 4 |
| ENSG00000163527 | STT3B | 0,076829675 | 1,054697792 | 0,8576526 | 0,988893234 | STT3 oligosaccharyltransferase complex catalytic subunit B |
| ENSG00000089057 | SLC23A2 | 0,076799207 | 1,054675518 | 0,74364908 | 0,975310206 | solute carrier family 23 member 2 |
| ENSG00000115211 | EIF2B4 | 0,076756848 | 1,054644552 | 0,72142458 | 0,972142678 | eukaryotic translation initiation factor 2B subunit delta |
| ENSG00000131389 | SLC6A6 | 0,076730814 | 1,054625521 | 0,74516892 | 0,975666349 | solute carrier family 6 member 6 |
| ENSG00000140988 | RPS2 | 0,076715368 | 1,05461423 | 0,74988981 | 0,975901808 | ribosomal protein S2 |
| ENSG00000163517 | HDAC11 | 0,076697508 | 1,054601174 | 0,85309059 | 0,987815708 | histone deacetylase 11 |
| ENSG00000197535 | MYO5A | 0,076664654 | 1,054577158 | 0,77525788 | 0,978587255 | myosin VA |
| ENSG00000254419 | AL139349,1 | 0,076609403 | 1,054536772 | 0,93500686 | 0,994479239 | novel transcript |
| ENSG00000256751 | PLBD1-AS1 | 0,076582677 | 1,054517237 | 0,89730982 | 0,991644439 | PLBD1 antisense RNA 1 |
| ENSG00000125878 | TCF15 | 0,076497889 | 1,054455264 | 0,938808 | 0,994709504 | transcription factor 15 |
| ENSG00000277517 | AC212493,1 | 0,076470443 | 1,054435204 | 0,85843452 | 0,989067345 | novel transcript, antisense to MAMDC2 |

| | | | | | | |
|---|---|---|---|---|---|---|
| ENSG00000143436 | MRPL9 | 0,076464143 | 1,054430599 | 0,70747748 | 0,969647757 | mitochondrial ribosomal protein L9 |
| ENSG00000230314 | ELOVL2-AS1 | 0,076461741 | 1,054428844 | 0,87756795 | 0,990109814 | ELOVL2 antisense RNA 1 |
| ENSG00000287892 | AC090907,3 | 0,0763939 | 1,054379262 | 0,949876 | 0,996432861 | novel transcript, antisense to LRRK1 |
| ENSG00000272574 | AL596325,2 | 0,076360728 | 1,054355019 | 0,92969049 | 0,994293848 | novel transcript, sense intronic to UAP1 |
| ENSG00000239268 | AC092691,1 | 0,076347807 | 1,054345576 | 0,9262118 | 0,993915537 | novel transcript |
| ENSG00000143398 | PIP5K1A | 0,076337549 | 1,054338079 | 0,73650609 | 0,974353474 | phosphatidylinositol-4-phosphate 5-kinase type 1 alpha |
| ENSG00000167550 | RHEBL1 | 0,076299099 | 1,05430998 | 0,85875251 | 0,989074806 | RHEB like 1 |
| ENSG00000186010 | NDUFA13 | 0,076281712 | 1,054297273 | 0,7493948 | 0,975901808 | NADH:ubiquinone oxidoreductase subunit A13 |
| ENSG00000287611 | AC007922,4 | 0,076280688 | 1,054296525 | 0,93063097 | 0,994344567 | novel transcript |
| ENSG00000085433 | WDR47 | 0,07624887 | 1,054273273 | 0,8360964 | 0,985159643 | WD repeat domain 47 |
| ENSG00000253857 | AC022679,1 | 0,076238217 | 1,054265489 | 0,94413812 | 0,995542144 | novel transcript |
| ENSG00000205423 | CNEP1R1 | 0,076193259 | 1,054232636 | 0,84993656 | 0,987036295 | CTD nuclear envelope phosphatase 1 regulatory subunit 1 |
| ENSG00000175643 | RMI2 | 0,076175229 | 1,054219461 | 0,79866976 | 0,980161849 | RecQ mediated genome instability 2 |
| ENSG00000112972 | HMGCS1 | 0,076164034 | 1,05421128 | 0,86566295 | 0,989074806 | 3-hydroxy-3-methylglutaryl-CoA synthase 1 |
| ENSG00000106723 | SPIN1 | 0,076138231 | 1,054192425 | 0,84254402 | 0,986175464 | spindlin 1 |
| ENSG00000160746 | ANO10 | 0,076128715 | 1,054185472 | 0,79671747 | 0,980146131 | anoctamin 10 |
| ENSG00000139971 | ARMH4 | 0,076099685 | 1,05416426 | 0,78132485 | 0,978597905 | armadillo like helical domain containing 4 |
| ENSG00000267892 | AC022144,1 | 0,076066979 | 1,054140362 | 0,9591122 | 0,996608711 | novel transcript, antisense to CAPN12 |
| ENSG00000163029 | SMC6 | 0,076061936 | 1,054136677 | 0,82940886 | 0,984081562 | structural maintenance of chromosomes 6 |
| ENSG00000127054 | INTS11 | 0,076025183 | 1,054109823 | 0,79277508 | 0,979471509 | integrator complex subunit 11 |
| ENSG00000181220 | ZNF746 | 0,076012245 | 1,05410037 | 0,81045502 | 0,981213877 | zinc finger protein 746 |
| ENSG00000131470 | PSMC3IP | 0,07598848 | 1,054083006 | 0,74571578 | 0,975707496 | PSMC3 interacting protein |
| ENSG00000156374 | PCGF6 | 0,075978977 | 1,054076063 | 0,73536108 | 0,974353474 | polycomb group ring finger 6 |
| ENSG00000049769 | PPP1R3F | 0,075917308 | 1,054031007 | 0,85525534 | 0,988016959 | protein phosphatase 1 regulatory subunit 3F |
| ENSG00000235286 | AP000925,1 | 0,075912948 | 1,054027822 | 0,92354921 | 0,99390845 | keratin 8 (KRT8) pseudogene |
| ENSG00000275621 | LRRC37A3 | 0,075903044 | 1,054020585 | 0,83425081 | 0,984905093 | leucine rich repeat containing 37 member A3 |
| ENSG00000223972 | DDX11L1 | 0,0758399 | 1,053974454 | 0,89463406 | 0,99120559 | DEAD/H-box helicase 11 like 1 (pseudogene) |
| ENSG00000130669 | PAK4 | 0,075712403 | 1,053881314 | 0,8166578 | 0,981961827 | p21 (RAC1) activated kinase 4 |
| ENSG00000176894 | PXMP2 | 0,075700924 | 1,053872929 | 0,81031618 | 0,981213877 | peroxisomal membrane protein 2 |
| ENSG00000171121 | KCNMB3 | 0,075657939 | 1,053841529 | 0,84777032 | 0,987035044 | potassium calcium-activated channel subfamily M regulatory beta subunit 3 |

| | | | | | | |
|---|---|---|---|---|---|---|
| ENSG00000277501 | AC243571,2 | 0,07562525 | 1,053817652 | 0,92036538 | 0,993787755 | novel transcript, antisense DDX52 |
| ENSG00000143643 | TTC13 | 0,075620915 | 1,053814485 | 0,8390256 | 0,985672131 | tetratricopeptide repeat domain 13 |
| ENSG00000229207 | SERPINH1P1 | 0,075620607 | 1,05381426 | 0,92066567 | 0,993787755 | serpin family H member 1 pseudogene 1 |
| ENSG00000162384 | CZIB | 0,075598176 | 1,053797875 | 0,79151208 | 0,979471509 | CXXC motif containing zinc binding protein |
| ENSG00000267510 | AC011451,1 | 0,075562298 | 1,053771669 | 0,94886033 | 0,996432861 | novel transcript, antisense to ZNF699 |
| ENSG00000067840 | PDZD4 | 0,075559465 | 1,0537696 | 0,8847636 | 0,990717729 | PDZ domain containing 4 |
| ENSG00000158711 | ELK4 | 0,075504739 | 1,053729628 | 0,81063171 | 0,981213877 | ETS transcription factor ELK4 |
| ENSG00000280798 | LINC00294 | 0,075467269 | 1,05370226 | 0,79991952 | 0,980161849 | long intergenic non-protein coding RNA 294 |
| ENSG00000148291 | SURF2 | 0,075463855 | 1,053699767 | 0,8124577 | 0,981213877 | surfeit 2 |
| ENSG00000281024 | SURF2 | 0,075463855 | 1,053699767 | 0,8124577 | 0,981213877 | surfeit 2 |
| ENSG00000254105 | VENTXP6 | 0,075456701 | 1,053694542 | 0,95982865 | 0,996608711 | VENT homeobox pseudogene 6 |
| ENSG00000162408 | NOL9 | 0,07544785 | 1,053688077 | 0,70962114 | 0,970482408 | nucleolar protein 9 |
| ENSG00000128245 | YWHAH | 0,075373415 | 1,053633714 | 0,73001144 | 0,974178338 | tyrosine 3-monooxygenase/tryptophan 5-monooxygenase activation protein eta |
| ENSG00000230358 | SPDYE21 | 0,075240979 | 1,053536998 | 0,92642244 | 0,993915537 | speedy/RINGO cell cycle regulator family member E21 |
| ENSG00000287629 | AC006059,5 | 0,075240792 | 1,053536861 | 0,88705774 | 0,990717729 | novel transcript, antisense to ZBTB47 |
| ENSG00000125319 | HROB | 0,075218156 | 1,053520331 | 0,78032081 | 0,978587255 | homologous recombination factor with OB-fold |
| ENSG00000232328 | AC084030,1 | 0,075107291 | 1,053439376 | 0,91717353 | 0,993361299 | novel transcript |
| ENSG00000101400 | SNTA1 | 0,075105959 | 1,053438403 | 0,84056952 | 0,985672131 | syntrophin alpha 1 |
| ENSG00000084463 | WBP11 | 0,075081566 | 1,053420592 | 0,79099992 | 0,979471509 | WW domain binding protein 11 |
| ENSG00000112242 | E2F3 | 0,075050191 | 1,053397683 | 0,80651731 | 0,981213877 | E2F transcription factor 3 |
| ENSG00000266913 | LINC01841 | 0,074838067 | 1,053242809 | 0,96223893 | 0,996608711 | long intergenic non-protein coding RNA 1841 |
| ENSG00000215162 | LINC00269 | 0,074789324 | 1,053207225 | 0,94637539 | 0,996027077 | long intergenic non-protein coding RNA 269 |
| ENSG00000285870 | AC012673,1 | 0,074759073 | 1,053185141 | 0,93085506 | 0,994433362 | novel transcript |
| ENSG00000114978 | MOB1A | 0,074741114 | 1,053172031 | 0,8591115 | 0,989074806 | MOB kinase activator 1A |
| ENSG00000091127 | PUS7 | 0,074718702 | 1,053155671 | 0,79827401 | 0,980161849 | pseudouridine synthase 7 |
| ENSG00000253708 | AC026979,1 | 0,074710542 | 1,053149714 | 0,94589767 | 0,996027077 | novel transcript |
| ENSG00000288068 | AC027088,5 | 0,074701673 | 1,05314324 | 0,9242975 | 0,993915537 | novel transcript, sense intronic to NOX5 |
| ENSG00000287216 | AC106865,2 | 0,074656868 | 1,053110533 | 0,93870313 | 0,994699525 | novel transcript, antisense to KIAA0922 |
| ENSG00000169919 | GUSB | 0,074610971 | 1,053077031 | 0,77107064 | 0,978033855 | glucuronidase beta |
| ENSG00000140632 | GLYR1 | 0,074436511 | 1,052949694 | 0,70983866 | 0,970482408 | glyoxylate reductase 1 homolog |
| ENSG00000284809 | AL365394,1 | 0,074406566 | 1,052927839 | 0,96244916 | 0,996608711 | novel transcript |

| | | | | | | |
|---|---|---|---|---|---|---|
| ENSG00000260852 | FBXL19-AS1 | 0,074346526 | 1,05288402 | 0,85448139 | 0,987861659 | FBXL19 antisense RNA 1 |
| ENSG00000158941 | CCAR2 | 0,074304701 | 1,052853496 | 0,79516126 | 0,97981137 | cell cycle and apoptosis regulator 2 |
| ENSG00000250116 | RHOQ-AS1 | 0,074249568 | 1,052813262 | 0,87548089 | 0,990078774 | RHOQ antisense RNA 1 |
| ENSG00000100335 | MIEF1 | 0,074219719 | 1,05279148 | 0,71402451 | 0,970867497 | mitochondrial elongation factor 1 |
| ENSG00000130826 | DKC1 | 0,074215927 | 1,052788713 | 0,78713638 | 0,979424965 | dyskerin pseudouridine synthase 1 |
| ENSG00000138614 | INTS14 | 0,07418877 | 1,052768896 | 0,81527208 | 0,981521413 | integrator complex subunit 14 |
| ENSG00000146282 | RARS2 | 0,074129932 | 1,052725961 | 0,83986862 | 0,985672131 | arginyl-tRNA synthetase 2, mitochondrial |
| ENSG00000138386 | NAB1 | 0,074129909 | 1,052725944 | 0,82088341 | 0,982719623 | NGFI-A binding protein 1 |
| ENSG00000113558 | SKP1 | 0,074069006 | 1,052681505 | 0,82940197 | 0,984081562 | S-phase kinase associated protein 1 |
| ENSG00000258551 | CRAT37 | 0,074036374 | 1,052657695 | 0,922009 | 0,993797669 | cervical cancer-associated transcript 37 |
| ENSG00000163597 | SNHG16 | 0,074015919 | 1,052642769 | 0,79840798 | 0,980161849 | small nucleolar RNA host gene 16 |
| ENSG00000106780 | MEGF9 | 0,073978765 | 1,052615661 | 0,83986075 | 0,985672131 | multiple EGF like domains 9 |
| ENSG00000204438 | GPANK1 | 0,07390375 | 1,05256093 | 0,86335381 | 0,989074806 | G-patch domain and ankyrin repeats 1 |
| ENSG00000222375 | RN7SKP127 | 0,07385603 | 1,052526115 | 0,94927273 | 0,996432861 | RN7SK pseudogene 127 |
| ENSG00000138018 | SELENOI | 0,073819342 | 1,05249935 | 0,86156023 | 0,989074806 | selenoprotein I |
| ENSG00000241269 | AC093620,1 | 0,073733461 | 1,052436699 | 0,92806136 | 0,994108531 | novel transcript |
| ENSG00000169213 | RAB3B | 0,073713817 | 1,052422368 | 0,83167468 | 0,984456953 | RAB3B, member RAS oncogene family |
| ENSG00000110955 | ATP5F1B | 0,0736593 | 1,052382599 | 0,75760697 | 0,976120207 | ATP synthase F1 subunit beta |
| ENSG00000108578 | BLMH | 0,0735968 | 1,05233701 | 0,76607532 | 0,977102081 | bleomycin hydrolase |
| ENSG00000264635 | AP001020,2 | 0,073586728 | 1,052329663 | 0,85669645 | 0,988312374 | novel transcript, sense intronic toYES1 |
| ENSG00000173960 | UBXN2A | 0,073535512 | 1,052292306 | 0,72394174 | 0,973141049 | UBX domain protein 2A |
| ENSG00000073614 | KDM5A | 0,073502029 | 1,052267883 | 0,74603104 | 0,975707496 | lysine demethylase 5A |
| ENSG00000288353 | ARHGAP17 | 0,073478266 | 1,052250552 | 0,75300388 | 0,975901808 | Rho GTPase activating protein 17 |
| ENSG00000120925 | RNF170 | 0,073392069 | 1,052187685 | 0,82735385 | 0,983595244 | ring finger protein 170 |
| ENSG00000149657 | LSM14B | 0,073377585 | 1,052177121 | 0,72061781 | 0,972142678 | LSM family member 14B |
| ENSG00000228549 | BX284668,2 | 0,073367901 | 1,052170059 | 0,80139952 | 0,980557464 | novel transcript |
| ENSG00000124214 | STAU1 | 0,0733481 | 1,052155617 | 0,78845365 | 0,979462318 | staufen double-stranded RNA binding protein 1 |
| ENSG00000163938 | GNL3 | 0,073312453 | 1,052129621 | 0,78325491 | 0,978919905 | G protein nucleolar 3 |
| ENSG00000274012 | RN7SL2 | 0,073299751 | 1,052120357 | 0,80768793 | 0,981213877 | RNA component of signal recognition particle 7SL2 |
| ENSG00000184388 | PABPC1L2B | 0,073269238 | 1,052098105 | 0,95294605 | 0,99652169 | poly(A) binding protein cytoplasmic 1 like 2B |
| ENSG00000188725 | SMIM15 | 0,073261205 | 1,052092247 | 0,85781289 | 0,988925186 | small integral membrane protein 15 |
| ENSG00000151498 | ACAD8 | 0,073256214 | 1,052088607 | 0,73375162 | 0,974353474 | acyl-CoA dehydrogenase family member 8 |

| | | | | | | |
|---|---|---|---|---|---|---|
| ENSG00000168067 | MAP4K2 | 0,073232189 | 1,052071087 | 0,8218206 | 0,982952211 | mitogen-activated protein kinase kinase kinase kinase 2 |
| ENSG00000137413 | TAF8 | 0,073215754 | 1,052059102 | 0,79560048 | 0,979879951 | TATA-box binding protein associated factor 8 |
| ENSG00000130818 | ZNF426 | 0,07313453 | 1,051999873 | 0,77718858 | 0,978587255 | zinc finger protein 426 |
| ENSG00000174891 | RSRC1 | 0,0730916 | 1,051968569 | 0,84845402 | 0,987036295 | arginine and serine rich coiled-coil 1 |
| ENSG00000217527 | RPS16P5 | 0,073054791 | 1,051941729 | 0,91441345 | 0,993361299 | ribosomal protein S16 pseudogene 5 |
| ENSG00000120334 | CENPL | 0,07297443 | 1,051883136 | 0,76396958 | 0,976640468 | centromere protein L |
| ENSG00000256897 | AC018410,1 | 0,072953904 | 1,05186817 | 0,90425821 | 0,992590661 | novel transcript |
| ENSG00000276550 | HERC2P2 | 0,072935251 | 1,05185457 | 0,74750068 | 0,975901808 | hect domain and RLD 2 pseudogene 2 |
| ENSG00000110218 | PANX1 | 0,072934889 | 1,051854306 | 0,79968413 | 0,980161849 | pannexin 1 |
| ENSG00000172939 | OXSR1 | 0,07293192 | 1,051852142 | 0,76326021 | 0,976622569 | oxidative stress responsive kinase 1 |
| ENSG00000176903 | PNMA1 | 0,072806795 | 1,051760918 | 0,71412229 | 0,970867497 | PNMA family member 1 |
| ENSG00000246548 | LINC02288 | 0,072675989 | 1,051665562 | 0,9080099 | 0,992590661 | long intergenic non-protein coding RNA 2288 |
| ENSG00000125386 | FAM193A | 0,072563156 | 1,051583315 | 0,73146361 | 0,974353474 | family with sequence similarity 193 member A |
| ENSG00000282947 | HNRNPL | 0,072552276 | 1,051575384 | 0,7386919 | 0,974353474 | heterogeneous nuclear ribonucleoprotein L |
| ENSG00000104824 | HNRNPL | 0,072552276 | 1,051575384 | 0,7386919 | 0,974353474 | heterogeneous nuclear ribonucleoprotein L |
| ENSG00000221933 | OR2A25 | 0,072422662 | 1,051480913 | 0,93396625 | 0,994479239 | olfactory receptor family 2 subfamily A member 25 |
| ENSG00000171763 | SPATA5L1 | 0,072329583 | 1,051413077 | 0,78048173 | 0,978597905 | spermatogenesis associated 5 like 1 |
| ENSG00000147604 | RPL7 | 0,072285746 | 1,051381129 | 0,82438128 | 0,983374097 | ribosomal protein L7 |
| ENSG00000151694 | ADAM17 | 0,072249504 | 1,051354718 | 0,78831581 | 0,979462318 | ADAM metallopeptidase domain 17 |
| ENSG00000244119 | PDCL3P4 | 0,072141616 | 1,051276098 | 0,92117363 | 0,993787755 | PDCL3 pseudogene 4 |
| ENSG00000160208 | RRP1B | 0,072095405 | 1,051242425 | 0,73017912 | 0,974303405 | ribosomal RNA processing 1B |
| ENSG00000105372 | RPS19 | 0,072081354 | 1,051232187 | 0,79248052 | 0,979471509 | ribosomal protein S19 |
| ENSG00000186615 | KTN1-AS1 | 0,072079352 | 1,051230728 | 0,87215244 | 0,989217334 | KTN1 antisense RNA 1 |
| ENSG00000196456 | ZNF775 | 0,072037309 | 1,051200093 | 0,86083044 | 0,989074806 | zinc finger protein 775 |
| ENSG00000154309 | DISP1 | 0,071993371 | 1,051168079 | 0,82843834 | 0,983717437 | dispatched RND transporter family member 1 |
| ENSG00000103449 | SALL1 | 0,071937921 | 1,051127679 | 0,81804415 | 0,982323352 | spalt like transcription factor 1 |
| ENSG00000283486 | FAM95C | 0,071920542 | 1,051115016 | 0,93151966 | 0,994433362 | family with sequence similarity 95 member C |
| ENSG00000267497 | NFE2L3P1 | 0,071829553 | 1,051048726 | 0,9268135 | 0,993915537 | nuclear factor, erythroid 2 like 3 pseudogene 1 |
| ENSG00000285451 | ADCK5 | 0,071785638 | 1,051016733 | 0,8975606 | 0,991708791 | aarF domain containing kinase 5 |
| ENSG00000173137 | ADCK5 | 0,071785638 | 1,051016733 | 0,8975606 | 0,991708791 | aarF domain containing kinase 5 |
| ENSG00000142920 | AZIN2 | 0,071716127 | 1,050966094 | 0,83731655 | 0,985577601 | antizyme inhibitor 2 |
| ENSG00000130653 | PNPLA7 | 0,071714647 | 1,050965016 | 0,88690222 | 0,990717729 | patatin like phospholipase domain containing 7 |

| | | | | | | |
|---|---|---|---|---|---|---|
| ENSG00000091483 | FH | 0,071652211 | 1,050919534 | 0,8283154 | 0,983646162 | fumarate hydratase |
| ENSG00000214265 | AC124312,1 | 0,071614184 | 1,050891835 | 0,90685688 | 0,992590661 | novel protein |
| ENSG00000156050 | FAM161B | 0,071603161 | 1,050883805 | 0,76965129 | 0,977708263 | FAM161 centrosomal protein B |
| ENSG00000149273 | RPS3 | 0,071600354 | 1,05088176 | 0,74448289 | 0,975649186 | ribosomal protein S3 |
| ENSG00000178922 | HYI | 0,071561679 | 1,050853589 | 0,82479246 | 0,983374097 | hydroxypyruvate isomerase (putative) |
| ENSG00000140090 | SLC24A4 | 0,071551709 | 1,050846327 | 0,93007937 | 0,994331958 | solute carrier family 24 member 4 |
| ENSG00000137860 | SLC28A2 | 0,071460684 | 1,050780028 | 0,8776186 | 0,990109814 | solute carrier family 28 member 2 |
| ENSG00000162772 | ATF3 | 0,071406566 | 1,050740612 | 0,83366751 | 0,984905093 | activating transcription factor 3 |
| ENSG00000112357 | PEX7 | 0,071361764 | 1,050707982 | 0,80345088 | 0,980869547 | peroxisomal biogenesis factor 7 |
| ENSG00000274577 | LHX1 | 0,071342338 | 1,050693834 | 0,86684262 | 0,989074806 | LIM homeobox 1 |
| ENSG00000262587 | AC133552,2 | 0,071330993 | 1,050685572 | 0,89748361 | 0,991708791 | leucine carboxyl methyltransferase 1 (LCMT1) pseudogene |
| ENSG00000176953 | NFATC2IP | 0,07121881 | 1,050603875 | 0,75806382 | 0,976120207 | nuclear factor of activated T cells 2 interacting protein |
| ENSG00000184436 | THAP7 | 0,071203725 | 1,050592889 | 0,79040532 | 0,979462318 | THAP domain containing 7 |
| ENSG00000155975 | VPS37A | 0,071193807 | 1,050585667 | 0,83472826 | 0,984905093 | VPS37A subunit of ESCRT-I |
| ENSG00000154813 | DPH3 | 0,071169905 | 1,050568261 | 0,87486827 | 0,990078774 | diphthamide biosynthesis 3 |
| ENSG00000171163 | ZNF692 | 0,071043737 | 1,05047639 | 0,79478 | 0,979776981 | zinc finger protein 692 |
| ENSG00000171208 | NETO2 | 0,071032742 | 1,050468384 | 0,84808221 | 0,987036295 | neuropilin and tolloid like 2 |
| ENSG00000283515 | AC020915,4 | 0,071032549 | 1,050468243 | 0,82949628 | 0,984081562 | novel protein |
| ENSG00000150656 | CNDP1 | 0,070987257 | 1,050435266 | 0,92908006 | 0,994293848 | carnosine dipeptidase 1 |
| ENSG00000248643 | RBM14-RBM4 | 0,070980891 | 1,05043063 | 0,80638556 | 0,981213877 | RBM14-RBM4 readthrough |
| ENSG00000136854 | STXBP1 | 0,070910914 | 1,050379682 | 0,75350841 | 0,975901808 | syntaxin binding protein 1 |
| ENSG00000134444 | RELCH | 0,070909173 | 1,050378414 | 0,81154464 | 0,981213877 | RAB11 binding and LisH domain, coiled-coil and HEAT repeat containing |
| ENSG00000212993 | POU5F1B | 0,070901364 | 1,050372728 | 0,92328944 | 0,993825008 | POU class 5 homeobox 1B |
| ENSG00000280618 | CYFIP1 | 0,070722545 | 1,050242545 | 0,77030073 | 0,977872051 | cytoplasmic FMR1 interacting protein 1 |
| ENSG00000204070 | SYS1 | 0,070659903 | 1,050196944 | 0,79099449 | 0,979471509 | SYS1 golgi trafficking protein |
| ENSG00000133028 | SCO1 | 0,070625567 | 1,05017195 | 0,72515519 | 0,97326334 | synthesis of cytochrome C oxidase 1 |
| ENSG00000184708 | EIF4ENIF1 | 0,070548751 | 1,050116036 | 0,7309228 | 0,974353474 | eukaryotic translation initiation factor 4E nuclear import factor 1 |
| ENSG00000111052 | LIN7A | 0,070540836 | 1,050110274 | 0,88558969 | 0,990717729 | lin-7 homolog A, crumbs cell polarity complex component |

| | | | | | | |
|---|---|---|---|---|---|---|
| ENSG00000226853 | AC010894,1 | 0,070488875 | 1,050072453 | 0,90760262 | 0,992590661 | novel transcript |
| ENSG00000198218 | QRICH1 | 0,070379126 | 1,049992575 | 0,78646121 | 0,979320406 | glutamine rich 1 |
| ENSG00000255438 | AL354813,1 | 0,070363125 | 1,049980929 | 0,91267276 | 0,993361299 | novel transcript, antisense to SULF2 |
| ENSG00000237846 | AL773545,3 | 0,070355405 | 1,049975311 | 0,89570241 | 0,991377987 | novel pseudogene |
| ENSG00000125870 | SNRPB2 | 0,070350894 | 1,049972028 | 0,850569 | 0,987147801 | small nuclear ribonucleoprotein polypeptide B2 |
| ENSG00000229474 | PATL2 | 0,070348715 | 1,049970442 | 0,9321534 | 0,994479239 | PAT1 homolog 2 |
| ENSG00000268205 | AC005261,1 | 0,070339262 | 1,049963563 | 0,75726792 | 0,975901808 | novel transcript |
| ENSG00000070770 | CSNK2A2 | 0,070318029 | 1,04994811 | 0,81865973 | 0,9824772 | casein kinase 2 alpha 2 |
| ENSG00000070366 | SMG6 | 0,070304832 | 1,049938506 | 0,78502324 | 0,979320406 | SMG6 nonsense mediated mRNA decay factor |
| ENSG00000125149 | C16orf70 | 0,070191866 | 1,049856296 | 0,78093163 | 0,978597905 | chromosome 16 open reading frame 70 |
| ENSG00000232893 | IQCA1-AS1 | 0,07017985 | 1,049847552 | 0,96306786 | 0,996608711 | IQCA1 antisense RNA 1 |
| ENSG00000258199 | AC073896,4 | 0,070166592 | 1,049837904 | 0,95913597 | 0,996608711 | novel transcript, sense overlapping to SMARCC2 |
| ENSG00000122507 | BBS9 | 0,070157646 | 1,049831394 | 0,83002104 | 0,984156028 | Bardet-Biedl syndrome 9 |
| ENSG00000100902 | PSMA6 | 0,070132797 | 1,049813312 | 0,85227235 | 0,987416817 | proteasome 20S subunit alpha 6 |
| ENSG00000188242 | PP7080 | 0,070112648 | 1,049798651 | 0,81098067 | 0,981213877 | uncharacterized LOC25845 |
| ENSG00000155508 | CNOT8 | 0,07001037 | 1,049724229 | 0,84693379 | 0,986939308 | CCR4-NOT transcription complex subunit 8 |
| ENSG00000180776 | ZDHHC20 | 0,069998887 | 1,049715874 | 0,85814833 | 0,989067345 | zinc finger DHHC-type palmitoyltransferase 20 |
| ENSG00000171606 | ZNF274 | 0,069898975 | 1,04964318 | 0,81415176 | 0,981521413 | zinc finger protein 274 |
| ENSG00000167526 | RPL13 | 0,06989731 | 1,049641968 | 0,77293548 | 0,978310326 | ribosomal protein L13 |
| ENSG00000104897 | SF3A2 | 0,069788953 | 1,049563135 | 0,77385053 | 0,978415098 | splicing factor 3a subunit 2 |
| ENSG00000107679 | PLEKHA1 | 0,06973654 | 1,049525005 | 0,8198217 | 0,98249617 | pleckstrin homology domain containing A1 |
| ENSG00000186660 | ZFP91 | 0,069713985 | 1,049508598 | 0,8190271 | 0,98249617 | ZFP91 zinc finger protein, atypical E3 ubiquitin ligase |
| ENSG00000088682 | COQ9 | 0,069696077 | 1,04949557 | 0,74271619 | 0,975201495 | coenzyme Q9 |
| ENSG00000267365 | KCNJ2-AS1 | 0,069678889 | 1,049483067 | 0,87880315 | 0,990109814 | KCNJ2 antisense RNA 1 |
| ENSG00000213533 | STIMATE | 0,069672215 | 1,049478211 | 0,751045 | 0,975901808 | STIM activating enhancer |
| ENSG00000015285 | WAS | 0,069594275 | 1,049421516 | 0,89639728 | 0,991638165 | WASP actin nucleation promoting factor |
| ENSG00000269904 | MAP2K4P1 | 0,069528096 | 1,049373378 | 0,94374304 | 0,995483785 | mitogen-activated protein kinase kinase 4 pseudogene 1 |
| ENSG00000277268 | LHX1-DT | 0,069440681 | 1,049309797 | 0,90511308 | 0,992590661 | LHX1 divergent transcript |
| ENSG00000234056 | LINC00463 | 0,069407702 | 1,049285811 | 0,88069682 | 0,990533002 | long intergenic non-protein coding RNA 463 |
| ENSG00000010256 | UQCRC1 | 0,069388661 | 1,049271963 | 0,76122972 | 0,97624106 | ubiquinol-cytochrome c reductase core protein 1 |
| ENSG00000188001 | TPRG1 | 0,069374175 | 1,049261427 | 0,89033636 | 0,990717729 | tumor protein p63 regulated 1 |
| ENSG00000273702 | AC091271,1 | 0,069371527 | 1,049259501 | 0,84253049 | 0,986175464 | novel transcript |

| | | | | | | |
|---|---|---|---|---|---|---|
| ENSG00000223797 | ENTPD3-AS1 | 0,069342832 | 1,049238632 | 0,85519962 | 0,988016959 | ENTPD3 antisense RNA 1 |
| ENSG00000171530 | TBCA | 0,069285447 | 1,049196898 | 0,84484907 | 0,986557476 | tubulin folding cofactor A |
| ENSG00000285497 | USP17L7 | 0,069251703 | 1,049172358 | 0,90196632 | 0,992182254 | ubiquitin specific peptidase 17 like family member 7 |
| ENSG00000283447 | NDUFS1 | 0,069195177 | 1,049131251 | 0,86596742 | 0,989074806 | NADH:ubiquinone oxidoreductase core subunit S1 |
| ENSG00000287726 | AC005740,5 | 0,06912533 | 1,049080459 | 0,89211185 | 0,99102448 | novel transcript |
| ENSG00000261467 | AC099398,1 | 0,069124882 | 1,049080134 | 0,91928457 | 0,993598006 | novel transcript |
| ENSG00000145781 | COMMD10 | 0,069095274 | 1,049058604 | 0,88996341 | 0,990717729 | COMM domain containing 10 |
| ENSG00000258938 | AL162311,3 | 0,069084099 | 1,049050478 | 0,89744196 | 0,991708791 | novel transcript, antisense to BRMS1L |
| ENSG00000213753 | CENPBD1P1 | 0,069082663 | 1,049049434 | 0,74386905 | 0,975394113 | CENPB DNA-binding domains containing 1 pseudogene 1 |
| ENSG00000232973 | CYP1B1-AS1 | 0,068928068 | 1,048937027 | 0,90067901 | 0,991865079 | CYP1B1 antisense RNA 1 |
| ENSG00000233288 | AC092447,5 | 0,068907365 | 1,048921975 | 0,93261772 | 0,994479239 | novel transcript |
| ENSG00000227171 | RNF39 | 0,068770713 | 1,048822626 | 0,94013877 | 0,994949796 | ring finger protein 39 |
| ENSG00000273179 | AC092535,5 | 0,068758061 | 1,048813428 | 0,9649309 | 0,997113804 | novel transcript, antisense to SPON2 |
| ENSG00000112977 | DAP | 0,068735147 | 1,048796769 | 0,76063721 | 0,97624106 | death associated protein |
| ENSG00000181652 | ATG9B | 0,068652868 | 1,048736957 | 0,89317234 | 0,991135494 | autophagy related 9B |
| ENSG00000259475 | AC036108,1 | 0,068628644 | 1,048719348 | 0,94044576 | 0,995028467 | novel transcript, antisense to IGF1R and SYNM |
| ENSG00000255141 | HNRNPA1P76 | 0,068551134 | 1,048663006 | 0,94594741 | 0,996027077 | heterogeneous nuclear ribonucleoprotein A1 pseudogene 76 |
| ENSG00000166507 | NDST2 | 0,068522394 | 1,048642116 | 0,77421856 | 0,978503783 | N-deacetylase and N-sulfotransferase 2 |
| ENSG00000286488 | AC103858,3 | 0,068481206 | 1,048612179 | 0,91207014 | 0,993361299 | novel transcript, antisense to STARD5 |
| ENSG00000205930 | C21orf62-AS1 | 0,06838417 | 1,04854165 | 0,8379456 | 0,985672131 | C21orf62 antisense RNA 1 |
| ENSG00000121753 | ADGRB2 | 0,068372088 | 1,04853287 | 0,8377206 | 0,98566906 | adhesion G protein-coupled receptor B2 |
| ENSG00000255384 | AP001267,2 | 0,06833487 | 1,04850582 | 0,9352043 | 0,994479239 | novel transcript |
| ENSG00000263938 | AC100832,2 | 0,06830126 | 1,048481394 | 0,93827364 | 0,994547792 | novel transcript, sense intronic to PPM1E |
| ENSG00000152439 | ZNF773 | 0,068298338 | 1,048479271 | 0,81180028 | 0,981213877 | zinc finger protein 773 |
| ENSG00000105677 | TMEM147 | 0,068230878 | 1,048430245 | 0,78731353 | 0,979462318 | transmembrane protein 147 |
| ENSG00000113658 | SMAD5 | 0,068221367 | 1,048423333 | 0,86578768 | 0,989074806 | SMAD family member 5 |
| ENSG00000221986 | MYBPHL | 0,068201515 | 1,048408907 | 0,95124409 | 0,996432861 | myosin binding protein H like |
| ENSG00000102145 | GATA1 | 0,068113916 | 1,04834525 | 0,95700294 | 0,996608711 | GATA binding protein 1 |
| ENSG00000083812 | ZNF324 | 0,068109645 | 1,048342147 | 0,8291679 | 0,983957145 | zinc finger protein 324 |
| ENSG00000279643 | TAMM41 | 0,068097389 | 1,048333241 | 0,87163509 | 0,989217334 | TAM41 mitochondrial translocator assembly and maintenance homolog |

| | | | | | | |
|---|---|---|---|---|---|---|
| ENSG00000273218 | AC005776,2 | 0,067995458 | 1,048259175 | 0,90554277 | 0,992590661 | novel transcript |
| ENSG00000230423 | AL672310,1 | 0,067909401 | 1,048196648 | 0,9236136 | 0,99390845 | novel transcript |
| ENSG00000177383 | MAGEF1 | 0,067895887 | 1,04818683 | 0,77327608 | 0,978373116 | MAGE family member F1 |
| ENSG00000013583 | HEBP1 | 0,06786384 | 1,048163547 | 0,79061256 | 0,979462318 | heme binding protein 1 |
| ENSG00000280927 | CTBP1-AS | 0,067840567 | 1,048146638 | 0,91124727 | 0,993222655 | CTBP1 antisense RNA |
| ENSG00000124092 | CTCFL | 0,067814495 | 1,048127696 | 0,89885785 | 0,991832796 | CCCTC-binding factor like |
| ENSG00000184992 | BRI3BP | 0,067799897 | 1,048117091 | 0,76972229 | 0,977719248 | BRI3 binding protein |
| ENSG00000164062 | APEH | 0,067767126 | 1,048093283 | 0,79519056 | 0,97981137 | acylaminoacyl-peptide hydrolase |
| ENSG00000232656 | IDI2-AS1 | 0,067721116 | 1,048059858 | 0,84911928 | 0,987036295 | IDI2 antisense RNA 1 |
| ENSG00000143321 | HDGF | 0,067642665 | 1,048002868 | 0,80799915 | 0,981213877 | heparin binding growth factor |
| ENSG00000164211 | STARD4 | 0,067627391 | 1,047991773 | 0,89847055 | 0,991832796 | StAR related lipid transfer domain containing 4 |
| ENSG00000152061 | RABGAP1L | 0,067610344 | 1,04797939 | 0,82636782 | 0,983471137 | RAB GTPase activating protein 1 like |
| ENSG00000120860 | WASHC3 | 0,067574609 | 1,047953432 | 0,85032473 | 0,987036295 | WASH complex subunit 3 |
| ENSG00000121879 | PIK3CA | 0,067557942 | 1,047941326 | 0,85143327 | 0,987259878 | phosphatidylinositol-4,5-bisphosphate 3-kinase catalytic subunit alpha |
| ENSG00000266469 | AC005288,1 | 0,067487176 | 1,047889924 | 0,94791736 | 0,99619108 | novel transcript, antisense to MED1 & FBXL20 |
| ENSG00000137100 | DCTN3 | 0,067433726 | 1,047851102 | 0,76130822 | 0,97624106 | dynactin subunit 3 |
| ENSG00000138375 | SMARCAL1 | 0,067408222 | 1,047832578 | 0,75313191 | 0,975901808 | SWI/SNF related, matrix associated, actin dependent regulator of chromatin, subfamily a like 1 |
| ENSG00000132716 | DCAF8 | 0,067330101 | 1,04777584 | 0,75075918 | 0,975901808 | DDB1 and CUL4 associated factor 8 |
| ENSG00000236227 | BRD2 | 0,067294871 | 1,047750254 | 0,97142256 | 0,998088605 | bromodomain containing 2 |
| ENSG00000234507 | BRD2 | 0,067294871 | 1,047750254 | 0,97142256 | 0,998088605 | bromodomain containing 2 |
| ENSG00000036257 | CUL3 | 0,067292969 | 1,047748872 | 0,8464718 | 0,986939308 | cullin 3 |
| ENSG00000287733 | AC083862,2 | 0,06728634 | 1,047744058 | 0,88579973 | 0,990717729 | novel transcript |
| ENSG00000116138 | DNAJC16 | 0,067145936 | 1,047642096 | 0,80946679 | 0,981213877 | DnaJ heat shock protein family (Hsp40) member C16 |
| ENSG00000139546 | TARBP2 | 0,066904157 | 1,047466538 | 0,78534945 | 0,979320406 | TARBP2 subunit of RISC loading complex |
| ENSG00000160336 | ZNF761 | 0,066860665 | 1,047434961 | 0,84365531 | 0,986273696 | zinc finger protein 761 |
| ENSG00000233276 | GPX1 | 0,06679927 | 1,047390388 | 0,82583373 | 0,983383773 | glutathione peroxidase 1 |
| ENSG00000226476 | LINC01748 | 0,06673514 | 1,04734383 | 0,86761989 | 0,989074806 | long intergenic non-protein coding RNA 1748 |
| ENSG00000288559 | AC016394,2 | 0,066734084 | 1,047343065 | 0,89324801 | 0,991135494 | novel transcript |
| ENSG00000158019 | BABAM2 | 0,066730148 | 1,047340207 | 0,80665946 | 0,981213877 | BRISC and BRCA1 A complex member 2 |
| ENSG00000231952 | DPY19L1P2 | 0,066701626 | 1,047319501 | 0,92444745 | 0,993915537 | DPY19L1 pseudogene 2 |
| ENSG00000231965 | AF131215,1 | 0,066656286 | 1,047286587 | 0,92719127 | 0,993915537 | ribosomal protein L17 (RPL17) pseudogene |

| | | | | | | |
|---|---|---|---|---|---|---|
| ENSG00000285026 | AC270301,5 | 0,066656286 | 1,047286587 | 0,92719127 | 0,993915537 | ribosomal protein L17 (RPL17) pseudogene |
| ENSG00000125735 | TNFSF14 | 0,066653448 | 1,047284527 | 0,93978911 | 0,994943406 | TNF superfamily member 14 |
| ENSG00000196335 | STK31 | 0,066612103 | 1,047254514 | 0,87072523 | 0,98911248 | serine/threonine kinase 31 |
| ENSG00000177302 | TOP3A | 0,066509635 | 1,047180135 | 0,8668489 | 0,989074806 | DNA topoisomerase III alpha |
| ENSG00000115207 | GTF3C2 | 0,066443947 | 1,047132457 | 0,76825175 | 0,977381742 | general transcription factor IIIC subunit 2 |
| ENSG00000123091 | RNF11 | 0,066427199 | 1,047120301 | 0,86238231 | 0,989074806 | ring finger protein 11 |
| ENSG00000256628 | ZBTB11-AS1 | 0,066409965 | 1,047107792 | 0,8613386 | 0,989074806 | ZBTB11 antisense RNA 1 |
| ENSG00000182484 | WASH6P | 0,066360249 | 1,047071709 | 0,82573651 | 0,983383773 | WASP family homolog 6, pseudogene |
| ENSG00000267385 | AC011498,4 | 0,066321936 | 1,047043902 | 0,94813725 | 0,996321687 | novel protein |
| ENSG00000106628 | POLD2 | 0,066281055 | 1,047014233 | 0,79513146 | 0,97981137 | DNA polymerase delta 2, accessory subunit |
| ENSG00000102931 | ARL2BP | 0,066239332 | 1,046983955 | 0,8789892 | 0,990109814 | ADP ribosylation factor like GTPase 2 binding protein |
| ENSG00000272690 | LINC02018 | 0,066217534 | 1,046968135 | 0,85580902 | 0,988145402 | long intergenic non-protein coding RNA 2018 |
| ENSG00000260095 | AC106820,3 | 0,066214031 | 1,046965593 | 0,9376941 | 0,994479239 | novel transcript, antisense to CCNF |
| ENSG00000223770 | CACNA2D1-AS1 | 0,066208225 | 1,04696138 | 0,90320365 | 0,992311172 | CACNA2D1 antisense RNA 1 |
| ENSG00000276111 | SDCCAG8 | 0,066202163 | 1,04695698 | 0,88151481 | 0,990717729 | SHH signaling and ciliogenesis regulator SDCCAG8 |
| ENSG00000267515 | AP001029,3 | 0,066181617 | 1,04694207 | 0,93640541 | 0,994479239 | high mobility group box 3 (HMGB3) pseudogene |
| ENSG00000242034 | AC097493,1 | 0,066086511 | 1,046873056 | 0,96275572 | 0,996608711 | ribosomal protein S24 (RPS24) pseudogene |
| ENSG00000157578 | LCA5L | 0,066083874 | 1,046871142 | 0,84402996 | 0,986273696 | lebercilin LCA5 like |
| ENSG00000090621 | PABPC4 | 0,065846394 | 1,046698832 | 0,7399763 | 0,974715504 | poly(A) binding protein cytoplasmic 4 |
| ENSG00000272758 | AC083798,2 | 0,065838679 | 1,046693235 | 0,90087881 | 0,991865079 | novel transcript, antisense to KPNA1 |
| ENSG00000107020 | PLGRKT | 0,065810301 | 1,046672646 | 0,84969847 | 0,987036295 | plasminogen receptor with a C-terminal lysine |
| ENSG00000142453 | CARM1 | 0,065614272 | 1,046530438 | 0,76714933 | 0,977201891 | coactivator associated arginine methyltransferase 1 |
| ENSG00000005889 | ZFX | 0,065592141 | 1,046514384 | 0,8466424 | 0,986939308 | zinc finger protein X-linked |
| ENSG00000283914 | AC087645,4 | 0,065589138 | 1,046512206 | 0,96651359 | 0,997263201 | novel transcript |
| ENSG00000164506 | STXBP5 | 0,065513411 | 1,046457276 | 0,86085592 | 0,989074806 | syntaxin binding protein 5 |
| ENSG00000137404 | NRM | 0,065481854 | 1,046434386 | 0,81241939 | 0,981213877 | nurim |
| ENSG00000228867 | NRM | 0,065481854 | 1,046434386 | 0,81241939 | 0,981213877 | nurim |
| ENSG00000235773 | NRM | 0,065481854 | 1,046434386 | 0,81241939 | 0,981213877 | nurim |
| ENSG00000234809 | NRM | 0,065481854 | 1,046434386 | 0,81241939 | 0,981213877 | nurim |
| ENSG00000236843 | NRM | 0,065481854 | 1,046434386 | 0,81241939 | 0,981213877 | nurim |
| ENSG00000228854 | NRM | 0,065481854 | 1,046434386 | 0,81241939 | 0,981213877 | nurim |
| ENSG00000206484 | NRM | 0,065481854 | 1,046434386 | 0,81241939 | 0,981213877 | nurim |
| ENSG00000165264 | NDUFB6 | 0,065446342 | 1,046408628 | 0,83511186 | 0,984905093 | NADH:ubiquinone oxidoreductase subunit B6 |

| | | | | | | |
|---|---|---|---|---|---|---|
| ENSG00000288438 | MG910338,3 | 0,065422317 | 1,046391203 | 0,96169273 | 0,996608711 | uncharacterized LOC102723360 |
| ENSG00000120137 | PANK3 | 0,065385945 | 1,046364822 | 0,88167993 | 0,990717729 | pantothenate kinase 3 |
| ENSG00000137055 | PLAA | 0,065345382 | 1,046335403 | 0,85394077 | 0,987861659 | phospholipase A2 activating protein |
| ENSG00000153037 | SRP19 | 0,065298724 | 1,046301564 | 0,81953988 | 0,98249617 | signal recognition particle 19 |
| ENSG00000267787 | AC027097,2 | 0,065262874 | 1,046275565 | 0,86334355 | 0,989074806 | novel transcript, antisense to ATP8B1 |
| ENSG00000182397 | DNM1P46 | 0,065192061 | 1,04622421 | 0,86619733 | 0,989074806 | dynamin 1 pseudogene 46 |
| ENSG00000113811 | SELENOK | 0,06518356 | 1,046218046 | 0,82685738 | 0,983548538 | selenoprotein K |
| ENSG00000137563 | GGH | 0,065095423 | 1,046154133 | 0,88030511 | 0,990315537 | gamma-glutamyl hydrolase |
| ENSG00000225721 | AL592166,1 | 0,065088913 | 1,046149411 | 0,93843431 | 0,994549499 | novel transcript |
| ENSG00000204256 | BRD2 | 0,065083927 | 1,046145796 | 0,75339425 | 0,975901808 | bromodomain containing 2 |
| ENSG00000255639 | AC005833,1 | 0,065079911 | 1,046142884 | 0,82536401 | 0,983383773 | novel protein |
| ENSG00000177125 | ZBTB34 | 0,065014148 | 1,046095199 | 0,84815745 | 0,987036295 | zinc finger and BTB domain containing 34 |
| ENSG00000277450 | AC002094,4 | 0,064971529 | 1,046064296 | 0,93316241 | 0,994479239 | novel transcript, sense intronic to transcript readthrough between TMEM199 and SARM1 |
| ENSG00000267491 | AC100788,1 | 0,064889728 | 1,046004986 | 0,94753255 | 0,996107107 | novel transcript, antisense to TIMP2 |
| ENSG00000138138 | ATAD1 | 0,064868442 | 1,045989553 | 0,87616213 | 0,990109814 | ATPase family AAA domain containing 1 |
| ENSG00000283024 | ATAD1 | 0,064868442 | 1,045989553 | 0,87616213 | 0,990109814 | ATPase family AAA domain containing 1 |
| ENSG00000215883 | CYB5RL | 0,064860963 | 1,045984131 | 0,79193336 | 0,979471509 | cytochrome b5 reductase like |
| ENSG00000172530 | BANP | 0,064831524 | 1,045962787 | 0,80189892 | 0,980710009 | BTG3 associated nuclear protein |
| ENSG00000008853 | RHOBTB2 | 0,064803116 | 1,045942191 | 0,8629094 | 0,989074806 | Rho related BTB domain containing 2 |
| ENSG00000271314 | AL161729,2 | 0,064731367 | 1,045890175 | 0,91902568 | 0,993585187 | novel transcript, antisense to PTCH1 |
| ENSG00000254812 | AC067930,2 | 0,064551747 | 1,045759966 | 0,94353166 | 0,995402494 | novel transcript |
| ENSG00000145220 | LYAR | 0,064309819 | 1,045584615 | 0,78963305 | 0,979462318 | Ly1 antibody reactive |
| ENSG00000041988 | THAP3 | 0,064272958 | 1,045557901 | 0,83998363 | 0,985672131 | THAP domain containing 3 |
| ENSG00000224414 | AC010886,1 | 0,064266882 | 1,045553498 | 0,93562021 | 0,994479239 | ATPase, H+/K+ transporting, nongastric, alpha polypeptide (ATP12A) pseudogene |
| ENSG00000204084 | INPP5B | 0,064207622 | 1,045510551 | 0,78521359 | 0,979320406 | inositol polyphosphate-5-phosphatase B |
| ENSG00000115282 | TTC31 | 0,064176518 | 1,045488011 | 0,77592003 | 0,978587255 | tetratricopeptide repeat domain 31 |
| ENSG00000168393 | DTYMK | 0,064121386 | 1,045448059 | 0,82574321 | 0,983383773 | deoxythymidylate kinase |
| ENSG00000162227 | TAF6L | 0,064098584 | 1,045431535 | 0,84623991 | 0,986939308 | TATA-box binding protein associated factor 6 like |
| ENSG00000286307 | AC017028,1 | 0,064050302 | 1,045396549 | 0,94233956 | 0,995373376 | novel transcript, antisense to HDAC4 |
| ENSG00000133393 | CEP20 | 0,064042411 | 1,045390831 | 0,86488408 | 0,989074806 | centrosomal protein 20 |
| ENSG00000178921 | PFAS | 0,06400837 | 1,045366165 | 0,87340375 | 0,989633545 | phosphoribosylformylglycinamidine synthase |

| | | | | | | |
|---|---|---|---|---|---|---|
| ENSG00000176716 | OR10AB1P | 0,063943219 | 1,045318958 | 0,96186157 | 0,996608711 | olfactory receptor family 10 subfamily AB member 1 pseudogene |
| ENSG00000131067 | GGT7 | 0,063928728 | 1,045308458 | 0,80071097 | 0,980505307 | gamma-glutamyltransferase 7 |
| ENSG00000286497 | AP001924,2 | 0,063914971 | 1,045298491 | 0,96011786 | 0,996608711 | novel transcript |
| ENSG00000180992 | MRPL14 | 0,06388956 | 1,04528008 | 0,78760163 | 0,979462318 | mitochondrial ribosomal protein L14 |
| ENSG00000082213 | C5orf22 | 0,063887064 | 1,045278271 | 0,88559162 | 0,990717729 | chromosome 5 open reading frame 22 |
| ENSG00000232846 | SLC25A6P3 | 0,063867689 | 1,045264233 | 0,95355498 | 0,99652169 | solute carrier family 25 member 6 pseudogene 3 |
| ENSG00000283201 | AC092329,3 | 0,063846259 | 1,045248707 | 0,96043679 | 0,996608711 | zinc finger protein 724 |
| ENSG00000259458 | MGC15885 | 0,063810047 | 1,045222472 | 0,93825464 | 0,994547792 | uncharacterized protein MGC15885 |
| ENSG00000277072 | STAG3L2 | 0,063781054 | 1,045201466 | 0,8129931 | 0,981246822 | stromal antigen 3-like 2 (pseudogene) |
| ENSG00000116161 | CACYBP | 0,063757391 | 1,045184323 | 0,86590788 | 0,989074806 | calcyclin binding protein |
| ENSG00000174939 | ASPHD1 | 0,063710466 | 1,045150329 | 0,92110692 | 0,993787755 | aspartate beta-hydroxylase domain containing 1 |
| ENSG00000172115 | CYCS | 0,063707942 | 1,0451485 | 0,87863945 | 0,990109814 | cytochrome c, somatic |
| ENSG00000154274 | C4orf19 | 0,063701477 | 1,045143816 | 0,89856007 | 0,991832796 | chromosome 4 open reading frame 19 |
| ENSG00000048342 | CC2D2A | 0,063587309 | 1,045061112 | 0,75454639 | 0,975901808 | coiled-coil and C2 domain containing 2A |
| ENSG00000185033 | SEMA4B | 0,063439131 | 1,04495378 | 0,81939589 | 0,98249617 | semaphorin 4B |
| ENSG00000126070 | AGO3 | 0,063390101 | 1,044918268 | 0,76747362 | 0,977201891 | argonaute RISC catalytic component 3 |
| ENSG00000223646 | AC002463,1 | 0,063385841 | 1,044915182 | 0,95434583 | 0,996608711 | novel transcript |
| ENSG00000270808 | AC022400,3 | 0,063370336 | 1,044903953 | 0,96105583 | 0,996608711 | ataxin 1-like (ATXN1L) pseudogene |
| ENSG00000103356 | EARS2 | 0,063350986 | 1,044889938 | 0,78149173 | 0,978597905 | glutamyl-tRNA synthetase 2, mitochondrial |
| ENSG00000123570 | RAB9B | 0,063300003 | 1,044853014 | 0,91089584 | 0,993222655 | RAB9B, member RAS oncogene family |
| ENSG00000272808 | AC015712,6 | 0,063268653 | 1,044830309 | 0,92603315 | 0,993915537 | uncharacterized LOC105369201 |
| ENSG00000141027 | NCOR1 | 0,063257844 | 1,044822481 | 0,81457218 | 0,981521413 | nuclear receptor corepressor 1 |
| ENSG00000229146 | SNX18P4 | 0,063204454 | 1,044783816 | 0,95731107 | 0,996608711 | sorting nexin 18 pseudogene 4 |
| ENSG00000156052 | GNAQ | 0,06319735 | 1,044778671 | 0,87117349 | 0,989217334 | G protein subunit alpha q |
| ENSG00000166225 | FRS2 | 0,063099385 | 1,044707729 | 0,89601493 | 0,991528124 | fibroblast growth factor receptor substrate 2 |
| ENSG00000227502 | MROCKI | 0,06306517 | 1,044682953 | 0,87658765 | 0,990109814 | MARCKS cis regulating lncRNA promoter of cytokines and inflammation |
| ENSG00000120049 | KCNIP2 | 0,063025002 | 1,044653867 | 0,84984997 | 0,987036295 | potassium voltage-gated channel interacting protein 2 |
| ENSG00000139687 | RB1 | 0,063014979 | 1,044646609 | 0,88707899 | 0,990717729 | RB transcriptional corepressor 1 |
| ENSG00000115998 | C2orf42 | 0,063014304 | 1,04464612 | 0,81268494 | 0,981246822 | chromosome 2 open reading frame 42 |
| ENSG00000108523 | RNF167 | 0,063002794 | 1,044637786 | 0,76704456 | 0,977201891 | ring finger protein 167 |

| | | | | | | |
|---|---|---|---|---|---|---|
| ENSG00000124181 | PLCG1 | 0,062985126 | 1,044624993 | 0,76957223 | 0,977683747 | phospholipase C gamma 1 |
| ENSG00000277005 | AC006449,4 | 0,06277731 | 1,044474529 | 0,91477984 | 0,993361299 | |
| ENSG00000095906 | NUBP2 | 0,062726325 | 1,044437617 | 0,86683979 | 0,989074806 | nucleotide binding protein 2 |
| ENSG00000244694 | PTCHD4 | 0,062659665 | 1,04438936 | 0,87014775 | 0,989074806 | patched domain containing 4 |
| ENSG00000131375 | CAPN7 | 0,062642332 | 1,044376813 | 0,86944011 | 0,989074806 | calpain 7 |
| ENSG00000148814 | LRRC27 | 0,06256844 | 1,044323323 | 0,84936815 | 0,987036295 | leucine rich repeat containing 27 |
| ENSG00000249245 | NCOA4P3 | 0,062490413 | 1,044266843 | 0,92276644 | 0,993797669 | nuclear receptor coactivator 4 pseudogene 3 |
| ENSG00000220804 | LINC01881 | 0,062432894 | 1,04422521 | 0,85785149 | 0,98893322 | long intergenic non-protein coding RNA 1881 |
| ENSG00000258692 | SALL4P7 | 0,062416402 | 1,044213273 | 0,96257061 | 0,996608711 | spalt like transcription factor 4 pseudogene 7 |
| ENSG00000235171 | HNRNPA1P2 | 0,062416402 | 1,044213273 | 0,96257061 | 0,996608711 | heterogeneous nuclear ribonucleoprotein A1 pseudogene 2 |
| ENSG00000231275 | HNRNPA1P2 | 0,062416402 | 1,044213273 | 0,96257061 | 0,996608711 | heterogeneous nuclear ribonucleoprotein A1 pseudogene 2 |
| ENSG00000231863 | AL139393,1 | 0,062403374 | 1,044203844 | 0,91828647 | 0,993367295 | novel transcript |
| ENSG00000101448 | EPPIN | 0,062191368 | 1,044050408 | 0,89989502 | 0,991845106 | epididymal peptidase inhibitor |
| ENSG00000166326 | TRIM44 | 0,062178384 | 1,044041012 | 0,78220962 | 0,978597905 | tripartite motif containing 44 |
| ENSG00000249787 | AC113385,1 | 0,061945748 | 1,043872672 | 0,95560564 | 0,996608711 | novel transcript |
| ENSG00000059758 | CDK17 | 0,061927506 | 1,043859473 | 0,86621486 | 0,989074806 | cyclin dependent kinase 17 |
| ENSG00000165661 | QSOX2 | 0,06186733 | 1,043815934 | 0,78171519 | 0,978597905 | quiescin sulfhydryl oxidase 2 |
| ENSG00000115970 | THADA | 0,061808071 | 1,04377306 | 0,81298654 | 0,981246822 | THADA armadillo repeat containing |
| ENSG00000279762 | AC005899,8 | 0,061804939 | 1,043770794 | 0,85265114 | 0,98761642 | novel transcript, antisense to PSMD11 |
| ENSG00000272325 | NUDT3 | 0,061709621 | 1,043701835 | 0,82104383 | 0,982753437 | nudix hydrolase 3 |
| ENSG00000086827 | ZW10 | 0,061657772 | 1,043664326 | 0,87149921 | 0,989217334 | zw10 kinetochore protein |
| ENSG00000126453 | BCL2L12 | 0,061634125 | 1,04364722 | 0,84917848 | 0,987036295 | BCL2 like 12 |
| ENSG00000156990 | RPUSD3 | 0,061632038 | 1,04364571 | 0,78427121 | 0,979320406 | RNA pseudouridine synthase D3 |
| ENSG00000137414 | FAM8A1 | 0,061627092 | 1,043642132 | 0,8940359 | 0,991135494 | family with sequence similarity 8 member A1 |
| ENSG00000259100 | COILP1 | 0,061621859 | 1,043638346 | 0,91797119 | 0,993361299 | coilin pseudogene 1 |
| ENSG00000102572 | STK24 | 0,061591156 | 1,043616136 | 0,83872761 | 0,985672131 | serine/threonine kinase 24 |
| ENSG00000232815 | DUX4L50 | 0,061589275 | 1,043614775 | 0,90855002 | 0,992603769 | double homeobox 4 like 50 (pseudogene) |
| ENSG00000090889 | KIF4A | 0,061542559 | 1,043580982 | 0,76835385 | 0,977381742 | kinesin family member 4A |
| ENSG00000085832 | EPS15 | 0,06146305 | 1,043523471 | 0,86102916 | 0,989074806 | epidermal growth factor receptor pathway substrate 15 |
| ENSG00000285813 | AP000813,1 | 0,061428613 | 1,043498562 | 0,8808768 | 0,990566588 | novel transcript, antisense to MSANTD4 |

| | | | | | | |
|---|---|---|---|---|---|---|
| ENSG00000256591 | AP003108,2 | 0,061417955 | 1,043490854 | 0,81566559 | 0,981590958 | novel transcript |
| ENSG00000172006 | ZNF554 | 0,061415477 | 1,043489061 | 0,85660292 | 0,988299298 | zinc finger protein 554 |
| ENSG00000233930 | KRTAP5-AS1 | 0,061252127 | 1,043370918 | 0,90591643 | 0,992590661 | KRTAP5-1/KRTAP5-2 antisense RNA 1 |
| ENSG00000198793 | MTOR | 0,061240883 | 1,043362787 | 0,76231932 | 0,976315588 | mechanistic target of rapamycin kinase |
| ENSG00000167283 | ATP5MG | 0,061138372 | 1,043288653 | 0,83054045 | 0,984156028 | ATP synthase membrane subunit g |
| ENSG00000288218 | AL358787,2 | 0,061085401 | 1,043250347 | 0,96697709 | 0,997263201 | SET domain and mariner transposase fusion gene (SETMAR) pseudogene |
| ENSG00000110844 | PRPF40B | 0,060934868 | 1,043141499 | 0,82871519 | 0,983784613 | pre-mRNA processing factor 40 homolog B |
| ENSG00000189233 | NUGGC | 0,060909466 | 1,043123132 | 0,94155034 | 0,99535504 | nuclear GTPase, germinal center associated |
| ENSG00000246366 | LACTB2-AS1 | 0,06085793 | 1,04308587 | 0,87397916 | 0,989711917 | LACTB2 antisense RNA 1 |
| ENSG00000288093 | AL592295,5 | 0,060775688 | 1,04302641 | 0,89953018 | 0,991845106 | novel transcript |
| ENSG00000238196 | NOTCH4 | 0,060769505 | 1,04302194 | 0,90638133 | 0,992590661 | notch receptor 4 |
| ENSG00000281207 | SLFNL1-AS1 | 0,06072546 | 1,042990097 | 0,90822484 | 0,992595473 | SLFNL1 antisense RNA 1 |
| ENSG00000254589 | EIF4A2P3 | 0,060711332 | 1,042979884 | 0,9535056 | 0,99652169 | eukaryotic translation initiation factor 4A2 pseudogene 3 |
| ENSG00000144410 | CPO | 0,060695625 | 1,042968528 | 0,96413499 | 0,996980762 | carboxypeptidase O |
| ENSG00000085449 | WDFY1 | 0,060669633 | 1,042949738 | 0,87004824 | 0,989074806 | WD repeat and FYVE domain containing 1 |
| ENSG00000140836 | ZFHX3 | 0,060650007 | 1,04293555 | 0,84543599 | 0,986639902 | zinc finger homeobox 3 |
| ENSG00000135900 | MRPL44 | 0,060643393 | 1,042930769 | 0,80418408 | 0,980875302 | mitochondrial ribosomal protein L44 |
| ENSG00000129749 | CHRNA10 | 0,060581616 | 1,042886111 | 0,90993286 | 0,992951496 | cholinergic receptor nicotinic alpha 10 subunit |
| ENSG00000183458 | PKD1P3 | 0,060523799 | 1,042844318 | 0,90836521 | 0,992595616 | polycystin 1, transient receptor potential channel interacting pseudogene 3 |
| ENSG00000186310 | NAP1L3 | 0,060521703 | 1,042842802 | 0,85389151 | 0,987861659 | nucleosome assembly protein 1 like 3 |
| ENSG00000143612 | C1orf43 | 0,060456584 | 1,042795733 | 0,83281468 | 0,984905093 | chromosome 1 open reading frame 43 |
| ENSG00000111450 | STX2 | 0,060446647 | 1,04278855 | 0,83248775 | 0,98489473 | syntaxin 2 |
| ENSG00000203879 | GDI1 | 0,060374682 | 1,042736535 | 0,8023164 | 0,980710009 | GDP dissociation inhibitor 1 |
| ENSG00000173276 | ZBTB21 | 0,060338021 | 1,042710038 | 0,86005798 | 0,989074806 | zinc finger and BTB domain containing 21 |
| ENSG00000258539 | AC068896,1 | 0,060268941 | 1,042660111 | 0,89079946 | 0,990717729 | novel transcript, METTL10-FAM53B readthrough |
| ENSG00000241258 | CRCP | 0,060257718 | 1,042652 | 0,80023769 | 0,980161849 | CGRP receptor component |
| ENSG00000169126 | ARMC4 | 0,06022982 | 1,042631838 | 0,84044214 | 0,985672131 | armadillo repeat containing 4 |
| ENSG00000185730 | ZNF696 | 0,060188907 | 1,04260227 | 0,86781016 | 0,989074806 | zinc finger protein 696 |
| ENSG00000235033 | DAAM2-AS1 | 0,06008892 | 1,042530015 | 0,90221884 | 0,992230188 | DAAM2 antisense RNA 1 |
| ENSG00000213695 | RPS7P14 | 0,060046284 | 1,042499206 | 0,96603949 | 0,997263201 | ribosomal protein S7 pseudogene 14 |

| | | | | | | |
|---|---|---|---|---|---|---|
| ENSG00000174996 | KLC2 | 0,060026177 | 1,042484676 | 0,86899185 | 0,989074806 | kinesin light chain 2 |
| ENSG00000121851 | POLR3GL | 0,0599882 | 1,042457234 | 0,81351425 | 0,981428111 | RNA polymerase III subunit GL |
| ENSG00000213853 | EMP2 | 0,059952731 | 1,042431605 | 0,80849409 | 0,981213877 | epithelial membrane protein 2 |
| ENSG00000239696 | AC092104,1 | 0,059713789 | 1,04225897 | 0,96035517 | 0,996608711 | zinc finger protein 655 (ZNF655) pseudogene |
| ENSG00000105732 | ZNF574 | 0,059690166 | 1,042241905 | 0,85679013 | 0,988312374 | zinc finger protein 574 |
| ENSG00000240005 | STIM2-AS1 | 0,05968676 | 1,042239444 | 0,95247098 | 0,996432861 | STIM2 antisense RNA 1 |
| ENSG00000113387 | SUB1 | 0,059680183 | 1,042234692 | 0,87885763 | 0,990109814 | SUB1 regulator of transcription |
| ENSG00000146540 | C7orf50 | 0,05964751 | 1,042211089 | 0,8740387 | 0,989711917 | chromosome 7 open reading frame 50 |
| ENSG00000001167 | NFYA | 0,05961174 | 1,042185249 | 0,85887905 | 0,989074806 | nuclear transcription factor Y subunit alpha |
| ENSG00000133731 | IMPA1 | 0,059595116 | 1,04217324 | 0,8843486 | 0,990717729 | inositol monophosphatase 1 |
| ENSG00000142168 | SOD1 | 0,059574673 | 1,042158473 | 0,84313786 | 0,986218419 | superoxide dismutase 1 |
| ENSG00000138231 | DBR1 | 0,059571697 | 1,042156323 | 0,83873876 | 0,985672131 | debranching RNA lariats 1 |
| ENSG00000130349 | MTRES1 | 0,059526826 | 1,04212391 | 0,83586717 | 0,985102385 | mitochondrial transcription rescue factor 1 |
| ENSG00000029725 | RABEP1 | 0,059348374 | 1,041995014 | 0,84601051 | 0,98689126 | rabaptin, RAB GTPase binding effector protein 1 |
| ENSG00000181773 | GPR3 | 0,059279372 | 1,041945178 | 0,83514055 | 0,984905093 | G protein-coupled receptor 3 |
| ENSG00000188613 | NANOS1 | 0,059204523 | 1,041891122 | 0,83912009 | 0,985672131 | nanos C2HC-type zinc finger 1 |
| ENSG00000273199 | AP000692,2 | 0,059196531 | 1,04188535 | 0,91571702 | 0,993361299 | novel transcript, antisense to MORC3 |
| ENSG00000179456 | ZBTB18 | 0,059184509 | 1,041876668 | 0,82049727 | 0,982609114 | zinc finger and BTB domain containing 18 |
| ENSG00000143333 | RGS16 | 0,059019138 | 1,041757248 | 0,86695407 | 0,989074806 | regulator of G protein signaling 16 |
| ENSG00000116830 | TTF2 | 0,058813825 | 1,041609004 | 0,79074827 | 0,979471509 | transcription termination factor 2 |
| ENSG00000258951 | KRT18P7 | 0,05879395 | 1,041594654 | 0,91359194 | 0,993361299 | keratin 18 pseudogene 7 |
| ENSG00000105607 | GCDH | 0,058744554 | 1,041558992 | 0,8409028 | 0,985672131 | glutaryl-CoA dehydrogenase |
| ENSG00000263934 | SNORD3A | 0,058709165 | 1,041533443 | 0,8956514 | 0,991377987 | small nucleolar RNA, C/D box 3A |
| ENSG00000063177 | RPL18 | 0,058685886 | 1,041516638 | 0,8163089 | 0,981806631 | ribosomal protein L18 |
| ENSG00000147533 | GOLGA7 | 0,058677498 | 1,041510582 | 0,89127874 | 0,990820775 | golgin A7 |
| ENSG00000283463 | HSFX4 | 0,058677389 | 1,041510504 | 0,95335575 | 0,99652169 | heat shock transcription factor family, X-linked member 4 |
| ENSG00000256139 | AC007637,1 | 0,05863334 | 1,041478704 | 0,90312376 | 0,992311172 | novel transcript |
| ENSG00000109670 | FBXW7 | 0,058557515 | 1,041423968 | 0,83328653 | 0,984905093 | F-box and WD repeat domain containing 7 |
| ENSG00000090263 | MRPS33 | 0,058490926 | 1,041375901 | 0,85770717 | 0,988897417 | mitochondrial ribosomal protein S33 |
| ENSG00000058056 | USP13 | 0,058486543 | 1,041372737 | 0,82786171 | 0,983595244 | ubiquitin specific peptidase 13 |
| ENSG00000247373 | TMED2-DT | 0,058448191 | 1,041345054 | 0,90946902 | 0,992864002 | TMED2 divergent transcript |
| ENSG00000132661 | NXT1 | 0,058414646 | 1,041320841 | 0,81119394 | 0,981213877 | nuclear transport factor 2 like export factor 1 |

| | | | | | | |
|---|---|---|---|---|---|---|
| ENSG00000198881 | ASB12 | 0,058364564 | 1,041284693 | 0,95043375 | 0,996432861 | ankyrin repeat and SOCS box containing 12 |
| ENSG00000164031 | DNAJB14 | 0,058363062 | 1,041283609 | 0,90003912 | 0,991845106 | DnaJ heat shock protein family (Hsp40) member B14 |
| ENSG00000197818 | SLC9A8 | 0,058334982 | 1,041263342 | 0,80433158 | 0,980875302 | solute carrier family 9 member A8 |
| ENSG00000237840 | FAM21FP | 0,058323774 | 1,041255253 | 0,92524564 | 0,993915537 | family with sequence similarity 21 member F, pseudogene |
| ENSG00000275468 | AC004678,1 | 0,058272491 | 1,041218241 | 0,94945664 | 0,996432861 | novel pseudogene |
| ENSG00000215421 | ZNF407 | 0,05827076 | 1,041216991 | 0,80254862 | 0,980798762 | zinc finger protein 407 |
| ENSG00000211584 | SLC48A1 | 0,058230107 | 1,041187652 | 0,83146706 | 0,984440275 | solute carrier family 48 member 1 |
| ENSG00000230732 | AC016949,1 | 0,058222204 | 1,041181948 | 0,85346762 | 0,987861659 | novel transcript |
| ENSG00000120008 | WDR11 | 0,058109323 | 1,041100486 | 0,84767229 | 0,987035044 | WD repeat domain 11 |
| ENSG00000275903 | PSMB3 | 0,05810112 | 1,041094566 | 0,87528105 | 0,990078774 | proteasome 20S subunit beta 3 |
| ENSG00000282770 | DLG5-AS1 | 0,058007662 | 1,041027126 | 0,90793347 | 0,992590661 | DLG5 antisense RNA 1 |
| ENSG00000150637 | CD226 | 0,057957721 | 1,040991091 | 0,92133701 | 0,993787755 | CD226 molecule |
| ENSG00000122386 | ZNF205 | 0,057905182 | 1,040953181 | 0,87721506 | 0,990109814 | zinc finger protein 205 |
| ENSG00000142676 | RPL11 | 0,05783589 | 1,040903186 | 0,87397261 | 0,989711917 | ribosomal protein L11 |
| ENSG00000171469 | ZNF561 | 0,057810123 | 1,040884595 | 0,87185925 | 0,989217334 | zinc finger protein 561 |
| ENSG00000167258 | CDK12 | 0,057782688 | 1,040864801 | 0,80339245 | 0,980866933 | cyclin dependent kinase 12 |
| ENSG00000269956 | MKNK1-AS1 | 0,057769161 | 1,040855042 | 0,91934059 | 0,993598006 | MKNK1 antisense RNA 1 |
| ENSG00000083099 | LYRM2 | 0,057671281 | 1,040784427 | 0,82965945 | 0,984081562 | LYR motif containing 2 |
| ENSG00000099968 | BCL2L13 | 0,057658557 | 1,040775248 | 0,7752978 | 0,978587255 | BCL2 like 13 |
| ENSG00000269855 | RNF225 | 0,0575802 | 1,040718722 | 0,96801708 | 0,997263201 | ring finger protein 225 |
| ENSG00000250999 | AC136604,2 | 0,057503443 | 1,040663353 | 0,93301075 | 0,994479239 | novel transcript |
| ENSG00000177084 | POLE | 0,057461593 | 1,040633166 | 0,79585541 | 0,979879951 | DNA polymerase epsilon, catalytic subunit |
| ENSG00000160221 | GATD3A | 0,057417812 | 1,040601586 | 0,85337504 | 0,987861659 | glutamine amidotransferase like class 1 domain containing 3A |
| ENSG00000278214 | LINC02139 | 0,057272858 | 1,040497038 | 0,96649428 | 0,997263201 | long intergenic non-protein coding RNA 2139 |
| ENSG00000119402 | FBXW2 | 0,057200603 | 1,040444928 | 0,82140144 | 0,982950156 | F-box and WD repeat domain containing 2 |
| ENSG00000197756 | RPL37A | 0,057196117 | 1,040441692 | 0,81960535 | 0,98249617 | ribosomal protein L37a |
| ENSG00000123505 | AMD1 | 0,057174548 | 1,040426137 | 0,86584625 | 0,989074806 | adenosylmethionine decarboxylase 1 |
| ENSG00000268686 | AC010643,1 | 0,05714751 | 1,040406638 | 0,91408451 | 0,993361299 | novel transcript, antisense to DKKL1 |
| ENSG00000003756 | RBM5 | 0,057137471 | 1,040399399 | 0,80094168 | 0,980524734 | RNA binding motif protein 5 |
| ENSG00000086589 | RBM22 | 0,057131445 | 1,040395053 | 0,86324902 | 0,989074806 | RNA binding motif protein 22 |
| ENSG00000163154 | TNFAIP8L2 | 0,057119011 | 1,040386086 | 0,94019254 | 0,994949796 | TNF alpha induced protein 8 like 2 |

| | | | | | | |
|---|---|---|---|---|---|---|
| ENSG00000232586 | KIAA1614-AS1 | 0,057091001 | 1,040365888 | 0,91837278 | 0,99340194 | KIAA1614 antisense RNA 1 |
| ENSG00000228661 | STIM1-AS1 | 0,057058802 | 1,040342669 | 0,9521097 | 0,996432861 | STIM1 antisense RNA 1 |
| ENSG00000179981 | TSHZ1 | 0,057050002 | 1,040336323 | 0,85837539 | 0,989067345 | teashirt zinc finger homeobox 1 |
| ENSG00000105341 | DMAC2 | 0,056876597 | 1,040211287 | 0,7949745 | 0,979776981 | distal membrane arm assembly complex 2 |
| ENSG00000271933 | AL603756,1 | 0,056857499 | 1,040197517 | 0,88769982 | 0,990717729 | novel transcript, antisense to GHITM |
| ENSG00000101391 | CDK5RAP1 | 0,056800813 | 1,040156646 | 0,80341434 | 0,980866933 | CDK5 regulatory subunit associated protein 1 |
| ENSG00000263290 | SCAMP3 | 0,056642194 | 1,040042292 | 0,84912053 | 0,987036295 | secretory carrier membrane protein 3 |
| ENSG00000005471 | ABCB4 | 0,056581791 | 1,039998747 | 0,89040955 | 0,990717729 | ATP binding cassette subfamily B member 4 |
| ENSG00000122335 | SERAC1 | 0,056561045 | 1,039983793 | 0,86064682 | 0,989074806 | serine active site containing 1 |
| ENSG00000197771 | MCMBP | 0,05655347 | 1,039978332 | 0,87436609 | 0,98990147 | minichromosome maintenance complex binding protein |
| ENSG00000249962 | AL157400,5 | 0,056535078 | 1,039965074 | 0,91561971 | 0,993361299 | pantothenate kinase 1 (PANK1) pseudogene |
| ENSG00000234704 | BRD2 | 0,05648095 | 1,039926057 | 0,79183893 | 0,979471509 | bromodomain containing 2 |
| ENSG00000119820 | YIPF4 | 0,056478292 | 1,039924141 | 0,79043475 | 0,979462318 | Yip1 domain family member 4 |
| ENSG00000183495 | EP400 | 0,056378222 | 1,039852011 | 0,7917682 | 0,979471509 | E1A binding protein p400 |
| ENSG00000152332 | UHMK1 | 0,056371228 | 1,03984697 | 0,88694041 | 0,990717729 | U2AF homology motif kinase 1 |
| ENSG00000258745 | AL358334,3 | 0,05630479 | 1,039799085 | 0,96663113 | 0,997263201 | novel transcript, antisense to PYGL |
| ENSG00000101654 | RNMT | 0,056290864 | 1,039789047 | 0,84766297 | 0,987035044 | RNA guanine-7 methyltransferase |
| ENSG00000096384 | HSP90AB1 | 0,05628631 | 1,039785766 | 0,801551 | 0,980623173 | heat shock protein 90 alpha family class B member 1 |
| ENSG00000196862 | RGPD4 | 0,056263141 | 1,039769067 | 0,88653416 | 0,990717729 | RANBP2 like and GRIP domain containing 4 |
| ENSG00000144452 | ABCA12 | 0,056240383 | 1,039752665 | 0,91671152 | 0,993361299 | ATP binding cassette subfamily A member 12 |
| ENSG00000268392 | AC003682,1 | 0,056222644 | 1,03973988 | 0,95814838 | 0,996608711 | novel transcript |
| ENSG00000136521 | NDUFB5 | 0,05612794 | 1,03967163 | 0,86748826 | 0,989074806 | NADH:ubiquinone oxidoreductase subunit B5 |
| ENSG00000224086 | PPM1F-AS1 | 0,055981789 | 1,039566313 | 0,89439672 | 0,991135494 | PPM1F antisense RNA 1 |
| ENSG00000047315 | POLR2B | 0,055937548 | 1,039534434 | 0,88103175 | 0,990598309 | RNA polymerase II subunit B |
| ENSG00000087053 | MTMR2 | 0,055867357 | 1,039483859 | 0,88882089 | 0,990717729 | myotubularin related protein 2 |
| ENSG00000160803 | UBQLN4 | 0,055814895 | 1,03944606 | 0,80601287 | 0,981213877 | ubiquilin 4 |
| ENSG00000184209 | SNRNP35 | 0,055786685 | 1,039425736 | 0,84627615 | 0,986939308 | small nuclear ribonucleoprotein U11/U12 subunit 35 |
| ENSG00000010361 | FUZ | 0,055772446 | 1,039415477 | 0,83461035 | 0,984905093 | fuzzy planar cell polarity protein |
| ENSG00000121957 | GPSM2 | 0,055768518 | 1,039412647 | 0,8456688 | 0,986639902 | G protein signaling modulator 2 |
| ENSG00000156603 | MED19 | 0,0557603 | 1,039406726 | 0,8288299 | 0,983795764 | mediator complex subunit 19 |
| ENSG00000278743 | AC087239,1 | 0,055735548 | 1,039388893 | 0,93474224 | 0,994479239 | novel transcript |
| ENSG00000138593 | SECISBP2L | 0,055694429 | 1,039359269 | 0,88082989 | 0,990566588 | SECIS binding protein 2 like |

| | | | | | | |
|---|---|---|---|---|---|---|
| ENSG00000280356 | AL021877,2 | 0,055662062 | 1,039335952 | 0,96089946 | 0,996608711 | novel transcript |
| ENSG00000138134 | STAMBPL1 | 0,055656592 | 1,039332011 | 0,8493919 | 0,987036295 | STAM binding protein like 1 |
| ENSG00000274750 | H3C6 | 0,055623761 | 1,03930836 | 0,88863167 | 0,990717729 | H3 clustered histone 6 |
| ENSG00000114544 | SLC41A3 | 0,055543534 | 1,039250566 | 0,81347344 | 0,981416747 | solute carrier family 41 member 3 |
| ENSG00000167272 | POP5 | 0,055477298 | 1,039202854 | 0,82213102 | 0,982952211 | POP5 homolog, ribonuclease P/MRP subunit |
| ENSG00000197457 | STMN3 | 0,055474824 | 1,039201072 | 0,86401536 | 0,989074806 | stathmin 3 |
| ENSG00000241468 | ATP5MF | 0,055465964 | 1,03919469 | 0,86119437 | 0,989074806 | ATP synthase membrane subunit f |
| ENSG00000198730 | CTR9 | 0,055424485 | 1,039164813 | 0,84737918 | 0,987035044 | CTR9 homolog, Paf1/RNA polymerase II complex component |
| ENSG00000163900 | TMEM41A | 0,055410698 | 1,039154882 | 0,80738015 | 0,981213877 | transmembrane protein 41A |
| ENSG00000207145 | SNORA18 | 0,055406662 | 1,039151974 | 0,96286161 | 0,996608711 | small nucleolar RNA, H/ACA box 18 |
| ENSG00000108591 | DRG2 | 0,055316905 | 1,039087326 | 0,81308004 | 0,981246822 | developmentally regulated GTP binding protein 2 |
| ENSG00000128463 | EMC4 | 0,05519721 | 1,03900112 | 0,82856759 | 0,983721455 | ER membrane protein complex subunit 4 |
| ENSG00000288150 | AC090826,4 | 0,05516177 | 1,038975597 | 0,98332152 | 0,99837368 | novel transcript, antisense to CYP11A1 |
| ENSG00000120742 | SERP1 | 0,055089939 | 1,038923869 | 0,88085223 | 0,990566588 | stress associated endoplasmic reticulum protein 1 |
| ENSG00000258584 | FAM181A-AS1 | 0,055078856 | 1,038915888 | 0,89524545 | 0,991303328 | FAM181A antisense RNA 1 |
| ENSG00000251410 | AC007106,1 | 0,055049103 | 1,038894462 | 0,90861828 | 0,992603769 | novel transcript |
| ENSG00000154767 | XPC | 0,055045238 | 1,038891679 | 0,83509308 | 0,984905093 | XPC complex subunit, DNA damage recognition and repair factor |
| ENSG00000239335 | LLPH-DT | 0,055036529 | 1,038885407 | 0,94216276 | 0,995373376 | LLPH divergent transcript |
| ENSG00000051596 | THOC3 | 0,055025198 | 1,038877248 | 0,83776453 | 0,98566906 | THO complex 3 |
| ENSG00000104643 | MTMR9 | 0,054952083 | 1,0388246 | 0,88189629 | 0,990717729 | myotubularin related protein 9 |
| ENSG00000133103 | COG6 | 0,054902721 | 1,038789057 | 0,90052809 | 0,991865079 | component of oligomeric golgi complex 6 |
| ENSG00000100567 | PSMA3 | 0,054849414 | 1,038750675 | 0,9006746 | 0,991865079 | proteasome 20S subunit alpha 3 |
| ENSG00000130414 | NDUFA10 | 0,054834808 | 1,038740159 | 0,80584412 | 0,981213877 | NADH:ubiquinone oxidoreductase subunit A10 |
| ENSG00000205937 | RNPS1 | 0,054736356 | 1,038669275 | 0,79402484 | 0,979776981 | RNA binding protein with serine rich domain 1 |
| ENSG00000111640 | GAPDH | 0,05472142 | 1,038658522 | 0,82568354 | 0,983383773 | glyceraldehyde-3-phosphate dehydrogenase |
| ENSG00000262312 | AC004494,1 | 0,054638635 | 1,038598924 | 0,93821935 | 0,994547792 | novel transcript, antisense to NLRC3 |
| ENSG00000154429 | CCSAP | 0,05457572 | 1,038553632 | 0,8768313 | 0,990109814 | centriole, cilia and spindle associated protein |
| ENSG00000103657 | HERC1 | 0,054549114 | 1,038534479 | 0,81024645 | 0,981213877 | HECT and RLD domain containing E3 ubiquitin protein ligase family member 1 |
| ENSG00000226954 | AL023802,1 | 0,054509008 | 1,038505609 | 0,95578647 | 0,996608711 | novel transcript |
| ENSG00000083223 | TUT7 | 0,054507275 | 1,038504361 | 0,84733686 | 0,987035044 | terminal uridylyl transferase 7 |

| | | | | | | |
|---|---|---|---|---|---|---|
| ENSG00000156873 | PHKG2 | 0,054472224 | 1,038479131 | 0,83403286 | 0,984905093 | phosphorylase kinase catalytic subunit gamma 2 |
| ENSG00000116874 | WARS2 | 0,054374124 | 1,038408519 | 0,88394624 | 0,990717729 | tryptophanyl tRNA synthetase 2, mitochondrial |
| ENSG00000056586 | RC3H2 | 0,054306869 | 1,038360112 | 0,88445463 | 0,990717729 | ring finger and CCCH-type domains 2 |
| ENSG00000176473 | WDR25 | 0,054305564 | 1,038359172 | 0,8304334 | 0,984156028 | WD repeat domain 25 |
| ENSG00000149948 | HMGA2 | 0,054226191 | 1,038302047 | 0,91141205 | 0,993222655 | high mobility group AT-hook 2 |
| ENSG00000147684 | NDUFB9 | 0,054215422 | 1,038294296 | 0,78819774 | 0,979462318 | NADH:ubiquinone oxidoreductase subunit B9 |
| ENSG00000258301 | VASH1-AS1 | 0,054158026 | 1,038252989 | 0,91346483 | 0,993361299 | VASH1 antisense RNA 1 |
| ENSG00000115226 | FNDC4 | 0,054123641 | 1,038228244 | 0,87527565 | 0,990078774 | fibronectin type III domain containing 4 |
| ENSG00000286826 | AL136129,1 | 0,053918221 | 1,038080425 | 0,94621146 | 0,996027077 | novel transcript |
| ENSG00000058673 | ZC3H11A | 0,053914833 | 1,038077987 | 0,88007843 | 0,990268593 | zinc finger CCCH-type containing 11A |
| ENSG00000277311 | AC012354,7 | 0,053900598 | 1,038067745 | 0,95396772 | 0,996608711 | |
| ENSG00000005339 | CREBBP | 0,053894546 | 1,038063391 | 0,80311237 | 0,980866933 | CREB binding protein |
| ENSG00000263731 | AC145207,5 | 0,053870619 | 1,038046174 | 0,86411003 | 0,989074806 | novel transcript |
| ENSG00000170385 | SLC30A1 | 0,053831193 | 1,038017807 | 0,890402 | 0,990717729 | solute carrier family 30 member 1 |
| ENSG00000005189 | REXO5 | 0,053830071 | 1,038017 | 0,84526938 | 0,986639902 | RNA exonuclease 5 |
| ENSG00000143947 | RPS27A | 0,053796114 | 1,037992568 | 0,81982465 | 0,98249617 | ribosomal protein S27a |
| ENSG00000258643 | BCL2L2-PABPN | 0,053785598 | 1,037985002 | 0,91855835 | 0,993481057 | BCL2L2-PABPN1 readthrough |
| ENSG00000176896 | TCEANC | 0,053723395 | 1,037940249 | 0,88875105 | 0,990717729 | transcription elongation factor A N-terminal and central domain containing |
| ENSG00000255517 | AP002748,4 | 0,053701458 | 1,037924467 | 0,89356985 | 0,991135494 | novel transcript |
| ENSG00000137992 | DBT | 0,053630497 | 1,037873417 | 0,8511928 | 0,987259878 | dihydrolipoamide branched chain transacylase E2 |
| ENSG00000100577 | GSTZ1 | 0,053613561 | 1,037861233 | 0,83310983 | 0,984905093 | glutathione S-transferase zeta 1 |
| ENSG00000149716 | LTO1 | 0,053504157 | 1,037782532 | 0,83361683 | 0,984905093 | LTO1 maturation factor of ABCE1 |
| ENSG00000249502 | AC006160,1 | 0,053465587 | 1,037754787 | 0,90415451 | 0,992538156 | novel transcript, antisense to LAP3 |
| ENSG00000257920 | LINC02409 | 0,053413747 | 1,037717498 | 0,95336979 | 0,99652169 | long intergenic non-protein coding RNA 2409 |
| ENSG00000205746 | PKD1P4 | 0,053327506 | 1,037655468 | 0,90862241 | 0,992603769 | polycystin 1, transient receptor potential channel interacting pseudogene 4 |
| ENSG00000099194 | SCD | 0,053315297 | 1,037646687 | 0,85867317 | 0,989074806 | stearoyl-CoA desaturase |
| ENSG00000136875 | PRPF4 | 0,05316798 | 1,037540735 | 0,8108833 | 0,981213877 | pre-mRNA processing factor 4 |
| ENSG00000100664 | EIF5 | 0,053141082 | 1,037521391 | 0,87634862 | 0,990109814 | eukaryotic translation initiation factor 5 |
| ENSG00000285794 | AL031281,1 | 0,053099817 | 1,037491716 | 0,96805034 | 0,997263201 | SCAN domain containing 1 (SCAND1) pseudogene |
| ENSG00000106993 | CDC37L1 | 0,05307127 | 1,037471187 | 0,88410719 | 0,990717729 | cell division cycle 37 like 1 |

| | | | | | | |
|---|---|---|---|---|---|---|
| ENSG00000134318 | ROCK2 | 0,053067661 | 1,037468592 | 0,87757667 | 0,990109814 | Rho associated coiled-coil containing protein kinase 2 |
| ENSG00000101265 | RASSF2 | 0,053056113 | 1,037460287 | 0,88250344 | 0,990717729 | Ras association domain family member 2 |
| ENSG00000111481 | COPZ1 | 0,053055849 | 1,037460097 | 0,85417239 | 0,987861659 | COPI coat complex subunit zeta 1 |
| ENSG00000138696 | BMPR1B | 0,053047817 | 1,037454321 | 0,91141219 | 0,993222655 | bone morphogenetic protein receptor type 1B |
| ENSG00000184787 | UBE2G2 | 0,053028071 | 1,037440122 | 0,82568476 | 0,983383773 | ubiquitin conjugating enzyme E2 G2 |
| ENSG00000142657 | PGD | 0,05301828 | 1,037433082 | 0,79906334 | 0,980161849 | phosphogluconate dehydrogenase |
| ENSG00000108106 | UBE2S | 0,053004036 | 1,037422839 | 0,8443937 | 0,986340494 | ubiquitin conjugating enzyme E2 S |
| ENSG00000236447 | ATG10-IT1 | 0,052971859 | 1,037399701 | 0,95444626 | 0,996608711 | ATG10 intronic transcript 1 |
| ENSG00000135945 | REV1 | 0,05292255 | 1,037364245 | 0,88279907 | 0,990717729 | REV1 DNA directed polymerase |
| ENSG00000235535 | TRDN-AS1 | 0,052834687 | 1,03730107 | 0,94176695 | 0,995373376 | TRDN antisense RNA 1 |
| ENSG00000287576 | AC067930,9 | 0,052804451 | 1,03727933 | 0,89791142 | 0,991711719 | novel transcript |
| ENSG00000272419 | LINC01145 | 0,052798842 | 1,037275297 | 0,84477008 | 0,986557476 | long intergenic non-protein coding RNA 1145 |
| ENSG00000245112 | SMARCA5-AS1 | 0,052742922 | 1,037235092 | 0,87505821 | 0,990078774 | SMARCA5 antisense RNA 1 |
| ENSG00000057704 | TMCC3 | 0,052644433 | 1,037164285 | 0,91446207 | 0,993361299 | transmembrane and coiled-coil domain family 3 |
| ENSG00000225548 | LINC01980 | 0,052628857 | 1,037153088 | 0,9017949 | 0,992136926 | long intergenic non-protein coding RNA 1980 |
| ENSG00000262873 | AC127496,5 | 0,052604536 | 1,037135604 | 0,94056211 | 0,995084223 | novel transcript |
| ENSG00000224837 | GCSHP5 | 0,052587572 | 1,037123408 | 0,83205558 | 0,984653014 | glycine cleavage system protein H pseudogene 5 |
| ENSG00000278616 | BEND3P3 | 0,052580536 | 1,03711835 | 0,91098839 | 0,993222655 | BEN domain containing 3 pseudogene 3 |
| ENSG00000182070 | OR52A1 | 0,052560383 | 1,037103863 | 0,89518473 | 0,991303328 | olfactory receptor family 52 subfamily A member 1 |
| ENSG00000236430 | KRT8P29 | 0,052548028 | 1,037094981 | 0,95689487 | 0,996608711 | keratin 8 pseudogene 29 |
| ENSG00000167552 | TUBA1A | 0,052467168 | 1,037036856 | 0,81072908 | 0,981213877 | tubulin alpha 1a |
| ENSG00000236510 | AC011284,1 | 0,052425018 | 1,037006558 | 0,93454699 | 0,994479239 | novel transcript |
| ENSG00000230337 | EXOSC10-AS1 | 0,052382756 | 1,036976181 | 0,94800517 | 0,996216385 | EXOSC10 antisense RNA 1 |
| ENSG00000124380 | SNRNP27 | 0,052372092 | 1,036968516 | 0,8755539 | 0,990078774 | small nuclear ribonucleoprotein U4/U6,U5 subunit 27 |
| ENSG00000115204 | MPV17 | 0,052351108 | 1,036953433 | 0,8244158 | 0,983374097 | mitochondrial inner membrane protein MPV17 |
| ENSG00000185000 | DGAT1 | 0,052294073 | 1,03691244 | 0,86699143 | 0,989074806 | diacylglycerol O-acyltransferase 1 |
| ENSG00000285482 | DGAT1 | 0,052294073 | 1,03691244 | 0,86699143 | 0,989074806 | diacylglycerol O-acyltransferase 1 |
| ENSG00000127527 | EPS15L1 | 0,052275367 | 1,036898995 | 0,80892996 | 0,981213877 | epidermal growth factor receptor pathway substrate 15 like 1 |
| ENSG00000130227 | XPO7 | 0,052261512 | 1,036889037 | 0,87529714 | 0,990078774 | exportin 7 |
| ENSG00000175105 | ZNF654 | 0,052254275 | 1,036883835 | 0,91502245 | 0,993361299 | zinc finger protein 654 |

| | | | | | | |
|---|---|---|---|---|---|---|
| ENSG00000075399 | VPS9D1 | 0,052233507 | 1,036868909 | 0,87059277 | 0,989074806 | VPS9 domain containing 1 |
| ENSG00000130255 | RPL36 | 0,052179969 | 1,036830433 | 0,8190952 | 0,98249617 | ribosomal protein L36 |
| ENSG00000276336 | SCARF1 | 0,052162151 | 1,036817627 | 0,97276346 | 0,99837368 | scavenger receptor class F member 1 |
| ENSG00000213689 | TREX1 | 0,052085116 | 1,036762266 | 0,88203099 | 0,990717729 | three prime repair exonuclease 1 |
| ENSG00000092853 | CLSPN | 0,052043578 | 1,036732416 | 0,86535938 | 0,989074806 | claspin |
| ENSG00000179859 | RNF227 | 0,051984243 | 1,036689778 | 0,90015465 | 0,991845106 | ring finger protein 227 |
| ENSG00000102882 | MAPK3 | 0,051975536 | 1,036683521 | 0,84538294 | 0,986639902 | mitogen-activated protein kinase 3 |
| ENSG00000166851 | PLK1 | 0,051974195 | 1,036682558 | 0,83816665 | 0,985672131 | polo like kinase 1 |
| ENSG00000009780 | FAM76A | 0,051731911 | 1,036508474 | 0,86563785 | 0,989074806 | family with sequence similarity 76 member A |
| ENSG00000106554 | CHCHD3 | 0,051647246 | 1,036447648 | 0,84242931 | 0,986175464 | coiled-coil-helix-coiled-coil-helix domain containing 3 |
| ENSG00000105829 | BET1 | 0,051640186 | 1,036442576 | 0,90303355 | 0,992311172 | Bet1 golgi vesicular membrane trafficking protein |
| ENSG00000177000 | MTHFR | 0,051630931 | 1,036435927 | 0,84560532 | 0,986639902 | methylenetetrahydrofolate reductase |
| ENSG00000205531 | NAP1L4 | 0,051519103 | 1,036355592 | 0,88478784 | 0,990717729 | nucleosome assembly protein 1 like 4 |
| ENSG00000237649 | KIFC1 | 0,051435864 | 1,0362958 | 0,81107773 | 0,981213877 | kinesin family member C1 |
| ENSG00000114744 | COMMD2 | 0,051351256 | 1,036235027 | 0,89293371 | 0,991120066 | COMM domain containing 2 |
| ENSG00000121005 | CRISPLD1 | 0,05132824 | 1,036218496 | 0,896279 | 0,991636805 | cysteine rich secretory protein LCCL domain containing 1 |
| ENSG00000110395 | CBL | 0,051274198 | 1,036179681 | 0,83617567 | 0,985159643 | Cbl proto-oncogene |
| ENSG00000270629 | NBPF14 | 0,051222313 | 1,036142416 | 0,84360571 | 0,986273696 | NBPF member 14 |
| ENSG00000189091 | SF3B3 | 0,051204732 | 1,03612979 | 0,79947504 | 0,980161849 | splicing factor 3b subunit 3 |
| ENSG00000198755 | RPL10A | 0,051203341 | 1,036128791 | 0,85148896 | 0,987259878 | ribosomal protein L10a |
| ENSG00000204536 | CCHCR1 | 0,051197417 | 1,036124536 | 0,92934808 | 0,994293848 | coiled-coil alpha-helical rod protein 1 |
| ENSG00000224895 | VPS26BP1 | 0,051081169 | 1,036041052 | 0,92796806 | 0,994108531 | VPS26B pseudogene 1 |
| ENSG00000120437 | ACAT2 | 0,050965362 | 1,035957891 | 0,87094738 | 0,989217334 | acetyl-CoA acetyltransferase 2 |
| ENSG00000105364 | MRPL4 | 0,050963535 | 1,035956579 | 0,8433549 | 0,986260952 | mitochondrial ribosomal protein L4 |
| ENSG00000124608 | AARS2 | 0,050958425 | 1,035952909 | 0,89348969 | 0,991135494 | alanyl-tRNA synthetase 2, mitochondrial |
| ENSG00000252139 | AC090227,1 | 0,050916587 | 1,035922868 | 0,97387186 | 0,99837368 | |
| ENSG00000259994 | AL353796,1 | 0,050915226 | 1,03592189 | 0,89793447 | 0,991711719 | novel transcript, overlapping to MTPAP |
| ENSG00000066739 | ATG2B | 0,050828461 | 1,035859591 | 0,88266484 | 0,990717729 | autophagy related 2B |
| ENSG00000262188 | LINC01978 | 0,050821435 | 1,035854546 | 0,94125779 | 0,995251853 | long intergenic non-protein coding RNA 1978 |
| ENSG00000117399 | CDC20 | 0,050799808 | 1,035839018 | 0,83345756 | 0,984905093 | cell division cycle 20 |
| ENSG00000279213 | SNX18P10 | 0,050781331 | 1,035825752 | 0,95836874 | 0,996608711 | sorting nexin 18 pseudogene 10 |

| | | | | | | |
|---|---|---|---|---|---|---|
| ENSG00000159885 | ZNF222 | 0,050777848 | 1,035823251 | 0,88519133 | 0,990717729 | zinc finger protein 222 |
| ENSG00000179523 | EIF3J-DT | 0,050710736 | 1,035775067 | 0,87777119 | 0,990109814 | EIF3J divergent transcript |
| ENSG00000231560 | CLEC12A-AS1 | 0,050603523 | 1,035698097 | 0,92976525 | 0,994293848 | CLEC12A antisense RNA 1 |
| ENSG00000145191 | EIF2B5 | 0,050498139 | 1,035622446 | 0,80559882 | 0,981213877 | eukaryotic translation initiation factor 2B subunit epsilon |
| ENSG00000149305 | HTR3B | 0,050452187 | 1,03558946 | 0,94235045 | 0,995373376 | 5-hydroxytryptamine receptor 3B |
| ENSG00000204713 | TRIM27 | 0,05036509 | 1,035526942 | 0,90182149 | 0,992136926 | tripartite motif containing 27 |
| ENSG00000145147 | SLIT2 | 0,050322103 | 1,035496088 | 0,91742156 | 0,993361299 | slit guidance ligand 2 |
| ENSG00000112339 | HBS1L | 0,050260237 | 1,035451684 | 0,87061445 | 0,989074806 | HBS1 like translational GTPase |
| ENSG00000113013 | HSPA9 | 0,050220192 | 1,035422943 | 0,86981962 | 0,989074806 | heat shock protein family A (Hsp70) member 9 |
| ENSG00000139318 | DUSP6 | 0,050216657 | 1,035420406 | 0,88459699 | 0,990717729 | dual specificity phosphatase 6 |
| ENSG00000135070 | ISCA1 | 0,05019059 | 1,035401698 | 0,90447429 | 0,992590661 | iron-sulfur cluster assembly 1 |
| ENSG00000164604 | GPR85 | 0,050119051 | 1,035350357 | 0,91564416 | 0,993361299 | G protein-coupled receptor 85 |
| ENSG00000090266 | NDUFB2 | 0,050107075 | 1,035341763 | 0,81541028 | 0,981521413 | NADH:ubiquinone oxidoreductase subunit B2 |
| ENSG00000119906 | SLF2 | 0,050078671 | 1,035321379 | 0,8832191 | 0,990717729 | SMC5-SMC6 complex localization factor 2 |
| ENSG00000088340 | FER1L4 | 0,049971223 | 1,035244274 | 0,93727562 | 0,994479239 | fer-1 like family member 4 (pseudogene) |
| ENSG00000213071 | LPAL2 | 0,04991765 | 1,035205832 | 0,89306769 | 0,991135494 | lipoprotein(a) like 2, pseudogene |
| ENSG00000109814 | UGDH | 0,049883625 | 1,035181418 | 0,88415808 | 0,990717729 | UDP-glucose 6-dehydrogenase |
| ENSG00000107882 | SUFU | 0,049877589 | 1,035177087 | 0,84801205 | 0,987036295 | SUFU negative regulator of hedgehog signaling |
| ENSG00000183281 | PLGLB1 | 0,049872481 | 1,035173421 | 0,92109973 | 0,993787755 | plasminogen like B1 |
| ENSG00000251661 | AC136475,1 | 0,049851034 | 1,035158033 | 0,88987457 | 0,990717729 | novel transcript, antisense to IFITM3 |
| ENSG00000159214 | CCDC24 | 0,049831619 | 1,035144102 | 0,88559384 | 0,990717729 | coiled-coil domain containing 24 |
| ENSG00000259431 | THTPA | 0,04981807 | 1,035134381 | 0,86142437 | 0,989074806 | thiamine triphosphatase |
| ENSG00000153201 | RANBP2 | 0,049797468 | 1,035119599 | 0,8804782 | 0,990367704 | RAN binding protein 2 |
| ENSG00000168273 | SMIM4 | 0,049777183 | 1,035105044 | 0,85603648 | 0,988185425 | small integral membrane protein 4 |
| ENSG00000215458 | AATBC | 0,049746637 | 1,035083129 | 0,93251081 | 0,994479239 | apoptosis associated transcript in bladder cancer |
| ENSG00000089009 | RPL6 | 0,049700178 | 1,035049797 | 0,87679966 | 0,990109814 | ribosomal protein L6 |
| ENSG00000099804 | CDC34 | 0,049681109 | 1,035036116 | 0,835574 | 0,985034063 | cell division cycle 34, ubiqiutin conjugating enzyme |
| ENSG00000270890 | AL049844,2 | 0,049645831 | 1,035010806 | 0,93532989 | 0,994479239 | synaptophysin-like 1 (SYPL1) pseudogene |
| ENSG00000273108 | AL121929,2 | 0,049614587 | 1,034988392 | 0,93616607 | 0,994479239 | novel transcript, antisense to SH3PXD2A |
| ENSG00000124224 | PPP4R1L | 0,049597367 | 1,034976038 | 0,86078216 | 0,989074806 | protein phosphatase 4 regulatory subunit 1 like (pseudogene) |
| ENSG00000116266 | STXBP3 | 0,049538058 | 1,034933492 | 0,91007089 | 0,992965454 | syntaxin binding protein 3 |

| | | | | | | |
|---|---|---|---|---|---|---|
| ENSG00000187048 | CYP4A11 | 0,049506933 | 1,034911164 | 0,94274142 | 0,995402494 | cytochrome P450 family 4 subfamily A member 11 |
| ENSG00000274220 | AC009163,7 | 0,049414224 | 1,034844662 | 0,88966566 | 0,990717729 | novel transcript |
| ENSG00000163320 | CGGBP1 | 0,049323807 | 1,034779807 | 0,90994734 | 0,992951496 | CGG triplet repeat binding protein 1 |
| ENSG00000213371 | NAP1L1P3 | 0,049226001 | 1,034709658 | 0,97041383 | 0,998013218 | nucleosome assembly protein 1 like 1 pseudogene 3 |
| ENSG00000120129 | DUSP1 | 0,04921892 | 1,03470458 | 0,8400013 | 0,985672131 | dual specificity phosphatase 1 |
| ENSG00000214401 | KANSL1-AS1 | 0,049201849 | 1,034692336 | 0,92175849 | 0,993787755 | KANSL1 antisense RNA 1 |
| ENSG00000282799 | KANSL1-AS1 | 0,049201849 | 1,034692336 | 0,92175849 | 0,993787755 | KANSL1 antisense RNA 1 |
| ENSG00000179136 | LINC00670 | 0,049179277 | 1,034676148 | 0,94648572 | 0,996027077 | long intergenic non-protein coding RNA 670 |
| ENSG00000135763 | URB2 | 0,049162325 | 1,034663991 | 0,8405434 | 0,985672131 | URB2 ribosome biogenesis homolog |
| ENSG00000100324 | TAB1 | 0,049138726 | 1,034647066 | 0,87655143 | 0,990109814 | TGF-beta activated kinase 1 (MAP3K7) binding protein 1 |
| ENSG00000127948 | POR | 0,04911539 | 1,03463033 | 0,84994694 | 0,987036295 | cytochrome p450 oxidoreductase |
| ENSG00000164556 | FAM183BP | 0,049028076 | 1,034567715 | 0,96492023 | 0,997113804 | family with sequence similarity 183 member B, pseudogene |
| ENSG00000260790 | AC092338,2 | 0,048915173 | 1,034486754 | 0,9534591 | 0,99652169 | novel transcript, antisense to CDR2 |
| ENSG00000206562 | METTL6 | 0,048828003 | 1,034424251 | 0,82739282 | 0,983595244 | methyltransferase like 6 |
| ENSG00000180878 | C11orf42 | 0,048823764 | 1,034421211 | 0,94681142 | 0,996027077 | chromosome 11 open reading frame 42 |
| ENSG00000174695 | TMEM167A | 0,048761555 | 1,034376608 | 0,92260132 | 0,993797669 | transmembrane protein 167A |
| ENSG00000230074 | AL162231,2 | 0,048729194 | 1,034353406 | 0,89288947 | 0,991120066 | novel transcript |
| ENSG00000167106 | FAM102A | 0,048688821 | 1,034324461 | 0,86450909 | 0,989074806 | family with sequence similarity 102 member A |
| ENSG00000175854 | SWI5 | 0,048597144 | 1,034258737 | 0,86639071 | 0,989074806 | SWI5 homologous recombination repair protein |
| ENSG00000111707 | SUDS3 | 0,048533974 | 1,034213451 | 0,85110732 | 0,987259878 | SDS3 homolog, SIN3A corepressor complex component |
| ENSG00000260120 | AC104066,3 | 0,048448656 | 1,034152292 | 0,95009834 | 0,996432861 | novel transcript |
| ENSG00000164919 | COX6C | 0,048409683 | 1,034124355 | 0,89693776 | 0,991644439 | cytochrome c oxidase subunit 6C |
| ENSG00000237057 | LINC02087 | 0,048264445 | 1,034020254 | 0,95901885 | 0,996608711 | long intergenic non-protein coding RNA 2087 |
| ENSG00000122674 | CCZ1 | 0,048244164 | 1,034005718 | 0,90169429 | 0,992136926 | CCZ1 homolog, vacuolar protein trafficking and biogenesis associated |
| ENSG00000230325 | AL359921,1 | 0,04814283 | 1,033933093 | 0,93762209 | 0,994479239 | novel transcript |
| ENSG00000224393 | MPIG6B | 0,048086609 | 1,033892802 | 0,98506899 | 0,99837368 | megakaryocyte and platelet inhibitory receptor G6b |
| ENSG00000120318 | ARAP3 | 0,048085298 | 1,033891862 | 0,86035676 | 0,989074806 | ArfGAP with RhoGAP domain, ankyrin repeat and PH domain 3 |
| ENSG00000053524 | MCF2L2 | 0,048078809 | 1,033887212 | 0,91793228 | 0,993361299 | MCF,2 cell line derived transforming sequence-like 2 |

| | | | | | | |
|---|---|---|---|---|---|---|
| ENSG00000176624 | MEX3C | 0,048069272 | 1,033880378 | 0,89178434 | 0,990937593 | mex-3 RNA binding family member C |
| ENSG00000139190 | VAMP1 | 0,048066274 | 1,033878229 | 0,8707635 | 0,989120015 | vesicle associated membrane protein 1 |
| ENSG00000132846 | ZBED3 | 0,047999082 | 1,033830078 | 0,84164377 | 0,98571999 | zinc finger BED-type containing 3 |
| ENSG00000223503 | AL353807,1 | 0,047968333 | 1,033808044 | 0,96328747 | 0,996608711 | claudin 6 (CLDN6) pseudogene |
| ENSG00000105072 | C19orf44 | 0,047935712 | 1,033784669 | 0,83931649 | 0,985672131 | chromosome 19 open reading frame 44 |
| ENSG00000266313 | AC026254,2 | 0,047888587 | 1,033750901 | 0,92589654 | 0,993915537 | novel transcript |
| ENSG00000087206 | UIMC1 | 0,047857818 | 1,033728854 | 0,82306995 | 0,983204348 | ubiquitin interaction motif containing 1 |
| ENSG00000115484 | CCT4 | 0,047839939 | 1,033716043 | 0,87059185 | 0,989074806 | chaperonin containing TCP1 subunit 4 |
| ENSG00000130713 | EXOSC2 | 0,047819729 | 1,033701563 | 0,87005455 | 0,989074806 | exosome component 2 |
| ENSG00000163625 | WDFY3 | 0,047784518 | 1,033676334 | 0,87760349 | 0,990109814 | WD repeat and FYVE domain containing 3 |
| ENSG00000133706 | LARS1 | 0,04768075 | 1,033601988 | 0,84544205 | 0,986639902 | leucyl-tRNA synthetase 1 |
| ENSG00000205592 | MUC19 | 0,047653298 | 1,033582321 | 0,96039377 | 0,996608711 | mucin 19, oligomeric |
| ENSG00000131508 | UBE2D2 | 0,047570943 | 1,033523321 | 0,89298933 | 0,991120066 | ubiquitin conjugating enzyme E2 D2 |
| ENSG00000225693 | LAGE3P1 | 0,047534817 | 1,033497441 | 0,95287181 | 0,99652169 | L antigen family member 3 pseudogene 1 |
| ENSG00000008394 | MGST1 | 0,047375276 | 1,033383158 | 0,92247168 | 0,993797669 | microsomal glutathione S-transferase 1 |
| ENSG00000286348 | AC084834,1 | 0,047367757 | 1,033377772 | 0,96817431 | 0,997263201 | novel transcript |
| ENSG00000172007 | RAB33B | 0,047365049 | 1,033375832 | 0,85355434 | 0,987861659 | RAB33B, member RAS oncogene family |
| ENSG00000272599 | AC016394,1 | 0,04733484 | 1,033354195 | 0,91653943 | 0,993361299 | novel transcript to NUDT13 |
| ENSG00000286475 | AL592156,1 | 0,047312228 | 1,033337998 | 0,9677101 | 0,997263201 | novel transcript |
| ENSG00000102221 | JADE3 | 0,047100047 | 1,033186034 | 0,88282085 | 0,990717729 | jade family PHD finger 3 |
| ENSG00000090534 | THPO | 0,047098648 | 1,033185032 | 0,94472608 | 0,995741983 | thrombopoietin |
| ENSG00000083520 | DIS3 | 0,047051852 | 1,03315152 | 0,89413327 | 0,991135494 | DIS3 homolog, exosome endoribonuclease and 3'-5' exoribonuclease |
| ENSG00000234741 | GAS5 | 0,046914761 | 1,03305335 | 0,86968824 | 0,989074806 | growth arrest specific 5 |
| ENSG00000123297 | TSFM | 0,046893267 | 1,033037959 | 0,8547985 | 0,987890746 | Ts translation elongation factor, mitochondrial |
| ENSG00000168434 | COG7 | 0,04686459 | 1,033017425 | 0,83366402 | 0,984905093 | component of oligomeric golgi complex 7 |
| ENSG00000134962 | KLB | 0,046841474 | 1,033000874 | 0,9059992 | 0,992590661 | klotho beta |
| ENSG00000237331 | XIAP-AS1 | 0,046824082 | 1,03298842 | 0,94642967 | 0,996027077 | XIAP antisense RNA 1 |
| ENSG00000257743 | MGAM2 | 0,046741388 | 1,032929212 | 0,96783035 | 0,997263201 | maltase-glucoamylase 2 (putative) |
| ENSG00000278563 | MGAM2 | 0,046741388 | 1,032929212 | 0,96783035 | 0,997263201 | maltase-glucoamylase 2 (putative) |
| ENSG00000028528 | SNX1 | 0,046703521 | 1,032902101 | 0,85178348 | 0,987289464 | sorting nexin 1 |
| ENSG00000165819 | METTL3 | 0,04669897 | 1,032898843 | 0,83510769 | 0,984905093 | methyltransferase like 3 |
| ENSG00000144476 | ACKR3 | 0,046667274 | 1,03287615 | 0,89215026 | 0,99102448 | atypical chemokine receptor 3 |

| | | | | | | |
|---|---|---|---|---|---|---|
| ENSG00000115128 | SF3B6 | 0,0465957 | 1,032824909 | 0,93035498 | 0,994336072 | splicing factor 3b subunit 6 |
| ENSG00000185697 | MYBL1 | 0,046595445 | 1,032824726 | 0,89648353 | 0,991638165 | MYB proto-oncogene like 1 |
| ENSG00000141570 | CBX8 | 0,046524952 | 1,032774261 | 0,86240768 | 0,989074806 | chromobox 8 |
| ENSG00000130173 | ANGPTL8 | 0,046508276 | 1,032762324 | 0,97169273 | 0,998088605 | angiopoietin like 8 |
| ENSG00000102103 | PQBP1 | 0,046506374 | 1,032760963 | 0,83143088 | 0,984440275 | polyglutamine binding protein 1 |
| ENSG00000112164 | GLP1R | 0,046504594 | 1,032759689 | 0,94436312 | 0,995542144 | glucagon like peptide 1 receptor |
| ENSG00000144306 | SCRN3 | 0,046483694 | 1,032744727 | 0,86105331 | 0,989074806 | secernin 3 |
| ENSG00000177600 | RPLP2 | 0,046435702 | 1,032710373 | 0,83252212 | 0,98489473 | ribosomal protein lateral stalk subunit P2 |
| ENSG00000106330 | MOSPD3 | 0,046431551 | 1,032707401 | 0,87897111 | 0,990109814 | motile sperm domain containing 3 |
| ENSG00000177453 | NIM1K | 0,046427762 | 1,032704689 | 0,92462276 | 0,993915537 | NIM1 serine/threonine protein kinase |
| ENSG00000166398 | GARRE1 | 0,046426791 | 1,032703994 | 0,83147166 | 0,984440275 | granule associated Rac and RHOG effector 1 |
| ENSG00000132604 | TERF2 | 0,046389761 | 1,032677488 | 0,83969368 | 0,985672131 | telomeric repeat binding factor 2 |
| ENSG00000166226 | CCT2 | 0,046380805 | 1,032671077 | 0,86874241 | 0,989074806 | chaperonin containing TCP1 subunit 2 |
| ENSG00000223802 | CERS1 | 0,046335413 | 1,032638586 | 0,91355801 | 0,993361299 | ceramide synthase 1 |
| ENSG00000113272 | THG1L | 0,046324625 | 1,032630865 | 0,87020399 | 0,989074806 | tRNA-histidine guanylyltransferase 1 like |
| ENSG00000272764 | AL596094,1 | 0,046316436 | 1,032625003 | 0,95953974 | 0,996608711 | novel transcript |
| ENSG00000149483 | TMEM138 | 0,046213095 | 1,032551038 | 0,87209053 | 0,989217334 | transmembrane protein 138 |
| ENSG00000258881 | AC007040,2 | 0,046139346 | 1,032498257 | 0,96182333 | 0,996608711 | novel protein |
| ENSG00000184497 | TMEM255B | 0,046124964 | 1,032487964 | 0,90259975 | 0,992230188 | transmembrane protein 255B |
| ENSG00000141736 | ERBB2 | 0,046077925 | 1,0324543 | 0,87939124 | 0,990178196 | erb-b2 receptor tyrosine kinase 2 |
| ENSG00000121766 | ZCCHC17 | 0,045984098 | 1,032387156 | 0,89367837 | 0,991135494 | zinc finger CCHC-type containing 17 |
| ENSG00000184809 | B3GALT5-AS1 | 0,045970654 | 1,032377536 | 0,8893013 | 0,990717729 | B3GALT5 antisense RNA 1 |
| ENSG00000136986 | DERL1 | 0,045865597 | 1,03230236 | 0,8972636 | 0,991644439 | derlin 1 |
| ENSG00000255459 | AC105233,2 | 0,045854709 | 1,03229457 | 0,93569736 | 0,994479239 | ribosomal protein S24 pseudogene |
| ENSG00000140025 | EFCAB11 | 0,045846915 | 1,032288993 | 0,89080802 | 0,990717729 | EF-hand calcium binding domain 11 |
| ENSG00000284024 | AC069544,2 | 0,045787559 | 1,032246523 | 0,87826262 | 0,990109814 | zinc finger and SCAN domain containing 29 |
| ENSG00000205488 | CALML3-AS1 | 0,045783875 | 1,032243887 | 0,96511719 | 0,997208095 | CALML3 antisense RNA 1 |
| ENSG00000286182 | AC112172,2 | 0,045762667 | 1,032228713 | 0,9149311 | 0,993361299 | novel transcript |
| ENSG00000146729 | NIPSNAP2 | 0,045710924 | 1,032191692 | 0,91529907 | 0,993361299 | nipsnap homolog 2 |
| ENSG00000163634 | THOC7 | 0,045698108 | 1,032182523 | 0,90784966 | 0,992590661 | THO complex 7 |
| ENSG00000129003 | VPS13C | 0,045669142 | 1,032161799 | 0,90234971 | 0,992230188 | vacuolar protein sorting 13 homolog C |
| ENSG00000184515 | BEX5 | 0,045614934 | 1,032123017 | 0,91282044 | 0,993361299 | brain expressed X-linked 5 |
| ENSG00000279085 | AL022323,2 | 0,045580834 | 1,032098622 | 0,9450123 | 0,995797387 | novel transcript |

| | | | | | | |
|---|---|---|---|---|---|---|
| ENSG00000141391 | PRELID3A | 0,045534221 | 1,032065276 | 0,87030141 | 0,989074806 | PRELI domain containing 3A |
| ENSG00000264739 | AC093484,2 | 0,045532354 | 1,03206394 | 0,96730741 | 0,997263201 | novel transcript |
| ENSG00000061987 | MON2 | 0,045448515 | 1,032003966 | 0,89427625 | 0,991135494 | MON2 homolog, regulator of endosome-to-Golgi trafficking |
| ENSG00000189190 | ZNF600 | 0,045447072 | 1,032002934 | 0,91551508 | 0,993361299 | zinc finger protein 600 |
| ENSG00000213064 | SFT2D2 | 0,045367648 | 1,031946121 | 0,82829849 | 0,983646162 | SFT2 domain containing 2 |
| ENSG00000071189 | SNX13 | 0,045331912 | 1,03192056 | 0,9159247 | 0,993361299 | sorting nexin 13 |
| ENSG00000173145 | NOC3L | 0,045287374 | 1,031888703 | 0,88683522 | 0,990717729 | NOC3 like DNA replication regulator |
| ENSG00000188825 | LINC00910 | 0,045267003 | 1,031874133 | 0,88047742 | 0,990367704 | long intergenic non-protein coding RNA 910 |
| ENSG00000196465 | MYL6B | 0,045232577 | 1,031849511 | 0,83506175 | 0,984905093 | myosin light chain 6B |
| ENSG00000228116 | PPT2 | 0,045183894 | 1,031814692 | 0,94614009 | 0,996027077 | palmitoyl-protein thioesterase 2 |
| ENSG00000226520 | KIRREL1-IT1 | 0,045119726 | 1,0317688 | 0,92672724 | 0,993915537 | KIRREL1 intronic transcript 1 |
| ENSG00000198945 | L3MBTL3 | 0,045103029 | 1,031756859 | 0,91079992 | 0,993222655 | L3MBTL histone methyl-lysine binding protein 3 |
| ENSG00000170779 | CDCA4 | 0,045069234 | 1,031732691 | 0,83470896 | 0,984905093 | cell division cycle associated 4 |
| ENSG00000273476 | AL133215,3 | 0,045064299 | 1,031729161 | 0,97106161 | 0,998088605 | novel transcript |
| ENSG00000163866 | SMIM12 | 0,045035163 | 1,031708325 | 0,84251992 | 0,986175464 | small integral membrane protein 12 |
| ENSG00000155621 | C9orf85 | 0,044992724 | 1,031677977 | 0,8601663 | 0,989074806 | chromosome 9 open reading frame 85 |
| ENSG00000258102 | MAP1LC3B2 | 0,044956116 | 1,031651798 | 0,95520969 | 0,996608711 | microtubule associated protein 1 light chain 3 beta 2 |
| ENSG00000249853 | HS3ST5 | 0,044948034 | 1,031646018 | 0,91282616 | 0,993361299 | heparan sulfate-glucosamine 3-sulfotransferase 5 |
| ENSG00000122008 | POLK | 0,044846841 | 1,03157366 | 0,90262874 | 0,992230188 | DNA polymerase kappa |
| ENSG00000172014 | ANKRD20A4P | 0,044786279 | 1,031530357 | 0,91881239 | 0,993489338 | ankyrin repeat domain 20 family member A4, pseudogene |
| ENSG00000142661 | MYOM3 | 0,044777452 | 1,031524046 | 0,94348323 | 0,995402494 | myomesin 3 |
| ENSG00000121897 | LIAS | 0,044769921 | 1,031518661 | 0,856284 | 0,988229014 | lipoic acid synthetase |
| ENSG00000197406 | DIO3 | 0,044685807 | 1,031458522 | 0,91877733 | 0,993485783 | iodothyronine deiodinase 3 |
| ENSG00000266733 | TBC1D29P | 0,044681415 | 1,031455382 | 0,94297393 | 0,995402494 | TBC1 domain family member 29, pseudogene |
| ENSG00000143458 | GABPB2 | 0,044676402 | 1,031451797 | 0,89358201 | 0,991135494 | GA binding protein transcription factor subunit beta 2 |
| ENSG00000259804 | AC027682,1 | 0,044551159 | 1,031362259 | 0,95583201 | 0,996608711 | novel transcript, antisense to CTCF |
| ENSG00000168062 | BATF2 | 0,044441784 | 1,031284071 | 0,96152155 | 0,996608711 | basic leucine zipper ATF-like transcription factor 2 |
| ENSG00000164823 | OSGIN2 | 0,04433352 | 1,031206684 | 0,92475581 | 0,993915537 | oxidative stress induced growth inhibitor family member 2 |
| ENSG00000285803 | AL442003,1 | 0,044321564 | 1,031198138 | 0,89797745 | 0,991711719 | novel transcript, antisense to AGAP6 |

| | | | | | | |
|---|---|---|---|---|---|---|
| ENSG00000139684 | ESD | 0,04421201 | 1,031119835 | 0,91285933 | 0,993361299 | esterase D |
| ENSG00000180155 | LYNX1 | 0,044173326 | 1,031092187 | 0,91589454 | 0,993361299 | Ly6/neurotoxin 1 |
| ENSG00000232545 | AC253536,3 | 0,044150836 | 1,031076114 | 0,95987218 | 0,996608711 | novel transcript |
| ENSG00000260231 | KDM7A-DT | 0,044113722 | 1,031049589 | 0,90715697 | 0,992590661 | KDM7A divergent transcript |
| ENSG00000132635 | PCED1A | 0,044106608 | 1,031044505 | 0,87438662 | 0,98990147 | PC-esterase domain containing 1A |
| ENSG00000229676 | ZNF492 | 0,044006542 | 1,030972994 | 0,95750252 | 0,996608711 | zinc finger protein 492 |
| ENSG00000278741 | ABR | 0,043987491 | 1,03095938 | 0,89774475 | 0,991708791 | ABR activator of RhoGEF and GTPase |
| ENSG00000111880 | RNGTT | 0,043927489 | 1,030916503 | 0,88455017 | 0,990717729 | RNA guanylyltransferase and 5'-phosphatase |
| ENSG00000287626 | AL021978,1 | 0,043898608 | 1,030895866 | 0,94974206 | 0,996432861 | novel transcript, antisense to DTNBP1 |
| ENSG00000070718 | AP3M2 | 0,043844189 | 1,03085698 | 0,87145771 | 0,989217334 | adaptor related protein complex 3 subunit mu 2 |
| ENSG00000250473 | DUTP7 | 0,043816223 | 1,030836998 | 0,93951605 | 0,994943406 | deoxyuridine triphosphatase pseudogene 7 |
| ENSG00000106588 | PSMA2 | 0,043749425 | 1,03078927 | 0,90703009 | 0,992590661 | proteasome 20S subunit alpha 2 |
| ENSG00000212123 | PRR22 | 0,043747348 | 1,030787786 | 0,92857444 | 0,994108531 | proline rich 22 |
| ENSG00000267534 | S1PR2 | 0,043726077 | 1,030772589 | 0,87372644 | 0,989633545 | sphingosine-1-phosphate receptor 2 |
| ENSG00000262585 | LINC01979 | 0,043673385 | 1,030734942 | 0,9735318 | 0,99837368 | long intergenic non-protein coding RNA 1979 |
| ENSG00000138381 | ASNSD1 | 0,043662084 | 1,030726868 | 0,90172122 | 0,992136926 | asparagine synthetase domain containing 1 |
| ENSG00000160678 | S100A1 | 0,043565452 | 1,030657832 | 0,94981714 | 0,996432861 | S100 calcium binding protein A1 |
| ENSG00000105767 | CADM4 | 0,043546988 | 1,030644642 | 0,91436097 | 0,993361299 | cell adhesion molecule 4 |
| ENSG00000196152 | ZNF79 | 0,04353214 | 1,030634034 | 0,85420875 | 0,987861659 | zinc finger protein 79 |
| ENSG00000240898 | AC132942,1 | 0,043525081 | 1,030628991 | 0,95494656 | 0,996608711 | ribosomal protein L21 (RPL21)pseudogene |
| ENSG00000102096 | PIM2 | 0,043492672 | 1,030605839 | 0,88576462 | 0,990717729 | Pim-2 proto-oncogene, serine/threonine kinase |
| ENSG00000172339 | ALG14 | 0,04343005 | 1,030561106 | 0,87724867 | 0,990109814 | ALG14 UDP-N-acetylglucosaminyltransferase subunit |
| ENSG00000167562 | ZNF701 | 0,043401286 | 1,030540559 | 0,87102594 | 0,989217334 | zinc finger protein 701 |
| ENSG00000272356 | AL080317,2 | 0,043363599 | 1,030513639 | 0,88872203 | 0,990717729 | novel transcript, antisense to REV3L |
| ENSG00000108654 | DDX5 | 0,04335632 | 1,030508439 | 0,87541968 | 0,990078774 | DEAD-box helicase 5 |
| ENSG00000112053 | SLC26A8 | 0,043206715 | 1,030401583 | 0,95308526 | 0,99652169 | solute carrier family 26 member 8 |
| ENSG00000203760 | CENPW | 0,043205694 | 1,030400854 | 0,89646735 | 0,991638165 | centromere protein W |
| ENSG00000109775 | UFSP2 | 0,043198485 | 1,030395705 | 0,92171773 | 0,993787755 | UFM1 specific peptidase 2 |
| ENSG00000136861 | CDK5RAP2 | 0,043175652 | 1,030379397 | 0,84142891 | 0,985672131 | CDK5 regulatory subunit associated protein 2 |
| ENSG00000114956 | DGUOK | 0,04314687 | 1,030358841 | 0,86405303 | 0,989074806 | deoxyguanosine kinase |
| ENSG00000148297 | MED22 | 0,043084363 | 1,0303142 | 0,88893712 | 0,990717729 | mediator complex subunit 22 |
| ENSG00000004455 | AK2 | 0,043080983 | 1,030311786 | 0,83906285 | 0,985672131 | adenylate kinase 2 |
| ENSG00000113812 | ACTR8 | 0,043078682 | 1,030310144 | 0,86189827 | 0,989074806 | actin related protein 8 |

| | | | | | | |
|---|---|---|---|---|---|---|
| ENSG00000108465 | CDK5RAP3 | 0,043060881 | 1,030297431 | 0,87351724 | 0,989633545 | CDK5 regulatory subunit associated protein 3 |
| ENSG00000149557 | FEZ1 | 0,043046648 | 1,030287266 | 0,87677992 | 0,990109814 | fasciculation and elongation protein zeta 1 |
| ENSG00000233937 | CTC-338M12,4 | 0,042987612 | 1,030245107 | 0,91663712 | 0,993361299 | uncharacterized LOC101928649 |
| ENSG00000092148 | HECTD1 | 0,042944567 | 1,030214368 | 0,87401007 | 0,989711917 | HECT domain E3 ubiquitin protein ligase 1 |
| ENSG00000140854 | KATNB1 | 0,042866422 | 1,030158567 | 0,88906738 | 0,990717729 | katanin regulatory subunit B1 |
| ENSG00000285041 | AL035412,1 | 0,042779351 | 1,030096397 | 0,96111632 | 0,996608711 | novel transcript |
| ENSG00000187514 | PTMA | 0,042738456 | 1,030067197 | 0,82780023 | 0,983595244 | prothymosin alpha |
| ENSG00000033627 | ATP6V0A1 | 0,042736948 | 1,030066121 | 0,83729939 | 0,985577601 | ATPase H+ transporting V0 subunit a1 |
| ENSG00000131379 | C3orf20 | 0,042674619 | 1,030021619 | 0,95224015 | 0,996432861 | chromosome 3 open reading frame 20 |
| ENSG00000229180 | AC006001,3 | 0,042639538 | 1,029996574 | 0,87871883 | 0,990109814 | RAB guanine nucleotide exchange factor (GEF) 1 (RABGEF1) pseudogene |
| ENSG00000122958 | VPS26A | 0,042575965 | 1,029951187 | 0,92372419 | 0,993915537 | VPS26, retromer complex component A |
| ENSG00000174684 | B4GAT1 | 0,042545169 | 1,029929202 | 0,85357394 | 0,987861659 | beta-1,4-glucuronyltransferase 1 |
| ENSG00000094631 | HDAC6 | 0,04253854 | 1,02992447 | 0,84132723 | 0,985672131 | histone deacetylase 6 |
| ENSG00000122952 | ZWINT | 0,042427956 | 1,029845528 | 0,85456009 | 0,987861659 | ZW10 interacting kinetochore protein |
| ENSG00000237978 | KCNMB2-AS1 | 0,042426459 | 1,029844459 | 0,86523468 | 0,989074806 | KCNMB2 antisense RNA 1 |
| ENSG00000106479 | ZNF862 | 0,042396513 | 1,029823083 | 0,86482596 | 0,989074806 | zinc finger protein 862 |
| ENSG00000285581 | AC091196,1 | 0,042396179 | 1,029822844 | 0,9655323 | 0,997208095 | novel transcript |
| ENSG00000260000 | AL133338,1 | 0,042347629 | 1,029788189 | 0,93452909 | 0,994479239 | novel transcript, antisense to ASCC3 |
| ENSG00000083817 | ZNF416 | 0,042321705 | 1,029769685 | 0,89088531 | 0,990717729 | zinc finger protein 416 |
| ENSG00000272688 | AP005329,3 | 0,042311249 | 1,029762222 | 0,96272454 | 0,996608711 | novel transcript |
| ENSG00000261766 | AC133550,3 | 0,0422324 | 1,029705943 | 0,95344781 | 0,99652169 | novel transcript, antisense to SH2B1 |
| ENSG00000248994 | AC124852,1 | 0,042228264 | 1,029702991 | 0,90447104 | 0,992590661 | novel transcript |
| ENSG00000284543 | LINC01226 | 0,042200838 | 1,029683416 | 0,94942072 | 0,996432861 | long intergenic non-protein coding RNA 1226 |
| ENSG00000101363 | MANBAL | 0,042192839 | 1,029677707 | 0,86541433 | 0,989074806 | mannosidase beta like |
| ENSG00000168806 | LCMT2 | 0,042073099 | 1,02959225 | 0,87045111 | 0,989074806 | leucine carboxyl methyltransferase 2 |
| ENSG00000227248 | FAM155A-IT1 | 0,041964796 | 1,029514962 | 0,94983909 | 0,996432861 | FAM155A intronic transcript 1 |
| ENSG00000197329 | PELI1 | 0,041948186 | 1,029503109 | 0,92588077 | 0,993915537 | pellino E3 ubiquitin protein ligase 1 |
| ENSG00000103061 | SLC7A6OS | 0,041901505 | 1,029469797 | 0,85675272 | 0,988312374 | solute carrier family 7 member 6 opposite strand |
| ENSG00000154305 | MIA3 | 0,041830714 | 1,029419284 | 0,87357923 | 0,989633545 | MIA SH3 domain ER export factor 3 |
| ENSG00000169432 | SCN9A | 0,041683591 | 1,029314312 | 0,91508893 | 0,993361299 | sodium voltage-gated channel alpha subunit 9 |
| ENSG00000120519 | SLC10A7 | 0,041653889 | 1,02929312 | 0,92164756 | 0,993787755 | solute carrier family 10 member 7 |
| ENSG00000148690 | FRA10AC1 | 0,041575381 | 1,02923711 | 0,9077541 | 0,992590661 | FRA10A associated CGG repeat 1 |

| | | | | | | |
|---|---|---|---|---|---|---|
| ENSG00000185722 | ANKFY1 | 0,041425955 | 1,029130513 | 0,86473057 | 0,989074806 | ankyrin repeat and FYVE domain containing 1 |
| ENSG00000146733 | PSPH | 0,041390428 | 1,029105171 | 0,87618042 | 0,990109814 | phosphoserine phosphatase |
| ENSG00000164631 | ZNF12 | 0,04136137 | 1,029084443 | 0,91876322 | 0,993485783 | zinc finger protein 12 |
| ENSG00000169894 | MUC3A | 0,041355056 | 1,029079939 | 0,95794358 | 0,996608711 | mucin 3A, cell surface associated |
| ENSG00000084731 | KIF3C | 0,041325001 | 1,029058502 | 0,89375108 | 0,991135494 | kinesin family member 3C |
| ENSG00000174886 | NDUFA11 | 0,041320289 | 1,029055141 | 0,85960756 | 0,989074806 | NADH:ubiquinone oxidoreductase subunit A11 |
| ENSG00000281930 | SMIM10L1 | 0,040994328 | 1,028822663 | 0,92238639 | 0,993797669 | small integral membrane protein 10 like 1 |
| ENSG00000213281 | NRAS | 0,040908286 | 1,028761306 | 0,92035551 | 0,993787755 | NRAS proto-oncogene, GTPase |
| ENSG00000163428 | LRRC58 | 0,040902394 | 1,028757105 | 0,85682426 | 0,988312374 | leucine rich repeat containing 58 |
| ENSG00000131626 | PPFIA1 | 0,040810648 | 1,028691684 | 0,86914356 | 0,989074806 | PTPRF interacting protein alpha 1 |
| ENSG00000281289 | LSS | 0,040804377 | 1,028687213 | 0,89368682 | 0,991135494 | lanosterol synthase |
| ENSG00000182308 | DCAF4L1 | 0,040800005 | 1,028684096 | 0,9272497 | 0,993915537 | DDB1 and CUL4 associated factor 4 like 1 |
| ENSG00000282458 | WASH5P | 0,040738867 | 1,028640503 | 0,90995405 | 0,992951496 | WASP family homolog 5, pseudogene |
| ENSG00000214676 | RPL9P16 | 0,040731 | 1,028634894 | 0,95007966 | 0,996432861 | ribosomal protein L9 pseudogene 16 |
| ENSG00000180901 | KCTD2 | 0,040654008 | 1,028580001 | 0,85523236 | 0,988016959 | potassium channel tetramerization domain containing 2 |
| ENSG00000128789 | PSMG2 | 0,040631442 | 1,028563913 | 0,9122478 | 0,993361299 | proteasome assembly chaperone 2 |
| ENSG00000217733 | CCT7P1 | 0,040623695 | 1,028558389 | 0,96168985 | 0,996608711 | chaperonin containing TCP1 subunit 7 pseudogene 1 |
| ENSG00000133597 | ADCK2 | 0,04054872 | 1,028504938 | 0,90742047 | 0,992590661 | aarF domain containing kinase 2 |
| ENSG00000176383 | B3GNT4 | 0,04054492 | 1,028502229 | 0,91522902 | 0,993361299 | UDP-GlcNAc:betaGal beta-1,3-N-acetylglucosaminyltransferase 4 |
| ENSG00000186166 | CENATAC | 0,040448041 | 1,028433165 | 0,86143497 | 0,989074806 | centrosomal AT-AC splicing factor |
| ENSG00000288453 | DCAF15 | 0,040443585 | 1,028429989 | 0,88101383 | 0,990598309 | DDB1 and CUL4 associated factor 15 |
| ENSG00000132017 | DCAF15 | 0,040443585 | 1,028429989 | 0,88101383 | 0,990598309 | DDB1 and CUL4 associated factor 15 |
| ENSG00000153317 | ASAP1 | 0,040417715 | 1,028411548 | 0,89544087 | 0,991303328 | ArfGAP with SH3 domain, ankyrin repeat and PH domain 1 |
| ENSG00000205763 | RP9P | 0,040400606 | 1,028399351 | 0,87463992 | 0,990039342 | RP9 pseudogene |
| ENSG00000272677 | AC124016,1 | 0,040333716 | 1,028351671 | 0,92677132 | 0,993915537 | novel transcript, antisense to HNRNPD |
| ENSG00000288121 | MDN1 | 0,040312016 | 1,028336204 | 0,86919543 | 0,989074806 | midasin AAA ATPase 1 |
| ENSG00000112159 | MDN1 | 0,040312016 | 1,028336204 | 0,86919543 | 0,989074806 | midasin AAA ATPase 1 |
| ENSG00000089157 | RPLP0 | 0,040228457 | 1,028276646 | 0,87042396 | 0,989074806 | ribosomal protein lateral stalk subunit P0 |
| ENSG00000155636 | RBM45 | 0,040227993 | 1,028276315 | 0,91428086 | 0,993361299 | RNA binding motif protein 45 |
| ENSG00000100418 | DESI1 | 0,040175184 | 1,028238676 | 0,85721122 | 0,988665109 | desumoylating isopeptidase 1 |

| | | | | | | |
|---|---|---|---|---|---|---|
| ENSG00000175390 | EIF3F | 0,040148988 | 1,028220006 | 0,89912049 | 0,991832796 | eukaryotic translation initiation factor 3 subunit F |
| ENSG00000258636 | AL121821,2 | 0,040118185 | 1,028198053 | 0,93977216 | 0,994943406 | novel transcript |
| ENSG00000172831 | CES2 | 0,040104657 | 1,028188412 | 0,87668694 | 0,990109814 | carboxylesterase 2 |
| ENSG00000287743 | AL031289,2 | 0,040094838 | 1,028181414 | 0,97576138 | 0,99837368 | novel transcript, antisense to RIMS3 |
| ENSG00000276384 | AC016876,3 | 0,040074045 | 1,028166595 | 0,97225972 | 0,998214946 | novel transcript, antisense to TNFSF12 |
| ENSG00000125633 | CCDC93 | 0,040056426 | 1,028154039 | 0,87981026 | 0,990261566 | coiled-coil domain containing 93 |
| ENSG00000169567 | HINT1 | 0,039975094 | 1,028096078 | 0,88901902 | 0,990717729 | histidine triad nucleotide binding protein 1 |
| ENSG00000100485 | SOS2 | 0,039939808 | 1,028070933 | 0,92007976 | 0,993787755 | SOS Ras/Rho guanine nucleotide exchange factor 2 |
| ENSG00000159708 | LRRC36 | 0,039843206 | 1,028002096 | 0,97434824 | 0,99837368 | leucine rich repeat containing 36 |
| ENSG00000008283 | CYB561 | 0,039816894 | 1,027983347 | 0,87860944 | 0,990109814 | cytochrome b561 |
| ENSG00000270039 | AC025165,5 | 0,039748601 | 1,027934687 | 0,94838963 | 0,99635783 | novel transcript |
| ENSG00000146830 | GIGYF1 | 0,039703632 | 1,027902647 | 0,92833013 | 0,994108531 | GRB10 interacting GYF protein 1 |
| ENSG00000274021 | AC024909,1 | 0,039697999 | 1,027898633 | 0,92215447 | 0,993797669 | novel transcript, antisense to DUSP6 |
| ENSG00000147121 | KRBOX4 | 0,039662499 | 1,02787334 | 0,89483854 | 0,991296984 | KRAB box domain containing 4 |
| ENSG00000186526 | CYP4F8 | 0,039617159 | 1,027841038 | 0,96371039 | 0,996772476 | cytochrome P450 family 4 subfamily F member 8 |
| ENSG00000248550 | OTX2-AS1 | 0,039603789 | 1,027831512 | 0,91082006 | 0,993222655 | OTX2 antisense RNA 1 (head to head) |
| ENSG00000284906 | AC091057,5 | 0,039543324 | 1,027788435 | 0,95168647 | 0,996432861 | Rho GTPase-activating protein 11B |
| ENSG00000262576 | PCDHGA4 | 0,039503419 | 1,027760007 | 0,93408936 | 0,994479239 | protocadherin gamma subfamily A, 4 |
| ENSG00000135974 | C2orf49 | 0,039470344 | 1,027736445 | 0,92059561 | 0,993787755 | chromosome 2 open reading frame 49 |
| ENSG00000286585 | AC104472,4 | 0,039455075 | 1,027725568 | 0,96073195 | 0,996608711 | novel transcript |
| ENSG00000100422 | CERK | 0,03938995 | 1,027679176 | 0,85189103 | 0,987322912 | ceramide kinase |
| ENSG00000152749 | GPR180 | 0,039230506 | 1,027565605 | 0,93038509 | 0,994336072 | G protein-coupled receptor 180 |
| ENSG00000244952 | AC123768,1 | 0,039157008 | 1,027513257 | 0,9597779 | 0,996608711 | novel transcript |
| ENSG00000136715 | SAP130 | 0,039134725 | 1,027497387 | 0,89041386 | 0,990717729 | Sin3A associated protein 130 |
| ENSG00000157765 | SLC34A2 | 0,03913246 | 1,027495774 | 0,96390656 | 0,996942396 | solute carrier family 34 member 2 |
| ENSG00000144713 | RPL32 | 0,039063978 | 1,027447002 | 0,91122061 | 0,993222655 | ribosomal protein L32 |
| ENSG00000117528 | ABCD3 | 0,03905716 | 1,027442146 | 0,91781322 | 0,993361299 | ATP binding cassette subfamily D member 3 |
| ENSG00000124767 | GLO1 | 0,038999634 | 1,027401179 | 0,93500949 | 0,994479239 | glyoxalase I |
| ENSG00000118402 | ELOVL4 | 0,038985384 | 1,027391031 | 0,92665967 | 0,993915537 | ELOVL fatty acid elongase 4 |
| ENSG00000088986 | DYNLL1 | 0,038974502 | 1,027383281 | 0,92605528 | 0,993915537 | dynein light chain LC8-type 1 |
| ENSG00000236409 | NRADDP | 0,038943949 | 1,027361524 | 0,96165298 | 0,996608711 | neurotrophin receptor associated death domain, pseudogene |
| ENSG00000163684 | RPP14 | 0,038939076 | 1,027358054 | 0,86799423 | 0,989074806 | ribonuclease P/MRP subunit p14 |

| | | | | | | |
|---|---|---|---|---|---|---|
| ENSG00000286724 | AL732509,1 | 0,038866006 | 1,027306021 | 0,91568211 | 0,993361299 | novel transcript, antisense to ADGRG2and PHKA2 |
| ENSG00000284691 | AC073111,3 | 0,038819999 | 1,027273262 | 0,91551442 | 0,993361299 | novel zinc finger protein |
| ENSG00000068001 | HYAL2 | 0,038804307 | 1,027262088 | 0,86583006 | 0,989074806 | hyaluronidase 2 |
| ENSG00000257859 | CASC18 | 0,038755473 | 1,027227316 | 0,97035995 | 0,998004261 | cancer susceptibility 18 |
| ENSG00000214940 | NPIPA8 | 0,038529919 | 1,027066731 | 0,97749312 | 0,99837368 | nuclear pore complex interacting protein family member A8 |
| ENSG00000198718 | TOGARAM1 | 0,038406133 | 1,02697861 | 0,89912883 | 0,991832796 | TOG array regulator of axonemal microtubules 1 |
| ENSG00000239218 | RPS20P22 | 0,038314157 | 1,026913139 | 0,94341706 | 0,995402494 | ribosomal protein S20 pseudogene 22 |
| ENSG00000165678 | GHITM | 0,03828895 | 1,026895197 | 0,90677673 | 0,992590661 | growth hormone inducible transmembrane protein |
| ENSG00000246922 | UBAP1L | 0,038231023 | 1,026853966 | 0,90560444 | 0,992590661 | ubiquitin associated protein 1 like |
| ENSG00000075975 | MKRN2 | 0,038202625 | 1,026833753 | 0,88645918 | 0,990717729 | makorin ring finger protein 2 |
| ENSG00000139263 | LRIG3 | 0,038110179 | 1,026767957 | 0,89069974 | 0,990717729 | leucine rich repeats and immunoglobulin like domains 3 |
| ENSG00000150687 | PRSS23 | 0,038047489 | 1,026723342 | 0,90921129 | 0,992864002 | serine protease 23 |
| ENSG00000149262 | INTS4 | 0,037988351 | 1,026681256 | 0,9014697 | 0,992041918 | integrator complex subunit 4 |
| ENSG00000287978 | AC245407,2 | 0,037939294 | 1,026646345 | 0,92778858 | 0,994108531 | novel transcript |
| ENSG00000177854 | TMEM187 | 0,037936784 | 1,02664456 | 0,91420426 | 0,993361299 | transmembrane protein 187 |
| ENSG00000111775 | COX6A1 | 0,037804863 | 1,026550686 | 0,87298607 | 0,989308263 | cytochrome c oxidase subunit 6A1 |
| ENSG00000186487 | MYT1L | 0,037664095 | 1,026450528 | 0,93338452 | 0,994479239 | myelin transcription factor 1 like |
| ENSG00000198431 | TXNRD1 | 0,037649472 | 1,026440124 | 0,8895527 | 0,990717729 | thioredoxin reductase 1 |
| ENSG00000198198 | SZT2 | 0,037635099 | 1,026429898 | 0,89126767 | 0,990820775 | SZT2 subunit of KICSTOR complex |
| ENSG00000066084 | DIP2B | 0,037614234 | 1,026415053 | 0,89362618 | 0,991135494 | disco interacting protein 2 homolog B |
| ENSG00000269958 | AL049840,5 | 0,0376011 | 1,026405709 | 0,90720081 | 0,992590661 | novel transcript, sense intronic to KLC1 |
| ENSG00000135211 | TMEM60 | 0,037579211 | 1,026390136 | 0,92224597 | 0,993797669 | transmembrane protein 60 |
| ENSG00000235530 | AC087294,1 | 0,037554698 | 1,026372697 | 0,9265029 | 0,993915537 | novel transcript, antisense to TMEM11 |
| ENSG00000100813 | ACIN1 | 0,037548852 | 1,026368538 | 0,90911865 | 0,992828195 | apoptotic chromatin condensation inducer 1 |
| ENSG00000112592 | TBP | 0,037536737 | 1,026359919 | 0,89917514 | 0,991832796 | TATA-box binding protein |
| ENSG00000111783 | RFX4 | 0,037493802 | 1,026329375 | 0,91447799 | 0,993361299 | regulatory factor X4 |
| ENSG00000152102 | FAM168B | 0,03738889 | 1,026254743 | 0,86854827 | 0,989074806 | family with sequence similarity 168 member B |
| ENSG00000151466 | SCLT1 | 0,037387232 | 1,026253564 | 0,89935606 | 0,991832796 | sodium channel and clathrin linker 1 |
| ENSG00000260288 | AC019294,2 | 0,037261635 | 1,026164225 | 0,93753559 | 0,994479239 | novel transcript |
| ENSG00000155096 | AZIN1 | 0,037192287 | 1,0261149 | 0,91451033 | 0,993361299 | antizyme inhibitor 1 |
| ENSG00000138802 | SEC24B | 0,037183417 | 1,026108591 | 0,92976166 | 0,994293848 | SEC24 homolog B, COPII coat complex component |

| | | | | | | |
|---|---|---|---|---|---|---|
| ENSG00000134371 | CDC73 | 0,037160872 | 1,026092557 | 0,91718383 | 0,993361299 | cell division cycle 73 |
| ENSG00000226604 | PAPPA-AS2 | 0,03715832 | 1,026090741 | 0,94637737 | 0,996027077 | PAPPA antisense RNA 2 |
| ENSG00000117174 | ZNHIT6 | 0,037153023 | 1,026086974 | 0,90319006 | 0,992311172 | zinc finger HIT-type containing 6 |
| ENSG00000141030 | COPS3 | 0,03710047 | 1,026049598 | 0,92108339 | 0,993787755 | COP9 signalosome subunit 3 |
| ENSG00000160972 | PPP1R16A | 0,037093466 | 1,026044616 | 0,92271581 | 0,993797669 | protein phosphatase 1 regulatory subunit 16A |
| ENSG00000203258 | AC013549,1 | 0,037065901 | 1,026025012 | 0,96402133 | 0,996980762 | novel transcript |
| ENSG00000139350 | NEDD1 | 0,037012659 | 1,025987148 | 0,92722038 | 0,993915537 | NEDD1 gamma-tubulin ring complex targeting factor |
| ENSG00000078142 | PIK3C3 | 0,0370116 | 1,025986394 | 0,91656888 | 0,993361299 | phosphatidylinositol 3-kinase catalytic subunit type 3 |
| ENSG00000182831 | C16orf72 | 0,037002201 | 1,02597971 | 0,88688182 | 0,990717729 | chromosome 16 open reading frame 72 |
| ENSG00000089723 | OTUB2 | 0,036994855 | 1,025974487 | 0,89616371 | 0,991600147 | OTU deubiquitinase, ubiquitin aldehyde binding 2 |
| ENSG00000101901 | ALG13 | 0,036964117 | 1,025952627 | 0,91649596 | 0,993361299 | ALG13 UDP-N-acetylglucosaminyltransferase subunit |
| ENSG00000140326 | CDAN1 | 0,036940879 | 1,025936102 | 0,89984983 | 0,991845106 | codanin 1 |
| ENSG00000205629 | LCMT1 | 0,036935298 | 1,025932133 | 0,86979411 | 0,989074806 | leucine carboxyl methyltransferase 1 |
| ENSG00000196646 | ZNF136 | 0,036927527 | 1,025926607 | 0,90259393 | 0,992230188 | zinc finger protein 136 |
| ENSG00000165934 | CPSF2 | 0,036899387 | 1,025906597 | 0,90542223 | 0,992590661 | cleavage and polyadenylation specific factor 2 |
| ENSG00000285999 | AC025442,2 | 0,036791629 | 1,025829972 | 0,94181577 | 0,995373376 | novel transcipt |
| ENSG00000283883 | VPS53 | 0,03666717 | 1,02574148 | 0,94264699 | 0,995390305 | VPS53 subunit of GARP complex |
| ENSG00000254224 | AP003465,1 | 0,036654437 | 1,025732426 | 0,97108028 | 0,998088605 | novel transcript |
| ENSG00000164346 | NSA2 | 0,036571923 | 1,025673762 | 0,93937547 | 0,99490615 | NSA2 ribosome biogenesis factor |
| ENSG00000105443 | CYTH2 | 0,036547092 | 1,025656109 | 0,92386646 | 0,993915537 | cytohesin 2 |
| ENSG00000277945 | AC107308,1 | 0,036447108 | 1,025585029 | 0,976235 | 0,99837368 | novel transcript, sense intronic to HMGA2 |
| ENSG00000118894 | EEF2KMT | 0,036412112 | 1,025560152 | 0,8912818 | 0,990820775 | eukaryotic elongation factor 2 lysine methyltransferase |
| ENSG00000177058 | SLC38A9 | 0,036335915 | 1,025505987 | 0,9215809 | 0,993787755 | solute carrier family 38 member 9 |
| ENSG00000254672 | AL136146,1 | 0,036314472 | 1,025490746 | 0,95058216 | 0,996432861 | WEE1 homolog (S, pombe) (WEE1) pseudogene |
| ENSG00000183317 | EPHA10 | 0,03630249 | 1,025482228 | 0,95408999 | 0,996608711 | EPH receptor A10 |
| ENSG00000156256 | USP16 | 0,036300585 | 1,025480874 | 0,92761318 | 0,994108531 | ubiquitin specific peptidase 16 |
| ENSG00000255269 | LINC02710 | 0,036249412 | 1,025444501 | 0,97891678 | 0,99837368 | long intergenic non-protein coding RNA 2710 |
| ENSG00000064419 | TNPO3 | 0,036230367 | 1,025430964 | 0,87810033 | 0,990109814 | transportin 3 |
| ENSG00000145715 | RASA1 | 0,036218551 | 1,025422565 | 0,91456233 | 0,993361299 | RAS p21 protein activator 1 |
| ENSG00000240132 | ETF1P2 | 0,036204528 | 1,025412598 | 0,94002281 | 0,994949796 | eukaryotic translation termination factor 1 pseudogene 2 |

| | | | | | | |
|---|---|---|---|---|---|---|
| ENSG00000206312 | NOTCH4 | 0,036164396 | 1,025384074 | 0,94565713 | 0,995958379 | notch receptor 4 |
| ENSG00000172888 | ZNF621 | 0,036126082 | 1,025356844 | 0,88504395 | 0,990717729 | zinc finger protein 621 |
| ENSG00000247416 | AP000802,1 | 0,03611042 | 1,025345712 | 0,92055173 | 0,993787755 | novel transcript, antisense to NCAM1 |
| ENSG00000124784 | RIOK1 | 0,03609603 | 1,025335485 | 0,89746229 | 0,991708791 | RIO kinase 1 |
| ENSG00000158470 | B4GALT5 | 0,036063662 | 1,025312481 | 0,8840819 | 0,990717729 | beta-1,4-galactosyltransferase 5 |
| ENSG00000163754 | GYG1 | 0,036015086 | 1,025277959 | 0,91624531 | 0,993361299 | glycogenin 1 |
| ENSG00000228450 | AL049734,1 | 0,036001998 | 1,025268658 | 0,97597829 | 0,99837368 | pseudogene similar to part of PYRIN-containing Apaf1-like protein 3 (PYPAF3) |
| ENSG00000130921 | C12orf65 | 0,035935751 | 1,025221579 | 0,904988 | 0,992590661 | chromosome 12 open reading frame 65 |
| ENSG00000143178 | TBX19 | 0,035832273 | 1,025148047 | 0,9292567 | 0,994293848 | T-box transcription factor 19 |
| ENSG00000262202 | AC007952,4 | 0,03575695 | 1,025094526 | 0,88632685 | 0,990717729 | novel transcript |
| ENSG00000139344 | AMDHD1 | 0,035735519 | 1,025079299 | 0,95672907 | 0,996608711 | amidohydrolase domain containing 1 |
| ENSG00000145736 | GTF2H2 | 0,035582236 | 1,024970393 | 0,89809002 | 0,991796521 | general transcription factor IIH subunit 2 |
| ENSG00000100523 | DDHD1 | 0,035558481 | 1,024953516 | 0,92162349 | 0,993787755 | DDHD domain containing 1 |
| ENSG00000259494 | MRPL46 | 0,035500424 | 1,02491227 | 0,90015074 | 0,991845106 | mitochondrial ribosomal protein L46 |
| ENSG00000237051 | GABBR1 | 0,035491104 | 1,024905649 | 0,97430357 | 0,99837368 | gamma-aminobutyric acid type B receptor subunit 1 |
| ENSG00000160050 | CCDC28B | 0,035461321 | 1,024884491 | 0,90826896 | 0,992595473 | coiled-coil domain containing 28B |
| ENSG00000223450 | AL590632,1 | 0,035447884 | 1,024874946 | 0,97635519 | 0,99837368 | ribosomal protein S24 (RPS24) pseudogene |
| ENSG00000163833 | FBXO40 | 0,035445441 | 1,02487321 | 0,98198272 | 0,99837368 | F-box protein 40 |
| ENSG00000165105 | RASEF | 0,035420586 | 1,024855554 | 0,93156359 | 0,994433362 | RAS and EF-hand domain containing |
| ENSG00000160957 | RECQL4 | 0,035360754 | 1,024813051 | 0,9268662 | 0,993915537 | RecQ like helicase 4 |
| ENSG00000197465 | GYPE | 0,035352973 | 1,024807524 | 0,95457455 | 0,996608711 | glycophorin E (MNS blood group) |
| ENSG00000276529 | AP001505,1 | 0,035303241 | 1,024772198 | 0,95580955 | 0,996608711 | novel transcript |
| ENSG00000102383 | ZDHHC15 | 0,035280136 | 1,024755786 | 0,92377305 | 0,993915537 | zinc finger DHHC-type palmitoyltransferase 15 |
| ENSG00000144535 | DIS3L2 | 0,035228481 | 1,024719096 | 0,88386258 | 0,990717729 | DIS3 like 3'-5' exoribonuclease 2 |
| ENSG00000134291 | TMEM106C | 0,035052981 | 1,024594449 | 0,91672945 | 0,993361299 | transmembrane protein 106C |
| ENSG00000135250 | SRPK2 | 0,035039967 | 1,024585207 | 0,90908813 | 0,992828195 | SRSF protein kinase 2 |
| ENSG00000258471 | AL161668,3 | 0,035022092 | 1,024572512 | 0,97692699 | 0,99837368 | novel transcript, antisense to SLC39A2 |
| ENSG00000225791 | TRAM2-AS1 | 0,034957161 | 1,024526401 | 0,89511683 | 0,991303328 | TRAM2 antisense RNA 1 (head to head) |
| ENSG00000240579 | AC006529,1 | 0,034923117 | 1,024502225 | 0,96676676 | 0,997263201 | pseudogene similar to part of CD2 (cytoplasmic tail) binding protein 2 (CD2BP2) |
| ENSG00000164934 | DCAF13 | 0,034909516 | 1,024492566 | 0,91748338 | 0,993361299 | DDB1 and CUL4 associated factor 13 |
| ENSG00000253309 | SERPINE3 | 0,034884153 | 1,024474555 | 0,96746563 | 0,997263201 | serpin family E member 3 |

| | | | | | | |
|---|---|---|---|---|---|---|
| ENSG00000117360 | PRPF3 | 0,034882725 | 1,024473541 | 0,86867991 | 0,989074806 | pre-mRNA processing factor 3 |
| ENSG00000280604 | AJ239328,1 | 0,034833429 | 1,024438536 | 0,94509512 | 0,99579981 | PCBP3 overlapping transcript 1 |
| ENSG00000285830 | AL109628,2 | 0,034810266 | 1,024422089 | 0,94106098 | 0,995251853 | novel transcript, antisense to TRAPPC6B |
| ENSG00000154380 | ENAH | 0,034729704 | 1,024364885 | 0,87031048 | 0,989074806 | ENAH actin regulator |
| ENSG00000227165 | WDR11-AS1 | 0,034721485 | 1,02435905 | 0,95577597 | 0,996608711 | WDR11 antisense RNA 1 |
| ENSG00000156261 | CCT8 | 0,034718044 | 1,024356606 | 0,93246247 | 0,994479239 | chaperonin containing TCP1 subunit 8 |
| ENSG00000267680 | ZNF224 | 0,034704179 | 1,024346762 | 0,92591675 | 0,993915537 | zinc finger protein 224 |
| ENSG00000101546 | RBFA | 0,034632461 | 1,024295841 | 0,90827058 | 0,992595473 | ribosome binding factor A |
| ENSG00000261546 | AC135782,3 | 0,034570852 | 1,024252101 | 0,9575155 | 0,996608711 | novel transcript, antisense to ACSF3 |
| ENSG00000227855 | DPY19L2P3 | 0,034559731 | 1,024244205 | 0,93796941 | 0,994488324 | DPY19L2 pseudogene 3 |
| ENSG00000287550 | AL450345,2 | 0,0345583 | 1,024243189 | 0,95317683 | 0,99652169 | novel transcript |
| ENSG00000173818 | ENDOV | 0,034541082 | 1,024230966 | 0,89990983 | 0,991845106 | endonuclease V |
| ENSG00000185630 | PBX1 | 0,034508969 | 1,024208168 | 0,92220234 | 0,993797669 | PBX homeobox 1 |
| ENSG00000230989 | HSBP1 | 0,034431538 | 1,024153199 | 0,94283273 | 0,995402494 | heat shock factor binding protein 1 |
| ENSG00000223960 | CHROMR | 0,03437251 | 1,024111296 | 0,88277284 | 0,990717729 | cholesterol induced regulator of metabolism RNA |
| ENSG00000218305 | CDC14C | 0,034356506 | 1,024099936 | 0,96248778 | 0,996608711 | cell division cycle 14C |
| ENSG00000111642 | CHD4 | 0,034322075 | 1,024075495 | 0,88317547 | 0,990717729 | chromodomain helicase DNA binding protein 4 |
| ENSG00000184182 | UBE2F | 0,034288879 | 1,024051931 | 0,91415149 | 0,993361299 | ubiquitin conjugating enzyme E2 F (putative) |
| ENSG00000251161 | AC020661,1 | 0,034280703 | 1,024046128 | 0,9500359 | 0,996432861 | novel transcript |
| ENSG00000130714 | POMT1 | 0,034279613 | 1,024045354 | 0,90227926 | 0,992230188 | protein O-mannosyltransferase 1 |
| ENSG00000141380 | SS18 | 0,034247303 | 1,024022421 | 0,93229504 | 0,994479239 | SS18 subunit of BAF chromatin remodeling complex |
| ENSG00000169612 | RAMAC | 0,034151559 | 1,023954464 | 0,93547147 | 0,994479239 | RNA guanine-7 methyltransferase activating subunit |
| ENSG00000180198 | RCC1 | 0,034119791 | 1,023931917 | 0,90214106 | 0,992230188 | regulator of chromosome condensation 1 |
| ENSG00000168092 | PAFAH1B2 | 0,034034664 | 1,023871501 | 0,92070945 | 0,993787755 | platelet activating factor acetylhydrolase 1b catalytic subunit 2 |
| ENSG00000069329 | VPS35 | 0,034007341 | 1,023852111 | 0,92954312 | 0,994293848 | VPS35 retromer complex component |
| ENSG00000276107 | AC037198,1 | 0,034000735 | 1,023847422 | 0,95944128 | 0,996608711 | novel transcript, sense intronic to THBS1 |
| ENSG00000149231 | CCDC82 | 0,033975776 | 1,023829709 | 0,92787058 | 0,994108531 | coiled-coil domain containing 82 |
| ENSG00000261526 | AC012615,1 | 0,033887866 | 1,023767325 | 0,93221774 | 0,994479239 | novel transcript |
| ENSG00000175581 | MRPL48 | 0,033775139 | 1,023687334 | 0,90439412 | 0,992590661 | mitochondrial ribosomal protein L48 |
| ENSG00000266236 | NARF-IT1 | 0,033759618 | 1,023676321 | 0,96147001 | 0,996608711 | NARF intronic transcript 1 |
| ENSG00000079435 | LIPE | 0,033699614 | 1,023633746 | 0,91053521 | 0,99320976 | lipase E, hormone sensitive type |
| ENSG00000271119 | AC026412,3 | 0,033677128 | 1,023617791 | 0,93888268 | 0,994713928 | novel transcript |

| | | | | | | |
|---|---|---|---|---|---|---|
| ENSG00000103343 | ZNF174 | 0,0336545 | 1,023601736 | 0,91233432 | 0,993361299 | zinc finger protein 174 |
| ENSG00000101574 | METTL4 | 0,033646034 | 1,02359573 | 0,93932087 | 0,994882028 | methyltransferase like 4 |
| ENSG00000232434 | AJM1 | 0,033558896 | 1,023533907 | 0,95637583 | 0,996608711 | apical junction component 1 homolog |
| ENSG00000224826 | USP21P1 | 0,033537183 | 1,023518503 | 0,97932828 | 0,99837368 | USP21 pseudogene 1 |
| ENSG00000166848 | TERF2IP | 0,033516858 | 1,023504083 | 0,87804127 | 0,990109814 | TERF2 interacting protein |
| ENSG00000111664 | GNB3 | 0,033506683 | 1,023496865 | 0,93416305 | 0,994479239 | G protein subunit beta 3 |
| ENSG00000124177 | CHD6 | 0,033501858 | 1,023493442 | 0,88100027 | 0,990598309 | chromodomain helicase DNA binding protein 6 |
| ENSG00000269473 | AC012313,8 | 0,033490878 | 1,023485652 | 0,93353702 | 0,994479239 | novel transcript |
| ENSG00000236516 | KLF2P4 | 0,033391835 | 1,023415391 | 0,95443587 | 0,996608711 | Kruppel like factor 2 pseudogene 4 |
| ENSG00000254838 | GVINP1 | 0,033380439 | 1,023407307 | 0,9181545 | 0,993361299 | GTPase, very large interferon inducible pseudogene 1 |
| ENSG00000163348 | PYGO2 | 0,033351296 | 1,023386634 | 0,87312657 | 0,989395767 | pygopus family PHD finger 2 |
| ENSG00000169413 | RNASE6 | 0,033299014 | 1,023349548 | 0,96797669 | 0,997263201 | ribonuclease A family member k6 |
| ENSG00000225934 | FAM133FP | 0,03327276 | 1,023330925 | 0,95865515 | 0,996608711 | family with sequence similarity 133 member F, pseudogene |
| ENSG00000117899 | MESD | 0,033232586 | 1,02330243 | 0,88560583 | 0,990717729 | mesoderm development LRP chaperone |
| ENSG00000178974 | FBXO34 | 0,033197598 | 1,023277613 | 0,92467902 | 0,993915537 | F-box protein 34 |
| ENSG00000161981 | SNRNP25 | 0,033184222 | 1,023268126 | 0,89147924 | 0,990820775 | small nuclear ribonucleoprotein U11/U12 subunit 25 |
| ENSG00000253210 | AC040970,1 | 0,033171927 | 1,023259405 | 0,9582916 | 0,996608711 | novel transcript |
| ENSG00000139714 | MORN3 | 0,03314432 | 1,023239824 | 0,95015339 | 0,996432861 | MORN repeat containing 3 |
| ENSG00000125484 | GTF3C4 | 0,03307438 | 1,023190221 | 0,90928082 | 0,992864002 | general transcription factor IIIC subunit 4 |
| ENSG00000258424 | AL512791,1 | 0,033057153 | 1,023178002 | 0,96756644 | 0,997263201 | novel transcript, antisense to CALM1 |
| ENSG00000136877 | FPGS | 0,033018756 | 1,023150771 | 0,92438041 | 0,993915537 | folylpolyglutamate synthase |
| ENSG00000206315 | PBX2 | 0,032925438 | 1,023084593 | 0,95889243 | 0,996608711 | PBX homeobox 2 |
| ENSG00000159082 | SYNJ1 | 0,032895588 | 1,023063425 | 0,91613868 | 0,993361299 | synaptojanin 1 |
| ENSG00000119705 | SLIRP | 0,032876999 | 1,023050243 | 0,90943356 | 0,992864002 | SRA stem-loop interacting RNA binding protein |
| ENSG00000146476 | ARMT1 | 0,032804743 | 1,022999006 | 0,94600827 | 0,996027077 | acidic residue methyltransferase 1 |
| ENSG00000213139 | CRYGS | 0,032799195 | 1,022995072 | 0,94294541 | 0,995402494 | crystallin gamma S |
| ENSG00000137073 | UBAP2 | 0,032787044 | 1,022986456 | 0,90319617 | 0,992311172 | ubiquitin associated protein 2 |
| ENSG00000143771 | CNIH4 | 0,032773414 | 1,022976791 | 0,90843888 | 0,992595616 | cornichon family AMPA receptor auxiliary protein 4 |
| ENSG00000159961 | OR3A3 | 0,032743942 | 1,022955893 | 0,97466065 | 0,99837368 | olfactory receptor family 3 subfamily A member 3 |
| ENSG00000215218 | UBE2QL1 | 0,032729847 | 1,0229459 | 0,94207601 | 0,995373376 | ubiquitin conjugating enzyme E2 Q family like 1 |
| ENSG00000176155 | CCDC57 | 0,0327105 | 1,022932181 | 0,94469951 | 0,995741983 | coiled-coil domain containing 57 |

| | | | | | | |
|---|---|---|---|---|---|---|
| ENSG00000198258 | UBL5 | 0,032676444 | 1,022908035 | 0,88967096 | 0,990717729 | ubiquitin like 5 |
| ENSG00000276248 | AL442125,1 | 0,032659023 | 1,022895683 | 0,94606398 | 0,996027077 | novel transcript, sense intronic to TMCO3 |
| ENSG00000156381 | ANKRD9 | 0,032620571 | 1,02286842 | 0,93638942 | 0,994479239 | ankyrin repeat domain 9 |
| ENSG00000198685 | LINC01565 | 0,0325856 | 1,022843625 | 0,97604173 | 0,99837368 | long intergenic non-protein coding RNA 1565 |
| ENSG00000118292 | C1orf54 | 0,032536977 | 1,022809153 | 0,93183819 | 0,994433362 | chromosome 1 open reading frame 54 |
| ENSG00000124614 | RPS10 | 0,032525325 | 1,022800893 | 0,89141036 | 0,990820775 | ribosomal protein S10 |
| ENSG00000110075 | PPP6R3 | 0,032518524 | 1,022796071 | 0,92127184 | 0,993787755 | protein phosphatase 6 regulatory subunit 3 |
| ENSG00000177303 | CASKIN2 | 0,032502761 | 1,022784896 | 0,9198945 | 0,993758187 | CASK interacting protein 2 |
| ENSG00000213033 | AURKAP1 | 0,032475035 | 1,02276524 | 0,94770378 | 0,996113938 | aurora kinase A pseudogene 1 |
| ENSG00000111276 | CDKN1B | 0,032446766 | 1,0227452 | 0,91810602 | 0,993361299 | cyclin dependent kinase inhibitor 1B |
| ENSG00000167394 | ZNF668 | 0,032367574 | 1,022689061 | 0,93164902 | 0,994433362 | zinc finger protein 668 |
| ENSG00000052126 | PLEKHA5 | 0,032351127 | 1,022677402 | 0,93099093 | 0,994433362 | pleckstrin homology domain containing A5 |
| ENSG00000180574 | EIF2S3B | 0,032316431 | 1,022652808 | 0,93600959 | 0,994479239 | eukaryotic translation initiation factor 2 subunit gamma B |
| ENSG00000204178 | MACO1 | 0,032275703 | 1,022623938 | 0,93332203 | 0,994479239 | macoilin 1 |
| ENSG00000171863 | RPS7 | 0,032231048 | 1,022592286 | 0,92144898 | 0,993787755 | ribosomal protein S7 |
| ENSG00000101146 | RAE1 | 0,032224354 | 1,022587541 | 0,89975938 | 0,991845106 | ribonucleic acid export 1 |
| ENSG00000010072 | SPRTN | 0,032196202 | 1,022567587 | 0,92310483 | 0,993825008 | SprT-like N-terminal domain |
| ENSG00000126217 | MCF2L | 0,032194462 | 1,022566354 | 0,9306396 | 0,994344567 | MCF,2 cell line derived transforming sequence like |
| ENSG00000187676 | B3GLCT | 0,032161449 | 1,022542955 | 0,94103332 | 0,995251853 | beta 3-glucosyltransferase |
| ENSG00000176595 | KBTBD11 | 0,032091394 | 1,022493303 | 0,96462385 | 0,997113804 | kelch repeat and BTB domain containing 11 |
| ENSG00000099942 | CRKL | 0,032022355 | 1,022444374 | 0,88579006 | 0,990717729 | CRK like proto-oncogene, adaptor protein |
| ENSG00000164096 | C4orf3 | 0,031974361 | 1,02241036 | 0,92754364 | 0,994094286 | chromosome 4 open reading frame 3 |
| ENSG00000072134 | EPN2 | 0,031972316 | 1,022408912 | 0,89287104 | 0,991120066 | epsin 2 |
| ENSG00000104320 | NBN | 0,031952332 | 1,022394749 | 0,93616794 | 0,994479239 | nibrin |
| ENSG00000114127 | XRN1 | 0,031950281 | 1,022393296 | 0,93565125 | 0,994479239 | 5'-3' exoribonuclease 1 |
| ENSG00000229061 | ATAT1 | 0,031940415 | 1,022386304 | 0,94550639 | 0,995917699 | alpha tubulin acetyltransferase 1 |
| ENSG00000206488 | ATAT1 | 0,031940415 | 1,022386304 | 0,94550639 | 0,995917699 | alpha tubulin acetyltransferase 1 |
| ENSG00000234549 | ATAT1 | 0,031940415 | 1,022386304 | 0,94550639 | 0,995917699 | alpha tubulin acetyltransferase 1 |
| ENSG00000138378 | STAT4 | 0,031924986 | 1,02237537 | 0,94232074 | 0,995373376 | signal transducer and activator of transcription 4 |
| ENSG00000109466 | KLHL2 | 0,03192407 | 1,022374721 | 0,92067208 | 0,993787755 | kelch like family member 2 |
| ENSG00000122550 | KLHL7 | 0,031842759 | 1,022317101 | 0,91459451 | 0,993361299 | kelch like family member 7 |
| ENSG00000226752 | CUTALP | 0,031838031 | 1,022313751 | 0,91367819 | 0,993361299 | cutA divalent cation tolerance like, pseudogene |

| | | | | | | |
|---|---|---|---|---|---|---|
| ENSG00000196814 | MVB12B | 0,031806664 | 1,022291524 | 0,89286876 | 0,991120066 | multivesicular body subunit 12B |
| ENSG00000163528 | CHCHD4 | 0,031714789 | 1,022226423 | 0,92584343 | 0,993915537 | coiled-coil-helix-coiled-coil-helix domain containing 4 |
| ENSG00000109381 | ELF2 | 0,031710826 | 1,022223615 | 0,93901445 | 0,994784787 | E74 like ETS transcription factor 2 |
| ENSG00000113448 | PDE4D | 0,03169041 | 1,02220915 | 0,93413633 | 0,994479239 | phosphodiesterase 4D |
| ENSG00000113657 | DPYSL3 | 0,031655619 | 1,022184499 | 0,8820375 | 0,990717729 | dihydropyrimidinase like 3 |
| ENSG00000084090 | STARD7 | 0,031647434 | 1,0221787 | 0,91373805 | 0,993361299 | StAR related lipid transfer domain containing 7 |
| ENSG00000267575 | AC006504,5 | 0,031559823 | 1,022116628 | 0,91802182 | 0,993361299 | novel transcript |
| ENSG00000230913 | NPM1P51 | 0,031390487 | 1,021996664 | 0,95823988 | 0,996608711 | nucleophosmin 1 pseudogene 51 |
| ENSG00000141378 | PTRH2 | 0,031389297 | 1,021995821 | 0,91938354 | 0,993598006 | peptidyl-tRNA hydrolase 2 |
| ENSG00000129292 | PHF20L1 | 0,031328598 | 1,021952823 | 0,92658941 | 0,993915537 | PHD finger protein 20 like 1 |
| ENSG00000100823 | APEX1 | 0,031316819 | 1,02194448 | 0,90684817 | 0,992590661 | apurinic/apyrimidinic endodeoxyribonuclease 1 |
| ENSG00000160584 | SIK3 | 0,03104348 | 1,021750876 | 0,88436629 | 0,990717729 | SIK family kinase 3 |
| ENSG00000111364 | DDX55 | 0,030995331 | 1,021716776 | 0,88367971 | 0,990717729 | DEAD-box helicase 55 |
| ENSG00000225589 | MDC1 | 0,030876202 | 1,021632413 | 0,95507486 | 0,996608711 | mediator of DNA damage checkpoint 1 |
| ENSG00000234012 | MDC1 | 0,030876202 | 1,021632413 | 0,95507486 | 0,996608711 | mediator of DNA damage checkpoint 1 |
| ENSG00000231135 | MDC1 | 0,030876202 | 1,021632413 | 0,95507486 | 0,996608711 | mediator of DNA damage checkpoint 1 |
| ENSG00000178988 | MRFAP1L1 | 0,030852607 | 1,021615704 | 0,91470025 | 0,993361299 | Morf4 family associated protein 1 like 1 |
| ENSG00000280873 | DGKD | 0,030736485 | 1,021533478 | 0,90194865 | 0,992182254 | diacylglycerol kinase delta |
| ENSG00000147400 | CETN2 | 0,030604069 | 1,021439722 | 0,90999019 | 0,992951496 | centrin 2 |
| ENSG00000277785 | SNORD116-21 | 0,030595272 | 1,021433493 | 0,97172294 | 0,998088605 | small nucleolar RNA, C/D box 116-21 |
| ENSG00000183765 | CHEK2 | 0,030557175 | 1,021406521 | 0,90581497 | 0,992590661 | checkpoint kinase 2 |
| ENSG00000240230 | COX19 | 0,030517273 | 1,021378271 | 0,89254059 | 0,991048031 | cytochrome c oxidase assembly factor COX19 |
| ENSG00000149485 | FADS1 | 0,030424888 | 1,021312868 | 0,9267926 | 0,993915537 | fatty acid desaturase 1 |
| ENSG00000164403 | SHROOM1 | 0,030391625 | 1,021289321 | 0,96211046 | 0,996608711 | shroom family member 1 |
| ENSG00000213020 | ZNF611 | 0,030325865 | 1,02124277 | 0,9099938 | 0,992951496 | zinc finger protein 611 |
| ENSG00000168671 | UGT3A2 | 0,030270932 | 1,021203885 | 0,96825779 | 0,997263201 | UDP glycosyltransferase family 3 member A2 |
| ENSG00000186462 | NAP1L2 | 0,030253416 | 1,021191487 | 0,92931587 | 0,994293848 | nucleosome assembly protein 1 like 2 |
| ENSG00000253200 | AC037459,2 | 0,030187149 | 1,021144582 | 0,92078347 | 0,993787755 | novel transcript, antisense to KIAA1967 |
| ENSG00000146731 | CCT6A | 0,030176253 | 1,02113687 | 0,92101235 | 0,993787755 | chaperonin containing TCP1 subunit 6A |
| ENSG00000273424 | AL008582,1 | 0,03010737 | 1,021088116 | 0,97428041 | 0,99837368 | novel transcript |
| ENSG00000197093 | GAL3ST4 | 0,030101461 | 1,021083933 | 0,91450498 | 0,993361299 | galactose-3-O-sulfotransferase 4 |
| ENSG00000123933 | MXD4 | 0,030100127 | 1,021082989 | 0,92909446 | 0,994293848 | MAX dimerization protein 4 |

| | | | | | | |
|---|---|---|---|---|---|---|
| ENSG00000234912 | SNHG20 | 0,030077426 | 1,021066922 | 0,93695793 | 0,994479239 | small nucleolar RNA host gene 20 |
| ENSG00000181896 | ZNF101 | 0,030044617 | 1,021043702 | 0,92472527 | 0,993915537 | zinc finger protein 101 |
| ENSG00000115468 | EFHD1 | 0,030042158 | 1,021041962 | 0,9279361 | 0,994108531 | EF-hand domain family member D1 |
| ENSG00000032389 | EIPR1 | 0,030007539 | 1,021017462 | 0,89162266 | 0,990889051 | EARP complex and GARP complex interacting protein 1 |
| ENSG00000075151 | EIF4G3 | 0,02995635 | 1,020981235 | 0,91442615 | 0,993361299 | eukaryotic translation initiation factor 4 gamma 3 |
| ENSG00000230734 | RPL10P3 | 0,029829044 | 1,020891146 | 0,96256868 | 0,996608711 | ribosomal protein L10 pseudogene 3 |
| ENSG00000054523 | KIF1B | 0,02976163 | 1,020843442 | 0,91789032 | 0,993361299 | kinesin family member 1B |
| ENSG00000253910 | PCDHGB2 | 0,029741253 | 1,020829024 | 0,91428852 | 0,993361299 | protocadherin gamma subfamily B, 2 |
| ENSG00000171045 | TSNARE1 | 0,029705896 | 1,020804006 | 0,94377365 | 0,995483785 | t-SNARE domain containing 1 |
| ENSG00000239672 | NME1 | 0,029686424 | 1,020790228 | 0,90017969 | 0,991845106 | NME/NM23 nucleoside diphosphate kinase 1 |
| ENSG00000278505 | C17orf78 | 0,029672525 | 1,020780394 | 0,9817178 | 0,99837368 | chromosome 17 open reading frame 78 |
| ENSG00000225215 | SMARCE1P1 | 0,029661923 | 1,020772893 | 0,96781945 | 0,997263201 | SWI/SNF related, matrix associated, actin dependent regulator of chromatin, subfamily e, member 1 pseudogene 1 |
| ENSG00000126950 | TMEM35A | 0,029644996 | 1,020760916 | 0,94649447 | 0,996027077 | transmembrane protein 35A |
| ENSG00000167186 | COQ7 | 0,029615564 | 1,020740092 | 0,91731481 | 0,993361299 | coenzyme Q7, hydroxylase |
| ENSG00000187980 | PLA2G2C | 0,029611683 | 1,020737346 | 0,95940738 | 0,996608711 | phospholipase A2 group IIC |
| ENSG00000287142 | AL355053,1 | 0,029573445 | 1,020710292 | 0,97892987 | 0,99837368 | novel transcript, antisense to PRKD1 |
| ENSG00000260853 | AC109460,2 | 0,029530897 | 1,02068019 | 0,93601104 | 0,994479239 | novel transcript, antisense to NFATC2IP |
| ENSG00000244528 | SEPTIN14P2 | 0,029451208 | 1,020623813 | 0,98447102 | 0,99837368 | septin 14 pseudogene 2 |
| ENSG00000105193 | RPS16 | 0,029424947 | 1,020605235 | 0,92523077 | 0,993915537 | ribosomal protein S16 |
| ENSG00000077232 | DNAJC10 | 0,029371899 | 1,020567708 | 0,93388005 | 0,994479239 | DnaJ heat shock protein family (Hsp40) member C10 |
| ENSG00000166847 | DCTN5 | 0,029361286 | 1,0205602 | 0,90852079 | 0,992603769 | dynactin subunit 5 |
| ENSG00000134202 | GSTM3 | 0,029347648 | 1,020550553 | 0,89276782 | 0,991120066 | glutathione S-transferase mu 3 |
| ENSG00000095015 | MAP3K1 | 0,029310814 | 1,020524497 | 0,93529909 | 0,994479239 | mitogen-activated protein kinase kinase kinase 1 |
| ENSG00000277104 | TADA2A | 0,029270221 | 1,020495783 | 0,93127329 | 0,994433362 | transcriptional adaptor 2A |
| ENSG00000151835 | SACS | 0,029261655 | 1,020489724 | 0,9298462 | 0,994331958 | sacsin molecular chaperone |
| ENSG00000272741 | AC069257,3 | 0,029239477 | 1,020474037 | 0,96526765 | 0,997208095 | novel protein |
| ENSG00000273174 | AC108673,2 | 0,029160521 | 1,02041819 | 0,9793609 | 0,99837368 | novel transcript, antisense to RAB43 |
| ENSG00000218537 | MIF-AS1 | 0,029118463 | 1,020388442 | 0,95317956 | 0,99652169 | MIF antisense RNA 1 |
| ENSG00000281335 | MIF-AS1 | 0,029118463 | 1,020388442 | 0,95317956 | 0,99652169 | MIF antisense RNA 1 |
| ENSG00000181704 | YIPF6 | 0,029103219 | 1,020377661 | 0,89597813 | 0,991528124 | Yip1 domain family member 6 |

| | | | | | | |
|---|---|---|---|---|---|---|
| ENSG00000260064 | AC092332,1 | 0,029100525 | 1,020375755 | 0,97618042 | 0,99837368 | novel transcript, sense intronic to CDYL2 |
| ENSG00000187735 | TCEA1 | 0,029035171 | 1,020329533 | 0,93471134 | 0,994479239 | transcription elongation factor A1 |
| ENSG00000128973 | CLN6 | 0,028960307 | 1,020276588 | 0,91065836 | 0,993222655 | CLN6 transmembrane ER protein |
| ENSG00000235060 | VDAC1P4 | 0,028932895 | 1,020257202 | 0,97959437 | 0,99837368 | voltage dependent anion channel 1 pseudogene 4 |
| ENSG00000146067 | FAM193B | 0,028858588 | 1,020204655 | 0,92847162 | 0,994108531 | family with sequence similarity 193 member B |
| ENSG00000055950 | MRPL43 | 0,028757473 | 1,020133153 | 0,90634444 | 0,992590661 | mitochondrial ribosomal protein L43 |
| ENSG00000183508 | TENT5C | 0,028716208 | 1,020103975 | 0,94838213 | 0,99635783 | terminal nucleotidyltransferase 5C |
| ENSG00000135775 | COG2 | 0,028620868 | 1,020036565 | 0,92563937 | 0,993915537 | component of oligomeric golgi complex 2 |
| ENSG00000258711 | AL358334,2 | 0,028619776 | 1,020035792 | 0,95868286 | 0,996608711 | novel transcript |
| ENSG00000164187 | LMBRD2 | 0,028577109 | 1,020005626 | 0,94977083 | 0,996432861 | LMBR1 domain containing 2 |
| ENSG00000128000 | ZNF780B | 0,028538996 | 1,01997868 | 0,96606443 | 0,997263201 | zinc finger protein 780B |
| ENSG00000080986 | NDC80 | 0,028535396 | 1,019976134 | 0,93379278 | 0,994479239 | NDC80 kinetochore complex component |
| ENSG00000196605 | ZNF846 | 0,028339834 | 1,019837883 | 0,92341424 | 0,993825008 | zinc finger protein 846 |
| ENSG00000165629 | ATP5F1C | 0,028300566 | 1,019810125 | 0,94552294 | 0,995917699 | ATP synthase F1 subunit gamma |
| ENSG00000249425 | LINC02477 | 0,028196093 | 1,019736278 | 0,98769083 | 0,99837368 | long intergenic non-protein coding RNA 2477 |
| ENSG00000213918 | DNASE1 | 0,0281862 | 1,019729285 | 0,90787392 | 0,992590661 | deoxyribonuclease 1 |
| ENSG00000231665 | OGFOD1P1 | 0,028107152 | 1,019673414 | 0,95935042 | 0,996608711 | 2-oxoglutarate and iron dependent oxygenase domain containing 1 pseudogene 1 |
| ENSG00000043355 | ZIC2 | 0,028084508 | 1,01965741 | 0,92736044 | 0,994000163 | Zic family member 2 |
| ENSG00000159111 | MRPL10 | 0,028032056 | 1,019620338 | 0,92692432 | 0,993915537 | mitochondrial ribosomal protein L10 |
| ENSG00000133878 | DUSP26 | 0,028008222 | 1,019603494 | 0,92710819 | 0,993915537 | dual specificity phosphatase 26 |
| ENSG00000084072 | PPIE | 0,027933261 | 1,019550518 | 0,90880005 | 0,992688304 | peptidylprolyl isomerase E |
| ENSG00000227220 | AL133346,1 | 0,027885247 | 1,019516587 | 0,96929001 | 0,997412278 | novel transcript |
| ENSG00000280975 | CENATAC | 0,027828515 | 1,019476496 | 0,90408569 | 0,992531054 | centrosomal AT-AC splicing factor |
| ENSG00000197301 | HMGA2-AS1 | 0,027736874 | 1,01941174 | 0,91112778 | 0,993222655 | HMGA2 antisense RNA 1 |
| ENSG00000136243 | NUP42 | 0,027719793 | 1,019399671 | 0,92629732 | 0,993915537 | nucleoporin 42 |
| ENSG00000162757 | C1orf74 | 0,027696989 | 1,019383558 | 0,93746896 | 0,994479239 | chromosome 1 open reading frame 74 |
| ENSG00000137496 | IL18BP | 0,027592678 | 1,019309856 | 0,92452405 | 0,993915537 | interleukin 18 binding protein |
| ENSG00000169016 | E2F6 | 0,027591369 | 1,019308932 | 0,9369226 | 0,994479239 | E2F transcription factor 6 |
| ENSG00000133226 | SRRM1 | 0,027443041 | 1,019204139 | 0,90944145 | 0,992864002 | serine and arginine repetitive matrix 1 |
| ENSG00000274570 | SPDYE10P | 0,027433946 | 1,019197714 | 0,96652937 | 0,997263201 | speedy/RINGO cell cycle regulator family member E10, pseudogene |
| ENSG00000251434 | AC104071,1 | 0,027378517 | 1,019158556 | 0,98016123 | 0,99837368 | novel transcript |

| | | | | | | |
|---|---|---|---|---|---|---|
| ENSG00000161939 | RNASEK-C17or[illegible] | 0,02735099 | 1,01913911 | 0,94692102 | 0,996027077 | RNASEK-C17orf49 readthrough |
| ENSG00000214367 | HAUS3 | 0,02720249 | 1,019034213 | 0,92538572 | 0,993915537 | HAUS augmin like complex subunit 3 |
| ENSG00000171204 | TMEM126B | 0,027166717 | 1,019008946 | 0,95789389 | 0,996608711 | transmembrane protein 126B |
| ENSG00000130299 | GTPBP3 | 0,027147098 | 1,018995089 | 0,93378984 | 0,994479239 | GTP binding protein 3, mitochondrial |
| ENSG00000254746 | AC103855,2 | 0,027118122 | 1,018974623 | 0,97008961 | 0,997890354 | novel transcript |
| ENSG00000276799 | WNT9B | 0,027059283 | 1,018933066 | 0,97988736 | 0,99837368 | Wnt family member 9B |
| ENSG00000177971 | IMP3 | 0,027013652 | 1,018900838 | 0,90714578 | 0,992590661 | IMP U3 small nucleolar ribonucleoprotein 3 |
| ENSG00000231652 | AL590428,1 | 0,027013387 | 1,018900651 | 0,96550498 | 0,997208095 | novel transcript |
| ENSG00000176148 | TCP11L1 | 0,026959287 | 1,018862444 | 0,90980493 | 0,992951496 | t-complex 11 like 1 |
| ENSG00000253741 | LNCOC1 | 0,02694356 | 1,018851337 | 0,96566728 | 0,997212942 | lncRNA associated with ovarian cancer 1 |
| ENSG00000111832 | RWDD1 | 0,026927218 | 1,018839796 | 0,93603997 | 0,994479239 | RWD domain containing 1 |
| ENSG00000249429 | AC016598,1 | 0,026924215 | 1,018837675 | 0,98732099 | 0,99837368 | novel transcript |
| ENSG00000152683 | SLC30A6 | 0,026855612 | 1,018789229 | 0,94909217 | 0,996432861 | solute carrier family 30 member 6 |
| ENSG00000064703 | DDX20 | 0,026778406 | 1,01873471 | 0,93957721 | 0,994943406 | DEAD-box helicase 20 |
| ENSG00000137332 | DDR1 | 0,026757864 | 1,018720204 | 0,92338451 | 0,993825008 | discoidin domain receptor tyrosine kinase 1 |
| ENSG00000132768 | DPH2 | 0,026685513 | 1,018669117 | 0,93116417 | 0,994433362 | diphthamide biosynthesis 2 |
| ENSG00000277879 | AL391988,1 | 0,026662194 | 1,018652652 | 0,95857364 | 0,996608711 | novel transcript, antisense to SLC18A2 |
| ENSG00000182180 | MRPS16 | 0,026636679 | 1,018634637 | 0,89425838 | 0,991135494 | mitochondrial ribosomal protein S16 |
| ENSG00000236485 | LINC01945 | 0,026546093 | 1,018570679 | 0,98670531 | 0,99837368 | long intergenic non-protein coding RNA 1945 |
| ENSG00000202363 | SNORA62 | 0,026520216 | 1,01855241 | 0,98740469 | 0,99837368 | small nucleolar RNA, H/ACA box 62 |
| ENSG00000271889 | AC016747,2 | 0,026458534 | 1,018508862 | 0,9813983 | 0,99837368 | novel transcript |
| ENSG00000058729 | RIOK2 | 0,026458162 | 1,0185086 | 0,94211355 | 0,995373376 | RIO kinase 2 |
| ENSG00000135423 | GLS2 | 0,026443926 | 1,01849855 | 0,97593709 | 0,99837368 | glutaminase 2 |
| ENSG00000103018 | CYB5B | 0,026359531 | 1,018438971 | 0,93717125 | 0,994479239 | cytochrome b5 type B |
| ENSG00000175792 | RUVBL1 | 0,026312925 | 1,018406071 | 0,91115422 | 0,993222655 | RuvB like AAA ATPase 1 |
| ENSG00000173744 | AGFG1 | 0,026279572 | 1,018382527 | 0,95219493 | 0,996432861 | ArfGAP with FG repeats 1 |
| ENSG00000169733 | RFNG | 0,026274352 | 1,018378842 | 0,92156673 | 0,993787755 | RFNG O-fucosylpeptide 3-beta-N-acetylglucosaminyltransferase |
| ENSG00000149300 | C11orf52 | 0,026247211 | 1,018359684 | 0,97971041 | 0,99837368 | chromosome 11 open reading frame 52 |
| ENSG00000239908 | RN7SL75P | 0,026224767 | 1,018343842 | 0,97914746 | 0,99837368 | RNA, 7SL, cytoplasmic 75, pseudogene |
| ENSG00000213563 | C8orf82 | 0,02619031 | 1,01831952 | 0,95199458 | 0,996432861 | chromosome 8 open reading frame 82 |
| ENSG00000111725 | PRKAB1 | 0,026130234 | 1,018277117 | 0,91399233 | 0,993361299 | protein kinase AMP-activated non-catalytic subunit beta 1 |

| | | | | | | |
|---|---|---|---|---|---|---|
| ENSG00000165525 | NEMF | 0,026071822 | 1,01823589 | 0,93538912 | 0,994479239 | nuclear export mediator factor |
| ENSG00000119318 | RAD23B | 0,025941383 | 1,018143831 | 0,93774232 | 0,994479239 | RAD23 homolog B, nucleotide excision repair protein |
| ENSG00000180182 | MED14 | 0,025906774 | 1,018119408 | 0,94188484 | 0,995373376 | mediator complex subunit 14 |
| ENSG00000100146 | SOX10 | 0,025901641 | 1,018115785 | 0,97787542 | 0,99837368 | SRY-box transcription factor 10 |
| ENSG00000284977 | AL160272,1 | 0,025898152 | 1,018113323 | 0,94705716 | 0,996027077 | novel transcript |
| ENSG00000198088 | NUP62CL | 0,025885347 | 1,018104286 | 0,96590616 | 0,997263201 | nucleoporin 62 C-terminal like |
| ENSG00000140403 | DNAJA4 | 0,02579757 | 1,018042345 | 0,92948256 | 0,994293848 | DnaJ heat shock protein family (Hsp40) member A4 |
| ENSG00000182307 | C8orf33 | 0,025774777 | 1,01802626 | 0,9194137 | 0,993598006 | chromosome 8 open reading frame 33 |
| ENSG00000102967 | DHODH | 0,025760474 | 1,018016168 | 0,9293574 | 0,994293848 | dihydroorotate dehydrogenase (quinone) |
| ENSG00000040933 | INPP4A | 0,025750668 | 1,018009248 | 0,90840053 | 0,992595616 | inositol polyphosphate-4-phosphatase type I A |
| ENSG00000129951 | PLPPR3 | 0,025718717 | 1,017986703 | 0,95616624 | 0,996608711 | phospholipid phosphatase related 3 |
| ENSG00000149735 | GPHA2 | 0,025668341 | 1,017951158 | 0,98424264 | 0,99837368 | glycoprotein hormone subunit alpha 2 |
| ENSG00000273008 | AC010864,1 | 0,025665401 | 1,017949083 | 0,94709209 | 0,996027077 | novel transcript |
| ENSG00000241863 | RPP21 | 0,02566434 | 1,017948334 | 0,91328296 | 0,993361299 | ribonuclease P/MRP subunit p21 |
| ENSG00000267665 | AC021683,3 | 0,025657776 | 1,017943703 | 0,97136868 | 0,998088605 | novel transcript |
| ENSG00000166529 | ZSCAN21 | 0,025610603 | 1,017910419 | 0,92275802 | 0,993797669 | zinc finger and SCAN domain containing 21 |
| ENSG00000153339 | TRAPPC8 | 0,025515788 | 1,017843524 | 0,95285323 | 0,99652169 | trafficking protein particle complex 8 |
| ENSG00000080815 | PSEN1 | 0,025415623 | 1,017772858 | 0,93424863 | 0,994479239 | presenilin 1 |
| ENSG00000230224 | PHBP9 | 0,025387904 | 1,017753304 | 0,96111817 | 0,996608711 | prohibitin pseudogene 9 |
| ENSG00000226797 | AC005828,1 | 0,025344436 | 1,017722639 | 0,9761223 | 0,99837368 | novel transcript, antisense to TANC2 |
| ENSG00000280733 | AC093642,2 | 0,025244023 | 1,017651807 | 0,97339346 | 0,99837368 | novel transcript |
| ENSG00000281213 | AC093642,2 | 0,025244023 | 1,017651807 | 0,97339346 | 0,99837368 | novel transcript |
| ENSG00000102978 | POLR2C | 0,02522481 | 1,017638255 | 0,92802015 | 0,994108531 | RNA polymerase II subunit C |
| ENSG00000115073 | ACTR1B | 0,025202991 | 1,017622864 | 0,91191656 | 0,993361299 | actin related protein 1B |
| ENSG00000173207 | CKS1B | 0,025194431 | 1,017616827 | 0,93738236 | 0,994479239 | CDC28 protein kinase regulatory subunit 1B |
| ENSG00000287503 | AC027243,3 | 0,025191239 | 1,017614575 | 0,9712785 | 0,998088605 | novel transcript, antisense to ETFA |
| ENSG00000139197 | PEX5 | 0,025160933 | 1,017593199 | 0,91151367 | 0,993292552 | peroxisomal biogenesis factor 5 |
| ENSG00000257267 | ZNF271P | 0,025136604 | 1,017576038 | 0,95170466 | 0,996432861 | zinc finger protein 271, pseudogene |
| ENSG00000100271 | TTLL1 | 0,02510674 | 1,017554975 | 0,91707718 | 0,993361299 | tubulin tyrosine ligase like 1 |
| ENSG00000273759 | AL117379,1 | 0,025060693 | 1,017522498 | 0,95778165 | 0,996608711 | novel transcript |
| ENSG00000175832 | ETV4 | 0,025023254 | 1,017496093 | 0,91054818 | 0,99320976 | ETS variant transcription factor 4 |
| ENSG00000257279 | AC127164,1 | 0,025014625 | 1,017490006 | 0,95355468 | 0,99652169 | novel transcript |

| | | | | | | |
|---|---|---|---|---|---|---|
| ENSG00000129055 | ANAPC13 | 0,024986489 | 1,017470164 | 0,94758257 | 0,996107107 | anaphase promoting complex subunit 13 |
| ENSG00000235374 | SSR4P1 | 0,024966747 | 1,01745624 | 0,95832212 | 0,996608711 | signal sequence receptor subunit 4 pseudogene 1 |
| ENSG00000155304 | HSPA13 | 0,02492067 | 1,017423745 | 0,95557813 | 0,996608711 | heat shock protein family A (Hsp70) member 13 |
| ENSG00000254876 | SUGT1P4-STRA | 0,024911434 | 1,017417232 | 0,94613212 | 0,996027077 | SUGT1P4-STRA6LP readthrough |
| ENSG00000287140 | AC066595,2 | 0,024831201 | 1,017360651 | 0,97402705 | 0,99837368 | novel transcript |
| ENSG00000160055 | TMEM234 | 0,024807556 | 1,017343977 | 0,91821738 | 0,993361299 | transmembrane protein 234 |
| ENSG00000185787 | MORF4L1 | 0,024728598 | 1,0172883 | 0,94210947 | 0,995373376 | mortality factor 4 like 1 |
| ENSG00000066926 | FECH | 0,024709208 | 1,017274628 | 0,92940679 | 0,994293848 | ferrochelatase |
| ENSG00000178035 | IMPDH2 | 0,024678647 | 1,017253079 | 0,92105604 | 0,993787755 | inosine monophosphate dehydrogenase 2 |
| ENSG00000254996 | ANKHD1-EIF4EI | 0,024678285 | 1,017252824 | 0,92615964 | 0,993915537 | ANKHD1-EIF4EBP3 readthrough |
| ENSG00000172336 | POP7 | 0,024667103 | 1,017244939 | 0,92795648 | 0,994108531 | POP7 homolog, ribonuclease P/MRP subunit |
| ENSG00000119314 | PTBP3 | 0,02455332 | 1,017164714 | 0,95492891 | 0,996608711 | polypyrimidine tract binding protein 3 |
| ENSG00000183311 | TUBB | 0,024453663 | 1,017094454 | 0,9426662 | 0,995390305 | tubulin beta class I |
| ENSG00000271551 | AL355297,3 | 0,024447267 | 1,017089944 | 0,96133433 | 0,996608711 | novel transcript, antisense to ARID1B |
| ENSG00000114062 | UBE3A | 0,024430994 | 1,017078472 | 0,93855374 | 0,994608643 | ubiquitin protein ligase E3A |
| ENSG00000165349 | SLC7A3 | 0,024411341 | 1,017064617 | 0,95191547 | 0,996432861 | solute carrier family 7 member 3 |
| ENSG00000110031 | LPXN | 0,024306845 | 1,016990953 | 0,93947103 | 0,994943406 | leupaxin |
| ENSG00000156017 | CARNMT1 | 0,024287798 | 1,016977526 | 0,9506523 | 0,996432861 | carnosine N-methyltransferase 1 |
| ENSG00000151353 | TMEM18 | 0,024267546 | 1,01696325 | 0,91700427 | 0,993361299 | transmembrane protein 18 |
| ENSG00000282405 | AC019257,7 | 0,024256894 | 1,016955742 | 0,95852117 | 0,996608711 | novel transcript, antisense to ARHGEF10 |
| ENSG00000117523 | PRRC2C | 0,024168392 | 1,016893358 | 0,92403023 | 0,993915537 | proline rich coiled-coil 2C |
| ENSG00000164576 | SAP30L | 0,024073618 | 1,016826558 | 0,92269297 | 0,993797669 | SAP30 like |
| ENSG00000255302 | EID1 | 0,024048013 | 1,016808512 | 0,94137288 | 0,99529668 | EP300 interacting inhibitor of differentiation 1 |
| ENSG00000147174 | GCNA | 0,024024327 | 1,016791818 | 0,95246171 | 0,996432861 | germ cell nuclear acidic peptidase |
| ENSG00000252481 | SCARNA13 | 0,024019804 | 1,016788631 | 0,92642561 | 0,993915537 | small Cajal body-specific RNA 13 |
| ENSG00000174007 | CEP19 | 0,023937798 | 1,016730836 | 0,94658149 | 0,996027077 | centrosomal protein 19 |
| ENSG00000165533 | TTC8 | 0,023871784 | 1,016684314 | 0,95011928 | 0,996432861 | tetratricopeptide repeat domain 8 |
| ENSG00000227372 | TP73-AS1 | 0,023867726 | 1,016681454 | 0,93973947 | 0,994943406 | TP73 antisense RNA 1 |
| ENSG00000163832 | ELP6 | 0,023862194 | 1,016677556 | 0,91437296 | 0,993361299 | elongator acetyltransferase complex subunit 6 |
| ENSG00000270072 | AC090559,2 | 0,023738249 | 1,016590215 | 0,97969752 | 0,99837368 | novel transcript |
| ENSG00000173227 | SYT12 | 0,023539037 | 1,016449851 | 0,96877591 | 0,997302342 | synaptotagmin 12 |
| ENSG00000198205 | ZXDA | 0,023352384 | 1,016318353 | 0,93626102 | 0,994479239 | zinc finger X-linked duplicated A |

| | | | | | | |
|---|---|---|---|---|---|---|
| ENSG00000066056 | TIE1 | 0,023329639 | 1,016302329 | 0,96220876 | 0,996608711 | tyrosine kinase with immunoglobulin like and EGF like domains 1 |
| ENSG00000204138 | PHACTR4 | 0,023252448 | 1,016247954 | 0,91133281 | 0,993222655 | phosphatase and actin regulator 4 |
| ENSG00000256612 | CYP2B7P | 0,023251332 | 1,016247168 | 0,97838772 | 0,99837368 | cytochrome P450 family 2 subfamily B member 7, pseudogene |
| ENSG00000187266 | EPOR | 0,023176079 | 1,01619416 | 0,95892921 | 0,996608711 | erythropoietin receptor |
| ENSG00000138326 | RPS24 | 0,023148 | 1,016174383 | 0,95133313 | 0,996432861 | ribosomal protein S24 |
| ENSG00000241228 | RPL12P31 | 0,023120821 | 1,016155239 | 0,97542782 | 0,99837368 | ribosomal protein L12 pseudogene 31 |
| ENSG00000197444 | OGDHL | 0,023119277 | 1,016154152 | 0,9339063 | 0,994479239 | oxoglutarate dehydrogenase L |
| ENSG00000116641 | DOCK7 | 0,023118802 | 1,016153817 | 0,95143272 | 0,996432861 | dedicator of cytokinesis 7 |
| ENSG00000236636 | AL627308,2 | 0,023088723 | 1,016132632 | 0,98102362 | 0,99837368 | novel pseudogene |
| ENSG00000172059 | KLF11 | 0,023068912 | 1,016118678 | 0,93825371 | 0,994547792 | Kruppel like factor 11 |
| ENSG00000229358 | DPY19L1P1 | 0,022997357 | 1,016068282 | 0,94710395 | 0,996027077 | DPY19L1 pseudogene 1 |
| ENSG00000009844 | VTA1 | 0,022994029 | 1,016065938 | 0,95849374 | 0,996608711 | vesicle trafficking 1 |
| ENSG00000114735 | HEMK1 | 0,022991971 | 1,016064488 | 0,9387777 | 0,994709504 | HemK methyltransferase family member 1 |
| ENSG00000021300 | PLEKHB1 | 0,022963062 | 1,016044128 | 0,94458912 | 0,99570566 | pleckstrin homology domain containing B1 |
| ENSG00000162594 | IL23R | 0,022958381 | 1,016040832 | 0,98614445 | 0,99837368 | interleukin 23 receptor |
| ENSG00000287047 | AL133395,1 | 0,022890718 | 1,01599318 | 0,97551952 | 0,99837368 | novel transcript, antisense to SORCS1 |
| ENSG00000137106 | GRHPR | 0,022856016 | 1,015968742 | 0,92286815 | 0,99381888 | glyoxylate and hydroxypyruvate reductase |
| ENSG00000272056 | AC013472,2 | 0,022828529 | 1,015949386 | 0,95953706 | 0,996608711 | novel transcript, antisense to AGBL5 |
| ENSG00000273489 | AC008264,2 | 0,022817011 | 1,015941274 | 0,95111504 | 0,996432861 | novel transcript, antisense to MKLN1 |
| ENSG00000284052 | AC006460,2 | 0,022791412 | 1,015923248 | 0,95233293 | 0,996432861 | novel transcript |
| ENSG00000145425 | RPS3A | 0,022772947 | 1,015910245 | 0,94872908 | 0,996432861 | ribosomal protein S3A |
| ENSG00000177465 | ACOT4 | 0,022726262 | 1,015877371 | 0,9558204 | 0,996608711 | acyl-CoA thioesterase 4 |
| ENSG00000177202 | SPACA4 | 0,022695211 | 1,015855507 | 0,98679165 | 0,99837368 | sperm acrosome associated 4 |
| ENSG00000285933 | AP003498,2 | 0,022643827 | 1,015819326 | 0,96921456 | 0,997412278 | novel transcript |
| ENSG00000179363 | TMEM31 | 0,022580337 | 1,015774623 | 0,97608511 | 0,99837368 | transmembrane protein 31 |
| ENSG00000174233 | ADCY6 | 0,022565792 | 1,015764382 | 0,94426583 | 0,995542144 | adenylate cyclase 6 |
| ENSG00000127540 | UQCR11 | 0,022525318 | 1,015735886 | 0,92612928 | 0,993915537 | ubiquinol-cytochrome c reductase, complex III subunit XI |
| ENSG00000050130 | JKAMP | 0,022464787 | 1,01569327 | 0,95519039 | 0,996608711 | JNK1/MAPK8 associated membrane protein |
| ENSG00000104140 | RHOV | 0,02245443 | 1,015685978 | 0,98144916 | 0,99837368 | ras homolog family member V |
| ENSG00000120694 | HSPH1 | 0,022418546 | 1,015660716 | 0,94122456 | 0,995251853 | heat shock protein family H (Hsp110) member 1 |

| | | | | | | |
|---|---|---|---|---|---|---|
| ENSG00000125629 | INSIG2 | 0,022363654 | 1,015622072 | 0,96020346 | 0,996608711 | insulin induced gene 2 |
| ENSG00000240996 | AL157400,4 | 0,022321 | 1,015592045 | 0,98080813 | 0,99837368 | novel transcript |
| ENSG00000285331 | AC090517,5 | 0,02230804 | 1,015582922 | 0,92183033 | 0,993797669 | novel transcript |
| ENSG00000139990 | DCAF5 | 0,022302391 | 1,015578945 | 0,93409754 | 0,994479239 | DDB1 and CUL4 associated factor 5 |
| ENSG00000132153 | DHX30 | 0,022297896 | 1,015575781 | 0,920273 | 0,993787755 | DExH-box helicase 30 |
| ENSG00000127947 | PTPN12 | 0,022275147 | 1,015559767 | 0,95106162 | 0,996432861 | protein tyrosine phosphatase non-receptor type 12 |
| ENSG00000060762 | MPC1 | 0,022224328 | 1,015523995 | 0,96214455 | 0,996608711 | mitochondrial pyruvate carrier 1 |
| ENSG00000282086 | PKD1P6 | 0,022180101 | 1,015492864 | 0,94313873 | 0,995402494 | polycystin 1, transient receptor potential channel interacting pseudogene 6 |
| ENSG00000105486 | LIG1 | 0,022171257 | 1,015486638 | 0,94754311 | 0,996107107 | DNA ligase 1 |
| ENSG00000092108 | SCFD1 | 0,022159631 | 1,015478455 | 0,95251383 | 0,996432861 | sec1 family domain containing 1 |
| ENSG00000213041 | AC093422,1 | 0,022115604 | 1,015447466 | 0,98505242 | 0,99837368 | ribosomal protein L17 (RPL17) pseudogene |
| ENSG00000157450 | RNF111 | 0,022056875 | 1,01540613 | 0,94726883 | 0,996063635 | ring finger protein 111 |
| ENSG00000215154 | AC141586,1 | 0,021971791 | 1,015346247 | 0,93591549 | 0,994479239 | potassium channel tetramerisation domain containing 5 (KCTD5) pseudogene |
| ENSG00000258634 | AL160006,1 | 0,021912711 | 1,015304669 | 0,93789088 | 0,994479239 | novel transcript, antisense to ALX3 |
| ENSG00000181513 | ACBD4 | 0,021910645 | 1,015303215 | 0,95190538 | 0,996432861 | acyl-CoA binding domain containing 4 |
| ENSG00000158201 | ABHD3 | 0,021891581 | 1,015289798 | 0,95880654 | 0,996608711 | abhydrolase domain containing 3, phospholipase |
| ENSG00000132510 | KDM6B | 0,021872527 | 1,01527639 | 0,93785069 | 0,994479239 | lysine demethylase 6B |
| ENSG00000187624 | C17orf97 | 0,021808094 | 1,015231047 | 0,94207797 | 0,995373376 | chromosome 17 open reading frame 97 |
| ENSG00000283985 | C17orf97 | 0,021808094 | 1,015231047 | 0,94207797 | 0,995373376 | chromosome 17 open reading frame 97 |
| ENSG00000225568 | AC093155,1 | 0,021803826 | 1,015228044 | 0,9759663 | 0,99837368 | pseudogene similar to part of SUMO-1 activating enzyme subunit 2 (UBA2) |
| ENSG00000156795 | NTAQ1 | 0,021779473 | 1,015210906 | 0,95318603 | 0,99652169 | N-terminal glutamine amidase 1 |
| ENSG00000107779 | BMPR1A | 0,021750888 | 1,015190791 | 0,96212063 | 0,996608711 | bone morphogenetic protein receptor type 1A |
| ENSG00000088325 | TPX2 | 0,021739654 | 1,015182886 | 0,92877681 | 0,994173587 | TPX2 microtubule nucleation factor |
| ENSG00000111142 | METAP2 | 0,021668968 | 1,015133148 | 0,94234978 | 0,995373376 | methionyl aminopeptidase 2 |
| ENSG00000196455 | PIK3R4 | 0,021615176 | 1,015095299 | 0,95030529 | 0,996432861 | phosphoinositide-3-kinase regulatory subunit 4 |
| ENSG00000087586 | AURKA | 0,021581612 | 1,015071683 | 0,94560729 | 0,995939435 | aurora kinase A |
| ENSG00000157045 | NTAN1 | 0,021558504 | 1,015055425 | 0,96455832 | 0,997113804 | N-terminal asparagine amidase |
| ENSG00000163046 | ANKRD30BL | 0,021445 | 1,014975568 | 0,98518971 | 0,99837368 | ankyrin repeat domain 30B like |
| ENSG00000158793 | NIT1 | 0,021433297 | 1,014967334 | 0,92933525 | 0,994293848 | nitrilase 1 |
| ENSG00000285390 | ATP5IF1 | 0,021425117 | 1,01496158 | 0,93449933 | 0,994479239 | ATP synthase inhibitory factor subunit 1 |

| | | | | | | |
|---|---|---|---|---|---|---|
| ENSG00000130770 | ATP5IF1 | 0,021425117 | 1,01496158 | 0,93449933 | 0,994479239 | ATP synthase inhibitory factor subunit 1 |
| ENSG00000204628 | RACK1 | 0,021411991 | 1,014952346 | 0,93136805 | 0,994433362 | receptor for activated C kinase 1 |
| ENSG00000088247 | KHSRP | 0,021336155 | 1,014898996 | 0,91838205 | 0,99340194 | KH-type splicing regulatory protein |
| ENSG00000224078 | SNHG14 | 0,021315227 | 1,014884273 | 0,93320077 | 0,994479239 | small nucleolar RNA host gene 14 |
| ENSG00000055130 | CUL1 | 0,021231457 | 1,014825345 | 0,95187593 | 0,996432861 | cullin 1 |
| ENSG00000010292 | NCAPD2 | 0,021143548 | 1,014763511 | 0,92469341 | 0,993915537 | non-SMC condensin I complex subunit D2 |
| ENSG00000265272 | RN7SL693P | 0,021078623 | 1,014717844 | 0,98178903 | 0,99837368 | RNA, 7SL, cytoplasmic 693, pseudogene |
| ENSG00000126759 | CFP | 0,021056577 | 1,014702339 | 0,97551341 | 0,99837368 | complement factor properdin |
| ENSG00000245060 | LINC00847 | 0,021043578 | 1,014693196 | 0,93764329 | 0,994479239 | long intergenic non-protein coding RNA 847 |
| ENSG00000104221 | BRF2 | 0,0210393 | 1,014690187 | 0,93006446 | 0,994331958 | BRF2 RNA polymerase III transcription initiation factor subunit |
| ENSG00000159256 | MORC3 | 0,020901749 | 1,014593448 | 0,95467465 | 0,996608711 | MORC family CW-type zinc finger 3 |
| ENSG00000176024 | ZNF613 | 0,020899871 | 1,014592127 | 0,95518022 | 0,996608711 | zinc finger protein 613 |
| ENSG00000083845 | RPS5 | 0,020843693 | 1,01455262 | 0,93481338 | 0,994479239 | ribosomal protein S5 |
| ENSG00000151366 | NDUFC2 | 0,020814205 | 1,014531884 | 0,9409903 | 0,995251853 | NADH:ubiquinone oxidoreductase subunit C2 |
| ENSG00000224668 | IPO8P1 | 0,020758108 | 1,014492435 | 0,98430368 | 0,99837368 | importin 8 pseudogene 1 |
| ENSG00000277903 | U2 | 0,020726267 | 1,014470046 | 0,96098589 | 0,996608711 | U2 spliceosomal RNA |
| ENSG00000274862 | U2 | 0,020726267 | 1,014470046 | 0,96098589 | 0,996608711 | U2 spliceosomal RNA |
| ENSG00000274062 | U2 | 0,020726267 | 1,014470046 | 0,96098589 | 0,996608711 | U2 spliceosomal RNA |
| ENSG00000273709 | U2 | 0,020726267 | 1,014470046 | 0,96098589 | 0,996608711 | U2 spliceosomal RNA |
| ENSG00000278048 | U2 | 0,020726267 | 1,014470046 | 0,96098589 | 0,996608711 | U2 spliceosomal RNA |
| ENSG00000275219 | U2 | 0,020726267 | 1,014470046 | 0,96098589 | 0,996608711 | U2 spliceosomal RNA |
| ENSG00000275616 | U2 | 0,020726267 | 1,014470046 | 0,96098589 | 0,996608711 | U2 spliceosomal RNA |
| ENSG00000278591 | U2 | 0,020726267 | 1,014470046 | 0,96098589 | 0,996608711 | U2 spliceosomal RNA |
| ENSG00000274432 | U2 | 0,020726267 | 1,014470046 | 0,96098589 | 0,996608711 | U2 spliceosomal RNA |
| ENSG00000276596 | U2 | 0,020726267 | 1,014470046 | 0,96098589 | 0,996608711 | U2 spliceosomal RNA |
| ENSG00000274585 | RNU2-1 | 0,020726267 | 1,014470046 | 0,96098589 | 0,996608711 | RNA, U2 small nuclear 1 |
| ENSG00000257218 | GATC | 0,020705176 | 1,014455215 | 0,93346288 | 0,994479239 | glutamyl-tRNA amidotransferase subunit C |
| ENSG00000269463 | AP001160,4 | 0,020698959 | 1,014450843 | 0,98716317 | 0,99837368 | novel transcript, sense intronic to STX5 |
| ENSG00000225335 | AC016027,1 | 0,020612988 | 1,014390393 | 0,96886189 | 0,997302342 | novel transcript |
| ENSG00000085721 | RRN3 | 0,020606271 | 1,01438567 | 0,95603775 | 0,996608711 | RRN3 homolog, RNA polymerase I transcription factor |
| ENSG00000101079 | NDRG3 | 0,020576546 | 1,014364771 | 0,9578916 | 0,996608711 | NDRG family member 3 |

| | | | | | | |
|---|---|---|---|---|---|---|
| ENSG00000100288 | CHKB | 0,020567755 | 1,01435859 | 0,93172398 | 0,994433362 | choline kinase beta |
| ENSG00000261460 | AC009690,2 | 0,020491722 | 1,014305132 | 0,96595357 | 0,997263201 | novel transcript, antisense to HEXA and CELF6 |
| ENSG00000168137 | SETD5 | 0,020486425 | 1,014301408 | 0,9357634 | 0,994479239 | SET domain containing 5 |
| ENSG00000141076 | UTP4 | 0,020485836 | 1,014300994 | 0,93098094 | 0,994433362 | UTP4 small subunit processome component |
| ENSG00000218336 | TENM3 | 0,02046012 | 1,014282914 | 0,92952816 | 0,994293848 | teneurin transmembrane protein 3 |
| ENSG00000143303 | RRNAD1 | 0,020420193 | 1,014254844 | 0,94164659 | 0,995373376 | ribosomal RNA adenine dimethylase domain containing 1 |
| ENSG00000254682 | AP002387,1 | 0,020404969 | 1,014244142 | 0,96490669 | 0,997113804 | novel transcript |
| ENSG00000155827 | RNF20 | 0,020373158 | 1,014221778 | 0,94242396 | 0,995373376 | ring finger protein 20 |
| ENSG00000109536 | FRG1 | 0,020290348 | 1,014163563 | 0,96493341 | 0,997113804 | FSHD region gene 1 |
| ENSG00000283153 | FRG1 | 0,020290348 | 1,014163563 | 0,96493341 | 0,997113804 | FSHD region gene 1 |
| ENSG00000283630 | FRG1 | 0,020290348 | 1,014163563 | 0,96493341 | 0,997113804 | FSHD region gene 1 |
| ENSG00000177943 | MAMDC4 | 0,020272834 | 1,014151252 | 0,96709972 | 0,997263201 | MAM domain containing 4 |
| ENSG00000224790 | AP000704,1 | 0,020184836 | 1,014089395 | 0,96948271 | 0,997495804 | novel transcript |
| ENSG00000120664 | SPART-AS1 | 0,020135775 | 1,01405491 | 0,96515037 | 0,997208095 | SPART antisense RNA 1 |
| ENSG00000272202 | AC097358,2 | 0,020119051 | 1,014043155 | 0,97340393 | 0,99837368 | novel transcript, antisense to ARHGEF3 |
| ENSG00000146006 | LRRTM2 | 0,020111888 | 1,01403812 | 0,96176596 | 0,996608711 | leucine rich repeat transmembrane neuronal 2 |
| ENSG00000146477 | SLC22A3 | 0,020111839 | 1,014038086 | 0,98011149 | 0,99837368 | solute carrier family 22 member 3 |
| ENSG00000259820 | AC083843,2 | 0,020084321 | 1,014018744 | 0,96691888 | 0,997263201 | novel transcript |
| ENSG00000197111 | PCBP2 | 0,020082493 | 1,014017459 | 0,95056004 | 0,996432861 | poly(rC) binding protein 2 |
| ENSG00000246323 | AC113382,1 | 0,020075757 | 1,014012725 | 0,9805848 | 0,99837368 | novel transcript, antisense to FAM13B |
| ENSG00000180008 | SOCS4 | 0,020058359 | 1,014000496 | 0,9655844 | 0,997208095 | suppressor of cytokine signaling 4 |
| ENSG00000161281 | COX7A1 | 0,020050569 | 1,013995021 | 0,98894623 | 0,99837368 | cytochrome c oxidase subunit 7A1 |
| ENSG00000286994 | AC069243,1 | 0,020045437 | 1,013991415 | 0,96563069 | 0,997208095 | novel transcript, sense intronic to GOLIM4 |
| ENSG00000169962 | TAS1R3 | 0,01996927 | 1,013937882 | 0,98053727 | 0,99837368 | taste 1 receptor member 3 |
| ENSG00000213920 | MDP1 | 0,019940099 | 1,013917381 | 0,94668266 | 0,996027077 | magnesium dependent phosphatase 1 |
| ENSG00000285200 | MDP1 | 0,019940099 | 1,013917381 | 0,94668266 | 0,996027077 | magnesium dependent phosphatase 1 |
| ENSG00000237523 | LINC00857 | 0,019938998 | 1,013916607 | 0,97889825 | 0,99837368 | long intergenic non-protein coding RNA 857 |
| ENSG00000103091 | WDR59 | 0,019912458 | 1,013897955 | 0,93628042 | 0,994479239 | WD repeat domain 59 |
| ENSG00000182901 | RGS7 | 0,019878834 | 1,013874325 | 0,96227556 | 0,996608711 | regulator of G protein signaling 7 |
| ENSG00000054148 | PHPT1 | 0,019837882 | 1,013845546 | 0,94018847 | 0,994949796 | phosphohistidine phosphatase 1 |
| ENSG00000258777 | HIF1A-AS1 | 0,019833731 | 1,013842629 | 0,9834399 | 0,99837368 | HIF1A antisense RNA 1 |
| ENSG00000198018 | ENTPD7 | 0,019813721 | 1,013828567 | 0,95407156 | 0,996608711 | ectonucleoside triphosphate diphosphohydrolase 7 |

| | | | | | | |
|---|---|---|---|---|---|---|
| ENSG00000187954 | CYHR1 | 0,01975494 | 1,01378726 | 0,95786867 | 0,996608711 | cysteine and histidine rich 1 |
| ENSG00000159714 | ZDHHC1 | 0,019703231 | 1,013750925 | 0,94897106 | 0,996432861 | zinc finger DHHC-type containing 1 |
| ENSG00000180098 | TRNAU1AP | 0,019686544 | 1,0137392 | 0,93034395 | 0,994336072 | tRNA selenocysteine 1 associated protein 1 |
| ENSG00000083635 | NUFIP1 | 0,01961735 | 1,01369058 | 0,9547303 | 0,996608711 | nuclear FMR1 interacting protein 1 |
| ENSG00000110713 | NUP98 | 0,019576409 | 1,013661814 | 0,9490089 | 0,996432861 | nucleoporin 98 and 96 precursor |
| ENSG00000184905 | TCEAL2 | 0,019575124 | 1,013660911 | 0,96591342 | 0,997263201 | transcription elongation factor A like 2 |
| ENSG00000188846 | RPL14 | 0,019548219 | 1,013642007 | 0,95235859 | 0,996432861 | ribosomal protein L14 |
| ENSG00000137177 | KIF13A | 0,019546084 | 1,013640507 | 0,93116642 | 0,994433362 | kinesin family member 13A |
| ENSG00000143376 | SNX27 | 0,019499605 | 1,013607851 | 0,93637654 | 0,994479239 | sorting nexin 27 |
| ENSG00000108924 | HLF | 0,019414367 | 1,013547967 | 0,97712755 | 0,99837368 | HLF transcription factor, PAR bZIP family member |
| ENSG00000223576 | AL355001,1 | 0,019413901 | 1,013547639 | 0,98821846 | 0,99837368 | novel transcript |
| ENSG00000242818 | RN7SL846P | 0,019403781 | 1,01354053 | 0,990455 | 0,998672296 | RNA, 7SL, cytoplasmic 846, pseudogene |
| ENSG00000228127 | LINC01649 | 0,019185608 | 1,013387268 | 0,98695781 | 0,99837368 | long intergenic non-protein coding RNA 1649 |
| ENSG00000170606 | HSPA4 | 0,019184233 | 1,013386302 | 0,95300189 | 0,99652169 | heat shock protein family A (Hsp70) member 4 |
| ENSG00000140307 | GTF2A2 | 0,019138137 | 1,013353923 | 0,96909323 | 0,997408593 | general transcription factor IIA subunit 2 |
| ENSG00000285723 | AL034430,2 | 0,019089568 | 1,013319809 | 0,95853508 | 0,996608711 | novel protein |
| ENSG00000146859 | TMEM140 | 0,019011457 | 1,013264947 | 0,93428855 | 0,994479239 | transmembrane protein 140 |
| ENSG00000115760 | BIRC6 | 0,018963157 | 1,013231024 | 0,9573409 | 0,996608711 | baculoviral IAP repeat containing 6 |
| ENSG00000169689 | CENPX | 0,018943755 | 1,013217398 | 0,93978521 | 0,994943406 | centromere protein X |
| ENSG00000197557 | TTC30A | 0,018910721 | 1,013194198 | 0,96684819 | 0,997263201 | tetratricopeptide repeat domain 30A |
| ENSG00000157911 | PEX10 | 0,018903462 | 1,0131891 | 0,94495168 | 0,99578567 | peroxisomal biogenesis factor 10 |
| ENSG00000258033 | SNX5P2 | 0,018808617 | 1,013122494 | 0,97862054 | 0,99837368 | sorting nexin 5 pseudogene 2 |
| ENSG00000186184 | POLR1D | 0,018779223 | 1,013101852 | 0,94755702 | 0,996107107 | RNA polymerase I and III subunit D |
| ENSG00000186625 | KATNA1 | 0,01876592 | 1,01309251 | 0,95149943 | 0,996432861 | katanin catalytic subunit A1 |
| ENSG00000167549 | CORO6 | 0,018750054 | 1,013081369 | 0,96332892 | 0,996608711 | coronin 6 |
| ENSG00000159761 | C16orf86 | 0,018635749 | 1,013001105 | 0,9684726 | 0,997302342 | chromosome 16 open reading frame 86 |
| ENSG00000282024 | AL009031,1 | 0,018400469 | 1,012835915 | 0,97868864 | 0,99837368 | novel transcript, antisense to GMPR |
| ENSG00000126653 | NSRP1 | 0,01839656 | 1,012833171 | 0,96104503 | 0,996608711 | nuclear speckle splicing regulatory protein 1 |
| ENSG00000285203 | LTB4R2 | 0,018386757 | 1,012826289 | 0,97124561 | 0,998088605 | leukotriene B4 receptor 2 |
| ENSG00000213906 | LTB4R2 | 0,018386757 | 1,012826289 | 0,97124561 | 0,998088605 | leukotriene B4 receptor 2 |
| ENSG00000186532 | SMYD4 | 0,018353276 | 1,012802785 | 0,93313305 | 0,994479239 | SET and MYND domain containing 4 |
| ENSG00000168301 | KCTD6 | 0,018346332 | 1,012797909 | 0,97092384 | 0,998088605 | potassium channel tetramerization domain containing 6 |

| | | | | | | |
|---|---|---|---|---|---|---|
| ENSG00000149743 | TRPT1 | 0,018331536 | 1,012787522 | 0,94718732 | 0,996060023 | tRNA phosphotransferase 1 |
| ENSG00000101440 | ASIP | 0,018283963 | 1,012754126 | 0,98753288 | 0,99837368 | agouti signaling protein |
| ENSG00000265800 | AC022211,3 | 0,018136011 | 1,012650271 | 0,97382391 | 0,99837368 | novel transcript, sense intronic to HN1 |
| ENSG00000132207 | SLX1A | 0,018087994 | 1,012616568 | 0,94142681 | 0,99529668 | SLX1 homolog A, structure-specific endonuclease subunit |
| ENSG00000172602 | RND1 | 0,018015411 | 1,012565624 | 0,96351434 | 0,996733015 | Rho family GTPase 1 |
| ENSG00000165501 | LRR1 | 0,017982021 | 1,012542189 | 0,95516463 | 0,996608711 | leucine rich repeat protein 1 |
| ENSG00000084110 | HAL | 0,017963739 | 1,012529358 | 0,98849534 | 0,99837368 | histidine ammonia-lyase |
| ENSG00000242082 | SLC5A4-AS1 | 0,01790494 | 1,012488092 | 0,95285063 | 0,99652169 | SLC5A4 antisense RNA 1 |
| ENSG00000131374 | TBC1D5 | 0,017787978 | 1,012406011 | 0,95645945 | 0,996608711 | TBC1 domain family member 5 |
| ENSG00000281935 | DNAH10 | 0,017737989 | 1,012370931 | 0,96224327 | 0,996608711 | dynein axonemal heavy chain 10 |
| ENSG00000166170 | BAG5 | 0,017721134 | 1,012359104 | 0,95402255 | 0,996608711 | BAG cochaperone 5 |
| ENSG00000170473 | PYM1 | 0,017715608 | 1,012355227 | 0,94608905 | 0,996027077 | PYM homolog 1, exon junction complex associated factor |
| ENSG00000262049 | AC139530,1 | 0,017707312 | 1,012349406 | 0,94963648 | 0,996432861 | novel transcript, antisense to HGS |
| ENSG00000146085 | MMUT | 0,017626499 | 1,012292699 | 0,95594812 | 0,996608711 | methylmalonyl-CoA mutase |
| ENSG00000149136 | SSRP1 | 0,017624845 | 1,012291539 | 0,93156473 | 0,994433362 | structure specific recognition protein 1 |
| ENSG00000210127 | MT-TA | 0,017608918 | 1,012280364 | 0,97876945 | 0,99837368 | mitochondrially encoded tRNA-Ala (GCN) |
| ENSG00000286607 | AL135818,3 | 0,017595528 | 1,012270969 | 0,98479912 | 0,99837368 | novel transcript |
| ENSG00000125970 | RALY | 0,017499316 | 1,012203464 | 0,93497419 | 0,994479239 | RALY heterogeneous nuclear ribonucleoprotein |
| ENSG00000230783 | RPS3AP13 | 0,017450457 | 1,012169184 | 0,98788497 | 0,99837368 | RPS3A pseudogene 13 |
| ENSG00000181045 | SLC26A11 | 0,017418546 | 1,012146796 | 0,95292689 | 0,99652169 | solute carrier family 26 member 11 |
| ENSG00000162927 | PUS10 | 0,017416758 | 1,012145542 | 0,96788379 | 0,997263201 | pseudouridine synthase 10 |
| ENSG00000173673 | HES3 | 0,017370135 | 1,012112833 | 0,98045356 | 0,99837368 | hes family bHLH transcription factor 3 |
| ENSG00000114391 | RPL24 | 0,017256287 | 1,012032967 | 0,95875734 | 0,996608711 | ribosomal protein L24 |
| ENSG00000100038 | TOP3B | 0,017254918 | 1,012032007 | 0,9451562 | 0,99579981 | DNA topoisomerase III beta |
| ENSG00000145414 | NAF1 | 0,016957693 | 1,011823529 | 0,94707195 | 0,996027077 | nuclear assembly factor 1 ribonucleoprotein |
| ENSG00000267100 | ILF3-DT | 0,016954595 | 1,011821356 | 0,94207676 | 0,995373376 | ILF3 divergent transcript |
| ENSG00000197713 | RPE | 0,016928591 | 1,011803118 | 0,97091874 | 0,998088605 | ribulose-5-phosphate-3-epimerase |
| ENSG00000198252 | STYX | 0,016872162 | 1,011763544 | 0,95172293 | 0,996432861 | serine/threonine/tyrosine interacting protein |
| ENSG00000136937 | NCBP1 | 0,01679171 | 1,011707125 | 0,96191927 | 0,996608711 | nuclear cap binding protein subunit 1 |
| ENSG00000112984 | KIF20A | 0,016643979 | 1,011603532 | 0,9571059 | 0,996608711 | kinesin family member 20A |
| ENSG00000067533 | RRP15 | 0,016579044 | 1,011558001 | 0,9558238 | 0,996608711 | ribosomal RNA processing 15 homolog |

| | | | | | | |
|---|---|---|---|---|---|---|
| ENSG00000247134 | AC090204,1 | 0,016561573 | 1,011545751 | 0,9726278 | 0,998267798 | novel transcript |
| ENSG00000171988 | JMJD1C | 0,016548661 | 1,011536698 | 0,96883243 | 0,997302342 | jumonji domain containing 1C |
| ENSG00000270055 | AC127502,2 | 0,016514797 | 1,011512954 | 0,95195871 | 0,996432861 | novel transcript |
| ENSG00000151422 | FER | 0,016508771 | 1,01150873 | 0,96868396 | 0,997302342 | FER tyrosine kinase |
| ENSG00000272391 | POM121C | 0,016490834 | 1,011496154 | 0,93982784 | 0,994943406 | POM121 transmembrane nucleoporin C |
| ENSG00000136381 | IREB2 | 0,01644049 | 1,011460857 | 0,9667506 | 0,997263201 | iron responsive element binding protein 2 |
| ENSG00000216475 | AL024474,1 | 0,01641457 | 1,011442685 | 0,98744355 | 0,99837368 | single-stranded DNA binding protein 1 (SSBP1) pseudogene |
| ENSG00000051382 | PIK3CB | 0,01638652 | 1,01142302 | 0,96301386 | 0,996608711 | phosphatidylinositol-4,5-bisphosphate 3-kinase catalytic subunit beta |
| ENSG00000108219 | TSPAN14 | 0,016386064 | 1,011422701 | 0,94420292 | 0,995542144 | tetraspanin 14 |
| ENSG00000285554 | AC242988,2 | 0,016371549 | 1,011412524 | 0,9815856 | 0,99837368 | Novel transcript |
| ENSG00000166313 | APBB1 | 0,016367562 | 1,01140973 | 0,94834231 | 0,99635783 | amyloid beta precursor protein binding family B member 1 |
| ENSG00000205639 | MFSD2B | 0,016332096 | 1,011384866 | 0,97888708 | 0,99837368 | major facilitator superfamily domain containing 2B |
| ENSG00000235316 | DUSP8P5 | 0,016309608 | 1,011369102 | 0,96015382 | 0,996608711 | dual specificity phosphatase 8 pseudogene 5 |
| ENSG00000196531 | NACA | 0,016232162 | 1,011314811 | 0,96814013 | 0,997263201 | nascent polypeptide associated complex subunit alpha |
| ENSG00000180815 | MAP3K15 | 0,01622026 | 1,011306468 | 0,96827428 | 0,997263201 | mitogen-activated protein kinase kinase kinase 15 |
| ENSG00000167654 | ATCAY | 0,016188046 | 1,011283887 | 0,9446768 | 0,995741983 | ATCAY kinesin light chain interacting caytaxin |
| ENSG00000143258 | USP21 | 0,016165393 | 1,011268007 | 0,94451998 | 0,995666356 | ubiquitin specific peptidase 21 |
| ENSG00000144021 | CIAO1 | 0,016147811 | 1,011255684 | 0,94771651 | 0,996113938 | cytosolic iron-sulfur assembly component 1 |
| ENSG00000213339 | QTRT1 | 0,016130224 | 1,011243356 | 0,96125195 | 0,996608711 | queuine tRNA-ribosyltransferase catalytic subunit 1 |
| ENSG00000270136 | MICOS10-NBL1 | 0,016123814 | 1,011238863 | 0,99675026 | 0,999708243 | MICOS10-NBL1 readthrough |
| ENSG00000101544 | ADNP2 | 0,016110707 | 1,011229676 | 0,95837998 | 0,996608711 | ADNP homeobox 2 |
| ENSG00000115524 | SF3B1 | 0,016092774 | 1,011217106 | 0,96799266 | 0,997263201 | splicing factor 3b subunit 1 |
| ENSG00000114166 | KAT2B | 0,01604973 | 1,011186936 | 0,96777868 | 0,997263201 | lysine acetyltransferase 2B |
| ENSG00000227963 | RBM15-AS1 | 0,01597636 | 1,011135512 | 0,98527132 | 0,99837368 | RBM15 antisense RNA 1 |
| ENSG00000270870 | AC004805,2 | 0,015917099 | 1,011093979 | 0,9884349 | 0,99837368 | dynactin 6 (DCTN6) pseudogene |
| ENSG00000164144 | ARFIP1 | 0,01589061 | 1,011075415 | 0,97052827 | 0,998013218 | ADP ribosylation factor interacting protein 1 |
| ENSG00000108559 | NUP88 | 0,015796231 | 1,011009274 | 0,95844959 | 0,996608711 | nucleoporin 88 |
| ENSG00000096717 | SIRT1 | 0,015782206 | 1,010999446 | 0,96439417 | 0,997113804 | sirtuin 1 |
| ENSG00000152284 | TCF7L1 | 0,015777074 | 1,010995849 | 0,93805425 | 0,994488324 | transcription factor 7 like 1 |

| | | | | | | |
|---|---|---|---|---|---|---|
| ENSG00000100387 | RBX1 | 0,015761297 | 1,010984793 | 0,96930582 | 0,997412278 | ring-box 1 |
| ENSG00000101138 | CSTF1 | 0,015754666 | 1,010980147 | 0,96297213 | 0,996608711 | cleavage stimulation factor subunit 1 |
| ENSG00000172260 | NEGR1 | 0,01568833 | 1,010933662 | 0,94242731 | 0,995373376 | neuronal growth regulator 1 |
| ENSG00000144909 | OSBPL11 | 0,015665483 | 1,010917653 | 0,9566325 | 0,996608711 | oxysterol binding protein like 11 |
| ENSG00000205923 | CEMP1 | 0,015657625 | 1,010912146 | 0,99533358 | 0,999650137 | cementum protein 1 |
| ENSG00000256053 | COA8 | 0,015656016 | 1,010911019 | 0,95068638 | 0,996432861 | cytochrome c oxidase assembly factor 8 |
| ENSG00000138032 | PPM1B | 0,015601292 | 1,010872674 | 0,96116449 | 0,996608711 | protein phosphatase, Mg2+/Mn2+ dependent 1B |
| ENSG00000166246 | C16orf71 | 0,015599966 | 1,010871745 | 0,9761514 | 0,99837368 | chromosome 16 open reading frame 71 |
| ENSG00000256625 | AC092747,2 | 0,015540972 | 1,01083041 | 0,98494755 | 0,99837368 | liprin-beta-1-like (PPFIBP1) pseudogene |
| ENSG00000089916 | GPATCH2L | 0,015522291 | 1,010817321 | 0,95809017 | 0,996608711 | G-patch domain containing 2 like |
| ENSG00000186130 | ZBTB6 | 0,015511455 | 1,010809729 | 0,97488108 | 0,99837368 | zinc finger and BTB domain containing 6 |
| ENSG00000118804 | STBD1 | 0,015479593 | 1,010787405 | 0,96819842 | 0,997263201 | starch binding domain 1 |
| ENSG00000259877 | AC009113,1 | 0,015363657 | 1,010706181 | 0,95256922 | 0,996448973 | novel transcript, antisense to ZNF778 |
| ENSG00000152944 | MED21 | 0,015300205 | 1,010661729 | 0,97669934 | 0,99837368 | mediator complex subunit 21 |
| ENSG00000109534 | GAR1 | 0,01527705 | 1,010645509 | 0,96639546 | 0,997263201 | GAR1 ribonucleoprotein |
| ENSG00000276166 | AC092118,2 | 0,015255702 | 1,010630554 | 0,98610158 | 0,99837368 | novel transcript, antisense to KIFC3 |
| ENSG00000182795 | C1orf116 | 0,015232294 | 1,010614156 | 0,98930991 | 0,99837368 | chromosome 1 open reading frame 116 |
| ENSG00000188766 | SPRED3 | 0,015227242 | 1,010610617 | 0,96406871 | 0,996980762 | sprouty related EVH1 domain containing 3 |
| ENSG00000123106 | CCDC91 | 0,015184149 | 1,010580431 | 0,96799841 | 0,997263201 | coiled-coil domain containing 91 |
| ENSG00000075914 | EXOSC7 | 0,015172101 | 1,010571992 | 0,95680991 | 0,996608711 | exosome component 7 |
| ENSG00000287872 | AC026956,2 | 0,015157766 | 1,01056195 | 0,98159681 | 0,99837368 | novel transcript |
| ENSG00000166261 | ZNF202 | 0,015040984 | 1,010480152 | 0,94909937 | 0,996432861 | zinc finger protein 202 |
| ENSG00000066185 | ZMYND12 | 0,015040579 | 1,010479868 | 0,98594863 | 0,99837368 | zinc finger MYND-type containing 12 |
| ENSG00000279028 | AC010223,1 | 0,015020859 | 1,010466056 | 0,99124874 | 0,998884788 | novel transcript, antisense to PCDHAC2 |
| ENSG00000121274 | TENT4B | 0,014935303 | 1,010406134 | 0,96819939 | 0,997263201 | terminal nucleotidyltransferase 4B |
| ENSG00000240303 | ACAD11 | 0,014934287 | 1,010405423 | 0,96891636 | 0,997302342 | acyl-CoA dehydrogenase family member 11 |
| ENSG00000100281 | HMGXB4 | 0,014900236 | 1,010381575 | 0,96160664 | 0,996608711 | HMG-box containing 4 |
| ENSG00000162620 | LRRIQ3 | 0,014823651 | 1,010327941 | 0,97969333 | 0,99837368 | leucine rich repeats and IQ motif containing 3 |
| ENSG00000132763 | MMACHC | 0,014650316 | 1,01020656 | 0,95229613 | 0,996432861 | metabolism of cobalamin associated C |
| ENSG00000143363 | PRUNE1 | 0,014620928 | 1,010185982 | 0,96143195 | 0,996608711 | prune exopolyphosphatase 1 |
| ENSG00000099622 | CIRBP | 0,014519261 | 1,010114797 | 0,95395165 | 0,996608711 | cold inducible RNA binding protein |
| ENSG00000196793 | ZNF239 | 0,014514616 | 1,010111545 | 0,95585235 | 0,996608711 | zinc finger protein 239 |
| ENSG00000262919 | CCNQ | 0,014499734 | 1,010101125 | 0,95443351 | 0,996608711 | cyclin Q |

| | | | | | | |
|---|---|---|---|---|---|---|
| ENSG00000269910 | AL049840,3 | 0,014490452 | 1,010094626 | 0,9777117 | 0,99837368 | novel transcript, antisense to KLC1 |
| ENSG00000100320 | RBFOX2 | 0,01440171 | 1,010032496 | 0,96523431 | 0,997208095 | RNA binding fox-1 homolog 2 |
| ENSG00000267698 | AC002116,2 | 0,014371745 | 1,010011518 | 0,9855742 | 0,99837368 | uncharacterized LOC101927572 |
| ENSG00000145907 | G3BP1 | 0,014318006 | 1,009973897 | 0,96279159 | 0,996608711 | G3BP stress granule assembly factor 1 |
| ENSG00000276633 | AJ011931,1 | 0,014271379 | 1,009941255 | 0,99046289 | 0,998672296 | novel transcript |
| ENSG00000100084 | HIRA | 0,014267897 | 1,009938818 | 0,94753469 | 0,996107107 | histone cell cycle regulator |
| ENSG00000231409 | AC018868,1 | 0,014224342 | 1,009908328 | 0,98853667 | 0,99837368 | ribosomal protein L36a (RPL36A) pseudogene |
| ENSG00000196704 | AMZ2 | 0,014208576 | 1,009897292 | 0,96472002 | 0,997113804 | archaelysin family metallopeptidase 2 |
| ENSG00000224609 | HSD52 | 0,014099635 | 1,009821035 | 0,97445454 | 0,99837368 | uncharacterized LOC729467 |
| ENSG00000180979 | LRRC57 | 0,014092622 | 1,009816126 | 0,94780209 | 0,996170391 | leucine rich repeat containing 57 |
| ENSG00000260719 | AC009133,3 | 0,01409213 | 1,009815782 | 0,97806443 | 0,99837368 | novel transcript, antisense to C16orf54 |
| ENSG00000033030 | ZCCHC8 | 0,014068926 | 1,009799541 | 0,95659872 | 0,996608711 | zinc finger CCHC-type containing 8 |
| ENSG00000235501 | AC105942,1 | 0,013986077 | 1,009741553 | 0,96419944 | 0,997009661 | novel transcript |
| ENSG00000242880 | AC026341,1 | 0,013939844 | 1,009709195 | 0,98957641 | 0,99837368 | novel transcript |
| ENSG00000170145 | SIK2 | 0,013928177 | 1,00970103 | 0,94915981 | 0,996432861 | salt inducible kinase 2 |
| ENSG00000078246 | TULP3 | 0,013897543 | 1,00967959 | 0,96523534 | 0,997208095 | TUB like protein 3 |
| ENSG00000249328 | AC036214,1 | 0,013896756 | 1,009679039 | 0,98541352 | 0,99837368 | novel transcript |
| ENSG00000124733 | MEA1 | 0,013896754 | 1,009679038 | 0,95265999 | 0,996485725 | male-enhanced antigen 1 |
| ENSG00000120875 | DUSP4 | 0,013599823 | 1,00947125 | 0,95909001 | 0,996608711 | dual specificity phosphatase 4 |
| ENSG00000225793 | TMEM30A-DT | 0,013596347 | 1,009468818 | 0,97467064 | 0,99837368 | TMEM30A divergent transcript |
| ENSG00000184677 | ZBTB40 | 0,013554463 | 1,009439512 | 0,94980697 | 0,996432861 | zinc finger and BTB domain containing 40 |
| ENSG00000182578 | CSF1R | 0,013531427 | 1,009423394 | 0,98337151 | 0,99837368 | colony stimulating factor 1 receptor |
| ENSG00000214814 | FER1L6 | 0,013524027 | 1,009418216 | 0,97876962 | 0,99837368 | fer-1 like family member 6 |
| ENSG00000171552 | BCL2L1 | 0,013521833 | 1,009416681 | 0,95088969 | 0,996432861 | BCL2 like 1 |
| ENSG00000285387 | NDUFS3 | 0,013475889 | 1,009384536 | 0,95721997 | 0,996608711 | NADH:ubiquinone oxidoreductase core subunit S3 |
| ENSG00000221978 | CCNL2 | 0,013452905 | 1,009368455 | 0,96103761 | 0,996608711 | cyclin L2 |
| ENSG00000178149 | DALRD3 | 0,013437342 | 1,009357566 | 0,95386594 | 0,996608711 | DALR anticodon binding domain containing 3 |
| ENSG00000198826 | ARHGAP11A | 0,013422841 | 1,009347421 | 0,96162589 | 0,996608711 | Rho GTPase activating protein 11A |
| ENSG00000261537 | AC027279,2 | 0,013334595 | 1,009285684 | 0,98876539 | 0,99837368 | novel transcript, sense intronic to WWOX |
| ENSG00000108187 | PBLD | 0,013306971 | 1,009266359 | 0,95542109 | 0,996608711 | phenazine biosynthesis like protein domain containing |
| ENSG00000253695 | AC011008,1 | 0,013269316 | 1,009240017 | 0,99195529 | 0,999049649 | novel transcript |
| ENSG00000161955 | TNFSF13 | 0,01320388 | 1,009194242 | 0,97043528 | 0,998013218 | TNF superfamily member 13 |

| | | | | | | |
|---|---|---|---|---|---|---|
| ENSG00000188312 | CENPP | 0,013153466 | 1,009158977 | 0,96366805 | 0,996761656 | centromere protein P |
| ENSG00000149480 | MTA2 | 0,013128051 | 1,009141199 | 0,95147437 | 0,996432861 | metastasis associated 1 family member 2 |
| ENSG00000286539 | AC073973,1 | 0,013099872 | 1,009121489 | 0,98901033 | 0,99837368 | novel transcript |
| ENSG00000141543 | EIF4A3 | 0,013075091 | 1,009104156 | 0,95165435 | 0,996432861 | eukaryotic translation initiation factor 4A3 |
| ENSG00000129696 | TTI2 | 0,013071141 | 1,009101393 | 0,95900052 | 0,996608711 | TELO2 interacting protein 2 |
| ENSG00000180189 | HMGB1P14 | 0,013028215 | 1,009071368 | 0,9799104 | 0,99837368 | high mobility group box 1 pseudogene 14 |
| ENSG00000161914 | ZNF653 | 0,01301307 | 1,009060775 | 0,97592709 | 0,99837368 | zinc finger protein 653 |
| ENSG00000186185 | KIF18B | 0,012999917 | 1,009051575 | 0,96774406 | 0,997263201 | kinesin family member 18B |
| ENSG00000167113 | COQ4 | 0,012986695 | 1,009042328 | 0,95975002 | 0,996608711 | coenzyme Q4 |
| ENSG00000177669 | MBOAT4 | 0,012984077 | 1,009040497 | 0,98607889 | 0,99837368 | membrane bound O-acyltransferase domain containing 4 |
| ENSG00000159140 | SON | 0,012840708 | 1,008940228 | 0,95951346 | 0,996608711 | SON DNA and RNA binding protein |
| ENSG00000167081 | PBX3 | 0,012797676 | 1,008910134 | 0,97454604 | 0,99837368 | PBX homeobox 3 |
| ENSG00000100092 | SH3BP1 | 0,012793755 | 1,008907392 | 0,96412826 | 0,996980762 | SH3 domain binding protein 1 |
| ENSG00000135912 | TTLL4 | 0,012666516 | 1,008818415 | 0,96122936 | 0,996608711 | tubulin tyrosine ligase like 4 |
| ENSG00000242628 | AC009228,1 | 0,012658841 | 1,008813048 | 0,97990997 | 0,99837368 | novel transcript |
| ENSG00000112964 | GHR | 0,012609022 | 1,008778213 | 0,97763609 | 0,99837368 | growth hormone receptor |
| ENSG00000100296 | THOC5 | 0,012586657 | 1,008762574 | 0,95626911 | 0,996608711 | THO complex 5 |
| ENSG00000214736 | TOMM6 | 0,01249493 | 1,008698439 | 0,96422672 | 0,997009661 | translocase of outer mitochondrial membrane 6 |
| ENSG00000181798 | LINC00471 | 0,012479344 | 1,008687541 | 0,98362172 | 0,99837368 | long intergenic non-protein coding RNA 471 |
| ENSG00000161905 | ALOX15 | 0,012452957 | 1,008669093 | 0,98097903 | 0,99837368 | arachidonate 15-lipoxygenase |
| ENSG00000156171 | DRAM2 | 0,012442007 | 1,008661437 | 0,97682616 | 0,99837368 | DNA damage regulated autophagy modulator 2 |
| ENSG00000242085 | RPS20P33 | 0,012440609 | 1,00866046 | 0,98733344 | 0,99837368 | ribosomal protein S20 pseudogene 33 |
| ENSG00000260645 | AL359715,2 | 0,01243503 | 1,008656559 | 0,98104437 | 0,99837368 | novel transcript |
| ENSG00000243749 | TMEM35B | 0,012433104 | 1,008655212 | 0,98307935 | 0,99837368 | transmembrane protein 35B |
| ENSG00000148925 | BTBD10 | 0,012377354 | 1,008616236 | 0,972873 | 0,99837368 | BTB domain containing 10 |
| ENSG00000206052 | DOK6 | 0,012355742 | 1,008601127 | 0,97970552 | 0,99837368 | docking protein 6 |
| ENSG00000175265 | GOLGA8A | 0,012348609 | 1,00859614 | 0,9681218 | 0,997263201 | golgin A8 family member A |
| ENSG00000215105 | TTC3P1 | 0,012317095 | 1,008574109 | 0,97374915 | 0,99837368 | tetratricopeptide repeat domain 3 pseudogene 1 |
| ENSG00000165417 | GTF2A1 | 0,012289905 | 1,008555101 | 0,97859552 | 0,99837368 | general transcription factor IIA subunit 1 |
| ENSG00000269352 | PTOV1-AS2 | 0,012265775 | 1,008538232 | 0,98264753 | 0,99837368 | PTOV1 antisense RNA 2 |
| ENSG00000090530 | P3H2 | 0,01223723 | 1,008518278 | 0,97639207 | 0,99837368 | prolyl 3-hydroxylase 2 |
| ENSG00000246273 | SBF2-AS1 | 0,012231104 | 1,008513995 | 0,9633055 | 0,996608711 | SBF2 antisense RNA 1 |

| | | | | | | |
|---|---|---|---|---|---|---|
| ENSG00000217801 | AL390719,1 | 0,012162661 | 1,008466151 | 0,98367826 | 0,99837368 | protein tyrosine phosphatase family pseudogene |
| ENSG00000231050 | AL109917,1 | 0,0121491 | 1,008456672 | 0,9906085 | 0,998754686 | novel transcript |
| ENSG00000233056 | ERVH48-1 | 0,012089424 | 1,008414959 | 0,97681943 | 0,99837368 | endogenous retrovirus group 48 member 1 |
| ENSG00000228925 | AC016722,2 | 0,011998063 | 1,008351101 | 0,97426665 | 0,99837368 | novel transcript, antisense to MCFD2 |
| ENSG00000186517 | ARHGAP30 | 0,011980627 | 1,008338914 | 0,96743477 | 0,997263201 | Rho GTPase activating protein 30 |
| ENSG00000001630 | CYP51A1 | 0,011974172 | 1,008334403 | 0,97532745 | 0,99837368 | cytochrome P450 family 51 subfamily A member 1 |
| ENSG00000261334 | AL353803,4 | 0,011949312 | 1,008317028 | 0,98840406 | 0,99837368 | novel transcript |
| ENSG00000260211 | AC139426,1 | 0,01194397 | 1,008313294 | 0,98584862 | 0,99837368 | hyperpolarization activated cyclic nucleotide-gated potassium channel 2 (HCN2) pseudogene |
| ENSG00000240288 | GHRLOS | 0,011940328 | 1,008310749 | 0,97554699 | 0,99837368 | ghrelin opposite strand/antisense RNA |
| ENSG00000197045 | GMFB | 0,011884524 | 1,008271747 | 0,9820592 | 0,99837368 | glia maturation factor beta |
| ENSG00000145864 | GABRB2 | 0,011835833 | 1,008237719 | 0,98166553 | 0,99837368 | gamma-aminobutyric acid type A receptor subunit beta2 |
| ENSG00000283859 | NLGN2 | 0,011807063 | 1,008217613 | 0,97163712 | 0,998088605 | neuroligin 2 |
| ENSG00000169992 | NLGN2 | 0,011807063 | 1,008217613 | 0,97163712 | 0,998088605 | neuroligin 2 |
| ENSG00000128641 | MYO1B | 0,011805949 | 1,008216835 | 0,96828293 | 0,997263201 | myosin IB |
| ENSG00000264964 | AP001033,1 | 0,011750546 | 1,008178117 | 0,97701466 | 0,99837368 | novel transcript |
| ENSG00000225264 | ZNRF2P2 | 0,011628984 | 1,008093171 | 0,97553509 | 0,99837368 | zinc and ring finger 2 pseudogene 2 |
| ENSG00000198815 | FOXJ3 | 0,011607801 | 1,00807837 | 0,97125239 | 0,998088605 | forkhead box J3 |
| ENSG00000263513 | FAM72C | 0,011572054 | 1,008053392 | 0,97238473 | 0,998214946 | family with sequence similarity 72 member C |
| ENSG00000115540 | MOB4 | 0,011528702 | 1,008023101 | 0,98453579 | 0,99837368 | MOB family member 4, phocein |
| ENSG00000122035 | RASL11A | 0,01152287 | 1,008019026 | 0,98663901 | 0,99837368 | RAS like family 11 member A |
| ENSG00000167037 | SGSM1 | 0,011468437 | 1,007980994 | 0,98298787 | 0,99837368 | small G protein signaling modulator 1 |
| ENSG00000141098 | GFOD2 | 0,011420505 | 1,007947506 | 0,96992466 | 0,997819165 | glucose-fructose oxidoreductase domain containing 2 |
| ENSG00000273397 | NDUFA6 | 0,011401649 | 1,007934332 | 0,97511909 | 0,99837368 | NADH:ubiquinone oxidoreductase subunit A6 |
| ENSG00000277365 | NDUFA6 | 0,011401649 | 1,007934332 | 0,97511909 | 0,99837368 | NADH:ubiquinone oxidoreductase subunit A6 |
| ENSG00000287663 | AL122020,2 | 0,011395162 | 1,0079298 | 0,98280021 | 0,99837368 | novel transcript, antisense to TTC7B |
| ENSG00000269526 | ERVV-1 | 0,011362521 | 1,007906996 | 0,98466671 | 0,99837368 | endogenous retrovirus group V member 1, envelope |
| ENSG00000120942 | UBIAD1 | 0,011358778 | 1,007904381 | 0,95960799 | 0,996608711 | UbiA prenyltransferase domain containing 1 |
| ENSG00000140740 | UQCRC2 | 0,011291951 | 1,007857695 | 0,9747596 | 0,99837368 | ubiquinol-cytochrome c reductase core protein 2 |
| ENSG00000143889 | HNRNPLL | 0,011288131 | 1,007855026 | 0,96847547 | 0,997302342 | heterogeneous nuclear ribonucleoprotein L like |

| | | | | | | |
|---|---|---|---|---|---|---|
| ENSG00000204253 | HNRNPCP2 | 0,011270381 | 1,007842627 | 0,99331347 | 0,999049649 | heterogeneous nuclear ribonucleoprotein C pseudogene 2 |
| ENSG00000287671 | AC068418,3 | 0,011237235 | 1,007819472 | 0,98383872 | 0,99837368 | novel transcript |
| ENSG00000183250 | LINC01547 | 0,01122251 | 1,007809185 | 0,97123481 | 0,998088605 | long intergenic non-protein coding RNA 1547 |
| ENSG00000278730 | AC005332,6 | 0,011062288 | 1,007697267 | 0,96643239 | 0,997263201 | novel transcript |
| ENSG00000171055 | FEZ2 | 0,011057979 | 1,007694257 | 0,96780366 | 0,997263201 | fasciculation and elongation protein zeta 2 |
| ENSG00000275111 | ZNF2 | 0,011048432 | 1,007687589 | 0,96449698 | 0,997113804 | zinc finger protein 2 |
| ENSG00000101343 | CRNKL1 | 0,011005586 | 1,007657662 | 0,97236064 | 0,998214946 | crooked neck pre-mRNA splicing factor 1 |
| ENSG00000154079 | SDHAF4 | 0,010981553 | 1,007640876 | 0,97623257 | 0,99837368 | succinate dehydrogenase complex assembly factor 4 |
| ENSG00000073464 | CLCN4 | 0,010980308 | 1,007640007 | 0,97092486 | 0,998088605 | chloride voltage-gated channel 4 |
| ENSG00000198393 | ZNF26 | 0,010900193 | 1,007584052 | 0,96603029 | 0,997263201 | zinc finger protein 26 |
| ENSG00000158301 | GPRASP2 | 0,010818378 | 1,007526914 | 0,95897024 | 0,996608711 | G protein-coupled receptor associated sorting protein 2 |
| ENSG00000269534 | AC011466,3 | 0,010811818 | 1,007522333 | 0,98972177 | 0,99837368 | novel transcript, antisense to LIG1 |
| ENSG00000226688 | ENTPD1-AS1 | 0,010718724 | 1,007457321 | 0,97506041 | 0,99837368 | ENTPD1 antisense RNA 1 |
| ENSG00000112031 | MTRF1L | 0,010647863 | 1,00740784 | 0,97992803 | 0,99837368 | mitochondrial translational release factor 1 like |
| ENSG00000162600 | OMA1 | 0,010541543 | 1,007333601 | 0,98131315 | 0,99837368 | OMA1 zinc metallopeptidase |
| ENSG00000270362 | HMGN3-AS1 | 0,010497124 | 1,007302587 | 0,97828984 | 0,99837368 | HMGN3 antisense RNA 1 |
| ENSG00000189376 | C8orf76 | 0,010467444 | 1,007281864 | 0,96697932 | 0,997263201 | chromosome 8 open reading frame 76 |
| ENSG00000170638 | TRABD | 0,010447194 | 1,007267726 | 0,97523458 | 0,99837368 | TraB domain containing |
| ENSG00000287713 | AC006517,2 | 0,010349603 | 1,007199592 | 0,98829264 | 0,99837368 | novel transcript, antisense to SLC2A14 |
| ENSG00000278002 | AL627171,1 | 0,010300592 | 1,007165376 | 0,98630878 | 0,99837368 | novel transcript |
| ENSG00000168096 | ANKS3 | 0,010258883 | 1,007136259 | 0,97798646 | 0,99837368 | ankyrin repeat and sterile alpha motif domain containing 3 |
| ENSG00000184047 | DIABLO | 0,010219891 | 1,007109039 | 0,96887107 | 0,997302342 | diablo IAP-binding mitochondrial protein |
| ENSG00000127564 | PKMYT1 | 0,010198796 | 1,007094313 | 0,98151982 | 0,99837368 | protein kinase, membrane associated tyrosine/threonine 1 |
| ENSG00000161048 | NAPEPLD | 0,010096005 | 1,007022561 | 0,97292494 | 0,99837368 | N-acyl phosphatidylethanolamine phospholipase D |
| ENSG00000057757 | PITHD1 | 0,010082087 | 1,007012846 | 0,96409185 | 0,996980762 | PITH domain containing 1 |
| ENSG00000204685 | STARD7-AS1 | 0,010033125 | 1,00697867 | 0,96875295 | 0,997302342 | STARD7 antisense RNA 1 |
| ENSG00000286018 | AF129075,3 | 0,009985684 | 1,006945558 | 0,98460194 | 0,99837368 | novel transcript, antisense to CCT8 |
| ENSG00000109270 | LAMTOR3 | 0,009981168 | 1,006942406 | 0,98488953 | 0,99837368 | late endosomal/lysosomal adaptor, MAPK and MTOR activator 3 |

| | | | | | | |
|---|---|---|---|---|---|---|
| ENSG00000273541 | AL137060,1 | 0,009908197 | 1,006891477 | 0,99265427 | 0,999049649 | |
| ENSG00000270419 | CAHM | 0,00988598 | 1,006875971 | 0,97918364 | 0,99837368 | colon adenocarcinoma hypermethylated |
| ENSG00000284512 | AC092718,8 | 0,00986216 | 1,006859347 | 0,97740555 | 0,99837368 | novel protein |
| ENSG00000285600 | AC023593,1 | 0,009845597 | 1,006847787 | 0,98310729 | 0,99837368 | Novel transcript, antisense to MCM2 |
| ENSG00000237989 | LINC01679 | 0,009808035 | 1,006821573 | 0,98955176 | 0,99837368 | long intergenic non-protein coding RNA 1679 |
| ENSG00000267096 | AC008735,1 | 0,009796087 | 1,006813236 | 0,99137584 | 0,998916278 | novel transcript |
| ENSG00000274425 | AC114271,1 | 0,009792904 | 1,006811014 | 0,97060867 | 0,998030243 | novel transcript, antisense to ICAM3 |
| ENSG00000068305 | MEF2A | 0,00973421 | 1,006770054 | 0,98106641 | 0,99837368 | myocyte enhancer factor 2A |
| ENSG00000263740 | RN7SL4P | 0,009722708 | 1,006762027 | 0,98053477 | 0,99837368 | RNA, 7SL, cytoplasmic 4, pseudogene |
| ENSG00000229496 | DDX39B | 0,009698834 | 1,006745368 | 0,96611827 | 0,997263201 | DExD-box helicase 39B |
| ENSG00000141574 | SECTM1 | 0,009681799 | 1,00673348 | 0,99275147 | 0,999049649 | secreted and transmembrane 1 |
| ENSG00000099814 | CEP170B | 0,009389431 | 1,006529482 | 0,98631267 | 0,99837368 | centrosomal protein 170B |
| ENSG00000164117 | FBXO8 | 0,009376053 | 1,006520149 | 0,98192675 | 0,99837368 | F-box protein 8 |
| ENSG00000261807 | LINC02141 | 0,009368685 | 1,006515009 | 0,98940582 | 0,99837368 | long intergenic non-protein coding RNA 2141 |
| ENSG00000257966 | OLA1P3 | 0,009328156 | 1,006486734 | 0,98885112 | 0,99837368 | OLA1 pseudogene 3 |
| ENSG00000157837 | SPPL3 | 0,009270363 | 1,006446415 | 0,97262274 | 0,998267798 | signal peptide peptidase like 3 |
| ENSG00000228434 | AC004951,1 | 0,009241889 | 1,006426551 | 0,98691231 | 0,99837368 | novel transcript |
| ENSG00000260257 | AL035071,1 | 0,009214181 | 1,006407223 | 0,97509805 | 0,99837368 | novel transcript |
| ENSG00000170260 | ZNF212 | 0,009146728 | 1,00636017 | 0,97729499 | 0,99837368 | zinc finger protein 212 |
| ENSG00000001036 | FUCA2 | 0,009127145 | 1,006346509 | 0,97782131 | 0,99837368 | alpha-L-fucosidase 2 |
| ENSG00000234358 | AC003080,1 | 0,009042676 | 1,00628759 | 0,9901415 | 0,998477042 | ribosomal protein L8 (RPL8) pseudogene |
| ENSG00000217330 | SSXP10 | 0,009009211 | 1,006264248 | 0,99178894 | 0,999049649 | SSX family pseudogene 10 |
| ENSG00000237950 | AL357079,1 | 0,00896732 | 1,00623503 | 0,98902565 | 0,99837368 | novel transcript |
| ENSG00000179632 | MAF1 | 0,008950073 | 1,006223 | 0,97229867 | 0,998214946 | MAF1 homolog, negative regulator of RNA polymerase III |
| ENSG00000141510 | TP53 | 0,008942111 | 1,006217447 | 0,96776769 | 0,997263201 | tumor protein p53 |
| ENSG00000271771 | AC139792,1 | 0,008938508 | 1,006214935 | 0,98993319 | 0,99837368 | novel transcript |
| ENSG00000111615 | KRR1 | 0,00889201 | 1,006182505 | 0,98122152 | 0,99837368 | KRR1 small subunit processome component homolog |
| ENSG00000178397 | FAM220A | 0,008870522 | 1,006167519 | 0,97483088 | 0,99837368 | family with sequence similarity 220 member A |
| ENSG00000237071 | TRIM27 | 0,008791303 | 1,006112271 | 0,97014695 | 0,997916506 | tripartite motif containing 27 |
| ENSG00000226144 | RPS27AP3 | 0,008783407 | 1,006106764 | 0,99190593 | 0,999049649 | RPS27A pseudogene 3 |
| ENSG00000124835 | AC105760,1 | 0,008764925 | 1,006093876 | 0,99405422 | 0,999279985 | novel transcript |

| | | | | | | |
|---|---|---|---|---|---|---|
| ENSG00000288067 | AC105180,2 | 0,008760301 | 1,006090651 | 0,97945805 | 0,99837368 | novel transcript |
| ENSG00000281131 | SCHLAP1 | 0,008735834 | 1,006073589 | 0,98842668 | 0,99837368 | SWI/SNF complex antagonist associated with prostate cancer 1 |
| ENSG00000131263 | RLIM | 0,008711145 | 1,006056371 | 0,98094966 | 0,99837368 | ring finger protein, LIM domain interacting |
| ENSG00000233718 | MYCNOS | 0,008659975 | 1,006020689 | 0,98496091 | 0,99837368 | MYCN opposite strand |
| ENSG00000170571 | EMB | 0,008653021 | 1,00601584 | 0,98712989 | 0,99837368 | embigin |
| ENSG00000254536 | AL360181,3 | 0,008628859 | 1,005998992 | 0,98135491 | 0,99837368 | novel protein |
| ENSG00000204055 | AL158151,1 | 0,008446932 | 1,005872141 | 0,99109721 | 0,998857746 | novel transcript, antisense to IER5L |
| ENSG00000134686 | PHC2 | 0,008425718 | 1,00585735 | 0,9724955 | 0,998263101 | polyhomeotic homolog 2 |
| ENSG00000227761 | BAG6 | 0,008393458 | 1,005834858 | 0,98101202 | 0,99837368 | BAG cochaperone 6 |
| ENSG00000276809 | AL138955,1 | 0,008381503 | 1,005826523 | 0,99141485 | 0,998923388 | novel transcript, sense intronic to DNAJC3 |
| ENSG00000110958 | PTGES3 | 0,008377619 | 1,005823816 | 0,98050156 | 0,99837368 | prostaglandin E synthase 3 |
| ENSG00000111206 | FOXM1 | 0,008352251 | 1,00580613 | 0,97173571 | 0,998088605 | forkhead box M1 |
| ENSG00000282619 | AF186192,6 | 0,008337307 | 1,005795711 | 0,98899804 | 0,99837368 | zinc finger protein 154 (ZNF154) pseudogene |
| ENSG00000255085 | AF186192,2 | 0,008337307 | 1,005795711 | 0,98899804 | 0,99837368 | zinc finger protein 154 (ZNF154) pseudogene |
| ENSG00000136167 | LCP1 | 0,008192834 | 1,005694995 | 0,9842261 | 0,99837368 | lymphocyte cytosolic protein 1 |
| ENSG00000164674 | SYTL3 | 0,008131256 | 1,00565207 | 0,9863295 | 0,99837368 | synaptotagmin like 3 |
| ENSG00000259146 | AC005476,2 | 0,008114157 | 1,005640151 | 0,99172108 | 0,999049649 | novel transcript, antisense to SIPA1L1 |
| ENSG00000104679 | R3HCC1 | 0,008102507 | 1,00563203 | 0,97854474 | 0,99837368 | R3H domain and coiled-coil containing 1 |
| ENSG00000234965 | SHISA8 | 0,007997816 | 1,005559058 | 0,99439877 | 0,999385471 | shisa family member 8 |
| ENSG00000262410 | AC024361,1 | 0,007953978 | 1,005528504 | 0,98835032 | 0,99837368 | novel transcript, sense intronic FN3K |
| ENSG00000132434 | LANCL2 | 0,007935467 | 1,005515602 | 0,97744397 | 0,99837368 | LanC like 2 |
| ENSG00000137692 | DCUN1D5 | 0,007881927 | 1,005478286 | 0,97553684 | 0,99837368 | defective in cullin neddylation 1 domain containing 5 |
| ENSG00000132953 | XPO4 | 0,007875038 | 1,005473485 | 0,98411872 | 0,99837368 | exportin 4 |
| ENSG00000227110 | LMCD1-AS1 | 0,0078475 | 1,005454293 | 0,97896116 | 0,99837368 | LMCD1 antisense RNA 1 |
| ENSG00000078140 | UBE2K | 0,007829791 | 1,005441952 | 0,98727663 | 0,99837368 | ubiquitin conjugating enzyme E2 K |
| ENSG00000274064 | AC040978,1 | 0,007749988 | 1,005386337 | 0,99649415 | 0,999664158 | glyceraldehyde-3-phosphate dehydrogenase (GAPDH) pseudogene |
| ENSG00000214941 | ZSWIM7 | 0,007715302 | 1,005362165 | 0,97417464 | 0,99837368 | zinc finger SWIM-type containing 7 |
| ENSG00000285646 | AL021155,4 | 0,007712661 | 1,005360325 | 0,98629703 | 0,99837368 | novel transcript |
| ENSG00000283852 | BCAP29 | 0,007676089 | 1,00533484 | 0,98895517 | 0,99837368 | B cell receptor associated protein 29 |
| ENSG00000152234 | ATP5F1A | 0,007563361 | 1,005256288 | 0,9773813 | 0,99837368 | ATP synthase F1 subunit alpha |

| | | | | | | |
|---|---|---|---|---|---|---|
| ENSG00000014123 | UFL1 | 0,007559095 | 1,005253316 | 0,98544363 | 0,99837368 | UFM1 specific ligase 1 |
| ENSG00000176532 | PRR15 | 0,007542594 | 1,005241818 | 0,99348994 | 0,999082296 | proline rich 15 |
| ENSG00000151092 | NGLY1 | 0,007507705 | 1,005217509 | 0,97845586 | 0,99837368 | N-glycanase 1 |
| ENSG00000151461 | UPF2 | 0,007470333 | 1,00519147 | 0,97853734 | 0,99837368 | UPF2 regulator of nonsense mediated mRNA decay |
| ENSG00000149084 | HSD17B12 | 0,007411566 | 1,005150525 | 0,9860417 | 0,99837368 | hydroxysteroid 17-beta dehydrogenase 12 |
| ENSG00000204120 | GIGYF2 | 0,007366697 | 1,005119264 | 0,97789535 | 0,99837368 | GRB10 interacting GYF protein 2 |
| ENSG00000080839 | RBL1 | 0,00735439 | 1,00511069 | 0,97987847 | 0,99837368 | RB transcriptional corepressor like 1 |
| ENSG00000116199 | FAM20B | 0,007299853 | 1,005072695 | 0,98019168 | 0,99837368 | FAM20B glycosaminoglycan xylosylkinase |
| ENSG00000148444 | COMMD3 | 0,007271018 | 1,005052607 | 0,98622304 | 0,99837368 | COMM domain containing 3 |
| ENSG00000166971 | AKTIP | 0,007199474 | 1,005002767 | 0,98588679 | 0,99837368 | AKT interacting protein |
| ENSG00000225880 | LINC00115 | 0,007184779 | 1,004992531 | 0,98497844 | 0,99837368 | long intergenic non-protein coding RNA 115 |
| ENSG00000100354 | TNRC6B | 0,00716602 | 1,004979463 | 0,9771822 | 0,99837368 | trinucleotide repeat containing adaptor 6B |
| ENSG00000183354 | KIAA2026 | 0,006998057 | 1,004862467 | 0,97938117 | 0,99837368 | KIAA2026 |
| ENSG00000273314 | AC005229,4 | 0,006914059 | 1,004803962 | 0,98332777 | 0,99837368 | novel transcript, antisense to CUL1 |
| ENSG00000100206 | DMC1 | 0,006901976 | 1,004795547 | 0,98394878 | 0,99837368 | DNA meiotic recombinase 1 |
| ENSG00000140750 | ARHGAP17 | 0,006901001 | 1,004794868 | 0,97453304 | 0,99837368 | Rho GTPase activating protein 17 |
| ENSG00000164291 | ARSK | 0,006840494 | 1,004752727 | 0,98325522 | 0,99837368 | arylsulfatase family member K |
| ENSG00000040275 | SPDL1 | 0,006797501 | 1,004722786 | 0,98440562 | 0,99837368 | spindle apparatus coiled-coil protein 1 |
| ENSG00000230124 | ACBD6 | 0,006693051 | 1,004650048 | 0,97461874 | 0,99837368 | acyl-CoA binding domain containing 6 |
| ENSG00000022976 | ZNF839 | 0,006692268 | 1,004649502 | 0,98011859 | 0,99837368 | zinc finger protein 839 |
| ENSG00000067704 | IARS2 | 0,006674557 | 1,004637169 | 0,98397915 | 0,99837368 | isoleucyl-tRNA synthetase 2, mitochondrial |
| ENSG00000266717 | AC134407,1 | 0,006581318 | 1,004572243 | 0,99225808 | 0,999049649 | novel transcript, antisense to BPTF |
| ENSG00000132463 | GRSF1 | 0,006480743 | 1,004502214 | 0,98495752 | 0,99837368 | G-rich RNA sequence binding factor 1 |
| ENSG00000261884 | AC040162,1 | 0,006478428 | 1,004500601 | 0,98674267 | 0,99837368 | novel transcript |
| ENSG00000148450 | MSRB2 | 0,006429933 | 1,004466836 | 0,98266536 | 0,99837368 | methionine sulfoxide reductase B2 |
| ENSG00000196182 | STK40 | 0,006424801 | 1,004463264 | 0,9794851 | 0,99837368 | serine/threonine kinase 40 |
| ENSG00000197776 | KLHDC1 | 0,006350934 | 1,004411836 | 0,98981465 | 0,99837368 | kelch domain containing 1 |
| ENSG00000109180 | OCIAD1 | 0,006329569 | 1,004396962 | 0,98657209 | 0,99837368 | OCIA domain containing 1 |
| ENSG00000197568 | HHLA3 | 0,00626793 | 1,00435405 | 0,98053107 | 0,99837368 | HERV-H LTR-associating 3 |
| ENSG00000099785 | MARCHF2 | 0,006249891 | 1,004341492 | 0,98601393 | 0,99837368 | membrane associated ring-CH-type finger 2 |
| ENSG00000101412 | E2F1 | 0,006237543 | 1,004332896 | 0,98082869 | 0,99837368 | E2F transcription factor 1 |
| ENSG00000269556 | TMEM185A | 0,006211015 | 1,004314428 | 0,97869732 | 0,99837368 | transmembrane protein 185A |
| ENSG00000123200 | ZC3H13 | 0,006175876 | 1,004289967 | 0,98497581 | 0,99837368 | zinc finger CCCH-type containing 13 |

| | | | | | | |
|---|---|---|---|---|---|---|
| ENSG00000232110 | AL353751,1 | 0,006122886 | 1,00425308 | 0,99235194 | 0,999049649 | novel transcript, antisense to LIPA |
| ENSG00000205726 | ITSN1 | 0,006116374 | 1,004248547 | 0,97980911 | 0,99837368 | intersectin 1 |
| ENSG00000119953 | SMNDC1 | 0,006097157 | 1,00423517 | 0,98629189 | 0,99837368 | survival motor neuron domain containing 1 |
| ENSG00000273192 | AL671710,1 | 0,006048226 | 1,004201111 | 0,98598891 | 0,99837368 | novel transcript, antisense to ALG12 |
| ENSG00000100592 | DAAM1 | 0,006041638 | 1,004196525 | 0,98668097 | 0,99837368 | dishevelled associated activator of morphogenesis 1 |
| ENSG00000171566 | PLRG1 | 0,006012602 | 1,004176315 | 0,98755674 | 0,99837368 | pleiotropic regulator 1 |
| ENSG00000148943 | LIN7C | 0,005994576 | 1,004163768 | 0,98998401 | 0,99838264 | lin-7 homolog C, crumbs cell polarity complex component |
| ENSG00000062725 | APPBP2 | 0,005992173 | 1,004162096 | 0,98883194 | 0,99837368 | amyloid beta precursor protein binding protein 2 |
| ENSG00000146670 | CDCA5 | 0,005982156 | 1,004155123 | 0,97877728 | 0,99837368 | cell division cycle associated 5 |
| ENSG00000165752 | STK32C | 0,005906456 | 1,004102435 | 0,98851198 | 0,99837368 | serine/threonine kinase 32C |
| ENSG00000277868 | CCDC125 | 0,00589827 | 1,004096738 | 0,98346834 | 0,99837368 | coiled-coil domain containing 125 |
| ENSG00000102870 | ZNF629 | 0,005887235 | 1,004089058 | 0,97837822 | 0,99837368 | zinc finger protein 629 |
| ENSG00000225434 | LINC01504 | 0,005868164 | 1,004075785 | 0,98937881 | 0,99837368 | long intergenic non-protein coding RNA 1504 |
| ENSG00000172508 | CARNS1 | 0,00581075 | 1,004035827 | 0,99404664 | 0,999279985 | carnosine synthase 1 |
| ENSG00000234771 | SLC25A25-AS1 | 0,00576769 | 1,00400586 | 0,98693895 | 0,99837368 | SLC25A25 antisense RNA 1 |
| ENSG00000072135 | PTPN18 | 0,005753683 | 1,003996112 | 0,98220112 | 0,99837368 | protein tyrosine phosphatase non-receptor type 18 |
| ENSG00000186235 | LINC02610 | 0,005744606 | 1,003989796 | 0,98836253 | 0,99837368 | long intergenic non-protein coding RNA 2610 |
| ENSG00000257599 | OVCH1-AS1 | 0,005731996 | 1,00398102 | 0,98687051 | 0,99837368 | OVCH1 antisense RNA 1 |
| ENSG00000261067 | AC109460,3 | 0,005688817 | 1,003950972 | 0,99096725 | 0,998800453 | novel transcript |
| ENSG00000141349 | G6PC3 | 0,005687805 | 1,003950268 | 0,98092093 | 0,99837368 | glucose-6-phosphatase catalytic subunit 3 |
| ENSG00000113460 | BRIX1 | 0,005677556 | 1,003943136 | 0,98670258 | 0,99837368 | biogenesis of ribosomes BRX1 |
| ENSG00000137270 | GCM1 | 0,005661819 | 1,003932185 | 0,99752547 | 0,999899141 | glial cells missing transcription factor 1 |
| ENSG00000121417 | ZNF211 | 0,005657285 | 1,003929029 | 0,98302433 | 0,99837368 | zinc finger protein 211 |
| ENSG00000083168 | KAT6A | 0,005647498 | 1,003922219 | 0,98071063 | 0,99837368 | lysine acetyltransferase 6A |
| ENSG00000188283 | ZNF383 | 0,00563232 | 1,003911658 | 0,98294524 | 0,99837368 | zinc finger protein 383 |
| ENSG00000233851 | LATS2-AS1 | 0,005630541 | 1,003910419 | 0,99450035 | 0,999391175 | LATS2 antisense RNA 1 |
| ENSG00000166595 | CIAO2B | 0,005602714 | 1,003891056 | 0,98196827 | 0,99837368 | cytosolic iron-sulfur assembly component 2B |
| ENSG00000080819 | CPOX | 0,005597779 | 1,003887622 | 0,98000903 | 0,99837368 | coproporphyrinogen oxidase |
| ENSG00000234709 | UPF3AP3 | 0,005590089 | 1,003882271 | 0,99461327 | 0,999426291 | UPF3A pseudogene 3 |
| ENSG00000256806 | C17orf100 | 0,005492239 | 1,003814185 | 0,98379435 | 0,99837368 | chromosome 17 open reading frame 100 |
| ENSG00000133104 | SPART | 0,005427396 | 1,00376907 | 0,98747241 | 0,99837368 | spartin |
| ENSG00000124641 | MED20 | 0,005366697 | 1,003726839 | 0,98468776 | 0,99837368 | mediator complex subunit 20 |

| | | | | | | |
|---|---|---|---|---|---|---|
| ENSG00000214188 | ST7-OT4 | 0,005318817 | 1,003693527 | 0,98918071 | 0,99837368 | ST7 overlapping transcript 4 |
| ENSG00000170242 | USP47 | 0,005278319 | 1,003665353 | 0,98761563 | 0,99837368 | ubiquitin specific peptidase 47 |
| ENSG00000151025 | GPR158 | 0,005252192 | 1,003647177 | 0,98673641 | 0,99837368 | G protein-coupled receptor 158 |
| ENSG00000011485 | PPP5C | 0,005179032 | 1,003596283 | 0,97940159 | 0,99837368 | protein phosphatase 5 catalytic subunit |
| ENSG00000261349 | AL031432,2 | 0,005124515 | 1,003558359 | 0,99653417 | 0,999664158 | novel pseudogene |
| ENSG00000162378 | ZYG11B | 0,005116157 | 1,003552545 | 0,98826661 | 0,99837368 | zyg-11 family member B, cell cycle regulator |
| ENSG00000137171 | KLC4 | 0,005101805 | 1,003542562 | 0,98482865 | 0,99837368 | kinesin light chain 4 |
| ENSG00000206427 | PRRC2A | 0,00505428 | 1,003509504 | 0,99086933 | 0,998800453 | proline rich coiled-coil 2A |
| ENSG00000121236 | TRIM6 | 0,005034262 | 1,00349558 | 0,98686984 | 0,99837368 | tripartite motif containing 6 |
| ENSG00000100311 | PDGFB | 0,005012548 | 1,003480477 | 0,99298113 | 0,999049649 | platelet derived growth factor subunit B |
| ENSG00000185619 | PCGF3 | 0,004926567 | 1,003420673 | 0,98575441 | 0,99837368 | polycomb group ring finger 3 |
| ENSG00000082068 | WDR70 | 0,004892829 | 1,003397208 | 0,98586047 | 0,99837368 | WD repeat domain 70 |
| ENSG00000230679 | ENO1-AS1 | 0,004847617 | 1,003365764 | 0,99641389 | 0,999664158 | ENO1 antisense RNA 1 |
| ENSG00000196781 | TLE1 | 0,004845588 | 1,003364352 | 0,98709531 | 0,99837368 | TLE family member 1, transcriptional corepressor |
| ENSG00000135446 | CDK4 | 0,004828539 | 1,003352495 | 0,9833102 | 0,99837368 | cyclin dependent kinase 4 |
| ENSG00000100836 | PABPN1 | 0,004818458 | 1,003345484 | 0,98124089 | 0,99837368 | poly(A) binding protein nuclear 1 |
| ENSG00000272711 | AC019069,1 | 0,004762031 | 1,003306242 | 0,98916231 | 0,99837368 | novel transcript |
| ENSG00000285812 | AL390719,3 | 0,004759949 | 1,003304794 | 0,99414934 | 0,999327539 | novel transcript, antisense to C1orf159 |
| ENSG00000276618 | RAD17 | 0,004737803 | 1,003289393 | 0,99179345 | 0,999049649 | RAD17 checkpoint clamp loader component |
| ENSG00000169398 | PTK2 | 0,004571825 | 1,003173974 | 0,98611715 | 0,99837368 | protein tyrosine kinase 2 |
| ENSG00000126814 | TRMT5 | 0,004528077 | 1,003143554 | 0,98988706 | 0,99837368 | tRNA methyltransferase 5 |
| ENSG00000118939 | UCHL3 | 0,004480785 | 1,003110672 | 0,99088715 | 0,998800453 | ubiquitin C-terminal hydrolase L3 |
| ENSG00000262943 | ALOX12P2 | 0,004435659 | 1,003079296 | 0,99489594 | 0,999515206 | arachidonate 12-lipoxygenase pseudogene 2 |
| ENSG00000131732 | ZCCHC9 | 0,004383978 | 1,003043363 | 0,98958897 | 0,99837368 | zinc finger CCHC-type containing 9 |
| ENSG00000188636 | RTL6 | 0,004373387 | 1,003036 | 0,98231592 | 0,99837368 | retrotransposon Gag like 6 |
| ENSG00000205628 | LINC01446 | 0,00436991 | 1,003033583 | 0,99626214 | 0,999664158 | long intergenic non-protein coding RNA 1446 |
| ENSG00000177728 | TMEM94 | 0,004294002 | 1,002980809 | 0,98650984 | 0,99837368 | transmembrane protein 94 |
| ENSG00000251356 | RAB5CP2 | 0,004129618 | 1,002866534 | 0,99771017 | 0,999899141 | RAB5C, member RAS oncogene family pseudogene 2 |
| ENSG00000205765 | C5orf51 | 0,004121978 | 1,002861223 | 0,99155304 | 0,999030431 | chromosome 5 open reading frame 51 |
| ENSG00000152475 | ZNF837 | 0,004112471 | 1,002854614 | 0,99250788 | 0,999049649 | zinc finger protein 837 |
| ENSG00000205250 | E2F4 | 0,004104852 | 1,002849318 | 0,98688027 | 0,99837368 | E2F transcription factor 4 |
| ENSG00000138180 | CEP55 | 0,004099837 | 1,002845832 | 0,99019511 | 0,998498905 | centrosomal protein 55 |
| ENSG00000242396 | AC096536,2 | 0,004042462 | 1,002805951 | 0,9983087 | 0,999899141 | novel transcript |

| | | | | | | |
|---|---|---|---|---|---|---|
| ENSG00000121067 | SPOP | 0,004001466 | 1,002777455 | 0,9875266 | 0,99837368 | speckle type BTB/POZ protein |
| ENSG00000163703 | CRELD1 | 0,003905447 | 1,002710717 | 0,98692186 | 0,99837368 | cysteine rich with EGF like domains 1 |
| ENSG00000125166 | GOT2 | 0,003690264 | 1,00256117 | 0,98939854 | 0,99837368 | glutamic-oxaloacetic transaminase 2 |
| ENSG00000268231 | AC008743,1 | 0,003682134 | 1,002555521 | 0,99715954 | 0,999813856 | novel transcript, sense intronic to LRRC4B |
| ENSG00000167986 | DDB1 | 0,003641839 | 1,002527519 | 0,98751199 | 0,99837368 | damage specific DNA binding protein 1 |
| ENSG00000029639 | TFB1M | 0,003635389 | 1,002523037 | 0,99131611 | 0,998888288 | transcription factor B1, mitochondrial |
| ENSG00000109920 | FNBP4 | 0,003612866 | 1,002507386 | 0,98946614 | 0,99837368 | formin binding protein 4 |
| ENSG00000285182 | FNBP4 | 0,003612866 | 1,002507386 | 0,98946614 | 0,99837368 | formin binding protein 4 |
| ENSG00000130939 | UBE4B | 0,003612747 | 1,002507303 | 0,98723844 | 0,99837368 | ubiquitination factor E4B |
| ENSG00000165716 | DIPK1B | 0,003610046 | 1,002505427 | 0,98801657 | 0,99837368 | divergent protein kinase domain 1B |
| ENSG00000213693 | SEC14L1P1 | 0,003578636 | 1,0024836 | 0,9920783 | 0,999049649 | SEC14 like 1 pseudogene 1 |
| ENSG00000080822 | CLDND1 | 0,003398036 | 1,002358115 | 0,99372832 | 0,999193459 | claudin domain containing 1 |
| ENSG00000171103 | TRMT61B | 0,003390894 | 1,002353153 | 0,99223437 | 0,999049649 | tRNA methyltransferase 61B |
| ENSG00000108953 | YWHAE | 0,003352442 | 1,002326438 | 0,99283558 | 0,999049649 | tyrosine 3-monooxygenase/tryptophan 5-monooxygenase activation protein epsilon |
| ENSG00000101911 | PRPS2 | 0,003338948 | 1,002317063 | 0,9928183 | 0,999049649 | phosphoribosyl pyrophosphate synthetase 2 |
| ENSG00000196422 | PPP1R26 | 0,003245057 | 1,002251833 | 0,99380566 | 0,999197865 | protein phosphatase 1 regulatory subunit 26 |
| ENSG00000172053 | QARS1 | 0,003204062 | 1,002223355 | 0,98913256 | 0,99837368 | glutaminyl-tRNA synthetase 1 |
| ENSG00000258940 | AL132639,2 | 0,00316214 | 1,002194232 | 0,99630508 | 0,999664158 | novel transcript, antisense to CTAGE5 |
| ENSG00000124160 | NCOA5 | 0,00309864 | 1,002150122 | 0,98981533 | 0,99837368 | nuclear receptor coactivator 5 |
| ENSG00000168813 | ZNF507 | 0,003066937 | 1,0021281 | 0,99329134 | 0,999049649 | zinc finger protein 507 |
| ENSG00000152223 | EPG5 | 0,003064741 | 1,002126574 | 0,98990627 | 0,99837368 | ectopic P-granules autophagy protein 5 homolog |
| ENSG00000147155 | EBP | 0,003061328 | 1,002124204 | 0,99070033 | 0,998800453 | EBP cholestenol delta-isomerase |
| ENSG00000025796 | SEC63 | 0,003045669 | 1,002113327 | 0,99336501 | 0,999049649 | SEC63 homolog, protein translocation regulator |
| ENSG00000272767 | JMJD1C-AS1 | 0,003018468 | 1,002094433 | 0,99503658 | 0,99957662 | JMJD1C antisense RNA 1 |
| ENSG00000187860 | CCDC157 | 0,002978031 | 1,002066346 | 0,99468161 | 0,999444832 | coiled-coil domain containing 157 |
| ENSG00000274349 | ZNF658 | 0,002940629 | 1,002040368 | 0,9945351 | 0,999393973 | zinc finger protein 658 |
| ENSG00000125835 | SNRPB | 0,002933921 | 1,002035708 | 0,98958851 | 0,99837368 | small nuclear ribonucleoprotein polypeptides B and B1 |
| ENSG00000103275 | UBE2I | 0,002892825 | 1,002007165 | 0,98956572 | 0,99837368 | ubiquitin conjugating enzyme E2 I |
| ENSG00000142252 | GEMIN7 | 0,002868187 | 1,001990054 | 0,99191026 | 0,999049649 | gem nuclear organelle associated protein 7 |
| ENSG00000285121 | MTCH2 | 0,002810918 | 1,001950279 | 0,99290858 | 0,999049649 | mitochondrial carrier 2 |
| ENSG00000109919 | MTCH2 | 0,002810918 | 1,001950279 | 0,99290858 | 0,999049649 | mitochondrial carrier 2 |

| | | | | | | |
|---|---|---|---|---|---|---|
| ENSG00000095794 | CREM | 0,002802456 | 1,001944403 | 0,99335158 | 0,999049649 | cAMP responsive element modulator |
| ENSG00000258469 | CHMP4BP1 | 0,002760333 | 1,001915148 | 0,99679946 | 0,999708243 | charged multivesicular body protein 4B pseudogene 1 |
| ENSG00000169683 | LRRC45 | 0,002736166 | 1,001898365 | 0,99617937 | 0,999664158 | leucine rich repeat containing 45 |
| ENSG00000282958 | HNRNPR | 0,002712278 | 1,001881776 | 0,99207916 | 0,999049649 | heterogeneous nuclear ribonucleoprotein R |
| ENSG00000093000 | NUP50 | 0,002665354 | 1,001849191 | 0,99291633 | 0,999049649 | nucleoporin 50 |
| ENSG00000147687 | TATDN1 | 0,002560014 | 1,001776042 | 0,99559689 | 0,999664158 | TatD DNase domain containing 1 |
| ENSG00000287565 | AC108168,1 | 0,002472 | 1,001714929 | 0,9956975 | 0,999664158 | novel transcript, antisense to RP11-526A4,1 |
| ENSG00000198712 | MT-CO2 | 0,002409762 | 1,001671716 | 0,99447946 | 0,999391175 | mitochondrially encoded cytochrome c oxidase II |
| ENSG00000111679 | PTPN6 | 0,002406595 | 1,001669517 | 0,99696144 | 0,999768593 | protein tyrosine phosphatase non-receptor type 6 |
| ENSG00000214753 | HNRNPUL2 | 0,002401618 | 1,001666061 | 0,99317611 | 0,999049649 | heterogeneous nuclear ribonucleoprotein U like 2 |
| ENSG00000271461 | CKAP2LP1 | 0,002365564 | 1,001641029 | 0,99856973 | 0,999899141 | CKAP2L pseudogene 1 |
| ENSG00000100106 | TRIOBP | 0,002362274 | 1,001638745 | 0,99511911 | 0,99962741 | TRIO and F-actin binding protein |
| ENSG00000253641 | LINCR-0001 | 0,002359792 | 1,001637021 | 0,99825846 | 0,999899141 | uncharacterized LINCR-0001 |
| ENSG00000277007 | AC096642,1 | 0,002322611 | 1,001611208 | 0,99563985 | 0,999664158 | novel transcript |
| ENSG00000188243 | COMMD6 | 0,00229646 | 1,001593052 | 0,99574227 | 0,999664158 | COMM domain containing 6 |
| ENSG00000243005 | RN7SL16P | 0,002292398 | 1,001590232 | 0,99764488 | 0,999899141 | RNA, 7SL, cytoplasmic 16, pseudogene |
| ENSG00000137338 | PGBD1 | 0,002278689 | 1,001580715 | 0,99315512 | 0,999049649 | piggyBac transposable element derived 1 |
| ENSG00000108256 | NUFIP2 | 0,002260216 | 1,00156789 | 0,99518042 | 0,999650137 | nuclear FMR1 interacting protein 2 |
| ENSG00000180712 | LINC02363 | 0,002218624 | 1,001539016 | 0,99712355 | 0,999813217 | long intergenic non-protein coding RNA 2363 |
| ENSG00000092421 | SEMA6A | 0,002203527 | 1,001528536 | 0,99425463 | 0,999385471 | semaphorin 6A |
| ENSG00000131013 | PPIL4 | 0,002183946 | 1,001514942 | 0,99526118 | 0,999650137 | peptidylprolyl isomerase like 4 |
| ENSG00000094880 | CDC23 | 0,002134083 | 1,001480329 | 0,9949115 | 0,999515206 | cell division cycle 23 |
| ENSG00000218510 | LINC00339 | 0,002099115 | 1,001456054 | 0,9932701 | 0,999049649 | long intergenic non-protein coding RNA 339 |
| ENSG00000216901 | ZNF603P | 0,002064864 | 1,00143228 | 0,9988819 | 0,999899141 | zinc finger protein 603, pseudogene |
| ENSG00000152620 | NADK2 | 0,002029907 | 1,001408014 | 0,99541219 | 0,999664158 | NAD kinase 2, mitochondrial |
| ENSG00000142230 | SAE1 | 0,001999628 | 1,001386998 | 0,99222129 | 0,999049649 | SUMO1 activating enzyme subunit 1 |
| ENSG00000160813 | PPP1R35 | 0,001998934 | 1,001386516 | 0,99635827 | 0,999664158 | protein phosphatase 1 regulatory subunit 35 |
| ENSG00000160446 | ZDHHC12 | 0,001908471 | 1,001323727 | 0,9961363 | 0,999664158 | zinc finger DHHC-type palmitoyltransferase 12 |
| ENSG00000270231 | NBPF8 | 0,001903979 | 1,001320609 | 0,99436183 | 0,999385471 | NBPF member 8 |
| ENSG00000213799 | ZNF845 | 0,001880757 | 1,001304492 | 0,99546099 | 0,999664158 | zinc finger protein 845 |
| ENSG00000287847 | AC018678,1 | 0,001821255 | 1,001263195 | 0,99663794 | 0,999675593 | novel transcript |
| ENSG00000261063 | AC009139,2 | 0,001812919 | 1,00125741 | 0,99896045 | 0,999899141 | novel transcript, antisense to MON1B & SYCE1L |

| | | | | | | |
|---|---|---|---|---|---|---|
| ENSG00000253121 | AC120042,2 | 0,0017919 | 1,001242822 | 0,99894118 | 0,999899141 | novel transcript |
| ENSG00000216636 | RPL7P25 | 0,001748432 | 1,001212655 | 0,99903143 | 0,999899141 | ribosomal protein L7 pseudogene 25 |
| ENSG00000150787 | PTS | 0,001660602 | 1,001151704 | 0,99655583 | 0,999664158 | 6-pyruvoyltetrahydropterin synthase |
| ENSG00000271788 | AC008875,1 | 0,001637241 | 1,001135493 | 0,99871887 | 0,999899141 | novel transcript |
| ENSG00000260917 | AL158212,3 | 0,001605088 | 1,001113182 | 0,99604906 | 0,999664158 | novel transcript, overlapping to VTI1A |
| ENSG00000102921 | N4BP1 | 0,001601134 | 1,001110438 | 0,99485506 | 0,999515206 | NEDD4 binding protein 1 |
| ENSG00000149474 | KAT14 | 0,001589065 | 1,001102062 | 0,99435144 | 0,999385471 | lysine acetyltransferase 14 |
| ENSG00000164329 | TENT2 | 0,001454619 | 1,001008774 | 0,99698922 | 0,999768593 | terminal nucleotidyltransferase 2 |
| ENSG00000139726 | DENR | 0,001440444 | 1,000998938 | 0,99650388 | 0,999664158 | density regulated re-initiation and release factor |
| ENSG00000048545 | GUCA1A | 0,00142328 | 1,000987029 | 0,99807308 | 0,999899141 | guanylate cyclase activator 1A |
| ENSG00000132305 | IMMT | 0,001406931 | 1,000975686 | 0,99608623 | 0,999664158 | inner membrane mitochondrial protein |
| ENSG00000107249 | GLIS3 | 0,00134723 | 1,000934265 | 0,99625921 | 0,999664158 | GLIS family zinc finger 3 |
| ENSG00000080910 | CFHR2 | 0,00130153 | 1,000902559 | 0,99839627 | 0,999899141 | complement factor H related 2 |
| ENSG00000168883 | USP39 | 0,001166518 | 1,000808896 | 0,99637108 | 0,999664158 | ubiquitin specific peptidase 39 |
| ENSG00000175066 | GK5 | 0,001149461 | 1,000797063 | 0,99640698 | 0,999664158 | glycerol kinase 5 |
| ENSG00000197261 | C6orf141 | 0,001120942 | 1,00077728 | 0,99784795 | 0,999899141 | chromosome 6 open reading frame 141 |
| ENSG00000232892 | RGS5-AS1 | 0,001066702 | 1,000739655 | 0,99934666 | 0,999899141 | RGS5 antisense RNA 1 |
| ENSG00000219491 | TPT1P8 | 0,001040665 | 1,000721594 | 0,99942315 | 0,999899141 | tumor protein, translationally-controlled 1 pseudogene 8 |
| ENSG00000176485 | PLAAT3 | 0,001029078 | 1,000713557 | 0,99878875 | 0,999899141 | phospholipase A and acyltransferase 3 |
| ENSG00000106078 | COBL | 0,000983116 | 1,000681676 | 0,99847556 | 0,999899141 | cordon-bleu WH2 repeat protein |
| ENSG00000169668 | BCRP2 | 0,000970209 | 1,000672724 | 0,99770676 | 0,999899141 | BCR pseudogene 2 |
| ENSG00000137040 | RANBP6 | 0,000956603 | 1,000663287 | 0,99811955 | 0,999899141 | RAN binding protein 6 |
| ENSG00000089335 | ZNF302 | 0,000947409 | 1,000656909 | 0,99832023 | 0,999899141 | zinc finger protein 302 |
| ENSG00000004534 | RBM6 | 0,000939517 | 1,000651435 | 0,99656259 | 0,999664158 | RNA binding motif protein 6 |
| ENSG00000080371 | RAB21 | 0,000935535 | 1,000648674 | 0,99793405 | 0,999899141 | RAB21, member RAS oncogene family |
| ENSG00000104517 | UBR5 | 0,000883313 | 1,000612453 | 0,99768078 | 0,999899141 | ubiquitin protein ligase E3 component n-recognin 5 |
| ENSG00000010322 | NISCH | 0,000880847 | 1,000610743 | 0,99773489 | 0,999899141 | nischarin |
| ENSG00000142892 | PIGK | 0,00087374 | 1,000605814 | 0,99855973 | 0,999899141 | phosphatidylinositol glycan anchor biosynthesis class K |
| ENSG00000198890 | PRMT6 | 0,000772359 | 1,000535501 | 0,99741691 | 0,999899141 | protein arginine methyltransferase 6 |
| ENSG00000163170 | BOLA3 | 0,000706915 | 1,000490116 | 0,9984178 | 0,999899141 | bolA family member 3 |
| ENSG00000140830 | TXNL4B | 0,000688172 | 1,000477118 | 0,99784839 | 0,999899141 | thioredoxin like 4B |

| | | | | | | |
|---|---|---|---|---|---|---|
| ENSG00000122512 | PMS2 | 0,000609672 | 1,000422682 | 0,99770047 | 0,999899141 | PMS1 homolog 2, mismatch repair system component |
| ENSG00000226045 | AC234644,1 | 0,000603482 | 1,000418389 | 0,99948337 | 0,999899141 | glutamyl-tRNA(Gln) amidotransferase, subunit C homolog (bacterial)(GATC) pseudogene |
| ENSG00000197860 | SGTB | 0,000553472 | 1,000383711 | 0,99858168 | 0,999899141 | small glutamine rich tetratricopeptide repeat containing beta |
| ENSG00000168002 | POLR2G | 0,000544129 | 1,000377232 | 0,99836703 | 0,999899141 | RNA polymerase II subunit G |
| ENSG00000134057 | CCNB1 | 0,00047906 | 1,000332114 | 0,99883958 | 0,999899141 | cyclin B1 |
| ENSG00000265479 | DTX2P1-UPK3B | 0,000437959 | 1,000303616 | 0,9990846 | 0,999899141 | DTX2P1-UPK3BP1-PMS2P11 readthrough, transcribed pseudogene |
| ENSG00000163491 | NEK10 | 0,00041182 | 1,000285493 | 0,99902646 | 0,999899141 | NIMA related kinase 10 |
| ENSG00000143799 | PARP1 | 0,000402387 | 1,000278953 | 0,99854897 | 0,999899141 | poly(ADP-ribose) polymerase 1 |
| ENSG00000183864 | TOB2 | 0,000328047 | 1,000227411 | 0,99902124 | 0,999899141 | transducer of ERBB2, 2 |
| ENSG00000183230 | CTNNA3 | 0,000301988 | 1,000209344 | 0,99944729 | 0,999899141 | catenin alpha 3 |
| ENSG00000184675 | AMER1 | 0,000300858 | 1,00020856 | 0,99911964 | 0,999899141 | APC membrane recruitment protein 1 |
| ENSG00000186666 | BCDIN3D | 0,000239669 | 1,000166139 | 0,99924028 | 0,999899141 | BCDIN3 domain containing RNA methyltransferase |
| ENSG00000224138 | AC000123,1 | 0,000226469 | 1,000156989 | 0,99971362 | 0,999950872 | novel transcript |
| ENSG00000266714 | MYO15B | 0,000219386 | 1,000152078 | 0,99968558 | 0,999950872 | myosin XVB |
| ENSG00000198934 | MAGEE1 | 0,000175812 | 1,000121871 | 0,99946336 | 0,999899141 | MAGE family member E1 |
| ENSG00000201201 | RN7SKP118 | 0,000111075 | 1,000076994 | 0,99992099 | 0,999952973 | RN7SK pseudogene 118 |
| ENSG00000163558 | PRKCI | 0,000105734 | 1,000073292 | 0,99977462 | 0,999950872 | protein kinase C iota |
| ENSG00000119912 | IDE | 7,46E-05 | 1,000051711 | 0,99983117 | 0,999950872 | insulin degrading enzyme |
| ENSG00000254303 | AC037486,1 | -4,60E-05 | 0,999968111 | 0,99995341 | 0,999953412 | novel transcript |
| ENSG00000100162 | CENPM | -5,27E-05 | 0,999963487 | 0,99986584 | 0,999950872 | centromere protein M |
| ENSG00000231431 | FAR2P4 | -7,99E-05 | 0,999944633 | 0,9998869 | 0,999950872 | fatty acyl-CoA reductase 2 pseudogene 4 |
| ENSG00000135643 | KCNMB4 | -0,000187675 | 0,999869922 | 0,99952804 | 0,999911834 | potassium calcium-activated channel subfamily M regulatory beta subunit 4 |
| ENSG00000265204 | AC090772,2 | -0,00019248 | 0,999866592 | 0,99987753 | 0,999950872 | novel transcript, antisense to TTC39C |
| ENSG00000258515 | AL355075,2 | -0,000224679 | 0,999844277 | 0,99965135 | 0,999950872 | novel transcript, antisense to OSGEP |
| ENSG00000253174 | AC009630,2 | -0,000239654 | 0,999833898 | 0,99982313 | 0,999950872 | novel transcript, antisense to GINS4 |
| ENSG00000259891 | AC107375,1 | -0,000247447 | 0,999828498 | 0,99956872 | 0,999920538 | novel transcript, antisense to CHRAC1 |
| ENSG00000111300 | NAA25 | -0,000324056 | 0,999775407 | 0,999024 | 0,999899141 | N-alpha-acetyltransferase 25, NatB auxiliary subunit |
| ENSG00000011451 | WIZ | -0,000469475 | 0,999674638 | 0,99875379 | 0,999899141 | WIZ zinc finger |

| | | | | | | |
|---|---|---|---|---|---|---|
| ENSG00000235439 | DDX39B | -0,000525446 | 0,999635855 | 0,99907388 | 0,999899141 | DExD-box helicase 39B |
| ENSG00000257259 | LINC02388 | -0,000535166 | 0,99962912 | 0,99903646 | 0,999899141 | long intergenic non-protein coding RNA 2388 |
| ENSG00000258472 | AC005726,1 | -0,000558126 | 0,999613211 | 0,99930417 | 0,999899141 | novel protein |
| ENSG00000144224 | UBXN4 | -0,000583683 | 0,999595504 | 0,99873839 | 0,999899141 | UBX domain protein 4 |
| ENSG00000130338 | TULP4 | -0,000601797 | 0,999582953 | 0,99774616 | 0,999899141 | TUB like protein 4 |
| ENSG00000146425 | DYNLT1 | -0,000634649 | 0,999560191 | 0,99849704 | 0,999899141 | dynein light chain Tctex-type 1 |
| ENSG00000275632 | AL035461,2 | -0,000650802 | 0,999549 | 0,99861923 | 0,999899141 | novel transcript |
| ENSG00000275575 | PTP4A3 | -0,000706057 | 0,999510718 | 0,99799342 | 0,999899141 | protein tyrosine phosphatase 4A3 |
| ENSG00000286244 | AC073046,3 | -0,000711251 | 0,99950712 | 0,99946805 | 0,999899141 | novel transcript, antisense to STAMBP |
| ENSG00000039600 | SOX30 | -0,000803103 | 0,999443486 | 0,99893989 | 0,999899141 | SRY-box transcription factor 30 |
| ENSG00000196436 | NPIPB15 | -0,000811876 | 0,999437409 | 0,99832948 | 0,999899141 | nuclear pore complex interacting protein family member B15 |
| ENSG00000187736 | NHEJ1 | -0,000859116 | 0,999404684 | 0,99712692 | 0,999813217 | non-homologous end joining factor 1 |
| ENSG00000004660 | CAMKK1 | -0,000879752 | 0,999390388 | 0,99847923 | 0,999899141 | calcium/calmodulin dependent protein kinase kinase 1 |
| ENSG00000281010 | AC009065,9 | -0,001012399 | 0,999298505 | 0,99938269 | 0,999899141 | |
| ENSG00000174473 | GALNTL6 | -0,001031746 | 0,999285104 | 0,99843158 | 0,999899141 | polypeptide N-acetylgalactosaminyltransferase like 6 |
| ENSG00000128585 | MKLN1 | -0,001036351 | 0,999281914 | 0,99795053 | 0,999899141 | muskelin 1 |
| ENSG00000287551 | AL445526,1 | -0,001038355 | 0,999280526 | 0,99944987 | 0,999899141 | novel transcript, antisense to LINGO2 |
| ENSG00000264304 | AC024267,3 | -0,00106827 | 0,999259806 | 0,99839777 | 0,999899141 | novel transcript, antisense to FLOT2 |
| ENSG00000229203 | AC103564,1 | -0,001161292 | 0,999195378 | 0,99916818 | 0,999899141 | novel transcript |
| ENSG00000285918 | AC092376,3 | -0,001189856 | 0,999175595 | 0,99856854 | 0,999899141 | novel transcript, antisense to WWOX |
| ENSG00000262692 | AC116914,2 | -0,00135747 | 0,999059516 | 0,99905271 | 0,999899141 | novel transcript, antisense to ITGAE |
| ENSG00000135298 | ADGRB3 | -0,001433439 | 0,999006909 | 0,99681308 | 0,999708243 | adhesion G protein-coupled receptor B3 |
| ENSG00000154518 | ATP5MC3 | -0,001464486 | 0,998985411 | 0,99633319 | 0,999664158 | ATP synthase membrane subunit c locus 3 |
| ENSG00000120868 | APAF1 | -0,001533276 | 0,998937778 | 0,99614772 | 0,999664158 | apoptotic peptidase activating factor 1 |
| ENSG00000197283 | SYNGAP1 | -0,00154565 | 0,99892921 | 0,99437391 | 0,999385471 | synaptic Ras GTPase activating protein 1 |
| ENSG00000284801 | AC270301,2 | -0,001573462 | 0,998909954 | 0,99683037 | 0,999708243 | novel transcript |
| ENSG00000111726 | CMAS | -0,001581753 | 0,998904213 | 0,99591458 | 0,999664158 | cytidine monophosphate N-acetylneuraminic acid synthetase |
| ENSG00000133997 | MED6 | -0,00162477 | 0,998874429 | 0,99654465 | 0,999664158 | mediator complex subunit 6 |
| ENSG00000287625 | AC022210,1 | -0,001674448 | 0,998840035 | 0,99660556 | 0,99967519 | novel transcript |

| | | | | | | |
|---|---|---|---|---|---|---|
| ENSG00000269145 | AC007192,2 | -0,001689234 | 0,998829797 | 0,99925459 | 0,999899141 | novel transcript, antisense to MAST3 |
| ENSG00000140950 | MEAK7 | -0,00194603 | 0,998652024 | 0,99260422 | 0,999049649 | MTOR associated protein, eak-7 homolog |
| ENSG00000232613 | LINC02576 | -0,001947862 | 0,998650756 | 0,99869804 | 0,999899141 | long intergenic non-protein coding RNA 2576 |
| ENSG00000171843 | MLLT3 | -0,001976247 | 0,998631108 | 0,99613524 | 0,999664158 | MLLT3 super elongation complex subunit |
| ENSG00000088205 | DDX18 | -0,002025053 | 0,998597325 | 0,9952979 | 0,999650137 | DEAD-box helicase 18 |
| ENSG00000237641 | RPL28P2 | -0,002061806 | 0,998571886 | 0,99764973 | 0,999899141 | ribosomal protein L28 pseudogene 2 |
| ENSG00000127774 | EMC6 | -0,002144929 | 0,998514353 | 0,99358949 | 0,99911813 | ER membrane protein complex subunit 6 |
| ENSG00000176261 | ZBTB8OS | -0,002158167 | 0,998505191 | 0,99463119 | 0,999426291 | zinc finger and BTB domain containing 8 opposite strand |
| ENSG00000108799 | EZH1 | -0,002167714 | 0,998498583 | 0,99240566 | 0,999049649 | enhancer of zeste 1 polycomb repressive complex 2 subunit |
| ENSG00000024862 | CCDC28A | -0,002175653 | 0,998493089 | 0,99585457 | 0,999664158 | coiled-coil domain containing 28A |
| ENSG00000265491 | RNF115 | -0,002187796 | 0,998484685 | 0,99245683 | 0,999049649 | ring finger protein 115 |
| ENSG00000179094 | PER1 | -0,002199811 | 0,998476369 | 0,9932458 | 0,999049649 | period circadian regulator 1 |
| ENSG00000276023 | DUSP14 | -0,002219508 | 0,998462737 | 0,99701846 | 0,999768593 | dual specificity phosphatase 14 |
| ENSG00000288182 | AC011509,4 | -0,002243141 | 0,998446382 | 0,99823908 | 0,999899141 | novel transcript |
| ENSG00000261211 | AL031123,2 | -0,002267291 | 0,998429668 | 0,99690877 | 0,999754796 | novel transcript |
| ENSG00000157827 | FMNL2 | -0,002308141 | 0,998401398 | 0,99435455 | 0,999385471 | formin like 2 |
| ENSG00000112237 | CCNC | -0,002456424 | 0,998298785 | 0,9960274 | 0,999664158 | cyclin C |
| ENSG00000081052 | COL4A4 | -0,002516922 | 0,998256923 | 0,99557065 | 0,999664158 | collagen type IV alpha 4 chain |
| ENSG00000211643 | IGLV5-52 | -0,002609315 | 0,998192996 | 0,99824501 | 0,999899141 | immunoglobulin lambda variable 5-52 |
| ENSG00000143633 | C1orf131 | -0,002621996 | 0,998184222 | 0,99253447 | 0,999049649 | chromosome 1 open reading frame 131 |
| ENSG00000163975 | MELTF | -0,002635533 | 0,998174855 | 0,99619789 | 0,999664158 | melanotransferrin |
| ENSG00000148187 | MRRF | -0,002695212 | 0,998133565 | 0,99310149 | 0,999049649 | mitochondrial ribosome recycling factor |
| ENSG00000166974 | MAPRE2 | -0,002731161 | 0,998108694 | 0,99274941 | 0,999049649 | microtubule associated protein RP/EB family member 2 |
| ENSG00000152242 | C18orf25 | -0,002759431 | 0,998089136 | 0,99339357 | 0,999049649 | chromosome 18 open reading frame 25 |
| ENSG00000196419 | XRCC6 | -0,002812111 | 0,998052692 | 0,9908501 | 0,998800453 | X-ray repair cross complementing 6 |
| ENSG00000065457 | ADAT1 | -0,002863236 | 0,998017324 | 0,98978828 | 0,99837368 | adenosine deaminase tRNA specific 1 |
| ENSG00000091536 | MYO15A | -0,002923529 | 0,997975616 | 0,99497867 | 0,999550566 | myosin XVA |
| ENSG00000283243 | FO681507,1 | -0,002937914 | 0,997965666 | 0,99853241 | 0,999899141 | |
| ENSG00000262468 | LINC01569 | -0,0030332 | 0,997899754 | 0,9930463 | 0,999049649 | long intergenic non-protein coding RNA 1569 |
| ENSG00000224429 | LINC00539 | -0,003129773 | 0,997832958 | 0,99328282 | 0,999049649 | long intergenic non-protein coding RNA 539 |

| | | | | | | |
|---|---|---|---|---|---|---|
| ENSG00000283780 | MAML1 | -0,003135055 | 0,997829305 | 0,98826048 | 0,99837368 | mastermind like transcriptional coactivator 1 |
| ENSG00000250790 | AC127070,2 | -0,003182129 | 0,997796747 | 0,99382858 | 0,999197865 | novel transcript |
| ENSG00000235222 | MSH5 | -0,003185621 | 0,997794332 | 0,99362296 | 0,999119652 | mutS homolog 5 |
| ENSG00000099783 | HNRNPM | -0,003428057 | 0,997626673 | 0,99112606 | 0,998857746 | heterogeneous nuclear ribonucleoprotein M |
| ENSG00000131969 | ABHD12B | -0,00357978 | 0,997521761 | 0,99380028 | 0,999197865 | abhydrolase domain containing 12B |
| ENSG00000142937 | RPS8 | -0,003652764 | 0,997471299 | 0,99002371 | 0,998390465 | ribosomal protein S8 |
| ENSG00000214827 | MTCP1 | -0,00368841 | 0,997446655 | 0,99278762 | 0,999049649 | mature T cell proliferation 1 |
| ENSG00000100294 | MCAT | -0,00368943 | 0,997445949 | 0,98865141 | 0,99837368 | malonyl-CoA-acyl carrier protein transacylase |
| ENSG00000196678 | ERI2 | -0,00371096 | 0,997431064 | 0,99286013 | 0,999049649 | ERI1 exoribonuclease family member 2 |
| ENSG00000280080 | TBC1D22A-AS1 | -0,003720887 | 0,997424201 | 0,99592965 | 0,999664158 | TBC1D22A antisense RNA 1 |
| ENSG00000276920 | NKD2 | -0,003762773 | 0,997395243 | 0,99559055 | 0,999664158 | NKD inhibitor of WNT signaling pathway 2 |
| ENSG00000259221 | AC023905,1 | -0,003786283 | 0,99737899 | 0,99598043 | 0,999664158 | novel transcript, antisense to SEMA6D |
| ENSG00000118496 | FBXO30 | -0,003799539 | 0,997369826 | 0,99287481 | 0,999049649 | F-box protein 30 |
| ENSG00000156467 | UQCRB | -0,00384583 | 0,997337824 | 0,99245925 | 0,999049649 | ubiquinol-cytochrome c reductase binding protein |
| ENSG00000119414 | PPP6C | -0,003871664 | 0,997319965 | 0,99338282 | 0,999049649 | protein phosphatase 6 catalytic subunit |
| ENSG00000251192 | ZNF674 | -0,003945456 | 0,997268954 | 0,98978834 | 0,99837368 | zinc finger protein 674 |
| ENSG00000271538 | LINC02427 | -0,004013813 | 0,997221704 | 0,99481611 | 0,999515206 | long intergenic non-protein coding RNA 2427 |
| ENSG00000257365 | FNTB | -0,004027281 | 0,997212394 | 0,98790617 | 0,99837368 | farnesyltransferase, CAAX box, beta |
| ENSG00000112234 | FBXL4 | -0,004054693 | 0,997193447 | 0,99246467 | 0,999049649 | F-box and leucine rich repeat protein 4 |
| ENSG00000103510 | KAT8 | -0,004083086 | 0,997173822 | 0,98526238 | 0,99837368 | lysine acetyltransferase 8 |
| ENSG00000112992 | NNT | -0,004154498 | 0,997124464 | 0,98979002 | 0,99837368 | nicotinamide nucleotide transhydrogenase |
| ENSG00000152465 | NMT2 | -0,004172524 | 0,997112005 | 0,98461269 | 0,99837368 | N-myristoyltransferase 2 |
| ENSG00000109472 | CPE | -0,004210036 | 0,997086079 | 0,99046239 | 0,998672296 | carboxypeptidase E |
| ENSG00000125648 | SLC25A23 | -0,00425629 | 0,997054112 | 0,98563075 | 0,99837368 | solute carrier family 25 member 23 |
| ENSG00000183780 | SLC35F3 | -0,004334913 | 0,996999777 | 0,99347958 | 0,999082296 | solute carrier family 35 member F3 |
| ENSG00000104133 | SPG11 | -0,004383399 | 0,996966271 | 0,98890549 | 0,99837368 | SPG11 vesicle trafficking associated, spatacsin |
| ENSG00000115548 | KDM3A | -0,004419569 | 0,996941275 | 0,98912376 | 0,99837368 | lysine demethylase 3A |
| ENSG00000235288 | AC099329,1 | -0,004435348 | 0,996930372 | 0,98700996 | 0,99837368 | novel transcript, antisense to CCBP2 |
| ENSG00000274229 | SOCS7 | -0,004439154 | 0,996927742 | 0,99223918 | 0,999049649 | suppressor of cytokine signaling 7 |
| ENSG00000118298 | CA14 | -0,00445586 | 0,996916198 | 0,99097336 | 0,998800453 | carbonic anhydrase 14 |
| ENSG00000078304 | PPP2R5C | -0,004473055 | 0,996904316 | 0,99057538 | 0,998753504 | protein phosphatase 2 regulatory subunit B'gamma |
| ENSG00000285049 | CPSF1 | -0,00451677 | 0,99687411 | 0,98767889 | 0,99837368 | cleavage and polyadenylation specific factor 1 |
| ENSG00000071894 | CPSF1 | -0,00451677 | 0,99687411 | 0,98767889 | 0,99837368 | cleavage and polyadenylation specific factor 1 |

| | | | | | | |
|---|---|---|---|---|---|---|
| ENSG00000172922 | RNASEH2C | -0,004522986 | 0,996869814 | 0,9868192 | 0,99837368 | ribonuclease H2 subunit C |
| ENSG00000280758 | AL356481,3 | -0,004609835 | 0,996809806 | 0,99554895 | 0,999664158 | novel transcript, antisense to SPTAN1 |
| ENSG00000121775 | TMEM39B | -0,004633447 | 0,996793491 | 0,98484362 | 0,99837368 | transmembrane protein 39B |
| ENSG00000240775 | AC021205,1 | -0,004662144 | 0,996773664 | 0,99644465 | 0,999664158 | ribosomal protein S26 (RPS26) pseudogene |
| ENSG00000226239 | AL031658,1 | -0,004715419 | 0,996736856 | 0,99229597 | 0,999049649 | novel transcript |
| ENSG00000273367 | AL355472,3 | -0,004761502 | 0,996705019 | 0,99719206 | 0,999814398 | novel transcript |
| ENSG00000181090 | EHMT1 | -0,004790743 | 0,996684818 | 0,98635871 | 0,99837368 | euchromatic histone lysine methyltransferase 1 |
| ENSG00000261740 | BOLA2-SMG1P( | -0,004804278 | 0,996675467 | 0,98693532 | 0,99837368 | BOLA2-SMG1P6 readthrough |
| ENSG00000137267 | TUBB2A | -0,004947864 | 0,996576276 | 0,98369384 | 0,99837368 | tubulin beta 2A class IIa |
| ENSG00000141295 | SCRN2 | -0,004974388 | 0,996557955 | 0,98848752 | 0,99837368 | secernin 2 |
| ENSG00000248685 | LINC02484 | -0,004981857 | 0,996552795 | 0,9878981 | 0,99837368 | long intergenic non-protein coding RNA 2484 |
| ENSG00000235974 | VN2R19P | -0,005025166 | 0,99652288 | 0,99672072 | 0,999708243 | vomeronasal 2 receptor 19, pseudogene |
| ENSG00000170949 | ZNF160 | -0,005026124 | 0,996522218 | 0,98237202 | 0,99837368 | zinc finger protein 160 |
| ENSG00000163328 | GPR155 | -0,005030097 | 0,996519473 | 0,98494248 | 0,99837368 | G protein-coupled receptor 155 |
| ENSG00000197312 | DDI2 | -0,005055537 | 0,996501901 | 0,98485623 | 0,99837368 | DNA damage inducible 1 homolog 2 |
| ENSG00000254585 | MAGEL2 | -0,005059134 | 0,996499417 | 0,9863225 | 0,99837368 | MAGE family member L2 |
| ENSG00000287862 | AC114971,1 | -0,005076956 | 0,996487107 | 0,99471837 | 0,999449647 | novel transcript |
| ENSG00000137221 | TJAP1 | -0,005093799 | 0,996475474 | 0,98545966 | 0,99837368 | tight junction associated protein 1 |
| ENSG00000100519 | PSMC6 | -0,00509551 | 0,996474291 | 0,98921539 | 0,99837368 | proteasome 26S subunit, ATPase 6 |
| ENSG00000171202 | TMEM126A | -0,005106025 | 0,996467029 | 0,99076399 | 0,998800453 | transmembrane protein 126A |
| ENSG00000114902 | SPCS1 | -0,005134105 | 0,996447634 | 0,98977158 | 0,99837368 | signal peptidase complex subunit 1 |
| ENSG00000238045 | AC009133,1 | -0,005263743 | 0,996358099 | 0,98243996 | 0,99837368 | novel transcript, antisense to C16orf53 and MVP |
| ENSG00000276709 | AC245049,1 | -0,005273625 | 0,996351274 | 0,98256487 | 0,99837368 | Pyridoxal-dependent decarboxylase domain-containing protein 1 |
| ENSG00000267083 | KRT18P61 | -0,005344603 | 0,996302257 | 0,9960805 | 0,999664158 | keratin 18 pseudogene 61 |
| ENSG00000196372 | ASB13 | -0,005386132 | 0,996273578 | 0,98599256 | 0,99837368 | ankyrin repeat and SOCS box containing 13 |
| ENSG00000257803 | PIGAP1 | -0,005395675 | 0,996266988 | 0,98983522 | 0,99837368 | phosphatidylinositol glycan anchor biosynthesis class A pseudogene 1 |
| ENSG00000080824 | HSP90AA1 | -0,005415809 | 0,996253085 | 0,98381523 | 0,99837368 | heat shock protein 90 alpha family class A member 1 |
| ENSG00000072736 | NFATC3 | -0,005467817 | 0,996217171 | 0,98510981 | 0,99837368 | nuclear factor of activated T cells 3 |
| ENSG00000167191 | GPRC5B | -0,005529815 | 0,996174361 | 0,98701255 | 0,99837368 | G protein-coupled receptor class C group 5 member B |
| ENSG00000286216 | AC104335,1 | -0,005645447 | 0,996094521 | 0,99443142 | 0,999386159 | novel transcript |

| | | | | | | |
|---|---|---|---|---|---|---|
| ENSG00000125691 | RPL23 | -0,005655212 | 0,996087779 | 0,98817225 | 0,99837368 | ribosomal protein L23 |
| ENSG00000119326 | CTNNAL1 | -0,005656119 | 0,996087152 | 0,98805709 | 0,99837368 | catenin alpha like 1 |
| ENSG00000107537 | PHYH | -0,005676677 | 0,996072959 | 0,98832519 | 0,99837368 | phytanoyl-CoA 2-hydroxylase |
| ENSG00000170633 | RNF34 | -0,005677712 | 0,996072244 | 0,97991699 | 0,99837368 | ring finger protein 34 |
| ENSG00000051009 | FAM160A2 | -0,005744316 | 0,99602626 | 0,98360439 | 0,99837368 | family with sequence similarity 160 member A2 |
| ENSG00000168710 | AHCYL1 | -0,005870945 | 0,99593884 | 0,98281187 | 0,99837368 | adenosylhomocysteinase like 1 |
| ENSG00000240096 | AL132838,1 | -0,005974295 | 0,995867497 | 0,99532465 | 0,999650137 | ribosomal protein L18a (RPL18A) pseudogene |
| ENSG00000280562 | AL132838,3 | -0,005974295 | 0,995867497 | 0,99532465 | 0,999650137 | ribosomal protein L18a (RPL18A) pseudogene |
| ENSG00000145779 | TNFAIP8 | -0,006073413 | 0,995799079 | 0,9860671 | 0,99837368 | TNF alpha induced protein 8 |
| ENSG00000107372 | ZFAND5 | -0,006125294 | 0,99576327 | 0,9878047 | 0,99837368 | zinc finger AN1-type containing 5 |
| ENSG00000270823 | AC007938,2 | -0,006185331 | 0,995721833 | 0,98717035 | 0,99837368 | novel transcript, antisense to MEST |
| ENSG00000182944 | EWSR1 | -0,00630035 | 0,995642452 | 0,97480972 | 0,99837368 | EWS RNA binding protein 1 |
| ENSG00000188158 | NHS | -0,00631528 | 0,995632148 | 0,98603298 | 0,99837368 | NHS actin remodeling regulator |
| ENSG00000263563 | UBBP4 | -0,006356021 | 0,995604033 | 0,9909286 | 0,998800453 | ubiquitin B pseudogene 4 |
| ENSG00000013392 | RWDD2A | -0,00637002 | 0,995594372 | 0,98389496 | 0,99837368 | RWD domain containing 2A |
| ENSG00000185958 | FAM186A | -0,006448876 | 0,995539955 | 0,9935227 | 0,999083101 | family with sequence similarity 186 member A |
| ENSG00000173930 | SLCO4C1 | -0,006506309 | 0,995500324 | 0,99078112 | 0,998800453 | solute carrier organic anion transporter family member 4C1 |
| ENSG00000135698 | MPHOSPH6 | -0,006529877 | 0,995484062 | 0,9830598 | 0,99837368 | M-phase phosphoprotein 6 |
| ENSG00000284966 | AL138689,2 | -0,006548906 | 0,995470932 | 0,98686003 | 0,99837368 | novel transcript, antisense to EFNB2 |
| ENSG00000198743 | SLC5A3 | -0,006649068 | 0,995401822 | 0,9864537 | 0,99837368 | solute carrier family 5 member 3 |
| ENSG00000143207 | COP1 | -0,006808071 | 0,995292122 | 0,98372764 | 0,99837368 | COP1 E3 ubiquitin ligase |
| ENSG00000251442 | LINC01094 | -0,006849654 | 0,995263435 | 0,98971343 | 0,99837368 | long intergenic non-protein coding RNA 1094 |
| ENSG00000271151 | AC016737,1 | -0,006977705 | 0,995175101 | 0,98899326 | 0,99837368 | novel transcript |
| ENSG00000286921 | AC004522,4 | -0,007023175 | 0,995143736 | 0,98936096 | 0,99837368 | novel transcript, antisense to AZGP1 |
| ENSG00000071994 | PDCD2 | -0,007171715 | 0,995041281 | 0,98447293 | 0,99837368 | programmed cell death 2 |
| ENSG00000185414 | MRPL30 | -0,007187653 | 0,995030288 | 0,97513845 | 0,99837368 | mitochondrial ribosomal protein L30 |
| ENSG00000115042 | FAHD2A | -0,00718925 | 0,995029187 | 0,97548601 | 0,99837368 | fumarylacetoacetate hydrolase domain containing 2A |
| ENSG00000275120 | AC048382,5 | -0,007192619 | 0,995026863 | 0,98992417 | 0,99837368 | novel transcript, antisense to ZSCAN2 |
| ENSG00000184361 | SPATA32 | -0,007284736 | 0,994963332 | 0,9927238 | 0,999049649 | spermatogenesis associated 32 |
| ENSG00000107611 | CUBN | -0,007305342 | 0,994949122 | 0,98414478 | 0,99837368 | cubilin |
| ENSG00000137814 | HAUS2 | -0,007309663 | 0,994946141 | 0,97907211 | 0,99837368 | HAUS augmin like complex subunit 2 |

| | | | | | | |
|---|---|---|---|---|---|---|
| ENSG00000188295 | ZNF669 | -0,007314527 | 0,994942787 | 0,97577387 | 0,99837368 | zinc finger protein 669 |
| ENSG00000164808 | SPIDR | -0,007364309 | 0,994908456 | 0,97248504 | 0,998263101 | scaffold protein involved in DNA repair |
| ENSG00000132950 | ZMYM5 | -0,007415179 | 0,994873376 | 0,97894944 | 0,99837368 | zinc finger MYM-type containing 5 |
| ENSG00000228295 | LINC00392 | -0,007425462 | 0,994866285 | 0,99123855 | 0,998884788 | long intergenic non-protein coding RNA 392 |
| ENSG00000129518 | EAPP | -0,007456749 | 0,99484471 | 0,98441431 | 0,99837368 | E2F associated phosphoprotein |
| ENSG00000182923 | CEP63 | -0,007470631 | 0,994835137 | 0,97980477 | 0,99837368 | centrosomal protein 63 |
| ENSG00000250479 | CHCHD10 | -0,007574625 | 0,994763429 | 0,98439678 | 0,99837368 | coiled-coil-helix-coiled-coil-helix domain containing 10 |
| ENSG00000101278 | RPS10P5 | -0,007649293 | 0,994711945 | 0,99391534 | 0,99925296 | ribosomal protein S10 pseudogene 5 |
| ENSG00000146707 | POMZP3 | -0,007688627 | 0,994684826 | 0,97549373 | 0,99837368 | POM121 and ZP3 fusion |
| ENSG00000065518 | NDUFB4 | -0,007693526 | 0,994681448 | 0,98617844 | 0,99837368 | NADH:ubiquinone oxidoreductase subunit B4 |
| ENSG00000169402 | RSPH10B2 | -0,007752668 | 0,994640673 | 0,99128295 | 0,998887065 | radial spoke head 10 homolog B2 |
| ENSG00000253119 | HNRNPA3P7 | -0,007760678 | 0,99463515 | 0,98991866 | 0,99837368 | heterogeneous nuclear ribonucleoprotein A3 pseudogene 7 |
| ENSG00000096092 | TMEM14A | -0,007817926 | 0,994595683 | 0,98088413 | 0,99837368 | transmembrane protein 14A |
| ENSG00000230461 | PROX1-AS1 | -0,007835666 | 0,994583453 | 0,98772183 | 0,99837368 | PROX1 antisense RNA 1 |
| ENSG00000111788 | AC009533,1 | -0,00788812 | 0,994547292 | 0,97982697 | 0,99837368 | DEAD/H (Asp-Glu-Ala-Asp/His) box polypeptide like pseudogene |
| ENSG00000132466 | ANKRD17 | -0,00788857 | 0,994546982 | 0,97706182 | 0,99837368 | ankyrin repeat domain 17 |
| ENSG00000244389 | RN7SL242P | -0,007918773 | 0,994526161 | 0,99394814 | 0,999253795 | RNA, 7SL, cytoplasmic 242, pseudogene |
| ENSG00000047410 | TPR | -0,007961408 | 0,994496771 | 0,97726106 | 0,99837368 | translocated promoter region, nuclear basket protein |
| ENSG00000169955 | ZNF747 | -0,007993339 | 0,99447476 | 0,97569625 | 0,99837368 | zinc finger protein 747 |
| ENSG00000262470 | TVP23CP2 | -0,008005441 | 0,994466418 | 0,99296889 | 0,999049649 | TVP23C pseudogene 2 |
| ENSG00000104960 | PTOV1 | -0,008074363 | 0,994418911 | 0,97074813 | 0,998088605 | PTOV1 extended AT-hook containing adaptor protein |
| ENSG00000261315 | LARP4P | -0,008138666 | 0,994374589 | 0,98841952 | 0,99837368 | LARP4 pseudogene |
| ENSG00000100154 | TTC28 | -0,008139138 | 0,994374264 | 0,97072544 | 0,998088605 | tetratricopeptide repeat domain 28 |
| ENSG00000116212 | LRRC42 | -0,008200239 | 0,994332151 | 0,97819355 | 0,99837368 | leucine rich repeat containing 42 |
| ENSG00000124212 | PTGIS | -0,008233045 | 0,994309541 | 0,97854078 | 0,99837368 | prostaglandin I2 synthase |
| ENSG00000112282 | MED23 | -0,008276858 | 0,994279345 | 0,98304206 | 0,99837368 | mediator complex subunit 23 |
| ENSG00000196504 | PRPF40A | -0,008327018 | 0,994244776 | 0,98091684 | 0,99837368 | pre-mRNA processing factor 40 homolog A |
| ENSG00000231439 | WASIR2 | -0,00836425 | 0,994219117 | 0,99261655 | 0,999049649 | WASH and IL9R antisense RNA 2 |

| | | | | | | |
|---|---|---|---|---|---|---|
| ENSG00000180229 | HERC2P3 | -0,008435135 | 0,994170269 | 0,96930133 | 0,997412278 | hect domain and RLD 2 pseudogene 3 |
| ENSG00000134986 | NREP | -0,008454827 | 0,994156699 | 0,98314439 | 0,99837368 | neuronal regeneration related protein |
| ENSG00000224109 | CENPVL3 | -0,008489434 | 0,994132852 | 0,97653265 | 0,99837368 | centromere protein V like 3 |
| ENSG00000187837 | H1-2 | -0,008496168 | 0,994128212 | 0,97409144 | 0,99837368 | H1,2 linker histone, cluster member |
| ENSG00000241112 | RPL29P14 | -0,008502728 | 0,994123692 | 0,98822263 | 0,99837368 | ribosomal protein L29 pseudogene 14 |
| ENSG00000180233 | ZNRF2 | -0,008503143 | 0,994123405 | 0,97820169 | 0,99837368 | zinc and ring finger 2 |
| ENSG00000186118 | TEX38 | -0,008543976 | 0,994095269 | 0,99106625 | 0,998857746 | testis expressed 38 |
| ENSG00000112304 | ACOT13 | -0,008557044 | 0,994086264 | 0,98014266 | 0,99837368 | acyl-CoA thioesterase 13 |
| ENSG00000100425 | BRD1 | -0,008586764 | 0,994065786 | 0,97117428 | 0,998088605 | bromodomain containing 1 |
| ENSG00000269001 | AC092070,2 | -0,008599735 | 0,994056849 | 0,97952265 | 0,99837368 | zinc finger protein 818, pseudogene |
| ENSG00000123600 | METTL8 | -0,008607506 | 0,994051495 | 0,96785174 | 0,997263201 | methyltransferase like 8 |
| ENSG00000160688 | FLAD1 | -0,008623172 | 0,9940407 | 0,97034784 | 0,998004261 | flavin adenine dinucleotide synthetase 1 |
| ENSG00000164163 | ABCE1 | -0,008667885 | 0,994009893 | 0,98345538 | 0,99837368 | ATP binding cassette subfamily E member 1 |
| ENSG00000251463 | FKBP4P1 | -0,008746797 | 0,993955524 | 0,99305813 | 0,999049649 | FKBP prolyl isomerase 4 pseudogene 1 |
| ENSG00000227345 | PARG | -0,008748975 | 0,993954024 | 0,97566189 | 0,99837368 | poly(ADP-ribose) glycohydrolase |
| ENSG00000104756 | KCTD9 | -0,00886826 | 0,993871845 | 0,98043367 | 0,99837368 | potassium channel tetramerization domain containing 9 |
| ENSG00000169131 | ZNF354A | -0,008874757 | 0,993867369 | 0,98170504 | 0,99837368 | zinc finger protein 354A |
| ENSG00000137815 | RTF1 | -0,00890732 | 0,993844937 | 0,97235622 | 0,998214946 | RTF1 homolog, Paf1/RNA polymerase II complex component |
| ENSG00000261765 | AC009127,1 | -0,008944985 | 0,99381899 | 0,99334077 | 0,999049649 | novel transcript |
| ENSG00000227739 | TUBB | -0,008974494 | 0,993798663 | 0,97301374 | 0,99837368 | tubulin beta class I |
| ENSG00000236063 | ABHD16A | -0,009037483 | 0,993755274 | 0,97174388 | 0,998088605 | abhydrolase domain containing 16A, phospholipase |
| ENSG00000231488 | ABHD16A | -0,009037483 | 0,993755274 | 0,97174388 | 0,998088605 | abhydrolase domain containing 16A, phospholipase |
| ENSG00000235676 | ABHD16A | -0,009037483 | 0,993755274 | 0,97174388 | 0,998088605 | abhydrolase domain containing 16A, phospholipase |
| ENSG00000171848 | RRM2 | -0,009058381 | 0,993740879 | 0,97640797 | 0,99837368 | ribonucleotide reductase regulatory subunit M2 |
| ENSG00000119041 | GTF3C3 | -0,00913791 | 0,993686101 | 0,98069439 | 0,99837368 | general transcription factor IIIC subunit 3 |
| ENSG00000135473 | PAN2 | -0,00915185 | 0,993676499 | 0,97350461 | 0,99837368 | poly(A) specific ribonuclease subunit PAN2 |
| ENSG00000139287 | TPH2 | -0,009183444 | 0,993654738 | 0,98034129 | 0,99837368 | tryptophan hydroxylase 2 |
| ENSG00000173889 | PHC3 | -0,009277214 | 0,993590157 | 0,97664549 | 0,99837368 | polyhomeotic homolog 3 |
| ENSG00000270964 | AC016355,1 | -0,009338188 | 0,993548164 | 0,98185745 | 0,99837368 | novel transcript |
| ENSG00000265948 | AC015878,2 | -0,009345951 | 0,993542818 | 0,99236116 | 0,999049649 | novel transcript |
| ENSG00000255154 | HTD2 | -0,009363246 | 0,993530908 | 0,97999584 | 0,99837368 | hydroxyacyl-thioester dehydratase type 2 |

| | | | | | | |
|---|---|---|---|---|---|---|
| ENSG00000105185 | PDCD5 | -0,009368363 | 0,993527384 | 0,98079901 | 0,99837368 | programmed cell death 5 |
| ENSG00000108179 | PPIF | -0,009406054 | 0,993501428 | 0,96814679 | 0,997263201 | peptidylprolyl isomerase F |
| ENSG00000188786 | MTF1 | -0,009461998 | 0,993462903 | 0,96557916 | 0,997208095 | metal regulatory transcription factor 1 |
| ENSG00000118096 | IFT46 | -0,00948992 | 0,993443676 | 0,97720399 | 0,99837368 | intraflagellar transport 46 |
| ENSG00000197343 | ZNF655 | -0,009552873 | 0,993400327 | 0,97609724 | 0,99837368 | zinc finger protein 655 |
| ENSG00000004897 | CDC27 | -0,009620666 | 0,993353648 | 0,98259448 | 0,99837368 | cell division cycle 27 |
| ENSG00000100321 | SYNGR1 | -0,009632194 | 0,993345711 | 0,96957902 | 0,997562062 | synaptogyrin 1 |
| ENSG00000173436 | MICOS10 | -0,009654786 | 0,993330155 | 0,97640919 | 0,99837368 | mitochondrial contact site and cristae organizing system subunit 10 |
| ENSG00000244230 | RN7SL151P | -0,009661524 | 0,993325516 | 0,9705225 | 0,998013218 | RNA, 7SL, cytoplasmic 151, pseudogene |
| ENSG00000226833 | AC092164,1 | -0,009684382 | 0,993309778 | 0,97309768 | 0,99837368 | novel transcript |
| ENSG00000117614 | SYF2 | -0,009843774 | 0,993200041 | 0,96844833 | 0,997302342 | SYF2 pre-mRNA splicing factor |
| ENSG00000117592 | PRDX6 | -0,009955045 | 0,993123441 | 0,97288306 | 0,99837368 | peroxiredoxin 6 |
| ENSG00000183718 | TRIM52 | -0,009958716 | 0,993120914 | 0,9668599 | 0,997263201 | tripartite motif containing 52 |
| ENSG00000164366 | CCDC127 | -0,010027676 | 0,993073445 | 0,96252706 | 0,996608711 | coiled-coil domain containing 127 |
| ENSG00000260714 | AC133552,1 | -0,010076096 | 0,993040116 | 0,98984581 | 0,99837368 | thyroid hormone receptor interactor 11 (TRIP11) pseudogene |
| ENSG00000176222 | ZNF404 | -0,010147676 | 0,992990847 | 0,98394789 | 0,99837368 | zinc finger protein 404 |
| ENSG00000274336 | CDIP1 | -0,010245976 | 0,99292319 | 0,976158 | 0,99837368 | cell death inducing p53 target 1 |
| ENSG00000172113 | NME6 | -0,010255723 | 0,992916481 | 0,96151442 | 0,996608711 | NME/NM23 nucleoside diphosphate kinase 6 |
| ENSG00000237438 | CECR7 | -0,010475988 | 0,992764898 | 0,97840581 | 0,99837368 | cat eye syndrome chromosome region, candidate 7 |
| ENSG00000267681 | AC135721,1 | -0,010488924 | 0,992755997 | 0,98589508 | 0,99837368 | high mobility group nucleosome binding domain 1 (HMGN1) pseudogene |
| ENSG00000166046 | TCP11L2 | -0,010590385 | 0,992686181 | 0,9777874 | 0,99837368 | t-complex 11 like 2 |
| ENSG00000131669 | NINJ1 | -0,010620618 | 0,992665379 | 0,97229094 | 0,998214946 | ninjurin 1 |
| ENSG00000102753 | KPNA3 | -0,010654054 | 0,992642373 | 0,98134778 | 0,99837368 | karyopherin subunit alpha 3 |
| ENSG00000055609 | KMT2C | -0,0106604 | 0,992638007 | 0,96840396 | 0,997302342 | lysine methyltransferase 2C |
| ENSG00000115808 | STRN | -0,010696882 | 0,992612906 | 0,97768442 | 0,99837368 | striatin |
| ENSG00000243770 | RN7SL65P | -0,010724881 | 0,992593642 | 0,99160425 | 0,999049649 | RNA, 7SL, cytoplasmic 65, pseudogene |
| ENSG00000196547 | MAN2A2 | -0,010756503 | 0,992571886 | 0,966702 | 0,997263201 | mannosidase alpha class 2A member 2 |
| ENSG00000147679 | UTP23 | -0,010778061 | 0,992557054 | 0,97205876 | 0,998214946 | UTP23 small subunit processome component |
| ENSG00000163155 | LYSMD1 | -0,010799936 | 0,992542004 | 0,96493075 | 0,997113804 | LysM domain containing 1 |
| ENSG00000231707 | PABPC1P1 | -0,01080104 | 0,992541245 | 0,98263844 | 0,99837368 | poly(A) binding protein cytoplasmic 1 pseudogene 1 |

| | | | | | | |
|---|---|---|---|---|---|---|
| ENSG00000165794 | SLC39A2 | -0,01081481 | 0,992531772 | 0,99407008 | 0,999279985 | solute carrier family 39 member 2 |
| ENSG00000172869 | DMXL1 | -0,010882996 | 0,992484863 | 0,97713039 | 0,99837368 | Dmx like 1 |
| ENSG00000165572 | KBTBD6 | -0,010929736 | 0,992452709 | 0,97121696 | 0,998088605 | kelch repeat and BTB domain containing 6 |
| ENSG00000235360 | C6orf47 | -0,010983219 | 0,992415918 | 0,97828121 | 0,99837368 | chromosome 6 open reading frame 47 |
| ENSG00000164654 | MIOS | -0,011005335 | 0,992400705 | 0,9772563 | 0,99837368 | meiosis regulator for oocyte development |
| ENSG00000232693 | AC012370,1 | -0,011031337 | 0,992382818 | 0,99327777 | 0,999049649 | novel transcript, sense intronic to SPRED2 |
| ENSG00000236581 | STARD13-AS | -0,011038096 | 0,99237817 | 0,97896051 | 0,99837368 | STARD13 antisense RNA |
| ENSG00000197279 | ZNF165 | -0,011067478 | 0,992357959 | 0,98030173 | 0,99837368 | zinc finger protein 165 |
| ENSG00000096401 | CDC5L | -0,01111711 | 0,99232382 | 0,97483978 | 0,99837368 | cell division cycle 5 like |
| ENSG00000136052 | SLC41A2 | -0,011152514 | 0,992299469 | 0,97578459 | 0,99837368 | solute carrier family 41 member 2 |
| ENSG00000212978 | AC016747,1 | -0,011182237 | 0,992279025 | 0,96272952 | 0,996608711 | novel transcript |
| ENSG00000257185 | LINC02293 | -0,011211191 | 0,992259111 | 0,99087222 | 0,998800453 | long intergenic non-protein coding RNA 2293 |
| ENSG00000237361 | TUSC8 | -0,011230136 | 0,992246081 | 0,98784749 | 0,99837368 | tumor suppressor candidate 8 |
| ENSG00000095787 | WAC | -0,011428851 | 0,99210942 | 0,97434305 | 0,99837368 | WW domain containing adaptor with coiled-coil |
| ENSG00000272205 | AL451050,2 | -0,011502627 | 0,992058687 | 0,98803973 | 0,99837368 | novel transcript |
| ENSG00000103194 | USP10 | -0,011602161 | 0,991990245 | 0,97027944 | 0,997987113 | ubiquitin specific peptidase 10 |
| ENSG00000166164 | BRD7 | -0,011704182 | 0,991920099 | 0,96816108 | 0,997263201 | bromodomain containing 7 |
| ENSG00000277475 | AC213203,2 | -0,011724328 | 0,991906247 | 0,99319118 | 0,999049649 | |
| ENSG00000106460 | TMEM106B | -0,011779539 | 0,991868289 | 0,98183847 | 0,99837368 | transmembrane protein 106B |
| ENSG00000162777 | DENND2D | -0,011802153 | 0,991852741 | 0,98078947 | 0,99837368 | DENN domain containing 2D |
| ENSG00000166839 | ANKDD1A | -0,011807918 | 0,991848778 | 0,96612163 | 0,997263201 | ankyrin repeat and death domain containing 1A |
| ENSG00000139908 | TSSK4 | -0,011822965 | 0,991838433 | 0,97540165 | 0,99837368 | testis specific serine kinase 4 |
| ENSG00000285140 | TSSK4 | -0,011822965 | 0,991838433 | 0,97540165 | 0,99837368 | testis specific serine kinase 4 |
| ENSG00000181038 | METTL23 | -0,011846901 | 0,991821977 | 0,97280808 | 0,99837368 | methyltransferase like 23 |
| ENSG00000118363 | SPCS2 | -0,011856271 | 0,991815536 | 0,97526581 | 0,99837368 | signal peptidase complex subunit 2 |
| ENSG00000088930 | XRN2 | -0,011898985 | 0,991786171 | 0,97410509 | 0,99837368 | 5'-3' exoribonuclease 2 |
| ENSG00000118816 | CCNI | -0,011905132 | 0,991781946 | 0,96625594 | 0,997263201 | cyclin I |
| ENSG00000083093 | PALB2 | -0,011940486 | 0,991757642 | 0,97178279 | 0,998088605 | partner and localizer of BRCA2 |
| ENSG00000260276 | AC022167,2 | -0,01196855 | 0,99173835 | 0,97876432 | 0,99837368 | novel transcript, antisense to CARHSP1 |
| ENSG00000284819 | EIF2S2P1 | -0,011978119 | 0,991731772 | 0,99336218 | 0,999049649 | eukaryotic translation initiation factor 2 subunit 2 beta pseudogene 1 |
| ENSG00000234534 | CSNK1G2P1 | -0,012029649 | 0,99169635 | 0,98965683 | 0,99837368 | casein kinase 1 gamma 2 pseudogene 1 |

| | | | | | | |
|---|---|---|---|---|---|---|
| ENSG00000169282 | KCNAB1 | -0,01203926 | 0,991689743 | 0,98123021 | 0,99837368 | potassium voltage-gated channel subfamily A member regulatory beta subunit 1 |
| ENSG00000188033 | ZNF490 | -0,012048792 | 0,991683192 | 0,95526449 | 0,996608711 | zinc finger protein 490 |
| ENSG00000286888 | AC006427,2 | -0,012062103 | 0,991674042 | 0,98814976 | 0,99837368 | novel transcript |
| ENSG00000182379 | NXPH4 | -0,012189792 | 0,991586275 | 0,98082991 | 0,99837368 | neurexophilin 4 |
| ENSG00000071082 | RPL31 | -0,012214102 | 0,991569567 | 0,97229962 | 0,998214946 | ribosomal protein L31 |
| ENSG00000116985 | BMP8B | -0,012249556 | 0,991545199 | 0,97392311 | 0,99837368 | bone morphogenetic protein 8b |
| ENSG00000169057 | MECP2 | -0,012253785 | 0,991542293 | 0,95172849 | 0,996432861 | methyl-CpG binding protein 2 |
| ENSG00000093183 | SEC22C | -0,012418496 | 0,991429096 | 0,96270383 | 0,996608711 | SEC22 homolog C, vesicle trafficking protein |
| ENSG00000100330 | MTMR3 | -0,0125848 | 0,991314818 | 0,95408859 | 0,996608711 | myotubularin related protein 3 |
| ENSG00000120733 | KDM3B | -0,012621909 | 0,991289319 | 0,95041421 | 0,996432861 | lysine demethylase 3B |
| ENSG00000145494 | NDUFS6 | -0,012624887 | 0,991287273 | 0,95080825 | 0,996432861 | NADH:ubiquinone oxidoreductase subunit S6 |
| ENSG00000237036 | ZEB1-AS1 | -0,012639328 | 0,99127735 | 0,95768984 | 0,996608711 | ZEB1 antisense RNA 1 |
| ENSG00000068784 | SRBD1 | -0,012678226 | 0,991250624 | 0,96983681 | 0,997761621 | S1 RNA binding domain 1 |
| ENSG00000142794 | NBPF3 | -0,012686188 | 0,991245153 | 0,96213947 | 0,996608711 | NBPF member 3 |
| ENSG00000271383 | NBPF19 | -0,012702855 | 0,991233702 | 0,96682553 | 0,997263201 | NBPF member 19 |
| ENSG00000273137 | AL022328,2 | -0,012731448 | 0,991214056 | 0,98848348 | 0,99837368 | novel transcript, antisense to SELO |
| ENSG00000166451 | CENPN | -0,012737255 | 0,991210067 | 0,96792295 | 0,997263201 | centromere protein N |
| ENSG00000183527 | PSMG1 | -0,012758671 | 0,991195353 | 0,97500146 | 0,99837368 | proteasome assembly chaperone 1 |
| ENSG00000240857 | RDH14 | -0,012785998 | 0,991176578 | 0,9704816 | 0,998013218 | retinol dehydrogenase 14 |
| ENSG00000196151 | WDSUB1 | -0,012811774 | 0,99115887 | 0,97350251 | 0,99837368 | WD repeat, sterile alpha motif and U-box domain containing 1 |
| ENSG00000179476 | C14orf28 | -0,012824928 | 0,991149832 | 0,9795325 | 0,99837368 | chromosome 14 open reading frame 28 |
| ENSG00000223899 | SEC13P1 | -0,012881592 | 0,991110904 | 0,98664185 | 0,99837368 | SEC13 homolog, nuclear pore and COPII coat complex component pseudogene 1 |
| ENSG00000132155 | RAF1 | -0,012905006 | 0,99109482 | 0,96001708 | 0,996608711 | Raf-1 proto-oncogene, serine/threonine kinase |
| ENSG00000064313 | TAF2 | -0,012924096 | 0,991081706 | 0,96878269 | 0,997302342 | TATA-box binding protein associated factor 2 |
| ENSG00000059769 | DNAJC25 | -0,01292424 | 0,991081606 | 0,97001276 | 0,997844133 | DnaJ heat shock protein family (Hsp40) member C25 |
| ENSG00000171574 | ZNF584 | -0,012951098 | 0,991063156 | 0,95672321 | 0,996608711 | zinc finger protein 584 |
| ENSG00000111596 | CNOT2 | -0,012992352 | 0,991034817 | 0,97686461 | 0,99837368 | CCR4-NOT transcription complex subunit 2 |
| ENSG00000258744 | AL132800,1 | -0,012994009 | 0,991033679 | 0,97803361 | 0,99837368 | novel transcript, antisense to CMA1 |
| ENSG00000274112 | SMDT1 | -0,013154624 | 0,990923353 | 0,97454766 | 0,99837368 | single-pass membrane protein with aspartate rich tail 1 |

| | | | | | | |
|---|---|---|---|---|---|---|
| ENSG00000272835 | SMDT1 | -0,013154624 | 0,990923353 | 0,97454766 | 0,99837368 | single-pass membrane protein with aspartate rich tail 1 |
| ENSG00000001497 | LAS1L | -0,013258422 | 0,990852062 | 0,96042309 | 0,996608711 | LAS1 like ribosome biogenesis factor |
| ENSG00000252690 | AC105339,2 | -0,013312377 | 0,990815006 | 0,97257981 | 0,998267798 | TEC |
| ENSG00000117133 | RPF1 | -0,013352443 | 0,990787489 | 0,97361446 | 0,99837368 | ribosome production factor 1 homolog |
| ENSG00000256862 | PARP11-AS1 | -0,013463153 | 0,990711461 | 0,97324302 | 0,99837368 | PARP11 antisense RNA 1 |
| ENSG00000124091 | GCNT7 | -0,013469139 | 0,990707351 | 0,97965858 | 0,99837368 | glucosaminyl (N-acetyl) transferase family member 7 |
| ENSG00000181924 | COA4 | -0,013531052 | 0,990664836 | 0,95521183 | 0,996608711 | cytochrome c oxidase assembly factor 4 homolog |
| ENSG00000198690 | FAN1 | -0,013621241 | 0,990602907 | 0,94839472 | 0,99635783 | FANCD2 and FANCI associated nuclease 1 |
| ENSG00000102043 | MTMR8 | -0,013632696 | 0,990595041 | 0,97474688 | 0,99837368 | myotubularin related protein 8 |
| ENSG00000109771 | LRP2BP | -0,013659819 | 0,990576418 | 0,96538928 | 0,997208095 | LRP2 binding protein |
| ENSG00000244151 | AC010973,2 | -0,013690928 | 0,990555058 | 0,98430886 | 0,99837368 | novel transcript |
| ENSG00000265458 | AC132938,3 | -0,013705281 | 0,990545204 | 0,98098363 | 0,99837368 | novel transcript, antisense NARF |
| ENSG00000174010 | KLHL15 | -0,013752752 | 0,990512611 | 0,97489328 | 0,99837368 | kelch like family member 15 |
| ENSG00000232811 | AL360270,1 | -0,013798979 | 0,990480873 | 0,97576347 | 0,99837368 | novel transcript |
| ENSG00000170542 | SERPINB9 | -0,013937465 | 0,9903858 | 0,9723313 | 0,998214946 | serpin family B member 9 |
| ENSG00000154114 | TBCEL | -0,013960842 | 0,990369752 | 0,96737438 | 0,997263201 | tubulin folding cofactor E like |
| ENSG00000152954 | NRSN1 | -0,01400953 | 0,99033633 | 0,98220592 | 0,99837368 | neurensin 1 |
| ENSG00000067601 | PMS2P4 | -0,014016087 | 0,990331829 | 0,95990627 | 0,996608711 | PMS1 homolog 2, mismatch repair system component pseudogene 4 |
| ENSG00000157150 | TIMP4 | -0,014038321 | 0,990316566 | 0,97500853 | 0,99837368 | TIMP metallopeptidase inhibitor 4 |
| ENSG00000177212 | OR2T33 | -0,014050602 | 0,990308136 | 0,98073398 | 0,99837368 | olfactory receptor family 2 subfamily T member 33 |
| ENSG00000277304 | AC142086,6 | -0,0140609 | 0,990301068 | 0,99326835 | 0,999049649 | Rho GTPase activating protein 23 (ARHGAP23) pseudogene |
| ENSG00000259397 | AC021231,2 | -0,0140732 | 0,990292625 | 0,9894474 | 0,99837368 | U2 small nuclear RNA auxiliary factor 1 (U2AF1) pseudogene |
| ENSG00000288217 | PEX5 | -0,014111504 | 0,990266333 | 0,96638335 | 0,997263201 | peroxisomal biogenesis factor 5 |
| ENSG00000128039 | SRD5A3 | -0,014128864 | 0,990254416 | 0,96756332 | 0,997263201 | steroid 5 alpha-reductase 3 |
| ENSG00000115825 | PRKD3 | -0,01419973 | 0,990205776 | 0,97171711 | 0,998088605 | protein kinase D3 |
| ENSG00000165516 | KLHDC2 | -0,014219297 | 0,990192346 | 0,97128745 | 0,998088605 | kelch domain containing 2 |
| ENSG00000166881 | NEMP1 | -0,014281492 | 0,99014966 | 0,96879307 | 0,997302342 | nuclear envelope integral membrane protein 1 |
| ENSG00000278644 | PCGF2 | -0,014322062 | 0,990121816 | 0,94956491 | 0,996432861 | polycomb group ring finger 2 |

| | | | | | | |
|---|---|---|---|---|---|---|
| ENSG00000177576 | C18orf32 | -0,01433902 | 0,990110178 | 0,96570431 | 0,997218242 | chromosome 18 open reading frame 32 |
| ENSG00000164494 | PDSS2 | -0,01436873 | 0,990089789 | 0,95961592 | 0,996608711 | decaprenyl diphosphate synthase subunit 2 |
| ENSG00000237689 | AC007064,2 | -0,014378465 | 0,990083108 | 0,98669 | 0,99837368 | novel transcript |
| ENSG00000162852 | CNST | -0,014389301 | 0,990075671 | 0,96295075 | 0,996608711 | consortin, connexin sorting protein |
| ENSG00000048405 | ZNF800 | -0,01441051 | 0,990061116 | 0,97133843 | 0,998088605 | zinc finger protein 800 |
| ENSG00000123500 | COL10A1 | -0,014431904 | 0,990046434 | 0,97213569 | 0,998214946 | collagen type X alpha 1 chain |
| ENSG00000230092 | AL669831,4 | -0,014463404 | 0,990024818 | 0,97441877 | 0,99837368 | F-box only protein 25 (FBXO25) pseudogene |
| ENSG00000141577 | CEP131 | -0,014537382 | 0,989974053 | 0,98220157 | 0,99837368 | centrosomal protein 131 |
| ENSG00000157077 | ZFYVE9 | -0,014638967 | 0,989904348 | 0,95971175 | 0,996608711 | zinc finger FYVE-type containing 9 |
| ENSG00000125611 | CHCHD5 | -0,014697491 | 0,989864193 | 0,96675106 | 0,997263201 | coiled-coil-helix-coiled-coil-helix domain containing 5 |
| ENSG00000079974 | RABL2B | -0,014705118 | 0,98985896 | 0,94669844 | 0,996027077 | RAB, member of RAS oncogene family like 2B |
| ENSG00000187720 | THSD4 | -0,014791554 | 0,989799656 | 0,95958152 | 0,996608711 | thrombospondin type 1 domain containing 4 |
| ENSG00000285282 | DHRS4-AS1 | -0,014830911 | 0,989772655 | 0,96649444 | 0,997263201 | DHRS4 antisense RNA 1 |
| ENSG00000188763 | FZD9 | -0,014847527 | 0,989761255 | 0,97662357 | 0,99837368 | frizzled class receptor 9 |
| ENSG00000253431 | SRPK2P | -0,014898135 | 0,989726536 | 0,98516594 | 0,99837368 | SRSF protein kinase 2 pseudogene |
| ENSG00000134987 | WDR36 | -0,015039042 | 0,989629875 | 0,97123118 | 0,998088605 | WD repeat domain 36 |
| ENSG00000124383 | MPHOSPH10 | -0,015081187 | 0,989600966 | 0,95627519 | 0,996608711 | M-phase phosphoprotein 10 |
| ENSG00000215021 | PHB2 | -0,015088936 | 0,98959565 | 0,9459267 | 0,996027077 | prohibitin 2 |
| ENSG00000178127 | NDUFV2 | -0,015112395 | 0,989579559 | 0,97175792 | 0,998088605 | NADH:ubiquinone oxidoreductase core subunit V2 |
| ENSG00000259642 | ST20-AS1 | -0,015123884 | 0,989571678 | 0,9573419 | 0,996608711 | ST20 antisense RNA 1 |
| ENSG00000277130 | AC073569,3 | -0,015125885 | 0,989570306 | 0,98031613 | 0,99837368 | novel transcript, antisense to PPP1R12A |
| ENSG00000243335 | KCTD7 | -0,015211791 | 0,989511383 | 0,9534714 | 0,99652169 | potassium channel tetramerization domain containing 7 |
| ENSG00000101132 | PFDN4 | -0,015227336 | 0,989500721 | 0,97253355 | 0,998267798 | prefoldin subunit 4 |
| ENSG00000124275 | MTRR | -0,015319098 | 0,989437787 | 0,9654994 | 0,997208095 | 5-methyltetrahydrofolate-homocysteine methyltransferase reductase |
| ENSG00000143756 | FBXO28 | -0,015419643 | 0,989368833 | 0,96980045 | 0,99775705 | F-box protein 28 |
| ENSG00000101003 | GINS1 | -0,015490262 | 0,989320404 | 0,93733146 | 0,994479239 | GINS complex subunit 1 |
| ENSG00000270528 | FAM133EP | -0,01573838 | 0,989150274 | 0,98842371 | 0,99837368 | family with sequence similarity 133 member E, pseudogene |
| ENSG00000125457 | MIF4GD | -0,015759188 | 0,989136008 | 0,95644172 | 0,996608711 | MIF4G domain containing |
| ENSG00000226913 | BSN-DT | -0,015788462 | 0,989115937 | 0,9775812 | 0,99837368 | BSN divergent transcript |

| | | | | | | |
|---|---|---|---|---|---|---|
| ENSG00000165097 | KDM1B | -0,015825332 | 0,989090659 | 0,9591044 | 0,996608711 | lysine demethylase 1B |
| ENSG00000185591 | SP1 | -0,015859792 | 0,989067034 | 0,96269297 | 0,996608711 | Sp1 transcription factor |
| ENSG00000281915 | MUC20-OT1 | -0,015860187 | 0,989066763 | 0,9631481 | 0,996608711 | MUC20 overlapping transcript |
| ENSG00000132911 | NMUR2 | -0,015863005 | 0,989064832 | 0,98774354 | 0,99837368 | neuromedin U receptor 2 |
| ENSG00000287721 | AC061975,8 | -0,015867184 | 0,989061966 | 0,98023041 | 0,99837368 | novel transcript |
| ENSG00000119965 | C10orf88 | -0,015956892 | 0,989000467 | 0,95898153 | 0,996608711 | chromosome 10 open reading frame 88 |
| ENSG00000260417 | AC092127,1 | -0,016060077 | 0,988929734 | 0,98242841 | 0,99837368 | novel transcript |
| ENSG00000282965 | AP001005,3 | -0,016104727 | 0,988899128 | 0,98940432 | 0,99837368 | novel transcript |
| ENSG00000161395 | PGAP3 | -0,016133393 | 0,98887948 | 0,96110211 | 0,996608711 | post-GPI attachment to proteins phospholipase 3 |
| ENSG00000170631 | ZNF16 | -0,016160018 | 0,98886123 | 0,94353739 | 0,995402494 | zinc finger protein 16 |
| ENSG00000235106 | BRD3OS | -0,016200652 | 0,988833378 | 0,94712067 | 0,996027077 | BRD3 opposite strand |
| ENSG00000179988 | PSTK | -0,016298533 | 0,988766292 | 0,95092092 | 0,996432861 | phosphoseryl-tRNA kinase |
| ENSG00000163512 | AZI2 | -0,016345108 | 0,988734372 | 0,96307486 | 0,996608711 | 5-azacytidine induced 2 |
| ENSG00000100784 | RPS6KA5 | -0,016365997 | 0,988720056 | 0,9549732 | 0,996608711 | ribosomal protein S6 kinase A5 |
| ENSG00000240429 | LRRFIP1P1 | -0,016406054 | 0,988692605 | 0,95544821 | 0,996608711 | LRR binding FLII interacting protein 1 pseudogene 1 |
| ENSG00000178028 | DMAP1 | -0,016427627 | 0,988677821 | 0,94043651 | 0,995028467 | DNA methyltransferase 1 associated protein 1 |
| ENSG00000198604 | BAZ1A | -0,016437804 | 0,988670846 | 0,95334118 | 0,99652169 | bromodomain adjacent to zinc finger domain 1A |
| ENSG00000185344 | ATP6V0A2 | -0,016489608 | 0,988635346 | 0,9371526 | 0,994479239 | ATPase H+ transporting V0 subunit a2 |
| ENSG00000174748 | RPL15 | -0,01649737 | 0,988630027 | 0,96275866 | 0,996608711 | ribosomal protein L15 |
| ENSG00000239899 | RN7SL674P | -0,016518411 | 0,988615608 | 0,98506759 | 0,99837368 | RNA, 7SL, cytoplasmic 674, pseudogene |
| ENSG00000100380 | ST13 | -0,016605881 | 0,988555671 | 0,96360848 | 0,996733015 | ST13 Hsp70 interacting protein |
| ENSG00000145388 | METTL14 | -0,016613566 | 0,988550405 | 0,96029044 | 0,996608711 | methyltransferase like 14 |
| ENSG00000072501 | SMC1A | -0,016613796 | 0,988550248 | 0,96095271 | 0,996608711 | structural maintenance of chromosomes 1A |
| ENSG00000201901 | RN7SKP48 | -0,016629281 | 0,988539637 | 0,98389635 | 0,99837368 | RN7SK pseudogene 48 |
| ENSG00000102606 | ARHGEF7 | -0,016631528 | 0,988538097 | 0,94745035 | 0,996107107 | Rho guanine nucleotide exchange factor 7 |
| ENSG00000078319 | PMS2P1 | -0,016717889 | 0,988478924 | 0,94790904 | 0,99619108 | PMS1 homolog 2, mismatch repair system component pseudogene 1 |
| ENSG00000124688 | MAD2L1BP | -0,01674106 | 0,988463049 | 0,94112644 | 0,995251853 | MAD2L1 binding protein |
| ENSG00000148429 | USP6NL | -0,016780143 | 0,988436271 | 0,95271558 | 0,996510537 | USP6 N-terminal like |
| ENSG00000198646 | NCOA6 | -0,01683461 | 0,988398954 | 0,95030058 | 0,996432861 | nuclear receptor coactivator 6 |
| ENSG00000198728 | LDB1 | -0,016847467 | 0,988390146 | 0,93322811 | 0,994479239 | LIM domain binding 1 |
| ENSG00000128607 | KLHDC10 | -0,016851991 | 0,988387047 | 0,95833285 | 0,996608711 | kelch domain containing 10 |
| ENSG00000261235 | AC092142,1 | -0,016861705 | 0,988380392 | 0,9899432 | 0,99837368 | novel transcript, antisense to HSD17B2 |

| | | | | | | |
|---|---|---|---|---|---|---|
| ENSG00000112787 | FBRSL1 | -0,01697823 | 0,988300565 | 0,95803455 | 0,996608711 | fibrosin like 1 |
| ENSG00000253352 | TUG1 | -0,01704152 | 0,988257209 | 0,95964422 | 0,996608711 | taurine up-regulated 1 |
| ENSG00000238103 | RPL9P7 | -0,017067621 | 0,98823933 | 0,98504444 | 0,99837368 | ribosomal protein L9 pseudogene 7 |
| ENSG00000164347 | GFM2 | -0,01714001 | 0,988189745 | 0,9646105 | 0,997113804 | GTP dependent ribosome recycling factor mitochondrial 2 |
| ENSG00000100596 | SPTLC2 | -0,017178276 | 0,988163535 | 0,95565446 | 0,996608711 | serine palmitoyltransferase long chain base subunit 2 |
| ENSG00000134779 | TPGS2 | -0,017253073 | 0,988112305 | 0,95728192 | 0,996608711 | tubulin polyglutamylase complex subunit 2 |
| ENSG00000244383 | FAM3D-AS1 | -0,01728714 | 0,988088972 | 0,98373442 | 0,99837368 | FAM3D antisense RNA 1 |
| ENSG00000188467 | SLC24A5 | -0,017520912 | 0,987928877 | 0,98374796 | 0,99837368 | solute carrier family 24 member 5 |
| ENSG00000170088 | TMEM192 | -0,017535865 | 0,987918637 | 0,94941285 | 0,996432861 | transmembrane protein 192 |
| ENSG00000174938 | SEZ6L2 | -0,017548441 | 0,987910026 | 0,93950344 | 0,994943406 | seizure related 6 homolog like 2 |
| ENSG00000196290 | NIF3L1 | -0,017551143 | 0,987908175 | 0,95629353 | 0,996608711 | NGG1 interacting factor 3 like 1 |
| ENSG00000185049 | NELFA | -0,017590847 | 0,987880988 | 0,94346082 | 0,995402494 | negative elongation factor complex member A |
| ENSG00000247877 | AC021086,1 | -0,017596367 | 0,987877208 | 0,9829045 | 0,99837368 | novel transcript |
| ENSG00000159131 | GART | -0,017597733 | 0,987876273 | 0,94080248 | 0,995154793 | phosphoribosylglycinamide formyltransferase, phosphoribosylglycinamide synthetase, phosphoribosylaminoimidazole synthetase |
| ENSG00000132485 | ZRANB2 | -0,017668975 | 0,987827492 | 0,96609395 | 0,997263201 | zinc finger RANBP2-type containing 2 |
| ENSG00000012983 | MAP4K5 | -0,017681086 | 0,987819199 | 0,96279632 | 0,996608711 | mitogen-activated protein kinase kinase kinase kinase 5 |
| ENSG00000229853 | AL034418,1 | -0,01773429 | 0,987782771 | 0,98114595 | 0,99837368 | ribosomal protein S3A (RPS3A) pseudogene |
| ENSG00000274105 | AC084824,3 | -0,017799676 | 0,987738003 | 0,9682997 | 0,997263201 | novel transcript, sense intronic to YARS2 |
| ENSG00000276734 | KIF5C | -0,017862497 | 0,987694994 | 0,95930349 | 0,996608711 | kinesin family member 5C |
| ENSG00000237807 | AC022034,1 | -0,017881165 | 0,987682214 | 0,95958766 | 0,996608711 | novel transcript |
| ENSG00000145743 | FBXL17 | -0,017905229 | 0,987665739 | 0,95707472 | 0,996608711 | F-box and leucine rich repeat protein 17 |
| ENSG00000019995 | ZRANB1 | -0,017920723 | 0,987655132 | 0,95353585 | 0,99652169 | zinc finger RANBP2-type containing 1 |
| ENSG00000127533 | F2RL3 | -0,017991458 | 0,987606709 | 0,98682378 | 0,99837368 | F2R like thrombin or trypsin receptor 3 |
| ENSG00000274211 | SOCS7 | -0,018015574 | 0,9875902 | 0,93182179 | 0,994433362 | suppressor of cytokine signaling 7 |
| ENSG00000163605 | PPP4R2 | -0,018051283 | 0,987565756 | 0,95737917 | 0,996608711 | protein phosphatase 4 regulatory subunit 2 |
| ENSG00000078043 | PIAS2 | -0,018118158 | 0,987519979 | 0,96124443 | 0,996608711 | protein inhibitor of activated STAT 2 |
| ENSG00000149527 | PLCH2 | -0,018138576 | 0,987506003 | 0,98630539 | 0,99837368 | phospholipase C eta 2 |
| ENSG00000089063 | TMEM230 | -0,018145809 | 0,987501053 | 0,96893358 | 0,997302342 | transmembrane protein 230 |

| | | | | | | |
|---|---|---|---|---|---|---|
| ENSG00000198876 | DCAF12 | -0,018251608 | 0,987428638 | 0,95583427 | 0,996608711 | DDB1 and CUL4 associated factor 12 |
| ENSG00000255506 | AP003718,1 | -0,018270935 | 0,98741541 | 0,989525 | 0,99837368 | novel transcript |
| ENSG00000197584 | KCNMB2 | -0,018317985 | 0,987383208 | 0,98298022 | 0,99837368 | potassium calcium-activated channel subfamily M regulatory beta subunit 2 |
| ENSG00000168118 | RAB4A | -0,018388375 | 0,987335034 | 0,9632679 | 0,996608711 | RAB4A, member RAS oncogene family |
| ENSG00000251614 | HSPA8P19 | -0,018417976 | 0,987314776 | 0,98292206 | 0,99837368 | heat shock protein family A (Hsp70) member pseudogene 19 |
| ENSG00000120899 | PTK2B | -0,018454465 | 0,987289805 | 0,95092777 | 0,996432861 | protein tyrosine kinase 2 beta |
| ENSG00000177889 | UBE2N | -0,018596434 | 0,987192655 | 0,96877037 | 0,997302342 | ubiquitin conjugating enzyme E2 N |
| ENSG00000035928 | RFC1 | -0,018601168 | 0,987189416 | 0,95449574 | 0,996608711 | replication factor C subunit 1 |
| ENSG00000149531 | FRG1BP | -0,018699428 | 0,987122182 | 0,94929881 | 0,996432861 | FSHD region gene 1 family member B, pseudogene |
| ENSG00000260093 | AC034111,1 | -0,018709995 | 0,987114952 | 0,96583497 | 0,997263201 | novel transcript, antisense to MSRA |
| ENSG00000275211 | CYP2D6 | -0,018740381 | 0,987094162 | 0,98858977 | 0,99837368 | cytochrome P450 family 2 subfamily D member 6 |
| ENSG00000284070 | AP000356,2 | -0,018757718 | 0,9870823 | 0,95970982 | 0,996608711 | novel transcript |
| ENSG00000226472 | AC008013,1 | -0,018797084 | 0,987055366 | 0,96561107 | 0,997208095 | tetraspanin family pseudogene |
| ENSG00000158296 | SLC13A3 | -0,01879851 | 0,98705439 | 0,94980805 | 0,996432861 | solute carrier family 13 member 3 |
| ENSG00000261215 | AL162231,4 | -0,018825767 | 0,987035742 | 0,96780042 | 0,997263201 | novel transcript (novel gene-CCL27 readthrough) |
| ENSG00000126883 | NUP214 | -0,018844953 | 0,987022616 | 0,92998732 | 0,994331958 | nucleoporin 214 |
| ENSG00000142528 | ZNF473 | -0,018880809 | 0,986998085 | 0,94410128 | 0,995542144 | zinc finger protein 473 |
| ENSG00000172586 | CHCHD1 | -0,018998779 | 0,986917381 | 0,95899682 | 0,996608711 | coiled-coil-helix-coiled-coil-helix domain containing 1 |
| ENSG00000164172 | MOCS2 | -0,019003715 | 0,986914005 | 0,96232563 | 0,996608711 | molybdenum cofactor synthesis 2 |
| ENSG00000156858 | PRR14 | -0,019005564 | 0,98691274 | 0,94050508 | 0,995057557 | proline rich 14 |
| ENSG00000079785 | DDX1 | -0,019055283 | 0,986878729 | 0,95200486 | 0,996432861 | DEAD-box helicase 1 |
| ENSG00000039319 | ZFYVE16 | -0,01919664 | 0,986782038 | 0,96356472 | 0,996733015 | zinc finger FYVE-type containing 16 |
| ENSG00000144674 | GOLGA4 | -0,019236154 | 0,986755012 | 0,93703262 | 0,994479239 | golgin A4 |
| ENSG00000197714 | ZNF460 | -0,019241246 | 0,986751529 | 0,93559012 | 0,994479239 | zinc finger protein 460 |
| ENSG00000198695 | MT-ND6 | -0,019254813 | 0,986742249 | 0,93740491 | 0,994479239 | mitochondrially encoded NADH:ubiquinone oxidoreductase core subunit 6 |
| ENSG00000196313 | POM121 | -0,019295309 | 0,986714552 | 0,9268876 | 0,993915537 | POM121 transmembrane nucleoporin |
| ENSG00000205339 | IPO7 | -0,019342987 | 0,986681944 | 0,96129101 | 0,996608711 | importin 7 |
| ENSG00000286692 | AC118469,2 | -0,01956495 | 0,986530152 | 0,98648948 | 0,99837368 | novel transcript |
| ENSG00000149532 | CPSF7 | -0,019607265 | 0,986501216 | 0,93126971 | 0,994433362 | cleavage and polyadenylation specific factor 7 |

| | | | | | | |
|---|---|---|---|---|---|---|
| ENSG00000168275 | COA6 | -0,019634206 | 0,986482795 | 0,95053732 | 0,996432861 | cytochrome c oxidase assembly factor 6 |
| ENSG00000162643 | DNAI3 | -0,019657813 | 0,986466653 | 0,9669421 | 0,997263201 | dynein axonemal intermediate chain 3 |
| ENSG00000110696 | C11orf58 | -0,019666748 | 0,986460543 | 0,93239388 | 0,994479239 | chromosome 11 open reading frame 58 |
| ENSG00000100575 | TIMM9 | -0,019694081 | 0,986441855 | 0,96312901 | 0,996608711 | translocase of inner mitochondrial membrane 9 |
| ENSG00000120948 | TARDBP | -0,019755544 | 0,98639983 | 0,94066834 | 0,995129256 | TAR DNA binding protein |
| ENSG00000200463 | SNORD118 | -0,019761992 | 0,986395421 | 0,99566894 | 0,999664158 | small nucleolar RNA, C/D box 118 |
| ENSG00000225265 | TAF1A-AS1 | -0,019844938 | 0,986338712 | 0,9718677 | 0,998110538 | TAF1A antisense RNA 1 |
| ENSG00000070081 | NUCB2 | -0,019865766 | 0,986324472 | 0,94979811 | 0,996432861 | nucleobindin 2 |
| ENSG00000280993 | MUC20-OT1 | -0,019876293 | 0,986317275 | 0,95166155 | 0,996432861 | MUC20 overlapping transcript |
| ENSG00000247934 | AC022364,1 | -0,019887948 | 0,986309307 | 0,96816675 | 0,997263201 | novel transcript, antisense to CCDC91 |
| ENSG00000072849 | DERL2 | -0,019942566 | 0,986271967 | 0,95852623 | 0,996608711 | derlin 2 |
| ENSG00000104093 | DMXL2 | -0,019974854 | 0,986249894 | 0,95966647 | 0,996608711 | Dmx like 2 |
| ENSG00000116586 | LAMTOR2 | -0,020059576 | 0,986191979 | 0,94180065 | 0,995373376 | late endosomal/lysosomal adaptor, MAPK and MTOR activator 2 |
| ENSG00000206559 | ZCWPW2 | -0,020259845 | 0,986055089 | 0,96180626 | 0,996608711 | zinc finger CW-type and PWWP domain containing 2 |
| ENSG00000138379 | MSTN | -0,020270038 | 0,986048123 | 0,98624184 | 0,99837368 | myostatin |
| ENSG00000264808 | AC068025,1 | -0,020281077 | 0,986040578 | 0,98404656 | 0,99837368 | novel transcript, antisense to TAOK1 |
| ENSG00000213871 | TAF9BP1 | -0,020303304 | 0,986025387 | 0,97695679 | 0,99837368 | TATA-box binding protein associated factor 9b pseudogene 1 |
| ENSG00000153207 | AHCTF1 | -0,020335663 | 0,986003271 | 0,9513781 | 0,996432861 | AT-hook containing transcription factor 1 |
| ENSG00000182873 | PRKCZ-AS1 | -0,020422912 | 0,985943643 | 0,94996708 | 0,996432861 | PRKCZ antisense RNA 1 |
| ENSG00000141556 | TBCD | -0,020522339 | 0,985875696 | 0,9358642 | 0,994479239 | tubulin folding cofactor D |
| ENSG00000118922 | KLF12 | -0,020539852 | 0,985863728 | 0,9646805 | 0,997113804 | Kruppel like factor 12 |
| ENSG00000284784 | AC270301,1 | -0,020545763 | 0,985859689 | 0,98677975 | 0,99837368 | novel transcript |
| ENSG00000270076 | AF131215,7 | -0,020545763 | 0,985859689 | 0,98677975 | 0,99837368 | novel transcript |
| ENSG00000198839 | ZNF277 | -0,020591336 | 0,985828547 | 0,96065443 | 0,996608711 | zinc finger protein 277 |
| ENSG00000107829 | FBXW4 | -0,020606556 | 0,985818147 | 0,95967586 | 0,996608711 | F-box and WD repeat domain containing 4 |
| ENSG00000078328 | RBFOX1 | -0,020661391 | 0,985780678 | 0,96616265 | 0,997263201 | RNA binding fox-1 homolog 1 |
| ENSG00000249623 | PALLD-AS1 | -0,020770903 | 0,985705852 | 0,98591284 | 0,99837368 | PALLD antisense RNA 1 |
| ENSG00000176907 | TCIM | -0,020826311 | 0,985667996 | 0,94424661 | 0,995542144 | transcriptional and immune response regulator |
| ENSG00000198862 | LTN1 | -0,020828739 | 0,985666338 | 0,96192494 | 0,996608711 | listerin E3 ubiquitin protein ligase 1 |
| ENSG00000173875 | ZNF791 | -0,020853738 | 0,985649258 | 0,94505835 | 0,995797387 | zinc finger protein 791 |
| ENSG00000206262 | FOXL2NB | -0,020862148 | 0,985643513 | 0,97164355 | 0,998088605 | FOXL2 neighbor |

| | | | | | | |
|---|---|---|---|---|---|---|
| ENSG00000125630 | POLR1B | -0,020870478 | 0,985637821 | 0,94431081 | 0,995542144 | RNA polymerase I subunit B |
| ENSG00000143512 | HHIPL2 | -0,020934284 | 0,985594231 | 0,98200582 | 0,99837368 | HHIP like 2 |
| ENSG00000184900 | SUMO3 | -0,02095589 | 0,985579471 | 0,94304873 | 0,995402494 | small ubiquitin like modifier 3 |
| ENSG00000184220 | CMSS1 | -0,020973839 | 0,985567208 | 0,93201244 | 0,994479239 | cms1 ribosomal small subunit homolog |
| ENSG00000143514 | TP53BP2 | -0,021009265 | 0,985543008 | 0,94277871 | 0,995402494 | tumor protein p53 binding protein 2 |
| ENSG00000214517 | PPME1 | -0,021023892 | 0,985533015 | 0,9271192 | 0,993915537 | protein phosphatase methylesterase 1 |
| ENSG00000234272 | RPL30P2 | -0,021034462 | 0,985525795 | 0,98979936 | 0,99837368 | ribosomal protein L30 pseudogene 2 |
| ENSG00000269929 | MIRLET7A1HG | -0,021098146 | 0,985482293 | 0,96142908 | 0,996608711 | miRlet-7a-1/let-7f-1/let-7d cluster host gene |
| ENSG00000234509 | AP000253,1 | -0,021109159 | 0,98547477 | 0,96287658 | 0,996608711 | novel transcript |
| ENSG00000182774 | RPS17 | -0,021132912 | 0,985458545 | 0,94959644 | 0,996432861 | ribosomal protein S17 |
| ENSG00000124789 | NUP153 | -0,021185916 | 0,98542234 | 0,95229277 | 0,996432861 | nucleoporin 153 |
| ENSG00000149179 | C11orf49 | -0,021228315 | 0,98539338 | 0,92816555 | 0,994108531 | chromosome 11 open reading frame 49 |
| ENSG00000250462 | LRRC37BP1 | -0,021373193 | 0,98529443 | 0,9371953 | 0,994479239 | leucine rich repeat containing 37B pseudogene 1 |
| ENSG00000227835 | CARM1P1 | -0,02138064 | 0,985289344 | 0,98447107 | 0,99837368 | coactivator associated arginine methyltransferase 1 pseudogene 1 |
| ENSG00000138363 | ATIC | -0,02141976 | 0,985262628 | 0,92631973 | 0,993915537 | 5-aminoimidazole-4-carboxamide ribonucleotide formyltransferase/IMP cyclohydrolase |
| ENSG00000273673 | AC011295,1 | -0,021537766 | 0,985182041 | 0,95183903 | 0,996432861 | ribosomal protein L9 pseudogene 9 |
| ENSG00000267731 | AC005332,2 | -0,02158543 | 0,985149493 | 0,97553917 | 0,99837368 | novel transcript |
| ENSG00000261501 | AC079341,1 | -0,021653765 | 0,985102831 | 0,97614468 | 0,99837368 | novel transcript |
| ENSG00000260229 | PPIAP51 | -0,021654437 | 0,985102372 | 0,97021814 | 0,997956894 | peptidylprolyl isomerase A pseudogene 51 |
| ENSG00000076242 | MLH1 | -0,021709096 | 0,985065051 | 0,91793019 | 0,993361299 | mutL homolog 1 |
| ENSG00000158321 | AUTS2 | -0,021724375 | 0,985054619 | 0,91421528 | 0,993361299 | activator of transcription and developmental regulator AUTS2 |
| ENSG00000132823 | OSER1 | -0,021821411 | 0,984988365 | 0,9518046 | 0,996432861 | oxidative stress responsive serine rich 1 |
| ENSG00000188010 | MORN2 | -0,021929107 | 0,98491484 | 0,96230716 | 0,996608711 | MORN repeat containing 2 |
| ENSG00000146278 | PNRC1 | -0,021935049 | 0,984910783 | 0,95245425 | 0,996432861 | proline rich nuclear receptor coactivator 1 |
| ENSG00000133657 | ATP13A3 | -0,021963915 | 0,984891077 | 0,95200606 | 0,996432861 | ATPase 13A3 |
| ENSG00000141034 | GID4 | -0,022039199 | 0,984839684 | 0,93985618 | 0,994943406 | GID complex subunit 4 homolog |
| ENSG00000172005 | MAL | -0,022073888 | 0,984816004 | 0,97984713 | 0,99837368 | mal, T cell differentiation protein |
| ENSG00000141431 | ASXL3 | -0,022081474 | 0,984810826 | 0,96076191 | 0,996608711 | ASXL transcriptional regulator 3 |
| ENSG00000185477 | GPRIN3 | -0,022164371 | 0,98475424 | 0,93769958 | 0,994479239 | GPRIN family member 3 |
| ENSG00000168566 | SNRNP48 | -0,022173051 | 0,984748315 | 0,96107614 | 0,996608711 | small nuclear ribonucleoprotein U11/U12 subunit 48 |

| | | | | | | |
|---|---|---|---|---|---|---|
| ENSG00000142541 | RPL13A | -0,022179423 | 0,984743966 | 0,93016066 | 0,994336072 | ribosomal protein L13a |
| ENSG00000187904 | AC097382,1 | -0,022258632 | 0,984689901 | 0,98477236 | 0,99837368 | novel transcript, sense overlapping TBC1D14 |
| ENSG00000261220 | AC103706,1 | -0,022289421 | 0,984668887 | 0,97392246 | 0,99837368 | novel transcript |
| ENSG00000105583 | WDR83OS | -0,022497671 | 0,984526763 | 0,92705925 | 0,993915537 | WD repeat domain 83 opposite strand |
| ENSG00000146242 | TPBG | -0,022548863 | 0,984491829 | 0,94127814 | 0,995251853 | trophoblast glycoprotein |
| ENSG00000082701 | GSK3B | -0,022567559 | 0,984479071 | 0,94351544 | 0,995402494 | glycogen synthase kinase 3 beta |
| ENSG00000145216 | FIP1L1 | -0,022604468 | 0,984453885 | 0,92777905 | 0,994108531 | factor interacting with PAPOLA and CPSF1 |
| ENSG00000066654 | THUMPD1 | -0,022623331 | 0,984441013 | 0,95199691 | 0,996432861 | THUMP domain containing 1 |
| ENSG00000272008 | AL139274,2 | -0,022674266 | 0,984406257 | 0,95416757 | 0,996608711 | novel transcript, antisense to ZNF292 |
| ENSG00000121864 | ZNF639 | -0,022706143 | 0,984384507 | 0,94712414 | 0,996027077 | zinc finger protein 639 |
| ENSG00000272248 | AL138831,2 | -0,022720286 | 0,984374857 | 0,97986299 | 0,99837368 | novel transcript |
| ENSG00000072756 | TRNT1 | -0,022777577 | 0,984335767 | 0,94605383 | 0,996027077 | tRNA nucleotidyl transferase 1 |
| ENSG00000227159 | DDX11L16 | -0,022784163 | 0,984331273 | 0,97706389 | 0,99837368 | DEAD/H-box helicase 11 like 16 (pseudogene) |
| ENSG00000135720 | DYNC1LI2 | -0,022915333 | 0,984241782 | 0,93751538 | 0,994479239 | dynein cytoplasmic 1 light intermediate chain 2 |
| ENSG00000157212 | PAXIP1 | -0,023087646 | 0,984124233 | 0,94165126 | 0,995373376 | PAX interacting protein 1 |
| ENSG00000254226 | LINC01933 | -0,02311246 | 0,984107307 | 0,97522104 | 0,99837368 | long intergenic non-protein coding RNA 1933 |
| ENSG00000228794 | LINC01128 | -0,023113569 | 0,98410655 | 0,91999891 | 0,993787755 | long intergenic non-protein coding RNA 1128 |
| ENSG00000235673 | AL929472,4 | -0,023136942 | 0,984090606 | 0,98261753 | 0,99837368 | ribosomal protein S2 (RPS2) pseudogene |
| ENSG00000185739 | SRL | -0,023141914 | 0,984087216 | 0,96426041 | 0,997011523 | sarcalumenin |
| ENSG00000225890 | HLA-DQA1 | -0,023227373 | 0,984028924 | 0,96858836 | 0,997302342 | major histocompatibility complex, class II, DQ alpha 1 |
| ENSG00000280241 | AC079298,3 | -0,023262592 | 0,984004902 | 0,96264211 | 0,996608711 | novel transcript, antisense to DCHS2 |
| ENSG00000283917 | AC104657,1 | -0,023262592 | 0,984004902 | 0,96264211 | 0,996608711 | novel transcript, antisense to DCHS2 |
| ENSG00000256683 | ZNF350 | -0,023442128 | 0,983882456 | 0,9414633 | 0,99529668 | zinc finger protein 350 |
| ENSG00000233223 | AC016876,1 | -0,023443286 | 0,983881666 | 0,95644892 | 0,996608711 | novel transcript, antisense to MPDU1 and CD68 |
| ENSG00000184990 | SIVA1 | -0,023524173 | 0,983826505 | 0,94294685 | 0,995402494 | SIVA1 apoptosis inducing factor |
| ENSG00000086570 | FAT2 | -0,023536252 | 0,983818267 | 0,94704557 | 0,996027077 | FAT atypical cadherin 2 |
| ENSG00000267659 | LINC01482 | -0,023650254 | 0,983740529 | 0,98298724 | 0,99837368 | long intergenic non-protein coding RNA 1482 |
| ENSG00000272901 | SMDT1 | -0,023696493 | 0,983709 | 0,95348602 | 0,99652169 | single-pass membrane protein with aspartate rich tail 1 |
| ENSG00000213891 | RPL3P6 | -0,023738495 | 0,983680361 | 0,95774929 | 0,996608711 | ribosomal protein L3 pseudogene 6 |
| ENSG00000175216 | CKAP5 | -0,023749356 | 0,983672956 | 0,93047262 | 0,994336072 | cytoskeleton associated protein 5 |
| ENSG00000113597 | TRAPPC13 | -0,023767906 | 0,983660308 | 0,95770001 | 0,996608711 | trafficking protein particle complex 13 |

| | | | | | | |
|---|---|---|---|---|---|---|
| ENSG00000139372 | TDG | -0,023813436 | 0,983629265 | 0,94882969 | 0,996432861 | thymine DNA glycosylase |
| ENSG00000110060 | PUS3 | -0,023928346 | 0,983550923 | 0,94662287 | 0,996027077 | pseudouridine synthase 3 |
| ENSG00000233334 | FAM53B-AS1 | -0,023956608 | 0,983531655 | 0,9628203 | 0,996608711 | FAM53B antisense RNA 1 |
| ENSG00000244625 | MIATNB | -0,024016528 | 0,983490807 | 0,93689067 | 0,994479239 | MIAT neighbor |
| ENSG00000223914 | LINC02471 | -0,024062478 | 0,983459483 | 0,98679766 | 0,99837368 | long intergenic non-protein coding RNA 2471 |
| ENSG00000287584 | AC004837,4 | -0,024079468 | 0,983447901 | 0,96640786 | 0,997263201 | novel transcript |
| ENSG00000101040 | ZMYND8 | -0,024080456 | 0,983447228 | 0,9187081 | 0,993485783 | zinc finger MYND-type containing 8 |
| ENSG00000269758 | AC092070,4 | -0,024229535 | 0,98334561 | 0,97716996 | 0,99837368 | novel transcript |
| ENSG00000081237 | PTPRC | -0,024233297 | 0,983343046 | 0,98456828 | 0,99837368 | protein tyrosine phosphatase receptor type C |
| ENSG00000182541 | LIMK2 | -0,024238738 | 0,983339337 | 0,92778994 | 0,994108531 | LIM domain kinase 2 |
| ENSG00000117593 | DARS2 | -0,024250335 | 0,983331432 | 0,93384561 | 0,994479239 | aspartyl-tRNA synthetase 2, mitochondrial |
| ENSG00000102996 | MMP15 | -0,024296558 | 0,983299928 | 0,95242418 | 0,996432861 | matrix metallopeptidase 15 |
| ENSG00000139826 | ABHD13 | -0,024296575 | 0,983299916 | 0,96080387 | 0,996608711 | abhydrolase domain containing 13 |
| ENSG00000287879 | AC114341,2 | -0,024433279 | 0,983206747 | 0,94486615 | 0,995762666 | novel transcript |
| ENSG00000287678 | AL031778,1 | -0,024488072 | 0,983169406 | 0,95394025 | 0,996608711 | novel transcript, antisense to NFYA |
| ENSG00000013588 | GPRC5A | -0,024498787 | 0,983162104 | 0,97389619 | 0,99837368 | G protein-coupled receptor class C group 5 member A |
| ENSG00000139218 | SCAF11 | -0,024528859 | 0,98314161 | 0,94018794 | 0,994949796 | SR-related CTD associated factor 11 |
| ENSG00000273300 | AC000068,3 | -0,024597276 | 0,983094988 | 0,98713312 | 0,99837368 | novel transcript, antisense to UFD1L |
| ENSG00000205177 | C11orf91 | -0,024632371 | 0,983071074 | 0,98645389 | 0,99837368 | chromosome 11 open reading frame 91 |
| ENSG00000111012 | CYP27B1 | -0,024696218 | 0,983027569 | 0,95963034 | 0,996608711 | cytochrome P450 family 27 subfamily B member 1 |
| ENSG00000227527 | AC096540,1 | -0,024784098 | 0,982967691 | 0,98300815 | 0,99837368 | novel transcript |
| ENSG00000197943 | PLCG2 | -0,024899586 | 0,982889007 | 0,94823651 | 0,99635783 | phospholipase C gamma 2 |
| ENSG00000117533 | VAMP4 | -0,024932593 | 0,98286652 | 0,96077698 | 0,996608711 | vesicle associated membrane protein 4 |
| ENSG00000085415 | SEH1L | -0,025023045 | 0,982804899 | 0,9288645 | 0,994214789 | SEH1 like nucleoporin |
| ENSG00000261599 | HERC2P8 | -0,025063681 | 0,982777218 | 0,93026489 | 0,994336072 | hect domain and RLD 2 pseudogene 8 |
| ENSG00000093144 | ECHDC1 | -0,025087087 | 0,982761273 | 0,94676263 | 0,996027077 | ethylmalonyl-CoA decarboxylase 1 |
| ENSG00000144381 | HSPD1 | -0,025142598 | 0,98272346 | 0,92157921 | 0,993787755 | heat shock protein family D (Hsp60) member 1 |
| ENSG00000168216 | LMBRD1 | -0,02522082 | 0,982670179 | 0,95216799 | 0,996432861 | LMBR1 domain containing 1 |
| ENSG00000231744 | AL669818,1 | -0,025233262 | 0,982661704 | 0,98583344 | 0,99837368 | RAN binding protein 1 (RANBP1) pseudogene |
| ENSG00000214022 | REPIN1 | -0,02529529 | 0,982619456 | 0,93766379 | 0,994479239 | replication initiator 1 |
| ENSG00000119684 | MLH3 | -0,025361188 | 0,982574574 | 0,94255956 | 0,995390305 | mutL homolog 3 |
| ENSG00000176396 | EID2 | -0,02538158 | 0,982560685 | 0,91948179 | 0,993617097 | EP300 interacting inhibitor of differentiation 2 |

| | | | | | | |
|---|---|---|---|---|---|---|
| ENSG00000145833 | DDX46 | -0,025396985 | 0,982550194 | 0,92011122 | 0,993787755 | DEAD-box helicase 46 |
| ENSG00000156136 | DCK | -0,025430301 | 0,982527504 | 0,95598003 | 0,996608711 | deoxycytidine kinase |
| ENSG00000159917 | ZNF235 | -0,025470693 | 0,982499997 | 0,9333091 | 0,994479239 | zinc finger protein 235 |
| ENSG00000268858 | AL118506,1 | -0,025609948 | 0,982405166 | 0,93263647 | 0,994479239 | uncharacterized LOC112268269 |
| ENSG00000183628 | DGCR6 | -0,025710193 | 0,982336906 | 0,95246944 | 0,996432861 | DiGeorge syndrome critical region gene 6 |
| ENSG00000011295 | TTC19 | -0,025742304 | 0,982315042 | 0,92255139 | 0,993797669 | tetratricopeptide repeat domain 19 |
| ENSG00000100147 | CCDC134 | -0,025768335 | 0,982297318 | 0,90597814 | 0,992590661 | coiled-coil domain containing 134 |
| ENSG00000239200 | RPL22P20 | -0,025863589 | 0,982232464 | 0,98339706 | 0,99837368 | ribosomal protein L22 pseudogene 20 |
| ENSG00000167747 | C19orf48 | -0,025867271 | 0,982229957 | 0,92629546 | 0,993915537 | chromosome 19 open reading frame 48 |
| ENSG00000119283 | TRIM67 | -0,025915005 | 0,982197459 | 0,95136929 | 0,996432861 | tripartite motif containing 67 |
| ENSG00000100490 | CDKL1 | -0,0259763 | 0,982155729 | 0,93551592 | 0,994479239 | cyclin dependent kinase like 1 |
| ENSG00000105926 | MPP6 | -0,0260056 | 0,982135783 | 0,95227584 | 0,996432861 | membrane palmitoylated protein 6 |
| ENSG00000248309 | MEF2C-AS1 | -0,02604996 | 0,982105585 | 0,96872801 | 0,997302342 | MEF2C antisense RNA 1 |
| ENSG00000226792 | C13orf42 | -0,026109907 | 0,982064777 | 0,9846448 | 0,99837368 | chromosome 13 open reading frame 42 |
| ENSG00000119938 | PPP1R3C | -0,026269347 | 0,98195625 | 0,95023474 | 0,996432861 | protein phosphatase 1 regulatory subunit 3C |
| ENSG00000114790 | ARHGEF26 | -0,026309371 | 0,981929008 | 0,95485305 | 0,996608711 | Rho guanine nucleotide exchange factor 26 |
| ENSG00000132383 | RPA1 | -0,026338035 | 0,981909499 | 0,91902816 | 0,993585187 | replication protein A1 |
| ENSG00000182782 | HCAR2 | -0,026376437 | 0,981883363 | 0,98285823 | 0,99837368 | hydroxycarboxylic acid receptor 2 |
| ENSG00000122406 | RPL5 | -0,026387265 | 0,981875993 | 0,94699579 | 0,996027077 | ribosomal protein L5 |
| ENSG00000182512 | GLRX5 | -0,026536 | 0,981774772 | 0,93428324 | 0,994479239 | glutaredoxin 5 |
| ENSG00000155970 | MICU3 | -0,026561723 | 0,981757267 | 0,9555241 | 0,996608711 | mitochondrial calcium uptake family member 3 |
| ENSG00000174004 | NRROS | -0,026601581 | 0,981730144 | 0,9653791 | 0,997208095 | negative regulator of reactive oxygen species |
| ENSG00000253550 | AC016065,2 | -0,026602829 | 0,981729295 | 0,98285659 | 0,99837368 | TAF9 RNA polymerase II (TAF9) pseudogene |
| ENSG00000175787 | ZNF169 | -0,026722465 | 0,981647888 | 0,92879421 | 0,994173587 | zinc finger protein 169 |
| ENSG00000271851 | AC087501,4 | -0,026806606 | 0,981590638 | 0,93619363 | 0,994479239 | novel transcript, sense intronic to STX8 |
| ENSG00000196236 | XPNPEP3 | -0,026871021 | 0,981546812 | 0,90730449 | 0,992590661 | X-prolyl aminopeptidase 3 |
| ENSG00000267710 | EDDM13 | -0,026880065 | 0,981540658 | 0,97165228 | 0,998088605 | epididymal protein 13 |
| ENSG00000213304 | AC008481,2 | -0,026889134 | 0,981534488 | 0,97717477 | 0,99837368 | ribosomal protein L18a (RPL18A) pseudogene |
| ENSG00000256537 | SMIM10L1 | -0,026906441 | 0,981522713 | 0,93929214 | 0,994882028 | small integral membrane protein 10 like 1 |
| ENSG00000166197 | NOLC1 | -0,026931362 | 0,981505759 | 0,9123389 | 0,993361299 | nucleolar and coiled-body phosphoprotein 1 |
| ENSG00000267587 | AC090241,1 | -0,02704216 | 0,981430383 | 0,97859196 | 0,99837368 | Ubiquitin carboxyl-terminal hydrolase 4, pseudogene |
| ENSG00000176444 | CLK2 | -0,027065739 | 0,981414343 | 0,91514029 | 0,993361299 | CDC like kinase 2 |

| | | | | | | |
|---|---|---|---|---|---|---|
| ENSG00000014138 | POLA2 | -0,027074321 | 0,981408505 | 0,89320979 | 0,991135494 | DNA polymerase alpha 2, accessory subunit |
| ENSG00000188760 | TMEM198 | -0,027117236 | 0,981379312 | 0,94550205 | 0,995917699 | transmembrane protein 198 |
| ENSG00000118762 | PKD2 | -0,027157245 | 0,981352097 | 0,92361034 | 0,99390845 | polycystin 2, transient receptor potential cation channel |
| ENSG00000177688 | SUMO4 | -0,027240056 | 0,981295768 | 0,94622704 | 0,996027077 | small ubiquitin like modifier 4 |
| ENSG00000237130 | MRPS6P2 | -0,027320325 | 0,981241172 | 0,98512789 | 0,99837368 | mitochondrial ribosomal protein S6 pseudogene 2 |
| ENSG00000167770 | OTUB1 | -0,027470662 | 0,981138927 | 0,90075335 | 0,991865079 | OTU deubiquitinase, ubiquitin aldehyde binding 1 |
| ENSG00000137494 | ANKRD42 | -0,02751005 | 0,981112141 | 0,91712542 | 0,993361299 | ankyrin repeat domain 42 |
| ENSG00000099251 | HSD17B7P2 | -0,027630054 | 0,981030534 | 0,9408198 | 0,995154793 | hydroxysteroid 17-beta dehydrogenase 7 pseudogene 2 |
| ENSG00000288374 | HSD17B7P2 | -0,027630054 | 0,981030534 | 0,9408198 | 0,995154793 | hydroxysteroid 17-beta dehydrogenase 7 pseudogene 2 |
| ENSG00000288538 | AP002985,1 | -0,027638418 | 0,981024847 | 0,94129357 | 0,995251853 | novel transcript, antisense to KCTD21 |
| ENSG00000234078 | DDR1 | -0,02765842 | 0,981011246 | 0,915479 | 0,993361299 | discoidin domain receptor tyrosine kinase 1 |
| ENSG00000215522 | DDR1 | -0,02765842 | 0,981011246 | 0,915479 | 0,993361299 | discoidin domain receptor tyrosine kinase 1 |
| ENSG00000106819 | ASPN | -0,027763635 | 0,980939704 | 0,96678379 | 0,997263201 | asporin |
| ENSG00000177283 | FZD8 | -0,027768488 | 0,980936404 | 0,94927109 | 0,996432861 | frizzled class receptor 8 |
| ENSG00000150768 | DLAT | -0,027882493 | 0,980858892 | 0,94237774 | 0,995373376 | dihydrolipoamide S-acetyltransferase |
| ENSG00000095303 | PTGS1 | -0,027915163 | 0,98083668 | 0,96209229 | 0,996608711 | prostaglandin-endoperoxide synthase 1 |
| ENSG00000262777 | AC032044,1 | -0,027962334 | 0,980804611 | 0,97422322 | 0,99837368 | novel transcript, antisense to CRK |
| ENSG00000132286 | TIMM10B | -0,027981233 | 0,980791762 | 0,93430152 | 0,994479239 | translocase of inner mitochondrial membrane 10B |
| ENSG00000127329 | PTPRB | -0,028011859 | 0,980770942 | 0,91850747 | 0,993468867 | protein tyrosine phosphatase receptor type B |
| ENSG00000136485 | DCAF7 | -0,028082885 | 0,980722658 | 0,91721629 | 0,993361299 | DDB1 and CUL4 associated factor 7 |
| ENSG00000139977 | NAA30 | -0,028088982 | 0,980718514 | 0,93749164 | 0,994479239 | N-alpha-acetyltransferase 30, NatC catalytic subunit |
| ENSG00000023572 | GLRX2 | -0,028180436 | 0,980656347 | 0,95401779 | 0,996608711 | glutaredoxin 2 |
| ENSG00000160199 | PKNOX1 | -0,028221926 | 0,980628145 | 0,9038612 | 0,992462326 | PBX/knotted 1 homeobox 1 |
| ENSG00000101210 | EEF1A2 | -0,028280903 | 0,980588058 | 0,94478282 | 0,995741983 | eukaryotic translation elongation factor 1 alpha 2 |
| ENSG00000167528 | ZNF641 | -0,028341734 | 0,980546712 | 0,92687951 | 0,993915537 | zinc finger protein 641 |
| ENSG00000186806 | VSIG10L | -0,028407919 | 0,98050173 | 0,9431895 | 0,995402494 | V-set and immunoglobulin domain containing 10 like |
| ENSG00000269289 | AC011503,1 | -0,028436707 | 0,980482165 | 0,95810616 | 0,996608711 | novel transcript, antisense to ZNF726 |
| ENSG00000079616 | KIF22 | -0,028477947 | 0,980454138 | 0,89325137 | 0,991135494 | kinesin family member 22 |
| ENSG00000231249 | ITPR1-DT | -0,028479034 | 0,980453399 | 0,97682052 | 0,99837368 | ITPR1 divergent transcript |

| | | | | | | |
|---|---|---|---|---|---|---|
| ENSG00000124787 | RPP40 | -0,028511617 | 0,980431256 | 0,94513163 | 0,99579981 | ribonuclease P/MRP subunit p40 |
| ENSG00000247157 | LINC01252 | -0,028512408 | 0,980430718 | 0,95778152 | 0,996608711 | long intergenic non-protein coding RNA 1252 |
| ENSG00000288060 | AL355338,1 | -0,028555859 | 0,980401191 | 0,9461627 | 0,996027077 | novel transcript |
| ENSG00000223569 | USP17L15 | -0,028667743 | 0,980325161 | 0,94912779 | 0,996432861 | ubiquitin specific peptidase 17 like family member 15 |
| ENSG00000253797 | UTP14C | -0,028709434 | 0,980296832 | 0,94145443 | 0,99529668 | UTP14C small subunit processome component |
| ENSG00000162522 | KIAA1522 | -0,028753654 | 0,980266786 | 0,91658659 | 0,993361299 | KIAA1522 |
| ENSG00000287769 | AL353748,3 | -0,028838867 | 0,980208887 | 0,92427692 | 0,993915537 | novel transcript, antisense to SPIN1 |
| ENSG00000138002 | IFT172 | -0,028909608 | 0,980160825 | 0,92513551 | 0,993915537 | intraflagellar transport 172 |
| ENSG00000167604 | NFKBID | -0,028933327 | 0,980144711 | 0,93571844 | 0,994479239 | NFKB inhibitor delta |
| ENSG00000121542 | SEC22A | -0,028978682 | 0,980113898 | 0,92692216 | 0,993915537 | SEC22 homolog A, vesicle trafficking protein |
| ENSG00000140350 | ANP32A | -0,028990528 | 0,98010585 | 0,90301799 | 0,992311172 | acidic nuclear phosphoprotein 32 family member A |
| ENSG00000154451 | GBP5 | -0,029019524 | 0,980086152 | 0,95706489 | 0,996608711 | guanylate binding protein 5 |
| ENSG00000104522 | GFUS | -0,029099775 | 0,980031635 | 0,92025278 | 0,993787755 | GDP-L-fucose synthase |
| ENSG00000163319 | MRPS18C | -0,029111047 | 0,980023978 | 0,93929953 | 0,994882028 | mitochondrial ribosomal protein S18C |
| ENSG00000188092 | GPR89B | -0,029128045 | 0,980012431 | 0,94311706 | 0,995402494 | G protein-coupled receptor 89B |
| ENSG00000171612 | SLC25A33 | -0,029128192 | 0,980012332 | 0,91116007 | 0,993222655 | solute carrier family 25 member 33 |
| ENSG00000243147 | MRPL33 | -0,029132932 | 0,980009112 | 0,93142538 | 0,994433362 | mitochondrial ribosomal protein L33 |
| ENSG00000230995 | PPP1R10 | -0,029221215 | 0,979949144 | 0,93501878 | 0,994479239 | protein phosphatase 1 regulatory subunit 10 |
| ENSG00000227804 | PPP1R10 | -0,029221215 | 0,979949144 | 0,93501878 | 0,994479239 | protein phosphatase 1 regulatory subunit 10 |
| ENSG00000238104 | PPP1R10 | -0,029221215 | 0,979949144 | 0,93501878 | 0,994479239 | protein phosphatase 1 regulatory subunit 10 |
| ENSG00000281022 | MED22 | -0,02923533 | 0,979939557 | 0,92319744 | 0,993825008 | mediator complex subunit 22 |
| ENSG00000145287 | PLAC8 | -0,02926749 | 0,979917712 | 0,96672987 | 0,997263201 | placenta associated 8 |
| ENSG00000168395 | ING5 | -0,029303285 | 0,979893399 | 0,92372974 | 0,993915537 | inhibitor of growth family member 5 |
| ENSG00000273391 | AC083880,1 | -0,02936355 | 0,979852468 | 0,95797981 | 0,996608711 | novel transcript, antisense to LUC7L2 |
| ENSG00000083642 | PDS5B | -0,029399953 | 0,979827744 | 0,9447614 | 0,995741983 | PDS5 cohesin associated factor B |
| ENSG00000232838 | PET117 | -0,029403432 | 0,979825381 | 0,91091405 | 0,993222655 | PET117 cytochrome c oxidase chaperone |
| ENSG00000151006 | PRSS53 | -0,029420317 | 0,979813913 | 0,94061193 | 0,995103256 | serine protease 53 |
| ENSG00000122741 | DCAF10 | -0,029444354 | 0,979797588 | 0,91913378 | 0,993598006 | DDB1 and CUL4 associated factor 10 |
| ENSG00000101843 | PSMD10 | -0,029458226 | 0,979788168 | 0,94359453 | 0,995406323 | proteasome 26S subunit, non-ATPase 10 |
| ENSG00000189298 | ZKSCAN3 | -0,029473783 | 0,979777602 | 0,90186756 | 0,992152698 | zinc finger with KRAB and SCAN domains 3 |
| ENSG00000179841 | AKAP5 | -0,02947472 | 0,979776966 | 0,9059739 | 0,992590661 | A-kinase anchoring protein 5 |
| ENSG00000089177 | KIF16B | -0,029480846 | 0,979772806 | 0,90621378 | 0,992590661 | kinesin family member 16B |

| | | | | | | |
|---|---|---|---|---|---|---|
| ENSG00000101442 | ACTR5 | -0,0294947 | 0,979763397 | 0,90379364 | 0,992462326 | actin related protein 5 |
| ENSG00000234520 | HRAT17 | -0,029616987 | 0,979680353 | 0,96037587 | 0,996608711 | heart tissue-associated transcript 17 |
| ENSG00000264743 | DPRXP4 | -0,029629459 | 0,979671884 | 0,97722516 | 0,99837368 | divergent-paired related homeobox pseudogene 4 |
| ENSG00000261061 | AC092718,4 | -0,029645912 | 0,979660711 | 0,90931201 | 0,992864002 | novel transcript |
| ENSG00000158158 | CNNM4 | -0,029675373 | 0,979640706 | 0,90246918 | 0,992230188 | cyclin and CBS domain divalent metal cation transport mediator 4 |
| ENSG00000177565 | TBL1XR1 | -0,029714842 | 0,979613905 | 0,95162817 | 0,996432861 | TBL1X receptor 1 |
| ENSG00000276649 | AL117335,1 | -0,029733852 | 0,979600997 | 0,96660824 | 0,997263201 | novel transcript, antisense to SIRPA |
| ENSG00000134056 | MRPS36 | -0,029793534 | 0,979560474 | 0,95484773 | 0,996608711 | mitochondrial ribosomal protein S36 |
| ENSG00000164080 | RAD54L2 | -0,029856648 | 0,979517621 | 0,9013416 | 0,992041918 | RAD54 like 2 |
| ENSG00000260357 | DNM1P34 | -0,029876702 | 0,979504006 | 0,97224693 | 0,998214946 | dynamin 1 pseudogene 34 |
| ENSG00000083896 | YTHDC1 | -0,029910405 | 0,979481124 | 0,90834391 | 0,992595616 | YTH domain containing 1 |
| ENSG00000237505 | PKN2-AS1 | -0,029978636 | 0,979434801 | 0,94826046 | 0,99635783 | PKN2 antisense RNA 1 |
| ENSG00000225377 | NRSN2-AS1 | -0,030043748 | 0,979390598 | 0,9363648 | 0,994479239 | NRSN2 antisense RNA 1 |
| ENSG00000286204 | AC025475,2 | -0,030088478 | 0,979360234 | 0,95622359 | 0,996608711 | novel transcript |
| ENSG00000225963 | AC009950,1 | -0,030096141 | 0,979355031 | 0,91454061 | 0,993361299 | novel transcript, antisense to SP110 |
| ENSG00000261243 | AC022165,1 | -0,030238218 | 0,979258589 | 0,96894379 | 0,997302342 | novel transcript, antisense to KIAA1609 |
| ENSG00000168060 | NAALADL1 | -0,030331253 | 0,979195442 | 0,96543255 | 0,997208095 | N-acetylated alpha-linked acidic dipeptidase like 1 |
| ENSG00000057468 | MSH4 | -0,030412912 | 0,979140019 | 0,98173924 | 0,99837368 | mutS homolog 4 |
| ENSG00000255198 | SNHG9 | -0,030425557 | 0,979131437 | 0,92598359 | 0,993915537 | small nucleolar RNA host gene 9 |
| ENSG00000138382 | METTL5 | -0,030498335 | 0,979082045 | 0,94676324 | 0,996027077 | methyltransferase like 5 |
| ENSG00000259398 | AC090970,1 | -0,030504961 | 0,979077548 | 0,97832589 | 0,99837368 | novel transcript, intronic to MAPK6 |
| ENSG00000277283 | AC004812,2 | -0,030515224 | 0,979070583 | 0,93226272 | 0,994479239 | novel transcript, antisense to RAB35 |
| ENSG00000140323 | DISP2 | -0,030645469 | 0,978982198 | 0,9328194 | 0,994479239 | dispatched RND transporter family member 2 |
| ENSG00000165259 | HDX | -0,030658781 | 0,978973165 | 0,94071076 | 0,995140459 | highly divergent homeobox |
| ENSG00000116350 | SRSF4 | -0,030677169 | 0,978960687 | 0,89042873 | 0,990717729 | serine and arginine rich splicing factor 4 |
| ENSG00000284946 | AC068831,6 | -0,03069146 | 0,97895099 | 0,92334283 | 0,993825008 | novel protein |
| ENSG00000060709 | RIMBP2 | -0,030697627 | 0,978946805 | 0,91635092 | 0,993361299 | RIMS binding protein 2 |
| ENSG00000139613 | SMARCC2 | -0,030772346 | 0,978896106 | 0,88638792 | 0,990717729 | SWI/SNF related, matrix associated, actin dependent regulator of chromatin subfamily c member 2 |
| ENSG00000068745 | IP6K2 | -0,030908998 | 0,978803389 | 0,90517715 | 0,992590661 | inositol hexakisphosphate kinase 2 |
| ENSG00000082515 | MRPL22 | -0,030950894 | 0,978774965 | 0,93734142 | 0,994479239 | mitochondrial ribosomal protein L22 |

| | | | | | | |
|---|---|---|---|---|---|---|
| ENSG00000154237 | LRRK1 | -0,030985931 | 0,978751195 | 0,95386044 | 0,996608711 | leucine rich repeat kinase 1 |
| ENSG00000101198 | NKAIN4 | -0,030986239 | 0,978750986 | 0,91156653 | 0,99331553 | sodium/potassium transporting ATPase interacting 4 |
| ENSG00000144635 | DYNC1LI1 | -0,031072139 | 0,978692711 | 0,91564636 | 0,993361299 | dynein cytoplasmic 1 light intermediate chain 1 |
| ENSG00000232150 | ST13P4 | -0,031136986 | 0,978648722 | 0,9720995 | 0,998214946 | ST13, Hsp70 interacting protein pseudogene 4 |
| ENSG00000147996 | CBWD5 | -0,031158671 | 0,978634012 | 0,89524251 | 0,991303328 | COBW domain containing 5 |
| ENSG00000185728 | YTHDF3 | -0,031165102 | 0,97862965 | 0,94221412 | 0,995373376 | YTH N6-methyladenosine RNA binding protein 3 |
| ENSG00000177463 | NR2C2 | -0,031341149 | 0,978510238 | 0,91553239 | 0,993361299 | nuclear receptor subfamily 2 group C member 2 |
| ENSG00000235291 | PPP1R10 | -0,031345634 | 0,978507196 | 0,93045277 | 0,994336072 | protein phosphatase 1 regulatory subunit 10 |
| ENSG00000231737 | PPP1R10 | -0,031345634 | 0,978507196 | 0,93045277 | 0,994336072 | protein phosphatase 1 regulatory subunit 10 |
| ENSG00000224404 | Z68323,1 | -0,031434806 | 0,978446717 | 0,98034005 | 0,99837368 | novel transcript |
| ENSG00000270733 | AL355877,1 | -0,031451121 | 0,978435652 | 0,97735798 | 0,99837368 | mitochondrial ribosomal protein S6 (MRPS6) pseudogene |
| ENSG00000267309 | AC092295,2 | -0,031508363 | 0,978396832 | 0,95039029 | 0,996432861 | novel transcript, antisense to ZNF566 |
| ENSG00000241634 | AC069499,1 | -0,031525045 | 0,978385518 | 0,93685267 | 0,994479239 | ribosomal protein L13 (RPL13) pseudogene |
| ENSG00000286290 | AC011593,1 | -0,031597412 | 0,978336443 | 0,97108542 | 0,998088605 | novel transcript |
| ENSG00000161547 | SRSF2 | -0,031625774 | 0,97831721 | 0,90390593 | 0,992474358 | serine and arginine rich splicing factor 2 |
| ENSG00000124486 | USP9X | -0,031634275 | 0,978311445 | 0,92492664 | 0,993915537 | ubiquitin specific peptidase 9 X-linked |
| ENSG00000239974 | EGFL8 | -0,031637507 | 0,978309254 | 0,9855941 | 0,99837368 | EGF like domain multiple 8 |
| ENSG00000288012 | AP003307,1 | -0,031660756 | 0,978293488 | 0,97473404 | 0,99837368 | novel transcript, antisense to CUL5 |
| ENSG00000182979 | MTA1 | -0,031687306 | 0,978275485 | 0,88916875 | 0,990717729 | metastasis associated 1 |
| ENSG00000251536 | AC055717,1 | -0,031725068 | 0,978249879 | 0,96558788 | 0,997208095 | novel transcript |
| ENSG00000170791 | CHCHD7 | -0,031759556 | 0,978226494 | 0,92184602 | 0,993797669 | coiled-coil-helix-coiled-coil-helix domain containing 7 |
| ENSG00000181555 | SETD2 | -0,0318291 | 0,978179341 | 0,91715456 | 0,993361299 | SET domain containing 2, histone lysine methyltransferase |
| ENSG00000112245 | PTP4A1 | -0,031846441 | 0,978167583 | 0,94323678 | 0,995402494 | protein tyrosine phosphatase 4A1 |
| ENSG00000106144 | CASP2 | -0,031859256 | 0,978158895 | 0,88197372 | 0,990717729 | caspase 2 |
| ENSG00000213735 | ANAPC10P1 | -0,031867327 | 0,978153422 | 0,97511445 | 0,99837368 | anaphase promoting complex subunit 10 pseudogene 1 |
| ENSG00000197070 | ARRDC1 | -0,031883713 | 0,978142312 | 0,92496528 | 0,993915537 | arrestin domain containing 1 |
| ENSG00000287044 | AL139106,1 | -0,031905408 | 0,978127603 | 0,95213844 | 0,996432861 | novel transcript, antisense to LACE1 |
| ENSG00000164175 | SLC45A2 | -0,0319972 | 0,978065372 | 0,971868 | 0,998110538 | solute carrier family 45 member 2 |
| ENSG00000147123 | NDUFB11 | -0,032045173 | 0,978032849 | 0,88748822 | 0,990717729 | NADH:ubiquinone oxidoreductase subunit B11 |

| | | | | | | |
|---|---|---|---|---|---|---|
| ENSG00000112695 | COX7A2 | -0,032126855 | 0,977977477 | 0,93509653 | 0,994479239 | cytochrome c oxidase subunit 7A2 |
| ENSG00000164284 | GRPEL2 | -0,032144548 | 0,977965483 | 0,93002806 | 0,994331958 | GrpE like 2, mitochondrial |
| ENSG00000256898 | ARPC3P4 | -0,032225457 | 0,977910638 | 0,97797377 | 0,99837368 | actin related protein 2/3 complex subunit 3 pseudogene 4 |
| ENSG00000173141 | MRPL57 | -0,032301991 | 0,977858763 | 0,89041365 | 0,990717729 | mitochondrial ribosomal protein L57 |
| ENSG00000237767 | LINC01370 | -0,032366834 | 0,977814813 | 0,97311256 | 0,99837368 | long intergenic non-protein coding RNA 1370 |
| ENSG00000215386 | MIR99AHG | -0,032385606 | 0,97780209 | 0,93520877 | 0,994479239 | mir-99a-let-7c cluster host gene |
| ENSG00000255476 | AC011092,2 | -0,03238983 | 0,977799227 | 0,97502029 | 0,99837368 | novel transcript |
| ENSG00000143493 | INTS7 | -0,032626699 | 0,9776387 | 0,92397507 | 0,993915537 | integrator complex subunit 7 |
| ENSG00000214174 | AMZ2P1 | -0,032647836 | 0,977624377 | 0,92303142 | 0,993825008 | archaelysin family metallopeptidase 2 pseudogene 1 |
| ENSG00000138081 | FBXO11 | -0,03265076 | 0,977622395 | 0,93381603 | 0,994479239 | F-box protein 11 |
| ENSG00000133027 | PEMT | -0,032662412 | 0,9776145 | 0,8877275 | 0,990717729 | phosphatidylethanolamine N-methyltransferase |
| ENSG00000165283 | STOML2 | -0,032664382 | 0,977613164 | 0,89454481 | 0,99116563 | stomatin like 2 |
| ENSG00000272402 | AL031775,2 | -0,032709501 | 0,977582591 | 0,96622606 | 0,997263201 | novel transcript, antisense to C6orf62 |
| ENSG00000278766 | AC092683,2 | -0,032783415 | 0,977532508 | 0,95416503 | 0,996608711 | novel transcript |
| ENSG00000273328 | AC099329,2 | -0,032916076 | 0,977442624 | 0,95940678 | 0,996608711 | novel transcript |
| ENSG00000085274 | MYNN | -0,03294618 | 0,977422229 | 0,9382246 | 0,994547792 | myoneurin |
| ENSG00000167619 | TMEM145 | -0,032947203 | 0,977421536 | 0,91118746 | 0,993222655 | transmembrane protein 145 |
| ENSG00000111786 | SRSF9 | -0,032959475 | 0,977413222 | 0,88296305 | 0,990717729 | serine and arginine rich splicing factor 9 |
| ENSG00000230624 | DDX39B | -0,033040329 | 0,977358445 | 0,87981405 | 0,990261566 | DExD-box helicase 39B |
| ENSG00000143450 | OAZ3 | -0,033139548 | 0,977291231 | 0,94326212 | 0,995402494 | ornithine decarboxylase antizyme 3 |
| ENSG00000155666 | KDM8 | -0,033226223 | 0,977232518 | 0,91809981 | 0,993361299 | lysine demethylase 8 |
| ENSG00000177542 | SLC25A22 | -0,033273836 | 0,977200268 | 0,91816596 | 0,993361299 | solute carrier family 25 member 22 |
| ENSG00000196470 | SIAH1 | -0,033274286 | 0,977199963 | 0,92707219 | 0,993915537 | siah E3 ubiquitin protein ligase 1 |
| ENSG00000166183 | ASPG | -0,033298008 | 0,977183895 | 0,98181732 | 0,99837368 | asparaginase |
| ENSG00000114859 | CLCN2 | -0,03334307 | 0,977153374 | 0,92944871 | 0,994293848 | chloride voltage-gated channel 2 |
| ENSG00000127589 | TUBBP1 | -0,033369218 | 0,977135664 | 0,94369209 | 0,995464929 | tubulin beta pseudogene 1 |
| ENSG00000108825 | PTGES3L-AARSI | -0,03338611 | 0,977124223 | 0,95250311 | 0,996432861 | PTGES3L-AARSD1 readthrough |
| ENSG00000233766 | CAVIN2-AS1 | -0,033404907 | 0,977111492 | 0,96401789 | 0,996980762 | CAVIN2 and TMEFF2 antisense RNA 1 |
| ENSG00000142864 | SERBP1 | -0,033427587 | 0,977096131 | 0,88872267 | 0,990717729 | SERPINE1 mRNA binding protein 1 |
| ENSG00000023608 | SNAPC1 | -0,033547749 | 0,977014752 | 0,92418887 | 0,993915537 | small nuclear RNA activating complex polypeptide 1 |
| ENSG00000178386 | ZNF223 | -0,033569368 | 0,977000111 | 0,93159941 | 0,994433362 | zinc finger protein 223 |
| ENSG00000140153 | WDR20 | -0,033594602 | 0,976983023 | 0,90971966 | 0,992951496 | WD repeat domain 20 |

| | | | | | | |
|---|---|---|---|---|---|---|
| ENSG00000260751 | AC008870,2 | -0,033603433 | 0,976977043 | 0,94001582 | 0,994949796 | novel transcript |
| ENSG00000145331 | TRMT10A | -0,033614144 | 0,97696979 | 0,94543425 | 0,995917699 | tRNA methyltransferase 10A |
| ENSG00000285485 | NFATC4 | -0,033661978 | 0,976937397 | 0,92595894 | 0,993915537 | nuclear factor of activated T cells 4 |
| ENSG00000100968 | NFATC4 | -0,033661978 | 0,976937397 | 0,92595894 | 0,993915537 | nuclear factor of activated T cells 4 |
| ENSG00000204584 | AC027801,1 | -0,03376185 | 0,976869771 | 0,96752989 | 0,997263201 | novel transcript, antisense to TAC4 |
| ENSG00000150459 | SAP18 | -0,033769065 | 0,976864885 | 0,91654316 | 0,993361299 | Sin3A associated protein 18 |
| ENSG00000162702 | ZNF281 | -0,033821436 | 0,976829425 | 0,93733294 | 0,994479239 | zinc finger protein 281 |
| ENSG00000278613 | NAIP | -0,033883124 | 0,976787658 | 0,94035243 | 0,995028467 | NLR family apoptosis inhibitory protein |
| ENSG00000146918 | NCAPG2 | -0,03394518 | 0,976745643 | 0,90790769 | 0,992590661 | non-SMC condensin II complex subunit G2 |
| ENSG00000088899 | LZTS3 | -0,034064532 | 0,976664842 | 0,94600307 | 0,996027077 | leucine zipper tumor suppressor family member 3 |
| ENSG00000235437 | LINC01278 | -0,034066559 | 0,976663469 | 0,88263194 | 0,990717729 | long intergenic non-protein coding RNA 1278 |
| ENSG00000141504 | SAT2 | -0,034125286 | 0,976623713 | 0,88332063 | 0,990717729 | spermidine/spermine N1-acetyltransferase family member 2 |
| ENSG00000232233 | LINC02043 | -0,034156876 | 0,976602329 | 0,93689447 | 0,994479239 | long intergenic non-protein coding RNA 2043 |
| ENSG00000127125 | PPCS | -0,03421725 | 0,976561461 | 0,92970358 | 0,994293848 | phosphopantothenoylcysteine synthetase |
| ENSG00000271730 | AL390208,1 | -0,034266738 | 0,976527963 | 0,97854519 | 0,99837368 | novel transcript, antisense to SESN1 |
| ENSG00000130856 | ZNF236 | -0,034291829 | 0,97651098 | 0,89673947 | 0,991644439 | zinc finger protein 236 |
| ENSG00000266472 | MRPS21 | -0,034418394 | 0,976425316 | 0,91500229 | 0,993361299 | mitochondrial ribosomal protein S21 |
| ENSG00000223367 | RPS18 | -0,034457291 | 0,976398991 | 0,93483234 | 0,994479239 | ribosomal protein S18 |
| ENSG00000226225 | RPS18 | -0,034457291 | 0,976398991 | 0,93483234 | 0,994479239 | ribosomal protein S18 |
| ENSG00000175782 | SLC35E3 | -0,03448452 | 0,976380563 | 0,92593435 | 0,993915537 | solute carrier family 35 member E3 |
| ENSG00000069275 | NUCKS1 | -0,034500155 | 0,976369981 | 0,92154748 | 0,993787755 | nuclear casein kinase and cyclin dependent kinase substrate 1 |
| ENSG00000137807 | KIF23 | -0,034518998 | 0,976357229 | 0,92164486 | 0,993787755 | kinesin family member 23 |
| ENSG00000259531 | AC022306,1 | -0,034532588 | 0,976348032 | 0,95703935 | 0,996608711 | signal peptide peptidase-like 2A (SPPL2A) pseudogene |
| ENSG00000262074 | SNORD3B-2 | -0,034569522 | 0,976323037 | 0,89264645 | 0,991048031 | small nucleolar RNA, C/D box 3B-2 |
| ENSG00000239665 | AL157392,3 | -0,034596471 | 0,9763048 | 0,86934792 | 0,989074806 | novel transcript |
| ENSG00000276517 | AL133243,3 | -0,034625085 | 0,976285436 | 0,91499951 | 0,993361299 | novel transcript, sense intronic to BIRC6 |
| ENSG00000100426 | ZBED4 | -0,03470315 | 0,97623261 | 0,87176865 | 0,989217334 | zinc finger BED-type containing 4 |
| ENSG00000103550 | KNOP1 | -0,034705365 | 0,976231111 | 0,86126987 | 0,989074806 | lysine rich nucleolar protein 1 |
| ENSG00000169139 | UBE2V2 | -0,034721033 | 0,97622051 | 0,94184033 | 0,995373376 | ubiquitin conjugating enzyme E2 V2 |
| ENSG00000122644 | ARL4A | -0,034731094 | 0,976213702 | 0,93642792 | 0,994479239 | ADP ribosylation factor like GTPase 4A |

| | | | | | | |
|---|---|---|---|---|---|---|
| ENSG00000065328 | MCM10 | -0,034779192 | 0,976181157 | 0,88264881 | 0,990717729 | minichromosome maintenance 10 replication initiation factor |
| ENSG00000260661 | AC116903,2 | -0,034845409 | 0,976136352 | 0,97689977 | 0,99837368 | novel transcript, antisense to SLCO3A1 |
| ENSG00000069431 | ABCC9 | -0,034874346 | 0,976116774 | 0,916986 | 0,993361299 | ATP binding cassette subfamily C member 9 |
| ENSG00000139697 | SBNO1 | -0,03489561 | 0,976102387 | 0,915422 | 0,993361299 | strawberry notch homolog 1 |
| ENSG00000126226 | PCID2 | -0,034921505 | 0,976084867 | 0,90149637 | 0,992041918 | PCI domain containing 2 |
| ENSG00000162601 | MYSM1 | -0,034925296 | 0,976082302 | 0,92812403 | 0,994108531 | Myb like, SWIRM and MPN domains 1 |
| ENSG00000279873 | LINC01126 | -0,034934452 | 0,976076107 | 0,93334297 | 0,994479239 | long intergenic non-protein coding RNA 1126 |
| ENSG00000203880 | PCMTD2 | -0,035078948 | 0,975978351 | 0,95053932 | 0,996432861 | protein-L-isoaspartate (D-aspartate) O-methyltransferase domain containing 2 |
| ENSG00000148773 | MKI67 | -0,035104039 | 0,975961377 | 0,88748008 | 0,990717729 | marker of proliferation Ki-67 |
| ENSG00000125430 | HS3ST3B1 | -0,035132002 | 0,975942461 | 0,92668542 | 0,993915537 | heparan sulfate-glucosamine 3-sulfotransferase 3B1 |
| ENSG00000261893 | CLK2 | -0,035318965 | 0,975815994 | 0,9031584 | 0,992311172 | CDC like kinase 2 |
| ENSG00000144481 | TRPM8 | -0,035421763 | 0,975746466 | 0,94938053 | 0,996432861 | transient receptor potential cation channel subfamily M member 8 |
| ENSG00000168000 | BSCL2 | -0,035449601 | 0,975727638 | 0,92699317 | 0,993915537 | BSCL2 lipid droplet biogenesis associated, seipin |
| ENSG00000248926 | AC114801,2 | -0,035550053 | 0,975659703 | 0,97378614 | 0,99837368 | FUS interacting protein (serine/arginine-rich) 1 (FUSIP1) pseudogene |
| ENSG00000173575 | CHD2 | -0,035552194 | 0,975658254 | 0,87364769 | 0,989633545 | chromodomain helicase DNA binding protein 2 |
| ENSG00000120159 | CAAP1 | -0,035594852 | 0,975629407 | 0,92163087 | 0,993787755 | caspase activity and apoptosis inhibitor 1 |
| ENSG00000197619 | ZNF615 | -0,035624464 | 0,975609382 | 0,91873805 | 0,993485783 | zinc finger protein 615 |
| ENSG00000196684 | HSH2D | -0,035643399 | 0,975596577 | 0,9787911 | 0,99837368 | hematopoietic SH2 domain containing |
| ENSG00000155545 | MIER3 | -0,035714556 | 0,97554846 | 0,92795705 | 0,994108531 | MIER family member 3 |
| ENSG00000259396 | AC087721,1 | -0,035769985 | 0,975510979 | 0,97306833 | 0,99837368 | novel transcript, sense intronic to NUSAP1 |
| ENSG00000119900 | OGFRL1 | -0,035851656 | 0,975455757 | 0,87736824 | 0,990109814 | opioid growth factor receptor like 1 |
| ENSG00000087258 | GNAO1 | -0,035869855 | 0,975443452 | 0,92557649 | 0,993915537 | G protein subunit alpha o1 |
| ENSG00000171806 | METTL18 | -0,035891173 | 0,975429038 | 0,93172609 | 0,994433362 | methyltransferase like 18 |
| ENSG00000156802 | ATAD2 | -0,035907468 | 0,975418021 | 0,91213876 | 0,993361299 | ATPase family AAA domain containing 2 |
| ENSG00000100568 | VTI1B | -0,035936786 | 0,975398199 | 0,90755405 | 0,992590661 | vesicle transport through interaction with t-SNAREs 1B |
| ENSG00000146776 | ATXN7L1 | -0,035962245 | 0,975380987 | 0,89884187 | 0,991832796 | ataxin 7 like 1 |
| ENSG00000213762 | ZNF134 | -0,03607467 | 0,975304981 | 0,90588102 | 0,992590661 | zinc finger protein 134 |
| ENSG00000266053 | NDUFV2-AS1 | -0,036087104 | 0,975296575 | 0,92869947 | 0,994173587 | NDUFV2 antisense RNA 1 |

| | | | | | | |
|---|---|---|---|---|---|---|
| ENSG00000152942 | RAD17 | -0,0361225 | 0,975272647 | 0,92661987 | 0,993915537 | RAD17 checkpoint clamp loader component |
| ENSG00000259425 | AC011767,1 | -0,036188912 | 0,975227753 | 0,97000104 | 0,997844133 | Golgin subfamily A member 6 pseudogene |
| ENSG00000241772 | AC092620,1 | -0,036238314 | 0,975194359 | 0,96784035 | 0,997263201 | novel transcript |
| ENSG00000080546 | SESN1 | -0,036243895 | 0,975190587 | 0,92638537 | 0,993915537 | sestrin 1 |
| ENSG00000234616 | JRK | -0,036246477 | 0,975188841 | 0,91873844 | 0,993485783 | Jrk helix-turn-helix protein |
| ENSG00000248103 | AC008443,2 | -0,036287665 | 0,975161001 | 0,93468534 | 0,994479239 | novel transcript |
| ENSG00000011454 | RABGAP1 | -0,036320372 | 0,975138893 | 0,90334469 | 0,992311172 | RAB GTPase activating protein 1 |
| ENSG00000137574 | TGS1 | -0,036373284 | 0,97510313 | 0,89520203 | 0,991303328 | trimethylguanosine synthase 1 |
| ENSG00000255097 | AP006259,1 | -0,036429108 | 0,9750654 | 0,96208219 | 0,996608711 | novel transcript |
| ENSG00000163660 | CCNL1 | -0,036533881 | 0,97499459 | 0,93183773 | 0,994433362 | cyclin L1 |
| ENSG00000078668 | VDAC3 | -0,036628996 | 0,974930312 | 0,91400838 | 0,993361299 | voltage dependent anion channel 3 |
| ENSG00000176476 | SGF29 | -0,036664046 | 0,974906627 | 0,87052803 | 0,989074806 | SAGA complex associated factor 29 |
| ENSG00000260179 | AL162741,1 | -0,036718 | 0,974870168 | 0,93833804 | 0,994549499 | novel transcript |
| ENSG00000241983 | RN7SL566P | -0,036719284 | 0,9748693 | 0,97384916 | 0,99837368 | RNA, 7SL, cytoplasmic 566, pseudogene |
| ENSG00000173272 | MZT2A | -0,036864008 | 0,974771511 | 0,92195599 | 0,993797669 | mitotic spindle organizing protein 2A |
| ENSG00000281912 | LINC01144 | -0,036887236 | 0,974755817 | 0,93639045 | 0,994479239 | long intergenic non-protein coding RNA 1144 |
| ENSG00000125676 | THOC2 | -0,036906559 | 0,974742761 | 0,91679232 | 0,993361299 | THO complex 2 |
| ENSG00000122565 | CBX3 | -0,036941854 | 0,974718915 | 0,9204251 | 0,993787755 | chromobox 3 |
| ENSG00000276691 | AC004466,3 | -0,036952868 | 0,974711474 | 0,95879857 | 0,996608711 | novel transcript, antisense to HDAC7 |
| ENSG00000279956 | AC073082,1 | -0,037083395 | 0,974623291 | 0,95831727 | 0,996608711 | novel transcript |
| ENSG00000101447 | FAM83D | -0,037125806 | 0,974594641 | 0,89551887 | 0,991303328 | family with sequence similarity 83 member D |
| ENSG00000140262 | TCF12 | -0,037153892 | 0,974575667 | 0,92969636 | 0,994293848 | transcription factor 12 |
| ENSG00000260008 | AP001527,1 | -0,037204868 | 0,974541232 | 0,9650006 | 0,99715028 | novel transcript |
| ENSG00000082153 | BZW1 | -0,037212676 | 0,974535958 | 0,93158361 | 0,994433362 | basic leucine zipper and W2 domains 1 |
| ENSG00000287884 | AC005594,1 | -0,037257548 | 0,974505648 | 0,95857215 | 0,996608711 | novel transcript, antisense to DPP9 |
| ENSG00000189164 | ZNF527 | -0,037329797 | 0,974456847 | 0,87125706 | 0,989217334 | zinc finger protein 527 |
| ENSG00000137474 | MYO7A | -0,037484426 | 0,974352409 | 0,94437031 | 0,995542144 | myosin VIIA |
| ENSG00000250654 | AC023794,3 | -0,037544922 | 0,974311553 | 0,9685551 | 0,997302342 | novel pseudogene |
| ENSG00000260084 | AC126773,3 | -0,037633647 | 0,974251635 | 0,97397421 | 0,99837368 | novel transcript, antisense to ZFP90 |
| ENSG00000065882 | TBC1D1 | -0,037691832 | 0,974212344 | 0,8856614 | 0,990717729 | TBC1 domain family member 1 |
| ENSG00000245293 | CYP2U1-AS1 | -0,037771864 | 0,974158302 | 0,94340011 | 0,995402494 | CYP2U1 and SGMS2 antisense RNA 1 |
| ENSG00000205978 | NYNRIN | -0,037793253 | 0,974143859 | 0,8686483 | 0,989074806 | NYN domain and retroviral integrase containing |
| ENSG00000177888 | ZBTB41 | -0,037800681 | 0,974138844 | 0,93365259 | 0,994479239 | zinc finger and BTB domain containing 41 |

| | | | | | | |
|---|---|---|---|---|---|---|
| ENSG00000245832 | MIR4300HG | -0,037801078 | 0,974138576 | 0,97420942 | 0,99837368 | MIR4300 host gene |
| ENSG00000172840 | PDP2 | -0,037827195 | 0,974120941 | 0,89104992 | 0,990780829 | pyruvate dehyrogenase phosphatase catalytic subunit 2 |
| ENSG00000068976 | PYGM | -0,037835923 | 0,974115048 | 0,95250166 | 0,996432861 | glycogen phosphorylase, muscle associated |
| ENSG00000168066 | SF1 | -0,037878231 | 0,974086482 | 0,85104829 | 0,987259878 | splicing factor 1 |
| ENSG00000162231 | NXF1 | -0,037928351 | 0,974052642 | 0,85838742 | 0,989067345 | nuclear RNA export factor 1 |
| ENSG00000260495 | AC009148,1 | -0,037930388 | 0,974051266 | 0,93512982 | 0,994479239 | novel transcript, antisense to GAN |
| ENSG00000272370 | AC092354,2 | -0,037943678 | 0,974042294 | 0,96672319 | 0,997263201 | novel transcript |
| ENSG00000104626 | ERI1 | -0,037968316 | 0,974025659 | 0,9338791 | 0,994479239 | exoribonuclease 1 |
| ENSG00000269161 | AC010618,2 | -0,037969603 | 0,974024791 | 0,97501341 | 0,99837368 | novel transcript, antisense PGLS |
| ENSG00000249637 | AC008438,1 | -0,038046734 | 0,973972717 | 0,93281378 | 0,994479239 | novel transcript |
| ENSG00000143952 | VPS54 | -0,038128712 | 0,973917376 | 0,91967979 | 0,993699236 | VPS54 subunit of GARP complex |
| ENSG00000128581 | IFT22 | -0,038156959 | 0,973898307 | 0,85143366 | 0,987259878 | intraflagellar transport 22 |
| ENSG00000144597 | EAF1 | -0,038165415 | 0,973892599 | 0,89730644 | 0,991644439 | ELL associated factor 1 |
| ENSG00000213619 | NDUFS3 | -0,038211423 | 0,973861541 | 0,87963256 | 0,990261566 | NADH:ubiquinone oxidoreductase core subunit S3 |
| ENSG00000121058 | COIL | -0,038235222 | 0,973845477 | 0,91957804 | 0,993639397 | coilin |
| ENSG00000134489 | HRH4 | -0,038366198 | 0,973757069 | 0,97563145 | 0,99837368 | histamine receptor H4 |
| ENSG00000084207 | GSTP1 | -0,038392545 | 0,973739287 | 0,89702349 | 0,991644439 | glutathione S-transferase pi 1 |
| ENSG00000056736 | IL17RB | -0,038410352 | 0,973727268 | 0,91268764 | 0,993361299 | interleukin 17 receptor B |
| ENSG00000154839 | SKA1 | -0,03859311 | 0,973603925 | 0,88527241 | 0,990717729 | spindle and kinetochore associated complex subunit 1 |
| ENSG00000198816 | ZNF358 | -0,038597725 | 0,973600811 | 0,9067454 | 0,992590661 | zinc finger protein 358 |
| ENSG00000234651 | BAG6 | -0,038603036 | 0,973597227 | 0,85951477 | 0,989074806 | BAG cochaperone 6 |
| ENSG00000181982 | CCDC149 | -0,038669105 | 0,973552642 | 0,8613567 | 0,989074806 | coiled-coil domain containing 149 |
| ENSG00000236489 | AC133473,1 | -0,0386842 | 0,973542455 | 0,9765085 | 0,99837368 | ribosomal protein L34 (RPL34) pseudogene |
| ENSG00000219395 | HSPA8P15 | -0,038701524 | 0,973530765 | 0,94314354 | 0,995402494 | heat shock protein family A (Hsp70) member 8 pseudogene 15 |
| ENSG00000163050 | COQ8A | -0,038715259 | 0,973521497 | 0,90641474 | 0,992590661 | coenzyme Q8A |
| ENSG00000169087 | HSPBAP1 | -0,038893405 | 0,973401292 | 0,88768312 | 0,990717729 | HSPB1 associated protein 1 |
| ENSG00000100211 | CBY1 | -0,038944279 | 0,973366968 | 0,85281903 | 0,987647582 | chibby family member 1, beta catenin antagonist |
| ENSG00000176890 | TYMS | -0,038978845 | 0,973343647 | 0,89609195 | 0,991555839 | thymidylate synthetase |
| ENSG00000243554 | AC004967,1 | -0,03901833 | 0,973317008 | 0,97965788 | 0,99837368 | CCZ1 homolog, vacuolar protein trafficking and biogenesis associated (CCZ1) pseudogene |

| | | | | | | |
|---|---|---|---|---|---|---|
| ENSG00000254618 | TMED10P1 | -0,039119669 | 0,973248641 | 0,9473537 | 0,996100967 | transmembrane p24 trafficking protein 10 pseudogene 1 |
| ENSG00000248734 | AC008906,1 | -0,039149125 | 0,973228771 | 0,94490671 | 0,99577185 | novel transcript |
| ENSG00000006652 | IFRD1 | -0,039178298 | 0,973209091 | 0,91680923 | 0,993361299 | interferon related developmental regulator 1 |
| ENSG00000165966 | PDZRN4 | -0,039338821 | 0,973100812 | 0,95723041 | 0,996608711 | PDZ domain containing ring finger 4 |
| ENSG00000132405 | TBC1D14 | -0,039358145 | 0,973087778 | 0,85746467 | 0,988888398 | TBC1 domain family member 14 |
| ENSG00000287620 | AC092053,4 | -0,039414573 | 0,973049718 | 0,94683482 | 0,996027077 | novel transcript |
| ENSG00000135018 | UBQLN1 | -0,039476904 | 0,973007679 | 0,90980422 | 0,992951496 | ubiquilin 1 |
| ENSG00000215440 | NPEPL1 | -0,039508289 | 0,972986512 | 0,9077233 | 0,992590661 | aminopeptidase like 1 |
| ENSG00000229766 | AL021396,1 | -0,039562243 | 0,972950125 | 0,95618152 | 0,996608711 | novel transcript, antisense to TMX4 |
| ENSG00000258885 | AL450267,1 | -0,039571222 | 0,97294407 | 0,97058709 | 0,998030243 | novel transcript, antisense to PAX5 |
| ENSG00000234327 | ZNF232-AS1 | -0,039581396 | 0,972937208 | 0,91223924 | 0,993361299 | ZNF232 antisense RNA 1 |
| ENSG00000107938 | EDRF1 | -0,039660588 | 0,972883804 | 0,90624217 | 0,992590661 | erythroid differentiation regulatory factor 1 |
| ENSG00000271369 | AC087783,1 | -0,039668791 | 0,972878272 | 0,92589303 | 0,993915537 | NADH dehydrogenase (ubiquinone) 1 beta subcomplex, 8, 19kDa, pseudogene |
| ENSG00000206489 | PPP1R10 | -0,039740819 | 0,972829701 | 0,91657197 | 0,993361299 | protein phosphatase 1 regulatory subunit 10 |
| ENSG00000285151 | AC140134,1 | -0,039993654 | 0,972659226 | 0,93490288 | 0,994479239 | novel transcript, antisense to SMN2 |
| ENSG00000285204 | AC139834,1 | -0,039993654 | 0,972659226 | 0,93490288 | 0,994479239 | novel transcript, antisense to SMN1 |
| ENSG00000197808 | ZNF461 | -0,039994298 | 0,972658792 | 0,88960276 | 0,990717729 | zinc finger protein 461 |
| ENSG00000189308 | LIN54 | -0,040026205 | 0,972637281 | 0,91165441 | 0,993342046 | lin-54 DREAM MuvB core complex component |
| ENSG00000198727 | MT-CYB | -0,040030993 | 0,972634052 | 0,90261728 | 0,992230188 | mitochondrially encoded cytochrome b |
| ENSG00000161179 | YDJC | -0,040070062 | 0,972607713 | 0,93288787 | 0,994479239 | YdjC chitooligosaccharide deacetylase homolog |
| ENSG00000197579 | TOPORS | -0,040120808 | 0,972573503 | 0,92326143 | 0,993825008 | TOP1 binding arginine/serine rich protein, E3 ubiquitin ligase |
| ENSG00000273784 | AL137058,2 | -0,040149112 | 0,972554422 | 0,89947378 | 0,991845106 | novel transcript |
| ENSG00000287414 | AC093787,2 | -0,040199915 | 0,972520176 | 0,94543433 | 0,995917699 | novel transcript |
| ENSG00000286563 | AC078983,1 | -0,040232514 | 0,972498201 | 0,94726718 | 0,996063635 | novel transcript |
| ENSG00000170222 | ADPRM | -0,040265956 | 0,972475658 | 0,92818954 | 0,994108531 | ADP-ribose/CDP-alcohol diphosphatase, manganese dependent |
| ENSG00000101019 | UQCC1 | -0,040283464 | 0,972463857 | 0,88808079 | 0,990717729 | ubiquinol-cytochrome c reductase complex assembly factor 1 |
| ENSG00000137337 | MDC1 | -0,040341103 | 0,972425005 | 0,85879803 | 0,989074806 | mediator of DNA damage checkpoint 1 |
| ENSG00000287682 | AC020651,2 | -0,040377829 | 0,972400251 | 0,95485415 | 0,996608711 | novel transcript |

| | | | | | | |
|---|---|---|---|---|---|---|
| ENSG00000134308 | YWHAQ | -0,040420584 | 0,972371434 | 0,91991926 | 0,993758187 | tyrosine 3-monooxygenase/tryptophan 5-monooxygenase activation protein theta |
| ENSG00000251687 | AC104791,1 | -0,040452561 | 0,972349882 | 0,95236791 | 0,996432861 | novel transcript |
| ENSG00000012963 | UBR7 | -0,04045591 | 0,972347625 | 0,89579332 | 0,991435932 | ubiquitin protein ligase E3 component n-recognin 7 |
| ENSG00000159063 | ALG8 | -0,040483791 | 0,972328833 | 0,90147931 | 0,992041918 | ALG8 alpha-1,3-glucosyltransferase |
| ENSG00000112306 | RPS12 | -0,040494286 | 0,97232176 | 0,90823435 | 0,992595473 | ribosomal protein S12 |
| ENSG00000093167 | LRRFIP2 | -0,040505267 | 0,97231436 | 0,88982213 | 0,990717729 | LRR binding FLII interacting protein 2 |
| ENSG00000165030 | NFIL3 | -0,040524751 | 0,972301228 | 0,92230685 | 0,993797669 | nuclear factor, interleukin 3 regulated |
| ENSG00000158773 | USF1 | -0,040535955 | 0,972293678 | 0,86468184 | 0,989074806 | upstream transcription factor 1 |
| ENSG00000164442 | CITED2 | -0,040559027 | 0,972278129 | 0,91347012 | 0,993361299 | Cbp/p300 interacting transactivator with Glu/Asp rich carboxy-terminal domain 2 |
| ENSG00000120029 | ARMH3 | -0,040573174 | 0,972268594 | 0,88729516 | 0,990717729 | armadillo like helical domain containing 3 |
| ENSG00000272221 | AL645933,2 | -0,040727989 | 0,972164266 | 0,93474607 | 0,994479239 | novel transcript |
| ENSG00000164978 | NUDT2 | -0,040765476 | 0,972139006 | 0,89147653 | 0,990820775 | nudix hydrolase 2 |
| ENSG00000154277 | UCHL1 | -0,04078038 | 0,972128963 | 0,86157883 | 0,989074806 | ubiquitin C-terminal hydrolase L1 |
| ENSG00000134014 | ELP3 | -0,040933444 | 0,972025829 | 0,89144728 | 0,990820775 | elongator acetyltransferase complex subunit 3 |
| ENSG00000110700 | RPS13 | -0,040974978 | 0,971997846 | 0,91958682 | 0,993639397 | ribosomal protein S13 |
| ENSG00000269502 | DMRTC1 | -0,04102481 | 0,971964273 | 0,96519167 | 0,997208095 | DMRT like family C1 |
| ENSG00000168569 | TMEM223 | -0,041070577 | 0,97193344 | 0,86937432 | 0,989074806 | transmembrane protein 223 |
| ENSG00000254959 | INMT-MINDY4 | -0,04107379 | 0,971931275 | 0,99121023 | 0,998884788 | INMT-MINDY4 readthrough (NMD candidate) |
| ENSG00000078579 | FGF20 | -0,041158793 | 0,971874011 | 0,97939665 | 0,99837368 | fibroblast growth factor 20 |
| ENSG00000286138 | AC010336,7 | -0,041180328 | 0,971859504 | 0,95190901 | 0,996432861 | novel transcript |
| ENSG00000168101 | NUDT16L1 | -0,041190363 | 0,971852744 | 0,92172476 | 0,993787755 | nudix hydrolase 16 like 1 |
| ENSG00000168795 | ZBTB5 | -0,041213564 | 0,971837115 | 0,9010335 | 0,991933219 | zinc finger and BTB domain containing 5 |
| ENSG00000260648 | AC020658,3 | -0,041247789 | 0,97181406 | 0,92662017 | 0,993915537 | novel transcript, sense intronic to PAK6 |
| ENSG00000163536 | SERPINI1 | -0,041254703 | 0,971809403 | 0,93866102 | 0,994688607 | serpin family I member 1 |
| ENSG00000157823 | AP3S2 | -0,041334398 | 0,971755722 | 0,85150513 | 0,987259878 | adaptor related protein complex 3 subunit sigma 2 |
| ENSG00000119333 | DYNC2I2 | -0,041384864 | 0,97172173 | 0,89640392 | 0,991638165 | dynein 2 intermediate chain 2 |
| ENSG00000116819 | TFAP2E | -0,041533006 | 0,971621954 | 0,94543264 | 0,995917699 | transcription factor AP-2 epsilon |
| ENSG00000146409 | SLC18B1 | -0,041538128 | 0,971618505 | 0,9233707 | 0,993825008 | solute carrier family 18 member B1 |
| ENSG00000100796 | PPP4R3A | -0,041573343 | 0,971594789 | 0,90311896 | 0,992311172 | protein phosphatase 4 regulatory subunit 3A |
| ENSG00000286561 | AC011092,3 | -0,041645354 | 0,971546293 | 0,94111725 | 0,995251853 | novel transcript, antisense to SBF2 |

| | | | | | | |
|---|---|---|---|---|---|---|
| ENSG00000275837 | SMARCB1 | -0,041649057 | 0,9715438 | 0,84842082 | 0,987036295 | SWI/SNF related, matrix associated, actin dependent regulator of chromatin, subfamily b, member 1 |
| ENSG00000097096 | SYDE2 | -0,04168809 | 0,971517514 | 0,90636879 | 0,992590661 | synapse defective Rho GTPase homolog 2 |
| ENSG00000258316 | KLF17P1 | -0,041716308 | 0,971498512 | 0,9673409 | 0,997263201 | Kruppel like factor 17 pseudogene 1 |
| ENSG00000169155 | ZBTB43 | -0,041744952 | 0,971479224 | 0,87282439 | 0,989268483 | zinc finger and BTB domain containing 43 |
| ENSG00000247809 | NR2F2-AS1 | -0,041799268 | 0,97144265 | 0,92651994 | 0,993915537 | NR2F2 antisense RNA 1 |
| ENSG00000254461 | AP001107,4 | -0,041813408 | 0,971433129 | 0,95938695 | 0,996608711 | novel transcript |
| ENSG00000242616 | GNG10 | -0,04182399 | 0,971426003 | 0,92944511 | 0,994293848 | G protein subunit gamma 10 |
| ENSG00000165609 | NUDT5 | -0,041839101 | 0,971415828 | 0,90655867 | 0,992590661 | nudix hydrolase 5 |
| ENSG00000238134 | EHMT2 | -0,042015649 | 0,97129696 | 0,90071731 | 0,991865079 | euchromatic histone lysine methyltransferase 2 |
| ENSG00000043093 | DCUN1D1 | -0,042057913 | 0,971268506 | 0,93112185 | 0,994433362 | defective in cullin neddylation 1 domain containing 1 |
| ENSG00000237351 | ITPK1P1 | -0,042073928 | 0,971257724 | 0,95157468 | 0,996432861 | ITPK1 pseudogene 1 |
| ENSG00000087470 | DNM1L | -0,042077902 | 0,971255049 | 0,91470304 | 0,993361299 | dynamin 1 like |
| ENSG00000239883 | PARGP1 | -0,042094537 | 0,97124385 | 0,87393125 | 0,989711917 | poly(ADP-ribose) glycohydrolase pseudogene 1 |
| ENSG00000124571 | XPO5 | -0,042119203 | 0,971227244 | 0,8549132 | 0,987986781 | exportin 5 |
| ENSG00000109189 | USP46 | -0,042218276 | 0,97116055 | 0,91751935 | 0,993361299 | ubiquitin specific peptidase 46 |
| ENSG00000235712 | RXRB | -0,0422462 | 0,971141753 | 0,9692351 | 0,997412278 | retinoid X receptor beta |
| ENSG00000185104 | FAF1 | -0,042293704 | 0,971109777 | 0,85207521 | 0,98732978 | Fas associated factor 1 |
| ENSG00000158717 | RNF166 | -0,042429536 | 0,97101835 | 0,89033021 | 0,990717729 | ring finger protein 166 |
| ENSG00000055044 | NOP58 | -0,042452246 | 0,971003065 | 0,88819632 | 0,990717729 | NOP58 ribonucleoprotein |
| ENSG00000261556 | SMG1P7 | -0,042495574 | 0,970973903 | 0,85438613 | 0,987861659 | SMG1 pseudogene 7 |
| ENSG00000238113 | LINC01410 | -0,042643932 | 0,970874059 | 0,90892609 | 0,992756615 | long intergenic non-protein coding RNA 1410 |
| ENSG00000156973 | PDE6D | -0,042670111 | 0,970856442 | 0,88797535 | 0,990717729 | phosphodiesterase 6D |
| ENSG00000139160 | ETFBKMT | -0,042712392 | 0,970827989 | 0,89848944 | 0,991832796 | electron transfer flavoprotein subunit beta lysine methyltransferase |
| ENSG00000250366 | TUNAR | -0,042718002 | 0,970824215 | 0,9167256 | 0,993361299 | TCL1 upstream neural differentiation-associated RNA |
| ENSG00000143443 | C1orf56 | -0,042728313 | 0,970817276 | 0,87959194 | 0,990261566 | chromosome 1 open reading frame 56 |
| ENSG00000182150 | ERCC6L2 | -0,042742546 | 0,970807698 | 0,90074168 | 0,991865079 | ERCC excision repair 6 like 2 |
| ENSG00000174173 | TRMT10C | -0,042783755 | 0,970779969 | 0,92843975 | 0,994108531 | tRNA methyltransferase 10C, mitochondrial RNase P subunit |

| | | | | | | |
|---|---|---|---|---|---|---|
| ENSG00000228709 | LINC02575 | -0,042853707 | 0,9707329 | 0,93749562 | 0,994479239 | long intergenic non-protein coding RNA 2575 |
| ENSG00000261141 | AC092718,5 | -0,042983039 | 0,970645881 | 0,95211857 | 0,996432861 | novel transcript, antisense to C16orf46 |
| ENSG00000217702 | AC073263,1 | -0,043020902 | 0,970620407 | 0,95574615 | 0,996608711 | novel transcript, antisense to BOLA3 |
| ENSG00000051128 | HOMER3 | -0,043043344 | 0,970605308 | 0,92990902 | 0,994331958 | homer scaffold protein 3 |
| ENSG00000125827 | TMX4 | -0,043077562 | 0,970582288 | 0,89076965 | 0,990717729 | thioredoxin related transmembrane protein 4 |
| ENSG00000225733 | FGD5-AS1 | -0,04315141 | 0,970532608 | 0,89915189 | 0,991832796 | FGD5 antisense RNA 1 |
| ENSG00000159239 | AC005041,1 | -0,043225242 | 0,97048294 | 0,95368739 | 0,996608711 | novel protein |
| ENSG00000130312 | MRPL34 | -0,043298225 | 0,970433847 | 0,88256393 | 0,990717729 | mitochondrial ribosomal protein L34 |
| ENSG00000100888 | CHD8 | -0,043417463 | 0,970353645 | 0,85151288 | 0,987259878 | chromodomain helicase DNA binding protein 8 |
| ENSG00000178338 | ZNF354B | -0,043519001 | 0,970285352 | 0,90291878 | 0,992311172 | zinc finger protein 354B |
| ENSG00000278133 | AC135050,6 | -0,043543212 | 0,97026907 | 0,90654706 | 0,992590661 | novel transcript, sense intronic to KAT8 |
| ENSG00000145912 | NHP2 | -0,043543676 | 0,970268758 | 0,85406297 | 0,987861659 | NHP2 ribonucleoprotein |
| ENSG00000136536 | MARCHF7 | -0,043558354 | 0,970258886 | 0,92367217 | 0,993915537 | membrane associated ring-CH-type finger 7 |
| ENSG00000131652 | THOC6 | -0,043721477 | 0,970149187 | 0,87987674 | 0,990261566 | THO complex 6 |
| ENSG00000198522 | GPN1 | -0,043763068 | 0,970121219 | 0,8909412 | 0,990730422 | GPN-loop GTPase 1 |
| ENSG00000099256 | PRTFDC1 | -0,043806751 | 0,970091845 | 0,91716922 | 0,993361299 | phosphoribosyl transferase domain containing 1 |
| ENSG00000247746 | USP51 | -0,043853041 | 0,97006072 | 0,91396489 | 0,993361299 | ubiquitin specific peptidase 51 |
| ENSG00000188693 | CYP51A1-AS1 | -0,043891973 | 0,970034542 | 0,92234609 | 0,993797669 | CYP51A1 antisense RNA 1 |
| ENSG00000285190 | AC018754,1 | -0,043967047 | 0,969984066 | 0,96082193 | 0,996608711 | novel transcript, antisense to CCNH |
| ENSG00000132664 | POLR3F | -0,044003467 | 0,96995958 | 0,92335746 | 0,993825008 | RNA polymerase III subunit F |
| ENSG00000073050 | XRCC1 | -0,044020179 | 0,969948344 | 0,85560717 | 0,988075747 | X-ray repair cross complementing 1 |
| ENSG00000152520 | PAN3 | -0,044025562 | 0,969944724 | 0,90507736 | 0,992590661 | poly(A) specific ribonuclease subunit PAN3 |
| ENSG00000166343 | MSS51 | -0,044049838 | 0,969928404 | 0,89236373 | 0,991048031 | MSS51 mitochondrial translational activator |
| ENSG00000119650 | IFT43 | -0,044066095 | 0,969917474 | 0,84735321 | 0,987035044 | intraflagellar transport 43 |
| ENSG00000164535 | DAGLB | -0,044109862 | 0,96988805 | 0,86816384 | 0,989074806 | diacylglycerol lipase beta |
| ENSG00000259543 | AC060809,1 | -0,044160035 | 0,969854321 | 0,97966654 | 0,99837368 | novel transcript |
| ENSG00000263028 | TAS2R19 | -0,044200011 | 0,969827447 | 0,95014269 | 0,996432861 | taste 2 receptor member 19 |
| ENSG00000212124 | TAS2R19 | -0,044200011 | 0,969827447 | 0,95014269 | 0,996432861 | taste 2 receptor member 19 |
| ENSG00000096070 | BRPF3 | -0,044204026 | 0,969824748 | 0,84718277 | 0,987035044 | bromodomain and PHD finger containing 3 |
| ENSG00000174106 | LEMD3 | -0,044262593 | 0,969785378 | 0,89715571 | 0,991644439 | LEM domain containing 3 |
| ENSG00000173914 | RBM4B | -0,044268826 | 0,969781188 | 0,89646295 | 0,991638165 | RNA binding motif protein 4B |
| ENSG00000233672 | RNASEH2B-AS1 | -0,04428323 | 0,969771506 | 0,93495377 | 0,994479239 | RNASEH2B antisense RNA 1 |
| ENSG00000173726 | TOMM20 | -0,044408635 | 0,969687213 | 0,90563458 | 0,992590661 | translocase of outer mitochondrial membrane 20 |

| | | | | | | |
|---|---|---|---|---|---|---|
| ENSG00000053702 | NRIP2 | -0,044507085 | 0,969621043 | 0,91637768 | 0,993361299 | nuclear receptor interacting protein 2 |
| ENSG00000104472 | CHRAC1 | -0,044517893 | 0,969613779 | 0,90300289 | 0,992311172 | chromatin accessibility complex subunit 1 |
| ENSG00000132698 | RAB25 | -0,044519164 | 0,969612926 | 0,95616974 | 0,996608711 | RAB25, member RAS oncogene family |
| ENSG00000089775 | ZBTB25 | -0,044561699 | 0,969584339 | 0,83094994 | 0,984269865 | zinc finger and BTB domain containing 25 |
| ENSG00000075234 | TTC38 | -0,044680252 | 0,969504667 | 0,87290261 | 0,989308263 | tetratricopeptide repeat domain 38 |
| ENSG00000068650 | ATP11A | -0,044697801 | 0,969492874 | 0,84979341 | 0,987036295 | ATPase phospholipid transporting 11A |
| ENSG00000239282 | CASTOR1 | -0,044745081 | 0,969461102 | 0,91669872 | 0,993361299 | cytosolic arginine sensor for mTORC1 subunit 1 |
| ENSG00000105011 | ASF1B | -0,044816465 | 0,969413135 | 0,86487495 | 0,989074806 | anti-silencing function 1B histone chaperone |
| ENSG00000288210 | ASF1B | -0,044816465 | 0,969413135 | 0,86487495 | 0,989074806 | anti-silencing function 1B histone chaperone |
| ENSG00000269937 | AC093525,7 | -0,044822717 | 0,969408934 | 0,89391576 | 0,991135494 | novel transcript, antisense to PDPK1 |
| ENSG00000138757 | G3BP2 | -0,044887442 | 0,969365443 | 0,88961911 | 0,990717729 | G3BP stress granule assembly factor 2 |
| ENSG00000241163 | LINC00877 | -0,045069694 | 0,969242994 | 0,9104696 | 0,99320976 | long intergenic non-protein coding RNA 877 |
| ENSG00000261502 | AC040174,1 | -0,045078084 | 0,969237357 | 0,96483251 | 0,997113804 | novel transcript |
| ENSG00000001084 | GCLC | -0,045209414 | 0,96914913 | 0,89522329 | 0,991303328 | glutamate-cysteine ligase catalytic subunit |
| ENSG00000160783 | PMF1 | -0,045419495 | 0,969008016 | 0,83552732 | 0,985034063 | polyamine modulated factor 1 |
| ENSG00000187953 | PMS2CL | -0,045449477 | 0,968987878 | 0,8387902 | 0,985672131 | PMS2 C-terminal like pseudogene |
| ENSG00000105497 | ZNF175 | -0,045472582 | 0,96897236 | 0,83963655 | 0,985672131 | zinc finger protein 175 |
| ENSG00000181061 | HIGD1A | -0,045548574 | 0,968921322 | 0,92835642 | 0,994108531 | HIG1 hypoxia inducible domain family member 1A |
| ENSG00000103528 | SYT17 | -0,04556723 | 0,968908793 | 0,89016622 | 0,990717729 | synaptotagmin 17 |
| ENSG00000152422 | XRCC4 | -0,045584349 | 0,968897296 | 0,89679581 | 0,991644439 | X-ray repair cross complementing 4 |
| ENSG00000230445 | LRRC37A6P | -0,045788448 | 0,968760235 | 0,88667782 | 0,990717729 | leucine rich repeat containing 37 member A6, pseudogene |
| ENSG00000033327 | GAB2 | -0,045886822 | 0,96869418 | 0,89629199 | 0,991636805 | GRB2 associated binding protein 2 |
| ENSG00000130449 | ZSWIM6 | -0,045908964 | 0,968679313 | 0,90394125 | 0,992478289 | zinc finger SWIM-type containing 6 |
| ENSG00000266710 | RN7SL48P | -0,045920352 | 0,968671666 | 0,9680901 | 0,997263201 | RNA, 7SL, cytoplasmic 48, pseudogene |
| ENSG00000171217 | CLDN20 | -0,045926142 | 0,968667778 | 0,90571161 | 0,992590661 | claudin 20 |
| ENSG00000131876 | SNRPA1 | -0,045951314 | 0,968650877 | 0,86434288 | 0,989074806 | small nuclear ribonucleoprotein polypeptide A' |
| ENSG00000203446 | SUGCT-AS1 | -0,045965874 | 0,968641101 | 0,95632637 | 0,996608711 | SUGCT antisense RNA 1 |
| ENSG00000137168 | PPIL1 | -0,046018501 | 0,968605768 | 0,90733893 | 0,992590661 | peptidylprolyl isomerase like 1 |
| ENSG00000245275 | SAP30L-AS1 | -0,046067817 | 0,968572658 | 0,86658524 | 0,989074806 | SAP30L antisense RNA 1 (head to head) |
| ENSG00000214046 | SMIM7 | -0,046167674 | 0,96850562 | 0,82442096 | 0,983374097 | small integral membrane protein 7 |
| ENSG00000132718 | SYT11 | -0,046170351 | 0,968503823 | 0,82968929 | 0,984081562 | synaptotagmin 11 |
| ENSG00000156502 | SUPV3L1 | -0,04618941 | 0,968491029 | 0,86990703 | 0,989074806 | Suv3 like RNA helicase |

| | | | | | | |
|---|---|---|---|---|---|---|
| ENSG00000169174 | PCSK9 | -0,046251139 | 0,96844959 | 0,90520842 | 0,992590661 | proprotein convertase subtilisin/kexin type 9 |
| ENSG00000223872 | AC006372,1 | -0,046261757 | 0,968442463 | 0,96944906 | 0,997494004 | novel transcript |
| ENSG00000142875 | PRKACB | -0,046271063 | 0,968436216 | 0,91018175 | 0,993017942 | protein kinase cAMP-activated catalytic subunit beta |
| ENSG00000176386 | CDC26 | -0,046277425 | 0,968431946 | 0,88462319 | 0,990717729 | cell division cycle 26 |
| ENSG00000013573 | DDX11 | -0,046293235 | 0,968421333 | 0,87904549 | 0,990109814 | DEAD/H-box helicase 11 |
| ENSG00000154719 | MRPL39 | -0,046336644 | 0,968392195 | 0,92251293 | 0,993797669 | mitochondrial ribosomal protein L39 |
| ENSG00000166435 | XRRA1 | -0,046374698 | 0,968366652 | 0,83851634 | 0,985672131 | X-ray radiation resistance associated 1 |
| ENSG00000153391 | INO80C | -0,046402803 | 0,968347787 | 0,8547288 | 0,987861659 | INO80 complex subunit C |
| ENSG00000183763 | TRAIP | -0,046444663 | 0,96831969 | 0,84956909 | 0,987036295 | TRAF interacting protein |
| ENSG00000169599 | NFU1 | -0,046460867 | 0,968308815 | 0,89773361 | 0,991708791 | NFU1 iron-sulfur cluster scaffold |
| ENSG00000247130 | AC138781,1 | -0,046489796 | 0,968289398 | 0,9566264 | 0,996608711 | novel transcript |
| ENSG00000236088 | COX10-AS1 | -0,046560235 | 0,968242123 | 0,88277374 | 0,990717729 | COX10 antisense RNA 1 |
| ENSG00000107341 | UBE2R2 | -0,046582412 | 0,968227239 | 0,83186923 | 0,984596888 | ubiquitin conjugating enzyme E2 R2 |
| ENSG00000197933 | ZNF823 | -0,046602195 | 0,968213963 | 0,89874782 | 0,991832796 | zinc finger protein 823 |
| ENSG00000102786 | INTS6 | -0,046718476 | 0,968135928 | 0,88744295 | 0,990717729 | integrator complex subunit 6 |
| ENSG00000121152 | NCAPH | -0,046812571 | 0,968072787 | 0,87632915 | 0,990109814 | non-SMC condensin I complex subunit H |
| ENSG00000164182 | NDUFAF2 | -0,046886575 | 0,96802313 | 0,86582208 | 0,989074806 | NADH:ubiquinone oxidoreductase complex assembly factor 2 |
| ENSG00000124766 | SOX4 | -0,046887468 | 0,968022531 | 0,83901495 | 0,985672131 | SRY-box transcription factor 4 |
| ENSG00000287574 | AC004938,2 | -0,046937434 | 0,967989005 | 0,94257833 | 0,995390305 | novel transcript |
| ENSG00000084774 | CAD | -0,046968171 | 0,967968382 | 0,88544885 | 0,990717729 | carbamoyl-phosphate synthetase 2, aspartate transcarbamylase, and dihydroorotase |
| ENSG00000236539 | HNRNPA1P54 | -0,047038871 | 0,967920947 | 0,92919574 | 0,994293848 | heterogeneous nuclear ribonucleoprotein A1 pseudogene 54 |
| ENSG00000213540 | TXN2P1 | -0,047054374 | 0,967910546 | 0,93662086 | 0,994479239 | TXN2 pseudogene 1 |
| ENSG00000181744 | DIPK2A | -0,047056016 | 0,967909445 | 0,89553508 | 0,991303328 | divergent protein kinase domain 2A |
| ENSG00000132004 | FBXW9 | -0,047095691 | 0,967882827 | 0,87870859 | 0,990109814 | F-box and WD repeat domain containing 9 |
| ENSG00000282386 | AL358472,4 | -0,047182468 | 0,967824612 | 0,93471625 | 0,994479239 | novel transcript, antisense to SLC39A1 |
| ENSG00000130427 | EPO | -0,047202008 | 0,967811503 | 0,97448039 | 0,99837368 | erythropoietin |
| ENSG00000119285 | HEATR1 | -0,047266179 | 0,967768456 | 0,89206905 | 0,99102448 | HEAT repeat containing 1 |
| ENSG00000164323 | CFAP97 | -0,047320935 | 0,967731726 | 0,91914417 | 0,993598006 | cilia and flagella associated protein 97 |
| ENSG00000272158 | AL139022,2 | -0,04740251 | 0,967677009 | 0,96049066 | 0,996608711 | novel transcript, antisense to CHURC1-FNTB |
| ENSG00000035687 | ADSS2 | -0,047411049 | 0,967671281 | 0,88301699 | 0,990717729 | adenylosuccinate synthase 2 |

| | | | | | | |
|---|---|---|---|---|---|---|
| ENSG00000244380 | AC134772,1 | -0,047431954 | 0,967657259 | 0,95765607 | 0,996608711 | novel transcript, antisense to TMA7 |
| ENSG00000259657 | PIGHP1 | -0,047442263 | 0,967650345 | 0,93276322 | 0,994479239 | phosphatidylinositol glycan anchor biosynthesis class H pseudogene 1 |
| ENSG00000137812 | KNL1 | -0,047608261 | 0,967539013 | 0,8898532 | 0,990717729 | kinetochore scaffold 1 |
| ENSG00000236993 | GAPDHP21 | -0,047626035 | 0,967527093 | 0,95140155 | 0,996432861 | glyceraldehyde 3 phosphate dehydrogenase pseudogene 21 |
| ENSG00000212170 | RNU1-77P | -0,047656731 | 0,967506507 | 0,95455535 | 0,996608711 | RNA, U1 small nuclear 77, pseudogene |
| ENSG00000272699 | AC007620,3 | -0,047704716 | 0,967474327 | 0,949727 | 0,996432861 | novel transcript, antisense to GNB4 |
| ENSG00000181192 | DHTKD1 | -0,047782762 | 0,967421991 | 0,84337607 | 0,986260952 | dehydrogenase E1 and transketolase domain containing 1 |
| ENSG00000174780 | SRP72 | -0,047783835 | 0,967421272 | 0,90680391 | 0,992590661 | signal recognition particle 72 |
| ENSG00000285675 | AC023385,1 | -0,047786809 | 0,967419278 | 0,96331659 | 0,996608711 | novel transcript |
| ENSG00000114115 | RBP1 | -0,047910975 | 0,96733602 | 0,84551035 | 0,986639902 | retinol binding protein 1 |
| ENSG00000131941 | RHPN2 | -0,047948919 | 0,967310579 | 0,88883475 | 0,990717729 | rhophilin Rho GTPase binding protein 2 |
| ENSG00000173065 | FAM222B | -0,047968472 | 0,967297468 | 0,84640139 | 0,986939308 | family with sequence similarity 222 member B |
| ENSG00000286859 | AL158214,2 | -0,048078395 | 0,96722377 | 0,92741235 | 0,994021729 | novel transcript, antisense to COG2 |
| ENSG00000238249 | HMGN2P17 | -0,048111336 | 0,967201686 | 0,9495459 | 0,996432861 | high mobility group nucleosomal binding domain 2 pseudogene 17 |
| ENSG00000175485 | OR52W1 | -0,048164248 | 0,967166213 | 0,95282839 | 0,99652169 | olfactory receptor family 52 subfamily W member 1 |
| ENSG00000278249 | SCARNA2 | -0,048195167 | 0,967145486 | 0,89878067 | 0,991832796 | small Cajal body-specific RNA 2 |
| ENSG00000282393 | AC016588,2 | -0,048250751 | 0,967108225 | 0,83167088 | 0,984456953 | novel transcript |
| ENSG00000152128 | TMEM163 | -0,048306143 | 0,967071094 | 0,86405839 | 0,989074806 | transmembrane protein 163 |
| ENSG00000144401 | METTL21A | -0,048386582 | 0,967017175 | 0,89404672 | 0,991135494 | methyltransferase like 21A |
| ENSG00000111231 | GPN3 | -0,048396648 | 0,967010427 | 0,88280521 | 0,990717729 | GPN-loop GTPase 3 |
| ENSG00000237037 | NDUFA6-DT | -0,048396812 | 0,967010317 | 0,87614862 | 0,990109814 | NDUFA6 divergent transcript |
| ENSG00000082996 | RNF13 | -0,048405367 | 0,967004583 | 0,91743111 | 0,993361299 | ring finger protein 13 |
| ENSG00000072803 | FBXW11 | -0,048535475 | 0,966917379 | 0,87926994 | 0,990148595 | F-box and WD repeat domain containing 11 |
| ENSG00000161929 | SCIMP | -0,04860167 | 0,966873015 | 0,94645231 | 0,996027077 | SLP adaptor and CSK interacting membrane protein |
| ENSG00000248503 | AL356235,1 | -0,048669401 | 0,966827624 | 0,90733049 | 0,992590661 | chromosome 4 open reading frame 46 (C4orf46) pseudogene |
| ENSG00000182378 | PLCXD1 | -0,048726526 | 0,966789342 | 0,86124068 | 0,989074806 | phosphatidylinositol specific phospholipase C X domain containing 1 |

| | | | | | | |
|---|---|---|---|---|---|---|
| ENSG00000279855 | AC005618,2 | -0,048744606 | 0,966777226 | 0,96779101 | 0,997263201 | novel transcript, antisense to protocadherin gamma cluster |
| ENSG00000113569 | NUP155 | -0,048784548 | 0,966750461 | 0,8807202 | 0,990533002 | nucleoporin 155 |
| ENSG00000108961 | RANGRF | -0,048790181 | 0,966746686 | 0,85270617 | 0,98761642 | RAN guanine nucleotide release factor |
| ENSG00000177595 | PIDD1 | -0,048793384 | 0,96674454 | 0,91338944 | 0,993361299 | p53-induced death domain protein 1 |
| ENSG00000165091 | TMC1 | -0,048919643 | 0,966659938 | 0,94557927 | 0,995939435 | transmembrane channel like 1 |
| ENSG00000130758 | MAP3K10 | -0,04898585 | 0,966615577 | 0,91353414 | 0,993361299 | mitogen-activated protein kinase kinase kinase 10 |
| ENSG00000196405 | EVL | -0,04902242 | 0,966591076 | 0,81544387 | 0,981521413 | Enah/Vasp-like |
| ENSG00000100150 | DEPDC5 | -0,049034518 | 0,966582971 | 0,81585088 | 0,981708551 | DEP domain containing 5, GATOR1 subcomplex subunit |
| ENSG00000245680 | ZNF585B | -0,049179845 | 0,966485608 | 0,8392774 | 0,985672131 | zinc finger protein 585B |
| ENSG00000129657 | SEC14L1 | -0,049186042 | 0,966481457 | 0,878056 | 0,990109814 | SEC14 like lipid binding 1 |
| ENSG00000133134 | BEX2 | -0,049207784 | 0,966466892 | 0,92097992 | 0,993787755 | brain expressed X-linked 2 |
| ENSG00000258377 | AL139099,1 | -0,049244064 | 0,966442588 | 0,98990667 | 0,99837368 | novel transcript |
| ENSG00000227827 | PKD1P2 | -0,049251834 | 0,966437383 | 0,91570469 | 0,993361299 | polycystin 1, transient receptor potential channel interacting pseudogene 2 |
| ENSG00000125375 | DMAC2L | -0,049289258 | 0,966412313 | 0,8266281 | 0,983510329 | distal membrane arm assembly complex 2 like |
| ENSG00000182173 | TSEN54 | -0,049353672 | 0,966369166 | 0,87039162 | 0,989074806 | tRNA splicing endonuclease subunit 54 |
| ENSG00000225393 | BX571846,1 | -0,049360906 | 0,96636432 | 0,93928726 | 0,994882028 | novel transcript |
| ENSG00000002016 | RAD52 | -0,049361324 | 0,96636404 | 0,83419213 | 0,984905093 | RAD52 homolog, DNA repair protein |
| ENSG00000229018 | PMS2P7 | -0,049449415 | 0,966305036 | 0,88816576 | 0,990717729 | PMS1 homolog 2, mismatch repair system component pseudogene 7 |
| ENSG00000182732 | RGS6 | -0,049500814 | 0,96627061 | 0,91924267 | 0,993598006 | regulator of G protein signaling 6 |
| ENSG00000186026 | ZNF284 | -0,049588484 | 0,966211893 | 0,89362023 | 0,991135494 | zinc finger protein 284 |
| ENSG00000284773 | AC114490,3 | -0,049596432 | 0,96620657 | 0,91690437 | 0,993361299 | novel protein |
| ENSG00000164620 | RELL2 | -0,049643939 | 0,966174754 | 0,87865626 | 0,990109814 | RELT like 2 |
| ENSG00000115233 | PSMD14 | -0,049650487 | 0,966170369 | 0,88662172 | 0,990717729 | proteasome 26S subunit, non-ATPase 14 |
| ENSG00000049449 | RCN1 | -0,049739806 | 0,966110554 | 0,8783087 | 0,990109814 | reticulocalbin 1 |
| ENSG00000117616 | RSRP1 | -0,049764356 | 0,966094114 | 0,86094376 | 0,989074806 | arginine and serine rich protein 1 |
| ENSG00000082516 | GEMIN5 | -0,049792659 | 0,966075161 | 0,8331243 | 0,984905093 | gem nuclear organelle associated protein 5 |
| ENSG00000285000 | AC008581,2 | -0,049796579 | 0,966072536 | 0,90179962 | 0,992136926 | novel protein, AGGF1-ZBED3-AS1 readthrough |
| ENSG00000066136 | NFYC | -0,049807809 | 0,966065016 | 0,86639214 | 0,989074806 | nuclear transcription factor Y subunit gamma |
| ENSG00000231187 | AL356056,2 | -0,049812277 | 0,966062025 | 0,90088455 | 0,991865079 | uncharacterized LOC102724593 |

| | | | | | | |
|---|---|---|---|---|---|---|
| ENSG00000120688 | WBP4 | -0,049821616 | 0,966055771 | 0,87734364 | 0,990109814 | WW domain binding protein 4 |
| ENSG00000135482 | ZC3H10 | -0,049875997 | 0,966019357 | 0,81874238 | 0,98249617 | zinc finger CCCH-type containing 10 |
| ENSG00000226124 | FTCDNL1 | -0,049894097 | 0,966007237 | 0,94912549 | 0,996432861 | formiminotransferase cyclodeaminase N-terminal like |
| ENSG00000062822 | POLD1 | -0,050027731 | 0,965917762 | 0,89395765 | 0,991135494 | DNA polymerase delta 1, catalytic subunit |
| ENSG00000241935 | HOGA1 | -0,050057812 | 0,965897623 | 0,93032839 | 0,994336072 | 4-hydroxy-2-oxoglutarate aldolase 1 |
| ENSG00000231407 | GORAB-AS1 | -0,050061632 | 0,965895065 | 0,91738901 | 0,993361299 | GORAB antisense RNA 1 |
| ENSG00000286885 | AL138824,1 | -0,050088544 | 0,965877047 | 0,94627674 | 0,996027077 | novel transcript |
| ENSG00000254315 | AC023632,5 | -0,050138476 | 0,965843619 | 0,97314677 | 0,99837368 | novel transcript |
| ENSG00000177042 | TMEM80 | -0,050277765 | 0,965750373 | 0,84111463 | 0,985672131 | transmembrane protein 80 |
| ENSG00000203705 | TATDN3 | -0,050377988 | 0,965683286 | 0,88399336 | 0,990717729 | TatD DNase domain containing 3 |
| ENSG00000264112 | AC015813,1 | -0,050413149 | 0,965659751 | 0,87798656 | 0,990109814 | novel transcript |
| ENSG00000264920 | AC018521,5 | -0,050659136 | 0,965495114 | 0,88582125 | 0,990717729 | novel transcript |
| ENSG00000230373 | GOLGA6L5P | -0,050660517 | 0,96549419 | 0,90332899 | 0,992311172 | golgin A6 family like 5, pseudogene |
| ENSG00000180304 | OAZ2 | -0,050665616 | 0,965490778 | 0,8186271 | 0,9824772 | ornithine decarboxylase antizyme 2 |
| ENSG00000158901 | WFDC8 | -0,05070647 | 0,965463438 | 0,94678697 | 0,996027077 | WAP four-disulfide core domain 8 |
| ENSG00000029363 | BCLAF1 | -0,0507557 | 0,965430493 | 0,89014616 | 0,990717729 | BCL2 associated transcription factor 1 |
| ENSG00000240489 | SETP14 | -0,050786697 | 0,965409751 | 0,96528375 | 0,997208095 | SET pseudogene 14 |
| ENSG00000172943 | PHF8 | -0,050897018 | 0,96533593 | 0,80562582 | 0,981213877 | PHD finger protein 8 |
| ENSG00000092439 | TRPM7 | -0,051070651 | 0,965219755 | 0,88657098 | 0,990717729 | transient receptor potential cation channel subfamily M member 7 |
| ENSG00000108306 | FBXL20 | -0,051075569 | 0,965216465 | 0,8325581 | 0,98489473 | F-box and leucine rich repeat protein 20 |
| ENSG00000244480 | AC005154,2 | -0,05110371 | 0,965197638 | 0,9276408 | 0,994108531 | pseudogene similar golgi autoantigen, golgin subfamily a, 8A GOLGA8A |
| ENSG00000178385 | PLEKHM3 | -0,051108186 | 0,965194643 | 0,96654853 | 0,997263201 | pleckstrin homology domain containing M3 |
| ENSG00000115750 | TAF1B | -0,051177702 | 0,965148137 | 0,89373863 | 0,991135494 | TATA-box binding protein associated factor, RNA polymerase I subunit B |
| ENSG00000147654 | EBAG9 | -0,05119475 | 0,965136732 | 0,91130321 | 0,993222655 | estrogen receptor binding site associated antigen 9 |
| ENSG00000214465 | SMARCE1P6 | -0,051221438 | 0,965118879 | 0,95117113 | 0,996432861 | SWI/SNF related, matrix associated, actin dependent regulator of chromatin, subfamily e, member 1 pseudogene 6 |
| ENSG00000136891 | TEX10 | -0,051267383 | 0,965088143 | 0,88781616 | 0,990717729 | testis expressed 10 |
| ENSG00000143499 | SMYD2 | -0,051274361 | 0,965083475 | 0,8339185 | 0,984905093 | SET and MYND domain containing 2 |

| | | | | | | |
|---|---|---|---|---|---|---|
| ENSG00000204659 | CBY3 | -0,051277229 | 0,965081557 | 0,9583641 | 0,996608711 | chibby family member 3 |
| ENSG00000283718 | CBY3 | -0,051277229 | 0,965081557 | 0,9583641 | 0,996608711 | chibby family member 3 |
| ENSG00000050426 | LETMD1 | -0,051322356 | 0,965051369 | 0,8307449 | 0,984156028 | LETM1 domain containing 1 |
| ENSG00000163362 | INAVA | -0,051392306 | 0,96500458 | 0,82748224 | 0,983595244 | innate immunity activator |
| ENSG00000225579 | EDNRB-AS1 | -0,051425069 | 0,964982665 | 0,92305892 | 0,993825008 | EDNRB antisense RNA 1 |
| ENSG00000196466 | ZNF799 | -0,051437803 | 0,964974147 | 0,87987694 | 0,990261566 | zinc finger protein 799 |
| ENSG00000109832 | DDX25 | -0,051441688 | 0,964971549 | 0,89727329 | 0,991644439 | DEAD-box helicase 25 |
| ENSG00000278921 | EPB41L4A-DT | -0,051493176 | 0,964937111 | 0,91141776 | 0,993222655 | EPB41L4A divergent transcript |
| ENSG00000229800 | ATP8A2P2 | -0,05154459 | 0,964902724 | 0,93959458 | 0,994943406 | ATPase phospholipid transporting 8A2 pseudogene 2 |
| ENSG00000172331 | BPGM | -0,051575081 | 0,964882331 | 0,88142608 | 0,990717729 | bisphosphoglycerate mutase |
| ENSG00000197062 | ZSCAN26 | -0,051595158 | 0,964868903 | 0,88719945 | 0,990717729 | zinc finger and SCAN domain containing 26 |
| ENSG00000125812 | GZF1 | -0,051731921 | 0,964777441 | 0,85593396 | 0,988180177 | GDNF inducible zinc finger protein 1 |
| ENSG00000182481 | KPNA2 | -0,051884423 | 0,964675463 | 0,88733269 | 0,990717729 | karyopherin subunit alpha 2 |
| ENSG00000130559 | CAMSAP1 | -0,051910244 | 0,964658198 | 0,79999255 | 0,980161849 | calmodulin regulated spectrin associated protein 1 |
| ENSG00000261513 | AC009097,4 | -0,051958455 | 0,964625962 | 0,97692272 | 0,99837368 | novel transcript, antisense to PHLPP1 |
| ENSG00000223834 | AL161935,1 | -0,05196045 | 0,964624628 | 0,90432954 | 0,992590661 | novel transcript |
| ENSG00000126215 | XRCC3 | -0,052030745 | 0,964577629 | 0,86829494 | 0,989074806 | X-ray repair cross complementing 3 |
| ENSG00000099810 | MTAP | -0,052042693 | 0,96456964 | 0,87652899 | 0,990109814 | methylthioadenosine phosphorylase |
| ENSG00000141298 | SSH2 | -0,052068496 | 0,964552389 | 0,84271535 | 0,986183927 | slingshot protein phosphatase 2 |
| ENSG00000143797 | MBOAT2 | -0,052104883 | 0,964528061 | 0,9023008 | 0,992230188 | membrane bound O-acyltransferase domain containing 2 |
| ENSG00000263883 | EEF1DP7 | -0,052105074 | 0,964527934 | 0,94481722 | 0,995744664 | eukaryotic translation elongation factor 1 delta pseudogene 7 |
| ENSG00000127957 | PMS2P3 | -0,052134085 | 0,964508538 | 0,84600266 | 0,98689126 | PMS1 homolog 2, mismatch repair system component pseudogene 3 |
| ENSG00000266601 | AC018521,6 | -0,052178723 | 0,964478696 | 0,96915075 | 0,997412278 | novel transcript, antisense to SP2 |
| ENSG00000116030 | SUMO1 | -0,052295297 | 0,964400766 | 0,89359249 | 0,991135494 | small ubiquitin like modifier 1 |
| ENSG00000100478 | AP4S1 | -0,05231935 | 0,964384688 | 0,85495912 | 0,98800332 | adaptor related protein complex 4 subunit sigma 1 |
| ENSG00000243927 | MRPS6 | -0,052325349 | 0,964380678 | 0,89265329 | 0,991048031 | mitochondrial ribosomal protein S6 |
| ENSG00000138092 | CENPO | -0,052355101 | 0,964360791 | 0,8192538 | 0,98249617 | centromere protein O |
| ENSG00000154727 | GABPA | -0,052375907 | 0,964346883 | 0,90992669 | 0,992951496 | GA binding protein transcription factor subunit alpha |
| ENSG00000225859 | DDX39B | -0,05243988 | 0,964304122 | 0,835739 | 0,985034063 | DExD-box helicase 39B |
| ENSG00000168876 | ANKRD49 | -0,052525595 | 0,964246831 | 0,90549903 | 0,992590661 | ankyrin repeat domain 49 |

| | | | | | | |
|---|---|---|---|---|---|---|
| ENSG00000126878 | AIF1L | -0,05257536 | 0,964213571 | 0,82090282 | 0,982719623 | allograft inflammatory factor 1 like |
| ENSG00000254984 | FTLP6 | -0,052638847 | 0,964171141 | 0,96354315 | 0,996733015 | ferritin light chain pseudogene 6 |
| ENSG00000128656 | CHN1 | -0,052687835 | 0,964138402 | 0,8884755 | 0,990717729 | chimerin 1 |
| ENSG00000183856 | IQGAP3 | -0,052703626 | 0,964127849 | 0,84909924 | 0,987036295 | IQ motif containing GTPase activating protein 3 |
| ENSG00000114473 | IQCG | -0,052720173 | 0,964116791 | 0,84865485 | 0,987036295 | IQ motif containing G |
| ENSG00000172123 | SLFN12 | -0,052729517 | 0,964110547 | 0,8621045 | 0,989074806 | schlafen family member 12 |
| ENSG00000140986 | RPL3L | -0,052761413 | 0,964089232 | 0,946361 | 0,996027077 | ribosomal protein L3 like |
| ENSG00000119685 | TTLL5 | -0,052772558 | 0,964081784 | 0,83343944 | 0,984905093 | tubulin tyrosine ligase like 5 |
| ENSG00000117411 | B4GALT2 | -0,052778208 | 0,964078008 | 0,823788 | 0,983374097 | beta-1,4-galactosyltransferase 2 |
| ENSG00000230262 | LINC02603 | -0,052790695 | 0,964069664 | 0,87201228 | 0,989217334 | long intergenic non-protein coding RNA 2603 |
| ENSG00000128268 | MGAT3 | -0,052836806 | 0,964038851 | 0,87840518 | 0,990109814 | beta-1,4-mannosyl-glycoprotein 4-beta-N-acetylglucosaminyltransferase |
| ENSG00000231806 | PCAT7 | -0,052853498 | 0,964027697 | 0,87464453 | 0,990039342 | prostate cancer associated transcript 7 |
| ENSG00000176641 | RNF152 | -0,052969788 | 0,963949994 | 0,88011892 | 0,990268593 | ring finger protein 152 |
| ENSG00000177614 | PGBD5 | -0,052982734 | 0,963941344 | 0,85069639 | 0,987147801 | piggyBac transposable element derived 5 |
| ENSG00000115816 | CEBPZ | -0,053076244 | 0,963878867 | 0,87032225 | 0,989074806 | CCAAT enhancer binding protein zeta |
| ENSG00000135829 | DHX9 | -0,053139869 | 0,963836359 | 0,86280333 | 0,989074806 | DExH-box helicase 9 |
| ENSG00000087302 | RTRAF | -0,053151336 | 0,963828699 | 0,89863471 | 0,991832796 | RNA transcription, translation and transport factor |
| ENSG00000138380 | CARF | -0,053154235 | 0,963826762 | 0,87744184 | 0,990109814 | calcium responsive transcription factor |
| ENSG00000253485 | PCDHGA5 | -0,053179088 | 0,963810158 | 0,92081632 | 0,993787755 | protocadherin gamma subfamily A, 5 |
| ENSG00000267808 | AC018755,1 | -0,053242309 | 0,963767924 | 0,95777143 | 0,996608711 | novel transcript |
| ENSG00000115840 | SLC25A12 | -0,053304625 | 0,963726295 | 0,88016836 | 0,990268593 | solute carrier family 25 member 12 |
| ENSG00000264956 | AC138761,1 | -0,053327998 | 0,963710683 | 0,92602525 | 0,993915537 | novel transcript |
| ENSG00000140795 | MYLK3 | -0,053333486 | 0,963707016 | 0,91004457 | 0,992965454 | myosin light chain kinase 3 |
| ENSG00000124562 | SNRPC | -0,053435975 | 0,963638557 | 0,8601167 | 0,989074806 | small nuclear ribonucleoprotein polypeptide C |
| ENSG00000151332 | MBIP | -0,053499934 | 0,963595837 | 0,90904239 | 0,992814265 | MAP3K12 binding inhibitory protein 1 |
| ENSG00000235194 | PPP1R3E | -0,053515245 | 0,963585611 | 0,84172331 | 0,98571999 | protein phosphatase 1 regulatory subunit 3E |
| ENSG00000123374 | CDK2 | -0,053554789 | 0,963559199 | 0,85826511 | 0,989067345 | cyclin dependent kinase 2 |
| ENSG00000215049 | PRDX2P1 | -0,053558151 | 0,963556954 | 0,95553462 | 0,996608711 | peroxiredoxin 2 pseudogene 1 |
| ENSG00000069956 | MAPK6 | -0,053579836 | 0,963542471 | 0,90449503 | 0,992590661 | mitogen-activated protein kinase 6 |
| ENSG00000185420 | SMYD3 | -0,053674029 | 0,963479564 | 0,87561566 | 0,990078774 | SET and MYND domain containing 3 |
| ENSG00000253385 | AP003696,1 | -0,053720186 | 0,963448739 | 0,93493738 | 0,994479239 | novel transcript |
| ENSG00000104691 | UBXN8 | -0,05374541 | 0,963431895 | 0,84389803 | 0,986273696 | UBX domain protein 8 |

| | | | | | | |
|---|---|---|---|---|---|---|
| ENSG00000269343 | ZNF587B | -0,05377227 | 0,963413957 | 0,81464756 | 0,981521413 | zinc finger protein 587B |
| ENSG00000139651 | ZNF740 | -0,053776132 | 0,963411378 | 0,81386155 | 0,981521413 | zinc finger protein 740 |
| ENSG00000152484 | USP12 | -0,053806159 | 0,963391327 | 0,90996805 | 0,992951496 | ubiquitin specific peptidase 12 |
| ENSG00000167257 | RNF214 | -0,053850918 | 0,963361438 | 0,80448611 | 0,980875302 | ring finger protein 214 |
| ENSG00000135457 | TFCP2 | -0,053859346 | 0,963355811 | 0,82454554 | 0,983374097 | transcription factor CP2 |
| ENSG00000224430 | MKRN5P | -0,053892452 | 0,963333704 | 0,92274149 | 0,993797669 | makorin ring finger protein 5, pseudogene |
| ENSG00000163293 | NIPAL1 | -0,053970807 | 0,963281386 | 0,91942139 | 0,993598006 | NIPA like domain containing 1 |
| ENSG00000232442 | MHENCR | -0,053998307 | 0,963263024 | 0,90365425 | 0,992441997 | melanoma highly expressed competing endogenous lncRNA for miR-425 and miR-489 |
| ENSG00000255507 | UVRAG-DT | -0,054040366 | 0,963234943 | 0,96911082 | 0,997408593 | UVRAG divergent transcript |
| ENSG00000144580 | CNOT9 | -0,054043861 | 0,963232609 | 0,86080216 | 0,989074806 | CCR4-NOT transcription complex subunit 9 |
| ENSG00000076685 | NT5C2 | -0,054076584 | 0,963210762 | 0,89493079 | 0,991303328 | 5'-nucleotidase, cytosolic II |
| ENSG00000214826 | DDX12P | -0,054285102 | 0,963071555 | 0,89299663 | 0,991120066 | DEAD/H-box helicase 12, pseudogene |
| ENSG00000232956 | SNHG15 | -0,054318901 | 0,963048993 | 0,85588007 | 0,988180177 | small nucleolar RNA host gene 15 |
| ENSG00000287311 | AC140121,1 | -0,054444585 | 0,962965098 | 0,93682816 | 0,994479239 | novel transcript |
| ENSG00000287513 | AL133390,1 | -0,054454525 | 0,962958464 | 0,95421474 | 0,996608711 | novel transcript |
| ENSG00000226715 | LINC01709 | -0,054466309 | 0,962950598 | 0,89965695 | 0,991845106 | long intergenic non-protein coding RNA 1709 |
| ENSG00000143429 | LSP1P4 | -0,054563131 | 0,962885975 | 0,8766826 | 0,990109814 | LSP1 pseudogene 4 |
| ENSG00000166024 | R3HCC1L | -0,05465887 | 0,962822078 | 0,87987705 | 0,990261566 | R3H domain and coiled-coil containing 1 like |
| ENSG00000066044 | ELAVL1 | -0,054680462 | 0,962807669 | 0,82989824 | 0,984156028 | ELAV like RNA binding protein 1 |
| ENSG00000267724 | AC012254,3 | -0,054838767 | 0,962702027 | 0,89239844 | 0,991048031 | novel transcript, antisense to HDHD2 |
| ENSG00000283643 | GPR1-AS | -0,054846615 | 0,96269679 | 0,96167088 | 0,996608711 | GPR1 antisense RNA |
| ENSG00000172901 | LVRN | -0,054850007 | 0,962694527 | 0,94419146 | 0,995542144 | laeverin |
| ENSG00000108395 | TRIM37 | -0,05485505 | 0,962691161 | 0,85683876 | 0,988312374 | tripartite motif containing 37 |
| ENSG00000100028 | SNRPD3 | -0,054885228 | 0,962671024 | 0,86185033 | 0,989074806 | small nuclear ribonucleoprotein D3 polypeptide |
| ENSG00000117000 | RLF | -0,054913724 | 0,96265201 | 0,89867871 | 0,991832796 | RLF zinc finger |
| ENSG00000180251 | SLC9A4 | -0,054936248 | 0,96263698 | 0,88760752 | 0,990717729 | solute carrier family 9 member A4 |
| ENSG00000173653 | RCE1 | -0,054936487 | 0,962636821 | 0,86709554 | 0,989074806 | Ras converting CAAX endopeptidase 1 |
| ENSG00000187847 | OR7E25P | -0,055011036 | 0,96258708 | 0,94392011 | 0,995537502 | olfactory receptor family 7 subfamily E member 25 pseudogene |
| ENSG00000176700 | SCAND2P | -0,055047804 | 0,962562548 | 0,80065239 | 0,980477724 | SCAN domain containing 2 pseudogene |
| ENSG00000188958 | UTS2B | -0,055066782 | 0,962549886 | 0,91327013 | 0,993361299 | urotensin 2B |
| ENSG00000172671 | ZFAND4 | -0,055073174 | 0,962545621 | 0,88249522 | 0,990717729 | zinc finger AN1-type containing 4 |

| | | | | | | |
|---|---|---|---|---|---|---|
| ENSG00000281178 | MLXIP | -0,055076321 | 0,962543521 | 0,98844018 | 0,99837368 | MLX interacting protein |
| ENSG00000234630 | AC245060,2 | -0,055194919 | 0,962464398 | 0,95613581 | 0,996608711 | novel transcript |
| ENSG00000150630 | VEGFC | -0,055198404 | 0,962462073 | 0,89264339 | 0,991048031 | vascular endothelial growth factor C |
| ENSG00000131778 | CHD1L | -0,055200286 | 0,962460818 | 0,83186551 | 0,984596888 | chromodomain helicase DNA binding protein 1 like |
| ENSG00000276615 | KEL | -0,055206393 | 0,962456743 | 0,92262327 | 0,993797669 | Kell metallo-endopeptidase (Kell blood group) |
| ENSG00000286730 | AL365295,2 | -0,055238456 | 0,962435353 | 0,89775276 | 0,991708791 | novel transcript |
| ENSG00000188895 | MSL1 | -0,05527584 | 0,962410415 | 0,80847916 | 0,981213877 | MSL complex subunit 1 |
| ENSG00000286108 | AL359916,1 | -0,055301693 | 0,962393168 | 0,88651755 | 0,990717729 | novel transcript, antisense to STK35 |
| ENSG00000181915 | ADO | -0,055305963 | 0,96239032 | 0,85653741 | 0,988292939 | 2-aminoethanethiol dioxygenase |
| ENSG00000157106 | SMG1 | -0,05557054 | 0,962213843 | 0,81726424 | 0,982048938 | SMG1 nonsense mediated mRNA decay associated PI3K related kinase |
| ENSG00000280807 | TUBGCP5 | -0,055638098 | 0,962168786 | 0,95586776 | 0,996608711 | tubulin gamma complex associated protein 5 |
| ENSG00000220201 | ZGLP1 | -0,055658339 | 0,962155287 | 0,89424217 | 0,991135494 | zinc finger GATA like protein 1 |
| ENSG00000142945 | KIF2C | -0,055753378 | 0,962091906 | 0,83051942 | 0,984156028 | kinesin family member 2C |
| ENSG00000237009 | GLIS3-AS1 | -0,055869634 | 0,962014381 | 0,96035591 | 0,996608711 | GLIS3 antisense RNA 1 |
| ENSG00000287935 | AC136297,1 | -0,055923031 | 0,961978776 | 0,90662813 | 0,992590661 | novel transcript |
| ENSG00000154920 | EME1 | -0,055974615 | 0,961944381 | 0,82800347 | 0,983595244 | essential meiotic structure-specific endonuclease 1 |
| ENSG00000213777 | AC011487,1 | -0,056021708 | 0,961912981 | 0,95987124 | 0,996608711 | vomeronasal 1 receptor (VN1R) pseudogene |
| ENSG00000176153 | GPX2 | -0,056129881 | 0,96184086 | 0,91204699 | 0,993361299 | glutathione peroxidase 2 |
| ENSG00000232467 | TMA16P2 | -0,056150892 | 0,961826852 | 0,96180396 | 0,996608711 | translation machinery associated 16 homolog pseudogene 2 |
| ENSG00000095564 | BTAF1 | -0,056153245 | 0,961825283 | 0,89182646 | 0,990939422 | B-TFIID TATA-box binding protein associated factor 1 |
| ENSG00000287561 | AL121899,4 | -0,056205061 | 0,961790738 | 0,96211122 | 0,996608711 | novel transcript |
| ENSG00000102974 | CTCF | -0,056273481 | 0,961745126 | 0,82435331 | 0,983374097 | CCCTC-binding factor |
| ENSG00000101493 | ZNF516 | -0,056274633 | 0,961744358 | 0,81846955 | 0,9824772 | zinc finger protein 516 |
| ENSG00000153253 | SCN3A | -0,056309069 | 0,961721403 | 0,90837409 | 0,992595616 | sodium voltage-gated channel alpha subunit 3 |
| ENSG00000256101 | AC092745,1 | -0,056316172 | 0,961716668 | 0,92085188 | 0,993787755 | novel transcript |
| ENSG00000259865 | AL390728,6 | -0,056327132 | 0,961709361 | 0,9372819 | 0,994479239 | novel transcript |
| ENSG00000135341 | MAP3K7 | -0,05635739 | 0,961689192 | 0,88685778 | 0,990717729 | mitogen-activated protein kinase kinase kinase 7 |
| ENSG00000198538 | ZNF28 | -0,056406839 | 0,96165623 | 0,86409086 | 0,989074806 | zinc finger protein 28 |
| ENSG00000173638 | SLC19A1 | -0,056461839 | 0,961619569 | 0,86422026 | 0,989074806 | solute carrier family 19 member 1 |
| ENSG00000287005 | AC090018,2 | -0,05648723 | 0,961602645 | 0,89834457 | 0,991832796 | novel transcript, antisense to XXYLT1 |
| ENSG00000130159 | ECSIT | -0,05651308 | 0,961585415 | 0,83976157 | 0,985672131 | ECSIT signaling integrator |

| | | | | | | |
|---|---|---|---|---|---|---|
| ENSG00000114416 | FXR1 | -0,056594766 | 0,961530972 | 0,85682372 | 0,988312374 | FMR1 autosomal homolog 1 |
| ENSG00000274677 | AC040169,3 | -0,056702002 | 0,961459504 | 0,94305176 | 0,995402494 | novel transcript, sense intronic to MBTPS1 |
| ENSG00000166471 | TMEM41B | -0,056703854 | 0,96145827 | 0,84021671 | 0,985672131 | transmembrane protein 41B |
| ENSG00000186687 | LYRM7 | -0,056856137 | 0,961356788 | 0,79060206 | 0,979462318 | LYR motif containing 7 |
| ENSG00000205037 | AC134312,1 | -0,056901526 | 0,961326543 | 0,94315231 | 0,995402494 | novel transcript |
| ENSG00000255495 | AC145124,1 | -0,056910826 | 0,961320347 | 0,85557187 | 0,988075747 | novel transcript |
| ENSG00000144504 | ANKMY1 | -0,056919085 | 0,961314843 | 0,90129585 | 0,992041918 | ankyrin repeat and MYND domain containing 1 |
| ENSG00000243004 | AC005062,1 | -0,056949206 | 0,961294773 | 0,95491461 | 0,996608711 | novel transcript, sense overlapping to MACC1 |
| ENSG00000063169 | BICRA | -0,056957997 | 0,961288915 | 0,86922754 | 0,989074806 | BRD4 interacting chromatin remodeling complex associated protein |
| ENSG00000214559 | AC019077,1 | -0,056984431 | 0,961271302 | 0,96469581 | 0,997113804 | novel transcript |
| ENSG00000277127 | Metazoa_SRP | -0,057052882 | 0,961225694 | 0,9590591 | 0,996608711 | Metazoan signal recognition particle RNA |
| ENSG00000187244 | BCAM | -0,057080949 | 0,961206994 | 0,85963269 | 0,989074806 | basal cell adhesion molecule (Lutheran blood group) |
| ENSG00000253772 | AC112191,1 | -0,05710206 | 0,961192929 | 0,93090852 | 0,994433362 | acid phosphatase 1, soluble (ACP1) pseudogene |
| ENSG00000166454 | ATMIN | -0,057208878 | 0,961121764 | 0,87678112 | 0,990109814 | ATM interactor |
| ENSG00000181722 | ZBTB20 | -0,057279984 | 0,961074394 | 0,85901786 | 0,989074806 | zinc finger and BTB domain containing 20 |
| ENSG00000241859 | ANOS2P | -0,057370638 | 0,961014006 | 0,89115589 | 0,990795338 | anosmin 2, pseudogene |
| ENSG00000260777 | AC061975,1 | -0,057375961 | 0,96101046 | 0,92016861 | 0,993787755 | novel transcript |
| ENSG00000198492 | YTHDF2 | -0,057467349 | 0,960949586 | 0,86729771 | 0,989074806 | YTH N6-methyladenosine RNA binding protein 2 |
| ENSG00000164751 | PEX2 | -0,057547703 | 0,960896066 | 0,86502633 | 0,989074806 | peroxisomal biogenesis factor 2 |
| ENSG00000186566 | GPATCH8 | -0,057579341 | 0,960874994 | 0,8247265 | 0,983374097 | G-patch domain containing 8 |
| ENSG00000175575 | PAAF1 | -0,057639966 | 0,960834617 | 0,81487597 | 0,981521413 | proteasomal ATPase associated factor 1 |
| ENSG00000224415 | AC007683,1 | -0,057695753 | 0,960797463 | 0,95773104 | 0,996608711 | ribosomal protein L19 (RPL19) pseudogene |
| ENSG00000282463 | AC216130,3 | -0,057695753 | 0,960797463 | 0,95773104 | 0,996608711 | ribosomal protein L19 (RPL19) pseudogene |
| ENSG00000284218 | NPIPB5 | -0,057717644 | 0,960782884 | 0,97569485 | 0,99837368 | nuclear pore complex interacting protein family member B5 |
| ENSG00000203739 | PRDX6-AS1 | -0,057770466 | 0,960747707 | 0,89329157 | 0,991135494 | PRDX6 antisense RNA 1 |
| ENSG00000251229 | AL645924,2 | -0,057779975 | 0,960741375 | 0,88152774 | 0,990717729 | ribosomal protein S3A (RPS3A) pseudogene |
| ENSG00000114021 | NIT2 | -0,057812599 | 0,96071965 | 0,82590885 | 0,983383773 | nitrilase family member 2 |
| ENSG00000179021 | C3orf38 | -0,057826915 | 0,960710116 | 0,90218537 | 0,992230188 | chromosome 3 open reading frame 38 |
| ENSG00000101236 | RNF24 | -0,057897899 | 0,960662848 | 0,79701421 | 0,980161849 | ring finger protein 24 |
| ENSG00000115266 | APC2 | -0,057912905 | 0,960652856 | 0,91790704 | 0,993361299 | APC regulator of WNT signaling pathway 2 |
| ENSG00000286299 | AL121938,1 | -0,05798004 | 0,960608154 | 0,91842825 | 0,99341755 | novel transcript, antisense to HDDC2 |

| | | | | | | |
|---|---|---|---|---|---|---|
| ENSG00000131477 | RAMP2 | -0,057994134 | 0,96059877 | 0,88943969 | 0,990717729 | receptor activity modifying protein 2 |
| ENSG00000206596 | RNU1-27P | -0,058063657 | 0,96055248 | 0,89073377 | 0,990717729 | RNA, U1 small nuclear 27, pseudogene |
| ENSG00000207389 | RNU1-4 | -0,058063657 | 0,96055248 | 0,89073377 | 0,990717729 | RNA, U1 small nuclear 4 |
| ENSG00000206652 | RNU1-1 | -0,058063657 | 0,96055248 | 0,89073377 | 0,990717729 | RNA, U1 small nuclear 1 |
| ENSG00000273768 | RNVU1-29 | -0,058063657 | 0,96055248 | 0,89073377 | 0,990717729 | RNA, variant U1 small nuclear 29 |
| ENSG00000206737 | RNVU1-18 | -0,058063657 | 0,96055248 | 0,89073377 | 0,990717729 | RNA, variant U1 small nuclear 18 |
| ENSG00000206588 | RNU1-28P | -0,058063657 | 0,96055248 | 0,89073377 | 0,990717729 | RNA, U1 small nuclear 28, pseudogene |
| ENSG00000275405 | U1 | -0,058063657 | 0,96055248 | 0,89073377 | 0,990717729 | U1 spliceosomal RNA |
| ENSG00000207005 | RNU1-2 | -0,058063657 | 0,96055248 | 0,89073377 | 0,990717729 | RNA, U1 small nuclear 2 |
| ENSG00000207513 | RNU1-3 | -0,058063657 | 0,96055248 | 0,89073377 | 0,990717729 | RNA, U1 small nuclear 3 |
| ENSG00000287431 | AC027601,5 | -0,05808815 | 0,960536172 | 0,86330115 | 0,989074806 | novel transcript |
| ENSG00000109475 | RPL34 | -0,058103202 | 0,960526151 | 0,87512893 | 0,990078774 | ribosomal protein L34 |
| ENSG00000111665 | CDCA3 | -0,058123583 | 0,960512582 | 0,81255924 | 0,981224866 | cell division cycle associated 3 |
| ENSG00000235908 | RHOA-IT1 | -0,058141415 | 0,96050071 | 0,92723292 | 0,993915537 | RHOA intronic transcript 1 |
| ENSG00000144935 | TRPC1 | -0,058283184 | 0,960406329 | 0,8813839 | 0,990717729 | transient receptor potential cation channel subfamily C member 1 |
| ENSG00000149187 | CELF1 | -0,058308716 | 0,960389332 | 0,77122635 | 0,978033855 | CUGBP Elav-like family member 1 |
| ENSG00000229427 | ANKRD26P4 | -0,058373548 | 0,960346175 | 0,93133118 | 0,994433362 | ankyrin repeat domain 26 pseudogene 4 |
| ENSG00000164134 | NAA15 | -0,058386304 | 0,960337684 | 0,86561464 | 0,989074806 | N-alpha-acetyltransferase 15, NatA auxiliary subunit |
| ENSG00000272655 | POLR2J4 | -0,058392394 | 0,96033363 | 0,82041114 | 0,982609114 | RNA polymerase II subunit J4, pseudogene |
| ENSG00000152049 | KCNE4 | -0,058403294 | 0,960326375 | 0,82605464 | 0,983464293 | potassium voltage-gated channel subfamily E regulatory subunit 4 |
| ENSG00000221829 | FANCG | -0,058414958 | 0,960318611 | 0,82449816 | 0,983374097 | FA complementation group G |
| ENSG00000230715 | AC018638,2 | -0,058443761 | 0,960299438 | 0,91761491 | 0,993361299 | novel pseudogene |
| ENSG00000100564 | PIGH | -0,058465481 | 0,960284981 | 0,86856796 | 0,989074806 | phosphatidylinositol glycan anchor biosynthesis class H |
| ENSG00000227517 | LINC01483 | -0,05846644 | 0,960284343 | 0,93965706 | 0,994943406 | long intergenic non-protein coding RNA 1483 |
| ENSG00000268006 | PTOV1-AS1 | -0,058515369 | 0,960251775 | 0,88001638 | 0,990268593 | PTOV1 antisense RNA 1 |
| ENSG00000240889 | NDUFB2-AS1 | -0,058540186 | 0,960235257 | 0,92361893 | 0,99390845 | NDUFB2 antisense RNA 1 |
| ENSG00000276565 | AP006285,3 | -0,058553373 | 0,96022648 | 0,95529067 | 0,996608711 | |
| ENSG00000131558 | EXOC4 | -0,058563418 | 0,960219794 | 0,84437969 | 0,986340494 | exocyst complex component 4 |
| ENSG00000116251 | RPL22 | -0,058606103 | 0,960191385 | 0,85619103 | 0,988221518 | ribosomal protein L22 |
| ENSG00000283101 | BMS1P21 | -0,0586471 | 0,960164099 | 0,93836134 | 0,994549499 | BMS1 pseudogene 21 |

| | | | | | | |
|---|---|---|---|---|---|---|
| ENSG00000237934 | AL590729,1 | -0,058689382 | 0,96013596 | 0,9450419 | 0,995797387 | novel transcript |
| ENSG00000183807 | FAM162B | -0,058706203 | 0,960124765 | 0,94383948 | 0,995486053 | family with sequence similarity 162 member B |
| ENSG00000254616 | AP001775,1 | -0,058771749 | 0,960081145 | 0,95691122 | 0,996608711 | Zinc finger protein 628 (ZNF628) pseudogene |
| ENSG00000166275 | BORCS7 | -0,058783045 | 0,960073627 | 0,83932515 | 0,985672131 | BLOC-1 related complex subunit 7 |
| ENSG00000204923 | FBXO48 | -0,058813537 | 0,960053336 | 0,88588741 | 0,990717729 | F-box protein 48 |
| ENSG00000104883 | PEX11G | -0,058866358 | 0,960018187 | 0,89983015 | 0,991845106 | peroxisomal biogenesis factor 11 gamma |
| ENSG00000234161 | PABPC5-AS1 | -0,058967807 | 0,959950681 | 0,93194035 | 0,994476862 | PABPC5 antisense RNA 1 |
| ENSG00000123213 | NLN | -0,058968056 | 0,959950516 | 0,86981717 | 0,989074806 | neurolysin |
| ENSG00000226798 | AL138749,1 | -0,059043194 | 0,959900521 | 0,95467954 | 0,996608711 | putatitive novel transcript |
| ENSG00000272078 | AL139423,1 | -0,05908394 | 0,959873411 | 0,8793183 | 0,990167381 | novel transcript, antisense to CASZ1 |
| ENSG00000220205 | VAMP2 | -0,059101857 | 0,95986149 | 0,78766727 | 0,979462318 | vesicle associated membrane protein 2 |
| ENSG00000234607 | AL355994,4 | -0,059275418 | 0,959746023 | 0,95122734 | 0,996432861 | chromosome 16 open reading frame 52 (C16orf52) pseudogene |
| ENSG00000123124 | WWP1 | -0,059326011 | 0,959712367 | 0,87557224 | 0,990078774 | WW domain containing E3 ubiquitin protein ligase 1 |
| ENSG00000157500 | APPL1 | -0,059411374 | 0,959655583 | 0,89494741 | 0,991303328 | adaptor protein, phosphotyrosine interacting with PH domain and leucine zipper 1 |
| ENSG00000138463 | SLC49A4 | -0,059467235 | 0,959618426 | 0,87670446 | 0,990109814 | solute carrier family 49 member 4 |
| ENSG00000110911 | SLC11A2 | -0,059501501 | 0,959595634 | 0,87258942 | 0,989217334 | solute carrier family 11 member 2 |
| ENSG00000260518 | BMS1P8 | -0,059571629 | 0,959548991 | 0,87402206 | 0,989711917 | BMS1 pseudogene 8 |
| ENSG00000123191 | ATP7B | -0,059577853 | 0,959544851 | 0,78883635 | 0,979462318 | ATPase copper transporting beta |
| ENSG00000285229 | LINCR-0001 | -0,059592458 | 0,959535137 | 0,92026793 | 0,993787755 | uncharacterized LINCR-0001 |
| ENSG00000243024 | AC012158,1 | -0,059642231 | 0,959502033 | 0,95207654 | 0,996432861 | ribosomal protein S11 pseudogene 6 |
| ENSG00000116698 | SMG7 | -0,059711188 | 0,959456173 | 0,84548782 | 0,986639902 | SMG7 nonsense mediated mRNA decay factor |
| ENSG00000206395 | DDAH2 | -0,059772491 | 0,959415404 | 0,9072323 | 0,992590661 | dimethylarginine dimethylaminohydrolase 2 |
| ENSG00000226634 | DDAH2 | -0,059772491 | 0,959415404 | 0,9072323 | 0,992590661 | dimethylarginine dimethylaminohydrolase 2 |
| ENSG00000139517 | LNX2 | -0,059783982 | 0,959407763 | 0,83397671 | 0,984905093 | ligand of numb-protein X 2 |
| ENSG00000205744 | DENND1C | -0,059829724 | 0,959377344 | 0,96116275 | 0,996608711 | DENN domain containing 1C |
| ENSG00000214199 | EEF1A1P12 | -0,059889374 | 0,959337679 | 0,94868353 | 0,996432861 | eukaryotic translation elongation factor 1 alpha 1 pseudogene 12 |
| ENSG00000269386 | RAB11B-AS1 | -0,059931934 | 0,959309378 | 0,84487187 | 0,986557476 | RAB11B antisense RNA 1 |
| ENSG00000213341 | CHUK | -0,059976452 | 0,959279777 | 0,88777984 | 0,990717729 | component of inhibitor of nuclear factor kappa B kinase complex |
| ENSG00000251666 | ZNF346-IT1 | -0,059979876 | 0,9592775 | 0,90282767 | 0,992311172 | ZNF346 intronic transcript 1 |

| | | | | | | |
|---|---|---|---|---|---|---|
| ENSG00000158715 | SLC45A3 | -0,059986318 | 0,959273217 | 0,88876184 | 0,990717729 | solute carrier family 45 member 3 |
| ENSG00000185115 | NSMCE3 | -0,059996155 | 0,959266676 | 0,88383785 | 0,990717729 | NSE3 homolog, SMC5-SMC6 complex component |
| ENSG00000145780 | FEM1C | -0,060212232 | 0,959123015 | 0,89027739 | 0,990717729 | fem-1 homolog C |
| ENSG00000170860 | LSM3 | -0,060254523 | 0,959094899 | 0,88418965 | 0,990717729 | LSM3 homolog, U6 small nuclear RNA and mRNA degradation associated |
| ENSG00000275902 | AC139530,3 | -0,060313164 | 0,959055915 | 0,9714086 | 0,998088605 | novel transcript, antisense to HGS |
| ENSG00000285680 | AL355481,1 | -0,060356654 | 0,959027005 | 0,91901861 | 0,993585187 | novel transcript, antisense to SLITRK1 |
| ENSG00000237788 | AL162615,1 | -0,060439098 | 0,958972203 | 0,89487509 | 0,991296984 | cytokine-like nuclear factor n-pac (N-PAC) pseudogene |
| ENSG00000147118 | ZNF182 | -0,060478407 | 0,958946074 | 0,85827445 | 0,989067345 | zinc finger protein 182 |
| ENSG00000263603 | AC127024,2 | -0,060482132 | 0,958943598 | 0,94019263 | 0,994949796 | novel transcript |
| ENSG00000231268 | AGER | -0,060482352 | 0,958943452 | 0,9378102 | 0,994479239 | advanced glycosylation end-product specific receptor |
| ENSG00000230514 | AGER | -0,060482352 | 0,958943452 | 0,9378102 | 0,994479239 | advanced glycosylation end-product specific receptor |
| ENSG00000275645 | AC068338,3 | -0,060551011 | 0,958897816 | 0,93890763 | 0,994713928 | novel transcript, antisense to NEIL1 |
| ENSG00000109332 | UBE2D3 | -0,060556992 | 0,958893841 | 0,87186724 | 0,989217334 | ubiquitin conjugating enzyme E2 D3 |
| ENSG00000122566 | HNRNPA2B1 | -0,06058376 | 0,958876049 | 0,84445095 | 0,986340494 | heterogeneous nuclear ribonucleoprotein A2/B1 |
| ENSG00000177200 | CHD9 | -0,060591536 | 0,958870881 | 0,86937331 | 0,989074806 | chromodomain helicase DNA binding protein 9 |
| ENSG00000125885 | MCM8 | -0,060672198 | 0,958817271 | 0,82190414 | 0,982952211 | minichromosome maintenance 8 homologous recombination repair factor |
| ENSG00000276880 | IFI27L1 | -0,060774225 | 0,958749467 | 0,88439078 | 0,990717729 | interferon alpha inducible protein 27 like 1 |
| ENSG00000009307 | CSDE1 | -0,0608284 | 0,958713465 | 0,85942534 | 0,989074806 | cold shock domain containing E1 |
| ENSG00000275582 | AL031670,1 | -0,060845709 | 0,958701963 | 0,88184062 | 0,990717729 | novel transcript, antisense to PANK2 |
| ENSG00000276612 | FP565260,2 | -0,060870012 | 0,958685813 | 0,95637915 | 0,996608711 | novel protein |
| ENSG00000230815 | AL807757,1 | -0,0608913 | 0,958671667 | 0,9533788 | 0,99652169 | novel transcript |
| ENSG00000250539 | KRT8P33 | -0,060900524 | 0,958665538 | 0,88210078 | 0,990717729 | keratin 8 pseudogene 33 |
| ENSG00000265778 | AC018413,1 | -0,06093619 | 0,958641838 | 0,88170646 | 0,990717729 | novel transcript, antisense to ZFN156 |
| ENSG00000171928 | TVP23B | -0,060964977 | 0,95862271 | 0,86191025 | 0,989074806 | trans-golgi network vesicle protein 23 homolog B |
| ENSG00000186468 | RPS23 | -0,061113064 | 0,958524316 | 0,85643381 | 0,988229014 | ribosomal protein S23 |
| ENSG00000269967 | AL136115,2 | -0,061197164 | 0,958468442 | 0,9248469 | 0,993915537 | novel transcript |
| ENSG00000269921 | AC068620,1 | -0,061204335 | 0,958463677 | 0,91482199 | 0,993361299 | novel transcript |
| ENSG00000280479 | CACFD1 | -0,061223809 | 0,95845074 | 0,86462568 | 0,989074806 | calcium channel flower domain containing 1 |

| | | | | | | |
|---|---|---|---|---|---|---|
| ENSG00000250240 | AC008840,1 | -0,061295007 | 0,958403441 | 0,90048521 | 0,991865079 | novel transcript |
| ENSG00000172345 | STARD5 | -0,061302637 | 0,958398373 | 0,89773234 | 0,991708791 | StAR related lipid transfer domain containing 5 |
| ENSG00000259456 | ADNP-AS1 | -0,061317939 | 0,958388207 | 0,88846615 | 0,990717729 | ADNP antisense RNA 1 |
| ENSG00000261728 | AL138690,1 | -0,061339342 | 0,958373989 | 0,9534557 | 0,99652169 | novel transcript |
| ENSG00000119242 | CCDC92 | -0,061361178 | 0,958359484 | 0,79187679 | 0,979471509 | coiled-coil domain containing 92 |
| ENSG00000226609 | AL390067,1 | -0,061426646 | 0,958315995 | 0,91928169 | 0,993598006 | novel transcript |
| ENSG00000245532 | NEAT1 | -0,061427608 | 0,958315357 | 0,82730168 | 0,983595244 | nuclear paraspeckle assembly transcript 1 |
| ENSG00000271601 | LIX1L | -0,061510847 | 0,958260066 | 0,81319709 | 0,981246822 | limb and CNS expressed 1 like |
| ENSG00000052344 | PRSS8 | -0,061527742 | 0,958248844 | 0,93258424 | 0,994479239 | serine protease 8 |
| ENSG00000153721 | CNKSR3 | -0,06156773 | 0,958222284 | 0,84359651 | 0,986273696 | CNKSR family member 3 |
| ENSG00000178935 | ZNF552 | -0,061588182 | 0,9582087 | 0,83125311 | 0,98438942 | zinc finger protein 552 |
| ENSG00000134852 | CLOCK | -0,061593659 | 0,958205063 | 0,89546435 | 0,991303328 | clock circadian regulator |
| ENSG00000275272 | YTHDC1 | -0,061635302 | 0,958177404 | 0,812279 | 0,981213877 | YTH domain containing 1 |
| ENSG00000231859 | AC079781,1 | -0,06170475 | 0,958131281 | 0,9171192 | 0,993361299 | novel pseudogene |
| ENSG00000182230 | FAM153B | -0,061818909 | 0,958055468 | 0,95510908 | 0,996608711 | family with sequence similarity 153 member B |
| ENSG00000009413 | REV3L | -0,061835559 | 0,958044411 | 0,83968512 | 0,985672131 | REV3 like, DNA directed polymerase zeta catalytic subunit |
| ENSG00000246334 | PRR7-AS1 | -0,061868441 | 0,958022576 | 0,90129322 | 0,992041918 | PRR7 antisense RNA 1 |
| ENSG00000287855 | AL023882,1 | -0,061972602 | 0,957953411 | 0,93211143 | 0,994479239 | novel transcript, antisense to LMF1 |
| ENSG00000255395 | AP001972,4 | -0,062096622 | 0,957871064 | 0,9072622 | 0,992590661 | novel transcript, antisense to SLCO2B1 |
| ENSG00000284534 | AC092634,7 | -0,062108702 | 0,957863044 | 0,95572849 | 0,996608711 | novel transcript |
| ENSG00000075131 | TIPIN | -0,062117738 | 0,957857044 | 0,85749912 | 0,988891647 | TIMELESS interacting protein |
| ENSG00000184752 | NDUFA12 | -0,062132572 | 0,957847196 | 0,87277212 | 0,989262536 | NADH:ubiquinone oxidoreductase subunit A12 |
| ENSG00000157985 | AGAP1 | -0,06215621 | 0,957831502 | 0,76773238 | 0,977309816 | ArfGAP with GTPase domain, ankyrin repeat and PH domain 1 |
| ENSG00000048649 | RSF1 | -0,062276182 | 0,957751853 | 0,82260835 | 0,98299282 | remodeling and spacing factor 1 |
| ENSG00000065559 | MAP2K4 | -0,062328009 | 0,957717448 | 0,88556424 | 0,990717729 | mitogen-activated protein kinase kinase 4 |
| ENSG00000231969 | MMADHC-DT | -0,062340709 | 0,957709018 | 0,89444399 | 0,991135494 | MMADHC divergent transcript |
| ENSG00000174231 | PRPF8 | -0,062407378 | 0,957664761 | 0,78170917 | 0,978597905 | pre-mRNA processing factor 8 |
| ENSG00000006118 | TMEM132A | -0,062483881 | 0,95761398 | 0,8770707 | 0,990109814 | transmembrane protein 132A |
| ENSG00000081154 | PCNP | -0,062591556 | 0,957542511 | 0,88596987 | 0,990717729 | PEST proteolytic signal containing nuclear protein |
| ENSG00000214226 | C17orf67 | -0,062619396 | 0,957524033 | 0,88593895 | 0,990717729 | chromosome 17 open reading frame 67 |
| ENSG00000073792 | IGF2BP2 | -0,06262974 | 0,957517169 | 0,80056316 | 0,980406861 | insulin like growth factor 2 mRNA binding protein 2 |

| | | | | | | |
|---|---|---|---|---|---|---|
| ENSG00000273184 | AC010655,4 | -0,062723968 | 0,957454631 | 0,91298076 | 0,993361299 | novel transcript, readthrough between HILPDA-METTL2B |
| ENSG00000078687 | TNRC6C | -0,062748103 | 0,957438614 | 0,7923979 | 0,979471509 | trinucleotide repeat containing adaptor 6C |
| ENSG00000241534 | CFB | -0,062754548 | 0,957434337 | 0,92827498 | 0,994108531 | complement factor B |
| ENSG00000272047 | GTF2H5 | -0,062774852 | 0,957420862 | 0,8759783 | 0,990109814 | general transcription factor IIH subunit 5 |
| ENSG00000057608 | GDI2 | -0,062805624 | 0,957400441 | 0,88421743 | 0,990717729 | GDP dissociation inhibitor 2 |
| ENSG00000257390 | AC023055,1 | -0,062864659 | 0,957361265 | 0,90932451 | 0,992864002 | novel protein |
| ENSG00000178074 | C2orf69 | -0,06286762 | 0,9573593 | 0,89846577 | 0,991832796 | chromosome 2 open reading frame 69 |
| ENSG00000155957 | TMBIM4 | -0,062917786 | 0,957326011 | 0,8882946 | 0,990717729 | transmembrane BAX inhibitor motif containing 4 |
| ENSG00000132475 | H3-3B | -0,062923627 | 0,957322135 | 0,83603333 | 0,98514044 | H3,3 histone B |
| ENSG00000259899 | AC010333,2 | -0,062982037 | 0,957283377 | 0,964858 | 0,997113804 | novel transcript, antisense to a novel protein |
| ENSG00000088992 | TESC | -0,063026745 | 0,957253712 | 0,94009465 | 0,994949796 | tescalcin |
| ENSG00000234456 | MAGI2-AS3 | -0,063126568 | 0,95718748 | 0,8706288 | 0,989074806 | MAGI2 antisense RNA 3 |
| ENSG00000188996 | HUS1B | -0,063147825 | 0,957173377 | 0,93060563 | 0,994344567 | HUS1 checkpoint clamp component B |
| ENSG00000223815 | DIAPH3-AS2 | -0,063158953 | 0,957165994 | 0,94784023 | 0,996176992 | DIAPH3 antisense RNA 2 |
| ENSG00000215425 | DDX39B | -0,063231367 | 0,957117951 | 0,7783663 | 0,978587255 | DExD-box helicase 39B |
| ENSG00000257017 | HP | -0,063248258 | 0,957106746 | 0,91275603 | 0,993361299 | haptoglobin |
| ENSG00000110871 | COQ5 | -0,063254129 | 0,957102851 | 0,82373163 | 0,983374097 | coenzyme Q5, methyltransferase |
| ENSG00000143921 | ABCG8 | -0,063307463 | 0,957067469 | 0,90351458 | 0,992361703 | ATP binding cassette subfamily G member 8 |
| ENSG00000237211 | SETP4 | -0,063310898 | 0,95706519 | 0,95177268 | 0,996432861 | SET pseudogene 4 |
| ENSG00000183426 | NPIPA1 | -0,063347611 | 0,957040836 | 0,79148729 | 0,979471509 | nuclear pore complex interacting protein family member A1 |
| ENSG00000170540 | ARL6IP1 | -0,063442135 | 0,956978133 | 0,86565116 | 0,989074806 | ADP ribosylation factor like GTPase 6 interacting protein 1 |
| ENSG00000183186 | C2CD4C | -0,0636152 | 0,956863341 | 0,88541154 | 0,990717729 | C2 calcium dependent domain containing 4C |
| ENSG00000141956 | PRDM15 | -0,063620094 | 0,956860096 | 0,79806512 | 0,980161849 | PR/SET domain 15 |
| ENSG00000285936 | AL392024,1 | -0,063646144 | 0,956842819 | 0,94654573 | 0,996027077 | novel transcript, antisense to DPF3 |
| ENSG00000225648 | SBDSP1 | -0,063679762 | 0,956820522 | 0,8109216 | 0,981213877 | SBDS pseudogene 1 |
| ENSG00000266420 | RN7SL118P | -0,06375709 | 0,956769238 | 0,9572784 | 0,996608711 | RNA, 7SL, cytoplasmic 118, pseudogene |
| ENSG00000188368 | PRR19 | -0,063916816 | 0,956663317 | 0,89190033 | 0,990986278 | proline rich 19 |
| ENSG00000237887 | RPL23AP32 | -0,063933637 | 0,956652163 | 0,93112743 | 0,994433362 | ribosomal protein L23a pseudogene 32 |
| ENSG00000273792 | AC022898,2 | -0,064039069 | 0,956582253 | 0,93805849 | 0,994488324 | novel transcript, sense intronic to RORA |
| ENSG00000282744 | AC246789,2 | -0,064153053 | 0,956506679 | 0,87472266 | 0,990078774 | novel transcript |

| | | | | | | |
|---|---|---|---|---|---|---|
| ENSG00000274010 | ZFP91P1 | -0,064187521 | 0,956483827 | 0,915214 | 0,993361299 | ZFP91 pseudogene 1 |
| ENSG00000179059 | ZFP42 | -0,064230435 | 0,956455376 | 0,93563727 | 0,994479239 | ZFP42 zinc finger protein |
| ENSG00000259407 | AC021739,2 | -0,064277378 | 0,956424255 | 0,87790285 | 0,990109814 | novel transcript, antisense to AKAP13 |
| ENSG00000183273 | CCDC60 | -0,064282911 | 0,956420587 | 0,92198067 | 0,993797669 | coiled-coil domain containing 60 |
| ENSG00000204394 | VARS1 | -0,064312296 | 0,956401106 | 0,86460829 | 0,989074806 | valyl-tRNA synthetase 1 |
| ENSG00000085511 | MAP3K4 | -0,064349333 | 0,956376554 | 0,8259241 | 0,983383773 | mitogen-activated protein kinase kinase kinase 4 |
| ENSG00000196757 | ZNF700 | -0,064494347 | 0,956280427 | 0,84561233 | 0,986639902 | zinc finger protein 700 |
| ENSG00000256975 | AC092821,2 | -0,064532749 | 0,956254974 | 0,95181778 | 0,996432861 | RAN binding protein (RANBP) pseudogene |
| ENSG00000242419 | PCDHGC4 | -0,064541825 | 0,956248957 | 0,77718253 | 0,978587255 | protocadherin gamma subfamily C, 4 |
| ENSG00000269066 | AC020908,3 | -0,064544902 | 0,956246918 | 0,9534505 | 0,99652169 | novel transcript, antisense MYO9B |
| ENSG00000255072 | PIGY | -0,064622009 | 0,956195811 | 0,86031638 | 0,989074806 | phosphatidylinositol glycan anchor biosynthesis class Y |
| ENSG00000145337 | PYURF | -0,064622009 | 0,956195811 | 0,86031638 | 0,989074806 | PIGY upstream reading frame |
| ENSG00000207445 | SNORD15B | -0,064626722 | 0,956192688 | 0,91648891 | 0,993361299 | small nucleolar RNA, C/D box 15B |
| ENSG00000176371 | ZSCAN2 | -0,064630639 | 0,956190092 | 0,76245354 | 0,976315588 | zinc finger and SCAN domain containing 2 |
| ENSG00000233668 | AL353662,1 | -0,064641007 | 0,95618322 | 0,93672687 | 0,994479239 | ribosomal protein S8 (RPS8) pseudogene |
| ENSG00000164683 | HEY1 | -0,064652884 | 0,956175348 | 0,86512236 | 0,989074806 | hes related family bHLH transcription factor with YRPW motif 1 |
| ENSG00000277806 | AC006213,4 | -0,064678612 | 0,956158296 | 0,92895596 | 0,994278638 | novel transcript, sense intronic to ZNF155 |
| ENSG00000237940 | LINC01238 | -0,06472545 | 0,956127254 | 0,92571986 | 0,993915537 | long intergenic non-protein coding RNA 1238 |
| ENSG00000269489 | AL589765,6 | -0,06477732 | 0,956092879 | 0,96699353 | 0,997263201 | novel transcript |
| ENSG00000251484 | AL356488,1 | -0,064779841 | 0,956091209 | 0,89444076 | 0,991135494 | poly(rC) binding protein 1 (PCBP1) pseudogene |
| ENSG00000198786 | MT-ND5 | -0,064835761 | 0,95605415 | 0,79991618 | 0,980161849 | mitochondrially encoded NADH:ubiquinone oxidoreductase core subunit 5 |
| ENSG00000235299 | MRPL53P1 | -0,064861372 | 0,956037179 | 0,94251767 | 0,995390305 | mitochondrial ribosomal protein L53 pseudogene 1 |
| ENSG00000272498 | AC090948,2 | -0,064969906 | 0,955965259 | 0,89771442 | 0,991708791 | novel transcript, antisense to RFTN1 |
| ENSG00000189339 | SLC35E2B | -0,064995769 | 0,955948121 | 0,74417215 | 0,975627885 | solute carrier family 35 member E2B |
| ENSG00000117859 | OSBPL9 | -0,065102817 | 0,955877192 | 0,84338813 | 0,986260952 | oxysterol binding protein like 9 |
| ENSG00000237373 | BRWD1-IT1 | -0,065130319 | 0,955858971 | 0,93486092 | 0,994479239 | BRWD1 intronic transcript 1 |
| ENSG00000236432 | MFF-DT | -0,065158289 | 0,955840439 | 0,8438657 | 0,986273696 | MFF divergent transcript |
| ENSG00000184831 | APOO | -0,065275874 | 0,955762538 | 0,88218807 | 0,990717729 | apolipoprotein O |
| ENSG00000267022 | AC067968,1 | -0,065303174 | 0,955744453 | 0,86821439 | 0,989074806 | novel protein |
| ENSG00000287766 | AC006130,3 | -0,065323344 | 0,95573109 | 0,92204942 | 0,993797669 | novel transcript |

| | | | | | | |
|---|---|---|---|---|---|---|
| ENSG00000122557 | HERPUD2 | -0,065391634 | 0,955685852 | 0,84300202 | 0,986218419 | HERPUD family member 2 |
| ENSG00000258653 | AC005520,1 | -0,065419192 | 0,955667597 | 0,912944 | 0,993361299 | novel protein |
| ENSG00000135316 | SYNCRIP | -0,065425193 | 0,955663621 | 0,83722593 | 0,985577601 | synaptotagmin binding cytoplasmic RNA interacting protein |
| ENSG00000149679 | CABLES2 | -0,065475869 | 0,955630054 | 0,81023729 | 0,981213877 | Cdk5 and Abl enzyme substrate 2 |
| ENSG00000279243 | LINC02736 | -0,06547635 | 0,955629735 | 0,93129932 | 0,994433362 | long intergenic non-protein coding RNA 2736 |
| ENSG00000224843 | LINC00240 | -0,065511707 | 0,955606315 | 0,89387892 | 0,991135494 | long intergenic non-protein coding RNA 240 |
| ENSG00000196214 | ZNF766 | -0,065535312 | 0,95559068 | 0,82219616 | 0,982952211 | zinc finger protein 766 |
| ENSG00000117153 | KLHL12 | -0,065560815 | 0,955573787 | 0,82141324 | 0,982950156 | kelch like family member 12 |
| ENSG00000233450 | KIFC1 | -0,065566045 | 0,955570324 | 0,90708267 | 0,992590661 | kinesin family member C1 |
| ENSG00000106346 | USP42 | -0,065601994 | 0,955546513 | 0,80526688 | 0,981209129 | ubiquitin specific peptidase 42 |
| ENSG00000108423 | TUBD1 | -0,065613231 | 0,95553907 | 0,82589158 | 0,983383773 | tubulin delta 1 |
| ENSG00000261609 | GAN | -0,06562057 | 0,955534209 | 0,81326225 | 0,981275558 | gigaxonin |
| ENSG00000225300 | AL591623,1 | -0,065627928 | 0,955529336 | 0,86157526 | 0,989074806 | novel transcript |
| ENSG00000160949 | TONSL | -0,065763228 | 0,955439728 | 0,87669612 | 0,990109814 | tonsoku like, DNA repair protein |
| ENSG00000141499 | WRAP53 | -0,06586108 | 0,955374927 | 0,82011149 | 0,982609114 | WD repeat containing antisense to TP53 |
| ENSG00000139636 | LMBR1L | -0,065925003 | 0,955332597 | 0,77782678 | 0,978587255 | limb development membrane protein 1 like |
| ENSG00000285350 | FO681525,1 | -0,065974159 | 0,955300047 | 0,96085519 | 0,996608711 | chymotrypsinogen B (CTRB) pseudogene |
| ENSG00000274333 | CU633967,1 | -0,065999622 | 0,955283187 | 0,8968122 | 0,991644439 | uncharacterized LOC102724219 |
| ENSG00000143457 | GOLPH3L | -0,066033125 | 0,955261003 | 0,86844325 | 0,989074806 | golgi phosphoprotein 3 like |
| ENSG00000145723 | GIN1 | -0,066068844 | 0,955237352 | 0,87860468 | 0,990109814 | gypsy retrotransposon integrase 1 |
| ENSG00000105640 | RPL18A | -0,066097278 | 0,955218526 | 0,81651288 | 0,981900869 | ribosomal protein L18a |
| ENSG00000287153 | AL161851,1 | -0,066155573 | 0,955179929 | 0,92213395 | 0,993797669 | novel transcript, antisense to NPAS3 |
| ENSG00000012660 | ELOVL5 | -0,066156021 | 0,955179633 | 0,88708452 | 0,990717729 | ELOVL fatty acid elongase 5 |
| ENSG00000235272 | RAMACL | -0,06616065 | 0,955176568 | 0,8533645 | 0,987861659 | RNA guanine-7 methyltransferase activating subunit like (pseudogene) |
| ENSG00000184207 | PGP | -0,066216336 | 0,9551397 | 0,7885565 | 0,979462318 | phosphoglycolate phosphatase |
| ENSG00000128346 | C22orf23 | -0,066240196 | 0,955123904 | 0,80875912 | 0,981213877 | chromosome 22 open reading frame 23 |
| ENSG00000131023 | LATS1 | -0,066381777 | 0,955030176 | 0,85565388 | 0,988075747 | large tumor suppressor kinase 1 |
| ENSG00000062582 | MRPS24 | -0,066392678 | 0,955022959 | 0,78275588 | 0,978687331 | mitochondrial ribosomal protein S24 |
| ENSG00000253327 | RAD21-AS1 | -0,066403154 | 0,955016024 | 0,87701258 | 0,990109814 | RAD21 antisense RNA 1 |
| ENSG00000155330 | C16orf87 | -0,066430215 | 0,954998111 | 0,83310496 | 0,984905093 | chromosome 16 open reading frame 87 |
| ENSG00000196951 | SCOC-AS1 | -0,06645481 | 0,954981831 | 0,90749277 | 0,992590661 | SCOC antisense RNA 1 |

| | | | | | | |
|---|---|---|---|---|---|---|
| ENSG00000234171 | RNASEH1-AS1 | -0,066525706 | 0,954934903 | 0,7798466 | 0,978587255 | RNASEH1 antisense RNA 1 |
| ENSG00000237973 | MTCO1P12 | -0,066677202 | 0,954834631 | 0,8884382 | 0,990717729 | MT-CO1 pseudogene 12 |
| ENSG00000049192 | ADAMTS6 | -0,066691822 | 0,954824955 | 0,87705511 | 0,990109814 | ADAM metallopeptidase with thrombospondin type 1 motif 6 |
| ENSG00000137601 | NEK1 | -0,066743342 | 0,954790858 | 0,84555046 | 0,986639902 | NIMA related kinase 1 |
| ENSG00000144228 | SPOPL | -0,066749333 | 0,954786893 | 0,89360581 | 0,991135494 | speckle type BTB/POZ protein like |
| ENSG00000257277 | AC092652,1 | -0,066881358 | 0,954699522 | 0,96017821 | 0,996608711 | novel transcript, antisense to ARHGAP15 |
| ENSG00000108510 | MED13 | -0,066928568 | 0,954668281 | 0,86074482 | 0,989074806 | mediator complex subunit 13 |
| ENSG00000159873 | CCDC117 | -0,067087499 | 0,954563118 | 0,85838094 | 0,989067345 | coiled-coil domain containing 117 |
| ENSG00000280831 | RPS25 | -0,067155081 | 0,954518404 | 0,83050135 | 0,984156028 | ribosomal protein S25 |
| ENSG00000118181 | RPS25 | -0,067155081 | 0,954518404 | 0,83050135 | 0,984156028 | ribosomal protein S25 |
| ENSG00000108468 | CBX1 | -0,067165385 | 0,954511586 | 0,81706598 | 0,981979397 | chromobox 1 |
| ENSG00000273026 | AL358472,3 | -0,067199199 | 0,954489215 | 0,90476093 | 0,992590661 | novel transcript, antisense to SLC39A1 |
| ENSG00000231066 | NPM1P9 | -0,06724728 | 0,954457405 | 0,90943334 | 0,992864002 | nucleophosmin 1 pseudogene 9 |
| ENSG00000276681 | LENG8 | -0,067276489 | 0,954438081 | 0,83731853 | 0,985577601 | leukocyte receptor cluster member 8 |
| ENSG00000171793 | CTPS1 | -0,067313923 | 0,954413316 | 0,76730296 | 0,977201891 | CTP synthase 1 |
| ENSG00000225683 | PACRG-AS3 | -0,067349176 | 0,954389995 | 0,91549498 | 0,993361299 | PACRG antisense RNA 3 |
| ENSG00000278486 | YRDCP1 | -0,067467706 | 0,954311587 | 0,93186778 | 0,994433362 | yrdC N(6)-threonylcarbamoyltransferase domain containing pseudogene 1 |
| ENSG00000198496 | NBR2 | -0,067523705 | 0,954274545 | 0,81717717 | 0,9820198 | neighbor of BRCA1 lncRNA 2 |
| ENSG00000185710 | SMG1P4 | -0,067652292 | 0,954189494 | 0,79401029 | 0,979776981 | SMG1 pseudogene 4 |
| ENSG00000173334 | TRIB1 | -0,067684613 | 0,954168118 | 0,77293791 | 0,978310326 | tribbles pseudokinase 1 |
| ENSG00000271380 | AL451085,2 | -0,067705038 | 0,95415461 | 0,85777266 | 0,988915281 | uncharacterized LOC101928120 |
| ENSG00000277741 | GOLGA6L17P | -0,06771535 | 0,954147789 | 0,96319992 | 0,996608711 | golgin A6 family like 17, pseudogene |
| ENSG00000229043 | ZFAND2A-DT | -0,067724765 | 0,954141563 | 0,81135318 | 0,981213877 | ZFAND2A divergent transcript |
| ENSG00000230650 | AC140479,2 | -0,06773679 | 0,95413361 | 0,93907523 | 0,994790361 | zinc finger protein 532 (ZNF532) pseudogene |
| ENSG00000105186 | ANKRD27 | -0,067796402 | 0,954094186 | 0,77228702 | 0,97820249 | ankyrin repeat domain 27 |
| ENSG00000253326 | AL606534,4 | -0,067853122 | 0,954056676 | 0,92248887 | 0,993797669 | novel pseudogene |
| ENSG00000272267 | AC021242,3 | -0,06787071 | 0,954045045 | 0,88456868 | 0,990717729 | novel transcript, antisense to TNKS |
| ENSG00000067191 | CACNB1 | -0,067961767 | 0,953984832 | 0,79121044 | 0,979471509 | calcium voltage-gated channel auxiliary subunit beta 1 |
| ENSG00000099901 | RANBP1 | -0,067968681 | 0,95398026 | 0,75263705 | 0,975901808 | RAN binding protein 1 |
| ENSG00000275482 | EPHB6 | -0,068140488 | 0,953866659 | 0,86836025 | 0,989074806 | EPH receptor B6 |

| | | | | | | |
|---|---|---|---|---|---|---|
| ENSG00000279247 | AK6 | -0,068143652 | 0,953864567 | 0,89570925 | 0,991377987 | adenylate kinase 6 |
| ENSG00000272473 | AC006273,1 | -0,068235708 | 0,953803705 | 0,93008712 | 0,994331958 | novel transcript |
| ENSG00000140463 | BBS4 | -0,068261377 | 0,953786735 | 0,85105416 | 0,987259878 | Bardet-Biedl syndrome 4 |
| ENSG00000129595 | EPB41L4A | -0,068301538 | 0,953760184 | 0,86028644 | 0,989074806 | erythrocyte membrane protein band 4,1 like 4A |
| ENSG00000108840 | HDAC5 | -0,068311299 | 0,953753731 | 0,80867673 | 0,981213877 | histone deacetylase 5 |
| ENSG00000185483 | ROR1 | -0,068337302 | 0,953736541 | 0,83057375 | 0,984156028 | receptor tyrosine kinase like orphan receptor 1 |
| ENSG00000275846 | AL513548,3 | -0,068345075 | 0,953731402 | 0,91097567 | 0,993222655 | novel transcript |
| ENSG00000155313 | USP25 | -0,06834632 | 0,953730579 | 0,86855351 | 0,989074806 | ubiquitin specific peptidase 25 |
| ENSG00000167637 | ZNF283 | -0,068615586 | 0,953552591 | 0,75260592 | 0,975901808 | zinc finger protein 283 |
| ENSG00000277946 | RN7SL478P | -0,068675243 | 0,953513161 | 0,93178782 | 0,994433362 | RNA, 7SL, cytoplasmic 478, pseudogene |
| ENSG00000171827 | ZNF570 | -0,068794127 | 0,953434591 | 0,77745608 | 0,978587255 | zinc finger protein 570 |
| ENSG00000226334 | AL359182,1 | -0,068794914 | 0,953434071 | 0,93702156 | 0,994479239 | novel transcript |
| ENSG00000140612 | SEC11A | -0,068828372 | 0,95341196 | 0,83881236 | 0,985672131 | SEC11 homolog A, signal peptidase complex subunit |
| ENSG00000135776 | ABCB10 | -0,068891878 | 0,953369992 | 0,78549474 | 0,979320406 | ATP binding cassette subfamily B member 10 |
| ENSG00000163257 | DCAF16 | -0,068915385 | 0,953354459 | 0,76810953 | 0,977377956 | DDB1 and CUL4 associated factor 16 |
| ENSG00000149196 | HIKESHI | -0,068931662 | 0,953343703 | 0,85116432 | 0,987259878 | heat shock protein nuclear import factor hikeshi |
| ENSG00000163686 | ABHD6 | -0,069023853 | 0,953282784 | 0,80330084 | 0,980866933 | abhydrolase domain containing 6, acylglycerol lipase |
| ENSG00000276390 | AC004241,3 | -0,069060715 | 0,953258427 | 0,94986276 | 0,996432861 | novel transcript, sense intronic to RPAP3 |
| ENSG00000166946 | CCNDBP1 | -0,069063557 | 0,953256549 | 0,75697104 | 0,975901808 | cyclin D1 binding protein 1 |
| ENSG00000286590 | AL132775,1 | -0,06913876 | 0,953206861 | 0,95584702 | 0,996608711 | novel transcript |
| ENSG00000189159 | JPT1 | -0,069175073 | 0,953182869 | 0,76161661 | 0,976285211 | Jupiter microtubule associated homolog 1 |
| ENSG00000204427 | ABHD16A | -0,069175335 | 0,953182695 | 0,88847213 | 0,990717729 | abhydrolase domain containing 16A, phospholipase |
| ENSG00000233013 | FAM157B | -0,069241822 | 0,953138768 | 0,91670562 | 0,993361299 | family with sequence similarity 157 member B |
| ENSG00000066422 | ZBTB11 | -0,069283687 | 0,95311111 | 0,85251727 | 0,98761642 | zinc finger and BTB domain containing 11 |
| ENSG00000260572 | AC069224,1 | -0,06930987 | 0,953093812 | 0,87892639 | 0,990109814 | novel transcript, antisense to PPM1L |
| ENSG00000156928 | MALSU1 | -0,069349763 | 0,953067458 | 0,85639275 | 0,988229014 | mitochondrial assembly of ribosomal large subunit 1 |
| ENSG00000163510 | CWC22 | -0,069358157 | 0,953061913 | 0,85520118 | 0,988016959 | CWC22 spliceosome associated protein homolog |
| ENSG00000130332 | LSM7 | -0,069393397 | 0,953038633 | 0,76855644 | 0,977381742 | LSM7 homolog, U6 small nuclear RNA and mRNA degradation associated |
| ENSG00000007516 | BAIAP3 | -0,069494492 | 0,952971853 | 0,85263032 | 0,98761642 | BAI1 associated protein 3 |
| ENSG00000204228 | HSD17B8 | -0,069506529 | 0,952963902 | 0,86577457 | 0,989074806 | hydroxysteroid 17-beta dehydrogenase 8 |
| ENSG00000135387 | CAPRIN1 | -0,069526494 | 0,952950714 | 0,84393055 | 0,986273696 | cell cycle associated protein 1 |
| ENSG00000092820 | EZR | -0,069538652 | 0,952942684 | 0,84188508 | 0,98571999 | ezrin |

| | | | | | | |
|---|---|---|---|---|---|---|
| ENSG00000214184 | GCC2-AS1 | -0,069590599 | 0,952908371 | 0,93333762 | 0,994479239 | GCC2 antisense RNA 1 |
| ENSG00000257337 | AC068888,1 | -0,069611348 | 0,952894667 | 0,84379318 | 0,986273696 | novel transcript, antisense to TENC1 & EIF4B |
| ENSG00000260796 | AC145285,3 | -0,069620675 | 0,952888506 | 0,93351583 | 0,994479239 | novel transcript, antisense to ATXN2L |
| ENSG00000182093 | GET1 | -0,06971171 | 0,95282838 | 0,86079947 | 0,989074806 | guided entry of tail-anchored proteins factor 1 |
| ENSG00000136918 | WDR38 | -0,06974388 | 0,952807134 | 0,93251307 | 0,994479239 | WD repeat domain 38 |
| ENSG00000133641 | C12orf29 | -0,069748284 | 0,952804225 | 0,8819248 | 0,990717729 | chromosome 12 open reading frame 29 |
| ENSG00000170871 | KIAA0232 | -0,069958636 | 0,952665312 | 0,83658841 | 0,985423106 | KIAA0232 |
| ENSG00000075239 | ACAT1 | -0,069976954 | 0,952653216 | 0,858181 | 0,989067345 | acetyl-CoA acetyltransferase 1 |
| ENSG00000269893 | SNHG8 | -0,070064504 | 0,952595406 | 0,81414034 | 0,981521413 | small nucleolar RNA host gene 8 |
| ENSG00000162244 | RPL29 | -0,070069042 | 0,95259241 | 0,85193866 | 0,987322912 | ribosomal protein L29 |
| ENSG00000262860 | LSM14A | -0,070069379 | 0,952592187 | 0,86431888 | 0,989074806 | LSM14A mRNA processing body assembly factor |
| ENSG00000162714 | ZNF496 | -0,070078793 | 0,952585971 | 0,79479442 | 0,979776981 | zinc finger protein 496 |
| ENSG00000171357 | LURAP1 | -0,070113117 | 0,952563308 | 0,84258953 | 0,986175464 | leucine rich adaptor protein 1 |
| ENSG00000127554 | GFER | -0,070165083 | 0,952528997 | 0,83790067 | 0,98566906 | growth factor, augmenter of liver regeneration |
| ENSG00000278540 | ACACA | -0,070218981 | 0,952493412 | 0,77812859 | 0,978587255 | acetyl-CoA carboxylase alpha |
| ENSG00000011198 | ABHD5 | -0,070222773 | 0,952490909 | 0,85473954 | 0,987861659 | abhydrolase domain containing 5, lysophosphatidic acid acyltransferase |
| ENSG00000143702 | CEP170 | -0,070281458 | 0,952452165 | 0,80615394 | 0,981213877 | centrosomal protein 170 |
| ENSG00000124444 | ZNF576 | -0,070301438 | 0,952438974 | 0,78325471 | 0,978919905 | zinc finger protein 576 |
| ENSG00000125743 | SNRPD2 | -0,070313212 | 0,952431201 | 0,83446406 | 0,984905093 | small nuclear ribonucleoprotein D2 polypeptide |
| ENSG00000100503 | NIN | -0,070356346 | 0,952402725 | 0,79254683 | 0,979471509 | ninein |
| ENSG00000238086 | PPP1R26P1 | -0,070365105 | 0,952396943 | 0,88755595 | 0,990717729 | protein phosphatase 1 regulatory subunit 26 pseudogene 1 |
| ENSG00000204620 | AC115618,1 | -0,070374887 | 0,952390485 | 0,90484469 | 0,992590661 | novel transcript, antisense to RBM3 |
| ENSG00000107186 | MPDZ | -0,070385609 | 0,952383407 | 0,82534046 | 0,983383773 | multiple PDZ domain crumbs cell polarity complex component |
| ENSG00000155511 | GRIA1 | -0,070397219 | 0,952375743 | 0,81366621 | 0,981497816 | glutamate ionotropic receptor AMPA type subunit 1 |
| ENSG00000099822 | HCN2 | -0,070449375 | 0,952341314 | 0,89889946 | 0,991832796 | hyperpolarization activated cyclic nucleotide gated potassium and sodium channel 2 |
| ENSG00000177932 | ZNF354C | -0,070513299 | 0,952299118 | 0,86096393 | 0,989074806 | zinc finger protein 354C |
| ENSG00000114268 | PFKFB4 | -0,07053214 | 0,952286681 | 0,78384665 | 0,979239248 | 6-phosphofructo-2-kinase/fructose-2,6-biphosphatase 4 |
| ENSG00000123179 | EBPL | -0,070605326 | 0,952238374 | 0,76832465 | 0,977381742 | EBP like |

| | | | | | | |
|---|---|---|---|---|---|---|
| ENSG00000134255 | CEPT1 | -0,070628583 | 0,952223023 | 0,87167677 | 0,989217334 | choline/ethanolamine phosphotransferase 1 |
| ENSG00000237159 | CNTFR-AS1 | -0,07065687 | 0,952204354 | 0,90245083 | 0,992230188 | CNTFR antisense RNA 1 |
| ENSG00000198455 | ZXDB | -0,070662046 | 0,952200938 | 0,81754818 | 0,982201377 | zinc finger X-linked duplicated B |
| ENSG00000263412 | NFE2L1-DT | -0,070714811 | 0,952166112 | 0,85519044 | 0,988016959 | NFE2L1 divergent transcript |
| ENSG00000024526 | DEPDC1 | -0,070739501 | 0,952149817 | 0,8853518 | 0,990717729 | DEP domain containing 1 |
| ENSG00000113360 | DROSHA | -0,070747114 | 0,952144793 | 0,80672331 | 0,981213877 | drosha ribonuclease III |
| ENSG00000271670 | AC010998,2 | -0,070807142 | 0,952105177 | 0,93266365 | 0,994479239 | novel transcript, antisense to WDR11 |
| ENSG00000162775 | RBM15 | -0,070814228 | 0,9521005 | 0,72428118 | 0,973178901 | RNA binding motif protein 15 |
| ENSG00000242412 | DBIL5P2 | -0,070840945 | 0,952082868 | 0,92131676 | 0,993787755 | diazepam binding inhibitor-like 5 pseudogene 2 |
| ENSG00000086189 | DIMT1 | -0,070848026 | 0,952078196 | 0,88492886 | 0,990717729 | DIMT1 rRNA methyltransferase and ribosome maturation factor |
| ENSG00000262833 | AC016245,1 | -0,070907737 | 0,952038791 | 0,90345646 | 0,992361703 | novel transcript, antisense to RPTOR |
| ENSG00000146872 | TLK2 | -0,070969153 | 0,951998264 | 0,81372795 | 0,981521413 | tousled like kinase 2 |
| ENSG00000197763 | TXNRD3 | -0,071052879 | 0,951943016 | 0,8084945 | 0,981213877 | thioredoxin reductase 3 |
| ENSG00000234284 | ZNF879 | -0,071120367 | 0,951898487 | 0,86631599 | 0,989074806 | zinc finger protein 879 |
| ENSG00000202058 | RN7SKP80 | -0,07120513 | 0,951842561 | 0,90385249 | 0,992462326 | RN7SK pseudogene 80 |
| ENSG00000214787 | MS4A4E | -0,071230708 | 0,951825686 | 0,83654275 | 0,985423106 | membrane spanning 4-domains A4E |
| ENSG00000062650 | WAPL | -0,071243924 | 0,951816966 | 0,84205215 | 0,985804848 | WAPL cohesin release factor |
| ENSG00000165512 | ZNF22 | -0,071282091 | 0,951791786 | 0,88174758 | 0,990717729 | zinc finger protein 22 |
| ENSG00000232374 | GPR79 | -0,071287937 | 0,951787929 | 0,9539806 | 0,996608711 | G protein-coupled receptor 79, pseudogene |
| ENSG00000153485 | TMEM251 | -0,071401812 | 0,951712805 | 0,83043157 | 0,984156028 | transmembrane protein 251 |
| ENSG00000276412 | AL591926,2 | -0,071443465 | 0,951685329 | 0,9104855 | 0,99320976 | novel transcript |
| ENSG00000164219 | PGGT1B | -0,071451093 | 0,951680296 | 0,87389086 | 0,989711917 | protein geranylgeranyltransferase type I subunit beta |
| ENSG00000263874 | LINC00672 | -0,071476499 | 0,951663537 | 0,79817922 | 0,980161849 | long intergenic non-protein coding RNA 672 |
| ENSG00000122378 | PRXL2A | -0,071494015 | 0,951651983 | 0,73994901 | 0,974715504 | peroxiredoxin like 2A |
| ENSG00000119048 | UBE2B | -0,071507704 | 0,951642954 | 0,8987318 | 0,991832796 | ubiquitin conjugating enzyme E2 B |
| ENSG00000112118 | MCM3 | -0,071535083 | 0,951624894 | 0,73129939 | 0,974353474 | minichromosome maintenance complex component 3 |
| ENSG00000116147 | TNR | -0,071541382 | 0,951620739 | 0,93161058 | 0,994433362 | tenascin R |
| ENSG00000160124 | CCDC58 | -0,071554987 | 0,951611765 | 0,86221465 | 0,989074806 | coiled-coil domain containing 58 |
| ENSG00000011347 | SYT7 | -0,071609911 | 0,951575537 | 0,78642337 | 0,979320406 | synaptotagmin 7 |
| ENSG00000254132 | MTND6P3 | -0,071621259 | 0,951568052 | 0,87756097 | 0,990109814 | MT-ND6 pseudogene 3 |

| | | | | | | |
|---|---|---|---|---|---|---|
| ENSG00000241651 | AC099340,1 | -0,071665381 | 0,951538951 | 0,93671115 | 0,994479239 | ribosomal protein L15 (RPL15) pseudogene |
| ENSG00000150977 | RILPL2 | -0,07180196 | 0,951448874 | 0,81677346 | 0,981977498 | Rab interacting lysosomal protein like 2 |
| ENSG00000072274 | TFRC | -0,071819652 | 0,951437206 | 0,87223642 | 0,989217334 | transferrin receptor |
| ENSG00000232855 | AF165147,1 | -0,071831854 | 0,951429159 | 0,82878376 | 0,983795764 | novel transcript |
| ENSG00000121486 | TRMT1L | -0,071832017 | 0,951429051 | 0,83958467 | 0,985672131 | tRNA methyltransferase 1 like |
| ENSG00000215251 | FASTKD5 | -0,071865082 | 0,951407246 | 0,81503452 | 0,981521413 | FAST kinase domains 5 |
| ENSG00000226754 | AL606760,1 | -0,071896827 | 0,951386311 | 0,86149017 | 0,989074806 | novel transcript, antisense to MAGOH |
| ENSG00000125124 | BBS2 | -0,071977197 | 0,951333313 | 0,80160789 | 0,980623173 | Bardet-Biedl syndrome 2 |
| ENSG00000248874 | C5orf17 | -0,071984559 | 0,951328458 | 0,92508256 | 0,993915537 | chromosome 5 putative open reading frame 17 |
| ENSG00000237765 | FAM200B | -0,071986617 | 0,951327101 | 0,88063373 | 0,990507013 | family with sequence similarity 200 member B |
| ENSG00000204899 | MZT1 | -0,072020916 | 0,951304484 | 0,9116442 | 0,993342046 | mitotic spindle organizing protein 1 |
| ENSG00000276342 | TMEM179 | -0,072021892 | 0,951303841 | 0,93783482 | 0,994479239 | transmembrane protein 179 |
| ENSG00000169255 | B3GALNT1 | -0,07202979 | 0,951298633 | 0,82225076 | 0,982952211 | beta-1,3-N-acetylgalactosaminyltransferase 1 (globoside blood group) |
| ENSG00000259612 | EEF1A1P22 | -0,07205477 | 0,951282161 | 0,87535839 | 0,990078774 | eukaryotic translation elongation factor 1 alpha 1 pseudogene 22 |
| ENSG00000040199 | PHLPP2 | -0,072123537 | 0,951236819 | 0,75185775 | 0,975901808 | PH domain and leucine rich repeat protein phosphatase 2 |
| ENSG00000065883 | CDK13 | -0,0721498 | 0,951219503 | 0,74487495 | 0,975649186 | cyclin dependent kinase 13 |
| ENSG00000185942 | NKAIN3 | -0,072230466 | 0,951166318 | 0,83034133 | 0,984156028 | sodium/potassium transporting ATPase interacting 3 |
| ENSG00000256294 | ZNF225 | -0,072288522 | 0,951128043 | 0,85172505 | 0,987289464 | zinc finger protein 225 |
| ENSG00000171453 | POLR1C | -0,072300875 | 0,951119899 | 0,76115854 | 0,97624106 | RNA polymerase I and III subunit C |
| ENSG00000267390 | AC036176,1 | -0,072321244 | 0,951106471 | 0,85545139 | 0,988075747 | novel transcript, antisense to KDSR |
| ENSG00000239756 | PHF1 | -0,072321426 | 0,951106351 | 0,89224495 | 0,991039277 | PHD finger protein 1 |
| ENSG00000225855 | RUSC1-AS1 | -0,072371746 | 0,951073177 | 0,83066362 | 0,984156028 | RUSC1 antisense RNA 1 |
| ENSG00000244115 | DNAJC25-GNG1 | -0,072401232 | 0,951053739 | 0,82005981 | 0,982609114 | DNAJC25-GNG10 readthrough |
| ENSG00000135069 | PSAT1 | -0,072405139 | 0,951051164 | 0,82444049 | 0,983374097 | phosphoserine aminotransferase 1 |
| ENSG00000144231 | POLR2D | -0,072485692 | 0,950998063 | 0,73462361 | 0,974353474 | RNA polymerase II subunit D |
| ENSG00000148459 | PDSS1 | -0,072525347 | 0,950971923 | 0,83253295 | 0,98489473 | decaprenyl diphosphate synthase subunit 1 |
| ENSG00000244560 | AC004890,2 | -0,072550701 | 0,950955212 | 0,88140841 | 0,990717729 | zinc finger protein pseudogene |
| ENSG00000187741 | FANCA | -0,072575793 | 0,950938672 | 0,76695811 | 0,977201891 | FA complementation group A |
| ENSG00000135002 | RFK | -0,072603467 | 0,950920431 | 0,88126331 | 0,990717729 | riboflavin kinase |
| ENSG00000214067 | AL360182,1 | -0,072655362 | 0,950886226 | 0,9412714 | 0,995251853 | ribosomal protein L21 (RPL21) pseudogene |

| | | | | | | |
|---|---|---|---|---|---|---|
| ENSG00000274460 | AC092119,2 | -0,072677811 | 0,95087143 | 0,89456629 | 0,99116563 | novel transcript |
| ENSG00000131044 | TTLL9 | -0,072683713 | 0,95086754 | 0,87024859 | 0,989074806 | tubulin tyrosine ligase like 9 |
| ENSG00000270842 | AC073413,2 | -0,072685103 | 0,950866624 | 0,95515172 | 0,996608711 | ribosomal protein S26 (RPS26) pseudogene |
| ENSG00000249115 | HAUS5 | -0,072712822 | 0,950848355 | 0,8065152 | 0,981213877 | HAUS augmin like complex subunit 5 |
| ENSG00000261613 | AC093525,6 | -0,072717484 | 0,950845282 | 0,80942592 | 0,981213877 | novel transcript, antisense to PDPK1 |
| ENSG00000182586 | LINC00334 | -0,072727108 | 0,950838939 | 0,94711845 | 0,996027077 | long intergenic non-protein coding RNA 334 |
| ENSG00000135372 | NAT10 | -0,072728145 | 0,950838256 | 0,75829943 | 0,976120207 | N-acetyltransferase 10 |
| ENSG00000121903 | ZSCAN20 | -0,072844481 | 0,950761585 | 0,74784802 | 0,975901808 | zinc finger and SCAN domain containing 20 |
| ENSG00000132388 | UBE2G1 | -0,072862085 | 0,950749985 | 0,84075348 | 0,985672131 | ubiquitin conjugating enzyme E2 G1 |
| ENSG00000169964 | TMEM42 | -0,072872977 | 0,950742806 | 0,81159662 | 0,981213877 | transmembrane protein 42 |
| ENSG00000280516 | TMEM42 | -0,072872977 | 0,950742806 | 0,81159662 | 0,981213877 | transmembrane protein 42 |
| ENSG00000157625 | TAB3 | -0,072874059 | 0,950742093 | 0,84708031 | 0,987034354 | TGF-beta activated kinase 1 (MAP3K7) binding protein 3 |
| ENSG00000248092 | NNT-AS1 | -0,072879684 | 0,950738387 | 0,81154604 | 0,981213877 | NNT antisense RNA 1 |
| ENSG00000287592 | AC073348,2 | -0,073048295 | 0,950627278 | 0,89720164 | 0,991644439 | novel transcript |
| ENSG00000129480 | DTD2 | -0,073058434 | 0,950620597 | 0,79855185 | 0,980161849 | D-aminoacyl-tRNA deacylase 2 |
| ENSG00000245648 | KLRK1-AS1 | -0,073060129 | 0,95061948 | 0,79043931 | 0,979462318 | KLRK1 antisense RNA 1 |
| ENSG00000166266 | CUL5 | -0,073078086 | 0,950607648 | 0,86885512 | 0,989074806 | cullin 5 |
| ENSG00000165832 | TRUB1 | -0,073093123 | 0,95059774 | 0,85108497 | 0,987259878 | TruB pseudouridine synthase family member 1 |
| ENSG00000101187 | SLCO4A1 | -0,073111914 | 0,950585359 | 0,86536185 | 0,989074806 | solute carrier organic anion transporter family member 4A1 |
| ENSG00000240751 | AC026348,1 | -0,073290158 | 0,950467922 | 0,92615977 | 0,993915537 | Rab geranylgeranyltransferase, beta subunit (RABGGTB) pseudogene |
| ENSG00000258670 | AL049874,3 | -0,073297462 | 0,95046311 | 0,86468243 | 0,989074806 | novel transcript, antisense to C14orf39 |
| ENSG00000116120 | FARSB | -0,0733483 | 0,950429618 | 0,83499271 | 0,984905093 | phenylalanyl-tRNA synthetase subunit beta |
| ENSG00000114841 | DNAH1 | -0,073400768 | 0,950395053 | 0,80705021 | 0,981213877 | dynein axonemal heavy chain 1 |
| ENSG00000168958 | MFF | -0,073422881 | 0,950380486 | 0,8461029 | 0,986939308 | mitochondrial fission factor |
| ENSG00000171497 | PPID | -0,073426617 | 0,950378025 | 0,81943529 | 0,98249617 | peptidylprolyl isomerase D |
| ENSG00000087448 | KLHL42 | -0,073446521 | 0,950364913 | 0,81656473 | 0,981919736 | kelch like family member 42 |
| ENSG00000135931 | ARMC9 | -0,073622634 | 0,950248907 | 0,72674373 | 0,973344786 | armadillo repeat containing 9 |
| ENSG00000247315 | ZCCHC3 | -0,073671441 | 0,95021676 | 0,7720566 | 0,978061414 | zinc finger CCHC-type containing 3 |
| ENSG00000074657 | ZNF532 | -0,073684903 | 0,950207894 | 0,79437124 | 0,979776981 | zinc finger protein 532 |
| ENSG00000213397 | HAUS7 | -0,073716576 | 0,950187033 | 0,80020995 | 0,980161849 | HAUS augmin like complex subunit 7 |

| | | | | | | |
|---|---|---|---|---|---|---|
| ENSG00000121671 | CRY2 | -0,073766034 | 0,95015446 | 0,74952613 | 0,975901808 | cryptochrome circadian regulator 2 |
| ENSG00000131051 | RBM39 | -0,073791381 | 0,950137766 | 0,83672861 | 0,985426146 | RNA binding motif protein 39 |
| ENSG00000101596 | SMCHD1 | -0,07379347 | 0,950136391 | 0,86087501 | 0,989074806 | structural maintenance of chromosomes flexible hinge domain containing 1 |
| ENSG00000155115 | GTF3C6 | -0,073808794 | 0,950126298 | 0,84082168 | 0,985672131 | general transcription factor IIIC subunit 6 |
| ENSG00000006695 | COX10 | -0,073888313 | 0,95007393 | 0,8101635 | 0,981213877 | cytochrome c oxidase assembly factor heme A:farnesyltransferase COX10 |
| ENSG00000172172 | MRPL13 | -0,073891898 | 0,95007157 | 0,85269082 | 0,98761642 | mitochondrial ribosomal protein L13 |
| ENSG00000142556 | ZNF614 | -0,073944515 | 0,95003692 | 0,86373715 | 0,989074806 | zinc finger protein 614 |
| ENSG00000263247 | PRR4 | -0,074005522 | 0,949996747 | 0,87644942 | 0,990109814 | proline rich 4 |
| ENSG00000173093 | CCDC63 | -0,074016202 | 0,949989714 | 0,9516875 | 0,996432861 | coiled-coil domain containing 63 |
| ENSG00000234750 | AC012618,2 | -0,074017653 | 0,949988759 | 0,92223472 | 0,993797669 | ribosomal protein S2 (RPS2) pseudogene |
| ENSG00000103942 | HOMER2 | -0,074032969 | 0,949978673 | 0,74173762 | 0,975176043 | homer scaffold protein 2 |
| ENSG00000273828 | AL133227,1 | -0,074037234 | 0,949975865 | 0,76284571 | 0,97647393 | novel transcript, antisense to ELMO2 |
| ENSG00000106049 | HIBADH | -0,074075467 | 0,94995069 | 0,81235519 | 0,981213877 | 3-hydroxyisobutyrate dehydrogenase |
| ENSG00000259244 | AC048382,2 | -0,074085549 | 0,949944051 | 0,88805122 | 0,990717729 | novel transcript |
| ENSG00000035115 | SH3YL1 | -0,074154213 | 0,94989884 | 0,7327907 | 0,974353474 | SH3 and SYLF domain containing 1 |
| ENSG00000248487 | ABHD14A | -0,074167482 | 0,949890104 | 0,79486272 | 0,979776981 | abhydrolase domain containing 14A |
| ENSG00000176533 | GNG7 | -0,07418435 | 0,949878998 | 0,80943629 | 0,981213877 | G protein subunit gamma 7 |
| ENSG00000174348 | PODN | -0,074290553 | 0,949809076 | 0,9230577 | 0,993825008 | podocan |
| ENSG00000175279 | CENPS | -0,07431828 | 0,949790822 | 0,82581783 | 0,983383773 | centromere protein S |
| ENSG00000196636 | SDHAF3 | -0,07433665 | 0,949778728 | 0,87427943 | 0,98990147 | succinate dehydrogenase complex assembly factor 3 |
| ENSG00000232045 | EHMT2 | -0,074366426 | 0,949759126 | 0,83027325 | 0,984156028 | euchromatic histone lysine methyltransferase 2 |
| ENSG00000268509 | AC026202,3 | -0,074367305 | 0,949758547 | 0,91443375 | 0,993361299 | novel transcript, antisense to EDEM1 |
| ENSG00000285269 | AL160269,1 | -0,074373953 | 0,949754171 | 0,96304459 | 0,996608711 | solute carrier family 35 member D2 |
| ENSG00000277232 | GTSE1-DT | -0,074471192 | 0,949690158 | 0,88548604 | 0,990717729 | GTSE1 divergent transcript |
| ENSG00000162650 | ATXN7L2 | -0,074598345 | 0,94960646 | 0,78580054 | 0,979320406 | ataxin 7 like 2 |
| ENSG00000163354 | DCST2 | -0,074713131 | 0,949530909 | 0,92798817 | 0,994108531 | DC-STAMP domain containing 2 |
| ENSG00000148842 | CNNM2 | -0,074801518 | 0,949472738 | 0,79388492 | 0,979776981 | cyclin and CBS domain divalent metal cation transport mediator 2 |
| ENSG00000181552 | EDDM3B | -0,074857037 | 0,9494362 | 0,95882986 | 0,996608711 | epididymal protein 3B |
| ENSG00000003509 | NDUFAF7 | -0,074884891 | 0,949417869 | 0,82061311 | 0,982609114 | NADH:ubiquinone oxidoreductase complex assembly factor 7 |

| | | | | | | |
|---|---|---|---|---|---|---|
| ENSG00000101213 | PTK6 | -0,074894938 | 0,949411258 | 0,88039617 | 0,990367704 | protein tyrosine kinase 6 |
| ENSG00000104885 | DOT1L | -0,074915364 | 0,949397816 | 0,81626439 | 0,981790866 | DOT1 like histone lysine methyltransferase |
| ENSG00000260060 | AC009088,1 | -0,074927079 | 0,949390107 | 0,86845996 | 0,989074806 | novel transcript, antisense to PYCARD |
| ENSG00000235262 | KDM5C-IT1 | -0,074962094 | 0,949367065 | 0,90430246 | 0,992590661 | KDM5C intronic transcript 1 |
| ENSG00000124164 | VAPB | -0,07497151 | 0,949360869 | 0,78716798 | 0,979425304 | VAMP associated protein B and C |
| ENSG00000115368 | WDR75 | -0,075019636 | 0,9493292 | 0,84398776 | 0,986273696 | WD repeat domain 75 |
| ENSG00000117713 | ARID1A | -0,075081629 | 0,949288408 | 0,70839138 | 0,970174703 | AT-rich interaction domain 1A |
| ENSG00000268095 | ZNF649-AS1 | -0,075196704 | 0,949212692 | 0,9521659 | 0,996432861 | ZNF649 antisense RNA 1 |
| ENSG00000132912 | DCTN4 | -0,075198335 | 0,949211619 | 0,84869401 | 0,987036295 | dynactin subunit 4 |
| ENSG00000166233 | ARIH1 | -0,075198697 | 0,949211381 | 0,80886384 | 0,981213877 | ariadne RBR E3 ubiquitin protein ligase 1 |
| ENSG00000153107 | ANAPC1 | -0,075222017 | 0,949196037 | 0,81969245 | 0,98249617 | anaphase promoting complex subunit 1 |
| ENSG00000155099 | PIP4P2 | -0,075228689 | 0,949191648 | 0,88555282 | 0,990717729 | phosphatidylinositol-4,5-bisphosphate 4-phosphatase 2 |
| ENSG00000163159 | VPS72 | -0,07522956 | 0,949191075 | 0,74437002 | 0,975641819 | vacuolar protein sorting 72 homolog |
| ENSG00000235695 | HIGD2AP2 | -0,075448317 | 0,949047159 | 0,92853294 | 0,994108531 | HIGD2A pseudogene 2 |
| ENSG00000138160 | KIF11 | -0,075511641 | 0,949005504 | 0,84319741 | 0,986218419 | kinesin family member 11 |
| ENSG00000143390 | RFX5 | -0,075581552 | 0,948959517 | 0,79478663 | 0,979776981 | regulatory factor X5 |
| ENSG00000104408 | EIF3E | -0,075639473 | 0,948921419 | 0,84271065 | 0,986183927 | eukaryotic translation initiation factor 3 subunit E |
| ENSG00000126016 | AMOT | -0,075686971 | 0,948890178 | 0,79974128 | 0,980161849 | angiomotin |
| ENSG00000143315 | PIGM | -0,07572926 | 0,948862365 | 0,81035614 | 0,981213877 | phosphatidylinositol glycan anchor biosynthesis class M |
| ENSG00000176623 | RMDN1 | -0,075757884 | 0,948843539 | 0,79987845 | 0,980161849 | regulator of microtubule dynamics 1 |
| ENSG00000105856 | HBP1 | -0,075767986 | 0,948836895 | 0,86409445 | 0,989074806 | HMG-box transcription factor 1 |
| ENSG00000100151 | PICK1 | -0,075858975 | 0,948777055 | 0,79616804 | 0,980015022 | protein interacting with PRKCA 1 |
| ENSG00000171298 | GAA | -0,075876938 | 0,948765242 | 0,80797149 | 0,981213877 | glucosidase alpha, acid |
| ENSG00000169951 | ZNF764 | -0,075994076 | 0,948688211 | 0,79542752 | 0,979879951 | zinc finger protein 764 |
| ENSG00000277795 | AC244517,3 | -0,076076828 | 0,948633796 | 0,94118926 | 0,995251853 | novel zinc finger pseudogene |
| ENSG00000287532 | AL353689,3 | -0,076138728 | 0,948593095 | 0,92210398 | 0,993797669 | novel transcript, antisense to CDC42BPA |
| ENSG00000185808 | PIGP | -0,076188912 | 0,948560099 | 0,830029 | 0,984156028 | phosphatidylinositol glycan anchor biosynthesis class P |
| ENSG00000234261 | AL138720,1 | -0,076289199 | 0,948494163 | 0,84321492 | 0,986218419 | novel transcript |
| ENSG00000131848 | ZSCAN5A | -0,076296014 | 0,948489683 | 0,77793924 | 0,978587255 | zinc finger and SCAN domain containing 5A |
| ENSG00000284900 | KBTBD4 | -0,076297221 | 0,948488889 | 0,79627065 | 0,980015022 | kelch repeat and BTB domain containing 4 |

| | | | | | | |
|---|---|---|---|---|---|---|
| ENSG00000123444 | KBTBD4 | -0,076297221 | 0,948488889 | 0,79627065 | 0,980015022 | kelch repeat and BTB domain containing 4 |
| ENSG00000162642 | C1orf52 | -0,076418651 | 0,94840906 | 0,80953686 | 0,981213877 | chromosome 1 open reading frame 52 |
| ENSG00000130023 | ERMARD | -0,076429461 | 0,948401953 | 0,86404229 | 0,989074806 | ER membrane associated RNA degradation |
| ENSG00000284887 | AC270254,1 | -0,07653496 | 0,948332602 | 0,9629918 | 0,996608711 | |
| ENSG00000005206 | SPPL2B | -0,076547985 | 0,948324041 | 0,83287257 | 0,984905093 | signal peptide peptidase like 2B |
| ENSG00000275017 | AL353748,2 | -0,076592502 | 0,948294779 | 0,88535998 | 0,990717729 | developmental pluripotency associated 3 (DPPA3) pseudogene |
| ENSG00000147421 | HMBOX1 | -0,076603031 | 0,948287858 | 0,816183 | 0,981730742 | homeobox containing 1 |
| ENSG00000183889 | AC138969,1 | -0,076614298 | 0,948280452 | 0,73666932 | 0,974353474 | novel member of the nuclear pore complex interacting protein NPIP gene family |
| ENSG00000284128 | AP000356,3 | -0,07671422 | 0,948214776 | 0,90468069 | 0,992590661 | breakpoint cluster region pseudogene 3 |
| ENSG00000198886 | MT-ND4 | -0,07671644 | 0,948213317 | 0,80444589 | 0,980875302 | mitochondrially encoded NADH:ubiquinone oxidoreductase core subunit 4 |
| ENSG00000156411 | ATP5MPL | -0,076746808 | 0,948193358 | 0,85765397 | 0,988893234 | ATP synthase membrane subunit 6,8PL |
| ENSG00000174839 | DENND6A | -0,076782162 | 0,948170122 | 0,75700647 | 0,975901808 | DENN domain containing 6A |
| ENSG00000288512 | C1R | -0,076830374 | 0,948138437 | 0,8431005 | 0,986218419 | complement C1r |
| ENSG00000183474 | GTF2H2C | -0,076835433 | 0,948135112 | 0,72047593 | 0,972142678 | GTF2H2 family member C |
| ENSG00000091164 | TXNL1 | -0,076871114 | 0,948111663 | 0,8441566 | 0,98632753 | thioredoxin like 1 |
| ENSG00000251330 | AC114939,1 | -0,076872021 | 0,948111067 | 0,9167088 | 0,993361299 | novel transcript, antisense to FBXO38 |
| ENSG00000140319 | SRP14 | -0,076914025 | 0,948083463 | 0,70361164 | 0,968967751 | signal recognition particle 14 |
| ENSG00000131591 | C1orf159 | -0,076987001 | 0,948035507 | 0,80861815 | 0,981213877 | chromosome 1 open reading frame 159 |
| ENSG00000287868 | AL355377,4 | -0,077027026 | 0,948009206 | 0,87251521 | 0,989217334 | novel transcript |
| ENSG00000173585 | CCR9 | -0,07706264 | 0,947985804 | 0,92659946 | 0,993915537 | C-C motif chemokine receptor 9 |
| ENSG00000143621 | ILF2 | -0,077080043 | 0,947974368 | 0,77181455 | 0,978033855 | interleukin enhancer binding factor 2 |
| ENSG00000196118 | CCDC189 | -0,077117389 | 0,947949829 | 0,82989644 | 0,984156028 | coiled-coil domain containing 189 |
| ENSG00000136146 | MED4 | -0,077130403 | 0,947941278 | 0,87607243 | 0,990109814 | mediator complex subunit 4 |
| ENSG00000171450 | CDK5R2 | -0,077152184 | 0,947926967 | 0,85664122 | 0,988303301 | cyclin dependent kinase 5 regulatory subunit 2 |
| ENSG00000134900 | TPP2 | -0,077179846 | 0,947908791 | 0,84735396 | 0,987035044 | tripeptidyl peptidase 2 |
| ENSG00000203761 | MSTO2P | -0,077185953 | 0,947904779 | 0,86704559 | 0,989074806 | misato family member 2, pseudogene |
| ENSG00000184277 | TM2D3 | -0,077288397 | 0,947837472 | 0,78936282 | 0,979462318 | TM2 domain containing 3 |
| ENSG00000276710 | CSPG4P10 | -0,077318063 | 0,947817982 | 0,83765854 | 0,98566906 | chondroitin sulfate proteoglycan 4 pseudogene 10 |
| ENSG00000287630 | AC011370,1 | -0,077345785 | 0,947799769 | 0,81028669 | 0,981213877 | novel transcript, antisense to JAKMIP2 |
| ENSG00000282269 | PRR4 | -0,077384603 | 0,947774268 | 0,81434344 | 0,981521413 | proline rich 4 |

| | | | | | | |
|---|---|---|---|---|---|---|
| ENSG00000104497 | SNX16 | -0,077530234 | 0,9476786 | 0,8625678 | 0,989074806 | sorting nexin 16 |
| ENSG00000185418 | TARS3 | -0,077574676 | 0,947649408 | 0,76450725 | 0,976640468 | threonyl-tRNA synthetase 3 |
| ENSG00000114383 | TUSC2 | -0,077577633 | 0,947647466 | 0,76206455 | 0,976315588 | tumor suppressor 2, mitochondrial calcium regulator |
| ENSG00000281398 | SNHG4 | -0,077614023 | 0,947623563 | 0,81883607 | 0,98249617 | small nucleolar RNA host gene 4 |
| ENSG00000172171 | TEFM | -0,077698014 | 0,947568395 | 0,82102467 | 0,982753437 | transcription elongation factor, mitochondrial |
| ENSG00000010318 | PHF7 | -0,077775316 | 0,947517625 | 0,83306813 | 0,984905093 | PHD finger protein 7 |
| ENSG00000126214 | KLC1 | -0,077777278 | 0,947516336 | 0,70652049 | 0,969503793 | kinesin light chain 1 |
| ENSG00000163960 | UBXN7 | -0,077813768 | 0,947492371 | 0,82897177 | 0,983844005 | UBX domain protein 7 |
| ENSG00000170160 | CCDC144A | -0,077853667 | 0,947466167 | 0,7485164 | 0,975901808 | coiled-coil domain containing 144A |
| ENSG00000122203 | KIAA1191 | -0,077919472 | 0,947422952 | 0,77756352 | 0,978587255 | KIAA1191 |
| ENSG00000287697 | Z99127,3 | -0,078010102 | 0,947363437 | 0,8875581 | 0,990717729 | novel transcript, antisense to CACYBP |
| ENSG00000183918 | SH2D1A | -0,078023429 | 0,947354686 | 0,92520836 | 0,993915537 | SH2 domain containing 1A |
| ENSG00000196628 | TCF4 | -0,078025246 | 0,947353492 | 0,84667233 | 0,986939308 | transcription factor 4 |
| ENSG00000144036 | EXOC6B | -0,078084894 | 0,947314325 | 0,81562118 | 0,981590958 | exocyst complex component 6B |
| ENSG00000237889 | DDX39B | -0,078129767 | 0,947284861 | 0,79048086 | 0,979462318 | DExD-box helicase 39B |
| ENSG00000273117 | AC144652,1 | -0,078139171 | 0,947278686 | 0,80749347 | 0,981213877 | novel transcript |
| ENSG00000154642 | C21orf91 | -0,078147631 | 0,947273131 | 0,88718353 | 0,990717729 | chromosome 21 open reading frame 91 |
| ENSG00000165300 | SLITRK5 | -0,078415805 | 0,947097064 | 0,77716036 | 0,978587255 | SLIT and NTRK like family member 5 |
| ENSG00000196668 | LINC00173 | -0,078437718 | 0,947082679 | 0,84928201 | 0,987036295 | long intergenic non-protein coding RNA 173 |
| ENSG00000119688 | ABCD4 | -0,078541987 | 0,947014233 | 0,7626503 | 0,97639149 | ATP binding cassette subfamily D member 4 |
| ENSG00000135365 | PHF21A | -0,078548705 | 0,947009822 | 0,76529281 | 0,976830764 | PHD finger protein 21A |
| ENSG00000103150 | MLYCD | -0,078552636 | 0,947007242 | 0,82550107 | 0,983383773 | malonyl-CoA decarboxylase |
| ENSG00000102893 | PHKB | -0,078602878 | 0,946974263 | 0,83471389 | 0,984905093 | phosphorylase kinase regulatory subunit beta |
| ENSG00000235961 | PNMA6A | -0,078681578 | 0,946922607 | 0,87784882 | 0,990109814 | PNMA family member 6A |
| ENSG00000262473 | GART | -0,078729346 | 0,946891254 | 0,83360609 | 0,984905093 | phosphoribosylglycinamide formyltransferase, phosphoribosylglycinamide synthetase, phosphoribosylaminoimidazole synthetase |
| ENSG00000136824 | SMC2 | -0,078744898 | 0,946881047 | 0,82233019 | 0,982960472 | structural maintenance of chromosomes 2 |
| ENSG00000213516 | RBMXL1 | -0,078850519 | 0,946811728 | 0,82443187 | 0,983374097 | RBMX like 1 |
| ENSG00000105146 | AURKC | -0,078880633 | 0,946791964 | 0,92126246 | 0,993787755 | aurora kinase C |
| ENSG00000184863 | RBM33 | -0,078931128 | 0,946758827 | 0,73543615 | 0,974353474 | RNA binding motif protein 33 |
| ENSG00000175518 | UBQLNL | -0,079079855 | 0,946661231 | 0,91533427 | 0,993361299 | ubiquilin like |
| ENSG00000176731 | RBIS | -0,079108933 | 0,946642151 | 0,86780436 | 0,989074806 | ribosomal biogenesis factor |

| | | | | | | |
|---|---|---|---|---|---|---|
| ENSG00000116497 | S100PBP | -0,079179433 | 0,946595893 | 0,75837527 | 0,976120207 | S100P binding protein |
| ENSG00000230797 | YY2 | -0,07923894 | 0,946556849 | 0,86924293 | 0,989074806 | YY2 transcription factor |
| ENSG00000198576 | ARC | -0,079406223 | 0,9464471 | 0,88878167 | 0,990717729 | activity regulated cytoskeleton associated protein |
| ENSG00000171723 | GPHN | -0,079414688 | 0,946441547 | 0,82171082 | 0,982952211 | gephyrin |
| ENSG00000273066 | AL355987,4 | -0,079454523 | 0,946415415 | 0,86603343 | 0,989074806 | novel transcript |
| ENSG00000092201 | SUPT16H | -0,07950292 | 0,946383667 | 0,74168731 | 0,975176043 | SPT16 homolog, facilitates chromatin remodeling subunit |
| ENSG00000182545 | RNASE10 | -0,079505272 | 0,946382124 | 0,88834561 | 0,990717729 | ribonuclease A family member 10 (inactive) |
| ENSG00000234882 | EIF3EP1 | -0,079575898 | 0,946335796 | 0,94382616 | 0,995486053 | eukaryotic translation initiation factor 3 subunit E pseudogene 1 |
| ENSG00000266921 | AC006213,1 | -0,079593164 | 0,94632447 | 0,89897618 | 0,991832796 | novel transcript, antisense to ZNF155 |
| ENSG00000137074 | APTX | -0,079674021 | 0,946271434 | 0,74673301 | 0,975796253 | aprataxin |
| ENSG00000274675 | GTF2H2C_2 | -0,079819518 | 0,946176006 | 0,8727875 | 0,989262536 | GTF2H2 family member C, copy 2 |
| ENSG00000149927 | DOC2A | -0,07984613 | 0,946158553 | 0,85961338 | 0,989074806 | double C2 domain alpha |
| ENSG00000212443 | SNORA53 | -0,079920514 | 0,946109772 | 0,81026623 | 0,981213877 | small nucleolar RNA, H/ACA box 53 |
| ENSG00000133243 | BTBD2 | -0,08000617 | 0,946053601 | 0,74372167 | 0,975343483 | BTB domain containing 2 |
| ENSG00000285632 | AC084024,4 | -0,080020002 | 0,94604453 | 0,9367513 | 0,994479239 | novel transcript |
| ENSG00000274904 | AC093512,1 | -0,080068065 | 0,946013013 | 0,92069916 | 0,993787755 | novel transcript, antisense to ALDOA |
| ENSG00000229833 | PET100 | -0,080097111 | 0,945993967 | 0,70066788 | 0,968522643 | PET100 cytochrome c oxidase chaperone |
| ENSG00000255423 | EBLN2 | -0,08009714 | 0,945993949 | 0,83124903 | 0,98438942 | endogenous Bornavirus like nucleoprotein 2 |
| ENSG00000183624 | HMCES | -0,080103808 | 0,945989576 | 0,70389079 | 0,968977218 | 5-hydroxymethylcytosine binding, ES cell specific |
| ENSG00000261654 | AL360270,2 | -0,080217166 | 0,945915249 | 0,85403024 | 0,987861659 | novel transcript, sense overlapping C1orf103 |
| ENSG00000277406 | SEC22B4P | -0,080236317 | 0,945902693 | 0,90479685 | 0,992590661 | SEC22 homolog B4, pseudogene |
| ENSG00000196715 | VKORC1L1 | -0,080274438 | 0,945877699 | 0,83357912 | 0,984905093 | vitamin K epoxide reductase complex subunit 1 like 1 |
| ENSG00000251600 | AC107223,1 | -0,080355092 | 0,945824821 | 0,88144195 | 0,990717729 | novel transcript, antisense to GYPA |
| ENSG00000103121 | CMC2 | -0,080359961 | 0,945821629 | 0,79801217 | 0,980161849 | C-X9-C motif containing 2 |
| ENSG00000278834 | AC073508,3 | -0,080365863 | 0,94581776 | 0,75442826 | 0,975901808 | novel transcript, antisense to SMARCE1 |
| ENSG00000138190 | EXOC6 | -0,080382842 | 0,945806628 | 0,84557953 | 0,986639902 | exocyst complex component 6 |
| ENSG00000115425 | PECR | -0,080555342 | 0,945693547 | 0,74126277 | 0,975176043 | peroxisomal trans-2-enoyl-CoA reductase |
| ENSG00000159164 | SV2A | -0,080607364 | 0,945659447 | 0,71230171 | 0,970812834 | synaptic vesicle glycoprotein 2A |
| ENSG00000265912 | AC037487,1 | -0,080726494 | 0,945581363 | 0,9161286 | 0,993361299 | novel transcript |
| ENSG00000170325 | PRDM10 | -0,080740693 | 0,945572057 | 0,73647338 | 0,974353474 | PR/SET domain 10 |

| | | | | | | |
|---|---|---|---|---|---|---|
| ENSG00000136870 | ZNF189 | -0,080754997 | 0,945562682 | 0,84087028 | 0,985672131 | zinc finger protein 189 |
| ENSG00000224552 | ABHD16A | -0,080755578 | 0,945562301 | 0,89255775 | 0,991048031 | abhydrolase domain containing 16A, phospholipase |
| ENSG00000157184 | CPT2 | -0,080776542 | 0,945548561 | 0,6957544 | 0,96776072 | carnitine palmitoyltransferase 2 |
| ENSG00000172239 | PAIP1 | -0,080782654 | 0,945544555 | 0,82029929 | 0,982609114 | poly(A) binding protein interacting protein 1 |
| ENSG00000254363 | AC011379,2 | -0,080808595 | 0,945527553 | 0,91555481 | 0,993361299 | novel transcript |
| ENSG00000129355 | CDKN2D | -0,080883507 | 0,945478458 | 0,76954813 | 0,977683747 | cyclin dependent kinase inhibitor 2D |
| ENSG00000125944 | HNRNPR | -0,08100068 | 0,945401671 | 0,82495444 | 0,983374097 | heterogeneous nuclear ribonucleoprotein R |
| ENSG00000234072 | AC074117,1 | -0,081037461 | 0,945377569 | 0,82769909 | 0,983595244 | novel transcript, antisense to GTF3C2 and EIF2B4 |
| ENSG00000287878 | AC022382,2 | -0,081071839 | 0,945355041 | 0,836884 | 0,985437134 | novel transcript, antisense to CPNE9and MTMR14 |
| ENSG00000136098 | NEK3 | -0,081081906 | 0,945348445 | 0,8049353 | 0,98109454 | NIMA related kinase 3 |
| ENSG00000265069 | AP002409,1 | -0,081109036 | 0,945330668 | 0,90610081 | 0,992590661 | novel transcript, antisense to LAMA1 |
| ENSG00000233671 | CENPNP1 | -0,081141177 | 0,945309608 | 0,94360469 | 0,995406323 | CENPN pseudogene 1 |
| ENSG00000188807 | TMEM201 | -0,081205323 | 0,945267578 | 0,7934493 | 0,979776981 | transmembrane protein 201 |
| ENSG00000162073 | PAQR4 | -0,081234969 | 0,945248154 | 0,76687243 | 0,977201891 | progestin and adipoQ receptor family member 4 |
| ENSG00000070061 | ELP1 | -0,081241152 | 0,945244102 | 0,73579304 | 0,974353474 | elongator complex protein 1 |
| ENSG00000247950 | SEC24B-AS1 | -0,081277452 | 0,945220319 | 0,80970768 | 0,981213877 | SEC24B antisense RNA 1 |
| ENSG00000275506 | H3P25 | -0,081299878 | 0,945205626 | 0,93718682 | 0,994479239 | H3 histone pseudogene 25 |
| ENSG00000100191 | SLC5A4 | -0,081301304 | 0,945204692 | 0,92702749 | 0,993915537 | solute carrier family 5 member 4 |
| ENSG00000233098 | CCDC144NL-AS | -0,081352113 | 0,945171405 | 0,75124179 | 0,975901808 | CCDC144NL antisense RNA 1 |
| ENSG00000134765 | DSC1 | -0,081431485 | 0,945119406 | 0,93738419 | 0,994479239 | desmocollin 1 |
| ENSG00000163389 | POGLUT1 | -0,081447828 | 0,945108699 | 0,84600555 | 0,98689126 | protein O-glucosyltransferase 1 |
| ENSG00000083097 | DOP1A | -0,081461395 | 0,945099812 | 0,85153691 | 0,987259878 | DOP1 leucine zipper like protein A |
| ENSG00000145916 | RMND5B | -0,081495812 | 0,945077266 | 0,72075038 | 0,972142678 | required for meiotic nuclear division 5 homolog B |
| ENSG00000141219 | C17orf80 | -0,081498928 | 0,945075224 | 0,8369222 | 0,985444996 | chromosome 17 open reading frame 80 |
| ENSG00000277936 | MRPL45 | -0,08151984 | 0,945061526 | 0,84009087 | 0,985672131 | mitochondrial ribosomal protein L45 |
| ENSG00000175387 | SMAD2 | -0,08158584 | 0,945018292 | 0,78614231 | 0,979320406 | SMAD family member 2 |
| ENSG00000263002 | ZNF234 | -0,081621313 | 0,944995056 | 0,78917584 | 0,979462318 | zinc finger protein 234 |
| ENSG00000259969 | AL049838,1 | -0,08165234 | 0,944974733 | 0,89516581 | 0,991303328 | novel transcript, overlapping C14orf37 |
| ENSG00000118246 | FASTKD2 | -0,081889919 | 0,94481913 | 0,83522012 | 0,984924633 | FAST kinase domains 2 |
| ENSG00000228486 | C2orf92 | -0,081923144 | 0,944797371 | 0,82544058 | 0,983383773 | chromosome 2 open reading frame 92 |
| ENSG00000132300 | PTCD3 | -0,082028458 | 0,944728405 | 0,81470681 | 0,981521413 | pentatricopeptide repeat domain 3 |
| ENSG00000136709 | WDR33 | -0,082161501 | 0,944641288 | 0,74783666 | 0,975901808 | WD repeat domain 33 |
| ENSG00000139531 | SUOX | -0,082162657 | 0,944640531 | 0,7609098 | 0,97624106 | sulfite oxidase |

| | | | | | | |
|---|---|---|---|---|---|---|
| ENSG00000171970 | ZNF57 | -0,082189088 | 0,944623225 | 0,84100953 | 0,985672131 | zinc finger protein 57 |
| ENSG00000284882 | AL359762,1 | -0,082199865 | 0,944616169 | 0,90709206 | 0,992590661 | novel transcript |
| ENSG00000228672 | PROB1 | -0,082240889 | 0,944589308 | 0,88045951 | 0,990367704 | proline rich basic protein 1 |
| ENSG00000213300 | HNRNPA3P6 | -0,082249384 | 0,944583746 | 0,95934251 | 0,996608711 | heterogeneous nuclear ribonucleoprotein A3 pseudogene 6 |
| ENSG00000148688 | RPP30 | -0,082263307 | 0,94457463 | 0,8411444 | 0,985672131 | ribonuclease P/MRP subunit p30 |
| ENSG00000065491 | TBC1D22B | -0,082274563 | 0,944567261 | 0,7445611 | 0,975649186 | TBC1 domain family member 22B |
| ENSG00000174547 | MRPL11 | -0,082310616 | 0,944543657 | 0,73427905 | 0,974353474 | mitochondrial ribosomal protein L11 |
| ENSG00000069020 | MAST4 | -0,082394892 | 0,944488482 | 0,69671782 | 0,968300554 | microtubule associated serine/threonine kinase family member 4 |
| ENSG00000128254 | C22orf24 | -0,082412682 | 0,944476836 | 0,7902085 | 0,979462318 | chromosome 22 open reading frame 24 |
| ENSG00000109819 | PPARGC1A | -0,082568019 | 0,944375148 | 0,84650283 | 0,986939308 | PPARG coactivator 1 alpha |
| ENSG00000275759 | AC026367,3 | -0,082622864 | 0,944339247 | 0,94175256 | 0,995373376 | novel transcript |
| ENSG00000260213 | AC092718,2 | -0,082658135 | 0,94431616 | 0,85219627 | 0,987401848 | novel transcript, antisense to CENPN |
| ENSG00000231825 | PRRC2A | -0,082674457 | 0,944305477 | 0,83807229 | 0,985672131 | proline rich coiled-coil 2A |
| ENSG00000068781 | STON1-GTF2A1 | -0,082678434 | 0,944302874 | 0,82815658 | 0,983644374 | STON1-GTF2A1L readthrough |
| ENSG00000107789 | MINPP1 | -0,082749001 | 0,944256686 | 0,84315579 | 0,986218419 | multiple inositol-polyphosphate phosphatase 1 |
| ENSG00000111716 | LDHB | -0,082759333 | 0,944249924 | 0,76039354 | 0,97624106 | lactate dehydrogenase B |
| ENSG00000130348 | QRSL1 | -0,0828135 | 0,944214472 | 0,81179909 | 0,981213877 | glutaminyl-tRNA amidotransferase subunit QRSL1 |
| ENSG00000224451 | ATP5PBP1 | -0,082943542 | 0,944129366 | 0,91406856 | 0,993361299 | ATP synthase peripheral stalk-membrane subunit b pseudogene 1 |
| ENSG00000118518 | RNF146 | -0,082957636 | 0,944120142 | 0,81588175 | 0,981708551 | ring finger protein 146 |
| ENSG00000198081 | ZBTB14 | -0,082972982 | 0,9441101 | 0,80174209 | 0,980659812 | zinc finger and BTB domain containing 14 |
| ENSG00000112983 | BRD8 | -0,082993133 | 0,944096913 | 0,73078379 | 0,974336839 | bromodomain containing 8 |
| ENSG00000267576 | AC011472,3 | -0,082996989 | 0,94409439 | 0,96456469 | 0,997113804 | novel transcript, sense intronic to TSPAN16 |
| ENSG00000178026 | LRRC75B | -0,083173227 | 0,943979067 | 0,78933151 | 0,979462318 | leucine rich repeat containing 75B |
| ENSG00000270194 | AC097359,2 | -0,083190024 | 0,943968077 | 0,79369333 | 0,979776981 | novel transcript, antisense to GOLGA4 |
| ENSG00000006042 | TMEM98 | -0,083209126 | 0,943955578 | 0,72995188 | 0,974178338 | transmembrane protein 98 |
| ENSG00000174720 | LARP7 | -0,083219087 | 0,943949061 | 0,78898848 | 0,979462318 | La ribonucleoprotein 7, transcriptional regulator |
| ENSG00000165490 | DDIAS | -0,083228177 | 0,943943113 | 0,85975618 | 0,989074806 | DNA damage induced apoptosis suppressor |
| ENSG00000249799 | SNRPCP13 | -0,083231888 | 0,943940685 | 0,94798586 | 0,996216385 | small nuclear ribonucleoprotein polypeptide C pseudogene 13 |
| ENSG00000155792 | DEPTOR | -0,083286369 | 0,943905039 | 0,8848769 | 0,990717729 | DEP domain containing MTOR interacting protein |

| | | | | | | |
|---|---|---|---|---|---|---|
| ENSG00000101464 | PIGU | -0,083302469 | 0,943894505 | 0,74465685 | 0,975649186 | phosphatidylinositol glycan anchor biosynthesis class U |
| ENSG00000127423 | AUNIP | -0,083307488 | 0,943891222 | 0,76410852 | 0,976640468 | aurora kinase A and ninein interacting protein |
| ENSG00000110074 | FOXRED1 | -0,083310751 | 0,943889087 | 0,77395687 | 0,978415098 | FAD dependent oxidoreductase domain containing 1 |
| ENSG00000151849 | CENPJ | -0,083340488 | 0,943869632 | 0,81488789 | 0,981521413 | centromere protein J |
| ENSG00000109606 | DHX15 | -0,083343994 | 0,943867338 | 0,80913666 | 0,981213877 | DEAH-box helicase 15 |
| ENSG00000276772 | AC025271,4 | -0,083370758 | 0,943849828 | 0,95033063 | 0,996432861 | novel transcript |
| ENSG00000166446 | CDYL2 | -0,083384402 | 0,943840902 | 0,77197488 | 0,978033855 | chromodomain Y like 2 |
| ENSG00000235016 | SEMA3F-AS1 | -0,08352159 | 0,943751155 | 0,80856023 | 0,981213877 | SEMA3F antisense RNA 1 |
| ENSG00000134864 | GGACT | -0,083531808 | 0,943744471 | 0,81501587 | 0,981521413 | gamma-glutamylamine cyclotransferase |
| ENSG00000100297 | MCM5 | -0,083563436 | 0,943723782 | 0,68491936 | 0,965415049 | minichromosome maintenance complex component 5 |
| ENSG00000203730 | TEDDM1 | -0,08358205 | 0,943711606 | 0,94030681 | 0,995016146 | transmembrane epididymal protein 1 |
| ENSG00000248441 | LINC01197 | -0,083606727 | 0,943695463 | 0,90432856 | 0,992590661 | long intergenic non-protein coding RNA 1197 |
| ENSG00000227540 | DNAJC9-AS1 | -0,083656498 | 0,943662908 | 0,83964603 | 0,985672131 | DNAJC9 antisense RNA 1 |
| ENSG00000234718 | AC007161,1 | -0,083688304 | 0,943642104 | 0,93036502 | 0,994336072 | ribosomal protein L23a (RPL23A) pseudogene |
| ENSG00000287839 | AL353807,5 | -0,083698391 | 0,943635506 | 0,74964552 | 0,975901808 | novel transcript |
| ENSG00000149346 | SLX4IP | -0,083808031 | 0,943563796 | 0,82206052 | 0,982952211 | SLX4 interacting protein |
| ENSG00000138095 | LRPPRC | -0,083812062 | 0,943561159 | 0,81412872 | 0,981521413 | leucine rich pentatricopeptide repeat containing |
| ENSG00000254153 | AC103957,2 | -0,083816542 | 0,943558229 | 0,88898938 | 0,990717729 | novel transcript |
| ENSG00000285450 | AC270259,1 | -0,083816542 | 0,943558229 | 0,88898938 | 0,990717729 | novel transcript |
| ENSG00000126858 | RHOT1 | -0,083867866 | 0,943524663 | 0,84616805 | 0,986939308 | ras homolog family member T1 |
| ENSG00000285139 | KF459542,2 | -0,083938512 | 0,943478461 | 0,90772082 | 0,992590661 | novel transcript, antisense to KBTBD4 |
| ENSG00000231880 | KF459542,1 | -0,083938512 | 0,943478461 | 0,90772082 | 0,992590661 | novel transcript, antisense to KBTBD4 |
| ENSG00000129353 | SLC44A2 | -0,083958347 | 0,94346549 | 0,69957535 | 0,968522643 | solute carrier family 44 member 2 |
| ENSG00000042429 | MED17 | -0,083977176 | 0,943453177 | 0,78437656 | 0,979320406 | mediator complex subunit 17 |
| ENSG00000259098 | AC025884,2 | -0,083983348 | 0,94344914 | 0,87602796 | 0,990109814 | zinc finger, CCHC domain containing 2 (ZCCHC2) pseudogene |
| ENSG00000171960 | PPIH | -0,083987557 | 0,943446388 | 0,74255551 | 0,975176043 | peptidylprolyl isomerase H |
| ENSG00000129933 | MAU2 | -0,084003185 | 0,943436168 | 0,69691066 | 0,968322071 | MAU2 sister chromatid cohesion factor |
| ENSG00000123219 | CENPK | -0,084021575 | 0,943424142 | 0,86216499 | 0,989074806 | centromere protein K |
| ENSG00000164815 | ORC5 | -0,084045922 | 0,943408221 | 0,82508085 | 0,983383773 | origin recognition complex subunit 5 |
| ENSG00000130881 | LRP3 | -0,084127401 | 0,943354942 | 0,83122574 | 0,98438942 | LDL receptor related protein 3 |

| | | | | | | |
|---|---|---|---|---|---|---|
| ENSG00000243317 | STMP1 | -0,084158883 | 0,943334357 | 0,82033502 | 0,982609114 | short transmembrane mitochondrial protein 1 |
| ENSG00000221874 | ZNF816-ZNF32 | -0,084160553 | 0,943333265 | 0,92077705 | 0,993787755 | ZNF816-ZNF321P readthrough |
| ENSG00000163214 | DHX57 | -0,084187438 | 0,943315686 | 0,77031798 | 0,977872051 | DExH-box helicase 57 |
| ENSG00000134461 | ANKRD16 | -0,084239669 | 0,943281534 | 0,74492538 | 0,975666349 | ankyrin repeat domain 16 |
| ENSG00000136319 | TTC5 | -0,084275521 | 0,943258094 | 0,78211264 | 0,978597905 | tetratricopeptide repeat domain 5 |
| ENSG00000260644 | HERC2P5 | -0,084302086 | 0,943240725 | 0,76794542 | 0,977368032 | hect domain and RLD 2 pseudogene 5 |
| ENSG00000285442 | FBXL6 | -0,084305448 | 0,943238527 | 0,82651285 | 0,983471137 | F-box and leucine rich repeat protein 6 |
| ENSG00000182325 | FBXL6 | -0,084305448 | 0,943238527 | 0,82651285 | 0,983471137 | F-box and leucine rich repeat protein 6 |
| ENSG00000225073 | DDX39B | -0,084325464 | 0,943225441 | 0,67726498 | 0,963580269 | DExD-box helicase 39B |
| ENSG00000129235 | TXNDC17 | -0,084654808 | 0,943010142 | 0,8137844 | 0,981521413 | thioredoxin domain containing 17 |
| ENSG00000155959 | VBP1 | -0,084683835 | 0,942991169 | 0,84981884 | 0,987036295 | VHL binding protein 1 |
| ENSG00000100749 | VRK1 | -0,084741888 | 0,942953224 | 0,85410607 | 0,987861659 | VRK serine/threonine kinase 1 |
| ENSG00000224083 | MTCO1P11 | -0,084750269 | 0,942947747 | 0,90497684 | 0,992590661 | MT-CO1 pseudogene 11 |
| ENSG00000166167 | BTRC | -0,084758157 | 0,942942591 | 0,80618925 | 0,981213877 | beta-transducin repeat containing E3 ubiquitin protein ligase |
| ENSG00000138709 | LARP1B | -0,084785462 | 0,942924744 | 0,79167851 | 0,979471509 | La ribonucleoprotein 1B |
| ENSG00000271382 | AC012464,3 | -0,084798959 | 0,942915923 | 0,93425852 | 0,994479239 | novel transcript |
| ENSG00000269246 | AC011445,1 | -0,084803066 | 0,942913239 | 0,9375878 | 0,994479239 | novel transcript |
| ENSG00000272941 | AC083862,1 | -0,084866161 | 0,942872002 | 0,83432154 | 0,984905093 | novel transcript, antisense to C7orf49 |
| ENSG00000187857 | OR6C75 | -0,084957357 | 0,942812403 | 0,94728634 | 0,996063635 | olfactory receptor family 6 subfamily C member 75 |
| ENSG00000178252 | WDR6 | -0,084961617 | 0,942809619 | 0,78388628 | 0,979239248 | WD repeat domain 6 |
| ENSG00000164053 | ATRIP | -0,085054193 | 0,942749122 | 0,70733495 | 0,969565008 | ATR interacting protein |
| ENSG00000100129 | EIF3L | -0,085078443 | 0,942733276 | 0,83814135 | 0,985672131 | eukaryotic translation initiation factor 3 subunit L |
| ENSG00000283078 | AL137077,2 | -0,085089418 | 0,942726104 | 0,86910724 | 0,989074806 | novel transcript |
| ENSG00000163867 | ZMYM6 | -0,085094921 | 0,942722508 | 0,85019201 | 0,987036295 | zinc finger MYM-type containing 6 |
| ENSG00000237883 | DGUOK-AS1 | -0,085101873 | 0,942717965 | 0,86120082 | 0,989074806 | DGUOK antisense RNA 1 |
| ENSG00000135537 | AFG1L | -0,085127414 | 0,942701276 | 0,80994022 | 0,981213877 | AFG1 like ATPase |
| ENSG00000225400 | RAB28P5 | -0,085141237 | 0,942692244 | 0,86143407 | 0,989074806 | RAB28, member RAS oncogene family pseudogene 5 |
| ENSG00000100014 | SPECC1L | -0,085154642 | 0,942683485 | 0,71600324 | 0,971360663 | sperm antigen with calponin homology and coiled-coil domains 1 like |
| ENSG00000169609 | C15orf40 | -0,085166054 | 0,942676028 | 0,82185776 | 0,982952211 | chromosome 15 open reading frame 40 |
| ENSG00000117620 | SLC35A3 | -0,08525092 | 0,942620577 | 0,70989613 | 0,970482408 | solute carrier family 35 member A3 |
| ENSG00000164638 | SLC29A4 | -0,085305209 | 0,942585106 | 0,78171872 | 0,978597905 | solute carrier family 29 member 4 |

| | | | | | | |
|---|---|---|---|---|---|---|
| ENSG00000112739 | PRPF4B | -0,085309794 | 0,942582111 | 0,82069465 | 0,982609114 | pre-mRNA processing factor 4B |
| ENSG00000241837 | ATP5PO | -0,085336457 | 0,942564691 | 0,78529575 | 0,979320406 | ATP synthase peripheral stalk subunit OSCP |
| ENSG00000115446 | UNC50 | -0,085338037 | 0,942563659 | 0,80442522 | 0,980875302 | unc-50 inner nuclear membrane RNA binding protein |
| ENSG00000231713 | AF064860,2 | -0,085396354 | 0,942525559 | 0,90072149 | 0,991865079 | novel transcript |
| ENSG00000120262 | CCDC170 | -0,085520555 | 0,942444421 | 0,92126349 | 0,993787755 | coiled-coil domain containing 170 |
| ENSG00000052723 | SIKE1 | -0,085523105 | 0,942442755 | 0,85612815 | 0,988185425 | suppressor of IKBKE 1 |
| ENSG00000175606 | TMEM70 | -0,08553728 | 0,942433495 | 0,85474174 | 0,987861659 | transmembrane protein 70 |
| ENSG00000114650 | SCAP | -0,085591084 | 0,942398348 | 0,75637058 | 0,975901808 | SREBF chaperone |
| ENSG00000244513 | AC109587,1 | -0,085613087 | 0,942383976 | 0,84032428 | 0,985672131 | novel transcript |
| ENSG00000112081 | SRSF3 | -0,085659 | 0,942353986 | 0,79035425 | 0,979462318 | serine and arginine rich splicing factor 3 |
| ENSG00000182568 | SATB1 | -0,085717482 | 0,942315786 | 0,79004246 | 0,979462318 | SATB homeobox 1 |
| ENSG00000051523 | CYBA | -0,085739004 | 0,942301729 | 0,84433693 | 0,986340494 | cytochrome b-245 alpha chain |
| ENSG00000235453 | SMIM27 | -0,085850995 | 0,942228585 | 0,75386951 | 0,975901808 | small integral membrane protein 27 |
| ENSG00000138794 | CASP6 | -0,085898954 | 0,942197263 | 0,79269244 | 0,979471509 | caspase 6 |
| ENSG00000205056 | LINC02397 | -0,085917763 | 0,942184979 | 0,89103762 | 0,990780829 | long intergenic non-protein coding RNA 2397 |
| ENSG00000101220 | C20orf27 | -0,086019132 | 0,94211878 | 0,82828658 | 0,983646162 | chromosome 20 open reading frame 27 |
| ENSG00000259814 | AC102941,1 | -0,086070668 | 0,942085126 | 0,92828795 | 0,994108531 | novel transcript |
| ENSG00000076382 | SPAG5 | -0,086088463 | 0,942073506 | 0,69828935 | 0,968522643 | sperm associated antigen 5 |
| ENSG00000132549 | VPS13B | -0,086126041 | 0,942048968 | 0,81596902 | 0,981708551 | vacuolar protein sorting 13 homolog B |
| ENSG00000235314 | LINC00957 | -0,086164058 | 0,942024144 | 0,88802191 | 0,990717729 | long intergenic non-protein coding RNA 957 |
| ENSG00000127483 | HP1BP3 | -0,086171346 | 0,942019385 | 0,75720416 | 0,975901808 | heterochromatin protein 1 binding protein 3 |
| ENSG00000100852 | ARHGAP5 | -0,086257528 | 0,941963114 | 0,85260037 | 0,98761642 | Rho GTPase activating protein 5 |
| ENSG00000095464 | PDE6C | -0,08625831 | 0,941962603 | 0,91418782 | 0,993361299 | phosphodiesterase 6C |
| ENSG00000187522 | HSPA14 | -0,086264169 | 0,941958778 | 0,82255393 | 0,98299282 | heat shock protein family A (Hsp70) member 14 |
| ENSG00000241360 | PDXP | -0,086331168 | 0,941915035 | 0,73376429 | 0,974353474 | pyridoxal phosphatase |
| ENSG00000259956 | RBM15B | -0,08633124 | 0,941914987 | 0,66719504 | 0,9611116 | RNA binding motif protein 15B |
| ENSG00000245261 | AL133375,1 | -0,086357261 | 0,941897999 | 0,96943234 | 0,997494004 | novel transcript |
| ENSG00000219410 | AC125494,1 | -0,086379318 | 0,941883599 | 0,83155922 | 0,984456953 | novel transcript |
| ENSG00000270585 | SNX2P1 | -0,086389784 | 0,941876766 | 0,87782271 | 0,990109814 | sorting nexin 2 pseudogene 1 |
| ENSG00000116183 | PAPPA2 | -0,086400481 | 0,941869782 | 0,87272778 | 0,989262536 | pappalysin 2 |
| ENSG00000196696 | PDXDC2P-NPIP | -0,086414198 | 0,941860827 | 0,69876476 | 0,968522643 | nuclear pore complex-interacting protein |
| ENSG00000206448 | PPIAP30 | -0,086472859 | 0,94182253 | 0,94021228 | 0,994949796 | peptidylprolyl isomerase A pseudogene 30 |

| | | | | | | |
|---|---|---|---|---|---|---|
| ENSG00000137770 | CTDSPL2 | -0,086512894 | 0,941796395 | 0,84743722 | 0,987035044 | CTD small phosphatase like 2 |
| ENSG00000151893 | CACUL1 | -0,086524438 | 0,941788859 | 0,79586074 | 0,979879951 | CDK2 associated cullin domain 1 |
| ENSG00000182054 | IDH2 | -0,086589512 | 0,94174638 | 0,68710194 | 0,965769168 | isocitrate dehydrogenase (NADP(+)) 2 |
| ENSG00000223838 | AC007091,1 | -0,086705309 | 0,941670795 | 0,93397098 | 0,994479239 | novel transcript |
| ENSG00000008294 | SPAG9 | -0,086802066 | 0,941607642 | 0,76841305 | 0,977381742 | sperm associated antigen 9 |
| ENSG00000162402 | USP24 | -0,086877316 | 0,941558529 | 0,77434911 | 0,978503783 | ubiquitin specific peptidase 24 |
| ENSG00000165185 | KIAA1958 | -0,086891151 | 0,9415495 | 0,81009837 | 0,981213877 | KIAA1958 |
| ENSG00000250934 | AC016924,1 | -0,086927127 | 0,941526022 | 0,8969263 | 0,991644439 | novel transcript |
| ENSG00000116194 | ANGPTL1 | -0,087037839 | 0,941453772 | 0,86378658 | 0,989074806 | angiopoietin like 1 |
| ENSG00000267289 | AC008752,2 | -0,087063103 | 0,941437286 | 0,94651945 | 0,996027077 | novel transcript |
| ENSG00000275481 | AC025031,4 | -0,087144861 | 0,941383935 | 0,86781078 | 0,989074806 | novel transcript |
| ENSG00000162572 | SCNN1D | -0,087154825 | 0,941377434 | 0,84653952 | 0,986939308 | sodium channel epithelial 1 subunit delta |
| ENSG00000111269 | CREBL2 | -0,087191783 | 0,941353318 | 0,80926622 | 0,981213877 | cAMP responsive element binding protein like 2 |
| ENSG00000136002 | ARHGEF4 | -0,087271836 | 0,941301086 | 0,81158547 | 0,981213877 | Rho guanine nucleotide exchange factor 4 |
| ENSG00000114738 | MAPKAPK3 | -0,087432577 | 0,941196214 | 0,75527869 | 0,975901808 | MAPK activated protein kinase 3 |
| ENSG00000172845 | SP3 | -0,0874674 | 0,941173496 | 0,83248534 | 0,98489473 | Sp3 transcription factor |
| ENSG00000091844 | RGS17 | -0,087504051 | 0,941149587 | 0,84829684 | 0,987036295 | regulator of G protein signaling 17 |
| ENSG00000275723 | NAPEPLD | -0,087606741 | 0,941082599 | 0,80860365 | 0,981213877 | N-acyl phosphatidylethanolamine phospholipase D |
| ENSG00000137218 | FRS3 | -0,087641143 | 0,941060159 | 0,77063081 | 0,977966587 | fibroblast growth factor receptor substrate 3 |
| ENSG00000122986 | HVCN1 | -0,087656566 | 0,941050098 | 0,82144123 | 0,982950156 | hydrogen voltage gated channel 1 |
| ENSG00000100505 | TRIM9 | -0,087660565 | 0,94104749 | 0,75715676 | 0,975901808 | tripartite motif containing 9 |
| ENSG00000225706 | PTPRD-AS1 | -0,087765169 | 0,940979261 | 0,84166513 | 0,98571999 | PTPRD antisense RNA 1 |
| ENSG00000151883 | PARP8 | -0,087779156 | 0,940970138 | 0,84508074 | 0,986639902 | poly(ADP-ribose) polymerase family member 8 |
| ENSG00000232112 | TMA7 | -0,087817865 | 0,940944891 | 0,77335448 | 0,97839731 | translation machinery associated 7 homolog |
| ENSG00000258521 | AL157871,2 | -0,087839399 | 0,940930846 | 0,92906491 | 0,994293848 | novel transcript, antisense to WARS |
| ENSG00000260625 | AC026471,2 | -0,087862744 | 0,940915621 | 0,8780363 | 0,990109814 | novel transcript, antisense to C16orf58 |
| ENSG00000286279 | AC108721,2 | -0,087866593 | 0,940913111 | 0,93337711 | 0,994479239 | novel transcript |
| ENSG00000147144 | CCDC120 | -0,088012523 | 0,940817941 | 0,73239425 | 0,974353474 | coiled-coil domain containing 120 |
| ENSG00000163728 | TTC14 | -0,08802626 | 0,940808983 | 0,84852715 | 0,987036295 | tetratricopeptide repeat domain 14 |
| ENSG00000261485 | PAN3-AS1 | -0,088045187 | 0,94079664 | 0,84954488 | 0,987036295 | PAN3 antisense RNA 1 |
| ENSG00000166135 | HIF1AN | -0,088057095 | 0,940788875 | 0,67015422 | 0,962026636 | hypoxia inducible factor 1 subunit alpha inhibitor |
| ENSG00000275650 | TTYH1 | -0,088090351 | 0,940767189 | 0,8664439 | 0,989074806 | tweety family member 1 |
| ENSG00000271828 | AC008937,3 | -0,088112121 | 0,940752993 | 0,80102661 | 0,980555343 | novel transcript, antisense to MIER3 |

| | | | | | | |
|---|---|---|---|---|---|---|
| ENSG00000183091 | NEB | -0,088204712 | 0,940692618 | 0,78490155 | 0,979320406 | nebulin |
| ENSG00000256060 | TRAPPC2B | -0,088222434 | 0,940681062 | 0,82044771 | 0,982609114 | trafficking protein particle complex 2B |
| ENSG00000170100 | ZNF778 | -0,088269064 | 0,940650659 | 0,70012384 | 0,968522643 | zinc finger protein 778 |
| ENSG00000104524 | PYCR3 | -0,088272454 | 0,940648448 | 0,73303837 | 0,974353474 | pyrroline-5-carboxylate reductase 3 |
| ENSG00000287188 | AC068989,1 | -0,088308439 | 0,940624987 | 0,92445402 | 0,993915537 | novel transcript, antisense to ANXA10 |
| ENSG00000280670 | CCDC163 | -0,088332541 | 0,940609272 | 0,81812102 | 0,982323352 | coiled-coil domain containing 163 |
| ENSG00000198342 | ZNF442 | -0,088340563 | 0,940604042 | 0,72246793 | 0,972594421 | zinc finger protein 442 |
| ENSG00000241769 | LINC00893 | -0,088343193 | 0,940602327 | 0,84802031 | 0,987036295 | long intergenic non-protein coding RNA 893 |
| ENSG00000109072 | VTN | -0,088408145 | 0,940559981 | 0,91229733 | 0,993361299 | vitronectin |
| ENSG00000166136 | NDUFB8 | -0,088464642 | 0,940523149 | 0,72671306 | 0,973344786 | NADH:ubiquinone oxidoreductase subunit B8 |
| ENSG00000136021 | SCYL2 | -0,088480963 | 0,940512509 | 0,84770011 | 0,987035044 | SCY1 like pseudokinase 2 |
| ENSG00000273611 | ZNHIT3 | -0,088485802 | 0,940509354 | 0,76351938 | 0,976640468 | zinc finger HIT-type containing 3 |
| ENSG00000184068 | SREBF2-AS1 | -0,088519963 | 0,940487084 | 0,86148543 | 0,989074806 | SREBF2 antisense RNA 1 |
| ENSG00000157216 | SSBP3 | -0,088660565 | 0,940395431 | 0,73255312 | 0,974353474 | single stranded DNA binding protein 3 |
| ENSG00000119707 | RBM25 | -0,088686714 | 0,940378387 | 0,74829523 | 0,975901808 | RNA binding motif protein 25 |
| ENSG00000166478 | ZNF143 | -0,088827642 | 0,940286531 | 0,80490482 | 0,98109454 | zinc finger protein 143 |
| ENSG00000181827 | RFX7 | -0,088846383 | 0,940274317 | 0,84378637 | 0,986273696 | regulatory factor X7 |
| ENSG00000052841 | TTC17 | -0,088989989 | 0,940180726 | 0,78495605 | 0,979320406 | tetratricopeptide repeat domain 17 |
| ENSG00000139620 | KANSL2 | -0,089017837 | 0,940162579 | 0,72526444 | 0,97326334 | KAT8 regulatory NSL complex subunit 2 |
| ENSG00000169288 | MRPL1 | -0,089121528 | 0,940095009 | 0,80188883 | 0,980710009 | mitochondrial ribosomal protein L1 |
| ENSG00000144354 | CDCA7 | -0,089131068 | 0,940088792 | 0,77667986 | 0,978587255 | cell division cycle associated 7 |
| ENSG00000103811 | CTSH | -0,089168272 | 0,94006455 | 0,81959805 | 0,98249617 | cathepsin H |
| ENSG00000228451 | SDAD1P1 | -0,089174684 | 0,940060371 | 0,84555282 | 0,986639902 | SDA1 domain containing 1 pseudogene 1 |
| ENSG00000274419 | TBC1D3D | -0,089199694 | 0,940044075 | 0,82678521 | 0,983548538 | TBC1 domain family member 3D |
| ENSG00000147586 | MRPS28 | -0,089459091 | 0,93987507 | 0,780004 | 0,978587255 | mitochondrial ribosomal protein S28 |
| ENSG00000145979 | TBC1D7 | -0,089511045 | 0,939841224 | 0,78552219 | 0,979320406 | TBC1 domain family member 7 |
| ENSG00000138376 | BARD1 | -0,089551879 | 0,939814623 | 0,82758365 | 0,983595244 | BRCA1 associated RING domain 1 |
| ENSG00000174529 | TMEM81 | -0,089575605 | 0,939799168 | 0,78468888 | 0,979320406 | transmembrane protein 81 |
| ENSG00000127804 | METTL16 | -0,089577996 | 0,93979761 | 0,65518638 | 0,957444211 | methyltransferase like 16 |
| ENSG00000285636 | AL450163,1 | -0,089627904 | 0,9397651 | 0,90151302 | 0,992041918 | Novel transcript, antisense to NOS1AP |
| ENSG00000167635 | ZNF146 | -0,089716694 | 0,939707264 | 0,79361961 | 0,979776981 | zinc finger protein 146 |
| ENSG00000160679 | CHTOP | -0,089722383 | 0,939703558 | 0,71659179 | 0,971438758 | chromatin target of PRMT1 |
| ENSG00000237401 | LINC01304 | -0,089795069 | 0,939656215 | 0,94264356 | 0,995390305 | long intergenic non-protein coding RNA 1304 |

| | | | | | | |
|---|---|---|---|---|---|---|
| ENSG00000229644 | NAMPTP1 | -0,089830117 | 0,939633388 | 0,86274892 | 0,989074806 | nicotinamide phosphoribosyltransferase pseudogene 1 |
| ENSG00000089234 | BRAP | -0,089934697 | 0,939565277 | 0,792006 | 0,979471509 | BRCA1 associated protein |
| ENSG00000153879 | CEBPG | -0,08996408 | 0,939546142 | 0,80912712 | 0,981213877 | CCAAT enhancer binding protein gamma |
| ENSG00000235586 | AC011247,1 | -0,08996874 | 0,939543107 | 0,93903815 | 0,994784787 | novel transcript |
| ENSG00000145626 | UGT3A1 | -0,08999095 | 0,939528643 | 0,94039971 | 0,995028467 | UDP glycosyltransferase family 3 member A1 |
| ENSG00000260733 | AC046158,1 | -0,090026079 | 0,939505766 | 0,91742465 | 0,993361299 | novel transcript, antisense to WWOX |
| ENSG00000273306 | AC018690,1 | -0,090086025 | 0,939466729 | 0,85304454 | 0,98780934 | novel transcript |
| ENSG00000257120 | AL356756,1 | -0,090111105 | 0,939450397 | 0,91218335 | 0,993361299 | novel transcript, antisense to PRKD1 |
| ENSG00000269293 | ZSCAN16-AS1 | -0,090129197 | 0,939438617 | 0,71510398 | 0,971041579 | ZSCAN16 antisense RNA 1 |
| ENSG00000226029 | LINC01772 | -0,09024302 | 0,939364501 | 0,79740261 | 0,980161849 | long intergenic non-protein coding RNA 1772 |
| ENSG00000112473 | SLC39A7 | -0,090355041 | 0,939291565 | 0,88471534 | 0,990717729 | solute carrier family 39 member 7 |
| ENSG00000272831 | AC027644,3 | -0,090404334 | 0,939259473 | 0,7246743 | 0,97326334 | novel transcript, antisense to RABGEF1 |
| ENSG00000260701 | AC025284,1 | -0,090425894 | 0,939245436 | 0,94771228 | 0,996113938 | novel transcript, antisense to ADAMTS18 |
| ENSG00000122515 | ZMIZ2 | -0,090426856 | 0,93924481 | 0,72889805 | 0,974009551 | zinc finger MIZ-type containing 2 |
| ENSG00000236296 | GUSBP5 | -0,090547957 | 0,939165972 | 0,8514274 | 0,987259878 | GUSB pseudogene 5 |
| ENSG00000168438 | CDC40 | -0,090588379 | 0,939139659 | 0,81179893 | 0,981213877 | cell division cycle 40 |
| ENSG00000143320 | CRABP2 | -0,090614257 | 0,939122813 | 0,84285269 | 0,986208256 | cellular retinoic acid binding protein 2 |
| ENSG00000282202 | AC015849,10 | -0,090621064 | 0,939118383 | 0,91786587 | 0,993361299 | novel transcript, antisense TAF15 |
| ENSG00000270871 | AC015849,3 | -0,090621064 | 0,939118383 | 0,91786587 | 0,993361299 | novel transcript, antisense TAF15 |
| ENSG00000182836 | PLCXD3 | -0,090688029 | 0,939074793 | 0,89731388 | 0,991644439 | phosphatidylinositol specific phospholipase C X domain containing 3 |
| ENSG00000287261 | AL139243,1 | -0,090691804 | 0,939072336 | 0,91590575 | 0,993361299 | novel transcript, antisense to HPS1 |
| ENSG00000186431 | FCAR | -0,090707788 | 0,939061932 | 0,9736865 | 0,99837368 | Fc fragment of IgA receptor |
| ENSG00000263006 | ROCK1P1 | -0,090866602 | 0,938958564 | 0,86566588 | 0,989074806 | Rho associated coiled-coil containing protein kinase 1 pseudogene 1 |
| ENSG00000285416 | AC137630,5 | -0,090883149 | 0,938947794 | 0,93244728 | 0,994479239 | novel pseudogene, ortholog of avian secreted frizzled-related protein 5 |
| ENSG00000100410 | PHF5A | -0,090940015 | 0,938910785 | 0,74244485 | 0,975176043 | PHD finger protein 5A |
| ENSG00000214193 | SH3D21 | -0,091057798 | 0,938834134 | 0,86046991 | 0,989074806 | SH3 domain containing 21 |
| ENSG00000131368 | MRPS25 | -0,091107654 | 0,938801691 | 0,68104937 | 0,964654539 | mitochondrial ribosomal protein S25 |
| ENSG00000182973 | CNOT10 | -0,091329657 | 0,938657239 | 0,75690345 | 0,975901808 | CCR4-NOT transcription complex subunit 10 |
| ENSG00000268743 | AC008737,1 | -0,091417055 | 0,938600377 | 0,94344721 | 0,995402494 | novel transcript, sense intronic SIN3B |

| | | | | | | |
|---|---|---|---|---|---|---|
| ENSG00000168944 | CEP120 | -0,091614787 | 0,938471744 | 0,80294971 | 0,980866933 | centrosomal protein 120 |
| ENSG00000078070 | MCCC1 | -0,091643229 | 0,938453242 | 0,7468199 | 0,975796253 | methylcrotonoyl-CoA carboxylase 1 |
| ENSG00000120729 | MYOT | -0,091692585 | 0,938421138 | 0,9226704 | 0,993797669 | myotilin |
| ENSG00000213862 | AC044787,1 | -0,09176843 | 0,938371805 | 0,90329441 | 0,992311172 | ribosomal protein L7a (RPL7A) pseudogene |
| ENSG00000238057 | ZEB2-AS1 | -0,091772158 | 0,938369379 | 0,84029772 | 0,985672131 | ZEB2 antisense RNA 1 |
| ENSG00000262848 | AC136624,2 | -0,091805848 | 0,938347467 | 0,89903806 | 0,991832796 | novel transcript, antisense to NOMO3 |
| ENSG00000282332 | AC244489,4 | -0,091805848 | 0,938347467 | 0,89903806 | 0,991832796 | novel transcript, antisense to NOMO3 |
| ENSG00000166797 | CIAO2A | -0,091854706 | 0,938315689 | 0,80153268 | 0,980623173 | cytosolic iron-sulfur assembly component 2A |
| ENSG00000241959 | RN7SL76P | -0,091866444 | 0,938308056 | 0,94583366 | 0,996027077 | RNA, 7SL, cytoplasmic 76, pseudogene |
| ENSG00000113119 | TMCO6 | -0,091894815 | 0,938289604 | 0,76988212 | 0,977742359 | transmembrane and coiled-coil domains 6 |
| ENSG00000073711 | PPP2R3A | -0,091981268 | 0,938233379 | 0,81198098 | 0,981213877 | protein phosphatase 2 regulatory subunit B''alpha |
| ENSG00000167904 | TMEM68 | -0,091996677 | 0,938223357 | 0,72905279 | 0,974009551 | transmembrane protein 68 |
| ENSG00000241882 | AC124893,1 | -0,091997833 | 0,938222606 | 0,89531891 | 0,991303328 | novel transcript |
| ENSG00000119720 | NRDE2 | -0,092008443 | 0,938215706 | 0,67107613 | 0,962026636 | NRDE-2, necessary for RNA interference, domain containing |
| ENSG00000287425 | AP003717,4 | -0,092021253 | 0,938207375 | 0,90032151 | 0,991865079 | novel transcript |
| ENSG00000162972 | MAIP1 | -0,092023875 | 0,93820567 | 0,81333047 | 0,981282128 | matrix AAA peptidase interacting protein 1 |
| ENSG00000124207 | CSE1L | -0,092034014 | 0,938199076 | 0,83285788 | 0,984905093 | chromosome segregation 1 like |
| ENSG00000273308 | AC024560,3 | -0,092076653 | 0,938171349 | 0,92337015 | 0,993825008 | novel transcript |
| ENSG00000265943 | AC090912,1 | -0,092138407 | 0,938131191 | 0,93805101 | 0,994488324 | novel transcript, antisense to RBBP8 |
| ENSG00000169375 | SIN3A | -0,092144222 | 0,93812741 | 0,73789389 | 0,974353474 | SIN3 transcription regulator family member A |
| ENSG00000147813 | NAPRT | -0,092169641 | 0,938110881 | 0,84683849 | 0,986939308 | nicotinate phosphoribosyltransferase |
| ENSG00000256222 | MTRNR2L3 | -0,092210933 | 0,938084031 | 0,90536092 | 0,992590661 | MT-RNR2 like 3 |
| ENSG00000006451 | RALA | -0,092294461 | 0,938029721 | 0,79359469 | 0,979776981 | RAS like proto-oncogene A |
| ENSG00000161682 | FAM171A2 | -0,092394868 | 0,937964439 | 0,82004525 | 0,982609114 | family with sequence similarity 171 member A2 |
| ENSG00000274849 | AC023043,4 | -0,092398422 | 0,937962129 | 0,9232062 | 0,993825008 | novel transcript, sense intronic to MOCOS |
| ENSG00000140391 | TSPAN3 | -0,092401042 | 0,937960425 | 0,78422 | 0,979320406 | tetraspanin 3 |
| ENSG00000286347 | AC114802,1 | -0,092458678 | 0,937922954 | 0,87741138 | 0,990109814 | novel transcript, antisense to ANTXR1 |
| ENSG00000275947 | TMEM251 | -0,092476692 | 0,937911243 | 0,8589816 | 0,989074806 | transmembrane protein 251 |
| ENSG00000257103 | LSM14A | -0,092549121 | 0,937864157 | 0,71807576 | 0,97178302 | LSM14A mRNA processing body assembly factor |
| ENSG00000119335 | SET | -0,092569609 | 0,937850838 | 0,72740076 | 0,973441576 | SET nuclear proto-oncogene |
| ENSG00000070761 | CFAP20 | -0,092754186 | 0,937730858 | 0,81529397 | 0,981521413 | cilia and flagella associated protein 20 |
| ENSG00000135597 | REPS1 | -0,092825602 | 0,93768444 | 0,77172101 | 0,978033855 | RALBP1 associated Eps domain containing 1 |

| | | | | | | |
|---|---|---|---|---|---|---|
| ENSG00000251497 | PITPNM2-AS1 | -0,09285715 | 0,937663936 | 0,91476545 | 0,993361299 | PITPNM2 antisense RNA 1 |
| ENSG00000229585 | RPL21P44 | -0,092867384 | 0,937657284 | 0,92640918 | 0,993915537 | ribosomal protein L21 pseudogene 44 |
| ENSG00000156875 | MFSD14A | -0,092876376 | 0,93765144 | 0,82432897 | 0,983374097 | major facilitator superfamily domain containing 14A |
| ENSG00000147257 | GPC3 | -0,093014159 | 0,937561895 | 0,84279992 | 0,986200189 | glypican 3 |
| ENSG00000166123 | GPT2 | -0,093071203 | 0,937524824 | 0,71815191 | 0,97178302 | glutamic--pyruvic transaminase 2 |
| ENSG00000144028 | SNRNP200 | -0,093088346 | 0,937513684 | 0,66481751 | 0,960551345 | small nuclear ribonucleoprotein U5 subunit 200 |
| ENSG00000164327 | RICTOR | -0,093125371 | 0,937489624 | 0,80521788 | 0,981187666 | RPTOR independent companion of MTOR complex 2 |
| ENSG00000018236 | CNTN1 | -0,093165607 | 0,937463479 | 0,80686717 | 0,981213877 | contactin 1 |
| ENSG00000135441 | BLOC1S1 | -0,093206038 | 0,937437207 | 0,70561533 | 0,969485452 | biogenesis of lysosomal organelles complex 1 subunit 1 |
| ENSG00000265293 | ARGFXP2 | -0,093276336 | 0,93739153 | 0,89222603 | 0,991039277 | arginine-fifty homeobox pseudogene 2 |
| ENSG00000138621 | PPCDC | -0,093389201 | 0,937318199 | 0,78701017 | 0,979424965 | phosphopantothenoylcysteine decarboxylase |
| ENSG00000133114 | GPALPP1 | -0,093391748 | 0,937316544 | 0,75417706 | 0,975901808 | GPALPP motifs containing 1 |
| ENSG00000181938 | GINS3 | -0,093393345 | 0,937315506 | 0,76532612 | 0,976830764 | GINS complex subunit 3 |
| ENSG00000113552 | GNPDA1 | -0,093399694 | 0,937311381 | 0,78079262 | 0,978597905 | glucosamine-6-phosphate deaminase 1 |
| ENSG00000107518 | ATRNL1 | -0,093426689 | 0,937293843 | 0,8249542 | 0,983374097 | attractin like 1 |
| ENSG00000189046 | ALKBH2 | -0,093468181 | 0,937266887 | 0,68783687 | 0,966105026 | alkB homolog 2, alpha-ketoglutarate dependent dioxygenase |
| ENSG00000137492 | THAP12 | -0,093490928 | 0,937252109 | 0,82309625 | 0,983204348 | THAP domain containing 12 |
| ENSG00000089094 | KDM2B | -0,093566714 | 0,937202876 | 0,72471877 | 0,97326334 | lysine demethylase 2B |
| ENSG00000151952 | TMEM132D | -0,093644783 | 0,937152162 | 0,84127456 | 0,985672131 | transmembrane protein 132D |
| ENSG00000086232 | EIF2AK1 | -0,093718215 | 0,937104463 | 0,78195468 | 0,978597905 | eukaryotic translation initiation factor 2 alpha kinase 1 |
| ENSG00000158161 | EYA3 | -0,093718216 | 0,937104462 | 0,72448709 | 0,97326334 | EYA transcriptional coactivator and phosphatase 3 |
| ENSG00000258168 | AC025569,1 | -0,093740498 | 0,937089989 | 0,88933062 | 0,990717729 | novel transcript, antisense to PTPRB |
| ENSG00000108733 | PEX12 | -0,093808396 | 0,937045887 | 0,78565532 | 0,979320406 | peroxisomal biogenesis factor 12 |
| ENSG00000207263 | SNORD116-16 | -0,093895307 | 0,93698944 | 0,86357655 | 0,989074806 | small nucleolar RNA, C/D box 116-16 |
| ENSG00000134874 | DZIP1 | -0,093895983 | 0,936989 | 0,7646666 | 0,976640468 | DAZ interacting zinc finger protein 1 |
| ENSG00000285261 | AC270285,6 | -0,093991086 | 0,936927236 | 0,89978351 | 0,991845106 | novel transcript |
| ENSG00000261329 | AC016597,1 | -0,094055269 | 0,936885555 | 0,76830123 | 0,977381742 | novel transcript, antisense to a novel protein |
| ENSG00000135837 | CEP350 | -0,094058164 | 0,936883675 | 0,78992323 | 0,979462318 | centrosomal protein 350 |
| ENSG00000126602 | TRAP1 | -0,09414301 | 0,936828577 | 0,7180184 | 0,971780402 | TNF receptor associated protein 1 |
| ENSG00000138083 | SIX3 | -0,094179063 | 0,936805166 | 0,81180204 | 0,981213877 | SIX homeobox 3 |

| | | | | | | |
|---|---|---|---|---|---|---|
| ENSG00000089486 | CDIP1 | -0,094253016 | 0,936757146 | 0,73248181 | 0,974353474 | cell death inducing p53 target 1 |
| ENSG00000102384 | CENPI | -0,094322083 | 0,936712302 | 0,78751641 | 0,979462318 | centromere protein I |
| ENSG00000243193 | PRKAR2B-AS1 | -0,094443025 | 0,93663378 | 0,9041163 | 0,992531054 | PRKAR2B antisense RNA 1 |
| ENSG00000260101 | AC008074,2 | -0,09450569 | 0,936593097 | 0,8812989 | 0,990717729 | novel transcript |
| ENSG00000102081 | FMR1 | -0,09476088 | 0,936427443 | 0,81030574 | 0,981213877 | FMRP translational regulator 1 |
| ENSG00000160191 | PDE9A | -0,094772074 | 0,936420177 | 0,68598033 | 0,965740648 | phosphodiesterase 9A |
| ENSG00000176834 | VSIG10 | -0,094829809 | 0,936382704 | 0,68445697 | 0,965415049 | V-set and immunoglobulin domain containing 10 |
| ENSG00000135968 | GCC2 | -0,094916533 | 0,936326417 | 0,77286407 | 0,978310326 | GRIP and coiled-coil domain containing 2 |
| ENSG00000168065 | SLC22A11 | -0,094960314 | 0,936298003 | 0,93775915 | 0,994479239 | solute carrier family 22 member 11 |
| ENSG00000105983 | LMBR1 | -0,095063426 | 0,936231086 | 0,80359254 | 0,980875302 | limb development membrane protein 1 |
| ENSG00000173581 | CCDC106 | -0,095119825 | 0,936194488 | 0,79628514 | 0,980015022 | coiled-coil domain containing 106 |
| ENSG00000134884 | ARGLU1 | -0,095130327 | 0,936187672 | 0,7868801 | 0,979417924 | arginine and glutamate rich 1 |
| ENSG00000128708 | HAT1 | -0,095201378 | 0,936141567 | 0,82870966 | 0,983784613 | histone acetyltransferase 1 |
| ENSG00000151247 | EIF4E | -0,095235546 | 0,936119397 | 0,8265637 | 0,983471137 | eukaryotic translation initiation factor 4E |
| ENSG00000146276 | GABRR1 | -0,09535161 | 0,93604409 | 0,93851471 | 0,994600991 | gamma-aminobutyric acid type A receptor subunit rho1 |
| ENSG00000261019 | AC010132,4 | -0,095396212 | 0,936015151 | 0,90233252 | 0,992230188 | novel transcript |
| ENSG00000171659 | GPR34 | -0,095405409 | 0,936009184 | 0,84848621 | 0,987036295 | G protein-coupled receptor 34 |
| ENSG00000144468 | RHBDD1 | -0,095435595 | 0,9359896 | 0,79912196 | 0,980161849 | rhomboid domain containing 1 |
| ENSG00000269176 | AP001160,3 | -0,095461381 | 0,935972871 | 0,81022349 | 0,981213877 | novel transcript, antisense to TAF6L |
| ENSG00000183647 | ZNF530 | -0,095507106 | 0,935943206 | 0,70975012 | 0,970482408 | zinc finger protein 530 |
| ENSG00000123473 | STIL | -0,095543647 | 0,935919501 | 0,80341736 | 0,980866933 | STIL centriolar assembly protein |
| ENSG00000126768 | TIMM17B | -0,095652489 | 0,935848894 | 0,69116951 | 0,967179456 | translocase of inner mitochondrial membrane 17B |
| ENSG00000120256 | LRP11 | -0,09566367 | 0,935841642 | 0,7304796 | 0,974321178 | LDL receptor related protein 11 |
| ENSG00000242553 | AP001432,1 | -0,095687102 | 0,935826442 | 0,90679681 | 0,992590661 | novel transcript, antisense to DSCR3 |
| ENSG00000165792 | METTL17 | -0,09575152 | 0,935784657 | 0,70349185 | 0,968940012 | methyltransferase like 17 |
| ENSG00000285437 | POLR2J3 | -0,09585714 | 0,935716151 | 0,79003348 | 0,979462318 | RNA polymerase II subunit J3 |
| ENSG00000157540 | DYRK1A | -0,09593327 | 0,935666775 | 0,76030095 | 0,97624106 | dual specificity tyrosine phosphorylation regulated kinase 1A |
| ENSG00000283235 | AC139493,2 | -0,095960869 | 0,935648875 | 0,9485833 | 0,996432861 | novel transcript |
| ENSG00000132199 | ENOSF1 | -0,095974681 | 0,935639918 | 0,64960677 | 0,956899963 | enolase superfamily member 1 |
| ENSG00000203489 | HMGB1P39 | -0,095986338 | 0,935632358 | 0,94339256 | 0,995402494 | high mobility group box 1 pseudogene 39 |
| ENSG00000272529 | AC090948,3 | -0,095992481 | 0,935628374 | 0,87310263 | 0,989395767 | novel transcript, antisense to RFTN1 |

| | | | | | | |
|---|---|---|---|---|---|---|
| ENSG00000228384 | AC007040,1 | -0,096006377 | 0,935619363 | 0,94399277 | 0,995542144 | novel transcript |
| ENSG00000112130 | RNF8 | -0,096092107 | 0,935563766 | 0,72451881 | 0,97326334 | ring finger protein 8 |
| ENSG00000162365 | CYP4A22 | -0,09610814 | 0,935553369 | 0,910661 | 0,993222655 | cytochrome P450 family 4 subfamily A member 22 |
| ENSG00000260884 | AC009120,3 | -0,096154344 | 0,935523407 | 0,87373242 | 0,989633545 | novel transcript |
| ENSG00000177875 | CCDC184 | -0,096171439 | 0,935512322 | 0,77597503 | 0,978587255 | coiled-coil domain containing 184 |
| ENSG00000259726 | CSPG4P11 | -0,096210461 | 0,935487019 | 0,74023672 | 0,974833668 | chondroitin sulfate proteoglycan 4 pseudogene 11 |
| ENSG00000132879 | FBXO44 | -0,096299973 | 0,935428978 | 0,7773764 | 0,978587255 | F-box protein 44 |
| ENSG00000269399 | AC008764,6 | -0,096415681 | 0,935353957 | 0,79413276 | 0,979776981 | novel transcript |
| ENSG00000266709 | AC005224,3 | -0,09642948 | 0,935345011 | 0,82387059 | 0,983374097 | novel transcript |
| ENSG00000284620 | AF228730,5 | -0,096475962 | 0,935314875 | 0,76753357 | 0,977201891 | RNA binding protein, fox-1 homolog (C, elegans) 1 (RBFOX1) pseudogene |
| ENSG00000216657 | GLRX3P2 | -0,096495845 | 0,935301985 | 0,92665636 | 0,993915537 | glutaredoxin 3 pseudogene 2 |
| ENSG00000007545 | CRAMP1 | -0,096529283 | 0,935280308 | 0,72240833 | 0,972594421 | cramped chromatin regulator homolog 1 |
| ENSG00000189362 | NEMP2 | -0,096535671 | 0,935276166 | 0,79958644 | 0,980161849 | nuclear envelope integral membrane protein 2 |
| ENSG00000147592 | LACTB2 | -0,096680407 | 0,935182341 | 0,84661317 | 0,986939308 | lactamase beta 2 |
| ENSG00000167065 | DUSP18 | -0,096685337 | 0,935179146 | 0,66836706 | 0,961351573 | dual specificity phosphatase 18 |
| ENSG00000242199 | AC022336,1 | -0,096686697 | 0,935178264 | 0,93123791 | 0,994433362 | ribosomal protein S26 (RPS26) pseudogene |
| ENSG00000205085 | FAM71F2 | -0,096687923 | 0,935177469 | 0,84920632 | 0,987036295 | family with sequence similarity 71 member F2 |
| ENSG00000267253 | WHSC1L2P | -0,09672867 | 0,935151056 | 0,82808198 | 0,983595244 | Wolf-Hirschhorn syndrome candidate 1-like 2, pseudogene |
| ENSG00000275333 | FAM120B | -0,096732058 | 0,935148861 | 0,74598047 | 0,975707496 | family with sequence similarity 120B |
| ENSG00000112584 | FAM120B | -0,096732058 | 0,935148861 | 0,74598047 | 0,975707496 | family with sequence similarity 120B |
| ENSG00000268350 | FAM156A | -0,096771949 | 0,935123004 | 0,8481643 | 0,987036295 | family with sequence similarity 156 member A |
| ENSG00000104765 | BNIP3L | -0,096778363 | 0,935118846 | 0,81444744 | 0,981521413 | BCL2 interacting protein 3 like |
| ENSG00000238151 | MLLT10P1 | -0,096779206 | 0,9351183 | 0,90766546 | 0,992590661 | MLLT10 pseudogene 1 |
| ENSG00000285230 | RALY-AS1 | -0,096788169 | 0,93511249 | 0,77878792 | 0,978587255 | RALY antisense RNA 1 |
| ENSG00000185163 | DDX51 | -0,096827409 | 0,935087056 | 0,89110918 | 0,990795338 | DEAD-box helicase 51 |
| ENSG00000169379 | ARL13B | -0,096851496 | 0,935071445 | 0,74587624 | 0,975707496 | ADP ribosylation factor like GTPase 13B |
| ENSG00000103740 | ACSBG1 | -0,096980627 | 0,934987753 | 0,85069114 | 0,987147801 | acyl-CoA synthetase bubblegum family member 1 |
| ENSG00000198899 | MT-ATP6 | -0,097006563 | 0,934970945 | 0,79519195 | 0,97981137 | mitochondrially encoded ATP synthase membrane subunit 6 |
| ENSG00000147874 | HAUS6 | -0,097024472 | 0,934959338 | 0,80357545 | 0,980875302 | HAUS augmin like complex subunit 6 |
| ENSG00000215241 | LINC02449 | -0,097105663 | 0,934906723 | 0,82378525 | 0,983374097 | long intergenic non-protein coding RNA 2449 |

| | | | | | | |
|---|---|---|---|---|---|---|
| ENSG00000213199 | ASIC3 | -0,097354997 | 0,934745161 | 0,87184707 | 0,989217334 | acid sensing ion channel subunit 3 |
| ENSG00000287908 | AC025165,6 | -0,097374971 | 0,93473222 | 0,86672015 | 0,989074806 | novel protein |
| ENSG00000250910 | AC097467,3 | -0,097432418 | 0,934695 | 0,87043983 | 0,989074806 | novel transcript, antisense to MAP9 |
| ENSG00000138101 | DTNB | -0,097475019 | 0,9346674 | 0,69085282 | 0,967179044 | dystrobrevin beta |
| ENSG00000114942 | EEF1B2 | -0,097606171 | 0,934582436 | 0,77635096 | 0,978587255 | eukaryotic translation elongation factor 1 beta 2 |
| ENSG00000283391 | EEF1B2 | -0,097606171 | 0,934582436 | 0,77635096 | 0,978587255 | eukaryotic translation elongation factor 1 beta 2 |
| ENSG00000160404 | TOR2A | -0,097637642 | 0,934562049 | 0,70470469 | 0,969219816 | torsin family 2 member A |
| ENSG00000261188 | Z95115,1 | -0,097681973 | 0,934533332 | 0,83680288 | 0,985426146 | novel transcript, antisense to TFIP11 |
| ENSG00000227199 | ST7-AS1 | -0,097736993 | 0,934497693 | 0,78554834 | 0,979320406 | ST7 antisense RNA 1 |
| ENSG00000198420 | TCAF1 | -0,097765797 | 0,934479035 | 0,71557762 | 0,971209849 | TRPM8 channel associated factor 1 |
| ENSG00000204947 | ZNF425 | -0,09779649 | 0,934459154 | 0,70282112 | 0,968742626 | zinc finger protein 425 |
| ENSG00000287866 | AC009102,4 | -0,097938452 | 0,934367208 | 0,92671909 | 0,993915537 | novel transcript, antisense to GNAO1 |
| ENSG00000119596 | YLPM1 | -0,09795005 | 0,934359696 | 0,69582501 | 0,96776072 | YLP motif containing 1 |
| ENSG00000156482 | RPL30 | -0,097966211 | 0,93434923 | 0,78882071 | 0,979462318 | ribosomal protein L30 |
| ENSG00000151611 | MMAA | -0,098009623 | 0,934321114 | 0,80131362 | 0,980557464 | metabolism of cobalamin associated A |
| ENSG00000147050 | KDM6A | -0,098025032 | 0,934311136 | 0,74405045 | 0,975509237 | lysine demethylase 6A |
| ENSG00000268364 | SMC5-AS1 | -0,098046854 | 0,934297003 | 0,74833465 | 0,975901808 | SMC5 antisense RNA 1 (head to head) |
| ENSG00000205808 | PLPP6 | -0,098049952 | 0,934294997 | 0,80985036 | 0,981213877 | phospholipid phosphatase 6 |
| ENSG00000158604 | TMED4 | -0,098065393 | 0,934284998 | 0,69083271 | 0,967179044 | transmembrane p24 trafficking protein 4 |
| ENSG00000219027 | RPS3AP2 | -0,098072819 | 0,934280188 | 0,92807509 | 0,994108531 | RPS3A pseudogene 2 |
| ENSG00000196534 | OR9K1P | -0,098082687 | 0,934273798 | 0,91934346 | 0,993598006 | olfactory receptor family 9 subfamily K member 1 pseudogene |
| ENSG00000113161 | HMGCR | -0,098111268 | 0,93425529 | 0,80776067 | 0,981213877 | 3-hydroxy-3-methylglutaryl-CoA reductase |
| ENSG00000107159 | CA9 | -0,098114528 | 0,934253179 | 0,92540751 | 0,993915537 | carbonic anhydrase 9 |
| ENSG00000198830 | HMGN2 | -0,098304829 | 0,934129953 | 0,70225601 | 0,968742626 | high mobility group nucleosomal binding domain 2 |
| ENSG00000232305 | CLCP1 | -0,098336863 | 0,934109211 | 0,9128669 | 0,993361299 | Charcot-Leyden crystal protein pseudogene 1 |
| ENSG00000250067 | YJEFN3 | -0,098355704 | 0,934097012 | 0,85837541 | 0,989067345 | YjeF N-terminal domain containing 3 |
| ENSG00000141279 | NPEPPS | -0,098402884 | 0,934066465 | 0,73336605 | 0,974353474 | aminopeptidase puromycin sensitive |
| ENSG00000233323 | TNXB | -0,098460167 | 0,934029378 | 0,9095146 | 0,992879102 | tenascin XB |
| ENSG00000276910 | GTF2H2 | -0,098479083 | 0,934017132 | 0,75343786 | 0,975901808 | general transcription factor IIH subunit 2 |
| ENSG00000198780 | FAM169A | -0,098488276 | 0,93401118 | 0,79894222 | 0,980161849 | family with sequence similarity 169 member A |
| ENSG00000259120 | SMIM6 | -0,098504546 | 0,934000647 | 0,90463011 | 0,992590661 | small integral membrane protein 6 |

| | | | | | | |
|---|---|---|---|---|---|---|
| ENSG00000223509 | AC135983,2 | -0,09855892 | 0,933965445 | 0,79879811 | 0,980161849 | WAS protein homolog associated with actin, golgi membranes and microtubules (WHAMM) pseudogene |
| ENSG00000205060 | SLC35B4 | -0,098594091 | 0,933942677 | 0,76618966 | 0,977110125 | solute carrier family 35 member B4 |
| ENSG00000274461 | AC023282,1 | -0,098705592 | 0,933870498 | 0,84356233 | 0,986273696 | novel transcript |
| ENSG00000168769 | TET2 | -0,098761319 | 0,933834426 | 0,77785554 | 0,978587255 | tet methylcytosine dioxygenase 2 |
| ENSG00000268896 | AC009955,3 | -0,098771551 | 0,933827804 | 0,92286644 | 0,99381888 | novel transcript, antisense to TMEM198 |
| ENSG00000138650 | PCDH10 | -0,098795262 | 0,933812456 | 0,8153625 | 0,981521413 | protocadherin 10 |
| ENSG00000110492 | MDK | -0,098808123 | 0,933804132 | 0,66290784 | 0,959897678 | midkine |
| ENSG00000226220 | CICP22 | -0,098868509 | 0,933765047 | 0,91194979 | 0,993361299 | capicua transcriptional repressor pseudogene 22 |
| ENSG00000104613 | INTS10 | -0,098890112 | 0,933751065 | 0,71700259 | 0,971583263 | integrator complex subunit 10 |
| ENSG00000133624 | ZNF767P | -0,09889801 | 0,933745953 | 0,65860233 | 0,95814533 | zinc finger family member 767, pseudogene |
| ENSG00000160808 | MYL3 | -0,098930998 | 0,933724602 | 0,86995247 | 0,989074806 | myosin light chain 3 |
| ENSG00000213366 | GSTM2 | -0,098943569 | 0,933716466 | 0,75039834 | 0,975901808 | glutathione S-transferase mu 2 |
| ENSG00000117834 | SLC5A9 | -0,09896806 | 0,933700616 | 0,80704945 | 0,981213877 | solute carrier family 5 member 9 |
| ENSG00000107262 | BAG1 | -0,099025772 | 0,933663266 | 0,74848819 | 0,975901808 | BAG cochaperone 1 |
| ENSG00000169252 | ADRB2 | -0,099046721 | 0,933649708 | 0,92429714 | 0,993915537 | adrenoceptor beta 2 |
| ENSG00000049541 | RFC2 | -0,099067685 | 0,933636142 | 0,64171957 | 0,95651945 | replication factor C subunit 2 |
| ENSG00000153487 | ING1 | -0,099082075 | 0,933626829 | 0,70888646 | 0,970282889 | inhibitor of growth family member 1 |
| ENSG00000271366 | AC002128,2 | -0,099099111 | 0,933615805 | 0,93349191 | 0,994479239 | novel transcript |
| ENSG00000014824 | SLC30A9 | -0,099103297 | 0,933613096 | 0,83454969 | 0,984905093 | solute carrier family 30 member 9 |
| ENSG00000178038 | ALS2CL | -0,099108173 | 0,933609941 | 0,90238235 | 0,992230188 | ALS2 C-terminal like |
| ENSG00000106477 | CEP41 | -0,099221768 | 0,933536432 | 0,73604591 | 0,974353474 | centrosomal protein 41 |
| ENSG00000250950 | AC093752,2 | -0,099241359 | 0,933523756 | 0,84514635 | 0,986639902 | novel transcript |
| ENSG00000111790 | FGFR1OP2 | -0,09926078 | 0,933511189 | 0,82807014 | 0,983595244 | FGFR1 oncogene partner 2 |
| ENSG00000039123 | MTREX | -0,099267009 | 0,933507159 | 0,79099546 | 0,979471509 | Mtr4 exosome RNA helicase |
| ENSG00000130649 | CYP2E1 | -0,099309506 | 0,933479661 | 0,87518542 | 0,990078774 | cytochrome P450 family 2 subfamily E member 1 |
| ENSG00000259719 | LINC02284 | -0,099326013 | 0,93346898 | 0,88839266 | 0,990717729 | long intergenic non-protein coding RNA 2284 |
| ENSG00000186272 | ZNF17 | -0,09941887 | 0,933408901 | 0,71347689 | 0,970812834 | zinc finger protein 17 |
| ENSG00000187984 | ANKRD19P | -0,0994564 | 0,93338462 | 0,87099557 | 0,989217334 | ankyrin repeat domain 19, pseudogene |
| ENSG00000275329 | AL138781,1 | -0,099515314 | 0,933346505 | 0,8921566 | 0,99102448 | novel transcript |
| ENSG00000254480 | LINC02749 | -0,09956469 | 0,933314562 | 0,90807568 | 0,992590661 | long intergenic non-protein coding RNA 2749 |
| ENSG00000204052 | LRRC73 | -0,099569057 | 0,933311737 | 0,82447839 | 0,983374097 | leucine rich repeat containing 73 |

| | | | | | | |
|---|---|---|---|---|---|---|
| ENSG00000089685 | BIRC5 | -0,099713411 | 0,933218355 | 0,67233911 | 0,962290553 | baculoviral IAP repeat containing 5 |
| ENSG00000186792 | HYAL3 | -0,099728685 | 0,933208475 | 0,80696049 | 0,981213877 | hyaluronidase 3 |
| ENSG00000168175 | MAPK1IP1L | -0,099760241 | 0,933188064 | 0,79149932 | 0,979471509 | mitogen-activated protein kinase 1 interacting protein 1 like |
| ENSG00000139800 | ZIC5 | -0,099763828 | 0,933185744 | 0,83614799 | 0,985159643 | Zic family member 5 |
| ENSG00000213062 | Z99572,1 | -0,099779964 | 0,933175306 | 0,67804084 | 0,963963232 | novel transcript |
| ENSG00000125652 | ALKBH7 | -0,099818194 | 0,933150578 | 0,75708752 | 0,975901808 | alkB homolog 7 |
| ENSG00000230091 | TMEM254-AS1 | -0,099828937 | 0,93314363 | 0,80551276 | 0,981213877 | TMEM254 antisense RNA 1 |
| ENSG00000124523 | SIRT5 | -0,099934935 | 0,933075072 | 0,68343484 | 0,965068292 | sirtuin 5 |
| ENSG00000250634 | LINC01182 | -0,099944757 | 0,93306872 | 0,80035399 | 0,980237962 | long intergenic non-protein coding RNA 1182 |
| ENSG00000153363 | LINC00467 | -0,099956447 | 0,933061159 | 0,67571486 | 0,963449137 | long intergenic non-protein coding RNA 467 |
| ENSG00000232554 | RSU1P2 | -0,100004598 | 0,933030018 | 0,93838867 | 0,994549499 | Ras suppressor protein 1 pseudogene 2 |
| ENSG00000287220 | AL590138,1 | -0,100019493 | 0,933020385 | 0,88394117 | 0,990717729 | novel transcript, antisense to PLXNA2 |
| ENSG00000260388 | LINC00562 | -0,100157274 | 0,932931283 | 0,82043317 | 0,982609114 | long intergenic non-protein coding RNA 562 |
| ENSG00000238961 | SNORA47 | -0,100186657 | 0,932912283 | 0,89255144 | 0,991048031 | small nucleolar RNA, H/ACA box 47 |
| ENSG00000186814 | ZSCAN30 | -0,100209684 | 0,932897393 | 0,75184842 | 0,975901808 | zinc finger and SCAN domain containing 30 |
| ENSG00000196653 | ZNF502 | -0,100210338 | 0,93289697 | 0,79868616 | 0,980161849 | zinc finger protein 502 |
| ENSG00000281448 | ZNF502 | -0,100210338 | 0,93289697 | 0,79868616 | 0,980161849 | zinc finger protein 502 |
| ENSG00000249691 | AC026117,1 | -0,100273792 | 0,932855939 | 0,89206359 | 0,99102448 | developmental pluripotency associated 4 (DPPA4) pseudogene |
| ENSG00000286219 | NOTCH2NLC | -0,100276805 | 0,932853991 | 0,7398227 | 0,974715504 | notch 2 N-terminal like C |
| ENSG00000286534 | AC021683,6 | -0,100292005 | 0,932844163 | 0,83779887 | 0,98566906 | novel transcript |
| ENSG00000134419 | RPS15A | -0,100309003 | 0,932833171 | 0,81826981 | 0,982426552 | ribosomal protein S15a |
| ENSG00000137038 | DMAC1 | -0,100333367 | 0,932817419 | 0,72039648 | 0,972142678 | distal membrane arm assembly complex 1 |
| ENSG00000233539 | AC011294,1 | -0,100338509 | 0,932814094 | 0,93788142 | 0,994479239 | novel transcript |
| ENSG00000196458 | ZNF605 | -0,100379211 | 0,932787777 | 0,74373684 | 0,975343483 | zinc finger protein 605 |
| ENSG00000163946 | TASOR | -0,100424567 | 0,932758452 | 0,82055642 | 0,982609114 | transcription activation suppressor |
| ENSG00000271754 | AL355802,3 | -0,100426271 | 0,93275735 | 0,86503403 | 0,989074806 | novel transcript, antisense to POLR1C |
| ENSG00000091409 | ITGA6 | -0,100440639 | 0,932748061 | 0,76326752 | 0,976622569 | integrin subunit alpha 6 |
| ENSG00000248323 | LUCAT1 | -0,10046911 | 0,932729654 | 0,82033513 | 0,982609114 | lung cancer associated transcript 1 |
| ENSG00000104442 | ARMC1 | -0,100646892 | 0,932614721 | 0,83266657 | 0,984905093 | armadillo repeat containing 1 |
| ENSG00000103160 | HSDL1 | -0,100649758 | 0,932612868 | 0,79846448 | 0,980161849 | hydroxysteroid dehydrogenase like 1 |
| ENSG00000198648 | STK39 | -0,100681965 | 0,932592049 | 0,71466565 | 0,971041579 | serine/threonine kinase 39 |

| | | | | | | |
|---|---|---|---|---|---|---|
| ENSG00000227587 | ATP6V1G2 | -0,10078571 | 0,932524989 | 0,82651696 | 0,983471137 | ATPase H+ transporting V1 subunit G2 |
| ENSG00000153130 | SCOC | -0,10086773 | 0,932471974 | 0,85169067 | 0,987289464 | short coiled-coil protein |
| ENSG00000287013 | AC091646,2 | -0,100926364 | 0,932434077 | 0,89056676 | 0,990717729 | novel transcript |
| ENSG00000182986 | ZNF320 | -0,101023987 | 0,932370984 | 0,69201188 | 0,967225449 | zinc finger protein 320 |
| ENSG00000189184 | PCDH18 | -0,101084966 | 0,932331576 | 0,84479672 | 0,986557476 | protocadherin 18 |
| ENSG00000281244 | ADAMTS13 | -0,101092883 | 0,93232646 | 0,83768783 | 0,98566906 | ADAM metallopeptidase with thrombospondin type 1 motif 13 |
| ENSG00000121764 | HCRTR1 | -0,101102346 | 0,932320344 | 0,87926895 | 0,990148595 | hypocretin receptor 1 |
| ENSG00000167720 | SRR | -0,101116436 | 0,932311239 | 0,80276178 | 0,980866933 | serine racemase |
| ENSG00000259964 | THSD4-AS1 | -0,101160059 | 0,932283049 | 0,87190871 | 0,989217334 | THSD4 antisense RNA 1 |
| ENSG00000275851 | MLLT6 | -0,101168103 | 0,932277851 | 0,77769208 | 0,978587255 | MLLT6, PHD finger containing |
| ENSG00000274070 | CASTOR2 | -0,10117767 | 0,932271669 | 0,68694777 | 0,965769168 | cytosolic arginine sensor for mTORC1 subunit 2 |
| ENSG00000249646 | OR7E94P | -0,101178492 | 0,932271138 | 0,87776031 | 0,990109814 | olfactory receptor family 7 subfamily E member 94 pseudogene |
| ENSG00000225675 | LINC01771 | -0,10119272 | 0,932261943 | 0,92804581 | 0,994108531 | long intergenic non-protein coding RNA 1771 |
| ENSG00000111666 | CHPT1 | -0,101214426 | 0,932247917 | 0,76214599 | 0,976315588 | choline phosphotransferase 1 |
| ENSG00000128805 | ARHGAP22 | -0,101270188 | 0,932211885 | 0,72861131 | 0,973955059 | Rho GTPase activating protein 22 |
| ENSG00000234608 | MAPKAPK5-AS1 | -0,101345806 | 0,932163025 | 0,75435259 | 0,975901808 | MAPKAPK5 antisense RNA 1 |
| ENSG00000132680 | KHDC4 | -0,101367599 | 0,932148944 | 0,75949527 | 0,976120207 | KH domain containing 4, pre-mRNA splicing factor |
| ENSG00000256557 | AC009509,3 | -0,10142834 | 0,932109699 | 0,93408723 | 0,994479239 | high-mobility group nucleosome binding domain 1 (HMGN1) pseudogene |
| ENSG00000065534 | MYLK | -0,101429485 | 0,93210896 | 0,75185747 | 0,975901808 | myosin light chain kinase |
| ENSG00000123737 | EXOSC9 | -0,101479248 | 0,932076809 | 0,76075626 | 0,97624106 | exosome component 9 |
| ENSG00000213551 | DNAJC9 | -0,101526269 | 0,932046431 | 0,68620389 | 0,965740648 | DnaJ heat shock protein family (Hsp40) member C9 |
| ENSG00000229124 | VIM-AS1 | -0,101640966 | 0,931972334 | 0,8161037 | 0,981708551 | VIM antisense RNA 1 |
| ENSG00000167513 | CDT1 | -0,101667044 | 0,931955488 | 0,7181743 | 0,97178302 | chromatin licensing and DNA replication factor 1 |
| ENSG00000120215 | MLANA | -0,101693837 | 0,931938181 | 0,83398753 | 0,984905093 | melan-A |
| ENSG00000217930 | PAM16 | -0,101710824 | 0,931927208 | 0,73167343 | 0,974353474 | presequence translocase associated motor 16 |
| ENSG00000242999 | RN7SL239P | -0,101728172 | 0,931916001 | 0,91174532 | 0,993357417 | RNA, 7SL, cytoplasmic 239, pseudogene |
| ENSG00000130638 | ATXN10 | -0,10174679 | 0,931903975 | 0,74835847 | 0,975901808 | ataxin 10 |
| ENSG00000140396 | NCOA2 | -0,10175506 | 0,931898633 | 0,77321498 | 0,978363975 | nuclear receptor coactivator 2 |
| ENSG00000231770 | TMEM44-AS1 | -0,101771004 | 0,931888334 | 0,78540298 | 0,979320406 | TMEM44 antisense RNA 1 |
| ENSG00000175354 | PTPN2 | -0,101875464 | 0,931820863 | 0,7533212 | 0,975901808 | protein tyrosine phosphatase non-receptor type 2 |

| | | | | | | |
|---|---|---|---|---|---|---|
| ENSG00000286548 | AL596214,1 | -0,101876142 | 0,931820424 | 0,80576724 | 0,981213877 | novel transcript, antisense to LRRC8B |
| ENSG00000151779 | NBAS | -0,101910488 | 0,931798241 | 0,73320619 | 0,974353474 | NBAS subunit of NRZ tethering complex |
| ENSG00000277399 | GPR179 | -0,10196365 | 0,931763906 | 0,86218376 | 0,989074806 | G protein-coupled receptor 179 |
| ENSG00000152556 | PFKM | -0,101993075 | 0,931744902 | 0,62457833 | 0,952793934 | phosphofructokinase, muscle |
| ENSG00000262904 | TMPOP2 | -0,102048249 | 0,931709269 | 0,84524557 | 0,986639902 | thymopoietin pseudogene 2 |
| ENSG00000260698 | AL591848,2 | -0,102133582 | 0,931654162 | 0,8868455 | 0,990717729 | novel transcript |
| ENSG00000213801 | ZNF321P | -0,10217473 | 0,93162759 | 0,70116314 | 0,968677171 | zinc finger protein 321, pseudogene |
| ENSG00000124193 | SRSF6 | -0,102210631 | 0,931604407 | 0,76494111 | 0,976725297 | serine and arginine rich splicing factor 6 |
| ENSG00000164611 | PTTG1 | -0,102213696 | 0,931602427 | 0,66760475 | 0,9611116 | PTTG1 regulator of sister chromatid separation, securin |
| ENSG00000245598 | DACT3-AS1 | -0,102225866 | 0,931594569 | 0,91219552 | 0,993361299 | DACT3 antisense RNA 1 |
| ENSG00000284850 | DCP1B | -0,102243935 | 0,931582901 | 0,74671614 | 0,975796253 | decapping mRNA 1B |
| ENSG00000151065 | DCP1B | -0,102243935 | 0,931582901 | 0,74671614 | 0,975796253 | decapping mRNA 1B |
| ENSG00000248319 | LINC02275 | -0,102321018 | 0,931533128 | 0,84732055 | 0,987035044 | long intergenic non-protein coding RNA 2275 |
| ENSG00000265494 | AC005304,2 | -0,102333869 | 0,931524831 | 0,79909055 | 0,980161849 | novel transcript |
| ENSG00000231908 | IDH1-AS1 | -0,102391564 | 0,931487579 | 0,83853157 | 0,985672131 | IDH1 antisense RNA 1 |
| ENSG00000267102 | AC060766,1 | -0,102411156 | 0,931474929 | 0,75173791 | 0,975901808 | RAB32, member RAS oncogene family (RAB32) pseudogene |
| ENSG00000189403 | HMGB1 | -0,102561053 | 0,931378153 | 0,78656603 | 0,979320406 | high mobility group box 1 |
| ENSG00000091140 | DLD | -0,102576702 | 0,93136805 | 0,81434725 | 0,981521413 | dihydrolipoamide dehydrogenase |
| ENSG00000226618 | PRRC2A | -0,102591751 | 0,931358335 | 0,82067587 | 0,982609114 | proline rich coiled-coil 2A |
| ENSG00000170881 | RNF139 | -0,102618866 | 0,931340831 | 0,82635479 | 0,983471137 | ring finger protein 139 |
| ENSG00000255769 | GOLGA2P10 | -0,102654307 | 0,931317952 | 0,82598476 | 0,983418544 | GOLGA2 pseudogene 10 |
| ENSG00000149016 | TUT1 | -0,102667442 | 0,931309473 | 0,769124 | 0,977475319 | terminal uridylyl transferase 1, U6 snRNA-specific |
| ENSG00000121579 | NAA50 | -0,102695063 | 0,931291643 | 0,80844391 | 0,981213877 | N-alpha-acetyltransferase 50, NatE catalytic subunit |
| ENSG00000197937 | ZNF347 | -0,102784384 | 0,931233986 | 0,61962537 | 0,952793934 | zinc finger protein 347 |
| ENSG00000161958 | FGF11 | -0,102819699 | 0,931211191 | 0,78763545 | 0,979462318 | fibroblast growth factor 11 |
| ENSG00000283903 | FGF11 | -0,102819699 | 0,931211191 | 0,78763545 | 0,979462318 | fibroblast growth factor 11 |
| ENSG00000180340 | FZD2 | -0,102844782 | 0,931195001 | 0,63172609 | 0,953915351 | frizzled class receptor 2 |
| ENSG00000105278 | ZFR2 | -0,102865561 | 0,931181589 | 0,77717872 | 0,978587255 | zinc finger RNA binding protein 2 |
| ENSG00000101745 | ANKRD12 | -0,102973757 | 0,931111757 | 0,76605258 | 0,977102081 | ankyrin repeat domain 12 |
| ENSG00000101444 | AHCY | -0,103071116 | 0,931048924 | 0,69408257 | 0,967359949 | adenosylhomocysteinase |
| ENSG00000178691 | SUZ12 | -0,103132828 | 0,931009098 | 0,80519382 | 0,981187666 | SUZ12 polycomb repressive complex 2 subunit |

| | | | | | | |
|---|---|---|---|---|---|---|
| ENSG00000184154 | LRTOMT | -0,103235887 | 0,930942594 | 0,67179936 | 0,962161102 | leucine rich transmembrane and O-methyltransferase domain containing |
| ENSG00000151093 | OXSM | -0,103260739 | 0,930926558 | 0,74673021 | 0,975796253 | 3-oxoacyl-ACP synthase, mitochondrial |
| ENSG00000224142 | AMMECR1-IT1 | -0,103270102 | 0,930920516 | 0,92513385 | 0,993915537 | AMMECR1 intronic transcript 1 |
| ENSG00000142396 | ERVK3-1 | -0,103387371 | 0,93084485 | 0,74459126 | 0,975649186 | endogenous retrovirus group K3 member 1 |
| ENSG00000255115 | AP002812,4 | -0,103406052 | 0,930832797 | 0,93965285 | 0,994943406 | family with sequence similarity 162, member A (FAM162A) pseudogene |
| ENSG00000124279 | FASTKD3 | -0,103423767 | 0,930821367 | 0,81613316 | 0,981708551 | FAST kinase domains 3 |
| ENSG00000133119 | RFC3 | -0,103484755 | 0,930782018 | 0,82449916 | 0,983374097 | replication factor C subunit 3 |
| ENSG00000139780 | METTL21C | -0,103490435 | 0,930778354 | 0,92961059 | 0,994293848 | methyltransferase like 21C |
| ENSG00000165119 | HNRNPK | -0,10355462 | 0,930736945 | 0,77544958 | 0,978587255 | heterogeneous nuclear ribonucleoprotein K |
| ENSG00000101052 | IFT52 | -0,103568894 | 0,930727736 | 0,76240093 | 0,976315588 | intraflagellar transport 52 |
| ENSG00000278200 | LINC01971 | -0,103597937 | 0,930709 | 0,85512348 | 0,988016959 | long intergenic non-protein coding RNA 1971 |
| ENSG00000004864 | SLC25A13 | -0,10361721 | 0,930696566 | 0,77807044 | 0,978587255 | solute carrier family 25 member 13 |
| ENSG00000156006 | NAT2 | -0,103647162 | 0,930677244 | 0,93586339 | 0,994479239 | N-acetyltransferase 2 |
| ENSG00000107949 | BCCIP | -0,103653795 | 0,930672965 | 0,73256104 | 0,974353474 | BRCA2 and CDKN1A interacting protein |
| ENSG00000188372 | ZP3 | -0,103656461 | 0,930671245 | 0,73120856 | 0,974353474 | zona pellucida glycoprotein 3 |
| ENSG00000113595 | TRIM23 | -0,103758743 | 0,930605267 | 0,79644576 | 0,980015022 | tripartite motif containing 23 |
| ENSG00000120709 | FAM53C | -0,103765984 | 0,930600596 | 0,65020923 | 0,956899963 | family with sequence similarity 53 member C |
| ENSG00000139154 | AEBP2 | -0,103860773 | 0,930539455 | 0,75218148 | 0,975901808 | AE binding protein 2 |
| ENSG00000128951 | DUT | -0,103998618 | 0,930450549 | 0,74775986 | 0,975901808 | deoxyuridine triphosphatase |
| ENSG00000282712 | DEAF1 | -0,104016073 | 0,930439292 | 0,88743951 | 0,990717729 | DEAF1 transcription factor |
| ENSG00000223965 | ZNF587P1 | -0,104152835 | 0,930351094 | 0,92701806 | 0,993915537 | zinc finger protein 587 pseudogene 1 |
| ENSG00000106344 | RBM28 | -0,104247526 | 0,930290032 | 0,62585944 | 0,952793934 | RNA binding motif protein 28 |
| ENSG00000272084 | AL137127,1 | -0,104262164 | 0,930280594 | 0,78787195 | 0,979462318 | novel transcript |
| ENSG00000113593 | PPWD1 | -0,104281673 | 0,930268014 | 0,81836977 | 0,9824772 | peptidylprolyl isomerase domain and WD repeat containing 1 |
| ENSG00000060140 | STYK1 | -0,104304457 | 0,930253322 | 0,88735232 | 0,990717729 | serine/threonine/tyrosine kinase 1 |
| ENSG00000242516 | LINC00960 | -0,104349032 | 0,930224581 | 0,79622868 | 0,980015022 | long intergenic non-protein coding RNA 960 |
| ENSG00000227848 | SUCLA2-AS1 | -0,104417729 | 0,930180287 | 0,92590423 | 0,993915537 | SUCLA2 antisense RNA 1 |
| ENSG00000237344 | PBX2 | -0,104437006 | 0,930167858 | 0,69169218 | 0,967204896 | PBX homeobox 2 |
| ENSG00000232098 | AC012313,1 | -0,104496635 | 0,930129414 | 0,76751739 | 0,977201891 | novel transcript |
| ENSG00000204104 | TRAF3IP1 | -0,104551886 | 0,930093793 | 0,77903765 | 0,978587255 | TRAF3 interacting protein 1 |

| | | | | | | |
|---|---|---|---|---|---|---|
| ENSG00000164615 | CAMLG | -0,104567638 | 0,930083638 | 0,71791202 | 0,971762632 | calcium modulating ligand |
| ENSG00000264343 | NOTCH2NLA | -0,104610365 | 0,930056093 | 0,70203541 | 0,968742626 | notch 2 N-terminal like A |
| ENSG00000146457 | WTAP | -0,104626195 | 0,930045888 | 0,74997186 | 0,975901808 | WT1 associated protein |
| ENSG00000278372 | MYO19 | -0,104646171 | 0,93003301 | 0,71243517 | 0,970812834 | myosin XIX |
| ENSG00000283001 | AL450426,1 | -0,104708606 | 0,929992762 | 0,8306848 | 0,984156028 | novel transcript, antisense to OR13C3 |
| ENSG00000170271 | FAXDC2 | -0,104729901 | 0,929979036 | 0,76240976 | 0,976315588 | fatty acid hydroxylase domain containing 2 |
| ENSG00000215790 | SLC35E2A | -0,104779876 | 0,929946822 | 0,70688328 | 0,969565008 | solute carrier family 35 member E2A |
| ENSG00000274554 | AC083806,2 | -0,104819223 | 0,929921459 | 0,76664617 | 0,977201891 | novel transcript, antisense to FBXW8 |
| ENSG00000287229 | AC009731,1 | -0,104826907 | 0,929916506 | 0,83380344 | 0,984905093 | novel transcript, antisense to RBM19 |
| ENSG00000180219 | FAM71C | -0,104847172 | 0,929903444 | 0,90668011 | 0,992590661 | family with sequence similarity 71 member C |
| ENSG00000112624 | BICRAL | -0,104892532 | 0,929874207 | 0,78013545 | 0,978587255 | BRD4 interacting chromatin remodeling complex associated protein like |
| ENSG00000227460 | SYNGAP1 | -0,104900549 | 0,92986904 | 0,69906016 | 0,968522643 | synaptic Ras GTPase activating protein 1 |
| ENSG00000168887 | C2orf68 | -0,104940658 | 0,929843189 | 0,60940115 | 0,951793902 | chromosome 2 open reading frame 68 |
| ENSG00000272953 | AC092171,4 | -0,104962154 | 0,929829334 | 0,86448068 | 0,989074806 | novel transcript |
| ENSG00000239822 | RN7SL754P | -0,105036235 | 0,92978159 | 0,9355206 | 0,994479239 | RNA, 7SL, cytoplasmic 754, pseudogene |
| ENSG00000175264 | CHST1 | -0,105063534 | 0,929763996 | 0,85896696 | 0,989074806 | carbohydrate sulfotransferase 1 |
| ENSG00000204934 | ATP6V0E2-AS1 | -0,105074674 | 0,929756817 | 0,70446081 | 0,969132783 | ATP6V0E2 antisense RNA 1 |
| ENSG00000230392 | AC004835,1 | -0,105157423 | 0,92970349 | 0,76715999 | 0,977201891 | novel transcript |
| ENSG00000244757 | AL137024,1 | -0,10528743 | 0,929619715 | 0,92915727 | 0,994293848 | novel transcript |
| ENSG00000115919 | KYNU | -0,105416289 | 0,929536687 | 0,65636173 | 0,957444211 | kynureninase |
| ENSG00000164032 | H2AZ1 | -0,105416485 | 0,92953656 | 0,74810819 | 0,975901808 | H2A,Z variant histone 1 |
| ENSG00000266936 | AC011442,1 | -0,105431002 | 0,929527207 | 0,87653327 | 0,990109814 | novel transcript, antisense to SMARCA4 |
| ENSG00000164929 | BAALC | -0,105462763 | 0,929506744 | 0,81305298 | 0,981246822 | BAALC binder of MAP3K1 and KLF4 |
| ENSG00000143157 | POGK | -0,105530621 | 0,929463025 | 0,68170321 | 0,964654539 | pogo transposable element derived with KRAB domain |
| ENSG00000177954 | RPS27 | -0,105533115 | 0,929461418 | 0,81803862 | 0,982323352 | ribosomal protein S27 |
| ENSG00000138780 | GSTCD | -0,105583258 | 0,929429114 | 0,74610006 | 0,975707496 | glutathione S-transferase C-terminal domain containing |
| ENSG00000161999 | JMJD8 | -0,105755077 | 0,929318429 | 0,67255202 | 0,962290553 | jumonji domain containing 8 |
| ENSG00000164414 | SLC35A1 | -0,10576459 | 0,929312302 | 0,83217377 | 0,984749455 | solute carrier family 35 member A1 |
| ENSG00000230319 | TTLL1-AS1 | -0,105802231 | 0,929288055 | 0,85077717 | 0,987147801 | TTLL1 antisense RNA 1 |
| ENSG00000224043 | CCNT2-AS1 | -0,105838493 | 0,929264698 | 0,73327924 | 0,974353474 | CCNT2 antisense RNA 1 |

| | | | | | | |
|---|---|---|---|---|---|---|
| ENSG00000262155 | LINC02175 | -0,105883747 | 0,929235549 | 0,76324521 | 0,976622569 | long intergenic non-protein coding RNA 2175 |
| ENSG00000271937 | AC104187,1 | -0,105937989 | 0,929200613 | 0,90879656 | 0,992688304 | novel transcript, antisense to TCAIM |
| ENSG00000159592 | GPBP1L1 | -0,106076272 | 0,929111553 | 0,74726963 | 0,97584218 | GC-rich promoter binding protein 1 like 1 |
| ENSG00000222020 | HDAC4-AS1 | -0,106091335 | 0,929101852 | 0,89824283 | 0,9918188 | HDAC4 antisense RNA 1 |
| ENSG00000271973 | AC141002,1 | -0,10615979 | 0,929057768 | 0,91624145 | 0,993361299 | novel transcript |
| ENSG00000255804 | OR6J1 | -0,10621167 | 0,929024359 | 0,83290823 | 0,984905093 | olfactory receptor family 6 subfamily J member 1 |
| ENSG00000178999 | AURKB | -0,106305553 | 0,928963905 | 0,65292734 | 0,95732962 | aurora kinase B |
| ENSG00000239207 | GAPDHP39 | -0,106328621 | 0,928949052 | 0,89625251 | 0,991636805 | glyceraldehyde 3 phosphate dehydrogenase pseudogene 39 |
| ENSG00000245571 | FAM111A-DT | -0,106341204 | 0,928940949 | 0,79904833 | 0,980161849 | FAM111A divergent transcript |
| ENSG00000198546 | ZNF511 | -0,106462686 | 0,928862732 | 0,67485399 | 0,963268551 | zinc finger protein 511 |
| ENSG00000260689 | HNRNPA3P11 | -0,106485007 | 0,92884836 | 0,81809689 | 0,982323352 | heterogeneous nuclear ribonucleoprotein A3 pseudogene 11 |
| ENSG00000206282 | RGL2 | -0,106491471 | 0,928844199 | 0,70229248 | 0,968742626 | ral guanine nucleotide dissociation stimulator like 2 |
| ENSG00000104064 | GABPB1 | -0,106520144 | 0,928825738 | 0,79585739 | 0,979879951 | GA binding protein transcription factor subunit beta 1 |
| ENSG00000198700 | IPO9 | -0,106600623 | 0,928773927 | 0,70471804 | 0,969219816 | importin 9 |
| ENSG00000265967 | AC100830,3 | -0,106656868 | 0,928737718 | 0,93251196 | 0,994479239 | novel transcript, antisense to ornithine decarboxylase antizyme 2 OAZ2 |
| ENSG00000189319 | FAM53B | -0,106671066 | 0,928728578 | 0,70921033 | 0,970282889 | family with sequence similarity 53 member B |
| ENSG00000221821 | C6orf226 | -0,106741516 | 0,928683227 | 0,77894929 | 0,978587255 | chromosome 6 open reading frame 226 |
| ENSG00000271414 | AC093107,2 | -0,106836317 | 0,928622204 | 0,93782808 | 0,994479239 | MYC associated factor X, pseudogene |
| ENSG00000261451 | AC104964,2 | -0,106851014 | 0,928612744 | 0,77398502 | 0,978415098 | novel transcript, overlapping to MSRA |
| ENSG00000116128 | BCL9 | -0,106862055 | 0,928605638 | 0,67567515 | 0,963449137 | BCL9 transcription coactivator |
| ENSG00000188493 | C19orf54 | -0,106877552 | 0,928595663 | 0,68822269 | 0,966426338 | chromosome 19 open reading frame 54 |
| ENSG00000167371 | PRRT2 | -0,106899326 | 0,928581648 | 0,65804965 | 0,957883832 | proline rich transmembrane protein 2 |
| ENSG00000249421 | ADAMTS19-AS1 | -0,106906037 | 0,928577328 | 0,85903313 | 0,989074806 | ADAMTS19 antisense RNA 1 |
| ENSG00000216863 | LY86-AS1 | -0,106944888 | 0,928552323 | 0,865729 | 0,989074806 | LY86 antisense RNA 1 |
| ENSG00000248213 | CICP16 | -0,106995107 | 0,928520001 | 0,77250146 | 0,978288572 | capicua transcriptional repressor pseudogene 16 |
| ENSG00000006704 | GTF2IRD1 | -0,107012422 | 0,928508857 | 0,66210975 | 0,959874876 | GTF2I repeat domain containing 1 |
| ENSG00000168228 | ZCCHC4 | -0,107013928 | 0,928507888 | 0,6392557 | 0,956018518 | zinc finger CCHC-type containing 4 |
| ENSG00000256552 | AC092745,2 | -0,107026914 | 0,928499531 | 0,90596872 | 0,992590661 | ribosomal protein S24 (RPS24) pseudogene |
| ENSG00000109133 | TMEM33 | -0,107153526 | 0,928418048 | 0,77232365 | 0,97820249 | transmembrane protein 33 |

| | | | | | | |
|---|---|---|---|---|---|---|
| ENSG00000259683 | AC243562,2 | -0,107154947 | 0,928417134 | 0,90484733 | 0,992590661 | golgin A6 family-like 9 (GOLGA6L9) pseudogene |
| ENSG00000239494 | RN7SL333P | -0,107173732 | 0,928405045 | 0,91543811 | 0,993361299 | RNA, 7SL, cytoplasmic 333, pseudogene |
| ENSG00000281245 | STKLD1 | -0,107275135 | 0,928339793 | 0,84656904 | 0,986939308 | serine/threonine kinase like domain containing 1 |
| ENSG00000186187 | ZNRF1 | -0,107309248 | 0,928317842 | 0,62441026 | 0,952793934 | zinc and ring finger 1 |
| ENSG00000143147 | GPR161 | -0,107377153 | 0,928274149 | 0,62437598 | 0,952793934 | G protein-coupled receptor 161 |
| ENSG00000170903 | MSANTD4 | -0,107395321 | 0,928262459 | 0,72630816 | 0,973344786 | Myb/SANT DNA binding domain containing 4 with coiled-coils |
| ENSG00000267075 | AC015818,8 | -0,107508767 | 0,928189468 | 0,85765628 | 0,988893234 | novel transcript, piRNA host |
| ENSG00000100650 | SRSF5 | -0,107528597 | 0,92817671 | 0,7066475 | 0,969503793 | serine and arginine rich splicing factor 5 |
| ENSG00000286760 | AL022069,3 | -0,107590748 | 0,928136726 | 0,78119262 | 0,978597905 | novel transcript |
| ENSG00000130024 | PHF10 | -0,107591135 | 0,928136476 | 0,78553222 | 0,979320406 | PHD finger protein 10 |
| ENSG00000215388 | ACTG1P3 | -0,107605598 | 0,928127172 | 0,86058176 | 0,989074806 | actin gamma 1 pseudogene 3 |
| ENSG00000100532 | CGRRF1 | -0,107653729 | 0,928096208 | 0,77793865 | 0,978587255 | cell growth regulator with ring finger domain 1 |
| ENSG00000275495 | AC114936,2 | -0,107664923 | 0,928089008 | 0,79691279 | 0,980161849 | THO complex 3 (THOC3) pseudogene |
| ENSG00000267058 | AC006213,2 | -0,107756298 | 0,928030228 | 0,75363544 | 0,975901808 | novel transcript |
| ENSG00000100941 | PNN | -0,10785559 | 0,927966359 | 0,75059615 | 0,975901808 | pinin, desmosome associated protein |
| ENSG00000217120 | Z98755,1 | -0,107868009 | 0,927958371 | 0,79898679 | 0,980161849 | ClpX caseinolytic protease X homolog (E, coli) (CLPX) pseudogene |
| ENSG00000205707 | ETFRF1 | -0,107911986 | 0,927930085 | 0,84761041 | 0,987035044 | electron transfer flavoprotein regulatory factor 1 |
| ENSG00000138398 | PPIG | -0,107946494 | 0,92790789 | 0,6847216 | 0,965415049 | peptidylprolyl isomerase G |
| ENSG00000111843 | TMEM14C | -0,107986503 | 0,927882157 | 0,74853547 | 0,975901808 | transmembrane protein 14C |
| ENSG00000284936 | TMEM14C | -0,107986503 | 0,927882157 | 0,74853547 | 0,975901808 | transmembrane protein 14C |
| ENSG00000232807 | AL137186,2 | -0,10801751 | 0,927862215 | 0,80962158 | 0,981213877 | novel transcript, antisense to FBXO18 |
| ENSG00000181201 | H2BU2P | -0,108060704 | 0,927834436 | 0,87538539 | 0,990078774 | H2B,U histone 2, pseudogene |
| ENSG00000285234 | H2BU2P | -0,108060704 | 0,927834436 | 0,87538539 | 0,990078774 | H2B,U histone 2, pseudogene |
| ENSG00000228242 | XPC-AS1 | -0,108092097 | 0,927814246 | 0,7667583 | 0,977201891 | XPC antisense RNA 1 |
| ENSG00000187626 | ZKSCAN4 | -0,108154252 | 0,927774274 | 0,72946661 | 0,974009551 | zinc finger with KRAB and SCAN domains 4 |
| ENSG00000101624 | CEP76 | -0,108164633 | 0,927767598 | 0,739823 | 0,974715504 | centrosomal protein 76 |
| ENSG00000205583 | STAG3L1 | -0,108359501 | 0,927642291 | 0,79427046 | 0,979776981 | stromal antigen 3-like 1 (pseudogene) |
| ENSG00000101230 | ISM1 | -0,108374235 | 0,927632817 | 0,89720834 | 0,991644439 | isthmin 1 |
| ENSG00000113368 | LMNB1 | -0,108480979 | 0,927564185 | 0,69156208 | 0,967201016 | lamin B1 |
| ENSG00000113163 | CERT1 | -0,108485379 | 0,927561356 | 0,69320867 | 0,967340633 | ceramide transporter 1 |
| ENSG00000138039 | LHCGR | -0,108488642 | 0,927559258 | 0,91574779 | 0,993361299 | luteinizing hormone/choriogonadotropin receptor |

| | | | | | | |
|---|---|---|---|---|---|---|
| ENSG00000147677 | EIF3H | -0,108494122 | 0,927555735 | 0,77932357 | 0,978587255 | eukaryotic translation initiation factor 3 subunit H |
| ENSG00000144357 | UBR3 | -0,108528688 | 0,927533512 | 0,76745329 | 0,977201891 | ubiquitin protein ligase E3 component n-recognin 3 |
| ENSG00000111678 | C12orf57 | -0,108564771 | 0,927510313 | 0,62540762 | 0,952793934 | chromosome 12 open reading frame 57 |
| ENSG00000178977 | LINC00324 | -0,108706554 | 0,927419165 | 0,78889625 | 0,979462318 | long intergenic non-protein coding RNA 324 |
| ENSG00000147853 | AK3 | -0,108739445 | 0,927398022 | 0,77937513 | 0,978587255 | adenylate kinase 3 |
| ENSG00000234694 | AL139289,1 | -0,108752827 | 0,92738942 | 0,92025934 | 0,993787755 | novel transcript |
| ENSG00000164039 | BDH2 | -0,108775402 | 0,927374908 | 0,80403434 | 0,980875302 | 3-hydroxybutyrate dehydrogenase 2 |
| ENSG00000164867 | NOS3 | -0,108902297 | 0,927293343 | 0,82534598 | 0,983383773 | nitric oxide synthase 3 |
| ENSG00000137504 | CREBZF | -0,108946417 | 0,927264985 | 0,68109283 | 0,964654539 | CREB/ATF bZIP transcription factor |
| ENSG00000008869 | HEATR5B | -0,108948287 | 0,927263783 | 0,73497973 | 0,974353474 | HEAT repeat containing 5B |
| ENSG00000156313 | RPGR | -0,108949408 | 0,927263063 | 0,63436118 | 0,955097925 | retinitis pigmentosa GTPase regulator |
| ENSG00000172340 | SUCLG2 | -0,108964953 | 0,927253072 | 0,74875786 | 0,975901808 | succinate-CoA ligase GDP-forming subunit beta |
| ENSG00000254231 | AC103760,1 | -0,108980912 | 0,927242815 | 0,88491167 | 0,990717729 | novel transcript |
| ENSG00000250327 | RPSAP70 | -0,10898726 | 0,927238735 | 0,9080512 | 0,992590661 | ribosomal protein SA pseudogene 70 |
| ENSG00000160551 | TAOK1 | -0,109036029 | 0,927207391 | 0,75941558 | 0,976120207 | TAO kinase 1 |
| ENSG00000121774 | KHDRBS1 | -0,109059618 | 0,927192231 | 0,63774597 | 0,956018518 | KH RNA binding domain containing, signal transduction associated 1 |
| ENSG00000198585 | NUDT16 | -0,10906463 | 0,927189009 | 0,73788319 | 0,974353474 | nudix hydrolase 16 |
| ENSG00000125901 | MRPS26 | -0,109089172 | 0,927173237 | 0,73798344 | 0,974353474 | mitochondrial ribosomal protein S26 |
| ENSG00000254273 | AC018620,1 | -0,109095818 | 0,927168965 | 0,91302542 | 0,993361299 | chromatin accessibility complex protein 1 (CHRAC1) pseudogene |
| ENSG00000240344 | PPIL3 | -0,109121026 | 0,927152766 | 0,81453555 | 0,981521413 | peptidylprolyl isomerase like 3 |
| ENSG00000068885 | IFT80 | -0,109209555 | 0,927095874 | 0,82805173 | 0,983595244 | intraflagellar transport 80 |
| ENSG00000157110 | RBPMS | -0,1093052 | 0,927034413 | 0,63617939 | 0,955399772 | RNA binding protein, mRNA processing factor |
| ENSG00000248288 | AC008868,1 | -0,109461139 | 0,926934217 | 0,92752601 | 0,994094286 | ribosomal protein L5 (RPL5) pseudogene |
| ENSG00000122824 | NUDT10 | -0,109564491 | 0,926867815 | 0,78887386 | 0,979462318 | nudix hydrolase 10 |
| ENSG00000276724 | AC123768,3 | -0,109619787 | 0,926832291 | 0,81096874 | 0,981213877 | novel transcript |
| ENSG00000257890 | AC011595,2 | -0,109623739 | 0,926829752 | 0,92454267 | 0,993915537 | novel transcript |
| ENSG00000216802 | AL590617,1 | -0,109638299 | 0,926820398 | 0,86703291 | 0,989074806 | mitochondrial carrier homolog 1 (MTCH1) pseudogene |
| ENSG00000101126 | ADNP | -0,109687658 | 0,926788689 | 0,74966664 | 0,975901808 | activity dependent neuroprotector homeobox |
| ENSG00000111802 | TDP2 | -0,109715703 | 0,926770673 | 0,68847557 | 0,966481084 | tyrosyl-DNA phosphodiesterase 2 |
| ENSG00000113391 | FAM172A | -0,109734778 | 0,926758419 | 0,76884513 | 0,977381742 | family with sequence similarity 172 member A |

| | | | | | | |
|---|---|---|---|---|---|---|
| ENSG00000287808 | AL355999,1 | -0,109750802 | 0,926748126 | 0,76362252 | 0,976640468 | novel transcript, antisense to DHX9 |
| ENSG00000261118 | AC092123,1 | -0,109914023 | 0,926643283 | 0,87696378 | 0,990109814 | novel transcript, antisense to SPG7 |
| ENSG00000247624 | CPEB2-DT | -0,110049949 | 0,926555982 | 0,92970407 | 0,994293848 | CPEB2 divergent transcript |
| ENSG00000282962 | CPEB2-DT | -0,110049949 | 0,926555982 | 0,92970407 | 0,994293848 | CPEB2 divergent transcript |
| ENSG00000175928 | LRRN1 | -0,110075842 | 0,926539353 | 0,76740755 | 0,977201891 | leucine rich repeat neuronal 1 |
| ENSG00000071794 | HLTF | -0,110115663 | 0,926513779 | 0,79824714 | 0,980161849 | helicase like transcription factor |
| ENSG00000051180 | RAD51 | -0,110163652 | 0,92648296 | 0,64333099 | 0,956633898 | RAD51 recombinase |
| ENSG00000175395 | ZNF25 | -0,110177045 | 0,92647436 | 0,80488123 | 0,98109454 | zinc finger protein 25 |
| ENSG00000068796 | KIF2A | -0,110219683 | 0,926446979 | 0,79281301 | 0,979471509 | kinesin family member 2A |
| ENSG00000107951 | MTPAP | -0,110248047 | 0,926428765 | 0,67892258 | 0,964151912 | mitochondrial poly(A) polymerase |
| ENSG00000187866 | FAM122A | -0,110326647 | 0,926378293 | 0,74824262 | 0,975901808 | family with sequence similarity 122A |
| ENSG00000251270 | AC108727,2 | -0,110352545 | 0,926361663 | 0,92659305 | 0,993915537 | novel transcript |
| ENSG00000153012 | LGI2 | -0,110352596 | 0,926361631 | 0,80357567 | 0,980875302 | leucine rich repeat LGI family member 2 |
| ENSG00000288077 | AC027129,1 | -0,110360481 | 0,926356568 | 0,92272938 | 0,993797669 | novel transcript, antisense to COLQ |
| ENSG00000260059 | AC092620,2 | -0,110368645 | 0,926351326 | 0,84955815 | 0,987036295 | novel transcript |
| ENSG00000011590 | ZBTB32 | -0,110432414 | 0,926310381 | 0,85407633 | 0,987861659 | zinc finger and BTB domain containing 32 |
| ENSG00000116906 | GNPAT | -0,110471409 | 0,926285344 | 0,76468724 | 0,976640468 | glyceronephosphate O-acyltransferase |
| ENSG00000142046 | TMEM91 | -0,110564852 | 0,92622535 | 0,83384858 | 0,984905093 | transmembrane protein 91 |
| ENSG00000102595 | UGGT2 | -0,110576843 | 0,926217651 | 0,74854857 | 0,975901808 | UDP-glucose glycoprotein glucosyltransferase 2 |
| ENSG00000259932 | AC051619,6 | -0,110691777 | 0,926143867 | 0,75011613 | 0,975901808 | novel transcript |
| ENSG00000287817 | AL096701,4 | -0,110794775 | 0,926077748 | 0,85413043 | 0,987861659 | novel transcript, antisense to EIF4ENIF1 |
| ENSG00000276537 | TTYH1 | -0,110821386 | 0,926060667 | 0,75498414 | 0,975901808 | tweety family member 1 |
| ENSG00000287828 | AL354743,2 | -0,111086796 | 0,925890317 | 0,90682697 | 0,992590661 | novel transcript, antisense to PRDM16 |
| ENSG00000117697 | NSL1 | -0,11113049 | 0,925862276 | 0,74514587 | 0,975666349 | NSL1 component of MIS12 kinetochore complex |
| ENSG00000256705 | AL137779,1 | -0,111231061 | 0,925797736 | 0,92477303 | 0,993915537 | novel transcript, antisense to PPP2R5C |
| ENSG00000225930 | LINC02249 | -0,111239265 | 0,925792471 | 0,68104231 | 0,964654539 | long intergenic non-protein coding RNA 2249 |
| ENSG00000273473 | BX649601,1 | -0,111243451 | 0,925789785 | 0,87442774 | 0,98990147 | novel transcript |
| ENSG00000287373 | AL122014,1 | -0,111335912 | 0,925730454 | 0,93175385 | 0,994433362 | novel transcript, antisense to DCDC1 |
| ENSG00000074356 | NCBP3 | -0,11136202 | 0,925713701 | 0,64965717 | 0,956899963 | nuclear cap binding subunit 3 |
| ENSG00000250569 | NTAN1P2 | -0,111370487 | 0,925708268 | 0,84738636 | 0,987035044 | N-terminal asparagine amidase pseudogene 2 |
| ENSG00000277448 | LINC02338 | -0,111371156 | 0,925707839 | 0,90134259 | 0,992041918 | long intergenic non-protein coding RNA 2338 |
| ENSG00000273890 | AC034102,8 | -0,111396101 | 0,925691833 | 0,89603519 | 0,991528124 | novel transcript, antisence to IKZF4 |
| ENSG00000056097 | ZFR | -0,111421119 | 0,925675781 | 0,76424143 | 0,976640468 | zinc finger RNA binding protein |

| | | | | | | |
|---|---|---|---|---|---|---|
| ENSG00000183826 | BTBD9 | -0,111459915 | 0,925650888 | 0,61695604 | 0,952793934 | BTB domain containing 9 |
| ENSG00000149269 | PAK1 | -0,111537221 | 0,925601289 | 0,62420129 | 0,952793934 | p21 (RAC1) activated kinase 1 |
| ENSG00000177410 | ZFAS1 | -0,111546744 | 0,92559518 | 0,67277254 | 0,962371283 | ZNFX1 antisense RNA 1 |
| ENSG00000168078 | PBK | -0,111670612 | 0,925515713 | 0,80883521 | 0,981213877 | PDZ binding kinase |
| ENSG00000272904 | AL390726,2 | -0,111671266 | 0,925515293 | 0,89300349 | 0,991120066 | novel transcript |
| ENSG00000271918 | AC034236,2 | -0,111719558 | 0,925484314 | 0,78343399 | 0,979026314 | novel transcript, antisense to COMMD10 |
| ENSG00000196295 | GARS1-DT | -0,111720709 | 0,925483575 | 0,61877146 | 0,952793934 | GARS1 divergent transcript |
| ENSG00000223522 | AC093690,1 | -0,111735318 | 0,925474203 | 0,78235535 | 0,978628303 | novel transcript |
| ENSG00000248802 | AC078850,2 | -0,111795019 | 0,925435907 | 0,95193361 | 0,996432861 | novel transcript |
| ENSG00000146205 | ANO7 | -0,11181329 | 0,925424186 | 0,8377356 | 0,98566906 | anoctamin 7 |
| ENSG00000120798 | NR2C1 | -0,111817286 | 0,925421623 | 0,80948185 | 0,981213877 | nuclear receptor subfamily 2 group C member 1 |
| ENSG00000104889 | RNASEH2A | -0,111818496 | 0,925420847 | 0,61805577 | 0,952793934 | ribonuclease H2 subunit A |
| ENSG00000245213 | AC105285,1 | -0,11183724 | 0,925408823 | 0,75161059 | 0,975901808 | novel transcript, antisense to GALNT7 |
| ENSG00000206597 | SNORA57 | -0,111869977 | 0,925387825 | 0,81464765 | 0,981521413 | small nucleolar RNA, H/ACA box 57 |
| ENSG00000205269 | TMEM170B | -0,111937895 | 0,925344261 | 0,82841959 | 0,983717437 | transmembrane protein 170B |
| ENSG00000125846 | ZNF133 | -0,112027941 | 0,925286508 | 0,64398303 | 0,956633898 | zinc finger protein 133 |
| ENSG00000274305 | LENG8 | -0,112032752 | 0,925283422 | 0,79626388 | 0,980015022 | leukocyte receptor cluster member 8 |
| ENSG00000164162 | ANAPC10 | -0,112039979 | 0,925278787 | 0,76175478 | 0,976285211 | anaphase promoting complex subunit 10 |
| ENSG00000189369 | GSPT2 | -0,112089624 | 0,925246948 | 0,70258198 | 0,968742626 | G1 to S phase transition 2 |
| ENSG00000157890 | MEGF11 | -0,112119313 | 0,925227907 | 0,69954212 | 0,968522643 | multiple EGF like domains 11 |
| ENSG00000133246 | PRAM1 | -0,112127175 | 0,925222865 | 0,89207086 | 0,99102448 | PML-RARA regulated adaptor molecule 1 |
| ENSG00000228506 | AL513550,1 | -0,112186111 | 0,925185069 | 0,77464731 | 0,978553937 | novel transcript |
| ENSG00000251143 | AP002490,1 | -0,112228802 | 0,925157692 | 0,82807254 | 0,983595244 | novel transcript |
| ENSG00000265519 | AC015922,2 | -0,11230994 | 0,925105663 | 0,82592227 | 0,983383773 | novel transcript |
| ENSG00000109572 | CLCN3 | -0,11231521 | 0,925102283 | 0,7571266 | 0,975901808 | chloride voltage-gated channel 3 |
| ENSG00000268520 | AC008750,4 | -0,11237031 | 0,925066952 | 0,91908436 | 0,993598006 | novel transcript, sense intronic to ETFB |
| ENSG00000198625 | MDM4 | -0,112439039 | 0,925022883 | 0,61356423 | 0,952413269 | MDM4 regulator of p53 |
| ENSG00000259668 | AC066613,1 | -0,112499109 | 0,924984369 | 0,76739846 | 0,977201891 | novel transcript, antisense to DMXL2 |
| ENSG00000231119 | AL031666,1 | -0,112546348 | 0,924954082 | 0,75584074 | 0,975901808 | novel transcript, antisense to ZMYND8 |
| ENSG00000183208 | GDPGP1 | -0,112575665 | 0,924935286 | 0,7432428 | 0,975310206 | GDP-D-glucose phosphorylase 1 |
| ENSG00000215571 | GRK6P1 | -0,112586555 | 0,924928305 | 0,86435644 | 0,989074806 | G protein-coupled receptor kinase 6 pseudogene 1 |
| ENSG00000274528 | AC090970,2 | -0,112617593 | 0,924908406 | 0,85068504 | 0,987147801 | novel transcript |
| ENSG00000237758 | BANF1P3 | -0,112634871 | 0,924897329 | 0,88776025 | 0,990717729 | BANF1 pseudogene 3 |

| | | | | | | |
|---|---|---|---|---|---|---|
| ENSG00000253880 | AC009435,1 | -0,112734915 | 0,924833194 | 0,87250052 | 0,989217334 | novel transcript |
| ENSG00000131844 | MCCC2 | -0,112743264 | 0,924827842 | 0,71501464 | 0,971041579 | methylcrotonoyl-CoA carboxylase 2 |
| ENSG00000146587 | RBAK | -0,112797877 | 0,924792833 | 0,80191253 | 0,980710009 | RB associated KRAB zinc finger |
| ENSG00000131495 | NDUFA2 | -0,112800307 | 0,924791276 | 0,70027157 | 0,968522643 | NADH:ubiquinone oxidoreductase subunit A2 |
| ENSG00000100058 | CRYBB2P1 | -0,112828831 | 0,924772992 | 0,65069692 | 0,956899963 | crystallin beta B2 pseudogene 1 |
| ENSG00000106771 | TMEM245 | -0,112832914 | 0,924770374 | 0,71950083 | 0,971869926 | transmembrane protein 245 |
| ENSG00000266794 | RN7SL7P | -0,112859162 | 0,924753549 | 0,81801489 | 0,982323352 | RNA, 7SL, cytoplasmic 7, pseudogene |
| ENSG00000270112 | AC090241,2 | -0,112914691 | 0,924717957 | 0,82853516 | 0,983721455 | novel transcript, antisense to ST8SIA5 |
| ENSG00000145808 | ADAMTS19 | -0,113038077 | 0,924638874 | 0,8542746 | 0,987861659 | ADAM metallopeptidase with thrombospondin type 1 motif 19 |
| ENSG00000205659 | LIN52 | -0,113078587 | 0,924612911 | 0,74447915 | 0,975649186 | lin-52 DREAM MuvB core complex component |
| ENSG00000053371 | AKR7A2 | -0,113121578 | 0,924585359 | 0,62606998 | 0,952793934 | aldo-keto reductase family 7 member A2 |
| ENSG00000218418 | AL591135,1 | -0,113380151 | 0,924419661 | 0,88246929 | 0,990717729 | Rho GTPase activating protein 21 (ARHGAP21) pseudogene |
| ENSG00000136931 | NR5A1 | -0,113414258 | 0,924397807 | 0,91337812 | 0,993361299 | nuclear receptor subfamily 5 group A member 1 |
| ENSG00000105605 | CACNG7 | -0,113457497 | 0,924370102 | 0,71781769 | 0,97176116 | calcium voltage-gated channel auxiliary subunit gamma 7 |
| ENSG00000137876 | RSL24D1 | -0,113472184 | 0,924360692 | 0,81001159 | 0,981213877 | ribosomal L24 domain containing 1 |
| ENSG00000150455 | TIRAP | -0,113503998 | 0,924340308 | 0,66926616 | 0,961754789 | TIR domain containing adaptor protein |
| ENSG00000118418 | HMGN3 | -0,113528187 | 0,92432481 | 0,71346133 | 0,970812834 | high mobility group nucleosomal binding domain 3 |
| ENSG00000284776 | AL121900,1 | -0,113537311 | 0,924318965 | 0,68140344 | 0,964654539 | novel protein |
| ENSG00000150776 | NKAPD1 | -0,113660551 | 0,924240009 | 0,74997425 | 0,975901808 | NKAP domain containing 1 |
| ENSG00000080802 | CNOT4 | -0,113714297 | 0,924205579 | 0,71522745 | 0,971041579 | CCR4-NOT transcription complex subunit 4 |
| ENSG00000225313 | AL513327,1 | -0,113785096 | 0,924160225 | 0,64851699 | 0,956899963 | novel transcript |
| ENSG00000101353 | MROH8 | -0,113799845 | 0,924150778 | 0,74249353 | 0,975176043 | maestro heat like repeat family member 8 |
| ENSG00000288427 | CLSTN3 | -0,113816013 | 0,924140421 | 0,61682965 | 0,952793934 | calsyntenin 3 |
| ENSG00000139182 | CLSTN3 | -0,113816013 | 0,924140421 | 0,61682965 | 0,952793934 | calsyntenin 3 |
| ENSG00000099625 | CBARP | -0,113817439 | 0,924139507 | 0,81297697 | 0,981246822 | CACN subunit beta associated regulatory protein |
| ENSG00000241429 | EEF1A1P25 | -0,11405627 | 0,923986533 | 0,81848048 | 0,9824772 | eukaryotic translation elongation factor 1 alpha 1 pseudogene 25 |
| ENSG00000228315 | GUSBP11 | -0,114176543 | 0,923909506 | 0,62139614 | 0,952793934 | GUSB pseudogene 11 |
| ENSG00000185158 | LRRC37B | -0,114181161 | 0,923906549 | 0,71302016 | 0,970812834 | leucine rich repeat containing 37B |
| ENSG00000168781 | PPIP5K1 | -0,114303242 | 0,923828372 | 0,63724224 | 0,955985674 | diphosphoinositol pentakisphosphate kinase 1 |

| | | | | | | |
|---|---|---|---|---|---|---|
| ENSG00000172493 | AFF1 | -0,114317144 | 0,923819469 | 0,71754051 | 0,971697135 | AF4/FMR2 family member 1 |
| ENSG00000255920 | CCND2-AS1 | -0,114365084 | 0,923788772 | 0,77291074 | 0,978310326 | CCND2 antisense RNA 1 |
| ENSG00000288040 | AC097713,2 | -0,114366511 | 0,923787858 | 0,70294996 | 0,968742626 | novel transcript |
| ENSG00000254843 | XIAPP2 | -0,114377598 | 0,923780759 | 0,82346926 | 0,983374097 | X-linked inhibitor of apoptosis pseudogene 2 |
| ENSG00000163482 | STK36 | -0,114427257 | 0,923748962 | 0,65457302 | 0,957444211 | serine/threonine kinase 36 |
| ENSG00000286190 | AC055839,2 | -0,114439919 | 0,923740855 | 0,59396555 | 0,947469769 | uncharacterized LOC728392 |
| ENSG00000113048 | MRPS27 | -0,114453005 | 0,923732475 | 0,71108443 | 0,970482408 | mitochondrial ribosomal protein S27 |
| ENSG00000284526 | AC015802,6 | -0,114459862 | 0,923728086 | 0,83080496 | 0,984156632 | novel protein |
| ENSG00000285976 | AL135905,2 | -0,114484729 | 0,923712164 | 0,82161499 | 0,982952211 | novel protein |
| ENSG00000213842 | SUGT1P2 | -0,114543908 | 0,923674274 | 0,83782672 | 0,98566906 | SUGT1 pseudogene 2 |
| ENSG00000263276 | AC020978,7 | -0,114548766 | 0,923671164 | 0,77199269 | 0,978033855 | novel transcript |
| ENSG00000105173 | CCNE1 | -0,114563469 | 0,92366175 | 0,7029403 | 0,968742626 | cyclin E1 |
| ENSG00000087995 | METTL2A | -0,114593143 | 0,923642752 | 0,5907144 | 0,947416795 | methyltransferase like 2A |
| ENSG00000105819 | PMPCB | -0,114599768 | 0,923638511 | 0,79386438 | 0,979776981 | peptidase, mitochondrial processing subunit beta |
| ENSG00000259033 | AL356804,1 | -0,114676574 | 0,92358934 | 0,87184511 | 0,989217334 | novel transcript |
| ENSG00000107669 | ATE1 | -0,114693978 | 0,923578198 | 0,71298165 | 0,970812834 | arginyltransferase 1 |
| ENSG00000160207 | HSF2BP | -0,114749157 | 0,923542874 | 0,77620545 | 0,978587255 | heat shock transcription factor 2 binding protein |
| ENSG00000163682 | RPL9 | -0,114793023 | 0,923514794 | 0,75965566 | 0,976120207 | ribosomal protein L9 |
| ENSG00000148153 | INIP | -0,114832162 | 0,92348974 | 0,6752607 | 0,963390425 | INTS3 and NABP interacting protein |
| ENSG00000254995 | STX16-NPEPL1 | -0,114850328 | 0,923478112 | 0,75286614 | 0,975901808 | STX16-NPEPL1 readthrough (NMD candidate) |
| ENSG00000047188 | YTHDC2 | -0,114877948 | 0,923460432 | 0,73565987 | 0,974353474 | YTH domain containing 2 |
| ENSG00000250920 | AC105460,1 | -0,114879835 | 0,923459224 | 0,85152787 | 0,987259878 | novel transcript |
| ENSG00000159596 | TMEM69 | -0,114922558 | 0,923431878 | 0,79420694 | 0,979776981 | transmembrane protein 69 |
| ENSG00000149313 | AASDHPPT | -0,115012347 | 0,923374408 | 0,77234416 | 0,97820249 | aminoadipate-semialdehyde dehydrogenase-phosphopantetheinyl transferase |
| ENSG00000277128 | AL589743,3 | -0,115024271 | 0,923366777 | 0,81902019 | 0,98249617 | novel transcript |
| ENSG00000145041 | DCAF1 | -0,115055042 | 0,923347082 | 0,64604963 | 0,956899963 | DDB1 and CUL4 associated factor 1 |
| ENSG00000168389 | MFSD2A | -0,115124137 | 0,923302862 | 0,60338087 | 0,950352286 | major facilitator superfamily domain containing 2A |
| ENSG00000180998 | GPR137C | -0,115145679 | 0,923289075 | 0,82488845 | 0,983374097 | G protein-coupled receptor 137C |
| ENSG00000116095 | PLEKHA3 | -0,115202637 | 0,923252624 | 0,72369064 | 0,972971932 | pleckstrin homology domain containing A3 |
| ENSG00000168906 | MAT2A | -0,115207712 | 0,923249376 | 0,71830953 | 0,971789526 | methionine adenosyltransferase 2A |
| ENSG00000187609 | EXD3 | -0,11520852 | 0,923248859 | 0,79583243 | 0,979879951 | exonuclease 3'-5' domain containing 3 |
| ENSG00000230426 | LINC01036 | -0,115218946 | 0,923242187 | 0,88431051 | 0,990717729 | long intergenic non-protein coding RNA 1036 |

| | | | | | | |
|---|---|---|---|---|---|---|
| ENSG00000138074 | SLC5A6 | -0,115286453 | 0,923198988 | 0,64628003 | 0,956899963 | solute carrier family 5 member 6 |
| ENSG00000100528 | CNIH1 | -0,115347423 | 0,923159973 | 0,82920672 | 0,983957145 | cornichon family AMPA receptor auxiliary protein 1 |
| ENSG00000135476 | ESPL1 | -0,115370754 | 0,923145044 | 0,68712647 | 0,965769168 | extra spindle pole bodies like 1, separase |
| ENSG00000271092 | TLCD4-RWDD3 | -0,115400494 | 0,923126014 | 0,81476713 | 0,981521413 | TLCD4-RWDD3 readthrough |
| ENSG00000233058 | LINC00884 | -0,115426984 | 0,923109064 | 0,68890012 | 0,966556275 | long intergenic non-protein coding RNA 884 |
| ENSG00000196072 | BLOC1S2 | -0,115447549 | 0,923095906 | 0,7780869 | 0,978587255 | biogenesis of lysosomal organelles complex 1 subunit 2 |
| ENSG00000185155 | MIXL1 | -0,115594699 | 0,923001758 | 0,84013998 | 0,985672131 | Mix paired-like homeobox |
| ENSG00000285886 | AC211476,6 | -0,115891995 | 0,922811575 | 0,82694335 | 0,983548538 | novel transcript |
| ENSG00000140905 | GCSH | -0,115946945 | 0,922776427 | 0,81695352 | 0,981977498 | glycine cleavage system protein H |
| ENSG00000273993 | PTPRK | -0,11596732 | 0,922763395 | 0,7314509 | 0,974353474 | protein tyrosine phosphatase receptor type K |
| ENSG00000077092 | RARB | -0,116039796 | 0,92271704 | 0,80089445 | 0,980505307 | retinoic acid receptor beta |
| ENSG00000181163 | NPM1 | -0,116060267 | 0,922703947 | 0,73594423 | 0,974353474 | nucleophosmin 1 |
| ENSG00000287031 | AC093307,1 | -0,116091795 | 0,922683783 | 0,89214731 | 0,99102448 | novel transcript |
| ENSG00000128266 | GNAZ | -0,116100984 | 0,922677906 | 0,60356991 | 0,950352286 | G protein subunit alpha z |
| ENSG00000196873 | CBWD3 | -0,116164242 | 0,92263745 | 0,62300084 | 0,952793934 | COBW domain containing 3 |
| ENSG00000228915 | OR7E128P | -0,116235887 | 0,922591633 | 0,79575731 | 0,979879951 | olfactory receptor family 7 subfamily E member 128 pseudogene |
| ENSG00000118961 | LDAH | -0,116278216 | 0,922564564 | 0,78184611 | 0,978597905 | lipid droplet associated hydrolase |
| ENSG00000166181 | API5 | -0,116363947 | 0,922509743 | 0,76486169 | 0,976722333 | apoptosis inhibitor 5 |
| ENSG00000102804 | TSC22D1 | -0,116452884 | 0,922452875 | 0,73028489 | 0,974321178 | TSC22 domain family member 1 |
| ENSG00000233024 | NPIPA9 | -0,116458593 | 0,922449225 | 0,73522651 | 0,974353474 | nuclear pore complex interacting protein family, member A9 |
| ENSG00000093217 | XYLB | -0,116477903 | 0,922436878 | 0,64703092 | 0,956899963 | xylulokinase |
| ENSG00000265962 | GACAT2 | -0,116575703 | 0,922374349 | 0,84951526 | 0,987036295 | gastric cancer associated transcript 2 |
| ENSG00000162735 | PEX19 | -0,116653396 | 0,922324677 | 0,64872091 | 0,956899963 | peroxisomal biogenesis factor 19 |
| ENSG00000185379 | RAD51D | -0,116654129 | 0,922324209 | 0,61144146 | 0,952139548 | RAD51 paralog D |
| ENSG00000270116 | AP001429,1 | -0,116719364 | 0,922282504 | 0,79928677 | 0,980161849 | novel transcript, sense intronic to TTC3 |
| ENSG00000115904 | SOS1 | -0,116757792 | 0,922257939 | 0,78369038 | 0,979210284 | SOS Ras/Rac guanine nucleotide exchange factor 1 |
| ENSG00000275383 | AC126773,6 | -0,116805427 | 0,922227488 | 0,83433274 | 0,984905093 | novel transcript |
| ENSG00000277149 | TYW1B | -0,116844848 | 0,922202289 | 0,71666219 | 0,971438758 | tRNA-yW synthesizing protein 1 homolog B |
| ENSG00000076003 | MCM6 | -0,116879901 | 0,922179883 | 0,68501167 | 0,965415049 | minichromosome maintenance complex component 6 |

| | | | | | | |
|---|---|---|---|---|---|---|
| ENSG00000228203 | GRASLND | -0,117019496 | 0,922090657 | 0,65166741 | 0,957089277 | glycosaminoglycan regulatory associated long non-coding RNA |
| ENSG00000228681 | AC008072,1 | -0,117029791 | 0,922084077 | 0,78769887 | 0,979462318 | ribosomal protein S27a (RPS27A) pseudogene |
| ENSG00000114388 | NPRL2 | -0,117099358 | 0,922039615 | 0,64968973 | 0,956899963 | NPR2 like, GATOR1 complex subunit |
| ENSG00000149295 | DRD2 | -0,11714171 | 0,922012548 | 0,83678369 | 0,985426146 | dopamine receptor D2 |
| ENSG00000121940 | CLCC1 | -0,11719095 | 0,92198108 | 0,6641452 | 0,960088568 | chloride channel CLIC like 1 |
| ENSG00000010270 | STARD3NL | -0,117191685 | 0,92198061 | 0,75949405 | 0,976120207 | STARD3 N-terminal like |
| ENSG00000259723 | AL354993,2 | -0,117197283 | 0,921977033 | 0,88240094 | 0,990717729 | novel transcript, antisense to TSHZ2 |
| ENSG00000051341 | POLQ | -0,117232749 | 0,921954368 | 0,72345109 | 0,972913663 | DNA polymerase theta |
| ENSG00000116213 | WRAP73 | -0,117249291 | 0,921943797 | 0,64776116 | 0,956899963 | WD repeat containing, antisense to TP73 |
| ENSG00000162961 | DPY30 | -0,117285864 | 0,921920425 | 0,78915024 | 0,979462318 | dpy-30 histone methyltransferase complex regulatory subunit |
| ENSG00000264058 | AC073508,2 | -0,117310297 | 0,921904812 | 0,86714025 | 0,989074806 | novel transcript |
| ENSG00000166004 | CEP295 | -0,117383854 | 0,921857809 | 0,76422622 | 0,976640468 | centrosomal protein 295 |
| ENSG00000232273 | FTH1P1 | -0,117477602 | 0,921797907 | 0,88356492 | 0,990717729 | ferritin heavy chain 1 pseudogene 1 |
| ENSG00000259632 | AC090971,4 | -0,117589181 | 0,921726618 | 0,87631962 | 0,990109814 | UTP6, small subunit (SSU) processome component, homolog (yeast) (UTP6) pseudogene |
| ENSG00000143409 | MINDY1 | -0,117687596 | 0,921663743 | 0,65211059 | 0,957241719 | MINDY lysine 48 deubiquitinase 1 |
| ENSG00000196275 | GTF2IRD2 | -0,117759145 | 0,921618036 | 0,70906164 | 0,970282889 | GTF2I repeat domain containing 2 |
| ENSG00000213757 | RPS3AP20 | -0,117768892 | 0,921611809 | 0,90178936 | 0,992136926 | RPS3A pseudogene 20 |
| ENSG00000075790 | BCAP29 | -0,117773275 | 0,921609009 | 0,75056298 | 0,975901808 | B cell receptor associated protein 29 |
| ENSG00000102805 | CLN5 | -0,117827179 | 0,921574576 | 0,71862912 | 0,971869926 | CLN5 intracellular trafficking protein |
| ENSG00000279298 | Z99916,3 | -0,117898457 | 0,921529045 | 0,88899733 | 0,990717729 | novel transcript |
| ENSG00000284319 | AC275454,2 | -0,117934829 | 0,921505812 | 0,83681163 | 0,985426146 | novel transcript, antisense to SLC25A24 |
| ENSG00000260879 | AL359258,2 | -0,117934829 | 0,921505812 | 0,83681163 | 0,985426146 | novel transcript, antisense to SLC25A24 |
| ENSG00000110697 | PITPNM1 | -0,11794076 | 0,921502024 | 0,72697593 | 0,973344786 | phosphatidylinositol transfer protein membrane associated 1 |
| ENSG00000142959 | BEST4 | -0,118048956 | 0,921432918 | 0,85800224 | 0,98899761 | bestrophin 4 |
| ENSG00000172469 | MANEA | -0,11806693 | 0,921421439 | 0,8350727 | 0,984905093 | mannosidase endo-alpha |
| ENSG00000172262 | ZNF131 | -0,118155986 | 0,921364562 | 0,70314149 | 0,968742626 | zinc finger protein 131 |
| ENSG00000257341 | AL928654,3 | -0,118191248 | 0,921342042 | 0,8941036 | 0,991135494 | novel protein |
| ENSG00000261364 | AC063923,2 | -0,118287219 | 0,921280755 | 0,89958796 | 0,991845106 | novel transcript |

| | | | | | | |
|---|---|---|---|---|---|---|
| ENSG00000260097 | SPDYE6 | -0,118341518 | 0,921246081 | 0,73096464 | 0,974353474 | speedy/RINGO cell cycle regulator family member E6 |
| ENSG00000114686 | MRPL3 | -0,11837269 | 0,921226176 | 0,75093892 | 0,975901808 | mitochondrial ribosomal protein L3 |
| ENSG00000215032 | GNL3LP1 | -0,118389788 | 0,921215258 | 0,74218911 | 0,975176043 | G protein nucleolar 3 like pseudogene 1 |
| ENSG00000275152 | CCL16 | -0,1184731 | 0,921162062 | 0,92851274 | 0,994108531 | C-C motif chemokine ligand 16 |
| ENSG00000119865 | CNRIP1 | -0,118488427 | 0,921152276 | 0,72035332 | 0,972142678 | cannabinoid receptor interacting protein 1 |
| ENSG00000137942 | FNBP1L | -0,118488594 | 0,921152169 | 0,77185365 | 0,978033855 | formin binding protein 1 like |
| ENSG00000249863 | AC021106,1 | -0,118581364 | 0,921092938 | 0,84194836 | 0,98572121 | synaptotagmin binding, cytoplasmic RNA interacting protein (SYNCRIP) pseudogene |
| ENSG00000273576 | AC009283,1 | -0,11859554 | 0,921083887 | 0,78513313 | 0,979320406 | novel transcript |
| ENSG00000184083 | FAM120C | -0,118705104 | 0,921013939 | 0,58061743 | 0,943832501 | family with sequence similarity 120C |
| ENSG00000147180 | ZNF711 | -0,118707218 | 0,92101259 | 0,8068524 | 0,981213877 | zinc finger protein 711 |
| ENSG00000283480 | AL512380,2 | -0,118740154 | 0,920991563 | 0,94101044 | 0,995251853 | novel transcript, antisense to SOX4 |
| ENSG00000135338 | LCA5 | -0,118792628 | 0,920958066 | 0,76542989 | 0,976830764 | lebercilin LCA5 |
| ENSG00000225718 | AC092100,1 | -0,118799373 | 0,92095376 | 0,91694178 | 0,993361299 | novel transcript |
| ENSG00000166987 | MBD6 | -0,118821902 | 0,920939379 | 0,64958188 | 0,956899963 | methyl-CpG binding domain protein 6 |
| ENSG00000224660 | SH3BP5-AS1 | -0,118846555 | 0,920923642 | 0,6685358 | 0,961351573 | SH3BP5 antisense RNA 1 |
| ENSG00000214106 | PAXIP1-AS2 | -0,118878649 | 0,920903155 | 0,70056839 | 0,968522643 | PAXIP1 antisense RNA 2 |
| ENSG00000287725 | AP003071,5 | -0,118902644 | 0,920887838 | 0,74842205 | 0,975901808 | novel protein, TPCN2 - SMIM38 readthrough |
| ENSG00000243660 | ZNF487 | -0,118960068 | 0,920851185 | 0,72227626 | 0,972594421 | zinc finger protein 487 |
| ENSG00000180044 | C3orf80 | -0,118964254 | 0,920848513 | 0,8680994 | 0,989074806 | chromosome 3 open reading frame 80 |
| ENSG00000232117 | LINC00384 | -0,119022925 | 0,920811065 | 0,89811769 | 0,991796521 | long intergenic non-protein coding RNA 384 |
| ENSG00000204160 | ZDHHC18 | -0,119080705 | 0,920774187 | 0,58555743 | 0,945213113 | zinc finger DHHC-type palmitoyltransferase 18 |
| ENSG00000267980 | AC007292,1 | -0,119205959 | 0,92069425 | 0,86538343 | 0,989074806 | novel transcript, antisense to SH3GL1 |
| ENSG00000228730 | LINC02013 | -0,119239348 | 0,920672942 | 0,86779569 | 0,989074806 | long intergenic non-protein coding RNA 2013 |
| ENSG00000213190 | MLLT11 | -0,119258561 | 0,920660681 | 0,73585364 | 0,974353474 | MLLT11 transcription factor 7 cofactor |
| ENSG00000160326 | SLC2A6 | -0,119347037 | 0,920604221 | 0,83876962 | 0,985672131 | solute carrier family 2 member 6 |
| ENSG00000065978 | YBX1 | -0,119351119 | 0,920601617 | 0,70022964 | 0,968522643 | Y-box binding protein 1 |
| ENSG00000114491 | UMPS | -0,119395849 | 0,920573074 | 0,69039351 | 0,967179044 | uridine monophosphate synthetase |
| ENSG00000246174 | KCTD21-AS1 | -0,119497659 | 0,920508112 | 0,71460437 | 0,971041579 | KCTD21 antisense RNA 1 |
| ENSG00000230928 | AL139241,1 | -0,11949796 | 0,92050792 | 0,84886844 | 0,987036295 | novel transcript |
| ENSG00000148399 | DPH7 | -0,11951917 | 0,920494387 | 0,60613604 | 0,950927937 | diphthamide biosynthesis 7 |
| ENSG00000138175 | ARL3 | -0,119529833 | 0,920487584 | 0,71833221 | 0,971789526 | ADP ribosylation factor like GTPase 3 |

| | | | | | | |
|---|---|---|---|---|---|---|
| ENSG00000167535 | CACNB3 | -0,119547082 | 0,920476579 | 0,59488427 | 0,947469769 | calcium voltage-gated channel auxiliary subunit beta 3 |
| ENSG00000225761 | AL596247,1 | -0,11972009 | 0,920366202 | 0,93594375 | 0,994479239 | novel transcript |
| ENSG00000288115 | AC008731,2 | -0,119812788 | 0,920307067 | 0,87923831 | 0,990148595 | novel transcript, antisense to TNRC6A |
| ENSG00000261669 | AC008731,1 | -0,119812788 | 0,920307067 | 0,87923831 | 0,990148595 | novel transcript, antisense to TNRC6A |
| ENSG00000154917 | RAB6B | -0,119824956 | 0,920299305 | 0,62708572 | 0,952793934 | RAB6B, member RAS oncogene family |
| ENSG00000188342 | GTF2F2 | -0,119840485 | 0,920289399 | 0,75569308 | 0,975901808 | general transcription factor IIF subunit 2 |
| ENSG00000233435 | AGGF1P2 | -0,119861936 | 0,920275715 | 0,84890221 | 0,987036295 | angiogenic factor with G-patch and FHA domains 1 pseudogene 2 |
| ENSG00000131269 | ABCB7 | -0,119902694 | 0,920249717 | 0,70917785 | 0,970282889 | ATP binding cassette subfamily B member 7 |
| ENSG00000143401 | ANP32E | -0,120095181 | 0,920126944 | 0,67061713 | 0,962026636 | acidic nuclear phosphoprotein 32 family member E |
| ENSG00000065060 | UHRF1BP1 | -0,120161341 | 0,920084749 | 0,62411607 | 0,952793934 | UHRF1 binding protein 1 |
| ENSG00000245573 | BDNF-AS | -0,12018441 | 0,920070037 | 0,69925536 | 0,968522643 | BDNF antisense RNA |
| ENSG00000119509 | INVS | -0,120300179 | 0,919996209 | 0,65377976 | 0,957444211 | inversin |
| ENSG00000270802 | AC005776,1 | -0,120326587 | 0,919979369 | 0,9142339 | 0,993361299 | ribosomal protein L21 (RPL21) pseudogene |
| ENSG00000148229 | POLE3 | -0,120355392 | 0,919961 | 0,61708017 | 0,952793934 | DNA polymerase epsilon 3, accessory subunit |
| ENSG00000287539 | AC022966,2 | -0,120470636 | 0,919887516 | 0,86839962 | 0,989074806 | novel transcript |
| ENSG00000258057 | BCDIN3D-AS1 | -0,120530268 | 0,919849494 | 0,76567343 | 0,976862445 | BCDIN3D antisense RNA 1 |
| ENSG00000063601 | MTMR1 | -0,1205437 | 0,91984093 | 0,73855516 | 0,974353474 | myotubularin related protein 1 |
| ENSG00000156162 | DPY19L4 | -0,120583661 | 0,919815452 | 0,74483186 | 0,975649186 | dpy-19 like 4 |
| ENSG00000206356 | AC114324,1 | -0,120640482 | 0,919779225 | 0,87142911 | 0,989217334 | putative POM121-like protein 1-like |
| ENSG00000232070 | TMEM253 | -0,120678 | 0,919755307 | 0,82693519 | 0,983548538 | transmembrane protein 253 |
| ENSG00000134748 | PRPF38A | -0,120743212 | 0,919713733 | 0,66581539 | 0,961016016 | pre-mRNA processing factor 38A |
| ENSG00000266947 | AC022916,1 | -0,120752444 | 0,919707848 | 0,70565513 | 0,969485452 | novel transcript, antisense SLFN5 |
| ENSG00000265261 | AC027575,1 | -0,120788069 | 0,919685137 | 0,82654228 | 0,983471137 | exportin, tRNA (nuclear export receptor for tRNAs) (XPOT) pseudogene |
| ENSG00000169427 | KCNK9 | -0,120811457 | 0,919670228 | 0,90867201 | 0,992617809 | potassium two pore domain channel subfamily K member 9 |
| ENSG00000149054 | ZNF215 | -0,120813028 | 0,919669227 | 0,79074525 | 0,979471509 | zinc finger protein 215 |
| ENSG00000149571 | KIRREL3 | -0,120897093 | 0,91961564 | 0,82066259 | 0,982609114 | kirre like nephrin family adhesion molecule 3 |
| ENSG00000281954 | AC208069,1 | -0,120912375 | 0,919605898 | 0,95206895 | 0,996432861 | novel transcript, antisense to DNAH10 |
| ENSG00000168298 | H1-4 | -0,120930285 | 0,919594482 | 0,57145523 | 0,942436428 | H1,4 linker histone, cluster member |
| ENSG00000225399 | AC121247,1 | -0,120950274 | 0,919581741 | 0,89929969 | 0,991832796 | novel transcript |

| | | | | | | |
|---|---|---|---|---|---|---|
| ENSG00000164011 | ZNF691 | -0,121028386 | 0,919531954 | 0,65130468 | 0,957089277 | zinc finger protein 691 |
| ENSG00000260436 | AC093525,3 | -0,121128591 | 0,919468088 | 0,93890297 | 0,994713928 | novel transcript, antisense to PDPK1 |
| ENSG00000121741 | ZMYM2 | -0,121200795 | 0,919422072 | 0,78072034 | 0,978597905 | zinc finger MYM-type containing 2 |
| ENSG00000286568 | AC007879,5 | -0,121211346 | 0,919415348 | 0,91359933 | 0,993361299 | novel transcript |
| ENSG00000259001 | AL355075,4 | -0,121237391 | 0,919398749 | 0,64652681 | 0,956899963 | ribonuclease P RNA component H1 |
| ENSG00000169246 | NPIPB3 | -0,12123755 | 0,919398648 | 0,75138344 | 0,975901808 | nuclear pore complex interacting protein family member B3 |
| ENSG00000120963 | ZNF706 | -0,121338225 | 0,919334492 | 0,7714516 | 0,978033855 | zinc finger protein 706 |
| ENSG00000152443 | ZNF776 | -0,121377768 | 0,919309295 | 0,73251207 | 0,974353474 | zinc finger protein 776 |
| ENSG00000287126 | AC108488,3 | -0,121449368 | 0,919263671 | 0,63860291 | 0,956018518 | novel transcript |
| ENSG00000287171 | AC006062,1 | -0,121473893 | 0,919248044 | 0,92495147 | 0,993915537 | novel transcript |
| ENSG00000235066 | AC009303,2 | -0,121505324 | 0,919228017 | 0,77478593 | 0,978553937 | novel transcript, antisense to CCDC93 |
| ENSG00000267672 | AC010632,2 | -0,121514204 | 0,91922236 | 0,80637889 | 0,981213877 | novel transcript |
| ENSG00000130935 | NOL11 | -0,121539581 | 0,91920619 | 0,78636793 | 0,979320406 | nucleolar protein 11 |
| ENSG00000178802 | MPI | -0,121547878 | 0,919200904 | 0,59482541 | 0,947469769 | mannose phosphate isomerase |
| ENSG00000281697 | AP003392,10 | -0,121616899 | 0,919156929 | 0,89438514 | 0,991135494 | potassium channel tetramerisation domain containing 6 (KCTD6) pseudogene |
| ENSG00000254909 | AP003392,2 | -0,121616899 | 0,919156929 | 0,89438514 | 0,991135494 | potassium channel tetramerisation domain containing 6 (KCTD6) pseudogene |
| ENSG00000265728 | AP001099,1 | -0,121717646 | 0,919092744 | 0,79200731 | 0,979471509 | novel transcript |
| ENSG00000103351 | CLUAP1 | -0,121766468 | 0,919061642 | 0,60344873 | 0,950352286 | clusterin associated protein 1 |
| ENSG00000185650 | ZFP36L1 | -0,121825835 | 0,919023823 | 0,62668912 | 0,952793934 | ZFP36 ring finger protein like 1 |
| ENSG00000215283 | HMGB3P24 | -0,121841199 | 0,919014036 | 0,86512134 | 0,989074806 | high mobility group box 3 pseudogene 24 |
| ENSG00000261087 | ZNNT1 | -0,121847062 | 0,919010302 | 0,79674494 | 0,980146131 | ZNF706 neighboring transcript 1 |
| ENSG00000123485 | HJURP | -0,12186653 | 0,9189979 | 0,69028383 | 0,967152173 | Holliday junction recognition protein |
| ENSG00000139946 | PELI2 | -0,12189875 | 0,918977376 | 0,77598698 | 0,978587255 | pellino E3 ubiquitin protein ligase family member 2 |
| ENSG00000284294 | AC007326,5 | -0,121906048 | 0,918972727 | 0,63840029 | 0,956018518 | novel transcript |
| ENSG00000250546 | AC079160,1 | -0,121912945 | 0,918968334 | 0,90463229 | 0,992590661 | novel transcript |
| ENSG00000129158 | SERGEF | -0,121920546 | 0,918963492 | 0,63590782 | 0,955399772 | secretion regulating guanine nucleotide exchange factor |
| ENSG00000274803 | BDP1 | -0,121975862 | 0,918928258 | 0,66231675 | 0,959897678 | B double prime 1, subunit of RNA polymerase III transcription initiation factor IIIB |
| ENSG00000090060 | PAPOLA | -0,122133398 | 0,918827921 | 0,77972646 | 0,978587255 | poly(A) polymerase alpha |

| | | | | | | |
|---|---|---|---|---|---|---|
| ENSG00000218358 | RAET1K | -0,122134651 | 0,918827123 | 0,86275689 | 0,989074806 | retinoic acid early transcript 1K pseudogene |
| ENSG00000089048 | ESF1 | -0,12215251 | 0,918815749 | 0,72029186 | 0,972142678 | ESF1 nucleolar pre-rRNA processing protein homolog |
| ENSG00000105514 | RAB3D | -0,122214569 | 0,918776225 | 0,56651915 | 0,941311563 | RAB3D, member RAS oncogene family |
| ENSG00000260278 | AC098818,2 | -0,122226789 | 0,918768443 | 0,8926121 | 0,991048031 | novel transcript, antisense to BMP2K |
| ENSG00000074054 | CLASP1 | -0,122298048 | 0,918723064 | 0,6626527 | 0,959897678 | cytoplasmic linker associated protein 1 |
| ENSG00000172878 | METAP1D | -0,12230043 | 0,918721547 | 0,67543435 | 0,963400586 | methionyl aminopeptidase type 1D, mitochondrial |
| ENSG00000244926 | ALKBH3-AS1 | -0,122355691 | 0,918686357 | 0,8243746 | 0,983374097 | ALKBH3 antisense RNA 1 |
| ENSG00000278703 | AC100847,1 | -0,122367541 | 0,918678811 | 0,93642989 | 0,994479239 | novel transcript |
| ENSG00000089123 | TASP1 | -0,122390321 | 0,918664305 | 0,62013767 | 0,952793934 | taspase 1 |
| ENSG00000282233 | AL450306,2 | -0,122541272 | 0,918568189 | 0,83058721 | 0,984156028 | novel transcript |
| ENSG00000182899 | RPL35A | -0,122625431 | 0,918514606 | 0,76046202 | 0,97624106 | ribosomal protein L35a |
| ENSG00000158623 | COPG2 | -0,122642021 | 0,918504044 | 0,69913304 | 0,968522643 | COPI coat complex subunit gamma 2 |
| ENSG00000074527 | NTN4 | -0,122652218 | 0,918497552 | 0,75444432 | 0,975901808 | netrin 4 |
| ENSG00000104432 | IL7 | -0,122671967 | 0,918484979 | 0,8186891 | 0,9824772 | interleukin 7 |
| ENSG00000276136 | AC016957,2 | -0,122678044 | 0,918481111 | 0,82373317 | 0,983374097 | novel transcript |
| ENSG00000255366 | AC120036,4 | -0,122737513 | 0,918443251 | 0,84892378 | 0,987036295 | novel transcript |
| ENSG00000270091 | AC015726,1 | -0,122743666 | 0,918439334 | 0,85128966 | 0,987259878 | novel transcript |
| ENSG00000102897 | LYRM1 | -0,122844005 | 0,918375459 | 0,74246342 | 0,975176043 | LYR motif containing 1 |
| ENSG00000143748 | NVL | -0,122954209 | 0,918305309 | 0,66580783 | 0,961016016 | nuclear VCP like |
| ENSG00000075188 | NUP37 | -0,122970943 | 0,918294657 | 0,77173221 | 0,978033855 | nucleoporin 37 |
| ENSG00000072315 | TRPC5 | -0,123171365 | 0,918167095 | 0,82391209 | 0,983374097 | transient receptor potential cation channel subfamily C member 5 |
| ENSG00000207375 | SNORD116-23 | -0,123171809 | 0,918166812 | 0,84417068 | 0,98632753 | small nucleolar RNA, C/D box 116-23 |
| ENSG00000180884 | ZNF792 | -0,123198457 | 0,918149853 | 0,60732945 | 0,950993594 | zinc finger protein 792 |
| ENSG00000126261 | UBA2 | -0,123217912 | 0,918137471 | 0,75729911 | 0,975901808 | ubiquitin like modifier activating enzyme 2 |
| ENSG00000251620 | STPG2-AS1 | -0,123228833 | 0,918130522 | 0,70218163 | 0,968742626 | STPG2 antisense RNA 1 |
| ENSG00000133460 | SLC2A11 | -0,123294161 | 0,918088948 | 0,74543706 | 0,975707496 | solute carrier family 2 member 11 |
| ENSG00000148082 | SHC3 | -0,123321887 | 0,918071304 | 0,64397715 | 0,956633898 | SHC adaptor protein 3 |
| ENSG00000150873 | C2orf50 | -0,123323205 | 0,918070465 | 0,75566095 | 0,975901808 | chromosome 2 open reading frame 50 |
| ENSG00000100105 | PATZ1 | -0,123359331 | 0,918047477 | 0,65743527 | 0,957853146 | POZ/BTB and AT hook containing zinc finger 1 |
| ENSG00000157036 | EXOG | -0,12341818 | 0,918010029 | 0,6415598 | 0,956465487 | exo/endonuclease G |
| ENSG00000163535 | SGO2 | -0,123462108 | 0,917982077 | 0,78158611 | 0,978597905 | shugoshin 2 |

| | | | | | | |
|---|---|---|---|---|---|---|
| ENSG00000132406 | TMEM128 | -0,123489736 | 0,917964498 | 0,73364967 | 0,974353474 | transmembrane protein 128 |
| ENSG00000278129 | ZNF8 | -0,123505412 | 0,917954524 | 0,57265189 | 0,942475308 | zinc finger protein 8 |
| ENSG00000223791 | UBE2E1-AS1 | -0,123520568 | 0,91794488 | 0,89925051 | 0,991832796 | UBE2E1 antisense RNA 1 |
| ENSG00000085999 | RAD54L | -0,123522203 | 0,91794384 | 0,5673094 | 0,941425553 | RAD54 like |
| ENSG00000152433 | ZNF547 | -0,123553375 | 0,917924007 | 0,69157348 | 0,967201016 | zinc finger protein 547 |
| ENSG00000234494 | SP2-AS1 | -0,123553903 | 0,917923671 | 0,75672181 | 0,975901808 | SP2 antisense RNA 1 |
| ENSG00000254400 | AC091564,2 | -0,123559797 | 0,91791992 | 0,92555834 | 0,993915537 | novel transcript |
| ENSG00000221817 | PPP3CB-AS1 | -0,123653196 | 0,917860497 | 0,71092988 | 0,970482408 | PPP3CB antisense RNA 1 (head to head) |
| ENSG00000273399 | AL159169,3 | -0,123776302 | 0,917782178 | 0,87901785 | 0,990109814 | novel transcript |
| ENSG00000128944 | KNSTRN | -0,12383337 | 0,917745875 | 0,72925826 | 0,974009551 | kinetochore localized astrin (SPAG5) binding protein |
| ENSG00000278666 | AC106795,4 | -0,123837678 | 0,917743134 | 0,77330406 | 0,978373116 | THO complex 3 (THOC3) pseudogene |
| ENSG00000206053 | JPT2 | -0,12389763 | 0,917704998 | 0,62775459 | 0,952793934 | Jupiter microtubule associated homolog 2 |
| ENSG00000280661 | ZNF660 | -0,12393629 | 0,917680406 | 0,86500392 | 0,989074806 | zinc finger protein 660 |
| ENSG00000152404 | CWF19L2 | -0,123964477 | 0,917662478 | 0,66540132 | 0,960882541 | CWF19 like cell cycle control factor 2 |
| ENSG00000258968 | AL049830,4 | -0,124039777 | 0,917614582 | 0,89825439 | 0,9918188 | intraflagellar transport 20 homolog (Chlamydomonas) (IFT20) pseudogene |
| ENSG00000196591 | HDAC2 | -0,124097818 | 0,917577666 | 0,75814871 | 0,976120207 | histone deacetylase 2 |
| ENSG00000272086 | AC025181,2 | -0,124146981 | 0,917546398 | 0,7737751 | 0,978415098 | novel transcript |
| ENSG00000273167 | AL359736,1 | -0,124155749 | 0,917540822 | 0,85024108 | 0,987036295 | novel transcript |
| ENSG00000182636 | NDN | -0,124171115 | 0,917531049 | 0,6848293 | 0,965415049 | necdin, MAGE family member |
| ENSG00000106692 | FKTN | -0,124171868 | 0,917530571 | 0,7568991 | 0,975901808 | fukutin |
| ENSG00000100364 | KIAA0930 | -0,124217888 | 0,917501303 | 0,62617516 | 0,952793934 | KIAA0930 |
| ENSG00000158571 | PFKFB1 | -0,124266351 | 0,917470483 | 0,84593997 | 0,98689126 | 6-phosphofructo-2-kinase/fructose-2,6-biphosphatase 1 |
| ENSG00000179941 | BBS10 | -0,124306959 | 0,917444659 | 0,80239846 | 0,980742323 | Bardet-Biedl syndrome 10 |
| ENSG00000181690 | PLAG1 | -0,124429047 | 0,917367024 | 0,80986413 | 0,981213877 | PLAG1 zinc finger |
| ENSG00000179029 | TMEM107 | -0,124473783 | 0,917338577 | 0,67650668 | 0,963580269 | transmembrane protein 107 |
| ENSG00000121406 | ZNF549 | -0,124585727 | 0,9172674 | 0,60709932 | 0,950993594 | zinc finger protein 549 |
| ENSG00000272288 | AL451165,2 | -0,124648374 | 0,91722757 | 0,77143043 | 0,978033855 | novel transcript, antisense to C6orf106 |
| ENSG00000278175 | GLIDR | -0,124669529 | 0,917214121 | 0,65033086 | 0,956899963 | glioblastoma down-regulated RNA |
| ENSG00000086712 | TXLNG | -0,124702331 | 0,917193267 | 0,70694492 | 0,969565008 | taxilin gamma |
| ENSG00000177791 | MYOZ1 | -0,124702335 | 0,917193264 | 0,897955 | 0,991711719 | myozenin 1 |
| ENSG00000262877 | AC110285,2 | -0,124738812 | 0,917170074 | 0,76091796 | 0,97624106 | novel transcript |

| | | | | | | |
|---|---|---|---|---|---|---|
| ENSG00000272501 | AL662844,4 | -0,12474844 | 0,917163954 | 0,69409988 | 0,967359949 | novel transcript, antisense to HCG27 |
| ENSG00000255843 | AP000593,3 | -0,124759143 | 0,917157149 | 0,91401661 | 0,993361299 | novel transcript |
| ENSG00000167074 | TEF | -0,124793992 | 0,917134995 | 0,63761756 | 0,956017934 | TEF transcription factor, PAR bZIP family member |
| ENSG00000214338 | SOGA3 | -0,124836684 | 0,917107856 | 0,77528855 | 0,978587255 | SOGA family member 3 |
| ENSG00000286230 | AC010327,7 | -0,124879761 | 0,917080473 | 0,85954464 | 0,989074806 | novel transcript |
| ENSG00000231697 | NANOGP5 | -0,124901377 | 0,917066732 | 0,83489936 | 0,984905093 | Nanog homeobox pseudogene 5 |
| ENSG00000204463 | BAG6 | -0,124915984 | 0,917057447 | 0,63459802 | 0,955149453 | BAG cochaperone 6 |
| ENSG00000276805 | AL133216,2 | -0,124995852 | 0,91700668 | 0,76861804 | 0,977381742 | general transcription factor IIi (GTF2I) pseudogene |
| ENSG00000164252 | AGGF1 | -0,125250701 | 0,916844707 | 0,72955796 | 0,974009551 | angiogenic factor with G-patch and FHA domains 1 |
| ENSG00000230568 | SF3A3P1 | -0,125335344 | 0,916790917 | 0,88292838 | 0,990717729 | splicing factor 3a, subunit 3 pseudogene 1 |
| ENSG00000223547 | ZNF844 | -0,125548793 | 0,916655286 | 0,60234234 | 0,950180678 | zinc finger protein 844 |
| ENSG00000163714 | U2SURP | -0,125555375 | 0,916651105 | 0,73451238 | 0,974353474 | U2 snRNP associated SURP domain containing |
| ENSG00000145555 | MYO10 | -0,125616351 | 0,916612363 | 0,64738434 | 0,956899963 | myosin X |
| ENSG00000103034 | NDRG4 | -0,125651958 | 0,91658974 | 0,61441643 | 0,952793934 | NDRG family member 4 |
| ENSG00000253607 | AC104316,1 | -0,125660076 | 0,916584582 | 0,90683424 | 0,992590661 | novel transcript, antisense to DERL1 |
| ENSG00000267426 | AC087289,3 | -0,125704773 | 0,916556186 | 0,91794945 | 0,993361299 | novel protein |
| ENSG00000197429 | IPP | -0,1257247 | 0,916543526 | 0,58489313 | 0,944857905 | intracisternal A particle-promoted polypeptide |
| ENSG00000258457 | AL132780,2 | -0,125729662 | 0,916540373 | 0,83667277 | 0,985426146 | novel transcript, antisense to C14orf93 |
| ENSG00000165672 | PRDX3 | -0,125822215 | 0,916481577 | 0,76564994 | 0,976862445 | peroxiredoxin 3 |
| ENSG00000140386 | SCAPER | -0,125826482 | 0,916478866 | 0,71529617 | 0,971041579 | S-phase cyclin A associated protein in the ER |
| ENSG00000197122 | SRC | -0,125847912 | 0,916465253 | 0,63964087 | 0,956018518 | SRC proto-oncogene, non-receptor tyrosine kinase |
| ENSG00000261286 | ATP2C2-AS1 | -0,125848546 | 0,91646485 | 0,81707166 | 0,981979397 | ATP2C2 antisense RNA 1 |
| ENSG00000100246 | DNAL4 | -0,125854505 | 0,916461065 | 0,60628115 | 0,950927937 | dynein axonemal light chain 4 |
| ENSG00000165409 | TSHR | -0,125907019 | 0,916427706 | 0,81279158 | 0,981246822 | thyroid stimulating hormone receptor |
| ENSG00000167755 | KLK6 | -0,125939288 | 0,916407209 | 0,91471446 | 0,993361299 | kallikrein related peptidase 6 |
| ENSG00000280663 | PCMTD2 | -0,125943236 | 0,9164047 | 0,64527278 | 0,956899963 | protein-L-isoaspartate (D-aspartate) O-methyltransferase domain containing 2 |
| ENSG00000282464 | AC137894,1 | -0,126082288 | 0,916316379 | 0,86338833 | 0,989074806 | novel transcript |
| ENSG00000010244 | ZNF207 | -0,126124078 | 0,916289836 | 0,7439938 | 0,975509237 | zinc finger protein 207 |
| ENSG00000196387 | ZNF140 | -0,126283753 | 0,916188429 | 0,74826379 | 0,975901808 | zinc finger protein 140 |
| ENSG00000280594 | BTG3-AS1 | -0,126286664 | 0,91618658 | 0,73930683 | 0,974715504 | BTG3 antisense RNA 1 |
| ENSG00000281317 | BTG3-AS1 | -0,126286664 | 0,91618658 | 0,73930683 | 0,974715504 | BTG3 antisense RNA 1 |

| | | | | | | |
|---|---|---|---|---|---|---|
| ENSG00000152219 | ARL14EP | -0,126293666 | 0,916182133 | 0,76802117 | 0,977377956 | ADP ribosylation factor like GTPase 14 effector protein |
| ENSG00000064393 | HIPK2 | -0,126424197 | 0,916099244 | 0,59033392 | 0,947297632 | homeodomain interacting protein kinase 2 |
| ENSG00000232458 | LINC01450 | -0,126431957 | 0,916094316 | 0,92173389 | 0,993787755 | long intergenic non-protein coding RNA 1450 |
| ENSG00000186451 | SPATA12 | -0,126464939 | 0,916073373 | 0,67524756 | 0,963390425 | spermatogenesis associated 12 |
| ENSG00000100379 | KCTD17 | -0,126528843 | 0,916032797 | 0,70506701 | 0,969413904 | potassium channel tetramerization domain containing 17 |
| ENSG00000000457 | SCYL3 | -0,126653572 | 0,915953604 | 0,67650501 | 0,963580269 | SCY1 like pseudokinase 3 |
| ENSG00000286873 | AC012306,3 | -0,126696626 | 0,91592627 | 0,67662607 | 0,963580269 | novel transcript |
| ENSG00000171345 | KRT19 | -0,126910316 | 0,915790614 | 0,82327846 | 0,983304474 | keratin 19 |
| ENSG00000213023 | SYT3 | -0,126941406 | 0,915770879 | 0,70666109 | 0,969503793 | synaptotagmin 3 |
| ENSG00000253729 | PRKDC | -0,12694934 | 0,915765843 | 0,67394947 | 0,963037987 | protein kinase, DNA-activated, catalytic subunit |
| ENSG00000081014 | AP4E1 | -0,126966791 | 0,915754766 | 0,66583188 | 0,961016016 | adaptor related protein complex 4 subunit epsilon 1 |
| ENSG00000242948 | EPS15P1 | -0,127053567 | 0,915699686 | 0,91826667 | 0,993367295 | epidermal growth factor receptor pathway substrate 15 pseudogene 1 |
| ENSG00000128408 | RIBC2 | -0,127060981 | 0,91569498 | 0,67409116 | 0,96304746 | RIB43A domain with coiled-coils 2 |
| ENSG00000272371 | AL591167,1 | -0,127130813 | 0,915650658 | 0,86655136 | 0,989074806 | novel transcript |
| ENSG00000166439 | RNF169 | -0,127280794 | 0,915555473 | 0,71197173 | 0,970812834 | ring finger protein 169 |
| ENSG00000118733 | OLFM3 | -0,127297966 | 0,915544576 | 0,83348041 | 0,984905093 | olfactomedin 3 |
| ENSG00000096060 | FKBP5 | -0,127357896 | 0,915506545 | 0,71264814 | 0,970812834 | FKBP prolyl isomerase 5 |
| ENSG00000092208 | GEMIN2 | -0,127358964 | 0,915505867 | 0,6673936 | 0,9611116 | gem nuclear organelle associated protein 2 |
| ENSG00000155974 | GRIP1 | -0,127453483 | 0,915445888 | 0,65741241 | 0,957853146 | glutamate receptor interacting protein 1 |
| ENSG00000122477 | LRRC39 | -0,127507177 | 0,915411818 | 0,80878394 | 0,981213877 | leucine rich repeat containing 39 |
| ENSG00000074696 | HACD3 | -0,127622953 | 0,91533836 | 0,74432833 | 0,975641819 | 3-hydroxyacyl-CoA dehydratase 3 |
| ENSG00000181315 | ZNF322 | -0,127626176 | 0,915336315 | 0,76091265 | 0,97624106 | zinc finger protein 322 |
| ENSG00000110429 | FBXO3 | -0,127694543 | 0,91529294 | 0,75487767 | 0,975901808 | F-box protein 3 |
| ENSG00000021776 | AQR | -0,127802293 | 0,915224582 | 0,74694501 | 0,975796253 | aquarius intron-binding spliceosomal factor |
| ENSG00000229240 | LINC00710 | -0,127874196 | 0,915178969 | 0,78486843 | 0,979320406 | long intergenic non-protein coding RNA 710 |
| ENSG00000276545 | PCDHGB9P | -0,127937125 | 0,915139051 | 0,73438704 | 0,974353474 | protocadherin gamma subfamily B, 9 pseudogene |
| ENSG00000187189 | TSPYL4 | -0,127989096 | 0,915106084 | 0,64873689 | 0,956899963 | TSPY like 4 |
| ENSG00000111796 | KLRB1 | -0,128064424 | 0,915058305 | 0,84687324 | 0,986939308 | killer cell lectin like receptor B1 |
| ENSG00000102710 | SUPT20H | -0,128070272 | 0,915054596 | 0,75539381 | 0,975901808 | SPT20 homolog, SAGA complex component |
| ENSG00000088882 | CPXM1 | -0,128115786 | 0,915025728 | 0,62491746 | 0,952793934 | carboxypeptidase X, M14 family member 1 |

| | | | | | | |
|---|---|---|---|---|---|---|
| ENSG00000162928 | PEX13 | -0,128116127 | 0,915025512 | 0,69206862 | 0,967225449 | peroxisomal biogenesis factor 13 |
| ENSG00000236144 | TMEM147-AS1 | -0,128136359 | 0,91501268 | 0,62941469 | 0,952812183 | TMEM147 antisense RNA 1 |
| ENSG00000286741 | AC008766,1 | -0,128225384 | 0,914956218 | 0,77291515 | 0,978310326 | novel transcript |
| ENSG00000253719 | ATXN7L3B | -0,128452954 | 0,914811905 | 0,7213607 | 0,972142678 | ataxin 7 like 3B |
| ENSG00000249767 | ENPP7P10 | -0,128502176 | 0,914780694 | 0,88367644 | 0,990717729 | ectonucleotide pyrophosphatase/phosphodiesterase 7 pseudogene 10 |
| ENSG00000187243 | MAGED4B | -0,128584281 | 0,914728635 | 0,5802831 | 0,943828882 | MAGE family member D4B |
| ENSG00000215057 | PKMP5 | -0,128622443 | 0,914704439 | 0,87580761 | 0,990109814 | pyruvate kinase M1/2 pseudogene 5 |
| ENSG00000223901 | AP001469,1 | -0,128748942 | 0,914624238 | 0,85029675 | 0,987036295 | novel transcript |
| ENSG00000088448 | ANKRD10 | -0,128837755 | 0,914567935 | 0,67044133 | 0,962026636 | ankyrin repeat domain 10 |
| ENSG00000235278 | ZNF652P1 | -0,128911354 | 0,914521281 | 0,9211758 | 0,993787755 | zinc finger protein 652 pseudogene 1 |
| ENSG00000116205 | TCEANC2 | -0,128943009 | 0,914501215 | 0,61622745 | 0,952793934 | transcription elongation factor A N-terminal and central domain containing 2 |
| ENSG00000163788 | SNRK | -0,128989513 | 0,914471737 | 0,73989955 | 0,974715504 | SNF related kinase |
| ENSG00000229989 | MIR181A1HG | -0,128997811 | 0,914466477 | 0,73766503 | 0,974353474 | MIR181A1 host gene |
| ENSG00000162068 | NTN3 | -0,129025134 | 0,914449158 | 0,85580121 | 0,988145402 | netrin 3 |
| ENSG00000104518 | GSDMD | -0,129063901 | 0,914424586 | 0,87188002 | 0,989217334 | gasdermin D |
| ENSG00000198589 | LRBA | -0,129099516 | 0,914402013 | 0,70725947 | 0,969565008 | LPS responsive beige-like anchor protein |
| ENSG00000085231 | AK6 | -0,12911714 | 0,914390842 | 0,73921711 | 0,974715504 | adenylate kinase 6 |
| ENSG00000135315 | CEP162 | -0,129122883 | 0,914387202 | 0,73817071 | 0,974353474 | centrosomal protein 162 |
| ENSG00000253738 | OTUD6B-AS1 | -0,129130731 | 0,914382228 | 0,79669051 | 0,980146131 | OTUD6B antisense RNA 1 (head to head) |
| ENSG00000182459 | TEX19 | -0,129146293 | 0,914372365 | 0,89455191 | 0,99116563 | testis expressed 19 |
| ENSG00000249084 | AC027627,1 | -0,129224691 | 0,914322678 | 0,80888481 | 0,981213877 | novel transcript |
| ENSG00000165868 | HSPA12A | -0,12935858 | 0,914237829 | 0,62363847 | 0,952793934 | heat shock protein family A (Hsp70) member 12A |
| ENSG00000189419 | SPATA41 | -0,129480349 | 0,914160667 | 0,80723646 | 0,981213877 | spermatogenesis associated 41 |
| ENSG00000199785 | SNORA52 | -0,12948794 | 0,914155857 | 0,83346428 | 0,984905093 | small nucleolar RNA, H/ACA box 52 |
| ENSG00000163626 | COX18 | -0,129510213 | 0,914141744 | 0,5854774 | 0,945213113 | cytochrome c oxidase assembly factor COX18 |
| ENSG00000215039 | CD27-AS1 | -0,129636779 | 0,91406155 | 0,75026115 | 0,975901808 | CD27 antisense RNA 1 |
| ENSG00000099338 | CATSPERG | -0,129719973 | 0,914008842 | 0,82059828 | 0,982609114 | cation channel sperm associated auxiliary subunit gamma |
| ENSG00000149656 | LINC00266-1 | -0,129737079 | 0,913998005 | 0,93057128 | 0,994344567 | long intergenic non-protein coding RNA 266-1 |
| ENSG00000233653 | CICP7 | -0,129754242 | 0,913987132 | 0,92302093 | 0,993825008 | capicua transcriptional repressor pseudogene 7 |

| | | | | | | |
|---|---|---|---|---|---|---|
| ENSG00000207357 | RNU6-2 | -0,129796518 | 0,913960349 | 0,90770172 | 0,992590661 | RNA, U6 small nuclear 2 |
| ENSG00000199562 | RNU6-37P | -0,129796518 | 0,913960349 | 0,90770172 | 0,992590661 | RNA, U6 small nuclear 37, pseudogene |
| ENSG00000206965 | RNU6-5P | -0,129796518 | 0,913960349 | 0,90770172 | 0,992590661 | RNA, U6 small nuclear 5, pseudogene |
| ENSG00000206932 | RNU6-4P | -0,129796518 | 0,913960349 | 0,90770172 | 0,992590661 | RNA, U6 small nuclear 4, pseudogene |
| ENSG00000207041 | RNU6-3P | -0,129796518 | 0,913960349 | 0,90770172 | 0,992590661 | RNA, U6 small nuclear 3, pseudogene |
| ENSG00000272055 | RNU6-6P | -0,129796518 | 0,913960349 | 0,90770172 | 0,992590661 | RNA, U6 small nuclear 6, pseudogene |
| ENSG00000207507 | RNU6-9 | -0,129796518 | 0,913960349 | 0,90770172 | 0,992590661 | RNA, U6 small nuclear 9 |
| ENSG00000201104 | RNU6-11P | -0,129796518 | 0,913960349 | 0,90770172 | 0,992590661 | RNA, U6 small nuclear 11, pseudogene |
| ENSG00000206625 | RNU6-1 | -0,129796518 | 0,913960349 | 0,90770172 | 0,992590661 | RNA, U6 small nuclear 1 |
| ENSG00000202337 | RNU6-8 | -0,129796518 | 0,913960349 | 0,90770172 | 0,992590661 | RNA, U6 small nuclear 8 |
| ENSG00000201654 | RNU6-7 | -0,129796518 | 0,913960349 | 0,90770172 | 0,992590661 | RNA, U6 small nuclear 7 |
| ENSG00000187231 | SESTD1 | -0,129808601 | 0,913952694 | 0,76960072 | 0,977683747 | SEC14 and spectrin domain containing 1 |
| ENSG00000136874 | STX17 | -0,12986263 | 0,913918467 | 0,71929467 | 0,971869926 | syntaxin 17 |
| ENSG00000165548 | TMEM63C | -0,12987255 | 0,913912183 | 0,74317766 | 0,975305642 | transmembrane protein 63C |
| ENSG00000047617 | ANO2 | -0,129983884 | 0,913841658 | 0,7670048 | 0,977201891 | anoctamin 2 |
| ENSG00000177191 | B3GNT8 | -0,129993921 | 0,913835301 | 0,79136161 | 0,979471509 | UDP-GlcNAc:betaGal beta-1,3-N-acetylglucosaminyltransferase 8 |
| ENSG00000271395 | LINC02336 | -0,130001476 | 0,913830515 | 0,9052508 | 0,992590661 | long intergenic non-protein coding RNA 2336 |
| ENSG00000283392 | AC090952,2 | -0,130065575 | 0,913789915 | 0,91784298 | 0,993361299 | novel transcript |
| ENSG00000136247 | ZDHHC4 | -0,13006921 | 0,913787612 | 0,64342574 | 0,956633898 | zinc finger DHHC-type palmitoyltransferase 4 |
| ENSG00000138468 | SENP7 | -0,130170605 | 0,913723392 | 0,76456079 | 0,976640468 | SUMO specific peptidase 7 |
| ENSG00000166295 | ANAPC16 | -0,130188508 | 0,913712053 | 0,62679363 | 0,952793934 | anaphase promoting complex subunit 16 |
| ENSG00000171044 | XKR6 | -0,130213918 | 0,91369596 | 0,62903314 | 0,952793934 | XK related 6 |
| ENSG00000274624 | AC007671,1 | -0,13021525 | 0,913695117 | 0,89089664 | 0,990717729 | novel transcript, sense intronic to ST8SIA1 |
| ENSG00000104738 | MCM4 | -0,130231519 | 0,913684813 | 0,61102284 | 0,951873283 | minichromosome maintenance complex component 4 |
| ENSG00000215908 | CROCCP2 | -0,130254971 | 0,913669961 | 0,79618481 | 0,980015022 | CROCC pseudogene 2 |
| ENSG00000169914 | OTUD3 | -0,130330679 | 0,913622016 | 0,62723202 | 0,952793934 | OTU deubiquitinase 3 |
| ENSG00000109881 | CCDC34 | -0,130374545 | 0,913594237 | 0,60193476 | 0,949939978 | coiled-coil domain containing 34 |
| ENSG00000274853 | MRM1 | -0,130378775 | 0,913591558 | 0,83061484 | 0,984156028 | mitochondrial rRNA methyltransferase 1 |
| ENSG00000135679 | MDM2 | -0,130379837 | 0,913590886 | 0,64253471 | 0,956633898 | MDM2 proto-oncogene |
| ENSG00000203667 | COX20 | -0,130424572 | 0,913562557 | 0,76133802 | 0,97624106 | cytochrome c oxidase assembly factor COX20 |

| | | | | | | |
|---|---|---|---|---|---|---|
| ENSG00000101247 | NDUFAF5 | -0,130533156 | 0,913493801 | 0,66778346 | 0,961113267 | NADH:ubiquinone oxidoreductase complex assembly factor 5 |
| ENSG00000165874 | SHLD2P1 | -0,130592527 | 0,913456209 | 0,83491382 | 0,984905093 | shieldin complex subunit 2 pseudogene 1 |
| ENSG00000288062 | AL136981,3 | -0,130592833 | 0,913456015 | 0,89692621 | 0,991644439 | uncharacterized LOC101929748 |
| ENSG00000079387 | SENP1 | -0,130602565 | 0,913449853 | 0,73822745 | 0,974353474 | SUMO specific peptidase 1 |
| ENSG00000238287 | AL603839,3 | -0,130614186 | 0,913442496 | 0,87362822 | 0,989633545 | novel transcript |
| ENSG00000256115 | LINC02443 | -0,13067315 | 0,913405163 | 0,74893766 | 0,975901808 | long intergenic non-protein coding RNA 2443 |
| ENSG00000138796 | HADH | -0,130743852 | 0,913360401 | 0,66367104 | 0,959920886 | hydroxyacyl-CoA dehydrogenase |
| ENSG00000178538 | CA8 | -0,130748918 | 0,913357194 | 0,79041407 | 0,979462318 | carbonic anhydrase 8 |
| ENSG00000108055 | SMC3 | -0,130918005 | 0,913250153 | 0,67185392 | 0,962161102 | structural maintenance of chromosomes 3 |
| ENSG00000285637 | AL161449,2 | -0,130918314 | 0,913249957 | 0,86781745 | 0,989074806 | Novel transcript |
| ENSG00000113648 | MACROH2A1 | -0,130938842 | 0,913236962 | 0,57463367 | 0,942518604 | macroH2A,1 histone |
| ENSG00000231304 | SGO1-AS1 | -0,130963177 | 0,913221558 | 0,81485687 | 0,981521413 | SGO1 antisense RNA 1 |
| ENSG00000231742 | LINC01273 | -0,131026713 | 0,913181341 | 0,84127689 | 0,985672131 | long intergenic non-protein coding RNA 1273 |
| ENSG00000053900 | ANAPC4 | -0,131066537 | 0,913156134 | 0,72842927 | 0,973955059 | anaphase promoting complex subunit 4 |
| ENSG00000111203 | ITFG2 | -0,13112757 | 0,913117504 | 0,60955437 | 0,951793902 | integrin alpha FG-GAP repeat containing 2 |
| ENSG00000259084 | AL133467,3 | -0,131155177 | 0,913100031 | 0,88724671 | 0,990717729 | novel transcript |
| ENSG00000113456 | RAD1 | -0,131170136 | 0,913090564 | 0,66923875 | 0,961754789 | RAD1 checkpoint DNA exonuclease |
| ENSG00000271122 | AC018647,2 | -0,131465547 | 0,912903615 | 0,64375336 | 0,956633898 | novel transcript |
| ENSG00000136522 | MRPL47 | -0,131466135 | 0,912903243 | 0,75497746 | 0,975901808 | mitochondrial ribosomal protein L47 |
| ENSG00000260725 | AC005307,1 | -0,131474525 | 0,912897934 | 0,89786276 | 0,991711719 | novel transcript |
| ENSG00000199631 | SNORD33 | -0,131503978 | 0,912879298 | 0,77931147 | 0,978587255 | small nucleolar RNA, C/D box 33 |
| ENSG00000259134 | LINC00924 | -0,131612157 | 0,912810849 | 0,86976292 | 0,989074806 | long intergenic non-protein coding RNA 924 |
| ENSG00000253420 | AC103853,1 | -0,131615806 | 0,91280854 | 0,86947885 | 0,989074806 | novel transcript |
| ENSG00000140104 | CLBA1 | -0,131751876 | 0,912722451 | 0,62047749 | 0,952793934 | clathrin binding box of aftiphilin containing 1 |
| ENSG00000085788 | DDHD2 | -0,131829473 | 0,91267336 | 0,7270961 | 0,973344786 | DDHD domain containing 2 |
| ENSG00000128050 | PAICS | -0,131860213 | 0,912653914 | 0,63934316 | 0,956018518 | phosphoribosylaminoimidazole carboxylase and phosphoribosylaminoimidazolesuccinocarboxamide synthase |
| ENSG00000286450 | AC021573,1 | -0,131874561 | 0,912644837 | 0,74530628 | 0,9756851 | novel transcript |
| ENSG00000226756 | AC007365,1 | -0,131875048 | 0,91264453 | 0,87841315 | 0,990109814 | novel transcript |
| ENSG00000123684 | LPGAT1 | -0,13188523 | 0,912638089 | 0,69990075 | 0,968522643 | lysophosphatidylglycerol acyltransferase 1 |
| ENSG00000224506 | LINC02523 | -0,131916622 | 0,91261823 | 0,94937195 | 0,996432861 | long intergenic non-protein coding RNA 2523 |

| | | | | | | |
|---|---|---|---|---|---|---|
| ENSG00000252947 | SCARNA1 | -0,131928486 | 0,912610725 | 0,86377995 | 0,989074806 | small Cajal body-specific RNA 1 |
| ENSG00000260293 | AC106820,4 | -0,131968848 | 0,912585193 | 0,7588916 | 0,976120207 | novel transcript, intronic to TBC1D24 |
| ENSG00000276701 | MIF | -0,131975905 | 0,91258073 | 0,74675927 | 0,975796253 | macrophage migration inhibitory factor |
| ENSG00000213089 | PDCL3P5 | -0,132003642 | 0,912563185 | 0,76308052 | 0,976583139 | PDCL3 pseudogene 5 |
| ENSG00000259527 | LINC00052 | -0,132029706 | 0,912546699 | 0,86864813 | 0,989074806 | long intergenic non-protein coding RNA 52 |
| ENSG00000261527 | AC026464,5 | -0,1320319 | 0,912545311 | 0,8562779 | 0,988229014 | transmembrane protein 111 (TMEM111) pseudogene |
| ENSG00000286998 | AC103974,2 | -0,132053156 | 0,912531866 | 0,82192184 | 0,982952211 | novel transcript, antisense to PTPN5 |
| ENSG00000150456 | EEF1AKMT1 | -0,132079339 | 0,912515305 | 0,74713936 | 0,975807355 | EEF1A lysine methyltransferase 1 |
| ENSG00000182957 | SPATA13 | -0,132085392 | 0,912511476 | 0,61746749 | 0,952793934 | spermatogenesis associated 13 |
| ENSG00000267001 | AC006538,2 | -0,132099106 | 0,912502802 | 0,85392385 | 0,987861659 | novel protein |
| ENSG00000175054 | ATR | -0,13212352 | 0,912487361 | 0,74984838 | 0,975901808 | ATR serine/threonine kinase |
| ENSG00000068097 | HEATR6 | -0,13217678 | 0,912453674 | 0,62697155 | 0,952793934 | HEAT repeat containing 6 |
| ENSG00000143379 | SETDB1 | -0,132297006 | 0,912377639 | 0,56924632 | 0,94219857 | SET domain bifurcated histone lysine methyltransferase 1 |
| ENSG00000141759 | TXNL4A | -0,132307214 | 0,912371183 | 0,61913603 | 0,952793934 | thioredoxin like 4A |
| ENSG00000197147 | LRRC8B | -0,132379794 | 0,912325284 | 0,77347374 | 0,978415098 | leucine rich repeat containing 8 VRAC subunit B |
| ENSG00000138777 | PPA2 | -0,132383169 | 0,91232315 | 0,71730823 | 0,971697135 | inorganic pyrophosphatase 2 |
| ENSG00000164898 | FMC1 | -0,132402863 | 0,912310696 | 0,58556094 | 0,945213113 | formation of mitochondrial complex V assembly factor 1 homolog |
| ENSG00000162688 | AGL | -0,132433375 | 0,912291402 | 0,77066795 | 0,977966587 | amylo-alpha-1, 6-glucosidase, 4-alpha-glucanotransferase |
| ENSG00000168214 | RBPJ | -0,132441637 | 0,912286177 | 0,77485504 | 0,978553937 | recombination signal binding protein for immunoglobulin kappa J region |
| ENSG00000132773 | TOE1 | -0,132501915 | 0,912248061 | 0,61756054 | 0,952793934 | target of EGR1, exonuclease |
| ENSG00000215548 | FRG1JP | -0,132535764 | 0,912226658 | 0,85860779 | 0,989074806 | FSHD region gene 1 family member J, pseudogene |
| ENSG00000178229 | ZNF543 | -0,132555385 | 0,912214252 | 0,54923446 | 0,936040289 | zinc finger protein 543 |
| ENSG00000143126 | CELSR2 | -0,132572855 | 0,912203205 | 0,64626367 | 0,956899963 | cadherin EGF LAG seven-pass G-type receptor 2 |
| ENSG00000271815 | AC008897,3 | -0,132593005 | 0,912190465 | 0,82754029 | 0,983595244 | novel transcript |
| ENSG00000161692 | DBF4B | -0,132598334 | 0,912187096 | 0,65580065 | 0,957444211 | DBF4 zinc finger B |
| ENSG00000088766 | CRLS1 | -0,132607872 | 0,912181065 | 0,67817039 | 0,964004688 | cardiolipin synthase 1 |
| ENSG00000141447 | OSBPL1A | -0,132608501 | 0,912180667 | 0,67544224 | 0,963400586 | oxysterol binding protein like 1A |
| ENSG00000130734 | ATG4D | -0,132610448 | 0,912179436 | 0,64325522 | 0,956633898 | autophagy related 4D cysteine peptidase |
| ENSG00000149308 | NPAT | -0,132689774 | 0,912129282 | 0,76808268 | 0,977377956 | nuclear protein, coactivator of histone transcription |

| | | | | | | |
|---|---|---|---|---|---|---|
| ENSG00000271605 | MILR1 | -0,132735656 | 0,912100274 | 0,88634027 | 0,990717729 | mast cell immunoglobulin like receptor 1 |
| ENSG00000228544 | CCDC183-AS1 | -0,132821855 | 0,912045779 | 0,7698998 | 0,977742359 | CCDC183 antisense RNA 1 |
| ENSG00000265749 | AC135178,4 | -0,132883697 | 0,912006684 | 0,76122594 | 0,97624106 | novel transcript, antisense to KRBA2 and RPL26 |
| ENSG00000281731 | METTL14-DT | -0,132911942 | 0,911988829 | 0,78424091 | 0,979320406 | METTL14 divergent transcript |
| ENSG00000049618 | ARID1B | -0,132931559 | 0,911976428 | 0,55684109 | 0,938264532 | AT-rich interaction domain 1B |
| ENSG00000083544 | TDRD3 | -0,13296787 | 0,911953475 | 0,73805948 | 0,974353474 | tudor domain containing 3 |
| ENSG00000273483 | AL354760,1 | -0,132969557 | 0,911952409 | 0,86107444 | 0,989074806 | novel transcript, antisense to WNT2B |
| ENSG00000173273 | TNKS | -0,13298875 | 0,911940277 | 0,74244578 | 0,975176043 | tankyrase |
| ENSG00000077254 | USP33 | -0,132998815 | 0,911933915 | 0,73004172 | 0,974178338 | ubiquitin specific peptidase 33 |
| ENSG00000154059 | IMPACT | -0,133099498 | 0,911870275 | 0,75319075 | 0,975901808 | impact RWD domain protein |
| ENSG00000165244 | ZNF367 | -0,133123345 | 0,911855202 | 0,68298931 | 0,964931888 | zinc finger protein 367 |
| ENSG00000117215 | PLA2G2D | -0,133142688 | 0,911842977 | 0,90328848 | 0,992311172 | phospholipase A2 group IID |
| ENSG00000163872 | YEATS2 | -0,133186525 | 0,91181527 | 0,5553596 | 0,937793851 | YEATS domain containing 2 |
| ENSG00000183054 | RGPD6 | -0,13324816 | 0,911776316 | 0,8630281 | 0,989074806 | RANBP2 like and GRIP domain containing 6 |
| ENSG00000272335 | AC093297,2 | -0,133302094 | 0,911742231 | 0,72200238 | 0,972516173 | novel transcript |
| ENSG00000226332 | AL354836,1 | -0,133373962 | 0,911696813 | 0,84658505 | 0,986939308 | novel transcript |
| ENSG00000153187 | HNRNPU | -0,133400842 | 0,911679827 | 0,62741508 | 0,952793934 | heterogeneous nuclear ribonucleoprotein U |
| ENSG00000129315 | CCNT1 | -0,13340361 | 0,911678078 | 0,73041775 | 0,974321178 | cyclin T1 |
| ENSG00000111530 | CAND1 | -0,133441402 | 0,911654196 | 0,75833896 | 0,976120207 | cullin associated and neddylation dissociated 1 |
| ENSG00000224121 | ATG12P2 | -0,133501002 | 0,911616535 | 0,86264661 | 0,989074806 | autophagy related 12 pseudogene 2 |
| ENSG00000168734 | PKIG | -0,133553265 | 0,911583512 | 0,66051514 | 0,959226315 | cAMP-dependent protein kinase inhibitor gamma |
| ENSG00000119927 | GPAM | -0,133621457 | 0,911540425 | 0,71591591 | 0,971360663 | glycerol-3-phosphate acyltransferase, mitochondrial |
| ENSG00000241131 | LINC02032 | -0,13366367 | 0,911513754 | 0,86840951 | 0,989074806 | long intergenic non-protein coding RNA 2032 |
| ENSG00000281013 | NDUFA6 | -0,133689659 | 0,911497333 | 0,82786919 | 0,983595244 | NADH:ubiquinone oxidoreductase subunit A6 |
| ENSG00000177679 | SRRM3 | -0,133705409 | 0,911487382 | 0,79268069 | 0,979471509 | serine/arginine repetitive matrix 3 |
| ENSG00000252010 | SCARNA5 | -0,133819071 | 0,911415574 | 0,62125281 | 0,952793934 | small Cajal body-specific RNA 5 |
| ENSG00000280652 | SCARNA5 | -0,133819071 | 0,911415574 | 0,62125281 | 0,952793934 | small Cajal body-specific RNA 5 |
| ENSG00000137817 | PARP6 | -0,133882436 | 0,911375545 | 0,58271781 | 0,944174844 | poly(ADP-ribose) polymerase family member 6 |
| ENSG00000112701 | SENP6 | -0,133906667 | 0,911360238 | 0,7011382 | 0,968677171 | SUMO specific peptidase 6 |
| ENSG00000259363 | AC090825,1 | -0,13397876 | 0,911314697 | 0,62107708 | 0,952793934 | novel transcript |
| ENSG00000198938 | MT-CO3 | -0,134060586 | 0,911263011 | 0,76200279 | 0,976315588 | mitochondrially encoded cytochrome c oxidase III |
| ENSG00000092199 | HNRNPC | -0,134062352 | 0,911261896 | 0,64523565 | 0,956899963 | heterogeneous nuclear ribonucleoprotein C |
| ENSG00000286565 | AC006348,1 | -0,134078386 | 0,911251768 | 0,90494182 | 0,992590661 | novel transcript |

| | | | | | | |
|---|---|---|---|---|---|---|
| ENSG00000132341 | RAN | -0,134241844 | 0,911148529 | 0,6645613 | 0,96046247 | RAN, member RAS oncogene family |
| ENSG00000197417 | SHPK | -0,134333254 | 0,9110908 | 0,58339356 | 0,944174844 | sedoheptulokinase |
| ENSG00000086475 | SEPHS1 | -0,134384399 | 0,911058501 | 0,72593466 | 0,973344786 | selenophosphate synthetase 1 |
| ENSG00000239887 | C1orf226 | -0,134488599 | 0,910992701 | 0,70148745 | 0,968742626 | chromosome 1 open reading frame 226 |
| ENSG00000004487 | KDM1A | -0,134535993 | 0,910962775 | 0,58612326 | 0,945246954 | lysine demethylase 1A |
| ENSG00000270048 | AC068790,4 | -0,134551823 | 0,910952779 | 0,85981049 | 0,989074806 | novel transcript, sense intronic to CCDC92 |
| ENSG00000158234 | FAIM | -0,134555434 | 0,910950499 | 0,6298041 | 0,952812183 | Fas apoptotic inhibitory molecule |
| ENSG00000153015 | CWC27 | -0,134595764 | 0,910925035 | 0,64405181 | 0,956633898 | CWC27 spliceosome associated cyclophilin |
| ENSG00000133138 | TBC1D8B | -0,134611302 | 0,910915224 | 0,62745598 | 0,952793934 | TBC1 domain family member 8B |
| ENSG00000203565 | AL450313,1 | -0,134697202 | 0,910860989 | 0,89554667 | 0,991303328 | novel transcript |
| ENSG00000288597 | AC234782,2 | -0,13474546 | 0,910830521 | 0,76028365 | 0,97624106 | novel transcript |
| ENSG00000178502 | KLHL11 | -0,134791657 | 0,910801355 | 0,63547645 | 0,955314044 | kelch like family member 11 |
| ENSG00000188785 | ZNF548 | -0,134910558 | 0,910726294 | 0,61034529 | 0,951841373 | zinc finger protein 548 |
| ENSG00000286966 | AL137800,1 | -0,134967906 | 0,910690093 | 0,8378552 | 0,98566906 | novel transcript |
| ENSG00000070476 | ZXDC | -0,135019761 | 0,91065736 | 0,5326704 | 0,931291085 | ZXD family zinc finger C |
| ENSG00000263050 | AC090617,5 | -0,135020205 | 0,91065708 | 0,86801664 | 0,989074806 | novel transcript |
| ENSG00000152413 | HOMER1 | -0,135100382 | 0,910606472 | 0,73513714 | 0,974353474 | homer scaffold protein 1 |
| ENSG00000090470 | PDCD7 | -0,135144333 | 0,910578731 | 0,5795846 | 0,943828882 | programmed cell death 7 |
| ENSG00000171791 | BCL2 | -0,13517927 | 0,91055668 | 0,69543339 | 0,967602563 | BCL2 apoptosis regulator |
| ENSG00000261471 | AC092145,1 | -0,135228535 | 0,910525587 | 0,83942339 | 0,985672131 | novel transcript, antisense to COTL1 |
| ENSG00000182628 | SKA2 | -0,135237946 | 0,910519648 | 0,75281161 | 0,975901808 | spindle and kinetochore associated complex subunit 2 |
| ENSG00000014919 | COX15 | -0,135246268 | 0,910514395 | 0,66139703 | 0,959582738 | cytochrome c oxidase assembly homolog COX15 |
| ENSG00000089022 | MAPKAPK5 | -0,135250899 | 0,910511473 | 0,49409321 | 0,917572329 | MAPK activated protein kinase 5 |
| ENSG00000198586 | TLK1 | -0,135365734 | 0,910439001 | 0,74096539 | 0,975146552 | tousled like kinase 1 |
| ENSG00000073536 | NLE1 | -0,135390015 | 0,910423679 | 0,61997327 | 0,952793934 | notchless homolog 1 |
| ENSG00000168522 | FNTA | -0,135578022 | 0,910305043 | 0,70834831 | 0,970158209 | farnesyltransferase, CAAX box, alpha |
| ENSG00000258725 | PRC1-AS1 | -0,135582446 | 0,910302251 | 0,63824441 | 0,956018518 | PRC1 antisense RNA 1 |
| ENSG00000162433 | AK4 | -0,135765065 | 0,910187031 | 0,7918835 | 0,979471509 | adenylate kinase 4 |
| ENSG00000143630 | HCN3 | -0,135780813 | 0,910177095 | 0,62646427 | 0,952793934 | hyperpolarization activated cyclic nucleotide gated potassium channel 3 |
| ENSG00000100483 | VCPKMT | -0,135815003 | 0,910155526 | 0,73244599 | 0,974353474 | valosin containing protein lysine methyltransferase |
| ENSG00000187323 | DCC | -0,135852525 | 0,910131855 | 0,72310798 | 0,972856862 | DCC netrin 1 receptor |

| | | | | | | |
|---|---|---|---|---|---|---|
| ENSG00000196876 | SCN8A | -0,13590236 | 0,910100417 | 0,62100621 | 0,952793934 | sodium voltage-gated channel alpha subunit 8 |
| ENSG00000176681 | LRRC37A | -0,13594729 | 0,910072074 | 0,88451264 | 0,990717729 | leucine rich repeat containing 37A |
| ENSG00000118596 | SLC16A7 | -0,135951052 | 0,9100697 | 0,72684817 | 0,973344786 | solute carrier family 16 member 7 |
| ENSG00000100201 | DDX17 | -0,135952553 | 0,910068753 | 0,69157673 | 0,967201016 | DEAD-box helicase 17 |
| ENSG00000206612 | SNORA2A | -0,135961262 | 0,91006326 | 0,79318318 | 0,979659177 | small nucleolar RNA, H/ACA box 2A |
| ENSG00000241868 | RN7SL434P | -0,136011587 | 0,910031515 | 0,90016885 | 0,991845106 | RNA, 7SL, cytoplasmic 434, pseudogene |
| ENSG00000111144 | LTA4H | -0,136029254 | 0,910020371 | 0,71575518 | 0,971282169 | leukotriene A4 hydrolase |
| ENSG00000229372 | SZT2-AS1 | -0,136109169 | 0,909969964 | 0,9103039 | 0,993116557 | SZT2 antisense RNA 1 |
| ENSG00000267582 | AC020916,2 | -0,136143052 | 0,909948593 | 0,888292 | 0,990717729 | novel transcript, antisense to ZSWIM4 |
| ENSG00000288519 | AC020916,6 | -0,136143052 | 0,909948593 | 0,888292 | 0,990717729 | novel transcript, antisense to ZSWIM4 |
| ENSG00000127946 | HIP1 | -0,136144914 | 0,909947418 | 0,55016753 | 0,93612918 | huntingtin interacting protein 1 |
| ENSG00000282417 | HERC2P9 | -0,136192017 | 0,90991771 | 0,54775357 | 0,935404974 | hect domain and RLD 2 pseudogene 9 |
| ENSG00000205133 | TRIQK | -0,136217111 | 0,909901883 | 0,78365985 | 0,979210284 | triple QxxK/R motif containing |
| ENSG00000231160 | KLF3-AS1 | -0,136230069 | 0,90989371 | 0,55619573 | 0,938046927 | KLF3 antisense RNA 1 |
| ENSG00000244515 | KRT18P34 | -0,136253875 | 0,909878696 | 0,82211715 | 0,982952211 | keratin 18 pseudogene 34 |
| ENSG00000214455 | RCN1P2 | -0,136256118 | 0,909877282 | 0,72491489 | 0,97326334 | reticulocalbin 1 pseudogene 2 |
| ENSG00000282932 | PTPRD | -0,136263252 | 0,909872782 | 0,73890926 | 0,974448853 | protein tyrosine phosphatase receptor type D |
| ENSG00000163235 | TGFA | -0,136326258 | 0,909833047 | 0,8836608 | 0,990717729 | transforming growth factor alpha |
| ENSG00000214439 | FAM185BP | -0,136382567 | 0,909797536 | 0,79823618 | 0,980161849 | family with sequence similarity 185 member B, pseudogene |
| ENSG00000130520 | LSM4 | -0,136450311 | 0,909754817 | 0,57382206 | 0,942475308 | LSM4 homolog, U6 small nuclear RNA and mRNA degradation associated |
| ENSG00000286550 | AL591002,1 | -0,136481712 | 0,909735015 | 0,81676776 | 0,981977498 | novel transcript, antisense to ANKS1A |
| ENSG00000165275 | TRMT10B | -0,136599643 | 0,909660654 | 0,54473119 | 0,934867206 | tRNA methyltransferase 10B |
| ENSG00000180822 | PSMG4 | -0,13661552 | 0,909650643 | 0,60684745 | 0,950927937 | proteasome assembly chaperone 4 |
| ENSG00000065615 | CYB5R4 | -0,136778792 | 0,909547702 | 0,64326489 | 0,956633898 | cytochrome b5 reductase 4 |
| ENSG00000086159 | AQP6 | -0,136786916 | 0,90954258 | 0,84916041 | 0,987036295 | aquaporin 6 |
| ENSG00000101292 | PROKR2 | -0,136787433 | 0,909542254 | 0,94007872 | 0,994949796 | prokineticin receptor 2 |
| ENSG00000250575 | AL732372,3 | -0,136841538 | 0,909508145 | 0,7461054 | 0,975707496 | novel pseudogene |
| ENSG00000196126 | HLA-DRB1 | -0,136856294 | 0,909498842 | 0,85636157 | 0,988229014 | major histocompatibility complex, class II, DR beta 1 |
| ENSG00000228363 | AC015971,1 | -0,136865683 | 0,909492923 | 0,82850839 | 0,983721455 | novel transcript |
| ENSG00000277758 | FO681492,1 | -0,136884224 | 0,909481234 | 0,77133197 | 0,978033855 | synaptotagmin-15 |

| | | | | | | |
|---|---|---|---|---|---|---|
| ENSG00000253975 | AC018861,1 | -0,136905334 | 0,909467927 | 0,81486093 | 0,981521413 | ubiquitin-conjugating enzyme E2L 3 (UBE2L3) pseudogene |
| ENSG00000271936 | AC012073,1 | -0,136914357 | 0,909462239 | 0,63947112 | 0,956018518 | novel transcript, antisense to ADCY3 |
| ENSG00000259316 | AC087632,2 | -0,137017284 | 0,909397357 | 0,5395676 | 0,933350549 | novel protein |
| ENSG00000162980 | ARL5A | -0,137122524 | 0,909331022 | 0,77455922 | 0,978553937 | ADP ribosylation factor like GTPase 5A |
| ENSG00000108406 | DHX40 | -0,137156513 | 0,909309598 | 0,73988656 | 0,974715504 | DEAH-box helicase 40 |
| ENSG00000184508 | HDDC3 | -0,137226711 | 0,909265355 | 0,59277487 | 0,947416795 | HD domain containing 3 |
| ENSG00000229807 | XIST | -0,137255695 | 0,909247088 | 0,89518597 | 0,991303328 | X inactive specific transcript |
| ENSG00000269356 | AL356740,2 | -0,137331008 | 0,909199624 | 0,87773657 | 0,990109814 | novel transcript |
| ENSG00000236287 | ZBED5 | -0,137402904 | 0,909154315 | 0,75971484 | 0,976120207 | zinc finger BED-type containing 5 |
| ENSG00000249241 | LINC02265 | -0,137461783 | 0,909117212 | 0,89372455 | 0,991135494 | long intergenic non-protein coding RNA 2265 |
| ENSG00000261455 | LINC01003 | -0,137482102 | 0,909104408 | 0,63614412 | 0,955399772 | long intergenic non-protein coding RNA 1003 |
| ENSG00000196922 | ZNF252P | -0,137501598 | 0,909092123 | 0,70606216 | 0,969503793 | zinc finger protein 252, pseudogene |
| ENSG00000284664 | AL161756,3 | -0,137504427 | 0,90909034 | 0,84963377 | 0,987036295 | novel transcript |
| ENSG00000162999 | DUSP19 | -0,137522433 | 0,909078994 | 0,71896245 | 0,971869926 | dual specificity phosphatase 19 |
| ENSG00000102054 | RBBP7 | -0,137559602 | 0,909055573 | 0,67814038 | 0,964004688 | RB binding protein 7, chromatin remodeling factor |
| ENSG00000092978 | GPATCH2 | -0,137593937 | 0,909033939 | 0,67774923 | 0,963712734 | G-patch domain containing 2 |
| ENSG00000160767 | FAM189B | -0,137599044 | 0,909030721 | 0,66761965 | 0,9611116 | family with sequence similarity 189 member B |
| ENSG00000236204 | LINC01376 | -0,13767362 | 0,908983732 | 0,77115815 | 0,978033855 | long intergenic non-protein coding RNA 1376 |
| ENSG00000178700 | DHFR2 | -0,137784648 | 0,908913781 | 0,61904224 | 0,952793934 | dihydrofolate reductase 2 |
| ENSG00000175611 | LINC00476 | -0,137807463 | 0,908899407 | 0,63070858 | 0,953375088 | long intergenic non-protein coding RNA 476 |
| ENSG00000268992 | CDC42EP3P1 | -0,137932136 | 0,908820866 | 0,78370649 | 0,979210284 | CDC42 effector protein 3 pseudogene 1 |
| ENSG00000110367 | DDX6 | -0,137971961 | 0,908795779 | 0,68291937 | 0,964931888 | DEAD-box helicase 6 |
| ENSG00000110315 | RNF141 | -0,137983537 | 0,908788487 | 0,74590102 | 0,975707496 | ring finger protein 141 |
| ENSG00000270574 | AC010680,2 | -0,137995914 | 0,90878069 | 0,72093536 | 0,972142678 | novel transcript, antisense to titin |
| ENSG00000125871 | MGME1 | -0,138023453 | 0,908763343 | 0,71611079 | 0,971360663 | mitochondrial genome maintenance exonuclease 1 |
| ENSG00000111011 | RSRC2 | -0,138072243 | 0,908732611 | 0,65620202 | 0,957444211 | arginine and serine rich coiled-coil 2 |
| ENSG00000260274 | AC068338,2 | -0,138080791 | 0,908727227 | 0,74302977 | 0,975240824 | novel transcript |
| ENSG00000251555 | AC096711,2 | -0,138082734 | 0,908726003 | 0,85204876 | 0,98732978 | novel transcript |
| ENSG00000224992 | AL445645,1 | -0,138124638 | 0,908699608 | 0,91378229 | 0,993361299 | novel transcript |
| ENSG00000233611 | AC019068,1 | -0,138169877 | 0,908671114 | 0,87503265 | 0,990078774 | novel transcript, antisense to ASB18 and GBX2 |
| ENSG00000143013 | LMO4 | -0,1382525 | 0,908619076 | 0,70102383 | 0,968677171 | LIM domain only 4 |
| ENSG00000013441 | CLK1 | -0,13828953 | 0,908595755 | 0,7383352 | 0,974353474 | CDC like kinase 1 |

| | | | | | | |
|---|---|---|---|---|---|---|
| ENSG00000121481 | RNF2 | -0,138313777 | 0,908580485 | 0,7968555 | 0,980161849 | ring finger protein 2 |
| ENSG00000122435 | TRMT13 | -0,138376497 | 0,908540986 | 0,73169082 | 0,974353474 | tRNA methyltransferase 13 homolog |
| ENSG00000055917 | PUM2 | -0,138415042 | 0,908516712 | 0,73856628 | 0,974353474 | pumilio RNA binding family member 2 |
| ENSG00000079246 | XRCC5 | -0,138495499 | 0,908466047 | 0,64980874 | 0,956899963 | X-ray repair cross complementing 5 |
| ENSG00000148143 | ZNF462 | -0,138524679 | 0,908447672 | 0,56667652 | 0,941311563 | zinc finger protein 462 |
| ENSG00000255100 | AP003119,2 | -0,138560406 | 0,908425176 | 0,86391367 | 0,989074806 | novel transcript |
| ENSG00000214021 | TTLL3 | -0,138604816 | 0,908397213 | 0,55941471 | 0,939727587 | tubulin tyrosine ligase like 3 |
| ENSG00000288596 | C8orf44 | -0,138658382 | 0,908363485 | 0,66823485 | 0,961289023 | chromosome 8 open reading frame 44 |
| ENSG00000226457 | RPL22P3 | -0,138696923 | 0,908339219 | 0,85366453 | 0,987861659 | ribosomal protein L22 pseudogene 3 |
| ENSG00000260329 | AC007541,1 | -0,138771525 | 0,908292249 | 0,77384814 | 0,978415098 | novel transcript, antisense to C12orf23 |
| ENSG00000271862 | AC104118,1 | -0,138805196 | 0,908271051 | 0,70419823 | 0,969101743 | novel transcript |
| ENSG00000258920 | FOXN3-AS1 | -0,138850166 | 0,90824274 | 0,72095923 | 0,972142678 | FOXN3 antisense RNA 1 |
| ENSG00000036549 | ZZZ3 | -0,13885814 | 0,90823772 | 0,73276067 | 0,974353474 | zinc finger ZZ-type containing 3 |
| ENSG00000212866 | HSPA1B | -0,138911051 | 0,908204411 | 0,63613473 | 0,955399772 | heat shock protein family A (Hsp70) member 1B |
| ENSG00000215498 | FAM230B | -0,138982084 | 0,908159696 | 0,88860936 | 0,990717729 | family with sequence similarity 230 member B |
| ENSG00000134825 | TMEM258 | -0,139015036 | 0,908138953 | 0,72845461 | 0,973955059 | transmembrane protein 258 |
| ENSG00000284644 | AC074386,1 | -0,139017284 | 0,908137538 | 0,85387057 | 0,987861659 | novel transcript, antisense to OR2A1 and ARHGEF5 |
| ENSG00000184924 | PTRHD1 | -0,139039281 | 0,908123691 | 0,70427342 | 0,969119904 | peptidyl-tRNA hydrolase domain containing 1 |
| ENSG00000116857 | TMEM9 | -0,139154499 | 0,908051169 | 0,53005181 | 0,931037682 | transmembrane protein 9 |
| ENSG00000248527 | MTATP6P1 | -0,139252901 | 0,907989235 | 0,58160556 | 0,944139372 | MT-ATP6 pseudogene 1 |
| ENSG00000260643 | AC092718,3 | -0,139269599 | 0,907978726 | 0,69186668 | 0,967204896 | novel protein |
| ENSG00000136518 | ACTL6A | -0,139329977 | 0,907940728 | 0,74434269 | 0,975641819 | actin like 6A |
| ENSG00000139835 | GRTP1 | -0,139350619 | 0,907927737 | 0,68186228 | 0,964654539 | growth hormone regulated TBC protein 1 |
| ENSG00000132781 | MUTYH | -0,139356155 | 0,907924253 | 0,64129715 | 0,956465487 | mutY DNA glycosylase |
| ENSG00000128654 | MTX2 | -0,139363647 | 0,907919538 | 0,70257398 | 0,968742626 | metaxin 2 |
| ENSG00000287160 | Z82195,3 | -0,139463403 | 0,907856761 | 0,54213923 | 0,933491609 | novel transcript |
| ENSG00000001631 | KRIT1 | -0,139513123 | 0,907825474 | 0,67493252 | 0,963268551 | KRIT1 ankyrin repeat containing |
| ENSG00000180921 | FAM83H | -0,139572751 | 0,907787954 | 0,84663906 | 0,986939308 | family with sequence similarity 83 member H |
| ENSG00000115568 | ZNF142 | -0,139593891 | 0,907774652 | 0,5245437 | 0,928616707 | zinc finger protein 142 |
| ENSG00000183513 | COA5 | -0,13963577 | 0,907748301 | 0,73786032 | 0,974353474 | cytochrome c oxidase assembly factor 5 |
| ENSG00000110427 | KIAA1549L | -0,139658194 | 0,907734192 | 0,53848886 | 0,933082867 | KIAA1549 like |
| ENSG00000213058 | AL365357,1 | -0,139672093 | 0,907725447 | 0,88961298 | 0,990717729 | ribosomal protein S14 (RPS14) pseudogene |
| ENSG00000099875 | MKNK2 | -0,139817228 | 0,907634134 | 0,60552668 | 0,950927937 | MAPK interacting serine/threonine kinase 2 |

| | | | | | | |
|---|---|---|---|---|---|---|
| ENSG00000169629 | RGPD8 | -0,139826912 | 0,907628042 | 0,70751038 | 0,969647757 | RANBP2 like and GRIP domain containing 8 |
| ENSG00000284639 | AP006294,2 | -0,139829279 | 0,907626553 | 0,92616421 | 0,993915537 | RNA binding protein, fox-1 homolog (C, elegans) 1 (RBFOX1) pseudogene |
| ENSG00000169679 | BUB1 | -0,139831633 | 0,907625072 | 0,71522374 | 0,971041579 | BUB1 mitotic checkpoint serine/threonine kinase |
| ENSG00000134644 | PUM1 | -0,139885414 | 0,907591238 | 0,63966501 | 0,956018518 | pumilio RNA binding family member 1 |
| ENSG00000248564 | AC079140,1 | -0,140017428 | 0,907508192 | 0,83767499 | 0,98566906 | novel pseudogene |
| ENSG00000151229 | SLC2A13 | -0,140048261 | 0,907488798 | 0,74231725 | 0,975176043 | solute carrier family 2 member 13 |
| ENSG00000128513 | POT1 | -0,140157772 | 0,907419915 | 0,76786842 | 0,977309816 | protection of telomeres 1 |
| ENSG00000085365 | SCAMP1 | -0,140232789 | 0,907372733 | 0,78494661 | 0,979320406 | secretory carrier membrane protein 1 |
| ENSG00000284484 | AC025287,4 | -0,140351618 | 0,907297999 | 0,82486663 | 0,983374097 | novel protein similar to cytoplasmic polyadenylated homeobox like CPHXL |
| ENSG00000081853 | PCDHGA2 | -0,140476816 | 0,907219267 | 0,76095364 | 0,97624106 | protocadherin gamma subfamily A, 2 |
| ENSG00000226706 | AL161452,1 | -0,140476885 | 0,907219223 | 0,88490709 | 0,990717729 | novel transcript |
| ENSG00000165775 | FUNDC2 | -0,140711634 | 0,907071616 | 0,64182198 | 0,95651945 | FUN14 domain containing 2 |
| ENSG00000258644 | SYNJ2BP-COX1 | -0,140737635 | 0,907055269 | 0,75090527 | 0,975901808 | SYNJ2BP-COX16 readthrough |
| ENSG00000282954 | FBXO17 | -0,140762254 | 0,90703979 | 0,6861441 | 0,965740648 | F-box protein 17 |
| ENSG00000269190 | FBXO17 | -0,140762254 | 0,90703979 | 0,6861441 | 0,965740648 | F-box protein 17 |
| ENSG00000286376 | AC006041,2 | -0,140788348 | 0,907023385 | 0,89048465 | 0,990717729 | novel transcript |
| ENSG00000260077 | AC104794,3 | -0,140814822 | 0,907006741 | 0,70183397 | 0,968742626 | novel transcript |
| ENSG00000064652 | SNX24 | -0,140835908 | 0,906993485 | 0,71842864 | 0,971830783 | sorting nexin 24 |
| ENSG00000277677 | Metazoa_SRP | -0,14084102 | 0,906990271 | 0,92724863 | 0,993915537 | Metazoan signal recognition particle RNA |
| ENSG00000261204 | AC004449,1 | -0,140929177 | 0,90693485 | 0,9012113 | 0,992041918 | novel transcript |
| ENSG00000155906 | RMND1 | -0,141000693 | 0,906889893 | 0,66787513 | 0,961113267 | required for meiotic nuclear division 1 homolog |
| ENSG00000069188 | SDK2 | -0,141025651 | 0,906874205 | 0,6135872 | 0,952413269 | sidekick cell adhesion molecule 2 |
| ENSG00000197162 | ZNF785 | -0,14102602 | 0,906873973 | 0,61604021 | 0,952793934 | zinc finger protein 785 |
| ENSG00000258405 | ZNF578 | -0,141115984 | 0,906817423 | 0,59115057 | 0,947416795 | zinc finger protein 578 |
| ENSG00000066629 | EML1 | -0,141134134 | 0,906806015 | 0,6638186 | 0,959971539 | EMAP like 1 |
| ENSG00000188647 | PTAR1 | -0,141240476 | 0,906739176 | 0,75833681 | 0,976120207 | protein prenyltransferase alpha subunit repeat containing 1 |
| ENSG00000272293 | AC083964,1 | -0,141296603 | 0,906703901 | 0,88293491 | 0,990717729 | novel transcript, antisense to FBXO25 |
| ENSG00000133943 | DGLUCY | -0,141332415 | 0,906681394 | 0,6005612 | 0,949439306 | D-glutamate cyclase |
| ENSG00000125534 | PPDPF | -0,141389533 | 0,906645498 | 0,56959971 | 0,94221458 | pancreatic progenitor cell differentiation and proliferation factor |

| | | | | | | |
|---|---|---|---|---|---|---|
| ENSG00000113318 | MSH3 | -0,141429007 | 0,906620691 | 0,61009823 | 0,951801946 | mutS homolog 3 |
| ENSG00000198736 | MSRB1 | -0,141437565 | 0,906615314 | 0,59622661 | 0,948282297 | methionine sulfoxide reductase B1 |
| ENSG00000135437 | RDH5 | -0,14155624 | 0,906540739 | 0,76492613 | 0,976725297 | retinol dehydrogenase 5 |
| ENSG00000174384 | PMS2P6 | -0,141621059 | 0,90650001 | 0,69665101 | 0,968300554 | PMS1 homolog 2, mismatch repair system component pseudogene 6 |
| ENSG00000123560 | PLP1 | -0,141626374 | 0,906496671 | 0,69085429 | 0,967179044 | proteolipid protein 1 |
| ENSG00000253958 | CLDN23 | -0,141637305 | 0,906489802 | 0,83887036 | 0,985672131 | claudin 23 |
| ENSG00000115041 | KCNIP3 | -0,141705739 | 0,906446804 | 0,76743585 | 0,977201891 | potassium voltage-gated channel interacting protein 3 |
| ENSG00000139168 | ZCRB1 | -0,141714416 | 0,906441352 | 0,71036477 | 0,970482408 | zinc finger CCHC-type and RNA binding motif containing 1 |
| ENSG00000272744 | AC107214,2 | -0,141752556 | 0,906417389 | 0,84375181 | 0,986273696 | novel transcript |
| ENSG00000152056 | AP1S3 | -0,141756117 | 0,906415152 | 0,76457065 | 0,976640468 | adaptor related protein complex 1 subunit sigma 3 |
| ENSG00000167487 | KLHL26 | -0,1417619 | 0,906411519 | 0,65462998 | 0,957444211 | kelch like family member 26 |
| ENSG00000235990 | RPL23AP20 | -0,141806171 | 0,906383704 | 0,86985127 | 0,989074806 | ribosomal protein L23a pseudogene 20 |
| ENSG00000101452 | DHX35 | -0,141831403 | 0,906367853 | 0,53797678 | 0,932883277 | DEAH-box helicase 35 |
| ENSG00000274594 | AL391669,1 | -0,141837797 | 0,906363836 | 0,86551293 | 0,989074806 | transmembrane protein 80 (TMEM80) pseudogene |
| ENSG00000141446 | ESCO1 | -0,14185611 | 0,906352331 | 0,71600364 | 0,971360663 | establishment of sister chromatid cohesion N-acetyltransferase 1 |
| ENSG00000272602 | ZNF595 | -0,141861968 | 0,906348651 | 0,70716732 | 0,969565008 | zinc finger protein 595 |
| ENSG00000206384 | COL6A6 | -0,141863174 | 0,906347892 | 0,73491717 | 0,974353474 | collagen type VI alpha 6 chain |
| ENSG00000272455 | MRPL20-DT | -0,141871298 | 0,906342789 | 0,62499446 | 0,952793934 | MRPL20 divergent transcript |
| ENSG00000116754 | SRSF11 | -0,141912485 | 0,906316915 | 0,72151326 | 0,972142678 | serine and arginine rich splicing factor 11 |
| ENSG00000104723 | TUSC3 | -0,1419842 | 0,906271864 | 0,71091975 | 0,970482408 | tumor suppressor candidate 3 |
| ENSG00000111860 | CEP85L | -0,142040293 | 0,906236628 | 0,69076018 | 0,967179044 | centrosomal protein 85 like |
| ENSG00000099956 | SMARCB1 | -0,142051632 | 0,906229505 | 0,63291712 | 0,954397372 | SWI/SNF related, matrix associated, actin dependent regulator of chromatin, subfamily b, member 1 |
| ENSG00000031691 | CENPQ | -0,142095743 | 0,906201797 | 0,7761821 | 0,978587255 | centromere protein Q |
| ENSG00000166503 | HDGFL3 | -0,142121701 | 0,906185492 | 0,71574178 | 0,971282169 | HDGF like 3 |
| ENSG00000162461 | SLC25A34 | -0,1421329 | 0,906178458 | 0,7510829 | 0,975901808 | solute carrier family 25 member 34 |
| ENSG00000232065 | LINC01063 | -0,142144686 | 0,906171055 | 0,90775197 | 0,992590661 | long intergenic non-protein coding RNA 1063 |
| ENSG00000183655 | KLHL25 | -0,142187342 | 0,906144263 | 0,64991063 | 0,956899963 | kelch like family member 25 |

| | | | | | | |
|---|---|---|---|---|---|---|
| ENSG00000267583 | AC007998,3 | -0,142321417 | 0,906060055 | 0,90801931 | 0,992590661 | novel transcript |
| ENSG00000250222 | AC008443,4 | -0,142329568 | 0,906054936 | 0,81927523 | 0,98249617 | novel transcript |
| ENSG00000175356 | SCUBE2 | -0,142335855 | 0,906050988 | 0,79587038 | 0,979879951 | signal peptide, CUB domain and EGF like domain containing 2 |
| ENSG00000165695 | AK8 | -0,142366432 | 0,906031785 | 0,74185903 | 0,975176043 | adenylate kinase 8 |
| ENSG00000151773 | CCDC122 | -0,142389141 | 0,906017524 | 0,65485966 | 0,957444211 | coiled-coil domain containing 122 |
| ENSG00000206149 | HERC2P9 | -0,142528155 | 0,905930227 | 0,5139952 | 0,924227101 | hect domain and RLD 2 pseudogene 9 |
| ENSG00000174680 | GRIK1-AS1 | -0,142559889 | 0,9059103 | 0,84741371 | 0,987035044 | GRIK1 antisense RNA 1 |
| ENSG00000251432 | LINC02615 | -0,142618529 | 0,905873478 | 0,75433746 | 0,975901808 | long intergenic non-protein coding RNA 2615 |
| ENSG00000268575 | AL031282,2 | -0,142647184 | 0,905855486 | 0,59437034 | 0,947469769 | novel transcript, readthrough between SLC35E2 and CDK11A |
| ENSG00000285679 | AC079142,1 | -0,142682581 | 0,905833261 | 0,83242511 | 0,98489473 | novel transcript |
| ENSG00000161813 | LARP4 | -0,142727053 | 0,905805339 | 0,74579564 | 0,975707496 | La ribonucleoprotein 4 |
| ENSG00000182264 | IZUMO1 | -0,142731128 | 0,90580278 | 0,85570274 | 0,988095675 | izumo sperm-egg fusion 1 |
| ENSG00000196792 | STRN3 | -0,142779607 | 0,905772343 | 0,64742347 | 0,956899963 | striatin 3 |
| ENSG00000224264 | VARS1 | -0,142942535 | 0,905670057 | 0,7595328 | 0,976120207 | valyl-tRNA synthetase 1 |
| ENSG00000231116 | VARS1 | -0,142942535 | 0,905670057 | 0,7595328 | 0,976120207 | valyl-tRNA synthetase 1 |
| ENSG00000196227 | FAM217B | -0,142962049 | 0,905657806 | 0,67426952 | 0,963103112 | family with sequence similarity 217 member B |
| ENSG00000282254 | AC004980,5 | -0,143028948 | 0,905615812 | 0,6541043 | 0,957444211 | pseudogene similar to part of postmeiotic segregation increased 2-like 3 PMS2L3 |
| ENSG00000169519 | METTL15 | -0,143033616 | 0,905612881 | 0,693781 | 0,967340633 | methyltransferase like 15 |
| ENSG00000162607 | USP1 | -0,143141822 | 0,90554496 | 0,73834227 | 0,974353474 | ubiquitin specific peptidase 1 |
| ENSG00000230433 | AL031770,1 | -0,143214215 | 0,905499522 | 0,84999962 | 0,987036295 | novel transcript |
| ENSG00000186017 | ZNF566 | -0,143227754 | 0,905491025 | 0,67762394 | 0,9635902 | zinc finger protein 566 |
| ENSG00000204387 | SNHG32 | -0,143244299 | 0,905480641 | 0,69468617 | 0,967417311 | small nucleolar RNA host gene 32 |
| ENSG00000144182 | LIPT1 | -0,143296671 | 0,905447771 | 0,617236 | 0,952793934 | lipoyltransferase 1 |
| ENSG00000175063 | UBE2C | -0,143389504 | 0,90538951 | 0,60914939 | 0,951793902 | ubiquitin conjugating enzyme E2 C |
| ENSG00000146950 | SHROOM2 | -0,143400234 | 0,905382776 | 0,51869311 | 0,926765093 | shroom family member 2 |
| ENSG00000254791 | FAR1-IT1 | -0,143469855 | 0,905339085 | 0,87578131 | 0,990109814 | FAR1 intronic transcript 1 |
| ENSG00000170144 | HNRNPA3 | -0,143531333 | 0,905300507 | 0,61951985 | 0,952793934 | heterogeneous nuclear ribonucleoprotein A3 |
| ENSG00000029153 | ARNTL2 | -0,143636442 | 0,905234552 | 0,55242234 | 0,936703121 | aryl hydrocarbon receptor nuclear translocator like 2 |
| ENSG00000266340 | AC138207,7 | -0,143651381 | 0,905225179 | 0,51623805 | 0,925176589 | novel transcript, antisense RNF135 |
| ENSG00000259581 | TYRO3P | -0,143700224 | 0,905194533 | 0,73973797 | 0,974715504 | TYRO3P protein tyrosine kinase pseudogene |

| | | | | | | |
|---|---|---|---|---|---|---|
| ENSG00000082898 | XPO1 | -0,143729389 | 0,905176234 | 0,72469607 | 0,97326334 | exportin 1 |
| ENSG00000185742 | C11orf87 | -0,14377068 | 0,905150327 | 0,78184822 | 0,978597905 | chromosome 11 open reading frame 87 |
| ENSG00000226009 | KCNIP2-AS1 | -0,143798598 | 0,905132812 | 0,85157821 | 0,987271154 | KCNIP2 antisense RNA 1 |
| ENSG00000270021 | AC026691,1 | -0,143894328 | 0,905072754 | 0,79482324 | 0,979776981 | novel transcript, antisense to H2AFY |
| ENSG00000255568 | BRWD1-AS2 | -0,143979807 | 0,90501913 | 0,72862916 | 0,973955059 | BRWD1 antisense RNA 2 |
| ENSG00000219926 | OR7E104P | -0,143979906 | 0,905019068 | 0,87361341 | 0,989633545 | olfactory receptor family 7 subfamily E member 104 pseudogene |
| ENSG00000205740 | AL359878,1 | -0,143986846 | 0,905014714 | 0,60932658 | 0,951793902 | uncharacterized LOC107984285 |
| ENSG00000112874 | NUDT12 | -0,144042498 | 0,904979804 | 0,76875494 | 0,977381742 | nudix hydrolase 12 |
| ENSG00000280795 | MPRIPP1 | -0,144096999 | 0,904945617 | 0,79425035 | 0,979776981 | myosin phosphatase Rho interacting protein pseudogene 1 |
| ENSG00000214820 | MPRIPP1 | -0,144096999 | 0,904945617 | 0,79425035 | 0,979776981 | myosin phosphatase Rho interacting protein pseudogene 1 |
| ENSG00000077152 | UBE2T | -0,144141132 | 0,904917935 | 0,73982064 | 0,974715504 | ubiquitin conjugating enzyme E2 T |
| ENSG00000141084 | RANBP10 | -0,1441727 | 0,904898134 | 0,52193354 | 0,927599011 | RAN binding protein 10 |
| ENSG00000237212 | AL449403,1 | -0,144314312 | 0,904809315 | 0,78779542 | 0,979462318 | novel transcript |
| ENSG00000172348 | RCAN2 | -0,144350938 | 0,904786345 | 0,73570798 | 0,974353474 | regulator of calcineurin 2 |
| ENSG00000217648 | AL136116,3 | -0,14437256 | 0,904772785 | 0,78153572 | 0,978597905 | pseudogene similar to presenilin associated, rhomboid-like PARL |
| ENSG00000129317 | PUS7L | -0,144374301 | 0,904771694 | 0,74162039 | 0,975176043 | pseudouridine synthase 7 like |
| ENSG00000120616 | EPC1 | -0,144396835 | 0,904757561 | 0,62577355 | 0,952793934 | enhancer of polycomb homolog 1 |
| ENSG00000232671 | ZNF687-AS1 | -0,14453138 | 0,904673188 | 0,75871726 | 0,976120207 | ZNF687 antisense RNA 1 |
| ENSG00000143228 | NUF2 | -0,144538284 | 0,904668859 | 0,72585998 | 0,973344786 | NUF2 component of NDC80 kinetochore complex |
| ENSG00000065150 | IPO5 | -0,144612488 | 0,904622329 | 0,67246585 | 0,962290553 | importin 5 |
| ENSG00000163145 | C1QTNF7 | -0,144649076 | 0,904599387 | 0,73605177 | 0,974353474 | C1q and TNF related 7 |
| ENSG00000156687 | UNC5D | -0,144672891 | 0,904584455 | 0,77903711 | 0,978587255 | unc-5 netrin receptor D |
| ENSG00000287937 | AC079753,2 | -0,144682961 | 0,904578141 | 0,79645028 | 0,980015022 | novel transcript |
| ENSG00000155868 | MED7 | -0,144701453 | 0,904566547 | 0,70311702 | 0,968742626 | mediator complex subunit 7 |
| ENSG00000198435 | NRARP | -0,144787765 | 0,904512431 | 0,65093226 | 0,957027739 | NOTCH regulated ankyrin repeat protein |
| ENSG00000152454 | ZNF256 | -0,14484038 | 0,904479444 | 0,67689662 | 0,963580269 | zinc finger protein 256 |
| ENSG00000230550 | ERLNC1 | -0,144923781 | 0,904427158 | 0,91241621 | 0,993361299 | estrogen receptor responsive lncRNA 1 |
| ENSG00000120802 | TMPO | -0,144928981 | 0,904423898 | 0,64945308 | 0,956899963 | thymopoietin |
| ENSG00000186298 | PPP1CC | -0,144961036 | 0,904403803 | 0,74628731 | 0,975796253 | protein phosphatase 1 catalytic subunit gamma |

| | | | | | | |
|---|---|---|---|---|---|---|
| ENSG00000237101 | AC092809,4 | -0,145008525 | 0,904374034 | 0,87118921 | 0,989217334 | novel transcript |
| ENSG00000071243 | ING3 | -0,145009814 | 0,904373225 | 0,75572009 | 0,975901808 | inhibitor of growth family member 3 |
| ENSG00000184347 | SLIT3 | -0,145106581 | 0,904312568 | 0,64681703 | 0,956899963 | slit guidance ligand 3 |
| ENSG00000260742 | AC009962,1 | -0,145122584 | 0,904302537 | 0,73346089 | 0,974353474 | novel transcript, antisense to SSFA2 |
| ENSG00000183604 | SMG1P5 | -0,145248721 | 0,904223476 | 0,55934537 | 0,939727587 | SMG1 pseudogene 5 |
| ENSG00000205485 | AC004980,1 | -0,145292872 | 0,904195804 | 0,64886983 | 0,956899963 | pseudogene similar to part of postmeiotic segregation increased 2-like 3 PMS2L3 |
| ENSG00000104164 | BLOC1S6 | -0,14531554 | 0,904181597 | 0,58096431 | 0,943914064 | biogenesis of lysosomal organelles complex 1 subunit 6 |
| ENSG00000136159 | NUDT15 | -0,145319521 | 0,904179102 | 0,61250153 | 0,952413269 | nudix hydrolase 15 |
| ENSG00000232814 | COL4A2-AS1 | -0,145333838 | 0,904170129 | 0,85723944 | 0,988665109 | COL4A2 antisense RNA 1 |
| ENSG00000235725 | AC007389,2 | -0,145398034 | 0,904129897 | 0,78868409 | 0,979462318 | novel transcript |
| ENSG00000112877 | CEP72 | -0,145418805 | 0,90411688 | 0,70962132 | 0,970482408 | centrosomal protein 72 |
| ENSG00000176809 | LRRC37A3 | -0,145425692 | 0,904112564 | 0,57446968 | 0,942475308 | leucine rich repeat containing 37 member A3 |
| ENSG00000136044 | APPL2 | -0,145448585 | 0,904098217 | 0,64696986 | 0,956899963 | adaptor protein, phosphotyrosine interacting with PH domain and leucine zipper 2 |
| ENSG00000223945 | AC244033,1 | -0,145449962 | 0,904097355 | 0,86760774 | 0,989074806 | ribosomal protein L6 (RPL6) pseudogene |
| ENSG00000275029 | HMGB1P24 | -0,145473198 | 0,904082793 | 0,80844749 | 0,981213877 | high mobility group box 1 pseudogene 24 |
| ENSG00000282695 | HMGB1P24 | -0,145473198 | 0,904082793 | 0,80844749 | 0,981213877 | high mobility group box 1 pseudogene 24 |
| ENSG00000197563 | PIGN | -0,145484672 | 0,904075603 | 0,49238495 | 0,917008021 | phosphatidylinositol glycan anchor biosynthesis class N |
| ENSG00000105879 | CBLL1 | -0,145497308 | 0,904067685 | 0,73909857 | 0,974657403 | Cbl proto-oncogene like 1 |
| ENSG00000163516 | ANKZF1 | -0,145519451 | 0,904053809 | 0,54372634 | 0,934259495 | ankyrin repeat and zinc finger peptidyl tRNA hydrolase 1 |
| ENSG00000228510 | AL359091,2 | -0,145537125 | 0,904042734 | 0,90143048 | 0,992041918 | novel pseudogene |
| ENSG00000166783 | MARF1 | -0,145672129 | 0,903958139 | 0,61978385 | 0,952793934 | meiosis regulator and mRNA stability factor 1 |
| ENSG00000213213 | CCDC183 | -0,1458337 | 0,903856908 | 0,77774675 | 0,978587255 | coiled-coil domain containing 183 |
| ENSG00000270562 | AC097634,1 | -0,145850752 | 0,903846226 | 0,84787103 | 0,987036295 | novel transcript |
| ENSG00000261840 | AC093249,6 | -0,145913462 | 0,903806938 | 0,7286906 | 0,973955059 | uncharacterized LOC730183 |
| ENSG00000153048 | CARHSP1 | -0,145941277 | 0,903789514 | 0,48587044 | 0,913997663 | calcium regulated heat stable protein 1 |
| ENSG00000285979 | AC009090,6 | -0,145946654 | 0,903786145 | 0,70261797 | 0,968742626 | novel transcript, antisense to FAM192A |
| ENSG00000267065 | LINC02080 | -0,146007785 | 0,90374785 | 0,55136204 | 0,936366211 | long intergenic non-protein coding RNA 2080 |
| ENSG00000260331 | AC079148,1 | -0,146035567 | 0,903730446 | 0,85554421 | 0,988075747 | novel transcript |

| | | | | | | |
|---|---|---|---|---|---|---|
| ENSG00000285994 | AL731559,1 | -0,146069152 | 0,903709409 | 0,72927182 | 0,974009551 | novel transcript, sense intronic to CAMK1D |
| ENSG00000072818 | ACAP1 | -0,146163333 | 0,903650415 | 0,75470576 | 0,975901808 | ArfGAP with coiled-coil, ankyrin repeat and PH domains 1 |
| ENSG00000223668 | EEF1A1P24 | -0,146193817 | 0,903631321 | 0,84441748 | 0,986340494 | eukaryotic translation elongation factor 1 alpha 1 pseudogene 24 |
| ENSG00000228808 | HMGB3P4 | -0,146259425 | 0,903590229 | 0,8370633 | 0,985462684 | high mobility group box 3 pseudogene 4 |
| ENSG00000069248 | NUP133 | -0,146287331 | 0,903572751 | 0,64364622 | 0,956633898 | nucleoporin 133 |
| ENSG00000227811 | INKA2-AS1 | -0,146305411 | 0,903561427 | 0,79914337 | 0,980161849 | INKA2 antisense RNA 1 |
| ENSG00000277600 | CNOT3 | -0,146366378 | 0,903523244 | 0,82427146 | 0,983374097 | CCR4-NOT transcription complex subunit 3 |
| ENSG00000154473 | BUB3 | -0,146404807 | 0,903499178 | 0,69725472 | 0,968522643 | BUB3 mitotic checkpoint protein |
| ENSG00000174428 | GTF2IRD2B | -0,146426758 | 0,903485431 | 0,53542093 | 0,932037868 | GTF2I repeat domain containing 2B |
| ENSG00000162032 | SPSB3 | -0,146459143 | 0,90346515 | 0,54536679 | 0,935085411 | splA/ryanodine receptor domain and SOCS box containing 3 |
| ENSG00000284548 | METTL9 | -0,146497784 | 0,903440952 | 0,79932139 | 0,980161849 | methyltransferase like 9 |
| ENSG00000263189 | AC023934,1 | -0,146529404 | 0,903421151 | 0,9084612 | 0,992595616 | Zinc finger protein pseudogene |
| ENSG00000263004 | AC007114,1 | -0,146603528 | 0,903374736 | 0,78883433 | 0,979462318 | novel transcript |
| ENSG00000236618 | PITPNA-AS1 | -0,14663847 | 0,903352856 | 0,63151865 | 0,953845566 | PITPNA antisense RNA 1 |
| ENSG00000255585 | AL590627,1 | -0,146653436 | 0,903343485 | 0,80452231 | 0,980875302 | forkhead box H1 (FOXH1) pseudogene |
| ENSG00000226571 | AL592429,2 | -0,146702214 | 0,903312943 | 0,85409575 | 0,987861659 | novel transcript |
| ENSG00000198740 | ZNF652 | -0,146761626 | 0,903275744 | 0,68023795 | 0,964580199 | zinc finger protein 652 |
| ENSG00000284938 | FAM180B | -0,146824159 | 0,903236594 | 0,89952461 | 0,991845106 | family with sequence similarity 180 member B |
| ENSG00000213742 | ZNF337-AS1 | -0,146860786 | 0,903213662 | 0,69269864 | 0,967271667 | ZNF337 antisense RNA 1 |
| ENSG00000261442 | AC023830,1 | -0,146964949 | 0,903148452 | 0,83397467 | 0,984905093 | novel transcript, antisense to MGRN1 |
| ENSG00000273703 | H2BC14 | -0,146966063 | 0,903147755 | 0,76158482 | 0,976285211 | H2B clustered histone 14 |
| ENSG00000205771 | CATSPER2P1 | -0,147032171 | 0,903106371 | 0,73150348 | 0,974353474 | cation channel sperm associated 2 pseudogene 1 |
| ENSG00000135336 | ORC3 | -0,147056411 | 0,903091197 | 0,74648064 | 0,975796253 | origin recognition complex subunit 3 |
| ENSG00000261504 | LINC01686 | -0,147056478 | 0,903091155 | 0,78463942 | 0,979320406 | long intergenic non-protein coding RNA 1686 |
| ENSG00000198668 | CALM1 | -0,14718425 | 0,903011177 | 0,66706217 | 0,9611116 | calmodulin 1 |
| ENSG00000198799 | LRIG2 | -0,147209989 | 0,902995067 | 0,59395584 | 0,947469769 | leucine rich repeats and immunoglobulin like domains 2 |
| ENSG00000122026 | RPL21 | -0,147270354 | 0,902957285 | 0,63387114 | 0,954929949 | ribosomal protein L21 |
| ENSG00000233327 | USP32P2 | -0,147318304 | 0,902927274 | 0,81038264 | 0,981213877 | ubiquitin specific peptidase 32 pseudogene 2 |
| ENSG00000129654 | FOXJ1 | -0,147337989 | 0,902914954 | 0,66484408 | 0,960551345 | forkhead box J1 |

| | | | | | | |
|---|---|---|---|---|---|---|
| ENSG00000188375 | H3-5 | -0,147343707 | 0,902911375 | 0,82589652 | 0,983383773 | H3,5 histone |
| ENSG00000059145 | UNKL | -0,147421886 | 0,902862448 | 0,67971403 | 0,964459885 | unk like zinc finger |
| ENSG00000214413 | BBIP1 | -0,147549161 | 0,902782801 | 0,72095647 | 0,972142678 | BBSome interacting protein 1 |
| ENSG00000241322 | CDRT1 | -0,147582922 | 0,902761675 | 0,69351441 | 0,967340633 | CMT1A duplicated region transcript 1 |
| ENSG00000255389 | Z97989,1 | -0,147614541 | 0,90274189 | 0,76087314 | 0,97624106 | chromosome 6 open reading frame 3 |
| ENSG00000205913 | SRRM2-AS1 | -0,147632152 | 0,90273087 | 0,71623717 | 0,97143026 | SRRM2 antisense RNA 1 |
| ENSG00000137656 | BUD13 | -0,147637802 | 0,902727335 | 0,52525496 | 0,928752911 | BUD13 homolog |
| ENSG00000140474 | ULK3 | -0,147664739 | 0,90271048 | 0,55179197 | 0,936366211 | unc-51 like kinase 3 |
| ENSG00000185305 | ARL15 | -0,147682373 | 0,902699446 | 0,78204971 | 0,978597905 | ADP ribosylation factor like GTPase 15 |
| ENSG00000256984 | AC008013,2 | -0,147733137 | 0,902667683 | 0,85293164 | 0,987741407 | mitochondrially encoded cytochrome c oxidase I (MT-CO1) pseudogene |
| ENSG00000267122 | AC004490,1 | -0,147737158 | 0,902665168 | 0,81186197 | 0,981213877 | novel transcript, antisense to DOT1L |
| ENSG00000092470 | WDR76 | -0,14775188 | 0,902655956 | 0,66297012 | 0,959920886 | WD repeat domain 76 |
| ENSG00000172215 | CXCR6 | -0,147843968 | 0,902598341 | 0,72669033 | 0,973344786 | C-X-C motif chemokine receptor 6 |
| ENSG00000224812 | TMEM72-AS1 | -0,147848071 | 0,902595774 | 0,75125695 | 0,975901808 | TMEM72 antisense RNA 1 |
| ENSG00000231925 | TAPBP | -0,14794881 | 0,902532751 | 0,87045863 | 0,989074806 | TAP binding protein |
| ENSG00000287248 | AC009271,2 | -0,148041186 | 0,902474963 | 0,84818062 | 0,987036295 | novel transcript |
| ENSG00000228412 | LNC-LBCS | -0,148053542 | 0,902467234 | 0,6580395 | 0,957883832 | lncRNA bladder and prostate cancer suppressor, hnRNPK interacting |
| ENSG00000145014 | TMEM44 | -0,148056025 | 0,902465681 | 0,61112602 | 0,951932839 | transmembrane protein 44 |
| ENSG00000179292 | TMEM151A | -0,148058233 | 0,9024643 | 0,81468985 | 0,981521413 | transmembrane protein 151A |
| ENSG00000119203 | CPSF3 | -0,14810317 | 0,90243619 | 0,67346974 | 0,962780262 | cleavage and polyadenylation specific factor 3 |
| ENSG00000083312 | TNPO1 | -0,148128278 | 0,902420485 | 0,74237368 | 0,975176043 | transportin 1 |
| ENSG00000165138 | ANKS6 | -0,14815206 | 0,902405609 | 0,62320128 | 0,952793934 | ankyrin repeat and sterile alpha motif domain containing 6 |
| ENSG00000213214 | ARHGEF35 | -0,148178925 | 0,902388805 | 0,79004099 | 0,979462318 | Rho guanine nucleotide exchange factor 35 |
| ENSG00000234268 | AP000936,3 | -0,14820554 | 0,902372158 | 0,91295611 | 0,993361299 | ribosomal protein S27 (metallopanstimulin 1) (RPS27) pseudogene |
| ENSG00000101773 | RBBP8 | -0,148276127 | 0,902328008 | 0,70277859 | 0,968742626 | RB binding protein 8, endonuclease |
| ENSG00000214113 | LYRM4 | -0,14834032 | 0,90228786 | 0,64344673 | 0,956633898 | LYR motif containing 4 |
| ENSG00000258342 | AL133304,3 | -0,148396206 | 0,902252909 | 0,89815496 | 0,991802654 | novel transcript, sense overlapping BRMS1L |
| ENSG00000205334 | LINC01460 | -0,148410618 | 0,902243895 | 0,81973083 | 0,98249617 | long intergenic non-protein coding RNA 1460 |
| ENSG00000183506 | PI4KAP2 | -0,148491982 | 0,902193013 | 0,63179137 | 0,953915351 | phosphatidylinositol 4-kinase alpha pseudogene 2 |

| | | | | | | |
|---|---|---|---|---|---|---|
| ENSG00000236021 | AL591043,2 | -0,148507892 | 0,902183064 | 0,87359567 | 0,989633545 | novel transcript |
| ENSG00000088808 | PPP1R13B | -0,148613896 | 0,902116777 | 0,5382426 | 0,932900071 | protein phosphatase 1 regulatory subunit 13B |
| ENSG00000274047 | SYNRG | -0,148614517 | 0,902116388 | 0,61854958 | 0,952793934 | synergin gamma |
| ENSG00000275867 | KANSL1 | -0,148631049 | 0,902106051 | 0,59747072 | 0,948282297 | KAT8 regulatory NSL complex subunit 1 |
| ENSG00000173480 | ZNF417 | -0,148646324 | 0,9020965 | 0,50129342 | 0,920520138 | zinc finger protein 417 |
| ENSG00000138050 | THUMPD2 | -0,148810837 | 0,901993638 | 0,69627264 | 0,96810276 | THUMP domain containing 2 |
| ENSG00000180817 | PPA1 | -0,148852067 | 0,901967861 | 0,69351155 | 0,967340633 | inorganic pyrophosphatase 1 |
| ENSG00000182362 | YBEY | -0,148858789 | 0,901963658 | 0,62776832 | 0,952793934 | ybeY metalloendoribonuclease |
| ENSG00000094975 | SUCO | -0,148923441 | 0,901923239 | 0,71073066 | 0,970482408 | SUN domain containing ossification factor |
| ENSG00000254860 | TMEM9B-AS1 | -0,148962578 | 0,901898772 | 0,68695291 | 0,965769168 | TMEM9B antisense RNA 1 |
| ENSG00000123545 | NDUFAF4 | -0,148990799 | 0,90188113 | 0,74246805 | 0,975176043 | NADH:ubiquinone oxidoreductase complex assembly factor 4 |
| ENSG00000275052 | PPP4R3B | -0,149068228 | 0,901832728 | 0,65534337 | 0,957444211 | protein phosphatase 4 regulatory subunit 3B |
| ENSG00000051620 | HEBP2 | -0,14908128 | 0,901824569 | 0,56484589 | 0,940865625 | heme binding protein 2 |
| ENSG00000269984 | AC078795,1 | -0,149181354 | 0,901762015 | 0,701925 | 0,968742626 | novel transcript, antisense to MYNN |
| ENSG00000114446 | IFT57 | -0,149223302 | 0,901735796 | 0,51930676 | 0,926800715 | intraflagellar transport 57 |
| ENSG00000277103 | AL445670,1 | -0,149295514 | 0,901690662 | 0,83837025 | 0,985672131 | sorting nexin 18 (SNX18) pseudogene |
| ENSG00000176978 | DPP7 | -0,149480306 | 0,901575174 | 0,7529205 | 0,975901808 | dipeptidyl peptidase 7 |
| ENSG00000237289 | CKMT1B | -0,149568143 | 0,901520284 | 0,78035261 | 0,978587255 | creatine kinase, mitochondrial 1B |
| ENSG00000102796 | DHRS12 | -0,149744576 | 0,90141004 | 0,61063275 | 0,951841373 | dehydrogenase/reductase 12 |
| ENSG00000166068 | SPRED1 | -0,149759988 | 0,90140041 | 0,65636287 | 0,957444211 | sprouty related EVH1 domain containing 1 |
| ENSG00000166268 | MYRFL | -0,149818577 | 0,901363804 | 0,75413604 | 0,975901808 | myelin regulatory factor like |
| ENSG00000286665 | AP000648,9 | -0,149853833 | 0,901341778 | 0,87738351 | 0,990109814 | novel transcript |
| ENSG00000288058 | AP000648,10 | -0,149853833 | 0,901341778 | 0,87738351 | 0,990109814 | novel transcript |
| ENSG00000285871 | AL359844,1 | -0,149865263 | 0,901334637 | 0,84114716 | 0,985672131 | novel transcript |
| ENSG00000173933 | RBM4 | -0,149900582 | 0,901312571 | 0,54503047 | 0,934923038 | RNA binding motif protein 4 |
| ENSG00000287068 | AP004606,1 | -0,149902212 | 0,901311553 | 0,8792386 | 0,990148595 | novel transcript, antisense to OPCML |
| ENSG00000285901 | AC008012,1 | -0,150033522 | 0,901229522 | 0,65409078 | 0,957444211 | novel transcript |
| ENSG00000196323 | ZBTB44 | -0,150038368 | 0,901226495 | 0,74183924 | 0,975176043 | zinc finger and BTB domain containing 44 |
| ENSG00000107625 | DDX50 | -0,1500869 | 0,901196178 | 0,67627291 | 0,963580269 | DExD-box helicase 50 |
| ENSG00000286729 | AC137695,3 | -0,150104101 | 0,901185433 | 0,78468884 | 0,979320406 | novel transcript, antisense to COPG1 |
| ENSG00000177683 | THAP5 | -0,150107886 | 0,901183069 | 0,72700752 | 0,973344786 | THAP domain containing 5 |
| ENSG00000239474 | KLHL41 | -0,150420657 | 0,900987717 | 0,76553044 | 0,97683946 | kelch like family member 41 |

| | | | | | | |
|---|---|---|---|---|---|---|
| ENSG00000167525 | PROCA1 | -0,150441204 | 0,900974885 | 0,6251675 | 0,952793934 | protein interacting with cyclin A1 |
| ENSG00000223773 | CD99P1 | -0,150655743 | 0,900840914 | 0,58179804 | 0,944145586 | CD99 molecule pseudogene 1 |
| ENSG00000166788 | SAAL1 | -0,150672162 | 0,900830661 | 0,6558521 | 0,957444211 | serum amyloid A like 1 |
| ENSG00000228748 | AL450306,1 | -0,150722643 | 0,900799141 | 0,79141115 | 0,979471509 | novel transcript |
| ENSG00000254221 | PCDHGB1 | -0,150741654 | 0,900787271 | 0,69957475 | 0,968522643 | protocadherin gamma subfamily B, 1 |
| ENSG00000101266 | CSNK2A1 | -0,150803643 | 0,900748567 | 0,62887508 | 0,952793934 | casein kinase 2 alpha 1 |
| ENSG00000125170 | DOK4 | -0,150948446 | 0,900658164 | 0,51619385 | 0,925176589 | docking protein 4 |
| ENSG00000288632 | AC133555,6 | -0,151016606 | 0,900615613 | 0,72287261 | 0,972707638 | novel transcript |
| ENSG00000237939 | IDI1P1 | -0,151044191 | 0,900598393 | 0,88612263 | 0,990717729 | IDI1 pseudogene 1 |
| ENSG00000175768 | TOMM5 | -0,151080591 | 0,900575671 | 0,6994155 | 0,968522643 | translocase of outer mitochondrial membrane 5 |
| ENSG00000235034 | C19orf81 | -0,151121195 | 0,900550325 | 0,77807794 | 0,978587255 | chromosome 19 open reading frame 81 |
| ENSG00000128602 | SMO | -0,15112959 | 0,900545085 | 0,60554831 | 0,950927937 | smoothened, frizzled class receptor |
| ENSG00000171858 | RPS21 | -0,151208305 | 0,900495952 | 0,5732606 | 0,942475308 | ribosomal protein S21 |
| ENSG00000285007 | REC8 | -0,151217179 | 0,900490413 | 0,56622733 | 0,941123407 | REC8 meiotic recombination protein |
| ENSG00000100918 | REC8 | -0,151217179 | 0,900490413 | 0,56622733 | 0,941123407 | REC8 meiotic recombination protein |
| ENSG00000251288 | AC018797,2 | -0,151235489 | 0,900478984 | 0,85414315 | 0,987861659 | WD repeat domain 12 (WDR12) pseudogene |
| ENSG00000127328 | RAB3IP | -0,151245737 | 0,900472588 | 0,7381013 | 0,974353474 | RAB3A interacting protein |
| ENSG00000280810 | NSFP1 | -0,151258098 | 0,900464872 | 0,74605771 | 0,975707496 | N-ethylmaleimide-sensitive factor pseudogene 1 |
| ENSG00000138286 | FAM149B1 | -0,151297723 | 0,900440141 | 0,69754131 | 0,968522643 | family with sequence similarity 149 member B1 |
| ENSG00000102781 | KATNAL1 | -0,151311823 | 0,90043134 | 0,67570879 | 0,963449137 | katanin catalytic subunit A1 like 1 |
| ENSG00000288360 | ZSWIM4 | -0,15133038 | 0,900419759 | 0,65770165 | 0,957864535 | zinc finger SWIM-type containing 4 |
| ENSG00000132003 | ZSWIM4 | -0,15133038 | 0,900419759 | 0,65770165 | 0,957864535 | zinc finger SWIM-type containing 4 |
| ENSG00000241547 | ACTG1P20 | -0,151368802 | 0,900395778 | 0,77721003 | 0,978587255 | actin gamma 1 pseudogene 20 |
| ENSG00000185238 | PRMT3 | -0,151376063 | 0,900391247 | 0,68841425 | 0,966481084 | protein arginine methyltransferase 3 |
| ENSG00000111046 | MYF6 | -0,151385589 | 0,900385302 | 0,84873219 | 0,987036295 | myogenic factor 6 |
| ENSG00000267496 | FAM215A | -0,151404621 | 0,900373424 | 0,85602576 | 0,988185425 | family with sequence similarity 215 member A |
| ENSG00000259673 | IQCH-AS1 | -0,151427731 | 0,900359001 | 0,59476802 | 0,947469769 | IQCH antisense RNA 1 |
| ENSG00000271267 | AL353807,4 | -0,151495653 | 0,900316613 | 0,91332447 | 0,993361299 | ATP synthase, H+ transporting, mitochondrial Fo complex, subunit F2 (ATP5J2) pseudogene |
| ENSG00000249459 | ZNF286B | -0,151513058 | 0,900305752 | 0,62395183 | 0,952793934 | zinc finger protein 286B (pseudogene) |
| ENSG00000269945 | AC004817,4 | -0,151548944 | 0,900283358 | 0,6947135 | 0,967417311 | novel transcript |
| ENSG00000272944 | AC079834,1 | -0,151554595 | 0,900279831 | 0,73744282 | 0,974353474 | novel transcript |
| ENSG00000106459 | NRF1 | -0,15158675 | 0,900259766 | 0,51590661 | 0,925152007 | nuclear respiratory factor 1 |

| | | | | | | |
|---|---|---|---|---|---|---|
| ENSG00000109805 | NCAPG | -0,151589422 | 0,900258099 | 0,69272417 | 0,967271667 | non-SMC condensin I complex subunit G |
| ENSG00000166526 | ZNF3 | -0,151609107 | 0,900245815 | 0,51426994 | 0,924319732 | zinc finger protein 3 |
| ENSG00000118058 | KMT2A | -0,151628858 | 0,900233491 | 0,55279218 | 0,936894267 | lysine methyltransferase 2A |
| ENSG00000169714 | CNBP | -0,151679549 | 0,90020186 | 0,6876996 | 0,966064784 | CCHC-type zinc finger nucleic acid binding protein |
| ENSG00000261770 | AC006504,1 | -0,151696189 | 0,900191477 | 0,65512063 | 0,957444211 | novel transcript |
| ENSG00000287706 | AC092924,2 | -0,151700172 | 0,900188992 | 0,88354587 | 0,990717729 | novel transcript |
| ENSG00000272886 | DCP1A | -0,151796854 | 0,900128668 | 0,63759047 | 0,956017934 | decapping mRNA 1A |
| ENSG00000115274 | INO80B | -0,15180823 | 0,900121571 | 0,6011687 | 0,949882873 | INO80 complex subunit B |
| ENSG00000023287 | RB1CC1 | -0,151825503 | 0,900110794 | 0,68706168 | 0,965769168 | RB1 inducible coiled-coil 1 |
| ENSG00000281569 | KANSL1-AS1 | -0,151830706 | 0,900107547 | 0,78243933 | 0,978628303 | KANSL1 antisense RNA 1 |
| ENSG00000282944 | AC140481,3 | -0,151921921 | 0,900050639 | 0,823123 | 0,983204348 | novel transcript |
| ENSG00000146232 | NFKBIE | -0,152007555 | 0,899997217 | 0,67071846 | 0,962026636 | NFKB inhibitor epsilon |
| ENSG00000226882 | GNL1 | -0,152083223 | 0,899950014 | 0,73244411 | 0,974353474 | G protein nucleolar 1 (putative) |
| ENSG00000238917 | SNORD10 | -0,152085793 | 0,899948411 | 0,79791883 | 0,980161849 | small nucleolar RNA, C/D box 10 |
| ENSG00000020256 | ZFP64 | -0,152086461 | 0,899947994 | 0,48057064 | 0,912847708 | ZFP64 zinc finger protein |
| ENSG00000269194 | AC006942,1 | -0,152102316 | 0,899938104 | 0,88317506 | 0,990717729 | novel transcript, antisense to FUZ |
| ENSG00000286490 | AC067942,4 | -0,152107247 | 0,899935028 | 0,91498192 | 0,993361299 | novel transcript |
| ENSG00000131019 | ULBP3 | -0,152200877 | 0,899876625 | 0,51502879 | 0,924720046 | UL16 binding protein 3 |
| ENSG00000206567 | AC022007,1 | -0,15221572 | 0,899867366 | 0,57118658 | 0,942436428 | novel transcript |
| ENSG00000182963 | GJC1 | -0,152247023 | 0,899847842 | 0,52383621 | 0,928308318 | gap junction protein gamma 1 |
| ENSG00000167632 | TRAPPC9 | -0,152288985 | 0,899821669 | 0,49286162 | 0,917090312 | trafficking protein particle complex 9 |
| ENSG00000159496 | RGL4 | -0,152340987 | 0,899789236 | 0,79026134 | 0,979462318 | ral guanine nucleotide dissociation stimulator like 4 |
| ENSG00000095574 | IKZF5 | -0,152403992 | 0,899749942 | 0,74200548 | 0,975176043 | IKAROS family zinc finger 5 |
| ENSG00000181035 | SLC25A42 | -0,152478574 | 0,899703429 | 0,67232554 | 0,962290553 | solute carrier family 25 member 42 |
| ENSG00000143416 | SELENBP1 | -0,152506978 | 0,899685715 | 0,62905285 | 0,952793934 | selenium binding protein 1 |
| ENSG00000272240 | AC004908,1 | -0,152562978 | 0,899650794 | 0,69779589 | 0,968522643 | novel transcript, antisense to ZNF596 |
| ENSG00000159905 | ZNF221 | -0,152684405 | 0,899575076 | 0,55728782 | 0,938438347 | zinc finger protein 221 |
| ENSG00000279529 | AC008764,8 | -0,152741338 | 0,899539577 | 0,65640756 | 0,957444211 | novel transcript, antisense to MED26 |
| ENSG00000224893 | LINC02840 | -0,152795665 | 0,899505704 | 0,72262018 | 0,972594421 | long intergenic non-protein coding RNA 2840 |
| ENSG00000185219 | ZNF445 | -0,152850762 | 0,899471352 | 0,69788393 | 0,968522643 | zinc finger protein 445 |
| ENSG00000261716 | H2BC20P | -0,152993435 | 0,899382405 | 0,71368533 | 0,970812834 | H2B clustered histone 20, pseudogene |
| ENSG00000134321 | RSAD2 | -0,153022402 | 0,899364347 | 0,77993445 | 0,978587255 | radical S-adenosyl methionine domain containing 2 |
| ENSG00000271327 | AC010201,2 | -0,15309553 | 0,89931876 | 0,71557582 | 0,971209849 | novel transcript |

| | | | | | | |
|---|---|---|---|---|---|---|
| ENSG00000175130 | MARCKSL1 | -0,153127791 | 0,899298651 | 0,46355492 | 0,907101775 | MARCKS like 1 |
| ENSG00000273749 | CYFIP1 | -0,153213888 | 0,899244984 | 0,68393538 | 0,965395984 | cytoplasmic FMR1 interacting protein 1 |
| ENSG00000114120 | SLC25A36 | -0,153318789 | 0,8991796 | 0,75411197 | 0,975901808 | solute carrier family 25 member 36 |
| ENSG00000130005 | GAMT | -0,153445556 | 0,899100594 | 0,68961211 | 0,966868992 | guanidinoacetate N-methyltransferase |
| ENSG00000257954 | AC125611,2 | -0,153473671 | 0,899083073 | 0,81268958 | 0,981246822 | FGFR1 oncogene partner 2 (FGFR1OP2) pseudogene |
| ENSG00000251032 | CUL1P1 | -0,15351062 | 0,899060047 | 0,90401008 | 0,992484155 | cullin 1 pseudogene 1 |
| ENSG00000140534 | TICRR | -0,153511935 | 0,899059228 | 0,50446447 | 0,921400408 | TOPBP1 interacting checkpoint and replication regulator |
| ENSG00000105339 | DENND3 | -0,153514293 | 0,899057758 | 0,63104739 | 0,953647802 | DENN domain containing 3 |
| ENSG00000251595 | ABCA11P | -0,153557934 | 0,899030562 | 0,66219346 | 0,959874876 | ATP binding cassette subfamily A member 11, pseudogene |
| ENSG00000251615 | AC104825,1 | -0,153593096 | 0,899008651 | 0,63757026 | 0,956017934 | novel transcript |
| ENSG00000274322 | AL136531,2 | -0,153629676 | 0,898985857 | 0,8860287 | 0,990717729 | novel protein |
| ENSG00000179604 | CDC42EP4 | -0,153638873 | 0,898980126 | 0,52486851 | 0,928616707 | CDC42 effector protein 4 |
| ENSG00000259705 | AC084757,3 | -0,153648171 | 0,898974332 | 0,74141257 | 0,975176043 | novel transcript |
| ENSG00000287467 | AC037459,4 | -0,153713492 | 0,89893363 | 0,91511381 | 0,993361299 | novel transcript, sense intronic to SORBS3 |
| ENSG00000166912 | MTMR10 | -0,153733799 | 0,898920977 | 0,68241161 | 0,964769679 | myotubularin related protein 10 |
| ENSG00000196811 | CHRNG | -0,153756829 | 0,898906627 | 0,83599665 | 0,985134338 | cholinergic receptor nicotinic gamma subunit |
| ENSG00000101868 | POLA1 | -0,153768755 | 0,898899196 | 0,58586608 | 0,945246954 | DNA polymerase alpha 1, catalytic subunit |
| ENSG00000226254 | PTP4A1P3 | -0,153769245 | 0,898898892 | 0,88122285 | 0,990717729 | PTP4A1 pseudogene 3 |
| ENSG00000277209 | RPPH1 | -0,153784662 | 0,898889285 | 0,50908294 | 0,922427526 | ribonuclease P RNA component H1 |
| ENSG00000267682 | AC016590,2 | -0,153882487 | 0,898828336 | 0,8072232 | 0,981213877 | novel transcript, sense intronic to HKR1 |
| ENSG00000185760 | KCNQ5 | -0,153889349 | 0,898824061 | 0,72089339 | 0,972142678 | potassium voltage-gated channel subfamily Q member 5 |
| ENSG00000253477 | AC012213,1 | -0,153911697 | 0,898810138 | 0,7383298 | 0,974353474 | novel transcript |
| ENSG00000237595 | LINC01275 | -0,153924645 | 0,898802072 | 0,88141426 | 0,990717729 | long intergenic non-protein coding RNA 1275 |
| ENSG00000271709 | AC017033,1 | -0,154010153 | 0,898748801 | 0,85251115 | 0,98761642 | novel transcript |
| ENSG00000183723 | CMTM4 | -0,154031634 | 0,89873542 | 0,57488942 | 0,942533938 | CKLF like MARVEL transmembrane domain containing 4 |
| ENSG00000242474 | AC093627,5 | -0,154189805 | 0,898636891 | 0,73248722 | 0,974353474 | novel transcript |
| ENSG00000288472 | AC093627,29 | -0,154189805 | 0,898636891 | 0,73248722 | 0,974353474 | novel transcript |
| ENSG00000286302 | AC084724,1 | -0,154190315 | 0,898636574 | 0,83008922 | 0,984156028 | novel transcript, antisense to NEK1 |

| | | | | | | |
|---|---|---|---|---|---|---|
| ENSG00000165195 | PIGA | -0,154256263 | 0,898595497 | 0,73700642 | 0,974353474 | phosphatidylinositol glycan anchor biosynthesis class A |
| ENSG00000260219 | CD2BP2-DT | -0,154279061 | 0,898581297 | 0,73662693 | 0,974353474 | CD2BP2 divergent transcript |
| ENSG00000275221 | H2AC15 | -0,154372174 | 0,898523304 | 0,56269152 | 0,940865625 | H2A clustered histone 15 |
| ENSG00000167380 | ZNF226 | -0,154375221 | 0,898521405 | 0,65039573 | 0,956899963 | zinc finger protein 226 |
| ENSG00000286811 | AL353138,1 | -0,154432816 | 0,898485536 | 0,89482385 | 0,991296984 | novel transcript |
| ENSG00000184489 | PTP4A3 | -0,154591818 | 0,898386518 | 0,66954059 | 0,961928077 | protein tyrosine phosphatase 4A3 |
| ENSG00000151729 | SLC25A4 | -0,154613455 | 0,898373044 | 0,6163407 | 0,952793934 | solute carrier family 25 member 4 |
| ENSG00000113851 | CRBN | -0,15463077 | 0,898362262 | 0,7173009 | 0,971697135 | cereblon |
| ENSG00000166448 | TMEM130 | -0,154643071 | 0,898354602 | 0,69900437 | 0,968522643 | transmembrane protein 130 |
| ENSG00000277575 | AL162233,1 | -0,154648531 | 0,898351203 | 0,79090171 | 0,979471509 | CAS1 domain containing 1 (CASD1) pseudogene |
| ENSG00000228274 | AL021707,2 | -0,154654672 | 0,898347379 | 0,51782782 | 0,926006346 | novel transcript, antisense to CBY1 |
| ENSG00000251537 | AC005324,2 | -0,154737604 | 0,898295739 | 0,73760243 | 0,974353474 | novel tripartite motif-containing 16 (TRIM16) and CMT1A duplicated region transcript 1 (CDRT1) protein |
| ENSG00000196172 | ZNF681 | -0,154768518 | 0,89827649 | 0,70375867 | 0,968976619 | zinc finger protein 681 |
| ENSG00000239306 | RBM14 | -0,154791361 | 0,898262268 | 0,49632717 | 0,918284563 | RNA binding motif protein 14 |
| ENSG00000143198 | MGST3 | -0,15481441 | 0,898247917 | 0,67495018 | 0,963268551 | microsomal glutathione S-transferase 3 |
| ENSG00000248711 | THUMPD3P1 | -0,154986292 | 0,898140907 | 0,85713293 | 0,988615208 | THUMP domain containing 3 pseudogene 1 |
| ENSG00000282044 | AC083849,3 | -0,15500686 | 0,898128102 | 0,88912281 | 0,990717729 | novel transcript |
| ENSG00000284990 | USP17L2 | -0,155056341 | 0,898097299 | 0,71173911 | 0,97080996 | ubiquitin specific peptidase 17 like family member 2 |
| ENSG00000161610 | HCRT | -0,155121033 | 0,898057028 | 0,91129695 | 0,993222655 | hypocretin neuropeptide precursor |
| ENSG00000153558 | FBXL2 | -0,155129423 | 0,898051806 | 0,5673307 | 0,941425553 | F-box and leucine rich repeat protein 2 |
| ENSG00000170632 | ARMC10 | -0,155141551 | 0,898044257 | 0,57457713 | 0,942475308 | armadillo repeat containing 10 |
| ENSG00000023041 | ZDHHC6 | -0,155280207 | 0,89795795 | 0,70068712 | 0,968522643 | zinc finger DHHC-type palmitoyltransferase 6 |
| ENSG00000151576 | QTRT2 | -0,155294395 | 0,89794912 | 0,50060433 | 0,920520138 | queuine tRNA-ribosyltransferase accessory subunit 2 |
| ENSG00000273783 | AL136040,1 | -0,155304852 | 0,897942611 | 0,78598933 | 0,979320406 | novel transcript, antisense to GTF2A1 |
| ENSG00000131153 | GINS2 | -0,15556905 | 0,897778188 | 0,48526824 | 0,913831238 | GINS complex subunit 2 |
| ENSG00000248971 | KRT8P46 | -0,155802215 | 0,897633102 | 0,71259389 | 0,970812834 | keratin 8 pseudogene 46 |
| ENSG00000260708 | AL118516,1 | -0,155835677 | 0,897612283 | 0,67441447 | 0,963103112 | novel transcript, antisense to TBC1D22A |
| ENSG00000133606 | MKRN1 | -0,155858532 | 0,897598063 | 0,58849702 | 0,946292796 | makorin ring finger protein 1 |
| ENSG00000147419 | CCDC25 | -0,155887313 | 0,897580157 | 0,64093831 | 0,956410077 | coiled-coil domain containing 25 |

| | | | | | | |
|---|---|---|---|---|---|---|
| ENSG00000170142 | UBE2E1 | -0,155897913 | 0,897573562 | 0,75927979 | 0,976120207 | ubiquitin conjugating enzyme E2 E1 |
| ENSG00000153531 | ADPRHL1 | -0,155935781 | 0,897550003 | 0,74135371 | 0,975176043 | ADP-ribosylhydrolase like 1 |
| ENSG00000111728 | ST8SIA1 | -0,155949713 | 0,897541335 | 0,6607333 | 0,959226315 | ST8 alpha-N-acetyl-neuraminide alpha-2,8-sialyltransferase 1 |
| ENSG00000187091 | PLCD1 | -0,155954163 | 0,897538566 | 0,58290625 | 0,944174844 | phospholipase C delta 1 |
| ENSG00000153896 | ZNF599 | -0,155956783 | 0,897536937 | 0,6200735 | 0,952793934 | zinc finger protein 599 |
| ENSG00000116062 | MSH6 | -0,155984311 | 0,897519811 | 0,57456305 | 0,942475308 | mutS homolog 6 |
| ENSG00000285923 | AL160171,1 | -0,15602531 | 0,897494305 | 0,90938691 | 0,992864002 | Novel transcript |
| ENSG00000185607 | ACTBP7 | -0,156041448 | 0,897484266 | 0,85772554 | 0,988897417 | ACTB pseudogene 7 |
| ENSG00000197782 | ZNF780A | -0,156103334 | 0,897445768 | 0,69755216 | 0,968522643 | zinc finger protein 780A |
| ENSG00000280568 | ZNF780A | -0,156103334 | 0,897445768 | 0,69755216 | 0,968522643 | zinc finger protein 780A |
| ENSG00000254842 | LINC02551 | -0,156250475 | 0,897354242 | 0,86644917 | 0,989074806 | long intergenic non-protein coding RNA 2551 |
| ENSG00000231551 | AC245100,4 | -0,15632705 | 0,897306614 | 0,68365083 | 0,965175217 | phosphodiesterase 4D interacting protein (PDE4DIP) pseudogene |
| ENSG00000239213 | NCK1-DT | -0,156338047 | 0,897299774 | 0,58010684 | 0,943828882 | NCK1 divergent transcript |
| ENSG00000173757 | STAT5B | -0,156384743 | 0,897270731 | 0,46990554 | 0,908378389 | signal transducer and activator of transcription 5B |
| ENSG00000286172 | RNVU1-8 | -0,156443181 | 0,897234387 | 0,86719248 | 0,989074806 | RNA, variant U1 small nuclear 8 |
| ENSG00000123607 | TTC21B | -0,156450928 | 0,897229569 | 0,6545454 | 0,957444211 | tetratricopeptide repeat domain 21B |
| ENSG00000245322 | H2AZ1-DT | -0,156469406 | 0,897218078 | 0,72351627 | 0,972913663 | H2AZ1 divergent transcript |
| ENSG00000137103 | TMEM8B | -0,156478683 | 0,897212308 | 0,48226368 | 0,913067988 | transmembrane protein 8B |
| ENSG00000258674 | AC011448,1 | -0,156510599 | 0,89719246 | 0,83819126 | 0,985672131 | novel protein |
| ENSG00000177733 | HNRNPA0 | -0,156511178 | 0,8971921 | 0,46755176 | 0,907940653 | heterogeneous nuclear ribonucleoprotein A0 |
| ENSG00000129071 | MBD4 | -0,156515536 | 0,89718939 | 0,65002977 | 0,956899963 | methyl-CpG binding domain 4, DNA glycosylase |
| ENSG00000242398 | RN7SL800P | -0,156619914 | 0,897124481 | 0,83919087 | 0,985672131 | RNA, 7SL, cytoplasmic 800, pseudogene |
| ENSG00000232912 | RERE-AS1 | -0,15664297 | 0,897110144 | 0,81877011 | 0,98249617 | RERE antisense RNA 1 |
| ENSG00000283050 | GTF2IP12 | -0,156669524 | 0,897093632 | 0,53093293 | 0,931265667 | general transcription factor IIi pseudogene 12 |
| ENSG00000134698 | AGO4 | -0,156669583 | 0,897093596 | 0,65402791 | 0,957444211 | argonaute RISC component 4 |
| ENSG00000135048 | CEMIP2 | -0,156683304 | 0,897085063 | 0,57963604 | 0,943828882 | cell migration inducing hyaluronidase 2 |
| ENSG00000285528 | AC097104,1 | -0,156687307 | 0,897082574 | 0,94406823 | 0,995542144 | novel protein |
| ENSG00000258420 | LONRF2P3 | -0,156714985 | 0,897065364 | 0,90725778 | 0,992590661 | LONRF2 pseudogene 3 |
| ENSG00000198346 | ZNF813 | -0,156796917 | 0,89701442 | 0,65623209 | 0,957444211 | zinc finger protein 813 |
| ENSG00000285872 | AC007240,3 | -0,157069908 | 0,896844701 | 0,8349052 | 0,984905093 | novel transcript |
| ENSG00000149089 | APIP | -0,157090196 | 0,896832089 | 0,65591022 | 0,957444211 | APAF1 interacting protein |

| | | | | | | |
|---|---|---|---|---|---|---|
| ENSG00000197013 | ZNF429 | -0,15714037 | 0,896800899 | 0,58104404 | 0,943919685 | zinc finger protein 429 |
| ENSG00000145348 | TBCK | -0,157161823 | 0,896787564 | 0,60801561 | 0,951163381 | TBC1 domain containing kinase |
| ENSG00000197894 | ADH5 | -0,15717174 | 0,896781399 | 0,63659807 | 0,95557086 | alcohol dehydrogenase 5 (class III), chi polypeptide |
| ENSG00000090612 | ZNF268 | -0,157195575 | 0,896766584 | 0,63945828 | 0,956018518 | zinc finger protein 268 |
| ENSG00000197863 | ZNF790 | -0,157242265 | 0,896737562 | 0,59599799 | 0,948258585 | zinc finger protein 790 |
| ENSG00000207233 | SNORA37 | -0,157289769 | 0,896708035 | 0,78922614 | 0,979462318 | small nucleolar RNA, H/ACA box 37 |
| ENSG00000075089 | ACTR6 | -0,157305061 | 0,896698531 | 0,73135971 | 0,974353474 | actin related protein 6 |
| ENSG00000136425 | CIB2 | -0,157431843 | 0,896619734 | 0,53174255 | 0,931291085 | calcium and integrin binding family member 2 |
| ENSG00000136104 | RNASEH2B | -0,157457751 | 0,896603632 | 0,54162333 | 0,933491609 | ribonuclease H2 subunit B |
| ENSG00000287264 | AP003778,1 | -0,157461606 | 0,896601237 | 0,83500794 | 0,984905093 | novel transcript |
| ENSG00000281801 | KRTAP5-AS1 | -0,157465825 | 0,896598614 | 0,78943173 | 0,979462318 | KRTAP5-1/KRTAP5-2 antisense RNA 1 |
| ENSG00000171720 | HDAC3 | -0,157475145 | 0,896592822 | 0,49279174 | 0,917090312 | histone deacetylase 3 |
| ENSG00000236178 | VARS2 | -0,157495673 | 0,896580065 | 0,53438556 | 0,931584152 | valyl-tRNA synthetase 2, mitochondrial |
| ENSG00000234707 | SEC61G-DT | -0,157501723 | 0,896576305 | 0,83484179 | 0,984905093 | SEC61G divergent transcript |
| ENSG00000213599 | SLX1A-SULT1A3 | -0,157598987 | 0,896515861 | 0,66919198 | 0,961736598 | SLX1A-SULT1A3 readthrough (NMD candidate) |
| ENSG00000227233 | CICP17 | -0,157624968 | 0,896499717 | 0,8915611 | 0,990855862 | capicua transcriptional repressor pseudogene 17 |
| ENSG00000233029 | AC244453,2 | -0,157763792 | 0,896413454 | 0,92706506 | 0,993915537 | novel transcript |
| ENSG00000136450 | SRSF1 | -0,157769182 | 0,896410105 | 0,6849223 | 0,965415049 | serine and arginine rich splicing factor 1 |
| ENSG00000167178 | ISLR2 | -0,157910701 | 0,896322178 | 0,64205443 | 0,956548736 | immunoglobulin superfamily containing leucine rich repeat 2 |
| ENSG00000272995 | AC097505,1 | -0,157940389 | 0,896303733 | 0,92249154 | 0,993797669 | novel transcript, antisense to KCNIP4 |
| ENSG00000174899 | SLC66A1L | -0,158038493 | 0,896242787 | 0,85593215 | 0,988180177 | solute carrier family 66 member 1 like |
| ENSG00000170790 | OR10A2 | -0,15805054 | 0,896235303 | 0,779326 | 0,978587255 | olfactory receptor family 10 subfamily A member 2 |
| ENSG00000157349 | DDX19B | -0,158063023 | 0,896227548 | 0,48495083 | 0,913690171 | DEAD-box helicase 19B |
| ENSG00000136925 | TSTD2 | -0,158090746 | 0,896210326 | 0,60591288 | 0,950927937 | thiosulfate sulfurtransferase like domain containing 2 |
| ENSG00000180628 | PCGF5 | -0,158107878 | 0,896199683 | 0,71139794 | 0,970635643 | polycomb group ring finger 5 |
| ENSG00000065665 | SEC61A2 | -0,15811211 | 0,896197054 | 0,6709363 | 0,962026636 | SEC61 translocon subunit alpha 2 |
| ENSG00000167842 | MIS12 | -0,158119083 | 0,896192723 | 0,71376546 | 0,970812834 | MIS12 kinetochore complex component |
| ENSG00000138767 | CNOT6L | -0,158192206 | 0,896147301 | 0,73420511 | 0,974353474 | CCR4-NOT transcription complex subunit 6 like |
| ENSG00000174796 | THAP6 | -0,158247176 | 0,896113156 | 0,66743815 | 0,9611116 | THAP domain containing 6 |
| ENSG00000283064 | AL353759,1 | -0,158337368 | 0,896057136 | 0,77158357 | 0,978033855 | novel transcript, antisense to HIST1H2BD |
| ENSG00000277969 | AC006449,6 | -0,158391315 | 0,89602363 | 0,72128545 | 0,972142678 | novel transcript |

| | | | | | | |
|---|---|---|---|---|---|---|
| ENSG00000282494 | AC006449,9 | -0,158391315 | 0,89602363 | 0,72128545 | 0,972142678 | novel transcript |
| ENSG00000253636 | AC022893,1 | -0,158417853 | 0,896007148 | 0,62738194 | 0,952793934 | novel transcript |
| ENSG00000156853 | ZNF689 | -0,158437416 | 0,895994998 | 0,50764357 | 0,922140988 | zinc finger protein 689 |
| ENSG00000240991 | RPL23AP67 | -0,158443486 | 0,895991229 | 0,84752328 | 0,987035044 | ribosomal protein L23a pseudogene 67 |
| ENSG00000277687 | AL139407,1 | -0,158495112 | 0,895959167 | 0,74351752 | 0,975310206 | novel transcript, sense intronic to CACUL1 |
| ENSG00000258422 | AL160191,1 | -0,158585473 | 0,895903051 | 0,76379908 | 0,976640468 | novel transcript, antisense to SLC8A3 |
| ENSG00000266173 | STRADA | -0,158614659 | 0,895884927 | 0,43009864 | 0,894889114 | STE20 related adaptor alpha |
| ENSG00000223865 | HLA-DPB1 | -0,158830926 | 0,89575064 | 0,74696155 | 0,975796253 | major histocompatibility complex, class II, DP beta 1 |
| ENSG00000181227 | DLSTP1 | -0,158850457 | 0,895738513 | 0,67664218 | 0,963580269 | dihydrolipoamide S-succinyltransferase pseudogene 1 |
| ENSG00000188993 | LRRC66 | -0,158856955 | 0,895734479 | 0,75226278 | 0,975901808 | leucine rich repeat containing 66 |
| ENSG00000076248 | UNG | -0,158870495 | 0,895726072 | 0,57666669 | 0,943261034 | uracil DNA glycosylase |
| ENSG00000278017 | AC064801,1 | -0,158884931 | 0,895717109 | 0,84531599 | 0,986639902 | novel transcript, antisense to ZCCHC2 |
| ENSG00000188352 | FOCAD | -0,158930557 | 0,895688782 | 0,60372208 | 0,950391097 | focadhesin |
| ENSG00000179399 | GPC5 | -0,159068613 | 0,895603075 | 0,8629033 | 0,989074806 | glypican 5 |
| ENSG00000114503 | NCBP2 | -0,159333488 | 0,89543866 | 0,62656544 | 0,952793934 | nuclear cap binding protein subunit 2 |
| ENSG00000164597 | COG5 | -0,159378197 | 0,895410911 | 0,73187089 | 0,974353474 | component of oligomeric golgi complex 5 |
| ENSG00000260260 | SNHG19 | -0,159408231 | 0,895392271 | 0,6763731 | 0,963580269 | small nucleolar RNA host gene 19 |
| ENSG00000169607 | CKAP2L | -0,159461138 | 0,895359435 | 0,69218283 | 0,967236299 | cytoskeleton associated protein 2 like |
| ENSG00000205643 | CDPF1 | -0,159535413 | 0,895313339 | 0,58766106 | 0,946111354 | cysteine rich DPF motif domain containing 1 |
| ENSG00000263072 | ZNF213-AS1 | -0,159571441 | 0,895290981 | 0,60390317 | 0,950551453 | ZNF213 antisense RNA 1 (head to head) |
| ENSG00000134343 | ANO3 | -0,159578465 | 0,895286623 | 0,80720632 | 0,981213877 | anoctamin 3 |
| ENSG00000137764 | MAP2K5 | -0,15959078 | 0,895278981 | 0,5684458 | 0,941656837 | mitogen-activated protein kinase kinase 5 |
| ENSG00000133627 | ACTR3B | -0,159636127 | 0,89525084 | 0,54763562 | 0,935404974 | actin related protein 3B |
| ENSG00000229992 | HMGB3P9 | -0,159636267 | 0,895250754 | 0,77663807 | 0,978587255 | high mobility group box 3 pseudogene 9 |
| ENSG00000183444 | OR7E38P | -0,159732192 | 0,89519123 | 0,66728879 | 0,9611116 | olfactory receptor family 7 subfamily E member 38 pseudogene |
| ENSG00000197837 | H4-16 | -0,159750271 | 0,895180012 | 0,61588951 | 0,952793934 | H4 histone 16 |
| ENSG00000130695 | CEP85 | -0,159750942 | 0,895179596 | 0,50427898 | 0,921400408 | centrosomal protein 85 |
| ENSG00000085224 | ATRX | -0,159754629 | 0,895177308 | 0,62481443 | 0,952793934 | ATRX chromatin remodeler |
| ENSG00000268987 | AC011462,3 | -0,159760651 | 0,895173572 | 0,8677578 | 0,989074806 | novel transcript, sense intronic to BCKDHA |
| ENSG00000254064 | AC105206,2 | -0,159858598 | 0,895112799 | 0,81683085 | 0,981977498 | novel transcript |
| ENSG00000265574 | WDR45BP1 | -0,159865327 | 0,895108624 | 0,88678703 | 0,990717729 | WD repeat domain 45B pseudogene 1 |

| | | | | | | |
|---|---|---|---|---|---|---|
| ENSG00000284627 | AC079781,4 | -0,159875956 | 0,895102029 | 0,88897728 | 0,990717729 | RNA binding protein, fox-1 homolog 1 (RBFOX1) pseudogene |
| ENSG00000259609 | AC027807,1 | -0,159914357 | 0,895078204 | 0,90095701 | 0,991883933 | family with sequence similarity 154, member B (Fam154b) pseudogene |
| ENSG00000275401 | AL391095,3 | -0,159963895 | 0,89504747 | 0,74622236 | 0,975778823 | novel transcript |
| ENSG00000136169 | SETDB2 | -0,159970416 | 0,895043424 | 0,6758141 | 0,963546698 | SET domain bifurcated histone lysine methyltransferase 2 |
| ENSG00000110583 | NAA40 | -0,159975033 | 0,89504056 | 0,44239039 | 0,89933631 | N-alpha-acetyltransferase 40, NatD catalytic subunit |
| ENSG00000155542 | SETD9 | -0,159988309 | 0,895032324 | 0,73179801 | 0,974353474 | SET domain containing 9 |
| ENSG00000270846 | MYG1P1 | -0,159992931 | 0,895029457 | 0,85378181 | 0,987861659 | MYG1 pseudogene 1 |
| ENSG00000197134 | ZNF257 | -0,160013708 | 0,895016567 | 0,88327943 | 0,990717729 | zinc finger protein 257 |
| ENSG00000183605 | SFXN4 | -0,160132848 | 0,894942658 | 0,5806471 | 0,943832501 | sideroflexin 4 |
| ENSG00000278434 | AC023830,3 | -0,160132919 | 0,894942614 | 0,72623305 | 0,973344786 | novel transcript, sense intronic to CDIP1 |
| ENSG00000223704 | LINC01422 | -0,160159977 | 0,894925829 | 0,65446795 | 0,957444211 | long intergenic non-protein coding RNA 1422 |
| ENSG00000223393 | AL118511,1 | -0,160187963 | 0,894908469 | 0,7494132 | 0,975901808 | novel transcript |
| ENSG00000204304 | PBX2 | -0,160224249 | 0,894885961 | 0,60606967 | 0,950927937 | PBX homeobox 2 |
| ENSG00000146963 | LUC7L2 | -0,160236415 | 0,894878415 | 0,55368524 | 0,937338325 | LUC7 like 2, pre-mRNA splicing factor |
| ENSG00000228801 | AC064807,1 | -0,160338888 | 0,894814855 | 0,5061944 | 0,921780819 | novel transcript |
| ENSG00000245248 | USP2-AS1 | -0,160381378 | 0,894788501 | 0,79574617 | 0,979879951 | USP2 antisense RNA 1 (head to head) |
| ENSG00000176933 | TOB2P1 | -0,160445927 | 0,894748467 | 0,78669632 | 0,979320406 | transducer of ERBB2, 2 pseudogene 1 |
| ENSG00000204574 | ABCF1 | -0,16048725 | 0,89472284 | 0,65530919 | 0,957444211 | ATP binding cassette subfamily F member 1 |
| ENSG00000233296 | TMEM18-DT | -0,160516101 | 0,894704947 | 0,78708582 | 0,979424965 | TMEM18 divergent transcript |
| ENSG00000178150 | ZNF114 | -0,160538874 | 0,894690825 | 0,59167138 | 0,947416795 | zinc finger protein 114 |
| ENSG00000197889 | MEIG1 | -0,160590127 | 0,89465904 | 0,79128009 | 0,979471509 | meiosis/spermiogenesis associated 1 |
| ENSG00000244314 | RN7SL36P | -0,160596883 | 0,894654851 | 0,87808326 | 0,990109814 | RNA, 7SL, cytoplasmic 36, pseudogene |
| ENSG00000285210 | LINC02795 | -0,160640942 | 0,894627529 | 0,76677911 | 0,977201891 | long intergenic non-protein coding RNA 2795 |
| ENSG00000268266 | AC003005,2 | -0,160687682 | 0,894598546 | 0,87712573 | 0,990109814 | novel transcript |
| ENSG00000269713 | NBPF9 | -0,160874326 | 0,894482817 | 0,47164147 | 0,908896785 | NBPF member 9 |
| ENSG00000116478 | HDAC1 | -0,160901753 | 0,894465812 | 0,50378069 | 0,921200643 | histone deacetylase 1 |
| ENSG00000266368 | AC005410,2 | -0,160965965 | 0,894426002 | 0,88900586 | 0,990717729 | novel transcript, antisense to DNAH9 |
| ENSG00000120699 | EXOSC8 | -0,160995389 | 0,89440776 | 0,65505349 | 0,957444211 | exosome component 8 |
| ENSG00000046651 | OFD1 | -0,16100765 | 0,894400159 | 0,61100855 | 0,951873283 | OFD1 centriole and centriolar satellite protein |
| ENSG00000215256 | DHRS4-AS1 | -0,161043559 | 0,894377898 | 0,62865112 | 0,952793934 | DHRS4 antisense RNA 1 |

| | | | | | | |
|---|---|---|---|---|---|---|
| ENSG00000197363 | ZNF517 | -0,161140361 | 0,894317889 | 0,67247205 | 0,962290553 | zinc finger protein 517 |
| ENSG00000157992 | KRTCAP3 | -0,161262119 | 0,894242415 | 0,75337372 | 0,975901808 | keratinocyte associated protein 3 |
| ENSG00000282971 | AC044810,9 | -0,1613224 | 0,894205051 | 0,87119976 | 0,989217334 | novel transcript, sense overlapping OR5P2, OR5P3 and OR5P1P |
| ENSG00000147642 | SYBU | -0,161335398 | 0,894196994 | 0,61430322 | 0,952793934 | syntabulin |
| ENSG00000138459 | SLC35A5 | -0,161389031 | 0,894163753 | 0,74165966 | 0,975176043 | solute carrier family 35 member A5 |
| ENSG00000120647 | CCDC77 | -0,161401726 | 0,894155885 | 0,56818394 | 0,94163744 | coiled-coil domain containing 77 |
| ENSG00000232864 | NUCKS1P1 | -0,161414705 | 0,89414784 | 0,71817575 | 0,97178302 | nuclear casein kinase and cyclin dependent kinase substrate 1 pseudogene 1 |
| ENSG00000138182 | KIF20B | -0,161514436 | 0,894086032 | 0,67408239 | 0,96304746 | kinesin family member 20B |
| ENSG00000178567 | EPM2AIP1 | -0,161535455 | 0,894073006 | 0,71647538 | 0,971438758 | EPM2A interacting protein 1 |
| ENSG00000100285 | NEFH | -0,161607405 | 0,894028418 | 0,75290571 | 0,975901808 | neurofilament heavy |
| ENSG00000250848 | AC021087,2 | -0,161655312 | 0,893998731 | 0,75527209 | 0,975901808 | protein phosphatase 4, regulatory subunit 2 (PPP4R2) pseudogene |
| ENSG00000164695 | CHMP4C | -0,161674344 | 0,893986937 | 0,7362537 | 0,974353474 | charged multivesicular body protein 4C |
| ENSG00000198860 | TSEN15 | -0,161686477 | 0,893979419 | 0,69436288 | 0,967417311 | tRNA splicing endonuclease subunit 15 |
| ENSG00000034713 | GABARAPL2 | -0,161722456 | 0,893957124 | 0,66257805 | 0,959897678 | GABA type A receptor associated protein like 2 |
| ENSG00000158246 | TENT5B | -0,161736382 | 0,893948495 | 0,53746709 | 0,932883277 | terminal nucleotidyltransferase 5B |
| ENSG00000226259 | GTF2H2B | -0,161755962 | 0,893936363 | 0,69747367 | 0,968522643 | general transcription factor IIH subunit 2B (pseudogene) |
| ENSG00000275246 | TLCD2 | -0,161786785 | 0,893917264 | 0,80745368 | 0,981213877 | TLC domain containing 2 |
| ENSG00000234197 | ETV5-AS1 | -0,161809718 | 0,893903055 | 0,74663779 | 0,975796253 | ETV5 antisense RNA 1 |
| ENSG00000213079 | SCAF8 | -0,161811549 | 0,89390192 | 0,60486172 | 0,950927937 | SR-related CTD associated factor 8 |
| ENSG00000100429 | HDAC10 | -0,161958942 | 0,893810599 | 0,69180306 | 0,967204896 | histone deacetylase 10 |
| ENSG00000119787 | ATL2 | -0,162003058 | 0,893783268 | 0,67889994 | 0,964151912 | atlastin GTPase 2 |
| ENSG00000162065 | TBC1D24 | -0,162012008 | 0,893777723 | 0,63136268 | 0,953801536 | TBC1 domain family member 24 |
| ENSG00000177839 | PCDHB9 | -0,162028888 | 0,893767266 | 0,65304387 | 0,95732962 | protocadherin beta 9 |
| ENSG00000270277 | AC009948,1 | -0,162125697 | 0,893707294 | 0,73482306 | 0,974353474 | novel transcript, antisense to titin |
| ENSG00000163882 | POLR2H | -0,1621512 | 0,893691495 | 0,52597361 | 0,929094237 | RNA polymerase II, I and III subunit H |
| ENSG00000213204 | AL049697,1 | -0,16217435 | 0,893677155 | 0,78106198 | 0,978597905 | novel transcript, C6orf165-SLC35A1 readthrough |
| ENSG00000270236 | AL162730,1 | -0,162250115 | 0,893630224 | 0,90819136 | 0,992595473 | adenylate kinase 3 (AK3) pseudogene |
| ENSG00000274810 | NPHP3-ACAD1: | -0,162260789 | 0,893623612 | 0,67401443 | 0,963044976 | NPHP3-ACAD11 readthrough (NMD candidate) |
| ENSG00000249731 | AC126768,2 | -0,162269448 | 0,893618248 | 0,52489732 | 0,928616707 | novel transcript |

| | | | | | | |
|---|---|---|---|---|---|---|
| ENSG00000159263 | SIM2 | -0,16227448 | 0,893615131 | 0,81017176 | 0,981213877 | SIM bHLH transcription factor 2 |
| ENSG00000111602 | TIMELESS | -0,162333402 | 0,893578636 | 0,45654664 | 0,903545728 | timeless circadian regulator |
| ENSG00000226266 | AC009961,1 | -0,162343621 | 0,893572306 | 0,7409915 | 0,975146552 | novel transcript |
| ENSG00000254481 | PTP4A2P2 | -0,162370045 | 0,89355594 | 0,79749865 | 0,980161849 | PTP4A2 pseudogene 2 |
| ENSG00000137094 | DNAJB5 | -0,162445381 | 0,89350928 | 0,55363401 | 0,937334216 | DnaJ heat shock protein family (Hsp40) member B5 |
| ENSG00000214942 | AC113167,1 | -0,16254827 | 0,89344556 | 0,75273329 | 0,975901808 | novel transcript |
| ENSG00000130755 | GMFG | -0,162549872 | 0,893444568 | 0,81983624 | 0,98249617 | glia maturation factor gamma |
| ENSG00000072954 | TMEM38A | -0,162606188 | 0,893409693 | 0,80413726 | 0,980875302 | transmembrane protein 38A |
| ENSG00000155561 | NUP205 | -0,162614076 | 0,893404808 | 0,62872946 | 0,952793934 | nucleoporin 205 |
| ENSG00000271894 | AC007744,1 | -0,162626692 | 0,893396995 | 0,85464552 | 0,987861659 | novel transcript |
| ENSG00000018510 | AGPS | -0,162627 | 0,893396805 | 0,70433671 | 0,969132783 | alkylglycerone phosphate synthase |
| ENSG00000008988 | RPS20 | -0,162635986 | 0,89339124 | 0,59934051 | 0,949084827 | ribosomal protein S20 |
| ENSG00000124532 | MRS2 | -0,162755356 | 0,893317323 | 0,58602852 | 0,945246954 | magnesium transporter MRS2 |
| ENSG00000258738 | AL121603,2 | -0,162806976 | 0,893285361 | 0,64151355 | 0,956465487 | novel transcript, antisense to BAZ1A |
| ENSG00000215695 | RSC1A1 | -0,162807478 | 0,89328505 | 0,9335975 | 0,994479239 | regulator of solute carriers 1 |
| ENSG00000237135 | DDX10P1 | -0,162815873 | 0,893279852 | 0,80044423 | 0,980299611 | DEAD-box helicase 10 pseudogene 1 |
| ENSG00000102977 | ACD | -0,162849576 | 0,893258984 | 0,55540988 | 0,937793851 | ACD shelterin complex subunit and telomerase recruitment factor |
| ENSG00000196511 | TPK1 | -0,162861426 | 0,893251647 | 0,66848026 | 0,961351573 | thiamin pyrophosphokinase 1 |
| ENSG00000279714 | Z95114,2 | -0,162897798 | 0,893229127 | 0,84652784 | 0,986939308 | novel transcript |
| ENSG00000255021 | AC093496,1 | -0,163085357 | 0,89311301 | 0,83044973 | 0,984156028 | novel transcript |
| ENSG00000163918 | RFC4 | -0,163125441 | 0,893088196 | 0,6853936 | 0,965475755 | replication factor C subunit 4 |
| ENSG00000243970 | PPIEL | -0,163149484 | 0,893073312 | 0,68746772 | 0,965965731 | peptidylprolyl isomerase E like pseudogene |
| ENSG00000275079 | LPCAT1 | -0,163184987 | 0,893051335 | 0,50012738 | 0,92041337 | lysophosphatidylcholine acyltransferase 1 |
| ENSG00000259548 | AC021483,1 | -0,163197727 | 0,893043449 | 0,89115804 | 0,990795338 | novel transcript |
| ENSG00000226002 | GTF2IP14 | -0,16331272 | 0,89297227 | 0,90263622 | 0,992230188 | general transcription factor IIi pseudogene 14 |
| ENSG00000275714 | H3C1 | -0,163320466 | 0,892967476 | 0,70692984 | 0,969565008 | H3 clustered histone 1 |
| ENSG00000104205 | SGK3 | -0,163376206 | 0,892932975 | 0,64633436 | 0,956899963 | serum/glucocorticoid regulated kinase family member 3 |
| ENSG00000060339 | CCAR1 | -0,163404291 | 0,892915593 | 0,52763717 | 0,929546857 | cell division cycle and apoptosis regulator 1 |
| ENSG00000104973 | MED25 | -0,163418251 | 0,892906953 | 0,52209959 | 0,927599011 | mediator complex subunit 25 |
| ENSG00000164754 | RAD21 | -0,163430529 | 0,892899354 | 0,68689875 | 0,965769168 | RAD21 cohesin complex component |
| ENSG00000136682 | CBWD2 | -0,163543539 | 0,892829413 | 0,56720605 | 0,941425553 | COBW domain containing 2 |

| | | | | | | |
|---|---|---|---|---|---|---|
| ENSG00000227885 | AL590652,1 | -0,163634651 | 0,892773029 | 0,73419581 | 0,974353474 | novel transcript |
| ENSG00000286834 | AL360020,1 | -0,163693232 | 0,892736779 | 0,4951427 | 0,917777383 | novel transcript |
| ENSG00000104835 | SARS2 | -0,163742184 | 0,892706488 | 0,51707918 | 0,925746338 | seryl-tRNA synthetase 2, mitochondrial |
| ENSG00000228933 | AC107419,1 | -0,163748206 | 0,892702762 | 0,75708716 | 0,975901808 | novel transcript |
| ENSG00000136897 | MRPL50 | -0,163748944 | 0,892702305 | 0,76487631 | 0,976722333 | mitochondrial ribosomal protein L50 |
| ENSG00000242110 | AMACR | -0,16385274 | 0,892638081 | 0,66286612 | 0,959897678 | alpha-methylacyl-CoA racemase |
| ENSG00000198408 | OGA | -0,16389332 | 0,892612973 | 0,55627527 | 0,938046927 | O-GlcNAcase |
| ENSG00000189423 | USP32P3 | -0,163902282 | 0,892607428 | 0,48633498 | 0,914091794 | ubiquitin specific peptidase 32 pseudogene 3 |
| ENSG00000224525 | AL591686,1 | -0,163953183 | 0,892575936 | 0,75967108 | 0,976120207 | novel transcript, antisense to PLD5 |
| ENSG00000188060 | RAB42 | -0,164026584 | 0,892530525 | 0,6808034 | 0,964654539 | RAB42, member RAS oncogene family |
| ENSG00000248098 | BCKDHA | -0,164046271 | 0,892518346 | 0,5396064 | 0,933350549 | branched chain keto acid dehydrogenase E1 subunit alpha |
| ENSG00000238083 | LRRC37A2 | -0,164106898 | 0,89248084 | 0,43696344 | 0,896876641 | leucine rich repeat containing 37 member A2 |
| ENSG00000235251 | AC093155,2 | -0,164214595 | 0,892414219 | 0,86115091 | 0,989074806 | pseudogene similar to part of SUMO-1 activating enzyme subunit 2 (UBA2) |
| ENSG00000135914 | HTR2B | -0,164241402 | 0,892397637 | 0,70589404 | 0,969503793 | 5-hydroxytryptamine receptor 2B |
| ENSG00000131668 | BARX1 | -0,164287597 | 0,892369063 | 0,82218318 | 0,982952211 | BARX homeobox 1 |
| ENSG00000156574 | NODAL | -0,164328176 | 0,892343963 | 0,71068353 | 0,970482408 | nodal growth differentiation factor |
| ENSG00000157214 | STEAP2 | -0,164342699 | 0,89233498 | 0,6330189 | 0,954408281 | STEAP2 metalloreductase |
| ENSG00000213860 | RPL21P75 | -0,16439529 | 0,892302453 | 0,92161628 | 0,993787755 | ribosomal protein L21 pseudogene 75 |
| ENSG00000283341 | AC068205,2 | -0,164506136 | 0,892233897 | 0,57735162 | 0,94337651 | novel transcript |
| ENSG00000123815 | COQ8B | -0,164575033 | 0,892191288 | 0,59015318 | 0,947262972 | coenzyme Q8B |
| ENSG00000166938 | DIS3L | -0,164609379 | 0,892170049 | 0,52729558 | 0,929454507 | DIS3 like exosome 3'-5' exoribonuclease |
| ENSG00000196510 | ANAPC7 | -0,164611619 | 0,892168664 | 0,57560502 | 0,942700009 | anaphase promoting complex subunit 7 |
| ENSG00000180929 | GPR62 | -0,164636031 | 0,892153567 | 0,81179341 | 0,981213877 | G protein-coupled receptor 62 |
| ENSG00000250317 | SMIM20 | -0,164718504 | 0,892102568 | 0,71015722 | 0,970482408 | small integral membrane protein 20 |
| ENSG00000185515 | BRCC3 | -0,164767374 | 0,892072349 | 0,63455732 | 0,955149453 | BRCA1/BRCA2-containing complex subunit 3 |
| ENSG00000270558 | AC025449,1 | -0,164929135 | 0,891972332 | 0,7821509 | 0,978597905 | chromobox homolog 3 (CBX3) pseudogene |
| ENSG00000168538 | TRAPPC11 | -0,164971311 | 0,891946256 | 0,66683111 | 0,9611116 | trafficking protein particle complex 11 |
| ENSG00000215875 | ST13P20 | -0,165029137 | 0,891910506 | 0,86439511 | 0,989074806 | ST13, Hsp70 interacting protein pseudogene 20 |
| ENSG00000188234 | AGAP4 | -0,165211773 | 0,891797603 | 0,48091604 | 0,912907448 | ArfGAP with GTPase domain, ankyrin repeat and PH domain 4 |
| ENSG00000134690 | CDCA8 | -0,165219119 | 0,891793062 | 0,58441419 | 0,944741266 | cell division cycle associated 8 |

| | | | | | | |
|---|---|---|---|---|---|---|
| ENSG00000282514 | AC010724,6 | -0,165256389 | 0,891770024 | 0,83789372 | 0,98566906 | dynamin 1 (DNM1) pseudogene |
| ENSG00000113805 | CNTN3 | -0,165256537 | 0,891769932 | 0,68432534 | 0,965415049 | contactin 3 |
| ENSG00000281649 | EBLN3P | -0,165295388 | 0,891745918 | 0,69286668 | 0,967271667 | endogenous Bornavirus like nucleoprotein 3, pseudogene |
| ENSG00000054967 | RELT | -0,165330976 | 0,891723921 | 0,63074496 | 0,953375088 | RELT TNF receptor |
| ENSG00000132952 | USPL1 | -0,165334704 | 0,891721617 | 0,67618873 | 0,963580269 | ubiquitin specific peptidase like 1 |
| ENSG00000279278 | AC245060,6 | -0,165356666 | 0,891708043 | 0,49384061 | 0,917303403 | novel transcript |
| ENSG00000198824 | CHAMP1 | -0,165411035 | 0,891674438 | 0,50708739 | 0,921951873 | chromosome alignment maintaining phosphoprotein 1 |
| ENSG00000111581 | NUP107 | -0,165448063 | 0,891651553 | 0,60623919 | 0,950927937 | nucleoporin 107 |
| ENSG00000285825 | CCDC15-DT | -0,165465777 | 0,891640605 | 0,73231326 | 0,974353474 | CCDC15 divergent transcript |
| ENSG00000204792 | LINC01291 | -0,165517739 | 0,891608491 | 0,66630629 | 0,961079583 | long intergenic non-protein coding RNA 1291 |
| ENSG00000286579 | AC023483,1 | -0,165593727 | 0,891561531 | 0,8843245 | 0,990717729 | novel transcript, antisense to SETMAR |
| ENSG00000184588 | PDE4B | -0,165626026 | 0,891541571 | 0,69340439 | 0,967340633 | phosphodiesterase 4B |
| ENSG00000108474 | PIGL | -0,165742017 | 0,891469894 | 0,64397533 | 0,956633898 | phosphatidylinositol glycan anchor biosynthesis class L |
| ENSG00000127989 | MTERF1 | -0,165783952 | 0,891443983 | 0,70049994 | 0,968522643 | mitochondrial transcription termination factor 1 |
| ENSG00000230316 | FEZF1-AS1 | -0,165815313 | 0,891424605 | 0,66819723 | 0,961289023 | FEZF1 antisense RNA 1 |
| ENSG00000237515 | SHISA9 | -0,165829162 | 0,891416047 | 0,4745163 | 0,91069468 | shisa family member 9 |
| ENSG00000135999 | EPC2 | -0,165855125 | 0,891400006 | 0,66816629 | 0,961289023 | enhancer of polycomb homolog 2 |
| ENSG00000235919 | ASH1L-AS1 | -0,165885168 | 0,891381443 | 0,6748454 | 0,963268551 | ASH1L antisense RNA 1 |
| ENSG00000172366 | MCRIP2 | -0,165981699 | 0,891321802 | 0,67397873 | 0,963037987 | MAPK regulated corepressor interacting protein 2 |
| ENSG00000101407 | TTI1 | -0,166011896 | 0,891303146 | 0,59487409 | 0,947469769 | TELO2 interacting protein 1 |
| ENSG00000178741 | COX5A | -0,166016912 | 0,891300047 | 0,70196219 | 0,968742626 | cytochrome c oxidase subunit 5A |
| ENSG00000253553 | AC090578,1 | -0,166063365 | 0,891271349 | 0,68006406 | 0,964580199 | novel transcript, antisense to MMP16 |
| ENSG00000280591 | AC237191,1 | -0,166136702 | 0,891226044 | 0,84999469 | 0,987036295 | novel transcript |
| ENSG00000281099 | AC244167,1 | -0,166136702 | 0,891226044 | 0,84999469 | 0,987036295 | novel transcript |
| ENSG00000171116 | HSFX1 | -0,166170606 | 0,8912051 | 0,7065408 | 0,969503793 | heat shock transcription factor family, X-linked 1 |
| ENSG00000178904 | DPY19L3 | -0,166441136 | 0,891038 | 0,7126449 | 0,970812834 | dpy-19 like C-mannosyltransferase 3 |
| ENSG00000185271 | KLHL33 | -0,166460093 | 0,891026291 | 0,47689825 | 0,91143003 | kelch like family member 33 |
| ENSG00000156384 | SFR1 | -0,166529001 | 0,890983734 | 0,68313582 | 0,964965151 | SWI5 dependent homologous recombination repair protein 1 |
| ENSG00000282905 | NFKBIB | -0,166529922 | 0,890983165 | 0,81153897 | 0,981213877 | NFKB inhibitor beta |

| | | | | | | |
|---|---|---|---|---|---|---|
| ENSG00000100583 | SAMD15 | -0,166544524 | 0,890974148 | 0,61248987 | 0,952413269 | sterile alpha motif domain containing 15 |
| ENSG00000168038 | ULK4 | -0,166545142 | 0,890973765 | 0,58532781 | 0,945213113 | unc-51 like kinase 4 |
| ENSG00000165494 | PCF11 | -0,166553013 | 0,890968905 | 0,53337166 | 0,931381467 | PCF11 cleavage and polyadenylation factor subunit |
| ENSG00000164904 | ALDH7A1 | -0,166575893 | 0,890954775 | 0,44659461 | 0,901644586 | aldehyde dehydrogenase 7 family member A1 |
| ENSG00000271966 | AC021321,1 | -0,166622944 | 0,890925718 | 0,69884989 | 0,968522643 | novel transcript |
| ENSG00000015153 | YAF2 | -0,166687828 | 0,890885651 | 0,63988055 | 0,956018518 | YY1 associated factor 2 |
| ENSG00000131503 | ANKHD1 | -0,16677505 | 0,890831791 | 0,61599004 | 0,952793934 | ankyrin repeat and KH domain containing 1 |
| ENSG00000246089 | AC016065,1 | -0,166817201 | 0,890805765 | 0,59656693 | 0,948282297 | novel transcript |
| ENSG00000238035 | AC138035,1 | -0,166822719 | 0,890802357 | 0,58818761 | 0,946176848 | novel transcript |
| ENSG00000198804 | MT-CO1 | -0,167032793 | 0,890672655 | 0,68777731 | 0,966064784 | mitochondrially encoded cytochrome c oxidase I |
| ENSG00000057935 | MTA3 | -0,167044332 | 0,890665531 | 0,56543497 | 0,940865625 | metastasis associated 1 family member 3 |
| ENSG00000282377 | AC011295,3 | -0,167045415 | 0,890664862 | 0,75893256 | 0,976120207 | chondroitin sulfate proteoglycan 4 (CSPG4) pseudogene |
| ENSG00000253973 | AC079296,1 | -0,167123169 | 0,890616862 | 0,78055846 | 0,978597905 | novel transcript |
| ENSG00000126838 | PZP | -0,16713138 | 0,890611793 | 0,7602111 | 0,97624106 | PZP alpha-2-macroglobulin like |
| ENSG00000138756 | BMP2K | -0,16713714 | 0,890608237 | 0,7180079 | 0,971780402 | BMP2 inducible kinase |
| ENSG00000062370 | ZNF112 | -0,16714739 | 0,890601909 | 0,63996185 | 0,956074313 | zinc finger protein 112 |
| ENSG00000256407 | AL357673,1 | -0,167165789 | 0,890590552 | 0,89703328 | 0,991644439 | novel transcript |
| ENSG00000196104 | SPOCK3 | -0,167193079 | 0,890573705 | 0,81162113 | 0,981213877 | SPARC (osteonectin), cwcv and kazal like domains proteoglycan 3 |
| ENSG00000132330 | SCLY | -0,167204887 | 0,890566416 | 0,46760911 | 0,907940653 | selenocysteine lyase |
| ENSG00000114346 | ECT2 | -0,16721899 | 0,890557711 | 0,69880838 | 0,968522643 | epithelial cell transforming 2 |
| ENSG00000170089 | AC106795,1 | -0,167273788 | 0,890523885 | 0,66703325 | 0,9611116 | THO complex 3 (THOC3) pseudogene |
| ENSG00000224859 | POLR1H | -0,167363579 | 0,890468462 | 0,57381131 | 0,942475308 | RNA polymerase I subunit H |
| ENSG00000128699 | ORMDL1 | -0,167413699 | 0,890437527 | 0,75772305 | 0,976120207 | ORMDL sphingolipid biosynthesis regulator 1 |
| ENSG00000163001 | CFAP36 | -0,16744767 | 0,89041656 | 0,47723069 | 0,91143003 | cilia and flagella associated protein 36 |
| ENSG00000160753 | RUSC1 | -0,167599629 | 0,890322778 | 0,5489717 | 0,936040289 | RUN and SH3 domain containing 1 |
| ENSG00000273254 | AF129075,2 | -0,167649673 | 0,890291895 | 0,79942104 | 0,980161849 | novel transcript, antisense to USP16 |
| ENSG00000227999 | MTND5P1 | -0,167650578 | 0,890291336 | 0,75865555 | 0,976120207 | MT-ND5 pseudogene 1 |
| ENSG00000162385 | MAGOH | -0,167663968 | 0,890283074 | 0,68525295 | 0,965425099 | mago homolog, exon junction complex subunit |
| ENSG00000225822 | UBXN7-AS1 | -0,167679855 | 0,89027327 | 0,80222791 | 0,980710009 | UBXN7 antisense RNA 1 |
| ENSG00000122482 | ZNF644 | -0,167743164 | 0,890234203 | 0,71023922 | 0,970482408 | zinc finger protein 644 |
| ENSG00000138385 | SSB | -0,16774338 | 0,89023407 | 0,63186587 | 0,953915351 | small RNA binding exonuclease protection factor La |

| | | | | | | |
|---|---|---|---|---|---|---|
| ENSG00000169203 | NPIPB12 | -0,167804716 | 0,890196223 | 0,58785063 | 0,946129322 | nuclear pore complex interacting protein family member B12 |
| ENSG00000264364 | DYNLL2 | -0,167946901 | 0,890108493 | 0,62618248 | 0,952793934 | dynein light chain LC8-type 2 |
| ENSG00000163521 | GLB1L | -0,167962947 | 0,890098593 | 0,51982577 | 0,926822772 | galactosidase beta 1 like |
| ENSG00000230267 | HERC2P4 | -0,167968937 | 0,890094898 | 0,7045628 | 0,969176848 | hect domain and RLD 2 pseudogene 4 |
| ENSG00000105968 | H2AZ2 | -0,168003612 | 0,890073505 | 0,54517021 | 0,934923038 | H2A,Z variant histone 2 |
| ENSG00000231148 | HMGB1P7 | -0,168040834 | 0,890050541 | 0,8301064 | 0,984156028 | high mobility group box 1 pseudogene 7 |
| ENSG00000135972 | MRPS9 | -0,168060581 | 0,890038358 | 0,65496914 | 0,957444211 | mitochondrial ribosomal protein S9 |
| ENSG00000267302 | RNFT1-DT | -0,168081593 | 0,890025395 | 0,76192359 | 0,976315588 | RNFT1 divergent transcript |
| ENSG00000223705 | NSUN5P1 | -0,168083355 | 0,890024309 | 0,52783825 | 0,929604268 | NSUN5 pseudogene 1 |
| ENSG00000066923 | STAG3 | -0,168084452 | 0,890023632 | 0,52740188 | 0,929454507 | stromal antigen 3 |
| ENSG00000275560 | AC008115,3 | -0,168092584 | 0,890018615 | 0,71600438 | 0,971360663 | novel transcript |
| ENSG00000233388 | AL022334,1 | -0,168155981 | 0,889979505 | 0,88784749 | 0,990717729 | tRNA methyltransferase 11 homolog (S, cerevisiae) (TRMT11) pseudogene |
| ENSG00000240853 | RN7SL328P | -0,168329111 | 0,88987271 | 0,91807969 | 0,993361299 | RNA, 7SL, cytoplasmic 328, pseudogene |
| ENSG00000272256 | AC044849,1 | -0,168329111 | 0,88987271 | 0,91807969 | 0,993361299 | novel transcript, antisense to TMEM66 |
| ENSG00000232388 | SMIM26 | -0,168339907 | 0,889866051 | 0,65407005 | 0,957444211 | small integral membrane protein 26 |
| ENSG00000274987 | AC092794,1 | -0,168406236 | 0,88982514 | 0,8643875 | 0,989074806 | novel transcript, antisense to KRAS |
| ENSG00000198089 | SFI1 | -0,168470551 | 0,889785473 | 0,63748324 | 0,956017934 | SFI1 centrin binding protein |
| ENSG00000103769 | RAB11A | -0,168531127 | 0,889748113 | 0,6979764 | 0,968522643 | RAB11A, member RAS oncogene family |
| ENSG00000140265 | ZSCAN29 | -0,16853465 | 0,88974594 | 0,50094191 | 0,920520138 | zinc finger and SCAN domain containing 29 |
| ENSG00000250318 | AC003072,1 | -0,168634426 | 0,889684408 | 0,7540454 | 0,975901808 | ribosomal protein L13a (RPL13A) pseudogene |
| ENSG00000142731 | PLK4 | -0,168654876 | 0,889671797 | 0,68774977 | 0,966064784 | polo like kinase 4 |
| ENSG00000264589 | MAPT-AS1 | -0,168677847 | 0,889657631 | 0,81227431 | 0,981213877 | MAPT antisense RNA 1 |
| ENSG00000137726 | FXYD6 | -0,168691103 | 0,889649457 | 0,44961959 | 0,902062594 | FXYD domain containing ion transport regulator 6 |
| ENSG00000005100 | DHX33 | -0,168707564 | 0,889639306 | 0,55324648 | 0,936992468 | DEAH-box helicase 33 |
| ENSG00000043514 | TRIT1 | -0,168741327 | 0,889618486 | 0,61699232 | 0,952793934 | tRNA isopentenyltransferase 1 |
| ENSG00000157657 | ZNF618 | -0,168865445 | 0,889541954 | 0,42385678 | 0,890949676 | zinc finger protein 618 |
| ENSG00000144115 | THNSL2 | -0,168866611 | 0,889541235 | 0,53208518 | 0,931291085 | threonine synthase like 2 |
| ENSG00000177082 | WDR73 | -0,168873079 | 0,889537247 | 0,49280905 | 0,917090312 | WD repeat domain 73 |
| ENSG00000239754 | CFB | -0,168943692 | 0,88949371 | 0,76624577 | 0,977114158 | complement factor B |
| ENSG00000124406 | ATP8A1 | -0,168948717 | 0,889490611 | 0,70977191 | 0,970482408 | ATPase phospholipid transporting 8A1 |
| ENSG00000215866 | LINC01356 | -0,169076417 | 0,889411882 | 0,73803522 | 0,974353474 | long intergenic non-protein coding RNA 1356 |

| | | | | | | |
|---|---|---|---|---|---|---|
| ENSG00000277675 | AC211486,5 | -0,16908722 | 0,889405221 | 0,72965732 | 0,974009551 | novel transcript |
| ENSG00000100068 | LRP5L | -0,169128007 | 0,889380077 | 0,72517965 | 0,97326334 | LDL receptor related protein 5 like |
| ENSG00000152193 | OBI1 | -0,169164509 | 0,889357575 | 0,66128482 | 0,959582738 | ORC ubiquitin ligase 1 |
| ENSG00000183011 | NAA38 | -0,169165271 | 0,889357106 | 0,50145135 | 0,920520138 | N-alpha-acetyltransferase 38, NatC auxiliary subunit |
| ENSG00000272462 | U91328,1 | -0,169177068 | 0,889349834 | 0,47435992 | 0,910676852 | novel transcript |
| ENSG00000179826 | MRGPRX3 | -0,16917722 | 0,889349739 | 0,82752265 | 0,983595244 | MAS related GPR family member X3 |
| ENSG00000132563 | REEP2 | -0,169337117 | 0,889251176 | 0,48423989 | 0,913690171 | receptor accessory protein 2 |
| ENSG00000104343 | UBE2W | -0,169445321 | 0,889184484 | 0,76133732 | 0,97624106 | ubiquitin conjugating enzyme E2 W |
| ENSG00000270344 | POC1B-AS1 | -0,169545325 | 0,88912285 | 0,63539311 | 0,955314044 | POC1B antisense RNA 1 |
| ENSG00000269973 | AC010969,2 | -0,169604938 | 0,889086112 | 0,50568584 | 0,921780819 | novel transcript |
| ENSG00000198742 | SMURF1 | -0,16961629 | 0,889079116 | 0,82273874 | 0,983107285 | SMAD specific E3 ubiquitin protein ligase 1 |
| ENSG00000104299 | INTS9 | -0,169694454 | 0,889030948 | 0,55917426 | 0,939727587 | integrator complex subunit 9 |
| ENSG00000185829 | ARL17A | -0,169739148 | 0,889003406 | 0,48543968 | 0,913907266 | ADP ribosylation factor like GTPase 17A |
| ENSG00000130347 | RTN4IP1 | -0,169814327 | 0,888957082 | 0,58811042 | 0,94615778 | reticulon 4 interacting protein 1 |
| ENSG00000230002 | ALMS1-IT1 | -0,169863637 | 0,888926698 | 0,70921536 | 0,970282889 | ALMS1 intronic transcript 1 |
| ENSG00000146802 | TMEM168 | -0,169881581 | 0,888915642 | 0,69972365 | 0,968522643 | transmembrane protein 168 |
| ENSG00000189043 | NDUFA4 | -0,169909636 | 0,888898356 | 0,73346975 | 0,974353474 | NDUFA4 mitochondrial complex associated |
| ENSG00000139496 | NUP58 | -0,169947188 | 0,888875219 | 0,65554358 | 0,957444211 | nucleoporin 58 |
| ENSG00000146416 | AIG1 | -0,170037528 | 0,888819561 | 0,57151328 | 0,942436428 | androgen induced 1 |
| ENSG00000287979 | AC253572,1 | -0,170082612 | 0,888791786 | 0,69540612 | 0,967602563 | novel transcript |
| ENSG00000165943 | MOAP1 | -0,170094087 | 0,888784716 | 0,555154 | 0,937793851 | modulator of apoptosis 1 |
| ENSG00000145996 | CDKAL1 | -0,170171238 | 0,888737188 | 0,60634562 | 0,950927937 | CDK5 regulatory subunit associated protein 1 like 1 |
| ENSG00000167355 | OR51B5 | -0,17018202 | 0,888730546 | 0,85860906 | 0,989074806 | olfactory receptor family 51 subfamily B member 5 |
| ENSG00000269906 | AL606834,1 | -0,170242122 | 0,888693523 | 0,70060722 | 0,968522643 | novel transcript, sense intronic to SAV1 |
| ENSG00000106261 | ZKSCAN1 | -0,170278527 | 0,888671098 | 0,52530603 | 0,928752911 | zinc finger with KRAB and SCAN domains 1 |
| ENSG00000258957 | AL359317,2 | -0,1703413 | 0,888632431 | 0,78462587 | 0,979320406 | novel transcript |
| ENSG00000267078 | AC015802,1 | -0,170344071 | 0,888630725 | 0,91228503 | 0,993361299 | novel transcript, antisense to ST6GALNAC2 |
| ENSG00000230806 | SRGAP2-AS1 | -0,170347664 | 0,888628512 | 0,65222994 | 0,957257009 | SRGAP2 antisense RNA 1 |
| ENSG00000130717 | UCK1 | -0,17035475 | 0,888624147 | 0,46595989 | 0,907940653 | uridine-cytidine kinase 1 |
| ENSG00000101639 | CEP192 | -0,170372546 | 0,888613185 | 0,6252451 | 0,952793934 | centrosomal protein 192 |
| ENSG00000069869 | NEDD4 | -0,170525128 | 0,888519209 | 0,51967244 | 0,926822772 | NEDD4 E3 ubiquitin protein ligase |
| ENSG00000121892 | PDS5A | -0,170534484 | 0,888513447 | 0,666685 | 0,9611116 | PDS5 cohesin associated factor A |
| ENSG00000161664 | ASB16 | -0,170572958 | 0,888489752 | 0,72150722 | 0,972142678 | ankyrin repeat and SOCS box containing 16 |

| | | | | | | |
|---|---|---|---|---|---|---|
| ENSG00000144559 | TAMM41 | -0,170573282 | 0,888489553 | 0,62929286 | 0,952793934 | TAM41 mitochondrial translocator assembly and maintenance homolog |
| ENSG00000070087 | PFN2 | -0,170582513 | 0,888483868 | 0,68195991 | 0,964654539 | profilin 2 |
| ENSG00000214491 | SEC14L6 | -0,170586719 | 0,888481278 | 0,64685921 | 0,956899963 | SEC14 like lipid binding 6 |
| ENSG00000229728 | AL136531,1 | -0,170633143 | 0,888452688 | 0,78029901 | 0,978587255 | novel transcript |
| ENSG00000234841 | AC017007,2 | -0,170637295 | 0,888450131 | 0,80300681 | 0,980866933 | TAF4b RNA polymerase II, TATA box binding protein (TBP)-associated factor, 105kDa (TAF4B) pseudogene |
| ENSG00000284626 | AC091167,7 | -0,17069092 | 0,888417108 | 0,89174921 | 0,990937593 | novel protein |
| ENSG00000259158 | ADAM20P1 | -0,170732278 | 0,88839164 | 0,7119205 | 0,970812834 | ADAM metallopeptidase domain 20 pseudogene 1 |
| ENSG00000124251 | TP53TG5 | -0,170820192 | 0,888337505 | 0,74467937 | 0,975649186 | TP53 target 5 |
| ENSG00000224975 | INE1 | -0,170822365 | 0,888336167 | 0,66116988 | 0,959558771 | inactivation escape 1 |
| ENSG00000247311 | AC010255,1 | -0,170906363 | 0,888284448 | 0,71397912 | 0,970867497 | novel transcript |
| ENSG00000167100 | SAMD14 | -0,170914812 | 0,888279246 | 0,59523228 | 0,947715519 | sterile alpha motif domain containing 14 |
| ENSG00000129351 | ILF3 | -0,171232606 | 0,888083598 | 0,45127798 | 0,902062594 | interleukin enhancer binding factor 3 |
| ENSG00000279345 | Z98885,3 | -0,171286113 | 0,888050662 | 0,80889313 | 0,981213877 | novel transcript, antisense to BRD1 |
| ENSG00000196150 | ZNF250 | -0,171300677 | 0,888041697 | 0,47932413 | 0,912714933 | zinc finger protein 250 |
| ENSG00000258230 | AC063950,1 | -0,171334199 | 0,888021063 | 0,8288708 | 0,983795764 | suppression of tumorigenicity 13 (colon carcinoma) (Hsp70 interacting protein) (ST13) pseudogene |
| ENSG00000224418 | STK24-AS1 | -0,171360963 | 0,888004589 | 0,71745625 | 0,971697135 | STK24 antisense RNA 1 |
| ENSG00000184619 | KRBA2 | -0,171393836 | 0,887984355 | 0,39378365 | 0,871635019 | KRAB-A domain containing 2 |
| ENSG00000113555 | PCDH12 | -0,17141601 | 0,887970707 | 0,58482159 | 0,944857905 | protocadherin 12 |
| ENSG00000104147 | OIP5 | -0,171536433 | 0,887896591 | 0,63527425 | 0,955314044 | Opa interacting protein 5 |
| ENSG00000205832 | C16orf96 | -0,171551093 | 0,887887568 | 0,71978528 | 0,972014805 | chromosome 16 open reading frame 96 |
| ENSG00000267498 | AC007786,1 | -0,171566553 | 0,887878054 | 0,84051368 | 0,985672131 | uncharacterized LOC105372352 |
| ENSG00000286112 | AL441992,2 | -0,171573835 | 0,887873572 | 0,79624368 | 0,980015022 | kynurenine aminotransferase 1 |
| ENSG00000164466 | SFXN1 | -0,17164304 | 0,887830982 | 0,57825987 | 0,943425812 | sideroflexin 1 |
| ENSG00000181544 | FANCB | -0,171727069 | 0,887779273 | 0,71943655 | 0,971869926 | FA complementation group B |
| ENSG00000188095 | MESP2 | -0,171771473 | 0,887751948 | 0,78136579 | 0,978597905 | mesoderm posterior bHLH transcription factor 2 |
| ENSG00000236583 | AP004290,1 | -0,171790629 | 0,887740161 | 0,76766516 | 0,977309816 | insulin-like growth factor 2 mRNA binding protein 2 (IGF2BP2) pseudogene |
| ENSG00000133805 | AMPD3 | -0,171797036 | 0,887736219 | 0,75635835 | 0,975901808 | adenosine monophosphate deaminase 3 |
| ENSG00000101104 | PABPC1L | -0,171803957 | 0,88773196 | 0,54048202 | 0,933385624 | poly(A) binding protein cytoplasmic 1 like |
| ENSG00000132535 | DLG4 | -0,17182552 | 0,887718692 | 0,45041144 | 0,902062594 | discs large MAGUK scaffold protein 4 |

| | | | | | | |
|---|---|---|---|---|---|---|
| ENSG00000268471 | MIR4453HG | -0,171900588 | 0,887672502 | 0,59944018 | 0,949137642 | MIR4453 host gene |
| ENSG00000241560 | ZBTB20-AS1 | -0,171920049 | 0,887660528 | 0,62011179 | 0,952793934 | ZBTB20 antisense RNA 1 |
| ENSG00000243943 | ZNF512 | -0,1719676 | 0,887631272 | 0,63882336 | 0,956018518 | zinc finger protein 512 |
| ENSG00000244754 | N4BP2L2 | -0,172066773 | 0,887570257 | 0,62771344 | 0,952793934 | NEDD4 binding protein 2 like 2 |
| ENSG00000128989 | ARPP19 | -0,172088253 | 0,887557042 | 0,7072788 | 0,969565008 | cAMP regulated phosphoprotein 19 |
| ENSG00000145592 | RPL37 | -0,172115119 | 0,887540514 | 0,59060895 | 0,947416795 | ribosomal protein L37 |
| ENSG00000137513 | NARS2 | -0,172117157 | 0,88753926 | 0,54597965 | 0,935201756 | asparaginyl-tRNA synthetase 2, mitochondrial |
| ENSG00000021826 | CPS1 | -0,172197498 | 0,887489836 | 0,66789439 | 0,961113267 | carbamoyl-phosphate synthase 1 |
| ENSG00000287243 | THCAT155 | -0,172253647 | 0,887455296 | 0,73888113 | 0,974448853 | thyroid cancer-associated transcript 155 |
| ENSG00000137714 | FDX1 | -0,172349548 | 0,887396306 | 0,65350936 | 0,957444211 | ferredoxin 1 |
| ENSG00000227097 | RPS28P7 | -0,172375082 | 0,8873806 | 0,61914902 | 0,952793934 | ribosomal protein S28 pseudogene 7 |
| ENSG00000286672 | AL096712,2 | -0,172390155 | 0,887371329 | 0,67666612 | 0,963580269 | novel transcript, antisense to TBC1D22B |
| ENSG00000143324 | XPR1 | -0,172479059 | 0,887316648 | 0,65559013 | 0,957444211 | xenotropic and polytropic retrovirus receptor 1 |
| ENSG00000267060 | PTGES3L | -0,17252832 | 0,88728635 | 0,6319262 | 0,953915351 | prostaglandin E synthase 3 like |
| ENSG00000156735 | BAG4 | -0,172550901 | 0,887272463 | 0,70553329 | 0,969485452 | BAG cochaperone 4 |
| ENSG00000060749 | QSER1 | -0,172559323 | 0,887267283 | 0,70304598 | 0,968742626 | glutamine and serine rich 1 |
| ENSG00000123552 | USP45 | -0,172620848 | 0,887229445 | 0,66873299 | 0,961386309 | ubiquitin specific peptidase 45 |
| ENSG00000265533 | AC114689,3 | -0,172626608 | 0,887225903 | 0,87225417 | 0,989217334 | novel transcript |
| ENSG00000155755 | TMEM237 | -0,17266055 | 0,88720503 | 0,66788072 | 0,961113267 | transmembrane protein 237 |
| ENSG00000277290 | AC136475,10 | -0,172693479 | 0,88718478 | 0,76008453 | 0,97624106 | signal recognition particle 19kDa (SRP19) pseudogene |
| ENSG00000272841 | MAP3K4-AS1 | -0,172720607 | 0,887168098 | 0,57651894 | 0,943231961 | MAP3K4 antisense RNA 1 |
| ENSG00000090905 | TNRC6A | -0,172796098 | 0,887121677 | 0,56970127 | 0,94221458 | trinucleotide repeat containing adaptor 6A |
| ENSG00000236953 | ZDHHC20-IT1 | -0,172848354 | 0,887089545 | 0,9057958 | 0,992590661 | ZDHHC20 intronic transcript 1 |
| ENSG00000229007 | EXOSC3P1 | -0,172872315 | 0,887074812 | 0,91497871 | 0,993361299 | exosome component 3 pseudogene 1 |
| ENSG00000242779 | ZNF702P | -0,172985481 | 0,887005232 | 0,7034253 | 0,968940012 | zinc finger protein 702, pseudogene |
| ENSG00000233597 | AC133435,1 | -0,173011734 | 0,886989091 | 0,84895617 | 0,987036295 | ribosomal protein L19 (RPL19) pseudogene |
| ENSG00000136108 | CKAP2 | -0,173011775 | 0,886989066 | 0,71646653 | 0,971438758 | cytoskeleton associated protein 2 |
| ENSG00000118162 | KPTN | -0,173070534 | 0,886952941 | 0,6193754 | 0,952793934 | kaptin, actin binding protein |
| ENSG00000114784 | EIF1B | -0,173130692 | 0,886915957 | 0,7116645 | 0,97080996 | eukaryotic translation initiation factor 1B |
| ENSG00000287164 | AC112254,1 | -0,173135965 | 0,886912715 | 0,82210133 | 0,982952211 | novel transcript, antisense to AFAP1 |
| ENSG00000234623 | MOG | -0,173179115 | 0,886886189 | 0,90612347 | 0,992590661 | myelin oligodendrocyte glycoprotein |
| ENSG00000163608 | NEPRO | -0,173187187 | 0,886881227 | 0,66704959 | 0,9611116 | nucleolus and neural progenitor protein |

| | | | | | | |
|---|---|---|---|---|---|---|
| ENSG00000105793 | GTPBP10 | -0,173240215 | 0,886848628 | 0,62145866 | 0,952793934 | GTP binding protein 10 |
| ENSG00000163607 | GTPBP8 | -0,17330461 | 0,886809045 | 0,68978521 | 0,966868992 | GTP binding protein 8 (putative) |
| ENSG00000165060 | FXN | -0,173462952 | 0,886711719 | 0,41177774 | 0,883966506 | frataxin |
| ENSG00000259485 | LINC02253 | -0,173470723 | 0,886706943 | 0,74042916 | 0,974833668 | long intergenic non-protein coding RNA 2253 |
| ENSG00000206511 | GABBR1 | -0,173483473 | 0,886699107 | 0,5849486 | 0,944857905 | gamma-aminobutyric acid type B receptor subunit 1 |
| ENSG00000235602 | POU5F1P3 | -0,173487227 | 0,886696799 | 0,89055293 | 0,990717729 | POU class 5 homeobox 1 pseudogene 3 |
| ENSG00000178053 | MLF1 | -0,17351643 | 0,886678851 | 0,61087089 | 0,951873283 | myeloid leukemia factor 1 |
| ENSG00000188306 | LRRIQ4 | -0,173626814 | 0,886611012 | 0,8503577 | 0,987036295 | leucine rich repeats and IQ motif containing 4 |
| ENSG00000172139 | SLC9C1 | -0,173640271 | 0,886602741 | 0,85551291 | 0,988075747 | solute carrier family 9 member C1 |
| ENSG00000285044 | SLC9C1 | -0,173640271 | 0,886602741 | 0,85551291 | 0,988075747 | solute carrier family 9 member C1 |
| ENSG00000247572 | CKMT2-AS1 | -0,173669362 | 0,886584864 | 0,45541429 | 0,903545728 | CKMT2 antisense RNA 1 |
| ENSG00000245970 | AP003352,1 | -0,173801298 | 0,886503789 | 0,59193593 | 0,947416795 | novel transcript |
| ENSG00000268895 | A1BG-AS1 | -0,173957404 | 0,88640787 | 0,57305625 | 0,942475308 | A1BG antisense RNA 1 |
| ENSG00000284431 | AL022238,3 | -0,174008896 | 0,886376233 | 0,8130273 | 0,981246822 | novel protein |
| ENSG00000167377 | ZNF23 | -0,174056966 | 0,8863467 | 0,54632972 | 0,935401821 | zinc finger protein 23 |
| ENSG00000133318 | RTN3 | -0,174080554 | 0,886332209 | 0,60565672 | 0,950927937 | reticulon 3 |
| ENSG00000111224 | PARP11 | -0,174086523 | 0,886328542 | 0,62434291 | 0,952793934 | poly(ADP-ribose) polymerase family member 11 |
| ENSG00000281189 | GHET1 | -0,174097781 | 0,886321625 | 0,54399945 | 0,934259495 | gastric carcinoma proliferation enhancing transcript 1 |
| ENSG00000078699 | CBFA2T2 | -0,17414403 | 0,886293212 | 0,49129652 | 0,916995341 | CBFA2/RUNX1 partner transcriptional co-repressor 2 |
| ENSG00000284980 | AGPAT5 | -0,174174267 | 0,886274637 | 0,66527405 | 0,960743128 | 1-acylglycerol-3-phosphate O-acyltransferase 5 |
| ENSG00000155189 | AGPAT5 | -0,174174267 | 0,886274637 | 0,66527405 | 0,960743128 | 1-acylglycerol-3-phosphate O-acyltransferase 5 |
| ENSG00000283580 | AC098484,3 | -0,174261619 | 0,886220977 | 0,64502551 | 0,956899963 | novel protein |
| ENSG00000263753 | LINC00667 | -0,174388417 | 0,886143091 | 0,59154028 | 0,947416795 | long intergenic non-protein coding RNA 667 |
| ENSG00000272568 | AC005162,2 | -0,174469315 | 0,886093402 | 0,81028096 | 0,981213877 | novel transcript, antisense to CPVL |
| ENSG00000259805 | AC022558,2 | -0,174508707 | 0,886069209 | 0,84438443 | 0,986340494 | novel transcript, antisense to HOMER2, C15orf40 and BTBD1 |
| ENSG00000176209 | SMIM19 | -0,174527657 | 0,88605757 | 0,65360229 | 0,957444211 | small integral membrane protein 19 |
| ENSG00000143815 | LBR | -0,174540254 | 0,886049833 | 0,66498476 | 0,960635969 | lamin B receptor |
| ENSG00000157193 | LRP8 | -0,17456992 | 0,886031614 | 0,52682869 | 0,92927181 | LDL receptor related protein 8 |
| ENSG00000152795 | HNRNPDL | -0,174579282 | 0,886025864 | 0,65828034 | 0,957906876 | heterogeneous nuclear ribonucleoprotein D like |
| ENSG00000214617 | SLC6A10P | -0,174593991 | 0,886016831 | 0,75875565 | 0,976120207 | solute carrier family 6 member 10, pseudogene |
| ENSG00000184661 | CDCA2 | -0,174675947 | 0,885966499 | 0,51195977 | 0,923222596 | cell division cycle associated 2 |

| | | | | | | |
|---|---|---|---|---|---|---|
| ENSG00000279078 | SND1-IT1 | -0,174744916 | 0,885924146 | 0,58487448 | 0,944857905 | SND1 intronic transcript 1 |
| ENSG00000128886 | ELL3 | -0,17481145 | 0,88588329 | 0,77955855 | 0,978587255 | elongation factor for RNA polymerase II 3 |
| ENSG00000115942 | ORC2 | -0,174850573 | 0,885859267 | 0,59303599 | 0,947416795 | origin recognition complex subunit 2 |
| ENSG00000112167 | SAYSD1 | -0,174894237 | 0,885832457 | 0,58997575 | 0,947252221 | SAYSVFN motif domain containing 1 |
| ENSG00000229587 | AL158825,2 | -0,174951188 | 0,885797489 | 0,86759472 | 0,989074806 | novel transcript |
| ENSG00000007392 | LUC7L | -0,175059037 | 0,885731273 | 0,50129732 | 0,920520138 | LUC7 like |
| ENSG00000124615 | MOCS1 | -0,175097376 | 0,885707735 | 0,52936877 | 0,930491829 | molybdenum cofactor synthesis 1 |
| ENSG00000235859 | AC006978,1 | -0,175139384 | 0,885681946 | 0,60645564 | 0,950927937 | proteasome (prosome, macropain) 26S subunit, ATPase, 5 (PSMC5) pseudogene |
| ENSG00000100479 | POLE2 | -0,175159357 | 0,885669685 | 0,69859763 | 0,968522643 | DNA polymerase epsilon 2, accessory subunit |
| ENSG00000276340 | ZNF623 | -0,175209671 | 0,885638798 | 0,60187093 | 0,949939978 | zinc finger protein 623 |
| ENSG00000215067 | ALOX12-AS1 | -0,175271562 | 0,885600804 | 0,54135546 | 0,933429141 | ALOX12 antisense RNA 1 |
| ENSG00000151665 | PIGF | -0,175336562 | 0,885560905 | 0,73822493 | 0,974353474 | phosphatidylinositol glycan anchor biosynthesis class F |
| ENSG00000264546 | AC008026,3 | -0,175354601 | 0,885549833 | 0,73368374 | 0,974353474 | novel transcript |
| ENSG00000167965 | MLST8 | -0,175407248 | 0,885517517 | 0,51361961 | 0,92410971 | MTOR associated protein, LST8 homolog |
| ENSG00000285178 | ZFP1 | -0,175416114 | 0,885512076 | 0,69937104 | 0,968522643 | ZFP1 zinc finger protein |
| ENSG00000184517 | ZFP1 | -0,175416114 | 0,885512076 | 0,69937104 | 0,968522643 | ZFP1 zinc finger protein |
| ENSG00000141127 | PRPSAP2 | -0,175471075 | 0,885478342 | 0,53147322 | 0,931291085 | phosphoribosyl pyrophosphate synthetase associated protein 2 |
| ENSG00000179958 | DCTPP1 | -0,175486738 | 0,885468728 | 0,49113944 | 0,916995341 | dCTP pyrophosphatase 1 |
| ENSG00000188730 | VWC2 | -0,175507712 | 0,885455855 | 0,76875724 | 0,977381742 | von Willebrand factor C domain containing 2 |
| ENSG00000152409 | JMY | -0,175540186 | 0,885435925 | 0,61046121 | 0,951841373 | junction mediating and regulatory protein, p53 cofactor |
| ENSG00000269421 | AC011503,3 | -0,175646318 | 0,88537079 | 0,79815803 | 0,980161849 | zinc finger protein 92 pseudogene 3 |
| ENSG00000006837 | CDKL3 | -0,175649857 | 0,885368618 | 0,5396059 | 0,933350549 | cyclin dependent kinase like 3 |
| ENSG00000071564 | TCF3 | -0,175712229 | 0,885330342 | 0,50452393 | 0,921400408 | transcription factor 3 |
| ENSG00000149476 | TKFC | -0,175758397 | 0,88530201 | 0,59832334 | 0,948675374 | triokinase and FMN cyclase |
| ENSG00000120333 | MRPS14 | -0,175779765 | 0,885288898 | 0,67530848 | 0,963390425 | mitochondrial ribosomal protein S14 |
| ENSG00000167468 | GPX4 | -0,175823645 | 0,885261973 | 0,41927304 | 0,88768414 | glutathione peroxidase 4 |
| ENSG00000263956 | NBPF11 | -0,175839456 | 0,88525227 | 0,43372143 | 0,896272827 | NBPF member 11 |
| ENSG00000108506 | INTS2 | -0,175961633 | 0,885177304 | 0,61704538 | 0,952793934 | integrator complex subunit 2 |
| ENSG00000120539 | MASTL | -0,175986393 | 0,885162113 | 0,66604954 | 0,961020116 | microtubule associated serine/threonine kinase like |

| | | | | | | |
|---|---|---|---|---|---|---|
| ENSG00000273148 | LINC00653 | -0,176138529 | 0,885068775 | 0,76611129 | 0,977102081 | long intergenic non-protein coding RNA 653 |
| ENSG00000115159 | GPD2 | -0,176196438 | 0,88503325 | 0,64803122 | 0,956899963 | glycerol-3-phosphate dehydrogenase 2 |
| ENSG00000086061 | DNAJA1 | -0,176198649 | 0,885031894 | 0,61989837 | 0,952793934 | DnaJ heat shock protein family (Hsp40) member A1 |
| ENSG00000223959 | AFG3L1P | -0,176225744 | 0,885015272 | 0,42375764 | 0,890949676 | AFG3 like matrix AAA peptidase subunit 1, pseudogene |
| ENSG00000139921 | TMX1 | -0,176327801 | 0,884952668 | 0,64031637 | 0,956251063 | thioredoxin related transmembrane protein 1 |
| ENSG00000142534 | RPS11 | -0,17647073 | 0,884864999 | 0,63890938 | 0,956018518 | ribosomal protein S11 |
| ENSG00000278468 | MUC4 | -0,176477704 | 0,884860722 | 0,9197064 | 0,993699236 | mucin 4, cell surface associated |
| ENSG00000242268 | LINC02082 | -0,176626846 | 0,884769252 | 0,71767468 | 0,971697135 | long intergenic non-protein coding RNA 2082 |
| ENSG00000228057 | SEC63P1 | -0,176636098 | 0,884763578 | 0,72238322 | 0,972594421 | SEC63 homolog, protein translocation regulator pseudogene 1 |
| ENSG00000103024 | NME3 | -0,176710266 | 0,884718094 | 0,69938452 | 0,968522643 | NME/NM23 nucleoside diphosphate kinase 3 |
| ENSG00000114098 | ARMC8 | -0,176749737 | 0,884693889 | 0,63673033 | 0,95572216 | armadillo repeat containing 8 |
| ENSG00000246731 | MGC16275 | -0,17680308 | 0,884661178 | 0,70440787 | 0,969132783 | uncharacterized protein MGC16275 |
| ENSG00000276787 | FAN1 | -0,176845654 | 0,884635072 | 0,70592664 | 0,969503793 | FANCD2 and FANCI associated nuclease 1 |
| ENSG00000116752 | BCAS2 | -0,176847105 | 0,884634183 | 0,68663947 | 0,965769168 | BCAS2 pre-mRNA processing factor |
| ENSG00000135951 | TSGA10 | -0,176871844 | 0,884619014 | 0,59340825 | 0,947416795 | testis specific 10 |
| ENSG00000233613 | DCUN1D2-AS | -0,176906013 | 0,884598062 | 0,77003946 | 0,977778135 | DCUN1D2 antisense RNA |
| ENSG00000206561 | COLQ | -0,176917196 | 0,884591205 | 0,60090959 | 0,94964552 | collagen like tail subunit of asymmetric acetylcholinesterase |
| ENSG00000260361 | AC106028,3 | -0,176932503 | 0,88458182 | 0,76779513 | 0,977309816 | novel transcript |
| ENSG00000187775 | DNAH17 | -0,176944378 | 0,884574539 | 0,73350584 | 0,974353474 | dynein axonemal heavy chain 17 |
| ENSG00000164764 | SBSPON | -0,176973908 | 0,884556433 | 0,70946653 | 0,970414391 | somatomedin B and thrombospondin type 1 domain containing |
| ENSG00000138399 | FASTKD1 | -0,177040037 | 0,884515888 | 0,66582304 | 0,961016016 | FAST kinase domains 1 |
| ENSG00000167536 | DHRS13 | -0,177058956 | 0,884504289 | 0,64577309 | 0,956899963 | dehydrogenase/reductase 13 |
| ENSG00000111653 | ING4 | -0,17708148 | 0,88449048 | 0,49522993 | 0,917777383 | inhibitor of growth family member 4 |
| ENSG00000118193 | KIF14 | -0,177128856 | 0,884461435 | 0,63702257 | 0,955945807 | kinesin family member 14 |
| ENSG00000137843 | PAK6 | -0,177142515 | 0,884453061 | 0,68713438 | 0,965769168 | p21 (RAC1) activated kinase 6 |
| ENSG00000272693 | AC073107,1 | -0,177161425 | 0,884441469 | 0,79716315 | 0,980161849 | nuclear protein 2, transcriptional regulator (NUPR2) pseudogene |
| ENSG00000261379 | AC010735,1 | -0,177279241 | 0,884369244 | 0,86104652 | 0,989074806 | novel transcript |
| ENSG00000154814 | OXNAD1 | -0,177279716 | 0,884368953 | 0,58602085 | 0,945246954 | oxidoreductase NAD binding domain containing 1 |

| | | | | | | |
|---|---|---|---|---|---|---|
| ENSG00000185519 | FAM131C | -0,177285416 | 0,884365459 | 0,81960777 | 0,98249617 | family with sequence similarity 131 member C |
| ENSG00000185716 | MOSMO | -0,177291986 | 0,884361432 | 0,72549102 | 0,97326334 | modulator of smoothened |
| ENSG00000244556 | ODCP | -0,177308621 | 0,884351235 | 0,64094711 | 0,956410077 | ornithine decarboxylase pseudogene |
| ENSG00000124249 | KCNK15 | -0,177396434 | 0,884297409 | 0,84626723 | 0,986939308 | potassium two pore domain channel subfamily K member 15 |
| ENSG00000158483 | FAM86C1P | -0,177435307 | 0,884273582 | 0,47707609 | 0,91143003 | family with sequence similarity 86 member C1, pseudogene |
| ENSG00000168702 | LRP1B | -0,177456179 | 0,884260788 | 0,77042098 | 0,977891334 | LDL receptor related protein 1B |
| ENSG00000274818 | AC004825,2 | -0,177510063 | 0,884227762 | 0,81983407 | 0,98249617 | novel transcript |
| ENSG00000142949 | PTPRF | -0,177583251 | 0,884182907 | 0,62188323 | 0,952793934 | protein tyrosine phosphatase receptor type F |
| ENSG00000286964 | AL136169,1 | -0,177639085 | 0,884148688 | 0,73237122 | 0,974353474 | novel transcript |
| ENSG00000110195 | FOLR1 | -0,177656923 | 0,884137756 | 0,83943242 | 0,985672131 | folate receptor alpha |
| ENSG00000205581 | HMGN1 | -0,177668499 | 0,884130662 | 0,64065007 | 0,956348702 | high mobility group nucleosome binding domain 1 |
| ENSG00000168803 | ADAL | -0,177712143 | 0,884103916 | 0,63504088 | 0,955158181 | adenosine deaminase like |
| ENSG00000267855 | NDUFA7 | -0,177781554 | 0,884061381 | 0,45778324 | 0,90388627 | NADH:ubiquinone oxidoreductase subunit A7 |
| ENSG00000165046 | LETM2 | -0,177794781 | 0,884053276 | 0,60564968 | 0,950927937 | leucine zipper and EF-hand containing transmembrane protein 2 |
| ENSG00000223891 | OSER1-DT | -0,177944869 | 0,88396131 | 0,47202419 | 0,909171312 | OSER1 divergent transcript |
| ENSG00000182162 | P2RY8 | -0,178089209 | 0,883872875 | 0,90785668 | 0,992590661 | P2Y receptor family member 8 |
| ENSG00000137500 | CCDC90B | -0,178233887 | 0,883784242 | 0,59768258 | 0,9482846 | coiled-coil domain containing 90B |
| ENSG00000254907 | AC087276,2 | -0,178319011 | 0,883732097 | 0,73765182 | 0,974353474 | novel transcript, antisense to API5 and TTC17 |
| ENSG00000233297 | RASA4DP | -0,178363325 | 0,883704953 | 0,89486713 | 0,991296984 | RAS p21 protein activator 4CD, pseudogene |
| ENSG00000102924 | CBLN1 | -0,17837869 | 0,883695541 | 0,61633134 | 0,952793934 | cerebellin 1 precursor |
| ENSG00000260898 | ADPGK-AS1 | -0,178405731 | 0,883678978 | 0,72221406 | 0,972594421 | ADPGK antisense RNA 1 |
| ENSG00000277027 | RMRP | -0,178412052 | 0,883675106 | 0,43632773 | 0,896850871 | RNA component of mitochondrial RNA processing endoribonuclease |
| ENSG00000162552 | WNT4 | -0,178435592 | 0,883660688 | 0,7452805 | 0,9756851 | Wnt family member 4 |
| ENSG00000287012 | AC108667,2 | -0,178461937 | 0,883644551 | 0,82303128 | 0,983204348 | novel transcript |
| ENSG00000150756 | ATPSCKMT | -0,178498313 | 0,883622271 | 0,6300112 | 0,952910985 | ATP synthase c subunit lysine N-methyltransferase |
| ENSG00000151838 | CCDC175 | -0,17862624 | 0,883543922 | 0,73798198 | 0,974353474 | coiled-coil domain containing 175 |
| ENSG00000121410 | A1BG | -0,178672472 | 0,883515609 | 0,73842692 | 0,974353474 | alpha-1-B glycoprotein |
| ENSG00000197951 | ZNF71 | -0,178680835 | 0,883510487 | 0,42273975 | 0,890403187 | zinc finger protein 71 |
| ENSG00000130821 | SLC6A8 | -0,178811274 | 0,88343061 | 0,49009986 | 0,916416395 | solute carrier family 6 member 8 |

| | | | | | | |
|---|---|---|---|---|---|---|
| ENSG00000011009 | LYPLA2 | -0,179003042 | 0,883313189 | 0,45514495 | 0,903545728 | lysophospholipase 2 |
| ENSG00000268182 | SMIM17 | -0,179005222 | 0,883311854 | 0,72627282 | 0,973344786 | small integral membrane protein 17 |
| ENSG00000134152 | KATNBL1 | -0,179007514 | 0,883310451 | 0,70903994 | 0,970282889 | katanin regulatory subunit B1 like 1 |
| ENSG00000113810 | SMC4 | -0,179042827 | 0,88328883 | 0,65210131 | 0,957241719 | structural maintenance of chromosomes 4 |
| ENSG00000206579 | XKR4 | -0,179049945 | 0,883284472 | 0,62380491 | 0,952793934 | XK related 4 |
| ENSG00000175711 | B3GNTL1 | -0,179070494 | 0,883271891 | 0,48097937 | 0,912907448 | UDP-GlcNAc:betaGal beta-1,3-N-acetylglucosaminyltransferase like 1 |
| ENSG00000166801 | FAM111A | -0,179073093 | 0,8832703 | 0,60730241 | 0,950993594 | family with sequence similarity 111 member A |
| ENSG00000181004 | BBS12 | -0,179095341 | 0,883256679 | 0,72146442 | 0,972142678 | Bardet-Biedl syndrome 12 |
| ENSG00000176749 | CDK5R1 | -0,179163887 | 0,883214715 | 0,59222183 | 0,947416795 | cyclin dependent kinase 5 regulatory subunit 1 |
| ENSG00000168314 | MOBP | -0,179168762 | 0,88321173 | 0,83293271 | 0,984905093 | myelin associated oligodendrocyte basic protein |
| ENSG00000145725 | PPIP5K2 | -0,179170621 | 0,883210592 | 0,67149668 | 0,962085713 | diphosphoinositol pentakisphosphate kinase 2 |
| ENSG00000258789 | AL162171,1 | -0,179186206 | 0,883201051 | 0,76748689 | 0,977201891 | novel transcript, antisense to PTPN21 |
| ENSG00000283034 | ZNF445 | -0,179222648 | 0,883178742 | 0,57561461 | 0,942700009 | zinc finger protein 445 |
| ENSG00000288558 | DUS4L-BCAP29 | -0,179244902 | 0,883165119 | 0,62329288 | 0,952793934 | DUS4L-BCAP29 readthrough |
| ENSG00000154001 | PPP2R5E | -0,179252419 | 0,883160517 | 0,63253943 | 0,954288117 | protein phosphatase 2 regulatory subunit B'epsilon |
| ENSG00000287214 | AC021321,2 | -0,179273419 | 0,883147662 | 0,72623927 | 0,973344786 | novel transcript, sense intronic to ARFGEF1 |
| ENSG00000276232 | AC006064,5 | -0,179526697 | 0,882992631 | 0,56963297 | 0,94221458 | small Cajal body-specific RNA 10 |
| ENSG00000065243 | PKN2 | -0,179559735 | 0,882972411 | 0,66647193 | 0,9611116 | protein kinase N2 |
| ENSG00000285847 | AL713852,1 | -0,179613888 | 0,882939268 | 0,62204948 | 0,952793934 | novel transcript |
| ENSG00000132423 | COQ3 | -0,179680987 | 0,882898204 | 0,57703387 | 0,943288205 | coenzyme Q3, methyltransferase |
| ENSG00000163701 | IL17RE | -0,179700394 | 0,882886327 | 0,76412739 | 0,976640468 | interleukin 17 receptor E |
| ENSG00000125869 | LAMP5 | -0,17970428 | 0,882883949 | 0,81977733 | 0,98249617 | lysosomal associated membrane protein family member 5 |
| ENSG00000162825 | NBPF20 | -0,179745018 | 0,882859019 | 0,6178789 | 0,952793934 | NBPF member 20 |
| ENSG00000240869 | RN7SL128P | -0,179860991 | 0,882788052 | 0,79295161 | 0,979565591 | RNA, 7SL, cytoplasmic 128, pseudogene |
| ENSG00000132825 | PPP1R3D | -0,179898259 | 0,882765248 | 0,58493068 | 0,944857905 | protein phosphatase 1 regulatory subunit 3D |
| ENSG00000151632 | AKR1C2 | -0,179925535 | 0,882748558 | 0,72135263 | 0,972142678 | aldo-keto reductase family 1 member C2 |
| ENSG00000261559 | FSCN1P1 | -0,179936199 | 0,882742033 | 0,84330596 | 0,986260952 | fascin actin-bundling protein 1 pseudogene 1 |
| ENSG00000240625 | RN7SL403P | -0,179956322 | 0,882729721 | 0,88439064 | 0,990717729 | RNA, 7SL, cytoplasmic 403, pseudogene |
| ENSG00000277548 | AC018926,3 | -0,179971429 | 0,882720477 | 0,86677287 | 0,989074806 | novel transcript, sense intronic to CCPG1 |
| ENSG00000075292 | ZNF638 | -0,179987964 | 0,88271036 | 0,58909104 | 0,946663699 | zinc finger protein 638 |
| ENSG00000178460 | MCMDC2 | -0,179997816 | 0,882704333 | 0,64923829 | 0,956899963 | minichromosome maintenance domain containing 2 |

| | | | | | | |
|---|---|---|---|---|---|---|
| ENSG00000120662 | MTRF1 | -0,180026102 | 0,882687026 | 0,61941187 | 0,952793934 | mitochondrial translation release factor 1 |
| ENSG00000204856 | FAM216A | -0,180028243 | 0,882685716 | 0,63307699 | 0,954408281 | family with sequence similarity 216 member A |
| ENSG00000088854 | C20orf194 | -0,180182589 | 0,882591288 | 0,41035162 | 0,882614012 | chromosome 20 open reading frame 194 |
| ENSG00000231829 | AL157834,1 | -0,180306634 | 0,882515404 | 0,64814658 | 0,956899963 | novel transcript |
| ENSG00000165434 | PGM2L1 | -0,180350782 | 0,882488399 | 0,60262906 | 0,950352286 | phosphoglucomutase 2 like 1 |
| ENSG00000185480 | PARPBP | -0,180398584 | 0,882459159 | 0,7008758 | 0,968622631 | PARP1 binding protein |
| ENSG00000183978 | COA3 | -0,180483607 | 0,882407154 | 0,56615871 | 0,941123407 | cytochrome c oxidase assembly factor 3 |
| ENSG00000119638 | NEK9 | -0,180573703 | 0,88235205 | 0,42052263 | 0,888563911 | NIMA related kinase 9 |
| ENSG00000263001 | GTF2I | -0,180584865 | 0,882345223 | 0,57358686 | 0,942475308 | general transcription factor IIi |
| ENSG00000253273 | AC022826,1 | -0,180622365 | 0,882322289 | 0,80317051 | 0,980866933 | regulator of G-protein signaling 10 (RGS10) pseudogene |
| ENSG00000139746 | RBM26 | -0,180722276 | 0,882261187 | 0,64464077 | 0,956633898 | RNA binding motif protein 26 |
| ENSG00000103355 | PRSS33 | -0,180801118 | 0,882212974 | 0,65620776 | 0,957444211 | serine protease 33 |
| ENSG00000170703 | TTLL6 | -0,180845671 | 0,88218573 | 0,83473049 | 0,984905093 | tubulin tyrosine ligase like 6 |
| ENSG00000241288 | LINC02614 | -0,180852555 | 0,88218152 | 0,59645006 | 0,948282297 | long intergenic non-protein coding RNA 2614 |
| ENSG00000213713 | PIGCP1 | -0,181142885 | 0,882004007 | 0,62361076 | 0,952793934 | phosphatidylinositol glycan anchor biosynthesis class C pseudogene 1 |
| ENSG00000285796 | AL162458,1 | -0,181144034 | 0,882003304 | 0,53669254 | 0,932745314 | novel transcript |
| ENSG00000256193 | LINC00507 | -0,18118684 | 0,881977135 | 0,65345682 | 0,957444211 | long intergenic non-protein coding RNA 507 |
| ENSG00000276482 | SLC12A7 | -0,18126282 | 0,881930687 | 0,57466891 | 0,942526961 | solute carrier family 12 member 7 |
| ENSG00000149256 | TENM4 | -0,181277944 | 0,881921442 | 0,42881193 | 0,893639261 | teneurin transmembrane protein 4 |
| ENSG00000243081 | AC092894,1 | -0,181326016 | 0,881892055 | 0,78496746 | 0,979320406 | novel transcript |
| ENSG00000234969 | AARSD1P1 | -0,181336732 | 0,881885505 | 0,70977906 | 0,970482408 | AARSD1 pseudogene 1 |
| ENSG00000219445 | AC019155,2 | -0,181436203 | 0,881824702 | 0,86519687 | 0,989074806 | novel transcript |
| ENSG00000178971 | CTC1 | -0,18150069 | 0,881785287 | 0,44075135 | 0,898633772 | CST telomere replication complex component 1 |
| ENSG00000156304 | SCAF4 | -0,181527068 | 0,881769164 | 0,427417 | 0,892552417 | SR-related CTD associated factor 4 |
| ENSG00000015479 | MATR3 | -0,181532574 | 0,881765799 | 0,60189315 | 0,949939978 | matrin 3 |
| ENSG00000266126 | AC005730,3 | -0,181609513 | 0,881718776 | 0,88894755 | 0,990717729 | novel transcript |
| ENSG00000123739 | PLA2G12A | -0,181627647 | 0,881707693 | 0,59140838 | 0,947416795 | phospholipase A2 group XIIA |
| ENSG00000198556 | ZNF789 | -0,181776239 | 0,881616885 | 0,40106891 | 0,876659568 | zinc finger protein 789 |
| ENSG00000164190 | NIPBL | -0,181780561 | 0,881614244 | 0,62707545 | 0,952793934 | NIPBL cohesin loading factor |
| ENSG00000178425 | NT5DC1 | -0,181793461 | 0,881606361 | 0,64071285 | 0,956348702 | 5'-nucleotidase domain containing 1 |
| ENSG00000240002 | YBX1P3 | -0,181848767 | 0,881572565 | 0,89920932 | 0,991832796 | Y-box binding protein 1 pseudogene 3 |

| | | | | | | |
|---|---|---|---|---|---|---|
| ENSG00000116747 | RO60 | -0,181917396 | 0,88153063 | 0,72103791 | 0,972142678 | Ro60, Y RNA binding protein |
| ENSG00000185009 | AP3M1 | -0,181952755 | 0,881509025 | 0,68250836 | 0,964781531 | adaptor related protein complex 3 subunit mu 1 |
| ENSG00000214114 | MYCBP | -0,181994423 | 0,881483565 | 0,68700056 | 0,965769168 | MYC binding protein |
| ENSG00000196550 | FAM72A | -0,18199536 | 0,881482993 | 0,66749977 | 0,9611116 | family with sequence similarity 72 member A |
| ENSG00000196659 | TTC30B | -0,18202299 | 0,881466111 | 0,69943248 | 0,968522643 | tetratricopeptide repeat domain 30B |
| ENSG00000258875 | AL135818,1 | -0,182037793 | 0,881457067 | 0,79546598 | 0,979879951 | novel transcript, antisense to GPR68 |
| ENSG00000230590 | FTX | -0,182042816 | 0,881453998 | 0,49537028 | 0,917799558 | FTX transcript, XIST regulator |
| ENSG00000084444 | FAM234B | -0,182068073 | 0,881438566 | 0,4352565 | 0,896272827 | family with sequence similarity 234 member B |
| ENSG00000229852 | AC019205,1 | -0,182068354 | 0,881438395 | 0,60435257 | 0,950809109 | novel transcript, antisense to KHDC1 |
| ENSG00000225746 | MEG8 | -0,182089955 | 0,881425197 | 0,73523037 | 0,974353474 | maternally expressed 8, small nucleolar RNA host gene |
| ENSG00000249558 | RCC2P4 | -0,182210468 | 0,881351572 | 0,85009771 | 0,987036295 | regulator of chromosome condensation 2 pseudogene 4 |
| ENSG00000021574 | SPAST | -0,182272311 | 0,881313793 | 0,59307173 | 0,947416795 | spastin |
| ENSG00000286274 | AC008679,1 | -0,182326931 | 0,881280427 | 0,62541195 | 0,952793934 | novel transcript |
| ENSG00000249592 | AC139887,2 | -0,182426996 | 0,881219304 | 0,68043456 | 0,964654539 | novel transcript, antisense to PCGF3 |
| ENSG00000089280 | FUS | -0,182429136 | 0,881217996 | 0,53365374 | 0,931381467 | FUS RNA binding protein |
| ENSG00000279159 | AC003681,1 | -0,182467603 | 0,881194501 | 0,46795572 | 0,907940653 | novel transcript, antisense to MTMR3 |
| ENSG00000170677 | SOCS6 | -0,182584643 | 0,881123016 | 0,6586392 | 0,958154383 | suppressor of cytokine signaling 6 |
| ENSG00000198464 | ZNF480 | -0,182595952 | 0,881116109 | 0,68650217 | 0,965769168 | zinc finger protein 480 |
| ENSG00000100722 | ZC3H14 | -0,18263437 | 0,881092646 | 0,59023096 | 0,947278267 | zinc finger CCCH-type containing 14 |
| ENSG00000227375 | DLG1-AS1 | -0,182721899 | 0,881039191 | 0,799411 | 0,980161849 | DLG1 antisense RNA 1 |
| ENSG00000260634 | AC012508,2 | -0,182737124 | 0,881029894 | 0,84007903 | 0,985672131 | novel transcript |
| ENSG00000247556 | OIP5-AS1 | -0,182821252 | 0,880978519 | 0,55512223 | 0,937793851 | OIP5 antisense RNA 1 |
| ENSG00000053770 | AP5M1 | -0,182826127 | 0,880975542 | 0,68967802 | 0,966868992 | adaptor related protein complex 5 subunit mu 1 |
| ENSG00000269044 | AC024075,2 | -0,183004677 | 0,880866518 | 0,53457417 | 0,93162317 | novel transcript, sense intronic C19orf42 |
| ENSG00000287826 | AC090994,1 | -0,183049652 | 0,880839058 | 0,65704175 | 0,957612854 | novel transcript |
| ENSG00000257474 | AC027288,1 | -0,183113558 | 0,880800042 | 0,80598418 | 0,981213877 | novel transcript |
| ENSG00000234323 | LINC01505 | -0,183175091 | 0,880762475 | 0,67757217 | 0,9635902 | long intergenic non-protein coding RNA 1505 |
| ENSG00000107371 | EXOSC3 | -0,183250242 | 0,880716596 | 0,56118056 | 0,940567693 | exosome component 3 |
| ENSG00000062194 | GPBP1 | -0,18327767 | 0,880699853 | 0,64322617 | 0,956633898 | GC-rich promoter binding protein 1 |
| ENSG00000139737 | SLAIN1 | -0,183297502 | 0,880687746 | 0,61573153 | 0,952793934 | SLAIN motif family member 1 |
| ENSG00000163281 | GNPDA2 | -0,183388758 | 0,880632041 | 0,71916397 | 0,971869926 | glucosamine-6-phosphate deaminase 2 |

| | | | | | | |
|---|---|---|---|---|---|---|
| ENSG00000285327 | XKR5 | -0,183394569 | 0,880628494 | 0,81164608 | 0,981213877 | XK related 5 |
| ENSG00000271533 | Z83843,1 | -0,183441997 | 0,880599544 | 0,67017981 | 0,962026636 | novel transcript, sense intronic FTX |
| ENSG00000172466 | ZNF24 | -0,183514881 | 0,880555058 | 0,64760539 | 0,956899963 | zinc finger protein 24 |
| ENSG00000287983 | AL035251,1 | -0,183531605 | 0,88054485 | 0,75591684 | 0,975901808 | novel transcript |
| ENSG00000138668 | HNRNPD | -0,183567135 | 0,880523165 | 0,52412815 | 0,928404671 | heterogeneous nuclear ribonucleoprotein D |
| ENSG00000204954 | C12orf73 | -0,18357323 | 0,880519446 | 0,53229952 | 0,931291085 | chromosome 12 open reading frame 73 |
| ENSG00000091651 | ORC6 | -0,183585839 | 0,88051175 | 0,44451766 | 0,900210555 | origin recognition complex subunit 6 |
| ENSG00000254244 | PAICSP4 | -0,183601742 | 0,880502044 | 0,8823194 | 0,990717729 | phosphoribosylaminoimidazole carboxylase, phosphoribosylaminoimidazole succinocarboxamide synthetase pseudogene 4 |
| ENSG00000135587 | SMPD2 | -0,183635445 | 0,880481474 | 0,58896907 | 0,946604802 | sphingomyelin phosphodiesterase 2 |
| ENSG00000136378 | ADAMTS7 | -0,183656963 | 0,880468342 | 0,55911085 | 0,939686141 | ADAM metallopeptidase with thrombospondin type 1 motif 7 |
| ENSG00000245466 | AL357153,1 | -0,183721142 | 0,880429175 | 0,86437442 | 0,989074806 | novel transcript |
| ENSG00000184507 | NUTM1 | -0,183727756 | 0,880425139 | 0,83508831 | 0,984905093 | NUT midline carcinoma family member 1 |
| ENSG00000171634 | BPTF | -0,183771007 | 0,880398745 | 0,51300754 | 0,923725914 | bromodomain PHD finger transcription factor |
| ENSG00000109390 | NDUFC1 | -0,183780626 | 0,880392875 | 0,53917656 | 0,933350549 | NADH:ubiquinone oxidoreductase subunit C1 |
| ENSG00000182013 | PNMA8A | -0,183797677 | 0,880382469 | 0,42437609 | 0,890949676 | PNMA family member 8A |
| ENSG00000272986 | AC009570,1 | -0,183804568 | 0,880378264 | 0,75588687 | 0,975901808 | novel transcript, antisense to RUFY3 |
| ENSG00000101162 | TUBB1 | -0,183883922 | 0,880329841 | 0,61170608 | 0,952161039 | tubulin beta 1 class VI |
| ENSG00000066855 | MTFR1 | -0,183888243 | 0,880327205 | 0,58194193 | 0,944149815 | mitochondrial fission regulator 1 |
| ENSG00000237892 | KLF7-IT1 | -0,183908279 | 0,880314979 | 0,67364287 | 0,962830239 | KLF7 intronic transcript 1 |
| ENSG00000171951 | SCG2 | -0,183956646 | 0,880285467 | 0,56303852 | 0,940865625 | secretogranin II |
| ENSG00000120526 | NUDCD1 | -0,183971939 | 0,880276135 | 0,69349163 | 0,967340633 | NudC domain containing 1 |
| ENSG00000281500 | SLC37A4 | -0,18401215 | 0,8802516 | 0,53509305 | 0,931830289 | solute carrier family 37 member 4 |
| ENSG00000102738 | MRPS31 | -0,184018891 | 0,880247487 | 0,6543179 | 0,957444211 | mitochondrial ribosomal protein S31 |
| ENSG00000185621 | LMLN | -0,184049909 | 0,880228562 | 0,53320466 | 0,931291085 | leishmanolysin like peptidase |
| ENSG00000153310 | CYRIB | -0,184182811 | 0,880147479 | 0,62980585 | 0,952812183 | CYFIP related Rac1 interactor B |
| ENSG00000185130 | H2BC13 | -0,18422652 | 0,880120814 | 0,63676378 | 0,955726489 | H2B clustered histone 13 |
| ENSG00000163288 | GABRB1 | -0,184238165 | 0,880113709 | 0,84082183 | 0,985672131 | gamma-aminobutyric acid type A receptor subunit beta1 |
| ENSG00000288016 | AL162493,1 | -0,184242244 | 0,880111221 | 0,87846342 | 0,990109814 | novel transcript |
| ENSG00000211456 | SACM1L | -0,184288906 | 0,880082755 | 0,67513498 | 0,963390425 | SAC1 like phosphatidylinositide phosphatase |

| | | | | | | |
|---|---|---|---|---|---|---|
| ENSG00000205476 | CCDC85C | -0,184337645 | 0,880053024 | 0,55162249 | 0,936366211 | coiled-coil domain containing 85C |
| ENSG00000240695 | AC117382,1 | -0,184342916 | 0,880049809 | 0,70052043 | 0,968522643 | DnaJ (Hsp40) homolog, subfamily C, member 8 (DNAJC8) pseudogene |
| ENSG00000100614 | PPM1A | -0,184393021 | 0,880019245 | 0,65853436 | 0,958135616 | protein phosphatase, Mg2+/Mn2+ dependent 1A |
| ENSG00000230736 | AL021937,1 | -0,184434023 | 0,879994235 | 0,69365905 | 0,967340633 | novel transcript |
| ENSG00000086848 | ALG9 | -0,184483843 | 0,879963847 | 0,62985528 | 0,952812183 | ALG9 alpha-1,2-mannosyltransferase |
| ENSG00000268628 | AL121761,1 | -0,184611234 | 0,879886149 | 0,67264428 | 0,962355986 | novel transcript |
| ENSG00000161010 | MRNIP | -0,184688468 | 0,879839046 | 0,58118734 | 0,943919685 | MRN complex interacting protein |
| ENSG00000020922 | MRE11 | -0,184819526 | 0,879759122 | 0,58018399 | 0,943828882 | MRE11 homolog, double strand break repair nuclease |
| ENSG00000104894 | CD37 | -0,184873479 | 0,879726223 | 0,77485547 | 0,978553937 | CD37 molecule |
| ENSG00000274939 | TBC1D3B | -0,184892925 | 0,879714365 | 0,74043969 | 0,974833668 | TBC1 domain family member 3B |
| ENSG00000261335 | AC005837,1 | -0,184975796 | 0,879663834 | 0,7884653 | 0,979462318 | novel transcript, antisense to MXRA7 |
| ENSG00000214756 | CSKMT | -0,185053104 | 0,879616698 | 0,50649959 | 0,921824694 | citrate synthase lysine methyltransferase |
| ENSG00000109685 | NSD2 | -0,185092664 | 0,879592578 | 0,35990182 | 0,850280272 | nuclear receptor binding SET domain protein 2 |
| ENSG00000122367 | LDB3 | -0,18525524 | 0,879493463 | 0,54484708 | 0,93489368 | LIM domain binding 3 |
| ENSG00000116299 | ELAPOR1 | -0,18526186 | 0,879489427 | 0,66864924 | 0,961375797 | endosome-lysosome associated apoptosis and autophagy regulator 1 |
| ENSG00000237651 | C2orf74 | -0,185316721 | 0,879455984 | 0,60577068 | 0,950927937 | chromosome 2 open reading frame 74 |
| ENSG00000174038 | C9orf131 | -0,185368496 | 0,879424423 | 0,71828957 | 0,971789526 | chromosome 9 open reading frame 131 |
| ENSG00000081138 | CDH7 | -0,185549394 | 0,879314159 | 0,7267876 | 0,973344786 | cadherin 7 |
| ENSG00000114742 | WDR48 | -0,185612462 | 0,879275721 | 0,6505147 | 0,956899963 | WD repeat domain 48 |
| ENSG00000145386 | CCNA2 | -0,185671037 | 0,879240022 | 0,65798961 | 0,957883832 | cyclin A2 |
| ENSG00000254726 | MEX3A | -0,185691068 | 0,879227814 | 0,40705011 | 0,880078305 | mex-3 RNA binding family member A |
| ENSG00000150764 | DIXDC1 | -0,185701607 | 0,879221391 | 0,56833723 | 0,94163744 | DIX domain containing 1 |
| ENSG00000173214 | MFSD4B | -0,185712073 | 0,879215014 | 0,4051023 | 0,878176155 | major facilitator superfamily domain containing 4B |
| ENSG00000082684 | SEMA5B | -0,185736462 | 0,87920015 | 0,59999141 | 0,949340299 | semaphorin 5B |
| ENSG00000184492 | FOXD4L1 | -0,185745674 | 0,879194537 | 0,81860534 | 0,9824772 | forkhead box D4 like 1 |
| ENSG00000151552 | QDPR | -0,185917255 | 0,879089979 | 0,41186954 | 0,884023707 | quinoid dihydropteridine reductase |
| ENSG00000266777 | SH3GL1P1 | -0,185939376 | 0,8790765 | 0,58830926 | 0,946185547 | SH3 domain containing GRB2 like 1, endophilin A2 pseudogene 1 |
| ENSG00000238221 | AL139095,4 | -0,185985067 | 0,87904866 | 0,86611834 | 0,989074806 | novel transcript |
| ENSG00000275127 | SNORD116-22 | -0,186192313 | 0,878922392 | 0,77559505 | 0,978587255 | small nucleolar RNA, C/D box 116-22 |

| | | | | | | |
|---|---|---|---|---|---|---|
| ENSG00000224934 | AL391684,1 | -0,186226786 | 0,87890139 | 0,5807782 | 0,943832501 | novel transcript |
| ENSG00000095002 | MSH2 | -0,186243674 | 0,878891102 | 0,64873065 | 0,956899963 | mutS homolog 2 |
| ENSG00000274588 | DGKK | -0,186254408 | 0,878884563 | 0,82436269 | 0,983374097 | diacylglycerol kinase kappa |
| ENSG00000163655 | GMPS | -0,186274702 | 0,8788722 | 0,65604208 | 0,957444211 | guanine monophosphate synthase |
| ENSG00000162490 | DRAXIN | -0,186317198 | 0,878846312 | 0,4117245 | 0,883966506 | dorsal inhibitory axon guidance protein |
| ENSG00000256682 | AC006518,2 | -0,18664339 | 0,878647628 | 0,84672347 | 0,986939308 | taste receptor, type 2, member 12 pseudogene |
| ENSG00000282363 | AC006518,8 | -0,18664339 | 0,878647628 | 0,84672347 | 0,986939308 | taste receptor, type 2, member 12 pseudogene |
| ENSG00000185973 | TMLHE | -0,186647002 | 0,878645429 | 0,66513911 | 0,960681381 | trimethyllysine hydroxylase, epsilon |
| ENSG00000226519 | LINC00390 | -0,186687169 | 0,878620966 | 0,90298109 | 0,992311172 | long intergenic non-protein coding RNA 390 |
| ENSG00000276744 | AC105137,2 | -0,186704676 | 0,878610304 | 0,73719181 | 0,974353474 | novel transcript, sense intronic to SIN3A |
| ENSG00000257167 | TMPO-AS1 | -0,186717822 | 0,878602298 | 0,52658186 | 0,929261495 | TMPO antisense RNA 1 |
| ENSG00000250331 | LINC01340 | -0,186742109 | 0,878587507 | 0,81028282 | 0,981213877 | long intergenic non-protein coding RNA 1340 |
| ENSG00000183814 | LIN9 | -0,186746164 | 0,878585038 | 0,66315207 | 0,959920886 | lin-9 DREAM MuvB core complex component |
| ENSG00000105136 | ZNF419 | -0,186757999 | 0,87857783 | 0,45153141 | 0,902062594 | zinc finger protein 419 |
| ENSG00000232907 | DLGAP4-AS1 | -0,186762452 | 0,878575119 | 0,74691915 | 0,975796253 | DLGAP4 antisense RNA 1 |
| ENSG00000283041 | AC008038,1 | -0,186831659 | 0,878532974 | 0,78889532 | 0,979462318 | eukaryotic translation elongation factor 1 gamma ( |
| ENSG00000115504 | EHBP1 | -0,186844567 | 0,878525114 | 0,54719098 | 0,935404974 | EH domain binding protein 1 |
| ENSG00000143553 | SNAPIN | -0,186857828 | 0,878517039 | 0,6323999 | 0,954224236 | SNAP associated protein |
| ENSG00000123119 | NECAB1 | -0,186861747 | 0,878514652 | 0,77363657 | 0,978415098 | N-terminal EF-hand calcium binding protein 1 |
| ENSG00000174579 | MSL2 | -0,186865556 | 0,878512333 | 0,54818609 | 0,935404974 | MSL complex subunit 2 |
| ENSG00000267342 | AC087289,2 | -0,186881331 | 0,878502726 | 0,75521161 | 0,975901808 | novel transcript, antisense to MRPL38 |
| ENSG00000133739 | LRRCC1 | -0,18689526 | 0,878494245 | 0,62579673 | 0,952793934 | leucine rich repeat and coiled-coil centrosomal protein 1 |
| ENSG00000219392 | ZNF602P | -0,186912347 | 0,87848384 | 0,71946876 | 0,971869926 | zinc finger protein 602, pseudogene |
| ENSG00000268705 | BNIP3P26 | -0,186938058 | 0,878468184 | 0,83653807 | 0,985423106 | BCL2 interacting protein 3 pseudogene 26 |
| ENSG00000163666 | HESX1 | -0,18694798 | 0,878462143 | 0,73069232 | 0,974336839 | HESX homeobox 1 |
| ENSG00000141441 | GAREM1 | -0,187077656 | 0,878383186 | 0,58810518 | 0,94615778 | GRB2 associated regulator of MAPK1 subtype 1 |
| ENSG00000248049 | UBA6-AS1 | -0,187102166 | 0,878368263 | 0,51552387 | 0,924901769 | UBA6 antisense RNA 1 (head to head) |
| ENSG00000035141 | FAM136A | -0,187162797 | 0,87833135 | 0,58724633 | 0,945935868 | family with sequence similarity 136 member A |
| ENSG00000170846 | AC093323,1 | -0,187177883 | 0,878322165 | 0,43235961 | 0,895599994 | Putative MORF4 family-associated protein 1-like protein UPP |
| ENSG00000285669 | AC026979,4 | -0,187209467 | 0,878302937 | 0,61194162 | 0,952187506 | novel transcript |
| ENSG00000104369 | JPH1 | -0,187286741 | 0,878255894 | 0,63975635 | 0,956018518 | junctophilin 1 |

| | | | | | | |
|---|---|---|---|---|---|---|
| ENSG00000287917 | AP000944,7 | -0,187375107 | 0,878202102 | 0,71666016 | 0,971438758 | novel transcript, antisense to POLA2 |
| ENSG00000160948 | VPS28 | -0,187401501 | 0,878186036 | 0,55328446 | 0,936992468 | VPS28 subunit of ESCRT-I |
| ENSG00000184983 | NDUFA6 | -0,187499339 | 0,878126482 | 0,6551349 | 0,957444211 | NADH:ubiquinone oxidoreductase subunit A6 |
| ENSG00000274024 | AL590282,1 | -0,187504198 | 0,878123525 | 0,85543635 | 0,988075747 | tigger transposable element derived 7 (TIGD7) pseudogene |
| ENSG00000157306 | ZFHX2-AS1 | -0,187516718 | 0,878115905 | 0,68044786 | 0,964654539 | ZFHX2 antisense RNA 1 |
| ENSG00000143476 | DTL | -0,187575163 | 0,878080332 | 0,62721036 | 0,952793934 | denticleless E3 ubiquitin protein ligase homolog |
| ENSG00000224952 | PBX2 | -0,187577667 | 0,878078808 | 0,43322329 | 0,895845815 | PBX homeobox 2 |
| ENSG00000100263 | RHBDD3 | -0,18772332 | 0,877990162 | 0,58289043 | 0,944174844 | rhomboid domain containing 3 |
| ENSG00000229235 | RPL37P18 | -0,187925567 | 0,877867088 | 0,83346302 | 0,984905093 | ribosomal protein L37 pseudogene 18 |
| ENSG00000249087 | ZNF436-AS1 | -0,187950242 | 0,877852074 | 0,57595688 | 0,942955084 | ZNF436 antisense RNA 1 |
| ENSG00000283106 | ZNF436-AS1 | -0,187950242 | 0,877852074 | 0,57595688 | 0,942955084 | ZNF436 antisense RNA 1 |
| ENSG00000180481 | GLIPR1L2 | -0,188126264 | 0,877744974 | 0,5445918 | 0,934679287 | GLIPR1 like 2 |
| ENSG00000120457 | KCNJ5 | -0,188182985 | 0,877710466 | 0,77183395 | 0,978033855 | potassium inwardly rectifying channel subfamily J member 5 |
| ENSG00000262334 | RPH3AL | -0,188225671 | 0,877684496 | 0,55228343 | 0,936619966 | rabphilin 3A like (without C2 domains) |
| ENSG00000198553 | KCNRG | -0,188257889 | 0,877664896 | 0,69013886 | 0,967053457 | potassium channel regulator |
| ENSG00000102547 | CAB39L | -0,188334955 | 0,877618015 | 0,58234829 | 0,944174844 | calcium binding protein 39 like |
| ENSG00000141562 | NARF | -0,188391733 | 0,877583476 | 0,36330138 | 0,853554847 | nuclear prelamin A recognition factor |
| ENSG00000100360 | IFT27 | -0,188421514 | 0,877565361 | 0,37711561 | 0,862818448 | intraflagellar transport 27 |
| ENSG00000164684 | ZNF704 | -0,188425592 | 0,87756288 | 0,61250151 | 0,952413269 | zinc finger protein 704 |
| ENSG00000115875 | SRSF7 | -0,18844309 | 0,877552237 | 0,53698591 | 0,932883277 | serine and arginine rich splicing factor 7 |
| ENSG00000275888 | AC132872,3 | -0,188457711 | 0,877543343 | 0,81296541 | 0,981246822 | novel transcript, antisense to CSNK1D |
| ENSG00000179083 | FAM133A | -0,188490578 | 0,877523351 | 0,68161925 | 0,964654539 | family with sequence similarity 133 member A |
| ENSG00000258890 | CEP95 | -0,188559495 | 0,877481433 | 0,5654505 | 0,940865625 | centrosomal protein 95 |
| ENSG00000064999 | ANKS1A | -0,188568617 | 0,877475885 | 0,46767771 | 0,907940653 | ankyrin repeat and sterile alpha motif domain containing 1A |
| ENSG00000270170 | NCBP2AS2 | -0,188572014 | 0,877473819 | 0,50716156 | 0,921951873 | NCBP2 antisense 2 (head to head) |
| ENSG00000149311 | ATM | -0,188574791 | 0,87747213 | 0,60399153 | 0,950575944 | ATM serine/threonine kinase |
| ENSG00000231181 | AL954705,1 | -0,188589895 | 0,877462943 | 0,80877548 | 0,981213877 | ribosomal protein L9 (RPL9) pseudogene |
| ENSG00000274026 | FAM27E3 | -0,188640411 | 0,87743222 | 0,53938032 | 0,933350549 | family with sequence similarity 27 member E3 |
| ENSG00000284642 | AL139424,2 | -0,188703696 | 0,877393731 | 0,62831 | 0,952793934 | novel transcript |
| ENSG00000169031 | COL4A3 | -0,188729977 | 0,877377748 | 0,79943738 | 0,980161849 | collagen type IV alpha 3 chain |

| | | | | | | |
|---|---|---|---|---|---|---|
| ENSG00000187474 | FPR3 | -0,188795005 | 0,877338203 | 0,87000243 | 0,989074806 | formyl peptide receptor 3 |
| ENSG00000287603 | AC008403,3 | -0,188800507 | 0,877334857 | 0,87530674 | 0,990078774 | novel transcript, antisense to SULT2B1 |
| ENSG00000152078 | TLCD4 | -0,188905515 | 0,877271001 | 0,72350681 | 0,972913663 | TLC domain containing 4 |
| ENSG00000135451 | TROAP | -0,188912373 | 0,877266831 | 0,46200432 | 0,906277266 | trophinin associated protein |
| ENSG00000276766 | NDUFA3 | -0,188946528 | 0,877246063 | 0,57417087 | 0,942475308 | NADH:ubiquinone oxidoreductase subunit A3 |
| ENSG00000278365 | NDUFA3 | -0,188946528 | 0,877246063 | 0,57417087 | 0,942475308 | NADH:ubiquinone oxidoreductase subunit A3 |
| ENSG00000277722 | NDUFA3 | -0,188946528 | 0,877246063 | 0,57417087 | 0,942475308 | NADH:ubiquinone oxidoreductase subunit A3 |
| ENSG00000276220 | NDUFA3 | -0,188946528 | 0,877246063 | 0,57417087 | 0,942475308 | NADH:ubiquinone oxidoreductase subunit A3 |
| ENSG00000273642 | NDUFA3 | -0,188946528 | 0,877246063 | 0,57417087 | 0,942475308 | NADH:ubiquinone oxidoreductase subunit A3 |
| ENSG00000274359 | NDUFA3 | -0,188946528 | 0,877246063 | 0,57417087 | 0,942475308 | NADH:ubiquinone oxidoreductase subunit A3 |
| ENSG00000275724 | NDUFA3 | -0,188946528 | 0,877246063 | 0,57417087 | 0,942475308 | NADH:ubiquinone oxidoreductase subunit A3 |
| ENSG00000275605 | NDUFA3 | -0,188946528 | 0,877246063 | 0,57417087 | 0,942475308 | NADH:ubiquinone oxidoreductase subunit A3 |
| ENSG00000254579 | AC090720,1 | -0,188998225 | 0,877214628 | 0,80794728 | 0,981213877 | zinc finger protein pseudogene |
| ENSG00000111877 | MCM9 | -0,189051816 | 0,877182043 | 0,52354778 | 0,928038684 | minichromosome maintenance 9 homologous recombination repair factor |
| ENSG00000054267 | ARID4B | -0,189097195 | 0,877154452 | 0,54218535 | 0,933491609 | AT-rich interaction domain 4B |
| ENSG00000159267 | HLCS | -0,189284667 | 0,877040477 | 0,52085605 | 0,927046149 | holocarboxylase synthetase |
| ENSG00000083457 | ITGAE | -0,189333624 | 0,877010716 | 0,61875086 | 0,952793934 | integrin subunit alpha E |
| ENSG00000205670 | SMIM11A | -0,189393407 | 0,876974375 | 0,62671302 | 0,952793934 | small integral membrane protein 11A |
| ENSG00000102543 | CDADC1 | -0,189476043 | 0,876924144 | 0,5996907 | 0,949340299 | cytidine and dCMP deaminase domain containing 1 |
| ENSG00000145354 | CISD2 | -0,189483739 | 0,876919466 | 0,68164581 | 0,964654539 | CDGSH iron sulfur domain 2 |
| ENSG00000206346 | DXO | -0,189523907 | 0,876895051 | 0,64099444 | 0,956410077 | decapping exoribonuclease |
| ENSG00000261242 | AL136038,3 | -0,189562269 | 0,876871734 | 0,78035948 | 0,978587255 | novel transcript, antisense to PPP2R5E |
| ENSG00000176695 | OR4F17 | -0,189605106 | 0,876845698 | 0,50112056 | 0,920520138 | olfactory receptor family 4 subfamily F member 17 |
| ENSG00000137285 | TUBB2B | -0,189629887 | 0,876830637 | 0,36584413 | 0,855700097 | tubulin beta 2B class IIb |
| ENSG00000111752 | PHC1 | -0,18966683 | 0,876808184 | 0,50575854 | 0,921780819 | polyhomeotic homolog 1 |
| ENSG00000281852 | LINC00891 | -0,189673198 | 0,876804314 | 0,60465669 | 0,950878607 | long intergenic non-protein coding RNA 891 |
| ENSG00000163083 | INHBB | -0,189742174 | 0,876762395 | 0,66259727 | 0,959897678 | inhibin subunit beta B |
| ENSG00000145945 | FAM50B | -0,189770924 | 0,876744923 | 0,57196006 | 0,942475308 | family with sequence similarity 50 member B |
| ENSG00000124074 | ENKD1 | -0,189774297 | 0,876742873 | 0,59448553 | 0,947469769 | enkurin domain containing 1 |
| ENSG00000004779 | NDUFAB1 | -0,189776613 | 0,876741465 | 0,64942563 | 0,956899963 | NADH:ubiquinone oxidoreductase subunit AB1 |
| ENSG00000174599 | TRAM1L1 | -0,189851213 | 0,876696131 | 0,71104051 | 0,970482408 | translocation associated membrane protein 1 like 1 |
| ENSG00000255121 | CENATAC-DT | -0,189851872 | 0,876695731 | 0,6484671 | 0,956899963 | CENATAC divergent transcript |

| | | | | | | |
|---|---|---|---|---|---|---|
| ENSG00000267871 | ZNF460-AS1 | -0,189930031 | 0,876648237 | 0,67883342 | 0,964151912 | ZNF460 antisense RNA 1 |
| ENSG00000286953 | AC122697,1 | -0,190198053 | 0,876485389 | 0,84102652 | 0,985672131 | novel transcript |
| ENSG00000237406 | NDUFA9P1 | -0,190251748 | 0,876452768 | 0,71378782 | 0,970812834 | NADH:ubiquinone oxidoreductase subunit A9 pseudogene 1 |
| ENSG00000118707 | TGIF2 | -0,190269253 | 0,876442134 | 0,41074745 | 0,882835102 | TGFB induced factor homeobox 2 |
| ENSG00000234663 | LINC01934 | -0,190313704 | 0,87641513 | 0,68883578 | 0,966556275 | long intergenic non-protein coding RNA 1934 |
| ENSG00000235478 | LINC01664 | -0,190362996 | 0,876385187 | 0,81243311 | 0,981213877 | long intergenic non-protein coding RNA 1664 |
| ENSG00000260233 | ZNRD2-AS1 | -0,190371488 | 0,876380028 | 0,66573038 | 0,961016016 | ZNRD2 antisense RNA 1 (head to head) |
| ENSG00000261542 | AC011978,2 | -0,190373155 | 0,876379016 | 0,74956681 | 0,975901808 | novel transcript |
| ENSG00000152904 | GGPS1 | -0,190373374 | 0,876378883 | 0,61186069 | 0,952187506 | geranylgeranyl diphosphate synthase 1 |
| ENSG00000267890 | AC010624,2 | -0,190539387 | 0,876278042 | 0,82962331 | 0,984081562 | novel transcript, similar to SYT3 |
| ENSG00000268496 | AC245884,9 | -0,190578978 | 0,876253995 | 0,75859179 | 0,976120207 | novel transcript, sense intronic to TTYH1 |
| ENSG00000227518 | MIR1302-9HG | -0,190590008 | 0,876247296 | 0,58491891 | 0,944857905 | MIR1302-9 host gene |
| ENSG00000163635 | ATXN7 | -0,190614498 | 0,876232421 | 0,44976529 | 0,902062594 | ataxin 7 |
| ENSG00000077935 | SMC1B | -0,190652206 | 0,876209519 | 0,72053429 | 0,972142678 | structural maintenance of chromosomes 1B |
| ENSG00000274137 | MYOM2 | -0,19067402 | 0,876196271 | 0,5964641 | 0,948282297 | myomesin 2 |
| ENSG00000163507 | CIP2A | -0,190732325 | 0,876160861 | 0,61983392 | 0,952793934 | cellular inhibitor of PP2A |
| ENSG00000060688 | SNRNP40 | -0,190765051 | 0,876140987 | 0,47943901 | 0,912714933 | small nuclear ribonucleoprotein U5 subunit 40 |
| ENSG00000179044 | EXOC3L1 | -0,190787216 | 0,876127526 | 0,84364187 | 0,986273696 | exocyst complex component 3 like 1 |
| ENSG00000224238 | WARS2-IT1 | -0,19080959 | 0,876113939 | 0,60745295 | 0,950993594 | WARS2 intronic transcript 1 |
| ENSG00000158106 | RHPN1 | -0,190835463 | 0,876098227 | 0,65344646 | 0,957444211 | rhophilin Rho GTPase binding protein 1 |
| ENSG00000250039 | AC096719,1 | -0,190900635 | 0,876058651 | 0,85402982 | 0,987861659 | novel transcript |
| ENSG00000205918 | PDPK2P | -0,191006561 | 0,875994331 | 0,44201413 | 0,899273096 | 3-phosphoinositide dependent protein kinase 2, pseudogene |
| ENSG00000228400 | AC079154,1 | -0,19107635 | 0,875951957 | 0,84936132 | 0,987036295 | novel transcript |
| ENSG00000115541 | HSPE1 | -0,191175267 | 0,8758919 | 0,65685634 | 0,957476526 | heat shock protein family E (Hsp10) member 1 |
| ENSG00000182791 | CCDC87 | -0,191195643 | 0,875879529 | 0,6420615 | 0,956548736 | coiled-coil domain containing 87 |
| ENSG00000160179 | ABCG1 | -0,191237746 | 0,875853968 | 0,53256766 | 0,931291085 | ATP binding cassette subfamily G member 1 |
| ENSG00000181790 | ADGRB1 | -0,191306128 | 0,875812455 | 0,70448099 | 0,969132783 | adhesion G protein-coupled receptor B1 |
| ENSG00000123453 | SARDH | -0,191330063 | 0,875797925 | 0,54499996 | 0,934923038 | sarcosine dehydrogenase |
| ENSG00000230844 | ZNF674-AS1 | -0,191387559 | 0,875763023 | 0,57985615 | 0,943828882 | ZNF674 antisense RNA 1 (head to head) |
| ENSG00000105552 | BCAT2 | -0,191509317 | 0,875689115 | 0,60393257 | 0,950551453 | branched chain amino acid transaminase 2 |
| ENSG00000259118 | AL139022,1 | -0,191610153 | 0,875627911 | 0,77081541 | 0,978033855 | novel transcript, antisense to MAX |

| | | | | | | |
|---|---|---|---|---|---|---|
| ENSG00000103202 | NME4 | -0,191623069 | 0,875620072 | 0,47129591 | 0,908869685 | NME/NM23 nucleoside diphosphate kinase 4 |
| ENSG00000228405 | RNF5 | -0,191626184 | 0,875618181 | 0,43630965 | 0,896850871 | ring finger protein 5 |
| ENSG00000131089 | ARHGEF9 | -0,191637998 | 0,875611011 | 0,49464098 | 0,91758518 | Cdc42 guanine nucleotide exchange factor 9 |
| ENSG00000236209 | AC104135,1 | -0,191654043 | 0,875601273 | 0,84940659 | 0,987036295 | novel transcript |
| ENSG00000175087 | PDIK1L | -0,191807686 | 0,875508029 | 0,64281222 | 0,956633898 | PDLIM1 interacting kinase 1 like |
| ENSG00000180257 | ZNF816 | -0,191842083 | 0,875487155 | 0,62427556 | 0,952793934 | zinc finger protein 816 |
| ENSG00000164169 | PRMT9 | -0,191900748 | 0,875451556 | 0,59675135 | 0,948282297 | protein arginine methyltransferase 9 |
| ENSG00000167483 | NIBAN3 | -0,191931723 | 0,875432759 | 0,47315994 | 0,909714799 | niban apoptosis regulator 3 |
| ENSG00000156970 | BUB1B | -0,191961324 | 0,875414798 | 0,52547932 | 0,92884555 | BUB1 mitotic checkpoint serine/threonine kinase B |
| ENSG00000183891 | TTC32 | -0,192068974 | 0,875349479 | 0,65543253 | 0,957444211 | tetratricopeptide repeat domain 32 |
| ENSG00000281614 | INPP5D | -0,192097941 | 0,875331903 | 0,82280261 | 0,983107285 | inositol polyphosphate-5-phosphatase D |
| ENSG00000136451 | VEZF1 | -0,192134527 | 0,875309706 | 0,66687373 | 0,9611116 | vascular endothelial zinc finger 1 |
| ENSG00000247699 | FABP6-AS1 | -0,192215658 | 0,875260484 | 0,87998335 | 0,990268593 | FABP6 antisense RNA 1 |
| ENSG00000152256 | PDK1 | -0,192262894 | 0,875231827 | 0,58905435 | 0,946653391 | pyruvate dehydrogenase kinase 1 |
| ENSG00000198513 | ATL1 | -0,192274711 | 0,875224658 | 0,60010092 | 0,949340299 | atlastin GTPase 1 |
| ENSG00000213066 | CEP43 | -0,192360562 | 0,875172577 | 0,45839235 | 0,90420131 | centrosomal protein 43 |
| ENSG00000286451 | AL499602,1 | -0,192372383 | 0,875165406 | 0,90725239 | 0,992590661 | novel transcript |
| ENSG00000253878 | AC087752,3 | -0,192402981 | 0,875146845 | 0,77483578 | 0,978553937 | novel transcript |
| ENSG00000180011 | ZADH2 | -0,192415549 | 0,875139221 | 0,57344687 | 0,942475308 | zinc binding alcohol dehydrogenase domain containing 2 |
| ENSG00000165698 | SPACA9 | -0,192416703 | 0,875138521 | 0,49256185 | 0,917016235 | sperm acrosome associated 9 |
| ENSG00000125107 | CNOT1 | -0,192461618 | 0,875111276 | 0,58355418 | 0,944174844 | CCR4-NOT transcription complex subunit 1 |
| ENSG00000288147 | MG910336,1 | -0,19252649 | 0,875071927 | 0,87884403 | 0,990109814 | uncharacterized LOC389831 |
| ENSG00000164105 | SAP30 | -0,192537973 | 0,875064962 | 0,49143214 | 0,916995341 | Sin3A associated protein 30 |
| ENSG00000121210 | TMEM131L | -0,192642712 | 0,875001435 | 0,58768816 | 0,946111354 | transmembrane 131 like |
| ENSG00000286886 | AJ011931,2 | -0,192655618 | 0,874993608 | 0,80517302 | 0,981187666 | novel transcript, antisense to PCBP3 |
| ENSG00000277744 | AC011462,4 | -0,192678101 | 0,874979971 | 0,7318844 | 0,974353474 | novel transcript, sense intronic TMEM91 |
| ENSG00000134759 | ELP2 | -0,192714816 | 0,874957705 | 0,56301843 | 0,940865625 | elongator acetyltransferase complex subunit 2 |
| ENSG00000102109 | PCSK1N | -0,192754564 | 0,874933599 | 0,65639154 | 0,957444211 | proprotein convertase subtilisin/kexin type 1 inhibitor |
| ENSG00000181264 | TLCD5 | -0,192760164 | 0,874930203 | 0,46926234 | 0,907940653 | TLC domain containing 5 |
| ENSG00000059588 | TARBP1 | -0,192832121 | 0,874886565 | 0,4311723 | 0,89530663 | TAR (HIV-1) RNA binding protein 1 |
| ENSG00000182841 | RRP7BP | -0,192838258 | 0,874882843 | 0,61284084 | 0,952413269 | ribosomal RNA processing 7 homolog B, pseudogene |

| | | | | | | |
|---|---|---|---|---|---|---|
| ENSG00000172954 | LCLAT1 | -0,192871254 | 0,874862834 | 0,55658458 | 0,938062556 | lysocardiolipin acyltransferase 1 |
| ENSG00000287408 | AC027811,1 | -0,192953246 | 0,874813115 | 0,68314834 | 0,964965151 | novel transcript, antisense to TMED3 |
| ENSG00000121454 | LHX4 | -0,193092287 | 0,874728808 | 0,5277545 | 0,9295615 | LIM homeobox 4 |
| ENSG00000256238 | SUPT16HP1 | -0,193123762 | 0,874709724 | 0,80589516 | 0,981213877 | SPT16 homolog, facilitates chromatin remodeling subunit pseudogene 1 |
| ENSG00000160867 | FGFR4 | -0,193128189 | 0,87470704 | 0,73741078 | 0,974353474 | fibroblast growth factor receptor 4 |
| ENSG00000144554 | FANCD2 | -0,193149726 | 0,874693982 | 0,50299169 | 0,920831325 | FA complementation group D2 |
| ENSG00000092964 | DPYSL2 | -0,193166043 | 0,87468409 | 0,38044701 | 0,86586292 | dihydropyrimidinase like 2 |
| ENSG00000073605 | GSDMB | -0,193183505 | 0,874673503 | 0,59049332 | 0,947407528 | gasdermin B |
| ENSG00000228253 | MT-ATP8 | -0,193267311 | 0,874622694 | 0,52524847 | 0,928752911 | mitochondrially encoded ATP synthase membrane subunit 8 |
| ENSG00000206573 | THUMPD3-AS1 | -0,193287115 | 0,874610689 | 0,57724016 | 0,94337651 | THUMPD3 antisense RNA 1 |
| ENSG00000062282 | DGAT2 | -0,19330741 | 0,874598385 | 0,70383062 | 0,968976619 | diacylglycerol O-acyltransferase 2 |
| ENSG00000265845 | AC024267,4 | -0,193310119 | 0,874596743 | 0,76312507 | 0,976600177 | novel transcript, antisense to PHF12 |
| ENSG00000272277 | AL031963,3 | -0,193345305 | 0,874575413 | 0,74844032 | 0,975901808 | novel transcript, antisense to RIPK1 |
| ENSG00000273204 | AC104506,1 | -0,193461881 | 0,874504746 | 0,75814618 | 0,976120207 | novel transcript, antisense to EXTL2 |
| ENSG00000213390 | ARHGAP19 | -0,193465676 | 0,874502446 | 0,57044749 | 0,9423888 | Rho GTPase activating protein 19 |
| ENSG00000259826 | AC072061,1 | -0,193489771 | 0,87448784 | 0,60586831 | 0,950927937 | novel transcript, antisense to CDC2L5 |
| ENSG00000249684 | AC106795,2 | -0,193557077 | 0,874447044 | 0,78032619 | 0,978587255 | novel transcript, antisense to PROP1 |
| ENSG00000257181 | AC025423,4 | -0,193589185 | 0,874427583 | 0,68867086 | 0,966556275 | novel transcript |
| ENSG00000240877 | RN7SL521P | -0,193753312 | 0,87432811 | 0,87982993 | 0,990261566 | RNA, 7SL, cytoplasmic 521, pseudogene |
| ENSG00000283103 | AC010642,2 | -0,193880946 | 0,874250763 | 0,40489166 | 0,878176155 | novel transcript, antisense to ERVK3-1 |
| ENSG00000271888 | AL136162,1 | -0,193897756 | 0,874240575 | 0,75219158 | 0,975901808 | novel transcript |
| ENSG00000170748 | RBMXL2 | -0,194032137 | 0,874159148 | 0,8635705 | 0,989074806 | RBMX like 2 |
| ENSG00000229808 | AL391825,1 | -0,194034282 | 0,874157848 | 0,78581288 | 0,979320406 | proteasome (prosome, macropain) activator subunit 3 (PA28 gamma; Ki) pseudogene |
| ENSG00000004468 | CD38 | -0,194080977 | 0,874129555 | 0,8426298 | 0,986183927 | CD38 molecule |
| ENSG00000210194 | MT-TE | -0,194109146 | 0,874112488 | 0,75610927 | 0,975901808 | mitochondrially encoded tRNA-Glu (GAA/G) |
| ENSG00000162086 | ZNF75A | -0,194165697 | 0,874078225 | 0,43828034 | 0,897177404 | zinc finger protein 75a |
| ENSG00000156206 | CFAP161 | -0,194319879 | 0,873984816 | 0,82463272 | 0,983374097 | cilia and flagella associated protein 161 |
| ENSG00000246283 | AC090510,1 | -0,194329464 | 0,87397901 | 0,88167605 | 0,990717729 | novel transcript |
| ENSG00000224858 | RPL29P11 | -0,19434362 | 0,873970434 | 0,75781704 | 0,976120207 | ribosomal protein L29 pseudogene 11 |

| | | | | | | |
|---|---|---|---|---|---|---|
| ENSG00000255561 | FDXACB1 | -0,194437042 | 0,873913842 | 0,50701856 | 0,921951873 | ferredoxin-fold anticodon binding domain containing 1 |
| ENSG00000113327 | GABRG2 | -0,194445449 | 0,873908749 | 0,40098919 | 0,876659568 | gamma-aminobutyric acid type A receptor subunit gamma2 |
| ENSG00000198380 | GFPT1 | -0,194462121 | 0,87389865 | 0,50169348 | 0,920520138 | glutamine--fructose-6-phosphate transaminase 1 |
| ENSG00000184076 | UQCR10 | -0,194539407 | 0,873851836 | 0,62772425 | 0,952793934 | ubiquinol-cytochrome c reductase, complex III subunit X |
| ENSG00000212232 | SNORD17 | -0,194547309 | 0,87384705 | 0,62836892 | 0,952793934 | small nucleolar RNA, C/D box 17 |
| ENSG00000260565 | ERVK13-1 | -0,194572225 | 0,873831958 | 0,44864486 | 0,902062594 | endogenous retrovirus group K13 member 1 |
| ENSG00000250582 | SMAD1-AS2 | -0,194614725 | 0,873806217 | 0,75499825 | 0,975901808 | SMAD1 antisense RNA 2 |
| ENSG00000241158 | ADAMTS9-AS1 | -0,194730073 | 0,873736356 | 0,56695462 | 0,941425553 | ADAMTS9 antisense RNA 1 |
| ENSG00000153707 | PTPRD | -0,194868163 | 0,873652729 | 0,65472991 | 0,957444211 | protein tyrosine phosphatase receptor type D |
| ENSG00000085382 | HACE1 | -0,194891851 | 0,873638384 | 0,6123891 | 0,952413269 | HECT domain and ankyrin repeat containing E3 ubiquitin protein ligase 1 |
| ENSG00000147548 | NSD3 | -0,194975723 | 0,873587596 | 0,58776139 | 0,946124093 | nuclear receptor binding SET domain protein 3 |
| ENSG00000186628 | FSD2 | -0,194981091 | 0,873584346 | 0,57267378 | 0,942475308 | fibronectin type III and SPRY domain containing 2 |
| ENSG00000253919 | THAP12P7 | -0,195017851 | 0,873562087 | 0,81503071 | 0,981521413 | THAP domain containing 12 pseudogene 7 |
| ENSG00000224635 | AL391095,1 | -0,195132998 | 0,873492367 | 0,53081999 | 0,931265667 | novel transcript |
| ENSG00000197885 | NKIRAS1 | -0,195263993 | 0,873413059 | 0,56961151 | 0,94221458 | NFKB inhibitor interacting Ras like 1 |
| ENSG00000143442 | POGZ | -0,195297818 | 0,873392581 | 0,48039816 | 0,912847708 | pogo transposable element derived with ZNF domain |
| ENSG00000257896 | AC093012,1 | -0,195330209 | 0,873372972 | 0,79829349 | 0,980161849 | La ribonucleoprotein domain family, member 7 (LARP7) |
| ENSG00000167088 | SNRPD1 | -0,195347849 | 0,873362294 | 0,45708918 | 0,90378766 | small nuclear ribonucleoprotein D1 polypeptide |
| ENSG00000182796 | TMEM198B | -0,195363272 | 0,873352957 | 0,48879875 | 0,91557151 | transmembrane protein 198B (pseudogene) |
| ENSG00000014257 | ACP3 | -0,195437091 | 0,873308271 | 0,84884012 | 0,987036295 | acid phosphatase 3 |
| ENSG00000232871 | SEC1P | -0,195476651 | 0,873284324 | 0,84794214 | 0,987036295 | secretory blood group 1, pseudogene |
| ENSG00000123965 | PMS2P5 | -0,195547142 | 0,873241657 | 0,71024816 | 0,970482408 | PMS1 homolog 2, mismatch repair system component pseudogene 5 |
| ENSG00000242259 | C22orf39 | -0,19554978 | 0,87324006 | 0,49192355 | 0,916995341 | chromosome 22 open reading frame 39 |
| ENSG00000153561 | RMND5A | -0,195589927 | 0,873215759 | 0,6180878 | 0,952793934 | required for meiotic nuclear division 5 homolog A |
| ENSG00000070950 | RAD18 | -0,195610078 | 0,873203563 | 0,64146807 | 0,956465487 | RAD18 E3 ubiquitin protein ligase |
| ENSG00000135503 | ACVR1B | -0,195690378 | 0,873154962 | 0,35142777 | 0,84405766 | activin A receptor type 1B |

| | | | | | | |
|---|---|---|---|---|---|---|
| ENSG00000128872 | TMOD2 | -0,195814045 | 0,873080119 | 0,55652385 | 0,938061341 | tropomodulin 2 |
| ENSG00000269376 | AL356740,3 | -0,195833424 | 0,873068391 | 0,80388242 | 0,980875302 | novel transcript |
| ENSG00000235440 | RPS2P15 | -0,195842739 | 0,873062754 | 0,79384247 | 0,979776981 | ribosomal protein S2 pseudogene 15 |
| ENSG00000274628 | AL669942,1 | -0,195852066 | 0,87305711 | 0,78772325 | 0,979462318 | novel cytochrome P450 family pseudogene |
| ENSG00000287896 | AC073548,2 | -0,195856554 | 0,873054394 | 0,79005113 | 0,979462318 | novel transcript, antisense to KPTNand SLC8A2 |
| ENSG00000205758 | CRYZL1 | -0,195861341 | 0,873051497 | 0,47223299 | 0,909171312 | crystallin zeta like 1 |
| ENSG00000107672 | NSMCE4A | -0,196063143 | 0,872929384 | 0,54721751 | 0,935404974 | NSE4 homolog A, SMC5-SMC6 complex component |
| ENSG00000153086 | ACMSD | -0,196157421 | 0,872872342 | 0,77155972 | 0,978033855 | aminocarboxymuconate semialdehyde decarboxylase |
| ENSG00000285456 | LTB4R | -0,196233625 | 0,872826238 | 0,65528391 | 0,957444211 | leukotriene B4 receptor |
| ENSG00000213903 | LTB4R | -0,196233625 | 0,872826238 | 0,65528391 | 0,957444211 | leukotriene B4 receptor |
| ENSG00000135108 | FBXO21 | -0,196273704 | 0,87280199 | 0,44286925 | 0,899490955 | F-box protein 21 |
| ENSG00000113328 | CCNG1 | -0,196337692 | 0,872763279 | 0,70865606 | 0,970181083 | cyclin G1 |
| ENSG00000182256 | GABRG3 | -0,196414193 | 0,872717001 | 0,57683494 | 0,943261034 | gamma-aminobutyric acid type A receptor subunit gamma3 |
| ENSG00000276016 | ABR | -0,196439454 | 0,87270172 | 0,45298803 | 0,902428411 | ABR activator of RhoGEF and GTPase |
| ENSG00000167889 | MGAT5B | -0,196443362 | 0,872699356 | 0,61309395 | 0,952413269 | alpha-1,6-mannosylglycoprotein 6-beta-N-acetylglucosaminyltransferase B |
| ENSG00000204967 | PCDHA4 | -0,196654144 | 0,872571862 | 0,49297193 | 0,917090312 | protocadherin alpha 4 |
| ENSG00000255036 | STRA6LP | -0,196654509 | 0,872571641 | 0,52949581 | 0,930531584 | STRA6 like, pseudogene |
| ENSG00000268204 | AC008763,1 | -0,196663884 | 0,872565971 | 0,84063899 | 0,985672131 | novel transcript, antisense to STXBP2 |
| ENSG00000236125 | USP17L4 | -0,196725268 | 0,872528846 | 0,84977525 | 0,987036295 | ubiquitin specific peptidase 17 like family member 4 |
| ENSG00000172992 | DCAKD | -0,196729551 | 0,872526255 | 0,37484201 | 0,861771195 | dephospho-CoA kinase domain containing |
| ENSG00000166508 | MCM7 | -0,196734331 | 0,872523364 | 0,39262308 | 0,870909282 | minichromosome maintenance complex component 7 |
| ENSG00000130684 | ZNF337 | -0,19674806 | 0,872515061 | 0,40786187 | 0,88080912 | zinc finger protein 337 |
| ENSG00000033800 | PIAS1 | -0,19675809 | 0,872508995 | 0,6803977 | 0,964654539 | protein inhibitor of activated STAT 1 |
| ENSG00000165972 | CCDC38 | -0,196825606 | 0,872468164 | 0,75448451 | 0,975901808 | coiled-coil domain containing 38 |
| ENSG00000130731 | METTL26 | -0,196847338 | 0,872455022 | 0,5584208 | 0,939334338 | methyltransferase like 26 |
| ENSG00000153914 | SREK1 | -0,196870898 | 0,872440774 | 0,56716996 | 0,941425553 | splicing regulatory glutamic acid and lysine rich protein 1 |
| ENSG00000248449 | PCDHGB8P | -0,196889217 | 0,872429696 | 0,56452478 | 0,940865625 | protocadherin gamma subfamily B, 8 pseudogene |

| | | | | | | |
|---|---|---|---|---|---|---|
| ENSG00000288424 | AGAP12P | -0,196901592 | 0,872422213 | 0,84892844 | 0,987036295 | ArfGAP with GTPase domain, ankyrin repeat and PH domain 12, pseudogene |
| ENSG00000265018 | AGAP12P | -0,196901592 | 0,872422213 | 0,84892844 | 0,987036295 | ArfGAP with GTPase domain, ankyrin repeat and PH domain 12, pseudogene |
| ENSG00000163913 | IFT122 | -0,196928467 | 0,872405961 | 0,35007736 | 0,843789477 | intraflagellar transport 122 |
| ENSG00000139323 | POC1B | -0,196994938 | 0,872365767 | 0,65201359 | 0,957241719 | POC1 centriolar protein B |
| ENSG00000215068 | AC025171,2 | -0,197045159 | 0,8723354 | 0,45394002 | 0,902912642 | novel transcript, antisense to ANXA2R |
| ENSG00000287729 | AF241728,2 | -0,197057695 | 0,87232782 | 0,77875811 | 0,978587255 | novel transcript |
| ENSG00000262803 | AL354943,1 | -0,197062001 | 0,872325216 | 0,80710305 | 0,981213877 | transmembrane protein 192 (TMEM192) pseudogene |
| ENSG00000155592 | ZKSCAN2 | -0,197102142 | 0,872300945 | 0,45489622 | 0,903424009 | zinc finger with KRAB and SCAN domains 2 |
| ENSG00000160325 | CACFD1 | -0,197150387 | 0,872271775 | 0,5749228 | 0,942533938 | calcium channel flower domain containing 1 |
| ENSG00000273824 | AC008033,3 | -0,197164578 | 0,872263195 | 0,770943 | 0,978033855 | novel transcript |
| ENSG00000284752 | GMPR2 | -0,197204118 | 0,872239289 | 0,44848883 | 0,902062594 | guanosine monophosphate reductase 2 |
| ENSG00000105251 | SHD | -0,19731964 | 0,872169449 | 0,56225482 | 0,940865625 | Src homology 2 domain containing transforming protein D |
| ENSG00000251303 | CAB39P1 | -0,197361421 | 0,872144191 | 0,78521084 | 0,979320406 | calcium binding protein 39 pseudogene 1 |
| ENSG00000228736 | RGL2 | -0,197401529 | 0,872119944 | 0,44442559 | 0,900210555 | ral guanine nucleotide dissociation stimulator like 2 |
| ENSG00000224841 | RGL2 | -0,197401529 | 0,872119944 | 0,44442559 | 0,900210555 | ral guanine nucleotide dissociation stimulator like 2 |
| ENSG00000237825 | RGL2 | -0,197401529 | 0,872119944 | 0,44442559 | 0,900210555 | ral guanine nucleotide dissociation stimulator like 2 |
| ENSG00000080345 | RIF1 | -0,197443213 | 0,872094747 | 0,59058674 | 0,947416795 | replication timing regulatory factor 1 |
| ENSG00000287063 | AL356489,3 | -0,197461388 | 0,87208376 | 0,70562282 | 0,969485452 | novel transcript |
| ENSG00000070367 | EXOC5 | -0,197469562 | 0,87207882 | 0,67730223 | 0,963580269 | exocyst complex component 5 |
| ENSG00000232645 | LINC01431 | -0,197497147 | 0,872062145 | 0,78783536 | 0,979462318 | long intergenic non-protein coding RNA 1431 |
| ENSG00000246859 | STARD4-AS1 | -0,197674362 | 0,871955031 | 0,60863255 | 0,951557131 | STARD4 antisense RNA 1 |
| ENSG00000262516 | AC007151,1 | -0,197692929 | 0,871943809 | 0,77415586 | 0,978503783 | developmental pluripotency associated 3 (DPPA3) pseudogene |
| ENSG00000247092 | SNHG10 | -0,197700559 | 0,871939198 | 0,5105729 | 0,922693987 | small nucleolar RNA host gene 10 |
| ENSG00000126216 | TUBGCP3 | -0,19771623 | 0,871929727 | 0,51446296 | 0,924434989 | tubulin gamma complex associated protein 3 |
| ENSG00000234222 | LIX1L-AS1 | -0,197818005 | 0,871868218 | 0,73617008 | 0,974353474 | LIX1L antisense RNA 1 |
| ENSG00000132967 | HMGB1P5 | -0,197918939 | 0,871807222 | 0,71504139 | 0,971041579 | high mobility group box 1 pseudogene 5 |
| ENSG00000180357 | ZNF609 | -0,197984994 | 0,871767307 | 0,3843787 | 0,868358674 | zinc finger protein 609 |
| ENSG00000081148 | IMPG2 | -0,197985764 | 0,871766842 | 0,59910412 | 0,948894897 | interphotoreceptor matrix proteoglycan 2 |
| ENSG00000110318 | CEP126 | -0,197992631 | 0,871762692 | 0,56386955 | 0,940865625 | centrosomal protein 126 |

| | | | | | | |
|---|---|---|---|---|---|---|
| ENSG00000116922 | C1orf109 | -0,19800706 | 0,871753973 | 0,4148054 | 0,885579089 | chromosome 1 open reading frame 109 |
| ENSG00000231177 | LINC00852 | -0,19801465 | 0,871749387 | 0,57205447 | 0,942475308 | long intergenic non-protein coding RNA 852 |
| ENSG00000286552 | AL390839,1 | -0,198186073 | 0,871645811 | 0,82065763 | 0,982609114 | novel transcript |
| ENSG00000111911 | HINT3 | -0,198221522 | 0,871624394 | 0,53780239 | 0,932883277 | histidine triad nucleotide binding protein 3 |
| ENSG00000130844 | ZNF331 | -0,198255829 | 0,871603667 | 0,48713031 | 0,914593485 | zinc finger protein 331 |
| ENSG00000172733 | PURG | -0,198274489 | 0,871592394 | 0,5960859 | 0,948282297 | purine rich element binding protein G |
| ENSG00000253106 | AC090198,1 | -0,198277743 | 0,871590428 | 0,49936951 | 0,920135598 | novel transcript, antisense to TATDN1 |
| ENSG00000180228 | PRKRA | -0,198472661 | 0,871472678 | 0,50944822 | 0,922427526 | protein activator of interferon induced protein kinase EIF2AK2 |
| ENSG00000280616 | VPS11 | -0,198602594 | 0,871394194 | 0,56292013 | 0,940865625 | VPS11 core subunit of CORVET and HOPS complexes |
| ENSG00000272027 | AC073218,1 | -0,198637403 | 0,87137317 | 0,89926999 | 0,991832796 | novel transcript |
| ENSG00000236859 | NIFK-AS1 | -0,19868946 | 0,871341728 | 0,39633334 | 0,87298265 | NIFK antisense RNA 1 |
| ENSG00000250012 | AC079848,1 | -0,1987443 | 0,871308607 | 0,69236486 | 0,967236299 | novel transcript |
| ENSG00000108064 | TFAM | -0,198842679 | 0,871249194 | 0,64421027 | 0,956633898 | transcription factor A, mitochondrial |
| ENSG00000145879 | SPINK7 | -0,198971338 | 0,871171499 | 0,84564944 | 0,986639902 | serine peptidase inhibitor Kazal type 7 |
| ENSG00000138658 | ZGRF1 | -0,198985629 | 0,87116287 | 0,61740112 | 0,952793934 | zinc finger GRF-type containing 1 |
| ENSG00000143919 | CAMKMT | -0,199007749 | 0,871149513 | 0,58019272 | 0,943828882 | calmodulin-lysine N-methyltransferase |
| ENSG00000283934 | IGSF6 | -0,199145475 | 0,871066353 | 0,60204252 | 0,94999532 | immunoglobulin superfamily member 6 |
| ENSG00000255092 | AC010768,2 | -0,199149196 | 0,871064106 | 0,89179312 | 0,990937593 | novel transcript |
| ENSG00000196865 | NHLRC2 | -0,199153747 | 0,871061359 | 0,62509583 | 0,952793934 | NHL repeat containing 2 |
| ENSG00000245614 | DDX11-AS1 | -0,199351358 | 0,870942055 | 0,54578199 | 0,935146964 | DDX11 antisense RNA 1 |
| ENSG00000198498 | TMA16 | -0,19945563 | 0,870879109 | 0,61752825 | 0,952793934 | translation machinery associated 16 homolog |
| ENSG00000225484 | NUTM2B-AS1 | -0,19950069 | 0,870851909 | 0,56239648 | 0,940865625 | NUTM2B antisense RNA 1 |
| ENSG00000224324 | THAP5P1 | -0,199542044 | 0,870826947 | 0,84518101 | 0,986639902 | THAP domain containing 5 pseudogene 1 |
| ENSG00000197275 | RAD54B | -0,199615382 | 0,87078268 | 0,62095071 | 0,952793934 | RAD54 homolog B |
| ENSG00000196712 | NF1 | -0,199716596 | 0,870721591 | 0,55347737 | 0,93717052 | neurofibromin 1 |
| ENSG00000132436 | FIGNL1 | -0,19972799 | 0,870714715 | 0,68211116 | 0,964699524 | fidgetin like 1 |
| ENSG00000179532 | DNHD1 | -0,199760087 | 0,870695344 | 0,48804678 | 0,915188998 | dynein heavy chain domain 1 |
| ENSG00000128609 | NDUFA5 | -0,199808018 | 0,870666417 | 0,69346605 | 0,967340633 | NADH:ubiquinone oxidoreductase subunit A5 |
| ENSG00000049883 | PTCD2 | -0,199889676 | 0,870617137 | 0,57397231 | 0,942475308 | pentatricopeptide repeat domain 2 |
| ENSG00000048991 | R3HDM1 | -0,199891259 | 0,870616182 | 0,58195024 | 0,944149815 | R3H domain containing 1 |
| ENSG00000235169 | SMIM1 | -0,199936848 | 0,870588671 | 0,73511406 | 0,974353474 | small integral membrane protein 1 (Vel blood group) |

| | | | | | | |
|---|---|---|---|---|---|---|
| ENSG00000174672 | BRSK2 | -0,200144078 | 0,870463628 | 0,46696515 | 0,907940653 | BR serine/threonine kinase 2 |
| ENSG00000187105 | HEATR4 | -0,200225521 | 0,87041449 | 0,72387071 | 0,973129253 | HEAT repeat containing 4 |
| ENSG00000282923 | AC093567,2 | -0,200239065 | 0,870406319 | 0,77560942 | 0,978587255 | novel transcript, antisense to CTIF |
| ENSG00000267764 | AC093567,1 | -0,200239065 | 0,870406319 | 0,77560942 | 0,978587255 | novel transcript, antisense to CTIF |
| ENSG00000163467 | TSACC | -0,200325617 | 0,870354102 | 0,7480301 | 0,975901808 | TSSK6 activating cochaperone |
| ENSG00000163611 | SPICE1 | -0,200496851 | 0,870250805 | 0,60674712 | 0,950927937 | spindle and centriole associated protein 1 |
| ENSG00000163939 | PBRM1 | -0,200502885 | 0,870247166 | 0,57687113 | 0,943268782 | polybromo 1 |
| ENSG00000164045 | CDC25A | -0,200514291 | 0,870240285 | 0,42272087 | 0,890403187 | cell division cycle 25A |
| ENSG00000261663 | AC009065,8 | -0,20054295 | 0,870222998 | 0,69853149 | 0,968522643 | novel transcript, antisense to DCI |
| ENSG00000197409 | H3C4 | -0,200610206 | 0,870182431 | 0,45474837 | 0,903424009 | H3 clustered histone 4 |
| ENSG00000168014 | C2CD3 | -0,200687533 | 0,870135791 | 0,37113816 | 0,859119168 | C2 domain containing 3 centriole elongation regulator |
| ENSG00000224080 | UBE2FP1 | -0,200711232 | 0,870121498 | 0,79640522 | 0,980015022 | UBE2F pseudogene 1 |
| ENSG00000112312 | GMNN | -0,200737958 | 0,870105379 | 0,54950788 | 0,936083284 | geminin DNA replication inhibitor |
| ENSG00000272765 | NDUFA6 | -0,200769701 | 0,870086234 | 0,69993289 | 0,968522643 | NADH:ubiquinone oxidoreductase subunit A6 |
| ENSG00000244004 | RPS3AP19 | -0,200785632 | 0,870076626 | 0,80332398 | 0,980866933 | RPS3A pseudogene 19 |
| ENSG00000255491 | AC100858,2 | -0,200788143 | 0,870075112 | 0,76981152 | 0,977742359 | uncharacterized LOC105375743 |
| ENSG00000141428 | C18orf21 | -0,200864089 | 0,870029311 | 0,65314131 | 0,95732962 | chromosome 18 open reading frame 21 |
| ENSG00000228878 | SEPTIN7-DT | -0,20096624 | 0,86996771 | 0,72652657 | 0,973344786 | SEPTIN7 divergent transcript |
| ENSG00000255201 | AC087623,1 | -0,20097041 | 0,869965196 | 0,88322791 | 0,990717729 | novel transcript |
| ENSG00000269900 | RMRP | -0,201033742 | 0,869927006 | 0,46656419 | 0,907940653 | RNA component of mitochondrial RNA processing endoribonuclease |
| ENSG00000276187 | ERMARD | -0,201039242 | 0,86992369 | 0,53812258 | 0,932900071 | ER membrane associated RNA degradation |
| ENSG00000134077 | THUMPD3 | -0,201097789 | 0,869888388 | 0,52970469 | 0,930679791 | THUMP domain containing 3 |
| ENSG00000284656 | AL645937,4 | -0,201160522 | 0,869850563 | 0,87297165 | 0,989308263 | novel transcript |
| ENSG00000229222 | KRT18P4 | -0,201198713 | 0,869827537 | 0,8010896 | 0,980557464 | keratin 18 pseudogene 4 |
| ENSG00000176401 | EID2B | -0,201239545 | 0,869802918 | 0,42963011 | 0,894509566 | EP300 interacting inhibitor of differentiation 2B |
| ENSG00000065057 | NTHL1 | -0,201352832 | 0,86973462 | 0,56960449 | 0,94221458 | nth like DNA glycosylase 1 |
| ENSG00000270165 | AC010530,1 | -0,201382286 | 0,869716864 | 0,82428009 | 0,983374097 | novel transcript |
| ENSG00000261692 | AC092120,1 | -0,201393213 | 0,869710277 | 0,87742416 | 0,990109814 | novel transcript |
| ENSG00000135049 | AGTPBP1 | -0,201661588 | 0,869548505 | 0,65822073 | 0,957906876 | ATP/GTP binding protein 1 |
| ENSG00000145375 | SPATA5 | -0,201766473 | 0,869485291 | 0,45133931 | 0,902062594 | spermatogenesis associated 5 |
| ENSG00000111196 | MAGOHB | -0,201801268 | 0,869464321 | 0,66187987 | 0,959707806 | mago homolog B, exon junction complex subunit |

| | | | | | | |
|---|---|---|---|---|---|---|
| ENSG00000277789 | TMC4 | -0,201889777 | 0,869410981 | 0,77465438 | 0,978553937 | transmembrane channel like 4 |
| ENSG00000135469 | COQ10A | -0,20189522 | 0,869407701 | 0,52617205 | 0,929261495 | coenzyme Q10A |
| ENSG00000173409 | ARV1 | -0,202023985 | 0,869330107 | 0,6548017 | 0,957444211 | ARV1 homolog, fatty acid homeostasis modulator |
| ENSG00000246477 | AF131216,1 | -0,202112832 | 0,869276572 | 0,58676016 | 0,945395317 | uncharacterized LOC101929290 |
| ENSG00000285340 | AC270277,5 | -0,202112832 | 0,869276572 | 0,58676016 | 0,945395317 | uncharacterized LOC101929290 |
| ENSG00000281693 | AC217775,1 | -0,20214085 | 0,86925969 | 0,52530369 | 0,928752911 | novel transcript |
| ENSG00000142197 | DOP1B | -0,202147998 | 0,869255383 | 0,3901459 | 0,870057277 | DOP1 leucine zipper like protein B |
| ENSG00000261777 | AC012184,3 | -0,202150534 | 0,869253855 | 0,59296523 | 0,947416795 | novel transcript, antisense to DDX19B and DDX19A |
| ENSG00000116127 | ALMS1 | -0,202152899 | 0,86925243 | 0,56664656 | 0,941311563 | ALMS1 centrosome and basal body associated protein |
| ENSG00000257135 | ODC1-DT | -0,202169068 | 0,869242689 | 0,5325535 | 0,931291085 | ODC1 divergent transcript |
| ENSG00000157259 | GATAD1 | -0,202187833 | 0,869231382 | 0,44797403 | 0,902062594 | GATA zinc finger domain containing 1 |
| ENSG00000164040 | PGRMC2 | -0,20219385 | 0,869227757 | 0,59269404 | 0,947416795 | progesterone receptor membrane component 2 |
| ENSG00000177432 | NAP1L5 | -0,20223802 | 0,869201145 | 0,55007564 | 0,93612918 | nucleosome assembly protein 1 like 5 |
| ENSG00000197841 | ZNF181 | -0,202300955 | 0,869163229 | 0,62522269 | 0,952793934 | zinc finger protein 181 |
| ENSG00000273063 | AC007250,1 | -0,202337991 | 0,869140916 | 0,89858302 | 0,991832796 | novel transcript |
| ENSG00000172476 | RAB40A | -0,202398236 | 0,869104623 | 0,58461122 | 0,944857905 | RAB40A, member RAS oncogene family |
| ENSG00000110066 | KMT5B | -0,202443294 | 0,869077479 | 0,55567998 | 0,937852451 | lysine methyltransferase 5B |
| ENSG00000273014 | AC018645,3 | -0,202444637 | 0,869076671 | 0,80432982 | 0,980875302 | novel transcript |
| ENSG00000108852 | MPP2 | -0,202484798 | 0,869052478 | 0,41587122 | 0,885986046 | membrane palmitoylated protein 2 |
| ENSG00000273893 | AL133520,1 | -0,202543967 | 0,869016836 | 0,61389848 | 0,952494509 | novel transcript, sense intronic to SLC13A3 |
| ENSG00000152558 | TMEM123 | -0,202563344 | 0,869005164 | 0,68179667 | 0,964654539 | transmembrane protein 123 |
| ENSG00000077312 | SNRPA | -0,202611408 | 0,868976214 | 0,46867029 | 0,907940653 | small nuclear ribonucleoprotein polypeptide A |
| ENSG00000231113 | AL035587,1 | -0,202636805 | 0,868960916 | 0,68831292 | 0,966426338 | novel transcript |
| ENSG00000229980 | TOB1-AS1 | -0,202652769 | 0,868951301 | 0,74485916 | 0,975649186 | TOB1 antisense RNA 1 |
| ENSG00000274712 | AC005332,4 | -0,202658948 | 0,868947579 | 0,41678922 | 0,886368152 | bromodomain PHD finger transcription factor (BPTF) pseudogene |
| ENSG00000114982 | KANSL3 | -0,202793765 | 0,868866382 | 0,36220669 | 0,85231292 | KAT8 regulatory NSL complex subunit 3 |
| ENSG00000165480 | SKA3 | -0,20287858 | 0,868815303 | 0,62479518 | 0,952793934 | spindle and kinetochore associated complex subunit 3 |
| ENSG00000261603 | PRSS46P | -0,203003343 | 0,868740172 | 0,89019818 | 0,990717729 | serine protease 46, pseudogene |
| ENSG00000225891 | DHDDS-AS1 | -0,203028724 | 0,868724889 | 0,68068663 | 0,964654539 | DHDDS antisense RNA 1 |
| ENSG00000213669 | AL137074,1 | -0,203041205 | 0,868717373 | 0,89246295 | 0,991048031 | ribosomal protein L21 (RPL21) pseudogene |

| | | | | | | |
|---|---|---|---|---|---|---|
| ENSG00000176973 | FAM89B | -0,203188902 | 0,868628442 | 0,4712352 | 0,908869685 | family with sequence similarity 89 member B |
| ENSG00000144120 | TMEM177 | -0,203303602 | 0,868559385 | 0,44323918 | 0,899774672 | transmembrane protein 177 |
| ENSG00000138750 | NUP54 | -0,203385729 | 0,868509943 | 0,6700881 | 0,962026636 | nucleoporin 54 |
| ENSG00000170364 | SETMAR | -0,203416004 | 0,868491718 | 0,4814447 | 0,912981134 | SET domain and mariner transposase fusion gene |
| ENSG00000224914 | LINC00863 | -0,203499892 | 0,86844122 | 0,41302621 | 0,884663685 | long intergenic non-protein coding RNA 863 |
| ENSG00000262145 | ATP23 | -0,203528975 | 0,868423713 | 0,80119838 | 0,980557464 | ATP23 metallopeptidase and ATP synthase assembly factor homolog |
| ENSG00000227014 | FKBP14-AS1 | -0,203550393 | 0,86841082 | 0,65138234 | 0,957089277 | FKBP14 antisense RNA 1 |
| ENSG00000147536 | GINS4 | -0,203581133 | 0,868392317 | 0,37408978 | 0,861369706 | GINS complex subunit 4 |
| ENSG00000135870 | RC3H1 | -0,203589427 | 0,868387325 | 0,58075435 | 0,943832501 | ring finger and CCCH-type domains 1 |
| ENSG00000163041 | H3-3A | -0,203784493 | 0,868269918 | 0,6905122 | 0,967179044 | H3,3 histone A |
| ENSG00000269155 | AL009178,2 | -0,203784703 | 0,868269792 | 0,689058 | 0,966647634 | novel transcript |
| ENSG00000138688 | KIAA1109 | -0,203832709 | 0,8682409 | 0,59084392 | 0,947416795 | KIAA1109 |
| ENSG00000147117 | ZNF157 | -0,203833916 | 0,868240174 | 0,70359982 | 0,968967751 | zinc finger protein 157 |
| ENSG00000139537 | CCDC65 | -0,203896052 | 0,86820278 | 0,48987603 | 0,916063942 | coiled-coil domain containing 65 |
| ENSG00000188124 | OR2AG2 | -0,203938472 | 0,868177253 | 0,7385075 | 0,974353474 | olfactory receptor family 2 subfamily AG member 2 |
| ENSG00000166347 | CYB5A | -0,203944345 | 0,868173718 | 0,3719416 | 0,859937929 | cytochrome b5 type A |
| ENSG00000278719 | MCM8-AS1 | -0,203980647 | 0,868151873 | 0,79860834 | 0,980161849 | MCM8 antisense RNA 1 |
| ENSG00000224402 | OR6D1P | -0,20398636 | 0,868148435 | 0,61155531 | 0,952155617 | olfactory receptor family 6 subfamily D member 1 pseudogene |
| ENSG00000270133 | AC025766,1 | -0,204054863 | 0,868107214 | 0,86561649 | 0,989074806 | novel transcript, sense intronic to C5orf36 |
| ENSG00000170921 | TANC2 | -0,20406013 | 0,868104045 | 0,44844838 | 0,902062594 | tetratricopeptide repeat, ankyrin repeat and coiled-coil containing 2 |
| ENSG00000145730 | PAM | -0,20410098 | 0,868079465 | 0,52830102 | 0,930000172 | peptidylglycine alpha-amidating monooxygenase |
| ENSG00000276495 | APBA2 | -0,204150507 | 0,868049665 | 0,32372171 | 0,825794844 | amyloid beta precursor protein binding family A member 2 |
| ENSG00000188612 | SUMO2 | -0,204162271 | 0,868042587 | 0,61163451 | 0,952155617 | small ubiquitin like modifier 2 |
| ENSG00000100842 | EFS | -0,204225012 | 0,868004838 | 0,40130787 | 0,876659568 | embryonal Fyn-associated substrate |
| ENSG00000149582 | TMEM25 | -0,204229315 | 0,868002248 | 0,3638033 | 0,854140004 | transmembrane protein 25 |
| ENSG00000142549 | IGLON5 | -0,204338927 | 0,867936303 | 0,46237307 | 0,9064797 | IgLON family member 5 |
| ENSG00000258477 | PPIAP6 | -0,204347392 | 0,86793121 | 0,85420506 | 0,987861659 | peptidylprolyl isomerase A pseudogene 6 |
| ENSG00000089220 | PEBP1 | -0,204387724 | 0,867906946 | 0,45254849 | 0,902285042 | phosphatidylethanolamine binding protein 1 |
| ENSG00000007255 | TRAPPC6A | -0,204498209 | 0,867840483 | 0,4575208 | 0,903831777 | trafficking protein particle complex 6A |

| | | | | | | |
|---|---|---|---|---|---|---|
| ENSG00000235369 | RPL36AP15 | -0,204525293 | 0,867824191 | 0,78580708 | 0,979320406 | ribosomal protein L36a pseudogene 15 |
| ENSG00000137210 | TMEM14B | -0,204537507 | 0,867816843 | 0,56248934 | 0,940865625 | transmembrane protein 14B |
| ENSG00000237094 | AL732372,2 | -0,204553448 | 0,867807255 | 0,46923546 | 0,907940653 | pseudogene similar to part of septin 14 SEPT14 |
| ENSG00000257534 | AC023794,4 | -0,204628994 | 0,867761814 | 0,84020075 | 0,985672131 | novel transcript |
| ENSG00000273451 | AL031666,3 | -0,204745543 | 0,867691714 | 0,59697112 | 0,948282297 | novel transcript, sense intronic to ZMYND8 |
| ENSG00000288130 | TNRC6A | -0,204775101 | 0,867673937 | 0,49218044 | 0,916995341 | trinucleotide repeat containing adaptor 6A |
| ENSG00000272853 | AC069544,1 | -0,204924854 | 0,867583876 | 0,76878434 | 0,977381742 | novel transcript |
| ENSG00000158023 | CFAP251 | -0,204928346 | 0,867581776 | 0,50829957 | 0,922427526 | cilia and flagella associated protein 251 |
| ENSG00000117906 | RCN2 | -0,204970226 | 0,867556592 | 0,57840306 | 0,943425812 | reticulocalbin 2 |
| ENSG00000125821 | DTD1 | -0,205224747 | 0,86740355 | 0,40373362 | 0,877691457 | D-aminoacyl-tRNA deacylase 1 |
| ENSG00000122547 | EEPD1 | -0,205302778 | 0,867356636 | 0,50565229 | 0,921780819 | endonuclease/exonuclease/phosphatase family domain containing 1 |
| ENSG00000172728 | FUT10 | -0,205365447 | 0,86731896 | 0,51915164 | 0,926800715 | fucosyltransferase 10 |
| ENSG00000253948 | VPS13B-DT | -0,205497228 | 0,86723974 | 0,59443055 | 0,947469769 | VPS13B divergent transcript |
| ENSG00000249379 | AL033397,2 | -0,205512803 | 0,867230377 | 0,8020631 | 0,980710009 | novel transcript |
| ENSG00000166793 | YPEL4 | -0,205551246 | 0,867207269 | 0,56330604 | 0,940865625 | yippee like 4 |
| ENSG00000281761 | AL078605,3 | -0,205584927 | 0,867187023 | 0,86219133 | 0,989074806 | novel transcript |
| ENSG00000271234 | AL078605,1 | -0,205584927 | 0,867187023 | 0,86219133 | 0,989074806 | novel transcript |
| ENSG00000162924 | REL | -0,205602104 | 0,867176698 | 0,44095076 | 0,898675567 | REL proto-oncogene, NF-kB subunit |
| ENSG00000158985 | CDC42SE2 | -0,205744856 | 0,867090897 | 0,63796622 | 0,956018518 | CDC42 small effector 2 |
| ENSG00000112137 | PHACTR1 | -0,205755681 | 0,867084391 | 0,39092935 | 0,870057277 | phosphatase and actin regulator 1 |
| ENSG00000224315 | RPL7P7 | -0,205769737 | 0,867075943 | 0,70931087 | 0,970328821 | ribosomal protein L7 pseudogene 7 |
| ENSG00000230799 | RPS2P16 | -0,205835295 | 0,867036543 | 0,78090818 | 0,978597905 | ribosomal protein S2 pseudogene 16 |
| ENSG00000130193 | THEM6 | -0,205836874 | 0,867035594 | 0,5589649 | 0,939635125 | thioesterase superfamily member 6 |
| ENSG00000188199 | NUTM2B | -0,205852586 | 0,867026152 | 0,66166235 | 0,959603271 | NUT family member 2B |
| ENSG00000174456 | C12orf76 | -0,20591595 | 0,866988072 | 0,40215424 | 0,87718264 | chromosome 12 open reading frame 76 |
| ENSG00000066735 | KIF26A | -0,206031302 | 0,866918754 | 0,76383942 | 0,976640468 | kinesin family member 26A |
| ENSG00000166845 | C18orf54 | -0,206037023 | 0,866915316 | 0,6277779 | 0,952793934 | chromosome 18 open reading frame 54 |
| ENSG00000186665 | C17orf58 | -0,206083781 | 0,866887219 | 0,50670117 | 0,921824694 | chromosome 17 open reading frame 58 |
| ENSG00000258448 | AL442663,2 | -0,206172181 | 0,866834103 | 0,80165056 | 0,980623173 | required for meiotic nuclear division 1 homolog (S, cerevisiae) (RMND1) pseudogene |
| ENSG00000109323 | MANBA | -0,206252816 | 0,866785656 | 0,58029295 | 0,943828882 | mannosidase beta |
| ENSG00000136527 | TRA2B | -0,206287484 | 0,866764827 | 0,60274704 | 0,950352286 | transformer 2 beta homolog |

| | | | | | | |
|---|---|---|---|---|---|---|
| ENSG00000205041 | AC118344,1 | -0,206335114 | 0,866736212 | 0,73822124 | 0,974353474 | novel transcript, sense intronic to AKT2 |
| ENSG00000111271 | ACAD10 | -0,206402868 | 0,866695508 | 0,39130795 | 0,870117476 | acyl-CoA dehydrogenase family member 10 |
| ENSG00000288033 | AC137894,4 | -0,20644891 | 0,866667849 | 0,57188907 | 0,942475308 | novel transcript |
| ENSG00000117748 | RPA2 | -0,206481281 | 0,866648403 | 0,56925502 | 0,94219857 | replication protein A2 |
| ENSG00000134146 | DPH6 | -0,206543013 | 0,86661132 | 0,36311879 | 0,853318246 | diphthamine biosynthesis 6 |
| ENSG00000273597 | AL355377,1 | -0,206565893 | 0,866597577 | 0,7526956 | 0,975901808 | SPATA31 subfamily D, member 1 (SPATA31D1) pseudogene |
| ENSG00000076108 | BAZ2A | -0,206574355 | 0,866592494 | 0,35601405 | 0,847322116 | bromodomain adjacent to zinc finger domain 2A |
| ENSG00000112996 | MRPS30 | -0,206582234 | 0,866587761 | 0,57152351 | 0,942436428 | mitochondrial ribosomal protein S30 |
| ENSG00000153446 | C16orf89 | -0,206604875 | 0,866574161 | 0,89226502 | 0,991039277 | chromosome 16 open reading frame 89 |
| ENSG00000259460 | AC078909,1 | -0,206633787 | 0,866556795 | 0,87292881 | 0,989308263 | novel transcript, antisense to MEIS2 |
| ENSG00000270706 | PRMT1P1 | -0,206747879 | 0,866488268 | 0,63792835 | 0,956018518 | protein arginine methyltransferase 1 pseudogene 1 |
| ENSG00000244056 | RN7SL417P | -0,20687902 | 0,866409508 | 0,78194417 | 0,978597905 | RNA, 7SL, cytoplasmic 417, pseudogene |
| ENSG00000137941 | TTLL7 | -0,206975972 | 0,866351285 | 0,53632009 | 0,93228049 | tubulin tyrosine ligase like 7 |
| ENSG00000286863 | AL513166,1 | -0,207270893 | 0,866174201 | 0,68599945 | 0,965740648 | novel transcript |
| ENSG00000117419 | ERI3 | -0,207330518 | 0,866138403 | 0,38837866 | 0,869820053 | ERI1 exoribonuclease family member 3 |
| ENSG00000214135 | AC132008,2 | -0,207510532 | 0,866030337 | 0,50435001 | 0,921400408 | succinate dehydrogenase complex, subunit A, flavoprotein (Fp) (SDHA) pseudogene |
| ENSG00000286248 | AC069209,2 | -0,20754209 | 0,866011393 | 0,84923047 | 0,987036295 | novel transcript, antisense to P2RX7 |
| ENSG00000231357 | AC006028,1 | -0,207542462 | 0,86601117 | 0,90650403 | 0,992590661 | prefoldin subunit 4 (PFDN4) pseudogene |
| ENSG00000234740 | AL162386,2 | -0,207548279 | 0,866007678 | 0,7415035 | 0,975176043 | novel transcript |
| ENSG00000140948 | ZCCHC14 | -0,207549956 | 0,866006671 | 0,39288048 | 0,871121102 | zinc finger CCHC-type containing 14 |
| ENSG00000278259 | MYO19 | -0,207604467 | 0,865973951 | 0,50683603 | 0,921852439 | myosin XIX |
| ENSG00000236177 | HLA-DPA1 | -0,20770828 | 0,865911639 | 0,86142633 | 0,989074806 | major histocompatibility complex, class II, DP alpha 1 |
| ENSG00000175003 | SLC22A1 | -0,2077226 | 0,865903045 | 0,7909786 | 0,979471509 | solute carrier family 22 member 1 |
| ENSG00000136770 | DNAJC1 | -0,207873017 | 0,865812769 | 0,43205737 | 0,89558934 | DnaJ heat shock protein family (Hsp40) member C1 |
| ENSG00000167094 | TTC16 | -0,20788448 | 0,86580589 | 0,84069999 | 0,985672131 | tetratricopeptide repeat domain 16 |
| ENSG00000135040 | NAA35 | -0,207892867 | 0,865800856 | 0,50969207 | 0,922427526 | N-alpha-acetyltransferase 35, NatC auxiliary subunit |
| ENSG00000073331 | ALPK1 | -0,207897023 | 0,865798362 | 0,51749307 | 0,925831376 | alpha kinase 1 |
| ENSG00000260473 | AC068987,2 | -0,207901372 | 0,865795752 | 0,77697004 | 0,978587255 | novel transcript |
| ENSG00000235307 | BRD2 | -0,207953448 | 0,865764501 | 0,89145924 | 0,990820775 | bromodomain containing 2 |
| ENSG00000148204 | CRB2 | -0,207996444 | 0,865738699 | 0,64369638 | 0,956633898 | crumbs cell polarity complex component 2 |

| | | | | | | |
|---|---|---|---|---|---|---|
| ENSG00000168564 | CDKN2AIP | -0,208000693 | 0,86573615 | 0,59238147 | 0,947416795 | CDKN2A interacting protein |
| ENSG00000171735 | CAMTA1 | -0,208066944 | 0,865696394 | 0,57573919 | 0,942847946 | calmodulin binding transcription activator 1 |
| ENSG00000234545 | FAM133B | -0,208071046 | 0,865693933 | 0,55602604 | 0,938031417 | family with sequence similarity 133 member B |
| ENSG00000196376 | SLC35F1 | -0,208172812 | 0,86563287 | 0,52321765 | 0,927995032 | solute carrier family 35 member F1 |
| ENSG00000197006 | METTL9 | -0,208331979 | 0,865537374 | 0,57909905 | 0,943828882 | methyltransferase like 9 |
| ENSG00000104321 | TRPA1 | -0,208345799 | 0,865529082 | 0,67508713 | 0,963388897 | transient receptor potential cation channel subfamily A member 1 |
| ENSG00000167775 | CD320 | -0,208374713 | 0,865511736 | 0,5919812 | 0,947416795 | CD320 molecule |
| ENSG00000135439 | AGAP2 | -0,208426085 | 0,865480917 | 0,52081742 | 0,927046149 | ArfGAP with GTPase domain, ankyrin repeat and PH domain 2 |
| ENSG00000288105 | AC117461,1 | -0,208479366 | 0,865448954 | 0,63265787 | 0,954291097 | novel transcript, antisense to ROBO1 |
| ENSG00000281706 | LINC01012 | -0,2085183 | 0,865425599 | 0,55102416 | 0,936366211 | long intergenic non-protein coding RNA 1012 |
| ENSG00000276740 | AL445649,1 | -0,208776227 | 0,86527089 | 0,73345624 | 0,974353474 | novel transcript, sense intronic to FAM155A |
| ENSG00000253406 | AC012613,2 | -0,208848471 | 0,865227562 | 0,85853318 | 0,989074806 | novel transcript, antisense to ABLIM3 |
| ENSG00000261864 | AC130462,2 | -0,208912199 | 0,865189343 | 0,80436133 | 0,980875302 | novel transcript |
| ENSG00000267395 | DM1-AS | -0,20894234 | 0,865171268 | 0,78855993 | 0,979462318 | DM1 locus antisense RNA |
| ENSG00000134905 | CARS2 | -0,209233549 | 0,86499665 | 0,32247914 | 0,824955371 | cysteinyl-tRNA synthetase 2, mitochondrial |
| ENSG00000126870 | DYNC2I1 | -0,209264047 | 0,864978365 | 0,46363496 | 0,907133332 | dynein 2 intermediate chain 1 |
| ENSG00000227056 | RPL6P2 | -0,209359563 | 0,864921099 | 0,83406092 | 0,984905093 | ribosomal protein L6 pseudogene 2 |
| ENSG00000184828 | ZBTB7C | -0,209404807 | 0,864893975 | 0,69955702 | 0,968522643 | zinc finger and BTB domain containing 7C |
| ENSG00000198482 | ZNF808 | -0,209407177 | 0,864892554 | 0,54933685 | 0,936040289 | zinc finger protein 808 |
| ENSG00000131899 | LLGL1 | -0,209474699 | 0,864852076 | 0,62653424 | 0,952793934 | LLGL scribble cell polarity complex component 1 |
| ENSG00000113522 | RAD50 | -0,209530512 | 0,864818618 | 0,53624297 | 0,932274935 | RAD50 double strand break repair protein |
| ENSG00000154640 | BTG3 | -0,209533568 | 0,864816787 | 0,6210101 | 0,952793934 | BTG anti-proliferation factor 3 |
| ENSG00000134955 | SLC37A2 | -0,209604596 | 0,86477421 | 0,61832902 | 0,952793934 | solute carrier family 37 member 2 |
| ENSG00000243302 | AC018638,4 | -0,209604794 | 0,864774092 | 0,44056849 | 0,898495324 | novel pseudogene |
| ENSG00000253764 | AC019257,1 | -0,209646926 | 0,864748837 | 0,53701489 | 0,932883277 | novel transcript |
| ENSG00000278000 | AC139100,2 | -0,209672579 | 0,864733461 | 0,69854195 | 0,968522643 | novel transcript |
| ENSG00000131747 | TOP2A | -0,209712358 | 0,864709618 | 0,59813086 | 0,948582224 | DNA topoisomerase II alpha |
| ENSG00000273004 | AL078644,1 | -0,209855174 | 0,864624023 | 0,65049328 | 0,956899963 | novel transcript |
| ENSG00000138078 | PREPL | -0,209924361 | 0,864582559 | 0,62859365 | 0,952793934 | prolyl endopeptidase like |
| ENSG00000260942 | CAPN10-DT | -0,210133618 | 0,864457164 | 0,48936182 | 0,915936198 | CAPN10 divergent transcript |
| ENSG00000268043 | NBPF12 | -0,210137439 | 0,864454875 | 0,35395933 | 0,845346751 | NBPF member 12 |

| | | | | | | |
|---|---|---|---|---|---|---|
| ENSG00000263878 | DLGAP1-AS4 | -0,210139097 | 0,864453881 | 0,7723557 | 0,97820249 | DLGAP1 antisense RNA 4 |
| ENSG00000275764 | AC092747,4 | -0,210157816 | 0,864442665 | 0,65860146 | 0,95814533 | novel transcript |
| ENSG00000230551 | AC021078,1 | -0,210217246 | 0,864407056 | 0,47900157 | 0,912714933 | novel transcript |
| ENSG00000233232 | NPIPB7 | -0,21024208 | 0,864392177 | 0,72711202 | 0,973344786 | nuclear pore complex interacting protein family member B7 |
| ENSG00000285119 | FAM86B3P | -0,210326322 | 0,864341705 | 0,51111893 | 0,922893865 | family with sequence similarity 86 member B3, pseudogene |
| ENSG00000251369 | ZNF550 | -0,210345341 | 0,86433031 | 0,40901066 | 0,881822587 | zinc finger protein 550 |
| ENSG00000140993 | TIGD7 | -0,210405492 | 0,864294274 | 0,58536235 | 0,945213113 | tigger transposable element derived 7 |
| ENSG00000183648 | NDUFB1 | -0,210411109 | 0,864290908 | 0,52354262 | 0,928038684 | NADH:ubiquinone oxidoreductase subunit B1 |
| ENSG00000271803 | AL121906,1 | -0,210484887 | 0,864246711 | 0,87492552 | 0,990078774 | novel transcript, antisense to NECAB3 |
| ENSG00000081665 | ZNF506 | -0,210642601 | 0,864152237 | 0,49918065 | 0,919933362 | zinc finger protein 506 |
| ENSG00000288470 | LINC02429 | -0,2107006 | 0,864117498 | 0,80131625 | 0,980557464 | long intergenic non-protein coding RNA 2429 |
| ENSG00000267128 | RNF157-AS1 | -0,21080384 | 0,864055663 | 0,66703933 | 0,9611116 | RNF157 antisense RNA 1 |
| ENSG00000117625 | RCOR3 | -0,210836747 | 0,864035955 | 0,59199983 | 0,947416795 | REST corepressor 3 |
| ENSG00000277382 | AC005837,4 | -0,210837933 | 0,864035245 | 0,85330716 | 0,987861659 | novel transcript, antisense to MXRA7 |
| ENSG00000170291 | ELP5 | -0,210839935 | 0,864034045 | 0,40962022 | 0,882177108 | elongator acetyltransferase complex subunit 5 |
| ENSG00000288485 | ELP5 | -0,210839935 | 0,864034045 | 0,40962022 | 0,882177108 | elongator acetyltransferase complex subunit 5 |
| ENSG00000239556 | AC004951,2 | -0,210894954 | 0,864001095 | 0,82869616 | 0,983784613 | uroplakin-like protein (UPLP) pseudogene |
| ENSG00000253159 | PCDHGA12 | -0,210920424 | 0,863985842 | 0,63989392 | 0,956018518 | protocadherin gamma subfamily A, 12 |
| ENSG00000164967 | RPP25L | -0,210985015 | 0,863947161 | 0,45871995 | 0,904420115 | ribonuclease P/MRP subunit p25 like |
| ENSG00000250132 | AC004803,1 | -0,21099707 | 0,863939942 | 0,48113474 | 0,912911021 | novel transcript |
| ENSG00000081721 | DUSP12 | -0,211031245 | 0,863919477 | 0,50010032 | 0,92041337 | dual specificity phosphatase 12 |
| ENSG00000013523 | ANGEL1 | -0,211041944 | 0,86391307 | 0,47069976 | 0,908869097 | angel homolog 1 |
| ENSG00000285218 | AC026316,4 | -0,211090506 | 0,863883991 | 0,95753365 | 0,996608711 | novel protein |
| ENSG00000086200 | IPO11 | -0,211132377 | 0,863858919 | 0,61984902 | 0,952793934 | importin 11 |
| ENSG00000287452 | AL391477,1 | -0,211160618 | 0,863842009 | 0,63640229 | 0,955547532 | novel transcript |
| ENSG00000178163 | ZNF518B | -0,211239202 | 0,863794956 | 0,62105713 | 0,952793934 | zinc finger protein 518B |
| ENSG00000166368 | OR2D2 | -0,211251206 | 0,863787769 | 0,78672246 | 0,979320406 | olfactory receptor family 2 subfamily D member 2 |
| ENSG00000140598 | EFL1 | -0,211272222 | 0,863775187 | 0,53791853 | 0,932883277 | elongation factor like GTPase 1 |
| ENSG00000235297 | FAUP1 | -0,211274474 | 0,863773838 | 0,86403795 | 0,989074806 | FAU pseudogene 1 |
| ENSG00000196981 | WDR5B | -0,211394224 | 0,863702144 | 0,61291018 | 0,952413269 | WD repeat domain 5B |
| ENSG00000257621 | PSMA3-AS1 | -0,211440522 | 0,863674427 | 0,50378813 | 0,921200643 | PSMA3 antisense RNA 1 |

| | | | | | | |
|---|---|---|---|---|---|---|
| ENSG00000270433 | H3P37 | -0,211538652 | 0,863615683 | 0,72208873 | 0,972531855 | H3 histone pseudogene 37 |
| ENSG00000286817 | AC105339,6 | -0,211587892 | 0,863586208 | 0,67457378 | 0,963183892 | novel transcript |
| ENSG00000214070 | MANEALP1 | -0,211656921 | 0,863544889 | 0,84356568 | 0,986273696 | MANEAL pseudogene 1 |
| ENSG00000146143 | PRIM2 | -0,211673621 | 0,863534893 | 0,59016083 | 0,947262972 | DNA primase subunit 2 |
| ENSG00000144229 | THSD7B | -0,211738307 | 0,863496175 | 0,51507201 | 0,924720046 | thrombospondin type 1 domain containing 7B |
| ENSG00000082146 | STRADB | -0,211852164 | 0,863428032 | 0,62388997 | 0,952793934 | STE20 related adaptor beta |
| ENSG00000175155 | YPEL2 | -0,211910757 | 0,863392965 | 0,46668089 | 0,907940653 | yippee like 2 |
| ENSG00000230910 | AL391807,1 | -0,211925297 | 0,863384264 | 0,61104901 | 0,951873283 | novel transcript |
| ENSG00000148604 | RGR | -0,211932104 | 0,86338019 | 0,72960711 | 0,974009551 | retinal G protein coupled receptor |
| ENSG00000187999 | HNRNPA1P61 | -0,211933748 | 0,863379206 | 0,83939403 | 0,985672131 | heterogeneous nuclear ribonucleoprotein A1 pseudogene 61 |
| ENSG00000135299 | ANKRD6 | -0,211953668 | 0,863367285 | 0,48756312 | 0,914936533 | ankyrin repeat domain 6 |
| ENSG00000239511 | POM121L7P | -0,212001103 | 0,863338898 | 0,74282095 | 0,975201495 | POM121 transmembrane nucleoporin like 7 pseudogene |
| ENSG00000185220 | PGBD2 | -0,212039378 | 0,863315994 | 0,50118335 | 0,920520138 | piggyBac transposable element derived 2 |
| ENSG00000237704 | AP004289,2 | -0,212148407 | 0,863250753 | 0,7566775 | 0,975901808 | male-specific lethal 3-like 1 (Drosophila) (MSL3L1) pseudogene |
| ENSG00000231329 | AL031772,1 | -0,212291244 | 0,86316529 | 0,6967542 | 0,968300554 | novel transcript |
| ENSG00000206633 | SNORA80B | -0,212415036 | 0,863091228 | 0,77074098 | 0,977979869 | small nucleolar RNA, H/ACA box 80B |
| ENSG00000230393 | AC092667,1 | -0,212541021 | 0,863015861 | 0,65554781 | 0,957444211 | novel transcript |
| ENSG00000287222 | AC234917,3 | -0,212578719 | 0,86299331 | 0,51648362 | 0,925349219 | novel transcript, antisense to PUF60 |
| ENSG00000168246 | UBTD2 | -0,21259265 | 0,862984977 | 0,56811722 | 0,94163744 | ubiquitin domain containing 2 |
| ENSG00000262879 | AC005670,3 | -0,212627146 | 0,862964342 | 0,38071019 | 0,865900094 | novel transcript |
| ENSG00000122257 | RBBP6 | -0,212683364 | 0,862930716 | 0,44307658 | 0,899690238 | RB binding protein 6, ubiquitin ligase |
| ENSG00000120306 | CYSTM1 | -0,212830347 | 0,862842804 | 0,55225447 | 0,936619966 | cysteine rich transmembrane module containing 1 |
| ENSG00000124802 | EEF1E1 | -0,212963719 | 0,862763041 | 0,69962766 | 0,968522643 | eukaryotic translation elongation factor 1 epsilon 1 |
| ENSG00000247498 | GPRC5D-AS1 | -0,213016443 | 0,862731512 | 0,74724216 | 0,97584218 | GPRC5D and HEBP1 antisense RNA 1 |
| ENSG00000282641 | AC183084,1 | -0,21303036 | 0,862723189 | 0,70141604 | 0,968742626 | aminopeptidase puromycin sensitive (NPEPPS) pseudogene |
| ENSG00000164543 | STK17A | -0,213033889 | 0,862721079 | 0,47539762 | 0,91115935 | serine/threonine kinase 17a |
| ENSG00000176490 | DIRAS1 | -0,213051771 | 0,862710386 | 0,3518179 | 0,844406182 | DIRAS family GTPase 1 |
| ENSG00000267234 | AC092067,1 | -0,213175397 | 0,862636462 | 0,85645039 | 0,988229014 | novel transcript, antisense to GATAD2A |
| ENSG00000104450 | SPAG1 | -0,213344816 | 0,862535167 | 0,60354797 | 0,950352286 | sperm associated antigen 1 |

| | | | | | | |
|---|---|---|---|---|---|---|
| ENSG00000138430 | OLA1 | -0,213424602 | 0,862487467 | 0,62211681 | 0,952793934 | Obg like ATPase 1 |
| ENSG00000259125 | LRP1-AS | -0,213456137 | 0,862468614 | 0,83915763 | 0,985672131 | LRP1 antisense RNA |
| ENSG00000287105 | AC090577,1 | -0,213542712 | 0,86241686 | 0,66489546 | 0,960551345 | novel transcript, antisense to NKAIN3 |
| ENSG00000204514 | ZNF814 | -0,213571873 | 0,862399428 | 0,3766465 | 0,862818448 | zinc finger protein 814 |
| ENSG00000152457 | DCLRE1C | -0,213613683 | 0,862374436 | 0,50921633 | 0,922427526 | DNA cross-link repair 1C |
| ENSG00000049323 | LTBP1 | -0,213624796 | 0,862367793 | 0,43840938 | 0,897220559 | latent transforming growth factor beta binding protein 1 |
| ENSG00000256594 | AC010186,2 | -0,213686731 | 0,862330773 | 0,48662631 | 0,914296452 | C-type lectin domain family 2 (CLEC2) pseudogene |
| ENSG00000253961 | AC009563,1 | -0,213688312 | 0,862329828 | 0,834102 | 0,984905093 | novel transcript |
| ENSG00000227456 | LINC00310 | -0,213756626 | 0,862288996 | 0,66072394 | 0,959226315 | long intergenic non-protein coding RNA 310 |
| ENSG00000254206 | NPIPB11 | -0,213791976 | 0,862267868 | 0,65762628 | 0,957864535 | nuclear pore complex interacting protein family member B11 |
| ENSG00000188610 | FAM72B | -0,213892656 | 0,862207695 | 0,54069836 | 0,933385624 | family with sequence similarity 72 member B |
| ENSG00000188365 | AC092171,1 | -0,213898148 | 0,862204413 | 0,67439832 | 0,963103112 | novel transcript |
| ENSG00000203647 | AC106827,1 | -0,213993986 | 0,862147139 | 0,80250724 | 0,980789891 | novel pseudogene |
| ENSG00000134627 | PIWIL4 | -0,214013415 | 0,862135528 | 0,65619931 | 0,957444211 | piwi like RNA-mediated gene silencing 4 |
| ENSG00000005249 | PRKAR2B | -0,214045644 | 0,862116269 | 0,59288787 | 0,947416795 | protein kinase cAMP-dependent type II regulatory subunit beta |
| ENSG00000284096 | PRKAR2B | -0,214045644 | 0,862116269 | 0,59288787 | 0,947416795 | protein kinase cAMP-dependent type II regulatory subunit beta |
| ENSG00000115966 | ATF2 | -0,214140947 | 0,86205932 | 0,65173281 | 0,957103407 | activating transcription factor 2 |
| ENSG00000171320 | ESCO2 | -0,214145481 | 0,862056611 | 0,57699668 | 0,943288205 | establishment of sister chromatid cohesion N-acetyltransferase 2 |
| ENSG00000177425 | PAWR | -0,214151811 | 0,862052829 | 0,54405974 | 0,934259495 | pro-apoptotic WT1 regulator |
| ENSG00000272518 | AC036214,2 | -0,214291679 | 0,861969257 | 0,644646 | 0,956633898 | novel transcript, sense intronic to MRPS28 |
| ENSG00000260007 | AC107871,1 | -0,214292653 | 0,861968675 | 0,495901 | 0,918105073 | novel protein |
| ENSG00000171466 | ZNF562 | -0,214434804 | 0,861883748 | 0,34564796 | 0,841168721 | zinc finger protein 562 |
| ENSG00000267313 | KC6 | -0,214464655 | 0,861865915 | 0,55899683 | 0,939635125 | keratoconus gene 6 |
| ENSG00000250159 | AC106791,1 | -0,214479576 | 0,861857002 | 0,36477889 | 0,854906078 | novel transcript |
| ENSG00000278588 | H2BC10 | -0,214481842 | 0,861855648 | 0,60317912 | 0,950352286 | H2B clustered histone 10 |
| ENSG00000287965 | AC018557,3 | -0,214545977 | 0,861817335 | 0,78387314 | 0,979239248 | novel transcript, antisense to DYNC1LI2 |
| ENSG00000137135 | ARHGEF39 | -0,214742488 | 0,861699954 | 0,49894927 | 0,919706965 | Rho guanine nucleotide exchange factor 39 |
| ENSG00000286584 | AC023494,1 | -0,215189983 | 0,861432714 | 0,75606559 | 0,975901808 | novel transcript, antisense to COX17 |

| | | | | | | |
|---|---|---|---|---|---|---|
| ENSG00000167700 | MFSD3 | -0,215239645 | 0,861403061 | 0,65093003 | 0,957027739 | major facilitator superfamily domain containing 3 |
| ENSG00000072832 | CRMP1 | -0,215280283 | 0,861378797 | 0,29471376 | 0,804859966 | collapsin response mediator protein 1 |
| ENSG00000079257 | LXN | -0,215340929 | 0,861342588 | 0,62167442 | 0,952793934 | latexin |
| ENSG00000101680 | LAMA1 | -0,215436222 | 0,861285697 | 0,49343026 | 0,917289924 | laminin subunit alpha 1 |
| ENSG00000251474 | RPL32P3 | -0,215439102 | 0,861283977 | 0,33405996 | 0,834518413 | ribosomal protein L32 pseudogene 3 |
| ENSG00000260018 | AC040169,1 | -0,215476874 | 0,861261428 | 0,75362602 | 0,975901808 | novel transcript, antisense to MBTPS1 |
| ENSG00000241713 | LY6G5B | -0,215566459 | 0,861207949 | 0,55943223 | 0,939727587 | lymphocyte antigen 6 family member G5B |
| ENSG00000047230 | CTPS2 | -0,215668499 | 0,861147039 | 0,55006821 | 0,93612918 | CTP synthase 2 |
| ENSG00000173473 | SMARCC1 | -0,215773232 | 0,861084526 | 0,41724606 | 0,886651836 | SWI/SNF related, matrix associated, actin dependent regulator of chromatin subfamily c member 1 |
| ENSG00000179314 | WSCD1 | -0,215812635 | 0,861061008 | 0,35797682 | 0,849032846 | WSC domain containing 1 |
| ENSG00000186020 | ZNF529 | -0,215857223 | 0,861034397 | 0,56469916 | 0,940865625 | zinc finger protein 529 |
| ENSG00000221818 | EBF2 | -0,215964408 | 0,860970429 | 0,68525285 | 0,965425099 | EBF transcription factor 2 |
| ENSG00000283445 | AL136985,3 | -0,215966172 | 0,860969376 | 0,39193962 | 0,870625724 | novel transcript |
| ENSG00000129028 | THAP10 | -0,216026194 | 0,860933557 | 0,56060218 | 0,940170934 | THAP domain containing 10 |
| ENSG00000255693 | LINC02389 | -0,216095124 | 0,860892423 | 0,71925904 | 0,971869926 | long intergenic non-protein coding RNA 2389 |
| ENSG00000224360 | DDR1-DT | -0,216391468 | 0,860715606 | 0,76620154 | 0,977110125 | DDR1 divergent transcript |
| ENSG00000163875 | MEAF6 | -0,216494736 | 0,860653998 | 0,56702746 | 0,941425553 | MYST/Esa1 associated factor 6 |
| ENSG00000272148 | AC013403,2 | -0,216539753 | 0,860627143 | 0,72715539 | 0,973344786 | novel transcript, antisense to AGBL5 |
| ENSG00000261096 | AC073476,2 | -0,216684417 | 0,86054085 | 0,57795415 | 0,943425812 | novel transcript |
| ENSG00000272192 | AC100812,1 | -0,216700227 | 0,860531419 | 0,78223665 | 0,978597905 | novel transcript |
| ENSG00000229990 | AC022400,2 | -0,216724655 | 0,860516849 | 0,76648081 | 0,977174732 | novel transcript, antisense to CAMK2G |
| ENSG00000233684 | LINC01865 | -0,216793525 | 0,860475771 | 0,75860105 | 0,976120207 | long intergenic non-protein coding RNA 1865 |
| ENSG00000267412 | AC092068,2 | -0,216854877 | 0,860439179 | 0,86244081 | 0,989074806 | novel transcript, sense intronic to GNG7 |
| ENSG00000213090 | SCYL2P1 | -0,216980427 | 0,860364303 | 0,64300614 | 0,956633898 | SCYL2 pseudogene 1 |
| ENSG00000111445 | RFC5 | -0,216997444 | 0,860354155 | 0,55299297 | 0,936959262 | replication factor C subunit 5 |
| ENSG00000197653 | DNAH10 | -0,217034425 | 0,860332101 | 0,58743098 | 0,946068138 | dynein axonemal heavy chain 10 |
| ENSG00000102098 | SCML2 | -0,217037928 | 0,860330013 | 0,65629031 | 0,957444211 | Scm polycomb group protein like 2 |
| ENSG00000242588 | AC108010,1 | -0,21704184 | 0,86032768 | 0,37164494 | 0,859783004 | novel transcript |
| ENSG00000285584 | AL158800,1 | -0,217067539 | 0,860312355 | 0,83776977 | 0,98566906 | novel transcript |
| ENSG00000164548 | TRA2A | -0,217080058 | 0,860304889 | 0,51149794 | 0,922893865 | transformer 2 alpha homolog |
| ENSG00000170456 | DENND5B | -0,217146712 | 0,860265143 | 0,54729308 | 0,935404974 | DENN domain containing 5B |

| | | | | | | |
|---|---|---|---|---|---|---|
| ENSG00000277715 | AC079174,2 | -0,217229177 | 0,860215971 | 0,65805492 | 0,957883832 | novel transcript, antisense to CKAP4 |
| ENSG00000123908 | AGO2 | -0,217267448 | 0,860193153 | 0,43195091 | 0,89558934 | argonaute RISC catalytic component 2 |
| ENSG00000168303 | MPLKIP | -0,217285082 | 0,860182639 | 0,40345396 | 0,877691113 | M-phase specific PLK1 interacting protein |
| ENSG00000254644 | AC007751,1 | -0,217286156 | 0,860181998 | 0,87424372 | 0,98990147 | mediator complex subunit 21 (MED21) pseudogene |
| ENSG00000099866 | MADCAM1 | -0,217331422 | 0,860155009 | 0,73673692 | 0,974353474 | mucosal vascular addressin cell adhesion molecule 1 |
| ENSG00000166192 | SENP8 | -0,217363309 | 0,860135998 | 0,35457259 | 0,846034781 | SUMO peptidase family member, NEDD8 specific |
| ENSG00000249604 | AC096564,1 | -0,217430344 | 0,860096033 | 0,62990455 | 0,952812183 | novel transcript |
| ENSG00000273151 | AC073957,3 | -0,217432324 | 0,860094852 | 0,58495587 | 0,944857905 | novel transcript, antisense to GET4 |
| ENSG00000260528 | FAM157C | -0,217435507 | 0,860092955 | 0,49642799 | 0,918362406 | family with sequence similarity 157 member C |
| ENSG00000286789 | AL161645,2 | -0,217451948 | 0,860083153 | 0,59442936 | 0,947469769 | novel transcript, sense intronic to CYP2E1 |
| ENSG00000183137 | CEP57L1 | -0,217493886 | 0,860058151 | 0,50135761 | 0,920520138 | centrosomal protein 57 like 1 |
| ENSG00000260461 | AL133355,1 | -0,217608436 | 0,859989866 | 0,89933333 | 0,991832796 | novel transcript, overlapping to OBFC1 |
| ENSG00000230658 | KLHL7-DT | -0,217614327 | 0,859986354 | 0,68976522 | 0,966868992 | KLHL7 divergent transcript |
| ENSG00000202392 | RN7SKP292 | -0,217661717 | 0,859958106 | 0,86900079 | 0,989074806 | RN7SK pseudogene 292 |
| ENSG00000165416 | SUGT1 | -0,217700176 | 0,859935181 | 0,5500394 | 0,93612918 | SGT1 homolog, MIS12 kinetochore complex assembly cochaperone |
| ENSG00000105258 | POLR2I | -0,217707979 | 0,85993053 | 0,40861999 | 0,88153752 | RNA polymerase II subunit I |
| ENSG00000248830 | ZNF807P | -0,217777912 | 0,859888847 | 0,88462076 | 0,990717729 | zinc finger protein 807, pseudogene |
| ENSG00000178750 | STX19 | -0,217803672 | 0,859873493 | 0,75139971 | 0,975901808 | syntaxin 19 |
| ENSG00000102984 | ZNF821 | -0,217921753 | 0,859803118 | 0,32973183 | 0,831550849 | zinc finger protein 821 |
| ENSG00000073111 | MCM2 | -0,218093627 | 0,859700692 | 0,4215237 | 0,889538769 | minichromosome maintenance complex component 2 |
| ENSG00000286535 | AC139019,1 | -0,2181747 | 0,859652382 | 0,52229221 | 0,927622768 | novel transcript, antisense to ASAP1 |
| ENSG00000247903 | AC024896,1 | -0,218212084 | 0,859630107 | 0,64400567 | 0,956633898 | novel transcript, antisense to TM7SF3 |
| ENSG00000276276 | ARL17B | -0,218216802 | 0,859627295 | 0,5282578 | 0,92998848 | ADP ribosylation factor like GTPase 17B |
| ENSG00000250312 | ZNF718 | -0,218363722 | 0,859539758 | 0,41851678 | 0,887094957 | zinc finger protein 718 |
| ENSG00000156453 | PCDH1 | -0,2184484 | 0,859489309 | 0,56922436 | 0,94219857 | protocadherin 1 |
| ENSG00000274979 | AC020656,2 | -0,218467192 | 0,859478114 | 0,73394733 | 0,974353474 | novel transcript |
| ENSG00000125967 | NECAB3 | -0,218512391 | 0,859451188 | 0,55652368 | 0,938061341 | N-terminal EF-hand calcium binding protein 3 |
| ENSG00000198967 | OR10Z1 | -0,218529439 | 0,859441031 | 0,82154844 | 0,982952211 | olfactory receptor family 10 subfamily Z member 1 |
| ENSG00000149554 | CHEK1 | -0,218537768 | 0,85943607 | 0,53887128 | 0,933348247 | checkpoint kinase 1 |
| ENSG00000278472 | AC009268,2 | -0,218546277 | 0,859431001 | 0,80119589 | 0,980557464 | novel transcript, sense intronic to AVEN |
| ENSG00000242086 | MUC20-OT1 | -0,218551416 | 0,85942794 | 0,551107 | 0,936366211 | MUC20 overlapping transcript |

| | | | | | | |
|---|---|---|---|---|---|---|
| ENSG00000241489 | AC244197,3 | -0,218559688 | 0,859423012 | 0,54860217 | 0,935825387 | novel protein |
| ENSG00000171962 | DRC3 | -0,218586332 | 0,85940714 | 0,43408549 | 0,896272827 | dynein regulatory complex subunit 3 |
| ENSG00000028116 | VRK2 | -0,218673314 | 0,859355326 | 0,54909108 | 0,936040289 | VRK serine/threonine kinase 2 |
| ENSG00000135119 | RNFT2 | -0,218674583 | 0,859354571 | 0,32742654 | 0,829416898 | ring finger protein, transmembrane 2 |
| ENSG00000163629 | PTPN13 | -0,218729402 | 0,859321918 | 0,54218429 | 0,933491609 | protein tyrosine phosphatase non-receptor type 13 |
| ENSG00000237480 | AL355306,2 | -0,218746729 | 0,859311598 | 0,70774947 | 0,969847883 | novel transcript |
| ENSG00000204653 | ASPDH | -0,218774986 | 0,859294767 | 0,84993331 | 0,987036295 | aspartate dehydrogenase domain containing |
| ENSG00000197299 | BLM | -0,218879573 | 0,859232475 | 0,5024516 | 0,920585659 | BLM RecQ like helicase |
| ENSG00000100526 | CDKN3 | -0,218887668 | 0,859227654 | 0,66776785 | 0,961113267 | cyclin dependent kinase inhibitor 3 |
| ENSG00000213250 | RBMS2P1 | -0,219115706 | 0,859091852 | 0,79998361 | 0,980161849 | RNA binding motif single stranded interacting protein 2 pseudogene 1 |
| ENSG00000278839 | SERF1B | -0,219181388 | 0,85905274 | 0,62090307 | 0,952793934 | small EDRK-rich factor 1B |
| ENSG00000231424 | BX284613,2 | -0,219185358 | 0,859050377 | 0,73211641 | 0,974353474 | novel transcript, antisense to FMO1 |
| ENSG00000285067 | AC270277,1 | -0,219204573 | 0,859038935 | 0,73814963 | 0,974353474 | novel transcript, antisense to MTMR9 |
| ENSG00000280273 | AF131216,4 | -0,219204573 | 0,859038935 | 0,73814963 | 0,974353474 | novel transcript, antisense to MTMR9 |
| ENSG00000240622 | RPL7P15 | -0,219403613 | 0,858920427 | 0,76438775 | 0,976640468 | ribosomal protein L7 pseudogene 15 |
| ENSG00000286147 | AC092445,1 | -0,219411759 | 0,858915577 | 0,37658105 | 0,862818448 | novel transcript, antisense to DKK2 |
| ENSG00000240086 | AC092969,1 | -0,219702914 | 0,858742254 | 0,56047495 | 0,940170934 | novel transcript, antisense to EPHB1 |
| ENSG00000272094 | AC091614,1 | -0,219718792 | 0,858732803 | 0,59853968 | 0,948804147 | novel transcript |
| ENSG00000277728 | AC097641,2 | -0,219780474 | 0,858696089 | 0,71287865 | 0,970812834 | novel transcript, antisense to COG1 |
| ENSG00000163661 | PTX3 | -0,219835077 | 0,85866359 | 0,51769819 | 0,925915783 | pentraxin 3 |
| ENSG00000233436 | BTBD18 | -0,219854578 | 0,858651983 | 0,46416123 | 0,90737035 | BTB domain containing 18 |
| ENSG00000287760 | AC099521,3 | -0,219868428 | 0,85864374 | 0,71341869 | 0,970812834 | novel transcript |
| ENSG00000264207 | AC239868,1 | -0,219872948 | 0,85864105 | 0,47375312 | 0,91029547 | novel transcript, antisense HIST2H4A |
| ENSG00000164941 | INTS8 | -0,220047157 | 0,858537373 | 0,61134727 | 0,952062377 | integrator complex subunit 8 |
| ENSG00000236532 | LINC01695 | -0,220105048 | 0,858502924 | 0,70735697 | 0,969565008 | long intergenic non-protein coding RNA 1695 |
| ENSG00000270986 | HMGB1P51 | -0,220108996 | 0,858500574 | 0,68024131 | 0,964580199 | high mobility group box 1 pseudogene 51 |
| ENSG00000215472 | RPL17-C18orf3 | -0,220165913 | 0,858466705 | 0,67204922 | 0,962290553 | RPL17-C18orf32 readthrough |
| ENSG00000233421 | LINC01783 | -0,220211696 | 0,858439463 | 0,53140234 | 0,931291085 | long intergenic non-protein coding RNA 1783 |
| ENSG00000277958 | Metazoa_SRP | -0,220266287 | 0,85840698 | 0,85445294 | 0,987861659 | Metazoan signal recognition particle RNA |
| ENSG00000143971 | ETAA1 | -0,220361007 | 0,858350624 | 0,64209561 | 0,956548736 | ETAA1 activator of ATR kinase |
| ENSG00000010803 | SCMH1 | -0,220543067 | 0,858242311 | 0,36214984 | 0,852258387 | Scm polycomb group protein homolog 1 |
| ENSG00000184486 | POU3F2 | -0,220572027 | 0,858225084 | 0,51573712 | 0,925006591 | POU class 3 homeobox 2 |

| | | | | | | |
|---|---|---|---|---|---|---|
| ENSG00000174953 | DHX36 | -0,220615431 | 0,858199264 | 0,54810374 | 0,935404974 | DEAH-box helicase 36 |
| ENSG00000258026 | AC078955,1 | -0,220640272 | 0,858184487 | 0,82313174 | 0,983204348 | novel transcript, antisense to ACSS3 |
| ENSG00000261616 | AC036108,3 | -0,220704288 | 0,858146408 | 0,56367646 | 0,940865625 | novel transcript, antisense to TTC23 |
| ENSG00000275004 | ZNF280B | -0,220716837 | 0,858138944 | 0,58672999 | 0,945395317 | zinc finger protein 280B |
| ENSG00000260071 | AC012178,1 | -0,220765755 | 0,858109847 | 0,88722258 | 0,990717729 | novel transcript |
| ENSG00000109107 | ALDOC | -0,22077466 | 0,858104551 | 0,50624507 | 0,921780819 | aldolase, fructose-bisphosphate C |
| ENSG00000170312 | CDK1 | -0,220808593 | 0,858084368 | 0,64044719 | 0,956251063 | cyclin dependent kinase 1 |
| ENSG00000122971 | ACADS | -0,220949422 | 0,85800061 | 0,58073417 | 0,943832501 | acyl-CoA dehydrogenase short chain |
| ENSG00000054983 | GALC | -0,221055359 | 0,857937609 | 0,57976629 | 0,943828882 | galactosylceramidase |
| ENSG00000131115 | ZNF227 | -0,221102511 | 0,857909569 | 0,4886912 | 0,91557151 | zinc finger protein 227 |
| ENSG00000179899 | PHC1P1 | -0,221137835 | 0,857888564 | 0,44097306 | 0,898675567 | polyhomeotic homolog 1 pseudogene 1 |
| ENSG00000266897 | AC005546,1 | -0,221219058 | 0,857840267 | 0,88840598 | 0,990717729 | novel transcript |
| ENSG00000260035 | AC051619,7 | -0,221278645 | 0,857804836 | 0,4318514 | 0,895555998 | novel transcript |
| ENSG00000182261 | NLRP10 | -0,221329553 | 0,857774568 | 0,93285222 | 0,994479239 | NLR family pyrin domain containing 10 |
| ENSG00000205560 | CPT1B | -0,221347312 | 0,857764009 | 0,48054839 | 0,912847708 | carnitine palmitoyltransferase 1B |
| ENSG00000022567 | SLC45A4 | -0,221371359 | 0,857749712 | 0,46094997 | 0,906029574 | solute carrier family 45 member 4 |
| ENSG00000130377 | ACSBG2 | -0,221440669 | 0,857708504 | 0,61890694 | 0,952793934 | acyl-CoA synthetase bubblegum family member 2 |
| ENSG00000186106 | ANKRD46 | -0,221481581 | 0,857684182 | 0,60748988 | 0,950993594 | ankyrin repeat domain 46 |
| ENSG00000228779 | Z69666,1 | -0,221631173 | 0,857595254 | 0,79469673 | 0,979776981 | novel transcript |
| ENSG00000188070 | C11orf95 | -0,221690594 | 0,857559933 | 0,50035036 | 0,920485547 | chromosome 11 open reading frame 95 |
| ENSG00000185246 | PRPF39 | -0,221804753 | 0,857492077 | 0,64942867 | 0,956899963 | pre-mRNA processing factor 39 |
| ENSG00000272173 | U47924,2 | -0,22188202 | 0,857446153 | 0,55619539 | 0,938046927 | novel transcript, antisense to C12orf57 |
| ENSG00000196091 | MYBPC1 | -0,222087117 | 0,857324266 | 0,88749188 | 0,990717729 | myosin binding protein C1 |
| ENSG00000244462 | RBM12 | -0,222128175 | 0,857299867 | 0,56989779 | 0,94221458 | RNA binding motif protein 12 |
| ENSG00000225361 | PPP1R26-AS1 | -0,222176976 | 0,857270868 | 0,64921234 | 0,956899963 | PPP1R26 antisense RNA 1 |
| ENSG00000042088 | TDP1 | -0,222197329 | 0,857258774 | 0,48911533 | 0,915835287 | tyrosyl-DNA phosphodiesterase 1 |
| ENSG00000106125 | MINDY4 | -0,222269172 | 0,857216086 | 0,37693212 | 0,862818448 | MINDY lysine 48 deubiquitinase 4 |
| ENSG00000134138 | MEIS2 | -0,222292285 | 0,857202353 | 0,36543459 | 0,855323027 | Meis homeobox 2 |
| ENSG00000197044 | ZNF441 | -0,222299079 | 0,857198316 | 0,60329694 | 0,950352286 | zinc finger protein 441 |
| ENSG00000236494 | AC008080,4 | -0,222439298 | 0,857115007 | 0,85277928 | 0,987638133 | novel transcript |
| ENSG00000279833 | AL031846,2 | -0,222497286 | 0,857080557 | 0,69793832 | 0,968522643 | novel transcript, antisense to CBX7 |
| ENSG00000286403 | AC010378,1 | -0,222544858 | 0,857052295 | 0,39518833 | 0,871981986 | novel transcript |
| ENSG00000265799 | AC090844,3 | -0,222689843 | 0,856966169 | 0,76407627 | 0,976640468 | novel transcript |

| | | | | | | |
|---|---|---|---|---|---|---|
| ENSG00000258210 | AC144548,1 | -0,222740265 | 0,856936219 | 0,54402604 | 0,934259495 | novel transcript |
| ENSG00000250102 | LINC02377 | -0,222850596 | 0,856870687 | 0,81380694 | 0,981521413 | long intergenic non-protein coding RNA 2377 |
| ENSG00000265241 | RBM8A | -0,222885757 | 0,856849803 | 0,297416 | 0,806635068 | RNA binding motif protein 8A |
| ENSG00000260034 | LCMT1-AS2 | -0,222945287 | 0,856814448 | 0,43526453 | 0,896272827 | LCMT1 antisense RNA 2 |
| ENSG00000275600 | PIGW | -0,22297778 | 0,856795151 | 0,79495234 | 0,979776981 | phosphatidylinositol glycan anchor biosynthesis class W |
| ENSG00000249174 | AC106744,1 | -0,223171801 | 0,856679932 | 0,79946158 | 0,980161849 | novel transcript |
| ENSG00000253251 | SHLD3 | -0,223175901 | 0,856677498 | 0,731029 | 0,974353474 | shieldin complex subunit 3 |
| ENSG00000287011 | AC003077,1 | -0,223291008 | 0,85660915 | 0,67243959 | 0,962290553 | novel transcript, antisense to DOCK4 |
| ENSG00000143624 | INTS3 | -0,223361862 | 0,856567081 | 0,56029268 | 0,940070839 | integrator complex subunit 3 |
| ENSG00000281490 | CICP14 | -0,22340842 | 0,856539438 | 0,57944075 | 0,943828882 | capicua transcriptional repressor pseudogene 14 |
| ENSG00000246877 | DNM1P35 | -0,22340888 | 0,856539165 | 0,72130409 | 0,972142678 | dynamin 1 pseudogene 35 |
| ENSG00000283900 | TPTEP2-CSNK1I | -0,223554217 | 0,856452882 | 0,60966686 | 0,951793902 | TPTEP2-CSNK1E readthrough |
| ENSG00000173269 | MMRN2 | -0,223566369 | 0,856445668 | 0,53422036 | 0,93151582 | multimerin 2 |
| ENSG00000237082 | COX5BP6 | -0,223637452 | 0,856403471 | 0,90090816 | 0,991865079 | cytochrome c oxidase subunit 5B pseudogene 6 |
| ENSG00000048028 | USP28 | -0,223703769 | 0,856364105 | 0,51260123 | 0,923540891 | ubiquitin specific peptidase 28 |
| ENSG00000272844 | AC074044,1 | -0,22371145 | 0,856359546 | 0,86552665 | 0,989074806 | novel transcript, antisense to GTPBP8 |
| ENSG00000234846 | NEU1 | -0,223759441 | 0,85633106 | 0,81913931 | 0,98249617 | neuraminidase 1 |
| ENSG00000094916 | CBX5 | -0,223815892 | 0,856297553 | 0,52552683 | 0,92884555 | chromobox 5 |
| ENSG00000260296 | AC095057,3 | -0,223950728 | 0,856217526 | 0,69681333 | 0,968300554 | novel transcript, overlapping N4BP2 |
| ENSG00000185187 | SIGIRR | -0,223956093 | 0,856214342 | 0,53193744 | 0,931291085 | single Ig and TIR domain containing |
| ENSG00000186998 | EMID1 | -0,223963914 | 0,856209701 | 0,5503149 | 0,936146662 | EMI domain containing 1 |
| ENSG00000152763 | DNAI4 | -0,224121513 | 0,856116174 | 0,57162671 | 0,942436428 | dynein axonemal intermediate chain 4 |
| ENSG00000231390 | SNX18P8 | -0,224344314 | 0,855983971 | 0,72596586 | 0,973344786 | sorting nexin 18 pseudogene 8 |
| ENSG00000130751 | NPAS1 | -0,224519543 | 0,85588001 | 0,73363229 | 0,974353474 | neuronal PAS domain protein 1 |
| ENSG00000275932 | DUSP14 | -0,224732029 | 0,855753962 | 0,54175823 | 0,933491609 | dual specificity phosphatase 14 |
| ENSG00000276128 | AL591441,1 | -0,224836371 | 0,855692072 | 0,67721534 | 0,963580269 | glucoside xylosyltransferase 1 pseudogene 5 |
| ENSG00000288534 | AP001931,2 | -0,224934515 | 0,855633863 | 0,40480208 | 0,878176155 | TMX2-CTNND1 readthrough (NMD candidate) |
| ENSG00000274031 | AC092140,2 | -0,224990689 | 0,855600548 | 0,79211406 | 0,979471509 | novel transcript, antisense to OGFOD1 |
| ENSG00000168255 | POLR2J3 | -0,22504827 | 0,8555664 | 0,37254341 | 0,860420124 | RNA polymerase II subunit J3 |
| ENSG00000042781 | USH2A | -0,225049418 | 0,855565719 | 0,69398794 | 0,9673565 | usherin |
| ENSG00000196284 | SUPT3H | -0,225057078 | 0,855561176 | 0,60442797 | 0,950813598 | SPT3 homolog, SAGA and STAGA complex component |

| | | | | | | |
|---|---|---|---|---|---|---|
| ENSG00000213057 | C1orf220 | -0,225191981 | 0,855481178 | 0,55538059 | 0,937793851 | chromosome 1 putative open reading frame 220 |
| ENSG00000184270 | H2AC21 | -0,225289814 | 0,855423168 | 0,39921712 | 0,875433675 | H2A clustered histone 21 |
| ENSG00000205424 | PCBP3-AS1 | -0,225320627 | 0,855404898 | 0,76104048 | 0,97624106 | PCBP3 antisense RNA 1 |
| ENSG00000235303 | AL023495,1 | -0,22532501 | 0,855402299 | 0,8203839 | 0,982609114 | novel transcript |
| ENSG00000284796 | DCAF11 | -0,225360695 | 0,855381141 | 0,61604578 | 0,952793934 | DDB1 and CUL4 associated factor 11 |
| ENSG00000176597 | B3GNT5 | -0,225361544 | 0,855380637 | 0,63968482 | 0,956018518 | UDP-GlcNAc:betaGal beta-1,3-N-acetylglucosaminyltransferase 5 |
| ENSG00000151967 | SCHIP1 | -0,225410339 | 0,855351707 | 0,53969777 | 0,933350549 | schwannomin interacting protein 1 |
| ENSG00000118482 | PHF3 | -0,225435401 | 0,855336849 | 0,56122327 | 0,940567693 | PHD finger protein 3 |
| ENSG00000134539 | KLRD1 | -0,225504835 | 0,855295684 | 0,53400369 | 0,93148979 | killer cell lectin like receptor D1 |
| ENSG00000227253 | AL158834,2 | -0,225532594 | 0,855279227 | 0,75777786 | 0,976120207 | novel transcript |
| ENSG00000170264 | FAM161A | -0,225534695 | 0,855277982 | 0,44551698 | 0,901068127 | FAM161 centrosomal protein A |
| ENSG00000287110 | AC112484,5 | -0,22553846 | 0,85527575 | 0,52963108 | 0,930664682 | novel transcript |
| ENSG00000112029 | FBXO5 | -0,225560978 | 0,855262401 | 0,62869796 | 0,952793934 | F-box protein 5 |
| ENSG00000286129 | AC245060,7 | -0,225565374 | 0,855259794 | 0,34186493 | 0,840048641 | novel transcript |
| ENSG00000276966 | H4C5 | -0,22562474 | 0,855224602 | 0,38233251 | 0,866930927 | H4 clustered histone 5 |
| ENSG00000257875 | AC089998,3 | -0,225631586 | 0,855220543 | 0,86771985 | 0,989074806 | F-box and WD repeat domain containing 12 (FBXW12) pseudogene |
| ENSG00000184402 | SS18L1 | -0,22568138 | 0,855191026 | 0,44308248 | 0,899690238 | SS18L1 subunit of BAF chromatin remodeling complex |
| ENSG00000137269 | LRRC1 | -0,225681857 | 0,855190744 | 0,54746064 | 0,935404974 | leucine rich repeat containing 1 |
| ENSG00000250673 | REELD1 | -0,225702655 | 0,855178415 | 0,86394517 | 0,989074806 | reeler domain containing 1 |
| ENSG00000188153 | COL4A5 | -0,225763778 | 0,855142185 | 0,40855636 | 0,88153752 | collagen type IV alpha 5 chain |
| ENSG00000273374 | AC069222,1 | -0,225766057 | 0,855140834 | 0,6211837 | 0,952793934 | novel transcript |
| ENSG00000265222 | AC079336,2 | -0,225785226 | 0,855129471 | 0,85844237 | 0,989067345 | novel transcript, sense intronic MYO1D |
| ENSG00000100580 | TMED8 | -0,225818055 | 0,855110013 | 0,51094049 | 0,922715081 | transmembrane p24 trafficking protein family member 8 |
| ENSG00000080947 | CROCCP3 | -0,225820759 | 0,85510841 | 0,56298239 | 0,940865625 | CROCC pseudogene 3 |
| ENSG00000255857 | PXN-AS1 | -0,225855071 | 0,855088073 | 0,49499766 | 0,917691373 | PXN antisense RNA 1 |
| ENSG00000126457 | PRMT1 | -0,225856623 | 0,855087154 | 0,36949822 | 0,857991124 | protein arginine methyltransferase 1 |
| ENSG00000168356 | SCN11A | -0,225899008 | 0,855062032 | 0,59330555 | 0,947416795 | sodium voltage-gated channel alpha subunit 11 |
| ENSG00000005486 | RHBDD2 | -0,225905364 | 0,855058265 | 0,29378244 | 0,803641108 | rhomboid domain containing 2 |
| ENSG00000250299 | MRPS31P4 | -0,225967988 | 0,85502115 | 0,57611972 | 0,943026545 | mitochondrial ribosomal protein S31 pseudogene 4 |

| | | | | | | |
|---|---|---|---|---|---|---|
| ENSG00000260447 | AC009065,3 | -0,225969829 | 0,855020059 | 0,86622883 | 0,989074806 | novel transcript, antisense to PKD1 |
| ENSG00000144026 | ZNF514 | -0,225989408 | 0,855008455 | 0,34228556 | 0,840048641 | zinc finger protein 514 |
| ENSG00000225091 | SNORA71A | -0,226026764 | 0,854986316 | 0,62271947 | 0,952793934 | small nucleolar RNA, H/ACA box 71A |
| ENSG00000173175 | ADCY5 | -0,226040714 | 0,854978049 | 0,55803323 | 0,939078529 | adenylate cyclase 5 |
| ENSG00000173209 | AHSA2P | -0,226106699 | 0,854938946 | 0,41348708 | 0,884943878 | activator of HSP90 ATPase homolog 2, pseudogene |
| ENSG00000226644 | AL121899,1 | -0,226142503 | 0,854917729 | 0,67125068 | 0,962026636 | novel transcript |
| ENSG00000205208 | C4orf46 | -0,226152354 | 0,854911891 | 0,63632849 | 0,955547532 | chromosome 4 open reading frame 46 |
| ENSG00000000460 | C1orf112 | -0,226176509 | 0,854897578 | 0,57358339 | 0,942475308 | chromosome 1 open reading frame 112 |
| ENSG00000163125 | RPRD2 | -0,226404345 | 0,85476258 | 0,43169877 | 0,89541765 | regulation of nuclear pre-mRNA domain containing 2 |
| ENSG00000160917 | CPSF4 | -0,226494587 | 0,854709115 | 0,3115566 | 0,817751312 | cleavage and polyadenylation specific factor 4 |
| ENSG00000118260 | CREB1 | -0,226551629 | 0,854675322 | 0,59072391 | 0,947416795 | cAMP responsive element binding protein 1 |
| ENSG00000181616 | OR52H1 | -0,226711814 | 0,854580431 | 0,85193259 | 0,987322912 | olfactory receptor family 52 subfamily H member 1 |
| ENSG00000268401 | AC007785,3 | -0,226759247 | 0,854552334 | 0,6714844 | 0,962085713 | novel transcript |
| ENSG00000132906 | CASP9 | -0,226762069 | 0,854550663 | 0,3678571 | 0,857401877 | caspase 9 |
| ENSG00000272556 | GTF2IP13 | -0,226796803 | 0,854530089 | 0,47411392 | 0,910491353 | general transcription factor IIi pseudogene 13 |
| ENSG00000137145 | DENND4C | -0,227077793 | 0,854363671 | 0,53201563 | 0,931291085 | DENN domain containing 4C |
| ENSG00000267746 | AC104985,1 | -0,22710232 | 0,854349146 | 0,7626697 | 0,97639149 | novel transcript, antisense to NOL4 |
| ENSG00000198650 | TAT | -0,227173907 | 0,854306754 | 0,76021266 | 0,97624106 | tyrosine aminotransferase |
| ENSG00000286192 | AC069288,1 | -0,227186977 | 0,854299014 | 0,78607559 | 0,979320406 | Novel protein |
| ENSG00000232040 | ZBED9 | -0,227210247 | 0,854285235 | 0,55137528 | 0,936366211 | zinc finger BED-type containing 9 |
| ENSG00000144406 | UNC80 | -0,22721433 | 0,854282817 | 0,40655465 | 0,879715952 | unc-80 homolog, NALCN channel complex subunit |
| ENSG00000267274 | AC008770,3 | -0,227285275 | 0,854240809 | 0,6212213 | 0,952793934 | novel transcript, sense overlapping with ZNF700 |
| ENSG00000258752 | AL357093,2 | -0,227378378 | 0,854185683 | 0,65370026 | 0,957444211 | novel transcript, antisense to FOXN3 |
| ENSG00000176920 | FUT2 | -0,227651564 | 0,854023951 | 0,66188651 | 0,959707806 | fucosyltransferase 2 |
| ENSG00000267481 | AC011477,1 | -0,22766968 | 0,854013227 | 0,56398861 | 0,940865625 | novel transcript, sense intronic to ZNF506 |
| ENSG00000051825 | MPHOSPH9 | -0,227866158 | 0,853896928 | 0,53294791 | 0,931291085 | M-phase phosphoprotein 9 |
| ENSG00000087095 | NLK | -0,227913823 | 0,853868717 | 0,58358565 | 0,944174844 | nemo like kinase |
| ENSG00000066427 | ATXN3 | -0,228095559 | 0,853761162 | 0,39110677 | 0,870057277 | ataxin 3 |
| ENSG00000231701 | BMS1P13 | -0,228176033 | 0,85371354 | 0,71929792 | 0,971869926 | BMS1 pseudogene 13 |
| ENSG00000125703 | ATG4C | -0,228179304 | 0,853711605 | 0,64809766 | 0,956899963 | autophagy related 4C cysteine peptidase |
| ENSG00000168661 | ZNF30 | -0,228180591 | 0,853710843 | 0,57120451 | 0,942436428 | zinc finger protein 30 |
| ENSG00000280484 | AL732314,1 | -0,228183138 | 0,853709336 | 0,84146588 | 0,985672131 | novel transcript, antisense to PPP2R3B |
| ENSG00000281849 | AL732314,10 | -0,228183138 | 0,853709336 | 0,84146588 | 0,985672131 | novel transcript, antisense to PPP2R3B |

| | | | | | | |
|---|---|---|---|---|---|---|
| ENSG00000280593 | AL732314,3 | -0,228183138 | 0,853709336 | 0,84146588 | 0,985672131 | novel transcript, antisense to PPP2R3B |
| ENSG00000105520 | PLPPR2 | -0,228350448 | 0,853610337 | 0,44934691 | 0,902062594 | phospholipid phosphatase related 2 |
| ENSG00000127366 | TAS2R5 | -0,228376523 | 0,853594909 | 0,70057274 | 0,968522643 | taste 2 receptor member 5 |
| ENSG00000005156 | LIG3 | -0,22838358 | 0,853590734 | 0,36030916 | 0,850808572 | DNA ligase 3 |
| ENSG00000176956 | LY6H | -0,228411452 | 0,853574243 | 0,74044828 | 0,974833668 | lymphocyte antigen 6 family member H |
| ENSG00000275964 | AL355001,2 | -0,22847421 | 0,853537113 | 0,48051626 | 0,912847708 | novel transcript |
| ENSG00000176320 | AL138899,1 | -0,228488282 | 0,853528788 | 0,70203306 | 0,968742626 | novel transcript |
| ENSG00000126091 | ST3GAL3 | -0,228498004 | 0,853523036 | 0,39531859 | 0,871981986 | ST3 beta-galactoside alpha-2,3-sialyltransferase 3 |
| ENSG00000232079 | LINC01697 | -0,228524871 | 0,853507141 | 0,73094713 | 0,974353474 | long intergenic non-protein coding RNA 1697 |
| ENSG00000231881 | AL109615,2 | -0,228583317 | 0,853472565 | 0,65664034 | 0,957444211 | novel transcript |
| ENSG00000268107 | AC003005,1 | -0,22864722 | 0,853434762 | 0,83025452 | 0,984156028 | novel transcript |
| ENSG00000119328 | ABITRAM | -0,228709776 | 0,853397757 | 0,64601981 | 0,956899963 | actin binding transcription modulator |
| ENSG00000234450 | AC005534,1 | -0,228747584 | 0,853375393 | 0,84638029 | 0,986939308 | ribosomal protein L26-like 1 (RPL26L1) pseudogene |
| ENSG00000276214 | AC008833,1 | -0,22881584 | 0,853335019 | 0,84006886 | 0,985672131 | endogenous retrovirus group V, member pseudogene |
| ENSG00000172058 | SERF1A | -0,22890577 | 0,853281828 | 0,51015207 | 0,922427526 | small EDRK-rich factor 1A |
| ENSG00000052795 | FNIP2 | -0,228932431 | 0,85326606 | 0,48414801 | 0,913690171 | folliculin interacting protein 2 |
| ENSG00000271009 | AC116667,1 | -0,228967942 | 0,853245058 | 0,71954558 | 0,971869926 | novel transcript |
| ENSG00000132639 | SNAP25 | -0,229121217 | 0,853154412 | 0,60654533 | 0,950927937 | synaptosome associated protein 25 |
| ENSG00000122483 | CCDC18 | -0,229172801 | 0,853123908 | 0,4687501 | 0,907940653 | coiled-coil domain containing 18 |
| ENSG00000226352 | PSPC1-AS2 | -0,22921812 | 0,853097109 | 0,82822925 | 0,983646162 | PSPC1 antisense RNA 2 |
| ENSG00000164038 | SLC9B2 | -0,229273359 | 0,853064446 | 0,46732064 | 0,907940653 | solute carrier family 9 member B2 |
| ENSG00000286837 | AC087533,1 | -0,229464896 | 0,852951198 | 0,67428284 | 0,963103112 | novel transcript, antisense to THAP1 |
| ENSG00000239388 | ASB14 | -0,229529303 | 0,85291312 | 0,63266991 | 0,954291097 | ankyrin repeat and SOCS box containing 14 |
| ENSG00000023697 | DERA | -0,229567161 | 0,852890739 | 0,60030391 | 0,94936019 | deoxyribose-phosphate aldolase |
| ENSG00000276887 | TTYH1 | -0,229708427 | 0,852807229 | 0,45143754 | 0,902062594 | tweety family member 1 |
| ENSG00000258559 | AC005519,1 | -0,229732387 | 0,852793066 | 0,45003876 | 0,902062594 | novel transcript, sense overlapping ABCD4 |
| ENSG00000040731 | CDH10 | -0,229743616 | 0,852786428 | 0,64523212 | 0,956899963 | cadherin 10 |
| ENSG00000118473 | SGIP1 | -0,229775235 | 0,852767738 | 0,56704136 | 0,941425553 | SH3GL interacting endocytic adaptor 1 |
| ENSG00000006530 | AGK | -0,229824713 | 0,852738493 | 0,39872836 | 0,875433675 | acylglycerol kinase |
| ENSG00000135447 | PPP1R1A | -0,229834176 | 0,8527329 | 0,43134039 | 0,89530663 | protein phosphatase 1 regulatory inhibitor subunit 1A |
| ENSG00000243008 | AL359538,1 | -0,229892757 | 0,852698275 | 0,78956597 | 0,979462318 | novel transcript |

| | | | | | | |
|---|---|---|---|---|---|---|
| ENSG00000159086 | PAXBP1 | -0,229945877 | 0,852666879 | 0,5515691 | 0,936366211 | PAX3 and PAX7 binding protein 1 |
| ENSG00000142609 | CFAP74 | -0,229955993 | 0,8526609 | 0,72650951 | 0,973344786 | cilia and flagella associated protein 74 |
| ENSG00000230530 | LIMD1-AS1 | -0,230015919 | 0,852625484 | 0,58302484 | 0,944174844 | LIMD1 antisense RNA 1 |
| ENSG00000198060 | MARCHF5 | -0,230076363 | 0,852589762 | 0,53095744 | 0,931265667 | membrane associated ring-CH-type finger 5 |
| ENSG00000278715 | SNORD116-20 | -0,230173752 | 0,85253221 | 0,85075454 | 0,987147801 | small nucleolar RNA, C/D box 116-20 |
| ENSG00000066557 | LRRC40 | -0,230202229 | 0,852515382 | 0,57853632 | 0,943425812 | leucine rich repeat containing 40 |
| ENSG00000091009 | RBM27 | -0,230280179 | 0,852469322 | 0,60232645 | 0,950180678 | RNA binding motif protein 27 |
| ENSG00000229422 | AL512625,2 | -0,230289379 | 0,852463886 | 0,6237065 | 0,952793934 | novel transcript |
| ENSG00000267397 | AC090229,1 | -0,230293559 | 0,852461416 | 0,60784493 | 0,951015776 | novel transcript |
| ENSG00000145284 | SCD5 | -0,230309882 | 0,85245177 | 0,42296724 | 0,890541302 | stearoyl-CoA desaturase 5 |
| ENSG00000235248 | OR13C1P | -0,230321595 | 0,85244485 | 0,84129118 | 0,985672131 | olfactory receptor family 13 subfamily C member 1 pseudogene |
| ENSG00000134258 | VTCN1 | -0,230339466 | 0,85243429 | 0,73676508 | 0,974353474 | V-set domain containing T cell activation inhibitor 1 |
| ENSG00000259849 | VENTXP1 | -0,230395903 | 0,852400944 | 0,65235019 | 0,957275891 | VENT homeobox pseudogene 1 |
| ENSG00000231607 | DLEU2 | -0,230402042 | 0,852397318 | 0,55787027 | 0,938984031 | deleted in lymphocytic leukemia 2 |
| ENSG00000285025 | AL022312,1 | -0,230412893 | 0,852390906 | 0,95213652 | 0,996432861 | MIEF1 upstream open reading frame protein |
| ENSG00000178694 | NSUN3 | -0,23046728 | 0,852358773 | 0,51303662 | 0,923725914 | NOP2/Sun RNA methyltransferase 3 |
| ENSG00000248489 | LINC02062 | -0,230494073 | 0,852342944 | 0,47762818 | 0,911578505 | long intergenic non-protein coding RNA 2062 |
| ENSG00000072571 | HMMR | -0,230503851 | 0,852337167 | 0,56983613 | 0,94221458 | hyaluronan mediated motility receptor |
| ENSG00000228995 | MTND1P9 | -0,230527138 | 0,85232341 | 0,71654909 | 0,971438758 | MT-ND1 pseudogene 9 |
| ENSG00000173728 | C1orf100 | -0,230557334 | 0,85230557 | 0,83270553 | 0,984905093 | chromosome 1 open reading frame 100 |
| ENSG00000224216 | AC234781,1 | -0,230566645 | 0,85230007 | 0,84372606 | 0,986273696 | novel transcript |
| ENSG00000161653 | NAGS | -0,230567348 | 0,852299654 | 0,61498802 | 0,952793934 | N-acetylglutamate synthase |
| ENSG00000273507 | AL354809,1 | -0,230593272 | 0,852284339 | 0,47482749 | 0,91069468 | novel transcript |
| ENSG00000147316 | MCPH1 | -0,230622733 | 0,852266935 | 0,45557986 | 0,903545728 | microcephalin 1 |
| ENSG00000170293 | CMTM8 | -0,230701318 | 0,852220512 | 0,64241893 | 0,956633898 | CKLF like MARVEL transmembrane domain containing 8 |
| ENSG00000203737 | GPR52 | -0,230745024 | 0,852194695 | 0,68762167 | 0,966063455 | G protein-coupled receptor 52 |
| ENSG00000266290 | AC015813,3 | -0,230815921 | 0,852152817 | 0,74285436 | 0,975201495 | novel transcript |
| ENSG00000267216 | ZNF8-ERVK3-1 | -0,23082307 | 0,852148595 | 0,36229796 | 0,852348131 | ZNF8-ERVK3-1 readthrough |
| ENSG00000234504 | AL353801,2 | -0,230841508 | 0,852137704 | 0,848023 | 0,987036295 | novel transcript |
| ENSG00000159588 | CCDC17 | -0,230912279 | 0,852095904 | 0,64760652 | 0,956899963 | coiled-coil domain containing 17 |
| ENSG00000224420 | ADM5 | -0,230916265 | 0,85209355 | 0,58173119 | 0,944145586 | adrenomedullin 5 (putative) |

| | | | | | | |
|---|---|---|---|---|---|---|
| ENSG00000204311 | PJVK | -0,230942814 | 0,852077869 | 0,4590767 | 0,90476838 | pejvakin |
| ENSG00000286113 | AC022868,1 | -0,230953232 | 0,852071716 | 0,80963908 | 0,981213877 | long intergenic non-protein coding RNA 1617 |
| ENSG00000235370 | DNM1P51 | -0,230985917 | 0,852052412 | 0,51079133 | 0,922693987 | dynamin 1 pseudogene 51 |
| ENSG00000169045 | HNRNPH1 | -0,230999041 | 0,852044661 | 0,44896863 | 0,902062594 | heterogeneous nuclear ribonucleoprotein H1 |
| ENSG00000152117 | SMPD4BP | -0,231001734 | 0,852043071 | 0,31639732 | 0,821646322 | sphingomyelin phosphodiesterase 4B, pseudogene |
| ENSG00000264116 | AC116003,1 | -0,231080148 | 0,851996761 | 0,78319497 | 0,978919905 | novel transcript, antisense to SMIM21 |
| ENSG00000114904 | NEK4 | -0,23118658 | 0,85193391 | 0,43770002 | 0,897021798 | NIMA related kinase 4 |
| ENSG00000230561 | CCDC192 | -0,231254544 | 0,851893777 | 0,70247861 | 0,968742626 | coiled-coil domain containing 192 |
| ENSG00000157869 | RAB28 | -0,231373769 | 0,851823379 | 0,47171002 | 0,908896785 | RAB28, member RAS oncogene family |
| ENSG00000255441 | SIGLEC10-AS1 | -0,231390684 | 0,851813391 | 0,72917802 | 0,974009551 | SIGLEC10 antisense RNA 1 |
| ENSG00000242510 | NDUFB4P1 | -0,231464171 | 0,851770004 | 0,83812125 | 0,985672131 | NADH:ubiquinone oxidoreductase subunit B4 pseudogene 1 |
| ENSG00000236693 | HLA-DPB1 | -0,231588589 | 0,85169655 | 0,67323483 | 0,962605657 | major histocompatibility complex, class II, DP beta 1 |
| ENSG00000230763 | HLA-DPB1 | -0,231588589 | 0,85169655 | 0,67323483 | 0,962605657 | major histocompatibility complex, class II, DP beta 1 |
| ENSG00000223812 | AC073365,1 | -0,231629838 | 0,851672199 | 0,84290145 | 0,986208256 | novel transcript |
| ENSG00000108950 | FAM20A | -0,231634893 | 0,851669215 | 0,46729295 | 0,907940653 | FAM20A golgi associated secretory pathway pseudokinase |
| ENSG00000115827 | DCAF17 | -0,231645921 | 0,851662705 | 0,56554433 | 0,940865625 | DDB1 and CUL4 associated factor 17 |
| ENSG00000146463 | ZMYM4 | -0,231684932 | 0,851639676 | 0,54973165 | 0,936108121 | zinc finger MYM-type containing 4 |
| ENSG00000174371 | EXO1 | -0,231702185 | 0,851629491 | 0,50938319 | 0,922427526 | exonuclease 1 |
| ENSG00000117139 | KDM5B | -0,231767954 | 0,851590668 | 0,41476472 | 0,885579089 | lysine demethylase 5B |
| ENSG00000270049 | AC009061,2 | -0,231779639 | 0,851583771 | 0,51950305 | 0,926800715 | novel transcript, antisense to AGRP |
| ENSG00000213672 | NCKIPSD | -0,231849798 | 0,851542359 | 0,39156002 | 0,870333649 | NCK interacting protein with SH3 domain |
| ENSG00000277585 | MUC4 | -0,231854416 | 0,851539633 | 0,86777894 | 0,989074806 | mucin 4, cell surface associated |
| ENSG00000074219 | TEAD2 | -0,23187397 | 0,851528092 | 0,30603061 | 0,815073553 | TEA domain transcription factor 2 |
| ENSG00000145919 | BOD1 | -0,231891279 | 0,851517875 | 0,45970471 | 0,905107873 | biorientation of chromosomes in cell division 1 |
| ENSG00000100802 | C14orf93 | -0,231936456 | 0,851491211 | 0,35158893 | 0,844115834 | chromosome 14 open reading frame 93 |
| ENSG00000105755 | ETHE1 | -0,232035669 | 0,851432657 | 0,44025511 | 0,897973376 | ETHE1 persulfide dioxygenase |
| ENSG00000287643 | AC004825,3 | -0,232083794 | 0,851404255 | 0,50762525 | 0,922140988 | novel transcript, antisense to PCNX1 |
| ENSG00000204116 | CHIC1 | -0,232257696 | 0,851301633 | 0,65213317 | 0,957241719 | cysteine rich hydrophobic domain 1 |
| ENSG00000283162 | AL390726,5 | -0,232260772 | 0,851299819 | 0,68095231 | 0,964654539 | STRA6-like (Stra6l) pseudogene |
| ENSG00000143158 | MPC2 | -0,232267355 | 0,851295934 | 0,44365974 | 0,900090367 | mitochondrial pyruvate carrier 2 |
| ENSG00000092036 | HAUS4 | -0,232302795 | 0,851275022 | 0,36836608 | 0,857548638 | HAUS augmin like complex subunit 4 |

| | | | | | | |
|---|---|---|---|---|---|---|
| ENSG00000269481 | AC010319,4 | -0,23237533 | 0,851232224 | 0,7259744 | 0,973344786 | novel transcript, antisense FAM125A |
| ENSG00000079462 | PAFAH1B3 | -0,232408095 | 0,851212891 | 0,30709341 | 0,815892645 | platelet activating factor acetylhydrolase 1b catalytic subunit 3 |
| ENSG00000149212 | SESN3 | -0,23244643 | 0,851190273 | 0,62819106 | 0,952793934 | sestrin 3 |
| ENSG00000142512 | SIGLEC10 | -0,23247238 | 0,851174963 | 0,68916333 | 0,966685038 | sialic acid binding Ig like lectin 10 |
| ENSG00000121310 | ECHDC2 | -0,232497646 | 0,851160056 | 0,68389687 | 0,965385216 | enoyl-CoA hydratase domain containing 2 |
| ENSG00000286976 | AC084335,1 | -0,232521825 | 0,851145791 | 0,79172449 | 0,979471509 | novel transcript |
| ENSG00000175970 | UNC119B | -0,232529379 | 0,851141335 | 0,34404612 | 0,840938551 | unc-119 lipid binding chaperone B |
| ENSG00000171790 | SLFNL1 | -0,232560381 | 0,851123045 | 0,41144371 | 0,883655718 | schlafen like 1 |
| ENSG00000249242 | TMEM150C | -0,232562836 | 0,851121597 | 0,51237576 | 0,923445033 | transmembrane protein 150C |
| ENSG00000274437 | DERL3 | -0,232610447 | 0,851093509 | 0,62843707 | 0,952793934 | derlin 3 |
| ENSG00000188859 | FAM78B | -0,232651444 | 0,851069324 | 0,6730178 | 0,962500737 | family with sequence similarity 78 member B |
| ENSG00000273175 | BX323046,1 | -0,232656725 | 0,851066208 | 0,81115473 | 0,981213877 | novel transcript |
| ENSG00000137804 | NUSAP1 | -0,232660378 | 0,851064053 | 0,60618331 | 0,950927937 | nucleolar and spindle associated protein 1 |
| ENSG00000215712 | TMEM242 | -0,23272307 | 0,851027071 | 0,51484675 | 0,924720046 | transmembrane protein 242 |
| ENSG00000152931 | PART1 | -0,232796458 | 0,850983782 | 0,61704299 | 0,952793934 | prostate androgen-regulated transcript 1 |
| ENSG00000228172 | AL020996,1 | -0,23286844 | 0,850941324 | 0,69761583 | 0,968522643 | novel transcript |
| ENSG00000186496 | ZNF396 | -0,232896116 | 0,850925 | 0,44878025 | 0,902062594 | zinc finger protein 396 |
| ENSG00000240137 | ERICH6-AS1 | -0,233027555 | 0,850847479 | 0,47253803 | 0,909357939 | ERICH6 antisense RNA 1 |
| ENSG00000161551 | ZNF577 | -0,233123953 | 0,850790628 | 0,55164291 | 0,936366211 | zinc finger protein 577 |
| ENSG00000287672 | AC003086,1 | -0,233161451 | 0,850768515 | 0,85023772 | 0,987036295 | novel transcript |
| ENSG00000173681 | BCLAF3 | -0,233249609 | 0,85071653 | 0,39121263 | 0,870083449 | BCLAF1 and THRAP3 family member 3 |
| ENSG00000168779 | SHOX2 | -0,233290957 | 0,850692148 | 0,35891341 | 0,849814998 | short stature homeobox 2 |
| ENSG00000081791 | DELE1 | -0,233296386 | 0,850688947 | 0,34561849 | 0,841168721 | DAP3 binding cell death enhancer 1 |
| ENSG00000261459 | AC002310,4 | -0,233309629 | 0,850681138 | 0,90351765 | 0,992361703 | novel transcript |
| ENSG00000262370 | AC108134,3 | -0,233392932 | 0,85063202 | 0,65457657 | 0,957444211 | novel transcript |
| ENSG00000264575 | LINC00526 | -0,233405906 | 0,850624371 | 0,45272983 | 0,902285042 | long intergenic non-protein coding RNA 526 |
| ENSG00000204832 | ST8SIA6-AS1 | -0,233502579 | 0,850567373 | 0,81706264 | 0,981979397 | ST8SIA6 antisense RNA 1 |
| ENSG00000141294 | LRRC46 | -0,233532735 | 0,850549594 | 0,45962529 | 0,905088088 | leucine rich repeat containing 46 |
| ENSG00000169762 | TAPT1 | -0,233590712 | 0,850515415 | 0,46782484 | 0,907940653 | transmembrane anterior posterior transformation 1 |
| ENSG00000288670 | AL592295,6 | -0,233667235 | 0,850470303 | 0,38090653 | 0,865900094 | novel transcript |
| ENSG00000272631 | AC067750,1 | -0,233737405 | 0,850428938 | 0,3472714 | 0,841896177 | novel transcript, antisense to BMPR1A |
| ENSG00000285382 | FO393400,1 | -0,233826352 | 0,850376508 | 0,74703132 | 0,975796253 | novel protein |

| | | | | | | |
|---|---|---|---|---|---|---|
| ENSG00000215515 | IFIT1P1 | -0,233826564 | 0,850376383 | 0,74800564 | 0,975901808 | interferon induced protein with tetratricopeptide repeats 1 pseudogene 1 |
| ENSG00000148584 | A1CF | -0,233848406 | 0,850363509 | 0,71345613 | 0,970812834 | APOBEC1 complementation factor |
| ENSG00000285725 | AC004967,2 | -0,233886884 | 0,850340829 | 0,4443932 | 0,900210555 | novel transcript |
| ENSG00000129422 | MTUS1 | -0,234000943 | 0,850273604 | 0,52295102 | 0,92790906 | microtubule associated scaffold protein 1 |
| ENSG00000246067 | RAB30-DT | -0,23411855 | 0,850204293 | 0,33092913 | 0,832153798 | RAB30 divergent transcript |
| ENSG00000101413 | RPRD1B | -0,234126936 | 0,850199352 | 0,47088648 | 0,908869685 | regulation of nuclear pre-mRNA domain containing 1B |
| ENSG00000100626 | GALNT16 | -0,234139528 | 0,850191931 | 0,36696694 | 0,856374902 | polypeptide N-acetylgalactosaminyltransferase 16 |
| ENSG00000164626 | KCNK5 | -0,234155889 | 0,85018229 | 0,55130766 | 0,936366211 | potassium two pore domain channel subfamily K member 5 |
| ENSG00000189089 | RIMKLBP1 | -0,23417479 | 0,850171151 | 0,66995786 | 0,962026636 | ribosomal modification protein rimK like family member B pseudogene 1 |
| ENSG00000214293 | APTR | -0,234188192 | 0,850163254 | 0,33172098 | 0,832548086 | Alu-mediated CDKN1A/p21 transcriptional regulator |
| ENSG00000077458 | FAM76B | -0,234202324 | 0,850154926 | 0,59333716 | 0,947416795 | family with sequence similarity 76 member B |
| ENSG00000208772 | SNORD94 | -0,234212557 | 0,850148896 | 0,73998441 | 0,974715504 | small nucleolar RNA, C/D box 94 |
| ENSG00000255082 | GRM5-AS1 | -0,234315328 | 0,850088337 | 0,76243205 | 0,976315588 | GRM5 antisense RNA 1 |
| ENSG00000223546 | LINC00630 | -0,234408524 | 0,850033424 | 0,34220295 | 0,840048641 | long intergenic non-protein coding RNA 630 |
| ENSG00000229212 | AC044860,1 | -0,234557947 | 0,849945389 | 0,67327591 | 0,962605657 | golgin subfamily A pseudogene |
| ENSG00000239524 | RPL32P34 | -0,234586513 | 0,84992856 | 0,86466862 | 0,989074806 | ribosomal protein L32 pseudogene 34 |
| ENSG00000137185 | ZSCAN9 | -0,234693117 | 0,849865759 | 0,40239744 | 0,87718264 | zinc finger and SCAN domain containing 9 |
| ENSG00000123329 | ARHGAP9 | -0,234771483 | 0,849819597 | 0,74773546 | 0,975901808 | Rho GTPase activating protein 9 |
| ENSG00000143994 | ABHD1 | -0,234781416 | 0,849813746 | 0,66390779 | 0,959985685 | abhydrolase domain containing 1 |
| ENSG00000074855 | ANO8 | -0,234816112 | 0,849793308 | 0,65805308 | 0,957883832 | anoctamin 8 |
| ENSG00000240240 | BX664727,3 | -0,234858304 | 0,849768456 | 0,54547666 | 0,935119936 | novel transcript |
| ENSG00000159685 | CHCHD6 | -0,234887695 | 0,849751145 | 0,38979114 | 0,870057277 | coiled-coil-helix-coiled-coil-helix domain containing 6 |
| ENSG00000167595 | PROSER3 | -0,234891432 | 0,849748944 | 0,42702484 | 0,89229411 | proline and serine rich 3 |
| ENSG00000287690 | AL023775,2 | -0,234895425 | 0,849746592 | 0,74911574 | 0,975901808 | novel transcript, antisense to RPS6KA2 |
| ENSG00000148225 | WDR31 | -0,234926222 | 0,849728452 | 0,4619008 | 0,906277266 | WD repeat domain 31 |
| ENSG00000287221 | AL589794,2 | -0,234954726 | 0,849711664 | 0,68902211 | 0,966640663 | novel transcript, antisense to ASAH2B |
| ENSG00000254656 | RTL1 | -0,234971159 | 0,849701986 | 0,64043425 | 0,956251063 | retrotransposon Gag like 1 |
| ENSG00000113231 | PDE8B | -0,234998205 | 0,849686056 | 0,6422865 | 0,956633898 | phosphodiesterase 8B |

| | | | | | | |
|---|---|---|---|---|---|---|
| ENSG00000227398 | KIF9-AS1 | -0,235005877 | 0,849681538 | 0,40913127 | 0,882021797 | KIF9 antisense RNA 1 |
| ENSG00000205002 | AARD | -0,235054733 | 0,849652764 | 0,57844616 | 0,943425812 | alanine and arginine rich domain containing protein |
| ENSG00000271109 | AC008555,4 | -0,235059157 | 0,849650159 | 0,50630188 | 0,921780819 | novel transcript |
| ENSG00000264215 | AC126365,1 | -0,235072698 | 0,849642184 | 0,78978417 | 0,979462318 | novel transcript |
| ENSG00000149548 | CCDC15 | -0,235116224 | 0,849616551 | 0,52657362 | 0,929261495 | coiled-coil domain containing 15 |
| ENSG00000287860 | AC079804,3 | -0,235116438 | 0,849616425 | 0,7688628 | 0,977381742 | novel transcript |
| ENSG00000275911 | NDE1 | -0,235141032 | 0,849601941 | 0,49408717 | 0,917572329 | nudE neurodevelopment protein 1 |
| ENSG00000178104 | PDE4DIP | -0,2351411 | 0,849601902 | 0,23811525 | 0,747367171 | phosphodiesterase 4D interacting protein |
| ENSG00000165071 | TMEM71 | -0,235146565 | 0,849598683 | 0,72393991 | 0,973141049 | transmembrane protein 71 |
| ENSG00000207191 | SNORD116-5 | -0,235165597 | 0,849587475 | 0,79783352 | 0,980161849 | small nucleolar RNA, C/D box 116-5 |
| ENSG00000207133 | SNORD116-7 | -0,235165597 | 0,849587475 | 0,79783352 | 0,980161849 | small nucleolar RNA, C/D box 116-7 |
| ENSG00000143156 | NME7 | -0,235225979 | 0,849551918 | 0,44982204 | 0,902062594 | NME/NM23 family member 7 |
| ENSG00000119888 | EPCAM | -0,235238516 | 0,849544535 | 0,67349041 | 0,962780262 | epithelial cell adhesion molecule |
| ENSG00000168237 | GLYCTK | -0,235247167 | 0,849539441 | 0,40958313 | 0,882177108 | glycerate kinase |
| ENSG00000143552 | NUP210L | -0,235291232 | 0,849513493 | 0,67676622 | 0,963580269 | nucleoporin 210 like |
| ENSG00000214544 | GTF2IRD2P1 | -0,235298776 | 0,849509051 | 0,7836018 | 0,979196878 | GTF2I repeat domain containing 2 pseudogene 1 |
| ENSG00000040341 | STAU2 | -0,235346949 | 0,849480686 | 0,60164274 | 0,949939978 | staufen double-stranded RNA binding protein 2 |
| ENSG00000264577 | AC010761,1 | -0,235360056 | 0,849472968 | 0,41394368 | 0,885456174 | novel transcript, antisense to RPL23A |
| ENSG00000100290 | BIK | -0,235364976 | 0,849470071 | 0,78775448 | 0,979462318 | BCL2 interacting killer |
| ENSG00000285596 | AC017116,2 | -0,235407906 | 0,849444794 | 0,57727599 | 0,94337651 | novel transcript |
| ENSG00000172031 | EPHX4 | -0,235515484 | 0,849381456 | 0,54090289 | 0,933385624 | epoxide hydrolase 4 |
| ENSG00000107099 | DOCK8 | -0,235532803 | 0,849371259 | 0,67280167 | 0,962371283 | dedicator of cytokinesis 8 |
| ENSG00000144134 | RABL2A | -0,235608459 | 0,849326719 | 0,3277522 | 0,829743891 | RAB, member of RAS oncogene family like 2A |
| ENSG00000154065 | ANKRD29 | -0,23563131 | 0,849313266 | 0,56716443 | 0,941425553 | ankyrin repeat domain 29 |
| ENSG00000112667 | DNPH1 | -0,235701888 | 0,849271718 | 0,43473814 | 0,896272827 | 2'-deoxynucleoside 5'-phosphate N-hydrolase 1 |
| ENSG00000213152 | RPL7AP60 | -0,235732139 | 0,84925391 | 0,64786536 | 0,956899963 | ribosomal protein L7a pseudogene 60 |
| ENSG00000254432 | AC113143,1 | -0,235902016 | 0,849153917 | 0,86901849 | 0,989074806 | novel transcript |
| ENSG00000155016 | CYP2U1 | -0,235929075 | 0,84913799 | 0,53779666 | 0,932883277 | cytochrome P450 family 2 subfamily U member 1 |
| ENSG00000198915 | RASGEF1A | -0,236047254 | 0,849068435 | 0,67322684 | 0,962605657 | RasGEF domain family member 1A |
| ENSG00000165959 | CLMN | -0,236167627 | 0,848997595 | 0,45621163 | 0,903545728 | calmin |
| ENSG00000269896 | AL513477,1 | -0,236178762 | 0,848991043 | 0,51220253 | 0,923394062 | small nuclear ribonucleoprotein N (SNRPN) pseudogene |
| ENSG00000174279 | EVX2 | -0,236234828 | 0,84895805 | 0,83898841 | 0,985672131 | even-skipped homeobox 2 |

| | | | | | | |
|---|---|---|---|---|---|---|
| ENSG00000126003 | PLAGL2 | -0,236419907 | 0,848849146 | 0,34037548 | 0,839010589 | PLAG1 like zinc finger 2 |
| ENSG00000288066 | AC097448,1 | -0,236503773 | 0,848799803 | 0,5549302 | 0,937793851 | novel transcript, antisense to ARL5A |
| ENSG00000230699 | AL645608,2 | -0,236531433 | 0,84878353 | 0,59731273 | 0,948282297 | novel transcript |
| ENSG00000165669 | FAM204A | -0,236585938 | 0,848751464 | 0,4645641 | 0,907588071 | family with sequence similarity 204 member A |
| ENSG00000278768 | BACE1-AS | -0,236634251 | 0,848723041 | 0,30391219 | 0,812968394 | BACE1 antisense RNA |
| ENSG00000122966 | CIT | -0,236680635 | 0,848695754 | 0,2878718 | 0,799107075 | citron rho-interacting serine/threonine kinase |
| ENSG00000153993 | SEMA3D | -0,236743452 | 0,848658802 | 0,52738592 | 0,929454507 | semaphorin 3D |
| ENSG00000223599 | AL513523,1 | -0,236811182 | 0,84861896 | 0,68058239 | 0,964654539 | adipose differentiation-related protein (ADFP) pseudogene |
| ENSG00000262025 | AL513523,5 | -0,236811182 | 0,84861896 | 0,68058239 | 0,964654539 | adipose differentiation-related protein (ADFP) pseudogene |
| ENSG00000272817 | AL359198,1 | -0,236820352 | 0,848613567 | 0,70648151 | 0,969503793 | novel transcript, antisense to FGFBP3 |
| ENSG00000134398 | ERN2 | -0,236837745 | 0,848603336 | 0,71368714 | 0,970812834 | endoplasmic reticulum to nucleus signaling 2 |
| ENSG00000149294 | NCAM1 | -0,236862774 | 0,848588614 | 0,48617317 | 0,914091794 | neural cell adhesion molecule 1 |
| ENSG00000166260 | COX11 | -0,236936828 | 0,848545056 | 0,56081941 | 0,940368987 | cytochrome c oxidase copper chaperone COX11 |
| ENSG00000136531 | SCN2A | -0,237049754 | 0,84847864 | 0,68256758 | 0,964815657 | sodium voltage-gated channel alpha subunit 2 |
| ENSG00000185105 | MYADML2 | -0,237069014 | 0,848467313 | 0,78790985 | 0,979462318 | myeloid associated differentiation marker like 2 |
| ENSG00000109758 | HGFAC | -0,237107147 | 0,848444886 | 0,71153618 | 0,970739528 | HGF activator |
| ENSG00000227028 | SLC8A1-AS1 | -0,237121282 | 0,848436574 | 0,60328423 | 0,950352286 | SLC8A1 antisense RNA 1 |
| ENSG00000167325 | RRM1 | -0,237168024 | 0,848409086 | 0,48586383 | 0,913997663 | ribonucleotide reductase catalytic subunit M1 |
| ENSG00000285919 | AC087639,2 | -0,237211898 | 0,848383285 | 0,65441804 | 0,957444211 | novel transcript, antisense to SPESP1 |
| ENSG00000260456 | C16orf95 | -0,237294505 | 0,848334709 | 0,51547482 | 0,924901769 | chromosome 16 open reading frame 95 |
| ENSG00000081307 | UBA5 | -0,237297872 | 0,848332729 | 0,45582901 | 0,903545728 | ubiquitin like modifier activating enzyme 5 |
| ENSG00000225630 | MTND2P28 | -0,237411256 | 0,84826606 | 0,62496069 | 0,952793934 | MT-ND2 pseudogene 28 |
| ENSG00000128059 | PPAT | -0,237487398 | 0,848221291 | 0,56534529 | 0,940865625 | phosphoribosyl pyrophosphate amidotransferase |
| ENSG00000270084 | GAS5-AS1 | -0,237578629 | 0,848167655 | 0,68068592 | 0,964654539 | GAS5 antisense RNA 1 |
| ENSG00000065371 | ROPN1 | -0,237636354 | 0,848133718 | 0,85469552 | 0,987861659 | rhophilin associated tail protein 1 |
| ENSG00000260911 | AC135050,3 | -0,237669565 | 0,848114194 | 0,86807135 | 0,989074806 | novel transcript |
| ENSG00000257220 | PRXL2AP1 | -0,237690135 | 0,848102102 | 0,8599404 | 0,989074806 | peroxiredoxin like 2A pseudogene 1 |
| ENSG00000229832 | AL360091,1 | -0,237692561 | 0,848100676 | 0,75505797 | 0,975901808 | novel transcript, sense intronic to PPP2R5A |
| ENSG00000146938 | NLGN4X | -0,237708443 | 0,84809134 | 0,51385547 | 0,924227101 | neuroligin 4 X-linked |
| ENSG00000273598 | AC211486,3 | -0,237712898 | 0,848088721 | 0,7294335 | 0,974009551 | postmeiotic segregation increased 2 pseudogene 6 |
| ENSG00000132780 | NASP | -0,237789003 | 0,848043983 | 0,37898793 | 0,864362316 | nuclear autoantigenic sperm protein |

| | | | | | | |
|---|---|---|---|---|---|---|
| ENSG00000228509 | AC006460,1 | -0,23779441 | 0,848040805 | 0,56249938 | 0,940865625 | novel transcript, antisense to NAB1 |
| ENSG00000255893 | AP000786,1 | -0,237837247 | 0,848015626 | 0,69453509 | 0,967417311 | novel transcript |
| ENSG00000223776 | LGALS8-AS1 | -0,237853951 | 0,848005807 | 0,73324981 | 0,974353474 | LGALS8 antisense RNA 1 |
| ENSG00000254837 | AP001372,2 | -0,237860409 | 0,848002011 | 0,56341729 | 0,940865625 | uncharacterized LOC100287896 |
| ENSG00000171747 | LGALS4 | -0,237861547 | 0,848001342 | 0,81965915 | 0,98249617 | galectin 4 |
| ENSG00000226174 | TEX22 | -0,237904645 | 0,84797601 | 0,77425072 | 0,978503783 | testis expressed 22 |
| ENSG00000117262 | GPR89A | -0,237915069 | 0,847969883 | 0,51497336 | 0,924720046 | G protein-coupled receptor 89A |
| ENSG00000241313 | WWTR1-AS1 | -0,238005064 | 0,847916988 | 0,63694626 | 0,955908589 | WWTR1 antisense RNA 1 |
| ENSG00000253167 | WASHC5-AS1 | -0,238063199 | 0,847882821 | 0,77391844 | 0,978415098 | WASHC5 antisense RNA 1 |
| ENSG00000224310 | LINC01567 | -0,238075787 | 0,847875423 | 0,64604232 | 0,956899963 | long intergenic non-protein coding RNA 1567 |
| ENSG00000228876 | AC010745,2 | -0,238166875 | 0,847821892 | 0,47593025 | 0,911390632 | novel transcript |
| ENSG00000214530 | STARD10 | -0,238174927 | 0,84781716 | 0,53498887 | 0,931805607 | StAR related lipid transfer domain containing 10 |
| ENSG00000085662 | AKR1B1 | -0,238251901 | 0,847771927 | 0,39021269 | 0,870057277 | aldo-keto reductase family 1 member B |
| ENSG00000133818 | RRAS2 | -0,238356178 | 0,847710653 | 0,6047473 | 0,950927937 | RAS related 2 |
| ENSG00000277692 | AL121583,1 | -0,238364916 | 0,847705518 | 0,78004275 | 0,978587255 | novel transcript |
| ENSG00000205863 | C1QTNF9B | -0,238488467 | 0,847632925 | 0,83047654 | 0,984156028 | C1q and TNF related 9B |
| ENSG00000283156 | AC068620,3 | -0,238496613 | 0,847628139 | 0,53086677 | 0,931265667 | novel Myb/SANT-like DNA-binding domain containing pseudogene |
| ENSG00000243433 | AC010973,1 | -0,238686178 | 0,847516771 | 0,78510394 | 0,979320406 | novel transcript |
| ENSG00000164830 | OXR1 | -0,238723097 | 0,847495083 | 0,59424259 | 0,947469769 | oxidation resistance 1 |
| ENSG00000124140 | SLC12A5 | -0,238784564 | 0,847458975 | 0,43316294 | 0,895845815 | solute carrier family 12 member 5 |
| ENSG00000266258 | LINC01909 | -0,23881407 | 0,847441644 | 0,70751005 | 0,969647757 | long intergenic non-protein coding RNA 1909 |
| ENSG00000163104 | SMARCAD1 | -0,238863514 | 0,847412601 | 0,59974375 | 0,949340299 | SWI/SNF-related, matrix-associated actin-dependent regulator of chromatin, subfamily a, containing DEAD/H box 1 |
| ENSG00000182606 | TRAK1 | -0,238879271 | 0,847403345 | 0,29296561 | 0,803365244 | trafficking kinesin protein 1 |
| ENSG00000119844 | AFTPH | -0,238881938 | 0,847401779 | 0,53420945 | 0,93151582 | aftiphilin |
| ENSG00000073584 | SMARCE1 | -0,238897738 | 0,847392498 | 0,51033285 | 0,922523629 | SWI/SNF related, matrix associated, actin dependent regulator of chromatin, subfamily e, member 1 |
| ENSG00000205940 | HSP90AB2P | -0,239075999 | 0,8472878 | 0,65519884 | 0,957444211 | heat shock protein 90 alpha family class B member 2, pseudogene |

| | | | | | | |
|---|---|---|---|---|---|---|
| ENSG00000007402 | CACNA2D2 | -0,239107255 | 0,847269444 | 0,37958516 | 0,865157837 | calcium voltage-gated channel auxiliary subunit alpha2delta 2 |
| ENSG00000143653 | SCCPDH | -0,239113516 | 0,847265767 | 0,45465474 | 0,903414621 | saccharopine dehydrogenase (putative) |
| ENSG00000170584 | NUDCD2 | -0,239215179 | 0,847206064 | 0,50941675 | 0,922427526 | NudC domain containing 2 |
| ENSG00000274828 | AC068473,5 | -0,239251807 | 0,847184555 | 0,56759642 | 0,941596868 | novel transcript |
| ENSG00000282330 | AC068473,8 | -0,239251807 | 0,847184555 | 0,56759642 | 0,941596868 | novel transcript |
| ENSG00000180152 | XIAPP3 | -0,239408443 | 0,84709258 | 0,79839929 | 0,980161849 | X-linked inhibitor of apoptosis pseudogene 3 |
| ENSG00000102763 | VWA8 | -0,239430057 | 0,847079889 | 0,49530189 | 0,917777383 | von Willebrand factor A domain containing 8 |
| ENSG00000170836 | PPM1D | -0,239432532 | 0,847078436 | 0,35800921 | 0,849032846 | protein phosphatase, Mg2+/Mn2+ dependent 1D |
| ENSG00000262133 | AC087392,2 | -0,239562441 | 0,847002163 | 0,81079119 | 0,981213877 | novel transcript, antisense to NXN |
| ENSG00000157551 | KCNJ15 | -0,23961134 | 0,846973455 | 0,36305154 | 0,853318246 | potassium inwardly rectifying channel subfamily J member 15 |
| ENSG00000275318 | AL136981,2 | -0,239668732 | 0,846939762 | 0,66505594 | 0,960681381 | novel zinc finger protein pseudogene |
| ENSG00000273302 | AC016747,3 | -0,239744476 | 0,846895298 | 0,58213933 | 0,944174844 | novel transcript |
| ENSG00000254507 | DEFB131E | -0,239815752 | 0,846853458 | 0,8049904 | 0,98109454 | defensin beta 131E (pseudogene) |
| ENSG00000285431 | DEFB131E | -0,239815752 | 0,846853458 | 0,8049904 | 0,98109454 | defensin beta 131E (pseudogene) |
| ENSG00000288263 | GRK1 | -0,239829884 | 0,846845163 | 0,85461419 | 0,987861659 | G protein-coupled receptor kinase 1 |
| ENSG00000185974 | GRK1 | -0,239829884 | 0,846845163 | 0,85461419 | 0,987861659 | G protein-coupled receptor kinase 1 |
| ENSG00000165898 | ISCA2 | -0,23983041 | 0,846844854 | 0,40290339 | 0,877551313 | iron-sulfur cluster assembly 2 |
| ENSG00000260802 | SERTM2 | -0,239863898 | 0,846825197 | 0,75730114 | 0,975901808 | serine rich and transmembrane domain containing 2 |
| ENSG00000133111 | RFXAP | -0,240081063 | 0,846697736 | 0,60455161 | 0,950864972 | regulatory factor X associated protein |
| ENSG00000286524 | AC092567,2 | -0,240166239 | 0,846647749 | 0,68452157 | 0,965415049 | novel transcript, antisense to EHBP1 |
| ENSG00000148288 | GBGT1 | -0,240168889 | 0,846646194 | 0,43504422 | 0,896272827 | globoside alpha-1,3-N-acetylgalactosaminyltransferase 1 (FORS blood group) |
| ENSG00000187535 | IFT140 | -0,240172361 | 0,846644156 | 0,43671876 | 0,896866043 | intraflagellar transport 140 |
| ENSG00000188573 | FBLL1 | -0,240178799 | 0,846640378 | 0,57523084 | 0,942667371 | fibrillarin like 1 |
| ENSG00000272914 | AL359532,1 | -0,240262783 | 0,846591094 | 0,78198213 | 0,978597905 | novel transcript, sense intronic to ZNF438 |
| ENSG00000244482 | LILRA6 | -0,240276003 | 0,846583336 | 0,84902932 | 0,987036295 | leukocyte immunoglobulin like receptor A6 |
| ENSG00000162062 | TEDC2 | -0,240285938 | 0,846577506 | 0,37846325 | 0,864031013 | tubulin epsilon and delta complex 2 |
| ENSG00000157017 | GHRL | -0,240360189 | 0,846533937 | 0,64101256 | 0,956410077 | ghrelin and obestatin prepropeptide |
| ENSG00000277067 | CU634019,1 | -0,240418782 | 0,846499557 | 0,79256619 | 0,979471509 | uncharacterized LOC102724843 |
| ENSG00000140471 | LINS1 | -0,240448813 | 0,846481937 | 0,49429522 | 0,91758518 | lines homolog 1 |

| | | | | | | |
|---|---|---|---|---|---|---|
| ENSG00000275437 | AL121832,3 | -0,240463239 | 0,846473472 | 0,58250392 | 0,944174844 | novel transcript, sense intronic to CABLES2 |
| ENSG00000260805 | AC092803,1 | -0,240494501 | 0,84645513 | 0,56239875 | 0,940865625 | novel transcript |
| ENSG00000259802 | AC012640,2 | -0,240502173 | 0,846450629 | 0,52088739 | 0,927046149 | novel transcript, antisense to MARCH6 |
| ENSG00000267080 | ASB16-AS1 | -0,24061648 | 0,846383566 | 0,38452329 | 0,868434303 | ASB16 antisense RNA 1 |
| ENSG00000236698 | EIF1AXP1 | -0,240899171 | 0,846217736 | 0,72934385 | 0,974009551 | EIF1AX pseudogene 1 |
| ENSG00000196159 | FAT4 | -0,240957013 | 0,846183809 | 0,45855364 | 0,904322271 | FAT atypical cadherin 4 |
| ENSG00000197362 | ZNF786 | -0,240957734 | 0,846183386 | 0,27752984 | 0,789067728 | zinc finger protein 786 |
| ENSG00000121057 | AKAP1 | -0,240977632 | 0,846171716 | 0,33931767 | 0,838276153 | A-kinase anchoring protein 1 |
| ENSG00000075043 | KCNQ2 | -0,241034547 | 0,846138335 | 0,57843971 | 0,943425812 | potassium voltage-gated channel subfamily Q member 2 |
| ENSG00000229558 | SACS-AS1 | -0,241046581 | 0,846131277 | 0,59207832 | 0,947416795 | SACS antisense RNA 1 |
| ENSG00000285693 | AP002381,2 | -0,241165181 | 0,846061721 | 0,54654719 | 0,935401821 | Novel transcript |
| ENSG00000205682 | AC020741,1 | -0,241225672 | 0,846026247 | 0,65935562 | 0,958425175 | novel transcript |
| ENSG00000101974 | ATP11C | -0,241251635 | 0,846011023 | 0,52433394 | 0,928579043 | ATPase phospholipid transporting 11C |
| ENSG00000135205 | CCDC146 | -0,24127696 | 0,845996172 | 0,54795114 | 0,935404974 | coiled-coil domain containing 146 |
| ENSG00000183044 | ABAT | -0,2412999 | 0,84598272 | 0,39922701 | 0,875433675 | 4-aminobutyrate aminotransferase |
| ENSG00000258344 | AC078778,1 | -0,241315712 | 0,845973448 | 0,66686089 | 0,9611116 | novel transcript, sense overlapping to COPZ1 & HNRNPA1 |
| ENSG00000203734 | ECT2L | -0,241328269 | 0,845966085 | 0,61288103 | 0,952413269 | epithelial cell transforming 2 like |
| ENSG00000178761 | FAM219B | -0,241356165 | 0,845949727 | 0,33019301 | 0,831800372 | family with sequence similarity 219 member B |
| ENSG00000230366 | DSCR9 | -0,241359165 | 0,845947968 | 0,70212338 | 0,968742626 | Down syndrome critical region 9 |
| ENSG00000235660 | LINC00345 | -0,241404602 | 0,845921326 | 0,64250068 | 0,956633898 | long intergenic non-protein coding RNA 345 |
| ENSG00000198939 | ZFP2 | -0,24141514 | 0,845915147 | 0,522978 | 0,92790906 | ZFP2 zinc finger protein |
| ENSG00000155085 | AK9 | -0,241419855 | 0,845912382 | 0,39091291 | 0,870057277 | adenylate kinase 9 |
| ENSG00000204634 | TBC1D8 | -0,241491622 | 0,845870303 | 0,33484428 | 0,834814326 | TBC1 domain family member 8 |
| ENSG00000135540 | NHSL1 | -0,241611859 | 0,84579981 | 0,30962804 | 0,816986489 | NHS like 1 |
| ENSG00000185278 | ZBTB37 | -0,241629233 | 0,845789624 | 0,49165353 | 0,916995341 | zinc finger and BTB domain containing 37 |
| ENSG00000251322 | SHANK3 | -0,241632331 | 0,845787808 | 0,51161528 | 0,922920779 | SH3 and multiple ankyrin repeat domains 3 |
| ENSG00000167740 | CYB5D2 | -0,241639198 | 0,845783782 | 0,31446864 | 0,819877431 | cytochrome b5 domain containing 2 |
| ENSG00000248383 | PCDHAC1 | -0,24166273 | 0,845769986 | 0,41616321 | 0,885986046 | protocadherin alpha subfamily C, 1 |
| ENSG00000163026 | WDCP | -0,241672731 | 0,845764123 | 0,46538142 | 0,90786072 | WD repeat and coiled coil containing |
| ENSG00000170448 | NFXL1 | -0,241770959 | 0,84570654 | 0,45700092 | 0,903711377 | nuclear transcription factor, X-box binding like 1 |
| ENSG00000286676 | ACTBP4 | -0,241782115 | 0,8457 | 0,7243332 | 0,973206978 | ACTB pseudogene 4 |

| | | | | | | |
|---|---|---|---|---|---|---|
| ENSG00000272751 | AC006518,4 | -0,24181582 | 0,845680243 | 0,8032245 | 0,980866933 | taste receptor, type 2, member 12 pseudogene |
| ENSG00000278705 | H4C2 | -0,241851513 | 0,845659321 | 0,35994619 | 0,850280272 | H4 clustered histone 2 |
| ENSG00000185737 | NRG3 | -0,24188591 | 0,845639159 | 0,46088884 | 0,906029574 | neuregulin 3 |
| ENSG00000213904 | LIPE-AS1 | -0,241942008 | 0,845606278 | 0,52477084 | 0,928616707 | LIPE antisense RNA 1 |
| ENSG00000151806 | GUF1 | -0,24207573 | 0,845527903 | 0,50712695 | 0,921951873 | GUF1 homolog, GTPase |
| ENSG00000143942 | CHAC2 | -0,242099968 | 0,845513697 | 0,58938881 | 0,946844281 | ChaC glutathione specific gamma-glutamylcyclotransferase 2 |
| ENSG00000182389 | CACNB4 | -0,242123391 | 0,84549997 | 0,53363578 | 0,931381467 | calcium voltage-gated channel auxiliary subunit beta 4 |
| ENSG00000128594 | LRRC4 | -0,242172433 | 0,845471229 | 0,41428453 | 0,885579089 | leucine rich repeat containing 4 |
| ENSG00000276282 | AC022960,2 | -0,242191163 | 0,845460253 | 0,77815668 | 0,978587255 | novel transcript, sense intronic to TRAPPC8 |
| ENSG00000213714 | FAM209B | -0,242234065 | 0,845435112 | 0,84527631 | 0,986639902 | family with sequence similarity 209 member B |
| ENSG00000260118 | AL157700,1 | -0,242238059 | 0,845432771 | 0,79360235 | 0,979776981 | novel transcript |
| ENSG00000227954 | TARID | -0,242307265 | 0,845392217 | 0,85992036 | 0,989074806 | TCF21 antisense RNA inducing promoter demethylation |
| ENSG00000273419 | AC004877,1 | -0,242377446 | 0,845351093 | 0,6830803 | 0,964965151 | novel transcript, antisense to ZNF862 |
| ENSG00000287853 | AL031668,2 | -0,242549759 | 0,845250132 | 0,64608975 | 0,956899963 | novel transcript |
| ENSG00000105426 | PTPRS | -0,242600964 | 0,845220132 | 0,38369636 | 0,868000076 | protein tyrosine phosphatase receptor type S |
| ENSG00000106603 | COA1 | -0,242667891 | 0,845180923 | 0,35247945 | 0,844912682 | cytochrome c oxidase assembly factor 1 homolog |
| ENSG00000064687 | ABCA7 | -0,24268926 | 0,845168405 | 0,58668702 | 0,945395317 | ATP binding cassette subfamily A member 7 |
| ENSG00000134551 | PRH2 | -0,24288393 | 0,845054369 | 0,57116727 | 0,942436428 | proline rich protein HaeIII subfamily 2 |
| ENSG00000155744 | FAM126B | -0,242895506 | 0,845047589 | 0,51219821 | 0,923394062 | family with sequence similarity 126 member B |
| ENSG00000163935 | SFMBT1 | -0,242957278 | 0,845011407 | 0,40401521 | 0,877939445 | Scm like with four mbt domains 1 |
| ENSG00000285730 | Z94721,3 | -0,243154089 | 0,844896139 | 0,48713571 | 0,914593485 | novel transcript |
| ENSG00000188803 | SHISA6 | -0,243313968 | 0,844802513 | 0,37393947 | 0,861369706 | shisa family member 6 |
| ENSG00000253793 | AC008066,1 | -0,243325873 | 0,844795542 | 0,87374721 | 0,989633545 | lysophosphatidylglycerol acyltransferase 1 (LPGAT1) pseudogene |
| ENSG00000189227 | C15orf61 | -0,243326568 | 0,844795135 | 0,49483584 | 0,917622746 | chromosome 15 open reading frame 61 |
| ENSG00000186260 | MRTFB | -0,243376282 | 0,844766025 | 0,48176954 | 0,912981134 | myocardin related transcription factor B |
| ENSG00000242852 | ZNF709 | -0,243439849 | 0,844728804 | 0,52310273 | 0,92790906 | zinc finger protein 709 |
| ENSG00000104218 | CSPP1 | -0,243466378 | 0,844713271 | 0,32877679 | 0,830659639 | centrosome and spindle pole associated protein 1 |
| ENSG00000273275 | AC017083,2 | -0,243468897 | 0,844711796 | 0,9299124 | 0,994331958 | novel transcript |
| ENSG00000269867 | AC010326,3 | -0,243490152 | 0,844699351 | 0,64549498 | 0,956899963 | novel transcript, sense intronic to ZNF814 |

| | | | | | | |
|---|---|---|---|---|---|---|
| ENSG00000184669 | OR7E14P | -0,243605139 | 0,844632029 | 0,57275854 | 0,942475308 | olfactory receptor family 7 subfamily E member 14 pseudogene |
| ENSG00000156650 | KAT6B | -0,243642266 | 0,844610293 | 0,27666999 | 0,788353146 | lysine acetyltransferase 6B |
| ENSG00000125482 | TTF1 | -0,24364475 | 0,844608839 | 0,39563401 | 0,872227623 | transcription termination factor 1 |
| ENSG00000011143 | MKS1 | -0,24365922 | 0,844600367 | 0,29108405 | 0,801689099 | MKS transition zone complex subunit 1 |
| ENSG00000105662 | CRTC1 | -0,243735599 | 0,844555654 | 0,29555571 | 0,805426883 | CREB regulated transcription coactivator 1 |
| ENSG00000278637 | H4C1 | -0,243748178 | 0,84454829 | 0,30373249 | 0,812795736 | H4 clustered histone 1 |
| ENSG00000272871 | AL159169,2 | -0,243757468 | 0,844542852 | 0,64790594 | 0,956899963 | novel transcript |
| ENSG00000225376 | TMEM246-AS1 | -0,243766022 | 0,844537844 | 0,78457467 | 0,979320406 | TMEM246 antisense RNA 1 |
| ENSG00000160051 | IQCC | -0,243818781 | 0,84450696 | 0,36629704 | 0,856103393 | IQ motif containing C |
| ENSG00000256250 | AC073912,1 | -0,243839322 | 0,844494936 | 0,71953139 | 0,971869926 | novel transcript |
| ENSG00000183067 | IGSF5 | -0,243842659 | 0,844492983 | 0,67824928 | 0,9640473 | immunoglobulin superfamily member 5 |
| ENSG00000240401 | AC012358,2 | -0,243854095 | 0,844486289 | 0,53290904 | 0,931291085 | novel transcript, antisense to CCDC88A |
| ENSG00000223752 | ATAT1 | -0,243883414 | 0,844469127 | 0,47652375 | 0,91143003 | alpha tubulin acetyltransferase 1 |
| ENSG00000214077 | GNAQP1 | -0,243921967 | 0,844446561 | 0,5968406 | 0,948282297 | G protein subunit alpha q pseudogene 1 |
| ENSG00000175189 | INHBC | -0,243935542 | 0,844438615 | 0,75986992 | 0,976190543 | inhibin subunit beta C |
| ENSG00000260672 | AC100827,4 | -0,243946331 | 0,8444323 | 0,72840435 | 0,973955059 | novel transcript |
| ENSG00000135749 | PCNX2 | -0,244097057 | 0,844344082 | 0,36385468 | 0,854140004 | pecanex 2 |
| ENSG00000246451 | AL049840,2 | -0,244168322 | 0,844302375 | 0,28511979 | 0,796602794 | novel transcript, antisense to KLC1 |
| ENSG00000196776 | CD47 | -0,24420243 | 0,844282415 | 0,54023432 | 0,933385624 | CD47 molecule |
| ENSG00000138660 | AP1AR | -0,244311334 | 0,844218685 | 0,59649601 | 0,948282297 | adaptor related protein complex 1 associated regulatory protein |
| ENSG00000130787 | HIP1R | -0,244315728 | 0,844216114 | 0,60369849 | 0,950391097 | huntingtin interacting protein 1 related |
| ENSG00000163806 | SPDYA | -0,244322085 | 0,844212394 | 0,52480066 | 0,928616707 | speedy/RINGO cell cycle regulator family member A |
| ENSG00000260589 | STAM-AS1 | -0,244419898 | 0,844155159 | 0,57494393 | 0,942533938 | STAM antisense RNA 1 (head to head) |
| ENSG00000122481 | RWDD3 | -0,244484693 | 0,844117247 | 0,52697279 | 0,92927181 | RWD domain containing 3 |
| ENSG00000281742 | MCCC2 | -0,244510113 | 0,844102374 | 0,6347176 | 0,955149453 | methylcrotonoyl-CoA carboxylase 2 |
| ENSG00000115687 | PASK | -0,244577048 | 0,844063212 | 0,34931779 | 0,843141089 | PAS domain containing serine/threonine kinase |
| ENSG00000258727 | AL135999,1 | -0,244606066 | 0,844046235 | 0,58336899 | 0,944174844 | novel transcript, antisense to AP1G2 |
| ENSG00000110172 | CHORDC1 | -0,244625077 | 0,844035113 | 0,57238425 | 0,942475308 | cysteine and histidine rich domain containing 1 |
| ENSG00000160298 | C21orf58 | -0,244641441 | 0,844025539 | 0,41560086 | 0,885986046 | chromosome 21 open reading frame 58 |
| ENSG00000061656 | SPAG4 | -0,244676711 | 0,844004905 | 0,72135641 | 0,972142678 | sperm associated antigen 4 |
| ENSG00000286677 | AP006545,3 | -0,244682419 | 0,844001566 | 0,729666 | 0,974009551 | novel transcript |

| | | | | | | |
|---|---|---|---|---|---|---|
| ENSG00000105270 | CLIP3 | -0,244690987 | 0,843996554 | 0,24873258 | 0,760720365 | CAP-Gly domain containing linker protein 3 |
| ENSG00000259366 | AC108449,2 | -0,244719513 | 0,843979866 | 0,49693337 | 0,918645042 | novel transcript, sense intronic to HMBOX1 |
| ENSG00000250130 | GAPDHP43 | -0,244739978 | 0,843967894 | 0,71359143 | 0,970812834 | glyceraldehyde 3 phosphate dehydrogenase pseudogene 43 |
| ENSG00000254263 | AC022973,4 | -0,244758697 | 0,843956943 | 0,77366421 | 0,978415098 | novel transcript |
| ENSG00000083828 | ZNF586 | -0,244862076 | 0,84389647 | 0,25279183 | 0,765053653 | zinc finger protein 586 |
| ENSG00000275091 | AC022098,4 | -0,245184436 | 0,843707929 | 0,84256783 | 0,986175464 | novel transcript, antisense to RFX1 |
| ENSG00000288351 | AC022098,7 | -0,245184436 | 0,843707929 | 0,84256783 | 0,986175464 | novel transcript, antisense to RFX1 |
| ENSG00000044446 | PHKA2 | -0,245223092 | 0,843685322 | 0,32283123 | 0,824955371 | phosphorylase kinase regulatory subunit alpha 2 |
| ENSG00000287841 | AC096585,1 | -0,245235621 | 0,843677996 | 0,80834945 | 0,981213877 | novel transcript |
| ENSG00000278275 | AC103705,1 | -0,245387426 | 0,843589226 | 0,65580972 | 0,957444211 | protein tyrosine phosphatase type IVA, member 1 (PTP4A1) pseudogene |
| ENSG00000185917 | SETD4 | -0,245437698 | 0,843559831 | 0,29586131 | 0,80547842 | SET domain containing 4 |
| ENSG00000131797 | CLUHP3 | -0,245484577 | 0,84353242 | 0,39779579 | 0,874263835 | clustered mitochondria homolog pseudogene 3 |
| ENSG00000120071 | KANSL1 | -0,245554807 | 0,843491358 | 0,31217682 | 0,818547047 | KAT8 regulatory NSL complex subunit 1 |
| ENSG00000198093 | ZNF649 | -0,245559672 | 0,843488514 | 0,49943389 | 0,920166254 | zinc finger protein 649 |
| ENSG00000125863 | MKKS | -0,245591209 | 0,843470076 | 0,60261633 | 0,950352286 | McKusick-Kaufman syndrome |
| ENSG00000234597 | AC010096,1 | -0,245594634 | 0,843468073 | 0,83457758 | 0,984905093 | novel transcript |
| ENSG00000185261 | KIAA0825 | -0,245628446 | 0,843448305 | 0,53107765 | 0,931265667 | KIAA0825 |
| ENSG00000200091 | RN7SKP163 | -0,245686867 | 0,843414151 | 0,85565084 | 0,988075747 | RN7SK pseudogene 163 |
| ENSG00000215223 | CYP51A1P3 | -0,245702786 | 0,843404845 | 0,76896583 | 0,977406998 | cytochrome P450 family 51 subfamily A member 1 pseudogene 3 |
| ENSG00000277221 | LRRC37A2 | -0,245725206 | 0,843391738 | 0,42643675 | 0,892138574 | leucine rich repeat containing 37 member A2 |
| ENSG00000237886 | NALT1 | -0,245728505 | 0,843389809 | 0,71192881 | 0,970812834 | NOTCH1 associated lncRNA in T cell acute lymphoblastic leukemia 1 |
| ENSG00000147471 | PLPBP | -0,245732578 | 0,843387428 | 0,46593603 | 0,907940653 | pyridoxal phosphate binding protein |
| ENSG00000255321 | AC068389,3 | -0,245876389 | 0,843303362 | 0,75458799 | 0,975901808 | novel transcript |
| ENSG00000254469 | AP002495,1 | -0,245910234 | 0,843283579 | 0,36405764 | 0,854207909 | XRCC1 N-terminal domain containing 1-like |
| ENSG00000155093 | PTPRN2 | -0,245945074 | 0,843263214 | 0,25949014 | 0,772164235 | protein tyrosine phosphatase receptor type N2 |
| ENSG00000254934 | LINC00678 | -0,245951543 | 0,843259433 | 0,67802861 | 0,963963232 | long intergenic non-protein coding RNA 678 |
| ENSG00000163082 | SGPP2 | -0,245962971 | 0,843252754 | 0,43650971 | 0,896866043 | sphingosine-1-phosphate phosphatase 2 |
| ENSG00000204682 | MIR1915HG | -0,24603024 | 0,843213435 | 0,36061159 | 0,851271174 | MIR1915 host gene |
| ENSG00000235389 | AL583805,2 | -0,246121703 | 0,84315998 | 0,78656028 | 0,979320406 | novel transcript |

| | | | | | | |
|---|---|---|---|---|---|---|
| ENSG00000275457 | AL117332,1 | -0,246166252 | 0,843133944 | 0,55499641 | 0,937793851 | novel transcript, antisense to XRN2 |
| ENSG00000106028 | SSBP1 | -0,246218687 | 0,843103301 | 0,64412295 | 0,956633898 | single stranded DNA binding protein 1 |
| ENSG00000147601 | TERF1 | -0,246266443 | 0,843075393 | 0,42437655 | 0,890949676 | telomeric repeat binding factor 1 |
| ENSG00000270061 | AC068790,5 | -0,246446847 | 0,842969976 | 0,64844643 | 0,956899963 | novel transcript, sense intronic to CCDC92 |
| ENSG00000165566 | AMER2 | -0,246481474 | 0,842949744 | 0,4371902 | 0,897021798 | APC membrane recruitment protein 2 |
| ENSG00000129810 | SGO1 | -0,246486655 | 0,842946717 | 0,51623434 | 0,925176589 | shugoshin 1 |
| ENSG00000186812 | ZNF397 | -0,246795079 | 0,842766528 | 0,30331708 | 0,812669425 | zinc finger protein 397 |
| ENSG00000241127 | YAE1 | -0,246882923 | 0,842715214 | 0,62466932 | 0,952793934 | YAE1 maturation factor of ABCE1 |
| ENSG00000261056 | AC079416,2 | -0,246888198 | 0,842712133 | 0,4463918 | 0,901491705 | thyroid hormone receptor interactor 13 (TRIP13) pseudogene |
| ENSG00000186918 | ZNF395 | -0,24694464 | 0,842679165 | 0,36410487 | 0,854207909 | zinc finger protein 395 |
| ENSG00000197050 | ZNF420 | -0,247020248 | 0,842635003 | 0,56779622 | 0,94163744 | zinc finger protein 420 |
| ENSG00000144034 | TPRKB | -0,247042399 | 0,842622065 | 0,65259307 | 0,957275891 | TP53RK binding protein |
| ENSG00000231993 | EP300-AS1 | -0,247063633 | 0,842609664 | 0,75819468 | 0,976120207 | EP300 antisense RNA 1 |
| ENSG00000266896 | AL354892,3 | -0,247134593 | 0,84256822 | 0,93345759 | 0,994479239 | novel transcript |
| ENSG00000221676 | RNU6ATAC | -0,247146628 | 0,842561192 | 0,64385031 | 0,956633898 | RNA, U6atac small nuclear (U12-dependent splicing) |
| ENSG00000106868 | SUSD1 | -0,247232082 | 0,842511286 | 0,40484445 | 0,878176155 | sushi domain containing 1 |
| ENSG00000229298 | TUBB8P1 | -0,247238476 | 0,842507552 | 0,62362879 | 0,952793934 | tubulin beta 8 class VIII pseudogene 1 |
| ENSG00000066583 | ISOC1 | -0,247314565 | 0,842463119 | 0,53184235 | 0,931291085 | isochorismatase domain containing 1 |
| ENSG00000260488 | AC007216,1 | -0,247322008 | 0,842458772 | 0,84117574 | 0,985672131 | novel transcript, antisense to RUNDC2A |
| ENSG00000228224 | NACA4P | -0,247331815 | 0,842453046 | 0,794716 | 0,979776981 | NACA family member 4, pseudogene |
| ENSG00000179277 | MEIS3P1 | -0,247382821 | 0,842423262 | 0,44895391 | 0,902062594 | Meis homeobox 3 pseudogene 1 |
| ENSG00000110077 | MS4A6A | -0,247411556 | 0,842406483 | 0,69087521 | 0,967179044 | membrane spanning 4-domains A6A |
| ENSG00000278530 | CHMP1B2P | -0,247412892 | 0,842405703 | 0,60686138 | 0,950927937 | charged multivesicular body protein 1B2, pseudogene |
| ENSG00000136449 | MYCBPAP | -0,247544056 | 0,842329118 | 0,80112253 | 0,980557464 | MYCBP associated protein |
| ENSG00000286923 | AC004522,5 | -0,247661021 | 0,84226083 | 0,8599094 | 0,989074806 | novel transcript |
| ENSG00000141579 | ZNF750 | -0,247684029 | 0,842247398 | 0,65054534 | 0,956899963 | zinc finger protein 750 |
| ENSG00000074201 | CLNS1A | -0,247838651 | 0,842157134 | 0,45184806 | 0,902062594 | chloride nucleotide-sensitive channel 1A |
| ENSG00000189266 | PNRC2 | -0,247895175 | 0,842124139 | 0,6187993 | 0,952793934 | proline rich nuclear receptor coactivator 2 |
| ENSG00000249614 | LINC02503 | -0,247950976 | 0,842091568 | 0,32138571 | 0,824933727 | long intergenic non-protein coding RNA 2503 |
| ENSG00000178498 | DTX3 | -0,247968816 | 0,842081155 | 0,29353137 | 0,803461221 | deltex E3 ubiquitin ligase 3 |
| ENSG00000183734 | ASCL2 | -0,248029107 | 0,842045965 | 0,54687412 | 0,935404974 | achaete-scute family bHLH transcription factor 2 |

| | | | | | | |
|---|---|---|---|---|---|---|
| ENSG00000279548 | AL022323,4 | -0,248153519 | 0,841973354 | 0,79964316 | 0,980161849 | novel transcript |
| ENSG00000266537 | SPDYE22P | -0,248158622 | 0,841970375 | 0,79382082 | 0,979776981 | speedy/RINGO cell cycle regulator family member E22, pseudogene |
| ENSG00000079482 | OPHN1 | -0,248173335 | 0,841961789 | 0,31801394 | 0,822977232 | oligophrenin 1 |
| ENSG00000198763 | MT-ND2 | -0,248212683 | 0,841938826 | 0,42171146 | 0,889720796 | mitochondrially encoded NADH:ubiquinone oxidoreductase core subunit 2 |
| ENSG00000106610 | STAG3L4 | -0,248215926 | 0,841936933 | 0,44585867 | 0,901173082 | stromal antigen 3-like 4 (pseudogene) |
| ENSG00000180764 | PIPSL | -0,248263838 | 0,841908972 | 0,60230276 | 0,950180678 | PIP5K1A and PSMD4 like (pseudogene) |
| ENSG00000182912 | TSPEAR-AS2 | -0,248375854 | 0,841843606 | 0,68869978 | 0,966556275 | TSPEAR antisense RNA 2 |
| ENSG00000270935 | AL021808,1 | -0,248440886 | 0,841805659 | 0,83504506 | 0,984905093 | cell division cycle associated 7 (CDCA7) pseudogene |
| ENSG00000234917 | AC098484,2 | -0,248480994 | 0,841782257 | 0,75637719 | 0,975901808 | novel transcript |
| ENSG00000188817 | SNTN | -0,248515157 | 0,841762324 | 0,82444789 | 0,983374097 | sentan, cilia apical structure protein |
| ENSG00000171368 | TPPP | -0,248607088 | 0,841708687 | 0,44004004 | 0,897906863 | tubulin polymerization promoting protein |
| ENSG00000226746 | SMCR5 | -0,248621406 | 0,841700333 | 0,65781808 | 0,957864535 | Smith-Magenis syndrome chromosome region, candidate 5 |
| ENSG00000285094 | LINC01488 | -0,248660015 | 0,841677809 | 0,83910096 | 0,985672131 | long intergenic non-protein coding RNA 1488 |
| ENSG00000240445 | FOXO3B | -0,248676367 | 0,841668268 | 0,31842971 | 0,823270714 | forkhead box O3B |
| ENSG00000164114 | MAP9 | -0,248742781 | 0,841629524 | 0,54624247 | 0,93535572 | microtubule associated protein 9 |
| ENSG00000138613 | APH1B | -0,248753737 | 0,841623132 | 0,54734299 | 0,935404974 | aph-1 homolog B, gamma-secretase subunit |
| ENSG00000223478 | AL441992,1 | -0,248762254 | 0,841618164 | 0,62470777 | 0,952793934 | novel transcript |
| ENSG00000166436 | TRIM66 | -0,248771327 | 0,841612871 | 0,32500752 | 0,826983176 | tripartite motif containing 66 |
| ENSG00000122417 | ODF2L | -0,24883797 | 0,841573995 | 0,39297202 | 0,871121102 | outer dense fiber of sperm tails 2 like |
| ENSG00000113211 | PCDHB6 | -0,248863279 | 0,841559231 | 0,54457344 | 0,934679287 | protocadherin beta 6 |
| ENSG00000131697 | NPHP4 | -0,248933071 | 0,841518521 | 0,45153293 | 0,902062594 | nephrocystin 4 |
| ENSG00000232093 | DCST1-AS1 | -0,2490424 | 0,841454752 | 0,64026724 | 0,956251063 | DCST1 antisense RNA 1 |
| ENSG00000224699 | LAMTOR5-AS1 | -0,249128546 | 0,841404509 | 0,45794738 | 0,904135472 | LAMTOR5 antisense RNA 1 |
| ENSG00000164406 | LEAP2 | -0,24912949 | 0,841403958 | 0,50396056 | 0,921285401 | liver enriched antimicrobial peptide 2 |
| ENSG00000204044 | SLC12A5-AS1 | -0,24918078 | 0,841374045 | 0,63395743 | 0,954929949 | SLC12A5 and MMP9 antisense RNA 1 |
| ENSG00000285652 | AL591428,1 | -0,249194121 | 0,841366265 | 0,75654837 | 0,975901808 | novel transcript |
| ENSG00000171365 | CLCN5 | -0,249249325 | 0,841334071 | 0,53113425 | 0,931265667 | chloride voltage-gated channel 5 |
| ENSG00000165646 | SLC18A2 | -0,24926893 | 0,841322638 | 0,75465599 | 0,975901808 | solute carrier family 18 member A2 |
| ENSG00000178301 | AQP11 | -0,24929369 | 0,841308199 | 0,51509034 | 0,924720046 | aquaporin 11 |
| ENSG00000250041 | AC069360,1 | -0,249411287 | 0,841239626 | 0,61193538 | 0,952187506 | novel transcript |

| | | | | | | |
|---|---|---|---|---|---|---|
| ENSG00000205559 | CHKB-DT | -0,249429408 | 0,841229059 | 0,36746512 | 0,857089629 | CHKB divergent transcript |
| ENSG00000282207 | AC254944,1 | -0,249439113 | 0,8412234 | 0,8113666 | 0,981213877 | novel transcript, antisense to AC026150,11 |
| ENSG00000270016 | AC026150,3 | -0,249439113 | 0,8412234 | 0,8113666 | 0,981213877 | novel transcript, antisense to AC026150,11 |
| ENSG00000266335 | AC016383,1 | -0,249566518 | 0,841149115 | 0,83575402 | 0,985034063 | novel transcript |
| ENSG00000106133 | NSUN5P2 | -0,24958548 | 0,841138059 | 0,35228554 | 0,844912682 | NSUN5 pseudogene 2 |
| ENSG00000243716 | NPIPB5 | -0,249634507 | 0,841109476 | 0,23168248 | 0,739869375 | nuclear pore complex interacting protein family member B5 |
| ENSG00000028203 | VEZT | -0,249683443 | 0,841080945 | 0,54362019 | 0,934259495 | vezatin, adherens junctions transmembrane protein |
| ENSG00000137142 | IGFBPL1 | -0,249698683 | 0,841072061 | 0,48675184 | 0,914421915 | insulin like growth factor binding protein like 1 |
| ENSG00000186908 | ZDHHC17 | -0,249925391 | 0,840939903 | 0,62307422 | 0,952793934 | zinc finger DHHC-type palmitoyltransferase 17 |
| ENSG00000287180 | AC106818,2 | -0,249951004 | 0,840924974 | 0,69414512 | 0,967359949 | novel transcript, antisense to FAM172A |
| ENSG00000275949 | AC139677,1 | -0,250173602 | 0,840795235 | 0,33309239 | 0,833301711 | ADP-ribosylation factor-like protein 17 |
| ENSG00000253368 | TRNP1 | -0,25028609 | 0,84072968 | 0,40953536 | 0,882177108 | TMF1 regulated nuclear protein 1 |
| ENSG00000153237 | CCDC148 | -0,250302998 | 0,840719827 | 0,55398646 | 0,937524781 | coiled-coil domain containing 148 |
| ENSG00000072415 | MPP5 | -0,250407788 | 0,840658763 | 0,52872613 | 0,930033144 | membrane palmitoylated protein 5 |
| ENSG00000249006 | AL136360,1 | -0,250460972 | 0,840627774 | 0,83258005 | 0,98489473 | pseudogene similar to part of HBS1-like |
| ENSG00000260285 | AL133367,1 | -0,250513259 | 0,840597308 | 0,4205898 | 0,888569577 | novel transcript, antisense to TRMT61A |
| ENSG00000287959 | AC021236,1 | -0,250514881 | 0,840596363 | 0,53614958 | 0,932216256 | novel transcript, antisense to PRKDC |
| ENSG00000204406 | MBD5 | -0,250545734 | 0,840578386 | 0,45604452 | 0,903545728 | methyl-CpG binding domain protein 5 |
| ENSG00000274492 | AL121768,1 | -0,250637847 | 0,840524719 | 0,59241486 | 0,947416795 | novel transcript |
| ENSG00000268531 | AC060814,2 | -0,250706954 | 0,840484457 | 0,81078785 | 0,981213877 | novel transcript |
| ENSG00000120832 | MTERF2 | -0,250716809 | 0,840478716 | 0,52767038 | 0,929546857 | mitochondrial transcription termination factor 2 |
| ENSG00000272426 | BX284668,6 | -0,250757253 | 0,840455155 | 0,75360155 | 0,975901808 | novel transcript |
| ENSG00000140006 | WDR89 | -0,25075726 | 0,840455151 | 0,5679361 | 0,94163744 | WD repeat domain 89 |
| ENSG00000268041 | ERFL | -0,250883014 | 0,840381895 | 0,8615819 | 0,989074806 | ETS repressor factor like |
| ENSG00000247627 | MTND4P12 | -0,250928534 | 0,840355379 | 0,73078196 | 0,974336839 | MT-ND4 pseudogene 12 |
| ENSG00000178623 | GPR35 | -0,250954218 | 0,840340419 | 0,71240053 | 0,970812834 | G protein-coupled receptor 35 |
| ENSG00000153147 | SMARCA5 | -0,25105105 | 0,840284018 | 0,53019469 | 0,931131717 | SWI/SNF related, matrix associated, actin dependent regulator of chromatin, subfamily a, member 5 |
| ENSG00000248727 | LINC01948 | -0,25115282 | 0,840224745 | 0,48639949 | 0,914091794 | long intergenic non-protein coding RNA 1948 |
| ENSG00000196368 | NUDT11 | -0,251174053 | 0,840212379 | 0,53138693 | 0,931291085 | nudix hydrolase 11 |
| ENSG00000237413 | MGC27382 | -0,251208077 | 0,840192564 | 0,61042355 | 0,951841373 | uncharacterized MGC27382 |

| | | | | | | |
|---|---|---|---|---|---|---|
| ENSG00000251022 | THAP9-AS1 | -0,251240886 | 0,840173457 | 0,52144899 | 0,927389576 | THAP9 antisense RNA 1 |
| ENSG00000170113 | NIPA1 | -0,251338032 | 0,840116885 | 0,51998809 | 0,926822772 | NIPA magnesium transporter 1 |
| ENSG00000088543 | C3orf18 | -0,251381661 | 0,840091479 | 0,2974312 | 0,806635068 | chromosome 3 open reading frame 18 |
| ENSG00000228696 | ARL17B | -0,251445239 | 0,840054458 | 0,5291355 | 0,93021211 | ADP ribosylation factor like GTPase 17B |
| ENSG00000174951 | FUT1 | -0,251471986 | 0,840038883 | 0,68161995 | 0,964654539 | fucosyltransferase 1 (H blood group) |
| ENSG00000115008 | IL1A | -0,251559357 | 0,839988012 | 0,67185706 | 0,962161102 | interleukin 1 alpha |
| ENSG00000271335 | AL117336,2 | -0,251596188 | 0,839966568 | 0,51907862 | 0,926800715 | novel transcript, antisense to CCNY |
| ENSG00000228883 | LY6G5C | -0,251785315 | 0,839856462 | 0,73274258 | 0,974353474 | lymphocyte antigen 6 family member G5C |
| ENSG00000196233 | LCOR | -0,251831016 | 0,839829857 | 0,50559013 | 0,921780819 | ligand dependent nuclear receptor corepressor |
| ENSG00000285820 | AL157829,1 | -0,251837011 | 0,839826367 | 0,80746602 | 0,981213877 | Novel transcript, antisense to ASTN2 |
| ENSG00000226801 | OSTCP8 | -0,251888075 | 0,839796642 | 0,75414193 | 0,975901808 | oligosaccharyltransferase complex subunit pseudogene 8 |
| ENSG00000230950 | MTND2P21 | -0,251888629 | 0,83979632 | 0,80387938 | 0,980875302 | MT-ND2 pseudogene 21 |
| ENSG00000223930 | AC117453,1 | -0,251899525 | 0,839789977 | 0,70179507 | 0,968742626 | novel transcript, antisense to KCNMB2 |
| ENSG00000100629 | CEP128 | -0,251977208 | 0,839744759 | 0,32651784 | 0,828450693 | centrosomal protein 128 |
| ENSG00000167005 | NUDT21 | -0,251996975 | 0,839733254 | 0,58266247 | 0,944174844 | nudix hydrolase 21 |
| ENSG00000112200 | ZNF451 | -0,252021309 | 0,83971909 | 0,53759514 | 0,932883277 | zinc finger protein 451 |
| ENSG00000144451 | SPAG16 | -0,252056111 | 0,839698834 | 0,43951948 | 0,89764124 | sperm associated antigen 16 |
| ENSG00000172594 | SMPDL3A | -0,252120925 | 0,839661111 | 0,59942111 | 0,949137642 | sphingomyelin phosphodiesterase acid like 3A |
| ENSG00000204380 | PKP4-AS1 | -0,252220987 | 0,839602875 | 0,48308543 | 0,91363215 | PKP4 antisense RNA 1 |
| ENSG00000117448 | AKR1A1 | -0,252277813 | 0,839569805 | 0,30874936 | 0,816730513 | aldo-keto reductase family 1 member A1 |
| ENSG00000243103 | RN7SL452P | -0,252286131 | 0,839564964 | 0,86894531 | 0,989074806 | RNA, 7SL, cytoplasmic 452, pseudogene |
| ENSG00000287608 | AC022137,3 | -0,252397188 | 0,839500338 | 0,52300918 | 0,92790906 | novel transcript, antisense to ZNF761and ZNF813 |
| ENSG00000203727 | SAMD5 | -0,252414044 | 0,83949053 | 0,43023006 | 0,895058897 | sterile alpha motif domain containing 5 |
| ENSG00000259295 | CSPG4P12 | -0,252605534 | 0,839379111 | 0,42924737 | 0,894029049 | chondroitin sulfate proteoglycan 4 pseudogene 12 |
| ENSG00000254595 | AC084337,1 | -0,252614379 | 0,839373965 | 0,60145788 | 0,949939978 | voltage-dependent anion channel 1(VDAC1) pseudogene |
| ENSG00000096063 | SRPK1 | -0,252656237 | 0,839349612 | 0,51902836 | 0,926800715 | SRSF protein kinase 1 |
| ENSG00000178295 | GEN1 | -0,252676246 | 0,839337971 | 0,52655882 | 0,929261495 | GEN1 Holliday junction 5' flap endonuclease |
| ENSG00000170180 | GYPA | -0,252702419 | 0,839322744 | 0,77991573 | 0,978587255 | glycophorin A (MNS blood group) |
| ENSG00000078403 | MLLT10 | -0,252838009 | 0,839243865 | 0,49203259 | 0,916995341 | MLLT10 histone lysine methyltransferase DOT1L cofactor |
| ENSG00000004777 | ARHGAP33 | -0,252854011 | 0,839234556 | 0,42591824 | 0,892133432 | Rho GTPase activating protein 33 |

| | | | | | | |
|---|---|---|---|---|---|---|
| ENSG00000165392 | WRN | -0,25289891 | 0,839208438 | 0,45115481 | 0,902062594 | WRN RecQ like helicase |
| ENSG00000146521 | LINC01558 | -0,252969145 | 0,839167584 | 0,83701703 | 0,985462684 | long intergenic non-protein coding RNA 1558 |
| ENSG00000236065 | AL020995,1 | -0,2529751 | 0,839164121 | 0,65875867 | 0,958206085 | novel transcript |
| ENSG00000031003 | FAM13B | -0,252985714 | 0,839157947 | 0,5060075 | 0,921780819 | family with sequence similarity 13 member B |
| ENSG00000235100 | AL157400,3 | -0,253081205 | 0,839102405 | 0,63985162 | 0,956018518 | novel transcript |
| ENSG00000259321 | AL136295,2 | -0,253114109 | 0,839083268 | 0,69349591 | 0,967340633 | novel transcript |
| ENSG00000285298 | AL136295,16 | -0,253114109 | 0,839083268 | 0,69349591 | 0,967340633 | novel transcript |
| ENSG00000167524 | RSKR | -0,253203206 | 0,83903145 | 0,37348816 | 0,861319894 | ribosomal protein S6 kinase related |
| ENSG00000141424 | SLC39A6 | -0,25329119 | 0,838980282 | 0,49618516 | 0,918239188 | solute carrier family 39 member 6 |
| ENSG00000078674 | PCM1 | -0,253326907 | 0,838959512 | 0,4363904 | 0,896850871 | pericentriolar material 1 |
| ENSG00000231240 | KLF2P1 | -0,253468437 | 0,838877213 | 0,75929021 | 0,976120207 | Kruppel like factor 2 pseudogene 1 |
| ENSG00000285671 | AL139405,1 | -0,253468542 | 0,838877152 | 0,6122163 | 0,9523974 | novel transcript |
| ENSG00000102531 | FNDC3A | -0,253486919 | 0,838866466 | 0,56556167 | 0,940865625 | fibronectin type III domain containing 3A |
| ENSG00000250959 | GLUD1P3 | -0,253550382 | 0,838829566 | 0,49676583 | 0,918457927 | glutamate dehydrogenase 1 pseudogene 3 |
| ENSG00000249621 | AC113349,1 | -0,253580239 | 0,838812207 | 0,77198493 | 0,978033855 | novel transcript |
| ENSG00000187950 | OVCH1 | -0,25360012 | 0,838800647 | 0,77686727 | 0,978587255 | ovochymase 1 |
| ENSG00000162664 | ZNF326 | -0,253620717 | 0,838788672 | 0,37672009 | 0,862818448 | zinc finger protein 326 |
| ENSG00000135093 | USP30 | -0,253657459 | 0,838767311 | 0,51719051 | 0,925746338 | ubiquitin specific peptidase 30 |
| ENSG00000141258 | SGSM2 | -0,253696336 | 0,838744708 | 0,43902785 | 0,897402199 | small G protein signaling modulator 2 |
| ENSG00000176102 | CSTF3 | -0,253710422 | 0,838736519 | 0,50458251 | 0,921400408 | cleavage stimulation factor subunit 3 |
| ENSG00000261079 | AC009053,2 | -0,253748652 | 0,838714294 | 0,52938649 | 0,930491829 | novel transcript |
| ENSG00000213741 | RPS29 | -0,253765037 | 0,838704768 | 0,45458921 | 0,903383444 | ribosomal protein S29 |
| ENSG00000242882 | RPL5P11 | -0,253773086 | 0,838700089 | 0,81561917 | 0,981590958 | ribosomal protein L5 pseudogene 11 |
| ENSG00000170954 | ZNF415 | -0,253800883 | 0,838683929 | 0,52184742 | 0,927599011 | zinc finger protein 415 |
| ENSG00000148835 | TAF5 | -0,253902063 | 0,838625112 | 0,50183852 | 0,920520138 | TATA-box binding protein associated factor 5 |
| ENSG00000275426 | AC253576,2 | -0,253970253 | 0,838585475 | 0,82888005 | 0,983795764 | novel transcript |
| ENSG00000188933 | USP32P1 | -0,254153297 | 0,838479085 | 0,528135 | 0,929969741 | ubiquitin specific peptidase 32 pseudogene 1 |
| ENSG00000166822 | TMEM170A | -0,254218464 | 0,838441211 | 0,33717993 | 0,836323083 | transmembrane protein 170A |
| ENSG00000261308 | FIGNL2 | -0,254232373 | 0,838433129 | 0,53223645 | 0,931291085 | fidgetin like 2 |
| ENSG00000178217 | SH2D4B | -0,254307004 | 0,838389757 | 0,7084791 | 0,970181083 | SH2 domain containing 4B |
| ENSG00000162521 | RBBP4 | -0,254341324 | 0,838369813 | 0,50599765 | 0,921780819 | RB binding protein 4, chromatin remodeling factor |
| ENSG00000160284 | SPATC1L | -0,254388105 | 0,838342628 | 0,57283369 | 0,942475308 | spermatogenesis and centriole associated 1 like |
| ENSG00000137101 | CD72 | -0,25444499 | 0,838309573 | 0,55023764 | 0,936146588 | CD72 molecule |

| | | | | | | |
|---|---|---|---|---|---|---|
| ENSG00000281332 | LINC00997 | -0,254477867 | 0,83829047 | 0,32945011 | 0,831334931 | long intergenic non-protein coding RNA 997 |
| ENSG00000127081 | ZNF484 | -0,254501902 | 0,838276504 | 0,56735613 | 0,941425553 | zinc finger protein 484 |
| ENSG00000228701 | TNKS2-AS1 | -0,254605664 | 0,838216215 | 0,59212689 | 0,947416795 | TNKS2 antisense RNA 1 (head to head) |
| ENSG00000157315 | TMED6 | -0,254659951 | 0,838184675 | 0,60953506 | 0,951793902 | transmembrane p24 trafficking protein 6 |
| ENSG00000174718 | RESF1 | -0,254739011 | 0,838138743 | 0,58803825 | 0,946139044 | retroelement silencing factor 1 |
| ENSG00000257545 | AC079385,1 | -0,254755251 | 0,838129309 | 0,61688582 | 0,952793934 | novel transcript, antisense to POLR3B, RFX4 and RIC8B, |
| ENSG00000101222 | SPEF1 | -0,254813254 | 0,838095613 | 0,62462312 | 0,952793934 | sperm flagellar 1 |
| ENSG00000253731 | PCDHGA6 | -0,25482501 | 0,838088784 | 0,63410432 | 0,954940644 | protocadherin gamma subfamily A, 6 |
| ENSG00000203668 | CHML | -0,254884125 | 0,838054443 | 0,59731034 | 0,948282297 | CHM like Rab escort protein |
| ENSG00000187730 | GABRD | -0,255042141 | 0,837962658 | 0,7318414 | 0,974353474 | gamma-aminobutyric acid type A receptor subunit delta |
| ENSG00000196935 | SRGAP1 | -0,255070881 | 0,837945964 | 0,40233915 | 0,87718264 | SLIT-ROBO Rho GTPase activating protein 1 |
| ENSG00000275131 | AC241952,1 | -0,255124443 | 0,837914855 | 0,36368571 | 0,854008572 | phosphodiesterase 4D interacting protein (PDE4DIP) pseudogene |
| ENSG00000072401 | UBE2D1 | -0,255252046 | 0,837840747 | 0,62865363 | 0,952793934 | ubiquitin conjugating enzyme E2 D1 |
| ENSG00000219188 | CACYBPP3 | -0,255291496 | 0,837817837 | 0,82455419 | 0,983374097 | calcyclin binding protein pseudogene 3 |
| ENSG00000007952 | NOX1 | -0,255292989 | 0,83781697 | 0,82686482 | 0,983548538 | NADPH oxidase 1 |
| ENSG00000254701 | AC138866,1 | -0,255391758 | 0,837759613 | 0,58797931 | 0,946139044 | glucuronidase, beta (GUSB) pseudogene |
| ENSG00000239039 | SNORD13 | -0,255520676 | 0,837684755 | 0,55477959 | 0,937793851 | small nucleolar RNA, C/D box 13 |
| ENSG00000259479 | SORD2P | -0,255535794 | 0,837675977 | 0,49467942 | 0,91758518 | sorbitol dehydrogenase 2, pseudogene |
| ENSG00000070731 | ST6GALNAC2 | -0,255571967 | 0,837654974 | 0,6966056 | 0,968300554 | ST6 N-acetylgalactosaminide alpha-2,6-sialyltransferase 2 |
| ENSG00000223591 | CENPVL1 | -0,255599828 | 0,837638798 | 0,60903482 | 0,951793902 | centromere protein V like 1 |
| ENSG00000283093 | CENPVL2 | -0,255599828 | 0,837638798 | 0,60903482 | 0,951793902 | centromere protein V like 2 |
| ENSG00000109184 | DCUN1D4 | -0,255605471 | 0,837635522 | 0,60999158 | 0,951801946 | defective in cullin neddylation 1 domain containing 4 |
| ENSG00000154146 | NRGN | -0,25561261 | 0,837631377 | 0,5469727 | 0,935404974 | neurogranin |
| ENSG00000240053 | LY6G5B | -0,255757857 | 0,83754705 | 0,61700075 | 0,952793934 | lymphocyte antigen 6 family member G5B |
| ENSG00000140548 | ZNF710 | -0,25579254 | 0,837526915 | 0,42424521 | 0,890949676 | zinc finger protein 710 |
| ENSG00000049167 | ERCC8 | -0,255803503 | 0,837520551 | 0,44789227 | 0,902062594 | ERCC excision repair 8, CSA ubiquitin ligase complex subunit |
| ENSG00000113638 | TTC33 | -0,255895801 | 0,837466971 | 0,5556394 | 0,937834595 | tetratricopeptide repeat domain 33 |

| | | | | | | |
|---|---|---|---|---|---|---|
| ENSG00000273210 | AP001437,1 | -0,255915359 | 0,837455618 | 0,71418701 | 0,970867497 | novel transcript, antisense to DYRK1A |
| ENSG00000187045 | TMPRSS6 | -0,256094634 | 0,83735156 | 0,63925035 | 0,956018518 | transmembrane serine protease 6 |
| ENSG00000081026 | MAGI3 | -0,256105867 | 0,83734504 | 0,34564068 | 0,841168721 | membrane associated guanylate kinase, WW and PDZ domain containing 3 |
| ENSG00000065029 | ZNF76 | -0,256153105 | 0,837317623 | 0,26258781 | 0,774111884 | zinc finger protein 76 |
| ENSG00000250127 | LINC02108 | -0,256185332 | 0,837298919 | 0,5071378 | 0,921951873 | long intergenic non-protein coding RNA 2108 |
| ENSG00000126733 | DACH2 | -0,256220539 | 0,837278486 | 0,69296006 | 0,967271667 | dachshund family transcription factor 2 |
| ENSG00000286140 | DERPC | -0,256274483 | 0,83724718 | 0,29930927 | 0,809131439 | DERPC proline and glycine rich nuclear protein |
| ENSG00000066651 | TRMT11 | -0,256276736 | 0,837245873 | 0,52630992 | 0,929261495 | tRNA methyltransferase 11 homolog |
| ENSG00000171180 | OR2M4 | -0,25628055 | 0,837243659 | 0,85415797 | 0,987861659 | olfactory receptor family 2 subfamily M member 4 |
| ENSG00000259585 | RBM17P4 | -0,256291373 | 0,837237378 | 0,71174306 | 0,97080996 | RNA binding motif protein 17 pseudogene 4 |
| ENSG00000083123 | BCKDHB | -0,256375962 | 0,83718829 | 0,5027797 | 0,920687909 | branched chain keto acid dehydrogenase E1 subunit beta |
| ENSG00000223979 | SMCR2 | -0,256376799 | 0,837187805 | 0,77355833 | 0,978415098 | Smith-Magenis syndrome chromosome region, candidate 2 |
| ENSG00000249850 | KRT18P31 | -0,256398175 | 0,837175401 | 0,53222413 | 0,931291085 | keratin 18 pseudogene 31 |
| ENSG00000134602 | STK26 | -0,256411076 | 0,837167914 | 0,62662217 | 0,952793934 | serine/threonine kinase 26 |
| ENSG00000249383 | LINC02071 | -0,256431154 | 0,837156263 | 0,73280001 | 0,974353474 | long intergenic non-protein coding RNA 2071 |
| ENSG00000168778 | TCTN2 | -0,256435163 | 0,837153937 | 0,32815266 | 0,830076239 | tectonic family member 2 |
| ENSG00000229065 | AL354893,2 | -0,256473124 | 0,83713191 | 0,66806006 | 0,961258727 | novel transcript |
| ENSG00000231125 | AF129075,1 | -0,256501942 | 0,837115188 | 0,70486436 | 0,969335769 | novel transcript, sense intronic to CCT8 |
| ENSG00000275494 | AC133552,5 | -0,256504496 | 0,837113706 | 0,5629207 | 0,940865625 | novel transcript |
| ENSG00000253452 | AC124290,1 | -0,256543916 | 0,837090833 | 0,39962829 | 0,875433675 | novel transcript |
| ENSG00000283843 | AC113189,6 | -0,256575312 | 0,837072617 | 0,84820068 | 0,987036295 | novel transcript |
| ENSG00000262880 | AC113189,2 | -0,256575312 | 0,837072617 | 0,84820068 | 0,987036295 | novel transcript |
| ENSG00000254187 | AC008708,2 | -0,25659308 | 0,837062307 | 0,68899599 | 0,966640663 | novel transcript |
| ENSG00000230499 | AC108463,2 | -0,256741965 | 0,836975928 | 0,78936762 | 0,979462318 | novel transcript |
| ENSG00000158089 | GALNT14 | -0,256836515 | 0,836921076 | 0,6299154 | 0,952812183 | polypeptide N-acetylgalactosaminyltransferase 14 |
| ENSG00000267781 | SLC25A36P1 | -0,256886632 | 0,836892004 | 0,81866043 | 0,9824772 | SLC25A36 pseudogene 1 |
| ENSG00000284057 | AP001273,2 | -0,256902829 | 0,836882608 | 0,89169505 | 0,990934274 | novel protein, C11orf54-MED17 readthrough |
| ENSG00000087299 | L2HGDH | -0,256949839 | 0,836855339 | 0,21199444 | 0,718583346 | L-2-hydroxyglutarate dehydrogenase |
| ENSG00000287275 | AC011451,4 | -0,257000013 | 0,836826235 | 0,61324478 | 0,952413269 | novel transcript |
| ENSG00000266820 | KPNA2P3 | -0,257093472 | 0,836772026 | 0,72693926 | 0,973344786 | karyopherin subunit alpha 2 pseudogene 3 |

| | | | | | | |
|---|---|---|---|---|---|---|
| ENSG00000100156 | SLC16A8 | -0,25715171 | 0,836738249 | 0,74803152 | 0,975901808 | solute carrier family 16 member 8 |
| ENSG00000285857 | AC016727,3 | -0,25721653 | 0,836700655 | 0,58096202 | 0,943914064 | novel transcript, antisense to XPO1 |
| ENSG00000198721 | ECI2 | -0,257260445 | 0,836675187 | 0,39385017 | 0,871635019 | enoyl-CoA delta isomerase 2 |
| ENSG00000201998 | SNORA23 | -0,257260713 | 0,836675031 | 0,58176278 | 0,944145586 | small nucleolar RNA, H/ACA box 23 |
| ENSG00000112742 | TTK | -0,257304953 | 0,836649375 | 0,62442004 | 0,952793934 | TTK protein kinase |
| ENSG00000280623 | PCAT14 | -0,257322219 | 0,836639362 | 0,44675287 | 0,901650934 | prostate cancer associated transcript 14 |
| ENSG00000257563 | AC126755,2 | -0,257447948 | 0,836566454 | 0,87743213 | 0,990109814 | microRNA 3180-3 |
| ENSG00000270140 | AC005520,3 | -0,257568897 | 0,836496323 | 0,77073546 | 0,977979869 | novel transcript |
| ENSG00000274251 | AP003419,2 | -0,257657107 | 0,836445179 | 0,80083674 | 0,980505307 | novel transcript |
| ENSG00000168878 | SFTPB | -0,257670581 | 0,836437366 | 0,59184378 | 0,947416795 | surfactant protein B |
| ENSG00000288065 | AC007881,4 | -0,257742449 | 0,8363957 | 0,6604147 | 0,959163386 | novel transcript, antisense to MCEE |
| ENSG00000167220 | HDHD2 | -0,257752142 | 0,836390081 | 0,47270059 | 0,909446845 | haloacid dehalogenase like hydrolase domain containing 2 |
| ENSG00000248736 | AC094105,2 | -0,25779181 | 0,836367084 | 0,50429614 | 0,921400408 | novel transcript |
| ENSG00000186280 | KDM4D | -0,257965215 | 0,836266563 | 0,53772099 | 0,932883277 | lysine demethylase 4D |
| ENSG00000160117 | ANKLE1 | -0,257975515 | 0,836260592 | 0,52531345 | 0,928752911 | ankyrin repeat and LEM domain containing 1 |
| ENSG00000110536 | PTPMT1 | -0,258111588 | 0,836181721 | 0,44454807 | 0,900210555 | protein tyrosine phosphatase mitochondrial 1 |
| ENSG00000285206 | PTPMT1 | -0,258111588 | 0,836181721 | 0,44454807 | 0,900210555 | protein tyrosine phosphatase mitochondrial 1 |
| ENSG00000169918 | OTUD7A | -0,25812314 | 0,836175026 | 0,39815193 | 0,874759093 | OTU deubiquitinase 7A |
| ENSG00000213406 | ANXA2P1 | -0,258125505 | 0,836173655 | 0,61755832 | 0,952793934 | annexin A2 pseudogene 1 |
| ENSG00000106258 | CYP3A5 | -0,258202843 | 0,836128832 | 0,37264863 | 0,860513096 | cytochrome P450 family 3 subfamily A member 5 |
| ENSG00000228941 | UBE3AP2 | -0,258203225 | 0,83612861 | 0,8314436 | 0,984440275 | ubiquitin protein ligase E3A pseudogene 2 |
| ENSG00000225151 | GOLGA2P7 | -0,258448784 | 0,835986306 | 0,37480694 | 0,861771195 | GOLGA2 pseudogene 7 |
| ENSG00000172785 | CBWD1 | -0,258472156 | 0,835972763 | 0,36898416 | 0,857932205 | COBW domain containing 1 |
| ENSG00000112419 | PHACTR2 | -0,258479962 | 0,83596824 | 0,52019296 | 0,926865961 | phosphatase and actin regulator 2 |
| ENSG00000151746 | BICD1 | -0,258497504 | 0,835958075 | 0,33640958 | 0,835713011 | BICD cargo adaptor 1 |
| ENSG00000162769 | FLVCR1 | -0,258597986 | 0,835899854 | 0,42086343 | 0,888805335 | FLVCR heme transporter 1 |
| ENSG00000267405 | AC005180,1 | -0,258696891 | 0,83584255 | 0,75716519 | 0,975901808 | novel transcript |
| ENSG00000179921 | GPBAR1 | -0,258714619 | 0,835832279 | 0,74960953 | 0,975901808 | G protein-coupled bile acid receptor 1 |
| ENSG00000197961 | ZNF121 | -0,258742145 | 0,835816332 | 0,4445382 | 0,900210555 | zinc finger protein 121 |
| ENSG00000233384 | CNIH3-AS2 | -0,258772936 | 0,835798494 | 0,4599285 | 0,905341196 | CNIH3 antisense RNA 2 |
| ENSG00000286753 | AC091978,1 | -0,258858175 | 0,835749114 | 0,57507905 | 0,942655666 | novel transcript, antisense to ICE1 |
| ENSG00000196812 | ZSCAN16 | -0,258880704 | 0,835736063 | 0,54639162 | 0,935401821 | zinc finger and SCAN domain containing 16 |

| | | | | | | |
|---|---|---|---|---|---|---|
| ENSG00000112659 | CUL9 | -0,258884389 | 0,835733928 | 0,37183557 | 0,859898346 | cullin 9 |
| ENSG00000264736 | BDP1P | -0,2589329 | 0,835705827 | 0,63800948 | 0,956018518 | B double prime 1 pseudogene |
| ENSG00000242595 | RPL26P26 | -0,258942599 | 0,835700208 | 0,83852917 | 0,985672131 | ribosomal protein L26 pseudogene 26 |
| ENSG00000170092 | SPDYE5 | -0,259023533 | 0,835653328 | 0,55874504 | 0,939635125 | speedy/RINGO cell cycle regulator family member E5 |
| ENSG00000227888 | FAM66A | -0,259065534 | 0,835629 | 0,61552316 | 0,952793934 | family with sequence similarity 66 member A |
| ENSG00000111875 | ASF1A | -0,259131826 | 0,835590603 | 0,62115483 | 0,952793934 | anti-silencing function 1A histone chaperone |
| ENSG00000287319 | AC079822,1 | -0,259247424 | 0,835523653 | 0,78139281 | 0,978597905 | novel transcript, antisense to SERPINI1 |
| ENSG00000115364 | MRPL19 | -0,259275981 | 0,835507115 | 0,51518191 | 0,924808725 | mitochondrial ribosomal protein L19 |
| ENSG00000117335 | CD46 | -0,259284748 | 0,835502038 | 0,53968435 | 0,933350549 | CD46 molecule |
| ENSG00000230565 | ZNF32-AS2 | -0,259285111 | 0,835501827 | 0,64618966 | 0,956899963 | ZNF32 antisense RNA 2 |
| ENSG00000274944 | AL139260,3 | -0,259310612 | 0,83548706 | 0,54780664 | 0,935404974 | novel protein |
| ENSG00000163013 | FBXO41 | -0,25949499 | 0,83538029 | 0,46489666 | 0,907804479 | F-box protein 41 |
| ENSG00000198791 | CNOT7 | -0,259621842 | 0,835306841 | 0,58302643 | 0,944174844 | CCR4-NOT transcription complex subunit 7 |
| ENSG00000122085 | MTERF4 | -0,259679184 | 0,835273641 | 0,33702002 | 0,836223219 | mitochondrial transcription termination factor 4 |
| ENSG00000267904 | AC024075,1 | -0,259680025 | 0,835273154 | 0,50460783 | 0,921400408 | novel transcript, sense intronic C19orf42 |
| ENSG00000161558 | TMEM143 | -0,259771176 | 0,835220382 | 0,38515188 | 0,868895792 | transmembrane protein 143 |
| ENSG00000113649 | TCERG1 | -0,259804899 | 0,835200859 | 0,43116759 | 0,89530663 | transcription elongation regulator 1 |
| ENSG00000116117 | PARD3B | -0,259832703 | 0,835184763 | 0,30955545 | 0,816986489 | par-3 family cell polarity regulator beta |
| ENSG00000250909 | AC138956,1 | -0,259860279 | 0,835168799 | 0,74689936 | 0,975796253 | novel transcript |
| ENSG00000237263 | MAPK6P3 | -0,259917568 | 0,835135635 | 0,57167484 | 0,942436428 | mitogen-activated protein kinase 6 pseudogene 3 |
| ENSG00000281530 | AC004461,2 | -0,259924229 | 0,83513178 | 0,63208949 | 0,953935145 | DiGeorge syndrome critical region gene 12 (non-protein coding) |
| ENSG00000198929 | NOS1AP | -0,259925941 | 0,835130789 | 0,48099863 | 0,912907448 | nitric oxide synthase 1 adaptor protein |
| ENSG00000160352 | ZNF714 | -0,259933946 | 0,835126155 | 0,31024937 | 0,817687618 | zinc finger protein 714 |
| ENSG00000237584 | RAC1P4 | -0,260059358 | 0,835053562 | 0,81501239 | 0,981521413 | Rac family small GTPase 1 pseudogene 4 |
| ENSG00000099958 | DERL3 | -0,260073397 | 0,835045436 | 0,46997931 | 0,908378389 | derlin 3 |
| ENSG00000102886 | GDPD3 | -0,260157912 | 0,834996519 | 0,48507971 | 0,913690171 | glycerophosphodiester phosphodiesterase domain containing 3 |
| ENSG00000154263 | ABCA10 | -0,260193523 | 0,834975908 | 0,67527866 | 0,963390425 | ATP binding cassette subfamily A member 10 |
| ENSG00000213121 | AL590867,1 | -0,260196094 | 0,83497442 | 0,27707993 | 0,788874154 | novel transcript |
| ENSG00000236360 | RRAS2P1 | -0,260520503 | 0,834786686 | 0,85523535 | 0,988016959 | RRAS2 pseudogene 1 |
| ENSG00000151692 | RNF144A | -0,26054668 | 0,83477154 | 0,43261712 | 0,89560871 | ring finger protein 144A |

| | | | | | | |
|---|---|---|---|---|---|---|
| ENSG00000119778 | ATAD2B | -0,260704488 | 0,834680234 | 0,42025497 | 0,888457169 | ATPase family AAA domain containing 2B |
| ENSG00000228492 | RAB11FIP1P1 | -0,260751905 | 0,834652801 | 0,49761264 | 0,918759905 | RAB11 family interacting protein 1 pseudogene 1 |
| ENSG00000261832 | AC138894,1 | -0,260795555 | 0,834627548 | 0,68581132 | 0,965740648 | novel protein |
| ENSG00000173338 | KCNK7 | -0,260876611 | 0,834580657 | 0,85632325 | 0,988229014 | potassium two pore domain channel subfamily K member 7 |
| ENSG00000117016 | RIMS3 | -0,260959723 | 0,834532579 | 0,35096909 | 0,844035671 | regulating synaptic membrane exocytosis 3 |
| ENSG00000250017 | AC117529,1 | -0,260961323 | 0,834531654 | 0,7981064 | 0,980161849 | tigger transposable element derived 1 (TIGD1) pseudogene |
| ENSG00000273363 | AL353801,3 | -0,260966801 | 0,834528485 | 0,73687824 | 0,974353474 | novel transcript |
| ENSG00000229466 | LINC02627 | -0,261011099 | 0,834502861 | 0,72744322 | 0,973441576 | long intergenic non-protein coding RNA 2627 |
| ENSG00000231305 | AC112484,1 | -0,26107223 | 0,834467502 | 0,55334224 | 0,936992468 | novel transcript |
| ENSG00000196834 | POTEI | -0,261117608 | 0,834441255 | 0,29778313 | 0,806960585 | POTE ankyrin domain family member I |
| ENSG00000276791 | AC092117,1 | -0,261141258 | 0,834427576 | 0,4576088 | 0,903831777 | novel transcript |
| ENSG00000184939 | ZFP90 | -0,261158623 | 0,834417533 | 0,43763632 | 0,897021798 | ZFP90 zinc finger protein |
| ENSG00000288605 | AC007461,1 | -0,261163092 | 0,834414948 | 0,84127537 | 0,985672131 | novel transcript |
| ENSG00000211445 | GPX3 | -0,261196804 | 0,83439545 | 0,37589216 | 0,862406526 | glutathione peroxidase 3 |
| ENSG00000223734 | LAPTM4A-DT | -0,261270025 | 0,834353103 | 0,78171682 | 0,978597905 | LAPTM4A divergent transcript |
| ENSG00000286044 | AP001977,1 | -0,261319705 | 0,834324372 | 0,3824015 | 0,866961613 | novel transcript |
| ENSG00000130270 | ATP8B3 | -0,261388285 | 0,834284712 | 0,54079382 | 0,933385624 | ATPase phospholipid transporting 8B3 |
| ENSG00000224939 | LINC00184 | -0,2614026 | 0,834276434 | 0,8469356 | 0,986939308 | long intergenic non-protein coding RNA 184 |
| ENSG00000270421 | AC022022,2 | -0,261530254 | 0,834202619 | 0,77111566 | 0,978033855 | leucine rich repeat containing 40 (LRRC40) pseudogene |
| ENSG00000132141 | CCT6B | -0,261610152 | 0,834156421 | 0,43144778 | 0,89530663 | chaperonin containing TCP1 subunit 6B |
| ENSG00000113761 | ZNF346 | -0,261686964 | 0,83411201 | 0,27888451 | 0,789775666 | zinc finger protein 346 |
| ENSG00000226648 | PLCG1-AS1 | -0,261796186 | 0,834048864 | 0,31053688 | 0,817751312 | PLCG1 antisense RNA 1 |
| ENSG00000287029 | AL162713,2 | -0,261862347 | 0,834010616 | 0,71969817 | 0,971992039 | novel transcript, antisense to ENOX1 |
| ENSG00000117543 | DPH5 | -0,261910106 | 0,833983008 | 0,4575265 | 0,903831777 | diphthamide biosynthesis 5 |
| ENSG00000266803 | AC127540,1 | -0,26201091 | 0,833924737 | 0,80686054 | 0,981213877 | novel transcript |
| ENSG00000288531 | AL669831,7 | -0,262031888 | 0,833912612 | 0,48202822 | 0,912981134 | novel transcript |
| ENSG00000114026 | OGG1 | -0,26215543 | 0,833841204 | 0,26968594 | 0,781791089 | 8-oxoguanine DNA glycosylase |
| ENSG00000280543 | ASAP1-IT2 | -0,26216952 | 0,833833061 | 0,56579923 | 0,940866081 | ASAP1 intronic transcript 2 |
| ENSG00000171551 | ECEL1 | -0,26225369 | 0,833784415 | 0,57836818 | 0,943425812 | endothelin converting enzyme like 1 |
| ENSG00000225975 | LINC01534 | -0,262370107 | 0,833717136 | 0,42317539 | 0,890714198 | long intergenic non-protein coding RNA 1534 |

| | | | | | | |
|---|---|---|---|---|---|---|
| ENSG00000133069 | TMCC2 | -0,262411589 | 0,833693164 | 0,29728211 | 0,806635068 | transmembrane and coiled-coil domain family 2 |
| ENSG00000164056 | SPRY1 | -0,262453144 | 0,833669151 | 0,27859081 | 0,789775666 | sprouty RTK signaling antagonist 1 |
| ENSG00000258484 | SPESP1 | -0,262490119 | 0,833647785 | 0,75674623 | 0,975901808 | sperm equatorial segment protein 1 |
| ENSG00000254553 | AL033529,1 | -0,262696816 | 0,833528356 | 0,88950016 | 0,990717729 | novel transcript |
| ENSG00000259810 | AC002519,1 | -0,262727854 | 0,833510423 | 0,7547048 | 0,975901808 | novel transcript |
| ENSG00000119915 | ELOVL3 | -0,262730468 | 0,833508913 | 0,83754696 | 0,98566906 | ELOVL fatty acid elongase 3 |
| ENSG00000246596 | AC139795,1 | -0,26274607 | 0,833499899 | 0,35944308 | 0,849982153 | SUMO interacting motifs containing 1 (SIMC1) pseudogene |
| ENSG00000227417 | AC114402,1 | -0,262754764 | 0,833494877 | 0,74317699 | 0,975305642 | SNF7 domain containing 2 (SNF7DC20) pseudogene |
| ENSG00000231241 | RPS3AP3 | -0,262775588 | 0,833482846 | 0,82829947 | 0,983646162 | RPS3A pseudogene 3 |
| ENSG00000229739 | PDC-AS1 | -0,262844795 | 0,833442864 | 0,41739443 | 0,886750438 | PDC antisense RNA 1 |
| ENSG00000223745 | CCDC18-AS1 | -0,262854375 | 0,83343733 | 0,36016185 | 0,850589209 | CCDC18 antisense RNA 1 |
| ENSG00000242430 | RN7SL274P | -0,262897052 | 0,833412676 | 0,84455785 | 0,986411265 | RNA, 7SL, cytoplasmic 274, pseudogene |
| ENSG00000225230 | AC008937,1 | -0,262918174 | 0,833400474 | 0,73966116 | 0,974715504 | novel transcript |
| ENSG00000243449 | C4orf48 | -0,262969705 | 0,833370707 | 0,54118095 | 0,933385624 | chromosome 4 open reading frame 48 |
| ENSG00000198888 | MT-ND1 | -0,262982255 | 0,833363458 | 0,41718116 | 0,886651836 | mitochondrially encoded NADH:ubiquinone oxidoreductase core subunit 1 |
| ENSG00000232587 | EEF1A1P3 | -0,263001502 | 0,83335234 | 0,49372595 | 0,917289924 | eukaryotic translation elongation factor 1 alpha 1 pseudogene 3 |
| ENSG00000172324 | OR5A2 | -0,263112767 | 0,833288071 | 0,55508251 | 0,937793851 | olfactory receptor family 5 subfamily A member 2 |
| ENSG00000234324 | RPL9P2 | -0,263144843 | 0,833269545 | 0,78847496 | 0,979462318 | ribosomal protein L9 pseudogene 2 |
| ENSG00000259713 | AC013391,2 | -0,263305431 | 0,833176798 | 0,76369558 | 0,976640468 | novel transcript |
| ENSG00000229206 | AL162408,1 | -0,26343119 | 0,833104173 | 0,72914661 | 0,974009551 | novel transcript |
| ENSG00000249773 | AC092647,5 | -0,263436988 | 0,833100825 | 0,49674939 | 0,918457927 | novel zinc finger protein 713 (ZNF713) and mitochondrial ribosomal protein S17 (MRPS17) protein |
| ENSG00000164070 | HSPA4L | -0,263501701 | 0,833063457 | 0,39003318 | 0,870057277 | heat shock protein family A (Hsp70) member 4 like |
| ENSG00000182685 | BRICD5 | -0,263619701 | 0,832995322 | 0,45730684 | 0,903831777 | BRICHOS domain containing 5 |
| ENSG00000006747 | SCIN | -0,263632092 | 0,832988168 | 0,65501128 | 0,957444211 | scinderin |
| ENSG00000188343 | CIBAR1 | -0,263747514 | 0,832921528 | 0,58557499 | 0,945213113 | CBY1 interacting BAR domain containing 1 |
| ENSG00000141576 | RNF157 | -0,263951213 | 0,832803933 | 0,26380577 | 0,775013078 | ring finger protein 157 |
| ENSG00000225611 | LINC02158 | -0,264147732 | 0,832690499 | 0,67293898 | 0,962432041 | long intergenic non-protein coding RNA 2158 |
| ENSG00000127951 | FGL2 | -0,264148441 | 0,83269009 | 0,64626278 | 0,956899963 | fibrinogen like 2 |

| | | | | | | |
|---|---|---|---|---|---|---|
| ENSG00000287104 | AC097382,3 | -0,264157946 | 0,832684604 | 0,58896069 | 0,946604802 | novel transcript, antisense to TBC1D14 |
| ENSG00000282787 | AL157888,1 | -0,264291775 | 0,832607365 | 0,85395109 | 0,987861659 | novel transcript |
| ENSG00000183773 | AIFM3 | -0,264316318 | 0,832593201 | 0,72533123 | 0,97326334 | apoptosis inducing factor mitochondria associated 3 |
| ENSG00000012048 | BRCA1 | -0,264395313 | 0,832547613 | 0,41038567 | 0,882614012 | BRCA1 DNA repair associated |
| ENSG00000275180 | AC048341,2 | -0,264436862 | 0,832523636 | 0,65662961 | 0,957444211 | novel transcript |
| ENSG00000257467 | PPFIA2-AS1 | -0,264526792 | 0,832471743 | 0,66679188 | 0,9611116 | PPFIA2 antisense RNA 1 |
| ENSG00000118513 | MYB | -0,264547819 | 0,83245961 | 0,39398828 | 0,871635019 | MYB proto-oncogene, transcription factor |
| ENSG00000161996 | WDR90 | -0,264587622 | 0,832436644 | 0,50898585 | 0,922427526 | WD repeat domain 90 |
| ENSG00000273707 | CDKN1C | -0,26462533 | 0,832414886 | 0,43115079 | 0,89530663 | cyclin dependent kinase inhibitor 1C |
| ENSG00000147246 | HTR2C | -0,264701405 | 0,832370993 | 0,67714894 | 0,963580269 | 5-hydroxytryptamine receptor 2C |
| ENSG00000172167 | MTBP | -0,264823862 | 0,832300344 | 0,48937285 | 0,915936198 | MDM2 binding protein |
| ENSG00000164938 | TP53INP1 | -0,264955249 | 0,832224549 | 0,54150624 | 0,933491609 | tumor protein p53 inducible nuclear protein 1 |
| ENSG00000139220 | PPFIA2 | -0,264959996 | 0,832221811 | 0,41603266 | 0,885986046 | PTPRF interacting protein alpha 2 |
| ENSG00000272994 | AC012360,3 | -0,264982161 | 0,832209025 | 0,52388392 | 0,928328101 | novel transcript |
| ENSG00000033867 | SLC4A7 | -0,265083771 | 0,832150414 | 0,43830796 | 0,897177404 | solute carrier family 4 member 7 |
| ENSG00000272563 | AC016745,2 | -0,265090397 | 0,832146592 | 0,63772567 | 0,956018518 | novel transcript |
| ENSG00000186723 | OR10H1 | -0,265092183 | 0,832145562 | 0,31580552 | 0,821233148 | olfactory receptor family 10 subfamily H member 1 |
| ENSG00000167840 | ZNF232 | -0,265136106 | 0,832120228 | 0,42441509 | 0,890949676 | zinc finger protein 232 |
| ENSG00000231793 | DOC2GP | -0,265169093 | 0,832101202 | 0,6194396 | 0,952793934 | double C2 domain gamma, pseudogene |
| ENSG00000159423 | ALDH4A1 | -0,265173264 | 0,832098796 | 0,40489572 | 0,878176155 | aldehyde dehydrogenase 4 family member A1 |
| ENSG00000145700 | ANKRD31 | -0,265254411 | 0,832051994 | 0,65428791 | 0,957444211 | ankyrin repeat domain 31 |
| ENSG00000162971 | TYW5 | -0,265262306 | 0,832047441 | 0,48768198 | 0,914936533 | tRNA-yW synthesizing protein 5 |
| ENSG00000266111 | AC068025,2 | -0,265328764 | 0,832009114 | 0,79298316 | 0,979565591 | novel transcript |
| ENSG00000287867 | AC007402,2 | -0,265479811 | 0,831922008 | 0,53107971 | 0,931265667 | novel transcript |
| ENSG00000171295 | ZNF440 | -0,265484073 | 0,831919551 | 0,26214268 | 0,773902615 | zinc finger protein 440 |
| ENSG00000213347 | MXD3 | -0,265492531 | 0,831914674 | 0,3538673 | 0,845346751 | MAX dimerization protein 3 |
| ENSG00000279442 | AP000542,2 | -0,265504662 | 0,831907678 | 0,34846459 | 0,842757646 | novel transcript |
| ENSG00000256525 | POLG2 | -0,265516973 | 0,831900579 | 0,26975618 | 0,781791089 | DNA polymerase gamma 2, accessory subunit |
| ENSG00000077684 | JADE1 | -0,265523471 | 0,831896833 | 0,30347489 | 0,812669425 | jade family PHD finger 1 |
| ENSG00000084764 | MAPRE3 | -0,265682487 | 0,831805145 | 0,26683401 | 0,779144341 | microtubule associated protein RP/EB family member 3 |
| ENSG00000204130 | RUFY2 | -0,265756381 | 0,831762541 | 0,45411705 | 0,90318049 | RUN and FYVE domain containing 2 |
| ENSG00000135486 | HNRNPA1 | -0,265778426 | 0,831749832 | 0,29045383 | 0,801030501 | heterogeneous nuclear ribonucleoprotein A1 |

| | | | | | | |
|---|---|---|---|---|---|---|
| ENSG00000256667 | KLRA1P | -0,265813984 | 0,831729332 | 0,64670253 | 0,956899963 | killer cell lectin like receptor A1, pseudogene |
| ENSG00000146005 | PSD2 | -0,265861269 | 0,831702072 | 0,57913694 | 0,943828882 | pleckstrin and Sec7 domain containing 2 |
| ENSG00000227495 | KIF1C-AS1 | -0,265938613 | 0,831657485 | 0,77884141 | 0,978587255 | KIF1C antisense RNA 1 |
| ENSG00000100442 | FKBP3 | -0,266074669 | 0,831579057 | 0,53466974 | 0,931663962 | FKBP prolyl isomerase 3 |
| ENSG00000273590 | SMIM11B | -0,266129579 | 0,831547408 | 0,87748214 | 0,990109814 | small integral membrane protein 11B |
| ENSG00000132964 | CDK8 | -0,266189881 | 0,831512652 | 0,43359289 | 0,896108974 | cyclin dependent kinase 8 |
| ENSG00000126787 | DLGAP5 | -0,266207071 | 0,831502744 | 0,52577052 | 0,929056124 | DLG associated protein 5 |
| ENSG00000157456 | CCNB2 | -0,266246859 | 0,831479812 | 0,51263845 | 0,923540891 | cyclin B2 |
| ENSG00000288589 | AC226119,3 | -0,266265761 | 0,831468918 | 0,43325686 | 0,895855976 | novel transcript |
| ENSG00000271795 | AC011337,1 | -0,266334985 | 0,831429023 | 0,5740215 | 0,942475308 | novel transcript, antisense to FAT2 |
| ENSG00000244040 | IL12A-AS1 | -0,266364869 | 0,831411801 | 0,55194468 | 0,936552095 | IL12A antisense RNA 1 |
| ENSG00000089876 | DHX32 | -0,266440227 | 0,831368374 | 0,47196464 | 0,909171312 | DEAH-box helicase 32 (putative) |
| ENSG00000170946 | DNAJC24 | -0,266466754 | 0,831353088 | 0,36412579 | 0,854207909 | DnaJ heat shock protein family (Hsp40) member C24 |
| ENSG00000146722 | AC211486,1 | -0,266487153 | 0,831341333 | 0,71596972 | 0,971360663 | FK506 binding protein 6, 36kDa (FKBP6 ) pseudogene |
| ENSG00000260923 | LINC02193 | -0,266529339 | 0,831317024 | 0,37381213 | 0,861319894 | long intergenic non-protein coding RNA 2193 |
| ENSG00000172731 | LRRC20 | -0,266602599 | 0,831274811 | 0,2733311 | 0,785672647 | leucine rich repeat containing 20 |
| ENSG00000178440 | TIMM23B-AGA | -0,266647071 | 0,831249187 | 0,45766891 | 0,903831777 | TIMM23B-AGAP6 readthrough (NMD candidate) |
| ENSG00000106823 | ECM2 | -0,266649099 | 0,831248018 | 0,58013068 | 0,943828882 | extracellular matrix protein 2 |
| ENSG00000275917 | CHRFAM7A | -0,26665891 | 0,831242366 | 0,46186272 | 0,906277266 | CHRNA7 (exons 5-10) and FAM7A (exons A-E) fusion |
| ENSG00000243244 | STON1 | -0,266661994 | 0,831240588 | 0,41779204 | 0,886857227 | stonin 1 |
| ENSG00000229953 | AL590666,2 | -0,266710849 | 0,83121244 | 0,81125308 | 0,981213877 | novel transcript |
| ENSG00000143353 | LYPLAL1 | -0,266732308 | 0,831200077 | 0,61350155 | 0,952413269 | lysophospholipase like 1 |
| ENSG00000272379 | AL008729,2 | -0,266752698 | 0,831188329 | 0,64211516 | 0,956548736 | novel transcript |
| ENSG00000133256 | PDE6B | -0,266844834 | 0,831135248 | 0,60023033 | 0,949340299 | phosphodiesterase 6B |
| ENSG00000170899 | GSTA4 | -0,266947883 | 0,831075884 | 0,50462746 | 0,921400408 | glutathione S-transferase alpha 4 |
| ENSG00000277895 | AC135279,3 | -0,266959667 | 0,831069095 | 0,58634016 | 0,945352943 | novel transcript, antisense to TBK1 |
| ENSG00000082269 | FAM135A | -0,267129017 | 0,830971546 | 0,44934722 | 0,902062594 | family with sequence similarity 135 member A |
| ENSG00000076944 | STXBP2 | -0,267261448 | 0,830895272 | 0,69944716 | 0,968522643 | syntaxin binding protein 2 |
| ENSG00000149292 | TTC12 | -0,267353005 | 0,830842542 | 0,35245261 | 0,844912682 | tetratricopeptide repeat domain 12 |
| ENSG00000239779 | WBP1 | -0,267521161 | 0,830745708 | 0,25753697 | 0,769908546 | WW domain binding protein 1 |
| ENSG00000156172 | C8orf37 | -0,267770399 | 0,830602201 | 0,38198251 | 0,866514369 | chromosome 8 open reading frame 37 |
| ENSG00000271992 | AL354872,1 | -0,267829591 | 0,830568124 | 0,78535387 | 0,979320406 | novel transcript |
| ENSG00000141026 | MED9 | -0,267853947 | 0,830554102 | 0,24206077 | 0,752344083 | mediator complex subunit 9 |

| | | | | | | |
|---|---|---|---|---|---|---|
| ENSG00000250250 | CTD-2350J17,1 | -0,268086297 | 0,83042035 | 0,81596338 | 0,981708551 | uncharacterized LOC101929472 |
| ENSG00000168509 | HJV | -0,268140149 | 0,830389353 | 0,83159627 | 0,984456953 | hemojuvelin BMP co-receptor |
| ENSG00000254676 | AP000873,4 | -0,268189119 | 0,830361167 | 0,62744878 | 0,952793934 | novel transcript, sense overlapping PCF11 |
| ENSG00000100632 | ERH | -0,268208388 | 0,830350077 | 0,56380965 | 0,940865625 | ERH mRNA splicing and mitosis factor |
| ENSG00000270953 | AC007938,3 | -0,268411403 | 0,830233239 | 0,53490984 | 0,931774961 | novel transcript, antisense to COPG2 |
| ENSG00000204308 | RNF5 | -0,268418735 | 0,830229019 | 0,27752546 | 0,789067728 | ring finger protein 5 |
| ENSG00000168763 | CNNM3 | -0,26844477 | 0,830214037 | 0,38702715 | 0,869441674 | cyclin and CBS domain divalent metal cation transport mediator 3 |
| ENSG00000171488 | LRRC8C | -0,26861872 | 0,830113942 | 0,54764066 | 0,935404974 | leucine rich repeat containing 8 VRAC subunit C |
| ENSG00000275155 | AC027348,1 | -0,268673559 | 0,830082389 | 0,42519335 | 0,891853867 | novel transcript, sense intronic to ZNF423 |
| ENSG00000259511 | UBE2Q2L | -0,268821112 | 0,829997496 | 0,52567433 | 0,929038005 | ubiquitin conjugating enzyme E2 Q2 like |
| ENSG00000225465 | RFPL1S | -0,268843019 | 0,829984892 | 0,38817434 | 0,869820053 | RFPL1 antisense RNA 1 |
| ENSG00000231344 | AL020997,1 | -0,268909354 | 0,82994673 | 0,82962922 | 0,984081562 | novel pseudogene |
| ENSG00000116833 | NR5A2 | -0,268913943 | 0,829944091 | 0,79919193 | 0,980161849 | nuclear receptor subfamily 5 group A member 2 |
| ENSG00000137822 | TUBGCP4 | -0,268927788 | 0,829936126 | 0,39388359 | 0,871635019 | tubulin gamma complex associated protein 4 |
| ENSG00000164118 | CEP44 | -0,269064765 | 0,829857331 | 0,51920666 | 0,926800715 | centrosomal protein 44 |
| ENSG00000236053 | LINC01067 | -0,269080029 | 0,829848551 | 0,68651921 | 0,965769168 | long intergenic non-protein coding RNA 1067 |
| ENSG00000176732 | PFN4 | -0,269218177 | 0,829769091 | 0,59557544 | 0,947923966 | profilin family member 4 |
| ENSG00000158555 | GDPD5 | -0,269308283 | 0,829717268 | 0,34036496 | 0,839010589 | glycerophosphodiester phosphodiesterase domain containing 5 |
| ENSG00000185904 | LINC00839 | -0,269331638 | 0,829703837 | 0,5542489 | 0,937715111 | long intergenic non-protein coding RNA 839 |
| ENSG00000140525 | FANCI | -0,269341024 | 0,829698439 | 0,39098757 | 0,870057277 | FA complementation group I |
| ENSG00000147655 | RSPO2 | -0,2695381 | 0,829585107 | 0,71529835 | 0,971041579 | R-spondin 2 |
| ENSG00000160469 | BRSK1 | -0,269605385 | 0,829546417 | 0,27948485 | 0,790302992 | BR serine/threonine kinase 1 |
| ENSG00000008324 | SS18L2 | -0,269613398 | 0,82954181 | 0,52329695 | 0,927995032 | SS18 like 2 |
| ENSG00000163923 | RPL39L | -0,269645167 | 0,829523543 | 0,33945376 | 0,838364485 | ribosomal protein L39 like |
| ENSG00000172292 | CERS6 | -0,269699087 | 0,829492541 | 0,53950584 | 0,933350549 | ceramide synthase 6 |
| ENSG00000152990 | ADGRA3 | -0,269763203 | 0,829455677 | 0,41420615 | 0,885579089 | adhesion G protein-coupled receptor A3 |
| ENSG00000135185 | TMEM243 | -0,269767522 | 0,829453194 | 0,37486774 | 0,861771195 | transmembrane protein 243 |
| ENSG00000157796 | WDR19 | -0,269792482 | 0,829438844 | 0,48705252 | 0,914593485 | WD repeat domain 19 |
| ENSG00000220370 | AL078595,1 | -0,269820734 | 0,829422602 | 0,50540065 | 0,921762096 | ribosomal protein (RPL7A) pseudogene |
| ENSG00000154898 | CCDC144CP | -0,269872273 | 0,829392972 | 0,46775153 | 0,907940653 | coiled-coil domain containing 144C, pseudogene |
| ENSG00000241217 | RN7SL809P | -0,269949229 | 0,829348732 | 0,75127568 | 0,975901808 | RNA, 7SL, cytoplasmic 809, pseudogene |

| | | | | | | |
|---|---|---|---|---|---|---|
| ENSG00000157181 | ODR4 | -0,269978765 | 0,829331752 | 0,51361027 | 0,92410971 | odr-4 GPCR localization factor homolog |
| ENSG00000162426 | SLC45A1 | -0,269983701 | 0,829328915 | 0,4206196 | 0,888569577 | solute carrier family 45 member 1 |
| ENSG00000219547 | RPL17P25 | -0,270002134 | 0,829318319 | 0,8054774 | 0,981213877 | ribosomal protein L17 pseudogene 25 |
| ENSG00000143578 | CREB3L4 | -0,270026566 | 0,829304275 | 0,47263091 | 0,909368754 | cAMP responsive element binding protein 3 like 4 |
| ENSG00000273711 | AC005520,5 | -0,270233227 | 0,829185488 | 0,55075327 | 0,9363105 | novel transcript, antisense to PTGR2 |
| ENSG00000145241 | CENPC | -0,270298618 | 0,829147906 | 0,46547245 | 0,907899037 | centromere protein C |
| ENSG00000228251 | AC012442,1 | -0,270303597 | 0,829145044 | 0,78983148 | 0,979462318 | novel transcript |
| ENSG00000225830 | ERCC6 | -0,270308002 | 0,829142513 | 0,25670125 | 0,769103136 | ERCC excision repair 6, chromatin remodeling factor |
| ENSG00000133247 | KMT5C | -0,270553358 | 0,829001514 | 0,4851641 | 0,913690171 | lysine methyltransferase 5C |
| ENSG00000162997 | PRORSD1P | -0,270563215 | 0,82899585 | 0,48258726 | 0,913348753 | prolyl-tRNA synthetase associated domain containing 1, pseudogene |
| ENSG00000170522 | ELOVL6 | -0,270598549 | 0,828975547 | 0,5248297 | 0,928616707 | ELOVL fatty acid elongase 6 |
| ENSG00000211452 | DIO1 | -0,270599149 | 0,828975202 | 0,64158732 | 0,956465487 | iodothyronine deiodinase 1 |
| ENSG00000146842 | TMEM209 | -0,270690835 | 0,828922521 | 0,36589247 | 0,855700097 | transmembrane protein 209 |
| ENSG00000228677 | TTC3-AS1 | -0,270748807 | 0,828889213 | 0,73466764 | 0,974353474 | TTC3 antisense RNA 1 |
| ENSG00000233175 | AC008105,1 | -0,27077374 | 0,828874888 | 0,70503917 | 0,969413904 | novel transcript, antisense to FMNL1 |
| ENSG00000196730 | DAPK1 | -0,270835892 | 0,828839181 | 0,2020364 | 0,701751592 | death associated protein kinase 1 |
| ENSG00000278231 | AL133342,1 | -0,270899516 | 0,828802629 | 0,66198616 | 0,959779887 | novel transcript |
| ENSG00000167384 | ZNF180 | -0,270931254 | 0,828784396 | 0,46869978 | 0,907940653 | zinc finger protein 180 |
| ENSG00000234753 | FOXP4-AS1 | -0,270980555 | 0,828756075 | 0,66747094 | 0,9611116 | FOXP4 antisense RNA 1 |
| ENSG00000232850 | PTGES2-AS1 | -0,271119203 | 0,828676432 | 0,36232429 | 0,852348131 | PTGES2 antisense RNA 1 (head to head) |
| ENSG00000287562 | AL109615,4 | -0,271154114 | 0,82865638 | 0,47444679 | 0,910676852 | novel transcript |
| ENSG00000233048 | LINC01722 | -0,271182909 | 0,828639841 | 0,71987569 | 0,972021834 | long intergenic non-protein coding RNA 1722 |
| ENSG00000257759 | AL357153,2 | -0,271211275 | 0,828623548 | 0,80835528 | 0,981213877 | novel transcript |
| ENSG00000166803 | PCLAF | -0,271249486 | 0,828601601 | 0,44292221 | 0,899540076 | PCNA clamp associated factor |
| ENSG00000272916 | AC022400,5 | -0,27126109 | 0,828594937 | 0,30704724 | 0,815892645 | novel transcript |
| ENSG00000099974 | DDTL | -0,271293422 | 0,828576368 | 0,88177075 | 0,990717729 | D-dopachrome tautomerase like |
| ENSG00000213857 | AL080243,1 | -0,271301225 | 0,828571886 | 0,81180212 | 0,981213877 | actin, beta (ACTB) pseudogene |
| ENSG00000174928 | C3orf33 | -0,271406698 | 0,828511313 | 0,40976313 | 0,882209844 | chromosome 3 open reading frame 33 |
| ENSG00000198551 | ZNF627 | -0,271419676 | 0,82850386 | 0,4303969 | 0,895198614 | zinc finger protein 627 |
| ENSG00000239219 | AC008040,1 | -0,271430353 | 0,828497729 | 0,62895644 | 0,952793934 | novel transcript, antisense to SEC62 |
| ENSG00000154447 | SH3RF1 | -0,271457934 | 0,82848189 | 0,31198463 | 0,818279156 | SH3 domain containing ring finger 1 |
| ENSG00000258035 | AC123567,2 | -0,271579927 | 0,828411837 | 0,63047826 | 0,953294597 | novel transcript, antisense to PLXNC1 |

| | | | | | | |
|---|---|---|---|---|---|---|
| ENSG00000274403 | AC090510,2 | -0,271586611 | 0,828407999 | 0,54387084 | 0,934259495 | novel transcript, antisense to CDAN1 |
| ENSG00000287695 | AC008625,1 | -0,271606128 | 0,828396792 | 0,62056948 | 0,952793934 | novel transcript, sense intronic to GALNT10 |
| ENSG00000288362 | CYP11A1 | -0,271677541 | 0,828355788 | 0,69171488 | 0,967204896 | cytochrome P450 family 11 subfamily A member 1 |
| ENSG00000140459 | CYP11A1 | -0,271677541 | 0,828355788 | 0,69171488 | 0,967204896 | cytochrome P450 family 11 subfamily A member 1 |
| ENSG00000259163 | AL096869,2 | -0,271799244 | 0,828285912 | 0,61354522 | 0,952413269 | novel transcript, antisense to TTC7B |
| ENSG00000259165 | DDX18P1 | -0,271810211 | 0,828279616 | 0,5306699 | 0,931265667 | DEAD-box helicase 18 pseudogene 1 |
| ENSG00000249860 | MTRNR2L5 | -0,271842784 | 0,828260915 | 0,77122447 | 0,978033855 | MT-RNR2 like 5 |
| ENSG00000159079 | CFAP298 | -0,271862117 | 0,828249816 | 0,21588002 | 0,722999018 | cilia and flagella associated protein 298 |
| ENSG00000225470 | JPX | -0,271898067 | 0,828229178 | 0,36739867 | 0,856998584 | JPX transcript, XIST activator |
| ENSG00000185933 | CALHM1 | -0,271899224 | 0,828228513 | 0,80336369 | 0,980866933 | calcium homeostasis modulator 1 |
| ENSG00000275339 | Z99129,1 | -0,271938816 | 0,828205785 | 0,83414506 | 0,984905093 | mitogen-activated protein kinase 6 (MAPK6) pseudogene |
| ENSG00000286035 | AC073840,1 | -0,271947924 | 0,828200556 | 0,55131058 | 0,936366211 | novel transcript |
| ENSG00000233967 | AL359715,1 | -0,271974986 | 0,828185021 | 0,45446559 | 0,903383444 | novel transcript |
| ENSG00000259172 | AC023024,1 | -0,271995923 | 0,828173002 | 0,59055528 | 0,947416795 | novel transcript, antisense to SNRPA1 and PCSK6 |
| ENSG00000056050 | HPF1 | -0,272007769 | 0,828166201 | 0,51681949 | 0,925606197 | histone PARylation factor 1 |
| ENSG00000122778 | KIAA1549 | -0,272030229 | 0,828153309 | 0,32946008 | 0,831334931 | KIAA1549 |
| ENSG00000288049 | AC010889,2 | -0,272067816 | 0,828131733 | 0,65070372 | 0,956899963 | novel transcript, antisense to KDM5D |
| ENSG00000213579 | AL096829,1 | -0,272094317 | 0,828116521 | 0,79521922 | 0,97981137 | ribosomal protein L29 (RPL29) pseudogene |
| ENSG00000256229 | ZNF486 | -0,272137087 | 0,828091971 | 0,26970493 | 0,781791089 | zinc finger protein 486 |
| ENSG00000127220 | ABHD8 | -0,272292243 | 0,828002918 | 0,40132265 | 0,876659568 | abhydrolase domain containing 8 |
| ENSG00000101098 | RIMS4 | -0,272457633 | 0,827908002 | 0,53824371 | 0,932900071 | regulating synaptic membrane exocytosis 4 |
| ENSG00000120784 | ZFP30 | -0,27249468 | 0,827886742 | 0,41483527 | 0,885579089 | ZFP30 zinc finger protein |
| ENSG00000163697 | APBB2 | -0,272543071 | 0,827858973 | 0,40262945 | 0,877321369 | amyloid beta precursor protein binding family B member 2 |
| ENSG00000179454 | KLHL28 | -0,272604184 | 0,827823906 | 0,40851044 | 0,88153752 | kelch like family member 28 |
| ENSG00000179085 | DPM3 | -0,272625205 | 0,827811844 | 0,31239155 | 0,818574551 | dolichyl-phosphate mannosyltransferase subunit 3, regulatory |
| ENSG00000124588 | NQO2 | -0,272796619 | 0,827713493 | 0,37422503 | 0,861369706 | N-ribosyldihydronicotinamide:quinone reductase 2 |
| ENSG00000265678 | AC129510,1 | -0,272798915 | 0,827712176 | 0,5880349 | 0,946139044 | novel transcript, antisense CCDC57 |
| ENSG00000170917 | NUDT6 | -0,272810834 | 0,827705338 | 0,51637287 | 0,925256849 | nudix hydrolase 6 |
| ENSG00000177853 | ZNF518A | -0,272856579 | 0,827679093 | 0,55534503 | 0,937793851 | zinc finger protein 518A |
| ENSG00000228198 | OR2M3 | -0,273017248 | 0,827586922 | 0,4804399 | 0,912847708 | olfactory receptor family 2 subfamily M member 3 |

| | | | | | | |
|---|---|---|---|---|---|---|
| ENSG00000164647 | STEAP1 | -0,273095402 | 0,827542091 | 0,53937431 | 0,933350549 | STEAP family member 1 |
| ENSG00000011523 | CEP68 | -0,273202029 | 0,827480931 | 0,22824739 | 0,736631691 | centrosomal protein 68 |
| ENSG00000287121 | AC007220,2 | -0,273213178 | 0,827474536 | 0,83252239 | 0,98489473 | novel transcript, antisense to CLEC16A |
| ENSG00000160131 | VMA21 | -0,273313021 | 0,827417272 | 0,56932045 | 0,94221458 | vacuolar ATPase assembly factor VMA21 |
| ENSG00000271882 | AP001330,4 | -0,27332997 | 0,827407552 | 0,74554938 | 0,975707496 | novel transcript |
| ENSG00000188037 | CLCN1 | -0,273347518 | 0,827397487 | 0,78750964 | 0,979462318 | chloride voltage-gated channel 1 |
| ENSG00000154743 | TSEN2 | -0,273360284 | 0,827390166 | 0,36658671 | 0,856103393 | tRNA splicing endonuclease subunit 2 |
| ENSG00000221843 | C2orf16 | -0,273402887 | 0,827365734 | 0,34758769 | 0,842271095 | chromosome 2 open reading frame 16 |
| ENSG00000272654 | AL358472,2 | -0,273646281 | 0,827226162 | 0,61330422 | 0,952413269 | novel transcript |
| ENSG00000114656 | CFAP92 | -0,27364902 | 0,827224592 | 0,26754608 | 0,779912412 | cilia and flagella associated protein 92 (putative) |
| ENSG00000176040 | TMPRSS7 | -0,273680201 | 0,827206713 | 0,81117727 | 0,981213877 | transmembrane serine protease 7 |
| ENSG00000270480 | AC073413,1 | -0,273710181 | 0,827189524 | 0,60714874 | 0,950993594 | PQ loop repeat containing 1 (PQLC1) pseudogene |
| ENSG00000273747 | AC022558,3 | -0,273716895 | 0,827185674 | 0,57974375 | 0,943828882 | novel transcript, antisense to BTBD1 |
| ENSG00000197077 | KIAA1671 | -0,273853379 | 0,827107423 | 0,36977459 | 0,858104601 | KIAA1671 |
| ENSG00000264365 | AC023983,1 | -0,273951159 | 0,827051367 | 0,78005335 | 0,978587255 | novel transcript, antisense to OSBPL1A |
| ENSG00000143469 | SYT14 | -0,273983171 | 0,827033016 | 0,5697388 | 0,94221458 | synaptotagmin 14 |
| ENSG00000167984 | NLRC3 | -0,274014443 | 0,827015089 | 0,50923988 | 0,922427526 | NLR family CARD domain containing 3 |
| ENSG00000227333 | EHMT2 | -0,274015303 | 0,827014596 | 0,35894404 | 0,849814998 | euchromatic histone lysine methyltransferase 2 |
| ENSG00000279765 | AC013394,1 | -0,274015326 | 0,827014583 | 0,34100384 | 0,839658516 | novel protein |
| ENSG00000096746 | HNRNPH3 | -0,274020615 | 0,827011551 | 0,36286952 | 0,853138593 | heterogeneous nuclear ribonucleoprotein H3 |
| ENSG00000084693 | AGBL5 | -0,274164709 | 0,826928954 | 0,32562572 | 0,827465638 | ATP/GTP binding protein like 5 |
| ENSG00000198885 | ITPRIPL1 | -0,274214244 | 0,826900562 | 0,46467718 | 0,907638841 | ITPRIP like 1 |
| ENSG00000149488 | TMC2 | -0,274400744 | 0,826793674 | 0,70226017 | 0,968742626 | transmembrane channel like 2 |
| ENSG00000178467 | P4HTM | -0,274448337 | 0,8267664 | 0,30130969 | 0,810823388 | prolyl 4-hydroxylase, transmembrane |
| ENSG00000238058 | AL355574,1 | -0,274482847 | 0,826746623 | 0,48432441 | 0,913690171 | novel transcript |
| ENSG00000287609 | AL589787,2 | -0,274550928 | 0,82670761 | 0,51016304 | 0,922427526 | novel transcript |
| ENSG00000257335 | MGAM | -0,274688115 | 0,826629001 | 0,70023609 | 0,968522643 | maltase-glucoamylase |
| ENSG00000187109 | NAP1L1 | -0,274699896 | 0,826622251 | 0,43050008 | 0,895238505 | nucleosome assembly protein 1 like 1 |
| ENSG00000245910 | SNHG6 | -0,2747056 | 0,826618983 | 0,5350314 | 0,931805607 | small nucleolar RNA host gene 6 |
| ENSG00000188732 | FAM221A | -0,274706635 | 0,82661839 | 0,26108069 | 0,773103493 | family with sequence similarity 221 member A |
| ENSG00000144331 | ZNF385B | -0,274713394 | 0,826614517 | 0,71474728 | 0,971041579 | zinc finger protein 385B |
| ENSG00000141028 | CDRT15P1 | -0,274805526 | 0,82656173 | 0,82299076 | 0,983204348 | CDRT15 pseudogene 1 |
| ENSG00000225738 | NRG3-AS1 | -0,274821434 | 0,826552617 | 0,67757534 | 0,9635902 | NRG3 antisense RNA 1 |

| | | | | | | |
|---|---|---|---|---|---|---|
| ENSG00000272009 | AL121944,1 | -0,274863157 | 0,826528713 | 0,8320215 | 0,984653014 | novel transcript, antisense to ZNF165 |
| ENSG00000138443 | ABI2 | -0,274929841 | 0,82649051 | 0,43813046 | 0,897177404 | abl interactor 2 |
| ENSG00000163818 | LZTFL1 | -0,274969078 | 0,826468032 | 0,5769536 | 0,943288205 | leucine zipper transcription factor like 1 |
| ENSG00000251002 | AC244502,1 | -0,275002674 | 0,826448786 | 0,28074396 | 0,791510309 | novel transcript, antisense to IG_gene_cluster |
| ENSG00000241743 | XACT | -0,275042628 | 0,826425899 | 0,24297744 | 0,753092752 | X active specific transcript |
| ENSG00000279170 | TSTD3 | -0,275072933 | 0,82640854 | 0,35096103 | 0,844035671 | thiosulfate sulfurtransferase like domain containing 3 |
| ENSG00000186193 | SAPCD2 | -0,275125864 | 0,82637822 | 0,50491271 | 0,921400408 | suppressor APC domain containing 2 |
| ENSG00000171130 | ATP6V0E2 | -0,275152415 | 0,826363011 | 0,32074524 | 0,824366234 | ATPase H+ transporting V0 subunit e2 |
| ENSG00000134769 | DTNA | -0,275181547 | 0,826346325 | 0,52613209 | 0,929261495 | dystrobrevin alpha |
| ENSG00000005801 | ZNF195 | -0,275301733 | 0,826277488 | 0,38804263 | 0,869820053 | zinc finger protein 195 |
| ENSG00000244055 | AC007566,1 | -0,275335225 | 0,826258306 | 0,52533478 | 0,928752911 | novel transcript, antisense to ERVW-1 and PEX1 |
| ENSG00000188811 | NHLRC3 | -0,275365171 | 0,826241156 | 0,49866108 | 0,919683359 | NHL repeat containing 3 |
| ENSG00000285900 | AC079177,1 | -0,275419276 | 0,82621017 | 0,68485358 | 0,965415049 | novel transcript, antisense to DMD |
| ENSG00000283104 | SARS2 | -0,275453658 | 0,82619048 | 0,61810642 | 0,952793934 | seryl-tRNA synthetase 2, mitochondrial |
| ENSG00000240771 | ARHGEF25 | -0,275487301 | 0,826171214 | 0,33034032 | 0,831982503 | Rho guanine nucleotide exchange factor 25 |
| ENSG00000232160 | RAP2C-AS1 | -0,275497462 | 0,826165395 | 0,4312754 | 0,89530663 | RAP2C antisense RNA 1 |
| ENSG00000165626 | BEND7 | -0,275511656 | 0,826157267 | 0,32122119 | 0,824723604 | BEN domain containing 7 |
| ENSG00000153790 | C7orf31 | -0,27557227 | 0,826122557 | 0,46924276 | 0,907940653 | chromosome 7 open reading frame 31 |
| ENSG00000274679 | SPATC1L | -0,275577439 | 0,826119598 | 0,5369007 | 0,932883277 | spermatogenesis and centriole associated 1 like |
| ENSG00000287335 | AL139351,3 | -0,275583491 | 0,826116132 | 0,55955632 | 0,939727587 | novel transcript |
| ENSG00000117009 | KMO | -0,275637692 | 0,826085096 | 0,71836706 | 0,971789526 | kynurenine 3-monooxygenase |
| ENSG00000231340 | ACTG1P10 | -0,27565358 | 0,826075999 | 0,50638762 | 0,921824694 | actin gamma 1 pseudogene 10 |
| ENSG00000288598 | AL354833,1 | -0,275698801 | 0,826050106 | 0,49069738 | 0,916995341 | uncharacterized LOC105370174 |
| ENSG00000124795 | DEK | -0,2757226 | 0,826036479 | 0,45574535 | 0,903545728 | DEK proto-oncogene |
| ENSG00000238741 | SCARNA7 | -0,275872581 | 0,82595061 | 0,46396926 | 0,907333149 | small Cajal body-specific RNA 7 |
| ENSG00000258732 | AC025884,1 | -0,275887298 | 0,825942184 | 0,47432075 | 0,910676852 | arginine-glutamic acid dipeptide (RE) repeats (RERE) pseudogene |
| ENSG00000283828 | AL137002,2 | -0,27600202 | 0,825876509 | 0,68628822 | 0,965752124 | WAS/WASL-interacting protein family member 2-like |
| ENSG00000155719 | OTOA | -0,276054986 | 0,825846189 | 0,71646489 | 0,971438758 | otoancorin |
| ENSG00000260442 | ATP2A1-AS1 | -0,276077886 | 0,82583308 | 0,35437074 | 0,845793337 | ATP2A1 antisense RNA 1 |

| | | | | | | |
|---|---|---|---|---|---|---|
| ENSG00000183454 | GRIN2A | -0,276136986 | 0,82579925 | 0,57061182 | 0,9423888 | glutamate ionotropic receptor NMDA type subunit 2A |
| ENSG00000270022 | Z93241,1 | -0,276189015 | 0,825769469 | 0,32436726 | 0,82642149 | RNA, U12 small nuclear |
| ENSG00000276256 | AC011043,1 | -0,276300663 | 0,825705567 | 0,46981002 | 0,908378389 | uncharacterized LOC389831 |
| ENSG00000170085 | SIMC1 | -0,276387955 | 0,825655608 | 0,41752326 | 0,886843356 | SUMO interacting motifs containing 1 |
| ENSG00000182584 | ACTL10 | -0,276448705 | 0,825620842 | 0,70563148 | 0,969485452 | actin like 10 |
| ENSG00000135913 | USP37 | -0,276581842 | 0,825544654 | 0,49844571 | 0,91953777 | ubiquitin specific peptidase 37 |
| ENSG00000269609 | RPARP-AS1 | -0,27660354 | 0,825532238 | 0,34835968 | 0,842757646 | RPARP antisense RNA 1 |
| ENSG00000266027 | MSH5-SAPCD1 | -0,276628163 | 0,825518148 | 0,80221681 | 0,980710009 | MSH5-SAPCD1 readthrough (NMD candidate) |
| ENSG00000277455 | Metazoa_SRP | -0,276636638 | 0,825513299 | 0,83495279 | 0,984905093 | Metazoan signal recognition particle RNA |
| ENSG00000165118 | C9orf64 | -0,276693224 | 0,825480921 | 0,76722035 | 0,977201891 | chromosome 9 open reading frame 64 |
| ENSG00000257815 | PRANCR | -0,276696367 | 0,825479123 | 0,34877694 | 0,842864225 | progenitor renewal associated non-coding RNA |
| ENSG00000116885 | OSCP1 | -0,276753236 | 0,825446584 | 0,39762682 | 0,874263835 | organic solute carrier partner 1 |
| ENSG00000275709 | AC090527,3 | -0,276891643 | 0,825367397 | 0,77849038 | 0,978587255 | novel transcript |
| ENSG00000143977 | SNRPG | -0,276952253 | 0,825332723 | 0,57807077 | 0,943425812 | small nuclear ribonucleoprotein polypeptide G |
| ENSG00000282535 | AC092192,1 | -0,276958831 | 0,82532896 | 0,66595437 | 0,961020116 | novel transcript |
| ENSG00000196689 | TRPV1 | -0,277008364 | 0,825300624 | 0,3841923 | 0,868125769 | transient receptor potential cation channel subfamily V member 1 |
| ENSG00000180834 | MAP6D1 | -0,277128321 | 0,825232005 | 0,55225269 | 0,936619966 | MAP6 domain containing 1 |
| ENSG00000261240 | AC009065,6 | -0,277184296 | 0,825199987 | 0,69734586 | 0,968522643 | novel transcript, antisense to PKD1 |
| ENSG00000186019 | AC021092,1 | -0,277226495 | 0,82517585 | 0,26055002 | 0,772692192 | novel transcript, antisense to ZNF225 and ZNF224 |
| ENSG00000261872 | AC002558,2 | -0,277240944 | 0,825167586 | 0,60153312 | 0,949939978 | novel transcript, antisense to CDC27 |
| ENSG00000196604 | POTEF | -0,277504101 | 0,825017084 | 0,2280177 | 0,736341996 | POTE ankyrin domain family member F |
| ENSG00000179152 | TCAIM | -0,277604883 | 0,824959453 | 0,52312696 | 0,92790906 | T cell activation inhibitor, mitochondrial |
| ENSG00000152578 | GRIA4 | -0,277608426 | 0,824957427 | 0,64914716 | 0,956899963 | glutamate ionotropic receptor AMPA type subunit 4 |
| ENSG00000281677 | AC005393,2 | -0,277782264 | 0,824858029 | 0,78210778 | 0,978597905 | novel transcript |
| ENSG00000276445 | AC005393,1 | -0,277782264 | 0,824858029 | 0,78210778 | 0,978597905 | novel transcript |
| ENSG00000263717 | AC079336,1 | -0,277834127 | 0,824828377 | 0,81174091 | 0,981213877 | novel transcript, sense intronic MYO1D |
| ENSG00000286176 | AC116317,1 | -0,277906505 | 0,824786998 | 0,75815184 | 0,976120207 | novel transcript |
| ENSG00000229956 | ZRANB2-AS2 | -0,277928634 | 0,824774347 | 0,38997753 | 0,870057277 | ZRANB2 antisense RNA 2 (head to head) |
| ENSG00000276900 | AC023157,2 | -0,277949125 | 0,824762633 | 0,49884241 | 0,919683359 | novel transcript, antisense to AMN1 |
| ENSG00000058453 | CROCC | -0,277953676 | 0,82476003 | 0,65679484 | 0,957476526 | ciliary rootlet coiled-coil, rootletin |
| ENSG00000172748 | ZNF596 | -0,27797512 | 0,824747772 | 0,35036022 | 0,843857545 | zinc finger protein 596 |

| | | | | | | |
|---|---|---|---|---|---|---|
| ENSG00000102890 | ELMO3 | -0,277992266 | 0,82473797 | 0,5420389 | 0,933491609 | engulfment and cell motility 3 |
| ENSG00000254433 | AP001001,1 | -0,278005492 | 0,824730409 | 0,71085604 | 0,970482408 | novel transcript |
| ENSG00000006007 | GDE1 | -0,278144276 | 0,824651076 | 0,44892342 | 0,902062594 | glycerophosphodiester phosphodiesterase 1 |
| ENSG00000245975 | AC090515,2 | -0,278190703 | 0,824624538 | 0,48444609 | 0,913690171 | novel transcript |
| ENSG00000270638 | AL023806,1 | -0,278257498 | 0,82458636 | 0,5181355 | 0,926344623 | novel transcript |
| ENSG00000179743 | FLJ37453 | -0,278268862 | 0,824579865 | 0,28186437 | 0,792821207 | uncharacterized LOC729614 |
| ENSG00000125814 | NAPB | -0,278339024 | 0,824539764 | 0,30275456 | 0,812344697 | NSF attachment protein beta |
| ENSG00000064692 | SNCAIP | -0,278426938 | 0,82448952 | 0,43932882 | 0,897434993 | synuclein alpha interacting protein |
| ENSG00000157999 | ANKRD61 | -0,278526862 | 0,824432417 | 0,45017914 | 0,902062594 | ankyrin repeat domain 61 |
| ENSG00000164199 | ADGRV1 | -0,278538911 | 0,824425532 | 0,49661547 | 0,918455764 | adhesion G protein-coupled receptor V1 |
| ENSG00000065809 | FAM107B | -0,278570763 | 0,82440733 | 0,54227126 | 0,933491609 | family with sequence similarity 107 member B |
| ENSG00000272033 | AL136984,1 | -0,278753739 | 0,824302777 | 0,56409292 | 0,940865625 | novel transcript, sense intronic to POU2F1 |
| ENSG00000118965 | WDR35 | -0,278800171 | 0,824276248 | 0,42867162 | 0,893524604 | WD repeat domain 35 |
| ENSG00000261369 | AC007533,1 | -0,27882258 | 0,824263445 | 0,75664273 | 0,975901808 | novel transcript, antisense to ITFG1 |
| ENSG00000164008 | C1orf50 | -0,278834524 | 0,824256621 | 0,21388367 | 0,721188427 | chromosome 1 open reading frame 50 |
| ENSG00000070961 | ATP2B1 | -0,278877986 | 0,82423179 | 0,40437896 | 0,878176155 | ATPase plasma membrane Ca2+ transporting 1 |
| ENSG00000282457 | CSPG4P10 | -0,2788852 | 0,824227669 | 0,60523876 | 0,950927937 | chondroitin sulfate proteoglycan 4 pseudogene 10 |
| ENSG00000169750 | RAC3 | -0,278986915 | 0,82416956 | 0,36918009 | 0,857942508 | Rac family small GTPase 3 |
| ENSG00000286045 | AC005034,6 | -0,279062685 | 0,824126276 | 0,45419655 | 0,90318049 | novel transcript, antisense to GCFC2 |
| ENSG00000105204 | DYRK1B | -0,27921087 | 0,824041631 | 0,31368836 | 0,819846908 | dual specificity tyrosine phosphorylation regulated kinase 1B |
| ENSG00000281320 | DYRK1B | -0,27921087 | 0,824041631 | 0,31368836 | 0,819846908 | dual specificity tyrosine phosphorylation regulated kinase 1B |
| ENSG00000276757 | RN7SL192P | -0,2792109 | 0,824041614 | 0,71095487 | 0,970482408 | RNA, 7SL, cytoplasmic 192, pseudogene |
| ENSG00000224287 | MSL3P1 | -0,279212721 | 0,824040574 | 0,46658236 | 0,907940653 | MSL complex subunit 3 pseudogene 1 |
| ENSG00000269235 | ZNF350-AS1 | -0,279225882 | 0,824033056 | 0,4458593 | 0,901173082 | ZNF350 antisense RNA 1 |
| ENSG00000086730 | LAT2 | -0,279239409 | 0,82402533 | 0,51448886 | 0,924434989 | linker for activation of T cells family member 2 |
| ENSG00000226367 | ST7-AS2 | -0,27925451 | 0,824016705 | 0,4934855 | 0,917289924 | ST7 antisense RNA 2 |
| ENSG00000114933 | INO80D | -0,279354601 | 0,823959539 | 0,43490971 | 0,896272827 | INO80 complex subunit D |
| ENSG00000283510 | INO80D | -0,279354601 | 0,823959539 | 0,43490971 | 0,896272827 | INO80 complex subunit D |
| ENSG00000177917 | ARL6IP6 | -0,279381555 | 0,823944144 | 0,43217904 | 0,89558934 | ADP ribosylation factor like GTPase 6 interacting protein 6 |
| ENSG00000197847 | SLC22A20P | -0,279443143 | 0,823908971 | 0,67069378 | 0,962026636 | solute carrier family 22 member 20, pseudogene |

| | | | | | | |
|---|---|---|---|---|---|---|
| ENSG00000250838 | AC091133,3 | -0,279467249 | 0,823895205 | 0,62771568 | 0,952793934 | novel transcript |
| ENSG00000123892 | RAB38 | -0,279482182 | 0,823886677 | 0,61677321 | 0,952793934 | RAB38, member RAS oncogene family |
| ENSG00000168026 | TTC21A | -0,279572548 | 0,823835073 | 0,35393148 | 0,845346751 | tetratricopeptide repeat domain 21A |
| ENSG00000089050 | RBBP9 | -0,279581613 | 0,823829897 | 0,32553819 | 0,827465638 | RB binding protein 9, serine hydrolase |
| ENSG00000103460 | TOX3 | -0,27959013 | 0,823825033 | 0,37385207 | 0,861319894 | TOX high mobility group box family member 3 |
| ENSG00000244733 | AL132656,2 | -0,279619742 | 0,823808124 | 0,64626921 | 0,956899963 | novel transcript |
| ENSG00000232063 | AL691447,2 | -0,279649589 | 0,823791081 | 0,25710489 | 0,769358275 | novel transcript |
| ENSG00000153395 | LPCAT1 | -0,279654531 | 0,823788259 | 0,58344133 | 0,944174844 | lysophosphatidylcholine acyltransferase 1 |
| ENSG00000259867 | AC105411,1 | -0,279764747 | 0,823725327 | 0,52066694 | 0,927046149 | novel transcript, antisense DYNLRB2 |
| ENSG00000232884 | AF127936,1 | -0,279791754 | 0,823709907 | 0,81608718 | 0,981708551 | novel transcript |
| ENSG00000271980 | AC012640,4 | -0,2798578 | 0,823672199 | 0,70385513 | 0,968976619 | novel transcript, antisense to CCT5 |
| ENSG00000286634 | AC146507,3 | -0,279884572 | 0,823656914 | 0,65539096 | 0,957444211 | novel transcript, sense intronic to NSD1 |
| ENSG00000235023 | AP001626,1 | -0,279920605 | 0,823636343 | 0,70843335 | 0,970181083 | novel transcript |
| ENSG00000173200 | PARP15 | -0,279938223 | 0,823626285 | 0,25451895 | 0,765935007 | poly(ADP-ribose) polymerase family member 15 |
| ENSG00000260558 | AC018557,2 | -0,279941166 | 0,823624605 | 0,57977189 | 0,943828882 | novel transcript |
| ENSG00000139163 | ETNK1 | -0,280063348 | 0,823554854 | 0,57247728 | 0,942475308 | ethanolamine kinase 1 |
| ENSG00000072210 | ALDH3A2 | -0,280240837 | 0,823453542 | 0,32250668 | 0,824955371 | aldehyde dehydrogenase 3 family member A2 |
| ENSG00000083067 | TRPM3 | -0,280329386 | 0,823403002 | 0,32026155 | 0,824366234 | transient receptor potential cation channel subfamily M member 3 |
| ENSG00000241449 | AC092666,1 | -0,280346212 | 0,823393399 | 0,58713867 | 0,945859922 | novel transcript |
| ENSG00000223482 | NUTM2A-AS1 | -0,28036032 | 0,823385347 | 0,30167904 | 0,811326505 | NUTM2A antisense RNA 1 |
| ENSG00000271938 | AC103724,4 | -0,280361028 | 0,823384943 | 0,80448269 | 0,980875302 | novel transcript |
| ENSG00000235036 | AL035456,1 | -0,280396494 | 0,823364702 | 0,68514962 | 0,965415049 | pseudogene similar to DNAJ protein |
| ENSG00000260266 | PPIAP46 | -0,28039933 | 0,823363084 | 0,62881703 | 0,952793934 | peptidylprolyl isomerase A pseudogene 46 |
| ENSG00000288314 | PPIAP46 | -0,28039933 | 0,823363084 | 0,62881703 | 0,952793934 | peptidylprolyl isomerase A pseudogene 46 |
| ENSG00000158406 | H4C8 | -0,280516188 | 0,823296394 | 0,3531999 | 0,845063255 | H4 clustered histone 8 |
| ENSG00000133302 | SLF1 | -0,280530913 | 0,823287991 | 0,52276091 | 0,92790906 | SMC5-SMC6 complex localization factor 1 |
| ENSG00000187147 | RNF220 | -0,280546335 | 0,82327919 | 0,32241475 | 0,824955371 | ring finger protein 220 |
| ENSG00000287825 | AL035587,2 | -0,280570854 | 0,823265198 | 0,64956745 | 0,956899963 | novel transcript, antisense to CNPY3 |
| ENSG00000204271 | SPIN3 | -0,280642086 | 0,823224551 | 0,461318 | 0,906165604 | spindlin family member 3 |
| ENSG00000135917 | SLC19A3 | -0,280668585 | 0,823209431 | 0,66876732 | 0,961391466 | solute carrier family 19 member 3 |
| ENSG00000256092 | SBNO1-AS1 | -0,28076298 | 0,82315557 | 0,44228974 | 0,899331131 | SBNO1 antisense RNA 1 |

| | | | | | | |
|---|---|---|---|---|---|---|
| ENSG00000219384 | AL138880,1 | -0,280764992 | 0,823154422 | 0,7430178 | 0,975240824 | pseudogene similar to part of poly(rC) binding protein 3 (PCBP3) |
| ENSG00000144445 | KANSL1L | -0,280770196 | 0,823151453 | 0,53063471 | 0,931265667 | KAT8 regulatory NSL complex subunit 1 like |
| ENSG00000244451 | RPL34P21 | -0,280811082 | 0,823128125 | 0,85177278 | 0,987289464 | ribosomal protein L34 pseudogene 21 |
| ENSG00000225506 | CYP4A22-AS1 | -0,280816074 | 0,823125277 | 0,6593267 | 0,958425175 | CYP4A22 antisense RNA 1 |
| ENSG00000176390 | CRLF3 | -0,280823736 | 0,823120905 | 0,26621016 | 0,777733913 | cytokine receptor like factor 3 |
| ENSG00000215271 | HOMEZ | -0,280885126 | 0,82308588 | 0,30284842 | 0,812446633 | homeobox and leucine zipper encoding |
| ENSG00000227218 | AL157935,1 | -0,280907408 | 0,823073168 | 0,64918922 | 0,956899963 | novel transcript |
| ENSG00000277531 | PNMA8C | -0,280920358 | 0,82306578 | 0,48127867 | 0,912972284 | PNMA family member 8C |
| ENSG00000065923 | SLC9A7 | -0,281035239 | 0,823000242 | 0,21600768 | 0,723046035 | solute carrier family 9 member A7 |
| ENSG00000277114 | CNOT3 | -0,281080092 | 0,822974656 | 0,3380939 | 0,837263082 | CCR4-NOT transcription complex subunit 3 |
| ENSG00000204779 | FOXD4L5 | -0,281107901 | 0,822958793 | 0,57775173 | 0,94337651 | forkhead box D4 like 5 |
| ENSG00000234890 | HNRNPDLP2 | -0,281203042 | 0,822904523 | 0,85442455 | 0,987861659 | HNRNPDL pseudogene 2 |
| ENSG00000260633 | AC010207,1 | -0,281251569 | 0,822876844 | 0,69779418 | 0,968522643 | novel transcript |
| ENSG00000165863 | C10orf82 | -0,281349004 | 0,822821272 | 0,44864955 | 0,902062594 | chromosome 10 open reading frame 82 |
| ENSG00000171595 | DNAI2 | -0,281373553 | 0,822807271 | 0,83729888 | 0,985577601 | dynein axonemal intermediate chain 2 |
| ENSG00000243406 | MRPS31P5 | -0,281400635 | 0,822791825 | 0,36848823 | 0,857548638 | mitochondrial ribosomal protein S31 pseudogene 5 |
| ENSG00000131127 | ZNF141 | -0,281454942 | 0,822760854 | 0,32958416 | 0,831379626 | zinc finger protein 141 |
| ENSG00000231325 | LY6G5C | -0,281483807 | 0,822744392 | 0,67122307 | 0,962026636 | lymphocyte antigen 6 family member G5C |
| ENSG00000227467 | LINC01537 | -0,281564359 | 0,822698456 | 0,63734745 | 0,956017912 | long intergenic non-protein coding RNA 1537 |
| ENSG00000173715 | C11orf80 | -0,281606312 | 0,822674533 | 0,24909519 | 0,761104998 | chromosome 11 open reading frame 80 |
| ENSG00000163462 | TRIM46 | -0,281677167 | 0,82263413 | 0,37858691 | 0,864078042 | tripartite motif containing 46 |
| ENSG00000198440 | ZNF583 | -0,281694127 | 0,822624459 | 0,39014792 | 0,870057277 | zinc finger protein 583 |
| ENSG00000179195 | ZNF664 | -0,281751541 | 0,822591722 | 0,49180009 | 0,916995341 | zinc finger protein 664 |
| ENSG00000287474 | AC110614,1 | -0,281768957 | 0,822581792 | 0,28178652 | 0,792816227 | novel transcript |
| ENSG00000125255 | SLC10A2 | -0,281852637 | 0,822534081 | 0,69384436 | 0,967340633 | solute carrier family 10 member 2 |
| ENSG00000198015 | MRPL42 | -0,281882008 | 0,822517336 | 0,4604887 | 0,90579581 | mitochondrial ribosomal protein L42 |
| ENSG00000141646 | SMAD4 | -0,281885565 | 0,822515308 | 0,47705659 | 0,91143003 | SMAD family member 4 |
| ENSG00000226180 | AC010536,1 | -0,282466502 | 0,822184169 | 0,56726781 | 0,941425553 | novel transcript |
| ENSG00000268469 | BNIP3P38 | -0,282638825 | 0,822085968 | 0,70382613 | 0,968976619 | BCL2 interacting protein 3 pseudogene 38 |
| ENSG00000260799 | KRT8P50 | -0,282664371 | 0,822071412 | 0,73926207 | 0,974715504 | keratin 8 pseudogene 50 |
| ENSG00000226383 | LINC01876 | -0,282795068 | 0,821996942 | 0,51676304 | 0,925584632 | long intergenic non-protein coding RNA 1876 |
| ENSG00000273763 | AC007318,1 | -0,282897224 | 0,821938739 | 0,53814573 | 0,932900071 | marker of proliferation Ki-67 (MKI67) pseudogene |

| | | | | | | |
|---|---|---|---|---|---|---|
| ENSG00000287859 | AC008507,4 | -0,282927688 | 0,821921383 | 0,71443867 | 0,971029555 | novel transcript |
| ENSG00000282684 | ZC3H3 | -0,282929877 | 0,821920136 | 0,48011789 | 0,912847708 | zinc finger CCCH-type containing 3 |
| ENSG00000264538 | SUZ12P1 | -0,283075334 | 0,821837272 | 0,20489094 | 0,706486192 | SUZ12 pseudogene 1 |
| ENSG00000189366 | ALG1L | -0,283324853 | 0,821695145 | 0,7797762 | 0,978587255 | ALG1 chitobiosyldiphosphodolichol beta-mannosyltransferase like |
| ENSG00000271843 | AC012557,1 | -0,283504988 | 0,821592554 | 0,55905044 | 0,939635125 | novel transcript |
| ENSG00000243199 | AC115223,1 | -0,283566949 | 0,821557269 | 0,7682814 | 0,977381742 | ribosomal protein L6 (RPL6) pseudogene |
| ENSG00000175262 | C1orf127 | -0,28363322 | 0,821519531 | 0,78633325 | 0,979320406 | chromosome 1 open reading frame 127 |
| ENSG00000166153 | DEPDC4 | -0,283780115 | 0,821435888 | 0,33130467 | 0,832361728 | DEP domain containing 4 |
| ENSG00000228477 | AL663070,1 | -0,283861006 | 0,821389832 | 0,61136145 | 0,952062377 | ribosomal protein S2 (RPS2) pseudogene |
| ENSG00000163291 | PAQR3 | -0,283886662 | 0,821375226 | 0,59467205 | 0,947469769 | progestin and adipoQ receptor family member 3 |
| ENSG00000188486 | H2AX | -0,283928645 | 0,821351324 | 0,25684943 | 0,769107677 | H2A,X variant histone |
| ENSG00000274833 | AC110285,5 | -0,283944375 | 0,821342368 | 0,82781695 | 0,983595244 | novel transcript |
| ENSG00000243663 | RPS4XP14 | -0,283959442 | 0,82133379 | 0,77966861 | 0,978587255 | ribosomal protein S4X pseudogene 14 |
| ENSG00000284817 | RPS4XP14 | -0,283959442 | 0,82133379 | 0,77966861 | 0,978587255 | ribosomal protein S4X pseudogene 14 |
| ENSG00000206404 | LY6G5C | -0,284037041 | 0,821289614 | 0,83817336 | 0,985672131 | lymphocyte antigen 6 family member G5C |
| ENSG00000286681 | AF235103,3 | -0,284134307 | 0,821234245 | 0,25177058 | 0,763986755 | novel transcript |
| ENSG00000280688 | CEP72 | -0,284214752 | 0,821188454 | 0,86813897 | 0,989074806 | centrosomal protein 72 |
| ENSG00000286437 | AF111169,4 | -0,284286513 | 0,821147608 | 0,70293345 | 0,968742626 | novel transcript, antisense to ANGEL1 |
| ENSG00000243304 | AC008494,1 | -0,284338588 | 0,821117969 | 0,7326129 | 0,974353474 | ribosomal protein S2 (RPS2) pseudogene |
| ENSG00000088727 | KIF9 | -0,284564533 | 0,820989381 | 0,25916274 | 0,771989931 | kinesin family member 9 |
| ENSG00000126773 | PCNX4 | -0,284620668 | 0,820957437 | 0,39360741 | 0,8715972 | pecanex 4 |
| ENSG00000269038 | AP001462,1 | -0,284640473 | 0,820946168 | 0,74499046 | 0,975666349 | novel transcript |
| ENSG00000155961 | RAB39B | -0,284667476 | 0,820930802 | 0,52711445 | 0,92927181 | RAB39B, member RAS oncogene family |
| ENSG00000109756 | RAPGEF2 | -0,284716826 | 0,820902721 | 0,28387114 | 0,79531744 | Rap guanine nucleotide exchange factor 2 |
| ENSG00000251023 | AC114980,1 | -0,284728737 | 0,820895944 | 0,52873452 | 0,930033144 | novel transcript |
| ENSG00000123870 | ZNF137P | -0,284818299 | 0,820844984 | 0,47762458 | 0,911578505 | zinc finger protein 137, pseudogene |
| ENSG00000185875 | THNSL1 | -0,28499497 | 0,82074447 | 0,459968 | 0,905341196 | threonine synthase like 1 |
| ENSG00000148730 | EIF4EBP2 | -0,285147671 | 0,820657604 | 0,42196905 | 0,889720796 | eukaryotic translation initiation factor 4E binding protein 2 |
| ENSG00000118520 | ARG1 | -0,285153033 | 0,820654554 | 0,73542206 | 0,974353474 | arginase 1 |
| ENSG00000255043 | NAV2-AS5 | -0,285307247 | 0,820566836 | 0,56174688 | 0,940728834 | NAV2 antisense RNA 5 |
| ENSG00000263126 | AC040162,3 | -0,285328976 | 0,820554477 | 0,44007629 | 0,897906863 | novel transcript |

| | | | | | | |
|---|---|---|---|---|---|---|
| ENSG00000265907 | AP000919,1 | -0,285433317 | 0,820495134 | 0,48831617 | 0,915394579 | novel transcript, antisense to LPIN2 |
| ENSG00000157734 | SNX22 | -0,285498427 | 0,820458105 | 0,31466464 | 0,81991112 | sorting nexin 22 |
| ENSG00000197056 | ZMYM1 | -0,285517616 | 0,820447193 | 0,53979282 | 0,933385624 | zinc finger MYM-type containing 1 |
| ENSG00000259402 | AC090515,5 | -0,285631897 | 0,820382205 | 0,71656171 | 0,971438758 | novel transcript, sense intronic to FAM63B |
| ENSG00000269834 | ZNF528-AS1 | -0,285640077 | 0,820377553 | 0,59403736 | 0,947469769 | ZNF528 antisense RNA 1 |
| ENSG00000246084 | LINC02325 | -0,285708159 | 0,82033884 | 0,69247857 | 0,967236299 | long intergenic non-protein coding RNA 2325 |
| ENSG00000132297 | HHLA1 | -0,285726618 | 0,820328344 | 0,55985798 | 0,939880787 | HERV-H LTR-associating 1 |
| ENSG00000273002 | AL355388,1 | -0,285734289 | 0,820323982 | 0,64499001 | 0,956899963 | uncharacterized LOC107985209 |
| ENSG00000250988 | SNHG21 | -0,285797 | 0,820288325 | 0,65847431 | 0,958135616 | small nucleolar RNA host gene 21 |
| ENSG00000188219 | POTEE | -0,285815114 | 0,820278025 | 0,38313298 | 0,867698804 | POTE ankyrin domain family member E |
| ENSG00000263841 | RN7SL44P | -0,285842308 | 0,820262564 | 0,62482226 | 0,952793934 | RNA, 7SL, cytoplasmic 44, pseudogene |
| ENSG00000106524 | ANKMY2 | -0,2858553 | 0,820255177 | 0,40643103 | 0,879536215 | ankyrin repeat and MYND domain containing 2 |
| ENSG00000249207 | AC079921,1 | -0,285977784 | 0,820185541 | 0,57337715 | 0,942475308 | novel transcript |
| ENSG00000139675 | HNRNPA1L2 | -0,285980991 | 0,820183718 | 0,26501278 | 0,776509969 | heterogeneous nuclear ribonucleoprotein A1 like 2 |
| ENSG00000271991 | AC013400,1 | -0,286048178 | 0,820145523 | 0,67763223 | 0,9635902 | novel transcript |
| ENSG00000028839 | TBPL1 | -0,28608554 | 0,820124283 | 0,5910487 | 0,947416795 | TATA-box binding protein like 1 |
| ENSG00000243230 | AC011005,4 | -0,286091384 | 0,820120961 | 0,71090858 | 0,970482408 | novel transcript |
| ENSG00000151743 | AMN1 | -0,286124225 | 0,820102292 | 0,5574784 | 0,938578675 | antagonist of mitotic exit network 1 homolog |
| ENSG00000181754 | AMIGO1 | -0,286207437 | 0,820054992 | 0,33112587 | 0,832246888 | adhesion molecule with Ig like domain 1 |
| ENSG00000214765 | SEPTIN7P2 | -0,286237229 | 0,820038058 | 0,32482925 | 0,826857333 | septin 7 pseudogene 2 |
| ENSG00000255330 | AL096711,2 | -0,286239332 | 0,820036862 | 0,36102213 | 0,851783601 | novel protein |
| ENSG00000141642 | ELAC1 | -0,28629454 | 0,820005482 | 0,40463144 | 0,878176155 | elaC ribonuclease Z 1 |
| ENSG00000163539 | CLASP2 | -0,286318291 | 0,819991983 | 0,47069143 | 0,908869097 | cytoplasmic linker associated protein 2 |
| ENSG00000280512 | MUC20-OT1 | -0,286331114 | 0,819984694 | 0,388237 | 0,869820053 | MUC20 overlapping transcript |
| ENSG00000198920 | KIAA0753 | -0,286346771 | 0,819975796 | 0,32377793 | 0,825794844 | KIAA0753 |
| ENSG00000248871 | TNFSF12-TNFSI | -0,286354346 | 0,81997149 | 0,85796705 | 0,98899351 | TNFSF12-TNFSF13 readthrough |
| ENSG00000130234 | ACE2 | -0,286400624 | 0,819945188 | 0,58870828 | 0,946486458 | angiotensin I converting enzyme 2 |
| ENSG00000227627 | AL080276,2 | -0,286435649 | 0,819925282 | 0,35776688 | 0,848908914 | novel transcript |
| ENSG00000232303 | DFFBP1 | -0,286586051 | 0,819839808 | 0,66696606 | 0,9611116 | DNA fragmentation factor subunit beta pseudogene 1 |
| ENSG00000164073 | MFSD8 | -0,28672059 | 0,819763358 | 0,51014284 | 0,922427526 | major facilitator superfamily domain containing 8 |
| ENSG00000167751 | KLK2 | -0,286930003 | 0,819644374 | 0,84698249 | 0,986957162 | kallikrein related peptidase 2 |
| ENSG00000213297 | ZNF625-ZNF20 | -0,28695394 | 0,819630775 | 0,20567511 | 0,708174755 | ZNF625-ZNF20 readthrough (NMD candidate) |

| | | | | | | |
|---|---|---|---|---|---|---|
| ENSG00000258779 | LINC01568 | -0,286988034 | 0,819611406 | 0,75418923 | 0,975901808 | long intergenic non-protein coding RNA 1568 |
| ENSG00000281813 | KAT6B | -0,286991028 | 0,819609705 | 0,20921629 | 0,713678247 | lysine acetyltransferase 6B |
| ENSG00000108666 | C17orf75 | -0,287028678 | 0,819588316 | 0,31819193 | 0,823060685 | chromosome 17 open reading frame 75 |
| ENSG00000249140 | PRDX2P3 | -0,287255181 | 0,81945965 | 0,71612372 | 0,971360663 | peroxiredoxin 2 pseudogene 3 |
| ENSG00000284556 | PRDX2P3 | -0,287255181 | 0,81945965 | 0,71612372 | 0,971360663 | peroxiredoxin 2 pseudogene 3 |
| ENSG00000251287 | ALG1L2 | -0,287270823 | 0,819450766 | 0,55846596 | 0,939359747 | ALG1 chitobiosyldiphosphodolichol beta-mannosyltransferase like 2 |
| ENSG00000219433 | BTBD10P2 | -0,287278184 | 0,819446585 | 0,72957546 | 0,974009551 | BTB domain containing 10 pseudogene 2 |
| ENSG00000165985 | C1QL3 | -0,28734438 | 0,819408986 | 0,6989691 | 0,968522643 | complement C1q like 3 |
| ENSG00000104231 | ZFAND1 | -0,28739008 | 0,81938303 | 0,53414044 | 0,93151582 | zinc finger AN1-type containing 1 |
| ENSG00000227972 | THAP12P3 | -0,28745401 | 0,819346722 | 0,8351803 | 0,984914821 | THAP domain containing 12 pseudogene 3 |
| ENSG00000058804 | NDC1 | -0,287489504 | 0,819326564 | 0,56364014 | 0,940865625 | NDC1 transmembrane nucleoporin |
| ENSG00000232081 | LARGE-IT1 | -0,287511238 | 0,819314221 | 0,83082286 | 0,984156632 | LARGE intronic transcript 1 |
| ENSG00000272769 | AC097532,2 | -0,287530109 | 0,819303505 | 0,54265328 | 0,933663748 | novel transcript |
| ENSG00000116560 | SFPQ | -0,287537832 | 0,819299119 | 0,28636585 | 0,797518439 | splicing factor proline and glutamine rich |
| ENSG00000260378 | AC109597,1 | -0,287667168 | 0,819225673 | 0,85317571 | 0,987861659 | novel transcript, antisense to CPPED1 |
| ENSG00000228606 | AL139011,1 | -0,287825298 | 0,819135885 | 0,60789167 | 0,951017078 | novel transcript |
| ENSG00000284287 | AC270107,7 | -0,287838978 | 0,819128118 | 0,52699532 | 0,92927181 | |
| ENSG00000284036 | AC270107,3 | -0,287838978 | 0,819128118 | 0,52699532 | 0,92927181 | |
| ENSG00000284230 | AC270107,6 | -0,287838978 | 0,819128118 | 0,52699532 | 0,92927181 | |
| ENSG00000283963 | AC270107,1 | -0,287838978 | 0,819128118 | 0,52699532 | 0,92927181 | |
| ENSG00000284151 | AC270107,5 | -0,287838978 | 0,819128118 | 0,52699532 | 0,92927181 | |
| ENSG00000283989 | AC270107,2 | -0,287838978 | 0,819128118 | 0,52699532 | 0,92927181 | |
| ENSG00000108389 | MTMR4 | -0,287945868 | 0,81906743 | 0,24158581 | 0,751948055 | myotubularin related protein 4 |
| ENSG00000236778 | INTS6-AS1 | -0,288179595 | 0,818934746 | 0,34125511 | 0,840012589 | INTS6 antisense RNA 1 |
| ENSG00000115239 | ASB3 | -0,288253205 | 0,818892963 | 0,25381633 | 0,76531122 | ankyrin repeat and SOCS box containing 3 |
| ENSG00000265802 | RN7SL49P | -0,288258246 | 0,818890101 | 0,74561742 | 0,975707496 | RNA, 7SL, cytoplasmic 49, pseudogene |
| ENSG00000271919 | AC005828,7 | -0,288258852 | 0,818889757 | 0,68886965 | 0,966556275 | density-regulated protein (DENR) pseudogene |
| ENSG00000279669 | AC079801,1 | -0,288287222 | 0,818873654 | 0,79753164 | 0,980161849 | novel transcript |
| ENSG00000273032 | DGCR5 | -0,288303438 | 0,81886445 | 0,29778353 | 0,806960585 | DiGeorge syndrome critical region gene 5 |
| ENSG00000103494 | RPGRIP1L | -0,2883285 | 0,818850226 | 0,47852001 | 0,91222253 | RPGRIP1 like |
| ENSG00000278728 | CNTNAP2 | -0,288342114 | 0,818842498 | 0,55746589 | 0,938578675 | contactin associated protein 2 |
| ENSG00000227066 | Z98257,1 | -0,288405729 | 0,818806392 | 0,78455575 | 0,979320406 | novel transcript |

| | | | | | | |
|---|---|---|---|---|---|---|
| ENSG00000132122 | SPATA6 | -0,288487767 | 0,818759833 | 0,50245886 | 0,920585659 | spermatogenesis associated 6 |
| ENSG00000286810 | AL513128,3 | -0,288632548 | 0,818677671 | 0,6511873 | 0,957089277 | novel transcript |
| ENSG00000282638 | AC243742,3 | -0,288687273 | 0,818646617 | 0,49425522 | 0,91758518 | novel transcript |
| ENSG00000280206 | AC026401,3 | -0,288687273 | 0,818646617 | 0,49425522 | 0,91758518 | novel transcript |
| ENSG00000170236 | USP50 | -0,288714447 | 0,818631198 | 0,62975529 | 0,952812183 | ubiquitin specific peptidase 50 |
| ENSG00000165359 | INTS6L | -0,288734111 | 0,81862004 | 0,50925016 | 0,922427526 | integrator complex subunit 6 like |
| ENSG00000144824 | PHLDB2 | -0,288778965 | 0,818594589 | 0,40194735 | 0,87718264 | pleckstrin homology like domain family B member 2 |
| ENSG00000214562 | NUTM2D | -0,288786074 | 0,818590555 | 0,399451 | 0,875433675 | NUT family member 2D |
| ENSG00000169660 | HEXD | -0,288793751 | 0,818586199 | 0,40369749 | 0,877691113 | hexosaminidase D |
| ENSG00000173285 | OR10K1 | -0,288815964 | 0,818573596 | 0,2427165 | 0,752892335 | olfactory receptor family 10 subfamily K member 1 |
| ENSG00000186638 | KIF24 | -0,288874637 | 0,818540306 | 0,30157161 | 0,811179119 | kinesin family member 24 |
| ENSG00000274512 | TBC1D3L | -0,288923304 | 0,818512694 | 0,33296153 | 0,833301711 | TBC1 domain family member 3L |
| ENSG00000228988 | AL353771,1 | -0,288930945 | 0,818508359 | 0,83139924 | 0,984440275 | novel transcript |
| ENSG00000205090 | TMEM240 | -0,2890095 | 0,818463792 | 0,44491466 | 0,90042802 | transmembrane protein 240 |
| ENSG00000105327 | BBC3 | -0,289011091 | 0,818462889 | 0,46204444 | 0,906277266 | BCL2 binding component 3 |
| ENSG00000166377 | ATP9B | -0,289043175 | 0,818444688 | 0,19247283 | 0,689049663 | ATPase phospholipid transporting 9B (putative) |
| ENSG00000152495 | CAMK4 | -0,289125113 | 0,818398206 | 0,51867198 | 0,926765093 | calcium/calmodulin dependent protein kinase IV |
| ENSG00000171174 | RBKS | -0,289173321 | 0,818370859 | 0,53443384 | 0,931616373 | ribokinase |
| ENSG00000240993 | RN7SL459P | -0,289220695 | 0,818343986 | 0,84434157 | 0,986340494 | RNA, 7SL, cytoplasmic 459, pseudogene |
| ENSG00000253384 | AC124242,2 | -0,28924807 | 0,818328459 | 0,79929793 | 0,980161849 | family with sequence similarity 175, member A (FAM175A) pseudogene |
| ENSG00000242299 | AC073861,1 | -0,289279798 | 0,818310462 | 0,52872323 | 0,930033144 | ribosomal protein S18 (RPS18) pseudogene |
| ENSG00000269947 | AC135178,6 | -0,289290136 | 0,818304598 | 0,62736082 | 0,952793934 | novel transcript |
| ENSG00000261630 | AC007496,2 | -0,28958635 | 0,818136602 | 0,76217702 | 0,976315588 | novel transcript, antisense to FTO |
| ENSG00000270915 | H3P8 | -0,289685542 | 0,818080353 | 0,77141472 | 0,978033855 | H3 histone pseudogene 8 |
| ENSG00000237984 | PTENP1 | -0,289731755 | 0,818054148 | 0,59727327 | 0,948282297 | phosphatase and tensin homolog pseudogene 1 |
| ENSG00000218757 | AL121952,1 | -0,289835297 | 0,817995438 | 0,52438621 | 0,928598732 | 60S ribosomal protein L17 (RPL17) pseudogene |
| ENSG00000163322 | ABRAXAS1 | -0,289835426 | 0,817995365 | 0,44118982 | 0,898885877 | abraxas 1, BRCA1 A complex subunit |
| ENSG00000158062 | UBXN11 | -0,28985647 | 0,817983434 | 0,40027919 | 0,87601134 | UBX domain protein 11 |
| ENSG00000287708 | AL137017,1 | -0,289951163 | 0,817929746 | 0,81035245 | 0,981213877 | novel transcript, antisense to NFIB |
| ENSG00000074266 | EED | -0,289959409 | 0,817925071 | 0,40615154 | 0,879163888 | embryonic ectoderm development |
| ENSG00000105483 | CARD8 | -0,289970891 | 0,817918561 | 0,34496165 | 0,841168721 | caspase recruitment domain family member 8 |
| ENSG00000256712 | AC134349,1 | -0,290102338 | 0,817844042 | 0,73669291 | 0,974353474 | novel transcript |

| | | | | | | |
|---|---|---|---|---|---|---|
| ENSG00000262531 | AC134349,4 | -0,290102338 | 0,817844042 | 0,73669291 | 0,974353474 | novel transcript |
| ENSG00000282546 | AC245078,4 | -0,290102338 | 0,817844042 | 0,73669291 | 0,974353474 | novel transcript |
| ENSG00000162929 | KIAA1841 | -0,290120758 | 0,8178336 | 0,34994449 | 0,843677595 | KIAA1841 |
| ENSG00000067369 | TP53BP1 | -0,290217971 | 0,817778494 | 0,25227076 | 0,76432863 | tumor protein p53 binding protein 1 |
| ENSG00000196296 | ATP2A1 | -0,290277479 | 0,817744764 | 0,43692476 | 0,896866043 | ATPase sarcoplasmic/endoplasmic reticulum Ca2+ transporting 1 |
| ENSG00000277870 | FAM230A | -0,290305096 | 0,81772911 | 0,67566901 | 0,963449137 | family with sequence similarity 230 member A |
| ENSG00000119661 | DNAL1 | -0,290354912 | 0,817700874 | 0,33946441 | 0,838364485 | dynein axonemal light chain 1 |
| ENSG00000149646 | CNBD2 | -0,290422312 | 0,817662674 | 0,52294696 | 0,92790906 | cyclic nucleotide binding domain containing 2 |
| ENSG00000128610 | FEZF1 | -0,290582447 | 0,817571921 | 0,50990471 | 0,922427526 | FEZ family zinc finger 1 |
| ENSG00000228791 | THRB-AS1 | -0,29072776 | 0,817489576 | 0,77999034 | 0,978587255 | THRB antisense RNA 1 |
| ENSG00000101290 | CDS2 | -0,290735129 | 0,8174854 | 0,34426723 | 0,84105522 | CDP-diacylglycerol synthase 2 |
| ENSG00000239407 | Z68871,1 | -0,290833507 | 0,817429658 | 0,45832954 | 0,904165485 | novel transcript |
| ENSG00000229267 | SNHG31 | -0,290884549 | 0,817400738 | 0,46698395 | 0,907940653 | small nucleolar RNA host gene 31 |
| ENSG00000120727 | PAIP2 | -0,290918743 | 0,817381365 | 0,50900998 | 0,922427526 | poly(A) binding protein interacting protein 2 |
| ENSG00000288569 | LINC01949 | -0,290961873 | 0,817356929 | 0,4380994 | 0,897177404 | long intergenic non-protein coding RNA 1949 |
| ENSG00000164050 | PLXNB1 | -0,291050257 | 0,817306857 | 0,36241444 | 0,852431928 | plexin B1 |
| ENSG00000175182 | FAM131A | -0,29111531 | 0,817270004 | 0,37098028 | 0,858959913 | family with sequence similarity 131 member A |
| ENSG00000235109 | ZSCAN31 | -0,291158172 | 0,817245723 | 0,33591628 | 0,835660292 | zinc finger and SCAN domain containing 31 |
| ENSG00000118557 | PMFBP1 | -0,29116827 | 0,817240003 | 0,40320577 | 0,877589651 | polyamine modulated factor 1 binding protein 1 |
| ENSG00000261645 | DISC1FP1 | -0,291339936 | 0,817142766 | 0,73391748 | 0,974353474 | DISC1 fusion partner 1 |
| ENSG00000142856 | ITGB3BP | -0,291342095 | 0,817141543 | 0,42996868 | 0,894726544 | integrin subunit beta 3 binding protein |
| ENSG00000141519 | CCDC40 | -0,291382619 | 0,817118591 | 0,33555479 | 0,835320086 | coiled-coil domain containing 40 |
| ENSG00000259104 | PTCSC3 | -0,291421211 | 0,817096733 | 0,53499611 | 0,931805607 | papillary thyroid carcinoma susceptibility candidate 3 |
| ENSG00000189157 | FAM47E | -0,291431438 | 0,817090941 | 0,41640419 | 0,886152099 | family with sequence similarity 47 member E |
| ENSG00000231555 | HSPA1B | -0,291484954 | 0,817060632 | 0,41868923 | 0,887094957 | heat shock protein family A (Hsp70) member 1B |
| ENSG00000224707 | E2F3-IT1 | -0,291497095 | 0,817053756 | 0,72743337 | 0,973441576 | E2F3 intronic transcript 1 |
| ENSG00000204666 | AC010624,1 | -0,29150096 | 0,817051567 | 0,74276628 | 0,975201495 | novel transcript, sense overlapping ZNF473 |
| ENSG00000109424 | UCP1 | -0,291560204 | 0,817018016 | 0,76373272 | 0,976640468 | uncoupling protein 1 |
| ENSG00000197462 | AC003975,1 | -0,29161779 | 0,816985405 | 0,50205737 | 0,920520138 | novel transcript |
| ENSG00000184672 | RALYL | -0,291628314 | 0,816979445 | 0,6377478 | 0,956018518 | RALY RNA binding protein like |
| ENSG00000230536 | AL360268,1 | -0,291632922 | 0,816976835 | 0,73751039 | 0,974353474 | novel transcript |

| | | | | | | |
|---|---|---|---|---|---|---|
| ENSG00000275594 | ADAM32 | -0,291713252 | 0,816931347 | 0,42009172 | 0,888457169 | ADAM metallopeptidase domain 32 |
| ENSG00000253392 | AC119403,1 | -0,291713326 | 0,816931305 | 0,66950263 | 0,961917746 | novel transcript, antisense to ZNF57 |
| ENSG00000125247 | TMTC4 | -0,291725159 | 0,816924605 | 0,35297292 | 0,845031687 | transmembrane O-mannosyltransferase targeting cadherins 4 |
| ENSG00000171877 | FRMD5 | -0,291886982 | 0,816832978 | 0,24915115 | 0,761135572 | FERM domain containing 5 |
| ENSG00000120685 | PROSER1 | -0,291915353 | 0,816816915 | 0,37229059 | 0,860223901 | proline and serine rich 1 |
| ENSG00000180773 | SLC36A4 | -0,291981973 | 0,816779197 | 0,53889467 | 0,933348247 | solute carrier family 36 member 4 |
| ENSG00000231731 | AC010976,1 | -0,292046772 | 0,816742512 | 0,39476047 | 0,871888817 | novel transcript, antisense to IWS1 |
| ENSG00000277511 | AC116407,2 | -0,292067823 | 0,816730594 | 0,69680921 | 0,968300554 | novel transcript |
| ENSG00000281750 | AC213220,3 | -0,292191876 | 0,816660369 | 0,75975986 | 0,976129381 | novel transcript, sense intronic to NDUFA6 |
| ENSG00000183696 | UPP1 | -0,292224356 | 0,816641984 | 0,41522399 | 0,885840844 | uridine phosphorylase 1 |
| ENSG00000256083 | AC090673,1 | -0,292244469 | 0,816630599 | 0,76988003 | 0,977742359 | novel transcript, antisense to HMGA2 |
| ENSG00000223516 | AFF2-IT1 | -0,292253416 | 0,816625534 | 0,72128875 | 0,972142678 | AFF2 intronic transcript 1 |
| ENSG00000277848 | MEGF11 | -0,292263749 | 0,816619685 | 0,80078337 | 0,980505307 | multiple EGF like domains 11 |
| ENSG00000196505 | GDAP2 | -0,292285621 | 0,816607305 | 0,47956988 | 0,912721304 | ganglioside induced differentiation associated protein 2 |
| ENSG00000278903 | CU633906,2 | -0,29240408 | 0,816540257 | 0,48207655 | 0,912981134 | novel transcript |
| ENSG00000278022 | AC118658,1 | -0,292504352 | 0,816483506 | 0,70924625 | 0,970282889 | novel transcript, sense intronic to IGF1R |
| ENSG00000196778 | OR52K1 | -0,292558565 | 0,816452826 | 0,59138905 | 0,947416795 | olfactory receptor family 52 subfamily K member 1 |
| ENSG00000278932 | CR381653,1 | -0,292566722 | 0,81644821 | 0,59149043 | 0,947416795 | novel transcript |
| ENSG00000135454 | B4GALNT1 | -0,292639932 | 0,81640678 | 0,26680244 | 0,779124932 | beta-1,4-N-acetyl-galactosaminyltransferase 1 |
| ENSG00000283538 | AC005972,3 | -0,292774112 | 0,816330852 | 0,59368827 | 0,947469769 | novel transcript |
| ENSG00000282226 | AC093614,2 | -0,292826666 | 0,816301116 | 0,58127097 | 0,943919685 | novel transcript |
| ENSG00000223650 | UHRF2P1 | -0,292894119 | 0,816262951 | 0,49491283 | 0,917622746 | ubiquitin like with PHD and ring finger domains 2 pseudogene 1 |
| ENSG00000247728 | AC091057,2 | -0,292912084 | 0,816252786 | 0,50043593 | 0,920485547 | novel transcript, antisense to ARHGAP11B |
| ENSG00000273015 | AC008124,1 | -0,293039964 | 0,816180437 | 0,48787391 | 0,915155099 | novel transcript |
| ENSG00000134755 | DSC2 | -0,293119159 | 0,816135635 | 0,49388383 | 0,917303403 | desmocollin 2 |
| ENSG00000135127 | BICDL1 | -0,293264277 | 0,816053545 | 0,44793875 | 0,902062594 | BICD family like cargo adaptor 1 |
| ENSG00000196123 | KIAA0895L | -0,293296641 | 0,816035239 | 0,2747049 | 0,786383635 | KIAA0895 like |
| ENSG00000111785 | RIC8B | -0,293353987 | 0,816002803 | 0,45006132 | 0,902062594 | RIC8 guanine nucleotide exchange factor B |
| ENSG00000147905 | ZCCHC7 | -0,293380355 | 0,815987889 | 0,47128334 | 0,908869685 | zinc finger CCHC-type containing 7 |
| ENSG00000139133 | ALG10 | -0,293522877 | 0,815907283 | 0,55820852 | 0,939161457 | ALG10 alpha-1,2-glucosyltransferase |

| | | | | | | |
|---|---|---|---|---|---|---|
| ENSG00000059377 | TBXAS1 | -0,293530343 | 0,81590306 | 0,40555681 | 0,878494653 | thromboxane A synthase 1 |
| ENSG00000273951 | AL031667,3 | -0,293607327 | 0,815859524 | 0,72197675 | 0,972516173 | novel transcript, sense intronic to CHD6 |
| ENSG00000109654 | TRIM2 | -0,29363221 | 0,815845453 | 0,48249169 | 0,913223171 | tripartite motif containing 2 |
| ENSG00000171792 | RHNO1 | -0,293670942 | 0,81582355 | 0,3910622 | 0,870057277 | RAD9-HUS1-RAD1 interacting nuclear orphan 1 |
| ENSG00000285541 | AC005548,1 | -0,293723451 | 0,815793857 | 0,47992792 | 0,912847708 | novel transcript, antisense to SHISA6 |
| ENSG00000251332 | PSME2P4 | -0,293733821 | 0,815787993 | 0,71344587 | 0,970812834 | proteasome activator subunit 2 pseudogene 4 |
| ENSG00000198924 | DCLRE1A | -0,293792533 | 0,815754795 | 0,54931814 | 0,936040289 | DNA cross-link repair 1A |
| ENSG00000280475 | AC007537,2 | -0,29380166 | 0,815749634 | 0,75091734 | 0,975901808 | ribosomal protein L21 (RPL21) pseudogene |
| ENSG00000242405 | AC007537,1 | -0,29380166 | 0,815749634 | 0,75091734 | 0,975901808 | ribosomal protein L21 (RPL21) pseudogene |
| ENSG00000259318 | HMGN1P1 | -0,293969401 | 0,815654793 | 0,59408927 | 0,947469769 | high mobility group nucleosome binding domain 1 pseudogene 1 |
| ENSG00000170537 | TMC7 | -0,294037969 | 0,815616028 | 0,26854379 | 0,780709773 | transmembrane channel like 7 |
| ENSG00000272645 | GTF2IP20 | -0,294040729 | 0,815614467 | 0,33291943 | 0,833301711 | general transcription factor IIi pseudogene 20 |
| ENSG00000136842 | TMOD1 | -0,294069821 | 0,81559802 | 0,53923393 | 0,933350549 | tropomodulin 1 |
| ENSG00000167393 | PPP2R3B | -0,294099864 | 0,815581037 | 0,60748218 | 0,950993594 | protein phosphatase 2 regulatory subunit B''beta |
| ENSG00000046604 | DSG2 | -0,294104331 | 0,815578511 | 0,4564939 | 0,903545728 | desmoglein 2 |
| ENSG00000102128 | RAB40AL | -0,294121379 | 0,815568874 | 0,66898284 | 0,961612856 | RAB40A like |
| ENSG00000238358 | AC004969,1 | -0,294243814 | 0,815499663 | 0,59224502 | 0,947416795 | novel transcript |
| ENSG00000261586 | AC068987,3 | -0,294337914 | 0,815446474 | 0,86830582 | 0,989074806 | novel transcript |
| ENSG00000267368 | UPK3BL1 | -0,294424192 | 0,815397709 | 0,47844537 | 0,912212078 | uroplakin 3B like 1 |
| ENSG00000234495 | TRIM27 | -0,294480105 | 0,815366108 | 0,35227741 | 0,844912682 | tripartite motif containing 27 |
| ENSG00000229905 | AL669831,2 | -0,294496836 | 0,815356652 | 0,66178997 | 0,959701366 | novel transcript |
| ENSG00000234798 | DXO | -0,294567703 | 0,815316602 | 0,39409944 | 0,871635019 | decapping exoribonuclease |
| ENSG00000224313 | DXO | -0,294567703 | 0,815316602 | 0,39409944 | 0,871635019 | decapping exoribonuclease |
| ENSG00000225682 | DXO | -0,294567703 | 0,815316602 | 0,39409944 | 0,871635019 | decapping exoribonuclease |
| ENSG00000174373 | RALGAPA1 | -0,29463245 | 0,815280012 | 0,4053902 | 0,878367308 | Ral GTPase activating protein catalytic subunit alpha 1 |
| ENSG00000237737 | DCTN1-AS1 | -0,294645878 | 0,815272424 | 0,52544264 | 0,928838563 | DCTN1 antisense RNA 1 |
| ENSG00000143727 | ACP1 | -0,294693008 | 0,815245791 | 0,5015133 | 0,920520138 | acid phosphatase 1 |
| ENSG00000287653 | AL355597,1 | -0,294728773 | 0,81522558 | 0,48114035 | 0,912911021 | novel transcript |
| ENSG00000229692 | SOS1-IT1 | -0,294752185 | 0,815212351 | 0,50222566 | 0,920520138 | SOS1 intronic transcript 1 |
| ENSG00000233813 | MRPS18B | -0,294792839 | 0,81518938 | 0,78224604 | 0,978597905 | mitochondrial ribosomal protein S18B |
| ENSG00000221947 | XKR9 | -0,294851335 | 0,815156327 | 0,41854973 | 0,887094957 | XK related 9 |

| | | | | | | |
|---|---|---|---|---|---|---|
| ENSG00000253686 | LINC01484 | -0,294855542 | 0,81515395 | 0,64462934 | 0,956633898 | long intergenic non-protein coding RNA 1484 |
| ENSG00000202444 | RNU5E-6P | -0,294895947 | 0,815131121 | 0,76361467 | 0,976640468 | RNA, U5E small nuclear 6, pseudogene |
| ENSG00000267629 | AC138430,1 | -0,294927531 | 0,815113276 | 0,60302969 | 0,950352286 | novel transcript, readthrough between TM6SF2 and HAPLN4 |
| ENSG00000226856 | THORLNC | -0,295083968 | 0,815024895 | 0,58181643 | 0,944145586 | testis associated oncogenic lncRNA |
| ENSG00000127585 | FBXL16 | -0,295130164 | 0,814998797 | 0,34412925 | 0,840938551 | F-box and leucine rich repeat protein 16 |
| ENSG00000114646 | CSPG5 | -0,295151366 | 0,81498682 | 0,36842511 | 0,857548638 | chondroitin sulfate proteoglycan 5 |
| ENSG00000180747 | SMG1P3 | -0,295302769 | 0,814901297 | 0,24281456 | 0,752892335 | SMG1 pseudogene 3 |
| ENSG00000288010 | AL035587,3 | -0,295385834 | 0,814854379 | 0,375358 | 0,862022464 | novel transcript |
| ENSG00000243775 | OSTCP1 | -0,295583396 | 0,814742801 | 0,51131752 | 0,922893865 | oligosaccharyltransferase complex subunit pseudogene 1 |
| ENSG00000247240 | UBL7-AS1 | -0,295771723 | 0,814636452 | 0,23584414 | 0,744966003 | UBL7 antisense RNA 1 (head to head) |
| ENSG00000273877 | AC236972,3 | -0,295824554 | 0,814606621 | 0,79315766 | 0,979659177 | uncharacterized LOC105373373 |
| ENSG00000228192 | AL512353,1 | -0,295899882 | 0,814564089 | 0,3643652 | 0,854320801 | novel transcript, antisense to ERMAP |
| ENSG00000111371 | SLC38A1 | -0,295946753 | 0,814537626 | 0,48761396 | 0,914936533 | solute carrier family 38 member 1 |
| ENSG00000065600 | PACC1 | -0,295963295 | 0,814528286 | 0,34199192 | 0,840048641 | proton activated chloride channel 1 |
| ENSG00000272562 | AL512343,2 | -0,296031644 | 0,814489698 | 0,58349903 | 0,944174844 | novel transcript, antisense to H3F3A |
| ENSG00000058404 | CAMK2B | -0,296192444 | 0,814398922 | 0,47261115 | 0,909368754 | calcium/calmodulin dependent protein kinase II beta |
| ENSG00000184897 | H1-10 | -0,296434371 | 0,814262365 | 0,27406639 | 0,7862019 | H1,10 linker histone |
| ENSG00000103365 | GGA2 | -0,296455076 | 0,81425068 | 0,19426278 | 0,691291421 | golgi associated, gamma adaptin ear containing, ARF binding protein 2 |
| ENSG00000203321 | CARNMT1-AS1 | -0,296463159 | 0,814246118 | 0,83961105 | 0,985672131 | CARNMT1 antisense RNA 1 |
| ENSG00000030066 | NUP160 | -0,296583462 | 0,814178222 | 0,41307717 | 0,884663685 | nucleoporin 160 |
| ENSG00000203644 | AC083799,1 | -0,296591492 | 0,814173691 | 0,56645766 | 0,941311563 | novel transcript, sense intronic to TMCC1 |
| ENSG00000125744 | RTN2 | -0,296646627 | 0,814142576 | 0,22827234 | 0,736631691 | reticulon 2 |
| ENSG00000164975 | SNAPC3 | -0,296693559 | 0,814116092 | 0,3951896 | 0,871981986 | small nuclear RNA activating complex polypeptide 3 |
| ENSG00000166866 | MYO1A | -0,296723713 | 0,814099076 | 0,78634439 | 0,979320406 | myosin IA |
| ENSG00000117724 | CENPF | -0,296859293 | 0,814022573 | 0,29499377 | 0,805123121 | centromere protein F |
| ENSG00000112367 | FIG4 | -0,296864217 | 0,814019795 | 0,41019047 | 0,882601171 | FIG4 phosphoinositide 5-phosphatase |
| ENSG00000261635 | AC103988,1 | -0,296870369 | 0,814016324 | 0,70171211 | 0,968742626 | tropomyosin 3 (TPM3) pseudogene |
| ENSG00000130779 | CLIP1 | -0,296892007 | 0,814004115 | 0,20503488 | 0,706634751 | CAP-Gly domain containing linker protein 1 |
| ENSG00000073150 | PANX2 | -0,29702092 | 0,813931382 | 0,57306673 | 0,942475308 | pannexin 2 |

| | | | | | | |
|---|---|---|---|---|---|---|
| ENSG00000270956 | AC009948,2 | -0,297061269 | 0,813908619 | 0,60292339 | 0,950352286 | novel transcript, antisense to TTN |
| ENSG00000114857 | NKTR | -0,297155803 | 0,813855288 | 0,40630272 | 0,879319405 | natural killer cell triggering receptor |
| ENSG00000261187 | AC079322,1 | -0,29715612 | 0,81385511 | 0,69683124 | 0,968300554 | novel transcript |
| ENSG00000125848 | FLRT3 | -0,297202967 | 0,813828683 | 0,56672848 | 0,941311563 | fibronectin leucine rich transmembrane protein 3 |
| ENSG00000170322 | NFRKB | -0,297255706 | 0,813798933 | 0,21318102 | 0,720528806 | nuclear factor related to kappaB binding protein |
| ENSG00000166532 | RIMKLB | -0,297281815 | 0,813784205 | 0,3435477 | 0,840901506 | ribosomal modification protein rimK like family member B |
| ENSG00000225806 | AL121917,1 | -0,297308934 | 0,813768909 | 0,67439673 | 0,963103112 | novel transcript, antisense to GNAS |
| ENSG00000128578 | STRIP2 | -0,297355653 | 0,813742557 | 0,31274037 | 0,818683344 | striatin interacting protein 2 |
| ENSG00000199477 | SNORA31 | -0,29738912 | 0,81372368 | 0,77862083 | 0,978587255 | small nucleolar RNA, H/ACA box 31 |
| ENSG00000239483 | RPS15AP16 | -0,297539068 | 0,813639109 | 0,68433537 | 0,965415049 | ribosomal protein S15a pseudogene 16 |
| ENSG00000270361 | AL451085,1 | -0,29759376 | 0,813608265 | 0,77322681 | 0,978363975 | novel transcript |
| ENSG00000250568 | AC098591,3 | -0,297679732 | 0,813559782 | 0,8116972 | 0,981213877 | poly(A) binding protein, cytoplasmic 1 (PABPC1) pseudogene |
| ENSG00000280383 | Z95331,1 | -0,297906567 | 0,813431876 | 0,29862763 | 0,808197215 | novel transcript, sense overlapping ATXN10 |
| ENSG00000274808 | TBC1D3B | -0,297965967 | 0,813398386 | 0,68975762 | 0,966868992 | TBC1 domain family member 3B |
| ENSG00000241537 | AC134050,1 | -0,297967069 | 0,813397764 | 0,75580703 | 0,975901808 | U2-associated SR140 protein (SR140) pseudogene |
| ENSG00000175906 | ARL4D | -0,297996397 | 0,81338123 | 0,3349071 | 0,834858938 | ADP ribosylation factor like GTPase 4D |
| ENSG00000177868 | SVBP | -0,298006874 | 0,813375323 | 0,45004354 | 0,902062594 | small vasohibin binding protein |
| ENSG00000198270 | TMEM116 | -0,298021132 | 0,813367284 | 0,26337716 | 0,774637968 | transmembrane protein 116 |
| ENSG00000188529 | SRSF10 | -0,29806994 | 0,813339767 | 0,52191922 | 0,927599011 | serine and arginine rich splicing factor 10 |
| ENSG00000136936 | XPA | -0,298084233 | 0,813331709 | 0,38519358 | 0,868895792 | XPA, DNA damage recognition and repair factor |
| ENSG00000273680 | AC009318,2 | -0,298119 | 0,81331211 | 0,51946371 | 0,926800715 | novel transcript, antisense to FAR2 |
| ENSG00000272674 | PCDHB16 | -0,298143819 | 0,813298118 | 0,28757806 | 0,798972766 | protocadherin beta 16 |
| ENSG00000248161 | AC098487,1 | -0,298165058 | 0,813286145 | 0,45828813 | 0,904140862 | novel transcript |
| ENSG00000021461 | CYP3A43 | -0,298220274 | 0,813255019 | 0,84978512 | 0,987036295 | cytochrome P450 family 3 subfamily A member 43 |
| ENSG00000157426 | AASDH | -0,298244792 | 0,813241198 | 0,46998732 | 0,908378389 | aminoadipate-semialdehyde dehydrogenase |
| ENSG00000213188 | YBX1P4 | -0,29828073 | 0,81322094 | 0,77275274 | 0,978310326 | Y-box binding protein 1 pseudogene 4 |
| ENSG00000284640 | AL590434,1 | -0,298387921 | 0,813160521 | 0,7189587 | 0,971869926 | novel transcript |
| ENSG00000267723 | AC011509,2 | -0,29848414 | 0,81310629 | 0,83897677 | 0,985672131 | novel transcript |
| ENSG00000242986 | RPL21P99 | -0,298590725 | 0,81304622 | 0,80565619 | 0,981213877 | ribosomal protein L21 pseudogene 99 |
| ENSG00000236772 | AL034550,1 | -0,298719227 | 0,812973805 | 0,72549348 | 0,97326334 | novel transcript, antisense to NOL4L |
| ENSG00000251079 | BMS1P2 | -0,298762179 | 0,812949602 | 0,26325257 | 0,774637968 | BMS1 pseudogene 2 |

| | | | | | | |
|---|---|---|---|---|---|---|
| ENSG00000286242 | AC018797,3 | -0,298796063 | 0,812930508 | 0,39505506 | 0,871981986 | novel transcript, antisense to UBE2D3 |
| ENSG00000278915 | AC005609,3 | -0,298807795 | 0,812923898 | 0,64712045 | 0,956899963 | novel transcript, antisense to PCDHA12 |
| ENSG00000183161 | FANCF | -0,298827444 | 0,812912826 | 0,32859896 | 0,830659639 | FA complementation group F |
| ENSG00000179387 | ELMOD2 | -0,29889648 | 0,812873927 | 0,56442537 | 0,940865625 | ELMO domain containing 2 |
| ENSG00000241644 | INMT | -0,298917551 | 0,812862055 | 0,64625019 | 0,956899963 | indolethylamine N-methyltransferase |
| ENSG00000106443 | PHF14 | -0,298957254 | 0,812839685 | 0,32894551 | 0,830774096 | PHD finger protein 14 |
| ENSG00000270181 | BIVM-ERCC5 | -0,298968627 | 0,812833278 | 0,86414064 | 0,989074806 | BIVM-ERCC5 readthrough |
| ENSG00000146281 | PM20D2 | -0,298990244 | 0,812821099 | 0,40965767 | 0,882177108 | peptidase M20 domain containing 2 |
| ENSG00000203527 | SCUBE1-AS1 | -0,299046259 | 0,81278954 | 0,70414998 | 0,969077997 | SCUBE1 antisense RNA 1 |
| ENSG00000247137 | AP000873,2 | -0,299084618 | 0,81276793 | 0,35407111 | 0,845465446 | novel transcript |
| ENSG00000269925 | Z98884,2 | -0,299102247 | 0,812757998 | 0,63136198 | 0,953801536 | novel transcript, sense intronic to VAMP3 |
| ENSG00000159055 | MIS18A | -0,299124438 | 0,812745497 | 0,33468762 | 0,834757602 | MIS18 kinetochore protein A |
| ENSG00000158769 | F11R | -0,29912645 | 0,812744363 | 0,46102462 | 0,906029574 | F11 receptor |
| ENSG00000233820 | AL589843,2 | -0,299236177 | 0,812682551 | 0,72471809 | 0,97326334 | makorin, ring finger protein, 2 (MKRN2) pseudogene |
| ENSG00000258311 | AC009779,3 | -0,299346835 | 0,812620218 | 0,26046386 | 0,772692192 | novel protein |
| ENSG00000287300 | AL080275,1 | -0,299480072 | 0,812545174 | 0,69340456 | 0,967340633 | novel transcript |
| ENSG00000274487 | NPEPPSP1 | -0,2994889 | 0,812540202 | 0,33380443 | 0,834152569 | NPEPPS pseudogene 1 |
| ENSG00000196177 | ACADSB | -0,299495879 | 0,812536272 | 0,34603173 | 0,841168721 | acyl-CoA dehydrogenase short/branched chain |
| ENSG00000135164 | DMTF1 | -0,299585539 | 0,812485776 | 0,37683844 | 0,862818448 | cyclin D binding myb like transcription factor 1 |
| ENSG00000271824 | SMIM32 | -0,299586793 | 0,812485069 | 0,68440157 | 0,965415049 | small integral membrane protein 32 |
| ENSG00000159593 | NAE1 | -0,299866997 | 0,812327282 | 0,51874491 | 0,926798149 | NEDD8 activating enzyme E1 subunit 1 |
| ENSG00000224728 | IMPDH1P8 | -0,29990798 | 0,812304206 | 0,60632357 | 0,950927937 | inosine monophosphate dehydrogenase 1 pseudogene 8 |
| ENSG00000154258 | ABCA9 | -0,299909961 | 0,812303091 | 0,55071664 | 0,9363105 | ATP binding cassette subfamily A member 9 |
| ENSG00000141698 | NT5C3B | -0,299926028 | 0,812294044 | 0,24275824 | 0,752892335 | 5'-nucleotidase, cytosolic IIIB |
| ENSG00000189042 | ZNF567 | -0,299947607 | 0,812281895 | 0,46909434 | 0,907940653 | zinc finger protein 567 |
| ENSG00000276480 | MYH11 | -0,300022992 | 0,812239452 | 0,54284353 | 0,933663748 | myosin heavy chain 11 |
| ENSG00000251013 | GAPDHP62 | -0,300088446 | 0,812202602 | 0,7099937 | 0,970482408 | glyceraldehyde 3 phosphate dehydrogenase pseudogene 62 |
| ENSG00000279277 | AC012020,1 | -0,300131782 | 0,812178205 | 0,55638306 | 0,938061341 | novel transcript |
| ENSG00000233203 | DHCR24-DT | -0,300163597 | 0,812160295 | 0,81602882 | 0,981708551 | DHCR24 divergent transcript |
| ENSG00000153832 | FBXO36 | -0,300171834 | 0,812155658 | 0,31134324 | 0,817751312 | F-box protein 36 |
| ENSG00000228327 | AL669831,1 | -0,300181599 | 0,812150161 | 0,27458065 | 0,786383635 | general transcription factor IIi (GTF2I) pseudogene |

| | | | | | | |
|---|---|---|---|---|---|---|
| ENSG00000257556 | LINC02298 | -0,300214223 | 0,812131796 | 0,75033009 | 0,975901808 | long intergenic non-protein coding RNA 2298 |
| ENSG00000260265 | LINC02562 | -0,300231557 | 0,812122038 | 0,75642265 | 0,975901808 | long intergenic non-protein coding RNA 2562 |
| ENSG00000257261 | AC008014,1 | -0,300237764 | 0,812118544 | 0,42345096 | 0,890929178 | novel transcript |
| ENSG00000287600 | AC122719,3 | -0,300330461 | 0,812066365 | 0,45877682 | 0,904420115 | novel transcript, antisense to TRIP13 |
| ENSG00000187902 | SHISA7 | -0,300382726 | 0,812036946 | 0,53074555 | 0,931265667 | shisa family member 7 |
| ENSG00000235958 | UBOX5-AS1 | -0,300398496 | 0,81202807 | 0,69224057 | 0,967236299 | UBOX5 antisense RNA 1 |
| ENSG00000286941 | AC244250,4 | -0,300398786 | 0,812027907 | 0,68884946 | 0,966556275 | novel transcript |
| ENSG00000261652 | C15orf65 | -0,300442633 | 0,812003228 | 0,63192584 | 0,953915351 | chromosome 15 open reading frame 65 |
| ENSG00000196705 | ZNF431 | -0,300478731 | 0,81198291 | 0,15370743 | 0,622713345 | zinc finger protein 431 |
| ENSG00000229414 | KCNQ1-AS1 | -0,300586945 | 0,811922008 | 0,77623924 | 0,978587255 | KCNQ1 antisense RNA 1 |
| ENSG00000226589 | VARS1 | -0,300622359 | 0,811902077 | 0,59339812 | 0,947416795 | valyl-tRNA synthetase 1 |
| ENSG00000278206 | AL031320,2 | -0,300629491 | 0,811898064 | 0,51067313 | 0,922693987 | novel transcript |
| ENSG00000121621 | KIF18A | -0,300793493 | 0,811805774 | 0,50146768 | 0,920520138 | kinesin family member 18A |
| ENSG00000032742 | IFT88 | -0,30088108 | 0,81175649 | 0,45910025 | 0,90476838 | intraflagellar transport 88 |
| ENSG00000115137 | DNAJC27 | -0,300898875 | 0,811746478 | 0,4312219 | 0,89530663 | DnaJ heat shock protein family (Hsp40) member C27 |
| ENSG00000172183 | ISG20 | -0,300935461 | 0,811725893 | 0,57194136 | 0,942475308 | interferon stimulated exonuclease gene 20 |
| ENSG00000283293 | RN7SK | -0,300945336 | 0,811720337 | 0,33844445 | 0,837263082 | RNA component of 7SK nuclear ribonucleoprotein |
| ENSG00000275894 | AL021578,1 | -0,30095175 | 0,811716728 | 0,68920834 | 0,966685038 | novel transcript, sense intronic to SDC4 |
| ENSG00000222112 | RN7SKP16 | -0,301012607 | 0,811682488 | 0,53809124 | 0,932900071 | RN7SK pseudogene 16 |
| ENSG00000171159 | C9orf16 | -0,301088157 | 0,811639984 | 0,30898692 | 0,816845699 | chromosome 9 open reading frame 16 |
| ENSG00000273061 | CDC37L1-DT | -0,301201529 | 0,811576204 | 0,70828593 | 0,97011892 | CDC37L1 divergent transcript |
| ENSG00000147162 | OGT | -0,301226891 | 0,811561937 | 0,44854476 | 0,902062594 | O-linked N-acetylglucosamine (GlcNAc) transferase |
| ENSG00000271040 | AL390955,2 | -0,301376435 | 0,811477818 | 0,7656681 | 0,976862445 | novel transcript |
| ENSG00000243368 | MCCC1-AS1 | -0,301424399 | 0,811450841 | 0,42991666 | 0,894689112 | MCCC1 antisense RNA 1 |
| ENSG00000228463 | AP006222,1 | -0,30149946 | 0,811408623 | 0,32944662 | 0,831334931 | ribosomal protein L23a (RPL23A) pseudogene |
| ENSG00000230510 | PPP5D1 | -0,301529178 | 0,811391909 | 0,40124566 | 0,876659568 | PPP5 tetratricopeptide repeat domain containing 1 |
| ENSG00000234233 | KCNH1-IT1 | -0,301538092 | 0,811386896 | 0,82393999 | 0,983374097 | KCNH1 intronic transcript 1 |
| ENSG00000282091 | AC139256,1 | -0,301556473 | 0,811376558 | 0,24000978 | 0,749843134 | phospholipase A2 group X (PLA2G10) pseudogene |
| ENSG00000286072 | AC002072,1 | -0,301562752 | 0,811373027 | 0,28861657 | 0,79998466 | novel transcript |
| ENSG00000286223 | AC064836,3 | -0,301693937 | 0,811299252 | 0,56832138 | 0,94163744 | novel transcript, antisense to NOP58 |
| ENSG00000280906 | AC225099,1 | -0,301706882 | 0,811291972 | 0,61024349 | 0,951841373 | novel transcript |
| ENSG00000280739 | EIF1B-AS1 | -0,301816575 | 0,811230289 | 0,22223742 | 0,730141952 | EIF1B antisense RNA 1 |
| ENSG00000176422 | SPRYD4 | -0,301907402 | 0,811179219 | 0,17681153 | 0,662651108 | SPRY domain containing 4 |

| | | | | | | |
|---|---|---|---|---|---|---|
| ENSG00000165084 | C8orf34 | -0,301959617 | 0,811149861 | 0,53314725 | 0,931291085 | chromosome 8 open reading frame 34 |
| ENSG00000270799 | AC013467,1 | -0,301995633 | 0,811129611 | 0,65167959 | 0,957089277 | Y box binding protein 1 (YBX1) pseudogene |
| ENSG00000162407 | PLPP3 | -0,302046141 | 0,811101214 | 0,36253706 | 0,852656173 | phospholipid phosphatase 3 |
| ENSG00000096171 | VARS1 | -0,302081186 | 0,811081512 | 0,57499 | 0,942559111 | valyl-tRNA synthetase 1 |
| ENSG00000267323 | SLC25A1P5 | -0,302086591 | 0,811078473 | 0,83162061 | 0,984456953 | solute carrier family 25 member 1 pseudogene 5 |
| ENSG00000166860 | ZBTB39 | -0,302190015 | 0,81102033 | 0,30850305 | 0,816730513 | zinc finger and BTB domain containing 39 |
| ENSG00000246982 | Z84485,1 | -0,302229589 | 0,810998084 | 0,53216122 | 0,931291085 | novel transcript, antisense to BRPF3 |
| ENSG00000243323 | PTPRVP | -0,302328717 | 0,810942362 | 0,71569368 | 0,971282169 | protein tyrosine phosphatase receptor type V, pseudogene |
| ENSG00000254884 | PRR13P2 | -0,302333663 | 0,810939582 | 0,77473926 | 0,978553937 | proline rich 13 pseudogene 2 |
| ENSG00000227227 | AC017101,1 | -0,302472823 | 0,810861364 | 0,71058886 | 0,970482408 | novel transcript |
| ENSG00000232564 | MRTFA-AS1 | -0,302505285 | 0,810843119 | 0,61194784 | 0,952187506 | MRTFA antisense RNA 1 |
| ENSG00000204805 | AL391987,2 | -0,30258038 | 0,810800914 | 0,71093419 | 0,970482408 | |
| ENSG00000175198 | PCCA | -0,302597789 | 0,81079113 | 0,3127095 | 0,818683344 | propionyl-CoA carboxylase subunit alpha |
| ENSG00000027869 | SH2D2A | -0,302706205 | 0,810730203 | 0,73952486 | 0,974715504 | SH2 domain containing 2A |
| ENSG00000196967 | ZNF585A | -0,302759173 | 0,810700438 | 0,41982972 | 0,88836254 | zinc finger protein 585A |
| ENSG00000066279 | ASPM | -0,302785485 | 0,810685652 | 0,50861425 | 0,922427526 | assembly factor for spindle microtubules |
| ENSG00000091181 | IL5RA | -0,302835244 | 0,810657692 | 0,76957967 | 0,977683747 | interleukin 5 receptor subunit alpha |
| ENSG00000168939 | SPRY3 | -0,302842013 | 0,810653888 | 0,31928096 | 0,824211281 | sprouty RTK signaling antagonist 3 |
| ENSG00000271639 | PPFIA1P1 | -0,302943546 | 0,810596839 | 0,66182601 | 0,959707806 | PPFIA1 pseudogene 1 |
| ENSG00000066117 | SMARCD1 | -0,302946477 | 0,810595192 | 0,19544731 | 0,693661566 | SWI/SNF related, matrix associated, actin dependent regulator of chromatin, subfamily d, member 1 |
| ENSG00000182103 | FAM181B | -0,303044695 | 0,810540009 | 0,32734467 | 0,829276688 | family with sequence similarity 181 member B |
| ENSG00000255435 | AP001267,3 | -0,303154509 | 0,810478315 | 0,44400988 | 0,900210555 | novel transcript, antisense to MLL and TTC36 |
| ENSG00000230266 | XXYLT1-AS2 | -0,303301248 | 0,810395884 | 0,7002591 | 0,968522643 | XXYLT1 antisense RNA 2 |
| ENSG00000277224 | H2BC7 | -0,303336019 | 0,810376353 | 0,46861777 | 0,907940653 | H2B clustered histone 7 |
| ENSG00000203791 | EEF1AKMT2 | -0,30336249 | 0,810361484 | 0,38197794 | 0,866514369 | EEF1A lysine methyltransferase 2 |
| ENSG00000277938 | AL035252,3 | -0,303364545 | 0,810360329 | 0,46318441 | 0,906932891 | novel transcript |
| ENSG00000273786 | AC020658,4 | -0,303417076 | 0,810330823 | 0,62621565 | 0,952793934 | novel transcript, sense intronic to novel protein |
| ENSG00000219294 | PIP5K1P1 | -0,303430202 | 0,810323451 | 0,75912217 | 0,976120207 | phosphatidylinositol-4-phosphate 5-kinase type 1 pseudogene 1 |
| ENSG00000173811 | CCDC13-AS1 | -0,303479189 | 0,810295936 | 0,64279796 | 0,956633898 | CCDC13 antisense RNA 1 |

| | | | | | | |
|---|---|---|---|---|---|---|
| ENSG00000176208 | ATAD5 | -0,303497678 | 0,810285552 | 0,44598515 | 0,901252648 | ATPase family AAA domain containing 5 |
| ENSG00000231784 | DBIL5P | -0,303505845 | 0,810280965 | 0,46919887 | 0,907940653 | diazepam binding inhibitor-like 5, pseudogene |
| ENSG00000241991 | AC114982,1 | -0,303638448 | 0,810206493 | 0,79693346 | 0,980161849 | ribosomal protein L31 (RPL31) pseudogene |
| ENSG00000283453 | AC244258,1 | -0,303672127 | 0,810187579 | 0,70515242 | 0,969413904 | primase, DNA, polypeptide 2 (58kDa) (PRIM2) pseudogene |
| ENSG00000198286 | CARD11 | -0,303737592 | 0,810150817 | 0,63017209 | 0,95306213 | caspase recruitment domain family member 11 |
| ENSG00000127184 | COX7C | -0,303750958 | 0,810143311 | 0,55261719 | 0,936881074 | cytochrome c oxidase subunit 7C |
| ENSG00000100031 | GGT1 | -0,303770202 | 0,810132504 | 0,59046308 | 0,947407528 | gamma-glutamyltransferase 1 |
| ENSG00000198908 | BHLHB9 | -0,303859595 | 0,810082308 | 0,44883673 | 0,902062594 | basic helix-loop-helix family member b9 |
| ENSG00000153922 | CHD1 | -0,303876023 | 0,810073084 | 0,39153594 | 0,870333649 | chromodomain helicase DNA binding protein 1 |
| ENSG00000288390 | UBL7-AS1 | -0,303965521 | 0,810022832 | 0,20220062 | 0,702087984 | UBL7 antisense RNA 1 (head to head) |
| ENSG00000254101 | LINC02055 | -0,303989206 | 0,810009534 | 0,57526461 | 0,942667371 | long intergenic non-protein coding RNA 2055 |
| ENSG00000282183 | GOLGA2P10 | -0,30400757 | 0,809999224 | 0,45796729 | 0,904135472 | GOLGA2 pseudogene 10 |
| ENSG00000227775 | AL031282,1 | -0,304014687 | 0,809995228 | 0,23993444 | 0,749799162 | ribosomal protein S7 (RPS7) pseudogene |
| ENSG00000286330 | AL353660,1 | -0,304063859 | 0,809967621 | 0,66161609 | 0,959582738 | novel transcript |
| ENSG00000273456 | AC064836,2 | -0,30407961 | 0,809958778 | 0,5099193 | 0,922427526 | novel transcript |
| ENSG00000229998 | PPP1R18 | -0,304111667 | 0,80994078 | 0,76055901 | 0,97624106 | protein phosphatase 1 regulatory subunit 18 |
| ENSG00000225060 | PPP1R18 | -0,304111667 | 0,80994078 | 0,76055901 | 0,97624106 | protein phosphatase 1 regulatory subunit 18 |
| ENSG00000286288 | AL109809,4 | -0,304145093 | 0,809922015 | 0,62320715 | 0,952793934 | novel transcript |
| ENSG00000286327 | AL035427,2 | -0,304155355 | 0,809916254 | 0,3361655 | 0,835660292 | novel transcript |
| ENSG00000150433 | TMEM218 | -0,304201522 | 0,809890337 | 0,28905077 | 0,800166929 | transmembrane protein 218 |
| ENSG00000204086 | RPA4 | -0,304280502 | 0,809846001 | 0,70354731 | 0,968967751 | replication protein A4 |
| ENSG00000196302 | GUSBP15 | -0,304444452 | 0,809753974 | 0,3930216 | 0,871139821 | GUSB pseudogene 15 |
| ENSG00000286522 | H3C2 | -0,304462588 | 0,809743794 | 0,29264129 | 0,803119489 | H3 clustered histone 2 |
| ENSG00000204764 | RANBP17 | -0,304486254 | 0,809730511 | 0,46033658 | 0,905553561 | RAN binding protein 17 |
| ENSG00000258947 | TUBB3 | -0,304492768 | 0,809726855 | 0,23135337 | 0,739268552 | tubulin beta 3 class III |
| ENSG00000171448 | ZBTB26 | -0,304538276 | 0,809701314 | 0,5374074 | 0,932883277 | zinc finger and BTB domain containing 26 |
| ENSG00000198563 | DDX39B | -0,304549373 | 0,809695086 | 0,43716298 | 0,897021798 | DExD-box helicase 39B |
| ENSG00000256399 | AC079866,1 | -0,304624036 | 0,809653184 | 0,50977821 | 0,922427526 | CTAGE family, member 5 (CTAGE5) pseudogene |
| ENSG00000283662 | AC138904,3 | -0,304663823 | 0,809630855 | 0,44238273 | 0,89933631 | novel transcript |
| ENSG00000227262 | HCG4B | -0,30472513 | 0,809596451 | 0,86790306 | 0,989074806 | HLA complex group 4B |
| ENSG00000174611 | KY | -0,304761526 | 0,809576026 | 0,52749066 | 0,929463343 | kyphoscoliosis peptidase |
| ENSG00000154997 | SEPTIN14 | -0,304812115 | 0,809547639 | 0,50498852 | 0,921400408 | septin 14 |

| | | | | | | |
|---|---|---|---|---|---|---|
| ENSG00000275226 | LINC00547 | -0,304836067 | 0,809534198 | 0,73196778 | 0,974353474 | long intergenic non-protein coding RNA 547 |
| ENSG00000007341 | ST7L | -0,304911455 | 0,809491898 | 0,3642756 | 0,854302928 | suppression of tumorigenicity 7 like |
| ENSG00000250608 | AC010210,1 | -0,305030294 | 0,80942522 | 0,32232046 | 0,824955371 | novel transcript |
| ENSG00000130475 | FCHO1 | -0,305079053 | 0,809397864 | 0,34082743 | 0,839475584 | FCH and mu domain containing endocytic adaptor 1 |
| ENSG00000228703 | AL355310,2 | -0,305083944 | 0,80939512 | 0,70243774 | 0,968742626 | novel transcript |
| ENSG00000269416 | LINC01224 | -0,305285648 | 0,809281966 | 0,37258085 | 0,860420124 | long intergenic non-protein coding RNA 1224 |
| ENSG00000196670 | ZFP62 | -0,305295365 | 0,809276515 | 0,39177576 | 0,870476676 | ZFP62 zinc finger protein |
| ENSG00000156876 | SASS6 | -0,30531297 | 0,80926664 | 0,55415342 | 0,937655056 | SAS-6 centriolar assembly protein |
| ENSG00000165891 | E2F7 | -0,30538122 | 0,809228357 | 0,28471066 | 0,796099998 | E2F transcription factor 7 |
| ENSG00000287756 | AL031293,1 | -0,305412618 | 0,809210745 | 0,63613709 | 0,955399772 | novel transcript, antisense to KAZN |
| ENSG00000229741 | BTNL2 | -0,305442591 | 0,809193933 | 0,82649022 | 0,983471137 | butyrophilin like 2 |
| ENSG00000229816 | DDX50P1 | -0,305490277 | 0,809167187 | 0,31794144 | 0,822857715 | DEAD-box helicase 50 pseudogene 1 |
| ENSG00000112294 | ALDH5A1 | -0,30553051 | 0,809144622 | 0,34287087 | 0,840419133 | aldehyde dehydrogenase 5 family member A1 |
| ENSG00000038274 | MAT2B | -0,305562337 | 0,809126772 | 0,53779885 | 0,932883277 | methionine adenosyltransferase 2B |
| ENSG00000272894 | AC004982,2 | -0,305590797 | 0,80911081 | 0,35853953 | 0,849814998 | novel transcript |
| ENSG00000140557 | ST8SIA2 | -0,305648086 | 0,809078682 | 0,18306656 | 0,673904498 | ST8 alpha-N-acetyl-neuraminide alpha-2,8-sialyltransferase 2 |
| ENSG00000221944 | TIGD1 | -0,305665236 | 0,809069064 | 0,18434624 | 0,676376113 | tigger transposable element derived 1 |
| ENSG00000234465 | PINLYP | -0,305815093 | 0,808985028 | 0,47725886 | 0,91143003 | phospholipase A2 inhibitor and LY6/PLAUR domain containing |
| ENSG00000169621 | APLF | -0,305851946 | 0,808964362 | 0,41721666 | 0,886651836 | aprataxin and PNKP like factor |
| ENSG00000153789 | CIBAR2 | -0,305893216 | 0,808941221 | 0,63240552 | 0,954224236 | CBY1 interacting BAR domain containing 2 |
| ENSG00000230660 | HCG18 | -0,305925556 | 0,808923089 | 0,55463256 | 0,937793851 | HLA complex group 18 |
| ENSG00000229536 | LINC02572 | -0,306052077 | 0,808852151 | 0,58493708 | 0,944857905 | long intergenic non-protein coding RNA 2572 |
| ENSG00000198671 | AC135895,1 | -0,306118807 | 0,808814739 | 0,8223649 | 0,982964373 | novel transcript, antisense to WIF1 |
| ENSG00000213872 | AC092798,1 | -0,306256682 | 0,808737446 | 0,7081832 | 0,97011892 | ribosomal protein L32 (RPL32) pseudogene |
| ENSG00000188921 | HACD4 | -0,306507222 | 0,808597012 | 0,41423589 | 0,885579089 | 3-hydroxyacyl-CoA dehydratase 4 |
| ENSG00000110931 | CAMKK2 | -0,306604906 | 0,808542265 | 0,16535639 | 0,644399285 | calcium/calmodulin dependent protein kinase kinase 2 |
| ENSG00000196747 | H2AC13 | -0,306607252 | 0,808540949 | 0,23094546 | 0,739268552 | H2A clustered histone 13 |
| ENSG00000260400 | AL513534,2 | -0,306681809 | 0,808499166 | 0,44397748 | 0,900210555 | novel transcript, overlapping to TET1 |
| ENSG00000143162 | CREG1 | -0,306734175 | 0,80846982 | 0,45606144 | 0,903545728 | cellular repressor of E1A stimulated genes 1 |
| ENSG00000100652 | SLC10A1 | -0,306765133 | 0,808452472 | 0,50472346 | 0,921400408 | solute carrier family 10 member 1 |

| | | | | | | |
|---|---|---|---|---|---|---|
| ENSG00000164167 | LSM6 | -0,306848508 | 0,808405752 | 0,45815662 | 0,904140862 | LSM6 homolog, U6 small nuclear RNA and mRNA degradation associated |
| ENSG00000273723 | SUGT1-DT | -0,306999819 | 0,80832097 | 0,64701778 | 0,956899963 | SUGT1 divergent transcript |
| ENSG00000257243 | AC020612,1 | -0,307000256 | 0,808320725 | 0,67172444 | 0,962161102 | Protein DJ-1 (PARK7) pseudogene |
| ENSG00000251301 | LINC02384 | -0,30702988 | 0,808304128 | 0,66247836 | 0,959897678 | long intergenic non-protein coding RNA 2384 |
| ENSG00000040608 | RTN4R | -0,307099367 | 0,808265197 | 0,27419908 | 0,7862019 | reticulon 4 receptor |
| ENSG00000085719 | CPNE3 | -0,307109459 | 0,808259543 | 0,50606355 | 0,921780819 | copine 3 |
| ENSG00000220323 | H2BC19P | -0,30716281 | 0,808229654 | 0,52260131 | 0,927855489 | H2B clustered histone 19, pseudogene |
| ENSG00000179008 | C14orf39 | -0,30716656 | 0,808227553 | 0,66013637 | 0,958904584 | chromosome 14 open reading frame 39 |
| ENSG00000258274 | AC012085,2 | -0,307194912 | 0,80821167 | 0,62009317 | 0,952793934 | novel transcript |
| ENSG00000246777 | AC044802,1 | -0,307207601 | 0,808204561 | 0,59285744 | 0,947416795 | novel transcript |
| ENSG00000127980 | PEX1 | -0,307231277 | 0,808191298 | 0,42422543 | 0,890949676 | peroxisomal biogenesis factor 1 |
| ENSG00000172244 | C5orf34 | -0,307244202 | 0,808184057 | 0,48424418 | 0,913690171 | chromosome 5 open reading frame 34 |
| ENSG00000175564 | UCP3 | -0,307265246 | 0,808172269 | 0,38766192 | 0,869820053 | uncoupling protein 3 |
| ENSG00000271870 | AC024060,2 | -0,307301501 | 0,80815196 | 0,66259838 | 0,959897678 | novel transcript, antisense to CRBN |
| ENSG00000245556 | SCAMP1-AS1 | -0,307454419 | 0,808066305 | 0,31703135 | 0,82206753 | SCAMP1 antisense RNA 1 |
| ENSG00000179935 | LINC00652 | -0,307473802 | 0,808055448 | 0,31545804 | 0,82082095 | long intergenic non-protein coding RNA 652 |
| ENSG00000158691 | ZSCAN12 | -0,307498968 | 0,808041352 | 0,34496598 | 0,841168721 | zinc finger and SCAN domain containing 12 |
| ENSG00000163743 | RCHY1 | -0,307557654 | 0,808008484 | 0,56650202 | 0,941311563 | ring finger and CHY zinc finger domain containing 1 |
| ENSG00000168676 | KCTD19 | -0,307621306 | 0,807972835 | 0,65262113 | 0,957275891 | potassium channel tetramerization domain containing 19 |
| ENSG00000121413 | ZSCAN18 | -0,307655934 | 0,807953442 | 0,38354644 | 0,868000076 | zinc finger and SCAN domain containing 18 |
| ENSG00000170161 | AL512625,1 | -0,307680268 | 0,807939815 | 0,43026613 | 0,895058897 | uncharacterized LOC554249 |
| ENSG00000170044 | ZPLD1 | -0,307689405 | 0,807934697 | 0,749906 | 0,975901808 | zona pellucida like domain containing 1 |
| ENSG00000288446 | SNX18P23 | -0,307739849 | 0,807906449 | 0,78221243 | 0,978597905 | sorting nexin 18 pseudogene 23 |
| ENSG00000250016 | SNX18P23 | -0,307739849 | 0,807906449 | 0,78221243 | 0,978597905 | sorting nexin 18 pseudogene 23 |
| ENSG00000277985 | SNORA67 | -0,307747458 | 0,807902187 | 0,51904595 | 0,926800715 | small nucleolar RNA, H/ACA box 67 |
| ENSG00000109684 | CLNK | -0,307824034 | 0,807859306 | 0,71761682 | 0,971697135 | cytokine dependent hematopoietic cell linker |
| ENSG00000257666 | CBX3P5 | -0,307916551 | 0,807807502 | 0,67688742 | 0,963580269 | chromobox 3 pseudogene 5 |
| ENSG00000256087 | ZNF432 | -0,307943032 | 0,807792674 | 0,27883488 | 0,789775666 | zinc finger protein 432 |
| ENSG00000224786 | CETN4P | -0,308014547 | 0,807752633 | 0,50530914 | 0,921715363 | centrin 4, pseudogene |
| ENSG00000266916 | ZNF793-AS1 | -0,30807916 | 0,807716457 | 0,5191211 | 0,926800715 | ZNF793 antisense RNA 1 (head to head) |
| ENSG00000176658 | MYO1D | -0,308087637 | 0,807711711 | 0,29488837 | 0,805040011 | myosin ID |

| | | | | | | |
|---|---|---|---|---|---|---|
| ENSG00000259083 | AL132639,3 | -0,308093695 | 0,80770832 | 0,80366883 | 0,980875302 | novel transcript, antisense to PNN |
| ENSG00000248243 | LINC02014 | -0,308206065 | 0,807645411 | 0,6464069 | 0,956899963 | long intergenic non-protein coding RNA 2014 |
| ENSG00000284959 | AC007262,2 | -0,30824999 | 0,807620821 | 0,25777455 | 0,7701554 | novel transcript, antisense to TSHR |
| ENSG00000281354 | AP000350,15 | -0,308284041 | 0,807601759 | 0,78407996 | 0,97928555 | novel transcript, antisense to SLC2A11 |
| ENSG00000272973 | AP000350,5 | -0,308284041 | 0,807601759 | 0,78407996 | 0,97928555 | novel transcript, antisense to SLC2A11 |
| ENSG00000270147 | AC068620,2 | -0,308288964 | 0,807599004 | 0,67251942 | 0,962290553 | novel transcript |
| ENSG00000101204 | CHRNA4 | -0,308417228 | 0,807527207 | 0,4699374 | 0,908378389 | cholinergic receptor nicotinic alpha 4 subunit |
| ENSG00000225828 | FAM229A | -0,308420826 | 0,807525193 | 0,29801119 | 0,807367576 | family with sequence similarity 229 member A |
| ENSG00000259191 | AC027237,1 | -0,3085207 | 0,807469292 | 0,67077599 | 0,962026636 | fumarylacetoacetate hydrolase domain containing 1 (FAHD1) pseudogene |
| ENSG00000287182 | AC131953,2 | -0,308525944 | 0,807466356 | 0,7223216 | 0,972594421 | novel transcript, antisense to NSUN7 |
| ENSG00000239653 | PSMD6-AS2 | -0,308644972 | 0,80739974 | 0,38769732 | 0,869820053 | PSMD6 antisense RNA 2 |
| ENSG00000117122 | MFAP2 | -0,308681738 | 0,807379165 | 0,2313331 | 0,739268552 | microfibril associated protein 2 |
| ENSG00000283031 | AC009242,1 | -0,308873344 | 0,807271943 | 0,58594361 | 0,945246954 | novel transcript |
| ENSG00000285967 | NIPBL-DT | -0,308886996 | 0,807264304 | 0,42470879 | 0,891326808 | NIPBL divergent transcript |
| ENSG00000010319 | SEMA3G | -0,309013914 | 0,807193289 | 0,72484201 | 0,97326334 | semaphorin 3G |
| ENSG00000259347 | AC087482,1 | -0,309083932 | 0,807154115 | 0,55156999 | 0,936366211 | novel transcript, antisense to SMAD3 |
| ENSG00000176761 | ZNF285B | -0,309159693 | 0,80711173 | 0,60324066 | 0,950352286 | zinc finger protein 285B (pseudogene) |
| ENSG00000260922 | AC009139,1 | -0,309170456 | 0,807105708 | 0,74884395 | 0,975901808 | novel transcript, antisense to ADAMTS18 |
| ENSG00000075407 | ZNF37A | -0,309230124 | 0,807072328 | 0,42828493 | 0,893037474 | zinc finger protein 37A |
| ENSG00000088451 | TGDS | -0,309241817 | 0,807065787 | 0,51821804 | 0,926413949 | TDP-glucose 4,6-dehydratase |
| ENSG00000064309 | CDON | -0,309242291 | 0,807065522 | 0,54598879 | 0,935201756 | cell adhesion associated, oncogene regulated |
| ENSG00000273362 | AP000547,2 | -0,309264258 | 0,807053233 | 0,72357517 | 0,972941126 | novel transcript |
| ENSG00000228857 | AC104653,1 | -0,309298724 | 0,807033953 | 0,73841229 | 0,974353474 | novel transcript |
| ENSG00000285344 | TRIM17 | -0,309310094 | 0,807027593 | 0,47747654 | 0,911574108 | tripartite motif containing 17 |
| ENSG00000162931 | TRIM17 | -0,309310094 | 0,807027593 | 0,47747654 | 0,911574108 | tripartite motif containing 17 |
| ENSG00000271147 | ARMCX5-GPRA | -0,309313251 | 0,807025826 | 0,43476241 | 0,896272827 | ARMCX5-GPRASP2 readthrough |
| ENSG00000134480 | CCNH | -0,309333992 | 0,807014225 | 0,44551916 | 0,901068127 | cyclin H |
| ENSG00000087903 | RFX2 | -0,309386381 | 0,80698492 | 0,18845731 | 0,682805588 | regulatory factor X2 |
| ENSG00000164758 | MED30 | -0,309396163 | 0,806979448 | 0,41325764 | 0,884678298 | mediator complex subunit 30 |
| ENSG00000236670 | KRT18P5 | -0,309428461 | 0,806961382 | 0,54060248 | 0,933385624 | keratin 18 pseudogene 5 |
| ENSG00000266602 | AP005262,1 | -0,30944445 | 0,806952439 | 0,2532059 | 0,765172364 | novel transcript |
| ENSG00000109046 | WSB1 | -0,309468555 | 0,806938956 | 0,41760221 | 0,886857227 | WD repeat and SOCS box containing 1 |

| | | | | | | |
|---|---|---|---|---|---|---|
| ENSG00000235651 | G3BP1P1 | -0,309593647 | 0,806868992 | 0,42952818 | 0,894365375 | G3BP1 pseudogene 1 |
| ENSG00000100578 | KIAA0586 | -0,309615162 | 0,806856959 | 0,32475858 | 0,826793081 | KIAA0586 |
| ENSG00000175344 | CHRNA7 | -0,309665794 | 0,806828643 | 0,32072901 | 0,824366234 | cholinergic receptor nicotinic alpha 7 subunit |
| ENSG00000261618 | LINC02605 | -0,30971468 | 0,806801304 | 0,49242547 | 0,917008021 | long intergenic non-protein coding RNA 2605 |
| ENSG00000142235 | LMTK3 | -0,309951901 | 0,806668653 | 0,46107122 | 0,906029574 | lemur tyrosine kinase 3 |
| ENSG00000204049 | AL391421,1 | -0,309952093 | 0,806668546 | 0,54192808 | 0,933491609 | novel transcript, antisense to DLG5 |
| ENSG00000282690 | AL391421,2 | -0,309952093 | 0,806668546 | 0,54192808 | 0,933491609 | novel transcript, antisense to DLG5 |
| ENSG00000089250 | NOS1 | -0,310003351 | 0,806639886 | 0,60982304 | 0,951801946 | nitric oxide synthase 1 |
| ENSG00000106341 | PPP1R17 | -0,310020984 | 0,806630027 | 0,61710291 | 0,952793934 | protein phosphatase 1 regulatory subunit 17 |
| ENSG00000206120 | EGFEM1P | -0,310080362 | 0,806596828 | 0,42376541 | 0,890949676 | EGF like and EMI domain containing 1, pseudogene |
| ENSG00000281661 | ZNF501 | -0,310222652 | 0,806517279 | 0,52192265 | 0,927599011 | zinc finger protein 501 |
| ENSG00000163002 | NUP35 | -0,310392861 | 0,806422132 | 0,52680049 | 0,92927181 | nucleoporin 35 |
| ENSG00000272870 | SAP30-DT | -0,31040978 | 0,806412675 | 0,28426361 | 0,795731744 | SAP30 divergent transcript |
| ENSG00000249286 | AMD1P3 | -0,310416879 | 0,806408707 | 0,60920178 | 0,951793902 | adenosylmethionine decarboxylase 1 pseudogene 3 |
| ENSG00000156968 | MPV17L | -0,310460203 | 0,806384491 | 0,27489911 | 0,786383635 | MPV17 mitochondrial inner membrane protein like |
| ENSG00000134245 | WNT2B | -0,310612709 | 0,806299253 | 0,25238547 | 0,76432863 | Wnt family member 2B |
| ENSG00000232882 | PHKA1P1 | -0,310634273 | 0,806287202 | 0,46985673 | 0,908378389 | phosphorylase kinase regulatory subunit alpha 1 pseudogene |
| ENSG00000282899 | E2F2 | -0,310665309 | 0,806269856 | 0,81739922 | 0,98215965 | E2F transcription factor 2 |
| ENSG00000287619 | AL021308,1 | -0,310715551 | 0,806241779 | 0,57738523 | 0,94337651 | novel transcript |
| ENSG00000271849 | MAN2A1-DT | -0,310806479 | 0,806190966 | 0,76266705 | 0,97639149 | MAN2A1 divergent transcript |
| ENSG00000171649 | ZIK1 | -0,310833012 | 0,806176139 | 0,29714865 | 0,806635068 | zinc finger protein interacting with K protein 1 |
| ENSG00000267199 | AP001029,2 | -0,31083885 | 0,806172876 | 0,35128648 | 0,84405766 | novel transcript, antisense SPIRE1 |
| ENSG00000168412 | MTNR1A | -0,310846498 | 0,806168603 | 0,68884859 | 0,966556275 | melatonin receptor 1A |
| ENSG00000254088 | SLC2A3P4 | -0,310852593 | 0,806165197 | 0,58730182 | 0,945938196 | solute carrier family 2 member 3 pseudogene 4 |
| ENSG00000164171 | ITGA2 | -0,310895504 | 0,806141219 | 0,44218155 | 0,899331131 | integrin subunit alpha 2 |
| ENSG00000164603 | BMT2 | -0,31106996 | 0,806043744 | 0,54682796 | 0,935404974 | base methyltransferase of 25S rRNA 2 homolog |
| ENSG00000287078 | AL513218,2 | -0,311086872 | 0,806034295 | 0,73254066 | 0,974353474 | novel transcript, antisense to CC2D1B |
| ENSG00000148090 | AUH | -0,311131119 | 0,806009574 | 0,35225464 | 0,844912682 | AU RNA binding methylglutaconyl-CoA hydratase |
| ENSG00000235954 | TTC28-AS1 | -0,311147285 | 0,806000543 | 0,14208913 | 0,600058555 | TTC28 antisense RNA 1 |
| ENSG00000230513 | THAP7-AS1 | -0,31127677 | 0,805928205 | 0,2842687 | 0,795731744 | THAP7 antisense RNA 1 |
| ENSG00000273807 | AC244489,1 | -0,311383929 | 0,805868346 | 0,26845885 | 0,780709773 | novel member of the nuclear pore complex interacting protein NPIP gene family |

| | | | | | | |
|---|---|---|---|---|---|---|
| ENSG00000243144 | AC079760,2 | -0,311386512 | 0,805866903 | 0,73742985 | 0,974353474 | novel transcript |
| ENSG00000178531 | CTXN1 | -0,311508077 | 0,805799002 | 0,48724381 | 0,914686581 | cortexin 1 |
| ENSG00000167363 | FN3K | -0,311531102 | 0,805786141 | 0,3207255 | 0,824366234 | fructosamine 3 kinase |
| ENSG00000285848 | AC025280,3 | -0,311597554 | 0,805749027 | 0,62655429 | 0,952793934 | novel transcript, antisense to CRISPLD2 |
| ENSG00000180610 | ZBTB12BP | -0,311655599 | 0,805716609 | 0,73765324 | 0,974353474 | zinc finger and BTB domain containing 12B, pseudogene |
| ENSG00000083535 | PIBF1 | -0,311738192 | 0,805670484 | 0,44802001 | 0,902062594 | progesterone immunomodulatory binding factor 1 |
| ENSG00000152240 | HAUS1 | -0,311738467 | 0,80567033 | 0,47799483 | 0,911886158 | HAUS augmin like complex subunit 1 |
| ENSG00000118690 | ARMC2 | -0,311829809 | 0,805619322 | 0,45164089 | 0,902062594 | armadillo repeat containing 2 |
| ENSG00000138759 | FRAS1 | -0,311843256 | 0,805611813 | 0,25113921 | 0,763478818 | Fraser extracellular matrix complex subunit 1 |
| ENSG00000180667 | YOD1 | -0,311877262 | 0,805592824 | 0,55788218 | 0,938984031 | YOD1 deubiquitinase |
| ENSG00000167766 | ZNF83 | -0,311897377 | 0,805581592 | 0,35661545 | 0,84797881 | zinc finger protein 83 |
| ENSG00000183476 | SH2D7 | -0,311897692 | 0,805581416 | 0,40037451 | 0,876132752 | SH2 domain containing 7 |
| ENSG00000111247 | RAD51AP1 | -0,312127011 | 0,805453378 | 0,46519423 | 0,907804479 | RAD51 associated protein 1 |
| ENSG00000249709 | ZNF564 | -0,312219148 | 0,80540194 | 0,43604177 | 0,896526626 | zinc finger protein 564 |
| ENSG00000141425 | RPRD1A | -0,312317821 | 0,805346856 | 0,48329598 | 0,913690171 | regulation of nuclear pre-mRNA domain containing 1A |
| ENSG00000093072 | ADA2 | -0,312415916 | 0,805292099 | 0,20758945 | 0,71186537 | adenosine deaminase 2 |
| ENSG00000259408 | AC010809,2 | -0,312464331 | 0,805265075 | 0,49252054 | 0,917016235 | novel transcript, antisense to RYR3 |
| ENSG00000202354 | RNY3 | -0,312483994 | 0,8052541 | 0,45385831 | 0,902912642 | RNA, Ro60-associated Y3 |
| ENSG00000241532 | AGGF1P3 | -0,312524778 | 0,805231336 | 0,72903233 | 0,974009551 | angiogenic factor with G-patch and FHA domains 1 pseudogene 3 |
| ENSG00000176542 | USF3 | -0,312664562 | 0,80515332 | 0,42035633 | 0,888563911 | upstream transcription factor family member 3 |
| ENSG00000225871 | AC245100,2 | -0,312670114 | 0,805150222 | 0,48791556 | 0,915157336 | family with sequence similarity 91, member A1 (FAM91A1) pseudogene |
| ENSG00000169258 | GPRIN1 | -0,31270168 | 0,805132605 | 0,41738396 | 0,886750438 | G protein regulated inducer of neurite outgrowth 1 |
| ENSG00000196312 | MFSD14C | -0,312767261 | 0,805096007 | 0,18529528 | 0,677679985 | major facilitator superfamily domain containing 14C |
| ENSG00000232044 | SILC1 | -0,31278984 | 0,805083407 | 0,31274102 | 0,818683344 | sciatic injury induced lincRNA upregulator of SOX11 |
| ENSG00000006453 | BAIAP2L1 | -0,312884833 | 0,805030399 | 0,49888115 | 0,919683359 | BAR/IMD domain containing adaptor protein 2 like 1 |
| ENSG00000237118 | CYP2F2P | -0,312952438 | 0,804992676 | 0,78489933 | 0,979320406 | cytochrome P450 family 2 subfamily F member 2, pseudogene |
| ENSG00000260083 | MIR762HG | -0,312984173 | 0,804974969 | 0,30443826 | 0,813181541 | MIR762 host gene |

| | | | | | | |
|---|---|---|---|---|---|---|
| ENSG00000249855 | EEF1A1P19 | -0,313016141 | 0,804957132 | 0,41774356 | 0,886857227 | eukaryotic translation elongation factor 1 alpha 1 pseudogene 19 |
| ENSG00000105697 | HAMP | -0,313139058 | 0,804888552 | 0,73506847 | 0,974353474 | hepcidin antimicrobial peptide |
| ENSG00000136492 | BRIP1 | -0,313193849 | 0,804857985 | 0,22922907 | 0,73730633 | BRCA1 interacting protein C-terminal helicase 1 |
| ENSG00000174963 | ZIC4 | -0,313305815 | 0,804795523 | 0,44969598 | 0,902062594 | Zic family member 4 |
| ENSG00000076258 | FMO4 | -0,313455091 | 0,804712255 | 0,5528481 | 0,936894267 | flavin containing dimethylaniline monoxygenase 4 |
| ENSG00000159259 | CHAF1B | -0,313491816 | 0,804691771 | 0,19452116 | 0,691630774 | chromatin assembly factor 1 subunit B |
| ENSG00000277972 | CISD3 | -0,31353084 | 0,804670005 | 0,51929148 | 0,926800715 | CDGSH iron sulfur domain 3 |
| ENSG00000285849 | AL021918,4 | -0,313551222 | 0,804658636 | 0,55558526 | 0,937793851 | novel transcript, antisense to ZNF184 |
| ENSG00000196597 | ZNF782 | -0,313573419 | 0,804646256 | 0,41006495 | 0,882513285 | zinc finger protein 782 |
| ENSG00000198416 | ZNF658B | -0,31361448 | 0,804623355 | 0,37554439 | 0,86204256 | zinc finger protein 658B (pseudogene) |
| ENSG00000233189 | RPL12P29 | -0,313674999 | 0,804589603 | 0,713011 | 0,970812834 | ribosomal protein L12 pseudogene 29 |
| ENSG00000168010 | ATG16L2 | -0,313696639 | 0,804577535 | 0,37495474 | 0,861771195 | autophagy related 16 like 2 |
| ENSG00000151023 | ENKUR | -0,31379908 | 0,804520406 | 0,45090495 | 0,902062594 | enkurin, TRPC channel interacting protein |
| ENSG00000118579 | MED28 | -0,313849677 | 0,804492192 | 0,32053879 | 0,824366234 | mediator complex subunit 28 |
| ENSG00000183496 | MEX3B | -0,31385325 | 0,804490199 | 0,16583939 | 0,645116689 | mex-3 RNA binding family member B |
| ENSG00000267172 | AC022031,1 | -0,314013459 | 0,804400866 | 0,77861766 | 0,978587255 | novel transcript |
| ENSG00000231365 | WARS2-AS1 | -0,314070449 | 0,804369091 | 0,19349657 | 0,690424879 | WARS2 antisense RNA 1 |
| ENSG00000100116 | GCAT | -0,314073847 | 0,804367197 | 0,29171133 | 0,802285739 | glycine C-acetyltransferase |
| ENSG00000233783 | AP001442,1 | -0,314080265 | 0,804363619 | 0,45196135 | 0,902062594 | novel transcript |
| ENSG00000237310 | GS1-124K5,4 | -0,314137317 | 0,80433181 | 0,42184977 | 0,889720796 | uncharacterized LOC100289098 |
| ENSG00000260141 | AC145350,2 | -0,31415728 | 0,80432068 | 0,41773096 | 0,886857227 | novel transcript |
| ENSG00000223519 | KIF28P | -0,314229649 | 0,804280335 | 0,7689114 | 0,977403783 | kinesin family member 28, pseudogene |
| ENSG00000244371 | PFN1P8 | -0,314237591 | 0,804275908 | 0,43735432 | 0,897021798 | profilin 1 pseudogene 8 |
| ENSG00000105672 | ETV2 | -0,314243304 | 0,804272723 | 0,49215011 | 0,916995341 | ETS variant transcription factor 2 |
| ENSG00000230782 | AL807761,4 | -0,314366636 | 0,80420397 | 0,42824818 | 0,893037474 | novel transcript |
| ENSG00000239246 | AC008026,1 | -0,314407159 | 0,804181382 | 0,75463364 | 0,975901808 | ribosomal protein S10 (RPS10) pseudogene |
| ENSG00000204991 | SPIRE2 | -0,314491433 | 0,804134408 | 0,44136378 | 0,898885877 | spire type actin nucleation factor 2 |
| ENSG00000109674 | NEIL3 | -0,31451839 | 0,804119382 | 0,42984859 | 0,894607 | nei like DNA glycosylase 3 |
| ENSG00000174136 | RGMB | -0,314552618 | 0,804100305 | 0,16412323 | 0,642888389 | repulsive guidance molecule BMP co-receptor b |
| ENSG00000124839 | RAB17 | -0,314642705 | 0,804050095 | 0,7602473 | 0,97624106 | RAB17, member RAS oncogene family |
| ENSG00000272667 | AC012306,2 | -0,3146607 | 0,804040067 | 0,44910112 | 0,902062594 | novel transcript |
| ENSG00000166348 | USP54 | -0,314660899 | 0,804039956 | 0,24958122 | 0,761448016 | ubiquitin specific peptidase 54 |

| | | | | | | |
|---|---|---|---|---|---|---|
| ENSG00000265148 | TSPOAP1-AS1 | -0,314738805 | 0,803996539 | 0,39998061 | 0,875830109 | TSPOAP1, SUPT4H1 and RNF43 antisense RNA 1 |
| ENSG00000271576 | AL359504,1 | -0,31477874 | 0,803974283 | 0,54369846 | 0,934259495 | novel transcript |
| ENSG00000261288 | AC093525,5 | -0,314832836 | 0,803944138 | 0,86549651 | 0,989074806 | novel transcript, intronic to PDPK1 |
| ENSG00000260176 | AC141586,2 | -0,314832836 | 0,803944138 | 0,86549651 | 0,989074806 | novel transcript |
| ENSG00000233251 | AC007743,1 | -0,31500087 | 0,803850506 | 0,20932834 | 0,713678247 | novel transcript, antisense to CCDC85A |
| ENSG00000248265 | FLJ12825 | -0,315007724 | 0,803846687 | 0,39316672 | 0,871240371 | uncharacterized LOC440101 |
| ENSG00000113300 | CNOT6 | -0,315033413 | 0,803832374 | 0,42005696 | 0,888457169 | CCR4-NOT transcription complex subunit 6 |
| ENSG00000100228 | RAB36 | -0,315091883 | 0,803799796 | 0,19394271 | 0,690714036 | RAB36, member RAS oncogene family |
| ENSG00000214776 | AC092821,1 | -0,315227784 | 0,803724082 | 0,40409604 | 0,878008106 | alpha-2-macroglobulin like 1 pseudogene |
| ENSG00000103995 | CEP152 | -0,315282376 | 0,80369367 | 0,30973872 | 0,817030292 | centrosomal protein 152 |
| ENSG00000234893 | HCG18 | -0,315313102 | 0,803676553 | 0,46292535 | 0,906849208 | HLA complex group 18 |
| ENSG00000223355 | NOTCH4 | -0,315357434 | 0,803651858 | 0,50673056 | 0,921824694 | notch receptor 4 |
| ENSG00000275343 | AC010999,1 | -0,315449633 | 0,8036005 | 0,82704275 | 0,983567657 | novel transcript, sense intronic to ZNF280D |
| ENSG00000223849 | AL354893,1 | -0,315486564 | 0,803579929 | 0,76283276 | 0,97647393 | ribosomal protein S26 (RPS26) pseudogene |
| ENSG00000223969 | AC002456,1 | -0,315580883 | 0,803527395 | 0,51748102 | 0,925831376 | novel transcript |
| ENSG00000235493 | LINC01967 | -0,315594686 | 0,803519708 | 0,34764366 | 0,842276143 | long intergenic non-protein coding RNA 1967 |
| ENSG00000276255 | LINC02809 | -0,31566628 | 0,803479834 | 0,60187428 | 0,949939978 | long intergenic non-protein coding RNA 2809 |
| ENSG00000079841 | RIMS1 | -0,315824351 | 0,803391804 | 0,3105303 | 0,817751312 | regulating synaptic membrane exocytosis 1 |
| ENSG00000198056 | PRIM1 | -0,316031315 | 0,803276561 | 0,50985603 | 0,922427526 | DNA primase subunit 1 |
| ENSG00000232005 | PBX2 | -0,316084215 | 0,803247107 | 0,44860062 | 0,902062594 | PBX homeobox 2 |
| ENSG00000288076 | AC018398,2 | -0,31608486 | 0,803246748 | 0,50019574 | 0,920484994 | novel transcript, antisense to MCPH1 |
| ENSG00000181458 | TMEM45A | -0,316152098 | 0,803209313 | 0,16072559 | 0,636469282 | transmembrane protein 45A |
| ENSG00000197121 | PGAP1 | -0,316180437 | 0,803193536 | 0,52111735 | 0,927046149 | post-GPI attachment to proteins inositol deacylase 1 |
| ENSG00000272918 | AC005070,3 | -0,316371098 | 0,803087395 | 0,23980645 | 0,749655932 | novel transcript, antisense to MLL5 |
| ENSG00000075303 | SLC25A40 | -0,316389571 | 0,803077112 | 0,52988884 | 0,930856025 | solute carrier family 25 member 40 |
| ENSG00000286141 | AC097480,2 | -0,316422343 | 0,80305887 | 0,56152562 | 0,94065996 | novel transcript |
| ENSG00000270641 | TSIX | -0,316556005 | 0,802984472 | 0,45146111 | 0,902062594 | TSIX transcript, XIST antisense RNA |
| ENSG00000250138 | AC139495,2 | -0,316588158 | 0,802966577 | 0,60425173 | 0,950746147 | POM121 membrane glycoprotein-like 1 (POM121L1) pseudogene |
| ENSG00000217165 | ANKRD18EP | -0,316690489 | 0,802909624 | 0,54395253 | 0,934259495 | ankyrin repeat domain 18E, pseudogene |
| ENSG00000131437 | KIF3A | -0,316749174 | 0,802876964 | 0,37543949 | 0,862022464 | kinesin family member 3A |
| ENSG00000164879 | CA3 | -0,316832959 | 0,802830338 | 0,40453628 | 0,878176155 | carbonic anhydrase 3 |
| ENSG00000171681 | ATF7IP | -0,31687073 | 0,802809319 | 0,35901165 | 0,849814998 | activating transcription factor 7 interacting protein |

| | | | | | | |
|---|---|---|---|---|---|---|
| ENSG00000276863 | AC124319,2 | -0,316872819 | 0,802808157 | 0,74005512 | 0,974724313 | novel transcript, antisense to RNF213 |
| ENSG00000099284 | MACROH2A2 | -0,31689435 | 0,802796176 | 0,16163561 | 0,63813307 | macroH2A,2 histone |
| ENSG00000276651 | AC007950,2 | -0,316902859 | 0,802791441 | 0,64419645 | 0,956633898 | novel transcript |
| ENSG00000198358 | AL354714,1 | -0,316958982 | 0,802760212 | 0,30539914 | 0,814326218 | antisense to ITLN2 |
| ENSG00000288533 | AP001178,4 | -0,317031581 | 0,802719817 | 0,48316729 | 0,91363215 | novel transcript, antisense to TYMS and CLUL1 |
| ENSG00000170919 | TPT1-AS1 | -0,31704454 | 0,802712606 | 0,1305555 | 0,577136255 | TPT1 antisense RNA 1 |
| ENSG00000141040 | ZNF287 | -0,317050956 | 0,802709036 | 0,42983553 | 0,894607 | zinc finger protein 287 |
| ENSG00000285427 | SOD2-OT1 | -0,317053955 | 0,802707368 | 0,41874553 | 0,887094957 | SOD2 overlapping transcript 1 |
| ENSG00000238286 | SLC35E1P1 | -0,317069943 | 0,802698472 | 0,54093295 | 0,933385624 | solute carrier family 35 member E1 pseudogene 1 |
| ENSG00000171492 | LRRC8D | -0,3171025 | 0,802680358 | 0,41299101 | 0,884663685 | leucine rich repeat containing 8 VRAC subunit D |
| ENSG00000215302 | AC127502,1 | -0,317154311 | 0,802651532 | 0,50063024 | 0,920520138 | WAS protein homolog associated with actin, golgi membranes and microtubules (WHAMM) pseudogene |
| ENSG00000225439 | BOLA3-AS1 | -0,317246672 | 0,802600148 | 0,32945881 | 0,831334931 | BOLA3 divergent transcript |
| ENSG00000250903 | GMDS-DT | -0,317292852 | 0,802574458 | 0,17022926 | 0,653377651 | GMDS divergent transcript |
| ENSG00000128694 | OSGEPL1 | -0,317340111 | 0,802548168 | 0,46916006 | 0,907940653 | O-sialoglycoprotein endopeptidase like 1 |
| ENSG00000173678 | SPDYE2B | -0,317460191 | 0,802481372 | 0,62705996 | 0,952793934 | speedy/RINGO cell cycle regulator family member E2B |
| ENSG00000269907 | PTMAP11 | -0,317462691 | 0,802479982 | 0,68122993 | 0,964654539 | prothymosin alpha pseudogene 11 |
| ENSG00000258648 | UBE2CP1 | -0,31750765 | 0,802454974 | 0,5617924 | 0,940754029 | ubiquitin conjugating enzyme E2 C pseudogene 1 |
| ENSG00000231312 | MAP4K3-DT | -0,317763365 | 0,802312753 | 0,2888485 | 0,800014136 | MAP4K3 divergent transcript |
| ENSG00000162694 | EXTL2 | -0,317825108 | 0,802278417 | 0,52189707 | 0,927599011 | exostosin like glycosyltransferase 2 |
| ENSG00000220522 | AL590002,1 | -0,317832499 | 0,802274308 | 0,86144996 | 0,989074806 | cornichon homolog 4 (Drosophila) (CNIH4) pseudogene |
| ENSG00000099957 | P2RX6 | -0,317973704 | 0,802195788 | 0,56706647 | 0,941425553 | purinergic receptor P2X 6 |
| ENSG00000166578 | IQCD | -0,318232463 | 0,802051921 | 0,42765507 | 0,892656446 | IQ motif containing D |
| ENSG00000244159 | RPS27AP13 | -0,318283732 | 0,802023418 | 0,68152605 | 0,964654539 | RPS27A pseudogene 13 |
| ENSG00000256218 | AC007848,2 | -0,318419084 | 0,801948177 | 0,54493913 | 0,934923038 | novel transcript |
| ENSG00000228570 | NUTM2E | -0,318431852 | 0,80194108 | 0,58778686 | 0,946124093 | NUT family member 2E |
| ENSG00000171823 | FBXL14 | -0,318503925 | 0,801901018 | 0,30562402 | 0,814456808 | F-box and leucine rich repeat protein 14 |
| ENSG00000138115 | CYP2C8 | -0,318637728 | 0,801826649 | 0,46768283 | 0,907940653 | cytochrome P450 family 2 subfamily C member 8 |
| ENSG00000267745 | AC060766,7 | -0,318653897 | 0,801817662 | 0,48967066 | 0,916063942 | novel transcript |
| ENSG00000234840 | LINC01239 | -0,318819513 | 0,801725622 | 0,46166226 | 0,906165804 | long intergenic non-protein coding RNA 1239 |

| | | | | | | |
|---|---|---|---|---|---|---|
| ENSG00000283589 | GOLGA8N | -0,318840868 | 0,801713755 | 0,40308488 | 0,877589651 | golgin A8 family member N |
| ENSG00000138640 | FAM13A | -0,318891736 | 0,801685488 | 0,32087106 | 0,824366234 | family with sequence similarity 13 member A |
| ENSG00000177034 | MTX3 | -0,318965031 | 0,80164476 | 0,46554209 | 0,907910177 | metaxin 3 |
| ENSG00000236264 | RPL26P30 | -0,318976287 | 0,801638506 | 0,75349076 | 0,975901808 | ribosomal protein L26 pseudogene 30 |
| ENSG00000231134 | TCF7L1-IT1 | -0,319044789 | 0,801600443 | 0,74090134 | 0,975142691 | TCF7L1 intronic transcript 1 |
| ENSG00000258776 | AL161757,4 | -0,319091847 | 0,801574297 | 0,53102166 | 0,931265667 | novel transcript |
| ENSG00000164109 | MAD2L1 | -0,319098585 | 0,801570553 | 0,49449288 | 0,91758518 | mitotic arrest deficient 2 like 1 |
| ENSG00000176927 | EFCAB5 | -0,31911099 | 0,801563661 | 0,60825099 | 0,951332922 | EF-hand calcium binding domain 5 |
| ENSG00000168283 | BMI1 | -0,31924637 | 0,801488447 | 0,49388458 | 0,917303403 | BMI1 proto-oncogene, polycomb ring finger |
| ENSG00000228409 | CCT6P1 | -0,319337365 | 0,801437896 | 0,35459816 | 0,846034781 | chaperonin containing TCP1 subunit 6 pseudogene 1 |
| ENSG00000284902 | AC074008,1 | -0,319382861 | 0,801412623 | 0,69150354 | 0,967201016 | novel transcript, sense intronic to ALMS1 |
| ENSG00000100197 | CYP2D6 | -0,319410522 | 0,801397258 | 0,61213384 | 0,952316495 | cytochrome P450 family 2 subfamily D member 6 |
| ENSG00000258655 | ARHGAP5-AS1 | -0,319487275 | 0,801354623 | 0,34492992 | 0,841168721 | ARHGAP5 antisense RNA 1 (head to head) |
| ENSG00000235076 | GAPDHP52 | -0,319657075 | 0,801260313 | 0,83934354 | 0,985672131 | glyceraldehyde 3 phosphate dehydrogenase pseudogene 52 |
| ENSG00000088538 | DOCK3 | -0,319964918 | 0,801089357 | 0,28421137 | 0,795731744 | dedicator of cytokinesis 3 |
| ENSG00000272812 | AC004908,2 | -0,319992451 | 0,801074069 | 0,67858539 | 0,9640473 | novel transcript, sense intronic to ZNF596 |
| ENSG00000235979 | AC004448,2 | -0,320202662 | 0,800957355 | 0,80102941 | 0,980555343 | novel transcript |
| ENSG00000174606 | ANGEL2 | -0,320304469 | 0,800900836 | 0,38302562 | 0,867682628 | angel homolog 2 |
| ENSG00000121570 | DPPA4 | -0,32033854 | 0,800881922 | 0,29893363 | 0,808745339 | developmental pluripotency associated 4 |
| ENSG00000242294 | STAG3L5P | -0,320358465 | 0,800870861 | 0,27360039 | 0,785719894 | stromal antigen 3-like 5 pseudogene |
| ENSG00000216937 | CCDC7 | -0,320401372 | 0,800847043 | 0,4050458 | 0,878176155 | coiled-coil domain containing 7 |
| ENSG00000174749 | FAM241A | -0,320418414 | 0,800837583 | 0,17180825 | 0,655690085 | family with sequence similarity 241 member A |
| ENSG00000214081 | CYP4F30P | -0,32042205 | 0,800835564 | 0,74605922 | 0,975707496 | cytochrome P450 family 4 subfamily F member 30, pseudogene |
| ENSG00000176728 | TTTY14 | -0,320535335 | 0,800772683 | 0,48212409 | 0,912981134 | testis-specific transcript, Y-linked 14 |
| ENSG00000258578 | AC008015,1 | -0,32064548 | 0,800711549 | 0,76054207 | 0,97624106 | ubiquitin-conjugating enzyme E2L 3 (UBE2L3) pseudogene |
| ENSG00000251669 | FAM86EP | -0,320686408 | 0,800688834 | 0,21363022 | 0,721033693 | family with sequence similarity 86 member E, pseudogene |
| ENSG00000222011 | FAM185A | -0,32069324 | 0,800685042 | 0,31613256 | 0,821394376 | family with sequence similarity 185 member A |
| ENSG00000143951 | WDPCP | -0,32070462 | 0,800678726 | 0,27078156 | 0,783073289 | WD repeat containing planar cell polarity effector |
| ENSG00000288050 | AC010727,1 | -0,320740752 | 0,800658674 | 0,58145148 | 0,944043373 | novel transcript, antisense to RFTN1 |

| | | | | | | |
|---|---|---|---|---|---|---|
| ENSG00000287364 | AL353709,1 | -0,320772377 | 0,800641123 | 0,76107124 | 0,97624106 | uncharacterized LOC105371672 |
| ENSG00000260410 | AC040169,2 | -0,320835916 | 0,800605862 | 0,72953254 | 0,974009551 | polo-like kinase 1 (PLK1) pseudogene |
| ENSG00000179902 | C1orf194 | -0,320940669 | 0,800547733 | 0,49608332 | 0,918105073 | chromosome 1 open reading frame 194 |
| ENSG00000237088 | AL163193,1 | -0,320941364 | 0,800547347 | 0,69393235 | 0,967340633 | ribosomal protein L7a (RPL7A) pseudogene |
| ENSG00000214654 | B3GNT10 | -0,320960715 | 0,800536609 | 0,4414907 | 0,898921877 | UDP-GlcNAc:betaGal beta-1,3-N-acetylglucosaminyltransferase 10 (putative) |
| ENSG00000082438 | COBLL1 | -0,321022521 | 0,800502315 | 0,45125888 | 0,902062594 | cordon-bleu WH2 repeat protein like 1 |
| ENSG00000186889 | TMEM17 | -0,321141048 | 0,800436551 | 0,34208962 | 0,840048641 | transmembrane protein 17 |
| ENSG00000143199 | ADCY10 | -0,321178152 | 0,800415965 | 0,69258372 | 0,967253522 | adenylate cyclase 10 |
| ENSG00000187257 | RSBN1L | -0,321182413 | 0,800413601 | 0,34724934 | 0,841896177 | round spermatid basic protein 1 like |
| ENSG00000197603 | CPLANE1 | -0,321196617 | 0,80040572 | 0,33011685 | 0,831783782 | ciliogenesis and planar polarity effector 1 |
| ENSG00000122643 | NT5C3A | -0,321343276 | 0,800324358 | 0,52944342 | 0,930491829 | 5'-nucleotidase, cytosolic IIIA |
| ENSG00000113494 | PRLR | -0,321488741 | 0,800243666 | 0,56306728 | 0,940865625 | prolactin receptor |
| ENSG00000236759 | EHMT2 | -0,321541069 | 0,800214641 | 0,25312024 | 0,76515438 | euchromatic histone lysine methyltransferase 2 |
| ENSG00000161082 | CELF5 | -0,321697107 | 0,800128097 | 0,39158227 | 0,870333649 | CUGBP Elav-like family member 5 |
| ENSG00000187742 | SECISBP2 | -0,321794733 | 0,800073955 | 0,22912972 | 0,737173617 | SECIS binding protein 2 |
| ENSG00000272144 | AC025171,4 | -0,321818003 | 0,80006105 | 0,61528411 | 0,952793934 | novel transcript |
| ENSG00000270141 | TERC | -0,321911213 | 0,800009362 | 0,27697662 | 0,788874154 | telomerase RNA component |
| ENSG00000054277 | OPN3 | -0,321949118 | 0,799988342 | 0,4506769 | 0,902062594 | opsin 3 |
| ENSG00000174469 | CNTNAP2 | -0,321995428 | 0,799962664 | 0,27449041 | 0,786383635 | contactin associated protein 2 |
| ENSG00000178235 | SLITRK1 | -0,322195767 | 0,799851585 | 0,35559804 | 0,846892875 | SLIT and NTRK like family member 1 |
| ENSG00000270690 | AC105129,3 | -0,322229623 | 0,799832815 | 0,76853126 | 0,977381742 | peroxiredoxin 3 (PRDX3) pseudogene |
| ENSG00000287238 | AP002954,2 | -0,32226447 | 0,799813496 | 0,27818132 | 0,789586738 | novel transcript |
| ENSG00000087338 | GMCL1 | -0,322371823 | 0,799753983 | 0,4592806 | 0,904862549 | germ cell-less 1, spermatogenesis associated |
| ENSG00000235778 | AC130360,1 | -0,322374863 | 0,799752297 | 0,74196376 | 0,975176043 | ubiquitin-specific protease (USP) pseudogene |
| ENSG00000221909 | FAM200A | -0,322380236 | 0,799749319 | 0,48516269 | 0,913690171 | family with sequence similarity 200 member A |
| ENSG00000186399 | GOLGA8R | -0,322431748 | 0,799720764 | 0,29936432 | 0,809210285 | golgin A8 family member R |
| ENSG00000108352 | RAPGEFL1 | -0,322439503 | 0,799716465 | 0,32477757 | 0,826793081 | Rap guanine nucleotide exchange factor like 1 |
| ENSG00000115947 | ORC4 | -0,322581225 | 0,79963791 | 0,34960343 | 0,843364329 | origin recognition complex subunit 4 |
| ENSG00000172824 | CES4A | -0,322606642 | 0,799623822 | 0,34370557 | 0,840901506 | carboxylesterase 4A |
| ENSG00000164221 | CCDC112 | -0,322680808 | 0,799582716 | 0,38672096 | 0,869441674 | coiled-coil domain containing 112 |
| ENSG00000116668 | SWT1 | -0,322699338 | 0,799572446 | 0,48397236 | 0,913690171 | SWT1 RNA endoribonuclease homolog |
| ENSG00000275580 | AC022306,2 | -0,322750802 | 0,799543924 | 0,7319712 | 0,974353474 | novel transcript, sense intronic to GALK2 |

| | | | | | | |
|---|---|---|---|---|---|---|
| ENSG00000117479 | SLC19A2 | -0,322759616 | 0,799539039 | 0,47911397 | 0,912714933 | solute carrier family 19 member 2 |
| ENSG00000254402 | LRRC24 | -0,32276352 | 0,799536876 | 0,63638842 | 0,955547532 | leucine rich repeat containing 24 |
| ENSG00000224543 | SNRPGP15 | -0,322799276 | 0,79951706 | 0,6525263 | 0,957275891 | small nuclear ribonucleoprotein polypeptide G pseudogene 15 |
| ENSG00000137857 | DUOX1 | -0,322818529 | 0,799506391 | 0,47985788 | 0,912847708 | dual oxidase 1 |
| ENSG00000269296 | AC005614,1 | -0,32282844 | 0,799500899 | 0,37714857 | 0,862818448 | novel transcript, antisense to ZNF780A |
| ENSG00000280797 | AC005614,3 | -0,32282844 | 0,799500899 | 0,37714857 | 0,862818448 | novel transcript, antisense to ZNF780A |
| ENSG00000265443 | AC127024,3 | -0,322843601 | 0,799492496 | 0,74797187 | 0,975901808 | novel transcript |
| ENSG00000237387 | AL022329,2 | -0,32289727 | 0,799462756 | 0,78159515 | 0,978597905 | novel transcript |
| ENSG00000153140 | CETN3 | -0,322915161 | 0,799452842 | 0,43522536 | 0,896272827 | centrin 3 |
| ENSG00000245479 | LINC01585 | -0,322997964 | 0,799406959 | 0,75131191 | 0,975901808 | long intergenic non-protein coding RNA 1585 |
| ENSG00000286058 | AC018865,2 | -0,323386842 | 0,799191508 | 0,46669292 | 0,907940653 | novel transcript |
| ENSG00000072657 | TRHDE | -0,323425053 | 0,799170341 | 0,60241906 | 0,950205821 | thyrotropin releasing hormone degrading enzyme |
| ENSG00000248588 | AC008517,1 | -0,323453065 | 0,799154824 | 0,64661412 | 0,956899963 | novel transcript |
| ENSG00000138036 | DYNC2LI1 | -0,323532132 | 0,799111028 | 0,29575858 | 0,805426883 | dynein cytoplasmic 2 light intermediate chain 1 |
| ENSG00000269873 | AC245884,10 | -0,323544051 | 0,799104425 | 0,79784228 | 0,980161849 | novel transcript |
| ENSG00000274045 | AC008746,1 | -0,323544051 | 0,799104425 | 0,79784228 | 0,980161849 | novel transcript |
| ENSG00000241120 | HMGN1P8 | -0,323601467 | 0,799072624 | 0,67602019 | 0,963560994 | high mobility group nucleosome binding domain 1 pseudogene 8 |
| ENSG00000227051 | C14orf132 | -0,323616385 | 0,79906436 | 0,25629857 | 0,768515424 | chromosome 14 open reading frame 132 |
| ENSG00000247796 | AC008966,1 | -0,323642128 | 0,799050103 | 0,33986879 | 0,838611198 | novel transcript, antisense to MOCS2 |
| ENSG00000272143 | FGF14-AS2 | -0,323703987 | 0,799015842 | 0,24296938 | 0,753092752 | FGF14 antisense RNA 2 |
| ENSG00000129534 | MIS18BP1 | -0,323738199 | 0,798996894 | 0,47945223 | 0,912714933 | MIS18 binding protein 1 |
| ENSG00000138606 | SHF | -0,323782389 | 0,798972422 | 0,15210692 | 0,620332737 | Src homology 2 domain containing F |
| ENSG00000276188 | AC069234,4 | -0,323824657 | 0,798949013 | 0,77034323 | 0,977872051 | novel transcript |
| ENSG00000205238 | SPDYE2 | -0,32385887 | 0,798930067 | 0,53579658 | 0,932139691 | speedy/RINGO cell cycle regulator family member E2 |
| ENSG00000197372 | ZNF675 | -0,323860256 | 0,798929299 | 0,43889786 | 0,897402199 | zinc finger protein 675 |
| ENSG00000226620 | LINC00343 | -0,32389881 | 0,79890795 | 0,17679651 | 0,662651108 | long intergenic non-protein coding RNA 343 |
| ENSG00000138778 | CENPE | -0,323951741 | 0,798878639 | 0,31458436 | 0,819877431 | centromere protein E |
| ENSG00000203279 | AL590705,1 | -0,324326858 | 0,798670948 | 0,5786 | 0,94348043 | novel transcript |
| ENSG00000286532 | PARTICL | -0,324497365 | 0,798576562 | 0,25273566 | 0,764984776 | promoter of MAT2A antisense radiation-induced circulating long non-coding RNA |

| | | | | | | |
|---|---|---|---|---|---|---|
| ENSG00000234329 | AL604028,2 | -0,324497525 | 0,798576474 | 0,49360671 | 0,917289924 | ribosomal protein L7a (RPL7A) pseudogene |
| ENSG00000287030 | AC107032,3 | -0,32464416 | 0,79849531 | 0,77428535 | 0,978503783 | novel transcript |
| ENSG00000101752 | MIB1 | -0,3247123 | 0,798457598 | 0,40363822 | 0,877691113 | mindbomb E3 ubiquitin protein ligase 1 |
| ENSG00000286833 | AC097532,3 | -0,324807247 | 0,798405051 | 0,42053337 | 0,888563911 | novel transcript |
| ENSG00000244274 | DBNDD2 | -0,324825783 | 0,798394793 | 0,2277667 | 0,736053199 | dysbindin domain containing 2 |
| ENSG00000185666 | SYN3 | -0,324931836 | 0,798336105 | 0,37474607 | 0,861771195 | synapsin III |
| ENSG00000251566 | HMGB1P35 | -0,324965398 | 0,798317533 | 0,72411673 | 0,973167061 | high mobility group box 1 pseudogene 35 |
| ENSG00000069943 | PIGB | -0,325022084 | 0,798286167 | 0,46778859 | 0,907940653 | phosphatidylinositol glycan anchor biosynthesis class B |
| ENSG00000275139 | AL133492,1 | -0,325042845 | 0,798274679 | 0,65483779 | 0,957444211 | novel transcript, antisense to PCBP3 |
| ENSG00000230961 | MSH5 | -0,32508913 | 0,798249069 | 0,43302543 | 0,895777109 | mutS homolog 5 |
| ENSG00000101746 | NOL4 | -0,32527489 | 0,798146294 | 0,41726294 | 0,886651836 | nucleolar protein 4 |
| ENSG00000179133 | C10orf67 | -0,325472455 | 0,798037002 | 0,54684485 | 0,935404974 | chromosome 10 open reading frame 67 |
| ENSG00000236200 | KDM4A-AS1 | -0,325516199 | 0,798012805 | 0,20333209 | 0,703503738 | KDM4A antisense RNA 1 |
| ENSG00000286667 | AC010641,2 | -0,325526052 | 0,798007355 | 0,41664599 | 0,886368152 | novel transcript |
| ENSG00000178222 | RNF212 | -0,325526664 | 0,798007016 | 0,5360161 | 0,932139691 | ring finger protein 212 |
| ENSG00000209582 | SNORA48 | -0,325621661 | 0,797954472 | 0,43262484 | 0,89560871 | small nucleolar RNA, H/ACA box 48 |
| ENSG00000108848 | LUC7L3 | -0,325643342 | 0,79794248 | 0,26818916 | 0,780692104 | LUC7 like 3 pre-mRNA splicing factor |
| ENSG00000177570 | SAMD12 | -0,325655511 | 0,797935749 | 0,14942366 | 0,614794108 | sterile alpha motif domain containing 12 |
| ENSG00000139618 | BRCA2 | -0,325725048 | 0,79789729 | 0,4536782 | 0,902908855 | BRCA2 DNA repair associated |
| ENSG00000232082 | RPS6KA2-IT1 | -0,325734472 | 0,797892078 | 0,81362314 | 0,981483735 | RPS6KA2 intronic transcript 1 |
| ENSG00000092847 | AGO1 | -0,325779633 | 0,797867102 | 0,21595361 | 0,723046035 | argonaute RISC component 1 |
| ENSG00000182676 | PPP1R27 | -0,325929185 | 0,797784398 | 0,71917817 | 0,971869926 | protein phosphatase 1 regulatory subunit 27 |
| ENSG00000243422 | RPL23AP49 | -0,326041004 | 0,797722567 | 0,60581754 | 0,950927937 | ribosomal protein L23a pseudogene 49 |
| ENSG00000103540 | CCP110 | -0,326092801 | 0,797693927 | 0,45216894 | 0,902062594 | centriolar coiled-coil protein 110 |
| ENSG00000285134 | ITFG2-AS1 | -0,326136291 | 0,79766988 | 0,77840819 | 0,978587255 | ITFG2 antisense RNA 1 |
| ENSG00000142611 | PRDM16 | -0,326142527 | 0,797666433 | 0,31984438 | 0,824366234 | PR/SET domain 16 |
| ENSG00000287202 | AC008119,1 | -0,32635981 | 0,797546306 | 0,36791849 | 0,857401877 | novel transcript, antisense to SART3 |
| ENSG00000180336 | MEIOC | -0,326387622 | 0,797530931 | 0,32295364 | 0,825051003 | meiosis specific with coiled-coil domain |
| ENSG00000093100 | AC016026,1 | -0,326575196 | 0,797427246 | 0,38564721 | 0,869017264 | novel transcript |
| ENSG00000167232 | ZNF91 | -0,326620114 | 0,797402418 | 0,45417896 | 0,90318049 | zinc finger protein 91 |
| ENSG00000116679 | IVNS1ABP | -0,326658046 | 0,797381453 | 0,34512241 | 0,841168721 | influenza virus NS1A binding protein |
| ENSG00000283503 | LINC01902 | -0,326679923 | 0,797369361 | 0,33993474 | 0,838611198 | long intergenic non-protein coding RNA 1902 |

| | | | | | | |
|---|---|---|---|---|---|---|
| ENSG00000150477 | KIAA1328 | -0,326842989 | 0,797279241 | 0,12556237 | 0,566089307 | KIAA1328 |
| ENSG00000005961 | ITGA2B | -0,326843384 | 0,797279023 | 0,54068942 | 0,933385624 | integrin subunit alpha 2b |
| ENSG00000261360 | AC010491,1 | -0,326901853 | 0,797246712 | 0,49637306 | 0,918315123 | novel transcript, antisense to FAM105B |
| ENSG00000172458 | IL17D | -0,326962395 | 0,797213256 | 0,29492514 | 0,805040011 | interleukin 17D |
| ENSG00000281709 | ZNF197 | -0,32700302 | 0,797190808 | 0,404525 | 0,878176155 | zinc finger protein 197 |
| ENSG00000261684 | AC018362,1 | -0,327030337 | 0,797175713 | 0,61799438 | 0,952793934 | novel transcript, antisense to SNAP23 |
| ENSG00000187186 | AL162231,1 | -0,327039275 | 0,797170774 | 0,33047913 | 0,832114797 | uncharacterized LOC730098 |
| ENSG00000140015 | KCNH5 | -0,32706157 | 0,797158455 | 0,63943821 | 0,956018518 | potassium voltage-gated channel subfamily H member 5 |
| ENSG00000237827 | RPS15AP29 | -0,327062608 | 0,797157882 | 0,72602973 | 0,973344786 | ribosomal protein S15a pseudogene 29 |
| ENSG00000069424 | KCNAB2 | -0,327179743 | 0,797093162 | 0,4070405 | 0,880078305 | potassium voltage-gated channel subfamily A regulatory beta subunit 2 |
| ENSG00000276728 | AC142472,1 | -0,327261885 | 0,797047779 | 0,28614439 | 0,797420923 | novel transcript |
| ENSG00000186448 | ZNF197 | -0,327290191 | 0,797032142 | 0,40403636 | 0,877939445 | zinc finger protein 197 |
| ENSG00000275467 | AC083806,3 | -0,327378932 | 0,796983117 | 0,76167672 | 0,976285211 | novel transcript |
| ENSG00000170289 | CNGB3 | -0,327411707 | 0,796965011 | 0,78487325 | 0,979320406 | cyclic nucleotide gated channel subunit beta 3 |
| ENSG00000247982 | LINC00926 | -0,327511356 | 0,796909966 | 0,40819217 | 0,881221125 | long intergenic non-protein coding RNA 926 |
| ENSG00000167196 | FBXO22 | -0,32753022 | 0,796899546 | 0,30417025 | 0,813056239 | F-box protein 22 |
| ENSG00000203666 | EFCAB2 | -0,327530799 | 0,796899226 | 0,25059533 | 0,762789652 | EF-hand calcium binding domain 2 |
| ENSG00000269825 | AC022150,4 | -0,327574151 | 0,79687528 | 0,22962347 | 0,73804339 | novel transcript, sense intronic to ZNF83 |
| ENSG00000238243 | OR2W3 | -0,32758356 | 0,796870083 | 0,55640683 | 0,938061341 | olfactory receptor family 2 subfamily W member 3 |
| ENSG00000278576 | AL162171,3 | -0,327586289 | 0,796868575 | 0,3139086 | 0,819877431 | novel transcript, sense intronic to ZC3H14 |
| ENSG00000237719 | Z95152,1 | -0,327901372 | 0,79669456 | 0,53153784 | 0,931291085 | nudix (nucleoside diphosphate linked moiety X) type 5 motif (NUDT5) pseudogene |
| ENSG00000227409 | ZMYM4-AS1 | -0,327952235 | 0,796666472 | 0,69062582 | 0,967179044 | ZMYM4 antisense RNA 1 |
| ENSG00000285560 | AC103739,3 | -0,327958114 | 0,796663225 | 0,63531378 | 0,955314044 | novel transcript, antisense to CRTC3 |
| ENSG00000249661 | TNRC18P1 | -0,327978455 | 0,796651994 | 0,57556399 | 0,942700009 | trinucleotide repeat containing 18 pseudogene 1 |
| ENSG00000262583 | TMEM231P1 | -0,328002478 | 0,796638728 | 0,81288147 | 0,981246822 | TMEM231 pseudogene 1 |
| ENSG00000188580 | NKAIN2 | -0,328017055 | 0,796630678 | 0,5650761 | 0,940865625 | sodium/potassium transporting ATPase interacting 2 |
| ENSG00000274099 | ABCB10P1 | -0,328084045 | 0,796593689 | 0,78659281 | 0,979320406 | ABCB10 pseudogene 1 |
| ENSG00000280461 | ABCB10P1 | -0,328084045 | 0,796593689 | 0,78659281 | 0,979320406 | ABCB10 pseudogene 1 |
| ENSG00000281173 | ABCB10P1 | -0,328084045 | 0,796593689 | 0,78659281 | 0,979320406 | ABCB10 pseudogene 1 |
| ENSG00000236546 | MYCL-AS1 | -0,328085024 | 0,796593148 | 0,64706436 | 0,956899963 | MYCL antisense RNA 1 |

| | | | | | | |
|---|---|---|---|---|---|---|
| ENSG00000288186 | PPP1R27 | -0,328104775 | 0,796582243 | 0,71952791 | 0,971869926 | protein phosphatase 1 regulatory subunit 27 |
| ENSG00000054793 | ATP9A | -0,328137102 | 0,796564394 | 0,23586617 | 0,744966003 | ATPase phospholipid transporting 9A (putative) |
| ENSG00000187122 | SLIT1 | -0,328191871 | 0,796534154 | 0,48459344 | 0,913690171 | slit guidance ligand 1 |
| ENSG00000238098 | ABCA17P | -0,32826788 | 0,79649219 | 0,58241026 | 0,944174844 | ATP binding cassette subfamily A member 17, pseudogene |
| ENSG00000153208 | MERTK | -0,328291146 | 0,796479345 | 0,31117957 | 0,817751312 | MER proto-oncogene, tyrosine kinase |
| ENSG00000099365 | STX1B | -0,32843276 | 0,796401167 | 0,18662151 | 0,680015718 | syntaxin 1B |
| ENSG00000197753 | LHFPL5 | -0,32844905 | 0,796392174 | 0,23543062 | 0,744654651 | LHFPL tetraspan subfamily member 5 |
| ENSG00000146426 | TIAM2 | -0,328479247 | 0,796375505 | 0,24095542 | 0,751467261 | TIAM Rac1 associated GEF 2 |
| ENSG00000270062 | AL606834,2 | -0,328481511 | 0,796374255 | 0,71279503 | 0,970812834 | novel transcript, sense intronic to NIN |
| ENSG00000231310 | TBL1XR1-AS1 | -0,328601927 | 0,796307788 | 0,58683654 | 0,945421911 | TBL1XR1 antisense RNA 1 |
| ENSG00000114770 | ABCC5 | -0,328627151 | 0,796293866 | 0,21131127 | 0,717057897 | ATP binding cassette subfamily C member 5 |
| ENSG00000166206 | GABRB3 | -0,328664153 | 0,796273443 | 0,36510075 | 0,855173479 | gamma-aminobutyric acid type A receptor subunit beta3 |
| ENSG00000146677 | AC004453,1 | -0,328723766 | 0,796240541 | 0,73599298 | 0,974353474 | ribosomal protein L32 (RPL32) pseudogene |
| ENSG00000285016 | AC017002,3 | -0,32873906 | 0,7962321 | 0,79886933 | 0,980161849 | novel transcript |
| ENSG00000153006 | SREK1IP1 | -0,328854295 | 0,796168504 | 0,46073022 | 0,905949965 | SREK1 interacting protein 1 |
| ENSG00000154556 | SORBS2 | -0,328919856 | 0,796132324 | 0,29365377 | 0,803461221 | sorbin and SH3 domain containing 2 |
| ENSG00000243854 | RN7SL67P | -0,328960586 | 0,796109848 | 0,56905674 | 0,942177697 | RNA, 7SL, cytoplasmic 67, pseudogene |
| ENSG00000256007 | ARAP1-AS1 | -0,328995007 | 0,796090854 | 0,80740966 | 0,981213877 | ARAP1 antisense RNA 1 |
| ENSG00000101871 | MID1 | -0,329269764 | 0,795939255 | 0,34579818 | 0,841168721 | midline 1 |
| ENSG00000158220 | ESYT3 | -0,329281927 | 0,795932545 | 0,43139986 | 0,89530663 | extended synaptotagmin 3 |
| ENSG00000225914 | TSBP1-AS1 | -0,329309059 | 0,795917576 | 0,17972032 | 0,668424474 | TSBP1 and BTNL2 antisense RNA 1 |
| ENSG00000286337 | AC025370,2 | -0,329446015 | 0,795842023 | 0,54063757 | 0,933385624 | novel transcript, antisense to FZD6 |
| ENSG00000277526 | AC245123,1 | -0,329472414 | 0,79582746 | 0,72141351 | 0,972142678 | novel transcript |
| ENSG00000213988 | ZNF90 | -0,329593431 | 0,795760707 | 0,12259762 | 0,560036813 | zinc finger protein 90 |
| ENSG00000167555 | ZNF528 | -0,329752234 | 0,795673119 | 0,28614177 | 0,797420923 | zinc finger protein 528 |
| ENSG00000214279 | SCART1 | -0,329812356 | 0,795639962 | 0,42613649 | 0,892133432 | scavenger receptor family member expressed on T cells 1 |
| ENSG00000200913 | SNORD46 | -0,329921349 | 0,795579855 | 0,65060922 | 0,956899963 | small nucleolar RNA, C/D box 46 |
| ENSG00000129173 | E2F8 | -0,329952898 | 0,795562458 | 0,28905723 | 0,800166929 | E2F transcription factor 8 |
| ENSG00000164411 | GJB7 | -0,329996914 | 0,795538186 | 0,51460403 | 0,924577921 | gap junction protein beta 7 |
| ENSG00000170396 | ZNF804A | -0,330002821 | 0,795534928 | 0,50201068 | 0,920520138 | zinc finger protein 804A |

| | | | | | | |
|---|---|---|---|---|---|---|
| ENSG00000213385 | AC105052,2 | -0,330069511 | 0,795498154 | 0,49882412 | 0,919683359 | ribosomal protein L7a (RPL7A) pseudogene |
| ENSG00000180914 | OXTR | -0,33019004 | 0,795431698 | 0,23618861 | 0,745253314 | oxytocin receptor |
| ENSG00000258757 | AL133453,1 | -0,330221352 | 0,795414435 | 0,68930418 | 0,966732729 | novel transcript, antisense to ERO1L |
| ENSG00000227863 | AC002383,1 | -0,330435994 | 0,795296103 | 0,52826463 | 0,92998848 | novel transcript |
| ENSG00000276698 | AL136295,6 | -0,330442639 | 0,795292439 | 0,75394841 | 0,975901808 | novel transcript |
| ENSG00000258654 | AC026495,1 | -0,330498591 | 0,795261596 | 0,64706526 | 0,956899963 | novel transcript |
| ENSG00000187607 | ZNF286A | -0,330558964 | 0,795228317 | 0,25786411 | 0,770217203 | zinc finger protein 286A |
| ENSG00000233907 | LINC01761 | -0,330618384 | 0,795195565 | 0,4051565 | 0,878176155 | long intergenic non-protein coding RNA 1761 |
| ENSG00000251562 | MALAT1 | -0,330668415 | 0,795167989 | 0,21085054 | 0,716463754 | metastasis associated lung adenocarcinoma transcript 1 |
| ENSG00000255867 | DENND5B-AS1 | -0,33070045 | 0,795150333 | 0,6564143 | 0,957444211 | DENND5B antisense RNA 1 |
| ENSG00000163704 | PRRT3 | -0,330785263 | 0,795103589 | 0,2374594 | 0,746907875 | proline rich transmembrane protein 3 |
| ENSG00000128271 | ADORA2A | -0,330786707 | 0,795102793 | 0,76232385 | 0,976315588 | adenosine A2a receptor |
| ENSG00000278165 | DUSP8 | -0,330816562 | 0,795086339 | 0,61185082 | 0,952187506 | dual specificity phosphatase 8 |
| ENSG00000204711 | C9orf135 | -0,330847585 | 0,795069242 | 0,62095662 | 0,952793934 | chromosome 9 open reading frame 135 |
| ENSG00000170153 | RNF150 | -0,330864346 | 0,795060006 | 0,3302423 | 0,831831721 | ring finger protein 150 |
| ENSG00000288074 | AC026341,3 | -0,330887286 | 0,795047363 | 0,72248856 | 0,972594421 | novel transcript |
| ENSG00000263235 | AC006111,2 | -0,330892808 | 0,795044321 | 0,48343731 | 0,913690171 | novel transcript, sense overlapping TRAP1 |
| ENSG00000268903 | AL627309,6 | -0,330975293 | 0,794998866 | 0,6559289 | 0,957444211 | novel pseudogene |
| ENSG00000198155 | ZNF876P | -0,331122369 | 0,794917823 | 0,37493043 | 0,861771195 | zinc finger protein 876, pseudogene |
| ENSG00000198722 | UNC13B | -0,331194165 | 0,794878265 | 0,13037308 | 0,577035665 | unc-13 homolog B |
| ENSG00000249348 | UGDH-AS1 | -0,331224059 | 0,794861795 | 0,1605419 | 0,636125619 | UGDH antisense RNA 1 |
| ENSG00000185909 | KLHDC8B | -0,331260966 | 0,794841461 | 0,25284023 | 0,765079081 | kelch domain containing 8B |
| ENSG00000095539 | SEMA4G | -0,331367582 | 0,794782724 | 0,19563265 | 0,69375328 | semaphorin 4G |
| ENSG00000244270 | RPL32P29 | -0,331381465 | 0,794775075 | 0,6208408 | 0,952793934 | ribosomal protein L32 pseudogene 29 |
| ENSG00000268153 | AC005176,1 | -0,331406716 | 0,794761165 | 0,69877949 | 0,968522643 | lectin, galactoside-binding, soluble, 14 (LGALS14) pseudogene |
| ENSG00000286360 | AC107081,3 | -0,331442444 | 0,794741483 | 0,3277804 | 0,829743891 | novel transcript, antisense to CCT4 |
| ENSG00000206127 | GOLGA8O | -0,331544336 | 0,794685356 | 0,34227677 | 0,840048641 | golgin A8 family member O |
| ENSG00000235488 | JARID2-AS1 | -0,331599456 | 0,794654995 | 0,70857428 | 0,970181083 | JARID2 antisense RNA 1 |
| ENSG00000272686 | WASL-DT | -0,33162196 | 0,794642599 | 0,42503399 | 0,891853867 | WASL divergent transcript |
| ENSG00000185513 | L3MBTL1 | -0,331666526 | 0,794618052 | 0,20498138 | 0,706634751 | L3MBTL histone methyl-lysine binding protein 1 |
| ENSG00000214182 | PTMAP5 | -0,331843553 | 0,794520554 | 0,45587701 | 0,903545728 | prothymosin alpha pseudogene 5 |

| | | | | | | |
|---|---|---|---|---|---|---|
| ENSG00000288071 | AL591135,2 | -0,331887762 | 0,794496208 | 0,81532623 | 0,981521413 | novel transcript |
| ENSG00000222594 | RN7SKP235 | -0,331948889 | 0,794462545 | 0,82252001 | 0,98299282 | RN7SK pseudogene 235 |
| ENSG00000130363 | RSPH3 | -0,332180865 | 0,794334811 | 0,14782916 | 0,612372753 | radial spoke head 3 |
| ENSG00000163633 | C4orf36 | -0,332231963 | 0,794306678 | 0,65898511 | 0,95828296 | chromosome 4 open reading frame 36 |
| ENSG00000251667 | BRCC3P1 | -0,332259175 | 0,794291696 | 0,56992523 | 0,94221458 | BRCA1/BRCA2-containing complex subunit 3 pseudogene 1 |
| ENSG00000227638 | HNRNPA1P14 | -0,332321904 | 0,79425716 | 0,39889777 | 0,875433675 | heterogeneous nuclear ribonucleoprotein A1 pseudogene 14 |
| ENSG00000268218 | AC137932,3 | -0,332336742 | 0,794248991 | 0,43957547 | 0,89764124 | novel transcript, antisense to ANKRD11 |
| ENSG00000028137 | TNFRSF1B | -0,332365703 | 0,794233047 | 0,45320461 | 0,902540855 | TNF receptor superfamily member 1B |
| ENSG00000099326 | MZF1 | -0,332404138 | 0,794211889 | 0,21472537 | 0,722164151 | myeloid zinc finger 1 |
| ENSG00000239374 | AC090013,1 | -0,332439279 | 0,794192544 | 0,76784598 | 0,977309816 | ribosomal protein L21 (RPL21) pseudogene |
| ENSG00000099617 | EFNA2 | -0,332442167 | 0,794190954 | 0,19319754 | 0,689984903 | ephrin A2 |
| ENSG00000155903 | RASA2 | -0,332479564 | 0,794170368 | 0,46140217 | 0,906165604 | RAS p21 protein activator 2 |
| ENSG00000188820 | CALHM6 | -0,332659057 | 0,794071567 | 0,72021119 | 0,972142678 | calcium homeostasis modulator family member 6 |
| ENSG00000259953 | AL138756,1 | -0,332681546 | 0,794059189 | 0,25670981 | 0,769103136 | novel transcript, overlapping SUSD1 |
| ENSG00000224597 | SVIL-AS1 | -0,332719246 | 0,794038439 | 0,36720357 | 0,856607413 | SVIL antisense RNA 1 |
| ENSG00000118495 | PLAGL1 | -0,332767881 | 0,794011671 | 0,33166057 | 0,832548086 | PLAG1 like zinc finger 1 |
| ENSG00000285254 | AL513128,2 | -0,332782907 | 0,794003402 | 0,69004907 | 0,967053457 | novel pseudogene |
| ENSG00000176912 | TYMSOS | -0,333019737 | 0,79387307 | 0,23232189 | 0,741229884 | TYMS opposite strand |
| ENSG00000196408 | NOXO1 | -0,333079999 | 0,793839911 | 0,79432077 | 0,979776981 | NADPH oxidase organizer 1 |
| ENSG00000259242 | AC002306,1 | -0,333103753 | 0,79382684 | 0,67059554 | 0,962026636 | novel transcript, antisense to PBX4 |
| ENSG00000234193 | ACSL3-AS1 | -0,333141785 | 0,793805913 | 0,70313934 | 0,968742626 | ACSL3 antisense RNA 1 |
| ENSG00000224751 | SHMT1P1 | -0,3331527 | 0,793799908 | 0,59656259 | 0,948282297 | serine hydroxymethyltransferase 1 (soluble) pseudogene 1 |
| ENSG00000280832 | GSEC | -0,333154168 | 0,7937991 | 0,26519999 | 0,77662164 | G-quadruplex forming sequence containing lncRNA |
| ENSG00000244676 | AL109761,1 | -0,33324692 | 0,793748068 | 0,64714142 | 0,956899963 | novel transcript, antisense to C21orf91 |
| ENSG00000165805 | C12orf50 | -0,333291582 | 0,793723496 | 0,27557276 | 0,787284069 | chromosome 12 open reading frame 50 |
| ENSG00000134253 | TRIM45 | -0,333336171 | 0,793698965 | 0,24644354 | 0,758198269 | tripartite motif containing 45 |
| ENSG00000147274 | RBMX | -0,333344087 | 0,79369461 | 0,39741152 | 0,874053739 | RNA binding motif protein X-linked |
| ENSG00000286372 | AL449223,1 | -0,33341028 | 0,793658195 | 0,78140609 | 0,978597905 | novel transcript, antisense to NMNAT2 |
| ENSG00000198720 | ANKRD13B | -0,333564398 | 0,793573416 | 0,19581408 | 0,69375328 | ankyrin repeat domain 13B |
| ENSG00000278632 | TBC1D3B | -0,333628087 | 0,793538383 | 0,67734041 | 0,963580269 | TBC1 domain family member 3B |

| | | | | | | |
|---|---|---|---|---|---|---|
| ENSG00000285708 | AC097634,4 | -0,333677554 | 0,793511175 | 0,79841461 | 0,980161849 | novel protein |
| ENSG00000286257 | AL137100,3 | -0,333714904 | 0,793490632 | 0,4630797 | 0,906849208 | novel transcript |
| ENSG00000205649 | HTN3 | -0,333723533 | 0,793485886 | 0,75645048 | 0,975901808 | histatin 3 |
| ENSG00000282967 | HTN3 | -0,333723533 | 0,793485886 | 0,75645048 | 0,975901808 | histatin 3 |
| ENSG00000162873 | KLHDC8A | -0,333780516 | 0,793454546 | 0,3788231 | 0,864112469 | kelch domain containing 8A |
| ENSG00000175548 | ALG10B | -0,333889987 | 0,793394341 | 0,53081367 | 0,931265667 | ALG10 alpha-1,2-glucosyltransferase B |
| ENSG00000258101 | AC010173,1 | -0,333952621 | 0,793359897 | 0,58022875 | 0,943828882 | novel transcript, antisense to TUBA1C |
| ENSG00000116151 | MORN1 | -0,334152404 | 0,793250041 | 0,27334352 | 0,785672647 | MORN repeat containing 1 |
| ENSG00000225928 | CACYBPP1 | -0,334237652 | 0,79320317 | 0,72477735 | 0,97326334 | calcyclin binding protein pseudogene 1 |
| ENSG00000169085 | VXN | -0,334246868 | 0,793198103 | 0,49947787 | 0,920186067 | vexin |
| ENSG00000198077 | CYP2A7 | -0,334280141 | 0,79317981 | 0,83475134 | 0,984905093 | cytochrome P450 family 2 subfamily A member 7 |
| ENSG00000228010 | AC073343,2 | -0,334289858 | 0,793174467 | 0,44835862 | 0,902062594 | novel transcript, antisense to ZNF12 |
| ENSG00000187187 | ZNF546 | -0,334327982 | 0,793153507 | 0,6968642 | 0,968300554 | zinc finger protein 546 |
| ENSG00000079134 | THOC1 | -0,334381702 | 0,793123974 | 0,40061385 | 0,8763498 | THO complex 1 |
| ENSG00000267053 | AC092296,2 | -0,33465048 | 0,792976227 | 0,54611469 | 0,935290591 | novel transcript |
| ENSG00000166863 | TAC3 | -0,334691279 | 0,792953802 | 0,67125835 | 0,962026636 | tachykinin precursor 3 |
| ENSG00000242889 | RN7SL449P | -0,334750266 | 0,792921381 | 0,51412013 | 0,924239655 | RNA, 7SL, cytoplasmic 449, pseudogene |
| ENSG00000213983 | AP1G2 | -0,334770081 | 0,792910491 | 0,33526101 | 0,835120721 | adaptor related protein complex 1 subunit gamma 2 |
| ENSG00000232774 | AL355916,1 | -0,334786991 | 0,792901197 | 0,1384882 | 0,593533053 | novel transcript |
| ENSG00000257681 | AC025265,1 | -0,33479331 | 0,792897724 | 0,72425249 | 0,973178901 | novel transcript, antisense to STAB2 |
| ENSG00000213337 | ANKRD39 | -0,334816324 | 0,792885076 | 0,19384358 | 0,690637119 | ankyrin repeat domain 39 |
| ENSG00000188687 | SLC4A5 | -0,334945895 | 0,792813868 | 0,2862047 | 0,797420923 | solute carrier family 4 member 5 |
| ENSG00000278023 | RDM1 | -0,33495034 | 0,792811426 | 0,50677159 | 0,921824694 | RAD52 motif containing 1 |
| ENSG00000223379 | AL391987,3 | -0,33498274 | 0,792793621 | 0,68941167 | 0,966840099 | novel transcript |
| ENSG00000044459 | CNTLN | -0,335038416 | 0,792763026 | 0,3310077 | 0,832186595 | centlein |
| ENSG00000198039 | ZNF273 | -0,33506015 | 0,792751084 | 0,32656984 | 0,828450693 | zinc finger protein 273 |
| ENSG00000272465 | AL031768,1 | -0,335164601 | 0,792693691 | 0,71539788 | 0,971134535 | novel transcript |
| ENSG00000257285 | AL132780,1 | -0,335273804 | 0,792633691 | 0,31193855 | 0,818226927 | novel transcript, antisense to HAUS4 |
| ENSG00000128534 | LSM8 | -0,335296357 | 0,7926213 | 0,44317322 | 0,899727948 | LSM8 homolog, U6 small nuclear RNA associated |
| ENSG00000176909 | MAMSTR | -0,335486702 | 0,792516731 | 0,44012084 | 0,897906863 | MEF2 activating motif and SAP domain containing transcriptional regulator |
| ENSG00000226200 | SGMS1-AS1 | -0,335539166 | 0,792487912 | 0,23670943 | 0,746175744 | SGMS1 antisense RNA 1 |
| ENSG00000221886 | ZBED8 | -0,33555938 | 0,792476808 | 0,44334131 | 0,899865141 | zinc finger BED-type containing 8 |

| | | | | | | |
|---|---|---|---|---|---|---|
| ENSG00000092871 | RFFL | -0,335603701 | 0,792452462 | 0,14866178 | 0,613346323 | ring finger and FYVE like domain containing E3 ubiquitin protein ligase |
| ENSG00000249721 | AC112206,3 | -0,335639001 | 0,792433073 | 0,80393936 | 0,980875302 | ribosomal protein L23a (RPL23A) pseudogene |
| ENSG00000158987 | RAPGEF6 | -0,335703925 | 0,792397413 | 0,29153627 | 0,802058262 | Rap guanine nucleotide exchange factor 6 |
| ENSG00000124721 | DNAH8 | -0,335721698 | 0,792387651 | 0,68217187 | 0,964699524 | dynein axonemal heavy chain 8 |
| ENSG00000229236 | TTTY10 | -0,335735672 | 0,792379976 | 0,20154239 | 0,701094807 | testis-specific transcript, Y-linked 10 |
| ENSG00000188321 | ZNF559 | -0,336022541 | 0,792222433 | 0,2585959 | 0,771444876 | zinc finger protein 559 |
| ENSG00000185252 | ZNF74 | -0,336108802 | 0,792175066 | 0,179939 | 0,668839959 | zinc finger protein 74 |
| ENSG00000255107 | AC079089,2 | -0,33634047 | 0,792047869 | 0,81038802 | 0,981213877 | BAG family molecular chaperone regulator 4 (BAG4) pseudogene |
| ENSG00000135778 | NTPCR | -0,336410073 | 0,792009657 | 0,23838005 | 0,747813944 | nucleoside-triphosphatase, cancer-related |
| ENSG00000266256 | LINC00683 | -0,336524865 | 0,791946641 | 0,41071907 | 0,882835102 | long intergenic non-protein coding RNA 683 |
| ENSG00000267298 | AC006116,5 | -0,336532875 | 0,791942244 | 0,6588619 | 0,958240464 | novel transcript, sense intronic to ZSCAN5A |
| ENSG00000078177 | N4BP2 | -0,336536793 | 0,791940094 | 0,38582116 | 0,869112622 | NEDD4 binding protein 2 |
| ENSG00000173588 | CEP83 | -0,336558389 | 0,791928239 | 0,29009702 | 0,800682868 | centrosomal protein 83 |
| ENSG00000282486 | LRRC37A9P | -0,336637657 | 0,791884728 | 0,5599693 | 0,940017197 | leucine rich repeat containing 37 member A9, pseudogene |
| ENSG00000277662 | AL354696,1 | -0,336686147 | 0,791858113 | 0,70112191 | 0,968677171 | novel transcript, sense intronic to MTRF1 |
| ENSG00000276763 | AC069499,2 | -0,336777514 | 0,791807966 | 0,61615094 | 0,952793934 | ribosomal protein L10 (RPL10) pseudogene |
| ENSG00000082258 | CCNT2 | -0,336844256 | 0,791771336 | 0,48454668 | 0,913690171 | cyclin T2 |
| ENSG00000259732 | AC092757,2 | -0,336857462 | 0,791764088 | 0,56222557 | 0,940865625 | novel transcript, sense overlapping to CCNB2 |
| ENSG00000282789 | CRYZP1 | -0,336864796 | 0,791760063 | 0,6000486 | 0,949340299 | crystallin zeta pseudogene 1 |
| ENSG00000233025 | CRYZP1 | -0,336864796 | 0,791760063 | 0,6000486 | 0,949340299 | crystallin zeta pseudogene 1 |
| ENSG00000188185 | LINC00265 | -0,336957069 | 0,791709425 | 0,2387147 | 0,747998807 | long intergenic non-protein coding RNA 265 |
| ENSG00000249228 | AC068944,1 | -0,337068592 | 0,791648227 | 0,36305154 | 0,853318246 | novel transcript |
| ENSG00000182518 | FAM104B | -0,337109381 | 0,791625845 | 0,44023764 | 0,897973376 | family with sequence similarity 104 member B |
| ENSG00000025156 | HSF2 | -0,337123759 | 0,791617955 | 0,46918638 | 0,907940653 | heat shock transcription factor 2 |
| ENSG00000144214 | LYG1 | -0,337163482 | 0,79159616 | 0,57433889 | 0,942475308 | lysozyme g1 |
| ENSG00000196096 | SPAG16-DT | -0,337169736 | 0,791592728 | 0,19987084 | 0,699357802 | SPAG16 divergent transcript |
| ENSG00000100427 | MLC1 | -0,337252232 | 0,791547464 | 0,58163141 | 0,944139372 | modulator of VRAC current 1 |
| ENSG00000215769 | ARHGAP27P1-E | -0,337270946 | 0,791537197 | 0,32150175 | 0,824933727 | ARHGAP27P1-BPTFP1-KPNA2P3 readthrough, transcribed pseudogene |
| ENSG00000205176 | REXO1L1P | -0,337280073 | 0,79153219 | 0,4287506 | 0,893571003 | REXO1 like 1, pseudogene |

| | | | | | | |
|---|---|---|---|---|---|---|
| ENSG00000277837 | AC090340,1 | -0,337295079 | 0,791523956 | 0,80381702 | 0,980875302 | novel transcript |
| ENSG00000189007 | ADAT2 | -0,337418901 | 0,791456025 | 0,3892102 | 0,870057277 | adenosine deaminase tRNA specific 2 |
| ENSG00000255417 | MTCO2P15 | -0,337478186 | 0,791423502 | 0,79456217 | 0,979776981 | MT-CO2 pseudogene 15 |
| ENSG00000162545 | CAMK2N1 | -0,337497165 | 0,791413091 | 0,29007323 | 0,800682868 | calcium/calmodulin dependent protein kinase II inhibitor 1 |
| ENSG00000250220 | AC053527,2 | -0,337549947 | 0,791384137 | 0,66335416 | 0,959920886 | novel transcript |
| ENSG00000170035 | UBE2E3 | -0,337621397 | 0,791344945 | 0,40337012 | 0,877589651 | ubiquitin conjugating enzyme E2 E3 |
| ENSG00000007933 | FMO3 | -0,337667001 | 0,79131993 | 0,74883659 | 0,975901808 | flavin containing dimethylaniline monoxygenase 3 |
| ENSG00000266642 | AC024267,6 | -0,337775521 | 0,791260409 | 0,31768248 | 0,822459851 | novel transcript, antisense to DHRS13 |
| ENSG00000153044 | CENPH | -0,338000714 | 0,79113691 | 0,39021654 | 0,870057277 | centromere protein H |
| ENSG00000137274 | BPHL | -0,338044979 | 0,791112636 | 0,12819391 | 0,571733888 | biphenyl hydrolase like |
| ENSG00000184792 | OSBP2 | -0,338230373 | 0,791010981 | 0,27346927 | 0,785672647 | oxysterol binding protein 2 |
| ENSG00000262732 | AC010401,1 | -0,338359184 | 0,790940358 | 0,47917343 | 0,912714933 | novel transcript, antisense to ERCC4 |
| ENSG00000198840 | MT-ND3 | -0,338369061 | 0,790934944 | 0,40330956 | 0,877589651 | mitochondrially encoded NADH:ubiquinone oxidoreductase core subunit 3 |
| ENSG00000232377 | AC016910,1 | -0,33838876 | 0,790924144 | 0,76869936 | 0,977381742 | novel transcript |
| ENSG00000250406 | AC109811,1 | -0,338482384 | 0,790872818 | 0,77597664 | 0,978587255 | novel transcript |
| ENSG00000250091 | DNAH10OS | -0,338522898 | 0,790850609 | 0,13257162 | 0,581174167 | dynein axonemal heavy chain 10 opposite strand |
| ENSG00000233122 | CTAGE7P | -0,338708093 | 0,790749096 | 0,38502705 | 0,868764521 | CTAGE family member 7, pseudogene |
| ENSG00000215022 | AL008729,1 | -0,338819549 | 0,790688009 | 0,35098549 | 0,844035671 | novel transcript, antisense to PHACTR1 |
| ENSG00000273674 | AC021752,1 | -0,338922025 | 0,790631847 | 0,66373438 | 0,959920886 | novel transcript |
| ENSG00000285793 | ANAPC1P2 | -0,338939066 | 0,790622509 | 0,23288515 | 0,74196691 | ANAPC1 pseudogene 2 |
| ENSG00000240399 | AC004801,2 | -0,339033409 | 0,790570809 | 0,44092579 | 0,898675567 | ribosomal protein L37 (RPL37) pseudogene |
| ENSG00000285579 | AC015687,1 | -0,33921731 | 0,790470041 | 0,80009208 | 0,980161849 | novel transcript |
| ENSG00000287813 | AC016705,3 | -0,339239891 | 0,790457669 | 0,55888024 | 0,939635125 | novel transcript, antisense to ARNT2 |
| ENSG00000183808 | RBM12B | -0,339318345 | 0,790414685 | 0,23870796 | 0,747998807 | RNA binding motif protein 12B |
| ENSG00000287495 | AL356489,4 | -0,339333761 | 0,790406238 | 0,72700651 | 0,973344786 | novel transcript |
| ENSG00000143768 | LEFTY2 | -0,339506214 | 0,790311763 | 0,77185366 | 0,978033855 | left-right determination factor 2 |
| ENSG00000205981 | DNAJC19 | -0,339571056 | 0,790276243 | 0,47971019 | 0,912821637 | DnaJ heat shock protein family (Hsp40) member C19 |
| ENSG00000156218 | ADAMTSL3 | -0,339599705 | 0,79026055 | 0,22801694 | 0,736341996 | ADAMTS like 3 |
| ENSG00000127364 | TAS2R4 | -0,33961065 | 0,790254555 | 0,39524752 | 0,871981986 | taste 2 receptor member 4 |
| ENSG00000228729 | AL627308,1 | -0,339776191 | 0,790163883 | 0,65133173 | 0,957089277 | cathepsin B (CTSB) pseudogene |
| ENSG00000225590 | VPS52 | -0,339810404 | 0,790145145 | 0,49653348 | 0,918448871 | VPS52 subunit of GARP complex |

| | | | | | | |
|---|---|---|---|---|---|---|
| ENSG00000228425 | VPS52 | -0,339810404 | 0,790145145 | 0,49653348 | 0,918448871 | VPS52 subunit of GARP complex |
| ENSG00000287480 | AL158832,2 | -0,339866104 | 0,790114639 | 0,49999169 | 0,92041337 | novel transcript |
| ENSG00000224376 | AC017104,1 | -0,339965746 | 0,79006007 | 0,61801428 | 0,952793934 | novel transcript |
| ENSG00000114455 | HHLA2 | -0,339986937 | 0,790048465 | 0,58648145 | 0,945395317 | HERV-H LTR-associating 2 |
| ENSG00000151500 | THYN1 | -0,340130751 | 0,789969714 | 0,31922582 | 0,824203151 | thymocyte nuclear protein 1 |
| ENSG00000244041 | LINC01011 | -0,340257939 | 0,789900073 | 0,29942975 | 0,809252953 | long intergenic non-protein coding RNA 1011 |
| ENSG00000135390 | ATP5MC2 | -0,340304912 | 0,789874355 | 0,19155598 | 0,687341458 | ATP synthase membrane subunit c locus 2 |
| ENSG00000233750 | CICP27 | -0,340325516 | 0,789863075 | 0,31416303 | 0,819877431 | capicua transcriptional repressor pseudogene 27 |
| ENSG00000196562 | SULF2 | -0,34034846 | 0,789850513 | 0,386022 | 0,869112622 | sulfatase 2 |
| ENSG00000233337 | UBE2FP3 | -0,340466521 | 0,789785879 | 0,66553814 | 0,960946962 | UBE2F pseudogene 3 |
| ENSG00000135414 | GDF11 | -0,340473601 | 0,789782003 | 0,25502753 | 0,76642479 | growth differentiation factor 11 |
| ENSG00000240972 | MIF | -0,340490433 | 0,789772789 | 0,23641516 | 0,745615211 | macrophage migration inhibitory factor |
| ENSG00000231192 | OR5H1 | -0,340499367 | 0,789767898 | 0,54403327 | 0,934259495 | olfactory receptor family 5 subfamily H member 1 |
| ENSG00000236137 | AL445231,1 | -0,340619169 | 0,789702319 | 0,6287293 | 0,952793934 | novel transcript |
| ENSG00000228716 | DHFR | -0,340623893 | 0,789699733 | 0,30160886 | 0,811209515 | dihydrofolate reductase |
| ENSG00000277281 | MUC6 | -0,340671037 | 0,789673927 | 0,7788308 | 0,978587255 | mucin 6, oligomeric mucus/gel-forming |
| ENSG00000124613 | ZNF391 | -0,340807777 | 0,789599085 | 0,35628738 | 0,847740014 | zinc finger protein 391 |
| ENSG00000127084 | FGD3 | -0,340867124 | 0,789566604 | 0,59993723 | 0,949340299 | FYVE, RhoGEF and PH domain containing 3 |
| ENSG00000174738 | NR1D2 | -0,340964686 | 0,789513212 | 0,49230431 | 0,916995341 | nuclear receptor subfamily 1 group D member 2 |
| ENSG00000233406 | AL162430,1 | -0,340968792 | 0,789510965 | 0,66131398 | 0,959582738 | ribosomal protein S2 (RPS2) pseudogene |
| ENSG00000235848 | RMDN2-AS1 | -0,34097943 | 0,789505143 | 0,68272947 | 0,964826385 | RMDN2 antisense RNA 1 |
| ENSG00000233155 | HMGA1P8 | -0,34099649 | 0,789495808 | 0,53633121 | 0,93228049 | high mobility group AT-hook 1 pseudogene 8 |
| ENSG00000288018 | AP003120,1 | -0,341088068 | 0,789445694 | 0,76134562 | 0,97624106 | novel transcript |
| ENSG00000224186 | C5orf66 | -0,341192135 | 0,78938875 | 0,37632226 | 0,862624756 | chromosome 5 open reading frame 66 |
| ENSG00000288617 | AL049777,1 | -0,341265108 | 0,789348824 | 0,51338269 | 0,924029735 | FOXG1-C14orf23 readthrough |
| ENSG00000267454 | ZNF582-AS1 | -0,341304505 | 0,789327268 | 0,40447499 | 0,878176155 | ZNF582 antisense RNA 1 (head to head) |
| ENSG00000174502 | SLC26A9 | -0,341366664 | 0,789293261 | 0,82910491 | 0,983948376 | solute carrier family 26 member 9 |
| ENSG00000272146 | ARF4-AS1 | -0,341403168 | 0,78927329 | 0,48443757 | 0,913690171 | ARF4 antisense RNA 1 |
| ENSG00000272702 | AC010913,1 | -0,341646843 | 0,78913999 | 0,62527269 | 0,952793934 | novel transcript |
| ENSG00000236501 | AC074286,1 | -0,341666094 | 0,789129461 | 0,72722576 | 0,973344786 | novel transcript |
| ENSG00000246863 | GPR176-DT | -0,341840136 | 0,789034268 | 0,67671487 | 0,963580269 | GPR176-DT |
| ENSG00000179994 | SPDYE7P | -0,341870615 | 0,789017599 | 0,75597098 | 0,975901808 | speedy/RINGO cell cycle regulator family member E7, pseudogene |

| | | | | | | |
|---|---|---|---|---|---|---|
| ENSG00000237594 | AP000251,1 | -0,341957712 | 0,788969967 | 0,35314181 | 0,845063255 | uncharacterized LOC150051 |
| ENSG00000163435 | ELF3 | -0,342032369 | 0,78892914 | 0,61105734 | 0,951873283 | E74 like ETS transcription factor 3 |
| ENSG00000241860 | AL627309,5 | -0,34206504 | 0,788911274 | 0,1657672 | 0,645076634 | novel transcript |
| ENSG00000262663 | AC087222,1 | -0,342107726 | 0,788887932 | 0,41594806 | 0,885986046 | novel transcript, sense intronic TBCD |
| ENSG00000231680 | LINC02723 | -0,342163538 | 0,788857414 | 0,81182229 | 0,981213877 | long intergenic non-protein coding RNA 2723 |
| ENSG00000277957 | SENP3-EIF4A1 | -0,342235205 | 0,788818228 | 0,34649333 | 0,841183985 | SENP3-EIF4A1 readthrough (NMD candidate) |
| ENSG00000254258 | AC087354,1 | -0,342304577 | 0,788780299 | 0,22388904 | 0,732710917 | novel transcript |
| ENSG00000165506 | DNAAF2 | -0,342327376 | 0,788767834 | 0,31330272 | 0,819604737 | dynein axonemal assembly factor 2 |
| ENSG00000260971 | AC119674,1 | -0,342403222 | 0,788726367 | 0,58129 | 0,943919685 | novel transcript |
| ENSG00000077097 | TOP2B | -0,342416158 | 0,788719295 | 0,35362676 | 0,845346751 | DNA topoisomerase II beta |
| ENSG00000227388 | AL133410,1 | -0,342506164 | 0,788670091 | 0,47892725 | 0,912714933 | novel transcript |
| ENSG00000188177 | ZC3H6 | -0,342604169 | 0,788616516 | 0,36764965 | 0,857206868 | zinc finger CCCH-type containing 6 |
| ENSG00000256282 | AC112694,1 | -0,342625238 | 0,788605 | 0,41486027 | 0,885579089 | mitochondrial carrier homolog 1 (C, elegans) (MTCH1) |
| ENSG00000275512 | AC007998,4 | -0,342706166 | 0,788560764 | 0,11965952 | 0,552426954 | novel transcript |
| ENSG00000005483 | KMT2E | -0,342770962 | 0,788525348 | 0,33622949 | 0,835660292 | lysine methyltransferase 2E (inactive) |
| ENSG00000278496 | AL158062,1 | -0,342784937 | 0,78851771 | 0,7544813 | 0,975901808 | retinoblastoma binding protein 8 (RBBP8) pseudogene |
| ENSG00000223922 | ASS1P2 | -0,342920191 | 0,788443789 | 0,58197003 | 0,944149815 | argininosuccinate synthetase 1 pseudogene 2 |
| ENSG00000110881 | ASIC1 | -0,343037764 | 0,788379537 | 0,15530449 | 0,625190083 | acid sensing ion channel subunit 1 |
| ENSG00000235381 | AL596202,1 | -0,343087443 | 0,78835239 | 0,38140773 | 0,865900094 | novel transcript |
| ENSG00000273891 | AL731566,2 | -0,343121637 | 0,788333705 | 0,31122299 | 0,817751312 | novel transcript, antisense to NSMCE4A |
| ENSG00000253537 | PCDHGA7 | -0,343158264 | 0,788313691 | 0,50291141 | 0,920767287 | protocadherin gamma subfamily A, 7 |
| ENSG00000281026 | N4BP2L2-IT2 | -0,343167395 | 0,788308702 | 0,20301266 | 0,703345263 | N4BPL2 intronic transcript 2 |
| ENSG00000117691 | NENF | -0,343275292 | 0,788249747 | 0,24246445 | 0,752612359 | neudesin neurotrophic factor |
| ENSG00000196911 | KPNA5 | -0,343410209 | 0,788176036 | 0,23693292 | 0,746456145 | karyopherin subunit alpha 5 |
| ENSG00000134297 | PLEKHA8P1 | -0,343445386 | 0,788156818 | 0,28785844 | 0,799107075 | pleckstrin homology domain containing A8 pseudogene 1 |
| ENSG00000136982 | DSCC1 | -0,343492541 | 0,788131058 | 0,39520877 | 0,871981986 | DNA replication and sister chromatid cohesion 1 |
| ENSG00000271964 | AC090948,1 | -0,343609343 | 0,788067252 | 0,53381759 | 0,931381467 | novel transcript, antisense to RFTN1 |
| ENSG00000198237 | GUSBP13 | -0,343820165 | 0,7879521 | 0,144396 | 0,604689513 | GUSB pseudogene 13 |
| ENSG00000261115 | TMEM178B | -0,343957935 | 0,787876858 | 0,2079864 | 0,712494639 | transmembrane protein 178B |
| ENSG00000285868 | AC008676,3 | -0,344094049 | 0,787802528 | 0,34098238 | 0,839658516 | Novel protein |

| | | | | | | |
|---|---|---|---|---|---|---|
| ENSG00000188747 | NOXA1 | -0,344215509 | 0,787736205 | 0,56490407 | 0,940865625 | NADPH oxidase activator 1 |
| ENSG00000164985 | PSIP1 | -0,344240115 | 0,78772277 | 0,39002675 | 0,870057277 | PC4 and SFRS1 interacting protein 1 |
| ENSG00000186230 | ZNF749 | -0,344265271 | 0,787709035 | 0,154406 | 0,623607455 | zinc finger protein 749 |
| ENSG00000275099 | GGNBP2 | -0,344279564 | 0,787701231 | 0,34236579 | 0,840048641 | gametogenetin binding protein 2 |
| ENSG00000232542 | DPYD-IT1 | -0,344306965 | 0,787686271 | 0,75371247 | 0,975901808 | DPYD intronic transcript 1 |
| ENSG00000235098 | ANKRD65 | -0,344310382 | 0,787684405 | 0,5047292 | 0,921400408 | ankyrin repeat domain 65 |
| ENSG00000171885 | AQP4 | -0,344325569 | 0,787676113 | 0,71162298 | 0,97080996 | aquaporin 4 |
| ENSG00000237682 | PRKCIP1 | -0,34445004 | 0,787608158 | 0,63416817 | 0,954948151 | PRKCI pseudogene 1 |
| ENSG00000112541 | PDE10A | -0,34449604 | 0,787583046 | 0,41715203 | 0,886651836 | phosphodiesterase 10A |
| ENSG00000156469 | MTERF3 | -0,344544785 | 0,787556436 | 0,37728341 | 0,862818448 | mitochondrial transcription termination factor 3 |
| ENSG00000255185 | PDXDC2P | -0,344590651 | 0,787531399 | 0,32887676 | 0,830667565 | pyridoxal dependent decarboxylase domain containing 2, pseudogene |
| ENSG00000237298 | TTN-AS1 | -0,344709346 | 0,787466609 | 0,27371019 | 0,785719894 | TTN antisense RNA 1 |
| ENSG00000280750 | AL021878,9 | -0,344716907 | 0,787462481 | 0,71676497 | 0,971438758 | novel transcript, sense intronic to NDUFA6 |
| ENSG00000281542 | AL021878,11 | -0,344716907 | 0,787462481 | 0,71676497 | 0,971438758 | novel transcript, sense intronic to NDUFA6 |
| ENSG00000270083 | AL021878,4 | -0,344716907 | 0,787462481 | 0,71676497 | 0,971438758 | novel transcript, sense intronic to NDUFA6 |
| ENSG00000259294 | AC005096,1 | -0,344790311 | 0,787422417 | 0,67652026 | 0,963580269 | novel transcript, antisense to CUX1 |
| ENSG00000235529 | AGAP1-IT1 | -0,344807544 | 0,787413011 | 0,67360423 | 0,962830239 | AGAP1 intronic transcript 1 |
| ENSG00000272375 | AC026979,3 | -0,344888128 | 0,78736903 | 0,69239899 | 0,967236299 | novel transcript |
| ENSG00000187325 | TAF9B | -0,344911762 | 0,787356131 | 0,53236215 | 0,931291085 | TATA-box binding protein associated factor 9b |
| ENSG00000228427 | AL590764,1 | -0,344978901 | 0,787319491 | 0,7820173 | 0,978597905 | novel transcript |
| ENSG00000132622 | HSPA12B | -0,344995204 | 0,787310594 | 0,34632567 | 0,841168721 | heat shock protein family A (Hsp70) member 12B |
| ENSG00000226091 | LINC00937 | -0,345046539 | 0,78728258 | 0,35843436 | 0,849814998 | long intergenic non-protein coding RNA 937 |
| ENSG00000143742 | SRP9 | -0,345059833 | 0,787275325 | 0,52705173 | 0,92927181 | signal recognition particle 9 |
| ENSG00000204519 | ZNF551 | -0,345127573 | 0,78723836 | 0,16396664 | 0,642745357 | zinc finger protein 551 |
| ENSG00000130943 | PKDREJ | -0,345182393 | 0,787208447 | 0,38503493 | 0,868764521 | polycystin family receptor for egg jelly |
| ENSG00000116675 | DNAJC6 | -0,345211077 | 0,787192796 | 0,31021546 | 0,817687618 | DnaJ heat shock protein family (Hsp40) member C6 |
| ENSG00000100744 | GSKIP | -0,345363254 | 0,787109767 | 0,43195839 | 0,89558934 | GSK3B interacting protein |
| ENSG00000162738 | VANGL2 | -0,345444462 | 0,787065462 | 0,10366998 | 0,513406963 | VANGL planar cell polarity protein 2 |
| ENSG00000231888 | MTND5P15 | -0,34553342 | 0,787016932 | 0,77384793 | 0,978415098 | MT-ND5 pseudogene 15 |
| ENSG00000183598 | H3C13 | -0,345576719 | 0,786993312 | 0,2571224 | 0,769358275 | H3 clustered histone 13 |
| ENSG00000164074 | ABHD18 | -0,345648819 | 0,786953982 | 0,19285198 | 0,689775116 | abhydrolase domain containing 18 |
| ENSG00000172915 | NBEA | -0,345649439 | 0,786953644 | 0,38416055 | 0,868125769 | neurobeachin |

| | | | | | | |
|---|---|---|---|---|---|---|
| ENSG00000231871 | IPO9-AS1 | -0,345685126 | 0,786934178 | 0,55142315 | 0,936366211 | IPO9 antisense RNA 1 |
| ENSG00000074935 | TUBE1 | -0,345777047 | 0,78688404 | 0,51399034 | 0,924227101 | tubulin epsilon 1 |
| ENSG00000225559 | AC083867,2 | -0,345803893 | 0,786869398 | 0,70919016 | 0,970282889 | novel transcript |
| ENSG00000189079 | ARID2 | -0,345812109 | 0,786864917 | 0,43319587 | 0,895845815 | AT-rich interaction domain 2 |
| ENSG00000282814 | AC010724,9 | -0,345940538 | 0,786794873 | 0,53261606 | 0,931291085 | novel transcript |
| ENSG00000082196 | C1QTNF3 | -0,346074382 | 0,786721883 | 0,48901261 | 0,915720281 | C1q and TNF related 3 |
| ENSG00000162543 | UBXN10 | -0,346142583 | 0,786684693 | 0,35147431 | 0,84405766 | UBX domain protein 10 |
| ENSG00000104047 | DTWD1 | -0,346164367 | 0,786672814 | 0,40322989 | 0,877589651 | DTW domain containing 1 |
| ENSG00000263787 | SKAP1-AS1 | -0,346172812 | 0,786668209 | 0,79382647 | 0,979776981 | SKAP1 antisense RNA 1 |
| ENSG00000257591 | ZNF625 | -0,346301368 | 0,786598114 | 0,27341334 | 0,785672647 | zinc finger protein 625 |
| ENSG00000267278 | MAP3K14-AS1 | -0,346338499 | 0,786577869 | 0,301998 | 0,811449876 | MAP3K14 antisense RNA 1 |
| ENSG00000114631 | PODXL2 | -0,346343556 | 0,786575112 | 0,09141896 | 0,481302729 | podocalyxin like 2 |
| ENSG00000273329 | AC078846,1 | -0,346412967 | 0,786537269 | 0,16178343 | 0,63836332 | novel transcript |
| ENSG00000275897 | AC021491,4 | -0,346442366 | 0,786521242 | 0,77491978 | 0,97857211 | novel transcript |
| ENSG00000201772 | SNORA5C | -0,346554194 | 0,786460278 | 0,58007327 | 0,943828882 | small nucleolar RNA, H/ACA box 5C |
| ENSG00000218502 | H2AZP3 | -0,346577838 | 0,786447389 | 0,75642858 | 0,975901808 | H2A,Z histone pseudogene 3 |
| ENSG00000168772 | CXXC4 | -0,346590471 | 0,786440502 | 0,45271361 | 0,902285042 | CXXC finger protein 4 |
| ENSG00000150593 | PDCD4 | -0,346603332 | 0,786433492 | 0,27965836 | 0,790385 | programmed cell death 4 |
| ENSG00000146757 | ZNF92 | -0,346715306 | 0,786372455 | 0,52735143 | 0,929454507 | zinc finger protein 92 |
| ENSG00000013503 | POLR3B | -0,346776158 | 0,786339288 | 0,31348331 | 0,819801802 | RNA polymerase III subunit B |
| ENSG00000232339 | NOTCH4 | -0,34679022 | 0,786331623 | 0,45502792 | 0,903467092 | notch receptor 4 |
| ENSG00000273162 | AL133215,2 | -0,346846379 | 0,786301015 | 0,34925611 | 0,843141089 | novel transcript |
| ENSG00000277386 | AL138999,1 | -0,346915185 | 0,786263515 | 0,814919 | 0,981521413 | novel transcript, sense intronic to STARD13 |
| ENSG00000120533 | ENY2 | -0,347060383 | 0,786184386 | 0,36570235 | 0,85559319 | ENY2 transcription and export complex 2 subunit |
| ENSG00000263069 | RNF213-AS1 | -0,347151265 | 0,786134862 | 0,25250699 | 0,764588712 | RNF213 antisense RNA 1 |
| ENSG00000272281 | TPTE2P2 | -0,347183859 | 0,786117102 | 0,5919589 | 0,947416795 | transmembrane phosphoinositide 3-phosphatase and tensin homolog 2 pseudogene 2 |
| ENSG00000256124 | LINC01152 | -0,347383199 | 0,78600849 | 0,33478647 | 0,834792227 | long intergenic non-protein coding RNA 1152 |
| ENSG00000279652 | Z82217,1 | -0,347415193 | 0,78599106 | 0,29016786 | 0,800682868 | novel transcript |
| ENSG00000101751 | POLI | -0,347532011 | 0,785927419 | 0,42433851 | 0,890949676 | DNA polymerase iota |
| ENSG00000182004 | SNRPE | -0,347779672 | 0,785792514 | 0,42755047 | 0,892656446 | small nuclear ribonucleoprotein polypeptide E |
| ENSG00000287310 | AL133445,3 | -0,347781447 | 0,785791547 | 0,51592967 | 0,925152007 | novel transcript |
| ENSG00000129535 | NRL | -0,347802306 | 0,785780186 | 0,48069378 | 0,912907448 | neural retina leucine zipper |

| | | | | | | |
|---|---|---|---|---|---|---|
| ENSG00000082212 | ME2 | -0,347814346 | 0,785773628 | 0,39111361 | 0,870057277 | malic enzyme 2 |
| ENSG00000226492 | CUTA | -0,347900208 | 0,785726864 | 0,25491859 | 0,76642479 | cutA divalent cation tolerance homolog |
| ENSG00000231296 | AL050341,1 | -0,347967838 | 0,785690032 | 0,74514554 | 0,975666349 | pseudogene similar to part of RIO kinase 2 (yeast) (RIOK2) |
| ENSG00000227487 | NCAM1-AS1 | -0,348010201 | 0,785666961 | 0,39134407 | 0,870135911 | NCAM1 antisense RNA1 |
| ENSG00000167968 | DNASE1L2 | -0,34805021 | 0,785645173 | 0,64904375 | 0,956899963 | deoxyribonuclease 1 like 2 |
| ENSG00000185559 | DLK1 | -0,348053143 | 0,785643576 | 0,54929256 | 0,936040289 | delta like non-canonical Notch ligand 1 |
| ENSG00000274225 | AP001065,3 | -0,34807863 | 0,785629697 | 0,49342866 | 0,917289924 | novel transcript |
| ENSG00000223653 | AL078459,1 | -0,348165531 | 0,785582376 | 0,14866743 | 0,613346323 | novel transcript |
| ENSG00000232043 | AL133230,1 | -0,348224431 | 0,785550304 | 0,35994944 | 0,850280272 | novel transcript, antisense to PTPN1 |
| ENSG00000280511 | AL591030,1 | -0,348322403 | 0,78549696 | 0,54861207 | 0,935825387 | novel transcript |
| ENSG00000159915 | ZNF233 | -0,348343564 | 0,785485439 | 0,25298518 | 0,76515438 | zinc finger protein 233 |
| ENSG00000287559 | AL136305,1 | -0,348360201 | 0,78547638 | 0,41275885 | 0,884576962 | novel transcript, antisense to RBM24 |
| ENSG00000183562 | AC131971,1 | -0,348420223 | 0,785443702 | 0,34600483 | 0,841168721 | novel transcript, antisense to NAP1L4 |
| ENSG00000282438 | AC001228,2 | -0,348420223 | 0,785443702 | 0,34600483 | 0,841168721 | novel transcript, antisense to NAP1L4 |
| ENSG00000072133 | RPS6KA6 | -0,348694494 | 0,785294396 | 0,42743394 | 0,892552417 | ribosomal protein S6 kinase A6 |
| ENSG00000265777 | RN7SL624P | -0,3488236 | 0,785224123 | 0,80797058 | 0,981213877 | RNA, 7SL, cytoplasmic 624, pseudogene |
| ENSG00000277429 | SERF1B | -0,348920201 | 0,785171547 | 0,62021054 | 0,952793934 | small EDRK-rich factor 1B |
| ENSG00000123636 | BAZ2B | -0,349010771 | 0,785122257 | 0,2288859 | 0,737173617 | bromodomain adjacent to zinc finger domain 2B |
| ENSG00000102349 | KLF8 | -0,34917154 | 0,78503477 | 0,29270057 | 0,803141172 | Kruppel like factor 8 |
| ENSG00000281324 | LRP6 | -0,34922363 | 0,785006427 | 0,29708125 | 0,806635068 | LDL receptor related protein 6 |
| ENSG00000070018 | LRP6 | -0,34922363 | 0,785006427 | 0,29708125 | 0,806635068 | LDL receptor related protein 6 |
| ENSG00000262671 | AC004918,1 | -0,349398339 | 0,784911369 | 0,49368071 | 0,917289924 | novel transcript, antisense to AGK |
| ENSG00000244701 | AC004918,1 | -0,349398339 | 0,784911369 | 0,49368071 | 0,917289924 | novel transcript, antisense to AGK |
| ENSG00000157224 | CLDN12 | -0,349471988 | 0,784871301 | 0,49812634 | 0,919274055 | claudin 12 |
| ENSG00000246560 | UBE2D3-AS1 | -0,349473304 | 0,784870584 | 0,50491938 | 0,921400408 | UBE2D3 antisense RNA 1 |
| ENSG00000137309 | HMGA1 | -0,349590972 | 0,784806572 | 0,18738957 | 0,681464181 | high mobility group AT-hook 1 |
| ENSG00000267520 | AC010733,1 | -0,349723056 | 0,784734723 | 0,34970468 | 0,843534837 | novel transcript |
| ENSG00000287799 | AC005014,4 | -0,349827123 | 0,78467812 | 0,79048113 | 0,979462318 | novel transcript, antisense to ANKMY2 |
| ENSG00000225076 | WWC3-AS1 | -0,349896691 | 0,784640283 | 0,73683444 | 0,974353474 | WWC3 antisense RNA 1 |
| ENSG00000181450 | ZNF678 | -0,350088473 | 0,784535985 | 0,45846338 | 0,90420131 | zinc finger protein 678 |
| ENSG00000173198 | CYSLTR1 | -0,350118231 | 0,784519803 | 0,1488603 | 0,613515281 | cysteinyl leukotriene receptor 1 |
| ENSG00000286053 | ASDURF | -0,350151409 | 0,784501761 | 0,66759613 | 0,9611116 | ASNSD1 upstream reading frame |

| | | | | | | |
|---|---|---|---|---|---|---|
| ENSG00000282614 | DND1P1 | -0,350446827 | 0,784341137 | 0,6276311 | 0,952793934 | DND microRNA-mediated repression inhibitor 1 pseudogene 1 |
| ENSG00000264070 | DND1P1 | -0,350446827 | 0,784341137 | 0,6276311 | 0,952793934 | DND microRNA-mediated repression inhibitor 1 pseudogene 1 |
| ENSG00000173894 | CBX2 | -0,350450239 | 0,784339282 | 0,21412136 | 0,721518909 | chromobox 2 |
| ENSG00000186409 | CCDC30 | -0,350560533 | 0,784279321 | 0,20802364 | 0,712494639 | coiled-coil domain containing 30 |
| ENSG00000255524 | NPIPB8 | -0,350671455 | 0,784219024 | 0,64376394 | 0,956633898 | nuclear pore complex interacting protein family member B8 |
| ENSG00000263612 | AF186192,3 | -0,35068234 | 0,784213107 | 0,73224971 | 0,974353474 | novel transcript |
| ENSG00000198130 | HIBCH | -0,350783373 | 0,78415819 | 0,33819404 | 0,837263082 | 3-hydroxyisobutyryl-CoA hydrolase |
| ENSG00000173531 | MST1 | -0,350806357 | 0,784145697 | 0,27491789 | 0,786383635 | macrophage stimulating 1 |
| ENSG00000140263 | SORD | -0,35081641 | 0,784140234 | 0,2213707 | 0,729254784 | sorbitol dehydrogenase |
| ENSG00000114757 | PEX5L | -0,350858675 | 0,784117262 | 0,32979497 | 0,831610673 | peroxisomal biogenesis factor 5 like |
| ENSG00000282217 | AC006449,8 | -0,350949711 | 0,784067785 | 0,78760569 | 0,979462318 | novel transcript, antisense to SRCIN1 |
| ENSG00000274996 | AC006449,1 | -0,350949711 | 0,784067785 | 0,78760569 | 0,979462318 | novel transcript, antisense to SRCIN1 |
| ENSG00000250073 | AP000866,2 | -0,350992252 | 0,784044665 | 0,42413431 | 0,890949676 | novel transcript, antisense to ESAM |
| ENSG00000256159 | AC022080,2 | -0,351070024 | 0,7840024 | 0,7417472 | 0,975176043 | mitochondrial ribosomal protein L30 (MRPL30) pseudogene |
| ENSG00000115109 | EPB41L5 | -0,351080247 | 0,783996844 | 0,36540859 | 0,855323027 | erythrocyte membrane protein band 4,1 like 5 |
| ENSG00000204802 | AL590399,1 | -0,351085646 | 0,783993911 | 0,23960646 | 0,749431667 | novel transcript |
| ENSG00000129197 | RPAIN | -0,351087994 | 0,783992635 | 0,09513258 | 0,489669163 | RPA interacting protein |
| ENSG00000236804 | RPS3AP12 | -0,35112635 | 0,783971792 | 0,68531526 | 0,965434008 | RPS3A pseudogene 12 |
| ENSG00000249825 | THBS4-AS1 | -0,351188804 | 0,783937854 | 0,57900595 | 0,943828882 | THBS4 antisense RNA 1 |
| ENSG00000197857 | ZNF44 | -0,351206148 | 0,78392843 | 0,16088117 | 0,63684337 | zinc finger protein 44 |
| ENSG00000115421 | PAPOLG | -0,351246247 | 0,783906641 | 0,44671403 | 0,901650934 | poly(A) polymerase gamma |
| ENSG00000188559 | RALGAPA2 | -0,351299634 | 0,783877634 | 0,22295168 | 0,731402952 | Ral GTPase activating protein catalytic subunit alpha 2 |
| ENSG00000174842 | GLMN | -0,351308348 | 0,783872898 | 0,42577135 | 0,892133432 | glomulin, FKBP associated protein |
| ENSG00000281344 | HELLPAR | -0,351328176 | 0,783862125 | 0,12515378 | 0,565471028 | HELLP associated long non-coding RNA |
| ENSG00000213109 | AL365223,1 | -0,351342935 | 0,783854106 | 0,56064305 | 0,940189042 | ribosomal protein L7a (RPL7A) pseudogene |
| ENSG00000266865 | AC138207,8 | -0,351420287 | 0,78381208 | 0,332362 | 0,83293962 | SMAD specific E3 ubiquitin protein ligase 2 (SMURF2) pseudogene |
| ENSG00000225205 | AC078883,1 | -0,351511288 | 0,783762641 | 0,53261333 | 0,931291085 | novel transcript |

| | | | | | | |
|---|---|---|---|---|---|---|
| ENSG00000167680 | SEMA6B | -0,351557343 | 0,783737621 | 0,47506214 | 0,911019008 | semaphorin 6B |
| ENSG00000245148 | ARAP1-AS2 | -0,351602792 | 0,783712932 | 0,57413195 | 0,942475308 | ARAP1 antisense RNA 2 |
| ENSG00000172367 | PDZD3 | -0,3516681 | 0,783677455 | 0,74850038 | 0,975901808 | PDZ domain containing 3 |
| ENSG00000260782 | AC007225,1 | -0,351739233 | 0,783638817 | 0,57194806 | 0,942475308 | nuclear factor (erythroid-derived 2)-like 3 (NFE2L3) pseudogene |
| ENSG00000286031 | AC232271,2 | -0,351958069 | 0,783519959 | 0,65250392 | 0,957275891 | novel transcript, antisense to PPP1R3F |
| ENSG00000204650 | LINC02210 | -0,351968495 | 0,783514297 | 0,15331325 | 0,621845544 | long intergenic non-protein coding RNA 2210 |
| ENSG00000251680 | AC008591,1 | -0,351975682 | 0,783510394 | 0,52976168 | 0,930684931 | novel transcript |
| ENSG00000162630 | B3GALT2 | -0,352159064 | 0,783410807 | 0,44267542 | 0,899437949 | beta-1,3-galactosyltransferase 2 |
| ENSG00000255475 | AP001007,2 | -0,352193905 | 0,783391888 | 0,72864347 | 0,973955059 | novel transcript |
| ENSG00000240024 | LINC00888 | -0,352207971 | 0,78338425 | 0,29320016 | 0,803365244 | long intergenic non-protein coding RNA 888 |
| ENSG00000247363 | AC090061,1 | -0,352223982 | 0,783375557 | 0,46686974 | 0,907940653 | novel transcript |
| ENSG00000125966 | MMP24 | -0,352329765 | 0,783318119 | 0,2346517 | 0,744134035 | matrix metallopeptidase 24 |
| ENSG00000108947 | EFNB3 | -0,352492396 | 0,783229823 | 0,27487367 | 0,786383635 | ephrin B3 |
| ENSG00000203865 | ATP1A1-AS1 | -0,352508981 | 0,783220819 | 0,21773659 | 0,72540192 | ATP1A1 antisense RNA 1 |
| ENSG00000255355 | AP000640,2 | -0,352564344 | 0,783190763 | 0,69711552 | 0,968520631 | novel transcript |
| ENSG00000226599 | AL450344,2 | -0,352645527 | 0,783146693 | 0,76662215 | 0,977201891 | novel transcript |
| ENSG00000120458 | MSANTD2 | -0,352708641 | 0,783112433 | 0,2853075 | 0,796937466 | Myb/SANT DNA binding domain containing 2 |
| ENSG00000278722 | AL445288,1 | -0,352779741 | 0,783073841 | 0,77535519 | 0,978587255 | novel transcript, antisense to PCDH17 |
| ENSG00000198315 | ZKSCAN8 | -0,352789597 | 0,783068491 | 0,38543634 | 0,868895792 | zinc finger with KRAB and SCAN domains 8 |
| ENSG00000188316 | ENO4 | -0,352822699 | 0,783050524 | 0,32078547 | 0,824366234 | enolase 4 |
| ENSG00000184166 | OR1D2 | -0,352933022 | 0,782990646 | 0,30400982 | 0,813056239 | olfactory receptor family 1 subfamily D member 2 |
| ENSG00000287811 | AL391840,3 | -0,35295825 | 0,782976955 | 0,51620753 | 0,925176589 | novel transcript |
| ENSG00000258066 | AC138331,1 | -0,353201288 | 0,782845064 | 0,71856827 | 0,971869926 | novel transcript |
| ENSG00000108309 | RUNDC3A | -0,353329976 | 0,782775238 | 0,33311812 | 0,833301711 | RUN domain containing 3A |
| ENSG00000287921 | AL713868,1 | -0,35354035 | 0,782661102 | 0,74479239 | 0,975649186 | novel transcript, antisense to SLC35F3 |
| ENSG00000286831 | AC244472,1 | -0,35356182 | 0,782649455 | 0,46511597 | 0,907804479 | novel transcript |
| ENSG00000281228 | AL132640,3 | -0,353695163 | 0,78257712 | 0,80906096 | 0,981213877 | |
| ENSG00000135248 | FAM71F1 | -0,353724387 | 0,782561268 | 0,63898082 | 0,956018518 | family with sequence similarity 71 member F1 |
| ENSG00000269570 | AP001350,1 | -0,353833681 | 0,782501986 | 0,47052736 | 0,908805405 | novel transcript, antisense to ZFP91 |
| ENSG00000125551 | PLGLB2 | -0,353857722 | 0,782488946 | 0,34994918 | 0,843677595 | plasminogen like B2 |
| ENSG00000253854 | AC010834,3 | -0,354105783 | 0,782354415 | 0,25914251 | 0,771989931 | novel transcript |
| ENSG00000273449 | AC093788,1 | -0,354106346 | 0,78235411 | 0,50082849 | 0,920520138 | novel transcript |

| | | | | | | |
|---|---|---|---|---|---|---|
| ENSG00000272323 | AC026801,2 | -0,354115973 | 0,782348889 | 0,59283928 | 0,947416795 | novel transcript, antisense to TTC23L |
| ENSG00000203288 | TDRKH-AS1 | -0,354214182 | 0,782295634 | 0,55206367 | 0,936552095 | TDRKH antisense RNA 1 |
| ENSG00000105538 | RASIP1 | -0,354220222 | 0,782292359 | 0,48234636 | 0,913113923 | Ras interacting protein 1 |
| ENSG00000114805 | PLCH1 | -0,354294266 | 0,78225221 | 0,27540882 | 0,787054963 | phospholipase C eta 1 |
| ENSG00000261330 | PPIAP50 | -0,354552764 | 0,782112061 | 0,6434463 | 0,956633898 | peptidylprolyl isomerase A pseudogene 50 |
| ENSG00000126249 | PDCD2L | -0,354560526 | 0,782107853 | 0,29402338 | 0,803927308 | programmed cell death 2 like |
| ENSG00000112425 | EPM2A | -0,354594378 | 0,782089501 | 0,21427922 | 0,72158829 | EPM2A glucan phosphatase, laforin |
| ENSG00000282953 | MUC20-OT1 | -0,354638975 | 0,782065326 | 0,24649218 | 0,758273287 | MUC20 overlapping transcript |
| ENSG00000090097 | PCBP4 | -0,354671293 | 0,782047807 | 0,14490162 | 0,605968311 | poly(rC) binding protein 4 |
| ENSG00000089091 | DZANK1 | -0,354720071 | 0,782021365 | 0,15299056 | 0,621569237 | double zinc ribbon and ankyrin repeat domains 1 |
| ENSG00000266079 | SNORA59B | -0,354873305 | 0,781938309 | 0,71213924 | 0,970812834 | small nucleolar RNA, H/ACA box 59B |
| ENSG00000260948 | AL390195,2 | -0,354907004 | 0,781920044 | 0,32620877 | 0,828184588 | novel transcript, sense overlapping WDR77 & OVGP1 |
| ENSG00000273772 | SMN2 | -0,355111171 | 0,781809396 | 0,30780644 | 0,816424917 | survival of motor neuron 2, centromeric |
| ENSG00000228863 | AL121985,1 | -0,355117757 | 0,781805827 | 0,38853112 | 0,869820053 | novel transcript, antisense to CD48 |
| ENSG00000243176 | AC092944,1 | -0,355124675 | 0,781802078 | 0,31454622 | 0,819877431 | novel transcript |
| ENSG00000151725 | CENPU | -0,355196647 | 0,781763077 | 0,38760038 | 0,869820053 | centromere protein U |
| ENSG00000197608 | ZNF841 | -0,355198486 | 0,781762081 | 0,20778519 | 0,712302214 | zinc finger protein 841 |
| ENSG00000285129 | RPS3AP30 | -0,355355914 | 0,781676779 | 0,77856655 | 0,978587255 | RPS3A pseudogene 30 |
| ENSG00000286362 | AP005717,2 | -0,35536153 | 0,781673736 | 0,6128242 | 0,952413269 | novel transcript, antisense to DSCC1 |
| ENSG00000240591 | AL096701,3 | -0,355363637 | 0,781672595 | 0,36401705 | 0,854207909 | novel transcript |
| ENSG00000126970 | ZC4H2 | -0,355393141 | 0,781656609 | 0,28517589 | 0,796688336 | zinc finger C4H2-type containing |
| ENSG00000288396 | AC213033,3 | -0,355408831 | 0,781648108 | 0,37320664 | 0,861257555 | |
| ENSG00000115355 | CCDC88A | -0,355511145 | 0,781592677 | 0,26733845 | 0,779594822 | coiled-coil domain containing 88A |
| ENSG00000284734 | AC099063,4 | -0,355774239 | 0,781450156 | 0,27766235 | 0,789067728 | novel transcript, antisense to GBP4 |
| ENSG00000225226 | LINC01795 | -0,355827578 | 0,781421265 | 0,45164258 | 0,902062594 | long intergenic non-protein coding RNA 1795 |
| ENSG00000237296 | SMG1P1 | -0,356093461 | 0,781277266 | 0,24178378 | 0,752128472 | SMG1 pseudogene 1 |
| ENSG00000184368 | MAP7D2 | -0,356177412 | 0,781231804 | 0,28173828 | 0,792816227 | MAP7 domain containing 2 |
| ENSG00000234719 | NPIPB2 | -0,356197934 | 0,781220691 | 0,23069419 | 0,739234644 | nuclear pore complex interacting protein family member B2 |
| ENSG00000269949 | AC069307,1 | -0,356310921 | 0,781159511 | 0,65402627 | 0,957444211 | novel transcript |
| ENSG00000128881 | TTBK2 | -0,356370221 | 0,781127403 | 0,31129217 | 0,817751312 | tau tubulin kinase 2 |
| ENSG00000228157 | AC007952,2 | -0,356376369 | 0,781124075 | 0,53976005 | 0,933385624 | novel transcript |
| ENSG00000119640 | ACYP1 | -0,356467663 | 0,781074647 | 0,36184038 | 0,851979035 | acylphosphatase 1 |

| | | | | | | |
|---|---|---|---|---|---|---|
| ENSG00000287601 | BX323046,2 | -0,356537419 | 0,781036881 | 0,58487757 | 0,944857905 | novel transcript |
| ENSG00000158402 | CDC25C | -0,356598907 | 0,781003594 | 0,26494462 | 0,776509969 | cell division cycle 25C |
| ENSG00000270175 | AC023509,3 | -0,356618263 | 0,780993116 | 0,43661462 | 0,896866043 | novel transcript, antisense to TARBP2 |
| ENSG00000144712 | CAND2 | -0,356635161 | 0,780983968 | 0,20592472 | 0,708488009 | cullin associated and neddylation dissociated 2 (putative) |
| ENSG00000273521 | AL162274,1 | -0,356704991 | 0,780946168 | 0,51597694 | 0,925176589 | novel transcript, antisense to BNIP3 |
| ENSG00000188428 | BLOC1S5 | -0,356953482 | 0,780811669 | 0,34907447 | 0,843141089 | biogenesis of lysosomal organelles complex 1 subunit 5 |
| ENSG00000277053 | GTF2IP1 | -0,357063524 | 0,780752114 | 0,35317883 | 0,845063255 | general transcription factor IIi pseudogene 1 |
| ENSG00000243819 | RN7SL832P | -0,357166185 | 0,780696558 | 0,4512142 | 0,902062594 | RNA, 7SL, cytoplasmic 832, pseudogene |
| ENSG00000251209 | LINC00923 | -0,357335616 | 0,780604878 | 0,13919103 | 0,594862544 | long intergenic non-protein coding RNA 923 |
| ENSG00000236031 | AL591721,1 | -0,357460479 | 0,780537321 | 0,60187402 | 0,949939978 | novel transcript |
| ENSG00000130775 | THEMIS2 | -0,357490936 | 0,780520843 | 0,38620196 | 0,869312875 | thymocyte selection associated family member 2 |
| ENSG00000205129 | C4orf47 | -0,357507363 | 0,780511956 | 0,62694803 | 0,952793934 | chromosome 4 open reading frame 47 |
| ENSG00000235740 | PHACTR2-AS1 | -0,357998056 | 0,780246531 | 0,52218589 | 0,927617924 | PHACTR2 antisense RNA 1 |
| ENSG00000274964 | AC026356,1 | -0,357999625 | 0,780245683 | 0,34314705 | 0,84076406 | novel transcript, sense intronic to BICD1 |
| ENSG00000077514 | POLD3 | -0,358067609 | 0,780208917 | 0,31009215 | 0,817567979 | DNA polymerase delta 3, accessory subunit |
| ENSG00000143498 | TAF1A | -0,358324068 | 0,780070236 | 0,26247131 | 0,774068771 | TATA-box binding protein associated factor, RNA polymerase I subunit A |
| ENSG00000242748 | RPL23AP81 | -0,358347609 | 0,780057507 | 0,5978102 | 0,948390834 | ribosomal protein L23a pseudogene 81 |
| ENSG00000068615 | REEP1 | -0,358527148 | 0,779960437 | 0,46812647 | 0,907940653 | receptor accessory protein 1 |
| ENSG00000287277 | AL392086,3 | -0,358567043 | 0,779938869 | 0,11179556 | 0,534594124 | novel transcript |
| ENSG00000234469 | CLDN34 | -0,358601114 | 0,77992045 | 0,66003135 | 0,958859629 | claudin 34 |
| ENSG00000261202 | Z83847,1 | -0,358616852 | 0,779911943 | 0,71334573 | 0,970812834 | novel transcript, antisense to FAM83F |
| ENSG00000117650 | NEK2 | -0,358707523 | 0,779862928 | 0,33064467 | 0,832114797 | NIMA related kinase 2 |
| ENSG00000261366 | MANEA-DT | -0,3587077 | 0,779862832 | 0,54396894 | 0,934259495 | MANEA divergent transcript |
| ENSG00000285501 | AGBL2 | -0,35874316 | 0,779843664 | 0,65097722 | 0,957027739 | ATP/GTP binding protein like 2 |
| ENSG00000165923 | AGBL2 | -0,35874316 | 0,779843664 | 0,65097722 | 0,957027739 | ATP/GTP binding protein like 2 |
| ENSG00000255171 | LINC01499 | -0,358859216 | 0,779780933 | 0,74227412 | 0,975176043 | long intergenic non-protein coding RNA 1499 |
| ENSG00000204283 | LINC01973 | -0,359007081 | 0,779701015 | 0,79249486 | 0,979471509 | long intergenic non-protein coding RNA 1973 |
| ENSG00000207955 | AL359091,1 | -0,359015859 | 0,779696272 | 0,44307553 | 0,899690238 | novel transcript |
| ENSG00000282176 | FLJ13224 | -0,359019637 | 0,77969423 | 0,52632044 | 0,929261495 | uncharacterized LOC79857 |
| ENSG00000177340 | FLJ13224 | -0,359019637 | 0,77969423 | 0,52632044 | 0,929261495 | uncharacterized LOC79857 |

| | | | | | | |
|---|---|---|---|---|---|---|
| ENSG00000271781 | AC026740,1 | -0,359037027 | 0,779684832 | 0,77228907 | 0,97820249 | novel transcript, antisense to TPPP |
| ENSG00000134184 | GSTM1 | -0,359167266 | 0,779614449 | 0,20708092 | 0,710823213 | glutathione S-transferase mu 1 |
| ENSG00000238197 | PAXBP1-AS1 | -0,359465201 | 0,779453465 | 0,19427093 | 0,691291421 | PAXBP1 antisense RNA 1 |
| ENSG00000254708 | MMADHCP2 | -0,359520695 | 0,779423484 | 0,52118282 | 0,927046149 | MMADHC pseudogene 2 |
| ENSG00000274204 | AL138689,1 | -0,359603127 | 0,779378951 | 0,49187302 | 0,916995341 | novel transcript, sense intronic to EFNB2 |
| ENSG00000132570 | PCBD2 | -0,359616102 | 0,779371941 | 0,11253301 | 0,535800356 | pterin-4 alpha-carbinolamine dehydratase 2 |
| ENSG00000075539 | FRYL | -0,359629956 | 0,779364457 | 0,33696522 | 0,836211475 | FRY like transcription coactivator |
| ENSG00000273033 | LINC02035 | -0,359634811 | 0,779361834 | 0,29237121 | 0,802771257 | long intergenic non-protein coding RNA 2035 |
| ENSG00000279932 | AP006248,3 | -0,359978759 | 0,779176051 | 0,81258153 | 0,981224866 | novel transcript, sense intronic SLC7A2 |
| ENSG00000171819 | ANGPTL7 | -0,359981188 | 0,77917474 | 0,45686149 | 0,903699469 | angiopoietin like 7 |
| ENSG00000155962 | CLIC2 | -0,359996779 | 0,779166319 | 0,57454631 | 0,942475308 | chloride intracellular channel 2 |
| ENSG00000183023 | SLC8A1 | -0,36008674 | 0,779117735 | 0,36836145 | 0,857548638 | solute carrier family 8 member A1 |
| ENSG00000188522 | FAM83G | -0,360097688 | 0,779111823 | 0,56284722 | 0,940865625 | family with sequence similarity 83 member G |
| ENSG00000100697 | DICER1 | -0,360203166 | 0,779054862 | 0,38899541 | 0,870057277 | dicer 1, ribonuclease III |
| ENSG00000227953 | LINC01341 | -0,360258512 | 0,779024976 | 0,23113717 | 0,739268552 | long intergenic non-protein coding RNA 1341 |
| ENSG00000132646 | PCNA | -0,360278376 | 0,77901425 | 0,38674212 | 0,869441674 | proliferating cell nuclear antigen |
| ENSG00000253377 | AC068672,2 | -0,36036062 | 0,778969842 | 0,67110128 | 0,962026636 | novel transcript |
| ENSG00000196074 | SYCP2 | -0,360369261 | 0,778965176 | 0,37680716 | 0,862818448 | synaptonemal complex protein 2 |
| ENSG00000255020 | AF131216,3 | -0,360403666 | 0,7789466 | 0,79012615 | 0,979462318 | novel transcript, antisense to TDH |
| ENSG00000259232 | AC105129,1 | -0,360529454 | 0,778878687 | 0,70723791 | 0,969565008 | target of EGR1, member 1 (nuclear) (TOE1) pseudogene |
| ENSG00000206656 | SNORD116-17 | -0,360531863 | 0,778877386 | 0,53340721 | 0,931381467 | small nucleolar RNA, C/D box 116-17 |
| ENSG00000207460 | SNORD116-19 | -0,360531863 | 0,778877386 | 0,53340721 | 0,931381467 | small nucleolar RNA, C/D box 116-19 |
| ENSG00000130413 | STK33 | -0,360617127 | 0,778831356 | 0,2764145 | 0,788263115 | serine/threonine kinase 33 |
| ENSG00000237596 | AL138828,1 | -0,360627099 | 0,778825972 | 0,19063492 | 0,685949202 | novel transcript |
| ENSG00000243620 | AC092957,1 | -0,360871915 | 0,778693822 | 0,53522552 | 0,931905243 | novel transcript |
| ENSG00000261167 | AC107027,3 | -0,36092602 | 0,778664619 | 0,39067287 | 0,870057277 | novel transcript, overlapping to MRPL3 |
| ENSG00000273261 | AC092953,2 | -0,36095652 | 0,778648157 | 0,720419 | 0,972142678 | novel transcript, antisense to ATP11B |
| ENSG00000173226 | IQCB1 | -0,360979656 | 0,778635671 | 0,37373134 | 0,861319894 | IQ motif containing B1 |
| ENSG00000186675 | MAGEE2 | -0,360988477 | 0,77863091 | 0,57773497 | 0,94337651 | MAGE family member E2 |
| ENSG00000237638 | LINC02245 | -0,361107866 | 0,778566478 | 0,54812771 | 0,935404974 | long intergenic non-protein coding RNA 2245 |
| ENSG00000261135 | AL137802,2 | -0,361248658 | 0,778490502 | 0,48602953 | 0,914018932 | novel transcript |
| ENSG00000182903 | ZNF721 | -0,361252177 | 0,778488603 | 0,31114647 | 0,817751312 | zinc finger protein 721 |

| | | | | | | |
|---|---|---|---|---|---|---|
| ENSG00000145687 | SSBP2 | -0,361323646 | 0,778450038 | 0,31738802 | 0,822252078 | single stranded DNA binding protein 2 |
| ENSG00000251152 | AC025539,1 | -0,361378928 | 0,77842021 | 0,56346086 | 0,940865625 | novel transcript |
| ENSG00000161642 | ZNF385A | -0,361405371 | 0,778405943 | 0,23055593 | 0,739234644 | zinc finger protein 385A |
| ENSG00000111605 | CPSF6 | -0,361487356 | 0,778361709 | 0,34414466 | 0,840938551 | cleavage and polyadenylation specific factor 6 |
| ENSG00000272892 | AL133551,1 | -0,361518684 | 0,778344807 | 0,45978368 | 0,905206367 | novel transcript |
| ENSG00000272829 | AC002470,1 | -0,361531914 | 0,778337669 | 0,67990403 | 0,964539408 | novel transcript |
| ENSG00000276368 | H2AC14 | -0,361690952 | 0,778251873 | 0,36946741 | 0,857991124 | H2A clustered histone 14 |
| ENSG00000234617 | SNRK-AS1 | -0,361867039 | 0,778156889 | 0,40611333 | 0,879152963 | SNRK antisense RNA 1 |
| ENSG00000188505 | NCCRP1 | -0,361870905 | 0,778154804 | 0,71046848 | 0,970482408 | NCCRP1, F-box associated domain containing |
| ENSG00000234187 | AIMP1P1 | -0,36192583 | 0,778125179 | 0,70315189 | 0,968742626 | aminoacyl tRNA synthetase complex interacting multifunctional protein 1 pseudogene 1 |
| ENSG00000196081 | ZNF724 | -0,361933168 | 0,778121221 | 0,22256667 | 0,731070016 | zinc finger protein 724 |
| ENSG00000165028 | NIPSNAP3B | -0,361959913 | 0,778106797 | 0,4161358 | 0,885986046 | nipsnap homolog 3B |
| ENSG00000272209 | AL023583,1 | -0,361971021 | 0,778100806 | 0,74077577 | 0,975059523 | novel transcript |
| ENSG00000268798 | AC027307,3 | -0,361982211 | 0,77809477 | 0,47944574 | 0,912714933 | novel transcript |
| ENSG00000214745 | GOT2P6 | -0,362148189 | 0,778005258 | 0,53289155 | 0,931291085 | GOT2 pseudogene 6 |
| ENSG00000107643 | MAPK8 | -0,362166561 | 0,777995351 | 0,43828021 | 0,897177404 | mitogen-activated protein kinase 8 |
| ENSG00000241764 | AC002467,1 | -0,362292427 | 0,777927478 | 0,45220467 | 0,902062594 | novel transcript, antisense to CBLL1 |
| ENSG00000239608 | RUVBL1-AS1 | -0,36230865 | 0,777918731 | 0,68176214 | 0,964654539 | RUVBL1 antisense RNA 1 |
| ENSG00000196724 | ZNF418 | -0,362319683 | 0,777912782 | 0,11597175 | 0,544078746 | zinc finger protein 418 |
| ENSG00000236977 | ANKRD44-IT1 | -0,362372085 | 0,777884527 | 0,77063332 | 0,977966587 | ANKRD44 intronic transcript 1 |
| ENSG00000136161 | RCBTB2 | -0,362505863 | 0,777812398 | 0,41575953 | 0,885986046 | RCC1 and BTB domain containing protein 2 |
| ENSG00000287168 | AL512625,3 | -0,362616953 | 0,777752508 | 0,33150097 | 0,832548086 | novel transcript |
| ENSG00000165168 | CYBB | -0,362806129 | 0,777650531 | 0,61303682 | 0,952413269 | cytochrome b-245 beta chain |
| ENSG00000237357 | BX088651,4 | -0,362843683 | 0,777630288 | 0,40172332 | 0,87705851 | novel transcript |
| ENSG00000161149 | TUBA3FP | -0,362894148 | 0,777603087 | 0,36148935 | 0,851893059 | tubulin alpha 3f pseudogene |
| ENSG00000271895 | AL109811,2 | -0,362971141 | 0,77756159 | 0,30916338 | 0,816845699 | novel transcript, antisense to MASP2 |
| ENSG00000242671 | LINC02010 | -0,362983942 | 0,777554691 | 0,61932292 | 0,952793934 | long intergenic non-protein coding RNA 2010 |
| ENSG00000253133 | AC009630,1 | -0,363037375 | 0,777525893 | 0,36136064 | 0,851875965 | novel transcript, antisense to AGPAT6 and GINS4 |
| ENSG00000143801 | PSEN2 | -0,363096224 | 0,777494177 | 0,11256195 | 0,535800356 | presenilin 2 |
| ENSG00000230751 | AC007036,2 | -0,363118473 | 0,777482187 | 0,77013093 | 0,977840776 | novel transcript |
| ENSG00000187790 | FANCM | -0,363170686 | 0,77745405 | 0,41331523 | 0,884678298 | FA complementation group M |
| ENSG00000152977 | ZIC1 | -0,363189162 | 0,777444093 | 0,26991231 | 0,78199781 | Zic family member 1 |

| | | | | | | |
|---|---|---|---|---|---|---|
| ENSG00000104427 | ZC2HC1A | -0,363225866 | 0,777424314 | 0,48551799 | 0,913907266 | zinc finger C2HC-type containing 1A |
| ENSG00000268442 | AC073534,1 | -0,363346825 | 0,777359135 | 0,49508052 | 0,917770377 | hepatitis A virus cellular receptor 1 (HAVCR1) pseudogene |
| ENSG00000206838 | SNORA5A | -0,36335025 | 0,77735729 | 0,71914416 | 0,971869926 | small nucleolar RNA, H/ACA box 5A |
| ENSG00000229109 | AL137847,1 | -0,363385449 | 0,777338325 | 0,47128569 | 0,908869685 | novel transcript |
| ENSG00000285156 | GS1-24F4,2 | -0,36339514 | 0,777333103 | 0,57723438 | 0,94337651 | uncharacterized LOC100652791 |
| ENSG00000167523 | SPATA33 | -0,36376965 | 0,777131341 | 0,15205567 | 0,620332737 | spermatogenesis associated 33 |
| ENSG00000229278 | AL133353,1 | -0,363799992 | 0,777114997 | 0,8714639 | 0,989217334 | novel transcript |
| ENSG00000137720 | C11orf1 | -0,363926718 | 0,777046738 | 0,36061398 | 0,851271174 | chromosome 11 open reading frame 1 |
| ENSG00000188985 | DHFRP1 | -0,364104122 | 0,776951193 | 0,85368635 | 0,987861659 | dihydrofolate reductase pseudogene 1 |
| ENSG00000250938 | AC073475,1 | -0,364109731 | 0,776948172 | 0,59376413 | 0,947469769 | novel transcript, antisense to MAD2L1 |
| ENSG00000258819 | LINC02289 | -0,364117052 | 0,77694423 | 0,41833834 | 0,887094957 | long intergenic non-protein coding RNA 2289 |
| ENSG00000166821 | PEX11A | -0,364125156 | 0,776939865 | 0,33449313 | 0,834713776 | peroxisomal biogenesis factor 11 alpha |
| ENSG00000066379 | POLR1H | -0,364129998 | 0,776937257 | 0,30028691 | 0,810130728 | RNA polymerase I subunit H |
| ENSG00000265908 | AC024267,5 | -0,364185396 | 0,776907425 | 0,51055219 | 0,922693987 | novel transcript, antisense to PHF12 |
| ENSG00000281128 | PTENP1-AS | -0,364317359 | 0,776836364 | 0,45265965 | 0,902285042 | PTENP1 antisense RNA |
| ENSG00000239445 | ST3GAL6-AS1 | -0,364430025 | 0,7767757 | 0,39231612 | 0,870909282 | ST3GAL6 antisense RNA 1 |
| ENSG00000203814 | H2BC18 | -0,364596842 | 0,776685888 | 0,39263902 | 0,870909282 | H2B clustered histone 18 |
| ENSG00000271078 | AC139426,3 | -0,364598069 | 0,776685227 | 0,60943569 | 0,951793902 | NEDD4 binding protein 2 (N4BP2) pseudogene |
| ENSG00000233006 | MIR3936HG | -0,364649673 | 0,776657447 | 0,30871434 | 0,816730513 | MIR3936 host gene |
| ENSG00000111816 | FRK | -0,364665123 | 0,776649129 | 0,13318922 | 0,582952249 | fyn related Src family tyrosine kinase |
| ENSG00000286369 | AP002813,1 | -0,364779745 | 0,776587427 | 0,50260502 | 0,920607749 | novel transcript |
| ENSG00000261140 | AC093525,4 | -0,364836446 | 0,776556906 | 0,37355965 | 0,861319894 | novel transcript, antisense to PDPK1 |
| ENSG00000123810 | B9D2 | -0,364862324 | 0,776542977 | 0,17410568 | 0,659373116 | B9 domain containing 2 |
| ENSG00000119397 | CNTRL | -0,364881408 | 0,776532705 | 0,10943179 | 0,527649315 | centriolin |
| ENSG00000080823 | MOK | -0,36529446 | 0,776310411 | 0,11206356 | 0,535057308 | MOK protein kinase |
| ENSG00000249014 | HMGN2P4 | -0,365369245 | 0,77627017 | 0,50721458 | 0,921951873 | high mobility group nucleosomal binding domain 2 pseudogene 4 |
| ENSG00000287918 | AF274853,1 | -0,365441111 | 0,776231502 | 0,42671743 | 0,892261855 | novel transcript, antisense to VMA21 |
| ENSG00000180828 | BHLHE22 | -0,365513845 | 0,776192369 | 0,51211315 | 0,923339417 | basic helix-loop-helix family member e22 |
| ENSG00000257027 | AC010186,3 | -0,365549132 | 0,776173384 | 0,43644289 | 0,896850871 | novel transcript |
| ENSG00000167992 | VWCE | -0,36563396 | 0,776127748 | 0,54427706 | 0,934381824 | von Willebrand factor C and EGF domains |
| ENSG00000229017 | LINC01277 | -0,365651142 | 0,776118505 | 0,64883915 | 0,956899963 | long intergenic non-protein coding RNA 1277 |

| | | | | | | |
|---|---|---|---|---|---|---|
| ENSG00000162613 | FUBP1 | -0,365664975 | 0,776111063 | 0,42165986 | 0,889705998 | far upstream element binding protein 1 |
| ENSG00000003137 | CYP26B1 | -0,365665413 | 0,776110827 | 0,3685225 | 0,857548638 | cytochrome P450 family 26 subfamily B member 1 |
| ENSG00000258429 | PDF | -0,365812232 | 0,776031849 | 0,36172942 | 0,851974425 | peptide deformylase, mitochondrial |
| ENSG00000272103 | AC026741,1 | -0,365844465 | 0,776014511 | 0,50213963 | 0,920520138 | novel transcript |
| ENSG00000266844 | AC093330,1 | -0,365847027 | 0,776013133 | 0,70243738 | 0,968742626 | novel transcript, antisense to MBP |
| ENSG00000169169 | CPT1C | -0,365966411 | 0,77594892 | 0,15252963 | 0,620962584 | carnitine palmitoyltransferase 1C |
| ENSG00000144893 | MED12L | -0,366018916 | 0,775920681 | 0,30404093 | 0,813056239 | mediator complex subunit 12L |
| ENSG00000133216 | EPHB2 | -0,366091798 | 0,775881484 | 0,11481481 | 0,541181798 | EPH receptor B2 |
| ENSG00000277775 | H3C7 | -0,366125854 | 0,775863169 | 0,19819483 | 0,697003721 | H3 clustered histone 7 |
| ENSG00000274561 | AC005332,3 | -0,366180005 | 0,775834048 | 0,55277935 | 0,936894267 | novel transcript |
| ENSG00000143669 | LYST | -0,366214335 | 0,775815586 | 0,34317503 | 0,840766725 | lysosomal trafficking regulator |
| ENSG00000255182 | AC084125,2 | -0,366281475 | 0,775779482 | 0,56488982 | 0,940865625 | novel transcript, antisense to PPP1R16A |
| ENSG00000225619 | MYT1L-AS1 | -0,366468753 | 0,775678784 | 0,50519714 | 0,921605995 | MYT1L antisense RNA 1 |
| ENSG00000156689 | GLYATL2 | -0,366477953 | 0,775673838 | 0,49605773 | 0,918105073 | glycine-N-acyltransferase like 2 |
| ENSG00000256361 | AC027544,1 | -0,366519646 | 0,775651421 | 0,5927515 | 0,947416795 | glycine cleavage system protein H (aminomethyl carrier) (GCSH) |
| ENSG00000205593 | DENND6B | -0,366617311 | 0,775598914 | 0,17684877 | 0,662711251 | DENN domain containing 6B |
| ENSG00000267886 | AC074135,1 | -0,366715722 | 0,77554601 | 0,28238355 | 0,793424943 | novel transcript |
| ENSG00000130783 | CCDC62 | -0,366726824 | 0,775540042 | 0,38738985 | 0,869631393 | coiled-coil domain containing 62 |
| ENSG00000225964 | NRIR | -0,366732013 | 0,775537252 | 0,55647532 | 0,938061341 | negative regulator of interferon response |
| ENSG00000196437 | ZNF569 | -0,36685908 | 0,775468949 | 0,45112937 | 0,902062594 | zinc finger protein 569 |
| ENSG00000279208 | CR381653,2 | -0,36688787 | 0,775453474 | 0,83279695 | 0,984905093 | ankyrin repeat domain 18A (ANKRD18A) pseudogene |
| ENSG00000287437 | AL161662,1 | -0,366950011 | 0,775420074 | 0,34586813 | 0,841168721 | novel transcript, antisense to NRDE2 |
| ENSG00000249713 | AC026725,1 | -0,366991492 | 0,775397779 | 0,68879062 | 0,966556275 | novel transcript |
| ENSG00000257060 | AC091078,1 | -0,367057625 | 0,775362236 | 0,14694527 | 0,610674677 | novel transcript |
| ENSG00000260920 | AL031985,3 | -0,367068884 | 0,775356185 | 0,44843852 | 0,902062594 | novel transcript, sense overlapping with ZNF643 |
| ENSG00000282961 | PRNCR1 | -0,367135103 | 0,775320597 | 0,26347628 | 0,774637968 | prostate cancer associated non-coding RNA 1 |
| ENSG00000228629 | AC092295,1 | -0,367320485 | 0,775220977 | 0,80533301 | 0,981213877 | novel transcript, antisense to ZNF260 |
| ENSG00000176230 | OR4K17 | -0,367354355 | 0,775202777 | 0,2415661 | 0,751948055 | olfactory receptor family 4 subfamily K member 17 |
| ENSG00000224389 | C4B | -0,367493902 | 0,775127798 | 0,43749034 | 0,897021798 | complement C4B (Chido blood group) |
| ENSG00000271344 | AC018638,6 | -0,367494874 | 0,775127276 | 0,55173238 | 0,936366211 | novel transcript |

| | | | | | | |
|---|---|---|---|---|---|---|
| ENSG00000250483 | PPM1AP1 | -0,367506003 | 0,775121297 | 0,64972685 | 0,956899963 | protein phosphatase, Mg2+/Mn2+ dependent 1A pseudogene 1 |
| ENSG00000246662 | LINC00535 | -0,367635783 | 0,775051572 | 0,44618893 | 0,901348735 | long intergenic non-protein coding RNA 535 |
| ENSG00000269388 | AC018755,3 | -0,367774372 | 0,774977123 | 0,60594855 | 0,950927937 | putative ATP-binding domain-containing protein 3-like protein (ABP3L) pseudogene |
| ENSG00000219481 | NBPF1 | -0,36787074 | 0,774925358 | 0,17084343 | 0,654404445 | NBPF member 1 |
| ENSG00000282418 | AC092811,1 | -0,367935483 | 0,774890583 | 0,74347011 | 0,975310206 | novel transcript |
| ENSG00000286162 | AL162253,2 | -0,368038943 | 0,774835016 | 0,45461015 | 0,903383444 | novel transcript |
| ENSG00000212907 | MT-ND4L | -0,368478462 | 0,774598997 | 0,33914278 | 0,838276153 | mitochondrially encoded NADH:ubiquinone oxidoreductase core subunit 4L |
| ENSG00000282866 | CFL1P1 | -0,368570503 | 0,774549581 | 0,56995785 | 0,94221458 | cofilin 1 pseudogene 1 |
| ENSG00000223820 | CFL1P1 | -0,368570503 | 0,774549581 | 0,56995785 | 0,94221458 | cofilin 1 pseudogene 1 |
| ENSG00000147509 | RGS20 | -0,368610665 | 0,774528019 | 0,24943487 | 0,761381055 | regulator of G protein signaling 20 |
| ENSG00000258682 | AL132989,1 | -0,3686913 | 0,77448473 | 0,52250736 | 0,927794062 | novel transcript, sense overlapping PSMA3 |
| ENSG00000171724 | VAT1L | -0,368696154 | 0,774482125 | 0,30190171 | 0,811368958 | vesicle amine transport 1 like |
| ENSG00000285702 | AC211476,5 | -0,368747119 | 0,774454765 | 0,17264977 | 0,657085135 | novel transcript |
| ENSG00000245711 | NADK2-AS1 | -0,368838528 | 0,774405697 | 0,54657585 | 0,935401821 | NADK2 antisense RNA 1 |
| ENSG00000183439 | TRIM61 | -0,36888522 | 0,774380635 | 0,3515377 | 0,84405766 | tripartite motif containing 61 |
| ENSG00000197816 | CCDC180 | -0,368945752 | 0,774348144 | 0,37982554 | 0,865325783 | coiled-coil domain containing 180 |
| ENSG00000284137 | LLGL1 | -0,368945985 | 0,774348019 | 0,40989621 | 0,882393109 | LLGL scribble cell polarity complex component 1 |
| ENSG00000255556 | AC087203,2 | -0,369013639 | 0,774311707 | 0,63731194 | 0,956017912 | ubiquitin specific peptidase 17-like 2 (USP17L2) pseudogene |
| ENSG00000165156 | ZHX1 | -0,369017424 | 0,774309676 | 0,38992896 | 0,870057277 | zinc fingers and homeoboxes 1 |
| ENSG00000235945 | AC002543,1 | -0,369017685 | 0,774309536 | 0,47210622 | 0,909171312 | novel pseudogene |
| ENSG00000213793 | ZNF888 | -0,369112001 | 0,774258917 | 0,3466321 | 0,841390219 | zinc finger protein 888 |
| ENSG00000287757 | AC244636,3 | -0,369128659 | 0,774249978 | 0,24423519 | 0,755642658 | novel transcript, antisense to ZNF630 |
| ENSG00000275569 | AL355073,2 | -0,369186728 | 0,774218814 | 0,79741428 | 0,980161849 | novel transcript |
| ENSG00000234019 | AL591212,1 | -0,369206409 | 0,774208252 | 0,7317705 | 0,974353474 | novel transcript |
| ENSG00000163879 | DNALI1 | -0,369278756 | 0,774169429 | 0,21771632 | 0,72540192 | dynein axonemal light intermediate chain 1 |
| ENSG00000276241 | AC243829,2 | -0,369397718 | 0,774105595 | 0,60976351 | 0,951801946 | novel transcript, antisense TBC1D3B |
| ENSG00000282100 | HSP90AB4P | -0,369490332 | 0,774055903 | 0,49023247 | 0,916554718 | heat shock protein 90 alpha family class B member 4, pseudogene |
| ENSG00000268362 | AC092279,1 | -0,369493134 | 0,7740544 | 0,06930007 | 0,415287103 | novel transcript |

| | | | | | | |
|---|---|---|---|---|---|---|
| ENSG00000070269 | TMEM260 | -0,36959012 | 0,774002365 | 0,34268886 | 0,840299954 | transmembrane protein 260 |
| ENSG00000287074 | AC137561,1 | -0,369745076 | 0,773919236 | 0,43919482 | 0,897434993 | novel transcript, sense intronic to AEBP2 |
| ENSG00000160190 | SLC37A1 | -0,369781571 | 0,773899659 | 0,14856451 | 0,613326406 | solute carrier family 37 member 1 |
| ENSG00000196267 | ZNF836 | -0,369957244 | 0,773805429 | 0,35724111 | 0,848693232 | zinc finger protein 836 |
| ENSG00000231074 | HCG18 | -0,369996637 | 0,7737843 | 0,23639023 | 0,745611805 | HLA complex group 18 |
| ENSG00000182141 | ZNF708 | -0,37002177 | 0,773770821 | 0,30404177 | 0,813056239 | zinc finger protein 708 |
| ENSG00000109927 | TECTA | -0,370079924 | 0,773739631 | 0,17118842 | 0,654843372 | tectorin alpha |
| ENSG00000110324 | IL10RA | -0,370115986 | 0,773720291 | 0,5982917 | 0,948673307 | interleukin 10 receptor subunit alpha |
| ENSG00000266907 | AC006116,1 | -0,370169109 | 0,773691801 | 0,64166668 | 0,956518714 | glycerol-3-phosphate dehydrogenase 2 (mitochondrial), pseudogene |
| ENSG00000226742 | HSBP1L1 | -0,370214365 | 0,773667532 | 0,24218793 | 0,752479103 | heat shock factor binding protein 1 like 1 |
| ENSG00000284196 | AL158064,1 | -0,370231125 | 0,773658544 | 0,13881424 | 0,594163262 | novel transcript |
| ENSG00000287945 | AL603824,1 | -0,370319267 | 0,773611278 | 0,70243604 | 0,968742626 | novel transcript, antisense to CCNY |
| ENSG00000232439 | RPL18AP7 | -0,37036204 | 0,773588343 | 0,55694451 | 0,938264532 | ribosomal protein L18a pseudogene 7 |
| ENSG00000140600 | SH3GL3 | -0,370382473 | 0,773577386 | 0,38850365 | 0,869820053 | SH3 domain containing GRB2 like 3, endophilin A3 |
| ENSG00000138336 | TET1 | -0,370514951 | 0,773506354 | 0,31405541 | 0,819877431 | tet methylcytosine dioxygenase 1 |
| ENSG00000188611 | ASAH2 | -0,370592898 | 0,773464564 | 0,43142481 | 0,89530663 | N-acylsphingosine amidohydrolase 2 |
| ENSG00000269924 | AC024451,4 | -0,37059756 | 0,773462065 | 0,58122162 | 0,943919685 | novel transcript, sense intronic to KIAA0146 |
| ENSG00000227331 | RPL7AP22 | -0,370759133 | 0,773375446 | 0,5114134 | 0,922893865 | ribosomal protein L7a pseudogene 22 |
| ENSG00000283930 | AL117339,4 | -0,370768032 | 0,773370676 | 0,30265376 | 0,812334076 | novel transcript |
| ENSG00000153291 | SLC25A27 | -0,370828191 | 0,773338428 | 0,30347734 | 0,812669425 | solute carrier family 25 member 27 |
| ENSG00000288048 | AC104088,3 | -0,370845232 | 0,773329294 | 0,70247859 | 0,968742626 | novel transcript |
| ENSG00000127241 | MASP1 | -0,370944154 | 0,77327627 | 0,48865193 | 0,91557151 | mannan binding lectin serine peptidase 1 |
| ENSG00000254449 | SF3A3P2 | -0,37101209 | 0,773239857 | 0,63584753 | 0,955399772 | splicing factor 3a, subunit 3 pseudogene 2 |
| ENSG00000232528 | AL109809,1 | -0,371109826 | 0,773187476 | 0,66489117 | 0,960551345 | pseudogene similar to F830045P16Rik |
| ENSG00000285509 | TBCEL-TECTA | -0,371209117 | 0,773134265 | 0,52237257 | 0,927689738 | TBCEL-TECTA readthrough |
| ENSG00000236438 | FAM157A | -0,371248051 | 0,7731134 | 0,42221243 | 0,889850969 | family with sequence similarity 157 member A |
| ENSG00000236130 | PTCSC2 | -0,371470228 | 0,772994349 | 0,47584667 | 0,911349573 | papillary thyroid carcinoma susceptibility candidate 2 |
| ENSG00000186104 | CYP2R1 | -0,371629427 | 0,772909055 | 0,27338031 | 0,785672647 | cytochrome P450 family 2 subfamily R member 1 |
| ENSG00000281449 | BCL2L14 | -0,371686829 | 0,772878303 | 0,44686279 | 0,901798674 | BCL2 like 14 |
| ENSG00000237217 | AC008074,1 | -0,371764052 | 0,772836934 | 0,78599614 | 0,979320406 | novel pseudogene |
| ENSG00000273133 | AC116651,1 | -0,371808434 | 0,772813159 | 0,45874271 | 0,904420115 | novel transcript, antisense to CC2D2A |

| | | | | | | |
|---|---|---|---|---|---|---|
| ENSG00000264801 | ERVFRD-3 | -0,371835362 | 0,772798735 | 0,50213556 | 0,920520138 | endogenous retrovirus group FRD member 3 |
| ENSG00000285649 | AL357079,2 | -0,371925859 | 0,772750261 | 0,74585242 | 0,975707496 | novel transcript, antisense to DPH2 and ATP6V0B |
| ENSG00000172578 | KLHL6 | -0,371987552 | 0,772717217 | 0,2313307 | 0,739268552 | kelch like family member 6 |
| ENSG00000270820 | AC016727,1 | -0,372146034 | 0,772632337 | 0,33698274 | 0,836211475 | novel transcript, antisense to XPO1 |
| ENSG00000262480 | SAMD11P1 | -0,372206934 | 0,772599723 | 0,63809613 | 0,956018518 | sterile alpha motif domain containing 11 pseudogene 1 |
| ENSG00000269887 | AL391001,1 | -0,3722073 | 0,772599527 | 0,57770118 | 0,94337651 | novel transcript, sense intronic to PBX1 |
| ENSG00000081386 | ZNF510 | -0,372300751 | 0,772549484 | 0,37550908 | 0,862024808 | zinc finger protein 510 |
| ENSG00000143033 | MTF2 | -0,372385058 | 0,772504339 | 0,36572887 | 0,85559319 | metal response element binding transcription factor 2 |
| ENSG00000288028 | AL049839,3 | -0,372431598 | 0,772479419 | 0,21386428 | 0,721188427 | novel transcript |
| ENSG00000099308 | MAST3 | -0,372567796 | 0,772406497 | 0,20436695 | 0,705535354 | microtubule associated serine/threonine kinase 3 |
| ENSG00000281950 | SPDYE16 | -0,372607729 | 0,772385117 | 0,25339081 | 0,765172364 | speedy/RINGO cell cycle regulator family member E16 |
| ENSG00000216977 | RPL21P65 | -0,372654514 | 0,77236007 | 0,67352767 | 0,96278949 | ribosomal protein L21 pseudogene 65 |
| ENSG00000288383 | AC018653,7 | -0,372697595 | 0,772337007 | 0,70618796 | 0,969503793 | novel transcript |
| ENSG00000256967 | AC018653,3 | -0,372697595 | 0,772337007 | 0,70618796 | 0,969503793 | novel transcript |
| ENSG00000260552 | AC023043,1 | -0,372715722 | 0,772327302 | 0,77781893 | 0,978587255 | novel transcript, antisense to MOCOS |
| ENSG00000228886 | AL138963,1 | -0,372754466 | 0,772306562 | 0,61521014 | 0,952793934 | novel transcript |
| ENSG00000096093 | EFHC1 | -0,372817435 | 0,772272854 | 0,28600019 | 0,797420923 | EF-hand domain containing 1 |
| ENSG00000275331 | ABCC6 | -0,372821383 | 0,77227074 | 0,41928198 | 0,88768414 | ATP binding cassette subfamily C member 6 |
| ENSG00000222000 | AC092675,1 | -0,372952084 | 0,77220078 | 0,53673484 | 0,932766987 | novel transcript, antisense to CNGA3 |
| ENSG00000285728 | AC098484,4 | -0,372965653 | 0,772193517 | 0,30316319 | 0,812669425 | novel transcript, antisense to PPIH |
| ENSG00000180769 | WDFY3-AS2 | -0,373055873 | 0,772145229 | 0,38545825 | 0,868895792 | WDFY3 antisense RNA 2 |
| ENSG00000272106 | AL691432,2 | -0,373081437 | 0,772131547 | 0,10670367 | 0,521491876 | artifact |
| ENSG00000259948 | AC124068,1 | -0,373182268 | 0,772077584 | 0,73411395 | 0,974353474 | VAMP (vesicle-associated membrane protein)-associated protein A, 33kDa (VAPA) pseudogene |
| ENSG00000147082 | CCNB3 | -0,373198885 | 0,772068691 | 0,41821359 | 0,887094957 | cyclin B3 |
| ENSG00000248458 | AL139147,1 | -0,373225783 | 0,772054296 | 0,24244551 | 0,752612359 | novel transcript, antisense to SGIP1 |
| ENSG00000273675 | AL118556,2 | -0,373352686 | 0,771986388 | 0,67442478 | 0,963103112 | zinc finger protein 124 (ZNF124) pseudogene |
| ENSG00000285597 | AC051619,10 | -0,373396702 | 0,771962835 | 0,72409833 | 0,973167061 | novel transcript |
| ENSG00000260368 | AC027373,1 | -0,373416811 | 0,771952075 | 0,32176185 | 0,824933727 | novel transcript, overlapping YWHAZ |
| ENSG00000226984 | AL035410,1 | -0,373419371 | 0,771950705 | 0,62021911 | 0,952793934 | ribosomal protein L26 (RPL26) pseudogene |

| | | | | | | |
|---|---|---|---|---|---|---|
| ENSG00000229036 | VDAC1P8 | -0,373431815 | 0,771944047 | 0,36976854 | 0,858104601 | voltage dependent anion channel 1 pseudogene 8 |
| ENSG00000196950 | SLC39A10 | -0,373555675 | 0,771877776 | 0,47005327 | 0,908378389 | solute carrier family 39 member 10 |
| ENSG00000156787 | TBC1D31 | -0,373606945 | 0,771850345 | 0,33539485 | 0,835232738 | TBC1 domain family member 31 |
| ENSG00000006757 | PNPLA4 | -0,373608702 | 0,771849406 | 0,33706771 | 0,836223219 | patatin like phospholipase domain containing 4 |
| ENSG00000267416 | AC025048,4 | -0,373631962 | 0,771836961 | 0,77676908 | 0,978587255 | novel transcript |
| ENSG00000260404 | AC110079,1 | -0,37369545 | 0,771802996 | 0,10745928 | 0,523140272 | general transcription factor IIi (GTF2I) pseudogene |
| ENSG00000269806 | BICRA-AS1 | -0,373699724 | 0,77180071 | 0,67669223 | 0,963580269 | BICRA antisense RNA 1 |
| ENSG00000275199 | AKT3 | -0,373792248 | 0,771751214 | 0,56238917 | 0,940865625 | AKT serine/threonine kinase 3 |
| ENSG00000274574 | U1 | -0,373912466 | 0,771686907 | 0,40029101 | 0,87601134 | U1 spliceosomal RNA |
| ENSG00000277918 | RNVU1-28 | -0,373912466 | 0,771686907 | 0,40029101 | 0,87601134 | RNA, variant U1 small nuclear 28 |
| ENSG00000265791 | AC127024,4 | -0,373959008 | 0,771662013 | 0,53222221 | 0,931291085 | novel transcript, sense intronic CRLF3 |
| ENSG00000246379 | AC040168,1 | -0,374009977 | 0,771634751 | 0,41570322 | 0,885986046 | novel transcript |
| ENSG00000285591 | AC092617,1 | -0,37403206 | 0,77162294 | 0,75730405 | 0,975901808 | novel transcript |
| ENSG00000283879 | MTND6P35 | -0,37403206 | 0,77162294 | 0,75730405 | 0,975901808 | MT-ND6 pseudogene 35 |
| ENSG00000180354 | MTURN | -0,374054979 | 0,771610682 | 0,30419392 | 0,813056239 | maturin, neural progenitor differentiation regulator homolog |
| ENSG00000115365 | LANCL1 | -0,374124816 | 0,771573332 | 0,43750831 | 0,897021798 | LanC like 1 |
| ENSG00000146386 | ABRACL | -0,374128614 | 0,7715713 | 0,30965932 | 0,816986489 | ABRA C-terminal like |
| ENSG00000248124 | RRN3P1 | -0,374164188 | 0,771552275 | 0,1972832 | 0,695943345 | RRN3 pseudogene 1 |
| ENSG00000180953 | ST20 | -0,374185795 | 0,77154072 | 0,25086877 | 0,763027659 | suppressor of tumorigenicity 20 |
| ENSG00000185869 | ZNF829 | -0,374198274 | 0,771534046 | 0,46074093 | 0,905949965 | zinc finger protein 829 |
| ENSG00000259768 | AC004943,2 | -0,374509253 | 0,771367756 | 0,18238958 | 0,672670499 | novel transcript, antisense to ZFHX3 |
| ENSG00000253873 | PCDHGA11 | -0,374520666 | 0,771361654 | 0,26083186 | 0,772952333 | protocadherin gamma subfamily A, 11 |
| ENSG00000243708 | PLA2G4B | -0,374593972 | 0,771322461 | 0,570836 | 0,942436428 | phospholipase A2 group IVB |
| ENSG00000240828 | RPL21P4 | -0,374641314 | 0,771297151 | 0,66160086 | 0,959582738 | ribosomal protein L21 pseudogene 4 |
| ENSG00000286472 | AL035252,5 | -0,374725568 | 0,771252108 | 0,14847429 | 0,613326406 | novel transcript |
| ENSG00000223908 | AC068657,1 | -0,374996668 | 0,771107194 | 0,78245815 | 0,978628303 | CCHC domain zinc finger protein pseudogene |
| ENSG00000287969 | AL355474,1 | -0,375001579 | 0,771104569 | 0,74637807 | 0,975796253 | novel transcript, antisense to FAM13C |
| ENSG00000170091 | NSG2 | -0,375012815 | 0,771098563 | 0,403146 | 0,877589651 | neuronal vesicle trafficking associated 2 |
| ENSG00000184445 | KNTC1 | -0,375193713 | 0,771001882 | 0,33770928 | 0,837066292 | kinetochore associated 1 |
| ENSG00000142530 | FAM71E1 | -0,375296017 | 0,770947211 | 0,44675904 | 0,901650934 | family with sequence similarity 71 member E1 |
| ENSG00000272578 | AP000347,1 | -0,375311467 | 0,770938955 | 0,2030057 | 0,703345263 | glucuronidase, beta (GUSB) pseudogene |
| ENSG00000135976 | ANKRD36 | -0,37531671 | 0,770936153 | 0,15243129 | 0,620962584 | ankyrin repeat domain 36 |

| | | | | | | |
|---|---|---|---|---|---|---|
| ENSG00000232479 | KRT8P52 | -0,375348237 | 0,770919306 | 0,61054801 | 0,951841373 | keratin 8 pseudogene 52 |
| ENSG00000137700 | SLC37A4 | -0,37534849 | 0,770919171 | 0,27707746 | 0,788874154 | solute carrier family 37 member 4 |
| ENSG00000235548 | HEBP2P1 | -0,375603325 | 0,770783009 | 0,58672644 | 0,945395317 | HEBP2 pseudogene 1 |
| ENSG00000112280 | COL9A1 | -0,375654474 | 0,770755683 | 0,3218032 | 0,824933727 | collagen type IX alpha 1 chain |
| ENSG00000234084 | AL049552,1 | -0,376055169 | 0,770541642 | 0,6449606 | 0,956899963 | novel transcript |
| ENSG00000245149 | RNF139-AS1 | -0,376067333 | 0,770535145 | 0,20823146 | 0,712801795 | RNF139 antisense RNA 1 (head to head) |
| ENSG00000120329 | SLC25A2 | -0,376092947 | 0,770521465 | 0,47836673 | 0,912208437 | solute carrier family 25 member 2 |
| ENSG00000259563 | AC025430,1 | -0,376256528 | 0,770434104 | 0,68408826 | 0,965403332 | novel transcript, antisense to CTDSPL2 |
| ENSG00000262038 | AC007599,2 | -0,376297621 | 0,770412159 | 0,77822106 | 0,978587255 | novel transcript |
| ENSG00000069493 | CLEC2D | -0,376560601 | 0,770271739 | 0,38520031 | 0,868895792 | C-type lectin domain family 2 member D |
| ENSG00000240291 | AL450384,2 | -0,376630802 | 0,770234259 | 0,51869676 | 0,926765093 | novel transcript, antisense to CACNB2 |
| ENSG00000003147 | ICA1 | -0,376668332 | 0,770214222 | 0,37687907 | 0,862818448 | islet cell autoantigen 1 |
| ENSG00000270006 | AC010531,6 | -0,376713473 | 0,770190123 | 0,61964096 | 0,952793934 | novel transcript, antisense to FBXO31 and C16orf95 readthrough |
| ENSG00000259055 | AL591770,1 | -0,376718161 | 0,77018762 | 0,59846752 | 0,948793037 | novel transcript, antisense to TRIM9 |
| ENSG00000151150 | ANK3 | -0,376811216 | 0,770137944 | 0,3188416 | 0,823755375 | ankyrin 3 |
| ENSG00000153944 | MSI2 | -0,376914991 | 0,770082549 | 0,20192184 | 0,701509559 | musashi RNA binding protein 2 |
| ENSG00000204581 | ACOXL-AS1 | -0,377078618 | 0,769995213 | 0,41311344 | 0,884663685 | ACOXL antisense RNA 1 |
| ENSG00000271259 | AC010201,1 | -0,377199783 | 0,769930548 | 0,66199762 | 0,959779887 | novel transcript |
| ENSG00000157240 | FZD1 | -0,37724118 | 0,769908456 | 0,1032802 | 0,512532106 | frizzled class receptor 1 |
| ENSG00000272150 | NBPF25P | -0,377245192 | 0,769906314 | 0,27472271 | 0,786383635 | NBPF member 25, pseudogene |
| ENSG00000272398 | CD24 | -0,377390438 | 0,769828807 | 0,46653033 | 0,907940653 | CD24 molecule |
| ENSG00000158467 | AHCYL2 | -0,37751331 | 0,769763244 | 0,20881451 | 0,713420724 | adenosylhomocysteinase like 2 |
| ENSG00000229863 | AC097358,1 | -0,377626305 | 0,769702957 | 0,45224313 | 0,902062594 | ribosomal protein L19 (RPL19) pseudogene |
| ENSG00000276308 | AC078880,4 | -0,377742683 | 0,76964087 | 0,17242403 | 0,656917487 | novel transcript |
| ENSG00000281151 | KCNQ2 | -0,377849211 | 0,769584042 | 0,11972216 | 0,552559297 | potassium voltage-gated channel subfamily Q member 2 |
| ENSG00000183666 | GUSBP1 | -0,377877835 | 0,769568773 | 0,24821337 | 0,760574609 | GUSB pseudogene 1 |
| ENSG00000259887 | FIGNL2-DT | -0,377917074 | 0,769547843 | 0,77830064 | 0,978587255 | FIGNL2 divergent transcript |
| ENSG00000198795 | ZNF521 | -0,377921602 | 0,769545427 | 0,32213422 | 0,824955371 | zinc finger protein 521 |
| ENSG00000167733 | HSD11B1L | -0,377993987 | 0,769506817 | 0,1647869 | 0,643964432 | hydroxysteroid 11-beta dehydrogenase 1 like |
| ENSG00000205307 | SAP25 | -0,378054518 | 0,769474532 | 0,62989478 | 0,952812183 | Sin3A associated protein 25 |
| ENSG00000228446 | ANKRD49P1 | -0,378097364 | 0,76945168 | 0,41493242 | 0,885668156 | ANKRD49 pseudogene 1 |

| | | | | | | |
|---|---|---|---|---|---|---|
| ENSG00000213722 | DDAH2 | -0,378135378 | 0,769431406 | 0,21173131 | 0,718113208 | dimethylarginine dimethylaminohydrolase 2 |
| ENSG00000219626 | FAM228B | -0,378201799 | 0,769395982 | 0,0802522 | 0,449320363 | family with sequence similarity 228 member B |
| ENSG00000175274 | TP53I11 | -0,378220406 | 0,769386059 | 0,2043005 | 0,705535354 | tumor protein p53 inducible protein 11 |
| ENSG00000281521 | AC116351,4 | -0,378358645 | 0,76931234 | 0,63646116 | 0,955547532 | novel transcript |
| ENSG00000272347 | AC116351,2 | -0,378358645 | 0,76931234 | 0,63646116 | 0,955547532 | novel transcript |
| ENSG00000111254 | AKAP3 | -0,378398987 | 0,769290828 | 0,28933901 | 0,800260154 | A-kinase anchoring protein 3 |
| ENSG00000259495 | AC016705,2 | -0,378629341 | 0,769168006 | 0,16174086 | 0,638306753 | novel transcript |
| ENSG00000241782 | AP002812,1 | -0,378660176 | 0,769151567 | 0,66574574 | 0,961016016 | ribosomal protein L21 (RPL21) pseudogene |
| ENSG00000152154 | TMEM178A | -0,378689462 | 0,769135953 | 0,17658493 | 0,662599918 | transmembrane protein 178A |
| ENSG00000112096 | SOD2 | -0,378747182 | 0,769105182 | 0,10732837 | 0,522893717 | superoxide dismutase 2 |
| ENSG00000148019 | CEP78 | -0,37888768 | 0,769030285 | 0,29164636 | 0,802285739 | centrosomal protein 78 |
| ENSG00000165115 | KIF27 | -0,378947968 | 0,76899815 | 0,34572462 | 0,841168721 | kinesin family member 27 |
| ENSG00000162598 | C1orf87 | -0,379137251 | 0,768897263 | 0,8085598 | 0,981213877 | chromosome 1 open reading frame 87 |
| ENSG00000005436 | GCFC2 | -0,379154133 | 0,768888265 | 0,32366889 | 0,825794844 | GC-rich sequence DNA-binding factor 2 |
| ENSG00000224616 | RTCA-AS1 | -0,379168722 | 0,76888049 | 0,12526485 | 0,565527849 | RTCA antisense RNA 1 |
| ENSG00000184678 | H2BC21 | -0,3791836 | 0,768872561 | 0,28565974 | 0,797398763 | H2B clustered histone 21 |
| ENSG00000251136 | AF117829,1 | -0,37919378 | 0,768867136 | 0,10470017 | 0,515975114 | novel transcript |
| ENSG00000181001 | OR52N1 | -0,379257374 | 0,768833245 | 0,48080779 | 0,912907448 | olfactory receptor family 52 subfamily N member 1 |
| ENSG00000199568 | RNU5A-1 | -0,379349112 | 0,768784358 | 0,59212904 | 0,947416795 | RNA, U5A small nuclear 1 |
| ENSG00000224152 | AC009506,1 | -0,379367072 | 0,768774787 | 0,20189116 | 0,701480895 | novel transcript, antisense to BAZ2B |
| ENSG00000207051 | SNORA27 | -0,379485567 | 0,768711647 | 0,66036055 | 0,959163386 | small nucleolar RNA, H/ACA box 27 |
| ENSG00000250198 | LINC02199 | -0,379588102 | 0,768657015 | 0,59283779 | 0,947416795 | long intergenic non-protein coding RNA 2199 |
| ENSG00000253877 | LINC01608 | -0,379904453 | 0,768488485 | 0,19827534 | 0,697208456 | long intergenic non-protein coding RNA 1608 |
| ENSG00000283709 | FAM238C | -0,379905343 | 0,768488011 | 0,54054335 | 0,933385624 | family with sequence similarity 238 member C |
| ENSG00000149564 | ESAM | -0,379964523 | 0,768456487 | 0,52893962 | 0,930181595 | endothelial cell adhesion molecule |
| ENSG00000283422 | LINC02452 | -0,379972097 | 0,768452453 | 0,58917818 | 0,946705487 | long intergenic non-protein coding RNA 2452 |
| ENSG00000140043 | PTGR2 | -0,379982596 | 0,768446861 | 0,19176455 | 0,687672451 | prostaglandin reductase 2 |
| ENSG00000266904 | LINC00663 | -0,37998525 | 0,768445447 | 0,31099604 | 0,817751312 | long intergenic non-protein coding RNA 663 |
| ENSG00000162437 | RAVER2 | -0,380245526 | 0,768306825 | 0,37140679 | 0,859476158 | ribonucleoprotein, PTB binding 2 |
| ENSG00000258787 | SEPHS1P2 | -0,380331611 | 0,768260982 | 0,63392459 | 0,954929949 | selenophosphate synthetase 1 pseudogene 2 |
| ENSG00000231704 | AC004895,1 | -0,380549582 | 0,768144917 | 0,75379743 | 0,975901808 | novel transcript |
| ENSG00000240395 | RPL5P23 | -0,380691265 | 0,768069483 | 0,61340349 | 0,952413269 | ribosomal protein L5 pseudogene 23 |
| ENSG00000248729 | MTCO1P30 | -0,380774437 | 0,768025205 | 0,77376302 | 0,978415098 | MT-CO1 pseudogene 30 |

| | | | | | | |
|---|---|---|---|---|---|---|
| ENSG00000259892 | AC027139,1 | -0,380814891 | 0,768003669 | 0,82537518 | 0,983383773 | amyloid beta (A4) precursor protein-binding, family A, member 2 (APBA2) pseudogene |
| ENSG00000245017 | LINC02453 | -0,38101579 | 0,76789673 | 0,49627717 | 0,918259281 | long intergenic non-protein coding RNA 2453 |
| ENSG00000288075 | AC108519,1 | -0,381141653 | 0,767829741 | 0,10796242 | 0,523928283 | novel transcript |
| ENSG00000287055 | AL359813,1 | -0,381234541 | 0,767780306 | 0,47553538 | 0,911264385 | novel transcript |
| ENSG00000265194 | AL359922,2 | -0,381235226 | 0,767779941 | 0,21515766 | 0,722445395 | novel transcript, antisense MTAP |
| ENSG00000145882 | PCYOX1L | -0,381282455 | 0,767754807 | 0,11855344 | 0,550275639 | prenylcysteine oxidase 1 like |
| ENSG00000251682 | AC122718,2 | -0,381365811 | 0,767710449 | 0,29767794 | 0,806956721 | DEAD (Asp-Glu-Ala-Asp) box polypeptide 25 (DDX25) pseudogene |
| ENSG00000198146 | ZNF770 | -0,381659542 | 0,76755416 | 0,46681763 | 0,907940653 | zinc finger protein 770 |
| ENSG00000268555 | AC123912,4 | -0,381674617 | 0,767546139 | 0,4990525 | 0,919788805 | novel transcript |
| ENSG00000276672 | AL161891,1 | -0,381737167 | 0,767512862 | 0,10588075 | 0,519214495 | novel transcript, sense intronic to RFC3 |
| ENSG00000261794 | GOLGA8H | -0,381740621 | 0,767511025 | 0,4341229 | 0,896272827 | golgin A8 family member H |
| ENSG00000181666 | ZNF875 | -0,381793432 | 0,76748293 | 0,06648075 | 0,405279626 | zinc finger protein 875 |
| ENSG00000287232 | AC079848,2 | -0,381826154 | 0,767465522 | 0,30464036 | 0,813276091 | novel transcript |
| ENSG00000251637 | LINC02754 | -0,381990601 | 0,767378047 | 0,3632799 | 0,853554847 | long intergenic non-protein coding RNA 2754 |
| ENSG00000229524 | BAG6 | -0,382156369 | 0,767289879 | 0,20127799 | 0,701094807 | BAG cochaperone 6 |
| ENSG00000156042 | CFAP70 | -0,382204638 | 0,767264208 | 0,34811406 | 0,842566991 | cilia and flagella associated protein 70 |
| ENSG00000224003 | YES1P1 | -0,382217397 | 0,767257423 | 0,57230507 | 0,942475308 | YES1 pseudogene 1 |
| ENSG00000170629 | DPY19L2P2 | -0,382307767 | 0,767209363 | 0,13055257 | 0,577136255 | DPY19L2 pseudogene 2 |
| ENSG00000116001 | TIA1 | -0,382311628 | 0,76720731 | 0,43860716 | 0,897246228 | TIA1 cytotoxic granule associated RNA binding protein |
| ENSG00000174483 | BBS1 | -0,382346275 | 0,767188885 | 0,14142828 | 0,599297054 | Bardet-Biedl syndrome 1 |
| ENSG00000274093 | AC009032,1 | -0,382481058 | 0,767117215 | 0,58424247 | 0,944650222 | novel transcript, sense intronic to NFAT5 |
| ENSG00000268119 | AC010615,2 | -0,38253929 | 0,767086252 | 0,24957642 | 0,761448016 | novel transcript |
| ENSG00000286416 | AL442067,3 | -0,382541923 | 0,767084852 | 0,25313046 | 0,76515438 | novel transcript |
| ENSG00000273181 | AC131235,3 | -0,382547079 | 0,76708211 | 0,74723624 | 0,97584218 | novel transcript |
| ENSG00000214946 | TBC1D26 | -0,382551301 | 0,767079866 | 0,75131934 | 0,975901808 | TBC1 domain family member 26 |
| ENSG00000204172 | AGAP9 | -0,382595713 | 0,767056252 | 0,13789787 | 0,592546669 | ArfGAP with GTPase domain, ankyrin repeat and PH domain 9 |
| ENSG00000206258 | TNXB | -0,38262277 | 0,767041866 | 0,80922414 | 0,981213877 | tenascin XB |
| ENSG00000231608 | TNXB | -0,38262277 | 0,767041866 | 0,80922414 | 0,981213877 | tenascin XB |
| ENSG00000229214 | LINC00242 | -0,382688214 | 0,767007073 | 0,41743866 | 0,886770159 | long intergenic non-protein coding RNA 242 |

| | | | | | | |
|---|---|---|---|---|---|---|
| ENSG00000182134 | TDRKH | -0,382762304 | 0,766967684 | 0,2794843 | 0,790302992 | tudor and KH domain containing |
| ENSG00000184465 | WDR27 | -0,382845973 | 0,766923205 | 0,11857591 | 0,550275639 | WD repeat domain 27 |
| ENSG00000278899 | AL358852,1 | -0,382852089 | 0,766919953 | 0,48267057 | 0,913451138 | novel transcript, antisense to LATS1 |
| ENSG00000117054 | ACADM | -0,38296116 | 0,766861975 | 0,36164053 | 0,851893059 | acyl-CoA dehydrogenase medium chain |
| ENSG00000248592 | STIMATE-MUST | -0,383091482 | 0,766792706 | 0,20086847 | 0,700418461 | STIMATE-MUSTN1 readthrough |
| ENSG00000278274 | SNORA61 | -0,383111174 | 0,766782239 | 0,3781167 | 0,863761554 | small nucleolar RNA, H/ACA box 61 |
| ENSG00000179240 | GVQW3 | -0,383225571 | 0,766721441 | 0,0728771 | 0,425795099 | GVQW motif containing 3 |
| ENSG00000121988 | ZRANB3 | -0,383406983 | 0,766625035 | 0,30653442 | 0,815375074 | zinc finger RANBP2-type containing 3 |
| ENSG00000135314 | KHDC1 | -0,383444157 | 0,766605281 | 0,14589617 | 0,608741186 | KH domain containing 1 |
| ENSG00000105808 | RASA4 | -0,38363662 | 0,76650302 | 0,1620136 | 0,638979845 | RAS p21 protein activator 4 |
| ENSG00000250461 | AC122718,1 | -0,383671173 | 0,766484662 | 0,33850096 | 0,837263082 | family with sequence similarity 133, member B (FAM133B) pseudogene |
| ENSG00000223403 | MEG9 | -0,383703093 | 0,766467703 | 0,60442682 | 0,950813598 | maternally expressed 9 |
| ENSG00000100307 | CBX7 | -0,383873961 | 0,766376931 | 0,32527713 | 0,827324367 | chromobox 7 |
| ENSG00000272777 | AC019131,2 | -0,384107704 | 0,766252774 | 0,53887787 | 0,933348247 | novel transcript |
| ENSG00000165197 | VEGFD | -0,384166468 | 0,766221563 | 0,58308303 | 0,944174844 | vascular endothelial growth factor D |
| ENSG00000247735 | AC120114,1 | -0,384368345 | 0,766114353 | 0,25839945 | 0,771153141 | novel transcript, antisense to KCTD13 |
| ENSG00000226430 | USP17L7 | -0,384445287 | 0,766073495 | 0,74222938 | 0,975176043 | ubiquitin specific peptidase 17 like family member 7 |
| ENSG00000152782 | PANK1 | -0,384474658 | 0,766057899 | 0,30912022 | 0,816845699 | pantothenate kinase 1 |
| ENSG00000229353 | TNXB | -0,384598005 | 0,765992406 | 0,71230116 | 0,970812834 | tenascin XB |
| ENSG00000278760 | CORO7-PAM16 | -0,384638202 | 0,765971064 | 0,82162863 | 0,982952211 | CORO7-PAM16 readthrough |
| ENSG00000103426 | CORO7-PAM16 | -0,384638202 | 0,765971064 | 0,82162863 | 0,982952211 | CORO7-PAM16 readthrough |
| ENSG00000228317 | AL158070,1 | -0,384723305 | 0,765925882 | 0,68118999 | 0,964654539 | novel transcript |
| ENSG00000122484 | RPAP2 | -0,38475056 | 0,765911412 | 0,12491791 | 0,564940487 | RNA polymerase II associated protein 2 |
| ENSG00000127337 | YEATS4 | -0,384800819 | 0,765884731 | 0,41415617 | 0,885579089 | YEATS domain containing 4 |
| ENSG00000286673 | LO018686,4 | -0,384813182 | 0,765878168 | 0,10363359 | 0,513379913 | novel transcript |
| ENSG00000286931 | AL136419,3 | -0,384813182 | 0,765878168 | 0,10363359 | 0,513379913 | novel transcript |
| ENSG00000186523 | FAM86B1 | -0,384900437 | 0,765831848 | 0,14996708 | 0,615621177 | family with sequence similarity 86 member B1 |
| ENSG00000143224 | PPOX | -0,384945643 | 0,765807852 | 0,12744487 | 0,570245289 | protoporphyrinogen oxidase |
| ENSG00000180346 | TIGD2 | -0,384983607 | 0,7657877 | 0,41500073 | 0,885697506 | tigger transposable element derived 2 |
| ENSG00000221475 | SNORA11D | -0,385034979 | 0,765760432 | 0,60830656 | 0,951332922 | small nucleolar RNA, H/ACA box 11D |
| ENSG00000221705 | SNORA11E | -0,385034979 | 0,765760432 | 0,60830656 | 0,951332922 | small nucleolar RNA, H/ACA box 11E |
| ENSG00000275191 | AC007497,1 | -0,385095058 | 0,765728544 | 0,67364856 | 0,962830239 | novel transcript, sense intronic to RPGRIP1L |

| | | | | | | |
|---|---|---|---|---|---|---|
| ENSG00000228043 | AC114763,1 | -0,385272525 | 0,765634357 | 0,70426275 | 0,969119904 | novel transcript |
| ENSG00000267143 | AP001120,2 | -0,385299291 | 0,765620152 | 0,54743124 | 0,935404974 | novel transcript |
| ENSG00000105708 | ZNF14 | -0,38530781 | 0,765615631 | 0,31478356 | 0,81991112 | zinc finger protein 14 |
| ENSG00000116761 | CTH | -0,385357833 | 0,765589085 | 0,38149435 | 0,865900094 | cystathionine gamma-lyase |
| ENSG00000254004 | ZNF260 | -0,385451089 | 0,765539599 | 0,39768765 | 0,874263835 | zinc finger protein 260 |
| ENSG00000126860 | EVI2A | -0,38547291 | 0,76552802 | 0,38182662 | 0,866412204 | ecotropic viral integration site 2A |
| ENSG00000231172 | AC007099,1 | -0,385789542 | 0,765360026 | 0,38718685 | 0,869441674 | novel transcript, antisense to EVA1A |
| ENSG00000235099 | AL138731,1 | -0,385804685 | 0,765351993 | 0,71014428 | 0,970482408 | novel transcript |
| ENSG00000229653 | ZNRD1ASP | -0,385841415 | 0,765332508 | 0,67082437 | 0,962026636 | zinc ribbon domain containing 1 antisense, pseudogene |
| ENSG00000174586 | ZNF497 | -0,385865887 | 0,765319526 | 0,30657414 | 0,815375074 | zinc finger protein 497 |
| ENSG00000205609 | EIF3CL | -0,385903515 | 0,765299565 | 0,292234 | 0,802771257 | eukaryotic translation initiation factor 3 subunit C like |
| ENSG00000231588 | SHQ1P1 | -0,385925541 | 0,765287881 | 0,7336221 | 0,974353474 | SHQ1, H/ACA ribonucleoprotein assembly factor pseudogene 1 |
| ENSG00000234390 | USP27X-AS1 | -0,385927669 | 0,765286752 | 0,36657401 | 0,856103393 | USP27X antisense RNA 1 (head to head) |
| ENSG00000287061 | AC124248,2 | -0,38593962 | 0,765280413 | 0,50877916 | 0,922427526 | novel transcript, antisense to BLM |
| ENSG00000226435 | ANKRD18DP | -0,386035015 | 0,765229812 | 0,78921128 | 0,979462318 | ankyrin repeat domain 18D, pseudogene |
| ENSG00000197119 | SLC25A29 | -0,386094448 | 0,765198288 | 0,14102233 | 0,598214217 | solute carrier family 25 member 29 |
| ENSG00000223356 | AL590666,1 | -0,386119599 | 0,765184949 | 0,75430786 | 0,975901808 | novel transcript |
| ENSG00000249493 | ANKRD20A18P | -0,386132315 | 0,765178204 | 0,81395728 | 0,981521413 | ankyrin repeat domain 20 family member A18, pseudogene |
| ENSG00000092140 | G2E3 | -0,386296068 | 0,765091358 | 0,38903512 | 0,870057277 | G2/M-phase specific E3 ubiquitin protein ligase |
| ENSG00000259287 | AC010809,1 | -0,386329522 | 0,765073617 | 0,41014376 | 0,882597401 | novel transcript, antisense to RYR3 |
| ENSG00000230606 | AC092683,1 | -0,386361152 | 0,765056843 | 0,23362355 | 0,742832118 | novel transcript |
| ENSG00000261324 | AC010168,2 | -0,386383086 | 0,765045212 | 0,15382649 | 0,622898002 | novel transcript, overlapping HIST4H4 |
| ENSG00000257354 | AC048341,1 | -0,386398881 | 0,765036836 | 0,29208016 | 0,802771257 | novel transcript |
| ENSG00000231980 | AC011899,1 | -0,386527127 | 0,764968832 | 0,51842557 | 0,926651255 | novel transcript |
| ENSG00000100181 | TPTEP1 | -0,386540075 | 0,764961966 | 0,38114376 | 0,865900094 | TPTE pseudogene 1 |
| ENSG00000235777 | DPYD-AS2 | -0,386622501 | 0,764918263 | 0,75919564 | 0,976120207 | DPYD antisense RNA 2 |
| ENSG00000286246 | AC130404,2 | -0,386640169 | 0,764908896 | 0,76080663 | 0,97624106 | novel transcript, antisense to TMEM132D |
| ENSG00000185842 | DNAH14 | -0,386663103 | 0,764896736 | 0,27158554 | 0,784591608 | dynein axonemal heavy chain 14 |
| ENSG00000176236 | RPP38-DT | -0,386726261 | 0,764863251 | 0,33280929 | 0,833301711 | RPP38 divergent transcript |

| | | | | | | |
|---|---|---|---|---|---|---|
| ENSG00000124243 | BCAS4 | -0,387147865 | 0,764639766 | 0,29581587 | 0,805468341 | breast carcinoma amplified sequence 4 |
| ENSG00000102053 | ZC3H12B | -0,387149433 | 0,764638934 | 0,33508451 | 0,834858938 | zinc finger CCCH-type containing 12B |
| ENSG00000277075 | H2AC8 | -0,387152396 | 0,764637364 | 0,40564861 | 0,878572779 | H2A clustered histone 8 |
| ENSG00000286293 | AP005901,5 | -0,387203624 | 0,764610213 | 0,14141911 | 0,599297054 | novel transcript |
| ENSG00000262703 | AC009121,2 | -0,387233769 | 0,764594237 | 0,36225839 | 0,852321354 | novel transcript, antisense to RMI2 |
| ENSG00000176601 | MAP3K19 | -0,387297186 | 0,764560628 | 0,64836169 | 0,956899963 | mitogen-activated protein kinase kinase kinase 19 |
| ENSG00000230683 | DDTP1 | -0,387326866 | 0,764544899 | 0,67610151 | 0,963560994 | D-dopachrome tautomerase pseudogene 1 |
| ENSG00000270606 | PPIAP52 | -0,387327026 | 0,764544814 | 0,47146315 | 0,908896785 | peptidylprolyl isomerase A pseudogene 52 |
| ENSG00000255410 | AC091564,5 | -0,387351271 | 0,764531966 | 0,47360827 | 0,910190245 | novel transcript |
| ENSG00000270074 | AC087203,3 | -0,387366801 | 0,764523736 | 0,29731605 | 0,806635068 | novel transcript |
| ENSG00000251340 | MTCYBP35 | -0,38740528 | 0,764503345 | 0,71616422 | 0,971373454 | MT-CYB pseudogene 35 |
| ENSG00000227706 | AL713998,1 | -0,387543172 | 0,764430278 | 0,13854549 | 0,593533053 | novel transcript |
| ENSG00000120837 | NFYB | -0,387599581 | 0,76440039 | 0,39610081 | 0,872705836 | nuclear transcription factor Y subunit beta |
| ENSG00000231831 | MTHFD1P1 | -0,387837772 | 0,764274196 | 0,61357882 | 0,952413269 | methylenetetrahydrofolate dehydrogenase (NADP+ dependent) 1 pseudogene 1 |
| ENSG00000197558 | SSPOP | -0,387855206 | 0,764264961 | 0,32208155 | 0,824955371 | SCO-spondin, pseudogene |
| ENSG00000249846 | LINC02021 | -0,38785584 | 0,764264625 | 0,16485027 | 0,643964432 | long intergenic non-protein coding RNA 2021 |
| ENSG00000042062 | RIPOR3 | -0,387858807 | 0,764263053 | 0,47136337 | 0,908887649 | RIPOR family member 3 |
| ENSG00000280500 | ELMO2P1 | -0,387869637 | 0,764257316 | 0,79895872 | 0,980161849 | engulfment and cell motility 2 pseudogene 1 |
| ENSG00000232960 | CSNK2B | -0,387872252 | 0,764255931 | 0,44140607 | 0,898913458 | casein kinase 2 beta |
| ENSG00000156475 | PPP2R2B | -0,387885217 | 0,764249063 | 0,29353602 | 0,803461221 | protein phosphatase 2 regulatory subunit Bbeta |
| ENSG00000268516 | AC020915,2 | -0,38789484 | 0,764243965 | 0,11190284 | 0,534735697 | novel transcript, antisense to ZNF544 |
| ENSG00000280868 | LINC00242 | -0,387910855 | 0,764235482 | 0,47573253 | 0,911331229 | long intergenic non-protein coding RNA 242 |
| ENSG00000135245 | HILPDA | -0,387987585 | 0,764194836 | 0,11680652 | 0,545783738 | hypoxia inducible lipid droplet associated |
| ENSG00000106003 | LFNG | -0,388052969 | 0,764160203 | 0,34620173 | 0,841168721 | LFNG O-fucosylpeptide 3-beta-N-acetylglucosaminyltransferase |
| ENSG00000214376 | VSTM5 | -0,388090713 | 0,764140211 | 0,31402422 | 0,819877431 | V-set and transmembrane domain containing 5 |
| ENSG00000185482 | STAC3 | -0,38809544 | 0,764137708 | 0,46843993 | 0,907940653 | SH3 and cysteine rich domain 3 |
| ENSG00000178409 | BEND3 | -0,388191781 | 0,764086682 | 0,09647672 | 0,493334689 | BEN domain containing 3 |
| ENSG00000138615 | CILP | -0,388241253 | 0,764060481 | 0,62064302 | 0,952793934 | cartilage intermediate layer protein |
| ENSG00000274290 | H2BC6 | -0,388281446 | 0,764039194 | 0,40786443 | 0,88080912 | H2B clustered histone 6 |
| ENSG00000137944 | KYAT3 | -0,388298497 | 0,764030164 | 0,38832964 | 0,869820053 | kynurenine aminotransferase 3 |
| ENSG00000176593 | AC008969,1 | -0,388314631 | 0,76402162 | 0,09542772 | 0,490483883 | novel transcript, antisense to ZNF606 |

| | | | | | | |
|---|---|---|---|---|---|---|
| ENSG00000213226 | AC242426,1 | -0,38832977 | 0,764013603 | 0,80398171 | 0,980875302 | prostaglandin reductase 1 (PTGR1) pseudogene |
| ENSG00000198131 | ZNF544 | -0,388349943 | 0,76400292 | 0,07968788 | 0,447684098 | zinc finger protein 544 |
| ENSG00000267422 | AC016582,2 | -0,38880955 | 0,763759566 | 0,58370046 | 0,944213336 | increased sodium tolerance 1 homolog (yeast) (IST1) pseudogene |
| ENSG00000269514 | AC024257,3 | -0,388817689 | 0,763755257 | 0,13702179 | 0,590712806 | novel transcript, antisense to OR10AD1 |
| ENSG00000281234 | LOH12CR2 | -0,388883819 | 0,763720249 | 0,826536 | 0,983471137 | loss of heterozygosity on chromosome 12, region 2 |
| ENSG00000286847 | AC007036,5 | -0,388901567 | 0,763710854 | 0,55718538 | 0,938366937 | novel transcript, antisense to PLEKHA8 |
| ENSG00000166111 | SVOP | -0,388923346 | 0,763699325 | 0,37035388 | 0,858545694 | SV2 related protein |
| ENSG00000281060 | MUC20-OT1 | -0,388970024 | 0,763674616 | 0,30814256 | 0,816424917 | MUC20 overlapping transcript |
| ENSG00000259972 | AC009120,2 | -0,388985611 | 0,763666365 | 0,13791099 | 0,592546669 | novel transcript, antisense to PSMD7 |
| ENSG00000249196 | TMEM132D-AS | -0,388991875 | 0,76366305 | 0,74128396 | 0,975176043 | TMEM132D antisense RNA 1 |
| ENSG00000203778 | FAM229B | -0,389026972 | 0,763644472 | 0,23174639 | 0,73999787 | family with sequence similarity 229 member B |
| ENSG00000087301 | TXNDC16 | -0,3890477 | 0,763633501 | 0,45214298 | 0,902062594 | thioredoxin domain containing 16 |
| ENSG00000156076 | WIF1 | -0,389091082 | 0,763610538 | 0,7758833 | 0,978587255 | WNT inhibitory factor 1 |
| ENSG00000279064 | FP236315,1 | -0,389191155 | 0,763557572 | 0,68772319 | 0,966064784 | novel transcript |
| ENSG00000275713 | H2BC9 | -0,389250037 | 0,763526409 | 0,39126482 | 0,870083449 | H2B clustered histone 9 |
| ENSG00000180376 | CCDC66 | -0,389277062 | 0,763512106 | 0,33367219 | 0,834001599 | coiled-coil domain containing 66 |
| ENSG00000169064 | ZBBX | -0,389348022 | 0,763474553 | 0,44013634 | 0,897906863 | zinc finger B-box domain containing |
| ENSG00000247324 | AC010547,1 | -0,38939557 | 0,763449391 | 0,416754 | 0,886368152 | novel transcript, antisense to ZNF23 |
| ENSG00000236740 | AL033384,1 | -0,389404479 | 0,763444677 | 0,77660965 | 0,978587255 | novel transcript |
| ENSG00000105866 | SP4 | -0,389441376 | 0,763425152 | 0,37949102 | 0,865067964 | Sp4 transcription factor |
| ENSG00000144161 | ZC3H8 | -0,389549187 | 0,763368104 | 0,10952873 | 0,527842443 | zinc finger CCCH-type containing 8 |
| ENSG00000078725 | BRINP1 | -0,389670046 | 0,763304157 | 0,40192014 | 0,87718264 | BMP/retinoic acid inducible neural specific 1 |
| ENSG00000273492 | AP000229,1 | -0,389673375 | 0,763302396 | 0,39048064 | 0,870057277 | novel transcript |
| ENSG00000285805 | AC012378,2 | -0,389698926 | 0,763288877 | 0,47127918 | 0,908869685 | novel transcript, antisense to PYGO1 |
| ENSG00000231890 | DARS-AS1 | -0,389760039 | 0,763256545 | 0,25506433 | 0,76642479 | DARS antisense RNA 1 |
| ENSG00000136231 | IGF2BP3 | -0,389783552 | 0,763244106 | 0,29708981 | 0,806635068 | insulin like growth factor 2 mRNA binding protein 3 |
| ENSG00000266405 | CBX3P2 | -0,389838836 | 0,763214859 | 0,33721179 | 0,836323083 | chromobox 3 pseudogene 2 |
| ENSG00000249898 | MCPH1-AS1 | -0,389848811 | 0,763209582 | 0,16061821 | 0,636125619 | MCPH1 antisense RNA 1 |
| ENSG00000230370 | RPL23AP52 | -0,389965669 | 0,763147764 | 0,69433664 | 0,967417311 | ribosomal protein L23a pseudogene 52 |
| ENSG00000249492 | AC114956,2 | -0,389997033 | 0,763131174 | 0,4275233 | 0,892656446 | novel transcript |
| ENSG00000286311 | AL034550,3 | -0,390046462 | 0,763105028 | 0,41358617 | 0,885095345 | novel transcript, antisense to NOL4L |
| ENSG00000164309 | CMYA5 | -0,390079873 | 0,763087356 | 0,21723371 | 0,724978097 | cardiomyopathy associated 5 |

| | | | | | | |
|---|---|---|---|---|---|---|
| ENSG00000156011 | PSD3 | -0,390095298 | 0,763079197 | 0,27222677 | 0,785283053 | pleckstrin and Sec7 domain containing 3 |
| ENSG00000186868 | MAPT | -0,390102885 | 0,763075184 | 0,29083744 | 0,80167048 | microtubule associated protein tau |
| ENSG00000259429 | UBE2Q2P2 | -0,390141031 | 0,763055008 | 0,49754507 | 0,918759905 | ubiquitin conjugating enzyme E2 Q2 pseudogene 2 |
| ENSG00000272606 | AC015982,2 | -0,390148368 | 0,763051127 | 0,45618896 | 0,903545728 | novel transcript |
| ENSG00000273297 | AC009275,1 | -0,390392825 | 0,762921844 | 0,4898096 | 0,916063942 | novel transcript |
| ENSG00000165125 | TRPV6 | -0,390411355 | 0,762912045 | 0,6280522 | 0,952793934 | transient receptor potential cation channel subfamily V member 6 |
| ENSG00000186960 | LINC01551 | -0,390599494 | 0,762812561 | 0,35071469 | 0,844035671 | long intergenic non-protein coding RNA 1551 |
| ENSG00000260683 | AL591643,1 | -0,390682733 | 0,762768551 | 0,63363139 | 0,954875723 | novel transcript |
| ENSG00000105131 | EPHX3 | -0,390752496 | 0,762731667 | 0,50254319 | 0,920607749 | epoxide hydrolase 3 |
| ENSG00000163138 | PACRGL | -0,390787844 | 0,76271298 | 0,28073999 | 0,791510309 | parkin coregulated like |
| ENSG00000198554 | WDHD1 | -0,390864294 | 0,762672563 | 0,31140817 | 0,817751312 | WD repeat and HMG-box DNA binding protein 1 |
| ENSG00000287258 | AL512283,2 | -0,390973371 | 0,762614902 | 0,27720534 | 0,789067728 | novel transcript, antisense to PKIB |
| ENSG00000221962 | TMEM14EP | -0,391080281 | 0,762558392 | 0,25636935 | 0,768541051 | transmembrane protein 14E, pseudogene |
| ENSG00000288170 | DDX11L9 | -0,391088071 | 0,762554274 | 0,51408359 | 0,924227101 | DEAD/H-box helicase 11 like 9 (pseudogene) |
| ENSG00000248472 | DDX11L9 | -0,391088071 | 0,762554274 | 0,51408359 | 0,924227101 | DEAD/H-box helicase 11 like 9 (pseudogene) |
| ENSG00000136267 | DGKB | -0,391212964 | 0,762488263 | 0,28983904 | 0,800629762 | diacylglycerol kinase beta |
| ENSG00000228335 | AC073063,1 | -0,391323128 | 0,762430042 | 0,50437177 | 0,921400408 | cytokine induced protein 29 kDa (CIP29) pseudogene |
| ENSG00000270228 | AC079880,1 | -0,3914103 | 0,762383975 | 0,46301883 | 0,906849208 | HEAT repeat containing 2 (HEATR2) pseudogene |
| ENSG00000244607 | CCDC13 | -0,391621973 | 0,762272126 | 0,48682243 | 0,914444641 | coiled-coil domain containing 13 |
| ENSG00000080618 | CPB2 | -0,391725398 | 0,762217482 | 0,30927317 | 0,816845699 | carboxypeptidase B2 |
| ENSG00000131233 | GJA9 | -0,391737697 | 0,762210984 | 0,44903852 | 0,902062594 | gap junction protein alpha 9 |
| ENSG00000119878 | CRIPT | -0,391854605 | 0,762149221 | 0,35493917 | 0,846374661 | CXXC repeat containing interactor of PDZ3 domain |
| ENSG00000188338 | SLC38A3 | -0,391911893 | 0,762118957 | 0,34584965 | 0,841168721 | solute carrier family 38 member 3 |
| ENSG00000106571 | GLI3 | -0,391923506 | 0,762112823 | 0,26899363 | 0,781285125 | GLI family zinc finger 3 |
| ENSG00000243297 | RPL31P61 | -0,391975471 | 0,762085372 | 0,80781499 | 0,981213877 | ribosomal protein L31 pseudogene 61 |
| ENSG00000263393 | AC011825,2 | -0,392057531 | 0,762042026 | 0,60313564 | 0,950352286 | novel transcript, antisense to RNF138 |
| ENSG00000215493 | AC007731,1 | -0,392078669 | 0,762030861 | 0,39474769 | 0,871888817 | kelch-like 12 (Drosophila) (KLH12) pseudogene |
| ENSG00000272223 | AL136304,1 | -0,392105015 | 0,762016945 | 0,51151061 | 0,922893865 | novel transcript, antisense to RRP36 |
| ENSG00000236478 | AC012513,2 | -0,392109169 | 0,762014751 | 0,25366226 | 0,765248462 | heterogeneous nuclear ribonucleoprotein A3 (HNRNPA3) pseudogene |
| ENSG00000254604 | AP002336,2 | -0,392160156 | 0,761987821 | 0,6436387 | 0,956633898 | novel transcript |
| ENSG00000288343 | AP002336,6 | -0,392160156 | 0,761987821 | 0,6436387 | 0,956633898 | novel transcript |

| | | | | | | |
|---|---|---|---|---|---|---|
| ENSG00000214435 | AS3MT | -0,392171739 | 0,761981703 | 0,12807029 | 0,571345532 | arsenite methyltransferase |
| ENSG00000244053 | RPL13AP2 | -0,392214576 | 0,761959078 | 0,71105439 | 0,970482408 | ribosomal protein L13a pseudogene 2 |
| ENSG00000176020 | AMIGO3 | -0,392327419 | 0,761899483 | 0,91899772 | 0,993585187 | adhesion molecule with Ig like domain 3 |
| ENSG00000184194 | GPR173 | -0,392345119 | 0,761890135 | 0,0996637 | 0,500760119 | G protein-coupled receptor 173 |
| ENSG00000243601 | RPL12P24 | -0,392357735 | 0,761883473 | 0,67909804 | 0,964151912 | ribosomal protein L12 pseudogene 24 |
| ENSG00000271254 | AC240274,1 | -0,39240787 | 0,761856997 | 0,15005936 | 0,615757437 | neuroblastoma breakpoint family member 1 |
| ENSG00000239503 | MARK2P8 | -0,392445858 | 0,761836937 | 0,42288306 | 0,890484004 | microtubule affinity regulating kinase 2 pseudogene 8 |
| ENSG00000274286 | ADRA2B | -0,392567531 | 0,761772688 | 0,59405423 | 0,947469769 | adrenoceptor alpha 2B |
| ENSG00000238109 | RNF14P3 | -0,392584351 | 0,761763807 | 0,37470879 | 0,861771195 | RNF14 pseudogene 3 |
| ENSG00000132872 | SYT4 | -0,39259181 | 0,761759869 | 0,39451214 | 0,871872016 | synaptotagmin 4 |
| ENSG00000287998 | AC104232,3 | -0,392601046 | 0,761754992 | 0,25713781 | 0,769358275 | novel transcript |
| ENSG00000281526 | ZNF546 | -0,392649254 | 0,761729538 | 0,64752616 | 0,956899963 | zinc finger protein 546 |
| ENSG00000147364 | FBXO25 | -0,392657109 | 0,761725391 | 0,24911245 | 0,761104998 | F-box protein 25 |
| ENSG00000270095 | AC068790,6 | -0,392659427 | 0,761724167 | 0,6570289 | 0,957612854 | novel transcript, sense intronic to CCDC92 |
| ENSG00000260352 | AC092287,1 | -0,392673037 | 0,761716981 | 0,76808161 | 0,977377956 | novel transcript, antisense to SMG1 |
| ENSG00000273437 | AC108673,3 | -0,392769898 | 0,761665841 | 0,56482327 | 0,940865625 | novel transcript |
| ENSG00000287560 | AL731533,3 | -0,392772157 | 0,761664649 | 0,64533107 | 0,956899963 | novel transcript |
| ENSG00000225246 | RPS2P1 | -0,392810844 | 0,761644225 | 0,67092796 | 0,962026636 | ribosomal protein S2 pseudogene 1 |
| ENSG00000267605 | AC016590,1 | -0,392912354 | 0,761590636 | 0,42834562 | 0,893058343 | novel transcript, antisense to ZNF383 |
| ENSG00000276855 | AC015922,3 | -0,392925049 | 0,761583935 | 0,5425659 | 0,933663748 | novel transcript |
| ENSG00000136274 | NACAD | -0,393230724 | 0,761422589 | 0,21219711 | 0,718991056 | NAC alpha domain containing |
| ENSG00000072080 | SPP2 | -0,393279894 | 0,761396639 | 0,29903157 | 0,808877417 | secreted phosphoprotein 2 |
| ENSG00000166619 | BLCAP | -0,393291526 | 0,7613905 | 0,12255979 | 0,560027662 | BLCAP apoptosis inducing factor |
| ENSG00000268947 | AC002128,1 | -0,393380725 | 0,761343426 | 0,33243111 | 0,83293962 | novel transcript |
| ENSG00000059728 | MXD1 | -0,393386078 | 0,761340601 | 0,26035398 | 0,772692192 | MAX dimerization protein 1 |
| ENSG00000175697 | GPR156 | -0,393418112 | 0,761323696 | 0,27988544 | 0,790565637 | G protein-coupled receptor 156 |
| ENSG00000259706 | HSP90B2P | -0,393451003 | 0,761306339 | 0,24150606 | 0,751948055 | heat shock protein 90 beta family member 2, pseudogene |
| ENSG00000107890 | ANKRD26 | -0,393743952 | 0,761151767 | 0,18689982 | 0,680474679 | ankyrin repeat domain 26 |
| ENSG00000259677 | AC027176,2 | -0,393786703 | 0,761129212 | 0,53106243 | 0,931265667 | Arginine/serine-rich coiled-coil protein (RSRC2) pseudogene |
| ENSG00000261824 | LINC00662 | -0,393805229 | 0,761119438 | 0,19392685 | 0,690714036 | long intergenic non-protein coding RNA 662 |

| | | | | | | |
|---|---|---|---|---|---|---|
| ENSG00000224370 | AC093392,2 | -0,393807715 | 0,761118127 | 0,48141502 | 0,912981134 | calmodulin 3 (phosphorylase kinase, delta) (CALM3) pseudogene |
| ENSG00000183778 | B3GALT5 | -0,393821635 | 0,761110783 | 0,13993016 | 0,596260721 | beta-1,3-galactosyltransferase 5 |
| ENSG00000018189 | RUFY3 | -0,393836828 | 0,761102768 | 0,26703378 | 0,779386557 | RUN and FYVE domain containing 3 |
| ENSG00000196912 | ANKRD36B | -0,393885016 | 0,761077347 | 0,13944132 | 0,595721964 | ankyrin repeat domain 36B |
| ENSG00000235724 | AC009299,2 | -0,393924289 | 0,761056629 | 0,73738493 | 0,974353474 | novel transcript, antisense to TANK |
| ENSG00000287014 | AC087883,2 | -0,394013953 | 0,76100933 | 0,30081401 | 0,810466143 | novel transcript |
| ENSG00000287242 | AC073593,2 | -0,394122144 | 0,760952263 | 0,38033046 | 0,865730737 | novel transcript, antisense to SCARB1 |
| ENSG00000185986 | SDHAP3 | -0,394325621 | 0,760844946 | 0,09905162 | 0,499450473 | succinate dehydrogenase complex flavoprotein subunit A pseudogene 3 |
| ENSG00000100814 | CCNB1IP1 | -0,39436214 | 0,760825687 | 0,29812714 | 0,807554329 | cyclin B1 interacting protein 1 |
| ENSG00000036672 | USP2 | -0,394392265 | 0,7608098 | 0,34225107 | 0,840048641 | ubiquitin specific peptidase 2 |
| ENSG00000244627 | TPTEP2 | -0,394466434 | 0,760770688 | 0,14197618 | 0,600058555 | TPTE pseudogene 2 |
| ENSG00000152382 | TADA1 | -0,394622933 | 0,760688166 | 0,3969351 | 0,873609704 | transcriptional adaptor 1 |
| ENSG00000103723 | AP3B2 | -0,394638215 | 0,760680108 | 0,09023606 | 0,478889887 | adaptor related protein complex 3 subunit beta 2 |
| ENSG00000145495 | MARCHF6 | -0,394729686 | 0,760631881 | 0,30238365 | 0,812035949 | membrane associated ring-CH-type finger 6 |
| ENSG00000233705 | SLC26A4-AS1 | -0,39484198 | 0,760572679 | 0,09321979 | 0,48515108 | SLC26A4 antisense RNA 1 |
| ENSG00000259725 | AC106738,2 | -0,394925511 | 0,760528643 | 0,78067553 | 0,978597905 | novel transcript |
| ENSG00000232493 | RPL12P11 | -0,394973567 | 0,760503311 | 0,60079373 | 0,949558426 | ribosomal protein L12 pseudogene 11 |
| ENSG00000284523 | AC004834,1 | -0,395003694 | 0,76048743 | 0,46475074 | 0,907725819 | novel transcript |
| ENSG00000258527 | ASB9P1 | -0,395047933 | 0,76046411 | 0,606081 | 0,950927937 | ankyrin repeat and SOCS box containing 9 pseudogene 1 |
| ENSG00000250882 | AC010261,1 | -0,395266779 | 0,760348762 | 0,79162372 | 0,979471509 | novel transcript |
| ENSG00000182613 | OR2V2 | -0,395295232 | 0,760333767 | 0,75969944 | 0,976120207 | olfactory receptor family 2 subfamily V member 2 |
| ENSG00000263307 | AC007216,4 | -0,395340631 | 0,760309841 | 0,56356523 | 0,940865625 | novel transcript, antisense to GSPT1 |
| ENSG00000186469 | GNG2 | -0,395597634 | 0,76017441 | 0,3256505 | 0,827465638 | G protein subunit gamma 2 |
| ENSG00000237853 | NFIA-AS1 | -0,395645057 | 0,760149423 | 0,57432402 | 0,942475308 | NFIA antisense RNA 1 |
| ENSG00000225224 | RPS27AP12 | -0,395656947 | 0,760143158 | 0,38756771 | 0,869820053 | RPS27A pseudogene 12 |
| ENSG00000146263 | MMS22L | -0,395670861 | 0,760135827 | 0,26064776 | 0,772692192 | MMS22 like, DNA repair protein |
| ENSG00000278600 | AC015871,3 | -0,395678524 | 0,76013179 | 0,34215906 | 0,840048641 | novel transcript, sense intronic to ST20 |
| ENSG00000152229 | PSTPIP2 | -0,395691069 | 0,76012518 | 0,16943425 | 0,651962158 | proline-serine-threonine phosphatase interacting protein 2 |
| ENSG00000196660 | SLC30A10 | -0,395779656 | 0,760078507 | 0,45142199 | 0,902062594 | solute carrier family 30 member 10 |

| | | | | | | |
|---|---|---|---|---|---|---|
| ENSG00000184984 | CHRM5 | -0,396031398 | 0,759945889 | 0,45470687 | 0,903424009 | cholinergic receptor muscarinic 5 |
| ENSG00000232268 | OR52I1 | -0,396035676 | 0,759943635 | 0,1369385 | 0,590598053 | olfactory receptor family 52 subfamily I member 1 |
| ENSG00000171262 | FAM98B | -0,396082236 | 0,75991911 | 0,33688655 | 0,836162806 | family with sequence similarity 98 member B |
| ENSG00000198707 | CEP290 | -0,396087704 | 0,75991623 | 0,26929699 | 0,781544346 | centrosomal protein 290 |
| ENSG00000067992 | PDK3 | -0,396152748 | 0,75988197 | 0,25989255 | 0,772441975 | pyruvate dehydrogenase kinase 3 |
| ENSG00000286125 | AC006115,2 | -0,396205784 | 0,759854036 | 0,38909256 | 0,870057277 | novel transcript (contains ZIM2-AS1 and PEG3-AS1) |
| ENSG00000201229 | SNORA63D | -0,396357884 | 0,75977393 | 0,53266976 | 0,931291085 | small nucleolar RNA, H/ACA box 63D |
| ENSG00000197183 | NOL4L | -0,396731962 | 0,759576954 | 0,12223936 | 0,559316128 | nucleolar protein 4 like |
| ENSG00000242615 | AC022415,1 | -0,396742257 | 0,759571533 | 0,31187936 | 0,818208978 | ribosomal protein L17 (RPL17) pseudogene |
| ENSG00000067141 | NEO1 | -0,396746775 | 0,759569154 | 0,21775515 | 0,72540192 | neogenin 1 |
| ENSG00000287509 | AP001033,3 | -0,39677123 | 0,759556279 | 0,51621676 | 0,925176589 | novel transcript, sense intronic to TWSG1 |
| ENSG00000255145 | STX17-AS1 | -0,396814013 | 0,759533755 | 0,09278936 | 0,484139956 | STX17 antisense RNA 1 |
| ENSG00000286282 | AC021733,4 | -0,396828765 | 0,759525988 | 0,65448674 | 0,957444211 | novel transcript, antisense to NOV |
| ENSG00000248405 | PRR5-ARHGAP8 | -0,396836879 | 0,759521717 | 0,85836461 | 0,989067345 | PRR5-ARHGAP8 readthrough |
| ENSG00000196352 | CD55 | -0,396938442 | 0,75946825 | 0,31034869 | 0,817751312 | CD55 molecule (Cromer blood group) |
| ENSG00000261219 | AC090651,1 | -0,397215956 | 0,759322174 | 0,73052617 | 0,974321178 | novel transcript |
| ENSG00000180573 | H2AC6 | -0,397357692 | 0,759247579 | 0,25303795 | 0,76515438 | H2A clustered histone 6 |
| ENSG00000250333 | AC093725,2 | -0,397368707 | 0,759241782 | 0,15156492 | 0,61913273 | novel transcript |
| ENSG00000198400 | NTRK1 | -0,397471511 | 0,759187681 | 0,38929975 | 0,870057277 | neurotrophic receptor tyrosine kinase 1 |
| ENSG00000197928 | ZNF677 | -0,397622114 | 0,759108434 | 0,19214216 | 0,688538797 | zinc finger protein 677 |
| ENSG00000125245 | GPR18 | -0,397722477 | 0,759055628 | 0,41677682 | 0,886368152 | G protein-coupled receptor 18 |
| ENSG00000204837 | FGF7P3 | -0,39777143 | 0,759029872 | 0,45064128 | 0,902062594 | fibroblast growth factor 7 pseudogene 3 |
| ENSG00000081913 | PHLPP1 | -0,398219279 | 0,758794287 | 0,14346078 | 0,602438935 | PH domain and leucine rich repeat protein phosphatase 1 |
| ENSG00000105865 | DUS4L | -0,398242566 | 0,758782039 | 0,42679638 | 0,892261855 | dihydrouridine synthase 4 like |
| ENSG00000284103 | DUS4L | -0,398242566 | 0,758782039 | 0,42679638 | 0,892261855 | dihydrouridine synthase 4 like |
| ENSG00000128915 | ICE2 | -0,398285726 | 0,758759339 | 0,35255395 | 0,844912682 | interactor of little elongation complex ELL subunit 2 |
| ENSG00000263163 | SLC27A3 | -0,398357458 | 0,758721614 | 0,5019043 | 0,920520138 | solute carrier family 27 member 3 |
| ENSG00000186153 | WWOX | -0,398542702 | 0,758624199 | 0,17017062 | 0,653348187 | WW domain containing oxidoreductase |
| ENSG00000250802 | ZBED3-AS1 | -0,398588941 | 0,758599885 | 0,07069704 | 0,418829978 | ZBED3 antisense RNA 1 |
| ENSG00000275056 | AC020663,3 | -0,398641044 | 0,758572489 | 0,34947737 | 0,843190352 | novel transcript, sense intronic to GLYR1 |
| ENSG00000188629 | ZNF177 | -0,398653056 | 0,758566173 | 0,41077924 | 0,882835102 | zinc finger protein 177 |
| ENSG00000134717 | BTF3L4 | -0,398773203 | 0,758503003 | 0,37855662 | 0,864072007 | basic transcription factor 3 like 4 |

| | | | | | | |
|---|---|---|---|---|---|---|
| ENSG00000197323 | TRIM33 | -0,398879212 | 0,75844727 | 0,34814441 | 0,842575226 | tripartite motif containing 33 |
| ENSG00000261584 | AL513548,1 | -0,399039638 | 0,758362937 | 0,41559175 | 0,885986046 | novel transcript |
| ENSG00000111305 | GSG1 | -0,399086185 | 0,758338469 | 0,39388412 | 0,871635019 | germ cell associated 1 |
| ENSG00000159784 | FAM131B | -0,399122739 | 0,758319255 | 0,07579125 | 0,434777568 | family with sequence similarity 131 member B |
| ENSG00000258917 | ZMYND19P1 | -0,399136815 | 0,758311857 | 0,56052661 | 0,940170934 | zinc finger MYND-type containing 19 pseudogene 1 |
| ENSG00000168300 | PCMTD1 | -0,399280521 | 0,758236325 | 0,47712076 | 0,91143003 | protein-L-isoaspartate (D-aspartate) O-methyltransferase domain containing 1 |
| ENSG00000248843 | AL645949,2 | -0,399359115 | 0,75819502 | 0,60038489 | 0,949439306 | ribosomal protein L7a (RPL7A) pseudogene |
| ENSG00000267353 | AC020928,2 | -0,399373409 | 0,758187508 | 0,5698905 | 0,94221458 | novel transcript |
| ENSG00000249784 | SCARNA22 | -0,399499242 | 0,758121381 | 0,25052923 | 0,762662687 | small Cajal body-specific RNA 22 |
| ENSG00000283689 | AC018553,2 | -0,399509602 | 0,758115937 | 0,6152833 | 0,952793934 | novel transcript, antisense to IRX3 |
| ENSG00000011405 | PIK3C2A | -0,399531166 | 0,758104605 | 0,32304091 | 0,825150495 | phosphatidylinositol-4-phosphate 3-kinase catalytic subunit type 2 alpha |
| ENSG00000231533 | AL391840,1 | -0,399537562 | 0,758101244 | 0,33185926 | 0,83268442 | novel transcript |
| ENSG00000206621 | SNORD116-14 | -0,399988858 | 0,757864136 | 0,69382049 | 0,967340633 | small nucleolar RNA, C/D box 116-14 |
| ENSG00000272905 | AC018648,1 | -0,400045135 | 0,757834574 | 0,69814009 | 0,968522643 | novel transcript |
| ENSG00000135605 | TEC | -0,400130366 | 0,757789804 | 0,48604033 | 0,914018932 | tec protein tyrosine kinase |
| ENSG00000277701 | AC159540,2 | -0,400169978 | 0,757768998 | 0,20435867 | 0,705535354 | novel transcript |
| ENSG00000258708 | SLC25A21-AS1 | -0,400220259 | 0,757742588 | 0,20301024 | 0,703345263 | SLC25A21 antisense RNA 1 |
| ENSG00000109084 | TMEM97 | -0,400314505 | 0,757693089 | 0,31296448 | 0,818922744 | transmembrane protein 97 |
| ENSG00000230724 | LINC01001 | -0,400376271 | 0,757660651 | 0,11717992 | 0,547080491 | long intergenic non-protein coding RNA 1001 |
| ENSG00000267769 | AC011498,6 | -0,400572497 | 0,757557606 | 0,67748739 | 0,9635902 | novel transcript, antisense to UBX6 |
| ENSG00000272195 | AL356512,1 | -0,400579651 | 0,75755385 | 0,39258889 | 0,870909282 | novel transcript, antisense to EFCAB2 |
| ENSG00000259090 | SEPTIN7P1 | -0,400617404 | 0,757534026 | 0,60053574 | 0,949439306 | septin 7 pseudogene 1 |
| ENSG00000224795 | NTM-AS1 | -0,400618338 | 0,757533535 | 0,48858946 | 0,915563475 | NTM antisense RNA 1 |
| ENSG00000233021 | AL669841,1 | -0,40071849 | 0,757480949 | 0,67717308 | 0,963580269 | novel transcript |
| ENSG00000224810 | AL355482,1 | -0,400765493 | 0,757456271 | 0,34475455 | 0,841168721 | novel transcript (LOC149157) |
| ENSG00000188558 | OR2G6 | -0,400772362 | 0,757452665 | 0,32865238 | 0,830659639 | olfactory receptor family 2 subfamily G member 6 |
| ENSG00000133639 | BTG1 | -0,400781669 | 0,757447778 | 0,26039373 | 0,772692192 | BTG anti-proliferation factor 1 |
| ENSG00000176225 | RTTN | -0,40082257 | 0,757426305 | 0,17703235 | 0,66300183 | rotatin |
| ENSG00000281357 | ARRDC3-AS1 | -0,400863273 | 0,757404935 | 0,10708329 | 0,522448794 | ARRDC3 antisense RNA 1 |
| ENSG00000234420 | ZNF37BP | -0,401002313 | 0,757331943 | 0,22343021 | 0,731984265 | zinc finger protein 37B, pseudogene |
| ENSG00000265217 | AC090358,1 | -0,401060784 | 0,75730125 | 0,56255926 | 0,940865625 | novel transcript |

| | | | | | | |
|---|---|---|---|---|---|---|
| ENSG00000261251 | Z97055,2 | -0,401183586 | 0,757236792 | 0,38713929 | 0,869441674 | novel transcript, antisense to EFCAB6 |
| ENSG00000215114 | UBXN2B | -0,401191193 | 0,757232799 | 0,35295687 | 0,845031687 | UBX domain protein 2B |
| ENSG00000127914 | AKAP9 | -0,401217503 | 0,75721899 | 0,2089064 | 0,713420724 | A-kinase anchoring protein 9 |
| ENSG00000258531 | BANF1P1 | -0,401246763 | 0,757203632 | 0,74008779 | 0,974724313 | BANF1 pseudogene 1 |
| ENSG00000227730 | MTND6P5 | -0,401367787 | 0,757140115 | 0,6815566 | 0,964654539 | MT-ND6 pseudogene 5 |
| ENSG00000108602 | ALDH3A1 | -0,401389699 | 0,757128615 | 0,63709002 | 0,955945807 | aldehyde dehydrogenase 3 family member A1 |
| ENSG00000218766 | AL450338,1 | -0,401477257 | 0,757082667 | 0,7880682 | 0,979462318 | ras-related C3 botulinum toxin substrate 3 (rho family, small GTP binding protein Rac3) pseudogene |
| ENSG00000224595 | AC073626,1 | -0,401495721 | 0,757072977 | 0,78188659 | 0,978597905 | novel transcript |
| ENSG00000133477 | FAM83F | -0,401549035 | 0,757045 | 0,48118797 | 0,912911021 | family with sequence similarity 83 member F |
| ENSG00000182185 | RAD51B | -0,401634554 | 0,757000126 | 0,24387843 | 0,754912399 | RAD51 paralog B |
| ENSG00000211450 | SELENOH | -0,401740351 | 0,756944615 | 0,06391815 | 0,39585974 | selenoprotein H |
| ENSG00000198658 | ABHD17AP1 | -0,401783727 | 0,756921858 | 0,70991016 | 0,970482408 | ABHD17A pseudogene 1 |
| ENSG00000236992 | RPL12P12 | -0,401904664 | 0,756858409 | 0,60748177 | 0,950993594 | ribosomal protein L12 pseudogene 12 |
| ENSG00000278767 | SERPINA10 | -0,402014152 | 0,756800973 | 0,66778923 | 0,961113267 | serpin family A member 10 |
| ENSG00000280341 | AP000542,3 | -0,402027967 | 0,756793726 | 0,1879906 | 0,68218766 | novel transcript |
| ENSG00000203812 | H2AC18 | -0,402062942 | 0,756775379 | 0,20946548 | 0,713913541 | H2A clustered histone 18 |
| ENSG00000249388 | AC023794,2 | -0,40209983 | 0,756756029 | 0,58620387 | 0,945256674 | novel transcript |
| ENSG00000153933 | DGKE | -0,402294006 | 0,756654183 | 0,31648297 | 0,821646322 | diacylglycerol kinase epsilon |
| ENSG00000251396 | LINC01301 | -0,402302516 | 0,756649719 | 0,18218573 | 0,672503417 | long intergenic non-protein coding RNA 1301 |
| ENSG00000135517 | MIP | -0,402394486 | 0,756601485 | 0,68733865 | 0,965965731 | major intrinsic protein of lens fiber |
| ENSG00000088035 | ALG6 | -0,402407645 | 0,756594584 | 0,41282464 | 0,884576962 | ALG6 alpha-1,3-glucosyltransferase |
| ENSG00000253051 | SNORA31B | -0,402416843 | 0,756589761 | 0,36667719 | 0,856146056 | small nucleolar RNA, H/ACA box 31B |
| ENSG00000241956 | AC109466,1 | -0,402418754 | 0,756588759 | 0,13370867 | 0,583882708 | novel transcript |
| ENSG00000232936 | AL157400,2 | -0,402429026 | 0,756583372 | 0,61092436 | 0,951873283 | novel transcript |
| ENSG00000152455 | SUV39H2 | -0,402465614 | 0,756564184 | 0,34212667 | 0,840048641 | suppressor of variegation 3-9 homolog 2 |
| ENSG00000225808 | DNAJC19P5 | -0,402514059 | 0,756538779 | 0,68741065 | 0,965965731 | DnaJ heat shock protein family (Hsp40) member C19 pseudogene 5 |
| ENSG00000276531 | SNX19P4 | -0,402632058 | 0,756476904 | 0,74601528 | 0,975707496 | sorting nexin 19 pseudogene 4 |
| ENSG00000231752 | EMBP1 | -0,402645732 | 0,756469734 | 0,18129214 | 0,670997703 | embigin pseudogene 1 |
| ENSG00000147894 | C9orf72 | -0,402698795 | 0,756441912 | 0,50734256 | 0,921988479 | C9orf72-SMCR8 complex subunit |
| ENSG00000242888 | AL133368,1 | -0,402845076 | 0,756365217 | 0,45021372 | 0,902062594 | ribosomal protein S2 (RPS2) pseudogene |
| ENSG00000269097 | TPRG1LP1 | -0,402883945 | 0,756344839 | 0,75795963 | 0,976120207 | tumor protein p63 regulated 1 like pseudogene 1 |

| | | | | | | |
|---|---|---|---|---|---|---|
| ENSG00000148057 | IDNK | -0,402885366 | 0,756344094 | 0,23868735 | 0,747998807 | IDNK gluconokinase |
| ENSG00000261426 | AC144833,1 | -0,403016775 | 0,756275205 | 0,54108967 | 0,933385624 | novel transcript, sense intronic to GABRG3 |
| ENSG00000258733 | LINC02328 | -0,403089014 | 0,756237338 | 0,68506989 | 0,965415049 | long intergenic non-protein coding RNA 2328 |
| ENSG00000232611 | AL683813,1 | -0,403216019 | 0,756170766 | 0,17437249 | 0,65976679 | novel transcript |
| ENSG00000260808 | AP003096,1 | -0,403351733 | 0,756099637 | 0,58606993 | 0,945246954 | novel transcript |
| ENSG00000173928 | SWSAP1 | -0,403434444 | 0,75605629 | 0,29137591 | 0,802058262 | SWIM-type zinc finger 7 associated protein 1 |
| ENSG00000138622 | HCN4 | -0,403490775 | 0,75602677 | 0,30181047 | 0,811332971 | hyperpolarization activated cyclic nucleotide gated potassium channel 4 |
| ENSG00000172817 | CYP7B1 | -0,403540855 | 0,756000527 | 0,32214313 | 0,824955371 | cytochrome P450 family 7 subfamily B member 1 |
| ENSG00000285410 | GABPB1-IT1 | -0,403561228 | 0,755989851 | 0,07362345 | 0,428474243 | GABPB1 intronic transcript |
| ENSG00000178803 | ADORA2A-AS1 | -0,403614628 | 0,755961869 | 0,61491164 | 0,952793934 | ADORA2A antisense RNA 1 |
| ENSG00000240322 | RN7SL481P | -0,403819582 | 0,755854482 | 0,61312531 | 0,952413269 | RNA, 7SL, cytoplasmic 481, pseudogene |
| ENSG00000174776 | WDR49 | -0,403860314 | 0,755833143 | 0,4130953 | 0,884663685 | WD repeat domain 49 |
| ENSG00000005448 | WDR54 | -0,403892092 | 0,755816494 | 0,16805443 | 0,649530679 | WD repeat domain 54 |
| ENSG00000053747 | LAMA3 | -0,404101576 | 0,755706755 | 0,56461937 | 0,940865625 | laminin subunit alpha 3 |
| ENSG00000049246 | PER3 | -0,404153572 | 0,755679519 | 0,1836506 | 0,674853346 | period circadian regulator 3 |
| ENSG00000124374 | PAIP2B | -0,404192631 | 0,75565906 | 0,30503707 | 0,81377925 | poly(A) binding protein interacting protein 2B |
| ENSG00000164649 | CDCA7L | -0,404431442 | 0,755533986 | 0,11170901 | 0,534425477 | cell division cycle associated 7 like |
| ENSG00000264345 | LINC01894 | -0,404481887 | 0,755507568 | 0,53548541 | 0,932098208 | long intergenic non-protein coding RNA 1894 |
| ENSG00000259380 | AC087473,1 | -0,404561594 | 0,755465829 | 0,08056786 | 0,450103921 | novel transcript |
| ENSG00000196418 | ZNF124 | -0,40473207 | 0,755376564 | 0,38704431 | 0,869441674 | zinc finger protein 124 |
| ENSG00000213399 | RPS2P17 | -0,404817742 | 0,755331709 | 0,74253407 | 0,975176043 | ribosomal protein S2 pseudogene 17 |
| ENSG00000287447 | AC099501,1 | -0,404819844 | 0,755330609 | 0,61401674 | 0,952591274 | novel transcript, antisense to FAM172A |
| ENSG00000282739 | SLC25A26 | -0,404928595 | 0,755273674 | 0,57333949 | 0,942475308 | solute carrier family 25 member 26 |
| ENSG00000259774 | AC243562,3 | -0,40499269 | 0,75524012 | 0,40513847 | 0,878176155 | novel transcript |
| ENSG00000259959 | AC107068,1 | -0,405001128 | 0,755235702 | 0,31288345 | 0,818889268 | novel transcript |
| ENSG00000154358 | OBSCN | -0,405081923 | 0,755193408 | 0,48645938 | 0,914147146 | obscurin, cytoskeletal calmodulin and titin-interacting RhoGEF |
| ENSG00000187416 | LHFPL3 | -0,405147065 | 0,755159309 | 0,50580417 | 0,921780819 | LHFPL tetraspan subfamily member 3 |
| ENSG00000180867 | PDIA3P1 | -0,405277767 | 0,755090898 | 0,24126467 | 0,751659063 | protein disulfide isomerase family A member 3 pseudogene 1 |
| ENSG00000286616 | AL008730,1 | -0,405280936 | 0,75508924 | 0,08232386 | 0,456410098 | novel transcript |
| ENSG00000282220 | AC110597,5 | -0,405395474 | 0,755029295 | 0,81320705 | 0,981246822 | novel transcript, antisense to DSEL |

| | | | | | | |
|---|---|---|---|---|---|---|
| ENSG00000263424 | AC110597,3 | -0,405395474 | 0,755029295 | 0,81320705 | 0,981246822 | novel transcript, antisense to DSEL |
| ENSG00000144366 | GULP1 | -0,405569287 | 0,754938335 | 0,37888914 | 0,864200043 | GULP PTB domain containing engulfment adaptor 1 |
| ENSG00000203724 | C1orf53 | -0,405608065 | 0,754918044 | 0,5418 | 0,933491609 | chromosome 1 open reading frame 53 |
| ENSG00000127252 | PLAAT1 | -0,405767782 | 0,754834474 | 0,37652198 | 0,862818448 | phospholipase A and acyltransferase 1 |
| ENSG00000037280 | FLT4 | -0,405793729 | 0,754820898 | 0,19632705 | 0,694046542 | fms related receptor tyrosine kinase 4 |
| ENSG00000275769 | AC068792,1 | -0,405842913 | 0,754795165 | 0,64719642 | 0,956899963 | novel transcript, sense intronic to DENND5B |
| ENSG00000248455 | LINC02217 | -0,405901855 | 0,754764328 | 0,7367953 | 0,974353474 | long intergenic non-protein coding RNA 2217 |
| ENSG00000226235 | LEMD1-AS1 | -0,406034987 | 0,754694682 | 0,52238926 | 0,927689738 | LEMD1 antisense RNA 1 |
| ENSG00000249131 | PSD2-AS1 | -0,406059913 | 0,754681643 | 0,68069059 | 0,964654539 | PSD2 antisense RNA 1 |
| ENSG00000227012 | LINC02527 | -0,406277629 | 0,754567763 | 0,60501401 | 0,950927937 | long intergenic non-protein coding RNA 2527 |
| ENSG00000270207 | AC077690,1 | -0,406417959 | 0,75449437 | 0,32012259 | 0,824366234 | novel transcript, antisense to GRM7 |
| ENSG00000258667 | HIF1A-AS3 | -0,406564515 | 0,754417729 | 0,06669225 | 0,40565499 | HIF1A antisense RNA 3 |
| ENSG00000227782 | AC002553,1 | -0,406685169 | 0,754354639 | 0,32879392 | 0,830659639 | novel transcript, antisense to NCOR1 |
| ENSG00000159648 | TEPP | -0,406687717 | 0,754353307 | 0,49882532 | 0,919683359 | testis, prostate and placenta expressed |
| ENSG00000286655 | AL158011,1 | -0,406750147 | 0,754320664 | 0,47161188 | 0,908896785 | novel transcript, antisense to TSEN15 |
| ENSG00000184908 | CLCNKB | -0,406951392 | 0,75421545 | 0,54814462 | 0,935404974 | chloride voltage-gated channel Kb |
| ENSG00000170324 | FRMPD2 | -0,406972557 | 0,754204385 | 0,7547054 | 0,975901808 | FERM and PDZ domain containing 2 |
| ENSG00000141404 | GNAL | -0,407174285 | 0,754098934 | 0,21015159 | 0,715238331 | G protein subunit alpha L |
| ENSG00000207279 | SNORD116-24 | -0,407183062 | 0,754094346 | 0,70548567 | 0,969485452 | small nucleolar RNA, C/D box 116-24 |
| ENSG00000145604 | SKP2 | -0,407218012 | 0,754076078 | 0,27827217 | 0,789652735 | S-phase kinase associated protein 2 |
| ENSG00000284989 | AL451062,2 | -0,407235925 | 0,754066715 | 0,80881831 | 0,981213877 | novel transcript |
| ENSG00000276334 | AL133243,2 | -0,407326931 | 0,754019149 | 0,50950758 | 0,922427526 | novel transcript, sense intronic to BIRC6 |
| ENSG00000097046 | CDC7 | -0,407425559 | 0,753967604 | 0,41859797 | 0,887094957 | cell division cycle 7 |
| ENSG00000287093 | AC003093,1 | -0,407431753 | 0,753964367 | 0,69129531 | 0,967201016 | novel transcript |
| ENSG00000277534 | AC007996,1 | -0,407438389 | 0,753960899 | 0,41538445 | 0,885986046 | novel transcript, sense intronic to KCTD1 |
| ENSG00000269019 | HOMER3-AS1 | -0,407448979 | 0,753955364 | 0,46702943 | 0,907940653 | HOMER3 antisense RNA 1 |
| ENSG00000178917 | ZNF852 | -0,407473836 | 0,753942374 | 0,22016213 | 0,728529712 | zinc finger protein 852 |
| ENSG00000281626 | ZNF852 | -0,407473836 | 0,753942374 | 0,22016213 | 0,728529712 | zinc finger protein 852 |
| ENSG00000268744 | AC008758,4 | -0,407664663 | 0,753842656 | 0,43773123 | 0,897021798 | novel transcript |
| ENSG00000227436 | FCF1P1 | -0,407702814 | 0,753822722 | 0,63612071 | 0,955399772 | FCF1 pseudogene 1 |
| ENSG00000259591 | AC022898,1 | -0,407737521 | 0,753804587 | 0,68146755 | 0,964654539 | novel transcript, antisense to RORA |
| ENSG00000205078 | SYCE1L | -0,40777686 | 0,753784033 | 0,36838004 | 0,857548638 | synaptonemal complex central element protein 1 like |

| | | | | | | |
|---|---|---|---|---|---|---|
| ENSG00000280734 | LINC01232 | -0,407800286 | 0,753771793 | 0,09168829 | 0,481715471 | long intergenic non-protein coding RNA 1232 |
| ENSG00000239969 | AC091390,3 | -0,407841898 | 0,753750052 | 0,30035611 | 0,810130728 | uncharacterized LOC100289561 |
| ENSG00000226711 | FAM66C | -0,407984708 | 0,753675443 | 0,12331814 | 0,561682809 | family with sequence similarity 66 member C |
| ENSG00000215156 | AC138409,1 | -0,407988399 | 0,753673515 | 0,51862488 | 0,926765093 | POM121-like protein 1 pseudogene |
| ENSG00000139988 | RDH12 | -0,40808181 | 0,753624718 | 0,73420409 | 0,974353474 | retinol dehydrogenase 12 |
| ENSG00000279094 | LINC01670 | -0,408149953 | 0,753589123 | 0,30077406 | 0,810428356 | long intergenic non-protein coding RNA 1670 |
| ENSG00000231327 | LINC01816 | -0,40819462 | 0,753565791 | 0,37336968 | 0,861257555 | long intergenic non-protein coding RNA 1816 |
| ENSG00000287569 | AL078587,2 | -0,40823683 | 0,753543744 | 0,59562061 | 0,947947604 | novel transcript |
| ENSG00000125954 | CHURC1-FNTB | -0,40833446 | 0,753492752 | 0,62450813 | 0,952793934 | CHURC1-FNTB readthrough |
| ENSG00000235823 | OLMALINC | -0,408363997 | 0,753477326 | 0,06656254 | 0,40541812 | oligodendrocyte maturation-associated long intergenic non-coding RNA |
| ENSG00000228809 | AL034428,1 | -0,408436888 | 0,753439258 | 0,61975848 | 0,952793934 | novel transcript, antisense to SNRPB2 |
| ENSG00000244671 | RN7SL280P | -0,408450756 | 0,753432015 | 0,69354926 | 0,967340633 | RNA, 7SL, cytoplasmic 280, pseudogene |
| ENSG00000223697 | AF230666,1 | -0,408556601 | 0,753376741 | 0,4880969 | 0,915188998 | novel transcript, antisense to PHF20L1 |
| ENSG00000137766 | UNC13C | -0,408562149 | 0,753373844 | 0,57827926 | 0,943425812 | unc-13 homolog C |
| ENSG00000272750 | AL592148,3 | -0,408644462 | 0,753330861 | 0,45491032 | 0,903424009 | novel transcript, antisense to MIA3 |
| ENSG00000246422 | DIAPH1-AS1 | -0,408693554 | 0,753305228 | 0,42787818 | 0,89282532 | DIAPH1 antisense RNA 1 |
| ENSG00000267187 | AC026898,1 | -0,408706757 | 0,753298334 | 0,5236508 | 0,928137575 | sal-like 2 (Drosophila), pseudogene |
| ENSG00000139973 | SYT16 | -0,408822209 | 0,753238053 | 0,45713743 | 0,903809806 | synaptotagmin 16 |
| ENSG00000214029 | ZNF891 | -0,408946036 | 0,753173405 | 0,07453316 | 0,431519399 | zinc finger protein 891 |
| ENSG00000165997 | ARL5B | -0,408981308 | 0,753154992 | 0,46918445 | 0,907940653 | ADP ribosylation factor like GTPase 5B |
| ENSG00000245317 | AC008393,1 | -0,409115634 | 0,75308487 | 0,49746385 | 0,918759905 | novel transcript |
| ENSG00000284274 | AC008393,3 | -0,409115634 | 0,75308487 | 0,49746385 | 0,918759905 | novel transcript |
| ENSG00000112182 | BACH2 | -0,409268984 | 0,753004826 | 0,13396984 | 0,584034405 | BTB domain and CNC homolog 2 |
| ENSG00000229932 | YWHAZP3 | -0,409284522 | 0,752996716 | 0,72071459 | 0,972142678 | tyrosine 3-monooxygenase/tryptophan 5-monooxygenase activation protein zeta pseudogene 3 |
| ENSG00000251046 | ZNF969P | -0,409531339 | 0,752867904 | 0,20036878 | 0,700040616 | zinc finger protein 969, pseudogene |
| ENSG00000164330 | EBF1 | -0,409565799 | 0,752849921 | 0,25672852 | 0,769103136 | EBF transcription factor 1 |
| ENSG00000170962 | PDGFD | -0,409726683 | 0,752765971 | 0,3886791 | 0,870026727 | platelet derived growth factor D |
| ENSG00000226581 | LINC02848 | -0,409745422 | 0,752756193 | 0,58491157 | 0,944857905 | long intergenic non-protein coding RNA 2848 |
| ENSG00000176884 | GRIN1 | -0,409835603 | 0,752709141 | 0,48377654 | 0,913690171 | glutamate ionotropic receptor NMDA type subunit 1 |
| ENSG00000280529 | TAF4 | -0,40983668 | 0,752708579 | 0,41042793 | 0,88262613 | TATA-box binding protein associated factor 4 |

| | | | | | | |
|---|---|---|---|---|---|---|
| ENSG00000155966 | AFF2 | -0,41015941 | 0,752540218 | 0,26239037 | 0,774068771 | AF4/FMR2 family member 2 |
| ENSG00000105855 | ITGB8 | -0,410215796 | 0,752510806 | 0,2915085 | 0,802058262 | integrin subunit beta 8 |
| ENSG00000178665 | ZNF713 | -0,410366684 | 0,752432107 | 0,06614229 | 0,404197127 | zinc finger protein 713 |
| ENSG00000134758 | RNF138 | -0,410416323 | 0,752406218 | 0,42433476 | 0,890949676 | ring finger protein 138 |
| ENSG00000228653 | HNRNPCP7 | -0,410454906 | 0,752386096 | 0,46549996 | 0,907899037 | heterogeneous nuclear ribonucleoprotein C pseudogene 7 |
| ENSG00000242100 | RPL9P32 | -0,410465656 | 0,75238049 | 0,43524563 | 0,896272827 | ribosomal protein L9 pseudogene 32 |
| ENSG00000273226 | AL391834,2 | -0,410500199 | 0,752362476 | 0,36826218 | 0,857548638 | novel transcript, antisense to RPS6 |
| ENSG00000216775 | AL109918,1 | -0,410574706 | 0,752323621 | 0,25196933 | 0,764267906 | heterogeneous nuclear ribonucleoprotein A/B (HNRNPAB) pseudogene |
| ENSG00000166128 | RAB8B | -0,41061631 | 0,752301926 | 0,43244505 | 0,89560871 | RAB8B, member RAS oncogene family |
| ENSG00000175175 | PPM1E | -0,410617161 | 0,752301483 | 0,2344524 | 0,744090329 | protein phosphatase, Mg2+/Mn2+ dependent 1E |
| ENSG00000235298 | AL354733,3 | -0,410650748 | 0,752283969 | 0,39538858 | 0,871996943 | novel transcript |
| ENSG00000231789 | PIK3CD-AS2 | -0,410727002 | 0,752244208 | 0,41517201 | 0,885821181 | PIK3CD antisense RNA 2 |
| ENSG00000106462 | EZH2 | -0,410850034 | 0,752180059 | 0,24942118 | 0,761381055 | enhancer of zeste 2 polycomb repressive complex 2 subunit |
| ENSG00000122970 | IFT81 | -0,410944521 | 0,752130799 | 0,31154707 | 0,817751312 | intraflagellar transport 81 |
| ENSG00000241627 | UBQLN4P1 | -0,411029317 | 0,752086593 | 0,79236152 | 0,979471509 | ubiquilin 4 pseudogene 1 |
| ENSG00000073067 | CYP2W1 | -0,411179921 | 0,752008086 | 0,69914861 | 0,968522643 | cytochrome P450 family 2 subfamily W member 1 |
| ENSG00000121898 | CPXM2 | -0,41127007 | 0,751961097 | 0,50373043 | 0,921200643 | carboxypeptidase X, M14 family member 2 |
| ENSG00000228149 | RPL3P1 | -0,41130774 | 0,751941463 | 0,50181604 | 0,920520138 | ribosomal protein L3 pseudogene 1 |
| ENSG00000261136 | AC023908,3 | -0,411476277 | 0,751853625 | 0,29217022 | 0,802771257 | novel transcript, antisense to FSIP1 |
| ENSG00000004846 | ABCB5 | -0,411606971 | 0,751785518 | 0,46340713 | 0,907101775 | ATP binding cassette subfamily B member 5 |
| ENSG00000175449 | RFESD | -0,411659224 | 0,751758289 | 0,41587058 | 0,885986046 | Rieske Fe-S domain containing |
| ENSG00000137075 | RNF38 | -0,411708163 | 0,751732789 | 0,28616063 | 0,797420923 | ring finger protein 38 |
| ENSG00000244691 | RPL10AP1 | -0,411759421 | 0,751706081 | 0,51489223 | 0,924720046 | ribosomal protein L10a pseudogene 1 |
| ENSG00000270068 | AC125494,3 | -0,411820012 | 0,751674511 | 0,75085192 | 0,975901808 | novel transcript, antisense to ZNF384 |
| ENSG00000006634 | DBF4 | -0,411852953 | 0,751657348 | 0,35396732 | 0,845346751 | DBF4 zinc finger |
| ENSG00000286118 | AC234031,1 | -0,411863725 | 0,751651736 | 0,26784566 | 0,780494619 | novel transcript, antisense to FAM156A |
| ENSG00000272432 | AL031432,3 | -0,411884149 | 0,751641095 | 0,65128444 | 0,957089277 | novel transcript |
| ENSG00000278765 | AC004477,2 | -0,412171644 | 0,751491325 | 0,48181226 | 0,912981134 | novel transcript |
| ENSG00000251034 | AC087854,1 | -0,412193368 | 0,75148001 | 0,08314937 | 0,45864181 | novel transcript |
| ENSG00000263624 | AC055811,1 | -0,412200536 | 0,751476276 | 0,48762364 | 0,914936533 | novel transcript |

| | | | | | | |
|---|---|---|---|---|---|---|
| ENSG00000172650 | AGAP5 | -0,412324242 | 0,751411842 | 0,14573014 | 0,608454479 | ArfGAP with GTPase domain, ankyrin repeat and PH domain 5 |
| ENSG00000007062 | PROM1 | -0,412328116 | 0,751409824 | 0,2568249 | 0,769107677 | prominin 1 |
| ENSG00000242290 | ZBTB20-AS5 | -0,41237046 | 0,75138777 | 0,71020659 | 0,970482408 | ZBTB20 antisense RNA 5 |
| ENSG00000270269 | IMMP1LP1 | -0,412384203 | 0,751380613 | 0,63479066 | 0,955149453 | inner mitochondrial membrane peptidase subunit 1 pseudogene 1 |
| ENSG00000169684 | CHRNA5 | -0,412467504 | 0,751337229 | 0,38469375 | 0,868536368 | cholinergic receptor nicotinic alpha 5 subunit |
| ENSG00000107362 | ABHD17B | -0,412491724 | 0,751324616 | 0,30370903 | 0,812795736 | abhydrolase domain containing 17B, depalmitoylase |
| ENSG00000232790 | LINC01162 | -0,412511523 | 0,751314306 | 0,75870878 | 0,976120207 | long intergenic non-protein coding RNA 1162 |
| ENSG00000236519 | LINC01424 | -0,412511697 | 0,751314215 | 0,45989834 | 0,905341196 | long intergenic non-protein coding RNA 1424 |
| ENSG00000160766 | GBAP1 | -0,412525761 | 0,75130689 | 0,16816625 | 0,649802219 | glucosylceramidase beta pseudogene 1 |
| ENSG00000166037 | CEP57 | -0,412640144 | 0,751247326 | 0,32497361 | 0,826983176 | centrosomal protein 57 |
| ENSG00000258123 | LINC02444 | -0,413027706 | 0,75104554 | 0,30616254 | 0,815320753 | long intergenic non-protein coding RNA 2444 |
| ENSG00000222489 | SNORA79B | -0,413052719 | 0,751032519 | 0,34554045 | 0,841168721 | small nucleolar RNA, H/ACA box 79B |
| ENSG00000282508 | LINC01002 | -0,413073954 | 0,751021464 | 0,19764317 | 0,696048902 | long intergenic non-protein coding RNA 1002 |
| ENSG00000166321 | NUDT13 | -0,413214648 | 0,750948227 | 0,23367265 | 0,742886071 | nudix hydrolase 13 |
| ENSG00000255129 | AP000880,1 | -0,413362885 | 0,750871071 | 0,48889526 | 0,915617728 | novel transcript |
| ENSG00000233836 | AC139769,1 | -0,413626508 | 0,750733877 | 0,39192678 | 0,870625724 | zinc finger protein 726 pseudogene 1 |
| ENSG00000205189 | ZBTB10 | -0,413644528 | 0,7507245 | 0,30859636 | 0,816730513 | zinc finger and BTB domain containing 10 |
| ENSG00000253762 | AC069133,1 | -0,413796138 | 0,750645612 | 0,59694281 | 0,948282297 | novel transcript |
| ENSG00000285027 | ALG1L13P | -0,413807538 | 0,750639681 | 0,6555272 | 0,957444211 | ALG1 like 13, pseudogene |
| ENSG00000253981 | ALG1L13P | -0,413807538 | 0,750639681 | 0,6555272 | 0,957444211 | ALG1 like 13, pseudogene |
| ENSG00000273448 | AC006480,2 | -0,413821161 | 0,750632593 | 0,29604065 | 0,805869821 | novel transcript |
| ENSG00000271199 | AL353804,2 | -0,413919269 | 0,750581549 | 0,78720767 | 0,979435718 | novel transcript, sense overlapping FTX |
| ENSG00000135747 | ZNF670-ZNF69 | -0,414018784 | 0,750529777 | 0,28343151 | 0,79458592 | ZNF670-ZNF695 readthrough (NMD candidate) |
| ENSG00000232859 | LYRM9 | -0,4140265 | 0,750525762 | 0,15261269 | 0,621071504 | LYR motif containing 9 |
| ENSG00000256146 | AC006206,1 | -0,414048919 | 0,7505141 | 0,43285081 | 0,89560871 | novel transcript |
| ENSG00000025423 | HSD17B6 | -0,414146765 | 0,7504632 | 0,44931276 | 0,902062594 | hydroxysteroid 17-beta dehydrogenase 6 |
| ENSG00000244471 | AL137058,1 | -0,414205505 | 0,750432646 | 0,54652657 | 0,935401821 | transmembrane phosphatase with tensin homology (TPTE) pseudogene |
| ENSG00000273321 | AC023983,2 | -0,414215368 | 0,750427515 | 0,68791361 | 0,966127143 | novel transcript |
| ENSG00000250538 | AC104407,1 | -0,414546928 | 0,750255072 | 0,65079101 | 0,956933968 | novel transcript |
| ENSG00000187372 | PCDHB13 | -0,414589443 | 0,750232963 | 0,13817657 | 0,592936302 | protocadherin beta 13 |

| | | | | | | |
|---|---|---|---|---|---|---|
| ENSG00000259070 | LINC00639 | -0,414595475 | 0,750229826 | 0,18976477 | 0,684955795 | long intergenic non-protein coding RNA 639 |
| ENSG00000180596 | H2BC4 | -0,41463421 | 0,750209683 | 0,32324914 | 0,82538884 | H2B clustered histone 4 |
| ENSG00000224165 | DNAJC27-AS1 | -0,414667713 | 0,750192262 | 0,34937605 | 0,843141089 | DNAJC27 antisense RNA 1 |
| ENSG00000260339 | HEXA-AS1 | -0,414701101 | 0,750174901 | 0,49205756 | 0,916995341 | HEXA antisense RNA 1 |
| ENSG00000261610 | AP000265,1 | -0,414701501 | 0,750174693 | 0,59512399 | 0,947639622 | novel transcript |
| ENSG00000254538 | AC027018,1 | -0,414756928 | 0,750145872 | 0,50862003 | 0,922427526 | novel transcript |
| ENSG00000283085 | TPBG | -0,414835893 | 0,750104814 | 0,32196742 | 0,824955371 | trophoblast glycoprotein |
| ENSG00000109103 | UNC119 | -0,414906438 | 0,750068137 | 0,09130779 | 0,481302729 | unc-119 lipid binding chaperone |
| ENSG00000132938 | MTUS2 | -0,414967832 | 0,750036218 | 0,24582033 | 0,757752583 | microtubule associated scaffold protein 2 |
| ENSG00000279636 | LINC00216 | -0,414995927 | 0,750021612 | 0,23849458 | 0,747998807 | long intergenic non-protein coding RNA 216 |
| ENSG00000273271 | AP000254,2 | -0,415319869 | 0,749853222 | 0,60841139 | 0,951449272 | novel transcript, antisense to SOD1 |
| ENSG00000163393 | SLC22A15 | -0,415329275 | 0,749848333 | 0,30785933 | 0,816424917 | solute carrier family 22 member 15 |
| ENSG00000219470 | AL355802,1 | -0,415429846 | 0,749796062 | 0,57848354 | 0,943425812 | 40S ribosomal protein S2 (RPS2) pseudogene |
| ENSG00000156239 | N6AMT1 | -0,415453155 | 0,749783948 | 0,32264602 | 0,824955371 | N-6 adenine-specific DNA methyltransferase 1 |
| ENSG00000242103 | AC079841,1 | -0,415523147 | 0,749747574 | 0,68655261 | 0,965769168 | actin related protein 2/3 complex, subunit 1A, 41kDa (ARPC1A) pseudogene |
| ENSG00000270108 | AL049840,6 | -0,415550635 | 0,749733289 | 0,29838825 | 0,80782909 | novel transcript, sense intronic to KLC1 |
| ENSG00000183828 | NUDT14 | -0,415561982 | 0,749727392 | 0,32815025 | 0,830076239 | nudix hydrolase 14 |
| ENSG00000268472 | AP002884,3 | -0,415828683 | 0,749588808 | 0,72349153 | 0,972913663 | novel transcript, antisense to C11orf34 |
| ENSG00000088305 | DNMT3B | -0,415890583 | 0,749556646 | 0,12782549 | 0,570756819 | DNA methyltransferase 3 beta |
| ENSG00000287458 | AL033519,5 | -0,415933573 | 0,749534311 | 0,16764038 | 0,648566191 | novel transcript, antisense to FKBP5 |
| ENSG00000156587 | UBE2L6 | -0,415949904 | 0,749525827 | 0,22075774 | 0,728572782 | ubiquitin conjugating enzyme E2 L6 |
| ENSG00000256271 | CACNA1C-AS2 | -0,415959125 | 0,749521036 | 0,74342679 | 0,975310206 | CACNA1C antisense RNA 2 |
| ENSG00000285101 | CACNA1C-AS2 | -0,415959125 | 0,749521036 | 0,74342679 | 0,975310206 | CACNA1C antisense RNA 2 |
| ENSG00000214855 | APOC1P1 | -0,415971561 | 0,749514575 | 0,75811885 | 0,976120207 | apolipoprotein C1 pseudogene 1 |
| ENSG00000174804 | FZD4 | -0,416009178 | 0,749495033 | 0,2149354 | 0,722242094 | frizzled class receptor 4 |
| ENSG00000277182 | AC006449,5 | -0,416045255 | 0,749476291 | 0,32180437 | 0,824933727 | novel transcript, antisense to PCGF2 |
| ENSG00000282749 | AC006449,10 | -0,416045255 | 0,749476291 | 0,32180437 | 0,824933727 | novel transcript, antisense to PCGF2 |
| ENSG00000242866 | STRC | -0,416096524 | 0,749449657 | 0,5321038 | 0,931291085 | stereocilin |
| ENSG00000161298 | ZNF382 | -0,416144111 | 0,749424937 | 0,10785764 | 0,523928283 | zinc finger protein 382 |
| ENSG00000273138 | AC005021,1 | -0,4161779 | 0,749407385 | 0,47675814 | 0,91143003 | novel transcript, antisense to PON2 |
| ENSG00000188013 | MEIS3P2 | -0,416241683 | 0,749374254 | 0,16172333 | 0,638306753 | Meis homeobox 3 pseudogene 2 |
| ENSG00000210049 | MT-TF | -0,416281245 | 0,749353704 | 0,71353312 | 0,970812834 | mitochondrially encoded tRNA-Phe (UUU/C) |

| | | | | | | |
|---|---|---|---|---|---|---|
| ENSG00000111554 | MDM1 | -0,416544525 | 0,749216966 | 0,26840252 | 0,780709773 | Mdm1 nuclear protein |
| ENSG00000227197 | AC009365,3 | -0,416558181 | 0,749209874 | 0,60647273 | 0,950927937 | novel transcript |
| ENSG00000283649 | ZDBF2 | -0,416664719 | 0,74915455 | 0,49548641 | 0,917815326 | zinc finger DBF-type containing 2 |
| ENSG00000224323 | DPRXP1 | -0,416717914 | 0,749126928 | 0,59624846 | 0,948282297 | divergent-paired related homeobox pseudogene 1 |
| ENSG00000169314 | C22orf15 | -0,41683983 | 0,749063625 | 0,45269463 | 0,902285042 | chromosome 22 open reading frame 15 |
| ENSG00000143554 | SLC27A3 | -0,416894282 | 0,749035353 | 0,25979575 | 0,772371097 | solute carrier family 27 member 3 |
| ENSG00000122194 | PLG | -0,416940273 | 0,749011475 | 0,56551629 | 0,940865625 | plasminogen |
| ENSG00000286478 | AC007130,1 | -0,416993661 | 0,748983758 | 0,54016699 | 0,933385624 | novel transcript, antisense to HIBADH |
| ENSG00000090447 | TFAP4 | -0,417019445 | 0,748970372 | 0,17517965 | 0,660653403 | transcription factor AP-4 |
| ENSG00000185946 | RNPC3 | -0,417062752 | 0,74894789 | 0,24870396 | 0,760720365 | RNA binding region (RNP1, RRM) containing 3 |
| ENSG00000258765 | AC107958,2 | -0,417243568 | 0,748854029 | 0,45388224 | 0,902912642 | novel transcript |
| ENSG00000275445 | AC092119,3 | -0,417304469 | 0,748822418 | 0,73520477 | 0,974353474 | novel transcript, antisense to C16orf52 |
| ENSG00000264278 | ZNF236-DT | -0,417307089 | 0,748821058 | 0,54932572 | 0,936040289 | ZNF236 divergent transcript |
| ENSG00000251247 | ZNF345 | -0,417337244 | 0,748805406 | 0,09083406 | 0,480889225 | zinc finger protein 345 |
| ENSG00000233547 | AL158212,2 | -0,41735625 | 0,748795541 | 0,65873401 | 0,958206085 | novel transcript |
| ENSG00000164385 | LINC01600 | -0,417446282 | 0,748748814 | 0,3860232 | 0,869112622 | long intergenic non-protein coding RNA 1600 |
| ENSG00000146083 | RNF44 | -0,417472741 | 0,748735082 | 0,13971567 | 0,595809743 | ring finger protein 44 |
| ENSG00000286214 | AUXG0100005[illegible] | -0,41747539 | 0,748733708 | 0,45229883 | 0,90209238 | COPG2 imprinted transcript 1 |
| ENSG00000213277 | MARCKSL1P1 | -0,417506447 | 0,74871759 | 0,66259126 | 0,959897678 | MARCKS like 1 pseudogene 1 |
| ENSG00000139343 | SNRPF | -0,417526796 | 0,748707029 | 0,3240011 | 0,826094468 | small nuclear ribonucleoprotein polypeptide F |
| ENSG00000233369 | GTF2IP4 | -0,417567176 | 0,748686073 | 0,23017813 | 0,738687017 | general transcription factor IIi pseudogene 4 |
| ENSG00000285644 | AC108448,3 | -0,417586085 | 0,748676261 | 0,09092508 | 0,480995222 | novel transcript, antisense to CARS |
| ENSG00000167103 | PIP5KL1 | -0,417612338 | 0,748662637 | 0,30665934 | 0,81549341 | phosphatidylinositol-4-phosphate 5-kinase like 1 |
| ENSG00000236120 | AC110995,1 | -0,417653939 | 0,748641049 | 0,55288653 | 0,936894267 | novel transcript |
| ENSG00000288526 | AL157834,4 | -0,417674813 | 0,748630218 | 0,71019532 | 0,970482408 | novel transcript |
| ENSG00000283236 | AC074141,1 | -0,417758311 | 0,748586891 | 0,3433801 | 0,840873784 | novel zinc finger protein pseudogene |
| ENSG00000240038 | AMY2B | -0,417775347 | 0,748578051 | 0,33850452 | 0,837263082 | amylase alpha 2B |
| ENSG00000281327 | LINC01338 | -0,417800271 | 0,748565119 | 0,62821214 | 0,952793934 | long intergenic non-protein coding RNA 1338 |
| ENSG00000230042 | AK3P3 | -0,417821986 | 0,748553852 | 0,30499786 | 0,813769074 | adenylate kinase 3 pseudogene 3 |
| ENSG00000011677 | GABRA3 | -0,418024357 | 0,748448857 | 0,36995234 | 0,858216968 | gamma-aminobutyric acid type A receptor subunit alpha3 |
| ENSG00000220721 | OR1F12 | -0,418068327 | 0,748426047 | 0,61283176 | 0,952413269 | olfactory receptor family 1 subfamily F member 12 |
| ENSG00000286686 | AL136040,2 | -0,418132198 | 0,748392913 | 0,38121926 | 0,865900094 | novel transcript |

| | | | | | | |
|---|---|---|---|---|---|---|
| ENSG00000090006 | LTBP4 | -0,418247198 | 0,74833326 | 0,27012391 | 0,782393358 | latent transforming growth factor beta binding protein 4 |
| ENSG00000197061 | H4C3 | -0,418351165 | 0,748279333 | 0,30632745 | 0,815375074 | H4 clustered histone 3 |
| ENSG00000271976 | AC012467,2 | -0,418558109 | 0,748172006 | 0,43041919 | 0,895198614 | novel transcript, antisense to IL17RB |
| ENSG00000176124 | DLEU1 | -0,41876631 | 0,748064042 | 0,07829478 | 0,442692864 | deleted in lymphocytic leukemia 1 |
| ENSG00000106809 | OGN | -0,418781258 | 0,748056291 | 0,40558444 | 0,878494653 | osteoglycin |
| ENSG00000179818 | PCBP1-AS1 | -0,418787528 | 0,74805304 | 0,12112039 | 0,555784698 | PCBP1 antisense RNA 1 |
| ENSG00000263160 | TMEM50B | -0,418817207 | 0,748037651 | 0,53196614 | 0,931291085 | transmembrane protein 50B |
| ENSG00000286316 | Z93020,1 | -0,418866283 | 0,748012206 | 0,63951955 | 0,956018518 | novel transcript, antisense to RANBP9 |
| ENSG00000232344 | AC087163,1 | -0,418870288 | 0,748010129 | 0,47916558 | 0,912714933 | ribosomal protein S2 (RPS2) pseudogene |
| ENSG00000259943 | AL050341,2 | -0,418989084 | 0,747948539 | 0,32061555 | 0,824366234 | novel transcript, antisense to ZMPSTE24 |
| ENSG00000248885 | AC118465,1 | -0,419122664 | 0,747879289 | 0,70548171 | 0,969485452 | septin family pseudogene |
| ENSG00000246250 | AC087521,2 | -0,419147474 | 0,747866427 | 0,18346279 | 0,674559627 | novel transcript |
| ENSG00000129195 | PIMREG | -0,419167162 | 0,747856222 | 0,05389924 | 0,357953158 | PICALM interacting mitotic regulator |
| ENSG00000150625 | GPM6A | -0,419187542 | 0,747845657 | 0,39433424 | 0,871799523 | glycoprotein M6A |
| ENSG00000170417 | TMEM182 | -0,419361137 | 0,747755676 | 0,09944311 | 0,50040065 | transmembrane protein 182 |
| ENSG00000103037 | SETD6 | -0,419486275 | 0,74769082 | 0,0573315 | 0,370182715 | SET domain containing 6, protein lysine methyltransferase |
| ENSG00000247708 | STX18-AS1 | -0,419705298 | 0,747577318 | 0,09243875 | 0,483278451 | STX18 antisense RNA 1 (head to head) |
| ENSG00000261377 | PDCD6IPP2 | -0,419705501 | 0,747577213 | 0,30811081 | 0,816424917 | PDCD6IP pseudogene 2 |
| ENSG00000107614 | TRDMT1 | -0,419721104 | 0,747569127 | 0,28132215 | 0,792365382 | tRNA aspartic acid methyltransferase 1 |
| ENSG00000255234 | AP000446,1 | -0,419779212 | 0,747539018 | 0,4760884 | 0,911413793 | novel transcript, antisense to CCDC90B |
| ENSG00000197375 | SLC22A5 | -0,419815354 | 0,747520291 | 0,16122334 | 0,6373908 | solute carrier family 22 member 5 |
| ENSG00000257941 | AC011611,5 | -0,419981226 | 0,747434351 | 0,31127913 | 0,817751312 | density-regulated protein (DENR) pseudogene |
| ENSG00000198711 | SSBP3-AS1 | -0,420010873 | 0,747418991 | 0,27953601 | 0,790302992 | SSBP3 antisense RNA 1 |
| ENSG00000228723 | SRGAP3-AS2 | -0,420039597 | 0,747404111 | 0,36386802 | 0,854140004 | SRGAP3 antisense RNA 2 |
| ENSG00000228372 | SMIM10L2B-AS | -0,420079636 | 0,747383368 | 0,73989246 | 0,974715504 | SMIM10L2B antisense RNA 1 |
| ENSG00000251503 | CENPS-CORT | -0,420154968 | 0,747344344 | 0,30705031 | 0,815892645 | CENPS-CORT readthrough |
| ENSG00000246851 | AL157938,3 | -0,420181553 | 0,747330572 | 0,24222587 | 0,752479103 | novel transcript |
| ENSG00000254254 | AC012349,1 | -0,420255058 | 0,747292497 | 0,14167546 | 0,599581043 | novel transcript |
| ENSG00000158560 | DYNC1I1 | -0,420381996 | 0,747226748 | 0,27302068 | 0,785672647 | dynein cytoplasmic 1 intermediate chain 1 |
| ENSG00000184206 | GOLGA6L4 | -0,420547202 | 0,747141186 | 0,12632089 | 0,567753944 | golgin A6 family like 4 |
| ENSG00000019102 | VSIG2 | -0,420579154 | 0,747124639 | 0,57675661 | 0,943261034 | V-set and immunoglobulin domain containing 2 |

| | | | | | | |
|---|---|---|---|---|---|---|
| ENSG00000107560 | RAB11FIP2 | -0,420908618 | 0,74695404 | 0,37680771 | 0,862818448 | RAB11 family interacting protein 2 |
| ENSG00000139540 | SLC39A5 | -0,420929086 | 0,746943443 | 0,5712395 | 0,942436428 | solute carrier family 39 member 5 |
| ENSG00000272010 | AC100814,1 | -0,420937143 | 0,746939271 | 0,71370942 | 0,970812834 | novel transcript |
| ENSG00000287658 | AC109642,1 | -0,420940615 | 0,746937474 | 0,22125929 | 0,729227317 | novel transcript |
| ENSG00000224143 | EHMT2 | -0,420985304 | 0,746914337 | 0,61233183 | 0,952413269 | euchromatic histone lysine methyltransferase 2 |
| ENSG00000136866 | ZFP37 | -0,421082054 | 0,746864249 | 0,35982725 | 0,850238665 | ZFP37 zinc finger protein |
| ENSG00000163012 | ZSWIM2 | -0,42112089 | 0,746844145 | 0,6195491 | 0,952793934 | zinc finger SWIM-type containing 2 |
| ENSG00000083814 | ZNF671 | -0,421154737 | 0,746826623 | 0,10622519 | 0,520129118 | zinc finger protein 671 |
| ENSG00000266445 | NARF-AS1 | -0,421167923 | 0,746819797 | 0,63169232 | 0,953915351 | NARF antisense RNA 1 |
| ENSG00000116106 | EPHA4 | -0,421396244 | 0,746701615 | 0,24784541 | 0,760164991 | EPH receptor A4 |
| ENSG00000273983 | H3C8 | -0,421506785 | 0,746644404 | 0,1272737 | 0,569742977 | H3 clustered histone 8 |
| ENSG00000105290 | APLP1 | -0,421573334 | 0,746609963 | 0,10212084 | 0,508635661 | amyloid beta precursor like protein 1 |
| ENSG00000077498 | TYR | -0,42180235 | 0,746491454 | 0,75446447 | 0,975901808 | tyrosinase |
| ENSG00000120696 | KBTBD7 | -0,421858942 | 0,746462173 | 0,38376737 | 0,868000076 | kelch repeat and BTB domain containing 7 |
| ENSG00000259709 | AC023906,4 | -0,421981603 | 0,746398709 | 0,45747717 | 0,903831777 | novel transcript, antisense to GNB5 |
| ENSG00000276337 | AC105429,1 | -0,42220787 | 0,746281656 | 0,4021735 | 0,87718264 | novel transcript, sense intronic to ZCCHC14 |
| ENSG00000287089 | AL138889,3 | -0,422280001 | 0,746244345 | 0,67370298 | 0,962863995 | novel transcript, antisense to MLN |
| ENSG00000105255 | FSD1 | -0,422289264 | 0,746239554 | 0,10974301 | 0,528090728 | fibronectin type III and SPRY domain containing 1 |
| ENSG00000187240 | DYNC2H1 | -0,422318517 | 0,746224423 | 0,29373002 | 0,803568033 | dynein cytoplasmic 2 heavy chain 1 |
| ENSG00000165995 | CACNB2 | -0,42232038 | 0,746223459 | 0,15818909 | 0,631383073 | calcium voltage-gated channel auxiliary subunit beta 2 |
| ENSG00000272968 | RBAK-RBAKDN | -0,422543971 | 0,746107817 | 0,73543701 | 0,974353474 | RBAK-RBAKDN readthrough |
| ENSG00000273058 | AL359921,2 | -0,42259572 | 0,746081055 | 0,52633499 | 0,929261495 | novel transcript |
| ENSG00000286021 | LINC02822 | -0,422598207 | 0,746079769 | 0,09734481 | 0,495746584 | long intergenic non-protein coding RNA 2822 |
| ENSG00000285828 | AC254945,4 | -0,422635278 | 0,746060598 | 0,34142514 | 0,840048641 | novel transcript, antisense to ARHGAP11A |
| ENSG00000285948 | AC123768,4 | -0,422635278 | 0,746060598 | 0,34142514 | 0,840048641 | novel transcript, antisense to ARHGAP11A |
| ENSG00000279249 | AC007614,1 | -0,422657076 | 0,746049326 | 0,37045146 | 0,858545694 | novel transcript, antisense to CBLN1 |
| ENSG00000200156 | RNU5B-1 | -0,422713126 | 0,746020342 | 0,46467398 | 0,907638841 | RNA, U5B small nuclear 1 |
| ENSG00000187961 | KLHL17 | -0,422729709 | 0,746011767 | 0,27043506 | 0,782859427 | kelch like family member 17 |
| ENSG00000227948 | AC003989,1 | -0,422815466 | 0,745967423 | 0,64756321 | 0,956899963 | pseudogene similar to part of argininosuccinate synthetase 1 (ASS1) |
| ENSG00000285310 | AC005183,2 | -0,42288063 | 0,74593373 | 0,47303578 | 0,909532026 | SDA1 domain containing 1 (SDAD1) pseudogene |
| ENSG00000271500 | AC005183,1 | -0,42288063 | 0,74593373 | 0,47303578 | 0,909532026 | SDA1 domain containing 1 (SDAD1) pseudogene |

| | | | | | | |
|---|---|---|---|---|---|---|
| ENSG00000273533 | FAM181A | -0,423090786 | 0,745825078 | 0,18509213 | 0,677443261 | family with sequence similarity 181 member A |
| ENSG00000140067 | FAM181A | -0,423090786 | 0,745825078 | 0,18509213 | 0,677443261 | family with sequence similarity 181 member A |
| ENSG00000286340 | AL356776,2 | -0,423304041 | 0,745714841 | 0,33805907 | 0,837263082 | novel transcript, antisense to PHIP |
| ENSG00000239377 | ZNF775-AS1 | -0,423478609 | 0,745624614 | 0,66248236 | 0,959897678 | ZNF775 antisense RNA 1 |
| ENSG00000125355 | TMEM255A | -0,423506228 | 0,74561034 | 0,45145088 | 0,902062594 | transmembrane protein 255A |
| ENSG00000285309 | AL136295,18 | -0,423612424 | 0,745555458 | 0,21522052 | 0,722501279 | novel transcript |
| ENSG00000278784 | AL136295,7 | -0,423612424 | 0,745555458 | 0,21522052 | 0,722501279 | novel transcript |
| ENSG00000166352 | IFTAP | -0,423666226 | 0,745527655 | 0,07271268 | 0,425231804 | intraflagellar transport associated protein |
| ENSG00000132849 | PATJ | -0,42369154 | 0,745514573 | 0,1679227 | 0,649182045 | PATJ crumbs cell polarity complex component |
| ENSG00000224023 | EDRF1-DT | -0,423767254 | 0,745475449 | 0,27192393 | 0,784843944 | EDRF1 divergent transcript |
| ENSG00000187118 | CMC1 | -0,423865232 | 0,745424823 | 0,25385777 | 0,76531122 | C-X9-C motif containing 1 |
| ENSG00000239265 | CLRN1-AS1 | -0,42394086 | 0,745385748 | 0,72642532 | 0,973344786 | CLRN1 antisense RNA 1 |
| ENSG00000225178 | RPSAP58 | -0,4240277 | 0,745340882 | 0,43097011 | 0,89530663 | ribosomal protein SA pseudogene 58 |
| ENSG00000235834 | Z93022,1 | -0,424234275 | 0,745234167 | 0,38682916 | 0,869441674 | novel transcript |
| ENSG00000241168 | AC128685,1 | -0,424304125 | 0,745198087 | 0,13961679 | 0,595782615 | novel transcript |
| ENSG00000229657 | AL391822,1 | -0,424460125 | 0,745117512 | 0,64185074 | 0,95651945 | ribosomal protein L13a (RPL13A) pseudogene |
| ENSG00000168143 | FAM83B | -0,424462703 | 0,745116181 | 0,54882371 | 0,936029622 | family with sequence similarity 83 member B |
| ENSG00000128596 | CCDC136 | -0,424599961 | 0,745045294 | 0,20884355 | 0,713420724 | coiled-coil domain containing 136 |
| ENSG00000254165 | AC090739,1 | -0,424685216 | 0,745001267 | 0,31062211 | 0,817751312 | novel transcript |
| ENSG00000278156 | TSC22D1-AS1 | -0,424692948 | 0,744997274 | 0,3160055 | 0,821394376 | TSC22D1 antisense RNA 1 |
| ENSG00000145416 | MARCHF1 | -0,424755828 | 0,744964804 | 0,40782797 | 0,88080912 | membrane associated ring-CH-type finger 1 |
| ENSG00000248360 | LINC00504 | -0,424808241 | 0,74493774 | 0,06838534 | 0,412055338 | long intergenic non-protein coding RNA 504 |
| ENSG00000204909 | SPINK9 | -0,424885116 | 0,744898047 | 0,31741813 | 0,822252078 | serine peptidase inhibitor Kazal type 9 |
| ENSG00000272275 | AC092687,3 | -0,424923501 | 0,744878228 | 0,63478759 | 0,955149453 | novel transcript |
| ENSG00000281384 | AC093802,4 | -0,425107772 | 0,744783093 | 0,63304074 | 0,954408281 | novel transcript |
| ENSG00000168928 | CTRB2 | -0,425114662 | 0,744779536 | 0,77875855 | 0,978587255 | chymotrypsinogen B2 |
| ENSG00000064933 | PMS1 | -0,425136088 | 0,744768475 | 0,32561643 | 0,827465638 | PMS1 homolog 1, mismatch repair system component |
| ENSG00000272888 | CHASERR | -0,425202483 | 0,7447342 | 0,25317247 | 0,765172364 | CHD2 adjacent suppressive regulatory RNA |
| ENSG00000246465 | AC138904,1 | -0,425219266 | 0,744725537 | 0,3740873 | 0,861369706 | novel transcript LOC100506705 |
| ENSG00000213049 | HNRNPA1P34 | -0,425310174 | 0,744678611 | 0,68489367 | 0,965415049 | heterogeneous nuclear ribonucleoprotein A1 pseudogene 34 |
| ENSG00000238755 | LINC02006 | -0,425321744 | 0,744672639 | 0,64323843 | 0,956633898 | long intergenic non-protein coding RNA 2006 |

| | | | | | | |
|---|---|---|---|---|---|---|
| ENSG00000207344 | SNORA22C | -0,425335756 | 0,744665407 | 0,6886703 | 0,966556275 | small nucleolar RNA, H/ACA box 22C |
| ENSG00000286269 | AC092661,1 | -0,425394235 | 0,744635222 | 0,18129052 | 0,670997703 | novel transcript |
| ENSG00000230946 | HNRNPA1P68 | -0,425495931 | 0,744582735 | 0,38826101 | 0,869820053 | heterogeneous nuclear ribonucleoprotein A1 pseudogene 68 |
| ENSG00000259707 | AC105339,4 | -0,425600809 | 0,744528609 | 0,57369144 | 0,942475308 | serine/arginine-rich splicing factor 9 (SRSF9) pseudogene |
| ENSG00000198105 | ZNF248 | -0,425828422 | 0,744411154 | 0,32143195 | 0,824933727 | zinc finger protein 248 |
| ENSG00000146243 | IRAK1BP1 | -0,426067079 | 0,744288021 | 0,32259019 | 0,824955371 | interleukin 1 receptor associated kinase 1 binding protein 1 |
| ENSG00000228113 | AC003991,1 | -0,426150198 | 0,744245141 | 0,2025777 | 0,702616942 | novel transcript, antisense to STEAP4 |
| ENSG00000286207 | AC012443,2 | -0,426166465 | 0,744236749 | 0,11436903 | 0,54053412 | novel transcript, antisense to FMNL2 |
| ENSG00000265763 | ZNF488 | -0,426239575 | 0,744199035 | 0,31353092 | 0,819801802 | zinc finger protein 488 |
| ENSG00000241738 | ZNF90P1 | -0,426304057 | 0,744165773 | 0,37999593 | 0,865511891 | zinc finger protein 90 pseudogene 1 |
| ENSG00000008735 | MAPK8IP2 | -0,426370957 | 0,744131266 | 0,33153027 | 0,832548086 | mitogen-activated protein kinase 8 interacting protein 2 |
| ENSG00000238795 | SCARNA12 | -0,426419094 | 0,744106438 | 0,15075441 | 0,617373838 | small Cajal body-specific RNA 12 |
| ENSG00000272843 | AC211476,2 | -0,426421941 | 0,744104969 | 0,38115212 | 0,865900094 | novel transcript, antisense to POM121 |
| ENSG00000166263 | STXBP4 | -0,426434003 | 0,744098748 | 0,21291332 | 0,720091093 | syntaxin binding protein 4 |
| ENSG00000187486 | KCNJ11 | -0,426458065 | 0,744086338 | 0,49734572 | 0,918759905 | potassium inwardly rectifying channel subfamily J member 11 |
| ENSG00000250295 | RDH10-AS1 | -0,426521902 | 0,744053414 | 0,52210081 | 0,927599011 | RDH10 antisense RNA 1 |
| ENSG00000233527 | ZNF529-AS1 | -0,426590334 | 0,744018122 | 0,17506903 | 0,660486623 | ZNF529 antisense RNA 1 |
| ENSG00000142065 | ZFP14 | -0,426683661 | 0,743969993 | 0,13291305 | 0,582037931 | ZFP14 zinc finger protein |
| ENSG00000262668 | AJ003147,2 | -0,426694232 | 0,743964542 | 0,41721088 | 0,886651836 | olfactory receptor family 1 subfamily F member 1 |
| ENSG00000281356 | GOLGA8IP | -0,426721113 | 0,74395068 | 0,61923819 | 0,952793934 | golgin A8 family member I, pseudogene |
| ENSG00000218586 | AC006971,1 | -0,426749723 | 0,743935927 | 0,68515156 | 0,965415049 | Rho GTPase activating protein 5 (ARHGAP5) pseudogene |
| ENSG00000237352 | LINC01358 | -0,426766237 | 0,743927412 | 0,20657247 | 0,709779284 | long intergenic non-protein coding RNA 1358 |
| ENSG00000228852 | AC092802,2 | -0,426782487 | 0,743919032 | 0,39337736 | 0,87137239 | novel transcript, antisense to RWDD3 |
| ENSG00000275202 | AL161421,1 | -0,426792644 | 0,743913795 | 0,28177484 | 0,792816227 | novel transcript, antisense to FNDC3A |
| ENSG00000268049 | AC012313,2 | -0,42681834 | 0,743900545 | 0,37614707 | 0,862539384 | novel transcript, antisense to ZNF497 |
| ENSG00000251544 | MTND5P12 | -0,426890665 | 0,743863253 | 0,75467871 | 0,975901808 | MT-ND5 pseudogene 12 |
| ENSG00000215196 | BASP1-AS1 | -0,426958504 | 0,743828276 | 0,38718366 | 0,869441674 | BASP1 antisense RNA 1 |

| | | | | | | |
|---|---|---|---|---|---|---|
| ENSG00000285970 | AC106820,6 | -0,427093131 | 0,743758867 | 0,65550368 | 0,957444211 | novel transcript, antisense to CCNF |
| ENSG00000113269 | RNF130 | -0,427237079 | 0,743684661 | 0,14203975 | 0,600058555 | ring finger protein 130 |
| ENSG00000144619 | CNTN4 | -0,427297146 | 0,743653698 | 0,30036092 | 0,810130728 | contactin 4 |
| ENSG00000215374 | FAM66B | -0,427473602 | 0,743562748 | 0,14704417 | 0,610840954 | family with sequence similarity 66 member B |
| ENSG00000242622 | AC092910,3 | -0,42762541 | 0,74348451 | 0,14409749 | 0,603895972 | novel transcript |
| ENSG00000256422 | LINC02552 | -0,427638875 | 0,743477571 | 0,18451847 | 0,67674498 | long intergenic non-protein coding RNA 2552 |
| ENSG00000271002 | AC129492,5 | -0,427738963 | 0,743425993 | 0,76842598 | 0,977381742 | ribosomal protein L11 (RPL11) pseudogene |
| ENSG00000156345 | CDK20 | -0,427928286 | 0,743328441 | 0,09817127 | 0,497481654 | cyclin dependent kinase 20 |
| ENSG00000096872 | IFT74 | -0,428030702 | 0,743275674 | 0,33228881 | 0,83293962 | intraflagellar transport 74 |
| ENSG00000226245 | ZNF32-AS1 | -0,428033395 | 0,743274287 | 0,6978152 | 0,968522643 | ZNF32 antisense RNA 1 |
| ENSG00000230022 | FNTAP2 | -0,428146455 | 0,743216041 | 0,56437842 | 0,940865625 | farnesyltransferase, CAAX box, alpha pseudogene 2 |
| ENSG00000140527 | WDR93 | -0,428238937 | 0,7431684 | 0,51137411 | 0,922893865 | WD repeat domain 93 |
| ENSG00000272604 | AC073073,2 | -0,428305786 | 0,743133965 | 0,44283848 | 0,899486888 | novel transcript, antisense to EFCAB10 |
| ENSG00000272491 | AL109659,2 | -0,428330092 | 0,743121445 | 0,41674938 | 0,886368152 | novel transcript, antisense to SLC5A9 |
| ENSG00000178597 | PSAPL1 | -0,428424213 | 0,743072965 | 0,65987197 | 0,958837121 | prosaposin like 1 |
| ENSG00000133678 | TMEM254 | -0,428437821 | 0,743065957 | 0,27866394 | 0,789775666 | transmembrane protein 254 |
| ENSG00000273011 | AC092681,3 | -0,428482139 | 0,74304313 | 0,63933686 | 0,956018518 | novel transcript |
| ENSG00000280164 | CU638689,5 | -0,428527407 | 0,743019816 | 0,19903395 | 0,698148483 | novel transcript |
| ENSG00000267430 | AC036176,2 | -0,428549166 | 0,74300861 | 0,53453923 | 0,93162317 | proteasome (prosome, macropain) 26S subunit, ATPase, 5 (PSMC5) pseudogene |
| ENSG00000260271 | AL132996,1 | -0,428714297 | 0,74292357 | 0,68134567 | 0,964654539 | novel transcript |
| ENSG00000260369 | AC120024,1 | -0,428776805 | 0,742891382 | 0,48101089 | 0,912907448 | novel transcript |
| ENSG00000160999 | SH2B2 | -0,428803024 | 0,742877881 | 0,21115175 | 0,716925331 | SH2B adaptor protein 2 |
| ENSG00000167920 | TMEM99 | -0,428816138 | 0,742871129 | 0,24943137 | 0,761381055 | transmembrane protein 99 (putative) |
| ENSG00000259367 | AC087286,2 | -0,428836988 | 0,742860392 | 0,44521479 | 0,90091879 | novel transcript, antisense to AKAP13 |
| ENSG00000238246 | AC069549,1 | -0,428955097 | 0,742799579 | 0,74971935 | 0,975901808 | novel transcript |
| ENSG00000162551 | ALPL | -0,428963418 | 0,742795295 | 0,40784666 | 0,88080912 | alkaline phosphatase, biomineralization associated |
| ENSG00000270177 | AC104109,2 | -0,429003027 | 0,742774902 | 0,36579981 | 0,855695111 | novel transcript |
| ENSG00000092295 | TGM1 | -0,42915395 | 0,742697203 | 0,48210697 | 0,912981134 | transglutaminase 1 |
| ENSG00000285348 | TGM1 | -0,42915395 | 0,742697203 | 0,48210697 | 0,912981134 | transglutaminase 1 |
| ENSG00000232454 | AL138752,1 | -0,429280945 | 0,742631829 | 0,55936445 | 0,939727587 | mitochondrial ribosomal protein S10 (MRPS10) pseudogene |
| ENSG00000274292 | AC084018,2 | -0,429346452 | 0,74259811 | 0,22992798 | 0,738414837 | novel transcript |

| | | | | | | |
|---|---|---|---|---|---|---|
| ENSG00000154016 | GRAP | -0,429368511 | 0,742586755 | 0,3664392 | 0,856103393 | GRB2 related adaptor protein |
| ENSG00000009724 | MASP2 | -0,429368792 | 0,742586611 | 0,24434418 | 0,755755471 | mannan binding lectin serine peptidase 2 |
| ENSG00000105143 | SLC1A6 | -0,429404828 | 0,742568063 | 0,27382351 | 0,785900879 | solute carrier family 1 member 6 |
| ENSG00000240720 | LRRD1 | -0,429451196 | 0,742544197 | 0,41401538 | 0,885528954 | leucine rich repeats and death domain containing 1 |
| ENSG00000231257 | ATAT1 | -0,429493816 | 0,742522261 | 0,13375665 | 0,583882708 | alpha tubulin acetyltransferase 1 |
| ENSG00000235658 | ATAT1 | -0,429493816 | 0,742522261 | 0,13375665 | 0,583882708 | alpha tubulin acetyltransferase 1 |
| ENSG00000259556 | AC090971,3 | -0,429513437 | 0,742512163 | 0,62666404 | 0,952793934 | Leo1, Paf1/RNA polymerase II complex component, homolog (S, cerevisiae) (LEO1) pseudogene |
| ENSG00000286523 | AL022151,1 | -0,429583388 | 0,742476162 | 0,06537448 | 0,401164726 | novel transcript |
| ENSG00000263324 | HCN3 | -0,429619461 | 0,742457597 | 0,17146474 | 0,65533907 | hyperpolarization activated cyclic nucleotide gated potassium channel 3 |
| ENSG00000265702 | AC005821,1 | -0,429725345 | 0,742403108 | 0,51896377 | 0,926800715 | novel transcript, antisense to MARCH10 |
| ENSG00000105516 | DBP | -0,429747073 | 0,742391927 | 0,16237605 | 0,639602491 | D-box binding PAR bZIP transcription factor |
| ENSG00000273604 | EPOP | -0,429916919 | 0,742304531 | 0,38399246 | 0,868113415 | elongin BC and polycomb repressive complex 2 associated protein |
| ENSG00000250474 | WBP1LP2 | -0,429941601 | 0,742291832 | 0,72578324 | 0,973344786 | WBP1L pseudogene 2 |
| ENSG00000242154 | AC004884,2 | -0,430233739 | 0,742141537 | 0,66473799 | 0,960501386 | novel transcript |
| ENSG00000135519 | KCNH3 | -0,430251508 | 0,742132396 | 0,5501256 | 0,93612918 | potassium voltage-gated channel subfamily H member 3 |
| ENSG00000267259 | ERVE-1 | -0,43035366 | 0,742079851 | 0,35366405 | 0,845346751 | endogenous retrovirus group E member 1 |
| ENSG00000287286 | AC024560,5 | -0,430369302 | 0,742071805 | 0,67748338 | 0,9635902 | novel transcript, antisense to RUBCN |
| ENSG00000173567 | ADGRF3 | -0,430375634 | 0,742068548 | 0,42043585 | 0,888563911 | adhesion G protein-coupled receptor F3 |
| ENSG00000237054 | PRMT5-AS1 | -0,430450359 | 0,742030113 | 0,30463776 | 0,813276091 | PRMT5 antisense RNA 1 |
| ENSG00000229021 | AL450992,1 | -0,430451526 | 0,742029513 | 0,30996928 | 0,817411297 | novel transcript |
| ENSG00000211451 | GNRHR2 | -0,430526201 | 0,741991106 | 0,12703179 | 0,569312214 | gonadotropin releasing hormone receptor 2 (pseudogene) |
| ENSG00000187123 | LYPD6 | -0,430526337 | 0,741991036 | 0,19932213 | 0,698610648 | LY6/PLAUR domain containing 6 |
| ENSG00000171016 | PYGO1 | -0,430541878 | 0,741983043 | 0,34691572 | 0,841625668 | pygopus family PHD finger 1 |
| ENSG00000217495 | AL136116,2 | -0,430696294 | 0,741903631 | 0,72698717 | 0,973344786 | actin protein pseudogene |
| ENSG00000157741 | UBN2 | -0,43078182 | 0,74185965 | 0,23613442 | 0,745253314 | ubinuclein 2 |
| ENSG00000156298 | TSPAN7 | -0,430996142 | 0,74174945 | 0,42613345 | 0,892133432 | tetraspanin 7 |
| ENSG00000241318 | WDR82P2 | -0,431010813 | 0,741741908 | 0,70005393 | 0,968522643 | WD repeat domain 82 pseudogene 2 |
| ENSG00000285608 | AL161665,1 | -0,431101011 | 0,741695535 | 0,74474675 | 0,975649186 | novel transcript, antisense to ARHGAP5 |

| | | | | | | |
|---|---|---|---|---|---|---|
| ENSG00000268995 | VN1R82P | -0,431130883 | 0,741680178 | 0,74965117 | 0,975901808 | vomeronasal 1 receptor 82 pseudogene |
| ENSG00000262115 | AC027601,2 | -0,431173507 | 0,741658266 | 0,73534047 | 0,974353474 | novel transcript, antisense to AZI1 |
| ENSG00000286762 | AL096870,9 | -0,431251105 | 0,741618375 | 0,63065941 | 0,953375088 | novel transcript, antisense to RABGGTA |
| ENSG00000286825 | AL096870,10 | -0,431251105 | 0,741618375 | 0,63065941 | 0,953375088 | novel transcript, antisense to RABGGTA |
| ENSG00000286649 | AC006566,2 | -0,431362116 | 0,741561312 | 0,56214701 | 0,940865625 | novel transcript, antisense to PTPRM |
| ENSG00000237775 | DDR1-DT | -0,43138759 | 0,741548218 | 0,80002311 | 0,980161849 | DDR1 divergent transcript |
| ENSG00000256443 | AP003559,1 | -0,431485546 | 0,74149787 | 0,67178448 | 0,962161102 | novel transcript |
| ENSG00000281763 | DHX36 | -0,431568586 | 0,741455191 | 0,42658588 | 0,892152052 | DEAH-box helicase 36 |
| ENSG00000236256 | DIAPH2-AS1 | -0,431637694 | 0,741419676 | 0,61089223 | 0,951873283 | DIAPH2 antisense RNA 1 |
| ENSG00000227486 | AL050309,1 | -0,431648402 | 0,741414173 | 0,09943 | 0,50040065 | novel transcript |
| ENSG00000254274 | TDGF1P5 | -0,431663845 | 0,741406236 | 0,65064959 | 0,956899963 | teratocarcinoma-derived growth factor 1 pseudogene 5 |
| ENSG00000243679 | AC018638,5 | -0,431911051 | 0,741279207 | 0,33070061 | 0,832114797 | novel pseudogene |
| ENSG00000227799 | CDPF1P1 | -0,432037488 | 0,741214244 | 0,33430541 | 0,83453565 | CDPF1 pseudogene 1 |
| ENSG00000107443 | CCNJ | -0,432038 | 0,741213982 | 0,39075144 | 0,870057277 | cyclin J |
| ENSG00000154153 | RETREG1 | -0,432180477 | 0,741140784 | 0,25182081 | 0,764065015 | reticulophagy regulator 1 |
| ENSG00000183784 | DOCK8-AS1 | -0,432282196 | 0,741088531 | 0,62846768 | 0,952793934 | DOCK8 antisense RNA 1 |
| ENSG00000254285 | KRT8P3 | -0,432285446 | 0,741086862 | 0,73841883 | 0,974353474 | keratin 8 pseudogene 3 |
| ENSG00000130540 | SULT4A1 | -0,432367267 | 0,741044833 | 0,17485685 | 0,660232445 | sulfotransferase family 4A member 1 |
| ENSG00000123975 | CKS2 | -0,432375553 | 0,741040577 | 0,38480084 | 0,868536368 | CDC28 protein kinase regulatory subunit 2 |
| ENSG00000288140 | AC006144,3 | -0,432420596 | 0,741017441 | 0,6711225 | 0,962026636 | novel transcript |
| ENSG00000287379 | AC006144,2 | -0,432420596 | 0,741017441 | 0,6711225 | 0,962026636 | novel transcript |
| ENSG00000224687 | RASAL2-AS1 | -0,432474068 | 0,740989976 | 0,12734771 | 0,56991107 | RASAL2 antisense RNA 1 |
| ENSG00000286483 | AL121762,1 | -0,432477289 | 0,740988322 | 0,54308191 | 0,93383087 | novel transcript |
| ENSG00000270392 | PFN1P2 | -0,432605424 | 0,740922513 | 0,2964566 | 0,806142623 | profilin 1 pseudogene 2 |
| ENSG00000109576 | AADAT | -0,432676994 | 0,740885758 | 0,25207417 | 0,764267906 | aminoadipate aminotransferase |
| ENSG00000285077 | ARHGAP11B | -0,432773612 | 0,740836142 | 0,20010069 | 0,699646707 | Rho GTPase activating protein 11B |
| ENSG00000106066 | CPVL | -0,432937552 | 0,740751962 | 0,32808054 | 0,830076239 | carboxypeptidase vitellogenic like |
| ENSG00000106605 | BLVRA | -0,433058273 | 0,740689981 | 0,15521723 | 0,625027256 | biliverdin reductase A |
| ENSG00000140398 | NEIL1 | -0,43308957 | 0,740673913 | 0,28704803 | 0,798434649 | nei like DNA glycosylase 1 |
| ENSG00000235280 | MCF2L-AS1 | -0,433100789 | 0,740668153 | 0,5716102 | 0,942436428 | MCF2L antisense RNA 1 |
| ENSG00000123496 | IL13RA2 | -0,433129399 | 0,740653465 | 0,3992691 | 0,875433675 | interleukin 13 receptor subunit alpha 2 |
| ENSG00000172238 | ATOH1 | -0,433294211 | 0,740568858 | 0,79422509 | 0,979776981 | atonal bHLH transcription factor 1 |

| | | | | | | |
|---|---|---|---|---|---|---|
| ENSG00000143195 | ILDR2 | -0,433317269 | 0,740557022 | 0,15304631 | 0,621569237 | immunoglobulin like domain containing receptor 2 |
| ENSG00000166262 | FAM227B | -0,433322969 | 0,740554096 | 0,2501337 | 0,762275082 | family with sequence similarity 227 member B |
| ENSG00000180730 | SHISA2 | -0,433327409 | 0,740551817 | 0,37079506 | 0,858780045 | shisa family member 2 |
| ENSG00000224509 | MRPS9-AS2 | -0,433414272 | 0,740507231 | 0,74102904 | 0,975146552 | MRPS9 antisense RNA 2 |
| ENSG00000229419 | RALGAPA1P1 | -0,433473855 | 0,740476648 | 0,29023913 | 0,800682868 | RALGAPA1 pseudogene 1 |
| ENSG00000231804 | AL353748,1 | -0,433587658 | 0,740418241 | 0,69299891 | 0,967271667 | ribosomal protein L21 (RPL21) pseudogene |
| ENSG00000234585 | CCT6P3 | -0,433637524 | 0,740392649 | 0,16787109 | 0,649062781 | chaperonin containing TCP1 subunit 6 pseudogene 3 |
| ENSG00000227492 | AL138921,1 | -0,433654029 | 0,740384178 | 0,6516798 | 0,957089277 | novel transcript |
| ENSG00000247853 | AC006064,2 | -0,433680631 | 0,740370526 | 0,23656064 | 0,745923541 | novel transcript, antisense to CHD4 |
| ENSG00000271793 | AL589666,1 | -0,433777882 | 0,74032062 | 0,80227726 | 0,980710009 | novel protein, SYNCRIP-SNX14 readthrough |
| ENSG00000250486 | FAM218A | -0,433933458 | 0,740240791 | 0,38160009 | 0,866023897 | family with sequence similarity 218 member A |
| ENSG00000237943 | PRKCQ-AS1 | -0,433982451 | 0,740215653 | 0,32034582 | 0,824366234 | PRKCQ antisense RNA 1 |
| ENSG00000118420 | UBE3D | -0,434092546 | 0,740159168 | 0,17544051 | 0,661199837 | ubiquitin protein ligase E3D |
| ENSG00000254985 | RSF1-IT2 | -0,434212056 | 0,740097857 | 0,44444033 | 0,900210555 | RSF1 intronic transcript 2 |
| ENSG00000281392 | LINC00506 | -0,434398937 | 0,740001994 | 0,07745039 | 0,439776428 | long intergenic non-protein coding RNA 506 |
| ENSG00000226026 | AC092802,1 | -0,434774226 | 0,739809522 | 0,1149107 | 0,541330721 | novel transcript |
| ENSG00000283674 | AC068587,4 | -0,434825187 | 0,739783389 | 0,11073275 | 0,531381215 | novel transcript |
| ENSG00000115392 | FANCL | -0,434889044 | 0,739750646 | 0,33743919 | 0,836578328 | FA complementation group L |
| ENSG00000265683 | SYPL1P2 | -0,434963851 | 0,739712289 | 0,4848623 | 0,913690171 | synaptophysin like 1 pseudogene 2 |
| ENSG00000269821 | KCNQ1OT1 | -0,434995059 | 0,739696288 | 0,06272363 | 0,391337366 | KCNQ1 opposite strand/antisense transcript 1 |
| ENSG00000160570 | DEDD2 | -0,435117648 | 0,739633437 | 0,17478941 | 0,660057513 | death effector domain containing 2 |
| ENSG00000163694 | RBM47 | -0,435131466 | 0,739626353 | 0,48293159 | 0,91363215 | RNA binding motif protein 47 |
| ENSG00000168116 | KIAA1586 | -0,435156183 | 0,739613681 | 0,46533777 | 0,90786072 | KIAA1586 |
| ENSG00000072041 | SLC6A15 | -0,435286722 | 0,739546762 | 0,32642956 | 0,828450693 | solute carrier family 6 member 15 |
| ENSG00000224363 | FP700111,1 | -0,435334165 | 0,739522443 | 0,1915462 | 0,687341458 | novel transcript |
| ENSG00000179299 | NSUN7 | -0,435335164 | 0,73952193 | 0,25982835 | 0,772394558 | NOP2/Sun RNA methyltransferase family member 7 |
| ENSG00000159217 | IGF2BP1 | -0,435342598 | 0,73951812 | 0,20507139 | 0,706634751 | insulin like growth factor 2 mRNA binding protein 1 |
| ENSG00000270996 | AC005034,4 | -0,435362748 | 0,739507791 | 0,59354127 | 0,947469769 | novel transcript |
| ENSG00000241058 | NSUN6 | -0,435433827 | 0,739471358 | 0,27331283 | 0,785672647 | NOP2/Sun RNA methyltransferase 6 |
| ENSG00000236526 | AL035448,1 | -0,435442446 | 0,73946694 | 0,26462176 | 0,775800337 | novel transcript |
| ENSG00000286746 | AL590807,1 | -0,435580326 | 0,739396271 | 0,05910534 | 0,377578532 | novel transcript |
| ENSG00000287357 | AL512484,1 | -0,435602146 | 0,739385089 | 0,33480872 | 0,834792227 | novel transcript |
| ENSG00000196220 | SRGAP3 | -0,435607974 | 0,739382102 | 0,05762786 | 0,3710973 | SLIT-ROBO Rho GTPase activating protein 3 |

| | | | | | | |
|---|---|---|---|---|---|---|
| ENSG00000162817 | C1orf115 | -0,43565244 | 0,739359314 | 0,31976364 | 0,824366234 | chromosome 1 open reading frame 115 |
| ENSG00000101846 | STS | -0,435700872 | 0,739334493 | 0,1760157 | 0,661519708 | steroid sulfatase |
| ENSG00000266171 | AP001020,3 | -0,435780342 | 0,739293768 | 0,28873405 | 0,800014136 | novel transcript, antisense to YES1 |
| ENSG00000167969 | ECI1 | -0,435878373 | 0,739243535 | 0,13783839 | 0,592546669 | enoyl-CoA delta isomerase 1 |
| ENSG00000283549 | AC007161,3 | -0,435888783 | 0,739238201 | 0,57678377 | 0,943261034 | novel transcript |
| ENSG00000271141 | AC010680,4 | -0,435911423 | 0,7392266 | 0,48622046 | 0,914091794 | novel transcript |
| ENSG00000115339 | GALNT3 | -0,436062648 | 0,739149118 | 0,37054734 | 0,858545694 | polypeptide N-acetylgalactosaminyltransferase 3 |
| ENSG00000163808 | KIF15 | -0,436126197 | 0,73911656 | 0,19604125 | 0,694033976 | kinesin family member 15 |
| ENSG00000280610 | KIF15 | -0,436126197 | 0,73911656 | 0,19604125 | 0,694033976 | kinesin family member 15 |
| ENSG00000234684 | SDCBP2-AS1 | -0,436208328 | 0,739074484 | 0,16548508 | 0,644620672 | SDCBP2 antisense RNA 1 |
| ENSG00000287112 | AC068787,3 | -0,436222229 | 0,739067363 | 0,54917228 | 0,936040289 | novel transcript |
| ENSG00000069966 | GNB5 | -0,436303972 | 0,739025488 | 0,05389081 | 0,357953158 | G protein subunit beta 5 |
| ENSG00000286291 | AC004047,1 | -0,43636464 | 0,738994412 | 0,60964581 | 0,951793902 | novel transcript |
| ENSG00000237484 | LINC01684 | -0,436419601 | 0,73896626 | 0,09784061 | 0,496997855 | long intergenic non-protein coding RNA 1684 |
| ENSG00000259635 | AC100830,1 | -0,436452496 | 0,73894941 | 0,61801476 | 0,952793934 | novel transcript, antisense OAZ2 |
| ENSG00000276662 | AL713922,2 | -0,436482736 | 0,738933922 | 0,44414824 | 0,900210555 | DEAD (Asp-Glu-Ala-Asp) box polypeptide 20 (DDX20) pseudogene |
| ENSG00000229598 | PRDX3P1 | -0,436509453 | 0,738920237 | 0,42583627 | 0,892133432 | peroxiredoxin 3 pseudogene 1 |
| ENSG00000215126 | CBWD6 | -0,436641865 | 0,738852422 | 0,24882298 | 0,760802113 | COBW domain containing 6 |
| ENSG00000259442 | AC105339,3 | -0,4368849 | 0,738727966 | 0,84430805 | 0,986340494 | novel transcript, antisense FSD2 |
| ENSG00000274602 | PI4KAP1 | -0,436955343 | 0,738691897 | 0,11003232 | 0,528913064 | phosphatidylinositol 4-kinase alpha pseudogene 1 |
| ENSG00000246763 | RGMB-AS1 | -0,436996246 | 0,738670954 | 0,16022015 | 0,635433547 | RGMB antisense RNA 1 |
| ENSG00000230612 | AC004039,1 | -0,437048923 | 0,738643984 | 0,17033657 | 0,653594444 | novel transcript, antisense to KIF3A |
| ENSG00000218283 | MORF4L1P1 | -0,437094052 | 0,738620878 | 0,37527498 | 0,862022464 | mortality factor 4 like 1 pseudogene 1 |
| ENSG00000001460 | STPG1 | -0,437139539 | 0,738597591 | 0,07651502 | 0,437005066 | sperm tail PG-rich repeat containing 1 |
| ENSG00000248508 | SRP14-AS1 | -0,437284002 | 0,738523635 | 0,25565669 | 0,767141836 | SRP14 antisense RNA1 (head to head) |
| ENSG00000112290 | WASF1 | -0,437348579 | 0,738490579 | 0,31606798 | 0,821394376 | WASP family member 1 |
| ENSG00000286855 | AC074024,1 | -0,437611671 | 0,738355919 | 0,65916214 | 0,958379996 | novel transcript |
| ENSG00000286074 | AC110795,1 | -0,437649724 | 0,738336444 | 0,65903598 | 0,958285698 | novel transcript |
| ENSG00000237481 | AL117350,1 | -0,437719278 | 0,738300849 | 0,68838776 | 0,966481084 | novel transcript |
| ENSG00000163040 | CCDC74A | -0,437720298 | 0,738300327 | 0,12690051 | 0,568970141 | coiled-coil domain containing 74A |
| ENSG00000260989 | AL133297,2 | -0,437926408 | 0,738194857 | 0,66254823 | 0,959897678 | novel transcript, antisense to IFT140 |
| ENSG00000206535 | LNP1 | -0,437949918 | 0,738182828 | 0,23573434 | 0,744895724 | leukemia NUP98 fusion partner 1 |

| | | | | | | |
|---|---|---|---|---|---|---|
| ENSG00000250536 | ABHD17AP3 | -0,4379914 | 0,738161603 | 0,585914 | 0,945246954 | ABHD17A pseudogene 3 |
| ENSG00000132424 | PNISR | -0,438235476 | 0,738036731 | 0,26737181 | 0,779594822 | PNN interacting serine and arginine rich protein |
| ENSG00000230084 | AC006059,1 | -0,438237496 | 0,738035698 | 0,41906348 | 0,887527661 | novel transcript |
| ENSG00000276952 | AL121772,3 | -0,438343776 | 0,73798133 | 0,48362097 | 0,913690171 | novel transcript, antisense to PYGB |
| ENSG00000077063 | CTTNBP2 | -0,438355925 | 0,737975116 | 0,25172636 | 0,763986755 | cortactin binding protein 2 |
| ENSG00000229534 | HNRNPA1P53 | -0,438430011 | 0,73793722 | 0,63392955 | 0,954929949 | heterogeneous nuclear ribonucleoprotein A1 pseudogene 53 |
| ENSG00000281263 | AC136352,8 | -0,438465969 | 0,737918828 | 0,58818308 | 0,946176848 | novel transcript |
| ENSG00000007174 | DNAH9 | -0,438470973 | 0,737916268 | 0,09993804 | 0,501634237 | dynein axonemal heavy chain 9 |
| ENSG00000228775 | WEE2-AS1 | -0,438482561 | 0,737910341 | 0,35235823 | 0,844912682 | WEE2 antisense RNA 1 |
| ENSG00000229920 | RPS4XP5 | -0,438509708 | 0,737896456 | 0,63951976 | 0,956018518 | ribosomal protein S4X pseudogene 5 |
| ENSG00000222018 | FAM243A | -0,438628439 | 0,737835731 | 0,54781681 | 0,935404974 | family with sequence similarity 243 member A |
| ENSG00000129204 | USP6 | -0,438640508 | 0,737829559 | 0,22837047 | 0,736649232 | ubiquitin specific peptidase 6 |
| ENSG00000228889 | UBAC2-AS1 | -0,438702987 | 0,737797606 | 0,16669502 | 0,646915725 | UBAC2 antisense RNA 1 |
| ENSG00000125772 | GPCPD1 | -0,438712465 | 0,737792759 | 0,24517039 | 0,757187317 | glycerophosphocholine phosphodiesterase 1 |
| ENSG00000267522 | LINC01864 | -0,438774106 | 0,737761237 | 0,63980555 | 0,956018518 | long intergenic non-protein coding RNA 1864 |
| ENSG00000227543 | SPAG5-AS1 | -0,438777268 | 0,73775962 | 0,24484837 | 0,756790838 | SPAG5 antisense RNA 1 |
| ENSG00000260903 | XKR7 | -0,438840653 | 0,737727206 | 0,29813542 | 0,807554329 | XK related 7 |
| ENSG00000125631 | HTR5BP | -0,438886493 | 0,737703767 | 0,68700802 | 0,965769168 | 5-hydroxytryptamine receptor 5B, pseudogene |
| ENSG00000140451 | PIF1 | -0,438920345 | 0,737686457 | 0,23772429 | 0,746907875 | PIF1 5'-to-3' DNA helicase |
| ENSG00000263089 | AC007114,2 | -0,438952586 | 0,737669972 | 0,29945776 | 0,809252953 | novel transcript, antisense to AKAP1 |
| ENSG00000269843 | AC008537,2 | -0,438958972 | 0,737666706 | 0,72626882 | 0,973344786 | novel transcript |
| ENSG00000228919 | AC097381,1 | -0,439013938 | 0,737638602 | 0,65143684 | 0,957089277 | uncharacterized LOC389199 |
| ENSG00000144061 | NPHP1 | -0,439129147 | 0,737579699 | 0,11190157 | 0,534735697 | nephrocystin 1 |
| ENSG00000048740 | CELF2 | -0,439182069 | 0,737552643 | 0,22186719 | 0,729383401 | CUGBP Elav-like family member 2 |
| ENSG00000271161 | BOLA2P2 | -0,439186955 | 0,737550145 | 0,35108664 | 0,844035671 | bolA family member 2 pseudogene 2 |
| ENSG00000248015 | AC005329,1 | -0,439195899 | 0,737545573 | 0,39573861 | 0,872227623 | novel transcript, antisense to NDUFS7 |
| ENSG00000170458 | CD14 | -0,439276836 | 0,737504196 | 0,6952968 | 0,967599788 | CD14 molecule |
| ENSG00000234219 | CDCA4P4 | -0,439442184 | 0,737419676 | 0,39496706 | 0,871888817 | cell division cycle associated 4 pseudogene 4 |
| ENSG00000286957 | AC079776,5 | -0,439482062 | 0,737399293 | 0,66465964 | 0,96046247 | novel transcript |
| ENSG00000178177 | LCORL | -0,439487477 | 0,737396525 | 0,35895167 | 0,849814998 | ligand dependent nuclear receptor corepressor like |
| ENSG00000185658 | BRWD1 | -0,439601925 | 0,73733803 | 0,27938676 | 0,790260344 | bromodomain and WD repeat domain containing 1 |
| ENSG00000137098 | SPAG8 | -0,439744509 | 0,737265161 | 0,31420839 | 0,819877431 | sperm associated antigen 8 |

| | | | | | | |
|---|---|---|---|---|---|---|
| ENSG00000138771 | SHROOM3 | -0,439755684 | 0,73725945 | 0,11223956 | 0,535189241 | shroom family member 3 |
| ENSG00000254757 | AC127526,1 | -0,439838272 | 0,737217247 | 0,52771645 | 0,929546857 | novel transcript |
| ENSG00000285953 | AC000120,3 | -0,44005509 | 0,737106461 | 0,57054647 | 0,9423888 | novel protein |
| ENSG00000274827 | LINC01297 | -0,440074095 | 0,737096752 | 0,59240951 | 0,947416795 | long intergenic non-protein coding RNA 1297 |
| ENSG00000176273 | SLC35G1 | -0,440089589 | 0,737088835 | 0,35153335 | 0,84405766 | solute carrier family 35 member G1 |
| ENSG00000154485 | MMP21 | -0,440253796 | 0,737004945 | 0,30270253 | 0,812334076 | matrix metallopeptidase 21 |
| ENSG00000158517 | NCF1 | -0,440362299 | 0,736949518 | 0,76147586 | 0,976285211 | neutrophil cytosolic factor 1 |
| ENSG00000006128 | TAC1 | -0,440389223 | 0,736935765 | 0,4947029 | 0,91758518 | tachykinin precursor 1 |
| ENSG00000197168 | NEK5 | -0,440650876 | 0,736802123 | 0,11098071 | 0,531999544 | NIMA related kinase 5 |
| ENSG00000227861 | SAPCD1 | -0,44067443 | 0,736790094 | 0,53378569 | 0,931381467 | suppressor APC domain containing 1 |
| ENSG00000234951 | SAPCD1 | -0,44067443 | 0,736790094 | 0,53378569 | 0,931381467 | suppressor APC domain containing 1 |
| ENSG00000227074 | SAPCD1 | -0,44067443 | 0,736790094 | 0,53378569 | 0,931381467 | suppressor APC domain containing 1 |
| ENSG00000135541 | AHI1 | -0,440706392 | 0,736773771 | 0,23628741 | 0,745362681 | Abelson helper integration site 1 |
| ENSG00000173451 | THAP2 | -0,440846989 | 0,736701973 | 0,43183024 | 0,895555998 | THAP domain containing 2 |
| ENSG00000198040 | ZNF84 | -0,440965896 | 0,736641256 | 0,29507937 | 0,805254732 | zinc finger protein 84 |
| ENSG00000270751 | FBXW7-AS1 | -0,441083272 | 0,736581326 | 0,74116849 | 0,975176043 | FBXW7 antisense RNA 1 |
| ENSG00000262461 | SPDYE9 | -0,441213867 | 0,736514653 | 0,29318498 | 0,803365244 | speedy/RINGO cell cycle regulator family member E9 |
| ENSG00000163596 | ICA1L | -0,441234725 | 0,736504005 | 0,08662532 | 0,468880561 | islet cell autoantigen 1 like |
| ENSG00000258804 | LINC01148 | -0,441312964 | 0,736464064 | 0,68439089 | 0,965415049 | long intergenic non-protein coding RNA 1148 |
| ENSG00000188227 | ZNF793 | -0,441335309 | 0,736452658 | 0,13717605 | 0,591024932 | zinc finger protein 793 |
| ENSG00000132182 | NUP210 | -0,441339602 | 0,736450466 | 0,07628138 | 0,43651514 | nucleoporin 210 |
| ENSG00000188869 | TMC3 | -0,441340912 | 0,736449798 | 0,41075678 | 0,882835102 | transmembrane channel like 3 |
| ENSG00000178226 | PRSS36 | -0,441389869 | 0,736424807 | 0,55772786 | 0,938875925 | serine protease 36 |
| ENSG00000235795 | AC093157,2 | -0,441409906 | 0,736414579 | 0,63616093 | 0,955399772 | novel transcript |
| ENSG00000267345 | AC010632,1 | -0,441616066 | 0,736309354 | 0,73802852 | 0,974353474 | novel transcript, antisense to ZNF420 |
| ENSG00000286942 | AC017104,5 | -0,441730934 | 0,736250731 | 0,75173013 | 0,975901808 | novel transcript |
| ENSG00000288246 | AC140725,6 | -0,441755592 | 0,736238147 | 0,57244134 | 0,942475308 | novel transcript |
| ENSG00000259553 | AC140725,1 | -0,441755592 | 0,736238147 | 0,57244134 | 0,942475308 | novel transcript |
| ENSG00000272752 | STAG3L5P-PVR | -0,441764867 | 0,736233414 | 0,19574354 | 0,69375328 | STAG3L5P-PVRIG2P-PILRB readthrough |
| ENSG00000272343 | AC107952,2 | -0,441846519 | 0,736191747 | 0,55490337 | 0,937793851 | novel transcript |
| ENSG00000278458 | KANSL1 | -0,441862607 | 0,736183537 | 0,38310833 | 0,867698804 | KAT8 regulatory NSL complex subunit 1 |

| | | | | | | |
|---|---|---|---|---|---|---|
| ENSG00000265798 | AC138207,6 | -0,441919948 | 0,736154278 | 0,4846187 | 0,913690171 | leucine rich repeat containing 37B (LRRC37B) pseudogene |
| ENSG00000269106 | AC012313,7 | -0,441960191 | 0,736133743 | 0,64675424 | 0,956899963 | novel transcript, antisense to ZNF446 |
| ENSG00000204962 | PCDHA8 | -0,441973903 | 0,736126747 | 0,13067382 | 0,577136255 | protocadherin alpha 8 |
| ENSG00000256196 | AP003721,1 | -0,441994728 | 0,736116121 | 0,329559 | 0,831379626 | novel transcript |
| ENSG00000287322 | AC068189,2 | -0,442052811 | 0,736086486 | 0,45566935 | 0,903545728 | novel transcript, antisense to NDUFAF6 |
| ENSG00000275527 | AC100835,2 | -0,442070208 | 0,736077609 | 0,83457831 | 0,984905093 | novel transcript |
| ENSG00000223692 | DIP2A-IT1 | -0,442078522 | 0,736073367 | 0,69512611 | 0,967519493 | DIP2A intronic transcript 1 |
| ENSG00000076555 | ACACB | -0,442109388 | 0,736057619 | 0,07075865 | 0,418829978 | acetyl-CoA carboxylase beta |
| ENSG00000219163 | HMGB1P20 | -0,442141352 | 0,736041312 | 0,73949931 | 0,974715504 | high mobility group box 1 pseudogene 20 |
| ENSG00000224881 | AC068279,1 | -0,44223176 | 0,735995188 | 0,52333618 | 0,927995032 | novel pseudogene |
| ENSG00000233339 | AC018878,1 | -0,442293579 | 0,735963652 | 0,57802349 | 0,943425812 | novel transcript |
| ENSG00000196366 | C9orf163 | -0,442321572 | 0,735949372 | 0,35014898 | 0,843789477 | chromosome 9 open reading frame 163 |
| ENSG00000250345 | AC104090,1 | -0,442380581 | 0,735919271 | 0,8117453 | 0,981213877 | ethanolamine kinase 1 (ETNK1) pseudogene |
| ENSG00000260876 | LINC01229 | -0,442453798 | 0,735881924 | 0,44230154 | 0,899331131 | long intergenic non-protein coding RNA 1229 |
| ENSG00000231307 | RPS3P2 | -0,442466762 | 0,735875311 | 0,49267151 | 0,917090312 | ribosomal protein S3 pseudogene 2 |
| ENSG00000139351 | SYCP3 | -0,442527591 | 0,735844285 | 0,29241238 | 0,802773151 | synaptonemal complex protein 3 |
| ENSG00000286328 | AC079052,1 | -0,442590952 | 0,735811969 | 0,55233708 | 0,936660142 | novel transcript |
| ENSG00000255339 | AL133352,1 | -0,442673565 | 0,735769835 | 0,27452026 | 0,786383635 | NADH dehydrogenase (ubiquinone) 1 beta subcomplex, 8, 19kDa (NDUFB8) and SEC31 homolog B (S, cerevisiae) (SEC31B) readthrough |
| ENSG00000261296 | AC126323,6 | -0,442766808 | 0,735722283 | 0,55694133 | 0,938264532 | novel transcript |
| ENSG00000197959 | DNM3 | -0,442875042 | 0,73566709 | 0,28921262 | 0,800260154 | dynamin 3 |
| ENSG00000240522 | RPL7AP10 | -0,442977987 | 0,735614597 | 0,27642582 | 0,788263115 | ribosomal protein L7a pseudogene 10 |
| ENSG00000223039 | RN7SKP268 | -0,443271925 | 0,735464737 | 0,73548431 | 0,974353474 | RN7SK pseudogene 268 |
| ENSG00000174501 | ANKRD36C | -0,443401724 | 0,73539857 | 0,15316597 | 0,621651423 | ankyrin repeat domain 36C |
| ENSG00000266801 | AC009137,2 | -0,443423651 | 0,735387393 | 0,64555396 | 0,956899963 | novel transcript, sense intronic to transmembrane and coiled-coil domains 7 TMCO7 |
| ENSG00000260269 | AC105036,3 | -0,443464163 | 0,735366743 | 0,52337566 | 0,927995032 | novel transcript |
| ENSG00000267040 | AC027097,1 | -0,443673633 | 0,73525998 | 0,17672456 | 0,662651108 | novel transcript, antisense to ATP8B1 |
| ENSG00000036828 | CASR | -0,44369501 | 0,735249086 | 0,07452694 | 0,431519399 | calcium sensing receptor |
| ENSG00000247199 | AC011346,1 | -0,443713945 | 0,735239436 | 0,05648335 | 0,366903759 | novel transcript, antisense to SPINK9, SPINK7 and SPINK13 |

| | | | | | | |
|---|---|---|---|---|---|---|
| ENSG00000255270 | NAV2-IT1 | -0,443809546 | 0,735190716 | 0,71562884 | 0,971237203 | NAV2 intronic transcript 1 |
| ENSG00000189269 | DRICH1 | -0,44383847 | 0,735175977 | 0,65162358 | 0,957089277 | aspartate rich 1 |
| ENSG00000286927 | AC103955,1 | -0,44393966 | 0,735124414 | 0,09436578 | 0,4879788 | novel transcript |
| ENSG00000205084 | TMEM231 | -0,443947475 | 0,735120432 | 0,07649521 | 0,436971724 | transmembrane protein 231 |
| ENSG00000213186 | TRIM59 | -0,444182804 | 0,73500053 | 0,32359678 | 0,825794844 | tripartite motif containing 59 |
| ENSG00000168658 | VWA3B | -0,444205692 | 0,73498887 | 0,41315238 | 0,884663685 | von Willebrand factor A domain containing 3B |
| ENSG00000250365 | AL139353,2 | -0,444341187 | 0,734919844 | 0,31487046 | 0,820069145 | novel transcript, antisense to C14orf126 |
| ENSG00000248930 | CHP1P1 | -0,444388256 | 0,734895867 | 0,72284453 | 0,972707638 | CHP1 pseudogene 1 |
| ENSG00000166257 | SCN3B | -0,444409816 | 0,734884885 | 0,30433559 | 0,813156808 | sodium voltage-gated channel beta subunit 3 |
| ENSG00000145332 | KLHL8 | -0,444430495 | 0,734874352 | 0,28899241 | 0,800166929 | kelch like family member 8 |
| ENSG00000286529 | AC090398,2 | -0,444436101 | 0,734871496 | 0,18359158 | 0,674715795 | novel transcript |
| ENSG00000237301 | AL121992,1 | -0,444538251 | 0,734819466 | 0,50076916 | 0,920520138 | novel transcript |
| ENSG00000249931 | GOLGA8K | -0,444660346 | 0,73475728 | 0,26503768 | 0,776509969 | golgin A8 family member K |
| ENSG00000119431 | HDHD3 | -0,444675978 | 0,734749319 | 0,16571413 | 0,645029358 | haloacid dehalogenase like hydrolase domain containing 3 |
| ENSG00000268521 | VN1R83P | -0,444717618 | 0,734728112 | 0,4116894 | 0,883966506 | vomeronasal 1 receptor 83 pseudogene |
| ENSG00000179862 | CITED4 | -0,444823027 | 0,734674432 | 0,2479338 | 0,760164991 | Cbp/p300 interacting transactivator with Glu/Asp rich carboxy-terminal domain 4 |
| ENSG00000259081 | AF111169,3 | -0,444992823 | 0,734587971 | 0,43775146 | 0,897021798 | novel transcript |
| ENSG00000249222 | ATP5MGL | -0,445002024 | 0,734583286 | 0,64882043 | 0,956899963 | ATP synthase membrane subunit g like |
| ENSG00000242261 | AC018635,1 | -0,44540264 | 0,734379331 | 0,46872415 | 0,907940653 | aurora kinase A (AURKA) pseudogene |
| ENSG00000276216 | AC245014,3 | -0,445425915 | 0,734367483 | 0,3066708 | 0,81549341 | novel transcript |
| ENSG00000112319 | EYA4 | -0,445504211 | 0,73432763 | 0,40171956 | 0,87705851 | EYA transcriptional coactivator and phosphatase 4 |
| ENSG00000199977 | SNORA73 | -0,445518036 | 0,734320593 | 0,72599523 | 0,973344786 | Small nucleolar RNA SNORA73 family |
| ENSG00000287049 | AC124862,1 | -0,445560389 | 0,734299036 | 0,60353017 | 0,950352286 | novel transcript, antisense to ANKMY1 |
| ENSG00000284923 | SCRT1 | -0,446193866 | 0,733976681 | 0,21638101 | 0,723599962 | scratch family transcriptional repressor 1 |
| ENSG00000261678 | SCRT1 | -0,446193866 | 0,733976681 | 0,21638101 | 0,723599962 | scratch family transcriptional repressor 1 |
| ENSG00000154040 | CABYR | -0,446206426 | 0,733970291 | 0,17961702 | 0,668298509 | calcium binding tyrosine phosphorylation regulated |
| ENSG00000173611 | SCAI | -0,446257309 | 0,733944405 | 0,04183325 | 0,306796776 | suppressor of cancer cell invasion |
| ENSG00000015133 | CCDC88C | -0,446416829 | 0,733863257 | 0,26787385 | 0,780504009 | coiled-coil domain containing 88C |
| ENSG00000142233 | NTN5 | -0,446435954 | 0,733853528 | 0,59128952 | 0,947416795 | netrin 5 |
| ENSG00000257169 | AC125612,1 | -0,446483818 | 0,733829182 | 0,47186795 | 0,909131671 | small nuclear ribonucleoprotein D1 polypeptide 16kDa (SNRPD1) pseudogene |

| | | | | | | |
|---|---|---|---|---|---|---|
| ENSG00000236254 | MTND4P14 | -0,44652443 | 0,733808525 | 0,64421244 | 0,956633898 | MT-ND4 pseudogene 14 |
| ENSG00000181409 | AATK | -0,44659216 | 0,733774076 | 0,37204198 | 0,859937929 | apoptosis associated tyrosine kinase |
| ENSG00000261094 | AC007066,2 | -0,446596451 | 0,733771893 | 0,35095808 | 0,844035671 | novel transcript |
| ENSG00000228199 | PAIP1P1 | -0,446598927 | 0,733770634 | 0,65582937 | 0,957444211 | poly(A) binding protein interacting protein 1 pseudogene 1 |
| ENSG00000232642 | AC008073,2 | -0,446668474 | 0,733735262 | 0,67948065 | 0,964416861 | novel transcript |
| ENSG00000134744 | TUT4 | -0,446712171 | 0,733713039 | 0,26264632 | 0,774111884 | terminal uridylyl transferase 4 |
| ENSG00000136999 | CCN3 | -0,446729519 | 0,733704216 | 0,48838222 | 0,915394579 | cellular communication network factor 3 |
| ENSG00000085831 | TTC39A | -0,446737181 | 0,73370032 | 0,33203588 | 0,832927044 | tetratricopeptide repeat domain 39A |
| ENSG00000287591 | AL360169,3 | -0,446872018 | 0,73363175 | 0,18165468 | 0,67167359 | novel transcript |
| ENSG00000235652 | FBXO30-DT | -0,447029694 | 0,733551574 | 0,17952093 | 0,668298509 | FBXO30 divergent transcript |
| ENSG00000221923 | ZNF880 | -0,447073742 | 0,733529177 | 0,29172161 | 0,802285739 | zinc finger protein 880 |
| ENSG00000253390 | AC104561,1 | -0,447074995 | 0,73352854 | 0,62001654 | 0,952793934 | novel transcript |
| ENSG00000235489 | DBF4P1 | -0,447377222 | 0,733374891 | 0,4503174 | 0,902062594 | DBF4 zinc finger pseudogene 1 |
| ENSG00000258332 | LINC02394 | -0,44744669 | 0,733339579 | 0,54359328 | 0,934259495 | long intergenic non-protein coding RNA 2394 |
| ENSG00000265750 | AC090772,3 | -0,44760308 | 0,733260089 | 0,59477514 | 0,947469769 | novel transcript |
| ENSG00000214998 | CCNB2P1 | -0,447680375 | 0,733220804 | 0,7501684 | 0,975901808 | cyclin B2 pseudogene 1 |
| ENSG00000260091 | AC093752,3 | -0,447754219 | 0,733183275 | 0,68409498 | 0,965403332 | novel transcript |
| ENSG00000253720 | AC022973,3 | -0,447755034 | 0,733182861 | 0,46015181 | 0,905443183 | novel transcript |
| ENSG00000225171 | DUTP6 | -0,447993636 | 0,733061612 | 0,4844993 | 0,913690171 | deoxyuridine triphosphatase pseudogene 6 |
| ENSG00000181418 | DDN | -0,448037442 | 0,733039354 | 0,64587542 | 0,956899963 | dendrin |
| ENSG00000184224 | C11orf72 | -0,448203034 | 0,732955221 | 0,06077673 | 0,383761049 | chromosome 11 putative open reading frame 72 |
| ENSG00000204060 | FOXO6 | -0,448356233 | 0,732877393 | 0,46974219 | 0,908378389 | forkhead box O6 |
| ENSG00000278829 | AC099811,5 | -0,448362442 | 0,732874239 | 0,33868947 | 0,837449623 | novel transcript, sense intronic to STAT5B |
| ENSG00000242173 | ARHGDIG | -0,448437948 | 0,732835883 | 0,46684737 | 0,907940653 | Rho GDP dissociation inhibitor gamma |
| ENSG00000254895 | MTCO3P15 | -0,448450018 | 0,732829752 | 0,65371657 | 0,957444211 | MT-CO3 pseudogene 15 |
| ENSG00000204852 | TCTN1 | -0,448671022 | 0,7327175 | 0,06624682 | 0,404519674 | tectonic family member 1 |
| ENSG00000223916 | AC097638,1 | -0,448760658 | 0,732671977 | 0,70490637 | 0,969350908 | ribosomal protein L18a (RPL18A) pseudogene |
| ENSG00000232593 | KANTR | -0,448788852 | 0,732657659 | 0,13868627 | 0,593837907 | KDM5C adjacent transcript |
| ENSG00000256085 | AC130404,1 | -0,449028662 | 0,732535884 | 0,64680868 | 0,956899963 | novel transcript |
| ENSG00000144395 | CCDC150 | -0,449240473 | 0,732428344 | 0,12167843 | 0,557385264 | coiled-coil domain containing 150 |
| ENSG00000222460 | RN7SKP271 | -0,449387648 | 0,73235363 | 0,54975488 | 0,936108121 | RN7SK pseudogene 271 |
| ENSG00000269951 | AC090181,2 | -0,449449943 | 0,732322008 | 0,36983001 | 0,858104601 | novel transcript, sense intronic to TSPAN3 |

| | | | | | | |
|---|---|---|---|---|---|---|
| ENSG00000275591 | XKR5 | -0,449460499 | 0,732316649 | 0,24064772 | 0,750950074 | XK related 5 |
| ENSG00000270265 | AC104819,3 | -0,449524088 | 0,732284372 | 0,6703346 | 0,962026636 | novel transcript |
| ENSG00000288312 | LINC02018 | -0,449634701 | 0,732228229 | 0,11667724 | 0,545587385 | long intergenic non-protein coding RNA 2018 |
| ENSG00000286588 | AC104841,1 | -0,449640065 | 0,732225507 | 0,23213981 | 0,740855165 | novel transcript, antisense to SEPT2 |
| ENSG00000166704 | ZNF606 | -0,449642386 | 0,732224329 | 0,14686788 | 0,610674677 | zinc finger protein 606 |
| ENSG00000265366 | GLUD1P2 | -0,449718426 | 0,732185737 | 0,52720763 | 0,929383663 | glutamate dehydrogenase 1 pseudogene 2 |
| ENSG00000233115 | FAM90A11P | -0,449733392 | 0,732178141 | 0,54144011 | 0,933491609 | family with sequence similarity 90 member A11, pseudogene |
| ENSG00000245534 | RORA-AS1 | -0,449812361 | 0,732138065 | 0,43909762 | 0,897434993 | RORA antisense RNA 1 |
| ENSG00000219133 | AL592114,1 | -0,450125278 | 0,731979283 | 0,56343184 | 0,940865625 | ribosomal protein L21 (RPL21) pseudogene |
| ENSG00000070501 | POLB | -0,450242588 | 0,731919766 | 0,1721993 | 0,65638124 | DNA polymerase beta |
| ENSG00000226539 | MLXP1 | -0,450340643 | 0,731870022 | 0,28614007 | 0,797420923 | MLX pseudogene 1 |
| ENSG00000153982 | GDPD1 | -0,450346485 | 0,731867058 | 0,06495053 | 0,399805715 | glycerophosphodiester phosphodiesterase domain containing 1 |
| ENSG00000170469 | SPATA24 | -0,450408475 | 0,731835612 | 0,21530035 | 0,722575157 | spermatogenesis associated 24 |
| ENSG00000270130 | AC068790,7 | -0,450578007 | 0,731749618 | 0,41746044 | 0,886770159 | novel transcript, sense intronic to CCDC92 |
| ENSG00000116711 | PLA2G4A | -0,450597533 | 0,731739714 | 0,42596167 | 0,892133432 | phospholipase A2 group IVA |
| ENSG00000185495 | AC138393,1 | -0,450786477 | 0,731643888 | 0,3065381 | 0,815375074 | septin 7 (SEPT7) pseudogene |
| ENSG00000132541 | RIDA | -0,450805429 | 0,731634277 | 0,46919903 | 0,907940653 | reactive intermediate imine deaminase A homolog |
| ENSG00000175893 | ZDHHC21 | -0,450942251 | 0,731564893 | 0,40386459 | 0,877749224 | zinc finger DHHC-type palmitoyltransferase 21 |
| ENSG00000187686 | KRT18P59 | -0,451081109 | 0,731494484 | 0,24192062 | 0,752203493 | keratin 18 pseudogene 59 |
| ENSG00000163864 | NMNAT3 | -0,451082543 | 0,731493757 | 0,05569523 | 0,364179132 | nicotinamide nucleotide adenylyltransferase 3 |
| ENSG00000233954 | UQCRHL | -0,451311386 | 0,731377736 | 0,27766195 | 0,789067728 | ubiquinol-cytochrome c reductase hinge protein like |
| ENSG00000064655 | EYA2 | -0,451484868 | 0,731289794 | 0,39428156 | 0,871794663 | EYA transcriptional coactivator and phosphatase 2 |
| ENSG00000286613 | AC006213,7 | -0,451549299 | 0,731257135 | 0,37731748 | 0,862818448 | novel transcript, sense intronic to ZNF404 |
| ENSG00000132958 | TPTE2 | -0,451651307 | 0,731205432 | 0,69391677 | 0,967340633 | transmembrane phosphoinositide 3-phosphatase and tensin homolog 2 |
| ENSG00000249267 | LINC00939 | -0,451701821 | 0,73117983 | 0,12281875 | 0,560473734 | long intergenic non-protein coding RNA 939 |
| ENSG00000183793 | NPIPA5 | -0,451731457 | 0,73116481 | 0,50333227 | 0,920918059 | nuclear pore complex interacting protein family member A5 |
| ENSG00000271913 | AL035530,2 | -0,451798388 | 0,73113089 | 0,15320957 | 0,62169919 | novel transcript, antisense to TAGAP |
| ENSG00000231769 | AL035701,1 | -0,451972086 | 0,731042869 | 0,63503707 | 0,955158181 | novel transcript |
| ENSG00000151338 | MIPOL1 | -0,451997734 | 0,731029872 | 0,09115139 | 0,481283407 | mirror-image polydactyly 1 |

| | | | | | | |
|---|---|---|---|---|---|---|
| ENSG00000287099 | AL121750,1 | -0,452006357 | 0,731025503 | 0,55008055 | 0,93612918 | novel transcript, antisense to ADD1 |
| ENSG00000200673 | RN7SKP174 | -0,452010429 | 0,73102344 | 0,62638065 | 0,952793934 | RN7SK pseudogene 174 |
| ENSG00000166920 | C15orf48 | -0,452014828 | 0,731021211 | 0,7277639 | 0,97367385 | chromosome 15 open reading frame 48 |
| ENSG00000229887 | HNRNPA1P6 | -0,452049537 | 0,731003624 | 0,56511937 | 0,940865625 | heterogeneous nuclear ribonucleoprotein A1 pseudogene 6 |
| ENSG00000280579 | AC006359,2 | -0,45212563 | 0,730965069 | 0,41527462 | 0,885858739 | novel transcript |
| ENSG00000255717 | SNHG1 | -0,452197084 | 0,730928867 | 0,28945114 | 0,800406909 | small nucleolar RNA host gene 1 |
| ENSG00000214700 | C12orf71 | -0,452247056 | 0,730903549 | 0,75198477 | 0,975901808 | chromosome 12 open reading frame 71 |
| ENSG00000256482 | AC023796,2 | -0,452334169 | 0,730859417 | 0,66841821 | 0,961351573 | novel transcript, antisense to BCAT1 |
| ENSG00000260691 | ANKRD20A1 | -0,452356656 | 0,730848025 | 0,36765247 | 0,857206868 | ankyrin repeat domain 20 family member A1 |
| ENSG00000275349 | SMN1 | -0,452391766 | 0,730830239 | 0,60025111 | 0,949340299 | survival of motor neuron 1, telomeric |
| ENSG00000105825 | TFPI2 | -0,452415568 | 0,730818182 | 0,22601529 | 0,734959876 | tissue factor pathway inhibitor 2 |
| ENSG00000069011 | PITX1 | -0,452469848 | 0,730790686 | 0,7762923 | 0,978587255 | paired like homeodomain 1 |
| ENSG00000251193 | AC112203,1 | -0,452493093 | 0,730778912 | 0,76669008 | 0,977201891 | novel DHHC-type zinc finger pseudogene |
| ENSG00000117281 | CD160 | -0,452569293 | 0,730740314 | 0,22161871 | 0,729254784 | CD160 molecule |
| ENSG00000139547 | RDH16 | -0,452615153 | 0,730717086 | 0,30317507 | 0,812669425 | retinol dehydrogenase 16 |
| ENSG00000260744 | AC007494,2 | -0,452741141 | 0,730653277 | 0,54805374 | 0,935404974 | novel transcript, antisense to ITFG1 |
| ENSG00000281994 | NPIPA1 | -0,45276525 | 0,730641067 | 0,25501353 | 0,76642479 | nuclear pore complex interacting protein family member A1 |
| ENSG00000196787 | H2AC11 | -0,452767867 | 0,730639742 | 0,13005035 | 0,57623217 | H2A clustered histone 11 |
| ENSG00000266456 | AP001178,3 | -0,452919217 | 0,730563096 | 0,31579945 | 0,821233148 | novel transcript, antisense to C18orf56 |
| ENSG00000231856 | AL162377,1 | -0,452963561 | 0,730540641 | 0,20184072 | 0,701461582 | novel transcript |
| ENSG00000256001 | AC079949,1 | -0,453187737 | 0,730427133 | 0,54918454 | 0,936040289 | novel transcript |
| ENSG00000229282 | AL031963,2 | -0,453198033 | 0,73042192 | 0,69078498 | 0,967179044 | novel transcript |
| ENSG00000283696 | AL592295,4 | -0,453200801 | 0,730420519 | 0,27684462 | 0,788777434 | novel transcript |
| ENSG00000284829 | AL662884,2 | -0,453349825 | 0,730345074 | 0,83816501 | 0,985672131 | novel transcript |
| ENSG00000264044 | AC005726,2 | -0,453403701 | 0,7303178 | 0,43515371 | 0,896272827 | novel transcript, antisense to KIAA0100 |
| ENSG00000248696 | AC011406,1 | -0,453452467 | 0,730293114 | 0,59762701 | 0,9482846 | Transmembrane protein pseudogene |
| ENSG00000171316 | CHD7 | -0,453455226 | 0,730291718 | 0,10259133 | 0,510071005 | chromodomain helicase DNA binding protein 7 |
| ENSG00000091831 | ESR1 | -0,453500524 | 0,730268788 | 0,34925805 | 0,843141089 | estrogen receptor 1 |
| ENSG00000180385 | EMC3-AS1 | -0,453543121 | 0,730247227 | 0,07500035 | 0,432694299 | EMC3 antisense RNA 1 |
| ENSG00000226899 | AL731577,1 | -0,453553965 | 0,730241738 | 0,45817871 | 0,904140862 | novel transcript |
| ENSG00000224775 | BRAFP1 | -0,453615275 | 0,730210705 | 0,56427446 | 0,940865625 | BRAF pseudogene 1 |

| | | | | | | |
|---|---|---|---|---|---|---|
| ENSG00000226396 | AL031727,1 | -0,453759782 | 0,730137568 | 0,48670911 | 0,914396582 | ribosomal protein S14 (RPS14) pseudogene |
| ENSG00000147434 | CHRNA6 | -0,453843095 | 0,730095405 | 0,7118471 | 0,970812834 | cholinergic receptor nicotinic alpha 6 subunit |
| ENSG00000273831 | LRRC56 | -0,454042155 | 0,729994675 | 0,42399245 | 0,890949676 | leucine rich repeat containing 56 |
| ENSG00000275936 | AC004263,1 | -0,454061479 | 0,729984897 | 0,59526903 | 0,947725777 | novel transcript, antisense to PXN |
| ENSG00000231741 | AL353743,3 | -0,454122326 | 0,72995411 | 0,73632561 | 0,974353474 | novel pseudogene |
| ENSG00000133740 | E2F5 | -0,454270436 | 0,729879175 | 0,26175802 | 0,773497425 | E2F transcription factor 5 |
| ENSG00000270012 | AC232271,1 | -0,454305074 | 0,729861652 | 0,11676293 | 0,545661612 | novel transcript |
| ENSG00000282556 | AC068733,3 | -0,454512761 | 0,72975659 | 0,21533498 | 0,722575157 | novel transcript, antisense to PRKCDBP |
| ENSG00000231688 | RPL21P43 | -0,454546585 | 0,729739481 | 0,49877345 | 0,919683359 | ribosomal protein L21 pseudogene 43 |
| ENSG00000081059 | TCF7 | -0,454733084 | 0,729645153 | 0,03723754 | 0,284946761 | transcription factor 7 |
| ENSG00000145911 | N4BP3 | -0,454750728 | 0,729636229 | 0,36101828 | 0,851783601 | NEDD4 binding protein 3 |
| ENSG00000257083 | AC073530,1 | -0,454798129 | 0,729612257 | 0,5521683 | 0,936619966 | novel transcript |
| ENSG00000104219 | ZDHHC2 | -0,454844713 | 0,729588699 | 0,34855145 | 0,842757646 | zinc finger DHHC-type palmitoyltransferase 2 |
| ENSG00000011332 | DPF1 | -0,454990334 | 0,72951506 | 0,16012617 | 0,635191158 | double PHD fingers 1 |
| ENSG00000239272 | RPL21P10 | -0,455185615 | 0,72941632 | 0,57641149 | 0,943158459 | ribosomal protein L21 pseudogene 10 |
| ENSG00000186714 | CCDC73 | -0,455270076 | 0,729373619 | 0,60656528 | 0,950927937 | coiled-coil domain containing 73 |
| ENSG00000122877 | EGR2 | -0,455309776 | 0,729353548 | 0,40948417 | 0,882177108 | early growth response 2 |
| ENSG00000214761 | HNRNPA1P15 | -0,455314898 | 0,729350959 | 0,49716965 | 0,918759905 | heterogeneous nuclear ribonucleoprotein A1 pseudogene 15 |
| ENSG00000224514 | LINC00620 | -0,455415353 | 0,729300176 | 0,68206689 | 0,964699524 | long intergenic non-protein coding RNA 620 |
| ENSG00000267653 | AC002546,1 | -0,455466043 | 0,729274552 | 0,65293395 | 0,95732962 | novel transcript |
| ENSG00000240003 | RPL7P24 | -0,455522651 | 0,729245937 | 0,66943666 | 0,961867165 | ribosomal protein L7 pseudogene 24 |
| ENSG00000227600 | PPT2 | -0,455558283 | 0,729227927 | 0,31824392 | 0,823093059 | palmitoyl-protein thioesterase 2 |
| ENSG00000286757 | AL137139,2 | -0,455589078 | 0,729212361 | 0,26713805 | 0,779493925 | novel transcript, antisense to CLYBL |
| ENSG00000204194 | RPL12P1 | -0,455642786 | 0,729185215 | 0,49228561 | 0,916995341 | ribosomal protein L12 pseudogene 1 |
| ENSG00000232733 | RPL12P1 | -0,455642786 | 0,729185215 | 0,49228561 | 0,916995341 | ribosomal protein L12 pseudogene 1 |
| ENSG00000267370 | AC008752,3 | -0,455783057 | 0,72911432 | 0,44794365 | 0,902062594 | zinc finger protein 420 (ZNF420) pseudogene |
| ENSG00000221988 | PPT2 | -0,455823707 | 0,729093777 | 0,10943188 | 0,527649315 | palmitoyl-protein thioesterase 2 |
| ENSG00000248408 | LINC02469 | -0,456326856 | 0,728839546 | 0,52071649 | 0,927046149 | long intergenic non-protein coding RNA 2469 |
| ENSG00000272097 | AL024498,1 | -0,456479755 | 0,728762306 | 0,48031588 | 0,912847708 | novel transcript |
| ENSG00000198182 | ZNF607 | -0,456550824 | 0,728726407 | 0,18280921 | 0,673503078 | zinc finger protein 607 |
| ENSG00000164104 | HMGB2 | -0,456551487 | 0,728726072 | 0,08615487 | 0,467467779 | high mobility group box 2 |
| ENSG00000169783 | LINGO1 | -0,456623813 | 0,72868954 | 0,0772333 | 0,439261743 | leucine rich repeat and Ig domain containing 1 |

| | | | | | | |
|---|---|---|---|---|---|---|
| ENSG00000203900 | AL121827,1 | -0,456754538 | 0,728623516 | 0,64526211 | 0,956899963 | novel transcript, antisense to CHRNA4 |
| ENSG00000272168 | CASC15 | -0,456860904 | 0,728569798 | 0,06802868 | 0,410826451 | cancer susceptibility 15 |
| ENSG00000181785 | OR5AS1 | -0,456912551 | 0,728543717 | 0,11168076 | 0,534372115 | olfactory receptor family 5 subfamily AS member 1 |
| ENSG00000184786 | TCTE3 | -0,45722887 | 0,728383997 | 0,34002743 | 0,838707505 | t-complex-associated-testis-expressed 3 |
| ENSG00000122779 | TRIM24 | -0,457240401 | 0,728378175 | 0,22167836 | 0,729254784 | tripartite motif containing 24 |
| ENSG00000163531 | NFASC | -0,457377067 | 0,728309179 | 0,09268686 | 0,484089861 | neurofascin |
| ENSG00000273017 | AP000240,1 | -0,457393985 | 0,728300639 | 0,69103819 | 0,967179456 | novel transcript, sense intronic BACH1 |
| ENSG00000270659 | AC079610,1 | -0,457454484 | 0,728270098 | 0,67950917 | 0,964416861 | novel transcript |
| ENSG00000235904 | RBMS3-AS3 | -0,457488528 | 0,728252913 | 0,18953333 | 0,684878649 | RBMS3 antisense RNA 3 |
| ENSG00000260038 | AC009090,1 | -0,457599447 | 0,728196925 | 0,76911483 | 0,977475319 | novel transcript, sense to intronic to ARL2BP |
| ENSG00000272910 | AC090425,3 | -0,45764075 | 0,728176077 | 0,6204911 | 0,952793934 | novel transcript, antisense to ACTL6A |
| ENSG00000272235 | AL590438,1 | -0,457691355 | 0,728150536 | 0,41196371 | 0,884165125 | novel transcript |
| ENSG00000140479 | PCSK6 | -0,457701576 | 0,728145377 | 0,26268557 | 0,774111884 | proprotein convertase subtilisin/kexin type 6 |
| ENSG00000285663 | AC108457,1 | -0,457710024 | 0,728141113 | 0,39261811 | 0,870909282 | novel transcript |
| ENSG00000272805 | TAS2R10 | -0,457717938 | 0,728137119 | 0,66752803 | 0,9611116 | taste 2 receptor member 10 |
| ENSG00000121318 | TAS2R10 | -0,457717938 | 0,728137119 | 0,66752803 | 0,9611116 | taste 2 receptor member 10 |
| ENSG00000277238 | TAS2R10 | -0,457717938 | 0,728137119 | 0,66752803 | 0,9611116 | taste 2 receptor member 10 |
| ENSG00000255443 | CD44-AS1 | -0,457925975 | 0,728032129 | 0,5188732 | 0,926800715 | CD44 antisense RNA 1 |
| ENSG00000227071 | FOCAD-AS1 | -0,457976594 | 0,728006585 | 0,7193929 | 0,971869926 | FOCAD antisense RNA 1 |
| ENSG00000179909 | ZNF154 | -0,458138451 | 0,727924915 | 0,0948075 | 0,489192643 | zinc finger protein 154 |
| ENSG00000270541 | AC069545,1 | -0,458379231 | 0,727803437 | 0,79905234 | 0,980161849 | zinc finger, MYM-type 4 (ZMYM4) pseudogene |
| ENSG00000273274 | ZBTB8B | -0,458428063 | 0,727778803 | 0,20174847 | 0,701374862 | zinc finger and BTB domain containing 8B |
| ENSG00000285618 | AC074132,1 | -0,458432527 | 0,727776551 | 0,57835307 | 0,943425812 | novel transcript, antisense to ADGRV1 |
| ENSG00000256162 | SMLR1 | -0,458436261 | 0,727774667 | 0,49231813 | 0,916995341 | small leucine rich protein 1 |
| ENSG00000109339 | MAPK10 | -0,458456037 | 0,727764691 | 0,2507484 | 0,762971073 | mitogen-activated protein kinase 10 |
| ENSG00000138111 | MFSD13A | -0,458624131 | 0,727679901 | 0,11528123 | 0,54238563 | major facilitator superfamily domain containing 13A |
| ENSG00000177272 | KCNA3 | -0,458703492 | 0,727639874 | 0,24418042 | 0,755571155 | potassium voltage-gated channel subfamily A member 3 |
| ENSG00000275895 | U2AF1L5 | -0,458776843 | 0,727602879 | 0,22118773 | 0,729145225 | U2 small nuclear RNA auxiliary factor 1 like 5 |
| ENSG00000228657 | RPL23AP58 | -0,458806627 | 0,727587858 | 0,63816241 | 0,956018518 | ribosomal protein L23a pseudogene 58 |
| ENSG00000266644 | AC103810,5 | -0,458941861 | 0,72751966 | 0,28544747 | 0,796948543 | novel transcript, antisense to LRRC37A3 |
| ENSG00000105982 | RNF32 | -0,459153977 | 0,727412702 | 0,18829002 | 0,682674151 | ring finger protein 32 |
| ENSG00000274841 | Metazoa_SRP | -0,459224918 | 0,727376934 | 0,74895448 | 0,975901808 | Metazoan signal recognition particle RNA |

| | | | | | | |
|---|---|---|---|---|---|---|
| ENSG00000228073 | AC133106,1 | -0,459229004 | 0,727374874 | 0,7640787 | 0,976640468 | novel transcript |
| ENSG00000101883 | RHOXF1 | -0,459365067 | 0,727306277 | 0,69762887 | 0,968522643 | Rhox homeobox family member 1 |
| ENSG00000221491 | SNORA2C | -0,45944964 | 0,727263642 | 0,31347451 | 0,819801802 | small nucleolar RNA, H/ACA box 2C |
| ENSG00000198298 | ZNF485 | -0,459536622 | 0,727219796 | 0,31638738 | 0,821646322 | zinc finger protein 485 |
| ENSG00000286808 | AC092338,3 | -0,459547851 | 0,727214136 | 0,25123597 | 0,763550239 | novel transcript, antisense to POLR3E |
| ENSG00000213420 | GPC2 | -0,459572273 | 0,727201826 | 0,20047432 | 0,700064356 | glypican 2 |
| ENSG00000102312 | PORCN | -0,459710802 | 0,727132003 | 0,15811324 | 0,631322122 | porcupine O-acyltransferase |
| ENSG00000107872 | FBXL15 | -0,45981505 | 0,727079462 | 0,28498272 | 0,796433333 | F-box and leucine rich repeat protein 15 |
| ENSG00000260317 | AC009812,4 | -0,460130274 | 0,726920615 | 0,46911241 | 0,907940653 | novel transcript |
| ENSG00000259798 | AC138625,1 | -0,460145867 | 0,726912759 | 0,66245476 | 0,959897678 | novel transcript HYDIN |
| ENSG00000008300 | CELSR3 | -0,460242987 | 0,726863826 | 0,16678428 | 0,647101483 | cadherin EGF LAG seven-pass G-type receptor 3 |
| ENSG00000133056 | PIK3C2B | -0,46037032 | 0,726799675 | 0,02507007 | 0,219719433 | phosphatidylinositol-4-phosphate 3-kinase catalytic subunit type 2 beta |
| ENSG00000272927 | AC107464,2 | -0,460406545 | 0,726781426 | 0,47346791 | 0,910083046 | novel transcript, antisense to PDE6B |
| ENSG00000158636 | EMSY | -0,460539345 | 0,726714529 | 0,26125889 | 0,77325059 | EMSY transcriptional repressor, BRCA2 interacting |
| ENSG00000231049 | OR52B5P | -0,460686185 | 0,726640566 | 0,39124061 | 0,870083449 | olfactory receptor family 52 subfamily B member 5 pseudogene |
| ENSG00000263272 | AC004148,1 | -0,460735058 | 0,726615951 | 0,10102268 | 0,505178054 | novel transcript, antisense to RPAIN |
| ENSG00000064225 | ST3GAL6 | -0,46079156 | 0,726587494 | 0,19355526 | 0,690424879 | ST3 beta-galactoside alpha-2,3-sialyltransferase 6 |
| ENSG00000253505 | AC103957,1 | -0,460851565 | 0,726557274 | 0,10798385 | 0,523928283 | novel transcript |
| ENSG00000270285 | AC124916,2 | -0,460904907 | 0,726530411 | 0,78281839 | 0,978687331 | ribosomal protein S27-like, pseudogene |
| ENSG00000157764 | BRAF | -0,461192694 | 0,726385499 | 0,1782317 | 0,665463944 | B-Raf proto-oncogene, serine/threonine kinase |
| ENSG00000255328 | AC136475,5 | -0,461280403 | 0,726341339 | 0,72114277 | 0,972142678 | novel transcript |
| ENSG00000260163 | AC012508,1 | -0,46130727 | 0,726327813 | 0,36490851 | 0,855057583 | novel transcript |
| ENSG00000286899 | AL139286,2 | -0,461406022 | 0,726278097 | 0,46753185 | 0,907940653 | novel transcript, antisense to AGO1 |
| ENSG00000269997 | AC068790,3 | -0,46143182 | 0,72626511 | 0,21078284 | 0,716372938 | novel transcript, sense intronic to CCDC92 |
| ENSG00000242522 | KLHL6-AS1 | -0,461501511 | 0,726230028 | 0,24697627 | 0,759015651 | KLHL6 antisense RNA 1 |
| ENSG00000232901 | CYCSP10 | -0,461590766 | 0,7261851 | 0,47795722 | 0,911872483 | CYCS pseudogene 10 |
| ENSG00000261386 | AC027682,4 | -0,461834234 | 0,72606256 | 0,39528613 | 0,871981986 | novel transcript |
| ENSG00000245849 | RAD51-AS1 | -0,461930697 | 0,726014015 | 0,12553282 | 0,566089307 | RAD51 antisense RNA 1 |
| ENSG00000146247 | PHIP | -0,461932542 | 0,726013086 | 0,26317671 | 0,774637968 | pleckstrin homology domain interacting protein |
| ENSG00000219665 | ZNF433-AS1 | -0,461945181 | 0,726006726 | 0,12273665 | 0,560344415 | ZNF433 and ZNF878 antisense RNA 1 |
| ENSG00000121594 | CD80 | -0,461971905 | 0,725993278 | 0,56811481 | 0,94163744 | CD80 molecule |

| | | | | | | |
|---|---|---|---|---|---|---|
| ENSG00000054392 | HHAT | -0,462036617 | 0,725960714 | 0,19969372 | 0,698923433 | hedgehog acyltransferase |
| ENSG00000280680 | HHAT | -0,462036617 | 0,725960714 | 0,19969372 | 0,698923433 | hedgehog acyltransferase |
| ENSG00000135436 | FAM186B | -0,462198225 | 0,725879398 | 0,34830443 | 0,842757646 | family with sequence similarity 186 member B |
| ENSG00000230832 | SSBL4P | -0,462303215 | 0,725826575 | 0,69394562 | 0,967340633 | SSB like 4, pseudogene |
| ENSG00000274421 | AL162390,1 | -0,462337628 | 0,725809262 | 0,29552468 | 0,805426883 | novel transcript |
| ENSG00000251292 | ERVH-1 | -0,462371494 | 0,725792224 | 0,57526711 | 0,942667371 | endogenous retrovirus group H member 1 |
| ENSG00000263338 | AC027796,5 | -0,462417509 | 0,725769075 | 0,64884167 | 0,956899963 | novel transcript, sense intronic to TRPV1 |
| ENSG00000286577 | AC117528,1 | -0,462456013 | 0,725749706 | 0,67901706 | 0,964151912 | novel transcript, antisense to KIAA0825 |
| ENSG00000286951 | AL356113,1 | -0,462470627 | 0,725742354 | 0,46947861 | 0,908222209 | novel transcript, antisense to KIAA1217 |
| ENSG00000248925 | AC021087,1 | -0,462548161 | 0,725703352 | 0,53946842 | 0,933350549 | novel transcript |
| ENSG00000228727 | SAPCD1 | -0,462573815 | 0,725690448 | 0,51556902 | 0,924926785 | suppressor APC domain containing 1 |
| ENSG00000269600 | AC016629,2 | -0,462707052 | 0,725623431 | 0,42302292 | 0,890598551 | novel transcript, antisense to RPL23AP79 |
| ENSG00000271049 | AC020910,4 | -0,462799209 | 0,725577081 | 0,61184291 | 0,952187506 | small nuclear ribonucleoprotein polypeptide E (SNRPE) pseudogene |
| ENSG00000226284 | ARPC3P1 | -0,462832036 | 0,725560572 | 0,48434941 | 0,913690171 | actin related protein 2/3 complex subunit 3 pseudogene 1 |
| ENSG00000286623 | AC005041,4 | -0,462855549 | 0,725548746 | 0,53880781 | 0,933348247 | novel transcript, antisense to C2orf81 |
| ENSG00000187049 | TMEM216 | -0,463019921 | 0,725466087 | 0,25864169 | 0,771485596 | transmembrane protein 216 |
| ENSG00000231864 | AL807752,3 | -0,463249944 | 0,725350427 | 0,70551424 | 0,969485452 | novel transcript |
| ENSG00000156170 | NDUFAF6 | -0,463336594 | 0,725306863 | 0,17617457 | 0,66169169 | NADH:ubiquinone oxidoreductase complex assembly factor 6 |
| ENSG00000267002 | AC060780,1 | -0,46337761 | 0,725286243 | 0,07943568 | 0,447115945 | novel transcript |
| ENSG00000276509 | AC239799,1 | -0,46349864 | 0,7252254 | 0,39269963 | 0,870981931 | novel transcript |
| ENSG00000248394 | FOSL1P1 | -0,463502183 | 0,725223619 | 0,66096979 | 0,959312917 | FOSL1 pseudogene 1 |
| ENSG00000272077 | AC124045,1 | -0,463523093 | 0,725213108 | 0,19883871 | 0,697657684 | novel transcript |
| ENSG00000281102 | AC092046,2 | -0,463523093 | 0,725213108 | 0,19883871 | 0,697657684 | novel transcript |
| ENSG00000228843 | AL133410,2 | -0,463699811 | 0,725124281 | 0,74476379 | 0,975649186 | novel transcript |
| ENSG00000127831 | VIL1 | -0,463811845 | 0,725067973 | 0,36127225 | 0,851821025 | villin 1 |
| ENSG00000287762 | AC108050,1 | -0,463832942 | 0,72505737 | 0,48710491 | 0,914593485 | novel transcript, antisense to LIMCH1 |
| ENSG00000258813 | AL442663,3 | -0,463846171 | 0,725050722 | 0,27934021 | 0,790200177 | novel transcript, antisense to Rbm25 |
| ENSG00000235947 | EGOT | -0,463949544 | 0,724998772 | 0,64751514 | 0,956899963 | eosinophil granule ontogeny transcript |
| ENSG00000169495 | HTRA4 | -0,464001622 | 0,724972601 | 0,45062355 | 0,902062594 | HtrA serine peptidase 4 |
| ENSG00000134072 | CAMK1 | -0,464034904 | 0,724955877 | 0,23154984 | 0,73957278 | calcium/calmodulin dependent protein kinase I |

| | | | | | | |
|---|---|---|---|---|---|---|
| ENSG00000264015 | AC124254,2 | -0,464071068 | 0,724937704 | 0,53419999 | 0,93151582 | novel transcript |
| ENSG00000160229 | ZNF66 | -0,464122608 | 0,724911807 | 0,03053437 | 0,249967703 | zinc finger protein 66 |
| ENSG00000104660 | LEPROTL1 | -0,464259578 | 0,724842987 | 0,37140005 | 0,859476158 | leptin receptor overlapping transcript like 1 |
| ENSG00000257470 | AC007656,1 | -0,464314614 | 0,724815336 | 0,66796264 | 0,961162795 | novel transcript |
| ENSG00000278828 | H3C10 | -0,464329172 | 0,724808022 | 0,12521197 | 0,565471028 | H3 clustered histone 10 |
| ENSG00000281836 | AC136352,12 | -0,464375832 | 0,724784581 | 0,24184837 | 0,752128472 | novel transcript |
| ENSG00000231934 | FAM242A | -0,464432156 | 0,724756285 | 0,41243484 | 0,884523593 | family with sequence similarity 242 member A |
| ENSG00000253516 | HMGB1P41 | -0,464551454 | 0,724696356 | 0,45487696 | 0,903424009 | high mobility group box 1 pseudogene 41 |
| ENSG00000169856 | ONECUT1 | -0,464700604 | 0,724621439 | 0,38305257 | 0,867682628 | one cut homeobox 1 |
| ENSG00000273343 | AC007663,4 | -0,464751006 | 0,724596124 | 0,41220574 | 0,88438705 | novel transcript, sense intronic to DGCR6L |
| ENSG00000273265 | CNNM3-DT | -0,464848512 | 0,724547153 | 0,73342556 | 0,974353474 | CNNM3 divergent transcript |
| ENSG00000230010 | AL049646,1 | -0,464873254 | 0,724534728 | 0,30489329 | 0,813743624 | novel transcript |
| ENSG00000089639 | GMIP | -0,464954011 | 0,724494172 | 0,21258345 | 0,719437301 | GEM interacting protein |
| ENSG00000187801 | ZFP69B | -0,465197107 | 0,724372104 | 0,23538773 | 0,744654651 | ZFP69 zinc finger protein B |
| ENSG00000258944 | AC004846,2 | -0,46533156 | 0,724304599 | 0,49829139 | 0,919439944 | novel transcript, antisense to PAPLN |
| ENSG00000237080 | EHMT2-AS1 | -0,465361985 | 0,724289324 | 0,37403137 | 0,861369706 | EHMT2 and SLC44A4 antisense RNA 1 |
| ENSG00000273608 | SRCIN1 | -0,465389829 | 0,724275345 | 0,71773931 | 0,971697135 | SRC kinase signaling inhibitor 1 |
| ENSG00000287329 | FP326651,1 | -0,465390072 | 0,724275223 | 0,33059656 | 0,832114797 | novel transcript |
| ENSG00000183018 | SPNS2 | -0,465545107 | 0,724197395 | 0,39826077 | 0,874936729 | sphingolipid transporter 2 |
| ENSG00000286430 | AC020558,6 | -0,465840183 | 0,72404929 | 0,7821411 | 0,978597905 | novel transcript, antisense to PEMT |
| ENSG00000212694 | LINC01089 | -0,465865143 | 0,724036763 | 0,12024808 | 0,553427919 | long intergenic non-protein coding RNA 1089 |
| ENSG00000259205 | PRKXP1 | -0,465968598 | 0,723984844 | 0,56946116 | 0,94221458 | PRKX pseudogene 1 |
| ENSG00000162139 | NEU3 | -0,46597313 | 0,72398257 | 0,04632656 | 0,326794561 | neuraminidase 3 |
| ENSG00000256987 | AC023050,5 | -0,466128031 | 0,723904841 | 0,69256697 | 0,967253522 | family with sequence similarity 98, member B (FAM98B) pseudogene |
| ENSG00000232163 | RPLP1P13 | -0,466295188 | 0,723820971 | 0,68073487 | 0,964654539 | ribosomal protein lateral stalk subunit P1 pseudogene 13 |
| ENSG00000224479 | AC104162,1 | -0,466343038 | 0,723796964 | 0,61953665 | 0,952793934 | novel cadherin domain containing pseudogene |
| ENSG00000263394 | AC007952,7 | -0,466416641 | 0,723760039 | 0,57544204 | 0,942700009 | novel transcript |
| ENSG00000019505 | SYT13 | -0,466648791 | 0,723643585 | 0,39325851 | 0,871294303 | synaptotagmin 13 |
| ENSG00000270001 | AL121894,2 | -0,466710454 | 0,723612656 | 0,59498014 | 0,947555351 | novel transcript |
| ENSG00000215424 | MCM3AP-AS1 | -0,466850674 | 0,723542329 | 0,12897612 | 0,573749206 | MCM3AP antisense RNA 1 |
| ENSG00000223583 | AL513365,1 | -0,466870874 | 0,723532199 | 0,57316206 | 0,942475308 | ribosomal protein L17 (RPL17) pseudogene |

| | | | | | | |
|---|---|---|---|---|---|---|
| ENSG00000240074 | RPL9P30 | -0,466949194 | 0,723492921 | 0,66287345 | 0,959897678 | ribosomal protein L9 pseudogene 30 |
| ENSG00000287201 | AC009682,1 | -0,466985654 | 0,723474637 | 0,45438905 | 0,903383444 | novel transcript, antisense to ASAP1 |
| ENSG00000138030 | KHK | -0,467024369 | 0,723455223 | 0,23363192 | 0,742832118 | ketohexokinase |
| ENSG00000272037 | AP002907,1 | -0,467038739 | 0,723448017 | 0,34870235 | 0,842757646 | novel transcript, antisense to UBR5 |
| ENSG00000277498 | AC107081,2 | -0,467132812 | 0,723400845 | 0,62179873 | 0,952793934 | serine/arginine repetitive matrix 1 (SRRM1) pseudogene |
| ENSG00000101327 | PDYN | -0,467151245 | 0,723391602 | 0,49275424 | 0,917090312 | prodynorphin |
| ENSG00000276155 | MAPT | -0,467200732 | 0,723366789 | 0,19742847 | 0,695991044 | microtubule associated protein tau |
| ENSG00000108001 | EBF3 | -0,467249674 | 0,72334225 | 0,30392703 | 0,812968394 | EBF transcription factor 3 |
| ENSG00000228063 | LYPLAL1-DT | -0,467340411 | 0,723296757 | 0,53489199 | 0,931774961 | LYPLAL1 divergent transcript |
| ENSG00000272034 | SNORD14A | -0,467342816 | 0,723295551 | 0,67811865 | 0,964004688 | small nucleolar RNA, C/D box 14A |
| ENSG00000231189 | AC013448,1 | -0,467396137 | 0,723268819 | 0,32414003 | 0,826248637 | novel transcript |
| ENSG00000285458 | AC093827,5 | -0,467498868 | 0,723217319 | 0,12625839 | 0,567554588 | novel protein |
| ENSG00000149809 | TM7SF2 | -0,467512773 | 0,723210348 | 0,0585903 | 0,375131502 | transmembrane 7 superfamily member 2 |
| ENSG00000188263 | IL17REL | -0,467589939 | 0,723171667 | 0,7892377 | 0,979462318 | interleukin 17 receptor E like |
| ENSG00000229116 | AL137026,1 | -0,46786767 | 0,723032464 | 0,71961238 | 0,971918153 | novel transcript |
| ENSG00000173542 | MOB1B | -0,467886898 | 0,723022828 | 0,34789722 | 0,842433491 | MOB kinase activator 1B |
| ENSG00000248423 | AC100850,1 | -0,467996873 | 0,722967714 | 0,45364245 | 0,902908855 | clathrin, light chain (Lca) (CLTA) pseudogene |
| ENSG00000148219 | ASTN2 | -0,468036999 | 0,722947607 | 0,09001913 | 0,478514897 | astrotactin 2 |
| ENSG00000104941 | RSPH6A | -0,468042961 | 0,722944619 | 0,61920182 | 0,952793934 | radial spoke head 6 homolog A |
| ENSG00000221953 | C1orf229 | -0,468073261 | 0,722929435 | 0,4624356 | 0,906545444 | chromosome 1 open reading frame 229 |
| ENSG00000105699 | LSR | -0,468126623 | 0,722902696 | 0,28659236 | 0,797864972 | lipolysis stimulated lipoprotein receptor |
| ENSG00000213462 | ERV3-1 | -0,468242844 | 0,722844463 | 0,22269758 | 0,73126953 | endogenous retrovirus group 3 member 1, envelope |
| ENSG00000173597 | SULT1B1 | -0,468243542 | 0,722844113 | 0,08523431 | 0,464892813 | sulfotransferase family 1B member 1 |
| ENSG00000162148 | PPP1R32 | -0,468392512 | 0,722769478 | 0,18470229 | 0,67694849 | protein phosphatase 1 regulatory subunit 32 |
| ENSG00000196247 | ZNF107 | -0,468421208 | 0,722755102 | 0,27342101 | 0,785672647 | zinc finger protein 107 |
| ENSG00000287777 | AL161851,2 | -0,468553704 | 0,722688728 | 0,45098587 | 0,902062594 | novel transcript, antisense to NPAS3 |
| ENSG00000164180 | TMEM161B | -0,468659011 | 0,722635978 | 0,33795408 | 0,837226333 | transmembrane protein 161B |
| ENSG00000233690 | EBAG9P1 | -0,468683256 | 0,722623834 | 0,53590704 | 0,932139691 | EBAG9 pseudogene 1 |
| ENSG00000179038 | AP001885,1 | -0,468778429 | 0,722576164 | 0,31171516 | 0,817984116 | tetratricopeptide repeat domain 3 (TTC3) pseudogene |
| ENSG00000172404 | DNAJB7 | -0,46899414 | 0,722468134 | 0,2781889 | 0,789586738 | DnaJ heat shock protein family (Hsp40) member B7 |
| ENSG00000234141 | AC006042,3 | -0,469070641 | 0,722429824 | 0,67962922 | 0,964459885 | novel transcript |

| | | | | | | |
|---|---|---|---|---|---|---|
| ENSG00000173295 | FAM86B3P | -0,469091487 | 0,722419386 | 0,13794656 | 0,592546669 | family with sequence similarity 86 member B3, pseudogene |
| ENSG00000275543 | MPV17L | -0,469302484 | 0,722313738 | 0,22516797 | 0,733832103 | MPV17 mitochondrial inner membrane protein like |
| ENSG00000277215 | SPANXA2-OT1 | -0,469360625 | 0,72228463 | 0,09275722 | 0,484139956 | SPANXA2 overlapping transcript 1 |
| ENSG00000213999 | MEF2B | -0,469508778 | 0,722210461 | 0,36143254 | 0,851893059 | myocyte enhancer factor 2B |
| ENSG00000287417 | AC006059,4 | -0,469599063 | 0,722165266 | 0,27245495 | 0,785492261 | novel transcript, antisense to CCDC13 |
| ENSG00000267470 | ZNF571-AS1 | -0,469673234 | 0,722128139 | 0,11794265 | 0,548984873 | ZNF571 antisense RNA 1 |
| ENSG00000286482 | AL117344,2 | -0,469728341 | 0,722100556 | 0,24407711 | 0,755452615 | novel transcript |
| ENSG00000228782 | MRPL45P2 | -0,469786081 | 0,722071657 | 0,16322472 | 0,64157124 | mitochondrial ribosomal protein L45 pseudogene 2 |
| ENSG00000279072 | AC118758,2 | -0,469812621 | 0,722058373 | 0,43432793 | 0,896272827 | novel piRNA host transcript |
| ENSG00000257433 | AC004241,1 | -0,470327715 | 0,721800618 | 0,32098838 | 0,824532132 | novel transcript, antisense to ENDOU and RAPGEF3 |
| ENSG00000248483 | POU5F2 | -0,470376413 | 0,721776255 | 0,19808726 | 0,696782178 | POU domain class 5, transcription factor 2 |
| ENSG00000280828 | AC090114,3 | -0,470481572 | 0,721723646 | 0,27804772 | 0,789397971 | pseudogene similar to part of tousled-like kinase 2 (TLK2) |
| ENSG00000227471 | AKR1B15 | -0,470506162 | 0,721711344 | 0,12362604 | 0,56228403 | aldo-keto reductase family 1 member B15 |
| ENSG00000189144 | ZNF573 | -0,470573794 | 0,721677512 | 0,16061846 | 0,636125619 | zinc finger protein 573 |
| ENSG00000131746 | TNS4 | -0,470640618 | 0,721644086 | 0,76443134 | 0,976640468 | tensin 4 |
| ENSG00000287642 | AC093599,2 | -0,470683113 | 0,72162283 | 0,69928913 | 0,968522643 | novel transcript, antisense to FHDC1 |
| ENSG00000129214 | SHBG | -0,470730333 | 0,721599212 | 0,27639521 | 0,788263115 | sex hormone binding globulin |
| ENSG00000137731 | FXYD2 | -0,470917539 | 0,721505582 | 0,56534499 | 0,940865625 | FXYD domain containing ion transport regulator 2 |
| ENSG00000269560 | AC010422,4 | -0,471014497 | 0,721457094 | 0,61524216 | 0,952793934 | novel transcript, sense intronic to ZNF564 |
| ENSG00000106399 | RPA3 | -0,471231541 | 0,721348563 | 0,21884231 | 0,727009468 | replication protein A3 |
| ENSG00000157851 | DPYSL5 | -0,471295084 | 0,721316792 | 0,03264638 | 0,261468653 | dihydropyrimidinase like 5 |
| ENSG00000235989 | MORC2-AS1 | -0,471412278 | 0,7212582 | 0,3667939 | 0,856302171 | MORC2 antisense RNA 1 |
| ENSG00000285934 | AC091074,2 | -0,471501087 | 0,721213803 | 0,09112605 | 0,481283407 | novel transcript |
| ENSG00000235677 | NPM1P26 | -0,471520731 | 0,721203983 | 0,37498514 | 0,861771195 | nucleophosmin 1 pseudogene 26 |
| ENSG00000225078 | AL365338,1 | -0,471616296 | 0,721156211 | 0,6553594 | 0,957444211 | COP9 constitutive photomorphogenic homolog subunit 2 (Arabidopsis) (COPS2) pseudogene |
| ENSG00000239523 | MYLK-AS1 | -0,471709921 | 0,721109413 | 0,14848169 | 0,613326406 | MYLK antisense RNA 1 |
| ENSG00000223525 | RABGAP1L-IT1 | -0,471742973 | 0,721092892 | 0,10146344 | 0,506329296 | RABGAP1L intronic transcript 1 |
| ENSG00000180263 | FGD6 | -0,471743737 | 0,721092511 | 0,24732109 | 0,7594781 | FYVE, RhoGEF and PH domain containing 6 |
| ENSG00000155761 | SPAG17 | -0,471889226 | 0,721019796 | 0,29767886 | 0,806956721 | sperm associated antigen 17 |
| ENSG00000106415 | GLCCI1 | -0,471951891 | 0,720988478 | 0,32560268 | 0,827465638 | glucocorticoid induced 1 |

| | | | | | | |
|---|---|---|---|---|---|---|
| ENSG00000131095 | GFAP | -0,472073141 | 0,720927886 | 0,45192197 | 0,902062594 | glial fibrillary acidic protein |
| ENSG00000163421 | PROK2 | -0,472092391 | 0,720918266 | 0,4793324 | 0,912714933 | prokineticin 2 |
| ENSG00000189057 | FAM111B | -0,472136633 | 0,720896159 | 0,32558073 | 0,827465638 | family with sequence similarity 111 member B |
| ENSG00000274611 | TBC1D3 | -0,472239652 | 0,720844683 | 0,13213471 | 0,580694357 | TBC1 domain family member 3 |
| ENSG00000203811 | H3C14 | -0,472245195 | 0,720841914 | 0,10216422 | 0,508689653 | H3 clustered histone 14 |
| ENSG00000203852 | H3C15 | -0,472245195 | 0,720841914 | 0,10216422 | 0,508689653 | H3 clustered histone 15 |
| ENSG00000260647 | AC127537,1 | -0,472293744 | 0,720817657 | 0,55950437 | 0,939727587 | novel transcript, antisense to TOP3A |
| ENSG00000284449 | AC127537,5 | -0,472293744 | 0,720817657 | 0,55950437 | 0,939727587 | novel transcript, antisense to TOP3A |
| ENSG00000089692 | LAG3 | -0,472383908 | 0,720772609 | 0,29655343 | 0,806142623 | lymphocyte activating 3 |
| ENSG00000282956 | MAGI1 | -0,472419933 | 0,720754611 | 0,08187249 | 0,454808375 | membrane associated guanylate kinase, WW and PDZ domain containing 1 |
| ENSG00000151276 | MAGI1 | -0,472419933 | 0,720754611 | 0,08187249 | 0,454808375 | membrane associated guanylate kinase, WW and PDZ domain containing 1 |
| ENSG00000255553 | LINC02733 | -0,472455573 | 0,720736806 | 0,37439443 | 0,861553618 | long intergenic non-protein coding RNA 2733 |
| ENSG00000196584 | XRCC2 | -0,472473834 | 0,720727683 | 0,09947841 | 0,500446627 | X-ray repair cross complementing 2 |
| ENSG00000267632 | AC067852,3 | -0,472547764 | 0,720690751 | 0,22636333 | 0,735064413 | novel transcript |
| ENSG00000102001 | CACNA1F | -0,47257979 | 0,720674753 | 0,63596501 | 0,955399772 | calcium voltage-gated channel subunit alpha1 F |
| ENSG00000242634 | RPS24P16 | -0,472624755 | 0,720652292 | 0,64908205 | 0,956899963 | ribosomal protein S24 pseudogene 16 |
| ENSG00000234210 | AC006372,3 | -0,472628256 | 0,720650543 | 0,52443716 | 0,928616707 | novel transcript |
| ENSG00000066468 | FGFR2 | -0,472631966 | 0,720648689 | 0,22433509 | 0,73307493 | fibroblast growth factor receptor 2 |
| ENSG00000239988 | RPL31P60 | -0,472645428 | 0,720641965 | 0,73554207 | 0,974353474 | ribosomal protein L31 pseudogene 60 |
| ENSG00000188171 | ZNF626 | -0,472693789 | 0,720617809 | 0,22269287 | 0,73126953 | zinc finger protein 626 |
| ENSG00000287281 | AC012101,2 | -0,472824147 | 0,720552699 | 0,18212173 | 0,672396513 | novel transcript |
| ENSG00000176857 | GJA1P1 | -0,472841101 | 0,720544231 | 0,31427786 | 0,819877431 | gap junction protein alpha 1 pseudogene 1 |
| ENSG00000255449 | AP002812,5 | -0,472869166 | 0,720530214 | 0,44918759 | 0,902062594 | novel transcript, antisense to AAMDC |
| ENSG00000247679 | AC139795,2 | -0,47288159 | 0,720524009 | 0,14578821 | 0,608479758 | novel transcript |
| ENSG00000246339 | EXTL3-AS1 | -0,472894511 | 0,720517556 | 0,15625793 | 0,627117842 | EXTL3 antisense RNA 1 |
| ENSG00000258441 | LINC00641 | -0,472908812 | 0,720510414 | 0,19224487 | 0,688548947 | long intergenic non-protein coding RNA 641 |
| ENSG00000148935 | GAS2 | -0,473426053 | 0,72025214 | 0,27375897 | 0,785787783 | growth arrest specific 2 |
| ENSG00000273247 | AC097376,3 | -0,473563746 | 0,720183401 | 0,08582888 | 0,46650804 | novel transcript, antisense to RAB33B |
| ENSG00000267265 | AC011476,3 | -0,473629351 | 0,720150652 | 0,4048282 | 0,878176155 | novel transcript, antisense to RDH13 and GP6 |
| ENSG00000247081 | BAALC-AS1 | -0,473690931 | 0,720119914 | 0,09428005 | 0,487847001 | BAALC antisense RNA 1 |

| | | | | | | |
|---|---|---|---|---|---|---|
| ENSG00000243307 | POM121L6P | -0,473694938 | 0,720117914 | 0,60579062 | 0,950927937 | POM121 transmembrane nucleoporin like 6, pseudogene |
| ENSG00000232136 | DUXAP7 | -0,473778906 | 0,720076003 | 0,70622528 | 0,969503793 | double homeobox A pseudogene 7 |
| ENSG00000162877 | PM20D1 | -0,473809989 | 0,720060489 | 0,45728609 | 0,903831777 | peptidase M20 domain containing 1 |
| ENSG00000284624 | AC092902,4 | -0,473895799 | 0,720017662 | 0,07151282 | 0,421367686 | novel transcript |
| ENSG00000152582 | SPEF2 | -0,473953278 | 0,719988976 | 0,2546026 | 0,766037493 | sperm flagellar 2 |
| ENSG00000276997 | AL513314,2 | -0,474074645 | 0,719928409 | 0,12364709 | 0,56228403 | novel transcript |
| ENSG00000259039 | AL161804,1 | -0,474140299 | 0,719895647 | 0,60134112 | 0,949939978 | novel transcript, antisense to SLC35F4 |
| ENSG00000285492 | AL356417,3 | -0,474154789 | 0,719888417 | 0,54234716 | 0,933491609 | uncharacterized LOC112267968 |
| ENSG00000251126 | AC017007,5 | -0,474159701 | 0,719885966 | 0,6766432 | 0,963580269 | novel transcript, antisense to ANK2 |
| ENSG00000276071 | AC074138,1 | -0,474401469 | 0,719765337 | 0,26848202 | 0,780709773 | novel transcript |
| ENSG00000232788 | ITGA6-AS1 | -0,474430497 | 0,719750855 | 0,58853988 | 0,946306024 | ITGA6 antisense RNA 1 |
| ENSG00000104177 | MYEF2 | -0,474434829 | 0,719748694 | 0,15552064 | 0,625765413 | myelin expression factor 2 |
| ENSG00000233044 | AL049820,1 | -0,474540469 | 0,719695993 | 0,6424651 | 0,956633898 | novel transcript |
| ENSG00000176049 | JAKMIP2 | -0,474589532 | 0,719671518 | 0,16915517 | 0,651208888 | janus kinase and microtubule interacting protein 2 |
| ENSG00000206585 | RNVU1-7 | -0,474754896 | 0,719589033 | 0,24745617 | 0,759682112 | RNA, variant U1 small nuclear 7 |
| ENSG00000286459 | AP006287,3 | -0,474764333 | 0,719584326 | 0,48399201 | 0,913690171 | novel transcript, antisense to GAL3ST3 |
| ENSG00000257657 | AC079950,1 | -0,474854746 | 0,719539231 | 0,70348542 | 0,968940012 | novel transcript |
| ENSG00000228126 | FALEC | -0,475042197 | 0,719445747 | 0,71482905 | 0,971041579 | focally amplified long non-coding RNA in epithelial cancer |
| ENSG00000215441 | CTAGE12P | -0,475182194 | 0,719375936 | 0,79158426 | 0,979471509 | CTAGE family member 12, pseudogene |
| ENSG00000235221 | LINC00383 | -0,475234738 | 0,719349736 | 0,43539997 | 0,89634451 | long intergenic non-protein coding RNA 383 |
| ENSG00000232485 | RPL37A-DT | -0,475363617 | 0,719285478 | 0,42427308 | 0,890949676 | RPL37A divergent transcript |
| ENSG00000111850 | SMIM8 | -0,475383406 | 0,719275612 | 0,25032437 | 0,762441549 | small integral membrane protein 8 |
| ENSG00000249216 | AC105389,2 | -0,475426679 | 0,719254038 | 0,72266404 | 0,972594421 | novel transcript |
| ENSG00000154646 | TMPRSS15 | -0,475452377 | 0,719241226 | 0,73781401 | 0,974353474 | transmembrane serine protease 15 |
| ENSG00000167216 | KATNAL2 | -0,475568112 | 0,71918353 | 0,17778737 | 0,664396898 | katanin catalytic subunit A1 like 2 |
| ENSG00000258851 | AL139300,2 | -0,47557266 | 0,719181263 | 0,75333543 | 0,975901808 | novel transcript, antisense to BAG5 |
| ENSG00000180116 | C12orf40 | -0,475694134 | 0,719120711 | 0,55445545 | 0,937793851 | chromosome 12 open reading frame 40 |
| ENSG00000217767 | NDUFAB1P1 | -0,475738725 | 0,719098485 | 0,75969727 | 0,976120207 | NADH:ubiquinone oxidoreductase subunit AB1 pseudogene 1 |
| ENSG00000162994 | CLHC1 | -0,4757537 | 0,719091021 | 0,27317282 | 0,785672647 | clathrin heavy chain linker domain containing 1 |
| ENSG00000168970 | JMJD7-PLA2G4 | -0,475845888 | 0,719045072 | 0,20393938 | 0,704681782 | JMJD7-PLA2G4B readthrough |

| | | | | | | |
|---|---|---|---|---|---|---|
| ENSG00000204977 | TRIM13 | -0,475955646 | 0,718990371 | 0,23590023 | 0,744968148 | tripartite motif containing 13 |
| ENSG00000283208 | AC001226,2 | -0,475963538 | 0,718986437 | 0,32310301 | 0,825150495 | novel protein |
| ENSG00000254988 | AP002498,1 | -0,476024175 | 0,718956219 | 0,71740575 | 0,971697135 | novel transcript |
| ENSG00000105376 | ICAM5 | -0,4761105 | 0,718913201 | 0,72726888 | 0,973344786 | intercellular adhesion molecule 5 |
| ENSG00000172421 | EFCAB3 | -0,476137662 | 0,718899666 | 0,10620116 | 0,520092902 | EF-hand calcium binding domain 3 |
| ENSG00000256223 | ZNF10 | -0,476341703 | 0,718797999 | 0,29007521 | 0,800682868 | zinc finger protein 10 |
| ENSG00000268163 | AC004076,1 | -0,476549775 | 0,718694338 | 0,63898371 | 0,956018518 | novel transcript |
| ENSG00000276148 | AC084824,5 | -0,476884701 | 0,71852751 | 0,51102959 | 0,922822691 | novel transcript, antisense to DNM1L |
| ENSG00000255580 | AP000462,1 | -0,47706813 | 0,71843616 | 0,56920175 | 0,94219857 | novel transcript |
| ENSG00000168959 | GRM5 | -0,477232323 | 0,718354399 | 0,3041151 | 0,813056239 | glutamate metabotropic receptor 5 |
| ENSG00000278463 | H2AC4 | -0,477304027 | 0,718318697 | 0,15900944 | 0,633144409 | H2A clustered histone 4 |
| ENSG00000266910 | AC008507,1 | -0,477388226 | 0,718276776 | 0,49229037 | 0,916995341 | TAF9 RNA polymerase II, TATA box binding protein (TBP)-associated factor, 32kDa (TAF9) pseudogene |
| ENSG00000244630 | AC022493,1 | -0,477660763 | 0,7181411 | 0,76688771 | 0,977201891 | ribosomal protein S26 (RPS26) pseudogene |
| ENSG00000232778 | RPL23AP50 | -0,477709938 | 0,718116622 | 0,47847327 | 0,912212078 | ribosomal protein L23a pseudogene 50 |
| ENSG00000073849 | ST6GAL1 | -0,477899193 | 0,718022425 | 0,20825012 | 0,712801795 | ST6 beta-galactoside alpha-2,6-sialyltransferase 1 |
| ENSG00000145439 | CBR4 | -0,477986318 | 0,717979065 | 0,3213787 | 0,824933727 | carbonyl reductase 4 |
| ENSG00000254691 | AP002812,3 | -0,478010255 | 0,717967152 | 0,78993934 | 0,979462318 | novel transcript |
| ENSG00000129048 | ACKR4 | -0,478015683 | 0,71796445 | 0,37592472 | 0,862409046 | atypical chemokine receptor 4 |
| ENSG00000145491 | ROPN1L | -0,478122958 | 0,717911067 | 0,42458495 | 0,891157406 | rhophilin associated tail protein 1 like |
| ENSG00000287365 | AC068787,4 | -0,478155981 | 0,717894634 | 0,69546576 | 0,967604531 | novel transcript |
| ENSG00000157429 | ZNF19 | -0,478166936 | 0,717889183 | 0,12321979 | 0,561566116 | zinc finger protein 19 |
| ENSG00000254911 | SCARNA9 | -0,478299769 | 0,717823087 | 0,30864127 | 0,816730513 | small Cajal body-specific RNA 9 |
| ENSG00000138028 | CGREF1 | -0,478305311 | 0,71782033 | 0,36325397 | 0,853554847 | cell growth regulator with EF-hand domain 1 |
| ENSG00000203872 | C6orf163 | -0,47839784 | 0,717774293 | 0,09841619 | 0,498059993 | chromosome 6 open reading frame 163 |
| ENSG00000272137 | AL451064,1 | -0,478525517 | 0,717710774 | 0,69481629 | 0,967417311 | novel transcript |
| ENSG00000245205 | EEF1A1P4 | -0,478739962 | 0,7176041 | 0,42189141 | 0,889720796 | eukaryotic translation elongation factor 1 alpha 1 pseudogene 4 |
| ENSG00000278416 | PMS2P2 | -0,478748067 | 0,717600068 | 0,22732592 | 0,735964811 | PMS1 homolog 2, mismatch repair system component pseudogene 2 |
| ENSG00000264660 | AC107926,1 | -0,478749537 | 0,717599337 | 0,41956926 | 0,887991693 | novel transcript |
| ENSG00000198807 | PAX9 | -0,478771988 | 0,71758817 | 0,59293217 | 0,947416795 | paired box 9 |

| | | | | | | |
|---|---|---|---|---|---|---|
| ENSG00000267800 | AC012254,4 | -0,478803647 | 0,717572423 | 0,70397026 | 0,969001288 | SUMO2 SMT3 suppressor of mif two 3 homolog 2 (S, cerevisiae), pseudogene |
| ENSG00000131781 | FMO5 | -0,478871864 | 0,717538494 | 0,21951091 | 0,727838473 | flavin containing dimethylaniline monoxygenase 5 |
| ENSG00000172426 | RSPH9 | -0,478948493 | 0,717500383 | 0,09662312 | 0,493760245 | radial spoke head component 9 |
| ENSG00000239415 | AP001469,3 | -0,479227284 | 0,717361744 | 0,09940736 | 0,50040065 | novel transcript, antisense to MCM3APAS |
| ENSG00000113296 | THBS4 | -0,479261724 | 0,71734462 | 0,10361947 | 0,513379913 | thrombospondin 4 |
| ENSG00000135702 | CHST5 | -0,479266041 | 0,717342473 | 0,46379391 | 0,907251937 | carbohydrate sulfotransferase 5 |
| ENSG00000272696 | AL359091,3 | -0,479464833 | 0,717243636 | 0,43343056 | 0,895978116 | novel transcript, antisense to TRUB2 |
| ENSG00000272583 | AL592494,2 | -0,479495445 | 0,717228417 | 0,48368875 | 0,913690171 | novel transcript |
| ENSG00000254042 | SLIT3-AS2 | -0,479700494 | 0,717126485 | 0,54043424 | 0,933385624 | SLIT3 antisense RNA 2 |
| ENSG00000269086 | AC008555,2 | -0,479707144 | 0,71712318 | 0,3573697 | 0,848693232 | novel transcript |
| ENSG00000287424 | AC080079,2 | -0,479770765 | 0,717091556 | 0,29290511 | 0,803304991 | novel transcript |
| ENSG00000255220 | DDX18P5 | -0,479812912 | 0,717070607 | 0,52584576 | 0,929056124 | DEAD-box helicase 18 pseudogene 5 |
| ENSG00000286619 | AC119673,2 | -0,479916425 | 0,71701916 | 0,05735345 | 0,370182715 | novel transcript, antisense to PM20D1 |
| ENSG00000155066 | PROM2 | -0,480197129 | 0,716879663 | 0,27643669 | 0,788263115 | prominin 2 |
| ENSG00000268403 | AC132192,2 | -0,480218755 | 0,716868918 | 0,19501109 | 0,69283135 | novel transcript, antisense to ZNF143 |
| ENSG00000273384 | AL137796,1 | -0,480270745 | 0,716843084 | 0,73190659 | 0,974353474 | novel transcript |
| ENSG00000265282 | AC005828,4 | -0,480337517 | 0,716809907 | 0,69890358 | 0,968522643 | novel transcript |
| ENSG00000261799 | AC007406,4 | -0,480507311 | 0,71672555 | 0,24493584 | 0,756911548 | novel transcript |
| ENSG00000079101 | CLUL1 | -0,48056376 | 0,716697506 | 0,48476569 | 0,913690171 | clusterin like 1 |
| ENSG00000175161 | CADM2 | -0,480572399 | 0,716693215 | 0,36347196 | 0,85380707 | cell adhesion molecule 2 |
| ENSG00000224551 | HMGB3P21 | -0,480577323 | 0,716690769 | 0,58382992 | 0,944276183 | high mobility group box 3 pseudogene 21 |
| ENSG00000224032 | EPB41L4A-AS1 | -0,480727324 | 0,716616256 | 0,08578673 | 0,46635999 | EPB41L4A antisense RNA 1 |
| ENSG00000233578 | EIF4EP1 | -0,480985386 | 0,716488083 | 0,35607078 | 0,84732813 | eukaryotic translation initiation factor 4E pseudogene 1 |
| ENSG00000224014 | AL390728,3 | -0,481197574 | 0,716382712 | 0,71135957 | 0,970635643 | SMT3 suppressor of mif two 3 homolog 3 (yeast) (SUMO3) pseudogene |
| ENSG00000285570 | AL590666,4 | -0,481199793 | 0,71638161 | 0,57061665 | 0,9423888 | Novel transcript, antisense to CRABP2 |
| ENSG00000183979 | NPB | -0,481222683 | 0,716370243 | 0,49145384 | 0,916995341 | neuropeptide B |
| ENSG00000280809 | LINC00836 | -0,481274018 | 0,716344753 | 0,07319893 | 0,426957349 | long intergenic non-protein coding RNA 836 |
| ENSG00000267655 | AC125437,1 | -0,481307322 | 0,716328217 | 0,34295251 | 0,840419133 | novel transcript, sense intronic to ATP9B |
| ENSG00000283953 | FCAR | -0,481349307 | 0,716307371 | 0,82207453 | 0,982952211 | Fc fragment of IgA receptor |
| ENSG00000106018 | VIPR2 | -0,481446464 | 0,716259134 | 0,52162506 | 0,927472429 | vasoactive intestinal peptide receptor 2 |

| | | | | | | |
|---|---|---|---|---|---|---|
| ENSG00000273901 | AC012313,9 | -0,481496026 | 0,716234528 | 0,57281639 | 0,942475308 | novel transcript, antisense to SLC27A5 |
| ENSG00000278811 | LINC00624 | -0,48159455 | 0,716185617 | 0,39463475 | 0,871872016 | long intergenic non-protein coding RNA 624 |
| ENSG00000254389 | RHPN1-AS1 | -0,481659907 | 0,716153173 | 0,14196122 | 0,600058555 | RHPN1 antisense RNA 1 (head to head) |
| ENSG00000269054 | AC012313,6 | -0,481697513 | 0,716134505 | 0,23489195 | 0,744576287 | novel transcript, antisense to ZNF497 |
| ENSG00000166664 | CHRFAM7A | -0,481796306 | 0,716085467 | 0,35390754 | 0,845346751 | CHRNA7 (exons 5-10) and FAM7A (exons A-E) fusion |
| ENSG00000224584 | UBE2V1P13 | -0,482067222 | 0,71595101 | 0,51373751 | 0,924189506 | ubiquitin conjugating enzyme E2 V1 pseudogene 13 |
| ENSG00000215630 | GUSBP9 | -0,482089661 | 0,715939875 | 0,17187941 | 0,655736851 | GUSB pseudogene 9 |
| ENSG00000257242 | LINC01619 | -0,482218436 | 0,715875973 | 0,56513868 | 0,940865625 | long intergenic non-protein coding RNA 1619 |
| ENSG00000103184 | SEC14L5 | -0,48239628 | 0,715787731 | 0,55766173 | 0,938815153 | SEC14 like lipid binding 5 |
| ENSG00000229659 | RPL26P6 | -0,482409221 | 0,71578131 | 0,5745262 | 0,942475308 | ribosomal protein L26 pseudogene 6 |
| ENSG00000268889 | AC008750,6 | -0,482448523 | 0,715761811 | 0,52477396 | 0,928616707 | novel transcript |
| ENSG00000183323 | CCDC125 | -0,482485651 | 0,715743391 | 0,033988 | 0,268360931 | coiled-coil domain containing 125 |
| ENSG00000255445 | AP003072,2 | -0,482507198 | 0,715732702 | 0,42262747 | 0,890403187 | aminoacyl tRNA synthetase complex-interacting multifunctional protein 2 (AIMP2) pseudogene |
| ENSG00000257894 | AC027288,3 | -0,482598406 | 0,715687454 | 0,05275174 | 0,352776586 | novel transcript |
| ENSG00000204371 | EHMT2 | -0,482700697 | 0,715636711 | 0,13610243 | 0,587965501 | euchromatic histone lysine methyltransferase 2 |
| ENSG00000223584 | TVP23CP1 | -0,482783857 | 0,715595462 | 0,38061879 | 0,865900094 | TVP23C pseudogene 1 |
| ENSG00000232000 | CLCN3P1 | -0,482878873 | 0,715548334 | 0,19139837 | 0,687158413 | chloride voltage-gated channel 3 pseudogene 1 |
| ENSG00000254929 | AL591684,2 | -0,482899649 | 0,71553803 | 0,36494219 | 0,855057583 | novel transcript |
| ENSG00000181619 | GPR135 | -0,482973602 | 0,715501352 | 0,12055546 | 0,554027014 | G protein-coupled receptor 135 |
| ENSG00000147027 | TMEM47 | -0,483084543 | 0,715446333 | 0,32105054 | 0,824556331 | transmembrane protein 47 |
| ENSG00000286331 | AC011388,1 | -0,483134285 | 0,715421666 | 0,67869374 | 0,9640473 | novel transcript |
| ENSG00000171984 | SHLD1 | -0,483241122 | 0,715368689 | 0,2002893 | 0,699920362 | shieldin complex subunit 1 |
| ENSG00000092051 | JPH4 | -0,483286829 | 0,715346025 | 0,15062804 | 0,617119003 | junctophilin 4 |
| ENSG00000106328 | FSCN3 | -0,483288433 | 0,715345229 | 0,43687447 | 0,896866043 | fascin actin-bundling protein 3 |
| ENSG00000212195 | U3 | -0,483491551 | 0,715244522 | 0,38108906 | 0,865900094 | Small nucleolar RNA U3 |
| ENSG00000081019 | RSBN1 | -0,483721396 | 0,715130581 | 0,23741973 | 0,746907875 | round spermatid basic protein 1 |
| ENSG00000240616 | RPS6P25 | -0,483836976 | 0,715073291 | 0,3908011 | 0,870057277 | ribosomal protein S6 pseudogene 25 |
| ENSG00000241286 | RPL29P25 | -0,483874561 | 0,715054663 | 0,73638118 | 0,974353474 | ribosomal protein L29 pseudogene 25 |
| ENSG00000163738 | MTHFD2L | -0,483909153 | 0,715037518 | 0,36068491 | 0,851374334 | methylenetetrahydrofolate dehydrogenase (NADP+ dependent) 2 like |
| ENSG00000235701 | PCBP2P1 | -0,483983498 | 0,715000672 | 0,4490927 | 0,902062594 | poly(rC) binding protein 2 pseudogene 1 |
| ENSG00000188994 | ZNF292 | -0,484055754 | 0,714964862 | 0,2876648 | 0,799000743 | zinc finger protein 292 |

| | | | | | | |
|---|---|---|---|---|---|---|
| ENSG00000237491 | LINC01409 | -0,484093261 | 0,714946275 | 0,11020628 | 0,529586409 | long intergenic non-protein coding RNA 1409 |
| ENSG00000273645 | KBTBD11 | -0,484115948 | 0,714935032 | 0,69729026 | 0,968522643 | kelch repeat and BTB domain containing 11 |
| ENSG00000116852 | KIF21B | -0,48413751 | 0,714924347 | 0,24791646 | 0,760164991 | kinesin family member 21B |
| ENSG00000288078 | AL133553,2 | -0,484149884 | 0,714918215 | 0,66043725 | 0,959163386 | novel transcript |
| ENSG00000224358 | TMCO1-AS1 | -0,484323129 | 0,71483237 | 0,30277185 | 0,812344697 | TMCO1 antisense RNA 1 |
| ENSG00000254528 | AP000757,1 | -0,484512563 | 0,714738515 | 0,3423024 | 0,840048641 | novel transcript, antisense to FXYD6 |
| ENSG00000173250 | GPR151 | -0,484588706 | 0,714700793 | 0,41859951 | 0,887094957 | G protein-coupled receptor 151 |
| ENSG00000184923 | NUTM2A | -0,484924194 | 0,714534614 | 0,31550043 | 0,82082095 | NUT family member 2A |
| ENSG00000231274 | SBK3 | -0,484961575 | 0,7145161 | 0,62732233 | 0,952793934 | SH3 domain binding kinase family member 3 |
| ENSG00000235397 | EPN2-AS1 | -0,485235011 | 0,71438069 | 0,41553651 | 0,885986046 | EPN2 antisense RNA 1 |
| ENSG00000225872 | LINC01529 | -0,485331779 | 0,714332775 | 0,32158894 | 0,824933727 | long intergenic non-protein coding RNA 1529 |
| ENSG00000188315 | C3orf62 | -0,485361209 | 0,714318203 | 0,03836163 | 0,290314236 | chromosome 3 open reading frame 62 |
| ENSG00000220695 | Z86062,1 | -0,485388191 | 0,714304844 | 0,61269929 | 0,952413269 | RP42 homolog (RP42) pseudogene |
| ENSG00000140543 | DET1 | -0,48544138 | 0,714278509 | 0,15650403 | 0,627541618 | DET1 partner of COP1 E3 ubiquitin ligase |
| ENSG00000225764 | P3H2-AS1 | -0,485865451 | 0,714068582 | 0,40576598 | 0,87874173 | P3H2 antisense RNA 1 |
| ENSG00000263164 | AC087500,2 | -0,485904571 | 0,71404922 | 0,53792796 | 0,932883277 | novel transcript |
| ENSG00000225193 | RPS12P26 | -0,486149304 | 0,713928102 | 0,60985138 | 0,951801946 | ribosomal protein S12 pseudogene 26 |
| ENSG00000271614 | ATP2B1-AS1 | -0,486319259 | 0,713844003 | 0,09952948 | 0,500593598 | ATP2B1 antisense RNA 1 |
| ENSG00000188322 | SBK1 | -0,486506387 | 0,713751419 | 0,05302053 | 0,354119602 | SH3 domain binding kinase 1 |
| ENSG00000179331 | RAB39A | -0,486726643 | 0,713642459 | 0,20635359 | 0,709417043 | RAB39A, member RAS oncogene family |
| ENSG00000272980 | Z94721,2 | -0,486730776 | 0,713640414 | 0,14949565 | 0,614794108 | novel transcript |
| ENSG00000273056 | AL354694,1 | -0,486843897 | 0,71358446 | 0,65103808 | 0,957072209 | novel transcript |
| ENSG00000272514 | CFAP206 | -0,486906773 | 0,713553361 | 0,36984546 | 0,858104601 | cilia and flagella associated protein 206 |
| ENSG00000274332 | LRRC37A2 | -0,487012703 | 0,713500971 | 0,30852199 | 0,816730513 | leucine rich repeat containing 37 member A2 |
| ENSG00000261336 | EIF4BP5 | -0,487016687 | 0,713499 | 0,45206554 | 0,902062594 | eukaryotic translation initiation factor 4B pseudogene 5 |
| ENSG00000260279 | AC137932,1 | -0,487085409 | 0,713465014 | 0,23276913 | 0,741684508 | novel transcript, antisense to ANKRD11 |
| ENSG00000267198 | AC091132,4 | -0,487140348 | 0,713437845 | 0,27793965 | 0,789165846 | novel transcript |
| ENSG00000282491 | AC243807,6 | -0,487140348 | 0,713437845 | 0,27793965 | 0,789165846 | novel transcript |
| ENSG00000228336 | OR9H1P | -0,487185083 | 0,713415723 | 0,07497473 | 0,432645898 | olfactory receptor family 9 subfamily H member 1 pseudogene |
| ENSG00000259453 | AC087286,4 | -0,487218935 | 0,713398983 | 0,45544451 | 0,903545728 | novel transcript, sense intronic to AKAP13 |

| | | | | | | |
|---|---|---|---|---|---|---|
| ENSG00000250325 | IGBP1P4 | -0,487243387 | 0,713386892 | 0,75135598 | 0,975901808 | immunoglobulin (CD79A) binding protein 1 pseudogene 4 |
| ENSG00000273568 | AC131009,3 | -0,487312339 | 0,713352797 | 0,61676772 | 0,952793934 | novel transcript, sense intronic to PUS1 |
| ENSG00000143627 | PKLR | -0,487393758 | 0,71331254 | 0,64238276 | 0,956633898 | pyruvate kinase L/R |
| ENSG00000275695 | UBE2Q2P6 | -0,487659095 | 0,713181362 | 0,49373576 | 0,917289924 | ubiquitin conjugating enzyme E2 Q2 pseudogene 6 |
| ENSG00000224261 | RPSAP18 | -0,487692805 | 0,713164698 | 0,64149836 | 0,956465487 | ribosomal protein SA pseudogene 18 |
| ENSG00000026652 | AGPAT4 | -0,487724509 | 0,713149026 | 0,08138816 | 0,452922666 | 1-acylglycerol-3-phosphate O-acyltransferase 4 |
| ENSG00000172320 | OR5A1 | -0,487750532 | 0,713136162 | 0,16205004 | 0,639001241 | olfactory receptor family 5 subfamily A member 1 |
| ENSG00000277454 | AC245078,1 | -0,487969655 | 0,713027856 | 0,61367803 | 0,952413269 | PRH1-PRR4 readthrough |
| ENSG00000105549 | THEG | -0,488028196 | 0,712998924 | 0,76362198 | 0,976640468 | theg spermatid protein |
| ENSG00000232433 | GXYLT1P3 | -0,488213978 | 0,712907114 | 0,36474734 | 0,854896238 | GXYLT1 pseudogene 3 |
| ENSG00000151640 | DPYSL4 | -0,488299692 | 0,71286476 | 0,04165124 | 0,306108252 | dihydropyrimidinase like 4 |
| ENSG00000032219 | ARID4A | -0,488449911 | 0,712790537 | 0,09077283 | 0,480677819 | AT-rich interaction domain 4A |
| ENSG00000234478 | ACBD3-AS1 | -0,48869721 | 0,712668365 | 0,60360841 | 0,950352286 | ACBD3 antisense RNA 1 |
| ENSG00000174429 | ABRA | -0,489162891 | 0,712438363 | 0,6311416 | 0,953697971 | actin binding Rho activating protein |
| ENSG00000267570 | STK25P1 | -0,489771397 | 0,712137931 | 0,69502621 | 0,967509655 | serine/threonine kinase 25 pseudogene 1 |
| ENSG00000259177 | AC018946,1 | -0,489909718 | 0,712069657 | 0,71473503 | 0,971041579 | novel transcript, antisense to IQGAP1 |
| ENSG00000137713 | PPP2R1B | -0,489934616 | 0,712057368 | 0,23696831 | 0,746456145 | protein phosphatase 2 scaffold subunit Abeta |
| ENSG00000254688 | AC013549,3 | -0,490204706 | 0,711924075 | 0,61371738 | 0,952413269 | novel transcript |
| ENSG00000287525 | AL627308,3 | -0,490232288 | 0,711910464 | 0,5490359 | 0,936040289 | novel transcript, antisense to CDC42BPA |
| ENSG00000204611 | ZNF616 | -0,490233882 | 0,711909677 | 0,10991401 | 0,528750831 | zinc finger protein 616 |
| ENSG00000188828 | GLRA4 | -0,490247342 | 0,711903035 | 0,4010559 | 0,876659568 | glycine receptor alpha 4 (pseudogene) |
| ENSG00000276704 | AL442067,2 | -0,490318097 | 0,711868122 | 0,68486683 | 0,965415049 | novel transcript, sense intronic to RAP2A |
| ENSG00000257711 | AC079385,2 | -0,490331551 | 0,711861484 | 0,19222607 | 0,688548947 | novel transcript, antisense to RFX4 |
| ENSG00000229399 | AL592148,1 | -0,490392974 | 0,711831176 | 0,72956311 | 0,974009551 | prefoldin 5 (PFDN5) pseudogene |
| ENSG00000236991 | EDRF1-AS1 | -0,490471497 | 0,711792434 | 0,30384262 | 0,812957289 | EDRF1 antisense RNA 1 |
| ENSG00000273344 | PAXIP1-AS1 | -0,490473425 | 0,711791483 | 0,16594632 | 0,645193807 | PAXIP1 antisense RNA 1 (head to head) |
| ENSG00000214629 | RPSAP6 | -0,490541244 | 0,711758023 | 0,63185147 | 0,953915351 | ribosomal protein SA pseudogene 6 |
| ENSG00000223429 | AL139156,1 | -0,490599472 | 0,711729297 | 0,51237568 | 0,923445033 | transformer-2 alpha (TRA2A) pseudogene |
| ENSG00000241984 | RPL7AP2 | -0,490684556 | 0,711687324 | 0,46163016 | 0,906165604 | ribosomal protein L7a pseudogene 2 |
| ENSG00000226950 | DANCR | -0,490891457 | 0,711585266 | 0,22280095 | 0,731378504 | differentiation antagonizing non-protein coding RNA |
| ENSG00000244642 | RN7SL396P | -0,490895705 | 0,71158317 | 0,60592575 | 0,950927937 | RNA, 7SL, cytoplasmic 396, pseudogene |

| | | | | | | |
|---|---|---|---|---|---|---|
| ENSG00000182477 | OR2B8P | -0,490918404 | 0,711571975 | 0,74583476 | 0,975707496 | olfactory receptor family 2 subfamily B member 8 pseudogene |
| ENSG00000274281 | AC022929,2 | -0,490951378 | 0,711555711 | 0,05572649 | 0,364179132 | novel transcript |
| ENSG00000136206 | SPDYE1 | -0,491036263 | 0,711513846 | 0,17513763 | 0,660574619 | speedy/RINGO cell cycle regulator family member E1 |
| ENSG00000281877 | RPS6KA1 | -0,491085802 | 0,711489414 | 0,15182971 | 0,619929987 | ribosomal protein S6 kinase A1 |
| ENSG00000117676 | RPS6KA1 | -0,491085802 | 0,711489414 | 0,15182971 | 0,619929987 | ribosomal protein S6 kinase A1 |
| ENSG00000184260 | H2AC20 | -0,491139569 | 0,711462899 | 0,20575912 | 0,708191543 | H2A clustered histone 20 |
| ENSG00000286406 | AC138776,1 | -0,491147724 | 0,711458877 | 0,19059453 | 0,685949202 | novel ankyrin repeat domain family pseudogene |
| ENSG00000143190 | POU2F1 | -0,491201611 | 0,711432304 | 0,19617995 | 0,694046542 | POU class 2 homeobox 1 |
| ENSG00000225673 | AC104164,1 | -0,49126102 | 0,711403008 | 0,73646213 | 0,974353474 | transmembrane emp24 domain trafficking protein 2 (TMED2) pseudogene |
| ENSG00000168131 | OR2B2 | -0,491334508 | 0,711366772 | 0,72559079 | 0,97326334 | olfactory receptor family 2 subfamily B member 2 |
| ENSG00000283633 | AP000547,3 | -0,491391249 | 0,711338794 | 0,31590921 | 0,821327495 | novel transcript |
| ENSG00000236802 | B3GALT4 | -0,491396118 | 0,711336393 | 0,81689257 | 0,981977498 | beta-1,3-galactosyltransferase 4 |
| ENSG00000226936 | B3GALT4 | -0,491396118 | 0,711336393 | 0,81689257 | 0,981977498 | beta-1,3-galactosyltransferase 4 |
| ENSG00000235155 | B3GALT4 | -0,491396118 | 0,711336393 | 0,81689257 | 0,981977498 | beta-1,3-galactosyltransferase 4 |
| ENSG00000105750 | ZNF85 | -0,491472862 | 0,711298555 | 0,253017 | 0,76515438 | zinc finger protein 85 |
| ENSG00000203965 | EFCAB7 | -0,49169771 | 0,711187706 | 0,33298072 | 0,833301711 | EF-hand calcium binding domain 7 |
| ENSG00000230769 | Z98048,1 | -0,491717687 | 0,711177858 | 0,41241581 | 0,884523593 | riboflavin kinase (RFK) pseudogene |
| ENSG00000214783 | POLR2J4 | -0,491813485 | 0,711130636 | 0,07964193 | 0,447586774 | RNA polymerase II subunit J4, pseudogene |
| ENSG00000179938 | GOLGA8J | -0,491959304 | 0,711058763 | 0,29019408 | 0,800682868 | golgin A8 family member J |
| ENSG00000232036 | AL078645,2 | -0,492090882 | 0,710993915 | 0,38098864 | 0,865900094 | novel pseudogene |
| ENSG00000237976 | RFX5-AS1 | -0,49214235 | 0,710968551 | 0,35168411 | 0,844279508 | RFX5 antisense RNA 1 |
| ENSG00000184307 | ZDHHC23 | -0,49223153 | 0,710924604 | 0,22698036 | 0,735436234 | zinc finger DHHC-type palmitoyltransferase 23 |
| ENSG00000164061 | BSN | -0,492366954 | 0,710857873 | 0,1550604 | 0,624717575 | bassoon presynaptic cytomatrix protein |
| ENSG00000162913 | OBSCN-AS1 | -0,492409886 | 0,71083672 | 0,30312901 | 0,812669425 | OBSCN antisense RNA 1 |
| ENSG00000236213 | AC006369,1 | -0,49264449 | 0,710721136 | 0,62166043 | 0,952793934 | novel transcript |
| ENSG00000231584 | FAHD2CP | -0,49270253 | 0,710692544 | 0,1185775 | 0,550275639 | fumarylacetoacetate hydrolase domain containing 2C, pseudogene |
| ENSG00000137960 | GIPC2 | -0,492745922 | 0,710671169 | 0,60175682 | 0,949939978 | GIPC PDZ domain containing family member 2 |
| ENSG00000282675 | AC245078,5 | -0,492788659 | 0,710650117 | 0,22982333 | 0,738230413 | taste receptor, type 2, member 18 pseudogene |
| ENSG00000207174 | SNORD116-15 | -0,492802445 | 0,710643326 | 0,23071064 | 0,739234644 | small nucleolar RNA, C/D box 116-15 |

| | | | | | | |
|---|---|---|---|---|---|---|
| ENSG00000119973 | PRLHR | -0,492943093 | 0,71057405 | 0,73098478 | 0,974353474 | prolactin releasing hormone receptor |
| ENSG00000257494 | AC004217,1 | -0,49294813 | 0,710571569 | 0,66245183 | 0,959897678 | novel transcript |
| ENSG00000246214 | AC024588,1 | -0,492967937 | 0,710561813 | 0,27949663 | 0,790302992 | novel transcript, antisense to FAM134B |
| ENSG00000231999 | LRRC8C-DT | -0,49306606 | 0,710513487 | 0,09400698 | 0,487240447 | LRRC8C divergent transcript |
| ENSG00000286909 | AC024560,4 | -0,493134573 | 0,710479746 | 0,23875 | 0,747998807 | novel transcript |
| ENSG00000105948 | TTC26 | -0,493241071 | 0,710427301 | 0,23764647 | 0,746907875 | tetratricopeptide repeat domain 26 |
| ENSG00000168243 | GNG4 | -0,493247323 | 0,710424222 | 0,0493695 | 0,338781394 | G protein subunit gamma 4 |
| ENSG00000204695 | OR14J1 | -0,493251425 | 0,710422202 | 0,07541195 | 0,433690715 | olfactory receptor family 14 subfamily J member 1 |
| ENSG00000256436 | TAS2R31 | -0,493266844 | 0,71041461 | 0,36799756 | 0,857401877 | taste 2 receptor member 31 |
| ENSG00000263097 | TAS2R31 | -0,493266844 | 0,71041461 | 0,36799756 | 0,857401877 | taste 2 receptor member 31 |
| ENSG00000275368 | TCTE3 | -0,493305681 | 0,710395486 | 0,30527731 | 0,814211732 | t-complex-associated-testis-expressed 3 |
| ENSG00000134326 | CMPK2 | -0,493479009 | 0,710310143 | 0,27909712 | 0,790013761 | cytidine/uridine monophosphate kinase 2 |
| ENSG00000267547 | AC060766,4 | -0,49350528 | 0,710297208 | 0,19299042 | 0,689849199 | novel transcript |
| ENSG00000277157 | H4C4 | -0,493555931 | 0,710272271 | 0,17507205 | 0,660486623 | H4 clustered histone 4 |
| ENSG00000256021 | ELOCP31 | -0,493605043 | 0,710248092 | 0,6774095 | 0,963580269 | elongin C pseudogene 31 |
| ENSG00000175779 | LINC02694 | -0,493630362 | 0,710235628 | 0,48701452 | 0,914593485 | long intergenic non-protein coding RNA 2694 |
| ENSG00000146001 | PCDHB18P | -0,493663834 | 0,71021915 | 0,16627883 | 0,645941997 | protocadherin beta 18 pseudogene |
| ENSG00000244294 | RN7SL740P | -0,493667843 | 0,710217176 | 0,71947748 | 0,971869926 | RNA, 7SL, cytoplasmic 740, pseudogene |
| ENSG00000274315 | AC009318,3 | -0,493753117 | 0,710175198 | 0,34129274 | 0,84003907 | novel transcript, antisense to FAR2 |
| ENSG00000242697 | RPL5P12 | -0,493782951 | 0,710160513 | 0,33421382 | 0,83453565 | ribosomal protein L5 pseudogene 12 |
| ENSG00000205534 | SMG1P2 | -0,493799352 | 0,710152439 | 0,41506132 | 0,885766354 | SMG1 pseudogene 2 |
| ENSG00000250517 | LDHAL6CP | -0,49384471 | 0,710130113 | 0,4742702 | 0,910673352 | lactate dehydrogenase A like 6C, pseudogene |
| ENSG00000259605 | AC074212,1 | -0,493922117 | 0,710092012 | 0,18843964 | 0,682805588 | novel transcript, antisense to SIX5 |
| ENSG00000225351 | ZNF311 | -0,493980329 | 0,710063361 | 0,54765987 | 0,935404974 | zinc finger protein 311 |
| ENSG00000235178 | ZNF311 | -0,493980329 | 0,710063361 | 0,54765987 | 0,935404974 | zinc finger protein 311 |
| ENSG00000235589 | ZNF311 | -0,493980329 | 0,710063361 | 0,54765987 | 0,935404974 | zinc finger protein 311 |
| ENSG00000258526 | AL049828,1 | -0,494235367 | 0,709937848 | 0,0505437 | 0,343953009 | novel transcript |
| ENSG00000285294 | LINC00842 | -0,494272943 | 0,709919357 | 0,66620551 | 0,961022895 | long intergenic non-protein coding RNA 842 |
| ENSG00000234945 | GTF3C2-AS1 | -0,494317748 | 0,70989731 | 0,35019779 | 0,843789477 | GTF3C2 antisense RNA 1 |
| ENSG00000046889 | PREX2 | -0,494375142 | 0,709869069 | 0,24010902 | 0,749945225 | phosphatidylinositol-3,4,5-trisphosphate dependent Rac exchange factor 2 |
| ENSG00000215018 | COL28A1 | -0,494375311 | 0,709868986 | 0,19768967 | 0,696048902 | collagen type XXVIII alpha 1 chain |

| | | | | | | |
|---|---|---|---|---|---|---|
| ENSG00000287751 | AC015818,10 | -0,494441677 | 0,709836331 | 0,73779331 | 0,974353474 | novel family with sequence similarity 106 pseudogene |
| ENSG00000229591 | AC006017,1 | -0,494584889 | 0,709765872 | 0,41451855 | 0,885579089 | novel transcript |
| ENSG00000243364 | EFNA4 | -0,494611635 | 0,709752714 | 0,08658694 | 0,468880561 | ephrin A4 |
| ENSG00000237489 | C10orf143 | -0,494764456 | 0,709677536 | 0,24213672 | 0,752479103 | chromosome 10 open reading frame 143 |
| ENSG00000231884 | NDUFB1P1 | -0,49483183 | 0,709644394 | 0,7417238 | 0,975176043 | NADH:ubiquinone oxidoreductase subunit B1 pseudogene 1 |
| ENSG00000111731 | C2CD5 | -0,494898493 | 0,709611604 | 0,21325218 | 0,720652258 | C2 calcium dependent domain containing 5 |
| ENSG00000243314 | AC106707,2 | -0,494949961 | 0,709586289 | 0,62715377 | 0,952793934 | ribosomal protein S12 (RPS12) pseudogene |
| ENSG00000214198 | TTC41P | -0,495060646 | 0,709531851 | 0,35716686 | 0,848693232 | tetratricopeptide repeat domain 41, pseudogene |
| ENSG00000203663 | OR2L2 | -0,495109643 | 0,709507755 | 0,11191069 | 0,534735697 | olfactory receptor family 2 subfamily L member 2 |
| ENSG00000259986 | AC103876,1 | -0,495127208 | 0,709499116 | 0,52029333 | 0,926865961 | novel transcript, antisense to HDGFRP3 |
| ENSG00000165821 | SALL2 | -0,495191458 | 0,70946752 | 0,07034755 | 0,417680252 | spalt like transcription factor 2 |
| ENSG00000247271 | ZBED5-AS1 | -0,495295929 | 0,709416146 | 0,09382854 | 0,486996636 | ZBED5 antisense RNA 1 |
| ENSG00000214626 | POLR3DP1 | -0,495307172 | 0,709410618 | 0,62486752 | 0,952793934 | RNA polymerase III subunit D pseudogene 1 |
| ENSG00000236305 | SLC12A9-AS1 | -0,495478893 | 0,709326183 | 0,79899196 | 0,980161849 | SLC12A9 antisense RNA 1 |
| ENSG00000197302 | ZNF720 | -0,495489624 | 0,709320907 | 0,21486779 | 0,722170133 | zinc finger protein 720 |
| ENSG00000104967 | NOVA2 | -0,495567518 | 0,70928261 | 0,02044219 | 0,189588991 | NOVA alternative splicing regulator 2 |
| ENSG00000279566 | ZNF43 | -0,495609828 | 0,709261809 | 0,43331555 | 0,895877116 | zinc finger protein 43 |
| ENSG00000275263 | AC135048,3 | -0,495678452 | 0,709228073 | 0,62493869 | 0,952793934 | novel transcript |
| ENSG00000178947 | SMIM10L2A | -0,495686266 | 0,709224232 | 0,04836207 | 0,335083246 | small integral membrane protein 10 like 2A |
| ENSG00000105737 | GRIK5 | -0,495830464 | 0,709153348 | 0,05494241 | 0,362021772 | glutamate ionotropic receptor kainate type subunit 5 |
| ENSG00000255151 | GLYATL1B | -0,495848455 | 0,709144504 | 0,72300788 | 0,972825184 | glycine-N-acyltransferase like 1B |
| ENSG00000261216 | AC007216,2 | -0,495973895 | 0,709082848 | 0,63496186 | 0,955158181 | novel transcript, antisense to GSPT1 |
| ENSG00000186777 | ZNF732 | -0,496245572 | 0,708949332 | 0,19060196 | 0,685949202 | zinc finger protein 732 |
| ENSG00000172766 | NAA16 | -0,496261998 | 0,70894126 | 0,22641025 | 0,735064413 | N-alpha-acetyltransferase 16, NatA auxiliary subunit |
| ENSG00000274775 | Z82195,2 | -0,496393332 | 0,708876725 | 0,75685657 | 0,975901808 | fibrous sheath interacting protein 2 (FSIP2) pseudogene |
| ENSG00000184117 | NIPSNAP1 | -0,496468214 | 0,708839933 | 0,12545762 | 0,565907826 | nipsnap homolog 1 |
| ENSG00000175868 | CALCB | -0,496528646 | 0,708810242 | 0,54910667 | 0,936040289 | calcitonin related polypeptide beta |
| ENSG00000105717 | PBX4 | -0,496632928 | 0,708759008 | 0,17458312 | 0,660057513 | PBX homeobox 4 |
| ENSG00000273013 | AC117490,2 | -0,496640257 | 0,708755408 | 0,6090649 | 0,951793902 | novel transcript, antisense to RNF168 |

| | | | | | | |
|---|---|---|---|---|---|---|
| ENSG00000282509 | ULK4P3 | -0,496679417 | 0,70873617 | 0,29531673 | 0,805426883 | ULK4 pseudogene 3 |
| ENSG00000178081 | ULK4P3 | -0,496679417 | 0,70873617 | 0,29531673 | 0,805426883 | ULK4 pseudogene 3 |
| ENSG00000006459 | KDM7A | -0,496744729 | 0,708704086 | 0,26105874 | 0,773103493 | lysine demethylase 7A |
| ENSG00000227379 | PPIAP2 | -0,496911481 | 0,708622176 | 0,60065632 | 0,949485239 | peptidylprolyl isomerase A pseudogene 2 |
| ENSG00000215160 | OR7E122P | -0,496913658 | 0,708621107 | 0,06439373 | 0,397835213 | olfactory receptor family 7 subfamily E member 122 pseudogene |
| ENSG00000274893 | AL136317,2 | -0,49694894 | 0,708603777 | 0,32332977 | 0,825527324 | novel transcript |
| ENSG00000273485 | AL139339,2 | -0,496994414 | 0,708581442 | 0,56830142 | 0,94163744 | novel transcript, antisense to CALHM2 |
| ENSG00000164621 | SMAD5-AS1 | -0,497067715 | 0,708545441 | 0,34359129 | 0,840901506 | SMAD5 antisense RNA 1 |
| ENSG00000166922 | SCG5 | -0,497088037 | 0,70853546 | 0,2379003 | 0,747090777 | secretogranin V |
| ENSG00000260423 | LINC02367 | -0,4971684 | 0,708495994 | 0,24324325 | 0,753617768 | long intergenic non-protein coding RNA 2367 |
| ENSG00000198003 | CCDC151 | -0,497237786 | 0,70846192 | 0,26189381 | 0,773825557 | coiled-coil domain containing 151 |
| ENSG00000153404 | PLEKHG4B | -0,497426926 | 0,708369045 | 0,11583488 | 0,543681398 | pleckstrin homology and RhoGEF domain containing G4B |
| ENSG00000287195 | AC131568,1 | -0,497462463 | 0,708351596 | 0,41077746 | 0,882835102 | novel transcript, antisense to FAM49B |
| ENSG00000123080 | CDKN2C | -0,497529936 | 0,708318468 | 0,32501683 | 0,826983176 | cyclin dependent kinase inhibitor 2C |
| ENSG00000224802 | TUBB4BP2 | -0,497539346 | 0,708313848 | 0,63540271 | 0,955314044 | TUBB4B pseudogene 2 |
| ENSG00000286996 | AC099332,2 | -0,49758563 | 0,708291125 | 0,6504693 | 0,956899963 | novel transcript |
| ENSG00000224223 | VSTM2A-OT1 | -0,497628347 | 0,708270153 | 0,69483226 | 0,967417311 | VSTM2A overlapping transcript 1 |
| ENSG00000111490 | TBC1D30 | -0,497746336 | 0,70821223 | 0,18102732 | 0,670752303 | TBC1 domain family member 30 |
| ENSG00000215417 | MIR17HG | -0,497804441 | 0,708183708 | 0,3911821 | 0,870083449 | miR-17-92a-1 cluster host gene |
| ENSG00000180178 | FAR2P1 | -0,498007415 | 0,70808408 | 0,11425555 | 0,540242823 | fatty acyl-CoA reductase 2 pseudogene 1 |
| ENSG00000136504 | KAT7 | -0,49810958 | 0,708033938 | 0,07255855 | 0,424727653 | lysine acetyltransferase 7 |
| ENSG00000254141 | AC103853,2 | -0,498233363 | 0,707973192 | 0,44569224 | 0,901173082 | novel transcript |
| ENSG00000164756 | SLC30A8 | -0,498552587 | 0,707816557 | 0,63655584 | 0,95557086 | solute carrier family 30 member 8 |
| ENSG00000164675 | IQUB | -0,498729077 | 0,707729972 | 0,21195856 | 0,718572144 | IQ motif and ubiquitin domain containing |
| ENSG00000259712 | AC023906,5 | -0,498922627 | 0,70763503 | 0,4311884 | 0,89530663 | novel transcript, antisense to MAPK6 |
| ENSG00000286342 | AC073210,3 | -0,499070208 | 0,707562646 | 0,30544428 | 0,814326218 | novel transcript |
| ENSG00000260918 | AC107398,3 | -0,499095335 | 0,707550323 | 0,52164993 | 0,927472429 | novel transcript, overlapping GABRB1 |
| ENSG00000253875 | AC013643,2 | -0,499385917 | 0,707407825 | 0,26396269 | 0,775030955 | novel transcript |
| ENSG00000164707 | SLC13A4 | -0,499624499 | 0,707290849 | 0,14892656 | 0,613619689 | solute carrier family 13 member 4 |
| ENSG00000146267 | FAXC | -0,499758579 | 0,707225118 | 0,02985799 | 0,246365813 | failed axon connections homolog, metaxin like GST domain containing |

| | | | | | | |
|---|---|---|---|---|---|---|
| ENSG00000244313 | AC024293,1 | -0,499866043 | 0,70717244 | 0,60239056 | 0,950205821 | ribosomal protein S10 (RPS10) pseudogene |
| ENSG00000279897 | BIRC6-AS2 | -0,500002704 | 0,707105456 | 0,39025928 | 0,870057277 | BIRC6 antisense RNA 2 |
| ENSG00000270882 | H4C14 | -0,500002994 | 0,707105314 | 0,09496625 | 0,489386708 | H4 clustered histone 14 |
| ENSG00000205822 | TPTE2P6 | -0,500043453 | 0,707085484 | 0,36205421 | 0,852161624 | transmembrane phosphoinositide 3-phosphatase and tensin homolog 2 pseudogene 6 |
| ENSG00000085840 | ORC1 | -0,500069134 | 0,707072897 | 0,08598543 | 0,467107732 | origin recognition complex subunit 1 |
| ENSG00000185864 | NPIPB4 | -0,500113764 | 0,707051024 | 0,03442812 | 0,270646388 | nuclear pore complex interacting protein family member B4 |
| ENSG00000249175 | AC008534,1 | -0,500583339 | 0,706820928 | 0,58508758 | 0,944990341 | novel transcript |
| ENSG00000170500 | LONRF2 | -0,500625523 | 0,706800261 | 0,0404515 | 0,300358836 | LON peptidase N-terminal domain and ring finger 2 |
| ENSG00000275544 | BDH1 | -0,500703229 | 0,706762192 | 0,09524335 | 0,489967108 | 3-hydroxybutyrate dehydrogenase 1 |
| ENSG00000254900 | AC091053,2 | -0,500736259 | 0,706746011 | 0,57142124 | 0,942436428 | novel transcript |
| ENSG00000172687 | ZNF738 | -0,50075555 | 0,706736561 | 0,06124339 | 0,385254164 | zinc finger protein 738 |
| ENSG00000213612 | FAM220CP | -0,5008641 | 0,706683387 | 0,70943319 | 0,970411233 | family with sequence similarity 220 member C, pseudogene |
| ENSG00000276541 | TAS2R14 | -0,500968008 | 0,706632491 | 0,6324618 | 0,954263106 | taste 2 receptor member 14 |
| ENSG00000286830 | AC105052,5 | -0,501008948 | 0,706612439 | 0,48939621 | 0,915936198 | novel transcript |
| ENSG00000272374 | Z97832,2 | -0,501016557 | 0,706608712 | 0,0365409 | 0,281313676 | novel transcript, antisense to SCUBE3 |
| ENSG00000223861 | AL365436,1 | -0,501034451 | 0,706599948 | 0,67941478 | 0,964375386 | ribosomal protein S10 (RPS10) pseudogene |
| ENSG00000272232 | RN7SL726P | -0,501131684 | 0,706552327 | 0,65396509 | 0,957444211 | RNA, 7SL, cytoplasmic 726, pseudogene |
| ENSG00000288551 | AL023806,5 | -0,501152948 | 0,706541913 | 0,33981318 | 0,838611198 | novel transcript, antisense to SHPRH |
| ENSG00000270900 | NUDT19P4 | -0,501172922 | 0,706532132 | 0,62835646 | 0,952793934 | NUDT19 pseudogene 4 |
| ENSG00000231663 | COA6-AS1 | -0,501402462 | 0,706419728 | 0,4290822 | 0,893964132 | COA6 antisense RNA 1 |
| ENSG00000225083 | GRTP1-AS1 | -0,501470933 | 0,706386201 | 0,53270444 | 0,931291085 | GRTP1 antisense RNA 1 |
| ENSG00000203877 | RIPPLY2 | -0,50166837 | 0,706289537 | 0,53074197 | 0,931265667 | ripply transcriptional repressor 2 |
| ENSG00000286144 | Z97056,2 | -0,501710375 | 0,706268973 | 0,74327577 | 0,975310206 | novel transcript |
| ENSG00000164953 | TMEM67 | -0,501855899 | 0,706197736 | 0,212621 | 0,719437301 | transmembrane protein 67 |
| ENSG00000167815 | PRDX2 | -0,501969765 | 0,706142001 | 0,06386463 | 0,395770809 | peroxiredoxin 2 |
| ENSG00000228284 | HLA-DQA1 | -0,502023199 | 0,706115847 | 0,0902126 | 0,478874121 | major histocompatibility complex, class II, DQ alpha 1 |
| ENSG00000250497 | AC007126,1 | -0,502052238 | 0,706101635 | 0,03606401 | 0,27832763 | novel transcript |
| ENSG00000253551 | AC021915,1 | -0,502060539 | 0,706097572 | 0,69730517 | 0,968522643 | novel transcript |
| ENSG00000197140 | ADAM32 | -0,502149474 | 0,706054046 | 0,08398831 | 0,461230231 | ADAM metallopeptidase domain 32 |

| | | | | | | |
|---|---|---|---|---|---|---|
| ENSG00000184903 | IMMP2L | -0,502178974 | 0,706039609 | 0,09685221 | 0,494365317 | inner mitochondrial membrane peptidase subunit 2 |
| ENSG00000187134 | AKR1C1 | -0,502181036 | 0,7060386 | 0,15644019 | 0,627446596 | aldo-keto reductase family 1 member C1 |
| ENSG00000224549 | AC017067,1 | -0,502319531 | 0,705970825 | 0,67020004 | 0,962026636 | novel transcript |
| ENSG00000186994 | KANK3 | -0,50235373 | 0,70595409 | 0,58011175 | 0,943828882 | KN motif and ankyrin repeat domains 3 |
| ENSG00000269535 | AC010320,4 | -0,502362624 | 0,705949738 | 0,30233037 | 0,812035949 | novel transcript, antisense to ZNF480 and ZNF610 |
| ENSG00000224356 | AL356966,1 | -0,502417827 | 0,705922726 | 0,22073626 | 0,728572782 | novel transcript |
| ENSG00000287933 | AL357141,1 | -0,502491671 | 0,705886595 | 0,06268845 | 0,391195976 | novel transcript, antisense to FRK |
| ENSG00000255135 | AP002360,1 | -0,502814315 | 0,705728748 | 0,14028865 | 0,597018607 | novel transcript |
| ENSG00000161180 | CCDC116 | -0,502867381 | 0,70570279 | 0,37487743 | 0,861771195 | coiled-coil domain containing 116 |
| ENSG00000220785 | MTMR9LP | -0,503067067 | 0,705605119 | 0,23837918 | 0,747813944 | myotubularin related protein 9 like, pseudogene |
| ENSG00000285103 | AL451123,1 | -0,503083481 | 0,705597092 | 0,1271298 | 0,569506518 | novel transcript |
| ENSG00000182218 | HHIPL1 | -0,503096971 | 0,705590494 | 0,19215999 | 0,688538797 | HHIP like 1 |
| ENSG00000140285 | FGF7 | -0,503109098 | 0,705584563 | 0,38838879 | 0,869820053 | fibroblast growth factor 7 |
| ENSG00000196482 | ESRRG | -0,503198395 | 0,705540891 | 0,25027253 | 0,762441549 | estrogen related receptor gamma |
| ENSG00000272354 | AC092354,1 | -0,503205277 | 0,705537526 | 0,64806842 | 0,956899963 | novel transcript |
| ENSG00000232934 | AL157786,1 | -0,503206942 | 0,705536712 | 0,53281515 | 0,931291085 | novel transcript, antisense to ACSL5 |
| ENSG00000274641 | H2BC17 | -0,503311215 | 0,705485719 | 0,15026822 | 0,615957904 | H2B clustered histone 17 |
| ENSG00000273062 | AL449106,1 | -0,503369325 | 0,705457304 | 0,37595915 | 0,862424749 | novel transcript, antisense to RALGPS2 |
| ENSG00000269938 | AC068790,2 | -0,503528655 | 0,705379398 | 0,26299889 | 0,77453036 | novel transcript, sense intronic to CCDC92 |
| ENSG00000250456 | LINC02260 | -0,503631076 | 0,705329323 | 0,56361089 | 0,940865625 | long intergenic non-protein coding RNA 2260 |
| ENSG00000270314 | AC007161,2 | -0,503633744 | 0,705328019 | 0,54579154 | 0,935146964 | 5-azacytidine induced 2 (AZI2) pseudogene |
| ENSG00000067715 | SYT1 | -0,503751145 | 0,705270624 | 0,08231619 | 0,456410098 | synaptotagmin 1 |
| ENSG00000114126 | TFDP2 | -0,503758182 | 0,705267184 | 0,16013871 | 0,635191158 | transcription factor Dp-2 |
| ENSG00000263884 | AP000845,1 | -0,503948951 | 0,705173932 | 0,26113426 | 0,7731889 | novel transcript |
| ENSG00000197980 | LEKR1 | -0,504112934 | 0,705093783 | 0,35559889 | 0,846892875 | leucine, glutamate and lysine rich 1 |
| ENSG00000249307 | LINC01088 | -0,5041916 | 0,705055338 | 0,07455557 | 0,431569222 | long intergenic non-protein coding RNA 1088 |
| ENSG00000269985 | AL021328,1 | -0,504241183 | 0,705031107 | 0,44780985 | 0,902062594 | novel transcript, antisense to FARS2 |
| ENSG00000223572 | CKMT1A | -0,5043051 | 0,704999872 | 0,32283471 | 0,824955371 | creatine kinase, mitochondrial 1A |
| ENSG00000008311 | AASS | -0,504364579 | 0,704970807 | 0,19632027 | 0,694046542 | aminoadipate-semialdehyde synthase |
| ENSG00000267251 | AC139100,1 | -0,504366644 | 0,704969798 | 0,2601473 | 0,772692192 | novel transcript, antisense to PARD6G |
| ENSG00000257568 | AC079035,1 | -0,504379084 | 0,704963719 | 0,43417467 | 0,896272827 | novel transcript, antisense to MON2 |
| ENSG00000277299 | AC084876,1 | -0,504397613 | 0,704954665 | 0,64922584 | 0,956899963 | novel transcript, antisense to GIT2 |
| ENSG00000272656 | AC024933,1 | -0,504453016 | 0,704927593 | 0,65052988 | 0,956899963 | novel transcript, antisense to MRPS22 |

| | | | | | | |
|---|---|---|---|---|---|---|
| ENSG00000116793 | PHTF1 | -0,504515431 | 0,704897097 | 0,17377359 | 0,65876772 | putative homeodomain transcription factor 1 |
| ENSG00000143194 | MAEL | -0,504569326 | 0,704870764 | 0,36840555 | 0,857548638 | maelstrom spermatogenic transposon silencer |
| ENSG00000227354 | RBM26-AS1 | -0,504643989 | 0,704834287 | 0,09661762 | 0,493760245 | RBM26 antisense RNA 1 |
| ENSG00000278878 | CU638689,1 | -0,504848977 | 0,704734146 | 0,1551313 | 0,624800779 | novel transcript |
| ENSG00000176714 | CCDC121 | -0,504939592 | 0,704689883 | 0,33107526 | 0,832186595 | coiled-coil domain containing 121 |
| ENSG00000224382 | LINC00703 | -0,50496948 | 0,704675285 | 0,46988194 | 0,908378389 | long intergenic non-protein coding RNA 703 |
| ENSG00000170430 | MGMT | -0,504972065 | 0,704674022 | 0,12690087 | 0,568970141 | O-6-methylguanine-DNA methyltransferase |
| ENSG00000260804 | LINC01963 | -0,505103911 | 0,704609626 | 0,2023082 | 0,702383517 | long intergenic non-protein coding RNA 1963 |
| ENSG00000234377 | OBI1-AS1 | -0,505294944 | 0,704516332 | 0,44797814 | 0,902062594 | OBI1 antisense RNA 1 |
| ENSG00000162779 | AXDND1 | -0,50534023 | 0,704494218 | 0,60004929 | 0,949340299 | axonemal dynein light chain domain containing 1 |
| ENSG00000243746 | EEF1A1P10 | -0,505414887 | 0,704457762 | 0,64029888 | 0,956251063 | eukaryotic translation elongation factor 1 alpha 1 pseudogene 10 |
| ENSG00000241787 | MTND4P16 | -0,505478549 | 0,704426677 | 0,68721918 | 0,965844929 | MT-ND4 pseudogene 16 |
| ENSG00000104808 | DHDH | -0,505533614 | 0,704399791 | 0,67244066 | 0,962290553 | dihydrodiol dehydrogenase |
| ENSG00000213123 | TCTEX1D2 | -0,505548318 | 0,704392611 | 0,19061836 | 0,685949202 | Tctex1 domain containing 2 |
| ENSG00000128617 | OPN1SW | -0,505617367 | 0,7043589 | 0,64325681 | 0,956633898 | opsin 1, short wave sensitive |
| ENSG00000181322 | NME9 | -0,505735495 | 0,704301229 | 0,11803973 | 0,549146283 | NME/NM23 family member 9 |
| ENSG00000217227 | AL079341,1 | -0,505853646 | 0,704243552 | 0,56820013 | 0,94163744 | TRK-fused gene (TFG) pseudogene |
| ENSG00000135097 | MSI1 | -0,505919529 | 0,704211392 | 0,02995598 | 0,246718575 | musashi RNA binding protein 1 |
| ENSG00000285567 | AC074051,3 | -0,505973271 | 0,70418516 | 0,3180677 | 0,822980102 | novel transcript |
| ENSG00000108639 | SYNGR2 | -0,506098014 | 0,704124275 | 0,08719726 | 0,470802426 | synaptogyrin 2 |
| ENSG00000253103 | LINC01609 | -0,506357412 | 0,703997684 | 0,03559326 | 0,276263058 | long intergenic non-protein coding RNA 1609 |
| ENSG00000260372 | AQP4-AS1 | -0,50640342 | 0,703975234 | 0,45170658 | 0,902062594 | AQP4 antisense RNA 1 |
| ENSG00000257391 | AC126763,1 | -0,50650513 | 0,703925605 | 0,39921069 | 0,875433675 | microRNA 3180-4 |
| ENSG00000224769 | MUC20P1 | -0,506589639 | 0,703884373 | 0,65649379 | 0,957444211 | mucin 20, cell surface associated pseudogene 1 |
| ENSG00000259616 | AC104574,2 | -0,50689536 | 0,703735228 | 0,73870301 | 0,974353474 | novel transcript |
| ENSG00000123405 | NFE2 | -0,507150721 | 0,703610676 | 0,48299262 | 0,91363215 | nuclear factor, erythroid 2 |
| ENSG00000134532 | SOX5 | -0,50761056 | 0,703386446 | 0,21183817 | 0,718397708 | SRY-box transcription factor 5 |
| ENSG00000271714 | AC010501,2 | -0,507626374 | 0,703378736 | 0,35819033 | 0,849333518 | novel transcript, antisense to FAM169A |
| ENSG00000236914 | LINC01852 | -0,507696388 | 0,703344601 | 0,12444143 | 0,563764231 | long intergenic non-protein coding RNA 1852 |
| ENSG00000196741 | LINC01560 | -0,507715801 | 0,703335138 | 0,29969963 | 0,809696605 | long intergenic non-protein coding RNA 1560 |
| ENSG00000236018 | AC004898,1 | -0,507744096 | 0,703321344 | 0,19579008 | 0,69375328 | UBX domain protein 2B (UBXN2B) pseudogene |
| ENSG00000184788 | SATL1 | -0,507878297 | 0,703255923 | 0,06932092 | 0,415287103 | spermidine/spermine N1-acetyl transferase like 1 |

| | | | | | | |
|---|---|---|---|---|---|---|
| ENSG00000241933 | DENND6A-DT | -0,508259439 | 0,703070156 | 0,63921835 | 0,956018518 | DENND6A divergent transcript |
| ENSG00000261773 | AC244090,2 | -0,508273249 | 0,703063426 | 0,62974165 | 0,952812183 | novel transcript, overlapping to FAM3A |
| ENSG00000185433 | LINC00158 | -0,508300369 | 0,70305021 | 0,60753741 | 0,950993594 | long intergenic non-protein coding RNA 158 |
| ENSG00000285713 | AC098588,2 | -0,508362455 | 0,703019955 | 0,51077891 | 0,922693987 | novel transcript |
| ENSG00000254598 | CSNK2A3 | -0,50839427 | 0,703004452 | 0,48838055 | 0,915394579 | casein kinase 2 alpha 3 |
| ENSG00000158528 | PPP1R9A | -0,508397008 | 0,703003117 | 0,13885885 | 0,594163262 | protein phosphatase 1 regulatory subunit 9A |
| ENSG00000248816 | AC079226,2 | -0,508724046 | 0,702843775 | 0,57343519 | 0,942475308 | novel transcript |
| ENSG00000229339 | AC093414,1 | -0,508882141 | 0,702766759 | 0,54232512 | 0,933491609 | ribosomal protein L7a (RPL7A) pseudogene |
| ENSG00000259216 | AC084757,1 | -0,508932969 | 0,702742 | 0,56383639 | 0,940865625 | oxidation resistance 1 (OXR1) pseudogene |
| ENSG00000248496 | ZBED9 | -0,509032929 | 0,702693311 | 0,24167411 | 0,751960515 | zinc finger BED-type containing 9 |
| ENSG00000225180 | PVALEF | -0,509035593 | 0,702692014 | 0,75131882 | 0,975901808 | parvalbumin like EF-hand containing |
| ENSG00000225539 | LINC01821 | -0,509070712 | 0,702674909 | 0,08729361 | 0,470988771 | long intergenic non-protein coding RNA 1821 |
| ENSG00000276486 | AC022960,3 | -0,509100042 | 0,702660623 | 0,56334468 | 0,940865625 | family with sequence similarity 76, member A (FAM76A) pseudogene |
| ENSG00000225299 | AL139240,1 | -0,509412125 | 0,70250864 | 0,61353014 | 0,952413269 | novel transcript |
| ENSG00000261879 | AC087500,1 | -0,509432173 | 0,702498878 | 0,29140473 | 0,802058262 | novel transcript, antisense to C17orf87 |
| ENSG00000185250 | PPIL6 | -0,509514062 | 0,702459005 | 0,07537755 | 0,433690715 | peptidylprolyl isomerase like 6 |
| ENSG00000282715 | ANKRD20A9P | -0,509763607 | 0,70233751 | 0,5948961 | 0,947469769 | ankyrin repeat domain 20 family member A9, pseudogene |
| ENSG00000285329 | AC025283,2 | -0,509889268 | 0,702276338 | 0,72059304 | 0,972142678 | novel protein |
| ENSG00000197935 | ZNF311 | -0,509912431 | 0,702265063 | 0,52685654 | 0,92927181 | zinc finger protein 311 |
| ENSG00000184349 | EFNA5 | -0,509956824 | 0,702243454 | 0,25615703 | 0,768274511 | ephrin A5 |
| ENSG00000159958 | TNFRSF13C | -0,509979414 | 0,702232458 | 0,19758286 | 0,696048902 | TNF receptor superfamily member 13C |
| ENSG00000285696 | AP002433,2 | -0,510036595 | 0,702204626 | 0,37359477 | 0,861319894 | novel transcript, antisense to ACAT1 |
| ENSG00000260664 | AC004158,1 | -0,510151198 | 0,702148847 | 0,20921736 | 0,713678247 | novel transcript |
| ENSG00000253369 | AC022034,3 | -0,510253041 | 0,702099283 | 0,31289373 | 0,818889268 | novel transcript |
| ENSG00000281501 | SEPSECS-AS1 | -0,510372808 | 0,702040999 | 0,06475563 | 0,399077493 | SEPSECS antisense RNA 1 (head to head) |
| ENSG00000250850 | AL161781,2 | -0,51048 | 0,70198884 | 0,69149777 | 0,967201016 | novel transcript, antisense to PAX5 |
| ENSG00000287223 | Z94277,1 | -0,510512513 | 0,70197302 | 0,05377965 | 0,357812498 | novel transcript |
| ENSG00000270462 | AC005034,2 | -0,510676595 | 0,701893187 | 0,46626337 | 0,907940653 | novel transcript, antisense to C2orf3 |
| ENSG00000287679 | AL355347,1 | -0,510716696 | 0,701873677 | 0,02613912 | 0,225624883 | novel transcript, antisense to KHDRBS2 |
| ENSG00000273722 | TMC4 | -0,510921481 | 0,701774056 | 0,54920288 | 0,936040289 | transmembrane channel like 4 |
| ENSG00000106537 | TSPAN13 | -0,510948646 | 0,701760842 | 0,30921058 | 0,816845699 | tetraspanin 13 |

| | | | | | | |
|---|---|---|---|---|---|---|
| ENSG00000139915 | MDGA2 | -0,511017433 | 0,701727383 | 0,16822841 | 0,649838265 | MAM domain containing glycosylphosphatidylinositol anchor 2 |
| ENSG00000245498 | AP000866,1 | -0,511021476 | 0,701725417 | 0,18843847 | 0,682805588 | novel transcript, antisense to C11orf61 |
| ENSG00000287416 | AC093746,1 | -0,511061674 | 0,701705865 | 0,57174999 | 0,942436428 | novel transcript |
| ENSG00000279726 | AC005609,5 | -0,51125286 | 0,701612881 | 0,25338573 | 0,765172364 | novel transcript, antisense to PCDHA5, PCDHA4, PCDHA3, PCDHA2 and PCDHA1 |
| ENSG00000152932 | RAB3C | -0,51127426 | 0,701602474 | 0,28982456 | 0,800629762 | RAB3C, member RAS oncogene family |
| ENSG00000122432 | SPATA1 | -0,511336835 | 0,701572044 | 0,23662196 | 0,746041653 | spermatogenesis associated 1 |
| ENSG00000277368 | AL138966,2 | -0,511449701 | 0,70151716 | 0,62734396 | 0,952793934 | novel transcript, antisense to ring finger protein (C3H2C3 type) 6 |
| ENSG00000278019 | AC245690,1 | -0,511578108 | 0,701454724 | 0,69774682 | 0,968522643 | glucuronidase, beta (GUSB) pseudogene |
| ENSG00000288480 | AC270129,4 | -0,51165206 | 0,701418769 | 0,38377575 | 0,868000076 | |
| ENSG00000099954 | CECR2 | -0,511684005 | 0,701403237 | 0,06579877 | 0,402522813 | CECR2 histone acetyl-lysine reader |
| ENSG00000183831 | ANKRD45 | -0,511736025 | 0,701377947 | 0,12482409 | 0,564757949 | ankyrin repeat domain 45 |
| ENSG00000285957 | AC019257,9 | -0,512012752 | 0,701243427 | 0,48394341 | 0,913690171 | novel transcript, antisense to ARHGEF10 |
| ENSG00000246889 | AP000487,1 | -0,512055316 | 0,701222739 | 0,19577624 | 0,69375328 | novel transcript |
| ENSG00000288166 | AP000487,3 | -0,512055316 | 0,701222739 | 0,19577624 | 0,69375328 | novel transcript |
| ENSG00000230359 | TPI1P2 | -0,512348943 | 0,701080035 | 0,26046505 | 0,772692192 | triosephosphate isomerase 1 pseudogene 2 |
| ENSG00000182359 | KBTBD3 | -0,512403353 | 0,701053595 | 0,29188783 | 0,802531095 | kelch repeat and BTB domain containing 3 |
| ENSG00000237126 | AC073254,1 | -0,512560001 | 0,700977479 | 0,27494446 | 0,786383635 | novel transcript |
| ENSG00000270174 | AL022097,1 | -0,51256466 | 0,700975215 | 0,60160504 | 0,949939978 | novel transcript, antisense to FARS2 |
| ENSG00000113946 | CLDN16 | -0,512599317 | 0,700958376 | 0,65476166 | 0,957444211 | claudin 16 |
| ENSG00000253517 | XRCC6P4 | -0,512632035 | 0,70094248 | 0,30733101 | 0,816138329 | X-ray repair cross complementing 6 pseudogene 4 |
| ENSG00000204623 | ZNRD1ASP | -0,512645827 | 0,700935779 | 0,20885331 | 0,713420724 | zinc ribbon domain containing 1 antisense, pseudogene |
| ENSG00000226622 | AC092155,1 | -0,512650171 | 0,700933668 | 0,19608369 | 0,694033976 | novel transcript |
| ENSG00000255326 | AP001922,5 | -0,512695056 | 0,700911861 | 0,40772796 | 0,880762246 | novel transcript |
| ENSG00000283839 | AC096667,1 | -0,51270086 | 0,700909041 | 0,63803511 | 0,956018518 | proline-rich protein 18-like |
| ENSG00000284948 | AC107959,4 | -0,512701456 | 0,700908752 | 0,59437712 | 0,947469769 | novel transcript |
| ENSG00000271833 | AL445222,1 | -0,512906484 | 0,70080915 | 0,48169539 | 0,912981134 | novel transcript |
| ENSG00000276005 | AC138749,8 | -0,512956617 | 0,700784798 | 0,29123422 | 0,80193294 | hect domain and RLD 2 (HERC2) pseudogene |
| ENSG00000286415 | AC017050,1 | -0,512964275 | 0,700781077 | 0,2072909 | 0,711123838 | novel transcript |
| ENSG00000230001 | AL359095,1 | -0,512967619 | 0,700779453 | 0,53571256 | 0,932139691 | novel transcript |

| | | | | | | |
|---|---|---|---|---|---|---|
| ENSG00000274296 | DECR2 | -0,513066535 | 0,700731407 | 0,34797118 | 0,842451708 | 2,4-dienoyl-CoA reductase 2 |
| ENSG00000186301 | MST1P2 | -0,513226354 | 0,700653786 | 0,48319023 | 0,91363215 | macrophage stimulating 1 pseudogene 2 |
| ENSG00000228662 | MCCD1 | -0,513258125 | 0,700638356 | 0,55510131 | 0,937793851 | mitochondrial coiled-coil domain 1 |
| ENSG00000204428 | LY6G5C | -0,51331928 | 0,700608657 | 0,5602813 | 0,940070839 | lymphocyte antigen 6 family member G5C |
| ENSG00000184602 | SNN | -0,513375484 | 0,700581364 | 0,01653394 | 0,164518442 | stannin |
| ENSG00000275885 | PRPF31 | -0,513397726 | 0,700570563 | 0,25012244 | 0,762275082 | pre-mRNA processing factor 31 |
| ENSG00000157653 | C9orf43 | -0,513534768 | 0,700504018 | 0,3767543 | 0,862818448 | chromosome 9 open reading frame 43 |
| ENSG00000254602 | AP000662,1 | -0,513738639 | 0,700405035 | 0,05093359 | 0,345795394 | novel transcript, sense overlapping CLP1 |
| ENSG00000273353 | AL008718,3 | -0,513744466 | 0,700402207 | 0,6444231 | 0,956633898 | novel transcript, sense intronic to NUP50 |
| ENSG00000286973 | AC087490,1 | -0,513810012 | 0,700370386 | 0,62984443 | 0,952812183 | novel transcript |
| ENSG00000261617 | LINC02177 | -0,513834864 | 0,700358321 | 0,5697838 | 0,94221458 | long intergenic non-protein coding RNA 2177 |
| ENSG00000180089 | TMEM86B | -0,5139119 | 0,700320925 | 0,37819148 | 0,863806148 | transmembrane protein 86B |
| ENSG00000197647 | ZNF433 | -0,51392652 | 0,700313828 | 0,05420137 | 0,359091243 | zinc finger protein 433 |
| ENSG00000109851 | DBX1 | -0,513935798 | 0,700309324 | 0,21745223 | 0,725179881 | developing brain homeobox 1 |
| ENSG00000287934 | AP001885,3 | -0,513995517 | 0,700280336 | 0,39621582 | 0,872845252 | novel transcript, antisense to KDM2A |
| ENSG00000239519 | CADM2-AS1 | -0,513999596 | 0,700278357 | 0,5470973 | 0,935404974 | CADM2 antisense RNA 1 |
| ENSG00000021488 | SLC7A9 | -0,51430076 | 0,700132188 | 0,36439812 | 0,854333929 | solute carrier family 7 member 9 |
| ENSG00000112309 | B3GAT2 | -0,514315398 | 0,700125085 | 0,3313988 | 0,832531293 | beta-1,3-glucuronyltransferase 2 |
| ENSG00000242853 | RN7SL749P | -0,51433452 | 0,700115805 | 0,61640326 | 0,952793934 | RNA, 7SL, cytoplasmic 749, pseudogene |
| ENSG00000250906 | AC131953,1 | -0,514513889 | 0,700028765 | 0,56532248 | 0,940865625 | novel transcript |
| ENSG00000164796 | CSMD3 | -0,514561048 | 0,700005883 | 0,29991904 | 0,809916717 | CUB and Sushi multiple domains 3 |
| ENSG00000164404 | GDF9 | -0,514888031 | 0,699847246 | 0,31451194 | 0,819877431 | growth differentiation factor 9 |
| ENSG00000259305 | ZHX1-C8orf76 | -0,51494455 | 0,69981983 | 0,30388562 | 0,812968394 | ZHX1-C8orf76 readthrough |
| ENSG00000160539 | PLPP7 | -0,515121018 | 0,699734234 | 0,08193079 | 0,454970566 | phospholipid phosphatase 7 (inactive) |
| ENSG00000268416 | AC010329,1 | -0,515580943 | 0,699511198 | 0,07700443 | 0,438408786 | novel transcript |
| ENSG00000280030 | AC010329,4 | -0,515580943 | 0,699511198 | 0,07700443 | 0,438408786 | novel transcript |
| ENSG00000177511 | ST8SIA3 | -0,515598503 | 0,699502683 | 0,40964234 | 0,882177108 | ST8 alpha-N-acetyl-neuraminide alpha-2,8-sialyltransferase 3 |
| ENSG00000229376 | CICP3 | -0,515620228 | 0,69949215 | 0,52419613 | 0,928419877 | capicua transcriptional repressor pseudogene 3 |
| ENSG00000196263 | ZNF471 | -0,515627987 | 0,699488388 | 0,09253102 | 0,48351825 | zinc finger protein 471 |
| ENSG00000197153 | H3C12 | -0,515678912 | 0,699463697 | 0,16726301 | 0,647913628 | H3 clustered histone 12 |
| ENSG00000091656 | ZFHX4 | -0,515888449 | 0,699362115 | 0,22894506 | 0,737173617 | zinc finger homeobox 4 |
| ENSG00000250656 | ST3GAL1P1 | -0,515998063 | 0,69930898 | 0,5083197 | 0,922427526 | ST3GAL1 pseudogene 1 |

| | | | | | | |
|---|---|---|---|---|---|---|
| ENSG00000159374 | M1AP | -0,516108205 | 0,699255594 | 0,42673001 | 0,892261855 | meiosis 1 associated protein |
| ENSG00000175097 | RAG2 | -0,516452766 | 0,69908861 | 0,50805492 | 0,922427526 | recombination activating 2 |
| ENSG00000285658 | AC127035,1 | -0,516487164 | 0,699071942 | 0,29406237 | 0,803927308 | novel transcript, antisense to AMBRA1 |
| ENSG00000283683 | MYOCOS | -0,51668296 | 0,698977073 | 0,22479136 | 0,733203418 | myocilin opposite strand |
| ENSG00000235481 | UBE2R2-AS1 | -0,516758282 | 0,698940581 | 0,13397821 | 0,584034405 | UBE2R2 antisense RNA 1 |
| ENSG00000127083 | OMD | -0,516781734 | 0,698929219 | 0,22392439 | 0,732710917 | osteomodulin |
| ENSG00000237916 | RPL37P12 | -0,516906578 | 0,69886874 | 0,4088335 | 0,881578006 | ribosomal protein L37 pseudogene 12 |
| ENSG00000259116 | AL049869,2 | -0,517043394 | 0,698802467 | 0,33737017 | 0,83650892 | novel transcript, antisense to HSPA2 & ZBTB1 |
| ENSG00000271410 | AC020728,1 | -0,517049322 | 0,698799595 | 0,39238571 | 0,870909282 | protein serine kinase H1 (PSKH1) pseudogene |
| ENSG00000277767 | AL442128,2 | -0,517404149 | 0,698627749 | 0,23347098 | 0,74254697 | novel transcript |
| ENSG00000258443 | AC005225,1 | -0,517408151 | 0,698625811 | 0,6514723 | 0,957089277 | novel transcript, sense intronic to HEATR4 |
| ENSG00000121390 | PSPC1 | -0,517443407 | 0,698608738 | 0,02502453 | 0,219551856 | paraspeckle component 1 |
| ENSG00000272849 | AC084018,1 | -0,517486294 | 0,698587971 | 0,22691418 | 0,735383523 | novel transcript |
| ENSG00000236364 | AL358115,1 | -0,517508125 | 0,6985774 | 0,17268612 | 0,657107871 | novel transcript, antisense to UCK2 |
| ENSG00000160271 | RALGDS | -0,517534261 | 0,698564745 | 0,03310758 | 0,264235013 | ral guanine nucleotide dissociation stimulator |
| ENSG00000262096 | PCDHB19P | -0,517543377 | 0,698560331 | 0,11329798 | 0,537503475 | protocadherin beta 19 pseudogene |
| ENSG00000253771 | TPTE2P1 | -0,517568048 | 0,698548385 | 0,11583027 | 0,543681398 | transmembrane phosphoinositide 3-phosphatase and tensin homolog 2 pseudogene 1 |
| ENSG00000250869 | AC087359,1 | -0,517618675 | 0,698523872 | 0,46153075 | 0,906165604 | tigger transposable element derived 1 (TIGD1) pseudogene |
| ENSG00000279342 | AP000866,6 | -0,517681816 | 0,698493301 | 0,26865329 | 0,780823861 | novel transcript, sense intronic to MSANTD2 |
| ENSG00000222724 | RNU2-63P | -0,517738458 | 0,698465878 | 0,40212121 | 0,87718264 | RNA, U2 small nuclear 63, pseudogene |
| ENSG00000267160 | AC091152,2 | -0,51795733 | 0,698359921 | 0,43817005 | 0,897177404 | novel transcript, antisense to C17orf104 |
| ENSG00000277550 | AL031651,1 | -0,518099936 | 0,698290894 | 0,54609769 | 0,935290591 | novel transcript |
| ENSG00000259349 | AC011921,1 | -0,518144888 | 0,698269137 | 0,30337879 | 0,812669425 | novel transcript, antisense to APPBP2 |
| ENSG00000204347 | BTBD17 | -0,518238225 | 0,698223963 | 0,13849446 | 0,593533053 | BTB domain containing 17 |
| ENSG00000287245 | AP001981,2 | -0,518342073 | 0,698173705 | 0,59185253 | 0,947416795 | novel transcript, antisense to RDX |
| ENSG00000151687 | ANKAR | -0,518357962 | 0,698166015 | 0,26420471 | 0,775595866 | ankyrin and armadillo repeat containing |
| ENSG00000171757 | LRRC34 | -0,518390087 | 0,698150469 | 0,31760499 | 0,822395418 | leucine rich repeat containing 34 |
| ENSG00000164989 | CCDC171 | -0,518390729 | 0,698150159 | 0,17013639 | 0,653308467 | coiled-coil domain containing 171 |
| ENSG00000189180 | ZNF33A | -0,518441495 | 0,698125593 | 0,22175746 | 0,729254784 | zinc finger protein 33A |
| ENSG00000116035 | VAX2 | -0,518467843 | 0,698112842 | 0,38656836 | 0,869441674 | ventral anterior homeobox 2 |
| ENSG00000267666 | AC004156,1 | -0,518595227 | 0,698051205 | 0,13327929 | 0,58326479 | novel transcript |

| | | | | | | |
|---|---|---|---|---|---|---|
| ENSG00000251441 | RTEL1P1 | -0,518805341 | 0,697949548 | 0,55170186 | 0,936366211 | regulator of telomere elongation helicase 1 pseudogene 1 |
| ENSG00000108551 | RASD1 | -0,518815069 | 0,697944842 | 0,24065279 | 0,750950074 | ras related dexamethasone induced 1 |
| ENSG00000171224 | FAM241B | -0,518839906 | 0,697932827 | 0,0982129 | 0,497493223 | family with sequence similarity 241 member B |
| ENSG00000265688 | MAFG-DT | -0,518934373 | 0,697887128 | 0,19963977 | 0,698923433 | MAFG divergent transcript |
| ENSG00000136367 | ZFHX2 | -0,518949263 | 0,697879925 | 0,03255086 | 0,261251213 | zinc finger homeobox 2 |
| ENSG00000250579 | CTD-2297D10,2 | -0,519008064 | 0,697851481 | 0,30327125 | 0,812669425 | uncharacterized LOC101929176 |
| ENSG00000230332 | RNF39 | -0,519193294 | 0,697761889 | 0,59032785 | 0,947297632 | ring finger protein 39 |
| ENSG00000204618 | RNF39 | -0,519193294 | 0,697761889 | 0,59032785 | 0,947297632 | ring finger protein 39 |
| ENSG00000243609 | RPS2P44 | -0,519234106 | 0,69774215 | 0,51499161 | 0,924720046 | ribosomal protein S2 pseudogene 44 |
| ENSG00000148985 | PGAP2 | -0,519308889 | 0,697705984 | 0,05430112 | 0,359622485 | post-GPI attachment to proteins 2 |
| ENSG00000270571 | AC007681,1 | -0,519365261 | 0,697678722 | 0,47779177 | 0,911721977 | novel transcript, antisense to TACR1 |
| ENSG00000242753 | AL445430,2 | -0,519405796 | 0,697659119 | 0,27781411 | 0,789162936 | novel transcript |
| ENSG00000258718 | LINC02311 | -0,519469557 | 0,697628286 | 0,26578038 | 0,77713706 | long intergenic non-protein coding RNA 2311 |
| ENSG00000233871 | DLG5-AS1 | -0,519475062 | 0,697625624 | 0,43940853 | 0,89753485 | DLG5 antisense RNA 1 |
| ENSG00000286705 | AC079140,6 | -0,519524275 | 0,697601827 | 0,27528856 | 0,786927091 | novel transcript |
| ENSG00000180425 | C11orf71 | -0,519660153 | 0,697536128 | 0,1783134 | 0,66556711 | chromosome 11 open reading frame 71 |
| ENSG00000262911 | DNAJC28 | -0,519664181 | 0,697534181 | 0,33684149 | 0,836162806 | DnaJ heat shock protein family (Hsp40) member C28 |
| ENSG00000270127 | AC027020,2 | -0,519677359 | 0,697527809 | 0,38627477 | 0,869312875 | novel transcript |
| ENSG00000281582 | AP000350,16 | -0,519680027 | 0,697526519 | 0,07150555 | 0,421367686 | novel transcript |
| ENSG00000273295 | AP000350,6 | -0,519680027 | 0,697526519 | 0,07150555 | 0,421367686 | novel transcript |
| ENSG00000204176 | SYT15 | -0,519760201 | 0,697487757 | 0,08706952 | 0,470307784 | synaptotagmin 15 |
| ENSG00000285002 | AC270254,2 | -0,519807998 | 0,697464649 | 0,64618747 | 0,956899963 | novel transcript |
| ENSG00000254340 | AC022784,5 | -0,519807998 | 0,697464649 | 0,64618747 | 0,956899963 | novel transcript |
| ENSG00000186212 | SOWAHB | -0,520175367 | 0,697287069 | 0,57143561 | 0,942436428 | sosondowah ankyrin repeat domain family member B |
| ENSG00000237194 | SNAI1P1 | -0,520253465 | 0,697249323 | 0,63056355 | 0,953331345 | snail family zinc finger 1 pseudogene 1 |
| ENSG00000276411 | AC106875,2 | -0,520338749 | 0,697208107 | 0,56416249 | 0,940865625 | novel transcript |
| ENSG00000235545 | AC103923,1 | -0,520433915 | 0,697162118 | 0,20006261 | 0,699646707 | novel transcript |
| ENSG00000169116 | PARM1 | -0,520573526 | 0,697094656 | 0,17416906 | 0,659374935 | prostate androgen-regulated mucin-like protein 1 |
| ENSG00000166800 | LDHAL6A | -0,520610747 | 0,697076672 | 0,26815829 | 0,780692104 | lactate dehydrogenase A like 6A |
| ENSG00000286943 | AP001992,2 | -0,520730523 | 0,697018801 | 0,23587572 | 0,744966003 | novel transcript, antisense to XRRA1 |
| ENSG00000146021 | KLHL3 | -0,520826406 | 0,696972478 | 0,08404361 | 0,461230231 | kelch like family member 3 |

| | | | | | | |
|---|---|---|---|---|---|---|
| ENSG00000095209 | TMEM38B | -0,521105699 | 0,696837564 | 0,27345804 | 0,785672647 | transmembrane protein 38B |
| ENSG00000206181 | ELOA2 | -0,5211446 | 0,696818774 | 0,47684593 | 0,91143003 | elongin A2 |
| ENSG00000217094 | PPIAP31 | -0,521299212 | 0,696744101 | 0,59859772 | 0,948821689 | peptidylprolyl isomerase A pseudogene 31 |
| ENSG00000112812 | PRSS16 | -0,521409904 | 0,696690645 | 0,38942486 | 0,870057277 | serine protease 16 |
| ENSG00000119703 | ZC2HC1C | -0,521414539 | 0,696688406 | 0,14064196 | 0,597543965 | zinc finger C2HC-type containing 1C |
| ENSG00000273432 | AC004951,4 | -0,521427243 | 0,696682272 | 0,41689216 | 0,886526763 | novel transcript, POLR2J4 pseudogene-RASA4 pseudogene readthrough |
| ENSG00000175471 | MCTP1 | -0,521498375 | 0,696647923 | 0,19104057 | 0,686751133 | multiple C2 and transmembrane domain containing 1 |
| ENSG00000185090 | MANEAL | -0,521533227 | 0,696631094 | 0,041516 | 0,30576122 | mannosidase endo-alpha like |
| ENSG00000243696 | AC006254,1 | -0,521612656 | 0,696592741 | 0,16263304 | 0,639889157 | novel MUSTN1-ITIH4 readthrough |
| ENSG00000178585 | CTNNBIP1 | -0,521735339 | 0,696533507 | 0,01456416 | 0,150888551 | catenin beta interacting protein 1 |
| ENSG00000196693 | ZNF33B | -0,521770726 | 0,696516423 | 0,19631562 | 0,694046542 | zinc finger protein 33B |
| ENSG00000260198 | AC009102,1 | -0,521824426 | 0,696490497 | 0,40510593 | 0,878176155 | novel transcript, sense intronic to GNAO1 |
| ENSG00000222035 | KIAA2012-AS1 | -0,521867131 | 0,69646988 | 0,59740579 | 0,948282297 | KIAA2012 antisense RNA 1 |
| ENSG00000286195 | AC007666,2 | -0,521880343 | 0,696463503 | 0,06659315 | 0,405525577 | novel transcript, antisense to CECR2 and SLC25A18 |
| ENSG00000144227 | NXPH2 | -0,521910826 | 0,696448787 | 0,31804463 | 0,822980102 | neurexophilin 2 |
| ENSG00000185133 | INPP5J | -0,521999226 | 0,696406114 | 0,14210594 | 0,600058555 | inositol polyphosphate-5-phosphatase J |
| ENSG00000248019 | FAM13A-AS1 | -0,522015008 | 0,696398496 | 0,17381222 | 0,658834301 | FAM13A antisense RNA 1 |
| ENSG00000273139 | AC007663,3 | -0,522348082 | 0,696237737 | 0,45609364 | 0,903545728 | novel transcript |
| ENSG00000275026 | GXYLT1P4 | -0,522399487 | 0,69621293 | 0,36212466 | 0,852258387 | GXYLT1 pseudogene 4 |
| ENSG00000099769 | IGFALS | -0,522580483 | 0,696125591 | 0,67971024 | 0,964459885 | insulin like growth factor binding protein acid labile subunit |
| ENSG00000168263 | KCNV2 | -0,522625364 | 0,696103935 | 0,52346634 | 0,928021082 | potassium voltage-gated channel modifier subfamily V member 2 |
| ENSG00000229893 | TAX1BP1-AS1 | -0,522738185 | 0,696049501 | 0,59744653 | 0,948282297 | TAX1BP1 antisense RNA 1 |
| ENSG00000273188 | AL022328,3 | -0,522754754 | 0,696041507 | 0,59480209 | 0,947469769 | novel transcript, antisense to TRABD |
| ENSG00000167964 | RAB26 | -0,522886325 | 0,695978032 | 0,26974847 | 0,781791089 | RAB26, member RAS oncogene family |
| ENSG00000259436 | AC010247,2 | -0,522951292 | 0,695946692 | 0,43284097 | 0,89560871 | novel transcript, antisense to POU2F2 |
| ENSG00000286739 | AC007387,1 | -0,522975035 | 0,695935239 | 0,06364781 | 0,395210559 | novel transcript |
| ENSG00000259756 | AC100839,1 | -0,523318546 | 0,695769553 | 0,62498549 | 0,952793934 | novel transcript, antisense to TLN2 |
| ENSG00000245552 | AP000787,1 | -0,523381641 | 0,695739125 | 0,05203764 | 0,349872693 | novel transcript, antisense to SESN3 |
| ENSG00000215641 | TRIM27 | -0,523383327 | 0,695738312 | 0,07035262 | 0,417680252 | tripartite motif containing 27 |

| | | | | | | |
|---|---|---|---|---|---|---|
| ENSG00000197903 | H2BC12 | -0,523396703 | 0,695731862 | 0,10409988 | 0,514557881 | H2B clustered histone 12 |
| ENSG00000257121 | AC126615,2 | -0,523449654 | 0,695706327 | 0,70050949 | 0,968522643 | peroxiredoxin 6 (PRDX6) pseudogene |
| ENSG00000264270 | AC087749,1 | -0,523588078 | 0,695639578 | 0,43160628 | 0,895347413 | novel transcript, antisense to SAP30BP |
| ENSG00000238039 | AC244197,2 | -0,523671702 | 0,695599257 | 0,42439541 | 0,890949676 | novel transcript |
| ENSG00000283946 | CHRNB1 | -0,523676261 | 0,69559706 | 0,69450984 | 0,967417311 | cholinergic receptor nicotinic beta 1 subunit |
| ENSG00000286827 | AC072026,2 | -0,52386677 | 0,695505212 | 0,05919391 | 0,377938736 | novel transcript, antisense to STXBP5L |
| ENSG00000287893 | AC083864,5 | -0,523969323 | 0,695455774 | 0,16181153 | 0,63836332 | novel transcript |
| ENSG00000182632 | CCNYL2 | -0,524006377 | 0,695437912 | 0,69218077 | 0,967236299 | cyclin Y like 2 (pseudogene) |
| ENSG00000226530 | AL158055,1 | -0,524056894 | 0,695413561 | 0,1068745 | 0,521837326 | novel transcript |
| ENSG00000206340 | C4A | -0,524109491 | 0,695388208 | 0,64750994 | 0,956899963 | complement C4A (Rodgers blood group) |
| ENSG00000236625 | C4B | -0,524109491 | 0,695388208 | 0,64750994 | 0,956899963 | complement C4B (Chido blood group) |
| ENSG00000285651 | AL450992,2 | -0,524281269 | 0,695305415 | 0,3223678 | 0,824955371 | novel transcript, antisense to THEM4 |
| ENSG00000226314 | ZNF192P1 | -0,524322306 | 0,695285638 | 0,18979568 | 0,684955795 | zinc finger protein 192 pseudogene 1 |
| ENSG00000287623 | AC037193,1 | -0,524359229 | 0,695267844 | 0,10129571 | 0,506227112 | novel transcript, antisense to SRGAP3 |
| ENSG00000150667 | FSIP1 | -0,524443652 | 0,695227159 | 0,17999511 | 0,668905046 | fibrous sheath interacting protein 1 |
| ENSG00000261714 | AC105137,1 | -0,524504911 | 0,69519764 | 0,64296547 | 0,956633898 | high mobility group nucleosomal binding domain 2 (HMGN2) pseudogene |
| ENSG00000286970 | AC010491,2 | -0,524544262 | 0,695178678 | 0,15545672 | 0,625669263 | novel transcript, antisense to ANKH |
| ENSG00000231527 | FAM27C | -0,524587557 | 0,695157816 | 0,02671119 | 0,228844863 | family with sequence similarity 27 member C |
| ENSG00000270589 | AL158163,1 | -0,524722103 | 0,695092988 | 0,28210712 | 0,793203731 | novel transcript, antisense to BBIP1 |
| ENSG00000160796 | NBEAL2 | -0,52477265 | 0,695068635 | 0,15056532 | 0,616942884 | neurobeachin like 2 |
| ENSG00000162669 | HFM1 | -0,524867448 | 0,695022964 | 0,32366043 | 0,825794844 | helicase for meiosis 1 |
| ENSG00000273455 | AC072039,2 | -0,524916689 | 0,694999243 | 0,71760141 | 0,971697135 | novel transcript, antisense to PPP2R3A |
| ENSG00000249167 | SEMA6A-AS2 | -0,524955368 | 0,69498061 | 0,22452538 | 0,733203418 | SEMA6A antisense RNA 2 |
| ENSG00000278535 | DHRS11 | -0,525162176 | 0,694880993 | 0,08275465 | 0,457565724 | dehydrogenase/reductase 11 |
| ENSG00000140873 | ADAMTS18 | -0,525172201 | 0,694876164 | 0,32816474 | 0,830076239 | ADAM metallopeptidase with thrombospondin type 1 motif 18 |
| ENSG00000182504 | CEP97 | -0,52543038 | 0,694751823 | 0,14316369 | 0,601880115 | centrosomal protein 97 |
| ENSG00000168152 | THAP9 | -0,525528801 | 0,694704429 | 0,28635039 | 0,797518439 | THAP domain containing 9 |
| ENSG00000260238 | PMF1-BGLAP | -0,525667021 | 0,694637874 | 0,25358255 | 0,765172364 | PMF1-BGLAP readthrough |
| ENSG00000256315 | AP000462,2 | -0,525672415 | 0,694635277 | 0,34289445 | 0,840419133 | novel transcript |
| ENSG00000255087 | AP001993,1 | -0,525708026 | 0,694618131 | 0,56437811 | 0,940865625 | novel transcript |
| ENSG00000282689 | ULK4P2 | -0,525797761 | 0,694574927 | 0,56493644 | 0,940865625 | ULK4 pseudogene 2 |

| | | | | | | |
|---|---|---|---|---|---|---|
| ENSG00000260128 | ULK4P2 | -0,525797761 | 0,694574927 | 0,56493644 | 0,940865625 | ULK4 pseudogene 2 |
| ENSG00000272948 | AP001412,1 | -0,525825064 | 0,694561783 | 0,46011329 | 0,905443183 | novel transcript, antisense to DSCR3 |
| ENSG00000267751 | AC009005,1 | -0,525940438 | 0,69450624 | 0,17679987 | 0,662651108 | novel transcript, antisense to BSG |
| ENSG00000236564 | YWHAQP5 | -0,525950355 | 0,694501466 | 0,73561387 | 0,974353474 | YWHAQ pseudogene 5 |
| ENSG00000286527 | AL161665,2 | -0,525951858 | 0,694500743 | 0,63202149 | 0,953935145 | novel transcript |
| ENSG00000283393 | AC068205,3 | -0,526116388 | 0,694421544 | 0,44090813 | 0,898675567 | ribosomal protein L23a (RPL23A) pseudogene |
| ENSG00000261068 | AL512274,1 | -0,526143782 | 0,694408358 | 0,57285057 | 0,942475308 | novel transcript |
| ENSG00000273007 | AC021205,3 | -0,526181426 | 0,69439024 | 0,05516587 | 0,362637904 | novel transcript, antisense to FGF2 |
| ENSG00000231006 | RPL7P32 | -0,526537785 | 0,69421874 | 0,54334346 | 0,934188893 | ribosomal protein L7 pseudogene 32 |
| ENSG00000278467 | AC138393,3 | -0,52667483 | 0,694152797 | 0,33241824 | 0,83293962 | novel transcript |
| ENSG00000272977 | AL008721,2 | -0,526724857 | 0,694128727 | 0,04596203 | 0,325324163 | novel transcript |
| ENSG00000236008 | LINC01814 | -0,52675973 | 0,694111949 | 0,01946887 | 0,183667726 | long intergenic non-protein coding RNA 1814 |
| ENSG00000249947 | XBP1P1 | -0,526913937 | 0,694037761 | 0,39956254 | 0,875433675 | X-box binding protein 1 pseudogene 1 |
| ENSG00000116983 | HPCAL4 | -0,526927363 | 0,694031302 | 0,12359108 | 0,562274686 | hippocalcin like 4 |
| ENSG00000054803 | CBLN4 | -0,527202876 | 0,693898774 | 0,55518675 | 0,937793851 | cerebellin 4 precursor |
| ENSG00000225572 | DOCK4-AS1 | -0,527217909 | 0,693891544 | 0,17304441 | 0,657758111 | DOCK4 antisense RNA 1 |
| ENSG00000231063 | RPL6P6 | -0,527292984 | 0,693855436 | 0,39799099 | 0,874589897 | ribosomal protein L6 pseudogene 6 |
| ENSG00000145975 | FAM217A | -0,527615294 | 0,693700441 | 0,42078955 | 0,888709349 | family with sequence similarity 217 member A |
| ENSG00000248445 | SEMA6A-AS1 | -0,527674668 | 0,693671892 | 0,27400478 | 0,7862019 | SEMA6A antisense RNA 1 |
| ENSG00000253558 | AC024568,1 | -0,527875376 | 0,693575395 | 0,26716124 | 0,779493925 | heat shock protein 90kDa alpha (cytosolic), class B member 1 (HSP90AB1) pseudogene |
| ENSG00000235816 | PRELID1P3 | -0,527877686 | 0,693574285 | 0,41828795 | 0,887094957 | PRELID1 pseudogene 3 |
| ENSG00000223529 | EEF1A1P8 | -0,528012126 | 0,693509656 | 0,62136548 | 0,952793934 | eukaryotic translation elongation factor 1 alpha 1 pseudogene 8 |
| ENSG00000253667 | AC100821,1 | -0,528020188 | 0,69350578 | 0,1806557 | 0,670229021 | ESF1, nucleolar pre-rRNA processing protein, homolog (S, cerevisiae) (ESF1) pseudogene |
| ENSG00000239877 | IGSF11-AS1 | -0,528111988 | 0,693461653 | 0,67609797 | 0,963560994 | IGSF11 antisense RNA 1 |
| ENSG00000286349 | AC108471,3 | -0,528238419 | 0,693400884 | 0,58250022 | 0,944174844 | novel transcript, antisense to SMIM14 |
| ENSG00000233190 | RPS24P13 | -0,528333058 | 0,693355399 | 0,65011122 | 0,956899963 | ribosomal protein S24 pseudogene 13 |
| ENSG00000239332 | LINC01119 | -0,52838584 | 0,693330033 | 0,52295674 | 0,92790906 | long intergenic non-protein coding RNA 1119 |
| ENSG00000170835 | CEL | -0,528456276 | 0,693296184 | 0,205652 | 0,708173174 | carboxyl ester lipase |
| ENSG00000109618 | SEPSECS | -0,528504687 | 0,69327292 | 0,27413934 | 0,7862019 | Sep (O-phosphoserine) tRNA:Sec (selenocysteine) tRNA synthase |

| | | | | | | |
|---|---|---|---|---|---|---|
| ENSG00000198483 | ANKRD35 | -0,528581572 | 0,693235974 | 0,29629364 | 0,805997082 | ankyrin repeat domain 35 |
| ENSG00000282717 | AC243651,2 | -0,52869302 | 0,693182424 | 0,34860512 | 0,842757646 | novel transcript, antisense to PDXDC1 |
| ENSG00000275910 | AC138932,4 | -0,52869302 | 0,693182424 | 0,34860512 | 0,842757646 | novel transcript, antisense to PDXDC1 |
| ENSG00000243069 | ARHGEF26-AS1 | -0,528739599 | 0,693160045 | 0,04864518 | 0,335726918 | ARHGEF26 antisense RNA 1 |
| ENSG00000117461 | PIK3R3 | -0,528856803 | 0,693103735 | 0,06745724 | 0,40855933 | phosphoinositide-3-kinase regulatory subunit 3 |
| ENSG00000199363 | SNORA63E | -0,528955346 | 0,693056394 | 0,44445374 | 0,900210555 | small nucleolar RNA, H/ACA box 63E |
| ENSG00000272661 | AC021097,1 | -0,52911819 | 0,692978169 | 0,67133237 | 0,962026636 | novel transcript, antisense to SMARCD3 |
| ENSG00000256458 | AC080075,1 | -0,529158681 | 0,692958721 | 0,47750925 | 0,911574108 | novel transcript, antisense to abl interactor 2 (ABI2) |
| ENSG00000269845 | CCNYL6 | -0,529170035 | 0,692953267 | 0,19208804 | 0,688460451 | cyclin Y like 6 (pseudogene) |
| ENSG00000279112 | BRI3BPP1 | -0,529170035 | 0,692953267 | 0,19208804 | 0,688460451 | BRI3 binding protein pseudogene 1 |
| ENSG00000232677 | LINC00665 | -0,529178119 | 0,692949384 | 0,03014835 | 0,247911111 | long intergenic non-protein coding RNA 665 |
| ENSG00000198010 | DLGAP2 | -0,529248691 | 0,692915488 | 0,64692962 | 0,956899963 | DLG associated protein 2 |
| ENSG00000253816 | GUSBP14 | -0,529279041 | 0,692900912 | 0,080085 | 0,448625211 | GUSB pseudogene 14 |
| ENSG00000187987 | ZSCAN23 | -0,52933807 | 0,692872562 | 0,16086044 | 0,636841924 | zinc finger and SCAN domain containing 23 |
| ENSG00000280169 | AL136526,1 | -0,529365088 | 0,692859586 | 0,31914902 | 0,82414099 | novel transcript, antisense to GGACT |
| ENSG00000260947 | AL356489,2 | -0,529420269 | 0,692833085 | 0,2840268 | 0,795610948 | novel transcript |
| ENSG00000260735 | AC139256,1 | -0,52959531 | 0,69274903 | 0,02892954 | 0,241573992 | phospholipase A2 group X (PLA2G10) pseudogene |
| ENSG00000235914 | MACROD2-AS1 | -0,529759974 | 0,692669966 | 0,58188156 | 0,944149815 | MACROD2 antisense RNA 1 |
| ENSG00000281195 | AC007878,1 | -0,529777805 | 0,692661405 | 0,14105195 | 0,598214217 | novel transcript |
| ENSG00000213433 | RPLP1P6 | -0,529784954 | 0,692657973 | 0,47897204 | 0,912714933 | ribosomal protein lateral stalk subunit P1 pseudogene 6 |
| ENSG00000276971 | TRPV6 | -0,529893508 | 0,692605857 | 0,66789748 | 0,961113267 | transient receptor potential cation channel subfamily V member 6 |
| ENSG00000270429 | KNOP1P2 | -0,529987625 | 0,692560675 | 0,32213462 | 0,824955371 | lysine rich nucleolar protein 1 pseudogene 2 |
| ENSG00000251188 | AC079140,3 | -0,530031773 | 0,692539482 | 0,34543722 | 0,841168721 | novel zinc finger protein pseudogene |
| ENSG00000232729 | AC211433,1 | -0,530125875 | 0,692494311 | 0,09455947 | 0,488468697 | novel transcript, antisense to GTF2I |
| ENSG00000230006 | ANKRD36BP2 | -0,530148572 | 0,692483417 | 0,14691985 | 0,610674677 | ankyrin repeat domain 36B pseudogene 2 |
| ENSG00000258323 | AC073575,1 | -0,530196642 | 0,692460344 | 0,44546297 | 0,901068127 | novel transcript, antisense to NAA25 |
| ENSG00000157856 | DRC1 | -0,530236398 | 0,692441262 | 0,22163987 | 0,729254784 | dynein regulatory complex subunit 1 |
| ENSG00000146350 | TBC1D32 | -0,530340544 | 0,692391278 | 0,08690208 | 0,469646795 | TBC1 domain family member 32 |
| ENSG00000171811 | CFAP46 | -0,530353716 | 0,692384956 | 0,17877204 | 0,66664232 | cilia and flagella associated protein 46 |
| ENSG00000171160 | MORN4 | -0,530393121 | 0,692366045 | 0,10798746 | 0,523928283 | MORN repeat containing 4 |
| ENSG00000288631 | ELOA3 | -0,530439143 | 0,692343959 | 0,68192564 | 0,964654539 | elongin A3 |

| | | | | | | |
|---|---|---|---|---|---|---|
| ENSG00000287466 | AL355531,1 | -0,530569615 | 0,692281348 | 0,4238344 | 0,890949676 | novel transcript, antisense to MGMT |
| ENSG00000278390 | AL354696,2 | -0,530702037 | 0,692217808 | 0,18852087 | 0,682877586 | novel transcript, antisense to KBTBD7 and MTRF1 |
| ENSG00000056487 | PHF21B | -0,5307448 | 0,692197291 | 0,02131294 | 0,194921664 | PHD finger protein 21B |
| ENSG00000163705 | FANCD2OS | -0,530801947 | 0,692169873 | 0,53221078 | 0,931291085 | FANCD2 opposite strand |
| ENSG00000267098 | AC113137,1 | -0,530950969 | 0,692098379 | 0,11214413 | 0,535189241 | novel transcript |
| ENSG00000257698 | GIHCG | -0,531903402 | 0,691641623 | 0,15570422 | 0,626262282 | GIHCG inhibitor of miR-200b/200a/429 expression |
| ENSG00000268926 | AL354861,3 | -0,532102104 | 0,69154637 | 0,61585722 | 0,952793934 | novel transcript |
| ENSG00000214525 | AC130709,1 | -0,532142145 | 0,691527177 | 0,57486143 | 0,942533938 | chromosome 15 open reading frame 2 (C15orf2) pseudogene |
| ENSG00000258134 | AC016954,1 | -0,532181312 | 0,691508403 | 0,37038716 | 0,858545694 | WW domain binding protein 2 (WBP2) pseudogene |
| ENSG00000180855 | ZNF443 | -0,532280341 | 0,691460938 | 0,25027326 | 0,762441549 | zinc finger protein 443 |
| ENSG00000245848 | CEBPA | -0,532330686 | 0,691436809 | 0,48988207 | 0,916063942 | CCAAT enhancer binding protein alpha |
| ENSG00000114796 | KLHL24 | -0,53265015 | 0,691283718 | 0,23520595 | 0,744654651 | kelch like family member 24 |
| ENSG00000228897 | AC012441,2 | -0,53267025 | 0,691274086 | 0,48282818 | 0,91363215 | novel pseudogene |
| ENSG00000285704 | AC004765,1 | -0,532792727 | 0,691215404 | 0,28750365 | 0,798896538 | novel transcript, antisense to WNK1 |
| ENSG00000285355 | AC005342,5 | -0,532845227 | 0,69119025 | 0,57605717 | 0,943019411 | novel transcript |
| ENSG00000183145 | RIPPLY3 | -0,532922735 | 0,691153117 | 0,21085539 | 0,716463754 | ripply transcriptional repressor 3 |
| ENSG00000266378 | AC005224,2 | -0,532929747 | 0,691149758 | 0,30268912 | 0,812334076 | novel transcript |
| ENSG00000186047 | DLEU7 | -0,532951865 | 0,691139163 | 0,09267606 | 0,484089861 | deleted in lymphocytic leukemia 7 |
| ENSG00000214049 | UCA1 | -0,53303569 | 0,691099006 | 0,48769887 | 0,914936533 | urothelial cancer associated 1 |
| ENSG00000225684 | FAM225B | -0,533164387 | 0,691037359 | 0,57014879 | 0,94228111 | family with sequence similarity 225 member B |
| ENSG00000227598 | Z94721,1 | -0,533217194 | 0,691012065 | 0,52787292 | 0,929612959 | novel transcript |
| ENSG00000204177 | BMS1P1 | -0,533224047 | 0,691008783 | 0,05750171 | 0,370798119 | BMS1 pseudogene 1 |
| ENSG00000203780 | FANK1 | -0,533228448 | 0,691006675 | 0,07896359 | 0,445215081 | fibronectin type III and ankyrin repeat domains 1 |
| ENSG00000262728 | AC123768,2 | -0,533258833 | 0,690992122 | 0,33817195 | 0,837263082 | novel transcript, antisense to GOLGA8N |
| ENSG00000249409 | AC025741,1 | -0,533303287 | 0,690970831 | 0,51959644 | 0,926822772 | novel transcript |
| ENSG00000197487 | GALP | -0,533306064 | 0,6909695 | 0,56840397 | 0,94163744 | galanin like peptide |
| ENSG00000162373 | BEND5 | -0,533375679 | 0,690936159 | 0,02036061 | 0,189112948 | BEN domain containing 5 |
| ENSG00000201098 | RNY1 | -0,53338349 | 0,690932419 | 0,32240703 | 0,824955371 | RNA, Ro60-associated Y1 |
| ENSG00000287731 | AL035603,1 | -0,533422433 | 0,690913768 | 0,46357712 | 0,907101775 | novel transcript |
| ENSG00000204930 | FAM221B | -0,533673943 | 0,69079333 | 0,24682286 | 0,758842554 | family with sequence similarity 221 member B |
| ENSG00000214425 | LRRC37A4P | -0,533929876 | 0,690670794 | 0,03787388 | 0,287819387 | leucine rich repeat containing 37 member A4, pseudogene |

| | | | | | | |
|---|---|---|---|---|---|---|
| ENSG00000135502 | SLC26A10 | -0,534128909 | 0,690575516 | 0,2845523 | 0,795919345 | solute carrier family 26 member 10 |
| ENSG00000269737 | AL691432,1 | -0,534185313 | 0,690548518 | 0,03925543 | 0,294877861 | ribosomal protein S7 (RPS7) pseudogene |
| ENSG00000234946 | SDHCP3 | -0,534269622 | 0,690508164 | 0,598719 | 0,948854865 | succinate dehydrogenase complex subunit C pseudogene 3 |
| ENSG00000104112 | SCG3 | -0,534571294 | 0,690363792 | 0,28926067 | 0,800260154 | secretogranin III |
| ENSG00000119522 | DENND1A | -0,534587795 | 0,690355896 | 0,03154401 | 0,256020734 | DENN domain containing 1A |
| ENSG00000273045 | C2orf15 | -0,534596678 | 0,690351645 | 0,28024226 | 0,790779983 | chromosome 2 open reading frame 15 |
| ENSG00000243243 | AC073130,2 | -0,534610378 | 0,69034509 | 0,45143236 | 0,902062594 | novel transcript |
| ENSG00000259144 | RANBP20P | -0,534624079 | 0,690338534 | 0,5927885 | 0,947416795 | RAN binding protein 20 pseudogene |
| ENSG00000286122 | AC016074,2 | -0,534646509 | 0,690327801 | 0,03722244 | 0,284946761 | novel transcript |
| ENSG00000288025 | AC093849,4 | -0,5346741 | 0,690314599 | 0,23581455 | 0,744966003 | novel transcript, antisense to SCRG1 |
| ENSG00000106633 | GCK | -0,534678809 | 0,690312346 | 0,2778232 | 0,789162936 | glucokinase |
| ENSG00000179406 | LINC00174 | -0,534741527 | 0,690282337 | 0,11984876 | 0,552892394 | long intergenic non-protein coding RNA 174 |
| ENSG00000118407 | FILIP1 | -0,534832051 | 0,690239025 | 0,24583817 | 0,757752583 | filamin A interacting protein 1 |
| ENSG00000163157 | TMOD4 | -0,535107556 | 0,690107226 | 0,34372124 | 0,840901506 | tropomodulin 4 |
| ENSG00000162482 | AKR7A3 | -0,535130461 | 0,690096269 | 0,23539785 | 0,744654651 | aldo-keto reductase family 7 member A3 |
| ENSG00000228568 | RPS15AP13 | -0,535137385 | 0,690092957 | 0,67877882 | 0,964124343 | ribosomal protein S15a pseudogene 13 |
| ENSG00000285710 | AC012184,4 | -0,535168144 | 0,690078244 | 0,44694295 | 0,901798674 | novel transcript, antisense to DDX19A |
| ENSG00000230866 | LINC02558 | -0,535344939 | 0,689993684 | 0,04638349 | 0,327048609 | long intergenic non-protein coding RNA 2558 |
| ENSG00000157152 | SYN2 | -0,535519245 | 0,689910324 | 0,14894958 | 0,613619689 | synapsin II |
| ENSG00000233860 | SHROOM3-AS1 | -0,535556382 | 0,689892565 | 0,12142917 | 0,55673288 | SHROOM3 antisense RNA 1 |
| ENSG00000184933 | OR6A2 | -0,535599333 | 0,689872026 | 0,37673459 | 0,862818448 | olfactory receptor family 6 subfamily A member 2 |
| ENSG00000185065 | AC000068,1 | -0,535624784 | 0,689859856 | 0,15020616 | 0,615923738 | novel transcript, antisense to C22orf39 |
| ENSG00000158966 | CACHD1 | -0,535698741 | 0,689824493 | 0,04679003 | 0,328507386 | cache domain containing 1 |
| ENSG00000284634 | AC092821,3 | -0,535826482 | 0,689763416 | 0,23133084 | 0,739268552 | novel transcript |
| ENSG00000139880 | CDH24 | -0,53586298 | 0,689745967 | 0,03414701 | 0,26913287 | cadherin 24 |
| ENSG00000246263 | UBR5-AS1 | -0,535955027 | 0,689701961 | 0,30145651 | 0,811013605 | UBR5 antisense RNA 1 |
| ENSG00000176826 | FKBP9P1 | -0,536008523 | 0,689676386 | 0,26266577 | 0,774111884 | FKBP prolyl isomerase 9 pseudogene 1 |
| ENSG00000237637 | FRY-AS1 | -0,53601772 | 0,68967199 | 0,62089735 | 0,952793934 | FRY antisense RNA 1 |
| ENSG00000250896 | RNPS1P1 | -0,536141877 | 0,68961264 | 0,69841394 | 0,968522643 | RNA binding protein with serine rich domain 1 pseudogene 1 |
| ENSG00000213003 | BTF3P12 | -0,536184024 | 0,689592494 | 0,50175605 | 0,920520138 | basic transcription factor 3 pseudogene 12 |
| ENSG00000227376 | FTH1P16 | -0,53626057 | 0,689555906 | 0,40687312 | 0,880078305 | ferritin heavy chain 1 pseudogene 16 |

| | | | | | | |
|---|---|---|---|---|---|---|
| ENSG00000284491 | THSD8 | -0,536299569 | 0,689537267 | 0,59515482 | 0,947640448 | thrombospondin type 1 domain containing 8 |
| ENSG00000277151 | AL138820,1 | -0,536461405 | 0,689459921 | 0,30539075 | 0,814326218 | novel transcript |
| ENSG00000152086 | TUBA3E | -0,536997868 | 0,689203595 | 0,63097203 | 0,953626097 | tubulin alpha 3e |
| ENSG00000144035 | NAT8 | -0,537105286 | 0,689152281 | 0,54954173 | 0,936083284 | N-acetyltransferase 8 (putative) |
| ENSG00000260089 | ADAM3B | -0,537210839 | 0,689101862 | 0,29911397 | 0,808953227 | ADAM metallopeptidase domain 3B (pseudogene) |
| ENSG00000215184 | RPS12P16 | -0,537366407 | 0,689027559 | 0,31701039 | 0,82206753 | ribosomal protein S12 pseudogene 16 |
| ENSG00000241316 | SUCLG2-AS1 | -0,537476115 | 0,688975165 | 0,05092645 | 0,345795394 | SUCLG2 antisense RNA 1 (head to head) |
| ENSG00000244128 | LINC01322 | -0,537641486 | 0,688896195 | 0,23908682 | 0,748678935 | long intergenic non-protein coding RNA 1322 |
| ENSG00000285827 | AP001267,5 | -0,537661599 | 0,68888659 | 0,07449907 | 0,431481852 | novel protein |
| ENSG00000187391 | MAGI2 | -0,537743103 | 0,688847673 | 0,09580515 | 0,491508392 | membrane associated guanylate kinase, WW and PDZ domain containing 2 |
| ENSG00000275485 | AL512652,1 | -0,537886546 | 0,688779187 | 0,57433124 | 0,942475308 | novel transcript, sense intronic to N6AMT2 |
| ENSG00000250045 | CBX3P3 | -0,538074901 | 0,688689267 | 0,46028008 | 0,9054994 | chromobox 3 pseudogene 3 |
| ENSG00000253284 | AC092828,1 | -0,538085651 | 0,688684136 | 0,05249555 | 0,351770604 | novel transcript |
| ENSG00000275734 | AC010538,1 | -0,538273225 | 0,688594601 | 0,05557713 | 0,363892597 | novel transcript, sense overlapping CDK10 |
| ENSG00000196866 | H2AC7 | -0,538407233 | 0,688530642 | 0,1066456 | 0,521491876 | H2A clustered histone 7 |
| ENSG00000110328 | GALNT18 | -0,538408385 | 0,688530093 | 0,09732041 | 0,49573736 | polypeptide N-acetylgalactosaminyltransferase 18 |
| ENSG00000282318 | DLGAP2 | -0,538452269 | 0,68850915 | 0,62182386 | 0,952793934 | DLG associated protein 2 |
| ENSG00000248278 | SUMO2P17 | -0,538671448 | 0,688404557 | 0,49328277 | 0,91725463 | SUMO2 pseudogene 17 |
| ENSG00000254431 | AC084083,1 | -0,538683587 | 0,688398764 | 0,49956241 | 0,920190594 | novel transcript |
| ENSG00000275581 | SERF1A | -0,538710414 | 0,688385964 | 0,43390463 | 0,896272827 | small EDRK-rich factor 1A |
| ENSG00000163617 | CCDC191 | -0,538771367 | 0,68835688 | 0,03637977 | 0,280280196 | coiled-coil domain containing 191 |
| ENSG00000125804 | FAM182A | -0,538831113 | 0,688328374 | 0,1757523 | 0,661199837 | family with sequence similarity 182 member A |
| ENSG00000279047 | AC244517,5 | -0,538831505 | 0,688328187 | 0,09201381 | 0,48210489 | novel transcript, antisense to PCDHB1 and PCDHB2 |
| ENSG00000112139 | MDGA1 | -0,538884049 | 0,688303118 | 0,05900758 | 0,377339533 | MAM domain containing glycosylphosphatidylinositol anchor 1 |
| ENSG00000184608 | FAM167A-AS1 | -0,539082073 | 0,688208649 | 0,49703847 | 0,918742673 | FAM167A antisense RNA 1 |
| ENSG00000281931 | SCG5 | -0,539128211 | 0,68818664 | 0,24003481 | 0,749844941 | secretogranin V |
| ENSG00000272145 | NFYC-AS1 | -0,539165909 | 0,688168657 | 0,13523444 | 0,586240921 | NFYC antisense RNA 1 |
| ENSG00000226148 | SLC25A39P1 | -0,539293665 | 0,68810772 | 0,22175341 | 0,729254784 | solute carrier family 25 member 39 pseudogene 1 |
| ENSG00000254208 | AC011773,1 | -0,539364824 | 0,688073781 | 0,27609196 | 0,788074712 | novel transcript, antisense to E2F5 |
| ENSG00000259377 | AC026770,1 | -0,539451769 | 0,688032315 | 0,64368446 | 0,956633898 | novel transcript, intronic to TMOD2 |
| ENSG00000149599 | DUSP15 | -0,539486793 | 0,688015612 | 0,1843931 | 0,676468655 | dual specificity phosphatase 15 |

| | | | | | | |
|---|---|---|---|---|---|---|
| ENSG00000254556 | AF131215,2 | -0,539546711 | 0,687987038 | 0,47153493 | 0,908896785 | novel transcript |
| ENSG00000251867 | AC009812,1 | -0,539557809 | 0,687981746 | 0,22595056 | 0,734959876 | novel transcript, antisense to ZBTB10 |
| ENSG00000233382 | NKAPP1 | -0,539605591 | 0,68795896 | 0,20130355 | 0,701094807 | NFKB activating protein pseudogene 1 |
| ENSG00000234982 | AL441963,1 | -0,539787502 | 0,68787222 | 0,40269475 | 0,877341338 | ribosomal protein L7a (RPL7A) pseudogene |
| ENSG00000227640 | SOX21-AS1 | -0,540086226 | 0,687729804 | 0,06849872 | 0,412469927 | SOX21 antisense divergent transcript 1 |
| ENSG00000182674 | KCNB2 | -0,540274644 | 0,687639992 | 0,02553631 | 0,222200708 | potassium voltage-gated channel subfamily B member 2 |
| ENSG00000259448 | LINC02352 | -0,540275719 | 0,687639479 | 0,29151644 | 0,802058262 | long intergenic non-protein coding RNA 2352 |
| ENSG00000197008 | ZNF138 | -0,540421675 | 0,687569915 | 0,30410331 | 0,813056239 | zinc finger protein 138 |
| ENSG00000270681 | AC095055,1 | -0,540522325 | 0,687521948 | 0,35541745 | 0,846840543 | novel transcript, antisense to SH3D19 |
| ENSG00000178966 | RMI1 | -0,540627177 | 0,687471982 | 0,27497268 | 0,786383635 | RecQ mediated genome instability 1 |
| ENSG00000233885 | YEATS2-AS1 | -0,540641674 | 0,687465074 | 0,06683155 | 0,406265118 | YEATS2 antisense RNA 1 |
| ENSG00000224903 | RNF32-AS1 | -0,540670116 | 0,687451521 | 0,340849 | 0,839475584 | RNF32 antisense RNA 1 |
| ENSG00000232139 | LINC00867 | -0,540815675 | 0,687382165 | 0,37875218 | 0,864112469 | long intergenic non-protein coding RNA 867 |
| ENSG00000231162 | COX11P1 | -0,540908047 | 0,687338155 | 0,65818591 | 0,957906876 | COX11 pseudogene 1 |
| ENSG00000270332 | SMC2-AS1 | -0,540917309 | 0,687333742 | 0,25133851 | 0,763639174 | SMC2 antisense RNA 1 (head to head) |
| ENSG00000011258 | MBTD1 | -0,540943449 | 0,687321289 | 0,25301575 | 0,76515438 | mbt domain containing 1 |
| ENSG00000171105 | INSR | -0,540947619 | 0,687319302 | 0,0404966 | 0,300590158 | insulin receptor |
| ENSG00000135750 | KCNK1 | -0,540990875 | 0,687298695 | 0,45935185 | 0,904862549 | potassium two pore domain channel subfamily K member 1 |
| ENSG00000259536 | AC091045,1 | -0,541032309 | 0,687278956 | 0,310286 | 0,817687618 | novel transcript, antisense to C15orf57 |
| ENSG00000268105 | AC124856,1 | -0,541095478 | 0,687248864 | 0,20762738 | 0,711917344 | ribosomal protein L5 (RPL5) pseudogene |
| ENSG00000227659 | CLYBL-AS2 | -0,541144515 | 0,687225505 | 0,62127844 | 0,952793934 | CLYBL antisense RNA 2 |
| ENSG00000282773 | GOLGA8Q | -0,541231397 | 0,68718412 | 0,60808953 | 0,951231418 | golgin A8 family member Q |
| ENSG00000286189 | AC108021,1 | -0,54128371 | 0,687159203 | 0,65668367 | 0,957448044 | novel transcript |
| ENSG00000235145 | RPSAP16 | -0,541327259 | 0,687138461 | 0,39905925 | 0,875433675 | ribosomal protein SA pseudogene 16 |
| ENSG00000283022 | HYDIN | -0,54137848 | 0,687114065 | 0,07080199 | 0,418829978 | HYDIN axonemal central pair apparatus protein |
| ENSG00000157423 | HYDIN | -0,54137848 | 0,687114065 | 0,07080199 | 0,418829978 | HYDIN axonemal central pair apparatus protein |
| ENSG00000113504 | SLC12A7 | -0,541415724 | 0,687096327 | 0,07175987 | 0,422186784 | solute carrier family 12 member 7 |
| ENSG00000187627 | RGPD1 | -0,541425656 | 0,687091597 | 0,50740068 | 0,922021199 | RANBP2 like and GRIP domain containing 1 |
| ENSG00000168913 | ENHO | -0,541443385 | 0,687083153 | 0,06274953 | 0,391420848 | energy homeostasis associated |
| ENSG00000223949 | ROR1-AS1 | -0,5418609 | 0,686884341 | 0,01950913 | 0,183714864 | ROR1 antisense RNA 1 |
| ENSG00000125257 | ABCC4 | -0,541903523 | 0,686864048 | 0,11961532 | 0,552362303 | ATP binding cassette subfamily C member 4 |

| | | | | | | |
|---|---|---|---|---|---|---|
| ENSG00000274259 | SYNGAP1-AS1 | -0,541991581 | 0,686822125 | 0,46150835 | 0,906165604 | SYNGAP1 antisense RNA 1 |
| ENSG00000121989 | ACVR2A | -0,542025259 | 0,686806092 | 0,21785701 | 0,72555786 | activin A receptor type 2A |
| ENSG00000261978 | AC116025,1 | -0,542067596 | 0,686785937 | 0,77888612 | 0,978587255 | novel transcript, antisense to TBC1D16 |
| ENSG00000259381 | AC090695,1 | -0,542120841 | 0,686760591 | 0,41426749 | 0,885579089 | WD repeat domain 12 (WDR12) pseudogene |
| ENSG00000255492 | H3P33 | -0,542182368 | 0,686731303 | 0,55691533 | 0,938264532 | H3 histone pseudogene 33 |
| ENSG00000253330 | AC024451,2 | -0,542236865 | 0,686705363 | 0,3211322 | 0,824630562 | NSFL1 (p97) cofactor (p47) (NSFL1C) pseudogene |
| ENSG00000133315 | MACROD1 | -0,542359056 | 0,686647203 | 0,16954813 | 0,652045207 | mono-ADP ribosylhydrolase 1 |
| ENSG00000177599 | ZNF491 | -0,542457898 | 0,686600162 | 0,11306558 | 0,53713451 | zinc finger protein 491 |
| ENSG00000224825 | RORB-AS1 | -0,542530344 | 0,686565684 | 0,35501994 | 0,846502691 | RORB antisense RNA 1 |
| ENSG00000275367 | AC092111,1 | -0,542628926 | 0,686518772 | 0,02656911 | 0,228077075 | novel transcript |
| ENSG00000266900 | AC027514,1 | -0,54262974 | 0,686518384 | 0,42208265 | 0,889775006 | novel transcript, sense intronic to KIAA1468 |
| ENSG00000286989 | AL691432,4 | -0,542730151 | 0,686470604 | 0,55451323 | 0,937793851 | novel transcript |
| ENSG00000287759 | AL353151,2 | -0,542761192 | 0,686455834 | 0,29017191 | 0,800682868 | novel transcript, antisense to AK3 |
| ENSG00000198521 | ZNF43 | -0,542887669 | 0,686395657 | 0,11040424 | 0,530211684 | zinc finger protein 43 |
| ENSG00000187080 | OR2AK2 | -0,542932204 | 0,686374469 | 0,58188604 | 0,944149815 | olfactory receptor family 2 subfamily AK member 2 |
| ENSG00000090661 | CERS4 | -0,543060311 | 0,686313524 | 0,06373441 | 0,395512649 | ceramide synthase 4 |
| ENSG00000270195 | AC016773,1 | -0,543157129 | 0,686267468 | 0,25336057 | 0,765172364 | novel transcript |
| ENSG00000143590 | EFNA3 | -0,543165916 | 0,686263288 | 0,12984468 | 0,575973903 | ephrin A3 |
| ENSG00000203943 | SAMD13 | -0,543320688 | 0,68618967 | 0,29689222 | 0,806635068 | sterile alpha motif domain containing 13 |
| ENSG00000235138 | AL445931,1 | -0,543357949 | 0,686171948 | 0,38238694 | 0,866961613 | novel transcript, antisense to BRD3 |
| ENSG00000201183 | RNVU1-3 | -0,543512339 | 0,686098521 | 0,41784183 | 0,886857227 | RNA, variant U1 small nuclear 3 |
| ENSG00000276434 | AL136221,1 | -0,543577749 | 0,686067415 | 0,65155964 | 0,957089277 | novel transcript, sense intronic to GRTP1 |
| ENSG00000176293 | ZNF135 | -0,543754215 | 0,685983502 | 0,0437685 | 0,31589834 | zinc finger protein 135 |
| ENSG00000221116 | SNORD110 | -0,543757792 | 0,685981801 | 0,65635897 | 0,957444211 | small nucleolar RNA, C/D box 110 |
| ENSG00000259212 | AC103739,1 | -0,543811116 | 0,685956447 | 0,52884312 | 0,930077007 | novel transcript, antisense to CRTC3 |
| ENSG00000131650 | KREMEN2 | -0,543944159 | 0,685893192 | 0,43552881 | 0,896403735 | kringle containing transmembrane protein 2 |
| ENSG00000267523 | AC008735,2 | -0,543954249 | 0,685888395 | 0,22704707 | 0,735496246 | novel transcript, antisense to U2AF2A |
| ENSG00000145642 | SHISAL2B | -0,544103357 | 0,68581751 | 0,63973111 | 0,956018518 | shisa like 2B |
| ENSG00000133687 | TMTC1 | -0,544165821 | 0,685787816 | 0,06808397 | 0,411001592 | transmembrane O-mannosyltransferase targeting cadherins 1 |
| ENSG00000241549 | GUSBP2 | -0,544211523 | 0,685766092 | 0,0166395 | 0,165046811 | GUSB pseudogene 2 |
| ENSG00000282364 | AC004980,6 | -0,544264097 | 0,685741102 | 0,38710538 | 0,869441674 | pseudogene similar to part of RB-associated KRAB zinc finger RBAK |

| | | | | | | |
|---|---|---|---|---|---|---|
| ENSG00000214243 | AC004980,2 | -0,544264097 | 0,685741102 | 0,38710538 | 0,869441674 | pseudogene similar to part of RB-associated KRAB zinc finger RBAK |
| ENSG00000284606 | AC105233,4 | -0,544294862 | 0,685726479 | 0,04042204 | 0,300321946 | novel transcript |
| ENSG00000117155 | SSX2IP | -0,54435073 | 0,685699925 | 0,19876469 | 0,697657684 | SSX family member 2 interacting protein |
| ENSG00000272825 | AL844908,1 | -0,544515067 | 0,685621822 | 0,43166989 | 0,89541715 | novel transcript |
| ENSG00000266385 | AC005899,5 | -0,544690824 | 0,685538301 | 0,6953076 | 0,967599788 | novel transcript |
| ENSG00000248671 | ALG1L9P | -0,544713074 | 0,685527728 | 0,0365724 | 0,281486846 | ALG1 like 9, pseudogene |
| ENSG00000280018 | CU634019,5 | -0,544779191 | 0,685496312 | 0,02096055 | 0,192913017 | novel transcript |
| ENSG00000261528 | CCNYL7 | -0,544806251 | 0,685483455 | 0,48623345 | 0,914091794 | cyclin Y like 7 (pseudogene) |
| ENSG00000151690 | MFSD6 | -0,544977703 | 0,685401995 | 0,15765978 | 0,630235931 | major facilitator superfamily domain containing 6 |
| ENSG00000199719 | RN7SKP74 | -0,544983324 | 0,685399325 | 0,64469146 | 0,956633898 | RN7SK pseudogene 74 |
| ENSG00000232970 | POLHP1 | -0,545224449 | 0,68528478 | 0,38711231 | 0,869441674 | DNA polymerase eta pseudogene 1 |
| ENSG00000120645 | IQSEC3 | -0,545428912 | 0,685187667 | 0,26066094 | 0,772692192 | IQ motif and Sec7 domain ArfGEF 3 |
| ENSG00000270071 | AP001172,1 | -0,545618426 | 0,685097665 | 0,0978108 | 0,496997855 | novel transcript |
| ENSG00000233405 | LINC01046 | -0,545665375 | 0,685075371 | 0,63335544 | 0,954689961 | long intergenic non-protein coding RNA 1046 |
| ENSG00000133124 | IRS4 | -0,545796918 | 0,68501291 | 0,32024426 | 0,824366234 | insulin receptor substrate 4 |
| ENSG00000189056 | RELN | -0,545995036 | 0,684918847 | 0,34920107 | 0,843141089 | reelin |
| ENSG00000240175 | MAGI1-AS1 | -0,546167906 | 0,684836782 | 0,63552881 | 0,955314044 | MAGI1 antisense RNA 1 |
| ENSG00000182648 | LINC01006 | -0,546216587 | 0,684813673 | 0,05157946 | 0,347838733 | long intergenic non-protein coding RNA 1006 |
| ENSG00000116191 | RALGPS2 | -0,546324587 | 0,68476241 | 0,28577119 | 0,797420923 | Ral GEF with PH domain and SH3 binding motif 2 |
| ENSG00000169760 | NLGN1 | -0,546454109 | 0,684700936 | 0,28542415 | 0,796948543 | neuroligin 1 |
| ENSG00000198919 | DZIP3 | -0,546504989 | 0,68467679 | 0,18285021 | 0,673574747 | DAZ interacting zinc finger protein 3 |
| ENSG00000267383 | AC011447,3 | -0,546580562 | 0,684640925 | 0,05660963 | 0,36741615 | novel transcript, antisense to ZNF486 and ZNF90 |
| ENSG00000262003 | AC087392,1 | -0,546594255 | 0,684634427 | 0,39114173 | 0,870057277 | novel transcript, antisense to NXN |
| ENSG00000169635 | HIC2 | -0,546830368 | 0,684522388 | 0,13512057 | 0,586038109 | HIC ZBTB transcriptional repressor 2 |
| ENSG00000278624 | AZU1 | -0,547133751 | 0,684378455 | 0,84790874 | 0,987036295 | azurocidin 1 |
| ENSG00000157343 | ARMC12 | -0,547406651 | 0,684249011 | 0,27182209 | 0,784843944 | armadillo repeat containing 12 |
| ENSG00000271137 | AC006254,3 | -0,547466193 | 0,684220771 | 0,56122283 | 0,940567693 | AKT interacting protein, pseudogene |
| ENSG00000147689 | FAM83A | -0,54760355 | 0,684155631 | 0,68628417 | 0,965752124 | family with sequence similarity 83 member A |
| ENSG00000125538 | IL1B | -0,547609729 | 0,684152701 | 0,29641594 | 0,806119355 | interleukin 1 beta |
| ENSG00000170315 | UBB | -0,547837136 | 0,684044869 | 0,01606345 | 0,161325917 | ubiquitin B |
| ENSG00000269559 | AC093677,2 | -0,547845684 | 0,684040816 | 0,22486452 | 0,733203418 | novel transcript, antisense to MTHFD2L |

| | | | | | | |
|---|---|---|---|---|---|---|
| ENSG00000242950 | ERVW-1 | -0,547872096 | 0,684028293 | 0,02218495 | 0,200749686 | endogenous retrovirus group W member 1, envelope |
| ENSG00000241057 | AC004951,3 | -0,548064005 | 0,683937309 | 0,6018183 | 0,949939978 | RAS p21 protein activator 4 (RASA4) pseudogene |
| ENSG00000170579 | DLGAP1 | -0,548142158 | 0,68390026 | 0,08921857 | 0,476498052 | DLG associated protein 1 |
| ENSG00000237575 | PYY2 | -0,548175013 | 0,683884685 | 0,53268438 | 0,931291085 | peptide YY 2 (pseudogene) |
| ENSG00000233393 | AP000688,2 | -0,548269693 | 0,683839805 | 0,30974254 | 0,817030292 | novel transcript |
| ENSG00000232931 | LINC00342 | -0,548325884 | 0,683813171 | 0,04569958 | 0,324200544 | long intergenic non-protein coding RNA 342 |
| ENSG00000204147 | ASAH2B | -0,548341549 | 0,683805746 | 0,23939975 | 0,749122637 | N-acylsphingosine amidohydrolase 2B |
| ENSG00000243789 | JMJD7 | -0,548406912 | 0,683774766 | 0,12565864 | 0,566207813 | jumonji domain containing 7 |
| ENSG00000214432 | VPS33B-DT | -0,548458521 | 0,683750306 | 0,33219882 | 0,83293962 | VPS33B divergent transcript |
| ENSG00000269427 | AC024075,3 | -0,548490352 | 0,68373522 | 0,60984653 | 0,951801946 | novel transcript, antisense C19orf42 |
| ENSG00000272902 | TBC1D8-AS1 | -0,548511713 | 0,683725097 | 0,52284257 | 0,92790906 | TBC1D8 antisense RNA 1 |
| ENSG00000120440 | TTLL2 | -0,548663449 | 0,683653189 | 0,46443205 | 0,907500217 | tubulin tyrosine ligase like 2 |
| ENSG00000177054 | ZDHHC13 | -0,548673282 | 0,68364853 | 0,19053261 | 0,685949202 | zinc finger DHHC-type palmitoyltransferase 13 |
| ENSG00000121211 | MND1 | -0,548872524 | 0,683554122 | 0,41090045 | 0,882983885 | meiotic nuclear divisions 1 |
| ENSG00000250170 | RASA2-IT1 | -0,548990756 | 0,683498105 | 0,48140319 | 0,912981134 | RASA2 intronic transcript 1 |
| ENSG00000285312 | C6orf52 | -0,548997878 | 0,683494731 | 0,29711272 | 0,806635068 | chromosome 6 open reading frame 52 |
| ENSG00000137434 | C6orf52 | -0,548997878 | 0,683494731 | 0,29711272 | 0,806635068 | chromosome 6 open reading frame 52 |
| ENSG00000288573 | AL357079,3 | -0,549167276 | 0,683414482 | 0,5777985 | 0,943403616 | novel transcript |
| ENSG00000235244 | DANT2 | -0,549208691 | 0,683394863 | 0,14619009 | 0,609479529 | DXZ4 associated non-coding transcript 2, distal |
| ENSG00000232928 | DDX3P1 | -0,549291615 | 0,683355584 | 0,33602591 | 0,835660292 | DEAD-box helicase 3 pseudogene 1 |
| ENSG00000229368 | AC090587,1 | -0,549370311 | 0,683318309 | 0,45112314 | 0,902062594 | novel transcript, sense overlapping STIM1 |
| ENSG00000254634 | SMG1P6 | -0,549483991 | 0,683264468 | 0,11328139 | 0,537503475 | SMG1 pseudogene 6 |
| ENSG00000272505 | AC104964,3 | -0,549835691 | 0,683097922 | 0,20866488 | 0,713284351 | novel transcript |
| ENSG00000196632 | WNK3 | -0,549853365 | 0,683089554 | 0,2319582 | 0,740447339 | WNK lysine deficient protein kinase 3 |
| ENSG00000268970 | AC022150,2 | -0,549904102 | 0,683065531 | 0,52896978 | 0,930182312 | novel transcript, sense intronic to ZNF83 |
| ENSG00000230912 | AL021707,4 | -0,549975874 | 0,683031551 | 0,53793931 | 0,932883277 | novel transcript |
| ENSG00000267577 | AC010327,4 | -0,550132856 | 0,682957233 | 0,68844597 | 0,966481084 | novel transcript, antisense to C19orf51 and SYT5 |
| ENSG00000271789 | AL080317,1 | -0,550226187 | 0,682913052 | 0,20608679 | 0,708889568 | novel transcript |
| ENSG00000102981 | PARD6A | -0,550433482 | 0,682814934 | 0,06538944 | 0,401164726 | par-6 family cell polarity regulator alpha |
| ENSG00000286997 | AC005856,1 | -0,550565894 | 0,682752268 | 0,16749544 | 0,648316613 | novel transcript, antisense to BCAS3 |
| ENSG00000277948 | AC007435,1 | -0,550928261 | 0,6825808 | 0,64084101 | 0,956410077 | |

| | | | | | | |
|---|---|---|---|---|---|---|
| ENSG00000124194 | GDAP1L1 | -0,550967863 | 0,682562063 | 0,04864171 | 0,335726918 | ganglioside induced differentiation associated protein 1 like 1 |
| ENSG00000229321 | AC008269,1 | -0,551157678 | 0,682472264 | 0,1421012 | 0,600058555 | novel transcript |
| ENSG00000264294 | SNORD55 | -0,55145925 | 0,68232962 | 0,41220855 | 0,88438705 | small nucleolar RNA, C/D box 55 |
| ENSG00000115718 | PROC | -0,551516165 | 0,682302702 | 0,59780622 | 0,948390834 | protein C, inactivator of coagulation factors Va and VIIIa |
| ENSG00000274925 | ZKSCAN2-DT | -0,551553489 | 0,68228505 | 0,06822179 | 0,411581814 | ZKSCAN2 divergent transcript |
| ENSG00000266573 | AC018697,1 | -0,551596481 | 0,682264719 | 0,38832071 | 0,869820053 | novel transcript |
| ENSG00000235162 | C12orf75 | -0,551763285 | 0,68218584 | 0,10794239 | 0,523928283 | chromosome 12 open reading frame 75 |
| ENSG00000273802 | H2BC8 | -0,551774761 | 0,682180414 | 0,22889162 | 0,737173617 | H2B clustered histone 8 |
| ENSG00000188848 | BEND4 | -0,551813981 | 0,682161869 | 0,21383077 | 0,721188427 | BEN domain containing 4 |
| ENSG00000275050 | ZNF66 | -0,552018942 | 0,682064962 | 0,18770581 | 0,681741934 | zinc finger protein 66 |
| ENSG00000102879 | CORO1A | -0,552056272 | 0,682047314 | 0,04864186 | 0,335726918 | coronin 1A |
| ENSG00000277462 | ZNF670 | -0,552223773 | 0,681968131 | 0,24785697 | 0,760164991 | zinc finger protein 670 |
| ENSG00000089163 | SIRT4 | -0,55224886 | 0,681956272 | 0,27483371 | 0,786383635 | sirtuin 4 |
| ENSG00000237017 | AC245052,4 | -0,552317723 | 0,681923722 | 0,38321704 | 0,867740551 | novel transcript |
| ENSG00000256664 | AC025423,2 | -0,552425286 | 0,681872881 | 0,22629256 | 0,734968883 | ribosomal L24 domain containing 1 (RSL24D1) pseudogene |
| ENSG00000139146 | SINHCAF | -0,552595521 | 0,681792426 | 0,26204124 | 0,773902615 | SIN3-HDAC complex associated factor |
| ENSG00000250241 | AC105383,1 | -0,552743371 | 0,681722559 | 0,18182699 | 0,672010829 | novel transcript |
| ENSG00000167614 | TTYH1 | -0,552874871 | 0,681660423 | 0,09818843 | 0,497481654 | tweety family member 1 |
| ENSG00000263846 | CIAPIN1P | -0,553050567 | 0,681577414 | 0,55826252 | 0,93916914 | cytokine induced apoptosis inhibitor 1 pseudogene |
| ENSG00000140455 | USP3 | -0,553307044 | 0,681456256 | 0,17419778 | 0,659374935 | ubiquitin specific peptidase 3 |
| ENSG00000286107 | AC007655,2 | -0,553342127 | 0,681439685 | 0,66563016 | 0,961016016 | novel transcript |
| ENSG00000244490 | RWDD4P1 | -0,553354328 | 0,681433922 | 0,35742046 | 0,848693232 | RWD domain containing 4 pseudogene 1 |
| ENSG00000242493 | RN7SL37P | -0,553405255 | 0,681409868 | 0,64844351 | 0,956899963 | RNA, 7SL, cytoplasmic 37, pseudogene |
| ENSG00000152926 | ZNF117 | -0,553408882 | 0,681408155 | 0,22936117 | 0,737655337 | zinc finger protein 117 |
| ENSG00000286787 | AL110114,1 | -0,553437898 | 0,68139445 | 0,13340637 | 0,583647193 | novel transcript, antisense to PSMF1 |
| ENSG00000162591 | MEGF6 | -0,55371873 | 0,681261824 | 0,26596197 | 0,777251371 | multiple EGF like domains 6 |
| ENSG00000124602 | UNC5CL | -0,553765066 | 0,681239944 | 0,22624175 | 0,734959876 | unc-5 family C-terminal like |
| ENSG00000271207 | MTCO1P22 | -0,553996858 | 0,681130501 | 0,57654692 | 0,943231961 | MT-CO1 pseudogene 22 |
| ENSG00000214018 | RRM2P3 | -0,554176052 | 0,681045904 | 0,52003508 | 0,926822772 | ribonucleotide reductase M2 polypeptide pseudogene 3 |

| | | | | | | |
|---|---|---|---|---|---|---|
| ENSG00000275185 | AC130324,3 | -0,554261924 | 0,681005368 | 0,51042151 | 0,922630564 | novel transcript, antisense to TEFM |
| ENSG00000154874 | CCDC144B | -0,554279522 | 0,680997061 | 0,05989727 | 0,38077031 | coiled-coil domain containing 144B (pseudogene) |
| ENSG00000167702 | KIFC2 | -0,554429063 | 0,680926477 | 0,23521125 | 0,744654651 | kinesin family member C2 |
| ENSG00000101850 | GPR143 | -0,55447113 | 0,680906623 | 0,1846204 | 0,676844797 | G protein-coupled receptor 143 |
| ENSG00000255794 | RMST | -0,554621012 | 0,680835887 | 0,26779501 | 0,780440321 | rhabdomyosarcoma 2 associated transcript |
| ENSG00000205277 | MUC12 | -0,554840834 | 0,680732156 | 0,2520908 | 0,764267906 | mucin 12, cell surface associated |
| ENSG00000151778 | SERP2 | -0,554863766 | 0,680721336 | 0,08544528 | 0,465393939 | stress associated endoplasmic reticulum protein family member 2 |
| ENSG00000184659 | FOXD4L4 | -0,555226219 | 0,680550338 | 0,38106877 | 0,865900094 | forkhead box D4 like 4 |
| ENSG00000131400 | NAPSA | -0,55543791 | 0,680450486 | 0,29638744 | 0,806111953 | napsin A aspartic peptidase |
| ENSG00000143479 | DYRK3 | -0,555451003 | 0,68044431 | 0,0594401 | 0,378942783 | dual specificity tyrosine phosphorylation regulated kinase 3 |
| ENSG00000282813 | ARMC10 | -0,555488744 | 0,68042651 | 0,4167435 | 0,886368152 | armadillo repeat containing 10 |
| ENSG00000116032 | GRIN3B | -0,555529922 | 0,680407089 | 0,52473622 | 0,928616707 | glutamate ionotropic receptor NMDA type subunit 3B |
| ENSG00000270497 | BX322635,1 | -0,555547106 | 0,680398985 | 0,40026804 | 0,87601134 | ribosomal protein S3A (RPS3A) pseudogene |
| ENSG00000250616 | AC012645,1 | -0,555875176 | 0,680244279 | 0,13965422 | 0,595782615 | novel transcript, antisense to YPEL3 |
| ENSG00000204366 | ZBTB12 | -0,555882341 | 0,680240901 | 0,15980547 | 0,63459396 | zinc finger and BTB domain containing 12 |
| ENSG00000063180 | CA11 | -0,555947163 | 0,680210337 | 0,06903149 | 0,414510104 | carbonic anhydrase 11 |
| ENSG00000234690 | EPCAM-DT | -0,555952755 | 0,680207701 | 0,28343213 | 0,79458592 | EPCAM divergent transcript |
| ENSG00000118276 | B4GALT6 | -0,55610672 | 0,680135113 | 0,34044618 | 0,839052015 | beta-1,4-galactosyltransferase 6 |
| ENSG00000132970 | WASF3 | -0,556164123 | 0,680108052 | 0,1486371 | 0,613346323 | WASP family member 3 |
| ENSG00000279668 | AC024610,2 | -0,556428126 | 0,679983608 | 0,31184178 | 0,818208978 | novel transcript |
| ENSG00000287524 | AC007736,1 | -0,556489143 | 0,67995485 | 0,40617462 | 0,879163888 | novel transcript, antisense to PARD3B |
| ENSG00000236671 | PRKG1-AS1 | -0,55655573 | 0,679923467 | 0,027932 | 0,236266271 | PRKG1 antisense RNA 1 |
| ENSG00000113966 | ARL6 | -0,556555898 | 0,679923388 | 0,30928774 | 0,816845699 | ADP ribosylation factor like GTPase 6 |
| ENSG00000130045 | NXNL2 | -0,556580115 | 0,679911975 | 0,19034982 | 0,685926325 | nucleoredoxin like 2 |
| ENSG00000237757 | EEF1A1P30 | -0,556607164 | 0,679899228 | 0,37933678 | 0,864853196 | eukaryotic translation elongation factor 1 alpha 1 pseudogene 30 |
| ENSG00000286139 | ARHGAP11B | -0,556621637 | 0,679892407 | 0,18328073 | 0,674320815 | Rho GTPase activating protein 11B |
| ENSG00000244476 | ERVFRD-1 | -0,556736389 | 0,67983833 | 0,09153742 | 0,481302729 | endogenous retrovirus group FRD member 1, envelope |
| ENSG00000261008 | LINC01572 | -0,557007626 | 0,679710528 | 0,07531288 | 0,433624677 | long intergenic non-protein coding RNA 1572 |

| | | | | | | |
|---|---|---|---|---|---|---|
| ENSG00000278700 | Metazoa_SRP | -0,557177427 | 0,679630533 | 0,61311506 | 0,952413269 | Metazoan signal recognition particle RNA |
| ENSG00000182885 | ADGRG3 | -0,557246435 | 0,679598025 | 0,60673989 | 0,950927937 | adhesion G protein-coupled receptor G3 |
| ENSG00000253976 | AC022679,2 | -0,557322987 | 0,679561965 | 0,60677456 | 0,950927937 | novel transcript |
| ENSG00000166402 | TUB | -0,55733022 | 0,679558558 | 0,06434125 | 0,397835213 | TUB bipartite transcription factor |
| ENSG00000166896 | ATP23 | -0,557471149 | 0,679492179 | 0,04881408 | 0,336595397 | ATP23 metallopeptidase and ATP synthase assembly factor homolog |
| ENSG00000166856 | GPR182 | -0,557608168 | 0,679427648 | 0,16118181 | 0,637387804 | G protein-coupled receptor 182 |
| ENSG00000170667 | RASA4B | -0,557618732 | 0,679422673 | 0,07357543 | 0,428274511 | RAS p21 protein activator 4B |
| ENSG00000210825 | SNORA40 | -0,557673452 | 0,679396904 | 0,39947545 | 0,875433675 | small nucleolar RNA, H/ACA box 40 |
| ENSG00000134940 | ACRV1 | -0,557873994 | 0,67930247 | 0,10174081 | 0,50730857 | acrosomal vesicle protein 1 |
| ENSG00000269707 | AC018730,1 | -0,55790189 | 0,679289336 | 0,06212888 | 0,389024096 | novel transcript, sense overlapping POU3F3 |
| ENSG00000187951 | AC091057,1 | -0,557912109 | 0,679284524 | 0,03852143 | 0,291183243 | OTU deubiquitinase 7A pseudogene |
| ENSG00000258758 | AL137100,1 | -0,557915738 | 0,679282815 | 0,64468103 | 0,956633898 | zinc finger, CCHC domain containing 7 (ZCCHC7) pseudogene |
| ENSG00000278041 | AL133325,3 | -0,557946835 | 0,679268174 | 0,55627028 | 0,938046927 | novel transcript |
| ENSG00000184613 | NELL2 | -0,557994079 | 0,67924593 | 0,12049446 | 0,553925447 | neural EGFL like 2 |
| ENSG00000285634 | AL583827,1 | -0,558150302 | 0,679172382 | 0,05116927 | 0,346267575 | Novel transcript |
| ENSG00000287840 | AC009268,3 | -0,558503661 | 0,679006052 | 0,68316718 | 0,964965151 | novel transcript, antisense to AVEN |
| ENSG00000240758 | AC010655,2 | -0,558515478 | 0,679000491 | 0,49461005 | 0,91758518 | novel transcript |
| ENSG00000274110 | LAIR1 | -0,558581131 | 0,678969592 | 0,2716543 | 0,784717743 | leukocyte associated immunoglobulin like receptor 1 |
| ENSG00000272182 | AC135507,1 | -0,558611875 | 0,678955123 | 0,40474108 | 0,878176155 | novel transcript, antisense to PDHB |
| ENSG00000253955 | AC008663,2 | -0,558738346 | 0,678895606 | 0,33644272 | 0,835713011 | novel transcript |
| ENSG00000132855 | ANGPTL3 | -0,558847568 | 0,678844212 | 0,49388972 | 0,917303403 | angiopoietin like 3 |
| ENSG00000144278 | GALNT13 | -0,559053876 | 0,678747142 | 0,15862027 | 0,632402923 | polypeptide N-acetylgalactosaminyltransferase 13 |
| ENSG00000251349 | MSANTD3-TME | -0,559129283 | 0,678711667 | 0,42209728 | 0,889775006 | MSANTD3-TMEFF1 readthrough |
| ENSG00000285545 | AC124798,2 | -0,559437696 | 0,67856659 | 0,73113366 | 0,974353474 | novel transcript, antisense to NUCB2 |
| ENSG00000130830 | MPP1 | -0,559480117 | 0,678546637 | 0,098539 | 0,498418285 | membrane palmitoylated protein 1 |
| ENSG00000277273 | CDK7 | -0,559558701 | 0,678509678 | 0,21733064 | 0,725037995 | cyclin dependent kinase 7 |
| ENSG00000234065 | MTND4P26 | -0,559643231 | 0,678469924 | 0,60114363 | 0,949882873 | MT-ND4 pseudogene 26 |
| ENSG00000198208 | RPS6KL1 | -0,55965274 | 0,678465452 | 0,07375116 | 0,429054144 | ribosomal protein S6 kinase like 1 |
| ENSG00000274614 | AL390754,1 | -0,559652984 | 0,678465338 | 0,41474289 | 0,885579089 | spalt-like transcription factor 4 (SALL4) pseudogene |
| ENSG00000286892 | AC084212,1 | -0,559656909 | 0,678463492 | 0,21794245 | 0,72555786 | novel transcript |

| | | | | | | |
|---|---|---|---|---|---|---|
| ENSG00000268230 | AC012313,3 | -0,559657629 | 0,678463153 | 0,29205442 | 0,802771257 | novel transcript |
| ENSG00000254423 | AC087203,1 | -0,559718384 | 0,678434582 | 0,50702304 | 0,921951873 | ubiquitin specific peptidase 17-like 2 ( USP17L2) pseudogene |
| ENSG00000275763 | ZNF516-DT | -0,559771576 | 0,678409569 | 0,44066725 | 0,898579495 | ZNF516 divergent transcript |
| ENSG00000275428 | AC024940,6 | -0,559811106 | 0,678390981 | 0,35720388 | 0,848693232 | ovostatin 2 |
| ENSG00000087495 | PHACTR3 | -0,559931256 | 0,678334486 | 0,2575855 | 0,769908546 | phosphatase and actin regulator 3 |
| ENSG00000231381 | RNF2P1 | -0,560232048 | 0,678193072 | 0,47379551 | 0,910320967 | ring finger protein 2 pseudogene 1 |
| ENSG00000226632 | UBE2V1P1 | -0,560307914 | 0,678157409 | 0,47457447 | 0,91069468 | ubiquitin conjugating enzyme E2 V1 pseudogene 1 |
| ENSG00000145088 | EAF2 | -0,560425881 | 0,67810196 | 0,33105378 | 0,832186595 | ELL associated factor 2 |
| ENSG00000174292 | TNK1 | -0,560469883 | 0,678081278 | 0,21717664 | 0,724864986 | tyrosine kinase non receptor 1 |
| ENSG00000283781 | TNK1 | -0,560469883 | 0,678081278 | 0,21717664 | 0,724864986 | tyrosine kinase non receptor 1 |
| ENSG00000216813 | AL358178,1 | -0,560540381 | 0,678048144 | 0,55550947 | 0,937793851 | 40S ribosomal protein S12 (RPS12) pseudogene |
| ENSG00000227973 | PIN4P1 | -0,560550034 | 0,678043607 | 0,43888716 | 0,897402199 | peptidylprolyl cis/trans isomerase, NIMA-interacting 4 pseudogene 1 |
| ENSG00000227236 | AL583844,1 | -0,560645091 | 0,677998933 | 0,17245419 | 0,656926048 | solute carrier family 39 (zinc transporter), member 14 pseudogene (SLC39A14) |
| ENSG00000230071 | RPL4P6 | -0,560783537 | 0,677933874 | 0,27483087 | 0,786383635 | ribosomal protein L4 pseudogene 6 |
| ENSG00000175536 | LIPT2 | -0,560788127 | 0,677931717 | 0,2246481 | 0,733203418 | lipoyl(octanoyl) transferase 2 |
| ENSG00000184060 | ADAP2 | -0,560838602 | 0,677907999 | 0,33575116 | 0,835476302 | ArfGAP with dual PH domains 2 |
| ENSG00000232260 | BTF3L4P1 | -0,560852184 | 0,677901617 | 0,46828329 | 0,907940653 | basic transcription factor 3 like 4 pseudogene 1 |
| ENSG00000267150 | AC006557,1 | -0,560897498 | 0,677880324 | 0,19021384 | 0,685594362 | novel transcript, antisense ZNF519 |
| ENSG00000270276 | H4C15 | -0,560923088 | 0,677868301 | 0,25310029 | 0,76515438 | H4 clustered histone 15 |
| ENSG00000259505 | AC009268,1 | -0,561259538 | 0,677710233 | 0,62004123 | 0,952793934 | Sjogren syndrome antigen B (autoantigen La) (SSB) pseudogene |
| ENSG00000225140 | AL358216,1 | -0,561272795 | 0,677704006 | 0,26909965 | 0,781347689 | novel transcript |
| ENSG00000276075 | AC027682,6 | -0,561702088 | 0,677502376 | 0,48482605 | 0,913690171 | novel transcript, antisense to FAM65A |
| ENSG00000213096 | ZNF254 | -0,561715938 | 0,677495873 | 0,25563037 | 0,767141836 | zinc finger protein 254 |
| ENSG00000285335 | CACNA1C-AS1 | -0,561733352 | 0,677487695 | 0,66616046 | 0,961020116 | CACNA1C antisense RNA 1 |
| ENSG00000246627 | CACNA1C-AS1 | -0,561733352 | 0,677487695 | 0,66616046 | 0,961020116 | CACNA1C antisense RNA 1 |
| ENSG00000140876 | NUDT7 | -0,561869374 | 0,677423822 | 0,28542944 | 0,796948543 | nudix hydrolase 7 |
| ENSG00000249359 | AC093274,1 | -0,562187728 | 0,677274354 | 0,46815339 | 0,907940653 | novel transcript |
| ENSG00000255559 | ZNF252P-AS1 | -0,562256848 | 0,677241906 | 0,33086038 | 0,832114797 | ZNF252P antisense RNA 1 |
| ENSG00000285747 | AL133485,3 | -0,56227973 | 0,677231165 | 0,47102795 | 0,908869685 | novel transcript |

| | | | | | | |
|---|---|---|---|---|---|---|
| ENSG00000249693 | THEGL | -0,56228661 | 0,677227935 | 0,3747727 | 0,861771195 | theg spermatid protein like |
| ENSG00000286545 | AC092645,2 | -0,562416444 | 0,677166992 | 0,18203547 | 0,672311616 | novel transcript, antisense to ZC3H8 |
| ENSG00000283886 | BX664615,2 | -0,562426615 | 0,677162218 | 0,19807078 | 0,696782178 | Uncharacterized protein FLJ76381 |
| ENSG00000243053 | RPL31P58 | -0,562527485 | 0,677114874 | 0,60225439 | 0,950180678 | ribosomal protein L31 pseudogene 58 |
| ENSG00000253554 | LINC01414 | -0,562679556 | 0,677043504 | 0,12532392 | 0,565631173 | long intergenic non-protein coding RNA 1414 |
| ENSG00000255411 | LINC02548 | -0,562787844 | 0,676992688 | 0,06376084 | 0,395519632 | long intergenic non-protein coding RNA 2548 |
| ENSG00000265480 | KRT18P55 | -0,562789837 | 0,676991752 | 0,6284287 | 0,952793934 | keratin 18 pseudogene 55 |
| ENSG00000278677 | H2AC17 | -0,562844108 | 0,676966286 | 0,1184561 | 0,550038851 | H2A clustered histone 17 |
| ENSG00000257957 | QRSL1P3 | -0,562938898 | 0,676921808 | 0,30814715 | 0,816424917 | QRSL1 pseudogene 3 |
| ENSG00000254717 | GLYATL1P2 | -0,563014528 | 0,676886323 | 0,18113655 | 0,670965888 | glycine-N-acyltransferase like 1 pseudogene 2 |
| ENSG00000276030 | AC073534,2 | -0,563060195 | 0,676864897 | 0,52080402 | 0,927046149 | novel transcript |
| ENSG00000273406 | AC245008,1 | -0,563129973 | 0,676832161 | 0,61510838 | 0,952793934 | novel transcript |
| ENSG00000165121 | AL353743,1 | -0,563140236 | 0,676827346 | 0,1283778 | 0,572390704 | kinesin family member 27 (KIF27) pseudogene |
| ENSG00000141337 | ARSG | -0,563479426 | 0,676668237 | 0,06223132 | 0,389221888 | arylsulfatase G |
| ENSG00000217488 | AL356057,1 | -0,563509022 | 0,676654355 | 0,27436556 | 0,786239603 | LSM14A, SCD6 homolog A (S, cerevisiae) (LSM14A) pseudogene |
| ENSG00000129295 | LRRC6 | -0,563572475 | 0,676624595 | 0,22883653 | 0,737173617 | leucine rich repeat containing 6 |
| ENSG00000270105 | AC136475,8 | -0,563648694 | 0,67658885 | 0,6840117 | 0,965403332 | novel transcript |
| ENSG00000272979 | AC093388,1 | -0,563694534 | 0,676567352 | 0,5760852 | 0,943019411 | novel transcript, antisense to MFSD6 |
| ENSG00000145087 | STXBP5L | -0,563928735 | 0,67645753 | 0,26473649 | 0,776063906 | syntaxin binding protein 5 like |
| ENSG00000213558 | HMGN2P32 | -0,563941659 | 0,67645147 | 0,52069491 | 0,927046149 | high mobility group nucleosomal binding domain 2 pseudogene 32 |
| ENSG00000233928 | AL591501,1 | -0,56395691 | 0,676444319 | 0,04817195 | 0,334380043 | novel transcript |
| ENSG00000177736 | FXNP2 | -0,563958498 | 0,676443574 | 0,48178629 | 0,912981134 | frataxin pseudogene 2 |
| ENSG00000260729 | AC009690,1 | -0,563960043 | 0,67644285 | 0,47232885 | 0,909179229 | novel protein |
| ENSG00000152076 | CCDC74B | -0,563983603 | 0,676431804 | 0,07755312 | 0,440002866 | coiled-coil domain containing 74B |
| ENSG00000118514 | ALDH8A1 | -0,564027135 | 0,676411393 | 0,49902016 | 0,919783412 | aldehyde dehydrogenase 8 family member A1 |
| ENSG00000224892 | RPS4XP16 | -0,56407301 | 0,676389885 | 0,4615251 | 0,906165604 | ribosomal protein S4X pseudogene 16 |
| ENSG00000286237 | ARMCX5-GPRA | -0,564084063 | 0,676384703 | 0,07661432 | 0,437332509 | ARMCX5-GPRASP2 readthrough |
| ENSG00000226429 | AL590399,2 | -0,56420863 | 0,676326304 | 0,55886955 | 0,939635125 | novel pseudogene |
| ENSG00000259976 | AC093010,2 | -0,564225592 | 0,676318352 | 0,78563521 | 0,979320406 | novel transcript |
| ENSG00000138823 | MTTP | -0,56437168 | 0,676249872 | 0,23719337 | 0,746624292 | microsomal triglyceride transfer protein |
| ENSG00000108641 | B9D1 | -0,564690936 | 0,67610024 | 0,01214042 | 0,131911723 | B9 domain containing 1 |

| | | | | | | |
|---|---|---|---|---|---|---|
| ENSG00000259866 | AC007599,1 | -0,564796253 | 0,676050886 | 0,53949558 | 0,933350549 | Rho GTPase activating protein 20 (ARHGAP20) pseudogene |
| ENSG00000223935 | LGALSL-DT | -0,564995585 | 0,675957485 | 0,24504841 | 0,757071585 | LGALSL divergent transcript |
| ENSG00000196476 | C20orf96 | -0,565089959 | 0,675913269 | 0,04610127 | 0,325793402 | chromosome 20 open reading frame 96 |
| ENSG00000104879 | CKM | -0,565265897 | 0,675830846 | 0,60988707 | 0,951801946 | creatine kinase, M-type |
| ENSG00000235706 | DICER1-AS1 | -0,565308705 | 0,675810793 | 0,06870363 | 0,413067344 | DICER1 antisense RNA 1 |
| ENSG00000250444 | CCT5P1 | -0,565368034 | 0,675783001 | 0,57894716 | 0,943828882 | chaperonin containing TCP1 subunit 5 pseudogene 1 |
| ENSG00000122733 | PHF24 | -0,565438761 | 0,675749872 | 0,25049499 | 0,762632703 | PHD finger protein 24 |
| ENSG00000092200 | RPGRIP1 | -0,565440733 | 0,675748948 | 0,56782964 | 0,94163744 | RPGR interacting protein 1 |
| ENSG00000154548 | SRSF12 | -0,565537001 | 0,675703859 | 0,15433125 | 0,623607455 | serine and arginine rich splicing factor 12 |
| ENSG00000170681 | CAVIN4 | -0,565546575 | 0,675699375 | 0,10250982 | 0,509923155 | caveolae associated protein 4 |
| ENSG00000236901 | MIR600HG | -0,565625385 | 0,675662464 | 0,07577236 | 0,434777568 | MIR600 host gene |
| ENSG00000248710 | AC079594,1 | -0,565705697 | 0,675624853 | 0,06795533 | 0,410542108 | TRIM59 and iFT80 readthrough |
| ENSG00000279110 | AL022323,3 | -0,565708429 | 0,675623573 | 0,14684769 | 0,610674677 | novel transcript |
| ENSG00000271926 | AC008972,1 | -0,565979189 | 0,675496786 | 0,64100608 | 0,956410077 | novel transcript |
| ENSG00000235105 | AL356968,2 | -0,566128357 | 0,675426947 | 0,32168106 | 0,824933727 | amyotrophic lateral sclerosis 2 (juvenile) chromosome region, candidate 2 (ALS2CR2) pseudogene |
| ENSG00000049130 | KITLG | -0,566129604 | 0,675426363 | 0,27108602 | 0,783727885 | KIT ligand |
| ENSG00000144199 | FAHD2B | -0,566343526 | 0,675326219 | 0,02054576 | 0,190211035 | fumarylacetoacetate hydrolase domain containing 2B |
| ENSG00000173627 | APOBEC4 | -0,566393217 | 0,675302959 | 0,25018728 | 0,762364037 | apolipoprotein B mRNA editing enzyme catalytic polypeptide like 4 |
| ENSG00000272163 | AF106564,1 | -0,566743387 | 0,675139069 | 0,09295746 | 0,484612663 | novel transcript, antisense to NEFM |
| ENSG00000182010 | RTKN2 | -0,566825971 | 0,675100423 | 0,23410695 | 0,743586269 | rhotekin 2 |
| ENSG00000198794 | SCAMP5 | -0,566878135 | 0,675076014 | 0,05494477 | 0,362021772 | secretory carrier membrane protein 5 |
| ENSG00000233230 | AC079807,1 | -0,567041 | 0,674999809 | 0,38698027 | 0,869441674 | novel transcript |
| ENSG00000205108 | FAM205A | -0,567167625 | 0,674940567 | 0,56166487 | 0,940714121 | family with sequence similarity 205 member A |
| ENSG00000224861 | YBX1P1 | -0,56718045 | 0,674934568 | 0,65945268 | 0,958490401 | Y-box binding protein 1 pseudogene 1 |
| ENSG00000147535 | PLPP5 | -0,567313015 | 0,674872553 | 0,03493442 | 0,272842801 | phospholipid phosphatase 5 |
| ENSG00000156398 | SFXN2 | -0,567373196 | 0,674844401 | 0,02327166 | 0,20787579 | sideroflexin 2 |
| ENSG00000259554 | AC084882,1 | -0,567475378 | 0,674796606 | 0,51309278 | 0,923745908 | novel transcript, intronic to SEMA6D |
| ENSG00000200312 | RN7SKP255 | -0,5675107 | 0,674780085 | 0,24153205 | 0,751948055 | RN7SK pseudogene 255 |

| | | | | | | |
|---|---|---|---|---|---|---|
| ENSG00000150275 | PCDH15 | -0,567653279 | 0,674713401 | 0,36119756 | 0,851821025 | protocadherin related 15 |
| ENSG00000106804 | C5 | -0,567896975 | 0,674599439 | 0,20485894 | 0,706459522 | complement C5 |
| ENSG00000136895 | GARNL3 | -0,56790181 | 0,674597179 | 0,03977067 | 0,297461745 | GTPase activating Rap/RanGAP domain like 3 |
| ENSG00000253218 | KLF3P1 | -0,568092916 | 0,674507824 | 0,13955722 | 0,595782615 | Kruppel like factor 3 pseudogene 1 |
| ENSG00000258986 | TMEM179 | -0,568096265 | 0,674506259 | 0,39902424 | 0,875433675 | transmembrane protein 179 |
| ENSG00000213592 | PPIAP42 | -0,5682696 | 0,674425224 | 0,42773875 | 0,892705337 | peptidylprolyl isomerase A pseudogene 42 |
| ENSG00000143850 | PLEKHA6 | -0,568329516 | 0,674397215 | 0,05010911 | 0,342279058 | pleckstrin homology domain containing A6 |
| ENSG00000081377 | CDC14B | -0,568396815 | 0,674365757 | 0,06651509 | 0,405287049 | cell division cycle 14B |
| ENSG00000220793 | RPL21P119 | -0,568511347 | 0,674312223 | 0,87687551 | 0,990109814 | ribosomal protein L21 pseudogene 119 |
| ENSG00000139209 | SLC38A4 | -0,568516433 | 0,674309845 | 0,3709991 | 0,858959913 | solute carrier family 38 member 4 |
| ENSG00000138764 | CCNG2 | -0,568637077 | 0,674253459 | 0,18373885 | 0,675018961 | cyclin G2 |
| ENSG00000286092 | AC117503,5 | -0,568698148 | 0,674224918 | 0,2230038 | 0,731429997 | novel transcript, antisense to ATP6V0A2 |
| ENSG00000170827 | CELP | -0,568824265 | 0,674165981 | 0,43664796 | 0,896866043 | carboxyl ester lipase pseudogene |
| ENSG00000180479 | ZNF571 | -0,568832724 | 0,674162028 | 0,11302871 | 0,537122619 | zinc finger protein 571 |
| ENSG00000134369 | NAV1 | -0,568859683 | 0,674149431 | 0,11995251 | 0,553044579 | neuron navigator 1 |
| ENSG00000234810 | AL603840,1 | -0,569015524 | 0,674076612 | 0,04456421 | 0,318908336 | novel transcript |
| ENSG00000287456 | AL157369,1 | -0,569153958 | 0,674011934 | 0,34541503 | 0,841168721 | novel transcript, antisense to SUCLA2 |
| ENSG00000229458 | AC010157,1 | -0,569240918 | 0,673971309 | 0,61320137 | 0,952413269 | novel transcript |
| ENSG00000250334 | LINC00989 | -0,569667189 | 0,673772201 | 0,48955684 | 0,916063942 | long intergenic non-protein coding RNA 989 |
| ENSG00000231720 | AL353747,3 | -0,569689312 | 0,673761869 | 0,54383136 | 0,934259495 | novel transcript |
| ENSG00000264937 | AC100830,2 | -0,569766988 | 0,673725594 | 0,57936845 | 0,943828882 | novel transcript, antisense to ornithine decarboxylase antizyme 2 OAZ2 |
| ENSG00000259326 | AC116158,1 | -0,569861473 | 0,673681472 | 0,32695365 | 0,828689008 | novel transcript, antisense to RASGRP1 |
| ENSG00000229031 | MTCO1P25 | -0,569865596 | 0,673679546 | 0,54481885 | 0,93489368 | MT-CO1 pseudogene 25 |
| ENSG00000246082 | NUDT16P1 | -0,570016434 | 0,673609115 | 0,22950401 | 0,737904201 | nudix hydrolase 16 pseudogene 1 |
| ENSG00000285572 | AL136441,1 | -0,570175573 | 0,673534816 | 0,02834322 | 0,238461392 | novel transcript |
| ENSG00000287837 | AL590064,1 | -0,570191803 | 0,673527239 | 0,18306228 | 0,673904498 | novel transcript |
| ENSG00000102468 | HTR2A | -0,570295983 | 0,673478604 | 0,35751846 | 0,848693232 | 5-hydroxytryptamine receptor 2A |
| ENSG00000282171 | LINC02210 | -0,570334958 | 0,673460409 | 0,22408498 | 0,732823516 | long intergenic non-protein coding RNA 2210 |
| ENSG00000271749 | AC005828,6 | -0,570585629 | 0,673343405 | 0,51422405 | 0,924319732 | mitochondrially encoded cytochrome c oxidase III (MT-CO3) pseudogene |
| ENSG00000225321 | LINC01427 | -0,570588889 | 0,673341883 | 0,05640103 | 0,366903759 | long intergenic non-protein coding RNA 1427 |
| ENSG00000205212 | CCDC144NL | -0,570661478 | 0,673308005 | 0,10326928 | 0,512532106 | CCDC144A N-terminal pseudogene |

| | | | | | | |
|---|---|---|---|---|---|---|
| ENSG00000286145 | AC068282,1 | -0,570757085 | 0,673263386 | 0,12779873 | 0,570756819 | novel transcript, antisense to MAP3K2 |
| ENSG00000224096 | HCG25 | -0,570785277 | 0,67325023 | 0,52202644 | 0,927599011 | HLA complex group 25 |
| ENSG00000233772 | HCG25 | -0,570785277 | 0,67325023 | 0,52202644 | 0,927599011 | HLA complex group 25 |
| ENSG00000231037 | HCG25 | -0,570785277 | 0,67325023 | 0,52202644 | 0,927599011 | HLA complex group 25 |
| ENSG00000173114 | LRRN3 | -0,5708636 | 0,673213681 | 0,24180224 | 0,752128472 | leucine rich repeat neuronal 3 |
| ENSG00000257660 | ADCY6-DT | -0,570956531 | 0,673170317 | 0,62760892 | 0,952793934 | ADCY6 divergent transcript |
| ENSG00000233184 | AC093157,1 | -0,571004829 | 0,673147781 | 0,12272871 | 0,560344415 | novel transcript |
| ENSG00000167011 | NAT16 | -0,571032329 | 0,673134951 | 0,38098287 | 0,865900094 | N-acetyltransferase 16 (putative) |
| ENSG00000117069 | ST6GALNAC5 | -0,571038463 | 0,673132088 | 0,22064623 | 0,728572782 | ST6 N-acetylgalactosaminide alpha-2,6-sialyltransferase 5 |
| ENSG00000272576 | AC027271,1 | -0,571049192 | 0,673127082 | 0,41625952 | 0,886060654 | novel transcript |
| ENSG00000251379 | AC099550,1 | -0,571062749 | 0,673120757 | 0,0304501 | 0,24960459 | novel transcript |
| ENSG00000188596 | CFAP54 | -0,571131473 | 0,673088693 | 0,02611409 | 0,225509966 | cilia and flagella associated protein 54 |
| ENSG00000127995 | CASD1 | -0,571158617 | 0,673076029 | 0,25392499 | 0,76531122 | CAS1 domain containing 1 |
| ENSG00000140450 | ARRDC4 | -0,571245205 | 0,673035634 | 0,07017608 | 0,417549035 | arrestin domain containing 4 |
| ENSG00000235258 | NDUFB4P6 | -0,571284476 | 0,673017314 | 0,57955951 | 0,943828882 | NADH:ubiquinone oxidoreductase subunit B4 pseudogene 6 |
| ENSG00000168477 | TNXB | -0,571444419 | 0,672942704 | 0,36987646 | 0,858104601 | tenascin XB |
| ENSG00000250771 | AC106865,1 | -0,571498001 | 0,672917711 | 0,0417074 | 0,306376906 | toll-like receptor 2 pseudogene |
| ENSG00000285853 | AL137802,3 | -0,571780283 | 0,672786059 | 0,23299483 | 0,742171568 | novel transcript |
| ENSG00000177640 | CASC2 | -0,571981069 | 0,672692431 | 0,13417506 | 0,58432222 | cancer susceptibility 2 |
| ENSG00000156395 | SORCS3 | -0,572524406 | 0,672439135 | 0,24973786 | 0,761653149 | sortilin related VPS10 domain containing receptor 3 |
| ENSG00000231562 | AL512593,1 | -0,572904613 | 0,672261944 | 0,18676346 | 0,680285072 | ribosomal protein L7a (RPL7A) pseudogene |
| ENSG00000223533 | CCHCR1 | -0,572916616 | 0,672256351 | 0,50354025 | 0,921005254 | coiled-coil alpha-helical rod protein 1 |
| ENSG00000224180 | CCHCR1 | -0,572916616 | 0,672256351 | 0,50354025 | 0,921005254 | coiled-coil alpha-helical rod protein 1 |
| ENSG00000270874 | RPAP2P1 | -0,572962207 | 0,672235107 | 0,25475533 | 0,766202086 | RNA polymerase II associated protein 2 pseudogene 1 |
| ENSG00000176165 | FOXG1 | -0,572973772 | 0,672229718 | 0,19561745 | 0,69375328 | forkhead box G1 |
| ENSG00000254622 | NAV2-AS4 | -0,573029507 | 0,672203749 | 0,51124759 | 0,922893865 | NAV2 antisense RNA 4 |
| ENSG00000233757 | AC092835,1 | -0,573105352 | 0,67216841 | 0,06639938 | 0,40505568 | novel C2H2 type zinc finger protein |
| ENSG00000228906 | AL353804,1 | -0,573410893 | 0,672026071 | 0,26241601 | 0,774068771 | novel transcript |
| ENSG00000115255 | REEP6 | -0,573435871 | 0,672014436 | 0,12722598 | 0,569692485 | receptor accessory protein 6 |
| ENSG00000111186 | WNT5B | -0,573509406 | 0,671980184 | 0,21745654 | 0,725179881 | Wnt family member 5B |

| | | | | | | |
|---|---|---|---|---|---|---|
| ENSG00000288579 | AC025472,1 | -0,573577126 | 0,671948642 | 0,5290401 | 0,930201325 | tigger transposable element derived 1 (TIGD1) pseudogene |
| ENSG00000262075 | DKFZP434A062 | -0,573652761 | 0,671913414 | 0,36410329 | 0,854207909 | uncharacterized LOC26102 |
| ENSG00000257176 | AC009318,1 | -0,573730175 | 0,671877361 | 0,06866767 | 0,413010032 | novel transcript, antisense to FAR2 |
| ENSG00000263142 | LRRC37A17P | -0,573747997 | 0,671869062 | 0,27293646 | 0,785672647 | leucine rich repeat containing 37 member A17, pseudogene |
| ENSG00000197822 | OCLN | -0,573760872 | 0,671863066 | 0,10247433 | 0,509827718 | occludin |
| ENSG00000226469 | ADAM1B | -0,573773321 | 0,671857268 | 0,2163685 | 0,723599962 | ADAM metallopeptidase domain 1B (pseudogene) |
| ENSG00000258502 | LINC02290 | -0,57385688 | 0,671818356 | 0,61538006 | 0,952793934 | long intergenic non-protein coding RNA 2290 |
| ENSG00000286974 | AL662884,5 | -0,57392102 | 0,671788489 | 0,26042023 | 0,772692192 | novel transcript, antisense to TNXB |
| ENSG00000233828 | MIR4280HG | -0,574008554 | 0,67174773 | 0,05522301 | 0,362784685 | MIR4280 host gene |
| ENSG00000138439 | FAM117B | -0,574046796 | 0,671729924 | 0,14819228 | 0,612760682 | family with sequence similarity 117 member B |
| ENSG00000287158 | AC117464,1 | -0,574082295 | 0,671713396 | 0,50936494 | 0,922427526 | novel transcript |
| ENSG00000170967 | DDI1 | -0,574104492 | 0,671703061 | 0,35900416 | 0,849814998 | DNA damage inducible 1 homolog 1 |
| ENSG00000262349 | GIHCG | -0,574233383 | 0,671643053 | 0,157422 | 0,629849164 | GIHCG inhibitor of miR-200b/200a/429 expression |
| ENSG00000124596 | OARD1 | -0,574245119 | 0,67163759 | 0,17607326 | 0,661629135 | O-acyl-ADP-ribose deacylase 1 |
| ENSG00000116783 | TNNI3K | -0,574247677 | 0,671636399 | 0,60245856 | 0,950220175 | TNNI3 interacting kinase |
| ENSG00000184612 | RPL7P26 | -0,574266821 | 0,671627486 | 0,28508336 | 0,796602794 | ribosomal protein L7 pseudogene 26 |
| ENSG00000182919 | C11orf54 | -0,574394449 | 0,671568074 | 0,11284794 | 0,53680967 | chromosome 11 open reading frame 54 |
| ENSG00000272316 | AL021368,2 | -0,574423137 | 0,67155472 | 0,22900988 | 0,737173617 | novel transcript |
| ENSG00000223390 | AL445685,1 | -0,574534689 | 0,671502796 | 0,6065188 | 0,950927937 | novel transcript |
| ENSG00000234031 | RPS3AP44 | -0,574572361 | 0,671485261 | 0,39213595 | 0,870909282 | RPS3A pseudogene 44 |
| ENSG00000268201 | AC020915,1 | -0,574594091 | 0,671475147 | 0,30252367 | 0,812236791 | novel transcript, sense intronic to ZNF544 |
| ENSG00000231096 | NDUFB4P3 | -0,574648426 | 0,671449859 | 0,64984067 | 0,956899963 | NADH:ubiquinone oxidoreductase subunit B4 pseudogene 3 |
| ENSG00000179564 | LSMEM2 | -0,574665029 | 0,671442132 | 0,307108 | 0,815892645 | leucine rich single-pass membrane protein 2 |
| ENSG00000272672 | AL391497,1 | -0,574712254 | 0,671420153 | 0,5320994 | 0,931291085 | novel transcript |
| ENSG00000250326 | AC104596,1 | -0,574735884 | 0,671409156 | 0,11766781 | 0,548330959 | novel transcript |
| ENSG00000073910 | FRY | -0,574793829 | 0,67138219 | 0,04765941 | 0,331927802 | FRY microtubule binding protein |
| ENSG00000264229 | RNU4ATAC | -0,5751194 | 0,671230697 | 0,32162522 | 0,824933727 | RNA, U4atac small nuclear (U12-dependent splicing) |
| ENSG00000281333 | AC024941,2 | -0,575158769 | 0,67121238 | 0,47408478 | 0,910491353 | novel transcript, antisense to KITLG |
| ENSG00000272797 | AC092954,1 | -0,575276686 | 0,671157522 | 0,62533597 | 0,952793934 | novel transcript |

| | | | | | | |
|---|---|---|---|---|---|---|
| ENSG00000286813 | AC015871,6 | -0,575679139 | 0,670970322 | 0,17688048 | 0,662750629 | novel transcript, sense intronic to ST20-MTHFSand MTHFS |
| ENSG00000220868 | MRPL35P1 | -0,575688804 | 0,670965827 | 0,39063032 | 0,870057277 | mitochondrial ribosomal protein L35 pseudogene 1 |
| ENSG00000228186 | PRR3 | -0,575762685 | 0,670931468 | 0,04897768 | 0,337574582 | proline rich 3 |
| ENSG00000117154 | IGSF21 | -0,575802875 | 0,670912778 | 0,31883845 | 0,823755375 | immunoglobin superfamily member 21 |
| ENSG00000254662 | AP000781,1 | -0,57594533 | 0,670846533 | 0,62199505 | 0,952793934 | novel transcript |
| ENSG00000286314 | AC010359,3 | -0,576306609 | 0,670678561 | 0,20058345 | 0,700127386 | novel transcript, sense intronic to ERBIN |
| ENSG00000181524 | RPL24P4 | -0,576498731 | 0,670589254 | 0,50206607 | 0,920520138 | RPL24 pseudogene 4 |
| ENSG00000278635 | AC141557,2 | -0,576524066 | 0,670577478 | 0,6239513 | 0,952793934 | WAS protein family homolog 1 (WASH1) pseudogene |
| ENSG00000189410 | SH2D5 | -0,576890635 | 0,670407115 | 0,34851772 | 0,842757646 | SH2 domain containing 5 |
| ENSG00000271880 | AGAP11 | -0,577030871 | 0,670341951 | 0,63709336 | 0,955945807 | ArfGAP with GTPase domain, ankyrin repeat and PH domain 11 |
| ENSG00000185565 | LSAMP | -0,577065164 | 0,670326018 | 0,14342999 | 0,602438935 | limbic system associated membrane protein |
| ENSG00000198429 | ZNF69 | -0,577069746 | 0,670323889 | 0,0161634 | 0,162061834 | zinc finger protein 69 |
| ENSG00000288054 | AL360178,2 | -0,577373647 | 0,670182701 | 0,39840369 | 0,875127721 | novel transcript, antisense to PDE7B |
| ENSG00000230988 | RPL23AP11 | -0,577407089 | 0,670167166 | 0,58253713 | 0,944174844 | ribosomal protein L23a pseudogene 11 |
| ENSG00000283769 | CR392000,2 | -0,577410737 | 0,670165472 | 0,56237145 | 0,940865625 | novel transcript, antisense to a novel protein |
| ENSG00000274089 | PRIMA1 | -0,57756247 | 0,670094992 | 0,29320045 | 0,803365244 | proline rich membrane anchor 1 |
| ENSG00000175785 | PRIMA1 | -0,57756247 | 0,670094992 | 0,29320045 | 0,803365244 | proline rich membrane anchor 1 |
| ENSG00000204624 | DISP3 | -0,577756232 | 0,670005 | 0,04167508 | 0,306211471 | dispatched RND transporter family member 3 |
| ENSG00000197472 | ZNF695 | -0,577962155 | 0,669909374 | 0,13203084 | 0,580641112 | zinc finger protein 695 |
| ENSG00000278344 | AC063943,1 | -0,578026172 | 0,669879649 | 0,61766468 | 0,952793934 | novel transcript |
| ENSG00000231794 | AC009542,1 | -0,578145172 | 0,669824396 | 0,44862316 | 0,902062594 | novel transcript, antisense to WDR91 |
| ENSG00000186314 | PRELID2 | -0,57824073 | 0,669780032 | 0,01870237 | 0,178483188 | PRELI domain containing 2 |
| ENSG00000285081 | AC004593,1 | -0,578552526 | 0,669635294 | 0,36906138 | 0,857942508 | Novel transcript, sense intronic to CPVL |
| ENSG00000171033 | PKIA | -0,578709301 | 0,66956253 | 0,29621349 | 0,805971225 | cAMP-dependent protein kinase inhibitor alpha |
| ENSG00000148798 | INA | -0,578878336 | 0,669484084 | 0,17793759 | 0,664799324 | internexin neuronal intermediate filament protein alpha |
| ENSG00000114378 | HYAL1 | -0,578972955 | 0,669440178 | 0,11256099 | 0,535800356 | hyaluronidase 1 |
| ENSG00000172717 | FAM71D | -0,57899514 | 0,669429883 | 0,59474731 | 0,947469769 | family with sequence similarity 71 member D |
| ENSG00000267709 | AC024592,2 | -0,579198639 | 0,669335464 | 0,36804834 | 0,857426477 | novel transcript, antisense to FUT3 |
| ENSG00000175322 | ZNF519 | -0,579215693 | 0,669327552 | 0,0649108 | 0,399718599 | zinc finger protein 519 |

| | | | | | | |
|---|---|---|---|---|---|---|
| ENSG00000184635 | ZNF93 | -0,579239005 | 0,669316737 | 0,13077864 | 0,577413268 | zinc finger protein 93 |
| ENSG00000258561 | AL359232,1 | -0,579289042 | 0,669293523 | 0,22507414 | 0,733632361 | novel transcript |
| ENSG00000266805 | AP005432,1 | -0,579339465 | 0,669270131 | 0,297216 | 0,806635068 | novel transcript, sense intronic to RALBP1 |
| ENSG00000204815 | TTC25 | -0,579441574 | 0,669222764 | 0,11606098 | 0,544096306 | tetratricopeptide repeat domain 25 |
| ENSG00000274191 | AC026333,4 | -0,579466025 | 0,669211422 | 0,38964789 | 0,870057277 | novel transcript |
| ENSG00000226205 | RPS3AP27 | -0,579513771 | 0,669189275 | 0,34072182 | 0,839360724 | RPS3A pseudogene 27 |
| ENSG00000232875 | HMGN2P35 | -0,579692504 | 0,669106375 | 0,65283123 | 0,95732962 | high mobility group nucleosomal binding domain 2 pseudogene 35 |
| ENSG00000257642 | KCCAT198 | -0,579711847 | 0,669097404 | 0,45452824 | 0,903383444 | renal clear cell carcinoma-associated transcript 198 |
| ENSG00000222012 | AC005481,1 | -0,579816247 | 0,669048987 | 0,09236109 | 0,483114784 | novel transcript, antisense to PTPRN2 |
| ENSG00000248553 | OR52H2P | -0,579871481 | 0,669023373 | 0,6277256 | 0,952793934 | olfactory receptor family 52 subfamily H member 2 pseudogene |
| ENSG00000250422 | AC117529,2 | -0,579997393 | 0,668964986 | 0,62823903 | 0,952793934 | tigger transposable element derived 1 (TIGD1) pseudogene |
| ENSG00000266490 | AC127024,5 | -0,580070574 | 0,668931054 | 0,22756817 | 0,736053199 | novel transcript |
| ENSG00000206828 | RNVU1-30 | -0,580201456 | 0,668870371 | 0,32579893 | 0,827657237 | RNA, variant U1 small nuclear 30 |
| ENSG00000249572 | AC034231,1 | -0,580258417 | 0,668843963 | 0,23675935 | 0,746249059 | novel transcript |
| ENSG00000250227 | TRIM60P14 | -0,580648819 | 0,668662994 | 0,62929382 | 0,952793934 | tripartite motif containing 60 pseudogene 14 |
| ENSG00000281305 | LINC00574 | -0,580665659 | 0,668655189 | 0,35914138 | 0,849846046 | long intergenic non-protein coding RNA 574 |
| ENSG00000129167 | TPH1 | -0,580736664 | 0,668622281 | 0,15722587 | 0,629552485 | tryptophan hydroxylase 1 |
| ENSG00000184274 | LINC00315 | -0,580785042 | 0,66859986 | 0,5749445 | 0,942533938 | long intergenic non-protein coding RNA 315 |
| ENSG00000079102 | RUNX1T1 | -0,580800414 | 0,668592736 | 0,06864999 | 0,413010032 | RUNX1 partner transcriptional co-repressor 1 |
| ENSG00000283462 | AC008505,1 | -0,580838745 | 0,668574972 | 0,60600657 | 0,950927937 | novel transcript |
| ENSG00000198099 | ADH4 | -0,581013716 | 0,668493892 | 0,09849574 | 0,498280077 | alcohol dehydrogenase 4 (class II), pi polypeptide |
| ENSG00000224593 | AC092427,1 | -0,581117705 | 0,668445709 | 0,27764696 | 0,789067728 | ribosomal protein L21 (RPL21) pseudogene |
| ENSG00000164066 | INTU | -0,581148237 | 0,668431563 | 0,05557242 | 0,363892597 | inturned planar cell polarity protein |
| ENSG00000270154 | AC130352,1 | -0,581321465 | 0,668351307 | 0,16906852 | 0,651115803 | novel transcript |
| ENSG00000136122 | BORA | -0,581351856 | 0,668337229 | 0,26050252 | 0,772692192 | BORA aurora kinase A activator |
| ENSG00000213160 | KLHL23 | -0,581459946 | 0,668287157 | 0,22810492 | 0,736341996 | kelch like family member 23 |
| ENSG00000229894 | GK3P | -0,581465682 | 0,6682845 | 0,37724749 | 0,862818448 | glycerol kinase 3 pseudogene |
| ENSG00000181333 | HEPHL1 | -0,581508976 | 0,668264446 | 0,31205032 | 0,818382785 | hephaestin like 1 |
| ENSG00000205268 | PDE7A | -0,581535645 | 0,668252093 | 0,13908664 | 0,594531666 | phosphodiesterase 7A |
| ENSG00000130957 | FBP2 | -0,581681961 | 0,668184323 | 0,41512588 | 0,885783213 | fructose-bisphosphatase 2 |

| | | | | | | |
|---|---|---|---|---|---|---|
| ENSG00000285160 | THSD1P1 | -0,58168456 | 0,668183119 | 0,6317197 | 0,953915351 | thrombospondin type 1 domain containing 1 pseudogene 1 |
| ENSG00000259241 | AC020892,1 | -0,581714089 | 0,668169443 | 0,75054122 | 0,975901808 | novel transcript |
| ENSG00000227257 | AL158827,1 | -0,581775236 | 0,668141124 | 0,7932156 | 0,979659177 | proline-rich nuclear receptor coactivator 2 (PNRC2) pseudogene |
| ENSG00000175077 | RTP1 | -0,582016156 | 0,668029558 | 0,55121313 | 0,936366211 | receptor transporter protein 1 |
| ENSG00000239002 | SCARNA10 | -0,582066596 | 0,668006203 | 0,04823776 | 0,334607086 | small Cajal body-specific RNA 10 |
| ENSG00000146414 | SHPRH | -0,582222641 | 0,667933953 | 0,16692967 | 0,647344249 | SNF2 histone linker PHD RING helicase |
| ENSG00000227542 | AC092614,1 | -0,582231972 | 0,667929634 | 0,16950424 | 0,652045207 | novel transcript |
| ENSG00000248752 | AC116362,1 | -0,582503602 | 0,667803888 | 0,27751084 | 0,789067728 | novel transcript |
| ENSG00000197969 | VPS13A | -0,582575204 | 0,667770745 | 0,2302469 | 0,738831879 | vacuolar protein sorting 13 homolog A |
| ENSG00000196782 | MAML3 | -0,582641692 | 0,667739971 | 0,0128102 | 0,137392093 | mastermind like transcriptional coactivator 3 |
| ENSG00000151655 | ITIH2 | -0,582646774 | 0,667737619 | 0,51367063 | 0,924122372 | inter-alpha-trypsin inhibitor heavy chain 2 |
| ENSG00000270020 | AC009108,3 | -0,582690381 | 0,667717436 | 0,26078681 | 0,772952333 | novel transcript |
| ENSG00000275748 | ARL17B | -0,582705646 | 0,667710371 | 0,06382289 | 0,395590561 | ADP ribosylation factor like GTPase 17B |
| ENSG00000230922 | AL079301,1 | -0,58281942 | 0,667657716 | 0,3052281 | 0,814167717 | novel transcript |
| ENSG00000237975 | FLG-AS1 | -0,58301007 | 0,667569492 | 0,46021197 | 0,905443183 | FLG antisense RNA 1 |
| ENSG00000268438 | BNIP3P27 | -0,583065674 | 0,667543763 | 0,32777251 | 0,829743891 | BCL2 interacting protein 3 pseudogene 27 |
| ENSG00000276888 | AL512624,2 | -0,583196493 | 0,667483235 | 0,50672004 | 0,921824694 | novel transcript, antisense to POTEM |
| ENSG00000070190 | DAPP1 | -0,583242895 | 0,667461767 | 0,40710561 | 0,880082273 | dual adaptor of phosphotyrosine and 3-phosphoinositides 1 |
| ENSG00000213252 | AP001282,1 | -0,583243607 | 0,667461437 | 0,45676594 | 0,903699469 | pyridine nucleotide-disulphide oxidoreductase domain 1 (PYROXD1) pseudogene |
| ENSG00000202415 | RN7SKP269 | -0,5832472 | 0,667459775 | 0,60731489 | 0,950993594 | RN7SK pseudogene 269 |
| ENSG00000240729 | AC146507,2 | -0,583359329 | 0,667407901 | 0,3887795 | 0,870057277 | ribosomal protein L21 (RPL21) pseudogene |
| ENSG00000233003 | CICP26 | -0,583462627 | 0,667360116 | 0,61758784 | 0,952793934 | capicua transcriptional repressor pseudogene 26 |
| ENSG00000184113 | CLDN5 | -0,583508574 | 0,667338862 | 0,2802762 | 0,790779983 | claudin 5 |
| ENSG00000166924 | NYAP1 | -0,583784612 | 0,667211189 | 0,21864734 | 0,726516149 | neuronal tyrosine phosphorylated phosphoinositide-3-kinase adaptor 1 |
| ENSG00000250062 | MAPK10-AS1 | -0,583842742 | 0,667184306 | 0,42066653 | 0,888569577 | MAPK10 antisense RNA 1 |
| ENSG00000281396 | AL122023,2 | -0,583931635 | 0,667143198 | 0,56419671 | 0,940865625 | novel transcript, sense intronic UNC79 |
| ENSG00000278396 | AL122023,1 | -0,583931635 | 0,667143198 | 0,56419671 | 0,940865625 | novel transcript, sense intronic UNC79 |
| ENSG00000287306 | AC016821,1 | -0,583974521 | 0,667123367 | 0,04053027 | 0,300657756 | novel transcript |

| | | | | | | |
|---|---|---|---|---|---|---|
| ENSG00000074803 | SLC12A1 | -0,584005184 | 0,667109188 | 0,39899501 | 0,875433675 | solute carrier family 12 member 1 |
| ENSG00000270903 | HNRNPA3P9 | -0,584097164 | 0,667066657 | 0,22389015 | 0,732710917 | heterogeneous nuclear ribonucleoprotein A3 pseudogene 9 |
| ENSG00000129596 | CDO1 | -0,584135984 | 0,667048708 | 0,18841533 | 0,682805588 | cysteine dioxygenase type 1 |
| ENSG00000139266 | MARCHF9 | -0,584250469 | 0,666995776 | 0,04709702 | 0,329700266 | membrane associated ring-CH-type finger 9 |
| ENSG00000259438 | MAPK6-DT | -0,58426514 | 0,666988994 | 0,62464227 | 0,952793934 | MAPK6 divergent transcript |
| ENSG00000256134 | EGLN3P1 | -0,584348263 | 0,666950565 | 0,4218344 | 0,889720796 | egl-9 family hypoxia-inducible factor 3 pseudogene 1 |
| ENSG00000259661 | AC068831,3 | -0,584353553 | 0,66694812 | 0,58972585 | 0,947051012 | novel transcript, antisense to MAN2A2 |
| ENSG00000235072 | ARNILA | -0,584417143 | 0,666918723 | 0,39942741 | 0,875433675 | androgen receptor negatively regulated lncRNA |
| ENSG00000100078 | PLA2G3 | -0,584452431 | 0,66690241 | 0,15614623 | 0,627034861 | phospholipase A2 group III |
| ENSG00000138741 | TRPC3 | -0,584593801 | 0,666837064 | 0,0724489 | 0,424244694 | transient receptor potential cation channel subfamily C member 3 |
| ENSG00000237437 | ASS1P12 | -0,584815706 | 0,666734504 | 0,73954539 | 0,974715504 | argininosuccinate synthetase 1 pseudogene 12 |
| ENSG00000234936 | AC010883,1 | -0,584831666 | 0,666727128 | 0,42810098 | 0,892931413 | novel transcript |
| ENSG00000224680 | PLA2G12AP1 | -0,584878926 | 0,666705287 | 0,40478027 | 0,878176155 | phospholipase A2 group XIIA pseudogene 1 |
| ENSG00000259411 | HNRNPA1P45 | -0,584898158 | 0,6666964 | 0,4081212 | 0,881221125 | heterogeneous nuclear ribonucleoprotein A1 pseudogene 45 |
| ENSG00000207118 | SNORD14D | -0,585154733 | 0,666577842 | 0,62087368 | 0,952793934 | small nucleolar RNA, C/D box 14D |
| ENSG00000274698 | AC099521,1 | -0,585165928 | 0,66657267 | 0,49961766 | 0,920190594 | novel transcript |
| ENSG00000136881 | BAAT | -0,585213948 | 0,666550483 | 0,25938168 | 0,772164235 | bile acid-CoA:amino acid N-acyltransferase |
| ENSG00000222046 | DCDC2B | -0,585225754 | 0,666545029 | 0,09004843 | 0,478543935 | doublecortin domain containing 2B |
| ENSG00000134007 | ADAM20 | -0,585316869 | 0,666502934 | 0,2111972 | 0,716986998 | ADAM metallopeptidase domain 20 |
| ENSG00000273080 | AC009309,1 | -0,585598397 | 0,666372885 | 0,2718596 | 0,784843944 | novel transcript, antisense to IMMT |
| ENSG00000277925 | Telomerase-ve | -0,58589245 | 0,666237077 | 0,0354719 | 0,27578235 | Vertebrate telomerase RNA |
| ENSG00000142449 | FBN3 | -0,586144674 | 0,666120611 | 0,0810093 | 0,451779326 | fibrillin 3 |
| ENSG00000259499 | AC090519,1 | -0,586229921 | 0,666081251 | 0,50497532 | 0,921400408 | novel transcript |
| ENSG00000198128 | OR2L3 | -0,586557949 | 0,66592982 | 0,549433 | 0,936051084 | olfactory receptor family 2 subfamily L member 3 |
| ENSG00000224822 | THRB-IT1 | -0,586839911 | 0,665799683 | 0,17420794 | 0,659374935 | THRB intronic transcript 1 |
| ENSG00000105048 | TNNT1 | -0,58684826 | 0,66579583 | 0,4627052 | 0,906789659 | troponin T1, slow skeletal type |
| ENSG00000224799 | AL139397,1 | -0,586869231 | 0,665786152 | 0,30810001 | 0,816424917 | novel transcript |
| ENSG00000231548 | OR55B1P | -0,586947805 | 0,665749892 | 0,18566511 | 0,678587091 | olfactory receptor family 55 subfamily B member 1 pseudogene |

| | | | | | | |
|---|---|---|---|---|---|---|
| ENSG00000235020 | AL390783,1 | -0,587082875 | 0,665687565 | 0,35381332 | 0,845346751 | novel transcript |
| ENSG00000229786 | SNRPFP2 | -0,5871294 | 0,665666098 | 0,61405511 | 0,952603524 | small nuclear ribonucleoprotein polypeptide F pseudogene 2 |
| ENSG00000283046 | HTN1 | -0,587336328 | 0,665570628 | 0,3107582 | 0,817751312 | histatin 1 |
| ENSG00000126550 | HTN1 | -0,587336328 | 0,665570628 | 0,3107582 | 0,817751312 | histatin 1 |
| ENSG00000180113 | TDRD6 | -0,587446759 | 0,665519683 | 0,32705166 | 0,828735859 | tudor domain containing 6 |
| ENSG00000229677 | AC018644,1 | -0,58748789 | 0,66550071 | 0,70514711 | 0,969413904 | ribosomal protein L17 (RPL17) pseudogene |
| ENSG00000267191 | AC006213,3 | -0,587494325 | 0,665497741 | 0,14988519 | 0,61544661 | novel transcript, antisense to ZNF45 |
| ENSG00000047346 | FAM214A | -0,587661169 | 0,665420783 | 0,19165304 | 0,687610821 | family with sequence similarity 214 member A |
| ENSG00000101342 | TLDC2 | -0,588049603 | 0,665241647 | 0,33159031 | 0,832548086 | TBC/LysM-associated domain containing 2 |
| ENSG00000181378 | CFAP65 | -0,588096903 | 0,665219837 | 0,17627044 | 0,661892749 | cilia and flagella associated protein 65 |
| ENSG00000121653 | MAPK8IP1 | -0,588132219 | 0,665203554 | 0,01260421 | 0,135601519 | mitogen-activated protein kinase 8 interacting protein 1 |
| ENSG00000165204 | OR1K1 | -0,58822999 | 0,665158475 | 0,57277586 | 0,942475308 | olfactory receptor family 1 subfamily K member 1 |
| ENSG00000123411 | IKZF4 | -0,588280614 | 0,665135135 | 0,03367712 | 0,266716135 | IKAROS family zinc finger 4 |
| ENSG00000243694 | LINC02027 | -0,588330584 | 0,665112097 | 0,18291381 | 0,673729645 | long intergenic non-protein coding RNA 2027 |
| ENSG00000237636 | ANKRD26P3 | -0,588610935 | 0,664982862 | 0,6015956 | 0,949939978 | ankyrin repeat domain 26 pseudogene 3 |
| ENSG00000288107 | AL731769,2 | -0,588625782 | 0,664976018 | 0,13798369 | 0,592546669 | novel transcript, antisense to FRG2Band SYCE1 |
| ENSG00000227288 | ARID3BP1 | -0,588775973 | 0,664906795 | 0,44488349 | 0,900423226 | ARID3B pseudogene 1 |
| ENSG00000263309 | KRT23 | -0,588842969 | 0,664875919 | 0,54231504 | 0,933491609 | keratin 23 |
| ENSG00000264057 | AC103810,1 | -0,588944225 | 0,664829256 | 0,37719298 | 0,862818448 | bromodomain PHD finger transcription factor (BPTF) pseudogene |
| ENSG00000282084 | UBE2Q2P2 | -0,589004321 | 0,664801563 | 0,12455718 | 0,563880062 | ubiquitin conjugating enzyme E2 Q2 pseudogene 2 |
| ENSG00000265817 | FSBP | -0,589138514 | 0,664739729 | 0,14100203 | 0,598214217 | fibrinogen silencer binding protein |
| ENSG00000168234 | TTC39C | -0,589204052 | 0,664709532 | 0,0951334 | 0,489669163 | tetratricopeptide repeat domain 39C |
| ENSG00000253773 | C8orf37-AS1 | -0,589239558 | 0,664693173 | 0,55553337 | 0,937793851 | C8orf37 antisense RNA 1 |
| ENSG00000215481 | BCRP3 | -0,589272231 | 0,66467812 | 0,44893222 | 0,902062594 | BCR pseudogene 3 |
| ENSG00000286364 | AL512308,1 | -0,589481256 | 0,664581825 | 0,31029446 | 0,817687618 | novel transcript |
| ENSG00000288087 | AC112770,1 | -0,58972003 | 0,664471842 | 0,42195068 | 0,889720796 | novel transcript |
| ENSG00000015592 | STMN4 | -0,589892835 | 0,664392257 | 0,18966085 | 0,684955795 | stathmin 4 |
| ENSG00000224924 | LINC00320 | -0,590018082 | 0,664334581 | 0,21755814 | 0,725286594 | long intergenic non-protein coding RNA 320 |
| ENSG00000230542 | LINC00102 | -0,590262194 | 0,664222181 | 0,3972588 | 0,873963772 | long intergenic non-protein coding RNA 102 |
| ENSG00000145020 | AMT | -0,590460072 | 0,664131084 | 0,0919873 | 0,482073274 | aminomethyltransferase |

| | | | | | | |
|---|---|---|---|---|---|---|
| ENSG00000232775 | BMS1P22 | -0,590564738 | 0,664082903 | 0,45614858 | 0,903545728 | BMS1 pseudogene 22 |
| ENSG00000196993 | NPIPB9 | -0,590600102 | 0,664066625 | 0,17959117 | 0,668298509 | nuclear pore complex interacting protein family member B9 |
| ENSG00000168280 | KIF5C | -0,590875753 | 0,663939756 | 0,11606238 | 0,544096306 | kinesin family member 5C |
| ENSG00000215893 | RPL23AP17 | -0,590914062 | 0,663922126 | 0,54230235 | 0,933491609 | ribosomal protein L23a pseudogene 17 |
| ENSG00000170782 | OR10A4 | -0,591201149 | 0,663790023 | 0,45841784 | 0,90420131 | olfactory receptor family 10 subfamily A member 4 |
| ENSG00000139233 | LLPH | -0,591380369 | 0,663707568 | 0,01338855 | 0,141603444 | LLP homolog, long-term synaptic facilitation factor |
| ENSG00000285630 | AL590068,3 | -0,59140925 | 0,663694282 | 0,24289709 | 0,753067691 | novel transcript |
| ENSG00000286755 | AC135893,1 | -0,591464274 | 0,663668969 | 0,56867886 | 0,941793403 | novel transcript, antisense to LMLN |
| ENSG00000286694 | AL161935,3 | -0,59157491 | 0,663618076 | 0,40427504 | 0,878176155 | novel transcript |
| ENSG00000280279 | LINC02887 | -0,59161621 | 0,663599079 | 0,13048479 | 0,577136255 | long intergenic non-protein coding RNA 2887 |
| ENSG00000282235 | LINC02887 | -0,59161621 | 0,663599079 | 0,13048479 | 0,577136255 | long intergenic non-protein coding RNA 2887 |
| ENSG00000114405 | C3orf14 | -0,591679718 | 0,663569868 | 0,2706165 | 0,783073289 | chromosome 3 open reading frame 14 |
| ENSG00000163006 | CCDC138 | -0,591728101 | 0,663547615 | 0,18455839 | 0,676757375 | coiled-coil domain containing 138 |
| ENSG00000270157 | AC004918,3 | -0,591859905 | 0,663486996 | 0,0914044 | 0,481302729 | novel transcript |
| ENSG00000271032 | AC020907,4 | -0,591876896 | 0,663479182 | 0,55700412 | 0,938292968 | novel transcript |
| ENSG00000263958 | AC091138,1 | -0,592206748 | 0,663327504 | 0,4352052 | 0,896272827 | novel transcript |
| ENSG00000255247 | NECTIN1-AS1 | -0,592280511 | 0,66329359 | 0,67710645 | 0,963580269 | NECTIN1 antisense RNA 1 |
| ENSG00000151131 | C12orf45 | -0,592495321 | 0,663194836 | 0,01155549 | 0,127208092 | chromosome 12 open reading frame 45 |
| ENSG00000248557 | AC024558,1 | -0,592611211 | 0,663141565 | 0,49202878 | 0,916995341 | methyltransferase like 5 (METTL5) pseudogene |
| ENSG00000196616 | ADH1B | -0,592693949 | 0,663103535 | 0,32212226 | 0,824955371 | alcohol dehydrogenase 1B (class I), beta polypeptide |
| ENSG00000115758 | ODC1 | -0,592838928 | 0,663036902 | 0,07523442 | 0,433332501 | ornithine decarboxylase 1 |
| ENSG00000101331 | CCM2L | -0,592900675 | 0,663008525 | 0,61512798 | 0,952793934 | CCM2 like scaffold protein |
| ENSG00000112852 | PCDHB2 | -0,593033261 | 0,662947596 | 0,0059284 | 0,078105982 | protocadherin beta 2 |
| ENSG00000281894 | ZKSCAN7 | -0,593197018 | 0,66287235 | 0,03163111 | 0,256419452 | zinc finger with KRAB and SCAN domains 7 |
| ENSG00000282935 | DUX4L34 | -0,593312973 | 0,662819075 | 0,38885193 | 0,870057277 | double homeobox 4 like 34 (pseudogene) |
| ENSG00000168824 | NSG1 | -0,593333496 | 0,662809646 | 0,07497683 | 0,432645898 | neuronal vesicle trafficking associated 1 |
| ENSG00000187595 | ZNF385C | -0,593364742 | 0,662795291 | 0,11250809 | 0,535800356 | zinc finger protein 385C |
| ENSG00000148948 | LRRC4C | -0,593545165 | 0,662712407 | 0,07424312 | 0,430477915 | leucine rich repeat containing 4C |
| ENSG00000270218 | AC013553,2 | -0,593632901 | 0,662672106 | 0,4484937 | 0,902062594 | transmembrane protein 126A (TMEM126A) pseudogene |
| ENSG00000278842 | AC008147,4 | -0,593856391 | 0,662569459 | 0,62069769 | 0,952793934 | novel pseudogene |
| ENSG00000259736 | CRTC3-AS1 | -0,593997045 | 0,662504865 | 0,28531605 | 0,796937466 | CRTC3 antisense RNA 1 |

| | | | | | | |
|---|---|---|---|---|---|---|
| ENSG00000267336 | EIF4A2P1 | -0,594016054 | 0,662496136 | 0,44278562 | 0,899437949 | eukaryotic translation initiation factor 4A2 pseudogene 1 |
| ENSG00000227304 | AC067942,1 | -0,594048241 | 0,662481356 | 0,43282127 | 0,89560871 | ribosomal protein S6 (RPS6) pseudogene |
| ENSG00000197020 | ZNF100 | -0,594199069 | 0,6624121 | 0,1988494 | 0,697657684 | zinc finger protein 100 |
| ENSG00000276115 | AC026356,2 | -0,594412436 | 0,66231414 | 0,09900755 | 0,499376869 | novel transcript, sense intronic to BICD1 |
| ENSG00000188818 | ZDHHC11 | -0,594449193 | 0,662297266 | 0,0693993 | 0,415571685 | zinc finger DHHC-type containing 11 |
| ENSG00000260350 | AC012173,1 | -0,594488552 | 0,662279198 | 0,47528806 | 0,911082311 | novel transcript, antisense to PMM2 |
| ENSG00000100167 | SEPTIN3 | -0,594537294 | 0,662256822 | 0,02656264 | 0,228077075 | septin 3 |
| ENSG00000146411 | SLC2A12 | -0,594685148 | 0,662188955 | 0,19948076 | 0,698904803 | solute carrier family 2 member 12 |
| ENSG00000258038 | LINC02327 | -0,594720595 | 0,662172685 | 0,37329205 | 0,861257555 | long intergenic non-protein coding RNA 2327 |
| ENSG00000230817 | LINC01362 | -0,594730185 | 0,662168283 | 0,68517805 | 0,965415049 | long intergenic non-protein coding RNA 1362 |
| ENSG00000271855 | AC073195,1 | -0,594797264 | 0,662137496 | 0,15917971 | 0,633235092 | novel transcript |
| ENSG00000114204 | SERPINI2 | -0,59506718 | 0,662013627 | 0,38477901 | 0,868536368 | serpin family I member 2 |
| ENSG00000198478 | SH3BGRL2 | -0,595151454 | 0,661974958 | 0,28220676 | 0,793283115 | SH3 domain binding glutamate rich protein like 2 |
| ENSG00000188290 | HES4 | -0,595414327 | 0,66185435 | 0,1366986 | 0,589807464 | hes family bHLH transcription factor 4 |
| ENSG00000284830 | AL049557,1 | -0,595610134 | 0,661764527 | 0,14856302 | 0,613326406 | novel transcript |
| ENSG00000006740 | ARHGAP44 | -0,59561851 | 0,661760685 | 0,07852843 | 0,443561759 | Rho GTPase activating protein 44 |
| ENSG00000261992 | GATAD2B | -0,595712522 | 0,661717564 | 0,78112862 | 0,978597905 | GATA zinc finger domain containing 2B |
| ENSG00000139626 | ITGB7 | -0,595892593 | 0,661634976 | 0,5985563 | 0,948804147 | integrin subunit beta 7 |
| ENSG00000215482 | CALM2P3 | -0,595956363 | 0,661605731 | 0,60184638 | 0,949939978 | calmodulin 2 pseudogene 3 |
| ENSG00000248174 | LINC02268 | -0,596117819 | 0,661531693 | 0,0369105 | 0,283461062 | long intergenic non-protein coding RNA 2268 |
| ENSG00000226360 | RPL10AP6 | -0,596184234 | 0,66150124 | 0,47053586 | 0,908805405 | ribosomal protein L10a pseudogene 6 |
| ENSG00000268583 | AC011466,1 | -0,596224051 | 0,661482983 | 0,21792774 | 0,72555786 | novel transcript, antisense to CARD8 |
| ENSG00000234129 | AC073529,1 | -0,596454672 | 0,661377251 | 0,12648044 | 0,56789955 | novel transcript |
| ENSG00000135622 | SEMA4F | -0,596507692 | 0,661352945 | 0,02195336 | 0,199346446 | ssemaphorin 4F |
| ENSG00000182993 | C12orf60 | -0,596551349 | 0,661332933 | 0,1746779 | 0,660057513 | chromosome 12 open reading frame 60 |
| ENSG00000271379 | AC023513,1 | -0,596641371 | 0,661291668 | 0,67183578 | 0,962161102 | family with sequence similarity 173, member B (FAM173B) pseudogene |
| ENSG00000250031 | AC009927,1 | -0,596738649 | 0,66124708 | 0,22001604 | 0,728200468 | protein tyrosine phosphatase, non-receptor type 11 (PTPN11) pseudogene |
| ENSG00000172361 | CFAP53 | -0,596813702 | 0,661212681 | 0,12913677 | 0,574218739 | cilia and flagella associated protein 53 |
| ENSG00000148200 | NR6A1 | -0,597078283 | 0,66109143 | 0,02957158 | 0,244583576 | nuclear receptor subfamily 6 group A member 1 |

| | | | | | | |
|---|---|---|---|---|---|---|
| ENSG00000228076 | AL591719,1 | -0,59708784 | 0,66108705 | 0,37740392 | 0,862854083 | chromosome 20 open reading frame 147 (C20orf147) pseudogene |
| ENSG00000235351 | AC114730,3 | -0,597104797 | 0,66107928 | 0,63492989 | 0,955158181 | novel transcript |
| ENSG00000104267 | CA2 | -0,597331466 | 0,660975423 | 0,07578817 | 0,434777568 | carbonic anhydrase 2 |
| ENSG00000258510 | AC007954,1 | -0,597392606 | 0,660947412 | 0,51854546 | 0,926748138 | ATP-binding cassette, sub-family G (WHITE), member 2 (ABCG2) pseudogene |
| ENSG00000185689 | C6orf201 | -0,597469927 | 0,66091199 | 0,46391941 | 0,907292403 | chromosome 6 open reading frame 201 |
| ENSG00000229815 | CCNB1IP1P1 | -0,597779871 | 0,660770017 | 0,68326135 | 0,965010965 | cyclin B1 interacting protein 1 pseudogene 1 |
| ENSG00000205856 | C22orf42 | -0,597854843 | 0,66073568 | 0,52289824 | 0,92790906 | chromosome 22 open reading frame 42 |
| ENSG00000229623 | METTL21AP1 | -0,597876653 | 0,660725691 | 0,35901734 | 0,849814998 | methyltransferase like 21A pseudogene 1 |
| ENSG00000258359 | PCNPP1 | -0,598016998 | 0,660661419 | 0,26864732 | 0,780823861 | PEST containing nuclear protein pseudogene 1 |
| ENSG00000112759 | SLC29A1 | -0,598218893 | 0,66056897 | 0,05789476 | 0,372126156 | solute carrier family 29 member 1 (Augustine blood group) |
| ENSG00000269934 | AL353593,1 | -0,598335309 | 0,660515669 | 0,06535187 | 0,401164726 | novel transcript, antisense to OBSCN |
| ENSG00000242445 | RPL7AP11 | -0,598400404 | 0,660485867 | 0,26338676 | 0,774637968 | ribosomal protein L7a pseudogene 11 |
| ENSG00000286093 | AC022267,1 | -0,598485652 | 0,66044684 | 0,31604514 | 0,821394376 | novel transcript |
| ENSG00000262648 | PHBP15 | -0,598525749 | 0,660428485 | 0,61780042 | 0,952793934 | prohibitin pseudogene 15 |
| ENSG00000262514 | AC020978,6 | -0,598667765 | 0,660363477 | 0,2624392 | 0,774068771 | novel transcript |
| ENSG00000251405 | AC016571,1 | -0,598700804 | 0,660348354 | 0,17168693 | 0,655397237 | novel transcript, sense overlapping ADAM19 |
| ENSG00000265514 | LINC01892 | -0,598755622 | 0,660323263 | 0,23957123 | 0,749431667 | long intergenic non-protein coding RNA 1892 |
| ENSG00000235436 | DPY19L2P4 | -0,598764259 | 0,66031931 | 0,251575 | 0,763892102 | DPY19L2 pseudogene 4 |
| ENSG00000246528 | AC079089,1 | -0,598777659 | 0,660313177 | 0,35639285 | 0,847771893 | novel transcript, antisense to SLCO5A1 |
| ENSG00000169189 | NSMCE1 | -0,598867869 | 0,66027189 | 0,00802477 | 0,097849669 | NSE1 homolog, SMC5-SMC6 complex component |
| ENSG00000234717 | TMEM212-AS1 | -0,598888037 | 0,660262659 | 0,4983424 | 0,919455689 | TMEM212 antisense RNA 1 |
| ENSG00000255931 | AP003108,1 | -0,599137971 | 0,660148285 | 0,49759054 | 0,918759905 | novel transcript |
| ENSG00000282917 | AC107068,2 | -0,599139684 | 0,660147501 | 0,30523477 | 0,814167717 | novel transcript, antisense to CORIN and NFXL1 |
| ENSG00000150773 | PIH1D2 | -0,599343631 | 0,660054186 | 0,24308627 | 0,753280718 | PIH1 domain containing 2 |
| ENSG00000207751 | AP000553,1 | -0,599358731 | 0,660047277 | 0,43900749 | 0,897402199 | novel transcript |
| ENSG00000109163 | GNRHR | -0,599513962 | 0,659976261 | 0,27428763 | 0,786239603 | gonadotropin releasing hormone receptor |
| ENSG00000255557 | AP001266,2 | -0,599607544 | 0,659933452 | 0,54745721 | 0,935404974 | novel transcript |
| ENSG00000285571 | AL513548,4 | -0,599640481 | 0,659918386 | 0,09842038 | 0,498059993 | novel transcript |
| ENSG00000287888 | AC087463,3 | -0,59966639 | 0,659906535 | 0,56019414 | 0,940070839 | novel transcript |
| ENSG00000288448 | AC087463,4 | -0,59966639 | 0,659906535 | 0,56019414 | 0,940070839 | novel transcript |

| | | | | | | |
|---|---|---|---|---|---|---|
| ENSG00000233776 | LINC01251 | -0,599684341 | 0,659898324 | 0,5588938 | 0,939635125 | long intergenic non-protein coding RNA 1251 |
| ENSG00000205038 | PKHD1L1 | -0,600142522 | 0,659688782 | 0,22173659 | 0,729254784 | PKHD1 like 1 |
| ENSG00000278083 | AC013476,1 | -0,600249398 | 0,659639914 | 0,49518197 | 0,917777383 | zinc finger protein 354B (ZNF354B) pseudogene |
| ENSG00000148826 | NKX6-2 | -0,600272448 | 0,659629375 | 0,42409774 | 0,890949676 | NK6 homeobox 2 |
| ENSG00000287960 | AC008567,3 | -0,600289748 | 0,659621465 | 0,27691265 | 0,788828896 | novel transcript |
| ENSG00000255306 | AC004923,4 | -0,600398838 | 0,65957159 | 0,60489488 | 0,950927937 | novel transcript |
| ENSG00000144827 | ABHD10 | -0,600580414 | 0,659488582 | 0,11193194 | 0,534745332 | abhydrolase domain containing 10, depalmitoylase |
| ENSG00000242267 | SKINT1L | -0,600651101 | 0,65945627 | 0,28582084 | 0,797420923 | Skint1 like (pseudogene) |
| ENSG00000123472 | ATPAF1 | -0,600983136 | 0,659304514 | 0,14643999 | 0,609870763 | ATP synthase mitochondrial F1 complex assembly factor 1 |
| ENSG00000183914 | DNAH2 | -0,60098786 | 0,659302355 | 0,26427186 | 0,775647329 | dynein axonemal heavy chain 2 |
| ENSG00000233237 | LINC00472 | -0,601141364 | 0,659232208 | 0,17495576 | 0,66036665 | long intergenic non-protein coding RNA 472 |
| ENSG00000140488 | CELF6 | -0,601173791 | 0,659217391 | 0,21462317 | 0,722164151 | CUGBP Elav-like family member 6 |
| ENSG00000286165 | AC012488,2 | -0,601242733 | 0,65918589 | 0,69458896 | 0,967417311 | novel protein |
| ENSG00000141433 | ADCYAP1 | -0,601371513 | 0,659127051 | 0,27059661 | 0,783073289 | adenylate cyclase activating polypeptide 1 |
| ENSG00000276761 | CWC25 | -0,601512574 | 0,659062608 | 0,26012447 | 0,772692192 | CWC25 spliceosome associated protein homolog |
| ENSG00000279685 | MAPT-IT1 | -0,601588675 | 0,659027844 | 0,65047872 | 0,956899963 | MAPT intronic transcript 1 |
| ENSG00000113532 | ST8SIA4 | -0,601791021 | 0,658935418 | 0,21055597 | 0,716131719 | ST8 alpha-N-acetyl-neuraminide alpha-2,8-sialyltransferase 4 |
| ENSG00000163412 | EIF4E3 | -0,601797122 | 0,658932631 | 0,19137151 | 0,687152738 | eukaryotic translation initiation factor 4E family member 3 |
| ENSG00000261072 | AC084783,1 | -0,602009797 | 0,658835502 | 0,20125001 | 0,701094807 | hydroxyacyl-CoA dehydrogenase (HADH) pseudogene |
| ENSG00000006468 | ETV1 | -0,602024291 | 0,658828883 | 0,25401677 | 0,76531122 | ETS variant transcription factor 1 |
| ENSG00000174514 | MFSD4A | -0,60229232 | 0,658706495 | 0,06372827 | 0,395512649 | major facilitator superfamily domain containing 4A |
| ENSG00000261120 | AL133299,1 | -0,602296974 | 0,65870437 | 0,38057315 | 0,865900094 | novel transcript, antisense to LRRC9 |
| ENSG00000259248 | USP3-AS1 | -0,602322997 | 0,658692488 | 0,05000547 | 0,341944699 | USP3 antisense RNA 1 |
| ENSG00000151365 | THRSP | -0,602344335 | 0,658682746 | 0,38730973 | 0,869513953 | thyroid hormone responsive |
| ENSG00000239521 | CASTOR3 | -0,60237252 | 0,658669878 | 0,01793768 | 0,174004579 | CASTOR family member 3 |
| ENSG00000227382 | EIF4A2P2 | -0,602513076 | 0,65860571 | 0,43117733 | 0,89530663 | eukaryotic translation initiation factor 4A2 pseudogene 2 |
| ENSG00000229999 | SGSM3-AS1 | -0,602550718 | 0,658588526 | 0,63469268 | 0,955149453 | SGSM3 antisense RNA 1 |
| ENSG00000226982 | CENPCP1 | -0,602581013 | 0,658574697 | 0,39410676 | 0,871635019 | centromere protein C pseudogene 1 |

| | | | | | | |
|---|---|---|---|---|---|---|
| ENSG00000286405 | AL132799,1 | -0,602843591 | 0,658454843 | 0,04375351 | 0,31589834 | novel transcript |
| ENSG00000255566 | AC135279,2 | -0,602914042 | 0,658422689 | 0,48554247 | 0,913907266 | mitochondrial ribosomal protein S25 (MRPS25) pseudogene |
| ENSG00000269837 | IPO5P1 | -0,603142546 | 0,658318412 | 0,02380069 | 0,210973821 | importin 5 pseudogene 1 |
| ENSG00000135324 | MRAP2 | -0,603165231 | 0,658308061 | 0,33966783 | 0,838482052 | melanocortin 2 receptor accessory protein 2 |
| ENSG00000107984 | DKK1 | -0,603173684 | 0,658304204 | 0,2604786 | 0,772692192 | dickkopf WNT signaling pathway inhibitor 1 |
| ENSG00000267474 | AC008569,2 | -0,603192639 | 0,658295555 | 0,46696826 | 0,907940653 | novel transcript, sense intronic to PKN1 |
| ENSG00000138738 | PRDM5 | -0,603528193 | 0,658142461 | 0,10428486 | 0,514902364 | PR/SET domain 5 |
| ENSG00000230202 | AL450405,1 | -0,603645318 | 0,658089032 | 0,32885541 | 0,830667565 | ribosomal protein L29 (RPL29) pseudogene |
| ENSG00000212127 | TAS2R14 | -0,603925497 | 0,65796124 | 0,30773082 | 0,816424917 | taste 2 receptor member 14 |
| ENSG00000163071 | SPATA18 | -0,603959047 | 0,657945939 | 0,1877802 | 0,681853677 | spermatogenesis associated 18 |
| ENSG00000273257 | AC069200,1 | -0,604218317 | 0,657827709 | 0,45567659 | 0,903545728 | novel transcript |
| ENSG00000271011 | AC010680,3 | -0,60437468 | 0,657756415 | 0,57375511 | 0,942475308 | novel transcript, antisense to TTN |
| ENSG00000010030 | ETV7 | -0,604525297 | 0,65768775 | 0,62733068 | 0,952793934 | ETS variant transcription factor 7 |
| ENSG00000216990 | HSPD1P10 | -0,604809373 | 0,657558259 | 0,62857796 | 0,952793934 | heat shock protein family D (Hsp60) member 1 pseudogene 10 |
| ENSG00000091622 | PITPNM3 | -0,604824659 | 0,657551292 | 0,11794816 | 0,548984873 | PITPNM family member 3 |
| ENSG00000265788 | NOS2P1 | -0,604829496 | 0,657549088 | 0,61557782 | 0,952793934 | nitric oxide synthase 2 pseudogene 1 |
| ENSG00000186481 | ANKRD20A5P | -0,604835769 | 0,657546228 | 0,10297128 | 0,511404924 | ankyrin repeat domain 20 family member A5, pseudogene |
| ENSG00000272239 | AC011373,1 | -0,605001827 | 0,657470547 | 0,62468958 | 0,952793934 | novel transcript, antisense to DPYSL3 |
| ENSG00000274397 | AL357153,4 | -0,605051002 | 0,657448138 | 0,31147085 | 0,817751312 | ADAM metallopeptidase domain 20 (ADAM20) pseudogene |
| ENSG00000281917 | SLC16A1 | -0,605076804 | 0,65743638 | 0,06079916 | 0,383772443 | solute carrier family 16 member 1 |
| ENSG00000288661 | AL451106,1 | -0,605088728 | 0,657430946 | 0,64866449 | 0,956899963 | novel protein |
| ENSG00000152705 | CATSPER3 | -0,60510111 | 0,657425303 | 0,3854729 | 0,868895792 | cation channel sperm associated 3 |
| ENSG00000285530 | AL445928,2 | -0,605171275 | 0,657393331 | 0,33683412 | 0,836162806 | novel transcript, antisense to DBT |
| ENSG00000141198 | TOM1L1 | -0,605294053 | 0,657337387 | 0,10945025 | 0,527656531 | target of myb1 like 1 membrane trafficking protein |
| ENSG00000224072 | VEZTP1 | -0,605367225 | 0,657304048 | 0,14987031 | 0,61544661 | VEZT pseudogene 1 |
| ENSG00000283317 | AL831711,1 | -0,6053779 | 0,657299184 | 0,43526311 | 0,896272827 | novel transcript |
| ENSG00000276027 | RNU12 | -0,60554998 | 0,657220789 | 0,12718828 | 0,569656846 | RNA, U12 small nuclear |
| ENSG00000264985 | AC124804,1 | -0,6056867 | 0,657158509 | 0,19749987 | 0,696048902 | novel transcript, antisense to SDK2 |
| ENSG00000230322 | AL136320,1 | -0,605689522 | 0,657157223 | 0,41990191 | 0,888425645 | novel transcript |

| | | | | | | |
|---|---|---|---|---|---|---|
| ENSG00000287021 | AC097462,3 | -0,605939753 | 0,657043251 | 0,30092888 | 0,810566078 | novel transcript, antisense to INTU |
| ENSG00000286983 | AC109635,5 | -0,606233898 | 0,656909303 | 0,62110494 | 0,952793934 | novel transcript |
| ENSG00000249593 | AC011405,1 | -0,606253271 | 0,656900482 | 0,26696964 | 0,779386557 | novel transcript |
| ENSG00000270424 | AC145285,5 | -0,606400559 | 0,656833421 | 0,64458334 | 0,956633898 | PRKC, apoptosis, WT1, regulator (PAWR) pseudogene |
| ENSG00000276061 | NDUFA3 | -0,606458008 | 0,656807266 | 0,15825155 | 0,631551738 | NADH:ubiquinone oxidoreductase subunit A3 |
| ENSG00000187889 | FYB2 | -0,606466487 | 0,656803406 | 0,36649343 | 0,856103393 | FYN binding protein 2 |
| ENSG00000213471 | TTLL13P | -0,606609312 | 0,656738386 | 0,45753029 | 0,903831777 | tubulin tyrosine ligase like 13, pseudogene |
| ENSG00000116544 | DLGAP3 | -0,606749816 | 0,65667443 | 0,07676901 | 0,437815845 | DLG associated protein 3 |
| ENSG00000113369 | ARRDC3 | -0,606784686 | 0,656658558 | 0,18629026 | 0,679406361 | arrestin domain containing 3 |
| ENSG00000141480 | ARRB2 | -0,606834672 | 0,656635807 | 0,04302992 | 0,312712094 | arrestin beta 2 |
| ENSG00000260582 | TPST2P1 | -0,606923625 | 0,656595321 | 0,61625654 | 0,952793934 | tyrosylprotein sulfotransferase 2 pseudogene 1 |
| ENSG00000282002 | AC243807,1 | -0,607075957 | 0,656525996 | 0,36362658 | 0,853933856 | novel transcript |
| ENSG00000267246 | AC091132,5 | -0,607075957 | 0,656525996 | 0,36362658 | 0,853933856 | novel transcript |
| ENSG00000155265 | GOLGA7B | -0,607144468 | 0,65649482 | 0,15131277 | 0,618627212 | golgin A7 family member B |
| ENSG00000258302 | AC025034,1 | -0,607247043 | 0,656448145 | 0,265561 | 0,776824277 | novel transcript, antisense to ATP2B1 |
| ENSG00000227454 | MTND4P30 | -0,60726973 | 0,656437822 | 0,69623517 | 0,96810276 | MT-ND4 pseudogene 30 |
| ENSG00000276768 | AL121890,2 | -0,607548221 | 0,656311119 | 0,57288737 | 0,942475308 | novel transcript |
| ENSG00000244945 | RUFY1-AS1 | -0,607595795 | 0,656289477 | 0,44575101 | 0,901173082 | RUFY1 antisense RNA 1 |
| ENSG00000283808 | RUFY1-AS1 | -0,607595795 | 0,656289477 | 0,44575101 | 0,901173082 | RUFY1 antisense RNA 1 |
| ENSG00000261611 | AC010547,4 | -0,608002539 | 0,656104472 | 0,13854927 | 0,593533053 | novel transcript |
| ENSG00000256061 | DNAAF4 | -0,608049364 | 0,656083178 | 0,08244134 | 0,45675103 | dynein axonemal assembly factor 4 |
| ENSG00000078053 | AMPH | -0,608082743 | 0,656067999 | 0,07342172 | 0,427777983 | amphiphysin |
| ENSG00000283812 | GPR22 | -0,608123902 | 0,656049282 | 0,34243817 | 0,840048641 | G protein-coupled receptor 22 |
| ENSG00000172209 | GPR22 | -0,608123902 | 0,656049282 | 0,34243817 | 0,840048641 | G protein-coupled receptor 22 |
| ENSG00000055163 | CYFIP2 | -0,608293139 | 0,655972328 | 0,03616653 | 0,279049926 | cytoplasmic FMR1 interacting protein 2 |
| ENSG00000228686 | AL590723,1 | -0,60833899 | 0,65595148 | 0,47426543 | 0,910673352 | novel transcript |
| ENSG00000233974 | AC138951,1 | -0,608538706 | 0,655860682 | 0,29006034 | 0,800682868 | POM121 membrane glycoprotein (POM121) pseudogene |
| ENSG00000276702 | AC010809,3 | -0,608596496 | 0,65583441 | 0,51187314 | 0,923184606 | novel transcript, antisense to RYR3 |
| ENSG00000182575 | NXPH3 | -0,609174507 | 0,655571705 | 0,4035357 | 0,877691113 | neurexophilin 3 |
| ENSG00000105967 | TFEC | -0,609398435 | 0,655469958 | 0,50078304 | 0,920520138 | transcription factor EC |
| ENSG00000197128 | ZNF772 | -0,609444606 | 0,655448982 | 0,01738038 | 0,170445528 | zinc finger protein 772 |

| | | | | | | |
|---|---|---|---|---|---|---|
| ENSG00000267293 | AC012569,1 | -0,609493849 | 0,655426609 | 0,30086392 | 0,810530781 | U3 small nucleolar RNA-associated protein 18 homolog, pseudogene |
| ENSG00000255002 | LINC02324 | -0,609801901 | 0,655286675 | 0,19954018 | 0,698904803 | long intergenic non-protein coding RNA 2324 |
| ENSG00000277883 | NLRP3P1 | -0,609826567 | 0,655275471 | 0,63285118 | 0,954396141 | NLR family pyrin domain containing 3 pseudogene 1 |
| ENSG00000264217 | RPL35AP23 | -0,60995844 | 0,655215577 | 0,59423365 | 0,947469769 | ribosomal protein L35a pseudogene 23 |
| ENSG00000228205 | AC131235,1 | -0,609998057 | 0,655197584 | 0,37544218 | 0,862022464 | ribosomal protein S3 (RPS3) pseudogene |
| ENSG00000239835 | AC092910,1 | -0,610188342 | 0,655111173 | 0,20985371 | 0,714847053 | presenilin associated, rhomboid-like (PARL) pseudogene |
| ENSG00000285878 | AP002961,1 | -0,610232079 | 0,655091312 | 0,53633562 | 0,93228049 | novel transcript, antisense to DYNC2H1 |
| ENSG00000229649 | LINC02681 | -0,610289389 | 0,65506529 | 0,58175337 | 0,944145586 | long intergenic non-protein coding RNA 2681 |
| ENSG00000250519 | AP002784,1 | -0,610401594 | 0,655014344 | 0,18631864 | 0,679406361 | novel transcript |
| ENSG00000258365 | AC073655,2 | -0,610819393 | 0,654824682 | 0,24691364 | 0,759015651 | novel transcript, antisense to PLXNC1 |
| ENSG00000003249 | DBNDD1 | -0,610864932 | 0,654804013 | 0,18110707 | 0,670948158 | dysbindin domain containing 1 |
| ENSG00000272263 | AC034198,2 | -0,611016991 | 0,654735001 | 0,19611654 | 0,694033976 | novel transcript, antisense to CAND2 |
| ENSG00000220867 | HSPE1P26 | -0,611018767 | 0,654734195 | 0,54974338 | 0,936108121 | heat shock protein family E (Hsp10) member 1 pseudogene 26 |
| ENSG00000245958 | AC093752,1 | -0,611112394 | 0,654691705 | 0,09215652 | 0,482558005 | septin 4 (sept4) pseudogene |
| ENSG00000244184 | AC091153,3 | -0,611144238 | 0,654677255 | 0,56371104 | 0,940865625 | novel transcript |
| ENSG00000158373 | H2BC5 | -0,611226993 | 0,654639703 | 0,14963896 | 0,615000989 | H2B clustered histone 5 |
| ENSG00000277171 | AC010975,1 | -0,611278703 | 0,654616239 | 0,53048413 | 0,931265667 | novel zinc finger protein pseudogene |
| ENSG00000260840 | LINC01964 | -0,611321498 | 0,654596821 | 0,66221665 | 0,959874876 | long intergenic non-protein coding RNA 1964 |
| ENSG00000229447 | AC114495,2 | -0,611349967 | 0,654583904 | 0,28464965 | 0,796071802 | general transcription factor IIA, 2, 12kDa (GTF2A2) pseudogene |
| ENSG00000185040 | SPDYE16 | -0,611465621 | 0,654531431 | 0,05668051 | 0,367607368 | speedy/RINGO cell cycle regulator family member E16 |
| ENSG00000288056 | AL023806,4 | -0,611543429 | 0,654496132 | 0,05699309 | 0,368909302 | novel transcript |
| ENSG00000139797 | RNF113B | -0,611727257 | 0,654412741 | 0,2994367 | 0,809252953 | ring finger protein 113B |
| ENSG00000237495 | LY6G5C | -0,611813072 | 0,654373816 | 0,37334288 | 0,861257555 | lymphocyte antigen 6 family member G5C |
| ENSG00000231845 | HMGB3P14 | -0,611878184 | 0,654344284 | 0,37735449 | 0,86283995 | high mobility group box 3 pseudogene 14 |
| ENSG00000206077 | ZDHHC11B | -0,611931882 | 0,654319929 | 0,10931046 | 0,527552049 | zinc finger DHHC-type containing 11B |
| ENSG00000233030 | AC243772,2 | -0,611991106 | 0,654293069 | 0,29920654 | 0,809131439 | novel transcript, antisense to FCGR1A |
| ENSG00000284076 | GOLGA8T | -0,612003182 | 0,654287593 | 0,0796258 | 0,447576616 | golgin A8 family member T |
| ENSG00000243813 | AC078857,1 | -0,612008039 | 0,65428539 | 0,57152489 | 0,942436428 | zinc finger protein pseudogene |

| | | | | | | |
|---|---|---|---|---|---|---|
| ENSG00000238622 | SNORD97 | -0,612012563 | 0,654283338 | 0,23851215 | 0,747998807 | small nucleolar RNA, C/D box 97 |
| ENSG00000183977 | PP2D1 | -0,612122797 | 0,654233347 | 0,25924777 | 0,771989931 | protein phosphatase 2C like domain containing 1 |
| ENSG00000277909 | SPC25 | -0,612134928 | 0,654227846 | 0,24696899 | 0,759015651 | SPC25 component of NDC80 kinetochore complex |
| ENSG00000272129 | AL359715,3 | -0,612177523 | 0,654208531 | 0,32537054 | 0,827359468 | novel transcript |
| ENSG00000228680 | AC004692,2 | -0,61219629 | 0,654200021 | 0,70570064 | 0,969503793 | novel transcript |
| ENSG00000135821 | GLUL | -0,612221061 | 0,654188788 | 0,0438716 | 0,31589834 | glutamate-ammonia ligase |
| ENSG00000286661 | AC092933,2 | -0,612311381 | 0,654147834 | 0,09090647 | 0,480978137 | novel transcript |
| ENSG00000044524 | EPHA3 | -0,612339467 | 0,654135099 | 0,17397628 | 0,659185304 | EPH receptor A3 |
| ENSG00000288021 | AL450338,2 | -0,612352519 | 0,654129181 | 0,41440116 | 0,885579089 | novel transcript |
| ENSG00000273035 | AC007684,2 | -0,612436888 | 0,654090929 | 0,45552507 | 0,903545728 | novel transcript, sense intronic to MAP4K3 |
| ENSG00000275326 | NOSTRIN | -0,612495987 | 0,654064135 | 0,21603035 | 0,723046035 | nitric oxide synthase trafficking |
| ENSG00000241370 | RPP21 | -0,612623735 | 0,654006222 | 0,25963122 | 0,772176335 | ribonuclease P/MRP subunit p21 |
| ENSG00000284657 | AL031432,5 | -0,612751826 | 0,653948157 | 0,46232738 | 0,9064797 | novel transcript |
| ENSG00000114107 | CEP70 | -0,612892585 | 0,653884357 | 0,19788465 | 0,696383642 | centrosomal protein 70 |
| ENSG00000277421 | ZNF280B | -0,612903759 | 0,653879292 | 0,11449085 | 0,540617781 | zinc finger protein 280B |
| ENSG00000240225 | ZNF542P | -0,613024874 | 0,653824402 | 0,04297578 | 0,312463916 | zinc finger protein 542, pseudogene |
| ENSG00000260161 | MTCYBP28 | -0,613055793 | 0,653810389 | 0,55323852 | 0,936992468 | MT-CYB pseudogene 28 |
| ENSG00000180530 | NRIP1 | -0,613059084 | 0,653808898 | 0,13258264 | 0,581174167 | nuclear receptor interacting protein 1 |
| ENSG00000196872 | CRACDL | -0,613066897 | 0,653805357 | 0,3798789 | 0,865384295 | CRACD like |
| ENSG00000165209 | STRBP | -0,613254307 | 0,653720431 | 0,17743302 | 0,663776507 | spermatid perinuclear RNA binding protein |
| ENSG00000249655 | AC008434,1 | -0,613385106 | 0,653661166 | 0,34281849 | 0,840419133 | novel transcript |
| ENSG00000169418 | NPR1 | -0,613539606 | 0,653591168 | 0,25795848 | 0,770351899 | natriuretic peptide receptor 1 |
| ENSG00000163794 | UCN | -0,613631275 | 0,65354964 | 0,40967915 | 0,882177108 | urocortin |
| ENSG00000278574 | ZNHIT3 | -0,61383045 | 0,653459419 | 0,05381444 | 0,357892949 | zinc finger HIT-type containing 3 |
| ENSG00000175520 | UBQLN3 | -0,613921886 | 0,653418005 | 0,59482138 | 0,947469769 | ubiquilin 3 |
| ENSG00000229152 | ANKRD10-IT1 | -0,614039212 | 0,653364869 | 0,34252073 | 0,840113448 | ANKRD10 intronic transcript 1 |
| ENSG00000213976 | AC010615,1 | -0,614079864 | 0,653346458 | 0,26304898 | 0,77453036 | zinc finger protein pseudogene |
| ENSG00000229839 | AC018462,1 | -0,614179956 | 0,653301132 | 0,41871942 | 0,887094957 | novel transcript |
| ENSG00000119969 | HELLS | -0,614349603 | 0,653224314 | 0,08382907 | 0,461082665 | helicase, lymphoid specific |
| ENSG00000244183 | PPIAP71 | -0,614379572 | 0,653210745 | 0,66453814 | 0,96046247 | peptidylprolyl isomerase A pseudogene 71 |
| ENSG00000162174 | ASRGL1 | -0,614507003 | 0,653153051 | 0,03083492 | 0,251966306 | asparaginase and isoaspartyl peptidase 1 |
| ENSG00000139187 | KLRG1 | -0,614568875 | 0,65312504 | 0,157861 | 0,630637128 | killer cell lectin like receptor G1 |
| ENSG00000178075 | GRAMD1C | -0,614849245 | 0,652998125 | 0,15904054 | 0,633144409 | GRAM domain containing 1C |

| | | | | | | |
|---|---|---|---|---|---|---|
| ENSG00000267270 | PARD6G-AS1 | -0,615016964 | 0,652922216 | 0,07403312 | 0,4296801 | PARD6G antisense RNA 1 |
| ENSG00000104381 | GDAP1 | -0,615110166 | 0,652880037 | 0,06992403 | 0,417175017 | ganglioside induced differentiation associated protein 1 |
| ENSG00000205325 | AC005863,1 | -0,615168277 | 0,65285374 | 0,15364221 | 0,622683206 | novel transcript |
| ENSG00000139910 | NOVA1 | -0,615178803 | 0,652848976 | 0,13352073 | 0,583791005 | NOVA alternative splicing regulator 1 |
| ENSG00000231064 | THBS3-AS1 | -0,615211294 | 0,652834274 | 0,0211172 | 0,193836827 | THBS3 antisense RNA 1 |
| ENSG00000177947 | ODF3 | -0,61529323 | 0,652797198 | 0,31919346 | 0,824187668 | outer dense fiber of sperm tails 3 |
| ENSG00000276571 | AC002550,2 | -0,615637858 | 0,652641278 | 0,46660881 | 0,907940653 | novel transcript, antisense to C16orf62 |
| ENSG00000235890 | TSPEAR-AS1 | -0,615772984 | 0,652580153 | 0,15226072 | 0,620540856 | TSPEAR antisense RNA 1 |
| ENSG00000101890 | GUCY2F | -0,615821021 | 0,652558424 | 0,2479741 | 0,760187537 | guanylate cyclase 2F, retinal |
| ENSG00000275212 | LINC02826 | -0,61583072 | 0,652554038 | 0,2416394 | 0,751953584 | long intergenic non-protein coding RNA 2826 |
| ENSG00000286447 | AC109782,1 | -0,615922916 | 0,652512337 | 0,2205387 | 0,728572782 | novel transcript |
| ENSG00000119411 | BSPRY | -0,615933141 | 0,652507712 | 0,62283346 | 0,952793934 | B-box and SPRY domain containing |
| ENSG00000287351 | AC092120,3 | -0,616054822 | 0,652452681 | 0,48506867 | 0,913690171 | novel transcript, antisense to ANKRD11 |
| ENSG00000162706 | CADM3 | -0,616060862 | 0,652449949 | 0,01063258 | 0,119935439 | cell adhesion molecule 3 |
| ENSG00000226375 | AL022310,1 | -0,616228114 | 0,652374315 | 0,48844959 | 0,915411008 | novel transcript |
| ENSG00000234426 | AL139042,1 | -0,61628664 | 0,65234785 | 0,18103275 | 0,670752303 | novel transcript |
| ENSG00000255622 | PCDHB17P | -0,616489455 | 0,652256149 | 0,15617069 | 0,627034861 | protocadherin beta 17 pseudogene |
| ENSG00000111218 | PRMT8 | -0,616526212 | 0,652239531 | 0,11228657 | 0,53522294 | protein arginine methyltransferase 8 |
| ENSG00000182224 | CYB5D1 | -0,616562357 | 0,65222319 | 0,01771819 | 0,172621197 | cytochrome b5 domain containing 1 |
| ENSG00000131730 | CKMT2 | -0,616590087 | 0,652210654 | 0,11185725 | 0,534735697 | creatine kinase, mitochondrial 2 |
| ENSG00000228109 | MELTF-AS1 | -0,616839511 | 0,652097905 | 0,0498057 | 0,340876825 | MELTF antisense RNA 1 |
| ENSG00000234232 | AC243772,3 | -0,616892626 | 0,652073897 | 0,61043692 | 0,951841373 | phosphodiesterase 4D interacting protein (PDE4DIP) pseudogene |
| ENSG00000285887 | AL009176,1 | -0,616930643 | 0,652056714 | 0,40866868 | 0,88153752 | novel transcript |
| ENSG00000179361 | ARID3B | -0,616998868 | 0,652025879 | 0,07564919 | 0,434488912 | AT-rich interaction domain 3B |
| ENSG00000253539 | AC018442,2 | -0,617207048 | 0,651931799 | 0,52249713 | 0,927794062 | novel transcript |
| ENSG00000153347 | FAM81B | -0,617373022 | 0,651856802 | 0,63559488 | 0,955347999 | family with sequence similarity 81 member B |
| ENSG00000217442 | SYCE3 | -0,617566057 | 0,651769589 | 0,53954605 | 0,933350549 | synaptonemal complex central element protein 3 |
| ENSG00000170558 | CDH2 | -0,617589745 | 0,651758887 | 0,04760285 | 0,331735648 | cadherin 2 |
| ENSG00000230773 | AC092650,1 | -0,617607067 | 0,651751062 | 0,39039763 | 0,870057277 | novel transcript |
| ENSG00000235330 | RPL12P25 | -0,617657405 | 0,651728321 | 0,65891292 | 0,958240464 | ribosomal protein L12 pseudogene 25 |
| ENSG00000263680 | AC007639,1 | -0,617768158 | 0,651678291 | 0,7072941 | 0,969565008 | novel transcript |

| | | | | | | |
|---|---|---|---|---|---|---|
| ENSG00000203808 | BVES-AS1 | -0,617819682 | 0,651655018 | 0,186963 | 0,680546189 | BVES antisense RNA 1 |
| ENSG00000233483 | AC008105,2 | -0,617929962 | 0,651605207 | 0,53946887 | 0,933350549 | novel pseudogene, ortholog of 1700023F06Rik (M, musculus) |
| ENSG00000198814 | GK | -0,618134878 | 0,651512662 | 0,20078625 | 0,700325598 | glycerol kinase |
| ENSG00000286975 | AP005136,4 | -0,618143899 | 0,651508588 | 0,62903485 | 0,952793934 | novel transcript |
| ENSG00000260325 | HSPB9 | -0,618193669 | 0,651486113 | 0,63963333 | 0,956018518 | heat shock protein family B (small) member 9 |
| ENSG00000270720 | AC104785,1 | -0,618327228 | 0,651425803 | 0,53516846 | 0,931905243 | novel transcript |
| ENSG00000176563 | CNTD1 | -0,618362736 | 0,65140977 | 0,13497359 | 0,586038109 | cyclin N-terminal domain containing 1 |
| ENSG00000236576 | AC241520,1 | -0,618375683 | 0,651403925 | 0,26172814 | 0,773497425 | zinc finger protein pseudogene |
| ENSG00000006432 | MAP3K9 | -0,618459401 | 0,651366125 | 0,04760309 | 0,331735648 | mitogen-activated protein kinase kinase kinase 9 |
| ENSG00000165555 | NOXRED1 | -0,618482367 | 0,651355757 | 0,11439044 | 0,540543423 | NADP dependent oxidoreductase domain containing 1 |
| ENSG00000283269 | AC005524,1 | -0,618538747 | 0,651330302 | 0,28698755 | 0,798400609 | SHC SH2-domain binding protein 1 (SHCBP1) pseudogene |
| ENSG00000165804 | ZNF219 | -0,61883377 | 0,651197122 | 0,04680067 | 0,328508325 | zinc finger protein 219 |
| ENSG00000258620 | AL135838,1 | -0,618909942 | 0,651162741 | 0,27989928 | 0,790565637 | novel transcript, intronic to WDR25 |
| ENSG00000261098 | AP000766,1 | -0,619276527 | 0,650997304 | 0,04707551 | 0,32967633 | novel transcript |
| ENSG00000273909 | AC138827,2 | -0,619277577 | 0,65099683 | 0,65987524 | 0,958837121 | novel transcript |
| ENSG00000110723 | EXPH5 | -0,619289047 | 0,650991654 | 0,16218239 | 0,63932293 | exophilin 5 |
| ENSG00000127074 | RGS13 | -0,61934525 | 0,650966294 | 0,59787583 | 0,948446825 | regulator of G protein signaling 13 |
| ENSG00000233785 | AC131011,1 | -0,619575354 | 0,650862476 | 0,52023498 | 0,926865961 | novel transcript |
| ENSG00000071073 | MGAT4A | -0,619674689 | 0,650817663 | 0,03932422 | 0,295181837 | alpha-1,3-mannosyl-glycoprotein 4-beta-N-acetylglucosaminyltransferase A |
| ENSG00000158113 | LRRC43 | -0,619697747 | 0,650807262 | 0,18767858 | 0,681722211 | leucine rich repeat containing 43 |
| ENSG00000264932 | AC115989,1 | -0,6197604 | 0,650778999 | 0,45385653 | 0,902912642 | novel transcript, antisense to AL3A2 |
| ENSG00000258545 | RHOXF1-AS1 | -0,619779133 | 0,650770549 | 0,20049237 | 0,700064356 | RHOXF1 antisense RNA 1 |
| ENSG00000064199 | SPA17 | -0,619791014 | 0,650765189 | 0,11125266 | 0,532731376 | sperm autoantigenic protein 17 |
| ENSG00000175229 | GAL3ST3 | -0,619860444 | 0,650733872 | 0,07132095 | 0,420712881 | galactose-3-O-sulfotransferase 3 |
| ENSG00000113249 | HAVCR1 | -0,620161439 | 0,650598121 | 0,34428815 | 0,84105522 | hepatitis A virus cellular receptor 1 |
| ENSG00000278301 | GRAMD4P3 | -0,620322402 | 0,650525537 | 0,110849 | 0,531538341 | GRAM domain containing 4 pseudogene 3 |
| ENSG00000120594 | PLXDC2 | -0,620493721 | 0,650448292 | 0,07648757 | 0,436971724 | plexin domain containing 2 |
| ENSG00000135973 | GPR45 | -0,620661797 | 0,650372518 | 0,43901944 | 0,897402199 | G protein-coupled receptor 45 |
| ENSG00000287972 | AC073648,7 | -0,620783351 | 0,650317724 | 0,0939803 | 0,487240447 | novel transcript |

| | | | | | | |
|---|---|---|---|---|---|---|
| ENSG00000285400 | MCPH1-AS1 | -0,620896973 | 0,650266509 | 0,10081191 | 0,504527553 | MCPH1 antisense RNA 1 |
| ENSG00000174628 | IQCK | -0,621083736 | 0,650182335 | 0,07985299 | 0,44796769 | IQ motif containing K |
| ENSG00000105664 | COMP | -0,621168332 | 0,650144211 | 0,56785858 | 0,94163744 | cartilage oligomeric matrix protein |
| ENSG00000180638 | SLC47A2 | -0,6213247 | 0,650073748 | 0,31581653 | 0,821233148 | solute carrier family 47 member 2 |
| ENSG00000288543 | AL035409,2 | -0,621549057 | 0,649972661 | 0,17833972 | 0,665585904 | novel transcript |
| ENSG00000262681 | AC005722,3 | -0,621562466 | 0,64996662 | 0,6355015 | 0,955314044 | novel transcript |
| ENSG00000231817 | LINC01198 | -0,621777128 | 0,649869918 | 0,10822736 | 0,524537831 | long intergenic non-protein coding RNA 1198 |
| ENSG00000284837 | AC007846,1 | -0,621883679 | 0,649821923 | 0,36204847 | 0,852161624 | novel transcript |
| ENSG00000274320 | GOLGA6L9 | -0,622065428 | 0,649740064 | 0,14180645 | 0,599761485 | golgin A6 family like 9 |
| ENSG00000214772 | AC010168,1 | -0,622155638 | 0,649699438 | 0,58798249 | 0,946139044 | novel transcript |
| ENSG00000151303 | AL136982,1 | -0,622208783 | 0,649675505 | 0,01782257 | 0,173394196 | novel transcript |
| ENSG00000259664 | LINC02254 | -0,62236889 | 0,64960341 | 0,03506909 | 0,273605654 | long intergenic non-protein coding RNA 2254 |
| ENSG00000189099 | PRSS48 | -0,62236983 | 0,649602986 | 0,40012337 | 0,875889731 | serine protease 48 |
| ENSG00000212916 | MAP10 | -0,622461775 | 0,649561588 | 0,15911437 | 0,633144409 | microtubule associated protein 10 |
| ENSG00000273619 | AL121832,2 | -0,622482037 | 0,649552465 | 0,10351669 | 0,513180277 | novel transcript, antisense to RPS21 |
| ENSG00000167554 | ZNF610 | -0,622503186 | 0,649542943 | 0,11449146 | 0,540617781 | zinc finger protein 610 |
| ENSG00000160013 | PTGIR | -0,622573172 | 0,649511434 | 0,64302224 | 0,956633898 | prostaglandin I2 receptor |
| ENSG00000171680 | PLEKHG5 | -0,622811724 | 0,649404045 | 0,22088743 | 0,728635155 | pleckstrin homology and RhoGEF domain containing G5 |
| ENSG00000287411 | AL031259,1 | -0,622981241 | 0,649327744 | 0,54577059 | 0,935146964 | novel transcript |
| ENSG00000106086 | PLEKHA8 | -0,623041788 | 0,649300494 | 0,1596621 | 0,634185749 | pleckstrin homology domain containing A8 |
| ENSG00000286707 | AC010896,1 | -0,623090441 | 0,649278597 | 0,23705194 | 0,74656913 | novel transcript |
| ENSG00000259228 | HNRNPA1P62 | -0,623250939 | 0,64920637 | 0,30749449 | 0,816364446 | heterogeneous nuclear ribonucleoprotein A1 pseudogene 62 |
| ENSG00000198846 | TOX | -0,623360337 | 0,649157143 | 0,13528215 | 0,586275832 | thymocyte selection associated high mobility group box |
| ENSG00000271743 | AF287957,1 | -0,623373135 | 0,649151385 | 0,31959662 | 0,824366234 | novel transcript |
| ENSG00000285267 | AF287957,2 | -0,623373135 | 0,649151385 | 0,31959662 | 0,824366234 | novel transcript |
| ENSG00000150471 | ADGRL3 | -0,62342679 | 0,649127243 | 0,05102725 | 0,345820358 | adhesion G protein-coupled receptor L3 |
| ENSG00000106927 | AMBP | -0,623432784 | 0,649124546 | 0,38375886 | 0,868000076 | alpha-1-microglobulin/bikunin precursor |
| ENSG00000258498 | DIO3OS | -0,623552855 | 0,649070524 | 0,1766912 | 0,662651108 | DIO3 opposite strand upstream RNA |
| ENSG00000244617 | ASPRV1 | -0,623571789 | 0,649062005 | 0,15310793 | 0,621632863 | aspartic peptidase retroviral like 1 |
| ENSG00000214842 | RAD51AP2 | -0,6235904 | 0,649053632 | 0,45687928 | 0,903699469 | RAD51 associated protein 2 |

| | | | | | | |
|---|---|---|---|---|---|---|
| ENSG00000275765 | AC091982,3 | -0,623993025 | 0,64887252 | 0,38143116 | 0,865900094 | novel transcript |
| ENSG00000276833 | TAF15 | -0,624014843 | 0,648862708 | 0,08960108 | 0,477709449 | TATA-box binding protein associated factor 15 |
| ENSG00000234196 | ZBTB12 | -0,624179135 | 0,648788821 | 0,74291114 | 0,975201495 | zinc finger and BTB domain containing 12 |
| ENSG00000287778 | AC006230,1 | -0,624355 | 0,648709738 | 0,34400853 | 0,840938551 | novel transcript, antisense to HS3ST1 |
| ENSG00000164627 | KIF6 | -0,624372829 | 0,648701721 | 0,09036231 | 0,479478496 | kinesin family member 6 |
| ENSG00000238064 | HCG25 | -0,624458294 | 0,648663293 | 0,61714087 | 0,952793934 | HLA complex group 25 |
| ENSG00000231573 | HCG25 | -0,624458294 | 0,648663293 | 0,61714087 | 0,952793934 | HLA complex group 25 |
| ENSG00000260484 | AC131902,1 | -0,624884964 | 0,648471483 | 0,43692959 | 0,896866043 | novel transcript |
| ENSG00000253357 | AC008708,1 | -0,624911701 | 0,648459465 | 0,397384 | 0,874053739 | novel transcript |
| ENSG00000141314 | RHBDL3 | -0,624963724 | 0,648436082 | 0,04011782 | 0,299101235 | rhomboid like 3 |
| ENSG00000228663 | PSMD10P1 | -0,625016574 | 0,648412328 | 0,04739561 | 0,330875511 | proteasome 26S subunit, non-ATPase, 10 pseudogene 1 |
| ENSG00000264290 | AC104564,3 | -0,62510707 | 0,648371656 | 0,47262261 | 0,909368754 | novel transcript |
| ENSG00000273654 | AC020904,2 | -0,625260441 | 0,648302733 | 0,50887993 | 0,922427526 | citrate synthase (CS) pseudogene |
| ENSG00000240180 | RPL12P28 | -0,625304066 | 0,648283129 | 0,44883712 | 0,902062594 | ribosomal protein L12 pseudogene 28 |
| ENSG00000176887 | SOX11 | -0,625428835 | 0,648227066 | 0,06723984 | 0,407637454 | SRY-box transcription factor 11 |
| ENSG00000260619 | AC068870,1 | -0,625648304 | 0,648128462 | 0,53960907 | 0,933350549 | novel transcript |
| ENSG00000288061 | AP000926,2 | -0,625671543 | 0,648118022 | 0,40243329 | 0,877199637 | novel transcript, antisense to HSPA8and CLMP |
| ENSG00000138411 | HECW2 | -0,625675114 | 0,648116418 | 0,03163414 | 0,256419452 | HECT, C2 and WW domain containing E3 ubiquitin protein ligase 2 |
| ENSG00000236745 | YRDCP2 | -0,625701822 | 0,64810442 | 0,35250651 | 0,844912682 | yrdC N(6)-threonylcarbamoyltransferase domain containing pseudogene 2 |
| ENSG00000230327 | MTCO1P42 | -0,625704092 | 0,6481034 | 0,61884155 | 0,952793934 | MT-CO1 pseudogene 42 |
| ENSG00000273091 | AP000255,1 | -0,625793497 | 0,648063238 | 0,40331736 | 0,877589651 | novel transcript |
| ENSG00000261505 | AL031714,1 | -0,625830032 | 0,648046827 | 0,10961483 | 0,528090728 | novel transcript, antisense to UBE2I |
| ENSG00000145920 | CPLX2 | -0,625902072 | 0,648014467 | 0,02572008 | 0,223230867 | complexin 2 |
| ENSG00000261124 | AC135050,4 | -0,625984519 | 0,647977436 | 0,45284778 | 0,902361309 | novel transcript, antisense to ZNF668 |
| ENSG00000259075 | POC1B-GALNT4 | -0,626081984 | 0,647933661 | 0,20057838 | 0,700127386 | POC1B-GALNT4 readthrough |
| ENSG00000139445 | FOXN4 | -0,626161618 | 0,647897898 | 0,15382203 | 0,622898002 | forkhead box N4 |
| ENSG00000124701 | APOBEC2 | -0,626187179 | 0,647886419 | 0,32956845 | 0,831379626 | apolipoprotein B mRNA editing enzyme catalytic subunit 2 |
| ENSG00000274718 | AL136964,1 | -0,626198157 | 0,647881489 | 0,32779449 | 0,829743891 | novel transcript |
| ENSG00000244468 | AC093001,1 | -0,626456811 | 0,647765344 | 0,37605255 | 0,862512424 | novel transcript |

| | | | | | | |
|---|---|---|---|---|---|---|
| ENSG00000186472 | PCLO | -0,626470053 | 0,647759398 | 0,05669149 | 0,367607368 | piccolo presynaptic cytomatrix protein |
| ENSG00000273428 | AC004832,6 | -0,626494223 | 0,647748546 | 0,59198741 | 0,947416795 | novel transcript |
| ENSG00000234118 | RPL13AP6 | -0,626963659 | 0,647537811 | 0,41453698 | 0,885579089 | ribosomal protein L13a pseudogene 6 |
| ENSG00000281557 | LRIG2-DT | -0,627035976 | 0,647505353 | 0,58234277 | 0,944174844 | LRIG2 divergent transcript |
| ENSG00000117477 | CCDC181 | -0,627601164 | 0,647251736 | 0,16380097 | 0,642705016 | coiled-coil domain containing 181 |
| ENSG00000101282 | RSPO4 | -0,627911628 | 0,647112465 | 0,50334723 | 0,920918059 | R-spondin 4 |
| ENSG00000225507 | AC069282,1 | -0,628003198 | 0,647071393 | 0,29725167 | 0,806635068 | pseudogene similar to part of TTK protein kinase TTK |
| ENSG00000234933 | CDC42P1 | -0,628025758 | 0,647061275 | 0,5085449 | 0,922427526 | cell division cycle 42 pseudogene 1 |
| ENSG00000275854 | AC084824,4 | -0,628215117 | 0,646976351 | 0,28620046 | 0,797420923 | novel transcript, sense intronic to YARS2 |
| ENSG00000254373 | AC112191,2 | -0,628357056 | 0,646912702 | 0,55410732 | 0,937637409 | ubiquinol-cytochrome c reductase binding protein (UQCRB) pseudogene |
| ENSG00000285739 | AC010157,2 | -0,628360104 | 0,646911335 | 0,28972338 | 0,800522458 | novel transcript, antisense to NRG3 |
| ENSG00000179163 | FUCA1 | -0,62840125 | 0,646892885 | 0,04644632 | 0,327343948 | alpha-L-fucosidase 1 |
| ENSG00000270035 | AL359881,2 | -0,628475435 | 0,646859622 | 0,34535892 | 0,841168721 | novel transcript |
| ENSG00000254427 | LINC02696 | -0,62871867 | 0,646750572 | 0,1481284 | 0,612658572 | long intergenic non-protein coding RNA 2696 |
| ENSG00000258170 | AC089998,4 | -0,628735097 | 0,646743208 | 0,61531711 | 0,952793934 | novel transcript |
| ENSG00000254932 | AP001007,1 | -0,628776533 | 0,646724633 | 0,37864178 | 0,864112469 | novel transcript, antisense to PKNOX2 |
| ENSG00000147724 | FAM135B | -0,628819794 | 0,64670524 | 0,33510251 | 0,834858938 | family with sequence similarity 135 member B |
| ENSG00000272842 | AL391834,1 | -0,629028728 | 0,64661159 | 0,4212309 | 0,889040893 | novel transcript, antisense to DENND4C |
| ENSG00000187753 | C9orf153 | -0,629152766 | 0,646555999 | 0,51246866 | 0,92347658 | chromosome 9 open reading frame 153 |
| ENSG00000263590 | LINC02591 | -0,62962876 | 0,646342714 | 0,6289028 | 0,952793934 | long intergenic non-protein coding RNA 2591 |
| ENSG00000268912 | AC012313,5 | -0,629648955 | 0,646333666 | 0,03641806 | 0,280436969 | novel transcript |
| ENSG00000206140 | TMEM191C | -0,629781803 | 0,646274152 | 0,16314786 | 0,641511104 | transmembrane protein 191C |
| ENSG00000276170 | AC244153,1 | -0,629897045 | 0,64622253 | 0,2973609 | 0,806635068 | novel transcript, antisense to ARHGAP23 |
| ENSG00000229822 | RPS15AP39 | -0,630078816 | 0,646141115 | 0,56454555 | 0,940865625 | ribosomal protein S15a pseudogene 39 |
| ENSG00000223843 | EFCAB6-AS1 | -0,630134231 | 0,646116297 | 0,02677131 | 0,229120826 | EFCAB6 antisense RNA 1 |
| ENSG00000203497 | PDCD4-AS1 | -0,63017139 | 0,646099655 | 0,17209375 | 0,656211582 | PDCD4 antisense RNA 1 |
| ENSG00000175106 | TVP23C | -0,630277757 | 0,646052021 | 0,00749522 | 0,093173215 | trans-golgi network vesicle protein 23 homolog C |
| ENSG00000268621 | IGFL2-AS1 | -0,630325819 | 0,646030499 | 0,4419284 | 0,899273096 | IGFL2 antisense RNA 1 |
| ENSG00000230572 | AC027612,2 | -0,630425152 | 0,64598602 | 0,62556287 | 0,952793934 | pseudogene similar to part of myeloid/lymphoid or mixed-lineage leukemia 3 (MLL3) |
| ENSG00000233579 | KRT8P15 | -0,630619484 | 0,645899011 | 0,3544441 | 0,84590384 | keratin 8 pseudogene 15 |
| ENSG00000259211 | AC013356,2 | -0,631083122 | 0,645691472 | 0,67562805 | 0,963449137 | novel transcript, antisense to BAHD1 |

| | | | | | | |
|---|---|---|---|---|---|---|
| ENSG00000232995 | RGS5 | -0,631120547 | 0,645674722 | 0,15952272 | 0,633929525 | regulator of G protein signaling 5 |
| ENSG00000210144 | MT-TY | -0,631192182 | 0,645642663 | 0,56241774 | 0,940865625 | mitochondrially encoded tRNA-Tyr (UAU/C) |
| ENSG00000167774 | AC010323,1 | -0,631224402 | 0,645628244 | 0,28744592 | 0,798896538 | novel transcript |
| ENSG00000260253 | AC084262,1 | -0,631250965 | 0,645616357 | 0,2467731 | 0,758836117 | novel transcript, overlapping to DUSP4 |
| ENSG00000035499 | DEPDC1B | -0,631361703 | 0,645566802 | 0,10358714 | 0,513379913 | DEP domain containing 1B |
| ENSG00000213963 | AC019080,1 | -0,631625835 | 0,645448622 | 0,0124786 | 0,134667197 | novel transcript |
| ENSG00000287399 | AC011503,4 | -0,631660403 | 0,645433156 | 0,32533478 | 0,827335829 | novel transcript |
| ENSG00000254305 | MRPL9P1 | -0,631764563 | 0,645386559 | 0,45549792 | 0,903545728 | mitochondrial ribosomal protein L9 pseudogene 1 |
| ENSG00000256742 | KDM2B-DT | -0,631820824 | 0,645361391 | 0,33221949 | 0,83293962 | KDM2B divergent transcript |
| ENSG00000259291 | ZNF710-AS1 | -0,631834904 | 0,645355093 | 0,00996256 | 0,114721665 | ZNF710 antisense RNA 1 |
| ENSG00000271711 | AC069236,1 | -0,631851955 | 0,645347466 | 0,3580029 | 0,849032846 | Sin3A-associated protein, 30kDa, pseudogene, |
| ENSG00000200320 | SNORA63 | -0,632030481 | 0,645267612 | 0,07000067 | 0,417274281 | small nucleolar RNA, H/ACA box 63 |
| ENSG00000188051 | TMEM221 | -0,632132424 | 0,645222018 | 0,29073196 | 0,801555946 | transmembrane protein 221 |
| ENSG00000287858 | AC020550,2 | -0,632175259 | 0,645202861 | 0,33194061 | 0,832821707 | novel transcript, antisense to RFTN2 |
| ENSG00000189014 | SHLD2P3 | -0,632411085 | 0,645097403 | 0,36569644 | 0,85559319 | shieldin complex subunit 2 pseudogene 3 |
| ENSG00000228158 | TLE1P1 | -0,632728878 | 0,644955319 | 0,58749423 | 0,946068138 | TLE family member 1 pseudogene 1 |
| ENSG00000227740 | LINC02803 | -0,632765084 | 0,644939133 | 0,26458773 | 0,775800337 | long intergenic non-protein coding RNA 2803 |
| ENSG00000249212 | ATP1B1P1 | -0,632852537 | 0,64490004 | 0,22831247 | 0,736631691 | ATPase Na+/K+ transporting subunit beta 1 pseudogene 1 |
| ENSG00000142149 | HUNK | -0,632853508 | 0,644899605 | 0,03790376 | 0,287976435 | hormonally up-regulated Neu-associated kinase |
| ENSG00000246203 | AL353807,3 | -0,633038436 | 0,644816946 | 0,24115766 | 0,751551821 | novel pseudogene |
| ENSG00000284862 | CCDC39 | -0,633070007 | 0,644802835 | 0,13511736 | 0,586038109 | coiled-coil domain containing 39 |
| ENSG00000145692 | BHMT | -0,633083477 | 0,644796815 | 0,28083301 | 0,791510309 | betaine--homocysteine S-methyltransferase |
| ENSG00000286918 | AL035411,3 | -0,633132072 | 0,644775096 | 0,08739885 | 0,47120083 | novel transcript, antisense to OMA1 |
| ENSG00000254582 | PSMA2P1 | -0,633271811 | 0,644712647 | 0,41509061 | 0,885768407 | proteasome subunit alpha 2 pseudogene 1 |
| ENSG00000188525 | AC010969,1 | -0,633333364 | 0,644685141 | 0,60433632 | 0,950809109 | novel transcript |
| ENSG00000249741 | AC093890,1 | -0,633694782 | 0,644523657 | 0,41843763 | 0,887094957 | salvador homolog 1 (Drosophila) (SAV1) pseudogene |
| ENSG00000197580 | BCO2 | -0,633898039 | 0,644432858 | 0,12349309 | 0,561992438 | beta-carotene oxygenase 2 |
| ENSG00000177842 | ZNF620 | -0,633902631 | 0,644430807 | 0,05778129 | 0,371626042 | zinc finger protein 620 |
| ENSG00000232630 | PRPS1P2 | -0,63392585 | 0,644420436 | 0,1805813 | 0,670229021 | phosphoribosyl pyrophosphate synthetase 1 pseudogene 2 |
| ENSG00000288349 | AC139272,1 | -0,633928298 | 0,644419342 | 0,29492459 | 0,805040011 | POM121 transmembrane nucleoporin C (POM121C) pseudogene |

| | | | | | | |
|---|---|---|---|---|---|---|
| ENSG00000104237 | RP1 | -0,633930394 | 0,644418406 | 0,55293697 | 0,936915156 | RP1 axonemal microtubule associated |
| ENSG00000237863 | AL035425,1 | -0,634118751 | 0,644334277 | 0,31381604 | 0,819877431 | novel transcript, antisense to IRS4 |
| ENSG00000253476 | AC091185,1 | -0,634142928 | 0,644323479 | 0,35027488 | 0,843812136 | novel transcript |
| ENSG00000246308 | AC116535,1 | -0,634145029 | 0,64432254 | 0,17857194 | 0,666018763 | novel transcript, antisense to ZBED5 |
| ENSG00000228625 | AL451047,1 | -0,634242842 | 0,644278858 | 0,39410758 | 0,871635019 | novel transcript |
| ENSG00000064270 | ATP2C2 | -0,634304381 | 0,644251376 | 0,18745773 | 0,681553495 | ATPase secretory pathway Ca2+ transporting 2 |
| ENSG00000270270 | BNIP3P10 | -0,634336349 | 0,6442371 | 0,23001625 | 0,738551869 | BCL2 interacting protein 3 pseudogene 10 |
| ENSG00000267343 | ZNF833P | -0,634368296 | 0,644222835 | 0,55524261 | 0,937793851 | zinc finger protein 833, pseudogene |
| ENSG00000261790 | AC005606,2 | -0,634408791 | 0,644204752 | 0,20121276 | 0,701094807 | novel transcript |
| ENSG00000196586 | MYO6 | -0,634455836 | 0,644183746 | 0,04540688 | 0,322783251 | myosin VI |
| ENSG00000167131 | CCDC103 | -0,634521 | 0,64415465 | 0,27988935 | 0,790565637 | coiled-coil domain containing 103 |
| ENSG00000274184 | AC011815,2 | -0,634569637 | 0,644132934 | 0,23016391 | 0,738687017 | novel transcript, antisense to ZSCAN30 |
| ENSG00000161040 | FBXL13 | -0,634903871 | 0,643983723 | 0,10232667 | 0,509336277 | F-box and leucine rich repeat protein 13 |
| ENSG00000249098 | AC055733,1 | -0,635116153 | 0,643888972 | 0,49968346 | 0,920246448 | cysteine and glycine-rich protein 2 pseudogene |
| ENSG00000159399 | HK2 | -0,635257277 | 0,64382599 | 0,08043537 | 0,449985752 | hexokinase 2 |
| ENSG00000130643 | CALY | -0,635360259 | 0,643780035 | 0,19300508 | 0,689849199 | calcyon neuron specific vesicular protein |
| ENSG00000254450 | ALG9-IT1 | -0,635379977 | 0,643771236 | 0,50421528 | 0,921400408 | ALG9 intronic transcript 1 |
| ENSG00000283738 | AC241640,1 | -0,63556954 | 0,643686653 | 0,02719608 | 0,231741141 | |
| ENSG00000266268 | AP005203,1 | -0,635977728 | 0,643504558 | 0,58554123 | 0,945213113 | novel transcript |
| ENSG00000260277 | AC009019,1 | -0,636002814 | 0,643493369 | 0,62380837 | 0,952793934 | novel transcript, antisense to C16orf52 and VWA3A |
| ENSG00000213424 | KRT222 | -0,636056739 | 0,643469316 | 0,41411629 | 0,885579089 | keratin 222 |
| ENSG00000132010 | ZNF20 | -0,636060667 | 0,643467565 | 0,15449684 | 0,623626246 | zinc finger protein 20 |
| ENSG00000231961 | AL035461,1 | -0,636221383 | 0,643395886 | 0,70858772 | 0,970181083 | pseudogene similar to part of cytochrome c oxidase I |
| ENSG00000165730 | STOX1 | -0,636265215 | 0,643376339 | 0,12276831 | 0,560407143 | storkhead box 1 |
| ENSG00000138075 | ABCG5 | -0,636319879 | 0,643351962 | 0,49370839 | 0,917289924 | ATP binding cassette subfamily G member 5 |
| ENSG00000242094 | FOXP1-IT1 | -0,636359166 | 0,643334442 | 0,11224292 | 0,535189241 | FOXP1 intronic transcript 1 |
| ENSG00000284841 | H2AW | -0,636785768 | 0,643144238 | 0,08787368 | 0,472122805 | H2A,W histone |
| ENSG00000181218 | H2AW | -0,636785768 | 0,643144238 | 0,08787368 | 0,472122805 | H2A,W histone |
| ENSG00000288009 | AL353135,2 | -0,636838718 | 0,643120633 | 0,21225947 | 0,719057938 | novel transcript |
| ENSG00000175170 | FAM182B | -0,637048904 | 0,643026944 | 0,15832449 | 0,631749049 | family with sequence similarity 182 member B |
| ENSG00000198744 | MTCO3P12 | -0,637204796 | 0,642957465 | 0,37294349 | 0,86087605 | MT-CO3 pseudogene 12 |
| ENSG00000266875 | RN7SL408P | -0,637555687 | 0,642801104 | 0,59076558 | 0,947416795 | RNA, 7SL, cytoplasmic 408, pseudogene |
| ENSG00000258476 | LINC02207 | -0,637603171 | 0,642779948 | 0,43178538 | 0,895537893 | long intergenic non-protein coding RNA 2207 |

| | | | | | | |
|---|---|---|---|---|---|---|
| ENSG00000081800 | SLC13A1 | -0,637852546 | 0,642668851 | 0,71083953 | 0,970482408 | solute carrier family 13 member 1 |
| ENSG00000276068 | NAIP | -0,638151226 | 0,642535813 | 0,13737777 | 0,591519288 | NLR family apoptosis inhibitory protein |
| ENSG00000171847 | FAM90A1 | -0,638315752 | 0,642462542 | 0,24911679 | 0,761104998 | family with sequence similarity 90 member A1 |
| ENSG00000228137 | AP001469,2 | -0,638327256 | 0,64245742 | 0,36104901 | 0,851783601 | novel transcript |
| ENSG00000276007 | AC079414,3 | -0,638501122 | 0,642379999 | 0,14542545 | 0,607506834 | novel transcript, antisense WWOX |
| ENSG00000255595 | AC007368,1 | -0,638589766 | 0,64234053 | 0,67070791 | 0,962026636 | uncharacterized LOC101927531 |
| ENSG00000237105 | HCP5 | -0,638610138 | 0,642331459 | 0,77589814 | 0,978587255 | HLA complex P5 |
| ENSG00000181800 | CELF2-AS1 | -0,638761251 | 0,642264183 | 0,07555502 | 0,434139326 | CELF2 antisense RNA 1 |
| ENSG00000253869 | PIGFP1 | -0,638957581 | 0,642176786 | 0,55811854 | 0,939078529 | phosphatidylinositol glycan anchor biosynthesis class F pseudogene 1 |
| ENSG00000245888 | NSMCE1-DT | -0,639008487 | 0,642154127 | 0,35419533 | 0,84556089 | NSMCE1 divergent transcript |
| ENSG00000252982 | RN7SKP234 | -0,63906608 | 0,642128492 | 0,63035469 | 0,953166783 | RN7SK pseudogene 234 |
| ENSG00000240891 | PLCXD2 | -0,639528057 | 0,641922904 | 0,12644401 | 0,567868238 | phosphatidylinositol specific phospholipase C X domain containing 2 |
| ENSG00000267924 | AC139769,2 | -0,639587383 | 0,641896508 | 0,12496398 | 0,565067097 | novel transcript |
| ENSG00000288548 | AL035653,1 | -0,639621863 | 0,641881167 | 0,36544924 | 0,855323027 | novel transcript |
| ENSG00000129946 | SHC2 | -0,639629479 | 0,641877778 | 0,09732715 | 0,49573736 | SHC adaptor protein 2 |
| ENSG00000177201 | OR2T12 | -0,640111998 | 0,641663134 | 0,17262406 | 0,657085135 | olfactory receptor family 2 subfamily T member 12 |
| ENSG00000077264 | PAK3 | -0,64011907 | 0,641659988 | 0,09000283 | 0,478514897 | p21 (RAC1) activated kinase 3 |
| ENSG00000243508 | DNAJB6P7 | -0,640360057 | 0,641552815 | 0,19612942 | 0,694033976 | DNAJB6 pseudogene 7 |
| ENSG00000139193 | CD27 | -0,640494893 | 0,641492858 | 0,57914811 | 0,943828882 | CD27 molecule |
| ENSG00000197124 | ZNF682 | -0,640572259 | 0,641458457 | 0,1037478 | 0,513629639 | zinc finger protein 682 |
| ENSG00000107864 | CPEB3 | -0,64058094 | 0,641454598 | 0,16225419 | 0,639443053 | cytoplasmic polyadenylation element binding protein 3 |
| ENSG00000256278 | AC048382,1 | -0,640657916 | 0,641420373 | 0,31289826 | 0,818889268 | novel transcript, antisense to ZSCAN2 |
| ENSG00000287117 | AC105916,1 | -0,64074904 | 0,641379861 | 0,15348746 | 0,622309924 | novel transcript |
| ENSG00000281404 | LINC01176 | -0,641066692 | 0,641238658 | 0,2188708 | 0,72702684 | long intergenic non-protein coding RNA 1176 |
| ENSG00000287966 | AL590062,1 | -0,641280576 | 0,6411436 | 0,17248931 | 0,656926048 | novel transcript |
| ENSG00000102904 | TSNAXIP1 | -0,641397401 | 0,641091684 | 0,05033552 | 0,343300557 | translin associated factor X interacting protein 1 |
| ENSG00000225889 | AC012368,1 | -0,641454112 | 0,641066483 | 0,02014283 | 0,187703555 | novel transcript, antisense to PELI1 |
| ENSG00000236411 | NDUFAF4P3 | -0,641530175 | 0,641032686 | 0,39999407 | 0,875830109 | NADH:ubiquinone oxidoreductase complex assembly factor 4 pseudogene 3 |
| ENSG00000236474 | GCNT1P1 | -0,641537905 | 0,641029251 | 0,33291176 | 0,833301711 | glucosaminyl (N-acetyl) transferase 1 pseudogene 1 |

| | | | | | | |
|---|---|---|---|---|---|---|
| ENSG00000203326 | ZNF525 | -0,641556357 | 0,641021052 | 0,17580565 | 0,661199837 | zinc finger protein 525 |
| ENSG00000263181 | CYP2D7 | -0,641681334 | 0,640965524 | 0,36309892 | 0,853318246 | cytochrome P450 family 2 subfamily D member 7 (gene/pseudogene) |
| ENSG00000265559 | RN7SL652P | -0,641723499 | 0,640946792 | 0,63614882 | 0,955399772 | RNA, 7SL, cytoplasmic 652, pseudogene |
| ENSG00000138346 | DNA2 | -0,641740383 | 0,64093929 | 0,15971774 | 0,63432613 | DNA replication helicase/nuclease 2 |
| ENSG00000285932 | AL133353,2 | -0,64182301 | 0,640902583 | 0,25756624 | 0,769908546 | novel transcript |
| ENSG00000163440 | PDCL2 | -0,642096278 | 0,640781198 | 0,4721295 | 0,909171312 | phosducin like 2 |
| ENSG00000175455 | CCDC14 | -0,642583091 | 0,640565014 | 0,16807762 | 0,649540012 | coiled-coil domain containing 14 |
| ENSG00000273799 | AC146335,1 | -0,642663517 | 0,640529305 | 0,17453302 | 0,660046011 | glucuronidase, beta (GUSB) pseudogene |
| ENSG00000050438 | SLC4A8 | -0,642939666 | 0,640406712 | 0,00848234 | 0,101620975 | solute carrier family 4 member 8 |
| ENSG00000286721 | AC010623,1 | -0,643164984 | 0,640306702 | 0,05952584 | 0,379411978 | novel transcript |
| ENSG00000166762 | CATSPER2 | -0,643357058 | 0,64022146 | 0,03126593 | 0,254424262 | cation channel sperm associated 2 |
| ENSG00000287406 | AC098656,1 | -0,643398537 | 0,640203053 | 0,40884082 | 0,881578006 | novel transcript |
| ENSG00000272156 | AC008280,2 | -0,643654118 | 0,640089648 | 0,63319595 | 0,9545416 | novel transcript |
| ENSG00000100077 | GRK3 | -0,643776527 | 0,64003534 | 0,01716417 | 0,168907949 | G protein-coupled receptor kinase 3 |
| ENSG00000234062 | AL390879,1 | -0,643897457 | 0,639981693 | 0,02752448 | 0,233711414 | transmembrane 9 superfamily member 2 (TM9SF2) pseudogene |
| ENSG00000070808 | CAMK2A | -0,643996137 | 0,63993792 | 0,13148237 | 0,579293249 | calcium/calmodulin dependent protein kinase II alpha |
| ENSG00000115290 | GRB14 | -0,643998168 | 0,639937019 | 0,08439568 | 0,462254126 | growth factor receptor bound protein 14 |
| ENSG00000236816 | ANKRD20A7P | -0,644132639 | 0,639877375 | 0,27181692 | 0,784843944 | ankyrin repeat domain 20 family member A7, pseudogene |
| ENSG00000278356 | AC005911,1 | -0,644141302 | 0,639873533 | 0,35433784 | 0,845779366 | novel transcript |
| ENSG00000137473 | TTC29 | -0,644186145 | 0,639853644 | 0,29349905 | 0,803461221 | tetratricopeptide repeat domain 29 |
| ENSG00000284707 | AC079781,5 | -0,644186927 | 0,639853297 | 0,01787442 | 0,173764302 | novel transcript |
| ENSG00000225345 | SNX18P3 | -0,644207862 | 0,639844012 | 0,14943964 | 0,614794108 | sorting nexin 18 pseudogene 3 |
| ENSG00000119139 | TJP2 | -0,644258366 | 0,639821614 | 0,0121367 | 0,131911723 | tight junction protein 2 |
| ENSG00000103599 | IQCH | -0,644376295 | 0,639769316 | 0,12522063 | 0,565471028 | IQ motif containing H |
| ENSG00000255710 | AP003170,1 | -0,64438571 | 0,63976514 | 0,14354469 | 0,602629403 | interferon stimulated exonuclease gene 20kDa-like 2 (ISG20L2) pseudogene |
| ENSG00000230467 | RNF39 | -0,644903805 | 0,639535432 | 0,60819426 | 0,951332922 | ring finger protein 39 |
| ENSG00000148655 | LRMDA | -0,645610457 | 0,639222255 | 0,4490416 | 0,902062594 | leucine rich melanocyte differentiation associated |

| | | | | | | |
|---|---|---|---|---|---|---|
| ENSG00000287276 | BX664718,2 | -0,645634413 | 0,639211641 | 0,64865176 | 0,956899963 | contactin associated protein like 3B (CNTNAP3B) pseudogene |
| ENSG00000130518 | IQCN | -0,645636615 | 0,639210665 | 0,03041828 | 0,249540007 | IQ motif containing N |
| ENSG00000054690 | PLEKHH1 | -0,645759331 | 0,639156296 | 0,03972918 | 0,297222562 | pleckstrin homology, MyTH4 and FERM domain containing H1 |
| ENSG00000251580 | LINC02482 | -0,64582419 | 0,639127562 | 0,13470687 | 0,585577813 | long intergenic non-protein coding RNA 2482 |
| ENSG00000272054 | AC007390,1 | -0,646043622 | 0,639030359 | 0,15104737 | 0,618269842 | novel transcript |
| ENSG00000273420 | AC008738,7 | -0,646133903 | 0,638990371 | 0,05561956 | 0,36392023 | novel transcript |
| ENSG00000226393 | IFNA20P | -0,646216655 | 0,63895372 | 0,41994031 | 0,888425645 | interferon alpha 20, pseudogene |
| ENSG00000204344 | STK19 | -0,646310187 | 0,638912297 | 0,21808277 | 0,72573766 | serine/threonine kinase 19 |
| ENSG00000277033 | B3GNTL1 | -0,646630911 | 0,638770277 | 0,51934995 | 0,926800715 | UDP-GlcNAc:betaGal beta-1,3-N-acetylglucosaminyltransferase like 1 |
| ENSG00000284299 | AL590132,1 | -0,646723411 | 0,638729323 | 0,1975239 | 0,696048902 | novel protein |
| ENSG00000282005 | LMNTD2-AS1 | -0,646823467 | 0,638685026 | 0,63016681 | 0,95306213 | LMNTD2 antisense RNA 1 |
| ENSG00000286872 | AC024270,4 | -0,647071588 | 0,638575192 | 0,21432975 | 0,721680711 | novel transcript, antisense to TM6SF1 |
| ENSG00000271522 | AC016831,5 | -0,647402318 | 0,638428819 | 0,40375903 | 0,877691457 | novel transcript |
| ENSG00000237423 | LINC01522 | -0,647465071 | 0,63840105 | 0,37462463 | 0,861771195 | long intergenic non-protein coding RNA 1522 |
| ENSG00000258836 | AL121579,2 | -0,647512107 | 0,638380236 | 0,57831488 | 0,943425812 | polycomb group ring finger 3 (PCGF3) pseudogene |
| ENSG00000162951 | LRRTM1 | -0,647595704 | 0,638343247 | 0,05967761 | 0,380139452 | leucine rich repeat transmembrane neuronal 1 |
| ENSG00000267501 | AC104365,2 | -0,64802349 | 0,638153994 | 0,55600791 | 0,938031417 | novel transcript |
| ENSG00000203801 | LINC00222 | -0,648107326 | 0,638116911 | 0,65617874 | 0,957444211 | long intergenic non-protein coding RNA 222 |
| ENSG00000166669 | ATF7IP2 | -0,648402418 | 0,637986403 | 0,06600778 | 0,403499746 | activating transcription factor 7 interacting protein 2 |
| ENSG00000207501 | RNVU1-14 | -0,648671581 | 0,637867385 | 0,13599841 | 0,587841041 | RNA, variant U1 small nuclear 14 |
| ENSG00000084710 | EFR3B | -0,648732869 | 0,637840288 | 0,00689125 | 0,087723139 | EFR3 homolog B |
| ENSG00000134323 | MYCN | -0,648756651 | 0,637829774 | 0,0719901 | 0,422984135 | MYCN proto-oncogene, bHLH transcription factor |
| ENSG00000273291 | AC092042,3 | -0,648823572 | 0,637800188 | 0,73637695 | 0,974353474 | novel protein |
| ENSG00000172264 | MACROD2 | -0,648900251 | 0,63776629 | 0,04497378 | 0,320506112 | mono-ADP ribosylhydrolase 2 |
| ENSG00000267795 | SMIM22 | -0,648949444 | 0,637744544 | 0,47969806 | 0,912821637 | small integral membrane protein 22 |
| ENSG00000226824 | AC006001,2 | -0,649262469 | 0,637606186 | 0,13898603 | 0,594182875 | novel transcript |
| ENSG00000281601 | ZNF780B | -0,64935507 | 0,637565262 | 0,32622483 | 0,828184588 | zinc finger protein 780B |
| ENSG00000224383 | PRR29 | -0,649370962 | 0,637558239 | 0,13517875 | 0,586080761 | proline rich 29 |
| ENSG00000235674 | LDHAP2 | -0,64953127 | 0,637487399 | 0,3939157 | 0,871635019 | lactate dehydrogenase A pseudogene 2 |
| ENSG00000263585 | AC145207,4 | -0,649563337 | 0,637473229 | 0,8656104 | 0,989074806 | novel transcript, antisense PYCR1 |

| | | | | | | |
|---|---|---|---|---|---|---|
| ENSG00000235560 | AC002310,1 | -0,64978454 | 0,637375496 | 0,1744293 | 0,659880557 | novel transcript, antisense to ZNF747 |
| ENSG00000282174 | AC006518,7 | -0,649799508 | 0,637368883 | 0,49318401 | 0,917245972 | taste receptor, type 2 pseudogene |
| ENSG00000272718 | AC006518,3 | -0,649799508 | 0,637368883 | 0,49318401 | 0,917245972 | taste receptor, type 2 pseudogene |
| ENSG00000256651 | AC006518,1 | -0,649799508 | 0,637368883 | 0,49318401 | 0,917245972 | taste receptor, type 2 pseudogene |
| ENSG00000172828 | CES3 | -0,649812401 | 0,637363187 | 0,02741104 | 0,232982843 | carboxylesterase 3 |
| ENSG00000167912 | AC090152,1 | -0,6498732 | 0,637336327 | 0,0303224 | 0,249039213 | novel transcript |
| ENSG00000150627 | WDR17 | -0,650029452 | 0,637267304 | 0,11440557 | 0,540543423 | WD repeat domain 17 |
| ENSG00000104290 | FZD3 | -0,650125091 | 0,63722506 | 0,1572193 | 0,629552485 | frizzled class receptor 3 |
| ENSG00000105655 | ISYNA1 | -0,650188062 | 0,637197247 | 0,01792144 | 0,173950874 | inositol-3-phosphate synthase 1 |
| ENSG00000225163 | LINC00618 | -0,650266976 | 0,637162393 | 0,37688541 | 0,862818448 | long intergenic non-protein coding RNA 618 |
| ENSG00000251689 | AC079140,5 | -0,65033006 | 0,637134533 | 0,22369934 | 0,732251704 | novel zinc finger protein pseudogene |
| ENSG00000154639 | CXADR | -0,650344963 | 0,637127952 | 0,20897784 | 0,713496019 | CXADR Ig-like cell adhesion molecule |
| ENSG00000075340 | ADD2 | -0,650873563 | 0,636894552 | 0,00229091 | 0,037418119 | adducin 2 |
| ENSG00000138617 | PARP16 | -0,650955291 | 0,636858474 | 0,11081248 | 0,531538341 | poly(ADP-ribose) polymerase family member 16 |
| ENSG00000235888 | AF064858,1 | -0,650979016 | 0,636848001 | 0,1865845 | 0,679960113 | novel transcript |
| ENSG00000269950 | AP000962,1 | -0,650987172 | 0,6368444 | 0,300987 | 0,810652787 | novel transcript |
| ENSG00000197479 | PCDHB11 | -0,651010965 | 0,636833898 | 0,01854421 | 0,177678855 | protocadherin beta 11 |
| ENSG00000262899 | AC004232,1 | -0,651013134 | 0,63683294 | 0,86997511 | 0,989074806 | novel transcript |
| ENSG00000160097 | FNDC5 | -0,651022571 | 0,636828774 | 0,04806476 | 0,33400684 | fibronectin type III domain containing 5 |
| ENSG00000274210 | RNVU1-27 | -0,651076201 | 0,636805102 | 0,13584192 | 0,587733415 | RNA, variant U1 small nuclear 27 |
| ENSG00000281106 | TMEM272 | -0,651085827 | 0,636800853 | 0,38423919 | 0,868128461 | transmembrane protein 272 |
| ENSG00000269425 | AC104521,1 | -0,65117444 | 0,63676174 | 0,51800008 | 0,926208443 | novel transcript, antisense to STAP2 |
| ENSG00000244879 | GABPB1-AS1 | -0,651208104 | 0,636746882 | 0,00180133 | 0,030740647 | GABPB1 antisense RNA 1 |
| ENSG00000183775 | KCTD16 | -0,651218187 | 0,636742432 | 0,16772391 | 0,648680831 | potassium channel tetramerization domain containing 16 |
| ENSG00000245768 | AC106793,1 | -0,651251848 | 0,636727576 | 0,52488646 | 0,928616707 | novel transcript |
| ENSG00000255474 | GAU1 | -0,651336172 | 0,636690361 | 0,18751032 | 0,681556081 | GALNT8 antisense upstream 1 |
| ENSG00000259575 | AC012404,2 | -0,651774041 | 0,63649715 | 0,47667502 | 0,91143003 | novel transcript, antisense to RORA |
| ENSG00000265590 | CFAP298-TCP1( | -0,651792066 | 0,636489197 | 0,03772707 | 0,287183177 | CFAP298-TCP10L readthrough |
| ENSG00000235703 | LINC00894 | -0,651984868 | 0,636404143 | 0,06090683 | 0,383945238 | long intergenic non-protein coding RNA 894 |
| ENSG00000030419 | IKZF2 | -0,652091865 | 0,636356946 | 0,24221553 | 0,752479103 | IKAROS family zinc finger 2 |
| ENSG00000237806 | FAUP2 | -0,652131183 | 0,636339603 | 0,54373639 | 0,934259495 | FAU pseudogene 2 |
| ENSG00000271978 | AL359643,2 | -0,652196087 | 0,636310976 | 0,26938286 | 0,781544346 | novel transcript |

| | | | | | | |
|---|---|---|---|---|---|---|
| ENSG00000261064 | LINC02256 | -0,652250278 | 0,636287075 | 0,43160763 | 0,895347413 | long intergenic non-protein coding RNA 2256 |
| ENSG00000231002 | TRIM26 | -0,652261488 | 0,636282131 | 0,72561785 | 0,97326334 | tripartite motif containing 26 |
| ENSG00000232648 | AC107214,1 | -0,65233364 | 0,63625031 | 0,13377815 | 0,58389504 | novel transcript |
| ENSG00000255776 | VDAC2P2 | -0,652392667 | 0,636224279 | 0,49311071 | 0,917245972 | VDAC2 pseudogene 2 |
| ENSG00000254778 | AL662828,1 | -0,652490226 | 0,636181257 | 0,14804281 | 0,612582848 | palmitoyl-protein thioesterase 2 (PPT2) and EGF-like-domain, multiple 8 (EGFL8) readthrough |
| ENSG00000248249 | AC110794,1 | -0,652645771 | 0,636112671 | 0,43456666 | 0,896272827 | Rac GTPase activating protein 1 (RACGAP1) pseudogene |
| ENSG00000183389 | OR56A4 | -0,652713335 | 0,636082881 | 0,2032729 | 0,703503738 | olfactory receptor family 56 subfamily A member 4 |
| ENSG00000226359 | ACTG1P24 | -0,652853255 | 0,636021193 | 0,36755732 | 0,8571768 | actin gamma 1 pseudogene 24 |
| ENSG00000287191 | AC055807,1 | -0,652863954 | 0,636016477 | 0,30552729 | 0,814326218 | novel transcript, antisense to IGF1R |
| ENSG00000224533 | TMLHE-AS1 | -0,652916188 | 0,63599345 | 0,15761513 | 0,630235931 | TMLHE antisense RNA 1 |
| ENSG00000268565 | AC005339,1 | -0,652921152 | 0,635991262 | 0,57051556 | 0,9423888 | novel transcript, antisense to TNFAIP8L1 |
| ENSG00000286128 | AC022413,2 | -0,652968889 | 0,635970218 | 0,5889168 | 0,946604802 | novel transcript |
| ENSG00000281076 | ZDHHC8P1 | -0,653424104 | 0,635769581 | 0,34329218 | 0,840790189 | ZDHHC8 pseudogene 1 |
| ENSG00000166363 | OR10A5 | -0,653439707 | 0,635762705 | 0,43989414 | 0,89788146 | olfactory receptor family 10 subfamily A member 5 |
| ENSG00000277956 | MAPT | -0,653443419 | 0,635761069 | 0,14338112 | 0,602428097 | microtubule associated protein tau |
| ENSG00000007001 | UPP2 | -0,653470517 | 0,635749128 | 0,48117166 | 0,912911021 | uridine phosphorylase 2 |
| ENSG00000241939 | RN7SL517P | -0,653500864 | 0,635735755 | 0,41215317 | 0,88438705 | RNA, 7SL, cytoplasmic 517, pseudogene |
| ENSG00000236015 | AC011290,2 | -0,653712605 | 0,635642456 | 0,14108156 | 0,59823327 | thioredoxin domain containing 1 (TXNDC1) pseudogene |
| ENSG00000251250 | AC091951,3 | -0,653777158 | 0,635614015 | 0,35273584 | 0,845031687 | pseudogene similar to part of hect domain and RLD 2 HERC2 |
| ENSG00000277545 | AL356752,1 | -0,653794911 | 0,635606194 | 0,45053182 | 0,902062594 | novel transcript, sense intronic to ATP11A |
| ENSG00000211454 | AKR7L | -0,654074181 | 0,635483168 | 0,07476936 | 0,432166624 | aldo-keto reductase family 7 like (gene/pseudogene) |
| ENSG00000287895 | AL136379,1 | -0,654290436 | 0,635387919 | 0,34408244 | 0,840938551 | novel transcript |
| ENSG00000288227 | AC012435,4 | -0,654423827 | 0,635329174 | 0,54639527 | 0,935401821 | coiled-coil-helix-coiled-coil-helix domain-containing protein 7 (CHCHD7) pseudogene |
| ENSG00000229046 | HMGN1P2 | -0,654440399 | 0,635321876 | 0,47392033 | 0,910448879 | high mobility group nucleosome binding domain 1 pseudogene 2 |
| ENSG00000269486 | ERVK9-11 | -0,654466076 | 0,635310569 | 0,13772471 | 0,592433314 | endogenous retrovirus group K9, member 11 |
| ENSG00000282930 | ERVK9-11 | -0,654466076 | 0,635310569 | 0,13772471 | 0,592433314 | endogenous retrovirus group K9, member 11 |

| | | | | | | |
|---|---|---|---|---|---|---|
| ENSG00000133874 | RNF122 | -0,654557179 | 0,635270451 | 0,0348008 | 0,272139119 | ring finger protein 122 |
| ENSG00000227107 | AC104667,1 | -0,654716508 | 0,635200297 | 0,45203351 | 0,902062594 | novel transcript |
| ENSG00000164794 | KCNV1 | -0,654957904 | 0,635094022 | 0,04903716 | 0,337686719 | potassium voltage-gated channel modifier subfamily V member 1 |
| ENSG00000286850 | AL358232,2 | -0,654975892 | 0,635086104 | 0,42573981 | 0,892133432 | novel transcript |
| ENSG00000278399 | AC026120,1 | -0,655091549 | 0,635035193 | 0,6466389 | 0,956899963 | hydroxysteroid (17-beta) dehydrogenase 6 (HSD17B6) pseudogene |
| ENSG00000239528 | RPS14P8 | -0,655104408 | 0,635029532 | 0,29360688 | 0,803461221 | ribosomal protein S14 pseudogene 8 |
| ENSG00000243107 | AC000120,1 | -0,655207409 | 0,634984196 | 0,17760055 | 0,664043981 | novel transcript |
| ENSG00000178814 | OPLAH | -0,655234174 | 0,634972416 | 0,24364608 | 0,754491993 | 5-oxoprolinase, ATP-hydrolysing |
| ENSG00000200169 | RNU5D-1 | -0,655297724 | 0,634944447 | 0,24039616 | 0,750433792 | RNA, U5D small nuclear 1 |
| ENSG00000048462 | TNFRSF17 | -0,655461846 | 0,634872219 | 0,47232308 | 0,909179229 | TNF receptor superfamily member 17 |
| ENSG00000229337 | AC079305,2 | -0,655632212 | 0,634797252 | 0,17004233 | 0,653256728 | novel transcript |
| ENSG00000267115 | AC022148,2 | -0,655741383 | 0,634749218 | 0,47887322 | 0,912714933 | novel transcript |
| ENSG00000285529 | AC016816,1 | -0,65579931 | 0,634723732 | 0,31581734 | 0,821233148 | novel transcript |
| ENSG00000225447 | RPS15AP10 | -0,656023056 | 0,634625301 | 0,37525865 | 0,862022464 | ribosomal protein S15a pseudogene 10 |
| ENSG00000276876 | MYT1 | -0,656051264 | 0,634612892 | 0,09701535 | 0,494713385 | myelin transcription factor 1 |
| ENSG00000227215 | AL355613,1 | -0,656058603 | 0,634609664 | 0,64847203 | 0,956899963 | novel transcript |
| ENSG00000248769 | AC139495,1 | -0,65612169 | 0,634581914 | 0,16921727 | 0,651287551 | POM121 membrane glycoprotein-like 1 (POM121L1) pseudogene |
| ENSG00000285035 | AC091057,6 | -0,656232355 | 0,634533239 | 0,32986729 | 0,831624145 | Serine/threonine-protein kinase ULK4 (ULK4), pseudogene |
| ENSG00000287403 | AC016245,2 | -0,656405245 | 0,634457202 | 0,20325132 | 0,703503738 | novel transcript, antisense to RPTOR |
| ENSG00000226726 | TMEM256P2 | -0,656636536 | 0,634355495 | 0,43966471 | 0,897665148 | transmembrane protein 256 pseudogene 2 |
| ENSG00000236988 | CTDSPL2P1 | -0,656723463 | 0,634317274 | 0,53207689 | 0,931291085 | CTDSPL2 pseudogene 1 |
| ENSG00000198729 | PPP1R14C | -0,656788962 | 0,634288477 | 0,08912128 | 0,47641326 | protein phosphatase 1 regulatory inhibitor subunit 14C |
| ENSG00000249055 | TBCAP3 | -0,656806411 | 0,634280805 | 0,61307747 | 0,952413269 | tubulin folding cofactor A pseudogene 3 |
| ENSG00000180138 | CSNK1A1L | -0,656884856 | 0,634246318 | 0,61061929 | 0,951841373 | casein kinase 1 alpha 1 like |
| ENSG00000238169 | LINC01053 | -0,65691544 | 0,634232873 | 0,33775024 | 0,837066292 | long intergenic non-protein coding RNA 1053 |
| ENSG00000234444 | ZNF736 | -0,657237866 | 0,634091144 | 0,01808133 | 0,174744891 | zinc finger protein 736 |
| ENSG00000213703 | AL138847,1 | -0,657330766 | 0,634050314 | 0,31118963 | 0,817751312 | ribosomal protein L13a (RPL13A) pseudogene |
| ENSG00000254463 | PPIAP41 | -0,657721598 | 0,633878571 | 0,31861886 | 0,823434042 | peptidylprolyl isomerase A pseudogene 41 |

| | | | | | | |
|---|---|---|---|---|---|---|
| ENSG00000180190 | TDRP | -0,657928675 | 0,633787594 | 0,03050717 | 0,24981045 | testis development related protein |
| ENSG00000224490 | TTC21B-AS1 | -0,657930469 | 0,633786806 | 0,30008918 | 0,809942221 | TTC21B antisense RNA 1 |
| ENSG00000278091 | ZNF85 | -0,658168406 | 0,633682287 | 0,32157981 | 0,824933727 | zinc finger protein 85 |
| ENSG00000228487 | AL450263,1 | -0,658171515 | 0,633680921 | 0,33217428 | 0,83293962 | novel transcript, antisense to RALGPS1 |
| ENSG00000259209 | AC004943,1 | -0,658175845 | 0,633679019 | 0,12777161 | 0,570756819 | novel transcript, antisense to ZFHX3 |
| ENSG00000175707 | KDF1 | -0,658209654 | 0,633664169 | 0,62071884 | 0,952793934 | keratinocyte differentiation factor 1 |
| ENSG00000175164 | ABO | -0,658408033 | 0,633577043 | 0,59665306 | 0,948282297 | ABO, alpha 1-3-N-acetylgalactosaminyltransferase and alpha 1-3-galactosyltransferase |
| ENSG00000287990 | AL358334,4 | -0,658718347 | 0,633440779 | 0,3563493 | 0,847771893 | novel transcript, antisense to TRIM9 |
| ENSG00000105617 | LENG1 | -0,658752658 | 0,633425715 | 0,68644203 | 0,965769168 | leukocyte receptor cluster member 1 |
| ENSG00000236809 | SNX25P1 | -0,658982069 | 0,633324998 | 0,51019117 | 0,922427526 | sorting nexin 25 pseudogene 1 |
| ENSG00000236418 | HLA-DQA1 | -0,65902383 | 0,633306666 | 0,38192868 | 0,866514369 | major histocompatibility complex, class II, DQ alpha 1 |
| ENSG00000034239 | EFCAB1 | -0,659081826 | 0,633281208 | 0,13837299 | 0,59326564 | EF-hand calcium binding domain 1 |
| ENSG00000162188 | GNG3 | -0,659155167 | 0,633249015 | 0,0684485 | 0,412326349 | G protein subunit gamma 3 |
| ENSG00000223984 | HNRNPRP1 | -0,659253842 | 0,633205705 | 0,06145713 | 0,386127478 | heterogeneous nuclear ribonucleoprotein R pseudogene 1 |
| ENSG00000162814 | SPATA17 | -0,659522954 | 0,633087601 | 0,10679871 | 0,521711701 | spermatogenesis associated 17 |
| ENSG00000170439 | METTL7B | -0,659580046 | 0,633062548 | 0,45949069 | 0,904914462 | methyltransferase like 7B |
| ENSG00000172782 | FADS6 | -0,659595097 | 0,633055944 | 0,59806905 | 0,948582224 | fatty acid desaturase 6 |
| ENSG00000222036 | POTEM | -0,659605824 | 0,633051237 | 0,24673478 | 0,758795604 | POTE ankyrin domain family member M |
| ENSG00000172146 | OR1A1 | -0,659978139 | 0,632887887 | 0,01133343 | 0,125514797 | olfactory receptor family 1 subfamily A member 1 |
| ENSG00000117569 | PTBP2 | -0,660012552 | 0,632872791 | 0,16869526 | 0,650640109 | polypyrimidine tract binding protein 2 |
| ENSG00000226055 | PAICSP1 | -0,660076244 | 0,632844851 | 0,1472403 | 0,611181688 | phosphoribosylaminoimidazole carboxylase, phosphoribosylaminoimidazole succinocarboxamide synthetase pseudogene 1 |
| ENSG00000170049 | KCNAB3 | -0,660267944 | 0,632760767 | 0,07846656 | 0,443372235 | potassium voltage-gated channel subfamily A regulatory beta subunit 3 |
| ENSG00000272159 | AC087623,3 | -0,660496988 | 0,632660317 | 0,19960105 | 0,698923433 | novel transcript, antisense to LETM2 |
| ENSG00000230185 | C9orf147 | -0,660705749 | 0,632568776 | 0,05156287 | 0,347838733 | chromosome 9 open reading frame 147 |
| ENSG00000234618 | RPSAP9 | -0,660729709 | 0,632558271 | 0,2918822 | 0,802531095 | ribosomal protein SA pseudogene 9 |
| ENSG00000270139 | AP001172,2 | -0,660889455 | 0,632488233 | 0,4573841 | 0,903831777 | novel transcript |
| ENSG00000284406 | AC004492,3 | -0,661005979 | 0,63243715 | 0,18622892 | 0,679406361 | novel transcript, antisense to HBP1 |

| | | | | | | |
|---|---|---|---|---|---|---|
| ENSG00000272072 | AC004492,1 | -0,661005979 | 0,63243715 | 0,18622892 | 0,679406361 | novel transcript, antisense to HBP1 |
| ENSG00000118849 | RARRES1 | -0,661104182 | 0,632394102 | 0,17192856 | 0,655750391 | retinoic acid receptor responder 1 |
| ENSG00000268093 | AC022154,1 | -0,661388016 | 0,632269698 | 0,37190594 | 0,859937929 | novel transcript, antisense to SPHK2 |
| ENSG00000215158 | AC138409,2 | -0,661485629 | 0,63222692 | 0,02793873 | 0,236266271 | Putative beta-glucuronidase-like protein FLJ75287 pseudogene |
| ENSG00000273369 | AC096586,2 | -0,661524415 | 0,632209923 | 0,38588275 | 0,869112622 | novel transcript, antisense to GUF1 |
| ENSG00000225282 | AP000350,2 | -0,66174541 | 0,632113087 | 0,23072594 | 0,739234644 | kelch-like 5 (Drosophila) (KLHL5) pseudogene |
| ENSG00000281900 | AP000350,17 | -0,66174541 | 0,632113087 | 0,23072594 | 0,739234644 | kelch-like 5 (Drosophila) (KLHL5) pseudogene |
| ENSG00000286086 | AC099661,1 | -0,661787351 | 0,632094711 | 0,54733166 | 0,935404974 | novel transcript |
| ENSG00000181007 | ZFP82 | -0,662185607 | 0,631920246 | 0,02726459 | 0,232168734 | ZFP82 zinc finger protein |
| ENSG00000213608 | SLC25A14P1 | -0,662327064 | 0,631858289 | 0,23493722 | 0,744634749 | solute carrier family 25 member 14 pseudogene 1 |
| ENSG00000261572 | AC097639,1 | -0,662557919 | 0,631757189 | 0,59708157 | 0,948282297 | novel transcript, intronic to CMTM8 |
| ENSG00000239828 | AC063944,1 | -0,662672199 | 0,631707147 | 0,55122062 | 0,936366211 | novel transcript |
| ENSG00000177990 | DPY19L2 | -0,662787887 | 0,631656494 | 0,05393148 | 0,357985923 | dpy-19 like 2 |
| ENSG00000145192 | AHSG | -0,663175606 | 0,631486761 | 0,62243951 | 0,952793934 | alpha 2-HS glycoprotein |
| ENSG00000258520 | AL359317,1 | -0,663196411 | 0,631477655 | 0,44671373 | 0,901650934 | novel transcript |
| ENSG00000254673 | AC110275,1 | -0,663282719 | 0,631439878 | 0,7561273 | 0,975901808 | novel protein |
| ENSG00000279232 | AC008522,1 | -0,663627084 | 0,631289174 | 0,19317145 | 0,689970564 | novel transcript, antisense to RGMB |
| ENSG00000286068 | AC010974,2 | -0,663684218 | 0,631264174 | 0,45604272 | 0,903545728 | novel transcript, antisense to NCKAP5 |
| ENSG00000123999 | INHA | -0,663778828 | 0,631222778 | 0,31148893 | 0,817751312 | inhibin subunit alpha |
| ENSG00000254346 | MTND1P6 | -0,663841514 | 0,631195352 | 0,48074386 | 0,912907448 | MT-ND1 pseudogene 6 |
| ENSG00000112293 | GPLD1 | -0,664085709 | 0,631088523 | 0,05448907 | 0,360376004 | glycosylphosphatidylinositol specific phospholipase D1 |
| ENSG00000230597 | AL136164,1 | -0,66415161 | 0,631059696 | 0,59224067 | 0,947416795 | novel transcript |
| ENSG00000283352 | AL021368,4 | -0,664180596 | 0,631047017 | 0,31450029 | 0,819877431 | novel transcript |
| ENSG00000254377 | MIR124-2HG | -0,66421529 | 0,631031841 | 0,09383533 | 0,486996636 | MIR124-2 host gene |
| ENSG00000127362 | TAS2R3 | -0,664323776 | 0,630984392 | 0,22853993 | 0,737046989 | taste 2 receptor member 3 |
| ENSG00000205890 | AC108134,1 | -0,664383734 | 0,630958169 | 0,58089904 | 0,943914064 | novel transcript, antisense to CCDC64B |
| ENSG00000234361 | AL391863,2 | -0,664536719 | 0,630891265 | 0,31674137 | 0,821646322 | novel transcript |
| ENSG00000229140 | CCDC26 | -0,664558723 | 0,630881643 | 0,00732544 | 0,091572382 | CCDC26 long non-coding RNA |
| ENSG00000179141 | MTUS2-AS1 | -0,664658983 | 0,630837801 | 0,44368259 | 0,900090367 | MTUS2 antisense RNA 1 |
| ENSG00000267165 | CHMP1B-AS1 | -0,665139502 | 0,630627723 | 0,72059162 | 0,972142678 | CHMP1B antisense RNA 1 |
| ENSG00000229127 | KANSL1L-AS1 | -0,665284352 | 0,630564409 | 0,02917099 | 0,242424711 | KANSL1L antisense RNA 1 |

| | | | | | | |
|---|---|---|---|---|---|---|
| ENSG00000275430 | MUC20 | -0,665497737 | 0,630471151 | 0,26336061 | 0,774637968 | mucin 20, cell surface associated |
| ENSG00000275744 | SLC2A11 | -0,665653049 | 0,630403282 | 0,03147981 | 0,255694831 | solute carrier family 2 member 11 |
| ENSG00000242428 | CFAP20DC-AS1 | -0,665675418 | 0,630393508 | 0,31320983 | 0,819430294 | CFAP20DC antisense RNA 1 |
| ENSG00000255837 | TAS2R20 | -0,665803401 | 0,630337587 | 0,22315772 | 0,731704543 | taste 2 receptor member 20 |
| ENSG00000273092 | TAS2R20 | -0,665803401 | 0,630337587 | 0,22315772 | 0,731704543 | taste 2 receptor member 20 |
| ENSG00000177076 | ACER2 | -0,665906123 | 0,630292708 | 0,11291583 | 0,53680967 | alkaline ceramidase 2 |
| ENSG00000282972 | GNG4 | -0,666042746 | 0,630233022 | 0,57332597 | 0,942475308 | G protein subunit gamma 4 |
| ENSG00000103021 | CCDC113 | -0,666145509 | 0,630188132 | 0,02523097 | 0,220341093 | coiled-coil domain containing 113 |
| ENSG00000153551 | CMTM7 | -0,66621478 | 0,630157874 | 0,06481033 | 0,399335875 | CKLF like MARVEL transmembrane domain containing 7 |
| ENSG00000271716 | AC022001,2 | -0,666521088 | 0,630024096 | 0,31134222 | 0,817751312 | novel transcript, antisense to VGLL4 |
| ENSG00000213280 | AC090114,1 | -0,666547791 | 0,630012435 | 0,42736264 | 0,892552417 | ribosomal protein S10 (RPS10) pseudogene |
| ENSG00000177483 | RBM44 | -0,66662178 | 0,629980125 | 0,30738518 | 0,816143546 | RNA binding motif protein 44 |
| ENSG00000196167 | COLCA1 | -0,666682923 | 0,629953427 | 0,03578705 | 0,277216643 | colorectal cancer associated 1 |
| ENSG00000224578 | HNRNPA1P48 | -0,666698444 | 0,62994665 | 0,18960974 | 0,684917268 | heterogeneous nuclear ribonucleoprotein A1 pseudogene 48 |
| ENSG00000241472 | PTPRG-AS1 | -0,666738313 | 0,629929241 | 0,0283864 | 0,238652911 | PTPRG antisense RNA 1 |
| ENSG00000273656 | AC139494,1 | -0,666823056 | 0,62989224 | 0,00507734 | 0,069494347 | glucuronidase, beta (GUSB) pseudogene |
| ENSG00000255760 | LINC02422 | -0,667110354 | 0,629766816 | 0,56287629 | 0,940865625 | long intergenic non-protein coding RNA 2422 |
| ENSG00000140022 | STON2 | -0,667290411 | 0,629688222 | 0,06685198 | 0,406310314 | stonin 2 |
| ENSG00000261390 | MAFTRR | -0,667451826 | 0,629617774 | 0,28745506 | 0,798896538 | MAF transcriptional regulator RNA |
| ENSG00000248932 | AC046134,2 | -0,667487372 | 0,629602262 | 0,04762209 | 0,331741728 | novel transcript, antisense to RBP2, RBP1 & NMNAT3 |
| ENSG00000178358 | OR2D3 | -0,667502757 | 0,629595547 | 0,554681 | 0,937793851 | olfactory receptor family 2 subfamily D member 3 |
| ENSG00000263220 | AC015727,1 | -0,667605086 | 0,629550892 | 0,57380203 | 0,942475308 | novel transcript, antisense to RABEP1 |
| ENSG00000236986 | AL157938,2 | -0,667728709 | 0,629496949 | 0,19058579 | 0,685949202 | novel transcript, antisense to NUP214 |
| ENSG00000181215 | C4orf50 | -0,668094246 | 0,629337473 | 0,34156793 | 0,840048641 | chromosome 4 open reading frame 50 |
| ENSG00000196337 | CGB7 | -0,668347031 | 0,629227212 | 0,62460057 | 0,952793934 | chorionic gonadotropin subunit beta 7 |
| ENSG00000133958 | UNC79 | -0,668378493 | 0,62921349 | 0,08409524 | 0,461255021 | unc-79 homolog, NALCN channel complex subunit |
| ENSG00000276416 | UNC79 | -0,668378493 | 0,62921349 | 0,08409524 | 0,461255021 | unc-79 homolog, NALCN channel complex subunit |
| ENSG00000080854 | IGSF9B | -0,668513382 | 0,629154663 | 0,10997983 | 0,528823343 | immunoglobulin superfamily member 9B |
| ENSG00000253549 | CA3-AS1 | -0,668564171 | 0,629132514 | 0,11317633 | 0,537252449 | CA3 antisense RNA 1 |
| ENSG00000242593 | AC006148,1 | -0,668678035 | 0,629082862 | 0,17611837 | 0,661639583 | novel transcript |
| ENSG00000261823 | AC084782,2 | -0,668759749 | 0,629047232 | 0,45171599 | 0,902062594 | novel transcript |

| | | | | | | |
|---|---|---|---|---|---|---|
| ENSG00000180245 | RRH | -0,66885318 | 0,629006495 | 0,38900501 | 0,870057277 | retinal pigment epithelium-derived rhodopsin homolog |
| ENSG00000197085 | NPSR1-AS1 | -0,668979558 | 0,628951397 | 0,48437483 | 0,913690171 | NPSR1 antisense RNA 1 |
| ENSG00000261186 | AC093642,2 | -0,66907423 | 0,628910126 | 0,24581728 | 0,757752583 | novel transcript |
| ENSG00000224215 | AL606469,1 | -0,669096345 | 0,628900486 | 0,46129297 | 0,906165604 | novel transcript |
| ENSG00000287518 | AC100778,4 | -0,669212703 | 0,628849765 | 0,35513548 | 0,846584484 | novel transcript, antisense to DYM |
| ENSG00000229176 | SAPCD1 | -0,669294681 | 0,628814033 | 0,39329104 | 0,871304649 | suppressor APC domain containing 1 |
| ENSG00000288044 | AL096870,12 | -0,669334184 | 0,628796816 | 0,24874332 | 0,760720365 | novel transcript, antisense to RABGGTAand DHRS1 |
| ENSG00000287436 | AL096870,11 | -0,669334184 | 0,628796816 | 0,24874332 | 0,760720365 | novel transcript, antisense to RABGGTAand DHRS1 |
| ENSG00000260806 | AL163051,1 | -0,669469506 | 0,628737838 | 0,11176532 | 0,534531289 | novel transcript, antisense to PAPOLA |
| ENSG00000081818 | PCDHB4 | -0,669621553 | 0,628671578 | 0,01834328 | 0,176288964 | protocadherin beta 4 |
| ENSG00000173041 | ZNF680 | -0,669661949 | 0,628653976 | 0,18970578 | 0,684955795 | zinc finger protein 680 |
| ENSG00000228784 | LINC00954 | -0,669717411 | 0,628629809 | 0,11404326 | 0,53948382 | long intergenic non-protein coding RNA 954 |
| ENSG00000125898 | FAM110A | -0,669883427 | 0,628557474 | 0,05797407 | 0,372428727 | family with sequence similarity 110 member A |
| ENSG00000067606 | PRKCZ | -0,670163412 | 0,628435501 | 0,03918991 | 0,294869635 | protein kinase C zeta |
| ENSG00000261326 | LINC01355 | -0,670184605 | 0,62842627 | 0,05572354 | 0,364179132 | long intergenic non-protein coding RNA 1355 |
| ENSG00000283124 | LINC01355 | -0,670184605 | 0,62842627 | 0,05572354 | 0,364179132 | long intergenic non-protein coding RNA 1355 |
| ENSG00000145569 | OTULINL | -0,670309025 | 0,628372076 | 0,06807345 | 0,411001592 | OTU deubiquitinase with linear linkage specificity like |
| ENSG00000091129 | NRCAM | -0,670957004 | 0,628089909 | 0,17006569 | 0,653266233 | neuronal cell adhesion molecule |
| ENSG00000205838 | TTC23L | -0,670969651 | 0,628084403 | 0,1111562 | 0,532514145 | tetratricopeptide repeat domain 23 like |
| ENSG00000275311 | AC139494,3 | -0,670992359 | 0,628074517 | 0,22359422 | 0,731984265 | novel transcript |
| | | | | | | chromosome X open reading frame 56 (CXorf56) |
| ENSG00000233170 | AC138356,2 | -0,671051791 | 0,628048644 | 0,20292632 | 0,703345263 | pseudogene |
| ENSG00000249437 | NAIP | -0,671065511 | 0,628042671 | 0,06533854 | 0,401164726 | NLR family apoptosis inhibitory protein |
| ENSG00000234040 | RPL10P12 | -0,671260903 | 0,627957618 | 0,34065782 | 0,839282032 | ribosomal protein L10 pseudogene 12 |
| ENSG00000233061 | TTLL7-IT1 | -0,671300011 | 0,627940596 | 0,06439132 | 0,397835213 | TTLL7 intronic transcript 1 |
| ENSG00000188488 | SERPINA5 | -0,671573427 | 0,627821601 | 0,21464709 | 0,722164151 | serpin family A member 5 |
| ENSG00000277610 | RNVU1-4 | -0,671675561 | 0,627777157 | 0,48962642 | 0,916063942 | RNA, variant U1 small nuclear 4 |
| ENSG00000166575 | TMEM135 | -0,671836906 | 0,627706953 | 0,23536729 | 0,744654651 | transmembrane protein 135 |
| ENSG00000272461 | AP005328,1 | -0,672036303 | 0,627620203 | 0,58827359 | 0,946176848 | novel transcript |
| | | | | | | ectonucleoside triphosphate diphosphohydrolase 5 |
| ENSG00000267005 | AC002984,1 | -0,672302827 | 0,627504266 | 0,60700163 | 0,950993594 | (ENTPD5) pseudogene |

| | | | | | | |
|---|---|---|---|---|---|---|
| ENSG00000168916 | ZNF608 | -0,67248909 | 0,627423256 | 0,0398966 | 0,298118347 | zinc finger protein 608 |
| ENSG00000166869 | CHP2 | -0,672500863 | 0,627418136 | 0,15227704 | 0,620540856 | calcineurin like EF-hand protein 2 |
| ENSG00000275956 | AC104446,2 | -0,672518135 | 0,627410624 | 0,18141336 | 0,671207946 | zinc finger protein 610 (ZNF610) pseudogene |
| ENSG00000143869 | GDF7 | -0,672726683 | 0,627319936 | 0,02951705 | 0,244377517 | growth differentiation factor 7 |
| ENSG00000258824 | AL122035,1 | -0,672842712 | 0,627269486 | 0,18643605 | 0,679577549 | novel transcript |
| ENSG00000287732 | AL731574,1 | -0,673564669 | 0,626955665 | 0,49606863 | 0,918105073 | novel transcript |
| ENSG00000271553 | AC018638,7 | -0,673779923 | 0,626862128 | 0,17823782 | 0,665463944 | novel transcript |
| ENSG00000159445 | THEM4 | -0,67395229 | 0,626787238 | 0,01402129 | 0,146560258 | thioesterase superfamily member 4 |
| ENSG00000241697 | TMEFF1 | -0,674290025 | 0,626640524 | 0,02959736 | 0,244732069 | transmembrane protein with EGF like and two follistatin like domains 1 |
| ENSG00000284812 | AC270289,1 | -0,674850293 | 0,626397216 | 0,32797696 | 0,830008227 | regulatory solute carrier protein, family 1, member 1 (RSC1A1) pseudogene |
| ENSG00000167080 | B4GALNT2 | -0,675007233 | 0,626329079 | 0,09943113 | 0,50040065 | beta-1,4-N-acetyl-galactosaminyltransferase 2 |
| ENSG00000036530 | CYP46A1 | -0,675097513 | 0,626289887 | 0,12333523 | 0,561682809 | cytochrome P450 family 46 subfamily A member 1 |
| ENSG00000165186 | PTCHD1 | -0,675104743 | 0,626286748 | 0,10080385 | 0,504527553 | patched domain containing 1 |
| ENSG00000164142 | FAM160A1 | -0,675335745 | 0,626186476 | 0,10600429 | 0,519559347 | family with sequence similarity 160 member A1 |
| ENSG00000170231 | FABP6 | -0,675346767 | 0,626181692 | 0,39956146 | 0,875433675 | fatty acid binding protein 6 |
| ENSG00000179698 | WDR97 | -0,675407261 | 0,626155436 | 0,23684366 | 0,746439525 | WD repeat domain 97 |
| ENSG00000287727 | AL139423,2 | -0,675688389 | 0,626033433 | 0,21013178 | 0,715238331 | novel transcript, antisense to PEX14 |
| ENSG00000228218 | ATF4P3 | -0,675705014 | 0,626026219 | 0,39633814 | 0,87298265 | activating transcription factor 4 pseudogene 3 |
| ENSG00000100218 | RSPH14 | -0,675803179 | 0,625983624 | 0,19584321 | 0,69375328 | radial spoke head 14 homolog |
| ENSG00000284428 | AC092329,4 | -0,675836555 | 0,625969142 | 0,00470562 | 0,065938206 | novel transcript |
| ENSG00000281708 | ERC2-IT1 | -0,675851748 | 0,62596255 | 0,22739957 | 0,735964811 | ERC2 intronic transcript 1 |
| ENSG00000224747 | MTCYBP21 | -0,675970224 | 0,625911148 | 0,2957328 | 0,805426883 | MT-CYB pseudogene 21 |
| ENSG00000185306 | C12orf56 | -0,676163993 | 0,625827087 | 0,23809927 | 0,747367171 | chromosome 12 open reading frame 56 |
| ENSG00000273183 | AC093726,2 | -0,676272201 | 0,625780149 | 0,38921313 | 0,870057277 | novel transcript, antisense to PAXIP1 |
| ENSG00000255189 | GLYATL1P1 | -0,676349068 | 0,625746808 | 0,54153276 | 0,933491609 | glycine-N-acyltransferase like 1 pseudogene 1 |
| ENSG00000285587 | AC012470,1 | -0,676517373 | 0,625673813 | 0,39885003 | 0,875433675 | novel transcript |
| ENSG00000166432 | ZMAT1 | -0,676801143 | 0,625550758 | 0,0754296 | 0,433690715 | zinc finger matrin-type 1 |
| ENSG00000124743 | KLHL31 | -0,676873878 | 0,625519221 | 0,34390246 | 0,840901506 | kelch like family member 31 |
| ENSG00000266598 | AC037487,2 | -0,676932479 | 0,625493814 | 0,37581201 | 0,862406526 | novel transcript, antisense to GNA13 |
| ENSG00000284925 | GCSAM | -0,677412134 | 0,625285889 | 0,27078921 | 0,783073289 | germinal center associated signaling and motility |
| ENSG00000174500 | GCSAM | -0,677412134 | 0,625285889 | 0,27078921 | 0,783073289 | germinal center associated signaling and motility |

| | | | | | | |
|---|---|---|---|---|---|---|
| ENSG00000249921 | AC034207,1 | -0,67752761 | 0,625235842 | 0,47212758 | 0,909171312 | guanine nucleotide binding protein (G protein), beta polypeptide 2-like 1 (GNB2L1) pseudogene |
| ENSG00000253208 | PSME2P5 | -0,677574588 | 0,625215484 | 0,15202262 | 0,620332737 | proteasome activator subunit 2 pseudogene 5 |
| ENSG00000223842 | AL360093,1 | -0,67780863 | 0,625114065 | 0,34880301 | 0,842864225 | novel transcript |
| ENSG00000136404 | TM6SF1 | -0,677935841 | 0,625058948 | 0,44591511 | 0,901227615 | transmembrane 6 superfamily member 1 |
| ENSG00000105649 | RAB3A | -0,678024348 | 0,625020603 | 0,02198505 | 0,19951833 | RAB3A, member RAS oncogene family |
| ENSG00000270587 | AC046185,2 | -0,678058562 | 0,62500578 | 0,45420666 | 0,90318049 | family with sequence similarity 136, member A (FAM136A) pseudogene |
| ENSG00000234722 | LINC01287 | -0,678194386 | 0,624946942 | 0,33975773 | 0,838611198 | long intergenic non-protein coding RNA 1287 |
| ENSG00000266993 | AL050343,2 | -0,678357427 | 0,624876319 | 0,09805104 | 0,497317904 | novel transcript, antisense to NRD1 |
| ENSG00000150054 | MPP7 | -0,678843292 | 0,624665911 | 0,03703191 | 0,284044574 | membrane palmitoylated protein 7 |
| ENSG00000281654 | AC233280,28 | -0,678934777 | 0,624626301 | 0,45056555 | 0,902062594 | novel transcript |
| ENSG00000280482 | AC233280,8 | -0,678934777 | 0,624626301 | 0,45056555 | 0,902062594 | novel transcript |
| ENSG00000281564 | AC233280,25 | -0,678934777 | 0,624626301 | 0,45056555 | 0,902062594 | novel transcript |
| ENSG00000280629 | AC233280,11 | -0,678934777 | 0,624626301 | 0,45056555 | 0,902062594 | novel transcript |
| ENSG00000280631 | AC233280,12 | -0,678934777 | 0,624626301 | 0,45056555 | 0,902062594 | novel transcript |
| ENSG00000280748 | AC233280,13 | -0,678934777 | 0,624626301 | 0,45056555 | 0,902062594 | novel transcript |
| ENSG00000229178 | AC233280,1 | -0,678934777 | 0,624626301 | 0,45056555 | 0,902062594 | novel transcript |
| ENSG00000281094 | AC233280,19 | -0,678934777 | 0,624626301 | 0,45056555 | 0,902062594 | novel transcript |
| ENSG00000233338 | TLR8-AS1 | -0,679078782 | 0,624563956 | 0,5756187 | 0,942700009 | TLR8 antisense RNA 1 |
| ENSG00000197813 | AC011450,1 | -0,679217822 | 0,624503766 | 0,17389636 | 0,659015241 | novel transcript |
| ENSG00000186648 | CARMIL3 | -0,679375431 | 0,624435546 | 0,04470039 | 0,319358544 | capping protein regulator and myosin 1 linker 3 |
| ENSG00000284765 | CARMIL3 | -0,679375431 | 0,624435546 | 0,04470039 | 0,319358544 | capping protein regulator and myosin 1 linker 3 |
| ENSG00000248521 | AC092612,1 | -0,67949265 | 0,624384812 | 0,50611453 | 0,921780819 | nei like 2 (E, coli) (NEIL2) pseudogene |
| ENSG00000208892 | SNORA49 | -0,679554253 | 0,624358152 | 0,01028756 | 0,11746904 | small nucleolar RNA, H/ACA box 49 |
| ENSG00000267277 | AC024575,1 | -0,680012788 | 0,624159742 | 0,54003957 | 0,933385624 | novel transcript, antisense to SWSAP1 |
| ENSG00000218857 | AL137222,1 | -0,680394982 | 0,623994413 | 0,39247957 | 0,870909282 | D4, zinc and double PHD fingers family 2 (DPF2) pseudogene |
| ENSG00000232373 | MTCYBP3 | -0,680558482 | 0,6239237 | 0,37825479 | 0,863824523 | MT-CYB pseudogene 3 |
| ENSG00000137841 | PLCB2 | -0,680872 | 0,623788128 | 0,02933742 | 0,243477915 | phospholipase C beta 2 |
| ENSG00000180525 | PRR26 | -0,681311359 | 0,623598188 | 0,26299228 | 0,77453036 | proline rich 26 |
| ENSG00000228828 | TLK2P2 | -0,681327898 | 0,623591039 | 0,20862439 | 0,713284351 | tousled like kinase 2 pseudogene 2 |
| ENSG00000261804 | AC007342,4 | -0,681666246 | 0,623444809 | 0,12028767 | 0,553447196 | novel transcript |

| | | | | | | |
|---|---|---|---|---|---|---|
| ENSG00000077274 | CAPN6 | -0,681732166 | 0,623416323 | 0,41906612 | 0,887527661 | calpain 6 |
| ENSG00000277684 | AL392048,1 | -0,681811725 | 0,623381945 | 0,61382495 | 0,952443383 | novel transcript, antisense to RB1 |
| ENSG00000234036 | TXNP6 | -0,681907947 | 0,623340369 | 0,25517335 | 0,766521096 | thioredoxin pseudogene 6 |
| ENSG00000004139 | SARM1 | -0,682242905 | 0,623195661 | 0,00716478 | 0,09003199 | sterile alpha and TIR motif containing 1 |
| ENSG00000242607 | RPS3AP34 | -0,682278891 | 0,623180117 | 0,27616897 | 0,788074712 | RPS3A pseudogene 34 |
| ENSG00000260306 | AC092375,2 | -0,68229559 | 0,623172904 | 0,13726273 | 0,591262337 | novel transcript |
| ENSG00000268870 | AC008758,5 | -0,682363239 | 0,623143683 | 0,4264883 | 0,892138574 | novel zinc finger protein |
| ENSG00000223642 | AC008277,1 | -0,682986333 | 0,622874608 | 0,27651334 | 0,788265979 | novel transcript, antisense to BAZ2B |
| ENSG00000262265 | AC002558,3 | -0,683082192 | 0,622833223 | 0,07264847 | 0,424935759 | novel transcript, sense intronic to CDC27 |
| ENSG00000276250 | AC127024,6 | -0,683117628 | 0,622817925 | 0,32010961 | 0,824366234 | novel transcript, antisense to CRLF3 |
| ENSG00000225998 | LSM2 | -0,683578533 | 0,622618982 | 0,08137225 | 0,452914752 | LSM2 homolog, U6 small nuclear RNA and mRNA degradation associated |
| ENSG00000224979 | LSM2 | -0,683578533 | 0,622618982 | 0,08137225 | 0,452914752 | LSM2 homolog, U6 small nuclear RNA and mRNA degradation associated |
| ENSG00000231502 | LSM2 | -0,683578533 | 0,622618982 | 0,08137225 | 0,452914752 | LSM2 homolog, U6 small nuclear RNA and mRNA degradation associated |
| ENSG00000172850 | LSM2 | -0,683578533 | 0,622618982 | 0,08137225 | 0,452914752 | LSM2 homolog, U6 small nuclear RNA and mRNA degradation associated |
| ENSG00000165762 | OR4K2 | -0,68362427 | 0,622599244 | 0,63168483 | 0,953915351 | olfactory receptor family 4 subfamily K member 2 |
| ENSG00000287925 | AL353576,1 | -0,683703054 | 0,622565246 | 0,44973052 | 0,902062594 | novel transcript, antisense to C1QL3 |
| ENSG00000285627 | AC005343,4 | -0,683739615 | 0,622549469 | 0,52968907 | 0,930679791 | novel transcript, antisense to ADIPOR2 |
| ENSG00000163072 | NOSTRIN | -0,683740077 | 0,622549269 | 0,11956445 | 0,552362303 | nitric oxide synthase trafficking |
| ENSG00000215867 | KRT18P57 | -0,683900622 | 0,622479995 | 0,28496313 | 0,796433333 | keratin 18 pseudogene 57 |
| ENSG00000129007 | CALML4 | -0,683924436 | 0,62246972 | 0,08100235 | 0,451779326 | calmodulin like 4 |
| ENSG00000288382 | AC256223,4 | -0,683998121 | 0,622437928 | 0,49325345 | 0,91725463 | |
| ENSG00000227449 | FGF7P6 | -0,684183131 | 0,622358113 | 0,28450881 | 0,795919345 | fibroblast growth factor 7 pseudogene 6 |
| ENSG00000230090 | AC108025,1 | -0,684422528 | 0,622254849 | 0,4121835 | 0,88438705 | novel transcript |
| ENSG00000231205 | ZNF826P | -0,684554875 | 0,622197768 | 0,1186927 | 0,550565054 | zinc finger protein 826, pseudogene |
| ENSG00000256538 | AC046130,2 | -0,684587593 | 0,622183658 | 0,38691347 | 0,869441674 | novel transcript |
| ENSG00000158077 | NLRP14 | -0,68465691 | 0,622153765 | 0,09948426 | 0,500446627 | NLR family pyrin domain containing 14 |
| ENSG00000286221 | AC009070,1 | -0,684945706 | 0,622029236 | 0,42854568 | 0,893382155 | novel protein |
| ENSG00000288011 | AC024597,1 | -0,684947201 | 0,622028591 | 0,17102656 | 0,654709629 | novel transcript, antisense to ZNF365 |
| ENSG00000164236 | ANKRD33B | -0,685040797 | 0,621988238 | 0,08970221 | 0,47775341 | ankyrin repeat domain 33B |

| | | | | | | |
|---|---|---|---|---|---|---|
| ENSG00000234026 | AL157834,2 | -0,685067528 | 0,621976713 | 0,18280123 | 0,673503078 | novel transcript, antisense to C10orf129 |
| ENSG00000274227 | AC073575,2 | -0,685129646 | 0,621949933 | 0,44857586 | 0,902062594 | novel transcript, antisense to ERP29 |
| ENSG00000259257 | AC037479,1 | -0,685189497 | 0,621924132 | 0,45338414 | 0,902754541 | Coiled-coil domain-containing protein 117 (CC117) pseudogene |
| ENSG00000235772 | AP001625,2 | -0,685273853 | 0,621887768 | 0,49423319 | 0,91758518 | novel transcript |
| ENSG00000236915 | CLCA4-AS1 | -0,685368501 | 0,621846971 | 0,0826889 | 0,45739309 | CLCA4 antisense RNA 1 |
| ENSG00000265113 | AC026474,1 | -0,685457811 | 0,621808477 | 0,60721984 | 0,950993594 | novel transcript, sense intronic to SNTB2 |
| ENSG00000214875 | MED28P1 | -0,685504374 | 0,621788408 | 0,45018845 | 0,902062594 | mediator complex subunit 28 pseudogene 1 |
| ENSG00000259161 | NPM1P20 | -0,685529078 | 0,621777761 | 0,41613936 | 0,885986046 | nucleophosmin 1 pseudogene 20 |
| ENSG00000223956 | LINC01767 | -0,685655484 | 0,621723285 | 0,33082873 | 0,832114797 | long intergenic non-protein coding RNA 1767 |
| ENSG00000263786 | AC022211,1 | -0,685878544 | 0,621627165 | 0,18944251 | 0,684708745 | novel transcript, sense intronic to HN1 |
| ENSG00000282834 | AP000787,3 | -0,685929341 | 0,621605278 | 0,42886302 | 0,893686151 | ribosomal protein S16 (RPS16) pseudogene |
| ENSG00000268869 | ESPNP | -0,686006309 | 0,621572116 | 0,20928569 | 0,713678247 | espin pseudogene |
| ENSG00000253168 | AC124069,1 | -0,686036724 | 0,621559013 | 0,34323977 | 0,840790189 | novel transcript |
| ENSG00000119608 | PROX2 | -0,686125067 | 0,621520952 | 0,15784883 | 0,630637128 | prospero homeobox 2 |
| ENSG00000272572 | AL138762,1 | -0,686746926 | 0,62125311 | 0,19439678 | 0,691503117 | novel transcript, antisense to FAM178A |
| ENSG00000131951 | LRRC9 | -0,6867787 | 0,621239428 | 0,27116782 | 0,7838194 | leucine rich repeat containing 9 |
| ENSG00000225650 | EIF2S2P5 | -0,686800999 | 0,621229825 | 0,2989309 | 0,808745339 | eukaryotic translation initiation factor 2 subunit 2 beta pseudogene 5 |
| ENSG00000155629 | PIK3AP1 | -0,68684644 | 0,621210259 | 0,18819036 | 0,682491978 | phosphoinositide-3-kinase adaptor protein 1 |
| ENSG00000183032 | SLC25A21 | -0,686950506 | 0,62116545 | 0,08776816 | 0,471948586 | solute carrier family 25 member 21 |
| ENSG00000261279 | ULK4P1 | -0,68716374 | 0,621073648 | 0,24011488 | 0,749945225 | ULK4 pseudogene 1 |
| ENSG00000253559 | OSGEPL1-AS1 | -0,687207139 | 0,621054965 | 0,28632894 | 0,797518439 | OSGEPL1 antisense RNA 1 |
| ENSG00000250742 | LINC02381 | -0,687247888 | 0,621037423 | 0,40134431 | 0,876659568 | long intergenic non-protein coding RNA 2381 |
| ENSG00000273980 | AL592071,1 | -0,687248677 | 0,621037083 | 0,5738275 | 0,942475308 | novel transcript |
| ENSG00000160145 | KALRN | -0,687335871 | 0,620999551 | 0,00691036 | 0,087880377 | kalirin RhoGEF kinase |
| ENSG00000237015 | AL031186,1 | -0,6873876 | 0,620977284 | 0,4128141 | 0,884576962 | novel transcript |
| ENSG00000282676 | AC026369,8 | -0,687402651 | 0,620970806 | 0,26382088 | 0,775013078 | |
| ENSG00000142513 | ACP4 | -0,687479004 | 0,620937943 | 0,56479833 | 0,940865625 | acid phosphatase 4 |
| ENSG00000283312 | AC017104,3 | -0,68752664 | 0,62091744 | 0,46616388 | 0,907940653 | GTF2I repeat domain containing 2(GTF2IRD2) pseudogene |
| ENSG00000247775 | SNCA-AS1 | -0,687617944 | 0,620878145 | 0,43385977 | 0,896272827 | SNCA antisense RNA 1 |

| | | | | | | |
|---|---|---|---|---|---|---|
| ENSG00000145365 | TIFA | -0,68771722 | 0,620835423 | 0,14135228 | 0,599218673 | TRAF interacting protein with forkhead associated domain |
| ENSG00000169306 | IL1RAPL1 | -0,688065179 | 0,620685704 | 0,16111235 | 0,63719373 | interleukin 1 receptor accessory protein like 1 |
| ENSG00000091704 | CPA1 | -0,688267818 | 0,620598529 | 0,56529172 | 0,940865625 | carboxypeptidase A1 |
| ENSG00000288282 | AC256230,1 | -0,688380996 | 0,620549845 | 0,58407526 | 0,944526397 | |
| ENSG00000170837 | GPR27 | -0,68843221 | 0,620527817 | 0,07603325 | 0,435686153 | G protein-coupled receptor 27 |
| ENSG00000261535 | AC005086,2 | -0,688496613 | 0,620500117 | 0,34978013 | 0,843534837 | novel transcript, antisense to CUX1 |
| ENSG00000143847 | PPFIA4 | -0,688549116 | 0,620477536 | 0,08403324 | 0,461230231 | PTPRF interacting protein alpha 4 |
| ENSG00000237513 | AC007384,1 | -0,688696531 | 0,620414139 | 0,04465405 | 0,319246353 | novel transcript |
| ENSG00000272931 | AC099568,2 | -0,688730598 | 0,620399488 | 0,33993016 | 0,838611198 | novel transcript |
| ENSG00000231427 | LINC01445 | -0,688737632 | 0,620396464 | 0,04322777 | 0,313858131 | long intergenic non-protein coding RNA 1445 |
| ENSG00000100628 | ASB2 | -0,688890881 | 0,620330566 | 0,47409601 | 0,910491353 | ankyrin repeat and SOCS box containing 2 |
| ENSG00000260469 | INSYN1-AS1 | -0,688979504 | 0,620292461 | 0,24239193 | 0,752612359 | INSYN1 antisense RNA 1 |
| ENSG00000215097 | AL731733,1 | -0,689038248 | 0,620267204 | 0,33008435 | 0,831768943 | dual specificity phosphatase 8 pseudogene 2 |
| ENSG00000232457 | SLC16A6P1 | -0,689119254 | 0,620232378 | 0,51152864 | 0,922893865 | SLC16A6 pseudogene 1 |
| ENSG00000260954 | AL133297,1 | -0,689483692 | 0,620075721 | 0,59170013 | 0,947416795 | novel transcript, intronic to IFT140 |
| ENSG00000214041 | PGAM1P4 | -0,689556572 | 0,620044398 | 0,50267483 | 0,92065763 | phosphoglycerate mutase 1 pseudogene 4 |
| ENSG00000260196 | AC124798,1 | -0,689573853 | 0,620036971 | 0,19341024 | 0,69035025 | antisense to KCNJ11 and overlapping to a novel gene |
| ENSG00000275143 | SCARNA16 | -0,689907217 | 0,619893715 | 0,24768906 | 0,759936282 | small Cajal body-specific RNA 16 |
| ENSG00000237877 | LINC01473 | -0,690260315 | 0,619742016 | 0,20979661 | 0,71473044 | long intergenic non-protein coding RNA 1473 |
| ENSG00000148950 | IMMP1L | -0,690351963 | 0,619702648 | 0,0892367 | 0,476498052 | inner mitochondrial membrane peptidase subunit 1 |
| ENSG00000243402 | AC022973,2 | -0,690384713 | 0,61968858 | 0,32159039 | 0,824933727 | ribosomal protein L15 (RPL15) pseudogene |
| ENSG00000273486 | AC096992,2 | -0,690527343 | 0,619627319 | 0,1813547 | 0,671070342 | novel transcript, antisense to SLC35G2 |
| ENSG00000283897 | AC011416,3 | -0,690884959 | 0,619473744 | 0,49830445 | 0,919439944 | novel transcript |
| ENSG00000214708 | AC116407,1 | -0,691030383 | 0,619411304 | 0,38625102 | 0,869312875 | novel transcript, antisense to RHOT1 |
| ENSG00000270427 | NRBF2P5 | -0,691110841 | 0,619376761 | 0,06448763 | 0,397974975 | nuclear receptor binding factor 2 pseudogene 5 |
| ENSG00000227068 | AL162584,1 | -0,691113703 | 0,619375532 | 0,42197991 | 0,889720796 | novel transcript |
| ENSG00000138356 | AOX1 | -0,691132377 | 0,619367515 | 0,16695777 | 0,647372904 | aldehyde oxidase 1 |
| ENSG00000224046 | AC005076,1 | -0,691359188 | 0,61927015 | 0,1624239 | 0,639629762 | novel transcript, antisense to DMTF1 |
| ENSG00000183960 | KCNH8 | -0,691387241 | 0,619258108 | 0,16352239 | 0,64209545 | potassium voltage-gated channel subfamily H member 8 |
| ENSG00000185664 | PMEL | -0,691646153 | 0,619146984 | 0,11215542 | 0,535189241 | premelanosome protein |
| ENSG00000212695 | AL583805,1 | -0,691660156 | 0,619140974 | 0,37046603 | 0,858545694 | ribosomal protein 18 (RPL18) pseudogene |

| | | | | | | |
|---|---|---|---|---|---|---|
| ENSG00000214659 | KRT8P26 | -0,691734224 | 0,619109188 | 0,35420655 | 0,84556089 | keratin 8 pseudogene 26 |
| ENSG00000276903 | H2AC16 | -0,692016921 | 0,618987886 | 0,04445274 | 0,318464072 | H2A clustered histone 16 |
| ENSG00000283578 | MTND5P42 | -0,692045699 | 0,618975539 | 0,58361626 | 0,944174844 | MT-ND5 pseudogene 42 |
| ENSG00000234776 | C11orf94 | -0,692207009 | 0,618906334 | 0,38022722 | 0,865730737 | chromosome 11 open reading frame 94 |
| ENSG00000286642 | AC091646,1 | -0,692250973 | 0,618887474 | 0,35864285 | 0,849814998 | novel transcript |
| ENSG00000175773 | ZBTB44-DT | -0,692410908 | 0,618818869 | 0,59550038 | 0,947901021 | ZBTB44 divergent transcript |
| ENSG00000255240 | AP001636,3 | -0,692768731 | 0,618665405 | 0,19713768 | 0,695927373 | novel transcript |
| ENSG00000243492 | HLA-L | -0,6933249 | 0,618426952 | 0,0183484 | 0,176288964 | major histocompatibility complex, class I, L (pseudogene) |
| ENSG00000270469 | AC006557,5 | -0,693385335 | 0,618401046 | 0,31443508 | 0,819877431 | non-SMC element 4 homolog A (S, cerevisiae), pseudogene |
| ENSG00000214919 | AC104472,1 | -0,693618072 | 0,618301293 | 0,13971214 | 0,595809743 | novel transcript, sense intronic to PLCH1 |
| ENSG00000231345 | BEND3P1 | -0,693694619 | 0,618268488 | 0,16423257 | 0,643026568 | BEN domain containing 3 pseudogene 1 |
| ENSG00000271270 | TMCC1-AS1 | -0,693730662 | 0,618253042 | 0,09197687 | 0,482073274 | TMCC1 antisense RNA 1 (head to head) |
| ENSG00000141622 | RNF165 | -0,693731025 | 0,618252886 | 0,00321306 | 0,049265853 | ring finger protein 165 |
| ENSG00000206557 | TRIM71 | -0,693789459 | 0,618227846 | 0,02969387 | 0,245400285 | tripartite motif containing 71 |
| ENSG00000176531 | PHLDB3 | -0,69406697 | 0,618108937 | 0,2887123 | 0,800014136 | pleckstrin homology like domain family B member 3 |
| ENSG00000279968 | CCDC28A-AS1 | -0,694109806 | 0,618090585 | 0,17477618 | 0,660057513 | CCDC28A antisense RNA 1 |
| ENSG00000233966 | UBE2SP1 | -0,694288813 | 0,618013898 | 0,3318466 | 0,83268442 | ubiquitin conjugating enzyme E2 S pseudogene 1 |
| ENSG00000125246 | CLYBL | -0,69431201 | 0,618003961 | 0,07054728 | 0,418200624 | citramalyl-CoA lyase |
| ENSG00000185386 | MAPK11 | -0,69431994 | 0,618000564 | 0,07284063 | 0,425741182 | mitogen-activated protein kinase 11 |
| ENSG00000235313 | HM13-IT1 | -0,694674055 | 0,617848892 | 0,50159954 | 0,920520138 | HM13 intronic transcript 1 |
| ENSG00000272630 | AL731563,3 | -0,694813126 | 0,617789336 | 0,05903629 | 0,377379048 | novel transcript |
| ENSG00000260600 | AC092125,1 | -0,69500586 | 0,61770681 | 0,49119733 | 0,916995341 | novel transcript |
| ENSG00000150510 | FAM124A | -0,695436318 | 0,617522531 | 0,01290521 | 0,138203846 | family with sequence similarity 124 member A |
| ENSG00000272885 | AC092574,1 | -0,695445209 | 0,617518726 | 0,29138501 | 0,802058262 | novel transcript |
| ENSG00000233589 | AL138789,1 | -0,695563648 | 0,617468032 | 0,2743232 | 0,786239603 | novel transcript |
| ENSG00000288467 | AL513166,2 | -0,695750246 | 0,617388174 | 0,36956446 | 0,858037898 | novel transcript |
| ENSG00000196391 | ZNF774 | -0,695839705 | 0,617349892 | 0,01933331 | 0,182760901 | zinc finger protein 774 |
| ENSG00000266541 | AP001381,1 | -0,695862249 | 0,617340245 | 0,34031428 | 0,839010589 | novel transcript, antisense to PPP4R1 |
| ENSG00000234997 | AC016745,1 | -0,696137125 | 0,617222635 | 0,19412532 | 0,691087892 | novel transcript |
| ENSG00000164669 | INTS4P1 | -0,696145832 | 0,61721891 | 0,16857795 | 0,650506523 | integrator complex subunit 4 pseudogene 1 |
| ENSG00000214357 | NEURL1B | -0,696224774 | 0,617185138 | 0,00952823 | 0,110822426 | neuralized E3 ubiquitin protein ligase 1B |

| | | | | | | |
|---|---|---|---|---|---|---|
| ENSG00000253206 | AC090155,1 | -0,696253689 | 0,617172768 | 0,42829465 | 0,893037474 | novel transcript |
| ENSG00000173124 | ACSM6 | -0,696295885 | 0,617154717 | 0,21513996 | 0,722445395 | acyl-CoA synthetase medium chain family member 6 |
| ENSG00000277361 | OCA2 | -0,696729577 | 0,616969221 | 0,38944823 | 0,870057277 | OCA2 melanosomal transmembrane protein |
| ENSG00000186446 | ZNF501 | -0,696806893 | 0,616936157 | 0,15137413 | 0,618716275 | zinc finger protein 501 |
| ENSG00000120756 | PLS1 | -0,697021224 | 0,61684451 | 0,12965097 | 0,575441236 | plastin 1 |
| ENSG00000282256 | TAS2R31 | -0,697186085 | 0,616774025 | 0,58605768 | 0,945246954 | taste 2 receptor member 31 |
| ENSG00000258754 | LINC01579 | -0,697280944 | 0,616733473 | 0,02901504 | 0,241642594 | long intergenic non-protein coding RNA 1579 |
| ENSG00000232204 | TET1P1 | -0,697448667 | 0,616661778 | 0,23077438 | 0,739234644 | tet methylcytosine dioxygenase 1 pseudogene 1 |
| ENSG00000162415 | ZSWIM5 | -0,69745598 | 0,616658652 | 0,04191468 | 0,306990518 | zinc finger SWIM-type containing 5 |
| ENSG00000225900 | HSPE1P13 | -0,697762481 | 0,616527657 | 0,51564721 | 0,924926785 | heat shock protein family E (Hsp10) member 1 pseudogene 13 |
| ENSG00000272140 | AC022400,4 | -0,697794233 | 0,616514088 | 0,02291064 | 0,20541387 | novel transcript |
| ENSG00000099994 | SUSD2 | -0,697798037 | 0,616512462 | 0,13664137 | 0,58964193 | sushi domain containing 2 |
| ENSG00000250219 | LTV1P1 | -0,697977905 | 0,616435603 | 0,56783017 | 0,94163744 | LTV1 ribosome biogenesis factor pseudogene 1 |
| ENSG00000168126 | OR2W6P | -0,698068794 | 0,61639677 | 0,60620739 | 0,950927937 | olfactory receptor family 2 subfamily W member 6 pseudogene |
| ENSG00000240418 | AC020917,1 | -0,698076709 | 0,616393388 | 0,29193366 | 0,802586512 | ribosomal protein S2 (RPS2) pseudogene |
| ENSG00000166455 | C16orf46 | -0,698183268 | 0,616347862 | 0,08438595 | 0,462254126 | chromosome 16 open reading frame 46 |
| ENSG00000149050 | ZNF214 | -0,698234076 | 0,616326156 | 0,13390819 | 0,584034405 | zinc finger protein 214 |
| ENSG00000236556 | AC025038,1 | -0,698431129 | 0,61624198 | 0,64646855 | 0,956899963 | novel transcript |
| ENSG00000226642 | ACTG1P12 | -0,698609326 | 0,616165868 | 0,55059754 | 0,936249395 | actin gamma 1 pseudogene 12 |
| ENSG00000285650 | AL157827,2 | -0,698802089 | 0,616083546 | 0,1755158 | 0,661199837 | Novel transcript, antisense to CENPP |
| ENSG00000196361 | ELAVL3 | -0,699053528 | 0,615976182 | 0,01456569 | 0,150888551 | ELAV like RNA binding protein 3 |
| ENSG00000274997 | H2AC12 | -0,699306231 | 0,615868297 | 0,0310857 | 0,253617788 | H2A clustered histone 12 |
| ENSG00000225423 | TNPO1P1 | -0,699353113 | 0,615848284 | 0,10443642 | 0,515254902 | transportin 1 pseudogene 1 |
| ENSG00000175567 | UCP2 | -0,699521911 | 0,615776233 | 0,01142497 | 0,126376669 | uncoupling protein 2 |
| ENSG00000179397 | CATSPERE | -0,699629473 | 0,615730324 | 0,17183054 | 0,655695088 | catsper channel auxiliary subunit epsilon |
| ENSG00000236863 | RPL23AP23 | -0,699721655 | 0,615690983 | 0,43432464 | 0,896272827 | ribosomal protein L23a pseudogene 23 |
| ENSG00000171084 | FAM86JP | -0,699941647 | 0,615597105 | 0,029475 | 0,244172299 | family with sequence similarity 86 member J, pseudogene |
| ENSG00000233585 | AC231533,2 | -0,69999047 | 0,615576273 | 0,41616577 | 0,885986046 | acetyl-Coenzyme A acyltransferase 2 (mitochondrial 3-oxoacyl-Coenzyme A thiolase) (ACAA2) pseudogene |

| | | | | | | |
|---|---|---|---|---|---|---|
| ENSG00000263766 | AC025682,1 | -0,700066256 | 0,615543937 | 0,24757317 | 0,759804402 | novel transcript, antisense to NPEPPS |
| ENSG00000257509 | AC073487,1 | -0,700185954 | 0,615492868 | 0,09441943 | 0,488002825 | novel transcript, sense intronic to SARNP |
| ENSG00000108984 | MAP2K6 | -0,700866414 | 0,615202634 | 0,12394775 | 0,562842145 | mitogen-activated protein kinase kinase 6 |
| ENSG00000095585 | BLNK | -0,700887319 | 0,61519372 | 0,24580727 | 0,757752583 | B cell linker |
| ENSG00000276603 | AL109614,1 | -0,700930378 | 0,615175359 | 0,22914065 | 0,737173617 | novel transcript, sense intronic to BLCAP |
| ENSG00000239212 | RPL6P7 | -0,70112077 | 0,61509418 | 0,43052427 | 0,895238505 | ribosomal protein L6 pseudogene 7 |
| ENSG00000091972 | CD200 | -0,701194043 | 0,615062941 | 0,10614159 | 0,51988717 | CD200 molecule |
| ENSG00000018869 | ZNF582 | -0,70157716 | 0,614899628 | 0,0785772 | 0,443676655 | zinc finger protein 582 |
| ENSG00000273897 | PMS2P8 | -0,701632149 | 0,614876192 | 0,21097889 | 0,716555746 | PMS1 homolog 2, mismatch repair system component pseudogene 8 |
| ENSG00000154645 | CHODL | -0,702062998 | 0,614692592 | 0,01792066 | 0,173950874 | chondrolectin |
| ENSG00000287720 | AC023480,1 | -0,702065305 | 0,614691608 | 0,13229265 | 0,580797367 | novel transcript, antisense to SUMF1 |
| ENSG00000186329 | TMEM212 | -0,702091871 | 0,614680289 | 0,01869921 | 0,178483188 | transmembrane protein 212 |
| ENSG00000121380 | BCL2L14 | -0,702107964 | 0,614673433 | 0,10684334 | 0,521837326 | BCL2 like 14 |
| ENSG00000228252 | COL6A4P2 | -0,702493359 | 0,614509254 | 0,26349632 | 0,774637968 | collagen type VI alpha 4 pseudogene 2 |
| ENSG00000213600 | U73169,1 | -0,702500976 | 0,614506009 | 0,53763907 | 0,932883277 | novel transcript, sense intronic to GNAI2 |
| ENSG00000287957 | AC007695,1 | -0,702735991 | 0,614405914 | 0,18010101 | 0,669203474 | novel transcript, antisense to RIC8B |
| ENSG00000220685 | AL139094,1 | -0,702965306 | 0,614308263 | 0,29704596 | 0,806635068 | transcription factor B2, mitochondrial (TFB2M) pseudogene |
| ENSG00000241007 | SEPTIN7P6 | -0,703452674 | 0,614100774 | 0,28937247 | 0,800260154 | septin 7 pseudogene 6 |
| ENSG00000163909 | HEYL | -0,703517246 | 0,614073288 | 0,23540193 | 0,744654651 | hes related family bHLH transcription factor with YRPW motif like |
| ENSG00000237646 | HLCS-IT1 | -0,703734114 | 0,613980987 | 0,50624159 | 0,921780819 | HLCS intronic transcript 1 |
| ENSG00000226874 | AC005154,1 | -0,70375801 | 0,613970818 | 0,53293694 | 0,931291085 | pseudogene similar to part of transient receptor potential cation channel, subfamily C, member 6 TRPC6 |
| ENSG00000120210 | INSL6 | -0,703917329 | 0,613903019 | 0,34388273 | 0,840901506 | insulin like 6 |
| ENSG00000287122 | AC026495,2 | -0,703993809 | 0,613870476 | 0,22738019 | 0,735964811 | novel transcript |
| ENSG00000176406 | RIMS2 | -0,703997973 | 0,613868705 | 0,01506423 | 0,154671635 | regulating synaptic membrane exocytosis 2 |
| ENSG00000187699 | C2orf88 | -0,704122545 | 0,613815701 | 0,10167018 | 0,507111388 | chromosome 2 open reading frame 88 |
| ENSG00000230649 | AC024084,1 | -0,704251131 | 0,613760995 | 0,3343332 | 0,834538367 | novel transcript |
| ENSG00000166323 | C11orf65 | -0,70430627 | 0,613737538 | 0,11792413 | 0,548984873 | chromosome 11 open reading frame 65 |

| | | | | | | |
|---|---|---|---|---|---|---|
| ENSG00000115474 | KCNJ13 | -0,704650585 | 0,61359108 | 0,08913132 | 0,47641326 | potassium inwardly rectifying channel subfamily J member 13 |
| ENSG00000266968 | AC023421,1 | -0,704679698 | 0,613578698 | 0,59583869 | 0,948101624 | novel transcript |
| ENSG00000260194 | AC007496,1 | -0,704698717 | 0,613570609 | 0,37973173 | 0,865238161 | novel transcript, antisense to FTO |
| ENSG00000181449 | SOX2 | -0,704780406 | 0,613535868 | 0,03278414 | 0,262266996 | SRY-box transcription factor 2 |
| ENSG00000144362 | PHOSPHO2 | -0,704837304 | 0,613511672 | 0,12594111 | 0,566777174 | phosphatase, orphan 2 |
| ENSG00000255052 | FAM66D | -0,705075746 | 0,613410282 | 0,01099376 | 0,122840875 | family with sequence similarity 66 member D |
| ENSG00000234614 | C2CD4D-AS1 | -0,705157798 | 0,613375396 | 0,36939184 | 0,857991124 | C2CD4D and THEM5 antisense RNA 1 |
| ENSG00000259520 | AC051619,4 | -0,705179195 | 0,613366299 | 0,14868987 | 0,613357982 | novel transcript, antisense to SLC28A2 |
| ENSG00000119737 | GPR75 | -0,705410401 | 0,613268008 | 0,0230249 | 0,206320004 | G protein-coupled receptor 75 |
| ENSG00000285434 | CRYGD | -0,705419418 | 0,613264176 | 0,26119494 | 0,773222089 | crystallin gamma D |
| ENSG00000118231 | CRYGD | -0,705419418 | 0,613264176 | 0,26119494 | 0,773222089 | crystallin gamma D |
| ENSG00000270059 | AC121493,1 | -0,705577099 | 0,613197152 | 0,20056498 | 0,700127386 | novel transcript, intronic to MAGI1 |
| ENSG00000284320 | SHMT1 | -0,705623248 | 0,613177537 | 0,14301678 | 0,601880115 | serine hydroxymethyltransferase 1 |
| ENSG00000204789 | ZNF204P | -0,705707214 | 0,613141851 | 0,27956804 | 0,790302992 | zinc finger protein 204, pseudogene |
| ENSG00000261723 | AC008870,4 | -0,705771291 | 0,613114619 | 0,6393411 | 0,956018518 | novel transcript, antisense to PALB2 |
| ENSG00000274392 | AC093724,1 | -0,70580709 | 0,613099405 | 0,5545388 | 0,937793851 | neurobeachin (NBEA) pseudogene |
| ENSG00000273345 | AC104109,4 | -0,7061094 | 0,612970947 | 0,25394051 | 0,76531122 | novel transcript |
| ENSG00000274642 | AC244669,2 | -0,706118337 | 0,612967149 | 0,71117325 | 0,970540858 | phosphodiesterase 4D interacting protein (PDE4DIP) pseudogene |
| ENSG00000287897 | AP000465,1 | -0,706120953 | 0,612966038 | 0,43851277 | 0,897220559 | novel transcript, sense intronic to CADM1 |
| ENSG00000287439 | AC092078,3 | -0,706173348 | 0,612943777 | 0,22909151 | 0,737173617 | novel transcript |
| ENSG00000287719 | AP001531,1 | -0,706295087 | 0,612892057 | 0,0378147 | 0,287579371 | novel transcript |
| ENSG00000231079 | KIF5C-AS1 | -0,706394046 | 0,612850018 | 0,04668045 | 0,32799008 | KIF5C antisense RNA 1 |
| ENSG00000234848 | AC008758,2 | -0,706468014 | 0,612818598 | 0,53120554 | 0,931265667 | novel transcript, antisense to ZNF709 |
| ENSG00000269189 | AL445223,1 | -0,706477005 | 0,612814779 | 0,13444195 | 0,584751412 | novel transcript, sense intronic to FARP1 |
| ENSG00000188176 | SMTNL2 | -0,70664038 | 0,612745386 | 0,38804813 | 0,869820053 | smoothelin like 2 |
| ENSG00000265179 | AP000894,2 | -0,706778767 | 0,612686612 | 0,30289812 | 0,81247062 | novel transcript, antisense to ADCYAP1 |
| ENSG00000238228 | OR7E7P | -0,706965566 | 0,612607287 | 0,17719986 | 0,663325859 | olfactory receptor family 7 subfamily E member 7 pseudogene |
| ENSG00000173947 | PIFO | -0,707191091 | 0,612511531 | 0,22473469 | 0,733203418 | primary cilia formation |
| ENSG00000242689 | CNTF | -0,707198184 | 0,61250852 | 0,17572528 | 0,661199837 | ciliary neurotrophic factor |
| ENSG00000239482 | AC112487,1 | -0,707219008 | 0,612499679 | 0,09379176 | 0,486996636 | novel transcript |

| | | | | | | |
|---|---|---|---|---|---|---|
| ENSG00000273814 | OCLN | -0,707255877 | 0,612484026 | 0,01388208 | 0,145299451 | occludin |
| ENSG00000182366 | FAM87A | -0,7072623 | 0,612481299 | 0,21384379 | 0,721188427 | family with sequence similarity 87 member A |
| ENSG00000159023 | EPB41 | -0,707348585 | 0,612444669 | 0,03598691 | 0,278282126 | erythrocyte membrane protein band 4,1 |
| ENSG00000280907 | ZNF197-AS1 | -0,707526517 | 0,612369139 | 0,30937249 | 0,816845699 | ZNF197 antisense RNA 1 |
| ENSG00000233509 | ZNF197-AS1 | -0,707526517 | 0,612369139 | 0,30937249 | 0,816845699 | ZNF197 antisense RNA 1 |
| ENSG00000250508 | LINC02701 | -0,707649693 | 0,612316857 | 0,31238018 | 0,818574551 | long intergenic non-protein coding RNA 2701 |
| ENSG00000164542 | KIAA0895 | -0,707750724 | 0,612273979 | 0,15360194 | 0,622683206 | KIAA0895 |
| ENSG00000185056 | C5orf47 | -0,707829205 | 0,612240673 | 0,62221918 | 0,952793934 | chromosome 5 open reading frame 47 |
| ENSG00000171435 | KSR2 | -0,707907762 | 0,612207336 | 0,11061502 | 0,530979255 | kinase suppressor of ras 2 |
| ENSG00000236819 | LINC01563 | -0,708131017 | 0,612112605 | 0,16736667 | 0,648154527 | long intergenic non-protein coding RNA 1563 |
| ENSG00000287555 | AC023424,2 | -0,708190277 | 0,612087463 | 0,02605105 | 0,225113315 | novel transcript, antisense to FBXW7 |
| ENSG00000228513 | AC023271,2 | -0,708441942 | 0,611980699 | 0,26105486 | 0,773103493 | zinc finger protein 14 (ZNF14) pseudogene |
| ENSG00000282537 | TAS2R43 | -0,708647613 | 0,611893461 | 0,1337409 | 0,583882708 | taste 2 receptor member 43 |
| ENSG00000261936 | TAS2R45 | -0,708647613 | 0,611893461 | 0,1337409 | 0,583882708 | taste 2 receptor member 45 |
| ENSG00000267475 | AC008736,1 | -0,708768121 | 0,611842352 | 0,55087964 | 0,936366211 | novel transcript |
| ENSG00000004799 | PDK4 | -0,708914578 | 0,611780243 | 0,31475415 | 0,81991112 | pyruvate dehydrogenase kinase 4 |
| ENSG00000188712 | OR13D3P | -0,708961684 | 0,611760268 | 0,44347267 | 0,900014899 | olfactory receptor family 13 subfamily D member 3 pseudogene |
| ENSG00000265205 | AC010761,3 | -0,708975167 | 0,61175455 | 0,16168811 | 0,63825975 | novel transcript, antisense to SUPT6H |
| ENSG00000232915 | RNF14P1 | -0,708998708 | 0,611744568 | 0,59463956 | 0,947469769 | RNF14 pseudogene 1 |
| ENSG00000224007 | USP12P2 | -0,709296677 | 0,611618234 | 0,45221904 | 0,902062594 | USP12 pseudogene 2 |
| ENSG00000166265 | CYYR1 | -0,709458757 | 0,611549526 | 0,20242584 | 0,702433811 | cysteine and tyrosine rich 1 |
| ENSG00000223442 | TH2LCRR | -0,709477571 | 0,61154155 | 0,48038001 | 0,912847708 | T helper type 2 locus control region associated RNA |
| ENSG00000153132 | CLGN | -0,709632912 | 0,611475706 | 0,11622225 | 0,544242846 | calmegin |
| ENSG00000259335 | HNRNPMP1 | -0,710045892 | 0,611300693 | 0,14111626 | 0,59829928 | heterogeneous nuclear ribonucleoprotein M pseudogene 1 |
| ENSG00000246090 | AP002026,1 | -0,710274032 | 0,611204033 | 0,08460712 | 0,462844615 | novel transcript, antisense to ADH4, ADH6 & ADH1A |
| ENSG00000251011 | TMEM108-AS1 | -0,710360134 | 0,611167557 | 0,16039002 | 0,635945906 | TMEM108 antisense RNA 1 |
| ENSG00000287508 | AC009560,4 | -0,710413169 | 0,61114509 | 0,40947581 | 0,882177108 | novel transcript, antisense to RORA |
| ENSG00000248180 | GAPDHP60 | -0,711064309 | 0,61086932 | 0,17572825 | 0,661199837 | glyceraldehyde-3-phosphate dehydrogenase pseudogene 60 |
| ENSG00000184111 | RPL11P4 | -0,711149697 | 0,610833166 | 0,41839518 | 0,887094957 | ribosomal protein L11 pseudogene 4 |
| ENSG00000118307 | CFAP94 | -0,711520087 | 0,610676364 | 0,26141878 | 0,77325059 | cilia and flagella associated protein 94 |

| | | | | | | |
|---|---|---|---|---|---|---|
| ENSG00000235253 | AC010240,2 | -0,711619099 | 0,610634455 | 0,08293752 | 0,457958054 | eukaryotic translation elongation factor 1 alpha 1 (EEF1A1) pseudogene |
| ENSG00000272848 | AL135910,1 | -0,71207354 | 0,610442139 | 0,15110887 | 0,618336884 | novel transcript |
| ENSG00000152527 | PLEKHH2 | -0,712109178 | 0,610427059 | 0,03572455 | 0,27700703 | pleckstrin homology, MyTH4 and FERM domain containing H2 |
| ENSG00000241975 | ELOCP19 | -0,712133463 | 0,610416784 | 0,12812232 | 0,571496122 | elongin C pseudogene 19 |
| ENSG00000217447 | AL021331,1 | -0,712542034 | 0,610243939 | 0,48902462 | 0,915720281 | MAD2 mitotic arrest deficient-like 1 (yeast) (MAD2L1) pseudogene |
| ENSG00000260536 | AL035071,2 | -0,712923854 | 0,610082455 | 0,44814642 | 0,902062594 | novel transcript, antisense to MAPRE1 |
| ENSG00000286024 | AC026803,3 | -0,713201985 | 0,609964851 | 0,52419526 | 0,928419877 | novel transcript, antisense to BCAT2 |
| ENSG00000254488 | AC007876,1 | -0,713214353 | 0,609959622 | 0,24658994 | 0,758424774 | novel transcript |
| ENSG00000286158 | AL034548,2 | -0,713592607 | 0,60979972 | 0,33836052 | 0,837263082 | novel transcript |
| ENSG00000226877 | AL354733,1 | -0,713613653 | 0,609790824 | 0,63457975 | 0,955149453 | novel transcript, antisense to KIF27 |
| ENSG00000110987 | BCL7A | -0,713644577 | 0,609777754 | 0,05492478 | 0,362021772 | BAF chromatin remodeling complex subunit BCL7A |
| ENSG00000277218 | AL139123,1 | -0,713723476 | 0,609744407 | 0,51721585 | 0,925746338 | novel transcript |
| ENSG00000272732 | AC004982,1 | -0,713748678 | 0,609733755 | 0,32490079 | 0,826972094 | novel transcript |
| ENSG00000106069 | CHN2 | -0,713969776 | 0,609640319 | 0,14222153 | 0,600366679 | chimerin 2 |
| ENSG00000169302 | STK32A | -0,714045476 | 0,609608331 | 0,08127797 | 0,452873718 | serine/threonine kinase 32A |
| ENSG00000213917 | RPL5P8 | -0,71413195 | 0,609571793 | 0,29874973 | 0,808457673 | ribosomal protein L5 pseudogene 8 |
| ENSG00000272425 | AC009902,3 | -0,714185612 | 0,609549119 | 0,11672624 | 0,545653262 | novel transcript |
| ENSG00000274430 | AL353804,3 | -0,714283615 | 0,609507714 | 0,52901622 | 0,930201325 | |
| ENSG00000286412 | AC007682,1 | -0,714354058 | 0,609477954 | 0,05225501 | 0,350880739 | novel transcript, antisense to NRXN1 |
| ENSG00000285542 | AC013717,1 | -0,714378315 | 0,609467707 | 0,2983584 | 0,807818219 | novel protein, PPM1B-SLC3A1 readthrough |
| ENSG00000184949 | FAM227A | -0,714486972 | 0,609421806 | 0,01157228 | 0,127348002 | family with sequence similarity 227 member A |
| ENSG00000188906 | LRRK2 | -0,714512931 | 0,609410841 | 0,10521753 | 0,517691472 | leucine rich repeat kinase 2 |
| ENSG00000267606 | AC006116,8 | -0,71470848 | 0,609328244 | 0,57042928 | 0,9423888 | novel transcript |
| ENSG00000225618 | ZNRD1ASP | -0,714798307 | 0,609290306 | 0,2147358 | 0,722164151 | zinc ribbon domain containing 1 antisense, pseudogene |
| ENSG00000236598 | ZNRD1ASP | -0,714798307 | 0,609290306 | 0,2147358 | 0,722164151 | zinc ribbon domain containing 1 antisense, pseudogene |
| ENSG00000154721 | JAM2 | -0,71483204 | 0,60927606 | 0,06834612 | 0,411986588 | junctional adhesion molecule 2 |
| ENSG00000224736 | HLA-V | -0,714962173 | 0,609221105 | 0,53763938 | 0,932883277 | major histocompatibility complex, class I, V (pseudogene) |

| | | | | | | |
|---|---|---|---|---|---|---|
| ENSG00000280145 | CU638689,4 | -0,7149718 | 0,60921704 | 0,02178991 | 0,198207677 | novel transcript |
| ENSG00000286608 | AC010201,3 | -0,715058635 | 0,609180373 | 0,39160657 | 0,870333649 | novel transcript |
| ENSG00000269578 | AC008764,7 | -0,715190754 | 0,609124588 | 0,43247634 | 0,89560871 | novel transcript, sense intronic MED26 |
| ENSG00000259818 | AL606760,3 | -0,715256112 | 0,609096993 | 0,18059915 | 0,670229021 | novel transcript, antisense to C1orf123 & MAGOH |
| ENSG00000158423 | RIBC1 | -0,715258041 | 0,609096179 | 0,04776658 | 0,332526018 | RIB43A domain with coiled-coils 1 |
| ENSG00000230583 | GTF2IRD1P1 | -0,715271784 | 0,609090377 | 0,29304934 | 0,803365244 | GTF2I repeat domain containing 1 pseudogene 1 |
| ENSG00000263648 | AC005244,1 | -0,715296467 | 0,609079956 | 0,49693291 | 0,918645042 | dynein, light chain, Tctex-type 1 (DYNLT1) pseudogene |
| ENSG00000287177 | AL583825,1 | -0,715581788 | 0,60895951 | 0,28741611 | 0,798896538 | novel transcript |
| ENSG00000253730 | AC015909,2 | -0,71575317 | 0,608887175 | 0,56135356 | 0,940567693 | novel transcript |
| ENSG00000124635 | H2BC11 | -0,715895838 | 0,608826965 | 0,15442076 | 0,623607455 | H2B clustered histone 11 |
| ENSG00000236274 | AC004865,1 | -0,715963655 | 0,608798346 | 0,42608539 | 0,892133432 | zinc finger protein pseudogene |
| ENSG00000187210 | GCNT1 | -0,716004547 | 0,60878109 | 0,08397385 | 0,461230231 | glucosaminyl (N-acetyl) transferase 1 |
| ENSG00000197766 | CFD | -0,716127072 | 0,60872939 | 0,31670989 | 0,821646322 | complement factor D |
| ENSG00000186231 | KLHL32 | -0,716165767 | 0,608713064 | 0,16417672 | 0,642888389 | kelch like family member 32 |
| ENSG00000228740 | GABRG3-AS1 | -0,716210613 | 0,608694142 | 0,2935033 | 0,803461221 | GABRG3 antisense RNA 1 |
| ENSG00000274618 | H4C6 | -0,716256246 | 0,608674889 | 0,14885581 | 0,613515281 | H4 clustered histone 6 |
| ENSG00000200792 | SNORA80A | -0,716279815 | 0,608664945 | 0,3947845 | 0,871888817 | small nucleolar RNA, H/ACA box 80A |
| ENSG00000149243 | KLHL35 | -0,716304993 | 0,608654323 | 0,02385784 | 0,211420503 | kelch like family member 35 |
| ENSG00000187672 | ERC2 | -0,716461569 | 0,608588269 | 0,02799304 | 0,236395267 | ELKS/RAB6-interacting/CAST family member 2 |
| ENSG00000231857 | MORF4L1P5 | -0,716535056 | 0,60855727 | 0,41096144 | 0,882983885 | mortality factor 4 like 1 pseudogene 5 |
| ENSG00000254440 | PBOV1 | -0,716584162 | 0,608536557 | 0,27637146 | 0,788263115 | prostate and breast cancer overexpressed 1 |
| ENSG00000244346 | AC092953,1 | -0,716600067 | 0,608529848 | 0,53135219 | 0,931291085 | chromosome 9 open reading frame 85 (C9orf85) pseudogene |
| ENSG00000282727 | AC091132,3 | -0,716681074 | 0,60849568 | 0,00551033 | 0,074320583 | TEC |
| ENSG00000275282 | CFL1P5 | -0,716768324 | 0,608458881 | 0,46236611 | 0,9064797 | cofilin 1 pseudogene 5 |
| ENSG00000213830 | CFL1P5 | -0,716768324 | 0,608458881 | 0,46236611 | 0,9064797 | cofilin 1 pseudogene 5 |
| ENSG00000256981 | AC134349,2 | -0,716841806 | 0,608427891 | 0,57051829 | 0,9423888 | taste receptor, type 2, member 18 pseudogene |
| ENSG00000261853 | AC134349,3 | -0,716841806 | 0,608427891 | 0,57051829 | 0,9423888 | taste receptor, type 2, member 18 pseudogene |
| ENSG00000260677 | AL445531,1 | -0,71693028 | 0,60839058 | 0,34768475 | 0,842293776 | novel transcript |
| ENSG00000168334 | XIRP1 | -0,71698096 | 0,608369208 | 0,61025963 | 0,951841373 | xin actin binding repeat containing 1 |
| ENSG00000271524 | BNIP3P17 | -0,717358589 | 0,608209987 | 0,0855486 | 0,465670107 | BCL2 interacting protein 3 pseudogene 17 |
| ENSG00000287650 | AL390071,1 | -0,717410129 | 0,608188259 | 0,09362229 | 0,48637543 | novel transcript, antisense to NBEA |

| | | | | | | |
|---|---|---|---|---|---|---|
| ENSG00000285601 | AC084262,2 | -0,717600408 | 0,60810805 | 0,19705918 | 0,695927373 | novel transcript |
| ENSG00000232499 | AL391058,1 | -0,717702616 | 0,60806497 | 0,31964609 | 0,824366234 | heterogeneous nuclear ribonucleoprotein A3 (hnRNPA3) pseudogene |
| ENSG00000207274 | SNORA70I | -0,71777138 | 0,608035988 | 0,32662539 | 0,828461469 | small nucleolar RNA, H/ACA box 70I |
| ENSG00000286952 | AC093928,1 | -0,71781232 | 0,608018734 | 0,54758627 | 0,935404974 | novel transcript, antisense to DNAH12 |
| ENSG00000212588 | SNORA26 | -0,717880121 | 0,60799016 | 0,43790328 | 0,897156497 | small nucleolar RNA, H/ACA box 26 |
| ENSG00000114786 | ABHD14A-ACY1 | -0,717992241 | 0,607942911 | 0,25083336 | 0,763027659 | ABHD14A-ACY1 readthrough |
| ENSG00000286553 | AL357075,2 | -0,718138646 | 0,60788122 | 0,23189604 | 0,740375324 | novel transcript, antisense to CNKSR3 |
| ENSG00000284885 | GCNT2P1 | -0,718155753 | 0,607874012 | 0,60685118 | 0,950927937 | GCNT2 pseudogene 1 |
| ENSG00000205318 | GCNT2P1 | -0,718155753 | 0,607874012 | 0,60685118 | 0,950927937 | GCNT2 pseudogene 1 |
| ENSG00000267641 | BNIP3P16 | -0,718219197 | 0,607847281 | 0,34158487 | 0,840048641 | BCL2 interacting protein 3 pseudogene 16 |
| ENSG00000229327 | AL391839,1 | -0,718336494 | 0,607797862 | 0,60948213 | 0,951793902 | novel transcript |
| ENSG00000230756 | RHOQP3 | -0,718753941 | 0,60762202 | 0,52045097 | 0,927003219 | ras homolog family member Q pseudogene 3 |
| ENSG00000146250 | PRSS35 | -0,718847447 | 0,607582639 | 0,05465831 | 0,361009299 | serine protease 35 |
| ENSG00000230077 | MTAPP2 | -0,71892465 | 0,607550127 | 0,48150467 | 0,912981134 | methylthioadenosine phosphorylase pseudogene 2 |
| ENSG00000259985 | AC017100,1 | -0,719195664 | 0,607436007 | 0,16675616 | 0,647072696 | novel transcript, antisense to B4GALT6 |
| ENSG00000272468 | AL021807,1 | -0,719210225 | 0,607429877 | 0,38810719 | 0,869820053 | novel transcript |
| ENSG00000259659 | AC009996,1 | -0,719256574 | 0,607410362 | 0,31994473 | 0,824366234 | novel transcript |
| ENSG00000243417 | RPS3AP18 | -0,719267949 | 0,607405573 | 0,64813257 | 0,956899963 | RPS3A pseudogene 18 |
| ENSG00000265713 | AC023389,2 | -0,719303079 | 0,607390783 | 0,22855906 | 0,737046989 | RNA terminal phosphate cyclase-like 1 (RCL1) pseudogene |
| ENSG00000286186 | AC018445,5 | -0,719448495 | 0,607329564 | 0,55375636 | 0,937338325 | novel transcript, antisense to NFATC1 |
| ENSG00000213569 | GTF3C6P2 | -0,719497306 | 0,607309017 | 0,43762855 | 0,897021798 | GTF3C6 pseudogene 2 |
| ENSG00000197332 | AC008543,1 | -0,719582389 | 0,607273202 | 0,08319247 | 0,45879863 | novel transcript |
| ENSG00000154118 | JPH3 | -0,719593991 | 0,607268318 | 0,054476 | 0,360376004 | junctophilin 3 |
| ENSG00000284048 | AC073111,2 | -0,719724067 | 0,607213568 | 0,65612376 | 0,957444211 | novel transcript |
| ENSG00000117115 | PADI2 | -0,719817812 | 0,607174113 | 0,1437682 | 0,603159913 | peptidyl arginine deiminase 2 |
| ENSG00000284716 | AL034417,3 | -0,719876443 | 0,607149438 | 0,45444902 | 0,903383444 | novel transcript |
| ENSG00000136286 | MYO1G | -0,720251128 | 0,606991775 | 0,37726308 | 0,862818448 | myosin IG |
| ENSG00000137343 | ATAT1 | -0,720259301 | 0,606988336 | 0,01697786 | 0,167390656 | alpha tubulin acetyltransferase 1 |
| ENSG00000104722 | NEFM | -0,720278998 | 0,606980049 | 0,18545158 | 0,678044469 | neurofilament medium |
| ENSG00000277606 | GUSBP1 | -0,720336682 | 0,60695578 | 0,01920416 | 0,182079045 | GUSB pseudogene 1 |
| ENSG00000287189 | AL121956,7 | -0,720369825 | 0,606941837 | 0,01350542 | 0,14230992 | novel transcript |

| | | | | | | |
|---|---|---|---|---|---|---|
| ENSG00000203709 | MIR29B2CHG | -0,720449066 | 0,606908501 | 0,00633165 | 0,082213892 | MIR29B2 and MIR29C host gene |
| ENSG00000175920 | DOK7 | -0,720491242 | 0,606890759 | 0,42395875 | 0,890949676 | docking protein 7 |
| ENSG00000143171 | RXRG | -0,720544556 | 0,606868332 | 0,39567605 | 0,872227623 | retinoid X receptor gamma |
| ENSG00000255875 | PABPN1P2 | -0,720609547 | 0,606840994 | 0,39217747 | 0,870909282 | PABPN1 pseudogene 2 |
| ENSG00000123201 | GUCY1B2 | -0,720719292 | 0,606794834 | 0,44134705 | 0,898885877 | guanylate cyclase 1 soluble subunit beta 2 (pseudogene) |
| ENSG00000260912 | AL158206,1 | -0,72082123 | 0,606751961 | 0,22610477 | 0,734959876 | novel transcript, overlapping ACER2 |
| ENSG00000205644 | AC025186,1 | -0,720992712 | 0,606679845 | 0,62656149 | 0,952793934 | ribosomal protein S2 (RPS2) pseudogene |
| ENSG00000039139 | DNAH5 | -0,721056047 | 0,606653212 | 0,070841 | 0,418829978 | dynein axonemal heavy chain 5 |
| ENSG00000181631 | P2RY13 | -0,721087319 | 0,606640062 | 0,2522516 | 0,76432863 | purinergic receptor P2Y13 |
| ENSG00000288125 | AC023300,4 | -0,721228124 | 0,606580858 | 0,2873719 | 0,798896538 | novel transcript, antisense to CCDC33 |
| ENSG00000277749 | AC023300,2 | -0,721228124 | 0,606580858 | 0,2873719 | 0,798896538 | novel transcript, antisense to CCDC33 |
| ENSG00000281367 | AC044810,2 | -0,721230913 | 0,606579685 | 0,75952975 | 0,976120207 | novel transcript, sense overlapping OR5P2, OR5P3 and OR5P1P |
| ENSG00000273144 | TAS2R15P | -0,721483645 | 0,606473433 | 0,23551514 | 0,744654651 | taste 2 receptor member 15 pseudogene |
| ENSG00000276217 | TAS2R15P | -0,721483645 | 0,606473433 | 0,23551514 | 0,744654651 | taste 2 receptor member 15 pseudogene |
| ENSG00000212125 | TAS2R15P | -0,721483645 | 0,606473433 | 0,23551514 | 0,744654651 | taste 2 receptor member 15 pseudogene |
| ENSG00000255319 | ENPP7P8 | -0,721552306 | 0,606444571 | 0,408709 | 0,88153752 | ectonucleotide pyrophosphatase/phosphodiesterase 7 pseudogene 8 |
| ENSG00000228187 | AC093433,1 | -0,721593649 | 0,606427192 | 0,46699503 | 0,907940653 | novel pseudogene |
| ENSG00000174453 | VWC2L | -0,721634155 | 0,606410166 | 0,14035844 | 0,597084546 | von Willebrand factor C domain containing 2 like |
| ENSG00000118432 | CNR1 | -0,721742195 | 0,606364755 | 0,06437317 | 0,397835213 | cannabinoid receptor 1 |
| ENSG00000119547 | ONECUT2 | -0,721830561 | 0,606327616 | 0,24510871 | 0,757071585 | one cut homeobox 2 |
| ENSG00000269069 | AC007842,1 | -0,721989009 | 0,606261028 | 0,07475027 | 0,432166624 | protein associated with topoisomerase II homolog 1 (yeast) (PATL1) pseudogene |
| ENSG00000281233 | AC007842,3 | -0,721989009 | 0,606261028 | 0,07475027 | 0,432166624 | protein associated with topoisomerase II homolog 1 (yeast) (PATL1) pseudogene |
| ENSG00000284434 | AC270107,9 | -0,722057075 | 0,606232426 | 0,37965362 | 0,865238161 | |
| ENSG00000181444 | ZNF467 | -0,722133519 | 0,606200304 | 0,09509142 | 0,489669163 | zinc finger protein 467 |
| ENSG00000138735 | PDE5A | -0,722150467 | 0,606193183 | 0,14946856 | 0,614794108 | phosphodiesterase 5A |
| ENSG00000245281 | AC124242,1 | -0,722216564 | 0,606165411 | 0,14950997 | 0,614794108 | novel transcript |
| ENSG00000232295 | AL589935,1 | -0,722249211 | 0,606151694 | 0,1665004 | 0,646481436 | novel transcript, antisense OGFRL1 |

| | | | | | | |
|---|---|---|---|---|---|---|
| ENSG00000274253 | AC138649,1 | -0,722266088 | 0,606144603 | 0,028126 | 0,237080451 | novel transcript, sense overlapping NIPA1 |
| ENSG00000259351 | AC015914,1 | -0,722537311 | 0,60603066 | 0,5542173 | 0,937712393 | novel transcript, antisense to DAPK2 |
| ENSG00000235902 | AC108472,1 | -0,722663222 | 0,605977771 | 0,03926495 | 0,294878559 | novel transcript |
| ENSG00000238161 | OR7E117P | -0,722665211 | 0,605976936 | 0,38044904 | 0,86586292 | olfactory receptor family 7 subfamily E member 117 pseudogene |
| ENSG00000236398 | TAS2R39 | -0,722718497 | 0,605954554 | 0,52780347 | 0,929595391 | taste 2 receptor member 39 |
| ENSG00000286803 | AL109824,1 | -0,722773032 | 0,605931649 | 0,57453857 | 0,942475308 | novel transcript, antisense to NCOA6 |
| ENSG00000234632 | NECAP1P2 | -0,723114025 | 0,605788449 | 0,44736998 | 0,902062594 | NECAP endocytosis associated 1 pseudogene 2 |
| ENSG00000148408 | CACNA1B | -0,723374713 | 0,605678996 | 0,09110368 | 0,481283407 | calcium voltage-gated channel subunit alpha1 B |
| ENSG00000231057 | AC096637,2 | -0,723632209 | 0,605570902 | 0,07236247 | 0,424120881 | novel transcript |
| ENSG00000276542 | AC097478,2 | -0,723706581 | 0,605539686 | 0,51281185 | 0,923640376 | novel transcript, antisense to SNCA |
| ENSG00000283268 | TEX54 | -0,72385352 | 0,605478014 | 0,53728911 | 0,932883277 | testis expressed 54 |
| ENSG00000286575 | AL356608,3 | -0,723963064 | 0,605432042 | 0,24256054 | 0,752770777 | novel transcript, antisense to CNNM2and AS3MT |
| ENSG00000186976 | EFCAB6 | -0,72397325 | 0,605427768 | 0,0546293 | 0,361008363 | EF-hand calcium binding domain 6 |
| ENSG00000284610 | AC107918,4 | -0,723996898 | 0,605417844 | 0,01670829 | 0,165411048 | RNA binding protein, fox-1 homolog (C, elegans) 1 (RBFOX1) pseudogene |
| ENSG00000276161 | SNORA17B | -0,724005145 | 0,605414383 | 0,13663674 | 0,58964193 | small nucleolar RNA, H/ACA box 17B |
| ENSG00000273243 | Z82243,1 | -0,724079296 | 0,605383267 | 0,23772098 | 0,746907875 | novel transcript |
| ENSG00000116990 | MYCL | -0,724348795 | 0,60527019 | 0,00844109 | 0,101461832 | MYCL proto-oncogene, bHLH transcription factor |
| ENSG00000066405 | CLDN18 | -0,724414028 | 0,605242823 | 0,04826838 | 0,334607086 | claudin 18 |
| ENSG00000250377 | AC008525,1 | -0,724470326 | 0,605219205 | 0,23571792 | 0,744895724 | novel transcript |
| ENSG00000162687 | KCNT2 | -0,724656462 | 0,605141125 | 0,08782655 | 0,472113008 | potassium sodium-activated channel subfamily T member 2 |
| ENSG00000144834 | TAGLN3 | -0,724704462 | 0,605120992 | 0,11957974 | 0,552362303 | transgelin 3 |
| ENSG00000213889 | PPM1N | -0,724760288 | 0,605097576 | 0,30202898 | 0,811449876 | protein phosphatase, Mg2+/Mn2+ dependent 1N (putative) |
| ENSG00000236560 | PPP1R11 | -0,724772014 | 0,605092659 | 0,18119655 | 0,670965888 | protein phosphatase 1 regulatory inhibitor subunit 11 |
| ENSG00000274240 | AC136352,1 | -0,724877564 | 0,60504839 | 0,19416774 | 0,691160239 | p21 protein (Cdc42/Rac)-activated kinase 2 (PAK2) pseudogene |
| ENSG00000286485 | Z80897,1 | -0,724954273 | 0,605016221 | 0,25875002 | 0,771610117 | novel transcript, antisense to ANKRD54and IFT27 |
| ENSG00000178568 | ERBB4 | -0,725469925 | 0,604800013 | 0,08456931 | 0,462844615 | erb-b2 receptor tyrosine kinase 4 |
| ENSG00000260261 | AC124944,2 | -0,725731438 | 0,604690392 | 0,02799935 | 0,236395267 | programmed cell death 6 (PDCD6) pseudogene |

| | | | | | | |
|---|---|---|---|---|---|---|
| ENSG00000272219 | AC005072,1 | -0,725746804 | 0,604683952 | 0,15268924 | 0,621087238 | novel transcript, sense intronic to CUX1 |
| ENSG00000226876 | SMYD3-AS1 | -0,725874333 | 0,604630502 | 0,46770734 | 0,907940653 | SMYD3 antisense RNA 1 |
| ENSG00000255856 | AC069503,1 | -0,725877906 | 0,604629005 | 0,38684518 | 0,869441674 | novel transcript |
| ENSG00000248858 | FLJ46284 | -0,725906026 | 0,60461722 | 0,27332487 | 0,785672647 | uncharacterized LOC441369 |
| ENSG00000206417 | H1-10-AS1 | -0,726071045 | 0,604548066 | 0,06882296 | 0,413705223 | H1-10 antisense RNA 1 |
| ENSG00000129757 | CDKN1C | -0,726084117 | 0,604542589 | 0,09339678 | 0,485607362 | cyclin dependent kinase inhibitor 1C |
| ENSG00000147576 | ADHFE1 | -0,726237517 | 0,604478312 | 0,17039842 | 0,653594444 | alcohol dehydrogenase iron containing 1 |
| ENSG00000231468 | PRDX3P2 | -0,72633058 | 0,604439321 | 0,60955776 | 0,951793902 | peroxiredoxin 3 pseudogene 2 |
| ENSG00000283483 | AC068205,4 | -0,726544474 | 0,604349713 | 0,14157354 | 0,599428784 | prohibitin (PHB) pseudogene |
| ENSG00000261758 | AC117382,2 | -0,726608796 | 0,604322769 | 0,12265502 | 0,560135346 | novel transcript, antisense to STAG1 |
| ENSG00000286220 | AC007546,3 | -0,726768817 | 0,604255742 | 0,02324786 | 0,207781894 | purinergic receptor P2X, ligand-gated ion channel, 7 (P2RX7) pseudogene |
| ENSG00000287066 | AP000962,3 | -0,72692784 | 0,604189141 | 0,14807783 | 0,612595581 | novel transcript |
| ENSG00000233623 | PGAM1P11 | -0,72711353 | 0,604111381 | 0,61639752 | 0,952793934 | phosphoglycerate mutase 1 pseudogene 11 |
| ENSG00000261596 | AC005632,3 | -0,727138045 | 0,604101115 | 0,06560994 | 0,402080164 | novel transcript, sense intronic to METTL9 |
| ENSG00000250284 | AC109439,1 | -0,727251802 | 0,604053483 | 0,0896757 | 0,47775341 | novel transcript |
| ENSG00000253523 | AC104350,1 | -0,727360504 | 0,604007972 | 0,55237182 | 0,936668248 | novel transcript |
| ENSG00000278911 | AC015819,3 | -0,727407211 | 0,603988418 | 0,52539046 | 0,928798838 | novel transcript |
| ENSG00000176244 | ACBD7 | -0,727582561 | 0,603915011 | 0,00385364 | 0,0570725 | acyl-CoA binding domain containing 7 |
| ENSG00000258345 | AC034102,7 | -0,727621884 | 0,603898551 | 0,63291289 | 0,954397372 | novel transcript, antisense to ZC3H10 and ESYT1 |
| ENSG00000105696 | TMEM59L | -0,7276821 | 0,603873345 | 0,0662352 | 0,404519674 | transmembrane protein 59 like |
| ENSG00000187959 | CPSF4L | -0,727819734 | 0,603815738 | 0,47164379 | 0,908896785 | cleavage and polyadenylation specific factor 4 like |
| ENSG00000274295 | AC131902,2 | -0,72788934 | 0,603786606 | 0,68602275 | 0,965740648 | polymerase (DNA directed), epsilon 2, accessory subunit (POLE2) pseudogene |
| ENSG00000144649 | GASK1A | -0,728128901 | 0,603686356 | 0,08489111 | 0,463749042 | golgi associated kinase 1A |
| ENSG00000259834 | AL365361,1 | -0,728359888 | 0,603589708 | 0,25473897 | 0,766202086 | novel transcript |
| ENSG00000070886 | EPHA8 | -0,728566758 | 0,603503165 | 0,44275364 | 0,899437949 | EPH receptor A8 |
| ENSG00000229666 | MAST4-AS1 | -0,728608514 | 0,603485698 | 0,43566884 | 0,896422069 | MAST4 antisense RNA 1 |
| ENSG00000233264 | AC006042,2 | -0,728918397 | 0,603356086 | 0,16536641 | 0,644399285 | peptidylprolyl isomerase (cyclophilin)-like 4 (PPIL4) pseudogene |
| ENSG00000168350 | DEGS2 | -0,72906871 | 0,603293227 | 0,21700724 | 0,724609024 | delta 4-desaturase, sphingolipid 2 |
| ENSG00000239827 | SUGT1P3 | -0,729123141 | 0,603270466 | 0,19256698 | 0,689307774 | SUGT1 pseudogene 3 |
| ENSG00000257842 | LINC02588 | -0,729425442 | 0,60314407 | 0,22040524 | 0,728572782 | long intergenic non-protein coding RNA 2588 |

| | | | | | | |
|---|---|---|---|---|---|---|
| ENSG00000226963 | AC078883,2 | -0,729841927 | 0,602969976 | 0,54000784 | 0,933385624 | novel transcript |
| ENSG00000257511 | AC084824,1 | -0,730405212 | 0,602734599 | 0,05644663 | 0,366903759 | nucleosome assembly protein 1-like 1 (NAP1L1) pseudogene |
| ENSG00000282984 | AL357556,3 | -0,730416741 | 0,602729782 | 0,35874466 | 0,849814998 | Tubulin-specific chaperone E |
| ENSG00000196132 | MYT1 | -0,730502358 | 0,602694014 | 0,29259236 | 0,803059213 | myelin transcription factor 1 |
| ENSG00000151575 | TEX9 | -0,730505433 | 0,60269273 | 0,11928867 | 0,552100834 | testis expressed 9 |
| ENSG00000261302 | AC106779,1 | -0,730513317 | 0,602689436 | 0,38395302 | 0,868113415 | novel transcript, antisense to NUP93 |
| ENSG00000147896 | IFNK | -0,730580891 | 0,602661208 | 0,20881498 | 0,713420724 | interferon kappa |
| ENSG00000279718 | SNX18P12 | -0,730866139 | 0,602542062 | 0,2583475 | 0,771145341 | sorting nexin 18 pseudogene 12 |
| ENSG00000151224 | MAT1A | -0,730965664 | 0,602500497 | 0,59531846 | 0,947756196 | methionine adenosyltransferase 1A |
| ENSG00000257489 | AC010203,1 | -0,73117751 | 0,602412032 | 0,26434268 | 0,775697617 | TBC1 domain family, member 2B (TBC1D2B) pseudogene |
| ENSG00000229263 | AC004691,1 | -0,731196742 | 0,602404001 | 0,54200334 | 0,933491609 | novel transcript |
| ENSG00000285865 | AC010285,3 | -0,731314921 | 0,602354657 | 0,10713167 | 0,522599956 | Novel transcript, antisense to RNF130 |
| ENSG00000279486 | OR2AG1 | -0,731367285 | 0,602332795 | 0,14670744 | 0,610674677 | olfactory receptor family 2 subfamily AG member 1 |
| ENSG00000268061 | NAPA-AS1 | -0,731388491 | 0,602323941 | 0,08466243 | 0,462904176 | NAPA antisense RNA 1 |
| ENSG00000264859 | DSG2-AS1 | -0,731446242 | 0,60229983 | 0,45496708 | 0,903424009 | DSG2 antisense RNA 1 |
| ENSG00000282472 | AC026950,1 | -0,731501441 | 0,602276786 | 0,57086124 | 0,942436428 | ERICH1 antisense RNA 1 |
| ENSG00000248027 | ARHGAP42-AS1 | -0,731524544 | 0,602267142 | 0,30924272 | 0,816845699 | ARHGAP42 antisense RNA 1 |
| ENSG00000270554 | AC069431,2 | -0,731730449 | 0,602181191 | 0,32071313 | 0,824366234 | eukaryotic translation initiation factor 3, subunit E, pseudogene |
| ENSG00000151320 | AKAP6 | -0,731780001 | 0,602160508 | 0,06665588 | 0,40559155 | A-kinase anchoring protein 6 |
| ENSG00000276505 | AP000892,3 | -0,732284229 | 0,601950087 | 0,49061006 | 0,916995341 | novel transcript, sense intronic to BACE1 |
| ENSG00000183850 | ZNF730 | -0,73228634 | 0,601949206 | 0,1675798 | 0,648417701 | zinc finger protein 730 |
| ENSG00000270885 | RASL10B | -0,73234349 | 0,601925362 | 0,03964401 | 0,296656351 | RAS like family 10 member B |
| ENSG00000178055 | PRSS42P | -0,732486556 | 0,601865674 | 0,50488678 | 0,921400408 | serine protease 42, pseudogene |
| ENSG00000243207 | PPAN-P2RY11 | -0,732590631 | 0,601822257 | 0,49378939 | 0,917303403 | PPAN-P2RY11 readthrough |
| ENSG00000228623 | ZNF883 | -0,732592085 | 0,601821651 | 0,14601708 | 0,609001881 | zinc finger protein 883 |
| ENSG00000237325 | AP000563,1 | -0,732743532 | 0,601758478 | 0,26938222 | 0,781544346 | chromosome 6 open reading frame 203 (C6orf203) pseudogene |
| ENSG00000203809 | LIN28B-AS1 | -0,732928003 | 0,601681539 | 0,20152845 | 0,701094807 | LIN28B antisense RNA 1 |
| ENSG00000234183 | LINC01952 | -0,733107532 | 0,60160667 | 0,49792696 | 0,919274055 | long intergenic non-protein coding RNA 1952 |
| ENSG00000271698 | AC233992,2 | -0,733166737 | 0,601581982 | 0,31933442 | 0,824211281 | novel transcript |

| | | | | | | |
|---|---|---|---|---|---|---|
| ENSG00000285047 | AC233992,6 | -0,733166737 | 0,601581982 | 0,31933442 | 0,824211281 | novel transcript |
| ENSG00000272196 | H2AC19 | -0,733365172 | 0,601499243 | 0,04412211 | 0,317111216 | H2A clustered histone 19 |
| ENSG00000196268 | ZNF493 | -0,73342205 | 0,60147553 | 0,09885642 | 0,499135531 | zinc finger protein 493 |
| ENSG00000221845 | C7orf65 | -0,733496693 | 0,601444411 | 0,46301553 | 0,906849208 | chromosome 7 open reading frame 65 |
| ENSG00000260526 | AC109347,1 | -0,733715285 | 0,601353289 | 0,23467661 | 0,744134035 | novel transcript, antisense to AP1AR |
| ENSG00000148541 | FAM13C | -0,733808615 | 0,601314388 | 0,03684423 | 0,283021613 | family with sequence similarity 13 member C |
| ENSG00000124507 | PACSIN1 | -0,733850676 | 0,601296857 | 0,25776097 | 0,7701554 | protein kinase C and casein kinase substrate in neurons 1 |
| ENSG00000287676 | AC096642,2 | -0,734010375 | 0,601230301 | 0,44408835 | 0,900210555 | novel transcript |
| ENSG00000122188 | LAX1 | -0,734022539 | 0,601225232 | 0,4511449 | 0,902062594 | lymphocyte transmembrane adaptor 1 |
| ENSG00000232998 | VPS13A-AS1 | -0,734024792 | 0,601224293 | 0,58917086 | 0,946705487 | VPS13A antisense RNA 1 |
| ENSG00000174871 | CNIH2 | -0,734087676 | 0,601198087 | 0,0080851 | 0,098546769 | cornichon family AMPA receptor auxiliary protein 2 |
| ENSG00000233295 | FAM90A20P | -0,734132902 | 0,601179241 | 0,53606205 | 0,932162269 | family with sequence similarity 90 member A20, pseudogene |
| ENSG00000232732 | AC097717,1 | -0,734254351 | 0,601128635 | 0,37698574 | 0,862818448 | novel transcript |
| ENSG00000235413 | KRT18P63 | -0,734314687 | 0,601103495 | 0,26157556 | 0,773396846 | keratin 18 pseudogene 63 |
| ENSG00000146013 | GFRA3 | -0,734631519 | 0,6009715 | 0,48601305 | 0,914018932 | GDNF family receptor alpha 3 |
| ENSG00000276398 | AC018659,5 | -0,734758598 | 0,600918566 | 0,04547001 | 0,323145642 | |
| ENSG00000112232 | KHDRBS2 | -0,734762256 | 0,600917043 | 0,06251049 | 0,390552401 | KH RNA binding domain containing, signal transduction associated 2 |
| ENSG00000149418 | ST14 | -0,73493946 | 0,600843237 | 0,47984279 | 0,912847708 | ST14 transmembrane serine protease matriptase |
| ENSG00000275811 | HTR1DP1 | -0,73518428 | 0,600741285 | 0,32104056 | 0,824556331 | 5-hydroxytryptamine receptor 1D pseudogene 1 |
| ENSG00000257576 | HSPD1P4 | -0,735210131 | 0,600730521 | 0,10443866 | 0,515254902 | heat shock protein family D (Hsp60) member 1 pseudogene 4 |
| ENSG00000267282 | AC011481,2 | -0,735210592 | 0,600730329 | 0,63946102 | 0,956018518 | novel transcript, antisense to PVRL2 |
| ENSG00000271216 | LINC01050 | -0,735421927 | 0,600642337 | 0,47135548 | 0,908887649 | long intergenic non-protein coding RNA 1050 |
| ENSG00000225756 | DBH-AS1 | -0,735428577 | 0,600639568 | 0,09791106 | 0,49709182 | DBH antisense RNA 1 |
| ENSG00000124171 | PARD6B | -0,735433277 | 0,600637611 | 0,06214179 | 0,389026994 | par-6 family cell polarity regulator beta |
| ENSG00000281705 | CENATAC-DT | -0,735526296 | 0,600598886 | 0,30975755 | 0,817030292 | CENATAC divergent transcript |
| ENSG00000206331 | PRRT1 | -0,735596654 | 0,600569596 | 0,26516718 | 0,776598285 | proline rich transmembrane protein 1 |
| ENSG00000249170 | AC096713,1 | -0,735719897 | 0,600518294 | 0,43320948 | 0,895845815 | solute carrier family 35, member A4 (SLC35A4) pseudogene |
| ENSG00000157404 | KIT | -0,736088517 | 0,600364877 | 0,23699799 | 0,746474416 | KIT proto-oncogene, receptor tyrosine kinase |

| | | | | | | |
|---|---|---|---|---|---|---|
| ENSG00000287899 | AC055714,1 | -0,736174577 | 0,600329065 | 0,50276062 | 0,920687909 | novel transcript |
| ENSG00000226803 | ZNF451-AS1 | -0,736231041 | 0,60030557 | 0,05004148 | 0,342116111 | ZNF451 regulatory antisense RNA 1 |
| ENSG00000226953 | NCKAP5-AS2 | -0,736357345 | 0,600253017 | 0,16533238 | 0,644399285 | NCKAP5 antisense RNA 2 |
| ENSG00000001626 | CFTR | -0,736460131 | 0,600210253 | 0,25691748 | 0,769163921 | CF transmembrane conductance regulator |
| ENSG00000285468 | EPHA1-AS1 | -0,736549495 | 0,600173076 | 0,60295962 | 0,950352286 | EPHA1 antisense RNA 1 |
| ENSG00000124215 | CDH26 | -0,736667463 | 0,600124002 | 0,34284909 | 0,840419133 | cadherin 26 |
| ENSG00000196890 | H2BU1 | -0,736729449 | 0,600098218 | 0,21627069 | 0,723540588 | H2B,U histone 1 |
| ENSG00000214773 | AC112512,1 | -0,736864336 | 0,600042114 | 0,03007651 | 0,247515702 | novel transcript, sense intronic SMARCC1 |
| ENSG00000277425 | AL121890,4 | -0,737166224 | 0,599916566 | 0,46267618 | 0,906789619 | novel transcript |
| ENSG00000130590 | SAMD10 | -0,737388316 | 0,599824221 | 0,0070916 | 0,08947206 | sterile alpha motif domain containing 10 |
| ENSG00000271971 | AC120053,1 | -0,737401485 | 0,599818746 | 0,26944674 | 0,781544346 | novel transcript, antisense to FAM92A1 |
| ENSG00000184774 | MGAT4EP | -0,737450961 | 0,599798175 | 0,59896397 | 0,948894897 | MGAT4 family member E, pseudogene |
| ENSG00000226330 | AL606489,1 | -0,737469748 | 0,599790365 | 0,55537634 | 0,937793851 | novel transcript |
| ENSG00000246145 | RRS1-AS1 | -0,737558846 | 0,599753324 | 0,24945068 | 0,761381055 | RRS1 antisense RNA 1 (head to head) |
| ENSG00000265218 | AC103810,2 | -0,737746719 | 0,599675227 | 0,11400643 | 0,53948382 | novel transcript, antisense to LRRC37A3 |
| ENSG00000287715 | AC026120,3 | -0,737862216 | 0,599627221 | 0,20314713 | 0,703503738 | novel transcript |
| ENSG00000228175 | GEMIN8P4 | -0,738096871 | 0,5995297 | 0,20911015 | 0,71355079 | gem nuclear organelle associated protein 8 pseudogene 4 |
| ENSG00000141665 | FBXO15 | -0,73829076 | 0,599449132 | 0,04555808 | 0,32341683 | F-box protein 15 |
| ENSG00000206952 | SNORA50A | -0,738916791 | 0,599189068 | 0,41481527 | 0,885579089 | small nucleolar RNA, H/ACA box 50A |
| ENSG00000265142 | MIR133A1HG | -0,739142226 | 0,599095447 | 0,24614061 | 0,758198269 | MIR133A1 host gene |
| ENSG00000156925 | ZIC3 | -0,739148882 | 0,599092683 | 0,10916008 | 0,527070407 | Zic family member 3 |
| ENSG00000286784 | AC116611,1 | -0,73926056 | 0,599046309 | 0,17335717 | 0,658330849 | novel transcript |
| ENSG00000257838 | OTOAP1 | -0,739477747 | 0,598956134 | 0,26728599 | 0,779594822 | OTOA pseudogene 1 |
| ENSG00000136883 | KIF12 | -0,739607417 | 0,598902302 | 0,57837157 | 0,943425812 | kinesin family member 12 |
| ENSG00000231616 | AL354733,2 | -0,739616349 | 0,598898594 | 0,05695951 | 0,368768315 | novel transcript, antisense to KIF27 |
| ENSG00000258439 | AC007956,2 | -0,739665581 | 0,598878157 | 0,30444888 | 0,813181541 | ribosomal protein S2 (RPS2) pseudogene |
| ENSG00000132259 | CNGA4 | -0,739697836 | 0,598864768 | 0,4582747 | 0,904140862 | cyclic nucleotide gated channel subunit alpha 4 |
| ENSG00000163545 | NUAK2 | -0,739714489 | 0,598857855 | 0,02838889 | 0,238652911 | NUAK family kinase 2 |
| ENSG00000248967 | AC022493,2 | -0,739821453 | 0,598813456 | 0,47713565 | 0,91143003 | S-phase kinase-associated protein 1 (SKP1) pseudogene |
| ENSG00000268945 | AC010422,2 | -0,739856417 | 0,598798944 | 0,34464134 | 0,841168721 | novel transcript |

| | | | | | | |
|---|---|---|---|---|---|---|
| ENSG00000267879 | AC011483,1 | -0,739989287 | 0,598743798 | 0,36460721 | 0,854695949 | novel transcript, antisense to KLK6, KLK7, KLK8 and KLK9 |
| ENSG00000204113 | BMP2KL | -0,740025279 | 0,598728861 | 0,18551291 | 0,678110103 | BMP2 inducible kinase like (pseudogene) |
| ENSG00000270103 | AL360012,1 | -0,740267629 | 0,598628293 | 0,02460015 | 0,21668006 | RNA, U11 small nuclear |
| ENSG00000166415 | WDR72 | -0,740279207 | 0,598623489 | 0,28332077 | 0,794558711 | WD repeat domain 72 |
| ENSG00000256973 | AC053513,2 | -0,740286965 | 0,598620269 | 0,08374674 | 0,461014667 | novel transcript |
| ENSG00000153162 | BMP6 | -0,740567945 | 0,598503693 | 0,05189264 | 0,349498355 | bone morphogenetic protein 6 |
| ENSG00000239922 | AC092958,1 | -0,740567991 | 0,598503674 | 0,09133253 | 0,481302729 | novel transcript |
| ENSG00000115271 | GCA | -0,740638004 | 0,59847463 | 0,1507721 | 0,617373838 | grancalcin |
| ENSG00000204186 | ZDBF2 | -0,740669204 | 0,598461688 | 0,05796136 | 0,372428727 | zinc finger DBF-type containing 2 |
| ENSG00000277504 | AC010536,3 | -0,740862468 | 0,598381522 | 0,51548059 | 0,924901769 | novel transcript, antisense to JPH3 |
| ENSG00000232046 | LINC01798 | -0,740932167 | 0,598352614 | 0,33916361 | 0,838276153 | long intergenic non-protein coding RNA 1798 |
| ENSG00000244378 | RPS2P45 | -0,740970536 | 0,598336701 | 0,26853295 | 0,780709773 | ribosomal protein S2 pseudogene 45 |
| ENSG00000213707 | HMGB1P10 | -0,741060107 | 0,598299554 | 0,67549591 | 0,963400586 | high mobility group box 1 pseudogene 10 |
| ENSG00000288032 | AC104472,5 | -0,741094331 | 0,598285361 | 0,07720023 | 0,439233464 | novel transcript, sense intronic to GMPS |
| ENSG00000260382 | AC087481,1 | -0,741103999 | 0,598281352 | 0,52108757 | 0,927046149 | transmembrane protein 183A (TMEM183A) pseudogene |
| ENSG00000249102 | AC034223,1 | -0,74114948 | 0,598262492 | 0,01945261 | 0,183597917 | novel transcript |
| ENSG00000230269 | LINC02525 | -0,741167477 | 0,598255028 | 0,16864145 | 0,650559718 | long intergenic non-protein coding RNA 2525 |
| ENSG00000233379 | AL139002,1 | -0,741191546 | 0,598245048 | 0,29957836 | 0,80943893 | novel transcript |
| ENSG00000263499 | AC007431,1 | -0,741193595 | 0,598244198 | 0,06980377 | 0,417035202 | novel transcript, antisense to MSI2 |
| ENSG00000235768 | BRD7P5 | -0,741266765 | 0,598213857 | 0,28370526 | 0,79499517 | bromodomain containing 7 pseudogene 5 |
| ENSG00000256588 | AC027544,2 | -0,741486793 | 0,59812263 | 0,37802517 | 0,863678679 | novel transcript |
| ENSG00000281103 | TRG-AS1 | -0,741551163 | 0,598095943 | 0,38905972 | 0,870057277 | T cell receptor gamma locus antisense RNA 1 |
| ENSG00000275413 | AC002553,2 | -0,741568503 | 0,598088755 | 0,38029054 | 0,865730737 | novel transcript, antisense to TTC19 |
| ENSG00000229325 | ACAP2-IT1 | -0,741669896 | 0,598046722 | 0,39446184 | 0,87185967 | ACAP2 intronic transcript 1 |
| ENSG00000232335 | AL354864,1 | -0,741746333 | 0,598015037 | 0,38636295 | 0,869441674 | novel transcript |
| ENSG00000260871 | AC093510,2 | -0,74179428 | 0,597995163 | 0,59983644 | 0,949340299 | novel transcript, overlapping CETN3 |
| ENSG00000188771 | PLET1 | -0,741855812 | 0,597969659 | 0,48510688 | 0,913690171 | placenta expressed transcript 1 |
| ENSG00000138400 | MDH1B | -0,741962928 | 0,597925263 | 0,11479666 | 0,541181798 | malate dehydrogenase 1B |
| ENSG00000245080 | MIR3150BHG | -0,742181562 | 0,597834657 | 0,3110611 | 0,817751312 | MIR3150B host gene |
| ENSG00000102174 | PHEX | -0,742276639 | 0,597795259 | 0,26017894 | 0,772692192 | phosphate regulating endopeptidase homolog X-linked |

| | | | | | | |
|---|---|---|---|---|---|---|
| ENSG00000248409 | AP000344,2 | -0,742284513 | 0,597791997 | 0,39436646 | 0,871799523 | coiled-coil domain containing 188 (CCDC188) pseudogene |
| ENSG00000231003 | MPIG6B | -0,742611342 | 0,597656588 | 0,84396975 | 0,986273696 | megakaryocyte and platelet inhibitory receptor G6b |
| ENSG00000212464 | SNORA12 | -0,742788024 | 0,597583399 | 0,22689857 | 0,735383523 | small nucleolar RNA, H/ACA box 12 |
| ENSG00000280708 | AC006359,3 | -0,74284626 | 0,597559278 | 0,26780207 | 0,780440321 | novel transcript |
| ENSG00000272023 | AC010240,3 | -0,743392593 | 0,597333031 | 0,57182377 | 0,942475308 | novel transcript |
| ENSG00000273142 | LINC02604 | -0,743438714 | 0,597313936 | 0,05005493 | 0,342117936 | long intergenic non-protein coding RNA 2604 |
| ENSG00000276848 | KIR3DX1 | -0,743661003 | 0,597221909 | 0,77831811 | 0,978587255 | killer cell immunoglobulin like receptor, three Ig domains X1 (pseudogene) |
| ENSG00000188620 | HMX3 | -0,743769063 | 0,597177178 | 0,49186351 | 0,916995341 | H6 family homeobox 3 |
| ENSG00000286786 | AC116158,3 | -0,743777571 | 0,597173656 | 0,01874324 | 0,17881864 | novel transcript, antisense to RASGRP1 |
| ENSG00000251323 | LINC02728 | -0,743866785 | 0,597136729 | 0,09161442 | 0,481464668 | long intergenic non-protein coding RNA 2728 |
| ENSG00000285733 | AL031315,1 | -0,744111907 | 0,597035281 | 0,08759395 | 0,471714073 | novel protein |
| ENSG00000275239 | FAM242F | -0,744147653 | 0,597020488 | 0,51520578 | 0,924808725 | family with sequence similarity 242 member F |
| ENSG00000220392 | FCF1P5 | -0,74426047 | 0,596973804 | 0,31436281 | 0,819877431 | FCF1 pseudogene 5 |
| ENSG00000226232 | NPIPB14P | -0,744734742 | 0,596777587 | 0,29622524 | 0,805971225 | nuclear pore complex interacting protein family member B14, pseudogene |
| ENSG00000207442 | SNORD116-6 | -0,744879926 | 0,596717534 | 0,53961466 | 0,933350549 | small nucleolar RNA, C/D box 116-6 |
| ENSG00000270708 | KF455155,1 | -0,744919211 | 0,596701285 | 0,52161421 | 0,927472429 | GABA(A) receptor-associated protein-like 2 (GABARAPL2) pseudogene |
| ENSG00000090376 | IRAK3 | -0,745011273 | 0,596663209 | 0,07023723 | 0,417549035 | interleukin 1 receptor associated kinase 3 |
| ENSG00000170476 | MZB1 | -0,745129323 | 0,596614389 | 0,5768097 | 0,943261034 | marginal zone B and B1 cell specific protein |
| ENSG00000274615 | AC233968,1 | -0,745139521 | 0,596610172 | 0,04107199 | 0,303420267 | aminopeptidase puromycin sensitive (NPEPPS) pseudogene |
| ENSG00000188211 | NCR3LG1 | -0,745261071 | 0,596559908 | 0,10697467 | 0,52216331 | natural killer cell cytotoxicity receptor 3 ligand 1 |
| ENSG00000007314 | SCN4A | -0,745547593 | 0,596441442 | 0,34556484 | 0,841168721 | sodium voltage-gated channel alpha subunit 4 |
| ENSG00000217512 | AL356776,1 | -0,745599593 | 0,596419944 | 0,31963534 | 0,824366234 | nucleoporin 155kDa pseudogene |
| ENSG00000198168 | SVIP | -0,746180677 | 0,596179768 | 0,10930139 | 0,527552049 | small VCP interacting protein |
| ENSG00000114739 | ACVR2B | -0,746453491 | 0,596067041 | 0,03334337 | 0,26518622 | activin A receptor type 2B |
| ENSG00000117114 | ADGRL2 | -0,747023315 | 0,595831658 | 0,08465381 | 0,462904176 | adhesion G protein-coupled receptor L2 |
| ENSG00000228624 | HDAC2-AS2 | -0,747495716 | 0,595636589 | 0,05972765 | 0,38023321 | HDAC2 and HS3ST5 antisense RNA 2 |
| ENSG00000277342 | AC048344,4 | -0,747506244 | 0,595632242 | 0,21660288 | 0,723909983 | novel transcript, sense intronic to BICD1 |

| | | | | | | |
|---|---|---|---|---|---|---|
| ENSG00000001561 | ENPP4 | -0,747538392 | 0,59561897 | 0,19729228 | 0,695943345 | ectonucleotide pyrophosphatase/phosphodiesterase 4 |
| ENSG00000283125 | AC022726,2 | -0,747599665 | 0,595593674 | 0,5616783 | 0,940714121 | novel transcript, antisense to BCL2 |
| ENSG00000136872 | ALDOB | -0,748032665 | 0,595414943 | 0,20625127 | 0,709299291 | aldolase, fructose-bisphosphate B |
| ENSG00000253203 | GUSBP3 | -0,748234955 | 0,595331462 | 0,01863182 | 0,17801357 | GUSB pseudogene 3 |
| ENSG00000250749 | AC122714,1 | -0,748464627 | 0,595236695 | 0,34175685 | 0,840048641 | proteolipid protein 2 (colonic epithelium-enriched) (PLP2) pseudogene |
| ENSG00000262111 | TAS2R30 | -0,749132196 | 0,594961329 | 0,11363239 | 0,538655668 | taste 2 receptor member 30 |
| ENSG00000256188 | TAS2R30 | -0,749132196 | 0,594961329 | 0,11363239 | 0,538655668 | taste 2 receptor member 30 |
| ENSG00000164048 | ZNF589 | -0,749185367 | 0,594939401 | 0,00495667 | 0,068387168 | zinc finger protein 589 |
| ENSG00000265972 | TXNIP | -0,749264773 | 0,594906657 | 0,01593596 | 0,160613099 | thioredoxin interacting protein |
| ENSG00000115257 | PCSK4 | -0,749819847 | 0,594677812 | 0,05906416 | 0,377469722 | proprotein convertase subtilisin/kexin type 4 |
| ENSG00000288400 | EP400P1 | -0,74994614 | 0,594625756 | 0,12004775 | 0,553128623 | EP400 pseudogene 1 |
| ENSG00000253363 | AC090809,1 | -0,750056229 | 0,594580383 | 0,32084737 | 0,824366234 | novel transcript |
| ENSG00000123454 | DBH | -0,750457477 | 0,594415039 | 0,17974347 | 0,668431057 | dopamine beta-hydroxylase |
| ENSG00000254238 | AC100782,1 | -0,750703434 | 0,594313709 | 0,2518578 | 0,764103086 | novel transcript |
| ENSG00000133863 | TEX15 | -0,750800311 | 0,594273803 | 0,13995005 | 0,596260721 | testis expressed 15, meiosis and synapsis associated |
| ENSG00000181741 | FDX1P1 | -0,75081719 | 0,59426685 | 0,39489384 | 0,871888817 | ferredoxin 1 pseudogene 1 |
| ENSG00000286621 | AC064843,1 | -0,750832823 | 0,59426041 | 0,04817036 | 0,334380043 | novel transcript, antisense to DAPL1 |
| ENSG00000243256 | RPL30P14 | -0,75092596 | 0,594222048 | 0,49188816 | 0,916995341 | ribosomal protein L30 pseudogene 14 |
| ENSG00000276564 | AC130650,2 | -0,750950024 | 0,594212136 | 0,23067558 | 0,739234644 | novel transcript |
| ENSG00000105612 | DNASE2 | -0,751017225 | 0,594184458 | 0,05258882 | 0,352155381 | deoxyribonuclease 2, lysosomal |
| ENSG00000254685 | FPGT | -0,751025945 | 0,594180867 | 0,19533848 | 0,693577833 | fucose-1-phosphate guanylyltransferase |
| ENSG00000261159 | AC112484,3 | -0,751081526 | 0,594157976 | 0,12016051 | 0,553269269 | novel transcript |
| ENSG00000253734 | LINC01289 | -0,751512148 | 0,593980655 | 0,14170625 | 0,599581043 | long intergenic non-protein coding RNA 1289 |
| ENSG00000276968 | AL158196,1 | -0,751591615 | 0,593947939 | 0,45083234 | 0,902062594 | novel transcript, antisense to NUDT15 |
| ENSG00000270011 | ZNF559-ZNF17 | -0,751718556 | 0,59389568 | 0,02805067 | 0,23672966 | ZNF559-ZNF177 readthrough |
| ENSG00000157778 | PSMG3 | -0,751802355 | 0,593861185 | 0,01817739 | 0,17534645 | proteasome assembly chaperone 3 |
| ENSG00000215859 | PDZK1P1 | -0,751858856 | 0,593837927 | 0,12920311 | 0,574432025 | PDZ domain containing 1 pseudogene 1 |
| ENSG00000287016 | AC010996,1 | -0,752148323 | 0,59371879 | 0,15371463 | 0,622713345 | novel transcript |
| ENSG00000131484 | AC091132,1 | -0,752161444 | 0,59371339 | 0,27477232 | 0,786383635 | novel transcript |
| ENSG00000204305 | AGER | -0,752399375 | 0,593615482 | 0,04061352 | 0,301085663 | advanced glycosylation end-product specific receptor |

| | | | | | | |
|---|---|---|---|---|---|---|
| ENSG00000224717 | LEMD1-DT | -0,752402071 | 0,593614373 | 0,02936962 | 0,243557543 | LEMD1 divergent transcript |
| ENSG00000196345 | ZKSCAN7 | -0,752471683 | 0,593585731 | 0,03181048 | 0,257287766 | zinc finger with KRAB and SCAN domains 7 |
| ENSG00000233644 | ARHGEF7-IT1 | -0,752528736 | 0,593562258 | 0,27894489 | 0,789808253 | ARHGEF7 intronic transcript 1 |
| ENSG00000225411 | CR786580,1 | -0,752665225 | 0,593506105 | 0,24988123 | 0,761802706 | novel transcript |
| ENSG00000283667 | AC009802,1 | -0,752719485 | 0,593483783 | 0,13293023 | 0,582037931 | novel transcript, antisense to SERPINB8 |
| ENSG00000065675 | PRKCQ | -0,752775339 | 0,593460807 | 0,13594073 | 0,587841041 | protein kinase C theta |
| ENSG00000237065 | NANOGP4 | -0,752786473 | 0,593456227 | 0,27341126 | 0,785672647 | Nanog homeobox pseudogene 4 |
| ENSG00000273542 | H4C12 | -0,752818633 | 0,593442998 | 0,0458002 | 0,324471967 | H4 clustered histone 12 |
| ENSG00000111644 | ACRBP | -0,753041932 | 0,593351153 | 0,12554936 | 0,566089307 | acrosin binding protein |
| ENSG00000254865 | AC100763,1 | -0,753166754 | 0,593299818 | 0,42214849 | 0,889776204 | novel transcript |
| ENSG00000286828 | AC008883,3 | -0,753279678 | 0,593253381 | 0,00960213 | 0,111515951 | novel transcript |
| ENSG00000131721 | RHOXF2 | -0,753467154 | 0,593176293 | 0,27616665 | 0,788074712 | Rhox homeobox family member 2 |
| ENSG00000235121 | AL645504,1 | -0,753761711 | 0,593055196 | 0,05337588 | 0,355733025 | novel transcript |
| ENSG00000232466 | H3P31 | -0,753820593 | 0,593030992 | 0,34780083 | 0,842330556 | H3 histone pseudogene 31 |
| ENSG00000233877 | AL606517,1 | -0,753896475 | 0,592999801 | 0,45670871 | 0,90366076 | SIPL protein (SIPL) pseudogene |
| ENSG00000186715 | MST1L | -0,754003313 | 0,592955888 | 0,12752788 | 0,570260369 | macrophage stimulating 1 like |
| ENSG00000256771 | ZNF253 | -0,754247894 | 0,592855372 | 0,10269148 | 0,510258481 | zinc finger protein 253 |
| ENSG00000256274 | TAS2R64P | -0,754363929 | 0,592807691 | 0,10522375 | 0,517691472 | taste 2 receptor member 64 pseudogene |
| ENSG00000263083 | TAS2R64P | -0,754363929 | 0,592807691 | 0,10522375 | 0,517691472 | taste 2 receptor member 64 pseudogene |
| ENSG00000233396 | LINC01719 | -0,754549515 | 0,592731438 | 0,01302922 | 0,139032628 | long intergenic non-protein coding RNA 1719 |
| ENSG00000205089 | CCNI2 | -0,754911744 | 0,592582635 | 0,22910839 | 0,737173617 | cyclin I family member 2 |
| ENSG00000261762 | AC027228,2 | -0,754962913 | 0,592561618 | 0,11587846 | 0,543804306 | novel transcript, antisense to CHRNA5 |
| ENSG00000214694 | ARHGEF33 | -0,755302297 | 0,592422239 | 0,06148102 | 0,386127478 | Rho guanine nucleotide exchange factor 33 |
| ENSG00000271550 | BNIP3P11 | -0,755375897 | 0,592392017 | 0,22488126 | 0,733203418 | BCL2 interacting protein 3 pseudogene 11 |
| ENSG00000273113 | AC133528,1 | -0,755382314 | 0,592389382 | 0,35181489 | 0,844406182 | novel transcript |
| ENSG00000240553 | AL031428,1 | -0,755626093 | 0,592289291 | 0,21971595 | 0,727931767 | novel transcript |
| ENSG00000237972 | TUBG1P | -0,755686281 | 0,592264582 | 0,31563095 | 0,821023886 | tubulin gamma 1 pseudogene |
| ENSG00000248334 | WHAMMP2 | -0,755808099 | 0,592214574 | 0,32234204 | 0,824955371 | WAS protein homolog associated with actin, golgi membranes and microtubules pseudogene 2 |
| ENSG00000183196 | CHST6 | -0,755822637 | 0,592208607 | 0,05316721 | 0,354644712 | carbohydrate sulfotransferase 6 |
| ENSG00000178636 | AC092656,1 | -0,755996434 | 0,59213727 | 0,28878004 | 0,800014136 | mitochondrial ribosomal protein L42 (MRPL42) pseudogene |
| ENSG00000180592 | SKIDA1 | -0,756052592 | 0,592114221 | 0,03618698 | 0,279138825 | SKI/DACH domain containing 1 |

| | | | | | | |
|---|---|---|---|---|---|---|
| ENSG00000253263 | AP003354,1 | -0,756276994 | 0,592022128 | 0,44132201 | 0,898885877 | novel transcript |
| ENSG00000138685 | FGF2 | -0,756372957 | 0,59198275 | 0,08763775 | 0,471714073 | fibroblast growth factor 2 |
| ENSG00000085552 | IGSF9 | -0,756400813 | 0,59197132 | 0,00762619 | 0,094425865 | immunoglobulin superfamily member 9 |
| ENSG00000257359 | AC073863,1 | -0,756773213 | 0,591818535 | 0,51715756 | 0,925746338 | novel transcript |
| ENSG00000123307 | NEUROD4 | -0,756870914 | 0,591778458 | 0,11496189 | 0,541371975 | neuronal differentiation 4 |
| ENSG00000238140 | AC104170,1 | -0,756999502 | 0,591725715 | 0,52218082 | 0,927617924 | novel transcript |
| ENSG00000286315 | AC087175,1 | -0,757083833 | 0,591691127 | 0,31103124 | 0,817751312 | novel transcript |
| ENSG00000285756 | BX890604,1 | -0,757498225 | 0,591521198 | 0,00063377 | 0,012750424 | novel transcript |
| ENSG00000233998 | SETP5 | -0,757565818 | 0,591493485 | 0,23717659 | 0,746624292 | SET pseudogene 5 |
| ENSG00000276523 | AC025287,3 | -0,757674889 | 0,591448768 | 0,13353024 | 0,583791005 | novel transcript, antisense to GABARAPL2 |
| ENSG00000099204 | ABLIM1 | -0,757765545 | 0,591411604 | 0,04930044 | 0,338604781 | actin binding LIM protein 1 |
| ENSG00000102445 | RUBCNL | -0,757906788 | 0,591353706 | 0,1260073 | 0,566777174 | rubicon like autophagy enhancer |
| ENSG00000273177 | AC092954,2 | -0,758011575 | 0,591310756 | 0,62601153 | 0,952793934 | novel transcript |
| ENSG00000147465 | STAR | -0,758582064 | 0,591076979 | 0,15219426 | 0,620418906 | steroidogenic acute regulatory protein |
| ENSG00000287646 | AC021701,1 | -0,758641398 | 0,591052669 | 0,2998197 | 0,809881047 | novel transcript, antisense to RTTN |
| ENSG00000047597 | XK | -0,758650222 | 0,591049055 | 0,18664942 | 0,680025828 | X-linked Kx blood group |
| ENSG00000261764 | KRT18P18 | -0,758671192 | 0,591040464 | 0,35008915 | 0,843789477 | keratin 18 pseudogene 18 |
| ENSG00000236709 | DAPK1-IT1 | -0,758690476 | 0,591032563 | 0,40479664 | 0,878176155 | DAPK1 intronic transcript 1 |
| ENSG00000204904 | LINC01545 | -0,758742261 | 0,591011349 | 0,48516047 | 0,913690171 | long intergenic non-protein coding RNA 1545 |
| ENSG00000167642 | SPINT2 | -0,758817893 | 0,590980367 | 0,0259728 | 0,224747726 | serine peptidase inhibitor, Kunitz type 2 |
| ENSG00000261439 | DKFZP434H168 | -0,759247824 | 0,590804277 | 0,15602677 | 0,626944439 | uncharacterized LOC26077 |
| ENSG00000273853 | AC023158,2 | -0,759805647 | 0,590575885 | 0,63285525 | 0,954396141 | novel transcript, sense intronic to SYT10 |
| ENSG00000228128 | DDAH2 | -0,7598802 | 0,590545367 | 0,01145074 | 0,126436253 | dimethylarginine dimethylaminohydrolase 2 |
| ENSG00000225635 | DDAH2 | -0,7598802 | 0,590545367 | 0,01145074 | 0,126436253 | dimethylarginine dimethylaminohydrolase 2 |
| ENSG00000233076 | DDAH2 | -0,7598802 | 0,590545367 | 0,01145074 | 0,126436253 | dimethylarginine dimethylaminohydrolase 2 |
| ENSG00000273245 | AC092653,1 | -0,760144146 | 0,590437335 | 0,35957529 | 0,849982153 | novel transcript |
| ENSG00000274591 | AC025031,3 | -0,760231289 | 0,590401672 | 0,63458545 | 0,955149453 | novel transcript, sense intronic to SLC38A1 |
| ENSG00000135472 | FAIM2 | -0,760255128 | 0,590391916 | 0,02209497 | 0,200225258 | Fas apoptotic inhibitory molecule 2 |
| ENSG00000260452 | TPRKBP2 | -0,760509579 | 0,590287796 | 0,52397999 | 0,928352631 | TP53RK binding protein pseudogene 2 |
| ENSG00000232684 | ATP11A-AS1 | -0,760837774 | 0,590153529 | 0,24098872 | 0,751473296 | ATP11A antisense RNA 1 |
| ENSG00000226498 | RPSAP21 | -0,761026001 | 0,590076537 | 0,32450327 | 0,82667087 | ribosomal protein SA pseudogene 21 |
| ENSG00000260855 | AL591848,3 | -0,761036306 | 0,590072322 | 0,00720547 | 0,090434273 | novel transcript |
| ENSG00000224733 | AL034349,1 | -0,761069982 | 0,590058549 | 0,35236857 | 0,844912682 | novel transcript, antisense to PTPRK |

| | | | | | | |
|---|---|---|---|---|---|---|
| ENSG00000281853 | AL034349,2 | -0,761069982 | 0,590058549 | 0,35236857 | 0,844912682 | novel transcript, antisense to PTPRK |
| ENSG00000258900 | HNRNPCP1 | -0,761113343 | 0,590040814 | 0,16286207 | 0,640548464 | heterogeneous nuclear ribonucleoprotein C pseudogene 1 |
| ENSG00000198520 | ARMH1 | -0,761118954 | 0,59003852 | 0,03999096 | 0,298466754 | armadillo like helical domain containing 1 |
| ENSG00000101197 | BIRC7 | -0,761202875 | 0,590004198 | 0,64774763 | 0,956899963 | baculoviral IAP repeat containing 7 |
| ENSG00000182255 | KCNA4 | -0,761239778 | 0,589989107 | 0,35016796 | 0,843789477 | potassium voltage-gated channel subfamily A member 4 |
| ENSG00000137571 | SLCO5A1 | -0,761401596 | 0,589922935 | 0,02538203 | 0,221166029 | solute carrier organic anion transporter family member 5A1 |
| ENSG00000225972 | MTND1P23 | -0,761517831 | 0,589875408 | 0,26052361 | 0,772692192 | MT-ND1 pseudogene 23 |
| ENSG00000255358 | NDUFAF2P2 | -0,76166083 | 0,589816943 | 0,40729862 | 0,880314995 | NDUFAF2 pseudogene 2 |
| ENSG00000137842 | TMEM62 | -0,761699368 | 0,589801188 | 0,01743126 | 0,170676808 | transmembrane protein 62 |
| ENSG00000258504 | AL157871,1 | -0,761736084 | 0,589786177 | 0,1275352 | 0,570260369 | novel transcript |
| ENSG00000285247 | AC270301,6 | -0,761756388 | 0,589777877 | 0,591603 | 0,947416795 | novel transcript |
| ENSG00000232237 | ASCL5 | -0,761859778 | 0,589735612 | 0,34975982 | 0,843534837 | achaete-scute family bHLH transcription factor 5 |
| ENSG00000262829 | AC242308,2 | -0,76242262 | 0,589505582 | 0,04559208 | 0,323511284 | novel taste receptor, type 2 family pseudogene |
| ENSG00000258212 | ZNF75BP | -0,76263599 | 0,589418403 | 0,27276799 | 0,78561201 | zinc finger protein 75B, pseudogene |
| ENSG00000079335 | CDC14A | -0,763052558 | 0,589248237 | 0,08717202 | 0,470780115 | cell division cycle 14A |
| ENSG00000253180 | AC104986,1 | -0,763437653 | 0,589090971 | 0,24642096 | 0,758198269 | ubiquitin-conjugating enzyme E2 variant 1 (UBE2V1) pseudogene |
| ENSG00000288581 | AL137139,3 | -0,763520717 | 0,589057055 | 0,30548969 | 0,814326218 | novel transcript, antisense to CLYBL |
| ENSG00000200966 | RN7SKP87 | -0,763563292 | 0,589039672 | 0,52462762 | 0,928616707 | RN7SK pseudogene 87 |
| ENSG00000278611 | ZNF426-DT | -0,76359924 | 0,589024995 | 0,12396785 | 0,562842145 | ZNF426 divergent transcript |
| ENSG00000254737 | OR10G4 | -0,763685163 | 0,588989915 | 0,19585039 | 0,69375328 | olfactory receptor family 10 subfamily G member 4 |
| ENSG00000151715 | TMEM45B | -0,763745153 | 0,588965424 | 0,10833906 | 0,524645663 | transmembrane protein 45B |
| ENSG00000228008 | AC105935,1 | -0,764029739 | 0,588849256 | 0,61815302 | 0,952793934 | novel transcript, antisense to MON1A |
| ENSG00000267127 | AC090360,1 | -0,764080548 | 0,588828519 | 0,05975261 | 0,380314665 | novel protein |
| ENSG00000096150 | RPS18 | -0,764086689 | 0,588826012 | 0,11489554 | 0,541330721 | ribosomal protein S18 |
| ENSG00000232470 | AL136368,1 | -0,764102843 | 0,588819419 | 0,47683061 | 0,91143003 | novel transcript |
| ENSG00000161021 | MAML1 | -0,764185354 | 0,588785744 | 0,20936483 | 0,713678247 | mastermind like transcriptional coactivator 1 |
| ENSG00000130222 | GADD45G | -0,764337933 | 0,588723478 | 0,06209853 | 0,389024096 | growth arrest and DNA damage inducible gamma |
| ENSG00000176160 | HSF5 | -0,765088547 | 0,588417253 | 0,50726016 | 0,921980206 | heat shock transcription factor 5 |
| ENSG00000165323 | FAT3 | -0,765111349 | 0,588407953 | 0,02554414 | 0,222207024 | FAT atypical cadherin 3 |

| | | | | | | |
|---|---|---|---|---|---|---|
| ENSG00000078589 | P2RY10 | -0,765618448 | 0,588201167 | 0,09963133 | 0,500760119 | P2Y receptor family member 10 |
| ENSG00000288063 | AC099050,1 | -0,765711984 | 0,588163033 | 0,41455324 | 0,885579089 | novel transcript, antisense to RAD54L2and TEX264 |
| ENSG00000273342 | AP000553,3 | -0,765763513 | 0,588142026 | 0,55805451 | 0,939078529 | novel transcript |
| ENSG00000236580 | AC073626,2 | -0,765800586 | 0,588126912 | 0,63797006 | 0,956018518 | ribosomal protein L36 (RPL36) pseudogene |
| ENSG00000272372 | AC009102,2 | -0,765831805 | 0,588114186 | 0,31251618 | 0,818574551 | novel transcript, antisense to GNAO1 |
| ENSG00000258924 | AC002094,1 | -0,765838308 | 0,588111535 | 0,4894036 | 0,915936198 | novel transcript readthrough between TMEM199 and SARM1 |
| ENSG00000201600 | RN7SKP124 | -0,765849188 | 0,5881071 | 0,2068776 | 0,710515357 | RN7SK pseudogene 124 |
| ENSG00000287312 | AC127526,4 | -0,766111237 | 0,588000287 | 0,11612746 | 0,544096306 | novel transcript |
| ENSG00000231918 | AC007402,1 | -0,766756014 | 0,587737553 | 0,64371803 | 0,956633898 | novel transcript |
| ENSG00000168852 | TPTE2P5 | -0,766945828 | 0,58766023 | 0,27876069 | 0,789775666 | transmembrane phosphoinositide 3-phosphatase and tensin homolog 2 pseudogene 5 |
| ENSG00000255647 | AC093510,1 | -0,767010007 | 0,587634088 | 0,30786579 | 0,816424917 | novel transcript |
| ENSG00000101203 | COL20A1 | -0,767070393 | 0,587609492 | 0,37756534 | 0,862943393 | collagen type XX alpha 1 chain |
| ENSG00000129993 | CBFA2T3 | -0,767307444 | 0,587512949 | 0,03459665 | 0,271313423 | CBFA2/RUNX1 partner transcriptional co-repressor 3 |
| ENSG00000063127 | SLC6A16 | -0,76747687 | 0,587443958 | 0,00342817 | 0,051902339 | solute carrier family 6 member 16 |
| ENSG00000256827 | AC068790,1 | -0,767662784 | 0,587368261 | 0,41001085 | 0,882457617 | spindle and kinetochore associated complex subunit 2 (SKA2) |
| ENSG00000133083 | DCLK1 | -0,76825913 | 0,587125519 | 0,03623116 | 0,279341761 | doublecortin like kinase 1 |
| ENSG00000286626 | AC090469,1 | -0,768281157 | 0,587116555 | 0,23044328 | 0,739158571 | novel transcript, antisense to CAPRIN1 |
| ENSG00000254420 | AP003086,1 | -0,768327507 | 0,587097693 | 0,5418394 | 0,933491609 | novel transcript |
| ENSG00000224464 | PGAM1P6 | -0,768596183 | 0,586988367 | 0,28752507 | 0,798896538 | phosphoglycerate mutase 1 pseudogene 6 |
| ENSG00000270728 | AL035413,1 | -0,76884371 | 0,586887665 | 0,12699766 | 0,569312214 | synaptonemal complex protein 3 (SYCP3) pseudogene |
| ENSG00000228839 | PIK3IP1-DT | -0,769124706 | 0,586773366 | 0,35545958 | 0,846840543 | PIK3IP1 divergent transcript |
| ENSG00000267565 | AC011477,2 | -0,769198542 | 0,586743336 | 0,13513142 | 0,586038109 | novel transcript, antisense to ZNF253 |
| ENSG00000278916 | CEP83-DT | -0,769213325 | 0,586737324 | 0,09326247 | 0,48515108 | CEP83 divergent transcript |
| ENSG00000229492 | FO681548,1 | -0,769368982 | 0,586674023 | 0,19705466 | 0,695927373 | Telomeric repeat-binding factor 2-interacting protein 1 (TERF2IP) pseudogene |
| ENSG00000226545 | AL357552,1 | -0,769421634 | 0,586652612 | 0,45748757 | 0,903831777 | ribosomal protein L27 (RPL27) pseudogene |
| ENSG00000216809 | AL589993,1 | -0,769444122 | 0,586643468 | 0,15638713 | 0,627418665 | DEAD/H (Asp-Glu-Ala-Asp/His) box polypeptide 24 (DDX24) pseudogene |

| | | | | | | |
|---|---|---|---|---|---|---|
| ENSG00000205482 | SPDYE18 | -0,769516821 | 0,586613907 | 0,47682518 | 0,91143003 | speedy/RINGO cell cycle regulator family member E18 |
| ENSG00000134709 | HOOK1 | -0,76972462 | 0,58652942 | 0,31150394 | 0,817751312 | hook microtubule tethering protein 1 |
| ENSG00000238105 | GOLGA2P5 | -0,769869398 | 0,586470563 | 0,05639225 | 0,366903759 | GOLGA2 pseudogene 5 |
| ENSG00000246523 | FZD4-DT | -0,770115963 | 0,58637034 | 0,22034522 | 0,728572782 | FZD4 divergent transcript |
| ENSG00000287833 | AL157997,1 | -0,770157633 | 0,586353405 | 0,00577971 | 0,076696396 | novel transcript |
| ENSG00000255345 | AP002957,1 | -0,770420291 | 0,586246662 | 0,22165624 | 0,729254784 | novel transcript |
| ENSG00000180626 | ZNF594 | -0,770523644 | 0,586204666 | 0,02017474 | 0,187832988 | zinc finger protein 594 |
| ENSG00000165795 | NDRG2 | -0,770791411 | 0,586095875 | 0,00483522 | 0,067096455 | NDRG family member 2 |
| ENSG00000231165 | TRBV26OR9-2 | -0,770889213 | 0,586056144 | 0,38391422 | 0,868113415 | T cell receptor beta variable 26/OR9-2 (pseudogene) |
| ENSG00000264384 | RN7SL431P | -0,770920534 | 0,586043421 | 0,64191877 | 0,956529637 | RNA, 7SL, cytoplasmic 431, pseudogene |
| ENSG00000240919 | AC022034,2 | -0,770920964 | 0,586043246 | 0,50915868 | 0,922427526 | ribosomal protein L21 (RPL21) pseudogene |
| ENSG00000174669 | SLC29A2 | -0,771435479 | 0,58583428 | 0,05981924 | 0,380382656 | solute carrier family 29 member 2 |
| ENSG00000254931 | MTATP6P15 | -0,771509982 | 0,585804028 | 0,4455067 | 0,901068127 | MT-ATP6 pseudogene 15 |
| ENSG00000205022 | PABPN1L | -0,771525291 | 0,585797812 | 0,46988483 | 0,908378389 | PABPN1 like, cytoplasmic |
| ENSG00000227314 | MSH5 | -0,771724433 | 0,585716957 | 0,19374063 | 0,690594523 | mutS homolog 5 |
| ENSG00000082482 | KCNK2 | -0,77173248 | 0,58571369 | 0,0670002 | 0,406736732 | potassium two pore domain channel subfamily K member 2 |
| ENSG00000269895 | AP000654,1 | -0,771778688 | 0,58569493 | 0,19368121 | 0,690594523 | novel transcript |
| ENSG00000287165 | AL078621,2 | -0,771961896 | 0,585620558 | 0,5240363 | 0,928399782 | novel transcript |
| ENSG00000286625 | AC113370,1 | -0,772162915 | 0,585538966 | 0,53656938 | 0,932583111 | novel transcript |
| ENSG00000232682 | AL592430,1 | -0,772416928 | 0,585435879 | 0,34418659 | 0,840938551 | novel transcript, antisense to ANK3 |
| ENSG00000286877 | AC090510,4 | -0,77254287 | 0,585384775 | 0,53435732 | 0,931584152 | novel transcript, antisense to TTBK2 |
| ENSG00000243101 | RPS3P7 | -0,772875476 | 0,585249833 | 0,39774112 | 0,874263835 | ribosomal protein S3 pseudogene 7 |
| ENSG00000174844 | DNAH12 | -0,773141164 | 0,585142063 | 0,24026254 | 0,750234283 | dynein axonemal heavy chain 12 |
| ENSG00000261441 | AC124068,2 | -0,773166773 | 0,585131676 | 0,15284439 | 0,62133458 | novel transcript, antisense to POLG |
| ENSG00000166165 | CKB | -0,773186141 | 0,585123821 | 0,00142994 | 0,025271661 | creatine kinase B |
| ENSG00000179766 | ATP8B5P | -0,773375393 | 0,58504707 | 0,57885881 | 0,943828882 | ATPase phospholipid transporting 8B5, pseudogene |
| ENSG00000267506 | AC021683,2 | -0,773614256 | 0,584950213 | 0,00586248 | 0,07756432 | novel transcript |
| ENSG00000260152 | AC113208,2 | -0,773988238 | 0,5847986 | 0,57309244 | 0,942475308 | stromal cell-derived factor 2 (SDF2) pseudogene |
| ENSG00000224844 | AC107079,1 | -0,774012014 | 0,584788962 | 0,51148826 | 0,922893865 | novel transcript |
| ENSG00000225279 | AL121987,1 | -0,774708107 | 0,584506872 | 0,41495865 | 0,885668156 | novel transcript |
| ENSG00000264924 | AC090772,1 | -0,774756816 | 0,584487138 | 0,37413844 | 0,861369706 | novel transcript, antisense to TTC39C |

| | | | | | | |
|---|---|---|---|---|---|---|
| ENSG00000199436 | SNORD9 | -0,775106236 | 0,584345593 | 0,50861289 | 0,922427526 | small nucleolar RNA, C/D box 9 |
| ENSG00000273525 | SNORA54 | -0,775939337 | 0,584008253 | 0,05104461 | 0,345820358 | small nucleolar RNA, H/ACA box 54 |
| ENSG00000207008 | SNORA54 | -0,775939337 | 0,584008253 | 0,05104461 | 0,345820358 | small nucleolar RNA, H/ACA box 54 |
| ENSG00000149289 | ZC3H12C | -0,776084784 | 0,583949379 | 0,08179647 | 0,454749316 | zinc finger CCCH-type containing 12C |
| ENSG00000230293 | MSH5 | -0,77611009 | 0,583939136 | 0,11817706 | 0,549318284 | mutS homolog 5 |
| ENSG00000171017 | LRRC8E | -0,776395835 | 0,58382349 | 0,31155072 | 0,817751312 | leucine rich repeat containing 8 VRAC subunit E |
| ENSG00000284483 | TMEM256-PLS( | -0,776528112 | 0,583769964 | 0,59913281 | 0,948894897 | TMEM256-PLSCR3 readthrough (NMD candidate) |
| ENSG00000262481 | TMEM256-PLS( | -0,776528112 | 0,583769964 | 0,59913281 | 0,948894897 | TMEM256-PLSCR3 readthrough (NMD candidate) |
| ENSG00000237001 | WASF3-AS1 | -0,776592927 | 0,583743738 | 0,13244543 | 0,581009552 | WASF3 antisense RNA 1 |
| ENSG00000125285 | SOX21 | -0,776657729 | 0,583717518 | 0,01523197 | 0,155828638 | SRY-box transcription factor 21 |
| ENSG00000268170 | AC073342,1 | -0,776849415 | 0,583639967 | 0,31856562 | 0,823392614 | novel transcript, antisense to TAS2R40 |
| ENSG00000262599 | DENND11 | -0,777161858 | 0,583513582 | 0,02235693 | 0,202005507 | DENN domain containing 11 |
| ENSG00000199683 | RN7SKP185 | -0,777198438 | 0,583498787 | 0,37751601 | 0,862893739 | RN7SK pseudogene 185 |
| ENSG00000254781 | GVINP2 | -0,777328275 | 0,583446277 | 0,44625271 | 0,901348735 | GTPase, very large interferon inducible pseudogene 2 |
| ENSG00000124343 | XG | -0,777514227 | 0,58337108 | 0,46208621 | 0,906277266 | Xg glycoprotein (Xg blood group) |
| ENSG00000278119 | GOLGA8J | -0,777552784 | 0,583355489 | 0,05059059 | 0,34398963 | golgin A8 family member J |
| ENSG00000270894 | AC015849,4 | -0,777666485 | 0,583309516 | 0,35287185 | 0,845031687 | novel transcript, sense overlapping TAF15 |
| ENSG00000282083 | AC015849,7 | -0,777666485 | 0,583309516 | 0,35287185 | 0,845031687 | novel transcript, sense overlapping TAF15 |
| ENSG00000285831 | AC027329,1 | -0,778207417 | 0,583090848 | 0,10583181 | 0,519178686 | novel transcript |
| ENSG00000274666 | AL133370,2 | -0,778479198 | 0,582981013 | 0,58311061 | 0,944174844 | zinc finger protein 585B (ZNF585B) pseudogene |
| ENSG00000267649 | AC010327,5 | -0,778532766 | 0,582959367 | 0,18882355 | 0,683182437 | novel transcript |
| ENSG00000235621 | LINC00494 | -0,779135947 | 0,582715686 | 0,34866204 | 0,842757646 | long intergenic non-protein coding RNA 494 |
| ENSG00000236896 | AL354726,1 | -0,779301816 | 0,582648695 | 0,37230237 | 0,860223901 | novel transcript |
| ENSG00000258982 | AL133523,1 | -0,779510992 | 0,582564223 | 0,23962365 | 0,749431667 | novel transcript, antisense to YY1 |
| ENSG00000159842 | ABR | -0,779666607 | 0,582501388 | 0,10352572 | 0,513180277 | ABR activator of RhoGEF and GTPase |
| ENSG00000231419 | LINC00689 | -0,779767919 | 0,582460484 | 0,49089556 | 0,916995341 | long intergenic non-protein coding RNA 689 |
| ENSG00000271996 | AC019080,4 | -0,779771866 | 0,582458891 | 0,48463663 | 0,913690171 | novel transcript, antisense to NFE2L2 |
| ENSG00000285996 | AC006116,9 | -0,780010638 | 0,582362499 | 0,01745333 | 0,17078586 | novel transcript, antisense to ZNF582 |
| ENSG00000196208 | GREB1 | -0,78030518 | 0,582243616 | 0,00246895 | 0,039685942 | growth regulating estrogen receptor binding 1 |
| ENSG00000236194 | AC099811,1 | -0,780457685 | 0,582182071 | 0,06705771 | 0,406871024 | novel transcript, sense intronic to STAT5B |
| ENSG00000185681 | MORN5 | -0,780499103 | 0,582165357 | 0,6119913 | 0,952189515 | MORN repeat containing 5 |
| ENSG00000285516 | NAALAD2 | -0,780853669 | 0,582022298 | 0,13922112 | 0,594862544 | N-acetylated alpha-linked acidic dipeptidase 2 |

| | | | | | | |
|---|---|---|---|---|---|---|
| ENSG00000077616 | NAALAD2 | -0,780853669 | 0,582022298 | 0,13922112 | 0,594862544 | N-acetylated alpha-linked acidic dipeptidase 2 |
| ENSG00000277199 | LINC01632 | -0,780877941 | 0,582012506 | 0,30759901 | 0,816380056 | long intergenic non-protein coding RNA 1632 |
| ENSG00000100027 | YPEL1 | -0,780960831 | 0,581979068 | 0,00549901 | 0,074231934 | yippee like 1 |
| ENSG00000231615 | AL645568,1 | -0,781007643 | 0,581960184 | 0,50937494 | 0,922427526 | novel pseudogene |
| ENSG00000239412 | RPL21P71 | -0,781009697 | 0,581959356 | 0,31097909 | 0,817751312 | ribosomal protein L21 pseudogene 71 |
| ENSG00000146856 | AGBL3 | -0,781069149 | 0,581935374 | 0,09860712 | 0,498520872 | ATP/GTP binding protein like 3 |
| ENSG00000226864 | ATE1-AS1 | -0,781070218 | 0,581934943 | 0,0807717 | 0,450856364 | ATE1 antisense RNA 1 |
| ENSG00000235609 | AF127577,4 | -0,781115609 | 0,581916634 | 0,25692102 | 0,769163921 | novel transcript |
| ENSG00000218676 | BRD7P4 | -0,781189456 | 0,581886848 | 0,25890459 | 0,771850203 | bromodomain containing 7 pseudogene 4 |
| ENSG00000226010 | AL355852,1 | -0,781379154 | 0,581810342 | 0,39543719 | 0,872042617 | zinc finger protein pseudogene |
| ENSG00000224713 | AC025165,1 | -0,781396548 | 0,581803327 | 0,05511728 | 0,362604555 | novel transcript, antisense to ARHGEF25 |
| ENSG00000175426 | PCSK1 | -0,781458191 | 0,581778469 | 0,0266573 | 0,228583033 | proprotein convertase subtilisin/kexin type 1 |
| ENSG00000225206 | MIR137HG | -0,781686221 | 0,581686521 | 0,01185542 | 0,129642445 | MIR137 host gene |
| ENSG00000164112 | TMEM155 | -0,781754 | 0,581659194 | 0,37202504 | 0,859937929 | transmembrane protein 155 |
| ENSG00000260337 | AC091544,4 | -0,781943927 | 0,581582625 | 0,49287197 | 0,917090312 | novel transcript |
| ENSG00000178162 | FAR2P2 | -0,78206897 | 0,581532219 | 0,02262668 | 0,203744369 | fatty acyl-CoA reductase 2 pseudogene 2 |
| ENSG00000197978 | GOLGA6L9 | -0,782074684 | 0,581529916 | 0,00366345 | 0,054792423 | golgin A6 family like 9 |
| ENSG00000233693 | PBX1-AS1 | -0,782088422 | 0,581524378 | 0,09475145 | 0,488991285 | PBX1 antisense RNA 1 |
| ENSG00000249318 | AC010468,1 | -0,782237823 | 0,581464161 | 0,58280049 | 0,944174844 | novel transcript |
| ENSG00000053328 | METTL24 | -0,78226339 | 0,581453856 | 0,16379109 | 0,642705016 | methyltransferase like 24 |
| ENSG00000229686 | SNORD56 | -0,782275965 | 0,581448788 | 0,47808461 | 0,911892945 | small nucleolar RNA, C/D box 56 |
| ENSG00000275805 | AC090206,1 | -0,782372909 | 0,581409718 | 0,19146097 | 0,687237165 | novel transcript, sense intronic to KCTD1 |
| ENSG00000214283 | RAD51AP1P1 | -0,782392578 | 0,581401792 | 0,27926892 | 0,790141487 | RAD51AP1 pseudogene 1 |
| ENSG00000228554 | AC004837,2 | -0,782455815 | 0,581376308 | 0,35096441 | 0,844035671 | novel transcript |
| ENSG00000250267 | AC021237,1 | -0,782768923 | 0,581250145 | 0,21839766 | 0,72607235 | novel transcript |
| ENSG00000281650 | AC243922,3 | -0,783025582 | 0,581146749 | 0,00152433 | 0,026623912 | glucuronidase, beta (GUSB) pseudogene |
| ENSG00000232578 | AC093311,1 | -0,783215459 | 0,581070267 | 0,33844654 | 0,837263082 | FAST kinase domains 1 (FASTKD1) pseudogene |
| ENSG00000220694 | AL096711,1 | -0,783399296 | 0,580996229 | 0,49330602 | 0,91725463 | ribosomal protein L17 (RPL17) pseudogene |
| ENSG00000198739 | LRRTM3 | -0,783954922 | 0,580772512 | 0,18985985 | 0,685029239 | leucine rich repeat transmembrane neuronal 3 |
| ENSG00000283380 | AL807742,1 | -0,783962442 | 0,580769485 | 0,31157413 | 0,817751312 | novel transcript |
| ENSG00000251187 | AC010261,2 | -0,783974735 | 0,580764536 | 0,52375368 | 0,928267286 | novel transcript |
| ENSG00000265552 | AC104961,1 | -0,784005272 | 0,580752244 | 0,39678701 | 0,873609704 | novel transcript |
| ENSG00000162722 | TRIM58 | -0,784019314 | 0,580746591 | 0,02915573 | 0,24236233 | tripartite motif containing 58 |

| | | | | | | |
|---|---|---|---|---|---|---|
| ENSG00000263327 | TAPT1-AS1 | -0,78420396 | 0,580672268 | 0,01959364 | 0,184067141 | TAPT1 antisense RNA 1 (head to head) |
| ENSG00000246792 | AC106038,1 | -0,784949922 | 0,580372103 | 0,22545378 | 0,734457344 | novel transcript |
| ENSG00000264549 | SNORD95 | -0,784972502 | 0,580363019 | 0,39053253 | 0,870057277 | small nucleolar RNA, C/D box 95 |
| ENSG00000134775 | FHOD3 | -0,785027847 | 0,580340755 | 0,00231214 | 0,037688666 | formin homology 2 domain containing 3 |
| ENSG00000221164 | SNORA11F | -0,785176665 | 0,580280894 | 0,42050765 | 0,888563911 | small nucleolar RNA, H/ACA box 11F |
| ENSG00000206481 | MDC1 | -0,785450861 | 0,580170618 | 0,17551632 | 0,661199837 | mediator of DNA damage checkpoint 1 |
| ENSG00000258445 | MAD2L1P1 | -0,785465978 | 0,580164539 | 0,38001797 | 0,865511891 | mitotic arrest deficient 2 like 1 pseudogene 1 |
| ENSG00000188386 | PPP3R2 | -0,785543008 | 0,580133563 | 0,49961141 | 0,920190594 | protein phosphatase 3 regulatory subunit B, beta |
| ENSG00000236078 | LINC01447 | -0,785583267 | 0,580117374 | 0,12258148 | 0,560036813 | long intergenic non-protein coding RNA 1447 |
| ENSG00000064547 | LPAR2 | -0,785610489 | 0,580106428 | 0,00470746 | 0,065938206 | lysophosphatidic acid receptor 2 |
| ENSG00000206147 | RPL23AP57 | -0,785793345 | 0,580032907 | 0,54221111 | 0,933491609 | ribosomal protein L23a pseudogene 57 |
| ENSG00000231444 | DDR1-DT | -0,785807574 | 0,580027186 | 0,56103263 | 0,940489231 | DDR1 divergent transcript |
| ENSG00000258910 | LINC01956 | -0,786128541 | 0,579898157 | 0,28284252 | 0,794000953 | long intergenic non-protein coding RNA 1956 |
| ENSG00000065609 | SNAP91 | -0,786220576 | 0,579861165 | 0,07168532 | 0,421847873 | synaptosome associated protein 91 |
| ENSG00000139239 | RPL14P1 | -0,786439387 | 0,579773225 | 0,42380602 | 0,890949676 | ribosomal protein L14 pseudogene 1 |
| ENSG00000232450 | AL133517,1 | -0,786569068 | 0,579721112 | 0,05587156 | 0,364974593 | ribosomal protein S2 (RPS2) pseudogene |
| ENSG00000242797 | GLYCTK-AS1 | -0,787087797 | 0,579512708 | 0,05292812 | 0,353577924 | GLYCTK antisense RNA 1 |
| ENSG00000273791 | AC007204,1 | -0,787283797 | 0,579433982 | 0,17673596 | 0,662651108 | BCL2/adenovirus E1B 19kDa interacting protein 3 (BNIP3) pseudogene |
| ENSG00000235903 | CPB2-AS1 | -0,787307099 | 0,579424624 | 0,00925117 | 0,108284762 | CPB2 antisense RNA 1 |
| ENSG00000137821 | LRRC49 | -0,788132899 | 0,579093056 | 0,04649047 | 0,327581282 | leucine rich repeat containing 49 |
| ENSG00000278077 | AL591926,6 | -0,788166978 | 0,579079376 | 0,40622184 | 0,879205242 | zinc finger protein 829 (ZNF829) pseudogene |
| ENSG00000266980 | AC087289,1 | -0,788347665 | 0,579006856 | 0,29267063 | 0,803129526 | novel transcript, antisense to UNK |
| ENSG00000286411 | AC002458,1 | -0,788432192 | 0,578972933 | 0,04282821 | 0,311753441 | novel transcript, antisense to CDK14 |
| ENSG00000284922 | AP000812,4 | -0,788520757 | 0,578937391 | 0,01351748 | 0,142389001 | leucine rich transmembrane and O-methyltransferase domain containing |
| ENSG00000135773 | CAPN9 | -0,788564803 | 0,578919716 | 0,08882532 | 0,475558385 | calpain 9 |
| ENSG00000160172 | FAM86C2P | -0,788806234 | 0,578822844 | 0,00421645 | 0,061085754 | family with sequence similarity 86 member C2, pseudogene |
| ENSG00000235672 | AC090286,3 | -0,789308336 | 0,578621431 | 0,35333758 | 0,84514549 | RAN binding protein 1 (RANBP1) pseudogene |
| ENSG00000271387 | AL445228,2 | -0,78938852 | 0,578589273 | 0,1221172 | 0,558903822 | novel transcript |
| ENSG00000265625 | AC104564,5 | -0,78993013 | 0,578372102 | 0,23881591 | 0,748055366 | novel transcript, sense intronic to SSH2 |
| ENSG00000274449 | AC008463,2 | -0,79017189 | 0,578275189 | 0,01126994 | 0,125033138 | novel transcript |

| | | | | | | |
|---|---|---|---|---|---|---|
| ENSG00000076344 | RGS11 | -0,790331986 | 0,578211021 | 0,04101147 | 0,303331616 | regulator of G protein signaling 11 |
| ENSG00000116396 | KCNC4 | -0,790482972 | 0,578150512 | 0,0599887 | 0,380964201 | potassium voltage-gated channel subfamily C member 4 |
| ENSG00000104435 | STMN2 | -0,790523658 | 0,578134207 | 0,18453336 | 0,67674498 | stathmin 2 |
| ENSG00000271825 | AC019080,3 | -0,790532524 | 0,578130654 | 0,50630342 | 0,921780819 | novel transcript, antisense to NFE2L2 |
| ENSG00000168589 | DYNLRB2 | -0,790684831 | 0,578069624 | 0,42792579 | 0,89282532 | dynein light chain roadblock-type 2 |
| ENSG00000269688 | AC008982,2 | -0,790891703 | 0,577986739 | 0,47485634 | 0,91069468 | novel transcript, sense intronic to HNRNPL |
| ENSG00000278383 | AL031673,1 | -0,790989254 | 0,577947658 | 0,26211924 | 0,773902615 | novel transcript, antisense to ZNF337 |
| ENSG00000250687 | NAIPP1 | -0,791021698 | 0,577934661 | 0,07355831 | 0,42825458 | NAIP pseudogene 1 |
| ENSG00000253320 | MAILR | -0,791034431 | 0,57792956 | 0,01155256 | 0,127208092 | macrophage interferon regulatory lncRNA |
| ENSG00000227929 | SRGAP3-AS3 | -0,791114897 | 0,577897328 | 0,12412163 | 0,563164667 | SRGAP3 antisense RNA 3 |
| ENSG00000102104 | RS1 | -0,791211798 | 0,577858513 | 0,36001264 | 0,850365296 | retinoschisin 1 |
| ENSG00000254048 | AC105150,1 | -0,791225386 | 0,577853071 | 0,50457497 | 0,921400408 | novel transcript |
| ENSG00000272588 | AC139887,4 | -0,791342618 | 0,577806117 | 0,25207023 | 0,764267906 | novel transcript, antisense to PCGF3 |
| ENSG00000223963 | THAP12P8 | -0,791364148 | 0,577797494 | 0,41250138 | 0,884523593 | THAP domain containing 12 pseudogene 8 |
| ENSG00000260882 | AC026458,1 | -0,791374606 | 0,577793306 | 0,57580663 | 0,942908987 | novel pseudogene |
| ENSG00000260773 | AC055855,1 | -0,791512635 | 0,577738028 | 0,46699296 | 0,907940653 | novel transcript, antisense to DIS3L |
| ENSG00000054654 | SYNE2 | -0,791581533 | 0,577710438 | 0,01053774 | 0,119169172 | spectrin repeat containing nuclear envelope protein 2 |
| ENSG00000286814 | AC009152,5 | -0,791639923 | 0,577687057 | 0,47661429 | 0,91143003 | novel transcript, antisense to XYLT1 |
| ENSG00000284495 | MIEF2 | -0,791818093 | 0,577615718 | 0,09813009 | 0,497396091 | mitochondrial elongation factor 2 |
| ENSG00000227463 | AL390838,1 | -0,79189347 | 0,57758554 | 0,07292096 | 0,425892188 | novel transcript |
| ENSG00000184616 | SPDYE12P | -0,791974559 | 0,577553077 | 0,28901526 | 0,800166929 | speedy/RINGO cell cycle regulator family member E12, pseudogene |
| ENSG00000265393 | AC084125,4 | -0,792410394 | 0,577378626 | 0,45004277 | 0,902062594 | novel transcript, antisense to RECQL4 |
| ENSG00000266402 | SNHG25 | -0,792504271 | 0,577341057 | 0,11550913 | 0,543019233 | small nucleolar RNA host gene 25 |
| ENSG00000080493 | SLC4A4 | -0,792522708 | 0,577333679 | 0,1178853 | 0,548937448 | solute carrier family 4 member 4 |
| ENSG00000273785 | CYP2D8P | -0,792545258 | 0,577324655 | 0,40735405 | 0,880314995 | ccytochrome P450 family 2 subfamily D member 8, pseudogene |
| ENSG00000286424 | AL157996,2 | -0,792675343 | 0,577272601 | 0,41764711 | 0,886857227 | novel transcript |
| ENSG00000042317 | SPATA7 | -0,792714364 | 0,577256987 | 0,08239534 | 0,456577089 | spermatogenesis associated 7 |
| ENSG00000134317 | GRHL1 | -0,793093229 | 0,577105414 | 0,00365016 | 0,054628342 | grainyhead like transcription factor 1 |
| ENSG00000076351 | SLC46A1 | -0,793204451 | 0,577060925 | 0,01783205 | 0,173406251 | solute carrier family 46 member 1 |

| | | | | | | |
|---|---|---|---|---|---|---|
| ENSG00000237280 | AL136982,3 | -0,793270577 | 0,577034476 | 0,39028214 | 0,870057277 | novel transcript |
| ENSG00000251224 | CNOT10-AS1 | -0,79351626 | 0,576936219 | 0,51540724 | 0,924901769 | CNOT10 antisense RNA 1 |
| ENSG00000272338 | AC067838,1 | -0,793706312 | 0,576860221 | 0,21994013 | 0,728200468 | novel transcript |
| ENSG00000266965 | AC090220,1 | -0,794200783 | 0,576662541 | 0,06936575 | 0,415450351 | novel transcript |
| ENSG00000226403 | AL392089,1 | -0,794539319 | 0,576527241 | 0,35276298 | 0,845031687 | novel transcript |
| ENSG00000152910 | CNTNAP4 | -0,794555369 | 0,576520827 | 0,10145356 | 0,506329296 | contactin associated protein family member 4 |
| ENSG00000234876 | NOTCH4 | -0,794680827 | 0,576470694 | 0,11368801 | 0,538699894 | notch receptor 4 |
| ENSG00000272799 | AC006238,1 | -0,79468399 | 0,57646943 | 0,58237737 | 0,944174844 | novel transcript |
| ENSG00000229311 | BX088651,2 | -0,794824231 | 0,576413396 | 0,21203081 | 0,718583346 | novel transcript |
| ENSG00000177685 | CRACR2B | -0,794836074 | 0,576408664 | 0,22458164 | 0,733203418 | calcium release activated channel regulator 2B |
| ENSG00000185010 | F8 | -0,795088558 | 0,576307797 | 0,05058717 | 0,34398963 | coagulation factor VIII |
| ENSG00000152785 | BMP3 | -0,795443471 | 0,576166038 | 0,09192353 | 0,482073274 | bone morphogenetic protein 3 |
| ENSG00000261436 | AC009081,2 | -0,795504969 | 0,576141478 | 0,41607517 | 0,885986046 | novel transcript |
| ENSG00000113396 | SLC27A6 | -0,79555834 | 0,576120165 | 0,23718753 | 0,746624292 | solute carrier family 27 member 6 |
| ENSG00000277769 | RBM22P4 | -0,795639126 | 0,576087905 | 0,23710135 | 0,746624292 | RNA binding motif protein 22 pseudogene 4 |
| ENSG00000213529 | AL359073,1 | -0,795671423 | 0,576075008 | 0,50678866 | 0,921824694 | ribosomal protein L32 (RPL32) pseudogene |
| ENSG00000131242 | RAB11FIP4 | -0,795899431 | 0,575983971 | 0,04474816 | 0,31940789 | RAB11 family interacting protein 4 |
| ENSG00000169075 | BRD7P3 | -0,796032494 | 0,575930849 | 0,14877105 | 0,613515281 | bromodomain containing 7 pseudogene 3 |
| ENSG00000159625 | DRC7 | -0,796179673 | 0,575872098 | 0,26029749 | 0,772692192 | dynein regulatory complex subunit 7 |
| ENSG00000250328 | MGC32805 | -0,796256991 | 0,575841236 | 0,20245749 | 0,702433811 | uncharacterized LOC153163 |
| ENSG00000150672 | DLG2 | -0,796431805 | 0,575771464 | 0,03531734 | 0,274940552 | discs large MAGUK scaffold protein 2 |
| ENSG00000223756 | TSSC2 | -0,796544533 | 0,575726477 | 0,00201177 | 0,033778017 | tumor suppressing subtransferable candidate 2 (pseudogene) |
| ENSG00000233542 | AL391845,2 | -0,79662918 | 0,575692698 | 0,46822799 | 0,907940653 | novel transcript |
| ENSG00000122574 | WIPF3 | -0,796720678 | 0,575656188 | 0,04030353 | 0,299940355 | WAS/WASL interacting protein family member 3 |
| ENSG00000077080 | ACTL6B | -0,796826245 | 0,575614067 | 0,13775146 | 0,592433314 | actin like 6B |
| ENSG00000223343 | AC137630,1 | -0,797436777 | 0,575370526 | 0,26527978 | 0,77668189 | novel transcript |
| ENSG00000154760 | SLFN13 | -0,79761079 | 0,57530113 | 0,06726638 | 0,407719263 | schlafen family member 13 |
| ENSG00000272181 | AC012557,2 | -0,797807878 | 0,575222543 | 0,42455236 | 0,891157406 | novel transcript, antisense to PSMD6 |
| ENSG00000080644 | CHRNA3 | -0,798669161 | 0,57487924 | 0,06287941 | 0,391611013 | cholinergic receptor nicotinic alpha 3 subunit |
| ENSG00000283403 | AC005394,2 | -0,799155629 | 0,574685427 | 0,07015005 | 0,417549035 | novel transcript |
| ENSG00000261102 | ATP5MFP6 | -0,799191025 | 0,574671328 | 0,51575982 | 0,925006591 | ATP synthase membrane subunit f pseudogene 6 |
| ENSG00000201558 | RNVU1-6 | -0,799818052 | 0,574421617 | 0,18413515 | 0,675910428 | RNA, variant U1 small nuclear 6 |

| | | | | | | |
|---|---|---|---|---|---|---|
| ENSG00000182177 | ASB18 | -0,799826933 | 0,574418081 | 0,20155579 | 0,701094807 | ankyrin repeat and SOCS box containing 18 |
| ENSG00000177181 | RIMKLA | -0,79984607 | 0,574410462 | 0,01299281 | 0,138779332 | ribosomal modification protein rimK like family member A |
| ENSG00000259017 | AL133163,3 | -0,800190899 | 0,574273184 | 0,22354009 | 0,731984265 | novel transcript |
| ENSG00000187559 | FOXD4L3 | -0,800256923 | 0,574246903 | 0,64182647 | 0,95651945 | forkhead box D4 like 3 |
| ENSG00000264829 | AC087749,2 | -0,800437601 | 0,574174991 | 0,22778003 | 0,736053199 | novel transcript, antisense to SAP30BP |
| ENSG00000255983 | AC007848,1 | -0,800782861 | 0,574037598 | 0,24508703 | 0,757071585 | novel transcript |
| ENSG00000168828 | OR13J1 | -0,801008667 | 0,573947759 | 0,5520397 | 0,936552095 | olfactory receptor family 13 subfamily J member 1 |
| ENSG00000236549 | RPS27AP7 | -0,801129783 | 0,573899577 | 0,18586015 | 0,678777213 | RPS27A pseudogene 7 |
| ENSG00000287384 | AL096840,2 | -0,801247482 | 0,573852759 | 0,55558185 | 0,937793851 | novel transcript |
| ENSG00000171759 | PAH | -0,801494974 | 0,573754324 | 0,51303151 | 0,923725914 | phenylalanine hydroxylase |
| ENSG00000175894 | TSPEAR | -0,801499095 | 0,573752685 | 0,26966424 | 0,781791089 | thrombospondin type laminin G domain and EAR repeats |
| ENSG00000144485 | HES6 | -0,801569829 | 0,573724555 | 0,03853633 | 0,291225498 | hes family bHLH transcription factor 6 |
| ENSG00000089199 | CHGB | -0,801640175 | 0,573696581 | 0,11224593 | 0,535189241 | chromogranin B |
| ENSG00000186288 | PABPC1L2A | -0,801655445 | 0,573690509 | 0,63185179 | 0,953915351 | poly(A) binding protein cytoplasmic 1 like 2A |
| ENSG00000153157 | SYCP2L | -0,801731962 | 0,573660082 | 0,21833693 | 0,725947652 | synaptonemal complex protein 2 like |
| ENSG00000229299 | AL121845,1 | -0,80197398 | 0,573563856 | 0,32291692 | 0,825051003 | novel transcript |
| ENSG00000285958 | AC073174,1 | -0,801985408 | 0,573559313 | 0,06954722 | 0,416138637 | novel transcript |
| ENSG00000282266 | ATP9B | -0,802102288 | 0,573512848 | 0,54817993 | 0,935404974 | ATPase phospholipid transporting 9B (putative) |
| ENSG00000285130 | AL358113,1 | -0,802114806 | 0,573507872 | 0,09062682 | 0,480392819 | novel protein |
| ENSG00000152034 | MCHR2 | -0,802357258 | 0,573411499 | 0,35240334 | 0,844912682 | melanin concentrating hormone receptor 2 |
| ENSG00000269846 | AL136172,1 | -0,802369666 | 0,573406568 | 0,18120401 | 0,670965888 | novel transcript, antisense to RBL1 |
| ENSG00000256937 | KRT17P8 | -0,802499626 | 0,573354917 | 0,28698094 | 0,798400609 | keratin 17 pseudogene 8 |
| ENSG00000262870 | CYCSP40 | -0,80253291 | 0,573341689 | 0,48371651 | 0,913690171 | CYCS pseudogene 40 |
| ENSG00000273472 | AC096733,2 | -0,802609019 | 0,573311443 | 0,08973158 | 0,47775341 | novel transcript |
| ENSG00000287032 | AC018647,3 | -0,802654335 | 0,573293436 | 0,04577034 | 0,324471967 | novel transcript |
| ENSG00000261405 | AC136944,3 | -0,802735693 | 0,573261107 | 0,47734611 | 0,911540968 | CHK2 checkpoint homolog (S, pombe) (CHEK2) pseudogene |
| ENSG00000288612 | AL133351,4 | -0,80275922 | 0,573251758 | 0,12444005 | 0,563764231 | novel transcript |
| ENSG00000174417 | TRHR | -0,802954325 | 0,573174239 | 0,31812878 | 0,823060685 | thyrotropin releasing hormone receptor |
| ENSG00000189367 | KIAA0408 | -0,803285589 | 0,573042645 | 0,13067898 | 0,577136255 | KIAA0408 |
| ENSG00000227583 | RPS3AP37 | -0,803345569 | 0,573018821 | 0,15926863 | 0,633427594 | RPS3A pseudogene 37 |

| | | | | | | |
|---|---|---|---|---|---|---|
| ENSG00000224494 | HNRNPA3P14 | -0,803369331 | 0,573009383 | 0,40275086 | 0,877341338 | heterogeneous nuclear ribonucleoprotein A3 pseudogene 14 |
| ENSG00000284292 | AC004922,1 | -0,803445925 | 0,572978962 | 0,16692285 | 0,647344249 | novel protein, ARPC1A and ARPC1B readthrough |
| ENSG00000243795 | LINC02044 | -0,803684575 | 0,572884189 | 0,16910023 | 0,651157725 | long intergenic non-protein coding RNA 2044 |
| ENSG00000243431 | RPL5P30 | -0,80369854 | 0,572878643 | 0,12242258 | 0,559564268 | ribosomal protein L5 pseudogene 30 |
| ENSG00000170683 | OR10A3 | -0,803702771 | 0,572876963 | 0,82563503 | 0,983383773 | olfactory receptor family 10 subfamily A member 3 |
| ENSG00000233110 | AC093797,1 | -0,803845789 | 0,572820175 | 0,05807537 | 0,372903749 | novel transcript, antisense to SORBS2 |
| ENSG00000259378 | DCAF13P3 | -0,803932641 | 0,572785691 | 0,09350814 | 0,485956782 | DDB1 and CUL4 associated factor 13 pseudogene 3 |
| ENSG00000281380 | AC244635,2 | -0,803953612 | 0,572777365 | 0,65823443 | 0,957906876 | NLR family, apoptosis inhibitory protein pseudogene |
| ENSG00000179978 | NAIPP2 | -0,803953612 | 0,572777365 | 0,65823443 | 0,957906876 | NAIP pseudogene 2 |
| ENSG00000282124 | AC015849,5 | -0,804042213 | 0,57274219 | 0,14307546 | 0,601880115 | leucine rich repeat containing 37 member A3 (LRRC37A3) pseudogene |
| ENSG00000113205 | PCDHB3 | -0,804393374 | 0,572602798 | 0,00537688 | 0,072847486 | protocadherin beta 3 |
| ENSG00000288103 | AL359538,4 | -0,805000213 | 0,572361996 | 0,45486776 | 0,903424009 | novel transcript |
| ENSG00000164691 | TAGAP | -0,805068363 | 0,572334959 | 0,32571992 | 0,827574777 | T cell activation RhoGTPase activating protein |
| ENSG00000259551 | AC048382,4 | -0,805070078 | 0,572334279 | 0,46876616 | 0,907940653 | novel transcript |
| ENSG00000259694 | AC136698,1 | -0,805070078 | 0,572334279 | 0,46876616 | 0,907940653 | novel transcript |
| ENSG00000249803 | AC112178,1 | -0,80512842 | 0,572311134 | 0,00858518 | 0,102591349 | novel transcript |
| ENSG00000262652 | AC124283,3 | -0,805262377 | 0,572257997 | 0,1476027 | 0,611779236 | novel transcript, sense intronic FOXK2 |
| ENSG00000253138 | LINC00967 | -0,805288457 | 0,572247652 | 0,39964475 | 0,875433675 | long intergenic non-protein coding RNA 967 |
| ENSG00000225731 | PDE9A-AS1 | -0,805316477 | 0,572236538 | 0,2413792 | 0,751940958 | PDE9A antisense RNA 1 |
| ENSG00000288043 | AC006511,6 | -0,80573358 | 0,57207112 | 0,15009943 | 0,615828266 | novel transcript |
| ENSG00000275297 | AL162731,1 | -0,805812003 | 0,572040024 | 0,50545191 | 0,921780819 | novel transcript |
| ENSG00000077522 | ACTN2 | -0,80647856 | 0,57177579 | 0,12006984 | 0,553128623 | actinin alpha 2 |
| ENSG00000136828 | RALGPS1 | -0,806827273 | 0,571637603 | 0,01858642 | 0,177868046 | Ral GEF with PH domain and SH3 binding motif 1 |
| ENSG00000274819 | AC018659,3 | -0,806949821 | 0,571589048 | 0,7193772 | 0,971869926 | |
| ENSG00000256464 | YWHABP2 | -0,807119637 | 0,571521772 | 0,23764599 | 0,746907875 | tyrosine 3-monooxygenase/tryptophan 5-monooxygenase activation protein beta pseudogene 2 |
| ENSG00000219409 | AL590704,1 | -0,807405068 | 0,57140871 | 0,46892312 | 0,907940653 | rhomboid domain containing 1 (RHBDD1) pseudogene |
| ENSG00000257947 | AC025030,2 | -0,807541934 | 0,571354504 | 0,53295248 | 0,931291085 | novel transcript, antisense to TMEM117 |

| | | | | | | |
|---|---|---|---|---|---|---|
| ENSG00000275389 | AC068790,9 | -0,807571695 | 0,571342718 | 0,27761387 | 0,789067728 | novel transcript |
| ENSG00000186838 | SELENOV | -0,807707502 | 0,571288937 | 0,42509605 | 0,891853867 | selenoprotein V |
| ENSG00000106701 | FSD1L | -0,807744932 | 0,571274115 | 0,12828027 | 0,572037431 | fibronectin type III and SPRY domain containing 1 like |
| ENSG00000157927 | RADIL | -0,807793006 | 0,57125508 | 0,03868608 | 0,292074747 | Rap associating with DIL domain |
| ENSG00000251259 | AC004069,1 | -0,80799115 | 0,571176627 | 0,4948946 | 0,917622746 | novel transcript |
| ENSG00000258903 | AL390816,2 | -0,808012527 | 0,571168164 | 0,30110418 | 0,810679941 | novel transcript |
| ENSG00000233183 | AL138889,1 | -0,808195848 | 0,571095591 | 0,00667208 | 0,085630448 | novel transcript |
| ENSG00000278925 | AC005753,1 | -0,808297986 | 0,571055161 | 0,36557457 | 0,85548827 | novel transcript |
| ENSG00000283360 | AL592295,3 | -0,809161343 | 0,570713524 | 0,50705159 | 0,921951873 | novel transcript |
| ENSG00000264853 | AC011933,2 | -0,809397369 | 0,570620163 | 0,03456502 | 0,271177119 | novel transcript, sense intronic to GRB2 |
| ENSG00000187758 | ADH1A | -0,809502608 | 0,57057854 | 0,19870834 | 0,697657684 | alcohol dehydrogenase 1A (class I), alpha polypeptide |
| ENSG00000287348 | AC099535,2 | -0,809660755 | 0,570515997 | 0,1910544 | 0,686751133 | novel transcript, antisense to SLC4A7 |
| ENSG00000266401 | AP002478,1 | -0,809689885 | 0,570504478 | 0,26211875 | 0,773902615 | novel transcript, antisense to DLGAP1 |
| ENSG00000158164 | TMSB15A | -0,810392177 | 0,570226828 | 0,21931908 | 0,727742708 | thymosin beta 15a |
| ENSG00000158292 | GPR153 | -0,810423325 | 0,570214517 | 0,00260914 | 0,041407116 | G protein-coupled receptor 153 |
| ENSG00000151490 | PTPRO | -0,810442149 | 0,570207077 | 0,05219214 | 0,350608424 | protein tyrosine phosphatase receptor type O |
| ENSG00000152670 | DDX4 | -0,810988584 | 0,569991147 | 0,48501979 | 0,913690171 | DEAD-box helicase 4 |
| ENSG00000277039 | Metazoa_SRP | -0,811055169 | 0,56996484 | 0,48018374 | 0,912847708 | Metazoan signal recognition particle RNA |
| ENSG00000167771 | RCOR2 | -0,811281218 | 0,569875542 | 0,01296286 | 0,138601472 | REST corepressor 2 |
| ENSG00000241890 | RPL13P4 | -0,811338956 | 0,569852736 | 0,43926628 | 0,897434993 | ribosomal protein L13 pseudogene 4 |
| ENSG00000135960 | EDAR | -0,811499851 | 0,569789187 | 0,20452224 | 0,705837666 | ectodysplasin A receptor |
| ENSG00000167769 | ACER1 | -0,811560542 | 0,569765218 | 0,41956578 | 0,887991693 | alkaline ceramidase 1 |
| ENSG00000288181 | CTAGE8 | -0,811603638 | 0,569748198 | 0,75936982 | 0,976120207 | CTAGE family member 8 |
| ENSG00000165113 | GKAP1 | -0,811624703 | 0,569739879 | 0,05220324 | 0,350608424 | G kinase anchoring protein 1 |
| ENSG00000102678 | FGF9 | -0,811690441 | 0,569713919 | 0,12007689 | 0,553128623 | fibroblast growth factor 9 |
| ENSG00000224452 | RSL24D1P6 | -0,811896372 | 0,569632604 | 0,53358811 | 0,931381467 | ribosomal L24 domain containing 1 pseudogene 6 |
| ENSG00000230368 | FAM41C | -0,812106608 | 0,5695496 | 0,3363191 | 0,835660292 | family with sequence similarity 41 member C |
| ENSG00000223462 | AL353801,1 | -0,812131052 | 0,56953995 | 0,39691078 | 0,873609704 | novel transcript |
| ENSG00000275619 | GON7 | -0,812536252 | 0,56938001 | 0,65852993 | 0,958135616 | GON7 subunit of KEOPS complex |
| ENSG00000253966 | AC008514,2 | -0,813130643 | 0,569145473 | 0,25594246 | 0,767778267 | tubulin, epsilon 1 (TUBE1) pseudogene |
| ENSG00000280620 | SCAANT1 | -0,813230039 | 0,569106263 | 0,47085567 | 0,908869685 | SCA7/ATXN7 antisense RNA 1 |

| | | | | | | |
|---|---|---|---|---|---|---|
| ENSG00000286864 | AC012076,1 | -0,813282295 | 0,56908565 | 0,03369252 | 0,266741689 | novel transcript, antisense to LRRFIP1 |
| ENSG00000121207 | LRAT | -0,813385846 | 0,569044804 | 0,13698366 | 0,590711323 | lecithin retinol acyltransferase |
| ENSG00000272991 | AF129408,1 | -0,813407843 | 0,569036128 | 0,63742096 | 0,956017934 | novel transcript, antisense to BRWD1 |
| ENSG00000287133 | AC007279,1 | -0,813418985 | 0,569031733 | 0,26847273 | 0,780709773 | novel transcript, antisense to MPP4and TMEM237 |
| ENSG00000007264 | MATK | -0,813524664 | 0,568990053 | 0,55808655 | 0,939078529 | megakaryocyte-associated tyrosine kinase |
| ENSG00000285287 | AL354855,1 | -0,813719581 | 0,568913184 | 0,3611278 | 0,851821025 | novel pseudogene similar to avian transmembrane protein 180 |
| ENSG00000227141 | AL160286,1 | -0,814054817 | 0,568781002 | 0,34859411 | 0,842757646 | centaurin, gamma 2 (CENTG2) pseudogene |
| ENSG00000287434 | AC109133,1 | -0,814386512 | 0,568650247 | 0,33966634 | 0,838482052 | novel transcript, antisense to DEPDC1B |
| ENSG00000233370 | ACTR1AP1 | -0,814614389 | 0,568560434 | 0,32361882 | 0,825794844 | ACTR1A pseudogene 1 |
| ENSG00000145198 | VWA5B2 | -0,814677956 | 0,568535383 | 0,34196171 | 0,840048641 | von Willebrand factor A domain containing 5B2 |
| ENSG00000230116 | RPS3AP11 | -0,814714694 | 0,568520906 | 0,63718441 | 0,955985674 | RPS3A pseudogene 11 |
| ENSG00000286473 | AC133485,7 | -0,814730267 | 0,568514769 | 0,14065327 | 0,597543965 | novel transcript |
| ENSG00000213935 | RPL22P16 | -0,814730942 | 0,568514503 | 0,41141791 | 0,883655718 | ribosomal protein L22 pseudogene 16 |
| ENSG00000229635 | AL137159,1 | -0,814987442 | 0,568413435 | 0,3246261 | 0,826676737 | pseudogene similar to part of F-box and WD-40 domain protein 1B (FBXW1B) |
| ENSG00000215452 | ZNF663P | -0,815015283 | 0,568402465 | 0,17061825 | 0,654022685 | zinc finger protein 663, pseudogene |
| ENSG00000286819 | Z98745,1 | -0,815043096 | 0,568391507 | 0,23437627 | 0,744054861 | novel transcript |
| ENSG00000250027 | AC022272,1 | -0,815160474 | 0,568345265 | 0,22585418 | 0,734959876 | novel transcript |
| ENSG00000242385 | LINC00901 | -0,815297122 | 0,568291435 | 0,5477409 | 0,935404974 | long intergenic non-protein coding RNA 901 |
| ENSG00000249626 | AC024560,2 | -0,815499637 | 0,568211668 | 0,07627366 | 0,43651514 | protein phosphatase 4, regulatory subunit 2 (PPP4R2) pseudogene |
| ENSG00000249768 | HSPE1P10 | -0,815510726 | 0,568207301 | 0,51362184 | 0,92410971 | heat shock protein family E (Hsp10) member 1 pseudogene 10 |
| ENSG00000106829 | TLE4 | -0,81554043 | 0,568195602 | 0,02065059 | 0,190898872 | TLE family member 4, transcriptional corepressor |
| ENSG00000146166 | LGSN | -0,816041543 | 0,567998276 | 0,32669486 | 0,828503246 | lengsin, lens protein with glutamine synthetase domain |
| ENSG00000226854 | AL451105,1 | -0,816252172 | 0,567915357 | 0,12595969 | 0,566777174 | novel transcript |
| ENSG00000288504 | YBX2 | -0,816269079 | 0,567908701 | 0,2535928 | 0,765172364 | Y-box binding protein 2 |
| ENSG00000006047 | YBX2 | -0,816269079 | 0,567908701 | 0,2535928 | 0,765172364 | Y-box binding protein 2 |
| ENSG00000184261 | KCNK12 | -0,816377683 | 0,567865951 | 0,01644695 | 0,163860559 | potassium two pore domain channel subfamily K member 12 |
| ENSG00000287022 | AC079841,2 | -0,816608907 | 0,567774945 | 0,07275247 | 0,425305383 | novel transcript, antisense to STXBP5L |

| | | | | | | |
|---|---|---|---|---|---|---|
| ENSG00000207205 | RNVU1-15 | -0,816935367 | 0,567646481 | 0,15764169 | 0,630235931 | RNA, variant U1 small nuclear 15 |
| ENSG00000005059 | MCUB | -0,816983245 | 0,567627643 | 0,01215576 | 0,131911723 | mitochondrial calcium uniporter dominant negative subunit beta |
| ENSG00000238168 | IFITM3P5 | -0,817208383 | 0,56753907 | 0,28881627 | 0,800014136 | IFITM3 pseudogene 5 |
| ENSG00000237986 | CELF2-AS2 | -0,817433954 | 0,56745034 | 0,05772544 | 0,371372961 | CELF2 antisense RNA 2 |
| ENSG00000236259 | PPIAP64 | -0,817449145 | 0,567444365 | 0,37481376 | 0,861771195 | peptidylprolyl isomerase A pseudogene 64 |
| ENSG00000118976 | OTUD4P1 | -0,817453559 | 0,567442629 | 0,10734173 | 0,522893717 | OTUD4 pseudogene 1 |
| ENSG00000266783 | AP005136,3 | -0,817460153 | 0,567440035 | 0,14407797 | 0,603895134 | novel transcript, sense intronic METTL4 |
| ENSG00000175730 | BAK1P1 | -0,818189767 | 0,567153136 | 0,38102488 | 0,865900094 | BCL2 antagonist/killer 1 pseudogene 1 |
| ENSG00000259385 | AC066612,2 | -0,818673285 | 0,566963087 | 0,43129895 | 0,89530663 | novel transcript, antisense to CTXN2 |
| ENSG00000251476 | MTCO1P31 | -0,818810108 | 0,56690932 | 0,50510602 | 0,921493523 | MT-CO1 pseudogene 31 |
| ENSG00000141506 | PIK3R5 | -0,81909768 | 0,566796329 | 0,28153506 | 0,792608024 | phosphoinositide-3-kinase regulatory subunit 5 |
| ENSG00000254454 | RCC2P6 | -0,819133728 | 0,566782167 | 0,17154017 | 0,655397237 | regulator of chromosome condensation 2 pseudogene 6 |
| ENSG00000233084 | RPL23AP25 | -0,819238796 | 0,566740891 | 0,54748368 | 0,935404974 | ribosomal protein L23a pseudogene 25 |
| ENSG00000197532 | OR6Y1 | -0,819626778 | 0,566588499 | 0,05149736 | 0,347735102 | olfactory receptor family 6 subfamily Y member 1 |
| ENSG00000215840 | AL592295,1 | -0,82012264 | 0,566393793 | 0,41978468 | 0,888327348 | ribosomal protein S2 (RPS2) pseudogene |
| ENSG00000264529 | DNAJB6P8 | -0,820858479 | 0,56610498 | 0,27041802 | 0,782859427 | DNAJB6 pseudogene 8 |
| ENSG00000270955 | AC135731,2 | -0,821000275 | 0,566049343 | 0,38135236 | 0,865900094 | NEDD4 binding protein 2 (N4BP2) pseudogene |
| ENSG00000264895 | AC006141,1 | -0,821303792 | 0,565930269 | 0,0147861 | 0,152464587 | novel transcript, sense intronic to MBTD1 |
| ENSG00000287069 | AC061958,1 | -0,821322404 | 0,565922968 | 0,1635541 | 0,642139315 | novel transcript, antisense to KCNH8 |
| ENSG00000179869 | ABCA13 | -0,82134193 | 0,565915309 | 0,02537602 | 0,221166029 | ATP binding cassette subfamily A member 13 |
| ENSG00000262358 | AC116914,1 | -0,821488796 | 0,565857702 | 0,49136773 | 0,916995341 | Novel protein, antisense to ITGAE |
| ENSG00000163075 | CFAP221 | -0,822022966 | 0,565648227 | 0,02414474 | 0,213444732 | cilia and flagella associated protein 221 |
| ENSG00000222024 | AC004945,1 | -0,822286233 | 0,565545015 | 0,47747932 | 0,911574108 | pseudogene similar to part of transmembrane protein 198 TMEM198 |
| ENSG00000267372 | AC005330,1 | -0,822587461 | 0,565426944 | 0,0895728 | 0,477709449 | novel transcript |
| ENSG00000112394 | SLC16A10 | -0,822658377 | 0,565399151 | 0,09185039 | 0,481984693 | solute carrier family 16 member 10 |
| ENSG00000160345 | C9orf116 | -0,822856287 | 0,565321595 | 0,02756054 | 0,233890522 | chromosome 9 open reading frame 116 |
| ENSG00000287343 | AL022397,1 | -0,822902086 | 0,565303649 | 0,21235405 | 0,719211015 | novel transcript, antisense to SYT14 |
| ENSG00000153802 | TMPRSS11D | -0,82305934 | 0,565242034 | 0,20257057 | 0,702616942 | transmembrane serine protease 11D |
| ENSG00000235347 | Z84478,1 | -0,823284656 | 0,565153763 | 0,10936366 | 0,527645896 | tRNA nucleotidyl transferase, CCA-adding, 1 (TRNT1) pseudogene |

| | | | | | | |
|---|---|---|---|---|---|---|
| ENSG00000112796 | ENPP5 | -0,823397016 | 0,565109749 | 0,11687501 | 0,546022183 | ectonucleotide pyrophosphatase/phosphodiesterase family member 5 |
| ENSG00000273847 | AC139494,2 | -0,823421583 | 0,565100127 | 0,10798228 | 0,523928283 | novel transcript |
| ENSG00000075826 | SEC31B | -0,823712787 | 0,564986074 | 0,00169525 | 0,029169105 | SEC31 homolog B, COPII coat complex component |
| ENSG00000165181 | SHOC1 | -0,823873009 | 0,564923332 | 0,11464366 | 0,541095927 | shortage in chiasmata 1 |
| ENSG00000233647 | NENFP1 | -0,824055262 | 0,56485197 | 0,34951422 | 0,843214191 | neudesin neurotrophic factor pseudogene 1 |
| ENSG00000232640 | AL354892,2 | -0,824106988 | 0,564831719 | 0,22038154 | 0,728572782 | novel transcript |
| ENSG00000281625 | AL354892,5 | -0,824106988 | 0,564831719 | 0,22038154 | 0,728572782 | novel transcript |
| ENSG00000183578 | TNFAIP8L3 | -0,824424401 | 0,564707462 | 0,23126495 | 0,739268552 | TNF alpha induced protein 8 like 3 |
| ENSG00000259700 | AC012379,2 | -0,824841095 | 0,564544381 | 0,56839775 | 0,94163744 | novel transcript, antisense to CEP152 |
| ENSG00000260978 | AC126407,1 | -0,824890402 | 0,564525086 | 0,14897372 | 0,613638264 | MKRN3 antisense RNA 1 |
| ENSG00000282308 | DPRXP3 | -0,824890959 | 0,564524868 | 0,31672474 | 0,821646322 | divergent-paired related homeobox pseudogene 3 |
| ENSG00000108244 | KRT23 | -0,825107579 | 0,564440112 | 0,4281318 | 0,892936134 | keratin 23 |
| ENSG00000232027 | AL671986,1 | -0,825264013 | 0,564378912 | 0,2958711 | 0,80547842 | proteasome (prosome, macropain) 26S subunit, non-ATPase, 7 (Mov34 homolog) (PSMD7) pseudogene |
| ENSG00000243642 | RN7SL526P | -0,825346979 | 0,564346457 | 0,5960452 | 0,948282297 | RNA, 7SL, cytoplasmic 526, pseudogene |
| ENSG00000182722 | SEPHS1P1 | -0,825367972 | 0,564338245 | 0,31081939 | 0,817751312 | selenophosphate synthetase 1 pseudogene 1 |
| ENSG00000287422 | AL442071,2 | -0,825417009 | 0,564319063 | 0,17981302 | 0,668568618 | novel transcript, antisense to MACF1 |
| ENSG00000256690 | AP001160,1 | -0,825466988 | 0,564299514 | 0,29357194 | 0,803461221 | novel transcript |
| ENSG00000234835 | PHBP13 | -0,825765119 | 0,564182914 | 0,40171807 | 0,87705851 | prohibitin pseudogene 13 |
| ENSG00000232795 | SCAND3P1 | -0,82591448 | 0,564124508 | 0,30891902 | 0,816838423 | SCAN domain containing 3 pseudogene 1 |
| ENSG00000237353 | PATE4 | -0,826730293 | 0,563805598 | 0,36642886 | 0,856103393 | prostate and testis expressed 4 |
| ENSG00000227688 | HNRNPA3P2 | -0,827013261 | 0,563695024 | 0,54026419 | 0,933385624 | heterogeneous nuclear ribonucleoprotein A3 pseudogene 2 |
| ENSG00000237977 | EIF4HP2 | -0,827059697 | 0,563676882 | 0,23329423 | 0,742507431 | eukaryotic translation initiation factor 4H pseudogene 2 |
| ENSG00000169752 | NRG4 | -0,827519401 | 0,563497298 | 0,0611372 | 0,384884976 | neuregulin 4 |
| ENSG00000229221 | HNRNPA1P66 | -0,827918949 | 0,563341262 | 0,4534948 | 0,90277373 | heterogeneous nuclear ribonucleoprotein A1 pseudogene 66 |
| ENSG00000280518 | AC244260,1 | -0,827963088 | 0,563324027 | 0,00711457 | 0,089581081 | NLR family, apoptosis inhibitory protein pseudogene |

| | | | | | | |
|---|---|---|---|---|---|---|
| ENSG00000128262 | POM121L9P | -0,828006194 | 0,563307196 | 0,27349269 | 0,785672647 | POM121 transmembrane nucleoporin like 9, pseudogene |
| ENSG00000255062 | AP001318,2 | -0,828306548 | 0,563189933 | 0,31155068 | 0,817751312 | novel transcript, antisense to DCPS and TIRAP |
| ENSG00000272383 | AC006270,2 | -0,828384342 | 0,563159566 | 0,40071262 | 0,876443227 | zinc finger protein 329 (ZNF329) pseudogene |
| ENSG00000276272 | AC024884,2 | -0,828534661 | 0,563100891 | 0,20903805 | 0,71355079 | novel transcript |
| ENSG00000214289 | RPL39P5 | -0,828599762 | 0,563075482 | 0,51634369 | 0,925256849 | ribosomal protein L39 pseudogene 5 |
| ENSG00000227256 | MIS18A-AS1 | -0,828954554 | 0,562937026 | 0,39162787 | 0,870333649 | MIS18A antisense RNA 1 |
| ENSG00000187764 | SEMA4D | -0,82934812 | 0,562783478 | 0,01581444 | 0,159883663 | semaphorin 4D |
| ENSG00000186452 | TMPRSS12 | -0,829541037 | 0,562708228 | 0,64719055 | 0,956899963 | transmembrane serine protease 12 |
| ENSG00000144648 | ACKR2 | -0,829561898 | 0,562700091 | 0,44164281 | 0,899078209 | atypical chemokine receptor 2 |
| ENSG00000165591 | FAAH2 | -0,829685512 | 0,56265188 | 0,20127416 | 0,701094807 | fatty acid amide hydrolase 2 |
| ENSG00000187144 | SPATA21 | -0,829702006 | 0,562645447 | 0,524977 | 0,928645526 | spermatogenesis associated 21 |
| ENSG00000178171 | AMER3 | -0,829719017 | 0,562638813 | 0,07308131 | 0,426510021 | APC membrane recruitment protein 3 |
| ENSG00000139132 | FGD4 | -0,829733428 | 0,562633192 | 0,04781245 | 0,332623171 | FYVE, RhoGEF and PH domain containing 4 |
| ENSG00000235748 | SEPTIN14P12 | -0,829926718 | 0,562557817 | 0,06945922 | 0,415771228 | septin 14 pseudogene 12 |
| ENSG00000285841 | AC005692,3 | -0,829973689 | 0,562539502 | 0,15442582 | 0,623607455 | novel transcript |
| ENSG00000273156 | AC124016,2 | -0,82998423 | 0,562535391 | 0,11896095 | 0,551318589 | novel transcript |
| ENSG00000235008 | AL023775,1 | -0,830024553 | 0,562519669 | 0,16417301 | 0,642888389 | IGF-II mRNA-binding protein 3 (KOC1) pseudogene |
| ENSG00000286504 | AC092745,4 | -0,830364848 | 0,562387001 | 0,04305161 | 0,312797003 | novel transcript |
| ENSG00000112218 | GPR63 | -0,830401039 | 0,562372893 | 0,08862854 | 0,47503559 | G protein-coupled receptor 63 |
| ENSG00000257310 | AL589743,1 | -0,830595303 | 0,562297172 | 0,66575973 | 0,961016016 | novel transcript |
| ENSG00000230495 | AL132765,1 | -0,830737742 | 0,562241659 | 0,35951538 | 0,849982153 | ras-related pseudogene |
| ENSG00000228741 | AL445985,1 | -0,830926182 | 0,562168226 | 0,00817339 | 0,099236047 | novel transcript |
| ENSG00000285277 | AC254813,1 | -0,830937392 | 0,562163858 | 0,49918979 | 0,919933362 | family with sequence similarity 115, member C (FAM115C) pseudogene |
| ENSG00000254352 | AC100854,1 | -0,831046821 | 0,562121218 | 0,61683563 | 0,952793934 | chromosome 4 open reading frame 46 (C4orf46) pseudogene |
| ENSG00000279182 | CR559946,2 | -0,83123023 | 0,562049761 | 0,22540635 | 0,734455885 | novel transcript, antisense to DENND6B |
| ENSG00000174448 | STARD6 | -0,831280351 | 0,562030235 | 0,43070981 | 0,895267028 | StAR related lipid transfer domain containing 6 |
| ENSG00000139200 | PIANP | -0,83136445 | 0,561997473 | 0,00813521 | 0,098882383 | PILR alpha associated neural protein |
| ENSG00000261054 | AC036108,2 | -0,831628224 | 0,561894731 | 0,24300414 | 0,753100852 | novel transcript, antisense to SYNM |
| ENSG00000184566 | AC132216,1 | -0,831644474 | 0,561888402 | 0,4127555 | 0,884576962 | novel transcript, antisense to LMO2 |
| ENSG00000260160 | AC011468,1 | -0,831809067 | 0,561824301 | 0,26150595 | 0,77333732 | novel transcript |

| | | | | | | |
|---|---|---|---|---|---|---|
| ENSG00000132394 | EEFSEC | -0,831879161 | 0,561797005 | 0,26304344 | 0,77453036 | eukaryotic elongation factor, selenocysteine-tRNA specific |
| ENSG00000225420 | AC104134,1 | -0,832165318 | 0,561685584 | 0,34989517 | 0,843677595 | novel transcript |
| ENSG00000272442 | AL353588,1 | -0,832227613 | 0,561661331 | 0,10640745 | 0,520858399 | novel protein, TMEM151B-SPATS1 readtrough |
| ENSG00000258450 | AL139099,2 | -0,832332361 | 0,561620553 | 0,37170185 | 0,859817753 | novel transcript |
| ENSG00000136297 | MMD2 | -0,832353291 | 0,561612406 | 0,4729153 | 0,909532026 | monocyte to macrophage differentiation associated 2 |
| ENSG00000226031 | FGF13-AS1 | -0,832941659 | 0,561383412 | 0,18078551 | 0,670312865 | FGF13 antisense RNA 1 |
| ENSG00000207166 | SNORA68 | -0,833057343 | 0,561338399 | 0,00622519 | 0,0812289 | small nucleolar RNA, H/ACA box 68 |
| ENSG00000236011 | GPANK1 | -0,833416625 | 0,561198623 | 0,03282934 | 0,262500411 | G-patch domain and ankyrin repeats 1 |
| ENSG00000239149 | SNORA59A | -0,833469076 | 0,561178221 | 0,13414295 | 0,584263748 | small nucleolar RNA, H/ACA box 59A |
| ENSG00000258162 | AC069228,1 | -0,833551238 | 0,561146262 | 0,159356 | 0,633636845 | novel transcript, antisense to PPFIA2 |
| ENSG00000254034 | INTS9-AS1 | -0,833602285 | 0,561126408 | 0,08557046 | 0,465670107 | INTS9 antisense RNA 1 |
| ENSG00000223884 | AC068481,1 | -0,833646402 | 0,561109249 | 0,0898751 | 0,478293227 | novel transcript, antisense to RNF144A |
| ENSG00000205976 | AC091951,1 | -0,83367599 | 0,561097741 | 0,63026043 | 0,953103521 | golgi autoantigen, golgin subfamily a, 6-like 2 (GOLGA6L2) pseudogene |
| ENSG00000273691 | AC087284,1 | -0,833852066 | 0,561029265 | 0,06318158 | 0,393174573 | novel transcript, sense intronic to ZNF710 |
| ENSG00000231887 | PRH1 | -0,834070211 | 0,56094444 | 0,04073705 | 0,301587285 | proline rich protein HaeIII subfamily 1 |
| ENSG00000264707 | L3MBTL4-AS1 | -0,834246004 | 0,560876093 | 0,29739109 | 0,806635068 | L3MBTL4 antisense RNA 1 |
| ENSG00000104490 | NCALD | -0,834377657 | 0,560824913 | 0,03192469 | 0,257772047 | neurocalcin delta |
| ENSG00000145002 | FAM86B2 | -0,83449195 | 0,560780485 | 0,25499476 | 0,76642479 | family with sequence similarity 86 member B2 |
| ENSG00000227896 | AL731569,1 | -0,834529812 | 0,560765768 | 0,07457688 | 0,431593284 | novel transcript |
| ENSG00000160716 | CHRNB2 | -0,834751486 | 0,560679611 | 0,00404504 | 0,059268345 | cholinergic receptor nicotinic beta 2 subunit |
| ENSG00000080031 | PTPRH | -0,834939779 | 0,560606439 | 0,24590062 | 0,757752922 | protein tyrosine phosphatase receptor type H |
| ENSG00000276393 | AC105429,2 | -0,835187397 | 0,560510228 | 0,34436647 | 0,84111511 | nuclear receptor subfamily 3, group C, member 1 (glucocorticoid receptor) (NR3C1) pseudogene |
| ENSG00000119698 | PPP4R4 | -0,835283605 | 0,560472851 | 0,34224791 | 0,840048641 | protein phosphatase 4 regulatory subunit 4 |
| ENSG00000246820 | RIC3-DT | -0,835340218 | 0,560450857 | 0,1360627 | 0,587965501 | RIC3 divergent transcript |
| ENSG00000275450 | PTGER4P2-CDK | -0,835404176 | 0,560426012 | 0,19539306 | 0,693577833 | PTGER4P2-CDK2AP2P2 readthrough, transcribed pseudogene |
| ENSG00000112379 | ARFGEF3 | -0,835651274 | 0,560330033 | 0,03558339 | 0,276263058 | ARFGEF family member 3 |
| ENSG00000279068 | AC244517,6 | -0,835673054 | 0,560321574 | 0,14929505 | 0,614476095 | novel transcript |
| ENSG00000255236 | AP002992,1 | -0,835823002 | 0,560263339 | 0,31831076 | 0,823093059 | novel transcript |

| | | | | | | |
|---|---|---|---|---|---|---|
| ENSG00000166278 | C2 | -0,835987791 | 0,560199348 | 0,47443894 | 0,910676852 | complement C2 |
| ENSG00000269058 | CALR3 | -0,835996163 | 0,560196097 | 0,44080653 | 0,898675567 | calreticulin 3 |
| ENSG00000235731 | LINC02250 | -0,836089144 | 0,560159994 | 0,04131883 | 0,304452496 | long intergenic non-protein coding RNA 2250 |
| ENSG00000266893 | AC005394,1 | -0,836564599 | 0,559975418 | 0,46912947 | 0,907940653 | novel transcript |
| ENSG00000149926 | TLCD3B | -0,836605 | 0,559959736 | 0,01191123 | 0,129979951 | TLC domain containing 3B |
| ENSG00000282880 | B3GALNT2 | -0,836938992 | 0,559830118 | 0,0002973 | 0,006706242 | beta-1,3-N-acetylgalactosaminyltransferase 2 |
| ENSG00000249346 | LINC01016 | -0,837015513 | 0,559800425 | 0,21418939 | 0,721518909 | long intergenic non-protein coding RNA 1016 |
| ENSG00000198734 | F5 | -0,837367314 | 0,559663934 | 0,03555963 | 0,276207785 | coagulation factor V |
| ENSG00000186088 | GSAP | -0,83766917 | 0,559546848 | 0,08362036 | 0,460508555 | gamma-secretase activating protein |
| ENSG00000225882 | LINC01456 | -0,837763496 | 0,559510265 | 0,34903631 | 0,843141089 | long intergenic non-protein coding RNA 1456 |
| ENSG00000215093 | EEF1A1P29 | -0,837791236 | 0,559499507 | 0,46869381 | 0,907940653 | eukaryotic translation elongation factor 1 alpha 1 pseudogene 29 |
| ENSG00000119782 | FKBP1B | -0,837862121 | 0,559472017 | 0,00414309 | 0,06035854 | FKBP prolyl isomerase 1B |
| ENSG00000257596 | SCAT2 | -0,83793946 | 0,559442026 | 0,07430365 | 0,43058932 | S-phase cancer associated transcript 2 |
| ENSG00000199377 | RNU5F-1 | -0,837987188 | 0,559423519 | 0,21242455 | 0,719216058 | RNA, U5F small nuclear 1 |
| ENSG00000169862 | CTNND2 | -0,838050302 | 0,559399046 | 0,00503858 | 0,069151088 | catenin delta 2 |
| ENSG00000261868 | MFSD1P1 | -0,83810243 | 0,559378834 | 0,08866018 | 0,475123718 | major facilitator superfamily domain containing 1 pseudogene 1 |
| ENSG00000249345 | LINC02405 | -0,838105486 | 0,559377649 | 0,6690921 | 0,961684145 | long intergenic non-protein coding RNA 2405 |
| ENSG00000271580 | AL583832,1 | -0,838305144 | 0,559300241 | 0,11714033 | 0,547016492 | glycogenin 2 (GYG2) pseudogene |
| ENSG00000241255 | AL136126,1 | -0,838350032 | 0,559282839 | 0,59010678 | 0,947262972 | ribosomal protein S24 (RPS24) pseudogene |
| ENSG00000272217 | AL645940,1 | -0,838369353 | 0,559275349 | 0,48918046 | 0,915902374 | novel transcript |
| ENSG00000173258 | ZNF483 | -0,838391368 | 0,559266815 | 0,01399145 | 0,146346115 | zinc finger protein 483 |
| ENSG00000139116 | KIF21A | -0,838903165 | 0,559068449 | 0,03245037 | 0,26082374 | kinesin family member 21A |
| ENSG00000223974 | BNIP3P42 | -0,839011857 | 0,559026331 | 0,19244654 | 0,689034439 | BCL2 interacting protein 3 pseudogene 42 |
| ENSG00000260318 | COX6CP1 | -0,83938428 | 0,558882041 | 0,36759852 | 0,857206868 | cytochrome c oxidase subunit 6C pseudogene 1 |
| ENSG00000229855 | AC008568,1 | -0,839639164 | 0,55878331 | 0,31120214 | 0,817751312 | novel transcript |
| ENSG00000215023 | AC131097,1 | -0,83979413 | 0,558723292 | 0,38947239 | 0,870057277 | novel transcript |
| ENSG00000281082 | AC131097,4 | -0,83979413 | 0,558723292 | 0,38947239 | 0,870057277 | novel transcript |
| ENSG00000265739 | AC104984,3 | -0,839824178 | 0,558711655 | 0,3030402 | 0,812669425 | novel transcript, antisense to CCDC55 |
| ENSG00000259024 | TVP23C-CDRT4 | -0,84059514 | 0,558413165 | 0,02726854 | 0,232168734 | TVP23C-CDRT4 readthrough |
| ENSG00000267811 | AP001160,2 | -0,840852102 | 0,558313713 | 0,03590018 | 0,277817599 | novel transcript, antisense to TAF6L |
| ENSG00000143297 | FCRL5 | -0,84159858 | 0,558024906 | 0,17728623 | 0,663555205 | Fc receptor like 5 |

| | | | | | | |
|---|---|---|---|---|---|---|
| ENSG00000124256 | ZBP1 | -0,841622366 | 0,558015706 | 0,6568119 | 0,957476526 | Z-DNA binding protein 1 |
| ENSG00000173157 | ADAMTS20 | -0,84204543 | 0,557852094 | 0,009556 | 0,111104017 | ADAM metallopeptidase with thrombospondin type 1 motif 20 |
| ENSG00000248477 | NAIPP3 | -0,842218133 | 0,557785318 | 0,11821307 | 0,549318284 | NAIP pseudogene 3 |
| ENSG00000159904 | ZNF890P | -0,842697403 | 0,557600051 | 0,61561285 | 0,952793934 | zinc finger protein 890, pseudogene |
| ENSG00000259584 | AC021755,2 | -0,842701601 | 0,557598428 | 0,47229463 | 0,909179229 | novel transcript |
| ENSG00000283122 | HYMAI | -0,842743139 | 0,557582374 | 0,09195413 | 0,482073274 | hydatidiform mole associated and imprinted |
| ENSG00000048052 | HDAC9 | -0,84276809 | 0,557572731 | 0,0357651 | 0,277115226 | histone deacetylase 9 |
| ENSG00000225774 | SIRPAP1 | -0,843148257 | 0,557425823 | 0,25338609 | 0,765172364 | signal regulatory protein alpha pseudogene 1 |
| ENSG00000136928 | GABBR2 | -0,843166169 | 0,557418902 | 0,06390152 | 0,395842478 | gamma-aminobutyric acid type B receptor subunit 2 |
| ENSG00000141744 | PNMT | -0,843364005 | 0,557342469 | 0,27788814 | 0,789162936 | phenylethanolamine N-methyltransferase |
| ENSG00000207165 | SNORA70 | -0,843473506 | 0,557300168 | 0,00108423 | 0,020056204 | small nucleolar RNA, H/ACA box 70 |
| ENSG00000260455 | NBAT1 | -0,843613395 | 0,557246133 | 0,06375332 | 0,395519632 | neuroblastoma associated transcript 1 |
| ENSG00000287166 | AC104561,4 | -0,843671509 | 0,557223687 | 0,17144463 | 0,65533907 | novel transcript |
| ENSG00000117394 | SLC2A1 | -0,84416871 | 0,557031682 | 0,02741632 | 0,232982843 | solute carrier family 2 member 1 |
| ENSG00000213328 | AC104117,1 | -0,844280367 | 0,556988572 | 0,27253125 | 0,785492261 | DCN1, defective in cullin neddylation 1, domain containing 3 (S, cerevisiae) (DCUN1D3) pseudogene |
| ENSG00000171817 | ZNF540 | -0,844602214 | 0,556864329 | 0,0266718 | 0,228644721 | zinc finger protein 540 |
| ENSG00000235518 | AC011196,1 | -0,844790494 | 0,55679166 | 0,60573302 | 0,950927937 | novel transcript |
| ENSG00000255367 | AC127526,2 | -0,845034927 | 0,556697331 | 0,18891928 | 0,683291575 | novel transcript |
| ENSG00000271133 | AC004130,2 | -0,845048873 | 0,55669195 | 0,29563551 | 0,805426883 | novel transcript, antisense to ITGB8 |
| ENSG00000286939 | U82671,1 | -0,845101714 | 0,556671561 | 0,35557008 | 0,846892875 | novel transcript, antisense to ZNF185 |
| ENSG00000235760 | MSH2-OT1 | -0,845373886 | 0,556566552 | 0,32923754 | 0,831175897 | MSH2 overlapping transcript 1 |
| ENSG00000276180 | H4C9 | -0,845422032 | 0,556547978 | 0,00229595 | 0,037444257 | H4 clustered histone 9 |
| ENSG00000242307 | RPS26P52 | -0,845620913 | 0,556471261 | 0,38393068 | 0,868113415 | ribosomal protein S26 pseudogene 52 |
| ENSG00000253307 | AC011676,1 | -0,846039408 | 0,556309864 | 0,56197446 | 0,940865625 | novel transcript |
| ENSG00000227649 | MTND6P32 | -0,846076194 | 0,55629568 | 0,56835545 | 0,94163744 | MT-ND6 pseudogene 32 |
| ENSG00000226306 | NPY6R | -0,846378491 | 0,556179127 | 0,43335305 | 0,895877116 | neuropeptide Y receptor Y6 (pseudogene) |
| ENSG00000226658 | RPL23AP30 | -0,846437868 | 0,556156237 | 0,49201621 | 0,916995341 | ribosomal protein L23a pseudogene 30 |
| ENSG00000284991 | AC275401,4 | -0,846739675 | 0,556039904 | 0,19540153 | 0,693577833 | ubiquitin specific peptidase 17-like 2 ( USP17L2) pseudogene |
| ENSG00000057294 | PKP2 | -0,847055805 | 0,555918075 | 0,09788717 | 0,497051225 | plakophilin 2 |
| ENSG00000139597 | N4BP2L1 | -0,847429388 | 0,555774139 | 0,03551309 | 0,275914829 | NEDD4 binding protein 2 like 1 |

| | | | | | | |
|---|---|---|---|---|---|---|
| ENSG00000205837 | LINC00487 | -0,847558872 | 0,55572426 | 0,29425255 | 0,804089131 | long intergenic non-protein coding RNA 487 |
| ENSG00000197054 | ZNF763 | -0,848132187 | 0,555503464 | 0,00795894 | 0,097274568 | zinc finger protein 763 |
| ENSG00000228577 | AC010731,1 | -0,848523696 | 0,555352735 | 0,38127338 | 0,865900094 | novel transcript |
| ENSG00000227541 | SFR1P1 | -0,848712571 | 0,555280035 | 0,37742143 | 0,862854083 | SFR1 pseudogene 1 |
| ENSG00000116652 | DLEU2L | -0,849153706 | 0,555110272 | 0,09325302 | 0,48515108 | deleted in lymphocytic leukemia 2 like |
| ENSG00000282810 | MIR210HG | -0,84931752 | 0,555047244 | 0,06781984 | 0,410040516 | MIR210 host gene |
| ENSG00000258807 | AL359237,1 | -0,849464791 | 0,554990588 | 0,27650542 | 0,788265979 | novel transcript |
| ENSG00000198453 | ZNF568 | -0,849531733 | 0,554964836 | 0,00267816 | 0,042245222 | zinc finger protein 568 |
| ENSG00000258525 | AL049830,3 | -0,849721794 | 0,55489173 | 0,13412981 | 0,584263748 | novel transcript, antisense to COCH |
| ENSG00000260024 | MRPS21P7 | -0,849828269 | 0,554850779 | 0,47245635 | 0,90925672 | mitochondrial ribosomal protein S21 pseudogene 7 |
| ENSG00000281617 | AC073135,6 | -0,850310807 | 0,554665229 | 0,26893192 | 0,781256973 | Ankyrin repeat domain 18A (ANKRD18A) pseudogene |
| ENSG00000237941 | KCNQ1DN | -0,850333458 | 0,554656521 | 0,13869557 | 0,593837907 | KCNQ1 downstream neighbor |
| ENSG00000271265 | AL355297,2 | -0,850690218 | 0,554519378 | 0,29528369 | 0,805426883 | novel transcript |
| ENSG00000254806 | SYS1-DBNDD2 | -0,850870316 | 0,554450159 | 0,105417 | 0,518119329 | SYS1-DBNDD2 readthrough (NMD candidate) |
| ENSG00000232149 | FERP1 | -0,851461586 | 0,554222971 | 0,14524028 | 0,606976632 | FER tyrosine kinase pseudogene 1 |
| ENSG00000183098 | GPC6 | -0,851537899 | 0,554193656 | 0,06136711 | 0,38572203 | glypican 6 |
| ENSG00000120328 | PCDHB12 | -0,851691618 | 0,55413461 | 0,00177958 | 0,03041921 | protocadherin beta 12 |
| ENSG00000128604 | IRF5 | -0,851862271 | 0,554069067 | 0,12806585 | 0,571345532 | interferon regulatory factor 5 |
| ENSG00000250007 | AC087457,1 | -0,851919904 | 0,554046933 | 0,16624073 | 0,645874246 | novel transcript, antisense to ACTC1 |
| ENSG00000197635 | DPP4 | -0,852550878 | 0,553804669 | 0,0561321 | 0,366010789 | dipeptidyl peptidase 4 |
| ENSG00000106278 | PTPRZ1 | -0,852689736 | 0,553751369 | 0,03492186 | 0,272812858 | protein tyrosine phosphatase receptor type Z1 |
| ENSG00000187762 | HSPE1P11 | -0,852892433 | 0,553673573 | 0,37400699 | 0,861369706 | heat shock protein family E (Hsp10) member 1 pseudogene 11 |
| ENSG00000232692 | AP001596,1 | -0,852894638 | 0,553672726 | 0,27907239 | 0,790013761 | novel transcript |
| ENSG00000263829 | SINHCAFP1 | -0,8531237 | 0,553584825 | 0,40583192 | 0,878787169 | SINHCAF pseudogene 1 |
| ENSG00000272983 | AL117339,3 | -0,853220188 | 0,553547802 | 0,00175998 | 0,030137218 | novel transcript |
| ENSG00000223724 | RAD17P2 | -0,853481921 | 0,553447387 | 0,21481627 | 0,722164151 | RAD17 pseudogene 2 |
| ENSG00000272663 | AC093635,1 | -0,853488121 | 0,553445008 | 0,1937255 | 0,690594523 | novel transcript |
| ENSG00000240964 | RN7SL751P | -0,853932385 | 0,553274606 | 0,34488882 | 0,841168721 | RNA, 7SL, cytoplasmic 751, pseudogene |
| ENSG00000272489 | AL132656,3 | -0,854086045 | 0,553215681 | 0,31670982 | 0,821646322 | novel transcript |
| ENSG00000178201 | VN1R1 | -0,854093498 | 0,553212823 | 0,01782526 | 0,173394196 | vomeronasal 1 receptor 1 |
| ENSG00000225284 | AC018693,1 | -0,854722685 | 0,552971609 | 0,44172339 | 0,899091141 | novel transcript |

| | | | | | | |
|---|---|---|---|---|---|---|
| ENSG00000236086 | HMGN2P28 | -0,85506406 | 0,552840778 | 0,5131847 | 0,92377973 | high mobility group nucleosomal binding domain 2 pseudogene 28 |
| ENSG00000237149 | ZNF503-AS2 | -0,856078879 | 0,552452036 | 0,05642625 | 0,366903759 | ZNF503 antisense RNA 2 |
| ENSG00000287106 | AC106743,1 | -0,856094364 | 0,552446107 | 0,11226233 | 0,535189241 | novel transcript, antisense to WWOX |
| ENSG00000248677 | LINC02102 | -0,856856643 | 0,552154287 | 0,2824995 | 0,793528967 | long intergenic non-protein coding RNA 2102 |
| ENSG00000286658 | AC005153,1 | -0,857039334 | 0,552084371 | 0,06071234 | 0,38355752 | novel transcript, antisense to GRB10 |
| ENSG00000259940 | AC109449,1 | -0,857064893 | 0,552074591 | 0,14306703 | 0,601880115 | novel transcript, antisense to JMJD5 |
| ENSG00000245384 | CXXC4-AS1 | -0,85734664 | 0,551966785 | 0,14797133 | 0,612582848 | CXXC4 antisense RNA 1 |
| ENSG00000278238 | AL359513,1 | -0,858053781 | 0,551696303 | 0,15727276 | 0,629565097 | novel transcript |
| ENSG00000133640 | LRRIQ1 | -0,858259925 | 0,551617478 | 0,03835644 | 0,290314236 | leucine rich repeats and IQ motif containing 1 |
| ENSG00000268678 | AC005261,2 | -0,858379281 | 0,551571844 | 0,30960311 | 0,816986489 | novel transcript, antisense to ZNF304 |
| ENSG00000078295 | ADCY2 | -0,858394086 | 0,551566184 | 0,00062061 | 0,012533997 | adenylate cyclase 2 |
| ENSG00000231703 | AL354993,1 | -0,858420804 | 0,551555969 | 0,12559594 | 0,566123342 | novel transcript, antisense to TSHZ2 |
| ENSG00000278307 | TAS2R19 | -0,858795376 | 0,551412785 | 0,1020498 | 0,50860594 | taste 2 receptor member 19 |
| ENSG00000179542 | SLITRK4 | -0,858887967 | 0,551377397 | 0,11753466 | 0,548039322 | SLIT and NTRK like family member 4 |
| ENSG00000212607 | SNORA3B | -0,858923677 | 0,55136375 | 0,53474114 | 0,931714104 | small nucleolar RNA, H/ACA box 3B |
| ENSG00000243845 | RN7SL30P | -0,859311955 | 0,551215379 | 0,54279073 | 0,933663748 | RNA, 7SL, cytoplasmic 30, pseudogene |
| ENSG00000259006 | AC092143,2 | -0,859766268 | 0,551041825 | 0,63608847 | 0,955399772 | novel transcript, antisense to MC1R |
| ENSG00000277949 | RTP5 | -0,860184908 | 0,550881948 | 0,46519753 | 0,907804479 | receptor transporter protein 5 (putative) |
| ENSG00000188011 | RTP5 | -0,860184908 | 0,550881948 | 0,46519753 | 0,907804479 | receptor transporter protein 5 (putative) |
| ENSG00000286846 | Z93943,1 | -0,860198741 | 0,550876666 | 0,05809789 | 0,372971773 | novel transcript |
| ENSG00000288187 | CU633906,5 | -0,860219621 | 0,550868693 | 0,02122087 | 0,194445863 | novel transcript |
| ENSG00000152213 | ARL11 | -0,860625163 | 0,550713865 | 0,55465503 | 0,937793851 | ADP ribosylation factor like GTPase 11 |
| ENSG00000273793 | DUSP8 | -0,860757403 | 0,550663388 | 0,04510905 | 0,321177288 | dual specificity phosphatase 8 |
| ENSG00000274328 | AC007551,2 | -0,860853133 | 0,55062685 | 0,35536291 | 0,846840543 | gem (nuclear organelle) associated protein 7 (GEMIN7) pseudogene |
| ENSG00000222328 | RNU2-2P | -0,861012557 | 0,550566007 | 0,00636394 | 0,082420082 | RNA, U2 small nuclear 2, pseudogene |
| ENSG00000239686 | AL158801,1 | -0,861019151 | 0,550563491 | 0,25255621 | 0,76458964 | ribosomal protein L34 (RPL34) pseudogene |
| ENSG00000285780 | AC109129,1 | -0,861119181 | 0,550525318 | 0,27707637 | 0,788874154 | Novel transcript, antisense to EPHA3 |
| ENSG00000272824 | AC245100,6 | -0,861125897 | 0,550522755 | 0,1741231 | 0,659373116 | novel transcript |
| ENSG00000276788 | SNORD26 | -0,86116546 | 0,550507659 | 0,35402923 | 0,845430026 | small nucleolar RNA, C/D box 26 |
| ENSG00000232739 | AL607028,1 | -0,861683689 | 0,550309947 | 0,25662459 | 0,769018737 | novel transcript |
| ENSG00000227684 | CROCCP4 | -0,861706793 | 0,550301134 | 0,49474601 | 0,91758518 | CROCC pseudogene 4 |

| | | | | | | |
|---|---|---|---|---|---|---|
| ENSG00000255403 | HSPD1P13 | -0,861831097 | 0,550253721 | 0,25941689 | 0,772164235 | heat shock protein family D (Hsp60) member 1 pseudogene 13 |
| ENSG00000258590 | NBEAP1 | -0,86184177 | 0,550249651 | 0,14913631 | 0,614065274 | neurobeachin pseudogene 1 |
| ENSG00000237734 | AL163192,1 | -0,861950886 | 0,550208035 | 0,60138774 | 0,949939978 | novel transcript |
| ENSG00000219881 | GAPDHP42 | -0,862168663 | 0,550124987 | 0,31789371 | 0,822802293 | glyceraldehyde 3 phosphate dehydrogenase pseudogene 42 |
| ENSG00000272858 | Z93930,3 | -0,86217075 | 0,550124191 | 0,28139846 | 0,792425997 | novel transcript |
| ENSG00000261131 | AC012186,2 | -0,862236467 | 0,550099132 | 0,50999679 | 0,922427526 | novel transcript |
| ENSG00000166492 | FAM86GP | -0,862283892 | 0,55008105 | 0,43604733 | 0,896526626 | family with sequence similarity 86 member G, pseudogene |
| ENSG00000236867 | Z98885,1 | -0,862356621 | 0,55005332 | 0,20281284 | 0,703198476 | suppressor of cytokine signaling 2 (SOCS2) pseudogene |
| ENSG00000261195 | AC027130,1 | -0,862540316 | 0,549983287 | 0,16001051 | 0,635004887 | novel transcript, antisense to IQCK |
| ENSG00000158445 | KCNB1 | -0,862597116 | 0,549961634 | 0,02655238 | 0,228058653 | potassium voltage-gated channel subfamily B member 1 |
| ENSG00000271830 | AC012213,4 | -0,862631671 | 0,549948462 | 0,32545 | 0,827426913 | novel transcript |
| ENSG00000238160 | LINC02863 | -0,862755163 | 0,549901389 | 0,47473411 | 0,91069468 | long intergenic non-protein coding RNA 2863 |
| ENSG00000176435 | CLEC14A | -0,862806076 | 0,549881984 | 0,22690134 | 0,735383523 | C-type lectin domain containing 14A |
| ENSG00000163064 | EN1 | -0,862947142 | 0,549828219 | 0,04139637 | 0,304951943 | engrailed homeobox 1 |
| ENSG00000154265 | ABCA5 | -0,862981593 | 0,54981509 | 0,06739805 | 0,408279929 | ATP binding cassette subfamily A member 5 |
| ENSG00000286693 | AC105230,1 | -0,863173858 | 0,549741822 | 0,18520806 | 0,677629584 | novel transcript, antisense to UNC5D |
| ENSG00000188404 | SELL | -0,863581112 | 0,549586659 | 0,47560806 | 0,911264385 | selectin L |
| ENSG00000213055 | EEF1B2P7 | -0,86362554 | 0,549569734 | 0,22343196 | 0,731984265 | eukaryotic translation elongation factor 1 beta 2 pseudogene 7 |
| ENSG00000286339 | Z99496,1 | -0,863818803 | 0,549496119 | 0,17745151 | 0,663776507 | novel transcript, antisense to CEP85L |
| ENSG00000243960 | AL390195,1 | -0,86389615 | 0,54946666 | 0,23336355 | 0,742507431 | novel transcript |
| ENSG00000232119 | MCTS1 | -0,864570952 | 0,549209714 | 0,00973937 | 0,112876842 | MCTS1 re-initiation and release factor |
| ENSG00000248794 | AC026436,1 | -0,864576785 | 0,549207494 | 0,34027187 | 0,838979484 | ribosomal protein S12 (RPS12) pseudogene |
| ENSG00000261003 | AL008628,1 | -0,864695116 | 0,549162449 | 0,43805815 | 0,897177404 | novel transcript, overlapping to FAM120B |
| ENSG00000276417 | AC092111,2 | -0,864928733 | 0,54907353 | 0,00445791 | 0,063437482 | novel transcript |
| ENSG00000185862 | EVI2B | -0,864964274 | 0,549060003 | 0,018664 | 0,178225795 | ecotropic viral integration site 2B |
| ENSG00000213149 | CNN2P9 | -0,86510763 | 0,549005448 | 0,41212991 | 0,88438705 | calponin 2 pseudogene 9 |

| | | | | | | |
|---|---|---|---|---|---|---|
| ENSG00000273715 | AC138827,1 | -0,865480649 | 0,548863517 | 0,09018563 | 0,478874121 | NLR family, apoptosis inhibitory protein (NAIP) pseudogene |
| ENSG00000239291 | AC002558,1 | -0,865627743 | 0,548807559 | 0,17469283 | 0,660057513 | ribosomal protein S2 (RPS2) pseudogene |
| ENSG00000197993 | KEL | -0,865793057 | 0,548744676 | 0,5172499 | 0,925746338 | Kell metallo-endopeptidase (Kell blood group) |
| ENSG00000165621 | OXGR1 | -0,866326239 | 0,548541912 | 0,2216366 | 0,729254784 | oxoglutarate receptor 1 |
| ENSG00000271452 | AC005034,5 | -0,86695993 | 0,548301023 | 0,11038301 | 0,530191209 | novel transcript, sense intronic to MRPL19 |
| ENSG00000226348 | VN2R10P | -0,867023488 | 0,548276868 | 0,24875238 | 0,760720365 | vomeronasal 2 receptor 10, pseudogene |
| ENSG00000105792 | CFAP69 | -0,867035415 | 0,548272335 | 0,03605749 | 0,27832763 | cilia and flagella associated protein 69 |
| ENSG00000251579 | AC055733,3 | -0,867354314 | 0,548151156 | 0,36900634 | 0,857932205 | pseudogene similar to part of makorin ring finger protein 1 |
| ENSG00000187017 | ESPN | -0,867580826 | 0,5480651 | 0,21326832 | 0,720652258 | espin |
| ENSG00000137642 | SORL1 | -0,867798849 | 0,547982282 | 0,00965429 | 0,112038477 | sortilin related receptor 1 |
| ENSG00000120322 | PCDHB8 | -0,868074673 | 0,547877525 | 0,02643156 | 0,227383165 | protocadherin beta 8 |
| ENSG00000242600 | MBL1P | -0,868466433 | 0,54772877 | 0,04659824 | 0,327821686 | mannose binding lectin 1, pseudogene |
| ENSG00000110619 | CARS1 | -0,868600248 | 0,547677969 | 0,02211584 | 0,200298239 | cysteinyl-tRNA synthetase 1 |
| ENSG00000170745 | KCNS3 | -0,868631563 | 0,547666081 | 0,05053997 | 0,343953009 | potassium voltage-gated channel modifier subfamily S member 3 |
| ENSG00000287405 | AC017028,2 | -0,868633989 | 0,54766516 | 0,08763537 | 0,471714073 | novel transcript, antisense to HDAC4 |
| ENSG00000229202 | PRR3 | -0,869061028 | 0,547503075 | 0,00492181 | 0,068026349 | proline rich 3 |
| ENSG00000250853 | RNF138P1 | -0,86917509 | 0,54745979 | 0,29616788 | 0,805971225 | ring finger protein 138 pseudogene 1 |
| ENSG00000215788 | TNFRSF25 | -0,869203432 | 0,547449035 | 0,06231238 | 0,389548927 | TNF receptor superfamily member 25 |
| ENSG00000250069 | AC011379,1 | -0,869222872 | 0,547441658 | 0,01663983 | 0,165046811 | novel transcript |
| ENSG00000179031 | NSA2P3 | -0,869407948 | 0,547371434 | 0,61123027 | 0,952047794 | NSA2 pseudogene 3 |
| ENSG00000189212 | DPY19L2P1 | -0,86963568 | 0,547285038 | 0,04517042 | 0,321467761 | DPY19L2 pseudogene 1 |
| ENSG00000176771 | NCKAP5 | -0,869898397 | 0,547185385 | 0,00469711 | 0,065852182 | NCK associated protein 5 |
| ENSG00000274922 | AL139384,1 | -0,870308452 | 0,547029882 | 0,15157501 | 0,61913273 | novel transcript |
| ENSG00000258724 | AC105001,2 | -0,870359991 | 0,54701034 | 0,61349399 | 0,952413269 | novel protein |
| ENSG00000122025 | FLT3 | -0,870553831 | 0,546936849 | 0,26556736 | 0,776824277 | fms related receptor tyrosine kinase 3 |
| ENSG00000218596 | AL162578,1 | -0,870708534 | 0,546878203 | 0,4022353 | 0,87718264 | ribosomal protein S15a (RPS15A) pseudogene |
| ENSG00000180210 | F2 | -0,870722153 | 0,54687304 | 0,50242465 | 0,920585659 | coagulation factor II, thrombin |
| ENSG00000121101 | TEX14 | -0,870904393 | 0,546803964 | 0,14047325 | 0,597347645 | testis expressed 14, intercellular bridge forming factor |
| ENSG00000276794 | AC108108,1 | -0,87095448 | 0,546784981 | 0,00057583 | 0,011774291 | glucuronidase, beta (GUSB) pseudogene |

| | | | | | | |
|---|---|---|---|---|---|---|
| ENSG00000277858 | H2AB2 | -0,871139717 | 0,54671478 | 0,61691013 | 0,952793934 | H2A,B variant histone 2 |
| ENSG00000277745 | H2AB3 | -0,871139717 | 0,54671478 | 0,61691013 | 0,952793934 | H2A,B variant histone 3 |
| ENSG00000008118 | CAMK1G | -0,871140189 | 0,546714601 | 0,01721562 | 0,169201274 | calcium/calmodulin dependent protein kinase IG |
| ENSG00000237711 | MTCO3P11 | -0,871192702 | 0,546694701 | 0,35774134 | 0,848908914 | MT-CO3 pseudogene 11 |
| ENSG00000225726 | AC007000,2 | -0,871337247 | 0,54663993 | 0,14595373 | 0,608900104 | pseudogene similar to part of SWI/SNF related, matrix associated, actin dependent regulator of chromatin, subfamily a, member 1 SMARCA1 |
| ENSG00000204961 | PCDHA9 | -0,871542983 | 0,546561982 | 0,00157615 | 0,027391159 | protocadherin alpha 9 |
| ENSG00000273355 | AP000894,4 | -0,871571872 | 0,546551038 | 0,31510431 | 0,82040483 | novel transcript |
| ENSG00000259399 | TGIF2-RAB5IF | -0,871579879 | 0,546548004 | 0,45303167 | 0,902428411 | TGIF2-RAB5IF readthrough |
| ENSG00000259555 | AC067863,1 | -0,871826769 | 0,546454481 | 0,13394396 | 0,584034405 | novel transcript, antisense to CHRNB4 |
| ENSG00000162711 | NLRP3 | -0,871924066 | 0,546417628 | 0,29543924 | 0,805426883 | NLR family pyrin domain containing 3 |
| ENSG00000187690 | EZHIP | -0,872011889 | 0,546384367 | 0,35358921 | 0,845346751 | EZH inhibitory protein |
| ENSG00000204022 | LIPJ | -0,87207956 | 0,546358739 | 0,37873053 | 0,864112469 | lipase family member J |
| ENSG00000136944 | LMX1B | -0,872104331 | 0,546349358 | 0,03410402 | 0,269045724 | LIM homeobox transcription factor 1 beta |
| ENSG00000223730 | Z83838,1 | -0,87234356 | 0,546258769 | 0,05006364 | 0,342117936 | ribosomal protein L6 (RPL6) pseudogene |
| ENSG00000185046 | ANKS1B | -0,872502345 | 0,546198651 | 0,01753938 | 0,171492516 | ankyrin repeat and sterile alpha motif domain containing 1B |
| ENSG00000284974 | AC270301,4 | -0,872586744 | 0,546166698 | 0,12364485 | 0,56228403 | novel transcript |
| ENSG00000241362 | RPL36AP43 | -0,872610501 | 0,546157705 | 0,37702479 | 0,862818448 | ribosomal protein L36a pseudogene 43 |
| ENSG00000260206 | AC105020,2 | -0,872623852 | 0,546152651 | 0,5558996 | 0,938031417 | novel transcript, antisense to IMP3 |
| ENSG00000229955 | Z98749,1 | -0,872742512 | 0,546107732 | 0,19859908 | 0,697657684 | novel transcript, sense intronic to CSNK1E |
| ENSG00000019582 | CD74 | -0,872864503 | 0,546061556 | 0,02797363 | 0,236352093 | CD74 molecule |
| ENSG00000253223 | AC110998,1 | -0,872997194 | 0,546011335 | 0,430949 | 0,89530663 | PRP3 pre-mRNA processing factor 3 homolog (S, cerevisiae) (PRPF3) pseudogene |
| ENSG00000272906 | AL353708,3 | -0,873167204 | 0,545946996 | 0,34322164 | 0,840790189 | novel transcript, antisense to TOR1AIP1 |
| ENSG00000236924 | AL162411,1 | -0,873305816 | 0,545894544 | 0,22290399 | 0,731402952 | novel transcript |
| ENSG00000235829 | TBCAP2 | -0,873352667 | 0,545876817 | 0,27487339 | 0,786383635 | tubulin folding cofactor A pseudogene 2 |
| ENSG00000284838 | C1QTNF4 | -0,873360526 | 0,545873843 | 0,09870923 | 0,498721099 | C1q and TNF related 4 |
| ENSG00000172247 | C1QTNF4 | -0,873360526 | 0,545873843 | 0,09870923 | 0,498721099 | C1q and TNF related 4 |
| ENSG00000241170 | AP001992,1 | -0,873362287 | 0,545873177 | 0,05121017 | 0,346269002 | ribosomal protein L31 (RPL31) pseudogene |
| ENSG00000235429 | AC083875,1 | -0,87357891 | 0,545791219 | 0,401538 | 0,876898854 | ribosomal protein S15a(RPS15A)pseudogene |
| ENSG00000225133 | MORF4L1P4 | -0,874347299 | 0,545500605 | 0,33947408 | 0,838364485 | mortality factor 4 like 1 pseudogene 4 |

| | | | | | | |
|---|---|---|---|---|---|---|
| ENSG00000267016 | AC111170,1 | -0,874455273 | 0,54545978 | 0,23258649 | 0,741684508 | novel transcript, antisense to SEPT9 |
| ENSG00000234281 | LANCL1-AS1 | -0,874476256 | 0,545451847 | 0,27986701 | 0,790565637 | LANCL1 antisense RNA 1 |
| ENSG00000249685 | AC079921,2 | -0,874506462 | 0,545440427 | 0,38346218 | 0,867918747 | novel transcript |
| ENSG00000233948 | TRIM27 | -0,874518334 | 0,545435938 | 0,17522258 | 0,660735603 | tripartite motif containing 27 |
| ENSG00000282598 | AC138374,2 | -0,874747076 | 0,545349465 | 0,33077333 | 0,832114797 | novel transcript |
| ENSG00000144460 | NYAP2 | -0,874795379 | 0,545331206 | 0,03211376 | 0,258956228 | neuronal tyrosine-phosphorylated phosphoinositide-3-kinase adaptor 2 |
| ENSG00000251596 | HADHAP1 | -0,874873681 | 0,54530161 | 0,14078404 | 0,597646146 | HADHA pseudogene 1 |
| ENSG00000277840 | AC026368,1 | -0,874918037 | 0,545284845 | 0,24810147 | 0,760455336 | novel transcript, sense intronic to FBXW8 |
| ENSG00000281021 | AL078581,4 | -0,875204716 | 0,545176501 | 0,39317843 | 0,871240371 | novel transcript |
| ENSG00000132749 | TESMIN | -0,875212066 | 0,545173724 | 0,13183117 | 0,580175909 | testis expressed metallothionein like protein |
| ENSG00000226439 | AC108161,1 | -0,875292166 | 0,545143456 | 0,54101897 | 0,933385624 | ribosomal protein S12 (RPS12) pseudogene |
| ENSG00000227523 | RPS20P15 | -0,875345288 | 0,545123383 | 0,50444094 | 0,921400408 | ribosomal protein S20 pseudogene 15 |
| ENSG00000197238 | H4C11 | -0,875350116 | 0,545121559 | 0,0168251 | 0,16634097 | H4 clustered histone 11 |
| ENSG00000227122 | PRRT1 | -0,875372711 | 0,545113022 | 0,19740078 | 0,695991044 | proline rich transmembrane protein 1 |
| ENSG00000225141 | PRRT1 | -0,875372711 | 0,545113022 | 0,19740078 | 0,695991044 | proline rich transmembrane protein 1 |
| ENSG00000273320 | AC007032,1 | -0,875387205 | 0,545107545 | 0,51942306 | 0,926800715 | novel transcript, antisense to NAMPT |
| ENSG00000254057 | AC084346,1 | -0,875656531 | 0,545005793 | 0,38569868 | 0,869017264 | novel transcript |
| ENSG00000285893 | AL358777,4 | -0,875663582 | 0,545003129 | 0,46583078 | 0,907940653 | novel transcript, antisense to C6orf52 |
| ENSG00000250324 | MRPL22P1 | -0,875759628 | 0,544966847 | 0,36517317 | 0,85525313 | mitochondrial ribosomal protein L22 pseudogene 1 |
| ENSG00000121104 | FAM117A | -0,875812102 | 0,544947026 | 0,00997192 | 0,114787261 | family with sequence similarity 117 member A |
| ENSG00000288571 | AL442063,1 | -0,875993082 | 0,544878669 | 0,52152208 | 0,927466801 | novel transcript |
| ENSG00000164663 | USP49 | -0,876278605 | 0,544770843 | 0,00603953 | 0,079336024 | ubiquitin specific peptidase 49 |
| ENSG00000235522 | AC010978,1 | -0,876301486 | 0,544762203 | 0,06925922 | 0,415287103 | novel transcript |
| ENSG00000242992 | FTH1P4 | -0,876535374 | 0,544673894 | 0,17837563 | 0,665640458 | ferritin heavy chain 1 pseudogene 4 |
| ENSG00000248434 | AC107463,1 | -0,876685057 | 0,544617386 | 0,10341792 | 0,512889905 | novel transcript |
| ENSG00000273024 | INTS4P2 | -0,877019489 | 0,544491152 | 0,08585355 | 0,46656099 | integrator complex subunit 4 pseudogene 2 |
| ENSG00000118137 | APOA1 | -0,87741393 | 0,544342306 | 0,42100696 | 0,888848798 | apolipoprotein A1 |
| ENSG00000164181 | ELOVL7 | -0,877566462 | 0,544284757 | 0,05109852 | 0,345863615 | ELOVL fatty acid elongase 7 |
| ENSG00000287558 | AL109933,5 | -0,877633743 | 0,544259374 | 0,01132006 | 0,125455597 | novel transcript, antisense to PLG |
| ENSG00000217783 | LDHAL6FP | -0,877955068 | 0,544138168 | 0,39361825 | 0,8715972 | lactate dehydrogenase A like 6F, pseudogene |
| ENSG00000107105 | ELAVL2 | -0,877972299 | 0,544131669 | 0,08793427 | 0,472326951 | ELAV like RNA binding protein 2 |

| | | | | | | |
|---|---|---|---|---|---|---|
| ENSG00000232520 | GCSIR | -0,878216647 | 0,544039517 | 0,273629 | 0,785719894 | GPR55 cis regulatory suppressor of immune response RNA |
| ENSG00000250305 | TRMT9B | -0,878283564 | 0,544014284 | 0,00201719 | 0,033815247 | tRNA methyltransferase 9B (putative) |
| ENSG00000274515 | AC105020,5 | -0,87836718 | 0,543982754 | 0,52843093 | 0,930033144 | novel transcript, antisense to IMP3 |
| ENSG00000228835 | AC012123,1 | -0,878730431 | 0,543845804 | 0,12902534 | 0,573886522 | novel transcript, antisense to KLHL14 |
| ENSG00000177335 | C8orf31 | -0,878739199 | 0,543842499 | 0,00846458 | 0,101559908 | chromosome 8 open reading frame 31 |
| ENSG00000260865 | AC010287,1 | -0,87887307 | 0,543792037 | 0,47564621 | 0,911264385 | chromosome 10 open reading frame 57 (C10orf57) pseudogene |
| ENSG00000285852 | AL353147,1 | -0,878951198 | 0,543762589 | 0,14691269 | 0,610674677 | novel transcript |
| ENSG00000076864 | RAP1GAP | -0,879006748 | 0,543741652 | 0,0054301 | 0,073434973 | RAP1 GTPase activating protein |
| ENSG00000248415 | GAPDHP61 | -0,879081083 | 0,543713636 | 0,11784288 | 0,548903435 | glyceraldehyde 3 phosphate dehydrogenase pseudogene 61 |
| ENSG00000274423 | SEC22B2P | -0,87927316 | 0,543641253 | 0,37272249 | 0,860564851 | SEC22 homolog B2, pseudogene |
| ENSG00000259269 | AC109630,1 | -0,879677479 | 0,543488917 | 0,40125348 | 0,876659568 | novel transcript |
| ENSG00000115353 | TACR1 | -0,879679198 | 0,54348827 | 0,09736166 | 0,495751641 | tachykinin receptor 1 |
| ENSG00000163915 | IGF2BP2-AS1 | -0,879769629 | 0,543454204 | 0,01483644 | 0,152933228 | IGF2BP2 antisense RNA 1 |
| ENSG00000249635 | AC109361,1 | -0,879790602 | 0,543446303 | 0,3075669 | 0,816380056 | novel transcript |
| ENSG00000217897 | HSPE1P8 | -0,879956141 | 0,54338395 | 0,38314298 | 0,867698804 | heat shock protein family E (Hsp10) member 1 pseudogene 8 |
| ENSG00000278062 | ZNF251 | -0,880477124 | 0,54318776 | 0,00383572 | 0,056888064 | zinc finger protein 251 |
| ENSG00000183307 | TMEM121B | -0,880771807 | 0,543076821 | 0,00069856 | 0,013761766 | transmembrane protein 121B |
| ENSG00000224333 | GAPDHP20 | -0,8807911 | 0,543069558 | 0,36753625 | 0,8571768 | glyceraldehyde 3 phosphate dehydrogenase pseudogene 20 |
| ENSG00000163449 | TMEM169 | -0,880807528 | 0,543063374 | 0,02058739 | 0,190540057 | transmembrane protein 169 |
| ENSG00000229820 | CR391992,1 | -0,88105455 | 0,542970397 | 0,29555583 | 0,805426883 | DnaJ (Hsp40) homolog, subfamily C member 8 (DNAJC8) pseudogene |
| ENSG00000254090 | MTND2P32 | -0,881128023 | 0,542942746 | 0,22797698 | 0,736341996 | MT-ND2 pseudogene 32 |
| ENSG00000254459 | AP002812,2 | -0,881145772 | 0,542936066 | 0,33516548 | 0,834949294 | novel transcript, antisense to AAMDC |
| ENSG00000167971 | CASKIN1 | -0,881204357 | 0,542914019 | 0,01144424 | 0,126436253 | CASK interacting protein 1 |
| ENSG00000238121 | LINC00426 | -0,881558703 | 0,542780688 | 0,20154081 | 0,701094807 | long intergenic non-protein coding RNA 426 |
| ENSG00000138670 | RASGEF1B | -0,881641879 | 0,542749396 | 0,0338777 | 0,267747212 | RasGEF domain family member 1B |
| ENSG00000259030 | FPGT-TNNI3K | -0,881769664 | 0,542701325 | 0,07815901 | 0,44219384 | FPGT-TNNI3K readthrough |
| ENSG00000239381 | AC125613,1 | -0,881821411 | 0,54268186 | 0,1766767 | 0,662651108 | novel transcript |

| | | | | | | |
|---|---|---|---|---|---|---|
| ENSG00000229589 | ACVR2B-AS1 | -0,881902983 | 0,542651176 | 0,01255309 | 0,135289937 | ACVR2B antisense RNA 1 |
| ENSG00000230585 | PHBP12 | -0,881919449 | 0,542644983 | 0,11305646 | 0,53713451 | prohibitin pseudogene 12 |
| ENSG00000265511 | AC020558,2 | -0,882312142 | 0,542497298 | 0,2820937 | 0,793203731 | novel transcript |
| ENSG00000165443 | PHYHIPL | -0,88276672 | 0,54232639 | 0,10052384 | 0,503650199 | phytanoyl-CoA 2-hydroxylase interacting protein like |
| ENSG00000234683 | AL138799,2 | -0,882811665 | 0,542309495 | 0,51565615 | 0,924926785 | ribosomal protein S20 (RPS20) pseudogene |
| ENSG00000226015 | CCT8P1 | -0,882909471 | 0,542272731 | 0,26845937 | 0,780709773 | chaperonin containing TCP1 subunit 8 pseudogene 1 |
| ENSG00000237461 | AL359710,1 | -0,884035033 | 0,541849826 | 0,02369342 | 0,210380919 | novel transcript |
| ENSG00000261624 | NDUFB10P2 | -0,884075397 | 0,541834666 | 0,26246029 | 0,774068771 | NADH:ubiquinone oxidoreductase subunit B10 pseudogene 2 |
| ENSG00000257056 | LINC02282 | -0,884626319 | 0,541627795 | 0,1067631 | 0,521619225 | long intergenic non-protein coding RNA 2282 |
| ENSG00000128011 | LRFN1 | -0,88463613 | 0,541624112 | 0,00793493 | 0,097133046 | leucine rich repeat and fibronectin type III domain containing 1 |
| ENSG00000226056 | MTND4P32 | -0,88465795 | 0,54161592 | 0,32659464 | 0,828450693 | MT-ND4 pseudogene 32 |
| ENSG00000281123 | FCGBP | -0,884700464 | 0,541599959 | 0,40891104 | 0,881668607 | Fc fragment of IgG binding protein |
| ENSG00000270863 | DDX55P1 | -0,884858247 | 0,54154073 | 0,44167295 | 0,899078209 | DEAD-box helicase 55 pseudogene 1 |
| ENSG00000110693 | SOX6 | -0,885126191 | 0,541440162 | 0,03314635 | 0,264359088 | SRY-box transcription factor 6 |
| ENSG00000163016 | ALMS1P1 | -0,885188152 | 0,541416908 | 0,04223531 | 0,308497259 | ALMS1 pseudogene 1 |
| ENSG00000225446 | LINC01763 | -0,885237263 | 0,541398478 | 0,36660564 | 0,856103393 | long intergenic non-protein coding RNA 1763 |
| ENSG00000107954 | NEURL1 | -0,885379581 | 0,541345073 | 0,04363425 | 0,315563514 | neuralized E3 ubiquitin protein ligase 1 |
| ENSG00000267934 | AC010300,1 | -0,885545467 | 0,541282831 | 0,05102045 | 0,345820358 | novel transcript, antisense to ZNF91 |
| ENSG00000081248 | CACNA1S | -0,885595537 | 0,541264046 | 0,32643261 | 0,828450693 | calcium voltage-gated channel subunit alpha1 S |
| ENSG00000253521 | HPYR1 | -0,885914859 | 0,541144257 | 0,46072586 | 0,905949965 | Helicobacter pylori responsive 1 |
| ENSG00000200534 | SNORA33 | -0,886012436 | 0,541107658 | 0,49159265 | 0,916995341 | small nucleolar RNA, H/ACA box 33 |
| ENSG00000224905 | AP001347,1 | -0,886540392 | 0,540909675 | 0,0181179 | 0,174934575 | novel transcript, antisense to LIPI |
| ENSG00000223864 | NPM1P19 | -0,886559123 | 0,540902653 | 0,45546225 | 0,903545728 | nucleophosmin 1 pseudogene 19 |
| ENSG00000172575 | RASGRP1 | -0,886733382 | 0,540837322 | 0,15021809 | 0,615923738 | RAS guanyl releasing protein 1 |
| ENSG00000257126 | FOXG1-AS1 | -0,886875885 | 0,540783903 | 0,11403368 | 0,53948382 | FOXG1 antisense RNA 1 |
| ENSG00000196593 | ANKRD20A19P | -0,886929283 | 0,540763888 | 0,07238706 | 0,424120881 | ankyrin repeat domain 20 family member A19, pseudogene |
| ENSG00000259357 | AL590133,2 | -0,886976596 | 0,540746154 | 0,43110267 | 0,89530663 | novel transcript, antisense to CERS2 |
| ENSG00000260171 | AC008269,2 | -0,887227329 | 0,540652183 | 0,48984896 | 0,916063942 | novel transcript |
| ENSG00000247828 | TMEM161B-AS | -0,887493717 | 0,540552363 | 0,02131013 | 0,194921664 | TMEM161B antisense RNA 1 |

| | | | | | | |
|---|---|---|---|---|---|---|
| ENSG00000285804 | AC025774,1 | -0,887871113 | 0,540410978 | 0,10210261 | 0,508625897 | novel transcript |
| ENSG00000178445 | GLDC | -0,887877713 | 0,540408506 | 0,00423063 | 0,061234409 | glycine decarboxylase |
| ENSG00000249867 | LINC02742 | -0,887897802 | 0,540400981 | 0,06538585 | 0,401164726 | long intergenic non-protein coding RNA 2742 |
| ENSG00000005421 | PON1 | -0,88801517 | 0,540357019 | 0,17218396 | 0,65638124 | paraoxonase 1 |
| ENSG00000282685 | AC105219,8 | -0,888046496 | 0,540345286 | 0,16973391 | 0,652551597 | IQ motif and ankyrin repeat containing 1 |
| ENSG00000282725 | CORO7 | -0,888429525 | 0,540201846 | 0,26901658 | 0,781285125 | coronin 7 |
| ENSG00000265015 | AP005901,3 | -0,888453929 | 0,540192708 | 0,23689908 | 0,746456145 | kinase suppressor of ras 1 pseudogene |
| ENSG00000267526 | AC005702,2 | -0,889338222 | 0,539861701 | 0,14312219 | 0,601880115 | UDP-N-acetyl-alpha-D-galactosamine:polypeptide N-acetylgalactosaminyltransferase 1 (GalNAc-T1) (GALNT1) pseudogene |
| ENSG00000171533 | MAP6 | -0,889342911 | 0,539859947 | 0,01684791 | 0,166371804 | microtubule associated protein 6 |
| ENSG00000281216 | KIF26B | -0,88941643 | 0,539832436 | 0,260298 | 0,772692192 | kinesin family member 26B |
| ENSG00000287109 | AP000224,1 | -0,889467423 | 0,539813356 | 0,00493114 | 0,068095058 | novel transcript, antisense to JAM2 |
| ENSG00000261649 | GOLGA6L7 | -0,889532293 | 0,539789084 | 0,24717094 | 0,759323829 | golgin A6 family like 7 |
| ENSG00000226249 | AL365271,1 | -0,889747615 | 0,539708527 | 0,43057334 | 0,895267028 | novel transcript |
| ENSG00000200983 | SNORA3A | -0,889862016 | 0,539665731 | 0,22920447 | 0,73730307 | small nucleolar RNA, H/ACA box 3A |
| ENSG00000283413 | AC008703,1 | -0,889938895 | 0,539636974 | 0,46451448 | 0,907547847 | novel transcript |
| ENSG00000237307 | SRRM1P3 | -0,890058553 | 0,539592218 | 0,03459975 | 0,271313423 | serine/arginine repetitive matrix 1 pseudogene 3 |
| ENSG00000189195 | BTBD8 | -0,89007177 | 0,539587275 | 0,09389795 | 0,487159915 | BTB domain containing 8 |
| ENSG00000186642 | PDE2A | -0,890247823 | 0,539521433 | 0,14716828 | 0,611181688 | phosphodiesterase 2A |
| ENSG00000286518 | AL354743,1 | -0,890256944 | 0,539518021 | 0,18462553 | 0,676844797 | novel transcript, antisense to PRDM16 |
| ENSG00000104901 | DKKL1 | -0,890600489 | 0,539389563 | 0,01014643 | 0,11641031 | dickkopf like acrosomal protein 1 |
| ENSG00000282205 | AC115090,1 | -0,890785531 | 0,539320384 | 0,14652382 | 0,610138629 | novel transcript, antisense to ARHGAP23 |
| ENSG00000276907 | ARHGAP27 | -0,890945121 | 0,539260729 | 0,29466073 | 0,804859966 | Rho GTPase activating protein 27 |
| ENSG00000135094 | SDS | -0,890984545 | 0,539245993 | 0,22803424 | 0,736341996 | serine dehydratase |
| ENSG00000164082 | GRM2 | -0,891261094 | 0,539142635 | 0,04474151 | 0,31940789 | glutamate metabotropic receptor 2 |
| ENSG00000259931 | AC105036,1 | -0,89143971 | 0,539075889 | 0,49290615 | 0,917090312 | small nuclear ribonucleoprotein polypeptide F (SNRPF) pseudogene |
| ENSG00000214050 | FBXO16 | -0,892333329 | 0,538742084 | 0,02078882 | 0,191935899 | F-box protein 16 |
| ENSG00000234995 | RPS3AP28 | -0,892469575 | 0,538691208 | 0,09749274 | 0,496015136 | RPS3A pseudogene 28 |
| ENSG00000198185 | ZNF334 | -0,89275887 | 0,538583198 | 0,01701912 | 0,167691735 | zinc finger protein 334 |
| ENSG00000169508 | GPR183 | -0,89310861 | 0,53845265 | 0,04489685 | 0,320249835 | G protein-coupled receptor 183 |
| ENSG00000050730 | TNIP3 | -0,893190432 | 0,538422113 | 0,32609634 | 0,828127364 | TNFAIP3 interacting protein 3 |

| | | | | | | |
|---|---|---|---|---|---|---|
| ENSG00000267500 | ZNF887P | -0,893340837 | 0,538365984 | 0,09673828 | 0,494090395 | zinc finger protein 887, pseudogene |
| ENSG00000288091 | AC062022,2 | -0,89357803 | 0,538277479 | 0,00917562 | 0,107561558 | novel transcript, antisense to SLC4A10 |
| ENSG00000227355 | AL359644,1 | -0,893740569 | 0,538216838 | 0,45176536 | 0,902062594 | novel transcript, antisense STOM |
| ENSG00000213131 | YWHAZP4 | -0,893756633 | 0,538210845 | 0,06559909 | 0,402080164 | tyrosine 3-monooxygenase/tryptophan 5-monooxygenase activation protein zeta pseudogene 4 |
| ENSG00000249319 | AC068533,4 | -0,894398124 | 0,537971584 | 0,31438502 | 0,819877431 | novel protein |
| ENSG00000080224 | EPHA6 | -0,894450099 | 0,537952203 | 0,03573864 | 0,277047585 | EPH receptor A6 |
| ENSG00000283415 | AC087280,2 | -0,894552881 | 0,537913879 | 0,05262621 | 0,352163052 | novel transcript |
| ENSG00000255434 | AP001922,6 | -0,894837658 | 0,53780771 | 0,12891135 | 0,573624329 | novel transcript |
| ENSG00000256120 | SOX5-AS1 | -0,894891832 | 0,537787515 | 0,07303494 | 0,426318958 | SOX5 antisense RNA 1 |
| ENSG00000225051 | HMGB3P22 | -0,894947232 | 0,537766864 | 0,3948312 | 0,871888817 | high mobility group box 3 pseudogene 22 |
| ENSG00000286048 | AC008966,3 | -0,895083064 | 0,537716235 | 0,54573519 | 0,935146964 | novel transcript, antisense to ITGA2 |
| ENSG00000249068 | AC008417,1 | -0,895197013 | 0,537673766 | 0,2777928 | 0,789162936 | cancer/testis antigen family 45, member A5 pseudogene |
| ENSG00000237317 | AL022400,1 | -0,89522479 | 0,537663414 | 0,51823355 | 0,926413949 | forty-two-three domain containing 1 (FYTTD1) pseudogene |
| ENSG00000259588 | AC012050,1 | -0,895518003 | 0,537554151 | 0,34621247 | 0,841168721 | novel transcript, antisense to SEMA6D |
| ENSG00000180332 | KCTD4 | -0,895582689 | 0,537530049 | 0,06783336 | 0,41004296 | potassium channel tetramerization domain containing 4 |
| ENSG00000232615 | AC026412,2 | -0,895598709 | 0,53752408 | 0,40840591 | 0,881491356 | protein phosphatase 4, regulatory subunit 2 (PPP4R2) pseudogene |
| ENSG00000163053 | SLC16A14 | -0,895701068 | 0,537485944 | 0,01294353 | 0,138442226 | solute carrier family 16 member 14 |
| ENSG00000224598 | RPS5P2 | -0,895726819 | 0,53747635 | 0,44008767 | 0,897906863 | RPS5 pseudogene 2 |
| ENSG00000176399 | DMRTA1 | -0,895830404 | 0,537437761 | 0,20730457 | 0,711123838 | DMRT like family A1 |
| ENSG00000185008 | ROBO2 | -0,895865738 | 0,537424599 | 0,02877184 | 0,240839934 | roundabout guidance receptor 2 |
| ENSG00000006625 | GGCT | -0,896000095 | 0,537374551 | 0,02536055 | 0,221163837 | gamma-glutamylcyclotransferase |
| ENSG00000197748 | CFAP43 | -0,896146147 | 0,537320152 | 0,10570193 | 0,518867175 | cilia and flagella associated protein 43 |
| ENSG00000137252 | HCRTR2 | -0,896668679 | 0,537125575 | 0,3383746 | 0,837263082 | hypocretin receptor 2 |
| ENSG00000083454 | P2RX5 | -0,896855636 | 0,537055974 | 0,23738529 | 0,746907875 | purinergic receptor P2X 5 |
| ENSG00000275953 | AL359535,1 | -0,896966067 | 0,537014867 | 0,42999189 | 0,894726544 | Yes-associated protein 1 (YAP1) pseudogene |
| ENSG00000286735 | AC126182,3 | -0,896987879 | 0,537006748 | 0,31730231 | 0,822252078 | novel transcript, antisense to FSTL1 |
| ENSG00000285548 | AC068725,1 | -0,897208524 | 0,536924624 | 0,38424903 | 0,868128461 | Novel transcript, antisense to NRXN1 |

| | | | | | | |
|---|---|---|---|---|---|---|
| ENSG00000257611 | AC010182,1 | -0,8972865 | 0,536895605 | 0,47561401 | 0,911264385 | thyroid hormone receptor interactor 11 (TRIP11) pseudogene |
| ENSG00000261529 | AC100774,1 | -0,897345425 | 0,536873676 | 0,03962292 | 0,29656955 | novel transcript, sense intronic to UBE3A |
| ENSG00000228328 | PGK1P1 | -0,897485203 | 0,536821663 | 0,24575037 | 0,757752583 | phosphoglycerate kinase 1 pseudogene 1 |
| ENSG00000078114 | NEBL | -0,89799586 | 0,536631683 | 0,21064215 | 0,716131719 | nebulette |
| ENSG00000255545 | B3GAT1-DT | -0,898298431 | 0,536519149 | 0,0256327 | 0,22266762 | B3GAT1 diverget transcript |
| ENSG00000286167 | AC105402,1 | -0,898558051 | 0,536422609 | 0,18328881 | 0,674320815 | novel transcript, antisense to KIF5C |
| ENSG00000264016 | AC015908,3 | -0,898617737 | 0,536400417 | 0,28534798 | 0,796948543 | novel transcript |
| ENSG00000132623 | ANKEF1 | -0,89882857 | 0,536322034 | 0,07749853 | 0,439969864 | ankyrin repeat and EF-hand domain containing 1 |
| ENSG00000269469 | AC010619,1 | -0,898981936 | 0,536265023 | 0,61940553 | 0,952793934 | novel protein |
| ENSG00000267699 | AC091551,1 | -0,899018404 | 0,536251468 | 0,4715671 | 0,908896785 | novel protein |
| ENSG00000144792 | ZNF660 | -0,89905786 | 0,536236802 | 0,46633589 | 0,907940653 | zinc finger protein 660 |
| ENSG00000202408 | RNVU1-21 | -0,899234153 | 0,536171279 | 0,33494643 | 0,834858938 | RNA, variant U1 small nuclear 21 |
| ENSG00000235932 | AL359955,2 | -0,899260489 | 0,536161492 | 0,49809924 | 0,919274055 | pseudogene similar to tripartite motif-containing protein |
| ENSG00000083307 | GRHL2 | -0,899287237 | 0,536151551 | 0,05482613 | 0,361697451 | grainyhead like transcription factor 2 |
| ENSG00000272024 | AC064807,4 | -0,899348218 | 0,536128889 | 0,36688591 | 0,856313634 | novel transcript |
| ENSG00000269966 | AL136164,2 | -0,899605065 | 0,536033449 | 0,46768382 | 0,907940653 | novel transcript |
| ENSG00000227962 | AL391994,1 | -0,899652454 | 0,536015842 | 0,5763046 | 0,943158459 | ubiquitin-conjugating enzyme E2G 1 (UBC7 homolog, C, elegans) (UBE2G10 pseudogene |
| ENSG00000258631 | AC110023,1 | -0,899813104 | 0,535956158 | 0,33810824 | 0,837263082 | novel transcript |
| ENSG00000184357 | H1-5 | -0,900242931 | 0,535796503 | 8,69E-05 | 0,002323603 | H1,5 linker histone, cluster member |
| ENSG00000266853 | ITM2BP1 | -0,900519817 | 0,535693681 | 0,33584011 | 0,835631125 | integral membrane protein 2B pseudogene 1 |
| ENSG00000109911 | ELP4 | -0,901010705 | 0,535511438 | 0,03768048 | 0,287047407 | elongator acetyltransferase complex subunit 4 |
| ENSG00000261838 | AC092718,6 | -0,901017403 | 0,535508952 | 0,47848535 | 0,912212078 | novel transcript, antisense to C16orf46 |
| ENSG00000131849 | ZNF132 | -0,901120689 | 0,535470615 | 0,00266555 | 0,042088805 | zinc finger protein 132 |
| ENSG00000260398 | AC068700,1 | -0,901153583 | 0,535458406 | 0,05890537 | 0,37691718 | novel transcript, overlapping to PKIA |
| ENSG00000233593 | LINC02609 | -0,901203207 | 0,535439988 | 0,00144541 | 0,025458789 | long intergenic non-protein coding RNA 2609 |
| ENSG00000261024 | GS1-279B7,1 | -0,901388015 | 0,535371403 | 0,27539132 | 0,787054963 | microtubule associated protein 1 light chain 3 beta pseudogene |
| ENSG00000137968 | SLC44A5 | -0,9025148 | 0,534953427 | 0,05503446 | 0,362383581 | solute carrier family 44 member 5 |
| ENSG00000265478 | AC107982,3 | -0,902650768 | 0,534903012 | 0,08913846 | 0,47641326 | novel transcript |

| | | | | | | |
|---|---|---|---|---|---|---|
| ENSG00000270941 | AC006254,2 | -0,90265333 | 0,534902062 | 0,46160414 | 0,906165604 | Adenylate kinase isoenzyme 4, mitochondrial, pseudogene |
| ENSG00000229029 | CDCA4P1 | -0,902683298 | 0,534890951 | 0,23004742 | 0,7385708 | cell division cycle associated 4 pseudogene 1 |
| ENSG00000269892 | AC125494,2 | -0,902702926 | 0,534883674 | 0,26160854 | 0,77342121 | novel transcript |
| ENSG00000259467 | NDUFAF4P1 | -0,902813318 | 0,534842747 | 0,4748135 | 0,91069468 | NADH:ubiquinone oxidoreductase complex assembly factor 4 pseudogene 1 |
| ENSG00000281016 | AC092574,2 | -0,902996177 | 0,534774961 | 0,06748774 | 0,408664851 | novel transcript, antisense to ZNF721 |
| ENSG00000288475 | CASP8AP2 | -0,903307098 | 0,534659722 | 0,00072591 | 0,014228805 | caspase 8 associated protein 2 |
| ENSG00000267743 | AC107896,1 | -0,903671438 | 0,534524715 | 0,43562104 | 0,896422069 | uncharacterized LOC105372143 |
| ENSG00000204963 | PCDHA7 | -0,904104732 | 0,534364202 | 0,01621579 | 0,1624385 | protocadherin alpha 7 |
| ENSG00000267920 | SNX6P1 | -0,904340751 | 0,53427679 | 0,03960244 | 0,296512476 | sorting nexin 6 pseudogene 1 |
| ENSG00000238230 | LINC00391 | -0,904429822 | 0,534243805 | 0,00479379 | 0,066788288 | long intergenic non-protein coding RNA 391 |
| ENSG00000119862 | LGALSL | -0,904699948 | 0,534143784 | 0,05647861 | 0,366903759 | galectin like |
| ENSG00000286445 | AL355334,2 | -0,90502084 | 0,53402499 | 0,14696508 | 0,610674677 | novel transcript |
| ENSG00000231395 | ARL4AP4 | -0,905110952 | 0,533991635 | 0,21478007 | 0,722164151 | ADP ribosylation factor like GTPase 4A pseudogene 4 |
| ENSG00000250302 | LINC01618 | -0,905892399 | 0,533702473 | 0,42869975 | 0,893524604 | long intergenic non-protein coding RNA 1618 |
| ENSG00000214389 | RPS3AP26 | -0,906240219 | 0,533573818 | 0,6337688 | 0,954929949 | RPS3A pseudogene 26 |
| ENSG00000105613 | MAST1 | -0,906261569 | 0,533565921 | 0,00557984 | 0,07493472 | microtubule associated serine/threonine kinase 1 |
| ENSG00000239626 | RPSAP41 | -0,906539131 | 0,533463278 | 0,11951736 | 0,552362303 | ribosomal protein SA pseudogene 41 |
| ENSG00000225632 | AL161644,1 | -0,906913877 | 0,533324727 | 0,15310412 | 0,621632863 | novel transcript |
| ENSG00000231104 | AC022395,1 | -0,906949373 | 0,533311605 | 0,09308268 | 0,484790551 | novel transcript, antisense to RAB11FIP2 |
| ENSG00000241535 | CBX5P1 | -0,90706334 | 0,533269477 | 0,04910592 | 0,337936952 | chromobox 5 pseudogene 1 |
| ENSG00000112414 | ADGRG6 | -0,907218364 | 0,533212178 | 0,06392963 | 0,39585974 | adhesion G protein-coupled receptor G6 |
| ENSG00000185149 | NPY2R | -0,907263831 | 0,533195374 | 0,19917288 | 0,698400736 | neuropeptide Y receptor Y2 |
| ENSG00000160883 | HK3 | -0,907745749 | 0,533017295 | 0,52578237 | 0,929056124 | hexokinase 3 |
| ENSG00000226869 | LHFPL3-AS1 | -0,907808388 | 0,532994153 | 0,57051202 | 0,9423888 | LHFPL3 antisense RNA 1 |
| ENSG00000285272 | TMEM249 | -0,908104911 | 0,532884616 | 0,29792372 | 0,807200542 | transmembrane protein 249 |
| ENSG00000261587 | TMEM249 | -0,908104911 | 0,532884616 | 0,29792372 | 0,807200542 | transmembrane protein 249 |
| ENSG00000232486 | DEPDC1P2 | -0,908298858 | 0,532812983 | 0,10558639 | 0,5185443 | DEP domain containing 1 pseudogene 2 |
| ENSG00000229185 | OR2H2 | -0,908362798 | 0,532789369 | 0,27871953 | 0,789775666 | olfactory receptor family 2 subfamily H member 2 |
| ENSG00000141434 | MEP1B | -0,909056087 | 0,532533398 | 0,27504242 | 0,786439192 | meprin A subunit beta |
| ENSG00000112981 | NME5 | -0,909306355 | 0,532441026 | 0,1231601 | 0,561375905 | NME/NM23 family member 5 |
| ENSG00000183150 | GPR19 | -0,909343424 | 0,532427346 | 0,05391257 | 0,357953158 | G protein-coupled receptor 19 |

| | | | | | | |
|---|---|---|---|---|---|---|
| ENSG00000234886 | MTND5P26 | -0,90951021 | 0,532365797 | 0,10729627 | 0,522893717 | MT-ND5 pseudogene 26 |
| ENSG00000248401 | AC114781,1 | -0,909842048 | 0,53224336 | 0,1821937 | 0,672503417 | aminomethyltransferase (AMT) pseudogene |
| ENSG00000229994 | RPL5P4 | -0,90989938 | 0,53222221 | 0,08677779 | 0,469380749 | ribosomal protein L5 pseudogene 4 |
| ENSG00000286977 | Z83745,1 | -0,909917779 | 0,532215422 | 0,06436276 | 0,397835213 | novel transcript |
| ENSG00000172062 | SMN1 | -0,910400667 | 0,532037313 | 0,23723662 | 0,746624292 | survival of motor neuron 1, telomeric |
| ENSG00000111674 | ENO2 | -0,910460804 | 0,532015136 | 0,00024835 | 0,005725999 | enolase 2 |
| ENSG00000255084 | AP003110,1 | -0,910544667 | 0,531984211 | 0,46927498 | 0,907940653 | novel transcript |
| ENSG00000249069 | LINC01033 | -0,910651358 | 0,531944871 | 0,46139851 | 0,906165604 | long intergenic non-protein coding RNA 1033 |
| ENSG00000276203 | ANKRD20A3P | -0,910932909 | 0,531841069 | 0,03888567 | 0,293298341 | ankyrin repeat domain 20 family member A3, pseudogene |
| ENSG00000229105 | ASTN2-AS1 | -0,910962025 | 0,531830335 | 0,58988023 | 0,947250274 | ASTN2 antisense RNA 1 |
| ENSG00000249487 | LINC01586 | -0,91098576 | 0,531821586 | 0,26164438 | 0,773454051 | long intergenic non-protein coding RNA 1586 |
| ENSG00000206634 | SNORA22 | -0,911036952 | 0,531802715 | 0,18387558 | 0,675441919 | small nucleolar RNA, H/ACA box 22 |
| ENSG00000263400 | TMEM220-AS1 | -0,911039651 | 0,53180172 | 0,0975631 | 0,496292325 | TMEM220 antisense RNA 1 |
| ENSG00000274363 | AC087742,1 | -0,911111714 | 0,531775157 | 0,15548984 | 0,625722005 | novel transcript |
| ENSG00000164398 | ACSL6 | -0,911352786 | 0,531686306 | 0,07252484 | 0,424609868 | acyl-CoA synthetase long chain family member 6 |
| ENSG00000128482 | RNF112 | -0,911451691 | 0,531649857 | 0,25867867 | 0,771485596 | ring finger protein 112 |
| ENSG00000176428 | VPS37D | -0,911701287 | 0,531557886 | 0,0035174 | 0,053073291 | VPS37D subunit of ESCRT-I |
| ENSG00000228212 | OFD1P17 | -0,911826988 | 0,531511574 | 0,01554743 | 0,157881532 | OFD1 pseudogene 17 |
| ENSG00000265206 | AC004687,1 | -0,911938345 | 0,53147055 | 0,27546855 | 0,787153723 | novel transcript, MIR142 host |
| ENSG00000282752 | CTDP1 | -0,91197551 | 0,531456859 | 0,71912293 | 0,971869926 | CTD phosphatase subunit 1 |
| ENSG00000258571 | PTTG4P | -0,911981547 | 0,531454635 | 0,31907039 | 0,824074082 | pituitary tumor-transforming 4 pseudogene |
| ENSG00000231238 | LINC00349 | -0,911992508 | 0,531450597 | 0,45821149 | 0,904140862 | long intergenic non-protein coding RNA 349 |
| ENSG00000254388 | DUTP2 | -0,912146126 | 0,531394011 | 0,1659624 | 0,645193807 | deoxyuridine triphosphatase pseudogene 2 |
| ENSG00000282450 | AC012676,7 | -0,912566158 | 0,531239322 | 0,4689526 | 0,907940653 | novel transcript, sense intronic with DNAJA3 |
| ENSG00000101049 | SGK2 | -0,912807542 | 0,531150445 | 0,15096232 | 0,618002612 | serum/glucocorticoid regulated kinase 2 |
| ENSG00000276449 | AC004076,2 | -0,913063689 | 0,531056149 | 0,00767063 | 0,094825839 | novel transcript |
| ENSG00000128242 | GAL3ST1 | -0,913271832 | 0,530979537 | 0,07718838 | 0,439233464 | galactose-3-O-sulfotransferase 1 |
| ENSG00000132196 | HSD17B7 | -0,913630181 | 0,530847664 | 0,00781468 | 0,09611257 | hydroxysteroid 17-beta dehydrogenase 7 |
| ENSG00000262525 | TAS2R46 | -0,913898992 | 0,530748763 | 0,27770187 | 0,789067728 | taste 2 receptor member 46 |
| ENSG00000226761 | TAS2R46 | -0,913898992 | 0,530748763 | 0,27770187 | 0,789067728 | taste 2 receptor member 46 |
| ENSG00000198732 | SMOC1 | -0,913928499 | 0,530737908 | 0,00643489 | 0,083132372 | SPARC related modular calcium binding 1 |
| ENSG00000117971 | CHRNB4 | -0,914115662 | 0,530669059 | 0,02388712 | 0,211620024 | cholinergic receptor nicotinic beta 4 subunit |

| | | | | | | |
|---|---|---|---|---|---|---|
| ENSG00000129990 | SYT5 | -0,914546062 | 0,530510767 | 0,05756598 | 0,371004511 | synaptotagmin 5 |
| ENSG00000248735 | R3HDM2P1 | -0,914606748 | 0,530488452 | 0,10333445 | 0,512638569 | R3H domain containing 2 pseudogene 1 |
| ENSG00000165588 | OTX2 | -0,914672385 | 0,530464318 | 0,0826161 | 0,457152163 | orthodenticle homeobox 2 |
| ENSG00000188729 | OSTN | -0,914799925 | 0,530417425 | 0,40121852 | 0,876659568 | osteocrin |
| ENSG00000261172 | AC133919,2 | -0,914872259 | 0,530390831 | 0,39695 | 0,873609704 | novel transcript |
| ENSG00000175749 | EIF3KP1 | -0,914889692 | 0,530384422 | 0,33633295 | 0,835660292 | eukaryotic translation initiation factor 3 subunit K pseudogene 1 |
| ENSG00000168875 | SOX14 | -0,915159995 | 0,530285059 | 0,32998472 | 0,831651927 | SRY-box transcription factor 14 |
| ENSG00000225647 | AC005487,1 | -0,915519477 | 0,530152942 | 0,61567983 | 0,952793934 | novel transcript |
| ENSG00000257752 | AC091516,1 | -0,915632554 | 0,530111391 | 0,22350712 | 0,731984265 | transmembrane protein 38A (TMEM38A) |
| ENSG00000251613 | AC063979,1 | -0,915815038 | 0,530044342 | 0,43627839 | 0,896850871 | novel transcript |
| ENSG00000286924 | AL139815,1 | -0,915892035 | 0,530016054 | 0,44119323 | 0,898885877 | novel transcript, antisense to ARMC3and MSRB2 |
| ENSG00000285276 | AC270301,7 | -0,916176669 | 0,529911496 | 0,18166811 | 0,67167359 | novel transcript |
| ENSG00000198216 | CACNA1E | -0,916244448 | 0,529886601 | 0,19842003 | 0,697413278 | calcium voltage-gated channel subunit alpha1 E |
| ENSG00000180616 | SSTR2 | -0,916549576 | 0,529774542 | 0,00042089 | 0,009000416 | somatostatin receptor 2 |
| ENSG00000267039 | AC016493,1 | -0,917057091 | 0,52958821 | 0,43334534 | 0,895877116 | novel transcript, antisense to KIAA1328 |
| ENSG00000213996 | TM6SF2 | -0,917114387 | 0,529567178 | 0,12578245 | 0,566474006 | transmembrane 6 superfamily member 2 |
| ENSG00000251676 | SNHG27 | -0,917211398 | 0,52953157 | 0,12381257 | 0,562627332 | small nucleolar RNA host gene 27 |
| ENSG00000258529 | AP001781,2 | -0,917265326 | 0,529511776 | 0,80638602 | 0,981213877 | novel protein |
| ENSG00000260986 | AC126335,1 | -0,917365436 | 0,529475034 | 0,38325297 | 0,867740551 | novel transcript |
| ENSG00000109255 | NMU | -0,917523211 | 0,529417133 | 0,06037517 | 0,382021721 | neuromedin U |
| ENSG00000272138 | LINC01607 | -0,917607628 | 0,529386156 | 0,22075191 | 0,728572782 | long intergenic non-protein coding RNA 1607 |
| ENSG00000287234 | AL592211,2 | -0,917960496 | 0,52925669 | 0,21489408 | 0,722180847 | novel transcript |
| ENSG00000139767 | SRRM4 | -0,918038579 | 0,529228045 | 0,01561277 | 0,158325561 | serine/arginine repetitive matrix 4 |
| ENSG00000259040 | BLOC1S5-TXND | -0,918039048 | 0,529227873 | 0,01858835 | 0,177868046 | BLOC1S5-TXNDC5 readthrough (NMD candidate) |
| ENSG00000163630 | SYNPR | -0,918050099 | 0,529223819 | 0,26270877 | 0,774111884 | synaptoporin |
| ENSG00000226111 | MRPS18B | -0,918110383 | 0,529201706 | 0,43746818 | 0,897021798 | mitochondrial ribosomal protein S18B |
| ENSG00000223775 | MRPS18B | -0,918110383 | 0,529201706 | 0,43746818 | 0,897021798 | mitochondrial ribosomal protein S18B |
| ENSG00000227420 | MRPS18B | -0,918110383 | 0,529201706 | 0,43746818 | 0,897021798 | mitochondrial ribosomal protein S18B |
| ENSG00000005469 | CROT | -0,918293397 | 0,529134578 | 0,03899473 | 0,29390824 | carnitine O-octanoyltransferase |
| ENSG00000230869 | AGAP10P | -0,918540752 | 0,529043864 | 0,09120281 | 0,481302729 | ArfGAP with GTPase domain, ankyrin repeat and PH domain 10, pseudogene |
| ENSG00000179603 | GRM8 | -0,91856035 | 0,529036677 | 0,01988417 | 0,185791619 | glutamate metabotropic receptor 8 |

| | | | | | | |
|---|---|---|---|---|---|---|
| ENSG00000281909 | HERC2P7 | -0,918668426 | 0,528997047 | 0,14052695 | 0,597347645 | hect domain and RLD 2 pseudogene 7 |
| ENSG00000219755 | AL137784,1 | -0,918906556 | 0,528909738 | 0,50080522 | 0,920520138 | 40S ribososmal protein 3 (RPS3) pseudogene |
| ENSG00000105784 | RUNDC3B | -0,919070521 | 0,52884963 | 0,1809461 | 0,670669701 | RUN domain containing 3B |
| ENSG00000271858 | Z84492,1 | -0,919366383 | 0,528741187 | 0,28897481 | 0,800166929 | novel transcript, antisense to CACNA2D2 |
| ENSG00000175445 | LPL | -0,919571988 | 0,528665839 | 0,0926086 | 0,483842806 | lipoprotein lipase |
| ENSG00000222515 | RN7SKP240 | -0,919884481 | 0,528551341 | 0,2610318 | 0,773103493 | RN7SK pseudogene 240 |
| ENSG00000279082 | LINC01727 | -0,920621171 | 0,528281513 | 0,0620482 | 0,388908386 | long intergenic non-protein coding RNA 1727 |
| ENSG00000053918 | KCNQ1 | -0,920786535 | 0,528220964 | 0,28251733 | 0,793528967 | potassium voltage-gated channel subfamily Q member 1 |
| ENSG00000271856 | LINC01215 | -0,920928891 | 0,528168845 | 0,53332404 | 0,931381467 | long intergenic non-protein coding RNA 1215 |
| ENSG00000287380 | AL589935,3 | -0,921525559 | 0,527950451 | 0,09672437 | 0,494090395 | novel transcript |
| ENSG00000215187 | FAM166B | -0,921633322 | 0,527911017 | 0,14969656 | 0,615156862 | family with sequence similarity 166 member B |
| ENSG00000287617 | AC092902,6 | -0,921636673 | 0,52790979 | 0,13776998 | 0,592433314 | novel transcript |
| ENSG00000188801 | ZNF322P1 | -0,921646442 | 0,527906216 | 0,29076034 | 0,801555946 | zinc finger protein 322 pseudogene 1 |
| ENSG00000275976 | AC211476,3 | -0,921883564 | 0,527819456 | 0,38812629 | 0,869820053 | speedy/RINGO cell cycle regulator family member E1 (SPDYE1) pseudogene |
| ENSG00000270878 | GCATP1 | -0,921988728 | 0,527780982 | 0,0572834 | 0,370098825 | glycine C-acetyltransferase pseudogene 1 |
| ENSG00000250122 | AC122694,1 | -0,922016105 | 0,527770967 | 0,1906414 | 0,685949202 | novel transcript |
| ENSG00000284282 | POLR3E | -0,922979851 | 0,527418524 | 0,12026712 | 0,553434082 | RNA polymerase III subunit E |
| ENSG00000282805 | AC023315,3 | -0,922991723 | 0,527414184 | 0,35488563 | 0,846374661 | novel transcript, sense intronic AATF |
| ENSG00000277589 | AC244093,4 | -0,922991723 | 0,527414184 | 0,35488563 | 0,846374661 | novel transcript, sense intronic AATF |
| ENSG00000238207 | AC009312,1 | -0,923105274 | 0,527372675 | 0,21435878 | 0,721700728 | novel transcript |
| ENSG00000204301 | NOTCH4 | -0,923291419 | 0,527304634 | 0,05307409 | 0,354174701 | notch receptor 4 |
| ENSG00000233723 | LINC01122 | -0,923351885 | 0,527282534 | 0,03035274 | 0,249117099 | long intergenic non-protein coding RNA 1122 |
| ENSG00000258909 | AC091078,2 | -0,923420566 | 0,527257433 | 0,00493024 | 0,068095058 | novel transcript |
| ENSG00000162728 | KCNJ9 | -0,923443524 | 0,527249043 | 0,09690975 | 0,494497533 | potassium inwardly rectifying channel subfamily J member 9 |
| ENSG00000170482 | SLC23A1 | -0,923952537 | 0,527063051 | 0,04429831 | 0,31783836 | solute carrier family 23 member 1 |
| ENSG00000172186 | HMGN1P35 | -0,924132876 | 0,526997172 | 0,46301875 | 0,906849208 | high mobility group nucleosome binding domain 1 pseudogene 35 |
| ENSG00000236283 | AC019197,1 | -0,924332238 | 0,526924352 | 0,14186619 | 0,599899099 | novel transcript, antisense to SLC38A11 and SCN3A |
| ENSG00000286320 | AC108053,1 | -0,924390506 | 0,526903071 | 0,04161126 | 0,305958244 | novel transcript, antisense to MAML3 |

| | | | | | | |
|---|---|---|---|---|---|---|
| ENSG00000080709 | KCNN2 | -0,92467641 | 0,526798663 | 0,04338178 | 0,314319373 | potassium calcium-activated channel subfamily N member 2 |
| ENSG00000288674 | AL391628,1 | -0,92509569 | 0,526645586 | 0,18222828 | 0,672551648 | Atypical kinase COQ8A, mitochondrial |
| ENSG00000074276 | CDHR2 | -0,925122275 | 0,526635881 | 0,0538715 | 0,357953158 | cadherin related family member 2 |
| ENSG00000244018 | RPL35P6 | -0,925128653 | 0,526633553 | 0,39497701 | 0,871888817 | ribosomal protein L35 pseudogene 6 |
| ENSG00000269653 | AC011479,3 | -0,925185725 | 0,52661272 | 0,33436597 | 0,834553555 | novel transcript, sense intronic to SIPA1L3 |
| ENSG00000287385 | AL121820,3 | -0,925371102 | 0,526545058 | 0,07941479 | 0,447113975 | uncharacterized LOC107984016 |
| ENSG00000273447 | AC004067,1 | -0,925387223 | 0,526539175 | 0,03705204 | 0,284129266 | novel transcript, antisense to CASP6 |
| ENSG00000236333 | TRHDE-AS1 | -0,925594471 | 0,526463541 | 0,05851365 | 0,374871039 | TRHDE antisense RNA 1 |
| ENSG00000214244 | SETP21 | -0,925595604 | 0,526463127 | 0,44092121 | 0,898675567 | SET pseudogene 21 |
| ENSG00000227290 | LINC01364 | -0,925609156 | 0,526458182 | 0,45443643 | 0,903383444 | long intergenic non-protein coding RNA 1364 |
| ENSG00000182533 | CAV3 | -0,925751246 | 0,526406334 | 0,37497881 | 0,861771195 | caveolin 3 |
| ENSG00000187229 | AC100800,1 | -0,925984107 | 0,526321375 | 0,04240663 | 0,309296116 | novel transcript, antisense to PSD3 |
| ENSG00000262612 | TAS2R43 | -0,926053713 | 0,526295982 | 0,11472752 | 0,541181798 | taste 2 receptor member 43 |
| ENSG00000235018 | AL137077,1 | -0,926157535 | 0,526258109 | 0,34524725 | 0,841168721 | ribosomal protein L17 (RPL17) pseudogene |
| ENSG00000204860 | FAM201A | -0,92642828 | 0,526159358 | 0,19477118 | 0,692283541 | family with sequence similarity 201 member A |
| ENSG00000049283 | EPN3 | -0,926479146 | 0,526140807 | 0,26648564 | 0,778345195 | epsin 3 |
| ENSG00000182768 | NGRN | -0,926604836 | 0,526094971 | 1,40E-05 | 0,000456354 | neugrin, neurite outgrowth associated |
| ENSG00000112246 | SIM1 | -0,92692715 | 0,525977449 | 0,13270596 | 0,581488329 | SIM bHLH transcription factor 1 |
| ENSG00000120915 | EPHX2 | -0,927111624 | 0,525910198 | 0,02007035 | 0,187083879 | epoxide hydrolase 2 |
| ENSG00000120162 | MOB3B | -0,927118094 | 0,525907839 | 0,01901119 | 0,180768182 | MOB kinase activator 3B |
| ENSG00000267429 | AC006116,6 | -0,927261794 | 0,525855458 | 0,16591417 | 0,645166759 | novel transcript, sense intronic to ZSCAN5A |
| ENSG00000238231 | AL512288,2 | -0,927371514 | 0,525815468 | 0,42282553 | 0,890422842 | interferon-induced transmembrane protein pseudogene |
| ENSG00000269961 | AC010359,2 | -0,927605165 | 0,525730316 | 0,21590068 | 0,722999018 | novel transcript |
| ENSG00000275395 | FCGBP | -0,927628135 | 0,525721946 | 0,25164758 | 0,763981167 | Fc fragment of IgG binding protein |
| ENSG00000237031 | AC008067,1 | -0,927896556 | 0,525624141 | 0,49452833 | 0,91758518 | novel transcript |
| ENSG00000143434 | SEMA6C | -0,928063515 | 0,525563316 | 5,24E-05 | 0,001479106 | semaphorin 6C |
| ENSG00000257023 | AC087241,2 | -0,928152737 | 0,525530814 | 0,45079512 | 0,902062594 | novel transcript, antisense to ETNK1 |
| ENSG00000225527 | AL450384,1 | -0,928228666 | 0,525503156 | 0,35800314 | 0,849032846 | novel transcript |
| ENSG00000168421 | RHOH | -0,928286932 | 0,525481933 | 0,00797362 | 0,097415937 | ras homolog family member H |
| ENSG00000169548 | ZNF280A | -0,928312884 | 0,525472481 | 0,34868168 | 0,842757646 | zinc finger protein 280A |
| ENSG00000275499 | ZNF280A | -0,928312884 | 0,525472481 | 0,34868168 | 0,842757646 | zinc finger protein 280A |

| | | | | | | |
|---|---|---|---|---|---|---|
| ENSG00000275470 | LENG8-AS1 | -0,928337879 | 0,525463377 | 0,12875012 | 0,573336703 | LENG8 antisense RNA 1 |
| ENSG00000149970 | CNKSR2 | -0,928573167 | 0,525377686 | 0,04954877 | 0,33978781 | connector enhancer of kinase suppressor of Ras 2 |
| ENSG00000255408 | PCDHA3 | -0,928737711 | 0,525317769 | 0,02257484 | 0,203412591 | protocadherin alpha 3 |
| ENSG00000225084 | AL450226,1 | -0,928834619 | 0,525282483 | 0,25573926 | 0,767242317 | novel transcript, antisense to SMG6 |
| ENSG00000125414 | MYH2 | -0,92900948 | 0,525218821 | 0,29483754 | 0,805040011 | myosin heavy chain 2 |
| ENSG00000233680 | HNRNPA1P27 | -0,929059107 | 0,525200754 | 0,39377056 | 0,871635019 | heterogeneous nuclear ribonucleoprotein A1 pseudogene 27 |
| ENSG00000121905 | HPCA | -0,929652264 | 0,524984865 | 0,03464768 | 0,271577099 | hippocalcin |
| ENSG00000101180 | HRH3 | -0,929735215 | 0,524954681 | 0,16455781 | 0,643811841 | histamine receptor H3 |
| ENSG00000147127 | RAB41 | -0,930229092 | 0,524775004 | 0,19552945 | 0,693689284 | RAB41, member RAS oncogene family |
| ENSG00000241438 | TDGF1P6 | -0,930621281 | 0,524632366 | 0,48736853 | 0,914865783 | teratocarcinoma-derived growth factor 1 pseudogene 6 |
| ENSG00000286647 | AC008459,1 | -0,930748111 | 0,524586247 | 0,28918326 | 0,800260154 | novel transcript |
| ENSG00000263158 | TAS2R63P | -0,930859646 | 0,524545692 | 0,38224293 | 0,866916418 | taste 2 receptor member 63 pseudogene |
| ENSG00000256019 | TAS2R63P | -0,930859646 | 0,524545692 | 0,38224293 | 0,866916418 | taste 2 receptor member 63 pseudogene |
| ENSG00000260583 | AP000223,1 | -0,930897079 | 0,524532082 | 0,13251387 | 0,581053809 | novel transcript, antisense to MRPL39 |
| ENSG00000285750 | AL157385,2 | -0,931065383 | 0,524470894 | 0,3827613 | 0,867356268 | novel transcript |
| ENSG00000008277 | ADAM22 | -0,931127646 | 0,52444826 | 0,07370482 | 0,428867922 | ADAM metallopeptidase domain 22 |
| ENSG00000278291 | AL161772,1 | -0,931467995 | 0,524324551 | 0,0505892 | 0,34398963 | novel transcript, antisense to IL17D |
| ENSG00000260838 | AC022893,2 | -0,931893055 | 0,524170092 | 0,01050169 | 0,118829088 | novel transcript |
| ENSG00000286218 | AC005040,2 | -0,932043489 | 0,524115438 | 0,51301934 | 0,923725914 | novel transcript |
| ENSG00000213047 | DENND1B | -0,932079921 | 0,524102203 | 0,06254783 | 0,390552401 | DENN domain containing 1B |
| ENSG00000243926 | TIPARP-AS1 | -0,932829484 | 0,523829973 | 0,17715991 | 0,663320644 | TIPARP antisense RNA 1 |
| ENSG00000226903 | LINC00354 | -0,933017237 | 0,523761806 | 0,1238722 | 0,562734731 | long intergenic non-protein coding RNA 354 |
| ENSG00000255471 | AP001528,1 | -0,933050096 | 0,523749876 | 0,10498378 | 0,517046764 | novel transcript |
| ENSG00000248568 | KRT8P48 | -0,933318182 | 0,523652561 | 0,37254372 | 0,860420124 | keratin 8 pseudogene 48 |
| ENSG00000214975 | PPIAP29 | -0,933446755 | 0,523605895 | 0,42355433 | 0,890949676 | peptidylprolyl isomerase A pseudogene 29 |
| ENSG00000105289 | TJP3 | -0,933726874 | 0,523504239 | 0,42982641 | 0,894607 | tight junction protein 3 |
| ENSG00000160460 | SPTBN4 | -0,933792642 | 0,523480375 | 0,2536053 | 0,765172364 | spectrin beta, non-erythrocytic 4 |
| ENSG00000164972 | C9orf24 | -0,934059324 | 0,523383619 | 0,22871005 | 0,737173617 | chromosome 9 open reading frame 24 |
| ENSG00000081479 | LRP2 | -0,934543721 | 0,523207918 | 0,00418245 | 0,06076214 | LDL receptor related protein 2 |
| ENSG00000131152 | AC010531,1 | -0,934913736 | 0,523073746 | 0,45581917 | 0,903545728 | novel protein |
| ENSG00000092850 | TEKT2 | -0,93492362 | 0,523070162 | 0,02187643 | 0,198849969 | tektin 2 |

| | | | | | | |
|---|---|---|---|---|---|---|
| ENSG00000229727 | AC013460,1 | -0,93501229 | 0,523038014 | 0,08937782 | 0,477088647 | novel transcript |
| ENSG00000226977 | HMGN1P24 | -0,935091207 | 0,523009404 | 0,41181196 | 0,883966506 | high mobility group nucleosome binding domain 1 pseudogene 24 |
| ENSG00000278601 | AL158163,2 | -0,935327933 | 0,522923593 | 0,32146727 | 0,824933727 | novel transcript, antisense to BBIP1 |
| ENSG00000274553 | AC138035,3 | -0,935338999 | 0,522919582 | 0,06644765 | 0,405191993 | capicua transcriptional repressor (CIC) pseudogene |
| ENSG00000233716 | MTFR2P1 | -0,93561495 | 0,52281957 | 0,15113755 | 0,618336884 | MTFR2 pseudogene 1 |
| ENSG00000108387 | SEPTIN4 | -0,935666254 | 0,522800978 | 0,01812403 | 0,174939666 | septin 4 |
| ENSG00000188451 | SRP72P2 | -0,936064109 | 0,522656824 | 0,0137478 | 0,144425809 | signal recognition particle 72 pseudogene 2 |
| ENSG00000260351 | AC024270,1 | -0,936107498 | 0,522641105 | 0,45079593 | 0,902062594 | novel transcript, antisense to BTBD1 |
| ENSG00000157703 | SVOPL | -0,936135952 | 0,522630798 | 0,2462451 | 0,758198269 | SVOP like |
| ENSG00000237112 | GABBR1 | -0,936453399 | 0,522515812 | 0,01534926 | 0,156640518 | gamma-aminobutyric acid type B receptor subunit 1 |
| ENSG00000206466 | GABBR1 | -0,936453399 | 0,522515812 | 0,01534926 | 0,156640518 | gamma-aminobutyric acid type B receptor subunit 1 |
| ENSG00000214988 | RPL7AP26 | -0,936556612 | 0,522478432 | 0,38538086 | 0,868895792 | ribosomal protein L7a pseudogene 26 |
| ENSG00000177989 | ODF3B | -0,936568625 | 0,522474081 | 0,14235011 | 0,600436282 | outer dense fiber of sperm tails 3B |
| ENSG00000236044 | FABP5P2 | -0,936892553 | 0,522356783 | 0,34770551 | 0,842293776 | fatty acid binding protein 5 pseudogene 2 |
| ENSG00000228939 | AKT3-IT1 | -0,937153456 | 0,522262327 | 0,23538548 | 0,744654651 | AKT3 intronic transcript 1 |
| ENSG00000005075 | POLR2J | -0,937487548 | 0,522141398 | 0,00093232 | 0,017633484 | RNA polymerase II subunit J |
| ENSG00000261292 | AC110491,1 | -0,937987665 | 0,521960426 | 0,26390492 | 0,775013078 | novel transcript |
| ENSG00000230894 | AL133284,1 | -0,938250109 | 0,521865484 | 0,11083096 | 0,531538341 | novel transcript |
| ENSG00000284308 | C2orf81 | -0,93851303 | 0,521770386 | 0,03271646 | 0,261798703 | chromosome 2 open reading frame 81 |
| ENSG00000231169 | EEF1B2P1 | -0,938686469 | 0,521707664 | 0,26142716 | 0,77325059 | eukaryotic translation elongation factor 1 beta 2 pseudogene 1 |
| ENSG00000273454 | AC112503,2 | -0,938915258 | 0,521624936 | 0,43831524 | 0,897177404 | novel transcript |
| ENSG00000233786 | CDC27P1 | -0,93891931 | 0,52162347 | 0,62113674 | 0,952793934 | cell division cycle 27 pseudogene 1 |
| ENSG00000185674 | LYG2 | -0,93896403 | 0,521607301 | 0,37369625 | 0,861319894 | lysozyme g2 |
| ENSG00000259924 | AC011939,1 | -0,939111786 | 0,521553883 | 0,1825522 | 0,673032069 | DnaJ (Hsp40) homolog, subfamily B, member 14 pseudogene |
| ENSG00000224559 | LINC01087 | -0,939203745 | 0,52152064 | 0,30792666 | 0,816424917 | long intergenic non-protein coding RNA 1087 |
| ENSG00000273338 | AC103591,3 | -0,939268961 | 0,521497065 | 0,26499731 | 0,776509969 | novel transcript, antisense to DNAJB4 |
| ENSG00000270832 | AC092120,2 | -0,939514976 | 0,521408145 | 0,34372966 | 0,840901506 | cytidine monophosphate (UMP-CMP) kinase 1, cytosolic (CMPK1) pseudogene |
| ENSG00000262119 | AL079343,1 | -0,939516839 | 0,521407471 | 0,06288621 | 0,391611013 | novel transcript, sense intronic to NOVA1 |
| ENSG00000188931 | CFAP126 | -0,939879006 | 0,521276596 | 0,07403629 | 0,4296801 | cilia and flagella associated protein 126 |

| | | | | | | |
|---|---|---|---|---|---|---|
| ENSG00000267440 | LINC02594 | -0,940094968 | 0,52119857 | 0,49976516 | 0,920310866 | long intergenic non-protein coding RNA 2594 |
| ENSG00000239466 | RN7SL552P | -0,940567965 | 0,52102772 | 0,43688071 | 0,896866043 | RNA, 7SL, cytoplasmic 552, pseudogene |
| ENSG00000236806 | RPL7AP15 | -0,940657049 | 0,520995549 | 0,38795973 | 0,869820053 | ribosomal protein L7a pseudogene 15 |
| ENSG00000251495 | PPIAP11 | -0,940897582 | 0,520908693 | 0,36801036 | 0,857401877 | peptidylprolyl isomerase A pseudogene 11 |
| ENSG00000254979 | AP000781,2 | -0,941308248 | 0,520760436 | 0,45269604 | 0,902285042 | novel protein |
| ENSG00000286802 | AC104459,1 | -0,941333479 | 0,520751329 | 0,40048822 | 0,876197594 | novel transcript, antisense to GBP7and GBP2 |
| ENSG00000249624 | AP000295,1 | -0,941522503 | 0,520683104 | 0,13778092 | 0,592433314 | novel protein |
| ENSG00000258739 | ENO1P2 | -0,941549493 | 0,520673363 | 0,0602436 | 0,381629797 | enolase 1 pseudogene 2 |
| ENSG00000107242 | PIP5K1B | -0,941991957 | 0,520513701 | 0,03118276 | 0,253879661 | phosphatidylinositol-4-phosphate 5-kinase type 1 beta |
| ENSG00000260975 | AC007333,2 | -0,942024598 | 0,520501924 | 0,05430456 | 0,359622485 | novel transcript |
| ENSG00000188659 | SAXO2 | -0,942132052 | 0,520463158 | 0,01043049 | 0,118366144 | stabilizer of axonemal microtubules 2 |
| ENSG00000236279 | CLEC2L | -0,942149234 | 0,520456959 | 0,2660971 | 0,77755567 | C-type lectin domain family 2 member L |
| ENSG00000234925 | ATP5PDP4 | -0,942159203 | 0,520453363 | 0,05344243 | 0,355858082 | ATP synthase peripheral stalk subunit d pseudogene 4 |
| ENSG00000036448 | MYOM2 | -0,942435751 | 0,520353608 | 0,43251159 | 0,89560871 | myomesin 2 |
| ENSG00000262209 | PCDHGB3 | -0,94308712 | 0,520118724 | 0,0662858 | 0,404599624 | protocadherin gamma subfamily B, 3 |
| ENSG00000230793 | SMARCE1P5 | -0,943288813 | 0,520046015 | 0,30119672 | 0,810679941 | SWI/SNF related, matrix associated, actin dependent regulator of chromatin, subfamily e, member 1 pseudogene 5 |
| ENSG00000258704 | SRP54-AS1 | -0,943306322 | 0,520039703 | 0,16653738 | 0,646493505 | SRP54 antisense RNA 1 (head to head) |
| ENSG00000218896 | TUBB8P2 | -0,943533703 | 0,519957747 | 0,24368111 | 0,754507765 | tubulin beta 8 class VIII pseudogene 2 |
| ENSG00000130226 | DPP6 | -0,94365668 | 0,519913427 | 0,04396821 | 0,316175438 | dipeptidyl peptidase like 6 |
| ENSG00000183837 | PNMA3 | -0,943780301 | 0,519868879 | 0,02194403 | 0,199319602 | PNMA family member 3 |
| ENSG00000267549 | AC006116,7 | -0,943995664 | 0,51979128 | 0,02178022 | 0,198207677 | novel transcript, antisense to ZSCAN5A |
| ENSG00000226758 | DISC1-IT1 | -0,944149905 | 0,519735711 | 0,01411484 | 0,147301149 | DISC1 intronic transcript 1 |
| ENSG00000248200 | AC093770,1 | -0,944534422 | 0,519597206 | 0,24604125 | 0,75807958 | ribosomal protein L17 (RPL17) pseudogene |
| ENSG00000243836 | WDR86-AS1 | -0,944672918 | 0,519547328 | 0,42866056 | 0,893524604 | WDR86 antisense RNA 1 |
| ENSG00000281977 | AC244131,3 | -0,944700493 | 0,519537398 | 0,44771103 | 0,902062594 | taste receptor, type 2, member 7 pseudogene |
| ENSG00000256657 | AC244131,1 | -0,944700493 | 0,519537398 | 0,44771103 | 0,902062594 | taste receptor, type 2, member 7 pseudogene |
| ENSG00000275025 | AC002401,3 | -0,944815275 | 0,519496064 | 0,52861717 | 0,930033144 | novel transcript, antisense to PDK2 |
| ENSG00000283294 | AP005212,4 | -0,945498254 | 0,519250191 | 0,12374336 | 0,562483871 | novel transcript |
| ENSG00000164049 | FBXW12 | -0,945659472 | 0,519192169 | 0,07509578 | 0,43291799 | F-box and WD repeat domain containing 12 |

| | | | | | | |
|---|---|---|---|---|---|---|
| ENSG00000130032 | PRRG3 | -0,945666665 | 0,51918958 | 0,00039986 | 0,00862441 | proline rich and Gla domain 3 |
| ENSG00000230828 | AL583843,1 | -0,946123465 | 0,519025215 | 0,48824798 | 0,915394579 | novel pseudogene |
| ENSG00000232978 | AL161909,2 | -0,946685326 | 0,518823119 | 0,49324573 | 0,91725463 | novel transcript |
| ENSG00000273373 | AL355488,1 | -0,946862669 | 0,518759347 | 0,03851259 | 0,291183243 | novel transcript, antisense to SLC16A4 |
| ENSG00000271907 | SNORA35B | -0,94729165 | 0,518605118 | 0,16303057 | 0,64113052 | small nucleolar RNA, H/ACA box 35B |
| ENSG00000186645 | SPDYE17 | -0,947595259 | 0,518495992 | 0,00877998 | 0,104076013 | speedy/RINGO cell cycle regulator family member E17 |
| ENSG00000248283 | CCNL2P1 | -0,947759462 | 0,518436981 | 0,33549032 | 0,835320086 | cyclin L2 pseudogene 1 |
| ENSG00000267041 | ZNF850 | -0,948213479 | 0,518273855 | 2,19E-05 | 0,000683521 | zinc finger protein 850 |
| ENSG00000168811 | IL12A | -0,949263791 | 0,517896678 | 0,27831319 | 0,789652735 | interleukin 12A |
| ENSG00000268530 | AC008403,2 | -0,949528932 | 0,517801506 | 0,29108455 | 0,801689099 | novel transcript, antisense to KCNJ14 |
| ENSG00000114541 | FRMD4B | -0,949714842 | 0,517734785 | 0,02563056 | 0,22266762 | FERM domain containing 4B |
| ENSG00000204571 | KRTAP5-11 | -0,949824851 | 0,517695308 | 0,16713978 | 0,647757288 | keratin associated protein 5-11 |
| ENSG00000233073 | GRM3-AS1 | -0,949855748 | 0,517684221 | 0,0911618 | 0,481283407 | GRM3 antisense RNA 1 |
| ENSG00000254689 | LINC02235 | -0,949966445 | 0,517644501 | 0,20171397 | 0,701332924 | long intergenic non-protein coding RNA 2235 |
| ENSG00000188394 | GPR21 | -0,950252118 | 0,517542011 | 0,01722686 | 0,169258473 | G protein-coupled receptor 21 |
| ENSG00000285879 | AC018628,2 | -0,95038332 | 0,517494947 | 0,04353373 | 0,315096443 | novel transcript, antisense to MED13 |
| ENSG00000261826 | AC121333,1 | -0,950543863 | 0,517437363 | 0,15531766 | 0,625190083 | novel transcript |
| ENSG00000264204 | AGAP7P | -0,950611795 | 0,517412999 | 0,13958714 | 0,595782615 | ArfGAP with GTPase domain, ankyrin repeat and PH domain 7, pseudogene |
| ENSG00000278195 | SSTR3 | -0,950799947 | 0,517345524 | 0,16157786 | 0,63813307 | somatostatin receptor 3 |
| ENSG00000271737 | AC008608,2 | -0,95083307 | 0,517333646 | 0,09107913 | 0,481283407 | novel transcript |
| ENSG00000269405 | NXF2 | -0,951315837 | 0,517160561 | 0,54193164 | 0,933491609 | nuclear RNA export factor 2 |
| ENSG00000249937 | LINC02223 | -0,951530999 | 0,517083438 | 0,14617597 | 0,609479529 | long intergenic non-protein coding RNA 2223 |
| ENSG00000232265 | LINC02805 | -0,95280777 | 0,516626026 | 0,08508632 | 0,464352786 | long intergenic non-protein coding RNA 2805 |
| ENSG00000233524 | RANP8 | -0,952818715 | 0,516622107 | 0,3571557 | 0,848693232 | RAN pseudogene 8 |
| ENSG00000233583 | AL356273,1 | -0,952863796 | 0,516605964 | 0,44439455 | 0,900210555 | tyrosine aminotransferase (TAT) pseudogene |
| ENSG00000287370 | AL356317,1 | -0,953372259 | 0,516423924 | 0,40239348 | 0,87718264 | novel transcript |
| ENSG00000281249 | AL049612,2 | -0,953374213 | 0,516423224 | 0,46417143 | 0,90737035 | novel transcript |
| ENSG00000236173 | AL049612,1 | -0,953374213 | 0,516423224 | 0,46417143 | 0,90737035 | novel transcript |
| ENSG00000204010 | IFIT1B | -0,953380994 | 0,516420797 | 0,36259622 | 0,852667044 | interferon induced protein with tetratricopeptide repeats 1B |
| ENSG00000176945 | MUC20 | -0,953450756 | 0,516395826 | 0,04491189 | 0,320284066 | mucin 20, cell surface associated |

| | | | | | | |
|---|---|---|---|---|---|---|
| ENSG00000285188 | AC008397,2 | -0,953457687 | 0,516393345 | 0,13230786 | 0,580797367 | novel protein |
| ENSG00000248859 | LINC01574 | -0,953690745 | 0,516309932 | 0,49468815 | 0,91758518 | long intergenic non-protein coding RNA 1574 |
| ENSG00000231822 | SMC3P1 | -0,953849221 | 0,51625322 | 0,19336735 | 0,690313354 | SMC3 pseudogene 1 |
| ENSG00000237329 | AL356320,2 | -0,953867907 | 0,516246533 | 0,06755834 | 0,408854824 | TAF9 RNA polymerase II, TATA box binding protein (TBP)-associated factor, 32kDa (TAF9) pseudogene |
| ENSG00000215217 | C5orf49 | -0,953939592 | 0,516220882 | 0,00661762 | 0,085142822 | chromosome 5 open reading frame 49 |
| ENSG00000235635 | AC016644,1 | -0,954113182 | 0,516158772 | 0,34640846 | 0,841168721 | novel transcript |
| ENSG00000251364 | AC107884,1 | -0,954542276 | 0,516005277 | 0,01848879 | 0,177256534 | novel transcript, antisense to OLFML1 |
| ENSG00000284189 | AC141424,2 | -0,954655855 | 0,515964655 | 0,42025764 | 0,888457169 | novel transcript, antisense to C17orf97 |
| ENSG00000241525 | AC141424,1 | -0,954655855 | 0,515964655 | 0,42025764 | 0,888457169 | novel transcript, antisense to C17orf97 |
| ENSG00000162755 | KLHDC9 | -0,954937322 | 0,515864001 | 0,00878747 | 0,104076013 | kelch domain containing 9 |
| ENSG00000252355 | RN7SKP287 | -0,955078044 | 0,515813685 | 0,25078186 | 0,762986007 | RN7SK pseudogene 287 |
| ENSG00000074706 | IPCEF1 | -0,955593696 | 0,515629355 | 0,07086916 | 0,418838091 | interaction protein for cytohesin exchange factors 1 |
| ENSG00000257350 | AC027287,1 | -0,955622511 | 0,515619056 | 0,49105241 | 0,916995341 | heat shock transcription factor 2 (HSF2) pseudogene |
| ENSG00000251158 | AC131392,1 | -0,955777569 | 0,515563642 | 0,1609822 | 0,637001317 | POM121 membrane glycoprotein-like (POM121L1) pseudogene |
| ENSG00000259512 | HNRNPA1P5 | -0,95635588 | 0,515357017 | 0,3761447 | 0,862539384 | heterogeneous nuclear ribonucleoprotein A1 pseudogene 5 |
| ENSG00000288385 | MG910337,1 | -0,95648833 | 0,515309706 | 0,35459891 | 0,846034781 | |
| ENSG00000272800 | AC021851,1 | -0,956624434 | 0,515261093 | 0,05996404 | 0,380884973 | novel transcript |
| ENSG00000270292 | AC079880,2 | -0,957828802 | 0,514831131 | 0,17265719 | 0,657085135 | tubulin, alpha 4a (TUBA4A) pseudogene |
| ENSG00000218313 | AL139274,1 | -0,958080228 | 0,514741416 | 0,01971884 | 0,18474376 | novel TWIST neighbor (TWISTNB) pseudogene |
| ENSG00000263505 | SLC25A3P3 | -0,958257748 | 0,514678082 | 0,36159757 | 0,851893059 | solute carrier family 25 member 3 pseudogene 3 |
| ENSG00000164946 | FREM1 | -0,958365483 | 0,51463965 | 0,01639216 | 0,163524062 | FRAS1 related extracellular matrix 1 |
| ENSG00000225952 | AL512444,1 | -0,958582851 | 0,514562116 | 0,52869732 | 0,930033144 | novel transcript |
| ENSG00000266983 | AC011444,1 | -0,959149037 | 0,514360215 | 0,51286878 | 0,923689697 | novel transcript, antisense to RFX2 |
| ENSG00000124466 | LYPD3 | -0,959164126 | 0,514354836 | 0,15910763 | 0,633144409 | LY6/PLAUR domain containing 3 |
| ENSG00000261839 | AL358933,1 | -0,95923431 | 0,514329814 | 0,10828114 | 0,524537831 | novel transcript, overlapping to ZNF435 |
| ENSG00000271427 | AL358072,1 | -0,959770105 | 0,514138835 | 0,4032853 | 0,877589651 | novel transcript |
| ENSG00000099849 | RASSF7 | -0,960064041 | 0,514034095 | 0,01465273 | 0,151489059 | Ras association domain family member 7 |
| ENSG00000179455 | MKRN3 | -0,960294675 | 0,513951926 | 0,00220374 | 0,036319322 | makorin ring finger protein 3 |
| ENSG00000241962 | AC079447,1 | -0,960533084 | 0,513867001 | 0,04552418 | 0,323249559 | novel protein, C2orf15-MRPL30-novel lincRNA readthrough |

| | | | | | | |
|---|---|---|---|---|---|---|
| ENSG00000130822 | PNCK | -0,960567716 | 0,513854666 | 0,04914934 | 0,33801251 | pregnancy up-regulated nonubiquitous CaM kinase |
| ENSG00000253868 | FER1L6-AS2 | -0,960870911 | 0,513746686 | 0,33447461 | 0,834713776 | FER1L6 antisense RNA 2 |
| ENSG00000271428 | MPHOSPH6P1 | -0,960945885 | 0,513719989 | 0,1475848 | 0,611779236 | MPHOSPH6 pseudogene 1 |
| ENSG00000241261 | RPL17P19 | -0,961044838 | 0,513684754 | 0,10059187 | 0,503748876 | ribosomal protein L17 pseudogene 19 |
| ENSG00000140057 | AK7 | -0,961338433 | 0,513580228 | 0,00870177 | 0,103520575 | adenylate kinase 7 |
| ENSG00000075213 | SEMA3A | -0,961349919 | 0,513576139 | 0,05508049 | 0,362534017 | semaphorin 3A |
| ENSG00000226765 | RPSAP33 | -0,961495373 | 0,513524362 | 0,09454825 | 0,488468697 | ribosomal protein SA pseudogene 33 |
| ENSG00000270977 | AC015849,5 | -0,961508841 | 0,513519569 | 0,07501289 | 0,432694299 | leucine rich repeat containing 37 member A3 (LRRC37A3) pseudogene |
| ENSG00000255206 | AC011853,2 | -0,961680181 | 0,513458585 | 0,38527665 | 0,868895792 | novel transcript |
| ENSG00000231028 | LINC00271 | -0,961833421 | 0,513404049 | 0,0030096 | 0,046603377 | long intergenic non-protein coding RNA 271 |
| ENSG00000268322 | BNIP3P25 | -0,961860542 | 0,513394398 | 0,07096108 | 0,419143629 | BCL2 interacting protein 3 pseudogene 25 |
| ENSG00000223878 | PPIAP53 | -0,961962382 | 0,513358159 | 0,14069657 | 0,597573348 | peptidylprolyl isomerase A pseudogene 53 |
| ENSG00000130224 | LRCH2 | -0,962601942 | 0,513130632 | 0,0387025 | 0,292128222 | leucine rich repeats and calponin homology domain containing 2 |
| ENSG00000161647 | MPP3 | -0,96280323 | 0,513059044 | 0,00039568 | 0,008560972 | membrane palmitoylated protein 3 |
| ENSG00000166582 | CENPV | -0,963032776 | 0,512977419 | 0,00082619 | 0,01587587 | centromere protein V |
| ENSG00000286064 | AL035413,2 | -0,963514547 | 0,512806144 | 0,47719388 | 0,91143003 | aldo-keto reductase family pseudogene |
| ENSG00000268066 | FMR1-AS1 | -0,963968631 | 0,512644765 | 0,16158351 | 0,63813307 | FMR1 antisense RNA 1 |
| ENSG00000244604 | AC025518,1 | -0,964477453 | 0,512463993 | 0,32306652 | 0,825150495 | ribosomal protein S26 (RPS26) pseudogene |
| ENSG00000272610 | MAGI1-IT1 | -0,964525067 | 0,512447081 | 0,02601268 | 0,225030586 | MAGI1 intronic transcript 1 |
| ENSG00000202198 | 7SK | -0,964693604 | 0,512387219 | 0,02125993 | 0,194704806 | 7SK RNA |
| ENSG00000185055 | EFCAB10 | -0,964702942 | 0,512383903 | 0,11876415 | 0,550651408 | EF-hand calcium binding domain 10 |
| ENSG00000186051 | TAL2 | -0,96481746 | 0,512343233 | 0,09895407 | 0,499306007 | TAL bHLH transcription factor 2 |
| ENSG00000250049 | AC114316,1 | -0,964902789 | 0,512312931 | 0,13610077 | 0,587965501 | novel transcript |
| ENSG00000179055 | OR13D1 | -0,965124089 | 0,512234351 | 0,43272065 | 0,89560871 | olfactory receptor family 13 subfamily D member 1 |
| ENSG00000203397 | SHISA5P2 | -0,965424414 | 0,512127731 | 0,47613234 | 0,911413793 | SHISA5 pseudogene 2 |
| ENSG00000146285 | SCML4 | -0,965680973 | 0,512036666 | 0,07534535 | 0,433690715 | Scm polycomb group protein like 4 |
| ENSG00000287577 | AC009533,4 | -0,96573555 | 0,512017296 | 0,38923268 | 0,870057277 | novel transcript |
| ENSG00000258741 | H2AZ2P1 | -0,965848708 | 0,511977137 | 0,05812699 | 0,373081984 | H2AZ2 pseudogene 1 |
| ENSG00000267288 | AC138150,2 | -0,966477545 | 0,511754027 | 0,26096836 | 0,77309354 | novel transcript, antisense to HEXIM2 |
| ENSG00000267758 | AC099811,4 | -0,966568316 | 0,51172183 | 0,1898834 | 0,685035137 | novel transcript |
| ENSG00000283757 | AL031686,1 | -0,966657723 | 0,511690118 | 0,09285642 | 0,484408979 | novel transcript |

| | | | | | | |
|---|---|---|---|---|---|---|
| ENSG00000159314 | ARHGAP27 | -0,966936914 | 0,511591105 | 0,17364155 | 0,658552229 | Rho GTPase activating protein 27 |
| ENSG00000233270 | SNRPEP4 | -0,967266374 | 0,511474289 | 0,18939453 | 0,684693635 | SNRPE pseudogene 4 |
| ENSG00000235422 | KATNBL1P3 | -0,967594346 | 0,511358028 | 0,21692565 | 0,724544002 | katanin regulatory subunit B1 like 1 pseudogene 3 |
| ENSG00000234806 | RPL26P29 | -0,967733118 | 0,511308843 | 0,20434526 | 0,705535354 | ribosomal protein L26 pseudogene 29 |
| ENSG00000256616 | AP002414,2 | -0,967734556 | 0,511308333 | 0,0729745 | 0,426045723 | G protein-coupled receptor 125 (GPR125) pseudogene |
| ENSG00000225725 | FAM66E | -0,967759075 | 0,511299644 | 0,05185812 | 0,349341151 | family with sequence similarity 66 member E |
| ENSG00000162374 | ELAVL4 | -0,968154629 | 0,511159476 | 0,15324998 | 0,621750224 | ELAV like RNA binding protein 4 |
| ENSG00000220506 | AL136310,1 | -0,968360709 | 0,511086465 | 0,18474042 | 0,67694849 | novel pseudogene |
| ENSG00000267708 | RDM1P3 | -0,968961723 | 0,510873595 | 0,35557082 | 0,846892875 | RDM1 pseudogene 3 |
| ENSG00000253598 | SLC10A5 | -0,969046226 | 0,510843673 | 0,26097842 | 0,77309354 | solute carrier family 10 member 5 |
| ENSG00000287936 | AL391425,1 | -0,969117427 | 0,510818462 | 0,09635095 | 0,493094813 | novel transcript |
| ENSG00000287857 | AL592546,3 | -0,969687111 | 0,510616792 | 0,0825889 | 0,457152163 | novel transcript, antisense to NHLRC2 |
| ENSG00000257643 | AC084824,2 | -0,970015103 | 0,510500718 | 0,30641346 | 0,815375074 | mitochondrial assembly of ribosomal large subunit 1 (MALSU1) pseudogene |
| ENSG00000210107 | MT-TQ | -0,970354547 | 0,51038062 | 0,43231745 | 0,89558934 | mitochondrially encoded tRNA-Gln (CAA/G) |
| ENSG00000204657 | OR2H2 | -0,97035991 | 0,510378723 | 0,06963983 | 0,416512828 | olfactory receptor family 2 subfamily H member 2 |
| ENSG00000232623 | MRAP-AS1 | -0,970509332 | 0,510325864 | 0,3880233 | 0,869820053 | MRAP antisense RNA 1 |
| ENSG00000270739 | AC090953,1 | -0,970736603 | 0,510245478 | 0,3200972 | 0,824366234 | THAP domain containing 5, pseudogene |
| ENSG00000111837 | MAK | -0,971482734 | 0,509981658 | 0,01919353 | 0,182079045 | male germ cell associated kinase |
| ENSG00000260907 | AC015818,2 | -0,971636008 | 0,50992748 | 0,15449262 | 0,623626246 | novel transcript |
| ENSG00000080298 | RFX3 | -0,97168258 | 0,509911019 | 0,03604357 | 0,27830728 | regulatory factor X3 |
| ENSG00000250197 | HMGN1P15 | -0,971823715 | 0,509861138 | 0,18089126 | 0,67062545 | high mobility group nucleosome binding domain 1 pseudogene 15 |
| ENSG00000115138 | POMC | -0,971892345 | 0,509836885 | 0,25158869 | 0,763892102 | proopiomelanocortin |
| ENSG00000272180 | AC011306,1 | -0,972082919 | 0,509769541 | 0,43475005 | 0,896272827 | novel transcript |
| ENSG00000280336 | AC244517,12 | -0,972339205 | 0,509678992 | 0,30290934 | 0,81247062 | novel transcript |
| ENSG00000253300 | AC018541,1 | -0,972438462 | 0,509643928 | 0,24382167 | 0,754811426 | novel transcript |
| ENSG00000258343 | AC090001,1 | -0,97246156 | 0,509635768 | 0,33608327 | 0,835660292 | novel transcript |
| ENSG00000215796 | AL512637,1 | -0,972964408 | 0,509458167 | 0,10523462 | 0,517691472 | primase polypeptide 1 49kDa (PRIM1) pseudogene |
| ENSG00000283567 | C19orf85 | -0,973088615 | 0,509414308 | 0,520382 | 0,926968424 | chromosome 19 open reading frame 85 |
| ENSG00000214300 | SPDYE3 | -0,97311289 | 0,509405736 | 0,00038943 | 0,008455021 | speedy/RINGO cell cycle regulator family member E3 |

| | | | | | | |
|---|---|---|---|---|---|---|
| ENSG00000104332 | SFRP1 | -0,973175166 | 0,509383747 | 0,0058191 | 0,07715373 | secreted frizzled related protein 1 |
| ENSG00000278558 | TMEM191B | -0,973211607 | 0,509370881 | 0,36166762 | 0,851893059 | transmembrane protein 191B |
| ENSG00000275197 | AC092794,2 | -0,973403809 | 0,509303025 | 0,40701342 | 0,880078305 | novel transcript, antisense to KRAS |
| ENSG00000288393 | AP003086,3 | -0,973462285 | 0,509282382 | 0,03775791 | 0,287287228 | zinc finger protein 75C, pseudogene |
| ENSG00000260693 | AC026150,1 | -0,973607796 | 0,509231018 | 0,39101715 | 0,870057277 | novel transcript |
| ENSG00000281960 | AC254960,1 | -0,973607796 | 0,509231018 | 0,39101715 | 0,870057277 | novel transcript |
| ENSG00000236463 | LINC00427 | -0,973707694 | 0,509195758 | 0,36236774 | 0,852386203 | long intergenic non-protein coding RNA 427 |
| ENSG00000287000 | AC024651,3 | -0,973892543 | 0,50913052 | 0,19788486 | 0,696383642 | novel transcript |
| ENSG00000270772 | PDHA1P1 | -0,974190681 | 0,509025318 | 0,17093545 | 0,654596502 | PDHA1 pseudogene 1 |
| ENSG00000285844 | FO393414,3 | -0,97445636 | 0,508931587 | 0,28971621 | 0,800522458 | novel transcript |
| ENSG00000264007 | AC104564,2 | -0,974468991 | 0,508927131 | 0,46847797 | 0,907940653 | novel transcript |
| ENSG00000284672 | AC109583,4 | -0,974542238 | 0,508901293 | 0,10399704 | 0,514293503 | novel transcript |
| ENSG00000146216 | TTBK1 | -0,974662126 | 0,508859005 | 0,05237665 | 0,351288432 | tau tubulin kinase 1 |
| ENSG00000281379 | SEPTIN14P19 | -0,974746211 | 0,508829348 | 0,17168959 | 0,655397237 | septin 14 pseudogene 19 |
| ENSG00000265752 | AC010754,1 | -0,975178976 | 0,508676737 | 0,4754877 | 0,911264385 | novel transcript |
| ENSG00000121749 | TBC1D15 | -0,975302563 | 0,508633164 | 0,01465806 | 0,151494064 | TBC1 domain family member 15 |
| ENSG00000103154 | NECAB2 | -0,975357554 | 0,508613777 | 0,00653238 | 0,084252613 | N-terminal EF-hand calcium binding protein 2 |
| ENSG00000238276 | AL354863,1 | -0,976249762 | 0,508299331 | 0,21532215 | 0,722575157 | uncharacterized LOC101927760 |
| ENSG00000258535 | AL079307,1 | -0,976267428 | 0,508293107 | 0,00917014 | 0,107537595 | novel transcript |
| ENSG00000269431 | BNIP3P8 | -0,976986357 | 0,508039876 | 0,33558332 | 0,835324592 | BCL2 interacting protein 3 pseudogene 8 |
| ENSG00000239508 | PLCH1-AS1 | -0,977183201 | 0,507970563 | 0,05028512 | 0,343031651 | PLCH1 antisense RNA 1 |
| ENSG00000169989 | TIGD4 | -0,977276319 | 0,507937777 | 0,02573324 | 0,223230867 | tigger transposable element derived 4 |
| ENSG00000240922 | LSAMP-AS1 | -0,977313133 | 0,507924816 | 0,18737784 | 0,681464181 | LSAMP antisense RNA 1 |
| ENSG00000240207 | AC080013,1 | -0,9773333 | 0,507917716 | 0,00041314 | 0,008865126 | novel transcript, antisense to RARRES1 |
| ENSG00000261570 | AC073878,1 | -0,977508619 | 0,507855996 | 0,27522938 | 0,786829871 | novel transcript, antisense to AGK |
| ENSG00000286703 | AC005682,2 | -0,977836641 | 0,50774054 | 0,35331672 | 0,84514549 | novel transcript |
| ENSG00000096395 | MLN | -0,977975259 | 0,507691757 | 0,49988188 | 0,920340799 | motilin |
| ENSG00000255040 | MORF4L1P3 | -0,978104918 | 0,507646131 | 0,29332676 | 0,803461221 | mortality factor 4 like 1 pseudogene 3 |
| ENSG00000163590 | PPM1L | -0,978186231 | 0,50761752 | 0,01571194 | 0,159066362 | protein phosphatase, Mg2+/Mn2+ dependent 1L |
| ENSG00000174808 | BTC | -0,978726135 | 0,507427588 | 0,30671982 | 0,815554392 | betacellulin |
| ENSG00000282381 | AC104073,4 | -0,978942162 | 0,507351613 | 0,01315223 | 0,139782152 | novel transcript |
| ENSG00000260570 | AC133550,2 | -0,979272528 | 0,507235446 | 0,58641866 | 0,945395317 | novel transcript, antisense to ATXN2L |
| ENSG00000258640 | RPL21P5 | -0,979603878 | 0,507118961 | 0,38672703 | 0,869441674 | ribosomal protein L21 pseudogene 5 |

| | | | | | | |
|---|---|---|---|---|---|---|
| ENSG00000187772 | LIN28B | -0,979941646 | 0,507000246 | 0,18581438 | 0,678777213 | lin-28 homolog B |
| ENSG00000286273 | AC078816,1 | -0,980052701 | 0,50696122 | 0,17165915 | 0,655397237 | novel transcript, antisense to GPR87 |
| ENSG00000078900 | TP73 | -0,980195535 | 0,506911031 | 0,22981605 | 0,738230413 | tumor protein p73 |
| ENSG00000251629 | LINC02241 | -0,98117251 | 0,506567874 | 0,03542724 | 0,275521695 | long intergenic non-protein coding RNA 2241 |
| ENSG00000219451 | RPL23P8 | -0,981335536 | 0,506510634 | 0,2543331 | 0,765826923 | ribosomal protein L23 pseudogene 8 |
| ENSG00000230005 | SNAP47-AS1 | -0,981436978 | 0,506475021 | 0,11050693 | 0,530623376 | SNAP47 antisense RNA 1 |
| ENSG00000231212 | AL590399,3 | -0,981836145 | 0,506334908 | 0,19292987 | 0,689849199 | novel transcript |
| ENSG00000185272 | RBM11 | -0,982591868 | 0,506069745 | 0,08058504 | 0,450103921 | RNA binding motif protein 11 |
| ENSG00000232190 | LINC02181 | -0,983012164 | 0,505922335 | 0,39768226 | 0,874263835 | long intergenic non-protein coding RNA 2181 |
| ENSG00000271597 | AC112230,1 | -0,983118442 | 0,505885067 | 0,50674063 | 0,921824694 | golgi reassembly stacking protein 2, 55kDa (GORASP2) pseudogene |
| ENSG00000249412 | AC010285,1 | -0,98319996 | 0,505856483 | 0,05925753 | 0,378088799 | novel transcript |
| ENSG00000004838 | ZMYND10 | -0,983280156 | 0,505828365 | 0,02544553 | 0,221657538 | zinc finger MYND-type containing 10 |
| ENSG00000182175 | RGMA | -0,98396407 | 0,505588632 | 0,00256934 | 0,040920913 | repulsive guidance molecule BMP co-receptor a |
| ENSG00000269040 | BNIP3P24 | -0,984094692 | 0,505542858 | 0,21282009 | 0,719853652 | BCL2 interacting protein 3 pseudogene 24 |
| ENSG00000280428 | BNIP3P24 | -0,984094692 | 0,505542858 | 0,21282009 | 0,719853652 | BCL2 interacting protein 3 pseudogene 24 |
| ENSG00000233771 | CICP5 | -0,984264941 | 0,505483203 | 0,34280416 | 0,840419133 | capicua transcriptional repressor pseudogene 5 |
| ENSG00000168490 | PHYHIP | -0,984649573 | 0,505348456 | 0,01612918 | 0,161882053 | phytanoyl-CoA 2-hydroxylase interacting protein |
| ENSG00000146360 | GPR6 | -0,985101519 | 0,505190173 | 0,42092416 | 0,888848798 | G protein-coupled receptor 6 |
| ENSG00000144152 | FBLN7 | -0,985534502 | 0,505038577 | 0,0054489 | 0,073650815 | fibulin 7 |
| ENSG00000166676 | TVP23A | -0,985739359 | 0,504966869 | 0,00061467 | 0,012430211 | trans-golgi network vesicle protein 23 homolog A |
| ENSG00000086300 | SNX10 | -0,986121014 | 0,504833301 | 0,13246712 | 0,581009552 | sorting nexin 10 |
| ENSG00000268758 | ADGRE4P | -0,98613912 | 0,504826966 | 0,38327243 | 0,867740551 | adhesion G protein-coupled receptor E4, pseudogene |
| ENSG00000187726 | DNAJB13 | -0,987119737 | 0,504483945 | 0,251558 | 0,763892102 | DnaJ heat shock protein family (Hsp40) member B13 |
| ENSG00000285873 | AL445253,1 | -0,987590815 | 0,504319245 | 0,20706879 | 0,710823213 | novel transcript |
| ENSG00000215319 | EIF5P1 | -0,987766893 | 0,504257697 | 0,18861635 | 0,682944308 | eukaryotic translation initiation factor 5 pseudogene 1 |
| ENSG00000198580 | AC073343,1 | -0,988042188 | 0,504161484 | 0,20486061 | 0,706459522 | zinc finger protein pseudogene |
| ENSG00000264324 | AC006030,1 | -0,988129975 | 0,504130807 | 0,2633022 | 0,774637968 | novel protein |
| ENSG00000133665 | DYDC2 | -0,988294786 | 0,504073219 | 0,01992465 | 0,186058616 | DPY30 domain containing 2 |
| ENSG00000286512 | AL031768,2 | -0,989173474 | 0,503766302 | 0,0434721 | 0,314827799 | novel transcript |
| ENSG00000231905 | SETP10 | -0,989299366 | 0,503722345 | 0,11945721 | 0,552362303 | SET pseudogene 10 |

| | | | | | | |
|---|---|---|---|---|---|---|
| ENSG00000128573 | FOXP2 | -0,989479197 | 0,50365956 | 0,045719 | 0,324264732 | forkhead box P2 |
| ENSG00000181240 | SLC25A41 | -0,989717117 | 0,503576506 | 0,05705953 | 0,36911008 | solute carrier family 25 member 41 |
| ENSG00000271732 | AL137798,1 | -0,990496663 | 0,503304477 | 0,45728118 | 0,903831777 | novel transcript |
| ENSG00000238063 | LINC01685 | -0,990533259 | 0,503291711 | 0,56541719 | 0,940865625 | long intergenic non-protein coding RNA 1685 |
| ENSG00000214671 | RPL6P12 | -0,990636847 | 0,503255575 | 0,1943249 | 0,6913261 | ribosomal protein L6 pseudogene 12 |
| ENSG00000282390 | AL392083,1 | -0,990843594 | 0,50318346 | 0,09835811 | 0,49790609 | novel transcript, sense intronic to ADARB2 |
| ENSG00000204983 | PRSS1 | -0,990871982 | 0,503173559 | 0,41554284 | 0,885986046 | serine protease 1 |
| ENSG00000240804 | NPM1P28 | -0,991158147 | 0,503073762 | 0,33055545 | 0,832114797 | nucleophosmin 1 pseudogene 28 |
| ENSG00000181291 | TMEM132E | -0,991336798 | 0,50301147 | 0,13681046 | 0,590208676 | transmembrane protein 132E |
| ENSG00000229488 | PRRT1 | -0,991403814 | 0,502988105 | 0,17108127 | 0,654834089 | proline rich transmembrane protein 1 |
| ENSG00000232491 | SAPCD2P3 | -0,991582862 | 0,502925684 | 0,52512233 | 0,928752911 | suppressor APC domain containing 2 pseudogene 3 |
| ENSG00000284872 | AC270289,3 | -0,992249633 | 0,5026933 | 0,01085868 | 0,121796931 | tRNA methyltransferase 9B (putative) |
| ENSG00000228998 | AC091167,1 | -0,992506191 | 0,502603913 | 0,01947725 | 0,183691282 | PERP, TP53 apoptosis effector (PERP) pseudogene |
| ENSG00000178115 | GOLGA8Q | -0,99291994 | 0,502459793 | 0,29346256 | 0,803461221 | golgin A8 family member Q |
| ENSG00000117013 | KCNQ4 | -0,993139499 | 0,502383331 | 0,05723618 | 0,369932153 | potassium voltage-gated channel subfamily Q member 4 |
| ENSG00000265055 | AC145343,1 | -0,993535201 | 0,502245556 | 0,28862819 | 0,79998466 | novel transcript |
| ENSG00000143365 | RORC | -0,993546883 | 0,502241489 | 0,23793446 | 0,747090777 | RAR related orphan receptor C |
| ENSG00000279206 | AC004943,3 | -0,994032768 | 0,502072368 | 0,21303967 | 0,72028467 | novel transcript |
| ENSG00000259594 | AC023034,1 | -0,994243795 | 0,501998934 | 0,29607274 | 0,805887018 | novel transcript |
| ENSG00000146352 | CLVS2 | -0,994460638 | 0,501923487 | 0,02096736 | 0,192913017 | clavesin 2 |
| ENSG00000213578 | CPLX3 | -0,994579512 | 0,501882131 | 0,30884183 | 0,816831843 | complexin 3 |
| ENSG00000185615 | PDIA2 | -0,995275547 | 0,501640055 | 0,2160235 | 0,723046035 | protein disulfide isomerase family A member 2 |
| ENSG00000274769 | AC016747,4 | -0,995720648 | 0,501485312 | 0,06691534 | 0,406616339 | novel transcript |
| ENSG00000146197 | SCUBE3 | -0,995921203 | 0,501415604 | 0,00546417 | 0,073825264 | signal peptide, CUB domain and EGF like domain containing 3 |
| ENSG00000233993 | DTD1-AS1 | -0,996169314 | 0,501329379 | 0,15204141 | 0,620332737 | DTD1 antisense RNA 1 |
| ENSG00000237672 | KRR1P1 | -0,996327475 | 0,501274422 | 0,09431876 | 0,487885809 | KRR1 pseudogene 1 |
| ENSG00000106038 | EVX1 | -0,996381273 | 0,501255729 | 0,34264942 | 0,84026916 | even-skipped homeobox 1 |
| ENSG00000224097 | AC021148,1 | -0,997800202 | 0,500762974 | 0,29815761 | 0,807554329 | WD repeat domain 5 (WDR5) pseudogene |
| ENSG00000130021 | PUDP | -0,998500259 | 0,500520041 | 0,00624446 | 0,081378445 | pseudouridine 5'-phosphatase |
| ENSG00000173988 | LRRC63 | -0,998868552 | 0,500392284 | 0,13504928 | 0,586038109 | leucine rich repeat containing 63 |
| ENSG00000182165 | TP53TG1 | -0,99908598 | 0,500316876 | 0,00571872 | 0,076202714 | TP53 target 1 |

| | | | | | | |
|---|---|---|---|---|---|---|
| ENSG00000164185 | ZNF474 | -0,999092985 | 0,500314446 | 0,12410113 | 0,563164667 | zinc finger protein 474 |
| ENSG00000136546 | SCN7A | -0,999119353 | 0,500305302 | 0,04936446 | 0,338781394 | sodium voltage-gated channel alpha subunit 7 |
| ENSG00000242540 | AC010729,2 | -0,99922998 | 0,50026694 | 0,21416341 | 0,721518909 | novel transcript |
| ENSG00000118492 | ADGB | -0,999589098 | 0,500142428 | 0,32984311 | 0,831624145 | androglobin |
| ENSG00000168004 | PLAAT5 | -0,999758634 | 0,500083658 | 0,02912434 | 0,24236233 | phospholipase A and acyltransferase 5 |
| ENSG00000276853 | AC026124,2 | -0,999993106 | 0,500002389 | 0,28747721 | 0,798896538 | novel transcript, sense intronic to LEMD3 |
| ENSG00000242808 | SOX2-OT | -1,000274968 | 0,499904712 | 0,01280577 | 0,137391732 | SOX2 overlapping transcript |
| ENSG00000132837 | DMGDH | -1,000496543 | 0,499827941 | 0,17550067 | 0,661199837 | dimethylglycine dehydrogenase |
| ENSG00000182352 | C17orf77 | -1,000785168 | 0,499727956 | 0,49167387 | 0,916995341 | chromosome 17 open reading frame 77 |
| ENSG00000074211 | PPP2R2C | -1,001585381 | 0,499450851 | 0,11565201 | 0,543151268 | protein phosphatase 2 regulatory subunit Bgamma |
| ENSG00000224488 | SAMM50P1 | -1,001999786 | 0,499307407 | 0,10057186 | 0,503729332 | SAMM50 sorting and assembly machinery component pseudogene 1 |
| ENSG00000114547 | ROPN1B | -1,002283614 | 0,499209186 | 0,10030819 | 0,502892138 | rhophilin associated tail protein 1B |
| ENSG00000254664 | AC103681,2 | -1,00310864 | 0,498923787 | 0,45968882 | 0,905107873 | novel transcript |
| ENSG00000270893 | AC008277,2 | -1,003131867 | 0,498915755 | 0,21501872 | 0,722298056 | elongation factor Tu GTP binding domain containing 1 (EFTUD1) pseudogene |
| ENSG00000241597 | AC022121,1 | -1,004193238 | 0,498548844 | 0,19430634 | 0,6913261 | ribosomal protein S9 (RPS9) pseudogene |
| ENSG00000226448 | RPL7AP51 | -1,004263031 | 0,498524727 | 0,21537486 | 0,72263142 | ribosomal protein L7a pseudogene 51 |
| ENSG00000236133 | LINC01069 | -1,004554367 | 0,498424065 | 0,42012543 | 0,888457169 | long intergenic non-protein coding RNA 1069 |
| ENSG00000253661 | ZFHX4-AS1 | -1,004618068 | 0,498402059 | 0,0787764 | 0,444319927 | ZFHX4 antisense RNA 1 |
| ENSG00000219085 | NPM1P37 | -1,005101247 | 0,498235165 | 0,21192993 | 0,718553023 | nucleophosmin 1 pseudogene 37 |
| ENSG00000141401 | IMPA2 | -1,00553715 | 0,498084648 | 0,00702884 | 0,088859506 | inositol monophosphatase 2 |
| ENSG00000110042 | DTX4 | -1,006029342 | 0,49791475 | 1,54E-06 | 6,33E-05 | deltex E3 ubiquitin ligase 4 |
| ENSG00000003987 | MTMR7 | -1,006225381 | 0,497847096 | 0,0426375 | 0,310582197 | myotubularin related protein 7 |
| ENSG00000229948 | B3GALNT1P1 | -1,006241714 | 0,49784146 | 0,19299485 | 0,689849199 | beta-1,3-N-acetylgalactosaminyltransferase 1 pseudogene 1 |
| ENSG00000162409 | PRKAA2 | -1,006280374 | 0,497828119 | 0,06457138 | 0,398334582 | protein kinase AMP-activated catalytic subunit alpha 2 |
| ENSG00000275536 | AC118459,1 | -1,007248272 | 0,49749424 | 0,24570794 | 0,757752583 | novel transcript |
| ENSG00000258593 | AL583810,1 | -1,007418933 | 0,497435393 | 0,43976157 | 0,897728093 | novel transcript, antisense to KIAA0284 |
| ENSG00000227230 | AL606534,1 | -1,007457529 | 0,497422086 | 0,04757027 | 0,331676387 | novel transcript |
| ENSG00000111834 | RSPH4A | -1,007530183 | 0,497397036 | 0,0537909 | 0,357812498 | radial spoke head component 4A |
| ENSG00000170959 | DCDC1 | -1,007533711 | 0,49739582 | 0,0255928 | 0,222568404 | doublecortin domain containing 1 |

| | | | | | | |
|---|---|---|---|---|---|---|
| ENSG00000285701 | AL357140,4 | -1,007889393 | 0,497273207 | 0,14725117 | 0,611181688 | novel transcript, antisense to CTNNBIP1 |
| ENSG00000244124 | ATP1B3-AS1 | -1,007987294 | 0,497239464 | 0,17188343 | 0,655736851 | ATP1B3 antisense RNA 1 |
| ENSG00000265136 | AC124283,4 | -1,008292931 | 0,497134134 | 0,31145925 | 0,817751312 | ADP-ribosylation factor-like 2 binding protein (ARL2BP) pseudogene |
| ENSG00000287218 | AL359962,3 | -1,00831482 | 0,497126591 | 0,29321797 | 0,803365244 | novel transcript, antisense to CD247 |
| ENSG00000218521 | AL589655,1 | -1,008659894 | 0,497007699 | 0,18392959 | 0,675485923 | pseudogene similar to an integrin Alpha-E precursor fragment |
| ENSG00000228414 | LINC01185 | -1,008663442 | 0,497006477 | 0,43603239 | 0,896526626 | long intergenic non-protein coding RNA 1185 |
| ENSG00000228393 | LINC01004 | -1,008835248 | 0,496947293 | 0,03338883 | 0,265327213 | long intergenic non-protein coding RNA 1004 |
| ENSG00000278662 | GOLGA6L10 | -1,009497956 | 0,496719071 | 0,06964973 | 0,416512828 | golgin A6 family like 10 |
| ENSG00000175048 | ZDHHC14 | -1,009694834 | 0,496651291 | 0,02589154 | 0,224198135 | zinc finger DHHC-type palmitoyltransferase 14 |
| ENSG00000204175 | GPRIN2 | -1,009881656 | 0,496586981 | 0,03694885 | 0,283685844 | G protein regulated inducer of neurite outgrowth 2 |
| ENSG00000234772 | LINC00412 | -1,009972075 | 0,496555859 | 0,25500781 | 0,76642479 | long intergenic non-protein coding RNA 412 |
| ENSG00000276017 | AC007325,1 | -1,010076267 | 0,496519999 | 0,15193349 | 0,620191762 | |
| ENSG00000100314 | CABP7 | -1,010426948 | 0,496399323 | 0,12414163 | 0,563164667 | calcium binding protein 7 |
| ENSG00000232472 | EEF1B2P3 | -1,010458684 | 0,496388403 | 0,09096787 | 0,481140155 | eukaryotic translation elongation factor 1 beta 2 pseudogene 3 |
| ENSG00000224727 | FCF1P7 | -1,01063169 | 0,496328881 | 0,39588324 | 0,872308364 | FCF1 pseudogene 7 |
| ENSG00000212128 | TAS2R13 | -1,010690728 | 0,49630857 | 0,11482627 | 0,541181798 | taste 2 receptor member 13 |
| ENSG00000277254 | TAS2R13 | -1,010690728 | 0,49630857 | 0,11482627 | 0,541181798 | taste 2 receptor member 13 |
| ENSG00000273457 | TAS2R13 | -1,010690728 | 0,49630857 | 0,11482627 | 0,541181798 | taste 2 receptor member 13 |
| ENSG00000286268 | AF196969,1 | -1,01069154 | 0,496308291 | 0,33873501 | 0,837449623 | novel protein |
| ENSG00000154822 | PLCL2 | -1,010891954 | 0,49623935 | 0,01602395 | 0,161013527 | phospholipase C like 2 |
| ENSG00000284017 | PLCL2 | -1,010891954 | 0,49623935 | 0,01602395 | 0,161013527 | phospholipase C like 2 |
| ENSG00000241884 | AC114401,1 | -1,01101115 | 0,496198353 | 0,39403477 | 0,871635019 | novel transcript |
| ENSG00000075643 | MOCOS | -1,011416431 | 0,49605898 | 0,03176808 | 0,257126831 | molybdenum cofactor sulfurase |
| ENSG00000237336 | AL445489,1 | -1,011569478 | 0,496006359 | 0,26968304 | 0,781791089 | novel transcript |
| ENSG00000130812 | ANGPTL6 | -1,011695689 | 0,495962969 | 0,4821594 | 0,912981134 | angiopoietin like 6 |
| ENSG00000196277 | GRM7 | -1,011762357 | 0,495940051 | 0,04790727 | 0,333134539 | glutamate metabotropic receptor 7 |
| ENSG00000270100 | AC012065,3 | -1,011818665 | 0,495920695 | 0,34904153 | 0,843141089 | novel transcript |
| ENSG00000287843 | AL161628,1 | -1,012752578 | 0,49559977 | 0,37148482 | 0,859476158 | novel transcript |
| ENSG00000163618 | CADPS | -1,013290996 | 0,495414845 | 0,00016804 | 0,004085144 | calcium dependent secretion activator |

| | | | | | | |
|---|---|---|---|---|---|---|
| ENSG00000237731 | RNGTTP1 | -1,013574272 | 0,495317579 | 0,09697766 | 0,494601865 | RNA guanylyltransferase and 5'-phosphatase pseudogene 1 |
| ENSG00000256779 | AP002383,3 | -1,014212554 | 0,495098487 | 0,37659467 | 0,862818448 | serine/arginine-rich splicing factor 8 (SRSF8) pseudogene |
| ENSG00000230221 | AL157384,1 | -1,014344635 | 0,495053162 | 0,02320776 | 0,207542201 | ATM interactor (ATMIN) pseudogene |
| ENSG00000111339 | ART4 | -1,014657432 | 0,494945839 | 0,07092428 | 0,419005435 | ADP-ribosyltransferase 4 (Dombrock blood group) |
| ENSG00000268058 | BNIP3P40 | -1,015169946 | 0,494770042 | 0,0846014 | 0,462844615 | BCL2 interacting protein 3 pseudogene 40 |
| ENSG00000179915 | NRXN1 | -1,015302857 | 0,494724462 | 0,03258955 | 0,261251213 | neurexin 1 |
| ENSG00000258769 | RAP1AP | -1,015427422 | 0,494681749 | 0,40179349 | 0,877131444 | RAP1A, member of RAS oncogene family pseudogene |
| ENSG00000178175 | ZNF366 | -1,015808462 | 0,494551112 | 0,21694142 | 0,724544002 | zinc finger protein 366 |
| ENSG00000253213 | USP12P1 | -1,016065996 | 0,494462839 | 0,18867036 | 0,682944308 | USP12 pseudogene 1 |
| ENSG00000277823 | PRH1 | -1,016089024 | 0,494454946 | 0,29718305 | 0,806635068 | proline rich protein HaeIII subfamily 1 |
| ENSG00000215246 | AC116351,1 | -1,016134637 | 0,494439313 | 0,1500518 | 0,615757437 | novel transcript |
| ENSG00000287967 | AC004882,3 | -1,016736201 | 0,494233189 | 0,27245702 | 0,785492261 | novel transcript, antisense to UQCR10 |
| ENSG00000284697 | AC006511,5 | -1,016821613 | 0,494203929 | 0,31714526 | 0,822226508 | novel transcript |
| ENSG00000229927 | RHEBP1 | -1,017504206 | 0,493970159 | 0,21219333 | 0,718991056 | RHEB pseudogene 1 |
| ENSG00000270614 | AC011451,2 | -1,0176835 | 0,493908773 | 0,10828324 | 0,524537831 | UBX domain protein 2A (UBXN2A) pseudogene |
| ENSG00000276653 | RN7SL856P | -1,017752469 | 0,493885162 | 0,42715664 | 0,892343796 | RNA, 7SL, cytoplasmic 856, pseudogene |
| ENSG00000281406 | BLACAT1 | -1,017997325 | 0,493801346 | 0,01973014 | 0,184776117 | bladder cancer associated transcript 1 |
| ENSG00000141748 | ARL5C | -1,018331016 | 0,493687145 | 0,28208693 | 0,793203731 | ADP ribosylation factor like GTPase 5C |
| ENSG00000264176 | MAGOH2P | -1,018622132 | 0,493587536 | 0,2453932 | 0,757479368 | mago homolog 2, pseudogene |
| ENSG00000270179 | AP002840,2 | -1,018693908 | 0,49356298 | 0,16751805 | 0,648316613 | novel transcript |
| ENSG00000270402 | AC011458,1 | -1,019160497 | 0,49340338 | 0,29541491 | 0,805426883 | ankyrin repeat domain 26 (ANKRD26) pseudogene |
| ENSG00000204315 | FKBPL | -1,020132329 | 0,493071124 | 0,65545902 | 0,957444211 | FKBP prolyl isomerase like |
| ENSG00000239830 | RPS4XP22 | -1,020359413 | 0,492993519 | 0,4424855 | 0,899414493 | ribosomal protein S4X pseudogene 22 |
| ENSG00000287578 | AL034348,1 | -1,020985584 | 0,492779593 | 0,35750071 | 0,848693232 | novel transcript |
| ENSG00000214688 | C10orf105 | -1,02103321 | 0,492763325 | 0,20722801 | 0,711093909 | chromosome 10 open reading frame 105 |
| ENSG00000286722 | AC009244,1 | -1,021518267 | 0,492597678 | 0,03802193 | 0,288593771 | novel transcript, antisense to AHCYL2 |
| ENSG00000237179 | LINC01797 | -1,021977957 | 0,492440745 | 0,26087752 | 0,773014385 | long intergenic non-protein coding RNA 1797 |
| ENSG00000237248 | LINC00987 | -1,023003484 | 0,492090823 | 0,0567931 | 0,367995736 | long intergenic non-protein coding RNA 987 |
| ENSG00000122548 | KIAA0087 | -1,023108873 | 0,492054877 | 0,03943487 | 0,295799395 | KIAA0087 lncRNA |
| ENSG00000236172 | MIR7515HG | -1,023744404 | 0,491838166 | 0,48092164 | 0,912907448 | MIR7515 host gene |

| | | | | | | |
|---|---|---|---|---|---|---|
| ENSG00000002079 | MYH16 | -1,023885311 | 0,491790131 | 0,30786399 | 0,816424917 | myosin heavy chain 16 pseudogene |
| ENSG00000228016 | RAPGEF4-AS1 | -1,024069987 | 0,491727182 | 0,29282264 | 0,803295982 | RAPGEF4 antisense RNA 1 |
| ENSG00000184564 | SLITRK6 | -1,024142339 | 0,491702522 | 0,07596506 | 0,435455001 | SLIT and NTRK like family member 6 |
| ENSG00000185002 | RFX6 | -1,024488917 | 0,491584415 | 0,28904499 | 0,800166929 | regulatory factor X6 |
| ENSG00000286773 | AL353593,3 | -1,024542187 | 0,491566264 | 0,46105067 | 0,906029574 | novel transcript, sense intronic to OBSCN |
| ENSG00000275898 | AC125603,4 | -1,025594919 | 0,4912077 | 0,4241136 | 0,890949676 | novel transcript |
| ENSG00000177133 | PRDM16-DT | -1,02565947 | 0,491185722 | 0,00623998 | 0,081353878 | PRDM16 divergent transcript |
| ENSG00000256349 | AP002748,5 | -1,025715696 | 0,49116658 | 0,69246894 | 0,967236299 | novel protein |
| ENSG00000234003 | MTATP6P27 | -1,026035569 | 0,491057691 | 0,15909121 | 0,633144409 | MT-ATP6 pseudogene 27 |
| ENSG00000235421 | AC114501,2 | -1,026430257 | 0,490923367 | 0,2685303 | 0,780709773 | novel pseudogene similar to part of integrator complex subunit 4 INTS4 |
| ENSG00000078018 | MAP2 | -1,026649995 | 0,4908486 | 0,01524206 | 0,155858486 | microtubule associated protein 2 |
| ENSG00000256420 | OSBPL9P4 | -1,026674257 | 0,490840345 | 0,08376573 | 0,461014667 | oxysterol binding protein like 9 pseudogene 4 |
| ENSG00000165325 | DEUP1 | -1,026806329 | 0,490795413 | 0,04456597 | 0,318908336 | deuterosome assembly protein 1 |
| ENSG00000228084 | AC118553,1 | -1,026837739 | 0,490784728 | 0,3928047 | 0,871091361 | novel transcript |
| ENSG00000140009 | ESR2 | -1,026846719 | 0,490781673 | 0,00017099 | 0,004143974 | estrogen receptor 2 |
| ENSG00000267940 | AC022762,2 | -1,027738804 | 0,490478294 | 0,02389817 | 0,211657934 | novel transcript |
| ENSG00000253245 | MTND1P36 | -1,028101535 | 0,49035499 | 0,26423354 | 0,775607676 | MT-ND1 pseudogene 36 |
| ENSG00000282872 | C1orf232 | -1,028283167 | 0,49029326 | 0,27608398 | 0,788074712 | chromosome 1 open reading frame 230 |
| ENSG00000288184 | AC118282,7 | -1,028292804 | 0,490289985 | 0,2573056 | 0,769504717 | |
| ENSG00000288249 | AC270106,1 | -1,028292804 | 0,490289985 | 0,2573056 | 0,769504717 | |
| ENSG00000244232 | RN7SL698P | -1,028306074 | 0,490285475 | 0,45195393 | 0,902062594 | RNA, 7SL, cytoplasmic 698, pseudogene |
| ENSG00000224448 | AC005020,1 | -1,028417533 | 0,490247598 | 0,25610382 | 0,768188606 | ADP-ribosylation factor 6 (ARF6) pseudogene |
| ENSG00000226647 | AL365356,1 | -1,028434267 | 0,490241912 | 0,07722106 | 0,439261743 | novel transcript |
| ENSG00000173825 | TIGD3 | -1,028867063 | 0,490094865 | 0,00187561 | 0,031834377 | tigger transposable element derived 3 |
| ENSG00000228368 | AC106873,1 | -1,029067578 | 0,490026754 | 0,46752607 | 0,907940653 | novel transcript, antisense to ST7 |
| ENSG00000166405 | RIC3 | -1,029270714 | 0,489957761 | 0,01225975 | 0,13267178 | RIC3 acetylcholine receptor chaperone |
| ENSG00000237483 | RPS27AP8 | -1,029283887 | 0,489953288 | 0,27320403 | 0,785672647 | RPS27A pseudogene 8 |
| ENSG00000250860 | AC093225,1 | -1,029356317 | 0,48992869 | 0,17444706 | 0,659880557 | novel zinc finger protein pseudogene |
| ENSG00000008226 | DLEC1 | -1,029456767 | 0,489894579 | 0,00048546 | 0,010153285 | DLEC1 cilia and flagella associated protein |
| ENSG00000163568 | AIM2 | -1,030602807 | 0,489505574 | 0,01637557 | 0,163462876 | absent in melanoma 2 |
| ENSG00000237436 | CAMTA1-DT | -1,030777279 | 0,489446379 | 0,38126297 | 0,865900094 | CAMTA1 divergent transcript |
| ENSG00000185915 | KLHL34 | -1,030836608 | 0,489426252 | 0,10385217 | 0,513912548 | kelch like family member 34 |

| | | | | | | |
|---|---|---|---|---|---|---|
| ENSG00000241596 | AC092468,1 | -1,031686153 | 0,489138133 | 0,22776317 | 0,736053199 | novel transcript |
| ENSG00000244558 | KCNK15-AS1 | -1,031859203 | 0,489079465 | 0,30546316 | 0,814326218 | KCNK15 and WISP2 antisense RNA 1 |
| ENSG00000280374 | AC019080,5 | -1,031964656 | 0,489043717 | 0,05449779 | 0,360376004 | novel transcript |
| ENSG00000175664 | TEX26 | -1,032136403 | 0,488985502 | 0,15257775 | 0,621037742 | testis expressed 26 |
| ENSG00000283991 | AC017000,1 | -1,032212785 | 0,488959614 | 0,41854947 | 0,887094957 | novel transcript, antisense to SDK1 |
| ENSG00000120156 | TEK | -1,032311033 | 0,488926317 | 0,01313245 | 0,13975671 | TEK receptor tyrosine kinase |
| ENSG00000275991 | Metazoa_SRP | -1,032381951 | 0,488902284 | 0,64009888 | 0,956157624 | Metazoan signal recognition particle RNA |
| ENSG00000235079 | ZRANB2-AS1 | -1,032706001 | 0,488792481 | 0,14169279 | 0,599581043 | ZRANB2 antisense RNA 1 |
| ENSG00000198765 | SYCP1 | -1,032871428 | 0,488736437 | 0,21409297 | 0,72150508 | synaptonemal complex protein 1 |
| ENSG00000285293 | TBILA | -1,033018787 | 0,488686519 | 0,13829919 | 0,593030588 | TGF-beta induced lncRNA |
| ENSG00000261488 | TBILA | -1,033018787 | 0,488686519 | 0,13829919 | 0,593030588 | TGF-beta induced lncRNA |
| ENSG00000285892 | AC006043,1 | -1,033104811 | 0,488657381 | 0,30001739 | 0,809925194 | novel transcript |
| ENSG00000274933 | TBC1D3I | -1,033266074 | 0,488602763 | 0,6343678 | 0,955097925 | TBC1 domain family member 3I |
| ENSG00000256745 | ARPC3P3 | -1,033502872 | 0,488522572 | 0,28084135 | 0,791510309 | actin related protein 2/3 complex subunit 3 pseudogene 3 |
| ENSG00000186205 | MTARC1 | -1,033987955 | 0,488358342 | 0,00260818 | 0,041407116 | mitochondrial amidoxime reducing component 1 |
| ENSG00000249065 | PCNAP1 | -1,034140441 | 0,488306727 | 0,26582343 | 0,77713706 | proliferating cell nuclear antigen pseudogene 1 |
| ENSG00000101349 | PAK5 | -1,034204368 | 0,488285091 | 0,04085257 | 0,302299378 | p21 (RAC1) activated kinase 5 |
| ENSG00000128833 | MYO5C | -1,034392166 | 0,488221534 | 0,00846355 | 0,101559908 | myosin VC |
| ENSG00000230470 | AL078645,1 | -1,034437739 | 0,488206112 | 0,45495716 | 0,903424009 | novel transcript |
| ENSG00000269978 | AL359881,1 | -1,034438451 | 0,488205871 | 0,22343941 | 0,731984265 | novel transcript |
| ENSG00000130035 | GALNT8 | -1,034502703 | 0,488184128 | 0,00868385 | 0,103346713 | polypeptide N-acetylgalactosaminyltransferase 8 |
| ENSG00000233210 | GPANK1 | -1,034576313 | 0,488159221 | 0,36434538 | 0,854320801 | G-patch domain and ankyrin repeats 1 |
| ENSG00000184156 | KCNQ3 | -1,034715035 | 0,488112284 | 0,00290264 | 0,045148393 | potassium voltage-gated channel subfamily Q member 3 |
| ENSG00000282260 | AC019257,6 | -1,035157756 | 0,48796252 | 0,27481896 | 0,786383635 | novel transcript |
| ENSG00000270988 | AC019257,2 | -1,035157756 | 0,48796252 | 0,27481896 | 0,786383635 | novel transcript |
| ENSG00000285257 | AL157893,1 | -1,035370984 | 0,487890405 | 0,20393689 | 0,704681782 | synaptotagmin 15 (SYT15) pseudogene |
| ENSG00000231170 | AC002451,1 | -1,035545829 | 0,48783128 | 0,30123122 | 0,81068201 | novel transcript |
| ENSG00000165837 | ERICH6B | -1,035780525 | 0,487751926 | 0,28141971 | 0,792425997 | glutamate rich 6B |
| ENSG00000226088 | HHLA3-AS1 | -1,036038324 | 0,487664776 | 0,43591046 | 0,896422069 | HHLA3 antisense RNA 1 |
| ENSG00000276945 | AC008498,1 | -1,036112198 | 0,487639806 | 0,28209589 | 0,793203731 | protocadherin 19 (PCDH19) pseudogene |
| ENSG00000146090 | RASGEF1C | -1,036173933 | 0,487618939 | 0,01660835 | 0,164996341 | RasGEF domain family member 1C |

| | | | | | | |
|---|---|---|---|---|---|---|
| ENSG00000280722 | AL008628,2 | -1,036530975 | 0,487498277 | 0,35729481 | 0,848693232 | novel transcript, overlapping to FAM120B |
| ENSG00000262050 | AC005696,1 | -1,036702883 | 0,487440191 | 0,42925633 | 0,894029049 | novel transcript, antisense to CLUH |
| ENSG00000107742 | SPOCK2 | -1,036822791 | 0,48739968 | 0,0401405 | 0,299154351 | SPARC (osteonectin), cwcv and kazal like domains proteoglycan 2 |
| ENSG00000267466 | AC021683,1 | -1,036852816 | 0,487389536 | 0,00622288 | 0,0812289 | novel transcript |
| ENSG00000287244 | AL020997,5 | -1,036937398 | 0,487360963 | 0,31219588 | 0,818547047 | novel transcript |
| ENSG00000238056 | PRRT1 | -1,037000395 | 0,487339682 | 0,15424407 | 0,623607455 | proline rich transmembrane protein 1 |
| ENSG00000235956 | PRRT1 | -1,037000395 | 0,487339682 | 0,15424407 | 0,623607455 | proline rich transmembrane protein 1 |
| ENSG00000239351 | NPM1P29 | -1,037515734 | 0,487165633 | 0,07831801 | 0,442692864 | nucleophosmin 1 pseudogene 29 |
| ENSG00000273402 | AC004908,3 | -1,037612288 | 0,487133029 | 0,26439934 | 0,775730197 | novel transcript, antisense to ZNF596 |
| ENSG00000173838 | MARCHF10 | -1,03770262 | 0,487102529 | 0,12281034 | 0,560473734 | membrane associated ring-CH-type finger 10 |
| ENSG00000259564 | AC009554,1 | -1,038165541 | 0,486946257 | 0,43572767 | 0,896422069 | novel transcript, antisense to VPS13C |
| ENSG00000288174 | SLC2A4 | -1,039055274 | 0,486646042 | 0,11569132 | 0,543236711 | solute carrier family 2 member 4 |
| ENSG00000272361 | AC005014,2 | -1,039063959 | 0,486643112 | 0,20988805 | 0,714886155 | novel transcript |
| ENSG00000230991 | AC092625,1 | -1,039666739 | 0,486439828 | 0,53026879 | 0,931157231 | novel transcript |
| ENSG00000287827 | AC002529,1 | -1,039769399 | 0,486405215 | 0,51379988 | 0,924227101 | novel transcript |
| ENSG00000266976 | AC079466,1 | -1,040527982 | 0,486149525 | 0,00499008 | 0,068726827 | novel transcript |
| ENSG00000235833 | AC017099,1 | -1,041389617 | 0,485859264 | 0,01811741 | 0,174934575 | coiled-coil domain containing 144C (CCDC144C) pseudogene |
| ENSG00000286881 | AL358780,1 | -1,041951752 | 0,48566999 | 0,05305229 | 0,354145183 | novel transcript, sense intronic to MLLT10 |
| ENSG00000180458 | AC022148,1 | -1,042024128 | 0,485645626 | 0,10536158 | 0,518119329 | novel transcript, antisense to ZNF793 |
| ENSG00000258759 | AL049779,2 | -1,042373363 | 0,485528079 | 0,38408227 | 0,868125769 | DnaJ (Hsp40) homolog, subfamily C, member 9 (DNAJC9) pseudogene |
| ENSG00000237349 | BX679664,2 | -1,042380515 | 0,485525672 | 0,46905839 | 0,907940653 | NADH dehydrogenase (ubiquinone) 1 beta subcomplex, 3, 12kDa (NDUFB3) pseudogene |
| ENSG00000181856 | SLC2A4 | -1,044041249 | 0,484967089 | 0,23263239 | 0,741684508 | solute carrier family 2 member 4 |
| ENSG00000285535 | AC021683,5 | -1,044081015 | 0,484953722 | 0,02341581 | 0,20868642 | novel transcript |
| ENSG00000089101 | CFAP61 | -1,044134766 | 0,484935654 | 0,06907992 | 0,414636184 | cilia and flagella associated protein 61 |
| ENSG00000205704 | LINC00634 | -1,044197831 | 0,484914456 | 0,02900481 | 0,241642594 | long intergenic non-protein coding RNA 634 |
| ENSG00000235012 | JCADP1 | -1,044338964 | 0,484867021 | 0,3106651 | 0,817751312 | JCAD pseudogene 1 |
| ENSG00000142621 | FHAD1 | -1,045110916 | 0,48460765 | 0,01515208 | 0,155214504 | forkhead associated phosphopeptide binding domain 1 |

| | | | | | | |
|---|---|---|---|---|---|---|
| ENSG00000233063 | SAR1AP4 | -1,045383602 | 0,484516062 | 0,41238369 | 0,884523593 | secretion associated Ras related GTPase 1A pseudogene 4 |
| ENSG00000198719 | DLL1 | -1,045487857 | 0,48448105 | 0,00144493 | 0,025458789 | delta like canonical Notch ligand 1 |
| ENSG00000180662 | RPL21P8 | -1,045519972 | 0,484470266 | 0,17367461 | 0,658552229 | ribosomal protein L21 pseudogene 8 |
| ENSG00000260785 | CASC17 | -1,046039269 | 0,484295912 | 0,00561159 | 0,075200114 | cancer susceptibility 17 |
| ENSG00000272438 | AL645608,6 | -1,046319093 | 0,484201988 | 0,37148061 | 0,859476158 | novel transcript |
| ENSG00000184524 | CEND1 | -1,046391558 | 0,484177667 | 0,00462407 | 0,065237814 | cell cycle exit and neuronal differentiation 1 |
| ENSG00000134376 | CRB1 | -1,046947501 | 0,483991125 | 0,03190512 | 0,257747195 | crumbs cell polarity complex component 1 |
| ENSG00000225037 | EIF1AX-AS1 | -1,047113374 | 0,483935482 | 0,25922778 | 0,771989931 | EIF1AX antisense RNA 1 |
| ENSG00000259773 | AC012100,2 | -1,047629694 | 0,483762319 | 0,48209049 | 0,912981134 | novel transcript, antisense to SPPL2A |
| ENSG00000271993 | AC126118,1 | -1,04803485 | 0,483626482 | 0,08044328 | 0,449985752 | novel transcript, antisense to LRRFIP2 |
| ENSG00000248228 | SLIT2-IT1 | -1,048521813 | 0,483463268 | 0,17352868 | 0,658349911 | SLIT2 intronic transcript 1 |
| ENSG00000287129 | AC097500,1 | -1,048937171 | 0,483324097 | 0,30858277 | 0,816730513 | novel transcript |
| ENSG00000227317 | DDAH2 | -1,049068822 | 0,483279994 | 0,02106171 | 0,193441015 | dimethylarginine dimethylaminohydrolase 2 |
| ENSG00000262874 | C19orf84 | -1,049120158 | 0,483262797 | 0,14980165 | 0,615346044 | chromosome 19 open reading frame 84 |
| ENSG00000163995 | ABLIM2 | -1,049183908 | 0,483241443 | 0,00424678 | 0,061411368 | actin binding LIM protein family member 2 |
| ENSG00000124678 | TCP11 | -1,049446908 | 0,483153358 | 0,32147379 | 0,824933727 | t-complex 11 |
| ENSG00000213050 | TPM3P1 | -1,049541928 | 0,483121537 | 0,15199438 | 0,620332737 | tropomyosin 3 pseudogene 1 |
| ENSG00000227170 | AF178030,1 | -1,05011156 | 0,482930819 | 0,28698906 | 0,798400609 | novel transcript |
| ENSG00000213760 | ATP6V1G2 | -1,050516197 | 0,482795389 | 0,18527577 | 0,677679985 | ATPase H+ transporting V1 subunit G2 |
| ENSG00000163631 | ALB | -1,051112125 | 0,482596004 | 0,36876309 | 0,857685565 | albumin |
| ENSG00000242816 | AC092691,3 | -1,051145076 | 0,482584982 | 0,25914263 | 0,771989931 | novel transcript |
| ENSG00000286981 | AL731553,1 | -1,051266509 | 0,482544364 | 0,30630806 | 0,815375074 | novel transcript |
| ENSG00000286319 | AL135978,1 | -1,051337652 | 0,482520569 | 0,16765967 | 0,648566191 | novel transcript, antisense to MPP5 |
| ENSG00000283913 | AL512662,1 | -1,052110821 | 0,482262046 | 0,07958284 | 0,44749602 | novel transcript |
| ENSG00000230847 | OCLNP1 | -1,052397501 | 0,482166224 | 0,12417146 | 0,563164667 | OCLN pseudogene 1 |
| ENSG00000267152 | AC093227,1 | -1,052480228 | 0,482138577 | 0,33308831 | 0,833301711 | novel transcript |
| ENSG00000229961 | AL357143,1 | -1,052491362 | 0,482134856 | 0,42594384 | 0,892133432 | novel SLAM family member pseudogene |
| ENSG00000137878 | GCOM1 | -1,052853538 | 0,482013835 | 0,08497204 | 0,46411003 | GRINL1A complex locus 1 |
| ENSG00000208308 | SNORA40B | -1,05320266 | 0,481897205 | 0,28980008 | 0,800629762 | small nucleolar RNA, H/ACA box 40B |
| ENSG00000260259 | LINC02166 | -1,053915785 | 0,481659062 | 0,28851605 | 0,799943767 | long intergenic non-protein coding RNA 2166 |
| ENSG00000268174 | BNIP3P22 | -1,054012916 | 0,481626635 | 0,0948876 | 0,489386708 | BCL2 interacting protein 3 pseudogene 22 |
| ENSG00000270765 | GAS2L2 | -1,054103645 | 0,481596347 | 0,34946127 | 0,843190352 | growth arrest specific 2 like 2 |

| | | | | | | |
|---|---|---|---|---|---|---|
| ENSG00000277036 | AC139494,6 | -1,054124881 | 0,481589258 | 0,07262797 | 0,424935759 | novel transcript |
| ENSG00000280677 | FAM66B | -1,054133523 | 0,481586374 | 0,00857213 | 0,102485328 | family with sequence similarity 66 member B |
| ENSG00000248152 | ANK2-AS1 | -1,054306377 | 0,481528677 | 0,01048812 | 0,118761539 | ANK2 antisense RNA 1 |
| ENSG00000185053 | SGCZ | -1,054968811 | 0,481307627 | 0,28295238 | 0,794166737 | sarcoglycan zeta |
| ENSG00000178125 | PPP1R42 | -1,055230966 | 0,481220175 | 0,13960206 | 0,595782615 | protein phosphatase 1 regulatory subunit 42 |
| ENSG00000267868 | AL356740,1 | -1,055268914 | 0,481207517 | 0,09773884 | 0,496782149 | novel transcript |
| ENSG00000232872 | CTAGE3P | -1,055791325 | 0,4810333 | 0,02707274 | 0,230753051 | CTAGE family member 3, pseudogene |
| ENSG00000213434 | VTI1BP2 | -1,055816077 | 0,481025047 | 0,22681852 | 0,735383523 | vesicle transport through interaction with t-SNAREs 1B pseudogene 2 |
| ENSG00000287580 | AC073349,4 | -1,055844719 | 0,481015498 | 0,0427871 | 0,311526734 | novel transcript, antisense to ZNF138 |
| ENSG00000272601 | AC010655,3 | -1,055852245 | 0,481012988 | 0,02100888 | 0,193069221 | EF-hand calcium binding domain 3 (EFCAB3) pseudogene |
| ENSG00000285394 | C3orf52 | -1,056097827 | 0,480931115 | 0,20066544 | 0,70017905 | chromosome 3 open reading frame 52 |
| ENSG00000155980 | KIF5A | -1,056111058 | 0,480926705 | 0,00507915 | 0,069494347 | kinesin family member 5A |
| ENSG00000186369 | LINC00643 | -1,056222094 | 0,480889692 | 0,10474163 | 0,516098066 | long intergenic non-protein coding RNA 643 |
| ENSG00000278704 | BX004987,1 | -1,056267025 | 0,480874715 | 0,03711932 | 0,284435918 | |
| ENSG00000270343 | UNGP3 | -1,056317832 | 0,480857781 | 0,16090997 | 0,636876744 | uracil-DNA glycosylase pseudogene 3 |
| ENSG00000282022 | AC006581,2 | -1,056406199 | 0,480828328 | 0,36600238 | 0,85572078 | novel transcript, antisense to PHC1 and A2ML1 |
| ENSG00000112041 | TULP1 | -1,056496231 | 0,480798323 | 0,1456947 | 0,608454479 | TUB like protein 1 |
| ENSG00000115252 | PDE1A | -1,056518031 | 0,480791058 | 0,09797564 | 0,497096776 | phosphodiesterase 1A |
| ENSG00000225569 | CCT4P2 | -1,056531332 | 0,480786625 | 0,17965177 | 0,668298509 | chaperonin containing TCP1 subunit 4 pseudogene 2 |
| ENSG00000285772 | AC000120,2 | -1,057767869 | 0,480374719 | 0,3437029 | 0,840901506 | novel protein |
| ENSG00000235292 | AL109838,1 | -1,057814626 | 0,48035915 | 0,32236313 | 0,824955371 | novel transcript |
| ENSG00000113212 | PCDHB7 | -1,05794745 | 0,480314927 | 0,0037923 | 0,056324176 | protocadherin beta 7 |
| ENSG00000285795 | FAM90A23P | -1,058083296 | 0,480269702 | 0,40462452 | 0,878176155 | family with sequence similarity 90 member A23, pseudogene |
| ENSG00000188130 | MAPK12 | -1,058231358 | 0,480220415 | 0,01612274 | 0,161869455 | mitogen-activated protein kinase 12 |
| ENSG00000164123 | C4orf45 | -1,058498747 | 0,480131419 | 0,32119935 | 0,824723604 | chromosome 4 open reading frame 45 |
| ENSG00000253582 | AC090579,1 | -1,058615271 | 0,480092641 | 0,49538983 | 0,917799558 | novel transcript |
| ENSG00000251155 | SEPTIN14P4 | -1,059748093 | 0,479715815 | 0,43209458 | 0,89558934 | septin 14 pseudogene 4 |
| ENSG00000185101 | ANO9 | -1,059845088 | 0,479683564 | 0,3437874 | 0,840901506 | anoctamin 9 |
| ENSG00000233602 | ERI3-IT1 | -1,059914772 | 0,479660395 | 0,21137428 | 0,717057897 | ERI3 intronic transcript 1 |
| ENSG00000251552 | COQ10BP2 | -1,060457536 | 0,479479973 | 0,27267974 | 0,78557496 | coenzyme Q10B pseudogene 2 |

| | | | | | | |
|---|---|---|---|---|---|---|
| ENSG00000180316 | PNPLA1 | -1,06050373 | 0,479464621 | 0,44200692 | 0,899273096 | patatin like phospholipase domain containing 1 |
| ENSG00000233626 | AC092814,2 | -1,060963036 | 0,479312 | 0,43588403 | 0,896422069 | protein (peptidyl-prolyl cis\\/trans isomerase) NIMA-interacting, 4 (parvulin) (PIN4) pseudogene |
| ENSG00000287042 | AC090945,1 | -1,061157233 | 0,479247485 | 0,30696321 | 0,815854622 | novel transcript |
| ENSG00000259103 | AF107885,2 | -1,061609213 | 0,479097366 | 0,02929596 | 0,243294836 | novel transcript, antisense to TTLL5 |
| ENSG00000140835 | CHST4 | -1,061732036 | 0,47905658 | 0,13700866 | 0,590712806 | carbohydrate sulfotransferase 4 |
| ENSG00000242152 | AC108729,1 | -1,061879128 | 0,47900774 | 0,47442997 | 0,910676852 | mitochondrially encoded cytochrome c oxidase (MT-CO1) pseudogene |
| ENSG00000278831 | AL513190,1 | -1,06202289 | 0,47896001 | 0,13247434 | 0,581009552 | novel transcript, antisense to FAM53B |
| ENSG00000122735 | DNAI1 | -1,062272317 | 0,47887721 | 0,01153751 | 0,1270996 | dynein axonemal intermediate chain 1 |
| ENSG00000218574 | HNRNPA1P37 | -1,063288569 | 0,478540002 | 0,52101042 | 0,927046149 | heterogeneous nuclear ribonucleoprotein A1 pseudogene 37 |
| ENSG00000265942 | RN7SL577P | -1,063384069 | 0,478508325 | 0,34463359 | 0,841168721 | RNA, 7SL, cytoplasmic 577, pseudogene |
| ENSG00000241599 | AL627309,4 | -1,063493353 | 0,47847208 | 0,32073717 | 0,824366234 | novel transcript |
| ENSG00000118004 | COLEC11 | -1,063526366 | 0,478461131 | 0,31904382 | 0,824073525 | collectin subfamily member 11 |
| ENSG00000287618 | AC025576,3 | -1,063816883 | 0,478364793 | 0,1743324 | 0,659766151 | novel transcript, antisense to SRGAP1 |
| ENSG00000224307 | AL161785,1 | -1,063853516 | 0,478352646 | 0,34597656 | 0,841168721 | novel transcript |
| ENSG00000144191 | CNGA3 | -1,063941263 | 0,478323553 | 0,04939802 | 0,338828378 | cyclic nucleotide gated channel subunit alpha 3 |
| ENSG00000225125 | RANP4 | -1,064199024 | 0,4782381 | 0,11626154 | 0,544294818 | RAN pseudogene 4 |
| ENSG00000259926 | AC007342,1 | -1,065065835 | 0,477950848 | 0,25111441 | 0,76347766 | novel transcript |
| ENSG00000135333 | EPHA7 | -1,065244853 | 0,477891545 | 0,10115817 | 0,505774682 | EPH receptor A7 |
| ENSG00000206062 | AL121823,1 | -1,065478099 | 0,477814288 | 0,40862126 | 0,88153752 | novel transcript |
| ENSG00000185522 | LMNTD2 | -1,06557714 | 0,477781487 | 0,44411455 | 0,900210555 | lamin tail domain containing 2 |
| ENSG00000231464 | AC024937,2 | -1,06577398 | 0,477716304 | 0,17042291 | 0,653594444 | protein phosphatase 4, regulatory subunit 2 (PPP4R2) pseudogene |
| ENSG00000214548 | MEG3 | -1,065782081 | 0,477713621 | 0,04224273 | 0,308497259 | maternally expressed 3 |
| ENSG00000285595 | AC105114,2 | -1,066179185 | 0,477582148 | 0,027511 | 0,233660421 | novel transcript, antisense to ZNF521 |
| ENSG00000231043 | AC007238,1 | -1,066757345 | 0,477390795 | 0,28478132 | 0,796197593 | IK cytokine, down-regulator of HLA II (IK) pseudogene |
| ENSG00000261476 | AC009163,4 | -1,066879554 | 0,477350358 | 0,12113347 | 0,555784698 | novel transcript |
| ENSG00000104043 | ATP8B4 | -1,066968436 | 0,47732095 | 0,16892718 | 0,650892223 | ATPase phospholipid transporting 8B4 (putative) |
| ENSG00000204965 | PCDHA5 | -1,067248886 | 0,477228171 | 0,00066202 | 0,01315798 | protocadherin alpha 5 |
| ENSG00000283706 | PRSS50 | -1,067597097 | 0,477113001 | 0,21393307 | 0,721277214 | serine protease 50 |

| | | | | | | |
|---|---|---|---|---|---|---|
| ENSG00000118997 | DNAH7 | -1,067801956 | 0,477045256 | 0,02511711 | 0,219776434 | dynein axonemal heavy chain 7 |
| ENSG00000223905 | AC103592,1 | -1,067980434 | 0,476986244 | 0,21979516 | 0,727931767 | peroxiredoxin 5 (PRDX5) pseudogene |
| ENSG00000287794 | AC119150,2 | -1,069164521 | 0,47659492 | 0,37043792 | 0,858545694 | novel ancient vomeronasal 1 receptor pseudogene |
| ENSG00000253333 | AC146944,1 | -1,069274647 | 0,476558541 | 0,08339575 | 0,45983856 | glucuronidase, beta (GUSB) pseudogene |
| ENSG00000271086 | NAMA | -1,06942771 | 0,476507983 | 0,05042099 | 0,343470607 | non-protein coding RNA, associated with MAP kinase pathway and growth arrest |
| ENSG00000223476 | VN1R42P | -1,069565374 | 0,476462516 | 0,20261408 | 0,702665191 | vomeronasal 1 receptor 42 pseudogene |
| ENSG00000286633 | AL160400,1 | -1,069566142 | 0,476462263 | 0,48847882 | 0,915411008 | novel transcript, antisense to RIPOR2 |
| ENSG00000241511 | RPS15AP24 | -1,069687887 | 0,476422057 | 0,29231412 | 0,802771257 | ribosomal protein S15a pseudogene 24 |
| ENSG00000182463 | TSHZ2 | -1,069835376 | 0,476373354 | 0,01159157 | 0,127515405 | teashirt zinc finger homeobox 2 |
| ENSG00000144130 | NT5DC4 | -1,070265184 | 0,476231454 | 0,35306052 | 0,845053131 | 5'-nucleotidase domain containing 4 |
| ENSG00000254092 | AC021613,1 | -1,070869327 | 0,476032069 | 0,38693594 | 0,869441674 | novel transcript |
| ENSG00000230833 | RPEP3 | -1,071165419 | 0,475934381 | 0,16411182 | 0,642888389 | ribulose-5-phosphate-3-epimerase pseudogene 3 |
| ENSG00000259404 | EFL1P1 | -1,071230216 | 0,475913005 | 0,00197742 | 0,033381193 | elongation factor like GTPase 1 pseudogene 1 |
| ENSG00000251246 | AL691442,1 | -1,071504736 | 0,475822456 | 0,61807544 | 0,952793934 | novel ephrin-A4 (EFNA4) and ephrin-A3 (EFNA3) protein |
| ENSG00000232531 | KMT5AP2 | -1,071744079 | 0,475743523 | 0,35041774 | 0,843857545 | KMT5A pseudogene 2 |
| ENSG00000259314 | AC103739,2 | -1,072077062 | 0,475633731 | 0,21937809 | 0,727838473 | novel transcript, antisense to CRTC3 |
| ENSG00000272070 | AC005618,1 | -1,072333113 | 0,475549323 | 0,00634261 | 0,082213892 | novel transcript |
| ENSG00000215237 | AL592293,1 | -1,07252751 | 0,475485249 | 0,32189965 | 0,824955371 | chromosome 4 open reading frame 27 (C4orf27) pseudogene |
| ENSG00000236790 | LINC00299 | -1,072890144 | 0,475365747 | 0,3463004 | 0,841168721 | long intergenic non-protein coding RNA 299 |
| ENSG00000118160 | SLC8A2 | -1,073013088 | 0,475325238 | 0,10604236 | 0,519559347 | solute carrier family 8 member A2 |
| ENSG00000281665 | KIAA1143 | -1,073446065 | 0,475182607 | 0,06790709 | 0,410329987 | KIAA1143 |
| ENSG00000276432 | RDM1 | -1,073712344 | 0,47509491 | 0,0344178 | 0,27063333 | RAD52 motif containing 1 |
| ENSG00000260022 | AL031716,1 | -1,073814413 | 0,475061299 | 0,45216499 | 0,902062594 | novel transcript, antisense to LMF1 |
| ENSG00000183643 | C15orf32 | -1,074411019 | 0,474864885 | 0,16854237 | 0,650506523 | chromosome 15 putative open reading frame 32 |
| ENSG00000112539 | C6orf118 | -1,074413723 | 0,474863995 | 0,00015093 | 0,003729453 | chromosome 6 open reading frame 118 |
| ENSG00000279168 | AC105052,4 | -1,074802957 | 0,474735895 | 0,03103778 | 0,253292923 | novel transcript, sense overlapping POLR2J2, UPK3BL and RASA4 |
| ENSG00000244342 | LINC00698 | -1,075173263 | 0,474614057 | 0,37231174 | 0,860223901 | long intergenic non-protein coding RNA 698 |
| ENSG00000251294 | AC091885,2 | -1,075227751 | 0,474596132 | 0,06575218 | 0,402442454 | novel transcript |
| ENSG00000272123 | AC008966,2 | -1,075939415 | 0,474362078 | 0,42606142 | 0,892133432 | novel transcript, antisense to ITGA2 |

| | | | | | | |
|---|---|---|---|---|---|---|
| ENSG00000233290 | AL157944,1 | -1,076924537 | 0,474038278 | 0,40130729 | 0,876659568 | novel transcript |
| ENSG00000219355 | RPL31P52 | -1,077142862 | 0,473966546 | 0,17571813 | 0,661199837 | ribosomal protein L31 pseudogene 52 |
| ENSG00000263219 | RYKP1 | -1,077150041 | 0,473964188 | 0,02142032 | 0,195699842 | RYK pseudogene 1 |
| ENSG00000256596 | AC026358,1 | -1,077155865 | 0,473962274 | 0,18263466 | 0,673171032 | novel transcript |
| ENSG00000271401 | AC010680,5 | -1,077228406 | 0,473938443 | 0,1659848 | 0,645200651 | novel transcript |
| ENSG00000275613 | AC243830,1 | -1,077302354 | 0,473914151 | 0,13206257 | 0,580641112 | novel transcript |
| ENSG00000167861 | HID1 | -1,077362061 | 0,473894538 | 0,00372426 | 0,055577708 | HID1 domain containing |
| ENSG00000101445 | PPP1R16B | -1,07758074 | 0,473822712 | 0,00558675 | 0,074995323 | protein phosphatase 1 regulatory subunit 16B |
| ENSG00000155530 | LRGUK | -1,077618829 | 0,473810203 | 0,01433133 | 0,149152676 | leucine rich repeats and guanylate kinase domain containing |
| ENSG00000264273 | AC107982,2 | -1,077657559 | 0,473797483 | 0,15266788 | 0,621081138 | novel transcript, antisense to PRPSAP2 |
| ENSG00000131401 | NAPSB | -1,077687016 | 0,47378781 | 0,13098383 | 0,577911147 | napsin B aspartic peptidase, pseudogene |
| ENSG00000259113 | AL118556,1 | -1,078023535 | 0,473677308 | 0,16865361 | 0,650559718 | novel transcript |
| ENSG00000166840 | GLYATL1 | -1,078111959 | 0,473648277 | 0,01173583 | 0,12878521 | glycine-N-acyltransferase like 1 |
| ENSG00000249485 | RBBP4P1 | -1,078163037 | 0,473631508 | 0,17741342 | 0,663776507 | RBBP4 pseudogene 1 |
| ENSG00000163624 | CDS1 | -1,078173539 | 0,47362806 | 0,10779228 | 0,523928283 | CDP-diacylglycerol synthase 1 |
| ENSG00000281425 | AC243922,2 | -1,07831232 | 0,473582501 | 0,69065095 | 0,967179044 | glucuronidase, beta (GUSB) pseudogene |
| ENSG00000222005 | LINC01118 | -1,078976478 | 0,473364534 | 0,3096327 | 0,816986489 | long intergenic non-protein coding RNA 1118 |
| ENSG00000133661 | SFTPD | -1,079378533 | 0,473232633 | 0,01928723 | 0,182560125 | surfactant protein D |
| ENSG00000214870 | AC004540,1 | -1,079632637 | 0,473149289 | 0,00993583 | 0,114625025 | novel transcript |
| ENSG00000280770 | AC216281,1 | -1,079656498 | 0,473141464 | 0,13007346 | 0,576252873 | novel transcript sense intronic to TRAPPC12 |
| ENSG00000169860 | P2RY1 | -1,07984933 | 0,473078228 | 0,0288015 | 0,240931045 | purinergic receptor P2Y1 |
| ENSG00000102003 | SYP | -1,080103453 | 0,472994905 | 0,00327477 | 0,05008927 | synaptophysin |
| ENSG00000234338 | AC073349,2 | -1,080384981 | 0,472902613 | 0,04369238 | 0,315756651 | endogenous retroviral family W, env(C7), member 2 (ERVWE2) pseudogene |
| ENSG00000238021 | ARMC4P1 | -1,080550854 | 0,472848245 | 0,34661091 | 0,841390219 | armadillo repeat containing 4 pseudogene 1 |
| ENSG00000232631 | MTND2P23 | -1,080862595 | 0,472746082 | 0,16485578 | 0,643964432 | MT-ND2 pseudogene 23 |
| ENSG00000288518 | AC275572,1 | -1,081232349 | 0,472624935 | 0,53320415 | 0,931291085 | |
| ENSG00000170370 | EMX2 | -1,081407866 | 0,472567439 | 0,06359568 | 0,394965288 | empty spiracles homeobox 2 |
| ENSG00000236811 | GAPDHP2 | -1,081572643 | 0,472513468 | 0,20735713 | 0,711147468 | glyceraldehyde-3-phosphate dehydrogenase pseudogene 2 |
| ENSG00000227466 | PDE4B-AS1 | -1,082045494 | 0,472358625 | 0,40274904 | 0,877341338 | PDE4B antisense RNA 1 |
| ENSG00000197210 | AP000550,1 | -1,082271745 | 0,472284553 | 0,13894042 | 0,594174432 | novel transcript |

| | | | | | | |
|---|---|---|---|---|---|---|
| ENSG00000272954 | AP000553,2 | -1,083477773 | 0,471889909 | 0,33326321 | 0,833504026 | novel transcript, sense intronic to PPIL2 |
| ENSG00000258131 | AC007656,2 | -1,083873664 | 0,471760436 | 0,27745872 | 0,789067728 | novel transcript |
| ENSG00000275601 | AC011330,2 | -1,083910846 | 0,471748277 | 0,1521767 | 0,620418906 | novel transcript, sense intronic to CATSPER2 |
| ENSG00000132932 | ATP8A2 | -1,08424825 | 0,471637962 | 0,01146112 | 0,126436253 | ATPase phospholipid transporting 8A2 |
| ENSG00000229388 | LINC01715 | -1,084403266 | 0,471587288 | 0,31671012 | 0,821646322 | long intergenic non-protein coding RNA 1715 |
| ENSG00000204161 | TMEM273 | -1,084796982 | 0,471458608 | 0,3477587 | 0,842293776 | transmembrane protein 273 |
| ENSG00000278464 | AC068506,1 | -1,085448448 | 0,471245763 | 0,19108393 | 0,686751133 | novel transcript |
| ENSG00000274156 | AC008121,2 | -1,086022714 | 0,47105822 | 0,35745133 | 0,848693232 | novel transcript |
| ENSG00000125337 | KIF25 | -1,086262444 | 0,470979952 | 0,12451267 | 0,56384187 | kinesin family member 25 |
| ENSG00000198822 | GRM3 | -1,086488924 | 0,470906022 | 0,02052465 | 0,190071839 | glutamate metabotropic receptor 3 |
| ENSG00000258702 | AL137786,1 | -1,086728187 | 0,470827931 | 0,04693185 | 0,329207362 | novel transcript |
| ENSG00000237772 | LINC02631 | -1,086971405 | 0,470748563 | 0,47164148 | 0,908896785 | long intergenic non-protein coding RNA 2631 |
| ENSG00000233797 | UFL1-AS1 | -1,08845886 | 0,470263459 | 0,15339869 | 0,622030694 | UFL1 antisense RNA 1 |
| ENSG00000276167 | TAS2R50 | -1,088547898 | 0,470234437 | 0,16500911 | 0,644037807 | taste 2 receptor member 50 |
| ENSG00000273431 | TAS2R50 | -1,088547898 | 0,470234437 | 0,16500911 | 0,644037807 | taste 2 receptor member 50 |
| ENSG00000212126 | TAS2R50 | -1,088547898 | 0,470234437 | 0,16500911 | 0,644037807 | taste 2 receptor member 50 |
| ENSG00000259096 | SHLD2P2 | -1,08908803 | 0,470058418 | 0,17092747 | 0,654596502 | shieldin complex subunit 2 pseudogene 2 |
| ENSG00000131037 | EPS8L1 | -1,089494677 | 0,469925944 | 0,00564191 | 0,075356774 | EPS8 like 1 |
| ENSG00000235547 | NDUFB11P1 | -1,089508033 | 0,469921593 | 0,33564224 | 0,835387231 | NADH:ubiquinone oxidoreductase subunit B11 pseudogene 1 |
| ENSG00000254266 | PKIA-AS1 | -1,089611675 | 0,469887836 | 0,04043934 | 0,300358836 | PKIA antisense RNA 1 |
| ENSG00000226701 | RPL15P14 | -1,089844707 | 0,469811943 | 0,28666427 | 0,797994114 | ribosomal protein L15 pseudogene 14 |
| ENSG00000267178 | PHF5CP | -1,090236666 | 0,469684319 | 0,39680574 | 0,873609704 | PHD finger protein 5C pseudogene |
| ENSG00000170011 | MYRIP | -1,090242826 | 0,469682314 | 0,03475702 | 0,271925393 | myosin VIIA and Rab interacting protein |
| ENSG00000064205 | CCN5 | -1,090629741 | 0,469556367 | 0,10219137 | 0,508743777 | cellular communication network factor 5 |
| ENSG00000258611 | YBX2P2 | -1,090715143 | 0,469528572 | 0,00819108 | 0,099373674 | YBX2 pseudogene 2 |
| ENSG00000077279 | DCX | -1,091134095 | 0,469392243 | 0,05104827 | 0,345820358 | doublecortin |
| ENSG00000177873 | ZNF619 | -1,092205957 | 0,469043634 | 0,00013591 | 0,00341569 | zinc finger protein 619 |
| ENSG00000260621 | AC092140,1 | -1,092374393 | 0,468988875 | 0,4202723 | 0,888457169 | novel transcript, antisense to AMFR |
| ENSG00000261351 | AC116913,1 | -1,092768095 | 0,468860909 | 0,00619795 | 0,081031602 | novel transcript, antisense to MAP2K1 |
| ENSG00000253846 | PCDHGA10 | -1,092841944 | 0,468836909 | 1,18E-05 | 0,000393781 | protocadherin gamma subfamily A, 10 |
| ENSG00000288038 | AL445588,1 | -1,093034425 | 0,468774362 | 0,34704711 | 0,841678758 | novel transcript |
| ENSG00000225187 | AC073283,1 | -1,093288769 | 0,468691726 | 0,25636529 | 0,768541051 | novel transcript |

| | | | | | | |
|---|---|---|---|---|---|---|
| ENSG00000285677 | AL355388,2 | -1,093779617 | 0,46853229 | 0,10468989 | 0,515975114 | Novel transcript, antisense to SSR2 |
| ENSG00000272254 | AC022893,3 | -1,094573222 | 0,468274628 | 0,15299471 | 0,621569237 | novel transcript, antisense to TERF1 |
| ENSG00000261373 | VPS9D1-AS1 | -1,094603394 | 0,468264835 | 0,21299926 | 0,72028467 | VPS9D1 antisense RNA 1 |
| ENSG00000251414 | AC138956,2 | -1,094686542 | 0,468237848 | 0,08484205 | 0,463562014 | novel transcript |
| ENSG00000243373 | RN7SL173P | -1,095277302 | 0,468046152 | 0,35415157 | 0,84556089 | RNA, 7SL, cytoplasmic 173, pseudogene |
| ENSG00000081842 | PCDHA6 | -1,096455155 | 0,467664183 | 0,00097375 | 0,01825309 | protocadherin alpha 6 |
| ENSG00000163606 | CD200R1 | -1,096619772 | 0,467610823 | 0,39737971 | 0,874053739 | CD200 receptor 1 |
| ENSG00000074370 | ATP2A3 | -1,096839729 | 0,467539536 | 0,01519782 | 0,155530211 | ATPase sarcoplasmic/endoplasmic reticulum Ca2+ transporting 3 |
| ENSG00000255382 | LINC02734 | -1,096875289 | 0,467528012 | 0,43645511 | 0,896850871 | long intergenic non-protein coding RNA 2734 |
| ENSG00000245482 | AC046130,1 | -1,096923102 | 0,467512517 | 0,32372066 | 0,825794844 | novel transcript, antisense to ALG10 |
| ENSG00000287546 | AL163534,1 | -1,097031678 | 0,467477334 | 0,4115889 | 0,8838461 | novel transcript |
| ENSG00000225342 | LRRK2-DT | -1,097074067 | 0,467463599 | 0,22471552 | 0,733203418 | LRRK2 divergent transcript |
| ENSG00000287789 | AL158136,1 | -1,097320817 | 0,467383654 | 0,04262839 | 0,310582197 | novel transcript, antisense to BACH2 |
| ENSG00000186281 | GPAT2 | -1,097366808 | 0,467368754 | 0,05623324 | 0,366342158 | glycerol-3-phosphate acyltransferase 2, mitochondrial |
| ENSG00000117266 | CDK18 | -1,097653857 | 0,467275773 | 0,02985235 | 0,246365813 | cyclin dependent kinase 18 |
| ENSG00000175267 | VWA3A | -1,097657532 | 0,467274582 | 0,30047877 | 0,810227614 | von Willebrand factor A domain containing 3A |
| ENSG00000260874 | AC106820,5 | -1,098047915 | 0,467148158 | 0,38186837 | 0,866444071 | novel transcript |
| ENSG00000259343 | TMC3-AS1 | -1,098594639 | 0,466971161 | 0,02144427 | 0,195861468 | TMC3 antisense RNA 1 |
| ENSG00000055118 | KCNH2 | -1,098607835 | 0,46696689 | 0,00381071 | 0,056543916 | potassium voltage-gated channel subfamily H member 2 |
| ENSG00000254610 | AC004923,1 | -1,099405554 | 0,466708758 | 0,25453079 | 0,765935007 | RNA binding protein, fox-1 homolog (C, elegans) 1 (RBFOX1) pseudogene |
| ENSG00000165521 | EML5 | -1,100273772 | 0,466427976 | 0,01716066 | 0,168907949 | EMAP like 5 |
| ENSG00000261173 | AC018845,3 | -1,10030506 | 0,46641786 | 0,44596149 | 0,901252648 | novel transcript, antisense to DNAJA2 |
| ENSG00000273165 | AL121652,1 | -1,100451313 | 0,46637058 | 0,17124484 | 0,654898934 | novel transcript |
| ENSG00000278177 | AL354811,1 | -1,100976269 | 0,466200912 | 0,1521489 | 0,620418906 | novel transcript, antisense to GPC6 |
| ENSG00000286861 | AL049779,4 | -1,101061081 | 0,466173506 | 0,01696834 | 0,167349589 | novel transcript, antisense to ARG2 |
| ENSG00000241832 | CECR3 | -1,101213885 | 0,466124133 | 0,11852715 | 0,550275639 | cat eye syndrome chromosome region, candidate 3 |
| ENSG00000286215 | AL356534,1 | -1,101761326 | 0,465947293 | 0,00046263 | 0,009759623 | vimentin 2, pseudogene |
| ENSG00000229593 | SUCLA2P3 | -1,101974936 | 0,465878308 | 0,11002447 | 0,528913064 | SUCLA2 pseudogene 3 |
| ENSG00000232282 | MTND1P32 | -1,102017625 | 0,465864523 | 0,10856016 | 0,525236658 | MT-ND1 pseudogene 32 |

| | | | | | | |
|---|---|---|---|---|---|---|
| ENSG00000235236 | AC137630,2 | -1,102041565 | 0,465856793 | 0,09940445 | 0,50040065 | novel transcript |
| ENSG00000286799 | AC124066,1 | -1,102178305 | 0,465812641 | 0,40971122 | 0,882177108 | novel transcript, antisense to B9D1 |
| ENSG00000213150 | AL512430,1 | -1,102672241 | 0,465653187 | 0,35138899 | 0,84405766 | novel pseudogene |
| ENSG00000230826 | AL390774,2 | -1,103428796 | 0,465409061 | 0,05213286 | 0,350361504 | novel transcript |
| ENSG00000054938 | CHRDL2 | -1,103507625 | 0,465383632 | 0,27960406 | 0,790302992 | chordin like 2 |
| ENSG00000124713 | GNMT | -1,103711976 | 0,465317717 | 0,06891561 | 0,414023399 | glycine N-methyltransferase |
| ENSG00000187172 | BAGE2 | -1,10378977 | 0,465292627 | 0,17252267 | 0,656973055 | BAGE family member 2 |
| ENSG00000206199 | ANKUB1 | -1,10383393 | 0,465278384 | 0,34744012 | 0,842200946 | ankyrin repeat and ubiquitin domain containing 1 |
| ENSG00000254061 | AC091144,2 | -1,104080438 | 0,465198891 | 0,25313433 | 0,76515438 | RNA binding motif protein 4 (RBM4) pseudogene |
| ENSG00000151376 | ME3 | -1,104503348 | 0,465062543 | 0,00622378 | 0,0812289 | malic enzyme 3 |
| ENSG00000224437 | PIGUP1 | -1,104545779 | 0,465048865 | 0,21266389 | 0,719481009 | phosphatidylinositol glycan anchor biosynthesis class U pseudogene 1 |
| ENSG00000255203 | OR7E2P | -1,104638245 | 0,46501906 | 0,09590858 | 0,491877624 | olfactory receptor family 7 subfamily E member 2 pseudogene |
| ENSG00000260639 | AC025678,1 | -1,10475369 | 0,46498185 | 0,25654836 | 0,768931843 | mitochondrially encoded cytochrome c oxidase III (MT-CO3) pseudogene |
| ENSG00000231550 | PTCHD3P2 | -1,104792773 | 0,464969254 | 0,02316644 | 0,207231888 | patched domain containing 3 pseudogene 2 |
| ENSG00000273687 | AC004223,4 | -1,105090065 | 0,464873449 | 0,29834953 | 0,807818219 | novel transcript, antisense to RFFL |
| ENSG00000186510 | CLCNKA | -1,105499296 | 0,464741603 | 0,45634481 | 0,903545728 | chloride voltage-gated channel Ka |
| ENSG00000185028 | LRRC14B | -1,105613968 | 0,464704665 | 0,38478531 | 0,868536368 | leucine rich repeat containing 14B |
| ENSG00000261668 | AC093591,2 | -1,105620453 | 0,464702576 | 0,01776819 | 0,173000543 | novel transcript, antisense HSPA4L |
| ENSG00000198523 | PLN | -1,106373038 | 0,464460226 | 0,03827974 | 0,290057638 | phospholamban |
| ENSG00000254630 | AP001922,2 | -1,106375002 | 0,464459594 | 0,35897867 | 0,849814998 | novel transcript |
| ENSG00000277245 | AC084782,3 | -1,106385618 | 0,464456176 | 0,22813976 | 0,736378443 | novel transcript, antisense to MNS1 |
| ENSG00000252634 | RN7SKP219 | -1,106894816 | 0,464292275 | 0,18705251 | 0,680713483 | RN7SK pseudogene 219 |
| ENSG00000228547 | OR7E26P | -1,106975986 | 0,464266154 | 0,25159003 | 0,763892102 | olfactory receptor family 7 subfamily E member 26 pseudogene |
| ENSG00000255964 | HSPD1P12 | -1,107138234 | 0,464213945 | 0,42541749 | 0,892125569 | heat shock protein family D (Hsp60) member 1 pseudogene 12 |
| ENSG00000255081 | AP003168,2 | -1,107521403 | 0,464090669 | 0,45536485 | 0,903545728 | novel transcript |
| ENSG00000238825 | RNVU1-2 | -1,107956805 | 0,463950629 | 0,25678965 | 0,769107677 | RNA, variant U1 small nuclear 2 |
| ENSG00000159712 | ANKRD18CP | -1,108023087 | 0,463929314 | 0,05921726 | 0,37798459 | ankyrin repeat domain 18C, pseudogene |
| ENSG00000171714 | ANO5 | -1,108041528 | 0,463923384 | 0,01075636 | 0,120922944 | anoctamin 5 |

| | | | | | | |
|---|---|---|---|---|---|---|
| ENSG00000224765 | AL035443,1 | -1,108118835 | 0,463898525 | 0,06199121 | 0,388629081 | novel transcript |
| ENSG00000251623 | AC139491,5 | -1,108247254 | 0,463857234 | 0,46350155 | 0,907101775 | centrosomal protein 192 (CEP192), pseudogene |
| ENSG00000260816 | AC027279,1 | -1,108705233 | 0,463710007 | 0,02883601 | 0,241114969 | novel transcript, sense intronic to WWOX |
| ENSG00000286593 | AP006261,1 | -1,110673821 | 0,463077697 | 0,22690991 | 0,735383523 | novel transcript |
| ENSG00000232077 | LINC01031 | -1,110680856 | 0,463075439 | 0,24850149 | 0,760720365 | long intergenic non-protein coding RNA 1031 |
| ENSG00000105409 | ATP1A3 | -1,11073598 | 0,463057745 | 0,00056274 | 0,011544199 | ATPase Na+/K+ transporting subunit alpha 3 |
| ENSG00000259868 | AL163952,1 | -1,111073305 | 0,462949488 | 0,23255112 | 0,74165848 | novel transcript |
| ENSG00000233932 | CTXN2 | -1,111166833 | 0,462919476 | 0,22700101 | 0,735436234 | cortexin 2 |
| ENSG00000275381 | AC019206,1 | -1,11181819 | 0,462710522 | 0,25331469 | 0,765172364 | Zinc finger protein 607 (ZNF607) pseudogene |
| ENSG00000286651 | PRSS43P | -1,111921079 | 0,462677524 | 0,15328057 | 0,621793648 | serine protease 43, pseudogene |
| ENSG00000272406 | AC108102,1 | -1,11225542 | 0,462570312 | 0,2540185 | 0,76531122 | novel transcript |
| ENSG00000146151 | HMGCLL1 | -1,112647385 | 0,462444653 | 0,12202445 | 0,558561133 | 3-hydroxymethyl-3-methylglutaryl-CoA lyase like 1 |
| ENSG00000157570 | TSPAN18 | -1,112785569 | 0,462400362 | 0,00247785 | 0,039747347 | tetraspanin 18 |
| ENSG00000237628 | MTCO2P19 | -1,112797114 | 0,462396661 | 0,23759339 | 0,746907875 | MT-CO2 pseudogene 19 |
| ENSG00000198723 | TEX45 | -1,112998286 | 0,462332188 | 0,19177696 | 0,687672451 | testis expressed 45 |
| ENSG00000234336 | JAZF1-AS1 | -1,113940285 | 0,46203041 | 0,3591676 | 0,849846046 | JAZF1 antisense RNA 1 |
| ENSG00000227960 | AL138799,1 | -1,114239696 | 0,461934532 | 0,38083333 | 0,865900094 | novel transcript |
| ENSG00000259666 | LINGO1-AS1 | -1,114415878 | 0,461878124 | 0,26981698 | 0,781794063 | LINGO1 antisense RNA 1 |
| ENSG00000287377 | AC090948,4 | -1,11518686 | 0,46163136 | 0,13473709 | 0,585627728 | novel transcript, antisense to RFTN1 |
| ENSG00000245857 | GS1-24F4,2 | -1,115681786 | 0,461473022 | 0,36828201 | 0,857548638 | uncharacterized LOC100652791 |
| ENSG00000213413 | PVRIG | -1,116147283 | 0,461324148 | 0,06971473 | 0,416662464 | PVR related immunoglobulin domain containing |
| ENSG00000235843 | AL390961,2 | -1,116206413 | 0,461305241 | 0,44715117 | 0,901918334 | novel transcript |
| ENSG00000272619 | AC093673,2 | -1,116525047 | 0,461203368 | 0,25924287 | 0,771989931 | novel transcript |
| ENSG00000286682 | AC025262,3 | -1,116651439 | 0,461162964 | 0,40089617 | 0,876599365 | novel transcript, antisense to TBC1D30 |
| ENSG00000181847 | TIGIT | -1,11731103 | 0,460952172 | 0,05935986 | 0,378585591 | T cell immunoreceptor with Ig and ITIM domains |
| ENSG00000234332 | BCAS2P2 | -1,118411822 | 0,460600594 | 0,10010717 | 0,502206432 | BCAS2 pseudogene 2 |
| ENSG00000101898 | MCTS2P | -1,118692345 | 0,460511042 | 0,13501582 | 0,586038109 | MCTS family member 2, pseudogene |
| ENSG00000231519 | AC007285,1 | -1,11911591 | 0,460375859 | 0,14881253 | 0,613515281 | novel transcript |
| ENSG00000126583 | PRKCG | -1,119230644 | 0,460339248 | 0,06664645 | 0,40559155 | protein kinase C gamma |
| ENSG00000230154 | AIDAP1 | -1,119514271 | 0,460248757 | 0,08219356 | 0,456024762 | AIDA pseudogene 1 |
| ENSG00000104833 | TUBB4A | -1,119906521 | 0,460123638 | 0,00269277 | 0,042390137 | tubulin beta 4A class IVa |
| ENSG00000241546 | AC092910,2 | -1,120089958 | 0,460065137 | 0,26821442 | 0,780692104 | riboflavin kinase (RFK) pseudogene |
| ENSG00000205336 | ADGRG1 | -1,120211845 | 0,46002627 | 1,04E-05 | 0,000351863 | adhesion G protein-coupled receptor G1 |

| | | | | | | |
|---|---|---|---|---|---|---|
| ENSG00000232300 | FAM215B | -1,120567157 | 0,459912987 | 0,03575092 | 0,27707409 | family with sequence similarity 215 member B |
| ENSG00000165164 | CFAP47 | -1,12154419 | 0,459601627 | 0,13248519 | 0,581009552 | cilia and flagella associated protein 47 |
| ENSG00000269662 | BNIP3P39 | -1,121972112 | 0,459465323 | 0,18516693 | 0,677558375 | BCL2 interacting protein 3 pseudogene 39 |
| ENSG00000227801 | COL11A2 | -1,122399419 | 0,459329256 | 0,45300191 | 0,902428411 | collagen type XI alpha 2 chain |
| ENSG00000218428 | NIP7P3 | -1,1224853 | 0,459301914 | 0,28579835 | 0,797420923 | NIP7 pseudogene 3 |
| ENSG00000272154 | AC244517,2 | -1,122777586 | 0,45920887 | 0,02345139 | 0,208943965 | novel transcript, antisense to PCDHB3, PCDHB4, PCDHB5, PCDHB6 and PCDHB7 |
| ENSG00000255148 | AC239802,2 | -1,12289431 | 0,459171718 | 0,37043509 | 0,858545694 | novel transcript |
| ENSG00000132204 | LINC00470 | -1,12298838 | 0,459141779 | 0,0213411 | 0,195090148 | long intergenic non-protein coding RNA 470 |
| ENSG00000278668 | AC005899,7 | -1,123513892 | 0,458974564 | 0,15468552 | 0,623971362 | novel transcript |
| ENSG00000287205 | AL451054,4 | -1,123573349 | 0,458955649 | 0,35055652 | 0,843970352 | novel transcript |
| ENSG00000005379 | TSPOAP1 | -1,123839076 | 0,458871122 | 0,00280912 | 0,044000102 | TSPO associated protein 1 |
| ENSG00000271998 | AC034229,3 | -1,123973193 | 0,458828467 | 0,67967659 | 0,964459885 | novel transcript |
| ENSG00000253007 | SNORA50B | -1,124506388 | 0,458658923 | 0,26238094 | 0,774068771 | small nucleolar RNA, H/ACA box 50B |
| ENSG00000230321 | MTCO2P27 | -1,125130683 | 0,458460491 | 0,36099751 | 0,851783601 | MT-CO2 pseudogene 27 |
| ENSG00000282249 | AC254962,1 | -1,12518786 | 0,458442322 | 0,30025451 | 0,810130728 | hyperpolarization activated cyclic nucleotide-gated potassium channel 2 (HCN2) pseudogene |
| ENSG00000287037 | AC097709,1 | -1,125303992 | 0,45840542 | 0,3014582 | 0,811013605 | novel transcript, antisense to CXCL5 |
| ENSG00000135902 | CHRND | -1,125322849 | 0,458399429 | 0,10730269 | 0,522893717 | cholinergic receptor nicotinic delta subunit |
| ENSG00000234043 | NUDT9P1 | -1,125983508 | 0,45818956 | 0,22639768 | 0,735064413 | nudix hydrolase 9 pseudogene 1 |
| ENSG00000272172 | AC138696,2 | -1,126026908 | 0,458175777 | 0,32527001 | 0,827324367 | novel transcript, antisense to ZNF696 |
| ENSG00000285639 | AL133259,1 | -1,126658389 | 0,457975273 | 0,35907293 | 0,849814998 | novel transcript |
| ENSG00000260618 | AC025917,1 | -1,127328116 | 0,457762721 | 0,15895308 | 0,633144409 | novel transcript, antisense to KIAA1370 |
| ENSG00000147697 | GSDMC | -1,127413451 | 0,457735645 | 0,5429555 | 0,933716215 | gasdermin C |
| ENSG00000131979 | GCH1 | -1,127778031 | 0,457619987 | 0,00441916 | 0,0631158 | GTP cyclohydrolase 1 |
| ENSG00000196381 | ZNF781 | -1,12821856 | 0,457480273 | 0,03092539 | 0,252639467 | zinc finger protein 781 |
| ENSG00000271917 | AL357568,1 | -1,128399503 | 0,4574229 | 0,09169296 | 0,481715471 | novel transcript |
| ENSG00000108785 | HSD17B1P1 | -1,12844546 | 0,457408329 | 0,13567949 | 0,587376532 | hydroxysteroid 17-beta dehydrogenase 1 pseudogene 1 |
| ENSG00000164651 | SP8 | -1,128756256 | 0,457309801 | 0,01636818 | 0,163441334 | Sp8 transcription factor |
| ENSG00000173674 | EIF1AX | -1,128764237 | 0,457307271 | 0,01624038 | 0,16263267 | eukaryotic translation initiation factor 1A X-linked |
| ENSG00000126861 | OMG | -1,128884921 | 0,457269018 | 0,00211322 | 0,03516114 | oligodendrocyte myelin glycoprotein |
| ENSG00000287340 | AC007222,2 | -1,128930902 | 0,457254445 | 0,37423186 | 0,861369706 | novel transcript, antisense to RBFOX1 |

| | | | | | | |
|---|---|---|---|---|---|---|
| ENSG00000237283 | AL691515,1 | -1,129154712 | 0,457183515 | 0,35301059 | 0,845031687 | novel transcript |
| ENSG00000186007 | LEMD1 | -1,129199633 | 0,45716928 | 0,00208879 | 0,034847287 | LEM domain containing 1 |
| ENSG00000230487 | PSMG3-AS1 | -1,129391929 | 0,457108348 | 0,00049905 | 0,01040156 | PSMG3 antisense RNA 1 (head to head) |
| ENSG00000251634 | NAIPP4 | -1,129456895 | 0,457087764 | 0,77566738 | 0,978587255 | NAIP pseudogene 4 |
| ENSG00000152503 | TRIM36 | -1,129668107 | 0,457020851 | 0,02397464 | 0,212275005 | tripartite motif containing 36 |
| ENSG00000224126 | UBE2SP2 | -1,130383341 | 0,456794333 | 0,27853408 | 0,789775666 | ubiquitin conjugating enzyme E2 S pseudogene 2 |
| ENSG00000273762 | VN1R76P | -1,130456975 | 0,45677102 | 0,0234979 | 0,209239044 | vomeronasal 1 receptor 76 pseudogene |
| ENSG00000249505 | AC108673,1 | -1,130683259 | 0,456699381 | 0,38150748 | 0,865900094 | jumonji domain containing 4 (JMJD4) pseudogene |
| ENSG00000227470 | RPL39P16 | -1,130737778 | 0,456682123 | 0,40961679 | 0,882177108 | ribosomal protein L39 pseudogene 16 |
| ENSG00000286555 | AP003065,1 | -1,130884265 | 0,456635756 | 0,3908577 | 0,870057277 | novel transcript, antisense to ARAP1 |
| ENSG00000102239 | BRS3 | -1,131347482 | 0,456489163 | 0,29683006 | 0,806635068 | bombesin receptor subtype 3 |
| ENSG00000271396 | AC091167,4 | -1,131479705 | 0,456447328 | 0,05561723 | 0,36392023 | MKI67 (FHA domain) interacting nucleolar phosphoprotein (MKI67IP) pseudogene |
| ENSG00000090932 | DLL3 | -1,131604557 | 0,456407828 | 0,00703222 | 0,088866368 | delta like canonical Notch ligand 3 |
| ENSG00000107018 | RLN1 | -1,131687065 | 0,456381727 | 0,31635858 | 0,821646322 | relaxin 1 |
| ENSG00000152270 | PDE3B | -1,131929938 | 0,456304903 | 0,008797 | 0,10409891 | phosphodiesterase 3B |
| ENSG00000130762 | ARHGEF16 | -1,132850987 | 0,456013681 | 0,00708656 | 0,089444543 | Rho guanine nucleotide exchange factor 16 |
| ENSG00000169836 | TACR3 | -1,133984019 | 0,455655688 | 0,07756221 | 0,440002866 | tachykinin receptor 3 |
| ENSG00000240253 | FAR2P3 | -1,134026123 | 0,45564239 | 0,01735832 | 0,170318153 | fatty acyl-CoA reductase 2 pseudogene 3 |
| ENSG00000225331 | LINC01678 | -1,134121098 | 0,455612395 | 0,47364013 | 0,910190245 | long intergenic non-protein coding RNA 1678 |
| ENSG00000235584 | AC008268,1 | -1,134257606 | 0,455569287 | 0,27761958 | 0,789067728 | novel transcript |
| ENSG00000125046 | SSUH2 | -1,134984967 | 0,455339662 | 0,04355282 | 0,315120457 | ssu-2 homolog |
| ENSG00000214189 | ZNF788P | -1,13505881 | 0,455316356 | 8,01E-07 | 3,47E-05 | zinc finger family member 788, pseudogene |
| ENSG00000166979 | EVA1C | -1,135254451 | 0,455254616 | 0,06171581 | 0,387290891 | eva-1 homolog C |
| ENSG00000101115 | SALL4 | -1,136060272 | 0,455000403 | 0,00068147 | 0,01348449 | spalt like transcription factor 4 |
| ENSG00000288172 | AL356419,1 | -1,136181478 | 0,454962178 | 0,20588513 | 0,708429733 | novel transcript |
| ENSG00000157470 | FAM81A | -1,137012965 | 0,45470004 | 0,06434549 | 0,397835213 | family with sequence similarity 81 member A |
| ENSG00000237716 | AL136309,2 | -1,137210993 | 0,454637631 | 0,11676211 | 0,545661612 | novel transcript |
| ENSG00000231443 | AC124944,1 | -1,137978383 | 0,454395867 | 0,00843712 | 0,101453094 | protein phosphatase 4, regulatory subunit 2 (PPP4R2) pseudogene |
| ENSG00000161594 | KLHL10 | -1,138008261 | 0,454386456 | 0,1462893 | 0,609649241 | kelch like family member 10 |
| ENSG00000273212 | AC000068,2 | -1,138101357 | 0,454357136 | 0,13609292 | 0,587965501 | novel transcript, antisense to UFD1L |
| ENSG00000234653 | LRRTM4-AS1 | -1,139709803 | 0,45385086 | 0,22771883 | 0,736053199 | LRRTM4 antisense RNA 1 |

| | | | | | | |
|---|---|---|---|---|---|---|
| ENSG00000188064 | WNT7B | -1,139740031 | 0,453841351 | 1,85E-05 | 0,000586923 | Wnt family member 7B |
| ENSG00000278145 | C17orf78 | -1,140052907 | 0,453742937 | 0,09427676 | 0,487847001 | chromosome 17 open reading frame 78 |
| ENSG00000262243 | CES1 | -1,140280865 | 0,453671248 | 0,02913322 | 0,24236233 | carboxylesterase 1 |
| ENSG00000172461 | FUT9 | -1,141186886 | 0,453386429 | 0,0559572 | 0,365178946 | fucosyltransferase 9 |
| ENSG00000278522 | POTEB3 | -1,141199713 | 0,453382398 | 0,16925625 | 0,651357396 | POTE ankyrin domain family member B3 |
| ENSG00000147437 | GNRH1 | -1,141732303 | 0,453215057 | 0,01524486 | 0,155858486 | gonadotropin releasing hormone 1 |
| ENSG00000225792 | AC004540,2 | -1,142071398 | 0,453108545 | 0,0523903 | 0,351288432 | novel transcript |
| ENSG00000206144 | MAPK6P1 | -1,142076496 | 0,453106944 | 0,02675953 | 0,229082684 | mitogen-activated protein kinase 6 pseudogene 1 |
| ENSG00000269139 | AC010336,5 | -1,142168141 | 0,453078161 | 0,06358216 | 0,394965288 | novel transcript, antisense to CTXN1 and TIMM44 |
| ENSG00000115423 | DNAH6 | -1,142218027 | 0,453062495 | 0,0177102 | 0,172597119 | dynein axonemal heavy chain 6 |
| ENSG00000278627 | AC005962,1 | -1,142582428 | 0,452948073 | 0,12910527 | 0,574160337 | novel transcript, antisense to EPX |
| ENSG00000231942 | HNRNPA1P36 | -1,142899907 | 0,452848409 | 0,17410658 | 0,659373116 | heterogeneous nuclear ribonucleoprotein A1 pseudogene 36 |
| ENSG00000285758 | AC036214,4 | -1,142910336 | 0,452845135 | 0,05476037 | 0,361492462 | novel transcript, antisense to MRPS28 |
| ENSG00000229071 | PRRT1 | -1,142914248 | 0,452843907 | 0,07643115 | 0,436971724 | proline rich transmembrane protein 1 |
| ENSG00000186105 | LRRC70 | -1,143092717 | 0,452787891 | 0,14959891 | 0,614957829 | leucine rich repeat containing 70 |
| ENSG00000227678 | AL355581,1 | -1,143197809 | 0,45275491 | 0,01139389 | 0,126095056 | novel transcript, antisense to L3MBTL3 |
| ENSG00000275996 | SNORD27 | -1,143390733 | 0,452694369 | 0,38568242 | 0,869017264 | small nucleolar RNA, C/D box 27 |
| ENSG00000236939 | BAALC-AS2 | -1,143929265 | 0,452525418 | 0,03448338 | 0,27087651 | BAALC antisense RNA 2 |
| ENSG00000250933 | GAPDHP66 | -1,144273004 | 0,452417612 | 0,17192893 | 0,655750391 | glyceraldehyde-3-phosphate dehydrogenase pseudogene 66 |
| ENSG00000101489 | CELF4 | -1,14438876 | 0,452381313 | 0,02120543 | 0,194418308 | CUGBP Elav-like family member 4 |
| ENSG00000267006 | AC008507,2 | -1,145771413 | 0,451947966 | 0,10467647 | 0,515975114 | novel transcript |
| ENSG00000233523 | PHBP5 | -1,14592808 | 0,451898891 | 0,07945804 | 0,447115945 | prohibitin pseudogene 5 |
| ENSG00000113763 | UNC5A | -1,145964946 | 0,451887343 | 0,01177749 | 0,1291517 | unc-5 netrin receptor A |
| ENSG00000181656 | GPR88 | -1,14612736 | 0,451836474 | 0,42275924 | 0,890403187 | G protein-coupled receptor 88 |
| ENSG00000236197 | PPP1R9A-AS1 | -1,146482136 | 0,451725375 | 0,0079576 | 0,097274568 | PPP1R9A antisense RNA 1 |
| ENSG00000257195 | HNRNPA1P50 | -1,146658526 | 0,451670149 | 0,14343398 | 0,602438935 | heterogeneous nuclear ribonucleoprotein A1 pseudogene 50 |
| ENSG00000258375 | AC079363,1 | -1,146715446 | 0,451652329 | 0,15953821 | 0,633929525 | transmembrane protein 66 (TMEM66) pseudogene |
| ENSG00000287807 | AL390242,1 | -1,147048424 | 0,451548098 | 0,12893448 | 0,573645596 | novel transcript |
| ENSG00000286890 | AC055840,1 | -1,147672886 | 0,451352691 | 0,03360372 | 0,266408273 | novel transcript, antisense to FHOD3 |
| ENSG00000276644 | DACH1 | -1,147800494 | 0,45131277 | 0,0485008 | 0,335471046 | dachshund family transcription factor 1 |

| | | | | | | |
|---|---|---|---|---|---|---|
| ENSG00000286804 | AC025871,3 | -1,147888657 | 0,451285191 | 0,32745613 | 0,829424659 | novel transcript, sense intronic to FBXO16 |
| ENSG00000167194 | C16orf92 | -1,147921576 | 0,451274894 | 0,32544617 | 0,827426913 | chromosome 16 open reading frame 92 |
| ENSG00000188582 | PAQR9 | -1,148302252 | 0,451155834 | 0,02120313 | 0,194418308 | progestin and adipoQ receptor family member 9 |
| ENSG00000276231 | PIK3R6 | -1,148575383 | 0,45107043 | 0,4321572 | 0,89558934 | phosphoinositide-3-kinase regulatory subunit 6 |
| ENSG00000238272 | AL139142,1 | -1,148607343 | 0,451060437 | 0,17473888 | 0,660057513 | novel transcript |
| ENSG00000267707 | AC015961,2 | -1,148678772 | 0,451038105 | 0,08669387 | 0,469007996 | novel transcript, antisense to CELF4 |
| ENSG00000170743 | SYT9 | -1,149105556 | 0,450904697 | 0,01226674 | 0,132701486 | synaptotagmin 9 |
| ENSG00000198821 | CD247 | -1,149318985 | 0,450837996 | 0,09967399 | 0,500760119 | CD247 molecule |
| ENSG00000286429 | BX004807,1 | -1,149636684 | 0,450738727 | 0,1425504 | 0,600806836 | novel transcript, antisense to ITGB3BP |
| ENSG00000268070 | AC006539,2 | -1,150003262 | 0,450624212 | 0,03785729 | 0,28776324 | BCL2/adenovirus E1B 19kDa interacting protein 3 (BNIP3) pseudogene |
| ENSG00000286643 | AP000248,1 | -1,150193965 | 0,45056465 | 0,18144627 | 0,671250291 | novel transcript, antisense to TIAM1 |
| ENSG00000231691 | AL592114,3 | -1,150205057 | 0,450561186 | 0,17819512 | 0,665463944 | novel transcript |
| ENSG00000225493 | LINC01107 | -1,150452329 | 0,450483969 | 0,16227662 | 0,639443053 | long intergenic non-protein coding RNA 1107 |
| ENSG00000286812 | AP001120,5 | -1,150477298 | 0,450476172 | 0,02226538 | 0,201302765 | novel transcript, antisense to GNAL |
| ENSG00000225674 | IPO7P2 | -1,150746228 | 0,450392207 | 0,04036875 | 0,300178807 | importin 7 pseudogene 2 |
| ENSG00000151892 | GFRA1 | -1,151214416 | 0,450246069 | 0,00143495 | 0,02534597 | GDNF family receptor alpha 1 |
| ENSG00000224697 | NEFMP1 | -1,151649989 | 0,450110153 | 0,20150781 | 0,701094807 | neurofilament medium pseudogene 1 |
| ENSG00000259544 | AC021739,3 | -1,152132212 | 0,449959728 | 0,1562172 | 0,627034861 | novel transcript, antisense to AKAP13 |
| ENSG00000267872 | AC073539,1 | -1,152895359 | 0,449721774 | 0,30458916 | 0,813276091 | novel transcript |
| ENSG00000244244 | RPS3AP42 | -1,153048806 | 0,449673943 | 0,11938447 | 0,552217224 | RPS3A pseudogene 42 |
| ENSG00000288457 | LJII02000414,1 | -1,153094962 | 0,449659557 | 0,04547853 | 0,323145642 | |
| ENSG00000181101 | ENTR1P2 | -1,153571056 | 0,449511193 | 0,23125061 | 0,739268552 | ENTR1 pseudogene 2 |
| ENSG00000253457 | SMIM18 | -1,154557063 | 0,44920408 | 0,14579168 | 0,608479758 | small integral membrane protein 18 |
| ENSG00000178631 | ACTG1P1 | -1,155045965 | 0,449051879 | 0,01398561 | 0,146334069 | actin gamma 1 pseudogene 1 |
| ENSG00000287764 | AL136038,5 | -1,155702512 | 0,44884757 | 0,11821046 | 0,549318284 | novel transcript |
| ENSG00000274015 | AL136038,4 | -1,155702512 | 0,44884757 | 0,11821046 | 0,549318284 | novel transcript |
| ENSG00000248335 | AC096733,1 | -1,156049245 | 0,448739708 | 0,31065564 | 0,817751312 | novel transcript |
| ENSG00000230498 | AL035409,1 | -1,156125417 | 0,448716016 | 0,00177148 | 0,030303261 | novel transcript |
| ENSG00000260988 | AC090260,1 | -1,156415184 | 0,4486259 | 0,05154104 | 0,347838733 | novel transcript |
| ENSG00000249876 | AC010285,2 | -1,157310864 | 0,448347462 | 0,0576528 | 0,371181442 | DnaJ (Hsp40) homolog, subfamily B, member 6 (DNAJB6) pseudogene |
| ENSG00000173404 | INSM1 | -1,15769681 | 0,448227537 | 0,00320993 | 0,049241959 | INSM transcriptional repressor 1 |

| | | | | | | |
|---|---|---|---|---|---|---|
| ENSG00000230911 | PPIHP1 | -1,157817314 | 0,4481901 | 0,08608962 | 0,467330023 | peptidylprolyl isomerase H pseudogene 1 |
| ENSG00000172955 | ADH6 | -1,15796571 | 0,448144001 | 0,00543058 | 0,073434973 | alcohol dehydrogenase 6 (class V) |
| ENSG00000187918 | OR51I2 | -1,158288316 | 0,448043801 | 0,38151784 | 0,865900094 | olfactory receptor family 51 subfamily I member 2 |
| ENSG00000242457 | RBBP4P2 | -1,158297669 | 0,448040897 | 0,0329574 | 0,263322314 | RBBP4 pseudogene 2 |
| ENSG00000257543 | AC063948,1 | -1,158457001 | 0,447991418 | 0,27124132 | 0,783959387 | novel transcript |
| ENSG00000185823 | NPAP1 | -1,158549451 | 0,44796271 | 0,00047356 | 0,009937001 | nuclear pore associated protein 1 |
| ENSG00000225075 | AL603832,1 | -1,159337667 | 0,447718033 | 0,2120265 | 0,718583346 | novel transcript |
| ENSG00000271958 | AC021860,1 | -1,15974366 | 0,447592057 | 0,34930446 | 0,843141089 | novel transcript |
| ENSG00000262402 | MCUR1P1 | -1,160456579 | 0,447370931 | 0,07958063 | 0,44749602 | MCUR1 pseudogene 1 |
| ENSG00000229794 | MTCYBP32 | -1,160898634 | 0,447233873 | 0,04387252 | 0,31589834 | MT-CYB pseudogene 32 |
| ENSG00000277507 | AP000351,2 | -1,161450531 | 0,447062818 | 0,22273132 | 0,731303518 | D-dopachrome tautomerase (DDT) pseudogene |
| ENSG00000253947 | AC008705,2 | -1,161474606 | 0,447055358 | 0,33790984 | 0,837226333 | novel transcript |
| ENSG00000236545 | AP001619,2 | -1,161542207 | 0,447034411 | 0,34092133 | 0,839587593 | novel transcript |
| ENSG00000221937 | TAS2R40 | -1,161589643 | 0,447019712 | 0,27188402 | 0,784843944 | taste 2 receptor member 40 |
| ENSG00000251141 | MRPS30-DT | -1,161677976 | 0,446992343 | 0,06030752 | 0,381748162 | MRPS30 divergent transcript |
| ENSG00000229241 | PNPT1P1 | -1,161865532 | 0,446934236 | 0,05049388 | 0,343780608 | polyribonucleotide nucleotidyltransferase 1 pseudogene 1 |
| ENSG00000234996 | ACTG1P25 | -1,162097229 | 0,446862464 | 0,00096514 | 0,018133545 | actin gamma 1 pseudogene 25 |
| ENSG00000287806 | AL135784,1 | -1,162101944 | 0,446861004 | 0,33984467 | 0,838611198 | novel transcript, antisense to IGSF1 |
| ENSG00000143674 | MAP3K21 | -1,162359284 | 0,446781302 | 0,03707986 | 0,284203181 | mitogen-activated protein kinase kinase kinase 21 |
| ENSG00000250980 | NSA2P6 | -1,163178662 | 0,446527625 | 0,07264134 | 0,424935759 | NSA2 pseudogene 6 |
| ENSG00000274918 | AC116447,1 | -1,163252998 | 0,446504618 | 0,1610917 | 0,637192669 | novel transcript |
| ENSG00000146374 | RSPO3 | -1,163307975 | 0,446487604 | 0,33793614 | 0,837226333 | R-spondin 3 |
| ENSG00000285700 | AC104260,3 | -1,164101401 | 0,44624212 | 0,23532311 | 0,744654651 | novel transcript |
| ENSG00000183090 | FREM3 | -1,164673391 | 0,446065232 | 0,18836395 | 0,682783835 | FRAS1 related extracellular matrix 3 |
| ENSG00000259577 | CERNA1 | -1,165094375 | 0,445935088 | 0,03601837 | 0,278307135 | competing endogenous lncRNA 1 for miR-4707-5p and miR-4767 |
| ENSG00000285569 | AC099344,3 | -1,165552441 | 0,445793522 | 0,01788963 | 0,17385807 | novel transcript |
| ENSG00000069535 | MAOB | -1,165585538 | 0,445783296 | 0,00114678 | 0,020942079 | monoamine oxidase B |
| ENSG00000223837 | AL645941,1 | -1,165616568 | 0,445773708 | 0,37667865 | 0,862818448 | novel transcript |
| ENSG00000163357 | DCST1 | -1,165703639 | 0,445746805 | 0,29223041 | 0,802771257 | DC-STAMP domain containing 1 |
| ENSG00000196189 | SEMA4A | -1,166395998 | 0,445532939 | 0,00589522 | 0,077832837 | semaphorin 4A |
| ENSG00000242602 | RPL35AP13 | -1,166906113 | 0,445375433 | 0,31460206 | 0,819877431 | ribosomal protein L35a pseudogene 13 |

| | | | | | | |
|---|---|---|---|---|---|---|
| ENSG00000213088 | ACKR1 | -1,167408854 | 0,445220259 | 0,00338444 | 0,051389635 | atypical chemokine receptor 1 (Duffy blood group) |
| ENSG00000248485 | PCP4L1 | -1,167794163 | 0,445101367 | 0,10430789 | 0,514934779 | Purkinje cell protein 4 like 1 |
| ENSG00000254727 | PTP4A1P6 | -1,167855794 | 0,445082353 | 0,38009998 | 0,865635619 | PTP4A1 pseudogene 6 |
| ENSG00000261474 | AC026471,4 | -1,168582278 | 0,444858284 | 0,07187078 | 0,422642955 | novel transcript |
| ENSG00000265801 | AC069366,2 | -1,168660266 | 0,444834237 | 0,03601098 | 0,278307135 | novel transcript |
| ENSG00000188992 | LIPI | -1,168964685 | 0,444740383 | 0,42652237 | 0,892138574 | lipase I |
| ENSG00000169032 | MAP2K1 | -1,168999863 | 0,444729539 | 5,57E-06 | 0,000203179 | mitogen-activated protein kinase kinase 1 |
| ENSG00000272733 | AP000345,2 | -1,169056443 | 0,444712098 | 0,38674997 | 0,869441674 | novel transcript |
| ENSG00000125531 | FNDC11 | -1,169516979 | 0,44457016 | 0,0163477 | 0,163324813 | fibronectin type III domain containing 11 |
| ENSG00000287419 | AL121929,3 | -1,170736591 | 0,444194492 | 0,17583212 | 0,661199837 | novel transcript, antisense to NEURL1 |
| ENSG00000232906 | AC092685,1 | -1,171052559 | 0,444097219 | 0,01589068 | 0,160311761 | novel pseudogene containing a Zinc finger, C2H2 type domain |
| ENSG00000231738 | TSPAN19 | -1,171478386 | 0,443966158 | 0,07817579 | 0,442208757 | tetraspanin 19 |
| ENSG00000204754 | LINC01951 | -1,171806405 | 0,443865227 | 0,15601011 | 0,626944439 | long intergenic non-protein coding RNA 1951 |
| ENSG00000240305 | AC091152,1 | -1,172825588 | 0,443551772 | 0,04472787 | 0,31940789 | ribosomal protein L6 (RPL6) pseudogene |
| ENSG00000287378 | AC129803,1 | -1,173264864 | 0,443416738 | 0,10728574 | 0,522893717 | novel transcript, antisense to CMSS1 |
| ENSG00000277991 | FP236241,1 | -1,175151959 | 0,442837113 | 0,32710763 | 0,828810524 | uncharacterized LOC102723360 |
| ENSG00000270923 | TAS2R6P | -1,175435003 | 0,442750241 | 0,11546661 | 0,543019233 | taste 2 receptor member 6 pseudogene |
| ENSG00000134873 | CLDN10 | -1,176681951 | 0,442367729 | 0,08952572 | 0,477633485 | claudin 10 |
| ENSG00000161798 | AQP5 | -1,176940988 | 0,442288309 | 0,09041903 | 0,479698016 | aquaporin 5 |
| ENSG00000273141 | AP001269,4 | -1,177340643 | 0,442165803 | 0,50882941 | 0,922427526 | novel transcript, antisense to MPPE1 |
| ENSG00000236814 | AC046176,1 | -1,177431941 | 0,442137822 | 0,22416412 | 0,73292901 | proteasome (prosome, macropain) 26S subunit, ATPase, 6 (PSMC6) pseudogene |
| ENSG00000224762 | CR769776,1 | -1,177592445 | 0,442088636 | 0,44226005 | 0,899331131 | novel zinc finger pseudogene |
| ENSG00000236776 | RPL21P23 | -1,178097417 | 0,441933923 | 0,19141705 | 0,687158413 | ribosomal protein L21 pseudogene 23 |
| ENSG00000155052 | CNTNAP5 | -1,178255478 | 0,441885508 | 0,01316717 | 0,139782152 | contactin associated protein family member 5 |
| ENSG00000170074 | FAM153A | -1,178352181 | 0,441855889 | 0,20084403 | 0,700411363 | family with sequence similarity 153 member A |
| ENSG00000231460 | AL365204,1 | -1,178361715 | 0,441852969 | 0,03321217 | 0,264681419 | novel transcript |
| ENSG00000236641 | Z99916,1 | -1,179166517 | 0,441606552 | 0,38496844 | 0,868764521 | novel transcript |
| ENSG00000260646 | AL031705,1 | -1,179382662 | 0,441540396 | 0,35773439 | 0,848908914 | novel transcript |
| ENSG00000279006 | AL022323,1 | -1,179502361 | 0,441503763 | 0,00991705 | 0,11451092 | novel transcript |
| ENSG00000254317 | AC022973,5 | -1,179681022 | 0,441449091 | 0,1232876 | 0,561682809 | novel transcript |
| ENSG00000260440 | LINC01544 | -1,179848964 | 0,441397706 | 0,10317785 | 0,512268134 | long intergenic non-protein coding RNA 1544 |

| | | | | | | |
|---|---|---|---|---|---|---|
| ENSG00000105877 | DNAH11 | -1,180585837 | 0,441172314 | 0,00217764 | 0,035984022 | dynein axonemal heavy chain 11 |
| ENSG00000180720 | CHRM4 | -1,180586592 | 0,441172083 | 0,13221878 | 0,580737262 | cholinergic receptor muscarinic 4 |
| ENSG00000250056 | LINC01018 | -1,181061204 | 0,441026972 | 0,18070341 | 0,670312865 | long intergenic non-protein coding RNA 1018 |
| ENSG00000180777 | ANKRD30B | -1,181250523 | 0,440969102 | 0,04353943 | 0,315096443 | ankyrin repeat domain 30B |
| ENSG00000269794 | AC010642,1 | -1,181963234 | 0,440751311 | 0,08535129 | 0,465128991 | zinc finger protein family pseudogene |
| ENSG00000249244 | AC110373,1 | -1,182777945 | 0,440502482 | 0,04860903 | 0,335726918 | glycerol kinase 2 pseudogene |
| ENSG00000288124 | C1RL | -1,183029596 | 0,440425652 | 0,12039384 | 0,553609804 | complement C1r subcomponent like |
| ENSG00000213755 | AC093270,1 | -1,183421939 | 0,440305894 | 0,20932498 | 0,713678247 | ribosomal protein L29 (RPL29) pseudogene |
| ENSG00000269815 | AC010463,3 | -1,183878749 | 0,440166499 | 0,17550566 | 0,661199837 | novel transcript, sense intronic ABHD8 |
| ENSG00000233151 | CYP21A2 | -1,183924507 | 0,440152538 | 0,53818274 | 0,932900071 | cytochrome P450 family 21 subfamily A member 2 |
| ENSG00000268810 | AC007193,1 | -1,184335504 | 0,440027165 | 0,64467332 | 0,956633898 | novel transcript, antisense to PPP5C |
| ENSG00000286245 | AC125437,3 | -1,18447943 | 0,439983269 | 0,00692531 | 0,088013355 | novel transcript, antisense to ATP9B |
| ENSG00000286431 | BX842570,1 | -1,184927384 | 0,439846676 | 3,03E-05 | 0,000905188 | novel transcript |
| ENSG00000006071 | ABCC8 | -1,185064722 | 0,439804807 | 0,00797748 | 0,097425006 | ATP binding cassette subfamily C member 8 |
| ENSG00000232627 | AC021876,1 | -1,185844358 | 0,4395672 | 0,1727909 | 0,657273971 | ribosomal protein L31 (RPL31) pseudogene |
| ENSG00000256427 | AC010175,1 | -1,186341323 | 0,439415808 | 0,23922189 | 0,748876847 | novel transcript |
| ENSG00000286965 | AC108066,2 | -1,186759687 | 0,439288401 | 0,21633781 | 0,723599962 | novel transcript |
| ENSG00000270282 | AL096855,2 | -1,188503162 | 0,438757849 | 0,26095698 | 0,77309354 | ferritin, light polypeptide (FTL) pseudogene |
| ENSG00000229191 | AL358473,1 | -1,1886731 | 0,438706169 | 0,04730421 | 0,330573569 | novel transcript, antisense KIF21B |
| ENSG00000223723 | TFDP1P2 | -1,188704135 | 0,438696732 | 0,33683236 | 0,836162806 | TFDP1 pseudogene 2 |
| ENSG00000180221 | TPT1P10 | -1,188899898 | 0,438637208 | 0,17955759 | 0,668298509 | tumor protein, translationally-controlled 1 pseudogene 10 |
| ENSG00000251408 | AC113615,1 | -1,190014219 | 0,438298541 | 0,17332319 | 0,658330849 | novel transcript |
| ENSG00000285731 | AC090644,1 | -1,190148355 | 0,438257792 | 0,15953188 | 0,633929525 | Novel transcript, antisense to TBC1D5 |
| ENSG00000286944 | AL137018,1 | -1,190702487 | 0,438089491 | 0,03887079 | 0,293256884 | novel transcript, antisense to RORB |
| ENSG00000125409 | TEKT3 | -1,191236887 | 0,437927245 | 0,17909164 | 0,667515625 | tektin 3 |
| ENSG00000243710 | CFAP57 | -1,191344672 | 0,437894528 | 0,02159564 | 0,196898814 | cilia and flagella associated protein 57 |
| ENSG00000261210 | CLEC19A | -1,191376014 | 0,437885015 | 0,16129608 | 0,637597747 | C-type lectin domain containing 19A |
| ENSG00000276410 | H2BC3 | -1,191446778 | 0,437863538 | 0,17066176 | 0,654109246 | H2B clustered histone 3 |
| ENSG00000251300 | AC104596,2 | -1,192361667 | 0,437585953 | 0,2048022 | 0,706459522 | LysM, putative peptidoglycan-binding, domain containing 4 (LYSMD4) pseudogene |
| ENSG00000119946 | CNNM1 | -1,192484 | 0,43754885 | 0,01160471 | 0,127615124 | cyclin and CBS domain divalent metal cation transport mediator 1 |

| | | | | | | |
|---|---|---|---|---|---|---|
| ENSG00000213785 | AKR1B1P3 | -1,192565018 | 0,437524279 | 0,35390244 | 0,845346751 | aldo-keto reductase family 1 member B1 pseudogene 3 |
| ENSG00000287305 | AC010969,3 | -1,192665965 | 0,437493666 | 0,45216876 | 0,902062594 | novel transcript |
| ENSG00000278673 | AC136352,5 | -1,192949943 | 0,437407559 | 0,05181795 | 0,349145779 | |
| ENSG00000237971 | AL445523,1 | -1,19318155 | 0,437337344 | 0,14947873 | 0,614794108 | karyopherin alpha 2 (RAG cohort 1, importin alpha 1) (KPNA2) pseudogene |
| ENSG00000287346 | AL132712,2 | -1,193297863 | 0,437302086 | 0,32085534 | 0,824366234 | novel transcript |
| ENSG00000285993 | AC018931,1 | -1,193323449 | 0,437294331 | 0,12226363 | 0,559316128 | novel transcript, antisense to RNF165 |
| ENSG00000172974 | VDAC2P5 | -1,194814816 | 0,436842517 | 0,28596576 | 0,797420923 | VDAC2 pseudogene 5 |
| ENSG00000130829 | DUSP9 | -1,195312888 | 0,436691729 | 0,34350122 | 0,840901506 | dual specificity phosphatase 9 |
| ENSG00000250541 | AC092436,3 | -1,195324697 | 0,436688155 | 0,32458268 | 0,826676737 | novel transcript |
| ENSG00000199319 | RN7SKP25 | -1,195978695 | 0,436490242 | 0,27609873 | 0,788074712 | RN7SK pseudogene 25 |
| ENSG00000238111 | RPS13P4 | -1,196280617 | 0,436398904 | 0,39649909 | 0,873142625 | ribosomal protein S13 pseudogene 4 |
| ENSG00000251171 | AC096741,1 | -1,196459066 | 0,436344928 | 0,24507632 | 0,757071585 | novel transcript |
| ENSG00000229089 | ANKRD20A8P | -1,196753419 | 0,43625591 | 0,00291313 | 0,045289003 | ankyrin repeat domain 20 family member A8, pseudogene |
| ENSG00000224763 | FDPSP8 | -1,196875749 | 0,43621892 | 0,29976613 | 0,809806306 | farnesyl diphosphate synthase pseudogene 8 |
| ENSG00000259440 | NPM1P43 | -1,197864669 | 0,435920009 | 0,07259886 | 0,424884099 | nucleophosmin 1 pseudogene 43 |
| ENSG00000226345 | AC083822,1 | -1,198322155 | 0,435781798 | 0,447555 | 0,902062594 | zinc finger, DHHC-type containing 1 (ZDHHC1) pseudogene |
| ENSG00000248125 | AC008700,1 | -1,198543388 | 0,435714978 | 0,16160944 | 0,63813307 | novel transcript |
| ENSG00000284975 | AC275401,3 | -1,198760672 | 0,43564936 | 0,08117286 | 0,452385684 | ubiquitin specific peptidase 17-like 2 (USP17L2) pseudogene |
| ENSG00000197934 | CYYR1-AS1 | -1,199216588 | 0,435511709 | 0,04476433 | 0,319450339 | CYYR1 antisense RNA 1 |
| ENSG00000168967 | PMCHL1 | -1,200209254 | 0,435212152 | 0,46990396 | 0,908378389 | pro-melanin concentrating hormone like 1 (pseudogene) |
| ENSG00000089116 | LHX5 | -1,201201556 | 0,434912911 | 0,00862775 | 0,102939044 | LIM homeobox 5 |
| ENSG00000257818 | C1GALT1P1 | -1,201710123 | 0,434759626 | 0,03191448 | 0,257756249 | C1GALT1 pseudogene 1 |
| ENSG00000229425 | AJ009632,2 | -1,201853633 | 0,434716381 | 0,09855755 | 0,498431446 | novel transcript |
| ENSG00000229668 | AL133384,1 | -1,202169868 | 0,434621103 | 0,24115285 | 0,751551821 | pseudogene similar to part of small nuclear ribonucleoprotein polypeptide B (SNRPB2) |
| ENSG00000213542 | AC007000,1 | -1,202510144 | 0,434518605 | 0,0777339 | 0,440506191 | ribosomal protein L7a (RPL7A) pseudogene |
| ENSG00000273356 | LINC02019 | -1,202600227 | 0,434491474 | 0,08502512 | 0,464318833 | long intergenic non-protein coding RNA 2019 |

| | | | | | | |
|---|---|---|---|---|---|---|
| ENSG00000286470 | AL034427,1 | -1,203093094 | 0,434343064 | 0,13829425 | 0,593030588 | novel transcript, antisense to PAK7 |
| ENSG00000229696 | KARS1P1 | -1,203220486 | 0,434304713 | 0,15128097 | 0,618627212 | lysyl-tRNA synthetase 1 pseudogene 1 |
| ENSG00000287922 | AC021979,3 | -1,204512607 | 0,433915911 | 0,06866549 | 0,413010032 | novel transcript |
| ENSG00000234373 | SNX18P7 | -1,20517808 | 0,433715804 | 0,37354662 | 0,861319894 | sorting nexin 18 pseudogene 7 |
| ENSG00000285846 | AL358613,3 | -1,205216036 | 0,433704394 | 0,31368615 | 0,819846908 | novel transcript |
| ENSG00000226277 | AC105393,1 | -1,205281665 | 0,433684665 | 0,36644022 | 0,856103393 | novel transcript |
| ENSG00000249106 | AC097537,1 | -1,20583624 | 0,433517988 | 0,07033496 | 0,417680252 | novel transcript, antisense to GPM6A |
| ENSG00000141449 | GREB1L | -1,206388366 | 0,43335211 | 0,00095674 | 0,018008092 | GREB1 like retinoic acid receptor coactivator |
| ENSG00000250371 | AC007370,2 | -1,20747946 | 0,433024494 | 0,07286665 | 0,425795099 | novel transcript |
| ENSG00000214867 | SRSF9P1 | -1,209414007 | 0,43244423 | 0,34107439 | 0,839766083 | serine and arginine rich splicing factor 9 pseudogene 1 |
| ENSG00000286980 | AC020584,1 | -1,209648719 | 0,432373881 | 0,23011456 | 0,73863465 | novel transcript |
| ENSG00000287279 | AL121932,2 | -1,20970476 | 0,432357086 | 0,1608478 | 0,636841924 | novel transcript |
| ENSG00000262299 | AL513523,6 | -1,209919061 | 0,432292868 | 0,25758839 | 0,769908546 | immunoglobulin superfamily, DCC subclass, member 3 (IGDCC3) pseudogene |
| ENSG00000164853 | UNCX | -1,209991581 | 0,432271138 | 0,00023212 | 0,005415695 | UNC homeobox |
| ENSG00000073670 | ADAM11 | -1,210866158 | 0,43200917 | 0,01423905 | 0,148389909 | ADAM metallopeptidase domain 11 |
| ENSG00000170426 | SDR9C7 | -1,210888773 | 0,432002398 | 0,42521666 | 0,891853867 | short chain dehydrogenase/reductase family 9C member 7 |
| ENSG00000271200 | AC099791,2 | -1,21107424 | 0,431946866 | 0,34886165 | 0,842940762 | novel transcript |
| ENSG00000121691 | CAT | -1,21131601 | 0,431874485 | 0,00064461 | 0,012877331 | catalase |
| ENSG00000214711 | CAPN14 | -1,21183478 | 0,431719218 | 0,19962462 | 0,698923433 | calpain 14 |
| ENSG00000236340 | GRM8-AS1 | -1,212095576 | 0,431641183 | 0,21974902 | 0,727931767 | GRM8 antisense RNA 1 |
| ENSG00000226957 | AL358075,1 | -1,212241983 | 0,431597382 | 0,08230843 | 0,456410098 | novel transcript |
| ENSG00000228903 | RASA4CP | -1,212261509 | 0,43159154 | 0,05938721 | 0,378682816 | RAS p21 protein activator 4C, pseudogene |
| ENSG00000230046 | BIRC6-AS1 | -1,212280144 | 0,431585965 | 0,26565521 | 0,776863208 | BIRC6 antisense RNA 1 |
| ENSG00000226891 | LINC01359 | -1,212968793 | 0,431380003 | 0,00374586 | 0,055793453 | long intergenic non-protein coding RNA 1359 |
| ENSG00000165215 | CLDN3 | -1,213299201 | 0,43128122 | 0,07796806 | 0,441433051 | claudin 3 |
| ENSG00000260019 | LINC01992 | -1,21366587 | 0,431171621 | 0,39425346 | 0,871794663 | long intergenic non-protein coding RNA 1992 |
| ENSG00000266839 | AC008088,1 | -1,213871925 | 0,431110043 | 0,13718493 | 0,591024932 | novel transcript |
| ENSG00000267717 | SRSF10P1 | -1,213880804 | 0,431107389 | 0,09234856 | 0,483114784 | serine and arginine rich splicing factor 10 pseudogene 1 |
| ENSG00000223813 | AC007255,1 | -1,21409397 | 0,431043696 | 0,08884777 | 0,475558385 | novel transcript, antisense to CHN2 |

| | | | | | | |
|---|---|---|---|---|---|---|
| ENSG00000234159 | RBPMSLP | -1,214142656 | 0,43102915 | 0,14504318 | 0,606233954 | RNA binding protein with multiple splicing-like pseudogene |
| ENSG00000100433 | KCNK10 | -1,215021026 | 0,430766802 | 0,00229196 | 0,037418119 | potassium two pore domain channel subfamily K member 10 |
| ENSG00000180934 | OR56A1 | -1,215499296 | 0,430624021 | 0,0186042 | 0,177872097 | olfactory receptor family 56 subfamily A member 1 |
| ENSG00000176204 | LRRTM4 | -1,215758933 | 0,43054653 | 0,00019485 | 0,004671685 | leucine rich repeat transmembrane neuronal 4 |
| ENSG00000253567 | AC025871,1 | -1,215760785 | 0,430545978 | 0,41996876 | 0,888425645 | novel transcript |
| ENSG00000172201 | ID4 | -1,216251512 | 0,430399554 | 0,00359882 | 0,054041002 | inhibitor of DNA binding 4, HLH protein |
| ENSG00000287645 | AC092939,1 | -1,216608995 | 0,430292919 | 0,22899857 | 0,737173617 | novel transcript, antisense to PEX5L |
| ENSG00000200785 | SNORD8 | -1,216899019 | 0,430206426 | 0,25379058 | 0,76531122 | small nucleolar RNA, C/D box 8 |
| ENSG00000126218 | F10 | -1,217009837 | 0,430173382 | 0,01921419 | 0,182079045 | coagulation factor X |
| ENSG00000267651 | AC015961,1 | -1,217391585 | 0,43005957 | 0,00550225 | 0,074243597 | novel transcript |
| ENSG00000254539 | AC239804,1 | -1,217652602 | 0,429981769 | 0,01791358 | 0,173950874 | novel transcript |
| ENSG00000149328 | GLB1L2 | -1,219094634 | 0,4295522 | 0,00034167 | 0,007575816 | galactosidase beta 1 like 2 |
| ENSG00000225513 | AL158824,1 | -1,219101485 | 0,42955016 | 0,32580532 | 0,827657237 | ribosomal protein L29 (RPL29) pseudogene |
| ENSG00000226508 | LINC01918 | -1,219692562 | 0,429374208 | 0,02510648 | 0,219776434 | long intergenic non-protein coding RNA 1918 |
| ENSG00000236676 | AC117944,1 | -1,219721662 | 0,429365547 | 0,21970378 | 0,727931767 | novel transcript |
| ENSG00000276846 | AC016590,3 | -1,220238273 | 0,429211824 | 0,0286462 | 0,240299088 | novel transcript, antisense to HKR1 |
| ENSG00000254877 | AP001636,2 | -1,220342398 | 0,429180848 | 0,10908326 | 0,526943592 | cAMP-dependent protein kinase type II-alpha regulatory subunit (PRKAR2A) pseudogene |
| ENSG00000272506 | AL357078,2 | -1,220548278 | 0,429119606 | 0,24078094 | 0,751152776 | novel transcript |
| ENSG00000084628 | NKAIN1 | -1,220898413 | 0,429015473 | 0,00201298 | 0,033778017 | sodium/potassium transporting ATPase interacting 1 |
| ENSG00000224981 | AC000367,1 | -1,221140937 | 0,42894336 | 0,08831217 | 0,473946233 | tigger transposable element derived 1 (TIGD1) pseudogene |
| ENSG00000178662 | CSRNP3 | -1,221217237 | 0,428920675 | 0,0655625 | 0,402068629 | cysteine and serine rich nuclear protein 3 |
| ENSG00000272942 | AL022324,3 | -1,222582968 | 0,428514828 | 0,47636318 | 0,91143003 | novel transcript |
| ENSG00000269993 | KC877982,1 | -1,222594088 | 0,428511525 | 0,02421427 | 0,213851655 | novel transcript |
| ENSG00000227586 | RPS26P39 | -1,222734191 | 0,428469913 | 0,36275102 | 0,85296691 | ribosomal protein S26 pseudogene 39 |
| ENSG00000180875 | GREM2 | -1,223146447 | 0,428347494 | 0,00029485 | 0,00665579 | gremlin 2, DAN family BMP antagonist |
| ENSG00000244738 | AC026316,3 | -1,22338115 | 0,428277814 | 0,32336677 | 0,825554369 | novel transcript |
| ENSG00000287315 | AL670729,3 | -1,224049501 | 0,428079454 | 0,02175723 | 0,19819875 | novel transcript, antisense to OBSCN |
| ENSG00000257376 | AC079601,2 | -1,224188298 | 0,428038272 | 0,17353755 | 0,658349911 | fragile X mental retardation, autosomal homolog 1 (FXR1) pseudogene |

| | | | | | | |
|---|---|---|---|---|---|---|
| ENSG00000220920 | AL023807,1 | -1,224525526 | 0,42793823 | 0,33960511 | 0,838481794 | ribosomal protein S12 (RPS12) pseudogene |
| ENSG00000288148 | AC254979,6 | -1,22561973 | 0,427613786 | 0,52623131 | 0,929261495 | |
| ENSG00000180660 | MAB21L1 | -1,226079899 | 0,427477414 | 0,00696273 | 0,088341044 | mab-21 like 1 |
| ENSG00000243607 | RPL35AP26 | -1,22615631 | 0,427454773 | 0,30461986 | 0,813276091 | ribosomal protein L35a pseudogene 26 |
| ENSG00000197826 | CFAP299 | -1,226753791 | 0,427277783 | 0,29761564 | 0,806925272 | cilia and flagella associated protein 299 |
| ENSG00000136160 | EDNRB | -1,226882592 | 0,427239638 | 0,0169682 | 0,167349589 | endothelin receptor type B |
| ENSG00000248216 | KCTD9P5 | -1,227039217 | 0,427193258 | 0,14860479 | 0,613346323 | potassium channel tetramerization domain containing 9 pseudogene 5 |
| ENSG00000231683 | AL033397,1 | -1,227157313 | 0,42715829 | 0,05105909 | 0,345820358 | novel transcript |
| ENSG00000221571 | RNU6ATAC35P | -1,227724323 | 0,426990441 | 0,25539366 | 0,766913772 | RNA, U6atac small nuclear 35, pseudogene |
| ENSG00000277135 | AC012409,3 | -1,227991289 | 0,426911435 | 0,34245202 | 0,840048641 | novel transcript |
| ENSG00000129682 | FGF13 | -1,228315528 | 0,426815499 | 0,0025777 | 0,041012301 | fibroblast growth factor 13 |
| ENSG00000286698 | AC027612,3 | -1,228511088 | 0,426757647 | 0,0905278 | 0,479949322 | novel transcript |
| ENSG00000164418 | GRIK2 | -1,228527973 | 0,426752653 | 9,80E-06 | 0,000335711 | glutamate ionotropic receptor kainate type subunit 2 |
| ENSG00000139890 | REM2 | -1,228986301 | 0,4266171 | 0,00475698 | 0,066334677 | RRAD and GEM like GTPase 2 |
| ENSG00000277352 | AP001065,4 | -1,229147237 | 0,426569513 | 0,17672794 | 0,662651108 | novel transcript |
| ENSG00000204128 | C2orf72 | -1,229587025 | 0,426439498 | 0,02603851 | 0,225107295 | chromosome 2 open reading frame 72 |
| ENSG00000259170 | AC090985,2 | -1,229665941 | 0,426416172 | 0,33399963 | 0,83443858 | novel transcript, antisense to ST8SIA2 |
| ENSG00000236290 | EEF1GP7 | -1,229876976 | 0,426353801 | 0,10971733 | 0,528090728 | eukaryotic translation elongation factor 1 gamma pseudogene 7 |
| ENSG00000223145 | RN7SKP112 | -1,229926754 | 0,426339091 | 0,36839119 | 0,857548638 | RN7SK pseudogene 112 |
| ENSG00000178796 | RIIAD1 | -1,230076749 | 0,426294767 | 0,01963347 | 0,184275245 | regulatory subunit of type II PKA R-subunit domain containing 1 |
| ENSG00000267213 | AC007773,1 | -1,230257919 | 0,426241238 | 0,04328748 | 0,313927122 | novel transcript, antisense to DPY19L3 |
| ENSG00000283945 | LINC00032 | -1,230826305 | 0,426073342 | 0,04668487 | 0,32799008 | long intergenic non-protein coding RNA 32 |
| ENSG00000275559 | AL137060,5 | -1,231514637 | 0,425870105 | 0,04740599 | 0,330875511 | |
| ENSG00000203624 | MRPS18B | -1,232824148 | 0,425483725 | 0,19049006 | 0,685949202 | mitochondrial ribosomal protein S18B |
| ENSG00000229861 | MRPS18B | -1,232824148 | 0,425483725 | 0,19049006 | 0,685949202 | mitochondrial ribosomal protein S18B |
| ENSG00000270324 | AC005972,1 | -1,23308979 | 0,425405388 | 0,32918728 | 0,831116147 | ribosomal protein L29 (RPL29) pseudogene |
| ENSG00000252757 | RN7SKP96 | -1,233179519 | 0,42537893 | 0,12981221 | 0,575911596 | RN7SK pseudogene 96 |
| ENSG00000205642 | VCX3B | -1,234082316 | 0,425112824 | 0,40001209 | 0,875830109 | variable charge X-linked 3B |
| ENSG00000204390 | HSPA1L | -1,234137641 | 0,425096522 | 0,46366576 | 0,907136822 | heat shock protein family A (Hsp70) member 1 like |

| | | | | | | |
|---|---|---|---|---|---|---|
| ENSG00000186952 | TMEM232 | -1,234148735 | 0,425093253 | 0,00415183 | 0,060441045 | transmembrane protein 232 |
| ENSG00000237027 | AL159174,1 | -1,234792157 | 0,424903709 | 0,29512314 | 0,805254732 | novel transcript |
| ENSG00000213315 | AL122020,1 | -1,235176128 | 0,424790637 | 0,30064062 | 0,81031109 | ribosomal protein S18 (RPS18) pseudogene |
| ENSG00000179799 | OR7E22P | -1,23562418 | 0,424658732 | 0,24978345 | 0,761653149 | olfactory receptor family 7 subfamily E member 22 pseudogene |
| ENSG00000116574 | RHOU | -1,235979441 | 0,424554173 | 0,01769028 | 0,172510533 | ras homolog family member U |
| ENSG00000284984 | RHOU | -1,235979441 | 0,424554173 | 0,01769028 | 0,172510533 | ras homolog family member U |
| ENSG00000204882 | GPR20 | -1,236146297 | 0,424505074 | 0,4199913 | 0,888425645 | G protein-coupled receptor 20 |
| ENSG00000178233 | TMEM151B | -1,236467421 | 0,424410595 | 0,00010036 | 0,002623342 | transmembrane protein 151B |
| ENSG00000229847 | EMX2OS | -1,236786752 | 0,424316665 | 0,01779413 | 0,173199181 | EMX2 opposite strand/antisense RNA |
| ENSG00000102195 | GPR50 | -1,236951643 | 0,424268171 | 0,000911 | 0,017272007 | G protein-coupled receptor 50 |
| ENSG00000232909 | AL157823,2 | -1,237051706 | 0,424238746 | 0,16464339 | 0,643811841 | novel transcript |
| ENSG00000177803 | YWHAQP4 | -1,23760155 | 0,42407709 | 0,30933005 | 0,816845699 | YWHAQ pseudogene 4 |
| ENSG00000230188 | AL158156,1 | -1,23760958 | 0,424074729 | 0,20124235 | 0,701094807 | ribosomal protein L21 (RPL21) pseudogene |
| ENSG00000251111 | FCF1P8 | -1,237862234 | 0,424000469 | 0,1403714 | 0,597084546 | FCF1 pseudogene 8 |
| ENSG00000275825 | AC139494,4 | -1,237922167 | 0,423982855 | 0,01882603 | 0,17950267 | novel transcript |
| ENSG00000287330 | AL353697,1 | -1,238024549 | 0,423952768 | 0,0148953 | 0,153337712 | novel transcript, antisense to PCCA |
| ENSG00000259547 | CYCSP2 | -1,238135809 | 0,423920075 | 0,33747844 | 0,836578328 | CYCS pseudogene 2 |
| ENSG00000145358 | DDIT4L | -1,238243051 | 0,423888564 | 0,02901062 | 0,241642594 | DNA damage inducible transcript 4 like |
| ENSG00000231690 | LINC00574 | -1,23883574 | 0,423714457 | 0,28962902 | 0,800507706 | long intergenic non-protein coding RNA 574 |
| ENSG00000267091 | CTBP2P7 | -1,23911338 | 0,423632923 | 0,34541549 | 0,841168721 | CTBP2 pseudogene 7 |
| ENSG00000204970 | PCDHA1 | -1,239543118 | 0,423506754 | 0,04527973 | 0,322025826 | protocadherin alpha 1 |
| ENSG00000172771 | EFCAB12 | -1,239553018 | 0,423503848 | 0,00039999 | 0,00862441 | EF-hand calcium binding domain 12 |
| ENSG00000254229 | FAM90A12P | -1,23962525 | 0,423482645 | 0,0609125 | 0,383945238 | family with sequence similarity 90 member A12, pseudogene |
| ENSG00000272807 | AC007038,1 | -1,239994847 | 0,423374168 | 0,10152952 | 0,5065782 | novel transcript, antisense to KANSL1L |
| ENSG00000076826 | CAMSAP3 | -1,240305535 | 0,423283003 | 0,0016026 | 0,027773592 | calmodulin regulated spectrin associated protein family member 3 |
| ENSG00000285911 | AL391834,3 | -1,240611632 | 0,423193205 | 0,42593869 | 0,892133432 | Novel transcript |
| ENSG00000135363 | LMO2 | -1,240697233 | 0,423168096 | 0,01740276 | 0,170557995 | LIM domain only 2 |
| ENSG00000002587 | HS3ST1 | -1,241005096 | 0,423077804 | 0,00203185 | 0,033995352 | heparan sulfate-glucosamine 3-sulfotransferase 1 |
| ENSG00000143995 | MEIS1 | -1,241116981 | 0,423044994 | 0,00116471 | 0,021207607 | Meis homeobox 1 |
| ENSG00000261357 | AC099518,2 | -1,241179749 | 0,423026589 | 0,40212327 | 0,87718264 | novel transcript |

| | | | | | | |
|---|---|---|---|---|---|---|
| ENSG00000266932 | AC006116,2 | -1,241625195 | 0,422895996 | 0,15480446 | 0,624266783 | NADH dehydrogenase (ubiquinone) complex I, assembly factor 1, pseudogene |
| ENSG00000197629 | MPEG1 | -1,241800767 | 0,422844534 | 0,00030086 | 0,006771804 | macrophage expressed 1 |
| ENSG00000224289 | IFIT6P | -1,24224707 | 0,422713745 | 0,28264694 | 0,793528967 | interferon induced protein with tetratricopeptide repeats 6, pseudogene |
| ENSG00000242979 | RPS15AP18 | -1,243087972 | 0,42246743 | 0,23976096 | 0,749639658 | ribosomal protein S15a pseudogene 18 |
| ENSG00000234817 | ECI2-DT | -1,243431874 | 0,422366737 | 0,23107842 | 0,739268552 | ECI2 divergent transcript |
| ENSG00000260001 | TGFBR3L | -1,243937421 | 0,422218758 | 0,41033421 | 0,882614012 | transforming growth factor beta receptor 3 like |
| ENSG00000101251 | SEL1L2 | -1,243955143 | 0,422213571 | 0,39565854 | 0,872227623 | SEL1L2 adaptor subunit of ERAD E3 ligase |
| ENSG00000278232 | CRHR1 | -1,244350492 | 0,422097886 | 0,16470678 | 0,643915584 | corticotropin releasing hormone receptor 1 |
| ENSG00000283669 | AC112487,2 | -1,245086234 | 0,42188268 | 0,43125792 | 0,89530663 | novel pseudogene |
| ENSG00000238251 | NSA2P7 | -1,245127801 | 0,421870525 | 0,13185112 | 0,580182048 | NSA2 pseudogene 7 |
| ENSG00000162843 | WDR64 | -1,246009691 | 0,421612723 | 0,41940194 | 0,887822044 | WD repeat domain 64 |
| ENSG00000215045 | GRID2IP | -1,246266662 | 0,421537633 | 0,28217828 | 0,793275939 | Grid2 interacting protein |
| ENSG00000229954 | MTND2P2 | -1,246269407 | 0,42153683 | 0,14391693 | 0,603381927 | MT-ND2 pseudogene 2 |
| ENSG00000287784 | AC112482,2 | -1,246571753 | 0,421448498 | 0,346457 | 0,841168721 | novel transcript, antisense to C3orf56 |
| ENSG00000242276 | RPL5P3 | -1,246925575 | 0,42134515 | 0,28482398 | 0,796203237 | ribosomal protein L5 pseudogene 3 |
| ENSG00000174946 | GPR171 | -1,246964665 | 0,421333734 | 0,08057994 | 0,450103921 | G protein-coupled receptor 171 |
| ENSG00000283982 | AL445623,2 | -1,247574575 | 0,42115565 | 0,04384867 | 0,31589834 | novel transcript |
| ENSG00000174498 | IGDCC3 | -1,247600588 | 0,421148056 | 0,00133152 | 0,02378773 | immunoglobulin superfamily DCC subclass member 3 |
| ENSG00000257097 | CLIP1-AS1 | -1,248092155 | 0,421004583 | 0,00681953 | 0,087129511 | CLIP1 antisense RNA 1 |
| ENSG00000212743 | LINC02656 | -1,248238267 | 0,420961948 | 0,36865939 | 0,857635769 | long intergenic non-protein coding RNA 2656 |
| ENSG00000182489 | XKRX | -1,249104413 | 0,420709292 | 0,1478226 | 0,612372753 | XK related X-linked |
| ENSG00000213927 | CCL27 | -1,249417388 | 0,420618034 | 0,26590748 | 0,777251371 | C-C motif chemokine ligand 27 |
| ENSG00000238181 | AHCYP2 | -1,249641063 | 0,420552827 | 0,37825318 | 0,863824523 | adenosylhomocysteinase pseudogene 2 |
| ENSG00000223896 | CCNJP2 | -1,250055703 | 0,420431974 | 0,05642117 | 0,366903759 | cyclin J pseudogene 2 |
| ENSG00000230749 | MEIS1-AS2 | -1,250061079 | 0,420430408 | 0,00015176 | 0,003744887 | MEIS1 antisense RNA 2 |
| ENSG00000286310 | AC008443,6 | -1,250318253 | 0,420355468 | 0,26170985 | 0,773497425 | novel transcript |
| ENSG00000242551 | POU5F1P6 | -1,25126643 | 0,420079291 | 0,39439521 | 0,871799523 | POU class 5 homeobox 1 pseudogene 6 |
| ENSG00000259528 | AC068397,1 | -1,251345915 | 0,420056147 | 0,20284084 | 0,703217554 | novel transcript, antisense to NEO1 |
| ENSG00000253335 | AC009884,1 | -1,251712136 | 0,419949531 | 0,16321817 | 0,64157124 | novel transcript |

| | | | | | | |
|---|---|---|---|---|---|---|
| ENSG00000186676 | EEF1GP1 | -1,252190223 | 0,41981039 | 0,18077593 | 0,670312865 | eukaryotic translation elongation factor 1 gamma pseudogene 1 |
| ENSG00000233242 | AL512604,2 | -1,252801428 | 0,419632573 | 0,36405709 | 0,854207909 | novel transcript |
| ENSG00000272512 | AL645608,7 | -1,253091624 | 0,419548173 | 0,00094061 | 0,017747187 | novel transcript |
| ENSG00000229944 | EIF4EP2 | -1,2536389 | 0,41938905 | 0,23746491 | 0,746907875 | eukaryotic translation initiation factor 4E pseudogene 2 |
| ENSG00000235363 | SNRPGP10 | -1,253678471 | 0,419377547 | 0,10849417 | 0,525152765 | small nuclear ribonucleoprotein polypeptide G pseudogene 10 |
| ENSG00000235035 | AC005234,1 | -1,253915952 | 0,419308519 | 0,3507062 | 0,844035671 | novel transcript |
| ENSG00000261266 | AC008870,3 | -1,253936757 | 0,419302473 | 0,40690635 | 0,880078305 | novel transcript |
| ENSG00000225670 | CADM3-AS1 | -1,254110759 | 0,419251904 | 0,00568367 | 0,075828159 | CADM3 antisense RNA 1 |
| ENSG00000270316 | BORCS7-ASMT | -1,254485441 | 0,419143034 | 0,21067628 | 0,716131719 | BORCS7-ASMT readthrough (NMD candidate) |
| ENSG00000257607 | AC073957,2 | -1,25552373 | 0,418841491 | 0,47517878 | 0,911082311 | novel transcript, antisense to GPR146 |
| ENSG00000228366 | AL592437,1 | -1,255765606 | 0,418771276 | 0,13839281 | 0,593269285 | FUS interacting protein (serine-arginine rich) 1 (FUSIP1) pseudogene |
| ENSG00000234855 | SLIT1-AS1 | -1,256527449 | 0,418550194 | 0,07666224 | 0,437367108 | SLIT1 antisense RNA 1 |
| ENSG00000213492 | NT5C3AP1 | -1,256740086 | 0,418488509 | 0,19070743 | 0,685985838 | NT5C3A pseudogene 1 |
| ENSG00000110975 | SYT10 | -1,258326758 | 0,418028509 | 0,0461246 | 0,325797698 | synaptotagmin 10 |
| ENSG00000233072 | RPS15AP6 | -1,259581637 | 0,41766506 | 0,17083125 | 0,654404445 | ribosomal protein S15a pseudogene 6 |
| ENSG00000185038 | MROH2A | -1,259960703 | 0,417555333 | 0,20853947 | 0,713245584 | maestro heat like repeat family member 2A |
| ENSG00000258437 | AL139193,1 | -1,260107628 | 0,417512811 | 0,27765003 | 0,789067728 | novel transcript, antisense to TTC7B |
| ENSG00000263657 | AC023389,1 | -1,260474847 | 0,417406552 | 0,20465394 | 0,706214197 | novel transcript, antisense to SSH2 |
| ENSG00000267243 | AC005381,1 | -1,261782919 | 0,417028267 | 0,22958587 | 0,737998412 | novel transcript |
| ENSG00000164440 | TXLNB | -1,261798098 | 0,417023879 | 0,01106707 | 0,123337997 | taxilin beta |
| ENSG00000145423 | SFRP2 | -1,262400135 | 0,416849892 | 4,35E-06 | 0,000162701 | secreted frizzled related protein 2 |
| ENSG00000168309 | FAM107A | -1,263135315 | 0,416637524 | 0,01344128 | 0,141928098 | family with sequence similarity 107 member A |
| ENSG00000278570 | NR2E3 | -1,263517998 | 0,416527023 | 0,03951982 | 0,296223341 | nuclear receptor subfamily 2 group E member 3 |
| ENSG00000221887 | HMSD | -1,263524968 | 0,416525011 | 0,01546053 | 0,157394364 | histocompatibility minor serpin domain containing |
| ENSG00000214851 | LINC00612 | -1,265766416 | 0,415878378 | 0,00221734 | 0,03650494 | long intergenic non-protein coding RNA 612 |
| ENSG00000174944 | P2RY14 | -1,265882242 | 0,41584499 | 0,01823047 | 0,175695833 | purinergic receptor P2Y14 |
| ENSG00000234773 | AC012618,3 | -1,265888218 | 0,415843268 | 6,67E-05 | 0,001832763 | novel zinc finger protein pseudogene |
| ENSG00000224986 | PPP1R8P1 | -1,266260502 | 0,415735975 | 0,02502824 | 0,219551856 | protein phosphatase 1 regulatory subunit 8 pseudogene 1 |

| | | | | | | |
|---|---|---|---|---|---|---|
| ENSG00000229891 | LINC01315 | -1,266352473 | 0,415709472 | 0,00185215 | 0,031470438 | long intergenic non-protein coding RNA 1315 |
| ENSG00000285603 | AL035693,1 | -1,267167081 | 0,415474811 | 0,27066775 | 0,783073289 | novel transcript, antisense to GMDS |
| ENSG00000177306 | OR7E125P | -1,267725101 | 0,415314141 | 0,46707646 | 0,907940653 | olfactory receptor family 7 subfamily E member 125 pseudogene |
| ENSG00000158457 | TSPAN33 | -1,267747553 | 0,415307677 | 0,00110531 | 0,020319143 | tetraspanin 33 |
| ENSG00000236928 | AC008267,4 | -1,267782997 | 0,415297474 | 0,12106 | 0,555610668 | GrpE-like 1, mitochondrial (E, coli) (GRPEL1) pseudogene |
| ENSG00000279177 | CTBP2P9 | -1,268619318 | 0,415056799 | 0,01304765 | 0,139032628 | CTBP2 pseudogene 9 |
| ENSG00000280330 | CTBP2P10 | -1,268619318 | 0,415056799 | 0,01304765 | 0,139032628 | CTBP2 pseudogene 10 |
| ENSG00000206549 | AC109583,1 | -1,269465825 | 0,414813334 | 0,15920114 | 0,633239748 | Probable threonine protease PRSS50 |
| ENSG00000237414 | AL365204,2 | -1,269522161 | 0,414797136 | 0,01962595 | 0,184259987 | novel transcript |
| ENSG00000161940 | BCL6B | -1,269526272 | 0,414795954 | 0,08686195 | 0,469592252 | BCL6B transcription repressor |
| ENSG00000285553 | AL137182,1 | -1,269552403 | 0,414788441 | 0,25749415 | 0,769908546 | novel transcript |
| ENSG00000255132 | AC090138,1 | -1,269604147 | 0,414773565 | 0,35135462 | 0,84405766 | novel transcript |
| ENSG00000118702 | GHRH | -1,269666823 | 0,414755546 | 0,16465966 | 0,643811841 | growth hormone releasing hormone |
| ENSG00000235784 | HNRNPA1P29 | -1,270255492 | 0,414586346 | 0,02956927 | 0,244583576 | heterogeneous nuclear ribonucleoprotein A1 pseudogene 29 |
| ENSG00000249715 | FER1L5 | -1,270261687 | 0,414584566 | 0,00033492 | 0,007442069 | fer-1 like family member 5 |
| ENSG00000225192 | ZNF33BP1 | -1,270407223 | 0,414542745 | 0,06654178 | 0,405370643 | zinc finger protein 33B pseudogene 1 |
| ENSG00000167434 | CA4 | -1,270409346 | 0,414542135 | 0,23336188 | 0,742507431 | carbonic anhydrase 4 |
| ENSG00000237161 | AC068446,1 | -1,270501636 | 0,414515618 | 0,02509345 | 0,219753919 | p21 protein (Cdc42/Rac)-activated kinase 2 (PAK2) pseudogene |
| ENSG00000255480 | AL353699,1 | -1,270693632 | 0,414460457 | 0,3426448 | 0,84026916 | novel transcript |
| ENSG00000117407 | ARTN | -1,271652632 | 0,414185045 | 0,02618756 | 0,225980605 | artemin |
| ENSG00000251520 | AL162431,2 | -1,271743261 | 0,414159027 | 0,07390113 | 0,429530795 | pseudogene similar to part of major histocompatibility complex, class I-related MR1 |
| ENSG00000108878 | CACNG1 | -1,27183486 | 0,414132732 | 0,15448801 | 0,623626246 | calcium voltage-gated channel auxiliary subunit gamma 1 |
| ENSG00000206142 | FAM230H | -1,272020442 | 0,414079463 | 0,41321468 | 0,884663685 | family with sequence similarity 230 member H |
| ENSG00000261147 | AC091167,2 | -1,272234802 | 0,414017943 | 0,24885683 | 0,760831212 | novel protein |
| ENSG00000255350 | MTCO1P15 | -1,27224014 | 0,414016411 | 0,08755843 | 0,471644859 | MT-CO1 pseudogene 15 |
| ENSG00000214290 | COLCA2 | -1,272344017 | 0,413986602 | 0,06780013 | 0,410000651 | colorectal cancer associated 2 |
| ENSG00000236542 | MED28P7 | -1,272613222 | 0,41390936 | 0,15333826 | 0,621866315 | mediator complex subunit 28 pseudogene 7 |

| | | | | | | |
|---|---|---|---|---|---|---|
| ENSG00000285187 | AC270290,1 | -1,272863688 | 0,413837507 | 0,47280669 | 0,909509054 | novel transcript, antisense to SOX7 and PINX1 |
| ENSG00000227422 | DDR1-DT | -1,273288873 | 0,413715561 | 0,63441851 | 0,955097925 | DDR1 divergent transcript |
| ENSG00000284852 | YPEL5P2 | -1,273851641 | 0,41355421 | 0,25968085 | 0,772176335 | YPEL5 pseudogene 2 |
| ENSG00000271557 | YPEL5P2 | -1,273851641 | 0,41355421 | 0,25968085 | 0,772176335 | YPEL5 pseudogene 2 |
| ENSG00000287528 | AL121749,2 | -1,274144974 | 0,413470133 | 0,1170351 | 0,546606705 | novel transcript, antisense to FZD8 |
| ENSG00000268696 | ZNF723 | -1,274448506 | 0,413383152 | 0,47485218 | 0,91069468 | zinc finger protein 723 |
| ENSG00000153234 | NR4A2 | -1,27458191 | 0,413344928 | 0,00064607 | 0,012898268 | nuclear receptor subfamily 4 group A member 2 |
| ENSG00000159409 | CELF3 | -1,274661342 | 0,413322171 | 0,00252288 | 0,040345363 | CUGBP Elav-like family member 3 |
| ENSG00000229759 | MRPS18AP1 | -1,274784982 | 0,41328675 | 0,0917476 | 0,481764036 | mitochondrial ribosomal protein S18A pseudogene 1 |
| ENSG00000280114 | AL021920,2 | -1,274790199 | 0,413285256 | 0,16043859 | 0,636057831 | phosphodiesterase 4D interacting protein (myomegalin) (PDE4DIP) pseudogene |
| ENSG00000144681 | STAC | -1,275046232 | 0,413211917 | 0,0023364 | 0,037975216 | SH3 and cysteine rich domain |
| ENSG00000249839 | AC011330,1 | -1,275969986 | 0,412947423 | 0,66395392 | 0,959985685 | histidine acid phosphatase domain containing 2A (HISPPD2A) pseudogene |
| ENSG00000127561 | SYNGR3 | -1,276122212 | 0,412903854 | 0,00484283 | 0,067172286 | synaptogyrin 3 |
| ENSG00000244073 | RPS4XP6 | -1,276150984 | 0,412895619 | 0,04407153 | 0,316820511 | ribosomal protein S4X pseudogene 6 |
| ENSG00000218823 | PAPOLB | -1,276930268 | 0,41267265 | 0,02274573 | 0,204639564 | poly(A) polymerase beta |
| ENSG00000251365 | LINC02236 | -1,277980548 | 0,412372334 | 0,29283407 | 0,803295982 | long intergenic non-protein coding RNA 2236 |
| ENSG00000270948 | MTDHP1 | -1,278241195 | 0,412297839 | 0,11960958 | 0,552362303 | metadherin pseudogene 1 |
| ENSG00000160188 | RSPH1 | -1,278274504 | 0,41228832 | 0,04508022 | 0,321045067 | radial spoke head component 1 |
| ENSG00000213640 | EEF1DP4 | -1,278336204 | 0,412270688 | 0,2080007 | 0,712494639 | eukaryotic translation elongation factor 1 delta pseudogene 4 |
| ENSG00000249279 | LINC02057 | -1,278412366 | 0,412248924 | 0,38414167 | 0,868125769 | long intergenic non-protein coding RNA 2057 |
| ENSG00000263724 | DLGAP1-AS3 | -1,278769225 | 0,412146965 | 0,09961075 | 0,500760119 | DLGAP1 antisense RNA 3 |
| ENSG00000196503 | ARL9 | -1,279483696 | 0,411942906 | 0,00055454 | 0,011398522 | ADP ribosylation factor like GTPase 9 |
| ENSG00000268144 | NIFKP6 | -1,280214317 | 0,41173434 | 0,11623303 | 0,544242846 | nucleolar protein interacting with the FHA domain of MKI67 pseudogene 6 |
| ENSG00000266371 | AC079915,1 | -1,280324481 | 0,411702901 | 0,00483081 | 0,067065015 | novel transcript, sense intronic NF1 |
| ENSG00000237187 | NR2F1-AS1 | -1,280412921 | 0,411677664 | 2,33E-06 | 9,32E-05 | NR2F1 antisense RNA 1 |
| ENSG00000237990 | CNTN4-AS1 | -1,280866331 | 0,411548302 | 0,21625124 | 0,723540588 | CNTN4 antisense RNA 1 |
| ENSG00000245904 | AC025164,1 | -1,281068213 | 0,411490716 | 0,0490058 | 0,337619537 | novel transcript, antisense to BTG1 |
| ENSG00000180919 | OR56B4 | -1,281120009 | 0,411475943 | 0,20237767 | 0,702433811 | olfactory receptor family 56 subfamily B member 4 |
| ENSG00000255105 | PTMAP15 | -1,281129897 | 0,411473123 | 0,16501628 | 0,644037807 | prothymosin alpha pseudogene 15 |

| | | | | | | |
|---|---|---|---|---|---|---|
| ENSG00000258928 | AL358333,2 | -1,281361554 | 0,411407057 | 0,19013327 | 0,685541073 | novel transcript, antisense to GNG2 |
| ENSG00000260409 | AC012414,4 | -1,281438345 | 0,411385159 | 0,02586038 | 0,224147228 | novel transcript |
| ENSG00000286500 | AL512643,2 | -1,281678627 | 0,411316648 | 0,01152942 | 0,127055311 | novel transcript, antisense to FREM1 |
| ENSG00000230623 | AC104461,1 | -1,282976837 | 0,410946691 | 0,0551492 | 0,362604555 | novel transcript, antisense to ZNF281 |
| ENSG00000249695 | AC026369,1 | -1,283451352 | 0,41081155 | 0,28183767 | 0,792817421 | novel transcript, antisense to IQSEC3 |
| ENSG00000263137 | AC026369,1 | -1,283451352 | 0,41081155 | 0,28183767 | 0,792817421 | novel transcript, antisense to IQSEC3 |
| ENSG00000119514 | GALNT12 | -1,283615324 | 0,410764861 | 0,01980511 | 0,185108204 | polypeptide N-acetylgalactosaminyltransferase 12 |
| ENSG00000254186 | AC113414,1 | -1,283781581 | 0,410717527 | 0,01173038 | 0,128770524 | novel transcript |
| ENSG00000262999 | AC099489,3 | -1,283850721 | 0,410697844 | 0,09147364 | 0,481302729 | novel transcript, antisense to a novel protein gene |
| ENSG00000055813 | CCDC85A | -1,2840945 | 0,410628452 | 0,09188241 | 0,481984693 | coiled-coil domain containing 85A |
| ENSG00000168955 | TM4SF20 | -1,284665286 | 0,410466024 | 0,04473444 | 0,31940789 | transmembrane 4 L six family member 20 |
| ENSG00000225555 | AP000320,1 | -1,285357333 | 0,410269175 | 0,21342939 | 0,720944157 | novel transcript |
| ENSG00000139572 | GPR84 | -1,285412127 | 0,410253593 | 0,36863896 | 0,857635769 | G protein-coupled receptor 84 |
| ENSG00000218233 | NEPNP | -1,285472636 | 0,410236386 | 0,2471265 | 0,759323829 | nephrocan, pseudogene |
| ENSG00000129437 | KLK14 | -1,285622699 | 0,410193718 | 0,37880876 | 0,864112469 | kallikrein related peptidase 14 |
| ENSG00000223409 | RPL17P17 | -1,285934349 | 0,410105117 | 0,27752367 | 0,789067728 | ribosomal protein L17 pseudogene 17 |
| ENSG00000254692 | AL136295,1 | -1,286009781 | 0,410083675 | 0,13347564 | 0,583715556 | novel protein |
| ENSG00000285502 | AL136295,20 | -1,286009781 | 0,410083675 | 0,13347564 | 0,583715556 | novel protein |
| ENSG00000260036 | AC013355,1 | -1,286579206 | 0,409921849 | 0,28430751 | 0,795731744 | thioesterase superfamily member 4 (THEM4) pseudogene |
| ENSG00000249717 | PARM1-AS1 | -1,28690378 | 0,409829636 | 0,04381874 | 0,31589834 | PARM1 antisense RNA 1 |
| ENSG00000271013 | LRRC37A9P | -1,287618372 | 0,409626691 | 0,22184611 | 0,729383401 | leucine rich repeat containing 37 member A9, pseudogene |
| ENSG00000282015 | AC004099,2 | -1,288654455 | 0,40933262 | 0,26935306 | 0,781544346 | ribosomal protein L24 (RPL24) pseudogene |
| ENSG00000275839 | AC243654,2 | -1,288654455 | 0,40933262 | 0,26935306 | 0,781544346 | ribosomal protein L24 (RPL24) pseudogene |
| ENSG00000249210 | GAPDHP38 | -1,290222913 | 0,408887847 | 0,27475057 | 0,786383635 | glyceraldehyde 3 phosphate dehydrogenase pseudogene 38 |
| ENSG00000285907 | AL161910,1 | -1,290978961 | 0,408673624 | 0,1241296 | 0,563164667 | novel transcript |
| ENSG00000225311 | AL024474,2 | -1,291761921 | 0,408451894 | 0,22399769 | 0,732710917 | novel transcript, antisense to UTRN |
| ENSG00000138722 | MMRN1 | -1,291786174 | 0,408445028 | 0,06089458 | 0,383945238 | multimerin 1 |
| ENSG00000278351 | AC009248,2 | -1,292917233 | 0,408124937 | 0,34026943 | 0,838979484 | novel transcript, sense intronic to ANO6 |
| ENSG00000270679 | AC138473,1 | -1,29390943 | 0,40784435 | 0,41658291 | 0,886367338 | mitochondrial ribosomal protein S15 (MRPS15) pseudogene |

| | | | | | | |
|---|---|---|---|---|---|---|
| ENSG00000204252 | HLA-DOA | -1,294773096 | 0,407600268 | 0,25389805 | 0,76531122 | major histocompatibility complex, class II, DO alpha |
| ENSG00000231473 | RB1-DT | -1,294876375 | 0,407571089 | 0,00721899 | 0,090558361 | RB1 divergent transcript |
| ENSG00000286932 | AC013716,1 | -1,294986181 | 0,40754007 | 0,17467177 | 0,660057513 | novel transcript, sense intronic to LRRTM4 |
| ENSG00000231322 | RPL13AP17 | -1,295592314 | 0,407368882 | 0,03320721 | 0,264681419 | ribosomal protein L13a pseudogene 17 |
| ENSG00000270171 | AL359881,3 | -1,295673632 | 0,407345921 | 0,01246767 | 0,134595733 | novel transcript |
| ENSG00000257935 | LHX5-AS1 | -1,29606137 | 0,407236458 | 0,01794367 | 0,174004579 | LHX5 antisense RNA 1 |
| ENSG00000254590 | AP003396,2 | -1,296356363 | 0,407153197 | 0,29416392 | 0,804046662 | hippocampus abundant transcript 1 (HIAT1) pseudogene |
| ENSG00000186453 | FAM228A | -1,296378701 | 0,407146893 | 0,02895363 | 0,241642594 | family with sequence similarity 228 member A |
| ENSG00000196542 | SPTSSB | -1,297801356 | 0,4067456 | 0,0189988 | 0,18070531 | serine palmitoyltransferase small subunit B |
| ENSG00000116703 | PDC | -1,298196918 | 0,406634093 | 0,16686784 | 0,647265026 | phosducin |
| ENSG00000272627 | AC016542,1 | -1,298426457 | 0,4065694 | 0,28009076 | 0,790779983 | novel transcript, sense intronic with MCU |
| ENSG00000233360 | PDXP-DT | -1,298708049 | 0,406490052 | 0,27370246 | 0,785719894 | PDXP divergent transcript |
| ENSG00000171119 | NRTN | -1,298739524 | 0,406481184 | 0,11801917 | 0,549146283 | neurturin |
| ENSG00000273301 | AC016717,2 | -1,299416394 | 0,40629052 | 0,04909832 | 0,337936952 | novel transcript |
| ENSG00000263264 | AC119396,1 | -1,299875676 | 0,406161198 | 0,31536595 | 0,82082095 | novel protein |
| ENSG00000224117 | PTPN2P2 | -1,30034421 | 0,406029313 | 0,09295131 | 0,484612663 | PTPN2 pseudogene 2 |
| ENSG00000264346 | SNORA77B | -1,300563337 | 0,405967647 | 0,07018926 | 0,417549035 | small nucleolar RNA, H/ACA box 77B |
| ENSG00000170464 | DNAJC18 | -1,300915153 | 0,40586866 | 0,11076544 | 0,531456515 | DnaJ heat shock protein family (Hsp40) member C18 |
| ENSG00000233473 | RAD1P2 | -1,301239456 | 0,405777435 | 0,12353943 | 0,562121493 | RAD1 pseudogene 2 |
| ENSG00000266100 | AC007431,2 | -1,301789313 | 0,40562281 | 0,16486346 | 0,643964432 | novel transcript, antisense to MSI2 |
| ENSG00000251660 | AC007036,3 | -1,303960915 | 0,405012709 | 0,13208789 | 0,580641112 | novel transcript |
| ENSG00000287007 | AC087683,3 | -1,304159701 | 0,404956907 | 0,30908731 | 0,816845699 | novel transcript |
| ENSG00000287482 | AC124312,5 | -1,304615875 | 0,404828882 | 0,21613031 | 0,72322574 | novel transcript |
| ENSG00000239465 | AC090543,1 | -1,306745923 | 0,404231618 | 0,18271676 | 0,673321159 | ribosomal protein L21 (RPL21) pseudogene |
| ENSG00000286723 | AL021368,5 | -1,306782954 | 0,404221243 | 0,01892688 | 0,180076047 | novel transcript |
| ENSG00000223548 | ACSL6-AS1 | -1,307495317 | 0,404021699 | 0,36587157 | 0,855700097 | ACSL6 antisense RNA 1 |
| ENSG00000236007 | MTCO1P46 | -1,307545817 | 0,404007557 | 0,04106712 | 0,303420267 | MT-CO1 pseudogene 46 |
| ENSG00000235396 | NOTCH4 | -1,30792755 | 0,403900672 | 0,02042304 | 0,189467657 | notch receptor 4 |
| ENSG00000272108 | AC244517,1 | -1,307976643 | 0,403886928 | 1,48E-05 | 0,000478112 | novel transcript |
| ENSG00000230251 | PHBP4 | -1,308184893 | 0,403828632 | 0,30119502 | 0,810679941 | prohibitin pseudogene 4 |
| ENSG00000275963 | AC008115,4 | -1,30844343 | 0,40375627 | 0,01499348 | 0,154196081 | novel transcript |
| ENSG00000184434 | LRRC19 | -1,308748498 | 0,403670902 | 0,00455093 | 0,064409388 | leucine rich repeat containing 19 |

| | | | | | | |
|---|---|---|---|---|---|---|
| ENSG00000237273 | RSL24D1P8 | -1,309527385 | 0,403453026 | 0,13001725 | 0,576206833 | ribosomal L24 domain containing 1 pseudogene 8 |
| ENSG00000281818 | MANSC1 | -1,309625027 | 0,403425721 | 0,02929904 | 0,243294836 | MANSC domain containing 1 |
| ENSG00000243658 | MTND5P16 | -1,30991159 | 0,403345596 | 0,19057563 | 0,685949202 | MT-ND5 pseudogene 16 |
| ENSG00000287062 | AL121582,1 | -1,311393798 | 0,402931416 | 0,36975273 | 0,858104601 | novel transcript, antisense to MACROD2 |
| ENSG00000180974 | OR52E4 | -1,311932134 | 0,402781092 | 0,11886308 | 0,551028361 | olfactory receptor family 52 subfamily E member 4 |
| ENSG00000220744 | RPL5P18 | -1,312363384 | 0,402660711 | 0,02467724 | 0,217204179 | ribosomal protein L5 pseudogene 18 |
| ENSG00000283443 | AC018688,1 | -1,312482603 | 0,402627438 | 0,04413439 | 0,317117513 | zinc finger protein 285 (ZNF285) pseudogene |
| ENSG00000286132 | AC022415,2 | -1,312580404 | 0,402600145 | 0,03815067 | 0,289219826 | novel protein |
| ENSG00000277561 | GOLGA8IP | -1,313249883 | 0,402413362 | 0,08259873 | 0,457152163 | golgin A8 family member I, pseudogene |
| ENSG00000282989 | PDCL3P2 | -1,313675185 | 0,40229475 | 0,00759895 | 0,094200507 | PDCL3 pseudogene 2 |
| ENSG00000264475 | AP005062,1 | -1,314219622 | 0,402142963 | 0,07565755 | 0,434488912 | novel transcript, antisense to LAMA1 |
| ENSG00000165714 | BORCS5 | -1,314828164 | 0,401973371 | 0,06111409 | 0,384884976 | BLOC-1 related complex subunit 5 |
| ENSG00000249738 | AC008691,1 | -1,315346465 | 0,401828984 | 0,01040039 | 0,118280376 | novel transcript |
| ENSG00000224058 | AC006509,1 | -1,315544914 | 0,401773715 | 0,28877233 | 0,800014136 | novel pseudogene |
| ENSG00000218793 | AL049697,3 | -1,315642792 | 0,401746458 | 0,16336312 | 0,641873148 | S-adenosylmethionine decarboxylase 1 (AMD1) pseudogene |
| ENSG00000260899 | AC106886,2 | -1,316110754 | 0,401616166 | 0,09440121 | 0,487989313 | novel transcript |
| ENSG00000189398 | OR7E12P | -1,317643013 | 0,401189843 | 0,00093872 | 0,017722241 | olfactory receptor family 7 subfamily E member 12 pseudogene |
| ENSG00000271101 | AC005181,1 | -1,318774183 | 0,400875407 | 0,28572804 | 0,797420923 | SERPINE1 mRNA binding protein 1 (SERBP1) pseudogene |
| ENSG00000202512 | RN7SKP230 | -1,3188768 | 0,400846894 | 0,43933094 | 0,897434993 | RN7SK pseudogene 230 |
| ENSG00000267296 | CEBPA-DT | -1,31889885 | 0,400840767 | 0,10407883 | 0,514535205 | CEBPA divergent transcript |
| ENSG00000286934 | AC027251,1 | -1,32001587 | 0,400530533 | 0,19772295 | 0,696048902 | novel transcript, antisense to CSMD1 |
| ENSG00000267254 | ZNF790-AS1 | -1,320089 | 0,40051023 | 0,00065044 | 0,012968892 | ZNF790 antisense RNA 1 |
| ENSG00000257191 | AC090709,1 | -1,320773018 | 0,400320384 | 0,24631964 | 0,758198269 | novel transcript, antisense to SYT1 |
| ENSG00000179774 | ATOH7 | -1,321084291 | 0,400234021 | 0,06978133 | 0,416980797 | atonal bHLH transcription factor 7 |
| ENSG00000267909 | CCDC177 | -1,321635832 | 0,400081041 | 0,00209682 | 0,0349627 | coiled-coil domain containing 177 |
| ENSG00000215304 | AC135983,1 | -1,321895693 | 0,400008984 | 0,07931226 | 0,446745131 | novel transcript |
| ENSG00000228060 | PABPC4-AS1 | -1,322161946 | 0,399935168 | 0,30805125 | 0,816424917 | PABPC4 antisense RNA 1 |
| ENSG00000283529 | AL035685,1 | -1,325689449 | 0,398958489 | 0,24075148 | 0,751152776 | somatomedin B and thrombospondin type 1 domain containing (SBSPON) pseudogene |
| ENSG00000165655 | ZNF503 | -1,325868544 | 0,398908966 | 0,00036479 | 0,00796418 | zinc finger protein 503 |

| | | | | | | |
|---|---|---|---|---|---|---|
| ENSG00000264545 | AL359922,1 | -1,326488384 | 0,398737616 | 0,60406769 | 0,950647947 | novel transcript |
| ENSG00000124827 | GCM2 | -1,326698609 | 0,398679517 | 0,39942182 | 0,875433675 | glial cells missing transcription factor 2 |
| ENSG00000261697 | AC010531,4 | -1,327133079 | 0,398559472 | 0,17420438 | 0,659374935 | novel transcript, sense intronic to C16orf95 |
| ENSG00000242661 | RPS3AP43 | -1,327301916 | 0,398512832 | 0,2736002 | 0,785719894 | RPS3A pseudogene 43 |
| ENSG00000234566 | RPL7AP71 | -1,327437984 | 0,398475248 | 0,23272967 | 0,741684508 | ribosomal protein L7a pseudogene 71 |
| ENSG00000198963 | RORB | -1,328099943 | 0,398292455 | 0,0081168 | 0,098779094 | RAR related orphan receptor B |
| ENSG00000248780 | ARL4AP2 | -1,328372725 | 0,398217154 | 0,17952926 | 0,668298509 | ADP ribosylation factor like GTPase 4A pseudogene 2 |
| ENSG00000226900 | AL451069,1 | -1,328546253 | 0,398169259 | 0,32381146 | 0,825812991 | novel transcript |
| ENSG00000276680 | AL109755,2 | -1,329762211 | 0,397833809 | 0,4342175 | 0,896272827 | |
| ENSG00000115380 | EFEMP1 | -1,329772008 | 0,397831107 | 0,00664307 | 0,085363366 | EGF containing fibulin extracellular matrix protein 1 |
| ENSG00000125895 | TMEM74B | -1,329931963 | 0,397787001 | 0,02325624 | 0,207797418 | transmembrane protein 74B |
| ENSG00000269981 | AL627309,7 | -1,330550935 | 0,397616372 | 0,20501937 | 0,706634751 | novel pseudogene |
| ENSG00000198046 | ZNF667 | -1,330662367 | 0,397585661 | 0,12246295 | 0,559666977 | zinc finger protein 667 |
| ENSG00000227533 | SLC2A1-AS1 | -1,330692011 | 0,397577492 | 0,00034592 | 0,007653827 | SLC2A1 antisense RNA 1 |
| ENSG00000275409 | AC026367,2 | -1,330832195 | 0,397538862 | 0,3405765 | 0,839201107 | novel transcript |
| ENSG00000287268 | AL050350,1 | -1,330907392 | 0,397518142 | 0,01508973 | 0,154803784 | novel transcript, antisense to METTL24 |
| ENSG00000133101 | CCNA1 | -1,331573908 | 0,397334533 | 0,01129927 | 0,125269653 | cyclin A1 |
| ENSG00000225094 | SETP20 | -1,331862093 | 0,397255172 | 0,08844427 | 0,474454658 | SET pseudogene 20 |
| ENSG00000143632 | ACTA1 | -1,332001028 | 0,397216917 | 0,11521284 | 0,542308656 | actin alpha 1, skeletal muscle |
| ENSG00000259153 | AC004816,1 | -1,332298516 | 0,397135018 | 0,01124762 | 0,124829844 | novel transcript |
| ENSG00000283141 | LINC02666 | -1,332324788 | 0,397127786 | 0,10602173 | 0,519559347 | long intergenic non-protein coding RNA 2666 |
| ENSG00000249690 | AC110813,1 | -1,332552954 | 0,397064985 | 0,03287375 | 0,26278825 | novel transcript, antisense to LRBA |
| ENSG00000272002 | AC010904,2 | -1,332748797 | 0,397011087 | 0,07244097 | 0,424244694 | novel transcript |
| ENSG00000156966 | B3GNT7 | -1,332796673 | 0,396997913 | 0,00634276 | 0,082213892 | UDP-GlcNAc:betaGal beta-1,3-N-acetylglucosaminyltransferase 7 |
| ENSG00000215873 | FEN1P1 | -1,333022088 | 0,396935888 | 0,08740807 | 0,47120083 | flap structure-specific endonuclease 1 pseudogene 1 |
| ENSG00000285777 | AL139142,2 | -1,333307232 | 0,396857443 | 0,19869985 | 0,697657684 | novel protein |
| ENSG00000214456 | PLIN5 | -1,33417291 | 0,396619383 | 0,00607253 | 0,079736075 | perilipin 5 |
| ENSG00000249478 | AC010307,3 | -1,33419422 | 0,396613525 | 0,35952581 | 0,849982153 | novel transcript |
| ENSG00000270110 | AL353593,2 | -1,33455764 | 0,396513629 | 0,09783665 | 0,496997855 | novel transcript, antisense to OBSCN |
| ENSG00000145428 | RNF175 | -1,334899256 | 0,396419749 | 0,00730146 | 0,091345734 | ring finger protein 175 |
| ENSG00000271787 | AC104794,5 | -1,334962581 | 0,39640235 | 0,42273606 | 0,890403187 | novel transcript, antisense to KLF11 |

| | | | | | | |
|---|---|---|---|---|---|---|
| ENSG00000223866 | AC002486,1 | -1,335277914 | 0,396315717 | 0,10886592 | 0,526415221 | pseudogene similar to part of pituitary tumor-transforming 1 PTTG1 |
| ENSG00000187260 | WDR86 | -1,335314789 | 0,396305587 | 0,00071358 | 0,014022199 | WD repeat domain 86 |
| ENSG00000287081 | AC104656,1 | -1,335752254 | 0,396185434 | 0,07053134 | 0,418185447 | novel transcript, antisense to LDB2 |
| ENSG00000260092 | AC009163,3 | -1,336333458 | 0,396025859 | 0,23541141 | 0,744654651 | TMEM231-CHST5 readthrough protein |
| ENSG00000164841 | TMEM74 | -1,336699472 | 0,3959254 | 0,05533219 | 0,363130125 | transmembrane protein 74 |
| ENSG00000285279 | AL353622,5 | -1,337215431 | 0,395783828 | 0,62919498 | 0,952793934 | novel transcript |
| ENSG00000250020 | AC113430,1 | -1,337315604 | 0,395756348 | 0,11194691 | 0,534745332 | Spi-C transcription factor (Spi-1/PU,1 related) (SPIC) pseudogene |
| ENSG00000152092 | ASTN1 | -1,338322901 | 0,395480125 | 0,0021806 | 0,0360138 | astrotactin 1 |
| ENSG00000204701 | OR2J3 | -1,339650222 | 0,395116439 | 0,32209971 | 0,824955371 | olfactory receptor family 2 subfamily J member 3 |
| ENSG00000230383 | AC009245,1 | -1,340206696 | 0,394964065 | 0,33690964 | 0,836162806 | ribosomal protein L6 (RPL6) pseudogene |
| ENSG00000258827 | AL079304,1 | -1,340855983 | 0,394786351 | 0,28159393 | 0,792702377 | novel transcript, sense intronic to SLC25A21 |
| ENSG00000269901 | AC010531,5 | -1,341415993 | 0,394633137 | 0,26869284 | 0,780823861 | novel transcript, antisense to FBXO31 |
| ENSG00000243753 | HLA-L | -1,342386248 | 0,394367823 | 0,1992712 | 0,698588788 | major histocompatibility complex, class I, L (pseudogene) |
| ENSG00000179598 | PLD6 | -1,344309459 | 0,393842455 | 0,00531725 | 0,072277606 | phospholipase D family member 6 |
| ENSG00000245526 | LINC00461 | -1,345150646 | 0,393612885 | 0,00018198 | 0,004383318 | long intergenic non-protein coding RNA 461 |
| ENSG00000254485 | AC026191,1 | -1,345636751 | 0,393480283 | 0,16856328 | 0,650506523 | novel transcript, antisense to THUMPD3 |
| ENSG00000266036 | AC016888,1 | -1,346016306 | 0,393376776 | 0,36597678 | 0,85572078 | novel transcript, antisense to SLC9A3R1 |
| ENSG00000166596 | CFAP52 | -1,346028198 | 0,393373534 | 0,09151597 | 0,481302729 | cilia and flagella associated protein 52 |
| ENSG00000235294 | RPL7AP25 | -1,346030954 | 0,393372782 | 0,12934727 | 0,57474603 | ribosomal protein L7a pseudogene 25 |
| ENSG00000101916 | TLR8 | -1,34668413 | 0,393194724 | 0,37503488 | 0,861822152 | toll like receptor 8 |
| ENSG00000199883 | RN7SKP90 | -1,34720659 | 0,393052358 | 0,27341507 | 0,785672647 | RN7SK pseudogene 90 |
| ENSG00000272692 | AC010997,4 | -1,347714264 | 0,39291407 | 0,14794789 | 0,612582848 | novel transcript |
| ENSG00000213046 | EEF1A1P32 | -1,34787834 | 0,392869387 | 0,06562467 | 0,402080164 | eukaryotic translation elongation factor 1 alpha 1 pseudogene 32 |
| ENSG00000278012 | AL031658,2 | -1,348424733 | 0,392720623 | 0,265154 | 0,776598285 | novel transcript, sense intronic to XKR7 |
| ENSG00000225920 | RIMKLBP2 | -1,348488685 | 0,392703215 | 0,01730907 | 0,169906008 | ribosomal modification protein rimK like family member B pseudogene 2 |
| ENSG00000198156 | NPIPB6 | -1,349989452 | 0,392294917 | 0,02625794 | 0,226400506 | nuclear pore complex interacting protein family member B6 |

| | | | | | | |
|---|---|---|---|---|---|---|
| ENSG00000197882 | OR7E13P | -1,352221658 | 0,391688409 | 0,021591 | 0,196898814 | olfactory receptor family 7 subfamily E member 13 pseudogene |
| ENSG00000227189 | AC092535,2 | -1,352259183 | 0,391678221 | 0,11564775 | 0,543151268 | novel transcript |
| ENSG00000229259 | LRRC37A12P | -1,352500058 | 0,391612832 | 0,25467326 | 0,766102636 | leucine rich repeat containing 37 member A12, pseudogene |
| ENSG00000169313 | P2RY12 | -1,352827778 | 0,391523884 | 0,09181317 | 0,481984693 | purinergic receptor P2Y12 |
| ENSG00000260192 | LINC02240 | -1,353063224 | 0,391459993 | 0,16747654 | 0,648316613 | long intergenic non-protein coding RNA 2240 |
| ENSG00000076554 | TPD52 | -1,35344079 | 0,391357557 | 0,01597453 | 0,160898053 | tumor protein D52 |
| ENSG00000244050 | DEFB109F | -1,353751408 | 0,391273306 | 0,22067962 | 0,728572782 | defensin beta 109F (pseudogene) |
| ENSG00000286365 | AP000553,9 | -1,355601402 | 0,39077189 | 0,00506734 | 0,069393478 | novel transcript, antisense to YPEL1 |
| ENSG00000242686 | PDE6B-AS1 | -1,355619319 | 0,390767037 | 0,00464082 | 0,065415041 | PDE6B antisense RNA 1 |
| ENSG00000285836 | AC092896,1 | -1,355915703 | 0,390686767 | 0,12454178 | 0,563880062 | Novel trancript, antisense to CCDC191 |
| ENSG00000220771 | BOLA2P3 | -1,358031298 | 0,390114277 | 0,28928981 | 0,800260154 | bolA family member 2 pseudogene 3 |
| ENSG00000253079 | NRON | -1,358067204 | 0,390104568 | 0,17892468 | 0,667131928 | non-coding repressor of NFAT |
| ENSG00000164078 | MST1R | -1,358334878 | 0,390032196 | 0,04440404 | 0,318259511 | macrophage stimulating 1 receptor |
| ENSG00000226626 | NOP56P2 | -1,358339092 | 0,390031056 | 0,01557259 | 0,158020616 | NOP56 ribonucleoprotein pseudogene 2 |
| ENSG00000226887 | ERVMER34-1 | -1,358443212 | 0,390002909 | 0,05882791 | 0,376575703 | endogenous retrovirus group MER34 member 1, envelope |
| ENSG00000273240 | AC013468,1 | -1,358444492 | 0,390002562 | 0,05408532 | 0,358398336 | novel transcript |
| ENSG00000244131 | RPSAP51 | -1,358717844 | 0,389928674 | 0,11475308 | 0,541181798 | ribosomal protein SA pseudogene 51 |
| ENSG00000248801 | C8orf34-AS1 | -1,359013099 | 0,389848882 | 0,05825092 | 0,373570598 | C8orf34 antisense RNA 1 |
| ENSG00000144488 | ESPNL | -1,359761459 | 0,38964671 | 0,07020703 | 0,417549035 | espin like |
| ENSG00000273989 | AC022079,1 | -1,35992582 | 0,389602322 | 0,19624809 | 0,694046542 | novel transcript, sense intronic to CCDC91 |
| ENSG00000196169 | KIF19 | -1,36208139 | 0,389020641 | 0,00133888 | 0,023905709 | kinesin family member 19 |
| ENSG00000232827 | LINC01189 | -1,362094461 | 0,389017116 | 0,04386451 | 0,31589834 | long intergenic non-protein coding RNA 1189 |
| ENSG00000174521 | TTC9B | -1,362874563 | 0,388806822 | 0,04905334 | 0,3377238 | tetratricopeptide repeat domain 9B |
| ENSG00000284476 | AC044810,10 | -1,363963588 | 0,38851344 | 0,24503214 | 0,757071585 | |
| ENSG00000255618 | LINC02440 | -1,364064257 | 0,388486331 | 0,05214549 | 0,350371085 | long intergenic non-protein coding RNA 2440 |
| ENSG00000258984 | UBE2F-SCLY | -1,364432721 | 0,388387124 | 0,17697708 | 0,662915139 | UBE2F-SCLY readthrough (NMD candidate) |
| ENSG00000153714 | LURAP1L | -1,364505752 | 0,388367464 | 0,00639677 | 0,082776796 | leucine rich adaptor protein 1 like |
| ENSG00000271350 | AC103792,1 | -1,365418264 | 0,388121898 | 0,12367838 | 0,562344506 | arsA arsenite transporter, ATP-binding, homolog 1 (bacterial), pseudogene |

| | | | | | | |
|---|---|---|---|---|---|---|
| ENSG00000175985 | PLEKHD1 | -1,365863845 | 0,388002043 | 0,00468921 | 0,06577186 | pleckstrin homology and coiled-coil domain containing D1 |
| ENSG00000153930 | ANKFN1 | -1,366116024 | 0,387934228 | 0,00069061 | 0,013630919 | ankyrin repeat and fibronectin type III domain containing 1 |
| ENSG00000284997 | TUBAP2 | -1,36635456 | 0,387870092 | 0,22603807 | 0,734959876 | tubulin alpha pseudogene 2 |
| ENSG00000214391 | TUBAP2 | -1,36635456 | 0,387870092 | 0,22603807 | 0,734959876 | tubulin alpha pseudogene 2 |
| ENSG00000242087 | RPL36AP41 | -1,366577993 | 0,387810026 | 0,16200095 | 0,638979845 | ribosomal protein L36a pseudogene 41 |
| ENSG00000261646 | AC093849,1 | -1,367000102 | 0,387696576 | 0,20509224 | 0,706634751 | novel transcript |
| ENSG00000253226 | HAUS1P3 | -1,367568723 | 0,3875438 | 0,08830369 | 0,473946233 | HAUS augmin like complex subunit 1 pseudogene 3 |
| ENSG00000183943 | PRKX | -1,367744953 | 0,387496463 | 2,14E-08 | 1,26E-06 | protein kinase X-linked |
| ENSG00000270255 | AC009884,2 | -1,369419328 | 0,387047 | 0,18165291 | 0,67167359 | chromosome 8 open reading frame 46 (C8orf46) pseudogene |
| ENSG00000137561 | TTPA | -1,370202035 | 0,386837072 | 0,05190976 | 0,349538418 | alpha tocopherol transfer protein |
| ENSG00000165973 | NELL1 | -1,370658004 | 0,38671483 | 0,01554503 | 0,157881532 | neural EGFL like 1 |
| ENSG00000285612 | AC004974,1 | -1,371014681 | 0,386619234 | 0,03344691 | 0,265604304 | novel transcript, antisense to SIPA1L1 |
| ENSG00000181541 | MAB21L2 | -1,371478614 | 0,386494928 | 0,0348535 | 0,272414936 | mab-21 like 2 |
| ENSG00000261837 | AC046158,2 | -1,371870411 | 0,386389981 | 0,29389592 | 0,803761875 | novel transcript, sense intronic to WWOX |
| ENSG00000228316 | MTATP6P19 | -1,372041345 | 0,386344203 | 0,19059202 | 0,685949202 | MT-ATP6 pseudogene 19 |
| ENSG00000251869 | SCARNA23 | -1,372502468 | 0,386220737 | 0,02093114 | 0,192865038 | small Cajal body-specific RNA 23 |
| ENSG00000276127 | AL589935,2 | -1,372606413 | 0,386192911 | 0,22035357 | 0,728572782 | coiled-coil domain containing 112 (CCDC112) pseudogene |
| ENSG00000254551 | AP000873,3 | -1,373981392 | 0,38582502 | 0,27252246 | 0,785492261 | novel transcript |
| ENSG00000228638 | FCF1P2 | -1,374092421 | 0,385795328 | 0,30442677 | 0,813181541 | FCF1 pseudogene 2 |
| ENSG00000132702 | HAPLN2 | -1,374343853 | 0,385728098 | 0,16224861 | 0,639443053 | hyaluronan and proteoglycan link protein 2 |
| ENSG00000254793 | FDPSP4 | -1,375637551 | 0,385382362 | 0,07853314 | 0,443561759 | farnesyl diphosphate synthase pseudogene 4 |
| ENSG00000085514 | PILRA | -1,376333809 | 0,385196418 | 0,01127989 | 0,125099154 | paired immunoglobin like type 2 receptor alpha |
| ENSG00000132164 | SLC6A11 | -1,376453886 | 0,385164359 | 0,00556483 | 0,074829592 | solute carrier family 6 member 11 |
| ENSG00000174788 | PCP2 | -1,376537696 | 0,385141984 | 0,36849253 | 0,857548638 | Purkinje cell protein 2 |
| ENSG00000104826 | LHB | -1,376896779 | 0,385046135 | 0,26029812 | 0,772692192 | luteinizing hormone subunit beta |
| ENSG00000266987 | AC104984,5 | -1,377695224 | 0,384833094 | 0,28369694 | 0,79499517 | novel transcript, antisense to NSRP1 |
| ENSG00000248585 | AC084024,2 | -1,37818373 | 0,384702809 | 0,18207723 | 0,672311616 | origin recognition complex, subunit 3-like (yeast) (ORC3L) pseudogene |
| ENSG00000158258 | CLSTN2 | -1,378733418 | 0,38455626 | 0,00050293 | 0,010454364 | calsyntenin 2 |

| | | | | | | |
|---|---|---|---|---|---|---|
| ENSG00000151079 | KCNA6 | -1,379035472 | 0,384475754 | 0,00095339 | 0,01795591 | potassium voltage-gated channel subfamily A member 6 |
| ENSG00000273570 | AC007351,1 | -1,379413306 | 0,384375076 | 0,02602372 | 0,225063767 | |
| ENSG00000285755 | AC132153,1 | -1,379692071 | 0,384300812 | 0,07000395 | 0,417274281 | Novel transcript |
| ENSG00000164434 | FABP7 | -1,379928139 | 0,384237934 | 8,98E-05 | 0,002387263 | fatty acid binding protein 7 |
| ENSG00000169884 | WNT10B | -1,380471808 | 0,384093164 | 0,08989052 | 0,478293227 | Wnt family member 10B |
| ENSG00000261520 | DLGAP1-AS5 | -1,381128953 | 0,38391825 | 0,02246915 | 0,202734598 | DLGAP1 antisense RNA 5 |
| ENSG00000260685 | AC027104,1 | -1,381377697 | 0,383852062 | 0,17575382 | 0,661199837 | RWD domain containing 1 (RWDD1) pseudogene |
| ENSG00000284227 | DCHS2 | -1,38265541 | 0,383512256 | 0,00041946 | 0,008982246 | dachsous cadherin-related 2 |
| ENSG00000197410 | DCHS2 | -1,38265541 | 0,383512256 | 0,00041946 | 0,008982246 | dachsous cadherin-related 2 |
| ENSG00000188523 | CFAP77 | -1,384083316 | 0,383132863 | 0,08427902 | 0,461857898 | cilia and flagella associated protein 77 |
| ENSG00000277544 | AC104073,3 | -1,384377621 | 0,383054713 | 0,16153207 | 0,63813307 | novel zinc finger protein pseudogene |
| ENSG00000197415 | VEPH1 | -1,384585475 | 0,382999529 | 0,01921449 | 0,182079045 | ventricular zone expressed PH domain containing 1 |
| ENSG00000183248 | PRR36 | -1,385259688 | 0,382820585 | 0,00018347 | 0,004415809 | proline rich 36 |
| ENSG00000227733 | AC239809,3 | -1,38558565 | 0,3827341 | 8,05E-05 | 0,002171169 | novel transcript |
| ENSG00000225017 | AP004289,1 | -1,385613842 | 0,382726621 | 0,00974299 | 0,112876842 | aconitase 2, mitochondrial (ACO2) pseudogene |
| ENSG00000272076 | AC090186,1 | -1,386495843 | 0,38249271 | 0,15610318 | 0,626979676 | novel transcript |
| ENSG00000232445 | EMSLR | -1,387442053 | 0,38224193 | 0,00142346 | 0,025199849 | E2F1 mRNA stabilizing lncRNA |
| ENSG00000232342 | AC022540,1 | -1,388547389 | 0,381949183 | 0,16326292 | 0,641596686 | novel transcript |
| ENSG00000186732 | MPPED1 | -1,38881511 | 0,381878311 | 0,00262303 | 0,0415655 | metallophosphoesterase domain containing 1 |
| ENSG00000167874 | TMEM88 | -1,388908896 | 0,381853487 | 0,00287036 | 0,044824652 | transmembrane protein 88 |
| ENSG00000107014 | RLN2 | -1,389268065 | 0,381758434 | 0,12506922 | 0,565297686 | relaxin 2 |
| ENSG00000163885 | CFAP100 | -1,391148074 | 0,38126128 | 0,09544224 | 0,490483883 | cilia and flagella associated protein 100 |
| ENSG00000240531 | RPL21P123 | -1,392516918 | 0,380899707 | 0,30955641 | 0,816986489 | ribosomal protein L21 pseudogene 123 |
| ENSG00000235669 | CHN2-AS1 | -1,392552873 | 0,380890214 | 0,04045583 | 0,300358836 | CHN2 antisense RNA 1 |
| ENSG00000228196 | PTPN2P1 | -1,393578301 | 0,380619584 | 0,04213572 | 0,307859592 | PTPN2 pseudogene 1 |
| ENSG00000146215 | CRIP3 | -1,39459819 | 0,380350606 | 0,05260738 | 0,352155381 | cysteine rich protein 3 |
| ENSG00000159753 | CARMIL2 | -1,394638948 | 0,380339861 | 0,01537411 | 0,156675544 | capping protein regulator and myosin 1 linker 2 |
| ENSG00000085276 | MECOM | -1,395665384 | 0,380069356 | 0,05276616 | 0,352797524 | MDS1 and EVI1 complex locus |
| ENSG00000236106 | AC010729,1 | -1,396876426 | 0,379750449 | 0,05669703 | 0,367607368 | novel transcript |
| ENSG00000261734 | AC116096,1 | -1,397029324 | 0,379710204 | 0,04797666 | 0,333468718 | novel transcript |
| ENSG00000264673 | AC098850,1 | -1,397740024 | 0,379523198 | 0,27615377 | 0,788074712 | novel transcript |
| ENSG00000185352 | HS6ST3 | -1,398584954 | 0,379300991 | 0,00980851 | 0,113491261 | heparan sulfate 6-O-sulfotransferase 3 |

| | | | | | | |
|---|---|---|---|---|---|---|
| ENSG00000205309 | NT5M | -1,398641524 | 0,379286118 | 0,00034558 | 0,007651681 | 5',3'-nucleotidase, mitochondrial |
| ENSG00000110786 | PTPN5 | -1,398695178 | 0,379272013 | 0,00010604 | 0,002753455 | protein tyrosine phosphatase non-receptor type 5 |
| ENSG00000271963 | AC026786,2 | -1,399236571 | 0,379129712 | 0,2495195 | 0,761442412 | novel transcript, antisense to ZNF891 |
| ENSG00000256108 | AC135586,1 | -1,399450571 | 0,379073479 | 0,15157504 | 0,61913273 | tumor protein D52-like 2 (TPD52L2) pseudogene |
| ENSG00000254768 | AC104009,1 | -1,400476909 | 0,378803901 | 0,08830133 | 0,473946233 | novel transcript |
| ENSG00000183690 | EFHC2 | -1,400649086 | 0,378758695 | 0,02454005 | 0,216301092 | EF-hand domain containing 2 |
| ENSG00000283345 | AC127502,3 | -1,402161275 | 0,3783619 | 0,38103938 | 0,865900094 | M-phase phosphoprotein 10 (MPHOSPH10) pseudogene |
| ENSG00000241829 | RPL21P54 | -1,402772929 | 0,378201521 | 0,27169098 | 0,784751179 | ribosomal protein L21 pseudogene 54 |
| ENSG00000242071 | RPL7AP6 | -1,403371528 | 0,378044631 | 0,0793207 | 0,446745131 | ribosomal protein L7a pseudogene 6 |
| ENSG00000279954 | BX324167,2 | -1,403410393 | 0,378034447 | 0,00043548 | 0,00927602 | novel transcript |
| ENSG00000263326 | AC133552,4 | -1,403719524 | 0,377953453 | 0,11495702 | 0,541371975 | thyroid hormone receptor interactor 11 (TRIP11) pseudogene |
| ENSG00000225578 | NCBP2-AS1 | -1,404144938 | 0,377842021 | 0,26302633 | 0,77453036 | NCBP2 antisense RNA 1 |
| ENSG00000254815 | LMNTD2-AS1 | -1,404215137 | 0,377823636 | 0,51064746 | 0,922693987 | LMNTD2 antisense RNA 1 |
| ENSG00000231799 | PA2G4P6 | -1,404858366 | 0,37765522 | 0,19760206 | 0,696048902 | proliferation-associated 2G4 pseudogene 6 |
| ENSG00000236324 | SNX9-AS1 | -1,405397732 | 0,377514056 | 0,18984888 | 0,685029239 | SNX9 antisense RNA 1 |
| ENSG00000275736 | OSCAR | -1,405881202 | 0,377387567 | 0,20585226 | 0,708394615 | osteoclast associated Ig-like receptor |
| ENSG00000286233 | AC087885,1 | -1,406654104 | 0,377185441 | 0,08778087 | 0,471948586 | novel transcript, antisense to SRRM4 |
| ENSG00000261970 | AL391840,2 | -1,406825432 | 0,377140651 | 0,17082176 | 0,654404445 | solute carrier family 19, member 3 (SLC19A3) pseudogene |
| ENSG00000134198 | TSPAN2 | -1,40939011 | 0,376470804 | 0,00634192 | 0,082213892 | tetraspanin 2 |
| ENSG00000250071 | AC093214,1 | -1,411755102 | 0,375854166 | 0,13158764 | 0,579455365 | actin related protein 2/3 complex, subunit 1A, 41kDa (ARPC1A) pseudogene |
| ENSG00000234369 | TATDN1P1 | -1,412378236 | 0,37569186 | 0,08278771 | 0,457613669 | TatD DNase domain containing 1 pseudogene 1 |
| ENSG00000225849 | MKRN7P | -1,413628452 | 0,375366433 | 0,01469462 | 0,151771634 | makorin ring finger protein 7, pseudogene |
| ENSG00000253726 | AC013562,1 | -1,414631406 | 0,375105571 | 0,00851744 | 0,101909406 | novel transcript |
| ENSG00000283692 | Z96074,1 | -1,414743439 | 0,375076443 | 0,02002537 | 0,186831751 | novel transcript |
| ENSG00000164362 | TERT | -1,416112782 | 0,374720606 | 0,09012289 | 0,478721526 | telomerase reverse transcriptase |
| ENSG00000239570 | SETP11 | -1,418763803 | 0,374032671 | 0,10740288 | 0,523028605 | SET pseudogene 11 |
| ENSG00000281919 | SLC45A2 | -1,419057176 | 0,373956619 | 0,1997142 | 0,698923433 | solute carrier family 45 member 2 |
| ENSG00000213137 | ARF1P2 | -1,419477381 | 0,373847715 | 0,15737174 | 0,629809263 | ADP ribosylation factor 1 pseudogene 2 |
| ENSG00000255000 | AC139103,1 | -1,420201433 | 0,373660137 | 0,12602877 | 0,566777174 | zinc finger protein pseudogene |

| | | | | | | |
|---|---|---|---|---|---|---|
| ENSG00000261842 | AC073314,1 | -1,42120535 | 0,373400211 | 0,06440136 | 0,397835213 | novel transcript |
| ENSG00000259138 | AL049780,2 | -1,421287543 | 0,373378939 | 0,39227065 | 0,870909282 | novel transcript, antisense to NEK9 and TMED10 |
| ENSG00000233956 | BTF3P6 | -1,421512201 | 0,3733208 | 0,09420378 | 0,487613759 | basic transcription factor 3 pseudogene 6 |
| ENSG00000176299 | OR4M1 | -1,421598104 | 0,373298572 | 0,33119947 | 0,832301933 | olfactory receptor family 4 subfamily M member 1 |
| ENSG00000223561 | AC005165,1 | -1,422077024 | 0,373174672 | 0,04184391 | 0,306802968 | novel transcript |
| ENSG00000267952 | AC008878,1 | -1,422625926 | 0,373032717 | 0,25441311 | 0,765909623 | novel transcript |
| ENSG00000240673 | AC006539,1 | -1,422661853 | 0,373023428 | 0,28708042 | 0,798441631 | ribosomal protein S8 (RPS8) pseudogene |
| ENSG00000251402 | FAM90A25P | -1,422788226 | 0,372990754 | 0,07080551 | 0,418829978 | family with sequence similarity 90 member A25, pseudogene |
| ENSG00000254592 | AC123788,1 | -1,423216798 | 0,372879969 | 0,23127546 | 0,739268552 | novel transcript |
| ENSG00000256947 | AP002518,2 | -1,423397015 | 0,372833393 | 0,38902332 | 0,870057277 | novel transcript |
| ENSG00000222019 | URAHP | -1,423485304 | 0,372810577 | 0,08599898 | 0,467107732 | urate (hydroxyiso-) hydrolase, pseudogene |
| ENSG00000225057 | AC012485,1 | -1,423917112 | 0,372699009 | 0,04187739 | 0,306904544 | novel transcript |
| ENSG00000144290 | SLC4A10 | -1,424024225 | 0,372671339 | 0,00696557 | 0,088341044 | solute carrier family 4 member 10 |
| ENSG00000285593 | AC055872,2 | -1,42408564 | 0,372655475 | 0,12468745 | 0,564306384 | novel transcript |
| ENSG00000287997 | AL354950,3 | -1,424133669 | 0,372643069 | 0,18226258 | 0,672565494 | novel transcript, antisense to EBF3 |
| ENSG00000203711 | C6orf99 | -1,42591608 | 0,372182963 | 0,04878193 | 0,33644788 | chromosome 6 open reading frame 99 |
| ENSG00000275918 | KIR3DX1 | -1,425942704 | 0,372176094 | 0,1984344 | 0,697413278 | killer cell immunoglobulin like receptor, three Ig domains X1 (pseudogene) |
| ENSG00000231534 | FBXO36-IT1 | -1,425955461 | 0,372172803 | 0,144441 | 0,604748614 | FBXO36 intronic transcript 1 |
| ENSG00000285856 | AL353704,1 | -1,426120198 | 0,372130309 | 0,01924655 | 0,182285456 | novel transcript, antisense to ARHGEF7 |
| ENSG00000228407 | AL139156,2 | -1,426273088 | 0,372090874 | 0,18500542 | 0,677443132 | pseudogene similar to part of alcohol dehydrogenase 5 (class III), chi polypeptidem (ADH5) |
| ENSG00000260515 | AC034229,2 | -1,426739199 | 0,371970677 | 0,70715373 | 0,969565008 | novel transcript |
| ENSG00000224354 | MTND2P5 | -1,42702556 | 0,371896852 | 0,21344227 | 0,720944157 | MT-ND2 pseudogene 5 |
| ENSG00000211683 | AP000346,1 | -1,427378396 | 0,371805909 | 0,22427226 | 0,73307493 | novel transcript |
| ENSG00000251455 | AC092611,1 | -1,427523176 | 0,371768599 | 0,18941874 | 0,684701996 | novel transcript, antisense to PET112L |
| ENSG00000233729 | NCKAP5-AS1 | -1,427627123 | 0,371741814 | 0,00191991 | 0,032509055 | NCKAP5 antisense RNA 1 |
| ENSG00000205231 | TTLL10-AS1 | -1,428659841 | 0,371475807 | 0,19723843 | 0,695943345 | TTLL10 antisense RNA 1 |
| ENSG00000249965 | CDC42P4 | -1,428784547 | 0,371443698 | 0,31832197 | 0,823093059 | cell division cycle 42 pseudogene 4 |
| ENSG00000259772 | AC012236,1 | -1,429782593 | 0,371186824 | 0,29700604 | 0,806635068 | uncharacterized LOC283710 |
| ENSG00000255372 | AC107886,1 | -1,429954397 | 0,371142624 | 0,07291757 | 0,425892188 | novel transcript |

| | | | | | | |
|---|---|---|---|---|---|---|
| ENSG00000182824 | FAM230E | -1,430300281 | 0,371053654 | 0,1541678 | 0,623607455 | family with sequence similarity 230 member E |
| ENSG00000287094 | AL049552,2 | -1,430422445 | 0,371022235 | 0,08255061 | 0,45713804 | novel transcript, antisense to AHI1 |
| ENSG00000270716 | BNIP3P15 | -1,431689384 | 0,370696556 | 0,21481489 | 0,722164151 | BCL2 interacting protein 3 pseudogene 15 |
| ENSG00000244464 | AC107029,2 | -1,431799236 | 0,370668331 | 0,22357809 | 0,731984265 | novel transcript |
| ENSG00000284138 | ATP6V0CP4 | -1,43229867 | 0,370540034 | 0,0134562 | 0,141982678 | ATPase H+ transporting V0 subunit c pseudogene 4 |
| ENSG00000278907 | AC005609,2 | -1,432361341 | 0,370523939 | 0,24862772 | 0,760720365 | novel transcript, antisense to PCDHA9 |
| ENSG00000278780 | ZNF430 | -1,432498111 | 0,370488814 | 0,00167775 | 0,028899884 | zinc finger protein 430 |
| ENSG00000210077 | MT-TV | -1,433084656 | 0,370338218 | 0,22910379 | 0,737173617 | mitochondrially encoded tRNA-Val (GUN) |
| ENSG00000107295 | SH3GL2 | -1,433833441 | 0,370146055 | 0,03489331 | 0,272657954 | SH3 domain containing GRB2 like 2, endophilin A1 |
| ENSG00000215354 | FAM90A24P | -1,43393962 | 0,370118814 | 0,08377125 | 0,461014667 | family with sequence similarity 90 member A24, pseudogene |
| ENSG00000198670 | LPA | -1,434330851 | 0,370018459 | 0,06040463 | 0,382130814 | lipoprotein(a) |
| ENSG00000188831 | DPPA3P2 | -1,434402012 | 0,370000208 | 0,33662758 | 0,835927285 | developmental pluripotency associated 3 pseudogene 2 |
| ENSG00000276996 | SLC6A3 | -1,434908831 | 0,36987025 | 0,15210556 | 0,620332737 | solute carrier family 6 member 3 |
| ENSG00000226194 | LINC02519 | -1,436184324 | 0,369543391 | 0,20711636 | 0,710866812 | long intergenic non-protein coding RNA 2519 |
| ENSG00000256988 | AC005833,2 | -1,437062314 | 0,369318564 | 0,00583479 | 0,077296171 | new variant |
| ENSG00000224993 | RPL29P12 | -1,437235444 | 0,369274246 | 0,28577864 | 0,797420923 | ribosomal protein L29 pseudogene 12 |
| ENSG00000283855 | NPIPB13 | -1,437414848 | 0,369228329 | 0,45646102 | 0,903545728 | nuclear pore complex interacting protein family, member B13 |
| ENSG00000286747 | AL627095,2 | -1,438077831 | 0,369058691 | 0,0182964 | 0,176080387 | novel transcript |
| ENSG00000249494 | AC008629,1 | -1,438348218 | 0,368989529 | 0,06022204 | 0,381629797 | novel transcript |
| ENSG00000281641 | SAMD12-AS1 | -1,438621285 | 0,368919695 | 0,39227477 | 0,870909282 | SAMD12 antisense RNA 1 |
| ENSG00000250764 | AC025178,1 | -1,439097719 | 0,368797883 | 0,2467965 | 0,758836117 | novel transcript |
| ENSG00000226759 | DAB1-AS1 | -1,441824005 | 0,368101618 | 0,0019946 | 0,03359865 | DAB1 antisense RNA 1 |
| ENSG00000106541 | AGR2 | -1,44234427 | 0,367968897 | 0,06379765 | 0,395590561 | anterior gradient 2, protein disulphide isomerase family member |
| ENSG00000238194 | PHACTR3-AS1 | -1,443521205 | 0,367668834 | 0,16520593 | 0,644240891 | PHACTR3 antisense RNA 1 |
| ENSG00000173452 | TMEM196 | -1,443531757 | 0,367666145 | 0,02375707 | 0,210728318 | transmembrane protein 196 |
| ENSG00000235448 | LURAP1L-AS1 | -1,443586287 | 0,367652248 | 0,12802987 | 0,571345532 | LURAP1L antisense RNA 1 |
| ENSG00000248266 | AC108142,1 | -1,444826118 | 0,367336429 | 0,39169646 | 0,870362343 | novel transcript |
| ENSG00000138798 | EGF | -1,444931788 | 0,367309525 | 0,00360072 | 0,054043592 | epidermal growth factor |
| ENSG00000226410 | AC104058,1 | -1,445305278 | 0,367214447 | 0,26208467 | 0,773902615 | ribosomal protein L29 (RPL29) pseudogene |

| | | | | | | |
|---|---|---|---|---|---|---|
| ENSG00000278099 | RNVU1-2A | -1,44626671 | 0,366969811 | 0,24997216 | 0,761931313 | RNA, variant U1 small nuclear 2A |
| ENSG00000273757 | RNVU1-2A | -1,44626671 | 0,366969811 | 0,24997216 | 0,761931313 | RNA, variant U1 small nuclear 2A |
| ENSG00000059915 | PSD | -1,447308204 | 0,366704988 | 5,52E-05 | 0,001554209 | pleckstrin and Sec7 domain containing |
| ENSG00000263081 | AC211486,2 | -1,448648564 | 0,366364453 | 0,32429373 | 0,826368833 | novel transcript |
| ENSG00000287146 | AC000085,1 | -1,449008785 | 0,366272988 | 0,12637663 | 0,567767162 | novel transcript |
| ENSG00000230725 | AL035252,2 | -1,449194685 | 0,366225795 | 0,04729558 | 0,330573569 | novel transcript |
| ENSG00000178715 | AL450998,1 | -1,450055488 | 0,366007347 | 0,16163146 | 0,63813307 | ribosomal protein S16 (RPS16) pseudogene |
| ENSG00000226659 | AC021028,1 | -1,450959884 | 0,365777976 | 0,20799975 | 0,712494639 | novel transcript |
| ENSG00000212864 | RNF208 | -1,451547384 | 0,365629053 | 0,00101953 | 0,019006859 | ring finger protein 208 |
| ENSG00000204110 | LINC02520 | -1,451966642 | 0,365522814 | 0,23549505 | 0,744654651 | long intergenic non-protein coding RNA 2520 |
| ENSG00000234293 | BACH1-IT3 | -1,452369242 | 0,365420825 | 0,2534175 | 0,765172364 | BACH1 intronic transcript 3 |
| ENSG00000213950 | RPS10P2 | -1,452893547 | 0,365288047 | 0,35117788 | 0,84405766 | ribosomal protein S10 pseudogene 2 |
| ENSG00000165076 | PRSS37 | -1,454335456 | 0,364923141 | 0,2791985 | 0,790013761 | serine protease 37 |
| ENSG00000287288 | AL121762,2 | -1,455675838 | 0,364584255 | 0,19844513 | 0,697413278 | novel transcript |
| ENSG00000228061 | AC131571,1 | -1,45674825 | 0,364313345 | 0,00073364 | 0,014344387 | novel transcript, antisense to ELP4 |
| ENSG00000138653 | NDST4 | -1,456754694 | 0,364311718 | 0,03833591 | 0,290314236 | N-deacetylase and N-sulfotransferase 4 |
| ENSG00000277112 | ANKRD20A21P | -1,458083407 | 0,363976344 | 0,04445296 | 0,318464072 | ankyrin repeat domain 20 family member A21, pseudogene |
| ENSG00000240589 | RN7SL258P | -1,458768663 | 0,363803502 | 0,23717802 | 0,746624292 | RNA, 7SL, cytoplasmic 258, pseudogene |
| ENSG00000272088 | AL512413,1 | -1,459186952 | 0,363698038 | 0,23616833 | 0,745253314 | novel transcript |
| ENSG00000234562 | TPMTP2 | -1,460230447 | 0,363435072 | 0,04195191 | 0,307024639 | thiopurine S-methyltransferase pseudogene 2 |
| ENSG00000213409 | C1QBPP2 | -1,460458984 | 0,363377505 | 0,00557666 | 0,074924241 | complement C1q binding protein pseudogene 2 |
| ENSG00000168348 | INSM2 | -1,461794358 | 0,363041314 | 0,00580472 | 0,076995684 | INSM transcriptional repressor 2 |
| ENSG00000226965 | AC073114,1 | -1,462330778 | 0,362906354 | 0,13032484 | 0,576957777 | novel transcript |
| ENSG00000249274 | PDLIM1P4 | -1,462419715 | 0,362883983 | 0,14401834 | 0,603726134 | PDZ and LIM domain 1 pseudogene 4 |
| ENSG00000284748 | AL513220,1 | -1,463098963 | 0,362713171 | 0,37287297 | 0,86077683 | novel transcript |
| ENSG00000137261 | KIAA0319 | -1,463302969 | 0,362661884 | 0,00015376 | 0,003782125 | KIAA0319 |
| ENSG00000270540 | AC093915,1 | -1,463433763 | 0,362629007 | 0,11811602 | 0,549193844 | novel transcript, antisense to IQCA1 |
| ENSG00000141086 | CTRL | -1,463532193 | 0,362604267 | 0,01288649 | 0,138068235 | chymotrypsin like |
| ENSG00000249234 | AC108063,1 | -1,46518344 | 0,362189483 | 0,20552705 | 0,707976813 | novel transcript |
| ENSG00000163273 | NPPC | -1,465835994 | 0,362025696 | 0,23896398 | 0,748444182 | natriuretic peptide C |
| ENSG00000128285 | MCHR1 | -1,466203334 | 0,361933528 | 0,00353713 | 0,053345281 | melanin concentrating hormone receptor 1 |
| ENSG00000228929 | RPS13P2 | -1,466407219 | 0,361882383 | 0,32062661 | 0,824366234 | ribosomal protein S13 pseudogene 2 |

| | | | | | | |
|---|---|---|---|---|---|---|
| ENSG00000165449 | SLC16A9 | -1,467198665 | 0,361683913 | 0,00026357 | 0,006032482 | solute carrier family 16 member 9 |
| ENSG00000251609 | SETP12 | -1,468085665 | 0,36146161 | 0,06994727 | 0,417175017 | SET pseudogene 12 |
| ENSG00000228213 | NLGN1-AS1 | -1,469624004 | 0,361076391 | 0,02131644 | 0,194921664 | NLGN1 antisense RNA 1 |
| ENSG00000248863 | AC097376,2 | -1,469825545 | 0,361025953 | 0,03232403 | 0,260224035 | glucose-fructose oxidoreductase domain containing 2 (GFOD2) pseudogene |
| ENSG00000106113 | CRHR2 | -1,471785946 | 0,360535707 | 0,04454572 | 0,318908336 | corticotropin releasing hormone receptor 2 |
| ENSG00000287613 | AC069549,2 | -1,47193895 | 0,360497473 | 0,0032873 | 0,050255868 | novel transcript, antisense to PLXDC2 |
| ENSG00000271239 | AC007423,1 | -1,472162286 | 0,36044167 | 0,27613721 | 0,788074712 | novel transcript |
| ENSG00000233487 | RPSAP69 | -1,472176821 | 0,360438039 | 0,25280736 | 0,765053653 | ribosomal protein SA pseudogene 69 |
| ENSG00000199879 | RNVU1-22 | -1,472292817 | 0,36040906 | 0,03724779 | 0,284946761 | RNA, variant U1 small nuclear 22 |
| ENSG00000253336 | AC018992,1 | -1,472648485 | 0,360320219 | 0,10056823 | 0,503729332 | hydroxysteroid (17-beta) dehydrogenase 4 (HSD17B4) pseudogene |
| ENSG00000287354 | AL034351,4 | -1,472904478 | 0,360256289 | 0,09619557 | 0,492702847 | novel transcript, antisense to HHAT |
| ENSG00000287371 | AL034351,5 | -1,472904478 | 0,360256289 | 0,09619557 | 0,492702847 | novel transcript, antisense to HHAT |
| ENSG00000213305 | HNRNPCP6 | -1,473885814 | 0,360011323 | 0,00904708 | 0,106453868 | heterogeneous nuclear ribonucleoprotein C pseudogene 6 |
| ENSG00000015413 | DPEP1 | -1,474031651 | 0,359974932 | 0,3883177 | 0,869820053 | dipeptidase 1 |
| ENSG00000171798 | KNDC1 | -1,474297101 | 0,359908704 | 5,71E-05 | 0,001599821 | kinase non-catalytic C-lobe domain containing 1 |
| ENSG00000242318 | AC058823,1 | -1,474808632 | 0,359781116 | 0,17450936 | 0,660036356 | ribosomal protein L21 (RPL21) pseudogene |
| ENSG00000260969 | WWOX-AS1 | -1,476830908 | 0,359277151 | 0,25235885 | 0,76432863 | WWOX antisense RNA 1 |
| ENSG00000286938 | AC092849,2 | -1,478867197 | 0,358770408 | 0,29470709 | 0,804859966 | novel transcript |
| ENSG00000223518 | CSNK1A1P1 | -1,479829401 | 0,358531206 | 0,11337674 | 0,537795521 | casein kinase 1 alpha 1 pseudogene 1 |
| ENSG00000108255 | CRYBA1 | -1,480403254 | 0,358388623 | 0,06408785 | 0,396603857 | crystallin beta A1 |
| ENSG00000259540 | AC027020,1 | -1,480747394 | 0,358303143 | 0,17677186 | 0,662651108 | novel transcript, antisense to LINS1 |
| ENSG00000267402 | TCF4-AS2 | -1,481035491 | 0,3582316 | 0,12326503 | 0,561682809 | TCF4 antisense RNA 2 |
| ENSG00000196218 | RYR1 | -1,482059739 | 0,357977362 | 0,00173222 | 0,02973983 | ryanodine receptor 1 |
| ENSG00000283647 | AL512643,1 | -1,482688384 | 0,35782141 | 0,01049884 | 0,118829088 | novel transcript, antisense to FREM1 |
| ENSG00000231201 | AF127577,2 | -1,484115743 | 0,357467567 | 0,04190701 | 0,306990518 | novel transcript |
| ENSG00000282670 | AC254944,4 | -1,484970823 | 0,35725576 | 0,21790422 | 0,72555786 | novel transcript |
| ENSG00000269930 | AC091057,3 | -1,484970823 | 0,35725576 | 0,21790422 | 0,72555786 | novel transcript |
| ENSG00000256571 | AC079866,2 | -1,485564632 | 0,357108745 | 0,10648811 | 0,521171627 | novel transcript |
| ENSG00000177182 | CLVS1 | -1,486047773 | 0,356989173 | 0,00992533 | 0,114546111 | clavesin 1 |
| ENSG00000198570 | RD3 | -1,486359863 | 0,356911956 | 0,00704107 | 0,088942188 | retinal degeneration 3, GUCY2D regulator |

| | | | | | | |
|---|---|---|---|---|---|---|
| ENSG00000223500 | RNF14P4 | -1,486463777 | 0,35688625 | 0,04431232 | 0,31783836 | RNF14 pseudogene 4 |
| ENSG00000287045 | AC072028,1 | -1,488056107 | 0,356492565 | 0,00078203 | 0,015169893 | novel transcript, antisense to PLS1 |
| ENSG00000099960 | SLC7A4 | -1,488253835 | 0,356443709 | 0,27237855 | 0,785492261 | solute carrier family 7 member 4 |
| ENSG00000259592 | PRELID1P4 | -1,489681473 | 0,35609116 | 0,22000617 | 0,728200468 | PRELID1 pseudogene 4 |
| ENSG00000235994 | HGC6,3 | -1,489683639 | 0,356090626 | 0,19395402 | 0,690714036 | uncharacterized LOC100128124 |
| ENSG00000197149 | AC107956,1 | -1,492011153 | 0,355516604 | 0,19538662 | 0,693577833 | ribosomal protein S2 (RPS2) pseudogene |
| ENSG00000249574 | AC226118,1 | -1,492146309 | 0,3554833 | 0,11808823 | 0,549146283 | novel transcript |
| ENSG00000278198 | AC011043,2 | -1,492298231 | 0,355445868 | 0,10420188 | 0,514802081 | |
| ENSG00000273496 | AC011841,1 | -1,492298231 | 0,355445868 | 0,10420188 | 0,514802081 | |
| ENSG00000167311 | ART5 | -1,492993825 | 0,355274532 | 0,03641344 | 0,280436969 | ADP-ribosyltransferase 5 |
| ENSG00000276867 | AC074050,4 | -1,494209994 | 0,354975167 | 0,21920362 | 0,727587652 | novel transcript |
| ENSG00000235140 | AC026391,1 | -1,494457889 | 0,354914177 | 0,03361966 | 0,266434764 | uncharacterized LOC105378318 |
| ENSG00000214534 | ZNF705E | -1,494800614 | 0,354829874 | 0,03475269 | 0,271925393 | zinc finger protein 705E |
| ENSG00000276174 | AC087683,2 | -1,494996577 | 0,354781681 | 0,41179507 | 0,883966506 | novel transcript, sense intronic to PIK3C3 |
| ENSG00000125851 | PCSK2 | -1,495601409 | 0,354632974 | 1,93E-05 | 0,00061048 | proprotein convertase subtilisin/kexin type 2 |
| ENSG00000258171 | LINC02412 | -1,495970024 | 0,354542375 | 0,3045073 | 0,813200217 | long intergenic non-protein coding RNA 2412 |
| ENSG00000283559 | AC139491,6 | -1,496867536 | 0,35432188 | 0,30357714 | 0,812727837 | centrosomal protein 192kDa (CEP192) pseudogene |
| ENSG00000232880 | NDUFB4P10 | -1,496873752 | 0,354320354 | 0,26547413 | 0,776824277 | NADH:ubiquinone oxidoreductase subunit B4 pseudogene 10 |
| ENSG00000288568 | AC092720,3 | -1,497585449 | 0,354145607 | 0,05832666 | 0,373902954 | novel transcript |
| ENSG00000036565 | SLC18A1 | -1,497880684 | 0,354073141 | 0,00813793 | 0,098882383 | solute carrier family 18 member A1 |
| ENSG00000198785 | GRIN3A | -1,499447584 | 0,353688794 | 0,00036917 | 0,008044632 | glutamate ionotropic receptor NMDA type subunit 3A |
| ENSG00000231556 | RSL24D1P3 | -1,499647072 | 0,353639891 | 0,09596243 | 0,492073066 | ribosomal L24 domain containing 1 pseudogene 3 |
| ENSG00000265287 | AC005726,4 | -1,500240794 | 0,353494386 | 0,15747414 | 0,629899768 | novel transcript, antisense to novel protein |
| ENSG00000287080 | H3C3 | -1,501043682 | 0,353297714 | 0,13622014 | 0,588311436 | H3 clustered histone 3 |
| ENSG00000170893 | TRH | -1,504331889 | 0,35249339 | 0,0125635 | 0,135303249 | thyrotropin releasing hormone |
| ENSG00000181085 | MAPK15 | -1,505047149 | 0,352318674 | 0,01961 | 0,184165545 | mitogen-activated protein kinase 15 |
| ENSG00000175718 | RBMXL3 | -1,505513607 | 0,35220478 | 0,05399201 | 0,358091488 | RBMX like 3 |
| ENSG00000253057 | RN7SKP57 | -1,505984099 | 0,352089937 | 0,23967743 | 0,74947742 | RN7SK pseudogene 57 |
| ENSG00000285154 | AC243837,4 | -1,5072679 | 0,351776765 | 0,34758652 | 0,842271095 | olfactory receptor 2A1/2A42-like |
| ENSG00000263720 | AC023394,2 | -1,507797534 | 0,351647646 | 0,29321906 | 0,803365244 | novel transcript, sense intronic to CDH7 |

| | | | | | | |
|---|---|---|---|---|---|---|
| ENSG00000183423 | LRIT3 | -1,50788793 | 0,351625614 | 0,21096384 | 0,716555746 | leucine rich repeat, Ig-like and transmembrane domains 3 |
| ENSG00000140368 | PSTPIP1 | -1,507900811 | 0,351622474 | 0,06671684 | 0,405725609 | proline-serine-threonine phosphatase interacting protein 1 |
| ENSG00000164746 | C7orf57 | -1,508068 | 0,351581728 | 0,12226481 | 0,559316128 | chromosome 7 open reading frame 57 |
| ENSG00000129991 | TNNI3 | -1,508209 | 0,351547368 | 0,11902925 | 0,551553363 | troponin I3, cardiac type |
| ENSG00000238278 | ALG1L6P | -1,50834238 | 0,351514869 | 0,30175664 | 0,811332971 | ALG1 like 6, pseudogene |
| ENSG00000129159 | KCNC1 | -1,509693988 | 0,351185702 | 5,88E-06 | 0,000212764 | potassium voltage-gated channel subfamily C member 1 |
| ENSG00000228045 | DNAJA1P6 | -1,509892502 | 0,351137382 | 0,21064845 | 0,716131719 | DnaJ heat shock protein family (Hsp40) member A1 pseudogene 6 |
| ENSG00000166819 | PLIN1 | -1,510911037 | 0,350889568 | 0,19966872 | 0,698923433 | perilipin 1 |
| ENSG00000069206 | ADAM7 | -1,512937938 | 0,350396935 | 0,23963883 | 0,749431667 | ADAM metallopeptidase domain 7 |
| ENSG00000251031 | AC132803,1 | -1,514074893 | 0,350120904 | 0,02797912 | 0,236352093 | novel transcript |
| ENSG00000250746 | AC105250,1 | -1,515389726 | 0,349801959 | 0,14635595 | 0,609683201 | heat shock 70kDa protein 5 (glucose-regulated protein, 78kDa) (HSPA5) pseudogene |
| ENSG00000256206 | AC018523,2 | -1,515457443 | 0,34978554 | 0,25085371 | 0,763027659 | novel protein |
| ENSG00000254370 | AC025871,2 | -1,515967386 | 0,349661925 | 0,14747686 | 0,611575703 | novel transcript |
| ENSG00000249112 | AC112196,1 | -1,516459895 | 0,349542577 | 0,00922561 | 0,108066503 | novel transcript, antisense to ZNF608 |
| ENSG00000075884 | ARHGAP15 | -1,516585622 | 0,349512117 | 0,05514076 | 0,362604555 | Rho GTPase activating protein 15 |
| ENSG00000197565 | COL4A6 | -1,518064376 | 0,349154053 | 0,00407617 | 0,059577367 | collagen type IV alpha 6 chain |
| ENSG00000232085 | AL606534,2 | -1,519041774 | 0,348917588 | 0,26334909 | 0,774637968 | novel transcript, antisense to CEP170 |
| ENSG00000118972 | FGF23 | -1,520613585 | 0,34853765 | 0,06927989 | 0,415287103 | fibroblast growth factor 23 |
| ENSG00000274578 | AC006249,1 | -1,521272446 | 0,348378514 | 0,0133029 | 0,140888193 | novel transcript, sense intronic to CDH2 |
| ENSG00000240204 | SMKR1 | -1,521862122 | 0,348236149 | 0,03922063 | 0,294869635 | small lysine rich protein 1 |
| ENSG00000248196 | AC092658,1 | -1,5219274 | 0,348220393 | 0,04925312 | 0,338502855 | uracil-DNA glycosylase (UNG) pseudogene |
| ENSG00000243979 | AC087752,1 | -1,522208765 | 0,348152487 | 0,15024517 | 0,615953993 | ribosomal protein L17 (RPL17) pseudogene |
| ENSG00000258783 | KRT18P6 | -1,522854732 | 0,347996637 | 0,07435578 | 0,430811562 | keratin 18 pseudogene 6 |
| ENSG00000231115 | RING1 | -1,522929713 | 0,347978551 | 0,00330777 | 0,050445936 | ring finger protein 1 |
| ENSG00000228520 | RING1 | -1,522929713 | 0,347978551 | 0,00330777 | 0,050445936 | ring finger protein 1 |
| ENSG00000235885 | LINC01828 | -1,52310591 | 0,347936055 | 0,22776947 | 0,736053199 | long intergenic non-protein coding RNA 1828 |
| ENSG00000224194 | AC008278,1 | -1,524784157 | 0,347531546 | 0,33254793 | 0,833045148 | novel transcript |

| | | | | | | |
|---|---|---|---|---|---|---|
| ENSG00000278308 | NPIPA3 | -1,525627266 | 0,347328508 | 0,09155767 | 0,48132821 | nuclear pore complex interacting protein family member A3 |
| ENSG00000229196 | AC087071,1 | -1,52589321 | 0,347264488 | 0,29364712 | 0,803461221 | novel transcript |
| ENSG00000264404 | LINC02675 | -1,526834681 | 0,347037945 | 0,24548638 | 0,757479368 | long intergenic non-protein coding RNA 2675 |
| ENSG00000252233 | RN7SKP253 | -1,526961814 | 0,347007364 | 0,22620477 | 0,734959876 | RN7SK pseudogene 253 |
| ENSG00000214381 | LINC00488 | -1,528288597 | 0,346688384 | 0,00407231 | 0,059549493 | long intergenic non-protein coding RNA 488 |
| ENSG00000231849 | UBE2V2P4 | -1,528554073 | 0,346624594 | 0,14696304 | 0,610674677 | ubiquitin conjugating enzyme E2 V2 pseudogene 4 |
| ENSG00000236483 | MTND2P40 | -1,529046927 | 0,3465062 | 0,01910429 | 0,18137766 | MT-ND2 pseudogene 40 |
| ENSG00000235875 | ARHGEF7-AS2 | -1,529330694 | 0,346438052 | 0,00465474 | 0,065581757 | ARHGEF7 antisense RNA 2 |
| ENSG00000270098 | LDHAL6EP | -1,529966362 | 0,346285441 | 0,18814012 | 0,682491978 | lactate dehydrogenase A like 6E, pseudogene |
| ENSG00000233924 | RPSAP13 | -1,530790755 | 0,346087621 | 0,01509602 | 0,154803784 | ribosomal protein SA pseudogene 13 |
| ENSG00000240265 | AC099328,1 | -1,530988242 | 0,346040249 | 0,07912451 | 0,44596152 | H1 histone family, member O, oocyte-specific (H1FOO) pseudogene |
| ENSG00000255886 | AC020611,2 | -1,531066997 | 0,34602136 | 0,14049735 | 0,597347645 | novel transcript |
| ENSG00000267480 | AP001542,3 | -1,531654641 | 0,345880446 | 0,03881225 | 0,292885897 | novel transcript, antisense IMPA2 |
| ENSG00000239533 | GOLGA2P2Y | -1,532164505 | 0,345758229 | 0,2910064 | 0,801689099 | GOLGA2 pseudogene 2 Y-linked |
| ENSG00000274765 | AC018926,1 | -1,533010926 | 0,345555435 | 0,26461046 | 0,775800337 | novel transcript, sense intronic to CCPG1 |
| ENSG00000287875 | AC068193,1 | -1,533122688 | 0,345528666 | 0,07392865 | 0,429530795 | novel transcript, antisense to VRK2 |
| ENSG00000268836 | Z69706,1 | -1,533819287 | 0,34536187 | 0,18203404 | 0,672311616 | novel transcript |
| ENSG00000188086 | PRSS45P | -1,533939354 | 0,345333128 | 0,3513603 | 0,84405766 | serine protease 45, pseudogene |
| ENSG00000241030 | RPL29P24 | -1,534553685 | 0,345186109 | 0,20891518 | 0,713420724 | ribosomal protein L29 pseudogene 24 |
| ENSG00000200198 | RN7SKP211 | -1,535282259 | 0,345011831 | 0,27553891 | 0,787282819 | RN7SK pseudogene 211 |
| ENSG00000254712 | AC087280,1 | -1,536855634 | 0,344635773 | 0,09773348 | 0,496782149 | phosphorylated adaptor for RNA export (PHAX) pseudogene |
| ENSG00000213731 | RAB5CP1 | -1,536884642 | 0,344628844 | 0,13360673 | 0,583882708 | RAB5C, member RAS oncogene family pseudogene 1 |
| ENSG00000261267 | AC026470,2 | -1,536991702 | 0,34460327 | 0,1711123 | 0,654843372 | novel transcript, antisense to N4BP1 |
| ENSG00000286689 | AC090946,1 | -1,53716119 | 0,344562789 | 0,00108972 | 0,020109687 | novel transcript |
| ENSG00000198515 | CNGA1 | -1,53826386 | 0,344299536 | 0,20808555 | 0,712550568 | cyclic nucleotide gated channel subunit alpha 1 |
| ENSG00000187747 | OR52B6 | -1,53920859 | 0,344074149 | 0,24814985 | 0,760508446 | olfactory receptor family 52 subfamily B member 6 |
| ENSG00000203334 | AP003027,1 | -1,539475438 | 0,344010514 | 0,26344569 | 0,774637968 | novel transcript |
| ENSG00000286474 | AL355672,1 | -1,540454036 | 0,343777246 | 0,46911891 | 0,907940653 | novel transcript |
| ENSG00000176387 | HSD11B2 | -1,540928357 | 0,34366424 | 0,00575538 | 0,076470932 | hydroxysteroid 11-beta dehydrogenase 2 |
| ENSG00000237605 | DYRK3-AS1 | -1,540947952 | 0,343659572 | 0,249818 | 0,761684205 | DYRK3 antisense RNA 1 |

| | | | | | | |
|---|---|---|---|---|---|---|
| ENSG00000234793 | AC114730,2 | -1,541829395 | 0,34344967 | 0,35886905 | 0,849814998 | novel transcript |
| ENSG00000197322 | C17orf102 | -1,543221181 | 0,3431185 | 0,23771302 | 0,746907875 | chromosome 17 open reading frame 102 |
| ENSG00000120471 | TP53AIP1 | -1,543454217 | 0,343063081 | 0,1549144 | 0,624591393 | tumor protein p53 regulated apoptosis inducing protein 1 |
| ENSG00000278981 | AC079298,1 | -1,544033672 | 0,342925318 | 0,04660848 | 0,327821686 | novel transcript, antisense to DCHS2 |
| ENSG00000284467 | AC079298,5 | -1,544033672 | 0,342925318 | 0,04660848 | 0,327821686 | novel transcript, antisense to DCHS2 |
| ENSG00000212122 | TSSK1B | -1,544979029 | 0,342700683 | 0,35860794 | 0,849814998 | testis specific serine kinase 1B |
| ENSG00000124302 | CHST8 | -1,545267783 | 0,342632098 | 0,00810281 | 0,098656485 | carbohydrate sulfotransferase 8 |
| ENSG00000286671 | AC025253,2 | -1,546848887 | 0,342256801 | 0,03808897 | 0,288962258 | novel transcript |
| ENSG00000184305 | CCSER1 | -1,546929771 | 0,342237613 | 0,0027705 | 0,043460535 | coiled-coil serine rich protein 1 |
| ENSG00000259648 | AL132640,2 | -1,547324809 | 0,342143914 | 0,30027803 | 0,810130728 | high mobility group box 1 (HMGB1) pseudogene |
| ENSG00000285222 | GCNT2 | -1,551319018 | 0,341197973 | 0,00312823 | 0,048201641 | glucosaminyl (N-acetyl) transferase 2 (I blood group) |
| ENSG00000144891 | AGTR1 | -1,551621697 | 0,341126397 | 0,00320305 | 0,04916054 | angiotensin II receptor type 1 |
| ENSG00000261409 | AL035425,3 | -1,552505725 | 0,340917432 | 0,0341495 | 0,26913287 | novel transcript, |
| ENSG00000242791 | AC117395,1 | -1,552563037 | 0,340903889 | 0,19686738 | 0,695464607 | novel transcript |
| ENSG00000150337 | FCGR1A | -1,552591339 | 0,340897201 | 0,11931179 | 0,552110068 | Fc fragment of IgG receptor Ia |
| ENSG00000234173 | AL353784,1 | -1,553326247 | 0,340723593 | 0,05533367 | 0,363130125 | novel transcript, antisense to PCDH15 |
| ENSG00000248873 | SERBP1P6 | -1,554443882 | 0,340459741 | 0,08554656 | 0,465670107 | SERPINE1 mRNA binding protein 1 pseudogene 6 |
| ENSG00000241231 | AC068308,1 | -1,555572358 | 0,340193538 | 0,00110894 | 0,020346144 | novel transcript |
| ENSG00000226121 | AHCTF1P1 | -1,557374553 | 0,339768838 | 0,00481853 | 0,06695396 | AT-hook containing transcription factor 1 pseudogene 1 |
| ENSG00000272940 | U62317,2 | -1,559605153 | 0,339243916 | 0,11525196 | 0,542364351 | novel transcript |
| ENSG00000275345 | AC091167,5 | -1,559687563 | 0,339224538 | 0,19401144 | 0,690839823 | novel transcript |
| ENSG00000215492 | HNRNPA1P7 | -1,559707602 | 0,339219826 | 0,24206209 | 0,752344083 | heterogeneous nuclear ribonucleoprotein A1 pseudogene 7 |
| ENSG00000286843 | AL160159,1 | -1,562143154 | 0,33864764 | 0,05157531 | 0,347838733 | novel transcript, antisense to GPC6 |
| ENSG00000180176 | TH | -1,562979329 | 0,338451419 | 0,00490669 | 0,067907399 | tyrosine hydroxylase |
| ENSG00000147676 | MAL2 | -1,563020429 | 0,338441777 | 0,00674792 | 0,08635575 | mal, T cell differentiation protein 2 |
| ENSG00000287167 | AL133313,1 | -1,563040333 | 0,338437108 | 0,12542647 | 0,565907826 | novel transcript, antisense to RAD51B |
| ENSG00000285162 | AC004593,2 | -1,564532696 | 0,3380872 | 0,19663489 | 0,694800312 | chimerin 2 |
| ENSG00000163285 | GABRG1 | -1,564718527 | 0,338043655 | 0,22757926 | 0,736053199 | gamma-aminobutyric acid type A receptor subunit gamma1 |
| ENSG00000140284 | SLC27A2 | -1,564930315 | 0,337994034 | 0,03228939 | 0,260130339 | solute carrier family 27 member 2 |

| | | | | | | |
|---|---|---|---|---|---|---|
| ENSG00000272175 | AL050343,3 | -1,565239965 | 0,337921497 | 0,12421064 | 0,563196389 | novel transcript, antisense to NRD1 |
| ENSG00000259416 | LINC02883 | -1,565653448 | 0,337824661 | 0,1381653 | 0,592936302 | long intergenic non-protein coding RNA 2883 |
| ENSG00000138769 | CDKL2 | -1,566042949 | 0,337733467 | 0,00538021 | 0,072847486 | cyclin dependent kinase like 2 |
| ENSG00000238217 | LINC01877 | -1,567693209 | 0,337347363 | 0,25989372 | 0,772441975 | long intergenic non-protein coding RNA 1877 |
| ENSG00000184845 | DRD1 | -1,570497523 | 0,336692264 | 0,04072749 | 0,301587285 | dopamine receptor D1 |
| ENSG00000164089 | ETNPPL | -1,571695539 | 0,33641279 | 0,07666235 | 0,437367108 | ethanolamine-phosphate phospho-lyase |
| ENSG00000206344 | HCG27 | -1,573543346 | 0,335982187 | 0,1861003 | 0,67937802 | HLA complex group 27 |
| ENSG00000228333 | RXRB | -1,573605432 | 0,335967729 | 0,00739861 | 0,092339375 | retinoid X receptor beta |
| ENSG00000259881 | AC092384,2 | -1,573696468 | 0,33594653 | 0,17826447 | 0,665463944 | novel transcript, antisense to CBFA2T3 |
| ENSG00000241723 | AC074276,1 | -1,57420667 | 0,335827745 | 0,10462808 | 0,515975114 | ribosomal protein L15 (RPL15) pseudogene |
| ENSG00000041515 | MYO16 | -1,574236899 | 0,335820708 | 0,00023962 | 0,005560475 | myosin XVI |
| ENSG00000183615 | FAM167B | -1,574921311 | 0,335661433 | 0,03581528 | 0,277297887 | family with sequence similarity 167 member B |
| ENSG00000255396 | AP000857,1 | -1,575395037 | 0,335551233 | 0,11392962 | 0,53948382 | ras homolog gene family, member B (RHOB) pseudogene |
| ENSG00000260132 | AL032819,1 | -1,57975047 | 0,334539746 | 0,35638727 | 0,847771893 | novel transcript, antisense to UNKL |
| ENSG00000104899 | AMH | -1,58000576 | 0,334480553 | 0,00053635 | 0,011075529 | anti-Mullerian hormone |
| ENSG00000267778 | AC004221,1 | -1,580020174 | 0,334477212 | 0,277339 | 0,789067728 | novel transcript |
| ENSG00000145721 | LIX1 | -1,580482581 | 0,334370023 | 0,00698517 | 0,088486344 | limb and CNS expressed 1 |
| ENSG00000144771 | LRTM1 | -1,581096563 | 0,334227752 | 0,18758924 | 0,681556081 | leucine rich repeats and transmembrane domains 1 |
| ENSG00000226465 | AL390198,1 | -1,581239056 | 0,334194743 | 0,06088375 | 0,383945238 | novel transcript |
| ENSG00000127377 | CRYGN | -1,581347993 | 0,334169509 | 0,26060563 | 0,772692192 | crystallin gamma N |
| ENSG00000249278 | AC112249,1 | -1,582268443 | 0,333956374 | 0,00572885 | 0,076280623 | ribosomal protein L18 (RPL18) pseudogene |
| ENSG00000184470 | TXNRD2 | -1,582447816 | 0,333914856 | 0,00063865 | 0,012799522 | thioredoxin reductase 2 |
| ENSG00000117245 | KIF17 | -1,582743604 | 0,333846402 | 0,0002153 | 0,005076179 | kinesin family member 17 |
| ENSG00000183346 | CABCOCO1 | -1,582822144 | 0,333828228 | 0,01516641 | 0,155310364 | ciliary associated calcium binding coiled-coil 1 |
| ENSG00000269102 | AC010320,3 | -1,583685679 | 0,333628472 | 0,22169672 | 0,729254784 | novel transcript, antisense to ZNF766 and ZNF480 |
| ENSG00000154479 | CCDC173 | -1,583763003 | 0,333610591 | 0,00019915 | 0,004760077 | coiled-coil domain containing 173 |
| ENSG00000167617 | CDC42EP5 | -1,584105365 | 0,333531433 | 0,66335285 | 0,959920886 | CDC42 effector protein 5 |
| ENSG00000145309 | CABS1 | -1,584563801 | 0,333425465 | 0,07343776 | 0,427791692 | calcium binding protein, spermatid associated 1 |
| ENSG00000243055 | GK-AS1 | -1,584969705 | 0,333331669 | 0,14694796 | 0,610674677 | GK antisense RNA 1 |
| ENSG00000141668 | CBLN2 | -1,58593991 | 0,33310758 | 0,02807443 | 0,236837254 | cerebellin 2 precursor |
| ENSG00000203690 | TCP10L3 | -1,585948589 | 0,333105576 | 0,15076317 | 0,617373838 | t-complex 10 like 3, pseudogene |

| | | | | | | |
|---|---|---|---|---|---|---|
| ENSG00000105642 | KCNN1 | -1,586516488 | 0,332974479 | 0,00075457 | 0,014698276 | potassium calcium-activated channel subfamily N member 1 |
| ENSG00000240328 | AC091805,1 | -1,587716242 | 0,332697691 | 0,1620599 | 0,639001241 | ribosomal protein L34 (RPL34) pseudogene |
| ENSG00000228879 | RPL35AP6 | -1,588976689 | 0,332407148 | 0,14538513 | 0,607500779 | ribosomal protein L35a pseudogene 6 |
| ENSG00000183206 | POTEC | -1,589837293 | 0,332208918 | 0,0068734 | 0,087531521 | POTE ankyrin domain family member C |
| ENSG00000237816 | RPL21P109 | -1,590315126 | 0,332098906 | 0,22558334 | 0,734461165 | ribosomal protein L21 pseudogene 109 |
| ENSG00000267703 | AC020917,2 | -1,59095247 | 0,331952226 | 0,34399882 | 0,840938551 | novel transcript, antisense EPS15L1 |
| ENSG00000255280 | AP001922,4 | -1,591382194 | 0,331853364 | 0,21734442 | 0,725037995 | ribosomal protein L37 (RPL37) pseudogene |
| ENSG00000213287 | AP001102,1 | -1,59304415 | 0,331471296 | 0,16571034 | 0,645029358 | translocase of outer mitochondrial membrane 22 homolog (yeast) (TOMM22) pseudogene |
| ENSG00000257674 | AC020629,1 | -1,59353405 | 0,331358757 | 0,23942021 | 0,749122637 | insulin-like growth factor 2 mRNA binding protein 2 (IGF2BP2) pseudogene |
| ENSG00000281732 | Z84468,1 | -1,594166453 | 0,331213538 | 0,15907451 | 0,633144409 | novel transcript |
| ENSG00000102172 | SMS | -1,594280266 | 0,33118741 | 7,34E-07 | 3,20E-05 | spermine synthase |
| ENSG00000261554 | AC037471,2 | -1,595214027 | 0,330973123 | 0,17237251 | 0,656881307 | vomeronasal 1 receptor (VN1R) pseudogene |
| ENSG00000241020 | AP000873,1 | -1,597110311 | 0,330538377 | 0,28163691 | 0,792751997 | ribosomal protein L32 (RPL32) pseudogene |
| ENSG00000236188 | PRKX-AS1 | -1,597750231 | 0,330391796 | 0,13085227 | 0,577575225 | PRKX antisense RNA 1 |
| ENSG00000234881 | PIGFP2 | -1,597911784 | 0,330354801 | 0,07220923 | 0,42371424 | phosphatidylinositol glycan anchor biosynthesis class F pseudogene 2 |
| ENSG00000269177 | L34079,2 | -1,597967873 | 0,330341958 | 0,11093879 | 0,531880111 | novel transcript, antisense to XRCC1 |
| ENSG00000259180 | AC012378,1 | -1,598871788 | 0,330135048 | 0,1695552 | 0,652045207 | novel transcript, antisense to PRTG |
| ENSG00000167646 | DNAAF3 | -1,599460261 | 0,330000414 | 0,00577052 | 0,076607039 | dynein axonemal assembly factor 3 |
| ENSG00000236154 | AL450311,1 | -1,599571084 | 0,329975065 | 0,05198429 | 0,349829381 | novel transcript |
| ENSG00000091664 | SLC17A6 | -1,600482774 | 0,329766608 | 0,0287558 | 0,240839934 | solute carrier family 17 member 6 |
| ENSG00000177994 | C2orf73 | -1,600569092 | 0,329746879 | 0,08105279 | 0,451941233 | chromosome 2 open reading frame 73 |
| ENSG00000234225 | AC244394,3 | -1,600768654 | 0,329701269 | 0,13369172 | 0,583882708 | novel transcript |
| ENSG00000237459 | MPIG6B | -1,601009181 | 0,329646306 | 0,29147518 | 0,802058262 | megakaryocyte and platelet inhibitory receptor G6b |
| ENSG00000259905 | PWRN1 | -1,601751978 | 0,329476625 | 0,00020427 | 0,004867516 | Prader-Willi region non-protein coding RNA 1 |
| ENSG00000285809 | AC093295,1 | -1,602141872 | 0,329387595 | 0,09226374 | 0,482910203 | Novel transcript |
| ENSG00000286387 | AC007432,1 | -1,60417051 | 0,328924754 | 0,02416482 | 0,213535623 | novel transcript |
| ENSG00000018607 | ZNF806 | -1,604282025 | 0,32889933 | 0,00192055 | 0,032509055 | zinc finger protein 806 |
| ENSG00000257835 | AC073591,1 | -1,605307352 | 0,328665664 | 0,07945256 | 0,447115945 | novel transcript |
| ENSG00000249163 | AL121809,1 | -1,605537899 | 0,328613146 | 0,21958473 | 0,727931767 | novel transcript, antisense to PRPF39 |

| | | | | | | |
|---|---|---|---|---|---|---|
| ENSG00000271967 | AL583856,2 | -1,606836217 | 0,328317552 | 0,33647639 | 0,835713011 | novel transcript |
| ENSG00000259202 | AC012568,1 | -1,607481054 | 0,328170837 | 0,06211942 | 0,389024096 | novel transcript, antisense to SMAD3 |
| ENSG00000237337 | ELAVL4-AS1 | -1,607909483 | 0,328073397 | 0,15842147 | 0,631907243 | ELAVL4 antisense RNA 1 |
| ENSG00000236826 | LSM2 | -1,609282653 | 0,327761282 | 0,00463405 | 0,065349052 | LSM2 homolog, U6 small nuclear RNA and mRNA degradation associated |
| ENSG00000231671 | LINC01307 | -1,609394805 | 0,327735804 | 0,0578523 | 0,371961929 | long intergenic non-protein coding RNA 1307 |
| ENSG00000213067 | AC092447,1 | -1,61024702 | 0,327542264 | 0,0795139 | 0,447349744 | insulin-like growth factor 2 mRNA binding protein 3 (IGF2BP3) pseudogene |
| ENSG00000260012 | AC007494,1 | -1,610660717 | 0,327448354 | 0,12296482 | 0,5608947 | proteasome (prosome, macropain) subunit, beta type, 5 (PSMB5) pseudogene |
| ENSG00000281734 | AL139246,13 | -1,611122231 | 0,327343621 | 0,53055257 | 0,931265667 | novel transcript |
| ENSG00000284920 | RPS3AP34 | -1,612772824 | 0,32696932 | 0,03110224 | 0,253664123 | RPS3A pseudogene 34 |
| ENSG00000248616 | ANKRD49P3 | -1,613269487 | 0,326856777 | 0,10144909 | 0,506329296 | ANKRD49 pseudogene 3 |
| ENSG00000081181 | ARG2 | -1,614643969 | 0,326545522 | 0,00773794 | 0,095469263 | arginase 2 |
| ENSG00000246526 | LINC02481 | -1,615180898 | 0,326424014 | 0,04708306 | 0,32967633 | long intergenic non-protein coding RNA 2481 |
| ENSG00000236327 | AL831737,1 | -1,615372315 | 0,326380707 | 0,26196917 | 0,773902615 | ribosomal protein L30 (RPL30) pseudogene |
| ENSG00000257802 | MRS2P2 | -1,616641786 | 0,326093641 | 0,02811688 | 0,237067462 | MRS2 pseudogene 2 |
| ENSG00000183833 | CFAP91 | -1,617125164 | 0,325984401 | 0,00706221 | 0,089173264 | cilia and flagella associated protein 91 |
| ENSG00000146221 | TCTE1 | -1,617408575 | 0,325920369 | 0,01634462 | 0,163324813 | t-complex-associated-testis-expressed 1 |
| ENSG00000268545 | VN1R107P | -1,617556981 | 0,325886844 | 0,27894904 | 0,789808253 | vomeronasal 1 receptor 107 pseudogene |
| ENSG00000184617 | ZNF840P | -1,617962061 | 0,325795355 | 0,02083264 | 0,192183647 | zinc finger protein 840, pseudogene |
| ENSG00000259277 | AC127522,1 | -1,620781588 | 0,325159259 | 0,09899934 | 0,499376869 | novel transcript |
| ENSG00000166450 | PRTG | -1,621886341 | 0,324910362 | 0,00499475 | 0,06873368 | protogenin |
| ENSG00000225146 | AC073957,1 | -1,622450769 | 0,324783272 | 0,08786793 | 0,472122805 | novel transcript |
| ENSG00000270911 | AC114878,2 | -1,622871685 | 0,324688528 | 0,14029881 | 0,597018607 | CTAGE family, member 5 (CTAGE5) pseudogene |
| ENSG00000232862 | AL138787,1 | -1,623748191 | 0,324491324 | 0,05390325 | 0,357953158 | implantation-associated protein (DKFZp564K142) pseudogene |
| ENSG00000248596 | AC139491,2 | -1,623940575 | 0,324448055 | 0,27429509 | 0,786239603 | novel transcript |
| ENSG00000286894 | AC009141,1 | -1,624657727 | 0,324286815 | 0,03453249 | 0,271098283 | novel transcript, antisense to WWOX |
| ENSG00000229057 | RPS3AP54 | -1,625753787 | 0,324040538 | 0,0995948 | 0,500760119 | RPS3A pseudogene 54 |
| ENSG00000275192 | LINC02197 | -1,625987877 | 0,323987964 | 0,35973234 | 0,850152836 | long intergenic non-protein coding RNA 2197 |
| ENSG00000070729 | CNGB1 | -1,626130521 | 0,323955932 | 0,00450248 | 0,063926219 | cyclic nucleotide gated channel subunit beta 1 |
| ENSG00000230593 | PPIAP40 | -1,629912121 | 0,323107888 | 0,13625009 | 0,588359488 | peptidylprolyl isomerase A pseudogene 40 |

| | | | | | | |
|---|---|---|---|---|---|---|
| ENSG00000241462 | AC100832,1 | -1,63064295 | 0,322944253 | 0,12435482 | 0,563535178 | ribosomal protein S12 (RPS12) pseudogene |
| ENSG00000167858 | TEKT1 | -1,631831511 | 0,322678305 | 0,0508566 | 0,345422694 | tektin 1 |
| ENSG00000236289 | GACAT3 | -1,632034884 | 0,322632821 | 0,09722069 | 0,495356529 | gastric cancer associated transcript 3 |
| ENSG00000209702 | SNORD41 | -1,63253454 | 0,322521102 | 0,10069889 | 0,504204073 | small nucleolar RNA, C/D box 41 |
| ENSG00000265107 | GJA5 | -1,633478482 | 0,322310148 | 0,0147054 | 0,151832809 | gap junction protein alpha 5 |
| ENSG00000259937 | AC023158,1 | -1,634117159 | 0,322167494 | 0,33120101 | 0,832301933 | novel transcript, sense intronic to SYT10 |
| ENSG00000283973 | AC099795,1 | -1,634236686 | 0,322140803 | 0,16384211 | 0,642745357 | novel transcript |
| ENSG00000268650 | AC005759,1 | -1,635326588 | 0,32189753 | 0,17565095 | 0,661199837 | novel transcript, antisense to PDE4C |
| ENSG00000237158 | AL353718,1 | -1,63581888 | 0,321787707 | 0,01331422 | 0,140952253 | pseudogene similar to part of DCN1, defective in cullin neddylation 1, domain containing 1 (S, cerevisiae) DCUN1D1 |
| ENSG00000147488 | ST18 | -1,636219619 | 0,321698336 | 0,00022478 | 0,005273175 | ST18 C2H2C-type zinc finger transcription factor |
| ENSG00000251456 | AC113398,2 | -1,636615393 | 0,321610097 | 0,00380024 | 0,05641529 | novel transcript |
| ENSG00000160111 | CPAMD8 | -1,637219994 | 0,321475346 | 2,26E-06 | 9,09E-05 | C3 and PZP like alpha-2-macroglobulin domain containing 8 |
| ENSG00000236114 | AL450326,2 | -1,637299211 | 0,321457694 | 0,08997269 | 0,478514897 | zinc finger protein pseudogene |
| ENSG00000276776 | TC2N | -1,637311117 | 0,321455042 | 0,12707495 | 0,569347582 | tandem C2 domains, nuclear |
| ENSG00000164037 | SLC9B1 | -1,637555646 | 0,321400561 | 0,00425365 | 0,061482281 | solute carrier family 9 member B1 |
| ENSG00000222343 | RN7SKP139 | -1,639777321 | 0,320906002 | 0,24164783 | 0,751953584 | RN7SK pseudogene 139 |
| ENSG00000255624 | C10orf88B | -1,640069334 | 0,320841055 | 0,43023786 | 0,895058897 | C10orf88B (pseudogene) |
| ENSG00000268120 | AC010336,3 | -1,640433163 | 0,320760153 | 0,10888969 | 0,526415221 | novel transcript |
| ENSG00000287677 | AC025839,1 | -1,641276348 | 0,320572739 | 0,07129904 | 0,420662976 | novel transcript |
| ENSG00000272990 | AC084036,1 | -1,64133489 | 0,320559731 | 0,06822397 | 0,411581814 | novel transcript, antisense to KCNAB1 |
| ENSG00000258693 | AL133368,2 | -1,643463315 | 0,320087154 | 0,1930632 | 0,689931655 | novel transcript |
| ENSG00000186417 | GLDN | -1,644404128 | 0,319878486 | 0,00695285 | 0,088323639 | gliomedin |
| ENSG00000280787 | HERC2P2 | -1,645577807 | 0,319618361 | 0,57821393 | 0,943425812 | hect domain and RLD 2 pseudogene 2 |
| ENSG00000262020 | AC007014,1 | -1,646306332 | 0,319457002 | 0,17211281 | 0,656211582 | novel transcript, antisense to CLEC16A |
| ENSG00000286758 | AL445438,1 | -1,646616853 | 0,319388251 | 0,12561439 | 0,566124892 | novel transcript |
| ENSG00000227769 | AC072062,1 | -1,64685508 | 0,319335516 | 0,08640087 | 0,468476705 | novel transcript, antisense to ABCA12 |
| ENSG00000243197 | TUSC7 | -1,648481459 | 0,318975725 | 0,03043442 | 0,249541473 | tumor suppressor candidate 7 |
| ENSG00000263300 | AC015853,3 | -1,649519437 | 0,318746314 | 0,22278706 | 0,731378504 | novel transcript, antisense to VPS53 |
| ENSG00000249806 | AC139720,1 | -1,649724305 | 0,318701054 | 0,31370997 | 0,819846908 | novel transcript |
| ENSG00000281880 | PAUPAR | -1,650999326 | 0,318419518 | 0,07702092 | 0,438408786 | PAX6 upstream antisense RNA |

| | | | | | | |
|---|---|---|---|---|---|---|
| ENSG00000278492 | AC006213,5 | -1,653121152 | 0,31795155 | 0,18879679 | 0,683164698 | novel transcript, sense intronic to ZNF155 |
| ENSG00000204990 | AC005692,1 | -1,653147887 | 0,317945658 | 0,02208819 | 0,200221829 | novel transcript |
| ENSG00000111879 | FAM184A | -1,654853012 | 0,317570099 | 0,00083196 | 0,01594761 | family with sequence similarity 184 member A |
| ENSG00000278733 | AC022079,2 | -1,65519281 | 0,317495311 | 0,1245785 | 0,563894916 | novel transcript, antisense to CCDC91 |
| ENSG00000137078 | SIT1 | -1,656095954 | 0,317296617 | 0,05236426 | 0,351288432 | signaling threshold regulating transmembrane adaptor 1 |
| ENSG00000238259 | PHF5GP | -1,658446277 | 0,316780124 | 0,08460091 | 0,462844615 | PHD finger protein 5G pseudogene |
| ENSG00000287814 | AC011407,1 | -1,658490969 | 0,316770311 | 0,18975449 | 0,684955795 | novel transcript |
| ENSG00000229048 | DUTP1 | -1,659142653 | 0,316627254 | 0,08229126 | 0,456410098 | deoxyuridine triphosphatase pseudogene 1 |
| ENSG00000266261 | AC005324,4 | -1,65981786 | 0,316479101 | 0,22169043 | 0,729254784 | novel transcript |
| ENSG00000273529 | HES5 | -1,662196724 | 0,315957688 | 0,19122807 | 0,686939473 | hes family bHLH transcription factor 5 |
| ENSG00000105427 | CNFN | -1,665743536 | 0,315181872 | 0,01403442 | 0,146599474 | cornifelin |
| ENSG00000267088 | AC087683,1 | -1,665944783 | 0,315137909 | 0,21136545 | 0,717057897 | Nucleosome assembly protein 1-like 2, pseudogene |
| ENSG00000274213 | AC015912,3 | -1,668817551 | 0,314511015 | 0,20818765 | 0,71274406 | novel transcript |
| ENSG00000285791 | AC009879,3 | -1,671136941 | 0,314005788 | 0,11768668 | 0,548330959 | novel transcript |
| ENSG00000242220 | TCP10L | -1,672107236 | 0,313794672 | 0,00453381 | 0,06422514 | t-complex 10 like |
| ENSG00000162676 | GFI1 | -1,67446063 | 0,313283213 | 0,00243442 | 0,039252031 | growth factor independent 1 transcriptional repressor |
| ENSG00000244226 | ILF2P1 | -1,675568538 | 0,313042721 | 0,02099159 | 0,192967084 | interleukin enhancer binding factor 2 pseudogene 1 |
| ENSG00000286040 | AL158058,2 | -1,675884748 | 0,312974116 | 0,23229966 | 0,741229884 | novel transcript |
| ENSG00000073350 | LLGL2 | -1,676465702 | 0,312848111 | 0,00200913 | 0,033752576 | LLGL scribble cell polarity complex component 2 |
| ENSG00000259915 | AC017071,1 | -1,676555349 | 0,312828672 | 0,09491314 | 0,489386708 | novel transcript |
| ENSG00000267767 | LINC01801 | -1,677469344 | 0,312630547 | 0,020307 | 0,188783285 | long intergenic non-protein coding RNA 1801 |
| ENSG00000140287 | HDC | -1,678758111 | 0,312351397 | 0,18957819 | 0,684882433 | histidine decarboxylase |
| ENSG00000249460 | LINC02500 | -1,679324082 | 0,312228886 | 0,2265213 | 0,735240631 | long intergenic non-protein coding RNA 2500 |
| ENSG00000241213 | LINC02024 | -1,679989659 | 0,312084874 | 0,08705703 | 0,470307784 | long intergenic non-protein coding RNA 2024 |
| ENSG00000273765 | AC024257,4 | -1,680154876 | 0,312049136 | 0,0912548 | 0,481302729 | novel transcript |
| ENSG00000106689 | LHX2 | -1,680888733 | 0,311890446 | 0,00139524 | 0,024808062 | LIM homeobox 2 |
| ENSG00000146530 | VWDE | -1,683609457 | 0,311302818 | 0,06473352 | 0,399019869 | von Willebrand factor D and EGF domains |
| ENSG00000271537 | AC009220,3 | -1,684127679 | 0,311191017 | 0,04741307 | 0,330875511 | FCF1 small subunit (SSU) processome component homolog (S, cerevisiae), pseudogene |
| ENSG00000261168 | AL592424,1 | -1,684354239 | 0,311142152 | 0,00014548 | 0,003618394 | novel transcript, sense overlapping SEMA6C |
| ENSG00000284783 | AC091057,4 | -1,684377879 | 0,311137053 | 0,18877655 | 0,683164698 | OTU deubiquitinase 7A (OTUD7A), pseudogene |

| | | | | | | |
|---|---|---|---|---|---|---|
| ENSG00000286098 | AC008770,4 | -1,68560007 | 0,310873583 | 0,35604958 | 0,84732813 | zinc finger protein 844 |
| ENSG00000285660 | AC092958,3 | -1,685761663 | 0,310838764 | 0,33156258 | 0,832548086 | novel transcript |
| ENSG00000282859 | AC003992,1 | -1,685914597 | 0,310805815 | 0,27958038 | 0,790302992 | novel transcript, sense intronic to FOXP2 |
| ENSG00000144550 | CPNE9 | -1,688649772 | 0,310217123 | 0,0295245 | 0,244377517 | copine family member 9 |
| ENSG00000134207 | SYT6 | -1,688950792 | 0,310152403 | 0,00031979 | 0,007146482 | synaptotagmin 6 |
| ENSG00000286446 | AL353688,1 | -1,690353595 | 0,309850973 | 0,33626406 | 0,835660292 | novel transcript |
| ENSG00000233040 | FAM204BP | -1,691033934 | 0,30970489 | 0,02952205 | 0,244377517 | family with sequence similarity 204 member B, pseudogene |
| ENSG00000251050 | RBX1P2 | -1,69352324 | 0,309170968 | 0,13134286 | 0,578977575 | RBX1 pseudogene 2 |
| ENSG00000261462 | AC004023,1 | -1,694373956 | 0,308988713 | 0,05717647 | 0,369637122 | novel transcript |
| ENSG00000214725 | CDIPTOSP | -1,697412027 | 0,30833872 | 0,22433093 | 0,73307493 | CDIP transferase opposite strand, pseudogene |
| ENSG00000270441 | LAMB2P1 | -1,69750442 | 0,308318974 | 0,12962252 | 0,575441236 | laminin subunit beta 2 pseudogene 1 |
| ENSG00000255397 | NASPP1 | -1,69988326 | 0,30781101 | 0,09527555 | 0,489967108 | nuclear autoantigenic sperm protein pseudogene 1 |
| ENSG00000237824 | RPL23AP33 | -1,70498795 | 0,306723806 | 0,25202355 | 0,764267906 | ribosomal protein L23a pseudogene 33 |
| ENSG00000239344 | AC090686,1 | -1,705076988 | 0,306704877 | 0,01029646 | 0,117527789 | ribosomal protein S6 (RPS6) pseudogene |
| ENSG00000267062 | AC018761,2 | -1,706164946 | 0,306473673 | 0,01036916 | 0,118054008 | novel transcript, antisense to PRDX2 |
| ENSG00000230707 | AL589987,1 | -1,707015983 | 0,30629294 | 2,65E-05 | 0,000805576 | novel pseudogene |
| ENSG00000135577 | NMBR | -1,707268931 | 0,306239242 | 0,18306905 | 0,673904498 | neuromedin B receptor |
| ENSG00000263041 | AL513324,1 | -1,708936484 | 0,305885477 | 0,19860954 | 0,697657684 | novel transcript |
| ENSG00000133048 | CHI3L1 | -1,709042252 | 0,305863052 | 0,02814464 | 0,237109673 | chitinase 3 like 1 |
| ENSG00000112706 | IMPG1 | -1,710128646 | 0,305632815 | 0,02834128 | 0,238461392 | interphotoreceptor matrix proteoglycan 1 |
| ENSG00000100473 | COCH | -1,711666627 | 0,305307169 | 0,00016181 | 0,003952288 | cochlin |
| ENSG00000221737 | MIR548I1 | -1,712062379 | 0,305223431 | 0,2256429 | 0,734461165 | microRNA 548i-1 |
| ENSG00000271367 | AL034374,1 | -1,712119731 | 0,305211297 | 0,21030202 | 0,715586415 | novel transcript |
| ENSG00000136305 | CIDEB | -1,712829063 | 0,30506127 | 0,07648034 | 0,436971724 | cell death inducing DFFA like effector b |
| ENSG00000285199 | CIDEB | -1,712829063 | 0,30506127 | 0,07648034 | 0,436971724 | cell death inducing DFFA like effector b |
| ENSG00000284456 | LINC02076 | -1,714555099 | 0,304696514 | 0,15406462 | 0,623437276 | long intergenic non-protein coding RNA 2076 |
| ENSG00000272305 | AC096887,1 | -1,715699455 | 0,304454923 | 0,22559147 | 0,734461165 | novel transcript |
| ENSG00000260548 | AL035425,2 | -1,716468834 | 0,304292602 | 0,05613555 | 0,366010789 | insulin receptor substrate 4 |
| ENSG00000278338 | VWA8-AS1 | -1,718434442 | 0,303878299 | 0,03147054 | 0,255689988 | VWA8 antisense RNA 1 (head to head) |
| ENSG00000233670 | PIRT | -1,718625932 | 0,303837968 | 0,07971886 | 0,447697198 | phosphoinositide interacting regulator of transient receptor potential channels |
| ENSG00000227017 | AC007036,1 | -1,720278842 | 0,303490057 | 0,18743963 | 0,681553495 | novel transcript |

| | | | | | | |
|---|---|---|---|---|---|---|
| ENSG00000234921 | AC006450,2 | -1,720712301 | 0,303398887 | 0,07489193 | 0,432482457 | novel transcript, antisense to LHX2 |
| ENSG00000232104 | RFX3-AS1 | -1,721884168 | 0,303152544 | 1,39E-05 | 0,000454557 | RFX3 antisense RNA 1 |
| ENSG00000244332 | AL138759,1 | -1,722504695 | 0,303022181 | 0,05406875 | 0,35836452 | novel transcript |
| ENSG00000225798 | AC025918,1 | -1,723128522 | 0,302891181 | 0,06350975 | 0,394666843 | novel transcript |
| ENSG00000281284 | MAPT-IT1 | -1,723959841 | 0,302716697 | 0,03979576 | 0,297578246 | MAPT intronic transcript 1 |
| ENSG00000275146 | AP003499,3 | -1,724090557 | 0,302689271 | 0,20011261 | 0,699646707 | |
| ENSG00000233469 | ST6GALNAC4P1 | -1,724270043 | 0,302651616 | 0,1615124 | 0,63813307 | ST6 N-acetylgalactosaminide alpha-2,6-sialyltransferase 4 pseudogene 1 |
| ENSG00000224204 | PHEX-AS1 | -1,724303787 | 0,302644537 | 0,13551763 | 0,586971877 | PHEX antisense RNA 1 |
| ENSG00000100987 | VSX1 | -1,724554876 | 0,302591869 | 0,10245092 | 0,509792407 | visual system homeobox 1 |
| ENSG00000188162 | OTOG | -1,7264205 | 0,302200824 | 0,09461291 | 0,488518255 | otogelin |
| ENSG00000070748 | CHAT | -1,726926193 | 0,302094915 | 0,00064285 | 0,012858692 | choline O-acetyltransferase |
| ENSG00000127578 | WFIKKN1 | -1,726949672 | 0,302089999 | 0,00628036 | 0,081714521 | WAP, follistatin/kazal, immunoglobulin, kunitz and netrin domain containing 1 |
| ENSG00000214872 | SMTNL1 | -1,728139613 | 0,301840937 | 0,0275751 | 0,233950565 | smoothelin like 1 |
| ENSG00000259374 | NDUFB4P11 | -1,728148456 | 0,301839086 | 0,22838859 | 0,736649232 | NADH:ubiquinone oxidoreductase subunit B4 pseudogene 11 |
| ENSG00000165370 | GPR101 | -1,728528355 | 0,301759615 | 0,04378344 | 0,31589834 | G protein-coupled receptor 101 |
| ENSG00000276798 | LILRA4 | -1,729403261 | 0,301576672 | 0,41771497 | 0,886857227 | leukocyte immunoglobulin like receptor A4 |
| ENSG00000285889 | AL355312,4 | -1,730905917 | 0,301262725 | 0,12745887 | 0,570245289 | novel transcript |
| ENSG00000283997 | TMEM159 | -1,73223959 | 0,300984356 | 0,00618832 | 0,080950485 | transmembrane protein 159 |
| ENSG00000248329 | APELA | -1,738048209 | 0,299774961 | 0,01101912 | 0,123080268 | apelin receptor early endogenous ligand |
| ENSG00000171786 | NHLH1 | -1,740383503 | 0,299290107 | 4,65E-05 | 0,001326605 | nescient helix-loop-helix 1 |
| ENSG00000255817 | AC025576,2 | -1,742094589 | 0,298935349 | 0,09115559 | 0,481283407 | novel transcript |
| ENSG00000231378 | NDUFAF4P4 | -1,743164635 | 0,298713711 | 0,18227506 | 0,672565494 | NADH:ubiquinone oxidoreductase complex assembly factor 4 pseudogene 4 |
| ENSG00000255581 | AC006511,2 | -1,747960524 | 0,297722359 | 0,0575221 | 0,370798119 | heat shock 60kDa protein 1 (chaperonin) (HSPD1) pseudogene |
| ENSG00000233431 | LINC02596 | -1,749101754 | 0,297486941 | 0,22031452 | 0,728572782 | long intergenic non-protein coding RNA 2596 |
| ENSG00000235091 | BX324167,1 | -1,751230214 | 0,297048372 | 0,01511505 | 0,154885911 | novel transcript |
| ENSG00000250686 | KHDRBS2-OT1 | -1,751519615 | 0,296988791 | 0,0324308 | 0,260781002 | KHDRBS2 3'UTR overlapping transcript 1 |
| ENSG00000273727 | U1 | -1,753617861 | 0,296557166 | 0,12285042 | 0,560536436 | U1 spliceosomal RNA |
| ENSG00000286632 | AC008751,3 | -1,754265105 | 0,29642415 | 0,147379 | 0,611530401 | novel transcript |

| | | | | | | |
|---|---|---|---|---|---|---|
| ENSG00000236980 | C3orf84 | -1,756203961 | 0,296026049 | 0,14224171 | 0,600366679 | chromosome 3 open reading frame 84 |
| ENSG00000257150 | PGAM1P5 | -1,756501448 | 0,295965014 | 0,06771787 | 0,409582442 | phosphoglycerate mutase 1 pseudogene 5 |
| ENSG00000241404 | EGFL8 | -1,756557452 | 0,295953526 | 0,04370322 | 0,315756651 | EGF like domain multiple 8 |
| ENSG00000287536 | AC012500,1 | -1,756677286 | 0,295928944 | 0,07953084 | 0,447364571 | novel transcript, antisense to PDE1A |
| ENSG00000287527 | AC005753,3 | -1,756728787 | 0,29591838 | 0,0087884 | 0,104076013 | novel transcript |
| ENSG00000224493 | AC093156,1 | -1,757253301 | 0,295810814 | 0,00718484 | 0,090211569 | pseudogene similar to part of nucleostemin (NS) |
| ENSG00000130433 | CACNG6 | -1,759170478 | 0,295417976 | 0,01056171 | 0,119335457 | calcium voltage-gated channel auxiliary subunit gamma 6 |
| ENSG00000232083 | RPL31P7 | -1,761532782 | 0,294934647 | 0,15399757 | 0,623246599 | ribosomal protein L31 pseudogene 7 |
| ENSG00000286363 | AC087427,1 | -1,762191121 | 0,294800092 | 0,04843255 | 0,335106518 | novel transcript, sense intronic to CNTN4 |
| ENSG00000266605 | LONRF2P1 | -1,762398151 | 0,29475779 | 0,18886447 | 0,683246297 | LONRF2 pseudogene 1 |
| ENSG00000255222 | SETP17 | -1,762670922 | 0,294702066 | 0,24001024 | 0,749843134 | SET pseudogene 17 |
| ENSG00000225711 | AL512326,3 | -1,763528316 | 0,294526976 | 0,02406053 | 0,212914898 | UDP-Gal:betaGlcNAc beta 1,3-galactosyltransferase, polypeptide 3 (B3GALT3) pseudogene |
| ENSG00000235376 | RPEL1 | -1,764113632 | 0,294407508 | 0,19502833 | 0,69283135 | ribulose-5-phosphate-3-epimerase like 1 |
| ENSG00000258710 | LINC01193 | -1,764644094 | 0,294299278 | 0,23554899 | 0,744686378 | long intergenic non-protein coding RNA 1193 |
| ENSG00000157005 | SST | -1,765834119 | 0,294056621 | 0,03900731 | 0,29393213 | somatostatin |
| ENSG00000279623 | AL359697,1 | -1,767228989 | 0,29377245 | 0,1206329 | 0,554057131 | novel transcript, sense intronic to DNAJC1 |
| ENSG00000287376 | AL359538,3 | -1,769204808 | 0,293370394 | 0,21907335 | 0,727390525 | novel transcript |
| ENSG00000205236 | AC105052,1 | -1,77000419 | 0,293207886 | 0,01321781 | 0,140068402 | novel protein, UPK3BL-RASA4 readthrough |
| ENSG00000236998 | AL354794,2 | -1,772879065 | 0,292624189 | 0,01255337 | 0,135289937 | adaptor-related protein complex 3, mu 2 subunit (AP3M2) pseudogene |
| ENSG00000140465 | CYP1A1 | -1,775814652 | 0,292029364 | 0,22663491 | 0,735240631 | cytochrome P450 family 1 subfamily A member 1 |
| ENSG00000134595 | SOX3 | -1,776484958 | 0,291893713 | 2,90E-05 | 0,000872002 | SRY-box transcription factor 3 |
| ENSG00000213430 | HSPD1P1 | -1,776977963 | 0,291793983 | 0,10833543 | 0,524645663 | heat shock protein family D (Hsp60) member 1 pseudogene 1 |
| ENSG00000137462 | TLR2 | -1,777123 | 0,291764649 | 0,00131984 | 0,023633129 | toll like receptor 2 |
| ENSG00000214796 | TUBA5P | -1,778727427 | 0,291440357 | 5,62E-06 | 0,000204936 | tubulin alpha 5, pseudogene |
| ENSG00000287595 | AC097369,4 | -1,78000675 | 0,291182034 | 0,10168505 | 0,507111388 | novel transcript |
| ENSG00000249771 | AC108210,1 | -1,785541351 | 0,290067115 | 0,16159141 | 0,63813307 | novel transcript |
| ENSG00000215325 | ASS1P10 | -1,787672267 | 0,289638991 | 0,00019806 | 0,004737798 | argininosuccinate synthetase 1 pseudogene 10 |
| ENSG00000262633 | AC005670,2 | -1,789384933 | 0,289295356 | 0,62320418 | 0,952793934 | novel protein |
| ENSG00000225368 | HCG15 | -1,794263736 | 0,28831869 | 0,65198407 | 0,957241719 | HLA complex group 15 |

| | | | | | | |
|---|---|---|---|---|---|---|
| ENSG00000169218 | RSPO1 | -1,798235961 | 0,287525943 | 0,03446748 | 0,270819603 | R-spondin 1 |
| ENSG00000165553 | NGB | -1,798638272 | 0,287445774 | 0,19761359 | 0,696048902 | neuroglobin |
| ENSG00000275005 | AL354950,1 | -1,798819954 | 0,287409578 | 0,15859282 | 0,632402923 | novel transcript, antisense to EBF3 |
| ENSG00000236165 | PRADC1P1 | -1,799564552 | 0,28726128 | 0,16060501 | 0,636125619 | protease associated domain containing 1 pseudogene 1 |
| ENSG00000236104 | ZBTB22 | -1,800127498 | 0,287149211 | 0,23409113 | 0,743586269 | zinc finger and BTB domain containing 22 |
| ENSG00000205143 | ARID3C | -1,801387581 | 0,286898518 | 0,08748171 | 0,471475303 | AT-rich interaction domain 3C |
| ENSG00000131914 | LIN28A | -1,801854296 | 0,28680572 | 0,00026663 | 0,006089157 | lin-28 homolog A |
| ENSG00000256473 | AC087260,1 | -1,802570097 | 0,286663456 | 0,02033385 | 0,188976655 | novel transcript, antisense to SOX5 |
| ENSG00000255780 | AC020611,1 | -1,804384495 | 0,286303161 | 0,29109438 | 0,801689099 | chromosome 3 open reading frame 19 (C3orf19) pseudogene |
| ENSG00000231827 | AL513523,2 | -1,804832259 | 0,286214316 | 0,0356571 | 0,276621272 | immunoglobulin superfamily, DCC subclass, member 3 (IGDCC3) pseudogene |
| ENSG00000130612 | CYP2G1P | -1,804992703 | 0,286182488 | 0,04836971 | 0,335083246 | cytochrome P450 family 2 subfamily G member 1, pseudogene |
| ENSG00000286945 | AL513478,4 | -1,80745397 | 0,285694671 | 0,13529875 | 0,586275832 | novel transcript |
| ENSG00000159251 | ACTC1 | -1,809143688 | 0,285360254 | 0,00020965 | 0,004965468 | actin alpha cardiac muscle 1 |
| ENSG00000187475 | H1-6 | -1,815426005 | 0,284120335 | 0,0108772 | 0,121820431 | H1,6 linker histone, cluster member |
| ENSG00000234948 | LINC01524 | -1,815886919 | 0,284029578 | 0,1130027 | 0,537080617 | long intergenic non-protein coding RNA 1524 |
| ENSG00000261754 | AC008555,1 | -1,815924556 | 0,284022168 | 0,00356336 | 0,053585776 | novel transcript |
| ENSG00000130528 | HRC | -1,816711078 | 0,283867369 | 0,01303048 | 0,139032628 | histidine rich calcium binding protein |
| ENSG00000149573 | MPZL2 | -1,823078098 | 0,282617342 | 0,04515935 | 0,321462197 | myelin protein zero like 2 |
| ENSG00000160161 | CILP2 | -1,824966196 | 0,282247715 | 0,00172098 | 0,02956301 | cartilage intermediate layer protein 2 |
| ENSG00000260880 | HCCAT5 | -1,826001941 | 0,282045155 | 0,16773166 | 0,648680831 | hepatocellular carcinoma associated transcript 5 |
| ENSG00000164076 | CAMKV | -1,829769287 | 0,281309604 | 9,61E-05 | 0,002526223 | CaM kinase like vesicle associated |
| ENSG00000255555 | AP000857,2 | -1,831524733 | 0,28096752 | 0,18570735 | 0,678662134 | novel transcript |
| ENSG00000168952 | STXBP6 | -1,832478666 | 0,280781801 | 0,0019355 | 0,032726533 | syntaxin binding protein 6 |
| ENSG00000286191 | AL158195,1 | -1,832669403 | 0,280744682 | 0,10523005 | 0,517691472 | novel transcript |
| ENSG00000184611 | KCNH7 | -1,833416913 | 0,280599256 | 0,00475292 | 0,066307569 | potassium voltage-gated channel subfamily H member 7 |
| ENSG00000249835 | VCAN-AS1 | -1,834981087 | 0,280295194 | 0,14463097 | 0,605120065 | VCAN antisense RNA 1 |
| ENSG00000264434 | AC110603,1 | -1,836687975 | 0,279963766 | 0,04041545 | 0,300321946 | novel transcript, antisense to ZNF521 |
| ENSG00000273825 | FP700111,2 | -1,837137607 | 0,279876526 | 0,22686841 | 0,735383523 | otopetrin 1 (OTOP1) pseudogene |

| | | | | | | |
|---|---|---|---|---|---|---|
| ENSG00000100060 | MFNG | -1,83861376 | 0,279590305 | 0,00140491 | 0,024956231 | MFNG O-fucosylpeptide 3-beta-N-acetylglucosaminyltransferase |
| ENSG00000286394 | AC117373,2 | -1,838933763 | 0,279528296 | 0,2441701 | 0,755571155 | novel transcript, antisense to TMEM132D |
| ENSG00000259336 | AC021231,1 | -1,839540726 | 0,279410719 | 0,05231915 | 0,351235995 | novel transcript |
| ENSG00000162571 | TTLL10 | -1,840736605 | 0,279179206 | 0,08764672 | 0,471714073 | tubulin tyrosine ligase like 10 |
| ENSG00000188452 | CERKL | -1,844127876 | 0,278523724 | 0,00813027 | 0,098882383 | ceramide kinase like |
| ENSG00000266369 | AC090774,2 | -1,845824824 | 0,278196307 | 0,11559909 | 0,543151268 | novel transcript, antisense to CCDC144NL |
| ENSG00000237517 | DGCR5 | -1,84651528 | 0,278063198 | 0,26379695 | 0,775013078 | DiGeorge syndrome critical region gene 5 |
| ENSG00000218475 | ACTG1P18 | -1,848528971 | 0,277675353 | 0,00112739 | 0,020648404 | actin gamma 1 pseudogene 18 |
| ENSG00000155754 | C2CD6 | -1,848835293 | 0,277616401 | 0,01183665 | 0,129527851 | C2 calcium dependent domain containing 6 |
| ENSG00000268288 | AL589765,5 | -1,851623381 | 0,27708041 | 0,17559013 | 0,661199837 | novel transcript, antisense to OAZ3 |
| ENSG00000250698 | AC131182,1 | -1,851634256 | 0,277078321 | 0,00027249 | 0,006207726 | novel transcript |
| ENSG00000137204 | SLC22A7 | -1,857730059 | 0,275910055 | 0,02419977 | 0,213783975 | solute carrier family 22 member 7 |
| ENSG00000253236 | LINC02143 | -1,858670095 | 0,275730335 | 0,10762758 | 0,52347056 | long intergenic non-protein coding RNA 2143 |
| ENSG00000220739 | AL513475,2 | -1,859368637 | 0,275596861 | 0,03430746 | 0,270005245 | 5'-nucleotidase domain containing 1 (NT5DC1) pseudogene |
| ENSG00000184811 | TRARG1 | -1,86051207 | 0,275378519 | 0,05834377 | 0,373935936 | trafficking regulator of GLUT4 (SLC2A4) 1 |
| ENSG00000248165 | AC110760,1 | -1,86158435 | 0,27517392 | 0,05710904 | 0,369353938 | novel transcript |
| ENSG00000281803 | AC136352,11 | -1,862142269 | 0,275067526 | 0,0304718 | 0,249663376 | novel transcript |
| ENSG00000224159 | HMGB1P9 | -1,864729656 | 0,274574651 | 0,16607699 | 0,645398515 | high mobility group box 1 pseudogene 9 |
| ENSG00000273027 | AL844908,2 | -1,865435549 | 0,274440338 | 0,2224974 | 0,730919269 | novel transcript, sense intronic to ITGB2 |
| ENSG00000253454 | NDUFA5P2 | -1,865493346 | 0,274429343 | 0,07664615 | 0,437367108 | NADH:ubiquinone oxidoreductase subunit A5 pseudogene 2 |
| ENSG00000280987 | MATR3 | -1,869607559 | 0,273647853 | 0,0013155 | 0,023568931 | matrin 3 |
| ENSG00000048540 | LMO3 | -1,873611826 | 0,272889384 | 0,00327226 | 0,050075384 | LIM domain only 3 |
| ENSG00000259276 | AC087286,1 | -1,876685863 | 0,272308541 | 0,15943526 | 0,633848434 | novel transcript, antisense to AKAP13 |
| ENSG00000217334 | AL355615,1 | -1,876797308 | 0,272287506 | 0,08300477 | 0,458031678 | pseudogene similar to part of heat shock 90kD protein 1, beta HSPCB) |
| ENSG00000172568 | FNDC9 | -1,877533714 | 0,272148556 | 0,01767348 | 0,172454361 | fibronectin type III domain containing 9 |
| ENSG00000152969 | JAKMIP1 | -1,87787894 | 0,272083441 | 0,01142742 | 0,126376669 | janus kinase and microtubule interacting protein 1 |
| ENSG00000236132 | AL022331,1 | -1,879078948 | 0,271857221 | 0,078701 | 0,444135046 | ribosomal protein L17 (RPL17) pseudogene |
| ENSG00000260392 | AC090518,1 | -1,880503147 | 0,271588981 | 0,05646451 | 0,366903759 | novel transcript |
| ENSG00000254760 | AC008750,1 | -1,880773592 | 0,271538074 | 0,06737145 | 0,408200237 | novel transcript, antisense to SIGLEC10 |

| | | | | | | |
|---|---|---|---|---|---|---|
| ENSG00000124785 | NRN1 | -1,881859667 | 0,271333735 | 0,00027262 | 0,006207726 | neuritin 1 |
| ENSG00000226677 | IGBP1P1 | -1,883427574 | 0,271039012 | 0,07030588 | 0,417640698 | immunoglobulin (CD79A) binding protein 1 pseudogene 1 |
| ENSG00000278812 | C8orf31 | -1,883523937 | 0,271020909 | 0,01745017 | 0,17078586 | chromosome 8 open reading frame 31 |
| ENSG00000185203 | WASIR1 | -1,883531358 | 0,271019515 | 0,30903031 | 0,816845699 | WASH and IL9R antisense RNA 1 |
| ENSG00000085563 | ABCB1 | -1,88544046 | 0,270661115 | 0,00048682 | 0,010173747 | ATP binding cassette subfamily B member 1 |
| ENSG00000225929 | AC000036,1 | -1,887255508 | 0,270320812 | 0,21502211 | 0,722298056 | novel transcript |
| ENSG00000175745 | NR2F1 | -1,889292837 | 0,269939342 | 1,05E-11 | 1,02E-09 | nuclear receptor subfamily 2 group F member 1 |
| ENSG00000276089 | AL137060,6 | -1,892513723 | 0,269337362 | 0,02589494 | 0,224198135 | |
| ENSG00000103832 | GOLGA8UP | -1,892762088 | 0,269290999 | 0,19743937 | 0,695991044 | golgin A8 family member U, pseudogene |
| ENSG00000267225 | WDR7-OT1 | -1,892785953 | 0,269286544 | 0,23566877 | 0,744839106 | WDR7 overlapping transcript 1 |
| ENSG00000277276 | OTUB2 | -1,89369496 | 0,269116927 | 0,40010652 | 0,875889731 | OTU deubiquitinase, ubiquitin aldehyde binding 2 |
| ENSG00000234280 | TSBP1 | -1,894554753 | 0,268956591 | 0,20133997 | 0,701094807 | testis expressed basic protein 1 |
| ENSG00000270981 | AC092944,2 | -1,895896967 | 0,268706483 | 0,20370067 | 0,704215578 | poly(rC) binding protein 1, pseudogene |
| ENSG00000196553 | CCDC196 | -1,895939044 | 0,268698646 | 0,07571822 | 0,43475749 | coiled-coil domain containing 196 |
| ENSG00000288442 | AC256231,3 | -1,896123263 | 0,268664338 | 0,00166865 | 0,028758864 | |
| ENSG00000112212 | TSPO2 | -1,89794644 | 0,268325033 | 0,16553016 | 0,644635638 | translocator protein 2 |
| ENSG00000269043 | AC008554,1 | -1,898602957 | 0,268202956 | 0,02879102 | 0,240931045 | novel transcript |
| ENSG00000286581 | AL450992,3 | -1,899716394 | 0,267996043 | 0,02348369 | 0,209172103 | novel transcript |
| ENSG00000271778 | AC080013,2 | -1,900463787 | 0,267857243 | 0,1020942 | 0,508625897 | novel transcript, sense intronic to MFSD1 |
| ENSG00000180613 | GSX2 | -1,905657029 | 0,266894776 | 0,04028728 | 0,299940355 | GS homeobox 2 |
| ENSG00000134917 | ADAMTS8 | -1,905998381 | 0,266831634 | 0,02627931 | 0,226459876 | ADAM metallopeptidase with thrombospondin type 1 motif 8 |
| ENSG00000287985 | AC073270,2 | -1,907335886 | 0,266584372 | 0,02567531 | 0,222913865 | novel transcript |
| ENSG00000240698 | RPS2P39 | -1,907895122 | 0,266481055 | 0,17272622 | 0,657107871 | ribosomal protein S2 pseudogene 39 |
| ENSG00000275496 | CU633906,1 | -1,909948161 | 0,266102107 | 0,00945408 | 0,1100828 | uncharacterized LOC102724701 |
| ENSG00000287355 | AL451123,2 | -1,912068955 | 0,265711218 | 0,0001961 | 0,004697893 | novel transcript, sense intronic to MOB3B |
| ENSG00000198133 | TMEM229B | -1,915967492 | 0,264994167 | 0,00626624 | 0,081628243 | transmembrane protein 229B |
| ENSG00000226287 | TMEM191A | -1,921967106 | 0,263894446 | 0,01259367 | 0,135534738 | transmembrane protein 191A (pseudogene) |
| ENSG00000270492 | AC137695,1 | -1,922477247 | 0,263801149 | 0,02002147 | 0,186831751 | MAP/microtubule affinity-regulating kinase 2, pseudogene |
| ENSG00000268560 | AC010636,1 | -1,923324481 | 0,263646275 | 0,17565394 | 0,661199837 | novel transcript |
| ENSG00000260391 | AC022336,2 | -1,924261568 | 0,263475082 | 0,05772442 | 0,371372961 | novel transcript, overlapping to KALRN |

| | | | | | | |
|---|---|---|---|---|---|---|
| ENSG00000260232 | PWRN4 | -1,926433483 | 0,26307873 | 0,00020989 | 0,004967509 | Prader-Willi region non-protein coding RNA 4 |
| ENSG00000283402 | AC087821,2 | -1,926751961 | 0,263020661 | 0,00661774 | 0,085142822 | mitochondrially encoded cytochrome b (MT-CYB) pseudogene |
| ENSG00000259946 | BX005019,1 | -1,928746054 | 0,262657365 | 0,00467325 | 0,065714391 | novel transcript |
| ENSG00000285940 | AC009899,1 | -1,932137386 | 0,262040664 | 0,04334512 | 0,314126529 | novel transcript, antisense to CELF4 |
| ENSG00000215914 | MMP23A | -1,932638287 | 0,2619497 | 0,19945688 | 0,698904803 | matrix metallopeptidase 23A (pseudogene) |
| ENSG00000228115 | AL161725,2 | -1,933874906 | 0,261725263 | 0,2234553 | 0,731984265 | novel transcript |
| ENSG00000159713 | TPPP3 | -1,934494439 | 0,261612895 | 0,00600078 | 0,078887556 | tubulin polymerization promoting protein family member 3 |
| ENSG00000163673 | DCLK3 | -1,935478584 | 0,261434495 | 0,00050326 | 0,010454364 | doublecortin like kinase 3 |
| ENSG00000284700 | AL049637,1 | -1,935588384 | 0,261414599 | 0,06516567 | 0,400662695 | novel transcript |
| ENSG00000120055 | C10orf95 | -1,937015311 | 0,261156169 | 0,26254701 | 0,774111884 | chromosome 10 open reading frame 95 |
| ENSG00000286391 | AL592078,2 | -1,938118069 | 0,260956625 | 0,02248019 | 0,202767423 | novel transcript |
| ENSG00000143355 | LHX9 | -1,938732632 | 0,260845485 | 0,01096176 | 0,122571027 | LIM homeobox 9 |
| ENSG00000272170 | AL355385,1 | -1,939833719 | 0,26064648 | 0,01535141 | 0,156640518 | novel transcript, antisense to KLC4 |
| ENSG00000287498 | AC007749,2 | -1,942597981 | 0,260147549 | 0,0686978 | 0,413067344 | novel transcript |
| ENSG00000227660 | UST-AS1 | -1,944211517 | 0,259856758 | 0,10659444 | 0,521482353 | UST antisense RNA 1 |
| ENSG00000185031 | SLC2A3P2 | -1,94428081 | 0,259844277 | 0,01799458 | 0,17412026 | solute carrier family 2 member 3 pseudogene 2 |
| ENSG00000250767 | AC139783,1 | -1,944380958 | 0,25982624 | 0,11315847 | 0,53724925 | fumarate hydratase (FH) pseudogene |
| ENSG00000224669 | AC016769,2 | -1,945740534 | 0,259581499 | 0,00818242 | 0,099307159 | glycine cleavage system protein H (aminomethyl carrier) (GCSH) pseudogene |
| ENSG00000271424 | GLRX5P3 | -1,947287631 | 0,259303281 | 0,12156383 | 0,557102244 | GLRX5 pseudogene 3 |
| ENSG00000227966 | LINC01623 | -1,94844523 | 0,259095304 | 0,56291305 | 0,940865625 | long intergenic non-protein coding RNA 1623 |
| ENSG00000237992 | LINC01808 | -1,948709775 | 0,259047798 | 0,12578233 | 0,566474006 | long intergenic non-protein coding RNA 1808 |
| ENSG00000230453 | ANKRD18B | -1,949031332 | 0,258990066 | 0,00267029 | 0,042142344 | ankyrin repeat domain 18B |
| ENSG00000132554 | RGS22 | -1,951204699 | 0,258600201 | 5,96E-05 | 0,001657591 | regulator of G protein signaling 22 |
| ENSG00000237818 | RPS3AP29 | -1,954906763 | 0,257937464 | 0,06074111 | 0,383638369 | RPS3A pseudogene 29 |
| ENSG00000261787 | TCF24 | -1,958706378 | 0,25725903 | 0,17265081 | 0,657085135 | transcription factor 24 |
| ENSG00000165309 | ARMC3 | -1,960668242 | 0,256909431 | 0,00634012 | 0,082213892 | armadillo repeat containing 3 |
| ENSG00000205181 | LINC00654 | -1,962332535 | 0,256613231 | 7,45E-13 | 8,47E-11 | long intergenic non-protein coding RNA 654 |
| ENSG00000273579 | C22orf15 | -1,962367243 | 0,256607058 | 0,03011301 | 0,247750834 | chromosome 22 open reading frame 15 |
| ENSG00000162496 | DHRS3 | -1,966263211 | 0,255915031 | 0,01316016 | 0,139782152 | dehydrogenase/reductase 3 |
| ENSG00000273155 | AC092587,1 | -1,966370159 | 0,25589606 | 0,10912204 | 0,527049531 | novel LIPT1-MRPL30 readthrough |

| | | | | | | |
|---|---|---|---|---|---|---|
| ENSG00000150628 | SPATA4 | -1,970608097 | 0,255145465 | 0,05047977 | 0,343759427 | spermatogenesis associated 4 |
| ENSG00000276781 | CHGA | -1,971242525 | 0,255033289 | 0,01933767 | 0,182760901 | chromogranin A |
| ENSG00000172460 | PRSS30P | -1,97409114 | 0,25453022 | 0,08286098 | 0,457858895 | serine protease 30, pseudogene |
| ENSG00000251186 | AC105345,2 | -1,974608905 | 0,254438889 | 0,14187736 | 0,599899099 | novel transcript |
| ENSG00000108622 | ICAM2 | -1,975473426 | 0,254286464 | 0,06693174 | 0,406636991 | intercellular adhesion molecule 2 |
| ENSG00000152760 | TCTEX1D1 | -1,977393573 | 0,253948248 | 0,03663211 | 0,281811808 | Tctex1 domain containing 1 |
| ENSG00000213216 | AC007066,1 | -1,977904475 | 0,253858333 | 0,01825272 | 0,175856062 | ribosomal protein S25 (RPS25) pseudogene |
| ENSG00000287894 | AC174469,1 | -1,977947174 | 0,25385082 | 0,17808326 | 0,665264078 | novel transcript, antisense to APBA2 |
| ENSG00000164483 | SAMD3 | -1,977964714 | 0,253847734 | 0,00594075 | 0,078235723 | sterile alpha motif domain containing 3 |
| ENSG00000246575 | AC093162,1 | -1,979230088 | 0,253625184 | 0,13037935 | 0,577035665 | retinol saturase (all-trans-retinol 13,14-reductase) (RESAT) pseudogene |
| ENSG00000258150 | AC133555,3 | -1,984629373 | 0,252677766 | 0,24816745 | 0,760508446 | coiled-coil domain containing 101 (CCDC101) pseudogene |
| ENSG00000285447 | ZNF883 | -1,985179831 | 0,252581376 | 0,6167541 | 0,952793934 | zinc finger protein 883 |
| ENSG00000214695 | NPAP1P2 | -1,98531922 | 0,252556973 | 0,06766078 | 0,409359696 | nuclear pore associated protein 1 pseudogene 2 |
| ENSG00000286022 | AL121899,2 | -1,985676063 | 0,252494512 | 0,36148998 | 0,851893059 | novel protein |
| ENSG00000274898 | AC001226,1 | -1,985744913 | 0,252482463 | 0,08615498 | 0,467467779 | novel transcript |
| ENSG00000265254 | AC015917,2 | -1,986921843 | 0,252276575 | 0,10260777 | 0,510071005 | novel transcript, antisense to SLC46A1 |
| ENSG00000253298 | LINC01845 | -1,987650606 | 0,252149172 | 0,02371409 | 0,210504604 | long intergenic non-protein coding RNA 1845 |
| ENSG00000276475 | AC016769,4 | -1,988902728 | 0,251930426 | 0,08751913 | 0,471587419 | BCL2/adenovirus E1B 19kDa interacting protein 3 (BNIP3) pseudogene |
| ENSG00000238199 | UBE2V2P3 | -1,989302904 | 0,251860555 | 0,13188363 | 0,580243421 | ubiquitin conjugating enzyme E2 V2 pseudogene 3 |
| ENSG00000185013 | NT5C1B | -1,991982029 | 0,251393277 | 0,0690364 | 0,414510104 | 5'-nucleotidase, cytosolic IB |
| ENSG00000243404 | RPL35AP32 | -1,996960922 | 0,250527187 | 0,0154683 | 0,157422163 | ribosomal protein L35a pseudogene 32 |
| ENSG00000235975 | Z83313,1 | -1,999522802 | 0,250082706 | 0,01626275 | 0,162750485 | tropomyosin 2 (beta) (TPM2) pseudogene |
| ENSG00000234224 | TMEM229A | -2,000682504 | 0,249881759 | 0,00115446 | 0,021056164 | transmembrane protein 229A |
| ENSG00000224019 | RPL21P32 | -2,002800566 | 0,24951517 | 0,03500945 | 0,273292232 | ribosomal protein L21 pseudogene 32 |
| ENSG00000257472 | AL355112,1 | -2,003101475 | 0,249463133 | 0,12994617 | 0,576178863 | novel transcript |
| ENSG00000178821 | TMEM52 | -2,004360952 | 0,249245446 | 0,00285184 | 0,044602313 | transmembrane protein 52 |
| ENSG00000226562 | CYP4F26P | -2,00506148 | 0,249124449 | 0,04289105 | 0,312138214 | cytochrome P450 family 4 subfamily F member 26, pseudogene |
| ENSG00000188910 | GJB3 | -2,013862868 | 0,247609253 | 0,02256671 | 0,203412591 | gap junction protein beta 3 |
| ENSG00000257883 | AC125603,2 | -2,014258546 | 0,247541352 | 0,11725192 | 0,547082438 | novel transcript |

| | | | | | | |
|---|---|---|---|---|---|---|
| ENSG00000221819 | GAS8-AS1 | -2,014434231 | 0,247511209 | 0,06553838 | 0,401999604 | GAS8 antisense RNA 1 |
| ENSG00000151834 | GABRA2 | -2,014669921 | 0,247470777 | 0,00199969 | 0,033666261 | gamma-aminobutyric acid type A receptor subunit alpha2 |
| ENSG00000286807 | AC093639,2 | -2,014998396 | 0,247414439 | 0,00496342 | 0,068419936 | novel transcript |
| ENSG00000182583 | VCX | -2,0170839 | 0,247057045 | 0,22045834 | 0,728572782 | variable charge X-linked |
| ENSG00000259803 | SLC22A31 | -2,018408224 | 0,246830363 | 0,0648971 | 0,399712976 | solute carrier family 22 member 31 |
| ENSG00000259286 | AC087639,1 | -2,018424487 | 0,24682758 | 0,05762449 | 0,3710973 | cytokine induced apoptosis inhibitor 1 (CIAPIN1) |
| ENSG00000187714 | SLC18A3 | -2,020987422 | 0,246389482 | 0,00030958 | 0,006948054 | solute carrier family 18 member A3 |
| ENSG00000269437 | NXF2B | -2,022113227 | 0,246197288 | 0,16897981 | 0,650934541 | nuclear RNA export factor 2B |
| ENSG00000204352 | C9orf129 | -2,023408789 | 0,245976298 | 0,01555067 | 0,157881532 | chromosome 9 open reading frame 129 |
| ENSG00000116014 | KISS1R | -2,027199788 | 0,24533079 | 0,12227884 | 0,559316128 | KISS1 receptor |
| ENSG00000235149 | AC096659,1 | -2,027310693 | 0,245311931 | 0,03795734 | 0,288313463 | novel transcript |
| ENSG00000213272 | RPL7AP9 | -2,031785251 | 0,244552268 | 0,08425775 | 0,461822296 | ribosomal protein L7a pseudogene 9 |
| ENSG00000086506 | HBQ1 | -2,032283361 | 0,244467848 | 0,07596306 | 0,435455001 | hemoglobin subunit theta 1 |
| ENSG00000243518 | AC108729,2 | -2,033357155 | 0,244285958 | 0,08769578 | 0,471734492 | DTW domain containing 1 (DTWD1) pseudogene |
| ENSG00000136535 | TBR1 | -2,034752603 | 0,244049787 | 0,11218166 | 0,535189241 | T-box brain transcription factor 1 |
| ENSG00000148053 | NTRK2 | -2,035107916 | 0,243989689 | 7,42E-06 | 0,000262822 | neurotrophic receptor tyrosine kinase 2 |
| ENSG00000140522 | RLBP1 | -2,03576114 | 0,24387924 | 0,01510038 | 0,154803784 | retinaldehyde binding protein 1 |
| ENSG00000176092 | CRYBG2 | -2,038579638 | 0,243403254 | 0,02084489 | 0,192183647 | crystallin beta-gamma domain containing 2 |
| ENSG00000276825 | DRD4 | -2,039362884 | 0,243271145 | 0,01723703 | 0,169264441 | dopamine receptor D4 |
| ENSG00000225460 | AL590233,1 | -2,040426364 | 0,243091885 | 0,14797887 | 0,612582848 | ubiquitin-conjugating enzyme E2 pseudogene |
| ENSG00000237224 | AL157832,2 | -2,041256586 | 0,242952034 | 0,11907039 | 0,551662213 | novel transcript |
| ENSG00000267250 | AC011591,1 | -2,043967262 | 0,24249598 | 0,14080628 | 0,597646146 | novel transcript |
| ENSG00000274205 | MAPK15 | -2,044585787 | 0,242392038 | 0,00527156 | 0,071750111 | mitogen-activated protein kinase 15 |
| ENSG00000280813 | AC092902,1 | -2,044790257 | 0,242357686 | 0,0850558 | 0,464324191 | ribosomal protein S24 (RPS24) pseudogene |
| ENSG00000242229 | RPS3AP14 | -2,044790257 | 0,242357686 | 0,0850558 | 0,464324191 | RPS3A pseudogene 14 |
| ENSG00000239881 | RPS27P25 | -2,055574238 | 0,240552844 | 0,0075509 | 0,093783925 | ribosomal protein S27 pseudogene 25 |
| ENSG00000224731 | AC016716,1 | -2,059345597 | 0,239924835 | 0,12022116 | 0,553427919 | novel transcript |
| ENSG00000275538 | RNVU1-19 | -2,059637788 | 0,239876247 | 0,16881899 | 0,650760243 | RNA, variant U1 small nuclear 19 |
| ENSG00000147571 | CRH | -2,060060059 | 0,239806047 | 2,42E-05 | 0,000745761 | corticotropin releasing hormone |
| ENSG00000275327 | AL354950,2 | -2,063919968 | 0,239165307 | 0,34920909 | 0,843141089 | novel transcript, antisense to EBF3 |
| ENSG00000288564 | AL355385,2 | -2,064126562 | 0,239131061 | 0,31558165 | 0,820963951 | novel protein |

| | | | | | | |
|---|---|---|---|---|---|---|
| ENSG00000256817 | TPT1P12 | -2,068001656 | 0,238489614 | 0,1138326 | 0,539221598 | tumor protein, translationally-controlled 1 pseudogene 12 |
| ENSG00000145526 | CDH18 | -2,071957517 | 0,237836573 | 0,00050314 | 0,010454364 | cadherin 18 |
| ENSG00000175513 | TSGA10IP | -2,072075627 | 0,237817102 | 0,00023016 | 0,005382099 | testis specific 10 interacting protein |
| ENSG00000178522 | AMBN | -2,072241939 | 0,237789689 | 0,00615515 | 0,080617746 | ameloblastin |
| ENSG00000144596 | GRIP2 | -2,07280416 | 0,237697039 | 0,00010034 | 0,002623342 | glutamate receptor interacting protein 2 |
| ENSG00000254705 | AP000722,1 | -2,072825579 | 0,237693511 | 0,09742448 | 0,495829213 | novel transcript |
| ENSG00000282740 | AL021920,3 | -2,073770777 | 0,237537834 | 0,38795767 | 0,869820053 | novel transcript |
| ENSG00000258580 | AL136298,1 | -2,076767636 | 0,237044917 | 0,14012159 | 0,596671397 | novel transcript, antisense to AKAP6 |
| ENSG00000198797 | BRINP2 | -2,077131984 | 0,23698506 | 0,00278834 | 0,043718459 | BMP/retinoic acid inducible neural specific 2 |
| ENSG00000124772 | CPNE5 | -2,080689 | 0,236401485 | 0,0015067 | 0,026345347 | copine 5 |
| ENSG00000271318 | MOB4P2 | -2,080701296 | 0,23639947 | 0,09541928 | 0,490483883 | MOB4 pseudogene 2 |
| ENSG00000178965 | ERICH3 | -2,081956363 | 0,236193904 | 0,04666628 | 0,32799008 | glutamate rich 3 |
| ENSG00000259627 | AC079328,2 | -2,082028897 | 0,236182029 | 0,59580545 | 0,948096995 | novel transcript, antisense to TPM1 |
| ENSG00000286480 | AC009501,2 | -2,082831007 | 0,236050754 | 0,01655005 | 0,164573987 | novel transcript, antisense to WDPCP |
| ENSG00000276087 | AC008073,3 | -2,086013781 | 0,235530569 | 0,00964574 | 0,11198088 | novel transcript |
| ENSG00000160870 | CYP3A7 | -2,08760333 | 0,235271206 | 0,05491819 | 0,362021772 | cytochrome P450 family 3 subfamily A member 7 |
| ENSG00000165905 | LARGE2 | -2,092377079 | 0,234494001 | 0,00846549 | 0,101559908 | LARGE xylosyl- and glucuronyltransferase 2 |
| ENSG00000215297 | AL354941,1 | -2,092619105 | 0,234454666 | 0,03059819 | 0,250349254 | ribosomal protein L7 (RPL7) pseudogene |
| ENSG00000154162 | CDH12 | -2,093115835 | 0,234373955 | 0,00012276 | 0,003133101 | cadherin 12 |
| ENSG00000234860 | AL162394,1 | -2,094784886 | 0,234102965 | 0,02643737 | 0,227383165 | novel transcript |
| ENSG00000260551 | PWRN2 | -2,095164173 | 0,234041427 | 3,12E-07 | 1,47E-05 | Prader-Willi region non-protein coding RNA 2 |
| ENSG00000286583 | AC105101,2 | -2,095293143 | 0,234020506 | 0,28989826 | 0,800682868 | novel transcript, antisense to ZBTB7C |
| ENSG00000127412 | TRPV5 | -2,095330077 | 0,234014515 | 0,04609968 | 0,325793402 | transient receptor potential cation channel subfamily V member 5 |
| ENSG00000277303 | EPOP | -2,09534304 | 0,234012412 | 0,03749965 | 0,286018323 | elongin BC and polycomb repressive complex 2 associated protein |
| ENSG00000157303 | SUSD3 | -2,099029561 | 0,233415203 | 0,04914326 | 0,33801251 | sushi domain containing 3 |
| ENSG00000115009 | CCL20 | -2,102062166 | 0,23292507 | 0,05978864 | 0,380382656 | C-C motif chemokine ligand 20 |
| ENSG00000227254 | VDAC1P12 | -2,103956675 | 0,2326194 | 0,19482448 | 0,692394302 | voltage dependent anion channel 1 pseudogene 12 |
| ENSG00000262503 | AC027763,1 | -2,105868886 | 0,23231128 | 0,13243885 | 0,581009552 | Spi-C transcription factor (Spi-1/PU,1 related) (SPIC) pseudogene |
| ENSG00000256564 | AC007552,2 | -2,106013483 | 0,232287998 | 0,03095652 | 0,252761701 | novel transcript |

| | | | | | | |
|---|---|---|---|---|---|---|
| ENSG00000260780 | AC087463,1 | -2,106984039 | 0,232131781 | 0,10614252 | 0,51988717 | novel transcript |
| ENSG00000237359 | AL354977,2 | -2,10754127 | 0,232042139 | 0,02129831 | 0,194921664 | novel transcript |
| ENSG00000285505 | AC010616,1 | -2,110456738 | 0,231573691 | 0,56793981 | 0,94163744 | novel protein, ATP1A3-RABAC1 readthrough |
| ENSG00000183662 | TAFA1 | -2,110907321 | 0,231501377 | 0,00147551 | 0,025901289 | TAFA chemokine like family member 1 |
| ENSG00000254923 | AC130366,1 | -2,113772568 | 0,231042063 | 0,07510706 | 0,43291799 | deubiquitinating enzyme 3 pseudogene |
| ENSG00000262655 | SPON1 | -2,119171441 | 0,23017907 | 5,51E-09 | 3,57E-07 | spondin 1 |
| ENSG00000259884 | NR4A1AS | -2,119623755 | 0,230106915 | 0,1524237 | 0,620962584 | NR4A1 antisense RNA |
| ENSG00000288080 | AC005104,2 | -2,120227366 | 0,230010661 | 0,13796679 | 0,592546669 | novel transcript, antisense to FARP2 |
| ENSG00000229369 | AL450263,2 | -2,1215719 | 0,2297964 | 0,09214752 | 0,482558005 | density-regulated protein (DENR) pseudogene |
| ENSG00000275624 | PRB3 | -2,122519832 | 0,22964546 | 0,02278099 | 0,204838933 | proline rich protein BstNI subfamily 3 |
| ENSG00000141013 | GAS8 | -2,123650152 | 0,229465608 | 7,45E-05 | 0,002026074 | growth arrest specific 8 |
| ENSG00000261763 | AC110716,2 | -2,126509017 | 0,229011346 | 0,00608135 | 0,079818402 | novel transcript |
| ENSG00000120160 | EQTN | -2,128815705 | 0,228645479 | 0,04783755 | 0,332723723 | equatorin |
| ENSG00000274161 | DLGAP2 | -2,129203517 | 0,228584024 | 0,01228138 | 0,132813932 | DLG associated protein 2 |
| ENSG00000234394 | AL353608,3 | -2,146221674 | 0,225903468 | 0,0018164 | 0,030930244 | uncharacterized LOC440896 |
| ENSG00000226636 | AC006020,1 | -2,156397478 | 0,224315703 | 0,09544857 | 0,490483883 | tetratricopeptide repeat domain 14 (TTC14) pseudogene |
| ENSG00000236854 | LINC00486 | -2,156875735 | 0,224241354 | 0,02321766 | 0,207571363 | long intergenic non-protein coding RNA 486 |
| ENSG00000189372 | AL034369,1 | -2,158026073 | 0,224062626 | 0,05746464 | 0,370798119 | pancreatic lipase-related protein 2 (PNLIPRP2) pseudogene |
| ENSG00000243915 | THAP12P2 | -2,159816366 | 0,22378475 | 0,02583124 | 0,223956742 | THAP domain containing 12 pseudogene 2 |
| ENSG00000276462 | BX255923,2 | -2,160797725 | 0,223632578 | 0,00599508 | 0,078884758 | novel transcript |
| ENSG00000273540 | AGBL1 | -2,161146479 | 0,223578524 | 0,06086165 | 0,383945238 | ATP/GTP binding protein like 1 |
| ENSG00000254896 | OPCML-IT1 | -2,162744357 | 0,223331033 | 0,09324038 | 0,48515108 | OPCML intronic transcript 1 |
| ENSG00000272912 | AL356608,1 | -2,164151367 | 0,223113333 | 0,05593514 | 0,365161047 | novel transcript |
| ENSG00000148482 | SLC39A12 | -2,165384716 | 0,222922676 | 0,01293372 | 0,138384635 | solute carrier family 39 member 12 |
| ENSG00000259045 | MTCO1P2 | -2,171963433 | 0,221908459 | 0,16775155 | 0,648680831 | MT-CO1 pseudogene 2 |
| ENSG00000271286 | CYTH1P1 | -2,175978504 | 0,221291738 | 0,06536372 | 0,401164726 | CYTH1 pseudogene 1 |
| ENSG00000280650 | KCNIP4-IT1 | -2,17994868 | 0,220683599 | 0,01037277 | 0,118054008 | KCNIP4 intronic transcript 1 |
| ENSG00000228599 | RPL7P52 | -2,182076708 | 0,220358323 | 0,07319524 | 0,426957349 | ribosomal protein L7 pseudogene 52 |
| ENSG00000136383 | ALPK3 | -2,18317394 | 0,220190794 | 6,99E-08 | 3,84E-06 | alpha kinase 3 |
| ENSG00000217612 | AL031119,1 | -2,187840044 | 0,219479782 | 0,0748199 | 0,432378836 | BTB/POZ domain zinc finger protein pseudogene |
| ENSG00000285833 | AL590989,1 | -2,190232098 | 0,219116176 | 0,28655671 | 0,797836764 | novel transcript, antisense to DISP3 |

| | | | | | | |
|---|---|---|---|---|---|---|
| ENSG00000226702 | MIR217HG | -2,190527156 | 0,219071368 | 0,00502034 | 0,068931066 | MIR217 host gene |
| ENSG00000237645 | AC092967,1 | -2,192226562 | 0,218813467 | 0,08726387 | 0,470950713 | chromosome 14 open reading frame 83 (C14orf83) pseudogene |
| ENSG00000104888 | SLC17A7 | -2,193791871 | 0,218576186 | 2,40E-07 | 1,17E-05 | solute carrier family 17 member 7 |
| ENSG00000285619 | AL449214,1 | -2,195989548 | 0,218243479 | 0,03094874 | 0,252761701 | Novel transcript, antisense to EEFSEC |
| ENSG00000236295 | AC007557,3 | -2,197569878 | 0,218004546 | 0,01215164 | 0,131911723 | novel transcript |
| ENSG00000254447 | OR7E11P | -2,207695469 | 0,216479832 | 0,10085271 | 0,504639949 | olfactory receptor family 7 subfamily E member 11 pseudogene |
| ENSG00000007171 | NOS2 | -2,210542261 | 0,216053085 | 0,00040908 | 0,008784033 | nitric oxide synthase 2 |
| ENSG00000159307 | SCUBE1 | -2,211546277 | 0,21590278 | 8,09E-06 | 0,00028244 | signal peptide, CUB domain and EGF like domain containing 1 |
| ENSG00000250433 | CLSTN2-AS1 | -2,227830364 | 0,213479529 | 0,03937356 | 0,295481253 | CLSTN2 antisense RNA 1 |
| ENSG00000105852 | PON3 | -2,230295485 | 0,213115069 | 0,02027825 | 0,188628172 | paraoxonase 3 |
| ENSG00000239880 | RALBP1P1 | -2,231943796 | 0,21287172 | 0,09146801 | 0,481302729 | RALBP1 pseudogene 1 |
| ENSG00000244227 | LRRC77P | -2,232653085 | 0,212767089 | 0,05026552 | 0,342972789 | leucine rich repeat containing 77, pseudogene |
| ENSG00000152377 | SPOCK1 | -2,235315336 | 0,212374825 | 5,67E-12 | 5,72E-10 | SPARC (osteonectin), cwcv and kazal like domains proteoglycan 1 |
| ENSG00000287025 | AC079117,1 | -2,236820636 | 0,21215335 | 0,02361322 | 0,210026619 | novel transcript, sense intronic to LRRTM4 |
| ENSG00000100101 | Z83844,1 | -2,240274813 | 0,211646009 | 0,42516186 | 0,891853867 | novel transcript |
| ENSG00000163646 | CLRN1 | -2,243160612 | 0,21122308 | 0,15469556 | 0,623971362 | clarin 1 |
| ENSG00000232474 | NCKAP5-IT1 | -2,243448708 | 0,211180904 | 0,06696457 | 0,40667839 | NCKAP5 intronic transcript 1 |
| ENSG00000237550 | AC243919,1 | -2,247170479 | 0,210636816 | 0,02288698 | 0,205260654 | ribosomal protein L9 pseudogene 9 |
| ENSG00000278373 | G6PC2 | -2,248645201 | 0,210421613 | 0,06886618 | 0,413805931 | glucose-6-phosphatase catalytic subunit 2 |
| ENSG00000249255 | PGAM1P9 | -2,252537524 | 0,20985467 | 0,01379518 | 0,144680458 | phosphoglycerate mutase 1 pseudogene 9 |
| ENSG00000283062 | PGAM1P9 | -2,252537524 | 0,20985467 | 0,01379518 | 0,144680458 | phosphoglycerate mutase 1 pseudogene 9 |
| ENSG00000286456 | AC016769,5 | -2,255129522 | 0,209477976 | 2,26E-07 | 1,11E-05 | novel transcript |
| ENSG00000268731 | AC010615,3 | -2,257765173 | 0,209095632 | 0,0804718 | 0,449985752 | iron-sulfur cluster assembly 1 homolog (S, cerevisiae) (ISCA1) pseudogene |
| ENSG00000240393 | AC092691,2 | -2,268806443 | 0,207501484 | 0,00615795 | 0,080620668 | developmentally regulated GTP binding protein 1 (DRG1) pseudogene |
| ENSG00000123977 | DAW1 | -2,270632536 | 0,207239005 | 0,00378068 | 0,056258542 | dynein assembly factor with WD repeats 1 |
| ENSG00000228810 | PABPC1P11 | -2,277440486 | 0,206263366 | 0,02884569 | 0,241131487 | poly(A) binding protein cytoplasmic 1 pseudogene 11 |

| | | | | | | |
|---|---|---|---|---|---|---|
| ENSG00000239921 | LINC01471 | -2,278899363 | 0,206054894 | 0,10588893 | 0,519214495 | long intergenic non-protein coding RNA 1471 |
| ENSG00000153266 | FEZF2 | -2,280491819 | 0,205827575 | 3,41E-05 | 0,001012184 | FEZ family zinc finger 2 |
| ENSG00000154764 | WNT7A | -2,287276246 | 0,204861921 | 1,98E-05 | 0,000625542 | Wnt family member 7A |
| ENSG00000254568 | AP003501,1 | -2,291246076 | 0,204298983 | 0,1513383 | 0,618650693 | novel transcript |
| ENSG00000250732 | RPEP1 | -2,293144823 | 0,204030279 | 0,05604211 | 0,365554067 | ribulose-5-phosphate-3-epimerase pseudogene 1 |
| ENSG00000181965 | NEUROG1 | -2,294766916 | 0,203801007 | 4,40E-06 | 0,000164415 | neurogenin 1 |
| ENSG00000109906 | ZBTB16 | -2,30224368 | 0,202747541 | 4,06E-06 | 0,000153851 | zinc finger and BTB domain containing 16 |
| ENSG00000288110 | AC010941,1 | -2,302888428 | 0,202656952 | 0,01974594 | 0,184776117 | novel transcript, antisense to CSMD1 |
| ENSG00000217325 | PRELID1P1 | -2,305622274 | 0,202273289 | 0,14994558 | 0,615613756 | PRELID1 pseudogene 1 |
| ENSG00000244321 | LINC01326 | -2,305703979 | 0,202261834 | 0,0324835 | 0,260936342 | long intergenic non-protein coding RNA 1326 |
| ENSG00000184144 | CNTN2 | -2,308517781 | 0,201867731 | 4,28E-11 | 3,83E-09 | contactin 2 |
| ENSG00000162367 | TAL1 | -2,31091736 | 0,201532252 | 0,07273483 | 0,425281767 | TAL bHLH transcription factor 1, erythroid differentiation factor |
| ENSG00000250816 | DCAF13P2 | -2,312407135 | 0,20132425 | 0,06669067 | 0,40565499 | DDB1 and CUL4 associated factor 13 pseudogene 2 |
| ENSG00000162753 | SLC9C2 | -2,312520995 | 0,201308362 | 0,00234144 | 0,037987956 | solute carrier family 9 member C2 (putative) |
| ENSG00000216723 | NUDT19P3 | -2,314749214 | 0,200997684 | 0,08661455 | 0,468880561 | NUDT19 pseudogene 3 |
| ENSG00000275478 | AL671879,2 | -2,317517986 | 0,200612306 | 0,09019402 | 0,478874121 | novel transcript |
| ENSG00000152430 | BOLL | -2,318467827 | 0,200480271 | 0,06305216 | 0,392447267 | boule homolog, RNA binding protein |
| ENSG00000161132 | AC007663,1 | -2,323484286 | 0,199784382 | 0,09588962 | 0,491861057 | proline dehydrogenase (oxidase) 1 (PRODH) pseudogene |
| ENSG00000233558 | AL050331,1 | -2,326945473 | 0,199305652 | 0,17760876 | 0,664043981 | RPS5 (40S Ribosomal Protein S5) pseudogene |
| ENSG00000275097 | AC024940,5 | -2,32874402 | 0,199057341 | 0,03330073 | 0,264914501 | novel transcript, overlapping 3'UTR of FAM60A |
| ENSG00000011201 | ANOS1 | -2,328899673 | 0,199035865 | 1,78E-05 | 0,000567536 | anosmin 1 |
| ENSG00000276938 | AC073135,4 | -2,332658187 | 0,198518011 | 0,10096315 | 0,504961114 | family with sequence similarity 157, member A |
| ENSG00000258039 | AC117377,1 | -2,333246263 | 0,198437107 | 0,08288016 | 0,457872768 | novel transcript, antisense & intronic to ANKS1B |
| ENSG00000284800 | AC235565,2 | -2,333306389 | 0,198428837 | 0,01004213 | 0,115425382 | novel transcript |
| ENSG00000126259 | KIRREL2 | -2,337663255 | 0,197830496 | 0,00015768 | 0,003857462 | kirre like nephrin family adhesion molecule 2 |
| ENSG00000223703 | AC027612,1 | -2,339880218 | 0,197526727 | 0,01186809 | 0,129690286 | pseudogene similar to part of immunoglobulin superfamily, member 3 (IGSF3) |
| ENSG00000120088 | CRHR1 | -2,34059966 | 0,197428249 | 0,04395124 | 0,316175438 | corticotropin releasing hormone receptor 1 |
| ENSG00000233157 | Z97055,1 | -2,341967206 | 0,197241194 | 0,07015535 | 0,417549035 | ribosomal protein L4 (RPL4) pseudogene |
| ENSG00000162992 | NEUROD1 | -2,344104715 | 0,196949176 | 9,83E-06 | 0,0003359 | neuronal differentiation 1 |
| ENSG00000187513 | GJA4 | -2,345179434 | 0,196802516 | 0,05670964 | 0,367607368 | gap junction protein alpha 4 |

| | | | | | | |
|---|---|---|---|---|---|---|
| ENSG00000241202 | ZIC4-AS1 | -2,348963201 | 0,196287036 | 0,07064496 | 0,418620939 | ZIC4 antisense RNA 1 |
| ENSG00000233114 | AL358075,3 | -2,349352405 | 0,19623409 | 0,00428198 | 0,061777436 | cytochrome c oxidase subunit VIIb (COX7B) pseudogene |
| ENSG00000249158 | PCDHA11 | -2,350038545 | 0,196140784 | 2,77E-23 | 8,02E-21 | protocadherin alpha 11 |
| ENSG00000202347 | RNU1-16P | -2,352035547 | 0,195869471 | 0,0941577 | 0,487613759 | RNA, U1 small nuclear 16, pseudogene |
| ENSG00000239840 | RPL23AP72 | -2,352349508 | 0,19582685 | 0,05454667 | 0,360614755 | ribosomal protein L23a pseudogene 72 |
| ENSG00000231466 | AL022324,2 | -2,357887518 | 0,195076579 | 0,00600286 | 0,078887556 | PHD finger protein 10 (PHF10) pseudogene |
| ENSG00000213721 | HMGN2P30 | -2,358196758 | 0,195034769 | 0,03170791 | 0,256751118 | high mobility group nucleosomal binding domain 2 pseudogene 30 |
| ENSG00000276975 | HYDIN2 | -2,36033142 | 0,194746402 | 5,01E-05 | 0,001418637 | HYDIN axonemal central pair apparatus protein 2 (pseudogene) |
| ENSG00000267713 | AC012433,1 | -2,360432016 | 0,194732823 | 0,13242307 | 0,581009552 | NADH dehydrogenase pseudogene |
| ENSG00000130988 | RGN | -2,361443524 | 0,194596339 | 9,53E-05 | 0,002518276 | regucalcin |
| ENSG00000286592 | LINC02885 | -2,374162941 | 0,192888235 | 0,0269206 | 0,229957863 | long intergenic non-protein coding RNA 2885 |
| ENSG00000232792 | FTH1P25 | -2,377046561 | 0,192503081 | 0,01863608 | 0,17801357 | ferritin heavy chain 1 pseudogene 25 |
| ENSG00000261241 | LINC02128 | -2,378579603 | 0,192298631 | 0,12096683 | 0,555346102 | long intergenic non-protein coding RNA 2128 |
| ENSG00000136014 | USP44 | -2,381222393 | 0,191946693 | 0,00040045 | 0,008628294 | ubiquitin specific peptidase 44 |
| ENSG00000278631 | SOWAHCP5 | -2,38162079 | 0,191893694 | 0,10588576 | 0,519214495 | SOWAHC pseudogene 5 |
| ENSG00000007372 | PAX6 | -2,381953773 | 0,191849409 | 6,64E-07 | 2,93E-05 | paired box 6 |
| ENSG00000232957 | HLA-DOA | -2,38857344 | 0,190971143 | 0,02202792 | 0,199791344 | major histocompatibility complex, class II, DO alpha |
| ENSG00000240048 | DDX50P2 | -2,397006758 | 0,189858072 | 0,00743456 | 0,092677055 | DEAD-box helicase 50 pseudogene 2 |
| ENSG00000271218 | AL033384,2 | -2,399754511 | 0,189496813 | 0,10323026 | 0,512446948 | novel transcript, antisense to LRRC1 |
| ENSG00000152467 | ZSCAN1 | -2,400821364 | 0,189356734 | 4,46E-07 | 2,03E-05 | zinc finger and SCAN domain containing 1 |
| ENSG00000240833 | HCG17 | -2,403119476 | 0,189055343 | 0,12014656 | 0,553269269 | HLA complex group 17 |
| ENSG00000241701 | HCG17 | -2,403119476 | 0,189055343 | 0,12014656 | 0,553269269 | HLA complex group 17 |
| ENSG00000128917 | DLL4 | -2,405707721 | 0,188716475 | 0,0001319 | 0,003333748 | delta like canonical Notch ligand 4 |
| ENSG00000233460 | RPL35AP31 | -2,40971081 | 0,188193563 | 0,00201389 | 0,033778017 | ribosomal protein L35a pseudogene 31 |
| ENSG00000160868 | CYP3A4 | -2,414739635 | 0,187538716 | 0,07078349 | 0,418829978 | cytochrome P450 family 3 subfamily A member 4 |
| ENSG00000272703 | AP005137,2 | -2,417832037 | 0,187137159 | 0,08824611 | 0,473879487 | novel transcript |
| ENSG00000226525 | RPS7P10 | -2,423876304 | 0,186354776 | 0,12996963 | 0,576201143 | ribosomal protein S7 pseudogene 10 |
| ENSG00000225868 | AC016582,1 | -2,436878762 | 0,184682777 | 0,08764479 | 0,471714073 | WD repeat domain 87 (WDR87) pseudogene |
| ENSG00000145194 | ECE2 | -2,438663399 | 0,184454463 | 0,11986984 | 0,552908046 | endothelin converting enzyme 2 |
| ENSG00000257680 | AC016144,1 | -2,438918109 | 0,1844219 | 0,14693002 | 0,610674677 | methionyl aminopeptidase 1 (METAP1) pseudogene |

| | | | | | | |
|---|---|---|---|---|---|---|
| ENSG00000229006 | TRIM27 | -2,444089802 | 0,183761978 | 0,0021946 | 0,036206842 | tripartite motif containing 27 |
| ENSG00000278769 | AC090510,3 | -2,445870433 | 0,183535312 | 0,12707611 | 0,569347582 | novel transcript, antisense to CDAN1 |
| ENSG00000182040 | USH1G | -2,458348758 | 0,181954702 | 0,06237966 | 0,389891608 | USH1 protein network component sans |
| ENSG00000226567 | LINC00606 | -2,464006456 | 0,181242543 | 0,09300298 | 0,484688303 | long intergenic non-protein coding RNA 606 |
| ENSG00000243686 | RPLP1P11 | -2,464484352 | 0,181182516 | 0,21484334 | 0,722165588 | ribosomal protein lateral stalk subunit P1 pseudogene 11 |
| ENSG00000272282 | LINC02084 | -2,467859517 | 0,180759138 | 0,09227486 | 0,482910203 | long intergenic non-protein coding RNA 2084 |
| ENSG00000286417 | AL133334,1 | -2,470610814 | 0,180414749 | 0,12391595 | 0,562842145 | novel transcript, antisense to SUPT3H |
| ENSG00000204969 | PCDHA2 | -2,471068216 | 0,180357558 | 4,06E-13 | 4,76E-11 | protocadherin alpha 2 |
| ENSG00000227380 | HCG15 | -2,472732882 | 0,180149571 | 0,30800384 | 0,816424917 | HLA complex group 15 |
| ENSG00000134201 | GSTM5 | -2,481590521 | 0,179046904 | 7,62E-05 | 0,002064403 | glutathione S-transferase mu 5 |
| ENSG00000227700 | AC239809,2 | -2,484668983 | 0,178665256 | 0,1403177 | 0,597018607 | novel pseudogene |
| ENSG00000236888 | RPS20P5 | -2,484991142 | 0,178625364 | 0,05994792 | 0,380859955 | ribosomal protein S20 pseudogene 5 |
| ENSG00000267590 | NDUFA3P1 | -2,485979645 | 0,178503016 | 0,07324056 | 0,427120474 | NADH:ubiquinone oxidoreductase subunit A3 pseudogene 1 |
| ENSG00000242163 | AL121769,1 | -2,486556919 | 0,178431605 | 0,08387313 | 0,461088375 | ribosomal protein S24 (RPS24) pseudogene |
| ENSG00000277445 | CPEB1 | -2,48783472 | 0,178273637 | 0,23507435 | 0,744654651 | cytoplasmic polyadenylation element binding protein 1 |
| ENSG00000261963 | CCDC92B | -2,491208515 | 0,177857225 | 0,12128639 | 0,556159845 | coiled-coil domain containing 92B |
| ENSG00000204296 | TSBP1 | -2,496659828 | 0,177186448 | 0,13553448 | 0,586971877 | testis expressed basic protein 1 |
| ENSG00000160856 | FCRL3 | -2,496864738 | 0,177161284 | 0,00871685 | 0,103621111 | Fc receptor like 3 |
| ENSG00000287975 | AC113143,2 | -2,497037129 | 0,177140116 | 0,00776296 | 0,095647955 | novel transcript |
| ENSG00000224477 | AL109933,1 | -2,502130243 | 0,176515864 | 0,10839975 | 0,524858259 | novel transcript |
| ENSG00000286418 | AC018716,2 | -2,502614395 | 0,176456638 | 0,13818238 | 0,592936302 | novel transcript |
| ENSG00000154080 | CHST9 | -2,504187108 | 0,176264383 | 2,44E-05 | 0,000751938 | carbohydrate sulfotransferase 9 |
| ENSG00000285706 | AL807776,1 | -2,517254638 | 0,174675038 | 0,0505522 | 0,343953009 | Novel transcript |
| ENSG00000166091 | CMTM5 | -2,527157338 | 0,173480169 | 0,04416751 | 0,317145854 | CKLF like MARVEL transmembrane domain containing 5 |
| ENSG00000137869 | CYP19A1 | -2,529355082 | 0,173216098 | 0,00015632 | 0,00383316 | cytochrome P450 family 19 subfamily A member 1 |
| ENSG00000236668 | AC006450,3 | -2,540813129 | 0,171845845 | 0,00083022 | 0,015923883 | novel transcript, antisense to LHX2 |
| ENSG00000187955 | COL14A1 | -2,559581253 | 0,169624768 | 4,14E-06 | 0,000156372 | collagen type XIV alpha 1 chain |
| ENSG00000235708 | COL11A2 | -2,571506395 | 0,168228449 | 0,14352873 | 0,602629403 | collagen type XI alpha 2 chain |

| | | | | | | |
|---|---|---|---|---|---|---|
| ENSG00000139292 | LGR5 | -2,573763908 | 0,167965413 | 0,00121741 | 0,021965248 | leucine rich repeat containing G protein-coupled receptor 5 |
| ENSG00000287683 | AL591519,1 | -2,575668027 | 0,167743873 | 0,04108733 | 0,303461931 | novel transcript, antisense to EPHA7 |
| ENSG00000286463 | AP002755,1 | -2,575692431 | 0,167741035 | 0,0190186 | 0,180783685 | novel transcript, antisense to ZBTB16 |
| ENSG00000278475 | AC009464,1 | -2,57783851 | 0,167491698 | 0,04017202 | 0,299317946 | novel transcript, sense intronic to ARID2 |
| ENSG00000277439 | GP6 | -2,578625878 | 0,167400312 | 0,00859084 | 0,102591349 | glycoprotein VI platelet |
| ENSG00000088053 | GP6 | -2,578625878 | 0,167400312 | 0,00859084 | 0,102591349 | glycoprotein VI platelet |
| ENSG00000143032 | BARHL2 | -2,579866181 | 0,167256458 | 0,00472604 | 0,066050442 | BarH like homeobox 2 |
| ENSG00000275164 | MUC4 | -2,581741329 | 0,167039207 | 0,11028088 | 0,529863467 | mucin 4, cell surface associated |
| ENSG00000110148 | CCKBR | -2,591539601 | 0,16590858 | 0,00104353 | 0,019419646 | cholecystokinin B receptor |
| ENSG00000285779 | AL358472,7 | -2,593572235 | 0,165674993 | 0,51509722 | 0,924720046 | novel protein |
| ENSG00000231920 | NEBL-AS1 | -2,598119435 | 0,165153628 | 0,13163037 | 0,579455365 | NEBL antisense RNA 1 |
| ENSG00000244052 | RPL5P24 | -2,599188805 | 0,165031256 | 0,03982974 | 0,297761121 | ribosomal protein L5 pseudogene 24 |
| ENSG00000224842 | AL161908,1 | -2,60282946 | 0,164615323 | 0,00232839 | 0,037897375 | novel transcript, antisense to LIM1B |
| ENSG00000229921 | KIF25-AS1 | -2,605793502 | 0,164277465 | 0,00011543 | 0,002975223 | KIF25 antisense RNA 1 |
| ENSG00000276754 | AC006111,3 | -2,611539585 | 0,163624469 | 0,02460383 | 0,21668006 | novel transcript, antisense to TRAP1 |
| ENSG00000144852 | NR1I2 | -2,614817019 | 0,163253178 | 0,00083306 | 0,015958724 | nuclear receptor subfamily 1 group I member 2 |
| ENSG00000122584 | NXPH1 | -2,615568065 | 0,163168213 | 0,00023551 | 0,005486645 | neurexophilin 1 |
| ENSG00000163508 | EOMES | -2,618019493 | 0,162891193 | 0,00087641 | 0,01666675 | eomesodermin |
| ENSG00000241179 | AC010301,1 | -2,62684465 | 0,161897807 | 0,06379009 | 0,395590561 | ribosomal protein L34 (RPL34) pseudogene |
| ENSG00000215029 | TCP11X2 | -2,629021696 | 0,161653685 | 0,01109799 | 0,123533851 | t-complex 11 family, X-linked 2 |
| ENSG00000263916 | AC100778,2 | -2,62919899 | 0,161633821 | 0,02866296 | 0,240375171 | novel transcript, sense intronic to C18orf32 |
| ENSG00000285137 | MGAT4C | -2,629679513 | 0,161579994 | 0,00266294 | 0,042068851 | MGAT4 family member C |
| ENSG00000170989 | S1PR1 | -2,644248036 | 0,159956549 | 2,64E-06 | 0,000104451 | sphingosine-1-phosphate receptor 1 |
| ENSG00000287076 | AC092805,1 | -2,647363102 | 0,159611543 | 0,0163872 | 0,163524062 | novel transcript |
| ENSG00000242795 | AC007849,1 | -2,670083122 | 0,157117619 | 0,04659086 | 0,327821686 | novel transcript |
| ENSG00000251621 | AC009487,2 | -2,672784914 | 0,156823654 | 0,07352628 | 0,428147811 | novel transcript |
| ENSG00000280913 | CTSLP3 | -2,679275488 | 0,156119701 | 0,00205795 | 0,0343696 | cathepsin L pseudogene 3 |
| ENSG00000230371 | KARS1P2 | -2,686624006 | 0,155326511 | 0,05785731 | 0,371961929 | lysyl-tRNA synthetase 1 pseudogene 2 |
| ENSG00000256810 | AC078925,1 | -2,689696475 | 0,154996068 | 0,00054709 | 0,011260086 | novel transcript |
| ENSG00000184302 | SIX6 | -2,697127759 | 0,15419974 | 8,62E-07 | 3,71E-05 | SIX homeobox 6 |
| ENSG00000271321 | CTAGE6 | -2,69928191 | 0,15396967 | 0,05761697 | 0,3710973 | CTAGE family member 6 |
| ENSG00000285123 | CTAGE6 | -2,69928191 | 0,15396967 | 0,05761697 | 0,3710973 | CTAGE family member 6 |

| | | | | | | |
|---|---|---|---|---|---|---|
| ENSG00000180767 | CHST13 | -2,707519286 | 0,153093052 | 0,05558953 | 0,363892597 | carbohydrate sulfotransferase 13 |
| ENSG00000259200 | AC068722,1 | -2,71114247 | 0,152709057 | 0,02449704 | 0,215982952 | novel transcript |
| ENSG00000142700 | DMRTA2 | -2,716735901 | 0,152118139 | 0,00141405 | 0,025104349 | DMRT like family A2 |
| ENSG00000229111 | MED4-AS1 | -2,728148793 | 0,150919508 | 0,02492442 | 0,219004288 | MED4 antisense RNA 1 |
| ENSG00000274428 | RNVU1-25 | -2,728908545 | 0,150840052 | 0,00390018 | 0,057625384 | RNA, variant U1 small nuclear 25 |
| ENSG00000285982 | AC012213,5 | -2,748868942 | 0,148767476 | 0,38481851 | 0,868536368 | novel protein |
| ENSG00000250472 | TRIM36-IT1 | -2,752980619 | 0,148344093 | 0,02287492 | 0,205211335 | TRIM36 intronic transcript 1 |
| ENSG00000077327 | SPAG6 | -2,75605913 | 0,148027884 | 0,0005153 | 0,010690406 | sperm associated antigen 6 |
| ENSG00000144010 | TRIM43B | -2,770248998 | 0,146579068 | 0,04779924 | 0,332623171 | tripartite motif containing 43B |
| ENSG00000237750 | KCNH7-AS1 | -2,780686992 | 0,145522386 | 2,35E-09 | 1,64E-07 | KCNH7 antisense RNA 1 |
| ENSG00000133475 | GGT2 | -2,788025232 | 0,144784069 | 0,0943848 | 0,487985199 | gamma-glutamyltransferase 2 |
| ENSG00000225419 | RPL21P3 | -2,791209062 | 0,144464902 | 0,03801641 | 0,288593771 | ribosomal protein L21 pseudogene 3 |
| ENSG00000174948 | GPR149 | -2,792053888 | 0,14438033 | 0,00354349 | 0,053389744 | G protein-coupled receptor 149 |
| ENSG00000233651 | AL590399,5 | -2,799318663 | 0,143655122 | 0,251671 | 0,763981167 | novel transcript |
| ENSG00000205018 | AC092384,1 | -2,810837284 | 0,142512732 | 0,04372722 | 0,315797603 | novel transcript, antisense to CBFA2T3 |
| ENSG00000108231 | LGI1 | -2,815309772 | 0,142071613 | 7,58E-08 | 4,13E-06 | leucine rich glioma inactivated 1 |
| ENSG00000273990 | RNVU1-26 | -2,823336309 | 0,141283382 | 2,74E-06 | 0,000107989 | RNA, variant U1 small nuclear 26 |
| ENSG00000275291 | RNVU1-26 | -2,823336309 | 0,141283382 | 2,74E-06 | 0,000107989 | RNA, variant U1 small nuclear 26 |
| ENSG00000225174 | OSTM1-AS1 | -2,825213383 | 0,14109968 | 0,00264418 | 0,041878228 | OSTM1 antisense RNA 1 |
| ENSG00000197921 | HES5 | -2,829427961 | 0,140688083 | 5,83E-05 | 0,001620656 | hes family bHLH transcription factor 5 |
| ENSG00000258399 | AL117190,1 | -2,832163862 | 0,140421538 | 0,12262104 | 0,560061984 | novel transcript |
| ENSG00000135638 | EMX1 | -2,84216221 | 0,139451735 | 4,49E-05 | 0,001285165 | empty spiracles homeobox 1 |
| ENSG00000168907 | PLA2G4F | -2,842928904 | 0,139377646 | 0,08155573 | 0,453774372 | phospholipase A2 group IVF |
| ENSG00000231609 | AC007098,1 | -2,844128833 | 0,13926177 | 4,96E-07 | 2,25E-05 | novel transcript, antisense to EHBP1 |
| ENSG00000188897 | AC099489,1 | -2,847220819 | 0,138963623 | 1,89E-07 | 9,41E-06 | novel lipoprotein amino terminal region containing protein |
| ENSG00000225549 | SELENOKP3 | -2,857118621 | 0,138013508 | 0,04162879 | 0,306015175 | selenoprotein K pseudogene 3 |
| ENSG00000178343 | SHISA3 | -2,859214125 | 0,13781319 | 0,0009734 | 0,01825309 | shisa family member 3 |
| ENSG00000177710 | SLC35G5 | -2,876184368 | 0,136201607 | 0,13364798 | 0,583882708 | solute carrier family 35 member G5 |
| ENSG00000265766 | CXADRP3 | -2,878082108 | 0,136022564 | 0,04423199 | 0,3175092 | CXADR pseudogene 3 |
| ENSG00000281593 | AC006978,2 | -2,878300568 | 0,136001968 | 0,03730032 | 0,285054324 | novel protein, ZNRF2-LINC01176 readthrough |
| ENSG00000257522 | AL133166,1 | -2,893941048 | 0,134535513 | 0,00427394 | 0,061690033 | novel transcript |
| ENSG00000231227 | AC006450,1 | -2,902551536 | 0,133734951 | 0,02215991 | 0,200639268 | novel transcript |

| | | | | | | |
|---|---|---|---|---|---|---|
| ENSG00000225286 | AC005105,1 | -2,906601598 | 0,133360045 | 0,07514668 | 0,43306652 | ribosomal protein S2 (RPS2) pseudogene |
| ENSG00000243806 | RPL7P18 | -2,907816339 | 0,133247804 | 0,01803084 | 0,17436315 | ribosomal protein L7 pseudogene 18 |
| ENSG00000258028 | AL135878,1 | -2,92339632 | 0,131816574 | 0,01098004 | 0,122731541 | novel transcript |
| ENSG00000271761 | AL021368,1 | -2,926602774 | 0,131523931 | 0,05622396 | 0,366342158 | novel transcript |
| ENSG00000133433 | GSTT2B | -2,95997349 | 0,12851659 | 0,1717553 | 0,655568029 | glutathione S-transferase theta 2B |
| ENSG00000262986 | SMIM10L1 | -2,98069006 | 0,126684326 | 0,00791594 | 0,09705248 | small integral membrane protein 10 like 1 |
| ENSG00000249509 | NEUROG2-AS1 | -3,011158855 | 0,124036888 | 0,00714817 | 0,089895541 | NEUROG2 and ZGRF1 antisense RNA 1 |
| ENSG00000267319 | SELENOKP1 | -3,030481725 | 0,122386665 | 0,06816172 | 0,411391518 | selenoprotein K pseudogene 1 |
| ENSG00000287869 | AC073349,5 | -3,04051031 | 0,12153887 | 0,04963134 | 0,339981227 | novel transcript |
| ENSG00000250651 | PABPC1P7 | -3,04683608 | 0,121007126 | 0,02661547 | 0,228371442 | poly(A) binding protein cytoplasmic 1 pseudogene 7 |
| ENSG00000260314 | MRC1 | -3,050225187 | 0,120723196 | 0,00409585 | 0,059781846 | mannose receptor C-type 1 |
| ENSG00000267640 | AC016582,3 | -3,050676695 | 0,12068542 | 5,24E-19 | 1,09E-16 | novel zinc finger protein pseudogene |
| ENSG00000173376 | NDNF | -3,055330287 | 0,120296762 | 2,77E-07 | 1,33E-05 | neuron derived neurotrophic factor |
| ENSG00000234602 | MCIDAS | -3,060640096 | 0,119854826 | 0,19929411 | 0,698590752 | multiciliate differentiation and DNA synthesis associated cell cycle protein |
| ENSG00000227917 | AL449043,1 | -3,075526458 | 0,11862447 | 0,06509291 | 0,400524473 | novel transcript |
| ENSG00000286961 | AC091938,1 | -3,083925516 | 0,11793587 | 1,01E-08 | 6,23E-07 | novel transcript, antisense to CDH12 |
| ENSG00000234186 | C16orf82 | -3,091510899 | 0,117317416 | 0,00336715 | 0,051151909 | chromosome 16 open reading frame 82 |
| ENSG00000272449 | AL139246,5 | -3,105033262 | 0,116222938 | 0,12521036 | 0,565471028 | novel transcript |
| ENSG00000220749 | RPL21P28 | -3,120254175 | 0,115003193 | 0,00710716 | 0,089581081 | ribosomal protein L21 pseudogene 28 |
| ENSG00000236428 | PPP1R18 | -3,126065045 | 0,114540916 | 0,22752308 | 0,736053199 | protein phosphatase 1 regulatory subunit 18 |
| ENSG00000287088 | AP001066,1 | -3,126996931 | 0,114466954 | 0,0555575 | 0,363892597 | novel transcript, antisense to TSPEAR |
| ENSG00000281630 | MUC20 | -3,136118587 | 0,113745503 | 0,03804518 | 0,288700106 | mucin 20, cell surface associated |
| ENSG00000251056 | ANKRD20A17P | -3,152542459 | 0,112457949 | 0,07875307 | 0,4442685 | ankyrin repeat domain 20 family member A17, pseudogene |
| ENSG00000240083 | RPS3AP22 | -3,160845838 | 0,11181256 | 0,03249188 | 0,260936563 | RPS3A pseudogene 22 |
| ENSG00000260246 | AC000032,1 | -3,166863239 | 0,111347167 | 0,04031991 | 0,299990909 | novel transcript, antisense to GSTM1 |
| ENSG00000280660 | AL157396,1 | -3,170455421 | 0,111070268 | 1,84E-07 | 9,22E-06 | novel transcript, sense intronic to LGI1 |
| ENSG00000270773 | AC137695,2 | -3,197899026 | 0,108977407 | 0,02652445 | 0,227881399 | MAP/microtubule affinity-regulating kinase 2, pseudogene |
| ENSG00000276143 | AC138749,7 | -3,200934108 | 0,108748386 | 0,03431337 | 0,270005245 | family with sequence similarity 208, member A (FAM208A) pseudogene |
| ENSG00000265818 | EEF1E1-BLOC1S | -3,203432875 | 0,108560195 | 0,08849365 | 0,474566472 | EEF1E1-BLOC1S5 readthrough (NMD candidate) |

| | | | | | | |
|---|---|---|---|---|---|---|
| ENSG00000004848 | ARX | -3,242479615 | 0,105661404 | 1,57E-09 | 1,14E-07 | aristaless related homeobox |
| ENSG00000206245 | TSBP1 | -3,246463149 | 0,105370056 | 0,4152379 | 0,885840844 | testis expressed basic protein 1 |
| ENSG00000277831 | AL138960,1 | -3,251691551 | 0,104988881 | 0,00972012 | 0,112718772 | novel transcript |
| ENSG00000236534 | H3P36 | -3,271219879 | 0,103577326 | 0,03300156 | 0,263607792 | H3 histone pseudogene 36 |
| ENSG00000226669 | AL354977,1 | -3,274154808 | 0,103366829 | 0,00864966 | 0,103057511 | novel transcript |
| ENSG00000082556 | OPRK1 | -3,314960301 | 0,100484139 | 3,87E-09 | 2,58E-07 | opioid receptor kappa 1 |
| ENSG00000196299 | ZNRD1ASP | -3,317671583 | 0,100295475 | 0,38932006 | 0,870057277 | zinc ribbon domain containing 1 antisense, pseudogene |
| ENSG00000080166 | DCT | -3,32250639 | 0,099959924 | 1,21E-05 | 0,000399084 | dopachrome tautomerase |
| ENSG00000265579 | AC023301,1 | -3,358167464 | 0,097519365 | 0,1755945 | 0,661199837 | novel transcript , sense intronic to NETO1 |
| ENSG00000280938 | TRARG1 | -3,358167464 | 0,097519365 | 0,1755945 | 0,661199837 | trafficking regulator of GLUT4 (SLC2A4) 1 |
| ENSG00000178403 | NEUROG2 | -3,382426441 | 0,095893282 | 1,42E-12 | 1,56E-10 | neurogenin 2 |
| ENSG00000179073 | TAAR3P | -3,391540783 | 0,095289378 | 0,09525878 | 0,489967108 | trace amine associated receptor 3, pseudogene |
| ENSG00000225477 | AC099730,1 | -3,396095526 | 0,094989014 | 0,01886937 | 0,179692945 | isocitrate dehydrogenase 3 (NAD+) gamma (IDH3G) pseudogene |
| ENSG00000214132 | AC094103,1 | -3,444240685 | 0,09187138 | 0,00538247 | 0,072847486 | nucleoporin 50kDa (NUP50) pseudogene |
| ENSG00000270096 | AC078795,2 | -3,445156116 | 0,091813104 | 0,38653026 | 0,869441674 | novel transcript, antisense to LRRC34 |
| ENSG00000075290 | WNT8B | -3,446199018 | 0,091746757 | 9,69E-17 | 1,67E-14 | Wnt family member 8B |
| ENSG00000228999 | AC096570,1 | -3,466356467 | 0,090473778 | 5,68E-07 | 2,53E-05 | long intergenic non-protein coding RNA 1830 |
| ENSG00000265984 | AC022809,1 | -3,467285618 | 0,090415529 | 0,01102749 | 0,123085791 | novel transcript |
| ENSG00000213185 | FAM24B | -3,48014361 | 0,089613282 | 0,02505005 | 0,219619974 | family with sequence similarity 24 member B |
| ENSG00000285438 | SOX7 | -3,524021452 | 0,086928831 | 0,13870379 | 0,593837907 | SRY-box transcription factor 7 |
| ENSG00000280485 | AL358154,1 | -3,533267043 | 0,086373524 | 0,04576007 | 0,324471967 | novel transcript, antisense to LGI1 |
| ENSG00000261949 | GFY | -3,537896405 | 0,08609681 | 8,76E-05 | 0,002335344 | golgi associated olfactory signaling regulator |
| ENSG00000284731 | AC083906,5 | -3,541109995 | 0,085905244 | 0,01214468 | 0,131911723 | RNA binding protein, fox-1 homolog (C, elegans) 1 (RBFOX1) pseudogene |
| ENSG00000164708 | PGAM2 | -3,567913348 | 0,084323973 | 6,41E-09 | 4,11E-07 | phosphoglycerate mutase 2 |
| ENSG00000235590 | GNAS-AS1 | -3,6044135 | 0,08221734 | 1,53E-07 | 7,93E-06 | GNAS antisense RNA 1 |
| ENSG00000282706 | LINC01043 | -3,643295626 | 0,08003109 | 0,09631696 | 0,49300157 | long intergenic non-protein coding RNA 1043 |
| ENSG00000285515 | CR936875,2 | -3,737056546 | 0,074995272 | 0,28936554 | 0,800260154 | novel protein |
| ENSG00000285213 | AL773541,2 | -3,737056546 | 0,074995272 | 0,28936554 | 0,800260154 | novel protein |
| ENSG00000233587 | LINC01884 | -3,768131897 | 0,073397163 | 0,12803664 | 0,571345532 | long intergenic non-protein coding RNA 1884 |

| | | | | | | |
|---|---|---|---|---|---|---|
| ENSG00000223691 | AC068044,1 | -3,783174331 | 0,072635854 | 0,00834049 | 0,100717272 | Mdm2, transformed 3T3 cell double minute 2, p53 binding protein (mouse) (MDM2) pseudogene |
| ENSG00000093010 | COMT | -3,828966757 | 0,070366533 | 2,05E-20 | 4,90E-18 | catechol-O-methyltransferase |
| ENSG00000267260 | AC020928,1 | -3,855302796 | 0,069093663 | 6,61E-10 | 5,07E-08 | novel transcript |
| ENSG00000144015 | TRIM43 | -3,878346939 | 0,067998798 | 0,00836108 | 0,100849048 | tripartite motif containing 43 |
| ENSG00000223626 | LINC01044 | -3,908590111 | 0,066588179 | 0,00028707 | 0,006508339 | long intergenic non-protein coding RNA 1044 |
| ENSG00000164512 | ANKRD55 | -3,99237176 | 0,062831344 | 0,00254727 | 0,040714725 | ankyrin repeat domain 55 |
| ENSG00000249854 | CDH18-AS1 | -4,127456266 | 0,057215258 | 4,26E-05 | 0,001226323 | CDH18 antisense RNA 1 |
| ENSG00000214415 | GNAT3 | -4,143531303 | 0,056581283 | 0,01488674 | 0,153300037 | G protein subunit alpha transducin 3 |
| ENSG00000282737 | LINC01044 | -4,1662946 | 0,055695532 | 0,00014903 | 0,003691955 | long intergenic non-protein coding RNA 1044 |
| ENSG00000267374 | MIR924HG | -4,176416273 | 0,05530615 | 2,60E-24 | 8,04E-22 | MIR924 host gene |
| ENSG00000273992 | NLRP2 | -4,184255144 | 0,055006459 | 8,90E-07 | 3,82E-05 | NLR family pyrin domain containing 2 |
| ENSG00000115507 | OTX1 | -4,21458671 | 0,053862063 | 6,49E-11 | 5,68E-09 | orthodenticle homeobox 1 |
| ENSG00000234892 | Z82214,1 | -4,226535635 | 0,0534178 | 0,02363379 | 0,210149803 | novel transcript |
| ENSG00000267219 | AC010504,1 | -4,256076506 | 0,05233513 | 0,02438857 | 0,215149831 | novel transcript, antisense to KIAA0355 |
| ENSG00000069696 | DRD4 | -4,295863006 | 0,050911557 | 0,03474265 | 0,271925393 | dopamine receptor D4 |
| ENSG00000119614 | VSX2 | -4,426216775 | 0,046513175 | 0,00530947 | 0,072203249 | visual system homeobox 2 |
| ENSG00000165246 | NLGN4Y | -4,42772696 | 0,046464511 | 2,23E-10 | 1,85E-08 | neuroligin 4 Y-linked |
| ENSG00000159248 | GJD2 | -4,546820859 | 0,042782931 | 9,78E-13 | 1,10E-10 | gap junction protein delta 2 |
| ENSG00000175886 | RPL7AP66 | -4,552561729 | 0,042613025 | 0,0003273 | 0,007293371 | ribosomal protein L7a pseudogene 66 |
| ENSG00000286559 | AC099845,1 | -4,58343213 | 0,041710889 | 1,18E-14 | 1,64E-12 | novel transcript |
| ENSG00000249506 | ZEB2P1 | -4,656913103 | 0,039639619 | 0,00051656 | 0,010702297 | zinc finger E-box binding homeobox 2 pseudogene 1 |
| ENSG00000231641 | TRIM26 | -4,695618565 | 0,038590283 | 0,00287687 | 0,04490384 | tripartite motif containing 26 |
| ENSG00000232962 | HLA-DOA | -4,790873043 | 0,036124638 | 1,15E-05 | 0,00038535 | major histocompatibility complex, class II, DO alpha |
| ENSG00000224173 | LINC02831 | -5,102974767 | 0,029097222 | 0,00466256 | 0,065632751 | long intergenic non-protein coding RNA 2831 |
| ENSG00000270149 | AL591806,2 | -5,135007239 | 0,02845829 | 0,01148692 | 0,126676175 | novel transcript, TSTD1 - F11R readthrough |
| ENSG00000204396 | VWA7 | -5,171565243 | 0,027746214 | 1,41E-05 | 0,000459645 | von Willebrand factor A domain containing 7 |
| ENSG00000241166 | HLA-L | -5,384946366 | 0,023931483 | 0,18331156 | 0,674320815 | major histocompatibility complex, class I, L (pseudogene) |
| ENSG00000234123 | RHBDF1P1 | -5,484879944 | 0,022329892 | 0,00212666 | 0,03532839 | RHBDF1 pseudogene 1 |
| ENSG00000179886 | TIGD5 | -5,550098134 | 0,021342927 | 0,10908103 | 0,526943592 | tigger transposable element derived 5 |
| ENSG00000286663 | AL513524,1 | -5,562321647 | 0,021162859 | 0,0012333 | 0,022185215 | novel transcript |
| ENSG00000232229 | LINC00865 | -5,631983654 | 0,020165267 | 3,72E-27 | 1,35E-24 | long intergenic non-protein coding RNA 865 |

| | | | | | | |
|---|---|---|---|---|---|---|
| ENSG00000185985 | SLITRK2 | -5,70255189 | 0,019202635 | 9,33E-12 | 9,20E-10 | SLIT and NTRK like family member 2 |
| ENSG00000214076 | CPSF1P1 | -5,759650025 | 0,018457487 | 0,00040412 | 0,008689322 | CPSF1 pseudogene 1 |
| ENSG00000241345 | AC004690,2 | -5,781079695 | 0,018185348 | 0,00023893 | 0,005549706 | novel transcript |
| ENSG00000112333 | NR2E1 | -5,871748992 | 0,017077623 | 1,28E-14 | 1,75E-12 | nuclear receptor subfamily 2 group E member 1 |
| ENSG00000224006 | AL645935,3 | -5,974835772 | 0,01589993 | 0,02630406 | 0,22653539 | HLA complex group 4 pseudogene 3 |
| ENSG00000228695 | CES1P1 | -6,050262735 | 0,015090007 | 0,0022346 | 0,036693513 | carboxylesterase 1 pseudogene 1 |
| ENSG00000146833 | TRIM4 | -6,064612388 | 0,014940659 | 5,20E-32 | 2,71E-29 | tripartite motif containing 4 |
| ENSG00000236561 | MOG | -6,151780305 | 0,014064672 | 0,01842417 | 0,176745366 | myelin oligodendrocyte glycoprotein |
| ENSG00000281670 | CABIN1 | -6,23939525 | 0,013235943 | 0,00039811 | 0,008595634 | calcineurin binding protein 1 |
| ENSG00000268649 | AL132655,2 | -6,320794837 | 0,012509823 | 0,00035209 | 0,007735532 | novel transcript |
| ENSG00000285839 | AL445685,3 | -6,416799272 | 0,011704448 | 0,01000519 | 0,115085444 | novel transcript |
| ENSG00000284330 | AC256236,1 | -6,538949484 | 0,010754248 | 0,00777943 | 0,095754377 | |
| ENSG00000227124 | ZNF717 | -6,546946263 | 0,010694803 | 7,06E-42 | 7,88E-39 | zinc finger protein 717 |
| ENSG00000281948 | CES1P1 | -6,820224802 | 0,008849276 | 0,09511466 | 0,489669163 | carboxylesterase 1 pseudogene 1 |
| ENSG00000204197 | KIFC1 | -6,878457597 | 0,008499198 | 0,09230532 | 0,482984696 | kinesin family member C1 |
| ENSG00000268333 | AL132655,1 | -6,88739535 | 0,008446707 | 7,14E-06 | 0,000253546 | novel transcript |
| ENSG00000162623 | TYW3 | -6,964091828 | 0,00800939 | 3,55E-30 | 1,66E-27 | tRNA-yW synthesizing protein 3 homolog |
| ENSG00000116791 | CRYZ | -7,173397208 | 0,006927751 | 1,02E-31 | 5,16E-29 | crystallin zeta |
| ENSG00000230733 | AC092171,2 | -7,233603071 | 0,006644595 | 0,07659194 | 0,43728459 | novel transcript |
| ENSG00000125084 | WNT1 | -7,255561011 | 0,006544229 | 3,00E-08 | 1,72E-06 | Wnt family member 1 |
| ENSG00000247033 | AC099508,1 | -7,405801714 | 0,005897006 | 1,27E-07 | 6,66E-06 | novel transcript, antisense to ZNRF1 |
| ENSG00000022556 | NLRP2 | -8,12166892 | 0,003590327 | 1,47E-46 | 2,00E-43 | NLR family pyrin domain containing 2 |
| ENSG00000130283 | GDF1 | -8,310308808 | 0,00315027 | 0,04179477 | 0,306600121 | growth differentiation factor 1 |
| ENSG00000277060 | NLRP2 | -10,05510294 | 0,000939967 | 1,58E-17 | 2,91E-15 | NLR family pyrin domain containing 2 |
| ENSG00000180543 | TSPYL5 | -11,24210595 | 0,000412847 | 2,59E-25 | 8,70E-23 | TSPY like 5 |
| ENSG00000275379 | H3C11 | -12,33536193 | 0,000193502 | 3,42E-88 | 5,34E-84 | H3 clustered histone 11 |
| ENSG00000275126 | H4C13 | -14,22153293 | 5,23E-05 | 1,84E-39 | 1,65E-36 | H4 clustered histone 13 |
