## Supplementary Table 4 for "Huntingtin loss-of-function contributes to transcriptional deregulation in Huntington’s disease"

**Supplementary Table 4. GO enrichment analysis of DEGs in HD and KO-NSCs**

Terms presented in Fig. 2 f and g are marked in yellow.

**HD-NSCs**

| | |
|---|---|
| Analysis Type: | PANTHER Overrepresentation Test (Released 20221013) |
| Annotation Version and Release Date: | PANTHER version 17.0 Released 2022-02-22 |
| Analyzed List: | HD-NSCs (Homo sapiens) |
| Reference List: | Homo sapiens (all genes in database) |
| Test Type: | FISHER |
| Correction: | FDR |

| **PANTHER GO-Slim Molecular Function** | Homo sapiens - REFLIST (20589) | HD NSCs | expected | over/ under | fold Enrichment | raw P-value | FDR |
|---|---|---|---|---|---|---|---|
| Wnt-protein binding (GO:0017147) | 21 | 5 | 0,55 | + | 9,01 | 4,66E-04 | 1,21E-02 |
| protein kinase inhibitor activity (GO:0004860) | 18 | 4 | 0,48 | + | 8,41 | 2,20E-03 | 4,00E-02 |
| growth factor binding (GO:0019838) | 35 | 6 | 0,92 | + | 6,49 | 5,92E-04 | 1,35E-02 |
| transmembrane receptor protein tyrosine kinase activity (GO:0004714) | 52 | 8 | 1,37 | + | 5,82 | 1,44E-04 | 4,38E-03 |
| integrin binding (GO:0005178) | 49 | 7 | 1,29 | + | 5,41 | 5,61E-04 | 1,33E-02 |
| transmembrane receptor protein kinase activity (GO:0019199) | 64 | 8 | 1,69 | + | 4,73 | 5,16E-04 | 1,28E-02 |
| cell adhesion molecule binding (GO:0050839) | 141 | 17 | 3,73 | + | 4,56 | 7,18E-07 | 4,35E-05 |
| metalloendopeptidase activity (GO:0004222) | 84 | 10 | 2,22 | + | 4,51 | 1,53E-04 | 4,40E-03 |
| metallopeptidase activity (GO:0008237) | 116 | 13 | 3,06 | + | 4,24 | 2,92E-05 | 9,97E-04 |
| protein tyrosine kinase activity (GO:0004713) | 96 | 9 | 2,54 | + | 3,55 | 1,56E-03 | 3,05E-02 |
| receptor ligand activity (GO:0048018) | 245 | 16 | 6,47 | + | 2,47 | 1,27E-03 | 2,66E-02 |
| signaling receptor activator activity (GO:0030546) | 248 | 16 | 6,55 | + | 2,44 | 1,43E-03 | 2,88E-02 |
| DNA-binding transcription factor activity, RNA polymerase II-specific (GO:0000981) | 984 | 61 | 26 | + | 2,35 | 2,39E-09 | 1,30E-06 |
| signaling receptor regulator activity (GO:0030545) | 276 | 17 | 7,29 | + | 2,33 | 2,10E-03 | 3,95E-02 |
| DNA-binding transcription factor activity (GO:0003700) | 1054 | 62 | 27,85 | + | 2,23 | 1,44E-08 | 3,93E-06 |
| RNA polymerase II transcription regulatory region sequence-specific DNA binding (GO:0000977) | 1059 | 62 | 27,98 | + | 2,22 | 1,57E-08 | 2,85E-06 |
| transcription cis-regulatory region binding (GO:0000976) | 1099 | 62 | 29,04 | + | 2,14 | 5,23E-08 | 7,14E-06 |

| | | | | | | | |
|---|---|---|---|---|---|---|---|
| transcription regulatory region nucleic acid binding (GO:0001067) | 1099 | 62 | 29,04 | + | 2,14 | 5,23E-08 | 5,71E-06 |
| RNA polymerase II cis-regulatory region sequence-specific DNA binding (GO:0000978) | 798 | 45 | 21,08 | + | 2,13 | 3,84E-06 | 2,10E-04 |
| cis-regulatory region sequence-specific DNA binding (GO:0000987) | 810 | 45 | 21,4 | + | 2,1 | 6,87E-06 | 3,13E-04 |
| sequence-specific double-stranded DNA binding (GO:1990837) | 1117 | 62 | 29,51 | + | 2,1 | 1,11E-07 | 1,01E-05 |
| double-stranded DNA binding (GO:0003690) | 1164 | 63 | 30,76 | + | 2,05 | 1,98E-07 | 1,55E-05 |
| sequence-specific DNA binding (GO:0043565) | 1148 | 62 | 30,33 | + | 2,04 | 2,76E-07 | 1,88E-05 |
| signaling receptor binding (GO:0005102) | 698 | 36 | 18,44 | + | 1,95 | 3,01E-04 | 8,22E-03 |
| molecular transducer activity (GO:0060089) | 1085 | 55 | 28,67 | + | 1,92 | 7,50E-06 | 3,15E-04 |
| signaling receptor activity (GO:0038023) | 1085 | 55 | 28,67 | + | 1,92 | 7,50E-06 | 2,93E-04 |
| transcription regulator activity (GO:0140110) | 1265 | 62 | 33,42 | + | 1,85 | 5,82E-06 | 2,89E-04 |
| DNA binding (GO:0003677) | 1361 | 64 | 35,96 | + | 1,78 | 1,26E-05 | 4,59E-04 |
| transmembrane signaling receptor activity (GO:0004888) | 684 | 32 | 18,07 | + | 1,77 | 2,48E-03 | 4,36E-02 |
| binding (GO:0005488) | 5893 | 199 | 155,7 | + | 1,28 | 8,09E-05 | 2,60E-03 |
| RNA binding (GO:0003723) | 617 | 4 | 16,3 | - | 0,25 | 6,85E-04 | 1,50E-02 |

**KO-NSCs**

| | |
|---|---|
| Analysis Type: | PANTHER Overrepresentation Test (Released 20221013) |
| Annotation Version and Release Date: | PANTHER version 17.0 Released 2022-02-22 |
| Analyzed List: | KO-NSCs (Homo sapiens) |
| Reference List: | Homo sapiens (all genes in database) |
| Test Type: | FISHER |
| Correction: | FDR |

| **PANTHER GO-Slim Molecular Function** | Homo sapiens - REFLIST (20589) | KO NSCs | expected | over/ under | fold Enrichment | raw P-value | FDR |
|---|---|---|---|---|---|---|---|
| metalloendopeptidase activity (GO:0004222) | 84 | 26 | 6,2 | + | 4,19 | 2,03E-08 | 5,82E-07 |
| integrin binding (GO:0005178) | 49 | 15 | 3,62 | + | 4,15 | 2,18E-05 | 3,84E-04 |
| G protein-coupled amine receptor activity (GO:0008227) | 35 | 10 | 2,58 | + | 3,87 | 7,94E-04 | 9,63E-03 |
| cytokine binding (GO:0019955) | 67 | 18 | 4,95 | + | 3,64 | 1,53E-05 | 2,79E-04 |
| metallopeptidase activity (GO:0008237) | 116 | 31 | 8,56 | + | 3,62 | 1,68E-08 | 5,11E-07 |
| growth factor binding (GO:0019838) | 35 | 9 | 2,58 | + | 3,48 | 2,65E-03 | 2,58E-02 |
| growth factor activity (GO:0008083) | 40 | 10 | 2,95 | + | 3,39 | 1,87E-03 | 1,92E-02 |
| cytokine receptor activity (GO:0004896) | 49 | 12 | 3,62 | + | 3,32 | 7,95E-04 | 9,44E-03 |
| cell adhesion molecule binding (GO:0050839) | 141 | 32 | 10,41 | + | 3,07 | 2,51E-07 | 5,72E-06 |
| cytokine activity (GO:0005125) | 131 | 29 | 9,67 | + | 3 | 1,39E-06 | 2,82E-05 |
| cadherin binding (GO:0045296) | 50 | 11 | 3,69 | + | 2,98 | 2,70E-03 | 2,58E-02 |
| UDP-glycosyltransferase activity (GO:0008194) | 62 | 13 | 4,58 | + | 2,84 | 1,68E-03 | 1,76E-02 |
| G protein-coupled peptide receptor activity (GO:0008528) | 78 | 15 | 5,76 | + | 2,6 | 1,64E-03 | 1,75E-02 |
| receptor ligand activity (GO:0048018) | 245 | 47 | 18,09 | + | 2,6 | 6,03E-08 | 1,65E-06 |
| peptide receptor activity (GO:0001653) | 84 | 16 | 6,2 | + | 2,58 | 2,06E-03 | 2,09E-02 |
| signaling receptor activator activity (GO:0030546) | 248 | 47 | 18,31 | + | 2,57 | 7,27E-08 | 1,73E-06 |
| signaling receptor regulator activity (GO:0030545) | 276 | 52 | 20,38 | + | 2,55 | 1,52E-08 | 4,89E-07 |
| transmembrane receptor protein kinase activity (GO:0019199) | 64 | 12 | 4,72 | + | 2,54 | 5,48E-03 | 4,75E-02 |
| calcium ion binding (GO:0005509) | 168 | 30 | 12,4 | + | 2,42 | 4,80E-05 | 7,70E-04 |
| neurotransmitter receptor activity (GO:0030594) | 101 | 17 | 7,46 | + | 2,28 | 3,13E-03 | 2,95E-02 |
| neurotransmitter binding (GO:0042165) | 102 | 17 | 7,53 | + | 2,26 | 5,19E-03 | 4,57E-02 |

| | | | | | | | |
|---|---|---|---|---|---|---|---|
| G protein-coupled receptor activity (GO:0004930) | 269 | 42 | 19,86 | + | 2,11 | 3,55E-05 | 6,05E-04 |
| DNA-binding transcription factor activity, RNA polymerase II-specific (GO:0000981) | 984 | 152 | 72,64 | + | 2,09 | 1,10E-15 | 5,98E-13 |
| peptide binding (GO:0042277) | 182 | 28 | 13,44 | + | 2,08 | 8,10E-04 | 9,41E-03 |
| signaling receptor binding (GO:0005102) | 698 | 106 | 51,53 | + | 2,06 | 8,67E-11 | 3,16E-09 |
| gated channel activity (GO:0022836) | 209 | 31 | 15,43 | + | 2,01 | 7,12E-04 | 8,83E-03 |
| RNA polymerase II cis-regulatory region sequence-specific DNA binding (GO:0000978) | 798 | 118 | 58,91 | + | 2 | 3,39E-11 | 1,54E-09 |
| molecular transducer activity (GO:0060089) | 1085 | 160 | 80,1 | + | 2 | 7,41E-15 | 2,02E-12 |
| signaling receptor activity (GO:0038023) | 1085 | 160 | 80,1 | + | 2 | 7,41E-15 | 1,35E-12 |
| RNA polymerase II transcription regulatory region sequence-specific DNA binding (GO:0000977) | 1059 | 156 | 78,18 | + | 2 | 1,56E-14 | 2,13E-12 |
| DNA-binding transcription factor activity (GO:0003700) | 1054 | 155 | 77,81 | + | 1,99 | 2,17E-14 | 2,37E-12 |
| G protein-coupled receptor binding (GO:0001664) | 157 | 23 | 11,59 | + | 1,98 | 4,27E-03 | 3,82E-02 |
| cis-regulatory region sequence-specific DNA binding (GO:0000987) | 810 | 118 | 59,8 | + | 1,97 | 7,27E-11 | 3,05E-09 |
| endopeptidase activity (GO:0004175) | 324 | 47 | 23,92 | + | 1,96 | 4,60E-05 | 7,61E-04 |
| cation binding (GO:0043169) | 325 | 47 | 23,99 | + | 1,96 | 6,89E-05 | 1,08E-03 |
| amide binding (GO:0033218) | 201 | 29 | 14,84 | + | 1,95 | 1,47E-03 | 1,61E-02 |
| transcription cis-regulatory region binding (GO:0000976) | 1099 | 156 | 81,13 | + | 1,92 | 2,46E-13 | 1,68E-11 |
| transcription regulatory region nucleic acid binding (GO:0001067) | 1099 | 156 | 81,13 | + | 1,92 | 2,46E-13 | 1,49E-11 |
| sequence-specific double-stranded DNA binding (GO:1990837) | 1117 | 157 | 82,46 | + | 1,9 | 5,42E-13 | 2,96E-11 |
| double-stranded DNA binding (GO:0003690) | 1164 | 163 | 85,93 | + | 1,9 | 1,90E-13 | 1,48E-11 |
| metal ion binding (GO:0046872) | 268 | 37 | 19,79 | + | 1,87 | 8,24E-04 | 9,37E-03 |
| ion channel activity (GO:0005216) | 319 | 44 | 23,55 | + | 1,87 | 2,27E-04 | 3,34E-03 |
| cation channel activity (GO:0005261) | 261 | 36 | 19,27 | + | 1,87 | 1,06E-03 | 1,18E-02 |
| sequence-specific DNA binding (GO:0043565) | 1148 | 158 | 84,75 | + | 1,86 | 1,63E-12 | 8,09E-11 |
| passive transmembrane transporter activity (GO:0022803) | 338 | 45 | 24,95 | + | 1,8 | 4,67E-04 | 6,22E-03 |
| channel activity (GO:0015267) | 338 | 45 | 24,95 | + | 1,8 | 4,67E-04 | 6,07E-03 |
| transmembrane signaling receptor activity (GO:0004888) | 684 | 90 | 50,5 | + | 1,78 | 1,01E-06 | 2,12E-05 |
| transcription regulator activity (GO:0140110) | 1265 | 163 | 93,39 | + | 1,75 | 8,63E-11 | 3,37E-09 |
| peptidase activity (GO:0008233) | 433 | 55 | 31,97 | + | 1,72 | 2,77E-04 | 3,98E-03 |
| DNA binding (GO:0003677) | 1361 | 170 | 100,48 | + | 1,69 | 2,54E-10 | 8,67E-09 |
| molecular function regulator (GO:0098772) | 821 | 100 | 60,61 | + | 1,65 | 5,30E-06 | 1,03E-04 |

| | | | | | | | |
|---|---|---|---|---|---|---|---|
| ion binding (GO:0043167) | 741 | 86 | 54,7 | + | 1,57 | 1,11E-04 | 1,68E-03 |
| ion transmembrane transporter activity (GO:0015075) | 530 | 59 | 39,13 | + | 1,51 | 3,72E-03 | 3,39E-02 |
| protein binding (GO:0005515) | 2812 | 282 | 207,6 | + | 1,36 | 3,25E-07 | 7,11E-06 |
| binding (GO:0005488) | 5893 | 574 | 435,06 | + | 1,32 | 1,53E-13 | 1,39E-11 |
| nucleic acid binding (GO:0003676) | 1974 | 188 | 145,73 | + | 1,29 | 6,59E-04 | 8,37E-03 |
| heterocyclic compound binding (GO:1901363) | 2375 | 223 | 175,34 | + | 1,27 | 3,79E-04 | 5,17E-03 |
| organic cyclic compound binding (GO:0097159) | 2412 | 226 | 178,07 | + | 1,27 | 3,57E-04 | 5,00E-03 |
| molecular_function (GO:0003674) | 9609 | 819 | 709,39 | + | 1,15 | 6,25E-08 | 1,63E-06 |
| Unclassified (UNCLASSIFIED) | 10980 | 701 | 810,61 | - | 0,86 | 6,25E-08 | 1,55E-06 |
| RNA binding (GO:0003723) | 617 | 18 | 45,55 | - | 0,4 | 7,28E-06 | 1,37E-04 |
| catalytic activity, acting on a nucleic acid (GO:0140640) | 311 | 9 | 22,96 | - | 0,39 | 2,41E-03 | 2,39E-02 |
| ubiquitin-protein transferase activity (GO:0004842) | 225 | 6 | 16,61 | - | 0,36 | 5,81E-03 | 4,95E-02 |
| ubiquitin-like protein transferase activity (GO:0019787) | 239 | 6 | 17,64 | - | 0,34 | 3,22E-03 | 2,98E-02 |
