## Supplementary Table 5 for "Huntingtin loss-of-function contributes to transcriptional deregulation in Huntington’s disease"

**Supplementary Table 5. Lists of deregulated miRNAs in HD and KO-NSCs**

**HD-NSCs vs IC1-NSCs**

| Sequence | Name | log2FC | Fold_change | Adjusted_p-value |
|---|---|---|---|---|
| ACAGCAGGCACAGACAGGCAGT | hsa-miR-214-3p | 7,941153153 | 245,7679796 | 9,52E-17 |
| CCCAGTGTTCAGACTACCTGTTC | hsa-miR-199a-5p | 7,83398958 | 228,173839 | 7,39E-46 |
| TGCCTGTCTACACTTGCTGTGC | hsa-miR-214-5p | 7,47002219 | 177,2967389 | 0,000000665 |
| ACAGTAGTCTGCACATTGGTTA | hsa-miR-199a-3p#hsa-miR-199b-3p | 5,540193224 | 46,53335247 | 9,28E-21 |
| TGAGATGAAGCACTGTAGCTC | hsa-miR-143-3p | 5,189140553 | 36,48269885 | 2,91E-23 |
| CTGGACTGAGCCGTGCTACTGG | hsa-miR-1269a | 4,702871658 | 26,04386496 | 3,24E-26 |
| GTCCAGTTTTCCCAGGAATCCCT | hsa-miR-145-5p | 4,238871657 | 18,88110975 | 8,34E-13 |
| CCCAGTGTTTAGACTATCTGTTC | hsa-miR-199b-5p | 3,824050047 | 14,16295154 | 1,07E-08 |
| TCCTTCATTCCACCGGAGTCTG | hsa-miR-205-5p | 3,148567217 | 8,867744582 | 0,0000102 |
| CAAAACGTGAGGCGCTGCTAT | hsa-miR-424-3p | 2,707859089 | 6,533513754 | 9,52E-08 |
| TTCATTCGGCTGTCCAGATGTA | hsa-miR-1298-5p | 2,355441903 | 5,117509598 | 0,017767558 |
| CAGCAGCAATTCATGTTTTGAA | hsa-miR-424-5p | 2,171015418 | 4,503402473 | 0,00000142 |
| TTGCATATGTAGGATGTCCCAT | hsa-miR-448 | 2,010563674 | 4,029396215 | 0,020279128 |
| TCGGGGATCATCATGTCACGAGA | hsa-miR-542-5p | 1,674766031 | 3,192675731 | 0,000244137 |
| CGGGTGGATCACGATGCAATTT | hsa-miR-363-5p | 1,671664514 | 3,185819465 | 0,027074773 |
| TGGAGTGTGACAATGGTGTTTG | hsa-miR-122-5p | 1,665429827 | 3,172081474 | 0,328422226 |
| TGATATGTTTGATATATTAGGT | hsa-miR-190a-5p | 1,495138135 | 2,818911403 | 0,048873964 |
| TTTAGAGACGGGGTCTTGCTCT | hsa-miR-1303 | 1,475422108 | 2,780649891 | 0,025722883 |
| TTTTGCAATATGTTCCTGAATA | hsa-miR-450b-5p | 1,39255935 | 2,625440227 | 0,000340533 |
| TGAGTACCGCCATGTCTGTTGGG | hsa-miR-1911-5p | 1,368875263 | 2,582691389 | 0,532963607 |
| CATTATTACTTTTGGTACGCG | hsa-miR-126-5p | 1,361673456 | 2,569830946 | 0,013711767 |
| GCAGTCCATGGGCATATACAC | hsa-miR-455-3p | 1,336953945 | 2,52617388 | 0,094039558 |
| CAAAGTGCTCATAGTGCAGGTAG | hsa-miR-20b-5p | 1,316415355 | 2,490465378 | 0,00000142 |
| GGGGTTCCTGGGGATGGGATTT | hsa-miR-23a-5p | 1,296044749 | 2,455547532 | 0,196655153 |
| TTTGTTCGTTCGGCTCGCGTGA | hsa-miR-375-3p | 1,237211081 | 2,357423706 | 0,236237735 |
| TTTGGCAATGGTAGAACTCACACT | hsa-miR-182-5p | 1,208622693 | 2,311168896 | 0,004007851 |
| TAAGTGCTTCCATGTTTCAGTGG | hsa-miR-302c-3p | 1,153470226 | 2,224483235 | 0,402551647 |
| AAAAGTGCTTACAGTGCAGGTAG | hsa-miR-106a-5p | 1,094943232 | 2,136046764 | 0,0000913 |
| TGAGAACTGAATTCCATAGGCTG | hsa-miR-146b-5p | 1,083997101 | 2,119901307 | 0,001301312 |
| CTCCTGACTCCAGGTCCTGTGT | hsa-miR-378a-5p | 1,066065418 | 2,0937155 | 0,297249769 |

| | | | | |
|---|---|---|---|---|
| GCCCTGTGGACTCAGTTCTGGT | hsa-miR-146b-3p | 1,046558543 | 2,065596617 | 0,020279128 |
| TAAGGTGCATCTAGTGCAGTTAG | hsa-miR-18b-5p | 1,040300353 | 2,056655782 | 0,093403293 |
| TATGGCACTGGTAGAATTCACT | hsa-miR-183-5p | 0,974568862 | 1,965053869 | 0,274244792 |
| TGTGACAGATTGATAACTGAAA | hsa-miR-542-3p | 0,974454374 | 1,964897934 | 0,094039558 |
| TTTGGTCCCCTTCAACCAGCTG | hsa-miR-133a-3p | 0,96002109 | 1,945338333 | 0,563740504 |
| ACTGACAGGAGAGCATTTTGA | hsa-miR-3660 | 0,951409439 | 1,93376092 | 0,678132372 |
| TGGAATGTAAAGAAGTATGTAT | hsa-miR-1-3p | 0,940936279 | 1,919773728 | 0,317540786 |
| TATGTGCCTTTGGACTACATCG | hsa-miR-455-5p | 0,934356374 | 1,911037878 | 0,099105096 |
| CAAAACTGGCAATTACTTTTGC | hsa-miR-548a-3p | 0,902072468 | 1,868748561 | 0,41738564 |
| TGAAGGTCTACTGTGTGCCAGG | hsa-miR-493-3p | 0,873125036 | 1,83162611 | 0,632712806 |
| ACTGGACTTGGAGTCAGAAGGC | hsa-miR-378a-3p | 0,841966604 | 1,792491905 | 0,008238257 |
| TGAGGTAGTAGGTTGTGTGGTT | hsa-let-7b-5p | 0,828056039 | 1,775291631 | 0,845937371 |
| CTCCCACATGCAGGGTTTGCA | hsa-miR-188-3p | 0,810936243 | 1,754349566 | 0,317540786 |
| GTGACATCACATATACGGCAGC | hsa-miR-489-3p | 0,802515344 | 1,744139391 | 0,578660922 |
| AAAGTTCTGAGACACTCCGACT | hsa-miR-148a-5p | 0,76377588 | 1,697928702 | 0,060245312 |
| ACTGTAGTATGGGCACTTCCAG | hsa-miR-20b-3p | 0,733782642 | 1,66299363 | 0,585308588 |
| TGTTGTACTTTTTTTTTTGTTC | hsa-miR-3613-5p | 0,731500339 | 1,6603649 | 0,49395756 |
| GCAGGAACTTGTGAGTCTCCT | hsa-miR-873-5p | 0,729197942 | 1,657717237 | 0,512116215 |
| TGCCCTAAATGCCCCTTCTGGC | hsa-miR-18b-3p | 0,723096823 | 1,650721598 | 0,402551647 |
| CGGCCCCACGCACCAGGGTAAGA | hsa-miR-874-5p | 0,717422471 | 1,644241795 | 0,328422226 |
| ATCATAGAGGAAAATCCACGT | hsa-miR-376a-3p | 0,71629268 | 1,642954675 | 0,698597151 |
| TGGGTTTACGTTGGGAGAACT | hsa-miR-629-5p | 0,702575383 | 1,627407318 | 0,291670269 |
| CTGGGAGGTGGATGTTTACTTC | hsa-miR-30b-3p | 0,668311411 | 1,5892118 | 0,563740504 |
| ATGCACCTGGGCAAGGATTCTG | hsa-miR-500a-3p | 0,66784711 | 1,588700428 | 0,317540786 |
| CACGCTCATGCACACACCCACA | hsa-miR-574-3p | 0,663504268 | 1,583925267 | 0,317540786 |
| TATGTGGGATGGTAAACCGCTT | hsa-miR-299-3p | 0,65295527 | 1,572385833 | 0,846300755 |
| TTCACAGTGGCTAAGTTCCGC | hsa-miR-27a-3p | 0,643524213 | 1,562140496 | 0,12273694 |
| AATTGCACGGTATCCATCTGTA | hsa-miR-363-3p | 0,631972867 | 1,549682716 | 0,105753321 |
| TTTTGCGATGTGTTCCTAATAT | hsa-miR-450a-5p | 0,629731412 | 1,547276909 | 0,49395756 |
| TAGTAGACCGTATAGCGTACG | hsa-miR-411-5p | 0,621304805 | 1,538265794 | 0,539209501 |
| AGGCGGAGACTTGGGCAATTG | hsa-miR-25-5p | 0,605430589 | 1,521432775 | 0,766989224 |
| AGGGCTTAGCTGCTTGTGAGCA | hsa-miR-27a-5p | 0,603570316 | 1,519472238 | 0,549540111 |
| ACAGGTGAGGTTCTTGGGAGCC | hsa-miR-125a-3p | 0,585379687 | 1,50043382 | 0,483998812 |

| | | | | |
|---|---|---|---|---|
| TATTGCACTTGTCCCGGCCTGT | hsa-miR-92a-3p | 0,580865434 | 1,495746238 | 0,402551647 |
| CATGCCTTGAGTGTAGGACCGT | hsa-miR-532-5p | 0,577180138 | 1,491930301 | 0,143445685 |
| TGAGTGTGTGTGTGTGAGTGTGT | hsa-miR-574-5p | 0,568731384 | 1,483218747 | 0,529934047 |
| TGAGAACTGAATTCCATGGGTT | hsa-miR-146a-5p | 0,555282636 | 1,469456484 | 0,634026883 |
| CTGCCCTGGCCCGAGGGACCGA | hsa-miR-874-3p | 0,542682985 | 1,456678991 | 0,324725905 |
| TACCCATTGCATATCGGAGTTG | hsa-miR-660-5p | 0,533057731 | 1,446992788 | 0,317540786 |
| GGAGACTGATGAGTTCCCGGGA | hsa-miR-873-3p | 0,527044011 | 1,440973707 | 0,704500821 |
| TGTCTTGCAGGCCGTCATGCA | hsa-miR-431-5p | 0,526352985 | 1,44028367 | 0,902114121 |
| TTGTGCTTGATCTAACCATGT | hsa-miR-218-5p | 0,526311129 | 1,440241885 | 0,450756185 |
| AGTGCCTGAGGGAGTAAGAGCCC | hsa-miR-550a-5p | 0,522866035 | 1,436806752 | 0,442956856 |
| TGGACGGAGAACTGATAAGGGT | hsa-miR-184 | 0,522038595 | 1,435982926 | 0,678132372 |
| ATCACATTGCCAGGGATTTCC | hsa-miR-23a-3p | 0,514883503 | 1,428878754 | 0,360064457 |
| TGGGTCTTTGCGGGCGAGATGA | hsa-miR-193a-5p | 0,504823244 | 1,418949499 | 0,694349795 |
| AGGTTGGGATCGGTTGCAATGCT | hsa-miR-92a-1-5p | 0,504027409 | 1,418166978 | 0,669141799 |
| GTCATACACGGCTCTCCTCTCT | hsa-miR-485-3p | 0,499568152 | 1,413790303 | 0,930361142 |
| CCTTCACTGTGACTCTGCTGCAG | hsa-miR-6837-3p | 0,486024432 | 1,400580039 | 0,699202027 |
| AAACAAACATGGTGCACTTCTT | hsa-miR-495-3p | 0,479508632 | 1,394268711 | 0,810365248 |
| TGGCTCAGTTCAGCAGGAACAG | hsa-miR-24-3p | 0,458877412 | 1,3744719 | 0,532963607 |
| TGCACGGCACTGGGGACACGT | hsa-miR-3177-3p | 0,45353952 | 1,369395826 | 0,699202027 |
| AATCCTTGGAACCTAGGTGTGAGT | hsa-miR-362-5p | 0,448442205 | 1,364566028 | 0,619739175 |
| TTCTCGAGGAAAGAAGCACTTTC | hsa-miR-516a-5p | 0,445742392 | 1,362014811 | 0,894939919 |
| AATGCACCCGGGCAAGGATTCT | hsa-miR-501-3p | 0,439473485 | 1,356109322 | 0,398428398 |
| AAGCTGCCAGTTGAAGAACTGT | hsa-miR-22-3p | 0,431689804 | 1,348812493 | 0,400732344 |
| TCAGTGCACTACAGAACTTTGT | hsa-miR-148a-3p | 0,426729172 | 1,344182636 | 0,539209501 |
| TGTCCTCTAGGGCCTGCAGTCT | hsa-miR-3909 | 0,408810985 | 1,327591212 | 0,589535041 |
| ATCGGGAATGTCGTGTCCGCCC | hsa-miR-425-3p | 0,40314626 | 1,322388662 | 0,751898825 |
| CTGGAGATATGGAAGAGCTGTGT | hsa-miR-1270 | 0,398944341 | 1,318542744 | 0,559237537 |
| AATAATACATGGTTGATCTTT | hsa-miR-369-3p | 0,397890596 | 1,317580032 | 0,845937371 |
| AGGCTGTGATGCTCTCCTGAGCCC | hsa-miR-7974 | 0,355134071 | 1,279104444 | 0,780780131 |
| CAGTGGTTTTACCCTATGGTAG | hsa-miR-140-5p | 0,355000812 | 1,278986301 | 0,728021243 |
| CTGGGAGAGGGTTGTTTACTCC | hsa-miR-30c-1-3p | 0,352836926 | 1,277069398 | 0,738867108 |
| TATACAAGGGCAAGCTCTCTGT | hsa-miR-381-3p | 0,351111285 | 1,275542779 | 0,885318949 |
| AAAGACATAGGATAGAGTCACCTC | hsa-miR-641 | 0,343689943 | 1,26899813 | 0,845937371 |

| | | | | |
|---|---|---|---|---|
| AATATAACACAGATGGCCTGT | hsa-miR-410-3p | 0,343638382 | 1,268952777 | 0,874993347 |
| TGTGCAAATCCATGCAAAACTGA | hsa-miR-19b-3p | 0,339562128 | 1,265372483 | 0,736897829 |
| CTGGGCCCGCGGCGGGCGTGGGG | hsa-miR-6724-5p | 0,33343727 | 1,260011822 | 0,878939539 |
| TAAGTGCTTCCATGTTTGAGTGT | hsa-miR-302d-3p | 0,333354954 | 1,259939932 | 0,932974149 |
| TCCGGTTCTCAGGGCTCCACC | hsa-miR-671-3p | 0,330092132 | 1,257093651 | 0,626625979 |
| ATCATACAAGGACAATTTCTTT | hsa-miR-539-3p | 0,32100104 | 1,249197026 | 0,931755298 |
| AGGTTCTGTGATACACTCCGACT | hsa-miR-152-5p | 0,319916465 | 1,24825827 | 0,827028048 |
| TGAGGGGCAGAGAGCGAGACTTT | hsa-miR-423-5p | 0,304719814 | 1,23517873 | 0,917226903 |
| CTTTCAGTCGGATGTTTACAGC | hsa-miR-30e-3p | 0,29732746 | 1,228865874 | 0,678132372 |
| CGGGGCCGTAGCACTGTCTGAGA | hsa-miR-128-1-5p | 0,295063629 | 1,226939089 | 0,72639342 |
| CTAGACTGAAGCTCCTTGAGG | hsa-miR-151a-3p | 0,29386007 | 1,22591595 | 0,624506689 |
| ATTCTAATTTCTCCACGTCTTT | hsa-miR-576-5p | 0,293051199 | 1,225228813 | 0,846300755 |
| CAAAGTGCTTACAGTGCAGGTAG | hsa-miR-17-5p | 0,292678007 | 1,224911915 | 0,626625979 |
| TTGTACATGGTAGGCTTTCATT | hsa-miR-493-5p | 0,290512558 | 1,223074732 | 0,931755298 |
| TAGGAGCTCAACAGATGCCTGTT | hsa-miR-3139 | 0,278528025 | 1,212956679 | 0,931172169 |
| CCAGTCCTGTGCCTGCCGCCT | hsa-miR-1910-5p | 0,27385904 | 1,20903754 | 0,874993347 |
| CCTATTCTTGATTACTTGTTTC | hsa-miR-26a-2-3p | 0,267441689 | 1,20367148 | 0,704500821 |
| AGATCAGAAGGTGATTGTGGCT | hsa-miR-383-5p | 0,265402105 | 1,201971014 | 0,931172169 |
| CACATTACACGGTCGACCTCT | hsa-miR-323a-3p | 0,257951451 | 1,195779556 | 0,893644373 |
| CCAAAACTGCAGTTACTTTTGC | hsa-miR-548o-3p | 0,252234788 | 1,191050668 | 0,766989224 |
| AAAAGCTGGGTTGAGAGGGCGA | hsa-miR-320a-3p | 0,250459137 | 1,18958564 | 0,799093566 |
| CCACCTCCCCTGCAAACGTCCA | hsa-miR-1306-5p | 0,237489762 | 1,178939563 | 0,919777469 |
| TAAAGTGCTTATAGTGCAGGTAG | hsa-miR-20a-5p | 0,209483086 | 1,15627382 | 0,752477997 |
| CTTCCGGTCTGTGAGCCCCGTC | hsa-miR-4664-3p | 0,204298375 | 1,152125898 | 0,952638859 |
| CATCATCGTCTCAAATGAGTCT | hsa-miR-136-3p | 0,202081377 | 1,150356779 | 0,935917968 |
| AAGTTCTGTTATACACTCAGGC | hsa-miR-148b-5p | 0,200575422 | 1,149156607 | 0,932974149 |
| TGCTGGATCAGTGGTTCGAGTC | hsa-miR-1287-5p | 0,20047178 | 1,149074056 | 0,810365248 |
| TCCCTGAGACCCTTTAACCTGTGA | hsa-miR-125a-5p | 0,200323957 | 1,148956324 | 0,931172169 |
| AAGGGCTTCCTCTCTGCAGGAC | hsa-miR-3158-3p | 0,199261782 | 1,148110724 | 0,932974149 |
| TCTGGGCAACAAAGTGAGACCT | hsa-miR-1285-3p | 0,198856454 | 1,147788205 | 0,916906684 |
| TGCGGGGCTAGGGCTAACAGCA | hsa-miR-744-5p | 0,196028412 | 1,145540458 | 0,806173445 |
| TTAGCCAATTGTCCATCTTTAG | hsa-miR-4662a-5p | 0,190715787 | 1,141329841 | 0,931172169 |
| TGTCTTACTCCCTCAGGCACAT | hsa-miR-550a-3p | 0,189950848 | 1,140724851 | 0,846300755 |

| | | | | |
|---|---|---|---|---|
| CCGGTCCCAGGAGAACCTGCAGA | hsa-miR-4746-5p | 0,188423192 | 1,139517588 | 0,898227653 |
| AGGCAGTGTAGTTAGCTGATTGC | hsa-miR-34c-5p | 0,184330035 | 1,136289176 | 0,932822306 |
| TCCTGTCTTTCCTTGTTGGAGC | hsa-miR-5699-3p | 0,183812092 | 1,135881309 | 0,935917968 |
| ACCCGTCCCGTTCGTCCCCGGA | hsa-miR-1247-5p | 0,180344657 | 1,133154561 | 0,952638859 |
| GAATGTTGCTCGGTGAACCCCT | hsa-miR-409-3p | 0,175714642 | 1,129523778 | 0,952638859 |
| CTAGGTATGGTCCCAGGGATCC | hsa-miR-331-5p | 0,175065726 | 1,129015839 | 0,952638859 |
| CCAGTTACCGCTTCCGCTACCGC | hsa-miR-935 | 0,16753402 | 1,123137079 | 0,952638859 |
| CGGGCGTGGTGGTGGGGG | hsa-miR-1268a | 0,166434066 | 1,122281092 | 0,932974149 |
| TGGAGAGAAAGGCAGTTCCTGA | hsa-miR-185-5p | 0,165967169 | 1,121917949 | 0,932974149 |
| AATCATACAGGGACATCCAGTT | hsa-miR-487a-3p | 0,162494028 | 1,119220295 | 0,952638859 |
| CCTCCCACACCCAAGGCTTGCA | hsa-miR-532-3p | 0,162009724 | 1,118844642 | 0,952638859 |
| TTAGGGCCCTGGCTCCATCTCC | hsa-miR-1296-5p | 0,153060164 | 1,111925527 | 0,932822306 |
| TTAATATCGGACAACCATTGT | hsa-miR-889-3p | 0,151800286 | 1,110954927 | 0,952638859 |
| GCTCTGACGAGGTTGCACTACT | hsa-miR-301b-5p | 0,151237815 | 1,110521878 | 0,952638859 |
| CCGCACTGTGGGTACTTGCTGC | hsa-miR-106b-3p | 0,150960465 | 1,110308407 | 0,902114121 |
| CAACTAGACTGTGAGCTTCTAG | hsa-miR-708-3p | 0,14618743 | 1,106641115 | 0,952638859 |
| TAGCAGCACAGAAATATTGGC | hsa-miR-195-5p | 0,145208429 | 1,105890412 | 0,94993674 |
| ATGACCTATGAATTGACAGAC | hsa-miR-215-5p | 0,144438079 | 1,105300062 | 0,952638859 |
| TTTCCGGCTCGCGTGGGTGTGT | hsa-miR-1180-3p | 0,141306868 | 1,102903731 | 0,915097122 |
| AGCTCGGTCTGAGGCCCCTCAGT | hsa-miR-423-3p | 0,138427844 | 1,100704985 | 0,932974149 |
| CTTTCAGTCAGATGTTTGCTGC | hsa-miR-30d-3p | 0,134978293 | 1,098076292 | 0,931172169 |
| ACTCGGCGTGGCGTCGGTCGTG | hsa-miR-1307-3p | 0,131586232 | 1,09549753 | 0,928426431 |
| TTCAAGTAATCCAGGATAGGCT | hsa-miR-26a-5p | 0,131039804 | 1,095082684 | 0,931172169 |
| AGAGCTTAGCTGATTGGTGAAC | hsa-miR-27b-5p | 0,126904596 | 1,091948334 | 0,952638859 |
| GTGGGGGAGAGGCTGTC | hsa-miR-1275 | 0,126528721 | 1,091663878 | 0,952638859 |
| CAAAGAATTCTCCTTTTGGGCT | hsa-miR-186-5p | 0,121458086 | 1,087833746 | 0,917226903 |
| TCCTGTACTGAGCTGCCCCGAG | hsa-miR-486-5p | 0,111634133 | 1,080451365 | 0,952638859 |
| TCGAGGAGCTCACAGTCT | hsa-miR-151b | 0,10729603 | 1,077207388 | 0,952638859 |
| TGGTAGACTATGGAACGTAGG | hsa-miR-379-5p | 0,102335847 | 1,073510161 | 0,971445722 |
| TGGCAGTGTATTGTTAGCTGGT | hsa-miR-449a | 0,101814839 | 1,073122549 | 0,967918429 |
| CCTCCGTGTTACCTGTCCTCTAG | hsa-miR-3605-3p | 0,100528226 | 1,072165952 | 0,967918429 |
| AATATTATACAGTCAACCTCT | hsa-miR-656-3p | 0,097609431 | 1,069998987 | 0,971445722 |
| AGGACCTTCCCTGAACCAAGGA | hsa-miR-659-5p | 0,092625161 | 1,066308698 | 0,971445722 |

| | | | | |
|---|---|---|---|---|
| GCCTGCTGGGGTGGAACCTGGT | hsa-miR-370-3p | 0,091925833 | 1,065791944 | 0,967918429 |
| AATGACACGATCACTCCCGTTGA | hsa-miR-425-5p | 0,087117576 | 1,062245753 | 0,952638859 |
| TCGAGGAGCTCACAGTCTAGT | hsa-miR-151a-5p | 0,084681208 | 1,060453387 | 0,952638859 |
| CCAATATTACTGTGCTGCTTTA | hsa-miR-16-2-3p | 0,083599205 | 1,059658359 | 0,967918429 |
| TGAGTATTACATGGCCAATCTC | hsa-miR-496 | 0,082679364 | 1,058982952 | 0,97690947 |
| ACTGCAGTGAAGGCACTTGTAG | hsa-miR-17-3p | 0,080643422 | 1,057489561 | 0,967918429 |
| TGGAGACGCGGCCCTGTTGGAGT | hsa-miR-139-3p | 0,075360841 | 1,053624531 | 0,971445722 |
| CATTGCACTTGTCTCGGTCTGA | hsa-miR-25-3p | 0,075346198 | 1,053613837 | 0,952638859 |
| TGTGACTGGTTGACCAGAGGGG | hsa-miR-134-5p | 0,072598924 | 1,051609386 | 0,97690947 |
| AAACATTCGCGGTGCACTTCTT | hsa-miR-543 | 0,07131626 | 1,050674842 | 0,97690947 |
| ATCCCCAGATACAATGGACAA | hsa-miR-2355-5p | 0,065428204 | 1,046395473 | 0,97690947 |
| TCCCTGTCCTCCAGGAGCTCACG | hsa-miR-339-5p | 0,065237061 | 1,046256845 | 0,97690947 |
| TCGGATCCGTCTGAGCTTGGCT | hsa-miR-127-3p | 0,063693152 | 1,045137785 | 0,97690947 |
| CCCGCAGGTGAGATGAGGGCT | hsa-miR-6886-5p | 0,059813218 | 1,042330804 | 0,971445722 |
| AATCGTACAGGGTCATCCACTT | hsa-miR-487b-3p | 0,059270131 | 1,041938504 | 0,97690947 |
| TAGCAGCACATAATGGTTTGTG | hsa-miR-15a-5p | 0,057352887 | 1,040554758 | 0,967918429 |
| CGAATCATTATTTGCTGCTCTA | hsa-miR-15b-3p | 0,057162538 | 1,040417476 | 0,973097564 |
| TACAGTACTGTGATAACTGAA | hsa-miR-101-3p | 0,050093578 | 1,035332077 | 0,97690947 |
| TTAAGACTTGCAGTGATGTTT | hsa-miR-499a-5p | 0,049205723 | 1,034695115 | 0,988067158 |
| TGTAAACATCCTACACTCAGCT | hsa-miR-30b-5p | 0,048977796 | 1,034531659 | 0,967918429 |
| CAAAGTGCTGTTCGTGCAGGTAG | hsa-miR-93-5p | 0,04862793 | 1,034280807 | 0,971445722 |
| GACCGAGAGGGCCTCGGCTGT | hsa-miR-4523 | 0,046386897 | 1,032675438 | 0,979134231 |
| AAAAGTACTTGCGGATTTTGCT | hsa-miR-548k | 0,03659697 | 1,025691569 | 0,971445722 |
| TTCAAGTAATTCAGGATAGGT | hsa-miR-26b-5p | 0,025993115 | 1,018180341 | 0,97690947 |
| AGGCAGTGTATTGTTAGCTGGC | hsa-miR-449b-5p | 0,023471539 | 1,016402296 | 0,993639967 |
| ACTGCCCTAAGTGCTCCTTCTGG | hsa-miR-18a-3p | 0,020583132 | 1,014369401 | 0,993639967 |
| TTATCAGAATCTCCAGGGGTAC | hsa-miR-361-5p | 0,015421161 | 1,010746467 | 0,991002008 |
| CAAGCTTGTATCTATAGGTATG | hsa-miR-100-3p | 0,015173233 | 1,010572785 | 0,993639967 |
| TGTAAACATCCCCGACTGGAAG | hsa-miR-30d-5p | 0,014943558 | 1,010411916 | 0,993639967 |
| TGAGCGCCTCGACGACAGAGCCG | hsa-miR-339-3p | 0,014037076 | 1,009777248 | 0,993639967 |
| TTTGTGACCTGGTCCACTAACC | hsa-miR-758-3p | 0,009554682 | 1,00664478 | 0,99568213 |
| AATTTGGTTTCTGAGGCACTTAGT | hsa-miR-5002-5p | 0,008786699 | 1,00610906 | 0,99568213 |
| TCAGGCTCAGTCCCCTCCCGAT | hsa-miR-484 | 0,007676424 | 1,005335073 | 0,99568213 |

| | | | | |
|---|---|---|---|---|
| CAACGGAATCCCAAAAGCAGCTG | hsa-miR-191-5p | 0,004441517 | 1,003083369 | 0,99568213 |
| CAGGTCGTCTTGCAGGGCTTCT | hsa-miR-431-3p | 0,00159992 | 1,001109595 | 0,998822107 |
| CTCACTGAACAATGAATGCAA | hsa-miR-181b-3p | -0,000862313 | 0,999402469 | 0,998822107 |
| CTGACCTATGAATTGACAGCC | hsa-miR-192-5p | -0,001062005 | 0,999264145 | 0,998822107 |
| TGATATGTTTGATATTGGGTTG | hsa-miR-190b-5p | -0,005233902 | 0,996378708 | 0,998822107 |
| TTTTTCATTATTGCTCCTGACC | hsa-miR-335-3p | -0,010355744 | 0,992847646 | 0,99568213 |
| TAGCACCATTTGAAATCAGTGTT | hsa-miR-29b-3p | -0,011436336 | 0,992104272 | 0,99568213 |
| CAAGCTCGTGTCTGTGGGTCCG | hsa-miR-99b-3p | -0,019871805 | 0,986320343 | 0,993639967 |
| GCTGCGCTTGGATTTCGTCCCC | hsa-miR-191-3p | -0,020722539 | 0,985738897 | 0,99568213 |
| CACCCGTAGAACCGACCTTGCG | hsa-miR-99b-5p | -0,021661028 | 0,985097872 | 0,991002008 |
| AACATTCATTGCTGTCGGTGGGT | hsa-miR-181b-5p | -0,021956471 | 0,984896159 | 0,97690947 |
| CACCCGGCTGTGTGCACATGTGC | hsa-miR-941 | -0,023097267 | 0,98411767 | 0,97690947 |
| AACATAGAGGAAATTCCACGT | hsa-miR-376c-3p | -0,024975971 | 0,982836968 | 0,993639967 |
| ATGTAGGGCTAAAAGCCATGGG | hsa-miR-135b-3p | -0,039469505 | 0,973012669 | 0,987478179 |
| AATGGCGCCACTAGGGTTGTG | hsa-miR-652-3p | -0,041098064 | 0,971914922 | 0,97690947 |
| TGAAGCGCCTGTGCTCTGCCGAGA | hsa-miR-7706 | -0,046496373 | 0,968284984 | 0,967918429 |
| AGAATTGTGGCTGGACATCTGT | hsa-miR-219a-2-3p | -0,050771286 | 0,965420063 | 0,988726858 |
| CGGGCTGTCCGGAGGGGTCGGCT | hsa-miR-4741 | -0,055778233 | 0,962075331 | 0,97690947 |
| TAAGGTGCATCTAGTGCAGATAG | hsa-miR-18a-5p | -0,058644263 | 0,960165988 | 0,967918429 |
| TAATGCCCCTAAAAATCCTTAT | hsa-miR-365b-3p#hsa-miR-365a-3p | -0,05883723 | 0,96003757 | 0,971445722 |
| ATATACAGGGGGAGACTCTTAT | hsa-miR-1185-1-3p | -0,064855917 | 0,956040793 | 0,97690947 |
| CAGCCACAACTACCCTGCCACT | hsa-miR-449b-3p | -0,06594163 | 0,955321587 | 0,987478179 |
| TAGCAGCACGTAAATATTGGCG | hsa-miR-16-5p | -0,072269199 | 0,951140782 | 0,952638859 |
| TCGACCGGACCTCGACCGGCT | hsa-miR-1307-5p | -0,081809665 | 0,94487169 | 0,967918429 |
| AGGTTACCCGAGCAACTTTGCAT | hsa-miR-409-5p | -0,087340652 | 0,941256187 | 0,971445722 |
| CTGTTGCCACTAACCTCAACCT | hsa-miR-744-3p | -0,090238381 | 0,939367522 | 0,967918429 |
| AAGGAGCTCACAGTCTATTGAG | hsa-miR-28-5p | -0,090799634 | 0,93900215 | 0,952638859 |
| TAGCACCATCTGAAATCGGTTA | hsa-miR-29a-3p | -0,091610828 | 0,938474319 | 0,952638859 |
| CAGTGCAATGATGAAAGGGCAT | hsa-miR-130b-3p | -0,091884368 | 0,938296398 | 0,935917968 |
| TTCACAGTGGCTAAGTTCTGC | hsa-miR-27b-3p | -0,092096903 | 0,93815818 | 0,94993674 |
| TGCACCATGGTTGTCTGAGCATG | hsa-miR-767-5p | -0,095336449 | 0,936053926 | 0,95576515 |
| AGCAGCATTGTACAGGGCTATGA | hsa-miR-103a-3p | -0,095751611 | 0,935784598 | 0,931755298 |
| TAATTTTATGTATAAGCTAGT | hsa-miR-590-3p | -0,096323586 | 0,935413668 | 0,952638859 |

| | | | | |
|---|---|---|---|---|
| TGGCAGTGTCTTAGCTGGTTGT | hsa-miR-34a-5p | -0,097185095 | 0,93485525 | 0,952638859 |
| TAAAGTGCTGACAGTGCAGAT | hsa-miR-106b-5p | -0,097881966 | 0,934403792 | 0,932822306 |
| CTCCTGGGGCCCGCACTCTCGC | hsa-miR-1343-3p | -0,100040765 | 0,933006628 | 0,967918429 |
| TGGAAGACTAGTGATTTTGTTGTT | hsa-miR-7-5p | -0,100061931 | 0,93299294 | 0,952638859 |
| TCTCACACAGAAATCGCACCCGT | hsa-miR-342-3p | -0,104096153 | 0,930387647 | 0,94993674 |
| TGTGCAAATCTATGCAAAACTGA | hsa-miR-19a-3p | -0,109844931 | 0,926687662 | 0,952638859 |
| TAGGCAGTGTATTGCTAGCGGCTGT | hsa-miR-449c-5p | -0,113403066 | 0,924404978 | 0,967918429 |
| TGAGCACCACACAGGCCGGGCGC | hsa-miR-3663-3p | -0,114642524 | 0,923611138 | 0,967918429 |
| CCTCAGGGCTGTAGAACAGGGCT | hsa-miR-1266-5p | -0,11526707 | 0,923211391 | 0,952638859 |
| TAGTGAGTTAGAGATGCAGAGCC | hsa-miR-3174 | -0,115929288 | 0,922787721 | 0,967918429 |
| AACATTCAACGCTGTCGGTGAGT | hsa-miR-181a-5p | -0,118005749 | 0,921460514 | 0,940327702 |
| AAGGAGCTTACAATCTAGCTGGG | hsa-miR-708-5p | -0,129705928 | 0,91401774 | 0,94993674 |
| ACGCCCTTCCCCCCCTTCTTCA | hsa-miR-1249-3p | -0,131171481 | 0,913089712 | 0,952638859 |
| TAAGTGCTTCCATGTTTTAGTAG | hsa-miR-302b-3p | -0,13342513 | 0,911664479 | 0,967918429 |
| TAGTGCAATATTGCTTATAGGGT | hsa-miR-454-3p | -0,134661827 | 0,910883323 | 0,931172169 |
| AAACTCTACTTGTCCTTCTGAGT | hsa-miR-618 | -0,135623631 | 0,910276265 | 0,967918429 |
| CAACACCAGTCGATGGGCTGT | hsa-miR-21-3p | -0,135724747 | 0,910212468 | 0,935917968 |
| TCTTGGAGTAGGTCATTGGGTGG | hsa-miR-432-5p | -0,139863007 | 0,907605334 | 0,967918429 |
| ACTCTTTCCCTGTTGCACTAC | hsa-miR-130b-5p | -0,141352908 | 0,906668515 | 0,932974149 |
| ACCGTGGCTTTCGATTGTTACT | hsa-miR-132-5p | -0,143320513 | 0,905432807 | 0,952638859 |
| ATGTAGGGATGGAAGCCATGAA | hsa-miR-135a-2-3p | -0,146848775 | 0,903221182 | 0,932974149 |
| TGCAGGACCAAGATGAGCCCT | hsa-miR-1286 | -0,149168541 | 0,901770024 | 0,932974149 |
| CTTGGCACCTAGCAAGCACTCA | hsa-miR-1271-5p | -0,154089255 | 0,898699525 | 0,921686549 |
| ACCATCGACCGTTGATTGTACC | hsa-miR-181a-3p | -0,154797172 | 0,89825865 | 0,874993347 |
| TCAAGAGCAATAACGAAAAATGT | hsa-miR-335-5p | -0,155007897 | 0,898127457 | 0,931172169 |
| TATGGCTTTTCATTCCTATGTGA | hsa-miR-135b-5p | -0,156871383 | 0,896968121 | 0,937266994 |
| CGGCTCTGGGTCTGTGGGGA | hsa-miR-760 | -0,158819577 | 0,895757686 | 0,952638859 |
| AGAGGCTGGCCGTGATGAATTC | hsa-miR-485-5p | -0,164533177 | 0,892217174 | 0,967918429 |
| TAGGACACATGGTCTACTTCT | hsa-miR-1197 | -0,165810735 | 0,891427433 | 0,952638859 |
| TTCACCACCTTCTCCACCCAGC | hsa-miR-197-3p | -0,17375825 | 0,886530241 | 0,931172169 |
| TAGCCCCCAGGCTTCACTTGGCG | hsa-miR-3943 | -0,17646784 | 0,884866772 | 0,948384295 |
| ATCACATTGCCAGGGATTACCAC | hsa-miR-23b-3p | -0,177057518 | 0,884505171 | 0,878939539 |
| CTTTCAGTCGGATGTTTGCAGC | hsa-miR-30a-3p | -0,185413735 | 0,879396847 | 0,839694702 |

| | | | | |
|---|---|---|---|---|
| TCACAGTGAACCGGTCTCTTT | hsa-miR-128-3p | -0,188845554 | 0,877307463 | 0,845937371 |
| AGAGTTGAGTCTGGACGTCCCG | hsa-miR-219a-1-3p | -0,196419944 | 0,872713522 | 0,902114121 |
| CATCCCTTGCATGGTGGAGGG | hsa-miR-188-5p | -0,197740807 | 0,871914873 | 0,952638859 |
| TATTCATTTATCCCCAGCCTACA | hsa-miR-664a-3p | -0,200126974 | 0,870473948 | 0,94993674 |
| TGTAACAGCAACTCCATGTGGA | hsa-miR-194-5p | -0,201750815 | 0,869494728 | 0,931172169 |
| ATCCGCGCTCTGACTCTCTGCC | hsa-miR-937-3p | -0,202177837 | 0,869237405 | 0,932974149 |
| TTGCAGCTGCCTGGGAGTGACTTC | hsa-miR-1301-3p | -0,202771997 | 0,868879492 | 0,935917968 |
| TCCGTCTCAGTTACTTTATAGC | hsa-miR-340-3p | -0,214311246 | 0,861957567 | 0,791024805 |
| GATATCAGCTCAGTAGGCACCG | hsa-miR-3074-3p | -0,2220044 | 0,857373422 | 0,932974149 |
| CTCGTGGGCTCTGGCCACGGCC | hsa-miR-3677-3p | -0,226998575 | 0,854410585 | 0,781392374 |
| TTAATTTTTTGTTTCGGTCACT | hsa-miR-4775 | -0,227931801 | 0,853858077 | 0,898227653 |
| CAAGCTCGCTTCTATGGGTCTG | hsa-miR-99a-3p | -0,228995918 | 0,853228512 | 0,931755298 |
| GTGAACGGGCGCCATCCCGAGG | hsa-miR-887-3p | -0,239254801 | 0,847182797 | 0,875167309 |
| TAGCAGCACATCATGGTTTACA | hsa-miR-15b-5p | -0,241136738 | 0,846078402 | 0,699202027 |
| CAGGCAGTGACTGTTCAGACGTC | hsa-miR-2682-5p | -0,249497164 | 0,841189552 | 0,874993347 |
| TTCCCTTTGTCATCCTATGCCT | hsa-miR-204-5p | -0,249763077 | 0,841034521 | 0,810365248 |
| GCTGACTCCTAGTCCAGGGCTC | hsa-miR-345-5p | -0,250433355 | 0,840643866 | 0,845937371 |
| AGGGCCCCCCCTCAATCCTGT | hsa-miR-296-5p | -0,250666715 | 0,8405079 | 0,845937371 |
| TTGTGTCAATATGCGATGATGT | hsa-miR-592 | -0,252579934 | 0,839394006 | 0,780780131 |
| TGGTGGGCCGCAGAACATGTGC | hsa-miR-654-5p | -0,252657556 | 0,839348845 | 0,948384295 |
| CAGTGCAATGTTAAAAGGGCAT | hsa-miR-130a-3p | -0,259664668 | 0,835282045 | 0,759354533 |
| TGTAAACATCCTTGACTGGAAG | hsa-miR-30e-5p | -0,261634109 | 0,834142569 | 0,872309274 |
| AACCCGTAGATCCGAACTTGTG | hsa-miR-100-5p | -0,26599133 | 0,831627096 | 0,912361079 |
| CGGCGGGGACGGCGATTGGTC | hsa-miR-1908-5p | -0,274520563 | 0,82672501 | 0,787414452 |
| ATGCTGACATATTTACTAGAGG | hsa-miR-628-5p | -0,275490172 | 0,82616957 | 0,678132372 |
| ATCAAGGATCTTAAACTTTGCC | hsa-miR-561-5p | -0,277060547 | 0,825270773 | 0,733132978 |
| TGGAAACATTTCTGCACAAACT | hsa-miR-147b-5p | -0,283681646 | 0,821491956 | 0,931172169 |
| AATTCCCTTGTAGATAACCCGG | hsa-miR-3938 | -0,28470894 | 0,820907208 | 0,791024805 |
| AGGAAGCCCTGGAGGGGCTGGAG | hsa-miR-671-5p | -0,287730345 | 0,819189799 | 0,935917968 |
| AAGGCAGGGCCCCCGCTCCCC | hsa-miR-940 | -0,287948682 | 0,819065832 | 0,845937371 |
| CTTATCAGATTGTATTGTAATT | hsa-miR-374a-3p | -0,290098529 | 0,817846202 | 0,736897829 |
| TTTTCAACTCTAATGGGAGAGA | hsa-miR-1305 | -0,294502481 | 0,815353462 | 0,845937371 |
| TGGTCGACCAGTTGGAAAGTAAT | hsa-miR-412-5p | -0,298964802 | 0,812835433 | 0,874993347 |

| | | | | |
|---|---|---|---|---|
| CACCTTGCGCTACTCAGGTCTG | hsa-miR-3200-3p | -0,299753048 | 0,812391445 | 0,902114121 |
| TATGGAGGTCTCTGTCTGGC | hsa-miR-1843 | -0,30164721 | 0,811325529 | 0,931172169 |
| TTATAATACAACCTGATAAGTG | hsa-miR-374a-5p | -0,303313706 | 0,810388886 | 0,699202027 |
| CTTAGCAGGTTGTATTATCATT | hsa-miR-374b-3p | -0,303890229 | 0,810065107 | 0,684757038 |
| GCTCTGACTTTATTGCACTACT | hsa-miR-301a-5p | -0,305737203 | 0,809028705 | 0,734319545 |
| ACCCTATCAATATTGTCTCTGC | hsa-miR-454-5p | -0,30694888 | 0,808349511 | 0,724424064 |
| TTTAGGATAAGCTTGACTTTTG | hsa-miR-651-5p | -0,311343935 | 0,805890686 | 0,932974149 |
| TACGCGCAGACCACAGGATGTC | hsa-miR-3939 | -0,321844149 | 0,800046551 | 0,738867108 |
| AGCGCGGGCTGAGCGCTGCCAGTC | hsa-miR-2277-5p | -0,324399704 | 0,798630622 | 0,788940866 |
| TGAGACCTCTGGGTTCTGAGCT | hsa-miR-769-5p | -0,327582975 | 0,796870406 | 0,512096705 |
| AACATTCAACCTGTCGGTGAGT | hsa-miR-181c-5p | -0,331657804 | 0,794622856 | 0,896141844 |
| TGTAAACATCCTCGACTGGAAG | hsa-miR-30a-5p | -0,332407689 | 0,794209934 | 0,738867108 |
| AAAAGTAATTGCGGATTTTGCC | hsa-miR-548i | -0,333771914 | 0,793459277 | 0,827028048 |
| TTATAAAGCAATGAGACTGATT | hsa-miR-340-5p | -0,338710604 | 0,79074772 | 0,585090965 |
| TCAGAACAAATGCCGGTTCCCAGA | hsa-miR-589-3p | -0,339275293 | 0,790438272 | 0,872309274 |
| CACTAGATTGTGAGCTCCTGGA | hsa-miR-28-3p | -0,343191488 | 0,788295537 | 0,845937371 |
| TTAATGCTAATCGTGATAGGGGTT | hsa-miR-155-5p | -0,345678092 | 0,786938015 | 0,724424064 |
| TCTGTGAGACCAAAGAACTACT | hsa-miR-4677-3p | -0,356694322 | 0,780951943 | 0,691215547 |
| TCAAATGCTCAGACTCCTGTGGT | hsa-miR-105-5p | -0,356760572 | 0,780916082 | 0,733132978 |
| TACGTCATCGTTGTCATCGTCA | hsa-miR-598-3p | -0,357536054 | 0,780496434 | 0,66140515 |
| TCCCCCAGGTGTGATTCTGATTT | hsa-miR-361-3p | -0,362109627 | 0,778026054 | 0,84846502 |
| AAAAACTGAGACTACTTTTGCA | hsa-miR-548e-3p | -0,366569037 | 0,775624867 | 0,902114121 |
| ATAAGACGAACAAAAGGTTTGT | hsa-miR-208b-3p | -0,368726578 | 0,774465792 | 0,799093566 |
| ACTCCAGCCCCACAGCCTCAGC | hsa-miR-766-3p | -0,369456038 | 0,774074303 | 0,803181225 |
| ACGGGTTAGGCTCTTGGGAGCT | hsa-miR-125b-1-3p | -0,369850889 | 0,773862476 | 0,692505052 |
| TGAGAACCACGTCTGCTCTGAG | hsa-miR-589-5p | -0,381973492 | 0,767387148 | 0,738867108 |
| CTGGCCCTCTCTGCCCTTCCGT | hsa-miR-328-3p | -0,389322817 | 0,763487892 | 0,684577389 |
| CAGTGCAATGATATTGTCAAAGC | hsa-miR-301b-3p | -0,390122711 | 0,763064698 | 0,41463432 |
| CAAAAACTGCAGTTACTTTTGC | hsa-miR-548ah-3p | -0,390244922 | 0,763000061 | 0,59275089 |
| TCAGTGCATGACAGAACTTGG | hsa-miR-152-3p | -0,396834508 | 0,759522965 | 0,780780131 |
| GAAGTTGTTCGTGGTGGATTCG | hsa-miR-382-5p | -0,399658232 | 0,758037838 | 0,912361079 |
| TGACGCCCCTTCTGATTCTGCCT | hsa-miR-6786-3p | -0,400924177 | 0,757372962 | 0,913199343 |
| CAACAAATCACAGTCTGCCATA | hsa-miR-7-1-3p | -0,404211753 | 0,755649045 | 0,738867108 |

| | | | | |
|---|---|---|---|---|
| TATGGCTTTTTATTCCTATGTGA | hsa-miR-135a-5p | -0,410378746 | 0,752425816 | 0,559237537 |
| TGGGGCGGAGCTTCCGGAGGCC | hsa-miR-3180-3p | -0,411285978 | 0,751952805 | 0,845937371 |
| CCTGTTCTCCATTACTTGGCT | hsa-miR-26b-3p | -0,419781518 | 0,747537823 | 0,802507706 |
| ACCACTGACCGTTGACTGTACC | hsa-miR-181a-2-3p | -0,420357707 | 0,747239328 | 0,402551647 |
| TCAGTGCATCACAGAACTTTGT | hsa-miR-148b-3p | -0,424078469 | 0,745314654 | 0,526925062 |
| AACATTCATTGTTGTCGGTGGGT | hsa-miR-181d-5p | -0,426393579 | 0,744119598 | 0,559237537 |
| TAGTGGATGATGCACTCTGTGC | hsa-miR-3681-5p | -0,441332613 | 0,736454034 | 0,692505052 |
| CAGTGCAATAGTATTGTCAAAGC | hsa-miR-301a-3p | -0,444170129 | 0,735006988 | 0,398428398 |
| TAGCTTATCAGACTGATGTTGA | hsa-miR-21-5p | -0,450181893 | 0,731950559 | 0,604964084 |
| TATTGCACTCGTCCCGGCCTCC | hsa-miR-92b-3p | -0,451262215 | 0,731402663 | 0,626625979 |
| CTGGGATCTCCGGGGTCTTGGTT | hsa-miR-769-3p | -0,456967718 | 0,728515858 | 0,626625979 |
| CCATGGATCTCCAGGTGGGT | hsa-miR-490-5p | -0,45832 | 0,727833318 | 0,892990833 |
| CGCATCCCCTAGGGCATTGGTG | hsa-miR-324-5p | -0,459830865 | 0,727071492 | 0,695212154 |
| TGTAAACATCCTACACTCTCAGC | hsa-miR-30c-5p | -0,460199095 | 0,72688594 | 0,559237537 |
| ATCAACAGACATTAATTGGGCGC | hsa-miR-421 | -0,460407871 | 0,726780758 | 0,402551647 |
| CGTGTTCACAGCGGACCTTGAT | hsa-miR-124-5p | -0,461262835 | 0,726350184 | 0,585308588 |
| CTGGGAGAAGGCTGTTTACTCT | hsa-miR-30c-2-3p | -0,462282868 | 0,725836812 | 0,402551647 |
| TCTGGCTCCGTGTCTTCACTCCC | hsa-miR-149-5p | -0,465603155 | 0,724168257 | 0,49395756 |
| TAATCTCAGCTGGCAACTGTGA | hsa-miR-216a-5p | -0,469559498 | 0,722185071 | 0,72639342 |
| CTCTCACCACTGCCCTCCCACAG | hsa-miR-1229-3p | -0,473508158 | 0,720211151 | 0,846300755 |
| TAGCACCATTTGAAATCGGTTA | hsa-miR-29c-3p | -0,475042898 | 0,719445397 | 0,634026883 |
| AGCTACATTGTCTGCTGGGTTTC | hsa-miR-221-3p | -0,499935823 | 0,707138237 | 0,49395756 |
| TAAGGCACGCGGTGAATGCCAA | hsa-miR-124-3p | -0,50211888 | 0,706069019 | 0,398428398 |
| ATATGGGTTTACTAGTTGGT | hsa-miR-3115 | -0,507889239 | 0,703250589 | 0,704500821 |
| TGACCTGGGACTCGGACAGCTG | hsa-miR-3661 | -0,513320701 | 0,700607967 | 0,806173445 |
| TTGCATAGTCACAAAAGTGATC | hsa-miR-153-3p | -0,52550401 | 0,694716366 | 0,30789554 |
| ACTGCTGAGCTAGCACTTCCCG | hsa-miR-93-3p | -0,5257714 | 0,694587619 | 0,736897829 |
| AGCAGCATTGTACAGGGCTATCA | hsa-miR-107 | -0,528333029 | 0,693355413 | 0,549540111 |
| TCTACAGTGCACGTGTCTCCAGT | hsa-miR-139-5p | -0,551085199 | 0,682506552 | 0,688933922 |
| TAAAGAGCCCTGTGGAGACA | hsa-miR-1276 | -0,553755324 | 0,681244544 | 0,799093566 |
| TAATACTGCCGGGTAATGATGGA | hsa-miR-200c-3p | -0,559042918 | 0,678752298 | 0,691215547 |
| AAAGGATTCTGCTGTCGGTCCCACT | hsa-miR-541-5p | -0,589837933 | 0,664417541 | 0,810365248 |
| TCCGAACTCTCCATTCCTCTGC | hsa-miR-6716-3p | -0,590690377 | 0,664025073 | 0,733132978 |

| | | | | |
|---|---|---|---|---|
| TCCCTGAGACCCTAACTTGTGA | hsa-miR-125b-5p | -0,594752607 | 0,662157992 | 0,684757038 |
| AAAAGTAATTGTGGATTTTGCT | hsa-miR-548ab | -0,621273178 | 0,650096964 | 0,492245209 |
| GATGATGCTGCTGATGCTG | hsa-miR-1322 | -0,629036147 | 0,646608265 | 0,759354533 |
| AACCCGTAGATCCGATCTTGTG | hsa-miR-99a-5p | -0,642663901 | 0,640529135 | 0,402551647 |
| TGCAACGAACCTGAGCCACTGA | hsa-miR-891a-5p | -0,651866395 | 0,636456406 | 0,731674409 |
| AGCTACATCTGGCTACTGGGT | hsa-miR-222-3p | -0,654326645 | 0,635371972 | 0,492245209 |
| CTGTGCGTGTGACAGCGGCTGA | hsa-miR-210-3p | -0,666182066 | 0,630172164 | 0,395270609 |
| CGTCAACACTTGCTGGTTTCCT | hsa-miR-505-3p | -0,681936315 | 0,623328112 | 0,398428398 |
| GCTCTTTTCACATTGTGCTACT | hsa-miR-130a-5p | -0,688070001 | 0,620683629 | 0,568795286 |
| ACCCCACTCCTGGTACC | hsa-miR-4286 | -0,698386061 | 0,616261231 | 0,626625979 |
| TAGATAAAATATTGGTACCTG | hsa-miR-577 | -0,698948183 | 0,616021162 | 0,780780131 |
| ATATAATACAACCTGCTAAGTG | hsa-miR-374b-5p | -0,700069282 | 0,615542646 | 0,297249769 |
| ACCTTGGCTCTAGACTGCTTACT | hsa-miR-212-5p | -0,700197713 | 0,615487852 | 0,402551647 |
| TCTCTGGGCCTGTGTCTTAGGC | hsa-miR-330-5p | -0,706508918 | 0,612801223 | 0,692505052 |
| TGAGGTAGGAGGTTGTATAGTT | hsa-let-7e-5p | -0,709970666 | 0,611332569 | 0,673177989 |
| TAACAGTCTACAGCCATGGTCG | hsa-miR-132-3p | -0,71759185 | 0,608111657 | 0,44136437 |
| TGGTGGGCACAGAATCTGGACT | hsa-miR-541-3p | -0,719828909 | 0,607169443 | 0,398428398 |
| GTAGAGGAGATGGCGCAGGG | hsa-miR-877-5p | -0,734385978 | 0,601073792 | 0,400732344 |
| TGAGACCAGGACTGGATGCACC | hsa-miR-4786-5p | -0,742814313 | 0,59757251 | 0,684757038 |
| TTGAAAGGCTATTTCTTGGTC | hsa-miR-488-3p | -0,762289411 | 0,589560016 | 0,094039558 |
| GCCCCTGGGCCTATCCTAGAA | hsa-miR-331-3p | -0,781227557 | 0,581871481 | 0,526925062 |
| TTATTGCTTAAGAATACGCGTAG | hsa-miR-137-3p | -0,781735759 | 0,581666548 | 0,684577389 |
| CTATACGGCCTCCTAGCTTTCC | hsa-let-7e-3p | -0,783810008 | 0,580830852 | 0,619739175 |
| AACTGGCCCTCAAAGTCCCGCT | hsa-miR-193b-3p | -0,792163094 | 0,577477606 | 0,317540786 |
| TGAGGTAGTAGGTTGTATAGTT | hsa-let-7a-5p | -0,796933083 | 0,575571442 | 0,698597151 |
| TAACAGTCTCCAGTCACGGCC | hsa-miR-212-3p | -0,799337864 | 0,57461284 | 0,5536546 |
| AGACCCTGGTCTGCACTCTATC | hsa-miR-504-5p | -0,812776205 | 0,569285317 | 0,409422741 |
| AAATCTCTGCAGGCAAATGTGA | hsa-miR-216b-5p | -0,848458783 | 0,555377724 | 0,137386105 |
| TACTGCATCAGGAACTGATTGGA | hsa-miR-217-5p | -0,873503711 | 0,545819669 | 0,421923918 |
| TCACAAGTCAGGCTCTTGGGAC | hsa-miR-125b-2-3p | -0,87970219 | 0,543479608 | 0,328422226 |
| AGGCAAGATGCTGGCATAGCT | hsa-miR-31-5p | -0,900930648 | 0,535541155 | 0,539209501 |
| GAAAAAGTCATGGAGGCC | hsa-miR-12136 | -0,90282779 | 0,534837382 | 0,559237537 |
| TCAGCACCAGGATATTGTTGGAG | hsa-miR-3065-3p | -0,911965802 | 0,531460435 | 0,49395756 |

| | | | | |
|---|---|---|---|---|
| TTCAACGGGTATTTATTGAGCA | hsa-miR-95-3p | -0,912280834 | 0,531344396 | 0,454399633 |
| ACTCTAGCTGCCAAAGGCGCT | hsa-miR-1251-5p | -0,926073476 | 0,526288773 | 0,116086396 |
| ATCATGATGGGCTCCTCGGTGT | hsa-miR-433-3p | -0,993407347 | 0,502290068 | 0,402551647 |
| TAACTGGTTGAACAACTGAACC | hsa-miR-582-3p | -0,998079125 | 0,500666168 | 0,559237537 |
| AAGCCCTTACCCCAAAAAGCAT | hsa-miR-129-2-3p | -1,012480601 | 0,495693209 | 0,402551647 |
| CTTTTTGCGGTCTGGGCTTGC | hsa-miR-129-5p | -1,018318745 | 0,493691344 | 0,402551647 |
| TACAGATGCAGATTCTCTGACTTC | hsa-miR-5683 | -1,063765426 | 0,478381855 | 0,020279128 |
| TCACCAGCCCTGTGTTCCCTAG | hsa-miR-1226-3p | -1,082267639 | 0,472285897 | 0,402551647 |
| AGGGACGGGACGCGGTGCAGTG | hsa-miR-92b-5p | -1,138470765 | 0,454240811 | 0,530860193 |
| GACTATAGAACTTTCCCCCTCA | hsa-miR-625-3p | -1,141269148 | 0,453360578 | 0,221918461 |
| TTACAGTTGTTCAACCAGTTACT | hsa-miR-582-5p | -1,141953122 | 0,453145693 | 0,426837512 |
| GTGAGGGCATGCAGGCCTGGATGGGG | hsa-miR-1226-5p | -1,210767013 | 0,43203886 | 0,317540786 |
| CATCAGTTCCTAATGCATTGCC | hsa-miR-217-3p | -1,213725242 | 0,431153877 | 0,492245209 |
| AAAAGTAATCGCGGTTTTTGTC | hsa-miR-548h-5p | -1,227095552 | 0,427176577 | 0,224983992 |
| TGAGGTAGTAGATTGTATAGTT | hsa-let-7f-5p | -1,270410137 | 0,414541908 | 0,324725905 |
| AAAAGTAATTGTGGTTTTGGCC | hsa-miR-548b-5p | -1,324112094 | 0,399394925 | 0,213361222 |
| AGCTGGTGTTGTGAATCAGGCCG | hsa-miR-138-5p | -1,43847242 | 0,368957764 | 0,297249769 |
| ATAAAGCTAGATAACCGAAAGT | hsa-miR-9-3p | -1,470085525 | 0,3609609 | 0,000340533 |
| TGAGGTAGTAGTTTGTGCTGTT | hsa-let-7i-5p | -1,652599085 | 0,318066628 | 0,543409045 |
| TGAGGTAGTAGTTTGTACAGTT | hsa-let-7g-5p | -1,675938931 | 0,312962362 | 0,49395756 |
| TGAGGTAGTAGGTTGTATGGTT | hsa-let-7c-5p | -1,709094536 | 0,305851968 | 0,226006464 |
| TCTTTGGTTATCTAGCTGTATGA | hsa-miR-9-5p | -1,923966057 | 0,263529056 | 1,91E-09 |
| TGAGGTAGTAAGTTGTATTGTT | hsa-miR-98-5p | -1,949489242 | 0,258907876 | 0,416276553 |
| AGAGGTAGTAGGTTGCATAGTT | hsa-let-7d-5p | -2,026974729 | 0,245369064 | 0,402551647 |
| CTATACGACCTGCTGCCTTTCT | hsa-let-7d-3p | -2,159031517 | 0,223906526 | 0,328422226 |
| CTGTACAACCTTCTAGCTTTCC | hsa-let-7c-3p | -2,318579816 | 0,200464709 | 0,090829287 |

**KO-NSC vs IC1-NSC**

| Sequence | Name | log2FC | Fold_change | Adjusted_p-value |
|---|---|---|---|---|
| TACCCTGTAGATCCGAATTTGTG | hsa-miR-10a-5p | 7,5661858 | 189,5173022 | 2,02E-24 |
| GGGGTTCCTGGGGATGGGATTT | hsa-miR-23a-5p | 3,2030858 | 9,209263516 | 6,12E-13 |
| TGAGGTAGTAGGTTGTGTGGTT | hsa-let-7b-5p | 6,7105491 | 104,7313191 | 5,77E-12 |
| TCCTTCATTCCACCGGAGTCTG | hsa-miR-205-5p | 4,2071291 | 18,47021932 | 2,20E-10 |
| AGGGCCCCCCCTCAATCCTGT | hsa-miR-296-5p | -7,1955129 | 0,006822363 | 2,30E-09 |
| TTCACAGTGGCTAAGTTCCGC | hsa-miR-27a-3p | 2,0389913 | 4,109581024 | 5,83E-09 |
| CAACACCAGTCGATGGGCTGT | hsa-miR-21-3p | 2,5475768 | 5,846514584 | 2,98E-08 |
| TCGGGGATCATCATGTCACGAGA | hsa-miR-542-5p | 2,2621813 | 4,797162493 | 1,48E-06 |
| TTGCATAGTCACAAAAGTGATC | hsa-miR-153-3p | -1,3304993 | 0,397630615 | 2,03E-06 |
| TGAGAACTGAATTCCATGGGTT | hsa-miR-146a-5p | 3,2996148 | 9,846526095 | 2,95E-06 |
| TAGCACCATCTGAAATCGGTTA | hsa-miR-29a-3p | 2,1312565 | 4,380988645 | 3,23E-06 |
| TGAGGTAGTAGGTTGTATAGTT | hsa-let-7a-5p | 2,6175791 | 6,137193729 | 6,59E-06 |
| ATCACATTGCCAGGGATTTCC | hsa-miR-23a-3p | 1,770615 | 3,411993741 | 6,59E-06 |
| TGGGTCTTTGCGGGCGAGATGA | hsa-miR-193a-5p | 2,4861776 | 5,602915068 | 7,59E-06 |
| AACTGGCCCTCAAAGTCCCGCT | hsa-miR-193b-3p | 2,1051348 | 4,302379671 | 7,59E-06 |
| AAGACGGGAGGAAAGAAGGGAG | hsa-miR-483-5p | 6,0231933 | 65,03720454 | 9,35E-06 |
| ATTTGTGCTTGGCTCTGTCAC | hsa-miR-2113 | 3,6904077 | 12,9099162 | 1,03E-05 |
| CAAAACGTGAGGCGCTGCTAT | hsa-miR-424-3p | 2,8953947 | 7,440475059 | 2,06E-05 |
| AAGCTGCCAGTTGAAGAACTGT | hsa-miR-22-3p | 1,663362 | 3,167538116 | 2,06E-05 |
| TTAATGCTAATCGTGATAGGGGTT | hsa-miR-155-5p | 2,2659387 | 4,80967262 | 3,09E-05 |
| AGGGCTTAGCTGCTTGTGAGCA | hsa-miR-27a-5p | 2,1952698 | 4,579753141 | 3,33E-05 |
| TGAGTGTGTGTGTGTGAGTGTGT | hsa-miR-574-5p | 1,3869037 | 2,615168088 | 4,34E-05 |
| CATCAGTTCCTAATGCATTGCC | hsa-miR-217-3p | -3,168469 | 0,1112233 | 5,45E-05 |
| AGGCAGTGTAGTTAGCTGATTGC | hsa-miR-34c-5p | 1,5336284 | 2,895130586 | 7,07E-05 |
| TTGTGTCAATATGCGATGATGT | hsa-miR-592 | -1,3402881 | 0,394941787 | 8,98E-05 |
| AAAGTGCTGCGACATTTGAGCGT | hsa-miR-372-3p | 4,5183905 | 22,91770207 | 0,000130023 |
| TGGCAGTGTCTTAGCTGGTTGT | hsa-miR-34a-5p | 2,6586706 | 6,314509057 | 0,000130023 |
| TGAGCACCACACAGGCCGGGCGC | hsa-miR-3663-3p | 2,3852816 | 5,224458669 | 0,000176572 |
| CAGTGCAATAGTATTGTCAAAGC | hsa-miR-301a-3p | -1,1193253 | 0,46030904 | 0,000189255 |
| TGTGCAAATCTATGCAAAACTGA | hsa-miR-19a-3p | -1,6894394 | 0,310047373 | 0,000218857 |
| TGAGGTAGTAGTTTGTGCTGTT | hsa-let-7i-5p | 3,2017313 | 9,200621659 | 0,00023993 |

| | | | | |
|---|---|---|---|---|
| TGAGGTAGTAGATTGTATAGTT | hsa-let-7f-5p | 2,3358593 | 5,048515658 | 0,000338702 |
| TACAGATGCAGATTCTCTGACTTC | hsa-miR-5683 | -1,660891 | 0,31624378 | 0,000338702 |
| TGAGGTAGGAGGTTGTATAGTT | hsa-let-7e-5p | 1,6262837 | 3,087167301 | 0,000373311 |
| TGAGGTAGTAAGTTGTATTGTT | hsa-miR-98-5p | 3,3759993 | 10,3819052 | 0,00037741 |
| TAATGCCCCTAAAAATCCTTAT | hsa-miR-365b-3p#hsa | 1,5269779 | 2,881815301 | 0,00037741 |
| TAGCTTATCAGACTGATGTTGA | hsa-miR-21-5p | 1,590511 | 3,011560042 | 0,000387134 |
| AGCTCATCCATAGTTGTCACTG | hsa-miR-549a-5p | 5,1176245 | 34,7183016 | 0,000694737 |
| GCTCTGACTTTATTGCACTACT | hsa-miR-301a-5p | -1,1972602 | 0,43610269 | 0,00073146 |
| AAATCTCTGCAGGCAAATGTGA | hsa-miR-216b-5p | -1,9792702 | 0,253618129 | 0,000904367 |
| TTGAAAGGCTATTTCTTGGTC | hsa-miR-488-3p | -1,9829098 | 0,25297911 | 0,001076883 |
| GCTGGTTTCATATGGTGGTTTAGA | hsa-miR-29b-1-5p | 4,0290816 | 16,32579776 | 0,001248373 |
| AATCACTAACCACACGGCCAGG | hsa-miR-34c-3p | 5,5398141 | 46,52112422 | 0,001396951 |
| TCCGTCTCAGTTACTTTATAGC | hsa-miR-340-3p | -0,9641771 | 0,512570698 | 0,00169912 |
| AGAGGTAGTAGGTTGCATAGTT | hsa-let-7d-5p | 3,1517845 | 8,887542025 | 0,002034977 |
| GAGGGTTGGGTGGAGGCTCTCC | hsa-miR-296-3p | -4,9234162 | 0,032953689 | 0,002034977 |
| CACTAGATTGTGAGCTCCTGGA | hsa-miR-28-3p | 1,4511291 | 2,734219642 | 0,002210596 |
| CAGTGCAATGATATTGTCAAAGC | hsa-miR-301b-3p | -1,2605065 | 0,417397384 | 0,002847039 |
| CTATACGACCTGCTGCCTTTCT | hsa-let-7d-3p | 3,3544904 | 10,22827129 | 0,00357215 |
| TAAGGTGCATCTAGTGCAGATAG | hsa-miR-18a-5p | -1,2553619 | 0,418888472 | 0,004141913 |
| CCTGTTCTCCATTACTTGGCT | hsa-miR-26b-3p | -1,8127531 | 0,284647209 | 0,004141913 |
| TGAGGTAGTAGTTTGTACAGTT | hsa-let-7g-5p | 2,0910409 | 4,260553712 | 0,004200096 |
| TGGCTCAGTTCAGCAGGAACAG | hsa-miR-24-3p | 1,2042521 | 2,304177903 | 0,004200096 |
| ATCACATTGCCAGGGATTACCAC | hsa-miR-23b-3p | 0,8668915 | 1,823729113 | 0,004503483 |
| CAGCAGCAATTCATGTTTTGAA | hsa-miR-424-5p | 1,8233916 | 3,539122192 | 0,005556175 |
| TAAAGTGCTGACAGTGCAGAT | hsa-miR-106b-5p | -0,9113485 | 0,531687873 | 0,005812198 |
| ACTTAAACGTGGATGTACTTGCT | hsa-miR-302a-5p | 2,4267853 | 5,376939706 | 0,006924936 |
| TCACTCCTCTCCTCCCGTCTT | hsa-miR-483-3p | 3,7187469 | 13,16601534 | 0,007250559 |
| TGCAACGAACCTGAGCCACTGA | hsa-miR-891a-5p | -2,7919445 | 0,144391282 | 0,007713442 |
| CAAGAACCTCAGTTGCTTTTGT | hsa-miR-548b-3p | -3,1637792 | 0,111585446 | 0,00863242 |
| ACTGACAGGAGAGCATTTTGA | hsa-miR-3660 | 2,0812853 | 4,231840582 | 0,009191146 |
| TAGCACCATTTGAAATCGGTTA | hsa-miR-29c-3p | -1,4597968 | 0,363544319 | 0,009627057 |
| ACTCAAACTGTGGGGGCACT | hsa-miR-371a-5p | 3,822891 | 14,15157737 | 0,012638661 |
| TAAAGTGCTTATAGTGCAGGTAG | hsa-miR-20a-5p | -0,885251 | 0,541393326 | 0,012971228 |

| | | | | |
|---|---|---|---|---|
| TCGACCGGACCTCGACCGGCT | hsa-miR-1307-5p | -0,9240153 | 0,527040112 | 0,012980382 |
| TTATCAGAATCTCCAGGGGTAC | hsa-miR-361-5p | 0,7012261 | 1,625886017 | 0,013287136 |
| CACGCTCATGCACACACCCACA | hsa-miR-574-3p | 1,2471484 | 2,373717684 | 0,014007781 |
| GGTGCAGTGCTGCATCTCTGGT | hsa-miR-143-5p | 4,5996022 | 24,24477861 | 0,017189605 |
| ACCACTGACCGTTGACTGTACC | hsa-miR-181a-2-3p | 0,8016583 | 1,743103588 | 0,017850263 |
| TACTGCATCAGGAACTGATTGGA | hsa-miR-217-5p | -2,4593542 | 0,181827933 | 0,017850263 |
| TAATTTTATGTATAAGCTAGT | hsa-miR-590-3p | -1,0560238 | 0,480955807 | 0,017959689 |
| CGGGGTTTTGAGGGCGAGATGA | hsa-miR-193b-5p | 1,9093443 | 3,756383255 | 0,019312036 |
| TGGAATGTAAAGAAGTATGTAT | hsa-miR-1-3p | 1,475657 | 2,781102725 | 0,019891907 |
| CAAGCTCGTGTCTGTGGGTCCG | hsa-miR-99b-3p | 0,8454598 | 1,796837357 | 0,02045106 |
| AGATCAGAAGGTGATTGTGGCT | hsa-miR-383-5p | -1,9932795 | 0,251167287 | 0,020715993 |
| ACCTTGGCTCTAGACTGCTTACT | hsa-miR-212-5p | -1,0862577 | 0,470981497 | 0,021343665 |
| TGGGTTTACGTTGGGAGAACT | hsa-miR-629-5p | 1,1422134 | 2,207194007 | 0,024136729 |
| AGTTCTTCAGTGGCAAGCTTTA | hsa-miR-22-5p | 1,6983842 | 3,245372678 | 0,024416095 |
| AACTGGCCTACAAAGTCCCAGT | hsa-miR-193a-3p | 3,0385118 | 8,216430632 | 0,025782005 |
| TGTGCAAATCCATGCAAAACTGA | hsa-miR-19b-3p | -1,2237474 | 0,428169097 | 0,030409428 |
| TAAGGCACGCGGTGAATGCCAA | hsa-miR-124-3p | -0,6541203 | 0,63546285 | 0,031652497 |
| AATAATACATGGTTGATCTTT | hsa-miR-369-3p | -1,5341885 | 0,345273494 | 0,031652497 |
| CAAAGTGCTTACAGTGCAGGTAG | hsa-miR-17-5p | -0,8208151 | 0,566122013 | 0,031983799 |
| CTGCGCAAGCTACTGCCTTGCT | hsa-let-7i-3p | 5,1000557 | 34,29807483 | 0,03768106 |
| AGAGCTTAGCTGATTGGTGAAC | hsa-miR-27b-5p | 1,0592374 | 2,083829703 | 0,03768106 |
| AAAAGTAATTGTGGATTTTGCT | hsa-miR-548ab | -1,0690754 | 0,476624373 | 0,03768106 |
| ACTCTAGCTGCCAAAGGCGCT | hsa-miR-1251-5p | -1,1175323 | 0,46088147 | 0,039075329 |
| TGGTCTGTCTCTGCCCTGGCAC | hsa-miR-6753-3p | 4,3797647 | 20,81807371 | 0,043099886 |
| TGGAAGACTAGTGATTTTGTTGTT | hsa-miR-7-5p | -1,0181332 | 0,493754856 | 0,043099886 |
| TGAGGTAGTAGGTTGTATGGTT | hsa-let-7c-5p | 1,2322273 | 2,349294108 | 0,044812059 |
| TGGAGAGAAAGGCAGTTCCTGA | hsa-miR-185-5p | 1,1215912 | 2,175868202 | 0,044812059 |
| GAAGTGCTTCGATTTTGGGGTGT | hsa-miR-373-3p | 3,9764694 | 15,7411539 | 0,047007936 |
| TTTAACATGGGGGTACCTGCTG | hsa-miR-302c-5p | 3,9284102 | 15,22542134 | 0,047007936 |
| CAGTGCAATGTTAAAAGGGCAT | hsa-miR-130a-3p | -0,7149524 | 0,609225244 | 0,047007936 |
| TAATCTCAGCTGGCAACTGTGA | hsa-miR-216a-5p | -2,1414545 | 0,226651168 | 0,047007936 |
| CCAGTCCTGTGCCTGCCGCCT | hsa-miR-1910-5p | 1,0420248 | 2,059115562 | 0,048453666 |
| TGTAAACATCCTTGACTGGAAG | hsa-miR-30e-5p | -1,1036997 | 0,465321665 | 0,048568573 |

| | | | | |
|---|---|---|---|---|
| CAAAAACTGCAGTTACTTTTGC | hsa-miR-548ah-3p | -0,9088784 | 0,532598975 | 0,048945096 |
| AATGACACGATCACTCCCGTTGA | hsa-miR-425-5p | -0,804262 | 0,572654923 | 0,049619684 |
| GCGGTGATCCCGATGGTGTGAGC | hsa-miR-598-5p | 1,5645367 | 2,957825072 | 0,050540463 |
| ACTCCAGCCCCACAGCCTCAGC | hsa-miR-766-3p | 1,4711344 | 2,772398065 | 0,050540463 |
| AAAAGTAATCGCGGTTTTTGTC | hsa-miR-548h-5p | 1,2326737 | 2,350021089 | 0,050540463 |
| AAAGTTCTGAGACACTCCGACT | hsa-miR-148a-5p | 1,0265537 | 2,03715215 | 0,051709935 |
| TATGGCTTTTCATTCCTATGTGA | hsa-miR-135b-5p | -0,8782755 | 0,544017328 | 0,052269761 |
| GTTCTCCCAACGTAAGCCCAGC | hsa-miR-629-3p | 3,5360568 | 11,60003152 | 0,052608634 |
| TAAGTGCTTCCATGTTTGAGTGT | hsa-miR-302d-3p | 1,5234839 | 2,874844446 | 0,05375986 |
| TAGCCCCCAGGCTTCACTTGGCG | hsa-miR-3943 | 1,289088 | 2,443735199 | 0,055693611 |
| TACCCAGAGCATGCAGTGTGAA | hsa-miR-1912-3p | 2,6688209 | 6,359092332 | 0,056060173 |
| TGAGATGAAGCACTGTAGCTC | hsa-miR-143-3p | 3,4821598 | 11,17466569 | 0,05662077 |
| AATTCCCTTGTAGATAACCCGG | hsa-miR-3938 | -0,7939964 | 0,576744223 | 0,060514636 |
| ATGTAGGGCTAAAAGCCATGGG | hsa-miR-135b-3p | -1,0388455 | 0,486716823 | 0,06561142 |
| GGGGTATTGTTTCCGCTGCCAGG | hsa-miR-503-3p | 3,5338531 | 11,58232573 | 0,065840777 |
| GTCCAGTTTTCCCAGGAATCCCT | hsa-miR-145-5p | 3,2291789 | 9,377341347 | 0,065840777 |
| TATGGCTTTTTATTCCTATGTGA | hsa-miR-135a-5p | -0,8371186 | 0,559760429 | 0,066654936 |
| AGGATGAGCAAAGAAAGTAGATT | hsa-miR-1255a | 1,3781167 | 2,599288418 | 0,070411821 |
| ATCCCCAGATACAATGGACAA | hsa-miR-2355-5p | 1,1844178 | 2,2727166 | 0,070411821 |
| AGGTTACCCGAGCAACTTTGCAT | hsa-miR-409-5p | -1,3282405 | 0,398253664 | 0,070411821 |
| CAAGCTTGTATCTATAGGTATG | hsa-miR-100-3p | 0,937725 | 1,915505277 | 0,072543847 |
| ACAGGTGAGGTTCTTGGGAGCC | hsa-miR-125a-3p | 0,6834194 | 1,60594152 | 0,072543847 |
| CATTATTACTTTTGGTACGCG | hsa-miR-126-5p | -1,0700435 | 0,476304651 | 0,072543847 |
| TAGCACCATTTGAAATCAGTGTT | hsa-miR-29b-3p | 1,3264181 | 2,507792754 | 0,074682378 |
| TAAGTGCTTCCATGTTTTGGTGA | hsa-miR-302a-3p | 1,5723118 | 2,973808567 | 0,075609886 |
| TTTGGTCCCCTTCAACCAGCTG | hsa-miR-133a-3p | 1,3796319 | 2,602019779 | 0,075609886 |
| TTTTGCAATATGTTCCTGAATA | hsa-miR-450b-5p | 1,1502667 | 2,219549231 | 0,075609886 |
| TTTTGCGATGTGTTCCTAATAT | hsa-miR-450a-5p | 1,0491385 | 2,069293857 | 0,075609886 |
| AACCCGTAGATCCGAACTTGTG | hsa-miR-100-5p | 0,777694 | 1,714388391 | 0,075609886 |
| GTGAACGGGCGCCATCCCGAGG | hsa-miR-887-3p | -0,694817 | 0,617787698 | 0,075609886 |
| TGATATGTTTGATATTGGGTTG | hsa-miR-190b-5p | -1,728933 | 0,301674992 | 0,075609886 |
| TACCCTGTAGAACCGAATTTGTG | hsa-miR-10b-5p | 2,1565902 | 4,458598158 | 0,076147644 |
| TAAGTGCTTCCATGTTTCAGTGG | hsa-miR-302c-3p | 1,6075497 | 3,04733836 | 0,076147644 |

| | | | | |
|---|---|---|---|---|
| AGCGCGGGCTGAGCGCTGCCAGTC | hsa-miR-2277-5p | -0,9761266 | 0,508342714 | 0,076147644 |
| TCAGTGCACTACAGAACTTTGT | hsa-miR-148a-3p | 0,8534485 | 1,806814649 | 0,083337591 |
| AAAAGTAATTGTGGTTTTGGCC | hsa-miR-548b-5p | -1,1686043 | 0,444851497 | 0,083337591 |
| TCTCACACAGAAATCGCACCCGT | hsa-miR-342-3p | -0,6921844 | 0,618916032 | 0,086028881 |
| TCCCTGAGACCCTTTAACCTGTGA | hsa-miR-125a-5p | 0,8207005 | 1,766263434 | 0,092945212 |
| CTAGACTGAAGCTCCTTGAGG | hsa-miR-151a-3p | 0,7530082 | 1,685303211 | 0,093867694 |
| TAGCAGCACATCATGGTTTACA | hsa-miR-15b-5p | -0,4801127 | 0,716921625 | 0,095311865 |
| CGCGCCTGCAGGAACTGGTAGA | hsa-miR-6720-3p | 4,3614701 | 20,55575009 | 0,095536712 |
| CTCCTGACTCCAGGTCCTGTGT | hsa-miR-378a-5p | 1,0557675 | 2,078823884 | 0,09881933 |
| AGACCCTGGTCTGCACTCTATC | hsa-miR-504-5p | -1,2387255 | 0,423746839 | 0,101392326 |
| TTGCAGCTGCCTGGGAGTGACTTC | hsa-miR-1301-3p | 0,8170037 | 1,761743297 | 0,101541504 |
| AAAGGTAACTGTGATTTTTGCT | hsa-miR-548ba | 1,5382959 | 2,904512152 | 0,104342363 |
| ATCAAGGATCTTAAACTTTGCC | hsa-miR-561-5p | -0,6196022 | 0,650850374 | 0,108418837 |
| AACATTCATTGCTGTCGGTGGGT | hsa-miR-181b-5p | 0,494194 | 1,408533573 | 0,110631419 |
| CTATACGGCCTCCTAGCTTTCC | hsa-let-7e-3p | 1,00733 | 2,010187388 | 0,110971473 |
| TTCACCACCTTCTCCACCCAGC | hsa-miR-197-3p | 0,7965138 | 1,736898933 | 0,114704619 |
| CAAAAGCAATCGCGGTTTTTGC | hsa-miR-548e-5p | -1,7772335 | 0,291742305 | 0,120679074 |
| TGTCTGCCCGCATGCCTGCCTCT | hsa-miR-346 | 1,9209814 | 3,786805685 | 0,124792439 |
| CTGGGAGGTGGATGTTTACTTC | hsa-miR-30b-3p | 1,0787497 | 2,112204714 | 0,13109944 |
| TTTTCAACTCTAATGGGAGAGA | hsa-miR-1305 | -0,7508798 | 0,594241079 | 0,13109944 |
| TGCCCTAAATGCCCCTTCTGGC | hsa-miR-18b-3p | -1,1611967 | 0,447141477 | 0,137563432 |
| ACCCTATCAATATTGTCTCTGC | hsa-miR-454-5p | -0,4594699 | 0,727253406 | 0,138162723 |
| ACTCTTTCCCTGTTGCACTAC | hsa-miR-130b-5p | -0,5059348 | 0,704203919 | 0,138162723 |
| AATTGCACGGTATCCATCTGTA | hsa-miR-363-3p | -0,7905217 | 0,578135001 | 0,138162723 |
| TCCCCCAGGTGTGATTCTGATTT | hsa-miR-361-3p | 0,9825645 | 1,975974772 | 0,138174393 |
| GTGACATCACATATACGGCAGC | hsa-miR-489-3p | 1,3208886 | 2,498199334 | 0,140096843 |
| TCAAATGCTCAGACTCCTGTGGT | hsa-miR-105-5p | -0,7580208 | 0,591306978 | 0,141055607 |
| TGTGACAGATTGATAACTGAAA | hsa-miR-542-3p | 1,2471055 | 2,373647243 | 0,149416203 |
| TTATAATACAACCTGATAAGTG | hsa-miR-374a-5p | -0,62297 | 0,649332808 | 0,1524642 |
| CTATACAATCTACTGTCTTTC | hsa-let-7a-3p | 2,2423087 | 4,731536172 | 0,153194836 |
| ACAGGGCCGCAGATGGAGACT | hsa-miR-3918 | 2,9224442 | 7,581294229 | 0,157648618 |
| ACTGCAGTGAAGGCACTTGTAG | hsa-miR-17-3p | -0,9602977 | 0,513950854 | 0,162816773 |
| GGATTCCTGGAAATACTGTTCT | hsa-miR-145-3p | 2,6875879 | 6,442353995 | 0,166881541 |

| | | | | |
|---|---|---|---|---|
| AAGATGTGGAAAAATTGGAATC | hsa-miR-576-3p | 1,9287635 | 3,807287465 | 0,166881541 |
| GAAATCAAGCGTGGGTGAGACC | hsa-miR-551b-5p | 1,548241 | 2,924603356 | 0,173666405 |
| AAGGAGCTCACAGTCTATTGAG | hsa-miR-28-5p | 0,75475 | 1,687339141 | 0,174264524 |
| CAAAGTGCTGTTCGTGCAGGTAG | hsa-miR-93-5p | -0,6224053 | 0,649586998 | 0,174264524 |
| TAACTGGTTGAACAACTGAACC | hsa-miR-582-3p | 1,1321379 | 2,191833028 | 0,176623991 |
| TGGACGGAGAACTGATAAGGGT | hsa-miR-184 | 0,8438598 | 1,794845664 | 0,176623991 |
| AGGTTCTGTGATACACTCCGACT | hsa-miR-152-5p | 0,7877612 | 1,72639334 | 0,176623991 |
| TCTTCTCTGTTTTGGCCATGTG | hsa-miR-942-5p | -1,6021094 | 0,329395011 | 0,176623991 |
| AAGTGCTGTCATAGCTGAGGTC | hsa-miR-512-3p | 1,4032688 | 2,645002048 | 0,182105908 |
| TATTCATTTATCCCCAGCCTACA | hsa-miR-664a-3p | 0,9702955 | 1,959241836 | 0,182105908 |
| AGGCAAGATGCTGGCATAGCT | hsa-miR-31-5p | 1,1702183 | 2,25045749 | 0,182259125 |
| AAAAGTAATTGCGGTTTTTGCC | hsa-miR-548o-5p#hsa | -2,7072672 | 0,153119806 | 0,182259125 |
| CAGTGCAATGATGAAAGGGCAT | hsa-miR-130b-3p | -0,5008945 | 0,706668494 | 0,183222314 |
| TAGCAGCACGTAAATATTGGCG | hsa-miR-16-5p | -0,4066034 | 0,75439738 | 0,183264168 |
| TTCATTTGCCTCCCAGCCTACA | hsa-miR-664b-3p | 2,2062581 | 4,614767793 | 0,184590185 |
| CACCCGTAGAACCGACCTTGCG | hsa-miR-99b-5p | 0,5112666 | 1,425300955 | 0,186834279 |
| TGACTTCTACCTCTTCCAAAG | hsa-miR-6505-3p | -2,851029 | 0,138597291 | 0,190263854 |
| ACTCCAAGAAGAATCTAGACAG | hsa-miR-5695 | -2,4421587 | 0,184008112 | 0,19457327 |
| AACATTCAACCTGTCGGTGAGT | hsa-miR-181c-5p | 0,7288742 | 1,657345229 | 0,197358012 |
| CTGTTGCCACTAACCTCAACCT | hsa-miR-744-3p | -0,8278407 | 0,56337181 | 0,197999829 |
| TATGGCACTGGTAGAATTCACT | hsa-miR-183-5p | -0,8457855 | 0,556407787 | 0,198888258 |
| TAAGTGCTTCCATGTTTTAGTAG | hsa-miR-302b-3p | 1,3634745 | 2,57304106 | 0,20079168 |
| AGCTACATTGTCTGCTGGGTTTC | hsa-miR-221-3p | 0,7596846 | 1,693120412 | 0,20079168 |
| TTATAAAGCAATGAGACTGATT | hsa-miR-340-5p | -0,5407981 | 0,687390541 | 0,20079168 |
| TGACTCTGCCTGTAGGCCGGT | hsa-miR-4522 | -1,6646098 | 0,315429657 | 0,20079168 |
| TCACAGTGGTCTCTGGGATTAT | hsa-miR-216a-3p | -2,7315846 | 0,150560516 | 0,20079168 |
| TGAGGATGGATAGCAAGGAAGCC | hsa-miR-3605-5p | 1,5126798 | 2,853395546 | 0,205385386 |
| AACATAGAGGAAATTCCACGT | hsa-miR-376c-3p | -1,0115365 | 0,496017698 | 0,205385386 |
| TCGTACCGTGAGTAATAATGCG | hsa-miR-126-3p | -1,7165757 | 0,304270073 | 0,205385386 |
| AACATTCAACGCTGTCGGTGAGT | hsa-miR-181a-5p | 0,566438 | 1,480862818 | 0,205453205 |
| TGTCTTGCAGGCCGTCATGCA | hsa-miR-431-5p | -2,8917347 | 0,134741421 | 0,205453205 |
| CAACCTGGAGGACTCCATGCTG | hsa-miR-490-3p | -2,7006353 | 0,153825302 | 0,206015221 |
| TGTGACTGGTTGACCAGAGGGG | hsa-miR-134-5p | -0,9130894 | 0,531046675 | 0,206077937 |

| | | | | |
|---|---|---|---|---|
| ATATACAGGGGGAGACTCTTAT | hsa-miR-1185-1-3p | -1,3451848 | 0,393603562 | 0,206077937 |
| AAAAGTGCTTACAGTGCAGGTAG | hsa-miR-106a-5p | -0,7871139 | 0,579502242 | 0,208923743 |
| CCGCACTGTGGGTACTTGCTGC | hsa-miR-106b-3p | -0,5145219 | 0,700024868 | 0,210062142 |
| CCAGTTACCGCTTCCGCTACCGC | hsa-miR-935 | 0,8837836 | 1,845208213 | 0,211830867 |
| AAAATGGTTCCCTTTAGAGTGT | hsa-miR-522-3p | 2,5572999 | 5,886050304 | 0,212028059 |
| TAGTGAGTTAGAGATGCAGAGCC | hsa-miR-3174 | 1,0393737 | 2,055335219 | 0,212028059 |
| TTGTGTGAGTACAGAGAGCATC | hsa-miR-6818-5p | -1,4663544 | 0,361895635 | 0,212028059 |
| TCTGTGAGACCAAAGAACTACT | hsa-miR-4677-3p | 0,5799049 | 1,494750758 | 0,216517897 |
| TTGAGGGGAGAATGAGGTGGAGA | hsa-miR-6734-5p | 2,8445228 | 7,182682562 | 0,216572291 |
| TTCAACGGGTATTTATTGAGCA | hsa-miR-95-3p | -0,9068739 | 0,533339523 | 0,228233523 |
| GCCCTGTGGACTCAGTTCTGGT | hsa-miR-146b-3p | -0,8322774 | 0,561641967 | 0,228746608 |
| CTGGACTGAGCCGTGCTACTGG | hsa-miR-1269a | 2,3634115 | 5,145857636 | 0,229414261 |
| TGGGGATTTGGAGAAGTGGTGA | hsa-miR-4450 | -1,8844163 | 0,270853316 | 0,229881042 |
| ATCATACAAGGACAATTTCTTT | hsa-miR-539-3p | -1,3207626 | 0,400323267 | 0,238000817 |
| ATGCTGACATATTTACTAGAGG | hsa-miR-628-5p | -0,4862941 | 0,713856461 | 0,239780445 |
| AATCGTACAGGGTCATCCACTT | hsa-miR-487b-3p | -1,0817506 | 0,47245518 | 0,239780445 |
| AAAAGTAATTGCGGTTTTTGC | hsa-miR-548au-5p | -1,4190719 | 0,373952814 | 0,239780445 |
| GTGCATTGTAGTTGCATTGCA | hsa-miR-33a-5p | -2,615993 | 0,163120161 | 0,239780445 |
| TACCACAGGGTAGAACCACGG | hsa-miR-140-3p | -1,749996 | 0,2973026 | 0,248052103 |
| TTTGTTCGTTCGGCTCGCGTGA | hsa-miR-375-3p | 1,0435284 | 2,061262757 | 0,250069263 |
| GCAGGAACTTGTGAGTCTCCT | hsa-miR-873-5p | 0,7892482 | 1,72817367 | 0,25294674 |
| TAGGAGCTCAACAGATGCCTGTT | hsa-miR-3139 | 0,8502588 | 1,802824332 | 0,258141452 |
| TTTCTTCTTAGACATGGCAACG | hsa-miR-4659a-3p | 2,7666085 | 6,805062945 | 0,259367051 |
| TAGTGCAATATTGCTTATAGGGT | hsa-miR-454-3p | -0,4562648 | 0,728870898 | 0,260714075 |
| TCACAAGTCAGGCTCTTGGGAC | hsa-miR-125b-2-3p | 0,6920544 | 1,615582527 | 0,263245604 |
| ATGTAGGGATGGAAGCCATGAA | hsa-miR-135a-2-3p | -0,605973 | 0,657028099 | 0,273054917 |
| CCACCTCCCCTGCAAACGTCCA | hsa-miR-1306-5p | 0,647512 | 1,566464454 | 0,274367359 |
| GCGACCCATACTTGGTTTCAG | hsa-miR-551b-3p | -1,2520051 | 0,419864259 | 0,278416142 |
| CTAGCACACAGATACGCCCAGA | hsa-miR-585-5p | -2,6274142 | 0,1618339 | 0,282478743 |
| TACAGTACTGTGATAACTGAA | hsa-miR-101-3p | -0,6180023 | 0,65157255 | 0,288656384 |
| CCCGCAGGTGAGATGAGGGCT | hsa-miR-6886-5p | 0,6155391 | 1,532130396 | 0,291000264 |
| GCCTGCTGGGGTGGAACCTGGT | hsa-miR-370-3p | -0,9772414 | 0,507950064 | 0,291000264 |
| AACTGTTTGCAGAGGAAACTGA | hsa-miR-452-5p | 2,3370153 | 5,052562501 | 0,292273323 |

| | | | | |
|---|---|---|---|---|
| TGAGGGGCAGAGAGCGAGACTTT | hsa-miR-423-5p | 0,8270963 | 1,774111005 | 0,292273323 |
| TTAGCCAATTGTCCATCTTTAG | hsa-miR-4662a-5p | -0,5996945 | 0,659893692 | 0,292273323 |
| GTAGAGGAGATGGCGCAGGG | hsa-miR-877-5p | -0,7259379 | 0,604603869 | 0,292273323 |
| TTAAGACTTGCAGTGATGTTT | hsa-miR-499a-5p | -1,2923371 | 0,40828909 | 0,292273323 |
| TCTTGGAGTAGGTCATTGGGTGG | hsa-miR-432-5p | -0,9492071 | 0,517917018 | 0,297443941 |
| AAGGAGCTTACAATCTAGCTGGG | hsa-miR-708-5p | -0,5252287 | 0,69484893 | 0,301273492 |
| AACACACCTATTCAAGGATTCA | hsa-miR-362-3p | 2,5673762 | 5,927304456 | 0,301286625 |
| TGCCCCAACAAGGAAGGACAAG | hsa-miR-5699-5p | 1,5210362 | 2,86997101 | 0,301286625 |
| CTTTCAGTCAGATGTTTGCTGC | hsa-miR-30d-3p | -0,4426251 | 0,735794533 | 0,301286625 |
| TCTACAGTGCACGTGTCTCCAGT | hsa-miR-139-5p | -1,0565013 | 0,480796626 | 0,301286625 |
| TAGGCAGTGTATTGCTAGCGGCTGT | hsa-miR-449c-5p | -2,2983249 | 0,203299007 | 0,301286625 |
| CATAAAGTAGAAAGCACTACT | hsa-miR-142-5p | 1,9366966 | 3,82828076 | 0,301742948 |
| TGCACCATGGTTGTCTGAGCATG | hsa-miR-767-5p | -0,6526596 | 0,636106566 | 0,306776135 |
| TTAATATCGGACAACCATTGT | hsa-miR-889-3p | -0,7655346 | 0,588235341 | 0,310607336 |
| AATGGATTTTTGGAGCAGG | hsa-miR-1246 | -1,2642497 | 0,416315835 | 0,311280979 |
| CTTCCGGTCTGTGAGCCCCGTC | hsa-miR-4664-3p | -1,3908106 | 0,381350465 | 0,311280979 |
| CATTGCACTTGTCTCGGTCTGA | hsa-miR-25-3p | -0,4634295 | 0,725260152 | 0,31188335 |
| TAAGGTGCATCTAGTGCAGTTAG | hsa-miR-18b-5p | -0,8838155 | 0,541932279 | 0,315290308 |
| ATGACCTATGAATTGACAGAC | hsa-miR-215-5p | -0,7040635 | 0,613840832 | 0,325582818 |
| GCTGCACCGGAGACTGGGTAA | hsa-miR-3130-3p | 2,1330161 | 4,38633523 | 0,326207508 |
| TGGGAACGGGTTCCGGCAGACGCTG | hsa-miR-1292-5p | 1,1515485 | 2,221522074 | 0,327622029 |
| TTCACAGTGGCTAAGTTCTGC | hsa-miR-27b-3p | 0,4062308 | 1,325219011 | 0,327622029 |
| CTTATCAGATTGTATTGTAATT | hsa-miR-374a-3p | -0,4204103 | 0,747212106 | 0,327622029 |
| AAGCCCTTACCCCAAAAAGCAT | hsa-miR-129-2-3p | -0,9984713 | 0,500530104 | 0,327622029 |
| CCACCGGGGGATGAATGTCAC | hsa-miR-181d-3p | 2,2680568 | 4,816739139 | 0,328726906 |
| CGGGGCAGCTCAGTACAGGAT | hsa-miR-486-3p | 1,7898752 | 3,45784972 | 0,328726906 |
| AGCCCCTGCCCACCGCACACTG | hsa-miR-210-5p | 1,1905002 | 2,282318598 | 0,328726906 |
| CGGGGCCGTAGCACTGTCTGAGA | hsa-miR-128-1-5p | 0,5001167 | 1,414327919 | 0,328726906 |
| TAGGACACATGGTCTACTTCT | hsa-miR-1197 | -0,8605865 | 0,550728634 | 0,328726906 |
| TTCATTTGGTATAAACCGCGATT | hsa-miR-579-3p | -1,748218 | 0,297669233 | 0,328726906 |
| TTGTGCTTGATCTAACCATGT | hsa-miR-218-5p | -0,5685554 | 0,674291615 | 0,334242491 |
| AAAAGTAATTGCGGATTTTGCC | hsa-miR-548i | -0,6162669 | 0,652356786 | 0,344320235 |
| GCTGGTCTGCGTGGTGCTCGG | hsa-miR-3663-5p | 1,346796 | 2,543466391 | 0,344961949 |

| | | | | |
|---|---|---|---|---|
| TTAATTTTTTGTTTCGGTCACT | hsa-miR-4775 | 0,5257504 | 1,439682235 | 0,344961949 |
| TGACGCCCCTTCTGATTCTGCCT | hsa-miR-6786-3p | -1,1627801 | 0,446651 | 0,344961949 |
| CATGCCTTGAGTGTAGGACCGT | hsa-miR-532-5p | 0,5957265 | 1,511233424 | 0,347178443 |
| TGTCCTCTAGGGCCTGCAGTCT | hsa-miR-3909 | 0,4446296 | 1,360964663 | 0,347178443 |
| CTCATTTAAGTAGTCTGATGCC | hsa-miR-5696 | -2,5083756 | 0,175753393 | 0,350239327 |
| TCAGCACCAGGATATTGTTGGAG | hsa-miR-3065-3p | 0,6818129 | 1,604154295 | 0,350866605 |
| TGGTCGACCAGTTGGAAAGTAAT | hsa-miR-412-5p | -0,7834063 | 0,580993401 | 0,350866605 |
| TGGAAACATTTCTGCACAAACT | hsa-miR-147b-5p | -0,7867568 | 0,579645679 | 0,350866605 |
| AAGTTCTGTTATACACTCAGGC | hsa-miR-148b-5p | -0,7977756 | 0,575235409 | 0,350866605 |
| CATCATCGTCTCAAATGAGTCT | hsa-miR-136-3p | -0,8217049 | 0,565772929 | 0,350866605 |
| AGTTGCCTTTTTGTTCCCATGC | hsa-miR-4423-5p | -1,7531222 | 0,296659071 | 0,350866605 |
| ATCATGATGGGCTCCTCGGTGT | hsa-miR-433-3p | -0,7786517 | 0,582911297 | 0,35302127 |
| CAAAGTGCTCATAGTGCAGGTAG | hsa-miR-20b-5p | -0,6791777 | 0,624521134 | 0,356828724 |
| GAACGGCTTCATACAGGAGTT | hsa-miR-337-5p | -1,5571422 | 0,339823562 | 0,356828724 |
| GTGAGGGCATGCAGGCCTGGATGGGG | hsa-miR-1226-5p | -0,9798965 | 0,507016108 | 0,35893817 |
| CAAAGAATTCTCCTTTTGGGCT | hsa-miR-186-5p | -0,3188881 | 0,801687514 | 0,359705476 |
| TGGCAGTGTATTGTTAGCTGGT | hsa-miR-449a | -1,7859292 | 0,28998914 | 0,359705476 |
| TGTAAACATCCCCGACTGGAAG | hsa-miR-30d-5p | -0,4312358 | 0,741626249 | 0,364844661 |
| GAAGTTGTTCGTGGTGGATTCG | hsa-miR-382-5p | -0,8524682 | 0,553836404 | 0,369804258 |
| AGAGTTGAGTCTGGACGTCCCG | hsa-miR-219a-1-3p | -0,4654642 | 0,724238032 | 0,371289521 |
| AGGCGGAGACTTGGGCAATTG | hsa-miR-25-5p | -0,8293663 | 0,562776372 | 0,376629033 |
| GGAGACTGATGAGTTCCCGGGA | hsa-miR-873-3p | 0,6176873 | 1,534413486 | 0,382510431 |
| TCCTGTACTGAGCTGCCCCGAG | hsa-miR-486-5p | 0,6332576 | 1,551063346 | 0,38446381 |
| AATATTATACAGTCAACCTCT | hsa-miR-656-3p | -0,9721859 | 0,509733147 | 0,385332951 |
| TCTTTTCTTTGAGACTCACT | hsa-miR-627-3p | -1,2225957 | 0,428511055 | 0,389011869 |
| AAACTCTACTTGTCCTTCTGAGT | hsa-miR-618 | 0,6719766 | 1,593254356 | 0,390519179 |
| CGGGCGTGGTGGTGGGGG | hsa-miR-1268a | 0,5527577 | 1,466886913 | 0,390519179 |
| AGTGCCTGAGGGAGTAAGAGCCC | hsa-miR-550a-5p | 0,4568688 | 1,372559592 | 0,390519179 |
| TCCGGTTCTCAGGGCTCCACC | hsa-miR-671-3p | 0,3499895 | 1,274551353 | 0,390519179 |
| TGGAGACGCGGCCCTGTTGGAGT | hsa-miR-139-3p | -0,7225098 | 0,606042216 | 0,394918451 |
| TAGTGGATGATGCACTCTGTGC | hsa-miR-3681-5p | 0,3960734 | 1,315921519 | 0,397378629 |
| AGAGGCTGGCCGTGATGAATTC | hsa-miR-485-5p | -1,083001 | 0,472045872 | 0,397378629 |
| AAACTAATCTCTACACTGCTGC | hsa-miR-3129-3p | 1,4146825 | 2,666010483 | 0,397504641 |

| | | | | |
|---|---|---|---|---|
| GCAAAGCACACGGCCTGCAGAGA | hsa-miR-330-3p | -1,5147256 | 0,349963017 | 0,399052832 |
| AAAAGTAATTGTGGTTTTTGC | hsa-miR-548ay-5p | -1,8962349 | 0,268643558 | 0,399052832 |
| CTGTGCGTGTGACAGCGGCTGA | hsa-miR-210-3p | -0,4634824 | 0,725233569 | 0,403631203 |
| CGTCAACACTTGCTGGTTTCCT | hsa-miR-505-3p | -0,5712664 | 0,673025768 | 0,413153533 |
| CACCTTGCGCTACTCAGGTCTG | hsa-miR-3200-3p | -0,6035242 | 0,6581443 | 0,414380296 |
| CAAGCTCGCTTCTATGGGTCTG | hsa-miR-99a-3p | 0,678283 | 1,600234095 | 0,417284241 |
| TAGCAGCACATAATGGTTTGTG | hsa-miR-15a-5p | -0,4220874 | 0,746344001 | 0,417284241 |
| CTCGTGGGCTCTGGCCACGGCC | hsa-miR-3677-3p | -0,5491815 | 0,683407765 | 0,417284241 |
| CAACCTCGACGATCTCCTCAGC | hsa-miR-3150a-5p | -1,4015119 | 0,378532235 | 0,417284241 |
| TATGTCTGCTGACCATCACCTT | hsa-miR-654-3p | -1,7658361 | 0,29405621 | 0,417284241 |
| TATTGCACTCGTCCCGGCCTCC | hsa-miR-92b-3p | -0,378948 | 0,768998134 | 0,429594971 |
| CAGCCACAACTACCCTGCCACT | hsa-miR-449b-3p | -1,4037876 | 0,377935614 | 0,429594971 |
| CAATCAGCAAGTATACTGCCCT | hsa-miR-34a-3p | 1,210675 | 2,314459069 | 0,430917424 |
| ACCATCGACCGTTGATTGTACC | hsa-miR-181a-3p | 0,3586483 | 1,282224022 | 0,432447844 |
| ATCATAGAGGAAAATCCATGTT | hsa-miR-376b-3p | -1,6157622 | 0,326292511 | 0,440327553 |
| TCGGATCCGTCTGAGCTTGGCT | hsa-miR-127-3p | -0,6241381 | 0,648807269 | 0,449737817 |
| TTCGCGGGCGAAGGCAAAGTC | hsa-miR-3124-5p | 1,0093903 | 2,013060215 | 0,452251046 |
| TAACAGTCTCCAGTCACGGCC | hsa-miR-212-3p | -0,5624885 | 0,677133163 | 0,452251046 |
| TGGTGGGCCGCAGAACATGTGC | hsa-miR-654-5p | -0,9746717 | 0,508855615 | 0,452251046 |
| ACGGTGCTGGATGTGGCCTTT | hsa-miR-1250-5p | -0,9368198 | 0,522383117 | 0,45765683 |
| TTTCCCTTCAGAGCCTGGCTTT | hsa-miR-4755-5p | 2,2932235 | 4,901500572 | 0,457796726 |
| TTTGTGACCTGGTCCACTAACC | hsa-miR-758-3p | -0,7621485 | 0,589617622 | 0,472214823 |
| ACCGTGGCTTTCGATTGTTACT | hsa-miR-132-5p | -0,5363198 | 0,689527608 | 0,473117792 |
| CAGCAGCACACTGTGGTTTGT | hsa-miR-497-5p | -1,1872366 | 0,439143206 | 0,476304627 |
| ACACATGGGTGGCTGTGGCCT | hsa-miR-4717-3p | -0,9126198 | 0,531219556 | 0,484736974 |
| AATCATACAGGGACATCCAGTT | hsa-miR-487a-3p | -0,8888296 | 0,540052079 | 0,487470174 |
| GAAAAAGTCATGGAGGCC | hsa-miR-12136 | -0,6536592 | 0,635665992 | 0,490063452 |
| TGGGGCGGAGCTTCCGGAGGCC | hsa-miR-3180-3p | 0,5985643 | 1,514208972 | 0,492488082 |
| ACTCGGCGTGGCGTCGGTCGTG | hsa-miR-1307-3p | 0,3813719 | 1,302579886 | 0,492507042 |
| ATAAGACGAACAAAAGGTTTGT | hsa-miR-208b-3p | -0,5096103 | 0,702412161 | 0,495421985 |
| TTTCCTACCCTACCTGAAGACT | hsa-miR-3685 | -1,0695914 | 0,476453918 | 0,503291308 |
| ACTGTAGTATGGGCACTTCCAG | hsa-miR-20b-3p | -0,759764 | 0,590592946 | 0,507343855 |
| AAACATTCGCGGTGCACTTCTT | hsa-miR-543 | -0,8511618 | 0,554338165 | 0,507343855 |

| | | | | |
|---|---|---|---|---|
| TTTAGAGACGGGGTCTTGCTCT | hsa-miR-1303 | 0,6873356 | 1,610306868 | 0,511203533 |
| TTTCCGGCTCGCGTGGGTGTGT | hsa-miR-1180-3p | -0,3005012 | 0,81197029 | 0,511203533 |
| TCAGTGCATCACAGAACTTTGT | hsa-miR-148b-3p | -0,3224297 | 0,799721884 | 0,511203533 |
| CAGGCACGGGAGCTCAGGTGAG | hsa-miR-3622a-5p | -1,3818953 | 0,383714366 | 0,511565122 |
| TCAATCACTTGGTAATTGCTGT | hsa-miR-4705 | 1,6706624 | 3,183607244 | 0,517740504 |
| CGGGTAGAGAGGGCAGTGGGAGG | hsa-miR-197-5p | 0,9370184 | 1,914567304 | 0,517740504 |
| ATCATAGAGGAAAATCCACGT | hsa-miR-376a-3p | -0,975175 | 0,508678152 | 0,518952784 |
| AACATTCATTGTTGTCGGTGGGT | hsa-miR-181d-5p | 0,3027122 | 1,233461081 | 0,523111502 |
| AACACACCTGGTTAACCTCTTT | hsa-miR-329-3p | -1,1830272 | 0,440426397 | 0,523111502 |
| ACGGGTATTCTTGGGTGGATAAT | hsa-miR-137-5p | -0,9984243 | 0,50054639 | 0,523677942 |
| GCGACCCACTCTTGGTTTCCA | hsa-miR-551a | -1,2506082 | 0,42027099 | 0,523677942 |
| TGGCTGTTGGAGGGGGCAGGC | hsa-miR-4687-3p | -1,3535947 | 0,391315807 | 0,523677942 |
| CAAAACTGGCAATTACTTTTGC | hsa-miR-548a-3p | -0,694714 | 0,617831789 | 0,525726474 |
| TGGTAGACTATGGAACGTAGG | hsa-miR-379-5p | -0,6280106 | 0,647068063 | 0,526886238 |
| GCAGTCCATGGGCATATACAC | hsa-miR-455-3p | 0,7517819 | 1,683871336 | 0,541115558 |
| TGTAAACATCCTACACTCTCAGC | hsa-miR-30c-5p | -0,3120227 | 0,805511589 | 0,541115558 |
| AGGTTGGGATCGGTTGCAATGCT | hsa-miR-92a-1-5p | -0,4178237 | 0,748552968 | 0,541115558 |
| TTTAGGATAAGCTTGACTTTTG | hsa-miR-651-5p | -0,6832279 | 0,622770303 | 0,541115558 |
| TTATGGTTTGCCTGGGACTGAG | hsa-miR-584-5p | 1,0172708 | 2,024086366 | 0,543469929 |
| ACAGCAGGCACAGACAGGCAGT | hsa-miR-214-3p | 2,0588526 | 4,166548023 | 0,548838337 |
| GCTCTGACGAGGTTGCACTACT | hsa-miR-301b-5p | -0,6431309 | 0,64032181 | 0,548838337 |
| CTGGGCCCGCGGCGGGCGTGGGG | hsa-miR-6724-5p | -0,6924494 | 0,618802365 | 0,551395676 |
| CTGGATGGCTCCTCCATGTCT | hsa-miR-432-3p | -1,4163826 | 0,374650532 | 0,551395676 |
| TGAGGATATGGCAGGGAAGGGGA | hsa-miR-3679-5p | 0,6412251 | 1,559653027 | 0,553206991 |
| CCAATATTGGCTGTGCTGCTCC | hsa-miR-195-3p | 1,6916814 | 3,230329736 | 0,563917722 |
| GATATCAGCTCAGTAGGCACCG | hsa-miR-3074-3p | -0,5922687 | 0,663299011 | 0,563917722 |
| TGCTGGATCAGTGGTTCGAGTC | hsa-miR-1287-5p | 0,2590252 | 1,196669897 | 0,56559717 |
| ACCGAAGACTGTGCGCTAATCT | hsa-miR-4671-5p | -1,882452 | 0,271222354 | 0,56559717 |
| GCTGACTCCTAGTCCAGGGCTC | hsa-miR-345-5p | -0,2379609 | 0,847942922 | 0,571782469 |
| TTTGGCAATGGTAGAACTCACACT | hsa-miR-182-5p | -0,4284337 | 0,743068058 | 0,571782469 |
| AATATAACACAGATGGCCTGT | hsa-miR-410-3p | -0,5623338 | 0,677205773 | 0,571782469 |
| ATATACAGGGGGAGACTCTCAT | hsa-miR-1185-2-3p | -1,1142461 | 0,461932491 | 0,571782469 |
| TTCTCGAGGAAAGAAGCACTTTC | hsa-miR-516a-5p | -0,6744473 | 0,626572229 | 0,577259995 |

| | | | | |
|---|---|---|---|---|
| ACCTGAGGTTGTGCATTTCTAA | hsa-miR-544b | -1,2492378 | 0,420670394 | 0,577362249 |
| CAGGCAGTGACTGTTCAGACGTC | hsa-miR-2682-5p | 0,4409035 | 1,357454194 | 0,579024066 |
| TATAGGGATTGGAGCCGTGGCG | hsa-miR-135a-3p | -1,2332387 | 0,42536147 | 0,592104397 |
| AGGGACTTTTGGGGGCAGATGTG | hsa-miR-365a-5p | 1,025125 | 2,035135768 | 0,597992138 |
| TGTCGTGGGGCTTGCTGGCTTG | hsa-miR-4440 | 0,933346 | 1,909699943 | 0,597992138 |
| AAAACTGCAGTTACTTTTGC | hsa-miR-548av-3p | -1,7869271 | 0,28978864 | 0,597992138 |
| TTCAAGTAATCCAGGATAGGCT | hsa-miR-26a-5p | 0,2247648 | 1,168586726 | 0,602087317 |
| TAGGGTACTCAGAGCAAGTTGT | hsa-miR-6841-5p | 1,4141727 | 2,665068666 | 0,606191598 |
| TGAGGCTCTGTTAGCCTTGGCTC | hsa-miR-2467-5p | 0,6769529 | 1,598759428 | 0,606191598 |
| GATGATGCTGCTGATGCTG | hsa-miR-1322 | 0,6508732 | 1,570118213 | 0,606191598 |
| ACCCGTCCCGTTCGTCCCCGGA | hsa-miR-1247-5p | -0,5047865 | 0,704764646 | 0,606191598 |
| TATACAAGGGCAAGCTCTCTGT | hsa-miR-381-3p | -0,4534206 | 0,730309262 | 0,607576823 |
| TGGTTTACCGTCCCACATACAT | hsa-miR-299-5p | 1,4430641 | 2,71897731 | 0,609434103 |
| TCTAGTAAGAGTGGCAGTCGA | hsa-miR-628-3p | 1,0057544 | 2,007993157 | 0,609434103 |
| TAACAGTCTACAGCCATGGTCG | hsa-miR-132-3p | 0,3239594 | 1,251761231 | 0,609434103 |
| AGCAGCATTGTACAGGGCTATCA | hsa-miR-107 | -0,3340315 | 0,793316514 | 0,609434103 |
| GAGAAATGCTGGACTAATCTGC | hsa-miR-5680 | -1,3067138 | 0,404240623 | 0,609434103 |
| CTGGGAGAAGGCTGTTTACTCT | hsa-miR-30c-2-3p | -0,2478867 | 0,842129065 | 0,610502051 |
| TGAAGCGCCTGTGCTCTGCCGAGA | hsa-miR-7706 | -0,2868161 | 0,819709069 | 0,610502051 |
| AGGCTGTGATGCTCTCCTGAGCCC | hsa-miR-7974 | -0,3500008 | 0,784583658 | 0,610502051 |
| AAAGATAGACAATTGGCTAAAT | hsa-miR-4662a-3p | -1,1152609 | 0,46160766 | 0,616344275 |
| AGCCAGGCTCTGAAGGGAAAGT | hsa-miR-4755-3p | -1,1904375 | 0,43816995 | 0,616817779 |
| TTGGAATAGGGGATATCTCAGC | hsa-miR-6505-5p | 1,0979836 | 2,140553127 | 0,617602878 |
| AAGGGCTTCCTCTCTGCAGGAC | hsa-miR-3158-3p | 0,4591519 | 1,374733476 | 0,617602878 |
| TGCGGGGACAGGCCAGGGCATC | hsa-miR-4749-5p | -0,9741662 | 0,509033953 | 0,617602878 |
| AGCTTCTTTACAGTGCTGCCTTG | hsa-miR-103a-2-5p | -1,0952829 | 0,468044347 | 0,617602878 |
| TATTGCACTTGTCCCGGCCTGT | hsa-miR-92a-3p | -0,3052202 | 0,809318668 | 0,622629619 |
| AAAAGCTGGGTTGAGAGGGCGA | hsa-miR-320a-3p | 0,263962 | 1,200771818 | 0,623394841 |
| TCGAGGAGCTCACAGTCT | hsa-miR-151b | 0,3720365 | 1,294178425 | 0,624887336 |
| TATGGAGGTCTCTGTCTGGC | hsa-miR-1843 | -0,5017514 | 0,706248876 | 0,632499321 |
| TGAGAACTGAATTCCATAGGCTG | hsa-miR-146b-5p | -0,4167907 | 0,74908914 | 0,638023959 |
| CCTTCACTGTGACTCTGCTGCAG | hsa-miR-6837-3p | -0,4557814 | 0,729115144 | 0,638023959 |
| TCCCTGTCCTCCAGGAGCTCACG | hsa-miR-339-5p | -0,4645665 | 0,724688785 | 0,638023959 |

| | | | | |
|---|---|---|---|---|
| CATCCCTTGCATGGTGGAGGG | hsa-miR-188-5p | -0,5852981 | 0,666511624 | 0,638023959 |
| TTTGGCACTAGCACATTTTTGCT | hsa-miR-96-5p | -1,0484533 | 0,483486228 | 0,639464815 |
| CGAAAACAGCAATTACCTTTGC | hsa-miR-570-3p | -1,3262262 | 0,398810075 | 0,639464815 |
| CTAGGTATGGTCCCAGGGATCC | hsa-miR-331-5p | 0,3286404 | 1,255829344 | 0,64126431 |
| CATCTTCCAGTACAGTGTTGGA | hsa-miR-141-5p | 1,2725198 | 2,415831372 | 0,64523538 |
| CTGCCCTGGCCCGAGGGACCGA | hsa-miR-874-3p | -0,2349705 | 0,849702392 | 0,64596356 |
| TAGTAGACCGTATAGCGTACG | hsa-miR-411-5p | -0,4281826 | 0,743197408 | 0,64596356 |
| TAAAGAGCCCTGTGGAGACA | hsa-miR-1276 | 0,4368124 | 1,353610252 | 0,646264957 |
| TTCCCTTTGTCATCCTATGCCT | hsa-miR-204-5p | 0,3351068 | 1,261470777 | 0,646264957 |
| GGGTGGGGATTTGTTGCATTAC | hsa-miR-92a-2-5p | 0,8175334 | 1,762390232 | 0,648419177 |
| TGTGCTTGCTCGTCCCGCCCGCA | hsa-miR-636 | -1,1412971 | 0,453351807 | 0,649251689 |
| GAATGTTGCTCGGTGAACCCCT | hsa-miR-409-3p | -0,432949 | 0,740746093 | 0,650753215 |
| CAGGATGTGGTCAAGTGTTGTT | hsa-miR-1265 | -1,0871176 | 0,470700851 | 0,650753215 |
| CTGGGATCTCCGGGGTCTTGGTT | hsa-miR-769-3p | -0,2887983 | 0,818583595 | 0,653108689 |
| TGGAGTGTGACAATGGTGTTTG | hsa-miR-122-5p | -0,7814592 | 0,581778058 | 0,65336091 |
| AGCTCGGTCTGAGGCCCCTCAGT | hsa-miR-423-3p | -0,2432013 | 0,844868466 | 0,653748268 |
| ACGCCCTTCCCCCCCTTCTTCA | hsa-miR-1249-3p | 0,3494696 | 1,274092126 | 0,656541499 |
| AATCCTTGGAACCTAGGTGTGAGT | hsa-miR-362-5p | -0,28073 | 0,823174394 | 0,656541499 |
| ACCCCCGGGCAAAGACCTGCAGAT | hsa-miR-6840-5p | 0,776456 | 1,712917842 | 0,662344875 |
| AAAAGTAATCACTGTTTTTGCC | hsa-miR-548y | 0,7374458 | 1,667221462 | 0,662344875 |
| TTCATTCGGCTGTCCAGATGTA | hsa-miR-1298-5p | 0,365313 | 1,2881611 | 0,662344875 |
| TTATTGCTTAAGAATACGCGTAG | hsa-miR-137-3p | -0,4598176 | 0,727078177 | 0,662344875 |
| TAACGCATAATATGGACATGT | hsa-miR-3912-3p | -0,7660823 | 0,588012086 | 0,662344875 |
| AAGCATTCTTTCATTGGTTGG | hsa-miR-1179 | -1,043572 | 0,48512487 | 0,662344875 |
| TCCGAACTCTCCATTCCTCTGC | hsa-miR-6716-3p | 0,4375115 | 1,354266321 | 0,663009209 |
| TCAGAACAAATGCCGGTTCCCAGA | hsa-miR-589-3p | -0,3365201 | 0,791949264 | 0,664599414 |
| CTCACTGAACAATGAATGCAA | hsa-miR-181b-3p | -0,3503515 | 0,784392948 | 0,664599414 |
| ACTGCCCTAAGTGCTCCTTCTGG | hsa-miR-18a-3p | -0,2827237 | 0,822037594 | 0,666384702 |
| CTTCCCCCCAGTAATCTTCATC | hsa-miR-3679-3p | 1,1775262 | 2,26188595 | 0,668068758 |
| CTGACCTATGAATTGACAGCC | hsa-miR-192-5p | 0,255541 | 1,193783295 | 0,670865885 |
| CACATTACACGGTCGACCTCT | hsa-miR-323a-3p | -0,3598175 | 0,779263163 | 0,670865885 |
| GAGAGGAACATGGGCTCAGGACA | hsa-miR-6859-5p | 0,5470885 | 1,46113398 | 0,675707461 |
| CTCTAGCCACAGATGCAGTGAT | hsa-miR-1287-3p | 1,1643794 | 2,241367809 | 0,676498779 |

| | | | | |
|---|---|---|---|---|
| CCTATTCTTGATTACTTGTTTC | hsa-miR-26a-2-3p | -0,2282171 | 0,853689253 | 0,679690105 |
| TTCCATGCCTCCTAGAAGTTCC | hsa-miR-5581-3p | 0,9724123 | 1,962118636 | 0,685072744 |
| CCAATATTACTGTGCTGCTTTA | hsa-miR-16-2-3p | -0,330199 | 0,795426739 | 0,68599859 |
| CATCTGGCATCCGTCACACAGA | hsa-miR-3126-3p | 0,8482454 | 1,800310019 | 0,693832661 |
| TCAACAAAATCACTGATGCTGGA | hsa-miR-3065-5p | 0,8271759 | 1,774208945 | 0,693832661 |
| TGCACGGCACTGGGGACACGT | hsa-miR-3177-3p | 0,3419741 | 1,267489777 | 0,693832661 |
| ACTGGACTTGGAGTCAGAAGGC | hsa-miR-378a-3p | 0,3041891 | 1,23472441 | 0,693832661 |
| CAACAAATCACAGTCTGCCATA | hsa-miR-7-1-3p | -0,3061886 | 0,808775628 | 0,693832661 |
| TTGAGAATGATGAATCATTAGG | hsa-miR-580-3p | -0,951982 | 0,516921804 | 0,693832661 |
| AGCCCGCCCCAGCCGAGGTTCT | hsa-miR-4707-3p | 0,7206935 | 1,647974009 | 0,697474339 |
| ATGGCCAGAGCTCACACAGAGG | hsa-miR-4435 | 0,5214873 | 1,435434315 | 0,697474339 |
| TAGCAGCGGGAACAGTTCTGCAG | hsa-miR-503-5p | 1,0381106 | 2,053536467 | 0,700594206 |
| TCTGGGCAACAAAGTGAGACCT | hsa-miR-1285-3p | 0,2881918 | 1,221108821 | 0,700594206 |
| AATGGCTGTCCGTAGTATGGTC | hsa-miR-889-5p | -0,854433 | 0,553082661 | 0,700604496 |
| GTGTGCGGAAATGCTTCTGCT | hsa-miR-147b-3p | 1,0189688 | 2,026470005 | 0,702087654 |
| ACTGCATTATGAGCACTTAAAG | hsa-miR-20a-3p | -0,4949713 | 0,70957578 | 0,702087654 |
| CCAGCCACGGACTGAGAGTGCAT | hsa-miR-4691-3p | -0,8979168 | 0,536661103 | 0,702087654 |
| TTCCCTTTGTCATCCTTCGCCT | hsa-miR-211-5p | -0,6106282 | 0,654911488 | 0,703925025 |
| ATATAATACAACCTGCTAAGTG | hsa-miR-374b-5p | -0,2412582 | 0,846007193 | 0,704125445 |
| TGATATGTTTGATATATTAGGT | hsa-miR-190a-5p | 0,4132989 | 1,331727473 | 0,706723971 |
| TCGTGTCTTGTGTTGCAGCCGG | hsa-miR-187-3p | 1,2579568 | 2,391567914 | 0,707898234 |
| TCGGCTCTCTCCCTCACCCTAG | hsa-miR-6741-3p | 0,8845621 | 1,846204128 | 0,707898234 |
| TAGAGGAAGCTGTGGAGAGA | hsa-miR-3125 | 0,693691 | 1,617416214 | 0,707898234 |
| TATGTGCCTTTGGACTACATCG | hsa-miR-455-5p | 0,3381567 | 1,264140437 | 0,707898234 |
| AAAAGTACTTGCGGATTTTGCT | hsa-miR-548k | -0,1444708 | 0,904711198 | 0,707898234 |
| TGAGACCAGGACTGGATGCACC | hsa-miR-4786-5p | -0,4256981 | 0,744478382 | 0,707898234 |
| CTCAGTAGCCAGTGTAGATCCT | hsa-miR-222-5p | -0,4399894 | 0,737140047 | 0,707898234 |
| CCATGGATCTCCAGGTGGGT | hsa-miR-490-5p | -0,4446489 | 0,734763135 | 0,707898234 |
| AAAAGTAATTGTGGTTTTTGCC | hsa-miR-548d-5p | -0,6400945 | 0,641670911 | 0,707898234 |
| CCTCTGGGCCCTTCCTCCAG | hsa-miR-326 | -0,9938431 | 0,502138391 | 0,707898234 |
| AAGCAATACTGTTACCTGAAAT | hsa-miR-3942-5p | 0,8060224 | 1,748384351 | 0,710361784 |
| AGTGGGGAACCCTTCCATGAGG | hsa-miR-491-5p | 0,5569155 | 1,471120548 | 0,710361784 |
| ACGGGTTAGGCTCTTGGGAGCT | hsa-miR-125b-1-3p | 0,1741684 | 1,128313857 | 0,710361784 |

| | | | | |
|---|---|---|---|---|
| ATCACACAAAGGCAACTTTTGT | hsa-miR-377-3p | -0,9962326 | 0,501307394 | 0,710361784 |
| ATTCTCTCTGGATCCCATGGAT | hsa-miR-4768-5p | -0,6253274 | 0,648272636 | 0,713865786 |
| CGGCTCTGGGTCTGTGGGGA | hsa-miR-760 | -0,461503 | 0,726229267 | 0,717670559 |
| TGAGTATTACATGGCCAATCTC | hsa-miR-496 | -0,6473687 | 0,638443716 | 0,71915089 |
| CTTTCAGTCGGATGTTTGCAGC | hsa-miR-30a-3p | -0,1901713 | 0,876501648 | 0,72071796 |
| TGATTGTAGCCTTTTGGAGTAGA | hsa-miR-508-3p | 0,6563231 | 1,576060693 | 0,722168735 |
| TTGTTCTCAAACTGGCTGTCAGA | hsa-miR-7156-5p | -0,3710771 | 0,773205028 | 0,722168735 |
| AAAGACATAGGATAGAGTCACCTC | hsa-miR-641 | 0,2782053 | 1,212685377 | 0,724309138 |
| TAATACTGTCTGGTAAAACCGT | hsa-miR-429 | -0,5681915 | 0,674461727 | 0,72967846 |
| GTGAGTCTCTAAGAAAAGAGGA | hsa-miR-627-5p | -0,7086582 | 0,611888974 | 0,72967846 |
| AGGGGGAAAGTTCTATAGTCC | hsa-miR-625-5p | 0,8519187 | 1,804899766 | 0,733485904 |
| TGAGGGACAGATGCCAGAAGCA | hsa-miR-3126-5p | 0,6248673 | 1,542068936 | 0,733485904 |
| TTACAGTTGTTCAACCAGTTACT | hsa-miR-582-5p | 0,309804 | 1,239539296 | 0,733485904 |
| TGTAAACATCCTACACTCAGCT | hsa-miR-30b-5p | -0,1244752 | 0,917337667 | 0,733485904 |
| ACCCCACTCCTGGTACC | hsa-miR-4286 | -0,3180097 | 0,802175782 | 0,733485904 |
| AGAGGTTGCCCTTGGTGAATTC | hsa-miR-377-5p | -0,7540987 | 0,592916699 | 0,733485904 |
| ATCCCACCACTGCCACCAT | hsa-miR-1260b | 0,663198 | 1,583589089 | 0,737469098 |
| TGAAGGTCTACTGTGTGCCAGG | hsa-miR-493-3p | 0,4013534 | 1,320746317 | 0,737469098 |
| AATGGCGCCACTAGGGTTGTG | hsa-miR-652-3p | 0,1724139 | 1,126942494 | 0,737469098 |
| TCAAGAGCAATAACGAAAAATGT | hsa-miR-335-5p | -0,2117298 | 0,863501263 | 0,737469098 |
| CAGGTCGTCTTGCAGGGCTTCT | hsa-miR-431-3p | -0,4511043 | 0,731482701 | 0,751948895 |
| GATGATGATGGCAGCAAATTCTGAAA | hsa-miR-1272 | -0,5297182 | 0,692690026 | 0,758476123 |
| TCTGAGTTCCTGGAGCCTGGTCT | hsa-miR-4682 | 0,8581795 | 1,812749384 | 0,760383683 |
| GTGGGGGAGAGGCTGTC | hsa-miR-1275 | 0,2657683 | 1,202276114 | 0,760780111 |
| CGTGTTCACAGCGGACCTTGAT | hsa-miR-124-5p | -0,1965685 | 0,872623647 | 0,760780111 |
| ATATGGGTTTACTAGTTGGT | hsa-miR-3115 | -0,2987376 | 0,812963448 | 0,761390098 |
| TCTCTGGGCCTGTGTCTTAGGC | hsa-miR-330-5p | -0,3161522 | 0,803209239 | 0,761390098 |
| CTTATGCAAGATTCCCTTCTAC | hsa-miR-491-3p | 0,4769249 | 1,391773909 | 0,761734823 |
| TTGTACATGGTAGGCTTTCATT | hsa-miR-493-5p | -0,3107114 | 0,806244092 | 0,761734823 |
| CTGCAATGTAAGCACTTCTTAC | hsa-miR-106a-3p | -0,5469877 | 0,68444776 | 0,76232865 |
| AGGGACGGGACGCGGTGCAGTG | hsa-miR-92b-5p | 0,2630207 | 1,199988637 | 0,762571107 |
| CTAGTGCTCTCCGTTACAAGTA | hsa-miR-4473 | -0,5434351 | 0,686135243 | 0,762571107 |
| AGAATTGCGTTTGGACAATCAGT | hsa-miR-219b-3p | 0,5624324 | 1,476756928 | 0,763376827 |

| | | | | |
|---|---|---|---|---|
| TTAGGGCCCTGGCTCCATCTCC | hsa-miR-1296-5p | 0,1860682 | 1,137658991 | 0,766726962 |
| AATGCACCCGGGCAAGGATTCT | hsa-miR-501-3p | -0,1558246 | 0,897619184 | 0,766726962 |
| ATCAACAGACATTAATTGGGCGC | hsa-miR-421 | -0,1581491 | 0,896174106 | 0,767308325 |
| TTAGTGCATAGTCTTTGGTCT | hsa-miR-4671-3p | 0,5955795 | 1,511079428 | 0,768973571 |
| CTGGGGGACGCGTGAGCGCGAGC | hsa-miR-4665-5p | 0,6069148 | 1,522998816 | 0,770922739 |
| ACTGCTGAGCTAGCACTTCCCG | hsa-miR-93-3p | -0,3570957 | 0,780734716 | 0,770922739 |
| AAAGGATTCTGCTGTCGGTCCCACT | hsa-miR-541-5p | -0,4988313 | 0,707679844 | 0,770922739 |
| TGAGACCTCTGGGTTCTGAGCT | hsa-miR-769-5p | 0,2447048 | 1,184850279 | 0,77236302 |
| TGGACTGCCCTGATCTGGAGA | hsa-miR-1288-3p | 0,4593337 | 1,374906692 | 0,772755718 |
| GATGCGCCGCCCACTGCCCCGCGC | hsa-miR-4787-3p | 0,3980049 | 1,317684402 | 0,772755718 |
| AGCTACATCTGGCTACTGGGT | hsa-miR-222-3p | 0,2407794 | 1,181630866 | 0,772755718 |
| CGGCGGGGACGGCGATTGGTC | hsa-miR-1908-5p | 0,2079773 | 1,155067627 | 0,772755718 |
| TGGTGGTTTACAAAGTAATTCA | hsa-miR-876-3p | -0,8988321 | 0,536320726 | 0,772961003 |
| AGCAGCATTGTACAGGGCTATGA | hsa-miR-103a-3p | -0,1417913 | 0,906393076 | 0,774182693 |
| CCTCAGGGCTGTAGAACAGGGCT | hsa-miR-1266-5p | -0,2374752 | 0,848228448 | 0,774182693 |
| AGGGCTGGACTCAGCGGCGGAGCT | hsa-miR-5001-5p | 0,543109 | 1,457109158 | 0,776804207 |
| AACTCGTGTTCAAAGCCTTTAG | hsa-miR-4636 | 0,3772611 | 1,298873618 | 0,776804207 |
| CTTTTTGCGGTCTGGGCTTGC | hsa-miR-129-5p | -0,3022363 | 0,810994286 | 0,776804207 |
| TCCCTGAGACCCTAACTTGTGA | hsa-miR-125b-5p | 0,2260983 | 1,169667365 | 0,777243748 |
| TTTTTCATTATTGCTCCTGACC | hsa-miR-335-3p | 0,1343889 | 1,097627774 | 0,781267862 |
| AATCATTCACGGACAACACTT | hsa-miR-382-3p | -0,6090368 | 0,655634262 | 0,781267862 |
| AGTTTTGCAGGTTTGCATCCAGC | hsa-miR-19b-1-5p | 0,4522278 | 1,368151355 | 0,785009361 |
| TATGTGGGATGGTAAACCGCTT | hsa-miR-299-3p | -0,4135959 | 0,750749805 | 0,787531226 |
| CTGTACAGCCTCCTAGCTTTCC | hsa-let-7a-2-3p | 0,838276 | 1,787912329 | 0,788914117 |
| TGTTGTACTTTTTTTTTTGTTC | hsa-miR-3613-5p | 0,2676371 | 1,203834494 | 0,788914117 |
| ATAAAGCTAGATAACCGAAAGT | hsa-miR-9-3p | 0,2531745 | 1,191826757 | 0,788914117 |
| AATCAGTGAATGCCTTGAACCT | hsa-miR-5094 | -0,4627304 | 0,725611678 | 0,788914117 |
| CTGAAGCTCAGAGGGCTCTGAT | hsa-miR-127-5p | -0,6109318 | 0,654773668 | 0,788914117 |
| GACCGAGAGGGCCTCGGCTGT | hsa-miR-4523 | -0,2635262 | 0,833049309 | 0,789659126 |
| CAGTGGTTTTACCCTATGGTAG | hsa-miR-140-5p | -0,1829625 | 0,880892283 | 0,792538109 |
| CTGGCCCTCTCTGCCCTTCCGT | hsa-miR-328-3p | 0,1837219 | 1,135810308 | 0,796335946 |
| TGTGTCACTCGATGACCACTGT | hsa-miR-597-5p | -0,6055167 | 0,657235961 | 0,796335946 |
| ATTCTAATTTCTCCACGTCTTT | hsa-miR-576-5p | 0,1440293 | 1,104986924 | 0,803545216 |

| | | | | |
|---|---|---|---|---|
| TGCAGGACCAAGATGAGCCCT | hsa-miR-1286 | 0,1680252 | 1,123519566 | 0,809218498 |
| TAATACTGCCGGGTAATGATGGA | hsa-miR-200c-3p | -0,2241892 | 0,856076025 | 0,816626658 |
| TCTGCAAGTGTCAGAGGCGAGG | hsa-miR-2276-3p | -0,5251863 | 0,694869371 | 0,816626658 |
| GTGAAATGTTTAGGACCACTAG | hsa-miR-203a-3p | -0,588814 | 0,664889273 | 0,816626658 |
| TGTGTGGATCCTGGAGGAGGCA | hsa-miR-3911 | 0,4867559 | 1,401290365 | 0,818171966 |
| GCCCTGAACGAGGGGTCTGGAG | hsa-miR-345-3p | -0,5146461 | 0,69996464 | 0,818171966 |
| TGTAAACATCCTCGACTGGAAG | hsa-miR-30a-5p | -0,1646895 | 0,892120506 | 0,819181974 |
| TAATCCTTGCTACCTGGGTGAGA | hsa-miR-500a-5p | 0,3669882 | 1,289657709 | 0,820806589 |
| TTGGAGGGTGTGGAAGACATC | hsa-miR-6515-5p | 0,2736291 | 1,208844887 | 0,827292081 |
| TTGCATATGTAGGATGTCCCAT | hsa-miR-448 | 0,2294196 | 1,172363238 | 0,827292081 |
| TGAGAACCACGTCTGCTCTGAG | hsa-miR-589-5p | -0,1634501 | 0,892887246 | 0,827292081 |
| GCCCCTGGGCCTATCCTAGAA | hsa-miR-331-3p | -0,2104216 | 0,86428463 | 0,827292081 |
| AGGAAGCCCTGGAGGGGCTGGAG | hsa-miR-671-5p | -0,2436502 | 0,844605676 | 0,827292081 |
| CATCTGGGCAACTGACTGAAC | hsa-miR-1298-3p | -0,6243 | 0,64873448 | 0,827292081 |
| CCTAATTTGAACACCTTCGGTA | hsa-miR-4735-5p | 0,3364771 | 1,262669498 | 0,836303663 |
| CCTCACCATCCCTTCTGCCTGC | hsa-miR-6511a-3p | 0,3550356 | 1,279017138 | 0,842020552 |
| CAGCCCGGATCCCAGCCCACTT | hsa-miR-3940-3p | -0,4188388 | 0,748026467 | 0,843622155 |
| GAGCTTATTCATAAAAGTGCAG | hsa-miR-590-5p | 0,581068 | 1,49595629 | 0,847595832 |
| TGCCTGGAACATAGTAGGGACT | hsa-miR-3116 | 0,3455427 | 1,270628873 | 0,847595832 |
| ACCTGGCATACAATGTAGATTT | hsa-miR-221-5p | 0,2874764 | 1,220503482 | 0,847595832 |
| TGACCTGGGACTCGGACAGCTG | hsa-miR-3661 | 0,2199073 | 1,164658764 | 0,848220569 |
| CTTGGCACCTAGCAAGCACTCA | hsa-miR-1271-5p | 0,0946363 | 1,067796222 | 0,848220569 |
| TCAGTAAATGTTTATTAGATGA | hsa-miR-545-5p | -0,4669949 | 0,723469996 | 0,848275183 |
| ACAGAGGACAGTGGAGTGTGAGC | hsa-miR-6847-5p | 0,3273506 | 1,2547071 | 0,85117295 |
| AACTAGCTCTGTGGATCCTGAC | hsa-miR-4661-5p | -0,3501567 | 0,784498908 | 0,852306372 |
| TGGTTCTCTTGTGGCTCAAGCGT | hsa-miR-597-3p | -0,5074002 | 0,703489022 | 0,852306372 |
| CCTCCCACACCCAAGGCTTGCA | hsa-miR-532-3p | 0,1724627 | 1,126980609 | 0,855970484 |
| TGAGCGCCTCGACGACAGAGCCG | hsa-miR-339-3p | 0,1131365 | 1,08157711 | 0,858537927 |
| CAGGGGGACTGGGGGTGAGC | hsa-miR-6794-5p | 0,5304772 | 1,44440693 | 0,863069369 |
| TCGAGGAGCTCACAGTCTAGT | hsa-miR-151a-5p | 0,0804487 | 1,057346855 | 0,863069369 |
| CTGGGAGAGGGTTGTTTACTCC | hsa-miR-30c-1-3p | -0,1204355 | 0,919909945 | 0,863069369 |
| TACCCATTGCATATCGGAGTTG | hsa-miR-660-5p | -0,1297691 | 0,91397775 | 0,863069369 |
| TGCCCTGCCTGTTTTCTCCTTT | hsa-miR-3173-5p | 0,3716713 | 1,293850808 | 0,864008503 |

| | | | | |
|---|---|---|---|---|
| CTCTCACCACTGCCCTCCCACAG | hsa-miR-1229-3p | -0,2024275 | 0,869087004 | 0,864008503 |
| GGGGGCCGATACACTGTACGAGA | hsa-miR-128-2-5p | 0,3386864 | 1,264604604 | 0,865224168 |
| CGGGTGGATCACGATGCAATTT | hsa-miR-363-5p | -0,2131 | 0,862681537 | 0,865224168 |
| TAACAAACACCTGTAAAACAGC | hsa-miR-5688 | -0,4071836 | 0,754094085 | 0,865224168 |
| AAGGCAGGGCCCCCGCTCCCC | hsa-miR-940 | -0,1322285 | 0,91242099 | 0,867579573 |
| TGGCCCGGCGACGTCTCACGGTC | hsa-miR-4745-3p | -0,6206243 | 0,650389419 | 0,867579573 |
| TCAGTGCATGACAGAACTTGG | hsa-miR-152-3p | 0,1774426 | 1,130877452 | 0,867830541 |
| TAGCAGCACAGAAATATTGGC | hsa-miR-195-5p | -0,1298858 | 0,913903764 | 0,867830541 |
| TAATACTGCCTGGTAATGATGA | hsa-miR-200b-3p | -0,4410124 | 0,736617492 | 0,875058877 |
| CCCTGTGCCCGGCCCACTTCTG | hsa-miR-1914-5p | -0,412877 | 0,751124004 | 0,881682816 |
| TCAGGCTCAGTCCCCTCCCGAT | hsa-miR-484 | -0,0799752 | 0,946073878 | 0,883240604 |
| CGGCCCGGGCTGCTGCTGTTCCT | hsa-miR-1538 | -0,2513606 | 0,840103722 | 0,883240604 |
| CCGGTCCCAGGAGAACCTGCAGA | hsa-miR-4746-5p | 0,1060332 | 1,076264859 | 0,884394596 |
| TAGTACCAGTACCTTGTGTTCA | hsa-miR-624-5p | 0,3960179 | 1,315870809 | 0,888931032 |
| GGAGGAACCTTGGAGCTTCGGC | hsa-miR-3928-3p | 0,2926405 | 1,224880042 | 0,888931032 |
| ATGGGTGAATTTGTAGAAGGAT | hsa-miR-1262 | 0,256461 | 1,194544838 | 0,888931032 |
| ATCAGGGCTTGTGGAATGGGAAG | hsa-miR-3127-5p | 0,1827731 | 1,135063595 | 0,888931032 |
| GACTATAGAACTTTCCCCCTCA | hsa-miR-625-3p | -0,1483375 | 0,902289649 | 0,888931032 |
| CCGGCCGCCGGCTCCGCCCCG | hsa-miR-1908-3p | -0,2324229 | 0,851204169 | 0,888931032 |
| CAACAAATCCCAGTCTACCTAA | hsa-miR-7-2-3p | -0,4404101 | 0,736925093 | 0,888931032 |
| TCTCTCGGCTCCTCGCGGCTC | hsa-miR-3615 | 0,1873087 | 1,138637619 | 0,890240523 |
| TGGTGGGCACAGAATCTGGACT | hsa-miR-541-3p | -0,1677722 | 0,890216272 | 0,890240523 |
| GTTCCTGCTGAACTGAGCCAG | hsa-miR-3074-5p | -0,254379 | 0,838347907 | 0,895383552 |
| TTCACTGGAGTTTGTTTCAATA | hsa-miR-653-3p | 0,6619327 | 1,582200817 | 0,896840342 |
| TCTTTGGTTATCTAGCTGTATGA | hsa-miR-9-5p | 0,0954251 | 1,068380135 | 0,899887234 |
| ATCGGGAATGTCGTGTCCGCCC | hsa-miR-425-3p | 0,1025315 | 1,073655723 | 0,903104728 |
| TACGTCATCGTTGTCATCGTCA | hsa-miR-598-3p | -0,0941784 | 0,936805566 | 0,903104728 |
| CGAATCATTATTTGCTGCTCTA | hsa-miR-15b-3p | -0,1193642 | 0,920593262 | 0,903104728 |
| TGCCTACTGAGCTGATATCAGT | hsa-miR-24-1-5p | -0,2273685 | 0,854191507 | 0,903104728 |
| GGGAGAAGGGTCGGGGC | hsa-miR-4516 | -0,2918858 | 0,816833646 | 0,903104728 |
| TCAGGACACTTCTGAACTTGGA | hsa-miR-5000-3p | -0,3186445 | 0,801822872 | 0,903104728 |
| TTCCTGGGCTTCTCCTCTGTAG | hsa-miR-6783-3p | 0,3545905 | 1,278622607 | 0,903923352 |
| ACTCAAAACCCTTCAGTGACTT | hsa-miR-616-5p | 0,1811672 | 1,133800776 | 0,903923352 |

| | | | | |
|---|---|---|---|---|
| ACACACTTACCCGTAGAGATTCTA | hsa-miR-216b-3p | -0,5005156 | 0,706854101 | 0,907373001 |
| CCTATTCTTGGTTACTTGCACG | hsa-miR-26a-1-3p | -0,1938307 | 0,874281226 | 0,917277933 |
| CACCCGGCTGTGTGCACATGTGC | hsa-miR-941 | 0,0445064 | 1,031330238 | 0,917348043 |
| AACCCGTAGATCCGATCTTGTG | hsa-miR-99a-5p | -0,067432 | 0,954335219 | 0,917348043 |
| GAAAATGATGAGTAGTGACTGATG | hsa-miR-3662 | -0,33752 | 0,791400548 | 0,919342082 |
| CCTCCGTGTTACCTGTCCTCTAG | hsa-miR-3605-3p | 0,1094577 | 1,078822659 | 0,921090452 |
| TCACCAGCCCTGTGTTCCCTAG | hsa-miR-1226-3p | 0,0997776 | 1,071608219 | 0,921090452 |
| CTTAGCAGGTTGTATTATCATT | hsa-miR-374b-3p | 0,05838 | 1,041295867 | 0,921090452 |
| TGCGGGGCTAGGGCTAACAGCA | hsa-miR-744-5p | 0,0530543 | 1,037458999 | 0,921090452 |
| TGTAACAGCAACTCCATGTGGA | hsa-miR-194-5p | -0,06593 | 0,955329277 | 0,921090452 |
| ATCCGCGCTCTGACTCTCTGCC | hsa-miR-937-3p | -0,0962988 | 0,935429719 | 0,921090452 |
| GCTCTTTTCACATTGTGCTACT | hsa-miR-130a-5p | -0,1066115 | 0,928766928 | 0,921090452 |
| CAACTAGACTGTGAGCTTCTAG | hsa-miR-708-3p | -0,1148995 | 0,923446615 | 0,921090452 |
| AGAAGGAAATTGAATTCATTTA | hsa-miR-1252-5p | -0,1419938 | 0,906265858 | 0,921090452 |
| TGGGCCAGGGAGCAGCTGGTGGG | hsa-miR-4640-5p | -0,2112222 | 0,863805116 | 0,921090452 |
| TGCCCTTCTCTCCTCCTGCCT | hsa-miR-6886-3p | 0,3142398 | 1,243356292 | 0,923860024 |
| CTGTCCTAAGGTTGTTGAGTT | hsa-miR-676-3p | 0,1262289 | 1,091437029 | 0,926273702 |
| AGGACCTTCCCTGAACCAAGGA | hsa-miR-659-5p | 0,1035305 | 1,074399455 | 0,926273702 |
| AAAAGTTATTGCGGTTTTGGCT | hsa-miR-548at-5p | -0,1382913 | 0,908594652 | 0,926273702 |
| TGAGTGGGGCTCCCGGGACGGCG | hsa-miR-4745-5p | -0,1710299 | 0,888208405 | 0,926273702 |
| ATGCACCTGGGCAAGGATTCTG | hsa-miR-500a-3p | 0,0622633 | 1,044102436 | 0,926822354 |
| TGTGAGGTTGGCATTGTTGTCT | hsa-miR-1294 | 0,1330846 | 1,096635925 | 0,927596438 |
| CAGGGAAATGGGAAGAACTAGA | hsa-miR-5584-5p | -0,208307 | 0,865552347 | 0,928335316 |
| CTCACTGATCAATGAATGCA | hsa-miR-181b-2-3p | 0,1133472 | 1,081735087 | 0,929161782 |
| TTACGGACCAGCTAAGGGAGGC | hsa-miR-4788 | 0,133195 | 1,096719822 | 0,930355898 |
| AAAAACTGAGACTACTTTTGCA | hsa-miR-548e-3p | 0,112786 | 1,081314355 | 0,930355898 |
| TCTGGCTCCGTGTCTTCACTCCC | hsa-miR-149-5p | -0,0478758 | 0,967359581 | 0,930702984 |
| AAAGCAAATGTTGGGTGAACGGC | hsa-miR-10527-5p | -0,2610597 | 0,834474764 | 0,931636453 |
| AAAAGTATTTGCGGGTTTTGTC | hsa-miR-548l | 0,1216487 | 1,08797745 | 0,932022494 |
| AGAATTGTGGCTGGACATCTGT | hsa-miR-219a-2-3p | -0,0805722 | 0,9456825 | 0,932747329 |
| CAGCCCTCCTCCCGCACCCAAA | hsa-miR-4687-5p | 0,1192043 | 1,086135625 | 0,935560381 |
| GTGGGTTGGGGCGGGCTCTG | hsa-miR-3940-5p | -0,1674174 | 0,890435269 | 0,935560381 |
| TTATCCTCCAGTAGACTAGGGA | hsa-miR-9903 | -0,1003659 | 0,932796413 | 0,936567634 |

| | | | | |
|---|---|---|---|---|
| CCAAAACTGCAGTTACTTTTGC | hsa-miR-548o-3p | 0,0436418 | 1,030712377 | 0,936933942 |
| CAACGGAATCCCAAAAGCAGCTG | hsa-miR-191-5p | 0,0336509 | 1,02359917 | 0,93815184 |
| AGCTGGTGTTGTGAATCAGGCCG | hsa-miR-138-5p | 0,0652909 | 1,046295857 | 0,939923512 |
| CGGGCTGTCCGGAGGGGTCGGCT | hsa-miR-4741 | -0,0711214 | 0,951897826 | 0,945958847 |
| GAAAGTAATTGCTGTTTTTGCC | hsa-miR-548aq-5p | 0,1970544 | 1,146355398 | 0,952421037 |
| CTCCTATATGATGCCTTTCTTC | hsa-miR-337-3p | 0,1572719 | 1,115176343 | 0,954878059 |
| AAAAGCTGGGTTGAGAGGGT | hsa-miR-320c | 0,1082457 | 1,077916693 | 0,960720802 |
| GCTGCGCTTGGATTTCGTCCCC | hsa-miR-191-3p | -0,0633254 | 0,957055591 | 0,965893512 |
| AAACAAACATGGTGCACTTCTT | hsa-miR-495-3p | -0,0968205 | 0,935091547 | 0,965893512 |
| CTGTACAACCTTCTAGCTTTCC | hsa-let-7c-3p | 0,067578 | 1,047955861 | 0,966311643 |
| CTTTCAGTCGGATGTTTACAGC | hsa-miR-30e-3p | -0,0340315 | 0,976687224 | 0,966311643 |
| TGCAGCTCTGGTGGAAAATGGAG | hsa-miR-4660 | 0,1215891 | 1,087932502 | 0,967960788 |
| CGGCCCCACGCACCAGGGTAAGA | hsa-miR-874-5p | 0,0421126 | 1,029620474 | 0,967960788 |
| AAAAGCTGGGTTGAGAGGGCAA | hsa-miR-320b | 0,118033 | 1,085254202 | 0,968132256 |
| TGGGGAGCTGAGGCTCTGGGGGTG | hsa-miR-939-5p | 0,1464775 | 1,106863652 | 0,968736238 |
| CTCCCACATGCAGGGTTTGCA | hsa-miR-188-3p | -0,0381008 | 0,973936247 | 0,968736238 |
| TAGATAAAATATTGGTACCTG | hsa-miR-577 | -0,0519249 | 0,964648366 | 0,968736238 |
| CGGGGAGAGAACGCAGTGACGT | hsa-miR-3175 | -0,0657494 | 0,955448879 | 0,968736238 |
| TAACACTGTCTGGTAACGATGT | hsa-miR-200a-3p | -0,1321086 | 0,912496793 | 0,968736238 |
| TGGGGAGCGGCCCCCGGGTGGG | hsa-miR-1343-5p | -0,1530398 | 0,899353516 | 0,968736238 |
| CGCATCCCCTAGGGCATTGGTG | hsa-miR-324-5p | -0,033331 | 0,977161573 | 0,968980838 |
| AATGCACCTGGGCAAGGATTCA | hsa-miR-502-3p | 0,0525688 | 1,037109933 | 0,969754778 |
| TGGGAAAGACAAACTCAGAGTT | hsa-miR-6733-5p | -0,0722862 | 0,951129546 | 0,972530417 |
| CTCCTGGGGCCCGCACTCTCGC | hsa-miR-1343-3p | -0,027491 | 0,981125108 | 0,97584891 |
| TTTGGGACTGATCTTGATGTCT | hsa-miR-3913-5p | -0,0417079 | 0,971504205 | 0,97584891 |
| TCACAGTGAACCGGTCTCTTT | hsa-miR-128-3p | 0,015464 | 1,010776502 | 0,979583806 |
| AATTTGGTTTCTGAGGCACTTAGT | hsa-miR-5002-5p | -0,0279508 | 0,980812479 | 0,979583806 |
| TTCAAGTAATTCAGGATAGGT | hsa-miR-26b-5p | 0,0138078 | 1,009616809 | 0,982240211 |
| GTCATACACGGCTCTCCTCTCT | hsa-miR-485-3p | -0,0471905 | 0,967819237 | 0,982240211 |
| TATTGCACATTACTAAGTTGCA | hsa-miR-32-5p | 0,054292 | 1,038349427 | 0,982610283 |
| GCTGGGAAGGCAAAGGGACGT | hsa-miR-204-3p | -0,0403673 | 0,97240733 | 0,982610283 |
| TCAGCTACTACCTCTATTAGG | hsa-miR-5690 | 0,0411838 | 1,028957779 | 0,985999953 |
| ATTGTCCTTGCTGTTTGGAGAT | hsa-miR-2355-3p | 0,0351369 | 1,024654019 | 0,987435694 |

| | | | | |
|---|---|---|---|---|
| GGGGCTGGGCGCGCGCC | hsa-miR-4492 | -0,0463346 | 0,968393553 | 0,987435694 |
| TACGCGCAGACCACAGGATGTC | hsa-miR-3939 | -0,0126446 | 0,991273711 | 0,988069143 |
| TGAGTACCGCCATGTCTGTTGGG | hsa-miR-1911-5p | -0,0228709 | 0,984272083 | 0,98901559 |
| AGGGGTGCTATCTGTGATTGA | hsa-miR-342-5p | 0,0248068 | 1,017343432 | 0,990867774 |
| TATGGAAAGACTTTGCCACTCT | hsa-miR-3688-3p | 0,0149709 | 1,010431088 | 0,990867774 |
| CTGGAGATATGGAAGAGCTGTGT | hsa-miR-1270 | 0,0054838 | 1,003808317 | 0,990867774 |
| CCTCACCACCCCTTCTGCCTGCA | hsa-miR-6511b-3p | -0,0232856 | 0,983989185 | 0,990867774 |
| GCTCGGACTGAGCAGGTGGG | hsa-miR-3917 | -0,0277822 | 0,980927097 | 0,990867774 |
| TCCTGTCTTTCCTTGTTGGAGC | hsa-miR-5699-3p | 0,0053525 | 1,003716961 | 0,993194384 |
| TGTCTTACTCCCTCAGGCACAT | hsa-miR-550a-3p | -0,0024535 | 0,998300774 | 0,993194384 |
| CATGGGAGTTCGGGGTGGTTGC | hsa-miR-6871-5p | -0,0092167 | 0,993631805 | 0,993194384 |
