## Supplementary Table 6 for "Huntingtin loss-of-function contributes to transcriptional deregulation in Huntington’s disease"

**Supplementary Table 6. Analysis of miRNA biogenesis and function genes in HD-NSCs**

| Gene/protein ID (alias) | miRNA-related function | log2FC HD-NSCs vs IC1-NSCs | P_adjusted |
|---|---|---|---|
| **Nuclear steps of miRNA biogenesis** | | **DE in HD-NSCs** | |
| SMAD4 | Activates miRNA precursor transcription upon TGFB/BMP activation | 0,280837555 | 0,559448515 |
| FUS | Facilitates cotranscriptional DROSHA recruitment to pri-miRNAs; shown to bind a terminal loop of specific neuronal pri-miRNAs and to enhance their processing | -0,258757616 | 0,611235301 |
| SRSF3 (SRP20) | Enhances mammalian pri-miRNA processing upon binding to the CNNC motif in the 3p flanking sequence | NA | NA |
| DROSHA | RNase III; catalyzes pri-miRNA to pre-miRNA for processing/cleavage | 0,461335559 | 0,552820446 |
| DGCR8 | Cofactor of DROSHA; coordinates the recognition of pri-miRNA at the dsRNA-ssRNA junction; functions as a molecular anchor and direct DROSHA to cleave pri-miRNA ~11 bp before the junction | -0,079667911 | 0,83353389 |
| DDX5 (P68) | Plays a role in recognition/binding of pri-miRNAs by the DROSHA complex; recruits DROSHA/DGCR8 to pri-miRNA | NA | NA |
| DDX17 (P72) | Plays a role in recognition/binding of pri-miRNAs by the DROSHA complex | 0,276297836 | 0,557946299 |
| GSK3B | Facilitates pri-miRNA binding by DROSHA and enhances DROSHA association with cofactors DGCR8 and P72 (149); DROSHA phosphorylation/stabilization | 0,277252802 | 0,590378219 |
| SMAD2 | Accelerates pri-miRNA processing by DROSHA; in complex with SMAD4 activates miRNA precursor transcription upon TGFB/BMP signaling | 0,21405338 | 0,529053796 |
| **Export to cytoplasm** | | | |

| | | | | |
|---|---|---|---|---|
| XPO5 (EXP5) | Plays a role in the nuclear export of pre-miRNA to the cytoplasm; facilitates the nuclear cleavage of clustered pri-miRNAs | | 0,382861323 | 0,535050852 |
| RAN | Interacts with XPO5; plays a role in the export of pre-miRNA to the cytoplasm | | 0,34610446 | 0,563301276 |
| **Cytoplasmic steps of miRNA biogenesis** | | | | |
| DICER1 (DICER) | RNase III; catalyzes pre-miRNA to miRNA-duplex processing by cutting off the pre-miRNA terminal loop | | 0,452925932 | 0,478857489 |
| TARBP2 (TRBP) | Coordinates pre-miRNA recognition by DICER1 and the precision of DICER1 cleavage | | 0,335102277 | 0,51603779 |
| PRKRA (PACT) | Coordinates pre-miRNA cleavage by DICER1 and assures the precision of the cleavage; plays a role in miRISC assembly and thus participates in miRNA stabilization and accumulation in the cell | | -0,270650736 | 0,591156638 |
| ADAR | Double-stranded RNA-specific adenosine deaminase; plays a role in pri- and pre-miRNA stem editing, which makes miRNA precursors resistant to DICER1 cleavage | | 0,580124111 | 0,262513243 |
| KHSRP (FUBP2, KSRP) | Binds to the terminal loop sequence of a subset of miRNA precursors, promoting their maturation | | 0,271882414 | 0,489679316 |
| LIN28A and LIN28B | Bind to the terminal loops of specific pre-miRNAs (including pre-let-7 and pre-miR-9) and, upon recruitment of ZCCHC11 or ZCCHC6 that induces pre-miRNA uridylation, inhibit DICER1 processing | A | 1,812101534 | 0,040158676 |
| | | B | 1,324502527 | 0,082755166 |
| ZCCHC11 (TUT4) and ZCCHC6 (TUT7) | Play a role in pre-miRNA uridylation and thus inhibit DICER1 processing | TUT4 | 0,241084277 | 0,586498766 |
| | | TUT7 | 0,285549132 | 0,668563866 |
| DIS3L2 | Exoribonuclease; targets the uridylated let-7 precursors | | 0,071822129 | 0,820662051 |
| **miRNA functioning** | | | | |

| | | | | |
|---|---|---|---|---|
| AGO1, AGO2, AGO3, and AGO4 | Play a role in miRISC formation/loading; catalytically active AGO2 functions as an endonuclease upon complementary mRNA:miRNA interaction | AGO1 | -0,169165522 | 0,661042623 |
| | | AGO2 | 0,253816482 | 0,64134584 |
| | | AGO3 | -0,779118561 | 0,328658971 |
| | | AGO4 | 0,428140155 | 0,343742567 |
| GEMIN4 | Binds to the miRNA guide strand and facilitates the formation of a miRISC by unwinding the miRNA duplex | | 0,280908231 | 0,39178722 |
| MOV10 | Upon interaction with the miRNA-loaded AGO-protein complexes, plays a role in mRNA degradation; present in P-bodies | | 0,318822195 | 0,445042268 |
| FMR1 | Interacts with DICER1 and AGO1 during mRNA degradation; controls DROSHA expression | | 0,332973116 | 0,556575672 |
| TNRC6A (GW182) | Component of P-bodies; plays a role in mRNA degradation upon interaction with Argonaute proteins | | 0,335982834 | 0,58011711 |
