## Supplementary Table 7 for "Huntingtin loss-of-function contributes to transcriptional deregulation in Huntington’s disease"

**Supplementary Table 7. Correlation analysis of expression level of selected TFs with subsequent passages of HD and KO-NSCs**

**a. RNA-seq**

| | **HD-NSCs TPM** | | | | Correlation | | **KO-NSCs TPM** | | | | Correlation | |
|---|---|---|---|---|---|---|---|---|---|---|---|---|
| Gene | p4 | p5 | p6 | p7 | r | p value | p4 | p5 | p6 | p7 | r | p value |
| *TWIST1* | 2,82 | 9,29 | 15,52 | 26,87 | 0,9882 | 0,0059 (**) | 3,33 | 4,19 | 3,15 | 1,94 | -0,7256 | 0,1372 |
| *FOXD1* | 1,06 | 1,57 | 1,22 | 1,73 | 0,6944 | 0,1528 | 3,68 | 4,17 | 3,00 | 6,24 | 0,6018 | 0,1991 |
| *MSX2* | 0,89 | 1,59 | 2,46 | 7,40 | 0,8935 | 0,0532 | 6,79 | 8,61 | 7,30 | 5,94 | -0,4459 | 0,2771 |
| *SIX1* | 3,11 | 2,90 | 3,31 | 4,61 | 0,8231 | 0,0884 | 3,87 | 3,91 | 3,80 | 3,42 | -0,8390 | 0,0805 |
| *MEOX2* | 0,32 | 0,57 | 1,49 | 4,76 | 0,8985 | 0,0508 | 0,22 | 0,16 | 0,30 | 0,08 | -0,3883 | 0,3059 |
| *TBX1* | 3,11 | 4,10 | 5,06 | 6,40 | 0,9967 | 0,0017(**) | 0,19 | 0,37 | 0,39 | 0,72 | 0,9413 | 0,0294 (*) |
| *TBX15* | 0,79 | 1,17 | 1,61 | 4,05 | 0,8983 | 0,0508 | 1,67 | 3,87 | 4,84 | 4,76 | 0,8951 | 0,0525 |

**b. RT-qPCR**

| | **HD-NSCs Log2 FC** | | | | Correlation | | **KO-NSCs Log2 FC** | | | | Correlation | |
|---|---|---|---|---|---|---|---|---|---|---|---|---|
| Gene | p4 | p5 | p6 | p7 | r | p value | p4 | p5 | p6 | p7 | r | p value |
| *TWIST1* | 6,411262 | 8,290849 | 8,601905 | 9,197026 | 0,9303 | 0,0348(*) | 6,707766 | 6,394562 | 6,45617 | 6,585375 | -0,2830 | 0,3585 |
| *FOXD1* | 2,334676 | 2,0348 | 1,87803 | 1,912484 | -0,8849 | 0,0576 | 3,086765 | 3,328042 | 3,203475 | 3,698957 | 0,8328 | 0,0836 |
| *MSX2* | 3,4465 | 4,14452 | 4,898885 | 6,418962 | 0,9799 | 0,0101 (*) | 6,524709 | 6,714968 | 6,439334 | 6,23361 | -0,7430 | 0,1285 |
| *SIX1* | 6,478509 | 7,085191 | 7,033595 | 6,999492 | 0,6903 | 0,1549 | 7,157148 | 7,098113 | 7,071002 | 8,062421 | 0,7259 | 0,1370 |
| *MEOX2* | 3,929324 | 4,941405 | 5,989196 | 7,801708 | 0,9886 | 0,0057(**) | 3,588635 | 3,579065 | 3,376236 | 2,774422 | -0,8921 | 0,0540 |
| *TBX1* | 6,426056 | 7,146449 | 7,174399 | 7,919301 | 0,9544 | 0,0228 (*) | 3,66676 | 3,846049 | 3,989823 | 4,841315 | 0,9094 | 0,0453 (*) |
| *TBX15* | 5,079306 | 6,086971 | 6,552935 | 7,525179 | 0,9910 | 0,0045 (**) | 6,241198 | 7,978323 | 8,004343 | 8,457925 | 0,8813 | 0,0593 |

control sample for RT-qPCR: IC1

multiple t test

p value <0.05

p value <0.10

p4-7 - passage number

r-Pearson correlation
