## Supplementary Table 8 for "Huntingtin loss-of-function contributes to transcriptional deregulation in Huntington’s disease"

| KO-NSCs | | [TPM] | | | | |
|---|---|---|---|---|---|---|
| gene_ID | Gene_name | KO-NSC_1 | KO-NSC_2 | KO-NSC_3 | KO-NSC_4 | change_% |
| ENSG00000229327 | AL391839,1 | 1,15 | 0,63 | 0,48 | 0 | 100,0 |
| ENSG00000284057 | AP001273,2 | 1,17 | 0,53 | 0,35 | 0 | 100,0 |
| ENSG00000256040 | PAPPA-AS1 | 1,43 | 0,65 | 0,62 | 0 | 100,0 |
| ENSG00000277945 | AC107308,1 | 1,47 | 1,39 | 1,18 | 0 | 100,0 |
| ENSG00000270828 | | 1,5 | 0,27 | 0,17 | 0 | 100,0 |
| ENSG00000276390 | AC004241,3 | 1,74 | 1,59 | 1,13 | 0 | 100,0 |
| ENSG00000281468 | AC006504,7 | 2,24 | 2,19 | 1,77 | 0 | 100,0 |
| ENSG00000273275 | AC017083,2 | 2,39 | 0,67 | 0,32 | 0 | 100,0 |
| ENSG00000285323 | | 3,19 | 1,92 | 1,81 | 0 | 100,0 |
| ENSG00000215695 | RSC1A1 | 3,41 | 2,48 | 2,33 | 0 | 100,0 |
| ENSG00000212440 | | 3,57 | 2,93 | 1,53 | 0 | 100,0 |
| ENSG00000231884 | NDUFB1P1 | 4,91 | 4,12 | 0,53 | 0 | 100,0 |
| ENSG00000201806 | | 8,06 | 3,94 | 3,11 | 0 | 100,0 |
| ENSG00000273727 | U1 | 10,91 | 5,56 | 3,24 | 0 | 100,0 |
| ENSG00000274630 | AC125257,2 | 28,67 | 23,7 | 18,2 | 0 | 100,0 |
| ENSG00000079689 | SCGN | 1,8 | 0,76 | 0,28 | 0,01 | 99,4 |
| ENSG00000197487 | GALP | 2,41 | 0,33 | 0,27 | 0,04 | 98,3 |
| ENSG00000273645 | KBTBD11 | 3,75 | 3,64 | 2,92 | 0,11 | 97,1 |
| ENSG00000241131 | LINC02032 | 4,76 | 3,02 | 1,43 | 0,18 | 96,2 |
| ENSG00000238276 | AL354863,1 | 6,97 | 3,11 | 1,15 | 0,28 | 96,0 |
| ENSG00000260548 | AL035425,2 | 3,85 | 1,23 | 0,78 | 0,16 | 95,8 |
| ENSG00000254768 | AC104009,1 | 1,97 | 1,75 | 0,43 | 0,1 | 94,9 |
| ENSG00000136535 | TBR1 | 9,24 | 4,51 | 0,92 | 0,47 | 94,9 |
| ENSG00000279566 | ZNF43 | 6,8 | 6,09 | 5,59 | 0,37 | 94,6 |
| ENSG00000271133 | AC004130,2 | 1,89 | 1,24 | 1,09 | 0,11 | 94,2 |
| ENSG00000129991 | TNNI3 | 1,8 | 0,55 | 0,13 | 0,11 | 93,9 |
| ENSG00000188801 | ZNF322P1 | 1,3 | 0,87 | 0,54 | 0,08 | 93,8 |
| ENSG00000146374 | RSPO3 | 5,03 | 1,86 | 0,92 | 0,31 | 93,8 |
| ENSG00000163508 | EOMES | 9,66 | 3,61 | 1,48 | 0,61 | 93,7 |
| ENSG00000137766 | UNC13C | 1,22 | 0,64 | 0,26 | 0,08 | 93,4 |
| ENSG00000145808 | ADAMTS19 | 2,99 | 2,58 | 0,73 | 0,21 | 93,0 |
| ENSG00000207110 | | 3,44 | 2,34 | 1,37 | 0,25 | 92,7 |
| ENSG00000226702 | MIR217HG | 1,07 | 0,95 | 0,35 | 0,08 | 92,5 |
| ENSG00000272807 | AC007038,1 | 2,13 | 1,98 | 1,43 | 0,16 | 92,5 |
| ENSG00000200997 | RNVU1-34 | 11,14 | 8,93 | 6,16 | 0,88 | 92,1 |
| ENSG00000227449 | FGF7P6 | 1,94 | 1,76 | 0,66 | 0,16 | 91,8 |
| ENSG00000091664 | SLC17A6 | 4,82 | 3,22 | 1,28 | 0,4 | 91,7 |
| ENSG00000077274 | CAPN6 | 5,34 | 1,76 | 0,48 | 0,45 | 91,6 |
| ENSG00000138722 | MMRN1 | 7,28 | 5,67 | 3,83 | 0,62 | 91,5 |
| ENSG00000265179 | AP000894,2 | 2,79 | 1,54 | 0,27 | 0,24 | 91,4 |
| ENSG00000226352 | PSPC1-AS2 | 1,48 | 0,98 | 0,97 | 0,13 | 91,2 |
| ENSG00000261409 | AL035425,3 | 4,54 | 1,85 | 1 | 0,4 | 91,2 |
| ENSG00000251349 | MSANTD3-TMEI | 26,31 | 24,93 | 21,14 | 2,4 | 90,9 |
| ENSG00000275769 | AC068792,1 | 1,81 | 1,14 | 0,62 | 0,17 | 90,6 |
| ENSG00000236106 | AC010729,1 | 1,35 | 0,57 | 0,5 | 0,13 | 90,4 |

| | | | | | | |
|---|---|---|---|---|---|---|
| ENSG00000158164 | TMSB15A | 241,96 | 177,63 | 140,37 | 23,47 | 90,3 |
| ENSG00000147724 | FAM135B | 1,03 | 0,28 | 0,19 | 0,1 | 90,3 |
| ENSG00000176399 | DMRTA1 | 11,38 | 6,17 | 3,4 | 1,11 | 90,2 |
| ENSG00000225826 | LINC00626 | 1,02 | 0,39 | 0,13 | 0,1 | 90,2 |
| ENSG00000273284 | AP001033,2 | 1,6 | 1,48 | 0,96 | 0,16 | 90,0 |
| ENSG00000282033 | AC074387,1 | 1,58 | 1,1 | 0,44 | 0,16 | 89,9 |
| ENSG00000183793 | NPIPA5 | 2,34 | 1,43 | 0,79 | 0,24 | 89,7 |
| ENSG00000105048 | TNNT1 | 9,71 | 4,87 | 1,4 | 1,01 | 89,6 |
| ENSG00000144331 | ZNF385B | 3,07 | 0,98 | 0,76 | 0,32 | 89,6 |
| ENSG00000164841 | TMEM74 | 2,78 | 1,59 | 0,83 | 0,29 | 89,6 |
| ENSG00000139292 | LGR5 | 1,52 | 0,96 | 0,23 | 0,16 | 89,5 |
| ENSG00000188611 | ASAH2 | 1,8 | 1,47 | 1,05 | 0,19 | 89,4 |
| ENSG00000263958 | AC091138,1 | 2,83 | 1,59 | 1,57 | 0,3 | 89,4 |
| ENSG00000138653 | NDST4 | 3,96 | 2,85 | 1,3 | 0,42 | 89,4 |
| ENSG00000184611 | KCNH7 | 1,21 | 0,55 | 0,37 | 0,13 | 89,3 |
| ENSG00000227794 | RPS18 | 205,88 | 161,08 | 103,3 | 22,78 | 88,9 |
| ENSG00000259856 | RAB43P1 | 1,8 | 1,21 | 1,1 | 0,2 | 88,9 |
| ENSG00000225715 | PSMA1P1 | 1,06 | 0,45 | 0,37 | 0,12 | 88,7 |
| ENSG00000187475 | H1-6 | 1,85 | 0,7 | 0,59 | 0,21 | 88,6 |
| ENSG00000143355 | LHX9 | 6,31 | 3,32 | 1,09 | 0,73 | 88,4 |
| ENSG00000153822 | KCNJ16 | 1,71 | 0,89 | 0,54 | 0,2 | 88,3 |
| ENSG00000080031 | PTPRH | 1,28 | 0,86 | 0,19 | 0,15 | 88,3 |
| ENSG00000134709 | HOOK1 | 1,85 | 1,41 | 0,67 | 0,22 | 88,1 |
| ENSG00000197921 | HES5 | 9,21 | 3,2 | 2,14 | 1,1 | 88,1 |
| ENSG00000288658 | AC010980,1 | 2,93 | 1,74 | 1,35 | 0,35 | 88,1 |
| ENSG00000207205 | RNVU1-15 | 89,97 | 47,41 | 46,49 | 10,75 | 88,1 |
| ENSG00000271971 | AC120053,1 | 3,39 | 3,06 | 2,24 | 0,42 | 87,6 |
| ENSG00000157470 | FAM81A | 2,32 | 1,15 | 0,98 | 0,29 | 87,5 |
| ENSG00000198848 | CES1 | 1,17 | 1,1 | 0,22 | 0,15 | 87,2 |
| ENSG00000229152 | ANKRD10-IT1 | 6,44 | 5,92 | 4,78 | 0,83 | 87,1 |
| ENSG00000267340 | | 1,93 | 1,47 | 1,12 | 0,25 | 87,0 |
| ENSG00000288385 | MG910337,1 | 1,99 | 0,97 | 0,41 | 0,26 | 86,9 |
| ENSG00000231165 | TRBV26OR9-2 | 2,99 | 1,15 | 1 | 0,4 | 86,6 |
| ENSG00000222724 | RNU2-63P | 26,99 | 26,74 | 19,68 | 3,64 | 86,5 |
| ENSG00000273951 | AL031667,3 | 1,11 | 1,01 | 0,77 | 0,15 | 86,5 |
| ENSG00000281763 | DHX36 | 22,43 | 13,71 | 12,13 | 3,04 | 86,4 |
| ENSG00000234257 | SOD2P1 | 1,09 | 0,85 | 0,65 | 0,15 | 86,2 |
| ENSG00000186369 | LINC00643 | 2,31 | 0,98 | 0,48 | 0,32 | 86,1 |
| ENSG00000078114 | NEBL | 10,04 | 6,3 | 2,25 | 1,4 | 86,1 |
| ENSG00000204789 | ZNF204P | 1,49 | 1,31 | 0,87 | 0,21 | 85,9 |
| ENSG00000177511 | ST8SIA3 | 1,56 | 1,27 | 0,68 | 0,22 | 85,9 |
| ENSG00000198216 | CACNA1E | 2,12 | 1,18 | 0,37 | 0,3 | 85,8 |
| ENSG00000273711 | AC005520,5 | 3,79 | 2,94 | 2,75 | 0,54 | 85,8 |
| ENSG00000107295 | SH3GL2 | 1,12 | 0,43 | 0,3 | 0,16 | 85,7 |
| ENSG00000132749 | TESMIN | 2,34 | 1,86 | 1,73 | 0,34 | 85,5 |
| ENSG00000187772 | LIN28B | 13,31 | 7,74 | 3,83 | 1,94 | 85,4 |
| ENSG00000079101 | CLUL1 | 2,6 | 2,02 | 1,17 | 0,38 | 85,4 |

| | | | | | | |
|---|---|---|---|---|---|---|
| ENSG00000050628 | PTGER3 | 1,23 | 0,93 | 0,55 | 0,18 | 85,4 |
| ENSG00000121211 | MND1 | 15,47 | 15,06 | 14,03 | 2,28 | 85,3 |
| ENSG00000259436 | AC010247,2 | 2,37 | 1,41 | 0,78 | 0,35 | 85,2 |
| ENSG00000261335 | AC005837,1 | 1 | 0,73 | 0,34 | 0,15 | 85,0 |
| ENSG00000145721 | LIX1 | 6,24 | 3,63 | 3,09 | 0,94 | 84,9 |
| ENSG00000132541 | RIDA | 13,87 | 12,75 | 11,23 | 2,11 | 84,8 |
| ENSG00000131951 | LRRC9 | 3,13 | 1,43 | 0,84 | 0,48 | 84,7 |
| ENSG00000165973 | NELL1 | 2,57 | 1,33 | 1,11 | 0,4 | 84,4 |
| ENSG00000230847 | OCLNP1 | 1,22 | 0,82 | 0,72 | 0,19 | 84,4 |
| ENSG00000157404 | KIT | 6,26 | 2,56 | 2,02 | 0,99 | 84,2 |
| ENSG00000121898 | CPXM2 | 1,01 | 0,71 | 0,31 | 0,16 | 84,2 |
| ENSG00000235447 | TRAPPC13P1 | 2,55 | 1,54 | 1,3 | 0,41 | 83,9 |
| ENSG00000230623 | AC104461,1 | 2,28 | 1,95 | 1,72 | 0,37 | 83,8 |
| ENSG00000240204 | SMKR1 | 1,29 | 0,97 | 0,34 | 0,21 | 83,7 |
| ENSG00000260907 | AC015818,2 | 1,34 | 1,19 | 0,9 | 0,22 | 83,6 |
| ENSG00000135750 | KCNK1 | 1,08 | 0,75 | 0,36 | 0,18 | 83,3 |
| ENSG00000147676 | MAL2 | 1,5 | 1,17 | 0,82 | 0,25 | 83,3 |
| ENSG00000283649 | ZDBF2 | 2,94 | 1,89 | 1,24 | 0,49 | 83,3 |
| ENSG00000213413 | PVRIG | 1,92 | 1,81 | 0,94 | 0,32 | 83,3 |
| ENSG00000255618 | LINC02440 | 2,38 | 0,99 | 0,63 | 0,4 | 83,2 |
| ENSG00000272192 | AC100812,1 | 1,66 | 1,15 | 0,44 | 0,28 | 83,1 |
| ENSG00000162374 | ELAVL4 | 12,55 | 8,83 | 4,74 | 2,12 | 83,1 |
| ENSG00000266910 | AC008507,1 | 1,18 | 0,76 | 0,68 | 0,2 | 83,1 |
| ENSG00000260317 | AC009812,4 | 1,12 | 1,02 | 0,87 | 0,19 | 83,0 |
| ENSG00000198088 | NUP62CL | 10,53 | 8,93 | 5,24 | 1,79 | 83,0 |
| ENSG00000126733 | DACH2 | 1,94 | 1,34 | 0,81 | 0,33 | 83,0 |
| ENSG00000260912 | AL158206,1 | 1,52 | 0,95 | 0,91 | 0,26 | 82,9 |
| ENSG00000276334 | AL133243,2 | 1,52 | 1,26 | 1,19 | 0,26 | 82,9 |
| ENSG00000177551 | NHLH2 | 14,02 | 8,87 | 5,36 | 2,41 | 82,8 |
| ENSG00000181541 | MAB21L2 | 2,21 | 1,22 | 0,48 | 0,38 | 82,8 |
| ENSG00000101888 | NXT2 | 18,31 | 17,37 | 12,29 | 3,21 | 82,5 |
| ENSG00000136014 | USP44 | 4,14 | 2,43 | 1,35 | 0,75 | 81,9 |
| ENSG00000170989 | S1PR1 | 2,58 | 1,17 | 0,92 | 0,47 | 81,8 |
| ENSG00000165997 | ARL5B | 12,91 | 11,4 | 8,62 | 2,36 | 81,7 |
| ENSG00000278356 | AC005911,1 | 2,12 | 1,6 | 1,06 | 0,39 | 81,6 |
| ENSG00000198865 | CCDC152 | 2,37 | 1,61 | 1,38 | 0,44 | 81,4 |
| ENSG00000228857 | AC104653,1 | 1,02 | 0,77 | 0,5 | 0,19 | 81,4 |
| ENSG00000269600 | AC016629,2 | 2,35 | 1,62 | 1,22 | 0,44 | 81,3 |
| ENSG00000089101 | CFAP61 | 1,44 | 0,82 | 0,44 | 0,27 | 81,3 |
| ENSG00000252355 | RN7SKP287 | 2,61 | 2,07 | 0,95 | 0,49 | 81,2 |
| ENSG00000170893 | TRH | 17,99 | 12,52 | 5,92 | 3,4 | 81,1 |
| ENSG00000198478 | SH3BGRL2 | 5,12 | 3,93 | 3,09 | 0,97 | 81,1 |
| ENSG00000172575 | RASGRP1 | 4,42 | 2,37 | 1,45 | 0,84 | 81,0 |
| ENSG00000114405 | C3orf14 | 26,55 | 23,06 | 19,58 | 5,05 | 81,0 |
| ENSG00000198963 | RORB | 1,84 | 1,73 | 1,13 | 0,35 | 81,0 |
| ENSG00000258571 | PTTG4P | 1,15 | 0,67 | 0,42 | 0,22 | 80,9 |
| ENSG00000215417 | MIR17HG | 19,95 | 19,35 | 10,72 | 3,85 | 80,7 |

| | | | | | | |
|---|---|---|---|---|---|---|
| ENSG00000183036 | PCP4 | 5,74 | 4,55 | 2,91 | 1,11 | 80,7 |
| ENSG00000255240 | AP001636,3 | 1,24 | 1,13 | 0,73 | 0,24 | 80,6 |
| ENSG00000189056 | RELN | 7,07 | 4,92 | 2,57 | 1,37 | 80,6 |
| ENSG00000123570 | RAB9B | 4,64 | 4,38 | 3,54 | 0,9 | 80,6 |
| ENSG00000183054 | RGPD6 | 26,07 | 24,76 | 20,19 | 5,06 | 80,6 |
| ENSG00000170370 | EMX2 | 27,83 | 19,45 | 9,97 | 5,41 | 80,6 |
| ENSG00000225507 | AC069282,1 | 3,48 | 3,22 | 2,21 | 0,68 | 80,5 |
| ENSG00000257935 | LHX5-AS1 | 21,51 | 15,83 | 8,03 | 4,23 | 80,3 |
| ENSG00000125355 | TMEM255A | 4,33 | 3,87 | 2,24 | 0,86 | 80,1 |
| ENSG00000115423 | DNAH6 | 1,6 | 1,17 | 0,69 | 0,32 | 80,0 |
| ENSG00000118997 | DNAH7 | 1,39 | 1,29 | 1,18 | 0,28 | 79,9 |
| ENSG00000137720 | C11orf1 | 11,75 | 11 | 7,39 | 2,37 | 79,8 |
| ENSG00000278811 | LINC00624 | 1,33 | 1,22 | 1,03 | 0,27 | 79,7 |
| ENSG00000168952 | STXBP6 | 1,92 | 1,56 | 0,89 | 0,39 | 79,7 |
| ENSG00000162409 | PRKAA2 | 1,92 | 1,39 | 1,25 | 0,39 | 79,7 |
| ENSG00000161249 | DMKN | 3,19 | 2,58 | 0,76 | 0,65 | 79,6 |
| ENSG00000135333 | EPHA7 | 4,45 | 2,76 | 1,52 | 0,91 | 79,6 |
| ENSG00000276644 | DACH1 | 8,63 | 5,95 | 3,48 | 1,77 | 79,5 |
| ENSG00000286810 | AL513128,3 | 1,17 | 1,05 | 0,48 | 0,24 | 79,5 |
| ENSG00000163909 | HEYL | 1,56 | 0,84 | 0,52 | 0,32 | 79,5 |
| ENSG00000179300 | RTL3 | 1,26 | 1,16 | 0,68 | 0,26 | 79,4 |
| ENSG00000273990 | RNVU1-26 | 12,18 | 7,64 | 5,27 | 2,52 | 79,3 |
| ENSG00000275291 | RNVU1-26 | 12,18 | 7,64 | 5,27 | 2,52 | 79,3 |
| ENSG00000197705 | KLHL14 | 5,92 | 5,73 | 3,69 | 1,23 | 79,2 |
| ENSG00000165443 | PHYHIPL | 5,62 | 4,27 | 2,79 | 1,17 | 79,2 |
| ENSG00000137821 | LRRC49 | 15,09 | 13,01 | 9,44 | 3,15 | 79,1 |
| ENSG00000081181 | ARG2 | 6,44 | 4,35 | 3,16 | 1,35 | 79,0 |
| ENSG00000135363 | LMO2 | 4,96 | 2,62 | 1,76 | 1,04 | 79,0 |
| ENSG00000261804 | AC007342,4 | 6,52 | 5,79 | 4,17 | 1,37 | 79,0 |
| ENSG00000183850 | ZNF730 | 17,25 | 14,77 | 11,71 | 3,63 | 79,0 |
| ENSG00000173376 | NDNF | 1,14 | 0,59 | 0,38 | 0,24 | 78,9 |
| ENSG00000270084 | GAS5-AS1 | 3,17 | 2,71 | 2,2 | 0,67 | 78,9 |
| ENSG00000203965 | EFCAB7 | 6,95 | 6,39 | 5,15 | 1,47 | 78,8 |
| ENSG00000201558 | RNVU1-6 | 21,65 | 21,39 | 14,49 | 4,58 | 78,8 |
| ENSG00000109929 | SC5D | 38,62 | 37,25 | 33,92 | 8,17 | 78,8 |
| ENSG00000180667 | YOD1 | 6,79 | 5,84 | 4,67 | 1,44 | 78,8 |
| ENSG00000197415 | VEPH1 | 12,53 | 8,19 | 5,2 | 2,66 | 78,8 |
| ENSG00000269397 | AC011503,2 | 1,36 | 1,21 | 0,91 | 0,29 | 78,7 |
| ENSG00000127184 | COX7C | 235,97 | 232,82 | 218,21 | 50,45 | 78,6 |
| ENSG00000197008 | ZNF138 | 11,73 | 8,96 | 7,81 | 2,52 | 78,5 |
| ENSG00000106689 | LHX2 | 24,36 | 13,94 | 7,3 | 5,25 | 78,4 |
| ENSG00000118276 | B4GALT6 | 9,26 | 8,85 | 6,58 | 2,01 | 78,3 |
| ENSG00000180769 | WDFY3-AS2 | 1,79 | 1,29 | 1,04 | 0,39 | 78,2 |
| ENSG00000104435 | STMN2 | 72,46 | 52,58 | 30,94 | 15,79 | 78,2 |
| ENSG00000278765 | AC004477,2 | 3,16 | 2,62 | 1,61 | 0,69 | 78,2 |
| ENSG00000162992 | NEUROD1 | 1,46 | 0,89 | 0,68 | 0,32 | 78,1 |
| ENSG00000275854 | AC084824,4 | 2,04 | 1,79 | 1,68 | 0,45 | 77,9 |

| | | | | | | |
|---|---|---|---|---|---|---|
| ENSG00000257489 | AC010203,1 | 1,13 | 0,82 | 0,52 | 0,25 | 77,9 |
| ENSG00000168843 | FSTL5 | 4,79 | 3,99 | 2,79 | 1,06 | 77,9 |
| ENSG00000108231 | LGI1 | 9,84 | 5,12 | 5,11 | 2,18 | 77,8 |
| ENSG00000198146 | ZNF770 | 18,69 | 16,96 | 14,54 | 4,15 | 77,8 |
| ENSG00000122643 | NT5C3A | 21,09 | 18,66 | 15,39 | 4,69 | 77,8 |
| ENSG00000136897 | MRPL50 | 7,55 | 7,15 | 6,9 | 1,69 | 77,6 |
| ENSG00000184672 | RALYL | 1,91 | 1,87 | 1,28 | 0,43 | 77,5 |
| ENSG00000173947 | PIFO | 1,42 | 0,62 | 0,55 | 0,32 | 77,5 |
| ENSG00000124802 | EEF1E1 | 26,2 | 21,58 | 19,9 | 5,91 | 77,4 |
| ENSG00000168348 | INSM2 | 3,9 | 1,94 | 1,38 | 0,88 | 77,4 |
| ENSG00000251456 | AC113398,2 | 2,29 | 2,23 | 1,4 | 0,52 | 77,3 |
| ENSG00000178662 | CSRNP3 | 12,67 | 8,65 | 4,85 | 2,88 | 77,3 |
| ENSG00000147145 | LPAR4 | 8,71 | 7,72 | 4,85 | 1,98 | 77,3 |
| ENSG00000166265 | CYYR1 | 7,17 | 5,23 | 4,4 | 1,63 | 77,3 |
| ENSG00000213047 | DENND1B | 8,65 | 5,31 | 3,96 | 1,97 | 77,2 |
| ENSG00000271976 | AC012467,2 | 2,29 | 1,98 | 1,51 | 0,53 | 76,9 |
| ENSG00000112742 | TTK | 24,39 | 21,29 | 20,44 | 5,68 | 76,7 |
| ENSG00000077279 | DCX | 45,87 | 35,29 | 20,63 | 10,7 | 76,7 |
| ENSG00000151575 | TEX9 | 16,47 | 13,7 | 11,05 | 3,86 | 76,6 |
| ENSG00000275947 | TMEM251 | 6,23 | 6,1 | 5,68 | 1,47 | 76,4 |
| ENSG00000169684 | CHRNA5 | 9,63 | 8,18 | 6,19 | 2,28 | 76,3 |
| ENSG00000106341 | PPP1R17 | 3,87 | 2,59 | 1,24 | 0,92 | 76,2 |
| ENSG00000138769 | CDKL2 | 1,68 | 0,88 | 0,6 | 0,4 | 76,2 |
| ENSG00000111834 | RSPH4A | 1,84 | 1,68 | 0,94 | 0,44 | 76,1 |
| ENSG00000156876 | SASS6 | 6,81 | 6,09 | 5,17 | 1,63 | 76,1 |
| ENSG00000107560 | RAB11FIP2 | 16,04 | 9,4 | 8,06 | 3,84 | 76,1 |
| ENSG00000112981 | NME5 | 3,8 | 3,46 | 3,3 | 0,91 | 76,1 |
| ENSG00000077080 | ACTL6B | 2,12 | 1,7 | 1,1 | 0,51 | 75,9 |
| ENSG00000123892 | RAB38 | 5,64 | 4,32 | 3,65 | 1,36 | 75,9 |
| ENSG00000285222 | GCNT2 | 2,72 | 1,73 | 1,15 | 0,66 | 75,7 |
| ENSG00000109255 | NMU | 29,54 | 23,62 | 16,66 | 7,21 | 75,6 |
| ENSG00000224578 | HNRNPA1P48 | 11,02 | 9,79 | 6,45 | 2,69 | 75,6 |
| ENSG00000169064 | ZBBX | 1,84 | 1,78 | 1,25 | 0,45 | 75,5 |
| ENSG00000145692 | BHMT | 1,1 | 0,52 | 0,45 | 0,27 | 75,5 |
| ENSG00000177485 | ZBTB33 | 24,13 | 23,11 | 20,38 | 5,99 | 75,2 |
| ENSG00000170917 | NUDT6 | 6,18 | 4,05 | 3,57 | 1,54 | 75,1 |
| ENSG00000254266 | PKIA-AS1 | 1,12 | 0,73 | 0,6 | 0,28 | 75,0 |
| ENSG00000133134 | BEX2 | 25,3 | 20,27 | 13,51 | 6,33 | 75,0 |
| ENSG00000143977 | SNRPG | 149,36 | 138,21 | 129,26 | 37,47 | 74,9 |
| ENSG00000086300 | SNX10 | 5,1 | 4,8 | 3,04 | 1,28 | 74,9 |
| ENSG00000176018 | LYSMD3 | 6,69 | 6,43 | 5,9 | 1,68 | 74,9 |
| ENSG00000145526 | CDH18 | 2,07 | 1,04 | 0,79 | 0,52 | 74,9 |
| ENSG00000276048 | AC012354,4 | 1,87 | 0,99 | 0,75 | 0,47 | 74,9 |
| ENSG00000162998 | FRZB | 8,46 | 7,78 | 6,47 | 2,14 | 74,7 |
| ENSG00000171757 | LRRC34 | 4,7 | 4,5 | 4,1 | 1,19 | 74,7 |
| ENSG00000267302 | RNFT1-DT | 1,34 | 1,08 | 0,73 | 0,34 | 74,6 |
| ENSG00000184305 | CCSER1 | 3,42 | 2,3 | 1,35 | 0,87 | 74,6 |

| | | | | | | |
|---|---|---|---|---|---|---|
| ENSG00000106537 | TSPAN13 | 18,66 | 16,57 | 14,33 | 4,75 | 74,5 |
| ENSG00000274618 | H4C6 | 266,65 | 198,45 | 176,09 | 67,91 | 74,5 |
| ENSG00000169860 | P2RY1 | 1,06 | 0,79 | 0,58 | 0,27 | 74,5 |
| ENSG00000278728 | CNTNAP2 | 19,7 | 16,06 | 13,19 | 5,02 | 74,5 |
| ENSG00000146757 | ZNF92 | 17,21 | 15,11 | 13,83 | 4,4 | 74,4 |
| ENSG00000166575 | TMEM135 | 16,46 | 11,72 | 10,09 | 4,21 | 74,4 |
| ENSG00000107443 | CCNJ | 19,93 | 16,82 | 15,46 | 5,1 | 74,4 |
| ENSG00000133101 | CCNA1 | 1,21 | 0,76 | 0,56 | 0,31 | 74,4 |
| ENSG00000226508 | LINC01918 | 1,99 | 1,83 | 1,54 | 0,51 | 74,4 |
| ENSG00000028839 | TBPL1 | 24,44 | 22,61 | 20,82 | 6,31 | 74,2 |
| ENSG00000133665 | DYDC2 | 1,82 | 1,5 | 1,13 | 0,47 | 74,2 |
| ENSG00000259985 | AC017100,1 | 2,98 | 2,9 | 1,89 | 0,77 | 74,2 |
| ENSG00000145428 | RNF175 | 9,12 | 6,26 | 5,04 | 2,37 | 74,0 |
| ENSG00000058804 | NDC1 | 16,94 | 14,23 | 13,1 | 4,41 | 74,0 |
| ENSG00000124785 | NRN1 | 1,69 | 1,1 | 0,99 | 0,44 | 74,0 |
| ENSG00000254004 | ZNF260 | 17,49 | 15,55 | 13,05 | 4,56 | 73,9 |
| ENSG00000178966 | RMI1 | 12,26 | 11 | 10,38 | 3,2 | 73,9 |
| ENSG00000228623 | ZNF883 | 2,95 | 2,62 | 2,2 | 0,77 | 73,9 |
| ENSG00000274675 | GTF2H2C_2 | 10,15 | 10,02 | 8 | 2,66 | 73,8 |
| ENSG00000170396 | ZNF804A | 2,44 | 1,81 | 1,23 | 0,64 | 73,8 |
| ENSG00000112218 | GPR63 | 2,12 | 1,58 | 1,31 | 0,56 | 73,6 |
| ENSG00000198924 | DCLRE1A | 7,98 | 7,82 | 6,31 | 2,11 | 73,6 |
| ENSG00000257181 | AC025423,4 | 11,19 | 9,06 | 7,6 | 2,96 | 73,5 |
| ENSG00000158806 | NPM2 | 2,91 | 2,39 | 0,83 | 0,77 | 73,5 |
| ENSG00000214198 | TTC41P | 1,13 | 0,82 | 0,77 | 0,3 | 73,5 |
| ENSG00000256667 | KLRA1P | 2,26 | 1,98 | 1,42 | 0,6 | 73,5 |
| ENSG00000239827 | SUGT1P3 | 2,22 | 1,76 | 1,09 | 0,59 | 73,4 |
| ENSG00000279232 | AC008522,1 | 3,98 | 3,32 | 3,15 | 1,06 | 73,4 |
| ENSG00000117569 | PTBP2 | 66,54 | 61,74 | 52,21 | 17,74 | 73,3 |
| ENSG00000235298 | AL354733,3 | 15,32 | 13,89 | 8,64 | 4,1 | 73,2 |
| ENSG00000280870 | MIR325HG | 6,15 | 5,53 | 4,46 | 1,65 | 73,2 |
| ENSG00000104112 | SCG3 | 9,79 | 9,08 | 6,31 | 2,63 | 73,1 |
| ENSG00000158427 | TMSB15B | 47,27 | 44,72 | 40,82 | 12,7 | 73,1 |
| ENSG00000189057 | FAM111B | 15,21 | 14,68 | 12,88 | 4,09 | 73,1 |
| ENSG00000145147 | SLIT2 | 67,77 | 54,66 | 26,49 | 18,26 | 73,1 |
| ENSG00000116574 | RHOU | 6,34 | 4,87 | 3,61 | 1,71 | 73,0 |
| ENSG00000284984 | RHOU | 6,34 | 4,87 | 3,61 | 1,71 | 73,0 |
| ENSG00000165338 | HECTD2 | 19,57 | 14,38 | 11,18 | 5,28 | 73,0 |
| ENSG00000205208 | C4orf46 | 12,05 | 11,75 | 9,65 | 3,26 | 72,9 |
| ENSG00000064309 | CDON | 25,11 | 17,26 | 11,53 | 6,8 | 72,9 |
| ENSG00000179941 | BBS10 | 6,79 | 6,73 | 5,38 | 1,84 | 72,9 |
| ENSG00000174695 | TMEM167A | 38,09 | 33,71 | 32,46 | 10,33 | 72,9 |
| ENSG00000132436 | FIGNL1 | 12,94 | 12,24 | 9,78 | 3,51 | 72,9 |
| ENSG00000189195 | BTBD8 | 1,99 | 1,6 | 1,51 | 0,54 | 72,9 |
| ENSG00000120696 | KBTBD7 | 4,16 | 3,63 | 3,01 | 1,13 | 72,8 |
| ENSG00000031691 | CENPQ | 8,29 | 7,83 | 7,17 | 2,26 | 72,7 |
| ENSG00000164604 | GPR85 | 5,39 | 5,11 | 4,89 | 1,47 | 72,7 |

| | | | | | | |
|---|---|---|---|---|---|---|
| ENSG00000230989 | HSBP1 | 131,6 | 128,51 | 114,85 | 35,92 | 72,7 |
| ENSG00000147592 | LACTB2 | 7,65 | 6,8 | 6,73 | 2,09 | 72,7 |
| ENSG00000282619 | AF186192,6 | 1,72 | 1,28 | 0,83 | 0,47 | 72,7 |
| ENSG00000255085 | AF186192,2 | 1,72 | 1,28 | 0,83 | 0,47 | 72,7 |
| ENSG00000286964 | AL136169,1 | 1,35 | 1,12 | 0,87 | 0,37 | 72,6 |
| ENSG00000136122 | BORA | 9,74 | 8,92 | 7,27 | 2,67 | 72,6 |
| ENSG00000268058 | BNIP3P40 | 1,64 | 1,17 | 0,9 | 0,45 | 72,6 |
| ENSG00000181004 | BBS12 | 2,58 | 2,29 | 2,2 | 0,71 | 72,5 |
| ENSG00000158815 | FGF17 | 55,57 | 43,99 | 16,81 | 15,3 | 72,5 |
| ENSG00000205277 | MUC12 | 4,61 | 2,49 | 1,53 | 1,27 | 72,5 |
| ENSG00000112319 | EYA4 | 1,45 | 0,71 | 0,42 | 0,4 | 72,4 |
| ENSG00000087495 | PHACTR3 | 2,79 | 2,02 | 1,26 | 0,77 | 72,4 |
| ENSG00000167646 | DNAAF3 | 1,05 | 0,71 | 0,53 | 0,29 | 72,4 |
| ENSG00000115252 | PDE1A | 1,81 | 1,8 | 1,57 | 0,5 | 72,4 |
| ENSG00000260232 | PWRN4 | 1,52 | 0,99 | 0,79 | 0,42 | 72,4 |
| ENSG00000176714 | CCDC121 | 3,29 | 1,85 | 1,77 | 0,91 | 72,3 |
| ENSG00000139826 | ABHD13 | 4,3 | 4,1 | 3,68 | 1,19 | 72,3 |
| ENSG00000152926 | ZNF117 | 14,88 | 13,8 | 10,21 | 4,12 | 72,3 |
| ENSG00000151665 | PIGF | 17,57 | 16,56 | 16,53 | 4,88 | 72,2 |
| ENSG00000260920 | AL031985,3 | 3,59 | 3,16 | 2,59 | 1 | 72,1 |
| ENSG00000132623 | ANKEF1 | 1,47 | 1,05 | 0,64 | 0,41 | 72,1 |
| ENSG00000057294 | PKP2 | 3,26 | 1,91 | 1,17 | 0,91 | 72,1 |
| ENSG00000165695 | AK8 | 1,11 | 0,9 | 0,59 | 0,31 | 72,1 |
| ENSG00000123307 | NEUROD4 | 2,04 | 1,51 | 1,06 | 0,57 | 72,1 |
| ENSG00000168116 | KIAA1586 | 17,7 | 16,31 | 12,85 | 4,96 | 72,0 |
| ENSG00000150773 | PIH1D2 | 3,53 | 2,58 | 2,48 | 0,99 | 72,0 |
| ENSG00000274210 | RNVU1-27 | 56,62 | 48,38 | 37,49 | 15,92 | 71,9 |
| ENSG00000169139 | UBE2V2 | 62,36 | 58,08 | 54,74 | 17,57 | 71,8 |
| ENSG00000272255 | AC113361,1 | 1,1 | 0,74 | 0,65 | 0,31 | 71,8 |
| ENSG00000163041 | H3-3A | 395,2 | 389,97 | 352,15 | 111,38 | 71,8 |
| ENSG00000182010 | RTKN2 | 11,44 | 10,7 | 9,76 | 3,23 | 71,8 |
| ENSG00000082212 | ME2 | 31,93 | 29,79 | 24,61 | 9,03 | 71,7 |
| ENSG00000204837 | FGF7P3 | 3,04 | 1,34 | 1,31 | 0,86 | 71,7 |
| ENSG00000154719 | MRPL39 | 23,08 | 20,74 | 19,26 | 6,55 | 71,6 |
| ENSG00000101349 | PAK5 | 1,23 | 1,22 | 0,67 | 0,35 | 71,5 |
| ENSG00000074935 | TUBE1 | 11,48 | 10,32 | 8,66 | 3,27 | 71,5 |
| ENSG00000281661 | ZNF501 | 1,86 | 1,42 | 1,13 | 0,53 | 71,5 |
| ENSG00000124610 | H1-1 | 56,65 | 40,83 | 34,43 | 16,17 | 71,5 |
| ENSG00000257056 | LINC02282 | 4,99 | 3,18 | 1,88 | 1,43 | 71,3 |
| ENSG00000229644 | NAMPTP1 | 1,08 | 1,05 | 0,95 | 0,31 | 71,3 |
| ENSG00000119328 | ABITRAM | 11,52 | 10,96 | 9,6 | 3,31 | 71,3 |
| ENSG00000175893 | ZDHHC21 | 5,21 | 5,17 | 4,81 | 1,5 | 71,2 |
| ENSG00000287697 | Z99127,3 | 1,25 | 1,24 | 0,81 | 0,36 | 71,2 |
| ENSG00000187325 | TAF9B | 20,43 | 19,3 | 18,12 | 5,9 | 71,1 |
| ENSG00000205981 | DNAJC19 | 23,5 | 19,34 | 16,83 | 6,79 | 71,1 |
| ENSG00000204406 | MBD5 | 26,29 | 13,82 | 13,54 | 7,64 | 70,9 |
| ENSG00000275825 | AC139494,4 | 4,29 | 2,83 | 2,73 | 1,25 | 70,9 |

| | | | | | | |
|---|---|---|---|---|---|---|
| ENSG00000164070 | HSPA4L | 7,8 | 5,95 | 4,99 | 2,28 | 70,8 |
| ENSG00000182916 | TCEAL7 | 68,78 | 63,03 | 54,19 | 20,13 | 70,7 |
| ENSG00000261762 | AC027228,2 | 2,59 | 1,96 | 1,66 | 0,76 | 70,7 |
| ENSG00000221818 | EBF2 | 3,61 | 3,03 | 1,69 | 1,06 | 70,6 |
| ENSG00000272316 | AL021368,2 | 2,21 | 2,08 | 1,73 | 0,65 | 70,6 |
| ENSG00000196418 | ZNF124 | 41,61 | 36,78 | 31,84 | 12,24 | 70,6 |
| ENSG00000137968 | SLC44A5 | 4,79 | 4,26 | 3,41 | 1,41 | 70,6 |
| ENSG00000183960 | KCNH8 | 1,46 | 1,45 | 1 | 0,43 | 70,5 |
| ENSG00000139800 | ZIC5 | 15,31 | 14,11 | 12,34 | 4,51 | 70,5 |
| ENSG00000160131 | VMA21 | 18,51 | 17,42 | 15,74 | 5,46 | 70,5 |
| ENSG00000132437 | DDC | 2 | 1,37 | 0,94 | 0,59 | 70,5 |
| ENSG00000231104 | AC022395,1 | 1,83 | 1,27 | 0,88 | 0,54 | 70,5 |
| ENSG00000133740 | E2F5 | 23,72 | 17,7 | 17,4 | 7 | 70,5 |
| ENSG00000217527 | RPS16P5 | 2,19 | 1,2 | 1,11 | 0,65 | 70,3 |
| ENSG00000276216 | AC245014,3 | 67,15 | 50,91 | 30,8 | 19,94 | 70,3 |
| ENSG00000129295 | LRRC6 | 7,37 | 5,47 | 4,89 | 2,19 | 70,3 |
| ENSG00000197969 | VPS13A | 10,26 | 9,27 | 7,26 | 3,06 | 70,2 |
| ENSG00000183833 | CFAP91 | 4,99 | 2,92 | 1,63 | 1,49 | 70,1 |
| ENSG00000076351 | SLC46A1 | 5,29 | 5,1 | 3,65 | 1,58 | 70,1 |
| ENSG00000279342 | AP000866,6 | 4,82 | 4,33 | 3,87 | 1,44 | 70,1 |
| ENSG00000163291 | PAQR3 | 14,44 | 12,5 | 9,13 | 4,32 | 70,1 |
| ENSG00000147251 | DOCK11 | 10,43 | 9,42 | 8,34 | 3,13 | 70,0 |
| ENSG00000129084 | PSMA1 | 96,58 | 96,15 | 87,58 | 29,03 | 69,9 |
| ENSG00000256664 | AC025423,2 | 22,85 | 22,42 | 19,9 | 6,88 | 69,9 |
| ENSG00000120526 | NUDCD1 | 15,89 | 14,21 | 11,59 | 4,79 | 69,9 |
| ENSG00000278621 | AC037198,2 | 1,69 | 1,26 | 0,61 | 0,51 | 69,8 |
| ENSG00000165521 | EML5 | 1,06 | 0,82 | 0,51 | 0,32 | 69,8 |
| ENSG00000183150 | GPR19 | 5 | 4,59 | 3,98 | 1,51 | 69,8 |
| ENSG00000137876 | RSL24D1 | 51,31 | 48,31 | 44,04 | 15,52 | 69,8 |
| ENSG00000088451 | TGDS | 5,45 | 4,56 | 3,56 | 1,65 | 69,7 |
| ENSG00000272033 | AL136984,1 | 2,08 | 1,91 | 1,36 | 0,63 | 69,7 |
| ENSG00000171174 | RBKS | 2,54 | 1,95 | 1,4 | 0,77 | 69,7 |
| ENSG00000117477 | CCDC181 | 6,03 | 4,32 | 3,03 | 1,83 | 69,7 |
| ENSG00000130224 | LRCH2 | 5,76 | 5,6 | 4,19 | 1,75 | 69,6 |
| ENSG00000153132 | CLGN | 7,04 | 5,84 | 4,75 | 2,14 | 69,6 |
| ENSG00000285704 | AC004765,1 | 1,71 | 1,5 | 1,17 | 0,52 | 69,6 |
| ENSG00000285595 | AC105114,2 | 1,97 | 1,31 | 0,92 | 0,6 | 69,5 |
| ENSG00000124767 | GLO1 | 99,76 | 97,61 | 93,38 | 30,39 | 69,5 |
| ENSG00000278023 | RDM1 | 1,28 | 1,13 | 1 | 0,39 | 69,5 |
| ENSG00000286473 | AC133485,7 | 1,05 | 0,68 | 0,55 | 0,32 | 69,5 |
| ENSG00000147687 | TATDN1 | 23,85 | 23,37 | 19,32 | 7,27 | 69,5 |
| ENSG00000282171 | LINC02210 | 10,89 | 10,18 | 9,48 | 3,32 | 69,5 |
| ENSG00000152484 | USP12 | 10,88 | 10,5 | 9,49 | 3,32 | 69,5 |
| ENSG00000148154 | UGCG | 10,42 | 9,1 | 7,11 | 3,18 | 69,5 |
| ENSG00000183281 | PLGLB1 | 1,9 | 1,79 | 1,37 | 0,58 | 69,5 |
| ENSG00000144834 | TAGLN3 | 48,96 | 30,55 | 20,39 | 14,99 | 69,4 |
| ENSG00000088035 | ALG6 | 9,87 | 9,67 | 7,09 | 3,03 | 69,3 |

| | | | | | | |
|---|---|---|---|---|---|---|
| ENSG00000255794 | RMST | 134,92 | 129,51 | 102,64 | 41,43 | 69,3 |
| ENSG00000180008 | SOCS4 | 9,15 | 8,59 | 8,53 | 2,81 | 69,3 |
| ENSG00000147669 | POLR2K | 39,09 | 38,18 | 34,02 | 12,03 | 69,2 |
| ENSG00000134755 | DSC2 | 9,37 | 8,13 | 5,6 | 2,89 | 69,2 |
| ENSG00000129682 | FGF13 | 17,23 | 12,76 | 9,52 | 5,32 | 69,1 |
| ENSG00000196659 | TTC30B | 3,46 | 3,08 | 2,62 | 1,07 | 69,1 |
| ENSG00000158373 | H2BC5 | 805,88 | 641,34 | 637,49 | 249,74 | 69,0 |
| ENSG00000153266 | FEZF2 | 8,13 | 4,83 | 3,03 | 2,54 | 68,8 |
| ENSG00000115541 | HSPE1 | 221,29 | 190,24 | 169,8 | 69,21 | 68,7 |
| ENSG00000233836 | AC139769,1 | 9,74 | 9,49 | 8,6 | 3,05 | 68,7 |
| ENSG00000197061 | H4C3 | 904,9 | 663,05 | 633,42 | 284,25 | 68,6 |
| ENSG00000057704 | TMCC3 | 1,11 | 0,73 | 0,43 | 0,35 | 68,5 |
| ENSG00000146352 | CLVS2 | 2,41 | 2,13 | 2,01 | 0,76 | 68,5 |
| ENSG00000100852 | ARHGAP5 | 48,95 | 45,68 | 39,16 | 15,45 | 68,4 |
| ENSG00000100060 | MFNG | 3,04 | 1,34 | 0,97 | 0,96 | 68,4 |
| ENSG00000082258 | CCNT2 | 18,45 | 17,21 | 17 | 5,83 | 68,4 |
| ENSG00000170743 | SYT9 | 2,31 | 1,96 | 1,31 | 0,73 | 68,4 |
| ENSG00000267745 | AC060766,7 | 5,94 | 5,76 | 4,38 | 1,88 | 68,4 |
| ENSG00000107105 | ELAVL2 | 12,04 | 11,63 | 8,39 | 3,82 | 68,3 |
| ENSG00000226856 | THORLNC | 1,67 | 1,29 | 1,07 | 0,53 | 68,3 |
| ENSG00000173041 | ZNF680 | 11,81 | 11,46 | 10,09 | 3,75 | 68,2 |
| ENSG00000185272 | RBM11 | 1,29 | 1,01 | 0,76 | 0,41 | 68,2 |
| ENSG00000177182 | CLVS1 | 1,95 | 1,28 | 0,68 | 0,62 | 68,2 |
| ENSG00000106355 | LSM5 | 64,17 | 61,87 | 55,13 | 20,42 | 68,2 |
| ENSG00000235875 | ARHGEF7-AS2 | 1,19 | 0,57 | 0,4 | 0,38 | 68,1 |
| ENSG00000102753 | KPNA3 | 24,97 | 24,52 | 22,89 | 7,99 | 68,0 |
| ENSG00000111846 | GCNT2 | 1 | 0,91 | 0,88 | 0,32 | 68,0 |
| ENSG00000000460 | C1orf112 | 15,84 | 13,39 | 11,5 | 5,07 | 68,0 |
| ENSG00000139133 | ALG10 | 6,86 | 6,57 | 4,71 | 2,2 | 67,9 |
| ENSG00000176076 | KCNE5 | 6,89 | 5,57 | 5,18 | 2,21 | 67,9 |
| ENSG00000072133 | RPS6KA6 | 2,93 | 2,92 | 1,81 | 0,94 | 67,9 |
| ENSG00000152382 | TADA1 | 10,93 | 10,82 | 9,51 | 3,51 | 67,9 |
| ENSG00000143971 | ETAA1 | 6,04 | 5,95 | 5,14 | 1,94 | 67,9 |
| ENSG00000188488 | SERPINA5 | 1,12 | 0,69 | 0,58 | 0,36 | 67,9 |
| ENSG00000262402 | MCUR1P1 | 1,15 | 0,85 | 0,55 | 0,37 | 67,8 |
| ENSG00000272398 | CD24 | 145,76 | 128,92 | 106,4 | 46,91 | 67,8 |
| ENSG00000163584 | RPL22L1 | 48,86 | 46,06 | 43,99 | 15,73 | 67,8 |
| ENSG00000225447 | RPS15AP10 | 1,21 | 0,85 | 0,8 | 0,39 | 67,8 |
| ENSG00000171659 | GPR34 | 1,02 | 0,78 | 0,7 | 0,33 | 67,6 |
| ENSG00000115421 | PAPOLG | 11,28 | 10,48 | 8,85 | 3,65 | 67,6 |
| ENSG00000159593 | NAE1 | 57,58 | 53,96 | 45,67 | 18,64 | 67,6 |
| ENSG00000270277 | AC009948,1 | 2,13 | 1,83 | 1,8 | 0,69 | 67,6 |
| ENSG00000155959 | VBP1 | 47,51 | 42,47 | 40,94 | 15,4 | 67,6 |
| ENSG00000137944 | KYAT3 | 15,63 | 13,76 | 12,84 | 5,07 | 67,6 |
| ENSG00000041515 | MYO16 | 2,77 | 1,4 | 1,18 | 0,9 | 67,5 |
| ENSG00000227230 | AL606534,1 | 1,6 | 1,39 | 1,13 | 0,52 | 67,5 |
| ENSG00000285417 | BX571818,1 | 11,37 | 11,08 | 6,95 | 3,7 | 67,5 |

| | | | | | | |
|---|---|---|---|---|---|---|
| ENSG00000257027 | AC010186,3 | 3,62 | 3,53 | 3,31 | 1,18 | 67,4 |
| ENSG00000231607 | DLEU2 | 13,07 | 12,35 | 10,51 | 4,27 | 67,3 |
| ENSG00000151287 | TEX30 | 10,77 | 9,03 | 8,9 | 3,52 | 67,3 |
| ENSG00000276663 | AC009090,3 | 1,56 | 1,29 | 0,69 | 0,51 | 67,3 |
| ENSG00000154813 | DPH3 | 17,62 | 16,84 | 12,73 | 5,77 | 67,3 |
| ENSG00000260196 | AC124798,1 | 1,19 | 0,66 | 0,6 | 0,39 | 67,2 |
| ENSG00000091009 | RBM27 | 9,67 | 9,54 | 7,57 | 3,19 | 67,0 |
| ENSG00000149970 | CNKSR2 | 6,84 | 4,61 | 3,42 | 2,26 | 67,0 |
| ENSG00000198553 | KCNRG | 1,63 | 1,55 | 1,33 | 0,54 | 66,9 |
| ENSG00000116752 | BCAS2 | 32,94 | 29,16 | 25,98 | 10,92 | 66,8 |
| ENSG00000276787 | FAN1 | 2,35 | 2,23 | 1,93 | 0,78 | 66,8 |
| ENSG00000163738 | MTHFD2L | 8,24 | 8,13 | 7,12 | 2,74 | 66,7 |
| ENSG00000221909 | FAM200A | 4,81 | 4,32 | 4,25 | 1,6 | 66,7 |
| ENSG00000184005 | ST6GALNAC3 | 18,94 | 17,11 | 15,67 | 6,31 | 66,7 |
| ENSG00000180855 | ZNF443 | 6,21 | 5,87 | 5,67 | 2,07 | 66,7 |
| ENSG00000213467 | HMGB1P37 | 1,02 | 0,99 | 0,62 | 0,34 | 66,7 |
| ENSG00000146414 | SHPRH | 12,6 | 10,12 | 7,67 | 4,2 | 66,7 |
| ENSG00000187240 | DYNC2H1 | 10,54 | 8,82 | 7,48 | 3,52 | 66,6 |
| ENSG00000197557 | TTC30A | 2,93 | 2,75 | 2,37 | 0,98 | 66,6 |
| ENSG00000278932 | CR381653,1 | 2,54 | 1,3 | 1,2 | 0,85 | 66,5 |
| ENSG00000116489 | CAPZA1 | 84,65 | 83,89 | 76,3 | 28,41 | 66,4 |
| ENSG00000102098 | SCML2 | 4,61 | 3,78 | 3,38 | 1,55 | 66,4 |
| ENSG00000153975 | ZUP1 | 10,02 | 8,61 | 8,35 | 3,37 | 66,4 |
| ENSG00000259905 | PWRN1 | 8,64 | 8,08 | 6,1 | 2,91 | 66,3 |
| ENSG00000108001 | EBF3 | 5,4 | 4,44 | 2,98 | 1,82 | 66,3 |
| ENSG00000196890 | H2BU1 | 5,01 | 4,54 | 3,53 | 1,69 | 66,3 |
| ENSG00000076770 | MBNL3 | 15,79 | 13,31 | 11,05 | 5,33 | 66,2 |
| ENSG00000169446 | MMGT1 | 10,6 | 10,02 | 9,44 | 3,58 | 66,2 |
| ENSG00000181201 | H2BU2P | 2,72 | 1,16 | 1,15 | 0,92 | 66,2 |
| ENSG00000285234 | H2BU2P | 2,72 | 1,16 | 1,15 | 0,92 | 66,2 |
| ENSG00000138382 | METTL5 | 30,06 | 27,19 | 26,33 | 10,17 | 66,2 |
| ENSG00000126950 | TMEM35A | 7,49 | 7,27 | 5,58 | 2,54 | 66,1 |
| ENSG00000115507 | OTX1 | 1,15 | 0,74 | 0,54 | 0,39 | 66,1 |
| ENSG00000115392 | FANCL | 17,75 | 17,46 | 16,25 | 6,03 | 66,0 |
| ENSG00000254837 | AP001372,2 | 2,59 | 2,55 | 2,35 | 0,88 | 66,0 |
| ENSG00000205268 | PDE7A | 23,22 | 19,05 | 16,02 | 7,89 | 66,0 |
| ENSG00000175548 | ALG10B | 5,65 | 4,54 | 3,7 | 1,92 | 66,0 |
| ENSG00000180346 | TIGD2 | 4,97 | 4,24 | 4,16 | 1,69 | 66,0 |
| ENSG00000128708 | HAT1 | 45,43 | 42,72 | 39,43 | 15,46 | 66,0 |
| ENSG00000008394 | MGST1 | 24,74 | 21,53 | 16,7 | 8,42 | 66,0 |
| ENSG00000154582 | ELOC | 48,97 | 48,81 | 46,81 | 16,67 | 66,0 |
| ENSG00000179152 | TCAIM | 11,13 | 9,28 | 8,91 | 3,79 | 65,9 |
| ENSG00000111196 | MAGOHB | 31,53 | 28,9 | 26,51 | 10,74 | 65,9 |
| ENSG00000185842 | DNAH14 | 20,38 | 17,56 | 15 | 6,95 | 65,9 |
| ENSG00000100567 | PSMA3 | 100,86 | 91,87 | 83,95 | 34,4 | 65,9 |
| ENSG00000257126 | FOXG1-AS1 | 2,49 | 1,92 | 1,74 | 0,85 | 65,9 |
| ENSG00000085871 | MGST2 | 11,98 | 11,31 | 10,2 | 4,09 | 65,9 |

| | | | | | | |
|---|---|---|---|---|---|---|
| ENSG00000274191 | AC026333,4 | 1,23 | 0,69 | 0,44 | 0,42 | 65,9 |
| ENSG00000069956 | MAPK6 | 32,37 | 29,43 | 24,79 | 11,07 | 65,8 |
| ENSG00000133818 | RRAS2 | 16,52 | 15,46 | 13,19 | 5,65 | 65,8 |
| ENSG00000155304 | HSPA13 | 21,49 | 20,33 | 19,71 | 7,35 | 65,8 |
| ENSG00000080224 | EPHA6 | 3,42 | 3,06 | 2,32 | 1,17 | 65,8 |
| ENSG00000125246 | CLYBL | 10,17 | 6,78 | 5,24 | 3,48 | 65,8 |
| ENSG00000177468 | OLIG3 | 2,86 | 1,74 | 1,35 | 0,98 | 65,7 |
| ENSG00000214114 | MYCBP | 11,23 | 9,01 | 7,33 | 3,85 | 65,7 |
| ENSG00000124207 | CSE1L | 73,84 | 65,41 | 60,35 | 25,33 | 65,7 |
| ENSG00000148798 | INA | 32,61 | 25,57 | 15,84 | 11,21 | 65,6 |
| ENSG00000208892 | SNORA49 | 672,42 | 615,74 | 549,99 | 231,58 | 65,6 |
| ENSG00000141646 | SMAD4 | 58,26 | 57,89 | 54,13 | 20,07 | 65,6 |
| ENSG00000278238 | AL359513,1 | 1,19 | 0,97 | 0,74 | 0,41 | 65,5 |
| ENSG00000127328 | RAB3IP | 18,72 | 18,01 | 12,53 | 6,45 | 65,5 |
| ENSG00000143469 | SYT14 | 2,06 | 1,99 | 1,84 | 0,71 | 65,5 |
| ENSG00000274818 | AC004825,2 | 1,01 | 0,84 | 0,8 | 0,35 | 65,3 |
| ENSG00000287979 | AC253572,1 | 15,72 | 10,81 | 6,81 | 5,45 | 65,3 |
| ENSG00000053328 | METTL24 | 2,22 | 1,71 | 1,2 | 0,77 | 65,3 |
| ENSG00000152749 | GPR180 | 2,88 | 2,71 | 2,69 | 1 | 65,3 |
| ENSG00000125618 | PAX8 | 20,9 | 11,93 | 8,45 | 7,26 | 65,3 |
| ENSG00000144681 | STAC | 1,84 | 1,5 | 0,75 | 0,64 | 65,2 |
| ENSG00000130226 | DPP6 | 1,15 | 0,93 | 0,52 | 0,4 | 65,2 |
| ENSG00000127980 | PEX1 | 11,69 | 10,87 | 10,61 | 4,07 | 65,2 |
| ENSG00000215218 | UBE2QL1 | 1,78 | 1,16 | 0,72 | 0,62 | 65,2 |
| ENSG00000154639 | CXADR | 22,29 | 18,84 | 16,94 | 7,77 | 65,1 |
| ENSG00000023041 | ZDHHC6 | 24,21 | 22,5 | 21,62 | 8,45 | 65,1 |
| ENSG00000135049 | AGTPBP1 | 22,28 | 22,21 | 19,4 | 7,78 | 65,1 |
| ENSG00000276005 | AC138749,8 | 2,09 | 1,52 | 1,46 | 0,73 | 65,1 |
| ENSG00000104047 | DTWD1 | 14,63 | 12,95 | 11,96 | 5,11 | 65,1 |
| ENSG00000172292 | CERS6 | 12,79 | 12,67 | 10,86 | 4,47 | 65,1 |
| ENSG00000177034 | MTX3 | 8,01 | 7,6 | 5,9 | 2,8 | 65,0 |
| ENSG00000121621 | KIF18A | 16,17 | 15,82 | 14,42 | 5,66 | 65,0 |
| ENSG00000249626 | AC024560,2 | 1,97 | 1,93 | 1,25 | 0,69 | 65,0 |
| ENSG00000250317 | SMIM20 | 17,03 | 16 | 15,73 | 5,97 | 64,9 |
| ENSG00000174842 | GLMN | 10,61 | 9,35 | 8,93 | 3,72 | 64,9 |
| ENSG00000145908 | ZNF300 | 8,38 | 8,31 | 7,24 | 2,94 | 64,9 |
| ENSG00000177853 | ZNF518A | 16,64 | 14,77 | 12,81 | 5,84 | 64,9 |
| ENSG00000188211 | NCR3LG1 | 3,33 | 2,7 | 1,87 | 1,17 | 64,9 |
| ENSG00000230084 | AC006059,1 | 6,4 | 6,04 | 5,74 | 2,25 | 64,8 |
| ENSG00000133119 | RFC3 | 21,16 | 20,19 | 17,67 | 7,44 | 64,8 |
| ENSG00000008324 | SS18L2 | 10,55 | 10,15 | 9,24 | 3,71 | 64,8 |
| ENSG00000187672 | ERC2 | 6,91 | 6,06 | 4,16 | 2,43 | 64,8 |
| ENSG00000139343 | SNRPF | 46,93 | 45,36 | 37,21 | 16,51 | 64,8 |
| ENSG00000056277 | ZNF280C | 16,62 | 14,81 | 12,96 | 5,85 | 64,8 |
| ENSG00000067064 | IDI1 | 64,1 | 57,43 | 52,6 | 22,58 | 64,8 |
| ENSG00000276368 | H2AC14 | 555,61 | 453,07 | 443,7 | 195,82 | 64,8 |
| ENSG00000145725 | PPIP5K2 | 35,04 | 29,83 | 26,75 | 12,36 | 64,7 |

| | | | | | | |
|---|---|---|---|---|---|---|
| ENSG00000091164 | TXNL1 | 39,83 | 36,33 | 33,99 | 14,05 | 64,7 |
| ENSG00000140006 | WDR89 | 8,71 | 8,18 | 7,1 | 3,08 | 64,6 |
| ENSG00000153485 | TMEM251 | 6,08 | 5,81 | 5,55 | 2,15 | 64,6 |
| ENSG00000231205 | ZNF826P | 13,37 | 11,86 | 11,01 | 4,73 | 64,6 |
| ENSG00000127081 | ZNF484 | 5,99 | 5,93 | 5,86 | 2,12 | 64,6 |
| ENSG00000143742 | SRP9 | 254,99 | 242,6 | 214,69 | 90,36 | 64,6 |
| ENSG00000160201 | U2AF1 | 97,44 | 86,48 | 56,8 | 34,59 | 64,5 |
| ENSG00000254539 | AC239804,1 | 1,04 | 0,69 | 0,66 | 0,37 | 64,4 |
| ENSG00000160124 | CCDC58 | 33,73 | 29,94 | 25,22 | 12,02 | 64,4 |
| ENSG00000129596 | CDO1 | 17,98 | 14,05 | 12,6 | 6,41 | 64,3 |
| ENSG00000000003 | TSPAN6 | 42,83 | 41,27 | 38,29 | 15,27 | 64,3 |
| ENSG00000162385 | MAGOH | 52,36 | 49,36 | 45,79 | 18,67 | 64,3 |
| ENSG00000077152 | UBE2T | 33,16 | 30,87 | 29,92 | 11,84 | 64,3 |
| ENSG00000171262 | FAM98B | 12,63 | 10,31 | 10,14 | 4,51 | 64,3 |
| ENSG00000214182 | PTMAP5 | 5,32 | 5,27 | 4,52 | 1,9 | 64,3 |
| ENSG00000196591 | HDAC2 | 179,05 | 164,07 | 149,35 | 63,97 | 64,3 |
| ENSG00000250950 | AC093752,2 | 3,05 | 2,58 | 2,23 | 1,09 | 64,3 |
| ENSG00000286757 | AL137139,2 | 3,58 | 3,2 | 2,6 | 1,28 | 64,2 |
| ENSG00000213741 | RPS29 | 515,49 | 442,22 | 428,72 | 184,41 | 64,2 |
| ENSG00000255717 | SNHG1 | 49,92 | 46,63 | 37,25 | 17,88 | 64,2 |
| ENSG00000171714 | ANO5 | 1,34 | 1,13 | 0,93 | 0,48 | 64,2 |
| ENSG00000061918 | GUCY1B1 | 10,3 | 9,71 | 8,54 | 3,69 | 64,2 |
| ENSG00000162694 | EXTL2 | 16,46 | 16,27 | 15,6 | 5,9 | 64,2 |
| ENSG00000125703 | ATG4C | 6,36 | 5,75 | 5,43 | 2,28 | 64,2 |
| ENSG00000260089 | ADAM3B | 1,31 | 1,27 | 1,16 | 0,47 | 64,1 |
| ENSG00000100442 | FKBP3 | 91,47 | 88,81 | 83,55 | 32,85 | 64,1 |
| ENSG00000180787 | ZFP3 | 1,03 | 0,81 | 0,79 | 0,37 | 64,1 |
| ENSG00000011258 | MBTD1 | 13,44 | 12,48 | 11,43 | 4,83 | 64,1 |
| ENSG00000163946 | TASOR | 33,58 | 33,15 | 28,52 | 12,07 | 64,1 |
| ENSG00000164167 | LSM6 | 44,17 | 41,22 | 39,33 | 15,9 | 64,0 |
| ENSG00000172058 | SERF1A | 17,88 | 11,18 | 8,91 | 6,44 | 64,0 |
| ENSG00000134602 | STK26 | 28,88 | 21,1 | 16,88 | 10,42 | 63,9 |
| ENSG00000150456 | EEF1AKMT1 | 7,87 | 6,85 | 5,75 | 2,84 | 63,9 |
| ENSG00000030419 | IKZF2 | 4,57 | 3,76 | 3,34 | 1,65 | 63,9 |
| ENSG00000286723 | AL021368,5 | 1,08 | 0,82 | 0,58 | 0,39 | 63,9 |
| ENSG00000102531 | FNDC3A | 26,86 | 24,89 | 20,47 | 9,7 | 63,9 |
| ENSG00000088986 | DYNLL1 | 299,83 | 291,62 | 291,6 | 108,4 | 63,8 |
| ENSG00000155313 | USP25 | 16,66 | 14,51 | 13,66 | 6,03 | 63,8 |
| ENSG00000097046 | CDC7 | 20 | 19,48 | 17,91 | 7,24 | 63,8 |
| ENSG00000091140 | DLD | 40,86 | 39,36 | 36,75 | 14,8 | 63,8 |
| ENSG00000196172 | ZNF681 | 4,5 | 3,79 | 2,85 | 1,63 | 63,8 |
| ENSG00000183527 | PSMG1 | 40,91 | 36,21 | 30,25 | 14,83 | 63,7 |
| ENSG00000128590 | DNAJB9 | 8,44 | 8,12 | 5,93 | 3,06 | 63,7 |
| ENSG00000136783 | NIPSNAP3A | 12,71 | 12,49 | 11,02 | 4,61 | 63,7 |
| ENSG00000213160 | KLHL23 | 23,67 | 22,93 | 17,33 | 8,59 | 63,7 |
| ENSG00000281601 | ZNF780B | 5,4 | 4,42 | 3,83 | 1,96 | 63,7 |
| ENSG00000114126 | TFDP2 | 44,31 | 39,12 | 34,51 | 16,09 | 63,7 |

| | | | | | | |
|---|---|---|---|---|---|---|
| ENSG00000140488 | CELF6 | 1,1 | 0,8 | 0,69 | 0,4 | 63,6 |
| ENSG00000164076 | CAMKV | 6,05 | 3,28 | 2,62 | 2,2 | 63,6 |
| ENSG00000237440 | ZNF737 | 9,34 | 9,19 | 6,95 | 3,4 | 63,6 |
| ENSG00000134595 | SOX3 | 6,01 | 5,5 | 4,71 | 2,19 | 63,6 |
| ENSG00000162994 | CLHC1 | 10,7 | 10,4 | 8,99 | 3,9 | 63,6 |
| ENSG00000138400 | MDH1B | 3,51 | 2,98 | 2,11 | 1,28 | 63,5 |
| ENSG00000156411 | ATP5MPL | 95,8 | 95,21 | 88,93 | 34,95 | 63,5 |
| ENSG00000225548 | LINC01980 | 7,37 | 6,38 | 5,21 | 2,69 | 63,5 |
| ENSG00000271452 | AC005034,5 | 1,89 | 1,8 | 1,3 | 0,69 | 63,5 |
| ENSG00000163568 | AIM2 | 2,3 | 2,01 | 1,39 | 0,84 | 63,5 |
| ENSG00000166450 | PRTG | 23,12 | 18,92 | 12,32 | 8,45 | 63,5 |
| ENSG00000049130 | KITLG | 7,96 | 6,17 | 5,3 | 2,91 | 63,4 |
| ENSG00000101856 | PGRMC1 | 152,92 | 147,62 | 140,37 | 55,95 | 63,4 |
| ENSG00000135341 | MAP3K7 | 19,7 | 18 | 15,92 | 7,21 | 63,4 |
| ENSG00000148019 | CEP78 | 31,92 | 29,61 | 24,19 | 11,69 | 63,4 |
| ENSG00000007372 | PAX6 | 11,1 | 7,3 | 4,98 | 4,07 | 63,3 |
| ENSG00000198856 | OSTC | 84,65 | 79,75 | 71,76 | 31,04 | 63,3 |
| ENSG00000075188 | NUP37 | 18,44 | 15,89 | 15,25 | 6,77 | 63,3 |
| ENSG00000165359 | INTS6L | 6,56 | 6,27 | 5,39 | 2,41 | 63,3 |
| ENSG00000104427 | ZC2HC1A | 13,9 | 13,37 | 10,23 | 5,11 | 63,2 |
| ENSG00000106701 | FSD1L | 11,88 | 11,21 | 10,06 | 4,37 | 63,2 |
| ENSG00000163104 | SMARCAD1 | 27,55 | 25,9 | 21,02 | 10,14 | 63,2 |
| ENSG00000135336 | ORC3 | 20,94 | 20,6 | 17,29 | 7,71 | 63,2 |
| ENSG00000146247 | PHIP | 21,24 | 19,4 | 15,29 | 7,84 | 63,1 |
| ENSG00000189212 | DPY19L2P1 | 1,57 | 1,25 | 1,2 | 0,58 | 63,1 |
| ENSG00000136108 | CKAP2 | 59,08 | 58,01 | 56,9 | 21,83 | 63,1 |
| ENSG00000198015 | MRPL42 | 39,65 | 34,4 | 28,62 | 14,66 | 63,0 |
| ENSG00000152240 | HAUS1 | 50,62 | 49,29 | 44,22 | 18,73 | 63,0 |
| ENSG00000141428 | C18orf21 | 17,57 | 16,02 | 15,88 | 6,51 | 62,9 |
| ENSG00000175054 | ATR | 11,2 | 9,34 | 8,93 | 4,15 | 62,9 |
| ENSG00000235833 | AC017099,1 | 2,32 | 1,98 | 1,85 | 0,86 | 62,9 |
| ENSG00000165209 | STRBP | 23,41 | 20,76 | 16,86 | 8,68 | 62,9 |
| ENSG00000145780 | FEM1C | 7,74 | 7,67 | 7,42 | 2,87 | 62,9 |
| ENSG00000146476 | ARMT1 | 14,18 | 13,2 | 11,14 | 5,26 | 62,9 |
| ENSG00000111261 | MANSC1 | 1,4 | 0,97 | 0,8 | 0,52 | 62,9 |
| ENSG00000049167 | ERCC8 | 10,74 | 9,57 | 8,16 | 3,99 | 62,8 |
| ENSG00000171786 | NHLH1 | 6,94 | 3,65 | 3,02 | 2,58 | 62,8 |
| ENSG00000056050 | HPF1 | 30,2 | 28,49 | 25,52 | 11,23 | 62,8 |
| ENSG00000162607 | USP1 | 39,06 | 38,79 | 33,56 | 14,53 | 62,8 |
| ENSG00000121741 | ZMYM2 | 51,14 | 50,86 | 43,11 | 19,04 | 62,8 |
| ENSG00000256073 | URB1-AS1 | 2,47 | 1,79 | 1,53 | 0,92 | 62,8 |
| ENSG00000247746 | USP51 | 3,3 | 2,85 | 2,31 | 1,23 | 62,7 |
| ENSG00000170222 | ADPRM | 3,86 | 3,76 | 3,4 | 1,44 | 62,7 |
| ENSG00000286458 | AC083870,1 | 1,5 | 1,29 | 0,71 | 0,56 | 62,7 |
| ENSG00000266916 | ZNF793-AS1 | 2,33 | 2,28 | 1,9 | 0,87 | 62,7 |
| ENSG00000116675 | DNAJC6 | 3,96 | 3,92 | 2,71 | 1,48 | 62,6 |
| ENSG00000165588 | OTX2 | 39,02 | 27,86 | 24,67 | 14,59 | 62,6 |

| | | | | | | |
|---|---|---|---|---|---|---|
| ENSG00000117000 | RLF | 8,98 | 8,92 | 8,27 | 3,36 | 62,6 |
| ENSG00000167232 | ZNF91 | 41,87 | 40,73 | 32,14 | 15,67 | 62,6 |
| ENSG00000270130 | AC068790,7 | 2,35 | 2,21 | 1,94 | 0,88 | 62,6 |
| ENSG00000250284 | AC109439,1 | 3,87 | 2,91 | 2,31 | 1,45 | 62,5 |
| ENSG00000114107 | CEP70 | 10,86 | 10,77 | 9,93 | 4,07 | 62,5 |
| ENSG00000171448 | ZBTB26 | 5,87 | 5,67 | 4,98 | 2,2 | 62,5 |
| ENSG00000179387 | ELMOD2 | 19,71 | 18,46 | 17,15 | 7,39 | 62,5 |
| ENSG00000243927 | MRPS6 | 50,66 | 43,69 | 41,72 | 19,01 | 62,5 |
| ENSG00000108384 | RAD51C | 24,65 | 22,68 | 19,78 | 9,25 | 62,5 |
| ENSG00000169612 | RAMAC | 19,48 | 18,07 | 17,45 | 7,31 | 62,5 |
| ENSG00000110700 | RPS13 | 431,05 | 404,26 | 399,23 | 161,76 | 62,5 |
| ENSG00000079134 | THOC1 | 39,37 | 34,9 | 29,72 | 14,78 | 62,5 |
| ENSG00000198791 | CNOT7 | 67,97 | 61,8 | 58,86 | 25,54 | 62,4 |
| ENSG00000009844 | VTA1 | 29,71 | 29,55 | 25,76 | 11,18 | 62,4 |
| ENSG00000116191 | RALGPS2 | 18,49 | 15,23 | 12,07 | 6,96 | 62,4 |
| ENSG00000174718 | RESF1 | 19,82 | 16,94 | 15,43 | 7,47 | 62,3 |
| ENSG00000152503 | TRIM36 | 14,75 | 11,54 | 7,72 | 5,56 | 62,3 |
| ENSG00000021776 | AQR | 16,09 | 14,96 | 12,83 | 6,07 | 62,3 |
| ENSG00000234840 | LINC01239 | 2,25 | 1,36 | 1,05 | 0,85 | 62,2 |
| ENSG00000118939 | UCHL3 | 22,72 | 17,17 | 15,58 | 8,59 | 62,2 |
| ENSG00000164338 | UTP15 | 12,93 | 12,67 | 11,23 | 4,89 | 62,2 |
| ENSG00000164187 | LMBRD2 | 3,91 | 3,55 | 3,38 | 1,48 | 62,1 |
| ENSG00000185480 | PARPBP | 12,07 | 10,71 | 10,48 | 4,57 | 62,1 |
| ENSG00000286707 | AC010896,1 | 1,03 | 0,87 | 0,85 | 0,39 | 62,1 |
| ENSG00000136960 | ENPP2 | 19,57 | 18,55 | 17,95 | 7,41 | 62,1 |
| ENSG00000184752 | NDUFA12 | 68,62 | 61,63 | 57,83 | 26,03 | 62,1 |
| ENSG00000196715 | VKORC1L1 | 12,25 | 11,86 | 10,16 | 4,65 | 62,0 |
| ENSG00000171488 | LRRC8C | 1,58 | 1,15 | 0,9 | 0,6 | 62,0 |
| ENSG00000078177 | N4BP2 | 14,19 | 12,94 | 12,12 | 5,39 | 62,0 |
| ENSG00000172348 | RCAN2 | 6,63 | 6,42 | 5,9 | 2,52 | 62,0 |
| ENSG00000165449 | SLC16A9 | 5,86 | 4,68 | 3,22 | 2,23 | 61,9 |
| ENSG00000112031 | MTRF1L | 10,9 | 10,09 | 9,73 | 4,15 | 61,9 |
| ENSG00000076053 | RBM7 | 18,49 | 16,69 | 16,4 | 7,04 | 61,9 |
| ENSG00000172915 | NBEA | 7,77 | 7 | 5,92 | 2,96 | 61,9 |
| ENSG00000284468 | SLC25A24 | 12,12 | 10,36 | 9,25 | 4,62 | 61,9 |
| ENSG00000085491 | SLC25A24 | 12,12 | 10,36 | 9,25 | 4,62 | 61,9 |
| ENSG00000154822 | PLCL2 | 1,81 | 1,69 | 1,26 | 0,69 | 61,9 |
| ENSG00000284017 | PLCL2 | 1,81 | 1,69 | 1,26 | 0,69 | 61,9 |
| ENSG00000008277 | ADAM22 | 6,19 | 4,22 | 3,02 | 2,36 | 61,9 |
| ENSG00000004779 | NDUFAB1 | 50,52 | 50,34 | 46,15 | 19,27 | 61,9 |
| ENSG00000175768 | TOMM5 | 94,08 | 77,4 | 74,67 | 35,89 | 61,9 |
| ENSG00000198814 | GK | 3,77 | 3,34 | 2,73 | 1,44 | 61,8 |
| ENSG00000066279 | ASPM | 16,3 | 15,87 | 14,63 | 6,23 | 61,8 |
| ENSG00000101773 | RBBP8 | 24,04 | 21,73 | 17,63 | 9,19 | 61,8 |
| ENSG00000198919 | DZIP3 | 10,71 | 9,92 | 7,12 | 4,1 | 61,7 |
| ENSG00000118402 | ELOVL4 | 4,31 | 4,13 | 3,76 | 1,65 | 61,7 |
| ENSG00000170860 | LSM3 | 15,62 | 14,04 | 13,21 | 5,98 | 61,7 |

| | | | | | | |
|---|---|---|---|---|---|---|
| ENSG00000105185 | PDCD5 | 91,21 | 84,61 | 78,18 | 34,92 | 61,7 |
| ENSG00000171016 | PYGO1 | 7,39 | 7,24 | 5,78 | 2,83 | 61,7 |
| ENSG00000095002 | MSH2 | 41,3 | 39,88 | 35,5 | 15,82 | 61,7 |
| ENSG00000197238 | H4C11 | 886,68 | 779,81 | 715,19 | 339,84 | 61,7 |
| ENSG00000259943 | AL050341,2 | 4,59 | 4,24 | 3,68 | 1,76 | 61,7 |
| ENSG00000196981 | WDR5B | 2,29 | 2,1 | 1,77 | 0,88 | 61,6 |
| ENSG00000151233 | GXYLT1 | 9,68 | 8,17 | 7,39 | 3,72 | 61,6 |
| ENSG00000106803 | SEC61B | 150,31 | 147,5 | 145,01 | 57,79 | 61,6 |
| ENSG00000066557 | LRRC40 | 17,79 | 16,79 | 15,59 | 6,84 | 61,6 |
| ENSG00000249412 | AC010285,1 | 1,43 | 1,33 | 1,17 | 0,55 | 61,5 |
| ENSG00000100433 | KCNK10 | 2,78 | 1,45 | 1,19 | 1,07 | 61,5 |
| ENSG00000168566 | SNRNP48 | 7,95 | 7,58 | 7,3 | 3,06 | 61,5 |
| ENSG00000213186 | TRIM59 | 16,95 | 16,63 | 15,97 | 6,53 | 61,5 |
| ENSG00000275470 | LENG8-AS1 | 1,09 | 0,99 | 0,43 | 0,42 | 61,5 |
| ENSG00000254806 | SYS1-DBNDD2 | 1,66 | 1,07 | 1,02 | 0,64 | 61,4 |
| ENSG00000041357 | PSMA4 | 139,55 | 138,48 | 130,55 | 53,81 | 61,4 |
| ENSG00000160345 | C9orf116 | 11,33 | 9,28 | 7,82 | 4,37 | 61,4 |
| ENSG00000233237 | LINC00472 | 14,44 | 12,06 | 11,08 | 5,57 | 61,4 |
| ENSG00000145386 | CCNA2 | 36,13 | 34,15 | 31,06 | 13,94 | 61,4 |
| ENSG00000133640 | LRRIQ1 | 4,12 | 3,68 | 1,85 | 1,59 | 61,4 |
| ENSG00000165392 | WRN | 12,9 | 11,86 | 10,35 | 4,98 | 61,4 |
| ENSG00000186871 | ERCC6L | 8,57 | 7,44 | 7,12 | 3,31 | 61,4 |
| ENSG00000162814 | SPATA17 | 3,65 | 2,11 | 1,86 | 1,41 | 61,4 |
| ENSG00000257596 | SCAT2 | 1,76 | 1,52 | 1,35 | 0,68 | 61,4 |
| ENSG00000163006 | CCDC138 | 13,93 | 12,46 | 10 | 5,39 | 61,3 |
| ENSG00000052802 | MSMO1 | 96,73 | 80,82 | 76,1 | 37,44 | 61,3 |
| ENSG00000196368 | NUDT11 | 15,84 | 14,18 | 12,14 | 6,14 | 61,2 |
| ENSG00000176273 | SLC35G1 | 4,2 | 3,83 | 3,06 | 1,63 | 61,2 |
| ENSG00000288632 | AC133555,6 | 2,55 | 2,52 | 2,06 | 0,99 | 61,2 |
| ENSG00000174405 | LIG4 | 4,84 | 4,63 | 4,47 | 1,88 | 61,2 |
| ENSG00000165113 | GKAP1 | 7,23 | 6,38 | 5,29 | 2,81 | 61,1 |
| ENSG00000003987 | MTMR7 | 3,19 | 2,69 | 1,9 | 1,24 | 61,1 |
| ENSG00000053438 | NNAT | 303,05 | 197,09 | 143,06 | 117,8 | 61,1 |
| ENSG00000150627 | WDR17 | 3,47 | 3,25 | 3,19 | 1,35 | 61,1 |
| ENSG00000196693 | ZNF33B | 14,43 | 12,31 | 9,07 | 5,62 | 61,1 |
| ENSG00000171806 | METTL18 | 6,08 | 5,52 | 4,41 | 2,37 | 61,0 |
| ENSG00000250366 | TUNAR | 3,77 | 2,88 | 2,56 | 1,47 | 61,0 |
| ENSG00000231965 | AF131215,1 | 1 | 0,62 | 0,43 | 0,39 | 61,0 |
| ENSG00000285026 | AC270301,5 | 1 | 0,62 | 0,43 | 0,39 | 61,0 |
| ENSG00000139116 | KIF21A | 18,33 | 14,57 | 13,99 | 7,15 | 61,0 |
| ENSG00000090060 | PAPOLA | 84,85 | 81,4 | 80 | 33,1 | 61,0 |
| ENSG00000137713 | PPP2R1B | 20,89 | 17,02 | 13,69 | 8,15 | 61,0 |
| ENSG00000167080 | B4GALNT2 | 1,05 | 0,95 | 0,74 | 0,41 | 61,0 |
| ENSG00000112232 | KHDRBS2 | 4,43 | 3,1 | 2,69 | 1,73 | 60,9 |
| ENSG00000156875 | MFSD14A | 17,51 | 17,17 | 16,59 | 6,84 | 60,9 |
| ENSG00000119314 | PTBP3 | 20,54 | 19,35 | 14,74 | 8,03 | 60,9 |
| ENSG00000132938 | MTUS2 | 1,33 | 1,19 | 0,67 | 0,52 | 60,9 |

| | | | | | | |
|---|---|---|---|---|---|---|
| ENSG00000153147 | SMARCA5 | 47,28 | 46,53 | 37,6 | 18,49 | 60,9 |
| ENSG00000150787 | PTS | 24,65 | 23,26 | 21,68 | 9,64 | 60,9 |
| ENSG00000277157 | H4C4 | 1137,15 | 758,46 | 731,39 | 444,75 | 60,9 |
| ENSG00000125629 | INSIG2 | 9,43 | 8,06 | 7,24 | 3,69 | 60,9 |
| ENSG00000278212 | AC134878,2 | 1,48 | 1,22 | 0,95 | 0,58 | 60,8 |
| ENSG00000111371 | SLC38A1 | 50,15 | 40,35 | 33,01 | 19,68 | 60,8 |
| ENSG00000128694 | OSGEPL1 | 7,82 | 7,44 | 7,41 | 3,07 | 60,7 |
| ENSG00000112290 | WASF1 | 27,84 | 24,9 | 21,39 | 10,93 | 60,7 |
| ENSG00000134825 | TMEM258 | 59,99 | 57,36 | 55,95 | 23,56 | 60,7 |
| ENSG00000177868 | SVBP | 27,62 | 25,38 | 23,88 | 10,85 | 60,7 |
| ENSG00000076685 | NT5C2 | 31,69 | 29,84 | 28,27 | 12,45 | 60,7 |
| ENSG00000196247 | ZNF107 | 9,57 | 8,61 | 8,46 | 3,76 | 60,7 |
| ENSG00000132321 | IQCA1 | 2,62 | 2,32 | 1,32 | 1,03 | 60,7 |
| ENSG00000132424 | PNISR | 72,25 | 69,52 | 53,86 | 28,41 | 60,7 |
| ENSG00000103599 | IQCH | 2,95 | 2,05 | 1,3 | 1,16 | 60,7 |
| ENSG00000177889 | UBE2N | 77,49 | 77,39 | 75,66 | 30,55 | 60,6 |
| ENSG00000152932 | RAB3C | 6,11 | 5,29 | 4,13 | 2,41 | 60,6 |
| ENSG00000143228 | NUF2 | 26,67 | 26,63 | 25,77 | 10,53 | 60,5 |
| ENSG00000137168 | PPIL1 | 45,25 | 39,91 | 35,48 | 17,87 | 60,5 |
| ENSG00000164659 | ELAPOR2 | 21,09 | 18,77 | 15,08 | 8,33 | 60,5 |
| ENSG00000196636 | SDHAF3 | 10,15 | 9,79 | 7,96 | 4,01 | 60,5 |
| ENSG00000107679 | PLEKHA1 | 8,78 | 8,42 | 6,95 | 3,47 | 60,5 |
| ENSG00000153044 | CENPH | 28,26 | 25,68 | 24,03 | 11,17 | 60,5 |
| ENSG00000173674 | EIF1AX | 29 | 23,72 | 21,51 | 11,47 | 60,4 |
| ENSG00000177076 | ACER2 | 1,39 | 0,95 | 0,78 | 0,55 | 60,4 |
| ENSG00000213123 | TCTEX1D2 | 32,31 | 31,79 | 26,44 | 12,79 | 60,4 |
| ENSG00000261799 | AC007406,4 | 3,46 | 3,34 | 2,96 | 1,37 | 60,4 |
| ENSG00000124596 | OARD1 | 16,98 | 15,88 | 13,78 | 6,73 | 60,4 |
| ENSG00000138399 | FASTKD1 | 10,54 | 9 | 8,27 | 4,18 | 60,3 |
| ENSG00000213281 | NRAS | 34,58 | 33,43 | 30,78 | 13,72 | 60,3 |
| ENSG00000156261 | CCT8 | 103,83 | 95,45 | 86,28 | 41,29 | 60,2 |
| ENSG00000006625 | GGCT | 21,85 | 19,4 | 17,4 | 8,7 | 60,2 |
| ENSG00000164542 | KIAA0895 | 9,11 | 8,7 | 7,09 | 3,63 | 60,2 |
| ENSG00000102078 | SLC25A14 | 24 | 19,5 | 18,91 | 9,57 | 60,1 |
| ENSG00000145354 | CISD2 | 14,82 | 14,28 | 12,99 | 5,91 | 60,1 |
| ENSG00000189007 | ADAT2 | 2,58 | 2,32 | 1,54 | 1,03 | 60,1 |
| ENSG00000204147 | ASAH2B | 2,63 | 2,29 | 2,14 | 1,05 | 60,1 |
| ENSG00000024526 | DEPDC1 | 14,7 | 13,77 | 13,4 | 5,87 | 60,1 |
| ENSG00000136161 | RCBTB2 | 15,29 | 14,07 | 11,92 | 6,11 | 60,0 |
| ENSG00000156469 | MTERF3 | 12,56 | 10,75 | 9,23 | 5,02 | 60,0 |
| ENSG00000104093 | DMXL2 | 16,9 | 15,92 | 15,59 | 6,76 | 60,0 |
| ENSG00000153140 | CETN3 | 26,48 | 21,45 | 20,37 | 10,6 | 60,0 |
| ENSG00000188994 | ZNF292 | 23,51 | 20,06 | 18,61 | 9,42 | 59,9 |
| ENSG00000270728 | AL035413,1 | 1,92 | 1,39 | 1,28 | 0,77 | 59,9 |
| ENSG00000118246 | FASTKD2 | 15,85 | 13,14 | 12,19 | 6,36 | 59,9 |
| ENSG00000182899 | RPL35A | 644,08 | 641,71 | 617,54 | 258,69 | 59,8 |
| ENSG00000173226 | IQCB1 | 25,7 | 22,42 | 19,93 | 10,33 | 59,8 |

| | | | | | | |
|---|---|---|---|---|---|---|
| ENSG00000151690 | MFSD6 | 1,79 | 1,36 | 1,27 | 0,72 | 59,8 |
| ENSG00000139767 | SRRM4 | 2,56 | 1,87 | 1,31 | 1,03 | 59,8 |
| ENSG00000189223 | PAX8-AS1 | 6,65 | 4,74 | 3,17 | 2,68 | 59,7 |
| ENSG00000152092 | ASTN1 | 6,55 | 5,96 | 3,83 | 2,64 | 59,7 |
| ENSG00000164118 | CEP44 | 8,46 | 6,88 | 5,87 | 3,41 | 59,7 |
| ENSG00000162999 | DUSP19 | 1,24 | 0,94 | 0,74 | 0,5 | 59,7 |
| ENSG00000079387 | SENP1 | 18,18 | 15,45 | 14,84 | 7,34 | 59,6 |
| ENSG00000245498 | AP000866,1 | 2,97 | 2,27 | 1,83 | 1,2 | 59,6 |
| ENSG00000229267 | SNHG31 | 1,46 | 1,14 | 0,77 | 0,59 | 59,6 |
| ENSG00000180530 | NRIP1 | 12,61 | 9,77 | 9,35 | 5,1 | 59,6 |
| ENSG00000138617 | PARP16 | 5,46 | 4,42 | 3,96 | 2,21 | 59,5 |
| ENSG00000271369 | AC087783,1 | 6,1 | 5,97 | 4,84 | 2,47 | 59,5 |
| ENSG00000136810 | TXN | 136,18 | 126,52 | 117,9 | 55,18 | 59,5 |
| ENSG00000189367 | KIAA0408 | 11,57 | 10,93 | 8,33 | 4,69 | 59,5 |
| ENSG00000112972 | HMGCS1 | 318,44 | 311,46 | 263,92 | 129,24 | 59,4 |
| ENSG00000139187 | KLRG1 | 2,71 | 2,18 | 1,32 | 1,1 | 59,4 |
| ENSG00000182004 | SNRPE | 179,75 | 162,08 | 149,63 | 72,97 | 59,4 |
| ENSG00000152270 | PDE3B | 5,54 | 3,46 | 3,24 | 2,25 | 59,4 |
| ENSG00000065615 | CYB5R4 | 6,3 | 6,29 | 6,24 | 2,56 | 59,4 |
| ENSG00000138802 | SEC24B | 14,8 | 14,78 | 13,89 | 6,02 | 59,3 |
| ENSG00000146267 | FAXC | 10,3 | 8,43 | 7,2 | 4,19 | 59,3 |
| ENSG00000205189 | ZBTB10 | 18,63 | 13,2 | 10,72 | 7,58 | 59,3 |
| ENSG00000189043 | NDUFA4 | 59,86 | 52,84 | 52,6 | 24,37 | 59,3 |
| ENSG00000156925 | ZIC3 | 30,72 | 25,5 | 20,71 | 12,51 | 59,3 |
| ENSG00000186976 | EFCAB6 | 1,52 | 1,04 | 0,82 | 0,62 | 59,2 |
| ENSG00000122481 | RWDD3 | 16,94 | 15,21 | 15,19 | 6,91 | 59,2 |
| ENSG00000196437 | ZNF569 | 5,17 | 5,15 | 4,22 | 2,11 | 59,2 |
| ENSG00000145604 | SKP2 | 32,18 | 26,7 | 22,33 | 13,14 | 59,2 |
| ENSG00000241261 | RPL17P19 | 1,2 | 1,08 | 0,83 | 0,49 | 59,2 |
| ENSG00000175455 | CCDC14 | 43,36 | 39,11 | 31,8 | 17,71 | 59,2 |
| ENSG00000175110 | MRPS22 | 28,35 | 28,23 | 21,22 | 11,59 | 59,1 |
| ENSG00000138439 | FAM117B | 7,38 | 7,08 | 5,78 | 3,02 | 59,1 |
| ENSG00000184349 | EFNA5 | 22,99 | 19,54 | 13,76 | 9,41 | 59,1 |
| ENSG00000229419 | RALGAPA1P1 | 2,32 | 2,23 | 1,88 | 0,95 | 59,1 |
| ENSG00000154760 | SLFN13 | 12,35 | 11,93 | 9,15 | 5,06 | 59,0 |
| ENSG00000187801 | ZFP69B | 5,93 | 5,42 | 4,25 | 2,43 | 59,0 |
| ENSG00000259959 | AC107068,1 | 1,61 | 1,41 | 1,02 | 0,66 | 59,0 |
| ENSG00000136273 | HUS1 | 18,68 | 16,19 | 15,47 | 7,66 | 59,0 |
| ENSG00000136982 | DSCC1 | 11,7 | 10,47 | 8,84 | 4,8 | 59,0 |
| ENSG00000001630 | CYP51A1 | 71,52 | 66,12 | 59,96 | 29,35 | 59,0 |
| ENSG00000119616 | FCF1 | 31,93 | 31,75 | 26,9 | 13,11 | 58,9 |
| ENSG00000156482 | RPL30 | 579,92 | 556,88 | 540,93 | 238,11 | 58,9 |
| ENSG00000188428 | BLOC1S5 | 8,2 | 7,48 | 6,27 | 3,37 | 58,9 |
| ENSG00000070269 | TMEM260 | 9,94 | 8,59 | 7,15 | 4,09 | 58,9 |
| ENSG00000154640 | BTG3 | 25,25 | 24,73 | 24,67 | 10,39 | 58,9 |
| ENSG00000100902 | PSMA6 | 85,29 | 84,47 | 80,23 | 35,1 | 58,8 |
| ENSG00000164651 | SP8 | 16,05 | 12,17 | 8,58 | 6,61 | 58,8 |

| | | | | | | |
|---|---|---|---|---|---|---|
| ENSG00000139131 | YARS2 | 13,45 | 13,11 | 11,39 | 5,54 | 58,8 |
| ENSG00000234345 | ELF2P1 | 2,57 | 2,11 | 1,81 | 1,06 | 58,8 |
| ENSG00000115368 | WDR75 | 22,27 | 21,25 | 16,04 | 9,19 | 58,7 |
| ENSG00000234056 | LINC00463 | 1,09 | 0,9 | 0,82 | 0,45 | 58,7 |
| ENSG00000166415 | WDR72 | 2,18 | 1,52 | 1,22 | 0,9 | 58,7 |
| ENSG00000233723 | LINC01122 | 1,21 | 1,15 | 0,59 | 0,5 | 58,7 |
| ENSG00000066583 | ISOC1 | 9,89 | 9,66 | 7,95 | 4,09 | 58,6 |
| ENSG00000196092 | PAX5 | 8,69 | 8,53 | 3,69 | 3,6 | 58,6 |
| ENSG00000032742 | IFT88 | 12,79 | 11,66 | 11,13 | 5,3 | 58,6 |
| ENSG00000261740 | BOLA2-SMG1P6 | 20,53 | 17,17 | 16,29 | 8,51 | 58,5 |
| ENSG00000125691 | RPL23 | 853,93 | 837,05 | 802,88 | 354,16 | 58,5 |
| ENSG00000120699 | EXOSC8 | 44,94 | 38,13 | 35,45 | 18,65 | 58,5 |
| ENSG00000084628 | NKAIN1 | 16,76 | 10,42 | 7,22 | 6,96 | 58,5 |
| ENSG00000100697 | DICER1 | 32,93 | 29,85 | 20,58 | 13,68 | 58,5 |
| ENSG00000135338 | LCA5 | 2,72 | 2,55 | 2,33 | 1,13 | 58,5 |
| ENSG00000111530 | CAND1 | 85,68 | 75,77 | 72,56 | 35,61 | 58,4 |
| ENSG00000198060 | MARCHF5 | 17,97 | 15,98 | 15,16 | 7,48 | 58,4 |
| ENSG00000269893 | SNHG8 | 54,82 | 43,32 | 41,08 | 22,82 | 58,4 |
| ENSG00000155755 | TMEM237 | 26,05 | 25,39 | 19,67 | 10,85 | 58,3 |
| ENSG00000186280 | KDM4D | 1,32 | 1,14 | 0,97 | 0,55 | 58,3 |
| ENSG00000271789 | AL080317,1 | 1,44 | 1,36 | 1,17 | 0,6 | 58,3 |
| ENSG00000179331 | RAB39A | 4,41 | 3,64 | 3,14 | 1,84 | 58,3 |
| ENSG00000066651 | TRMT11 | 13,95 | 12 | 11,18 | 5,83 | 58,2 |
| ENSG00000165704 | HPRT1 | 27,92 | 22,77 | 21,04 | 11,67 | 58,2 |
| ENSG00000163607 | GTPBP8 | 10,03 | 9,23 | 9,17 | 4,2 | 58,1 |
| ENSG00000149554 | CHEK1 | 46,51 | 44,59 | 39,7 | 19,48 | 58,1 |
| ENSG00000277936 | MRPL45 | 7,09 | 6,72 | 5,59 | 2,97 | 58,1 |
| ENSG00000008311 | AASS | 20,36 | 17,04 | 13,94 | 8,53 | 58,1 |
| ENSG00000235677 | NPM1P26 | 1,05 | 1,02 | 0,92 | 0,44 | 58,1 |
| ENSG00000112282 | MED23 | 11,08 | 10,56 | 9,14 | 4,65 | 58,0 |
| ENSG00000154721 | JAM2 | 14,55 | 12,14 | 12,11 | 6,11 | 58,0 |
| ENSG00000133678 | TMEM254 | 14 | 11,69 | 10,02 | 5,88 | 58,0 |
| ENSG00000117906 | RCN2 | 106,02 | 96,58 | 88,03 | 44,54 | 58,0 |
| ENSG00000006652 | IFRD1 | 28,03 | 22,8 | 21,38 | 11,79 | 57,9 |
| ENSG00000148688 | RPP30 | 36,25 | 34,42 | 28,71 | 15,25 | 57,9 |
| ENSG00000083457 | ITGAE | 21,15 | 20,08 | 18,82 | 8,9 | 57,9 |
| ENSG00000270638 | AL023806,1 | 1,33 | 1,19 | 0,91 | 0,56 | 57,9 |
| ENSG00000235369 | RPL36AP15 | 2,61 | 1,66 | 1,42 | 1,1 | 57,9 |
| ENSG00000162971 | TYW5 | 5,48 | 4,97 | 4,68 | 2,31 | 57,8 |
| ENSG00000181450 | ZNF678 | 11,86 | 11,16 | 10,83 | 5 | 57,8 |
| ENSG00000177054 | ZDHHC13 | 7,13 | 6,13 | 5,56 | 3,01 | 57,8 |
| ENSG00000128915 | ICE2 | 24,08 | 19,56 | 17,94 | 10,17 | 57,8 |
| ENSG00000188342 | GTF2F2 | 16,21 | 14,68 | 13,53 | 6,85 | 57,7 |
| ENSG00000153922 | CHD1 | 29,29 | 28,55 | 25,48 | 12,38 | 57,7 |
| ENSG00000132196 | HSD17B7 | 10,17 | 8,83 | 7,87 | 4,3 | 57,7 |
| ENSG00000152782 | PANK1 | 11,65 | 10,49 | 9,64 | 4,93 | 57,7 |
| ENSG00000185875 | THNSL1 | 3,45 | 3,15 | 2,42 | 1,46 | 57,7 |

| | | | | | | |
|---|---|---|---|---|---|---|
| ENSG00000225400 | RAB28P5 | 1,89 | 1,71 | 1,6 | 0,8 | 57,7 |
| ENSG00000115109 | EPB41L5 | 20,79 | 20,35 | 16,97 | 8,8 | 57,7 |
| ENSG00000267920 | SNX6P1 | 1,11 | 0,91 | 0,8 | 0,47 | 57,7 |
| ENSG00000271147 | ARMCX5-GPRAS | 10,01 | 9,02 | 7,75 | 4,24 | 57,6 |
| ENSG00000145087 | STXBP5L | 3,61 | 2,92 | 2,83 | 1,53 | 57,6 |
| ENSG00000168803 | ADAL | 7,38 | 6,2 | 5,28 | 3,13 | 57,6 |
| ENSG00000163689 | CFAP20DC | 3,89 | 3,62 | 2,59 | 1,65 | 57,6 |
| ENSG00000170522 | ELOVL6 | 24,17 | 23,62 | 19,61 | 10,26 | 57,6 |
| ENSG00000107789 | MINPP1 | 8,55 | 8,26 | 7,78 | 3,63 | 57,5 |
| ENSG00000172172 | MRPL13 | 21,49 | 19,37 | 18,64 | 9,13 | 57,5 |
| ENSG00000176165 | FOXG1 | 26,9 | 21,7 | 15,2 | 11,43 | 57,5 |
| ENSG00000108094 | CUL2 | 21,45 | 20,21 | 19,71 | 9,12 | 57,5 |
| ENSG00000129028 | THAP10 | 3,9 | 2,96 | 2,75 | 1,66 | 57,4 |
| ENSG00000126787 | DLGAP5 | 38,69 | 35,75 | 32,42 | 16,47 | 57,4 |
| ENSG00000270820 | AC016727,1 | 2,23 | 2,2 | 2,09 | 0,95 | 57,4 |
| ENSG00000235381 | AL596202,1 | 1,69 | 1,24 | 1,15 | 0,72 | 57,4 |
| ENSG00000013503 | POLR3B | 6,22 | 4,74 | 3,72 | 2,65 | 57,4 |
| ENSG00000047346 | FAM214A | 10,32 | 8,94 | 8,35 | 4,4 | 57,4 |
| ENSG00000197372 | ZNF675 | 13,1 | 12,83 | 10,32 | 5,59 | 57,3 |
| ENSG00000152683 | SLC30A6 | 10,73 | 10,71 | 7,86 | 4,58 | 57,3 |
| ENSG00000237513 | AC007384,1 | 1,92 | 1,79 | 1,57 | 0,82 | 57,3 |
| ENSG00000147488 | ST18 | 4,94 | 4,79 | 2,39 | 2,11 | 57,3 |
| ENSG00000154174 | TOMM70 | 23,36 | 23,27 | 20,5 | 9,98 | 57,3 |
| ENSG00000082146 | STRADB | 19,98 | 19,28 | 18,86 | 8,54 | 57,3 |
| ENSG00000167842 | MIS12 | 15,86 | 13,81 | 13,79 | 6,78 | 57,3 |
| ENSG00000169116 | PARM1 | 10,08 | 7,33 | 6,18 | 4,31 | 57,2 |
| ENSG00000163075 | CFAP221 | 3,32 | 3,18 | 2,45 | 1,42 | 57,2 |
| ENSG00000148835 | TAF5 | 5,12 | 4,29 | 3,77 | 2,19 | 57,2 |
| ENSG00000168246 | UBTD2 | 27,33 | 24,57 | 19,38 | 11,69 | 57,2 |
| ENSG00000165156 | ZHX1 | 11,5 | 11,21 | 9,83 | 4,92 | 57,2 |
| ENSG00000117133 | RPF1 | 17,78 | 15,58 | 15,3 | 7,61 | 57,2 |
| ENSG00000171533 | MAP6 | 38,52 | 33,3 | 23,9 | 16,49 | 57,2 |
| ENSG00000138688 | KIAA1109 | 24,69 | 24,46 | 21,64 | 10,57 | 57,2 |
| ENSG00000163510 | CWC22 | 14,97 | 13,4 | 10,44 | 6,41 | 57,2 |
| ENSG00000157426 | AASDH | 5,09 | 4,97 | 4,54 | 2,18 | 57,2 |
| ENSG00000260804 | LINC01963 | 7,12 | 7,06 | 6,34 | 3,05 | 57,2 |
| ENSG00000187049 | TMEM216 | 6,58 | 6,29 | 5,63 | 2,82 | 57,1 |
| ENSG00000164040 | PGRMC2 | 20,21 | 17,38 | 12,25 | 8,67 | 57,1 |
| ENSG00000129990 | SYT5 | 2,68 | 1,97 | 1,24 | 1,15 | 57,1 |
| ENSG00000167554 | ZNF610 | 7,01 | 6,02 | 4,85 | 3,01 | 57,1 |
| ENSG00000133739 | LRRCC1 | 10,05 | 9,67 | 7,64 | 4,32 | 57,0 |
| ENSG00000185495 | AC138393,1 | 2,53 | 2,15 | 1,91 | 1,09 | 56,9 |
| ENSG00000163682 | RPL9 | 430,95 | 421,26 | 382,56 | 185,78 | 56,9 |
| ENSG00000121481 | RNF2 | 23,19 | 20,48 | 20,45 | 10,01 | 56,8 |
| ENSG00000142892 | PIGK | 10,4 | 10,2 | 9,15 | 4,49 | 56,8 |
| ENSG00000010270 | STARD3NL | 41,2 | 36,85 | 34,92 | 17,79 | 56,8 |
| ENSG00000143162 | CREG1 | 6,6 | 6,55 | 5,33 | 2,85 | 56,8 |

| | | | | | | |
|---|---|---|---|---|---|---|
| ENSG00000101166 | PRELID3B | 35,13 | 34,29 | 33,41 | 15,17 | 56,8 |
| ENSG00000113161 | HMGCR | 98,72 | 87,31 | 81,6 | 42,65 | 56,8 |
| ENSG00000259456 | ADNP-AS1 | 2,06 | 1,92 | 1,71 | 0,89 | 56,8 |
| ENSG00000272008 | AL139274,2 | 1,18 | 1,01 | 0,85 | 0,51 | 56,8 |
| ENSG00000164211 | STARD4 | 39,02 | 35,04 | 27,87 | 16,9 | 56,7 |
| ENSG00000108666 | C17orf75 | 20,94 | 19,58 | 16,03 | 9,07 | 56,7 |
| ENSG00000134255 | CEPT1 | 9,61 | 8,8 | 8,17 | 4,17 | 56,6 |
| ENSG00000176208 | ATAD5 | 10,83 | 9,96 | 9,17 | 4,7 | 56,6 |
| ENSG00000163655 | GMPS | 55,65 | 55,54 | 49,62 | 24,17 | 56,6 |
| ENSG00000152193 | OBI1 | 14,73 | 14,47 | 12,51 | 6,4 | 56,6 |
| ENSG00000242474 | AC093627,5 | 1,38 | 1,27 | 0,98 | 0,6 | 56,5 |
| ENSG00000288472 | AC093627,29 | 1,38 | 1,27 | 0,98 | 0,6 | 56,5 |
| ENSG00000172476 | RAB40A | 2,69 | 2,21 | 1,64 | 1,17 | 56,5 |
| ENSG00000273447 | AC004067,1 | 2,23 | 2,17 | 1,33 | 0,97 | 56,5 |
| ENSG00000184575 | XPOT | 27 | 26,01 | 25,95 | 11,75 | 56,5 |
| ENSG00000228335 | AC073063,1 | 1,24 | 0,99 | 0,55 | 0,54 | 56,5 |
| ENSG00000277463 | AC080038,2 | 1,01 | 0,87 | 0,54 | 0,44 | 56,4 |
| ENSG00000198185 | ZNF334 | 7,43 | 6,55 | 5,69 | 3,24 | 56,4 |
| ENSG00000255517 | AP002748,4 | 2,43 | 2,36 | 1,8 | 1,06 | 56,4 |
| ENSG00000174574 | AKIRIN1 | 47,65 | 46,71 | 43,93 | 20,79 | 56,4 |
| ENSG00000184515 | BEX5 | 4,19 | 3,22 | 2,21 | 1,83 | 56,3 |
| ENSG00000006740 | ARHGAP44 | 2,77 | 2,02 | 1,48 | 1,21 | 56,3 |
| ENSG00000177888 | ZBTB41 | 5,47 | 5,31 | 4,82 | 2,39 | 56,3 |
| ENSG00000272599 | AC016394,1 | 4,21 | 3,93 | 3,67 | 1,84 | 56,3 |
| ENSG00000011201 | ANOS1 | 3,59 | 2,71 | 2,39 | 1,57 | 56,3 |
| ENSG00000180817 | PPA1 | 59,15 | 51,54 | 49,67 | 25,88 | 56,2 |
| ENSG00000172167 | MTBP | 7,63 | 6,98 | 6,24 | 3,34 | 56,2 |
| ENSG00000233871 | DLG5-AS1 | 1,05 | 0,87 | 0,54 | 0,46 | 56,2 |
| ENSG00000176542 | USF3 | 5,91 | 5,71 | 4,75 | 2,59 | 56,2 |
| ENSG00000120533 | ENY2 | 75,64 | 65,13 | 58,82 | 33,15 | 56,2 |
| ENSG00000138772 | ANXA3 | 13,21 | 9,97 | 8,04 | 5,79 | 56,2 |
| ENSG00000166432 | ZMAT1 | 4,54 | 4,09 | 3,08 | 1,99 | 56,2 |
| ENSG00000118193 | KIF14 | 8,94 | 8,28 | 7,07 | 3,92 | 56,2 |
| ENSG00000203667 | COX20 | 26,51 | 24,99 | 22,13 | 11,66 | 56,0 |
| ENSG00000099204 | ABLIM1 | 21,37 | 15,55 | 12,6 | 9,4 | 56,0 |
| ENSG00000146263 | MMS22L | 17,05 | 13,62 | 12,11 | 7,5 | 56,0 |
| ENSG00000232864 | NUCKS1P1 | 2,25 | 1,83 | 1,52 | 0,99 | 56,0 |
| ENSG00000198040 | ZNF84 | 25,16 | 24,53 | 20,44 | 11,08 | 56,0 |
| ENSG00000179902 | C1orf194 | 2,27 | 1,82 | 1,64 | 1 | 55,9 |
| ENSG00000124613 | ZNF391 | 4,63 | 3,68 | 3,01 | 2,04 | 55,9 |
| ENSG00000170837 | GPR27 | 2,61 | 2,32 | 1,56 | 1,15 | 55,9 |
| ENSG00000171497 | PPID | 28,12 | 23,86 | 20,98 | 12,4 | 55,9 |
| ENSG00000163918 | RFC4 | 43,04 | 41,35 | 38,22 | 18,98 | 55,9 |
| ENSG00000164796 | CSMD3 | 1,02 | 0,99 | 0,47 | 0,45 | 55,9 |
| ENSG00000163539 | CLASP2 | 36,05 | 30,51 | 24,79 | 15,91 | 55,9 |
| ENSG00000143727 | ACP1 | 78 | 74,77 | 68,92 | 34,48 | 55,8 |
| ENSG00000134900 | TPP2 | 29,43 | 27 | 23,84 | 13,02 | 55,8 |

| | | | | | | |
|---|---|---|---|---|---|---|
| ENSG00000184007 | PTP4A2 | 113,63 | 110,94 | 101,87 | 50,29 | 55,7 |
| ENSG00000287978 | AC245407,2 | 3,93 | 2,98 | 2,62 | 1,74 | 55,7 |
| ENSG00000248866 | USP46-DT | 2,37 | 1,96 | 1,81 | 1,05 | 55,7 |
| ENSG00000096093 | EFHC1 | 28,81 | 27,54 | 21,56 | 12,77 | 55,7 |
| ENSG00000268129 | AC026304,1 | 1,94 | 1,61 | 1,1 | 0,86 | 55,7 |
| ENSG00000149054 | ZNF215 | 3,88 | 2,82 | 2,75 | 1,72 | 55,7 |
| ENSG00000156239 | N6AMT1 | 3,97 | 3,57 | 3,12 | 1,76 | 55,7 |
| ENSG00000149948 | HMGA2 | 107,15 | 95,3 | 64,47 | 47,54 | 55,6 |
| ENSG00000133997 | MED6 | 21,09 | 20,97 | 18,44 | 9,36 | 55,6 |
| ENSG00000120675 | DNAJC15 | 10,7 | 10,54 | 8,56 | 4,75 | 55,6 |
| ENSG00000136522 | MRPL47 | 36,92 | 34,06 | 31,88 | 16,39 | 55,6 |
| ENSG00000138083 | SIX3 | 15,33 | 12,69 | 7,63 | 6,81 | 55,6 |
| ENSG00000155868 | MED7 | 8,59 | 7,98 | 7,69 | 3,82 | 55,5 |
| ENSG00000277564 | RBFOX2 | 9,82 | 9,64 | 7,08 | 4,37 | 55,5 |
| ENSG00000141441 | GAREM1 | 5,12 | 4,76 | 4,34 | 2,28 | 55,5 |
| ENSG00000109805 | NCAPG | 36,71 | 35,71 | 29,63 | 16,35 | 55,5 |
| ENSG00000141431 | ASXL3 | 15,19 | 13,44 | 10,7 | 6,77 | 55,4 |
| ENSG00000112304 | ACOT13 | 10,5 | 10,21 | 8,06 | 4,68 | 55,4 |
| ENSG00000182141 | ZNF708 | 16,64 | 15,25 | 13,97 | 7,42 | 55,4 |
| ENSG00000166004 | CEP295 | 13,05 | 12,64 | 11,26 | 5,82 | 55,4 |
| ENSG00000155636 | RBM45 | 7,06 | 6,94 | 5,99 | 3,15 | 55,4 |
| ENSG00000107362 | ABHD17B | 10,39 | 9,9 | 9,07 | 4,64 | 55,3 |
| ENSG00000102053 | ZC3H12B | 2,55 | 2,5 | 1,9 | 1,14 | 55,3 |
| ENSG00000144559 | TAMM41 | 8,03 | 6,59 | 5,93 | 3,59 | 55,3 |
| ENSG00000083544 | TDRD3 | 14,98 | 14,86 | 12,89 | 6,7 | 55,3 |
| ENSG00000178852 | EFCAB13 | 2,66 | 2,47 | 1,97 | 1,19 | 55,3 |
| ENSG00000267751 | AC009005,1 | 17,77 | 13,81 | 12,76 | 7,95 | 55,3 |
| ENSG00000109674 | NEIL3 | 7,71 | 7,67 | 5,86 | 3,45 | 55,3 |
| ENSG00000249456 | AL731577,2 | 8,67 | 8,48 | 6,64 | 3,88 | 55,2 |
| ENSG00000163412 | EIF4E3 | 9,25 | 8,05 | 7,58 | 4,14 | 55,2 |
| ENSG00000188811 | NHLRC3 | 8,31 | 8,09 | 7,8 | 3,72 | 55,2 |
| ENSG00000274315 | AC009318,3 | 1,72 | 1,61 | 1,5 | 0,77 | 55,2 |
| ENSG00000083097 | DOP1A | 10,22 | 9,87 | 8,45 | 4,58 | 55,2 |
| ENSG00000146143 | PRIM2 | 16,51 | 16,05 | 15,28 | 7,4 | 55,2 |
| ENSG00000208308 | SNORA40B | 13,43 | 10,14 | 7,84 | 6,03 | 55,1 |
| ENSG00000146802 | TMEM168 | 8,53 | 8,14 | 7,19 | 3,83 | 55,1 |
| ENSG00000119326 | CTNNAL1 | 24,45 | 24,16 | 21,23 | 10,98 | 55,1 |
| ENSG00000229692 | SOS1-IT1 | 3,45 | 3,33 | 2,82 | 1,55 | 55,1 |
| ENSG00000164074 | ABHD18 | 4,38 | 3,92 | 3,37 | 1,97 | 55,0 |
| ENSG00000145439 | CBR4 | 13,14 | 11,44 | 10 | 5,91 | 55,0 |
| ENSG00000114742 | WDR48 | 24,1 | 21,26 | 19,98 | 10,84 | 55,0 |
| ENSG00000143553 | SNAPIN | 25,32 | 24,12 | 24,04 | 11,4 | 55,0 |
| ENSG00000074266 | EED | 13,9 | 12,7 | 11,33 | 6,26 | 55,0 |
| ENSG00000164305 | CASP3 | 31,46 | 29,44 | 24,54 | 14,17 | 55,0 |
| ENSG00000118690 | ARMC2 | 6,17 | 6,07 | 5,45 | 2,78 | 54,9 |
| ENSG00000198464 | ZNF480 | 6,7 | 6,67 | 5,8 | 3,02 | 54,9 |
| ENSG00000147905 | ZCCHC7 | 15,07 | 15,02 | 10,77 | 6,8 | 54,9 |

| | | | | | | |
|---|---|---|---|---|---|---|
| ENSG00000174963 | ZIC4 | 59,55 | 50,38 | 36,17 | 26,9 | 54,8 |
| ENSG00000269984 | AC078795,1 | 1,46 | 1,3 | 1,19 | 0,66 | 54,8 |
| ENSG00000134759 | ELP2 | 31,92 | 29,95 | 26,83 | 14,43 | 54,8 |
| ENSG00000198039 | ZNF273 | 8,99 | 7,68 | 7,05 | 4,07 | 54,7 |
| ENSG00000064933 | PMS1 | 15,77 | 13,69 | 11,89 | 7,14 | 54,7 |
| ENSG00000185246 | PRPF39 | 17,07 | 16,8 | 15,48 | 7,73 | 54,7 |
| ENSG00000154548 | SRSF12 | 8,59 | 8,07 | 7,93 | 3,89 | 54,7 |
| ENSG00000240891 | PLCXD2 | 1,39 | 1,27 | 1,13 | 0,63 | 54,7 |
| ENSG00000119705 | SLIRP | 148,91 | 143,09 | 128,18 | 67,52 | 54,7 |
| ENSG00000134744 | TUT4 | 39,03 | 37,68 | 32,61 | 17,7 | 54,7 |
| ENSG00000122779 | TRIM24 | 49,62 | 44,31 | 36,09 | 22,51 | 54,6 |
| ENSG00000125870 | SNRPB2 | 57,06 | 55,97 | 53,53 | 25,91 | 54,6 |
| ENSG00000099282 | TSPAN15 | 8,01 | 7,89 | 4,28 | 3,64 | 54,6 |
| ENSG00000057608 | GDI2 | 176,47 | 171,73 | 150,02 | 80,27 | 54,5 |
| ENSG00000164169 | PRMT9 | 4,66 | 3,66 | 3,24 | 2,12 | 54,5 |
| ENSG00000124374 | PAIP2B | 2,11 | 2,05 | 1,67 | 0,96 | 54,5 |
| ENSG00000123472 | ATPAF1 | 28,39 | 21,94 | 19,65 | 12,92 | 54,5 |
| ENSG00000111247 | RAD51AP1 | 25,68 | 25,11 | 20,18 | 11,69 | 54,5 |
| ENSG00000086061 | DNAJA1 | 141,62 | 123,39 | 95,61 | 64,47 | 54,5 |
| ENSG00000144048 | DUSP11 | 10,5 | 10,49 | 9,01 | 4,78 | 54,5 |
| ENSG00000116459 | ATP5PB | 125,19 | 123,67 | 111,61 | 57 | 54,5 |
| ENSG00000166845 | C18orf54 | 12,78 | 12,22 | 8,46 | 5,82 | 54,5 |
| ENSG00000136891 | TEX10 | 20,21 | 18,68 | 16,23 | 9,21 | 54,4 |
| ENSG00000270823 | AC007938,2 | 2,5 | 2,44 | 1,8 | 1,14 | 54,4 |
| ENSG00000198860 | TSEN15 | 20,7 | 20,25 | 18,06 | 9,44 | 54,4 |
| ENSG00000180776 | ZDHHC20 | 22,58 | 21,66 | 20,49 | 10,3 | 54,4 |
| ENSG00000131732 | ZCCHC9 | 15,59 | 13,71 | 11,68 | 7,12 | 54,3 |
| ENSG00000145416 | MARCHF1 | 3,7 | 3,43 | 2,76 | 1,69 | 54,3 |
| ENSG00000150768 | DLAT | 17,3 | 17,05 | 13,83 | 7,91 | 54,3 |
| ENSG00000102738 | MRPS31 | 17,19 | 15,37 | 13,54 | 7,86 | 54,3 |
| ENSG00000138778 | CENPE | 19,9 | 15,21 | 11,8 | 9,1 | 54,3 |
| ENSG00000127561 | SYNGR3 | 1,29 | 0,85 | 0,62 | 0,59 | 54,3 |
| ENSG00000230409 | TCEA1P2 | 2,23 | 2,04 | 1,48 | 1,02 | 54,3 |
| ENSG00000137714 | FDX1 | 5,05 | 5 | 4,64 | 2,31 | 54,3 |
| ENSG00000146386 | ABRACL | 36,9 | 33,48 | 31,29 | 16,89 | 54,2 |
| ENSG00000138160 | KIF11 | 37,75 | 37,28 | 35 | 17,28 | 54,2 |
| ENSG00000078018 | MAP2 | 78,3 | 62,35 | 53,85 | 35,85 | 54,2 |
| ENSG00000228716 | DHFR | 56,24 | 53,03 | 45,54 | 25,75 | 54,2 |
| ENSG00000189190 | ZNF600 | 1,07 | 0,9 | 0,7 | 0,49 | 54,2 |
| ENSG00000226383 | LINC01876 | 4,06 | 4,04 | 3,26 | 1,86 | 54,2 |
| ENSG00000106804 | C5 | 4,78 | 4,41 | 3,36 | 2,19 | 54,2 |
| ENSG00000119013 | NDUFB3 | 85,55 | 83,95 | 79,24 | 39,2 | 54,2 |
| ENSG00000117748 | RPA2 | 30,84 | 27,11 | 24,92 | 14,14 | 54,2 |
| ENSG00000102678 | FGF9 | 2,66 | 1,9 | 1,47 | 1,22 | 54,1 |
| ENSG00000113569 | NUP155 | 23,48 | 22,25 | 20,24 | 10,77 | 54,1 |
| ENSG00000177565 | TBL1XR1 | 57,58 | 55,2 | 49,69 | 26,42 | 54,1 |
| ENSG00000115966 | ATF2 | 28,21 | 27,27 | 24,32 | 12,95 | 54,1 |

| | | | | | | |
|---|---|---|---|---|---|---|
| ENSG00000186777 | ZNF732 | 5,38 | 4,72 | 4,44 | 2,47 | 54,1 |
| ENSG00000165349 | SLC7A3 | 5,77 | 3,91 | 3,41 | 2,65 | 54,1 |
| ENSG00000285427 | SOD2-OT1 | 1,85 | 1,3 | 1,07 | 0,85 | 54,1 |
| ENSG00000082515 | MRPL22 | 35,48 | 34,28 | 33,39 | 16,31 | 54,0 |
| ENSG00000214765 | SEPTIN7P2 | 8 | 7,8 | 7,69 | 3,68 | 54,0 |
| ENSG00000062194 | GPBP1 | 48 | 46,87 | 40,89 | 22,1 | 54,0 |
| ENSG00000100387 | RBX1 | 31,44 | 31,04 | 29,99 | 14,48 | 53,9 |
| ENSG00000111875 | ASF1A | 17,8 | 16,75 | 16,61 | 8,2 | 53,9 |
| ENSG00000139684 | ESD | 73,22 | 70,53 | 69,11 | 33,8 | 53,8 |
| ENSG00000106588 | PSMA2 | 63,22 | 58,26 | 53,09 | 29,22 | 53,8 |
| ENSG00000129055 | ANAPC13 | 38,07 | 35,24 | 32,13 | 17,62 | 53,7 |
| ENSG00000060749 | QSER1 | 26,31 | 26,01 | 22,47 | 12,18 | 53,7 |
| ENSG00000138385 | SSB | 67,92 | 64,38 | 50,69 | 31,47 | 53,7 |
| ENSG00000066032 | CTNNA2 | 9,03 | 8,37 | 7,24 | 4,19 | 53,6 |
| ENSG00000140386 | SCAPER | 9,61 | 9,5 | 7,8 | 4,46 | 53,6 |
| ENSG00000245958 | AC093752,1 | 12,17 | 11,75 | 10,79 | 5,65 | 53,6 |
| ENSG00000116120 | FARSB | 10,02 | 8,39 | 7,78 | 4,66 | 53,5 |
| ENSG00000155903 | RASA2 | 8,47 | 7,86 | 5,64 | 3,94 | 53,5 |
| ENSG00000120868 | APAF1 | 13,09 | 11,87 | 10,71 | 6,09 | 53,5 |
| ENSG00000130227 | XPO7 | 30,65 | 28,21 | 27,41 | 14,26 | 53,5 |
| ENSG00000112306 | RPS12 | 590,39 | 557,28 | 539,11 | 274,71 | 53,5 |
| ENSG00000196268 | ZNF493 | 13,49 | 10,88 | 10,32 | 6,28 | 53,4 |
| ENSG00000155016 | CYP2U1 | 3,2 | 2,91 | 2,61 | 1,49 | 53,4 |
| ENSG00000104490 | NCALD | 35,07 | 32,42 | 27,59 | 16,33 | 53,4 |
| ENSG00000149548 | CCDC15 | 3,5 | 3,23 | 2,51 | 1,63 | 53,4 |
| ENSG00000080298 | RFX3 | 15,3 | 14,7 | 13,69 | 7,13 | 53,4 |
| ENSG00000110429 | FBXO3 | 13,38 | 12,73 | 11,04 | 6,24 | 53,4 |
| ENSG00000110987 | BCL7A | 13,85 | 11,26 | 9,58 | 6,46 | 53,4 |
| ENSG00000135972 | MRPS9 | 20,55 | 20,5 | 16,78 | 9,59 | 53,3 |
| ENSG00000213390 | ARHGAP19 | 12,55 | 10,16 | 9,79 | 5,86 | 53,3 |
| ENSG00000153250 | RBMS1 | 25,35 | 25,02 | 21,13 | 11,84 | 53,3 |
| ENSG00000079257 | LXN | 4,11 | 3,61 | 3,58 | 1,92 | 53,3 |
| ENSG00000107951 | MTPAP | 10,85 | 9,84 | 9,61 | 5,07 | 53,3 |
| ENSG00000220785 | MTMR9LP | 2,14 | 1,86 | 1,13 | 1 | 53,3 |
| ENSG00000115947 | ORC4 | 20,43 | 19,71 | 17,86 | 9,55 | 53,3 |
| ENSG00000006757 | PNPLA4 | 5,25 | 5,01 | 3,95 | 2,46 | 53,1 |
| ENSG00000104442 | ARMC1 | 19,1 | 17,08 | 16,14 | 8,95 | 53,1 |
| ENSG00000240497 | AC092919,1 | 1,13 | 1,06 | 0,81 | 0,53 | 53,1 |
| ENSG00000117054 | ACADM | 35,71 | 33,93 | 28,81 | 16,76 | 53,1 |
| ENSG00000072041 | SLC6A15 | 4,92 | 3,9 | 3,21 | 2,31 | 53,0 |
| ENSG00000164944 | VIRMA | 24,76 | 22,43 | 22,15 | 11,65 | 52,9 |
| ENSG00000047230 | CTPS2 | 26,97 | 26,32 | 23,27 | 12,69 | 52,9 |
| ENSG00000253719 | ATXN7L3B | 22,1 | 21,29 | 18,3 | 10,4 | 52,9 |
| ENSG00000100485 | SOS2 | 15,25 | 14,82 | 14,09 | 7,18 | 52,9 |
| ENSG00000166483 | WEE1 | 43,66 | 42,35 | 40,89 | 20,56 | 52,9 |
| ENSG00000101557 | USP14 | 35,68 | 33,85 | 30,62 | 16,81 | 52,9 |
| ENSG00000108506 | INTS2 | 7,28 | 7,22 | 5,36 | 3,43 | 52,9 |

| | | | | | | |
|---|---|---|---|---|---|---|
| ENSG00000141219 | C17orf80 | 15,85 | 15,16 | 13,53 | 7,47 | 52,9 |
| ENSG00000214654 | B3GNT10 | 6,98 | 6,92 | 4,34 | 3,29 | 52,9 |
| ENSG00000104231 | ZFAND1 | 17,25 | 16,09 | 12,25 | 8,14 | 52,8 |
| ENSG00000143207 | COP1 | 25,23 | 23,12 | 22,11 | 11,91 | 52,8 |
| ENSG00000197299 | BLM | 16,29 | 14,28 | 13,49 | 7,69 | 52,8 |
| ENSG00000067248 | DHX29 | 12,92 | 12,58 | 11,51 | 6,1 | 52,8 |
| ENSG00000119927 | GPAM | 5,59 | 5,37 | 4,77 | 2,64 | 52,8 |
| ENSG00000109911 | ELP4 | 10,5 | 9,41 | 8,33 | 4,96 | 52,8 |
| ENSG00000109606 | DHX15 | 82,99 | 80,57 | 72,49 | 39,23 | 52,7 |
| ENSG00000001617 | SEMA3F | 4,95 | 3,56 | 2,55 | 2,34 | 52,7 |
| ENSG00000235299 | MRPL53P1 | 1,29 | 0,95 | 0,62 | 0,61 | 52,7 |
| ENSG00000100522 | GNPNAT1 | 13,49 | 12,15 | 10,4 | 6,38 | 52,7 |
| ENSG00000145868 | FBXO38 | 21,83 | 21,17 | 17,16 | 10,34 | 52,6 |
| ENSG00000138018 | SELENOI | 8,55 | 7,43 | 6,64 | 4,05 | 52,6 |
| ENSG00000168813 | ZNF507 | 10,26 | 9,95 | 8,34 | 4,86 | 52,6 |
| ENSG00000213462 | ERV3-1 | 16,16 | 13,27 | 13,05 | 7,66 | 52,6 |
| ENSG00000124486 | USP9X | 45,44 | 44,06 | 36,04 | 21,55 | 52,6 |
| ENSG00000163322 | ABRAXAS1 | 8,96 | 8,92 | 8,12 | 4,25 | 52,6 |
| ENSG00000172296 | SPTLC3 | 4,49 | 3,59 | 2,58 | 2,13 | 52,6 |
| ENSG00000161813 | LARP4 | 25,39 | 22,76 | 20,65 | 12,05 | 52,5 |
| ENSG00000111581 | NUP107 | 46,55 | 40,38 | 37,21 | 22,1 | 52,5 |
| ENSG00000145996 | CDKAL1 | 14,78 | 12,99 | 11,72 | 7,02 | 52,5 |
| ENSG00000147654 | EBAG9 | 12,33 | 11,82 | 10,62 | 5,86 | 52,5 |
| ENSG00000006459 | KDM7A | 3,45 | 3,34 | 3,15 | 1,64 | 52,5 |
| ENSG00000113522 | RAD50 | 15,13 | 13,09 | 10,76 | 7,2 | 52,4 |
| ENSG00000231107 | LINC01508 | 2,92 | 1,84 | 1,65 | 1,39 | 52,4 |
| ENSG00000138430 | OLA1 | 70,81 | 69,41 | 62,87 | 33,71 | 52,4 |
| ENSG00000259209 | AC004943,1 | 2,98 | 2,81 | 2,05 | 1,42 | 52,3 |
| ENSG00000105143 | SLC1A6 | 2,77 | 2,04 | 1,77 | 1,32 | 52,3 |
| ENSG00000129003 | VPS13C | 15,77 | 14,19 | 13,42 | 7,52 | 52,3 |
| ENSG00000243422 | RPL23AP49 | 1,09 | 0,91 | 0,69 | 0,52 | 52,3 |
| ENSG00000131469 | RPL27 | 601,63 | 596,5 | 595,81 | 287,06 | 52,3 |
| ENSG00000170959 | DCDC1 | 1,36 | 1,23 | 0,93 | 0,65 | 52,2 |
| ENSG00000137992 | DBT | 4,31 | 3,86 | 3,42 | 2,06 | 52,2 |
| ENSG00000178568 | ERBB4 | 7,07 | 6,16 | 5,51 | 3,38 | 52,2 |
| ENSG00000266839 | AC008088,1 | 1,15 | 1,1 | 0,7 | 0,55 | 52,2 |
| ENSG00000168234 | TTC39C | 7,92 | 6,8 | 4,81 | 3,79 | 52,1 |
| ENSG00000228506 | AL513550,1 | 2,57 | 2,54 | 2,51 | 1,23 | 52,1 |
| ENSG00000182481 | KPNA2 | 185,09 | 181,41 | 169,67 | 88,65 | 52,1 |
| ENSG00000122435 | TRMT13 | 7,82 | 7,31 | 6,62 | 3,75 | 52,0 |
| ENSG00000169288 | MRPL1 | 21,33 | 19,29 | 15,92 | 10,23 | 52,0 |
| ENSG00000224046 | AC005076,1 | 1,23 | 1,08 | 0,67 | 0,59 | 52,0 |
| ENSG00000198729 | PPP1R14C | 2,46 | 2,13 | 1,28 | 1,18 | 52,0 |
| ENSG00000174606 | ANGEL2 | 16,05 | 14,7 | 13,04 | 7,7 | 52,0 |
| ENSG00000116906 | GNPAT | 35,37 | 34,35 | 32,58 | 16,97 | 52,0 |
| ENSG00000096060 | FKBP5 | 17,15 | 16,56 | 11,77 | 8,23 | 52,0 |
| ENSG00000038274 | MAT2B | 27,65 | 27,47 | 23,11 | 13,27 | 52,0 |

| | | | | | | |
|---|---|---|---|---|---|---|
| ENSG00000140463 | BBS4 | 16,69 | 16,67 | 14,48 | 8,01 | 52,0 |
| ENSG00000168522 | FNTA | 28,09 | 26,9 | 25,98 | 13,49 | 52,0 |
| ENSG00000166012 | TAF1D | 67,08 | 65,59 | 56,76 | 32,22 | 52,0 |
| ENSG00000138078 | PREPL | 23,92 | 22,52 | 21,92 | 11,49 | 52,0 |
| ENSG00000213190 | MLLT11 | 76,85 | 71,73 | 64,77 | 36,92 | 52,0 |
| ENSG00000196284 | SUPT3H | 11,74 | 9,93 | 8,58 | 5,65 | 51,9 |
| ENSG00000283085 | TPBG | 5,07 | 3,74 | 3,01 | 2,44 | 51,9 |
| ENSG00000196865 | NHLRC2 | 12,55 | 9,48 | 8,93 | 6,04 | 51,9 |
| ENSG00000247137 | AP000873,2 | 5,13 | 4,35 | 3,39 | 2,47 | 51,9 |
| ENSG00000046604 | DSG2 | 13,27 | 10,34 | 6,82 | 6,39 | 51,8 |
| ENSG00000117620 | SLC35A3 | 6,25 | 5,66 | 4,55 | 3,01 | 51,8 |
| ENSG00000149311 | ATM | 20,42 | 18,09 | 15,51 | 9,84 | 51,8 |
| ENSG00000211456 | SACM1L | 18,48 | 15,79 | 14,55 | 8,91 | 51,8 |
| ENSG00000106066 | CPVL | 6,24 | 4,97 | 4,57 | 3,01 | 51,8 |
| ENSG00000097096 | SYDE2 | 2,84 | 2,23 | 1,72 | 1,37 | 51,8 |
| ENSG00000152413 | HOMER1 | 13,82 | 12,67 | 10,35 | 6,67 | 51,7 |
| ENSG00000084453 | SLCO1A2 | 4,1 | 3,15 | 2,44 | 1,98 | 51,7 |
| ENSG00000116679 | IVNS1ABP | 49,62 | 48,4 | 36,23 | 23,98 | 51,7 |
| ENSG00000173436 | MICOS10 | 62,74 | 61,65 | 55,18 | 30,33 | 51,7 |
| ENSG00000121274 | TENT4B | 3,93 | 3,49 | 3,42 | 1,9 | 51,7 |
| ENSG00000164347 | GFM2 | 17,91 | 15,37 | 14,26 | 8,66 | 51,6 |
| ENSG00000053770 | AP5M1 | 16,44 | 15 | 13,52 | 7,95 | 51,6 |
| ENSG00000028203 | VEZT | 38,71 | 37,7 | 33,75 | 18,73 | 51,6 |
| ENSG00000028116 | VRK2 | 13,59 | 12,15 | 8,86 | 6,58 | 51,6 |
| ENSG00000187134 | AKR1C1 | 4,17 | 3,22 | 2,94 | 2,02 | 51,6 |
| ENSG00000125851 | PCSK2 | 3,24 | 2,66 | 2,61 | 1,57 | 51,5 |
| ENSG00000138095 | LRPPRC | 44,81 | 35,19 | 34,55 | 21,72 | 51,5 |
| ENSG00000171365 | CLCN5 | 4,6 | 3,95 | 3,19 | 2,23 | 51,5 |
| ENSG00000105948 | TTC26 | 6,33 | 5,2 | 5,09 | 3,07 | 51,5 |
| ENSG00000146250 | PRSS35 | 1,34 | 1,15 | 0,84 | 0,65 | 51,5 |
| ENSG00000132970 | WASF3 | 12,28 | 11,78 | 10,85 | 5,96 | 51,5 |
| ENSG00000101752 | MIB1 | 22,83 | 19,61 | 15,36 | 11,11 | 51,3 |
| ENSG00000120156 | TEK | 2,73 | 2,23 | 1,5 | 1,33 | 51,3 |
| ENSG00000072571 | HMMR | 31,46 | 26,79 | 23,65 | 15,33 | 51,3 |
| ENSG00000246250 | AC087521,2 | 1,97 | 1,64 | 1,51 | 0,96 | 51,3 |
| ENSG00000115594 | IL1R1 | 3,3 | 2,79 | 2,27 | 1,61 | 51,2 |
| ENSG00000151490 | PTPRO | 4,96 | 4,1 | 2,83 | 2,42 | 51,2 |
| ENSG00000184785 | SMIM10 | 2,09 | 2,08 | 2,07 | 1,02 | 51,2 |
| ENSG00000068615 | REEP1 | 4,2 | 3,61 | 3,04 | 2,05 | 51,2 |
| ENSG00000180376 | CCDC66 | 22,16 | 19,27 | 17,31 | 10,82 | 51,2 |
| ENSG00000198087 | CD2AP | 7,88 | 6,49 | 6,2 | 3,85 | 51,1 |
| ENSG00000104147 | OIP5 | 10,09 | 9,18 | 8,99 | 4,93 | 51,1 |
| ENSG00000286918 | AL035411,3 | 6,16 | 6,14 | 5,43 | 3,01 | 51,1 |
| ENSG00000243317 | STMP1 | 32,21 | 31,96 | 31,07 | 15,74 | 51,1 |
| ENSG00000178538 | CA8 | 1,33 | 0,86 | 0,78 | 0,65 | 51,1 |
| ENSG00000101938 | CHRDL1 | 9,35 | 9,15 | 5,98 | 4,57 | 51,1 |
| ENSG00000233184 | AC093157,1 | 5,79 | 5,17 | 4,53 | 2,83 | 51,1 |

| | | | | | | |
|---|---|---|---|---|---|---|
| ENSG00000096401 | CDC5L | 20,23 | 19,57 | 16,85 | 9,89 | 51,1 |
| ENSG00000262860 | LSM14A | 49,09 | 48,35 | 43,42 | 24 | 51,1 |
| ENSG00000111785 | RIC8B | 8,58 | 7,89 | 7,13 | 4,2 | 51,0 |
| ENSG00000182150 | ERCC6L2 | 13,4 | 12,94 | 11,44 | 6,56 | 51,0 |
| ENSG00000203880 | PCMTD2 | 12,35 | 12,07 | 11,6 | 6,05 | 51,0 |
| ENSG00000115233 | PSMD14 | 61,69 | 55,16 | 50,61 | 30,23 | 51,0 |
| ENSG00000138115 | CYP2C8 | 1,02 | 0,88 | 0,58 | 0,5 | 51,0 |
| ENSG00000122884 | P4HA1 | 17,58 | 17,18 | 16,83 | 8,62 | 51,0 |
| ENSG00000120265 | PCMT1 | 31,87 | 31,74 | 30,49 | 15,63 | 51,0 |
| ENSG00000134987 | WDR36 | 12,64 | 12,55 | 10,67 | 6,2 | 50,9 |
| ENSG00000141424 | SLC39A6 | 43,17 | 42,85 | 38,71 | 21,19 | 50,9 |
| ENSG00000198818 | SFT2D1 | 14,48 | 14,31 | 14,17 | 7,11 | 50,9 |
| ENSG00000258311 | AC009779,3 | 3,36 | 2,06 | 1,91 | 1,65 | 50,9 |
| ENSG00000128059 | PPAT | 13,45 | 10,71 | 9,03 | 6,61 | 50,9 |
| ENSG00000034533 | ASTE1 | 3,52 | 3,29 | 3,04 | 1,73 | 50,9 |
| ENSG00000143919 | CAMKMT | 4,17 | 4,14 | 3,44 | 2,05 | 50,8 |
| ENSG00000145425 | RPS3A | 769,54 | 737,11 | 707,3 | 378,33 | 50,8 |
| ENSG00000137513 | NARS2 | 14,03 | 13,11 | 10,43 | 6,9 | 50,8 |
| ENSG00000030066 | NUP160 | 27,48 | 23,33 | 21,61 | 13,52 | 50,8 |
| ENSG00000080345 | RIF1 | 24,59 | 23,17 | 21,44 | 12,1 | 50,8 |
| ENSG00000168288 | MMADHC | 45,66 | 43,9 | 42,61 | 22,47 | 50,8 |
| ENSG00000281556 | AC243960,18 | 2,54 | 2,19 | 1,56 | 1,25 | 50,8 |
| ENSG00000166037 | CEP57 | 28,69 | 27,98 | 24,16 | 14,12 | 50,8 |
| ENSG00000175175 | PPM1E | 7,31 | 6,85 | 5,69 | 3,6 | 50,8 |
| ENSG00000087053 | MTMR2 | 26,65 | 25,72 | 23,51 | 13,13 | 50,7 |
| ENSG00000107672 | NSMCE4A | 34,72 | 32,66 | 27,6 | 17,11 | 50,7 |
| ENSG00000242797 | GLYCTK-AS1 | 3 | 2,46 | 1,68 | 1,48 | 50,7 |
| ENSG00000044459 | CNTLN | 7,5 | 7,17 | 5,39 | 3,7 | 50,7 |
| ENSG00000172771 | EFCAB12 | 1,56 | 0,97 | 0,8 | 0,77 | 50,6 |
| ENSG00000288558 | DUS4L-BCAP29 | 1,74 | 1,64 | 1,62 | 0,86 | 50,6 |
| ENSG00000203666 | EFCAB2 | 9,77 | 9,02 | 6,2 | 4,83 | 50,6 |
| ENSG00000124789 | NUP153 | 24,81 | 23,82 | 20,73 | 12,27 | 50,5 |
| ENSG00000117155 | SSX2IP | 12,05 | 10,6 | 9,67 | 5,96 | 50,5 |
| ENSG00000198707 | CEP290 | 12,53 | 9,93 | 6,62 | 6,2 | 50,5 |
| ENSG00000229989 | MIR181A1HG | 7,15 | 6,85 | 6 | 3,54 | 50,5 |
| ENSG00000155561 | NUP205 | 31,08 | 26,91 | 25,36 | 15,39 | 50,5 |
| ENSG00000164329 | TENT2 | 17,97 | 17,31 | 15,13 | 8,9 | 50,5 |
| ENSG00000251595 | ABCA11P | 6,56 | 5,02 | 4,27 | 3,25 | 50,5 |
| ENSG00000204909 | SPINK9 | 3,31 | 2,22 | 1,68 | 1,64 | 50,5 |
| ENSG00000261251 | Z97055,2 | 1,19 | 1,11 | 1,01 | 0,59 | 50,4 |
| ENSG00000120162 | MOB3B | 7,36 | 6,82 | 5,56 | 3,65 | 50,4 |
| ENSG00000109184 | DCUN1D4 | 31,85 | 29,94 | 24,54 | 15,8 | 50,4 |
| ENSG00000148730 | EIF4EBP2 | 51,07 | 47,65 | 44,19 | 25,34 | 50,4 |
| ENSG00000185065 | AC000068,1 | 1,31 | 1,1 | 0,88 | 0,65 | 50,4 |
| ENSG00000205325 | AC005863,1 | 1,33 | 1,3 | 1,1 | 0,66 | 50,4 |
| ENSG00000168824 | NSG1 | 9 | 8,28 | 5,57 | 4,47 | 50,3 |
| ENSG00000162415 | ZSWIM5 | 3,26 | 2,37 | 1,81 | 1,62 | 50,3 |

| | | | | | | |
|---|---|---|---|---|---|---|
| ENSG00000109339 | MAPK10 | 49,02 | 47,42 | 42,6 | 24,37 | 50,3 |
| ENSG00000078668 | VDAC3 | 85,19 | 80,68 | 75,06 | 42,36 | 50,3 |
| ENSG00000174460 | ZCCHC12 | 5,61 | 5,26 | 4,3 | 2,79 | 50,3 |
| ENSG00000143493 | INTS7 | 11,36 | 11,33 | 9,75 | 5,65 | 50,3 |
| ENSG00000108375 | RNF43 | 2,01 | 1,73 | 1,61 | 1 | 50,2 |
| ENSG00000129810 | SGO1 | 14,41 | 12,89 | 12,05 | 7,17 | 50,2 |
| ENSG00000169914 | OTUD3 | 6,47 | 6 | 4,53 | 3,22 | 50,2 |
| ENSG00000188732 | FAM221A | 9,14 | 7,55 | 6,54 | 4,55 | 50,2 |
| ENSG00000276276 | ARL17B | 2,45 | 2,18 | 2,17 | 1,22 | 50,2 |
| ENSG00000143190 | POU2F1 | 12,35 | 10,36 | 8,29 | 6,15 | 50,2 |
| ENSG00000141380 | SS18 | 44,03 | 43,81 | 38,38 | 21,93 | 50,2 |
| ENSG00000114480 | GBE1 | 7,83 | 7,53 | 6,28 | 3,9 | 50,2 |
| ENSG00000070669 | ASNS | 27,54 | 21,3 | 15,74 | 13,72 | 50,2 |
| ENSG00000136527 | TRA2B | 128,74 | 126,06 | 110,67 | 64,17 | 50,2 |
| ENSG00000114098 | ARMC8 | 17,49 | 16,86 | 14,74 | 8,73 | 50,1 |
| ENSG00000188612 | SUMO2 | 310,17 | 305,17 | 288,09 | 154,88 | 50,1 |
| ENSG00000132964 | CDK8 | 13,91 | 13,14 | 10,57 | 6,95 | 50,0 |
| ENSG00000244041 | LINC01011 | 1,58 | 1,16 | 1,15 | 0,79 | 50,0 |
| ENSG00000272849 | AC084018,1 | 2,54 | 2,31 | 1,77 | 1,27 | 50,0 |
| ENSG00000272173 | U47924,2 | 2,62 | 2,12 | 1,67 | 1,31 | 50,0 |
| ENSG00000052795 | FNIP2 | 10,1 | 9,73 | 7,1 | 5,05 | 50,0 |
| ENSG00000164654 | MIOS | 16,61 | 15,26 | 14,77 | 8,31 | 50,0 |
| ENSG00000127920 | GNG11 | 13,65 | 13,21 | 12,22 | 6,83 | 50,0 |
| ENSG00000153201 | RANBP2 | 28,8 | 27,31 | 22,04 | 14,43 | 49,9 |
| ENSG00000158406 | H4C8 | 699,87 | 498,6 | 468,61 | 350,68 | 49,9 |
| ENSG00000196632 | WNK3 | 11,65 | 10,94 | 7,13 | 5,84 | 49,9 |
| ENSG00000104626 | ERI1 | 16,12 | 14,31 | 11,12 | 8,09 | 49,8 |
| ENSG00000158092 | NCK1 | 15,02 | 11,52 | 10,55 | 7,54 | 49,8 |
| ENSG00000181163 | NPM1 | 606,04 | 540,23 | 446,93 | 304,41 | 49,8 |
| ENSG00000109390 | NDUFC1 | 25,48 | 25,3 | 24,82 | 12,8 | 49,8 |
| ENSG00000241058 | NSUN6 | 7,92 | 7,55 | 6,37 | 3,98 | 49,7 |
| ENSG00000130935 | NOL11 | 31,22 | 29,89 | 24,57 | 15,69 | 49,7 |
| ENSG00000197147 | LRRC8B | 3,7 | 3,3 | 2,95 | 1,86 | 49,7 |
| ENSG00000106086 | PLEKHA8 | 19,41 | 14,81 | 13,3 | 9,76 | 49,7 |
| ENSG00000186448 | ZNF197 | 7,17 | 6,74 | 5,52 | 3,61 | 49,7 |
| ENSG00000281709 | ZNF197 | 7,17 | 6,74 | 5,52 | 3,61 | 49,7 |
| ENSG00000170892 | TSEN34 | 1,41 | 1,23 | 1,22 | 0,71 | 49,6 |
| ENSG00000040275 | SPDL1 | 20,89 | 20,12 | 16,06 | 10,52 | 49,6 |
| ENSG00000170681 | CAVIN4 | 2,6 | 2,33 | 1,83 | 1,31 | 49,6 |
| ENSG00000120705 | ETF1 | 50,61 | 46,35 | 43,99 | 25,51 | 49,6 |
| ENSG00000183439 | TRIM61 | 1,13 | 1,06 | 0,87 | 0,57 | 49,6 |
| ENSG00000082068 | WDR70 | 19,17 | 18,76 | 14 | 9,67 | 49,6 |
| ENSG00000175279 | CENPS | 16,49 | 13,41 | 13,33 | 8,32 | 49,5 |
| ENSG00000128833 | MYO5C | 3,27 | 2,83 | 1,93 | 1,65 | 49,5 |
| ENSG00000183814 | LIN9 | 6,54 | 6,06 | 5,33 | 3,3 | 49,5 |
| ENSG00000284052 | AC006460,2 | 1,09 | 0,67 | 0,61 | 0,55 | 49,5 |
| ENSG00000106688 | SLC1A1 | 1,07 | 0,88 | 0,7 | 0,54 | 49,5 |

| | | | | | | |
|---|---|---|---|---|---|---|
| ENSG00000145723 | GIN1 | 4,14 | 3,87 | 3,18 | 2,09 | 49,5 |
| ENSG00000135766 | EGLN1 | 15,36 | 15,27 | 13,24 | 7,76 | 49,5 |
| ENSG00000100580 | TMED8 | 5,68 | 5,48 | 4,61 | 2,87 | 49,5 |
| ENSG00000114391 | RPL24 | 710,38 | 681,33 | 633,27 | 358,97 | 49,5 |
| ENSG00000065609 | SNAP91 | 3,34 | 3,25 | 2,37 | 1,69 | 49,4 |
| ENSG00000106462 | EZH2 | 43,06 | 39,29 | 32,22 | 21,8 | 49,4 |
| ENSG00000236526 | AL035448,1 | 1,54 | 1,51 | 1,37 | 0,78 | 49,4 |
| ENSG00000162441 | LZIC | 23,02 | 21,95 | 20,83 | 11,66 | 49,3 |
| ENSG00000203760 | CENPW | 30,32 | 27,17 | 25,79 | 15,37 | 49,3 |
| ENSG00000147854 | UHRF2 | 30,21 | 29,71 | 24,99 | 15,32 | 49,3 |
| ENSG00000021574 | SPAST | 22,42 | 21,76 | 17,38 | 11,37 | 49,3 |
| ENSG00000186094 | AGBL4 | 1,34 | 1,08 | 0,87 | 0,68 | 49,3 |
| ENSG00000285077 | ARHGAP11B | 3,86 | 3,85 | 3,83 | 1,96 | 49,2 |
| ENSG00000069702 | TGFBR3 | 1,87 | 1,66 | 1,39 | 0,95 | 49,2 |
| ENSG00000100473 | COCH | 1,79 | 1,49 | 1,33 | 0,91 | 49,2 |
| ENSG00000166024 | R3HCC1L | 7,43 | 7,34 | 6,06 | 3,78 | 49,1 |
| ENSG00000112378 | PERP | 3,32 | 3,08 | 2,74 | 1,69 | 49,1 |
| ENSG00000176624 | MEX3C | 26,77 | 26,16 | 23,25 | 13,63 | 49,1 |
| ENSG00000130741 | EIF2S3 | 133,11 | 129,65 | 126,77 | 67,79 | 49,1 |
| ENSG00000129518 | EAPP | 18,23 | 18 | 17,55 | 9,29 | 49,0 |
| ENSG00000282978 | AC110994,2 | 1,04 | 0,92 | 0,85 | 0,53 | 49,0 |
| ENSG00000277401 | TJP1 | 21,55 | 21,29 | 19,75 | 10,99 | 49,0 |
| ENSG00000079841 | RIMS1 | 4,47 | 3,96 | 3,59 | 2,28 | 49,0 |
| ENSG00000183520 | UTP11 | 38,59 | 38,03 | 30,87 | 19,69 | 49,0 |
| ENSG00000121892 | PDS5A | 34,65 | 33,84 | 29,52 | 17,68 | 49,0 |
| ENSG00000100095 | SEZ6L | 8,03 | 6,26 | 4,92 | 4,1 | 48,9 |
| ENSG00000112245 | PTP4A1 | 43,91 | 41,53 | 32,78 | 22,42 | 48,9 |
| ENSG00000207165 | SNORA70 | 361,56 | 358,02 | 340,24 | 184,63 | 48,9 |
| ENSG00000233860 | SHROOM3-AS1 | 4,6 | 3,54 | 3,08 | 2,35 | 48,9 |
| ENSG00000065135 | GNAI3 | 16,85 | 16,52 | 15,61 | 8,61 | 48,9 |
| ENSG00000151500 | THYN1 | 30,49 | 28,16 | 26,08 | 15,58 | 48,9 |
| ENSG00000100519 | PSMC6 | 63,56 | 57 | 53,94 | 32,49 | 48,9 |
| ENSG00000031003 | FAM13B | 17,37 | 16,22 | 14,63 | 8,88 | 48,9 |
| ENSG00000143179 | UCK2 | 27,32 | 21,56 | 16,71 | 13,97 | 48,9 |
| ENSG00000129317 | PUS7L | 10,48 | 8,27 | 7,13 | 5,37 | 48,8 |
| ENSG00000164032 | H2AZ1 | 429,53 | 380,81 | 367,04 | 220,18 | 48,7 |
| ENSG00000056097 | ZFR | 41,9 | 41,07 | 37,49 | 21,48 | 48,7 |
| ENSG00000115758 | ODC1 | 175,46 | 138,57 | 108,94 | 89,96 | 48,7 |
| ENSG00000170448 | NFXL1 | 7,54 | 5,24 | 4,54 | 3,87 | 48,7 |
| ENSG00000128656 | CHN1 | 43,06 | 41,05 | 34,73 | 22,11 | 48,7 |
| ENSG00000189180 | ZNF33A | 15,74 | 14,89 | 11,98 | 8,09 | 48,6 |
| ENSG00000177125 | ZBTB34 | 5,72 | 5,01 | 4,95 | 2,94 | 48,6 |
| ENSG00000175606 | TMEM70 | 10,66 | 10,01 | 9,55 | 5,48 | 48,6 |
| ENSG00000147679 | UTP23 | 8,51 | 6,45 | 6,44 | 4,38 | 48,5 |
| ENSG00000185420 | SMYD3 | 11,56 | 10,23 | 9,6 | 5,95 | 48,5 |
| ENSG00000174796 | THAP6 | 7,34 | 6,28 | 6,17 | 3,78 | 48,5 |
| ENSG00000166405 | RIC3 | 7,32 | 6,52 | 4,83 | 3,77 | 48,5 |

| | | | | | | |
|---|---|---|---|---|---|---|
| ENSG00000136169 | SETDB2 | 3,61 | 3,53 | 3,39 | 1,86 | 48,5 |
| ENSG00000286621 | AC064843,1 | 1,3 | 1 | 0,85 | 0,67 | 48,5 |
| ENSG00000144036 | EXOC6B | 16,49 | 15,46 | 13,85 | 8,5 | 48,5 |
| ENSG00000127252 | PLAAT1 | 1,92 | 1,49 | 1,04 | 0,99 | 48,4 |
| ENSG00000270903 | HNRNPA3P9 | 1,26 | 1,05 | 0,97 | 0,65 | 48,4 |
| ENSG00000183023 | SLC8A1 | 9,36 | 7,78 | 5,22 | 4,83 | 48,4 |
| ENSG00000137601 | NEK1 | 5,89 | 5,8 | 4,68 | 3,04 | 48,4 |
| ENSG00000180263 | FGD6 | 6,95 | 6,32 | 4,56 | 3,59 | 48,3 |
| ENSG00000187118 | CMC1 | 9,92 | 9,52 | 8,82 | 5,13 | 48,3 |
| ENSG00000186522 | SEPTIN10 | 52,36 | 51,74 | 45,87 | 27,08 | 48,3 |
| ENSG00000231562 | AL512593,1 | 1,45 | 1,24 | 1,15 | 0,75 | 48,3 |
| ENSG00000135045 | C9orf40 | 8,97 | 7,34 | 7,11 | 4,64 | 48,3 |
| ENSG00000215199 | YWHAZP6 | 2,3 | 2,16 | 1,95 | 1,19 | 48,3 |
| ENSG00000273807 | AC244489,1 | 18,26 | 13,63 | 9,66 | 9,45 | 48,2 |
| ENSG00000110318 | CEP126 | 1,14 | 1,07 | 0,96 | 0,59 | 48,2 |
| ENSG00000114942 | EEF1B2 | 128,46 | 120,72 | 103,99 | 66,53 | 48,2 |
| ENSG00000283391 | EEF1B2 | 128,46 | 120,72 | 103,99 | 66,53 | 48,2 |
| ENSG00000091483 | FH | 33,92 | 33,06 | 30,41 | 17,57 | 48,2 |
| ENSG00000077097 | TOP2B | 94,98 | 92,18 | 72 | 49,2 | 48,2 |
| ENSG00000159023 | EPB41 | 35,77 | 28,65 | 22,88 | 18,53 | 48,2 |
| ENSG00000148187 | MRRF | 15,77 | 13,95 | 12,19 | 8,17 | 48,2 |
| ENSG00000169714 | CNBP | 127,3 | 119,45 | 111,54 | 65,97 | 48,2 |
| ENSG00000136521 | NDUFB5 | 41,92 | 39,98 | 38,09 | 21,73 | 48,2 |
| ENSG00000142875 | PRKACB | 18,41 | 16,83 | 13,52 | 9,55 | 48,1 |
| ENSG00000109472 | CPE | 35,29 | 31,82 | 23,89 | 18,31 | 48,1 |
| ENSG00000125772 | GPCPD1 | 16,12 | 14,91 | 11,18 | 8,37 | 48,1 |
| ENSG00000285103 | AL451123,1 | 1,29 | 0,9 | 0,81 | 0,67 | 48,1 |
| ENSG00000108947 | EFNB3 | 9,8 | 7,56 | 5,95 | 5,09 | 48,1 |
| ENSG00000102763 | VWA8 | 6,18 | 5,63 | 5,04 | 3,21 | 48,1 |
| ENSG00000258441 | LINC00641 | 12,49 | 12,04 | 10,1 | 6,49 | 48,0 |
| ENSG00000175581 | MRPL48 | 23,94 | 20,06 | 19,62 | 12,44 | 48,0 |
| ENSG00000120437 | ACAT2 | 175,72 | 140,04 | 121,52 | 91,32 | 48,0 |
| ENSG00000107938 | EDRF1 | 13,29 | 12,15 | 9,91 | 6,92 | 47,9 |
| ENSG00000134594 | RAB33A | 9,79 | 8,43 | 7,09 | 5,1 | 47,9 |
| ENSG00000198298 | ZNF485 | 1,9 | 1,5 | 1,19 | 0,99 | 47,9 |
| ENSG00000152409 | JMY | 2,13 | 1,95 | 1,61 | 1,11 | 47,9 |
| ENSG00000020922 | MRE11 | 15,94 | 12,96 | 12,19 | 8,31 | 47,9 |
| ENSG00000169213 | RAB3B | 4,45 | 3,86 | 2,96 | 2,32 | 47,9 |
| ENSG00000083635 | NUFIP1 | 4,87 | 4,55 | 4,43 | 2,54 | 47,8 |
| ENSG00000113643 | RARS1 | 47,45 | 45,66 | 41,27 | 24,76 | 47,8 |
| ENSG00000100077 | GRK3 | 4,12 | 3,52 | 2,42 | 2,15 | 47,8 |
| ENSG00000137500 | CCDC90B | 56,35 | 53,16 | 49,03 | 29,42 | 47,8 |
| ENSG00000114857 | NKTR | 43,5 | 42,25 | 34,61 | 22,72 | 47,8 |
| ENSG00000131263 | RLIM | 15,46 | 15,14 | 13,5 | 8,08 | 47,7 |
| ENSG00000196670 | ZFP62 | 13,91 | 13,32 | 11,89 | 7,27 | 47,7 |
| ENSG00000108064 | TFAM | 25,81 | 24,21 | 21,19 | 13,49 | 47,7 |
| ENSG00000137269 | LRRC1 | 3,06 | 2,98 | 2,44 | 1,6 | 47,7 |

| | | | | | | |
|---|---|---|---|---|---|---|
| ENSG00000151725 | CENPU | 44,9 | 42,4 | 39,9 | 23,49 | 47,7 |
| ENSG00000273344 | PAXIP1-AS1 | 2,35 | 2,17 | 2,02 | 1,23 | 47,7 |
| ENSG00000272145 | NFYC-AS1 | 2,12 | 1,79 | 1,39 | 1,11 | 47,6 |
| ENSG00000111843 | TMEM14C | 23,05 | 22,02 | 18,67 | 12,07 | 47,6 |
| ENSG00000284936 | TMEM14C | 23,05 | 22,02 | 18,67 | 12,07 | 47,6 |
| ENSG00000080822 | CLDND1 | 40,73 | 39,78 | 33,83 | 21,33 | 47,6 |
| ENSG00000144161 | ZC3H8 | 7,69 | 7,11 | 4,52 | 4,03 | 47,6 |
| ENSG00000263465 | SRSF8 | 9,06 | 8,68 | 7,47 | 4,75 | 47,6 |
| ENSG00000162928 | PEX13 | 10,16 | 9,19 | 8,15 | 5,33 | 47,5 |
| ENSG00000088448 | ANKRD10 | 62 | 59,01 | 52,45 | 32,53 | 47,5 |
| ENSG00000053900 | ANAPC4 | 11,95 | 11,65 | 9,35 | 6,27 | 47,5 |
| ENSG00000088727 | KIF9 | 5,16 | 4,87 | 4,45 | 2,71 | 47,5 |
| ENSG00000135968 | GCC2 | 12,96 | 12,19 | 8,79 | 6,81 | 47,5 |
| ENSG00000198018 | ENTPD7 | 5,67 | 5,43 | 4,5 | 2,98 | 47,4 |
| ENSG00000156831 | NSMCE2 | 17,75 | 17,53 | 17,26 | 9,33 | 47,4 |
| ENSG00000281649 | EBLN3P | 17,84 | 17,43 | 15,16 | 9,38 | 47,4 |
| ENSG00000178177 | LCORL | 12,55 | 10,83 | 10,49 | 6,6 | 47,4 |
| ENSG00000152422 | XRCC4 | 6,73 | 5,65 | 5,11 | 3,54 | 47,4 |
| ENSG00000132953 | XPO4 | 8,84 | 8,57 | 7,24 | 4,65 | 47,4 |
| ENSG00000164941 | INTS8 | 35,7 | 31,71 | 31,41 | 18,78 | 47,4 |
| ENSG00000153207 | AHCTF1 | 16,88 | 15,77 | 13,7 | 8,88 | 47,4 |
| ENSG00000168438 | CDC40 | 7,73 | 6,58 | 6,3 | 4,07 | 47,3 |
| ENSG00000177432 | NAP1L5 | 3,74 | 3,26 | 3,04 | 1,97 | 47,3 |
| ENSG00000197170 | PSMD12 | 36,08 | 34,34 | 29,84 | 19,01 | 47,3 |
| ENSG00000106603 | COA1 | 43,46 | 42,44 | 38,27 | 22,9 | 47,3 |
| ENSG00000115904 | SOS1 | 18,01 | 14,2 | 13,37 | 9,49 | 47,3 |
| ENSG00000196323 | ZBTB44 | 10,34 | 10,04 | 9,83 | 5,45 | 47,3 |
| ENSG00000004864 | SLC25A13 | 26,84 | 25,85 | 22,44 | 14,15 | 47,3 |
| ENSG00000106484 | MEST | 216,75 | 176,27 | 170,12 | 114,28 | 47,3 |
| ENSG00000122483 | CCDC18 | 6,65 | 6,16 | 4,57 | 3,51 | 47,2 |
| ENSG00000091972 | CD200 | 20,23 | 19 | 15,72 | 10,68 | 47,2 |
| ENSG00000153879 | CEBPG | 12,89 | 12,08 | 11,07 | 6,81 | 47,2 |
| ENSG00000116161 | CACYBP | 58,02 | 53,87 | 48,53 | 30,66 | 47,2 |
| ENSG00000121579 | NAA50 | 53,04 | 49,65 | 48,7 | 28,03 | 47,2 |
| ENSG00000188021 | UBQLN2 | 15,89 | 15,81 | 14,09 | 8,4 | 47,1 |
| ENSG00000213585 | VDAC1 | 105,47 | 102,44 | 99,64 | 55,79 | 47,1 |
| ENSG00000138381 | ASNSD1 | 21,27 | 18,95 | 17,27 | 11,26 | 47,1 |
| ENSG00000049883 | PTCD2 | 5,25 | 3,98 | 3,79 | 2,78 | 47,0 |
| ENSG00000071994 | PDCD2 | 32,58 | 31,38 | 31,04 | 17,26 | 47,0 |
| ENSG00000138286 | FAM149B1 | 12,47 | 11,82 | 11,11 | 6,61 | 47,0 |
| ENSG00000006634 | DBF4 | 31,34 | 27,69 | 24,35 | 16,62 | 47,0 |
| ENSG00000121766 | ZCCHC17 | 26,34 | 25,69 | 24,73 | 13,97 | 47,0 |
| ENSG00000154814 | OXNAD1 | 7,52 | 6,43 | 5,14 | 3,99 | 46,9 |
| ENSG00000280594 | BTG3-AS1 | 1,96 | 1,24 | 1,09 | 1,04 | 46,9 |
| ENSG00000281317 | BTG3-AS1 | 1,96 | 1,24 | 1,09 | 1,04 | 46,9 |
| ENSG00000111850 | SMIM8 | 5,88 | 5,85 | 5,09 | 3,12 | 46,9 |
| ENSG00000166439 | RNF169 | 8,91 | 8,58 | 8,52 | 4,73 | 46,9 |

| | | | | | | |
|---|---|---|---|---|---|---|
| ENSG00000151806 | GUF1 | 10,86 | 9,7 | 8,34 | 5,77 | 46,9 |
| ENSG00000118482 | PHF3 | 23,46 | 22,04 | 18,77 | 12,47 | 46,8 |
| ENSG00000153956 | CACNA2D1 | 10,25 | 10,04 | 9,89 | 5,45 | 46,8 |
| ENSG00000038210 | PI4K2B | 5,83 | 5,58 | 4,97 | 3,1 | 46,8 |
| ENSG00000170855 | TRIAP1 | 12,41 | 11,92 | 11,43 | 6,61 | 46,7 |
| ENSG00000139146 | SINHCAF | 41,99 | 34,75 | 30,48 | 22,39 | 46,7 |
| ENSG00000182890 | GLUD2 | 1,5 | 1,23 | 1,17 | 0,8 | 46,7 |
| ENSG00000288118 | GLUD2 | 1,5 | 1,23 | 1,17 | 0,8 | 46,7 |
| ENSG00000129315 | CCNT1 | 14,34 | 13,16 | 11,51 | 7,65 | 46,7 |
| ENSG00000254377 | MIR124-2HG | 27,87 | 26,9 | 24,71 | 14,87 | 46,6 |
| ENSG00000138764 | CCNG2 | 35,28 | 31,48 | 30,45 | 18,83 | 46,6 |
| ENSG00000147274 | RBMX | 238,88 | 231,37 | 221,76 | 127,52 | 46,6 |
| ENSG00000082458 | DLG3 | 33,35 | 32,12 | 30,15 | 17,81 | 46,6 |
| ENSG00000134901 | POGLUT2 | 11,83 | 11,54 | 10,32 | 6,32 | 46,6 |
| ENSG00000130177 | CDC16 | 36,91 | 36,35 | 33,88 | 19,74 | 46,5 |
| ENSG00000288295 | MCTS1 | 29,78 | 23,84 | 23,51 | 15,93 | 46,5 |
| ENSG00000168906 | MAT2A | 108,56 | 101,92 | 90,46 | 58,08 | 46,5 |
| ENSG00000184445 | KNTC1 | 38,35 | 31,99 | 28,16 | 20,52 | 46,5 |
| ENSG00000269837 | IPO5P1 | 10,82 | 8,98 | 7,16 | 5,79 | 46,5 |
| ENSG00000143498 | TAF1A | 8,37 | 7,7 | 5,51 | 4,48 | 46,5 |
| ENSG00000145979 | TBC1D7 | 16,12 | 15,38 | 13,83 | 8,63 | 46,5 |
| ENSG00000138780 | GSTCD | 12,85 | 12,29 | 10,57 | 6,88 | 46,5 |
| ENSG00000206557 | TRIM71 | 7,97 | 6,69 | 5,06 | 4,27 | 46,4 |
| ENSG00000235652 | FBXO30-DT | 46,22 | 44,26 | 38,22 | 24,77 | 46,4 |
| ENSG00000126261 | UBA2 | 82,71 | 80,04 | 78,14 | 44,33 | 46,4 |
| ENSG00000100815 | TRIP11 | 14,94 | 14,52 | 11,27 | 8,01 | 46,4 |
| ENSG00000115365 | LANCL1 | 18,55 | 17,49 | 15,04 | 9,95 | 46,4 |
| ENSG00000130348 | QRSL1 | 6,02 | 5,29 | 5,21 | 3,23 | 46,3 |
| ENSG00000214029 | ZNF891 | 3,39 | 3,25 | 2,8 | 1,82 | 46,3 |
| ENSG00000112996 | MRPS30 | 18,44 | 17,59 | 17,01 | 9,9 | 46,3 |
| ENSG00000117262 | GPR89A | 9,12 | 8,34 | 7,13 | 4,9 | 46,3 |
| ENSG00000287202 | AC008119,1 | 1,47 | 1,37 | 1,22 | 0,79 | 46,3 |
| ENSG00000137807 | KIF23 | 29,91 | 29,55 | 28,19 | 16,08 | 46,2 |
| ENSG00000184613 | NELL2 | 84,68 | 64,71 | 48,7 | 45,55 | 46,2 |
| ENSG00000145348 | TBCK | 16,43 | 14,83 | 14,24 | 8,84 | 46,2 |
| ENSG00000143033 | MTF2 | 29,44 | 26,96 | 23,25 | 15,84 | 46,2 |
| ENSG00000197771 | MCMBP | 29,92 | 29,77 | 28,85 | 16,1 | 46,2 |
| ENSG00000175322 | ZNF519 | 17,02 | 15,89 | 13,41 | 9,16 | 46,2 |
| ENSG00000121988 | ZRANB3 | 7,84 | 6,37 | 4,6 | 4,22 | 46,2 |
| ENSG00000233369 | GTF2IP4 | 105,18 | 98,71 | 88,86 | 56,62 | 46,2 |
| ENSG00000165678 | GHITM | 36,57 | 31,83 | 30,01 | 19,69 | 46,2 |
| ENSG00000099251 | HSD17B7P2 | 1,82 | 1,54 | 1,38 | 0,98 | 46,2 |
| ENSG00000288374 | HSD17B7P2 | 1,82 | 1,54 | 1,38 | 0,98 | 46,2 |
| ENSG00000086712 | TXLNG | 10,53 | 10,41 | 8,92 | 5,67 | 46,2 |
| ENSG00000134202 | GSTM3 | 23,25 | 22,7 | 22,18 | 12,52 | 46,2 |
| ENSG00000081479 | LRP2 | 9,97 | 7,41 | 6,66 | 5,37 | 46,1 |
| ENSG00000076554 | TPD52 | 4,27 | 3,08 | 2,81 | 2,3 | 46,1 |

| | | | | | | |
|---|---|---|---|---|---|---|
| ENSG00000126773 | PCNX4 | 48,94 | 41,68 | 37,96 | 26,37 | 46,1 |
| ENSG00000120594 | PLXDC2 | 7,42 | 6,76 | 5,54 | 4 | 46,1 |
| ENSG00000228065 | LINC01515 | 4,73 | 3,95 | 3,24 | 2,55 | 46,1 |
| ENSG00000163558 | PRKCI | 13,15 | 12,05 | 11,2 | 7,09 | 46,1 |
| ENSG00000287014 | AC087883,2 | 1,91 | 1,35 | 1,22 | 1,03 | 46,1 |
| ENSG00000230392 | AC004835,1 | 1,91 | 1,17 | 1,06 | 1,03 | 46,1 |
| ENSG00000022567 | SLC45A4 | 4,19 | 4,18 | 2,52 | 2,26 | 46,1 |
| ENSG00000163449 | TMEM169 | 10,6 | 10,07 | 6,27 | 5,72 | 46,0 |
| ENSG00000139973 | SYT16 | 1 | 0,95 | 0,64 | 0,54 | 46,0 |
| ENSG00000165416 | SUGT1 | 34,69 | 34,63 | 32,45 | 18,75 | 45,9 |
| ENSG00000171208 | NETO2 | 31,79 | 31,41 | 23,86 | 17,19 | 45,9 |
| ENSG00000113460 | BRIX1 | 25,13 | 22,6 | 19,38 | 13,6 | 45,9 |
| ENSG00000198270 | TMEM116 | 6,19 | 5,04 | 3,93 | 3,35 | 45,9 |
| ENSG00000269911 | FAM226B | 3,64 | 3,07 | 2,28 | 1,97 | 45,9 |
| ENSG00000120685 | PROSER1 | 21,08 | 19,45 | 16,79 | 11,41 | 45,9 |
| ENSG00000168795 | ZBTB5 | 10,25 | 9,5 | 9,27 | 5,55 | 45,9 |
| ENSG00000117543 | DPH5 | 26,72 | 22,2 | 18,92 | 14,47 | 45,8 |
| ENSG00000182134 | TDRKH | 8,27 | 8,03 | 6,14 | 4,48 | 45,8 |
| ENSG00000111144 | LTA4H | 29,09 | 27,4 | 24,78 | 15,76 | 45,8 |
| ENSG00000173542 | MOB1B | 15,32 | 14,3 | 13,46 | 8,3 | 45,8 |
| ENSG00000181007 | ZFP82 | 3,82 | 3,78 | 3,15 | 2,07 | 45,8 |
| ENSG00000119787 | ATL2 | 24,47 | 22,79 | 19,94 | 13,26 | 45,8 |
| ENSG00000202354 | RNY3 | 6339,32 | 4797,96 | 4738,66 | 3435,54 | 45,8 |
| ENSG00000021826 | CPS1 | 5,83 | 4,53 | 3,26 | 3,16 | 45,8 |
| ENSG00000048405 | ZNF800 | 15,09 | 14,36 | 14,21 | 8,18 | 45,8 |
| ENSG00000156313 | RPGR | 4,98 | 4,11 | 3,21 | 2,7 | 45,8 |
| ENSG00000085760 | MTIF2 | 15,95 | 13,08 | 10,66 | 8,65 | 45,8 |
| ENSG00000169126 | ARMC4 | 5,07 | 4,63 | 3,52 | 2,75 | 45,8 |
| ENSG00000141401 | IMPA2 | 11,63 | 7,93 | 7,78 | 6,31 | 45,7 |
| ENSG00000249790 | AC092490,1 | 18,78 | 17,39 | 12,84 | 10,21 | 45,6 |
| ENSG00000130024 | PHF10 | 27,54 | 25,85 | 21,09 | 14,99 | 45,6 |
| ENSG00000240024 | LINC00888 | 9,48 | 9,12 | 7,89 | 5,16 | 45,6 |
| ENSG00000081377 | CDC14B | 7,75 | 5,4 | 4,73 | 4,22 | 45,5 |
| ENSG00000166471 | TMEM41B | 14 | 13,7 | 11,35 | 7,63 | 45,5 |
| ENSG00000273749 | CYFIP1 | 27,52 | 26,71 | 23,68 | 15 | 45,5 |
| ENSG00000164221 | CCDC112 | 13,04 | 11,46 | 9 | 7,11 | 45,5 |
| ENSG00000241837 | ATP5PO | 144,55 | 130,95 | 128,7 | 78,82 | 45,5 |
| ENSG00000196678 | ERI2 | 16,87 | 16,02 | 15,4 | 9,2 | 45,5 |
| ENSG00000119285 | HEATR1 | 18,19 | 15,47 | 15,22 | 9,93 | 45,4 |
| ENSG00000169519 | METTL15 | 4,89 | 4,42 | 3,98 | 2,67 | 45,4 |
| ENSG00000008869 | HEATR5B | 6,7 | 6,35 | 5,37 | 3,66 | 45,4 |
| ENSG00000267547 | AC060766,4 | 3,88 | 3,7 | 2,71 | 2,12 | 45,4 |
| ENSG00000139697 | SBNO1 | 10,41 | 10,12 | 9 | 5,69 | 45,3 |
| ENSG00000166181 | API5 | 53,61 | 51,13 | 45,06 | 29,32 | 45,3 |
| ENSG00000109814 | UGDH | 54 | 47,19 | 33,09 | 29,54 | 45,3 |
| ENSG00000124171 | PARD6B | 2,12 | 1,97 | 1,58 | 1,16 | 45,3 |
| ENSG00000268471 | MIR4453HG | 2,43 | 2,17 | 1,84 | 1,33 | 45,3 |

| | | | | | | |
|---|---|---|---|---|---|---|
| ENSG00000127995 | CASD1 | 16,4 | 14,05 | 13,5 | 8,98 | 45,2 |
| ENSG00000135913 | USP37 | 9,66 | 8,74 | 8,19 | 5,29 | 45,2 |
| ENSG00000205611 | LINC01597 | 1,68 | 1,52 | 1,17 | 0,92 | 45,2 |
| ENSG00000134326 | CMPK2 | 1,57 | 1,41 | 1,31 | 0,86 | 45,2 |
| ENSG00000234585 | CCT6P3 | 3,43 | 3,09 | 2,88 | 1,88 | 45,2 |
| ENSG00000087253 | LPCAT2 | 3,83 | 3,1 | 2,67 | 2,1 | 45,2 |
| ENSG00000131791 | PRKAB2 | 7,86 | 7,81 | 7,25 | 4,31 | 45,2 |
| ENSG00000178295 | GEN1 | 19,54 | 16,16 | 12,77 | 10,72 | 45,1 |
| ENSG00000261770 | AC006504,1 | 1,33 | 1,22 | 0,97 | 0,73 | 45,1 |
| ENSG00000213801 | ZNF321P | 2,35 | 2,13 | 1,91 | 1,29 | 45,1 |
| ENSG00000151743 | AMN1 | 5,81 | 5,58 | 4,2 | 3,19 | 45,1 |
| ENSG00000108055 | SMC3 | 28,92 | 27,64 | 21,72 | 15,88 | 45,1 |
| ENSG00000241697 | TMEFF1 | 9,01 | 7,53 | 7,09 | 4,95 | 45,1 |
| ENSG00000143315 | PIGM | 4,35 | 3,94 | 3,25 | 2,39 | 45,1 |
| ENSG00000134970 | TMED7 | 25,76 | 25,65 | 22,31 | 14,16 | 45,0 |
| ENSG00000035499 | DEPDC1B | 19,93 | 17,94 | 16,68 | 10,97 | 45,0 |
| ENSG00000151229 | SLC2A13 | 2,67 | 2,03 | 2,01 | 1,47 | 44,9 |
| ENSG00000100557 | CCDC198 | 1,58 | 1,09 | 0,96 | 0,87 | 44,9 |
| ENSG00000159055 | MIS18A | 15,12 | 14,34 | 11,84 | 8,33 | 44,9 |
| ENSG00000147118 | ZNF182 | 2,65 | 2,56 | 2,39 | 1,46 | 44,9 |
| ENSG00000132300 | PTCD3 | 58,39 | 54,85 | 52,82 | 32,17 | 44,9 |
| ENSG00000196159 | FAT4 | 9,02 | 8,92 | 8,02 | 4,97 | 44,9 |
| ENSG00000138686 | BBS7 | 11,94 | 10,54 | 9,54 | 6,58 | 44,9 |
| ENSG00000049449 | RCN1 | 116,18 | 114,8 | 98,32 | 64,04 | 44,9 |
| ENSG00000163661 | PTX3 | 60,17 | 50,43 | 37,93 | 33,17 | 44,9 |
| ENSG00000109618 | SEPSECS | 2,92 | 2,48 | 2,41 | 1,61 | 44,9 |
| ENSG00000067704 | IARS2 | 20,84 | 20,42 | 18,57 | 11,5 | 44,8 |
| ENSG00000138663 | COPS4 | 26,24 | 26,16 | 23,15 | 14,48 | 44,8 |
| ENSG00000266053 | NDUFV2-AS1 | 2,41 | 1,87 | 1,76 | 1,33 | 44,8 |
| ENSG00000143653 | SCCPDH | 24,43 | 21,93 | 20,88 | 13,49 | 44,8 |
| ENSG00000008988 | RPS20 | 1092,77 | 1086,9 | 1049,33 | 603,43 | 44,8 |
| ENSG00000138738 | PRDM5 | 6,79 | 5,39 | 5,03 | 3,75 | 44,8 |
| ENSG00000086589 | RBM22 | 26,82 | 25,7 | 25,03 | 14,82 | 44,7 |
| ENSG00000272606 | AC015982,2 | 1,61 | 1,35 | 1,29 | 0,89 | 44,7 |
| ENSG00000237335 | PFDN6 | 7,63 | 6,98 | 6,9 | 4,22 | 44,7 |
| ENSG00000224782 | PFDN6 | 7,63 | 6,98 | 6,9 | 4,22 | 44,7 |
| ENSG00000085382 | HACE1 | 10,34 | 10,23 | 8,42 | 5,72 | 44,7 |
| ENSG00000278077 | AL591926,6 | 1,88 | 1,59 | 1,21 | 1,04 | 44,7 |
| ENSG00000268119 | AC010615,2 | 17,64 | 14,7 | 11,84 | 9,76 | 44,7 |
| ENSG00000186468 | RPS23 | 1100,52 | 1054,43 | 1030,88 | 609,34 | 44,6 |
| ENSG00000228409 | CCT6P1 | 3,9 | 3,85 | 3,34 | 2,16 | 44,6 |
| ENSG00000119844 | AFTPH | 6,86 | 6,68 | 5,84 | 3,8 | 44,6 |
| ENSG00000178917 | ZNF852 | 2,67 | 2,52 | 2,04 | 1,48 | 44,6 |
| ENSG00000281626 | ZNF852 | 2,67 | 2,52 | 2,04 | 1,48 | 44,6 |
| ENSG00000145332 | KLHL8 | 8,46 | 7,19 | 6,44 | 4,69 | 44,6 |
| ENSG00000106069 | CHN2 | 7,25 | 6,3 | 4,39 | 4,02 | 44,6 |
| ENSG00000174371 | EXO1 | 16,14 | 15,51 | 13,08 | 8,95 | 44,5 |

| | | | | | | |
|---|---|---|---|---|---|---|
| ENSG00000117475 | BLZF1 | 9,6 | 9,19 | 7,28 | 5,33 | 44,5 |
| ENSG00000172845 | SP3 | 36,67 | 34,89 | 33,36 | 20,36 | 44,5 |
| ENSG00000082213 | C5orf22 | 19,27 | 19,16 | 17,08 | 10,7 | 44,5 |
| ENSG00000169679 | BUB1 | 36,99 | 34,4 | 34,07 | 20,55 | 44,4 |
| ENSG00000186063 | AIDA | 20,48 | 19,35 | 18,85 | 11,38 | 44,4 |
| ENSG00000078053 | AMPH | 22,02 | 19,86 | 15,84 | 12,25 | 44,4 |
| ENSG00000123505 | AMD1 | 30,57 | 27,17 | 26,71 | 17,02 | 44,3 |
| ENSG00000116885 | OSCP1 | 7,02 | 5,72 | 5,64 | 3,91 | 44,3 |
| ENSG00000059588 | TARBP1 | 12,24 | 9,24 | 7,55 | 6,82 | 44,3 |
| ENSG00000197798 | FAM118B | 13,71 | 13,44 | 13,14 | 7,64 | 44,3 |
| ENSG00000287222 | AC234917,3 | 2,53 | 2,16 | 2,12 | 1,41 | 44,3 |
| ENSG00000186416 | NKRF | 7,21 | 6,91 | 6,03 | 4,02 | 44,2 |
| ENSG00000146457 | WTAP | 56,86 | 55,49 | 52,55 | 31,74 | 44,2 |
| ENSG00000116983 | HPCAL4 | 1,2 | 1,12 | 0,72 | 0,67 | 44,2 |
| ENSG00000119862 | LGALSL | 8,47 | 8,46 | 7,02 | 4,73 | 44,2 |
| ENSG00000266472 | MRPS21 | 42,01 | 40,94 | 39,61 | 23,47 | 44,1 |
| ENSG00000137812 | KNL1 | 16,59 | 16,32 | 15,56 | 9,27 | 44,1 |
| ENSG00000143036 | SLC44A3 | 1,95 | 1,86 | 1,57 | 1,09 | 44,1 |
| ENSG00000157869 | RAB28 | 12,54 | 11,32 | 10,51 | 7,01 | 44,1 |
| ENSG00000126953 | TIMM8A | 9,4 | 8,29 | 6,69 | 5,26 | 44,0 |
| ENSG00000186432 | KPNA4 | 44,82 | 43,17 | 37,69 | 25,09 | 44,0 |
| ENSG00000185104 | FAF1 | 26,99 | 25,78 | 21,43 | 15,12 | 44,0 |
| ENSG00000224985 | AL590714,1 | 3,48 | 3,15 | 2,02 | 1,95 | 44,0 |
| ENSG00000165506 | DNAAF2 | 7,53 | 6,34 | 5,47 | 4,22 | 44,0 |
| ENSG00000120688 | WBP4 | 9,28 | 9 | 8,09 | 5,21 | 43,9 |
| ENSG00000144468 | RHBDD1 | 12,73 | 9,72 | 9,36 | 7,15 | 43,8 |
| ENSG00000122406 | RPL5 | 752,64 | 710,29 | 691,38 | 422,78 | 43,8 |
| ENSG00000179455 | MKRN3 | 5,82 | 4,97 | 4,66 | 3,27 | 43,8 |
| ENSG00000285309 | AL136295,18 | 3,45 | 2,86 | 2,53 | 1,94 | 43,8 |
| ENSG00000278784 | AL136295,7 | 3,45 | 2,86 | 2,53 | 1,94 | 43,8 |
| ENSG00000259802 | AC012640,2 | 2,88 | 1,96 | 1,65 | 1,62 | 43,8 |
| ENSG00000035687 | ADSS2 | 18,08 | 17,9 | 13,93 | 10,17 | 43,8 |
| ENSG00000162639 | HENMT1 | 4,55 | 4,52 | 2,89 | 2,56 | 43,7 |
| ENSG00000111224 | PARP11 | 9,81 | 9 | 8,66 | 5,52 | 43,7 |
| ENSG00000119865 | CNRIP1 | 8,97 | 8,96 | 8,76 | 5,05 | 43,7 |
| ENSG00000150756 | ATPSCKMT | 12,24 | 12,05 | 9,83 | 6,9 | 43,6 |
| ENSG00000107949 | BCCIP | 46,98 | 41,41 | 35,36 | 26,49 | 43,6 |
| ENSG00000120694 | HSPH1 | 53 | 44,01 | 34,48 | 29,89 | 43,6 |
| ENSG00000132963 | POMP | 52,06 | 51,08 | 49,4 | 29,36 | 43,6 |
| ENSG00000077232 | DNAJC10 | 55,32 | 52,05 | 47,12 | 31,21 | 43,6 |
| ENSG00000187790 | FANCM | 5,14 | 5,12 | 4,83 | 2,9 | 43,6 |
| ENSG00000277222 | BTBD7 | 13,02 | 11,01 | 10,8 | 7,35 | 43,5 |
| ENSG00000287110 | AC112484,5 | 1,47 | 0,93 | 0,84 | 0,83 | 43,5 |
| ENSG00000196345 | ZKSCAN7 | 2,94 | 2,65 | 1,77 | 1,66 | 43,5 |
| ENSG00000272077 | AC124045,1 | 1,31 | 1,15 | 0,97 | 0,74 | 43,5 |
| ENSG00000281102 | AC092046,2 | 1,31 | 1,15 | 0,97 | 0,74 | 43,5 |
| ENSG00000162757 | C1orf74 | 1,31 | 1,09 | 0,93 | 0,74 | 43,5 |

| | | | | | | |
|---|---|---|---|---|---|---|
| ENSG00000176386 | CDC26 | 23,5 | 21,33 | 19,58 | 13,28 | 43,5 |
| ENSG00000163535 | SGO2 | 20,08 | 19,17 | 16,54 | 11,36 | 43,4 |
| ENSG00000158691 | ZSCAN12 | 6,31 | 6,05 | 4,79 | 3,57 | 43,4 |
| ENSG00000129007 | CALML4 | 1,82 | 1,46 | 1,39 | 1,03 | 43,4 |
| ENSG00000168958 | MFF | 40,32 | 37,62 | 37,1 | 22,82 | 43,4 |
| ENSG00000283079 | LINC01515 | 4,38 | 3,64 | 3,26 | 2,48 | 43,4 |
| ENSG00000231427 | LINC01445 | 1,43 | 1,37 | 1,31 | 0,81 | 43,4 |
| ENSG00000285070 | ADIPOR2 | 9,76 | 9,68 | 8,8 | 5,53 | 43,3 |
| ENSG00000006831 | ADIPOR2 | 9,76 | 9,68 | 8,8 | 5,53 | 43,3 |
| ENSG00000157224 | CLDN12 | 10,73 | 9,04 | 8,48 | 6,08 | 43,3 |
| ENSG00000153310 | CYRIB | 44,52 | 40,66 | 37,02 | 25,24 | 43,3 |
| ENSG00000147050 | KDM6A | 10,07 | 9,89 | 8,92 | 5,71 | 43,3 |
| ENSG00000173692 | PSMD1 | 58,61 | 56,34 | 51,1 | 33,24 | 43,3 |
| ENSG00000112541 | PDE10A | 7,56 | 7,25 | 6 | 4,29 | 43,3 |
| ENSG00000282269 | PRR4 | 5,16 | 4,62 | 4,05 | 2,93 | 43,2 |
| ENSG00000196912 | ANKRD36B | 14,51 | 12,93 | 12,19 | 8,24 | 43,2 |
| ENSG00000144278 | GALNT13 | 2,5 | 1,78 | 1,77 | 1,42 | 43,2 |
| ENSG00000066136 | NFYC | 40,46 | 36 | 34,28 | 23 | 43,2 |
| ENSG00000114850 | SSR3 | 68,88 | 67,17 | 61,34 | 39,16 | 43,1 |
| ENSG00000163714 | U2SURP | 67,76 | 67,47 | 56,71 | 38,53 | 43,1 |
| ENSG00000145675 | PIK3R1 | 20,61 | 19 | 15,21 | 11,72 | 43,1 |
| ENSG00000159063 | ALG8 | 29,99 | 24,77 | 24,16 | 17,06 | 43,1 |
| ENSG00000112312 | GMNN | 55,46 | 53,36 | 45,71 | 31,56 | 43,1 |
| ENSG00000234444 | ZNF736 | 4,85 | 4,53 | 4,17 | 2,76 | 43,1 |
| ENSG00000112167 | SAYSD1 | 6,08 | 5,65 | 5,15 | 3,46 | 43,1 |
| ENSG00000171492 | LRRC8D | 10,08 | 9,6 | 9,21 | 5,74 | 43,1 |
| ENSG00000271964 | AC090948,1 | 1 | 0,99 | 0,96 | 0,57 | 43,0 |
| ENSG00000075407 | ZNF37A | 7,21 | 6,8 | 5,96 | 4,11 | 43,0 |
| ENSG00000100578 | KIAA0586 | 11,28 | 11,19 | 10,04 | 6,44 | 42,9 |
| ENSG00000112893 | MAN2A1 | 18,37 | 17,65 | 16,35 | 10,49 | 42,9 |
| ENSG00000166167 | BTRC | 12,15 | 10,64 | 10,21 | 6,94 | 42,9 |
| ENSG00000164442 | CITED2 | 28,65 | 22,34 | 18,4 | 16,37 | 42,9 |
| ENSG00000286786 | AC116158,3 | 1,05 | 0,9 | 0,74 | 0,6 | 42,9 |
| ENSG00000136709 | WDR33 | 21,56 | 21,53 | 20,28 | 12,34 | 42,8 |
| ENSG00000134480 | CCNH | 21,97 | 19,05 | 18,28 | 12,58 | 42,7 |
| ENSG00000088756 | ARHGAP28 | 16,37 | 15,87 | 12,59 | 9,38 | 42,7 |
| ENSG00000152977 | ZIC1 | 135,5 | 116,3 | 96,58 | 77,74 | 42,6 |
| ENSG00000174013 | FBXO45 | 7,77 | 7,07 | 6,36 | 4,46 | 42,6 |
| ENSG00000067177 | PHKA1 | 2,7 | 2,62 | 2,12 | 1,55 | 42,6 |
| ENSG00000124356 | STAMBP | 22,19 | 21,43 | 19,78 | 12,74 | 42,6 |
| ENSG00000104695 | PPP2CB | 43,8 | 41,61 | 37,6 | 25,16 | 42,6 |
| ENSG00000196449 | YRDC | 6,42 | 5,21 | 4,62 | 3,69 | 42,5 |
| ENSG00000156504 | FAM122B | 17,57 | 16,9 | 14,54 | 10,1 | 42,5 |
| ENSG00000092978 | GPATCH2 | 9,67 | 9,27 | 7,92 | 5,56 | 42,5 |
| ENSG00000144357 | UBR3 | 13,67 | 12,68 | 11,57 | 7,86 | 42,5 |
| ENSG00000064703 | DDX20 | 11,77 | 11,61 | 11,01 | 6,77 | 42,5 |
| ENSG00000198825 | INPP5F | 67,21 | 66,76 | 57,69 | 38,66 | 42,5 |

| | | | | | | |
|---|---|---|---|---|---|---|
| ENSG00000100592 | DAAM1 | 33,9 | 32,82 | 28,51 | 19,5 | 42,5 |
| ENSG00000284967 | FDFT1 | 178,54 | 162,94 | 147,14 | 102,74 | 42,5 |
| ENSG00000261824 | LINC00662 | 15,63 | 13,35 | 11,49 | 9 | 42,4 |
| ENSG00000111911 | HINT3 | 9,76 | 9,74 | 9,09 | 5,62 | 42,4 |
| ENSG00000112539 | C6orf118 | 4,79 | 3,69 | 2,92 | 2,76 | 42,4 |
| ENSG00000141030 | COPS3 | 45,22 | 45 | 42,63 | 26,06 | 42,4 |
| ENSG00000146676 | PURB | 7,04 | 6,89 | 6,05 | 4,06 | 42,3 |
| ENSG00000100583 | SAMD15 | 4,75 | 4,57 | 3,08 | 2,74 | 42,3 |
| ENSG00000135778 | NTPCR | 26,51 | 24,85 | 23,8 | 15,3 | 42,3 |
| ENSG00000029153 | ARNTL2 | 4,59 | 4,13 | 3,84 | 2,65 | 42,3 |
| ENSG00000157259 | GATAD1 | 12,07 | 10,87 | 10,16 | 6,97 | 42,3 |
| ENSG00000081154 | PCNP | 51,44 | 51,03 | 48,66 | 29,74 | 42,2 |
| ENSG00000183161 | FANCF | 8,89 | 7,67 | 7,06 | 5,14 | 42,2 |
| ENSG00000214870 | AC004540,1 | 5,43 | 4,66 | 4,63 | 3,14 | 42,2 |
| ENSG00000112305 | SMAP1 | 19,07 | 18,9 | 17,57 | 11,03 | 42,2 |
| ENSG00000268322 | BNIP3P25 | 1,02 | 0,86 | 0,77 | 0,59 | 42,2 |
| ENSG00000164134 | NAA15 | 30,4 | 28,18 | 23,4 | 17,6 | 42,1 |
| ENSG00000177842 | ZNF620 | 7,23 | 6,35 | 5,38 | 4,19 | 42,0 |
| ENSG00000122565 | CBX3 | 111,62 | 110,95 | 101,15 | 64,72 | 42,0 |
| ENSG00000251022 | THAP9-AS1 | 31,6 | 30,02 | 23,67 | 18,34 | 42,0 |
| ENSG00000224429 | LINC00539 | 2,05 | 1,59 | 1,42 | 1,19 | 42,0 |
| ENSG00000157764 | BRAF | 13,9 | 13,25 | 11,61 | 8,07 | 41,9 |
| ENSG00000120656 | TAF12 | 16,07 | 15,81 | 14,1 | 9,33 | 41,9 |
| ENSG00000096063 | SRPK1 | 41,2 | 37,94 | 32,78 | 23,95 | 41,9 |
| ENSG00000143815 | LBR | 60,06 | 54,37 | 51,56 | 34,92 | 41,9 |
| ENSG00000131023 | LATS1 | 11,35 | 10,73 | 10,01 | 6,6 | 41,9 |
| ENSG00000232044 | SILC1 | 9,46 | 8,49 | 7,24 | 5,51 | 41,8 |
| ENSG00000138029 | HADHB | 29,52 | 29,43 | 27,7 | 17,21 | 41,7 |
| ENSG00000177917 | ARL6IP6 | 16,74 | 16,26 | 14,28 | 9,76 | 41,7 |
| ENSG00000180917 | CMTR2 | 11,54 | 10,36 | 9,63 | 6,73 | 41,7 |
| ENSG00000196262 | PPIA | 1173,51 | 1148,49 | 1122,93 | 684,4 | 41,7 |
| ENSG00000172071 | EIF2AK3 | 8,47 | 8,46 | 6,9 | 4,94 | 41,7 |
| ENSG00000141622 | RNF165 | 9,79 | 7,62 | 6,89 | 5,71 | 41,7 |
| ENSG00000197302 | ZNF720 | 12,23 | 10,49 | 8,42 | 7,14 | 41,6 |
| ENSG00000103591 | AAGAB | 38,51 | 35,7 | 30,94 | 22,49 | 41,6 |
| ENSG00000259408 | AC010809,2 | 1,01 | 0,98 | 0,63 | 0,59 | 41,6 |
| ENSG00000118965 | WDR35 | 8,97 | 8,57 | 7,19 | 5,24 | 41,6 |
| ENSG00000197780 | TAF13 | 23,86 | 23,68 | 20,81 | 13,94 | 41,6 |
| ENSG00000118418 | HMGN3 | 79,49 | 79,42 | 73,21 | 46,45 | 41,6 |
| ENSG00000062650 | WAPL | 17,95 | 17,9 | 15,6 | 10,49 | 41,6 |
| ENSG00000102172 | SMS | 70,21 | 59,91 | 52,7 | 41,05 | 41,5 |
| ENSG00000255330 | AL096711,2 | 12,27 | 11,05 | 8,93 | 7,18 | 41,5 |
| ENSG00000148153 | INIP | 14,5 | 14,37 | 14,05 | 8,49 | 41,4 |
| ENSG00000103540 | CCP110 | 11,27 | 10,82 | 8,96 | 6,6 | 41,4 |
| ENSG00000132485 | ZRANB2 | 57,78 | 56,2 | 48,87 | 33,86 | 41,4 |
| ENSG00000170035 | UBE2E3 | 121,93 | 105,27 | 96,16 | 71,47 | 41,4 |
| ENSG00000125378 | BMP4 | 7,98 | 7,65 | 6,4 | 4,68 | 41,4 |

| | | | | | | |
|---|---|---|---|---|---|---|
| ENSG00000111726 | CMAS | 22,49 | 21,12 | 18,93 | 13,19 | 41,4 |
| ENSG00000147586 | MRPS28 | 39,16 | 34,09 | 26,09 | 22,97 | 41,3 |
| ENSG00000119318 | RAD23B | 45,61 | 44,43 | 43,48 | 26,77 | 41,3 |
| ENSG00000178162 | FAR2P2 | 5,09 | 5,04 | 3,46 | 2,99 | 41,3 |
| ENSG00000101052 | IFT52 | 30,79 | 28,46 | 28,44 | 18,09 | 41,2 |
| ENSG00000081721 | DUSP12 | 18,46 | 17,07 | 14,83 | 10,85 | 41,2 |
| ENSG00000174628 | IQCK | 8,98 | 8,27 | 7,93 | 5,28 | 41,2 |
| ENSG00000064102 | INTS13 | 11,88 | 10,06 | 9,43 | 6,99 | 41,2 |
| ENSG00000232859 | LYRM9 | 3,28 | 2,58 | 2,34 | 1,93 | 41,2 |
| ENSG00000133083 | DCLK1 | 21,45 | 18,65 | 15,61 | 12,63 | 41,1 |
| ENSG00000113615 | SEC24A | 9,69 | 9,54 | 8,55 | 5,71 | 41,1 |
| ENSG00000133104 | SPART | 30,13 | 29,41 | 29,09 | 17,76 | 41,1 |
| ENSG00000163848 | ZNF148 | 23,49 | 22,61 | 21,02 | 13,85 | 41,0 |
| ENSG00000109084 | TMEM97 | 61,35 | 53,14 | 44,75 | 36,18 | 41,0 |
| ENSG00000072736 | NFATC3 | 17,94 | 17,86 | 16,94 | 10,59 | 41,0 |
| ENSG00000117519 | CNN3 | 291,97 | 287,19 | 286,42 | 172,4 | 41,0 |
| ENSG00000144228 | SPOPL | 5,08 | 4,96 | 4,45 | 3 | 40,9 |
| ENSG00000180628 | PCGF5 | 7,06 | 6,68 | 5,92 | 4,17 | 40,9 |
| ENSG00000105258 | POLR2I | 53,19 | 44,49 | 43,56 | 31,44 | 40,9 |
| ENSG00000151748 | SAV1 | 11,57 | 10,6 | 10,54 | 6,84 | 40,9 |
| ENSG00000259768 | AC004943,2 | 4,21 | 3,92 | 3,28 | 2,49 | 40,9 |
| ENSG00000077514 | POLD3 | 16,89 | 16,71 | 13,04 | 9,99 | 40,9 |
| ENSG00000164024 | METAP1 | 19,76 | 18,23 | 15,63 | 11,69 | 40,8 |
| ENSG00000151778 | SERP2 | 5,24 | 5,12 | 3,92 | 3,1 | 40,8 |
| ENSG00000249102 | AC034223,1 | 3,82 | 3,1 | 2,67 | 2,26 | 40,8 |
| ENSG00000111266 | DUSP16 | 4,14 | 3,69 | 3,28 | 2,45 | 40,8 |
| ENSG00000280962 | DUSP16 | 4,14 | 3,69 | 3,28 | 2,45 | 40,8 |
| ENSG00000110172 | CHORDC1 | 13,4 | 12,15 | 8,72 | 7,93 | 40,8 |
| ENSG00000101413 | RPRD1B | 17,32 | 14,38 | 11,89 | 10,25 | 40,8 |
| ENSG00000205659 | LIN52 | 7,7 | 7,13 | 6,62 | 4,56 | 40,8 |
| ENSG00000173588 | CEP83 | 15,41 | 12,68 | 11,55 | 9,13 | 40,8 |
| ENSG00000138180 | CEP55 | 19,59 | 19,53 | 19,48 | 11,61 | 40,7 |
| ENSG00000101812 | H2BW2 | 1,13 | 1,04 | 0,93 | 0,67 | 40,7 |
| ENSG00000166226 | CCT2 | 122,85 | 106,98 | 98,35 | 72,85 | 40,7 |
| ENSG00000151465 | CDC123 | 57,85 | 55,17 | 50,71 | 34,33 | 40,7 |
| ENSG00000146731 | CCT6A | 94,44 | 93,68 | 82,49 | 56,05 | 40,7 |
| ENSG00000132825 | PPP1R3D | 1,55 | 1,43 | 1,21 | 0,92 | 40,6 |
| ENSG00000172115 | CYCS | 70,69 | 62,26 | 55,92 | 42,01 | 40,6 |
| ENSG00000197329 | PELI1 | 16,15 | 15,71 | 12,51 | 9,6 | 40,6 |
| ENSG00000184047 | DIABLO | 34,26 | 29,39 | 27,77 | 20,38 | 40,5 |
| ENSG00000164934 | DCAF13 | 39,03 | 35,92 | 29,17 | 23,22 | 40,5 |
| ENSG00000163788 | SNRK | 8,79 | 8,14 | 7,08 | 5,23 | 40,5 |
| ENSG00000082438 | COBLL1 | 3,51 | 2,57 | 2,55 | 2,09 | 40,5 |
| ENSG00000171858 | RPS21 | 496,68 | 461,38 | 437,23 | 295,76 | 40,5 |
| ENSG00000203875 | SNHG5 | 89,52 | 89,5 | 85,25 | 53,33 | 40,4 |
| ENSG00000203778 | FAM229B | 18,15 | 16,36 | 14,14 | 10,82 | 40,4 |
| ENSG00000206149 | HERC2P9 | 18,32 | 17,15 | 15,75 | 10,93 | 40,3 |

| | | | | | | |
|---|---|---|---|---|---|---|
| ENSG00000112242 | E2F3 | 14,83 | 14,6 | 13,17 | 8,85 | 40,3 |
| ENSG00000165525 | NEMF | 34,45 | 34,22 | 25,82 | 20,57 | 40,3 |
| ENSG00000065150 | IPO5 | 92,04 | 82,87 | 76,98 | 54,96 | 40,3 |
| ENSG00000087502 | ERGIC2 | 32,33 | 27,97 | 27,62 | 19,31 | 40,3 |
| ENSG00000185658 | BRWD1 | 18,02 | 17,99 | 14,48 | 10,77 | 40,2 |
| ENSG00000196227 | FAM217B | 8,83 | 7,95 | 7,59 | 5,28 | 40,2 |
| ENSG00000118873 | RAB3GAP2 | 14,72 | 14,32 | 13,88 | 8,81 | 40,1 |
| ENSG00000224287 | MSL3P1 | 2,84 | 2,58 | 2,48 | 1,7 | 40,1 |
| ENSG00000108953 | YWHAE | 386,61 | 380,54 | 366,81 | 231,45 | 40,1 |
| ENSG00000101247 | NDUFAF5 | 17 | 13,71 | 12,33 | 10,18 | 40,1 |
| ENSG00000115380 | EFEMP1 | 5,26 | 3,47 | 3,46 | 3,15 | 40,1 |
| ENSG00000137055 | PLAA | 23,69 | 22,85 | 20,86 | 14,19 | 40,1 |
| ENSG00000101109 | STK4 | 15,89 | 15,36 | 14,53 | 9,52 | 40,1 |
| ENSG00000164163 | ABCE1 | 37,65 | 33,21 | 30,61 | 22,57 | 40,1 |
| ENSG00000181915 | ADO | 11,56 | 10,85 | 10,04 | 6,93 | 40,1 |
| ENSG00000111445 | RFC5 | 18,96 | 17,3 | 15,19 | 11,37 | 40,0 |
| ENSG00000061987 | MON2 | 15,99 | 15,91 | 13,89 | 9,59 | 40,0 |
| ENSG00000198130 | HIBCH | 16,94 | 15,17 | 13,6 | 10,16 | 40,0 |
| ENSG00000106771 | TMEM245 | 19,14 | 18,87 | 16,85 | 11,48 | 40,0 |
| ENSG00000107020 | PLGRKT | 16,55 | 16,1 | 14,66 | 9,93 | 40,0 |
| ENSG00000166797 | CIAO2A | 36,28 | 33,27 | 29,96 | 21,77 | 40,0 |
| ENSG00000104549 | SQLE | 100,54 | 91,42 | 83,48 | 60,34 | 40,0 |
| ENSG00000096092 | TMEM14A | 10,61 | 10,53 | 9,38 | 6,37 | 40,0 |
| ENSG00000227345 | PARG | 22,64 | 21,35 | 19,34 | 13,6 | 39,9 |
| ENSG00000123485 | HJURP | 23,38 | 23,22 | 20,94 | 14,05 | 39,9 |
| ENSG00000261893 | CLK2 | 9,9 | 8,49 | 7,35 | 5,95 | 39,9 |
| ENSG00000136628 | EPRS1 | 38,76 | 37,39 | 30,22 | 23,31 | 39,9 |
| ENSG00000117450 | PRDX1 | 247,16 | 215,05 | 213,32 | 148,67 | 39,8 |
| ENSG00000115159 | GPD2 | 16,14 | 15,79 | 12,74 | 9,71 | 39,8 |
| ENSG00000114686 | MRPL3 | 73,98 | 69,2 | 52,84 | 44,52 | 39,8 |
| ENSG00000177189 | RPS6KA3 | 8,69 | 8,43 | 7,61 | 5,23 | 39,8 |
| ENSG00000196628 | TCF4 | 45,45 | 44,28 | 39,67 | 27,36 | 39,8 |
| ENSG00000005059 | MCUB | 10,71 | 9,54 | 9,43 | 6,45 | 39,8 |
| ENSG00000101367 | MAPRE1 | 70,56 | 68,04 | 55,13 | 42,53 | 39,7 |
| ENSG00000285312 | C6orf52 | 1,16 | 1,12 | 0,78 | 0,7 | 39,7 |
| ENSG00000137434 | C6orf52 | 1,16 | 1,12 | 0,78 | 0,7 | 39,7 |
| ENSG00000222328 | RNU2-2P | 5395,44 | 4844,67 | 4303,11 | 3256,25 | 39,6 |
| ENSG00000154518 | ATP5MC3 | 109,47 | 93,39 | 92 | 66,08 | 39,6 |
| ENSG00000108010 | GLRX3 | 61,58 | 55,13 | 54,16 | 37,19 | 39,6 |
| ENSG00000089682 | RBM41 | 15,58 | 14,52 | 10,67 | 9,41 | 39,6 |
| ENSG00000164815 | ORC5 | 11,65 | 10,6 | 9,54 | 7,04 | 39,6 |
| ENSG00000136159 | NUDT15 | 16,05 | 13,71 | 11,11 | 9,71 | 39,5 |
| ENSG00000133935 | ERG28 | 18,46 | 17 | 14,3 | 11,17 | 39,5 |
| ENSG00000183137 | CEP57L1 | 14,05 | 13,82 | 11,92 | 8,51 | 39,4 |
| ENSG00000214736 | TOMM6 | 110,88 | 99,51 | 91,66 | 67,2 | 39,4 |
| ENSG00000135018 | UBQLN1 | 50,53 | 46,5 | 42,74 | 30,63 | 39,4 |
| ENSG00000074201 | CLNS1A | 73,51 | 72,11 | 66,08 | 44,58 | 39,4 |

| | | | | | | |
|---|---|---|---|---|---|---|
| ENSG00000142166 | IFNAR1 | 15,48 | 15,32 | 14,48 | 9,39 | 39,3 |
| ENSG00000132031 | MATN3 | 5,95 | 5,63 | 5,03 | 3,61 | 39,3 |
| ENSG00000112419 | PHACTR2 | 16,43 | 15,08 | 14,43 | 9,97 | 39,3 |
| ENSG00000179195 | ZNF664 | 68,93 | 68,7 | 61,28 | 41,83 | 39,3 |
| ENSG00000143479 | DYRK3 | 3,18 | 2,76 | 2,59 | 1,93 | 39,3 |
| ENSG00000108651 | UTP6 | 28,7 | 25,52 | 23,39 | 17,44 | 39,2 |
| ENSG00000170085 | SIMC1 | 14,39 | 13,45 | 10,56 | 8,75 | 39,2 |
| ENSG00000174197 | MGA | 24,14 | 23,86 | 21,34 | 14,68 | 39,2 |
| ENSG00000243725 | TTC4 | 14,78 | 13,18 | 11,4 | 8,99 | 39,2 |
| ENSG00000285571 | AL513548,4 | 1,66 | 1,37 | 1,07 | 1,01 | 39,2 |
| ENSG00000264895 | AC006141,1 | 1,15 | 1,12 | 0,82 | 0,7 | 39,1 |
| ENSG00000283022 | HYDIN | 2,07 | 2,02 | 1,6 | 1,26 | 39,1 |
| ENSG00000157423 | HYDIN | 2,07 | 2,02 | 1,6 | 1,26 | 39,1 |
| ENSG00000121644 | DESI2 | 22,04 | 21,7 | 21,06 | 13,43 | 39,1 |
| ENSG00000156787 | TBC1D31 | 8,55 | 8,24 | 7,85 | 5,21 | 39,1 |
| ENSG00000254585 | MAGEL2 | 3,61 | 3,38 | 2,81 | 2,2 | 39,1 |
| ENSG00000004766 | VPS50 | 11,43 | 11,26 | 10,92 | 6,97 | 39,0 |
| ENSG00000197713 | RPE | 18,92 | 18,08 | 16,2 | 11,54 | 39,0 |
| ENSG00000155100 | OTUD6B | 7,41 | 6,85 | 5,62 | 4,52 | 39,0 |
| ENSG00000154114 | TBCEL | 11,54 | 10,54 | 9,64 | 7,04 | 39,0 |
| ENSG00000147601 | TERF1 | 21,34 | 21,13 | 17,15 | 13,02 | 39,0 |
| ENSG00000010244 | ZNF207 | 89,84 | 88,46 | 87,62 | 54,82 | 39,0 |
| ENSG00000091656 | ZFHX4 | 31,98 | 29,04 | 20,59 | 19,54 | 38,9 |
| ENSG00000100027 | YPEL1 | 12,65 | 12,01 | 8,78 | 7,73 | 38,9 |
| ENSG00000099194 | SCD | 212,59 | 199,04 | 197,94 | 129,95 | 38,9 |
| ENSG00000142856 | ITGB3BP | 22,3 | 16,69 | 16,16 | 13,64 | 38,8 |
| ENSG00000262473 | GART | 1,65 | 1,35 | 1,09 | 1,01 | 38,8 |
| ENSG00000005483 | KMT2E | 41,08 | 38,3 | 32,2 | 25,15 | 38,8 |
| ENSG00000180182 | MED14 | 21,98 | 20,83 | 18,7 | 13,46 | 38,8 |
| ENSG00000262771 | SSBP1 | 94,24 | 93,51 | 92,91 | 57,72 | 38,8 |
| ENSG00000149084 | HSD17B12 | 40,44 | 33,9 | 31,96 | 24,77 | 38,7 |
| ENSG00000269994 | AL513318,1 | 6,56 | 6,16 | 5,7 | 4,02 | 38,7 |
| ENSG00000198721 | ECI2 | 30,44 | 26,96 | 26,92 | 18,66 | 38,7 |
| ENSG00000166860 | ZBTB39 | 6,65 | 6,28 | 5,34 | 4,08 | 38,6 |
| ENSG00000213066 | CEP43 | 14,88 | 14,03 | 12,6 | 9,13 | 38,6 |
| ENSG00000119703 | ZC2HC1C | 1,45 | 1,18 | 1,06 | 0,89 | 38,6 |
| ENSG00000120800 | UTP20 | 9,97 | 8,55 | 7,25 | 6,12 | 38,6 |
| ENSG00000227051 | C14orf132 | 18,06 | 16,54 | 13,48 | 11,09 | 38,6 |
| ENSG00000055044 | NOP58 | 42,3 | 39,44 | 31,58 | 25,98 | 38,6 |
| ENSG00000165417 | GTF2A1 | 15,33 | 15,32 | 14,79 | 9,42 | 38,6 |
| ENSG00000123737 | EXOSC9 | 32,67 | 31,25 | 27,91 | 20,08 | 38,5 |
| ENSG00000260257 | AL035071,1 | 6,08 | 5,22 | 4,63 | 3,74 | 38,5 |
| ENSG00000152154 | TMEM178A | 9,85 | 7,62 | 7,03 | 6,06 | 38,5 |
| ENSG00000150753 | CCT5 | 196,67 | 176,4 | 156,4 | 121 | 38,5 |
| ENSG00000111639 | MRPL51 | 140,25 | 133,73 | 131,69 | 86,34 | 38,4 |
| ENSG00000088538 | DOCK3 | 2,55 | 2,13 | 1,98 | 1,57 | 38,4 |
| ENSG00000170558 | CDH2 | 112,47 | 109,69 | 105,11 | 69,26 | 38,4 |

| | | | | | | |
|---|---|---|---|---|---|---|
| ENSG00000166352 | IFTAP | 25,3 | 25,06 | 21,16 | 15,59 | 38,4 |
| ENSG00000125249 | RAP2A | 14,43 | 14,35 | 13,47 | 8,9 | 38,3 |
| ENSG00000109133 | TMEM33 | 25,26 | 24,81 | 22,84 | 15,58 | 38,3 |
| ENSG00000145736 | GTF2H2 | 5,48 | 4,93 | 4,01 | 3,38 | 38,3 |
| ENSG00000152404 | CWF19L2 | 7,83 | 7,7 | 6,05 | 4,83 | 38,3 |
| ENSG00000184117 | NIPSNAP1 | 62,02 | 59,41 | 45,03 | 38,27 | 38,3 |
| ENSG00000065883 | CDK13 | 27,82 | 26,31 | 21,67 | 17,17 | 38,3 |
| ENSG00000134297 | PLEKHA8P1 | 5,08 | 4,44 | 3,98 | 3,14 | 38,2 |
| ENSG00000116793 | PHTF1 | 26,27 | 22,1 | 21,53 | 16,24 | 38,2 |
| ENSG00000176749 | CDK5R1 | 10,74 | 10,51 | 8,09 | 6,64 | 38,2 |
| ENSG00000180423 | HARBI1 | 3,67 | 3,06 | 2,78 | 2,27 | 38,1 |
| ENSG00000196584 | XRCC2 | 8,9 | 7,65 | 7,25 | 5,51 | 38,1 |
| ENSG00000112759 | SLC29A1 | 15,47 | 14,74 | 11,57 | 9,58 | 38,1 |
| ENSG00000123607 | TTC21B | 16 | 15,71 | 15,04 | 9,91 | 38,1 |
| ENSG00000187189 | TSPYL4 | 24,82 | 23,01 | 19,36 | 15,38 | 38,0 |
| ENSG00000136450 | SRSF1 | 174,92 | 160,85 | 148,26 | 108,47 | 38,0 |
| ENSG00000110218 | PANX1 | 21,05 | 20,14 | 17,77 | 13,06 | 38,0 |
| ENSG00000144909 | OSBPL11 | 9,12 | 8,35 | 7,77 | 5,66 | 37,9 |
| ENSG00000106571 | GLI3 | 36,86 | 36,04 | 28,32 | 22,89 | 37,9 |
| ENSG00000102921 | N4BP1 | 13,96 | 12,46 | 11,92 | 8,67 | 37,9 |
| ENSG00000116698 | SMG7 | 24,95 | 24,3 | 23,78 | 15,5 | 37,9 |
| ENSG00000083312 | TNPO1 | 72,8 | 65,75 | 62,49 | 45,26 | 37,8 |
| ENSG00000132341 | RAN | 491,55 | 423,1 | 388,26 | 305,64 | 37,8 |
| ENSG00000100814 | CCNB1IP1 | 35,55 | 29,13 | 25,36 | 22,11 | 37,8 |
| ENSG00000116747 | RO60 | 36,49 | 34,9 | 31,86 | 22,71 | 37,8 |
| ENSG00000215424 | MCM3AP-AS1 | 7,39 | 6,5 | 6,13 | 4,6 | 37,8 |
| ENSG00000164548 | TRA2A | 62,17 | 57,27 | 56,01 | 38,71 | 37,7 |
| ENSG00000134690 | CDCA8 | 22,17 | 21,15 | 19,98 | 13,81 | 37,7 |
| ENSG00000145495 | MARCHF6 | 49,69 | 45,67 | 42,63 | 30,96 | 37,7 |
| ENSG00000284980 | AGPAT5 | 10,38 | 9,14 | 7,54 | 6,47 | 37,7 |
| ENSG00000155189 | AGPAT5 | 10,38 | 9,14 | 7,54 | 6,47 | 37,7 |
| ENSG00000204272 | NBDY | 31,01 | 29,5 | 28,51 | 19,33 | 37,7 |
| ENSG00000249709 | ZNF564 | 5,55 | 5,54 | 5,2 | 3,46 | 37,7 |
| ENSG00000167840 | ZNF232 | 10,42 | 9,47 | 8,64 | 6,5 | 37,6 |
| ENSG00000136937 | NCBP1 | 25,52 | 21,64 | 18,99 | 15,92 | 37,6 |
| ENSG00000260708 | AL118516,1 | 4,23 | 3,95 | 3,47 | 2,64 | 37,6 |
| ENSG00000198732 | SMOC1 | 19,47 | 18,24 | 12,38 | 12,16 | 37,5 |
| ENSG00000141425 | RPRD1A | 26,29 | 24,84 | 24 | 16,42 | 37,5 |
| ENSG00000149547 | EI24 | 48,55 | 42,97 | 40 | 30,34 | 37,5 |
| ENSG00000155906 | RMND1 | 12,54 | 11,85 | 10,67 | 7,84 | 37,5 |
| ENSG00000151779 | NBAS | 17,8 | 17,12 | 15,84 | 11,13 | 37,5 |
| ENSG00000163811 | WDR43 | 28,11 | 25,64 | 21,14 | 17,58 | 37,5 |
| ENSG00000174804 | FZD4 | 2,03 | 1,77 | 1,52 | 1,27 | 37,4 |
| ENSG00000080608 | PUM3 | 23,11 | 20,58 | 17,91 | 14,46 | 37,4 |
| ENSG00000120616 | EPC1 | 17,1 | 14,89 | 14,68 | 10,7 | 37,4 |
| ENSG00000163867 | ZMYM6 | 6,36 | 6,2 | 5,29 | 3,98 | 37,4 |
| ENSG00000026652 | AGPAT4 | 12,14 | 9,94 | 9,73 | 7,6 | 37,4 |

| | | | | | | |
|---|---|---|---|---|---|---|
| ENSG00000171960 | PPIH | 35,64 | 29,42 | 27,87 | 22,32 | 37,4 |
| ENSG00000158402 | CDC25C | 11,75 | 11,45 | 9,82 | 7,36 | 37,4 |
| ENSG00000118007 | STAG1 | 22,27 | 22,26 | 18,74 | 13,95 | 37,4 |
| ENSG00000179454 | KLHL28 | 5,95 | 5,28 | 4,92 | 3,73 | 37,3 |
| ENSG00000132434 | LANCL2 | 9,89 | 8,98 | 7,68 | 6,2 | 37,3 |
| ENSG00000187742 | SECISBP2 | 26,47 | 26,13 | 22,87 | 16,6 | 37,3 |
| ENSG00000142731 | PLK4 | 29,34 | 27,87 | 23,8 | 18,4 | 37,3 |
| ENSG00000165675 | ENOX2 | 13,5 | 12,81 | 10,72 | 8,47 | 37,3 |
| ENSG00000081019 | RSBN1 | 9,24 | 7,88 | 7,3 | 5,8 | 37,2 |
| ENSG00000129493 | HEATR5A | 10,25 | 10,05 | 9 | 6,44 | 37,2 |
| ENSG00000166348 | USP54 | 16,54 | 15,25 | 15,18 | 10,41 | 37,1 |
| ENSG00000165060 | FXN | 12,71 | 12,28 | 11,5 | 8 | 37,1 |
| ENSG00000234741 | GAS5 | 94,93 | 93,97 | 85,15 | 59,76 | 37,0 |
| ENSG00000172795 | DCP2 | 22,31 | 21,41 | 20,45 | 14,05 | 37,0 |
| ENSG00000186329 | TMEM212 | 13,46 | 12,88 | 10,82 | 8,48 | 37,0 |
| ENSG00000241343 | RPL36A | 448,7 | 397,2 | 371,59 | 282,71 | 37,0 |
| ENSG00000224281 | SLC25A5-AS1 | 1,19 | 1,13 | 1 | 0,75 | 37,0 |
| ENSG00000184220 | CMSS1 | 26,63 | 25,76 | 22 | 16,79 | 37,0 |
| ENSG00000101407 | TTI1 | 17,08 | 16,65 | 15,06 | 10,77 | 36,9 |
| ENSG00000131844 | MCCC2 | 9,38 | 8,78 | 7,46 | 5,92 | 36,9 |
| ENSG00000144366 | GULP1 | 28,25 | 23,78 | 21,1 | 17,83 | 36,9 |
| ENSG00000092208 | GEMIN2 | 16,05 | 13,79 | 12,08 | 10,13 | 36,9 |
| ENSG00000196476 | C20orf96 | 6,86 | 6,22 | 5,77 | 4,33 | 36,9 |
| ENSG00000136925 | TSTD2 | 12,94 | 12,47 | 10,83 | 8,18 | 36,8 |
| ENSG00000139517 | LNX2 | 3,29 | 3,12 | 2,45 | 2,08 | 36,8 |
| ENSG00000276180 | H4C9 | 108,36 | 73,51 | 71,6 | 68,51 | 36,8 |
| ENSG00000112079 | STK38 | 15,23 | 15,06 | 12,53 | 9,63 | 36,8 |
| ENSG00000151092 | NGLY1 | 14,09 | 13,61 | 11,87 | 8,91 | 36,8 |
| ENSG00000134444 | RELCH | 11,22 | 11,06 | 9,5 | 7,1 | 36,7 |
| ENSG00000285967 | NIPBL-DT | 7,71 | 6,95 | 5,65 | 4,88 | 36,7 |
| ENSG00000184319 | RPL23AP82 | 18,83 | 18,19 | 17,93 | 11,92 | 36,7 |
| ENSG00000094916 | CBX5 | 152,36 | 151,69 | 123,95 | 96,54 | 36,6 |
| ENSG00000131747 | TOP2A | 99,47 | 96,78 | 91,02 | 63,05 | 36,6 |
| ENSG00000214050 | FBXO16 | 4,7 | 4,41 | 3,64 | 2,98 | 36,6 |
| ENSG00000129595 | EPB41L4A | 13,33 | 12,21 | 9,08 | 8,46 | 36,5 |
| ENSG00000066933 | MYO9A | 13,73 | 12,66 | 11,24 | 8,72 | 36,5 |
| ENSG00000115484 | CCT4 | 100,03 | 88,52 | 81,59 | 63,56 | 36,5 |
| ENSG00000239900 | ADSL | 37,12 | 33,52 | 32,62 | 23,59 | 36,4 |
| ENSG00000125885 | MCM8 | 13,62 | 12,37 | 11,49 | 8,66 | 36,4 |
| ENSG00000065911 | MTHFD2 | 39,3 | 27,41 | 25,49 | 24,99 | 36,4 |
| ENSG00000227953 | LINC01341 | 1,54 | 1,41 | 1,38 | 0,98 | 36,4 |
| ENSG00000056736 | IL17RB | 1,54 | 1,47 | 1,37 | 0,98 | 36,4 |
| ENSG00000159111 | MRPL10 | 11,22 | 10,84 | 10,55 | 7,14 | 36,4 |
| ENSG00000196586 | MYO6 | 8,64 | 8,48 | 7,09 | 5,5 | 36,3 |
| ENSG00000261098 | AP000766,1 | 1,02 | 0,97 | 0,71 | 0,65 | 36,3 |
| ENSG00000170153 | RNF150 | 4,77 | 4,49 | 3,66 | 3,04 | 36,3 |
| ENSG00000072849 | DERL2 | 35,63 | 35,42 | 30,68 | 22,72 | 36,2 |

| | | | | | | |
|---|---|---|---|---|---|---|
| ENSG00000110958 | PTGES3 | 161,85 | 156,67 | 143,19 | 103,29 | 36,2 |
| ENSG00000116984 | MTR | 14,1 | 13,9 | 10,8 | 9 | 36,2 |
| ENSG00000155438 | NIFK | 46,57 | 42,95 | 37,08 | 29,73 | 36,2 |
| ENSG00000169155 | ZBTB43 | 5,2 | 5,06 | 4,96 | 3,32 | 36,2 |
| ENSG00000142149 | HUNK | 4,85 | 3,46 | 3,14 | 3,1 | 36,1 |
| ENSG00000188613 | NANOS1 | 2,55 | 2,3 | 1,72 | 1,63 | 36,1 |
| ENSG00000106853 | PTGR1 | 25,34 | 24,66 | 22,2 | 16,2 | 36,1 |
| ENSG00000227782 | AC002553,1 | 1,5 | 1,04 | 1,01 | 0,96 | 36,0 |
| ENSG00000173660 | UQCRH | 145,55 | 142,59 | 141,22 | 93,16 | 36,0 |
| ENSG00000072210 | ALDH3A2 | 27,37 | 23,44 | 22,58 | 17,52 | 36,0 |
| ENSG00000117016 | RIMS3 | 5,68 | 5,36 | 3,91 | 3,64 | 35,9 |
| ENSG00000171951 | SCG2 | 8,02 | 7,68 | 5,79 | 5,14 | 35,9 |
| ENSG00000164603 | BMT2 | 6,22 | 6,19 | 5,4 | 3,99 | 35,9 |
| ENSG00000112294 | ALDH5A1 | 3,91 | 3,76 | 3,33 | 2,51 | 35,8 |
| ENSG00000136492 | BRIP1 | 11,01 | 9,93 | 9,43 | 7,07 | 35,8 |
| ENSG00000233183 | AL138889,1 | 1,23 | 1,22 | 0,94 | 0,79 | 35,8 |
| ENSG00000131437 | KIF3A | 12,19 | 11,56 | 8,37 | 7,83 | 35,8 |
| ENSG00000116106 | EPHA4 | 26,09 | 25,67 | 23,69 | 16,76 | 35,8 |
| ENSG00000134057 | CCNB1 | 106,74 | 105,82 | 97,72 | 68,57 | 35,8 |
| ENSG00000141698 | NT5C3B | 34,57 | 32,56 | 31,29 | 22,22 | 35,7 |
| ENSG00000163082 | SGPP2 | 1,12 | 1,09 | 0,73 | 0,72 | 35,7 |
| ENSG00000134077 | THUMPD3 | 27,73 | 27,28 | 23,78 | 17,83 | 35,7 |
| ENSG00000261140 | AC093525,4 | 2,92 | 2,55 | 2,05 | 1,88 | 35,6 |
| ENSG00000127720 | METTL25 | 4,13 | 4 | 3,85 | 2,66 | 35,6 |
| ENSG00000122026 | RPL21 | 582,98 | 550,15 | 504,45 | 375,53 | 35,6 |
| ENSG00000245970 | AP003352,1 | 1,66 | 1,53 | 1,2 | 1,07 | 35,5 |
| ENSG00000214013 | GANC | 6,5 | 6,21 | 6,05 | 4,19 | 35,5 |
| ENSG00000271122 | AC018647,2 | 3,63 | 3,34 | 3,14 | 2,34 | 35,5 |
| ENSG00000163597 | SNHG16 | 59,36 | 55,63 | 49,82 | 38,27 | 35,5 |
| ENSG00000103035 | PSMD7 | 55,46 | 53,08 | 49,78 | 35,79 | 35,5 |
| ENSG00000163938 | GNL3 | 44,01 | 39,11 | 31,67 | 28,41 | 35,4 |
| ENSG00000141404 | GNAL | 8,24 | 8,18 | 7,14 | 5,32 | 35,4 |
| ENSG00000141198 | TOM1L1 | 9,37 | 8,91 | 7,95 | 6,05 | 35,4 |
| ENSG00000198252 | STYX | 6,78 | 6,4 | 4,93 | 4,38 | 35,4 |
| ENSG00000273314 | AC005229,4 | 1,67 | 1,55 | 1,33 | 1,08 | 35,3 |
| ENSG00000001460 | STPG1 | 3,51 | 2,88 | 2,78 | 2,27 | 35,3 |
| ENSG00000244055 | AC007566,1 | 1,7 | 1,35 | 1,3 | 1,1 | 35,3 |
| ENSG00000237298 | TTN-AS1 | 21,99 | 19,42 | 19,03 | 14,23 | 35,3 |
| ENSG00000129159 | KCNC1 | 2,16 | 1,62 | 1,45 | 1,4 | 35,2 |
| ENSG00000078674 | PCM1 | 96,47 | 81,37 | 75,34 | 62,53 | 35,2 |
| ENSG00000285023 | CHORDC1 | 13,4 | 12,15 | 8,72 | 8,69 | 35,1 |
| ENSG00000065548 | ZC3H15 | 42,31 | 39,67 | 34,7 | 27,46 | 35,1 |
| ENSG00000135597 | REPS1 | 22,68 | 21,27 | 20,29 | 14,72 | 35,1 |
| ENSG00000139726 | DENR | 35,98 | 35 | 30,52 | 23,36 | 35,1 |
| ENSG00000070061 | ELP1 | 26 | 21,88 | 18,44 | 16,91 | 35,0 |
| ENSG00000158528 | PPP1R9A | 10,54 | 9,54 | 8,11 | 6,86 | 34,9 |
| ENSG00000276966 | H4C5 | 259,74 | 215,95 | 200,32 | 169,1 | 34,9 |

| | | | | | | |
|---|---|---|---|---|---|---|
| ENSG00000163875 | MEAF6 | 45,31 | 42,5 | 41,62 | 29,5 | 34,9 |
| ENSG00000017260 | ATP2C1 | 30,48 | 30,09 | 25,2 | 19,86 | 34,8 |
| ENSG00000172954 | LCLAT1 | 11,26 | 10,71 | 8,94 | 7,34 | 34,8 |
| ENSG00000152990 | ADGRA3 | 26,14 | 25,79 | 24,71 | 17,04 | 34,8 |
| ENSG00000119203 | CPSF3 | 38,47 | 35,63 | 27,48 | 25,08 | 34,8 |
| ENSG00000072756 | TRNT1 | 13,85 | 13,28 | 10,39 | 9,03 | 34,8 |
| ENSG00000183340 | JRKL | 4,95 | 4,64 | 3,91 | 3,23 | 34,7 |
| ENSG00000163933 | RFT1 | 8,35 | 7,42 | 6,8 | 5,45 | 34,7 |
| ENSG00000204177 | BMS1P1 | 6,08 | 5,89 | 4,89 | 3,97 | 34,7 |
| ENSG00000125743 | SNRPD2 | 190,45 | 184,02 | 181,87 | 124,37 | 34,7 |
| ENSG00000132286 | TIMM10B | 13,52 | 12,74 | 12,32 | 8,83 | 34,7 |
| ENSG00000113811 | SELENOK | 33,04 | 30,92 | 30,61 | 21,58 | 34,7 |
| ENSG00000116898 | MRPS15 | 42,93 | 41,29 | 39,21 | 28,06 | 34,6 |
| ENSG00000051825 | MPHOSPH9 | 32,39 | 31,94 | 28,82 | 21,18 | 34,6 |
| ENSG00000137996 | RTCA | 20,79 | 20,1 | 16,63 | 13,6 | 34,6 |
| ENSG00000106772 | PRUNE2 | 6,55 | 5,85 | 5,49 | 4,29 | 34,5 |
| ENSG00000128050 | PAICS | 105,05 | 99,51 | 87,11 | 68,85 | 34,5 |
| ENSG00000114388 | NPRL2 | 8,07 | 7,72 | 6,93 | 5,29 | 34,4 |
| ENSG00000113013 | HSPA9 | 73,58 | 72,64 | 61,38 | 48,28 | 34,4 |
| ENSG00000187736 | NHEJ1 | 10,56 | 10,13 | 7,59 | 6,93 | 34,4 |
| ENSG00000101935 | AMMECR1 | 9,6 | 8,95 | 6,75 | 6,3 | 34,4 |
| ENSG00000188725 | SMIM15 | 29,17 | 26,3 | 24,26 | 19,16 | 34,3 |
| ENSG00000121022 | COPS5 | 35,72 | 34,8 | 33,05 | 23,47 | 34,3 |
| ENSG00000107566 | ERLIN1 | 10,18 | 9,66 | 9,48 | 6,69 | 34,3 |
| ENSG00000175161 | CADM2 | 2,6 | 2,4 | 2 | 1,71 | 34,2 |
| ENSG00000167377 | ZNF23 | 5,47 | 5,24 | 4,49 | 3,6 | 34,2 |
| ENSG00000115840 | SLC25A12 | 8,6 | 8,53 | 7,84 | 5,66 | 34,2 |
| ENSG00000184675 | AMER1 | 4,74 | 4,36 | 3,3 | 3,12 | 34,2 |
| ENSG00000165996 | HACD1 | 10,72 | 10,19 | 9,14 | 7,06 | 34,1 |
| ENSG00000115875 | SRSF7 | 124,89 | 117,97 | 90,58 | 82,27 | 34,1 |
| ENSG00000105649 | RAB3A | 3,4 | 2,81 | 2,52 | 2,24 | 34,1 |
| ENSG00000287306 | AC016821,1 | 11,23 | 11,03 | 9,75 | 7,4 | 34,1 |
| ENSG00000228284 | HLA-DQA1 | 8,04 | 7,71 | 7,2 | 5,3 | 34,1 |
| ENSG00000141279 | NPEPPS | 59,47 | 59,28 | 54,3 | 39,21 | 34,1 |
| ENSG00000135390 | ATP5MC2 | 203,53 | 188,14 | 183,91 | 134,22 | 34,1 |
| ENSG00000203872 | C6orf163 | 2,38 | 1,97 | 1,85 | 1,57 | 34,0 |
| ENSG00000158615 | PPP1R15B | 15,88 | 15,36 | 13,59 | 10,48 | 34,0 |
| ENSG00000186638 | KIF24 | 4,5 | 4,12 | 4 | 2,97 | 34,0 |
| ENSG00000077312 | SNRPA | 62,52 | 55,99 | 55,73 | 41,28 | 34,0 |
| ENSG00000012174 | MBTPS2 | 7,83 | 7,61 | 6,6 | 5,17 | 34,0 |
| ENSG00000274746 | ZNF100 | 4,74 | 3,9 | 3,41 | 3,13 | 34,0 |
| ENSG00000176695 | OR4F17 | 41,66 | 41,39 | 30,96 | 27,51 | 34,0 |
| ENSG00000172572 | PDE3A | 4,33 | 4,06 | 3,83 | 2,86 | 33,9 |
| ENSG00000278705 | H4C2 | 637,35 | 436,69 | 433,5 | 421,04 | 33,9 |
| ENSG00000071082 | RPL31 | 674,36 | 673,89 | 664,02 | 445,94 | 33,9 |
| ENSG00000158292 | GPR153 | 4,55 | 3,69 | 3,02 | 3,01 | 33,8 |
| ENSG00000164114 | MAP9 | 11,7 | 11,08 | 9,09 | 7,74 | 33,8 |

| | | | | | | |
|---|---|---|---|---|---|---|
| ENSG00000198554 | WDHD1 | 20,02 | 19,56 | 15,84 | 13,25 | 33,8 |
| ENSG00000260855 | AL591848,3 | 1,51 | 1,27 | 1,16 | 1 | 33,8 |
| ENSG00000103160 | HSDL1 | 19,69 | 17,97 | 16,93 | 13,04 | 33,8 |
| ENSG00000004866 | ST7 | 11,26 | 10,67 | 10,21 | 7,46 | 33,7 |
| ENSG00000269416 | LINC01224 | 10,62 | 10,39 | 9,8 | 7,04 | 33,7 |
| ENSG00000105173 | CCNE1 | 8,52 | 7,46 | 6,6 | 5,65 | 33,7 |
| ENSG00000278053 | DDX52 | 23,4 | 21,46 | 18,47 | 15,52 | 33,7 |
| ENSG00000146842 | TMEM209 | 14,53 | 14,33 | 12,79 | 9,64 | 33,7 |
| ENSG00000188321 | ZNF559 | 10,8 | 9,42 | 9,19 | 7,17 | 33,6 |
| ENSG00000280832 | GSEC | 2,59 | 2,26 | 1,83 | 1,72 | 33,6 |
| ENSG00000112078 | KCTD20 | 26,07 | 25,87 | 24,32 | 17,32 | 33,6 |
| ENSG00000138386 | NAB1 | 26,84 | 26,5 | 21,92 | 17,84 | 33,5 |
| ENSG00000196233 | LCOR | 25,95 | 23,87 | 21,09 | 17,25 | 33,5 |
| ENSG00000112592 | TBP | 9,14 | 7,64 | 7,14 | 6,08 | 33,5 |
| ENSG00000060688 | SNRNP40 | 43,17 | 38,7 | 36,66 | 28,73 | 33,4 |
| ENSG00000236018 | AC004898,1 | 1,71 | 1,32 | 1,22 | 1,14 | 33,3 |
| ENSG00000271855 | AC073195,1 | 1,02 | 0,9 | 0,74 | 0,68 | 33,3 |
| ENSG00000197603 | CPLANE1 | 15,68 | 15,52 | 14,07 | 10,46 | 33,3 |
| ENSG00000248049 | UBA6-AS1 | 6,82 | 6,61 | 5,75 | 4,55 | 33,3 |
| ENSG00000070501 | POLB | 25,09 | 22,43 | 17,41 | 16,74 | 33,3 |
| ENSG00000149313 | AASDHPPT | 30,93 | 30,9 | 25,93 | 20,64 | 33,3 |
| ENSG00000005801 | ZNF195 | 27,45 | 22,85 | 20,1 | 18,32 | 33,3 |
| ENSG00000117461 | PIK3R3 | 27,04 | 25,21 | 20,83 | 18,05 | 33,2 |
| ENSG00000247556 | OIP5-AS1 | 32,76 | 31,73 | 29,45 | 21,87 | 33,2 |
| ENSG00000001167 | NFYA | 15,68 | 15,62 | 12,76 | 10,47 | 33,2 |
| ENSG00000143756 | FBXO28 | 14,31 | 14,18 | 12,16 | 9,56 | 33,2 |
| ENSG00000163618 | CADPS | 10,83 | 10,43 | 7,93 | 7,24 | 33,1 |
| ENSG00000101464 | PIGU | 16,39 | 14,29 | 13,34 | 10,96 | 33,1 |
| ENSG00000135521 | LTV1 | 17,12 | 16,55 | 13,1 | 11,45 | 33,1 |
| ENSG00000143195 | ILDR2 | 11,95 | 11,81 | 8,39 | 8 | 33,1 |
| ENSG00000165868 | HSPA12A | 6,9 | 5,84 | 5,23 | 4,62 | 33,0 |
| ENSG00000144407 | PTH2R | 1,12 | 1,09 | 0,93 | 0,75 | 33,0 |
| ENSG00000178150 | ZNF114 | 6,72 | 6,03 | 5,02 | 4,5 | 33,0 |
| ENSG00000087995 | METTL2A | 17,08 | 14,01 | 12,26 | 11,44 | 33,0 |
| ENSG00000131849 | ZNF132 | 1,06 | 1,01 | 0,87 | 0,71 | 33,0 |
| ENSG00000274047 | SYNRG | 7,42 | 6,29 | 5,07 | 4,97 | 33,0 |
| ENSG00000144635 | DYNC1LI1 | 35,66 | 31,34 | 27,21 | 23,9 | 33,0 |
| ENSG00000100225 | FBXO7 | 35,24 | 34,84 | 33,73 | 23,62 | 33,0 |
| ENSG00000103018 | CYB5B | 78,66 | 73,07 | 69,01 | 52,73 | 33,0 |
| ENSG00000074696 | HACD3 | 76,27 | 72,91 | 63,92 | 51,15 | 32,9 |
| ENSG00000185621 | LMLN | 4,16 | 4,14 | 3,19 | 2,79 | 32,9 |
| ENSG00000128654 | MTX2 | 15,46 | 15,25 | 14,31 | 10,37 | 32,9 |
| ENSG00000125107 | CNOT1 | 55,84 | 51,66 | 48,57 | 37,46 | 32,9 |
| ENSG00000219481 | NBPF1 | 40,16 | 34,23 | 27,36 | 26,95 | 32,9 |
| ENSG00000172264 | MACROD2 | 4,41 | 3,39 | 3,33 | 2,96 | 32,9 |
| ENSG00000174720 | LARP7 | 33,3 | 31,16 | 24,52 | 22,36 | 32,9 |
| ENSG00000144730 | IL17RD | 99,51 | 97,98 | 78,44 | 66,86 | 32,8 |

| | | | | | | |
|---|---|---|---|---|---|---|
| ENSG00000151461 | UPF2 | 10,15 | 9,89 | 7,93 | 6,82 | 32,8 |
| ENSG00000167315 | ACAA2 | 51,21 | 45,43 | 41,85 | 34,41 | 32,8 |
| ENSG00000165821 | SALL2 | 60,57 | 60,25 | 46,39 | 40,7 | 32,8 |
| ENSG00000033867 | SLC4A7 | 23,29 | 21,09 | 15,76 | 15,65 | 32,8 |
| ENSG00000168916 | ZNF608 | 22,9 | 22,57 | 20 | 15,39 | 32,8 |
| ENSG00000170364 | SETMAR | 16,41 | 14,95 | 13,12 | 11,03 | 32,8 |
| ENSG00000145919 | BOD1 | 55,02 | 52,68 | 48,12 | 36,99 | 32,8 |
| ENSG00000138640 | FAM13A | 9,46 | 9,15 | 7,17 | 6,36 | 32,8 |
| ENSG00000196267 | ZNF836 | 5,59 | 4,59 | 4,51 | 3,76 | 32,7 |
| ENSG00000198876 | DCAF12 | 16,13 | 15,05 | 15,04 | 10,86 | 32,7 |
| ENSG00000198648 | STK39 | 18,74 | 18,54 | 16,12 | 12,62 | 32,7 |
| ENSG00000165152 | PGAP4 | 5,77 | 5,57 | 5,54 | 3,89 | 32,6 |
| ENSG00000264247 | LINC00909 | 5,68 | 4,92 | 3,84 | 3,83 | 32,6 |
| ENSG00000180233 | ZNRF2 | 2,8 | 2,42 | 1,96 | 1,89 | 32,5 |
| ENSG00000144120 | TMEM177 | 7,48 | 6,77 | 6,65 | 5,05 | 32,5 |
| ENSG00000227739 | TUBB | 226,41 | 212,44 | 191,72 | 152,87 | 32,5 |
| ENSG00000175344 | CHRNA7 | 4,9 | 4,53 | 3,56 | 3,31 | 32,4 |
| ENSG00000233757 | AC092835,1 | 5,99 | 4,99 | 4,46 | 4,05 | 32,4 |
| ENSG00000029364 | SLC39A9 | 15,67 | 15,35 | 15,32 | 10,6 | 32,4 |
| ENSG00000069329 | VPS35 | 50,43 | 47,8 | 43,86 | 34,12 | 32,3 |
| ENSG00000165732 | DDX21 | 26,54 | 23,87 | 18,73 | 17,96 | 32,3 |
| ENSG00000125445 | MRPS7 | 36,99 | 35,27 | 34,95 | 25,04 | 32,3 |
| ENSG00000198755 | RPL10A | 518,54 | 480,82 | 447,28 | 351,1 | 32,3 |
| ENSG00000102781 | KATNAL1 | 12,16 | 11,52 | 10,65 | 8,24 | 32,2 |
| ENSG00000117593 | DARS2 | 14,93 | 13,54 | 12,67 | 10,12 | 32,2 |
| ENSG00000285943 | AC112128,1 | 1,46 | 1,25 | 1,21 | 0,99 | 32,2 |
| ENSG00000175193 | PARL | 26,21 | 25,75 | 24,62 | 17,78 | 32,2 |
| ENSG00000187699 | C2orf88 | 1,68 | 1,27 | 1,15 | 1,14 | 32,1 |
| ENSG00000215712 | TMEM242 | 4,3 | 4 | 3,56 | 2,92 | 32,1 |
| ENSG00000154001 | PPP2R5E | 28,45 | 27,82 | 24,88 | 19,33 | 32,1 |
| ENSG00000160752 | FDPS | 260,84 | 224,6 | 195,96 | 177,25 | 32,0 |
| ENSG00000120334 | CENPL | 9,4 | 7,85 | 7,78 | 6,39 | 32,0 |
| ENSG00000085840 | ORC1 | 9,9 | 8,17 | 7,58 | 6,73 | 32,0 |
| ENSG00000162378 | ZYG11B | 12,32 | 12,14 | 11,96 | 8,38 | 32,0 |
| ENSG00000113318 | MSH3 | 9,61 | 9,26 | 7,8 | 6,54 | 31,9 |
| ENSG00000133687 | TMTC1 | 3,32 | 3,22 | 2,78 | 2,26 | 31,9 |
| ENSG00000166130 | IKBIP | 15,63 | 15,28 | 14,31 | 10,65 | 31,9 |
| ENSG00000112701 | SENP6 | 46,47 | 44,54 | 38,5 | 31,67 | 31,8 |
| ENSG00000283447 | NDUFS1 | 17,87 | 17,29 | 13,05 | 12,18 | 31,8 |
| ENSG00000127947 | PTPN12 | 38,19 | 35,39 | 28,9 | 26,04 | 31,8 |
| ENSG00000242110 | AMACR | 6,35 | 5,95 | 5,94 | 4,33 | 31,8 |
| ENSG00000114503 | NCBP2 | 45,11 | 44,49 | 37,31 | 30,77 | 31,8 |
| ENSG00000170584 | NUDCD2 | 15,75 | 13,79 | 13,41 | 10,75 | 31,7 |
| ENSG00000059728 | MXD1 | 1,67 | 1,21 | 1,18 | 1,14 | 31,7 |
| ENSG00000067798 | NAV3 | 18,45 | 15,34 | 13,05 | 12,6 | 31,7 |
| ENSG00000164164 | OTUD4 | 11,59 | 11,42 | 10,28 | 7,92 | 31,7 |
| ENSG00000138592 | USP8 | 27,3 | 27,18 | 22,16 | 18,66 | 31,6 |

| | | | | | | |
|---|---|---|---|---|---|---|
| ENSG00000158161 | EYA3 | 12,36 | 11,04 | 10,77 | 8,45 | 31,6 |
| ENSG00000131876 | SNRPA1 | 38,97 | 38,56 | 31,43 | 26,65 | 31,6 |
| ENSG00000156011 | PSD3 | 15,89 | 15,29 | 15,18 | 10,87 | 31,6 |
| ENSG00000120802 | TMPO | 145,37 | 144 | 120,72 | 99,53 | 31,5 |
| ENSG00000224109 | CENPVL3 | 3,05 | 2,76 | 2,2 | 2,09 | 31,5 |
| ENSG00000251201 | TMED7-TICAM2 | 2,13 | 1,85 | 1,83 | 1,46 | 31,5 |
| ENSG00000170185 | USP38 | 8,46 | 8,27 | 7,17 | 5,8 | 31,4 |
| ENSG00000166822 | TMEM170A | 10,88 | 10,71 | 10,33 | 7,46 | 31,4 |
| ENSG00000239039 | SNORD13 | 3868,6 | 3555,06 | 2822,5 | 2652,86 | 31,4 |
| ENSG00000095951 | HIVEP1 | 14,26 | 13,76 | 11,23 | 9,79 | 31,3 |
| ENSG00000086200 | IPO11 | 12,29 | 10,81 | 10,62 | 8,44 | 31,3 |
| ENSG00000196712 | NF1 | 23,72 | 23,62 | 22,38 | 16,29 | 31,3 |
| ENSG00000163389 | POGLUT1 | 8,16 | 6,8 | 6,41 | 5,61 | 31,3 |
| ENSG00000267811 | AP001160,2 | 2,21 | 2,06 | 1,77 | 1,52 | 31,2 |
| ENSG00000151729 | SLC25A4 | 8,05 | 7,57 | 7,38 | 5,54 | 31,2 |
| ENSG00000175155 | YPEL2 | 1,64 | 1,57 | 1,34 | 1,13 | 31,1 |
| ENSG00000185238 | PRMT3 | 7,21 | 7,13 | 6,36 | 4,97 | 31,1 |
| ENSG00000141337 | ARSG | 3,51 | 3,22 | 2,54 | 2,42 | 31,1 |
| ENSG00000119125 | GDA | 2,48 | 2,14 | 1,81 | 1,71 | 31,0 |
| ENSG00000144597 | EAF1 | 11,31 | 10,17 | 9,04 | 7,8 | 31,0 |
| ENSG00000115944 | COX7A2L | 46,98 | 45,99 | 44,7 | 32,4 | 31,0 |
| ENSG00000170954 | ZNF415 | 5,74 | 4,86 | 4,37 | 3,96 | 31,0 |
| ENSG00000064651 | SLC12A2 | 13,62 | 13,2 | 12,35 | 9,4 | 31,0 |
| ENSG00000198521 | ZNF43 | 24,6 | 23,87 | 23,26 | 17 | 30,9 |
| ENSG00000008018 | PSMB1 | 108,69 | 105,88 | 103,74 | 75,13 | 30,9 |
| ENSG00000140740 | UQCRC2 | 54,35 | 53,25 | 49,22 | 37,63 | 30,8 |
| ENSG00000126947 | ARMCX1 | 15,37 | 15,02 | 13,15 | 10,65 | 30,7 |
| ENSG00000107669 | ATE1 | 8,13 | 7,4 | 6,88 | 5,64 | 30,6 |
| ENSG00000168002 | POLR2G | 64,8 | 63,53 | 61,11 | 44,97 | 30,6 |
| ENSG00000135069 | PSAT1 | 47,58 | 39,4 | 38,03 | 33,02 | 30,6 |
| ENSG00000120256 | LRP11 | 10,82 | 10,18 | 8,58 | 7,51 | 30,6 |
| ENSG00000196305 | IARS1 | 54,3 | 48,21 | 38,8 | 37,69 | 30,6 |
| ENSG00000182923 | CEP63 | 14,13 | 14,02 | 10,55 | 9,81 | 30,6 |
| ENSG00000197498 | RPF2 | 14,63 | 10,99 | 10,96 | 10,16 | 30,6 |
| ENSG00000101391 | CDK5RAP1 | 13,13 | 12,5 | 10,78 | 9,12 | 30,5 |
| ENSG00000071539 | TRIP13 | 20,25 | 19,38 | 18,45 | 14,07 | 30,5 |
| ENSG00000143224 | PPOX | 10,11 | 9,06 | 7,21 | 7,03 | 30,5 |
| ENSG00000140694 | PARN | 11,07 | 9,75 | 9,26 | 7,7 | 30,4 |
| ENSG00000135698 | MPHOSPH6 | 30,32 | 28,8 | 26,32 | 21,09 | 30,4 |
| ENSG00000187372 | PCDHB13 | 1,38 | 1,37 | 1,14 | 0,96 | 30,4 |
| ENSG00000183889 | AC138969,1 | 25,83 | 25,17 | 18,79 | 17,97 | 30,4 |
| ENSG00000229117 | RPL41 | 2259,12 | 2104,61 | 2040,93 | 1572,29 | 30,4 |
| ENSG00000102780 | DGKH | 3,75 | 3,27 | 2,62 | 2,61 | 30,4 |
| ENSG00000280651 | AC156455,2 | 2,21 | 1,96 | 1,7 | 1,54 | 30,3 |
| ENSG00000256546 | AC156455,1 | 2,21 | 1,96 | 1,7 | 1,54 | 30,3 |
| ENSG00000123213 | NLN | 24,95 | 22,12 | 18,71 | 17,4 | 30,3 |
| ENSG00000109971 | HSPA8 | 573,3 | 527,57 | 401,78 | 400,08 | 30,2 |

| | | | | | | |
|---|---|---|---|---|---|---|
| ENSG00000050130 | JKAMP | 21,47 | 19,81 | 19,21 | 14,99 | 30,2 |
| ENSG00000274265 | AC245297,3 | 15,04 | 14,11 | 12,34 | 10,51 | 30,1 |
| ENSG00000243406 | MRPS31P5 | 5,69 | 5,06 | 4,49 | 3,98 | 30,1 |
| ENSG00000089220 | PEBP1 | 129,13 | 129,06 | 124,44 | 90,36 | 30,0 |
| ENSG00000162148 | PPP1R32 | 1,2 | 1,12 | 1,06 | 0,84 | 30,0 |
| ENSG00000234684 | SDCBP2-AS1 | 4,3 | 4,14 | 3,77 | 3,01 | 30,0 |
| ENSG00000173065 | FAM222B | 13,88 | 12,98 | 10,66 | 9,72 | 30,0 |
| ENSG00000147121 | KRBOX4 | 9,58 | 9,23 | 7,54 | 6,71 | 30,0 |
| ENSG00000112200 | ZNF451 | 20,45 | 18,9 | 18,61 | 14,33 | 29,9 |
| ENSG00000186814 | ZSCAN30 | 9,98 | 9,18 | 8,52 | 7 | 29,9 |
| ENSG00000165494 | PCF11 | 24,15 | 21,9 | 21,26 | 16,94 | 29,9 |
| ENSG00000198408 | OGA | 63,15 | 62,4 | 52,59 | 44,33 | 29,8 |
| ENSG00000116095 | PLEKHA3 | 9,2 | 7,49 | 7,03 | 6,46 | 29,8 |
| ENSG00000043355 | ZIC2 | 51,17 | 45,52 | 37,43 | 35,94 | 29,8 |
| ENSG00000255052 | FAM66D | 2,96 | 2,8 | 2,3 | 2,08 | 29,7 |
| ENSG00000114956 | DGUOK | 46,32 | 42,86 | 36,89 | 32,56 | 29,7 |
| ENSG00000135870 | RC3H1 | 12,09 | 11,17 | 9,05 | 8,5 | 29,7 |
| ENSG00000162851 | TFB2M | 6,68 | 5,79 | 5,59 | 4,7 | 29,6 |
| ENSG00000173473 | SMARCC1 | 67,55 | 67,2 | 48,95 | 47,54 | 29,6 |
| ENSG00000138495 | COX17 | 50,69 | 48,17 | 45,27 | 35,68 | 29,6 |
| ENSG00000284901 | RUVBL1 | 18,93 | 16,63 | 15,97 | 13,33 | 29,6 |
| ENSG00000198000 | NOL8 | 16,3 | 15,75 | 13,77 | 11,48 | 29,6 |
| ENSG00000101901 | ALG13 | 25,41 | 22,49 | 19,24 | 17,91 | 29,5 |
| ENSG00000046651 | OFD1 | 9,46 | 9,19 | 7,87 | 6,67 | 29,5 |
| ENSG00000198522 | GPN1 | 22,84 | 21,87 | 21,06 | 16,11 | 29,5 |
| ENSG00000196459 | TRAPPC2 | 7,75 | 7,56 | 6,1 | 5,47 | 29,4 |
| ENSG00000180425 | C11orf71 | 3,3 | 2,99 | 2,62 | 2,33 | 29,4 |
| ENSG00000164104 | HMGB2 | 203,49 | 201,66 | 170,54 | 143,68 | 29,4 |
| ENSG00000187987 | ZSCAN23 | 6,88 | 5,54 | 5,47 | 4,86 | 29,4 |
| ENSG00000237190 | CDKN2AIPNL | 19,92 | 19,63 | 18,59 | 14,08 | 29,3 |
| ENSG00000120458 | MSANTD2 | 10,89 | 10,83 | 9,41 | 7,7 | 29,3 |
| ENSG00000272143 | FGF14-AS2 | 5,09 | 4,72 | 3,89 | 3,6 | 29,3 |
| ENSG00000248174 | LINC02268 | 3,11 | 2,87 | 2,28 | 2,2 | 29,3 |
| ENSG00000164066 | INTU | 11,86 | 9,59 | 9,14 | 8,39 | 29,3 |
| ENSG00000125977 | EIF2S2 | 50,06 | 45,37 | 41,5 | 35,43 | 29,2 |
| ENSG00000165792 | METTL17 | 25,26 | 22,5 | 21,22 | 17,89 | 29,2 |
| ENSG00000231074 | HCG18 | 15,88 | 14,96 | 13,72 | 11,26 | 29,1 |
| ENSG00000157212 | PAXIP1 | 8,22 | 8,09 | 7,21 | 5,83 | 29,1 |
| ENSG00000119231 | SENP5 | 20,24 | 17,94 | 17,48 | 14,36 | 29,1 |
| ENSG00000135314 | KHDC1 | 4,96 | 3,78 | 3,7 | 3,52 | 29,0 |
| ENSG00000078687 | TNRC6C | 12,6 | 10,37 | 9,3 | 8,95 | 29,0 |
| ENSG00000145414 | NAF1 | 10,4 | 9,31 | 8,06 | 7,39 | 28,9 |
| ENSG00000140525 | FANCI | 46,62 | 42,39 | 39,17 | 33,13 | 28,9 |
| ENSG00000158169 | FANCC | 9,82 | 8,97 | 8,05 | 6,98 | 28,9 |
| ENSG00000083093 | PALB2 | 9,24 | 9,1 | 8,39 | 6,57 | 28,9 |
| ENSG00000167005 | NUDT21 | 60,54 | 58 | 57,3 | 43,06 | 28,9 |
| ENSG00000177640 | CASC2 | 1,7 | 1,67 | 1,38 | 1,21 | 28,8 |

| | | | | | | |
|---|---|---|---|---|---|---|
| ENSG00000273611 | ZNHIT3 | 18,25 | 17,61 | 15,85 | 12,99 | 28,8 |
| ENSG00000147471 | PLPBP | 20,79 | 16,75 | 15,88 | 14,8 | 28,8 |
| ENSG00000171612 | SLC25A33 | 4,34 | 3,69 | 3,59 | 3,09 | 28,8 |
| ENSG00000123560 | PLP1 | 31,65 | 26,94 | 26,27 | 22,55 | 28,8 |
| ENSG00000144580 | CNOT9 | 29,84 | 27,63 | 26,31 | 21,27 | 28,7 |
| ENSG00000135040 | NAA35 | 12,99 | 12,16 | 10,81 | 9,26 | 28,7 |
| ENSG00000276523 | AC025287,3 | 1,15 | 1,14 | 0,92 | 0,82 | 28,7 |
| ENSG00000198824 | CHAMP1 | 20,77 | 19,17 | 16,62 | 14,81 | 28,7 |
| ENSG00000126457 | PRMT1 | 114,32 | 97,23 | 93,51 | 81,53 | 28,7 |
| ENSG00000084764 | MAPRE3 | 7,81 | 6,99 | 6,31 | 5,57 | 28,7 |
| ENSG00000257800 | FNBP1P1 | 2,76 | 2,55 | 2,08 | 1,97 | 28,6 |
| ENSG00000135537 | AFG1L | 3,74 | 3,38 | 3,13 | 2,67 | 28,6 |
| ENSG00000196510 | ANAPC7 | 37,9 | 37,6 | 31,92 | 27,08 | 28,5 |
| ENSG00000119661 | DNAL1 | 14,75 | 11,6 | 10,92 | 10,54 | 28,5 |
| ENSG00000135829 | DHX9 | 101,55 | 97,71 | 83,91 | 72,58 | 28,5 |
| ENSG00000130363 | RSPH3 | 3,05 | 2,96 | 2,23 | 2,18 | 28,5 |
| ENSG00000143443 | C1orf56 | 3,09 | 2,85 | 2,64 | 2,21 | 28,5 |
| ENSG00000284959 | AC007262,2 | 10,54 | 10,04 | 8,81 | 7,54 | 28,5 |
| ENSG00000147535 | PLPP5 | 16,31 | 13,15 | 12,85 | 11,68 | 28,4 |
| ENSG00000197323 | TRIM33 | 20,54 | 19,55 | 14,89 | 14,71 | 28,4 |
| ENSG00000136682 | CBWD2 | 20,47 | 19,61 | 17,27 | 14,69 | 28,2 |
| ENSG00000176444 | CLK2 | 10,98 | 10,89 | 9,08 | 7,88 | 28,2 |
| ENSG00000151304 | SRFBP1 | 9,07 | 8,67 | 7,26 | 6,51 | 28,2 |
| ENSG00000164031 | DNAJB14 | 16,81 | 16,61 | 14,69 | 12,07 | 28,2 |
| ENSG00000144283 | PKP4 | 24,36 | 21,28 | 20,85 | 17,5 | 28,2 |
| ENSG00000259664 | LINC02254 | 1,92 | 1,58 | 1,55 | 1,38 | 28,1 |
| ENSG00000247271 | ZBED5-AS1 | 4,34 | 4,02 | 3,72 | 3,12 | 28,1 |
| ENSG00000198182 | ZNF607 | 4,91 | 4,16 | 3,99 | 3,53 | 28,1 |
| ENSG00000081026 | MAGI3 | 9,02 | 8,74 | 7,9 | 6,49 | 28,0 |
| ENSG00000149485 | FADS1 | 70,87 | 63,51 | 59,22 | 51 | 28,0 |
| ENSG00000047410 | TPR | 44,55 | 41,44 | 33,03 | 32,06 | 28,0 |
| ENSG00000145041 | DCAF1 | 9,08 | 8,42 | 7,64 | 6,54 | 28,0 |
| ENSG00000101290 | CDS2 | 21,86 | 20,48 | 19,06 | 15,76 | 27,9 |
| ENSG00000165898 | ISCA2 | 15,27 | 13,45 | 11,27 | 11,02 | 27,8 |
| ENSG00000143621 | ILF2 | 134,82 | 124,17 | 113,05 | 97,35 | 27,8 |
| ENSG00000171135 | JAGN1 | 9,16 | 8,83 | 8,49 | 6,62 | 27,7 |
| ENSG00000122203 | KIAA1191 | 41,43 | 36,97 | 36,35 | 29,95 | 27,7 |
| ENSG00000256092 | SBNO1-AS1 | 1,95 | 1,92 | 1,88 | 1,41 | 27,7 |
| ENSG00000076003 | MCM6 | 45,85 | 43,15 | 39,07 | 33,16 | 27,7 |
| ENSG00000146109 | ABT1 | 7,94 | 7,44 | 6,66 | 5,75 | 27,6 |
| ENSG00000137185 | ZSCAN9 | 13,21 | 12,36 | 11,19 | 9,57 | 27,6 |
| ENSG00000273117 | AC144652,1 | 1,96 | 1,71 | 1,44 | 1,42 | 27,6 |
| ENSG00000106554 | CHCHD3 | 39,66 | 38,03 | 37,39 | 28,74 | 27,5 |
| ENSG00000204650 | LINC02210 | 10,51 | 9,21 | 8,39 | 7,63 | 27,4 |
| ENSG00000213593 | TMX2 | 32,84 | 31,84 | 30,88 | 23,86 | 27,3 |
| ENSG00000196396 | PTPN1 | 16,39 | 14,71 | 13,6 | 11,91 | 27,3 |
| ENSG00000163629 | PTPN13 | 27,08 | 24,85 | 21,7 | 19,68 | 27,3 |

| | | | | | | |
|---|---|---|---|---|---|---|
| ENSG00000165650 | PDZD8 | 13,39 | 11,88 | 10,37 | 9,74 | 27,3 |
| ENSG00000128203 | ASPHD2 | 3,09 | 2,57 | 2,27 | 2,25 | 27,2 |
| ENSG00000182831 | C16orf72 | 30,6 | 29,62 | 22,95 | 22,29 | 27,2 |
| ENSG00000105821 | DNAJC2 | 16,38 | 14,91 | 12,8 | 11,94 | 27,1 |
| ENSG00000259129 | LINC00648 | 3,95 | 3,59 | 3,37 | 2,88 | 27,1 |
| ENSG00000163468 | CCT3 | 107,38 | 104,03 | 99,62 | 78,31 | 27,1 |
| ENSG00000250131 | AC078881,1 | 2,07 | 2,06 | 1,56 | 1,51 | 27,1 |
| ENSG00000156928 | MALSU1 | 18,95 | 18,75 | 18,5 | 13,83 | 27,0 |
| ENSG00000237928 | NFIA-AS2 | 1,26 | 1,21 | 1,09 | 0,92 | 27,0 |
| ENSG00000126003 | PLAGL2 | 11,6 | 10,24 | 9,87 | 8,47 | 27,0 |
| ENSG00000128581 | IFT22 | 33,39 | 33,01 | 26,73 | 24,39 | 27,0 |
| ENSG00000176396 | EID2 | 11,8 | 10,64 | 10,58 | 8,62 | 26,9 |
| ENSG00000197013 | ZNF429 | 7,98 | 6,78 | 6,05 | 5,83 | 26,9 |
| ENSG00000235481 | UBE2R2-AS1 | 1,04 | 0,95 | 0,77 | 0,76 | 26,9 |
| ENSG00000100823 | APEX1 | 119,55 | 116,73 | 109,73 | 87,37 | 26,9 |
| ENSG00000167325 | RRM1 | 60,46 | 55,9 | 49,45 | 44,2 | 26,9 |
| ENSG00000147383 | NSDHL | 23,22 | 22,86 | 21,66 | 16,98 | 26,9 |
| ENSG00000270071 | AP001172,1 | 2,16 | 2,11 | 2,02 | 1,58 | 26,9 |
| ENSG00000134504 | KCTD1 | 9,81 | 8,65 | 7,28 | 7,18 | 26,8 |
| ENSG00000185737 | NRG3 | 3,4 | 2,7 | 2,68 | 2,49 | 26,8 |
| ENSG00000203780 | FANK1 | 2,28 | 2,13 | 2,07 | 1,67 | 26,8 |
| ENSG00000169021 | UQCRFS1 | 16,89 | 16,24 | 14,75 | 12,38 | 26,7 |
| ENSG00000139620 | KANSL2 | 18,3 | 17,54 | 16,83 | 13,43 | 26,6 |
| ENSG00000164053 | ATRIP | 6,62 | 6,4 | 5,87 | 4,86 | 26,6 |
| ENSG00000093167 | LRRFIP2 | 29,17 | 24,88 | 22,54 | 21,42 | 26,6 |
| ENSG00000125821 | DTD1 | 7,42 | 7,27 | 5,99 | 5,45 | 26,5 |
| ENSG00000241316 | SUCLG2-AS1 | 2,64 | 2,57 | 2,2 | 1,94 | 26,5 |
| ENSG00000173320 | STOX2 | 12,41 | 11,84 | 11,36 | 9,12 | 26,5 |
| ENSG00000167635 | ZNF146 | 49,03 | 42,76 | 42,13 | 36,06 | 26,5 |
| ENSG00000079246 | XRCC5 | 192,5 | 174,15 | 155,32 | 141,6 | 26,4 |
| ENSG00000172262 | ZNF131 | 20,12 | 20,01 | 19,06 | 14,8 | 26,4 |
| ENSG00000173597 | SULT1B1 | 6,78 | 6,67 | 6,02 | 4,99 | 26,4 |
| ENSG00000144231 | POLR2D | 28,31 | 26,11 | 24,4 | 20,84 | 26,4 |
| ENSG00000254389 | RHPN1-AS1 | 1,26 | 1,1 | 1,08 | 0,93 | 26,2 |
| ENSG00000181191 | PJA1 | 58,52 | 57,59 | 51,16 | 43,23 | 26,1 |
| ENSG00000152495 | CAMK4 | 9,09 | 8,53 | 7,93 | 6,72 | 26,1 |
| ENSG00000156508 | EEF1A1 | 3436,14 | 3279,87 | 3215,61 | 2541,68 | 26,0 |
| ENSG00000136231 | IGF2BP3 | 58,57 | 54,56 | 46,52 | 43,34 | 26,0 |
| ENSG00000225648 | SBDSP1 | 9,31 | 9,07 | 8,3 | 6,89 | 26,0 |
| ENSG00000165671 | NSD1 | 38,23 | 37,81 | 33,24 | 28,3 | 26,0 |
| ENSG00000155858 | LSM11 | 3,39 | 3,1 | 3 | 2,51 | 26,0 |
| ENSG00000186106 | ANKRD46 | 11,18 | 10,02 | 9,28 | 8,28 | 25,9 |
| ENSG00000109332 | UBE2D3 | 130,47 | 129,99 | 123,59 | 96,63 | 25,9 |
| ENSG00000161654 | LSM12 | 24,54 | 23,78 | 21,97 | 18,18 | 25,9 |
| ENSG00000004487 | KDM1A | 70,94 | 68,41 | 61,37 | 52,59 | 25,9 |
| ENSG00000135900 | MRPL44 | 16,55 | 16,02 | 13,7 | 12,27 | 25,9 |
| ENSG00000085415 | SEH1L | 20,19 | 17,42 | 16,69 | 14,97 | 25,9 |

| | | | | | | |
|---|---|---|---|---|---|---|
| ENSG00000174839 | DENND6A | 15,99 | 14,39 | 13,71 | 11,86 | 25,8 |
| ENSG00000250312 | ZNF718 | 9,18 | 8,46 | 8,31 | 6,81 | 25,8 |
| ENSG00000133706 | LARS1 | 55,05 | 50,61 | 47,03 | 40,84 | 25,8 |
| ENSG00000113742 | CPEB4 | 6,32 | 6,08 | 5,51 | 4,69 | 25,8 |
| ENSG00000286293 | AP005901,5 | 15,57 | 15,24 | 11,93 | 11,56 | 25,8 |
| ENSG00000197894 | ADH5 | 94,6 | 93,03 | 87,35 | 70,27 | 25,7 |
| ENSG00000204519 | ZNF551 | 7,33 | 7,03 | 5,89 | 5,45 | 25,6 |
| ENSG00000167272 | POP5 | 18,18 | 15,74 | 15,02 | 13,52 | 25,6 |
| ENSG00000196419 | XRCC6 | 162,82 | 154,53 | 139,72 | 121,26 | 25,5 |
| ENSG00000132849 | PATJ | 5,11 | 4,41 | 4,39 | 3,81 | 25,4 |
| ENSG00000164975 | SNAPC3 | 19,97 | 19,09 | 18,11 | 14,89 | 25,4 |
| ENSG00000225206 | MIR137HG | 2,95 | 2,68 | 2,31 | 2,2 | 25,4 |
| ENSG00000143368 | SF3B4 | 38,86 | 37,9 | 37,12 | 28,99 | 25,4 |
| ENSG00000083845 | RPS5 | 284,45 | 257,67 | 256,4 | 212,25 | 25,4 |
| ENSG00000139734 | DIAPH3 | 13,34 | 11,64 | 10,35 | 9,96 | 25,3 |
| ENSG00000159200 | RCAN1 | 16,96 | 14,21 | 13,95 | 12,68 | 25,2 |
| ENSG00000145912 | NHP2 | 72,88 | 67,41 | 64,63 | 54,54 | 25,2 |
| ENSG00000124784 | RIOK1 | 9,2 | 9,02 | 7,07 | 6,89 | 25,1 |
| ENSG00000144026 | ZNF514 | 7,65 | 7,3 | 6,95 | 5,73 | 25,1 |
| ENSG00000272140 | AC022400,4 | 2,75 | 2,63 | 2,59 | 2,06 | 25,1 |
| ENSG00000114115 | RBP1 | 57,14 | 55,29 | 47,65 | 42,83 | 25,0 |
| ENSG00000110955 | ATP5F1B | 196,82 | 177,58 | 172,19 | 147,53 | 25,0 |
