## Supplementary Table 9 for "Huntingtin loss-of-function contributes to transcriptional deregulation in Huntington’s disease"

**Supplementary Table 9. List of genes with expression increasing with time in IC1, HD and KO-NSCs**

| IC-NSCs | | | [TPM] | | |
|---|---|---|---|---|---|
| **gene_ID** | **Gene_name** | **IC1-NSC_1** | **IC1-NSC_2** | **IC1-NSC_3** | **change_%** |
| ENSG00000207174 | SNORD116-15 | 0 | 168,23 | 2434,63 | |
| ENSG00000206656 | SNORD116-17 | 0 | 37,05 | 594,93 | |
| ENSG00000207460 | SNORD116-19 | 0 | 37,05 | 594,93 | |
| ENSG00000207263 | SNORD116-16 | 0 | 3,45 | 451,67 | |
| ENSG00000275127 | SNORD116-22 | 0 | 30,24 | 338,15 | |
| ENSG00000207375 | SNORD116-23 | 0 | 8,12 | 218,03 | |
| ENSG00000207279 | SNORD116-24 | 0 | 14,16 | 215,51 | |
| ENSG00000206621 | SNORD116-14 | 0 | 10,46 | 155,47 | |
| ENSG00000252326 | SNORD116-25 | 0 | 77,23 | 83,88 | |
| ENSG00000207093 | | 0 | 7,24 | 71,86 | |
| ENSG00000207063 | | 0 | 18,46 | 37,69 | |
| ENSG00000252431 | | 0 | 2,8 | 32,68 | |
| ENSG00000222881 | | 0 | 12,14 | 18,54 | |
| ENSG00000207197 | | 0 | 16,12 | 16,58 | |
| ENSG00000275129 | | 0 | 3,97 | 14,72 | |
| ENSG00000251939 | | 0 | 3,47 | 12,36 | |
| ENSG00000202430 | | 0 | 6,31 | 11,02 | |
| ENSG00000201297 | | 0 | 1,44 | 9,86 | |
| ENSG00000206739 | | 0 | 4,14 | 9,35 | |
| ENSG00000222108 | | 0 | 0,58 | 9,13 | |
| ENSG00000238405 | | 0 | 0,58 | 9,13 | |
| ENSG00000206779 | | 0 | 4,72 | 8,93 | |
| ENSG00000202417 | | 0 | 3,96 | 8,86 | |
| ENSG00000201448 | | 0 | 2,22 | 8,5 | |
| ENSG00000206920 | | 0 | 0,44 | 8,46 | |
| ENSG00000200579 | | 0 | 2,24 | 8,38 | |
| ENSG00000202412 | | 0 | 2,24 | 8,38 | |
| ENSG00000206816 | | 0 | 2,24 | 8,38 | |
| ENSG00000199201 | | 0 | 2,12 | 7,87 | |
| ENSG00000206870 | | 0 | 3,35 | 7,01 | |
| ENSG00000206627 | | 0 | 1,91 | 6,96 | |
| ENSG00000223300 | | 0 | 3,83 | 6,96 | |
| ENSG00000199535 | | 0 | 1,04 | 6,58 | |
| ENSG00000222383 | | 0 | 5,41 | 6,52 | |
| ENSG00000201113 | | 0 | 5,2 | 6,18 | |
| ENSG00000285522 | | 0 | 1,17 | 5,42 | |
| ENSG00000200261 | | 0 | 2,76 | 4,67 | |
| ENSG00000199676 | | 0 | 1,32 | 4,43 | |
| ENSG00000275227 | | 0 | 0,49 | 3,96 | |
| ENSG00000238835 | | 0 | 0,64 | 3,51 | |
| ENSG00000221808 | | 0 | 3,12 | 3,29 | |
| ENSG00000232101 | | 0 | 0,83 | 3,02 | |
| ENSG00000207635 | | 0 | 1,86 | 2,86 | |
| ENSG00000276884 | | 0 | 0,21 | 2,4 | |
| ENSG00000270381 | | 0 | 0,73 | 2,24 | |
| ENSG00000275327 | AL354950,2 | 0 | 0,82 | 2,06 | |

| | | | | | |
|---|---|---|---|---|---|
| ENSG00000199273 | | 0 | 0,4 | 1,99 | |
| ENSG00000253092 | | 0 | 0,39 | 1,84 | |
| ENSG00000272359 | | 0 | 1,31 | 1,82 | |
| ENSG00000284932 | | 0 | 0,27 | 1,68 | |
| ENSG00000238749 | | 0 | 0,46 | 1,56 | |
| ENSG00000238933 | | 0 | 0,46 | 1,56 | |
| ENSG00000275904 | | 0 | 0,46 | 1,56 | |
| ENSG00000228027 | | 0 | 0,63 | 1,42 | |
| ENSG00000200176 | | 0 | 0,63 | 1,4 | |
| ENSG00000288603 | | 0 | 0,19 | 1,4 | |
| ENSG00000239614 | HMGN1P7 | 0 | 0,97 | 1,38 | |
| ENSG00000239194 | | 0 | 0,93 | 1,37 | |
| ENSG00000278722 | AL445288,1 | 0 | 0,15 | 1,32 | |
| ENSG00000244451 | RPL34P21 | 0 | 0,56 | 1,25 | |
| ENSG00000236580 | AC073626,2 | 0 | 0,48 | 1,24 | |
| ENSG00000220793 | RPL21P119 | 0 | 0,12 | 1,24 | |
| ENSG00000225701 | | 0 | 0,14 | 1,18 | |
| ENSG00000229146 | SNX18P4 | 0 | 0,56 | 1,13 | |
| ENSG00000206687 | | 0 | 0,45 | 1,13 | |
| ENSG00000271159 | | 0 | 0,13 | 1,12 | |
| ENSG00000263370 | | 0 | 0,21 | 1,11 | |
| ENSG00000241625 | | 0 | 0,65 | 1,07 | |
| ENSG00000243267 | | 0 | 0,2 | 1,02 | |
| ENSG00000244630 | AC022493,1 | 0 | 0,22 | 1 | |
| ENSG00000283782 | AC116366,2 | 0 | 0,13 | 1 | |
| ENSG00000262607 | | 0,03 | 0,04 | 8,51 | 28267 |
| ENSG00000263585 | AC145207,4 | 0,01 | 1,77 | 2,72 | 27100 |
| ENSG00000176994 | SMCR8 | 0,04 | 1,58 | 3,87 | 9575 |
| ENSG00000285449 | H2BU1 | 0,06 | 2,24 | 4,83 | 7950 |
| ENSG00000164778 | EN2 | 0,4 | 7,72 | 16,7 | 4075 |
| ENSG00000114113 | | 0,06 | 0,48 | 1,31 | 2083 |
| ENSG00000267319 | SELENOKP1 | 0,13 | 0,85 | 2,67 | 1954 |
| ENSG00000146910 | CNPY1 | 1,1 | 5,23 | 20,29 | 1745 |
| ENSG00000260426 | AC008060,4 | 0,07 | 0,2 | 1,24 | 1671 |
| ENSG00000178343 | SHISA3 | 0,1 | 1 | 1,62 | 1520 |
| ENSG00000254622 | NAV2-AS4 | 0,07 | 0,4 | 1,06 | 1414 |
| ENSG00000283128 | AC009403,2 | 0,5 | 1,66 | 7,39 | 1378 |
| ENSG00000234812 | | 0,08 | 1 | 1,09 | 1263 |
| ENSG00000276257 | | 0,08 | 1 | 1,09 | 1263 |
| ENSG00000156096 | UGT2B4 | 0,09 | 0,79 | 1,17 | 1200 |
| ENSG00000227962 | AL391994,1 | 0,13 | 0,48 | 1,58 | 1115 |
| ENSG00000218872 | | 0,09 | 0,14 | 1,09 | 1111 |
| ENSG00000228116 | PPT2 | 0,59 | 5,04 | 7,13 | 1108 |
| ENSG00000218672 | AC008060,1 | 0,15 | 0,62 | 1,77 | 1080 |
| ENSG00000236556 | AC025038,1 | 0,09 | 0,87 | 1 | 1011 |
| ENSG00000277276 | OTUB2 | 0,11 | 0,18 | 1,09 | 891 |
| ENSG00000122584 | NXPH1 | 0,67 | 0,8 | 6,31 | 842 |
| ENSG00000255475 | AP001007,2 | 0,11 | 0,3 | 1 | 809 |
| ENSG00000270808 | AC022400,3 | 0,13 | 0,85 | 1,12 | 762 |
| ENSG00000231720 | AL353747,3 | 0,15 | 0,31 | 1,28 | 753 |

| | | | | | |
|---|---|---|---|---|---|
| ENSG00000137965 | IFI44 | 0,37 | 1 | 3,04 | 722 |
| ENSG00000281076 | ZDHHC8P1 | 0,49 | 1,72 | 3,87 | 690 |
| ENSG00000180921 | FAM83H | 0,15 | 0,68 | 1,13 | 653 |
| ENSG00000285642 | | 0,16 | 0,32 | 1,17 | 631 |
| ENSG00000137959 | IFI44L | 1,21 | 2,19 | 8,67 | 617 |
| ENSG00000152578 | GRIA4 | 1,86 | 8,25 | 12,17 | 554 |
| ENSG00000282367 | PTDSS2 | 4,87 | 6,49 | 31,27 | 542 |
| ENSG00000255817 | AC025576,2 | 0,35 | 0,6 | 2,21 | 531 |
| ENSG00000230306 | | 0,16 | 0,43 | 1,01 | 531 |
| ENSG00000102466 | FGF14 | 0,4 | 1,78 | 2,51 | 528 |
| ENSG00000168702 | LRP1B | 0,19 | 0,31 | 1,19 | 526 |
| ENSG00000258640 | RPL21P5 | 0,2 | 0,27 | 1,23 | 515 |
| ENSG00000207313 | | 1,57 | 2,34 | 9,43 | 501 |
| ENSG00000233547 | AL158212,2 | 0,24 | 1,24 | 1,44 | 500 |
| ENSG00000226352 | PSPC1-AS2 | 0,3 | 1,16 | 1,79 | 497 |
| ENSG00000243433 | AC010973,1 | 0,24 | 1,32 | 1,35 | 463 |
| ENSG00000263990 | AC004253,1 | 0,24 | 0,8 | 1,33 | 454 |
| ENSG00000239412 | RPL21P71 | 0,74 | 2,22 | 4,03 | 445 |
| ENSG00000255571 | MIR9-3HG | 0,69 | 2,45 | 3,69 | 435 |
| ENSG00000248266 | AC108142,1 | 0,26 | 1,32 | 1,39 | 435 |
| ENSG00000074211 | PPP2R2C | 0,34 | 0,87 | 1,79 | 426 |
| ENSG00000274987 | AC092794,1 | 0,22 | 0,24 | 1,15 | 423 |
| ENSG00000225890 | HLA-DQA1 | 0,49 | 0,54 | 2,52 | 414 |
| ENSG00000243620 | AC092957,1 | 5,31 | 13,43 | 27,06 | 410 |
| ENSG00000254273 | AC018620,1 | 0,22 | 0,97 | 1,12 | 409 |
| ENSG00000175445 | LPL | 10,69 | 16,46 | 53,01 | 396 |
| ENSG00000187391 | MAGI2 | 7,92 | 18,48 | 39,16 | 394 |
| ENSG00000171540 | OTP | 0,48 | 2,08 | 2,37 | 394 |
| ENSG00000283879 | MTND6P35 | 0,22 | 0,82 | 1,06 | 382 |
| ENSG00000078549 | ADCYAP1R1 | 1,26 | 1,56 | 5,66 | 349 |
| ENSG00000256571 | AC079866,2 | 0,35 | 0,93 | 1,57 | 349 |
| ENSG00000242737 | AC012170,1 | 0,24 | 0,46 | 1,06 | 342 |
| ENSG00000284657 | AL031432,5 | 0,23 | 0,74 | 1,01 | 339 |
| ENSG00000206680 | SNORD21 | 26,9 | 98,8 | 117,99 | 339 |
| ENSG00000236611 | LINC02556 | 0,26 | 0,55 | 1,14 | 338 |
| ENSG00000150275 | PCDH15 | 1,96 | 3,75 | 8,51 | 334 |
| ENSG00000206952 | SNORA50A | 10,53 | 13,36 | 45,57 | 333 |
| ENSG00000260422 | Z97205,2 | 0,36 | 0,58 | 1,55 | 331 |
| ENSG00000252212 | | 0,45 | 0,95 | 1,91 | 324 |
| ENSG00000276500 | BMS1P14 | 0,38 | 0,78 | 1,59 | 318 |
| ENSG00000126785 | RHOJ | 1,04 | 2,25 | 4,32 | 315 |
| ENSG00000172995 | ARPP21 | 0,33 | 0,65 | 1,37 | 315 |
| ENSG00000171385 | KCND3 | 0,28 | 0,55 | 1,16 | 314 |
| ENSG00000199436 | SNORD9 | 23,77 | 50,33 | 96,14 | 304 |
| ENSG00000277423 | AC069234,5 | 0,31 | 0,58 | 1,23 | 297 |
| ENSG00000009709 | PAX7 | 0,8 | 1,63 | 3,17 | 296 |
| ENSG00000236404 | VLDLR-AS1 | 0,26 | 0,54 | 1,02 | 292 |
| ENSG00000151834 | GABRA2 | 0,3 | 0,33 | 1,17 | 290 |
| ENSG00000166342 | NETO1 | 0,59 | 1,08 | 2,3 | 290 |
| ENSG00000146005 | PSD2 | 0,55 | 1,01 | 2,14 | 289 |

| | | | | | |
|---|---|---|---|---|---|
| ENSG00000112473 | SLC39A7 | 1,65 | 3,38 | 6,36 | 285 |
| ENSG00000254305 | MRPL9P1 | 0,31 | 0,7 | 1,18 | 281 |
| ENSG00000213018 | PABPN1P1 | 0,35 | 0,45 | 1,29 | 269 |
| ENSG00000101134 | DOK5 | 5,26 | 6,28 | 19,38 | 268 |
| ENSG00000081138 | CDH7 | 1,38 | 3,68 | 5,03 | 264 |
| ENSG00000229980 | TOB1-AS1 | 0,37 | 0,68 | 1,34 | 262 |
| ENSG00000234383 | CTBP2P8 | 0,33 | 0,77 | 1,19 | 261 |
| ENSG00000242087 | RPL36AP41 | 0,67 | 1,77 | 2,41 | 260 |
| ENSG00000176842 | IRX5 | 2,22 | 6,73 | 7,98 | 259 |
| ENSG00000266378 | AC005224,2 | 0,42 | 0,74 | 1,48 | 252 |
| ENSG00000235204 | AL162724,2 | 0,44 | 0,75 | 1,55 | 252 |
| ENSG00000120833 | SOCS2 | 3,9 | 9,34 | 13,72 | 252 |
| ENSG00000115252 | PDE1A | 1,7 | 3,08 | 5,93 | 249 |
| ENSG00000225216 | AC007362,1 | 0,85 | 1,49 | 2,93 | 245 |
| ENSG00000203688 | LINC02487 | 0,83 | 1,25 | 2,85 | 243 |
| ENSG00000237732 | CT75 | 1,09 | 3,37 | 3,74 | 243 |
| ENSG00000232599 | AL008707,1 | 1,13 | 1,93 | 3,86 | 242 |
| ENSG00000248801 | C8orf34-AS1 | 0,3 | 0,4 | 1,02 | 240 |
| ENSG00000070748 | CHAT | 0,68 | 1,64 | 2,3 | 238 |
| ENSG00000127241 | MASP1 | 0,78 | 1,53 | 2,63 | 237 |
| ENSG00000069702 | TGFBR3 | 0,43 | 1,33 | 1,44 | 235 |
| ENSG00000285701 | AL357140,4 | 0,54 | 0,93 | 1,8 | 233 |
| ENSG00000212135 | | 10,36 | 23,05 | 34,52 | 233 |
| ENSG00000226045 | AC234644,1 | 0,38 | 1,13 | 1,26 | 232 |
| ENSG00000182021 | AL591379,1 | 1,16 | 2,98 | 3,84 | 231 |
| ENSG00000273780 | ARHGAP23 | 1,63 | 3,44 | 5,33 | 227 |
| ENSG00000239528 | RPS14P8 | 2,15 | 4,59 | 6,99 | 225 |
| ENSG00000272232 | RN7SL726P | 0,47 | 0,96 | 1,52 | 223 |
| ENSG00000182168 | UNC5C | 1,06 | 1,49 | 3,42 | 223 |
| ENSG00000082556 | OPRK1 | 1,27 | 1,99 | 4,09 | 222 |
| ENSG00000235488 | JARID2-AS1 | 0,54 | 0,7 | 1,72 | 219 |
| ENSG00000235829 | TBCAP2 | 1,07 | 1,52 | 3,4 | 218 |
| ENSG00000107742 | SPOCK2 | 1,18 | 3,16 | 3,72 | 215 |
| ENSG00000168505 | GBX2 | 11,69 | 16,03 | 36,66 | 214 |
| ENSG00000267707 | AC015961,2 | 0,49 | 0,98 | 1,53 | 212 |
| ENSG00000259322 | AC090607,1 | 0,35 | 0,95 | 1,09 | 211 |
| ENSG00000114251 | WNT5A | 3,65 | 3,91 | 11,33 | 210 |
| ENSG00000283060 | ID3 | 14,53 | 22,28 | 45,01 | 210 |
| ENSG00000117318 | ID3 | 14,53 | 22,28 | 45,01 | 210 |
| ENSG00000196569 | LAMA2 | 0,43 | 0,45 | 1,32 | 207 |
| ENSG00000237333 | MSH5 | 0,59 | 0,94 | 1,81 | 207 |
| ENSG00000280661 | ZNF660 | 2,87 | 4,96 | 8,79 | 206 |
| ENSG00000242999 | RN7SL239P | 0,58 | 0,87 | 1,76 | 203 |
| ENSG00000285633 | AL132633,1 | 0,39 | 0,49 | 1,18 | 203 |
| ENSG00000282205 | AC115090,1 | 0,83 | 1,59 | 2,51 | 202 |
| ENSG00000276170 | AC244153,1 | 0,83 | 1,59 | 2,51 | 202 |
| ENSG00000249685 | AC079921,2 | 0,43 | 0,75 | 1,3 | 202 |
| ENSG00000163064 | EN1 | 5,75 | 12,79 | 17,25 | 200 |
| ENSG00000230424 | EMC1-AS1 | 0,39 | 1,03 | 1,17 | 200 |
| ENSG00000255780 | AC020611,1 | 0,68 | 0,7 | 2,02 | 197 |

| | | | | | |
|---|---|---|---|---|---|
| ENSG00000153253 | SCN3A | 2,11 | 2,37 | 6,21 | 194 |
| ENSG00000276017 | AC007325,1 | 0,34 | 0,87 | 1 | 194 |
| ENSG00000278475 | AC009464,1 | 0,46 | 0,76 | 1,35 | 193 |
| ENSG00000198822 | GRM3 | 12,23 | 18,54 | 35,83 | 193 |
| ENSG00000136099 | PCDH8 | 1,68 | 4,39 | 4,91 | 192 |
| ENSG00000257954 | AC125611,2 | 0,45 | 0,66 | 1,31 | 191 |
| ENSG00000198216 | CACNA1E | 0,93 | 1,24 | 2,7 | 190 |
| ENSG00000242818 | RN7SL846P | 0,37 | 0,82 | 1,07 | 189 |
| ENSG00000237493 | AC034102,1 | 0,45 | 0,76 | 1,3 | 189 |
| ENSG00000168994 | PXDC1 | 1,72 | 2,13 | 4,92 | 186 |
| ENSG00000234159 | RBPMSLP | 0,41 | 0,58 | 1,17 | 185 |
| ENSG00000241882 | AC124893,1 | 0,41 | 1,01 | 1,17 | 185 |
| ENSG00000232433 | GXYLT1P3 | 1,07 | 1,33 | 3,03 | 183 |
| ENSG00000136750 | GAD2 | 0,39 | 0,54 | 1,1 | 182 |
| ENSG00000135525 | MAP7 | 0,49 | 0,91 | 1,38 | 182 |
| ENSG00000205683 | DPF3 | 0,54 | 0,94 | 1,52 | 181 |
| ENSG00000102362 | SYTL4 | 0,59 | 1,02 | 1,66 | 181 |
| ENSG00000212440 | | 2,03 | 3,01 | 5,71 | 181 |
| ENSG00000268442 | AC073534,1 | 0,53 | 0,94 | 1,49 | 181 |
| ENSG00000275832 | ARHGAP23 | 3,2 | 6,18 | 8,98 | 181 |
| ENSG00000198576 | ARC | 0,38 | 0,48 | 1,05 | 176 |
| ENSG00000270343 | UNGP3 | 0,45 | 0,82 | 1,24 | 176 |
| ENSG00000095739 | BAMBI | 0,87 | 2,07 | 2,37 | 172 |
| ENSG00000240163 | AC087385,1 | 0,43 | 0,57 | 1,16 | 170 |
| ENSG00000223039 | RN7SKP268 | 0,42 | 0,88 | 1,13 | 169 |
| ENSG00000237887 | RPL23AP32 | 0,63 | 0,65 | 1,69 | 168 |
| ENSG00000263606 | CHORDC1P4 | 0,44 | 0,49 | 1,18 | 168 |
| ENSG00000132437 | DDC | 0,59 | 0,89 | 1,58 | 168 |
| ENSG00000155657 | TTN | 1,28 | 2,94 | 3,42 | 167 |
| ENSG00000185985 | SLITRK2 | 0,8 | 0,84 | 2,13 | 166 |
| ENSG00000064042 | LIMCH1 | 7,49 | 14,3 | 19,81 | 164 |
| ENSG00000266709 | AC005224,3 | 2,05 | 3,36 | 5,41 | 164 |
| ENSG00000251621 | AC009487,2 | 0,47 | 0,54 | 1,24 | 164 |
| ENSG00000241231 | AC068308,1 | 1,73 | 2,36 | 4,54 | 162 |
| ENSG00000187122 | SLIT1 | 3,9 | 9,73 | 10,23 | 162 |
| ENSG00000213600 | U73169,1 | 0,39 | 0,63 | 1,02 | 162 |
| ENSG00000176058 | TPRN | 3,27 | 7,04 | 8,55 | 161 |
| ENSG00000235299 | MRPL53P1 | 0,51 | 0,94 | 1,33 | 161 |
| ENSG00000134121 | CHL1 | 2,91 | 5,87 | 7,52 | 158 |
| ENSG00000277956 | MAPT | 1 | 1,11 | 2,57 | 157 |
| ENSG00000227014 | FKBP14-AS1 | 0,39 | 0,45 | 1 | 156 |
| ENSG00000138650 | PCDH10 | 4,04 | 8,57 | 10,33 | 156 |
| ENSG00000168502 | MTCL1 | 12,27 | 24,55 | 31,35 | 156 |
| ENSG00000162599 | NFIA | 6,57 | 10,98 | 16,75 | 155 |
| ENSG00000237977 | EIF4HP2 | 0,66 | 0,96 | 1,68 | 155 |
| ENSG00000214595 | EML6 | 1,02 | 1,45 | 2,59 | 154 |
| ENSG00000278613 | NAIP | 0,87 | 2,03 | 2,2 | 153 |
| ENSG00000087258 | GNAO1 | 4,85 | 10,07 | 12,23 | 152 |
| ENSG00000189337 | KAZN | 3,67 | 9,2 | 9,25 | 152 |
| ENSG00000199883 | RN7SKP90 | 0,56 | 1,02 | 1,41 | 152 |

| | | | | | |
|---|---|---|---|---|---|
| ENSG00000179915 | NRXN1 | 5,05 | 12,45 | 12,66 | 151 |
| ENSG00000063015 | SEZ6 | 1,7 | 2,52 | 4,25 | 150 |
| ENSG00000184672 | RALYL | 0,97 | 0,98 | 2,42 | 149 |
| ENSG00000252577 | | 3,95 | 4,17 | 9,82 | 149 |
| ENSG00000264273 | AC107982,2 | 0,63 | 0,69 | 1,56 | 148 |
| ENSG00000082438 | COBLL1 | 1,73 | 2,31 | 4,27 | 147 |
| ENSG00000257711 | AC079385,2 | 0,57 | 0,88 | 1,4 | 146 |
| ENSG00000244131 | RPSAP51 | 0,42 | 0,48 | 1,03 | 145 |
| ENSG00000224707 | E2F3-IT1 | 0,66 | 1,46 | 1,61 | 144 |
| ENSG00000197177 | ADGRA1 | 0,96 | 1,86 | 2,34 | 144 |
| ENSG00000005513 | SOX8 | 1,84 | 2,63 | 4,48 | 143 |
| ENSG00000226266 | AC009961,1 | 0,97 | 2,03 | 2,35 | 142 |
| ENSG00000170549 | IRX1 | 2,96 | 4,9 | 7,16 | 142 |
| ENSG00000185811 | IKZF1 | 0,93 | 1,93 | 2,24 | 141 |
| ENSG00000033122 | LRRC7 | 0,81 | 1,51 | 1,94 | 140 |
| ENSG00000237986 | CELF2-AS2 | 1,61 | 2,01 | 3,84 | 139 |
| ENSG00000240240 | BX664727,3 | 3,03 | 5,03 | 7,22 | 138 |
| ENSG00000231134 | TCF7L1-IT1 | 0,42 | 0,46 | 1 | 138 |
| ENSG00000101463 | SYNDIG1 | 0,56 | 0,98 | 1,33 | 138 |
| ENSG00000215492 | HNRNPA1P7 | 0,51 | 0,78 | 1,21 | 137 |
| ENSG00000164161 | HHIP | 0,88 | 1,38 | 2,08 | 136 |
| ENSG00000240589 | RN7SL258P | 0,7 | 1,56 | 1,64 | 134 |
| ENSG00000244356 | RN7SL398P | 0,71 | 0,79 | 1,66 | 134 |
| ENSG00000165714 | BORCS5 | 1,21 | 2,35 | 2,82 | 133 |
| ENSG00000196562 | SULF2 | 11,91 | 13,11 | 27,69 | 132 |
| ENSG00000281385 | AP2A2 | 4,04 | 7,79 | 9,37 | 132 |
| ENSG00000145623 | OSMR | 0,47 | 0,78 | 1,09 | 132 |
| ENSG00000268201 | AC020915,1 | 1,04 | 1,16 | 2,4 | 131 |
| ENSG00000267016 | AC111170,1 | 0,49 | 0,53 | 1,13 | 131 |
| ENSG00000237853 | NFIA-AS1 | 0,85 | 0,9 | 1,96 | 131 |
| ENSG00000250328 | MGC32805 | 1,15 | 1,51 | 2,65 | 130 |
| ENSG00000124191 | TOX2 | 2,51 | 2,79 | 5,75 | 129 |
| ENSG00000258162 | AC069228,1 | 0,45 | 0,48 | 1,03 | 129 |
| ENSG00000115232 | ITGA4 | 1,08 | 1,64 | 2,47 | 129 |
| ENSG00000139915 | MDGA2 | 1,88 | 2,04 | 4,29 | 128 |
| ENSG00000199363 | SNORA63E | 24,87 | 30,77 | 56,65 | 128 |
| ENSG00000106069 | CHN2 | 5,36 | 10,96 | 12,08 | 125 |
| ENSG00000147119 | CHST7 | 0,56 | 1 | 1,26 | 125 |
| ENSG00000274893 | AL136317,2 | 0,56 | 0,58 | 1,26 | 125 |
| ENSG00000207955 | AL359091,1 | 2,06 | 4,49 | 4,62 | 124 |
| ENSG00000234814 | SVIL2P | 0,46 | 0,75 | 1,03 | 124 |
| ENSG00000227487 | NCAM1-AS1 | 0,48 | 1,01 | 1,07 | 123 |
| ENSG00000286478 | AC007130,1 | 0,57 | 0,81 | 1,27 | 123 |
| ENSG00000204231 | RXRB | 1,55 | 2,41 | 3,42 | 121 |
| ENSG00000213058 | AL365357,1 | 0,49 | 0,6 | 1,08 | 120 |
| ENSG00000133121 | STARD13 | 2,04 | 3,29 | 4,49 | 120 |
| ENSG00000148053 | NTRK2 | 2,99 | 4,68 | 6,58 | 120 |
| ENSG00000162878 | PKDCC | 3,97 | 4,63 | 8,73 | 120 |
| ENSG00000169085 | VXN | 0,71 | 0,72 | 1,56 | 120 |
| ENSG00000170561 | IRX2 | 6,71 | 12,88 | 14,71 | 119 |

| | | | | | |
|---|---|---|---|---|---|
| ENSG00000232439 | RPL18AP7 | 0,9 | 0,97 | 1,96 | 118 |
| ENSG00000125430 | HS3ST3B1 | 5,59 | 6,71 | 12,16 | 118 |
| ENSG00000164621 | SMAD5-AS1 | 0,46 | 0,88 | 1 | 117 |
| ENSG00000015592 | STMN4 | 14,12 | 21,11 | 30,63 | 117 |
| ENSG00000286139 | ARHGAP11B | 0,95 | 1,06 | 2,06 | 117 |
| ENSG00000206538 | VGLL3 | 2,79 | 4,22 | 6,04 | 116 |
| ENSG00000134115 | CNTN6 | 1,64 | 1,78 | 3,55 | 116 |
| ENSG00000201098 | RNY1 | 2745,09 | 3298,73 | 5938,77 | 116 |
| ENSG00000267085 | AL512605,2 | 0,5 | 0,92 | 1,08 | 116 |
| ENSG00000059804 | SLC2A3 | 34,02 | 41,02 | 73,09 | 115 |
| ENSG00000244218 | RN7SL81P | 1,44 | 1,55 | 3,09 | 115 |
| ENSG00000177272 | KCNA3 | 0,79 | 0,81 | 1,69 | 114 |
| ENSG00000152402 | GUCY1A2 | 1,9 | 2,85 | 4,06 | 114 |
| ENSG00000111913 | RIPOR2 | 4,21 | 4,49 | 8,95 | 113 |
| ENSG00000082684 | SEMA5B | 16,13 | 22,04 | 34,17 | 112 |
| ENSG00000261386 | AC027682,4 | 0,85 | 1,21 | 1,8 | 112 |
| ENSG00000178385 | PLEKHM3 | 1,14 | 1,95 | 2,41 | 111 |
| ENSG00000104964 | TLE5 | 40,15 | 56,27 | 84,59 | 111 |
| ENSG00000121440 | PDZRN3 | 4,23 | 4,72 | 8,91 | 111 |
| ENSG00000275482 | EPHB6 | 1,41 | 2,92 | 2,97 | 111 |
| ENSG00000144285 | SCN1A | 6,8 | 8,61 | 14,32 | 111 |
| ENSG00000288455 | SEMA7A | 0,95 | 1,77 | 2 | 111 |
| ENSG00000138623 | SEMA7A | 0,95 | 1,77 | 2 | 111 |
| ENSG00000278220 | CLN8 | 0,83 | 1,43 | 1,74 | 110 |
| ENSG00000242267 | SKINT1L | 1,07 | 2,2 | 2,24 | 109 |
| ENSG00000231473 | RB1-DT | 0,67 | 1,29 | 1,4 | 109 |
| ENSG00000168874 | ATOH8 | 3,34 | 3,88 | 6,96 | 108 |
| ENSG00000166016 | ABTB2 | 4,91 | 6,74 | 10,23 | 108 |
| ENSG00000120210 | INSL6 | 0,49 | 0,91 | 1,02 | 108 |
| ENSG00000224185 | SNX18P9 | 0,63 | 1,29 | 1,31 | 108 |
| ENSG00000230267 | HERC2P4 | 0,53 | 1,08 | 1,1 | 108 |
| ENSG00000212695 | AL583805,1 | 0,59 | 0,84 | 1,22 | 107 |
| ENSG00000113356 | POLR3G | 0,89 | 1,56 | 1,84 | 107 |
| ENSG00000262265 | AC002558,3 | 1,34 | 2,22 | 2,77 | 107 |
| ENSG00000237758 | BANF1P3 | 1,5 | 2,92 | 3,1 | 107 |
| ENSG00000172638 | EFEMP2 | 5,73 | 6,59 | 11,82 | 106 |
| ENSG00000168461 | RAB31 | 5,17 | 7,07 | 10,64 | 106 |
| ENSG00000225706 | PTPRD-AS1 | 1,55 | 1,77 | 3,19 | 106 |
| ENSG00000286833 | AC097532,3 | 1,05 | 1,87 | 2,16 | 106 |
| ENSG00000147481 | SNTG1 | 0,54 | 0,75 | 1,11 | 106 |
| ENSG00000147526 | TACC1 | 10,39 | 14,85 | 21,32 | 105 |
| ENSG00000162882 | HAAO | 0,86 | 1,09 | 1,76 | 105 |
| ENSG00000114529 | C3orf52 | 0,5 | 0,7 | 1,02 | 104 |
| ENSG00000136052 | SLC41A2 | 1,35 | 1,99 | 2,75 | 104 |
| ENSG00000233665 | AC060234,2 | 0,57 | 0,95 | 1,16 | 104 |
| ENSG00000140488 | CELF6 | 0,86 | 1,09 | 1,75 | 103 |
| ENSG00000125968 | ID1 | 4,18 | 4,19 | 8,5 | 103 |
| ENSG00000265474 | AC010761,4 | 0,9 | 1,37 | 1,83 | 103 |
| ENSG00000205363 | INSYN1 | 1,52 | 2,49 | 3,09 | 103 |
| ENSG00000128394 | APOBEC3F | 0,51 | 0,99 | 1,03 | 102 |

| | | | | | |
|---|---|---|---|---|---|
| ENSG00000270606 | PPIAP52 | 1,61 | 1,98 | 3,25 | 102 |
| ENSG00000232260 | BTF3L4P1 | 0,62 | 0,77 | 1,25 | 102 |
| ENSG00000273079 | GRIN2B | 0,77 | 1,42 | 1,54 | 100 |
| ENSG00000259079 | AC005476,1 | 0,74 | 1,13 | 1,48 | 100 |
| ENSG00000234964 | FABP5P7 | 12,91 | 16,65 | 25,62 | 98 |
| ENSG00000163884 | KLF15 | 0,57 | 0,97 | 1,13 | 98 |
| ENSG00000080503 | SMARCA2 | 4,97 | 6,82 | 9,84 | 98 |
| ENSG00000180834 | MAP6D1 | 0,89 | 1,09 | 1,76 | 98 |
| ENSG00000169991 | IFFO2 | 2,09 | 3,29 | 4,13 | 98 |
| ENSG00000242808 | SOX2-OT | 25,16 | 39,91 | 49,71 | 98 |
| ENSG00000172197 | MBOAT1 | 0,95 | 1,6 | 1,87 | 97 |
| ENSG00000275885 | PRPF31 | 1,17 | 1,67 | 2,3 | 97 |
| ENSG00000130287 | NCAN | 3,53 | 6,32 | 6,92 | 96 |
| ENSG00000250122 | AC122694,1 | 0,69 | 1,11 | 1,35 | 96 |
| ENSG00000131969 | ABHD12B | 0,57 | 1,04 | 1,11 | 95 |
| ENSG00000147437 | GNRH1 | 3,93 | 6,43 | 7,64 | 94 |
| ENSG00000201793 | RN7SKP9 | 1,8 | 2,46 | 3,49 | 94 |
| ENSG00000128567 | PODXL | 7,04 | 9,88 | 13,64 | 94 |
| ENSG00000182253 | SYNM | 0,75 | 1,25 | 1,45 | 93 |
| ENSG00000072818 | ACAP1 | 0,72 | 1,25 | 1,39 | 93 |
| ENSG00000238057 | ZEB2-AS1 | 1,26 | 1,61 | 2,43 | 93 |
| ENSG00000162493 | PDPN | 11,5 | 20,11 | 22,16 | 93 |
| ENSG00000250365 | AL139353,2 | 0,94 | 1,02 | 1,81 | 93 |
| ENSG00000153814 | JAZF1 | 1,86 | 2,14 | 3,57 | 92 |
| ENSG00000113594 | LIFR | 19,58 | 33,95 | 37,48 | 91 |
| ENSG00000265908 | AC024267,5 | 1,03 | 1,67 | 1,97 | 91 |
| ENSG00000170091 | NSG2 | 6,77 | 8,67 | 12,92 | 91 |
| ENSG00000181381 | DDX60L | 0,54 | 0,7 | 1,03 | 91 |
| ENSG00000106341 | PPP1R17 | 1,82 | 2,37 | 3,47 | 91 |
| ENSG00000224490 | TTC21B-AS1 | 0,64 | 0,65 | 1,22 | 91 |
| ENSG00000233230 | AC079807,1 | 0,53 | 0,56 | 1,01 | 91 |
| ENSG00000118432 | CNR1 | 3,54 | 5,29 | 6,74 | 90 |
| ENSG00000228295 | LINC00392 | 2,59 | 4,66 | 4,89 | 89 |
| ENSG00000227331 | RPL7AP22 | 0,53 | 0,86 | 1 | 89 |
| ENSG00000164483 | SAMD3 | 4,1 | 5,73 | 7,73 | 89 |
| ENSG00000166257 | SCN3B | 2,28 | 3,77 | 4,29 | 88 |
| ENSG00000259712 | AC023906,5 | 0,72 | 0,85 | 1,35 | 88 |
| ENSG00000228589 | SPCS2P4 | 1,15 | 2,05 | 2,15 | 87 |
| ENSG00000236483 | MTND2P40 | 0,84 | 1,55 | 1,57 | 87 |
| ENSG00000284648 | AC097493,3 | 1,02 | 1,72 | 1,9 | 86 |
| ENSG00000119669 | IRF2BPL | 22,27 | 26,19 | 41,43 | 86 |
| ENSG00000282228 | PAM16 | 3,41 | 4,59 | 6,32 | 85 |
| ENSG00000276545 | PCDHGB9P | 0,85 | 0,99 | 1,57 | 85 |
| ENSG00000229955 | Z98749,1 | 0,64 | 0,82 | 1,18 | 84 |
| ENSG00000184347 | SLIT3 | 0,64 | 0,71 | 1,18 | 84 |
| ENSG00000136206 | SPDYE1 | 0,76 | 1,25 | 1,4 | 84 |
| ENSG00000068615 | REEP1 | 2,8 | 3,27 | 5,15 | 84 |
| ENSG00000226882 | GNL1 | 1,81 | 1,91 | 3,32 | 83 |
| ENSG00000157064 | NMNAT2 | 3,19 | 3,65 | 5,84 | 83 |
| ENSG00000258134 | AC016954,1 | 0,82 | 1,13 | 1,5 | 83 |

| | | | | | |
|---|---|---|---|---|---|
| ENSG00000253335 | AC009884,1 | 0,57 | 0,67 | 1,04 | 82 |
| ENSG00000253159 | PCDHGA12 | 1,07 | 1,13 | 1,95 | 82 |
| ENSG00000206579 | XKR4 | 1,18 | 1,35 | 2,14 | 81 |
| ENSG00000142188 | TMEM50B | 3,75 | 4,65 | 6,79 | 81 |
| ENSG00000258670 | AL049874,3 | 0,94 | 1,51 | 1,7 | 81 |
| ENSG00000237854 | LINC00674 | 3,27 | 4,32 | 5,89 | 80 |
| ENSG00000200320 | SNORA63 | 1087,61 | 1341,52 | 1958,5 | 80 |
| ENSG00000153976 | HS3ST3A1 | 2,57 | 2,84 | 4,62 | 80 |
| ENSG00000138641 | HERC3 | 1,74 | 2,72 | 3,12 | 79 |
| ENSG00000241462 | AC100832,1 | 0,57 | 0,83 | 1,02 | 79 |
| ENSG00000206394 | CLIC1 | 10,73 | 11,6 | 19,14 | 78 |
| ENSG00000226651 | CLIC1 | 10,73 | 11,6 | 19,14 | 78 |
| ENSG00000171388 | APLN | 1,43 | 1,65 | 2,55 | 78 |
| ENSG00000259330 | INAFM2 | 1,79 | 3,06 | 3,19 | 78 |
| ENSG00000129657 | SEC14L1 | 22,6 | 29,49 | 40,09 | 77 |
| ENSG00000273004 | AL078644,1 | 1,23 | 2,11 | 2,18 | 77 |
| ENSG00000164972 | C9orf24 | 1,05 | 1,7 | 1,85 | 76 |
| ENSG00000078295 | ADCY2 | 4,09 | 5,01 | 7,2 | 76 |
| ENSG00000114166 | KAT2B | 2,49 | 3,07 | 4,38 | 76 |
| ENSG00000223639 | CLIC1 | 11,12 | 11,6 | 19,49 | 75 |
| ENSG00000226248 | CLIC1 | 11,12 | 11,6 | 19,49 | 75 |
| ENSG00000226417 | CLIC1 | 11,12 | 11,6 | 19,49 | 75 |
| ENSG00000213719 | CLIC1 | 11,12 | 11,6 | 19,49 | 75 |
| ENSG00000169047 | IRS1 | 6,77 | 8,36 | 11,82 | 75 |
| ENSG00000221890 | NPTXR | 0,63 | 1,03 | 1,1 | 75 |
| ENSG00000086205 | FOLH1 | 0,62 | 1,06 | 1,08 | 74 |
| ENSG00000276740 | AL445649,1 | 0,58 | 0,79 | 1,01 | 74 |
| ENSG00000278413 | | 1,64 | 2,45 | 2,85 | 74 |
| ENSG00000236047 | RPL13AP12 | 0,91 | 0,94 | 1,58 | 74 |
| ENSG00000282899 | E2F2 | 3,08 | 4,66 | 5,34 | 73 |
| ENSG00000008226 | DLEC1 | 1,01 | 1,37 | 1,75 | 73 |
| ENSG00000141314 | RHBDL3 | 1,6 | 2,61 | 2,77 | 73 |
| ENSG00000153551 | CMTM7 | 7,16 | 8,88 | 12,39 | 73 |
| ENSG00000250366 | TUNAR | 2,09 | 3,44 | 3,61 | 73 |
| ENSG00000100207 | TCF20 | 2,8 | 3,73 | 4,82 | 72 |
| ENSG00000272788 | AP000864,1 | 0,82 | 0,83 | 1,41 | 72 |
| ENSG00000074527 | NTN4 | 1,57 | 1,73 | 2,69 | 71 |
| ENSG00000255769 | GOLGA2P10 | 5,63 | 8,06 | 9,64 | 71 |
| ENSG00000163995 | ABLIM2 | 1,27 | 1,36 | 2,17 | 71 |
| ENSG00000274726 | ARHGEF10 | 4,2 | 5,79 | 7,17 | 71 |
| ENSG00000111961 | SASH1 | 4,92 | 5,72 | 8,39 | 71 |
| ENSG00000166886 | NAB2 | 9,61 | 11,24 | 16,37 | 70 |
| ENSG00000272254 | AC022893,3 | 0,74 | 0,89 | 1,26 | 70 |
| ENSG00000130477 | UNC13A | 1,68 | 2,12 | 2,86 | 70 |
| ENSG00000110619 | CARS1 | 0,6 | 0,76 | 1,02 | 70 |
| ENSG00000257135 | ODC1-DT | 0,73 | 0,9 | 1,24 | 70 |
| ENSG00000232818 | RPS2P32 | 0,82 | 1,04 | 1,39 | 70 |
| ENSG00000267710 | EDDM13 | 1,01 | 1,06 | 1,71 | 69 |
| ENSG00000170017 | ALCAM | 1,87 | 2,4 | 3,15 | 68 |
| ENSG00000120738 | EGR1 | 4,36 | 6,33 | 7,33 | 68 |

| | | | | | |
|---|---|---|---|---|---|
| ENSG00000287214 | AC021321,2 | 0,69 | 0,98 | 1,16 | 68 |
| ENSG00000147852 | VLDLR | 3,26 | 4,27 | 5,47 | 68 |
| ENSG00000159588 | CCDC17 | 0,74 | 0,89 | 1,24 | 68 |
| ENSG00000185634 | SHC4 | 1,51 | 1,59 | 2,53 | 68 |
| ENSG00000069431 | ABCC9 | 4,71 | 6,69 | 7,88 | 67 |
| ENSG00000082781 | ITGB5 | 27,6 | 45,75 | 46,15 | 67 |
| ENSG00000286242 | AC018797,3 | 2,58 | 3,87 | 4,3 | 67 |
| ENSG00000215386 | MIR99AHG | 32,48 | 34,97 | 54,1 | 67 |
| ENSG00000163491 | NEK10 | 0,98 | 1,01 | 1,63 | 66 |
| ENSG00000169184 | MN1 | 5,52 | 7,24 | 9,14 | 66 |
| ENSG00000275045 | GTF2H2 | 3,07 | 3,21 | 5,08 | 65 |
| ENSG00000124067 | SLC12A4 | 3,17 | 5,04 | 5,19 | 64 |
| ENSG00000178031 | ADAMTSL1 | 0,63 | 0,91 | 1,03 | 63 |
| ENSG00000273451 | AL031666,3 | 1,23 | 1,67 | 2,01 | 63 |
| ENSG00000221676 | RNU6ATAC | 424,84 | 619,78 | 692,83 | 63 |
| ENSG00000256977 | LIMS3 | 0,65 | 1,02 | 1,06 | 63 |
| ENSG00000101489 | CELF4 | 3,4 | 3,77 | 5,54 | 63 |
| ENSG00000263163 | SLC27A3 | 1,36 | 2,05 | 2,21 | 63 |
| ENSG00000095370 | SH2D3C | 1,41 | 1,97 | 2,29 | 62 |
| ENSG00000170092 | SPDYE5 | 1,4 | 1,86 | 2,27 | 62 |
| ENSG00000112964 | GHR | 1,58 | 2,19 | 2,56 | 62 |
| ENSG00000204442 | FAM155A | 0,63 | 0,65 | 1,02 | 62 |
| ENSG00000162545 | CAMK2N1 | 17,57 | 27,88 | 28,42 | 62 |
| ENSG00000286529 | AC090398,2 | 2,43 | 3,82 | 3,93 | 62 |
| ENSG00000121361 | KCNJ8 | 1,12 | 1,21 | 1,81 | 62 |
| ENSG00000203727 | SAMD5 | 1,08 | 1,67 | 1,74 | 61 |
| ENSG00000196361 | ELAVL3 | 21,75 | 22,82 | 35,02 | 61 |
| ENSG00000149929 | HIRIP3 | 7,39 | 8,25 | 11,89 | 61 |
| ENSG00000276161 | SNORA17B | 199,28 | 268,77 | 320,53 | 61 |
| ENSG00000085231 | AK6 | 2,77 | 3,16 | 4,45 | 61 |
| ENSG00000178235 | SLITRK1 | 3,07 | 3,67 | 4,93 | 61 |
| ENSG00000269707 | AC018730,1 | 3,77 | 5,04 | 6,05 | 60 |
| ENSG00000225968 | ELFN1 | 5,03 | 5,43 | 8,02 | 59 |
| ENSG00000146426 | TIAM2 | 8,56 | 12,94 | 13,64 | 59 |
| ENSG00000212607 | SNORA3B | 9,88 | 14,59 | 15,71 | 59 |
| ENSG00000248458 | AL139147,1 | 3,44 | 4,33 | 5,46 | 59 |
| ENSG00000231738 | TSPAN19 | 0,64 | 1 | 1,01 | 58 |
| ENSG00000284796 | DCAF11 | 8,49 | 11,52 | 13,38 | 58 |
| ENSG00000185219 | ZNF445 | 4,15 | 4,55 | 6,54 | 58 |
| ENSG00000265790 | RNASEH1P1 | 1,65 | 2 | 2,6 | 58 |
| ENSG00000072134 | EPN2 | 12,77 | 16,66 | 20,11 | 57 |
| ENSG00000129595 | EPB41L4A | 8,26 | 10,72 | 13 | 57 |
| ENSG00000276005 | AC138749,8 | 1,66 | 1,91 | 2,61 | 57 |
| ENSG00000277114 | CNOT3 | 1,49 | 2,2 | 2,34 | 57 |
| ENSG00000224687 | RASAL2-AS1 | 3,06 | 3,98 | 4,8 | 57 |
| ENSG00000247081 | BAALC-AS1 | 4,73 | 7,15 | 7,41 | 57 |
| ENSG00000256101 | AC092745,1 | 0,92 | 1,25 | 1,44 | 57 |
| ENSG00000203666 | EFCAB2 | 7,76 | 11,07 | 12,13 | 56 |
| ENSG00000171723 | GPHN | 8,86 | 10,22 | 13,84 | 56 |
| ENSG00000170382 | LRRN2 | 1,8 | 2,51 | 2,8 | 56 |

| | | | | | |
|---|---|---|---|---|---|
| ENSG00000152377 | SPOCK1 | 25,87 | 27,95 | 40,24 | 56 |
| ENSG00000250950 | AC093752,2 | 1,91 | 2,75 | 2,97 | 55 |
| ENSG00000109686 | SH3D19 | 7,53 | 8,34 | 11,7 | 55 |
| ENSG00000169249 | ZRSR2 | 4,13 | 5,2 | 6,41 | 55 |
| ENSG00000251620 | STPG2-AS1 | 1,31 | 1,42 | 2,03 | 55 |
| ENSG00000084731 | KIF3C | 9,69 | 12,22 | 15,01 | 55 |
| ENSG00000144619 | CNTN4 | 2,85 | 4,26 | 4,4 | 54 |
| ENSG00000113721 | PDGFRB | 1,03 | 1,24 | 1,59 | 54 |
| ENSG00000105928 | GSDME | 7,01 | 7,86 | 10,82 | 54 |
| ENSG00000067082 | KLF6 | 12,42 | 15,61 | 19,17 | 54 |
| ENSG00000118513 | MYB | 2,36 | 2,77 | 3,64 | 54 |
| ENSG00000232472 | EEF1B2P3 | 0,98 | 1,3 | 1,51 | 54 |
| ENSG00000134569 | LRP4 | 4,46 | 4,62 | 6,87 | 54 |
| ENSG00000272620 | AFAP1-AS1 | 0,67 | 0,74 | 1,03 | 54 |
| ENSG00000123080 | CDKN2C | 1,49 | 1,51 | 2,29 | 54 |
| ENSG00000228549 | BX284668,2 | 2,67 | 3,03 | 4,1 | 54 |
| ENSG00000168824 | NSG1 | 8,75 | 10,6 | 13,41 | 53 |
| ENSG00000073417 | PDE8A | 2,61 | 3,8 | 4 | 53 |
| ENSG00000166426 | CRABP1 | 15,79 | 16,77 | 24,19 | 53 |
| ENSG00000273026 | AL358472,3 | 1,75 | 1,92 | 2,68 | 53 |
| ENSG00000274429 | DLG5 | 8,8 | 9,54 | 13,47 | 53 |
| ENSG00000103154 | NECAB2 | 7,13 | 10,51 | 10,89 | 53 |
| ENSG00000249242 | TMEM150C | 1,29 | 1,53 | 1,97 | 53 |
| ENSG00000108950 | FAM20A | 1,14 | 1,66 | 1,74 | 53 |
| ENSG00000270986 | HMGB1P51 | 0,78 | 1,12 | 1,19 | 53 |
| ENSG00000287001 | AC010624,5 | 11,68 | 15,34 | 17,8 | 52 |
| ENSG00000196302 | GUSBP15 | 15,94 | 23,4 | 24,29 | 52 |
| ENSG00000223843 | EFCAB6-AS1 | 5,19 | 6,14 | 7,9 | 52 |
| ENSG00000169981 | ZNF35 | 3,41 | 3,74 | 5,19 | 52 |
| ENSG00000281306 | ZNF35 | 3,41 | 3,74 | 5,19 | 52 |
| ENSG00000177551 | NHLH2 | 4,14 | 5,84 | 6,3 | 52 |
| ENSG00000147010 | SH3KBP1 | 3,45 | 3,86 | 5,25 | 52 |
| ENSG00000248019 | FAM13A-AS1 | 0,71 | 0,77 | 1,08 | 52 |
| ENSG00000206120 | EGFEM1P | 13,5 | 17,12 | 20,5 | 52 |
| ENSG00000167778 | SPRYD3 | 2,18 | 2,85 | 3,31 | 52 |
| ENSG00000207014 | SNORD116-3 | 93,71 | 140,43 | 142,28 | 52 |
| ENSG00000206727 | SNORD116-9 | 93,71 | 140,43 | 142,28 | 52 |
| ENSG00000266993 | AL050343,2 | 1,06 | 1,51 | 1,6 | 51 |
| ENSG00000287998 | AC104232,3 | 1,65 | 2,19 | 2,49 | 51 |
| ENSG00000282002 | AC243807,1 | 0,71 | 0,91 | 1,07 | 51 |
| ENSG00000235731 | LINC02250 | 0,81 | 0,99 | 1,22 | 51 |
| ENSG00000108592 | FTSJ3 | 9,11 | 13,05 | 13,72 | 51 |
| ENSG00000204682 | MIR1915HG | 6,99 | 9,15 | 10,52 | 51 |
| ENSG00000254788 | CKLF-CMTM1 | 2,62 | 3,27 | 3,93 | 50 |
| ENSG00000258571 | PTTG4P | 0,92 | 1,15 | 1,38 | 50 |
| ENSG00000151490 | PTPRO | 5,26 | 7,06 | 7,86 | 49 |
| ENSG00000259345 | AC013652,1 | 1,79 | 2,06 | 2,67 | 49 |
| ENSG00000261485 | PAN3-AS1 | 1,1 | 1,25 | 1,64 | 49 |
| ENSG00000089737 | DDX24 | 9,91 | 11,43 | 14,76 | 49 |
| ENSG00000081913 | PHLPP1 | 14,92 | 17,67 | 22,22 | 49 |

| | | | | | |
|---|---|---|---|---|---|
| ENSG00000154917 | RAB6B | 6,05 | 6,67 | 9 | 49 |
| ENSG00000187605 | TET3 | 6,49 | 7,76 | 9,65 | 49 |
| ENSG00000144668 | ITGA9 | 0,7 | 0,92 | 1,04 | 49 |
| ENSG00000234494 | SP2-AS1 | 1,22 | 1,74 | 1,81 | 48 |
| ENSG00000286689 | AC090946,1 | 1,02 | 1,17 | 1,51 | 48 |
| ENSG00000116141 | MARK1 | 7,59 | 8,99 | 11,22 | 48 |
| ENSG00000185737 | NRG3 | 2,77 | 2,98 | 4,08 | 47 |
| ENSG00000253554 | LINC01414 | 3,29 | 3,39 | 4,83 | 47 |
| ENSG00000214756 | CSKMT | 16,01 | 19,79 | 23,49 | 47 |
| ENSG00000231172 | AC007099,1 | 2,36 | 2,79 | 3,46 | 47 |
| ENSG00000268496 | AC245884,9 | 1,66 | 1,99 | 2,43 | 46 |
| ENSG00000245694 | CRNDE | 19,37 | 21,69 | 28,31 | 46 |
| ENSG00000139187 | KLRG1 | 2,44 | 2,89 | 3,56 | 46 |
| ENSG00000166450 | PRTG | 34,9 | 38,18 | 50,91 | 46 |
| ENSG00000106976 | DNM1 | 20,79 | 22,3 | 30,27 | 46 |
| ENSG00000164465 | DCBLD1 | 2,37 | 3,15 | 3,45 | 46 |
| ENSG00000197813 | AC011450,1 | 4,49 | 6,4 | 6,5 | 45 |
| ENSG00000223768 | LINC00205 | 3,71 | 5,24 | 5,37 | 45 |
| ENSG00000281950 | SPDYE16 | 2,55 | 2,93 | 3,69 | 45 |
| ENSG00000231859 | AC079781,1 | 1,98 | 2 | 2,85 | 44 |
| ENSG00000160188 | RSPH1 | 0,78 | 1,1 | 1,12 | 44 |
| ENSG00000206395 | DDAH2 | 6,27 | 7,17 | 9 | 44 |
| ENSG00000226634 | DDAH2 | 6,27 | 7,17 | 9 | 44 |
| ENSG00000155816 | FMN2 | 6,13 | 6,42 | 8,78 | 43 |
| ENSG00000135269 | TES | 12,55 | 14,91 | 17,96 | 43 |
| ENSG00000162692 | VCAM1 | 1,94 | 2,5 | 2,77 | 43 |
| ENSG00000157350 | ST3GAL2 | 10,72 | 13,45 | 15,3 | 43 |
| ENSG00000145147 | SLIT2 | 35,15 | 43,08 | 50,13 | 43 |
| ENSG00000196440 | ARMCX4 | 4,16 | 5,77 | 5,93 | 43 |
| ENSG00000258515 | AL355075,2 | 2,13 | 2,62 | 3,03 | 42 |
| ENSG00000179242 | CDH4 | 4,07 | 4,75 | 5,78 | 42 |
| ENSG00000277450 | AC002094,4 | 1,1 | 1,25 | 1,56 | 42 |
| ENSG00000170949 | ZNF160 | 8,75 | 9,28 | 12,39 | 42 |
| ENSG00000188818 | ZDHHC11 | 3,74 | 3,93 | 5,29 | 41 |
| ENSG00000139908 | TSSK4 | 1 | 1,24 | 1,41 | 41 |
| ENSG00000285140 | TSSK4 | 1 | 1,24 | 1,41 | 41 |
| ENSG00000144810 | COL8A1 | 8,54 | 9,3 | 12,04 | 41 |
| ENSG00000196550 | FAM72A | 2,92 | 3,35 | 4,11 | 41 |
| ENSG00000240370 | RPL13P5 | 2,26 | 2,83 | 3,18 | 41 |
| ENSG00000181234 | TMEM132C | 1,16 | 1,21 | 1,63 | 41 |
| ENSG00000255441 | SIGLEC10-AS1 | 0,79 | 0,98 | 1,11 | 41 |
| ENSG00000184489 | PTP4A3 | 1,73 | 2,25 | 2,43 | 40 |
| ENSG00000161082 | CELF5 | 4,6 | 6,44 | 6,46 | 40 |
| ENSG00000257896 | AC093012,1 | 0,92 | 1,24 | 1,29 | 40 |
| ENSG00000113205 | PCDHB3 | 2,29 | 2,3 | 3,21 | 40 |
| ENSG00000154511 | DIPK1A | 1,37 | 1,77 | 1,92 | 40 |
| ENSG00000282956 | MAGI1 | 16,21 | 16,47 | 22,62 | 40 |
| ENSG00000151276 | MAGI1 | 16,21 | 16,47 | 22,62 | 40 |
| ENSG00000181744 | DIPK2A | 13,25 | 14 | 18,48 | 39 |
| ENSG00000119866 | BCL11A | 11,76 | 12,93 | 16,37 | 39 |

| | | | | | |
|---|---|---|---|---|---|
| ENSG00000287169 | AC107050,1 | 0,92 | 1,17 | 1,28 | 39 |
| ENSG00000198597 | ZNF536 | 3,63 | 4,33 | 5,05 | 39 |
| ENSG00000169554 | ZEB2 | 27,43 | 28,45 | 38,06 | 39 |
| ENSG00000132881 | CPLANE2 | 1,68 | 2,25 | 2,33 | 39 |
| ENSG00000198695 | MT-ND6 | 816,28 | 963,63 | 1130,8 | 39 |
| ENSG00000196937 | FAM3C | 11,34 | 13,73 | 15,67 | 38 |
| ENSG00000100376 | FAM118A | 8,54 | 11,15 | 11,8 | 38 |
| ENSG00000288091 | AC062022,2 | 0,84 | 0,94 | 1,16 | 38 |
| ENSG00000262663 | AC087222,1 | 2,97 | 3,39 | 4,1 | 38 |
| ENSG00000268070 | AC006539,2 | 1,71 | 2,12 | 2,36 | 38 |
| ENSG00000168280 | KIF5C | 32,73 | 38 | 45,11 | 38 |
| ENSG00000146909 | NOM1 | 5,81 | 7,62 | 8 | 38 |
| ENSG00000270722 | RNVU1-31 | 41,76 | 55,17 | 57,5 | 38 |
| ENSG00000213676 | ATF6B | 4,31 | 5,35 | 5,92 | 37 |
| ENSG00000114450 | GNB4 | 12,81 | 14,49 | 17,59 | 37 |
| ENSG00000071575 | TRIB2 | 46,01 | 49,63 | 63,05 | 37 |
| ENSG00000118473 | SGIP1 | 3,47 | 4,1 | 4,75 | 37 |
| ENSG00000248415 | GAPDHP61 | 1,04 | 1,16 | 1,42 | 37 |
| ENSG00000209482 | | 29,58 | 30,15 | 40,31 | 36 |
| ENSG00000106367 | AP1S1 | 20,48 | 22,4 | 27,89 | 36 |
| ENSG00000262333 | HNRNPA1P16 | 2,02 | 2,08 | 2,75 | 36 |
| ENSG00000250608 | AC010210,1 | 1,11 | 1,46 | 1,51 | 36 |
| ENSG00000250506 | CDK3 | 1,2 | 1,37 | 1,63 | 36 |
| ENSG00000181007 | ZFP82 | 4,85 | 5,08 | 6,58 | 36 |
| ENSG00000273820 | USP27X | 1,29 | 1,41 | 1,75 | 36 |
| ENSG00000268403 | AC132192,2 | 2,34 | 2,42 | 3,17 | 35 |
| ENSG00000198826 | ARHGAP11A | 21,02 | 24,03 | 28,47 | 35 |
| ENSG00000241852 | C8orf58 | 9,29 | 12,4 | 12,58 | 35 |
| ENSG00000201183 | RNVU1-3 | 11,93 | 15,64 | 16,15 | 35 |
| ENSG00000280708 | AC006359,3 | 1,26 | 1,36 | 1,7 | 35 |
| ENSG00000121039 | RDH10 | 2,55 | 3,42 | 3,44 | 35 |
| ENSG00000260081 | AF274858,1 | 2,35 | 3,09 | 3,17 | 35 |
| ENSG00000100344 | PNPLA3 | 3,02 | 3,64 | 4,07 | 35 |
| ENSG00000223508 | RPL23AP53 | 4,37 | 4,39 | 5,88 | 35 |
| ENSG00000006210 | CX3CL1 | 2,52 | 3 | 3,39 | 35 |
| ENSG00000141522 | ARHGDIA | 79,57 | 80,53 | 106,94 | 34 |
| ENSG00000109339 | MAPK10 | 50,39 | 55,54 | 67,7 | 34 |
| ENSG00000239039 | SNORD13 | 3446,01 | 3819,26 | 4629,54 | 34 |
| ENSG00000115902 | SLC1A4 | 9,68 | 11,49 | 13 | 34 |
| ENSG00000275145 | FRG1 | 4,98 | 5,47 | 6,68 | 34 |
| ENSG00000152433 | ZNF547 | 1,44 | 1,66 | 1,93 | 34 |
| ENSG00000273032 | DGCR5 | 4,01 | 4,39 | 5,37 | 34 |
| ENSG00000274675 | GTF2H2C_2 | 7,42 | 8,97 | 9,93 | 34 |
| ENSG00000228242 | XPC-AS1 | 2,4 | 3,04 | 3,21 | 34 |
| ENSG00000157999 | ANKRD61 | 0,89 | 0,92 | 1,19 | 34 |
| ENSG00000181722 | ZBTB20 | 8,45 | 9,12 | 11,28 | 33 |
| ENSG00000090975 | PITPNM2 | 1,41 | 1,65 | 1,88 | 33 |
| ENSG00000156030 | MIDEAS | 4,47 | 4,98 | 5,95 | 33 |
| ENSG00000172456 | FGGY | 6,27 | 6,43 | 8,33 | 33 |
| ENSG00000233885 | YEATS2-AS1 | 1,98 | 2,46 | 2,63 | 33 |

| | | | | | |
|---|---|---|---|---|---|
| ENSG00000251379 | AC099550,1 | 27,26 | 30,47 | 36,14 | 33 |
| ENSG00000103150 | MLYCD | 1,69 | 2,04 | 2,24 | 33 |
| ENSG00000188573 | FBLL1 | 1,88 | 2,13 | 2,49 | 32 |
| ENSG00000270175 | AC023509,3 | 0,93 | 0,98 | 1,23 | 32 |
| ENSG00000148803 | FUOM | 3,02 | 3,26 | 3,99 | 32 |
| ENSG00000165632 | TAF3 | 3,15 | 3,94 | 4,15 | 32 |
| ENSG00000145916 | RMND5B | 9,88 | 12,38 | 13,01 | 32 |
| ENSG00000160953 | PWWP3A | 17,52 | 18,07 | 23,07 | 32 |
| ENSG00000198752 | CDC42BPB | 18,78 | 22,27 | 24,72 | 32 |
| ENSG00000242732 | RTL5 | 6,72 | 7,62 | 8,84 | 32 |
| ENSG00000152061 | RABGAP1L | 9,33 | 10,84 | 12,27 | 32 |
| ENSG00000284294 | AC007326,5 | 5,04 | 5,28 | 6,62 | 31 |
| ENSG00000173889 | PHC3 | 11,2 | 13,61 | 14,71 | 31 |
| ENSG00000286904 | AC093675,1 | 0,83 | 0,93 | 1,09 | 31 |
| ENSG00000268947 | AC002128,1 | 0,99 | 1,04 | 1,3 | 31 |
| ENSG00000158417 | EIF5B | 13,37 | 13,76 | 17,55 | 31 |
| ENSG00000274315 | AC009318,3 | 1,74 | 1,8 | 2,28 | 31 |
| ENSG00000100647 | SUSD6 | 2,84 | 3,22 | 3,72 | 31 |
| ENSG00000286360 | AC107081,3 | 1,42 | 1,65 | 1,86 | 31 |
| ENSG00000173875 | ZNF791 | 5,24 | 5,35 | 6,86 | 31 |
| ENSG00000242950 | ERVW-1 | 5,91 | 7,6 | 7,73 | 31 |
| ENSG00000182175 | RGMA | 44,24 | 50,95 | 57,84 | 31 |
| ENSG00000106278 | PTPRZ1 | 112,98 | 121,72 | 147,51 | 31 |
| ENSG00000287933 | AL357141,1 | 10,82 | 13,49 | 14,12 | 30 |
| ENSG00000107282 | APBA1 | 1,05 | 1,28 | 1,37 | 30 |
| ENSG00000251034 | AC087854,1 | 25,47 | 31,28 | 33,21 | 30 |
| ENSG00000121417 | ZNF211 | 7,49 | 9,23 | 9,76 | 30 |
| ENSG00000274403 | AC090510,2 | 1,53 | 1,64 | 1,99 | 30 |
| ENSG00000143614 | GATAD2B | 9,65 | 11,46 | 12,55 | 30 |
| ENSG00000148339 | SLC25A25 | 5,66 | 7,27 | 7,35 | 30 |
| ENSG00000099251 | HSD17B7P2 | 1,32 | 1,64 | 1,71 | 30 |
| ENSG00000288374 | HSD17B7P2 | 1,32 | 1,64 | 1,71 | 30 |
| ENSG00000206077 | ZDHHC11B | 1,8 | 2,11 | 2,33 | 29 |
| ENSG00000040608 | RTN4R | 4,23 | 4,48 | 5,47 | 29 |
| ENSG00000174740 | PABPC5 | 4,04 | 4,83 | 5,22 | 29 |
| ENSG00000257354 | AC048341,1 | 1,85 | 2,02 | 2,39 | 29 |
| ENSG00000236526 | AL035448,1 | 1,59 | 1,62 | 2,05 | 29 |
| ENSG00000272146 | ARF4-AS1 | 1,39 | 1,43 | 1,79 | 29 |
| ENSG00000150625 | GPM6A | 25,55 | 27,81 | 32,9 | 29 |
| ENSG00000286966 | AL137800,1 | 1,53 | 1,82 | 1,97 | 29 |
| ENSG00000092853 | CLSPN | 13,37 | 16,14 | 17,19 | 29 |
| ENSG00000108669 | CYTH1 | 4,34 | 5,09 | 5,58 | 29 |
| ENSG00000187980 | PLA2G2C | 0,84 | 0,97 | 1,08 | 29 |
| ENSG00000125931 | CITED1 | 1,37 | 1,62 | 1,76 | 28 |
| ENSG00000158006 | PAFAH2 | 2,71 | 2,82 | 3,48 | 28 |
| ENSG00000228794 | LINC01128 | 7,01 | 7,9 | 9 | 28 |
| ENSG00000119899 | SLC17A5 | 3,57 | 3,69 | 4,58 | 28 |
| ENSG00000126107 | HECTD3 | 6,05 | 6,87 | 7,76 | 28 |
| ENSG00000198042 | MAK16 | 7,45 | 7,67 | 9,55 | 28 |
| ENSG00000234773 | AC012618,3 | 4,41 | 5,16 | 5,65 | 28 |

| | | | | | |
|---|---|---|---|---|---|
| ENSG00000188372 | ZP3 | 5,93 | 6,33 | 7,59 | 28 |
| ENSG00000127418 | FGFRL1 | 5,88 | 6,24 | 7,52 | 28 |
| ENSG00000164082 | GRM2 | 1,04 | 1,05 | 1,33 | 28 |
| ENSG00000198937 | CCDC167 | 44,05 | 45,71 | 56,27 | 28 |
| ENSG00000171877 | FRMD5 | 2,5 | 2,62 | 3,19 | 28 |
| ENSG00000205559 | CHKB-DT | 4,46 | 4,99 | 5,69 | 28 |
| ENSG00000138660 | AP1AR | 7,99 | 9,45 | 10,19 | 28 |
| ENSG00000242600 | MBL1P | 1,2 | 1,22 | 1,53 | 28 |
| ENSG00000164221 | CCDC112 | 11,5 | 11,52 | 14,64 | 27 |
| ENSG00000106089 | STX1A | 2,36 | 2,91 | 3 | 27 |
| ENSG00000170873 | MTSS1 | 13,92 | 15,85 | 17,66 | 27 |
| ENSG00000246889 | AP000487,1 | 0,83 | 1 | 1,05 | 27 |
| ENSG00000288166 | AP000487,3 | 0,83 | 1 | 1,05 | 27 |
| ENSG00000232729 | AC211433,1 | 3,02 | 3,7 | 3,82 | 26 |
| ENSG00000177030 | DEAF1 | 11,91 | 13,18 | 15,06 | 26 |
| ENSG00000236204 | LINC01376 | 2,06 | 2,2 | 2,6 | 26 |
| ENSG00000107957 | SH3PXD2A | 2,71 | 2,77 | 3,41 | 26 |
| ENSG00000183354 | KIAA2026 | 7,11 | 8,75 | 8,94 | 26 |
| ENSG00000221475 | SNORA11D | 30,08 | 32,69 | 37,8 | 26 |
| ENSG00000221705 | SNORA11E | 30,08 | 32,69 | 37,8 | 26 |
| ENSG00000162512 | SDC3 | 33,13 | 38,83 | 41,61 | 26 |
| ENSG00000159200 | RCAN1 | 4,23 | 4,76 | 5,31 | 26 |
| ENSG00000064666 | CNN2 | 83,67 | 83,82 | 104,92 | 25 |
| ENSG00000178922 | HYI | 10,62 | 11,59 | 13,31 | 25 |
| ENSG00000275575 | PTP4A3 | 3,99 | 4,63 | 5 | 25 |
| ENSG00000236018 | AC004898,1 | 1,75 | 1,98 | 2,19 | 25 |

| HD-NSCs | | [TPM] | | | | |
|---|---|---|---|---|---|---|
| gene_ID | Gene_name | HD-NSC_1 | HD-NSC_2 | HD-NSC_3 | HD-NSC_4 | change_% |
| ENSG00000019186 | CYP24A1 | 0 | 0,45 | 1,78 | 14,28 | |
| ENSG00000252050 | | 0 | 4,6 | 5,33 | 6,59 | |
| ENSG00000252135 | | 0 | 1,89 | 4,41 | 5,39 | |
| ENSG00000260318 | COX6CP1 | 0 | 1,18 | 1,72 | 4,48 | |
| ENSG00000278444 | CDC42EP5 | 0 | 2,86 | 2,99 | 3,05 | |
| ENSG00000259519 | AC051619,3 | 0 | 0,53 | 1,29 | 2,53 | |
| ENSG00000211829 | TRDC | 0 | 0,14 | 0,42 | 2,19 | |
| ENSG00000243883 | RN7SL419P | 0 | 0,14 | 0,7 | 1,88 | |
| ENSG00000220517 | ASS1P1 | 0 | 0,12 | 0,52 | 1,44 | |
| ENSG00000225544 | | 0 | 0,26 | 0,32 | 1,22 | |
| ENSG00000239856 | RN7SL225P | 0 | 0,13 | 0,22 | 1,21 | |
| ENSG00000236808 | POLR1H | 0 | 0,1 | 0,49 | 1,2 | |
| ENSG00000233795 | POLR1H | 0 | 0,1 | 0,49 | 1,2 | |
| ENSG00000235176 | POLR1H | 0 | 0,1 | 0,49 | 1,2 | |
| ENSG00000235443 | POLR1H | 0 | 0,1 | 0,5 | 1,2 | |
| ENSG00000236949 | POLR1H | 0 | 0,1 | 0,5 | 1,2 | |
| ENSG00000125879 | | 0 | 0,02 | 0,08 | 1,12 | |
| ENSG00000226890 | Z97652,1 | 0 | 0,16 | 0,29 | 1,04 | |
| ENSG00000078098 | FAP | 0,01 | 0,15 | 0,32 | 1,44 | 14300 |
| ENSG00000237898 | HCG15 | 0,01 | 0,62 | 0,63 | 1,23 | 12200 |
| ENSG00000263360 | RN7SL134P | 0,01 | 0,67 | 1,05 | 1,16 | 11500 |
| ENSG00000138207 | RBP4 | 0,02 | 0,2 | 0,4 | 2,12 | 10500 |
| ENSG00000111341 | MGP | 0,18 | 0,7 | 1,52 | 12,76 | 6989 |
| ENSG00000253819 | LINC01151 | 0,03 | 0,17 | 0,64 | 2,13 | 7000 |
| ENSG00000267280 | TBX2-AS1 | 0,03 | 0,5 | 0,72 | 1,71 | 5600 |
| ENSG00000234210 | AC006372,3 | 0,04 | 0,26 | 0,33 | 1,85 | 4525 |
| ENSG00000127528 | KLF2 | 0,03 | 0,36 | 0,82 | 1,38 | 4500 |
| ENSG00000152583 | SPARCL1 | 0,69 | 4,25 | 6,69 | 29,7 | 4204 |
| ENSG00000177103 | DSCAML1 | 0,03 | 0,33 | 0,55 | 1,23 | 4000 |
| ENSG00000147481 | SNTG1 | 0,09 | 0,13 | 0,59 | 2,89 | 3111 |
| ENSG00000121068 | TBX2 | 0,08 | 0,49 | 1,26 | 2,48 | 3000 |
| ENSG00000135525 | MAP7 | 0,21 | 1,03 | 1,41 | 6,08 | 2795 |
| ENSG00000223867 | | 0,09 | 0,2 | 1,12 | 2,58 | 2767 |
| ENSG00000113924 | HGD | 0,18 | 0,94 | 1,05 | 4,98 | 2667 |
| ENSG00000238222 | MKRN4P | 0,04 | 0,15 | 0,32 | 1,07 | 2575 |
| ENSG00000215875 | ST13P20 | 0,05 | 0,18 | 0,39 | 1,25 | 2400 |
| ENSG00000147869 | CER1 | 0,06 | 0,8 | 1,05 | 1,45 | 2317 |
| ENSG00000108691 | CCL2 | 3,16 | 19,65 | 52,57 | 74,84 | 2268 |
| ENSG00000120708 | TGFBI | 2,87 | 6,79 | 18,01 | 67,32 | 2246 |
| ENSG00000261857 | MIA | 0,57 | 3,24 | 3,89 | 13,37 | 2246 |
| ENSG00000136099 | PCDH8 | 0,48 | 1,44 | 3,69 | 10,67 | 2123 |
| ENSG00000283128 | AC009403,2 | 1,12 | 6,26 | 8,47 | 23,57 | 2004 |
| ENSG00000184221 | OLIG1 | 0,4 | 1,25 | 2,89 | 8,34 | 1985 |
| ENSG00000137252 | HCRTR2 | 0,05 | 0,37 | 0,63 | 1,04 | 1980 |
| ENSG00000146910 | CNPY1 | 3,45 | 21,76 | 31,38 | 71,42 | 1970 |
| ENSG00000285446 | Z84488,1 | 0,07 | 0,48 | 0,68 | 1,45 | 1971 |
| ENSG00000075035 | WSCD2 | 0,15 | 1,05 | 2,12 | 3,02 | 1913 |
| ENSG00000177468 | OLIG3 | 0,37 | 1,16 | 2,61 | 7,06 | 1808 |

| | | | | | | |
|---|---|---|---|---|---|---|
| ENSG00000164778 | EN2 | 2,01 | 11,2 | 15,1 | 38,3 | 1805 |
| ENSG00000286546 | AL079338,1 | 0,16 | 0,4 | 0,7 | 3 | 1775 |
| ENSG00000064300 | NGFR | 1,41 | 4,99 | 9,66 | 26,39 | 1772 |
| ENSG00000189058 | APOD | 0,07 | 0,13 | 0,4 | 1,27 | 1714 |
| ENSG00000206412 | GNL1 | 0,29 | 0,51 | 4,62 | 5,08 | 1652 |
| ENSG00000170961 | HAS2 | 1,35 | 5,11 | 10,73 | 23,32 | 1627 |
| ENSG00000137965 | IFI44 | 0,28 | 1,55 | 3,3 | 4,68 | 1571 |
| ENSG00000285633 | AL132633,1 | 0,67 | 1,81 | 3,45 | 11,17 | 1567 |
| ENSG00000207955 | AL359091,1 | 0,4 | 1,43 | 2,07 | 6,6 | 1550 |
| ENSG00000205927 | OLIG2 | 0,19 | 0,35 | 1,02 | 3,06 | 1511 |
| ENSG00000167779 | IGFBP6 | 0,07 | 0,26 | 0,45 | 1,05 | 1400 |
| ENSG00000106511 | MEOX2 | 0,32 | 0,57 | 1,49 | 4,76 | 1388 |
| ENSG00000270105 | AC136475,8 | 0,1 | 0,34 | 0,52 | 1,46 | 1360 |
| ENSG00000150275 | PCDH15 | 0,19 | 0,76 | 1,04 | 2,74 | 1342 |
| ENSG00000275695 | UBE2Q2P6 | 0,19 | 0,41 | 2,15 | 2,74 | 1342 |
| ENSG00000205325 | AC005863,1 | 0,12 | 0,48 | 0,78 | 1,68 | 1300 |
| ENSG00000227288 | ARID3BP1 | 0,08 | 0,21 | 0,32 | 1,11 | 1288 |
| ENSG00000177301 | KCNA2 | 0,26 | 0,92 | 1,56 | 3,57 | 1273 |
| ENSG00000168993 | CPLX1 | 0,26 | 0,56 | 1,82 | 3,46 | 1231 |
| ENSG00000122176 | FMOD | 0,54 | 1,71 | 3,98 | 7,07 | 1209 |
| ENSG00000196767 | POU3F4 | 1,08 | 3,27 | 5,33 | 13,92 | 1189 |
| ENSG00000227245 | | 0,09 | 0,28 | 0,33 | 1,15 | 1178 |
| ENSG00000285780 | AC109129,1 | 0,15 | 0,4 | 0,5 | 1,88 | 1153 |
| ENSG00000184672 | RALYL | 0,52 | 2,87 | 4,19 | 6,5 | 1150 |
| ENSG00000125430 | HS3ST3B1 | 0,92 | 2,57 | 4,98 | 11,49 | 1149 |
| ENSG00000196569 | LAMA2 | 0,6 | 2,83 | 4,93 | 7,4 | 1133 |
| ENSG00000163359 | COL6A3 | 1,73 | 6,55 | 7,97 | 20,75 | 1099 |
| ENSG00000073756 | PTGS2 | 0,11 | 0,14 | 0,48 | 1,32 | 1100 |
| ENSG00000286139 | ARHGAP11B | 0,13 | 0,69 | 1,35 | 1,55 | 1092 |
| ENSG00000137959 | IFI44L | 0,7 | 5,21 | 6,85 | 8,33 | 1090 |
| ENSG00000237321 | | 0,09 | 0,2 | 0,24 | 1,07 | 1089 |
| ENSG00000255571 | MIR9-3HG | 0,27 | 1,38 | 1,84 | 3,15 | 1067 |
| ENSG00000282435 | AC246817,4 | 0,11 | 0,5 | 0,52 | 1,28 | 1064 |
| ENSG00000241357 | | 0,11 | 0,24 | 0,38 | 1,23 | 1018 |
| ENSG00000113356 | POLR3G | 1,55 | 3,4 | 7,27 | 17,26 | 1014 |
| ENSG00000152578 | GRIA4 | 0,46 | 1,36 | 3,03 | 5,12 | 1013 |
| ENSG00000213654 | GPSM3 | 0,18 | 0,53 | 0,72 | 2 | 1011 |
| ENSG00000050767 | COL23A1 | 0,36 | 0,84 | 1,06 | 3,95 | 997 |
| ENSG00000239494 | RN7SL333P | 0,23 | 0,25 | 0,44 | 2,43 | 957 |
| ENSG00000184254 | ALDH1A3 | 0,19 | 0,81 | 1,14 | 2 | 953 |
| ENSG00000171956 | FOXB1 | 0,54 | 1,97 | 3,05 | 5,58 | 933 |
| ENSG00000120251 | GRIA2 | 0,64 | 2,57 | 3,2 | 6,46 | 909 |
| ENSG00000266709 | AC005224,3 | 0,66 | 1,41 | 2,67 | 6,66 | 909 |
| ENSG00000266378 | AC005224,2 | 0,13 | 0,17 | 0,37 | 1,31 | 908 |
| ENSG00000227094 | | 0,1 | 0,26 | 0,34 | 1 | 900 |
| ENSG00000160307 | S100B | 1,73 | 3,55 | 8,66 | 17,19 | 894 |
| ENSG00000232821 | AC003986,2 | 0,24 | 0,74 | 0,98 | 2,34 | 875 |
| ENSG00000243621 | | 0,6 | 0,68 | 1,1 | 5,74 | 857 |
| ENSG00000165061 | ZMAT4 | 2,82 | 8,35 | 12,73 | 26,88 | 853 |
| ENSG00000122691 | TWIST1 | 2,82 | 9,29 | 15,52 | 26,87 | 853 |

| | | | | | | |
|---|---|---|---|---|---|---|
| ENSG00000154930 | ACSS1 | 0,23 | 0,52 | 0,7 | 2,17 | 843 |
| ENSG00000137558 | PI15 | 0,34 | 0,4 | 0,79 | 3,09 | 809 |
| ENSG00000100234 | TIMP3 | 4,12 | 7,54 | 17,18 | 37,2 | 803 |
| ENSG00000101134 | DOK5 | 1,51 | 4,75 | 9,4 | 13,54 | 797 |
| ENSG00000237773 | AC073332,1 | 0,34 | 0,41 | 0,91 | 3,03 | 791 |
| ENSG00000166250 | CLMP | 1,84 | 4,89 | 7,51 | 16,01 | 770 |
| ENSG00000232143 | GNL1 | 0,29 | 0,44 | 0,89 | 2,52 | 769 |
| ENSG00000228581 | GNL1 | 0,29 | 0,44 | 0,89 | 2,52 | 769 |
| ENSG00000204590 | GNL1 | 0,29 | 0,44 | 0,89 | 2,52 | 769 |
| ENSG00000168505 | GBX2 | 7,85 | 29,01 | 43,55 | 68,15 | 768 |
| ENSG00000187017 | ESPN | 0,3 | 0,78 | 0,96 | 2,6 | 767 |
| ENSG00000170624 | SGCD | 0,25 | 0,61 | 0,8 | 2,16 | 764 |
| ENSG00000257986 | LINC02306 | 0,53 | 0,97 | 1,38 | 4,56 | 760 |
| ENSG00000143387 | CTSK | 1,99 | 3,29 | 5,31 | 16,85 | 747 |
| ENSG00000170891 | CYTL1 | 0,33 | 0,54 | 0,81 | 2,79 | 745 |
| ENSG00000244056 | RN7SL417P | 0,76 | 1,07 | 1,55 | 6,38 | 739 |
| ENSG00000155980 | KIF5A | 5,98 | 9,53 | 27,18 | 50,13 | 738 |
| ENSG00000261864 | AC130462,2 | 0,12 | 0,25 | 0,46 | 1 | 733 |
| ENSG00000120149 | MSX2 | 0,89 | 1,59 | 2,46 | 7,4 | 731 |
| ENSG00000111348 | ARHGDIB | 2,96 | 7,12 | 13,65 | 24,47 | 727 |
| ENSG00000078596 | ITM2A | 1,24 | 2,2 | 4,47 | 10,17 | 720 |
| ENSG00000115252 | PDE1A | 0,48 | 0,83 | 2,48 | 3,93 | 719 |
| ENSG00000257918 | AC079385,3 | 0,3 | 0,72 | 0,95 | 2,45 | 717 |
| ENSG00000237457 | LINC01351 | 0,17 | 0,18 | 0,44 | 1,38 | 712 |
| ENSG00000174600 | CMKLR1 | 0,2 | 0,68 | 0,76 | 1,61 | 705 |
| ENSG00000041982 | TNC | 2,96 | 3,74 | 11,61 | 23,81 | 704 |
| ENSG00000176842 | IRX5 | 0,92 | 1,48 | 3,03 | 7,37 | 701 |
| ENSG00000158955 | WNT9B | 0,13 | 0,2 | 0,54 | 1,04 | 700 |
| ENSG00000111799 | COL12A1 | 9,78 | 20,49 | 39,23 | 76,83 | 686 |
| ENSG00000259571 | BLID | 0,13 | 0,22 | 0,6 | 1,02 | 685 |
| ENSG00000078549 | ADCYAP1R1 | 0,94 | 1,67 | 3,59 | 7,37 | 684 |
| ENSG00000159231 | CBR3 | 0,4 | 0,74 | 1,06 | 3,11 | 678 |
| ENSG00000068078 | FGFR3 | 1 | 1,77 | 2,71 | 7,67 | 667 |
| ENSG00000111432 | FZD10 | 0,97 | 2,32 | 3,86 | 7,33 | 656 |
| ENSG00000075651 | PLD1 | 0,31 | 1,14 | 1,23 | 2,31 | 645 |
| ENSG00000230630 | DNM3OS | 2,06 | 6,43 | 8,91 | 15,26 | 641 |
| ENSG00000011465 | DCN | 10,17 | 22,58 | 44,32 | 73,92 | 627 |
| ENSG00000115738 | ID2 | 14,78 | 28,41 | 47,27 | 106,92 | 623 |
| ENSG00000106034 | CPED1 | 1,75 | 5,91 | 7,9 | 12,51 | 615 |
| ENSG00000229273 | BX664615,1 | 0,36 | 1,26 | 1,8 | 2,56 | 611 |
| ENSG00000138823 | MTTP | 1,53 | 4,08 | 5,6 | 10,83 | 608 |
| ENSG00000179915 | NRXN1 | 3,48 | 6,64 | 23,04 | 24,52 | 605 |
| ENSG00000108379 | WNT3 | 0,47 | 1,87 | 1,91 | 3,31 | 604 |
| ENSG00000044524 | EPHA3 | 4,26 | 9,67 | 14,34 | 29,95 | 603 |
| ENSG00000103316 | CRYM | 0,86 | 2,03 | 3,84 | 6,04 | 602 |
| ENSG00000114654 | EFCC1 | 0,66 | 1,52 | 1,95 | 4,62 | 600 |
| ENSG00000255690 | TRIL | 7,3 | 14,09 | 20,4 | 51,06 | 599 |
| ENSG00000278091 | ZNF85 | 1,33 | 6,94 | 7,21 | 9,28 | 598 |
| ENSG00000242971 | RN7SL233P | 0,41 | 0,45 | 0,64 | 2,86 | 598 |
| ENSG00000173673 | HES3 | 1,01 | 1,52 | 1,88 | 6,98 | 591 |

| | | | | | | |
|---|---|---|---|---|---|---|
| ENSG00000006283 | CACNA1G | 0,9 | 1,82 | 2,62 | 6,21 | 590 |
| ENSG00000237207 | RBM17P3 | 0,18 | 0,33 | 0,46 | 1,24 | 589 |
| ENSG00000101445 | PPP1R16B | 0,15 | 0,3 | 0,63 | 1,03 | 587 |
| ENSG00000172819 | RARG | 1,23 | 1,96 | 3,59 | 8,44 | 586 |
| ENSG00000121039 | RDH10 | 1,88 | 4,34 | 8,07 | 12,88 | 585 |
| ENSG00000005108 | THSD7A | 1,56 | 4,34 | 5,22 | 10,67 | 584 |
| ENSG00000100427 | MLC1 | 0,18 | 0,63 | 0,69 | 1,23 | 583 |
| ENSG00000226440 | LAMA4-AS1 | 0,23 | 0,63 | 0,75 | 1,55 | 574 |
| ENSG00000145794 | MEGF10 | 4,39 | 13,13 | 22,82 | 29,52 | 572 |
| ENSG00000256637 | LINC01965 | 0,49 | 1 | 1,11 | 3,29 | 571 |
| ENSG00000231977 | AL096828,1 | 0,33 | 0,56 | 0,8 | 2,2 | 567 |
| ENSG00000150048 | CLEC1A | 0,36 | 0,98 | 1,99 | 2,37 | 558 |
| ENSG00000267177 | AP002505,1 | 0,72 | 1,51 | 2,59 | 4,73 | 557 |
| ENSG00000255043 | NAV2-AS5 | 0,17 | 0,4 | 0,44 | 1,11 | 553 |
| ENSG00000008441 | NFIX | 0,21 | 0,4 | 0,43 | 1,37 | 552 |
| ENSG00000162599 | NFIA | 4,72 | 15,53 | 22,62 | 30,49 | 546 |
| ENSG00000219747 | AL133260,1 | 0,26 | 0,28 | 0,3 | 1,68 | 546 |
| ENSG00000277626 | WNT3 | 1,41 | 1,58 | 3,6 | 9,08 | 544 |
| ENSG00000136267 | DGKB | 0,43 | 0,51 | 1,7 | 2,73 | 535 |
| ENSG00000117791 | MTARC2 | 0,51 | 1,31 | 1,74 | 3,23 | 533 |
| ENSG00000184111 | RPL11P4 | 0,17 | 0,24 | 0,74 | 1,07 | 529 |
| ENSG00000134817 | APLNR | 0,55 | 1,05 | 2,46 | 3,43 | 524 |
| ENSG00000287800 | | 0,31 | 0,35 | 1 | 1,93 | 523 |
| ENSG00000112769 | LAMA4 | 6,91 | 12,21 | 21,68 | 42,89 | 521 |
| ENSG00000255566 | AC135279,2 | 0,23 | 0,43 | 0,68 | 1,42 | 517 |
| ENSG00000226965 | AC073114,1 | 0,24 | 0,83 | 1,17 | 1,47 | 513 |
| ENSG00000111913 | RIPOR2 | 2,69 | 5,47 | 11,51 | 16,39 | 509 |
| ENSG00000249669 | CARMN | 0,95 | 2,05 | 2,78 | 5,78 | 508 |
| ENSG00000288508 | ISLR | 2,14 | 5,94 | 9,43 | 12,95 | 505 |
| ENSG00000129009 | ISLR | 2,14 | 5,94 | 9,43 | 12,95 | 505 |
| ENSG00000285479 | CACNA1C | 0,63 | 1,37 | 2,39 | 3,81 | 505 |
| ENSG00000151067 | CACNA1C | 0,63 | 1,37 | 2,39 | 3,81 | 505 |
| ENSG00000169594 | BNC1 | 0,4 | 0,66 | 1,18 | 2,41 | 503 |
| ENSG00000184906 | AMYH02020865,1 | 0,31 | 1,02 | 1,54 | 1,86 | 500 |
| ENSG00000184486 | POU3F2 | 8,46 | 16,81 | 20,14 | 50,38 | 496 |
| ENSG00000107242 | PIP5K1B | 2,86 | 7,66 | 11,24 | 16,92 | 492 |
| ENSG00000065717 | TLE2 | 0,31 | 0,67 | 0,85 | 1,83 | 490 |
| ENSG00000274594 | AL391669,1 | 0,26 | 0,28 | 0,84 | 1,53 | 488 |
| ENSG00000231185 | SPRY4-AS1 | 0,36 | 0,81 | 1,15 | 2,1 | 483 |
| ENSG00000241983 | RN7SL566P | 0,24 | 0,51 | 0,66 | 1,4 | 483 |
| ENSG00000271228 | | 0,22 | 0,58 | 0,91 | 1,28 | 482 |
| ENSG00000233639 | PANTR1 | 11,58 | 26,34 | 31,27 | 66,7 | 476 |
| ENSG00000256630 | AC090424,1 | 0,32 | 0,36 | 0,57 | 1,84 | 475 |
| ENSG00000184584 | STING1 | 0,23 | 0,36 | 0,59 | 1,31 | 470 |
| ENSG00000277400 | AC145212,1 | 0,26 | 0,56 | 0,66 | 1,48 | 469 |
| ENSG00000164199 | ADGRV1 | 9,77 | 16,48 | 21,4 | 55,51 | 468 |
| ENSG00000183908 | LRRC55 | 2,65 | 7,01 | 11,23 | 15,01 | 466 |
| ENSG00000276203 | ANKRD20A3P | 0,58 | 0,7 | 0,88 | 3,26 | 462 |
| ENSG00000268643 | AC006486,1 | 0,73 | 0,89 | 1,3 | 4,1 | 462 |
| ENSG00000111783 | RFX4 | 3,97 | 7,75 | 9,93 | 22,27 | 461 |

| | | | | | | |
|---|---|---|---|---|---|---|
| ENSG00000285417 | BX571818,1 | 0,45 | 1 | 1,21 | 2,51 | 458 |
| ENSG00000272255 | AC113361,1 | 0,28 | 0,6 | 1,18 | 1,56 | 457 |
| ENSG00000017483 | SLC38A5 | 1,83 | 8,12 | 9,27 | 10,17 | 456 |
| ENSG00000260807 | CEROX1 | 1,89 | 4,35 | 8,06 | 10,49 | 455 |
| ENSG00000228592 | D21S2088E | 0,57 | 1,47 | 1,73 | 3,16 | 454 |
| ENSG00000259203 | AC016044,1 | 0,6 | 0,65 | 1,29 | 3,32 | 453 |
| ENSG00000173546 | CSPG4 | 0,9 | 2,32 | 2,5 | 4,87 | 441 |
| ENSG00000079308 | TNS1 | 4,18 | 9,88 | 12,33 | 22,6 | 441 |
| ENSG00000288243 | STING1 | 0,23 | 0,36 | 0,52 | 1,24 | 439 |
| ENSG00000189223 | PAX8-AS1 | 0,95 | 2,21 | 2,34 | 5,12 | 439 |
| ENSG00000286615 | AC011416,4 | 0,25 | 0,48 | 0,55 | 1,34 | 436 |
| ENSG00000272023 | AC010240,3 | 0,2 | 0,43 | 0,44 | 1,07 | 435 |
| ENSG00000138639 | ARHGAP24 | 0,97 | 1,48 | 1,99 | 5,18 | 434 |
| ENSG00000125968 | ID1 | 4,77 | 7,73 | 16,18 | 25,44 | 433 |
| ENSG00000076356 | PLXNA2 | 3,1 | 7,63 | 9,16 | 16,52 | 433 |
| ENSG00000176170 | SPHK1 | 3,48 | 5,82 | 9,27 | 18,54 | 433 |
| ENSG00000133519 | ZDHHC8P1 | 0,48 | 0,75 | 1,88 | 2,54 | 429 |
| ENSG00000125618 | PAX8 | 2,73 | 7,32 | 8,78 | 14,42 | 428 |
| ENSG00000166825 | ANPEP | 0,22 | 0,36 | 0,66 | 1,16 | 427 |
| ENSG00000082684 | SEMA5B | 5,06 | 9,79 | 12,9 | 26,59 | 425 |
| ENSG00000169891 | REPS2 | 0,24 | 0,64 | 0,81 | 1,26 | 425 |
| ENSG00000198914 | POU3F3 | 6,5 | 13,91 | 15,35 | 33,99 | 423 |
| ENSG00000262815 | AC087501,2 | 0,34 | 0,47 | 0,85 | 1,77 | 421 |
| ENSG00000104327 | CALB1 | 2,41 | 3,99 | 8,25 | 12,53 | 420 |
| ENSG00000177508 | IRX3 | 0,68 | 0,77 | 1,95 | 3,51 | 416 |
| ENSG00000181234 | TMEM132C | 2,98 | 7,63 | 8,27 | 15,29 | 413 |
| ENSG00000092607 | TBX15 | 0,79 | 1,17 | 1,61 | 4,05 | 413 |
| ENSG00000170561 | IRX2 | 8,88 | 17,31 | 24,24 | 45,4 | 411 |
| ENSG00000226179 | LINC00685 | 0,53 | 0,62 | 0,82 | 2,71 | 411 |
| ENSG00000151726 | ACSL1 | 0,74 | 2,14 | 2,43 | 3,78 | 411 |
| ENSG00000180287 | PLD5 | 0,56 | 0,91 | 2,46 | 2,86 | 411 |
| ENSG00000288115 | AC008731,2 | 0,2 | 0,41 | 0,45 | 1,02 | 410 |
| ENSG00000261669 | AC008731,1 | 0,2 | 0,41 | 0,45 | 1,02 | 410 |
| ENSG00000134569 | LRP4 | 4,08 | 8,86 | 10,75 | 20,74 | 408 |
| ENSG00000031081 | ARHGAP31 | 1,51 | 3,36 | 3,86 | 7,66 | 407 |
| ENSG00000283060 | ID3 | 21,89 | 34,08 | 77,15 | 110,85 | 406 |
| ENSG00000117318 | ID3 | 21,89 | 34,08 | 77,15 | 110,85 | 406 |
| ENSG00000226539 | MLXP1 | 0,33 | 0,78 | 1,06 | 1,67 | 406 |
| ENSG00000236007 | MTCO1P46 | 0,24 | 0,31 | 0,53 | 1,21 | 404 |
| ENSG00000114251 | WNT5A | 3,55 | 6,5 | 9,95 | 17,88 | 404 |
| ENSG00000257711 | AC079385,2 | 0,29 | 0,6 | 0,62 | 1,46 | 403 |
| ENSG00000164761 | TNFRSF11B | 3,72 | 6,7 | 10,61 | 18,5 | 397 |
| ENSG00000102230 | PCYT1B | 2,22 | 4,93 | 7,93 | 11 | 395 |
| ENSG00000266524 | GDF10 | 2,18 | 4,07 | 6,11 | 10,8 | 395 |
| ENSG00000047457 | CP | 1,89 | 3,34 | 4,3 | 9,31 | 393 |
| ENSG00000228643 | AC079779,2 | 0,22 | 0,34 | 0,68 | 1,08 | 391 |
| ENSG00000236166 | AL021408,1 | 0,32 | 0,86 | 1,01 | 1,57 | 391 |
| ENSG00000138650 | PCDH10 | 3,93 | 8,91 | 12,95 | 19,19 | 388 |
| ENSG00000247809 | NR2F2-AS1 | 1,7 | 3,27 | 3,68 | 8,27 | 386 |
| ENSG00000142173 | COL6A2 | 4,89 | 7,47 | 8,35 | 23,76 | 386 |

| | | | | | | |
|---|---|---|---|---|---|---|
| ENSG00000080503 | SMARCA2 | 3,31 | 7,94 | 8,54 | 15,96 | 382 |
| ENSG00000185614 | INKA1 | 0,49 | 1,42 | 2,05 | 2,35 | 380 |
| ENSG00000128567 | PODXL | 3,18 | 4,86 | 8,03 | 15,19 | 378 |
| ENSG00000244586 | WNT5A-AS1 | 1,29 | 3,1 | 5,87 | 6,16 | 378 |
| ENSG00000108375 | RNF43 | 0,55 | 0,66 | 1,47 | 2,61 | 375 |
| ENSG00000250062 | MAPK10-AS1 | 0,23 | 0,42 | 0,43 | 1,09 | 374 |
| ENSG00000183160 | TMEM119 | 0,43 | 0,55 | 0,89 | 2,03 | 372 |
| ENSG00000127863 | TNFRSF19 | 4,69 | 11,23 | 14,98 | 21,98 | 369 |
| ENSG00000182771 | GRID1 | 0,25 | 0,59 | 0,63 | 1,17 | 368 |
| ENSG00000254319 | AC246817,2 | 1,11 | 2,7 | 2,9 | 5,19 | 368 |
| ENSG00000198121 | LPAR1 | 5,04 | 9,55 | 14,73 | 23,55 | 367 |
| ENSG00000171189 | GRIK1 | 0,39 | 1,08 | 1,34 | 1,82 | 367 |
| ENSG00000124205 | EDN3 | 0,36 | 0,94 | 1,02 | 1,66 | 361 |
| ENSG00000118257 | NRP2 | 4,88 | 7,49 | 13,82 | 22,48 | 361 |
| ENSG00000285072 | AC270272,2 | 1,8 | 4,57 | 4,72 | 8,28 | 360 |
| ENSG00000234383 | CTBP2P8 | 0,44 | 0,89 | 0,93 | 2,02 | 359 |
| ENSG00000144285 | SCN1A | 3,94 | 12,14 | 15,2 | 17,76 | 351 |
| ENSG00000187957 | DNER | 3,22 | 7,97 | 11,56 | 14,46 | 349 |
| ENSG00000275180 | AC048341,2 | 4,63 | 7,22 | 7,62 | 20,77 | 349 |
| ENSG00000120833 | SOCS2 | 2,07 | 4,36 | 5,9 | 9,27 | 348 |
| ENSG00000229146 | SNX18P4 | 0,23 | 0,24 | 0,45 | 1,03 | 348 |
| ENSG00000121440 | PDZRN3 | 1,77 | 2,51 | 3,77 | 7,87 | 345 |
| ENSG00000229743 | LINC01159 | 6,06 | 11,76 | 15,49 | 26,92 | 344 |
| ENSG00000206838 | SNORA5A | 5,09 | 11,78 | 18,31 | 22,42 | 340 |
| ENSG00000223486 | AC092198,1 | 0,37 | 1,03 | 1,09 | 1,63 | 341 |
| ENSG00000139329 | LUM | 30,02 | 63,09 | 94,87 | 131,2 | 337 |
| ENSG00000064042 | LIMCH1 | 5,53 | 8,75 | 12,98 | 23,93 | 333 |
| ENSG00000162493 | PDPN | 8,08 | 12,66 | 19,59 | 34,93 | 332 |
| ENSG00000111218 | PRMT8 | 0,36 | 0,38 | 1,05 | 1,55 | 331 |
| ENSG00000096433 | ITPR3 | 0,83 | 1,56 | 2,04 | 3,57 | 330 |
| ENSG00000224216 | AC234781,1 | 0,3 | 0,59 | 0,91 | 1,29 | 330 |
| ENSG00000139364 | TMEM132B | 2,68 | 4,74 | 5,66 | 11,52 | 330 |
| ENSG00000272158 | AL139022,2 | 0,28 | 0,53 | 0,67 | 1,2 | 329 |
| ENSG00000234257 | SOD2P1 | 0,27 | 0,37 | 0,43 | 1,15 | 326 |
| ENSG00000287220 | AL590138,1 | 0,24 | 0,32 | 0,45 | 1,02 | 325 |
| ENSG00000180801 | ARSJ | 0,57 | 0,74 | 0,85 | 2,42 | 325 |
| ENSG00000070882 | OSBPL3 | 4,13 | 5,1 | 8,57 | 17,43 | 322 |
| ENSG00000082556 | OPRK1 | 0,33 | 0,52 | 0,71 | 1,39 | 321 |
| ENSG00000124191 | TOX2 | 0,76 | 1,82 | 2,16 | 3,2 | 321 |
| ENSG00000198597 | ZNF536 | 1,62 | 5,39 | 5,84 | 6,77 | 318 |
| ENSG00000232977 | LINC00327 | 0,32 | 0,59 | 0,87 | 1,33 | 316 |
| ENSG00000129595 | EPB41L4A | 6,77 | 18,53 | 20,35 | 28,1 | 315 |
| ENSG00000103723 | AP3B2 | 1,9 | 4,07 | 6,66 | 7,87 | 314 |
| ENSG00000109099 | PMP22 | 11,65 | 22,79 | 37,06 | 48,08 | 313 |
| ENSG00000272699 | AC007620,3 | 0,32 | 0,59 | 0,82 | 1,32 | 313 |
| ENSG00000141086 | CTRL | 0,25 | 0,39 | 0,46 | 1,03 | 312 |
| ENSG00000237819 | CDK6-AS1 | 2,45 | 3,75 | 7,38 | 10,05 | 310 |
| ENSG00000182022 | CHST15 | 2,85 | 5,25 | 6,32 | 11,54 | 305 |
| ENSG00000147862 | NFIB | 4,79 | 15,17 | 15,28 | 19,29 | 303 |
| ENSG00000271474 | AC106881,1 | 0,51 | 1,07 | 1,19 | 2,05 | 302 |

| | | | | | | |
|---|---|---|---|---|---|---|
| ENSG00000196092 | PAX5 | 2,85 | 4,33 | 5,42 | 11,44 | 301 |
| ENSG00000235920 | THRAP3P3 | 0,71 | 0,93 | 1,8 | 2,85 | 301 |
| ENSG00000121297 | TSHZ3 | 0,91 | 1,42 | 2,21 | 3,65 | 301 |
| ENSG00000104313 | EYA1 | 2,86 | 6,02 | 9,26 | 11,45 | 300 |
| ENSG00000095739 | BAMBI | 0,59 | 1,03 | 1,97 | 2,36 | 300 |
| ENSG00000124942 | AHNAK | 4,79 | 10,31 | 11,16 | 19,12 | 299 |
| ENSG00000116132 | PRRX1 | 9,91 | 13,32 | 17,39 | 39,52 | 299 |
| ENSG00000163762 | TM4SF18 | 0,45 | 1,07 | 1,62 | 1,78 | 296 |
| ENSG00000236107 | SCN1A-AS1 | 1,14 | 2,37 | 3,12 | 4,5 | 295 |
| ENSG00000163661 | PTX3 | 18,59 | 29,97 | 42,54 | 72,95 | 292 |
| ENSG00000272855 | AC104458,1 | 0,36 | 0,49 | 0,63 | 1,41 | 292 |
| ENSG00000112773 | TENT5A | 0,85 | 1,45 | 2,22 | 3,31 | 289 |
| ENSG00000185532 | PRKG1 | 2,07 | 2,47 | 3,74 | 8,05 | 289 |
| ENSG00000111907 | TPD52L1 | 1,48 | 1,64 | 3,01 | 5,69 | 284 |
| ENSG00000140323 | DISP2 | 0,5 | 0,9 | 1,89 | 1,92 | 284 |
| ENSG00000283979 | RFLNB | 0,62 | 0,96 | 1,67 | 2,38 | 284 |
| ENSG00000132854 | KANK4 | 0,91 | 1,45 | 2,88 | 3,47 | 281 |
| ENSG00000234685 | NUS1P2 | 0,95 | 1,03 | 1,39 | 3,62 | 281 |
| ENSG00000203688 | LINC02487 | 0,36 | 0,69 | 0,91 | 1,37 | 281 |
| ENSG00000241785 | RN7SL390P | 0,91 | 1,54 | 1,68 | 3,46 | 280 |
| ENSG00000162733 | DDR2 | 2,78 | 4,08 | 6,34 | 10,52 | 278 |
| ENSG00000185551 | NR2F2 | 8,7 | 15,25 | 19,46 | 32,89 | 278 |
| ENSG00000183801 | OLFML1 | 0,85 | 1,13 | 2,27 | 3,21 | 278 |
| ENSG00000140848 | CPNE2 | 11,01 | 18,81 | 29,96 | 41,32 | 275 |
| ENSG00000175463 | TBC1D10C | 0,39 | 0,81 | 0,9 | 1,46 | 274 |
| ENSG00000139540 | SLC39A5 | 0,3 | 0,53 | 0,7 | 1,12 | 273 |
| ENSG00000275559 | AL137060,5 | 22,72 | 36,45 | 58,88 | 84,54 | 272 |
| ENSG00000189292 | ALKAL2 | 5,27 | 7,71 | 12,17 | 19,57 | 271 |
| ENSG00000185432 | METTL7A | 0,91 | 1,58 | 2,05 | 3,37 | 270 |
| ENSG00000147509 | RGS20 | 3,9 | 5,93 | 7,12 | 14,44 | 270 |
| ENSG00000243687 | | 0,37 | 0,9 | 0,96 | 1,37 | 270 |
| ENSG00000278921 | EPB41L4A-DT | 1,6 | 2,55 | 3,82 | 5,9 | 269 |
| ENSG00000079215 | SLC1A3 | 14,94 | 27,94 | 30,04 | 54,6 | 265 |
| ENSG00000140682 | TGFB1I1 | 4,14 | 9,16 | 12,95 | 15,08 | 264 |
| ENSG00000178568 | ERBB4 | 4,81 | 8,1 | 9,94 | 17,5 | 264 |
| ENSG00000277101 | ARHGEF26 | 0,3 | 0,58 | 0,68 | 1,09 | 263 |
| ENSG00000232454 | AL138752,1 | 0,44 | 0,55 | 0,81 | 1,59 | 261 |
| ENSG00000252947 | SCARNA1 | 5,14 | 7,32 | 8,28 | 18,52 | 260 |
| ENSG00000196562 | SULF2 | 15,24 | 27,64 | 34,98 | 54,68 | 259 |
| ENSG00000137273 | FOXF2 | 0,75 | 1,39 | 1,87 | 2,69 | 259 |
| ENSG00000136960 | ENPP2 | 8,1 | 13,62 | 16,58 | 29,01 | 258 |
| ENSG00000253368 | TRNP1 | 4,2 | 7,97 | 12,08 | 15,04 | 258 |
| ENSG00000235586 | AC011247,1 | 0,63 | 1,16 | 1,57 | 2,25 | 257 |
| ENSG00000066468 | FGFR2 | 2,01 | 2,75 | 3,41 | 7,16 | 256 |
| ENSG00000167123 | CERCAM | 3,19 | 6,22 | 6,54 | 11,33 | 255 |
| ENSG00000215808 | LINC01139 | 0,66 | 1,9 | 1,98 | 2,34 | 255 |
| ENSG00000277290 | AC136475,10 | 0,78 | 1,2 | 1,25 | 2,76 | 254 |
| ENSG00000170549 | IRX1 | 3,79 | 6,92 | 8,47 | 13,38 | 253 |
| ENSG00000113580 | NR3C1 | 1,73 | 3,32 | 4,46 | 6,1 | 253 |
| ENSG00000141934 | PLPP2 | 0,48 | 0,78 | 1,38 | 1,69 | 252 |

| | | | | | | |
|---|---|---|---|---|---|---|
| ENSG00000168874 | ATOH8 | 3,42 | 5,11 | 6,59 | 12,03 | 252 |
| ENSG00000118322 | ATP10B | 0,55 | 1,2 | 1,41 | 1,93 | 251 |
| ENSG00000163132 | MSX1 | 2,22 | 3,31 | 5,64 | 7,78 | 250 |
| ENSG00000270504 | AL391422,4 | 0,44 | 0,75 | 1,23 | 1,54 | 250 |
| ENSG00000197177 | ADGRA1 | 0,3 | 0,49 | 0,92 | 1,05 | 250 |
| ENSG00000171867 | PRNP | 2,46 | 4,2 | 5,31 | 8,6 | 250 |
| ENSG00000105894 | PTN | 163,1 | 260,65 | 357,67 | 569,91 | 249 |
| ENSG00000198846 | TOX | 2,94 | 4,56 | 6,21 | 10,27 | 249 |
| ENSG00000122870 | BICC1 | 1,18 | 1,58 | 1,77 | 4,11 | 248 |
| ENSG00000246898 | LINC00920 | 0,36 | 0,43 | 0,47 | 1,25 | 247 |
| ENSG00000277954 | AC092376,2 | 0,39 | 0,72 | 0,89 | 1,35 | 246 |
| ENSG00000279068 | AC244517,6 | 0,59 | 0,63 | 1,05 | 2,04 | 246 |
| ENSG00000189184 | PCDH18 | 13,46 | 19,65 | 31,54 | 46,41 | 245 |
| ENSG00000154319 | FAM167A | 5,55 | 8,39 | 11,27 | 19,11 | 244 |
| ENSG00000092068 | SLC7A8 | 20,24 | 25,76 | 37,05 | 69,43 | 243 |
| ENSG00000172572 | PDE3A | 2,33 | 2,48 | 3,51 | 7,96 | 242 |
| ENSG00000010610 | CD4 | 1,21 | 1,98 | 2,82 | 4,11 | 240 |
| ENSG00000124126 | PREX1 | 9,62 | 15,23 | 19,78 | 32,67 | 240 |
| ENSG00000240583 | AQP1 | 0,47 | 0,55 | 0,69 | 1,59 | 238 |
| ENSG00000136110 | CNMD | 0,37 | 0,45 | 0,89 | 1,25 | 238 |
| ENSG00000261216 | AC007216,2 | 0,43 | 0,53 | 0,87 | 1,45 | 237 |
| ENSG00000235092 | ID2-AS1 | 0,58 | 0,75 | 1,18 | 1,95 | 236 |
| ENSG00000153071 | DAB2 | 6,89 | 11,49 | 17,78 | 23,11 | 235 |
| ENSG00000165124 | SVEP1 | 0,48 | 0,54 | 0,91 | 1,61 | 235 |
| ENSG00000189337 | KAZN | 2,99 | 4,94 | 6,89 | 10,01 | 235 |
| ENSG00000173926 | MARCHF3 | 0,74 | 1,87 | 2,02 | 2,47 | 234 |
| ENSG00000153814 | JAZF1 | 1,34 | 2,41 | 2,64 | 4,47 | 234 |
| ENSG00000276054 | AC243654,3 | 0,34 | 0,45 | 1 | 1,13 | 232 |
| ENSG00000264982 | AC015563,2 | 0,77 | 0,96 | 1,81 | 2,55 | 231 |
| ENSG00000257354 | AC048341,1 | 1,53 | 1,75 | 2,14 | 5,06 | 231 |
| ENSG00000232284 | GNG12-AS1 | 0,75 | 0,77 | 1,68 | 2,48 | 231 |
| ENSG00000082397 | EPB41L3 | 5,24 | 12,88 | 17,05 | 17,31 | 230 |
| ENSG00000154229 | PRKCA | 2,48 | 2,7 | 3,38 | 8,19 | 230 |
| ENSG00000227082 | LINC02798 | 0,53 | 0,68 | 0,95 | 1,75 | 230 |
| ENSG00000175806 | MSRA | 0,4 | 0,45 | 0,57 | 1,32 | 230 |
| ENSG00000130558 | OLFM1 | 5,88 | 10,55 | 13,15 | 19,4 | 230 |
| ENSG00000163082 | SGPP2 | 0,81 | 1,63 | 2,03 | 2,67 | 230 |
| ENSG00000213694 | S1PR3 | 9,12 | 16,77 | 21,92 | 29,84 | 227 |
| ENSG00000281980 | AC246817,3 | 0,37 | 0,52 | 0,67 | 1,21 | 227 |
| ENSG00000121871 | SLITRK3 | 0,71 | 1,13 | 1,35 | 2,32 | 227 |
| ENSG00000106538 | RARRES2 | 7,06 | 13,69 | 20,33 | 23,06 | 227 |
| ENSG00000103056 | SMPD3 | 0,79 | 1,57 | 2,25 | 2,58 | 227 |
| ENSG00000188158 | NHS | 3,05 | 5,33 | 6,69 | 9,96 | 227 |
| ENSG00000273254 | AF129075,2 | 0,34 | 0,77 | 0,9 | 1,11 | 226 |
| ENSG00000167994 | RAB3IL1 | 0,46 | 0,61 | 1,04 | 1,5 | 226 |
| ENSG00000020181 | ADGRA2 | 0,82 | 1,69 | 1,95 | 2,67 | 226 |
| ENSG00000250295 | RDH10-AS1 | 0,32 | 0,39 | 0,7 | 1,04 | 225 |
| ENSG00000144355 | DLX1 | 5,57 | 6,25 | 10,4 | 18,04 | 224 |
| ENSG00000047365 | ARAP2 | 0,86 | 1,17 | 1,5 | 2,78 | 223 |
| ENSG00000125864 | BFSP1 | 0,56 | 0,65 | 1,08 | 1,81 | 223 |

| | | | | | | |
|---|---|---|---|---|---|---|
| ENSG00000139915 | MDGA2 | 1,32 | 2,39 | 2,45 | 4,26 | 223 |
| ENSG00000173068 | BNC2 | 2,25 | 3,01 | 3,77 | 7,26 | 223 |
| ENSG00000104964 | TLE5 | 37,54 | 61,87 | 75,54 | 121,06 | 222 |
| ENSG00000130477 | UNC13A | 1,2 | 2,08 | 2,25 | 3,87 | 223 |
| ENSG00000273729 | AC007686,3 | 0,32 | 0,57 | 0,77 | 1,03 | 222 |
| ENSG00000276914 | CEP20 | 1,01 | 1,84 | 1,87 | 3,25 | 222 |
| ENSG00000187678 | SPRY4 | 24,32 | 47,55 | 68,46 | 78,19 | 222 |
| ENSG00000164626 | KCNK5 | 2,5 | 4,27 | 5,1 | 8,03 | 221 |
| ENSG00000231114 | AC078842,2 | 2,75 | 3,32 | 4,13 | 8,83 | 221 |
| ENSG00000270492 | AC137695,1 | 0,38 | 0,4 | 1,06 | 1,22 | 221 |
| ENSG00000176771 | NCKAP5 | 2,83 | 4,98 | 5,01 | 9,08 | 221 |
| ENSG00000239437 | RN7SL752P | 0,44 | 0,6 | 0,85 | 1,41 | 220 |
| ENSG00000228107 | AP000692,1 | 0,65 | 1,42 | 1,52 | 2,08 | 220 |
| ENSG00000163520 | FBLN2 | 3,33 | 6,35 | 6,37 | 10,65 | 220 |
| ENSG00000102760 | RGCC | 3,86 | 4,75 | 9,79 | 12,34 | 220 |
| ENSG00000066032 | CTNNA2 | 5,48 | 7,41 | 10,08 | 17,41 | 218 |
| ENSG00000172985 | SH3RF3 | 0,33 | 0,53 | 0,74 | 1,04 | 215 |
| ENSG00000179104 | TMTC2 | 6,57 | 9,4 | 11,56 | 20,66 | 214 |
| ENSG00000286500 | AL512643,2 | 0,43 | 0,51 | 0,66 | 1,35 | 214 |
| ENSG00000179242 | CDH4 | 2,99 | 3,66 | 5,25 | 9,38 | 214 |
| ENSG00000263883 | EEF1DP7 | 0,44 | 0,51 | 0,54 | 1,38 | 214 |
| ENSG00000240240 | BX664727,3 | 2,32 | 3,72 | 4,43 | 7,25 | 213 |
| ENSG00000273456 | AC064836,2 | 0,57 | 0,72 | 1,32 | 1,78 | 212 |
| ENSG00000162951 | LRRTM1 | 0,36 | 0,37 | 0,59 | 1,12 | 211 |
| ENSG00000272426 | BX284668,6 | 0,68 | 0,84 | 1,11 | 2,11 | 210 |
| ENSG00000102290 | PCDH11X | 2,28 | 4,05 | 4,18 | 7,07 | 210 |
| ENSG00000114200 | BCHE | 8,79 | 16,14 | 20,58 | 27,17 | 209 |
| ENSG00000183762 | KREMEN1 | 2,04 | 3,24 | 3,51 | 6,29 | 208 |
| ENSG00000103154 | NECAB2 | 4,45 | 5,61 | 10,22 | 13,72 | 208 |
| ENSG00000182168 | UNC5C | 8,7 | 14 | 17,95 | 26,82 | 208 |
| ENSG00000140807 | NKD1 | 1,98 | 2,98 | 3,14 | 6,1 | 208 |
| ENSG00000238251 | NSA2P7 | 0,5 | 0,77 | 0,96 | 1,54 | 208 |
| ENSG00000273355 | AP000894,4 | 0,63 | 0,77 | 1,4 | 1,94 | 208 |
| ENSG00000147526 | TACC1 | 9,65 | 14,2 | 17,28 | 29,64 | 207 |
| ENSG00000005513 | SOX8 | 1,85 | 2,42 | 3,61 | 5,68 | 207 |
| ENSG00000218690 | H2AC10P | 0,33 | 0,34 | 0,64 | 1,01 | 206 |
| ENSG00000141338 | ABCA8 | 0,36 | 0,59 | 0,71 | 1,1 | 206 |
| ENSG00000184226 | PCDH9 | 2,75 | 4,94 | 5,84 | 8,38 | 205 |
| ENSG00000139173 | TMEM117 | 1,11 | 2,68 | 3,3 | 3,37 | 204 |
| ENSG00000171451 | DSEL | 2,53 | 2,96 | 3,82 | 7,68 | 204 |
| ENSG00000226937 | CEP164P1 | 0,65 | 0,95 | 1,02 | 1,97 | 203 |
| ENSG00000231310 | TBL1XR1-AS1 | 0,69 | 0,7 | 1,2 | 2,09 | 203 |
| ENSG00000178573 | MAF | 4 | 6,87 | 9,36 | 12,11 | 203 |
| ENSG00000167191 | GPRC5B | 14,03 | 20,42 | 24,41 | 42,4 | 202 |
| ENSG00000144668 | ITGA9 | 2,1 | 4,06 | 4,28 | 6,34 | 202 |
| ENSG00000162692 | VCAM1 | 0,56 | 0,61 | 0,94 | 1,69 | 202 |
| ENSG00000255248 | MIR100HG | 11,16 | 29,43 | 31,39 | 33,67 | 202 |
| ENSG00000175161 | CADM2 | 1,98 | 3,19 | 4,29 | 5,95 | 201 |
| ENSG00000231688 | RPL21P43 | 0,6 | 0,64 | 0,75 | 1,8 | 200 |
| ENSG00000118971 | CCND2 | 68,43 | 110,29 | 127,8 | 204,81 | 199 |

| | | | | | | |
|---|---|---|---|---|---|---|
| ENSG00000175899 | A2M | 22,04 | 45,84 | 50,27 | 65,95 | 199 |
| ENSG00000169129 | AFAP1L2 | 5,35 | 7,6 | 9,26 | 15,99 | 199 |
| ENSG00000147010 | SH3KBP1 | 3,91 | 6,43 | 6,87 | 11,64 | 198 |
| ENSG00000154096 | THY1 | 15,93 | 22,7 | 37,33 | 47,41 | 198 |
| ENSG00000259657 | PIGHP1 | 0,42 | 0,81 | 1,01 | 1,25 | 198 |
| ENSG00000177519 | RPRM | 2,3 | 3,08 | 4,35 | 6,84 | 197 |
| ENSG00000155760 | FZD7 | 6,2 | 11,06 | 11,52 | 18,43 | 197 |
| ENSG00000277450 | AC002094,4 | 0,87 | 1,41 | 1,52 | 2,58 | 197 |
| ENSG00000168502 | MTCL1 | 7,43 | 13,44 | 15,46 | 22,03 | 197 |
| ENSG00000112175 | BMP5 | 2,75 | 3,3 | 4,51 | 8,14 | 196 |
| ENSG00000242156 | AC000041,1 | 17,12 | 20,45 | 20,52 | 50,58 | 195 |
| ENSG00000144847 | IGSF11 | 0,65 | 1,52 | 1,6 | 1,92 | 195 |
| ENSG00000153253 | SCN3A | 1,22 | 2,24 | 2,91 | 3,6 | 195 |
| ENSG00000154856 | APCDD1 | 8,58 | 16,51 | 19,79 | 25,26 | 194 |
| ENSG00000121904 | CSMD2 | 0,68 | 1,12 | 1,53 | 2 | 194 |
| ENSG00000136237 | RAPGEF5 | 3,71 | 5,22 | 8,13 | 10,91 | 194 |
| ENSG00000170425 | ADORA2B | 0,5 | 0,62 | 0,94 | 1,47 | 194 |
| ENSG00000165566 | AMER2 | 4,46 | 6,49 | 6,57 | 13,09 | 193 |
| ENSG00000278765 | AC004477,2 | 1,55 | 2,72 | 2,9 | 4,54 | 193 |
| ENSG00000185585 | OLFML2A | 0,7 | 1,06 | 1,31 | 2,05 | 193 |
| ENSG00000177272 | KCNA3 | 0,36 | 0,81 | 0,89 | 1,05 | 192 |
| ENSG00000164929 | BAALC | 16,81 | 26,04 | 29,94 | 48,99 | 191 |
| ENSG00000115183 | TANC1 | 3,33 | 5,6 | 6,12 | 9,69 | 191 |
| ENSG00000282413 | AL133461,1 | 6,06 | 12,06 | 14,84 | 17,58 | 190 |
| ENSG00000140859 | KIFC3 | 1,81 | 2,55 | 3,13 | 5,24 | 190 |
| ENSG00000137142 | IGFBPL1 | 44,46 | 73,71 | 94,66 | 128,7 | 189 |
| ENSG00000225216 | AC007362,1 | 0,37 | 0,49 | 0,7 | 1,07 | 189 |
| ENSG00000233450 | KIFC1 | 1,65 | 2,6 | 2,61 | 4,76 | 188 |
| ENSG00000185924 | RTN4RL1 | 0,58 | 1,47 | 1,52 | 1,67 | 188 |
| ENSG00000053524 | MCF2L2 | 0,47 | 0,71 | 0,8 | 1,35 | 187 |
| ENSG00000256940 | PPP1R14B-AS1 | 0,78 | 1,51 | 1,76 | 2,24 | 187 |
| ENSG00000206120 | EGFEM1P | 9,21 | 13,48 | 14,31 | 26,39 | 187 |
| ENSG00000150907 | FOXO1 | 2,28 | 3,91 | 4,85 | 6,51 | 186 |
| ENSG00000288169 | ACAP1 | 0,41 | 0,8 | 0,96 | 1,17 | 185 |
| ENSG00000132821 | VSTM2L | 2,9 | 4,37 | 5,42 | 8,27 | 185 |
| ENSG00000154127 | UBASH3B | 3,05 | 5,18 | 7,66 | 8,68 | 185 |
| ENSG00000144218 | AFF3 | 12,52 | 19,03 | 22,32 | 35,63 | 185 |
| ENSG00000177614 | PGBD5 | 3,04 | 4,07 | 5,24 | 8,64 | 184 |
| ENSG00000276934 | AC009704,2 | 0,44 | 0,51 | 0,77 | 1,25 | 184 |
| ENSG00000259040 | BLOC1S5-TXNDC5 | 1,48 | 1,79 | 2,94 | 4,19 | 183 |
| ENSG00000168994 | PXDC1 | 2,32 | 3,27 | 5,47 | 6,54 | 182 |
| ENSG00000230700 | CSNK2B | 2,48 | 2,83 | 4,77 | 6,99 | 182 |
| ENSG00000008196 | TFAP2B | 5,89 | 10,09 | 14,26 | 16,53 | 181 |
| ENSG00000158270 | COLEC12 | 6,14 | 9,52 | 14,14 | 17,21 | 180 |
| ENSG00000105810 | CDK6 | 14,49 | 19,61 | 25,53 | 40,6 | 180 |
| ENSG00000153707 | PTPRD | 17,82 | 24,45 | 30,56 | 49,92 | 180 |
| ENSG00000111077 | TNS2 | 2,87 | 5,78 | 7,94 | 8,03 | 180 |
| ENSG00000079931 | MOXD1 | 4,92 | 9,57 | 11,14 | 13,75 | 179 |
| ENSG00000047634 | SCML1 | 1,86 | 2,46 | 2,9 | 5,19 | 179 |
| ENSG00000119669 | IRF2BPL | 11,2 | 24,22 | 29,88 | 31,23 | 179 |

| | | | | | | |
|---|---|---|---|---|---|---|
| ENSG00000123104 | ITPR2 | 1,91 | 3,5 | 3,75 | 5,32 | 179 |
| ENSG00000278200 | LINC01971 | 0,4 | 0,45 | 0,88 | 1,11 | 178 |
| ENSG00000185920 | PTCH1 | 6,57 | 8,41 | 9,48 | 18,21 | 177 |
| ENSG00000266598 | AC037487,2 | 0,7 | 0,78 | 1,04 | 1,94 | 177 |
| ENSG00000276128 | AL591441,1 | 0,48 | 0,84 | 0,89 | 1,33 | 177 |
| ENSG00000132749 | TESMIN | 1,25 | 2,24 | 2,67 | 3,46 | 177 |
| ENSG00000284838 | C1QTNF4 | 0,77 | 1,33 | 1,82 | 2,13 | 177 |
| ENSG00000172247 | C1QTNF4 | 0,77 | 1,33 | 1,82 | 2,13 | 177 |
| ENSG00000101265 | RASSF2 | 5,73 | 8,21 | 10,27 | 15,83 | 176 |
| ENSG00000113594 | LIFR | 10,15 | 16,85 | 20,38 | 28,01 | 176 |
| ENSG00000154654 | NCAM2 | 1,96 | 2,38 | 4,61 | 5,39 | 175 |
| ENSG00000148123 | PLPPR1 | 1,77 | 2,44 | 4,1 | 4,85 | 174 |
| ENSG00000126785 | RHOJ | 6,99 | 11,13 | 16,14 | 19,09 | 173 |
| ENSG00000111817 | DSE | 8,14 | 12,15 | 15,37 | 22,19 | 173 |
| ENSG00000129654 | FOXJ1 | 1,29 | 2,53 | 3,18 | 3,51 | 172 |
| ENSG00000125266 | EFNB2 | 14,81 | 22,72 | 29,47 | 40,24 | 172 |
| ENSG00000109686 | SH3D19 | 5,51 | 6,55 | 8,54 | 14,96 | 172 |
| ENSG00000138061 | CYP1B1 | 5,74 | 9,47 | 10,41 | 15,57 | 171 |
| ENSG00000255339 | AL133352,1 | 1,65 | 2,4 | 2,46 | 4,47 | 171 |
| ENSG00000081138 | CDH7 | 0,88 | 1,25 | 2,14 | 2,38 | 170 |
| ENSG00000188573 | FBLL1 | 0,57 | 0,89 | 1,44 | 1,54 | 170 |
| ENSG00000182021 | AL591379,1 | 2,01 | 3,18 | 3,37 | 5,43 | 170 |
| ENSG00000137809 | ITGA11 | 1,25 | 1,67 | 2,25 | 3,37 | 170 |
| ENSG00000050555 | LAMC3 | 1,93 | 3,8 | 5,09 | 5,2 | 169 |
| ENSG00000282932 | PTPRD | 18,79 | 24,55 | 30,53 | 50,51 | 169 |
| ENSG00000185811 | IKZF1 | 2,61 | 3,29 | 3,81 | 7,01 | 169 |
| ENSG00000156140 | ADAMTS3 | 1 | 1,28 | 1,39 | 2,68 | 168 |
| ENSG00000111961 | SASH1 | 6,59 | 8,8 | 9,52 | 17,66 | 168 |
| ENSG00000147852 | VLDLR | 1,56 | 2,96 | 3,16 | 4,17 | 167 |
| ENSG00000197256 | KANK2 | 10,24 | 11 | 12,22 | 27,31 | 167 |
| ENSG00000255139 | AP000442,1 | 0,39 | 0,51 | 0,88 | 1,04 | 167 |
| ENSG00000151322 | NPAS3 | 4,8 | 5,66 | 7,49 | 12,79 | 166 |
| ENSG00000164398 | ACSL6 | 0,59 | 0,97 | 1,07 | 1,57 | 166 |
| ENSG00000018408 | WWTR1 | 8,63 | 11,89 | 17,09 | 22,96 | 166 |
| ENSG00000067082 | KLF6 | 10,85 | 17,86 | 25,53 | 28,83 | 166 |
| ENSG00000237333 | MSH5 | 0,89 | 1,55 | 1,9 | 2,36 | 165 |
| ENSG00000166387 | PPFIBP2 | 2,52 | 4,1 | 4,25 | 6,68 | 165 |
| ENSG00000105889 | STEAP1B | 3,09 | 4,08 | 5,77 | 8,19 | 165 |
| ENSG00000172380 | GNG12 | 13,49 | 18,57 | 21,14 | 35,69 | 165 |
| ENSG00000107968 | MAP3K8 | 2,18 | 3,13 | 3,91 | 5,76 | 164 |
| ENSG00000185565 | LSAMP | 9,11 | 11,6 | 14,07 | 24,05 | 164 |
| ENSG00000236308 | AL138921,2 | 0,52 | 0,66 | 0,87 | 1,37 | 163 |
| ENSG00000165269 | AQP7 | 0,6 | 0,88 | 1,01 | 1,58 | 163 |
| ENSG00000205611 | LINC01597 | 0,7 | 0,91 | 0,95 | 1,84 | 163 |
| ENSG00000132622 | HSPA12B | 0,85 | 1,19 | 1,52 | 2,23 | 162 |
| ENSG00000186493 | C5orf38 | 5,82 | 7,64 | 10,33 | 15,23 | 162 |
| ENSG00000116704 | SLC35D1 | 4,31 | 6,11 | 6,52 | 11,27 | 161 |
| ENSG00000006210 | CX3CL1 | 1,42 | 1,56 | 2,16 | 3,71 | 161 |
| ENSG00000171992 | SYNPO | 0,98 | 1,93 | 2,42 | 2,56 | 161 |
| ENSG00000184232 | OAF | 7,16 | 9,58 | 13,91 | 18,7 | 161 |

| | | | | | | |
|---|---|---|---|---|---|---|
| ENSG00000166016 | ABTB2 | 2,49 | 5,55 | 6,44 | 6,5 | 161 |
| ENSG00000197457 | STMN3 | 18,1 | 29,41 | 35,27 | 47,18 | 161 |
| ENSG00000131370 | SH3BP5 | 1,93 | 3,3 | 3,5 | 5,03 | 161 |
| ENSG00000168938 | PPIC | 1,57 | 3,25 | 3,98 | 4,09 | 161 |
| ENSG00000197594 | ENPP1 | 0,48 | 0,53 | 0,7 | 1,25 | 160 |
| ENSG00000151136 | BTBD11 | 1,31 | 1,48 | 2,97 | 3,41 | 160 |
| ENSG00000189067 | LITAF | 9,03 | 9,18 | 13,48 | 23,48 | 160 |
| ENSG00000181800 | CELF2-AS1 | 0,7 | 1,05 | 1,11 | 1,82 | 160 |
| ENSG00000089505 | CMTM1 | 1,56 | 2,73 | 3,03 | 4,05 | 160 |
| ENSG00000145431 | PDGFC | 10,65 | 14,49 | 15,54 | 27,63 | 159 |
| ENSG00000167114 | SLC27A4 | 3,94 | 5,26 | 5,36 | 10,19 | 159 |
| ENSG00000272764 | AL596094,1 | 0,48 | 0,75 | 0,77 | 1,24 | 158 |
| ENSG00000104447 | TRPS1 | 6,06 | 10,19 | 10,72 | 15,65 | 158 |
| ENSG00000112320 | SOBP | 5,96 | 8,63 | 10,6 | 15,39 | 158 |
| ENSG00000166833 | NAV2 | 4,8 | 6,31 | 9,12 | 12,39 | 158 |
| ENSG00000171243 | SOSTDC1 | 1,25 | 1,31 | 2 | 3,22 | 158 |
| ENSG00000151702 | FLI1 | 1,57 | 2,95 | 4,03 | 4,04 | 157 |
| ENSG00000019549 | SNAI2 | 8,36 | 10,56 | 16,68 | 21,49 | 157 |
| ENSG00000272986 | AC009570,1 | 0,76 | 1,02 | 1,14 | 1,95 | 157 |
| ENSG00000227220 | AL133346,1 | 0,66 | 0,7 | 1,45 | 1,69 | 156 |
| ENSG00000137267 | TUBB2A | 35,84 | 50,53 | 79,55 | 91,73 | 156 |
| ENSG00000183722 | LHFPL6 | 15,06 | 19,98 | 28,11 | 38,53 | 156 |
| ENSG00000254274 | TDGF1P5 | 0,54 | 0,62 | 0,65 | 1,38 | 156 |
| ENSG00000141582 | CBX4 | 3,04 | 3,71 | 5,16 | 7,76 | 155 |
| ENSG00000231407 | GORAB-AS1 | 0,42 | 0,63 | 0,85 | 1,07 | 155 |
| ENSG00000162849 | KIF26B | 1,59 | 2,25 | 2,45 | 4,05 | 155 |
| ENSG00000169126 | ARMC4 | 4,04 | 5,92 | 7,23 | 10,27 | 154 |
| ENSG00000234196 | ZBTB12 | 2,35 | 2,76 | 2,83 | 5,95 | 153 |
| ENSG00000145908 | ZNF300 | 3,07 | 4,29 | 5,17 | 7,77 | 153 |
| ENSG00000179902 | C1orf194 | 1,34 | 1,99 | 2,59 | 3,39 | 153 |
| ENSG00000071205 | ARHGAP10 | 0,78 | 1,4 | 1,85 | 1,97 | 153 |
| ENSG00000144824 | PHLDB2 | 13,29 | 19,11 | 22,17 | 33,56 | 153 |
| ENSG00000164976 | MYORG | 2,31 | 4,93 | 5,15 | 5,83 | 152 |
| ENSG00000184838 | PRR16 | 1,96 | 2,14 | 2,87 | 4,94 | 152 |
| ENSG00000116774 | OLFML3 | 6,95 | 9,11 | 13,88 | 17,51 | 152 |
| ENSG00000179832 | MROH1 | 0,6 | 1,09 | 1,12 | 1,51 | 152 |
| ENSG00000117114 | ADGRL2 | 44,84 | 54,76 | 67,99 | 112,83 | 152 |
| ENSG00000238057 | ZEB2-AS1 | 0,68 | 1,3 | 1,52 | 1,71 | 151 |
| ENSG00000232439 | RPL18AP7 | 0,56 | 1,04 | 1,28 | 1,4 | 150 |
| ENSG00000263873 | THY1-AS1 | 6,88 | 8,57 | 15,44 | 17,16 | 149 |
| ENSG00000198053 | SIRPA | 6,08 | 8,64 | 11,27 | 15,15 | 149 |
| ENSG00000007944 | MYLIP | 1,89 | 3,3 | 3,7 | 4,7 | 149 |
| ENSG00000232970 | POLHP1 | 0,76 | 1,12 | 1,13 | 1,88 | 147 |
| ENSG00000196843 | ARID5A | 2,03 | 2,61 | 3,15 | 5,02 | 147 |
| ENSG00000104361 | NIPAL2 | 0,45 | 0,8 | 0,94 | 1,11 | 147 |
| ENSG00000226824 | AC006001,2 | 1,12 | 1,38 | 1,5 | 2,76 | 146 |
| ENSG00000144642 | RBMS3 | 4,29 | 5,55 | 8,46 | 10,56 | 146 |
| ENSG00000104219 | ZDHHC2 | 7,04 | 10,74 | 12,41 | 17,29 | 146 |
| ENSG00000088881 | EBF4 | 0,88 | 1,08 | 1,73 | 2,16 | 145 |
| ENSG00000198822 | GRM3 | 9,52 | 11,17 | 14,11 | 23,36 | 145 |

| | | | | | | |
|---|---|---|---|---|---|---|
| ENSG00000178209 | | 2,02 | 2,38 | 2,97 | 4,95 | 145 |
| ENSG00000261934 | PCDHGA9 | 0,67 | 0,95 | 1,2 | 1,64 | 145 |
| ENSG00000187676 | B3GLCT | 5,61 | 7,1 | 8,36 | 13,73 | 145 |
| ENSG00000130940 | CASZ1 | 2,17 | 2,74 | 3,15 | 5,31 | 145 |
| ENSG00000188760 | TMEM198 | 1,37 | 1,96 | 2,98 | 3,35 | 145 |
| ENSG00000164292 | RHOBTB3 | 16,81 | 27,07 | 31,7 | 41,05 | 144 |
| ENSG00000108669 | CYTH1 | 4,26 | 5,29 | 6,49 | 10,4 | 144 |
| ENSG00000198929 | NOS1AP | 2,11 | 4,33 | 4,58 | 5,14 | 144 |
| ENSG00000185129 | PURA | 0,86 | 1,37 | 1,49 | 2,09 | 143 |
| ENSG00000147642 | SYBU | 1,86 | 3,69 | 4,23 | 4,52 | 143 |
| ENSG00000046653 | GPM6B | 19,2 | 23,17 | 29,3 | 46,65 | 143 |
| ENSG00000148841 | ITPRIP | 3,36 | 4,1 | 5,49 | 8,16 | 143 |
| ENSG00000146648 | EGFR | 2,87 | 3,85 | 4,51 | 6,97 | 143 |
| ENSG00000026559 | KCNG1 | 1,28 | 1,6 | 1,91 | 3,1 | 142 |
| ENSG00000225163 | LINC00618 | 0,5 | 0,53 | 0,77 | 1,21 | 142 |
| ENSG00000273084 | AC092171,5 | 1,66 | 2,11 | 3,24 | 4,01 | 142 |
| ENSG00000141905 | NFIC | 1,73 | 2,25 | 2,44 | 4,17 | 141 |
| ENSG00000242732 | RTL5 | 3,82 | 6,18 | 6,41 | 9,2 | 141 |
| ENSG00000277978 | AC010542,5 | 1,03 | 1,31 | 1,48 | 2,48 | 141 |
| ENSG00000202347 | RNU1-16P | 1,35 | 2,31 | 3,1 | 3,25 | 141 |
| ENSG00000164946 | FREM1 | 5,62 | 7,71 | 8,87 | 13,52 | 141 |
| ENSG00000148411 | NACC2 | 0,74 | 0,94 | 1,04 | 1,78 | 141 |
| ENSG00000116991 | SIPA1L2 | 9,82 | 13,57 | 14,96 | 23,58 | 140 |
| ENSG00000143842 | SOX13 | 5,74 | 6,96 | 7,43 | 13,78 | 140 |
| ENSG00000250241 | AC105383,1 | 1,25 | 2,49 | 2,98 | 3 | 140 |
| ENSG00000154310 | TNIK | 7,98 | 13,87 | 13,94 | 19,15 | 140 |
| ENSG00000196935 | SRGAP1 | 6,36 | 9,13 | 10,97 | 15,26 | 140 |
| ENSG00000151617 | EDNRA | 3,13 | 3,83 | 4,95 | 7,51 | 140 |
| ENSG00000183208 | GDPGP1 | 1,12 | 1,71 | 2,46 | 2,68 | 139 |
| ENSG00000215014 | AL645728,1 | 0,56 | 0,87 | 1,02 | 1,34 | 139 |
| ENSG00000204131 | NHSL2 | 3,8 | 4,5 | 5,86 | 9,09 | 139 |
| ENSG00000136153 | LMO7 | 1,56 | 2,08 | 2,37 | 3,72 | 138 |
| ENSG00000080493 | SLC4A4 | 2,5 | 4,38 | 4,82 | 5,96 | 138 |
| ENSG00000260691 | ANKRD20A1 | 1,13 | 1,78 | 1,91 | 2,69 | 138 |
| ENSG00000184005 | ST6GALNAC3 | 14,85 | 21,51 | 26,28 | 35,33 | 138 |
| ENSG00000174740 | PABPC5 | 2,8 | 4,29 | 4,62 | 6,66 | 138 |
| ENSG00000072840 | EVC | 2,83 | 3,44 | 3,46 | 6,73 | 138 |
| ENSG00000185989 | RASA3 | 3,05 | 4,65 | 5,1 | 7,25 | 138 |
| ENSG00000280477 | RASA3 | 3,05 | 4,65 | 5,1 | 7,25 | 138 |
| ENSG00000184588 | PDE4B | 13,41 | 17,22 | 23,3 | 31,86 | 138 |
| ENSG00000122863 | CHST3 | 3,42 | 5,16 | 5,77 | 8,11 | 137 |
| ENSG00000272674 | PCDHB16 | 1,27 | 2,56 | 2,89 | 3,01 | 137 |
| ENSG00000154122 | ANKH | 3,25 | 6,09 | 6,42 | 7,7 | 137 |
| ENSG00000178695 | KCTD12 | 3,34 | 5,26 | 6,69 | 7,91 | 137 |
| ENSG00000112559 | MDFI | 16,37 | 25,15 | 32,62 | 38,76 | 137 |
| ENSG00000152952 | PLOD2 | 8,13 | 12,78 | 13,77 | 19,17 | 136 |
| ENSG00000119865 | CNRIP1 | 4,3 | 6,03 | 6,24 | 10,13 | 136 |
| ENSG00000176055 | MBLAC2 | 1,47 | 1,66 | 2,04 | 3,45 | 135 |
| ENSG00000221949 | LINC01465 | 1,3 | 1,73 | 2,05 | 3,05 | 135 |
| ENSG00000287431 | AC027601,5 | 1,06 | 1,27 | 1,55 | 2,48 | 134 |

| | | | | | | |
|---|---|---|---|---|---|---|
| ENSG00000122756 | CNTFR | 21,08 | 29,33 | 37,14 | 49,31 | 134 |
| ENSG00000139926 | FRMD6 | 9,2 | 11,74 | 13,37 | 21,52 | 134 |
| ENSG00000125246 | CLYBL | 8,54 | 10,13 | 15,09 | 19,97 | 134 |
| ENSG00000261765 | AC009127,1 | 0,51 | 0,7 | 0,87 | 1,19 | 133 |
| ENSG00000162512 | SDC3 | 26,25 | 39,49 | 46,32 | 61,12 | 133 |
| ENSG00000267419 | ZNF56 | 0,9 | 1,32 | 1,4 | 2,09 | 132 |
| ENSG00000145335 | SNCA | 9,73 | 15,27 | 16,96 | 22,58 | 132 |
| ENSG00000182118 | FAM89A | 1,31 | 1,74 | 2,06 | 3,04 | 132 |
| ENSG00000023572 | GLRX2 | 4,62 | 6,7 | 7,26 | 10,68 | 131 |
| ENSG00000231945 | VARS1 | 4,42 | 5,01 | 6,77 | 10,21 | 131 |
| ENSG00000077782 | FGFR1 | 51,53 | 64,63 | 68,07 | 118,74 | 130 |
| ENSG00000277459 | AP001527,2 | 1,25 | 1,31 | 1,5 | 2,88 | 130 |
| ENSG00000164125 | GASK1B | 1,54 | 2,53 | 2,64 | 3,54 | 130 |
| ENSG00000184384 | MAML2 | 5,11 | 7,35 | 8,45 | 11,74 | 130 |
| ENSG00000214286 | PDCL3P3 | 0,58 | 0,67 | 0,96 | 1,33 | 129 |
| ENSG00000100979 | PLTP | 16,92 | 26,62 | 34,05 | 38,79 | 129 |
| ENSG00000152518 | ZFP36L2 | 7,91 | 9,28 | 13,12 | 18,12 | 129 |
| ENSG00000206284 | WDR46 | 0,93 | 1,28 | 1,44 | 2,13 | 129 |
| ENSG00000261663 | AC009065,8 | 0,59 | 1,13 | 1,2 | 1,35 | 129 |
| ENSG00000162433 | AK4 | 3,14 | 5,57 | 6,22 | 7,18 | 129 |
| ENSG00000172602 | RND1 | 1,52 | 1,75 | 2,32 | 3,47 | 128 |
| ENSG00000145107 | TM4SF19 | 0,53 | 0,64 | 0,81 | 1,21 | 128 |
| ENSG00000091136 | LAMB1 | 29,91 | 42,51 | 50,33 | 68,14 | 128 |
| ENSG00000257594 | GALNT4 | 1,51 | 2,55 | 2,74 | 3,44 | 128 |
| ENSG00000266710 | RN7SL48P | 0,81 | 1,37 | 1,38 | 1,84 | 127 |
| ENSG00000116544 | DLGAP3 | 0,67 | 0,94 | 1,16 | 1,52 | 127 |
| ENSG00000180592 | SKIDA1 | 11,42 | 13,4 | 15,91 | 25,84 | 126 |
| ENSG00000271161 | BOLA2P2 | 1,07 | 1,85 | 2,03 | 2,42 | 126 |
| ENSG00000154721 | JAM2 | 10,61 | 13,28 | 15,02 | 23,97 | 126 |
| ENSG00000100599 | RIN3 | 2,16 | 2,62 | 4,25 | 4,87 | 125 |
| ENSG00000284512 | AC092718,8 | 0,63 | 0,73 | 1,02 | 1,42 | 125 |
| ENSG00000135547 | HEY2 | 0,56 | 1,03 | 1,07 | 1,26 | 125 |
| ENSG00000054598 | FOXC1 | 2,11 | 3,76 | 4,44 | 4,74 | 125 |
| ENSG00000237462 | TRIM27 | 2,3 | 3,39 | 5,1 | 5,16 | 124 |
| ENSG00000182621 | PLCB1 | 6,21 | 7,36 | 7,71 | 13,93 | 124 |
| ENSG00000047648 | ARHGAP6 | 1,2 | 1,72 | 1,78 | 2,69 | 124 |
| ENSG00000251363 | LINC02315 | 2,77 | 3,48 | 4,31 | 6,19 | 123 |
| ENSG00000116729 | WLS | 30,47 | 40,84 | 55,63 | 68,08 | 123 |
| ENSG00000082497 | SERTAD4 | 1,89 | 2,07 | 3,89 | 4,22 | 123 |
| ENSG00000281230 | SERTAD4 | 1,89 | 2,07 | 3,89 | 4,22 | 123 |
| ENSG00000054793 | ATP9A | 6,29 | 10,47 | 11,41 | 14,04 | 123 |
| ENSG00000224379 | TCF19 | 2,69 | 4,1 | 5,01 | 6 | 123 |
| ENSG00000197380 | DACT3 | 2,88 | 4,82 | 5,1 | 6,42 | 123 |
| ENSG00000163995 | ABLIM2 | 0,57 | 0,92 | 0,94 | 1,27 | 123 |
| ENSG00000273604 | EPOP | 2,86 | 5,21 | 5,84 | 6,37 | 123 |
| ENSG00000279413 | AC112497,2 | 1,19 | 1,43 | 1,66 | 2,64 | 122 |
| ENSG00000286156 | AC026273,1 | 1,54 | 1,69 | 2,21 | 3,41 | 121 |
| ENSG00000184675 | AMER1 | 4,53 | 5,86 | 6,42 | 10,03 | 121 |
| ENSG00000221164 | SNORA11F | 19,44 | 20 | 30,81 | 43,02 | 121 |
| ENSG00000106780 | MEGF9 | 5,45 | 8,43 | 9,2 | 12,06 | 121 |

| | | | | | | |
|---|---|---|---|---|---|---|
| ENSG00000203325 | AL445248,1 | 1,13 | 1,41 | 2,09 | 2,49 | 120 |
| ENSG00000101187 | SLCO4A1 | 0,89 | 1,01 | 1,25 | 1,96 | 120 |
| ENSG00000114656 | CFAP92 | 5,55 | 6,04 | 7,22 | 12,22 | 120 |
| ENSG00000171503 | ETFDH | 4,02 | 6,3 | 7,32 | 8,85 | 120 |
| ENSG00000117877 | POLR1G | 2,19 | 2,63 | 2,88 | 4,82 | 120 |
| ENSG00000242808 | SOX2-OT | 6,23 | 8,91 | 10,67 | 13,7 | 120 |
| ENSG00000160216 | AGPAT3 | 4,02 | 5,86 | 6,21 | 8,82 | 119 |
| ENSG00000109063 | MYH3 | 1,24 | 1,6 | 2,33 | 2,72 | 119 |
| ENSG00000183579 | ZNRF3 | 3,63 | 4,68 | 5,48 | 7,96 | 119 |
| ENSG00000148926 | ADM | 3,6 | 5,22 | 7,82 | 7,89 | 119 |
| ENSG00000213109 | AL365223,1 | 0,58 | 0,85 | 0,91 | 1,27 | 119 |
| ENSG00000167912 | AC090152,1 | 2,81 | 3,44 | 4,74 | 6,15 | 119 |
| ENSG00000254535 | PABPC4L | 1,33 | 1,7 | 1,92 | 2,91 | 119 |
| ENSG00000169071 | ROR2 | 8,49 | 10,86 | 11,79 | 18,57 | 119 |
| ENSG00000115844 | DLX2 | 2,67 | 3,17 | 3,98 | 5,84 | 119 |
| ENSG00000150471 | ADGRL3 | 12,49 | 17,82 | 20,44 | 27,22 | 118 |
| ENSG00000138696 | BMPR1B | 3,66 | 4,83 | 5,73 | 7,97 | 118 |
| ENSG00000104112 | SCG3 | 4,79 | 8,46 | 10,22 | 10,41 | 117 |
| ENSG00000181751 | MACIR | 5,37 | 6,14 | 8,1 | 11,67 | 117 |
| ENSG00000131196 | NFATC1 | 1,79 | 1,86 | 2,01 | 3,89 | 117 |
| ENSG00000147234 | FRMPD3 | 0,52 | 0,58 | 0,67 | 1,13 | 117 |
| ENSG00000220842 | RPL21P16 | 70,46 | 98,75 | 99,57 | 152,7 | 117 |
| ENSG00000271815 | AC008897,3 | 1,02 | 1,18 | 1,58 | 2,21 | 117 |
| ENSG00000103187 | COTL1 | 118,31 | 158,37 | 212,6 | 256,01 | 116 |
| ENSG00000244486 | SCARF2 | 2,16 | 3,23 | 3,68 | 4,67 | 116 |
| ENSG00000178878 | APOLD1 | 5,07 | 6,33 | 8,67 | 10,94 | 116 |
| ENSG00000160963 | COL26A1 | 21,41 | 30,77 | 41,78 | 46,15 | 116 |
| ENSG00000173918 | C1QTNF1 | 0,52 | 0,72 | 0,76 | 1,12 | 115 |
| ENSG00000091986 | CCDC80 | 26,18 | 37,5 | 47,72 | 56,37 | 115 |
| ENSG00000164061 | BSN | 1,24 | 2,19 | 2,25 | 2,67 | 115 |
| ENSG00000132561 | MATN2 | 1,99 | 3,01 | 3,64 | 4,27 | 115 |
| ENSG00000233924 | RPSAP13 | 0,49 | 0,56 | 0,58 | 1,05 | 114 |
| ENSG00000196376 | SLC35F1 | 9,95 | 13,84 | 15,22 | 21,3 | 114 |
| ENSG00000261072 | AC084783,1 | 0,71 | 0,96 | 1,09 | 1,52 | 114 |
| ENSG00000018236 | CNTN1 | 0,71 | 1,25 | 1,35 | 1,52 | 114 |
| ENSG00000237187 | NR2F1-AS1 | 6,63 | 7,92 | 8,69 | 14,19 | 114 |
| ENSG00000229006 | TRIM27 | 0,5 | 0,52 | 0,86 | 1,07 | 114 |
| ENSG00000258932 | AL390334,1 | 1,23 | 1,44 | 1,46 | 2,63 | 114 |
| ENSG00000266028 | SRGAP2 | 29,07 | 35,97 | 46,57 | 62,03 | 113 |
| ENSG00000149557 | FEZ1 | 71,14 | 103,6 | 124,19 | 151,55 | 113 |
| ENSG00000275031 | METRNL | 4,84 | 7,91 | 8,36 | 10,31 | 113 |
| ENSG00000107863 | ARHGAP21 | 34,64 | 48,86 | 57,85 | 73,75 | 113 |
| ENSG00000186469 | GNG2 | 25,73 | 43,08 | 52,24 | 54,71 | 113 |
| ENSG00000182749 | PAQR7 | 1,03 | 1,35 | 1,66 | 2,19 | 113 |
| ENSG00000119771 | KLHL29 | 3,72 | 5,24 | 5,46 | 7,9 | 112 |
| ENSG00000285302 | CHMP4A | 1,63 | 2,92 | 3,11 | 3,46 | 112 |
| ENSG00000125148 | MT2A | 7,39 | 8,15 | 13,2 | 15,68 | 112 |
| ENSG00000170921 | TANC2 | 14,53 | 17,31 | 21,39 | 30,79 | 112 |
| ENSG00000165633 | VSTM4 | 1,44 | 2,05 | 2,21 | 3,05 | 112 |
| ENSG00000275426 | AC253576,2 | 0,52 | 0,89 | 0,97 | 1,1 | 112 |

| | | | | | | |
|---|---|---|---|---|---|---|
| ENSG00000136213 | CHST12 | 2 | 2,68 | 3,65 | 4,23 | 112 |
| ENSG00000228824 | MIR4500HG | 0,96 | 1,11 | 1,35 | 2,03 | 111 |
| ENSG00000105605 | CACNG7 | 7,48 | 11,76 | 12,04 | 15,76 | 111 |
| ENSG00000278932 | CR381653,1 | 1,13 | 1,17 | 2,32 | 2,38 | 111 |
| ENSG00000142178 | SIK1 | 3,04 | 5,18 | 6,24 | 6,4 | 111 |
| ENSG00000038427 | VCAN | 121,06 | 151,37 | 161,15 | 254,71 | 110 |
| ENSG00000183287 | CCBE1 | 0,49 | 0,52 | 0,69 | 1,03 | 110 |
| ENSG00000204950 | LRRC10B | 1,8 | 2,15 | 2,62 | 3,78 | 110 |
| ENSG00000175745 | NR2F1 | 12,37 | 13,82 | 17,12 | 25,94 | 110 |
| ENSG00000154274 | C4orf19 | 0,93 | 1,41 | 1,5 | 1,95 | 110 |
| ENSG00000139220 | PPFIA2 | 2,87 | 4,51 | 4,95 | 6,01 | 109 |
| ENSG00000151689 | INPP1 | 1,98 | 3,76 | 4,01 | 4,14 | 109 |
| ENSG00000142694 | EVA1B | 3,61 | 3,97 | 4,97 | 7,54 | 109 |
| ENSG00000082641 | NFE2L1 | 28,49 | 41,81 | 52,53 | 59,45 | 109 |
| ENSG00000114450 | GNB4 | 11,8 | 14,3 | 16,79 | 24,59 | 108 |
| ENSG00000105767 | CADM4 | 10,53 | 16,72 | 16,76 | 21,94 | 108 |
| ENSG00000187187 | ZNF546 | 0,72 | 1,28 | 1,44 | 1,5 | 108 |
| ENSG00000111145 | ELK3 | 5,75 | 9,03 | 9,53 | 11,97 | 108 |
| ENSG00000170153 | RNF150 | 3,1 | 4,33 | 4,38 | 6,45 | 108 |
| ENSG00000231187 | AL356056,2 | 0,62 | 0,94 | 1,1 | 1,29 | 108 |
| ENSG00000280810 | NSFP1 | 1,27 | 1,77 | 2,16 | 2,64 | 108 |
| ENSG00000237916 | RPL37P12 | 2,34 | 2,4 | 2,57 | 4,86 | 108 |
| ENSG00000196526 | AFAP1 | 5,47 | 9,69 | 10 | 11,36 | 108 |
| ENSG00000008086 | CDKL5 | 3,71 | 5,6 | 6,27 | 7,69 | 107 |
| ENSG00000244691 | RPL10AP1 | 0,7 | 0,74 | 0,78 | 1,45 | 107 |
| ENSG00000137691 | CFAP300 | 1,56 | 2,08 | 2,45 | 3,23 | 107 |
| ENSG00000213199 | ASIC3 | 0,58 | 0,62 | 0,7 | 1,2 | 107 |
| ENSG00000127955 | GNAI1 | 9,41 | 12,41 | 14,86 | 19,46 | 107 |
| ENSG00000101000 | PROCR | 3,41 | 4,1 | 4,31 | 7,05 | 107 |
| ENSG00000187605 | TET3 | 5,05 | 8,21 | 8,38 | 10,43 | 107 |
| ENSG00000146070 | PLA2G7 | 0,67 | 0,88 | 0,95 | 1,38 | 106 |
| ENSG00000149260 | CAPN5 | 3,75 | 4,96 | 6,22 | 7,72 | 106 |
| ENSG00000184058 | TBX1 | 3,11 | 4,1 | 5,06 | 6,4 | 106 |
| ENSG00000184640 | SEPTIN9 | 93,22 | 112,01 | 132,86 | 191,67 | 106 |
| ENSG00000162545 | CAMK2N1 | 5,76 | 8,71 | 10,74 | 11,84 | 106 |
| ENSG00000179761 | PIPOX | 2,89 | 3,99 | 4,85 | 5,94 | 106 |
| ENSG00000114423 | CBLB | 15,95 | 23,97 | 24,6 | 32,76 | 105 |
| ENSG00000065485 | PDIA5 | 5,07 | 6,4 | 8,28 | 10,41 | 105 |
| ENSG00000080561 | MID2 | 0,95 | 1,43 | 1,73 | 1,95 | 105 |
| ENSG00000137824 | RMDN3 | 6,58 | 7,94 | 8,57 | 13,5 | 105 |
| ENSG00000237441 | RGL2 | 3,71 | 4,84 | 6,04 | 7,61 | 105 |
| ENSG00000197959 | DNM3 | 1 | 1,39 | 1,91 | 2,05 | 105 |
| ENSG00000204682 | MIR1915HG | 4,05 | 4,48 | 5,5 | 8,29 | 105 |
| ENSG00000163110 | PDLIM5 | 14,54 | 19,2 | 21,67 | 29,74 | 105 |
| ENSG00000060491 | OGFR | 1,55 | 1,64 | 1,7 | 3,17 | 105 |
| ENSG00000135269 | TES | 10,06 | 14,22 | 16,39 | 20,57 | 104 |
| ENSG00000141682 | PMAIP1 | 4,36 | 5,8 | 6,98 | 8,89 | 104 |
| ENSG00000166562 | SEC11C | 24,51 | 28,49 | 39,93 | 49,91 | 104 |
| ENSG00000170667 | RASA4B | 2,27 | 2,88 | 3,41 | 4,62 | 104 |
| ENSG00000100242 | SUN2 | 6,95 | 9,03 | 10,98 | 14,14 | 103 |

| | | | | | | |
|---|---|---|---|---|---|---|
| ENSG00000156515 | HK1 | 14,82 | 24,18 | 24,44 | 30,13 | 103 |
| ENSG00000187239 | FNBP1 | 12,91 | 19,6 | 19,79 | 26,23 | 103 |
| ENSG00000088387 | DOCK9 | 2,64 | 3,1 | 3,22 | 5,36 | 103 |
| ENSG00000008516 | MMP25 | 2,36 | 2,74 | 2,99 | 4,79 | 103 |
| ENSG00000204054 | LINC00963 | 3,12 | 3,46 | 4,04 | 6,33 | 103 |
| ENSG00000181274 | FRAT2 | 1,04 | 1,08 | 1,2 | 2,11 | 103 |
| ENSG00000119147 | ECRG4 | 0,7 | 0,88 | 1,29 | 1,42 | 103 |
| ENSG00000259330 | INAFM2 | 2,47 | 4,13 | 4,54 | 5,01 | 103 |
| ENSG00000228395 | AL356481,1 | 0,71 | 0,92 | 1,16 | 1,44 | 103 |
| ENSG00000171943 | SRGAP2C | 8,42 | 11,13 | 12,39 | 17,05 | 102 |
| ENSG00000231298 | MANCR | 0,87 | 0,93 | 1,09 | 1,76 | 102 |
| ENSG00000157693 | TMEM268 | 2,41 | 3,89 | 4,6 | 4,87 | 102 |
| ENSG00000264044 | AC005726,2 | 0,55 | 0,6 | 0,64 | 1,11 | 102 |
| ENSG00000272797 | AC092954,1 | 0,55 | 0,58 | 0,72 | 1,11 | 102 |
| ENSG00000273167 | AL359736,1 | 0,55 | 0,83 | 0,98 | 1,11 | 102 |
| ENSG00000162390 | ACOT11 | 1,22 | 1,24 | 1,28 | 2,46 | 102 |
| ENSG00000128731 | HERC2 | 11,06 | 15,5 | 15,7 | 22,29 | 102 |
| ENSG00000240694 | PNMA2 | 11,04 | 14,91 | 15,21 | 22,19 | 101 |
| ENSG00000142156 | COL6A1 | 7,09 | 8,04 | 8,81 | 14,25 | 101 |
| ENSG00000132313 | MRPL35 | 12,35 | 16,17 | 18,69 | 24,77 | 101 |
| ENSG00000266754 | | 3,27 | 3,53 | 4,96 | 6,55 | 100 |
| ENSG00000187068 | C3orf70 | 3,82 | 4,93 | 5,97 | 7,65 | 100 |
| ENSG00000169554 | ZEB2 | 18,9 | 24,49 | 25,39 | 37,83 | 100 |
| ENSG00000165046 | LETM2 | 2,62 | 3,38 | 3,89 | 5,24 | 100 |
| ENSG00000154783 | FGD5 | 0,55 | 0,56 | 0,84 | 1,1 | 100 |
| ENSG00000131773 | KHDRBS3 | 17,87 | 23,89 | 30,92 | 35,69 | 100 |
| ENSG00000150990 | DHX37 | 2,45 | 3,18 | 4,18 | 4,89 | 100 |
| ENSG00000248564 | AC079140,1 | 1,44 | 1,56 | 2,25 | 2,87 | 99 |
| ENSG00000093217 | XYLB | 2,63 | 2,76 | 3,44 | 5,24 | 99 |
| ENSG00000196083 | IL1RAP | 1,04 | 1,83 | 1,88 | 2,07 | 99 |
| ENSG00000132640 | BTBD3 | 16,52 | 21,07 | 24,54 | 32,84 | 99 |
| ENSG00000147119 | CHST7 | 0,72 | 0,97 | 1,25 | 1,43 | 99 |
| ENSG00000065413 | ANKRD44 | 2,11 | 3,07 | 3,44 | 4,19 | 99 |
| ENSG00000135916 | ITM2C | 45,92 | 65,91 | 82,83 | 91,11 | 98 |
| ENSG00000278535 | DHRS11 | 3 | 3,44 | 4,05 | 5,94 | 98 |
| ENSG00000103226 | NOMO3 | 3,78 | 4,47 | 4,69 | 7,48 | 98 |
| ENSG00000285395 | XYLT1 | 0,79 | 1,07 | 1,28 | 1,56 | 97 |
| ENSG00000170873 | MTSS1 | 11,54 | 18,61 | 20,41 | 22,78 | 97 |
| ENSG00000233766 | CAVIN2-AS1 | 0,67 | 0,9 | 1,26 | 1,32 | 97 |
| ENSG00000162878 | PKDCC | 27,42 | 34,31 | 40,96 | 53,96 | 97 |
| ENSG00000108947 | EFNB3 | 8,79 | 12,95 | 13,84 | 17,29 | 97 |
| ENSG00000175093 | SPSB4 | 9,05 | 9,47 | 10,67 | 17,79 | 97 |
| ENSG00000244733 | AL132656,2 | 0,86 | 0,89 | 1,04 | 1,69 | 97 |
| ENSG00000247315 | ZCCHC3 | 14,64 | 18,3 | 19,65 | 28,75 | 96 |
| ENSG00000236255 | AC009404,1 | 0,82 | 0,84 | 0,93 | 1,61 | 96 |
| ENSG00000087303 | NID2 | 4,07 | 4,31 | 6,93 | 7,99 | 96 |
| ENSG00000125398 | SOX9 | 34,28 | 51,34 | 58,27 | 67,29 | 96 |
| ENSG00000144369 | FAM171B | 7,22 | 9,3 | 10,01 | 14,16 | 96 |
| ENSG00000162783 | IER5 | 3,31 | 4,55 | 5,22 | 6,49 | 96 |
| ENSG00000197093 | GAL3ST4 | 2,01 | 2,62 | 3,22 | 3,94 | 96 |

| | | | | | | |
|---|---|---|---|---|---|---|
| ENSG00000139083 | ETV6 | 4,86 | 5,99 | 6,18 | 9,52 | 96 |
| ENSG00000170265 | ZNF282 | 5,68 | 6,29 | 6,39 | 11,12 | 96 |
| ENSG00000135540 | NHSL1 | 14,14 | 20,93 | 22,73 | 27,67 | 96 |
| ENSG00000164970 | FAM219A | 5,08 | 7,57 | 8,58 | 9,94 | 96 |
| ENSG00000095203 | EPB41L4B | 0,69 | 0,77 | 1 | 1,35 | 96 |
| ENSG00000164741 | DLC1 | 6,05 | 7,6 | 8,98 | 11,83 | 96 |
| ENSG00000273245 | AC092653,1 | 0,67 | 0,88 | 0,9 | 1,31 | 96 |
| ENSG00000139289 | PHLDA1 | 6,18 | 9,29 | 9,86 | 12,08 | 95 |
| ENSG00000173320 | STOX2 | 7,78 | 9,39 | 11,81 | 15,19 | 95 |
| ENSG00000110900 | TSPAN11 | 8,17 | 11,71 | 13,25 | 15,93 | 95 |
| ENSG00000136811 | ODF2 | 18,59 | 22,28 | 23,87 | 36,23 | 95 |
| ENSG00000261716 | H2BC20P | 2,47 | 2,89 | 3,55 | 4,81 | 95 |
| ENSG00000184178 | SCFD2 | 3,49 | 4,22 | 6,01 | 6,79 | 95 |
| ENSG00000169856 | ONECUT1 | 3,67 | 4,45 | 5,36 | 7,14 | 95 |
| ENSG00000237238 | BMS1P10 | 0,9 | 1,14 | 1,23 | 1,75 | 94 |
| ENSG00000285301 | BOP1 | 3,15 | 3,68 | 3,91 | 6,12 | 94 |
| ENSG00000261236 | BOP1 | 3,15 | 3,68 | 3,91 | 6,12 | 94 |
| ENSG00000143344 | RGL1 | 6,07 | 6,81 | 9,16 | 11,79 | 94 |
| ENSG00000048740 | CELF2 | 11,54 | 16,03 | 17,16 | 22,41 | 94 |
| ENSG00000269439 | AC010618,3 | 1,2 | 1,69 | 1,77 | 2,33 | 94 |
| ENSG00000111859 | NEDD9 | 8,88 | 11,84 | 13,04 | 17,23 | 94 |
| ENSG00000165424 | ZCCHC24 | 3,73 | 6,05 | 6,55 | 7,23 | 94 |
| ENSG00000186312 | CA5BP1 | 7,92 | 11,73 | 12,28 | 15,35 | 94 |
| ENSG00000075213 | SEMA3A | 15,02 | 16,24 | 20,6 | 29,07 | 94 |
| ENSG00000249014 | HMGN2P4 | 1,08 | 1,1 | 1,82 | 2,09 | 94 |
| ENSG00000182326 | C1S | 1,51 | 2,18 | 2,29 | 2,92 | 93 |
| ENSG00000205277 | MUC12 | 2,76 | 3 | 3,28 | 5,33 | 93 |
| ENSG00000180787 | ZFP3 | 0,58 | 0,74 | 0,81 | 1,12 | 93 |
| ENSG00000163754 | GYG1 | 7,91 | 8,4 | 9,67 | 15,27 | 93 |
| ENSG00000145087 | STXBP5L | 3,01 | 3,9 | 5,14 | 5,81 | 93 |
| ENSG00000148834 | GSTO1 | 19,67 | 27,54 | 27,85 | 37,95 | 93 |
| ENSG00000153885 | KCTD15 | 8,17 | 10,37 | 11,77 | 15,76 | 93 |
| ENSG00000068137 | PLEKHH3 | 2,15 | 2,25 | 2,65 | 4,14 | 93 |
| ENSG00000161217 | PCYT1A | 7,55 | 8,78 | 11,25 | 14,53 | 92 |
| ENSG00000241749 | RPSAP52 | 2,3 | 2,73 | 3,65 | 4,42 | 92 |
| ENSG00000109339 | MAPK10 | 50,41 | 56,49 | 58,8 | 96,85 | 92 |
| ENSG00000135090 | TAOK3 | 4,98 | 7,21 | 7,39 | 9,56 | 92 |
| ENSG00000272525 | AC099522,2 | 0,87 | 0,88 | 1,06 | 1,67 | 92 |
| ENSG00000130810 | PPAN | 6,93 | 8,51 | 8,76 | 13,3 | 92 |
| ENSG00000168056 | LTBP3 | 3,21 | 3,96 | 3,99 | 6,16 | 92 |
| ENSG00000011422 | PLAUR | 4,19 | 4,89 | 4,9 | 8,03 | 92 |
| ENSG00000159579 | RSPRY1 | 21,54 | 26,96 | 32,23 | 41,22 | 91 |
| ENSG00000129657 | SEC14L1 | 17,74 | 22,16 | 24,87 | 33,94 | 91 |
| ENSG00000109654 | TRIM2 | 21,53 | 27,5 | 29,33 | 41,19 | 91 |
| ENSG00000118495 | PLAGL1 | 19,99 | 21,29 | 24,94 | 38,23 | 91 |
| ENSG00000050405 | LIMA1 | 10,27 | 12,79 | 14,55 | 19,62 | 91 |
| ENSG00000154146 | NRGN | 5,08 | 5,46 | 8,26 | 9,69 | 91 |
| ENSG00000170734 | POLH | 3,13 | 3,65 | 3,93 | 5,97 | 91 |
| ENSG00000140548 | ZNF710 | 8,15 | 9,15 | 9,4 | 15,54 | 91 |
| ENSG00000181826 | RELL1 | 10,56 | 14,37 | 16,1 | 20,12 | 91 |

| | | | | | | |
|---|---|---|---|---|---|---|
| ENSG00000173559 | NABP1 | 2,11 | 2,34 | 2,4 | 4,02 | 91 |
| ENSG00000151023 | ENKUR | 1,37 | 2,09 | 2,47 | 2,61 | 91 |
| ENSG00000166068 | SPRED1 | 18,86 | 22,78 | 26,72 | 35,9 | 90 |
| ENSG00000148180 | GSN | 10,51 | 14,99 | 16,07 | 19,98 | 90 |
| ENSG00000135929 | CYP27A1 | 2,76 | 3,56 | 3,66 | 5,24 | 90 |
| ENSG00000235535 | TRDN-AS1 | 2,56 | 3,08 | 3,79 | 4,86 | 90 |
| ENSG00000114767 | RRP9 | 6,02 | 6,98 | 9,53 | 11,42 | 90 |
| ENSG00000264558 | AC015674,1 | 38,73 | 55,5 | 55,83 | 73,4 | 90 |
| ENSG00000136379 | ABHD17C | 6,67 | 8,3 | 9,07 | 12,62 | 89 |
| ENSG00000043355 | ZIC2 | 45,55 | 66,35 | 80,71 | 86,07 | 89 |
| ENSG00000268069 | AC004466,1 | 0,71 | 0,81 | 0,83 | 1,34 | 89 |
| ENSG00000234332 | BCAS2P2 | 1,06 | 1,13 | 1,21 | 2 | 89 |
| ENSG00000203362 | POLH-AS1 | 0,61 | 0,87 | 0,9 | 1,15 | 89 |
| ENSG00000153993 | SEMA3D | 1,04 | 1,32 | 1,56 | 1,96 | 88 |
| ENSG00000147679 | UTP23 | 6,16 | 6,28 | 7,14 | 11,59 | 88 |
| ENSG00000276002 | Metazoa_SRP | 1,34 | 1,37 | 2,02 | 2,52 | 88 |
| ENSG00000229520 | LINC00404 | 1,94 | 2,49 | 2,52 | 3,64 | 88 |
| ENSG00000178764 | ZHX2 | 4,68 | 6,37 | 6,88 | 8,78 | 88 |
| ENSG00000136279 | DBNL | 14,97 | 18,56 | 20,64 | 28,07 | 88 |
| ENSG00000144118 | RALB | 15,15 | 20,16 | 23,07 | 28,36 | 87 |
| ENSG00000187866 | FAM122A | 1,88 | 2,54 | 2,78 | 3,51 | 87 |
| ENSG00000232044 | SILC1 | 5,22 | 7,21 | 7,6 | 9,73 | 86 |
| ENSG00000286132 | AC022415,2 | 1,9 | 2,24 | 2,58 | 3,54 | 86 |
| ENSG00000242086 | MUC20-OT1 | 5,23 | 5,36 | 6,66 | 9,72 | 86 |
| ENSG00000160888 | IER2 | 13,67 | 19,05 | 21,96 | 25,38 | 86 |
| ENSG00000274779 | NOMO1 | 17,86 | 22,05 | 25,25 | 33,14 | 86 |
| ENSG00000132967 | HMGB1P5 | 52,23 | 57,71 | 59,61 | 96,91 | 86 |
| ENSG00000183337 | BCOR | 16,33 | 23,65 | 26,76 | 30,29 | 85 |
| ENSG00000187800 | PEAR1 | 0,62 | 0,81 | 0,95 | 1,15 | 85 |
| ENSG00000105325 | FZR1 | 9,47 | 12,03 | 13,79 | 17,55 | 85 |
| ENSG00000141519 | CCDC40 | 5,58 | 6,96 | 8,17 | 10,34 | 85 |
| ENSG00000136478 | TEX2 | 4,06 | 6,26 | 7,33 | 7,52 | 85 |
| ENSG00000278087 | NOMO3 | 7,36 | 8,48 | 12,02 | 13,58 | 85 |
| ENSG00000172831 | CES2 | 8,93 | 12,18 | 15,09 | 16,47 | 84 |
| ENSG00000179314 | WSCD1 | 11,74 | 12,2 | 15,26 | 21,62 | 84 |
| ENSG00000239887 | C1orf226 | 2,65 | 3,73 | 4 | 4,88 | 84 |
| ENSG00000131018 | SYNE1 | 6,91 | 8,91 | 10,77 | 12,72 | 84 |
| ENSG00000174498 | IGDCC3 | 51,92 | 57,14 | 66,24 | 95,43 | 84 |
| ENSG00000140743 | CDR2 | 4,74 | 5,31 | 5,84 | 8,71 | 84 |
| ENSG00000273447 | AC004067,1 | 1,71 | 2,36 | 2,37 | 3,14 | 84 |
| ENSG00000076716 | GPC4 | 19,87 | 29,29 | 34,78 | 36,48 | 84 |
| ENSG00000143341 | HMCN1 | 4,41 | 5,91 | 7,02 | 8,09 | 83 |
| ENSG00000156103 | MMP16 | 10,24 | 13,23 | 14,46 | 18,78 | 83 |
| ENSG00000053438 | NNAT | 23,49 | 34,86 | 41,87 | 43,06 | 83 |
| ENSG00000144724 | PTPRG | 18,7 | 28 | 31,37 | 34,27 | 83 |
| ENSG00000160392 | C19orf47 | 4,17 | 4,46 | 6,94 | 7,64 | 83 |
| ENSG00000227331 | RPL7AP22 | 0,77 | 0,78 | 1,1 | 1,41 | 83 |
| ENSG00000247416 | AP000802,1 | 0,77 | 0,97 | 1,01 | 1,41 | 83 |
| ENSG00000174306 | ZHX3 | 2,68 | 3,99 | 4,15 | 4,9 | 83 |
| ENSG00000240184 | PCDHGC3 | 14,07 | 19,4 | 22,27 | 25,71 | 83 |

| | | | | | | |
|---|---|---|---|---|---|---|
| ENSG00000035862 | TIMP2 | 23,7 | 34,81 | 39,32 | 43,3 | 83 |
| ENSG00000260914 | AC026464,4 | 9,18 | 12,9 | 16,23 | 16,77 | 83 |
| ENSG00000224243 | SOX1-OT | 4,42 | 4,99 | 5,15 | 8,06 | 82 |
| ENSG00000148948 | LRRC4C | 2,21 | 2,67 | 2,98 | 4,03 | 82 |
| ENSG00000165632 | TAF3 | 2,94 | 3,63 | 3,95 | 5,36 | 82 |
| ENSG00000196141 | SPATS2L | 20,73 | 26,62 | 32,03 | 37,75 | 82 |
| ENSG00000163964 | PIGX | 8,67 | 10,63 | 10,97 | 15,78 | 82 |
| ENSG00000171791 | BCL2 | 3,49 | 3,86 | 4,22 | 6,35 | 82 |
| ENSG00000165259 | HDX | 2,92 | 4,29 | 5,23 | 5,31 | 82 |
| ENSG00000185885 | IFITM1 | 27,15 | 36,24 | 44,82 | 49,37 | 82 |
| ENSG00000272288 | AL451165,2 | 1,74 | 3,06 | 3,12 | 3,16 | 82 |
| ENSG00000265882 | RN7SL73P | 3,74 | 3,86 | 5,57 | 6,79 | 82 |
| ENSG00000174010 | KLHL15 | 3,88 | 4,56 | 4,75 | 7,04 | 81 |
| ENSG00000146950 | SHROOM2 | 4,7 | 7,14 | 7,8 | 8,52 | 81 |
| ENSG00000165916 | PSMC3 | 69,31 | 82,5 | 97,32 | 125,58 | 81 |
| ENSG00000169862 | CTNND2 | 12,26 | 17,45 | 21,46 | 22,2 | 81 |
| ENSG00000243406 | MRPS31P5 | 4,7 | 5,75 | 6,09 | 8,51 | 81 |
| ENSG00000178921 | PFAS | 15,88 | 17,21 | 17,87 | 28,74 | 81 |
| ENSG00000198746 | GPATCH3 | 2,67 | 3,53 | 4,15 | 4,83 | 81 |
| ENSG00000103174 | NAGPA | 1,78 | 2,02 | 2,08 | 3,22 | 81 |
| ENSG00000267127 | AC090360,1 | 0,94 | 1,18 | 1,2 | 1,7 | 81 |
| ENSG00000100280 | AP1B1 | 16,86 | 22,81 | 25,37 | 30,49 | 81 |
| ENSG00000005810 | MYCBP2 | 13,37 | 14,26 | 18,16 | 24,13 | 80 |
| ENSG00000283192 | AC007383,6 | 1,02 | 1,33 | 1,37 | 1,84 | 80 |
| ENSG00000227946 | AC007383,1 | 1,02 | 1,33 | 1,37 | 1,84 | 80 |
| ENSG00000232000 | CLCN3P1 | 0,56 | 0,68 | 0,8 | 1,01 | 80 |
| ENSG00000261490 | AC005674,1 | 0,81 | 1 | 1,14 | 1,46 | 80 |
| ENSG00000214655 | ZSWIM8 | 11,74 | 11,97 | 15 | 21,13 | 80 |
| ENSG00000112531 | QKI | 99,11 | 108,68 | 131,85 | 178,37 | 80 |
| ENSG00000138835 | RGS3 | 12,72 | 15,82 | 16,78 | 22,87 | 80 |
| ENSG00000040531 | CTNS | 4,4 | 4,6 | 7,15 | 7,91 | 80 |
| ENSG00000272077 | AC124045,1 | 1,08 | 1,47 | 1,48 | 1,94 | 80 |
| ENSG00000281102 | AC092046,2 | 1,08 | 1,47 | 1,48 | 1,94 | 80 |
| ENSG00000278488 | NAPRT | 0,93 | 1,2 | 1,56 | 1,67 | 80 |
| ENSG00000141858 | SAMD1 | 18,17 | 20,49 | 22,86 | 32,62 | 80 |
| ENSG00000253764 | AC019257,1 | 1,25 | 1,43 | 1,71 | 2,24 | 79 |
| ENSG00000176896 | TCEANC | 1,2 | 1,63 | 1,9 | 2,15 | 79 |
| ENSG00000224975 | INE1 | 2,39 | 2,56 | 2,73 | 4,28 | 79 |
| ENSG00000111879 | FAM184A | 4,96 | 5,84 | 6,32 | 8,88 | 79 |
| ENSG00000108175 | ZMIZ1 | 22,16 | 28,05 | 35,07 | 39,65 | 79 |
| ENSG00000183955 | KMT5A | 16,94 | 22,84 | 26,34 | 30,31 | 79 |
| ENSG00000103966 | EHD4 | 2,75 | 2,81 | 3,52 | 4,92 | 79 |
| ENSG00000079819 | EPB41L2 | 28,81 | 29,84 | 30,61 | 51,51 | 79 |
| ENSG00000141696 | P3H4 | 8,92 | 11,01 | 14,14 | 15,94 | 79 |
| ENSG00000131409 | LRRC4B | 9,93 | 14,22 | 15,92 | 17,74 | 79 |
| ENSG00000008256 | CYTH3 | 6,14 | 7,64 | 9,82 | 10,96 | 79 |
| ENSG00000170390 | DCLK2 | 10,62 | 13,96 | 15,33 | 18,95 | 78 |
| ENSG00000141522 | ARHGDIA | 76,27 | 100,66 | 119,24 | 135,95 | 78 |
| ENSG00000285972 | CERNA2 | 1,23 | 1,25 | 1,38 | 2,19 | 78 |
| ENSG00000176014 | TUBB6 | 40,35 | 44,29 | 57,39 | 71,82 | 78 |

| | | | | | | |
|---|---|---|---|---|---|---|
| ENSG00000082438 | COBLL1 | 2,88 | 3,57 | 4,29 | 5,12 | 78 |
| ENSG00000073712 | FERMT2 | 29,59 | 33,89 | 41,99 | 52,6 | 78 |
| ENSG00000102385 | DRP2 | 0,85 | 0,99 | 1,04 | 1,51 | 78 |
| ENSG00000242028 | HYPK | 3,53 | 5,09 | 5,67 | 6,27 | 78 |
| ENSG00000076928 | ARHGEF1 | 5,44 | 5,94 | 6,68 | 9,65 | 77 |
| ENSG00000254481 | PTP4A2P2 | 1,1 | 1,3 | 1,79 | 1,95 | 77 |
| ENSG00000281917 | SLC16A1 | 4,89 | 5,29 | 5,51 | 8,66 | 77 |
| ENSG00000128536 | CDHR3 | 1,48 | 1,63 | 1,78 | 2,62 | 77 |
| ENSG00000198342 | ZNF442 | 3,48 | 3,83 | 5,3 | 6,16 | 77 |
| ENSG00000101680 | LAMA1 | 33,84 | 43,36 | 49,96 | 59,9 | 77 |
| ENSG00000174238 | PITPNA | 22,09 | 26,32 | 27,39 | 39,1 | 77 |
| ENSG00000147065 | MSN | 47,8 | 60,15 | 61,85 | 84,59 | 77 |
| ENSG00000176105 | YES1 | 26,6 | 29,04 | 34,1 | 46,99 | 77 |
| ENSG00000198142 | SOWAHC | 4,01 | 5,32 | 5,4 | 7,08 | 77 |
| ENSG00000235499 | AC073046,1 | 3,73 | 4,55 | 5,2 | 6,58 | 76 |
| ENSG00000100417 | PMM1 | 6,06 | 6,4 | 8,67 | 10,69 | 76 |
| ENSG00000145349 | CAMK2D | 15,32 | 17,59 | 21,08 | 26,99 | 76 |
| ENSG00000259488 | AC023355,1 | 1,13 | 1,6 | 1,61 | 1,99 | 76 |
| ENSG00000100336 | APOL4 | 1,2 | 1,21 | 1,64 | 2,11 | 76 |
| ENSG00000105248 | YJU2 | 5,74 | 5,94 | 7,4 | 10,09 | 76 |
| ENSG00000230269 | LINC02525 | 1,27 | 1,29 | 1,45 | 2,23 | 76 |
| ENSG00000230565 | ZNF32-AS2 | 0,86 | 1,05 | 1,2 | 1,51 | 76 |
| ENSG00000282033 | AC074387,1 | 1,02 | 1,28 | 1,68 | 1,79 | 75 |
| ENSG00000232804 | HSPA1B | 3,46 | 4,75 | 4,81 | 6,07 | 75 |
| ENSG00000149150 | SLC43A1 | 0,57 | 0,7 | 0,71 | 1 | 75 |
| ENSG00000142330 | CAPN10 | 3,64 | 4,38 | 4,58 | 6,37 | 75 |
| ENSG00000233622 | CYP2T1P | 0,72 | 0,85 | 0,94 | 1,26 | 75 |
| ENSG00000130816 | DNMT1 | 47,85 | 54,46 | 63,63 | 83,69 | 75 |
| ENSG00000205476 | CCDC85C | 8,8 | 9,74 | 10,24 | 15,39 | 75 |
| ENSG00000107164 | FUBP3 | 19,05 | 22,11 | 22,51 | 33,29 | 75 |
| ENSG00000113734 | BNIP1 | 4,71 | 6,08 | 6,5 | 8,23 | 75 |
| ENSG00000162946 | DISC1 | 1,82 | 2,39 | 2,63 | 3,18 | 75 |
| ENSG00000147576 | ADHFE1 | 0,75 | 0,82 | 0,92 | 1,31 | 75 |
| ENSG00000103227 | LMF1 | 4,73 | 6,15 | 6,32 | 8,26 | 75 |
| ENSG00000286789 | AL161645,2 | 2,2 | 2,7 | 2,8 | 3,84 | 75 |
| ENSG00000064393 | HIPK2 | 16,31 | 22,97 | 24,66 | 28,44 | 74 |
| ENSG00000015676 | NUDCD3 | 18,28 | 25,48 | 29,32 | 31,85 | 74 |
| ENSG00000162639 | HENMT1 | 2,86 | 3,58 | 3,94 | 4,98 | 74 |
| ENSG00000086619 | ERO1B | 1,77 | 2,38 | 2,87 | 3,08 | 74 |
| ENSG00000101400 | SNTA1 | 1,52 | 2,06 | 2,57 | 2,64 | 74 |
| ENSG00000271828 | AC008937,3 | 0,76 | 0,8 | 1,15 | 1,32 | 74 |
| ENSG00000213397 | HAUS7 | 12,34 | 14,05 | 15,54 | 21,43 | 74 |
| ENSG00000106266 | SNX8 | 4,82 | 5,58 | 6,55 | 8,37 | 74 |
| ENSG00000137266 | SLC22A23 | 7,31 | 9,85 | 9,89 | 12,69 | 74 |
| ENSG00000125952 | MAX | 19,6 | 19,79 | 27,67 | 34 | 73 |
| ENSG00000197128 | ZNF772 | 5,08 | 5,96 | 6,17 | 8,81 | 73 |
| ENSG00000277972 | CISD3 | 1,46 | 1,76 | 2,25 | 2,53 | 73 |
| ENSG00000259315 | ACTG1P17 | 0,86 | 0,88 | 0,92 | 1,49 | 73 |
| ENSG00000275911 | NDE1 | 12 | 14,06 | 18,02 | 20,78 | 73 |
| ENSG00000087266 | SH3BP2 | 7,83 | 9,09 | 9,95 | 13,55 | 73 |

| | | | | | | |
|---|---|---|---|---|---|---|
| ENSG00000217555 | CKLF | 14,95 | 18,08 | 22,9 | 25,87 | 73 |
| ENSG00000121210 | TMEM131L | 7,51 | 9,08 | 9,69 | 12,99 | 73 |
| ENSG00000130052 | STARD8 | 1,48 | 1,76 | 2,06 | 2,56 | 73 |
| ENSG00000196507 | TCEAL3 | 5,57 | 9,25 | 9,34 | 9,63 | 73 |
| ENSG00000034693 | PEX3 | 4,32 | 4,64 | 5,57 | 7,46 | 73 |
| ENSG00000151491 | EPS8 | 13,31 | 15,09 | 17,97 | 22,98 | 73 |
| ENSG00000164574 | GALNT10 | 6,87 | 9,61 | 11,06 | 11,86 | 73 |
| ENSG00000213859 | KCTD11 | 0,91 | 1,13 | 1,36 | 1,57 | 73 |
| ENSG00000288399 | KCTD11 | 0,91 | 1,13 | 1,36 | 1,57 | 73 |
| ENSG00000148481 | MINDY3 | 8,04 | 9,91 | 11,77 | 13,87 | 73 |
| ENSG00000149294 | NCAM1 | 16,69 | 23,77 | 25,85 | 28,79 | 72 |
| ENSG00000135862 | LAMC1 | 75,19 | 95,94 | 112,58 | 129,6 | 72 |
| ENSG00000275481 | AC025031,4 | 0,76 | 0,8 | 0,85 | 1,31 | 72 |
| ENSG00000204469 | PRRC2A | 1,66 | 1,7 | 1,89 | 2,86 | 72 |
| ENSG00000104368 | PLAT | 5,07 | 5,73 | 6,72 | 8,73 | 72 |
| ENSG00000139800 | ZIC5 | 14,41 | 19,93 | 21,22 | 24,81 | 72 |
| ENSG00000169946 | ZFPM2 | 5,08 | 6,9 | 7,71 | 8,74 | 72 |
| ENSG00000005189 | REXO5 | 3,97 | 4,09 | 5,34 | 6,83 | 72 |
| ENSG00000196371 | FUT4 | 1,07 | 1,12 | 1,35 | 1,84 | 72 |
| ENSG00000136141 | LRCH1 | 3,92 | 5,15 | 5,66 | 6,74 | 72 |
| ENSG00000074047 | GLI2 | 8,08 | 9,54 | 10,98 | 13,89 | 72 |
| ENSG00000164330 | EBF1 | 4,97 | 5,34 | 8,19 | 8,54 | 72 |
| ENSG00000173465 | ZNRD2 | 9,01 | 11,37 | 13,54 | 15,48 | 72 |
| ENSG00000145736 | GTF2H2 | 4,14 | 5,4 | 5,43 | 7,11 | 72 |
| ENSG00000175895 | PLEKHF2 | 2,32 | 2,93 | 3,54 | 3,98 | 72 |
| ENSG00000285460 | BCAR1 | 4,79 | 5,99 | 7,84 | 8,21 | 71 |
| ENSG00000083444 | PLOD1 | 11,7 | 13,81 | 16,59 | 20,04 | 71 |
| ENSG00000078487 | ZCWPW1 | 0,73 | 1,06 | 1,11 | 1,25 | 71 |
| ENSG00000144840 | RABL3 | 9,88 | 10,33 | 11,7 | 16,9 | 71 |
| ENSG00000168917 | SLC35G2 | 3,1 | 4,08 | 4,29 | 5,3 | 71 |
| ENSG00000064651 | SLC12A2 | 9,01 | 10,75 | 11,78 | 15,4 | 71 |
| ENSG00000135506 | OS9 | 26,42 | 37,15 | 37,16 | 45,11 | 71 |
| ENSG00000163064 | EN1 | 1,5 | 2,22 | 2,43 | 2,56 | 71 |
| ENSG00000136158 | SPRY2 | 49,98 | 64,86 | 71,37 | 85,25 | 71 |
| ENSG00000056972 | TRAF3IP2 | 2,54 | 3,26 | 4,19 | 4,33 | 70 |
| ENSG00000082512 | TRAF5 | 5,46 | 6,86 | 7,69 | 9,3 | 70 |
| ENSG00000213020 | ZNF611 | 6,89 | 7,46 | 7,51 | 11,73 | 70 |
| ENSG00000269743 | SLC25A53 | 3,83 | 4,58 | 5,21 | 6,52 | 70 |
| ENSG00000230911 | PPIHP1 | 0,67 | 0,89 | 0,92 | 1,14 | 70 |
| ENSG00000081913 | PHLPP1 | 9,84 | 11,62 | 11,68 | 16,74 | 70 |
| ENSG00000250366 | TUNAR | 2,23 | 2,79 | 3,07 | 3,79 | 70 |
| ENSG00000151353 | TMEM18 | 8,18 | 8,95 | 10,89 | 13,9 | 70 |
| ENSG00000170035 | UBE2E3 | 85,8 | 106,71 | 125,06 | 145,71 | 70 |
| ENSG00000198648 | STK39 | 12,85 | 16,33 | 19,18 | 21,82 | 70 |
| ENSG00000089057 | SLC23A2 | 3,86 | 4,75 | 4,91 | 6,55 | 70 |
| ENSG00000119471 | HSDL2 | 7,41 | 9 | 10,57 | 12,57 | 70 |
| ENSG00000034533 | ASTE1 | 2,46 | 2,88 | 3,73 | 4,17 | 70 |
| ENSG00000244479 | OR2A1-AS1 | 3,41 | 4,78 | 4,84 | 5,78 | 70 |
| ENSG00000146072 | TNFRSF21 | 12,61 | 14,86 | 20,49 | 21,33 | 69 |
| ENSG00000168779 | SHOX2 | 2,91 | 4,58 | 4,66 | 4,92 | 69 |

| | | | | | | |
|---|---|---|---|---|---|---|
| ENSG00000224897 | POT1-AS1 | 1,06 | 1,36 | 1,46 | 1,79 | 69 |
| ENSG00000288516 | PRKACA | 17,43 | 23,34 | 28,19 | 29,43 | 69 |
| ENSG00000287038 | AL162388,2 | 0,93 | 1,46 | 1,54 | 1,57 | 69 |
| ENSG00000144619 | CNTN4 | 3,61 | 4,51 | 4,66 | 6,09 | 69 |
| ENSG00000087074 | PPP1R15A | 8,05 | 12,13 | 12,15 | 13,58 | 69 |
| ENSG00000182551 | ADI1 | 11,96 | 14,05 | 16,9 | 20,17 | 69 |
| ENSG00000134352 | IL6ST | 11,05 | 16,14 | 16,6 | 18,63 | 69 |
| ENSG00000131725 | WDR44 | 4,41 | 5,26 | 5,97 | 7,43 | 68 |
| ENSG00000131381 | RBSN | 7,2 | 9,27 | 9,32 | 12,13 | 68 |
| ENSG00000148935 | GAS2 | 2,44 | 2,85 | 3,27 | 4,11 | 68 |
| ENSG00000133401 | PDZD2 | 1,51 | 2,12 | 2,16 | 2,54 | 68 |
| ENSG00000174028 | FAM3C2P | 11,29 | 16,59 | 18,28 | 18,99 | 68 |
| ENSG00000187837 | H1-2 | 784,5 | 935,72 | 1084,76 | 1318,74 | 68 |
| ENSG00000152217 | SETBP1 | 8,92 | 10,88 | 11,26 | 14,97 | 68 |
| ENSG00000125386 | FAM193A | 10,59 | 13,67 | 14,44 | 17,77 | 68 |
| ENSG00000226916 | WDR46 | 5,62 | 7,57 | 7,98 | 9,43 | 68 |
| ENSG00000204221 | WDR46 | 5,62 | 7,57 | 7,98 | 9,43 | 68 |
| ENSG00000260032 | NORAD | 56,56 | 75,65 | 76,96 | 94,9 | 68 |
| ENSG00000118922 | KLF12 | 11,39 | 15,24 | 16,56 | 19,11 | 68 |
| ENSG00000110080 | ST3GAL4 | 6,43 | 6,57 | 8,13 | 10,77 | 67 |
| ENSG00000155858 | LSM11 | 2,43 | 2,62 | 3,05 | 4,07 | 67 |
| ENSG00000124783 | SSR1 | 25,33 | 29,48 | 35,06 | 42,4 | 67 |
| ENSG00000130764 | LRRC47 | 11,67 | 12,62 | 13,44 | 19,53 | 67 |
| ENSG00000203952 | CCDC160 | 13,61 | 13,73 | 16,56 | 22,77 | 67 |
| ENSG00000277270 | AL160412,1 | 1,1 | 1,31 | 1,39 | 1,84 | 67 |
| ENSG00000206357 | NELFE | 2,93 | 3,95 | 4,41 | 4,9 | 67 |
| ENSG00000231044 | NELFE | 2,93 | 3,95 | 4,41 | 4,9 | 67 |
| ENSG00000229363 | NELFE | 2,93 | 3,95 | 4,41 | 4,9 | 67 |
| ENSG00000233801 | NELFE | 2,93 | 3,95 | 4,41 | 4,9 | 67 |
| ENSG00000206268 | NELFE | 2,93 | 3,95 | 4,41 | 4,9 | 67 |
| ENSG00000101871 | MID1 | 15,73 | 19,71 | 23,08 | 26,3 | 67 |
| ENSG00000019485 | PRDM11 | 2,74 | 3,19 | 3,71 | 4,58 | 67 |
| ENSG00000278175 | GLIDR | 8,36 | 11,55 | 12,09 | 13,97 | 67 |
| ENSG00000196646 | ZNF136 | 4,1 | 4,97 | 4,98 | 6,85 | 67 |
| ENSG00000103647 | CORO2B | 9,89 | 14,88 | 16,36 | 16,51 | 67 |
| ENSG00000100034 | PPM1F | 5,56 | 6,65 | 7,97 | 9,28 | 67 |
| ENSG00000197620 | EOLA1 | 3,77 | 4,48 | 4,65 | 6,29 | 67 |
| ENSG00000119720 | NRDE2 | 6,6 | 6,63 | 8,87 | 11,01 | 67 |
| ENSG00000196453 | ZNF777 | 3,88 | 5,13 | 5,3 | 6,47 | 67 |
| ENSG00000263238 | CTSO | 1,77 | 2,55 | 2,81 | 2,95 | 67 |
| ENSG00000164100 | NDST3 | 1,11 | 1,59 | 1,8 | 1,85 | 67 |
| ENSG00000198354 | DCAF12L2 | 0,6 | 0,76 | 0,97 | 1 | 67 |
| ENSG00000101670 | LIPG | 11,41 | 14,15 | 17,37 | 19,01 | 67 |
| ENSG00000140577 | CRTC3 | 8,47 | 11,32 | 11,58 | 14,1 | 66 |
| ENSG00000011052 | NME1-NME2 | 1,34 | 1,71 | 2,07 | 2,23 | 66 |
| ENSG00000174013 | FBXO45 | 4,64 | 5,9 | 5,94 | 7,72 | 66 |
| ENSG00000077235 | GTF3C1 | 24,81 | 30,92 | 31,31 | 41,27 | 66 |
| ENSG00000013619 | MAMLD1 | 0,86 | 1,33 | 1,42 | 1,43 | 66 |
| ENSG00000134278 | SPIRE1 | 6,64 | 8,29 | 9,94 | 11,04 | 66 |
| ENSG00000111237 | VPS29 | 40,13 | 53,66 | 55,78 | 66,72 | 66 |

| | | | | | | |
|---|---|---|---|---|---|---|
| ENSG00000227372 | TP73-AS1 | 6,43 | 8,22 | 9,99 | 10,69 | 66 |
| ENSG00000246334 | PRR7-AS1 | 2,45 | 2,74 | 3,35 | 4,07 | 66 |
| ENSG00000110013 | SIAE | 2,18 | 3 | 3,37 | 3,62 | 66 |
| ENSG00000206432 | TMEM200C | 4,21 | 6,34 | 6,53 | 6,99 | 66 |
| ENSG00000167460 | TPM4 | 146,09 | 164,45 | 190,56 | 242,29 | 66 |
| ENSG00000111052 | LIN7A | 4,09 | 5,99 | 6,44 | 6,78 | 66 |
| ENSG00000272325 | NUDT3 | 14,45 | 18,25 | 18,73 | 23,95 | 66 |
| ENSG00000061936 | SFSWAP | 13,72 | 15,53 | 16,72 | 22,74 | 66 |
| ENSG00000037474 | NSUN2 | 17,53 | 20,55 | 21,24 | 29,05 | 66 |
| ENSG00000267191 | AC006213,3 | 0,7 | 0,71 | 0,78 | 1,16 | 66 |
| ENSG00000114948 | ADAM23 | 10,33 | 12,04 | 13,2 | 17,11 | 66 |
| ENSG00000156521 | TYSND1 | 2,24 | 2,43 | 2,52 | 3,71 | 66 |
| ENSG00000185630 | PBX1 | 31,4 | 39,26 | 43,04 | 51,97 | 66 |
| ENSG00000095321 | CRAT | 5,15 | 7,62 | 8,01 | 8,52 | 65 |
| ENSG00000197774 | EME2 | 1,27 | 1,77 | 1,8 | 2,1 | 65 |
| ENSG00000166341 | DCHS1 | 13,23 | 17,85 | 18,97 | 21,87 | 65 |
| ENSG00000105379 | ETFB | 15,1 | 16,91 | 18,28 | 24,96 | 65 |
| ENSG00000101608 | MYL12A | 96,46 | 109,5 | 136,02 | 159,34 | 65 |
| ENSG00000172748 | ZNF596 | 4,79 | 6,64 | 7,13 | 7,91 | 65 |
| ENSG00000108312 | UBTF | 22,64 | 30,31 | 31,18 | 37,37 | 65 |
| ENSG00000184164 | CRELD2 | 8,97 | 10,31 | 14,3 | 14,8 | 65 |
| ENSG00000141994 | DUS3L | 11,64 | 13,55 | 14,36 | 19,2 | 65 |
| ENSG00000183597 | TANGO2 | 3,28 | 3,59 | 4,5 | 5,41 | 65 |
| ENSG00000257950 | P2RX5-TAX1BP3 | 0,74 | 0,89 | 0,98 | 1,22 | 65 |
| ENSG00000107862 | GBF1 | 15,06 | 20,56 | 21,07 | 24,81 | 65 |
| ENSG00000008083 | JARID2 | 17,29 | 26,61 | 26,85 | 28,48 | 65 |
| ENSG00000180867 | PDIA3P1 | 1,7 | 2,29 | 2,4 | 2,8 | 65 |
| ENSG00000166886 | NAB2 | 8,81 | 10,06 | 10,09 | 14,51 | 65 |
| ENSG00000172534 | HCFC1 | 25,3 | 28,83 | 32,99 | 41,66 | 65 |
| ENSG00000002822 | MAD1L1 | 2,57 | 3,21 | 3,44 | 4,23 | 65 |
| ENSG00000161835 | TAMALIN | 2,4 | 2,72 | 2,8 | 3,95 | 65 |
| ENSG00000107282 | APBA1 | 1,27 | 1,56 | 1,66 | 2,09 | 65 |
| ENSG00000145715 | RASA1 | 13,22 | 14,88 | 15,41 | 21,75 | 65 |
| ENSG00000110880 | CORO1C | 56,15 | 62,86 | 68,03 | 92,35 | 64 |
| ENSG00000101868 | POLA1 | 24,16 | 24,25 | 25,56 | 39,73 | 64 |
| ENSG00000075945 | KIFAP3 | 8,24 | 11,29 | 12,73 | 13,55 | 64 |
| ENSG00000147459 | DOCK5 | 2,86 | 3,72 | 4,24 | 4,7 | 64 |
| ENSG00000183621 | ZNF438 | 1,43 | 1,55 | 1,93 | 2,35 | 64 |
| ENSG00000130511 | SSBP4 | 36,58 | 38,03 | 40,45 | 60,1 | 64 |
| ENSG00000166025 | AMOTL1 | 19,38 | 23,33 | 24,44 | 31,84 | 64 |
| ENSG00000214717 | ZBED1 | 7,95 | 9,61 | 11,5 | 13,06 | 64 |
| ENSG00000080603 | SRCAP | 15,59 | 20,1 | 20,12 | 25,61 | 64 |
| ENSG00000111412 | SPRING1 | 10,5 | 13,31 | 14,11 | 17,22 | 64 |
| ENSG00000197928 | ZNF677 | 9,2 | 10,64 | 10,98 | 15,08 | 64 |
| ENSG00000060762 | MPC1 | 9,08 | 10,77 | 13,51 | 14,88 | 64 |
| ENSG00000274565 | AC080038,1 | 0,94 | 1,1 | 1,21 | 1,54 | 64 |
| ENSG00000140022 | STON2 | 15,86 | 21,23 | 21,6 | 25,98 | 64 |
| ENSG00000143641 | GALNT2 | 16,62 | 23,06 | 23,37 | 27,19 | 64 |
| ENSG00000169855 | ROBO1 | 30,06 | 30,49 | 31,2 | 49,16 | 64 |
| ENSG00000186575 | NF2 | 10,88 | 14,84 | 16,49 | 17,79 | 64 |

| | | | | | | |
|---|---|---|---|---|---|---|
| ENSG00000166086 | JAM3 | 35,65 | 49,39 | 50,75 | 58,27 | 63 |
| ENSG00000159082 | SYNJ1 | 2,16 | 3,09 | 3,38 | 3,53 | 63 |
| ENSG00000285250 | MSRA | 2,57 | 2,84 | 3,63 | 4,2 | 63 |
| ENSG00000151458 | ANKRD50 | 8,3 | 11,2 | 11,44 | 13,55 | 63 |
| ENSG00000002586 | CD99 | 69,46 | 77,4 | 98,4 | 113,32 | 63 |
| ENSG00000159840 | ZYX | 13,88 | 16,68 | 22,61 | 22,64 | 63 |
| ENSG00000285443 | ZYX | 13,88 | 16,68 | 22,61 | 22,64 | 63 |
| ENSG00000102935 | ZNF423 | 24,56 | 33,04 | 33,6 | 40,06 | 63 |
| ENSG00000137478 | FCHSD2 | 10,71 | 12,42 | 15,08 | 17,45 | 63 |
| ENSG00000176788 | BASP1 | 64,09 | 92,31 | 97,14 | 104,42 | 63 |
| ENSG00000066933 | MYO9A | 8,68 | 10,99 | 11,57 | 14,14 | 63 |
| ENSG00000108819 | PPP1R9B | 2,77 | 3,07 | 3,3 | 4,51 | 63 |
| ENSG00000258732 | AC025884,1 | 10,19 | 12,44 | 15,91 | 16,59 | 63 |
| ENSG00000162944 | RFTN2 | 7,47 | 10,83 | 11,04 | 12,16 | 63 |
| ENSG00000198373 | WWP2 | 8,92 | 13,23 | 13,3 | 14,52 | 63 |
| ENSG00000166444 | DENND2B | 22,48 | 30,89 | 31,32 | 36,59 | 63 |
| ENSG00000101040 | ZMYND8 | 17,64 | 20,27 | 21,18 | 28,7 | 63 |
| ENSG00000113141 | IK | 23,15 | 28,32 | 30,36 | 37,66 | 63 |
| ENSG00000150760 | DOCK1 | 17,33 | 19,09 | 20,37 | 28,19 | 63 |
| ENSG00000129680 | MAP7D3 | 4,52 | 5,5 | 5,71 | 7,35 | 63 |
| ENSG00000135521 | LTV1 | 12,27 | 13,71 | 15,39 | 19,95 | 63 |
| ENSG00000221923 | ZNF880 | 5,45 | 6,8 | 7,6 | 8,86 | 63 |
| ENSG00000087269 | NOP14 | 8,44 | 9,04 | 9,79 | 13,72 | 63 |
| ENSG00000136631 | VPS45 | 18,82 | 22,88 | 25,42 | 30,59 | 63 |
| ENSG00000161904 | LEMD2 | 13,93 | 16,11 | 18,07 | 22,63 | 62 |
| ENSG00000100109 | TFIP11 | 7,53 | 8,43 | 9,36 | 12,23 | 62 |
| ENSG00000196814 | MVB12B | 16,85 | 19,15 | 20,87 | 27,34 | 62 |
| ENSG00000136040 | PLXNC1 | 3,55 | 4,36 | 4,93 | 5,76 | 62 |
| ENSG00000111300 | NAA25 | 11,41 | 11,58 | 12,87 | 18,51 | 62 |
| ENSG00000136819 | C9orf78 | 12,52 | 17,65 | 19,26 | 20,31 | 62 |
| ENSG00000277292 | PIP4K2B | 11,53 | 15,36 | 16,8 | 18,7 | 62 |
| ENSG00000276293 | PIP4K2B | 11,53 | 15,36 | 16,8 | 18,7 | 62 |
| ENSG00000174775 | HRAS | 11,34 | 14,33 | 16,98 | 18,39 | 62 |
| ENSG00000277118 | Metazoa_SRP | 1,4 | 1,66 | 2,01 | 2,27 | 62 |
| ENSG00000198783 | ZNF830 | 5,07 | 5,78 | 6,95 | 8,22 | 62 |
| ENSG00000156671 | SAMD8 | 4,17 | 5,52 | 5,77 | 6,76 | 62 |
| ENSG00000112902 | SEMA5A | 10,14 | 10,73 | 11,72 | 16,43 | 62 |
| ENSG00000085644 | ZNF213 | 2,7 | 3,67 | 3,8 | 4,37 | 62 |
| ENSG00000108349 | CASC3 | 30,11 | 36,63 | 37,51 | 48,71 | 62 |
| ENSG00000123159 | GIPC1 | 18,77 | 22,19 | 26,89 | 30,34 | 62 |
| ENSG00000100605 | ITPK1 | 4,69 | 5,64 | 5,9 | 7,58 | 62 |
| ENSG00000272941 | AC083862,1 | 0,99 | 1,12 | 1,14 | 1,6 | 62 |
| ENSG00000056487 | PHF21B | 10,18 | 12,48 | 13,17 | 16,45 | 62 |
| ENSG00000166170 | BAG5 | 11,11 | 13,06 | 14,91 | 17,95 | 62 |
| ENSG00000101812 | H2BW2 | 0,78 | 0,89 | 0,96 | 1,26 | 62 |
| ENSG00000198961 | PJA2 | 21,78 | 28,29 | 28,51 | 35,18 | 62 |
| ENSG00000117758 | STX12 | 6,77 | 8,99 | 9,44 | 10,93 | 61 |
| ENSG00000259342 | AC025580,1 | 3,16 | 3,59 | 3,99 | 5,1 | 61 |
| ENSG00000139910 | NOVA1 | 23,76 | 29,73 | 32,38 | 38,33 | 61 |
| ENSG00000137285 | TUBB2B | 171,5 | 238,65 | 246,28 | 276,39 | 61 |

| | | | | | | |
|---|---|---|---|---|---|---|
| ENSG00000160799 | CCDC12 | 13,47 | 15,9 | 16,3 | 21,69 | 61 |
| ENSG00000124570 | SERPINB6 | 14,68 | 21,29 | 22,23 | 23,63 | 61 |
| ENSG00000177728 | TMEM94 | 16,83 | 21,11 | 25,38 | 27,09 | 61 |
| ENSG00000091436 | MAP3K20 | 7,72 | 7,89 | 9,14 | 12,42 | 61 |
| ENSG00000168646 | AXIN2 | 13,44 | 15,68 | 18,84 | 21,62 | 61 |
| ENSG00000123191 | ATP7B | 5,44 | 6,61 | 6,78 | 8,75 | 61 |
| ENSG00000151135 | TMEM263 | 10,92 | 14,5 | 14,59 | 17,56 | 61 |
| ENSG00000233024 | NPIPA9 | 11,29 | 13,8 | 17,11 | 18,15 | 61 |
| ENSG00000180957 | PITPNB | 26,75 | 31,46 | 31,91 | 43 | 61 |
| ENSG00000225234 | TRAPPC12-AS1 | 0,84 | 0,96 | 1,07 | 1,35 | 61 |
| ENSG00000196159 | FAT4 | 7,38 | 7,85 | 8,25 | 11,86 | 61 |
| ENSG00000189306 | RRP7A | 10,21 | 11,19 | 12,72 | 16,4 | 61 |
| ENSG00000179403 | VWA1 | 2,92 | 3,31 | 3,87 | 4,69 | 61 |
| ENSG00000118855 | MFSD1 | 5,15 | 5,67 | 7,2 | 8,27 | 61 |
| ENSG00000179115 | FARSA | 12,93 | 14,73 | 19,37 | 20,76 | 61 |
| ENSG00000146094 | DOK3 | 0,71 | 0,78 | 0,99 | 1,14 | 61 |
| ENSG00000153395 | LPCAT1 | 4,74 | 5,95 | 6,65 | 7,61 | 61 |
| ENSG00000117481 | NSUN4 | 5,22 | 5,79 | 6,52 | 8,38 | 61 |
| ENSG00000087191 | PSMC5 | 59,85 | 68,53 | 76,12 | 96,03 | 60 |
| ENSG00000113441 | LNPEP | 5,7 | 5,87 | 6,28 | 9,14 | 60 |
| ENSG00000155254 | MARVELD1 | 9,43 | 11,76 | 13,52 | 15,12 | 60 |
| ENSG00000104129 | DNAJC17 | 13,3 | 13,38 | 16,98 | 21,32 | 60 |
| ENSG00000198917 | SPOUT1 | 8,34 | 8,71 | 10,03 | 13,36 | 60 |
| ENSG00000105419 | MEIS3 | 18,36 | 19,95 | 21,47 | 29,39 | 60 |
| ENSG00000275079 | LPCAT1 | 24,07 | 27,77 | 35,62 | 38,53 | 60 |
| ENSG00000101236 | RNF24 | 7,75 | 8,38 | 9,07 | 12,4 | 60 |
| ENSG00000119042 | SATB2 | 3,35 | 3,83 | 4,38 | 5,36 | 60 |
| ENSG00000005700 | IBTK | 5,51 | 6,79 | 7,01 | 8,81 | 60 |
| ENSG00000283068 | IBTK | 5,51 | 6,79 | 7,01 | 8,81 | 60 |
| ENSG00000133835 | HSD17B4 | 53,64 | 55,55 | 55,95 | 85,76 | 60 |
| ENSG00000141873 | SLC39A3 | 7,4 | 9,16 | 11,38 | 11,83 | 60 |
| ENSG00000121350 | PYROXD1 | 4,16 | 5,78 | 5,99 | 6,65 | 60 |
| ENSG00000197043 | ANXA6 | 50,77 | 59,96 | 72,42 | 81,12 | 60 |
| ENSG00000055609 | KMT2C | 14,01 | 16,54 | 16,73 | 22,38 | 60 |
| ENSG00000196937 | FAM3C | 13,12 | 14,53 | 15,96 | 20,94 | 60 |
| ENSG00000141524 | TMC6 | 5,99 | 7,32 | 7,79 | 9,56 | 60 |
| ENSG00000083290 | ULK2 | 6,01 | 8,45 | 9,53 | 9,59 | 60 |
| ENSG00000101928 | MOSPD1 | 5,06 | 5,18 | 5,97 | 8,07 | 59 |
| ENSG00000147027 | TMEM47 | 8,71 | 9,58 | 9,66 | 13,89 | 59 |
| ENSG00000107937 | GTPBP4 | 19,51 | 23,02 | 23,89 | 31,1 | 59 |
| ENSG00000114779 | ABHD14B | 1,33 | 1,52 | 1,75 | 2,12 | 59 |
| ENSG00000080608 | PUM3 | 19,39 | 19,49 | 22,02 | 30,9 | 59 |
| ENSG00000003249 | DBNDD1 | 3,05 | 4,34 | 4,61 | 4,86 | 59 |
| ENSG00000108829 | LRRC59 | 19,56 | 21,76 | 22,72 | 31,16 | 59 |
| ENSG00000276005 | AC138749,8 | 1,72 | 1,79 | 1,95 | 2,74 | 59 |
| ENSG00000133612 | AGAP3 | 15,46 | 18,22 | 20,59 | 24,62 | 59 |
| ENSG00000253079 | NRON | 0,76 | 0,97 | 0,99 | 1,21 | 59 |
| ENSG00000250462 | LRRC37BP1 | 7,09 | 9,79 | 10,58 | 11,28 | 59 |
| ENSG00000145220 | LYAR | 8,2 | 8,97 | 11,6 | 13,04 | 59 |
| ENSG00000111652 | COPS7A | 9,26 | 10,39 | 12,44 | 14,72 | 59 |

| | | | | | | |
|---|---|---|---|---|---|---|
| ENSG00000250616 | AC012645,1 | 1,29 | 1,37 | 1,5 | 2,05 | 59 |
| ENSG00000230797 | YY2 | 0,73 | 0,84 | 0,96 | 1,16 | 59 |
| ENSG00000004866 | ST7 | 8,88 | 10,2 | 11,25 | 14,11 | 59 |
| ENSG00000160818 | GPATCH4 | 10,04 | 10,47 | 10,56 | 15,95 | 59 |
| ENSG00000121005 | CRISPLD1 | 31,09 | 31,72 | 41,77 | 49,37 | 59 |
| ENSG00000111647 | UHRF1BP1L | 7,45 | 9,24 | 9,76 | 11,83 | 59 |
| ENSG00000131873 | CHSY1 | 16,84 | 23,99 | 25,44 | 26,74 | 59 |
| ENSG00000162073 | PAQR4 | 9,9 | 12,01 | 14,1 | 15,72 | 59 |
| ENSG00000114030 | KPNA1 | 23,6 | 27,42 | 28,85 | 37,47 | 59 |
| ENSG00000106244 | PDAP1 | 36,23 | 48,79 | 51,73 | 57,52 | 59 |
| ENSG00000136444 | RSAD1 | 7,15 | 9,3 | 9,51 | 11,35 | 59 |
| ENSG00000204410 | MSH5 | 4,89 | 5,34 | 7,43 | 7,76 | 59 |
| ENSG00000231113 | AL035587,1 | 1,45 | 1,54 | 1,9 | 2,3 | 59 |
| ENSG00000137841 | PLCB2 | 2,24 | 2,25 | 3,3 | 3,55 | 58 |
| ENSG00000143845 | ETNK2 | 2,02 | 2,2 | 2,39 | 3,2 | 58 |
| ENSG00000154240 | CEP112 | 9,97 | 11,52 | 13,86 | 15,79 | 58 |
| ENSG00000198894 | CIPC | 9,32 | 11,03 | 12,63 | 14,76 | 58 |
| ENSG00000258410 | AC087386,1 | 2,33 | 2,6 | 2,92 | 3,69 | 58 |
| ENSG00000157654 | PALM2AKAP2 | 31,44 | 38,06 | 40,59 | 49,78 | 58 |
| ENSG00000174446 | SNAPC5 | 7,75 | 9,16 | 12,1 | 12,27 | 58 |
| ENSG00000100266 | PACSIN2 | 13,26 | 15,12 | 16,16 | 20,99 | 58 |
| ENSG00000100029 | PES1 | 19,18 | 20,76 | 21,3 | 30,36 | 58 |
| ENSG00000163075 | CFAP221 | 3,74 | 4,14 | 4,77 | 5,92 | 58 |
| ENSG00000140931 | CMTM3 | 22,41 | 24,19 | 24,31 | 35,45 | 58 |
| ENSG00000137693 | YAP1 | 39,59 | 46,25 | 48,93 | 62,6 | 58 |
| ENSG00000073111 | MCM2 | 52,39 | 55,06 | 63,41 | 82,82 | 58 |
| ENSG00000154832 | CXXC1 | 9,28 | 9,34 | 10,9 | 14,67 | 58 |
| ENSG00000072134 | EPN2 | 14,91 | 16,05 | 21,86 | 23,56 | 58 |
| ENSG00000196810 | CTBP1-DT | 3,78 | 4,29 | 4,77 | 5,97 | 58 |
| ENSG00000143870 | PDIA6 | 101,54 | 111,27 | 122,74 | 160,22 | 58 |
| ENSG00000180104 | EXOC3 | 10,06 | 11,68 | 13,67 | 15,87 | 58 |
| ENSG00000169047 | IRS1 | 6,1 | 7,34 | 7,48 | 9,62 | 58 |
| ENSG00000116473 | RAP1A | 10,45 | 13,1 | 14,23 | 16,48 | 58 |
| ENSG00000130479 | MAP1S | 12,41 | 12,61 | 18 | 19,57 | 58 |
| ENSG00000122884 | P4HA1 | 11,92 | 14,36 | 15,11 | 18,79 | 58 |
| ENSG00000179218 | CALR | 213,39 | 250,84 | 267,81 | 336,34 | 58 |
| ENSG00000095637 | SORBS1 | 7,09 | 10,04 | 10,84 | 11,17 | 58 |
| ENSG00000179010 | MRFAP1 | 88,27 | 108,21 | 117,66 | 138,96 | 57 |
| ENSG00000184743 | ATL3 | 4,88 | 5,7 | 6,71 | 7,68 | 57 |
| ENSG00000176619 | LMNB2 | 17,96 | 20,2 | 20,62 | 28,26 | 57 |
| ENSG00000197694 | SPTAN1 | 39,76 | 51,65 | 52,62 | 62,55 | 57 |
| ENSG00000283341 | AC068205,2 | 1,03 | 1,04 | 1,17 | 1,62 | 57 |
| ENSG00000111860 | CEP85L | 3,79 | 4,31 | 4,4 | 5,96 | 57 |
| ENSG00000275779 | NTAN1 | 11,58 | 14,23 | 16,93 | 18,21 | 57 |
| ENSG00000113761 | ZNF346 | 10,78 | 12,02 | 13,13 | 16,95 | 57 |
| ENSG00000138071 | ACTR2 | 64,43 | 77,27 | 83,57 | 101,3 | 57 |
| ENSG00000242852 | ZNF709 | 3,62 | 4,53 | 4,82 | 5,69 | 57 |
| ENSG00000110906 | KCTD10 | 28,26 | 35,89 | 41,57 | 44,41 | 57 |
| ENSG00000162373 | BEND5 | 5,92 | 6,1 | 6,79 | 9,3 | 57 |
| ENSG00000028310 | BRD9 | 15,03 | 16,93 | 18,04 | 23,61 | 57 |

| | | | | | | |
|---|---|---|---|---|---|---|
| ENSG00000071127 | WDR1 | 59,65 | 64,68 | 82,28 | 93,69 | 57 |
| ENSG00000178605 | GTPBP6 | 12,22 | 13,52 | 15,9 | 19,19 | 57 |
| ENSG00000165244 | ZNF367 | 4,39 | 4,66 | 5,63 | 6,89 | 57 |
| ENSG00000164465 | DCBLD1 | 3,46 | 4,01 | 4,31 | 5,43 | 57 |
| ENSG00000156860 | FBRS | 11,19 | 12,64 | 13,62 | 17,56 | 57 |
| ENSG00000108946 | PRKAR1A | 59,83 | 78,02 | 90,35 | 93,87 | 57 |
| ENSG00000099219 | ERMP1 | 4,08 | 4,51 | 4,65 | 6,4 | 57 |
| ENSG00000072062 | PRKACA | 13,95 | 16,1 | 16,4 | 21,88 | 57 |
| ENSG00000112658 | SRF | 9,38 | 10,47 | 11,8 | 14,71 | 57 |
| ENSG00000163297 | ANTXR2 | 18,55 | 21,81 | 24,8 | 29,09 | 57 |
| ENSG00000177045 | SIX5 | 2,77 | 3,36 | 4,11 | 4,34 | 57 |
| ENSG00000270147 | AC068620,2 | 0,76 | 0,84 | 0,95 | 1,19 | 57 |
| ENSG00000177732 | SOX12 | 14,59 | 16,48 | 17,19 | 22,84 | 57 |
| ENSG00000152133 | GPATCH11 | 3,47 | 4,67 | 4,8 | 5,43 | 56 |
| ENSG00000285540 | AC022023,2 | 1,1 | 1,12 | 1,13 | 1,72 | 56 |
| ENSG00000275055 | AC011468,5 | 1,35 | 1,37 | 1,76 | 2,11 | 56 |
| ENSG00000274349 | ZNF658 | 3,34 | 4,11 | 4,22 | 5,22 | 56 |
| ENSG00000165757 | JCAD | 0,96 | 1,43 | 1,44 | 1,5 | 56 |
| ENSG00000067208 | EVI5 | 8,17 | 10,72 | 12,59 | 12,76 | 56 |
| ENSG00000111726 | CMAS | 17,48 | 20,09 | 20,64 | 27,29 | 56 |
| ENSG00000079313 | REXO1 | 5,56 | 6,16 | 7,18 | 8,67 | 56 |
| ENSG00000167962 | ZNF598 | 7,08 | 7,85 | 8,77 | 11,04 | 56 |
| ENSG00000140718 | FTO | 46,2 | 53,29 | 58,34 | 72,03 | 56 |
| ENSG00000213839 | TMX2P1 | 5,39 | 6,58 | 6,9 | 8,4 | 56 |
| ENSG00000245711 | NADK2-AS1 | 0,77 | 0,92 | 0,93 | 1,2 | 56 |
| ENSG00000163866 | SMIM12 | 11,22 | 13,53 | 16,29 | 17,47 | 56 |
| ENSG00000152778 | IFIT5 | 3,16 | 3,93 | 4,19 | 4,92 | 56 |
| ENSG00000196730 | DAPK1 | 15,7 | 15,93 | 16,98 | 24,44 | 56 |
| ENSG00000136933 | RABEPK | 6,83 | 6,89 | 7,93 | 10,63 | 56 |
| ENSG00000276386 | CNTNAP3P2 | 2,25 | 2,36 | 2,38 | 3,5 | 56 |
| ENSG00000108963 | DPH1 | 6,86 | 6,94 | 8,21 | 10,67 | 56 |
| ENSG00000106608 | URGCP | 4,65 | 5,88 | 6,79 | 7,23 | 55 |
| ENSG00000259699 | HMGB1P8 | 1,93 | 2,33 | 2,7 | 3 | 55 |
| ENSG00000125912 | NCLN | 8,27 | 8,49 | 10,68 | 12,85 | 55 |
| ENSG00000174705 | SH3PXD2B | 20,19 | 23,13 | 23,98 | 31,36 | 55 |
| ENSG00000150455 | TIRAP | 3,76 | 3,93 | 4,34 | 5,84 | 55 |
| ENSG00000164889 | SLC4A2 | 17,02 | 20,61 | 25,87 | 26,43 | 55 |
| ENSG00000176533 | GNG7 | 2,08 | 2,18 | 2,44 | 3,23 | 55 |
| ENSG00000077157 | PPP1R12B | 6,88 | 7,91 | 8,26 | 10,68 | 55 |
| ENSG00000130881 | LRP3 | 8,27 | 8,92 | 9,25 | 12,83 | 55 |
| ENSG00000286305 | AC004893,2 | 0,78 | 0,92 | 0,94 | 1,21 | 55 |
| ENSG00000154309 | DISP1 | 7,95 | 8,37 | 10,08 | 12,33 | 55 |
| ENSG00000270757 | HSPE1-MOB4 | 4,63 | 5,27 | 5,46 | 7,18 | 55 |
| ENSG00000181192 | DHTKD1 | 11,07 | 12,48 | 14,16 | 17,16 | 55 |
| ENSG00000170832 | USP32 | 9,29 | 11,11 | 11,25 | 14,4 | 55 |
| ENSG00000164742 | ADCY1 | 4,17 | 4,43 | 5,19 | 6,46 | 55 |
| ENSG00000138698 | RAP1GDS1 | 16,51 | 20,08 | 20,49 | 25,57 | 55 |
| ENSG00000172037 | LAMB2 | 20,67 | 26,26 | 31,36 | 32,01 | 55 |
| ENSG00000070423 | RNF126 | 11,46 | 12,72 | 14,71 | 17,74 | 55 |
| ENSG00000137076 | TLN1 | 19,67 | 22,59 | 26,46 | 30,43 | 55 |

| | | | | | | |
|---|---|---|---|---|---|---|
| ENSG00000188976 | NOC2L | 18,71 | 21,26 | 22,65 | 28,94 | 55 |
| ENSG00000166261 | ZNF202 | 7,12 | 7,9 | 9,19 | 11,01 | 55 |
| ENSG00000213563 | C8orf82 | 4,54 | 5,15 | 5,33 | 7,02 | 55 |
| ENSG00000054118 | THRAP3 | 53,19 | 60,23 | 64 | 82,24 | 55 |
| ENSG00000254635 | WAC-AS1 | 7,2 | 8,59 | 10,09 | 11,13 | 55 |
| ENSG00000266967 | AARSD1 | 15,78 | 16,97 | 19,1 | 24,39 | 55 |
| ENSG00000197329 | PELI1 | 15,14 | 18,38 | 21,48 | 23,39 | 54 |
| ENSG00000162129 | CLPB | 7,38 | 8,65 | 9,96 | 11,4 | 54 |
| ENSG00000196741 | LINC01560 | 1,12 | 1,54 | 1,55 | 1,73 | 54 |
| ENSG00000149091 | DGKZ | 11,11 | 12,28 | 15,96 | 17,16 | 54 |
| ENSG00000197496 | SLC2A10 | 2,83 | 3,39 | 3,49 | 4,37 | 54 |
| ENSG00000213390 | ARHGAP19 | 10,87 | 11,86 | 12,69 | 16,78 | 54 |
| ENSG00000171067 | C11orf24 | 10,44 | 10,54 | 12,1 | 16,1 | 54 |
| ENSG00000144848 | ATG3 | 14,74 | 17,52 | 19,94 | 22,73 | 54 |
| ENSG00000161149 | TUBA3FP | 1,79 | 2,35 | 2,63 | 2,76 | 54 |
| ENSG00000035928 | RFC1 | 26,97 | 30,72 | 32,13 | 41,52 | 54 |
| ENSG00000164086 | DUSP7 | 3,84 | 4,41 | 4,42 | 5,91 | 54 |
| ENSG00000215305 | VPS16 | 7,31 | 9,03 | 10 | 11,25 | 54 |
| ENSG00000266933 | AC005775,1 | 1,04 | 1,08 | 1,17 | 1,6 | 54 |
| ENSG00000166831 | RBPMS2 | 14,64 | 14,96 | 17,53 | 22,52 | 54 |
| ENSG00000173020 | GRK2 | 9,65 | 11,98 | 13,48 | 14,84 | 54 |
| ENSG00000129422 | MTUS1 | 7,85 | 9,09 | 10,27 | 12,07 | 54 |
| ENSG00000175029 | CTBP2 | 33,01 | 35,6 | 42,91 | 50,74 | 54 |
| ENSG00000140575 | IQGAP1 | 18,09 | 22,13 | 25,2 | 27,8 | 54 |
| ENSG00000277283 | AC004812,2 | 1,25 | 1,42 | 1,73 | 1,92 | 54 |
| ENSG00000204859 | ZBTB48 | 5,3 | 7,69 | 7,77 | 8,14 | 54 |
| ENSG00000109775 | UFSP2 | 18,24 | 22,98 | 23,52 | 28,01 | 54 |
| ENSG00000232940 | HCG25 | 2,26 | 2,54 | 2,62 | 3,47 | 54 |
| ENSG00000005889 | ZFX | 16,97 | 20,85 | 22,02 | 26,05 | 54 |
| ENSG00000276141 | WHAMMP3 | 1,14 | 1,5 | 1,73 | 1,75 | 54 |
| ENSG00000156983 | BRPF1 | 6,47 | 7,55 | 9,12 | 9,93 | 53 |
| ENSG00000171443 | ZNF524 | 1,31 | 1,39 | 1,54 | 2,01 | 53 |
| ENSG00000171241 | SHCBP1 | 8,46 | 9,04 | 9,64 | 12,98 | 53 |
| ENSG00000254377 | MIR124-2HG | 22,46 | 27,97 | 31,36 | 34,45 | 53 |
| ENSG00000140950 | MEAK7 | 10,06 | 10,97 | 11,12 | 15,43 | 53 |
| ENSG00000197816 | CCDC180 | 2,68 | 2,77 | 3,23 | 4,11 | 53 |
| ENSG00000261971 | MMP25-AS1 | 1,69 | 1,7 | 1,88 | 2,59 | 53 |
| ENSG00000166598 | HSP90B1 | 211,88 | 256,61 | 261,06 | 324,67 | 53 |
| ENSG00000114867 | EIF4G1 | 75,13 | 85,44 | 90,56 | 115,12 | 53 |
| ENSG00000143786 | CNIH3 | 1,09 | 1,16 | 1,53 | 1,67 | 53 |
| ENSG00000169018 | FEM1B | 14,25 | 16,27 | 17,92 | 21,83 | 53 |
| ENSG00000087274 | ADD1 | 61,03 | 71,76 | 83,01 | 93,48 | 53 |
| ENSG00000158292 | GPR153 | 5,72 | 6,02 | 6,92 | 8,76 | 53 |
| ENSG00000105497 | ZNF175 | 3,35 | 3,93 | 4,03 | 5,13 | 53 |
| ENSG00000135698 | MPHOSPH6 | 25,98 | 30,8 | 33,01 | 39,77 | 53 |
| ENSG00000134001 | EIF2S1 | 49,71 | 54,93 | 67,85 | 76,05 | 53 |
| ENSG00000221823 | PPP3R1 | 21,99 | 24,9 | 27,47 | 33,64 | 53 |
| ENSG00000144711 | IQSEC1 | 6,91 | 7,69 | 8,97 | 10,57 | 53 |
| ENSG00000121073 | SLC35B1 | 28,18 | 29,92 | 34,27 | 43,1 | 53 |
| ENSG00000269604 | AC005523,2 | 5,12 | 5,35 | 6,86 | 7,83 | 53 |

| | | | | | | |
|---|---|---|---|---|---|---|
| ENSG00000105486 | LIG1 | 27,37 | 31,22 | 32,52 | 41,84 | 53 |
| ENSG00000186001 | LRCH3 | 21,38 | 23,62 | 28,58 | 32,68 | 53 |
| ENSG00000127948 | POR | 15,4 | 16,87 | 19,41 | 23,53 | 53 |
| ENSG00000106992 | AK1 | 10,88 | 13,5 | 16,02 | 16,62 | 53 |
| ENSG00000141431 | ASXL3 | 10,83 | 13,87 | 15,71 | 16,53 | 53 |
| ENSG00000127334 | DYRK2 | 10,42 | 11,99 | 12,62 | 15,9 | 53 |
| ENSG00000196576 | PLXNB2 | 17,26 | 22,5 | 24,87 | 26,33 | 53 |
| ENSG00000149231 | CCDC82 | 15,62 | 17,01 | 19,39 | 23,81 | 52 |
| ENSG00000109929 | SC5D | 29,37 | 31,35 | 37,74 | 44,75 | 52 |
| ENSG00000001167 | NFYA | 13,36 | 15,94 | 16,91 | 20,35 | 52 |
| ENSG00000156709 | AIFM1 | 12,29 | 12,98 | 13,81 | 18,72 | 52 |
| ENSG00000148337 | CIZ1 | 35,23 | 38,31 | 47,19 | 53,65 | 52 |
| ENSG00000105656 | ELL | 1,53 | 1,84 | 2,16 | 2,33 | 52 |
| ENSG00000169925 | BRD3 | 15,59 | 18,77 | 19,05 | 23,74 | 52 |
| ENSG00000055147 | FAM114A2 | 7,71 | 9,92 | 11,62 | 11,74 | 52 |
| ENSG00000148339 | SLC25A25 | 4,71 | 4,88 | 6,06 | 7,17 | 52 |
| ENSG00000138758 | SEPTIN11 | 74,43 | 76,51 | 97,85 | 113,18 | 52 |
| ENSG00000115526 | CHST10 | 9,79 | 11,99 | 13,48 | 14,88 | 52 |
| ENSG00000172977 | KAT5 | 16,01 | 16,61 | 21,58 | 24,33 | 52 |
| ENSG00000184110 | EIF3C | 147,7 | 178,4 | 184,08 | 224,42 | 52 |
| ENSG00000101935 | AMMECR1 | 5,93 | 7,11 | 7,49 | 9,01 | 52 |
| ENSG00000113013 | HSPA9 | 62,75 | 68,03 | 80,65 | 95,33 | 52 |
| ENSG00000002834 | LASP1 | 33,6 | 42,99 | 46,02 | 51,04 | 52 |
| ENSG00000129128 | SPCS3 | 28,18 | 30,79 | 33,85 | 42,8 | 52 |
| ENSG00000162298 | SYVN1 | 6,65 | 7,37 | 9,14 | 10,1 | 52 |
| ENSG00000127990 | SGCE | 23,51 | 29,74 | 29,9 | 35,7 | 52 |
| ENSG00000107566 | ERLIN1 | 9,17 | 10,65 | 10,96 | 13,92 | 52 |
| ENSG00000175395 | ZNF25 | 3,67 | 4,21 | 4,57 | 5,57 | 52 |
| ENSG00000156802 | ATAD2 | 21,33 | 23,18 | 23,4 | 32,37 | 52 |
| ENSG00000109956 | B3GAT1 | 14,22 | 16 | 16,03 | 21,58 | 52 |
| ENSG00000162434 | JAK1 | 12,56 | 16,94 | 18,43 | 19,06 | 52 |
| ENSG00000087365 | SF3B2 | 60,71 | 74,87 | 77,25 | 92,12 | 52 |
| ENSG00000068489 | PRR11 | 23,16 | 24,47 | 27,35 | 35,14 | 52 |
| ENSG00000142208 | AKT1 | 39,61 | 42,55 | 48,56 | 60,09 | 52 |
| ENSG00000262473 | GART | 1,22 | 1,3 | 1,35 | 1,85 | 52 |
| ENSG00000123607 | TTC21B | 16,06 | 19 | 20,74 | 24,35 | 52 |
| ENSG00000155660 | PDIA4 | 68,56 | 76,66 | 92,28 | 103,94 | 52 |
| ENSG00000103495 | MAZ | 77,6 | 84,71 | 98,9 | 117,64 | 52 |
| ENSG00000163867 | ZMYM6 | 5,04 | 5,61 | 6,49 | 7,64 | 52 |
| ENSG00000144406 | UNC80 | 4,81 | 6,01 | 6,28 | 7,29 | 52 |
| ENSG00000168724 | DNAJC21 | 14,38 | 18,76 | 19,62 | 21,78 | 51 |
| ENSG00000165416 | SUGT1 | 26,06 | 29,71 | 34,52 | 39,47 | 51 |
| ENSG00000276071 | AC074138,1 | 1,03 | 1,26 | 1,36 | 1,56 | 51 |
| ENSG00000113758 | DBN1 | 60,84 | 74,59 | 79,37 | 92,09 | 51 |
| ENSG00000151611 | MMAA | 2,59 | 3,29 | 3,41 | 3,92 | 51 |
| ENSG00000189403 | HMGB1 | 436,37 | 496,25 | 572,27 | 660,44 | 51 |
| ENSG00000143549 | TPM3 | 133,35 | 151,01 | 190,34 | 201,82 | 51 |
| ENSG00000115756 | HPCAL1 | 4,19 | 4,95 | 5,92 | 6,34 | 51 |
| ENSG00000197037 | ZSCAN25 | 4,58 | 5,39 | 5,46 | 6,93 | 51 |
| ENSG00000173692 | PSMD1 | 44,92 | 51,28 | 54,01 | 67,96 | 51 |

| | | | | | | |
|---|---|---|---|---|---|---|
| ENSG00000276045 | ORAI1 | 2,71 | 2,85 | 2,92 | 4,1 | 51 |
| ENSG00000263528 | IKBKE | 1,6 | 1,77 | 1,79 | 2,42 | 51 |
| ENSG00000205682 | AC020741,1 | 0,82 | 0,91 | 1,06 | 1,24 | 51 |
| ENSG00000135945 | REV1 | 20,02 | 23,11 | 25,13 | 30,26 | 51 |
| ENSG00000177946 | CENPBD1 | 2,29 | 3,19 | 3,23 | 3,46 | 51 |
| ENSG00000124357 | NAGK | 13,57 | 17,14 | 17,77 | 20,5 | 51 |
| ENSG00000085872 | CHERP | 16,78 | 21,02 | 23,71 | 25,33 | 51 |
| ENSG00000105221 | AKT2 | 30,82 | 32,09 | 33,96 | 46,48 | 51 |
| ENSG00000197724 | PHF2 | 9,08 | 11,76 | 12,01 | 13,69 | 51 |
| ENSG00000163932 | PRKCD | 1,46 | 1,47 | 1,62 | 2,2 | 51 |
| ENSG00000107798 | LIPA | 11,89 | 13,39 | 13,89 | 17,91 | 51 |
| ENSG00000254510 | AP001107,5 | 1,68 | 1,76 | 1,77 | 2,53 | 51 |
| ENSG00000166734 | GOLM2 | 22,59 | 25,69 | 33,84 | 34 | 51 |
| ENSG00000065457 | ADAT1 | 23,5 | 27,88 | 28,66 | 35,36 | 50 |
| ENSG00000075336 | TIMM21 | 16,13 | 16,83 | 22,82 | 24,26 | 50 |
| ENSG00000005022 | SLC25A5 | 69,53 | 71,21 | 81,4 | 104,57 | 50 |
| ENSG00000071575 | TRIB2 | 33,43 | 41,75 | 44,62 | 50,27 | 50 |
| ENSG00000130244 | FAM98C | 3,26 | 3,63 | 3,64 | 4,9 | 50 |
| ENSG00000166548 | TK2 | 5,8 | 6,89 | 7,55 | 8,71 | 50 |
| ENSG00000090889 | KIF4A | 21,33 | 22,79 | 25,11 | 32,03 | 50 |
| ENSG00000169371 | SNUPN | 11,27 | 12,31 | 13,49 | 16,92 | 50 |
| ENSG00000100211 | CBY1 | 16,98 | 21,58 | 23,2 | 25,49 | 50 |
| ENSG00000121691 | CAT | 13,13 | 16,26 | 16,46 | 19,7 | 50 |
| ENSG00000108021 | TASOR2 | 18,75 | 21,21 | 21,94 | 28,13 | 50 |
| ENSG00000143878 | RHOB | 23,95 | 25,7 | 27,32 | 35,93 | 50 |
| ENSG00000197170 | PSMD12 | 28,09 | 30,41 | 32,29 | 42,14 | 50 |
| ENSG00000280529 | TAF4 | 4,16 | 4,66 | 5,48 | 6,24 | 50 |
| ENSG00000106012 | IQCE | 3,6 | 3,96 | 4,05 | 5,4 | 50 |
| ENSG00000178722 | C5orf64 | 1 | 1,05 | 1,06 | 1,5 | 50 |
| ENSG00000087087 | SRRT | 35,66 | 37,53 | 39,31 | 53,46 | 50 |
| ENSG00000197566 | ZNF624 | 3,97 | 4,85 | 4,87 | 5,95 | 50 |
| ENSG00000109066 | TMEM104 | 3,83 | 4,56 | 5,23 | 5,74 | 50 |
| ENSG00000122545 | SEPTIN7 | 110,34 | 126,73 | 143,7 | 165,32 | 50 |
| ENSG00000215067 | ALOX12-AS1 | 2,91 | 2,95 | 3,57 | 4,36 | 50 |
| ENSG00000166452 | AKIP1 | 12,45 | 13,65 | 14,53 | 18,65 | 50 |
| ENSG00000204217 | BMPR2 | 8,18 | 11,68 | 11,71 | 12,25 | 50 |
| ENSG00000029725 | RABEP1 | 22,9 | 26,5 | 29,26 | 34,29 | 50 |
| ENSG00000186767 | SPIN4 | 5,68 | 6,71 | 8,24 | 8,5 | 50 |
| ENSG00000120029 | ARMH3 | 6,55 | 7,89 | 8,07 | 9,8 | 50 |
| ENSG00000166855 | CLPX | 16,53 | 17,33 | 19,13 | 24,73 | 50 |
| ENSG00000165795 | NDRG2 | 3,71 | 4,29 | 5,37 | 5,55 | 50 |
| ENSG00000116990 | MYCL | 7,04 | 8,77 | 9,24 | 10,53 | 50 |
| ENSG00000166225 | FRS2 | 9,04 | 11,29 | 12,29 | 13,52 | 50 |
| ENSG00000144320 | LNPK | 12,72 | 13,32 | 16,76 | 19,02 | 50 |
| ENSG00000267655 | AC125437,1 | 0,97 | 1,22 | 1,35 | 1,45 | 49 |
| ENSG00000075131 | TIPIN | 13,04 | 13,67 | 15,25 | 19,49 | 49 |
| ENSG00000167987 | VPS37C | 6,45 | 6,86 | 7,71 | 9,64 | 49 |
| ENSG00000135365 | PHF21A | 15,61 | 20,68 | 21,81 | 23,33 | 49 |
| ENSG00000230280 | HNRNPA1P59 | 0,89 | 1,01 | 1,21 | 1,33 | 49 |
| ENSG00000143751 | SDE2 | 4,88 | 6,33 | 6,7 | 7,29 | 49 |

| | | | | | | |
|---|---|---|---|---|---|---|
| ENSG00000186318 | BACE1 | 8,85 | 11,6 | 11,8 | 13,22 | 49 |
| ENSG00000171790 | SLFNL1 | 0,75 | 0,78 | 0,8 | 1,12 | 49 |
| ENSG00000213614 | HEXA | 17,56 | 22,32 | 23,56 | 26,22 | 49 |
| ENSG00000104154 | SLC30A4 | 0,69 | 0,93 | 0,94 | 1,03 | 49 |
| ENSG00000249992 | TMEM158 | 1,34 | 1,57 | 1,78 | 2 | 49 |
| ENSG00000211460 | TSN | 33,61 | 37,28 | 46,83 | 50,16 | 49 |
| ENSG00000074755 | ZZEF1 | 9,27 | 9,9 | 11,83 | 13,83 | 49 |
| ENSG00000109452 | INPP4B | 3,48 | 4,06 | 5,16 | 5,19 | 49 |
| ENSG00000239569 | KMT2E-AS1 | 1,16 | 1,43 | 1,48 | 1,73 | 49 |
| ENSG00000242802 | AP5Z1 | 3,36 | 3,69 | 4,71 | 5,01 | 49 |
| ENSG00000086475 | SEPHS1 | 45,44 | 49,93 | 53,38 | 67,72 | 49 |
| ENSG00000115145 | STAM2 | 7,78 | 8,82 | 10,48 | 11,59 | 49 |
| ENSG00000185189 | NRBP2 | 3,82 | 3,92 | 4,07 | 5,69 | 49 |
| ENSG00000130640 | TUBGCP2 | 15,35 | 15,98 | 21,16 | 22,86 | 49 |
| ENSG00000117523 | PRRC2C | 50,99 | 57,71 | 59,07 | 75,93 | 49 |
| ENSG00000151612 | ZNF827 | 11,97 | 15,47 | 15,78 | 17,82 | 49 |
| ENSG00000119392 | GLE1 | 7,94 | 8,83 | 9,85 | 11,82 | 49 |
| ENSG00000088812 | ATRN | 9,7 | 11,78 | 12,27 | 14,44 | 49 |
| ENSG00000156273 | BACH1 | 19,81 | 23,05 | 25,84 | 29,49 | 49 |
| ENSG00000131236 | CAP1 | 65,29 | 77,64 | 85,45 | 97,19 | 49 |
| ENSG00000157212 | PAXIP1 | 8,52 | 8,94 | 10,2 | 12,68 | 49 |
| ENSG00000212978 | AC016747,1 | 3,32 | 3,42 | 3,49 | 4,94 | 49 |
| ENSG00000123552 | USP45 | 4,8 | 4,98 | 5,48 | 7,14 | 49 |
| ENSG00000175866 | BAIAP2 | 6,71 | 7,8 | 8,32 | 9,98 | 49 |
| ENSG00000158352 | SHROOM4 | 2,34 | 2,9 | 2,96 | 3,48 | 49 |
| ENSG00000104812 | GYS1 | 7,72 | 9,8 | 10,57 | 11,48 | 49 |
| ENSG00000169957 | ZNF768 | 6,4 | 6,98 | 7,36 | 9,51 | 49 |
| ENSG00000111737 | RAB35 | 13,11 | 15,79 | 17,23 | 19,48 | 49 |
| ENSG00000134897 | BIVM | 9,06 | 9,58 | 10,07 | 13,46 | 49 |
| ENSG00000100711 | ZFYVE21 | 12,17 | 12,9 | 14,52 | 18,07 | 48 |
| ENSG00000125741 | OPA3 | 2,89 | 3,1 | 3,68 | 4,29 | 48 |
| ENSG00000166794 | PPIB | 195,17 | 217,39 | 257,43 | 289,71 | 48 |
| ENSG00000282034 | AC106886,5 | 2,23 | 2,32 | 2,42 | 3,31 | 48 |
| ENSG00000125826 | RBCK1 | 10,56 | 11,83 | 12,51 | 15,67 | 48 |
| ENSG00000160233 | LRRC3 | 2,42 | 2,83 | 3,03 | 3,59 | 48 |
| ENSG00000282825 | CTIF | 3,83 | 4,21 | 5,07 | 5,68 | 48 |
| ENSG00000148297 | MED22 | 5,53 | 6 | 6,47 | 8,2 | 48 |
| ENSG00000189114 | BLOC1S3 | 1,16 | 1,34 | 1,64 | 1,72 | 48 |
| ENSG00000269929 | MIRLET7A1HG | 4,6 | 4,97 | 5,39 | 6,82 | 48 |
| ENSG00000106080 | FKBP14 | 4,78 | 6,93 | 6,99 | 7,08 | 48 |
| ENSG00000087842 | PIR | 16,17 | 16,36 | 19,01 | 23,95 | 48 |
| ENSG00000288663 | AC073611,1 | 0,79 | 1,05 | 1,09 | 1,17 | 48 |
| ENSG00000113532 | ST8SIA4 | 8,05 | 9,9 | 11,26 | 11,92 | 48 |
| ENSG00000276410 | H2BC3 | 170,28 | 188,33 | 207,76 | 252,13 | 48 |
| ENSG00000178202 | POGLUT3 | 13,67 | 16,49 | 18,28 | 20,24 | 48 |
| ENSG00000225748 | PRRC2A | 19,8 | 22,55 | 23,58 | 29,3 | 48 |
| ENSG00000088038 | CNOT3 | 20,24 | 22,47 | 24,95 | 29,95 | 48 |
| ENSG00000141560 | FN3KRP | 11,01 | 12,5 | 12,96 | 16,29 | 48 |
| ENSG00000231889 | TRAF3IP2-AS1 | 8,01 | 9,31 | 9,85 | 11,85 | 48 |
| ENSG00000166716 | ZNF592 | 8,58 | 9,96 | 10,9 | 12,69 | 48 |

| | | | | | | |
|---|---|---|---|---|---|---|
| ENSG00000146373 | RNF217 | 3,53 | 3,82 | 4,13 | 5,22 | 48 |
| ENSG00000070010 | UFD1 | 29,83 | 31,93 | 39,06 | 44,11 | 48 |
| ENSG00000254132 | MTND6P3 | 21,33 | 21,89 | 23,84 | 31,54 | 48 |
| ENSG00000272410 | AC022384,1 | 2,55 | 2,69 | 2,92 | 3,77 | 48 |
| ENSG00000169826 | CSGALNACT2 | 5,61 | 6,66 | 6,79 | 8,29 | 48 |
| ENSG00000148719 | DNAJB12 | 8,3 | 9,16 | 10,09 | 12,26 | 48 |
| ENSG00000187630 | DHRS4L2 | 5,1 | 5,9 | 7,35 | 7,53 | 48 |
| ENSG00000169410 | PTPN9 | 13,16 | 16,45 | 17,4 | 19,43 | 48 |
| ENSG00000112149 | CD83 | 2,12 | 2,5 | 2,7 | 3,13 | 48 |
| ENSG00000135299 | ANKRD6 | 11,18 | 14,08 | 14,56 | 16,5 | 48 |
| ENSG00000173621 | LRFN4 | 4,75 | 6,02 | 6,59 | 7,01 | 48 |
| ENSG00000115091 | ACTR3 | 74,62 | 87,33 | 102,83 | 110,12 | 48 |
| ENSG00000168495 | POLR3D | 16,03 | 19,46 | 20,56 | 23,65 | 48 |
| ENSG00000078140 | UBE2K | 23,06 | 24,83 | 29,68 | 34,02 | 48 |
| ENSG00000183250 | LINC01547 | 1,41 | 1,69 | 2,03 | 2,08 | 48 |
| ENSG00000170606 | HSPA4 | 72,17 | 80,25 | 87,84 | 106,45 | 47 |
| ENSG00000156011 | PSD3 | 15,54 | 18,68 | 18,7 | 22,92 | 47 |
| ENSG00000284807 | DHRS4 | 2,59 | 3,31 | 3,59 | 3,82 | 47 |
| ENSG00000157326 | DHRS4 | 2,59 | 3,31 | 3,59 | 3,82 | 47 |
| ENSG00000157193 | LRP8 | 13,3 | 15,07 | 15,79 | 19,61 | 47 |
| ENSG00000151835 | SACS | 13,26 | 14,67 | 14,96 | 19,55 | 47 |
| ENSG00000143753 | DEGS1 | 13,06 | 14,25 | 16,23 | 19,25 | 47 |
| ENSG00000163513 | TGFBR2 | 3,57 | 5,05 | 5,19 | 5,26 | 47 |
| ENSG00000134684 | YARS1 | 33,65 | 34,84 | 44,78 | 49,57 | 47 |
| ENSG00000272886 | DCP1A | 9,26 | 11,15 | 11,94 | 13,64 | 47 |
| ENSG00000129292 | PHF20L1 | 18,88 | 23,37 | 25,06 | 27,81 | 47 |
| ENSG00000105993 | DNAJB6 | 56,75 | 67,53 | 68,83 | 83,58 | 47 |
| ENSG00000158615 | PPP1R15B | 11,36 | 12,7 | 13,4 | 16,73 | 47 |
| ENSG00000100764 | PSMC1 | 94,42 | 106,32 | 122,95 | 139,01 | 47 |
| ENSG00000182973 | CNOT10 | 18,43 | 21,11 | 21,93 | 27,13 | 47 |
| ENSG00000105879 | CBLL1 | 16,86 | 17,44 | 18,65 | 24,8 | 47 |
| ENSG00000149136 | SSRP1 | 93,55 | 107,3 | 113,17 | 137,58 | 47 |
| ENSG00000198862 | LTN1 | 8,01 | 8,88 | 8,89 | 11,78 | 47 |
| ENSG00000088882 | CPXM1 | 37,37 | 45,12 | 48,32 | 54,93 | 47 |
| ENSG00000169607 | CKAP2L | 10,29 | 11,28 | 12,14 | 15,12 | 47 |
| ENSG00000153179 | RASSF3 | 7,14 | 8,59 | 9,09 | 10,49 | 47 |
| ENSG00000164715 | LMTK2 | 5,59 | 6,73 | 6,74 | 8,21 | 47 |
| ENSG00000150893 | FREM2 | 6,18 | 6,51 | 7,02 | 9,07 | 47 |
| ENSG00000224945 | AL353150,1 | 4,17 | 4,34 | 5,77 | 6,12 | 47 |
| ENSG00000095319 | NUP188 | 22,52 | 22,61 | 24,09 | 33,05 | 47 |
| ENSG00000172175 | MALT1 | 9,86 | 10,76 | 11,6 | 14,47 | 47 |
| ENSG00000285796 | AL162458,1 | 1,35 | 1,55 | 1,56 | 1,98 | 47 |
| ENSG00000162419 | GMEB1 | 7,76 | 8,85 | 9,68 | 11,38 | 47 |
| ENSG00000031823 | RANBP3 | 32,7 | 34,57 | 41,14 | 47,94 | 47 |
| ENSG00000126522 | ASL | 5,71 | 6,05 | 6,67 | 8,37 | 47 |
| ENSG00000264247 | LINC00909 | 3,95 | 4,47 | 4,8 | 5,79 | 47 |
| ENSG00000005302 | MSL3 | 12,11 | 13,48 | 15,69 | 17,75 | 47 |
| ENSG00000115042 | FAHD2A | 15,19 | 17,07 | 19,11 | 22,26 | 47 |
| ENSG00000071794 | HLTF | 22,14 | 26 | 26,27 | 32,44 | 47 |
| ENSG00000171914 | TLN2 | 9,33 | 10,5 | 12,19 | 13,67 | 47 |

| | | | | | | |
|---|---|---|---|---|---|---|
| ENSG00000010810 | FYN | 82,46 | 99,85 | 116,41 | 120,81 | 47 |
| ENSG00000282254 | AC004980,5 | 1,79 | 2,33 | 2,44 | 2,62 | 46 |
| ENSG00000111328 | CDK2AP1 | 117,22 | 137,22 | 149,92 | 171,54 | 46 |
| ENSG00000055483 | USP36 | 8,18 | 9,6 | 9,66 | 11,97 | 46 |
| ENSG00000131116 | ZNF428 | 25,73 | 30,02 | 34,73 | 37,65 | 46 |
| ENSG00000162704 | ARPC5 | 70,42 | 79,29 | 85,77 | 103,03 | 46 |
| ENSG00000108788 | MLX | 11,49 | 11,81 | 13,8 | 16,81 | 46 |
| ENSG00000134313 | KIDINS220 | 21,61 | 30,11 | 30,61 | 31,61 | 46 |
| ENSG00000165661 | QSOX2 | 6,16 | 6,19 | 7,35 | 9,01 | 46 |
| ENSG00000125827 | TMX4 | 7,98 | 9,64 | 10,25 | 11,67 | 46 |
| ENSG00000011007 | ELOA | 15,23 | 17,74 | 19,28 | 22,27 | 46 |
| ENSG00000185100 | ADSS1 | 2,25 | 2,33 | 2,35 | 3,29 | 46 |
| ENSG00000161036 | LRWD1 | 9,96 | 11,2 | 12,18 | 14,56 | 46 |
| ENSG00000120327 | PCDHB14 | 2,62 | 2,98 | 3,45 | 3,83 | 46 |
| ENSG00000259781 | HMGB1P6 | 174,79 | 205,57 | 230,22 | 255,5 | 46 |
| ENSG00000022567 | SLC45A4 | 5,96 | 6,16 | 6,62 | 8,71 | 46 |
| ENSG00000112033 | PPARD | 5,83 | 6,79 | 7,33 | 8,52 | 46 |
| ENSG00000090372 | STRN4 | 28,31 | 37,47 | 39,59 | 41,37 | 46 |
| ENSG00000140367 | UBE2Q2 | 12,12 | 15,27 | 16,78 | 17,71 | 46 |
| ENSG00000260643 | AC092718,3 | 2,45 | 2,98 | 3,54 | 3,58 | 46 |
| ENSG00000178105 | DDX10 | 10,63 | 11,08 | 12,21 | 15,53 | 46 |
| ENSG00000123094 | RASSF8 | 15,04 | 17,82 | 19,81 | 21,97 | 46 |
| ENSG00000136485 | DCAF7 | 31,13 | 41,91 | 43,58 | 45,47 | 46 |
| ENSG00000157985 | AGAP1 | 14,7 | 15,92 | 16,93 | 21,47 | 46 |
| ENSG00000149823 | VPS51 | 16,45 | 16,56 | 17,99 | 24,02 | 46 |
| ENSG00000119684 | MLH3 | 9,83 | 10,9 | 11,05 | 14,35 | 46 |
| ENSG00000135392 | DNAJC14 | 8,57 | 10,09 | 10,49 | 12,51 | 46 |
| ENSG00000153046 | CDYL | 25,64 | 29,28 | 35,51 | 37,41 | 46 |
| ENSG00000265185 | SNORD3B-1 | 18,04 | 20,99 | 23,41 | 26,32 | 46 |
| ENSG00000063587 | ZNF275 | 4,38 | 5,02 | 5,15 | 6,39 | 46 |
| ENSG00000163781 | TOPBP1 | 21,14 | 23,67 | 23,9 | 30,84 | 46 |
| ENSG00000138604 | GLCE | 3,38 | 4,33 | 4,68 | 4,93 | 46 |
| ENSG00000245534 | RORA-AS1 | 0,72 | 0,76 | 0,91 | 1,05 | 46 |
| ENSG00000067992 | PDK3 | 5,48 | 5,78 | 6,12 | 7,99 | 46 |
| ENSG00000134996 | OSTF1 | 2,73 | 3 | 3,86 | 3,98 | 46 |
| ENSG00000215301 | DDX3X | 87,97 | 95,5 | 99,82 | 128,24 | 46 |
| ENSG00000131779 | PEX11B | 5,31 | 6,9 | 7,14 | 7,74 | 46 |
| ENSG00000164162 | ANAPC10 | 10,85 | 10,89 | 12,14 | 15,81 | 46 |
| ENSG00000131931 | THAP1 | 4,16 | 5,07 | 5,31 | 6,06 | 46 |
| ENSG00000164163 | ABCE1 | 32,85 | 35,17 | 38,48 | 47,84 | 46 |
| ENSG00000072518 | MARK2 | 12,78 | 15,29 | 17,1 | 18,61 | 46 |
| ENSG00000151240 | DIP2C | 6,6 | 7,38 | 7,63 | 9,61 | 46 |
| ENSG00000140943 | MBTPS1 | 21,03 | 25,14 | 27,68 | 30,62 | 46 |
| ENSG00000183751 | TBL3 | 7,61 | 8,45 | 9,05 | 11,08 | 46 |
| ENSG00000105887 | MTPN | 38,91 | 42,71 | 47,79 | 56,65 | 46 |
| ENSG00000090863 | GLG1 | 37,94 | 46,08 | 47,89 | 55,23 | 46 |
| ENSG00000124160 | NCOA5 | 20,56 | 22,43 | 26,44 | 29,92 | 46 |
| ENSG00000235173 | HGH1 | 4,68 | 4,76 | 6,28 | 6,81 | 46 |
| ENSG00000184887 | BTBD6 | 10,36 | 12,25 | 13,77 | 15,07 | 45 |
| ENSG00000131100 | ATP6V1E1 | 19,36 | 20,73 | 26,86 | 28,16 | 45 |

| | | | | | | |
|---|---|---|---|---|---|---|
| ENSG00000121957 | GPSM2 | 14,81 | 14,91 | 15,42 | 21,54 | 45 |
| ENSG00000102317 | RBM3 | 109,36 | 114,03 | 132,82 | 159,02 | 45 |
| ENSG00000132434 | LANCL2 | 7,67 | 9,18 | 10,92 | 11,15 | 45 |
| ENSG00000088833 | NSFL1C | 38,88 | 43,19 | 45,65 | 56,52 | 45 |
| ENSG00000104490 | NCALD | 36,62 | 38,83 | 42,73 | 53,23 | 45 |
| ENSG00000075234 | TTC38 | 6,55 | 8,14 | 9,1 | 9,52 | 45 |
| ENSG00000133056 | PIK3C2B | 12,11 | 13,77 | 14,32 | 17,6 | 45 |
| ENSG00000198818 | SFT2D1 | 10,13 | 11,71 | 13,35 | 14,72 | 45 |
| ENSG00000071889 | FAM3A | 6,81 | 7,51 | 9,01 | 9,89 | 45 |
| ENSG00000129480 | DTD2 | 8,23 | 9,68 | 10,58 | 11,95 | 45 |
| ENSG00000283777 | CANX | 142,23 | 160,44 | 166,41 | 206,41 | 45 |
| ENSG00000065665 | SEC61A2 | 12,59 | 13 | 15,19 | 18,27 | 45 |
| ENSG00000110917 | MLEC | 37,24 | 43,37 | 48,85 | 54,03 | 45 |
| ENSG00000164654 | MIOS | 15,01 | 15,58 | 16,54 | 21,77 | 45 |
| ENSG00000177628 | GBA | 4,96 | 5,67 | 6,62 | 7,19 | 45 |
| ENSG00000171227 | TMEM37 | 0,69 | 0,82 | 0,97 | 1 | 45 |
| ENSG00000100239 | PPP6R2 | 10,11 | 11,59 | 11,84 | 14,65 | 45 |
| ENSG00000121486 | TRMT1L | 9,28 | 11,49 | 11,65 | 13,44 | 45 |
| ENSG00000131797 | CLUHP3 | 2,9 | 3,5 | 3,57 | 4,2 | 45 |
| ENSG00000138709 | LARP1B | 8,2 | 9,36 | 9,63 | 11,87 | 45 |
| ENSG00000158941 | CCAR2 | 56,07 | 75,86 | 77,59 | 81,15 | 45 |
| ENSG00000144040 | SFXN5 | 6,71 | 7,17 | 7,94 | 9,71 | 45 |
| ENSG00000136381 | IREB2 | 22,64 | 24,71 | 27,11 | 32,76 | 45 |
| ENSG00000156453 | PCDH1 | 5,28 | 6,67 | 7,22 | 7,64 | 45 |
| ENSG00000095139 | ARCN1 | 34,34 | 39,08 | 45,98 | 49,67 | 45 |
| ENSG00000148737 | TCF7L2 | 17,05 | 20,44 | 21,35 | 24,66 | 45 |
| ENSG00000133393 | CEP20 | 11,34 | 13,29 | 13,48 | 16,4 | 45 |
| ENSG00000163541 | SUCLG1 | 24,78 | 28,23 | 32,95 | 35,83 | 45 |
| ENSG00000075420 | FNDC3B | 21,36 | 24,79 | 29,4 | 30,88 | 45 |
| ENSG00000277149 | TYW1B | 1,01 | 1,06 | 1,17 | 1,46 | 45 |
| ENSG00000124370 | MCEE | 3,01 | 3,36 | 4,34 | 4,35 | 45 |
| ENSG00000027697 | IFNGR1 | 10,3 | 11,26 | 12,05 | 14,88 | 44 |
| ENSG00000167635 | ZNF146 | 37,06 | 42,67 | 46,95 | 53,53 | 44 |
| ENSG00000026103 | FAS | 0,99 | 1 | 1,24 | 1,43 | 44 |
| ENSG00000122008 | POLK | 8,53 | 9,64 | 10,13 | 12,32 | 44 |
| ENSG00000070444 | MNT | 7,61 | 7,62 | 10,39 | 10,99 | 44 |
| ENSG00000120690 | ELF1 | 4,64 | 5,79 | 6,64 | 6,7 | 44 |
| ENSG00000177613 | CSTF2T | 9,69 | 11,67 | 11,89 | 13,99 | 44 |
| ENSG00000044574 | HSPA5 | 82,81 | 96 | 108,64 | 119,55 | 44 |
| ENSG00000135677 | GNS | 16,64 | 19,53 | 20,28 | 24,02 | 44 |
| ENSG00000129691 | ASH2L | 17,61 | 20,42 | 23,4 | 25,42 | 44 |
| ENSG00000128791 | TWSG1 | 11,12 | 12,54 | 13,26 | 16,05 | 44 |
| ENSG00000198466 | ZNF587 | 13,27 | 14,32 | 14,99 | 19,15 | 44 |
| ENSG00000187742 | SECISBP2 | 27,42 | 29,6 | 30,9 | 39,56 | 44 |
| ENSG00000170275 | CRTAP | 23,81 | 26,02 | 29,82 | 34,35 | 44 |
| ENSG00000101079 | NDRG3 | 9,56 | 12,33 | 12,9 | 13,79 | 44 |
| ENSG00000143319 | ISG20L2 | 15,03 | 16,22 | 18,56 | 21,68 | 44 |
| ENSG00000153827 | TRIP12 | 63,06 | 83,83 | 85,51 | 90,96 | 44 |
| ENSG00000164296 | TIGD6 | 2,17 | 2,72 | 2,99 | 3,13 | 44 |
| ENSG00000068878 | PSME4 | 19,17 | 19,88 | 19,89 | 27,65 | 44 |

| | | | | | | |
|---|---|---|---|---|---|---|
| ENSG00000203620 | AL354919,1 | 1,56 | 1,85 | 1,93 | 2,25 | 44 |
| ENSG00000065308 | TRAM2 | 6,92 | 7,77 | 9,28 | 9,98 | 44 |
| ENSG00000254221 | PCDHGB1 | 0,95 | 0,99 | 1,08 | 1,37 | 44 |
| ENSG00000116918 | TSNAX | 12,85 | 15,83 | 16,14 | 18,53 | 44 |
| ENSG00000006740 | ARHGAP44 | 3,01 | 3,31 | 4,25 | 4,34 | 44 |
| ENSG00000140632 | GLYR1 | 28,7 | 32,68 | 35,8 | 41,38 | 44 |
| ENSG00000145817 | YIPF5 | 11,26 | 13,84 | 15,6 | 16,23 | 44 |
| ENSG00000236609 | ZNF853 | 1,18 | 1,44 | 1,59 | 1,7 | 44 |
| ENSG00000234345 | ELF2P1 | 2,27 | 2,29 | 2,49 | 3,27 | 44 |
| ENSG00000172375 | C2CD2L | 1,68 | 1,96 | 2,3 | 2,42 | 44 |
| ENSG00000166166 | TRMT61A | 2,07 | 2,17 | 2,46 | 2,98 | 44 |
| ENSG00000166987 | MBD6 | 8,15 | 10,13 | 10,65 | 11,73 | 44 |
| ENSG00000060709 | RIMBP2 | 1,48 | 1,85 | 1,88 | 2,13 | 44 |
| ENSG00000160961 | ZNF333 | 5,18 | 5,63 | 5,85 | 7,45 | 44 |
| ENSG00000277782 | AC068870,2 | 1,53 | 1,62 | 1,93 | 2,2 | 44 |
| ENSG00000171302 | CANT1 | 14,89 | 16,46 | 19,08 | 21,41 | 44 |
| ENSG00000160972 | PPP1R16A | 6,7 | 6,79 | 7,61 | 9,63 | 44 |
| ENSG00000175582 | RAB6A | 30,35 | 39,48 | 43,04 | 43,62 | 44 |
| ENSG00000110768 | GTF2H1 | 7,39 | 8,38 | 8,64 | 10,62 | 44 |
| ENSG00000288114 | GTF2H1 | 7,39 | 7,62 | 8,59 | 10,62 | 44 |
| ENSG00000276358 | PLEKHM1 | 1,26 | 1,35 | 1,64 | 1,81 | 44 |
| ENSG00000132436 | FIGNL1 | 11,25 | 11,34 | 12,08 | 16,16 | 44 |
| ENSG00000253305 | PCDHGB6 | 24,38 | 31,58 | 34,56 | 35,02 | 44 |
| ENSG00000147649 | MTDH | 27,93 | 30,77 | 39,97 | 40,1 | 44 |
| ENSG00000233223 | AC016876,1 | 2,41 | 2,61 | 2,84 | 3,46 | 44 |
| ENSG00000182473 | EXOC7 | 25,66 | 28,29 | 32,07 | 36,83 | 44 |
| ENSG00000088367 | EPB41L1 | 13,19 | 13,58 | 16,82 | 18,93 | 44 |
| ENSG00000142002 | DPP9 | 8,48 | 9,86 | 10,56 | 12,17 | 44 |
| ENSG00000079459 | FDFT1 | 27,66 | 28,61 | 32,09 | 39,69 | 43 |
| ENSG00000100227 | POLDIP3 | 25,57 | 29,68 | 33,38 | 36,69 | 43 |
| ENSG00000215472 | RPL17-C18orf32 | 12,99 | 18,6 | 18,62 | 18,63 | 43 |
| ENSG00000160310 | PRMT2 | 31,32 | 38,54 | 43,11 | 44,9 | 43 |
| ENSG00000169255 | B3GALNT1 | 6,53 | 8,01 | 9,19 | 9,36 | 43 |
| ENSG00000113248 | PCDHB15 | 3,93 | 5,14 | 5,37 | 5,63 | 43 |
| ENSG00000119778 | ATAD2B | 9,81 | 11,17 | 11,78 | 14,05 | 43 |
| ENSG00000184281 | TSSC4 | 8,91 | 10,81 | 11,06 | 12,76 | 43 |
| ENSG00000236397 | DDX11L2 | 1,32 | 1,4 | 1,67 | 1,89 | 43 |
| ENSG00000156052 | GNAQ | 17,12 | 18,47 | 20,44 | 24,51 | 43 |
| ENSG00000173406 | DAB1 | 9,2 | 9,71 | 11,6 | 13,17 | 43 |
| ENSG00000182944 | EWSR1 | 120,38 | 128,08 | 141,27 | 172,27 | 43 |
| ENSG00000257176 | AC009318,1 | 1,16 | 1,19 | 1,31 | 1,66 | 43 |
| ENSG00000167513 | CDT1 | 12,58 | 13,14 | 15,58 | 18 | 43 |
| ENSG00000104381 | GDAP1 | 17,97 | 23,93 | 25,04 | 25,71 | 43 |
| ENSG00000100726 | TELO2 | 5,49 | 5,75 | 6,68 | 7,85 | 43 |
| ENSG00000165934 | CPSF2 | 25,94 | 30,13 | 33,39 | 37,09 | 43 |
| ENSG00000136238 | RAC1 | 81,14 | 93,39 | 114,3 | 116,01 | 43 |
| ENSG00000060971 | ACAA1 | 10,08 | 10,5 | 11,52 | 14,41 | 43 |
| ENSG00000181915 | ADO | 9,41 | 10,73 | 11,87 | 13,45 | 43 |
| ENSG00000058063 | ATP11B | 15,55 | 16,09 | 16,89 | 22,22 | 43 |
| ENSG00000187555 | USP7 | 45,18 | 50,73 | 53,69 | 64,55 | 43 |

| | | | | | | |
|---|---|---|---|---|---|---|
| ENSG00000070371 | CLTCL1 | 2,17 | 2,24 | 2,3 | 3,1 | 43 |
| ENSG00000184702 | SEPTIN5 | 16,46 | 19,13 | 20,35 | 23,51 | 43 |
| ENSG00000004766 | VPS50 | 8,46 | 9,53 | 11,26 | 12,08 | 43 |
| ENSG00000154845 | PPP4R1 | 54,22 | 64,67 | 68,72 | 77,42 | 43 |
| ENSG00000172264 | MACROD2 | 3,04 | 3,18 | 3,24 | 4,34 | 43 |
| ENSG00000166747 | AP1G1 | 25,72 | 28,19 | 31,66 | 36,71 | 43 |
| ENSG00000183741 | CBX6 | 9,83 | 10,38 | 10,76 | 14,03 | 43 |
| ENSG00000143569 | UBAP2L | 95,22 | 106,3 | 116,75 | 135,87 | 43 |
| ENSG00000124198 | ARFGEF2 | 4,31 | 4,83 | 4,9 | 6,15 | 43 |
| ENSG00000122390 | NAA60 | 18,98 | 22,3 | 24,53 | 27,08 | 43 |
| ENSG00000102181 | CD99L2 | 14,84 | 18,31 | 20,89 | 21,17 | 43 |
| ENSG00000112146 | FBXO9 | 11,19 | 13,47 | 14,28 | 15,96 | 43 |
| ENSG00000175137 | SH3BP5L | 5,49 | 6,28 | 6,97 | 7,83 | 43 |
| ENSG00000064601 | CTSA | 8,12 | 9,13 | 10,38 | 11,58 | 43 |
| ENSG00000203667 | COX20 | 19,07 | 23,4 | 25,33 | 27,19 | 43 |
| ENSG00000049769 | PPP1R3F | 2,56 | 3,34 | 3,44 | 3,65 | 43 |
| ENSG00000130734 | ATG4D | 8,62 | 10,65 | 11,6 | 12,29 | 43 |
| ENSG00000104131 | EIF3J | 44,2 | 45,57 | 54,39 | 63 | 43 |
| ENSG00000109787 | KLF3 | 16,3 | 16,53 | 18,05 | 23,23 | 43 |
| ENSG00000088247 | KHSRP | 150,81 | 155,8 | 163,24 | 214,92 | 43 |
| ENSG00000236991 | EDRF1-AS1 | 1,46 | 1,47 | 1,57 | 2,08 | 42 |
| ENSG00000173960 | UBXN2A | 12,02 | 13,77 | 14,42 | 17,12 | 42 |
| ENSG00000239789 | MRPS17 | 6,34 | 6,94 | 8,25 | 9,03 | 42 |
| ENSG00000087245 | MMP2 | 28,2 | 32,03 | 35,28 | 40,16 | 42 |
| ENSG00000117408 | IPO13 | 5,9 | 7,15 | 8,3 | 8,4 | 42 |
| ENSG00000115935 | WIPF1 | 8,78 | 9,19 | 10,43 | 12,5 | 42 |
| ENSG00000165175 | MID1IP1 | 9,56 | 9,84 | 12,08 | 13,61 | 42 |
| ENSG00000050748 | MAPK9 | 12,61 | 14,75 | 14,81 | 17,95 | 42 |
| ENSG00000141564 | RPTOR | 6,45 | 7,34 | 7,42 | 9,18 | 42 |
| ENSG00000138073 | PREB | 11,24 | 12,09 | 15,34 | 15,99 | 42 |
| ENSG00000115761 | NOL10 | 10,39 | 10,95 | 11,75 | 14,78 | 42 |
| ENSG00000288091 | AC062022,2 | 0,71 | 0,75 | 0,76 | 1,01 | 42 |
| ENSG00000131446 | MGAT1 | 13,78 | 14,61 | 15,05 | 19,6 | 42 |
| ENSG00000048828 | FAM120A | 19,74 | 20,95 | 23,82 | 28,07 | 42 |
| ENSG00000116560 | SFPQ | 254,1 | 278,24 | 297,91 | 361,27 | 42 |
| ENSG00000140937 | CDH11 | 50,19 | 58,9 | 60,56 | 71,35 | 42 |
| ENSG00000111276 | CDKN1B | 17,98 | 19,34 | 21,82 | 25,56 | 42 |
| ENSG00000165312 | OTUD1 | 1,97 | 2,11 | 2,18 | 2,8 | 42 |
| ENSG00000179833 | SERTAD2 | 4,89 | 5,26 | 5,58 | 6,95 | 42 |
| ENSG00000136240 | KDELR2 | 33,67 | 35,71 | 45,15 | 47,84 | 42 |
| ENSG00000128606 | LRRC17 | 18,44 | 22,96 | 24,48 | 26,2 | 42 |
| ENSG00000153487 | ING1 | 9,44 | 9,89 | 11,63 | 13,41 | 42 |
| ENSG00000152102 | FAM168B | 29,06 | 34,08 | 34,56 | 41,28 | 42 |
| ENSG00000164077 | MON1A | 2,19 | 2,6 | 2,96 | 3,11 | 42 |
| ENSG00000035403 | VCL | 42,92 | 46,97 | 55,41 | 60,93 | 42 |
| ENSG00000135318 | NT5E | 4,6 | 6,11 | 6,23 | 6,53 | 42 |
| ENSG00000140299 | BNIP2 | 18,61 | 22,16 | 23,14 | 26,41 | 42 |
| ENSG00000198901 | PRC1 | 43,15 | 45,67 | 48,1 | 61,23 | 42 |
| ENSG00000051382 | PIK3CB | 5,37 | 5,89 | 6,66 | 7,62 | 42 |
| ENSG00000167881 | SRP68 | 23,7 | 26,54 | 27,94 | 33,63 | 42 |

| | | | | | | |
|---|---|---|---|---|---|---|
| ENSG00000167491 | GATAD2A | 50,42 | 52,29 | 64,83 | 71,54 | 42 |
| ENSG00000152782 | PANK1 | 12,99 | 13,24 | 13,55 | 18,43 | 42 |
| ENSG00000100335 | MIEF1 | 11,92 | 13,82 | 15,62 | 16,91 | 42 |
| ENSG00000022277 | RTF2 | 28,97 | 31,52 | 39,8 | 41,09 | 42 |
| ENSG00000125686 | MED1 | 12,36 | 13,44 | 15,47 | 17,53 | 42 |
| ENSG00000164576 | SAP30L | 13,27 | 18,65 | 18,68 | 18,82 | 42 |
| ENSG00000141543 | EIF4A3 | 39,55 | 41,66 | 42,31 | 56,09 | 42 |
| ENSG00000173786 | CNP | 34,18 | 37,43 | 39,14 | 48,47 | 42 |
| ENSG00000138674 | SEC31A | 45,12 | 56,92 | 57,56 | 63,98 | 42 |
| ENSG00000174353 | STAG3L3 | 10,41 | 11,2 | 11,3 | 14,76 | 42 |
| ENSG00000125834 | STK35 | 7,3 | 7,32 | 7,73 | 10,35 | 42 |
| ENSG00000241852 | C8orf58 | 9,72 | 9,99 | 12,31 | 13,78 | 42 |
| ENSG00000151422 | FER | 8,55 | 9,1 | 10,07 | 12,12 | 42 |
| ENSG00000139687 | RB1 | 13,58 | 14,4 | 15,26 | 19,25 | 42 |
| ENSG00000171467 | ZNF318 | 8,58 | 9,72 | 9,77 | 12,16 | 42 |
| ENSG00000102221 | JADE3 | 6,09 | 6,38 | 8,35 | 8,63 | 42 |
| ENSG00000133773 | CCDC59 | 19,35 | 21,85 | 24,75 | 27,42 | 42 |
| ENSG00000103342 | GSPT1 | 47,39 | 50,49 | 60,09 | 67,15 | 42 |
| ENSG00000125354 | SEPTIN6 | 19,26 | 23,43 | 24,29 | 27,29 | 42 |
| ENSG00000128408 | RIBC2 | 3,43 | 3,97 | 4,38 | 4,86 | 42 |
| ENSG00000167693 | NXN | 28,09 | 35,91 | 37,44 | 39,8 | 42 |
| ENSG00000141367 | CLTC | 83,74 | 92,93 | 103,52 | 118,64 | 42 |
| ENSG00000136932 | TRMO | 3,36 | 3,74 | 4,72 | 4,76 | 42 |
| ENSG00000104671 | DCTN6 | 20,34 | 25,85 | 26,97 | 28,81 | 42 |
| ENSG00000151929 | BAG3 | 6,46 | 7,84 | 8,27 | 9,15 | 42 |
| ENSG00000197226 | TBC1D9B | 14,16 | 14,87 | 15,78 | 20,05 | 42 |
| ENSG00000173166 | RAPH1 | 8,61 | 9,44 | 10,39 | 12,19 | 42 |
| ENSG00000100023 | PPIL2 | 9,48 | 9,58 | 10,88 | 13,42 | 42 |
| ENSG00000165512 | ZNF22 | 22,69 | 24,85 | 25,23 | 32,12 | 42 |
| ENSG00000223496 | EXOSC6 | 9,27 | 10,75 | 10,85 | 13,12 | 42 |
| ENSG00000106459 | NRF1 | 7,13 | 9,15 | 9,25 | 10,09 | 42 |
| ENSG00000175376 | EIF1AD | 13,95 | 17,48 | 17,65 | 19,74 | 42 |
| ENSG00000131023 | LATS1 | 9,38 | 11,08 | 11,89 | 13,27 | 41 |
| ENSG00000112977 | DAP | 31,29 | 33,87 | 39,19 | 44,26 | 41 |
| ENSG00000134153 | EMC7 | 13,61 | 16,35 | 17,46 | 19,25 | 41 |
| ENSG00000170876 | TMEM43 | 16,8 | 19,52 | 21,88 | 23,76 | 41 |
| ENSG00000205542 | TMSB4X | 1035,65 | 1185,09 | 1359,61 | 1464,62 | 41 |
| ENSG00000173812 | EIF1 | 182,19 | 206,36 | 238,2 | 257,65 | 41 |
| ENSG00000090520 | DNAJB11 | 27,37 | 31,28 | 33,29 | 38,7 | 41 |
| ENSG00000278768 | BACE1-AS | 5,63 | 6,05 | 7,16 | 7,96 | 41 |
| ENSG00000141002 | TCF25 | 37,67 | 41,92 | 50,88 | 53,25 | 41 |
| ENSG00000101350 | KIF3B | 7,86 | 9,96 | 10,06 | 11,11 | 41 |
| ENSG00000125870 | SNRPB2 | 48,3 | 50,3 | 60,88 | 68,23 | 41 |
| ENSG00000282958 | HNRNPR | 82,7 | 93,01 | 96,79 | 116,81 | 41 |
| ENSG00000096384 | HSP90AB1 | 394,44 | 427,24 | 435,72 | 557,05 | 41 |
| ENSG00000110693 | SOX6 | 9,47 | 9,93 | 10,5 | 13,37 | 41 |
| ENSG00000179331 | RAB39A | 4,3 | 4,71 | 5,11 | 6,07 | 41 |
| ENSG00000124496 | TRERF1 | 3,96 | 4,01 | 4,35 | 5,59 | 41 |
| ENSG00000013503 | POLR3B | 5,71 | 5,86 | 7,82 | 8,06 | 41 |
| ENSG00000070785 | EIF2B3 | 10,89 | 11,8 | 13,63 | 15,37 | 41 |

| | | | | | | |
|---|---|---|---|---|---|---|
| ENSG00000105447 | GRWD1 | 11,79 | 13,09 | 16,25 | 16,64 | 41 |
| ENSG00000160563 | MED27 | 14,69 | 15,92 | 17,29 | 20,73 | 41 |
| ENSG00000119820 | YIPF4 | 11,4 | 12,43 | 12,55 | 16,08 | 41 |
| ENSG00000111790 | FGFR1OP2 | 15,51 | 17,35 | 17,46 | 21,87 | 41 |
| ENSG00000186272 | ZNF17 | 3,93 | 4,24 | 4,32 | 5,54 | 41 |
| ENSG00000169100 | SLC25A6 | 116,69 | 144,66 | 147,33 | 164,49 | 41 |
| ENSG00000113209 | PCDHB5 | 0,83 | 0,86 | 0,87 | 1,17 | 41 |
| ENSG00000071537 | SEL1L | 10,28 | 12,63 | 14,03 | 14,49 | 41 |
| ENSG00000167380 | ZNF226 | 11,75 | 13,31 | 13,82 | 16,56 | 41 |
| ENSG00000135655 | USP15 | 23,6 | 27,96 | 28,95 | 33,26 | 41 |
| ENSG00000114480 | GBE1 | 4,94 | 5,24 | 6,68 | 6,96 | 41 |
| ENSG00000184277 | TM2D3 | 13,6 | 16,3 | 17,12 | 19,16 | 41 |
| ENSG00000163655 | GMPS | 60,3 | 60,43 | 62,71 | 84,94 | 41 |
| ENSG00000165732 | DDX21 | 19,63 | 20,38 | 22,46 | 27,65 | 41 |
| ENSG00000100604 | CHGA | 0,71 | 0,8 | 0,89 | 1 | 41 |
| ENSG00000108883 | EFTUD2 | 54,77 | 58,31 | 63,69 | 77,13 | 41 |
| ENSG00000154839 | SKA1 | 9,88 | 12,51 | 12,53 | 13,91 | 41 |
| ENSG00000257591 | ZNF625 | 12,14 | 12,83 | 14,56 | 17,09 | 41 |
| ENSG00000110514 | MADD | 10,67 | 11,69 | 12,16 | 15,02 | 41 |
| ENSG00000123066 | MED13L | 27,6 | 31,87 | 34,85 | 38,85 | 41 |
| ENSG00000058262 | SEC61A1 | 38,55 | 44,08 | 48,6 | 54,24 | 41 |
| ENSG00000162998 | FRZB | 22,29 | 25,95 | 31,09 | 31,36 | 41 |
| ENSG00000173889 | PHC3 | 11,57 | 13,99 | 16,08 | 16,27 | 41 |
| ENSG00000197157 | SND1 | 41,43 | 42,57 | 53,21 | 58,24 | 41 |
| ENSG00000078018 | MAP2 | 91,82 | 110,05 | 113,33 | 129,06 | 41 |
| ENSG00000119777 | TMEM214 | 9,45 | 10,12 | 12,99 | 13,28 | 41 |
| ENSG00000136731 | UGGT1 | 22,98 | 28,07 | 30,18 | 32,29 | 41 |
| ENSG00000116731 | PRDM2 | 16,86 | 19,61 | 20,94 | 23,69 | 41 |
| ENSG00000160953 | PWWP3A | 19,29 | 20,15 | 20,16 | 27,1 | 40 |
| ENSG00000124784 | RIOK1 | 8,14 | 8,36 | 9,98 | 11,43 | 40 |
| ENSG00000144895 | EIF2A | 33,46 | 34,34 | 36,26 | 46,98 | 40 |
| ENSG00000128656 | CHN1 | 34,51 | 35,94 | 38,13 | 48,45 | 40 |
| ENSG00000240849 | PEDS1 | 9,31 | 10,86 | 12,55 | 13,07 | 40 |
| ENSG00000128016 | ZFP36 | 1,04 | 1,31 | 1,44 | 1,46 | 40 |
| ENSG00000238227 | TMEM250 | 7,81 | 7,96 | 8,91 | 10,96 | 40 |
| ENSG00000115514 | TXNDC9 | 16,84 | 17,78 | 20,02 | 23,63 | 40 |
| ENSG00000107185 | RGP1 | 4,74 | 5,47 | 5,91 | 6,65 | 40 |
| ENSG00000087152 | ATXN7L3 | 19,2 | 21,09 | 22,28 | 26,93 | 40 |
| ENSG00000143376 | SNX27 | 15,31 | 16,83 | 16,85 | 21,47 | 40 |
| ENSG00000087053 | MTMR2 | 21,43 | 24,5 | 24,64 | 30,05 | 40 |
| ENSG00000103111 | MON1B | 9,41 | 11,77 | 12,48 | 13,19 | 40 |
| ENSG00000106263 | EIF3B | 47,2 | 49,05 | 51,88 | 66,16 | 40 |
| ENSG00000155393 | HEATR3 | 5,13 | 5,34 | 5,9 | 7,19 | 40 |
| ENSG00000007080 | CCDC124 | 30,61 | 35,45 | 36,58 | 42,89 | 40 |
| ENSG00000175115 | PACS1 | 12,49 | 13,85 | 14,14 | 17,5 | 40 |
| ENSG00000067596 | DHX8 | 14,56 | 17,13 | 17,17 | 20,4 | 40 |
| ENSG00000281706 | LINC01012 | 2,17 | 2,3 | 2,47 | 3,04 | 40 |
| ENSG00000093000 | NUP50 | 35,8 | 38,29 | 42,79 | 50,15 | 40 |
| ENSG00000160710 | ADAR | 44,69 | 53,74 | 57,44 | 62,59 | 40 |
| ENSG00000106636 | YKT6 | 23,1 | 25,93 | 29,91 | 32,35 | 40 |

| | | | | | | |
|---|---|---|---|---|---|---|
| ENSG00000183655 | KLHL25 | 5,22 | 5,5 | 5,94 | 7,31 | 40 |
| ENSG00000163374 | YY1AP1 | 18,26 | 19,89 | 21,99 | 25,57 | 40 |
| ENSG00000108479 | GALK1 | 3,95 | 3,99 | 4,11 | 5,53 | 40 |
| ENSG00000273270 | AC090114,2 | 2,7 | 3,59 | 3,61 | 3,78 | 40 |
| ENSG00000127022 | CANX | 63,87 | 75,99 | 84,14 | 89,4 | 40 |
| ENSG00000146223 | RPL7L1 | 31,91 | 37,14 | 38,12 | 44,66 | 40 |
| ENSG00000055332 | EIF2AK2 | 13,94 | 14,45 | 15,33 | 19,5 | 40 |
| ENSG00000127527 | EPS15L1 | 15,3 | 17,21 | 18,35 | 21,4 | 40 |
| ENSG00000106799 | TGFBR1 | 57,5 | 70,66 | 73,32 | 80,42 | 40 |
| ENSG00000173875 | ZNF791 | 5,72 | 6,21 | 7,02 | 8 | 40 |
| ENSG00000084463 | WBP11 | 41,35 | 42,98 | 46,95 | 57,82 | 40 |
| ENSG00000127946 | HIP1 | 27,68 | 33,68 | 34,93 | 38,7 | 40 |
| ENSG00000170445 | HARS1 | 22,64 | 24,33 | 27,45 | 31,65 | 40 |
| ENSG00000173950 | XXYLT1 | 15,18 | 16,97 | 19,47 | 21,22 | 40 |
| ENSG00000117069 | ST6GALNAC5 | 18,13 | 19,4 | 24,92 | 25,34 | 40 |
| ENSG00000284832 | POLR2A | 32,69 | 35,82 | 38,01 | 45,68 | 40 |
| ENSG00000259363 | AC090825,1 | 4,81 | 5,02 | 5,06 | 6,72 | 40 |
| ENSG00000197696 | NMB | 2,67 | 2,87 | 2,93 | 3,73 | 40 |
| ENSG00000196428 | TSC22D2 | 11,77 | 12,57 | 13,21 | 16,43 | 40 |
| ENSG00000113719 | ERGIC1 | 33,67 | 37,51 | 39,18 | 46,99 | 40 |
| ENSG00000173848 | NET1 | 15,27 | 15,94 | 17,21 | 21,31 | 40 |
| ENSG00000120333 | MRPS14 | 10,65 | 11,97 | 13,64 | 14,86 | 40 |
| ENSG00000168297 | PXK | 3,09 | 3,45 | 3,69 | 4,31 | 39 |
| ENSG00000125945 | ZNF436 | 3,85 | 4,58 | 4,79 | 5,37 | 39 |
| ENSG00000283009 | ZNF436 | 3,85 | 4,58 | 4,79 | 5,37 | 39 |
| ENSG00000168259 | DNAJC7 | 43,57 | 49,68 | 54,31 | 60,77 | 39 |
| ENSG00000143761 | ARF1 | 79,7 | 86,05 | 100,17 | 111,15 | 39 |
| ENSG00000154582 | ELOC | 35,16 | 39,93 | 44,33 | 49,03 | 39 |
| ENSG00000186889 | TMEM17 | 2,13 | 2,34 | 2,57 | 2,97 | 39 |
| ENSG00000102580 | DNAJC3 | 6,62 | 7,07 | 7,61 | 9,23 | 39 |
| ENSG00000198108 | CHSY3 | 1,32 | 1,37 | 1,82 | 1,84 | 39 |
| ENSG00000169895 | SYAP1 | 14,32 | 16,52 | 18,97 | 19,96 | 39 |
| ENSG00000135334 | AKIRIN2 | 20,27 | 24,25 | 27 | 28,25 | 39 |
| ENSG00000152359 | POC5 | 6,15 | 7,37 | 7,63 | 8,57 | 39 |
| ENSG00000196566 | AL138767,1 | 0,89 | 0,97 | 1,16 | 1,24 | 39 |
| ENSG00000242419 | PCDHGC4 | 6,16 | 6,67 | 6,93 | 8,58 | 39 |
| ENSG00000181830 | SLC35C1 | 2,98 | 3,01 | 3,38 | 4,15 | 39 |
| ENSG00000172315 | TP53RK | 7,11 | 7,91 | 8,5 | 9,9 | 39 |
| ENSG00000173706 | HEG1 | 14,68 | 16,89 | 17,41 | 20,44 | 39 |
| ENSG00000114999 | TTL | 26,89 | 27,66 | 31,99 | 37,44 | 39 |
| ENSG00000166311 | SMPD1 | 4,64 | 4,75 | 5,68 | 6,46 | 39 |
| ENSG00000272142 | LYRM4-AS1 | 2,32 | 2,35 | 2,43 | 3,23 | 39 |
| ENSG00000055070 | SZRD1 | 35,39 | 40,54 | 47,31 | 49,27 | 39 |
| ENSG00000126524 | SBDS | 16,26 | 16,8 | 20,67 | 22,63 | 39 |
| ENSG00000187372 | PCDHB13 | 1,2 | 1,35 | 1,36 | 1,67 | 39 |
| ENSG00000221988 | PPT2 | 5,11 | 5,7 | 6,95 | 7,11 | 39 |
| ENSG00000112763 | BTN2A1 | 7,21 | 8,13 | 9,94 | 10,03 | 39 |
| ENSG00000120265 | PCMT1 | 22,81 | 26,68 | 28,52 | 31,73 | 39 |
| ENSG00000198355 | PIM3 | 5,97 | 6,39 | 7,28 | 8,3 | 39 |
| ENSG00000158201 | ABHD3 | 3,69 | 3,83 | 4,1 | 5,13 | 39 |

| | | | | | | |
|---|---|---|---|---|---|---|
| ENSG00000047644 | WWC3 | 7 | 8,21 | 8,47 | 9,73 | 39 |
| ENSG00000075391 | RASAL2 | 8,54 | 9,51 | 11,6 | 11,87 | 39 |
| ENSG00000159352 | PSMD4 | 64,98 | 67,27 | 82,88 | 90,3 | 39 |
| ENSG00000162430 | SELENON | 35,75 | 38,5 | 41,15 | 49,67 | 39 |
| ENSG00000138069 | RAB1A | 31,31 | 37,18 | 39,82 | 43,5 | 39 |
| ENSG00000175782 | SLC35E3 | 6,37 | 6,74 | 7,27 | 8,85 | 39 |
| ENSG00000120314 | WDR55 | 5,5 | 6,21 | 6,57 | 7,64 | 39 |
| ENSG00000198668 | CALM1 | 157,41 | 163,65 | 202,16 | 218,61 | 39 |
| ENSG00000149948 | HMGA2 | 106,2 | 106,86 | 110,15 | 147,46 | 39 |
| ENSG00000177879 | AP3S1 | 33,98 | 39,64 | 42,07 | 47,17 | 39 |
| ENSG00000076242 | MLH1 | 31,83 | 35,62 | 37,52 | 44,18 | 39 |
| ENSG00000168610 | STAT3 | 16,56 | 19,53 | 21,06 | 22,98 | 39 |
| ENSG00000167004 | PDIA3 | 152,8 | 164,95 | 190,02 | 212,03 | 39 |
| ENSG00000132763 | MMACHC | 3,07 | 3,84 | 3,95 | 4,26 | 39 |
| ENSG00000131051 | RBM39 | 106,63 | 127,6 | 130,15 | 147,94 | 39 |
| ENSG00000163636 | PSMD6 | 47,87 | 49,07 | 55,76 | 66,41 | 39 |
| ENSG00000277443 | MARCKS | 156,79 | 186,19 | 198,28 | 217,46 | 39 |
| ENSG00000112078 | KCTD20 | 20,17 | 23,4 | 24,13 | 27,97 | 39 |
| ENSG00000161542 | PRPSAP1 | 30,56 | 31,21 | 32,39 | 42,37 | 39 |
| ENSG00000167792 | NDUFV1 | 33,97 | 37,23 | 39,23 | 47,09 | 39 |
| ENSG00000103266 | STUB1 | 18,78 | 19,61 | 24,36 | 26,03 | 39 |
| ENSG00000141644 | MBD1 | 16,25 | 17,22 | 17,94 | 22,52 | 39 |
| ENSG00000130164 | LDLR | 24,63 | 28,79 | 30,77 | 34,13 | 39 |
| ENSG00000169564 | PCBP1 | 98,84 | 102,83 | 119,93 | 136,96 | 39 |
| ENSG00000138246 | DNAJC13 | 14,04 | 16,17 | 17,18 | 19,45 | 39 |
| ENSG00000153904 | DDAH1 | 18,25 | 20,08 | 22,42 | 25,28 | 39 |
| ENSG00000136271 | DDX56 | 18,8 | 19,36 | 21,42 | 26,04 | 39 |
| ENSG00000070476 | ZXDC | 8,31 | 9,27 | 10,05 | 11,51 | 39 |
| ENSG00000089195 | TRMT6 | 6,91 | 7,01 | 8,45 | 9,57 | 38 |
| ENSG00000141965 | FEM1A | 1,43 | 1,84 | 1,85 | 1,98 | 38 |
| ENSG00000232024 | LSM12P1 | 14,1 | 18,25 | 18,57 | 19,52 | 38 |
| ENSG00000197586 | ENTPD6 | 14,04 | 14,62 | 14,94 | 19,43 | 38 |
| ENSG00000158850 | B4GALT3 | 11,58 | 11,7 | 14,28 | 16,02 | 38 |
| ENSG00000096070 | BRPF3 | 13,23 | 15,87 | 16,99 | 18,3 | 38 |
| ENSG00000112578 | BYSL | 9,58 | 10,2 | 11,36 | 13,25 | 38 |
| ENSG00000176022 | B3GALT6 | 8,72 | 10,09 | 11,42 | 12,06 | 38 |
| ENSG00000126214 | KLC1 | 71,54 | 84,77 | 86,41 | 98,93 | 38 |
| ENSG00000281540 | RCC2 | 30,43 | 34,04 | 34,29 | 42,08 | 38 |
| ENSG00000179051 | RCC2 | 30,43 | 34,04 | 34,29 | 42,08 | 38 |
| ENSG00000108433 | GOSR2 | 17,37 | 20,14 | 21,29 | 24,02 | 38 |
| ENSG00000262860 | LSM14A | 46,86 | 51,69 | 52,52 | 64,78 | 38 |
| ENSG00000134330 | IAH1 | 16,09 | 18,12 | 18,44 | 22,24 | 38 |
| ENSG00000124831 | LRRFIP1 | 21,34 | 22,2 | 26,1 | 29,49 | 38 |
| ENSG00000119912 | IDE | 13,96 | 15,59 | 16,34 | 19,29 | 38 |
| ENSG00000101152 | DNAJC5 | 10,27 | 12,81 | 14,07 | 14,19 | 38 |
| ENSG00000014164 | ZC3H3 | 4,64 | 5,49 | 6,02 | 6,41 | 38 |
| ENSG00000196204 | RNF216P1 | 15,24 | 15,29 | 19,36 | 21,05 | 38 |
| ENSG00000128595 | CALU | 94,48 | 102,99 | 110,5 | 130,46 | 38 |
| ENSG00000136295 | TTYH3 | 36,61 | 43,16 | 47,45 | 50,55 | 38 |
| ENSG00000120158 | RCL1 | 4,95 | 5,16 | 6,11 | 6,83 | 38 |

| | | | | | | |
|---|---|---|---|---|---|---|
| ENSG00000067601 | PMS2P4 | 3,53 | 3,97 | 4,69 | 4,87 | 38 |
| ENSG00000138032 | PPM1B | 15,1 | 17,95 | 18,8 | 20,83 | 38 |
| ENSG00000183207 | RUVBL2 | 35,67 | 37,63 | 42,67 | 49,2 | 38 |
| ENSG00000198646 | NCOA6 | 14,5 | 16,72 | 18,7 | 20 | 38 |
| ENSG00000117010 | ZNF684 | 2,69 | 2,81 | 3,28 | 3,71 | 38 |
| ENSG00000150316 | CWC15 | 33,79 | 40,01 | 43,47 | 46,6 | 38 |
| ENSG00000122550 | KLHL7 | 31,13 | 36,85 | 41,29 | 42,93 | 38 |
| ENSG00000183431 | SF3A3 | 68,21 | 70,43 | 78,97 | 94,06 | 38 |
| ENSG00000087502 | ERGIC2 | 19,43 | 21,36 | 24 | 26,79 | 38 |
| ENSG00000234545 | FAM133B | 22,09 | 22,38 | 23,52 | 30,45 | 38 |
| ENSG00000232533 | AC093673,1 | 3,99 | 4,38 | 5,44 | 5,5 | 38 |
| ENSG00000169217 | CD2BP2 | 24,2 | 27,33 | 30,72 | 33,35 | 38 |
| ENSG00000134539 | KLRD1 | 2,09 | 2,38 | 2,45 | 2,88 | 38 |
| ENSG00000279457 | WASH9P | 3,68 | 4,52 | 4,54 | 5,07 | 38 |
| ENSG00000173545 | ZNF622 | 7,97 | 8,12 | 9,26 | 10,98 | 38 |
| ENSG00000196968 | FUT11 | 8,98 | 11,13 | 11,3 | 12,37 | 38 |
| ENSG00000100281 | HMGXB4 | 21,59 | 22,98 | 25,15 | 29,74 | 38 |
| ENSG00000135250 | SRPK2 | 21,94 | 23,78 | 25,06 | 30,22 | 38 |
| ENSG00000196313 | POM121 | 15,74 | 17,6 | 21,02 | 21,68 | 38 |
| ENSG00000110651 | CD81 | 63,69 | 69,75 | 77 | 87,72 | 38 |
| ENSG00000073614 | KDM5A | 18,61 | 22,01 | 23,04 | 25,63 | 38 |
| ENSG00000142864 | SERBP1 | 206,54 | 223,72 | 234,51 | 284,37 | 38 |
| ENSG00000048162 | NOP16 | 13,59 | 14,35 | 16,56 | 18,71 | 38 |
| ENSG00000113194 | FAF2 | 15,45 | 15,73 | 17,55 | 21,27 | 38 |
| ENSG00000166913 | YWHAB | 57,08 | 63,36 | 72,12 | 78,57 | 38 |
| ENSG00000131470 | PSMC3IP | 15,98 | 16,62 | 20,64 | 21,99 | 38 |
| ENSG00000186973 | FAM183A | 2,26 | 2,53 | 2,78 | 3,11 | 38 |
| ENSG00000112874 | NUDT12 | 5,93 | 6,31 | 6,54 | 8,16 | 38 |
| ENSG00000067248 | DHX29 | 11,02 | 12,11 | 12,82 | 15,16 | 38 |
| ENSG00000042429 | MED17 | 18,29 | 21,33 | 22,52 | 25,16 | 38 |
| ENSG00000174106 | LEMD3 | 8,76 | 9,84 | 10,06 | 12,05 | 38 |
| ENSG00000155876 | RRAGA | 16,51 | 18,96 | 22,08 | 22,71 | 38 |
| ENSG00000103544 | VPS35L | 20,2 | 20,93 | 22,11 | 27,78 | 38 |
| ENSG00000137275 | RIPK1 | 5,33 | 6,46 | 6,73 | 7,33 | 38 |
| ENSG00000126216 | TUBGCP3 | 13,65 | 13,87 | 14,6 | 18,76 | 37 |
| ENSG00000112941 | TENT4A | 14,99 | 16,27 | 17,26 | 20,6 | 37 |
| ENSG00000250021 | ARPIN-AP3S2 | 10,05 | 12,32 | 13,09 | 13,81 | 37 |
| ENSG00000101421 | CHMP4B | 23,88 | 29,31 | 30,02 | 32,81 | 37 |
| ENSG00000087448 | KLHL42 | 14,11 | 16,35 | 16,75 | 19,38 | 37 |
| ENSG00000083544 | TDRD3 | 13,71 | 15,49 | 16,04 | 18,83 | 37 |
| ENSG00000078902 | TOLLIP | 7,15 | 8,05 | 9,05 | 9,82 | 37 |
| ENSG00000110042 | DTX4 | 15,94 | 17,04 | 19,39 | 21,89 | 37 |
| ENSG00000141456 | PELP1 | 14,12 | 14,21 | 14,85 | 19,39 | 37 |
| ENSG00000145703 | IQGAP2 | 10,94 | 12,11 | 13,97 | 15,02 | 37 |
| ENSG00000123728 | RAP2C | 23,33 | 26,44 | 29,68 | 32,03 | 37 |
| ENSG00000123562 | MORF4L2 | 159,95 | 169,06 | 188,39 | 219,58 | 37 |
| ENSG00000131871 | SELENOS | 19,18 | 21,21 | 24,04 | 26,33 | 37 |
| ENSG00000172775 | PSME3IP1 | 53,07 | 59,68 | 70,24 | 72,84 | 37 |
| ENSG00000029363 | BCLAF1 | 119,1 | 135,78 | 142,81 | 163,46 | 37 |
| ENSG00000128928 | IVD | 11,07 | 11,19 | 13,74 | 15,19 | 37 |

| | | | | | | |
|---|---|---|---|---|---|---|
| ENSG00000116199 | FAM20B | 13,14 | 14,2 | 14,64 | 18,03 | 37 |
| ENSG00000067955 | CBFB | 28,19 | 30,87 | 32,63 | 38,68 | 37 |
| ENSG00000136026 | CKAP4 | 42 | 47,24 | 54,59 | 57,62 | 37 |
| ENSG00000134283 | PPHLN1 | 24,72 | 25,26 | 26,76 | 33,91 | 37 |
| ENSG00000076650 | GPATCH1 | 5,01 | 5,1 | 5,56 | 6,87 | 37 |
| ENSG00000103274 | NUBP1 | 14,6 | 15,86 | 18,34 | 20,02 | 37 |
| ENSG00000131263 | RLIM | 12,29 | 13,5 | 13,95 | 16,85 | 37 |
| ENSG00000174437 | ATP2A2 | 51,41 | 54,88 | 62,45 | 70,48 | 37 |
| ENSG00000119335 | SET | 385,5 | 410,02 | 431,06 | 528,47 | 37 |
| ENSG00000114503 | NCBP2 | 37,95 | 44,19 | 44,82 | 52,02 | 37 |
| ENSG00000103194 | USP10 | 38,89 | 40,48 | 44,52 | 53,3 | 37 |
| ENSG00000164091 | WDR82 | 28,43 | 35,09 | 35,42 | 38,96 | 37 |
| ENSG00000114850 | SSR3 | 43,67 | 47,32 | 54,14 | 59,84 | 37 |
| ENSG00000165410 | CFL2 | 31,44 | 35,53 | 35,58 | 43,08 | 37 |
| ENSG00000124486 | USP9X | 27,88 | 27,91 | 28,64 | 38,19 | 37 |
| ENSG00000110721 | CHKA | 20,27 | 23,61 | 26,49 | 27,75 | 37 |
| ENSG00000144381 | HSPD1 | 224,34 | 237,79 | 270,58 | 307,12 | 37 |
| ENSG00000178567 | EPM2AIP1 | 14,26 | 16,67 | 18,35 | 19,52 | 37 |
| ENSG00000166266 | CUL5 | 13,42 | 14,47 | 16,05 | 18,37 | 37 |
| ENSG00000161981 | SNRNP25 | 12,58 | 12,88 | 15,35 | 17,22 | 37 |
| ENSG00000157625 | TAB3 | 5,83 | 6,27 | 6,84 | 7,98 | 37 |
| ENSG00000128228 | SDF2L1 | 6,21 | 7,02 | 7,21 | 8,5 | 37 |
| ENSG00000089597 | GANAB | 84,57 | 92,35 | 97,07 | 115,67 | 37 |
| ENSG00000140993 | TIGD7 | 3,1 | 3,75 | 4,19 | 4,24 | 37 |
| ENSG00000212747 | RTL8B | 7,56 | 7,75 | 10,1 | 10,34 | 37 |
| ENSG00000149257 | SERPINH1 | 120 | 124,38 | 143,69 | 164,07 | 37 |
| ENSG00000168005 | SPINDOC | 22,12 | 24,41 | 25,27 | 30,24 | 37 |
| ENSG00000173409 | ARV1 | 7,63 | 8,95 | 9,65 | 10,43 | 37 |
| ENSG00000080371 | RAB21 | 20,81 | 25,7 | 27,44 | 28,44 | 37 |
| ENSG00000185651 | UBE2L3 | 32,93 | 35,45 | 41,33 | 45 | 37 |
| ENSG00000110422 | HIPK3 | 11,68 | 13,67 | 15,38 | 15,96 | 37 |
| ENSG00000276345 | AC004556,3 | 11,3 | 11,93 | 13,78 | 15,44 | 37 |
| ENSG00000165630 | PRPF18 | 7,37 | 8,95 | 9,47 | 10,07 | 37 |
| ENSG00000175550 | DRAP1 | 55,75 | 61,81 | 64,27 | 76,17 | 37 |
| ENSG00000008710 | PKD1 | 15,51 | 15,87 | 16,35 | 21,19 | 37 |
| ENSG00000120948 | TARDBP | 77,49 | 86,41 | 90,57 | 105,84 | 37 |
| ENSG00000103550 | KNOP1 | 34,55 | 38,25 | 39,02 | 47,19 | 37 |
| ENSG00000058056 | USP13 | 10,75 | 11,64 | 13,71 | 14,68 | 37 |
| ENSG00000113615 | SEC24A | 7,14 | 7,46 | 8,04 | 9,75 | 37 |
| ENSG00000133739 | LRRCC1 | 7,77 | 8,21 | 9,08 | 10,61 | 37 |
| ENSG00000163466 | ARPC2 | 88,96 | 95,22 | 113,69 | 121,45 | 37 |
| ENSG00000119396 | RAB14 | 17,88 | 20,29 | 22,49 | 24,41 | 37 |
| ENSG00000213699 | SLC35F6 | 6,63 | 7,56 | 7,94 | 9,05 | 37 |
| ENSG00000076604 | TRAF4 | 40,91 | 44,13 | 54,55 | 55,84 | 36 |
| ENSG00000120694 | HSPH1 | 39,85 | 47,1 | 49,36 | 54,39 | 36 |
| ENSG00000244038 | DDOST | 54,87 | 62,52 | 69,64 | 74,89 | 36 |
| ENSG00000141551 | CSNK1D | 30,91 | 34,73 | 38,74 | 42,18 | 36 |
| ENSG00000075785 | RAB7A | 35,6 | 37,66 | 47,97 | 48,57 | 36 |
| ENSG00000146555 | SDK1 | 4,75 | 6,06 | 6,24 | 6,48 | 36 |
| ENSG00000132612 | VPS4A | 20,9 | 23,7 | 25,01 | 28,51 | 36 |

| | | | | | | |
|---|---|---|---|---|---|---|
| ENSG00000082153 | BZW1 | 142,19 | 150,58 | 177,7 | 193,93 | 36 |
| ENSG00000148450 | MSRB2 | 10,61 | 11,55 | 12,27 | 14,47 | 36 |
| ENSG00000142634 | EFHD2 | 6,35 | 6,7 | 6,98 | 8,66 | 36 |
| ENSG00000121067 | SPOP | 19,89 | 21,93 | 22,27 | 27,11 | 36 |
| ENSG00000104549 | SQLE | 75,19 | 78,7 | 93,72 | 102,46 | 36 |
| ENSG00000102753 | KPNA3 | 18,75 | 22,08 | 23,25 | 25,55 | 36 |
| ENSG00000108039 | XPNPEP1 | 26,76 | 31,73 | 34,33 | 36,45 | 36 |
| ENSG00000088826 | SMOX | 3,26 | 3,64 | 3,88 | 4,44 | 36 |
| ENSG00000136193 | SCRN1 | 24,49 | 27,56 | 30,33 | 33,35 | 36 |
| ENSG00000242615 | AC022415,1 | 10,15 | 12,01 | 12,02 | 13,82 | 36 |
| ENSG00000039319 | ZFYVE16 | 29,99 | 30,31 | 31,2 | 40,83 | 36 |
| ENSG00000067225 | PKM | 251,31 | 265,64 | 291,93 | 342,14 | 36 |
| ENSG00000011198 | ABHD5 | 5,65 | 5,74 | 6,45 | 7,69 | 36 |
| ENSG00000138867 | GUCD1 | 13,86 | 14,32 | 17,86 | 18,86 | 36 |
| ENSG00000140464 | PML | 7,77 | 8,31 | 9,77 | 10,57 | 36 |
| ENSG00000132670 | PTPRA | 35,86 | 38,28 | 41,28 | 48,78 | 36 |
| ENSG00000104679 | R3HCC1 | 14,19 | 16,91 | 17,95 | 19,3 | 36 |
| ENSG00000168255 | POLR2J3 | 8,86 | 9,6 | 10,03 | 12,05 | 36 |
| ENSG00000179295 | PTPN11 | 30,94 | 34,77 | 36,04 | 42,07 | 36 |
| ENSG00000284406 | AC004492,3 | 0,89 | 0,95 | 1 | 1,21 | 36 |
| ENSG00000272072 | AC004492,1 | 0,89 | 0,95 | 1 | 1,21 | 36 |
| ENSG00000165417 | GTF2A1 | 13,97 | 16,17 | 16,39 | 18,99 | 36 |
| ENSG00000128607 | KLHDC10 | 10,19 | 13,04 | 13,16 | 13,85 | 36 |
| ENSG00000130066 | SAT1 | 37,87 | 49,86 | 51,28 | 51,46 | 36 |
| ENSG00000130717 | UCK1 | 14,02 | 14,58 | 16,48 | 19,05 | 36 |
| ENSG00000198720 | ANKRD13B | 7,47 | 7,57 | 8,21 | 10,15 | 36 |
| ENSG00000205339 | IPO7 | 42,6 | 43,94 | 46,07 | 57,88 | 36 |
| ENSG00000134461 | ANKRD16 | 3,74 | 3,93 | 4,95 | 5,08 | 36 |
| ENSG00000071462 | BUD23 | 18,67 | 19 | 22,9 | 25,35 | 36 |
| ENSG00000111196 | MAGOHB | 26,98 | 29,62 | 33,02 | 36,63 | 36 |
| ENSG00000151148 | UBE3B | 12,36 | 13,51 | 14,96 | 16,78 | 36 |
| ENSG00000108262 | GIT1 | 16,91 | 20,46 | 22,27 | 22,95 | 36 |
| ENSG00000108424 | KPNB1 | 150,87 | 154,82 | 168,31 | 204,74 | 36 |
| ENSG00000063601 | MTMR1 | 8,1 | 9 | 9,66 | 10,99 | 36 |
| ENSG00000198663 | C6orf89 | 9,67 | 11,32 | 12,67 | 13,12 | 36 |
| ENSG00000106479 | ZNF862 | 5,69 | 6,18 | 7,69 | 7,72 | 36 |
| ENSG00000009335 | UBE3C | 17,27 | 17,65 | 19,28 | 23,43 | 36 |
| ENSG00000282086 | PKD1P6 | 5,44 | 6,94 | 7,04 | 7,38 | 36 |
| ENSG00000171202 | TMEM126A | 15,2 | 18,76 | 19,01 | 20,62 | 36 |
| ENSG00000125755 | SYMPK | 14,65 | 15,77 | 16,85 | 19,87 | 36 |
| ENSG00000165240 | ATP7A | 3,06 | 3,45 | 3,47 | 4,15 | 36 |
| ENSG00000100395 | L3MBTL2 | 9,04 | 10,26 | 10,34 | 12,26 | 36 |
| ENSG00000115825 | PRKD3 | 22,18 | 22,61 | 24,12 | 30,08 | 36 |
| ENSG00000172992 | DCAKD | 17,38 | 20,28 | 23,28 | 23,57 | 36 |
| ENSG00000132463 | GRSF1 | 27,18 | 30,14 | 32,22 | 36,86 | 36 |
| ENSG00000092108 | SCFD1 | 26,36 | 28,21 | 29,28 | 35,73 | 36 |
| ENSG00000196367 | TRRAP | 17,81 | 19,7 | 21,6 | 24,14 | 36 |
| ENSG00000149218 | ENDOD1 | 2,45 | 2,7 | 2,95 | 3,32 | 36 |
| ENSG00000185624 | P4HB | 108,98 | 112,03 | 146,07 | 147,64 | 35 |
| ENSG00000086200 | IPO11 | 12,97 | 13,46 | 14,03 | 17,57 | 35 |

| | | | | | | |
|---|---|---|---|---|---|---|
| ENSG00000110921 | MVK | 13,95 | 16,12 | 18,45 | 18,89 | 35 |
| ENSG00000269951 | AC090181,2 | 4,49 | 4,54 | 5,15 | 6,08 | 35 |
| ENSG00000178188 | SH2B1 | 10,6 | 11,24 | 12,12 | 14,35 | 35 |
| ENSG00000136824 | SMC2 | 19,59 | 23,01 | 23,02 | 26,51 | 35 |
| ENSG00000169093 | ASMTL | 11,41 | 12,51 | 12,63 | 15,44 | 35 |
| ENSG00000149658 | YTHDF1 | 14,1 | 15,67 | 17,62 | 19,08 | 35 |
| ENSG00000101158 | NELFCD | 34,43 | 38,56 | 44,74 | 46,59 | 35 |
| ENSG00000126804 | ZBTB1 | 14,16 | 16,37 | 16,77 | 19,16 | 35 |
| ENSG00000100325 | ASCC2 | 11,84 | 13,58 | 14,85 | 16,02 | 35 |
| ENSG00000141664 | ZCCHC2 | 5,98 | 7,49 | 7,63 | 8,09 | 35 |
| ENSG00000176597 | B3GNT5 | 15,59 | 15,78 | 17,67 | 21,09 | 35 |
| ENSG00000100591 | AHSA1 | 53,37 | 55,16 | 66,57 | 72,18 | 35 |
| ENSG00000127804 | METTL16 | 10,3 | 10,44 | 11,71 | 13,93 | 35 |
| ENSG00000196591 | HDAC2 | 155,25 | 175,72 | 191,99 | 209,94 | 35 |
| ENSG00000155034 | FBXL18 | 5,34 | 5,77 | 5,89 | 7,22 | 35 |
| ENSG00000064999 | ANKS1A | 7,9 | 9,62 | 10,16 | 10,68 | 35 |
| ENSG00000131828 | PDHA1 | 58,72 | 68,75 | 73,88 | 79,38 | 35 |
| ENSG00000062650 | WAPL | 16,43 | 18,47 | 18,74 | 22,21 | 35 |
| ENSG00000145293 | ENOPH1 | 16,3 | 16,96 | 19,57 | 22,03 | 35 |
| ENSG00000122218 | COPA | 37,96 | 41,64 | 46,39 | 51,3 | 35 |
| ENSG00000149636 | DSN1 | 12,24 | 12,67 | 14,04 | 16,54 | 35 |
| ENSG00000173145 | NOC3L | 9,14 | 9,2 | 9,47 | 12,35 | 35 |
| ENSG00000213516 | RBMXL1 | 6,35 | 6,68 | 7,29 | 8,58 | 35 |
| ENSG00000132432 | SEC61G | 75,11 | 79,83 | 85,47 | 101,46 | 35 |
| ENSG00000176946 | THAP4 | 14,88 | 16,25 | 17,06 | 20,09 | 35 |
| ENSG00000100412 | ACO2 | 26,24 | 30,5 | 32,96 | 35,42 | 35 |
| ENSG00000139684 | ESD | 54,96 | 67,8 | 72,18 | 74,18 | 35 |
| ENSG00000240057 | AC078785,1 | 1,03 | 1,12 | 1,28 | 1,39 | 35 |
| ENSG00000110925 | CSRNP2 | 14,77 | 17,9 | 18,17 | 19,93 | 35 |
| ENSG00000284776 | AL121900,1 | 18,32 | 19,76 | 23,79 | 24,72 | 35 |
| ENSG00000099341 | PSMD8 | 49,28 | 53,01 | 65,01 | 66,47 | 35 |
| ENSG00000103005 | USB1 | 13,88 | 16,75 | 17,33 | 18,72 | 35 |
| ENSG00000242247 | ARFGAP3 | 9,85 | 12,18 | 12,37 | 13,28 | 35 |
| ENSG00000140153 | WDR20 | 8,31 | 10,31 | 10,43 | 11,2 | 35 |
| ENSG00000100603 | SNW1 | 33,14 | 36,5 | 38,78 | 44,66 | 35 |
| ENSG00000273562 | NAP1L4 | 60,78 | 65,97 | 68,83 | 81,9 | 35 |
| ENSG00000198874 | TYW1 | 5,85 | 5,89 | 6,09 | 7,88 | 35 |
| ENSG00000117751 | PPP1R8 | 24,82 | 29,42 | 30,48 | 33,43 | 35 |
| ENSG00000110344 | UBE4A | 12,6 | 13,19 | 14,38 | 16,97 | 35 |
| ENSG00000100568 | VTI1B | 25,51 | 27,54 | 29,96 | 34,35 | 35 |
| ENSG00000167193 | CRK | 17,69 | 19,28 | 22,73 | 23,82 | 35 |
| ENSG00000011275 | RNF216 | 11,43 | 12,44 | 13,37 | 15,39 | 35 |
| ENSG00000055044 | NOP58 | 38,05 | 40,44 | 44,39 | 51,22 | 35 |
| ENSG00000006715 | VPS41 | 23,09 | 28,31 | 28,5 | 31,07 | 35 |
| ENSG00000175470 | PPP2R2D | 17,42 | 18,72 | 20,79 | 23,44 | 35 |
| ENSG00000172379 | ARNT2 | 16,07 | 17,56 | 18,7 | 21,62 | 35 |
| ENSG00000282977 | PCBP2-OT1 | 2,78 | 3 | 3,42 | 3,74 | 35 |
| ENSG00000112335 | SNX3 | 23,4 | 24,58 | 28,63 | 31,46 | 34 |
| ENSG00000251602 | AL928654,1 | 1,8 | 1,94 | 2,21 | 2,42 | 34 |
| ENSG00000139579 | NABP2 | 22,27 | 24,26 | 28,45 | 29,94 | 34 |

| | | | | | | |
|---|---|---|---|---|---|---|
| ENSG00000149531 | FRG1BP | 5,58 | 6,22 | 6,53 | 7,5 | 34 |
| ENSG00000135801 | TAF5L | 9,13 | 9,62 | 11,03 | 12,27 | 34 |
| ENSG00000110107 | PRPF19 | 45,77 | 47,76 | 59,15 | 61,51 | 34 |
| ENSG00000109689 | STIM2 | 9,16 | 9,57 | 10,01 | 12,31 | 34 |
| ENSG00000140262 | TCF12 | 86,88 | 92,19 | 97,38 | 116,75 | 34 |
| ENSG00000142039 | CCDC97 | 11,03 | 11,48 | 12,23 | 14,82 | 34 |
| ENSG00000130449 | ZSWIM6 | 11,76 | 13,72 | 13,86 | 15,8 | 34 |
| ENSG00000128203 | ASPHD2 | 2,62 | 3 | 3,06 | 3,52 | 34 |
| ENSG00000189050 | RNFT1 | 6,99 | 8,01 | 8,1 | 9,39 | 34 |
| ENSG00000136940 | PDCL | 8,39 | 10,2 | 10,59 | 11,27 | 34 |
| ENSG00000124535 | WRNIP1 | 11,19 | 14,24 | 14,27 | 15,02 | 34 |
| ENSG00000143702 | CEP170 | 49 | 60,53 | 63,7 | 65,77 | 34 |
| ENSG00000153443 | UBALD1 | 5,29 | 6,07 | 6,71 | 7,1 | 34 |
| ENSG00000143776 | CDC42BPA | 12,86 | 15,84 | 16,22 | 17,26 | 34 |
| ENSG00000136709 | WDR33 | 20,8 | 22,37 | 23,66 | 27,91 | 34 |
| ENSG00000166847 | DCTN5 | 27,36 | 31,6 | 31,85 | 36,71 | 34 |
| ENSG00000103051 | COG4 | 15,64 | 18,35 | 19,87 | 20,98 | 34 |
| ENSG00000119953 | SMNDC1 | 16,6 | 19,81 | 20,47 | 22,26 | 34 |
| ENSG00000165406 | MARCHF8 | 6,6 | 7,79 | 8,13 | 8,85 | 34 |
| ENSG00000205485 | AC004980,1 | 1,79 | 1,82 | 2,28 | 2,4 | 34 |
| ENSG00000121775 | TMEM39B | 7,99 | 8,31 | 10,25 | 10,71 | 34 |
| ENSG00000184863 | RBM33 | 22,01 | 24,42 | 25,26 | 29,5 | 34 |
| ENSG00000107263 | RAPGEF1 | 17,05 | 19,18 | 22,33 | 22,85 | 34 |
| ENSG00000136950 | ARPC5L | 13,38 | 13,73 | 14,42 | 17,93 | 34 |
| ENSG00000131504 | DIAPH1 | 22,91 | 24,32 | 27,05 | 30,7 | 34 |
| ENSG00000105723 | GSK3A | 22,74 | 23,47 | 27,48 | 30,47 | 34 |
| ENSG00000111711 | GOLT1B | 12,87 | 13,24 | 13,87 | 17,24 | 34 |
| ENSG00000135624 | CCT7 | 109,22 | 110,53 | 138,33 | 146,29 | 34 |
| ENSG00000102038 | SMARCA1 | 30,94 | 35,54 | 36,16 | 41,44 | 34 |
| ENSG00000130204 | TOMM40 | 29,06 | 29,12 | 34,81 | 38,92 | 34 |
| ENSG00000173852 | DPY19L1 | 16,87 | 19,81 | 20,53 | 22,59 | 34 |
| ENSG00000136643 | RPS6KC1 | 8,15 | 9,56 | 9,76 | 10,91 | 34 |
| ENSG00000116750 | UCHL5 | 18,97 | 20,87 | 21,12 | 25,39 | 34 |
| ENSG00000091527 | CDV3 | 62,5 | 72,5 | 78,24 | 83,63 | 34 |
| ENSG00000074842 | MYDGF | 19,07 | 20,05 | 23,16 | 25,51 | 34 |
| ENSG00000165650 | PDZD8 | 9,51 | 10,89 | 11,95 | 12,72 | 34 |
| ENSG00000144021 | CIAO1 | 14,17 | 16,04 | 17,61 | 18,95 | 34 |
| ENSG00000108256 | NUFIP2 | 15,09 | 15,13 | 18,23 | 20,18 | 34 |
| ENSG00000182484 | WASH6P | 20,28 | 21,87 | 25,77 | 27,12 | 34 |
| ENSG00000198890 | PRMT6 | 11,06 | 11,91 | 13,41 | 14,79 | 34 |
| ENSG00000135148 | TRAFD1 | 16,32 | 19,22 | 20,75 | 21,82 | 34 |
| ENSG00000176894 | PXMP2 | 11,94 | 12,3 | 14,42 | 15,96 | 34 |
| ENSG00000082482 | KCNK2 | 2,05 | 2,15 | 2,41 | 2,74 | 34 |
| ENSG00000104320 | NBN | 18,04 | 18,11 | 21,02 | 24,11 | 34 |
| ENSG00000286129 | AC245060,7 | 2,41 | 3,06 | 3,11 | 3,22 | 34 |
| ENSG00000101557 | USP14 | 28,3 | 30,78 | 30,9 | 37,81 | 34 |
| ENSG00000112249 | ASCC3 | 13,26 | 14,64 | 15,26 | 17,71 | 34 |
| ENSG00000127837 | AAMP | 28,97 | 30,86 | 35,18 | 38,68 | 34 |
| ENSG00000136448 | NMT1 | 22,53 | 23,17 | 27,68 | 30,07 | 33 |
| ENSG00000151327 | FAM177A1 | 18,95 | 21,29 | 23,93 | 25,29 | 33 |

| | | | | | | |
|---|---|---|---|---|---|---|
| ENSG00000144567 | RETREG2 | 13,87 | 14,4 | 16,48 | 18,51 | 33 |
| ENSG00000167986 | DDB1 | 67,8 | 76 | 76,96 | 90,45 | 33 |
| ENSG00000070214 | SLC44A1 | 4,7 | 5,48 | 5,81 | 6,27 | 33 |
| ENSG00000176108 | CHMP6 | 9,53 | 9,69 | 10,79 | 12,71 | 33 |
| ENSG00000183520 | UTP11 | 30,76 | 31,14 | 37,52 | 41,02 | 33 |
| ENSG00000100426 | ZBED4 | 11,31 | 13,06 | 14 | 15,08 | 33 |
| ENSG00000114745 | GORASP1 | 11,16 | 13,56 | 13,81 | 14,88 | 33 |
| ENSG00000047849 | MAP4 | 49,1 | 57,46 | 58,52 | 65,46 | 33 |
| ENSG00000157538 | VPS26C | 15,95 | 16,79 | 17,83 | 21,26 | 33 |
| ENSG00000140443 | IGF1R | 22,66 | 25,93 | 26,54 | 30,2 | 33 |
| ENSG00000097033 | SH3GLB1 | 20,75 | 23,97 | 26,47 | 27,65 | 33 |
| ENSG00000152454 | ZNF256 | 6,17 | 6,57 | 6,91 | 8,22 | 33 |
| ENSG00000144791 | LIMD1 | 5,87 | 6,62 | 7,03 | 7,82 | 33 |
| ENSG00000211456 | SACM1L | 15,75 | 18,67 | 19,09 | 20,98 | 33 |
| ENSG00000198642 | KLHL9 | 12,23 | 14,28 | 15,61 | 16,29 | 33 |
| ENSG00000181450 | ZNF678 | 12,75 | 12,92 | 14,59 | 16,98 | 33 |
| ENSG00000168872 | DDX19A | 17,95 | 20,02 | 21,53 | 23,9 | 33 |
| ENSG00000187953 | PMS2CL | 6,19 | 6,67 | 6,79 | 8,24 | 33 |
| ENSG00000138162 | TACC2 | 21,58 | 26,49 | 27,84 | 28,72 | 33 |
| ENSG00000178694 | NSUN3 | 5,08 | 5,2 | 5,7 | 6,76 | 33 |
| ENSG00000168813 | ZNF507 | 9,24 | 9,97 | 10,29 | 12,29 | 33 |
| ENSG00000136875 | PRPF4 | 15,07 | 15,45 | 17,92 | 20,03 | 33 |
| ENSG00000146085 | MMUT | 7,9 | 7,96 | 8,95 | 10,5 | 33 |
| ENSG00000063660 | GPC1 | 16,96 | 18,12 | 22,27 | 22,54 | 33 |
| ENSG00000031698 | SARS1 | 37,4 | 42,29 | 43,42 | 49,7 | 33 |
| ENSG00000125630 | POLR1B | 10,62 | 11,75 | 11,93 | 14,11 | 33 |
| ENSG00000165006 | UBAP1 | 11,94 | 13,17 | 14,59 | 15,86 | 33 |
| ENSG00000183495 | EP400 | 16,79 | 17,84 | 17,99 | 22,3 | 33 |
| ENSG00000131669 | NINJ1 | 6,98 | 7,79 | 7,85 | 9,27 | 33 |
| ENSG00000177125 | ZBTB34 | 5,06 | 5,62 | 5,69 | 6,72 | 33 |
| ENSG00000176148 | TCP11L1 | 8,43 | 9,77 | 10,16 | 11,19 | 33 |
| ENSG00000138613 | APH1B | 3,82 | 4,52 | 4,99 | 5,07 | 33 |
| ENSG00000142082 | SIRT3 | 7,98 | 8,63 | 9,33 | 10,59 | 33 |
| ENSG00000110200 | ANAPC15 | 25,35 | 25,64 | 30,3 | 33,64 | 33 |
| ENSG00000241553 | ARPC4 | 49,63 | 52,87 | 53,14 | 65,86 | 33 |
| ENSG00000083642 | PDS5B | 29,25 | 32,16 | 33,53 | 38,81 | 33 |
| ENSG00000146007 | ZMAT2 | 24,05 | 27,61 | 27,63 | 31,91 | 33 |
| ENSG00000165097 | KDM1B | 7,29 | 7,62 | 9,07 | 9,67 | 33 |
| ENSG00000116539 | ASH1L | 12,13 | 13,4 | 13,93 | 16,09 | 33 |
| ENSG00000157978 | LDLRAP1 | 3,86 | 4,11 | 4,13 | 5,12 | 33 |
| ENSG00000090061 | CCNK | 25,43 | 27,33 | 29,83 | 33,73 | 33 |
| ENSG00000064313 | TAF2 | 15,29 | 15,5 | 19,04 | 20,27 | 33 |
| ENSG00000158882 | TOMM40L | 4,3 | 4,61 | 4,81 | 5,7 | 33 |
| ENSG00000236778 | INTS6-AS1 | 1,72 | 1,79 | 2,09 | 2,28 | 33 |
| ENSG00000276276 | ARL17B | 2,55 | 2,64 | 2,84 | 3,38 | 33 |
| ENSG00000184432 | COPB2 | 37,73 | 43,08 | 44,05 | 50,01 | 33 |
| ENSG00000109576 | AADAT | 13,71 | 14,21 | 15,89 | 18,17 | 33 |
| ENSG00000139726 | DENR | 30,26 | 34,23 | 35,75 | 40,1 | 33 |
| ENSG00000138495 | COX17 | 37,58 | 39,52 | 42,78 | 49,8 | 33 |
| ENSG00000117899 | MESD | 13,75 | 14,77 | 15,99 | 18,22 | 33 |

| | | | | | | |
|---|---|---|---|---|---|---|
| ENSG00000181894 | ZNF329 | 5,63 | 6,1 | 6,74 | 7,46 | 33 |
| ENSG00000083857 | FAT1 | 66,41 | 75,62 | 79,66 | 87,97 | 32 |
| ENSG00000196547 | MAN2A2 | 9,06 | 9,84 | 11,01 | 12 | 32 |
| ENSG00000067182 | TNFRSF1A | 19,79 | 22,16 | 25,7 | 26,21 | 32 |
| ENSG00000132305 | IMMT | 37,92 | 39,76 | 44,65 | 50,22 | 32 |
| ENSG00000076513 | ANKRD13A | 9,25 | 10,95 | 11,5 | 12,25 | 32 |
| ENSG00000148082 | SHC3 | 5,18 | 5,48 | 5,51 | 6,86 | 32 |
| ENSG00000100505 | TRIM9 | 2,9 | 3,27 | 3,33 | 3,84 | 32 |
| ENSG00000115652 | UXS1 | 27,8 | 28,85 | 30,39 | 36,81 | 32 |
| ENSG00000007376 | RPUSD1 | 4,54 | 4,75 | 5,8 | 6,01 | 32 |
| ENSG00000226479 | TMEM185B | 3,83 | 4,23 | 4,76 | 5,07 | 32 |
| ENSG00000083099 | LYRM2 | 19,07 | 20 | 22,82 | 25,24 | 32 |
| ENSG00000075239 | ACAT1 | 17,62 | 18,82 | 20,03 | 23,32 | 32 |
| ENSG00000061794 | MRPS35 | 14,9 | 16,28 | 18,04 | 19,72 | 32 |
| ENSG00000105854 | PON2 | 57,26 | 64,04 | 64,95 | 75,78 | 32 |
| ENSG00000136936 | XPA | 8,69 | 9,72 | 9,85 | 11,5 | 32 |
| ENSG00000180901 | KCTD2 | 10,98 | 12,56 | 14,17 | 14,53 | 32 |
| ENSG00000092201 | SUPT16H | 74,29 | 75,46 | 76,09 | 98,28 | 32 |
| ENSG00000185787 | MORF4L1 | 341 | 378,44 | 400,5 | 451,07 | 32 |
| ENSG00000078668 | VDAC3 | 69,63 | 76,95 | 89,23 | 92,1 | 32 |
| ENSG00000105323 | HNRNPUL1 | 104,9 | 113,58 | 124,95 | 138,74 | 32 |
| ENSG00000213585 | VDAC1 | 86,12 | 91,18 | 96,85 | 113,9 | 32 |
| ENSG00000066422 | ZBTB11 | 5,93 | 6,32 | 6,73 | 7,84 | 32 |
| ENSG00000119383 | PTPA | 35,79 | 36,26 | 39,27 | 47,31 | 32 |
| ENSG00000108588 | CCDC47 | 24,08 | 25,86 | 27,07 | 31,83 | 32 |
| ENSG00000203644 | AC083799,1 | 4,35 | 4,87 | 5,29 | 5,75 | 32 |
| ENSG00000072110 | ACTN1 | 47,86 | 49,17 | 54,63 | 63,25 | 32 |
| ENSG00000108510 | MED13 | 21,24 | 21,46 | 22,14 | 28,07 | 32 |
| ENSG00000185022 | MAFF | 1,4 | 1,47 | 1,54 | 1,85 | 32 |
| ENSG00000147324 | MFHAS1 | 4,67 | 5,02 | 5,81 | 6,17 | 32 |
| ENSG00000286219 | NOTCH2NLC | 7,1 | 8,66 | 9,09 | 9,38 | 32 |
| ENSG00000275066 | SYNRG | 9,38 | 11,8 | 12,26 | 12,39 | 32 |
| ENSG00000198937 | CCDC167 | 39,03 | 40,08 | 49,49 | 51,55 | 32 |
| ENSG00000115685 | PPP1R7 | 27,32 | 28,16 | 32,28 | 36,08 | 32 |
| ENSG00000171566 | PLRG1 | 36,49 | 36,73 | 39,24 | 48,19 | 32 |
| ENSG00000286873 | AC012306,3 | 1,84 | 2,18 | 2,21 | 2,43 | 32 |
| ENSG00000136813 | ECPAS | 25,35 | 29,01 | 32,35 | 33,47 | 32 |
| ENSG00000130349 | MTRES1 | 9,13 | 10,42 | 11,18 | 12,05 | 32 |
| ENSG00000255302 | EID1 | 92,68 | 104,58 | 105,85 | 122,32 | 32 |
| ENSG00000171853 | TRAPPC12 | 19,45 | 20,86 | 21,14 | 25,67 | 32 |
| ENSG00000156697 | UTP14A | 8,82 | 9,41 | 11,43 | 11,64 | 32 |
| ENSG00000196782 | MAML3 | 4,41 | 5,01 | 5,48 | 5,82 | 32 |
| ENSG00000120616 | EPC1 | 15,33 | 17,51 | 18,52 | 20,23 | 32 |
| ENSG00000013288 | MAN2B2 | 1,69 | 1,9 | 2,13 | 2,23 | 32 |
| ENSG00000039123 | MTREX | 36,59 | 41,06 | 42,84 | 48,28 | 32 |
| ENSG00000167005 | NUDT21 | 57,19 | 62,44 | 69,44 | 75,46 | 32 |
| ENSG00000102931 | ARL2BP | 30,12 | 34,29 | 38,31 | 39,73 | 32 |
| ENSG00000095059 | DHPS | 22,41 | 22,58 | 23,6 | 29,56 | 32 |
| ENSG00000103855 | CD276 | 54,11 | 59,78 | 69,54 | 71,37 | 32 |
| ENSG00000103994 | ZNF106 | 17,95 | 19,06 | 20,46 | 23,67 | 32 |

| | | | | | | |
|---|---|---|---|---|---|---|
| ENSG00000204138 | PHACTR4 | 25,16 | 28,93 | 29,05 | 33,17 | 32 |
| ENSG00000102710 | SUPT20H | 36,61 | 39,52 | 41,51 | 48,26 | 32 |
| ENSG00000122565 | CBX3 | 96,3 | 98,57 | 102,6 | 126,93 | 32 |
| ENSG00000124155 | PIGT | 23,06 | 23,55 | 30,09 | 30,39 | 32 |
| ENSG00000106261 | ZKSCAN1 | 27,47 | 35,61 | 35,76 | 36,2 | 32 |
| ENSG00000132963 | POMP | 37,76 | 40,69 | 44,27 | 49,76 | 32 |
| ENSG00000172661 | WASHC2C | 11,58 | 12,97 | 13,52 | 15,26 | 32 |
| ENSG00000104853 | CLPTM1 | 23,55 | 25,53 | 29,5 | 31,03 | 32 |
| ENSG00000126883 | NUP214 | 18,5 | 18,6 | 20,29 | 24,37 | 32 |
| ENSG00000176915 | ANKLE2 | 26,19 | 27,16 | 28,38 | 34,5 | 32 |
| ENSG00000168538 | TRAPPC11 | 15,17 | 16,98 | 18,34 | 19,98 | 32 |
| ENSG00000129187 | DCTD | 25,95 | 28,11 | 33,43 | 34,17 | 32 |
| ENSG00000170185 | USP38 | 5,18 | 6,51 | 6,59 | 6,82 | 32 |
| ENSG00000145476 | CYP4V2 | 3,76 | 4,35 | 4,64 | 4,95 | 32 |
| ENSG00000237649 | KIFC1 | 32,01 | 32,28 | 33,04 | 42,14 | 32 |
| ENSG00000174231 | PRPF8 | 74,83 | 79,98 | 86,8 | 98,51 | 32 |
| ENSG00000175792 | RUVBL1 | 17,9 | 18,35 | 19,38 | 23,56 | 32 |
| ENSG00000167081 | PBX3 | 28,53 | 29,34 | 31,08 | 37,55 | 32 |
| ENSG00000015171 | ZMYND11 | 17,62 | 19,01 | 22,75 | 23,19 | 32 |
| ENSG00000160785 | SLC25A44 | 5,29 | 6,08 | 6,4 | 6,96 | 32 |
| ENSG00000062485 | CS | 66,34 | 71,73 | 78,8 | 87,28 | 32 |
| ENSG00000134758 | RNF138 | 25,78 | 27,68 | 28,22 | 33,91 | 32 |
| ENSG00000165671 | NSD1 | 34,76 | 37,74 | 38,77 | 45,72 | 32 |
| ENSG00000166435 | XRRA1 | 11,02 | 11,68 | 12,66 | 14,49 | 31 |
| ENSG00000166333 | ILK | 41,07 | 44,12 | 52,74 | 54 | 31 |
| ENSG00000100380 | ST13 | 94,38 | 102,24 | 120,37 | 124,06 | 31 |
| ENSG00000143368 | SF3B4 | 33,7 | 36,97 | 42,11 | 44,29 | 31 |
| ENSG00000099992 | TBC1D10A | 2,26 | 2,61 | 2,96 | 2,97 | 31 |
| ENSG00000244045 | TMEM199 | 8,03 | 8,98 | 10,18 | 10,55 | 31 |
| ENSG00000114331 | ACAP2 | 20,81 | 23,41 | 25,42 | 27,34 | 31 |
| ENSG00000077721 | UBE2A | 22,56 | 22,84 | 26,71 | 29,63 | 31 |
| ENSG00000168439 | STIP1 | 92,83 | 93,63 | 99,89 | 121,92 | 31 |
| ENSG00000117262 | GPR89A | 9,1 | 9,72 | 11,66 | 11,95 | 31 |
| ENSG00000170027 | YWHAG | 62,68 | 72,8 | 75,09 | 82,31 | 31 |
| ENSG00000156256 | USP16 | 17,53 | 18,31 | 20,04 | 23,02 | 31 |
| ENSG00000023330 | ALAS1 | 13,1 | 13,22 | 15,83 | 17,2 | 31 |
| ENSG00000105401 | CDC37 | 46,64 | 48,28 | 54,04 | 61,23 | 31 |
| ENSG00000108443 | RPS6KB1 | 12,19 | 12,99 | 14,57 | 16 | 31 |
| ENSG00000110321 | EIF4G2 | 320,42 | 352,01 | 403 | 420,34 | 31 |
| ENSG00000124786 | SLC35B3 | 4,17 | 4,44 | 4,91 | 5,47 | 31 |
| ENSG00000227199 | ST7-AS1 | 1,54 | 1,62 | 1,98 | 2,02 | 31 |
| ENSG00000187123 | LYPD6 | 3,66 | 4,19 | 4,71 | 4,8 | 31 |
| ENSG00000071553 | ATP6AP1 | 19,88 | 22,23 | 26,03 | 26,07 | 31 |
| ENSG00000172757 | CFL1 | 505,07 | 530,01 | 636,44 | 662,19 | 31 |
| ENSG00000123473 | STIL | 12,93 | 13,83 | 14,73 | 16,95 | 31 |
| ENSG00000120802 | TMPO | 148,69 | 154,79 | 169,71 | 194,89 | 31 |
| ENSG00000164934 | DCAF13 | 30,94 | 31,49 | 37,98 | 40,54 | 31 |
| ENSG00000156599 | ZDHHC5 | 13,38 | 16,69 | 17,07 | 17,53 | 31 |
| ENSG00000129245 | FXR2 | 13,8 | 14,72 | 15,05 | 18,08 | 31 |
| ENSG00000101294 | HM13 | 33,97 | 36,47 | 39,64 | 44,5 | 31 |

| | | | | | | |
|---|---|---|---|---|---|---|
| ENSG00000101189 | MRGBP | 7,78 | 8,54 | 9,05 | 10,19 | 31 |
| ENSG00000157916 | RER1 | 34,74 | 35,68 | 36,47 | 45,5 | 31 |
| ENSG00000213930 | GALT | 11,69 | 12,58 | 13,5 | 15,31 | 31 |
| ENSG00000181929 | PRKAG1 | 16,43 | 19,04 | 20,01 | 21,51 | 31 |
| ENSG00000139826 | ABHD13 | 3,72 | 3,85 | 3,86 | 4,87 | 31 |
| ENSG00000135956 | TMEM127 | 6,6 | 7,93 | 8,5 | 8,64 | 31 |
| ENSG00000109118 | PHF12 | 19,16 | 19,75 | 23,87 | 25,08 | 31 |
| ENSG00000048405 | ZNF800 | 11,25 | 12,41 | 13,92 | 14,72 | 31 |
| ENSG00000184634 | MED12 | 11,78 | 12,35 | 13,64 | 15,41 | 31 |
| ENSG00000244026 | FAM86DP | 5,29 | 5,59 | 5,84 | 6,92 | 31 |
| ENSG00000288323 | FAM86DP | 5,29 | 5,59 | 5,84 | 6,92 | 31 |
| ENSG00000135597 | REPS1 | 23,45 | 25,24 | 25,48 | 30,67 | 31 |
| ENSG00000166889 | PATL1 | 10,17 | 10,36 | 11,59 | 13,3 | 31 |
| ENSG00000116062 | MSH6 | 53,48 | 56,24 | 62,68 | 69,92 | 31 |
| ENSG00000109971 | HSPA8 | 394,63 | 436,19 | 515,54 | 515,92 | 31 |
| ENSG00000001629 | ANKIB1 | 15,52 | 19,41 | 19,94 | 20,29 | 31 |
| ENSG00000033100 | CHPF2 | 6,41 | 7,12 | 7,64 | 8,38 | 31 |
| ENSG00000247626 | MARS2 | 2,67 | 2,72 | 2,84 | 3,49 | 31 |
| ENSG00000138594 | TMOD3 | 12,83 | 15,17 | 16,14 | 16,77 | 31 |
| ENSG00000073584 | SMARCE1 | 102 | 119,77 | 122,5 | 133,3 | 31 |
| ENSG00000119725 | ZNF410 | 18,22 | 18,3 | 21,07 | 23,81 | 31 |
| ENSG00000104325 | DECR1 | 51,82 | 60,27 | 60,94 | 67,7 | 31 |
| ENSG00000179335 | CLK3 | 22,59 | 23,5 | 24,57 | 29,51 | 31 |
| ENSG00000189376 | C8orf76 | 12,96 | 13,94 | 15,45 | 16,93 | 31 |
| ENSG00000116688 | MFN2 | 21,55 | 23,51 | 25,4 | 28,15 | 31 |
| ENSG00000251474 | RPL32P3 | 8 | 8,03 | 8,22 | 10,45 | 31 |
| ENSG00000049656 | CLPTM1L | 26,62 | 30,02 | 32,09 | 34,77 | 31 |
| ENSG00000139998 | RAB15 | 10,55 | 11,61 | 12,5 | 13,78 | 31 |
| ENSG00000178922 | HYI | 9,94 | 10,37 | 12,12 | 12,98 | 31 |
| ENSG00000159069 | FBXW5 | 12,17 | 13,45 | 15,07 | 15,89 | 31 |
| ENSG00000120053 | GOT1 | 13,13 | 13,9 | 14,96 | 17,14 | 31 |
| ENSG00000110801 | PSMD9 | 19,97 | 22,68 | 24,46 | 26,06 | 30 |
| ENSG00000260804 | LINC01963 | 7,48 | 8,02 | 8,65 | 9,76 | 30 |
| ENSG00000144028 | SNRNP200 | 68,12 | 70,3 | 74,46 | 88,88 | 30 |
| ENSG00000085377 | PREP | 14,22 | 15,78 | 16,81 | 18,55 | 30 |
| ENSG00000078319 | PMS2P1 | 26,93 | 28,04 | 30,52 | 35,13 | 30 |
| ENSG00000102743 | SLC25A15 | 9,43 | 9,98 | 10,95 | 12,3 | 30 |
| ENSG00000103150 | MLYCD | 1,61 | 1,7 | 1,82 | 2,1 | 30 |
| ENSG00000140395 | WDR61 | 26,24 | 27,1 | 32,18 | 34,22 | 30 |
| ENSG00000096401 | CDC5L | 17,08 | 18,29 | 19,43 | 22,27 | 30 |
| ENSG00000178127 | NDUFV2 | 78,51 | 79,76 | 87,27 | 102,36 | 30 |
| ENSG00000183684 | ALYREF | 62,15 | 64,64 | 80,53 | 81,02 | 30 |
| ENSG00000108344 | PSMD3 | 40,43 | 41,09 | 44,35 | 52,7 | 30 |
| ENSG00000196110 | ZNF699 | 2,34 | 2,46 | 2,52 | 3,05 | 30 |
| ENSG00000078674 | PCM1 | 88,04 | 91,45 | 96,68 | 114,75 | 30 |
| ENSG00000160194 | NDUFV3 | 10,75 | 11,34 | 12,15 | 14,01 | 30 |
| ENSG00000116157 | GPX7 | 15,31 | 16,72 | 19,09 | 19,95 | 30 |
| ENSG00000177119 | ANO6 | 12,18 | 12,43 | 12,73 | 15,87 | 30 |
| ENSG00000213024 | NUP62 | 27,58 | 29,42 | 32,5 | 35,93 | 30 |
| ENSG00000127957 | PMS2P3 | 7,08 | 7,62 | 7,76 | 9,22 | 30 |

| | | | | | | |
|---|---|---|---|---|---|---|
| ENSG00000265241 | RBM8A | 113,85 | 121,2 | 147,61 | 148,26 | 30 |
| ENSG00000152291 | TGOLN2 | 16,08 | 18,23 | 20,25 | 20,94 | 30 |
| ENSG00000168385 | SEPTIN2 | 170,27 | 176,26 | 179,17 | 221,64 | 30 |
| ENSG00000112294 | ALDH5A1 | 4,21 | 4,53 | 4,81 | 5,48 | 30 |
| ENSG00000165495 | PKNOX2 | 10,51 | 12,7 | 12,99 | 13,68 | 30 |
| ENSG00000176371 | ZSCAN2 | 7,54 | 8,02 | 8,91 | 9,81 | 30 |
| ENSG00000172354 | GNB2 | 61,08 | 61,96 | 70,02 | 79,45 | 30 |
| ENSG00000086061 | DNAJA1 | 115,04 | 139,41 | 146,48 | 149,63 | 30 |
| ENSG00000215154 | AC141586,1 | 4,59 | 5,29 | 5,48 | 5,97 | 30 |
| ENSG00000178896 | EXOSC4 | 8,79 | 10 | 11,13 | 11,43 | 30 |
| ENSG00000104915 | STX10 | 11,49 | 11,98 | 14,3 | 14,94 | 30 |
| ENSG00000261578 | AP003119,3 | 1,1 | 1,22 | 1,25 | 1,43 | 30 |
| ENSG00000090382 | LYZ | 1,1 | 1,18 | 1,25 | 1,43 | 30 |
| ENSG00000099246 | RAB18 | 32,7 | 37,77 | 38,19 | 42,5 | 30 |
| ENSG00000067560 | RHOA | 173,19 | 179,36 | 215,56 | 225,08 | 30 |
| ENSG00000151348 | EXT2 | 20,14 | 22,56 | 25,85 | 26,17 | 30 |
| ENSG00000119682 | AREL1 | 14,23 | 14,35 | 15,26 | 18,48 | 30 |
| ENSG00000196177 | ACADSB | 4,99 | 5,37 | 5,5 | 6,48 | 30 |
| ENSG00000116285 | ERRFI1 | 16,68 | 17,28 | 18,2 | 21,66 | 30 |
| ENSG00000241973 | PI4KA | 12,28 | 12,5 | 13,97 | 15,94 | 30 |
| ENSG00000068796 | KIF2A | 38,17 | 41,82 | 43,45 | 49,54 | 30 |
| ENSG00000223802 | CERS1 | 6,11 | 7,36 | 7,68 | 7,93 | 30 |
| ENSG00000117505 | DR1 | 17,87 | 19,23 | 19,91 | 23,19 | 30 |
| ENSG00000010017 | RANBP9 | 13,92 | 16,59 | 16,91 | 18,06 | 30 |
| ENSG00000145041 | DCAF1 | 8,58 | 9,19 | 9,35 | 11,13 | 30 |
| ENSG00000276111 | SDCCAG8 | 3,03 | 3,27 | 3,75 | 3,93 | 30 |
| ENSG00000101577 | LPIN2 | 8,96 | 9,73 | 10,06 | 11,62 | 30 |
| ENSG00000116685 | KIAA2013 | 7,11 | 7,16 | 8,09 | 9,22 | 30 |
| ENSG00000186665 | C17orf58 | 4,79 | 5,49 | 6,2 | 6,21 | 30 |
| ENSG00000198369 | SPRED2 | 19,33 | 20,18 | 23,08 | 25,06 | 30 |
| ENSG00000149187 | CELF1 | 43,13 | 46,63 | 50,9 | 55,91 | 30 |
| ENSG00000116489 | CAPZA1 | 69,43 | 73,48 | 78,74 | 89,98 | 30 |
| ENSG00000068323 | TFE3 | 8,05 | 8,7 | 9,06 | 10,43 | 30 |
| ENSG00000142186 | SCYL1 | 20,1 | 21,8 | 24,69 | 26,04 | 30 |
| ENSG00000134440 | NARS1 | 47,07 | 53,77 | 56,37 | 60,98 | 30 |
| ENSG00000198786 | MT-ND5 | 976,91 | 1121,76 | 1156,41 | 1265,58 | 30 |
| ENSG00000166197 | NOLC1 | 29,21 | 29,99 | 30,99 | 37,84 | 30 |
| ENSG00000125447 | GGA3 | 12,9 | 14,25 | 14,44 | 16,71 | 30 |
| ENSG00000182628 | SKA2 | 81,51 | 82,44 | 93,87 | 105,57 | 30 |
| ENSG00000162065 | TBC1D24 | 3,12 | 3,35 | 3,7 | 4,04 | 29 |
| ENSG00000140990 | NDUFB10 | 58,32 | 67,01 | 74,44 | 75,5 | 29 |
| ENSG00000134824 | FADS2 | 149,63 | 157,3 | 173,11 | 193,67 | 29 |
| ENSG00000165280 | VCP | 99,64 | 100,22 | 107,6 | 128,96 | 29 |
| ENSG00000106144 | CASP2 | 17,74 | 19,67 | 22,79 | 22,96 | 29 |
| ENSG00000135316 | SYNCRIP | 65,36 | 71,34 | 75,26 | 84,58 | 29 |
| ENSG00000107929 | LARP4B | 15,41 | 15,89 | 17,16 | 19,94 | 29 |
| ENSG00000107949 | BCCIP | 41,65 | 42,98 | 49,29 | 53,89 | 29 |
| ENSG00000113387 | SUB1 | 119,75 | 122,17 | 152,69 | 154,93 | 29 |
| ENSG00000028203 | VEZT | 33,1 | 38,28 | 39,16 | 42,82 | 29 |
| ENSG00000095906 | NUBP2 | 14,54 | 14,9 | 18,58 | 18,8 | 29 |

| | | | | | | |
|---|---|---|---|---|---|---|
| ENSG00000130726 | TRIM28 | 252,98 | 261,58 | 280,67 | 327,08 | 29 |
| ENSG00000126261 | UBA2 | 84,95 | 87,15 | 93,88 | 109,83 | 29 |
| ENSG00000116874 | WARS2 | 4,61 | 4,67 | 5,03 | 5,96 | 29 |
| ENSG00000083520 | DIS3 | 14,39 | 14,6 | 15,87 | 18,6 | 29 |
| ENSG00000170471 | RALGAPB | 14,17 | 15,02 | 16,16 | 18,3 | 29 |
| ENSG00000119280 | C1orf198 | 26,85 | 27,49 | 30,02 | 34,67 | 29 |
| ENSG00000176171 | BNIP3 | 7,28 | 7,81 | 8,65 | 9,4 | 29 |
| ENSG00000111615 | KRR1 | 20,82 | 22,85 | 25,35 | 26,88 | 29 |
| ENSG00000158623 | COPG2 | 22,93 | 24,33 | 25,6 | 29,6 | 29 |
| ENSG00000062716 | VMP1 | 28,66 | 34,13 | 35,95 | 36,99 | 29 |
| ENSG00000070831 | CDC42 | 83,44 | 97,91 | 98,91 | 107,69 | 29 |
| ENSG00000113742 | CPEB4 | 4,37 | 5,23 | 5,59 | 5,64 | 29 |
| ENSG00000169016 | E2F6 | 11,19 | 11,51 | 12,79 | 14,44 | 29 |
| ENSG00000000419 | DPM1 | 26,55 | 27,93 | 29,29 | 34,26 | 29 |
| ENSG00000100697 | DICER1 | 34,51 | 34,64 | 35,55 | 44,53 | 29 |
| ENSG00000100614 | PPM1A | 15,95 | 16,38 | 17,69 | 20,58 | 29 |
| ENSG00000153879 | CEBPG | 12,3 | 12,39 | 13,25 | 15,87 | 29 |
| ENSG00000010322 | NISCH | 23,69 | 26,09 | 26,14 | 30,56 | 29 |
| ENSG00000151718 | WWC2 | 12,12 | 12,57 | 14,17 | 15,63 | 29 |
| ENSG00000163161 | ERCC3 | 18,72 | 20,89 | 21 | 24,14 | 29 |
| ENSG00000136937 | NCBP1 | 20,24 | 23 | 24,08 | 26,1 | 29 |
| ENSG00000056097 | ZFR | 34 | 39,46 | 39,8 | 43,84 | 29 |
| ENSG00000066697 | MSANTD3 | 19,74 | 21,08 | 24,71 | 25,45 | 29 |
| ENSG00000141068 | KSR1 | 7,85 | 8,65 | 9,55 | 10,12 | 29 |
| ENSG00000148606 | POLR3A | 10,73 | 11,13 | 11,18 | 13,83 | 29 |
| ENSG00000101474 | APMAP | 25,27 | 26,99 | 29,53 | 32,57 | 29 |
| ENSG00000110395 | CBL | 15,41 | 17,31 | 17,57 | 19,86 | 29 |
| ENSG00000150779 | TIMM8B | 49,33 | 54,96 | 55,15 | 63,56 | 29 |
| ENSG00000185049 | NELFA | 14,53 | 15,83 | 17,22 | 18,72 | 29 |
| ENSG00000176542 | USF3 | 5,76 | 6,18 | 6,24 | 7,42 | 29 |
| ENSG00000213347 | MXD3 | 9,49 | 9,9 | 11,09 | 12,22 | 29 |
| ENSG00000262664 | OVCA2 | 8,35 | 9,08 | 10,13 | 10,75 | 29 |
| ENSG00000185753 | CXorf38 | 2,68 | 3,12 | 3,17 | 3,45 | 29 |
| ENSG00000164494 | PDSS2 | 3,9 | 4,71 | 4,83 | 5,02 | 29 |
| ENSG00000214293 | APTR | 3,24 | 3,28 | 3,3 | 4,17 | 29 |
| ENSG00000101442 | ACTR5 | 3,31 | 3,58 | 4,04 | 4,26 | 29 |
| ENSG00000124802 | EEF1E1 | 24,22 | 25,9 | 26,68 | 31,17 | 29 |
| ENSG00000141642 | ELAC1 | 6,98 | 7,52 | 8,67 | 8,98 | 29 |
| ENSG00000272391 | POM121C | 21,08 | 22,35 | 25,71 | 27,12 | 29 |
| ENSG00000107771 | CCSER2 | 18,85 | 21,83 | 23,47 | 24,25 | 29 |
| ENSG00000120686 | UFM1 | 17,14 | 17,76 | 19,67 | 22,05 | 29 |
| ENSG00000162607 | USP1 | 38,22 | 39,41 | 43,01 | 49,16 | 29 |
| ENSG00000160999 | SH2B2 | 2,76 | 2,8 | 3,39 | 3,55 | 29 |
| ENSG00000114520 | SNX4 | 16,98 | 18,84 | 21,46 | 21,84 | 29 |
| ENSG00000162909 | CAPN2 | 41,8 | 41,98 | 52,92 | 53,76 | 29 |
| ENSG00000066654 | THUMPD1 | 18,14 | 18,6 | 21,24 | 23,33 | 29 |
| ENSG00000162613 | FUBP1 | 88,45 | 95,94 | 96,65 | 113,75 | 29 |
| ENSG00000277147 | LINC00869 | 5,07 | 5,13 | 5,21 | 6,52 | 29 |
| ENSG00000110066 | KMT5B | 25,95 | 30,17 | 30,33 | 33,37 | 29 |
| ENSG00000182158 | CREB3L2 | 9,55 | 10,06 | 10,55 | 12,28 | 29 |

| | | | | | | |
|---|---|---|---|---|---|---|
| ENSG00000076924 | XAB2 | 9,97 | 10,86 | 11,96 | 12,82 | 29 |
| ENSG00000153140 | CETN3 | 23,86 | 25,03 | 26,01 | 30,68 | 29 |
| ENSG00000158470 | B4GALT5 | 17,88 | 19,56 | 20,74 | 22,99 | 29 |
| ENSG00000224109 | CENPVL3 | 2,38 | 2,47 | 2,88 | 3,06 | 29 |
| ENSG00000113407 | TARS1 | 49,41 | 51,37 | 57,64 | 63,52 | 29 |
| ENSG00000132676 | DAP3 | 37,96 | 40,3 | 44,72 | 48,8 | 29 |
| ENSG00000187735 | TCEA1 | 41,85 | 43,04 | 48,09 | 53,8 | 29 |
| ENSG00000142669 | SH3BGRL3 | 64,86 | 66,56 | 77,98 | 83,35 | 29 |
| ENSG00000197479 | PCDHB11 | 1,58 | 1,68 | 1,81 | 2,03 | 28 |
| ENSG00000149532 | CPSF7 | 37,4 | 38,51 | 43,14 | 48,05 | 28 |
| ENSG00000152795 | HNRNPDL | 243,2 | 253,34 | 295,27 | 312,43 | 28 |
| ENSG00000179134 | SAMD4B | 49,4 | 55,67 | 59,25 | 63,45 | 28 |
| ENSG00000136732 | GYPC | 27,15 | 29,24 | 29,33 | 34,87 | 28 |
| ENSG00000100226 | GTPBP1 | 21,95 | 22,53 | 26,92 | 28,19 | 28 |
| ENSG00000127511 | SIN3B | 12,7 | 12,75 | 13,79 | 16,31 | 28 |
| ENSG00000169131 | ZNF354A | 7,29 | 7,9 | 8,5 | 9,36 | 28 |
| ENSG00000105953 | OGDH | 29,8 | 32,61 | 36,22 | 38,26 | 28 |
| ENSG00000108219 | TSPAN14 | 31,38 | 34,02 | 37,69 | 40,28 | 28 |
| ENSG00000111676 | ATN1 | 17,89 | 19,25 | 20,47 | 22,96 | 28 |
| ENSG00000129562 | DAD1 | 56,6 | 58,12 | 70,84 | 72,64 | 28 |
| ENSG00000183496 | MEX3B | 12,29 | 12,46 | 13,83 | 15,77 | 28 |
| ENSG00000115816 | CEBPZ | 22,02 | 22,71 | 23,75 | 28,25 | 28 |
| ENSG00000111832 | RWDD1 | 22,34 | 25,08 | 28,41 | 28,66 | 28 |
| ENSG00000234224 | TMEM229A | 0,99 | 1,05 | 1,18 | 1,27 | 28 |
| ENSG00000150456 | EEF1AKMT1 | 6,86 | 7,83 | 8,03 | 8,8 | 28 |
| ENSG00000062598 | ELMO2 | 17,54 | 18,39 | 21,33 | 22,5 | 28 |
| ENSG00000104886 | PLEKHJ1 | 25,19 | 25,93 | 31,72 | 32,31 | 28 |
| ENSG00000143486 | EIF2D | 10,31 | 11,7 | 11,8 | 13,22 | 28 |
| ENSG00000143373 | ZNF687 | 8,15 | 9,21 | 9,5 | 10,45 | 28 |
| ENSG00000171811 | CFAP46 | 1,56 | 1,69 | 1,9 | 2 | 28 |
| ENSG00000173727 | AP000769,1 | 1,56 | 1,88 | 1,96 | 2 | 28 |
| ENSG00000198911 | SREBF2 | 58,48 | 69,33 | 72,07 | 74,96 | 28 |
| ENSG00000113648 | MACROH2A1 | 150,96 | 163,75 | 165,19 | 193,46 | 28 |
| ENSG00000118894 | EEF2KMT | 8,29 | 8,83 | 9,6 | 10,62 | 28 |
| ENSG00000065802 | ASB1 | 6,98 | 7,03 | 7,68 | 8,94 | 28 |
| ENSG00000288307 | CTDNEP1 | 18,56 | 19,89 | 23,18 | 23,77 | 28 |
| ENSG00000175826 | CTDNEP1 | 18,56 | 19,89 | 23,18 | 23,77 | 28 |
| ENSG00000282813 | ARMC10 | 6,77 | 7,75 | 8,02 | 8,67 | 28 |
| ENSG00000139746 | RBM26 | 28,33 | 28,7 | 29,78 | 36,28 | 28 |
| ENSG00000167088 | SNRPD1 | 73,68 | 84,01 | 89,6 | 94,35 | 28 |
| ENSG00000155959 | VBP1 | 37,77 | 41,63 | 44,58 | 48,36 | 28 |
| ENSG00000089723 | OTUB2 | 1,82 | 1,95 | 2,02 | 2,33 | 28 |
| ENSG00000125962 | ARMCX5 | 6,18 | 7,1 | 7,75 | 7,91 | 28 |
| ENSG00000131089 | ARHGEF9 | 15,97 | 18,6 | 19,7 | 20,44 | 28 |
| ENSG00000153560 | UBP1 | 23,12 | 23,31 | 24,43 | 29,59 | 28 |
| ENSG00000279047 | AC244517,5 | 10,15 | 11,43 | 12,79 | 12,99 | 28 |
| ENSG00000102103 | PQBP1 | 26,68 | 27,93 | 30,75 | 34,14 | 28 |
| ENSG00000175203 | DCTN2 | 61,38 | 68,97 | 75,94 | 78,54 | 28 |
| ENSG00000100823 | APEX1 | 119,7 | 122,59 | 133,07 | 153,16 | 28 |
| ENSG00000105647 | PIK3R2 | 14,92 | 14,99 | 16,41 | 19,09 | 28 |

| | | | | | | |
|---|---|---|---|---|---|---|
| ENSG00000201098 | RNY1 | 2073,89 | 2423,5 | 2474,98 | 2652,65 | 28 |
| ENSG00000185085 | INTS5 | 4,3 | 4,83 | 5,14 | 5,5 | 28 |
| ENSG00000122687 | MRM2 | 12,51 | 12,79 | 13,52 | 16 | 28 |
| ENSG00000074582 | BCS1L | 12,45 | 12,63 | 13,08 | 15,92 | 28 |
| ENSG00000176624 | MEX3C | 23,8 | 25,19 | 27,68 | 30,43 | 28 |
| ENSG00000131171 | SH3BGRL | 45,57 | 51,14 | 53,28 | 58,25 | 28 |
| ENSG00000204934 | ATP6V0E2-AS1 | 1,26 | 1,47 | 1,53 | 1,61 | 28 |
| ENSG00000120334 | CENPL | 8,03 | 8,51 | 8,54 | 10,26 | 28 |
| ENSG00000169375 | SIN3A | 29,24 | 30,15 | 30,57 | 37,36 | 28 |
| ENSG00000164823 | OSGIN2 | 4,18 | 4,39 | 4,86 | 5,34 | 28 |
| ENSG00000115649 | CNPPD1 | 12,11 | 12,51 | 13,69 | 15,47 | 28 |
| ENSG00000125841 | NRSN2 | 10,13 | 11,75 | 12,72 | 12,94 | 28 |
| ENSG00000118523 | CCN2 | 35,74 | 35,81 | 38,25 | 45,65 | 28 |
| ENSG00000160679 | CHTOP | 44,12 | 45,84 | 46,32 | 56,34 | 28 |
| ENSG00000112685 | EXOC2 | 9,79 | 10,11 | 12,08 | 12,5 | 28 |
| ENSG00000102225 | CDK16 | 80,88 | 89,79 | 96,81 | 103,26 | 28 |
| ENSG00000132275 | RRP8 | 5,82 | 6,7 | 7,01 | 7,43 | 28 |
| ENSG00000113272 | THG1L | 6,11 | 6,47 | 7,6 | 7,8 | 28 |
| ENSG00000066044 | ELAVL1 | 61,45 | 64,21 | 70,37 | 78,44 | 28 |
| ENSG00000146063 | TRIM41 | 3,51 | 3,78 | 3,82 | 4,48 | 28 |
| ENSG00000113360 | DROSHA | 37,04 | 39,21 | 40,48 | 47,27 | 28 |
| ENSG00000116863 | ADPRS | 7,97 | 8,47 | 9,39 | 10,17 | 28 |
| ENSG00000132589 | FLOT2 | 27,07 | 27,88 | 28,69 | 34,54 | 28 |
| ENSG00000130818 | ZNF426 | 8,59 | 9,65 | 10,22 | 10,96 | 28 |
| ENSG00000065135 | GNAI3 | 13,16 | 14,35 | 14,51 | 16,79 | 28 |
| ENSG00000188483 | IER5L | 5,84 | 6,29 | 6,93 | 7,45 | 28 |
| ENSG00000229337 | AC079305,2 | 0,98 | 1,01 | 1,2 | 1,25 | 28 |
| ENSG00000154305 | MIA3 | 18,81 | 22,73 | 23,74 | 23,99 | 28 |
| ENSG00000100722 | ZC3H14 | 37,39 | 38,37 | 41,27 | 47,68 | 28 |
| ENSG00000155957 | TMBIM4 | 24,39 | 26,57 | 29,38 | 31,1 | 28 |
| ENSG00000137337 | MDC1 | 18,53 | 19,01 | 19,67 | 23,62 | 27 |
| ENSG00000164081 | TEX264 | 5,39 | 5,99 | 6,01 | 6,87 | 27 |
| ENSG00000118402 | ELOVL4 | 3,57 | 4,3 | 4,32 | 4,55 | 27 |
| ENSG00000100567 | PSMA3 | 86,64 | 89,53 | 91,85 | 110,42 | 27 |
| ENSG00000075618 | FSCN1 | 115,39 | 123,41 | 139,73 | 147,04 | 27 |
| ENSG00000198561 | CTNND1 | 35,09 | 35,43 | 38,45 | 44,71 | 27 |
| ENSG00000154723 | ATP5PF | 61,33 | 67,47 | 75,29 | 78,14 | 27 |
| ENSG00000112640 | PPP2R5D | 26,12 | 28,23 | 29,39 | 33,27 | 27 |
| ENSG00000197857 | ZNF44 | 4,46 | 4,51 | 4,65 | 5,68 | 27 |
| ENSG00000247556 | OIP5-AS1 | 31,55 | 34,39 | 35,71 | 40,18 | 27 |
| ENSG00000233237 | LINC00472 | 11,85 | 12,44 | 13,44 | 15,09 | 27 |
| ENSG00000004142 | POLDIP2 | 20,82 | 21,21 | 25,08 | 26,51 | 27 |
| ENSG00000164258 | NDUFS4 | 48,39 | 51,89 | 52,4 | 61,61 | 27 |
| ENSG00000101367 | MAPRE1 | 64,29 | 64,46 | 65,87 | 81,85 | 27 |
| ENSG00000167315 | ACAA2 | 40,91 | 46,83 | 48,42 | 52,08 | 27 |
| ENSG00000154719 | MRPL39 | 20,64 | 21,19 | 21,47 | 26,27 | 27 |
| ENSG00000157881 | PANK4 | 2,57 | 2,78 | 2,83 | 3,27 | 27 |
| ENSG00000108528 | SLC25A11 | 19,41 | 20,48 | 23,55 | 24,69 | 27 |
| ENSG00000122729 | ACO1 | 11,51 | 12,47 | 13,07 | 14,64 | 27 |
| ENSG00000151923 | TIAL1 | 50,07 | 51,21 | 55,78 | 63,68 | 27 |

| | | | | | | |
|---|---|---|---|---|---|---|
| ENSG00000011451 | WIZ | 17,55 | 18,21 | 21,9 | 22,32 | 27 |
| ENSG00000165169 | DYNLT3 | 3,94 | 4,3 | 4,75 | 5,01 | 27 |
| ENSG00000136231 | IGF2BP3 | 53,05 | 57,13 | 60,21 | 67,45 | 27 |
| ENSG00000100784 | RPS6KA5 | 2,14 | 2,17 | 2,36 | 2,72 | 27 |
| ENSG00000099904 | ZDHHC8 | 10,15 | 10,98 | 12,59 | 12,9 | 27 |
| ENSG00000243927 | MRPS6 | 45,56 | 46,89 | 51,23 | 57,9 | 27 |
| ENSG00000104897 | SF3A2 | 45,05 | 45,48 | 51,66 | 57,25 | 27 |
| ENSG00000066455 | GOLGA5 | 7,76 | 8,84 | 9,23 | 9,86 | 27 |
| ENSG00000181852 | RNF41 | 11,94 | 12,68 | 13,89 | 15,17 | 27 |
| ENSG00000165629 | ATP5F1C | 90,7 | 92,7 | 103,51 | 115,23 | 27 |
| ENSG00000119906 | SLF2 | 18,16 | 19,52 | 19,87 | 23,07 | 27 |
| ENSG00000188807 | TMEM201 | 8,99 | 9,04 | 10,44 | 11,42 | 27 |
| ENSG00000025800 | KPNA6 | 13,47 | 14,51 | 16,02 | 17,11 | 27 |
| ENSG00000158864 | NDUFS2 | 34,69 | 36,77 | 43,49 | 44,05 | 27 |
| ENSG00000165675 | ENOX2 | 9,86 | 9,98 | 11,62 | 12,52 | 27 |
| ENSG00000086589 | RBM22 | 23,32 | 26,21 | 28,95 | 29,61 | 27 |
| ENSG00000105717 | PBX4 | 2,04 | 2,55 | 2,58 | 2,59 | 27 |
| ENSG00000132294 | EFR3A | 8,28 | 8,96 | 9,02 | 10,51 | 27 |
| ENSG00000145741 | BTF3 | 230,08 | 242,9 | 271,85 | 291,99 | 27 |
| ENSG00000073921 | PICALM | 61,83 | 72,24 | 72,69 | 78,45 | 27 |
| ENSG00000169783 | LINGO1 | 6,66 | 7,96 | 8,07 | 8,45 | 27 |
| ENSG00000165194 | PCDH19 | 10,87 | 11,04 | 11,57 | 13,79 | 27 |
| ENSG00000285226 | PAK1IP1 | 5,55 | 5,72 | 6,3 | 7,04 | 27 |
| ENSG00000111845 | PAK1IP1 | 5,55 | 5,72 | 6,3 | 7,04 | 27 |
| ENSG00000148429 | USP6NL | 16,54 | 17,73 | 18,99 | 20,97 | 27 |
| ENSG00000198925 | ATG9A | 9,28 | 9,38 | 9,73 | 11,76 | 27 |
| ENSG00000181789 | COPG1 | 33,76 | 36,86 | 40,21 | 42,78 | 27 |
| ENSG00000175606 | TMEM70 | 8,61 | 9,14 | 9,88 | 10,91 | 27 |
| ENSG00000111602 | TIMELESS | 30,85 | 31,51 | 33,46 | 39,09 | 27 |
| ENSG00000187801 | ZFP69B | 6,86 | 7,15 | 7,2 | 8,69 | 27 |
| ENSG00000065548 | ZC3H15 | 33,03 | 36,59 | 39,18 | 41,84 | 27 |
| ENSG00000138942 | RNF185 | 10,21 | 10,22 | 10,78 | 12,93 | 27 |
| ENSG00000187243 | MAGED4B | 46,37 | 50,65 | 51,27 | 58,72 | 27 |
| ENSG00000196981 | WDR5B | 1,99 | 2,18 | 2,27 | 2,52 | 27 |
| ENSG00000108587 | GOSR1 | 17,51 | 19,83 | 21,09 | 22,17 | 27 |
| ENSG00000105698 | USF2 | 24,63 | 26,6 | 30,66 | 31,18 | 27 |
| ENSG00000172292 | CERS6 | 12,57 | 14,23 | 14,49 | 15,91 | 27 |
| ENSG00000126247 | CAPNS1 | 117,85 | 127,23 | 147,75 | 149,15 | 27 |
| ENSG00000077585 | GPR137B | 4,64 | 5,1 | 5,33 | 5,87 | 27 |
| ENSG00000116815 | CD58 | 4,98 | 5,57 | 5,87 | 6,3 | 27 |
| ENSG00000101457 | DNTTIP1 | 16,64 | 17,72 | 20,4 | 21,05 | 27 |
| ENSG00000164548 | TRA2A | 59,22 | 63,92 | 68,27 | 74,89 | 26 |
| ENSG00000282269 | PRR4 | 4,8 | 4,99 | 5,48 | 6,07 | 26 |
| ENSG00000135912 | TTLL4 | 23,64 | 26,24 | 26,67 | 29,89 | 26 |
| ENSG00000124422 | USP22 | 85,2 | 90,88 | 107,44 | 107,72 | 26 |
| ENSG00000170954 | ZNF415 | 5,6 | 5,79 | 6,89 | 7,08 | 26 |
| ENSG00000100897 | DCAF11 | 11,05 | 11,1 | 12,17 | 13,97 | 26 |
| ENSG00000163939 | PBRM1 | 48,62 | 54,63 | 55,34 | 61,46 | 26 |
| ENSG00000134590 | RTL8C | 30,9 | 35,76 | 38,9 | 39,06 | 26 |
| ENSG00000122882 | ECD | 18,41 | 18,56 | 18,57 | 23,27 | 26 |

| | | | | | | |
|---|---|---|---|---|---|---|
| ENSG00000129636 | ITFG1 | 15,32 | 18,24 | 18,39 | 19,36 | 26 |
| ENSG00000100258 | LMF2 | 11,16 | 11,64 | 13,92 | 14,1 | 26 |
| ENSG00000114631 | PODXL2 | 19,02 | 21,26 | 23,74 | 24,03 | 26 |
| ENSG00000167996 | FTH1 | 506,08 | 528,41 | 557,37 | 639,37 | 26 |
| ENSG00000185627 | PSMD13 | 44,12 | 45,43 | 51,23 | 55,74 | 26 |
| ENSG00000143437 | ARNT | 14,32 | 16,19 | 17,74 | 18,09 | 26 |
| ENSG00000141084 | RANBP10 | 6,23 | 6,92 | 7,52 | 7,87 | 26 |
| ENSG00000274425 | AC114271,1 | 1,18 | 1,39 | 1,42 | 1,49 | 26 |
| ENSG00000104960 | PTOV1 | 86,72 | 93,87 | 102,91 | 109,5 | 26 |
| ENSG00000104979 | C19orf53 | 62,73 | 63,67 | 76,14 | 79,19 | 26 |
| ENSG00000127445 | PIN1 | 39,58 | 40,6 | 49,31 | 49,96 | 26 |
| ENSG00000106367 | AP1S1 | 18 | 19,89 | 21,42 | 22,72 | 26 |
| ENSG00000258441 | LINC00641 | 13,73 | 13,82 | 14,07 | 17,33 | 26 |
| ENSG00000099817 | POLR2E | 74,23 | 74,75 | 83,9 | 93,68 | 26 |
| ENSG00000150961 | SEC24D | 9,16 | 10,59 | 10,67 | 11,56 | 26 |
| ENSG00000163902 | RPN1 | 58,78 | 62,54 | 70,85 | 74,18 | 26 |
| ENSG00000183475 | ASB7 | 4,47 | 4,79 | 5,1 | 5,64 | 26 |
| ENSG00000170325 | PRDM10 | 3,63 | 3,66 | 4,18 | 4,58 | 26 |
| ENSG00000108559 | NUP88 | 41,88 | 44,55 | 44,9 | 52,84 | 26 |
| ENSG00000175215 | CTDSP2 | 42,8 | 46,4 | 51,78 | 54 | 26 |
| ENSG00000170525 | PFKFB3 | 11,81 | 11,85 | 13,55 | 14,9 | 26 |
| ENSG00000136802 | LRRC8A | 12,6 | 13,47 | 15,57 | 15,89 | 26 |
| ENSG00000165119 | HNRNPK | 376,77 | 392 | 424,53 | 475,09 | 26 |
| ENSG00000120253 | NUP43 | 17,02 | 17,67 | 17,74 | 21,46 | 26 |
| ENSG00000120656 | TAF12 | 11,32 | 11,48 | 12,19 | 14,27 | 26 |
| ENSG00000182986 | ZNF320 | 17,12 | 18,24 | 18,58 | 21,58 | 26 |
| ENSG00000100888 | CHD8 | 32,02 | 34,97 | 36,25 | 40,36 | 26 |
| ENSG00000284869 | EEFSEC | 6,26 | 6,39 | 7,22 | 7,89 | 26 |
| ENSG00000284792 | PTEN | 11,03 | 12,26 | 13,48 | 13,9 | 26 |
| ENSG00000137161 | CNPY3 | 24,27 | 26,46 | 28,21 | 30,58 | 26 |
| ENSG00000137802 | MAPKBP1 | 6,81 | 8,18 | 8,26 | 8,58 | 26 |
| ENSG00000175224 | ATG13 | 20,65 | 21,99 | 23,42 | 26,01 | 26 |
| ENSG00000164045 | CDC25A | 21,4 | 21,47 | 21,67 | 26,95 | 26 |
| ENSG00000110218 | PANX1 | 16,48 | 18,99 | 19,8 | 20,75 | 26 |
| ENSG00000170364 | SETMAR | 15,37 | 17,52 | 18,09 | 19,35 | 26 |
| ENSG00000100883 | SRP54 | 25,12 | 26,33 | 28,71 | 31,61 | 26 |
| ENSG00000105568 | PPP2R1A | 92,55 | 97,97 | 108,25 | 116,46 | 26 |
| ENSG00000148248 | SURF4 | 17,07 | 17,95 | 21,07 | 21,48 | 26 |
| ENSG00000280951 | SURF4 | 17,07 | 17,95 | 21,07 | 21,48 | 26 |
| ENSG00000134910 | STT3A | 53,22 | 53,37 | 56,59 | 66,95 | 26 |
| ENSG00000146535 | GNA12 | 18,15 | 18,74 | 21,8 | 22,82 | 26 |
| ENSG00000234585 | CCT6P3 | 3,19 | 3,34 | 3,4 | 4,01 | 26 |
| ENSG00000132953 | XPO4 | 7,9 | 8,49 | 8,96 | 9,93 | 26 |
| ENSG00000214960 | CRPPA | 1,48 | 1,56 | 1,82 | 1,86 | 26 |
| ENSG00000100664 | EIF5 | 121,93 | 128,03 | 149,55 | 153,23 | 26 |
| ENSG00000132603 | NIP7 | 11,54 | 12,56 | 13,86 | 14,5 | 26 |
| ENSG00000119421 | NDUFA8 | 32,42 | 34,64 | 36,55 | 40,73 | 26 |
| ENSG00000100056 | ESS2 | 5,97 | 6,36 | 7,07 | 7,5 | 26 |
| ENSG00000124541 | RRP36 | 29,54 | 30,75 | 36,63 | 37,11 | 26 |
| ENSG00000172943 | PHF8 | 9,99 | 10,73 | 10,76 | 12,55 | 26 |

| | | | | | | |
|---|---|---|---|---|---|---|
| ENSG00000120063 | GNA13 | 19,2 | 21,73 | 23,86 | 24,12 | 26 |
| ENSG00000102226 | USP11 | 63,65 | 68,37 | 75,73 | 79,96 | 26 |
| ENSG00000089335 | ZNF302 | 14,99 | 15,13 | 15,86 | 18,83 | 26 |
| ENSG00000065427 | KARS1 | 56,67 | 56,94 | 65,62 | 71,18 | 26 |
| ENSG00000175354 | PTPN2 | 15,96 | 16,49 | 17,69 | 20,04 | 26 |
| ENSG00000204220 | PFDN6 | 7,63 | 8,69 | 8,77 | 9,58 | 26 |
| ENSG00000121022 | COPS5 | 29,81 | 31,14 | 36,08 | 37,42 | 26 |
| ENSG00000178741 | COX5A | 116,03 | 120,74 | 125,77 | 145,63 | 26 |
| ENSG00000186625 | KATNA1 | 6,94 | 7,29 | 8,25 | 8,71 | 26 |
| ENSG00000128245 | YWHAH | 66,24 | 70,95 | 79,99 | 83,13 | 25 |
| ENSG00000100519 | PSMC6 | 55,14 | 58,54 | 61,57 | 69,18 | 25 |
| ENSG00000064309 | CDON | 37,44 | 41,79 | 44,05 | 46,97 | 25 |
| ENSG00000128185 | DGCR6L | 15,49 | 16,2 | 19,29 | 19,43 | 25 |
| ENSG00000145495 | MARCHF6 | 45,99 | 52,78 | 55,94 | 57,65 | 25 |
| ENSG00000124795 | DEK | 128 | 135,66 | 146,62 | 160,41 | 25 |
| ENSG00000124313 | IQSEC2 | 3,12 | 3,18 | 3,71 | 3,91 | 25 |
| ENSG00000213593 | TMX2 | 28,77 | 29,36 | 30,57 | 36,05 | 25 |
| ENSG00000079785 | DDX1 | 52,31 | 52,86 | 58,63 | 65,54 | 25 |
| ENSG00000273494 | PANK4 | 7,91 | 8,33 | 8,52 | 9,91 | 25 |
| ENSG00000163320 | CGGBP1 | 39,91 | 41,52 | 46,27 | 50 | 25 |
| ENSG00000147416 | ATP6V1B2 | 26,4 | 29,5 | 32,41 | 33,06 | 25 |
| ENSG00000123106 | CCDC91 | 10,23 | 10,83 | 10,9 | 12,81 | 25 |
| ENSG00000148110 | MFSD14B | 11,42 | 12,15 | 12,34 | 14,3 | 25 |
| ENSG00000159259 | CHAF1B | 10,59 | 10,77 | 11,67 | 13,26 | 25 |
| ENSG00000148396 | SEC16A | 14,44 | 16,2 | 17,56 | 18,08 | 25 |
| ENSG00000198727 | MT-CYB | 1858,83 | 2099,35 | 2127,07 | 2327,27 | 25 |
| ENSG00000134248 | LAMTOR5 | 60,16 | 69,98 | 70,86 | 75,3 | 25 |
| ENSG00000107341 | UBE2R2 | 27,4 | 31,14 | 31,52 | 34,29 | 25 |
| ENSG00000164828 | SUN1 | 25,27 | 26,34 | 28,18 | 31,62 | 25 |
| ENSG00000154813 | DPH3 | 12,7 | 13,27 | 14,53 | 15,89 | 25 |
| ENSG00000047315 | POLR2B | 51,16 | 51,36 | 57,34 | 64,01 | 25 |
| ENSG00000149269 | PAK1 | 31,26 | 34,85 | 36,19 | 39,1 | 25 |
| ENSG00000131845 | ZNF304 | 3,23 | 3,51 | 3,9 | 4,04 | 25 |
| ENSG00000154781 | CCDC174 | 9,77 | 11,56 | 11,66 | 12,22 | 25 |
| ENSG00000163812 | ZDHHC3 | 15,06 | 16,43 | 17,66 | 18,83 | 25 |
| ENSG00000129084 | PSMA1 | 87,38 | 88,98 | 89,14 | 109,25 | 25 |
| ENSG00000147419 | CCDC25 | 21,15 | 22,09 | 23,24 | 26,44 | 25 |

| KO-NSCs | | | [TPM] | | | |
|---|---|---|---|---|---|---|
| gene_ID | Gene_name | KO-NSC_1 | KO-NSC_2 | KO-NSC_3 | KO-NSC_4 | change_% |
| ENSG00000221716 | SNORA11 | 0 | 3,25 | 7,84 | 16,7 | |
| ENSG00000130772 | MED18 | 0 | 3,27 | 3,95 | 7,84 | |
| ENSG00000281758 | KCNIP4 | 0 | 0,96 | 5,62 | 6,13 | |
| ENSG00000244563 | RPS26P19 | 0 | 0,68 | 1 | 4,63 | |
| ENSG00000259121 | | 0 | 0,93 | 1,07 | 2,36 | |
| ENSG00000277677 | Metazoa_SRP | 0 | 0,36 | 0,41 | 2,03 | |
| ENSG00000282278 | AC058822,1 | 0 | 0,34 | 0,68 | 1,86 | |
| ENSG00000243437 | RN7SL370P | 0 | 0,16 | 0,37 | 1,8 | |
| ENSG00000281234 | LOH12CR2 | 0 | 0,87 | 1,07 | 1,56 | |
| ENSG00000270549 | | 0 | 0,28 | 0,85 | 1,55 | |
| ENSG00000264769 | AC145207,8 | 0 | 0,05 | 0,08 | 1,52 | |
| ENSG00000241959 | RN7SL76P | 0 | 0,51 | 0,9 | 1,45 | |
| ENSG00000182742 | HOXB4 | 0 | 0,16 | 0,47 | 1,35 | |
| ENSG00000203593 | AC005342,1 | 0 | 0,27 | 0,52 | 1,35 | |
| ENSG00000268545 | VN1R107P | 0 | 0,41 | 0,94 | 1,27 | |
| ENSG00000264809 | | 0 | 0,19 | 0,23 | 1,23 | |
| ENSG00000244677 | | 0 | 0,17 | 0,25 | 1,22 | |
| ENSG00000231726 | HMGN2P38 | 0 | 0,42 | 0,64 | 1,11 | |
| ENSG00000274848 | | 0 | 0,16 | 0,8 | 1,01 | |
| ENSG00000168081 | PNOC | 0,02 | 0,22 | 0,67 | 2,39 | 11850 |
| ENSG00000273686 | B2M | 0,51 | 1,27 | 1,64 | 52 | 10096 |
| ENSG00000253974 | NRG1-IT1 | 0,03 | 0,3 | 1,01 | 2,53 | 8333 |
| ENSG00000276849 | TRBC2 | 0,38 | 2,66 | 4,95 | 12,99 | 3318 |
| ENSG00000109846 | CRYAB | 0,08 | 0,09 | 0,66 | 2,31 | 2788 |
| ENSG00000288199 | LSP1 | 0,05 | 0,26 | 0,4 | 1,26 | 2420 |
| ENSG00000144785 | | 1,31 | 1,49 | 1,81 | 30,52 | 2230 |
| ENSG00000128422 | KRT17 | 0,12 | 0,18 | 0,61 | 2,74 | 2183 |
| ENSG00000257605 | MYG1-AS1 | 0,05 | 0,09 | 0,28 | 1,11 | 2120 |
| ENSG00000234745 | HLA-B | 0,07 | 0,19 | 0,73 | 1,55 | 2114 |
| ENSG00000172061 | LRRC15 | 0,07 | 0,45 | 0,61 | 1,52 | 2071 |
| ENSG00000263940 | RN7SL275P | 0,2 | 0,44 | 0,76 | 4,16 | 1980 |
| ENSG00000277101 | ARHGEF26 | 0,18 | 0,43 | 0,84 | 3,47 | 1828 |
| ENSG00000198768 | APCDD1L | 0,14 | 0,51 | 0,99 | 2,62 | 1771 |
| ENSG00000274334 | VASN | 0,2 | 1,3 | 1,32 | 3,7 | 1750 |
| ENSG00000282806 | AC068400,2 | 0,26 | 0,64 | 1,01 | 4,56 | 1654 |
| ENSG00000170498 | KISS1 | 0,06 | 0,14 | 0,19 | 1,04 | 1633 |
| ENSG00000281369 | AC093826,1 | 4,59 | 28,63 | 45,1 | 78,88 | 1619 |
| ENSG00000123689 | G0S2 | 0,13 | 0,25 | 0,65 | 2,2 | 1592 |
| ENSG00000276396 | Metazoa_SRP | 0,13 | 0,17 | 0,38 | 2,14 | 1546 |
| ENSG00000140563 | MCTP2 | 0,11 | 0,36 | 1,26 | 1,81 | 1545 |
| ENSG00000281644 | AC106795,9 | 0,35 | 0,7 | 0,83 | 5,66 | 1517 |
| ENSG00000274505 | Metazoa_SRP | 0,14 | 0,18 | 0,27 | 2,2 | 1471 |
| ENSG00000162595 | DIRAS3 | 0,87 | 2,3 | 6,84 | 13,5 | 1452 |
| ENSG00000231898 | MYO3B-AS1 | 0,12 | 0,34 | 0,36 | 1,84 | 1433 |
| ENSG00000136630 | HLX | 0,13 | 0,17 | 0,27 | 1,98 | 1423 |
| ENSG00000243509 | TNFRSF6B | 0,18 | 0,72 | 1,45 | 2,68 | 1389 |
| ENSG00000253507 | AC104257,1 | 0,09 | 0,25 | 0,29 | 1,33 | 1378 |
| ENSG00000261371 | PECAM1 | 0,1 | 0,18 | 1,27 | 1,4 | 1300 |

| | | | | | | |
|---|---|---|---|---|---|---|
| ENSG00000234136 | AC055764,1 | 0,11 | 0,46 | 0,61 | 1,47 | 1236 |
| ENSG00000164488 | DACT2 | 0,51 | 0,89 | 3,06 | 6,5 | 1175 |
| ENSG00000224608 | HLA-B | 0,46 | 2,25 | 2,99 | 5,8 | 1161 |
| ENSG00000008441 | NFIX | 1 | 2,88 | 4,8 | 12,53 | 1153 |
| ENSG00000163814 | CDCP1 | 0,12 | 0,36 | 0,78 | 1,46 | 1117 |
| ENSG00000249565 | SERBP1P5 | 0,11 | 0,23 | 0,4 | 1,29 | 1073 |
| ENSG00000231437 | LINC01750 | 0,16 | 0,4 | 0,66 | 1,82 | 1038 |
| ENSG00000196878 | LAMB3 | 0,22 | 0,43 | 0,49 | 2,49 | 1032 |
| ENSG00000177363 | LRRN4CL | 0,39 | 0,61 | 1,09 | 4,39 | 1026 |
| ENSG00000167895 | TMC8 | 0,16 | 0,24 | 0,29 | 1,79 | 1019 |
| ENSG00000120075 | HOXB5 | 0,17 | 0,76 | 1,45 | 1,88 | 1006 |
| ENSG00000255026 | AC136475,3 | 0,23 | 0,27 | 0,88 | 2,51 | 991 |
| ENSG00000120093 | HOXB3 | 0,29 | 1,25 | 1,57 | 3,13 | 979 |
| ENSG00000198753 | PLXNB3 | 0,18 | 0,26 | 0,88 | 1,94 | 978 |
| ENSG00000277965 | Metazoa_SRP | 0,45 | 0,55 | 1,43 | 4,84 | 976 |
| ENSG00000178860 | MSC | 0,69 | 1,35 | 1,99 | 7,4 | 972 |
| ENSG00000223532 | HLA-B | 3,44 | 15,88 | 19,23 | 36,42 | 959 |
| ENSG00000128274 | A4GALT | 0,22 | 0,86 | 1,4 | 2,32 | 955 |
| ENSG00000196581 | AJAP1 | 0,28 | 0,65 | 1,12 | 2,93 | 946 |
| ENSG00000228526 | MIR34AHG | 0,39 | 0,89 | 1,14 | 4,05 | 938 |
| ENSG00000160223 | ICOSLG | 0,21 | 0,41 | 0,6 | 2,13 | 914 |
| ENSG00000234551 | LINC01309 | 0,16 | 0,18 | 0,25 | 1,61 | 906 |
| ENSG00000281903 | LINC02246 | 0,72 | 0,88 | 0,92 | 7,11 | 888 |
| ENSG00000261327 | AC134312,5 | 0,6 | 1,28 | 2,13 | 5,9 | 883 |
| ENSG00000137198 | GMPR | 0,18 | 0,69 | 0,9 | 1,74 | 867 |
| ENSG00000172137 | CALB2 | 0,7 | 1,97 | 2,67 | 6,76 | 866 |
| ENSG00000282830 | CALB2 | 0,7 | 1,97 | 2,67 | 6,76 | 866 |
| ENSG00000234546 | LNCTAM34A | 0,13 | 0,22 | 0,35 | 1,24 | 854 |
| ENSG00000133110 | POSTN | 2,27 | 7,22 | 19,13 | 21,63 | 853 |
| ENSG00000179057 | IGSF22 | 0,26 | 0,42 | 0,79 | 2,42 | 831 |
| ENSG00000174640 | SLCO2A1 | 0,17 | 0,24 | 0,55 | 1,58 | 829 |
| ENSG00000204277 | LINC01993 | 0,36 | 0,4 | 0,84 | 3,34 | 828 |
| ENSG00000261653 | AC092337,1 | 0,36 | 0,38 | 0,52 | 3,34 | 828 |
| ENSG00000185332 | TMEM105 | 0,18 | 0,28 | 0,41 | 1,67 | 828 |
| ENSG00000184254 | ALDH1A3 | 0,18 | 0,34 | 0,84 | 1,61 | 794 |
| ENSG00000239268 | AC092691,1 | 0,25 | 0,71 | 0,94 | 2,22 | 788 |
| ENSG00000156219 | ART3 | 0,13 | 0,29 | 0,35 | 1,15 | 785 |
| ENSG00000197182 | MIRLET7BHG | 0,43 | 0,71 | 0,78 | 3,78 | 779 |
| ENSG00000131620 | ANO1 | 2,06 | 4,34 | 10,13 | 18,04 | 776 |
| ENSG00000125735 | TNFSF14 | 0,12 | 0,16 | 0,19 | 1,04 | 767 |
| ENSG00000124749 | COL21A1 | 0,14 | 0,33 | 0,37 | 1,2 | 757 |
| ENSG00000092758 | COL9A3 | 3,11 | 4,12 | 4,27 | 25,79 | 729 |
| ENSG00000112493 | TAPBP | 2,46 | 12,55 | 17,51 | 20,2 | 721 |
| ENSG00000273829 | | 0,14 | 0,17 | 0,18 | 1,15 | 721 |
| ENSG00000267491 | AC100788,1 | 0,18 | 0,23 | 0,31 | 1,46 | 711 |
| ENSG00000282873 | BCL7A | 0,54 | 0,66 | 0,74 | 4,32 | 700 |
| ENSG00000145708 | CRHBP | 0,48 | 0,65 | 1,28 | 3,77 | 685 |
| ENSG00000262097 | LINC02185 | 0,16 | 0,17 | 0,48 | 1,25 | 681 |
| ENSG00000007237 | GAS7 | 2,47 | 7,02 | 14,68 | 19,28 | 681 |
| ENSG00000196754 | S100A2 | 0,22 | 0,47 | 0,76 | 1,71 | 677 |

| | | | | | | |
|---|---|---|---|---|---|---|
| ENSG00000175356 | SCUBE2 | 0,7 | 2,58 | 3,9 | 5,43 | 676 |
| ENSG00000135722 | FBXL8 | 0,13 | 0,27 | 0,43 | 1 | 669 |
| ENSG00000264548 | AC132872,2 | 0,19 | 0,27 | 0,29 | 1,46 | 668 |
| ENSG00000285784 | AL353595,1 | 0,17 | 0,19 | 0,2 | 1,29 | 659 |
| ENSG00000255851 | Metazoa_SRP | 1,86 | 2,39 | 3,56 | 13,97 | 651 |
| ENSG00000138316 | ADAMTS14 | 1,03 | 2,55 | 2,58 | 7,68 | 646 |
| ENSG00000229981 | LINC01435 | 0,38 | 0,52 | 0,54 | 2,82 | 642 |
| ENSG00000128342 | LIF | 1,09 | 1,3 | 1,55 | 8,08 | 641 |
| ENSG00000183840 | GPR39 | 1,22 | 1,52 | 2,45 | 9,04 | 641 |
| ENSG00000222022 | AC112721,1 | 0,15 | 0,61 | 0,9 | 1,11 | 640 |
| ENSG00000274206 | Metazoa_SRP | 3,18 | 3,54 | 3,95 | 23,24 | 631 |
| ENSG00000134668 | SPOCD1 | 1,09 | 1,77 | 2,28 | 7,93 | 628 |
| ENSG00000087510 | TFAP2C | 0,58 | 1,32 | 2,62 | 4,19 | 622 |
| ENSG00000167992 | VWCE | 0,62 | 0,68 | 0,84 | 4,38 | 606 |
| ENSG00000137880 | GCHFR | 1,06 | 3,04 | 5,19 | 7,45 | 603 |
| ENSG00000237022 | HLA-C | 0,36 | 0,81 | 0,99 | 2,53 | 603 |
| ENSG00000230317 | LINC01284 | 0,16 | 0,21 | 0,25 | 1,1 | 588 |
| ENSG00000226179 | LINC00685 | 0,5 | 1,39 | 1,42 | 3,4 | 580 |
| ENSG00000159674 | SPON2 | 3,47 | 7,26 | 9,75 | 23,38 | 574 |
| ENSG00000163694 | RBM47 | 0,42 | 0,58 | 0,64 | 2,81 | 569 |
| ENSG00000197355 | UAP1L1 | 0,2 | 0,38 | 0,43 | 1,33 | 565 |
| ENSG00000186340 | THBS2 | 2,79 | 5,64 | 6,13 | 18,08 | 548 |
| ENSG00000286112 | AL441992,2 | 0,17 | 0,27 | 0,33 | 1,1 | 547 |
| ENSG00000225691 | HLA-C | 2,16 | 7,91 | 8,88 | 13,92 | 544 |
| ENSG00000107485 | GATA3 | 0,48 | 0,9 | 0,95 | 3,08 | 542 |
| ENSG00000082397 | EPB41L3 | 16,81 | 38,87 | 59,49 | 105,23 | 526 |
| ENSG00000235204 | AL162724,2 | 0,58 | 0,73 | 0,78 | 3,59 | 519 |
| ENSG00000147408 | CSGALNACT1 | 0,33 | 0,95 | 1,59 | 2,04 | 518 |
| ENSG00000115107 | STEAP3 | 0,52 | 0,65 | 1,38 | 3,2 | 515 |
| ENSG00000115363 | EVA1A | 1,45 | 2,91 | 3,78 | 8,83 | 509 |
| ENSG00000268996 | MAN1B1-DT | 0,35 | 0,55 | 0,7 | 2,1 | 500 |
| ENSG00000221946 | FXYD7 | 1,93 | 5,16 | 10,23 | 11,53 | 497 |
| ENSG00000153823 | PID1 | 0,85 | 2,62 | 3,51 | 5,06 | 495 |
| ENSG00000176485 | PLAAT3 | 1,37 | 2,11 | 3,04 | 8,14 | 494 |
| ENSG00000225953 | SATB2-AS1 | 0,17 | 0,22 | 0,41 | 1,01 | 494 |
| ENSG00000026508 | CD44 | 7,13 | 13,57 | 21,02 | 42,33 | 494 |
| ENSG00000275650 | TTYH1 | 2,59 | 2,62 | 3,81 | 15,36 | 493 |
| ENSG00000284649 | AC009093,8 | 0,23 | 0,33 | 0,47 | 1,36 | 491 |
| ENSG00000160712 | IL6R | 0,32 | 0,33 | 0,43 | 1,89 | 491 |
| ENSG00000159231 | CBR3 | 1,11 | 2,8 | 4,77 | 6,54 | 489 |
| ENSG00000287586 | Z98886,1 | 0,28 | 0,29 | 0,31 | 1,65 | 489 |
| ENSG00000272836 | AL022328,1 | 0,41 | 0,52 | 0,54 | 2,39 | 483 |
| ENSG00000111331 | OAS3 | 0,75 | 1,84 | 2,23 | 4,37 | 483 |
| ENSG00000255348 | AP001775,2 | 0,44 | 0,64 | 1,03 | 2,55 | 480 |
| ENSG00000259175 | AC108451,1 | 0,48 | 0,67 | 0,76 | 2,78 | 479 |
| ENSG00000082126 | MPP4 | 0,6 | 0,62 | 0,76 | 3,46 | 477 |
| ENSG00000185274 | GALNT17 | 2,84 | 4,68 | 7,36 | 16,35 | 476 |
| ENSG00000276626 | 7SK | 0,41 | 1,56 | 1,98 | 2,35 | 473 |
| ENSG00000140853 | NLRC5 | 1,43 | 2,23 | 3,42 | 8,18 | 472 |
| ENSG00000126561 | STAT5A | 0,21 | 0,64 | 0,69 | 1,2 | 471 |

| | | | | | | |
|---|---|---|---|---|---|---|
| ENSG00000065717 | TLE2 | 0,84 | 2,08 | 2,12 | 4,78 | 469 |
| ENSG00000284968 | AC093827,4 | 0,31 | 0,51 | 0,6 | 1,76 | 468 |
| ENSG00000264449 | AP001021,1 | 0,37 | 0,82 | 0,93 | 2,08 | 462 |
| ENSG00000155897 | ADCY8 | 0,32 | 0,48 | 1,06 | 1,77 | 453 |
| ENSG00000237582 | POU5F1 | 0,98 | 1,35 | 1,42 | 5,36 | 447 |
| ENSG00000206454 | POU5F1 | 0,98 | 1,35 | 1,42 | 5,36 | 447 |
| ENSG00000185567 | | 1,15 | 1,48 | 1,49 | 6,24 | 443 |
| ENSG00000090339 | ICAM1 | 1,69 | 3,75 | 4,43 | 9,15 | 441 |
| ENSG00000287897 | AP000465,1 | 0,23 | 0,58 | 0,89 | 1,24 | 439 |
| ENSG00000141013 | GAS8 | 0,49 | 0,78 | 0,81 | 2,64 | 439 |
| ENSG00000182601 | HS3ST4 | 1,57 | 4,37 | 6 | 8,41 | 436 |
| ENSG00000043143 | JADE2 | 0,7 | 1,15 | 1,43 | 3,74 | 434 |
| ENSG00000126709 | IFI6 | 6,66 | 22,1 | 34,19 | 35,43 | 432 |
| ENSG00000130813 | SHFL | 1,96 | 2,57 | 3,19 | 10,4 | 431 |
| ENSG00000172780 | RAB43 | 0,86 | 0,96 | 1,47 | 4,56 | 430 |
| ENSG00000171189 | GRIK1 | 0,52 | 0,75 | 2,02 | 2,75 | 429 |
| ENSG00000135842 | NIBAN1 | 0,61 | 0,91 | 1,27 | 3,22 | 428 |
| ENSG00000270504 | AL391422,4 | 1,62 | 4,21 | 5,09 | 8,51 | 425 |
| ENSG00000278910 | BANCR | 0,28 | 0,71 | 1,39 | 1,47 | 425 |
| ENSG00000273079 | GRIN2B | 1,29 | 2,15 | 3,13 | 6,76 | 424 |
| ENSG00000133392 | MYH11 | 0,21 | 0,23 | 0,3 | 1,1 | 424 |
| ENSG00000228299 | HLA-C | 1,44 | 3,46 | 3,92 | 7,52 | 422 |
| ENSG00000196611 | MMP1 | 0,47 | 0,72 | 1,18 | 2,45 | 421 |
| ENSG00000284721 | AL662907,3 | 0,21 | 0,28 | 0,3 | 1,09 | 419 |
| ENSG00000224746 | AC015987,1 | 0,26 | 0,27 | 0,34 | 1,34 | 415 |
| ENSG00000130592 | LSP1 | 0,38 | 0,57 | 0,58 | 1,95 | 413 |
| ENSG00000076356 | PLXNA2 | 5,65 | 12,07 | 13,75 | 28,56 | 405 |
| ENSG00000226339 | | 0,21 | 0,54 | 1,03 | 1,06 | 405 |
| ENSG00000188643 | S100A16 | 3,6 | 6,36 | 8,68 | 18,06 | 402 |
| ENSG00000166145 | SPINT1 | 0,25 | 0,39 | 0,68 | 1,25 | 400 |
| ENSG00000232406 | AL121895,1 | 0,51 | 0,59 | 0,68 | 2,54 | 398 |
| ENSG00000090554 | FLT3LG | 0,58 | 1,13 | 1,37 | 2,87 | 395 |
| ENSG00000136826 | KLF4 | 0,56 | 1,24 | 2,02 | 2,77 | 395 |
| ENSG00000168394 | TAP1 | 0,52 | 1,71 | 1,95 | 2,56 | 392 |
| ENSG00000127954 | STEAP4 | 0,57 | 0,59 | 0,62 | 2,79 | 389 |
| ENSG00000248510 | LINC02267 | 0,33 | 0,41 | 0,49 | 1,61 | 388 |
| ENSG00000263234 | AC010401,2 | 0,97 | 1,1 | 1,24 | 4,73 | 388 |
| ENSG00000235531 | MSC-AS1 | 0,46 | 0,54 | 0,97 | 2,24 | 387 |
| ENSG00000160180 | TFF3 | 0,6 | 1,28 | 1,96 | 2,92 | 387 |
| ENSG00000147119 | CHST7 | 1,41 | 2,91 | 3,96 | 6,86 | 387 |
| ENSG00000272904 | AL390726,2 | 0,33 | 0,38 | 0,45 | 1,6 | 385 |
| ENSG00000284543 | LINC01226 | 0,63 | 0,8 | 1,11 | 3,02 | 379 |
| ENSG00000276890 | Metazoa_SRP | 0,29 | 0,36 | 0,67 | 1,39 | 379 |
| ENSG00000113319 | RASGRF2 | 0,32 | 0,55 | 0,81 | 1,53 | 378 |
| ENSG00000236404 | VLDLR-AS1 | 0,59 | 1,26 | 2,51 | 2,8 | 375 |
| ENSG00000173917 | HOXB2 | 2,13 | 4,44 | 6,87 | 10,08 | 373 |
| ENSG00000143318 | CASQ1 | 0,48 | 1,42 | 1,59 | 2,26 | 371 |
| ENSG00000274825 | AL023803,2 | 0,89 | 0,91 | 1,05 | 4,19 | 371 |
| ENSG00000139044 | B4GALNT3 | 0,56 | 0,8 | 1,05 | 2,61 | 366 |
| ENSG00000271437 | AL356423,1 | 0,26 | 0,32 | 0,33 | 1,21 | 365 |

| | | | | | | |
|---|---|---|---|---|---|---|
| ENSG00000025708 | TYMP | 0,63 | 0,77 | 0,88 | 2,93 | 365 |
| ENSG00000069399 | BCL3 | 1,62 | 2,09 | 2,83 | 7,52 | 364 |
| ENSG00000163131 | CTSS | 0,72 | 2,38 | 3,09 | 3,34 | 364 |
| ENSG00000132874 | SLC14A2 | 0,36 | 0,4 | 0,42 | 1,67 | 364 |
| ENSG00000177993 | ZNRF3-AS1 | 1,43 | 1,52 | 1,66 | 6,6 | 362 |
| ENSG00000086696 | HSD17B2 | 0,43 | 0,75 | 1,73 | 1,98 | 360 |
| ENSG00000134531 | EMP1 | 9,7 | 21,94 | 24 | 44,62 | 360 |
| ENSG00000250748 | AC025419,1 | 0,25 | 0,28 | 0,38 | 1,15 | 360 |
| ENSG00000147889 | CDKN2A | 5,02 | 9,21 | 11,64 | 22,93 | 357 |
| ENSG00000206435 | HLA-C | 0,51 | 1,06 | 1,25 | 2,32 | 355 |
| ENSG00000120708 | TGFBI | 49,44 | 132,75 | 177,92 | 224,39 | 354 |
| ENSG00000105643 | ARRDC2 | 0,27 | 0,39 | 0,63 | 1,21 | 348 |
| ENSG00000227007 | LINC01247 | 0,26 | 0,29 | 0,35 | 1,16 | 346 |
| ENSG00000116016 | EPAS1 | 0,66 | 0,83 | 1,08 | 2,93 | 344 |
| ENSG00000154930 | ACSS1 | 0,3 | 0,84 | 0,97 | 1,33 | 343 |
| ENSG00000101463 | SYNDIG1 | 1,8 | 3,43 | 5,84 | 7,95 | 342 |
| ENSG00000259402 | AC090515,5 | 0,27 | 0,55 | 0,64 | 1,19 | 341 |
| ENSG00000151117 | TMEM86A | 1,73 | 3,71 | 7,29 | 7,62 | 340 |
| ENSG00000171444 | MCC | 4,29 | 7,26 | 8,49 | 18,89 | 340 |
| ENSG00000148411 | NACC2 | 3,07 | 4,21 | 5,68 | 13,39 | 336 |
| ENSG00000242798 | AC073842,2 | 0,97 | 1,3 | 1,37 | 4,23 | 336 |
| ENSG00000187479 | C11orf96 | 0,77 | 1,06 | 1,34 | 3,35 | 335 |
| ENSG00000168497 | CAVIN2 | 0,56 | 0,94 | 1,51 | 2,43 | 334 |
| ENSG00000179588 | ZFPM1 | 1,28 | 1,67 | 2,17 | 5,52 | 331 |
| ENSG00000068079 | IFI35 | 1,16 | 1,84 | 2,8 | 5 | 331 |
| ENSG00000103740 | ACSBG1 | 0,96 | 2,13 | 2,86 | 4,11 | 328 |
| ENSG00000185112 | FAM43A | 1,41 | 2,74 | 3,09 | 6,03 | 328 |
| ENSG00000258674 | AC011448,1 | 0,27 | 0,31 | 0,41 | 1,15 | 326 |
| ENSG00000159216 | RUNX1 | 2,91 | 4,96 | 6,27 | 12,35 | 324 |
| ENSG00000171385 | KCND3 | 1,35 | 3,21 | 4,92 | 5,71 | 323 |
| ENSG00000278032 | LY6E | 3,17 | 6,66 | 7,67 | 13,34 | 321 |
| ENSG00000215089 | KRT18P11 | 0,24 | 0,41 | 0,77 | 1,01 | 321 |
| ENSG00000243811 | APOBEC3D | 0,86 | 0,87 | 0,9 | 3,58 | 316 |
| ENSG00000065320 | NTN1 | 1,89 | 2,15 | 2,24 | 7,84 | 315 |
| ENSG00000118785 | SPP1 | 3,3 | 6,52 | 11,94 | 13,67 | 314 |
| ENSG00000271538 | LINC02427 | 0,25 | 0,29 | 0,31 | 1,03 | 312 |
| ENSG00000157168 | NRG1 | 4,02 | 7,64 | 8,68 | 16,54 | 311 |
| ENSG00000137857 | DUOX1 | 1,63 | 2,43 | 2,62 | 6,69 | 310 |
| ENSG00000146722 | AC211486,1 | 0,29 | 0,35 | 0,51 | 1,19 | 310 |
| ENSG00000268750 | AC010522,1 | 1,83 | 2,04 | 2,14 | 7,47 | 308 |
| ENSG00000187013 | LINC02875 | 0,26 | 0,32 | 0,43 | 1,06 | 308 |
| ENSG00000102755 | FLT1 | 0,71 | 1,15 | 1,53 | 2,87 | 304 |
| ENSG00000166482 | MFAP4 | 1,02 | 2,03 | 2,72 | 4,12 | 304 |
| ENSG00000118257 | NRP2 | 18,56 | 33,04 | 41,46 | 74,35 | 301 |
| ENSG00000260577 | AC126773,4 | 1,25 | 2,75 | 3,29 | 4,99 | 299 |
| ENSG00000101000 | PROCR | 3,63 | 4,87 | 8,72 | 14,47 | 299 |
| ENSG00000161714 | PLCD3 | 1,55 | 3,13 | 3,37 | 6,17 | 298 |
| ENSG00000288508 | ISLR | 12,28 | 26,32 | 34,89 | 48,66 | 296 |
| ENSG00000129009 | ISLR | 12,28 | 26,32 | 34,89 | 48,66 | 296 |
| ENSG00000228692 | AL445307,1 | 0,26 | 0,5 | 0,66 | 1,03 | 296 |

| | | | | | | |
|---|---|---|---|---|---|---|
| ENSG00000222032 | AC112721,2 | 0,51 | 1,49 | 1,63 | 2,02 | 296 |
| ENSG00000161011 | SQSTM1 | 18,45 | 24,22 | 35,35 | 73,01 | 296 |
| ENSG00000198838 | RYR3 | 2,15 | 3,98 | 7,87 | 8,5 | 295 |
| ENSG00000229625 | AC016877,1 | 10,74 | 11,27 | 12,7 | 42,03 | 291 |
| ENSG00000174600 | CMKLR1 | 2,26 | 5,52 | 6,47 | 8,84 | 291 |
| ENSG00000136542 | GALNT5 | 0,29 | 0,75 | 0,92 | 1,13 | 290 |
| ENSG00000187608 | ISG15 | 9,41 | 22,8 | 32,84 | 36,65 | 289 |
| ENSG00000072682 | P4HA2 | 4,08 | 6,09 | 6,75 | 15,89 | 289 |
| ENSG00000108439 | PNPO | 4,6 | 5,76 | 5,85 | 17,86 | 288 |
| ENSG00000206432 | TMEM200C | 5,47 | 7,5 | 8,35 | 21,2 | 288 |
| ENSG00000179673 | RPRML | 1,33 | 1,7 | 1,72 | 5,14 | 286 |
| ENSG00000267452 | LINC02073 | 0,57 | 0,8 | 0,9 | 2,2 | 286 |
| ENSG00000245248 | USP2-AS1 | 0,28 | 0,32 | 0,49 | 1,08 | 286 |
| ENSG00000263873 | THY1-AS1 | 20,96 | 28,65 | 34,29 | 80,55 | 284 |
| ENSG00000131435 | PDLIM4 | 4,63 | 7,3 | 9,11 | 17,79 | 284 |
| ENSG00000258654 | AC026495,1 | 0,4 | 0,59 | 0,61 | 1,53 | 283 |
| ENSG00000260088 | DDX59-AS1 | 0,57 | 0,59 | 0,69 | 2,18 | 282 |
| ENSG00000152208 | GRID2 | 0,49 | 1,14 | 1,59 | 1,87 | 282 |
| ENSG00000011105 | TSPAN9 | 8,93 | 14,98 | 15,33 | 34 | 281 |
| ENSG00000057657 | PRDM1 | 0,31 | 0,49 | 0,97 | 1,18 | 281 |
| ENSG00000114757 | PEX5L | 0,44 | 0,71 | 1,19 | 1,66 | 277 |
| ENSG00000005513 | SOX8 | 4,18 | 6,11 | 7,54 | 15,75 | 277 |
| ENSG00000237172 | B3GNT9 | 0,3 | 0,51 | 0,87 | 1,13 | 277 |
| ENSG00000143819 | EPHX1 | 4,29 | 7,69 | 8,55 | 16 | 273 |
| ENSG00000233841 | HLA-C | 10,91 | 16,1 | 20,66 | 40,57 | 272 |
| ENSG00000280641 | CDH4 | 1,2 | 2,37 | 3,33 | 4,46 | 272 |
| ENSG00000284099 | SQSTM1 | 5,91 | 7,19 | 8,95 | 21,93 | 271 |
| ENSG00000178031 | ADAMTSL1 | 1,33 | 2,84 | 4,11 | 4,87 | 266 |
| ENSG00000167994 | RAB3IL1 | 0,59 | 0,93 | 1,39 | 2,16 | 266 |
| ENSG00000240463 | RPS19P3 | 0,41 | 0,44 | 1,01 | 1,5 | 266 |
| ENSG00000096433 | ITPR3 | 0,81 | 1,64 | 1,79 | 2,95 | 264 |
| ENSG00000230615 | AL139220,2 | 1,35 | 1,43 | 1,73 | 4,87 | 261 |
| ENSG00000204264 | PSMB8 | 0,28 | 0,65 | 0,66 | 1,01 | 261 |
| ENSG00000198879 | SFMBT2 | 1,31 | 2,45 | 3,14 | 4,72 | 260 |
| ENSG00000274737 | AC004466,2 | 0,69 | 1,03 | 1,09 | 2,48 | 259 |
| ENSG00000178573 | MAF | 7,7 | 12,18 | 13,98 | 27,57 | 258 |
| ENSG00000260549 | MT1L | 0,53 | 0,83 | 1,59 | 1,89 | 257 |
| ENSG00000227218 | AL157935,1 | 1,2 | 1,21 | 1,28 | 4,27 | 256 |
| ENSG00000169758 | TMEM266 | 0,34 | 0,59 | 0,66 | 1,21 | 256 |
| ENSG00000261170 | AC009053,3 | 0,48 | 0,6 | 0,7 | 1,7 | 254 |
| ENSG00000147010 | SH3KBP1 | 7,95 | 14,81 | 17,76 | 28,13 | 254 |
| ENSG00000103888 | CEMIP | 0,81 | 1,26 | 1,64 | 2,86 | 253 |
| ENSG00000171094 | ALK | 1,99 | 3,65 | 4,3 | 6,98 | 251 |
| ENSG00000102032 | RENBP | 1,25 | 1,43 | 1,45 | 4,37 | 250 |
| ENSG00000171608 | PIK3CD | 1,4 | 1,7 | 1,78 | 4,88 | 249 |
| ENSG00000171310 | CHST11 | 4,66 | 6,34 | 7,36 | 16,24 | 248 |
| ENSG00000184371 | CSF1 | 3,56 | 6,2 | 8,87 | 12,38 | 248 |
| ENSG00000133466 | C1QTNF6 | 14,08 | 22,71 | 35,46 | 48,9 | 247 |
| ENSG00000196187 | TMEM63A | 2,07 | 2,58 | 4,11 | 7,17 | 246 |
| ENSG00000082781 | ITGB5 | 59,25 | 59,64 | 70,12 | 204,99 | 246 |

| | | | | | | |
|---|---|---|---|---|---|---|
| ENSG00000186197 | EDARADD | 0,37 | 0,58 | 0,72 | 1,28 | 246 |
| ENSG00000162458 | FBLIM1 | 17,8 | 20,37 | 20,75 | 61,04 | 243 |
| ENSG00000127528 | KLF2 | 1,61 | 1,89 | 2,07 | 5,51 | 242 |
| ENSG00000174343 | CHRNA9 | 0,38 | 0,56 | 0,84 | 1,3 | 242 |
| ENSG00000141485 | SLC13A5 | 1,03 | 1,49 | 1,55 | 3,51 | 241 |
| ENSG00000114654 | EFCC1 | 2,1 | 2,81 | 3,05 | 7,15 | 240 |
| ENSG00000063015 | SEZ6 | 4,7 | 9,11 | 12,73 | 15,9 | 238 |
| ENSG00000156113 | KCNMA1 | 1,6 | 2,57 | 2,76 | 5,41 | 238 |
| ENSG00000052850 | ALX4 | 0,48 | 0,74 | 0,86 | 1,62 | 238 |
| ENSG00000273387 | AC005005,3 | 0,32 | 0,48 | 0,59 | 1,08 | 238 |
| ENSG00000175040 | CHST2 | 5,37 | 5,94 | 6,9 | 18,12 | 237 |
| ENSG00000162804 | SNED1 | 0,37 | 0,61 | 0,65 | 1,24 | 235 |
| ENSG00000104368 | PLAT | 14,19 | 26,71 | 27,74 | 47,5 | 235 |
| ENSG00000147526 | TACC1 | 25,38 | 38,5 | 41,98 | 84,92 | 235 |
| ENSG00000197956 | S100A6 | 48,6 | 98,72 | 140,75 | 162,42 | 234 |
| ENSG00000170485 | NPAS2 | 0,38 | 0,86 | 1,01 | 1,27 | 234 |
| ENSG00000277586 | NEFL | 23,76 | 33,95 | 46,96 | 79,25 | 234 |
| ENSG00000163702 | IL17RC | 2,09 | 2,8 | 3,12 | 6,96 | 233 |
| ENSG00000179715 | PCED1B | 0,48 | 0,72 | 0,82 | 1,59 | 231 |
| ENSG00000182253 | SYNM | 1,59 | 1,64 | 1,78 | 5,26 | 231 |
| ENSG00000140961 | OSGIN1 | 0,56 | 0,69 | 0,71 | 1,85 | 230 |
| ENSG00000127666 | TICAM1 | 0,43 | 0,45 | 0,49 | 1,42 | 230 |
| ENSG00000103316 | CRYM | 0,83 | 1,31 | 2,45 | 2,74 | 230 |
| ENSG00000231721 | LINC-PINT | 2,64 | 3,07 | 3,44 | 8,67 | 228 |
| ENSG00000204267 | TAP2 | 0,89 | 1,31 | 1,47 | 2,92 | 228 |
| ENSG00000275016 | AC015574,1 | 0,37 | 0,41 | 0,7 | 1,21 | 227 |
| ENSG00000123342 | MMP19 | 0,63 | 0,74 | 0,88 | 2,06 | 227 |
| ENSG00000179627 | ZBTB42 | 0,38 | 0,49 | 0,65 | 1,24 | 226 |
| ENSG00000059378 | PARP12 | 0,88 | 2,01 | 2,3 | 2,87 | 226 |
| ENSG00000233538 | AC017104,2 | 0,55 | 0,66 | 0,74 | 1,79 | 225 |
| ENSG00000226085 | UQCRFS1P1 | 0,71 | 0,76 | 1,3 | 2,31 | 225 |
| ENSG00000222460 | RN7SKP271 | 0,75 | 1,49 | 1,65 | 2,41 | 221 |
| ENSG00000188897 | AC099489,1 | 0,48 | 0,55 | 0,6 | 1,54 | 221 |
| ENSG00000103490 | PYCARD | 0,36 | 0,5 | 0,61 | 1,15 | 219 |
| ENSG00000108679 | LGALS3BP | 20,54 | 32,68 | 45,22 | 65,59 | 219 |
| ENSG00000123989 | CHPF | 11,53 | 14,35 | 18,11 | 36,75 | 219 |
| ENSG00000087494 | PTHLH | 2,23 | 2,41 | 3,19 | 7,1 | 218 |
| ENSG00000172159 | FRMD3 | 0,46 | 0,84 | 0,86 | 1,46 | 217 |
| ENSG00000273622 | CDC42EP5 | 4,42 | 8,42 | 8,85 | 13,99 | 217 |
| ENSG00000278915 | AC005609,3 | 0,79 | 0,97 | 1,09 | 2,49 | 215 |
| ENSG00000164251 | F2RL1 | 5,1 | 8,57 | 8,84 | 15,99 | 214 |
| ENSG00000168675 | LDLRAD4 | 6,32 | 7,99 | 8,58 | 19,81 | 213 |
| ENSG00000141933 | TPGS1 | 4,74 | 5,64 | 6,48 | 14,8 | 212 |
| ENSG00000178718 | RPP25 | 1,72 | 2,25 | 2,96 | 5,37 | 212 |
| ENSG00000143127 | ITGA10 | 0,41 | 0,58 | 0,63 | 1,28 | 212 |
| ENSG00000179242 | CDH4 | 5,26 | 8,28 | 10,83 | 16,38 | 211 |
| ENSG00000169991 | IFFO2 | 3 | 3,05 | 3,59 | 9,3 | 210 |
| ENSG00000285106 | AC016831,6 | 0,42 | 0,54 | 0,56 | 1,3 | 210 |
| ENSG00000229206 | AL162408,1 | 0,67 | 0,84 | 0,99 | 2,07 | 209 |
| ENSG00000206412 | GNL1 | 2,71 | 3,67 | 4,44 | 8,37 | 209 |

| | | | | | | |
|---|---|---|---|---|---|---|
| ENSG00000105639 | JAK3 | 1,44 | 2,14 | 2,44 | 4,44 | 208 |
| ENSG00000137936 | BCAR3 | 4,88 | 6,76 | 8,3 | 15,04 | 208 |
| ENSG00000206281 | TAPBP | 2,38 | 3,3 | 4,24 | 7,33 | 208 |
| ENSG00000075426 | FOSL2 | 5,6 | 7,06 | 8,16 | 17,2 | 207 |
| ENSG00000117643 | MAN1C1 | 2,15 | 3,15 | 4,85 | 6,6 | 207 |
| ENSG00000206315 | PBX2 | 0,35 | 0,57 | 0,63 | 1,07 | 206 |
| ENSG00000260070 | AC006960,3 | 0,41 | 0,53 | 0,58 | 1,25 | 205 |
| ENSG00000130589 | HELZ2 | 1,75 | 1,84 | 1,87 | 5,33 | 205 |
| ENSG00000213398 | LCAT | 1,85 | 2,27 | 2,99 | 5,6 | 203 |
| ENSG00000165886 | UBTD1 | 3,05 | 3,78 | 4,42 | 9,22 | 202 |
| ENSG00000173918 | C1QTNF1 | 0,91 | 1,72 | 2,27 | 2,75 | 202 |
| ENSG00000006210 | CX3CL1 | 2,48 | 3,62 | 5,08 | 7,49 | 202 |
| ENSG00000224593 | AC092427,1 | 1,03 | 1,79 | 1,98 | 3,11 | 202 |
| ENSG00000176978 | DPP7 | 2,12 | 3,06 | 3,47 | 6,4 | 202 |
| ENSG00000126785 | RHOJ | 9,31 | 17,78 | 20,15 | 27,99 | 201 |
| ENSG00000278839 | SERF1B | 2,03 | 3,49 | 3,62 | 6,09 | 200 |
| ENSG00000152642 | GPD1L | 1,68 | 2,16 | 2,92 | 5,04 | 200 |
| ENSG00000196196 | HRCT1 | 0,66 | 0,94 | 1,84 | 1,98 | 200 |
| ENSG00000104870 | FCGRT | 4,52 | 5,47 | 7,59 | 13,52 | 199 |
| ENSG00000186174 | BCL9L | 8,44 | 8,97 | 9,67 | 25,16 | 198 |
| ENSG00000263310 | SALL3 | 0,39 | 0,51 | 0,6 | 1,16 | 197 |
| ENSG00000276250 | AC127024,6 | 0,53 | 0,86 | 1,2 | 1,57 | 196 |
| ENSG00000197324 | LRP10 | 6,78 | 8,37 | 10,68 | 20,08 | 196 |
| ENSG00000132274 | TRIM22 | 9,07 | 13,96 | 16,94 | 26,85 | 196 |
| ENSG00000149260 | CAPN5 | 8,8 | 13,98 | 16,66 | 26,02 | 196 |
| ENSG00000118503 | TNFAIP3 | 2,71 | 6,85 | 7,45 | 8,01 | 196 |
| ENSG00000186567 | CEACAM19 | 0,65 | 0,85 | 1,28 | 1,92 | 195 |
| ENSG00000143786 | CNIH3 | 1,89 | 4,13 | 4,51 | 5,58 | 195 |
| ENSG00000214944 | ARHGEF28 | 0,39 | 0,7 | 0,84 | 1,15 | 195 |
| ENSG00000066248 | NGEF | 0,39 | 0,57 | 0,82 | 1,15 | 195 |
| ENSG00000184557 | SOCS3 | 13,49 | 19,8 | 21,92 | 39,74 | 195 |
| ENSG00000233621 | LINC01137 | 0,51 | 0,77 | 0,99 | 1,5 | 194 |
| ENSG00000110057 | UNC93B1 | 0,66 | 0,76 | 1,08 | 1,94 | 194 |
| ENSG00000134070 | IRAK2 | 0,42 | 0,44 | 0,58 | 1,23 | 193 |
| ENSG00000161835 | TAMALIN | 1,69 | 2,31 | 2,61 | 4,94 | 192 |
| ENSG00000121068 | TBX2 | 3,74 | 4,86 | 5,17 | 10,84 | 190 |
| ENSG00000183258 | DDX41 | 53,21 | 62,97 | 80,38 | 154,1 | 190 |
| ENSG00000125148 | MT2A | 33,54 | 40,48 | 48,51 | 97,11 | 190 |
| ENSG00000158458 | NRG2 | 0,55 | 0,91 | 1,06 | 1,59 | 189 |
| ENSG00000154274 | C4orf19 | 0,82 | 0,99 | 1,6 | 2,37 | 189 |
| ENSG00000099715 | PCDH11Y | 0,79 | 1,14 | 1,27 | 2,28 | 189 |
| ENSG00000189431 | RASSF10 | 0,92 | 1,21 | 1,4 | 2,65 | 188 |
| ENSG00000185340 | GAS2L1 | 2,87 | 2,9 | 3,43 | 8,26 | 188 |
| ENSG00000106785 | TRIM14 | 0,4 | 0,78 | 0,99 | 1,15 | 188 |
| ENSG00000169499 | PLEKHA2 | 1,4 | 2,84 | 3,25 | 4,02 | 187 |
| ENSG00000106236 | NPTX2 | 7,46 | 10,04 | 13,51 | 21,34 | 186 |
| ENSG00000133121 | STARD13 | 3,18 | 3,77 | 5,62 | 9,09 | 186 |
| ENSG00000182771 | GRID1 | 0,63 | 0,91 | 1,35 | 1,8 | 186 |
| ENSG00000123500 | COL10A1 | 0,54 | 0,62 | 1,2 | 1,54 | 185 |
| ENSG00000124104 | SNX21 | 4 | 4,67 | 6,16 | 11,35 | 184 |

| | | | | | | |
|---|---|---|---|---|---|---|
| ENSG00000271646 | IRF2-DT | 0,72 | 0,87 | 0,96 | 2,04 | 183 |
| ENSG00000230254 | HLA-E | 1,91 | 2,64 | 2,86 | 5,4 | 183 |
| ENSG00000229252 | HLA-E | 1,91 | 2,65 | 2,86 | 5,4 | 183 |
| ENSG00000030582 | GRN | 18,79 | 24,1 | 27,93 | 52,83 | 181 |
| ENSG00000171223 | JUNB | 6,62 | 7,16 | 9,45 | 18,57 | 181 |
| ENSG00000165084 | C8orf34 | 1,67 | 2,37 | 2,71 | 4,68 | 180 |
| ENSG00000278243 | GFUS | 3,93 | 4,18 | 5,02 | 11 | 180 |
| ENSG00000115556 | PLCD4 | 0,51 | 0,7 | 0,82 | 1,42 | 178 |
| ENSG00000064545 | TMEM161A | 5,87 | 6,57 | 7,29 | 16,18 | 176 |
| ENSG00000188763 | FZD9 | 0,65 | 0,86 | 1,35 | 1,79 | 175 |
| ENSG00000105088 | OLFM2 | 6,29 | 8,08 | 8,83 | 17,31 | 175 |
| ENSG00000232599 | AL008707,1 | 2,31 | 4,07 | 6,22 | 6,35 | 175 |
| ENSG00000187688 | TRPV2 | 2,67 | 2,95 | 3,44 | 7,33 | 175 |
| ENSG00000271815 | AC008897,3 | 0,66 | 1,39 | 1,5 | 1,81 | 174 |
| ENSG00000128335 | APOL2 | 4,01 | 5,03 | 6,07 | 10,98 | 174 |
| ENSG00000229520 | LINC00404 | 2,23 | 2,63 | 3,31 | 6,1 | 174 |
| ENSG00000111801 | BTN3A3 | 1,72 | 3,02 | 3,06 | 4,7 | 173 |
| ENSG00000102934 | PLLP | 1,52 | 1,9 | 2,1 | 4,14 | 172 |
| ENSG00000128294 | TPST2 | 5,41 | 6,35 | 8,15 | 14,73 | 172 |
| ENSG00000223839 | FAM95B1 | 1,26 | 1,51 | 1,65 | 3,43 | 172 |
| ENSG00000100599 | RIN3 | 2,02 | 2,79 | 2,94 | 5,48 | 171 |
| ENSG00000196843 | ARID5A | 3,24 | 3,96 | 4,36 | 8,78 | 171 |
| ENSG00000249684 | AC106795,2 | 0,48 | 0,87 | 1,1 | 1,3 | 171 |
| ENSG00000130147 | SH3BP4 | 17,18 | 21,95 | 24,23 | 46,4 | 170 |
| ENSG00000235659 | AL391987,4 | 0,42 | 0,52 | 0,54 | 1,13 | 169 |
| ENSG00000262243 | CES1 | 0,58 | 1,22 | 1,29 | 1,56 | 169 |
| ENSG00000224945 | AL353150,1 | 4,2 | 5,17 | 7,66 | 11,25 | 168 |
| ENSG00000164742 | ADCY1 | 4,18 | 5,44 | 6,82 | 11,17 | 167 |
| ENSG00000185585 | OLFML2A | 2,39 | 2,73 | 3,14 | 6,38 | 167 |
| ENSG00000142327 | RNPEPL1 | 3,45 | 3,47 | 4,34 | 9,19 | 166 |
| ENSG00000275214 | IFI27 | 0,91 | 1,26 | 2,34 | 2,42 | 166 |
| ENSG00000280707 | HPAT5 | 0,87 | 0,9 | 1,31 | 2,31 | 166 |
| ENSG00000130513 | GDF15 | 0,55 | 0,61 | 1,04 | 1,46 | 165 |
| ENSG00000102265 | TIMP1 | 26 | 34,58 | 52,38 | 68,93 | 165 |
| ENSG00000148180 | GSN | 27,62 | 43,12 | 46,96 | 73,14 | 165 |
| ENSG00000143507 | DUSP10 | 1,14 | 1,99 | 2,11 | 3 | 163 |
| ENSG00000100336 | APOL4 | 1,85 | 3,55 | 3,61 | 4,85 | 162 |
| ENSG00000196954 | CASP4 | 1,47 | 3,19 | 3,84 | 3,85 | 162 |
| ENSG00000196154 | S100A4 | 5,65 | 11,64 | 13,78 | 14,79 | 162 |
| ENSG00000257767 | AC002996,1 | 2,04 | 2,56 | 4,27 | 5,34 | 162 |
| ENSG00000167306 | MYO5B | 1,07 | 2 | 2,69 | 2,8 | 162 |
| ENSG00000239467 | AC007405,3 | 0,69 | 0,89 | 0,96 | 1,8 | 161 |
| ENSG00000166289 | PLEKHF1 | 3,91 | 5,37 | 5,57 | 10,19 | 161 |
| ENSG00000165475 | CRYL1 | 0,74 | 1,42 | 1,7 | 1,92 | 159 |
| ENSG00000231631 | PSMB8 | 0,44 | 0,69 | 1,12 | 1,14 | 159 |
| ENSG00000284691 | AC073111,3 | 1,88 | 2,26 | 2,45 | 4,87 | 159 |
| ENSG00000062038 | CDH3 | 8 | 11,89 | 16,32 | 20,68 | 159 |
| ENSG00000287263 | AC008875,3 | 0,57 | 1,15 | 1,46 | 1,47 | 158 |
| ENSG00000228981 | AC097658,1 | 0,45 | 0,75 | 0,98 | 1,16 | 158 |
| ENSG00000152953 | STK32B | 2,4 | 3,44 | 3,79 | 6,12 | 155 |

| | | | | | | |
|---|---|---|---|---|---|---|
| ENSG00000162873 | KLHDC8A | 14,74 | 17,82 | 25,37 | 37,55 | 155 |
| ENSG00000168792 | ABHD15 | 0,76 | 1,08 | 1,24 | 1,93 | 154 |
| ENSG00000139211 | AMIGO2 | 4,42 | 7,26 | 10,93 | 11,22 | 154 |
| ENSG00000257303 | AC073896,2 | 1,05 | 1,27 | 1,38 | 2,65 | 152 |
| ENSG00000107796 | ACTA2 | 122,21 | 143,23 | 158,75 | 308,37 | 152 |
| ENSG00000004776 | HSPB6 | 0,48 | 0,54 | 0,63 | 1,21 | 152 |
| ENSG00000119699 | TGFB3 | 1,91 | 2,34 | 2,79 | 4,8 | 151 |
| ENSG00000103249 | CLCN7 | 8,31 | 9,29 | 11,33 | 20,8 | 150 |
| ENSG00000235695 | HIGD2AP2 | 0,81 | 1,02 | 1,44 | 2,02 | 149 |
| ENSG00000159128 | IFNGR2 | 2,9 | 3,51 | 3,66 | 7,23 | 149 |
| ENSG00000235439 | DDX39B | 2,23 | 2,75 | 3,23 | 5,55 | 149 |
| ENSG00000172216 | CEBPB | 1,39 | 1,4 | 1,9 | 3,45 | 148 |
| ENSG00000205929 | C21orf62 | 2,72 | 3,82 | 5,97 | 6,75 | 148 |
| ENSG00000235257 | ITGA9-AS1 | 5,35 | 7,32 | 8,25 | 13,27 | 148 |
| ENSG00000089060 | SLC8B1 | 2,46 | 3,25 | 3,49 | 6,09 | 148 |
| ENSG00000140464 | PML | 10,92 | 12,83 | 13,51 | 26,98 | 147 |
| ENSG00000125845 | BMP2 | 0,85 | 1,49 | 2,07 | 2,1 | 147 |
| ENSG00000163032 | VSNL1 | 0,94 | 1,71 | 2,31 | 2,32 | 147 |
| ENSG00000198832 | SELENOM | 12,2 | 13,22 | 13,5 | 30,07 | 146 |
| ENSG00000140832 | MARVELD3 | 0,65 | 1,04 | 1,07 | 1,6 | 146 |
| ENSG00000144668 | ITGA9 | 1,87 | 2,51 | 2,76 | 4,6 | 146 |
| ENSG00000178882 | RFLNA | 1,61 | 2,06 | 2,54 | 3,96 | 146 |
| ENSG00000188152 | NUTM2G | 0,78 | 0,87 | 0,9 | 1,91 | 145 |
| ENSG00000228818 | AL359918,1 | 0,65 | 0,7 | 0,85 | 1,59 | 145 |
| ENSG00000275397 | DHRS11 | 0,81 | 1,36 | 1,43 | 1,98 | 144 |
| ENSG00000020577 | SAMD4A | 9,81 | 11,5 | 13,49 | 23,84 | 143 |
| ENSG00000183762 | KREMEN1 | 8,89 | 15,37 | 20,05 | 21,59 | 143 |
| ENSG00000186470 | BTN3A2 | 1,19 | 1,92 | 2,05 | 2,89 | 143 |
| ENSG00000240583 | AQP1 | 2 | 2,31 | 2,55 | 4,85 | 143 |
| ENSG00000106333 | PCOLCE | 10,22 | 15,87 | 16,2 | 24,66 | 141 |
| ENSG00000163520 | FBLN2 | 2,42 | 3,23 | 3,48 | 5,83 | 141 |
| ENSG00000130707 | ASS1 | 8,33 | 10,68 | 10,77 | 20,03 | 140 |
| ENSG00000250602 | AC093535,1 | 0,42 | 0,67 | 0,94 | 1,01 | 140 |
| ENSG00000198354 | DCAF12L2 | 0,89 | 1,14 | 1,73 | 2,14 | 140 |
| ENSG00000121904 | CSMD2 | 0,94 | 1,34 | 1,82 | 2,26 | 140 |
| ENSG00000185432 | METTL7A | 2,04 | 2,59 | 4,39 | 4,89 | 140 |
| ENSG00000165233 | CARD19 | 6,21 | 7,12 | 8,74 | 14,88 | 140 |
| ENSG00000159403 | C1R | 0,76 | 1,25 | 1,45 | 1,82 | 139 |
| ENSG00000230555 | AL450326,1 | 0,64 | 0,92 | 0,96 | 1,53 | 139 |
| ENSG00000090539 | CHRD | 0,47 | 0,73 | 0,79 | 1,12 | 138 |
| ENSG00000168938 | PPIC | 5,34 | 7,9 | 8,95 | 12,71 | 138 |
| ENSG00000224243 | SOX1-OT | 7,4 | 11,09 | 14,68 | 17,59 | 138 |
| ENSG00000198929 | NOS1AP | 2,42 | 3,77 | 4,14 | 5,75 | 138 |
| ENSG00000137834 | SMAD6 | 3,15 | 3,21 | 3,73 | 7,48 | 137 |
| ENSG00000283859 | NLGN2 | 5,55 | 6,54 | 6,75 | 13,16 | 137 |
| ENSG00000169992 | NLGN2 | 5,55 | 6,54 | 6,75 | 13,16 | 137 |
| ENSG00000258301 | VASH1-AS1 | 0,87 | 1,32 | 1,9 | 2,06 | 137 |
| ENSG00000139364 | TMEM132B | 5,35 | 8,54 | 10,61 | 12,64 | 136 |
| ENSG00000102359 | SRPX2 | 13,71 | 21,67 | 21,7 | 32,38 | 136 |
| ENSG00000111961 | SASH1 | 8,72 | 12,38 | 12,75 | 20,56 | 136 |

| | | | | | | |
|---|---|---|---|---|---|---|
| ENSG00000197536 | IRF1-AS1 | 0,93 | 1,2 | 1,46 | 2,19 | 135 |
| ENSG00000136099 | PCDH8 | 8,44 | 12,5 | 18,8 | 19,85 | 135 |
| ENSG00000239282 | CASTOR1 | 0,86 | 1,33 | 1,8 | 2,02 | 135 |
| ENSG00000107819 | SFXN3 | 4,59 | 6,57 | 6,72 | 10,78 | 135 |
| ENSG00000130309 | COLGALT1 | 25,97 | 26,71 | 26,83 | 60,96 | 135 |
| ENSG00000168077 | SCARA3 | 6,2 | 9,43 | 10,06 | 14,55 | 135 |
| ENSG00000223687 | ZNF311 | 0,44 | 0,63 | 0,71 | 1,03 | 134 |
| ENSG00000090581 | GNPTG | 7,68 | 7,87 | 8,13 | 17,92 | 133 |
| ENSG00000079337 | RAPGEF3 | 0,66 | 0,75 | 0,93 | 1,54 | 133 |
| ENSG00000140511 | HAPLN3 | 1,37 | 1,64 | 2,09 | 3,19 | 133 |
| ENSG00000169946 | ZFPM2 | 4,38 | 6,71 | 7,81 | 10,19 | 133 |
| ENSG00000088826 | SMOX | 4,1 | 4,93 | 5,53 | 9,53 | 132 |
| ENSG00000160226 | CFAP410 | 4,17 | 4,58 | 4,87 | 9,66 | 132 |
| ENSG00000165379 | LRFN5 | 2,43 | 3,7 | 4,55 | 5,62 | 131 |
| ENSG00000198563 | DDX39B | 2,4 | 2,75 | 3,23 | 5,55 | 131 |
| ENSG00000147481 | SNTG1 | 0,9 | 1,62 | 2,06 | 2,08 | 131 |
| ENSG00000087245 | MMP2 | 45,74 | 67,3 | 68,97 | 105,55 | 131 |
| ENSG00000139722 | VPS37B | 12,54 | 14,62 | 17,69 | 28,89 | 130 |
| ENSG00000050165 | DKK3 | 17,33 | 20,14 | 25,57 | 39,82 | 130 |
| ENSG00000158220 | ESYT3 | 0,47 | 0,49 | 0,63 | 1,08 | 130 |
| ENSG00000122642 | FKBP9 | 28,79 | 35,3 | 36,76 | 66,02 | 129 |
| ENSG00000099250 | NRP1 | 37,57 | 59,26 | 67,83 | 86,12 | 129 |
| ENSG00000136352 | NKX2-1 | 0,69 | 0,88 | 1,19 | 1,58 | 129 |
| ENSG00000115641 | FHL2 | 6,86 | 6,92 | 9,58 | 15,7 | 129 |
| ENSG00000083444 | PLOD1 | 21,97 | 24,89 | 25,39 | 50,18 | 128 |
| ENSG00000132530 | XAF1 | 1,71 | 2,14 | 2,33 | 3,9 | 128 |
| ENSG00000172638 | EFEMP2 | 7,19 | 9,99 | 10,48 | 16,39 | 128 |
| ENSG00000239697 | TNFSF12 | 1,34 | 1,68 | 2,44 | 3,05 | 128 |
| ENSG00000154803 | FLCN | 6,93 | 7,18 | 8,86 | 15,77 | 128 |
| ENSG00000172379 | ARNT2 | 15,32 | 20,44 | 25,96 | 34,85 | 127 |
| ENSG00000268087 | AC008764,2 | 2,49 | 2,74 | 2,84 | 5,66 | 127 |
| ENSG00000162512 | SDC3 | 36,33 | 41,46 | 45,8 | 82,57 | 127 |
| ENSG00000068903 | SIRT2 | 5,14 | 5,23 | 5,62 | 11,66 | 127 |
| ENSG00000103024 | NME3 | 2,37 | 2,52 | 3,59 | 5,37 | 127 |
| ENSG00000128052 | KDR | 5,18 | 5,84 | 9,31 | 11,73 | 126 |
| ENSG00000189114 | BLOC1S3 | 1,68 | 1,73 | 2,16 | 3,8 | 126 |
| ENSG00000198925 | ATG9A | 9,68 | 11,35 | 11,51 | 21,88 | 126 |
| ENSG00000139567 | ACVRL1 | 1 | 1,03 | 1,06 | 2,26 | 126 |
| ENSG00000140538 | NTRK3 | 3,23 | 4,95 | 5,72 | 7,29 | 126 |
| ENSG00000179772 | FOXS1 | 1,56 | 2,21 | 2,23 | 3,52 | 126 |
| ENSG00000167136 | ENDOG | 2,62 | 2,66 | 3,25 | 5,91 | 126 |
| ENSG00000113916 | BCL6 | 2,32 | 3,54 | 4,1 | 5,23 | 125 |
| ENSG00000089820 | ARHGAP4 | 0,75 | 0,81 | 0,99 | 1,69 | 125 |
| ENSG00000150630 | VEGFC | 0,95 | 1,25 | 1,46 | 2,14 | 125 |
| ENSG00000257524 | AL157935,2 | 0,49 | 0,54 | 0,8 | 1,1 | 124 |
| ENSG00000284161 | EEF2K | 2,3 | 2,88 | 3,03 | 5,15 | 124 |
| ENSG00000103044 | HAS3 | 2,22 | 2,61 | 3,4 | 4,97 | 124 |
| ENSG00000119408 | NEK6 | 21,73 | 27,96 | 35,31 | 48,6 | 124 |
| ENSG00000148053 | NTRK2 | 0,68 | 0,87 | 1,35 | 1,52 | 124 |
| ENSG00000066468 | FGFR2 | 2,32 | 3,59 | 3,67 | 5,18 | 123 |

| | | | | | | |
|---|---|---|---|---|---|---|
| ENSG00000285132 | AC270285,3 | 9,93 | 15,61 | 19,3 | 22,1 | 123 |
| ENSG00000135144 | DTX1 | 2,08 | 2,41 | 2,42 | 4,61 | 122 |
| ENSG00000159840 | ZYX | 31,54 | 32,75 | 34,2 | 69,65 | 121 |
| ENSG00000285443 | ZYX | 31,54 | 32,75 | 34,2 | 69,65 | 121 |
| ENSG00000002586 | CD99 | 173,6 | 230,39 | 270,46 | 382,72 | 120 |
| ENSG00000262246 | CORO7 | 4,55 | 5,38 | 6,28 | 10,02 | 120 |
| ENSG00000139190 | VAMP1 | 2,15 | 3,27 | 3,52 | 4,73 | 120 |
| ENSG00000106123 | EPHB6 | 0,65 | 1,02 | 1,15 | 1,43 | 120 |
| ENSG00000182580 | EPHB3 | 9,69 | 12,71 | 12,93 | 21,27 | 120 |
| ENSG00000176974 | SHMT1 | 4,26 | 4,74 | 6,5 | 9,35 | 119 |
| ENSG00000154928 | EPHB1 | 3,82 | 5,76 | 6,97 | 8,37 | 119 |
| ENSG00000173581 | CCDC106 | 3,04 | 3,57 | 3,68 | 6,64 | 118 |
| ENSG00000128536 | CDHR3 | 1,09 | 1,57 | 1,94 | 2,38 | 118 |
| ENSG00000264964 | AP001033,1 | 0,83 | 0,88 | 0,93 | 1,81 | 118 |
| ENSG00000137573 | SULF1 | 11,26 | 14,49 | 17,6 | 24,55 | 118 |
| ENSG00000171132 | PRKCE | 3,12 | 3,99 | 4,06 | 6,8 | 118 |
| ENSG00000133065 | SLC41A1 | 4,02 | 5,61 | 6,8 | 8,76 | 118 |
| ENSG00000208772 | SNORD94 | 50,49 | 54,55 | 59,6 | 109,98 | 118 |
| ENSG00000139926 | FRMD6 | 15,13 | 18,38 | 19,69 | 32,93 | 118 |
| ENSG00000224320 | HLA-A | 9,75 | 14,4 | 16,49 | 21,2 | 117 |
| ENSG00000229215 | HLA-A | 9,75 | 14,4 | 16,5 | 21,2 | 117 |
| ENSG00000135063 | FAM189A2 | 0,72 | 0,94 | 1,24 | 1,56 | 117 |
| ENSG00000144040 | SFXN5 | 8,27 | 9,64 | 12,55 | 17,91 | 117 |
| ENSG00000239887 | C1orf226 | 2,42 | 3,54 | 4,11 | 5,24 | 117 |
| ENSG00000233904 | HLA-E | 1,91 | 2,65 | 2,86 | 4,13 | 116 |
| ENSG00000231859 | AC079781,1 | 1,36 | 1,71 | 2,74 | 2,94 | 116 |
| ENSG00000124788 | ATXN1 | 4,25 | 5,74 | 6,69 | 9,17 | 116 |
| ENSG00000112769 | LAMA4 | 18,43 | 28,99 | 33,2 | 39,74 | 116 |
| ENSG00000234383 | CTBP2P8 | 0,64 | 0,9 | 0,92 | 1,38 | 116 |
| ENSG00000206493 | HLA-E | 1,91 | 2,64 | 2,86 | 4,11 | 115 |
| ENSG00000117280 | RAB29 | 0,94 | 1,04 | 1,67 | 2,02 | 115 |
| ENSG00000134107 | BHLHE40 | 6,99 | 8,87 | 8,93 | 15,01 | 115 |
| ENSG00000154096 | THY1 | 42,49 | 74,67 | 83,85 | 90,98 | 114 |
| ENSG00000116729 | WLS | 52,56 | 56,95 | 60,81 | 112,39 | 114 |
| ENSG00000166925 | TSC22D4 | 5,92 | 6,31 | 6,96 | 12,65 | 114 |
| ENSG00000199568 | RNU5A-1 | 305,74 | 366,71 | 470,28 | 652,56 | 113 |
| ENSG00000104946 | TBC1D17 | 6,05 | 7,25 | 7,34 | 12,91 | 113 |
| ENSG00000103966 | EHD4 | 8 | 9,56 | 11,09 | 17,06 | 113 |
| ENSG00000013288 | MAN2B2 | 2,45 | 3,18 | 3,69 | 5,22 | 113 |
| ENSG00000274229 | SOCS7 | 1,15 | 1,19 | 1,22 | 2,45 | 113 |
| ENSG00000135899 | SP110 | 3,25 | 5,98 | 6,04 | 6,92 | 113 |
| ENSG00000132561 | MATN2 | 4,08 | 4,42 | 4,62 | 8,66 | 112 |
| ENSG00000123096 | SSPN | 1,51 | 2,44 | 2,51 | 3,2 | 112 |
| ENSG00000089159 | PXN | 16,46 | 17,59 | 19,22 | 34,79 | 111 |
| ENSG00000132386 | SERPINF1 | 12,05 | 20,39 | 20,9 | 25,41 | 111 |
| ENSG00000170525 | PFKFB3 | 12,43 | 12,75 | 12,92 | 26,21 | 111 |
| ENSG00000196547 | MAN2A2 | 7,87 | 8,67 | 9,69 | 16,59 | 111 |
| ENSG00000173193 | PARP14 | 4,93 | 9,04 | 10,14 | 10,37 | 110 |
| ENSG00000184584 | STING1 | 0,91 | 1,11 | 1,29 | 1,91 | 110 |
| ENSG00000197019 | SERTAD1 | 1,62 | 1,72 | 1,99 | 3,4 | 110 |

| | | | | | | |
|---|---|---|---|---|---|---|
| ENSG00000133103 | COG6 | 7,28 | 8,08 | 8,23 | 15,27 | 110 |
| ENSG00000138641 | HERC3 | 2,1 | 3,54 | 3,94 | 4,4 | 110 |
| ENSG00000206503 | HLA-A | 18,33 | 26,23 | 28,45 | 38,38 | 109 |
| ENSG00000166682 | TMPRSS5 | 1,4 | 1,84 | 2,33 | 2,92 | 109 |
| ENSG00000167614 | TTYH1 | 15,06 | 16,1 | 16,87 | 31,32 | 108 |
| ENSG00000173281 | PPP1R3B | 6,69 | 7,91 | 8,83 | 13,91 | 108 |
| ENSG00000079385 | CEACAM1 | 0,69 | 0,77 | 1,07 | 1,43 | 107 |
| ENSG00000105974 | CAV1 | 24,12 | 27,36 | 29,21 | 49,97 | 107 |
| ENSG00000166780 | BMERB1 | 14,53 | 19,06 | 21,11 | 30,09 | 107 |
| ENSG00000249992 | TMEM158 | 4,38 | 5,4 | 6,82 | 9,05 | 107 |
| ENSG00000148120 | AOPEP | 14 | 14,89 | 16,37 | 28,91 | 107 |
| ENSG00000153902 | LGI4 | 0,49 | 0,56 | 0,71 | 1,01 | 106 |
| ENSG00000221955 | SLC12A8 | 1,84 | 2,04 | 3,06 | 3,79 | 106 |
| ENSG00000248905 | FMN1 | 2,11 | 3,27 | 3,4 | 4,34 | 106 |
| ENSG00000229422 | AL512625,2 | 0,58 | 0,68 | 0,88 | 1,19 | 105 |
| ENSG00000175352 | NRIP3 | 1,91 | 2,36 | 2,41 | 3,91 | 105 |
| ENSG00000187621 | TCL6 | 1,07 | 1,2 | 1,25 | 2,19 | 105 |
| ENSG00000166340 | TPP1 | 9,71 | 13,88 | 14,41 | 19,85 | 104 |
| ENSG00000166250 | CLMP | 3,93 | 4,06 | 6,74 | 8,03 | 104 |
| ENSG00000103260 | METRN | 18,67 | 20,45 | 27,46 | 38,1 | 104 |
| ENSG00000164932 | CTHRC1 | 23,92 | 37,01 | 46,86 | 48,81 | 104 |
| ENSG00000244691 | RPL10AP1 | 0,5 | 0,54 | 0,6 | 1,02 | 104 |
| ENSG00000100097 | LGALS1 | 294,09 | 378,01 | 430,22 | 599,6 | 104 |
| ENSG00000163013 | FBXO41 | 0,8 | 0,81 | 0,86 | 1,63 | 104 |
| ENSG00000256897 | AC018410,1 | 0,57 | 0,85 | 1,13 | 1,16 | 104 |
| ENSG00000181444 | ZNF467 | 0,6 | 0,71 | 1,19 | 1,22 | 103 |
| ENSG00000243449 | C4orf48 | 11,59 | 11,98 | 14,29 | 23,55 | 103 |
| ENSG00000287562 | AL109615,4 | 0,63 | 0,72 | 0,74 | 1,28 | 103 |
| ENSG00000012232 | EXTL3 | 25,73 | 30,46 | 36,03 | 52,2 | 103 |
| ENSG00000243232 | PCDHAC2 | 2,55 | 4,19 | 4,62 | 5,17 | 103 |
| ENSG00000240373 | SEC62-AS1 | 0,97 | 1,15 | 1,31 | 1,96 | 102 |
| ENSG00000214176 | PLEKHM1P1 | 2,85 | 3,07 | 3,11 | 5,75 | 102 |
| ENSG00000244731 | C4A | 0,66 | 0,72 | 1,23 | 1,33 | 102 |
| ENSG00000268350 | FAM156A | 4,02 | 4,06 | 6,22 | 8,1 | 101 |
| ENSG00000257446 | ZNF878 | 0,78 | 0,91 | 1,55 | 1,57 | 101 |
| ENSG00000267198 | AC091132,4 | 1,12 | 1,56 | 2,13 | 2,25 | 101 |
| ENSG00000282491 | AC243807,6 | 1,12 | 1,56 | 2,13 | 2,25 | 101 |
| ENSG00000145934 | TENM2 | 8,18 | 8,4 | 9,24 | 16,43 | 101 |
| ENSG00000277702 | AC239859,5 | 1,26 | 1,32 | 1,37 | 2,53 | 101 |
| ENSG00000022840 | RNF10 | 45,59 | 46,16 | 47,06 | 91,53 | 101 |
| ENSG00000141505 | ASGR1 | 3,41 | 3,86 | 4,98 | 6,82 | 100 |
| ENSG00000161013 | MGAT4B | 12,25 | 14,16 | 14,44 | 24,46 | 100 |
| ENSG00000284501 | MGAT4B | 12,25 | 14,16 | 14,44 | 24,46 | 100 |
| ENSG00000143845 | ETNK2 | 2,55 | 3,06 | 4,17 | 5,09 | 100 |
| ENSG00000148158 | SNX30 | 4,39 | 4,95 | 5,6 | 8,76 | 100 |
| ENSG00000166833 | NAV2 | 7,03 | 8,37 | 11,83 | 14,02 | 99 |
| ENSG00000134569 | LRP4 | 14,18 | 21,84 | 26,31 | 28,27 | 99 |
| ENSG00000123933 | MXD4 | 8,3 | 8,45 | 8,92 | 16,53 | 99 |
| ENSG00000135926 | TMBIM1 | 5,04 | 5,09 | 5,37 | 10,03 | 99 |
| ENSG00000188785 | ZNF548 | 4,44 | 4,98 | 5,19 | 8,82 | 99 |

| | | | | | | |
|---|---|---|---|---|---|---|
| ENSG00000228049 | POLR2J2 | 1,4 | 1,52 | 1,53 | 2,78 | 99 |
| ENSG00000167703 | SLC43A2 | 3,11 | 3,73 | 3,76 | 6,17 | 98 |
| ENSG00000244509 | APOBEC3C | 2,42 | 3,83 | 3,87 | 4,79 | 98 |
| ENSG00000181788 | SIAH2 | 3,56 | 4,59 | 5,99 | 7,03 | 97 |
| ENSG00000267279 | AC090409,1 | 2,06 | 2,31 | 2,78 | 4,06 | 97 |
| ENSG00000106367 | AP1S1 | 26,04 | 27,71 | 28,57 | 51,27 | 97 |
| ENSG00000198131 | ZNF544 | 10,15 | 11,82 | 13,89 | 19,92 | 96 |
| ENSG00000086730 | LAT2 | 0,79 | 0,83 | 0,9 | 1,55 | 96 |
| ENSG00000164171 | ITGA2 | 1,27 | 1,76 | 2,25 | 2,49 | 96 |
| ENSG00000223802 | CERS1 | 5,51 | 5,68 | 6,81 | 10,79 | 96 |
| ENSG00000263843 | AC022211,2 | 0,7 | 0,77 | 0,79 | 1,37 | 96 |
| ENSG00000134222 | PSRC1 | 64,69 | 79,2 | 94,19 | 126,46 | 95 |
| ENSG00000073464 | CLCN4 | 3,98 | 4,03 | 4,72 | 7,78 | 95 |
| ENSG00000134072 | CAMK1 | 2,72 | 3,47 | 3,86 | 5,31 | 95 |
| ENSG00000147862 | NFIB | 14,22 | 22,04 | 23,53 | 27,75 | 95 |
| ENSG00000269918 | AF131215,6 | 0,62 | 0,8 | 1,12 | 1,21 | 95 |
| ENSG00000205795 | CYS1 | 0,61 | 0,89 | 1,16 | 1,19 | 95 |
| ENSG00000160691 | SHC1 | 67,05 | 87,1 | 88,26 | 130,77 | 95 |
| ENSG00000186907 | RTN4RL2 | 0,78 | 0,8 | 0,99 | 1,52 | 95 |
| ENSG00000117385 | P3H1 | 24,51 | 26,83 | 27,69 | 47,69 | 95 |
| ENSG00000257221 | AC007569,1 | 0,55 | 0,66 | 0,77 | 1,07 | 95 |
| ENSG00000141522 | ARHGDIA | 125,07 | 127,49 | 129,2 | 242,64 | 94 |
| ENSG00000115183 | TANC1 | 7,49 | 10,92 | 11,71 | 14,53 | 94 |
| ENSG00000275401 | AL391095,3 | 1,03 | 1,06 | 1,09 | 1,99 | 93 |
| ENSG00000189283 | FHIT | 1,46 | 1,64 | 2,05 | 2,82 | 93 |
| ENSG00000106266 | SNX8 | 5,93 | 7,34 | 10,09 | 11,44 | 93 |
| ENSG00000100417 | PMM1 | 11,4 | 11,79 | 12,67 | 21,97 | 93 |
| ENSG00000109956 | B3GAT1 | 15,58 | 19,03 | 20,94 | 30,02 | 93 |
| ENSG00000150551 | LYPD1 | 8,77 | 11,73 | 14,96 | 16,89 | 93 |
| ENSG00000010704 | HFE | 1,35 | 2,03 | 2,38 | 2,6 | 93 |
| ENSG00000164949 | GEM | 1,08 | 1,77 | 1,99 | 2,08 | 93 |
| ENSG00000119632 | IFI27L2 | 5,81 | 9,04 | 10,7 | 11,18 | 92 |
| ENSG00000168016 | TRANK1 | 2,31 | 3,02 | 3,22 | 4,44 | 92 |
| ENSG00000034152 | MAP2K3 | 5,86 | 6,09 | 6,69 | 11,26 | 92 |
| ENSG00000140945 | CDH13 | 2,56 | 3 | 4,35 | 4,91 | 92 |
| ENSG00000018625 | ATP1A2 | 14,31 | 22,86 | 25,44 | 27,42 | 92 |
| ENSG00000160801 | PTH1R | 0,7 | 1,1 | 1,32 | 1,34 | 91 |
| ENSG00000177108 | ZDHHC22 | 1,13 | 1,78 | 1,91 | 2,16 | 91 |
| ENSG00000134780 | DAGLA | 2,14 | 2,33 | 2,44 | 4,09 | 91 |
| ENSG00000103227 | LMF1 | 4,3 | 4,88 | 6,23 | 8,21 | 91 |
| ENSG00000172667 | ZMAT3 | 2,64 | 3,72 | 3,77 | 5,03 | 91 |
| ENSG00000183741 | CBX6 | 12,61 | 12,93 | 13,09 | 24,02 | 90 |
| ENSG00000197816 | CCDC180 | 2,01 | 2,05 | 2,29 | 3,82 | 90 |
| ENSG00000177409 | SAMD9L | 0,79 | 1,33 | 1,44 | 1,5 | 90 |
| ENSG00000255571 | MIR9-3HG | 9,46 | 12,4 | 13,89 | 17,96 | 90 |
| ENSG00000109113 | RAB34 | 36,72 | 41,37 | 41,61 | 69,71 | 90 |
| ENSG00000064961 | HMG20B | 21,54 | 21,94 | 23,54 | 40,87 | 90 |
| ENSG00000271121 | NUDT4P2 | 13,6 | 16,65 | 18,87 | 25,8 | 90 |
| ENSG00000275993 | SIK1B | 2,78 | 2,91 | 3,09 | 5,26 | 89 |
| ENSG00000227110 | LMCD1-AS1 | 1,29 | 1,72 | 2,05 | 2,44 | 89 |

| | | | | | | |
|---|---|---|---|---|---|---|
| ENSG00000002330 | BAD | 20,47 | 21,5 | 25,28 | 38,64 | 89 |
| ENSG00000008256 | CYTH3 | 8,09 | 8,65 | 8,93 | 15,27 | 89 |
| ENSG00000154146 | NRGN | 6,75 | 6,76 | 6,99 | 12,65 | 87 |
| ENSG00000100605 | ITPK1 | 6,1 | 6,56 | 7,63 | 11,43 | 87 |
| ENSG00000143369 | ECM1 | 11,09 | 14,94 | 16,52 | 20,77 | 87 |
| ENSG00000151474 | FRMD4A | 48,14 | 63,64 | 65,6 | 90,04 | 87 |
| ENSG00000203706 | SERTAD4-AS1 | 2,07 | 2,67 | 2,89 | 3,87 | 87 |
| ENSG00000280963 | SERTAD4-AS1 | 2,07 | 2,67 | 2,89 | 3,87 | 87 |
| ENSG00000132669 | RIN2 | 14,32 | 18,96 | 22,04 | 26,75 | 87 |
| ENSG00000137266 | SLC22A23 | 12,02 | 13,29 | 13,5 | 22,44 | 87 |
| ENSG00000183779 | ZNF703 | 3,95 | 4,54 | 4,89 | 7,35 | 86 |
| ENSG00000137507 | LRRC32 | 0,92 | 0,99 | 1,32 | 1,71 | 86 |
| ENSG00000100906 | NFKBIA | 11,12 | 14,07 | 16,43 | 20,66 | 86 |
| ENSG00000126351 | THRA | 5,29 | 6,29 | 6,78 | 9,81 | 85 |
| ENSG00000283100 | SIRT2 | 4,57 | 5,23 | 6,51 | 8,47 | 85 |
| ENSG00000197746 | PSAP | 188,62 | 217,58 | 223,2 | 349,42 | 85 |
| ENSG00000131196 | NFATC1 | 1,55 | 1,75 | 2,02 | 2,87 | 85 |
| ENSG00000188916 | INSYN2A | 0,66 | 0,8 | 0,95 | 1,22 | 85 |
| ENSG00000157833 | GAREM2 | 3,97 | 4,56 | 6,35 | 7,33 | 85 |
| ENSG00000168874 | ATOH8 | 5,58 | 6,3 | 7,25 | 10,3 | 85 |
| ENSG00000106538 | RARRES2 | 14,62 | 14,74 | 17,35 | 26,97 | 84 |
| ENSG00000172936 | MYD88 | 9,13 | 10,38 | 11,42 | 16,78 | 84 |
| ENSG00000166311 | SMPD1 | 8,63 | 10,68 | 10,74 | 15,86 | 84 |
| ENSG00000272696 | AL359091,3 | 0,55 | 0,59 | 0,82 | 1,01 | 84 |
| ENSG00000154229 | PRKCA | 2,9 | 3,69 | 4,03 | 5,31 | 83 |
| ENSG00000184381 | PLA2G6 | 3,4 | 4,3 | 4,33 | 6,21 | 83 |
| ENSG00000099381 | SETD1A | 5,42 | 5,45 | 5,84 | 9,86 | 82 |
| ENSG00000169598 | DFFB | 2,31 | 2,4 | 2,42 | 4,2 | 82 |
| ENSG00000147234 | FRMPD3 | 0,87 | 0,96 | 1,21 | 1,58 | 82 |
| ENSG00000163513 | TGFBR2 | 9,51 | 12,79 | 13,51 | 17,27 | 82 |
| ENSG00000175662 | TOM1L2 | 7,68 | 8,36 | 9,23 | 13,93 | 81 |
| ENSG00000116096 | SPR | 0,59 | 0,62 | 0,68 | 1,07 | 81 |
| ENSG00000188186 | LAMTOR4 | 40,58 | 40,69 | 44,91 | 73,56 | 81 |
| ENSG00000149761 | NUDT22 | 13,23 | 14,7 | 14,94 | 23,98 | 81 |
| ENSG00000105605 | CACNG7 | 7,49 | 12,92 | 13,1 | 13,57 | 81 |
| ENSG00000119599 | DCAF4 | 2,64 | 2,83 | 3,27 | 4,78 | 81 |
| ENSG00000115935 | WIPF1 | 8,94 | 11 | 11,79 | 16,18 | 81 |
| ENSG00000122863 | CHST3 | 5,01 | 5,63 | 6,08 | 9,06 | 81 |
| ENSG00000142546 | NOSIP | 18,26 | 19,38 | 19,48 | 33 | 81 |
| ENSG00000126062 | TMEM115 | 11,33 | 11,51 | 12,37 | 20,46 | 81 |
| ENSG00000231566 | LINC02595 | 2,15 | 2,18 | 2,31 | 3,88 | 80 |
| ENSG00000069188 | SDK2 | 10,25 | 11,53 | 11,54 | 18,46 | 80 |
| ENSG00000189171 | S100A13 | 30,3 | 35,98 | 36,8 | 54,51 | 80 |
| ENSG00000114251 | WNT5A | 10,83 | 15,64 | 16,72 | 19,47 | 80 |
| ENSG00000213699 | SLC35F6 | 11,65 | 12,74 | 15,97 | 20,88 | 79 |
| ENSG00000136802 | LRRC8A | 15,13 | 15,19 | 15,29 | 27,1 | 79 |
| ENSG00000276600 | RAB7B | 0,91 | 1,21 | 1,4 | 1,63 | 79 |
| ENSG00000018408 | WWTR1 | 19,7 | 25,79 | 30,15 | 35,27 | 79 |
| ENSG00000110651 | CD81 | 75,26 | 77,14 | 86,6 | 134,67 | 79 |
| ENSG00000105928 | GSDME | 7,69 | 9,87 | 10,09 | 13,74 | 79 |

| | | | | | | |
|---|---|---|---|---|---|---|
| ENSG00000111087 | GLI1 | 1,62 | 2,58 | 2,65 | 2,89 | 78 |
| ENSG00000160233 | LRRC3 | 3,56 | 3,68 | 4,13 | 6,35 | 78 |
| ENSG00000159433 | STARD9 | 4,97 | 5,71 | 5,75 | 8,85 | 78 |
| ENSG00000165424 | ZCCHC24 | 6,97 | 9,79 | 9,97 | 12,4 | 78 |
| ENSG00000166750 | SLFN5 | 5,55 | 8,99 | 9,55 | 9,87 | 78 |
| ENSG00000005243 | COPZ2 | 5,59 | 6 | 6,22 | 9,93 | 78 |
| ENSG00000173295 | FAM86B3P | 2,26 | 2,36 | 2,78 | 4,01 | 77 |
| ENSG00000102178 | UBL4A | 6,73 | 7,18 | 7,33 | 11,93 | 77 |
| ENSG00000283060 | ID3 | 89,8 | 123,98 | 153,12 | 159,12 | 77 |
| ENSG00000117318 | ID3 | 89,8 | 123,98 | 153,12 | 159,12 | 77 |
| ENSG00000281836 | AC136352,12 | 0,6 | 0,78 | 1,01 | 1,06 | 77 |
| ENSG00000145794 | MEGF10 | 14,93 | 20,61 | 23,92 | 26,37 | 77 |
| ENSG00000101605 | MYOM1 | 0,9 | 1,13 | 1,5 | 1,58 | 76 |
| ENSG00000224897 | POT1-AS1 | 1,34 | 2,12 | 2,28 | 2,35 | 75 |
| ENSG00000119771 | KLHL29 | 3,45 | 4,1 | 4,2 | 6,05 | 75 |
| ENSG00000103005 | USB1 | 15,8 | 16,94 | 17,35 | 27,62 | 75 |
| ENSG00000010810 | FYN | 90,18 | 113,91 | 127,01 | 157,57 | 75 |
| ENSG00000124201 | ZNFX1 | 3,83 | 4,58 | 4,69 | 6,68 | 74 |
| ENSG00000117984 | CTSD | 28,88 | 29,57 | 35,68 | 50,33 | 74 |
| ENSG00000131018 | SYNE1 | 12,79 | 15,36 | 15,9 | 22,26 | 74 |
| ENSG00000087088 | BAX | 26,68 | 28,29 | 34,54 | 46,41 | 74 |
| ENSG00000095321 | CRAT | 6,37 | 7,66 | 8,33 | 11,08 | 74 |
| ENSG00000170271 | FAXDC2 | 2,98 | 3,32 | 3,94 | 5,18 | 74 |
| ENSG00000087460 | GNAS | 518,95 | 555,58 | 578,2 | 899,95 | 73 |
| ENSG00000144560 | VGLL4 | 19,37 | 24,75 | 24,86 | 33,54 | 73 |
| ENSG00000176842 | IRX5 | 4,76 | 5,53 | 6,76 | 8,23 | 73 |
| ENSG00000100647 | SUSD6 | 3,6 | 4,67 | 5,23 | 6,22 | 73 |
| ENSG00000138193 | PLCE1 | 3,69 | 3,85 | 4,7 | 6,37 | 73 |
| ENSG00000145012 | LPP | 26,1 | 26,74 | 31,05 | 45,04 | 73 |
| ENSG00000170382 | LRRN2 | 2,76 | 2,86 | 3,27 | 4,76 | 72 |
| ENSG00000124570 | SERPINB6 | 26,3 | 33,37 | 40,44 | 45,35 | 72 |
| ENSG00000173786 | CNP | 33,33 | 40,79 | 45,33 | 57,47 | 72 |
| ENSG00000157214 | STEAP2 | 0,58 | 0,76 | 0,94 | 1 | 72 |
| ENSG00000142227 | EMP3 | 23,11 | 25,89 | 30,66 | 39,8 | 72 |
| ENSG00000259630 | FABP5P9 | 0,75 | 0,77 | 0,91 | 1,29 | 72 |
| ENSG00000067182 | TNFRSF1A | 33,72 | 34,38 | 39,38 | 57,95 | 72 |
| ENSG00000105419 | MEIS3 | 27,74 | 28,3 | 33,87 | 47,66 | 72 |
| ENSG00000197457 | STMN3 | 38,5 | 39,71 | 39,99 | 66,12 | 72 |
| ENSG00000206120 | EGFEM1P | 9,97 | 12,25 | 17,01 | 17,12 | 72 |
| ENSG00000159208 | CIART | 0,99 | 1,65 | 1,66 | 1,7 | 72 |
| ENSG00000150967 | ABCB9 | 2,28 | 2,32 | 3,43 | 3,91 | 71 |
| ENSG00000198373 | WWP2 | 10,87 | 11,39 | 11,4 | 18,6 | 71 |
| ENSG00000166851 | PLK1 | 37,06 | 39,17 | 39,59 | 63,24 | 71 |
| ENSG00000276500 | BMS1P14 | 1,49 | 1,89 | 2,07 | 2,54 | 70 |
| ENSG00000125388 | GRK4 | 3,07 | 3,68 | 4,26 | 5,23 | 70 |
| ENSG00000063854 | HAGH | 14,83 | 15,17 | 16,38 | 25,25 | 70 |
| ENSG00000196083 | IL1RAP | 2,15 | 2,4 | 2,52 | 3,66 | 70 |
| ENSG00000182704 | TSKU | 9,81 | 10,75 | 11,6 | 16,66 | 70 |
| ENSG00000230749 | MEIS1-AS2 | 1,32 | 1,75 | 2,01 | 2,24 | 70 |
| ENSG00000090013 | BLVRB | 6,09 | 6,79 | 8,98 | 10,31 | 69 |

| | | | | | | |
|---|---|---|---|---|---|---|
| ENSG00000198597 | ZNF536 | 5,59 | 7,82 | 9,28 | 9,46 | 69 |
| ENSG00000107521 | HPS1 | 7,71 | 9,35 | 10,65 | 13,04 | 69 |
| ENSG00000161671 | EMC10 | 36,13 | 36,39 | 37,22 | 61,09 | 69 |
| ENSG00000131759 | RARA | 7,24 | 7,29 | 8,48 | 12,23 | 69 |
| ENSG00000100100 | PIK3IP1 | 1,44 | 1,89 | 2,02 | 2,43 | 69 |
| ENSG00000165995 | CACNB2 | 2,13 | 2,85 | 3,53 | 3,59 | 69 |
| ENSG00000186684 | CYP27C1 | 0,98 | 1,21 | 1,34 | 1,65 | 68 |
| ENSG00000133424 | LARGE1 | 7,52 | 7,6 | 7,83 | 12,66 | 68 |
| ENSG00000144579 | CTDSP1 | 13,38 | 13,73 | 14,25 | 22,5 | 68 |
| ENSG00000160606 | TLCD1 | 2,39 | 2,82 | 3,16 | 4,01 | 68 |
| ENSG00000180385 | EMC3-AS1 | 2,75 | 2,76 | 3,03 | 4,61 | 68 |
| ENSG00000179104 | TMTC2 | 10,16 | 11,8 | 16,34 | 17,02 | 68 |
| ENSG00000176014 | TUBB6 | 76,18 | 77,53 | 78,18 | 127,41 | 67 |
| ENSG00000155254 | MARVELD1 | 10,62 | 13,73 | 14,85 | 17,76 | 67 |
| ENSG00000103528 | SYT17 | 2,01 | 2,1 | 2,55 | 3,36 | 67 |
| ENSG00000087258 | GNAO1 | 5,76 | 7,77 | 8,13 | 9,62 | 67 |
| ENSG00000167930 | FAM234A | 27,98 | 30,77 | 32,29 | 46,72 | 67 |
| ENSG00000213347 | MXD3 | 8,6 | 8,66 | 10,61 | 14,32 | 67 |
| ENSG00000103066 | PLA2G15 | 4,32 | 5,12 | 5,17 | 7,19 | 66 |
| ENSG00000140092 | FBLN5 | 2,41 | 2,57 | 2,82 | 4,01 | 66 |
| ENSG00000277270 | AL160412,1 | 1,65 | 1,99 | 2,2 | 2,74 | 66 |
| ENSG00000184979 | USP18 | 1,08 | 1,58 | 1,68 | 1,79 | 66 |
| ENSG00000121440 | PDZRN3 | 7,99 | 11,79 | 12,69 | 13,24 | 66 |
| ENSG00000226149 | AL356124,1 | 0,69 | 0,87 | 0,88 | 1,14 | 65 |
| ENSG00000158552 | ZFAND2B | 8,44 | 8,88 | 9,48 | 13,94 | 65 |
| ENSG00000267648 | AC060766,5 | 0,63 | 0,88 | 1 | 1,04 | 65 |
| ENSG00000131323 | TRAF3 | 7,5 | 10,42 | 11,07 | 12,38 | 65 |
| ENSG00000109079 | TNFAIP1 | 11,22 | 12,04 | 12,61 | 18,52 | 65 |
| ENSG00000185670 | ZBTB3 | 2,39 | 2,62 | 2,75 | 3,94 | 65 |
| ENSG00000144749 | LRIG1 | 27,35 | 37,13 | 39,8 | 45,04 | 65 |
| ENSG00000110660 | SLC35F2 | 1,13 | 1,18 | 1,37 | 1,86 | 65 |
| ENSG00000135929 | CYP27A1 | 2,49 | 2,77 | 2,95 | 4,09 | 64 |
| ENSG00000242193 | CRYZL2P | 1,49 | 1,77 | 1,84 | 2,44 | 64 |
| ENSG00000250510 | GPR162 | 4,49 | 5,24 | 5,29 | 7,34 | 63 |
| ENSG00000254416 | LINC02732 | 1,2 | 1,25 | 1,91 | 1,96 | 63 |
| ENSG00000227500 | SCAMP4 | 18,09 | 20,25 | 22,33 | 29,52 | 63 |
| ENSG00000144567 | RETREG2 | 16,73 | 17,96 | 19,14 | 27,29 | 63 |
| ENSG00000237854 | LINC00674 | 2,23 | 3,04 | 3,62 | 3,63 | 63 |
| ENSG00000185000 | DGAT1 | 2,47 | 2,82 | 2,89 | 4,02 | 63 |
| ENSG00000285482 | DGAT1 | 2,47 | 2,82 | 2,89 | 4,02 | 63 |
| ENSG00000026297 | RNASET2 | 4,94 | 5,31 | 6,04 | 8,03 | 63 |
| ENSG00000099991 | CABIN1 | 22,14 | 22,16 | 23,44 | 35,96 | 62 |
| ENSG00000040633 | PHF23 | 22,78 | 23,55 | 24,58 | 36,9 | 62 |
| ENSG00000160050 | CCDC28B | 7,84 | 9,75 | 9,85 | 12,68 | 62 |
| ENSG00000102007 | PLP2 | 9,32 | 11,37 | 13,76 | 15,05 | 61 |
| ENSG00000104964 | TLE5 | 83,51 | 110,43 | 133,15 | 134,67 | 61 |
| ENSG00000138760 | SCARB2 | 39,98 | 55,4 | 61,08 | 64,47 | 61 |
| ENSG00000185033 | SEMA4B | 4,77 | 4,93 | 5,65 | 7,69 | 61 |
| ENSG00000112699 | GMDS | 7,84 | 8,33 | 8,87 | 12,63 | 61 |
| ENSG00000100243 | CYB5R3 | 49,71 | 50,41 | 54,44 | 80,05 | 61 |

| | | | | | | |
|---|---|---|---|---|---|---|
| ENSG00000181830 | SLC35C1 | 3,84 | 4,18 | 4,49 | 6,18 | 61 |
| ENSG00000107249 | GLIS3 | 8,42 | 8,91 | 9 | 13,55 | 61 |
| ENSG00000197586 | ENTPD6 | 15,53 | 15,54 | 15,57 | 24,98 | 61 |
| ENSG00000068323 | TFE3 | 9,74 | 10,76 | 11,65 | 15,66 | 61 |
| ENSG00000104853 | CLPTM1 | 27,72 | 32,31 | 32,6 | 44,56 | 61 |
| ENSG00000048140 | TSPAN17 | 13,76 | 14,76 | 15,6 | 22,1 | 61 |
| ENSG00000226419 | SLC16A1-AS1 | 2,89 | 3,09 | 3,22 | 4,63 | 60 |
| ENSG00000048828 | FAM120A | 25,53 | 26,01 | 26,84 | 40,89 | 60 |
| ENSG00000138061 | CYP1B1 | 16,49 | 18,98 | 19,15 | 26,4 | 60 |
| ENSG00000131748 | STARD3 | 17,26 | 17,84 | 18,33 | 27,59 | 60 |
| ENSG00000176134 | AL445665,1 | 1,02 | 1,04 | 1,54 | 1,63 | 60 |
| ENSG00000168310 | IRF2 | 5,29 | 8,13 | 8,19 | 8,42 | 59 |
| ENSG00000170581 | STAT2 | 13,71 | 17,32 | 18,36 | 21,82 | 59 |
| ENSG00000065485 | PDIA5 | 12,95 | 13,25 | 14,07 | 20,61 | 59 |
| ENSG00000241973 | PI4KA | 11,99 | 12,82 | 13,38 | 19,08 | 59 |
| ENSG00000134363 | FST | 29,54 | 33,24 | 34,34 | 47 | 59 |
| ENSG00000021300 | PLEKHB1 | 6,86 | 7,66 | 9,05 | 10,91 | 59 |
| ENSG00000182022 | CHST15 | 7,69 | 11,2 | 11,62 | 12,23 | 59 |
| ENSG00000198561 | CTNND1 | 36,61 | 47,52 | 49,61 | 58,22 | 59 |
| ENSG00000278126 | AC139768,1 | 1,63 | 2,02 | 2,38 | 2,59 | 59 |
| ENSG00000129566 | TEP1 | 3,11 | 3,23 | 3,47 | 4,94 | 59 |
| ENSG00000204673 | AKT1S1 | 12,97 | 13,6 | 14,24 | 20,56 | 59 |
| ENSG00000013619 | MAMLD1 | 2,12 | 2,7 | 3,01 | 3,36 | 58 |
| ENSG00000134853 | PDGFRA | 2,19 | 3,11 | 3,36 | 3,47 | 58 |
| ENSG00000105722 | ERF | 19,7 | 19,81 | 20,05 | 31,11 | 58 |
| ENSG00000064490 | RFXANK | 30,66 | 32,35 | 33,72 | 48,4 | 58 |
| ENSG00000103647 | CORO2B | 20,77 | 25,37 | 30,16 | 32,73 | 58 |
| ENSG00000168610 | STAT3 | 35,14 | 44,67 | 47,74 | 55,34 | 57 |
| ENSG00000124782 | RREB1 | 7,66 | 9,06 | 9,56 | 11,99 | 57 |
| ENSG00000099290 | WASHC2A | 12,49 | 13,55 | 14,24 | 19,54 | 56 |
| ENSG00000181350 | LRRC75A | 3,12 | 4,43 | 4,81 | 4,86 | 56 |
| ENSG00000106268 | NUDT1 | 28,77 | 30,47 | 31,48 | 44,58 | 55 |
| ENSG00000135404 | CD63 | 220,47 | 285,12 | 315,06 | 341,34 | 55 |
| ENSG00000067715 | SYT1 | 14,93 | 16,87 | 17,09 | 23,07 | 55 |
| ENSG00000141858 | SAMD1 | 21,41 | 22,14 | 24,96 | 33,06 | 54 |
| ENSG00000042445 | RETSAT | 4,29 | 4,99 | 5,12 | 6,62 | 54 |
| ENSG00000065308 | TRAM2 | 10,09 | 11,55 | 12,7 | 15,55 | 54 |
| ENSG00000177628 | GBA | 7,8 | 9,07 | 9,98 | 11,99 | 54 |
| ENSG00000075234 | TTC38 | 6,96 | 7,02 | 7,27 | 10,66 | 53 |
| ENSG00000110917 | MLEC | 42,48 | 54,3 | 59,56 | 65,06 | 53 |
| ENSG00000160211 | G6PD | 26,05 | 26,61 | 27,69 | 39,89 | 53 |
| ENSG00000277150 | F8A3 | 3,39 | 3,87 | 4,28 | 5,19 | 53 |
| ENSG00000197177 | ADGRA1 | 1,85 | 2 | 2,44 | 2,83 | 53 |
| ENSG00000278771 | RN7SL3 | 2231,79 | 2540,06 | 2745,17 | 3413,9 | 53 |
| ENSG00000198715 | GLMP | 11,19 | 14,54 | 15,21 | 17,08 | 53 |
| ENSG00000272620 | AFAP1-AS1 | 0,78 | 1,01 | 1,02 | 1,19 | 53 |
| ENSG00000040487 | SLC66A1 | 6,4 | 6,48 | 6,71 | 9,74 | 52 |
| ENSG00000236753 | MKLN1-AS | 1,19 | 1,53 | 1,55 | 1,81 | 52 |
| ENSG00000058668 | ATP2B4 | 17,3 | 20,31 | 22,19 | 26,24 | 52 |
| ENSG00000175283 | DOLK | 5,28 | 5,67 | 6,12 | 8 | 52 |

| | | | | | | |
|---|---|---|---|---|---|---|
| ENSG00000124126 | PREX1 | 27,62 | 33,38 | 34,52 | 41,83 | 51 |
| ENSG00000167323 | STIM1 | 7,31 | 9,48 | 10,67 | 11,07 | 51 |
| ENSG00000172757 | CFL1 | 740,16 | 759,4 | 805,09 | 1117,68 | 51 |
| ENSG00000103449 | SALL1 | 14,29 | 16,26 | 19,31 | 21,54 | 51 |
| ENSG00000126767 | ELK1 | 6,07 | 7,16 | 8,13 | 9,14 | 51 |
| ENSG00000198108 | CHSY3 | 2,08 | 2,78 | 2,96 | 3,13 | 50 |
| ENSG00000182534 | MXRA7 | 37,42 | 40,44 | 44,72 | 56,23 | 50 |
| ENSG00000073060 | SCARB1 | 10,18 | 10,88 | 12,49 | 15,29 | 50 |
| ENSG00000163481 | RNF25 | 10,85 | 11,55 | 11,56 | 16,27 | 50 |
| ENSG00000142089 | IFITM3 | 172,91 | 202,4 | 232,48 | 259,19 | 50 |
| ENSG00000175416 | CLTB | 10,3 | 11,83 | 12,14 | 15,42 | 50 |
| ENSG00000263740 | RN7SL4P | 71,15 | 77,59 | 85,05 | 106,49 | 50 |
| ENSG00000151929 | BAG3 | 8,86 | 9,55 | 9,89 | 13,26 | 50 |
| ENSG00000147509 | RGS20 | 8,66 | 10,05 | 12,81 | 12,96 | 50 |
| ENSG00000091947 | TMEM101 | 6,2 | 6,56 | 6,69 | 9,27 | 50 |
| ENSG00000154447 | SH3RF1 | 7,34 | 8,08 | 9,07 | 10,97 | 49 |
| ENSG00000129925 | PGAP6 | 12,58 | 13,88 | 14,52 | 18,78 | 49 |
| ENSG00000006194 | ZNF263 | 10,41 | 10,96 | 11,35 | 15,52 | 49 |
| ENSG00000137207 | YIPF3 | 33,1 | 43,15 | 43,52 | 49,34 | 49 |
| ENSG00000119673 | ACOT2 | 7,24 | 7,27 | 8,15 | 10,78 | 49 |
| ENSG00000288033 | AC137894,4 | 2,32 | 2,47 | 2,61 | 3,45 | 49 |
| ENSG00000122550 | KLHL7 | 37,85 | 39,52 | 39,65 | 56,22 | 49 |
| ENSG00000103145 | HCFC1R1 | 15,38 | 17,06 | 18,69 | 22,8 | 48 |
| ENSG00000104957 | CCDC130 | 13,22 | 13,32 | 13,7 | 19,59 | 48 |
| ENSG00000262580 | AC087741,1 | 1,08 | 1,13 | 1,24 | 1,6 | 48 |
| ENSG00000183722 | LHFPL6 | 28,55 | 34,81 | 37,74 | 42,27 | 48 |
| ENSG00000107821 | KAZALD1 | 3,23 | 3,41 | 3,68 | 4,78 | 48 |
| ENSG00000159348 | CYB5R1 | 7,61 | 8,03 | 8,37 | 11,26 | 48 |
| ENSG00000073910 | FRY | 2,05 | 2,34 | 2,46 | 3,03 | 48 |
| ENSG00000077782 | FGFR1 | 68,26 | 87,47 | 88,06 | 100,8 | 48 |
| ENSG00000275903 | PSMB3 | 21,99 | 23,4 | 24,32 | 32,45 | 48 |
| ENSG00000213689 | TREX1 | 1,77 | 2,35 | 2,38 | 2,61 | 47 |
| ENSG00000251602 | AL928654,1 | 1,96 | 2,04 | 2,1 | 2,88 | 47 |
| ENSG00000182718 | ANXA2 | 377,24 | 388,97 | 391,89 | 552,84 | 47 |
| ENSG00000132109 | TRIM21 | 4 | 4,74 | 5,02 | 5,86 | 47 |
| ENSG00000131653 | TRAF7 | 15,96 | 17,28 | 19,96 | 23,36 | 46 |
| ENSG00000137216 | TMEM63B | 11,2 | 12,05 | 13,68 | 16,36 | 46 |
| ENSG00000237441 | RGL2 | 5,16 | 5,28 | 5,53 | 7,53 | 46 |
| ENSG00000274333 | CU633967,1 | 11,12 | 12,02 | 12,41 | 16,2 | 46 |
| ENSG00000135956 | TMEM127 | 8,77 | 10,63 | 10,64 | 12,76 | 45 |
| ENSG00000136868 | SLC31A1 | 8,59 | 9,05 | 9,12 | 12,49 | 45 |
| ENSG00000034510 | TMSB10 | 2237,87 | 2434,12 | 2538,69 | 3253,38 | 45 |
| ENSG00000197619 | ZNF615 | 4,1 | 4,61 | 5,56 | 5,96 | 45 |
| ENSG00000163638 | ADAMTS9 | 10,56 | 13,15 | 13,2 | 15,33 | 45 |
| ENSG00000103855 | CD276 | 60,21 | 66,57 | 70,58 | 87,32 | 45 |
| ENSG00000164951 | PDP1 | 13,8 | 17,87 | 18,84 | 20,01 | 45 |
| ENSG00000260916 | CCPG1 | 10 | 11,73 | 12,21 | 14,42 | 44 |
| ENSG00000259953 | AL138756,1 | 1,03 | 1,05 | 1,11 | 1,48 | 44 |
| ENSG00000263290 | SCAMP3 | 9,2 | 10,02 | 11,86 | 13,19 | 43 |
| ENSG00000155366 | RHOC | 72,96 | 76,29 | 86,24 | 104,55 | 43 |

| | | | | | | |
|---|---|---|---|---|---|---|
| ENSG00000048707 | VPS13D | 7,35 | 7,96 | 8,89 | 10,52 | 43 |
| ENSG00000204950 | LRRC10B | 3,3 | 4,07 | 4,52 | 4,72 | 43 |
| ENSG00000119487 | MAPKAP1 | 32,18 | 33,03 | 34,81 | 45,94 | 43 |
| ENSG00000106330 | MOSPD3 | 6,72 | 7,87 | 8,34 | 9,59 | 43 |
| ENSG00000237493 | AC034102,1 | 0,75 | 0,93 | 0,98 | 1,07 | 43 |
| ENSG00000072952 | IRAG1 | 4,25 | 4,65 | 5,23 | 6,06 | 43 |
| ENSG00000257169 | AC125612,1 | 1,01 | 1,27 | 1,34 | 1,44 | 43 |
| ENSG00000162430 | SELENON | 40,79 | 47,58 | 52,98 | 58,02 | 42 |
| ENSG00000153815 | CMIP | 16,82 | 17,9 | 17,98 | 23,89 | 42 |
| ENSG00000197043 | ANXA6 | 76,39 | 88,81 | 90,89 | 108,48 | 42 |
| ENSG00000226472 | AC008013,1 | 1,36 | 1,66 | 1,72 | 1,93 | 42 |
| ENSG00000182240 | BACE2 | 9,24 | 11,03 | 11,98 | 13,05 | 41 |
| ENSG00000154328 | NEIL2 | 7,97 | 8,14 | 8,81 | 11,18 | 40 |
| ENSG00000179899 | PHC1P1 | 4,46 | 5,14 | 5,36 | 6,25 | 40 |
| ENSG00000171877 | FRMD5 | 2,12 | 2,41 | 2,79 | 2,97 | 40 |
| ENSG00000155093 | PTPRN2 | 8,47 | 8,48 | 9,08 | 11,86 | 40 |
| ENSG00000064687 | ABCA7 | 1,1 | 1,17 | 1,24 | 1,54 | 40 |
| ENSG00000170549 | IRX1 | 10,19 | 10,79 | 11,37 | 14,26 | 40 |
| ENSG00000163902 | RPN1 | 75,25 | 76,24 | 79,98 | 104,87 | 39 |
| ENSG00000157593 | SLC35B2 | 15,39 | 15,99 | 18,42 | 21,41 | 39 |
| ENSG00000088298 | EDEM2 | 4,9 | 5,53 | 5,9 | 6,81 | 39 |
| ENSG00000026025 | VIM | 1312,06 | 1504,74 | 1641,76 | 1821,8 | 39 |
| ENSG00000185339 | TCN2 | 11,64 | 12,43 | 13,19 | 16,11 | 38 |
| ENSG00000099910 | KLHL22 | 11,78 | 12,28 | 12,55 | 16,28 | 38 |
| ENSG00000064666 | CNN2 | 165,1 | 181,86 | 185,36 | 227,55 | 38 |
| ENSG00000185043 | CIB1 | 9,69 | 9,9 | 10,06 | 13,35 | 38 |
| ENSG00000120709 | FAM53C | 15,41 | 15,78 | 16,19 | 21,21 | 38 |
| ENSG00000055070 | SZRD1 | 46,13 | 46,84 | 49,3 | 63,32 | 37 |
| ENSG00000103353 | UBFD1 | 30,45 | 34,24 | 35,61 | 41,62 | 37 |
| ENSG00000090857 | PDPR | 6,93 | 7,36 | 7,57 | 9,45 | 36 |
| ENSG00000162073 | PAQR4 | 9,82 | 10,08 | 11,42 | 13,38 | 36 |
| ENSG00000127948 | POR | 15,99 | 16,41 | 19,28 | 21,75 | 36 |
| ENSG00000099977 | DDT | 9,96 | 10,29 | 10,66 | 13,5 | 36 |
| ENSG00000149483 | TMEM138 | 15,93 | 16,57 | 16,72 | 21,57 | 35 |
| ENSG00000166173 | LARP6 | 9,03 | 9,73 | 10,7 | 12,22 | 35 |
| ENSG00000271380 | AL451085,2 | 2,14 | 2,21 | 2,45 | 2,89 | 35 |
| ENSG00000152217 | SETBP1 | 11,97 | 13,15 | 14,32 | 16,15 | 35 |
| ENSG00000145349 | CAMK2D | 29,28 | 32,51 | 35,79 | 39,49 | 35 |
| ENSG00000162616 | DNAJB4 | 10,67 | 11,17 | 11,87 | 14,39 | 35 |
| ENSG00000117305 | HMGCL | 7,66 | 7,93 | 8,03 | 10,33 | 35 |
| ENSG00000248008 | NRAV | 5,09 | 5,5 | 6,16 | 6,85 | 35 |
| ENSG00000178096 | BOLA1 | 3,42 | 4,05 | 4,55 | 4,6 | 35 |
| ENSG00000278576 | AL162171,3 | 0,87 | 0,95 | 0,98 | 1,17 | 34 |
| ENSG00000173039 | RELA | 15,71 | 17,72 | 17,82 | 21,12 | 34 |
| ENSG00000132128 | LRRC41 | 22,53 | 23,2 | 24,75 | 29,97 | 33 |
| ENSG00000184009 | ACTG1 | 2413,95 | 2572,56 | 2633,49 | 3210,08 | 33 |
| ENSG00000184743 | ATL3 | 7,97 | 9,98 | 10 | 10,59 | 33 |
| ENSG00000187678 | SPRY4 | 51,05 | 61,24 | 63,91 | 67,65 | 33 |
| ENSG00000138867 | GUCD1 | 15,96 | 16,42 | 16,85 | 21,14 | 32 |
| ENSG00000151353 | TMEM18 | 9,88 | 11,36 | 11,6 | 13,04 | 32 |

| | | | | | | |
|---|---|---|---|---|---|---|
| ENSG00000105778 | AVL9 | 14,06 | 15,92 | 17,95 | 18,53 | 32 |
| ENSG00000183579 | ZNRF3 | 4,28 | 4,37 | 4,57 | 5,64 | 32 |
| ENSG00000172301 | COPRS | 29,33 | 30,02 | 33,3 | 38,45 | 31 |
| ENSG00000173402 | DAG1 | 26,09 | 28,03 | 31,09 | 34,19 | 31 |
| ENSG00000122705 | CLTA | 95,94 | 101,02 | 108,44 | 125,29 | 31 |
| ENSG00000173457 | PPP1R14B | 100,23 | 102,02 | 104,94 | 130,85 | 31 |
| ENSG00000182873 | PRKCZ-AS1 | 1,69 | 1,86 | 1,9 | 2,2 | 30 |
| ENSG00000116962 | NID1 | 10,42 | 11,81 | 12,08 | 13,53 | 30 |
| ENSG00000151702 | FLI1 | 1,56 | 1,9 | 1,92 | 2,02 | 29 |
| ENSG00000154889 | MPPE1 | 8,99 | 9,73 | 11,45 | 11,64 | 29 |
| ENSG00000162889 | MAPKAPK2 | 24,12 | 24,65 | 26,19 | 31,19 | 29 |
| ENSG00000142197 | DOP1B | 4,91 | 5,32 | 5,8 | 6,34 | 29 |
| ENSG00000189164 | ZNF527 | 2,58 | 2,6 | 2,94 | 3,31 | 28 |
| ENSG00000101460 | MAP1LC3A | 6,72 | 6,88 | 7,33 | 8,62 | 28 |
| ENSG00000107175 | CREB3 | 19,29 | 19,51 | 19,83 | 24,71 | 28 |
| ENSG00000155252 | PI4K2A | 6,45 | 6,78 | 7,28 | 8,26 | 28 |
| ENSG00000174529 | TMEM81 | 1,39 | 1,52 | 1,73 | 1,78 | 28 |
| ENSG00000131944 | FAAP24 | 2,32 | 2,35 | 2,72 | 2,97 | 28 |
| ENSG00000075461 | CACNG4 | 30,86 | 32 | 36,54 | 39,44 | 28 |
| ENSG00000085063 | CD59 | 43,36 | 48,39 | 53,41 | 55,18 | 27 |
| ENSG00000087086 | FTL | 348,02 | 370,47 | 441,19 | 441,8 | 27 |
| ENSG00000108175 | ZMIZ1 | 26,7 | 29,13 | 31,17 | 33,85 | 27 |
| ENSG00000163132 | MSX1 | 6,47 | 6,49 | 7,64 | 8,2 | 27 |
| ENSG00000004059 | ARF5 | 60,12 | 63,42 | 69,79 | 76,13 | 27 |
| ENSG00000145391 | SETD7 | 6,28 | 6,55 | 6,82 | 7,95 | 27 |
| ENSG00000258315 | C17orf49 | 65,02 | 66,29 | 72,23 | 82,28 | 27 |
| ENSG00000107551 | RASSF4 | 5,35 | 5,47 | 5,67 | 6,76 | 26 |
| ENSG00000160218 | TRAPPC10 | 11,07 | 11,11 | 11,5 | 13,97 | 26 |
| ENSG00000164054 | SHISA5 | 45,72 | 49,3 | 52,7 | 57,58 | 26 |
| ENSG00000103187 | COTL1 | 157,78 | 175,97 | 196,18 | 198,64 | 26 |
| ENSG00000286833 | AC097532,3 | 1,2 | 1,23 | 1,32 | 1,51 | 26 |
| ENSG00000120693 | SMAD9 | 3,95 | 4,02 | 4,32 | 4,97 | 26 |
| ENSG00000145246 | ATP10D | 4,34 | 4,44 | 5,27 | 5,45 | 26 |
| ENSG00000240694 | PNMA2 | 24,42 | 28,95 | 29,16 | 30,66 | 26 |
| ENSG00000166407 | LMO1 | 5,05 | 5,76 | 5,79 | 6,33 | 25 |
| ENSG00000124496 | TRERF1 | 4,5 | 5,31 | 5,59 | 5,64 | 25 |
| ENSG00000130818 | ZNF426 | 8,49 | 8,66 | 8,89 | 10,63 | 25 |
| ENSG00000157191 | NECAP2 | 16,47 | 17,02 | 17,16 | 20,62 | 25 |
