## Supplementary Table 10 for "Huntingtin loss-of-function contributes to transcriptional deregulation in Huntington’s disease"

**Supplementary Table 10. GO analysis of genes increasing and decreasing in time in HD and KO-NSCs**

**increasing genes specific for HD; BP**

| ID | Description | GeneRatio | BgRatio | pvalue | p.adjust | qvalue | geneID | Count |
|---|---|---|---|---|---|---|---|---|
| GO:0043161 | proteasome-mediated ubiquitin-dependent protein catabolic process | 98/2001 | 489/20772 | 1,47271E-12 | 8,86569E-09 | 7,8379E-09 | FOXF2/DAB2/SH3RF3/RHOBTB3/ECRG4/HERC2/SPSB4/TRIM2/PABIR1/FZR1/KLHL15/PSMC3/ZSWIM8/HSPA1B/OS9/FBXO45/GIPC1/BAG5/PJA2/AXIN2/PSMC5/AGAP3/CALR/KCTD10/ZNF598/RNF126/PELI1/HSP90B1/FEM1B/KAT5/SYVN1/ERLIN1/AKT1/PSMD1/PSMD12/RBCK1/RNF217/UFD1/UBE2K/PHF20L1/PSMC1/LTN1/ANAPC10/NSFL1C/CANX/CCAR2/HSPA5/TRIP12/USP7/FBXO9/UBXN2A/MAPK9/HSP90AB1/SEL1L/CANX/AKIRIN2/PSMD4/PSMD6/STUB1/FEM1A/FAF2/KLHL42/SELENOS/USP9X/CUL5/TRAF4/CSNK1D/SPOP/KLHDC10/ASCC2/FBXL18/PSMD8/UBE4A/RNF216/GSK3A/UCHL5/USP14/ASCC3/DDB1/ANAPC15/TRIM9/KCTD2/CCDC47/ECPAS/UBE2A/ANKIB1/FBXW5/PSMD3/VCP/SH3BGRL/PSMA3/RNF185/DCAF11/PSMD13/GNA12/PSMC6/MARCHF6/PSMA1 | 98 |
| GO:0016197 | endosomal transport | 59/2001 | 268/20772 | 1,07584E-09 | 3,23827E-06 | 2,86286E-06 | REPS2/TBC1D10C/ZDHHC2/RHOBTB3/TRIM27/ATP9A/TRIM27/CHMP4A/RBSN/VPS29/GBF1/CORO1C/ANKRD50/STX12/BLTP3B/EVI5/RNF126/VPS16/VPS37C/AP5Z1/STAM2/RAB35/LMTK2/VPS51/ARHGAP44/RAB6A/USP7/CLTCL1/VPS50/CLTC/RGP1/SNX27/YKT6/EPS15L1/VPS35L/CHMP4B/RAB14/RAB7A/VPS4A/ZFYVE16/VTI1B/SNX3/GOLT1B/WASH6P/CHMP6/VPS26C/UBAP1/WASHC2C/TBC1D10A/ACAP2/ATP6AP1/REPS1/STX10/GGA3/VCP/SNX4/PLEKHJ1/PICALM/GOSR1 | 59 |

| | | | | | | | | |
|---|---|---|---|---|---|---|---|---|
| GO:0042254 | ribosome biogenesis | 66/2001 | 331/20772 | 8,22347E-09 | 1,65018E-05 | 1,45887E-05 | WDR46/DHX37/BOP1/BOP1/PPAN/RRP9/UTP23/WDR46/WDR46/LTV1/NOP14/RRP7A/NSUN4/GTPBP4/LYAR/PES1/ZNF658/NOC2L/MPHOSPH6/SDE2/USP36/DDX10/DDX3X/ABCE1/TBL3/ISG20L2/NOL10/EIF4A3/EXOSC6/GRWD1/DDX21/RIOK1/EIF2A/HEATR3/RPL7L1/WBP11/SBDS/WDR55/DDX56/BYSL/RCL1/ZNF622/NOP16/DHX29/PELP1/BUD23/METTL16/NOP58/UTP11/NSUN3/RPUSD1/UTP14A/MTREX/DCAF13/EXOSC4/NOLC1/DIS3/KRR1/MRM2/RRP8/DROSHA/PAK1IP1/PAK1IP1/NUP88/NIP7/RRP36 | 66 |
| GO:0001503 | ossification | 85/2001 | 477/20772 | 1,71534E-08 | 2,58158E-05 | 2,2823E-05 | CYP24A1/MGP/CER1/PTGS2/TWIST1/CTSK/MSX2/TNC/FGFR3/WNT3/WNT3/ADGRV1/GDF10/TMEM119/TENT5A/RFLNB/DDR2/PTN/SMPD3/BMP5/CDK6/PTCH1/RASSF2/ITGA11/ENPP1/SNAI2/GPM6B/EGFR/ANKH/FOXC1/ROR2/BMPR1B/VCAN/COL6A1/XYLT1/SOX9/LTBP3/BCOR/MMP16/ZHX3/BCL2/IFITM1/FERMT2/GLI2/SHOX2/PRKACA/IL6ST/TPM4/PBX1/DCHS1/AXIN2/SATB2/HSD17B4/GTPBP4/CHSY1/YAP1/PRKACA/LRP3/AKT1/CAT/BMPR2/OSTF1/TWSG1/FIGNL1/CDH11/CLTC/DDX21/SND1/ADAR/SBDS/DNAJC13/CBFB/PTPN11/GIT1/LIMD1/MESD/CCDC47/ILK/SNRNP200/ALYREF/ANO6/RHOA/EXT2/SYNCRIP/CCN2 | 85 |
| GO:0018205 | peptidyl-lysine modification | 75/2001 | 417/20772 | 7,99962E-08 | 7,13629E-05 | 6,30898E-05 | TWIST1/PAX5/EYA1/SNAI2/CBX4/PLOD2/SNCA/TRIM27/TRIM27/BCOR/NNAT/KMT5A/P3H4/H1-2/NELFE/NELFE/NELFE/NELFE/NELFE/HCFC1/ZBED1/KMT2C/CXXC1/NOC2L/BRPF1/NFYA/KAT5/TAF4/PAXIP1/PHF20L1/MSL3/ASH2L/GLYR1/NAA60/JADE3/DDX21/ATXN7L3/BRPF3/RUVBL2/NCOA6/TOLLIP/MORF4L2/SET/WDR82/ATP7A/TRRAP/HDAC2/SNW1/TAF5L/EP400/SIRT3/ASH1L/TAF2/MORF4L1/EPC1/DHPS/SUPT20H/RUVBL1/MRGBP/SKIC8/DR1/TRIM28/UBA2/NELFA/KMT5B/EEF1AKMT1/EEF2KMT/SIN3A/SLF2/WDR5B/USP22/ARNT/TAF12/SETMAR/DEK | 75 |

| | | | | | | | | |
|---|---|---|---|---|---|---|---|---|
| GO:0006364 | rRNA processing | 50/2001 | 237/20772 | 8,29801E-08 | 7,13629E-05 | 6,30898E-05 | WDR46/DHX37/BOP1/BOP1/PPAN/RRP9/UTP23/WDR46/WDR46/NOP14/RRP7A/NSUN4/GTPBP4/LYAR/PES1/MPHOSPH6/SDE2/USP36/DDX10/TBL3/NOL10/EIF4A3/EXOSC6/DDX21/RIOK1/RPL7L1/WBP11/SBDS/WDR55/DDX56/BYSL/RCL1/PELP1/BUD23/METTL16/NOP58/UTP11/NSUN3/RPUSD1/UTP14A/MTREX/DCAF13/EXOSC4/NOLC1/DIS3/KRR1/MRM2/RRP8/DROSHA/RRP36 | 50 |
| GO:0016482 | cytosolic transport | 44/2001 | 195/20772 | 6,44044E-08 | 7,13629E-05 | 6,30898E-05 | KIF5A/TBC1D10C/DAB2/RHOBTB3/TRIM27/ATP9A/TRIM27/TANC2/PPFIA2/MSN/RBSN/VPS29/GBF1/NF2/BLTP3B/ACTR2/EVI5/RNF126/LMTK2/VPS51/RAB6A/USP7/CLTCL1/AP1G1/CLTC/MAP2/RGP1/MTMR2/YKT6/PXK/DNAJC13/RAB21/RAB14/RAB7A/VTI1B/SNX3/GOLT1B/WASH6P/WASHC2C/TBC1D10A/STX10/PLEKHJ1/GOSR1/CCDC91 | 44 |
| GO:0034975 | protein folding in endoplasmic reticulum | 9/2001 | 12/20772 | 1,17958E-07 | 8,87635E-05 | 7,84732E-05 | ERO1B/CALR/HSP90B1/CANX/HSPA5/CANX/DNAJC3/PDIA3/P4HB | 9 |
| GO:0072594 | establishment of protein localization to organelle | 83/2001 | 487/20772 | 1,86943E-07 | 0,000125044 | 0,000110548 | APOD/PTGS2/CHMP4A/CBLB/HK1/NAGPA/PEX3/TRAF3IP2/KPNA1/PPP3R1/AIFM1/SPCS3/AKT1/PRKCD/TIMM21/SNUPN/VPS37C/STAM2/ZNF827/HSPA4/NUP188/USP36/LRWD1/SEC61A2/HSPA5/GDAP1/IPO13/ING1/MON1A/SRP68/MED1/SEC61A1/MON1B/HEATR3/NUP50/PXK/AKIRIN2/RUVBL2/POM121/CHMP4B/SSR3/USP9X/HSPD1/UBE2L3/TARDBP/RAB7A/VPS4A/KPNA3/ZFYVE16/IPO7/KPNB1/IPO11/SEC61G/VPS41/NABP2/GSK3A/CCT7/TOMM40/NMT1/SH3GLB1/UBAP1/TOMM40L/NUP214/HSPA8/MFN2/NUP62/DNAJA1/NOLC1/GGA3/TRIM28/TIMM8B/POM121C/UFM1/SREBF2/MACROH2A1/PIK3R2/BCS1L/ELAVL1/KPNA6/RBM22/NUP88/ATG13/SRP54 | 83 |

| | | | | | | | | |
|---|---|---|---|---|---|---|---|---|
| GO:0034470 | ncRNA processing | 80/2001 | 475/20772 | 5,09411E-07 | 0,000278787 | 0,000246467 | WDR46/DHX37/BOP1/BOP1/PPAN/RRP9/UTP23/HENMT1/WDR46/WDR46/NSUN2/DUS3L/NOP14/RRP7A/NSUN4/SPOUT1/GTPBP4/LYAR/PES1/MPHOSPH6/ADAT1/SRRT/SDE2/TSN/USP36/DDX10/DDX3X/TBL3/TRMT1L/TYW1B/TSNAX/TRMT61A/NOL10/EIF4A3/TRMO/EXOSC6/DDX21/RIOK1/ADAR/RPL7L1/WBP11/TP53RK/SBDS/WDR55/DDX56/TRMT6/BYSL/RCL1/PELP1/NCBP2/ANKRD16/BUD23/GRSF1/METTL16/TYW1/NOP58/UTP11/NSUN3/SARS1/RPUSD1/UTP14A/MTREX/DCAF13/EXOSC4/NOLC1/DIS3/KRR1/DICER1/NCBP1/ELAC1/THUMPD1/INTS5/MRM2/RRP8/THG1L/DROSHA/RRP36/KARS1/DDX1/DPH3 | 80 |
| GO:0016072 | rRNA metabolic process | 54/2001 | 279/20772 | 4,94059E-07 | 0,000278787 | 0,000246467 | WDR46/POLR1G/DHX37/BOP1/BOP1/PPAN/RRP9/UTP23/WDR46/WDR46/GTF3C1/NOP14/RRP7A/NSUN4/GTPBP4/LYAR/PES1/MPHOSPH6/SDE2/USP36/DDX10/TBL3/NOL10/EIF4A3/EXOSC6/DDX21/RIOK1/RPL7L1/WBP11/SBDS/WDR55/DDX56/BYSL/RCL1/PELP1/BUD23/METTL16/NOP58/UTP11/NSUN3/POLR1B/RPUSD1/UTP14A/MTREX/DCAF13/EXOSC4/NOLC1/DIS3/KRR1/MACROH2A1/MRM2/RRP8/DROSHA/RRP36 | 54 |
| GO:0016055 | Wnt signaling pathway | 81/2001 | 485/20772 | 6,15243E-07 | 0,000308647 | 0,000272866 | IGFBP6/WNT9B/FZD10/WNT3/RARG/WNT3/RNF43/TGFB1I1/BICC1/DAB2/ADGRA2/NKD1/GPRC5B/FZD7/APCDD1/AMER2/FOXO1/CSNK2B/SOSTDC1/SNAI2/RBMS3/TMEM198/EGFR/SOX13/TNIK/EDNRA/MDFI/DACT3/AMER1/ROR2/FRAT2/SOX9/GPC4/CTNND2/FERMT2/DISC1/TMEM131L/AMOTL1/SEMA5A/AXIN2/KPNA1/YAP1/TTC21B/CBY1/NDRG2/ANKRD6/DDX3X/MARK2/CCAR2/TCF7L2/GNAQ/RAC1/NXN/LATS1/FRZB/PPM1B/RUVBL2/HMGXB4/PKD1/CSNK1D/SNX3/GSK3A/DDB1/LIMD1/MESD/RUVBL1/RNF138/ILK/LYPD6/MED12/SKIC8/RHOA/VCP/CDC42/PPM1A/CTDNEP1/CTDNEP1/PIN1/CHD8/PTEN/PPP2R1A | 81 |

| | | | | | | | | |
|---|---|---|---|---|---|---|---|---|
| GO:0198738 | cell-cell signaling by wnt | 81/2001 | 487/20772 | 7,29593E-07 | 0,000337857 | 0,00029869 | IGFBP6/WNT9B/FZD10/WNT3/RARG/WNT3/RNF43/TGFB1I1/BICC1/DAB2/ADGRA2/NKD1/GPRC5B/FZD7/APCDD1/AMER2/FOXO1/CSNK2B/SOSTDC1/SNAI2/RBMS3/TMEM198/EGFR/SOX13/TNIK/EDNRA/MDFI/DACT3/AMER1/ROR2/FRAT2/SOX9/GPC4/CTNND2/FERMT2/DISC1/TMEM131L/AMOTL1/SEMA5A/AXIN2/KPNA1/YAP1/TTC21B/CBY1/NDRG2/ANKRD6/DDX3X/MARK2/CCAR2/TCF7L2/GNAQ/RAC1/NXN/LATS1/FRZB/PPM1B/RUVBL2/HMGXB4/PKD1/CSNK1D/SNX3/GSK3A/DDB1/LIMD1/MESD/RUVBL1/RNF138/ILK/LYPD6/MED12/SKIC8/RHOA/VCP/CDC42/PPM1A/CTDNEP1/CTDNEP1/PIN1/CHD8/PTEN/PPP2R1A | 81 |
| GO:0071364 | cellular response to epidermal growth factor stimulus | 16/2001 | 43/20772 | 1,09446E-06 | 0,000470616 | 0,000416058 | ERBB4/DAB2/SNAI2/EGFR/ZFP36L2/FOXC1/SOX9/PPP1R9B/IQGAP1/AKT1/BAIAP2/MED1/ZFP36/SYAP1/PTPN11/ERRFI1 | 16 |
| GO:0034504 | protein localization to nucleus | 60/2001 | 332/20772 | 1,31499E-06 | 0,000494766 | 0,000437408 | NGFR/APOD/PTGS2/CBLB/SUN2/SOX9/DCLK2/FERMT2/TRAF3IP2/NF2/RRP7A/KPNA1/YAP1/CALR/LMNB2/RAP1GDS1/PPP3R1/CIZ1/AKT1/PRKCD/SNUPN/DNAJB6/NUP188/TCF7L2/IPO13/ING1/MED1/LATS1/HEATR3/NUP50/AKIRIN2/BYSL/POM121/TARDBP/KPNA3/IPO7/KPNB1/IPO11/CCT7/XPA/MFHAS1/PLRG1/ARL2BP/NUP214/NUP62/NOLC1/TRIM28/POM121C/UFM1/CTDNEP1/CTDNEP1/PIK3R2/SIN3A/ELAVL1/KPNA6/RBM22/PIN1/NUP88/LAMTOR5/SUN1 | 60 |
| GO:0042147 | retrograde transport, endosome to Golgi | 28/2001 | 110/20772 | 1,24412E-06 | 0,000494766 | 0,000437408 | TBC1D10C/RHOBTB3/TRIM27/ATP9A/TRIM27/RBSN/VPS29/GBF1/BLTP3B/EVI5/RNF126/VPS51/RAB6A/USP7/CLTCL1/CLTC/RGP1/YKT6/RAB7A/VTI1B/SNX3/GOLT1B/WASH6P/WASHC2C/TBC1D10A/STX10/PLEKHJ1/GOSR1 | 28 |

| | | | | | | | | |
|---|---|---|---|---|---|---|---|---|
| GO:0051568 | histone H3-K4 methylation | 21/2001 | 71/20772 | 1,96154E-06 | 0,000694616 | 0,00061409 | BCOR/H1-2/NELFE/NELFE/NELFE/NELFE/NELFE/HCFC1/KMT2C/CXXC1/NFYA/PAXIP1/ASH2L/NCOA6/WDR82/SNW1/ASH1L/SKIC8/NELFA/WDR5B/SETMAR | 21 |
| GO:0021543 | pallium development | 43/2001 | 213/20772 | 2,23651E-06 | 0,000747989 | 0,000661275 | KCNA2/POU3F2/POU3F3/PAX5/DLX1/CDK6/TUBB2A/EGFR/LAMB1/PLCB1/DLX2/SRGAP2/FEZ1/SUN2/SRGAP2C/BTBD3/DCLK2/CCDC85C/DISC1/NDE1/FBXO45/ROBO1/NF2/PPP1R9B/TUBB2B/FAT4/SRF/GART/LRP8/DAB1/AKIRIN2/CRK/IGF1R/TACC2/NOTCH2NLC/RHOA/NARS1/BNIP3/XAB2/OGDH/PTEN/CDON/SUN1 | 43 |
| GO:0021987 | cerebral cortex development | 33/2001 | 146/20772 | 2,62536E-06 | 0,000831824 | 0,000735391 | KCNA2/POU3F2/POU3F3/PAX5/TUBB2A/EGFR/LAMB1/PLCB1/SRGAP2/SUN2/SRGAP2C/BTBD3/CCDC85C/DISC1/NDE1/FBXO45/ROBO1/PPP1R9B/TUBB2B/FAT4/GART/LRP8/DAB1/AKIRIN2/CRK/TACC2/NOTCH2NLC/RHOA/NARS1/BNIP3/XAB2/CDON/SUN1 | 33 |
| GO:0010975 | regulation of neuron projection development | 78/2001 | 480/20772 | 2,89845E-06 | 0,000872433 | 0,000771292 | WNT3/EPHA3/WNT3/POU3F2/LPAR1/PMP22/PRRX1/DDR2/ALKAL2/PTN/TOX/DAB2/CTNNA2/BMP5/RTN4RL1/PTPRD/EFNB2/PTPRD/MYLIP/TNIK/ROR2/CNTN1/FEZ1/TANC2/NR2F1/PPFIA2/CDKL5/DNM3/PDLIM5/ITM2C/EFNB3/SEMA3A/SEMA3D/PTPRG/LRRC4C/MYCBP2/DISC1/SHOX2/ROBO1/PLXNC1/SEMA5A/BAG5/TUBB2B/ULK2/RAP1A/SRF/PLXNB2/AKT1/DBN1/BMPR2/BAIAP2/PTPN9/LRP8/KIDINS220/MARK2/HSPA5/ARHGAP44/DAB1/MAP2/CHN1/DDX56/RAB21/YTHDF1/HDAC2/CRK/SNX3/RAPGEF1/GSK3A/GORASP1/IGF1R/RHOA/TBC1D24/CREB3L2/PQBP1/SF3A2/PTEN/YWHAH/PAK1 | 78 |
| GO:0002062 | chondrocyte differentiation | 28/2001 | 116/20772 | 3,84474E-06 | 0,001052061 | 0,000930096 | CYTL1/MSX2/FGFR3/RARG/RFLNB/SMPD3/TRPS1/SNAI2/BMPR1B/SOX9/LTBP3/SHOX2/AXIN2/CHSY1/BMPR2/LNPK/GLG1/TWSG1/RB1/SOX6/TGFBR1/HMGA2/SERPINH1/PTPN11/EXT2/CREB3L2/MEX3C/CCN2 | 28 |
| GO:0030166 | proteoglycan biosynthetic process | 20/2001 | 68/20772 | 3,76671E-06 | 0,001052061 | 0,000930096 | CYTL1/DSEL/DSE/BMPR1B/CHST12/XYLT1/GAL3ST4/NDST3/CHSY1/SLC2A10/CHST10/BMPR2/CSGALNACT2/GLCE/TCF7L2/CANT1/B3GALT6/FAM20B/CHPF2/EXT2 | 20 |

| | | | | | | | | |
|---|---|---|---|---|---|---|---|---|
| GO:0070849 | response to epidermal growth factor | 16/2001 | 47/20772 | 4,27117E-06 | 0,001117933 | 0,000988332 | ERBB4/DAB2/SNAI2/EGFR/ZFP36L2/FOXC1/SOX9/PPP1R9B/IQGAP1/AKT1/BAIAP2/MED1/ZFP36/SYAP1/PTPN11/ERRFI1 | 16 |
| GO:0006913 | nucleocytoplasmic transport | 62/2001 | 361/20772 | 5,0005E-06 | 0,00120412 | 0,001064527 | APOD/PTGS2/CBLB/FRAT2/TRAF3IP2/PRKACA/NRDE2/NSUN2/LTV1/CASC3/KPNA1/CALR/PRKACA/PPP3R1/HSPA9/AKT1/PRKCD/SNUPN/GLE1/NUP188/MALT1/RANBP3/ABCE1/POLDIP3/IPO13/CDKN1B/ING1/MED1/EIF4A3/HEATR3/NUP50/AKIRIN2/POM121/NCBP2/PKD1/TARDBP/KPNA3/PTPN11/IPO7/MAGOHB/KPNB1/IPO11/DDX19A/NUP214/PRKAG1/ALYREF/NUP62/RBM8A/NOLC1/TRIM28/PPM1A/NCBP1/POM121C/UFM1/PIK3R2/CHTOP/ELAVL1/KPNA6/RBM22/NUP88/NUP43/XPO4 | 62 |
| GO:0051169 | nuclear transport | 62/2001 | 361/20772 | 5,0005E-06 | 0,00120412 | 0,001064527 | APOD/PTGS2/CBLB/FRAT2/TRAF3IP2/PRKACA/NRDE2/NSUN2/LTV1/CASC3/KPNA1/CALR/PRKACA/PPP3R1/HSPA9/AKT1/PRKCD/SNUPN/GLE1/NUP188/MALT1/RANBP3/ABCE1/POLDIP3/IPO13/CDKN1B/ING1/MED1/EIF4A3/HEATR3/NUP50/AKIRIN2/POM121/NCBP2/PKD1/TARDBP/KPNA3/PTPN11/IPO7/MAGOHB/KPNB1/IPO11/DDX19A/NUP214/PRKAG1/ALYREF/NUP62/RBM8A/NOLC1/TRIM28/PPM1A/NCBP1/POM121C/UFM1/PIK3R2/CHTOP/ELAVL1/KPNA6/RBM22/NUP88/NUP43/XPO4 | 62 |
| GO:0021537 | telencephalon development | 54/2001 | 302/20772 | 6,03927E-06 | 0,00138086 | 0,001220778 | KCNA2/FOXB1/ID2/POU3F2/RFX4/POU3F3/LPAR1/PAX5/ERBB4/DLX1/RTN4RL1/CDK6/TUBB2A/EGFR/LAMB1/PLCB1/DLX2/SRGAP2/FEZ1/SUN2/SRGAP2C/BTBD3/SEMA3A/DCLK2/CCDC85C/DISC1/NDE1/FBXO45/CORO1C/ROBO1/NF2/PPP1R9B/TUBB2B/FAT4/SRF/GART/LRP8/SECISBP2/DAB1/AKIRIN2/CRK/ZSWIM6/UCHL5/IGF1R/TACC2/NOTCH2NLC/RHOA/NARS1/BNIP3/XAB2/OGDH/PTEN/CDON/SUN1 | 54 |

| | | | | | | | | |
|---|---|---|---|---|---|---|---|---|
| GO:0006457 | protein folding | 46/2001 | 243/20772 | 6,24979E-06 | 0,00138086 | 0,001220778 | HYPK/HSPA1B/NUDCD3/ERO1B/BAG5/PDIA6/CALR/HSP90B1/HSPA9/PDIA4/DNAJC21/CLPX/PPIB/DNAJB12/HSPA4/SACS/DNAJB6/QSOX2/CANX/HSPA5/CRTAP/KHSRP/PPIL2/DNAJB11/HSP90AB1/UGGT1/CANX/DNAJC7/DNAJC3/PDIA3/DNAJC5/RUVBL2/HSPD1/SDF2L1/HSPH1/P4HB/AHSA1/CCT7/MESD/CCDC47/ST13/CDC37/HSPA8/DNAJA1/VBP1/PFDN6 | 46 |
| GO:0006029 | proteoglycan metabolic process | 25/2001 | 100/20772 | 6,42261E-06 | 0,00138086 | 0,001220778 | CYTL1/DCN/CNMD/DSEL/DSE/BMPR1B/CHST12/XYLT1/GAL3ST4/NDST3/CHSY1/SLC2A10/CHST10/BMPR2/HEXA/CSGALNACT2/PPARD/GLCE/TCF7L2/CANT1/B3GALT6/FAM20B/GPC1/CHPF2/EXT2 | 25 |
| GO:0000375 | RNA splicing, via transesterification reactions | 63/2001 | 372/20772 | 6,70851E-06 | 0,001392594 | 0,001231151 | SLC39A5/KHDRBS3/CELF2/QKI/YJU2/SFSWAP/IK/TFIP11/C9orf78/CASC3/NOVA1/DNAJC17/THRAP3/SF3B2/SDE2/RBM3/TSSC4/KHSRP/SFPQ/EIF4A3/SNRPB2/HNRNPR/EFTUD2/DHX8/RBM39/CWC15/SF3A3/CD2BP2/SRPK2/GPATCH1/NCBP2/SNRNP25/PRPF18/MAGOHB/METTL16/RBMXL1/SNW1/PRPF19/SMNDC1/PRPF4/ZMAT2/PLRG1/MTREX/PRPF8/SF3B4/FXR2/HSPA8/SNRNP200/CDC5L/ALYREF/RBM8A/CELF1/SYNCRIP/NCBP1/XAB2/SNRPD1/PQBP1/SF3A2/RBM22/TRA2A/HNRNPK/ESS2/DDX1 | 63 |
| GO:0051216 | cartilage development | 42/2001 | 215/20772 | 6,94546E-06 | 0,001393723 | 0,001232149 | OTOR/MGP/CER1/CTSK/CYTL1/MSX2/FGFR3/RARG/PRRX1/RFLNB/CNMD/SMPD3/BMP5/TRPS1/SNAI2/EVC/DLX2/BMPR1B/SOX9/LTBP3/SHOX2/AXIN2/SATB2/CHSY1/SRF/BMPR2/LNPK/GLG1/TWSG1/RB1/SOX6/FRZB/TGFBR1/HMGA2/SERPINH1/PKD1/PTPN11/ATP7A/EXT2/CREB3L2/MEX3C/CCN2 | 42 |

| | | | | | | | | |
|---|---|---|---|---|---|---|---|---|
| GO:2001020 | regulation of response to DNA damage stimulus | 60/2001 | 350/20772 | 7,48712E-06 | 0,00145395 | 0,001285395 | TWIST1/EYA1/SNAI2/EGFR/PMAIP1/POLH/SPRED1/BCL2/KLHL15/KMT5A/YJU2/ZCWPW1/SPIRE1/TFIP11/MAP3K20/AXIN2/ACTR2/FEM1B/KAT5/HMGB1/PRKCD/TAF4/PAXIP1/CCAR2/TRIP12/FIGNL1/TELO2/ATXN7L3/HMGA2/RUVBL2/MORF4L2/BCLAF1/TRRAP/TAF5L/UCHL5/EP400/TAF2/MORF4L1/EPC1/SUPT20H/RUVBL1/FXR2/MRGBP/SMARCE1/SPRED2/TRIM28/CEBPG/ACTR5/EEF1E1/USP1/KMT5B/OTUB2/SLF2/TIMELESS/USP22/PBRM1/HNRNPK/TAF12/SETMAR/DEK | 60 |
| GO:2001022 | positive regulation of response to DNA damage stimulus | 37/2001 | 181/20772 | 8,23088E-06 | 0,001548434 | 0,001368925 | EYA1/EGFR/PMAIP1/SPRED1/ZCWPW1/SPIRE1/MAP3K20/ACTR2/KAT5/HMGB1/PRKCD/PAXIP1/CCAR2/TELO2/RUVBL2/MORF4L2/BCLAF1/TRRAP/UCHL5/EP400/MORF4L1/EPC1/RUVBL1/FXR2/MRGBP/SMARCE1/SPRED2/TRIM28/CEBPG/ACTR5/EEF1E1/USP1/KMT5B/SLF2/TIMELESS/PBRM1/SETMAR | 37 |
| GO:0007015 | actin filament organization | 76/2001 | 478/20772 | 8,66522E-06 | 0,001580747 | 0,001397492 | CDC42EP5/ESPN/LPAR1/RFLNB/KANK4/RGCC/CTNNA2/SYNPO/PHLDB2/PLEC/RND1/TRIM27/ARHGAP6/TRIM27/DLC1/NEDD9/LIMA1/HMCN1/BCL2/SHROOM2/FERMT2/EPS8/BCAR1/GAS2/PPM1F/SPIRE1/TPM4/RASA1/CORO1C/NF2/FCHSD2/PPP1R9B/SEMA5A/SORBS1/SPTAN1/ACTR2/WDR1/SRF/HSP90B1/ADD1/DBN1/TPM3/PRKCD/BAIAP2/SHROOM4/ACTR3/ARPC5/MTPN/RAC1/FER/LATS1/TMSB4X/TGFBR1/HIP1/ARF1/MARCKS/IQGAP2/CFL2/ARPC2/MSRB2/ARPC5L/DIAPH1/WASH6P/ARPC4/FAT1/ACTN1/WASHC2C/TMOD3/RHOA/CAPZA1/CDC42/ELMO2/PIK3R2/CCN2/FSCN1/PAK1 | 76 |

| | | | | | | | | |
|---|---|---|---|---|---|---|---|---|
| GO:1903311 | regulation of mRNA metabolic process | 60/2001 | 353/20772 | 9,81628E-06 | 0,001709561 | 0,001511373 | TENT5A/SLC39A5/ZFP36L2/KHDRBS3/CELF2/QKI/TRAF3IP2/TRAF5/SFSWAP/CASC3/NOVA1/AXIN2/FTO/TIRAP/THRAP3/AKT1/IKBKE/PRKCD/CNOT3/DCP1A/CNOT10/RBM3/SECISBP2/KHSRP/EIF4A3/ZFP36/MLH1/RBM39/ZC3H3/SRPK2/SERBP1/TENT4A/NCBP2/TARDBP/MAGOHB/YTHDF1/METTL16/RBMXL1/SNW1/PRPF19/NUDT21/SF3B4/FXR2/PATL1/HSPA8/RBM8A/CELF1/SYNCRIP/LARP4B/DIS3/NCBP1/CPSF7/SAMD4B/GTPBP1/APEX1/ELAVL1/ZC3H14/IGF2BP3/TRA2A/HNRNPK | 60 |
| GO:0034976 | response to endoplasmic reticulum stress | 52/2001 | 292/20772 | 1,00236E-05 | 0,001709561 | 0,001511373 | TMEM117/PMAIP1/BCL2/OS9/PPP1R15A/PDIA6/CALR/ERMP1/HSP90B1/EIF4G1/EIF2S1/AIFM1/SYVN1/ERLIN1/PDIA4/UFD1/DNAJB12/PPP1R15B/DDX3X/MBTPS1/CANX/HSPA5/SEL1L/UGGT1/CANX/EIF2AK2/DNAJC3/PDIA3/MARCKS/STUB1/FAF2/SELENOS/ATP2A2/SDF2L1/USP13/P4HB/UBE4A/RNFT1/USP14/CCDC47/ECPAS/HM13/VCP/UFM1/CREB3L2/PIK3R2/RNF185/PTPN2/COPS5/PSMC6/MARCHF6/SEC16A | 52 |
| GO:0016573 | histone acetylation | 34/2001 | 162/20772 | 1,0456E-05 | 0,001709561 | 0,001511373 | TWIST1/SNAI2/SNCA/HCFC1/NOC2L/BRPF1/NFYA/KAT5/TAF4/PHF20L1/MSL3/GLYR1/NAA60/JADE3/DDX21/ATXN7L3/BRPF3/RUVBL2/MORF4L2/SET/TRRAP/TAF5L/EP400/TAF2/MORF4L1/EPC1/SUPT20H/RUVBL1/MRGBP/DR1/SIN3A/USP22/TAF12/DEK | 34 |
| GO:0006413 | translational initiation | 30/2001 | 135/20772 | 1,05073E-05 | 0,001709561 | 0,001511373 | PPP1R15A/EIF4G1/EIF2S1/EIF3C/CTIF/PPP1R15B/DDX3X/ABCE1/EIF3J/EIF1AD/EIF1/EIF2B3/EIF2A/EIF3B/KLHL25/EIF2AK2/DNAJC3/DHX29/NCBP2/BZW1/YTHDF1/DENR/RPS6KB1/EIF4G2/NCBP1/EIF2D/SH3BGRL/EIF5/COPS5/DDX1 | 30 |

| | | | | | | | | |
|---|---|---|---|---|---|---|---|---|
| GO:0001649 | osteoblast differentiation | 51/2001 | 286/20772 | 1,16896E-05 | 0,001851873 | 0,001637187 | CYP24A1/TWIST1/MSX2/TNC/WNT3/WNT3/GDF10/TMEM119/TENT5A/DDR2/CDK6/PTCH1/RASSF2/ITGA11/SNAI2/BMPR1B/VCAN/COL6A1/SOX9/ZHX3/IFITM1/FERMT2/GLI2/SHOX2/PRKACA/IL6ST/TPM4/AXIN2/SATB2/HSD17B4/GTPBP4/YAP1/PRKACA/LRP3/AKT1/CAT/BMPR2/TWSG1/FIGNL1/CLTC/DDX21/SND1/ADAR/DNAJC13/CBFB/LIMD1/CCDC47/ILK/SNRNP200/ALYREF/SYNCRIP | 51 |
| GO:0060070 | canonical Wnt signaling pathway | 56/2001 | 325/20772 | 1,27549E-05 | 0,001968834 | 0,001740588 | IGFBP6/WNT9B/FZD10/WNT3/RARG/WNT3/BICC1/DAB2/ADGRA2/NKD1/GPRC5B/FZD7/AMER2/FOXO1/SOSTDC1/SNAI2/RBMS3/TMEM198/EGFR/SOX13/EDNRA/DACT3/AMER1/FRAT2/SOX9/CTNND2/DISC1/TMEM131L/SEMA5A/AXIN2/KPNA1/YAP1/TTC21B/CBY1/ANKRD6/DDX3X/CCAR2/TCF7L2/GNAQ/LATS1/FRZB/PPM1B/RUVBL2/CSNK1D/GSK3A/LIMD1/RUVBL1/ILK/LYPD6/VCP/PPM1A/CTDNEP1/CTDNEP1/PIN1/CHD8/PTEN | 56 |
| GO:0034446 | substrate adhesion-dependent cell spreading | 25/2001 | 104/20772 | 1,34496E-05 | 0,00202416 | 0,0017895 | HAS2/DAB2/FZD7/LAMB1/SRGAP2/NEDD9/FERMT2/LAMC1/CORO1C/DOCK5/DOCK1/CALR/AKIP1/RAC1/FER/RAB1A/RCC2/RCC2/ARPC2/P4HB/CRK/WASHC2C/ILK/RHOA/CDC42 | 25 |
| GO:0007265 | Ras protein signal transduction | 61/2001 | 367/20772 | 1,68516E-05 | 0,002474301 | 0,002187457 | NGFR/ARHGDIB/PLD1/ARHGAP24/LPAR1/RAPGEF5/KANK2/RASA3/RASA3/ARHGAP6/CADM4/RASA4/DLC1/RGL1/DBNL/RALB/ARHGEF1/STARD8/EPS8/RABL3/SPRY2/GBF1/DOCK5/ROBO1/HRAS/DOK3/RAP1A/KCTD10/IQSEC1/RHOB/RAB35/PSD3/GPSM2/ARHGAP44/RAC1/ARFGEF2/PIK3CB/RB1/RAB39A/MADD/NET1/HEG1/RAP2C/RAB21/GIT1/KPNB1/CRK/RAPGEF1/PHACTR4/MFN2/RAB15/NUP62/RHOA/CDC42/NISCH/KSR1/CBL/SH2B2/GNA12/GNA13/IQSEC2 | 61 |

| | | | | | | | | |
|---|---|---|---|---|---|---|---|---|
| GO:0000377 | RNA splicing, via transesterification reactions with bulged adenosine as nucleophile | 61/2001 | 368/20772 | 1,83404E-05 | 0,002567659 | 0,002269992 | SLC39A5/KHDRBS3/CELF2/QKI/YJU2/SFSWAP/IK/TFIP11/C9orf78/CASC3/NOVA1/DNAJC17/THRAP3/SF3B2/SDE2/RBM3/TSSC4/SFPQ/EIF4A3/SNRPB2/HNRNPR/EFTUD2/DHX8/RBM39/CWC15/SF3A3/CD2BP2/SRPK2/GPATCH1/NCBP2/SNRNP25/PRPF18/MAGOHB/METTL16/RBMXL1/SNW1/PRPF19/PRPF4/ZMAT2/PLRG1/MTREX/PRPF8/SF3B4/FXR2/HSPA8/SNRNP200/CDC5L/ALYREF/RBM8A/CELF1/SYNCRIP/NCBP1/XAB2/SNRPD1/PQBP1/SF3A2/RBM22/TRA2A/HNRNPK/ESS2/DDX1 | 61 |
| GO:0000398 | mRNA splicing, via spliceosome | 61/2001 | 368/20772 | 1,83404E-05 | 0,002567659 | 0,002269992 | SLC39A5/KHDRBS3/CELF2/QKI/YJU2/SFSWAP/IK/TFIP11/C9orf78/CASC3/NOVA1/DNAJC17/THRAP3/SF3B2/SDE2/RBM3/TSSC4/SFPQ/EIF4A3/SNRPB2/HNRNPR/EFTUD2/DHX8/RBM39/CWC15/SF3A3/CD2BP2/SRPK2/GPATCH1/NCBP2/SNRNP25/PRPF18/MAGOHB/METTL16/RBMXL1/SNW1/PRPF19/PRPF4/ZMAT2/PLRG1/MTREX/PRPF8/SF3B4/FXR2/HSPA8/SNRNP200/CDC5L/ALYREF/RBM8A/CELF1/SYNCRIP/NCBP1/XAB2/SNRPD1/PQBP1/SF3A2/RBM22/TRA2A/HNRNPK/ESS2/DDX1 | 61 |
| GO:0006379 | mRNA cleavage | 11/2001 | 27/20772 | 1,95061E-05 | 0,002668789 | 0,002359398 | POLR1H/POLR1H/POLR1H/POLR1H/POLR1H/CSTF2T/CPSF2/NCBP2/NUDT21/NCBP1/CPSF7 | 11 |
| GO:0031330 | negative regulation of cellular catabolic process | 50/2001 | 284/20772 | 2,02113E-05 | 0,002703826 | 0,002390373 | TIMP3/TENT5A/SNCA/FEZ1/CHMP4A/TIMP2/BCL2/HIPK2/TRAF3IP2/TRAF5/NRDE2/NSUN2/GIPC1/BAG5/AXIN2/TIRAP/THRAP3/EIF4G1/AKT1/IKBKE/NRBP2/PHF20L1/USP36/CCAR2/SECISBP2/USP7/CTSA/RPTOR/DAP/HSP90AB1/ZFP36/MTMR2/RRAGA/TENT4A/CHMP4B/USP9X/TAB3/TARDBP/SCFD1/METTL16/GSK3A/UCHL5/USP14/EIF4G2/SYNCRIP/LARP4B/ELAVL1/POLDIP2/IGF2BP3/RNF41 | 50 |

| | | | | | | | | |
|---|---|---|---|---|---|---|---|---|
| GO:0042176 | regulation of protein catabolic process | 67/2001 | 419/20772 | 2,40481E-05 | 0,003147169 | 0,00278232 | TIMP3/FOXF2/DAB2/SH3RF3/NKD1/FOXO1/MYLIP/ZDHHC2/EGFR/SNCA/AMER1/CBLB/SOX9/PABIR1/FZR1/TIMP2/PSMC3/MYCBP2/MSN/HSPA1B/PRKACA/GIPC1/BAG5/AXIN2/LPCAT1/PSMC5/LPCAT1/PRKACA/AKT1/PSMD1/RNF217/UBE2K/PHF20L1/PSMC1/CCAR2/USP7/CTSA/UBXN2A/MAPK9/CDKN1B/HSP90AB1/RGP1/STUB1/LDLR/IDE/PSME3IP1/USP9X/PKD1/USP13/CSNK1D/RAB7A/SNX3/RNFT1/GSK3A/UCHL5/USP14/DDB1/CHMP6/ANKIB1/PSMD3/GGA3/VCP/RNF41/RNF185/PIN1/GNA12/PSMC6 | 67 |
| GO:0061640 | cytoskeleton-dependent cytokinesis | 26/2001 | 115/20772 | 2,89822E-05 | 0,003634846 | 0,003213461 | PLEC/CHMP4A/SEPTIN9/SPIRE1/RASA1/EXOC3/ACTR2/SEPTIN11/KIF4A/RHOB/SEPTIN7/RAB35/ACTR3/EXOC7/SEPTIN5/SEPTIN6/TMEM250/ARF1/CHMP4B/VPS4A/CHMP6/NUP62/SEPTIN2/RHOA/EXOC2/POLDIP2 | 26 |
| GO:0000271 | polysaccharide biosynthetic process | 19/2001 | 71/20772 | 2,89814E-05 | 0,003634846 | 0,003213461 | HAS2/SMPD3/ENPP1/GYG1/NDST3/SORBS1/DYRK2/AKT1/AKT2/GYS1/CSGALNACT2/PPP1R3F/CLTC/GBE1/SELENOS/EPM2AIP1/GSK3A/EXT2/B4GALT5 | 19 |
| GO:0035520 | monoubiquitinated protein deubiquitination | 13/2001 | 38/20772 | 3,18056E-05 | 0,003832771 | 0,003388441 | TAF4/USP7/USP15/ATXN7L3/USP9X/TRRAP/TAF5L/TAF2/SUPT20H/USP16/USP1/USP22/TAF12 | 13 |
| GO:0002063 | chondrocyte development | 12/2001 | 33/20772 | 3,18337E-05 | 0,003832771 | 0,003388441 | MSX2/RARG/RFLNB/SMPD3/BMPR1B/SOX9/SHOX2/AXIN2/CHSY1/BMPR2/SERPINH1/MEX3C | 12 |
| GO:0061448 | connective tissue development | 50/2001 | 289/20772 | 3,25233E-05 | 0,003839019 | 0,003393964 | OTOR/MGP/CER1/CTSK/CYTL1/MSX2/FGFR3/RARG/PRRX1/RFLNB/CNMD/SMPD3/BMP5/TRPS1/SNAI2/EVC/AMER1/DLX2/BMPR1B/SOX9/LTBP3/SHOX2/HRAS/AXIN2/SATB2/CHSY1/SRF/FTO/SH3PXD2B/BMPR2/LNPK/PAXIP1/PPARD/GLG1/TWSG1/RB1/SOX6/FRZB/TGFBR1/RASAL2/HMGA2/SERPINH1/PKD1/PTPN11/ATP7A/ACAT1/EXT2/CREB3L2/MEX3C/CCN2 | 50 |

| | | | | | | | | |
|---|---|---|---|---|---|---|---|---|
| GO:0043543 | protein acylation | 48/2001 | 274/20772 | 3,34276E-05 | 0,003862822 | 0,003415007 | TWIST1/SPHK1/FOXO1/SNAI2/ZDHHC2/SNCA/HCFC1/NAA25/NOC2L/BRPF1/NFYA/KAT5/TAF4/PHF20L1/MSL3/CDYL/DDX3X/GLYR1/NAA60/JADE3/DDX21/ATXN7L3/BRPF3/PPM1B/RUVBL2/MORF4L2/SET/TRRAP/HDAC2/TAF5L/NMT1/EP400/TAF2/MORF4L1/EPC1/SUPT20H/RUVBL1/ZDHHC5/MRGBP/DR1/PPM1A/OGDH/SIN3A/ZDHHC8/USP22/TAF12/DEK/ZDHHC3 | 48 |
| GO:0060828 | regulation of canonical Wnt signaling pathway | 47/2001 | 267/20772 | 3,54462E-05 | 0,003862822 | 0,003415007 | IGFBP6/BICC1/DAB2/ADGRA2/NKD1/GPRC5B/FZD7/AMER2/FOXO1/SOSTDC1/SNAI2/RBMS3/TMEM198/EGFR/SOX13/DACT3/AMER1/SOX9/CTNND2/TMEM131L/SEMA5A/AXIN2/KPNA1/YAP1/TTC21B/CBY1/ANKRD6/DDX3X/CCAR2/TCF7L2/GNAQ/LATS1/FRZB/PPM1B/RUVBL2/CSNK1D/GSK3A/LIMD1/RUVBL1/ILK/LYPD6/VCP/PPM1A/CTDNEP1/CTDNEP1/PIN1/CHD8 | 47 |
| GO:0006282 | regulation of DNA repair | 43/2001 | 237/20772 | 3,59332E-05 | 0,003862822 | 0,003415007 | TWIST1/EYA1/EGFR/POLH/KLHL15/ZCWPW1/SPIRE1/TFIP11/AXIN2/ACTR2/KAT5/HMGB1/TAF4/TRIP12/FIGNL1/ATXN7L3/HMGA2/RUVBL2/MORF4L2/TRRAP/TAF5L/UCHL5/EP400/TAF2/MORF4L1/EPC1/SUPT20H/RUVBL1/MRGBP/SMARCE1/TRIM28/CEBPG/ACTR5/USP1/KMT5B/OTUB2/SLF2/TIMELESS/USP22/PBRM1/TAF12/SETMAR/DEK | 43 |
| GO:0036503 | ERAD pathway | 27/2001 | 123/20772 | 3,58483E-05 | 0,003862822 | 0,003415007 | OS9/CALR/HSP90B1/SYVN1/ERLIN1/UFD1/DNAJB12/CANX/HSPA5/SEL1L/UGGT1/CANX/STUB1/FAF2/SELENOS/SDF2L1/USP13/UBE4A/RNFT1/USP14/CCDC47/ECPAS/HM13/VCP/RNF185/PSMC6/MARCHF6 | 27 |
| GO:0007044 | cell-substrate junction assembly | 24/2001 | 103/20772 | 3,44106E-05 | 0,003862822 | 0,003415007 | APOD/EPHA3/PHLDB2/PLEC/GPM6B/ARHGAP6/DLC1/BCL2/FERMT2/LAMC1/PPM1F/CORO1C/SORBS1/TLN1/RAC1/CDH11/VCL/RCC2/RCC2/PTPRA/ACTN1/RHOA/POLDIP2/PTEN | 24 |
| GO:0150115 | cell-substrate junction organization | 25/2001 | 110/20772 | 3,7309E-05 | 0,003872421 | 0,003423494 | APOD/EPHA3/PHLDB2/PLEC/GPM6B/ARHGAP6/DLC1/BCL2/FERMT2/LAMC1/PPM1F/CORO1C/SORBS1/TLN1/IQSEC1/RAC1/CDH11/VCL/RCC2/RCC2/PTPRA/ACTN1/RHOA/POLDIP2/PTEN | 25 |

| | | | | | | | | |
|---|---|---|---|---|---|---|---|---|
| GO:0043968 | histone H2A acetylation | 10/2001 | 24/20772 | 3,71354E-05 | 0,003872421 | 0,003423494 | KAT5/MSL3/RUVBL2/MORF4L2/TRRAP/EP400/MORF4L1/EPC1/RUVBL1/MRGBP | 10 |
| GO:0030900 | forebrain development | 68/2001 | 434/20772 | 4,05821E-05 | 0,004140753 | 0,003660718 | OLIG2/KCNA2/FOXB1/ID2/POU3F2/RFX4/POU3F3/LPAR1/PAX5/PRKG1/NR2F2/ERBB4/TOX/DLX1/RTN4RL1/CDK6/TUBB2A/EGFR/LAMB1/PLCB1/DLX2/SRGAP2/FEZ1/SUN2/SRGAP2C/BTBD3/DLC1/SEMA3A/DCLK2/CCDC85C/DISC1/NDE1/GLI2/FBXO45/CORO1C/ROBO1/NF2/PPP1R9B/SEMA5A/TUBB2B/FAT4/SRF/GART/TTC21B/FRS2/LRP8/TWSG1/SECISBP2/DAB1/AKIRIN2/ATP7A/CRK/ZSWIM6/UCHL5/IGF1R/TACC2/NOTCH2NLC/STIL/PCM1/RHOA/NARS1/BNIP3/XAB2/OGDH/SIN3A/PTEN/CDON/SUN1 | 68 |
| GO:0051656 | establishment of organelle localization | 72/2001 | 467/20772 | 4,182E-05 | 0,004195936 | 0,003709504 | KIF5A/AP3B2/SMPD3/ADORA2B/KIFC1/SYBU/SNCA/ATP9A/FEZ1/ARHGAP21/CHMP4A/TANC2/PPFIA2/SUN2/SHROOM2/NDE1/SPRY2/SPIRE1/LIN7A/GBF1/MAD1L1/KIFAP3/SYNJ1/LTV1/EXOC3/MAP1S/LMNB2/KAT5/ABCE1/GPSM2/NSFL1C/RAB6A/EXOC7/CDT1/SEPTIN5/PREB/FER/KIF3B/CHGA/MAP2/YKT6/TTL/RAB1A/AP3S1/MLH1/CHMP4B/CSNK1D/RAB7A/VPS4A/KPNB1/ARFGAP3/CHMP6/MAP4/TRAPPC12/TRAPPC11/KIFC1/PCM1/NUP62/COPG2/CDC42/CBL/SNX4/DCTN2/EXOC2/MAPRE1/TMEM201/PICALM/COPG1/NUP88/COPS5/SEC16A/SUN1 | 72 |
| GO:1902850 | microtubule cytoskeleton organization involved in mitosis | 35/2001 | 181/20772 | 4,86438E-05 | 0,004800589 | 0,00424406 | KIFC1/CHMP4A/GNAI1/SUN2/HSPA1B/NDE1/SPRY2/MAP1S/KAT5/KIF4A/GPSM2/NSFL1C/PRC1/CLTC/DCTN6/KIF3B/SBDS/LSM14A/CHMP4B/PKD1/KPNB1/CHMP6/MAP4/TACC2/PTPA/KIFC1/STIL/NUP62/RHOA/KIF2A/VCP/BCCIP/DCTN2/POLDIP2/MAPRE1 | 35 |
| GO:0006900 | vesicle budding from membrane | 21/2001 | 86/20772 | 5,01282E-05 | 0,004867287 | 0,004303025 | AP3B2/CHMP4A/GBF1/PREB/SEC31A/RAB1A/AP3S1/CHMP4B/SEC24A/CSNK1D/RAB7A/VPS4A/ARFGAP3/SNX3/CHMP6/TRAPPC12/TRAPPC11/MIA3/PICALM/SEC24D/SEC16A | 21 |

| | | | | | | | | |
|---|---|---|---|---|---|---|---|---|
| GO:0150116 | regulation of cell-substrate junction organization | 19/2001 | 74/20772 | 5,40691E-05 | 0,005166607 | 0,004567645 | APOD/EPHA3/PHLDB2/GPM6B/ARHGAP6/DLC1/FERMT2/PPM1F/CORO1C/TLN1/IQSEC1/RAC1/VCL/RCC2/RCC2/PTPRA/RHOA/POLDIP2/PTEN | 19 |
| GO:0048041 | focal adhesion assembly | 22/2001 | 93/20772 | 5,63291E-05 | 0,005298453 | 0,004684206 | APOD/EPHA3/PHLDB2/GPM6B/ARHGAP6/DLC1/BCL2/FERMT2/PPM1F/CORO1C/SORBS1/TLN1/RAC1/CDH11/VCL/RCC2/RCC2/PTPRA/ACTN1/RHOA/POLDIP2/PTEN | 22 |
| GO:0030111 | regulation of Wnt signaling pathway | 57/2001 | 351/20772 | 6,12023E-05 | 0,005668271 | 0,005011152 | IGFBP6/WNT3/WNT3/RNF43/BICC1/DAB2/ADGRA2/NKD1/GPRC5B/FZD7/APCDD1/AMER2/FOXO1/SOSTDC1/SNAI2/RBMS3/TMEM198/EGFR/SOX13/MDFI/DACT3/AMER1/SOX9/CTNND2/DISC1/TMEM131L/SEMA5A/AXIN2/KPNA1/YAP1/TTC21B/CBY1/ANKRD6/DDX3X/CCAR2/TCF7L2/GNAQ/NXN/LATS1/FRZB/PPM1B/RUVBL2/HMGXB4/CSNK1D/SNX3/GSK3A/LIMD1/RUVBL1/ILK/LYPD6/VCP/PPM1A/CTDNEP1/CTDNEP1/PIN1/CHD8/PPP2R1A | 57 |
| GO:0007052 | mitotic spindle organization | 30/2001 | 148/20772 | 6,68244E-05 | 0,005786324 | 0,005115519 | KIFC1/CHMP4A/GNAI1/SUN2/HSPA1B/MAP1S/KIF4A/GPSM2/PRC1/CLTC/DCTN6/KIF3B/SBDS/LSM14A/CHMP4B/PKD1/KPNB1/CHMP6/MAP4/TACC2/PTPA/KIFC1/STIL/NUP62/RHOA/KIF2A/VCP/BCCIP/DCTN2/POLDIP2 | 30 |
| GO:0032874 | positive regulation of stress-activated MAPK cascade | 30/2001 | 148/20772 | 6,68244E-05 | 0,005786324 | 0,005115519 | IGFBP6/SPHK1/TNFRSF19/TPD52L1/SH3RF3/FZD7/RASSF2/TNIK/PLCB1/SEMA3A/TAOK3/RELL1/HIPK2/TRAF5/STK39/MID1/HRAS/PJA2/TIRAP/HMGB1/ANKRD6/EIF2AK2/ZNF622/RIPK1/TRAF4/KLHDC10/CRK/MFHAS1/CCN2/MAPKBP1 | 30 |
| GO:0051893 | regulation of focal adhesion assembly | 18/2001 | 69/20772 | 6,72828E-05 | 0,005786324 | 0,005115519 | APOD/EPHA3/PHLDB2/GPM6B/ARHGAP6/DLC1/FERMT2/PPM1F/CORO1C/TLN1/RAC1/VCL/RCC2/RCC2/PTPRA/RHOA/POLDIP2/PTEN | 18 |
| GO:0090109 | regulation of cell-substrate junction assembly | 18/2001 | 69/20772 | 6,72828E-05 | 0,005786324 | 0,005115519 | APOD/EPHA3/PHLDB2/GPM6B/ARHGAP6/DLC1/FERMT2/PPM1F/CORO1C/TLN1/RAC1/VCL/RCC2/RCC2/PTPRA/RHOA/POLDIP2/PTEN | 18 |

| | | | | | | | | |
|---|---|---|---|---|---|---|---|---|
| GO:1900024 | regulation of substrate adhesion-dependent cell spreading | 17/2001 | 63/20772 | 6,70708E-05 | 0,005786324 | 0,005115519 | HAS2/DAB2/NEDD9/FERMT2/CORO1C/DOCK5/DOCK1/CALR/RAC1/RCC2/RCC2/ARPC2/P4HB/CRK/WASHC2C/ILK/CDC42 | 17 |
| GO:0034329 | cell junction assembly | 73/2001 | 483/20772 | 7,13729E-05 | 0,005967571 | 0,005275754 | APOD/EPHA3/APLNR/IRX3/DNER/ERBB4/SLITRK3/LRRTM1/PTPRD/EFNB2/PTPRD/SNAI2/PHLDB2/PLEC/GPM6B/PCDHB16/SNCA/ARHGAP6/ADGRL3/BSN/SRGAP2/DNM3/PDLIM5/SRGAP2C/EFNB3/DLC1/DBNL/GPC4/BCL2/CTNND2/MYCBP2/LRRC4B/FERMT2/LAMC1/PRKACA/PPM1F/FBXO45/CORO1C/JAM3/MYO9A/RAP1A/SORBS1/WDR1/PRKACA/SRF/TLN1/EIF4G1/PLXNB2/LRFN4/TLN2/PCDHB14/RAC1/CDH11/VCL/FER/PCDHB5/PCDHB13/RCC2/RCC2/SDK1/PTPRA/CRK/RAPGEF1/ACTN1/RHOA/CAPZA1/VMP1/CDC42/PCDHB11/ARHGEF9/FSCN1/POLDIP2/PTEN | 73 |
| GO:0051170 | import into nucleus | 34/2001 | 177/20772 | 7,10913E-05 | 0,005967571 | 0,005275754 | APOD/PTGS2/CBLB/TRAF3IP2/KPNA1/PPP3R1/AKT1/PRKCD/SNUPN/NUP188/IPO13/ING1/MED1/HEATR3/NUP50/AKIRIN2/POM121/TARDBP/KPNA3/IPO7/KPNB1/IPO11/NUP214/PRKAG1/NUP62/NOLC1/TRIM28/POM121C/UFM1/PIK3R2/ELAVL1/KPNA6/RBM22/NUP88 | 34 |
| GO:0000910 | cytokinesis | 36/2001 | 192/20772 | 7,47276E-05 | 0,006162467 | 0,005448057 | PLEC/CHMP4A/SEPTIN9/SPIRE1/RASA1/GIPC1/EXOC3/ACTR2/SEPTIN11/KIF4A/RHOB/SEPTIN7/RAB35/ACTR3/EXOC7/SEPTIN5/PRC1/SEPTIN6/KIF3B/TMEM250/ARF1/CALM1/CHMP4B/VPS4A/GIT1/CHMP6/IGF1R/SH3GLB1/KLHL9/NUP62/SEPTIN2/RHOA/CDC42/EXOC2/POLDIP2/PIN1 | 36 |

| | | | | | | | | |
|---|---|---|---|---|---|---|---|---|
| GO:0048193 | Golgi vesicle transport | 53/2001 | 322/20772 | 7,6172E-05 | 0,006196699 | 0,00547832 | ATP9A/BNIP1/RBSN/SPIRE1/GBF1/PITPNB/BLTP3B/RABEP1/VPS51/ARCN1/YIPF5/RAB6A/AP1G1/ARFGEF2/PREB/KDELR2/SEC31A/YIPF4/YKT6/ERGIC1/RAB1A/AP3S1/GOSR2/ERGIC2/VPS35L/SEC24A/RAB14/CSNK1D/SCFD1/P4HB/COPA/ARFGAP3/VTI1B/SNX3/COG4/GOLT1B/COPB2/TRAPPC12/TRAPPC11/RER1/SCYL1/GGA3/VCP/COPG2/CREB3L2/EXOC2/MIA3/GOLGA5/COPG1/GOSR1/SEC24D/CCDC91/SEC16A | 53 |
| GO:0032872 | regulation of stress-activated MAPK cascade | 39/2001 | 215/20772 | 8,13979E-05 | 0,006385178 | 0,005644948 | IGFBP6/SPHK1/TNFRSF19/TPD52L1/SH3RF3/FZD7/FOXO1/RASSF2/SIRPA/EGFR/TNIK/PLCB1/SEMA3A/TAOK3/RELL1/HIPK2/TRAF5/STK39/MID1/HRAS/PJA2/TIRAP/HMGB1/ANKRD6/FAS/EIF2AK2/ZNF622/RIPK1/HIPK3/TRAF4/KLHDC10/CRK/IGF1R/MFHAS1/ZMYND11/DNAJA1/CCN2/MAPKBP1/COPS5 | 39 |
| GO:0061035 | regulation of cartilage development | 19/2001 | 76/20772 | 8,00739E-05 | 0,006385178 | 0,005644948 | CTSK/RARG/RFLNB/SMPD3/TRPS1/SNAI2/BMPR1B/SOX9/LTBP3/SHOX2/AXIN2/BMPR2/LNPK/GLG1/SOX6/FRZB/TGFBR1/PTPN11/CCN2 | 19 |
| GO:0006890 | retrograde vesicle-mediated transport, Golgi to endoplasmic reticulum | 16/2001 | 58/20772 | 8,16709E-05 | 0,006385178 | 0,005644948 | ATP9A/BNIP1/GBF1/PITPNB/ARCN1/RAB6A/KDELR2/ERGIC1/ERGIC2/SCFD1/COPA/COG4/COPB2/RER1/SCYL1/COPG2 | 16 |
| GO:0033692 | cellular polysaccharide biosynthetic process | 17/2001 | 64/20772 | 8,32148E-05 | 0,006422473 | 0,00567792 | HAS2/ENPP1/GYG1/NDST3/SORBS1/DYRK2/AKT1/AKT2/GYS1/CSGALNACT2/PPP1R3F/GBE1/SELENOS/EPM2AIP1/GSK3A/EXT2/B4GALT5 | 17 |
| GO:0070304 | positive regulation of stress-activated protein kinase signaling cascade | 30/2001 | 150/20772 | 8,6475E-05 | 0,006589613 | 0,005825684 | IGFBP6/SPHK1/TNFRSF19/TPD52L1/SH3RF3/FZD7/RASSF2/TNIK/PLCB1/SEMA3A/TAOK3/RELL1/HIPK2/TRAF5/STK39/MID1/HRAS/PJA2/TIRAP/HMGB1/ANKRD6/EIF2AK2/ZNF622/RIPK1/TRAF4/KLHDC10/CRK/MFHAS1/CCN2/MAPKBP1 | 30 |

| | | | | | | | | |
|---|---|---|---|---|---|---|---|---|
| GO:0006606 | protein import into nucleus | 33/2001 | 172/20772 | 9,23491E-05 | 0,006949267 | 0,006143643 | APOD/PTGS2/CBLB/TRAF3IP2/KPNA1/PPP3R1/AKT1/PRKCD/SNUPN/NUP188/IPO13/ING1/MED1/HEATR3/NUP50/AKIRIN2/POM121/TARDBP/KPNA3/IPO7/KPNB1/IPO11/NUP214/NUP62/NOLC1/TRIM28/POM121C/UFM1/PIK3R2/ELAVL1/KPNA6/RBM22/NUP88 | 33 |
| GO:0032456 | endocytic recycling | 22/2001 | 96/20772 | 9,36155E-05 | 0,006957597 | 0,006151008 | ZDHHC2/ATP9A/VPS29/ANKRD50/STX12/RAB35/LMTK2/VPS51/ARHGAP44/VPS50/SNX27/VPS35L/RAB14/RAB7A/SNX3/WASH6P/VPS26C/WASHC2C/ACAP2/ATP6AP1/GGA3/SNX4 | 22 |
| GO:0070302 | regulation of stress-activated protein kinase signaling cascade | 39/2001 | 218/20772 | 0,000110677 | 0,008081096 | 0,00714426 | IGFBP6/SPHK1/TNFRSF19/TPD52L1/SH3RF3/FZD7/FOXO1/RASSF2/SIRPA/EGFR/TNIK/PLCB1/SEMA3A/TAOK3/RELL1/HIPK2/TRAF5/STK39/MID1/HRAS/PJA2/TIRAP/HMGB1/ANKRD6/FAS/EIF2AK2/ZNF622/RIPK1/HIPK3/TRAF4/KLHDC10/CRK/IGF1R/MFHAS1/ZMYND11/DNAJA1/CCN2/MAPKBP1/COPS5 | 39 |
| GO:0008286 | insulin receptor signaling pathway | 27/2001 | 131/20772 | 0,000112759 | 0,008081096 | 0,00714426 | FOXO1/TNS2/ENPP1/SLC27A4/BCAR1/PIP4K2B/PIP4K2B/HRAS/SORBS1/AKT1/AKT2/PRKCD/BAIAP2/ZNF592/NCOA5/FER/AP3S1/IDE/PTPRA/PTPN11/GSK3A/IGF1R/ZNF106/RPS6KB1/SH2B2/PIK3R2/PTPN2 | 27 |
| GO:0018210 | peptidyl-threonine modification | 27/2001 | 131/20772 | 0,000112759 | 0,008081096 | 0,00714426 | SPHK1/CSNK2B/GALNT4/CADM4/SPRED1/BCL2/HIPK2/SPRY2/STK39/PRKACA/GALNT2/PRKACA/GRK2/EIF4G1/DYRK2/AKT1/PRKCD/UBE2K/LMTK2/MARK2/RPTOR/TGFBR1/CALM1/HIPK3/CDC42BPA/GSK3A/SPRED2 | 27 |
| GO:0048705 | skeletal system morphogenesis | 41/2001 | 234/20772 | 0,000120395 | 0,008526768 | 0,007538265 | OTOR/MGP/DSCAML1/CER1/HAS2/TWIST1/MSX2/WNT9B/FGFR3/RARG/TBX15/TMEM119/PAX5/EYA1/PRRX1/RFLNB/SMPD3/FREM1/MDFI/FOXC1/DLX2/BMPR1B/TBX1/SOX9/LTBP3/MMP16/SHOX2/AXIN2/SATB2/SLC39A3/CHSY1/BMPR2/GLG1/EIF4A3/SOX6/TGFBR1/SERPINH1/PKD1/MED12/EXT2/CCN2 | 41 |

| | | | | | | | | |
|---|---|---|---|---|---|---|---|---|
| GO:0018393 | internal peptidyl-lysine acetylation | 34/2001 | 182/20772 | 0,000125699 | 0,00879895 | 0,007778894 | TWIST1/SNAI2/SNCA/HCFC1/NOC2L/BRPF1/NFYA/KAT5/TAF4/PHF20L1/MSL3/GLYR1/NAA60/JADE3/DDX21/ATXN7L3/BRPF3/RUVBL2/MORF4L2/SET/TRRAP/TAF5L/EP400/TAF2/MORF4L1/EPC1/SUPT20H/RUVBL1/MRGBP/DR1/SIN3A/USP22/TAF12/DEK | 34 |
| GO:0006401 | RNA catabolic process | 57/2001 | 361/20772 | 0,000133336 | 0,009226206 | 0,008156618 | TENT5A/ZFP36L2/ZHX2/ZSWIM8/HSPA1B/TRAF3IP2/TRAF5/NRDE2/NSUN2/CASC3/AXIN2/FTO/TIRAP/THRAP3/AKT1/IKBKE/PRKCD/CTIF/CNOT3/DCP1A/CNOT10/SECISBP2/KHSRP/EIF4A3/GSPT1/EXOSC6/SND1/ZFP36/MLH1/SERBP1/NUDT12/TENT4A/NCBP2/WDR82/TARDBP/MAGOHB/GRSF1/YTHDF1/METTL16/PPP1R8/MTREX/FXR2/PATL1/RBM8A/EXOSC4/CELF1/SYNCRIP/LARP4B/DIS3/DICER1/NCBP1/SAMD4B/GTPBP1/APEX1/ELAVL1/ZC3H14/IGF2BP3 | 57 |
| GO:0006446 | regulation of translational initiation | 21/2001 | 92/20772 | 0,00014207 | 0,009718855 | 0,008592155 | PPP1R15A/EIF4G1/EIF2S1/CTIF/PPP1R15B/DDX3X/EIF1/EIF3B/KLHL25/EIF2AK2/DNAJC3/DHX29/NCBP2/BZW1/YTHDF1/RPS6KB1/EIF4G2/NCBP1/SH3BGRL/EIF5/DDX1 | 21 |
| GO:0006997 | nucleus organization | 29/2001 | 147/20772 | 0,00014466 | 0,009784859 | 0,008650506 | PLEC/CHMP4A/SUN2/HIPK2/LEMD2/PITPNB/LMNB2/KAT5/PHF2/USP36/NSFL1C/PSME4/UBXN2A/TMEM43/SRPK2/SERBP1/CHMP4B/TARDBP/VPS4A/CHMP6/POLR1B/EPC1/ANKLE2/NOLC1/CTDNEP1/CTDNEP1/RRP8/PHF8/SUN1 | 29 |
| GO:0006888 | endoplasmic reticulum to Golgi vesicle-mediated transport | 28/2001 | 140/20772 | 0,000146431 | 0,009794601 | 0,008659119 | GBF1/ARCN1/YIPF5/PREB/KDELR2/SEC31A/YIPF4/YKT6/ERGIC1/RAB1A/ERGIC2/SEC24A/CSNK1D/SCFD1/P4HB/COPA/GOLT1B/COPB2/TRAPPC12/TRAPPC11/VCP/COPG2/CREB3L2/MIA3/COPG1/GOSR1/SEC24D/SEC16A | 28 |
| GO:0051168 | nuclear export | 34/2001 | 185/20772 | 0,000174316 | 0,011266496 | 0,009960378 | FRAT2/PRKACA/NRDE2/NSUN2/LTV1/CASC3/CALR/PRKACA/HSPA9/GLE1/NUP188/MALT1/RANBP3/ABCE1/POLDIP3/CDKN1B/EIF4A3/POM121/NCBP2/PKD1/PTPN11/MAGOHB/DDX19A/NUP214/ALYREF/NUP62/RBM8A/PPM1A/NCBP1/POM121C/CHTOP/RBM22/NUP88/XPO4 | 34 |

| | | | | | | | | |
|---|---|---|---|---|---|---|---|---|
| GO:0031056 | regulation of histone modification | 31/2001 | 163/20772 | 0,000175109 | 0,011266496 | 0,009960378 | TWIST1/APLNR/PAX5/SNAI2/SNCA/BCOR/NELFE/NELFE/NELFE/NELFE/NELFE/HCFC1/NOC2L/NFYA/PAXIP1/TRIP12/GLYR1/DDX21/RUVBL2/SET/C6orf89/SNW1/PRMT6/SKIC8/DR1/NELFA/OTUB2/SIN3A/CHTOP/RRP8/DEK | 31 |
| GO:1900026 | positive regulation of substrate adhesion-dependent cell spreading | 13/2001 | 44/20772 | 0,000175922 | 0,011266496 | 0,009960378 | HAS2/DAB2/NEDD9/FERMT2/DOCK5/DOCK1/CALR/RAC1/ARPC2/P4HB/CRK/ILK/CDC42 | 13 |
| GO:0051571 | positive regulation of histone H3-K4 methylation | 10/2001 | 28/20772 | 0,000173373 | 0,011266496 | 0,009960378 | NELFE/NELFE/NELFE/NELFE/NELFE/HCFC1/PAXIP1/SNW1/SKIC8/NELFA | 10 |
| GO:0018394 | peptidyl-lysine acetylation | 35/2001 | 193/20772 | 0,000184855 | 0,011713977 | 0,010355983 | TWIST1/SNAI2/SNCA/HCFC1/NOC2L/BRPF1/NFYA/KAT5/TAF4/PHF20L1/MSL3/GLYR1/NAA60/JADE3/DDX21/ATXN7L3/BRPF3/RUVBL2/MORF4L2/SET/TRRAP/HDAC2/TAF5L/EP400/TAF2/MORF4L1/EPC1/SUPT20H/RUVBL1/MRGBP/DR1/SIN3A/USP22/TAF12/DEK | 35 |
| GO:0006475 | internal protein amino acid acetylation | 34/2001 | 186/20772 | 0,000193923 | 0,011792072 | 0,010425025 | TWIST1/SNAI2/SNCA/HCFC1/NOC2L/BRPF1/NFYA/KAT5/TAF4/PHF20L1/MSL3/GLYR1/NAA60/JADE3/DDX21/ATXN7L3/BRPF3/RUVBL2/MORF4L2/SET/TRRAP/TAF5L/EP400/TAF2/MORF4L1/EPC1/SUPT20H/RUVBL1/MRGBP/DR1/SIN3A/USP22/TAF12/DEK | 34 |
| GO:0045931 | positive regulation of mitotic cell cycle | 28/2001 | 142/20772 | 0,0001883 | 0,011792072 | 0,010425025 | SPHK1/DDR2/RGCC/CCND2/EGFR/PLCB1/LSM11/PBX1/EIF4G1/AKT1/DDX3X/CDT1/RPTOR/RB1/TTL/RCC2/RCC2/PTPN11/PLRG1/RPS6KB1/STIL/TMOD3/APEX1/SIN3A/POLDIP2/DYNLT3/USP22/CDC25A | 28 |
| GO:0018107 | peptidyl-threonine phosphorylation | 25/2001 | 121/20772 | 0,000190969 | 0,011792072 | 0,010425025 | SPHK1/CSNK2B/CADM4/SPRED1/BCL2/HIPK2/SPRY2/STK39/PRKACA/PRKACA/GRK2/EIF4G1/DYRK2/AKT1/PRKCD/UBE2K/LMTK2/MARK2/RPTOR/TGFBR1/CALM1/HIPK3/CDC42BPA/GSK3A/SPRED2 | 25 |
| GO:0010761 | fibroblast migration | 15/2001 | 56/20772 | 0,000193856 | 0,011792072 | 0,010425025 | TNS1/DDR2/PLEC/MACIR/CORO1C/IQGAP1/AKT1/RAC1/FER/RCC2/RCC2/ILK/TMEM201/GNA12/GNA13 | 15 |

| | | | | | | | | |
|---|---|---|---|---|---|---|---|---|
| GO:0032885 | regulation of polysaccharide biosynthetic process | 12/2001 | 39/20772 | 0,000203023 | 0,012221973 | 0,010805087 | HAS2/SMPD3/ENPP1/SORBS1/DYRK2/AKT1/AKT2/PPP1R3F/CLTC/SELENOS/EPM2AIP1/GSK3A | 12 |
| GO:0032330 | regulation of chondrocyte differentiation | 15/2001 | 57/20772 | 0,000239915 | 0,014299905 | 0,012642126 | RARG/RFLNB/TRPS1/SNAI2/BMPR1B/SOX9/LTBP3/SHOX2/AXIN2/LNPK/GLG1/SOX6/TGFBR1/PTPN11/CCN2 | 15 |
| GO:0007173 | epidermal growth factor receptor signaling pathway | 24/2001 | 116/20772 | 0,000248676 | 0,014676773 | 0,012975304 | REPS2/ERBB4/AFAP1L2/RASSF2/EGFR/CBLB/SOX9/PLAUR/RALB/BCAR1/MVB12B/RNF126/IQGAP1/AKT1/FER/HIP1/RAB7A/PTPN11/CHMP6/SHC3/NUP62/ERRFI1/CBL/PTPN2 | 24 |
| GO:0002183 | cytoplasmic translational initiation | 12/2001 | 40/20772 | 0,000264822 | 0,015477963 | 0,013683613 | EIF4G1/EIF3C/EIF3J/EIF2B3/EIF3B/DHX29/NCBP2/DENR/NCBP1/EIF2D/SH3BGRL/EIF5 | 12 |
| GO:0042063 | gliogenesis | 56/2001 | 363/20772 | 0,00027962 | 0,016185685 | 0,014309289 | CCL2/OLIG1/OLIG2/ID2/POU3F2/CSPG4/LPAR1/DNER/NR3C1/PTN/DLX1/APCDD1/CDK6/LAMC3/PLEC/GPM6B/SOX13/LAMB1/ROR2/DLX2/CNTN1/SRGAP2/SUN2/SRGAP2C/SOX9/DISC1/TNFRSF21/IL6ST/NF2/SYNJ1/WDR1/LAMB2/PPP3R1/AKT1/TTC21B/AKT2/LRP8/IFNGR1/DAB1/RB1/SOX6/EIF2B3/LDLR/PTPN11/HDAC2/PRPF19/GPC1/ILK/MED12/RHOA/BNIP3/DICER1/B4GALT5/NDUFS2/PTEN/SUN1 | 56 |
| GO:0046328 | regulation of JNK cascade | 30/2001 | 160/20772 | 0,000285575 | 0,016372982 | 0,014474873 | TNFRSF19/TPD52L1/SH3RF3/FZD7/RASSF2/SIRPA/EGFR/TNIK/PLCB1/SEMA3A/TAOK3/HIPK2/TRAF5/HRAS/PJA2/TIRAP/HMGB1/ANKRD6/ZNF622/RIPK1/HIPK3/TRAF4/CRK/IGF1R/MFHAS1/ZMYND11/DNAJA1/CCN2/MAPKBP1/COPS5 | 30 |

| | | | | | | | | |
|---|---|---|---|---|---|---|---|---|
| GO:0032956 | regulation of actin cytoskeleton organization | 57/2001 | 372/20772 | 0,000296061 | 0,016656874 | 0,014725854 | CDC42EP5/ARHGDIB/FZD10/EPHA3/LPAR1/KANK4/RGCC/CTNNA2/SYNPO/PHLDB2/GPM6B/TRIM27/ARHGAP6/TRIM27/DLC1/NEDD9/LIMA1/FERMT2/EPS8/PPM1F/RASA1/NF2/JAM3/FCHSD2/HRAS/SEMA5A/SPTAN1/WDR1/SLC4A2/IQGAP1/ADD1/PRKCD/BAIAP2/ARPC5/MTPN/ARHGAP44/RAC1/FER/LATS1/TMSB4X/TGFBR1/ARF1/IQGAP2/CFL2/ARPC2/CRK/ARPC5L/WASH6P/WASHC2C/TMOD3/RHOA/CAPZA1/CDC42/PIK3R2/CCN2/FSCN1/PAK1 | 57 |
| GO:0140014 | mitotic nuclear division | 50/2001 | 315/20772 | 0,000293805 | 0,016656874 | 0,014725854 | SPHK1/EDN3/SMPD3/RGCC/KIFC1/CHMP4A/KMT5A/HSPA1B/NDE1/EPS8/MAD1L1/PPP1R9B/IK/KAT5/KIF4A/NSFL1C/CDT1/CDKN1B/PRC1/RB1/KIF3B/LSM14A/TENT4A/CHMP4B/VPS4A/KPNB1/SH2B1/SMC2/WAPL/DSN1/PPP2R2D/CHMP6/ANAPC15/PDS5B/ANKLE2/KIFC1/USP16/FBXW5/NUP62/RHOA/KIF2A/BCCIP/CDC42/MACROH2A1/DCTN2/POLDIP2/MAPRE1/SLF2/PIN1/PPP2R1A | 50 |
| GO:0051895 | negative regulation of focal adhesion assembly | 8/2001 | 20/20772 | 0,000318653 | 0,016680788 | 0,014746995 | APOD/PHLDB2/ARHGAP6/DLC1/CORO1C/RCC2/RCC2/PTEN | 8 |
| GO:0150118 | negative regulation of cell-substrate junction organization | 8/2001 | 20/20772 | 0,000318653 | 0,016680788 | 0,014746995 | APOD/PHLDB2/ARHGAP6/DLC1/CORO1C/RCC2/RCC2/PTEN | 8 |
| GO:0061013 | regulation of mRNA catabolic process | 39/2001 | 229/20772 | 0,000317272 | 0,016680788 | 0,014746995 | TENT5A/ZFP36L2/TRAF3IP2/TRAF5/CASC3/AXIN2/FTO/TIRAP/THRAP3/AKT1/IKBKE/PRKCD/CNOT3/DCP1A/CNOT10/SECISBP2/KHSRP/EIF4A3/ZFP36/MLH1/SERBP1/TENT4A/TARDBP/MAGOHB/YTHDF1/METTL16/FXR2/PATL1/RBM8A/CELF1/SYNCRIP/LARP4B/DIS3/SAMD4B/GTPBP1/APEX1/ELAVL1/ZC3H14/IGF2BP3 | 39 |

| | | | | | | | | |
|---|---|---|---|---|---|---|---|---|
| GO:1903008 | organelle disassembly | 28/2001 | 146/20772 | 0,000305178 | 0,016680788 | 0,014746995 | CTSK/STING1/NEDD9/GBF1/RRP7A/ULK2/ATG3/KAT5/SLC25A5/USP36/MARK2/ATG4D/KLC1/ASCC2/GSK3A/ASCC3/DENR/VDAC1/CDC37/MFN2/VCP/BNIP3/UFM1/EIF2D/SREBF2/TEX264/RNF41/ATG13 | 28 |
| GO:1904375 | regulation of protein localization to cell periphery | 27/2001 | 139/20772 | 0,000313153 | 0,016680788 | 0,014746995 | CPLX1/EPHA3/PRNP/DAB2/ZDHHC2/EGFR/GNAI1/GPC4/EPB41L2/PPP1R9B/HRAS/RAP1A/SORBS1/AKT1/PTPN9/GPSM2/ARHGAP44/CLTC/VPS4A/VTI1B/LDLRAP1/ZDHHC5/RER1/PIK3R2/ZDHHC8/PICALM/IQSEC2 | 27 |
| GO:0030433 | ubiquitin-dependent ERAD pathway | 21/2001 | 97/20772 | 0,000310168 | 0,016680788 | 0,014746995 | OS9/CALR/HSP90B1/SYVN1/ERLIN1/UFD1/CANX/HSPA5/SEL1L/CANX/STUB1/FAF2/SELENOS/UBE4A/USP14/CCDC47/ECPAS/VCP/RNF185/PSMC6/MARCHF6 | 21 |
| GO:0034637 | cellular carbohydrate biosynthetic process | 18/2001 | 77/20772 | 0,000302995 | 0,016680788 | 0,014746995 | HAS2/ENPP1/GYG1/NDST3/SORBS1/DYRK2/AKT1/AKT2/GYS1/CSGALNACT2/PPP1R3F/GBE1/SELENOS/EPM2AIP1/GSK3A/GOT1/EXT2/B4GALT5 | 18 |
| GO:0051569 | regulation of histone H3-K4 methylation | 11/2001 | 35/20772 | 0,000302332 | 0,016680788 | 0,014746995 | BCOR/NELFE/NELFE/NELFE/NELFE/NELFE/HCFC1/PAXIP1/SNW1/SKIC8/NELFA | 11 |
| GO:1903649 | regulation of cytoplasmic transport | 10/2001 | 30/20772 | 0,000331739 | 0,017216104 | 0,015220252 | DAB2/ATP9A/MSN/NF2/USP7/MAP2/MTMR2/DNAJC13/RAB21/SNX3 | 10 |
| GO:0061162 | establishment of monopolar cell polarity | 9/2001 | 25/20772 | 0,000342319 | 0,017349106 | 0,015337835 | FOXJ1/LAMA1/MSN/GBF1/MYO9A/FAT1/RHOA/CDC42/FSCN1 | 9 |
| GO:0051098 | regulation of binding | 61/2001 | 407/20772 | 0,000338594 | 0,017349106 | 0,015337835 | TWIST1/MSX2/STING1/PRMT8/DAB2/UBASH3B/CSNK2B/MDFI/HEY2/FOXC1/CBLB/PLAUR/RALB/DISC1/HIPK2/PRKACA/TFIP11/GTPBP4/PRKACA/CTBP2/EIF4G1/ADD1/EIF2S1/EIF3C/AKT1/HMGB1/PRKCD/MARK2/TCF7L2/CDT1/EIF4A3/RB1/TMSB4X/MAP2/TGFBR1/CALM1/HMGA2/STUB1/IDE/PSME3IP1/ATP2A2/USP9X/PKD1/SYMPK/HDAC2/WAPL/CRK/LDLRAP1/NSD1/SUB1/TRIM28/CDC42/CEBPG/NCBP1/SIN3A/POLDIP2/MAPRE1/PIN1/CDON/LAMTOR5/DPH3 | 61 |

| | | | | | | | | |
|---|---|---|---|---|---|---|---|---|
| GO:0050684 | gliogenesis | 28/2001 | 147/20772 | 0,000342947 | 0,017349106 | 0,015337835 | SLC39A5/KHDRBS3/CELF2/QKI/SFSWAP/NOVA1/THRAP3/RBM3/RBM39/ZC3H3/SRPK2/NCBP2/MAGOHB/METTL16/RBMXL1/SNW1/PRPF19/NUDT21/SF3B4/FXR2/HSPA8/RBM8A/CELF1/NCBP1/CPSF7/ZC3H14/TRA2A/HNRNPK | 28 |
| GO:0038127 | ERBB signaling pathway | 26/2001 | 133/20772 | 0,000361306 | 0,018125539 | 0,016024257 | REPS2/ERBB4/AFAP1L2/RASSF2/EGFR/CBLB/SOX9/PLAUR/RALB/BCAR1/MVB12B/RNF126/IQGAP1/AKT1/FER/HIP1/STUB1/CUL5/RAB7A/PTPN11/CHMP6/SHC3/NUP62/ERRFI1/CBL/PTPN2 | 26 |
| GO:0006402 | mRNA catabolic process | 49/2001 | 310/20772 | 0,000370786 | 0,018225251 | 0,01611241 | TENT5A/ZFP36L2/ZHX2/HSPA1B/TRAF3IP2/TRAF5/CASC3/AXIN2/FTO/TIRAP/THRAP3/AKT1/IKBKE/PRKCD/CTIF/CNOT3/DCP1A/CNOT10/SECISBP2/KHSRP/EIF4A3/GSPT1/EXOSC6/SND1/ZFP36/MLH1/SERBP1/NUDT12/TENT4A/NCBP2/TARDBP/MAGOHB/YTHDF1/METTL16/FXR2/PATL1/RBM8A/EXOSC4/CELF1/SYNCRIP/LARP4B/DIS3/NCBP1/SAMD4B/GTPBP1/APEX1/ELAVL1/ZC3H14/IGF2BP3 | 49 |
| GO:0048762 | mesenchymal cell differentiation | 45/2001 | 278/20772 | 0,000372376 | 0,018225251 | 0,01611241 | HAS2/TWIST1/MSX2/EPHA3/EDN3/RFLNB/TGFB1I1/ERBB4/FOXF2/DAB2/OLFM1/RGCC/BMP5/SNAI2/PHLDB2/EDNRA/HEY2/FOXC1/DACT3/AMER1/TBX1/SOX9/SEMA3A/SPRED1/SEMA3D/BCL2/FERMT2/SPRY2/CORO1C/SEMA5A/AXIN2/PPP3R1/TCF7L2/FRZB/TGFBR1/HMGA2/HDAC2/POLR1B/PHACTR4/SPRED2/NOLC1/TRIM28/CDC42/USF3/PTEN | 45 |
| GO:0035307 | positive regulation of protein dephosphorylation | 14/2001 | 53/20772 | 0,000367451 | 0,018225251 | 0,01611241 | PPP1R16B/DLC1/PPP1R15A/PRKCD/PPP1R15B/HSP90AB1/CALM1/SYMPK/PTPA/PPP1R7/ANKLE2/PPP2R5D/PIN1/GNA12 | 14 |
| GO:0006473 | protein acetylation | 39/2001 | 231/20772 | 0,00037974 | 0,018435784 | 0,016298535 | TWIST1/SPHK1/FOXO1/SNAI2/SNCA/HCFC1/NAA25/NOC2L/BRPF1/NFYA/KAT5/TAF4/PHF20L1/MSL3/DDX3X/GLYR1/NAA60/JADE3/DDX21/ATXN7L3/BRPF3/RUVBL2/MORF4L2/SET/TRRAP/HDAC2/TAF5L/EP400/TAF2/MORF4L1/EPC1/SUPT20H/RUVBL1/MRGBP/DR1/SIN3A/USP22/TAF12/DEK | 39 |

| | | | | | | | | |
|---|---|---|---|---|---|---|---|---|
| GO:0010810 | regulation of cell-substrate adhesion | 40/2001 | 239/20772 | 0,000386615 | 0,018594096 | 0,016438494 | APOD/HAS2/EPHA3/DAB2/FZD7/CDK6/PHLDB2/GPM6B/ARHGAP6/COL26A1/CCDC80/DLC1/ONECUT1/NEDD9/BCL2/FERMT2/DISC1/PPM1F/RASA1/CORO1C/DOCK5/NF2/DOCK1/CALR/TLN1/RAC1/VCL/PIK3CB/RCC2/RCC2/ARPC2/PTPRA/P4HB/CRK/WASHC2C/ILK/RHOA/CDC42/POLDIP2/PTEN | 40 |
| GO:0022029 | telencephalon cell migration | 17/2001 | 72/20772 | 0,000389179 | 0,018594096 | 0,016438494 | FOXB1/POU3F2/POU3F3/EGFR/LAMB1/SRGAP2/SUN2/SRGAP2C/DISC1/FBXO45/ROBO1/SRF/LRP8/DAB1/RHOA/OGDH/SUN1 | 17 |
| GO:0097152 | mesenchymal cell apoptotic process | 7/2001 | 16/20772 | 0,000397876 | 0,01868164 | 0,01651589 | POU3F4/MSX2/PAX8/EDNRA/TBX1/SOX9/ETV6 | 7 |
| GO:0071453 | cellular response to oxygen levels | 32/2001 | 178/20772 | 0,000400321 | 0,01868164 | 0,01651589 | PTGS2/TWIST1/LPAR1/DDR2/RGCC/FOXO1/AK4/PMAIP1/BCL2/HIPK2/AIFM1/AKT1/PPARD/PDK3/FAS/ADO/RPTOR/CDKN1B/PIK3CB/MIEF1/STUB1/DDAH1/ATP7A/P4HB/KCNK2/TMEM199/ATP6AP1/BNIP3/CPEB4/CBL/ACAA2/NDUFS2 | 32 |
| GO:0030010 | establishment of cell polarity | 30/2001 | 163/20772 | 0,000397196 | 0,01868164 | 0,01651589 | FOXJ1/KIF26B/PHLDB2/FEZ1/LAMA1/MSN/NDE1/SPRY2/TRAF3IP2/GBF1/AMOTL1/JAM3/MYO9A/KAT5/MARK2/GPSM2/NSFL1C/HSP90AB1/SNX27/PKD1/CRK/MAP4/IGF1R/FAT1/RHOA/SDCCAG8/CDC42/FLOT2/FSCN1/MAPRE1 | 30 |
| GO:0060485 | mesenchyme development | 53/2001 | 344/20772 | 0,000414113 | 0,018949427 | 0,016752632 | CER1/HAS2/TWIST1/MSX2/EPHA3/APLNR/EDN3/RFLNB/TGFB1I1/ERBB4/FOXF2/DAB2/OLFM1/BNC2/RGCC/BMP5/SNAI2/PHLDB2/EDNRA/HEY2/FOXC1/DACT3/AMER1/TBX1/SOX9/SEMA3A/SPRED1/SEMA3D/BCL2/FERMT2/SPRY2/DCHS1/CORO1C/ROBO1/BASP1/SEMA5A/AXIN2/YAP1/PPP3R1/BMPR2/TCF7L2/FRZB/TGFBR1/HMGA2/HDAC2/POLR1B/PHACTR4/SPRED2/NOLC1/TRIM28/CDC42/USF3/PTEN | 53 |
| GO:0050000 | chromosome localization | 21/2001 | 99/20772 | 0,000415502 | 0,018949427 | 0,016752632 | KIFC1/CHMP4A/NDE1/MAD1L1/MAP1S/ACTR2/KAT5/ACTR3/CDT1/TTL/MLH1/CHMP4B/VPS4A/KPNB1/CHMP6/TRAPPC12/KIFC1/NUP62/DCTN2/MAPRE1/SUN1 | 21 |

| | | | | | | | | |
|---|---|---|---|---|---|---|---|---|
| GO:0060349 | bone morphogenesis | 21/2001 | 99/20772 | 0,000415502 | 0,018949427 | 0,016752632 | CER1/HAS2/TWIST1/MSX2/FGFR3/RARG/TMEM119/SMPD3/FREM1/FOXC1/BMPR1B/SOX9/LTBP3/MMP16/SHOX2/AXIN2/CHSY1/BMPR2/GLG1/SERPINH1/EXT2 | 21 |
| GO:0031647 | regulation of protein stability | 56/2001 | 369/20772 | 0,000424607 | 0,019075649 | 0,016864221 | ADGRV1/PRNP/RASSF2/MYLIP/SNCA/BCL2/HYPK/HSPA1B/RABL3/HCFC1/NF2/GIPC1/BAG5/STX12/MFSD1/GTPBP4/CALR/NCLN/SYVN1/SUGT1/PRKCD/PPIB/USP36/CCAR2/CRTAP/GNAQ/TELO2/USP7/CTSA/HSP90AB1/SEL1L/HIP1/STUB1/RUVBL2/USP9X/HSPD1/RAB21/TARDBP/USP13/DDOST/CCT7/SH3GLB1/RUVBL1/CDC37/HSPA8/GGA3/B4GALT5/DAD1/VBP1/FLOT2/WIZ/PIN1/POLR2E/PTEN/PFDN6/SEC16A | 56 |
| GO:0046330 | positive regulation of JNK cascade | 23/2001 | 113/20772 | 0,000424208 | 0,019075649 | 0,016864221 | TNFRSF19/TPD52L1/SH3RF3/FZD7/RASSF2/TNIK/PLCB1/SEMA3A/TAOK3/HIPK2/TRAF5/HRAS/PJA2/TIRAP/HMGB1/ANKRD6/ZNF622/RIPK1/TRAF4/CRK/MFHAS1/CCN2/MAPKBP1 | 23 |
| GO:0006998 | nuclear envelope organization | 15/2001 | 60/20772 | 0,000438892 | 0,01957135 | 0,017302456 | CHMP4A/SUN2/LEMD2/LMNB2/NSFL1C/UBXN2A/TMEM43/CHMP4B/TARDBP/VPS4A/CHMP6/ANKLE2/CTDNEP1/CTDNEP1/SUN1 | 15 |
| GO:0031345 | negative regulation of cell projection organization | 35/2001 | 202/20772 | 0,000451764 | 0,019997212 | 0,017678948 | WNT3/WNT3/ARHGAP24/LPAR1/PMP22/PRNP/DAB2/RTN4RL1/EFNB2/MYLIP/NR2F1/DNM3/SRGAP2C/ITM2C/EFNB3/SEMA3A/SEMA3D/PTPRG/SPRY2/SEMA5A/BAG5/ULK2/YAP1/PRKCD/PTPN9/ARHGAP44/DAB1/MAP2/HDAC2/GSK3A/GORASP1/MAP4/RHOA/PTEN/YWHAH | 35 |
| GO:0051651 | maintenance of location in cell | 40/2001 | 241/20772 | 0,000459526 | 0,020192321 | 0,017851438 | APLNR/CACNA1C/CACNA1C/CD4/UBASH3B/ITPR2/RASA3/RASA3/SNCA/PLCB1/ARHGAP21/SUN2/HK1/GSTO1/OS9/PRKACA/ATP7B/SPOUT1/IBTK/IBTK/PLCB2/CALR/PRKACA/HSP90B1/CIZ1/AKT1/DBN1/CHERP/HEXA/SLC30A4/GPSM2/HSPA5/KDELR2/TMSB4X/CALM1/DIAPH1/ARL2BP/RER1/FTH1/SUN1 | 40 |

| | | | | | | | | |
|---|---|---|---|---|---|---|---|---|
| GO:0010001 | glial cell differentiation | 42/2001 | 257/20772 | 0,00046846 | 0,020435729 | 0,018066628 | OLIG1/OLIG2/ID2/POU3F2/LPAR1/DNER/NR3C1/PTN/DLX1/CDK6/LAMC3/PLEC/GPM6B/SOX13/ROR2/DLX2/CNTN1/SOX9/TNFRSF21/IL6ST/WDR1/LAMB2/PPP3R1/AKT1/TTC21B/AKT2/IFNGR1/DAB1/SOX6/EIF2B3/LDLR/PTPN11/HDAC2/PRPF19/GPC1/ILK/MED12/RHOA/BNIP3/DICER1/B4GALT5/PTEN | 42 |
| GO:0071826 | ribonucleoprotein complex subunit organization | 44/2001 | 273/20772 | 0,000472328 | 0,020456212 | 0,018084736 | BOP1/BOP1/CELF2/PPAN/YJU2/SFSWAP/TFIP11/RRP7A/DNAJC17/EIF3C/SF3B2/TSSC4/EIF3J/SNRPB2/HSP90AB1/EIF3B/DHX8/ADAR/KLC1/RUVBL2/SF3A3/CD2BP2/SRPK2/DHX29/PRPF18/PRPF19/DENR/NUDT21/PRPF8/RUVBL1/SF3B4/SNRNP200/CELF1/VCP/DICER1/NCBP1/XAB2/CPSF7/EIF2D/SNRPD1/MRM2/SF3A2/EIF5/DDX1 | 44 |
| GO:0061339 | establishment or maintenance of monopolar cell polarity | 9/2001 | 26/20772 | 0,000479078 | 0,020527381 | 0,018147655 | FOXJ1/LAMA1/MSN/GBF1/MYO9A/FAT1/RHOA/CDC42/FSCN1 | 9 |
| GO:0043484 | regulation of RNA splicing | 34/2001 | 195/20772 | 0,000480791 | 0,020527381 | 0,018147655 | AHNAK/SLC39A5/KHDRBS3/CELF2/QKI/SFSWAP/NOVA1/THRAP3/RBM3/ATXN7L3/POLR2A/RBM39/SRPK2/GRSF1/TRRAP/METTL16/RBMXL1/SNW1/TAF5L/PRPF19/SUPT20H/SF3B4/FXR2/HSPA8/CLK3/RBM8A/CELF1/NCBP1/PQBP1/RBM22/TRA2A/USP22/HNRNPK/TAF12 | 34 |
| GO:0032984 | protein-containing complex disassembly | 42/2001 | 258/20772 | 0,000508531 | 0,021558853 | 0,019059549 | NCKAP5/CHMP4A/FRAT2/LIMA1/EPS8/MID1/SYNJ1/PPP1R9B/TFIP11/SEMA5A/DNAJC17/MAP1S/SPTAN1/WDR1/VPS16/ADD1/SSRP1/ABCE1/MTPN/MID1IP1/GSPT1/GRWD1/DHX8/CALM1/KLC1/CHMP4B/ATP2A2/SET/CFL2/VPS4A/CHMP6/IGF1R/SUPT16H/WASHC2C/HSPA8/TMOD3/SMARCE1/KIF2A/CAPZA1/VCP/BNIP3/VMP1 | 42 |

| | | | | | | | | |
|---|---|---|---|---|---|---|---|---|
| GO:0007059 | chromosome segregation | 71/2001 | 499/20772 | 0,000542867 | 0,022460652 | 0,019856803 | NR3C1/KIFC1/CHMP4A/KMT5A/P3H4/HSPA1B/HAUS7/NDE1/ZCWPW1/EME2/MAD1L1/IK/C9orf78/MAP3K20/MSH5/MAP1S/ACTR2/KAT5/SLC25A5/KIF4A/TUBGCP2/ACTR3/MLH3/DDX3X/GPSM2/CDT1/NAA60/PRC1/RB1/LATS1/KIF3B/SKA1/TTL/MLH1/RCC2/RCC2/LSM14A/PSMC3IP/TUBGCP3/TENT4A/CHMP4B/USP9X/CSNK1D/VPS4A/KPNB1/SMC2/WAPL/DSN1/CIAO1/CHMP6/ANAPC15/PDS5B/TRAPPC12/KIFC1/STIL/SMARCE1/NUP62/RHOA/KIF2A/SKA2/BCCIP/CDC42/MACROH2A1/DCTN2/POLDIP2/MAPRE1/SLF2/PBRM1/NUP43/PPP2R1A/SUN1 | 71 |
| GO:0009100 | glycoprotein metabolic process | 63/2001 | 431/20772 | 0,000538033 | 0,022460652 | 0,019856803 | CYTL1/ITM2A/DCN/PTX3/CNMD/DSEL/DSE/B3GLCT/ST6GALNAC3/GALNT4/PLCB1/BMPR1B/CHST12/ITM2C/XYLT1/GAL3ST4/PHLDA1/BCL2/NAGPA/GALNT10/FUT4/ST3GAL4/NDST3/GALNT2/CHSY1/SLC2A10/CHST10/SYVN1/BMPR2/HEXA/ST8SIA4/POGLUT3/CSGALNACT2/PPARD/GLCE/TCF7L2/CANT1/B3GALNT1/MGAT1/UGGT1/XXYLT1/ST6GALNAC5/RAB1A/B4GALT3/B3GALT6/FUT11/FAM20B/GANAB/DDOST/ATP7A/B3GNT5/DPY19L1/GORASP1/GPC1/CHPF2/EXT2/DPM1/B4GALT5/DAD1/RNF185/STT3A/CRPPA/MARCHF6 | 63 |
| GO:0006479 | protein methylation | 35/2001 | 204/20772 | 0,000544727 | 0,022460652 | 0,019856803 | PRMT8/PAX5/BCOR/KMT5A/H1-2/NELFE/NELFE/NELFE/NELFE/NELFE/HCFC1/KMT2C/CXXC1/NFYA/PAXIP1/ASH2L/RAB6A/PRMT2/GSPT1/PRDM2/PCMT1/NCOA6/WDR82/SNW1/PRMT6/ASH1L/SKIC8/NELFA/KMT5B/EEF1AKMT1/EEF2KMT/CHTOP/RRP8/WDR5B/SETMAR | 35 |
| GO:0008213 | protein alkylation | 35/2001 | 204/20772 | 0,000544727 | 0,022460652 | 0,019856803 | PRMT8/PAX5/BCOR/KMT5A/H1-2/NELFE/NELFE/NELFE/NELFE/NELFE/HCFC1/KMT2C/CXXC1/NFYA/PAXIP1/ASH2L/RAB6A/PRMT2/GSPT1/PRDM2/PCMT1/NCOA6/WDR82/SNW1/PRMT6/ASH1L/SKIC8/NELFA/KMT5B/EEF1AKMT1/EEF2KMT/CHTOP/RRP8/WDR5B/SETMAR | 35 |

| | | | | | | | | |
|---|---|---|---|---|---|---|---|---|
| GO:0016358 | dendrite development | 42/2001 | 259/20772 | 0,000551623 | 0,022513215 | 0,019903272 | LPAR1/PRKG1/PTN/CTNNA2/BMP5/PTPRD/PTPRD/TNIK/SRGAP2/TANC2/PPFIA2/CDKL5/DNM3/PDLIM5/SRGAP2C/BTBD3/SEMA3A/CTNND2/DISC1/PPP1R9B/MAP1S/IQSEC1/DBN1/BAIAP2/LRP8/KIDINS220/ARHGAP44/DAB1/MAP2/RAB21/SDK1/HDAC2/CRK/GSK3A/GORASP1/RHOA/TBC1D24/CDC42/PQBP1/PICALM/PTEN/YWHAH | 42 |
| GO:0099175 | regulation of postsynapse organization | 22/2001 | 108/20772 | 0,000553481 | 0,022513215 | 0,019903272 | PRNP/PTPRD/PTPRD/ROR2/TANC2/PPFIA2/CDKL5/DNM3/PDLIM5/NEDD9/LRRC4B/DISC1/AKT1/DBN1/BAIAP2/LRFN4/LRP8/ARHGAP44/HSPA8/CDC42/RPS6KA5/PTEN | 22 |
| GO:0032970 | regulation of actin filament-based process | 61/2001 | 415/20772 | 0,000565698 | 0,02284104 | 0,020193092 | CDC42EP5/ARHGDIB/FZD10/EPHA3/CACNA1C/CACNA1C/LPAR1/KANK4/RGCC/CTNNA2/SYNPO/PHLDB2/GPM6B/PDE4B/TRIM27/ARHGAP6/TRIM27/DLC1/NEDD9/LIMA1/FERMT2/EPS8/PPM1F/RASA1/NF2/JAM3/FCHSD2/HRAS/SEMA5A/SPTAN1/WDR1/SLC4A2/IQGAP1/ADD1/PRKCD/BAIAP2/ARPC5/MTPN/ARHGAP44/RAC1/FER/LATS1/TMSB4X/TGFBR1/ARF1/IQGAP2/ATP2A2/CFL2/ARPC2/CRK/ARPC5L/WASH6P/WASHC2C/TMOD3/RHOA/CAPZA1/CDC42/PIK3R2/CCN2/FSCN1/PAK1 | 61 |
| GO:0009895 | negative regulation of catabolic process | 58/2001 | 390/20772 | 0,000569129 | 0,02284104 | 0,020193092 | TIMP3/TENT5A/EGFR/SNCA/FEZ1/CHMP4A/TIMP2/BCL2/MYCBP2/HIPK2/TRAF3IP2/TRAF5/NRDE2/CRTC3/NSUN2/GIPC1/BAG5/AXIN2/TIRAP/THRAP3/EIF4G1/AKT1/IKBKE/NRBP2/PHF20L1/USP36/CCAR2/SECISBP2/USP7/CTSA/RPTOR/DAP/HSP90AB1/ZFP36/RGP1/MTMR2/RRAGA/TENT4A/CHMP4B/PSME3IP1/USP9X/TAB3/TARDBP/GIT1/SCFD1/METTL16/SNX3/GSK3A/UCHL5/USP14/EIF4G2/SYNCRIP/LARP4B/ELAVL1/POLDIP2/IGF2BP3/RNF41/PIN1 | 58 |

| | | | | | | | | |
|---|---|---|---|---|---|---|---|---|
| GO:0031098 | stress-activated protein kinase signaling cascade | 44/2001 | 276/20772 | 0,000598046 | 0,023555581 | 0,020824798 | CDC42EP5/IGFBP6/SPHK1/TNFRSF19/TPD52L1/SH3RF3/FZD7/FOXO1/RASSF2/SIRPA/EGFR/TNIK/PLCB1/SEMA3A/TAOK3/RELL1/HIPK2/TRAF5/STK39/MID1/HRAS/PJA2/MAP3K20/TIRAP/HMGB1/ANKRD6/FAS/MAPK9/ZFP36/EIF2AK2/ZNF622/RIPK1/HIPK3/TRAF4/KLHDC10/CRK/IGF1R/MFHAS1/ZMYND11/DNAJA1/RHOA/CCN2/MAPKBP1/COPS5 | 44 |
| GO:0051403 | stress-activated MAPK cascade | 43/2001 | 268/20772 | 0,000598672 | 0,023555581 | 0,020824798 | CDC42EP5/IGFBP6/SPHK1/TNFRSF19/TPD52L1/SH3RF3/FZD7/FOXO1/RASSF2/SIRPA/EGFR/TNIK/PLCB1/SEMA3A/TAOK3/RELL1/HIPK2/TRAF5/STK39/MID1/HRAS/PJA2/MAP3K20/TIRAP/HMGB1/ANKRD6/FAS/MAPK9/ZFP36/EIF2AK2/ZNF622/RIPK1/HIPK3/TRAF4/KLHDC10/CRK/IGF1R/MFHAS1/ZMYND11/DNAJA1/CCN2/MAPKBP1/COPS5 | 43 |
| GO:0048026 | positive regulation of mRNA splicing, via spliceosome | 10/2001 | 32/20772 | 0,000595538 | 0,023555581 | 0,020824798 | SLC39A5/THRAP3/RBM3/RBMXL1/SNW1/PRPF19/SF3B4/HSPA8/NCBP1/TRA2A | 10 |
| GO:0003188 | heart valve formation | 7/2001 | 17/20772 | 0,000620328 | 0,024249184 | 0,021437991 | TWIST1/OLFM1/HEY2/SOX9/DCHS1/CDH11/RHOA | 7 |
| GO:0021885 | forebrain cell migration | 17/2001 | 75/20772 | 0,000645107 | 0,025055133 | 0,022150508 | FOXB1/POU3F2/POU3F3/EGFR/LAMB1/SRGAP2/SUN2/SRGAP2C/DISC1/FBXO45/ROBO1/SRF/LRP8/DAB1/RHOA/OGDH/SUN1 | 17 |
| GO:0031589 | cell-substrate adhesion | 59/2001 | 401/20772 | 0,000673192 | 0,025682446 | 0,022705096 | APOD/HAS2/EPHA3/DAB2/FZD7/CDK6/ITGA11/PHLDB2/GPM6B/FREM1/LAMB1/ARHGAP6/COL26A1/CCDC80/SRGAP2/PPFIA2/NID2/DLC1/ONECUT1/NEDD9/BCL2/FERMT2/DISC1/LAMC1/PPM1F/RASA1/CORO1C/DOCK5/NF2/JAM3/DOCK1/CALR/SORBS1/SRF/TLN1/SGCE/AKIP1/PPARD/RAC1/CDH11/VCL/PIK3CB/FER/RAB1A/RCC2/RCC2/PKD1/ARPC2/PTPRA/P4HB/CRK/ACTN1/WASHC2C/ILK/RHOA/CDC42/CCN2/POLDIP2/PTEN | 59 |

| | | | | | | | | |
|---|---|---|---|---|---|---|---|---|
| GO:0016571 | histone methylation | 28/2001 | 153/20772 | 0,000668906 | 0,025682446 | 0,022705096 | PRMT8/PAX5/BCOR/H1-2/NELFE/NELFE/NELFE/NELFE/NELFE/HCFC1/KMT2C/CXXC1/NFYA/PAXIP1/ASH2L/PRDM2/NCOA6/WDR82/SNW1/PRMT6/ASH1L/SKIC8/NELFA/KMT5B/CHTOP/RRP8/WDR5B/SETMAR | 28 |
| GO:0034314 | Arp2/3 complex-mediated actin nucleation | 14/2001 | 56/20772 | 0,000674058 | 0,025682446 | 0,022705096 | CTNNA2/TRIM27/TRIM27/FCHSD2/ACTR2/ACTR3/ARPC5/ARF1/IQGAP2/ARPC2/ARPC5L/WASH6P/ARPC4/WASHC2C | 14 |
| GO:0006984 | ER-nucleus signaling pathway | 13/2001 | 50/20772 | 0,000695364 | 0,025841326 | 0,022845558 | PPP1R15A/SPRING1/EIF2S1/ERLIN1/PPP1R15B/MBTPS1/HSPA5/EIF2A/SELENOS/ATP2A2/CCDC47/SREBF2/PTPN2 | 13 |
| GO:0016578 | histone deubiquitination | 13/2001 | 50/20772 | 0,000695364 | 0,025841326 | 0,022845558 | EPOP/TAF4/USP36/USP7/USP15/ATXN7L3/TRRAP/TAF5L/TAF2/SUPT20H/USP16/USP22/TAF12 | 13 |
| GO:0000462 | maturation of SSU-rRNA from tricistronic rRNA transcript (SSU-rRNA, 5.8S rRNA, LSU-rRNA) | 12/2001 | 44/20772 | 0,000695398 | 0,025841326 | 0,022845558 | WDR46/DHX37/UTP23/WDR46/WDR46/NOP14/TBL3/NOL10/BYSL/RCL1/DCAF13/RRP36 | 12 |
| GO:0050685 | positive regulation of mRNA processing | 12/2001 | 44/20772 | 0,000695398 | 0,025841326 | 0,022845558 | SLC39A5/THRAP3/RBM3/NCBP2/RBMXL1/SNW1/PRPF19/NUDT21/SF3B4/HSPA8/NCBP1/TRA2A | 12 |
| GO:0035966 | response to topologically incorrect protein | 34/2001 | 199/20772 | 0,000699888 | 0,025848619 | 0,022852005 | KLHL15/HSPA1B/OS9/PPP1R15A/ERMP1/RNF126/EIF2S1/HSPA9/UFD1/DNAJB12/HSPA4/PPP1R15B/MBTPS1/HSPA5/HSP90AB1/UGGT1/EIF2AK2/DNAJC3/AKIRIN2/STUB1/FAF2/SELENOS/HSPD1/SDF2L1/SERPINH1/HSPH1/HSPA8/MFN2/DNAJA1/VCP/CREB3L2/RNF185/PTPN2/COPS5 | 34 |
| GO:2000779 | regulation of double-strand break repair | 26/2001 | 139/20772 | 0,000729327 | 0,026771636 | 0,023668017 | TWIST1/KLHL15/ZCWPW1/SPIRE1/TFIP11/ACTR2/KAT5/TRIP12/FIGNL1/HMGA2/RUVBL2/MORF4L2/TRRAP/EP400/MORF4L1/EPC1/RUVBL1/MRGBP/SMARCE1/KMT5B/OTUB2/SLF2/TIMELESS/PBRM1/SETMAR/DEK | 26 |

| | | | | | | | | |
|---|---|---|---|---|---|---|---|---|
| GO:0098693 | regulation of synaptic vesicle cycle | 6/2001 | 13/20772 | 0,000748989 | 0,027162134 | 0,024013245 | PLD1/LPAR1/BSN/PPP3R1/DNAJC5/SCRN1 | 6 |
| GO:0098789 | pre-mRNA cleavage required for polyadenylation | 6/2001 | 13/20772 | 0,000748989 | 0,027162134 | 0,024013245 | CSTF2T/CPSF2/NCBP2/NUDT21/NCBP1/CPSF7 | 6 |
| GO:0048255 | mRNA stabilization | 15/2001 | 63/20772 | 0,000764802 | 0,027569496 | 0,024373382 | TENT5A/TRAF3IP2/TRAF5/AXIN2/TIRAP/THRAP3/IKBKE/ZFP36/TENT4A/TARDBP/METTL16/SYNCRIP/LARP4B/ELAVL1/IGF2BP3 | 15 |
| GO:0009101 | glycoprotein biosynthetic process | 52/2001 | 345/20772 | 0,00078321 | 0,027898952 | 0,024664645 | CYTL1/ITM2A/DSEL/DSE/B3GLCT/ST6GALNAC3/GALNT4/PLCB1/BMPR1B/CHST12/ITM2C/XYLT1/GAL3ST4/PHLDA1/BCL2/NAGPA/GALNT10/FUT4/ST3GAL4/NDST3/GALNT2/CHSY1/SLC2A10/CHST10/BMPR2/ST8SIA4/POGLUT3/CSGALNACT2/GLCE/TCF7L2/CANT1/B3GALNT1/MGAT1/UGGT1/XXYLT1/ST6GALNAC5/B4GALT3/B3GALT6/FUT11/FAM20B/DDOST/ATP7A/B3GNT5/DPY19L1/GORASP1/CHPF2/EXT2/DPM1/B4GALT5/DAD1/STT3A/CRPPA | 52 |
| GO:0032388 | positive regulation of intracellular transport | 35/2001 | 208/20772 | 0,000782684 | 0,027898952 | 0,024664645 | PTGS2/MLC1/PRNP/DAB2/ZDHHC2/FEZ1/MSN/CAPN10/PRKACA/NRDE2/NF2/HRAS/PRKACA/AKT2/PRKCD/USP36/MIEF1/MAP2/MTMR2/NCBP2/RAB21/UBE2L3/TARDBP/GSK3A/NMT1/SH3GLB1/LDLRAP1/PCM1/TRIM28/PPM1A/SREBF2/PIK3R2/RBM22/ATG13/XPO4 | 35 |
| GO:0072665 | protein localization to vacuole | 18/2001 | 83/20772 | 0,000793122 | 0,028085838 | 0,024829865 | NAGPA/MFSD1/MEAK7/VPS37C/STAM2/MON1A/MON1B/PXK/RAB7A/VPS4A/ZFYVE16/VPS41/SH3GLB1/UBAP1/HSPA8/GGA3/GPR137B/LAMTOR5 | 18 |
| GO:0061136 | regulation of proteasomal protein catabolic process | 37/2001 | 224/20772 | 0,00080557 | 0,028359834 | 0,025072097 | FOXF2/DAB2/SH3RF3/PABIR1/FZR1/PSMC3/HSPA1B/PRKACA/GIPC1/BAG5/AXIN2/PSMC5/PRKACA/AKT1/RNF217/UBE2K/PHF20L1/PSMC1/CCAR2/USP7/MAPK9/HSP90AB1/STUB1/PSME3IP1/USP9X/PKD1/USP13/CSNK1D/RNFT1/GSK3A/UCHL5/USP14/ANKIB1/VCP/RNF185/GNA12/PSMC6 | 37 |

| | | | | | | | | |
|---|---|---|---|---|---|---|---|---|
| GO:0007409 | axonogenesis | 65/2001 | 456/20772 | 0,000867721 | 0,030370244 | 0,026849441 | DSCAML1/NGFR/FOXB1/S100B/KIF5A/WNT3/EPHA3/WNT3/POU3F2/EDN3/PRKG1/OLFM1/SLITRK3/CTNNA2/PTCH1/EFNB2/EDNRA/BMPR1B/CNTN1/FEZ1/CDKL5/EFNB3/SEMA3A/SEMA3D/LRRC4C/BCL2/MYCBP2/DISC1/NCAM1/GLI2/SHOX2/FBXO45/ROBO1/PLXNC1/SEMA5A/TUBB2B/ULK2/MAP1S/SRF/LAMB2/PLXNB2/BMPR2/BAIAP2/MARK2/DAB1/RAC1/CDH11/VCL/RAPH1/HSP90AB1/MAP2/CHN1/TTL/USP9X/RAB21/PTPN11/YTHDF1/IGF1R/B4GALT5/SIN3A/RPS6KA5/PICALM/PTEN/CRPPA/PAK1 | 65 |
| GO:0016236 | macroautophagy | 53/2001 | 355/20772 | 0,000876758 | 0,030509168 | 0,02697226 | DCN/STING1/FEZ1/CHMP4A/RALB/PRKACA/PIP4K2B/PIP4K2B/STX12/ULK2/CALR/PRKACA/ATG3/VPS16/AKT1/SLC25A5/VPS37C/STAM2/NRBP2/USP36/ATP6V1E1/NSFL1C/PIK3C2B/PLEKHM1/EXOC7/ATG4D/UBXN2A/RAB1A/CALM1/CHMP4B/ATP2A2/RAB7A/VPS4A/SCFD1/VTI1B/VPS41/CHMP6/SH3GLB1/VDAC1/CDC37/MFN2/VCP/BNIP3/VMP1/UFM1/SNX4/GNAI3/TEX264/POLDIP2/RNF41/CAPNS1/ATG13/ATP6V1B2 | 53 |
| GO:0042059 | negative regulation of epidermal growth factor receptor signaling pathway | 9/2001 | 28/20772 | 0,000887046 | 0,030514375 | 0,026976864 | EGFR/CBLB/RNF126/RAB7A/CHMP6/NUP62/ERRFI1/CBL/PTPN2 | 9 |
| GO:0006469 | negative regulation of protein kinase activity | 39/2001 | 241/20772 | 0,000887022 | 0,030514375 | 0,026976864 | NR2F2/UBASH3B/SH3BP5/PLEC/SNCA/TRIM27/TRIM27/CBLB/SPRED1/SPRY2/PPM1F/CORO1C/NF2/IBTK/IBTK/DBNDD1/PRKAR1A/DUSP7/AKT1/PRKCD/GNAQ/SH3BP5L/CDKN1B/PIK3CB/RB1/LATS1/ADAR/HEG1/HIPK3/IPO7/CHMP6/YWHAG/NUP62/DNAJA1/ERRFI1/CBL/MACROH2A1/PTEN/PTPN2 | 39 |

| | | | | | | | | |
|---|---|---|---|---|---|---|---|---|
| GO:0001837 | epithelial to mesenchymal transition | 33/2001 | 194/20772 | 0,000898962 | 0,030748591 | 0,027183927 | HAS2/TWIST1/MSX2/EPHA3/RFLNB/TGFB1I1/FOXF2/DAB2/OLFM1/RGCC/BMP5/SNAI2/PHLDB2/EDNRA/HEY2/FOXC1/DACT3/SOX9/SPRED1/FERMT2/SPRY2/AXIN2/PPP3R1/TCF7L2/TGFBR1/HMGA2/HDAC2/POLR1B/SPRED2/NOLC1/TRIM28/USF3/PTEN | 33 |
| GO:0016050 | vesicle organization | 57/2001 | 389/20772 | 0,000907944 | 0,03076444 | 0,027197938 | CPLX1/SPHK1/AP3B2/SNCA/CHMP4A/BCL2/SHROOM2/AP1B1/PLEKHF2/GBF1/CORO1C/SYNJ1/PIP4K2B/PIP4K2B/STX12/VPS16/AKT2/VPS37C/STAM2/ATP6V1E1/YIPF5/PLEKHM1/AP1G1/ARFGEF2/PREB/SEC31A/RAB39A/YIPF4/RAB1A/AP3S1/DNAJC13/GOSR2/CHMP4B/SEC24A/RAB14/CSNK1D/RAB7A/VPS4A/ZFYVE16/ARFGAP3/VTI1B/VPS41/SNX3/CHMP6/UBAP1/TRAPPC12/TRAPPC11/ATP6AP1/STX10/PLEKHJ1/GNAI3/MIA3/PICALM/GOSR1/SEC24D/ATP6V1B2/SEC16A | 57 |
| GO:0018022 | peptidyl-lysine methylation | 26/2001 | 141/20772 | 0,000909646 | 0,03076444 | 0,027197938 | PAX5/BCOR/KMT5A/H1-2/NELFE/NELFE/NELFE/NELFE/NELFE/HCFC1/KMT2C/CXXC1/NFYA/PAXIP1/ASH2L/NCOA6/WDR82/SNW1/ASH1L/SKIC8/NELFA/KMT5B/EEF1AKMT1/EEF2KMT/WDR5B/SETMAR | 26 |
| GO:0035089 | establishment of apical/basal cell polarity | 8/2001 | 23/20772 | 0,000953417 | 0,030923125 | 0,027338227 | FOXJ1/LAMA1/MSN/MYO9A/FAT1/RHOA/CDC42/FSCN1 | 8 |
| GO:0035988 | chondrocyte proliferation | 8/2001 | 23/20772 | 0,000953417 | 0,030923125 | 0,027338227 | FGFR3/DDR2/BMPR1B/SOX9/MMP16/BMPR2/HMGA2/CCN2 | 8 |
| GO:1902683 | regulation of receptor localization to synapse | 8/2001 | 23/20772 | 0,000953417 | 0,030923125 | 0,027338227 | CPLX1/GPC4/HRAS/RAP1A/DBN1/ARHGAP44/IQSEC2/ZDHHC3 | 8 |
| GO:0060644 | mammary gland epithelial cell differentiation | 7/2001 | 18/20772 | 0,000931089 | 0,030923125 | 0,027338227 | FOXB1/ID2/ERBB4/PTCH1/AKT1/AKT2/LATS1 | 7 |

| | | | | | | | | |
|---|---|---|---|---|---|---|---|---|
| GO:0033673 | negative regulation of kinase activity | 42/2001 | 266/20772 | 0,000955432 | 0,030923125 | 0,027338227 | DDR2/NR2F2/UBASH3B/SH3BP5/PLEC/SNCA/TRIM27/TRIM27/CBLB/SPRED1/SPRY2/PPM1F/CORO1C/NF2/PIP4K2B/PIP4K2B/IBTK/IBTK/DBNDD1/PRKAR1A/DUSP7/AKT1/PRKCD/GNAQ/SH3BP5L/CDKN1B/PIK3CB/RB1/LATS1/ADAR/HEG1/HIPK3/IPO7/CHMP6/YWHAG/NUP62/DNAJA1/ERRFI1/CBL/MACROH2A1/PTEN/PTPN2 | 42 |
| GO:0048024 | regulation of mRNA splicing, via spliceosome | 22/2001 | 112/20772 | 0,000925197 | 0,030923125 | 0,027338227 | SLC39A5/KHDRBS3/CELF2/QKI/SFSWAP/NOVA1/THRAP3/RBM3/RBM39/SRPK2/METTL16/RBMXL1/SNW1/PRPF19/SF3B4/FXR2/HSPA8/RBM8A/CELF1/NCBP1/TRA2A/HNRNPK | 22 |
| GO:1901992 | positive regulation of mitotic cell cycle phase transition | 21/2001 | 105/20772 | 0,000939017 | 0,030923125 | 0,027338227 | DDR2/RGCC/CCND2/EGFR/PLCB1/LSM11/PBX1/EIF4G1/AKT1/DDX3X/CDT1/RPTOR/RB1/RCC2/RCC2/PLRG1/STIL/TMOD3/APEX1/SIN3A/CDC25A | 21 |
| GO:0051303 | establishment of chromosome localization | 19/2001 | 91/20772 | 0,000938551 | 0,030923125 | 0,027338227 | KIFC1/CHMP4A/NDE1/MAD1L1/MAP1S/KAT5/CDT1/TTL/MLH1/CHMP4B/VPS4A/KPNB1/CHMP6/TRAPPC12/KIFC1/NUP62/DCTN2/MAPRE1/SUN1 | 19 |
| GO:0031346 | positive regulation of cell projection organization | 56/2001 | 382/20772 | 0,000990759 | 0,031874413 | 0,028179232 | CDC42EP5/WNT3/EPHA3/WNT3/DDR2/ALKAL2/ENPP2/PTN/TOX/BMP5/PTPRD/PTPRD/ROR2/CNTN1/FEZ1/CDKL5/SEPTIN9/DNM3/NEDD9/DISC1/EPS8/SHOX2/CORO1C/ROBO1/PLXNC1/HRAS/SEMA5A/TUBB2B/RAP1A/ACTR2/SRF/PLXNB2/DBN1/SEPTIN7/BMPR2/BAIAP2/ACTR3/LRP8/KIDINS220/MARK2/HSPA5/PLEKHM1/RAC1/TGFBR1/DDX56/RAB21/ARPC2/ATP7A/SNX3/RAPGEF1/IGF1R/TBC1D24/CDC42/CREB3L2/FSCN1/SF3A2 | 56 |
| GO:0007156 | homophilic cell adhesion via plasma membrane adhesion molecules | 30/2001 | 172/20772 | 0,000995414 | 0,031874413 | 0,028179232 | DSCAML1/PCDH18/PCDH11X/PCDH9/CADM2/IGSF11/PCDHGA9/PCDHB16/CADM4/HMCN1/PCDHGC3/DCHS1/ROBO1/FAT4/PLXNB2/PCDHB14/PCDH1/PCDHGB1/PCDHGB6/PCDHB15/CDH11/PIK3CB/PCDHB5/PCDHGC4/PCDHB13/PKD1/SDK1/FAT1/PCDHB11/PCDH19 | 30 |

| | | | | | | | | |
|---|---|---|---|---|---|---|---|---|
| GO:0043087 | regulation of GTPase activity | 59/2001 | 408/20772 | 0,001033888 | 0,032123962 | 0,028399852 | CCL2/FZD10/EPHA3/ARHGAP24/PRKG1/TBC1D10C/ARAP2/RAPGEF5/FOXJ1/SLC27A4/SIPA1L2/RASA3/RASA3/ARHGAP6/SRGAP2/CDKL5/CBLB/RASA4/DOCK9/FGD5/RGL1/NEDD9/FERMT2/LRCH1/SPRY2/RASA1/CORO1C/MYO9A/PLXNC1/HRAS/AGAP3/RAP1A/EVI5/RAP1GDS1/IQGAP1/EIF2S1/PLXNB2/AKT2/AGAP1/ARHGAP44/TBC1D9B/CHN1/RGP1/NET1/RASAL2/RCC2/RCC2/IQGAP2/CRK/RAPGEF1/MMUT/TBC1D10A/RALGAPB/POLDIP2/PICALM/USP6NL/ZC3H15/GPR137B/PIN1 | 59 |
| GO:0000209 | protein polyubiquitination | 43/2001 | 275/20772 | 0,001022864 | 0,032123962 | 0,028399852 | FOXF2/TRIM27/TRIM27/SPSB4/TRIM2/FZR1/BCL2/TRAF3IP2/TRAF5/UBE2E3/LNPEP/RNF126/PELI1/SYVN1/RBCK1/RNF217/UBE2K/UBE2Q2/DDX3X/ANAPC10/TRIP12/PPIL2/STUB1/KLHL42/CBFB/UBE2L3/TRAF4/SPOP/UBE3B/UBE3C/UBE4A/RNF216/PRPF19/MARCHF8/UBE2A/ANKIB1/AREL1/CBL/OTUB2/RNF41/RNF185/MARCHF6/UBE2R2 | 43 |
| GO:0016051 | carbohydrate biosynthetic process | 37/2001 | 227/20772 | 0,001035224 | 0,032123962 | 0,028399852 | RBP4/HAS2/NR3C1/SMPD3/DSEL/FOXO1/DSE/ENPP1/SNCA/CHST12/SIK1/GYG1/FUT4/ST3GAL4/NDST3/SORBS1/DYRK2/CHST10/AKT1/AKT2/GYS1/CSGALNACT2/B3GALNT1/USP7/PPP1R3F/CLTC/GBE1/ST6GALNAC5/SELENOS/EPM2AIP1/GSK3A/DDB1/GOT1/EXT2/B4GALT5/SLC25A11/PTPN2 | 37 |
| GO:0051028 | mRNA transport | 26/2001 | 142/20772 | 0,001013503 | 0,032123962 | 0,028399852 | QKI/NSUN2/CASC3/GLE1/NUP188/POLDIP3/KHSRP/EIF4A3/ZFP36/NUP50/ZC3H3/POM121/NCBP2/MAGOHB/DDX19A/NUP214/ALYREF/NUP62/RBM8A/NCBP1/POM121C/CETN3/CHTOP/IGF2BP3/NUP88/NUP43 | 26 |
| GO:0006023 | aminoglycan biosynthetic process | 17/2001 | 78/20772 | 0,001033359 | 0,032123962 | 0,028399852 | HAS2/SMPD3/DSEL/DSE/CHST12/XYLT1/ST3GAL4/NDST3/CHSY1/HEXA/CSGALNACT2/GLCE/CLTC/B3GALT6/CHPF2/EXT2/B4GALT5 | 17 |
| GO:0051310 | metaphase plate congression | 17/2001 | 78/20772 | 0,001033359 | 0,032123962 | 0,028399852 | KIFC1/CHMP4A/MAD1L1/MAP1S/KAT5/CDT1/TTL/MLH1/CHMP4B/VPS4A/KPNB1/CHMP6/TRAPPC12/KIFC1/NUP62/DCTN2/MAPRE1 | 17 |

| | | | | | | | | |
|---|---|---|---|---|---|---|---|---|
| GO:0034655 | nucleobase-containing compound catabolic process | 69/2001 | 494/20772 | 0,001064876 | 0,03224036 | 0,028502755 | TENT5A/ENPP1/PDE4B/ZFP36L2/ZHX2/ZSWIM8/HSPA1B/TRAF3IP2/TRAF5/NRDE2/NUDT3/NSUN2/CASC3/AXIN2/FTO/TIRAP/THRAP3/AIFM1/AKT1/IKBKE/PRKCD/CTIF/CNOT3/DCP1A/CNOT10/SUCLG1/SECISBP2/KHSRP/MGAT1/NT5E/EIF4A3/GSPT1/EXOSC6/SND1/ZFP36/MLH1/SERBP1/NUDT12/TENT4A/NCBP2/WDR82/TARDBP/MAGOHB/GRSF1/YTHDF1/METTL16/PPP1R8/GSK3A/ACAT1/MTREX/FXR2/GALT/PATL1/RBM8A/EXOSC4/CELF1/VCP/SYNCRIP/LARP4B/DIS3/DICER1/NCBP1/SAMD4B/GTPBP1/APEX1/ELAVL1/ZC3H14/IGF2BP3/SETMAR | 69 |
| GO:0048813 | dendrite morphogenesis | 27/2001 | 150/20772 | 0,001071108 | 0,03224036 | 0,028502755 | PTN/CTNNA2/PTPRD/PTPRD/TNIK/TANC2/PPFIA2/CDKL5/DNM3/PDLIM5/BTBD3/SEMA3A/CTNND2/DBN1/BAIAP2/LRP8/KIDINS220/ARHGAP44/MAP2/RAB21/GORASP1/TBC1D24/CDC42/PQBP1/PICALM/PTEN/YWHAH | 27 |
| GO:1901796 | regulation of signal transduction by p53 class mediator | 22/2001 | 113/20772 | 0,001046315 | 0,03224036 | 0,028502755 | TWIST1/SNAI2/PMAIP1/BOP1/BOP1/SPRED1/BCL2/KMT5A/YJU2/HIPK2/DYRK2/AKT1/USP7/USP15/TP53RK/PRMT6/SPRED2/EEF1E1/ARMC10/PAK1IP1/PAK1IP1/HNRNPK | 22 |
| GO:0005978 | glycogen biosynthetic process | 12/2001 | 46/20772 | 0,001070506 | 0,03224036 | 0,028502755 | ENPP1/GYG1/SORBS1/DYRK2/AKT1/AKT2/GYS1/PPP1R3F/GBE1/SELENOS/EPM2AIP1/GSK3A | 12 |
| GO:0009250 | glucan biosynthetic process | 12/2001 | 46/20772 | 0,001070506 | 0,03224036 | 0,028502755 | ENPP1/GYG1/SORBS1/DYRK2/AKT1/AKT2/GYS1/PPP1R3F/GBE1/SELENOS/EPM2AIP1/GSK3A | 12 |
| GO:0090148 | membrane fission | 12/2001 | 46/20772 | 0,001070506 | 0,03224036 | 0,028502755 | CHMP4A/CORO1C/EXOC3/VPS37C/STAM2/EXOC7/CHMP4B/VPS4A/CHMP6/SH3GLB1/UBAP1/EXOC2 | 12 |
| GO:0006470 | protein dephosphorylation | 45/2001 | 292/20772 | 0,001077003 | 0,0322565 | 0,028517025 | PPP1R16B/EYA1/UBASH3B/CSNK2B/PTPRD/TNS2/PTPRD/DLC1/PTPRG/BCL2/PPP1R15A/PPM1F/PPP1R9B/DUSP7/HSP90B1/PPP3R1/PRKCD/PTPN9/PPP1R15B/PPP6R2/PPP4R1/HSP90AB1/MTMR2/CALM1/PPM1B/YWHAB/PTPRA/PTPN11/SYMPK/PPP1R8/PTPA/MFHAS1/PPP1R7/ANKLE2/PPM1A/CTDNEP1/CTDNEP1/PPP2R5D/PIN1/CTDSP2/PTEN/CDC25A/PPP2R1A/GNA12/PTPN2 | 45 |

| | | | | | | | | |
|---|---|---|---|---|---|---|---|---|
| GO:0030970 | retrograde protein transport, ER to cytosol | 9/2001 | 29/20772 | 0,001177359 | 0,034406321 | 0,030417619 | OS9/HSP90B1/SYVN1/UFD1/SEL1L/FAF2/SELENOS/HM13/VCP | 9 |
| GO:0072207 | metanephric epithelium development | 9/2001 | 29/20772 | 0,001177359 | 0,034406321 | 0,030417619 | WNT9B/PAX8/POU3F3/CALB1/SOX9/YAP1/LAMB2/PKD1/ACAT1 | 9 |
| GO:1903513 | endoplasmic reticulum to cytosol transport | 9/2001 | 29/20772 | 0,001177359 | 0,034406321 | 0,030417619 | OS9/HSP90B1/SYVN1/UFD1/SEL1L/FAF2/SELENOS/HM13/VCP | 9 |
| GO:0043414 | macromolecule methylation | 52/2001 | 351/20772 | 0,001161488 | 0,034406321 | 0,030417619 | PRMT8/PAX5/BCOR/KMT5A/DNMT1/HENMT1/H1-2/NELFE/NELFE/NELFE/NELFE/NELFE/NSUN2/HCFC1/NSUN4/KMT2C/CXXC1/NFYA/PAXIP1/CBLL1/TRMT1L/ASH2L/TRMT61A/RAB6A/PRMT2/GATAD2A/GSPT1/TRMO/PRDM2/PCMT1/TRMT6/NCOA6/WDR82/BUD23/METTL16/SNW1/PRMT6/NSUN3/KDM1B/ASH1L/SKIC8/TRIM28/NELFA/KMT5B/EEF1AKMT1/EEF2KMT/SNRPD1/MRM2/CHTOP/RRP8/WDR5B/SETMAR | 52 |
| GO:0021795 | cerebral cortex cell migration | 14/2001 | 59/20772 | 0,001173856 | 0,034406321 | 0,030417619 | POU3F2/POU3F3/EGFR/LAMB1/SRGAP2/SUN2/SRGAP2C/DISC1/FBXO45/ROBO1/LRP8/DAB1/RHOA/SUN1 | 14 |
| GO:0090305 | nucleic acid phosphodiester bond hydrolysis | 43/2001 | 277/20772 | 0,001184901 | 0,034459453 | 0,030464591 | POLR1H/POLR1H/POLR1H/POLR1H/POLR1H/ENPP2/PGBD5/ENPP1/BOP1/BOP1/UTP23/REXO5/ASTE1/EME2/NOP14/REXO1/SDE2/TSN/DCP1A/TBL3/CSTF2T/ISG20L2/TSNAX/CPSF2/SND1/ZFP36/RCL1/NUDT12/NCBP2/ENDOD1/PPP1R8/NUDT21/EXOSC4/DIS3/DICER1/NCBP1/ELAC1/CPSF7/APEX1/DROSHA/SETMAR/RRP36/DDX1 | 43 |
| GO:0098787 | mRNA cleavage involved in mRNA processing | 6/2001 | 14/20772 | 0,00120425 | 0,034853778 | 0,030813202 | CSTF2T/CPSF2/NCBP2/NUDT21/NCBP1/CPSF7 | 6 |

| | | | | | | | | |
|---|---|---|---|---|---|---|---|---|
| GO:0032386 | regulation of intracellular transport | 52/2001 | 352/20772 | 0,001238216 | 0,035327293 | 0,031231823 | APOD/PTGS2/MLC1/PRNP/DAB2/ZDHHC2/ATP9A/FEZ1/FRAT2/MSN/CAPN10/OS9/PRKACA/NRDE2/NSUN2/NF2/HRAS/PRKACA/TTC21B/AKT2/PRKCD/USP36/ARHGAP44/YIPF5/USP7/MIEF1/MAP2/MTMR2/DNAJC13/NCBP2/ARV1/RAB21/UBE2L3/TARDBP/PTPN11/SCFD1/SNX3/GSK3A/NMT1/SH3GLB1/LDLRAP1/NUP214/PCM1/NOLC1/TRIM28/PPM1A/UFM1/SREBF2/PIK3R2/RBM22/ATG13/XPO4 | 52 |
| GO:0045185 | maintenance of protein location | 19/2001 | 93/20772 | 0,001234556 | 0,035327293 | 0,031231823 | CER1/CD4/IGSF11/MDFI/SUN2/HK1/OS9/CIZ1/AKT1/DBN1/HSPA5/KDELR2/LATS1/TMSB4X/YWHAB/PKD1/ARL2BP/RER1/SUN1 | 19 |
| GO:0070972 | protein localization to endoplasmic reticulum | 18/2001 | 86/20772 | 0,001226575 | 0,035327293 | 0,031231823 | CHMP4A/OS9/PPP1R15A/GBF1/SPCS3/SEC61A2/HSPA5/KDELR2/SRP68/SEC61A1/CHMP4B/SSR3/SEC61G/RER1/MIA3/BTF3/SRP54/SEC16A | 18 |
| GO:0051235 | maintenance of location | 54/2001 | 369/20772 | 0,001255653 | 0,035488403 | 0,031374255 | CER1/APLNR/CACNA1C/CACNA1C/CD4/IGSF11/UBASH3B/ITPR2/ENPP1/RASA3/RASA3/MDFI/SNCA/PLCB1/ARHGAP21/SUN2/HK1/GSTO1/OS9/PRKACA/ATP7B/SPOUT1/IBTK/IBTK/PLCB2/CALR/PRKACA/FTO/HSP90B1/CIZ1/AKT1/DBN1/IKBKE/CHERP/HEXA/SLC30A4/GPSM2/HSPA5/KDELR2/LATS1/TMSB4X/CALM1/YWHAB/PKD1/SQLE/ABHD5/DIAPH1/COX17/ARL2BP/RER1/SREBF2/FTH1/PTPN2/SUN1 | 54 |
| GO:0045739 | positive regulation of DNA repair | 26/2001 | 144/20772 | 0,00125239 | 0,035488403 | 0,031374255 | EYA1/EGFR/ZCWPW1/SPIRE1/ACTR2/KAT5/HMGB1/RUVBL2/MORF4L2/TRRAP/UCHL5/EP400/MORF4L1/EPC1/RUVBL1/MRGBP/SMARCE1/TRIM28/CEBPG/ACTR5/USP1/KMT5B/SLF2/TIMELESS/PBRM1/SETMAR | 26 |
| GO:0050650 | chondroitin sulfate proteoglycan biosynthetic process | 8/2001 | 24/20772 | 0,001310402 | 0,035709262 | 0,03156951 | CYTL1/DSE/CHST12/XYLT1/CHSY1/CSGALNACT2/B3GALT6/CHPF2 | 8 |

| | | | | | | | | |
|---|---|---|---|---|---|---|---|---|
| GO:0072243 | metanephric nephron epithelium development | 8/2001 | 24/20772 | 0,001310402 | 0,035709262 | 0,03156951 | PAX8/POU3F3/CALB1/SOX9/YAP1/LAMB2/PKD1/ACAT1 | 8 |
| GO:0000478 | endonucleolytic cleavage involved in rRNA processing | 7/2001 | 19/20772 | 0,001352444 | 0,035709262 | 0,03156951 | BOP1/BOP1/UTP23/NOP14/SDE2/TBL3/RCL1 | 7 |
| GO:0000479 | endonucleolytic cleavage of tricistronic rRNA transcript (SSU-rRNA, 5.8S rRNA, LSU-rRNA) | 7/2001 | 19/20772 | 0,001352444 | 0,035709262 | 0,03156951 | BOP1/BOP1/UTP23/NOP14/SDE2/TBL3/RCL1 | 7 |
| GO:0030206 | chondroitin sulfate biosynthetic process | 7/2001 | 19/20772 | 0,001352444 | 0,035709262 | 0,03156951 | DSE/CHST12/XYLT1/CHSY1/CSGALNACT2/B3GALT6/CHPF2 | 7 |
| GO:0007163 | establishment or maintenance of cell polarity | 38/2001 | 238/20772 | 0,001319793 | 0,035709262 | 0,03156951 | FOXJ1/KIF26B/PHLDB2/FEZ1/LAMA1/MSN/NDE1/SPRY2/TRAF3IP2/LIN7A/GBF1/AMOTL1/JAM3/MYO9A/ACTR2/WDR1/KAT5/CAP1/ACTR3/MARK2/GPSM2/NSFL1C/RAC1/HSP90AB1/SNX27/PKD1/CRK/MAP4/IGF1R/FAT1/ILK/RHOA/SDCCAG8/CDC42/ATN1/FLOT2/FSCN1/MAPRE1 | 38 |
| GO:0032869 | cellular response to insulin stimulus | 36/2001 | 222/20772 | 0,001317403 | 0,035709262 | 0,03156951 | FOXO1/TNS2/ENPP1/SLC27A4/CAPN10/BCAR1/PIP4K2B/PIP4K2B/HRAS/SORBS1/AKT1/AKT2/PRKCD/BAIAP2/ZNF592/NCOA5/FER/RB1/SYAP1/AP3S1/IDE/SELENOS/PKM/PTPRA/PTPN11/GSK3A/IGF1R/ZNF106/YWHAG/RPS6KB1/GOT1/ERRFI1/LPIN2/SH2B2/PIK3R2/PTPN2 | 36 |
| GO:0006368 | transcription elongation by RNA polymerase II | 23/2001 | 122/20772 | 0,001285495 | 0,035709262 | 0,03156951 | NELFE/NELFE/NELFE/NELFE/NELFE/ZMYND8/ELL/MED22/ELOA/MED1/MED27/MED17/NCBP2/NELFCD/CCNK/SUPT16H/ZMYND11/SKIC8/ERCC3/NCBP1/NELFA/TCEA1/INTS5 | 23 |

| | | | | | | | | |
|---|---|---|---|---|---|---|---|---|
| GO:0034968 | histone lysine methylation | 23/2001 | 122/20772 | 0,001285495 | 0,035709262 | 0,03156951 | PAX5/BCOR/H1-2/NELFE/NELFE/NELFE/NELFE/NELFE/HCFC1/KMT2C/CXXC1/NFYA/PAXIP1/ASH2L/NCOA6/WDR82/SNW1/ASH1L/SKIC8/NELFA/KMT5B/WDR5B/SETMAR | 23 |
| GO:1902373 | negative regulation of mRNA catabolic process | 16/2001 | 73/20772 | 0,001346955 | 0,035709262 | 0,03156951 | TENT5A/TRAF3IP2/TRAF5/AXIN2/TIRAP/THRAP3/IKBKE/SECISBP2/ZFP36/TENT4A/TARDBP/METTL16/SYNCRIP/LARP4B/ELAVL1/IGF2BP3 | 16 |
| GO:0032881 | regulation of polysaccharide metabolic process | 12/2001 | 47/20772 | 0,001313288 | 0,035709262 | 0,03156951 | HAS2/SMPD3/ENPP1/SORBS1/DYRK2/AKT1/AKT2/PPP1R3F/CLTC/SELENOS/EPM2AIP1/GSK3A | 12 |
| GO:0048701 | embryonic cranial skeleton morphogenesis | 12/2001 | 47/20772 | 0,001313288 | 0,035709262 | 0,03156951 | TWIST1/WNT9B/TBX15/PAX5/PRRX1/DLX2/TBX1/MMP16/SLC39A3/EIF4A3/TGFBR1/MED12 | 12 |
| GO:0046627 | negative regulation of insulin receptor signaling pathway | 11/2001 | 41/20772 | 0,001333791 | 0,035709262 | 0,03156951 | TNS2/ENPP1/SLC27A4/PIP4K2B/PIP4K2B/PRKCD/ZNF592/NCOA5/GSK3A/RPS6KB1/PTPN2 | 11 |
| GO:1905168 | positive regulation of double-strand break repair via homologous recombination | 11/2001 | 41/20772 | 0,001333791 | 0,035709262 | 0,03156951 | ACTR2/KAT5/RUVBL2/MORF4L2/TRRAP/EP400/MORF4L1/EPC1/RUVBL1/MRGBP/TIMELESS | 11 |
| GO:0006301 | postreplication repair | 10/2001 | 35/20772 | 0,0012965 | 0,035709262 | 0,03156951 | POLH/REV1/POLK/USP10/ZBTB1/WDR33/UBE2A/VCP/USP1/POLDIP2 | 10 |
| GO:2000643 | positive regulation of early endosome to late endosome transport | 5/2001 | 10/20772 | 0,001373845 | 0,036115919 | 0,031929024 | DAB2/MSN/NF2/MTMR2/RAB21 | 5 |
| GO:0090174 | organelle membrane fusion | 24/2001 | 130/20772 | 0,001380083 | 0,036122171 | 0,031934551 | CPLX1/SPHK1/SNCA/CHMP4A/BNIP1/PIP4K2B/PIP4K2B/STX12/VPS16/AKT2/YIPF5/PLEKHM1/RAB39A/YIPF4/GOSR2/CHMP4B/RAB7A/VPS4A/VTI1B/VPS41/CHMP6/STX10/GNAI3/GOSR1 | 24 |

| | | | | | | | | |
|---|---|---|---|---|---|---|---|---|
| GO:0032259 | methylation | 57/2001 | 396/20772 | 0,00138813 | 0,036124703 | 0,031936789 | PRMT8/PAX5/BCOR/KMT5A/DNMT1/HENMT1/H1-2/NELFE/NELFE/NELFE/NELFE/NELFE/PRDM11/NSUN2/HCFC1/NSUN4/SPOUT1/KMT2C/CXXC1/NFYA/PAXIP1/CBLL1/TRMT1L/ASH2L/TRMT61A/RAB6A/PRMT2/GATAD2A/GSPT1/TRMO/PRDM2/PCMT1/TRMT6/NCOA6/WDR82/BUD23/ASMTL/METTL16/SNW1/PRMT6/NSUN3/KDM1B/ASH1L/NSD1/SKIC8/TRIM28/NELFA/KMT5B/EEF1AKMT1/EEF2KMT/SNRPD1/MRM2/CHTOP/RRP8/WDR5B/PRDM10/SETMAR | 57 |
| GO:1903050 | regulation of proteolysis involved in protein catabolic process | 41/2001 | 263/20772 | 0,001399751 | 0,036124703 | 0,031936789 | FOXF2/DAB2/SH3RF3/PABIR1/FZR1/PSMC3/HSPA1B/DISC1/HIPK2/PRKACA/GIPC1/BAG5/AXIN2/PSMC5/PRKACA/AKT1/RNF217/UBE2K/PHF20L1/PSMC1/CCAR2/USP7/MAPK9/LATS1/HSP90AB1/STUB1/PSME3IP1/USP9X/PKD1/USP13/CSNK1D/RNFT1/GSK3A/UCHL5/USP14/ANKIB1/VCP/RNF185/PTEN/GNA12/PSMC6 | 41 |
| GO:0072384 | organelle transport along microtubule | 19/2001 | 94/20772 | 0,001410184 | 0,036124703 | 0,031936789 | KIF5A/AP3B2/SYBU/FEZ1/ARHGAP21/SUN2/NDE1/KIFAP3/MAP1S/RAB6A/KIF3B/MAP2/RAB1A/AP3S1/COPG2/CDC42/TMEM201/COPG1/SUN1 | 19 |
| GO:1902369 | negative regulation of RNA catabolic process | 18/2001 | 87/20772 | 0,001409835 | 0,036124703 | 0,031936789 | TENT5A/TRAF3IP2/TRAF5/NRDE2/NSUN2/AXIN2/TIRAP/THRAP3/IKBKE/SECISBP2/ZFP36/TENT4A/TARDBP/METTL16/SYNCRIP/LARP4B/ELAVL1/IGF2BP3 | 18 |
| GO:0060350 | endochondral bone morphogenesis | 14/2001 | 60/20772 | 0,001397541 | 0,036124703 | 0,031936789 | CER1/FGFR3/RARG/TMEM119/SMPD3/FOXC1/BMPR1B/SOX9/MMP16/SHOX2/AXIN2/BMPR2/SERPINH1/EXT2 | 14 |
| GO:0045010 | actin nucleation | 15/2001 | 67/20772 | 0,00150032 | 0,03827087 | 0,033834152 | CTNNA2/TRIM27/TRIM27/SPIRE1/FCHSD2/ACTR2/ACTR3/ARPC5/ARF1/IQGAP2/ARPC2/ARPC5L/WASH6P/ARPC4/WASHC2C | 15 |
| GO:0007254 | JNK cascade | 33/2001 | 200/20772 | 0,001524492 | 0,038723381 | 0,034234204 | CDC42EP5/TNFRSF19/TPD52L1/SH3RF3/FZD7/RASSF2/SIRPA/EGFR/TNIK/PLCB1/SEMA3A/TAOK3/HIPK2/TRAF5/HRAS/PJA2/MAP3K20/TIRAP/HMGB1/ANKRD6/MAPK9/ZNF622/RIPK1/HIPK3/TRAF4/CRK/IGF1R/MFHAS1/ZMYND11/DNAJA1/CCN2/MAPKBP1/COPS5 | 33 |

| | | | | | | | | |
|---|---|---|---|---|---|---|---|---|
| GO:0043489 | RNA stabilization | 16/2001 | 74/20772 | 0,001567561 | 0,039650084 | 0,035053475 | TENT5A/TRAF3IP2/TRAF5/NSUN2/AXIN2/TIRAP/THRAP3/IKBKE/ZFP36/TENT4A/TARDBP/METTL16/SYNCRIP/LARP4B/ELAVL1/IGF2BP3 | 16 |
| GO:0072073 | kidney epithelium development | 27/2001 | 154/20772 | 0,001602441 | 0,040362729 | 0,035683503 | CER1/WNT9B/PAX8/POU3F3/CALB1/IRX3/EYA1/TFAP2B/PTCH1/FOXJ1/EFNB2/KANK2/KIF26B/EDNRA/FOXC1/SOX9/BCL2/PBX1/DCHS1/BASP1/YAP1/LAMB2/IQGAP1/CAT/PKD1/ACAT1/ILK | 27 |
| GO:0072009 | nephron epithelium development | 23/2001 | 124/20772 | 0,00160995 | 0,040382903 | 0,035701339 | WNT9B/PAX8/POU3F3/CALB1/IRX3/EYA1/TFAP2B/PTCH1/FOXJ1/KIF26B/EDNRA/FOXC1/SOX9/BCL2/PBX1/DCHS1/BASP1/YAP1/LAMB2/IQGAP1/PKD1/ACAT1/ILK | 23 |
| GO:0001952 | regulation of cell-matrix adhesion | 25/2001 | 139/20772 | 0,001625561 | 0,040605288 | 0,035897943 | APOD/EPHA3/CDK6/PHLDB2/GPM6B/ARHGAP6/DLC1/ONECUT1/BCL2/FERMT2/DISC1/PPM1F/RASA1/CORO1C/NF2/TLN1/RAC1/VCL/PIK3CB/RCC2/RCC2/PTPRA/RHOA/POLDIP2/PTEN | 25 |
| GO:0071218 | cellular response to misfolded protein | 10/2001 | 36/20772 | 0,00164216 | 0,040850438 | 0,036114673 | KLHL15/RNF126/UFD1/DNAJB12/UGGT1/AKIRIN2/STUB1/SDF2L1/VCP/RNF185 | 10 |
| GO:0044380 | protein localization to cytoskeleton | 14/2001 | 61/20772 | 0,001655701 | 0,041017778 | 0,036262614 | MID2/ABHD17C/DISC1/NUDCD3/MID1/PPP1R9B/NSFL1C/CSNK1D/DIAPH1/STIL/PCM1/NUP62/DCTN2/MAPRE1 | 14 |
| GO:0019985 | translesion synthesis | 8/2001 | 25/20772 | 0,001765971 | 0,043392436 | 0,038361979 | POLH/REV1/POLK/USP10/ZBTB1/VCP/USP1/POLDIP2 | 8 |
| GO:0072170 | metanephric tubule development | 8/2001 | 25/20772 | 0,001765971 | 0,043392436 | 0,038361979 | WNT9B/PAX8/POU3F3/CALB1/SOX9/YAP1/PKD1/ACAT1 | 8 |
| GO:0045911 | positive regulation of DNA recombination | 16/2001 | 75/20772 | 0,001817719 | 0,04448238 | 0,039325567 | ZCWPW1/ACTR2/KAT5/PAXIP1/EXOSC6/MLH1/RUVBL2/MORF4L2/TRRAP/EP400/MORF4L1/EPC1/RUVBL1/MRGBP/KMT5B/TIMELESS | 16 |
| GO:1903651 | positive regulation of cytoplasmic transport | 6/2001 | 15/20772 | 0,001844372 | 0,044951905 | 0,03974066 | DAB2/MSN/NF2/MAP2/MTMR2/RAB21 | 6 |

| | | | | | | | | |
|---|---|---|---|---|---|---|---|---|
| GO:0010563 | negative regulation of phosphorus metabolic process | 68/2001 | 496/20772 | 0,001863782 | 0,045060103 | 0,039836314 | PPP1R16B/DDR2/NR2F2/PRNP/UBASH3B/RASSF2/SH3BP5/ENPP1/SIRPA/PLEC/SNCA/TRIM27/TRIM27/CADM4/CBLB/SPRED1/SPRY2/PPP1R15A/PPM1F/CORO1C/NF2/PPP1R9B/PIP4K2B/PIP4K2B/LPCAT1/LPCAT1/IBTK/IBTK/DBNDD1/PRKAR1A/DUSP7/IQGAP1/EIF4G1/AKT1/PRKCD/PPP1R15B/GNAQ/SH3BP5L/CDKN1B/PIK3CB/RB1/LATS1/ADAR/DNAJC3/HEG1/CALM1/YWHAB/HIPK3/TARDBP/IPO7/GIT1/PPP1R8/GSK3A/CHMP6/PTPA/MFHAS1/ANKLE2/YWHAG/NUP62/DNAJA1/ERRFI1/SPRED2/CBL/MACROH2A1/CTDSP2/PTEN/PPP2R1A/PTPN2 | 68 |
| GO:0098813 | nuclear chromosome segregation | 55/2001 | 384/20772 | 0,001862008 | 0,045060103 | 0,039836314 | KIFC1/CHMP4A/KMT5A/P3H4/HSPA1B/ZCWPW1/EME2/MAD1L1/IK/C9orf78/MAP3K20/MSH5/MAP1S/ACTR2/KAT5/KIF4A/ACTR3/MLH3/CDT1/PRC1/RB1/LATS1/KIF3B/TTL/MLH1/RCC2/RCC2/LSM14A/PSMC3IP/TENT4A/CHMP4B/VPS4A/KPNB1/SMC2/WAPL/DSN1/CHMP6/ANAPC15/PDS5B/TRAPPC12/KIFC1/SMARCE1/NUP62/RHOA/KIF2A/BCCIP/CDC42/MACROH2A1/DCTN2/POLDIP2/MAPRE1/SLF2/PBRM1/PPP2R1A/SUN1 | 55 |
| GO:0001502 | cartilage condensation | 7/2001 | 20/20772 | 0,0019091 | 0,045069743 | 0,039844837 | OTOR/MGP/BMPR1B/SOX9/SOX6/PKD1/CCN2 | 7 |
| GO:2000641 | regulation of early endosome to late endosome transport | 7/2001 | 20/20772 | 0,0019091 | 0,045069743 | 0,039844837 | DAB2/MSN/NF2/MTMR2/DNAJC13/RAB21/SNX3 | 7 |
| GO:0110053 | regulation of actin filament organization | 44/2001 | 292/20772 | 0,001908062 | 0,045069743 | 0,039844837 | CDC42EP5/LPAR1/KANK4/RGCC/CTNNA2/SYNPO/PHLDB2/TRIM27/ARHGAP6/TRIM27/DLC1/LIMA1/FERMT2/EPS8/PPM1F/RASA1/NF2/FCHSD2/SEMA5A/SPTAN1/WDR1/ADD1/PRKCD/BAIAP2/ARPC5/MTPN/RAC1/FER/LATS1/TMSB4X/TGFBR1/ARF1/CFL2/ARPC2/ARPC5L/WASH6P/WASHC2C/TMOD3/RHOA/CAPZA1/CDC42/PIK3R2/CCN2/PAK1 | 44 |

| | | | | | | | | |
|---|---|---|---|---|---|---|---|---|
| GO:0050657 | nucleic acid transport | 30/2001 | 179/20772 | 0,001900908 | 0,045069743 | 0,039844837 | QKI/NRDE2/NSUN2/RFTN2/CASC3/SNUPN/GLE1/NUP188/POLDIP3/KHSRP/EIF4A3/ZFP36/NUP50/ZC3H3/POM121/NCBP2/MAGOHB/KPNB1/DDX19A/NUP214/ALYREF/NUP62/RBM8A/NCBP1/POM121C/CETN3/CHTOP/IGF2BP3/NUP88/NUP43 | 30 |
| GO:0050658 | RNA transport | 30/2001 | 179/20772 | 0,001900908 | 0,045069743 | 0,039844837 | QKI/NRDE2/NSUN2/RFTN2/CASC3/SNUPN/GLE1/NUP188/POLDIP3/KHSRP/EIF4A3/ZFP36/NUP50/ZC3H3/POM121/NCBP2/MAGOHB/KPNB1/DDX19A/NUP214/ALYREF/NUP62/RBM8A/NCBP1/POM121C/CETN3/CHTOP/IGF2BP3/NUP88/NUP43 | 30 |
| GO:0006354 | DNA-templated transcription elongation | 26/2001 | 148/20772 | 0,001878865 | 0,045069743 | 0,039844837 | NELFE/NELFE/NELFE/NELFE/NELFE/ZMYND8/ELL/MED22/ELOA/CCAR2/MED1/MED27/MED17/NCBP2/WDR82/NELFCD/CCNK/SUPT16H/ZMYND11/SKIC8/ALYREF/ERCC3/NCBP1/NELFA/TCEA1/INTS5 | 26 |
| GO:0015931 | nucleobase-containing compound transport | 38/2001 | 243/20772 | 0,001934539 | 0,045491887 | 0,040218042 | SLC35D1/QKI/SLC25A53/NRDE2/NSUN2/RFTN2/CASC3/SLC35B1/SLC25A5/SNUPN/GLE1/NUP188/POLDIP3/KHSRP/EIF4A3/SLC25A6/ZFP36/NUP50/SLC35E3/ZC3H3/POM121/RIPK1/NCBP2/MAGOHB/KPNB1/DDX19A/NUP214/SLC35B3/ALYREF/NUP62/RBM8A/NCBP1/POM121C/CETN3/CHTOP/IGF2BP3/NUP88/NUP43 | 38 |
| GO:0007051 | spindle organization | 37/2001 | 235/20772 | 0,001949044 | 0,04565465 | 0,040361936 | KIFC1/CHMP4A/GNAI1/SUN2/HSPA1B/HAUS7/MAP1S/KIF4A/TUBGCP2/GPSM2/PRC1/CLTC/DCTN6/KIF3B/SBDS/MLH1/LSM14A/TUBGCP3/CHMP4B/PKD1/CSNK1D/KPNB1/CHMP6/MAP4/TACC2/PTPA/KIFC1/STIL/NUP62/RHOA/KIF2A/VCP/BCCIP/DCTN2/POLDIP2/MAPRE1/PPP2R1A | 37 |
| GO:0001945 | lymph vessel development | 9/2001 | 31/20772 | 0,001986501 | 0,045995131 | 0,040662945 | NR2F2/SVEP1/EFNB2/FOXC1/CCBE1/TBX1/BMPR2/HEG1/PKD1 | 9 |
| GO:0005979 | regulation of glycogen biosynthetic process | 9/2001 | 31/20772 | 0,001986501 | 0,045995131 | 0,040662945 | ENPP1/SORBS1/DYRK2/AKT1/AKT2/PPP1R3F/SELENOS/EPM2AIP1/GSK3A | 9 |

| | | | | | | | | |
|---|---|---|---|---|---|---|---|---|
| GO:0010962 | regulation of glucan biosynthetic process | 9/2001 | 31/20772 | 0,001986501 | 0,045995131 | 0,040662945 | ENPP1/SORBS1/DYRK2/AKT1/AKT2/PPP1R3F/SELENOS/EPM2AIP1/GSK3A | 9 |
| GO:0090068 | positive regulation of cell cycle process | 44/2001 | 293/20772 | 0,002040883 | 0,046799071 | 0,041373685 | MSX2/SPHK1/EDN3/DDR2/SMPD3/RGCC/CCND2/EGFR/PLCB1/LSM11/PBX1/MAD1L1/GIPC1/MAP3K20/EIF4G1/KAT5/AKT1/DDX3X/GPSM2/NSFL1C/CDT1/RPTOR/MED1/RB1/KIF3B/RCC2/RCC2/SH2B1/SMC2/IGF1R/PLRG1/STIL/TMOD3/NUP62/RHOA/CDC42/MACROH2A1/APEX1/SIN3A/CCN2/POLDIP2/WIZ/SLF2/CDC25A | 44 |
| GO:0061572 | actin filament bundle organization | 30/2001 | 180/20772 | 0,00207564 | 0,046799071 | 0,041373685 | ESPN/LPAR1/RFLNB/RGCC/SYNPO/PHLDB2/ARHGAP6/DLC1/NEDD9/LIMA1/FERMT2/EPS8/PPM1F/SPIRE1/NF2/SORBS1/SRF/HSP90B1/ADD1/BAIAP2/RAC1/TGFBR1/MARCKS/ACTN1/RHOA/CDC42/PIK3R2/CCN2/FSCN1/PAK1 | 30 |
| GO:0006405 | RNA export from nucleus | 19/2001 | 97/20772 | 0,00206947 | 0,046799071 | 0,041373685 | NRDE2/NSUN2/CASC3/GLE1/NUP188/POLDIP3/EIF4A3/POM121/NCBP2/MAGOHB/DDX19A/NUP214/ALYREF/NUP62/RBM8A/NCBP1/POM121C/CHTOP/NUP88 | 19 |
| GO:1903312 | negative regulation of mRNA metabolic process | 19/2001 | 97/20772 | 0,00206947 | 0,046799071 | 0,041373685 | TENT5A/TRAF3IP2/TRAF5/SFSWAP/AXIN2/TIRAP/THRAP3/IKBKE/SECISBP2/ZFP36/TENT4A/TARDBP/METTL16/SYNCRIP/LARP4B/ELAVL1/ZC3H14/IGF2BP3/HNRNPK | 19 |
| GO:1900077 | negative regulation of cellular response to insulin stimulus | 11/2001 | 43/20772 | 0,002029494 | 0,046799071 | 0,041373685 | TNS2/ENPP1/SLC27A4/PIP4K2B/PIP4K2B/PRKCD/ZNF592/NCOA5/GSK3A/RPS6KB1/PTPN2 | 11 |
| GO:0006270 | DNA replication initiation | 10/2001 | 37/20772 | 0,002058788 | 0,046799071 | 0,041373685 | PURA/POLA1/MCM2/CIZ1/LRWD1/TOPBP1/CDT1/NOC3L/WRNIP1/NBN | 10 |
| GO:0006891 | intra-Golgi vesicle-mediated transport | 10/2001 | 37/20772 | 0,002058788 | 0,046799071 | 0,041373685 | RAB6A/GOSR2/COPA/VTI1B/COG4/COPB2/COPG2/GOLGA5/COPG1/GOSR1 | 10 |

| | | | | | | | | |
|---|---|---|---|---|---|---|---|---|
| GO:0045814 | negative regulation of gene expression, epigenetic | 22/2001 | 119/20772 | 0,002098302 | 0,046921218 | 0,041481671 | TRIM27/TRIM27/DNMT1/JARID2/LMNB2/HMGB1/PHF2/TASOR2/USP7/RB1/HMGA2/KDM5A/PPHLN1/GSK3A/EPC1/CBX3/TRIM28/MACROH2A1/SIN3A/RRP8/PHF8/ZNF304 | 22 |
| GO:2000781 | positive regulation of double-strand break repair | 18/2001 | 90/20772 | 0,00210444 | 0,046921218 | 0,041481671 | ZCWPW1/SPIRE1/ACTR2/KAT5/RUVBL2/MORF4L2/TRRAP/EP400/MORF4L1/EPC1/RUVBL1/MRGBP/SMARCE1/KMT5B/SLF2/TIMELESS/PBRM1/SETMAR | 18 |
| GO:0006024 | glycosaminoglycan biosynthetic process | 16/2001 | 76/20772 | 0,002100425 | 0,046921218 | 0,041481671 | HAS2/SMPD3/DSEL/DSE/CHST12/XYLT1/ST3GAL4/NDST3/CHSY1/HEXA/CSGALNACT2/GLCE/CLTC/B3GALT6/CHPF2/EXT2 | 16 |
| GO:0099173 | postsynapse organization | 33/2001 | 204/20772 | 0,002126401 | 0,047235916 | 0,041759887 | PRNP/SLITRK3/PTPRD/PTPRD/ZDHHC2/ROR2/TANC2/PPFIA2/CDKL5/DNM3/PDLIM5/NEDD9/CTNND2/LRRC4B/DISC1/AKT1/DBN1/BAIAP2/LRFN4/LRP8/ARHGAP44/MTMR2/ARF1/CRK/IGF1R/MESD/ACTN1/RER1/HSPA8/CDC42/ARHGEF9/RPS6KA5/PTEN | 33 |
| GO:0072698 | protein localization to microtubule cytoskeleton | 13/2001 | 56/20772 | 0,002146984 | 0,047517811 | 0,042009101 | MID2/ABHD17C/DISC1/NUDCD3/MID1/NSFL1C/CSNK1D/DIAPH1/STIL/PCM1/NUP62/DCTN2/MAPRE1 | 13 |
| GO:0048863 | stem cell differentiation | 40/2001 | 261/20772 | 0,002186034 | 0,048124251 | 0,042545238 | TWIST1/MSX2/WNT3/WNT3/EDN3/ERBB4/PTN/A2M/CDK6/SNAI2/EDNRA/ZFP36L2/FOXC1/EPOP/TBX1/SOX9/SEMA3A/LTBP3/SEMA3D/BCL2/NSUN2/JARID2/CORO1C/SEMA5A/YAP1/SRF/KAT5/HSPA9/GATAD2A/MED1/SOX6/FRZB/ZFP36/EIF2AK2/HMGA2/HDAC2/NUDT21/PHACTR4/NOLC1/CDC42 | 40 |
| GO:0048284 | organelle fusion | 28/2001 | 165/20772 | 0,002190373 | 0,048124251 | 0,042545238 | CPLX1/SPHK1/SNCA/CHMP4A/BNIP1/PIP4K2B/PIP4K2B/STX12/VPS16/AKT2/YIPF5/PLEKHM1/GDAP1/RAB39A/YIPF4/GOSR2/CHMP4B/RAB7A/VPS4A/VTI1B/VPS41/CHMP6/MFN2/STX10/BNIP3/THG1L/GNAI3/GOSR1 | 28 |

**increasing genes specific for HD; MF**

| ID | Description | GeneRatio | BgRatio | pvalue | p.adjust | qvalue | geneID | Count |
|---|---|---|---|---|---|---|---|---|
| GO:0043021 | ribonucleoprotein complex binding | 45/2043 | 186/20696 | 1,00607E-08 | 1,13686E-05 | 1,0389E-05 | LETM2/NOMO3/BOP1/BOP1/NOMO1/NOMO3/MAP3K20/GTPBP4/PES1/GTPBP6/ZNF598/NCLN/EIF2S1/EIF3C/LTN1/DDX3X/ABCE1/RBM3/SEC61A2/HSPA5/SECISBP2/SRP68/EIF4A3/EIF1/SNRPB2/SEC61A1/SND1/EIF2A/SBDS/RBM39/CD2BP2/ZNF622/SERBP1/DHX29/LEMD3/YTHDF1/SEC61G/CCDC47/MTRES1/NOLC1/CPEB4/SNRPD1/PQBP1/EEFSEC/SRP54 | 45 |
| GO:0045296 | cadherin binding | 69/2043 | 357/20696 | 3,76484E-08 | 2,12714E-05 | 1,94385E-05 | AHNAK/CTNNA2/PHLDB2/PLEC/EGFR/SEPTIN9/PDLIM5/DOCK9/LIMA1/DBNL/CTNND2/DCHS1/HCFC1/GIPC1/LRRC59/PACSIN2/EXOC3/SPTAN1/TLN1/IQGAP1/ADD1/LASP1/DBN1/SEPTIN7/BMPR2/BAIAP2/DDX3X/MARK2/HSPA5/CDH11/VCL/FER/HSP90AB1/SND1/EIF2A/TXNDC9/YKT6/EPS15L1/PCMT1/RAB1A/PCBP1/LRRFIP1/SERBP1/YWHAB/DHX29/CHMP4B/CSNK1D/BZW1/EFHD2/PKM/PTPN11/AHSA1/SH3GLB1/RUVBL1/TBC1D10A/EIF4G2/TMPO/HSPA8/TMOD3/SEPTIN2/CAPZA1/SCYL1/CBL/FSCN1/MAPRE1/PICALM/ZC3H15/HNRNPK/EIF5 | 69 |
| GO:0140034 | methylation-dependent protein binding | 22/2043 | 82/20696 | 9,8778E-06 | 0,003720639 | 0,003400044 | CBX4/ZCWPW1/ZMYND8/CXXC1/PHF2/SPIN4/ZZEF1/PHF20L1/MSL3/LRWD1/CDYL/GLYR1/ING1/KDM5A/TDRD3/L3MBTL2/CBX3/ZMYND11/MSH6/RRP8/CHD8/PHF8 | 22 |
| GO:0035064 | methylated histone binding | 21/2043 | 80/20696 | 2,24226E-05 | 0,005188125 | 0,004741081 | CBX4/ZCWPW1/ZMYND8/CXXC1/PHF2/SPIN4/ZZEF1/MSL3/LRWD1/CDYL/GLYR1/ING1/KDM5A/TDRD3/L3MBTL2/CBX3/ZMYND11/MSH6/RRP8/CHD8/PHF8 | 21 |
| GO:0140030 | modification-dependent protein binding | 39/2043 | 200/20696 | 2,64287E-05 | 0,005188125 | 0,004741081 | CBX4/ZCWPW1/JARID2/ZMYND8/AGAP3/CXXC1/BRD9/BRD3/IKBKE/PHF2/SPIN4/ZZEF1/UFD1/PHF20L1/MSL3/LRWD1/CDYL/PSME4/GLYR1/ATAD2B/ING1/USP15/PSMD4/IDE/KDM5A/TDRD3/TAB3/ANKRD13B/L3MBTL2/ZBTB1/ANKRD13A/CBX3/PRPF8/ZMYND11/MSH6/VCP/RRP8/CHD8/PHF8 | 39 |

| | | | | | | | | |
|---|---|---|---|---|---|---|---|---|
| GO:0003713 | transcription coactivator activity | 55/2043 | 319/20696 | 2,95965E-05 | 0,005188125 | 0,004741081 | TGFB1I1/MAML2/TRIM27/TRIM27/MID2/HIPK2/HCFC1/ZBED1/SRCAP/KMT2C/YAP1/THRAP3/CTBP2/KAT5/HMGB1/PHF2/MTDH/PRMT2/SERTAD2/MED1/MED27/ATXN7L3/CCDC124/ZXDC/NCOA6/KDM5A/PSMC3IP/MED17/TDRD3/RAP2C/CBFB/UBE2L3/DRAP1/SNW1/TAF5L/ACTN1/MED13/MAML3/ARL2BP/RUVBL1/USP16/MED12/SMARCE1/PSMD9/LPIN2/SUB1/TRIM28/ATN1/CEBPZ/PQBP1/APEX1/SIN3A/USP22/TAF12/COPS5 | 55 |
| GO:0051015 | actin filament binding | 41/2043 | 216/20696 | 3,21388E-05 | 0,005188125 | 0,004741081 | ESPN/CTNNA2/PLEC/EGFR/MYH3/LIMA1/DBNL/SHROOM2/FERMT2/GAS2/TPM4/CORO1C/MYO9A/PPP1R9B/MAP1S/SPTAN1/ACTR2/WDR1/TLN1/VPS16/IQGAP1/ADD1/LASP1/TPM3/SHROOM4/ACTR3/TLN2/ARPC5/HIP1/MARCKS/IQGAP2/CFL2/ARPC2/ARPC5L/ARPC4/ACTN1/PKNOX2/CAPZA1/FSCN1/TMEM201/PANX1 | 41 |
| GO:0043022 | ribosome binding | 21/2043 | 84/20696 | 4,93647E-05 | 0,006972763 | 0,006371943 | LETM2/NOMO3/NOMO1/NOMO3/MAP3K20/GTPBP6/ZNF598/NCLN/EIF2S1/EIF3C/SEC61A2/HSPA5/SRP68/SEC61A1/EIF2A/SBDS/SERBP1/YTHDF1/SEC61G/CCDC47/CPEB4 | 21 |
| GO:0042393 | histone binding | 46/2043 | 258/20696 | 5,59474E-05 | 0,00702451 | 0,006419231 | CBX4/SNCA/ZCWPW1/JARID2/SRCAP/ZMYND8/KMT2C/MCM2/CXXC1/BRD9/NOC2L/BRD3/KAT5/ATAD2/PHF2/SPIN4/ZZEF1/SSRP1/MSL3/LRWD1/CDYL/PSME4/GLYR1/ATAD2B/ING1/SAP30L/GRWD1/USP15/KDM5A/TDRD3/SET/IPO7/L3MBTL2/NAP1L4/PRMT6/KDM1B/CBX3/ZMYND11/USP16/MSH6/RRP8/WDR5B/CHD8/PHF8/DEK/CHAF1B | 46 |
| GO:0046332 | SMAD binding | 20/2043 | 80/20696 | 7,39961E-05 | 0,008361556 | 0,007641068 | TGFB1I1/DAB2/RGCC/PURA/BMPR1B/FERMT2/HIPK2/AXIN2/RANBP3/USP15/TGFBR1/HMGA2/STUB1/ZC3H3/USP9X/IPO7/SNW1/TCF12/PPM1A/DROSHA | 20 |
| GO:0051539 | 4 iron, 4 sulfur cluster binding | 13/2043 | 41/20696 | 0,000101067 | 0,010382385 | 0,009487769 | ETFDH/RSAD1/DPH1/ABCE1/IREB2/TYW1B/NDUFV1/NUBP1/ACO2/TYW1/NUBP2/ACO1/NDUFS2 | 13 |
| GO:0003743 | translation initiation factor activity | 15/2043 | 53/20696 | 0,000129601 | 0,012142099 | 0,011095854 | EIF4G1/EIF2S1/EIF3C/EIF3J/EIF1AD/EIF1/EIF2B3/EIF2A/EIF3B/DHX29/DENR/EIF4G2/EIF2D/EIF5/COPS5 | 15 |

| | | | | | | | | |
|---|---|---|---|---|---|---|---|---|
| GO:0031625 | ubiquitin protein ligase binding | 57/2043 | 354/20696 | 0,000149227 | 0,012142099 | 0,011095854 | NGFR/STING1/FOXO1/UBASH3B/RHOBTB3/EGFR/HERC2/RALB/BCL2/HSPA1B/TRAF5/PRKACA/MID1/CASC3/BAG5/AXIN2/CALR/PRKAR1A/PRKACA/HSPA9/ERLIN1/JAK1/PSMD1/IKBKE/SLC25A5/UBE2K/HLTF/HSPA5/USP7/CDKN1B/VCL/RB1/HSP90AB1/POLR2A/STUB1/FAF2/RRAGA/RIPK1/TOLLIP/HSPD1/CUL5/UBE2L3/USP13/TRAF4/SPOP/CRK/MFHAS1/UBE2A/HM13/HSPA8/MFN2/DNAJA1/VCP/TRIM28/PSMA3/SLF2/NDUFS2 | 57 |
| GO:0046966 | nuclear thyroid hormone receptor binding | 11/2043 | 32/20696 | 0,000152787 | 0,012142099 | 0,011095854 | THRAP3/TRIP12/GTF2H1/GTF2H1/PRMT2/MED1/NCOA6/MED17/MED13/NSD1/MED12 | 11 |
| GO:0019208 | phosphatase regulator activity | 27/2043 | 131/20696 | 0,000169097 | 0,012142099 | 0,011095854 | PPP1R16B/PTN/BMPR1B/PABIR1/PPP1R15A/PPP1R9B/PPP1R12B/PPP3R1/BMPR2/FRS2/PPP1R15B/LMTK2/PPP6R2/PPP1R16A/PPP4R1/EIF2AK2/CALM1/SET/PPP1R8/PPP2R2D/PTPA/PPP1R7/PHACTR4/ANKLE2/PPP2R5D/PPP2R1A/GNA12 | 27 |
| GO:0051082 | unfolded protein binding | 29/2043 | 145/20696 | 0,000171924 | 0,012142099 | 0,011095854 | HSPA1B/NUDCD3/ERO1B/CALR/HSP90B1/HSPA9/SYVN1/CLPX/PPIB/DNAJB6/CANX/CCAR2/HSPA5/DNAJB11/HSP90AB1/UGGT1/HEATR3/CANX/RUVBL2/HSPD1/SERPINH1/NAP1L4/CCT7/CDC37/HSPA8/DNAJA1/VBP1/PFDN6/CHAF1B | 29 |
| GO:0019001 | guanyl nucleotide binding | 66/2043 | 431/20696 | 0,000211924 | 0,012844615 | 0,011737837 | GNL1/GNL1/GNL1/STING1/PRKG1/TUBB2A/AK4/RND1/GNAI1/SEPTIN9/DNM3/RALB/RABL3/PIP4K2B/PIP4K2B/HRAS/TUBB2B/GTPBP4/AGAP3/RAP1A/GTPBP6/DAPK1/URGCP/SEPTIN11/MMAA/RHOB/SEPTIN7/SEPHS1/RAB35/ADSS1/AGAP1/LANCL2/RAB6A/GNAQ/RAC1/SEPTIN5/MIEF1/GSPT1/SEPTIN6/RAB39A/EFTUD2/ARF1/RAB1A/RRAGA/RAP2C/RAB21/RAB14/RAB7A/ARL17A/MFHAS1/MFN2/RAB15/SEPTIN2/RAB18/RHOA/NOLC1/CDC42/DAP3/GTPBP1/THG1L/GNAI3/EEFSEC/SRP54/GNA12/EIF5/GNA13 | 66 |

| | | | | | | | | |
|---|---|---|---|---|---|---|---|---|
| GO:0032561 | guanyl ribonucleotide binding | 66/2043 | 431/20696 | 0,000211924 | 0,012844615 | 0,011737837 | GNL1/GNL1/GNL1/STING1/PRKG1/TUBB2A/AK4/RND1/GNAI1/SEPTIN9/DNM3/RALB/RABL3/PIP4K2B/PIP4K2B/HRAS/TUBB2B/GTPBP4/AGAP3/RAP1A/GTPBP6/DAPK1/URGCP/SEPTIN11/MMAA/RHOB/SEPTIN7/SEPHS1/RAB35/ADSS1/AGAP1/LANCL2/RAB6A/GNAQ/RAC1/SEPTIN5/MIEF1/GSPT1/SEPTIN6/RAB39A/EFTUD2/ARF1/RAB1A/RRAGA/RAP2C/RAB21/RAB14/RAB7A/ARL17A/MFHAS1/MFN2/RAB15/SEPTIN2/RAB18/RHOA/NOLC1/CDC42/DAP3/GTPBP1/THG1L/GNAI3/EEFSEC/SRP54/GNA12/EIF5/GNA13 | 66 |
| GO:0005525 | GTP binding | 63/2043 | 407/20696 | 0,000215971 | 0,012844615 | 0,011737837 | GNL1/GNL1/GNL1/TUBB2A/AK4/RND1/GNAI1/SEPTIN9/DNM3/RALB/RABL3/PIP4K2B/PIP4K2B/HRAS/TUBB2B/GTPBP4/AGAP3/RAP1A/GTPBP6/DAPK1/URGCP/SEPTIN11/MMAA/RHOB/SEPTIN7/SEPHS1/RAB35/ADSS1/AGAP1/LANCL2/RAB6A/GNAQ/RAC1/SEPTIN5/GSPT1/SEPTIN6/RAB39A/EFTUD2/ARF1/RAB1A/RRAGA/RAP2C/RAB21/RAB14/RAB7A/ARL17A/MFHAS1/MFN2/RAB15/SEPTIN2/RAB18/RHOA/NOLC1/CDC42/DAP3/GTPBP1/THG1L/GNAI3/EEFSEC/SRP54/GNA12/EIF5/GNA13 | 63 |
| GO:0030674 | protein-macromolecule adaptor activity | 60/2043 | 386/20696 | 0,000269259 | 0,014495569 | 0,013246533 | PRNP/DAB2/SUN2/SPSB4/KLHL15/ZSWIM8/BNIP1/PEX3/GAS2/HCFC1/STON2/HRAS/STX12/SORBS1/TIRAP/IQGAP1/FEM1B/SRRT/FRS2/ZZEF1/BAIAP2/DCAF7/CDYL/AP1G1/RPTOR/GATAD2A/YKT6/KLHL25/HIP1/ELOC/AKIRIN2/STUB1/FEM1A/GOSR2/CUL5/TARDBP/PTPN11/KLHDC10/SH2B1/VTI1B/CRK/DDB1/APH1B/ARPC4/LDLRAP1/EPC1/ST13/HSPA8/NUP62/GNB2/STX10/NOLC1/KSR1/SH2B2/SH3BGRL/FSCN1/TIAL1/GOSR1/PSMC6/SUN1 | 60 |

| | | | | | | | | |
|---|---|---|---|---|---|---|---|---|
| GO:0003924 | GTPase activity | 55/2043 | 346/20696 | 0,000269387 | 0,014495569 | 0,013246533 | GNL1/GNL1/GNL1/ARHGDIB/TUBB2A/RND1/GNAI1/SEPTIN9/DNM3/RALB/RGS3/RABL3/RASA1/HRAS/TUBB2B/GTPBP4/AGAP3/RAP1A/SEPTIN11/MMAA/RHOB/SEPTIN7/RAB35/AGAP1/DDX3X/RAB6A/GNAQ/RAC1/SEPTIN5/GSPT1/SEPTIN6/RAB39A/EFTUD2/TMEM250/ARF1/RAB1A/RRAGA/RAP2C/RAB21/RAB14/RAB7A/MMUT/ARL17A/MFN2/SEPTIN2/GNB2/RAB18/RHOA/CDC42/GTPBP1/GNAI3/EEFSEC/SRP54/GNA12/GNA13 | 55 |
| GO:0051536 | iron-sulfur cluster binding | 17/2043 | 70/20696 | 0,00036319 | 0,017187486 | 0,015706496 | GLRX2/ETFDH/CISD3/RSAD1/DPH1/KIF4A/ABCE1/IREB2/TYW1B/NDUFV1/NUBP1/ACO2/TYW1/NDUFV2/NUBP2/ACO1/NDUFS2 | 17 |
| GO:0051540 | metal cluster binding | 17/2043 | 70/20696 | 0,00036319 | 0,017187486 | 0,015706496 | GLRX2/ETFDH/CISD3/RSAD1/DPH1/KIF4A/ABCE1/IREB2/TYW1B/NDUFV1/NUBP1/ACO2/TYW1/NDUFV2/NUBP2/ACO1/NDUFS2 | 17 |
| GO:0003779 | actin binding | 69/2043 | 464/20696 | 0,000365044 | 0,017187486 | 0,015706496 | ESPN/TNS1/CTNNA2/SYNPO/PLEC/EGFR/SNCA/MYH3/AFAP1/PDLIM5/LIMA1/DBNL/SHROOM2/EPB41L2/FERMT2/MSN/SNTA1/EPS8/GAS2/SPIRE1/TPM4/CORO1C/NF2/MYO9A/PPP1R9B/GIPC1/MAP1S/SORBS1/SPTAN1/ACTR2/WDR1/TLN1/VPS16/IQGAP1/ADD1/LASP1/DBN1/TPM3/CAP1/SHROOM4/ACTR3/TLN2/ARPC5/NCALD/EPB41L1/VCL/FER/TMSB4X/HIP1/PXK/MARCKS/IQGAP2/CFL2/ARPC2/MSRB2/P4HB/ARPC5L/DIAPH1/WASH6P/ARPC4/ACTN1/PHACTR4/TMOD3/PKNOX2/CAPZA1/FSCN1/TMEM201/PANX1/YWHAH | 69 |
| GO:0005048 | signal sequence binding | 14/2043 | 52/20696 | 0,00038136 | 0,01723745 | 0,015752155 | KPNA1/SEC61A2/KDELR2/SRP68/SEC61A1/POM121/KPNA3/KPNB1/TOMM40L/NUP214/NOLC1/POM121C/KPNA6/SRP54 | 14 |
| GO:0051087 | chaperone binding | 24/2043 | 118/20696 | 0,000465247 | 0,020220349 | 0,018478028 | CP/PRNP/SLC12A2/BAG5/CALR/SYVN1/SUGT1/SACS/DNAJB6/HSPA5/CDKN1B/DNAJC3/STUB1/HSPD1/SDF2L1/USP13/ATP7A/AHSA1/CDC37/HSPA8/DNAJA1/OGDH/CDC25A/PFDN6 | 24 |

| | | | | | | | | |
|---|---|---|---|---|---|---|---|---|
| GO:0019888 | protein phosphatase regulator activity | 23/2043 | 112/20696 | 0,000524518 | 0,021952036 | 0,020060501 | PPP1R16B/PTN/PABIR1/PPP1R15A/PPP1R9B/PPP3R1/PPP1R15B/LMTK2/PPP6R2/PPP1R16A/PPP4R1/EIF2AK2/CALM1/SET/PPP1R8/PPP2R2D/PTPA/PPP1R7/PHACTR4/ANKLE2/PPP2R5D/PPP2R1A/GNA12 | 23 |
| GO:0044389 | ubiquitin-like protein ligase binding | 57/2043 | 373/20696 | 0,000581425 | 0,023464642 | 0,02144277 | NGFR/STING1/FOXO1/UBASH3B/RHOBTB3/EGFR/HERC2/RALB/BCL2/HSPA1B/TRAF5/PRKACA/MID1/CASC3/BAG5/AXIN2/CALR/PRKAR1A/PRKACA/HSPA9/ERLIN1/JAK1/PSMD1/IKBKE/SLC25A5/UBE2K/HLTF/HSPA5/USP7/CDKN1B/VCL/RB1/HSP90AB1/POLR2A/STUB1/FAF2/RRAGA/RIPK1/TOLLIP/HSPD1/CUL5/UBE2L3/USP13/TRAF4/SPOP/CRK/MFHAS1/UBE2A/HM13/HSPA8/MFN2/DNAJA1/VCP/TRIM28/PSMA3/SLF2/NDUFS2 | 57 |
| GO:0051219 | phosphoprotein binding | 20/2043 | 94/20696 | 0,000738222 | 0,028765183 | 0,026286581 | UBASH3B/CBX4/SNCA/CBLB/MID2/YES1/SH3BP2/MID1/RASA1/THRAP3/RB1/YWHAB/RRAGA/PTPN11/CRK/LDLRAP1/SHC3/CBL/PIK3R2/PIN1 | 20 |
| GO:0032182 | ubiquitin-like protein binding | 24/2043 | 122/20696 | 0,000769208 | 0,028973493 | 0,026476942 | SOBP/CBX4/HERC2/JARID2/STAM2/RBCK1/NSFL1C/UBXN2A/SERBP1/FAF2/TOLLIP/PELP1/TAB3/USP13/ASCC2/RNFT1/UBAP1/UBE2A/USP16/NUP62/GGA3/UBA2/OTUB2/RNF185 | 24 |
| GO:0034450 | ubiquitin-ubiquitin ligase activity | 6/2043 | 13/20696 | 0,000854152 | 0,03055129 | 0,027918784 | PELI1/UBE2K/PPIL2/STUB1/UBE4A/PRPF19 | 6 |
| GO:0003899 | DNA-directed 5'-3' RNA polymerase activity | 12/2043 | 44/20696 | 0,000865169 | 0,03055129 | 0,027918784 | POLR1H/POLR1H/POLR1H/POLR1H/POLR1H/POLR3D/POLR3B/POLR2A/POLR1B/POLR3A/POLR2E/POLR2B | 12 |
| GO:0003756 | protein disulfide isomerase activity | 7/2043 | 18/20696 | 0,001078147 | 0,035832526 | 0,032744954 | GLRX2/CRELD2/PDIA6/PDIA4/QSOX2/PDIA3/P4HB | 7 |
| GO:0016864 | intramolecular oxidoreductase activity, transposing S-S bonds | 7/2043 | 18/20696 | 0,001078147 | 0,035832526 | 0,032744954 | GLRX2/CRELD2/PDIA6/PDIA4/QSOX2/PDIA3/P4HB | 7 |

| | | | | | | | | |
|---|---|---|---|---|---|---|---|---|
| GO:0016667 | oxidoreductase activity, acting on a sulfur group of donors | 14/2043 | 58/20696 | 0,00124488 | 0,040191829 | 0,03672863 | MSRA/GLRX2/GSTO1/ERO1B/MSRA/PDIA6/PDIA4/QSOX2/TMX4/NXN/PDIA3/MSRB2/P4HB/TMX2 | 14 |
| GO:0140098 | catalytic activity, acting on RNA | 66/2043 | 462/20696 | 0,001400841 | 0,043970848 | 0,040182023 | POLR1H/POLR1H/POLR1H/POLR1H/POLR1H/VARS1/DHX37/HENMT1/LRRC47/NSUN2/DUS3L/FARSA/NSUN4/FTO/AARSD1/POLR3D/YARS1/DCP1A/DDX10/DDX3X/DTD2/TRMT1L/TYW1B/ISG20L2/TSNAX/TRMT61A/EIF4A3/TRMO/POLR3B/DDX21/SND1/DHX8/HARS1/POLR2A/DDX56/RCL1/NUDT12/DHX29/PRPF18/BUD23/METTL16/TYW1/PPP1R8/DDX19A/NSUN3/SARS1/POLR1B/MTREX/MARS2/SNRNP200/EXOSC4/NARS1/WARS2/DIS3/DICER1/POLR3A/ELAC1/TARS1/APEX1/MRM2/THG1L/DROSHA/POLR2E/KARS1/DDX1/POLR2B | 66 |
| GO:0003725 | double-stranded RNA binding | 16/2043 | 72/20696 | 0,00149219 | 0,045572291 | 0,041645475 | MSN/ZNF346/MTDH/CLTC/HSP90AB1/DDX21/ADAR/EIF2AK2/LSM14A/LRRFIP1/HSPD1/ACTN1/DICER1/ZFR/ELAVL1/DDX1 | 16 |
| GO:0001054 | RNA polymerase I activity | 7/2043 | 19/20696 | 0,001562627 | 0,046467594 | 0,042463633 | POLR1H/POLR1H/POLR1H/POLR1H/POLR1H/POLR1B/POLR2E | 7 |
| GO:0015036 | disulfide oxidoreductase activity | 11/2043 | 41/20696 | 0,001627947 | 0,047168726 | 0,04310435 | GLRX2/GSTO1/ERO1B/PDIA6/PDIA4/QSOX2/TMX4/NXN/PDIA3/P4HB/TMX2 | 11 |

**decreasing genes specific for KO; BP**

| ID | Description | GeneRatio | BgRatio | pvalue | p.adjust | qvalue | geneID | Count |
|---|---|---|---|---|---|---|---|---|
| GO:0034470 | ncRNA processing | 84/1673 | 475/20772 | 4,94855E-12 | 2,64005E-08 | 2,44615E-08 | PIH1D2/DPH3/METTL5/DTWD1/FAM98B/LSM6/INTS6L/UTP15/METTL18/OSGEPL1/ADAT2/RPL35A/FCF1/WDR75/EXOSC8/DICER1/TRMT11/RPP30/TYW5/RPF1/ELP2/TUT4/TSEN15/NUP155/SSB/INTS2/ELP4/CDKAL1/RPL27/TRMT13/TENT4B/WDR36/INTS7/ERI1/NSUN6/NOL11/TSEN34/UTP11/UTP23/INTS8/TRIM71/BRIX1/TDRKH/ERI2/HEATR1/METTL15/SEPSECS/TARBP1/RPL5/HENMT1/YRDC/BMP4/INTS13/DCAF13/RPS21/UTP6/UTP20/NOP58/EXOSC9/NCBP1/WDR43/NIFK/THUMPD3/ELP1/TRNT1/DDX52/METTL2A/LARP7/DDX21/NCBP2/RPF2/CDK5RAP1/PARN/MPHOSPH6/TFB2M/NOL8/NAF1/ZNHIT3/SRFBP1/ABT1/MRPL44/POP5/NHP2/RIOK1 | 84 |
| GO:0006302 | double-strand break repair | 63/1673 | 318/20772 | 1,8142E-11 | 4,83937E-08 | 4,48393E-08 | ACTL6B/RMI1/DCLRE1A/FIGNL1/UBE2V2/FAN1/MBTD1/CDC7/SMARCAD1/ATR/ERCC8/HPF1/USP51/RAD51C/PARPBP/RBBP8/MSH2/WRN/LIG4/UBE2N/HUS1/KDM4D/CHEK1/MMS22L/HMGA2/WDR48/RAD51AP1/RPA2/BCL7A/BLM/ABRAXAS1/RAD50/ATM/PSMD14/RIF1/NSMCE4A/MRE11/NSMCE2/XRCC4/RNF169/PHF10/GEN1/FANCM/INIP/XRCC2/EPC1/MCM8/BRIP1/MEAF6/NHEJ1/POLB/SETMAR/EYA3/TRIP13/SMARCC1/RUVBL1/HMGB2/PALB2/MCM6/ATRIP/XRCC5/KDM1A/XRCC6 | 63 |
| GO:0006403 | RNA localization | 49/1673 | 221/20772 | 5,35158E-11 | 9,51689E-08 | 8,8179E-08 | DHX36/NXT2/NDC1/MAGOHB/MAGOH/NUP37/ATR/THOC1/CCT8/CETN3/XPOT/ENY2/NUP155/SSB/NUP107/ATM/LRPPRC/NUP160/NUP153/NUP205/RANBP2/NPM1/AHCTF1/PARP1/CCT2/CCT6A/TERF1/NOP58/CCT5/SRSF1/SMG7/RAN/NCBP1/DCP2/CCT4/SRSF7/UPF2/NCBP2/PARN/RUVBL1/NAF1/ZNHIT3/DHX9/TPR/CCT3/POLR2D/IGF2BP3/SEH1L/NHP2 | 49 |

| | | | | | | | | |
|---|---|---|---|---|---|---|---|---|
| GO:0031123 | RNA 3'-end processing | 32/1673 | 110/20772 | 7,94887E-11 | 1,06018E-07 | 9,82314E-08 | DHX36/PAPOLG/INTS6L/PAPOLA/EXOSC8/TUT4/SSB/MTPAP/VIRMA/INTS2/TENT4B/TENT2/INTS7/ERI1/INTS8/ERI2/WDR33/RPRD1B/RPS21/EXOSC9/NCBP1/RPRD1A/CPSF3/TRNT1/SNRPA/LARP7/NCBP2/PARN/PCF11/NUDT21/POLR2D/LSM11 | 32 |
| GO:0007059 | chromosome segregation | 83/1673 | 499/20772 | 1,63285E-10 | 1,74225E-07 | 1,61428E-07 | MND1/TTK/SASS6/ACTL6B/NDC1/RMI1/CENPQ/KIF18A/NUP37/SMARCAD1/HAUS1/RAD51C/ASPM/SMARCA5/NUF2/CENPH/DSCC1/DLGAP5/MIS12/KIF14/CLASP2/NCAPG/CENPE/KIF11/BCL7A/USP9X/ABRAXAS1/ATM/OIP5/LSM14A/SGO1/SPDL1/CENPS/CENPW/PDS5A/SETDB2/TOP2B/MRE11/ANAPC4/NSMCE2/NAA50/CDC16/KNTC1/KIF23/PHF10/GEN1/SMC3/MIS18A/BUB1/KNL1/BCCIP/FANCM/CDC26/SGO2/SRPK1/LATS1/WAPL/CIAO2A/HJURP/MAPRE1/TERF1/ZNF207/RAN/CCNB1IP1/CDCA8/STAG1/TOP2A/BRIP1/CCNB1/MAP9/CCNE1/DYNC1LI1/BOD1/TUBB/SENP6/CEP63/TRIP13/SMARCC1/CHAMP1/MAPRE3/ANAPC7/TPR/SEH1L | 83 |
| GO:0034502 | protein localization to chromosome | 33/1673 | 123/20772 | 4,26042E-10 | 3,70228E-07 | 3,43036E-07 | TTK/CENPQ/H4C3/TASOR/ATR/MSH2/H4C11/H4C4/CCT8/MIS12/MTBP/RPA2/H4C8/SPDL1/EZH2/XRCC4/KNTC1/KNL1/WAPL/CCT2/CCT6A/TERF1/CCT5/H4C9/CCT4/MCM8/GNL3/H4C5/H4C2/BOD1/CHAMP1/CCT3/XRCC5 | 33 |
| GO:0140053 | mitochondrial gene expression | 41/1673 | 177/20772 | 4,85772E-10 | 3,70228E-07 | 3,43036E-07 | MRPL50/MRPL39/MRPL42/MRPS6/MTRF1L/FASTKD1/MTERF3/FASTKD2/MRPS22/YARS2/MRPL45/MRPL13/MRPL47/MRPS31/MRPL22/MRPS9/MRPL1/GFM2/LRPPRC/MRPL48/TFAM/QRSL1/MRPS30/MTIF2/PTCD3/IARS2/MRPS21/MRPS28/MRPL3/MRPL51/RMND1/MRPL10/TRNT1/MRPS15/MRPS7/DARS2/CDK5RAP1/TFB2M/SLC25A33/MALSU1/MRPL44 | 41 |

| | | | | | | | | |
|---|---|---|---|---|---|---|---|---|
| GO:0006310 | DNA recombination | 64/1673 | 354/20772 | 6,88641E-10 | 4,53783E-07 | 4,20454E-07 | H1-6/MND1/RMI1/FIGNL1/H1-1/FAN1/MBTD1/CDC7/SMARCAD1/USP51/RAD51C/THOC1/PARPBP/RBBP8/MSH2/WRN/LIG4/UBE2N/HUS1/KDM4D/CHEK1/ATAD5/MMS22L/WDR48/RAD51AP1/RPA2/BLM/RAD50/KPNA2/ATM/PSMD14/RIF1/NSMCE4A/CENPS/TOP2B/MRE11/NSMCE2/XRCC4/ZRANB3/GEN1/EXO1/FANCM/INIP/XRCC2/CCNB1IP1/EPC1/TOP2A/MCM8/BRIP1/MEAF6/NHEJ1/POLB/MSH3/TRIP13/RUVBL1/HMGB2/PAXIP1/PALB2/DCAF1/MCM6/APEX1/XRCC5/KDM1A/XRCC6 | 64 |
| GO:0006260 | DNA replication | 57/1673 | 299/20772 | 7,65519E-10 | 4,53783E-07 | 4,20454E-07 | RMI1/NPM2/FAM111B/ETAA1/RFC3/CDC7/ORC3/ATR/THOC1/RBBP8/CCNA2/WRN/LIG4/SMARCA5/DSCC1/CHEK1/ATAD5/OBI1/MMS22L/RFC4/PRIM2/RPA2/RBMS1/ORC4/BLM/RAD50/CENPS/PDS5A/MRE11/DBF4/MCMBP/ZRANB3/USP37/GEN1/SMC3/FAF1/FANCM/GMNN/WAPL/POLD3/RFC5/ORC5/TERF1/SSBP1/MCM8/MEAF6/WDHD1/CCNE1/POLB/SETMAR/ORC1/RUVBL1/DHX9/MCM6/DNAJC2/RRM1/DTD1 | 57 |
| GO:0033044 | regulation of chromosome organization | 55/1673 | 285/20772 | 9,51199E-10 | 5,07465E-07 | 4,70193E-07 | DHX36/TTK/ACTL6B/TASOR/ATR/RESF1/LIG4/SMARCA5/NUF2/CCT8/DLGAP5/NCAPG/CENPE/BCL7A/RAD50/ATM/TENT4B/NSMCE4A/SPDL1/SETDB2/MRE11/ANAPC4/NSMCE2/CDC16/KNTC1/PHF10/GEN1/BUB1/CDC26/WAPL/CCT2/CCT6A/TERF1/ZNF207/SSBP1/CCT5/GTF2H2/CDCA8/DCP2/TOP2A/CCT4/CCNB1/GNL3/DYNC1LI1/SETMAR/SENP6/TRIP13/PARN/SMARCC1/RUVBL1/NAF1/ANAPC7/TPR/CCT3/XRCC5 | 55 |

| | | | | | | | | |
|---|---|---|---|---|---|---|---|---|
| GO:0000819 | sister chromatid segregation | 53/1673 | 272/20772 | 1,33981E-09 | 6,49808E-07 | 6,02081E-07 | TTK/ACTL6B/KIF18A/RAD51C/SMARCA5/NUF2/DSCC1/DLGAP5/MIS12/KIF14/CLASP2/NCAPG/CENPE/KIF11/BCL7A/ABRAXAS1/ATM/LSM14A/SGO1/SPDL1/PDS5A/TOP2B/ANAPC4/NSMCE2/NAA50/CDC16/KNTC1/KIF23/PHF10/GEN1/SMC3/BUB1/BCCIP/CDC26/SGO2/LATS1/WAPL/MAPRE1/ZNF207/RAN/CDCA8/STAG1/TOP2A/CCNB1/MAP9/DYNC1LI1/BOD1/TRIP13/SMARCC1/CHAMP1/ANAPC7/TPR/SEH1L | 53 |
| GO:0032543 | mitochondrial translation | 35/1673 | 143/20772 | 1,88588E-09 | 8,38433E-07 | 7,76852E-07 | MRPL50/MRPL39/MRPL42/MRPS6/MTRF1L/FASTKD2/MRPS22/YARS2/MRPL45/MRPL13/MRPL47/MRPS31/MRPL22/MRPS9/MRPL1/GFM2/LRPPRC/MRPL48/QRSL1/MRPS30/MTIF2/PTCD3/IARS2/MRPS21/MRPS28/MRPL3/MRPL51/RMND1/MRPL10/MRPS15/MRPS7/DARS2/CDK5RAP1/MALSU1/MRPL44 | 35 |
| GO:0098813 | nuclear chromosome segregation | 66/1673 | 384/20772 | 3,20954E-09 | 1,31715E-06 | 1,22041E-06 | MND1/TTK/ACTL6B/NDC1/RMI1/CENPQ/KIF18A/RAD51C/ASPM/SMARCA5/NUF2/DSCC1/DLGAP5/MIS12/KIF14/CLASP2/NCAPG/CENPE/KIF11/BCL7A/ABRAXAS1/ATM/LSM14A/SGO1/SPDL1/CENPS/PDS5A/TOP2B/MRE11/ANAPC4/NSMCE2/NAA50/CDC16/KNTC1/KIF23/PHF10/GEN1/SMC3/BUB1/KNL1/BCCIP/FANCM/CDC26/SGO2/LATS1/WAPL/MAPRE1/TERF1/ZNF207/RAN/CCNB1IP1/CDCA8/STAG1/TOP2A/BRIP1/CCNB1/MAP9/CCNE1/DYNC1LI1/BOD1/TRIP13/SMARCC1/CHAMP1/ANAPC7/TPR/SEH1L | 66 |
| GO:0006261 | DNA-templated DNA replication | 37/1673 | 162/20772 | 5,07844E-09 | 1,86422E-06 | 1,7273E-06 | ETAA1/RFC3/CDC7/ORC3/ATR/THOC1/RBBP8/WRN/DSCC1/ATAD5/MMS22L/RFC4/PRIM2/ORC4/BLM/RAD50/CENPS/MRE11/DBF4/MCMBP/ZRANB3/GEN1/FANCM/GMNN/POLD3/RFC5/ORC5/TERF1/SSBP1/MCM8/WDHD1/CCNE1/POLB/SETMAR/ORC1/MCM6/RRM1 | 37 |

| | | | | | | | | |
|---|---|---|---|---|---|---|---|---|
| GO:0032200 | telomere organization | 44/1673 | 213/20772 | 5,24148E-09 | 1,86422E-06 | 1,7273E-06 | DHX36/DCLRE1A/H3-3A/H4C3/HAT1/ATR/RAD51C/H4C11/WRN/H4C4/CCT8/HUS1/RPA2/BLM/RAD50/ATM/TENT4B/RIF1/NSMCE4A/H4C8/EZH2/MRE11/NSMCE2/EXO1/CCT2/CCT6A/TERF1/CCT5/DCP2/H4C9/CCT4/PTGES3/GNL3/H4C5/H4C2/CCNE1/PARN/RUVBL1/NAF1/CCT3/APEX1/XRCC5/XRCC6/NHP2 | 44 |
| GO:0051054 | positive regulation of DNA metabolic process | 60/1673 | 340/20772 | 6,03331E-09 | 2,01173E-06 | 1,86397E-06 | DHX36/ACTL6B/NPM2/EYA4/UBE2V2/RFC3/MBTD1/CDC7/ATR/USP1/RBBP8/MSH2/WRN/SMARCA5/UBE2N/CCT8/DSCC1/KDM4D/PARM1/ATAD5/ANXA3/RFC4/PRIM2/WDR48/RAD51AP1/BCL7A/USP9X/ABRAXAS1/RAD50/ATM/RIF1/MRE11/CEBPG/DBF4/PHF10/FAF1/MARCHF6-DT/CCT2/CCT6A/RFC5/TERF1/SSBP1/CCT5/EPC1/CCT4/PTGES3/GNL3/STPG1/MEAF6/SETMAR/OTUD4/EYA3/PARN/SMARCC1/RUVBL1/PAXIP1/NAF1/DHX9/CCT3/XRCC5 | 60 |
| GO:0002181 | cytoplasmic translation | 39/1673 | 180/20772 | 9,58383E-09 | 3,00763E-06 | 2,78673E-06 | RPS18/DHX36/RPL22L1/DPH3/RPS29/RPS13/RPL35A/RPL30/RPL23/RPL9/RPS12/DHX29/RPL27/RPS3A/NCK1/ETF1/RPL24/EIF2S3/CNBP/MCTS1/DPH5/RPS20/RPS23/RPL5/RPS21/NEMF/NCBP1/RPL36A/RPL21/ZC3H15/DENR/RPL31/RPL10A/NCBP2/RPL41/EIF2S2/DHX9/CPEB4/RPS5 | 39 |
| GO:0042254 | ribosome biogenesis | 58/1673 | 331/20772 | 1,40305E-08 | 4,15849E-06 | 3,85306E-06 | PIH1D2/RSL24D1/METTL5/LSM6/UTP15/METTL18/MTERF3/FASTKD2/RPL35A/FCF1/WDR75/EXOSC8/RPP30/RPF1/MRPL22/DHX29/RPL27/TENT4B/WDR36/ERI1/NPM1/NOL11/UTP11/UTP23/BRIX1/ERI2/HEATR1/METTL15/RPL5/DCAF13/RPS21/ABCE1/UTP6/UTP20/NOP58/EXOSC9/RAN/WDR43/NIFK/DDX52/LTV1/DDX21/MRPS7/RPF2/MPHOSPH6/TFB2M/NOL8/METTL17/NAF1/ZNHIT3/SRFBP1/ABT1/MALSU1/XRCC5/MRPL44/RPS5/NHP2/RIOK1 | 58 |

| | | | | | | | | |
|---|---|---|---|---|---|---|---|---|
| GO:0000725 | recombinational repair | 38/1673 | 183/20772 | 4,92218E-08 | 1,3821E-05 | 1,28059E-05 | RMI1/FIGNL1/FAN1/MBTD1/CDC7/USP51/RAD51C/PARPBP/RBBP8/WRN/UBE2N/HUS1/KDM4D/CHEK1/MMS22L/WDR48/RAD51AP1/RPA2/BLM/ATM/PSMD14/RIF1/NSMCE4A/MRE11/NSMCE2/GEN1/FANCM/INIP/XRCC2/EPC1/MCM8/MEAF6/RUVBL1/PALB2/MCM6/XRCC5/KDM1A/XRCC6 | 38 |
| GO:0042073 | intraciliary transport | 17/1673 | 47/20772 | 6,17162E-08 | 1,57277E-05 | 1,45725E-05 | BBS12/IFT70B/DYNC2H1/IFT70A/DYNLL1/DYNLT2B/IFT88/LCA5/IFT56/SSX2IP/RPGR/WDR35/IFT52/TTC21B/PCM1/INTU/IFT22 | 17 |
| GO:0032259 | methylation | 64/1673 | 396/20772 | 6,19084E-08 | 1,57277E-05 | 1,45725E-05 | EOMES/BHMT/SNRPG/DYDC2/METTL5/SMAD4/SNRPF/FAM98B/EEF1AKMT1/ARMT1/RLF/RAMAC/WDR5B/METTL18/SMARCA5/SNRPE/PAX5/TRMT11/N6AMT1/PRMT9/VIRMA/TRMT13/PCMTD2/PCMT1/CAMKMT/RIF1/NSUN6/ETF1/EZH2/SMYD3/SETDB2/MTF2/DPH5/METTL15/MIS18A/PRDM5/TARBP1/WTAP/HENMT1/ATPSCKMT/KDM6A/NFYC/CMTR2/NDUFAF5/CLNS1A/KMT2E/MTR/BMT2/THUMPD3/METTL25/SNRPD2/NFYA/METTL2A/LARP7/SETMAR/BOD1/PRMT3/TFB2M/METTL17/PAXIP1/PRMT1/KANSL2/NSD1/KDM1A | 64 |
| GO:1904872 | regulation of telomerase RNA localization to Cajal body | 11/1673 | 20/20772 | 7,60315E-08 | 1,84376E-05 | 1,70834E-05 | CCT8/CCT2/CCT6A/CCT5/DCP2/CCT4/PARN/RUVBL1/NAF1/CCT3/NHP2 | 11 |
| GO:0000070 | mitotic sister chromatid segregation | 42/1673 | 217/20772 | 8,18172E-08 | 1,8978E-05 | 1,75841E-05 | TTK/KIF18A/SMARCA5/NUF2/DSCC1/DLGAP5/MIS12/KIF14/CLASP2/NCAPG/CENPE/KIF11/ABRAXAS1/ATM/LSM14A/SGO1/SPDL1/PDS5A/NSMCE2/NAA50/CDC16/KNTC1/KIF23/GEN1/SMC3/BUB1/BCCIP/WAPL/MAPRE1/ZNF207/RAN/CDCA8/STAG1/CCNB1/MAP9/DYNC1LI1/BOD1/TRIP13/CHAMP1/ANAPC7/TPR/SEH1L | 42 |

| | | | | | | | | |
|---|---|---|---|---|---|---|---|---|
| GO:0006364 | rRNA processing | 44/1673 | 237/20772 | 1,42335E-07 | 2,82004E-05 | 2,61291E-05 | PIH1D2/METTL5/LSM6/UTP15/METTL18/RPL35A/FCF1/WDR75/EXOSC8/RPP30/RPF1/RPL27/TENT4B/WDR36/ERI1/NOL11/UTP11/UTP23/BRIX1/ERI2/HEATR1/METTL15/RPL5/DCAF13/RPS21/UTP6/UTP20/NOP58/EXOSC9/WDR43/NIFK/DDX52/DDX21/RPF2/MPHOSPH6/TFB2M/NOL8/NAF1/ZNHIT3/SRFBP1/ABT1/MRPL44/NHP2/RIOK1 | 44 |
| GO:0090670 | RNA localization to Cajal body | 11/1673 | 21/20772 | 1,48006E-07 | 2,82004E-05 | 2,61291E-05 | CCT8/CCT2/CCT6A/CCT5/DCP2/CCT4/PARN/RUVBL1/NAF1/CCT3/NHP2 | 11 |
| GO:0090671 | telomerase RNA localization to Cajal body | 11/1673 | 21/20772 | 1,48006E-07 | 2,82004E-05 | 2,61291E-05 | CCT8/CCT2/CCT6A/CCT5/DCP2/CCT4/PARN/RUVBL1/NAF1/CCT3/NHP2 | 11 |
| GO:0090672 | telomerase RNA localization | 11/1673 | 21/20772 | 1,48006E-07 | 2,82004E-05 | 2,61291E-05 | CCT8/CCT2/CCT6A/CCT5/DCP2/CCT4/PARN/RUVBL1/NAF1/CCT3/NHP2 | 11 |
| GO:0090685 | RNA localization to nucleus | 11/1673 | 21/20772 | 1,48006E-07 | 2,82004E-05 | 2,61291E-05 | CCT8/CCT2/CCT6A/CCT5/DCP2/CCT4/PARN/RUVBL1/NAF1/CCT3/NHP2 | 11 |
| GO:0032392 | DNA geometric change | 23/1673 | 86/20772 | 1,95765E-07 | 3,53327E-05 | 3,27376E-05 | DHX36/RFC3/WRN/DSCC1/RFC4/BLM/RAD50/MRE11/ZRANB3/FANCM/RFC5/SSBP1/GTF2H2/TOP2A/MCM8/BRIP1/RUVBL1/HMGB2/DHX9/MCM6/DTD1/XRCC5/XRCC6 | 23 |
| GO:0071459 | protein localization to chromosome, centromeric region | 16/1673 | 45/20772 | 1,98685E-07 | 3,53327E-05 | 3,27376E-05 | TTK/CENPQ/H4C3/H4C11/H4C4/MIS12/MTBP/H4C8/SPDL1/KNTC1/KNL1/H4C9/H4C5/H4C2/BOD1/CHAMP1 | 16 |
| GO:0000724 | double-strand break repair via homologous recombination | 36/1673 | 179/20772 | 2,53387E-07 | 4,32163E-05 | 4,00421E-05 | RMI1/FIGNL1/FAN1/MBTD1/CDC7/USP51/RAD51C/PARPBP/RBBP8/WRN/UBE2N/HUS1/KDM4D/CHEK1/MMS22L/WDR48/RAD51AP1/RPA2/BLM/ATM/PSMD14/RIF1/NSMCE4A/MRE11/NSMCE2/GEN1/FANCM/INIP/XRCC2/EPC1/MCM8/MEAF6/RUVBL1/PALB2/MCM6/KDM1A | 36 |

| | | | | | | | | |
|---|---|---|---|---|---|---|---|---|
| GO:0043414 | macromolecule methylation | 57/1673 | 351/20772 | 2,73873E-07 | 4,32163E-05 | 4,00421E-05 | EOMES/BHMT/SNRPG/DYDC2/METTL5/SMAD4/SNRPF/FAM98B/EEF1AKMT1/ARMT1/RLF/RAMAC/WDR5B/METTL18/SMARCA5/SNRPE/PAX5/TRMT11/N6AMT1/PRMT9/VIRMA/TRMT13/PCMTD2/PCMT1/CAMKMT/RIF1/NSUN6/ETF1/EZH2/SMYD3/SETDB2/MTF2/METTL15/MIS18A/PRDM5/TARBP1/WTAP/HENMT1/ATPSCKMT/KDM6A/NFYC/CMTR2/CLNS1A/KMT2E/THUMPD3/SNRPD2/NFYA/METTL2A/LARP7/SETMAR/BOD1/PRMT3/TFB2M/PAXIP1/PRMT1/KANSL2/KDM1A | 57 |
| GO:0030071 | regulation of mitotic metaphase/anaphase transition | 24/1673 | 94/20772 | 2,75418E-07 | 4,32163E-05 | 4,00421E-05 | TTK/ACTL6B/NUF2/DLGAP5/CENPE/BCL7A/ATM/SPDL1/ANAPC4/NSMCE2/CDC16/KNTC1/PHF10/GEN1/BUB1/CDC26/ZNF207/CDCA8/CCNB1/DYNC1LI1/TRIP13/SMARCC1/ANAPC7/TPR | 24 |
| GO:0071103 | DNA conformation change | 24/1673 | 94/20772 | 2,75418E-07 | 4,32163E-05 | 4,00421E-05 | DHX36/RFC3/WRN/DSCC1/RFC4/BLM/RAD50/TOP2B/MRE11/ZRANB3/FANCM/RFC5/SSBP1/GTF2H2/TOP2A/MCM8/BRIP1/RUVBL1/HMGB2/DHX9/MCM6/DTD1/XRCC5/XRCC6 | 24 |
| GO:0006913 | nucleocytoplasmic transport | 58/1673 | 361/20772 | 3,12019E-07 | 4,62395E-05 | 4,28434E-05 | NXT2/RGPD6/EFCAB7/NEUROD1/NDC1/KPNA3/MAGOHB/CSE1L/MAGOH/NUP37/THOC1/TXN/RPL23/STRADB/XPOT/ENY2/NUP155/ATF2/SSB/XPO7/NUP107/KPNA2/NUP160/NUP153/NUP205/RANBP2/NPM1/XPO4/AHCTF1/RBM22/KPNA4/PIK3R1/ZIC1/BMP4/DUSP16/DUSP16/E2F3/NEMF/IPO5/YWHAE/STK4/ABCE1/GLI3/SMG7/TNPO1/RAN/NCBP1/STYX/HSPA9/LTV1/UPF2/NCBP2/IPO11/NF1/DHX9/TPR/POLR2D/SEH1L | 58 |
| GO:0051169 | nuclear transport | 58/1673 | 361/20772 | 3,12019E-07 | 4,62395E-05 | 4,28434E-05 | NXT2/RGPD6/EFCAB7/NEUROD1/NDC1/KPNA3/MAGOHB/CSE1L/MAGOH/NUP37/THOC1/TXN/RPL23/STRADB/XPOT/ENY2/NUP155/ATF2/SSB/XPO7/NUP107/KPNA2/NUP160/NUP153/NUP205/RANBP2/NPM1/XPO4/AHCTF1/RBM22/KPNA4/PIK3R1/ZIC1/BMP4/DUSP16/DUSP16/E2F3/NEMF/IPO5/YWHAE/STK4/ABCE1/GLI3/SMG7/TNPO1/RAN/NCBP1/STYX/HSPA9/LTV1/UPF2/NCBP2/IPO11/NF1/DHX9/TPR/POLR2D/SEH1L | 58 |

| | | | | | | | | |
|---|---|---|---|---|---|---|---|---|
| GO:0090305 | nucleic acid phosphodiester bond hydrolysis | 48/1673 | 277/20772 | 3,41819E-07 | 4,92866E-05 | 4,56666E-05 | RIDA/DCLRE1A/LACTB2/TATDN1/FAN1/CNOT7/ENPP2/N4BP2/RBBP8/WRN/FCF1/EXOSC8/DICER1/RPP30/ATAD5/ZC3H12B/RAD50/ANGEL2/ASTE1/ERI1/TSEN34/UTP23/MRE11/ZRANB3/ERI2/GEN1/EXO1/FANCM/POLR2I/RPS21/UTP20/EXOSC9/HARBI1/N4BP1/NCBP1/DCP2/CPSF3/CNOT1/SETMAR/NCBP2/PARN/PCF11/NUDT21/ABT1/APEX1/MRPL44/POP5/NHP2 | 48 |
| GO:0000956 | nuclear-transcribed mRNA catabolic process | 32/1673 | 152/20772 | 3,95852E-07 | 5,55756E-05 | 5,14937E-05 | DHX36/MAGOHB/MAGOH/CNOT7/EXOSC8/ZCCHC7/TUT4/SSB/MTPAP/ATM/TENT4B/TENT2/ETF1/EXOSC9/SMG7/NBDY/NCBP1/NBAS/SECISBP2/DCP2/NANOS1/NT5C3B/CNOT1/UPF2/NCBP2/POLR2G/PARN/RC3H1/TNRC6C/CNOT9/DHX9/POLR2D | 32 |
| GO:0007091 | metaphase/anaphase transition of mitotic cell cycle | 24/1673 | 98/20772 | 6,31259E-07 | 8,41942E-05 | 7,80103E-05 | TTK/ACTL6B/NUF2/DLGAP5/CENPE/BCL7A/ATM/SPDL1/ANAPC4/NSMCE2/CDC16/KNTC1/PHF10/GEN1/BUB1/CDC26/ZNF207/CDCA8/CCNB1/DYNC1LI1/TRIP13/SMARCC1/ANAPC7/TPR | 24 |
| GO:1902099 | regulation of metaphase/anaphase transition of cell cycle | 24/1673 | 98/20772 | 6,31259E-07 | 8,41942E-05 | 7,80103E-05 | TTK/ACTL6B/NUF2/DLGAP5/CENPE/BCL7A/ATM/SPDL1/ANAPC4/NSMCE2/CDC16/KNTC1/PHF10/GEN1/BUB1/CDC26/ZNF207/CDCA8/CCNB1/DYNC1LI1/TRIP13/SMARCC1/ANAPC7/TPR | 24 |
| GO:0016126 | sterol biosynthetic process | 19/1673 | 66/20772 | 6,47827E-07 | 8,42965E-05 | 7,81052E-05 | CES1/PRKAA2/SC5D/IDI1/MSMO1/INSIG2/HMGCS1/CYP51A1/HSD17B7/HMGCR/FDFT1/LBR/SQLE/ERG28/ERLIN1/MBTPS2/ACAA2/FDPS/NSDHL | 19 |
| GO:0009451 | RNA modification | 36/1673 | 186/20772 | 6,73537E-07 | 8,55552E-05 | 7,92714E-05 | SNRPG/DPH3/METTL5/DTWD1/SNRPF/RAMAC/OSGEPL1/ADAT2/SNRPE/TRMT11/TYW5/ELP2/SSB/VIRMA/ELP4/CDKAL1/TRMT13/NSUN6/PUS7L/METTL15/SEPSECS/TARBP1/WTAP/HENMT1/YRDC/CMTR2/THUMPD3/ELP1/SNRPD2/METTL2A/LARP7/CDK5RAP1/PARN/TFB2M/NAF1/NHP2 | 36 |

| | | | | | | | | |
|---|---|---|---|---|---|---|---|---|
| GO:0006399 | tRNA metabolic process | 41/1673 | 226/20772 | 7,05921E-07 | 8,75834E-05 | 8,11507E-05 | DPH3/DTWD1/FAM98B/LSM6/OSGEPL1/ADAT2/YARS2/EXOSC8/DICER1/TRMT11/RPP30/TYW5/ZCCHC7/ELP2/TSEN15/SSB/FARSB/ELP4/CDKAL1/TRMT13/NARS2/NSUN6/TSEN34/RARS1/QRSL1/SEPSECS/IARS2/TARBP1/YRDC/EPRS1/EXOSC9/THUMPD3/ELP1/TRNT1/METTL2A/DARS2/IARS1/CDK5RAP1/DTD1/LARS1/POP5 | 41 |
| GO:0032508 | DNA duplex unwinding | 21/1673 | 80/20772 | 9,21247E-07 | 0,000111701 | 0,000103497 | DHX36/RFC3/WRN/DSCC1/RFC4/BLM/RAD50/MRE11/FANCM/RFC5/SSBP1/GTF2H2/TOP2A/MCM8/BRIP1/RUVBL1/DHX9/MCM6/DTD1/XRCC5/XRCC6 | 21 |
| GO:1904874 | positive regulation of telomerase RNA localization to Cajal body | 9/1673 | 16/20772 | 9,48263E-07 | 0,000112422 | 0,000104165 | CCT8/CCT2/CCT6A/CCT5/CCT4/RUVBL1/NAF1/CCT3/NHP2 | 9 |
| GO:0008334 | histone mRNA metabolic process | 10/1673 | 20/20772 | 9,74173E-07 | 0,000112983 | 0,000104685 | SSB/MTPAP/ATM/TENT4B/TENT2/NCBP1/DCP2/CPSF3/NCBP2/LSM11 | 10 |
| GO:0051236 | establishment of RNA localization | 35/1673 | 182/20772 | 1,11984E-06 | 0,000127114 | 0,000117778 | NXT2/NDC1/MAGOHB/MAGOH/NUP37/ATR/THOC1/CETN3/XPOT/ENY2/NUP155/SSB/NUP107/ATM/LRPPRC/NUP160/NUP153/NUP205/RANBP2/NPM1/AHCTF1/PARP11/TERF1/SRSF1/SMG7/RAN/NCBP1/SRSF7/UPF2/NCBP2/DHX9/TPR/POLR2D/IGF2BP3/SEH1L | 35 |
| GO:0140014 | mitotic nuclear division | 51/1673 | 315/20772 | 1,2542E-06 | 0,000139399 | 0,000129161 | TTK/NPM2/BORA/KIF18A/PHIP/SMARCA5/NUF2/DSCC1/CHEK1/DLGAP5/MIS12/MTBP/KIF14/CLASP2/NCAPG/CENPE/KIF11/ABRAXAS1/ATM/LSM14A/SGO1/SPDL1/SPAST/PDS5A/NSMCE2/NAA50/CDC16/KNTC1/KIF23/GEN1/SMC3/BUB1/BCCIP/WAPL/BMP4/MAPRE1/ZNF207/RAN/CDCA8/CDC25C/STAG1/CCNB1/TOM1L1/MAP9/DYNC1LI1/BOD1/TRIP13/CHAMP1/ANAPC7/TPR/SEH1L | 51 |

| | | | | | | | | |
|---|---|---|---|---|---|---|---|---|
| GO:0071826 | ribonucleoprotein complex subunit organization | 46/1673 | 273/20772 | 1,33867E-06 | 0,000145574 | 0,000134882 | RNVU1-15/RNVU1-6/SNRPG/PTBP2/PIH1D2/SNRPF/CELF6/ATR/RAMAC/FASTKD2/SNRPE/DICER1/PRPF39/SRSF12/SNRPB2/ZFAND1/DHX29/ATM/EIF2S3/NUFIP1/MCTS1/BRIX1/RPL5/DDX20/SRPK1/CLNS1A/SRSF1/NCBP1/GEMIN2/PTGES3/DENR/SNRPD2/MRPS7/SNRPA1/RPF2/RUVBL1/EIF2S2/NAF1/NUDT21/ZNHIT3/DHX9/ABT1/XRCC5/POLR2D/SF3B4/RPS5 | 46 |
| GO:0044784 | metaphase/anaphase transition of cell cycle | 24/1673 | 102/20772 | 1,37331E-06 | 0,000145574 | 0,000134882 | TTK/ACTL6B/NUF2/DLGAP5/CENPE/BCL7A/ATM/SPDL1/ANAPC4/NSMCE2/CDC16/KNTC1/PHF10/GEN1/BUB1/CDC26/ZNF207/CDCA8/CCNB1/DYNC1LI1/TRIP13/SMARCC1/ANAPC7/TPR | 24 |
| GO:0022618 | ribonucleoprotein complex assembly | 45/1673 | 265/20772 | 1,3977E-06 | 0,000145574 | 0,000134882 | RNVU1-15/RNVU1-6/SNRPG/PTBP2/PIH1D2/SNRPF/CELF6/ATR/RAMAC/FASTKD2/SNRPE/DICER1/PRPF39/SRSF12/SNRPB2/DHX29/ATM/EIF2S3/NUFIP1/MCTS1/BRIX1/RPL5/DDX20/SRPK1/CLNS1A/SRSF1/NCBP1/GEMIN2/PTGES3/DENR/SNRPD2/MRPS7/SNRPA1/RPF2/RUVBL1/EIF2S2/NAF1/NUDT21/ZNHIT3/DHX9/ABT1/XRCC5/POLR2D/SF3B4/RPS5 | 45 |
| GO:0031124 | mRNA 3'-end processing | 18/1673 | 63/20772 | 1,43518E-06 | 0,000145574 | 0,000134882 | DHX36/PAPOLG/PAPOLA/MTPAP/VIRMA/TENT4B/TENT2/WDR33/RPRD1B/NCBP1/RPRD1A/CPSF3/SNRPA/NCBP2/PCF11/NUDT21/POLR2D/LSM11 | 18 |
| GO:0045839 | negative regulation of mitotic nuclear division | 17/1673 | 57/20772 | 1,44619E-06 | 0,000145574 | 0,000134882 | TTK/NUF2/CHEK1/MTBP/ATM/SPDL1/KNTC1/GEN1/BUB1/BMP4/ZNF207/CDCA8/CCNB1/TOM1L1/DYNC1LI1/TRIP13/TPR | 17 |

| | | | | | | | | |
|---|---|---|---|---|---|---|---|---|
| GO:0071824 | protein-DNA complex subunit organization | 48/1673 | 292/20772 | 1,64476E-06 | 0,000161244 | 0,000149401 | H1-6/H3-3A/H1-1/H4C3/SHPRH/HAT1/RAD51C/H4C11/SMARCA5/H4C4/CENPH/RPL23/CAND1/MED23/CHD1/DLGAP5/MIS12/TAF5/MED6/MED7/CENPE/ASF1A/OIP5/H4C8/NPM1/CENPW/PSMC6/SMYD3/NAP1L5/KNTC1/MIS18A/GMNN/TAF12/TAF13/HJURP/TCF4/TERF1/ITGB3BP/MED14/GTF2A1/TSPYL4/H4C9/H4C5/H4C2/TBP/SENP6/SMARCC1/HMGB2 | 48 |
| GO:0051168 | nuclear export | 35/1673 | 185/20772 | 1,66231E-06 | 0,000161244 | 0,000149401 | NXT2/MAGOHB/CSE1L/MAGOH/THOC1/TXN/STRADB/XPOT/ENY2/NUP155/SSB/XPO7/NUP107/NUP160/NUP153/RANBP2/NPM1/XPO4/RBM22/DUSP16/DUSP16/NEMF/YWHAE/ABCE1/SMG7/RAN/NCBP1/STYX/HSPA9/LTV1/UPF2/NCBP2/DHX9/TPR/POLR2D | 35 |
| GO:0044782 | cilium organization | 61/1673 | 410/20772 | 2,22662E-06 | 0,000212125 | 0,000196545 | CFAP61/DNAH7/PIFO/RSPH4A/NME5/SNX10/BBS10/BBS12/DNAAF3/DNAAF11/IFT70B/DYNC2H1/IFT70A/DYNLL1/CEP78/CEP70/PIERCE1/DYNLT2B/IQCB1/IFT88/LCA5/TMEM237/CFAP221/TMEM216/ABLIM1/ARMC2/GDI2/RFX3/BBS4/IFT56/CCDC66/SSX2IP/CEP290/NEK1/CEP126/FAM149B1/TBC1D7/RPGR/ODAD2/BBS7/KIAA0586/WDR35/CCP110/IFT52/CEP83/MAPRE1/HYDIN/HYDIN/TBC1D31/TTC21B/RO60/PLK4/KIF3A/PCM1/MPHOSPH9/KIF24/CPLANE1/OFD1/INTU/DNAL1/IFT22 | 61 |
| GO:0065004 | protein-DNA complex assembly | 45/1673 | 270/20772 | 2,35306E-06 | 0,00021884 | 0,000202767 | H1-6/H3-3A/H1-1/H4C3/SHPRH/HAT1/RAD51C/H4C11/SMARCA5/H4C4/CENPH/CAND1/MED23/DLGAP5/MIS12/TAF5/MED6/MED7/CENPE/ASF1A/OIP5/H4C8/NPM1/CENPW/PSMC6/SMYD3/NAP1L5/KNTC1/MIS18A/GMNN/TAF12/TAF13/HJURP/TCF4/TERF1/ITGB3BP/MED14/GTF2A1/TSPYL4/H4C9/H4C5/H4C2/TBP/SENP6/HMGB2 | 45 |
| GO:0006695 | cholesterol biosynthetic process | 17/1673 | 59/20772 | 2,46118E-06 | 0,00021884 | 0,000202767 | CES1/PRKAA2/SC5D/IDI1/MSMO1/INSIG2/HMGCS1/CYP51A1/HSD17B7/HMGCR/FDFT1/LBR/ERLIN1/MBTPS2/ACAA2/FDPS/NSDHL | 17 |

| | | | | | | | | |
|---|---|---|---|---|---|---|---|---|
| GO:0010965 | regulation of mitotic sister chromatid separation | 17/1673 | 59/20772 | 2,46118E-06 | 0,00021884 | 0,000202767 | TTK/NUF2/DLGAP5/ATM/SPDL1/NSMCE2/CDC16/KNTC1/GEN1/BUB1/ZNF207/CDCA8/CCNB1/DYNC1LI1/TRIP13/ANAPC7/TPR | 17 |
| GO:1902653 | secondary alcohol biosynthetic process | 17/1673 | 59/20772 | 2,46118E-06 | 0,00021884 | 0,000202767 | CES1/PRKAA2/SC5D/IDI1/MSMO1/INSIG2/HMGCS1/CYP51A1/HSD17B7/HMGCR/FDFT1/LBR/ERLIN1/MBTPS2/ACAA2/FDPS/NSDHL | 17 |
| GO:0000723 | telomere maintenance | 34/1673 | 181/20772 | 2,74646E-06 | 0,000240202 | 0,00022256 | DHX36/DCLRE1A/ATR/RAD51C/WRN/CCT8/HUS1/RPA2/BLM/RAD50/ATM/TENT4B/RIF1/NSMCE4A/MRE11/NSMCE2/EXO1/CCT2/CCT6A/TERF1/CCT5/DCP2/CCT4/PTGES3/GNL3/CCNE1/PARN/RUVBL1/NAF1/CCT3/APEX1/XRCC5/XRCC6/NHP2 | 34 |
| GO:0006999 | nuclear pore organization | 8/1673 | 14/20772 | 3,36345E-06 | 0,000284826 | 0,000263906 | NDC1/NUP107/NUP153/NUP205/AHCTF1/TMEM170A/TPR/SEH1L | 8 |
| GO:1904814 | regulation of protein localization to chromosome, telomeric region | 8/1673 | 14/20772 | 3,36345E-06 | 0,000284826 | 0,000263906 | CCT8/CCT2/CCT6A/TERF1/CCT5/CCT4/GNL3/CCT3 | 8 |
| GO:1903311 | regulation of mRNA metabolic process | 54/1673 | 353/20772 | 3,63713E-06 | 0,000303189 | 0,000280921 | DHX36/RIDA/ELAVL4/MAGOHB/MAGOH/CELF6/CNOT7/RBM7/PAPOLA/KHDRBS2/FASTKD1/FASTKD2/SRRM4/EXOSC8/CWC22/RBFOX2/SRSF12/TUT4/VIRMA/ANGEL2/TENT4B/TRA2B/NPM1/PTCD2/RBMX/TRIM71/WTAP/SRPK1/CLNS1A/EXOSC9/TRA2A/NCBP1/NBAS/SECISBP2/DCP2/NANOS1/NT5C3B/SRSF7/SNRPA/CNOT1/LARP7/NCBP2/POLR2G/PARN/HSPA8/RC3H1/TNRC6C/NUDT21/CNOT9/DHX9/APEX1/POLR2D/IGF2BP3/SF3B4 | 54 |
| GO:0033045 | regulation of sister chromatid segregation | 24/1673 | 108/20772 | 4,03603E-06 | 0,000331265 | 0,000306934 | TTK/ACTL6B/NUF2/DLGAP5/CENPE/BCL7A/ATM/SPDL1/ANAPC4/NSMCE2/CDC16/KNTC1/PHF10/GEN1/BUB1/CDC26/ZNF207/CDCA8/CCNB1/DYNC1LI1/TRIP13/SMARCC1/ANAPC7/TPR | 24 |

| | | | | | | | | |
|---|---|---|---|---|---|---|---|---|
| GO:0031503 | protein-containing complex localization | 35/1673 | 194/20772 | 5,09129E-06 | 0,000408731 | 0,000378711 | RELN/NDC1/BBS12/IFT70B/DYNC2H1/IFT70A/DYNLL1/ATR/DYNLT2B/IFT88/LCA5/ARHGAP44/SNAPIN/ERBB4/ATM/IFT56/SSX2IP/NSG1/MIOS/NPM1/SLC1A1/NETO2/RPGR/WDR35/IFT52/ABCE1/TERF1/TTC21B/RAN/PCM1/LTV1/VPS35/INTU/IFT22/SEH1L | 35 |
| GO:0051306 | mitotic sister chromatid separation | 17/1673 | 62/20772 | 5,19963E-06 | 0,000408731 | 0,000378711 | TTK/NUF2/DLGAP5/ATM/SPDL1/NSMCE2/CDC16/KNTC1/GEN1/BUB1/ZNF207/CDCA8/CCNB1/DYNC1LI1/TRIP13/ANAPC7/TPR | 17 |
| GO:0070203 | regulation of establishment of protein localization to telomere | 7/1673 | 11/20772 | 5,36292E-06 | 0,000408731 | 0,000378711 | CCT8/CCT2/CCT6A/TERF1/CCT5/CCT4/CCT3 | 7 |
| GO:1904869 | regulation of protein localization to Cajal body | 7/1673 | 11/20772 | 5,36292E-06 | 0,000408731 | 0,000378711 | CCT8/CCT2/CCT6A/CCT5/CCT4/LARP7/CCT3 | 7 |
| GO:1904871 | positive regulation of protein localization to Cajal body | 7/1673 | 11/20772 | 5,36292E-06 | 0,000408731 | 0,000378711 | CCT8/CCT2/CCT6A/CCT5/CCT4/LARP7/CCT3 | 7 |
| GO:0016072 | rRNA metabolic process | 45/1673 | 279/20772 | 5,74092E-06 | 0,000431378 | 0,000399694 | PIH1D2/METTL5/LSM6/UTP15/METTL18/RPL35A/FCF1/WDR75/EXOSC8/RPP30/RPF1/ZCCHC7/RPL27/TENT4B/WDR36/ERI1/NOL11/UTP11/UTP23/BRIX1/ERI2/HEATR1/METTL15/RPL5/DCAF13/RPS21/UTP6/UTP20/NOP58/EXOSC9/WDR43/NIFK/DDX52/DDX21/RPF2/MPHOSPH6/TFB2M/NOL8/NAF1/ZNHIT3/SRFBP1/ABT1/MRPL44/NHP2/RIOK1 | 45 |

| | | | | | | | | |
|---|---|---|---|---|---|---|---|---|
| GO:2001020 | regulation of response to DNA damage stimulus | 53/1673 | 350/20772 | 6,02632E-06 | 0,000446533 | 0,000413737 | EEF1E1/ACTL6B/FIGNL1/EYA4/UBE2V2/TAF9B/TADA1/ETAA1/MBTD1/ATR/ARMT1/ERCC8/USP1/USP51/THOC1/PARPBP/RBBP8/SMARCA5/UBE2N/KDM4D/CHEK1/TAF5/ATAD5/ENY2/HMGA2/WDR48/RAD51AP1/RPA2/BCL7A/ABRAXAS1/SUPT3H/ATM/RIF1/NSMCE4A/CEBPG/RNF169/TRIAP1/PHF10/MARCHF6-DT/COPS3/TAF12/PARG/EPC1/TTI1/MEAF6/SETMAR/EYA3/SMARCC1/RUVBL1/PAXIP1/DHX9/ATRIP/KDM1A | 53 |
| GO:0006282 | regulation of DNA repair | 40/1673 | 237/20772 | 6,14005E-06 | 0,000448728 | 0,00041577 | ACTL6B/FIGNL1/EYA4/UBE2V2/TADA1/MBTD1/ATR/ERCC8/USP1/USP51/PARPBP/RBBP8/UBE2N/KDM4D/CHEK1/TAF5/ENY2/HMGA2/WDR48/RAD51AP1/RPA2/BCL7A/ABRAXAS1/SUPT3H/RIF1/CEBPG/RNF169/PHF10/MARCHF6-DT/TAF12/PARG/EPC1/MEAF6/SETMAR/EYA3/SMARCC1/RUVBL1/DHX9/ATRIP/KDM1A | 40 |
| GO:0051028 | mRNA transport | 28/1673 | 142/20772 | 7,8152E-06 | 0,000558725 | 0,000517688 | NXT2/NDC1/MAGOHB/MAGOH/NUP37/THOC1/CETN3/ENY2/NUP155/NUP107/LRPPRC/NUP160/NUP153/NUP205/RANBP2/AHCTF1/PARP11/SRSF1/SMG7/NCBP1/SRSF7/UPF2/NCBP2/DHX9/TPR/POLR2D/IGF2BP3/SEH1L | 28 |
| GO:0018205 | peptidyl-lysine modification | 60/1673 | 417/20772 | 7,85461E-06 | 0,000558725 | 0,000517688 | EOMES/PRKAA2/ACTL6B/DYDC2/KIAA1586/TADA1/HAT1/SMAD4/HDAC2/MBTD1/EEF1AKMT1/MAP3K7/RLF/NDUFAB1/WDR5B/NNAT/METTL18/SMARCA5/SENP1/RWDD3/PAX5/CHEK1/TAF5/ENY2/N6AMT1/ATF2/SUPT3H/CAMKMT/RIF1/NSMCE4A/SUMO2/RANBP2/EZH2/UHRF2/SMYD3/SETDB2/NSMCE2/NAA50/UBA2/MTF2/PRDM5/ATPSCKMT/KDM6A/NFYC/TAF12/MSL3P1/EPC1/GNL3/MEAF6/NFYA/SETMAR/BOD1/DDX21/SENP6/ZNF451/RUVBL1/PAXIP1/SENP5/KANSL2/KDM1A | 60 |
| GO:0051784 | negative regulation of nuclear division | 17/1673 | 64/20772 | 8,30653E-06 | 0,000583097 | 0,00054027 | TTK/NUF2/CHEK1/MTBP/ATM/SPDL1/KNTC1/GEN1/BUB1/BMP4/ZNF207/CDCA8/CCNB1/TOM1L1/DYNC1LI1/TRIP13/TPR | 17 |
| GO:0036297 | interstrand cross-link repair | 13/1673 | 40/20772 | 8,71237E-06 | 0,000603643 | 0,000559307 | DCLRE1A/FAN1/FANCL/ATR/NEIL3/RAD51AP1/ERCC6L2/CENPS/FANCM/FANCF/MCM8/FANCI/FANCC | 13 |

| | | | | | | | | |
|---|---|---|---|---|---|---|---|---|
| GO:0031297 | replication fork processing | 14/1673 | 46/20772 | 9,35797E-06 | 0,000640061 | 0,00059305 | ETAA1/ATR/THOC1/RBBP8/WRN/MMS22L/BLM/RAD50/CENPS/MRE11/ZRANB3/GEN1/FANCM/SETMAR | 14 |
| GO:0015931 | nucleobase-containing compound transport | 40/1673 | 243/20772 | 1,13573E-05 | 0,0007514 | 0,000696212 | RSC1A1/NXT2/NDC1/MAGOHB/MAGOH/NUP37/THOC1/LRRC8C/SLC25A24/SLC25A24/CETN3/XPOT/ENY2/NUP155/SSB/NUP107/SLC35A3/LRPPRC/NUP160/NUP153/NUP205/RANBP2/NPM1/AHCTF1/PARP11/SLC29A1/SRSF1/SMG7/RAN/NCBP1/SRSF7/UPF2/NCBP2/SLC25A4/SLC25A33/DHX9/TPR/POLR2D/IGF2BP3/SEH1L | 40 |
| GO:0000018 | regulation of DNA recombination | 27/1673 | 137/20772 | 1,13872E-05 | 0,0007514 | 0,000696212 | H1-6/FIGNL1/H1-1/MBTD1/SMARCAD1/USP51/THOC1/PARPBP/RBBP8/MSH2/CHEK1/ATAD5/WDR48/RAD51AP1/RPA2/BLM/RAD50/KPNA2/RIF1/MRE11/ZRANB3/EPC1/MEAF6/MSH3/RUVBL1/PAXIP1/KDM1A | 27 |
| GO:0034508 | centromere complex assembly | 11/1673 | 30/20772 | 1,16079E-05 | 0,0007514 | 0,000696212 | CENPH/DLGAP5/MIS12/CENPE/OIP5/CENPW/KNTC1/MIS18A/HJURP/ITGB3BP/SENP6 | 11 |
| GO:0070202 | regulation of establishment of protein localization to chromosome | 7/1673 | 12/20772 | 1,19717E-05 | 0,0007514 | 0,000696212 | CCT8/CCT2/CCT6A/TERF1/CCT5/CCT4/CCT3 | 7 |
| GO:1903405 | protein localization to nuclear body | 7/1673 | 12/20772 | 1,19717E-05 | 0,0007514 | 0,000696212 | CCT8/CCT2/CCT6A/CCT5/CCT4/LARP7/CCT3 | 7 |
| GO:1904816 | positive regulation of protein localization to chromosome, telomeric region | 7/1673 | 12/20772 | 1,19717E-05 | 0,0007514 | 0,000696212 | CCT8/CCT2/CCT6A/CCT5/CCT4/GNL3/CCT3 | 7 |
| GO:1904867 | protein localization to Cajal body | 7/1673 | 12/20772 | 1,19717E-05 | 0,0007514 | 0,000696212 | CCT8/CCT2/CCT6A/CCT5/CCT4/LARP7/CCT3 | 7 |

| | | | | | | | | |
|---|---|---|---|---|---|---|---|---|
| GO:0043161 | proteasome-mediated ubiquitin-dependent protein catabolic process | 67/1673 | 489/20772 | 1,24851E-05 | 0,000757666 | 0,000702017 | YOD1/RNF175/KBTBD7/PSMA1/CLGN/SOCS4/PSMA3/GLMN/DNAJB9/FEM1C/ERCC8/SEC61B/PSMA4/SKP2/PSMA6/CUL2/KIF14/NHLRC3/TBL1XR1/RBX1/PSMA2/ANAPC13/FBXO3/USP14/COP1/FBXO38/USP9X/CD2AP/PSMD14/ARMC8/SUMO2/PSMC6/ANAPC4/PSMD12/UBQLN2/CDC16/TRIM71/AMN1/BBS7/FAF1/DNAJC10/CDC26/PSMD1/BTRC/FBXO45/PPP2CB/UBR3/PCNP/RAD23B/SPOPL/PELI1/NEMF/PLAA/UBQLN1/N4BP1/MARCHF6/DERL2/PSMD7/STYX/ERLIN1/DCAF12/ZYG11B/PSMB1/JKAMP/SMARCC1/ANAPC7/UBE2D3 | 67 |
| GO:0000377 | RNA splicing, via transesterification reactions with bulged adenosine as nucleophile | 54/1673 | 368/20772 | 1,24976E-05 | 0,000757666 | 0,000702017 | RNVU1-15/RNVU1-6/SNRPG/PTBP2/LSM5/BCAS2/MAGOHB/SNRPF/U2AF1/MAGOH/LSM6/CELF6/AQR/RBM7/LSM3/KHDRBS2/PPIL1/SRRM4/SNRPE/CWC22/RBFOX2/PRPF39/SRSF12/SNRPB2/CDC5L/TRA2B/SRSF8/CDC40/RBMX/RBM22/WTAP/WBP4/DDX20/SRPK1/RBM41/CLNS1A/CWF19L2/CDK13/SRSF1/TRA2A/NCBP1/PPIH/GEMIN2/SNRPD2/SRSF7/SNRPA/SNRNP40/LARP7/NCBP2/SNRPA1/HSPA8/DHX9/KDM1A/SF3B4 | 54 |
| GO:0000398 | mRNA splicing, via spliceosome | 54/1673 | 368/20772 | 1,24976E-05 | 0,000757666 | 0,000702017 | RNVU1-15/RNVU1-6/SNRPG/PTBP2/LSM5/BCAS2/MAGOHB/SNRPF/U2AF1/MAGOH/LSM6/CELF6/AQR/RBM7/LSM3/KHDRBS2/PPIL1/SRRM4/SNRPE/CWC22/RBFOX2/PRPF39/SRSF12/SNRPB2/CDC5L/TRA2B/SRSF8/CDC40/RBMX/RBM22/WTAP/WBP4/DDX20/SRPK1/RBM41/CLNS1A/CWF19L2/CDK13/SRSF1/TRA2A/NCBP1/PPIH/GEMIN2/SNRPD2/SRSF7/SNRPA/SNRNP40/LARP7/NCBP2/SNRPA1/HSPA8/DHX9/KDM1A/SF3B4 | 54 |
| GO:0050657 | nucleic acid transport | 32/1673 | 179/20772 | 1,54085E-05 | 0,000903639 | 0,000837269 | NXT2/NDC1/MAGOHB/MAGOH/NUP37/THOC1/CETN3/XPOT/ENY2/NUP155/SSB/NUP107/LRPPRC/NUP160/NUP153/NUP205/RANBP2/NPM1/AHCTF1/PARP11/SRSF1/SMG7/RAN/NCBP1/SRSF7/UPF2/NCBP2/DHX9/TPR/POLR2D/IGF2BP3/SEH1L | 32 |

| | | | | | | | | |
|---|---|---|---|---|---|---|---|---|
| GO:0050658 | RNA transport | 32/1673 | 179/20772 | 1,54085E-05 | 0,000903639 | 0,000837269 | NXT2/NDC1/MAGOHB/MAGOH/NUP37/THOC1/CETN3/XPOT/ENY2/NUP155/SSB/NUP107/LRPPRC/NUP160/NUP153/NUP205/RANBP2/NPM1/AHCTF1/PARP11/SRSF1/SMG7/RAN/NCBP1/SRSF7/UPF2/NCBP2/DHX9/TPR/POLR2D/IGF2BP3/SEH1L | 32 |
| GO:0050684 | regulation of mRNA processing | 28/1673 | 147/20772 | 1,54135E-05 | 0,000903639 | 0,000837269 | DHX36/MAGOHB/MAGOH/CELF6/RBM7/PAPOLA/KHDRBS2/SRRM4/CWC22/RBFOX2/SRSF12/VIRMA/TRA2B/PTCD2/RBMX/WTAP/SRPK1/CLNS1A/TRA2A/NCBP1/SRSF7/SNRPA/LARP7/NCBP2/HSPA8/NUDT21/DHX9/SF3B4 | 28 |
| GO:0006479 | protein methylation | 35/1673 | 204/20772 | 1,58802E-05 | 0,000910976 | 0,000844068 | BHMT/DYDC2/SMAD4/FAM98B/EEF1AKMT1/ARMT1/RLF/WDR5B/METTL18/SMARCA5/PAX5/N6AMT1/PRMT9/PCMTD2/PCMT1/CAMKMT/RIF1/ETF1/EZH2/SMYD3/SETDB2/MTF2/PRDM5/ATPSCKMT/KDM6A/NFYC/CLNS1A/NFYA/SETMAR/BOD1/PRMT3/PAXIP1/PRMT1/KANSL2/KDM1A | 35 |
| GO:0008213 | protein alkylation | 35/1673 | 204/20772 | 1,58802E-05 | 0,000910976 | 0,000844068 | BHMT/DYDC2/SMAD4/FAM98B/EEF1AKMT1/ARMT1/RLF/WDR5B/METTL18/SMARCA5/PAX5/N6AMT1/PRMT9/PCMTD2/PCMT1/CAMKMT/RIF1/ETF1/EZH2/SMYD3/SETDB2/MTF2/PRDM5/ATPSCKMT/KDM6A/NFYC/CLNS1A/NFYA/SETMAR/BOD1/PRMT3/PAXIP1/PRMT1/KANSL2/KDM1A | 35 |
| GO:0006303 | double-strand break repair via nonhomologous end joining | 17/1673 | 67/20772 | 1,60992E-05 | 0,000913715 | 0,000846605 | DCLRE1A/ERCC8/USP51/LIG4/KDM4D/HMGA2/ATM/PSMD14/RIF1/MRE11/XRCC4/NHEJ1/POLB/SETMAR/HMGB2/XRCC5/XRCC6 | 17 |
| GO:0000729 | DNA double-strand break processing | 9/1673 | 21/20772 | 1,68024E-05 | 0,000924826 | 0,0008569 | UBE2V2/SMARCAD1/RBBP8/UBE2N/BLM/RAD50/MRE11/BRIP1/SETMAR | 9 |
| GO:0036260 | RNA capping | 9/1673 | 21/20772 | 1,68024E-05 | 0,000924826 | 0,0008569 | SNRPG/SNRPF/RAMAC/SNRPE/CMTR2/UTP20/NCBP1/SNRPD2/ABT1 | 9 |

| | | | | | | | | |
|---|---|---|---|---|---|---|---|---|
| GO:0060271 | cilium assembly | 55/1673 | 381/20772 | 1,6815E-05 | 0,000924826 | 0,0008569 | DNAH7/RSPH4A/NME5/SNX10/BBS10/DNAAF3/DNAAF11/IFT70B/DYNC2H1/DYNLL1/CEP70/PIERCE1/DYNLT2B/IQCB1/IFT88/TMEM237/CFAP221/TMEM216/ABLIM1/ARMC2/GDI2/RFX3/BBS4/IFT56/CCDC66/SSX2IP/CEP290/NEK1/CEP126/FAM149B1/TBC1D7/RPGR/ODAD2/BBS7/KIAA0586/WDR35/CCP110/IFT52/CEP83/MAPRE1/HYDIN/HYDIN/TBC1D31/TTC21B/RO60/PLK4/KIF3A/PCM1/MPHOSPH9/KIF24/CPLANE1/OFD1/INTU/DNAL1/IFT22 | 55 |
| GO:0000375 | RNA splicing, via transesterification reactions | 54/1673 | 372/20772 | 1,70597E-05 | 0,000928712 | 0,0008605 | RNVU1-15/RNVU1-6/SNRPG/PTBP2/LSM5/BCAS2/MAGOHB/SNRPF/U2AF1/MAGOH/LSM6/CELF6/AQR/RBM7/LSM3/KHDRBS2/PPIL1/SRRM4/SNRPE/CWC22/RBFOX2/PRPF39/SRSF12/SNRPB2/CDC5L/TRA2B/SRSF8/CDC40/RBMX/RBM22/WTAP/WBP4/DDX20/SRPK1/RBM41/CLNS1A/CWF19L2/CDK13/SRSF1/TRA2A/NCBP1/PPIH/GEMIN2/SNRPD2/SRSF7/SNRPA/SNRNP40/LARP7/NCBP2/SNRPA1/HSPA8/DHX9/KDM1A/SF3B4 | 54 |
| GO:0007004 | telomere maintenance via telomerase | 18/1673 | 75/20772 | 2,08911E-05 | 0,001114539 | 0,001032679 | ATR/CCT8/RAD50/ATM/TENT4B/MRE11/CCT2/CCT6A/TERF1/CCT5/DCP2/CCT4/PTGES3/PARN/NAF1/CCT3/XRCC5/NHP2 | 18 |
| GO:1905818 | regulation of chromosome separation | 18/1673 | 75/20772 | 2,08911E-05 | 0,001114539 | 0,001032679 | TTK/NUF2/DLGAP5/NCAPG/ATM/SPDL1/NSMCE2/CDC16/KNTC1/GEN1/BUB1/ZNF207/CDCA8/CCNB1/DYNC1LI1/TRIP13/ANAPC7/TPR | 18 |
| GO:0051304 | chromosome separation | 19/1673 | 82/20772 | 2,12514E-05 | 0,001122535 | 0,001040088 | TTK/SMARCAD1/NUF2/DLGAP5/NCAPG/ATM/SPDL1/NSMCE2/CDC16/KNTC1/GEN1/BUB1/ZNF207/CDCA8/CCNB1/DYNC1LI1/TRIP13/ANAPC7/TPR | 19 |
| GO:0000075 | cell cycle checkpoint signaling | 37/1673 | 224/20772 | 2,20386E-05 | 0,001152707 | 0,001068044 | TTK/ETAA1/NAE1/ATR/THOC1/RBBP8/MSH2/NUF2/HUS1/CHEK1/RPA2/ATF2/BLM/ABRAXAS1/RAD50/ATM/CDC5L/INTS7/SPDL1/MRE11/TRIAP1/KNTC1/GEN1/BUB1/INIP/ZNF207/CDCA8/TTI1/BRIP1/CCNB1/DYNC1LI1/SETMAR/ORC1/CEP63/TRIP13/TPR/ATRIP | 37 |

| | | | | | | | | |
|---|---|---|---|---|---|---|---|---|
| GO:0010212 | response to ionizing radiation | 28/1673 | 150/20772 | 2,27289E-05 | 0,001177269 | 0,001090802 | FIGNL1/ATR/ERCC8/MSH2/WRN/LIG4/HUS1/KDM4D/CASP3/RAD51AP1/COP1/BLM/ABRAXAS1/ATM/INTS7/XRCC4/INIP/CLK2/XRCC2/NHEJ1/POLB/IKBIP/EYA3/PAXIP1/CLK2/XRCC5/KDM1A/XRCC6 | 28 |
| GO:0010165 | response to X-ray | 11/1673 | 32/20772 | 2,35818E-05 | 0,0012097 | 0,001120851 | ERCC8/MSH2/LIG4/CASP3/BLM/ATM/XRCC4/XRCC2/IKBIP/XRCC5/XRCC6 | 11 |
| GO:0045005 | DNA-templated DNA replication maintenance of fidelity | 15/1673 | 56/20772 | 2,50662E-05 | 0,001273601 | 0,001180058 | ETAA1/ATR/THOC1/RBBP8/WRN/MMS22L/BLM/RAD50/CENPS/MRE11/ZRANB3/GEN1/FANCM/MCM8/SETMAR | 15 |
| GO:0006278 | RNA-templated DNA biosynthetic process | 18/1673 | 76/20772 | 2,53347E-05 | 0,001275102 | 0,001181449 | ATR/CCT8/RAD50/ATM/TENT4B/MRE11/CCT2/CCT6A/TERF1/CCT5/DCP2/CCT4/PTGES3/PARN/NAF1/CCT3/XRCC5/NHP2 | 18 |
| GO:2001251 | negative regulation of chromosome organization | 21/1673 | 98/20772 | 2,84282E-05 | 0,00141443 | 0,001310543 | TTK/SMARCA5/NUF2/RAD50/ATM/TENT4B/SPDL1/KNTC1/GEN1/BUB1/WAPL/TERF1/ZNF207/CDCA8/DCP2/TOP2A/CCNB1/DYNC1LI1/SETMAR/TRIP13/TPR | 21 |
| GO:2001252 | positive regulation of chromosome organization | 26/1673 | 136/20772 | 2,86333E-05 | 0,00141443 | 0,001310543 | DHX36/TASOR/ATR/RESF1/LIG4/CCT8/NCAPG/RAD50/ATM/SETDB2/MRE11/NSMCE2/CCT2/CCT6A/TERF1/SSBP1/CCT5/GTF2H2/CCT4/GNL3/PARN/RUVBL1/NAF1/TPR/CCT3/XRCC5 | 26 |
| GO:0043484 | regulation of RNA splicing | 33/1673 | 195/20772 | 3,64469E-05 | 0,001783893 | 0,001652871 | PTBP2/TADA1/MBNL3/MAGOH/CELF6/RPS13/RBM7/KHDRBS2/PTBP3/SRRM4/CWC22/ENY2/RBFOX2/SRSF12/SUPT3H/TRA2B/RBMX/RBM22/WTAP/PIK3R1/TAF12/SRPK1/CLK2/CLNS1A/SRSF1/TRA2A/NCBP1/SRSF7/ILDR2/LARP7/HSPA8/CLK2/SF3B4 | 33 |

| | | | | | | | | |
|---|---|---|---|---|---|---|---|---|
| GO:0010639 | negative regulation of organelle organization | 56/1673 | 401/20772 | 3,68171E-05 | 0,001785631 | 0,001654481 | S1PR1/STMN2/TTK/TMSB15B/SLIT2/FGF13/CAPZA1/SMAD4/CKAP2/SMARCA5/NUF2/CHEK1/MTBP/CLASP2/GDI2/MAP2/CTNNA2/RAD50/BBS4/ATM/TENT4B/NPM1/SPDL1/TJP1/TRIAP1/KNTC1/TBC1D7/GHITM/GEN1/BUB1/CYRIB/PIK3R1/ARHGAP28/WAPL/CCP110/BMP4/TMEM14A/MAPRE1/TERF1/ZNF207/CDCA8/NBDY/FXN/DCP2/TOP2A/CCNB1/TOM1L1/MPHOSPH9/KIF24/DYNC1LI1/ACAA2/SETMAR/PARL/SLC25A4/TRIP13/TPR | 56 |
| GO:2000779 | regulation of double-strand break repair | 26/1673 | 139/20772 | 4,23295E-05 | 0,002034484 | 0,001885056 | ACTL6B/FIGNL1/UBE2V2/MBTD1/ATR/USP51/PARPBP/RBBP8/UBE2N/KDM4D/CHEK1/HMGA2/WDR48/RAD51AP1/RPA2/BCL7A/RIF1/RNF169/PHF10/EPC1/MEAF6/SETMAR/SMARCC1/RUVBL1/ATRIP/KDM1A | 26 |
| GO:1904851 | positive regulation of establishment of protein localization to telomere | 6/1673 | 10/20772 | 4,27749E-05 | 0,002037539 | 0,001887887 | CCT8/CCT2/CCT6A/CCT5/CCT4/CCT3 | 6 |
| GO:1990173 | protein localization to nucleoplasm | 7/1673 | 14/20772 | 4,48936E-05 | 0,002119536 | 0,001963861 | CCT8/CCT2/CCT6A/CCT5/CCT4/LARP7/CCT3 | 7 |
| GO:0008608 | attachment of spindle microtubules to kinetochore | 13/1673 | 46/20772 | 4,66609E-05 | 0,002183648 | 0,002023264 | NUF2/MIS12/CENPE/ABRAXAS1/SGO1/KNL1/MAPRE1/ZNF207/CDCA8/CCNB1/BOD1/CHAMP1/SEH1L | 13 |
| GO:0006275 | regulation of DNA replication | 28/1673 | 156/20772 | 4,74821E-05 | 0,002202757 | 0,00204097 | NPM2/RFC3/CDC7/ORC3/ATR/CCNA2/SMARCA5/DSCC1/ATAD5/OBI1/RFC4/BLM/PDS5A/DBF4/ZRANB3/USP37/SMC3/FAF1/GMNN/WAPL/RFC5/ORC5/TERF1/SSBP1/MEAF6/RUVBL1/DHX9/MCM6 | 28 |
| GO:0007088 | regulation of mitotic nuclear division | 24/1673 | 125/20772 | 5,32447E-05 | 0,002448795 | 0,002268938 | TTK/BORA/PHIP/NUF2/CHEK1/DLGAP5/MTBP/ATM/SPDL1/NSMCE2/CDC16/KNTC1/GEN1/BUB1/BMP4/ZNF207/CDCA8/CDC25C/CCNB1/TOM1L1/DYNC1LI1/TRIP13/ANAPC7/TPR | 24 |

| | | | | | | | | |
|---|---|---|---|---|---|---|---|---|
| GO:0090501 | RNA phosphodiester bond hydrolysis | 31/1673 | 182/20772 | 5,45649E-05 | 0,002488065 | 0,002305323 | RIDA/LACTB2/CNOT7/FCF1/EXOSC8/DICER1/RPP30/ZC3H12B/ANGEL2/ERI1/TSEN34/UTP23/ERI2/EXO1/POLR2I/RPS21/UTP20/EXOSC9/N4BP1/NCBP1/DCP2/CPSF3/CNOT1/NCBP2/PARN/PCF11/NUDT21/ABT1/APEX1/POP5/NHP2 | 31 |
| GO:0007094 | mitotic spindle assembly checkpoint signaling | 13/1673 | 47/20772 | 5,97814E-05 | 0,002657782 | 0,002462575 | TTK/NUF2/ATM/SPDL1/KNTC1/GEN1/BUB1/ZNF207/CDCA8/CCNB1/DYNC1LI1/TRIP13/TPR | 13 |
| GO:0071173 | spindle assembly checkpoint signaling | 13/1673 | 47/20772 | 5,97814E-05 | 0,002657782 | 0,002462575 | TTK/NUF2/ATM/SPDL1/KNTC1/GEN1/BUB1/ZNF207/CDCA8/CCNB1/DYNC1LI1/TRIP13/TPR | 13 |
| GO:0071174 | mitotic spindle checkpoint signaling | 13/1673 | 47/20772 | 5,97814E-05 | 0,002657782 | 0,002462575 | TTK/NUF2/ATM/SPDL1/KNTC1/GEN1/BUB1/ZNF207/CDCA8/CCNB1/DYNC1LI1/TRIP13/TPR | 13 |
| GO:0010833 | telomere maintenance via telomere lengthening | 19/1673 | 88/20772 | 6,03252E-05 | 0,002659795 | 0,00246444 | DHX36/ATR/CCT8/RAD50/ATM/TENT4B/MRE11/CCT2/CCT6A/TERF1/CCT5/DCP2/CCT4/PTGES3/PARN/NAF1/CCT3/XRCC5/NHP2 | 19 |
| GO:0007018 | microtubule-based movement | 59/1673 | 437/20772 | 6,16428E-05 | 0,002695608 | 0,002497622 | CFAP61/RASGRP1/DNAH6/DNAH7/RSPH4A/NME5/HSBP1/BBS12/DNAAF11/VPS13A/IFT70B/DYNC2H1/IFT70A/DNAH14/AGTPBP1/KIF18A/DYNLL1/PIERCE1/KIF21A/WASF1/DYNLT2B/BLOC1S5/IFT88/LCA5/DLGAP5/CFAP221/KIF14/SNAPIN/ARMC2/SLIRP/DNAJA1/CENPE/KIF11/MAP2/RFX3/BBS4/LRPPRC/IFT56/SSX2IP/SPAST/AGBL4/KIF9/KIF23/RPGR/ODAD2/WDR35/IFT52/INTS13/HYDIN/HYDIN/TTC21B/KIF3A/PCM1/KIF24/DYNC1LI1/HSPA8/OFD1/INTU/IFT22 | 59 |
| GO:1904358 | positive regulation of telomere maintenance via telomere lengthening | 12/1673 | 41/20772 | 6,2281E-05 | 0,002701377 | 0,002502968 | DHX36/ATR/CCT8/ATM/CCT2/CCT6A/CCT5/CCT4/PARN/NAF1/CCT3/XRCC5 | 12 |

| | | | | | | | | |
|---|---|---|---|---|---|---|---|---|
| GO:0061077 | chaperone-mediated protein folding | 18/1673 | 82/20772 | 7,43924E-05 | 0,003200673 | 0,002965592 | HSPE1/HSPA13/PDCD5/CCT8/PPID/FKBP5/HSPH1/CHORDC1/CCT2/CCT6A/CCT5/CCT4/PTGES3/CHORDC1/HSPA9/HSPA8/DNAJB14/CCT3 | 18 |
| GO:0031577 | spindle checkpoint signaling | 13/1673 | 48/20772 | 7,59892E-05 | 0,003243219 | 0,003005013 | TTK/NUF2/ATM/SPDL1/KNTC1/GEN1/BUB1/ZNF207/CDCA8/CCNB1/DYNC1LI1/TRIP13/TPR | 13 |
| GO:0006405 | RNA export from nucleus | 20/1673 | 97/20772 | 7,77015E-05 | 0,003264076 | 0,003024338 | NXT2/MAGOHB/MAGOH/THOC1/XPOT/ENY2/NUP155/SSB/NUP107/NUP160/NUP153/NPM1/SMG7/RAN/NCBP1/UPF2/NCBP2/DHX9/TPR/POLR2D | 20 |
| GO:0042274 | ribosomal small subunit biogenesis | 20/1673 | 97/20772 | 7,77015E-05 | 0,003264076 | 0,003024338 | LSM6/FCF1/NPM1/NOL11/UTP23/HEATR1/DCAF13/RPS21/UTP6/UTP20/WDR43/DDX52/LTV1/MRPS7/METTL17/SRFBP1/ABT1/XRCC5/RPS5/RIOK1 | 20 |
| GO:0051983 | regulation of chromosome segregation | 25/1673 | 136/20772 | 7,92168E-05 | 0,003301731 | 0,003059227 | TTK/ACTL6B/NUF2/DLGAP5/NCAPG/CENPE/BCL7A/ATM/SPDL1/ANAPC4/NSMCE2/CDC16/KNTC1/PHF10/GEN1/BUB1/CDC26/ZNF207/CDCA8/CCNB1/DYNC1LI1/TRIP13/SMARCC1/ANAPC7/TPR | 25 |
| GO:0071044 | histone mRNA catabolic process | 6/1673 | 11/20772 | 8,76694E-05 | 0,003625706 | 0,003359407 | SSB/MTPAP/ATM/TENT4B/TENT2/DCP2 | 6 |
| GO:0034501 | protein localization to kinetochore | 8/1673 | 20/20772 | 9,08766E-05 | 0,003672929 | 0,003403162 | TTK/CENPQ/MIS12/MTBP/SPDL1/KNTC1/KNL1/CHAMP1 | 8 |
| GO:0045653 | negative regulation of megakaryocyte differentiation | 8/1673 | 20/20772 | 9,08766E-05 | 0,003672929 | 0,003403162 | H4C3/H4C11/H4C4/H4C8/H4C9/H4C5/H4C2/PRMT1 | 8 |
| GO:1903083 | protein localization to condensed chromosome | 8/1673 | 20/20772 | 9,08766E-05 | 0,003672929 | 0,003403162 | TTK/CENPQ/MIS12/MTBP/SPDL1/KNTC1/KNL1/CHAMP1 | 8 |
| GO:0033046 | negative regulation of sister chromatid segregation | 13/1673 | 49/20772 | 9,58686E-05 | 0,003760726 | 0,00348451 | TTK/NUF2/ATM/SPDL1/KNTC1/GEN1/BUB1/ZNF207/CDCA8/CCNB1/DYNC1LI1/TRIP13/TPR | 13 |

| | | | | | | | | |
|---|---|---|---|---|---|---|---|---|
| GO:0033048 | negative regulation of mitotic sister chromatid segregation | 13/1673 | 49/20772 | 9,58686E-05 | 0,003760726 | 0,00348451 | TTK/NUF2/ATM/SPDL1/KNTC1/GEN1/BUB1/ZNF207/CDCA8/CCNB1/DYNC1LI1/TRIP13/TPR | 13 |
| GO:0045841 | negative regulation of mitotic metaphase/anaphase transition | 13/1673 | 49/20772 | 9,58686E-05 | 0,003760726 | 0,00348451 | TTK/NUF2/ATM/SPDL1/KNTC1/GEN1/BUB1/ZNF207/CDCA8/CCNB1/DYNC1LI1/TRIP13/TPR | 13 |
| GO:2000816 | negative regulation of mitotic sister chromatid separation | 13/1673 | 49/20772 | 9,58686E-05 | 0,003760726 | 0,00348451 | TTK/NUF2/ATM/SPDL1/KNTC1/GEN1/BUB1/ZNF207/CDCA8/CCNB1/DYNC1LI1/TRIP13/TPR | 13 |
| GO:0000469 | cleavage involved in rRNA processing | 10/1673 | 31/20772 | 0,00010269 | 0,003998895 | 0,003705187 | FCF1/EXOSC8/ERI1/UTP23/ERI2/RPS21/UTP20/EXOSC9/ABT1/NHP2 | 10 |
| GO:0032212 | positive regulation of telomere maintenance via telomerase | 11/1673 | 37/20772 | 0,000107315 | 0,004148729 | 0,003844016 | ATR/CCT8/ATM/CCT2/CCT6A/CCT5/CCT4/PARN/NAF1/CCT3/XRCC5 | 11 |
| GO:0051783 | regulation of nuclear division | 27/1673 | 155/20772 | 0,000108987 | 0,004183057 | 0,003875822 | TTK/NPM2/BORA/PHIP/NUF2/CHEK1/DLGAP5/MTBP/RAD51AP1/ATM/SPDL1/NSMCE2/CDC16/KNTC1/GEN1/BUB1/BMP4/ZNF207/CDCA8/CDC25C/CCNB1/TOM1L1/PDE3A/DYNC1LI1/TRIP13/ANAPC7/TPR | 27 |
| GO:0006401 | RNA catabolic process | 50/1673 | 361/20772 | 0,000115589 | 0,004404756 | 0,004081238 | DHX36/RIDA/ELAVL4/LSM5/MAGOHB/MAGOH/LSM6/CNOT7/RBM7/FASTKD1/FASTKD2/EXOSC8/DICER1/ZCCHC7/SLIRP/TUT4/SSB/MTPAP/ANGEL2/ATM/TENT4B/LRPPRC/TENT2/NPM1/ETF1/ZCCHC17/TRIM71/EXOSC9/SMG7/NBDY/NCBP1/NBAS/SECISBP2/GAS5/DCP2/NANOS1/NT5C3B/CNOT1/UPF2/NCBP2/POLR2G/PARN/RC3H1/TNRC6C/NAF1/CNOT9/DHX9/APEX1/POLR2D/IGF2BP3 | 50 |

| | | | | | | | | |
|---|---|---|---|---|---|---|---|---|
| GO:0006457 | protein folding | 37/1673 | 243/20772 | 0,000129222 | 0,004889359 | 0,004530248 | DNAJC19/HSPA4L/CLGN/HSPE1/VBP1/HSPA13/PDCD5/PPIL1/CCT8/PPID/DNAJA1/FKBP5/RANBP2/NKTR/PFDN6/PFDN6/HSPH1/DNAJC10/NFYC/PPIA/CHORDC1/CCT2/CCT6A/TTC4/TBCEL/CCT5/PPIH/NGLY1/CCT4/PTGES3/CHORDC1/HSPA9/NUDCD2/HSPA8/DNAJB14/DNAJC2/CCT3 | 37 |
| GO:0001510 | RNA methylation | 19/1673 | 93/20772 | 0,000132155 | 0,00491904 | 0,004557749 | SNRPG/METTL5/SNRPF/RAMAC/SNRPE/TRMT11/VIRMA/TRMT13/NSUN6/METTL15/TARBP1/WTAP/HENMT1/CMTR2/THUMPD3/SNRPD2/METTL2A/LARP7/TFB2M | 19 |
| GO:0043631 | RNA polyadenylation | 12/1673 | 44/20772 | 0,00013259 | 0,00491904 | 0,004557749 | PAPOLG/PAPOLA/ZCCHC7/MTPAP/VIRMA/TENT4B/TENT2/WDR33/CPSF3/SNRPA/PCF11/NUDT21 | 12 |
| GO:0071897 | DNA biosynthetic process | 33/1673 | 208/20772 | 0,000132938 | 0,00491904 | 0,004557749 | RFC3/ATR/USP1/WRN/LIG4/CCT8/DSCC1/CHEK1/PARM1/RFC4/RAD50/ATM/TENT4B/LIN9/MRE11/POLD3/CCT2/CCT6A/RFC5/TERF1/CCT5/DCP2/CCT4/PTGES3/MEAF6/POLB/PARN/NAF1/DNAJC2/CCT3/RRM1/XRCC5/NHP2 | 33 |
| GO:0099111 | microtubule-based transport | 34/1673 | 217/20772 | 0,000135973 | 0,00491904 | 0,004557749 | RASGRP1/NME5/HSBP1/BBS12/DNAAF11/IFT70B/DYNC2H1/IFT70A/AGTPBP1/DYNLL1/WASF1/DYNLT2B/BLOC1S5/IFT88/LCA5/CFAP221/SNAPIN/MAP2/RFX3/LRPPRC/IFT56/SSX2IP/SPAST/AGBL4/RPGR/WDR35/IFT52/TTC21B/KIF3A/PCM1/HSPA8/OFD1/INTU/IFT22 | 34 |
| GO:0043628 | regulatory ncRNA 3'-end processing | 8/1673 | 21/20772 | 0,000136461 | 0,00491904 | 0,004557749 | EXOSC8/TENT4B/ERI1/ERI2/RPS21/EXOSC9/LARP7/PARN | 8 |
| GO:0046931 | pore complex assembly | 8/1673 | 21/20772 | 0,000136461 | 0,00491904 | 0,004557749 | NDC1/CCT8/NUP107/NUP153/NUP205/AHCTF1/TMEM170A/CCT3 | 8 |
| GO:0051315 | attachment of mitotic spindle microtubules to kinetochore | 8/1673 | 21/20772 | 0,000136461 | 0,00491904 | 0,004557749 | NUF2/MIS12/CENPE/MAPRE1/CDCA8/BOD1/CHAMP1/SEH1L | 8 |
| GO:2000573 | positive regulation of DNA biosynthetic process | 18/1673 | 86/20772 | 0,000142441 | 0,005100138 | 0,004725546 | RFC3/ATR/USP1/CCT8/DSCC1/PARM1/RFC4/ATM/CCT2/CCT6A/RFC5/CCT5/CCT4/PTGES3/PARN/NAF1/CCT3/XRCC5 | 18 |

| | | | | | | | | |
|---|---|---|---|---|---|---|---|---|
| GO:0051985 | negative regulation of chromosome segregation | 13/1673 | 51/20772 | 0,000149399 | 0,005243719 | 0,004858581 | TTK/NUF2/ATM/SPDL1/KNTC1/GEN1/BUB1/ZNF207/CDCA8/CCNB1/DYNC1LI1/TRIP13/TPR | 13 |
| GO:1902100 | negative regulation of metaphase/anaphase transition of cell cycle | 13/1673 | 51/20772 | 0,000149399 | 0,005243719 | 0,004858581 | TTK/NUF2/ATM/SPDL1/KNTC1/GEN1/BUB1/ZNF207/CDCA8/CCNB1/DYNC1LI1/TRIP13/TPR | 13 |
| GO:1905819 | negative regulation of chromosome separation | 13/1673 | 51/20772 | 0,000149399 | 0,005243719 | 0,004858581 | TTK/NUF2/ATM/SPDL1/KNTC1/GEN1/BUB1/ZNF207/CDCA8/CCNB1/DYNC1LI1/TRIP13/TPR | 13 |
| GO:1901990 | regulation of mitotic cell cycle phase transition | 53/1673 | 394/20772 | 0,000156498 | 0,00545697 | 0,005056169 | TTK/SASS6/ACTL6B/NPM2/NAE1/CDC7/RAD51C/RBBP8/NUF2/HUS1/CHEK1/DLGAP5/ATAD5/MTBP/KIF14/CENPE/RPA2/BCL7A/WEE1/BLM/ABRAXAS1/RAD50/ATM/SPDL1/EZH2/MRE11/ANAPC4/NSMCE2/TRIAP1/CDC16/KNTC1/PHF10/GEN1/BUB1/CDC26/CCNH/INIP/ZNF207/KMT2E/CDCA8/CDC25C/CCNB1/CPSF3/DYNC1LI1/FBXO7/ORC1/TRIP13/SMARCC1/ANAPC7/TPR/APEX1/RRM1/LSM11 | 53 |
| GO:0070646 | protein modification by small protein removal | 30/1673 | 185/20772 | 0,000177369 | 0,006144568 | 0,005693265 | YOD1/USP12/TADA1/USP25/USP1/USP51/UCHL3/SENP1/TAF5/ENY2/USP14/USP9X/ABRAXAS1/SUPT3H/PSMD14/OTUD3/USP37/COPS4/STAMBP/COPS3/TAF12/DESI2/OTUD6B/USP54/COPS5/SENP6/OTUD4/USP8/USP38/SENP5 | 30 |
| GO:0007093 | mitotic cell cycle checkpoint signaling | 28/1673 | 168/20772 | 0,000179256 | 0,00616988 | 0,005716718 | TTK/NAE1/RBBP8/MSH2/NUF2/HUS1/CHEK1/RPA2/ATF2/BLM/ABRAXAS1/RAD50/ATM/SPDL1/MRE11/TRIAP1/KNTC1/GEN1/BUB1/INIP/ZNF207/CDCA8/CCNB1/DYNC1LI1/SETMAR/ORC1/TRIP13/TPR | 28 |

| | | | | | | | | |
|---|---|---|---|---|---|---|---|---|
| GO:0006413 | translational initiation | 24/1673 | 135/20772 | 0,000186287 | 0,006370779 | 0,005902861 | EIF1AX/EIF4E3/PAIP2B/DHX29/RPS3A/EIF4EBP2/NCK1/NPM1/EIF2S3/MCTS1/MTIF2/EIF2AK3/CDC123/ABCE1/NCBP1/DENR/COPS5/PPP1R15B/NCBP2/POLR2G/EIF2S2/TPR/POLR2D/RPS5 | 24 |
| GO:0010970 | transport along microtubule | 29/1673 | 177/20772 | 0,000188079 | 0,006391087 | 0,005921678 | RASGRP1/HSBP1/BBS12/IFT70B/DYNC2H1/IFT70A/AGTPBP1/DYNLL1/WASF1/DYNLT2B/BLOC1S5/IFT88/LCA5/SNAPIN/MAP2/LRPPRC/IFT56/SSX2IP/SPAST/AGBL4/RPGR/WDR35/IFT52/TTC21B/KIF3A/PCM1/HSPA8/INTU/IFT22 | 29 |
| GO:0007062 | sister chromatid cohesion | 14/1673 | 59/20772 | 0,000192298 | 0,006411934 | 0,005940994 | RAD51C/DSCC1/SGO1/PDS5A/MRE11/NSMCE2/NAA50/MCMBP/SMC3/BUB1/SGO2/WAPL/STAG1/BOD1 | 14 |
| GO:0016073 | snRNA metabolic process | 14/1673 | 59/20772 | 0,000192298 | 0,006411934 | 0,005940994 | INTS6L/RBM7/EXOSC8/ICE2/ZCCHC7/INTS2/INTS7/ZC3H8/INTS8/INTS13/EXOSC9/LARP7/SNAPC3/NHP2 | 14 |
| GO:0032210 | regulation of telomere maintenance via telomerase | 14/1673 | 59/20772 | 0,000192298 | 0,006411934 | 0,005940994 | ATR/CCT8/ATM/TENT4B/CCT2/CCT6A/TERF1/CCT5/DCP2/CCT4/PARN/NAF1/CCT3/XRCC5 | 14 |
| GO:0033151 | V(D)J recombination | 7/1673 | 17/20772 | 0,000204942 | 0,006749186 | 0,006253475 | LIG4/ATM/XRCC4/NHEJ1/POLB/HMGB2/DCAF1 | 7 |
| GO:0070200 | establishment of protein localization to telomere | 7/1673 | 17/20772 | 0,000204942 | 0,006749186 | 0,006253475 | CCT8/CCT2/CCT6A/TERF1/CCT5/CCT4/CCT3 | 7 |
| GO:0031023 | microtubule organizing center organization | 27/1673 | 161/20772 | 0,000208872 | 0,006806376 | 0,006306465 | SASS6/TUBE1/HAUS1/CETN3/CEP44/CHEK1/CLASP2/CEP295/KIF11/BBS4/CNTLN/SSX2IP/SGO1/NPM1/GCC2/PARD6B/GEN1/CCP110/CHORDC1/XRCC2/PLK4/KIF3A/PCM1/CHORDC1/MAP9/CEP63/PATJ | 27 |
| GO:2000278 | regulation of DNA biosynthetic process | 24/1673 | 136/20772 | 0,000209231 | 0,006806376 | 0,006306465 | RFC3/ATR/USP1/CCT8/DSCC1/CHEK1/PARM1/RFC4/ATM/TENT4B/CCT2/CCT6A/RFC5/TERF1/CCT5/DCP2/CCT4/PTGES3/MEAF6/PARN/NAF1/DNAJC2/CCT3/XRCC5 | 24 |

| | | | | | | | | |
|---|---|---|---|---|---|---|---|---|
| GO:0070198 | protein localization to chromosome, telomeric region | 10/1673 | 34/20772 | 0,000242865 | 0,007852623 | 0,007275868 | ATR/CCT8/CCT2/CCT6A/TERF1/CCT5/CCT4/GNL3/CCT3/XRCC5 | 10 |
| GO:0051321 | meiotic cell cycle | 44/1673 | 316/20772 | 0,000256318 | 0,008237677 | 0,007632641 | TESMIN/MND1/TTK/CCNA1/NDC1/RMI1/NPM2/FIGNL1/KIF18A/RAD51C/RBM7/ASPM/RBBP8/SPATA17/NUF2/HUS1/RAD51AP1/ANAPC13/ORC4/RAD50/ATM/SGO1/CENPS/TOP2B/MRE11/ANAPC4/CDC16/TDRKH/SMC3/EXO1/BUB1/FANCM/CDC26/SGO2/TERF1/XRCC2/CCNB1IP1/CDC25C/TOP2A/BRIP1/PDE3A/CCNE1/TRIP13/ANAPC7 | 44 |
| GO:1904356 | regulation of telomere maintenance via telomere lengthening | 15/1673 | 68/20772 | 0,000273104 | 0,008713457 | 0,008073476 | DHX36/ATR/CCT8/ATM/TENT4B/CCT2/CCT6A/TERF1/CCT5/DCP2/CCT4/PARN/NAF1/CCT3/XRCC5 | 15 |
| GO:0006368 | transcription elongation by RNA polymerase II | 22/1673 | 122/20772 | 0,000275315 | 0,008713457 | 0,008073476 | MED23/GTF2F2/ENY2/MED6/MED7/ELP2/INTS2/ELP4/INTS7/EZH2/EAPP/INTS8/CCNT1/INTS13/POLR2I/MED14/CDK13/NCBP1/WDR43/LARP7/NCBP2/EAF1 | 22 |
| GO:0033047 | regulation of mitotic sister chromatid segregation | 13/1673 | 54/20772 | 0,00027697 | 0,008713457 | 0,008073476 | TTK/NUF2/ATM/SPDL1/KNTC1/GEN1/BUB1/ZNF207/CDCA8/CCNB1/DYNC1LI1/TRIP13/TPR | 13 |
| GO:2001022 | positive regulation of response to DNA damage stimulus | 29/1673 | 181/20772 | 0,000278068 | 0,008713457 | 0,008073476 | EEF1E1/ACTL6B/EYA4/UBE2V2/MBTD1/ATR/USP1/RBBP8/UBE2N/KDM4D/WDR48/RAD51AP1/BCL7A/ABRAXAS1/ATM/RIF1/NSMCE4A/CEBPG/PHF10/MARCHF6-DT/EPC1/TTI1/MEAF6/SETMAR/EYA3/SMARCC1/RUVBL1/PAXIP1/DHX9 | 29 |
| GO:0030490 | maturation of SSU-rRNA | 14/1673 | 61/20772 | 0,000279602 | 0,008713457 | 0,008073476 | LSM6/FCF1/NOL11/UTP23/HEATR1/DCAF13/RPS21/UTP6/UTP20/WDR43/DDX52/SRFBP1/ABT1/RIOK1 | 14 |

| | | | | | | | | |
|---|---|---|---|---|---|---|---|---|
| GO:0000288 | nuclear-transcribed mRNA catabolic process, deadenylation-dependent decay | 17/1673 | 83/20772 | 0,000283629 | 0,008713457 | 0,008073476 | DHX36/CNOT7/EXOSC8/TUT4/TENT4B/EXOSC9/DCP2/NANOS1/NT5C3B/CNOT1/POLR2G/PARN/RC3H1/TNRC6C/CNOT9/DHX9/POLR2D | 17 |
| GO:0009127 | purine nucleoside monophosphate biosynthetic process | 8/1673 | 23/20772 | 0,000284188 | 0,008713457 | 0,008073476 | HPRT1/GMPS/PPAT/ADSS2/GART/ADSL/PAICS/DGUOK | 8 |
| GO:0051383 | kinetochore organization | 8/1673 | 23/20772 | 0,000284188 | 0,008713457 | 0,008073476 | NUF2/CENPH/DLGAP5/MIS12/CENPE/CENPW/KNTC1/SENP6 | 8 |
| GO:0008033 | tRNA processing | 26/1673 | 156/20772 | 0,000299785 | 0,009139169 | 0,00846792 | DPH3/DTWD1/FAM98B/LSM6/OSGEPL1/ADAT2/TRMT11/RPP30/TYW5/ELP2/TSEN15/SSB/ELP4/CDKAL1/TRMT13/NSUN6/TSEN34/SEPSECS/TARBP1/YRDC/THUMPD3/ELP1/TRNT1/METTL2A/CDK5RAP1/POP5 | 26 |
| GO:0051382 | kinetochore assembly | 7/1673 | 18/20772 | 0,000312107 | 0,009407282 | 0,008716341 | CENPH/DLGAP5/MIS12/CENPE/CENPW/KNTC1/SENP6 | 7 |
| GO:0061644 | protein localization to CENP-A containing chromatin | 7/1673 | 18/20772 | 0,000312107 | 0,009407282 | 0,008716341 | H4C3/H4C11/H4C4/H4C8/H4C9/H4C5/H4C2 | 7 |
| GO:0007052 | mitotic spindle organization | 25/1673 | 148/20772 | 0,000314472 | 0,009425324 | 0,008733058 | TTK/SASS6/BORA/NUF2/CENPH/DLGAP5/CLASP2/EFHC1/CENPE/KIF11/ABRAXAS1/LSM14A/SPAST/CEP126/KIF23/SMC3/BCCIP/INTS13/ZNF207/RAN/CDCA8/STAG1/CCNB1/MAP9/TPR | 25 |
| GO:0046037 | GMP metabolic process | 9/1673 | 29/20772 | 0,000317021 | 0,00944865 | 0,008754671 | NT5C2/HPRT1/GMPS/PPAT/GART/ADSL/PAICS/GDA/MAGI3 | 9 |

| | | | | | | | | |
|---|---|---|---|---|---|---|---|---|
| GO:0006402 | mRNA catabolic process | 43/1673 | 310/20772 | 0,000325234 | 0,009639568 | 0,008931566 | DHX36/RIDA/ELAVL4/LSM5/MAGOHB/MAGOH/LSM6/CNOT7/FASTKD1/FASTKD2/EXOSC8/ZCCHC7/TUT4/SSB/MTPAP/ANGEL2/ATM/TENT4B/TENT2/NPM1/ETF1/TRIM71/EXOSC9/SMG7/NBDY/NCBP1/NBAS/SECISBP2/DCP2/NANOS1/NT5C3B/CNOT1/UPF2/NCBP2/POLR2G/PARN/RC3H1/TNRC6C/CNOT9/DHX9/APEX1/POLR2D/IGF2BP3 | 43 |
| GO:0006400 | tRNA modification | 19/1673 | 100/20772 | 0,000354236 | 0,010441169 | 0,009674292 | DPH3/DTWD1/OSGEPL1/ADAT2/TRMT11/TYW5/ELP2/SSB/ELP4/CDKAL1/TRMT13/NSUN6/SEPSECS/TARBP1/YRDC/THUMPD3/ELP1/METTL2A/CDK5RAP1 | 19 |
| GO:0000380 | alternative mRNA splicing, via spliceosome | 16/1673 | 77/20772 | 0,00035713 | 0,010468628 | 0,009699734 | MAGOH/CELF6/RBM7/KHDRBS2/SRRM4/RBFOX2/SRSF12/TRA2B/RBMX/WTAP/CDK13/SRSF1/NCBP1/NCBP2/DHX9/KDM1A | 16 |
| GO:0018022 | peptidyl-lysine methylation | 24/1673 | 141/20772 | 0,000365594 | 0,010658173 | 0,009875357 | DYDC2/SMAD4/EEF1AKMT1/RLF/WDR5B/METTL18/PAX5/N6AMT1/CAMKMT/RIF1/EZH2/SMYD3/SETDB2/MTF2/PRDM5/ATPSCKMT/KDM6A/NFYC/NFYA/SETMAR/BOD1/PAXIP1/KANSL2/KDM1A | 24 |
| GO:0006378 | mRNA polyadenylation | 11/1673 | 42/20772 | 0,000369466 | 0,010712498 | 0,009925693 | PAPOLG/PAPOLA/MTPAP/VIRMA/TENT4B/TENT2/WDR33/CPSF3/SNRPA/PCF11/NUDT21 | 11 |
| GO:0072594 | establishment of protein localization to organelle | 61/1673 | 487/20772 | 0,000376943 | 0,010870211 | 0,010071821 | NXT2/RGPD6/EFCAB7/RHOU/RHOU/DNAJC19/VPS13A/KPNA3/CSE1L/PEX1/SRP9/TOMM5/PDCD5/SEC61B/CCT8/RPL23/TOMM70/DNAJC15/DNAJA1/NUP155/ATF2/USP9X/NUP107/VPS13C/KPNA2/NUP153/RANBP2/PEX13/GCC2/RBM22/TIMM8A/KPNA4/MFF/SSR3/PIK3R1/ZIC1/BMP4/CCT2/CCT6A/E2F3/IPO5/RAB3GAP2/STK4/TOMM6/TERF1/CCT5/GLI3/TNPO1/RAN/CCT4/TIMM10B/NPEPPS/FBXO7/MTX2/PARL/IPO11/NF1/HSPA8/TPR/CCT3/UBE2D3 | 61 |
| GO:0009066 | aspartate family amino acid metabolic process | 12/1673 | 49/20772 | 0,000398016 | 0,011416204 | 0,010577713 | RIDA/BHMT/MTHFD2L/AASS/ASNS/SLC25A13/ASNSD1/ADSS2/SMS/MTR/SLC25A12/AASDHPPT | 12 |

| | | | | | | | | |
|---|---|---|---|---|---|---|---|---|
| GO:0034504 | protein localization to nucleus | 45/1673 | 332/20772 | 0,00040068 | 0,011431174 | 0,010591584 | NXT2/RGPD6/EFCAB7/KPNA3/CSE1L/PYGO1/WRN/CCT8/TXN/RPL23/NUP155/ATF2/NUP107/KPNA2/BBS4/CD2AP/NUP153/RANBP2/RBM22/KPNA4/PIK3R1/ZIC1/LATS1/EIF2AK3/BMP4/INTS13/DCLK1/CCT2/CCT6A/E2F3/IPO5/YWHAE/STK4/CCT5/GLI3/TNPO1/RAN/CCT4/LARP7/IPO11/NF1/RPF2/NOL8/TPR/CCT3 | 45 |
| GO:0021543 | pallium development | 32/1673 | 213/20772 | 0,000453714 | 0,012875333 | 0,011929673 | TBR1/EOMES/RELN/EMX2/NEUROD1/SLIT2/CDON/FGF13/ASPM/ZIC3/PAX5/ANXA3/KIF14/EFHC1/CASP3/BBS4/TENT2/TRA2B/EZH2/ARHGAP11B/PEX13/FAT4/ZIC1/FBXO45/YWHAE/GART/CDH2/CDK5R1/GLI3/RAN/NF1/KDM1A | 32 |
| GO:0046040 | IMP metabolic process | 7/1673 | 19/20772 | 0,000459949 | 0,012983222 | 0,012029637 | NT5C2/HPRT1/PPAT/ADSS2/GART/ADSL/PAICS | 7 |
| GO:0006119 | oxidative phosphorylation | 27/1673 | 169/20772 | 0,000464478 | 0,013042047 | 0,012084142 | COX7C/TMEM135/DLD/NDUFA12/NDUFAB1/MSH2/NDUFA4/DNAJC15/ATP5PB/NDUFB3/NDUFC1/NDUFB5/GHITM/ATP5PO/ATPSCKMT/CYCS/FXN/UQCRH/CCNB1/NDUFS1/COX7A2L/UQCRC2/DGUOK/SLC25A33/AFG1L/UQCRFS1/ATP5F1B | 27 |
| GO:0098727 | maintenance of cell number | 31/1673 | 205/20772 | 0,000492924 | 0,013768315 | 0,012757068 | EOMES/KIT/ACTL6B/PAX8/HDAC2/ASPM/LIG4/ZIC3/HMGA2/MED6/MED7/TUT4/BCL7A/RIF1/SS18/EZH2/SINHCAF/MTF2/PHF10/SMC3/ZIC1/BRAF/SAV1/MED14/CDH2/WDR43/GNL3/PCM1/CNOT1/SMARCC1/FANCC | 31 |
| GO:0046165 | alcohol biosynthetic process | 24/1673 | 144/20772 | 0,000502255 | 0,013955881 | 0,012930857 | ASAH2/CES1/PRKAA2/SC5D/P2RY1/IDI1/PPIP5K2/MSMO1/PTS/INSIG2/ACER2/HMGCS1/CYP51A1/HSD17B7/HMGCR/DHFR/IMPA2/FDFT1/LBR/ERLIN1/MBTPS2/ACAA2/FDPS/NSDHL | 24 |
| GO:1903046 | meiotic cell cycle process | 35/1673 | 242/20772 | 0,000520573 | 0,014389925 | 0,013333022 | TESMIN/MND1/TTK/CCNA1/NDC1/RMI1/NPM2/FIGNL1/KIF18A/RAD51C/ASPM/SPATA17/NUF2/HUS1/RAD51AP1/ORC4/RAD50/ATM/SGO1/CENPS/TOP2B/MRE11/TDRKH/SMC3/BUB1/FANCM/SGO2/TERF1/CCNB1IP1/CDC25C/TOP2A/BRIP1/PDE3A/CCNE1/TRIP13 | 35 |

| | | | | | | | | |
|---|---|---|---|---|---|---|---|---|
| GO:0034655 | nucleobase-containing compound catabolic process | 61/1673 | 494/20772 | 0,000546353 | 0,015024703 | 0,013921177 | DHX36/RIDA/ELAVL4/LSM5/PDE7A/MAGOHB/MAGOH/LSM6/CNOT7/RBM7/NUDT11/NT5C2/FASTKD1/FASTKD2/EXOSC8/DICER1/HPRT1/ADAL/ZCCHC7/SLIRP/TUT4/SSB/APAF1/MTPAP/ANGEL2/ATM/TENT4B/LRPPRC/TENT2/NPM1/ETF1/ENTPD7/ZCCHC17/TRIM71/PDE10A/NUDT15/EXOSC9/SMG7/NBDY/NCBP1/NBAS/SECISBP2/GAS5/DCP2/NANOS1/NT5C3B/CNOT1/UPF2/SETMAR/NCBP2/GDA/POLR2G/PARN/RC3H1/TNRC6C/NAF1/CNOT9/DHX9/APEX1/POLR2D/IGF2BP3 | 61 |
| GO:0010948 | negative regulation of cell cycle process | 48/1673 | 366/20772 | 0,000556176 | 0,015156042 | 0,014042869 | TTK/ETAA1/NAE1/ATR/THOC1/RBBP8/MSH2/SMARCA5/NUF2/HUS1/CHEK1/MTBP/RPA2/ATF2/WEE1/BLM/ABRAXAS1/RAD50/ATM/CDC5L/INTS7/NPM1/SPDL1/EZH2/MRE11/TRIAP1/KNTC1/GEN1/BUB1/WAPL/INIP/CCP110/BMP4/TERF1/ZNF207/CDCA8/TTI1/BRIP1/CCNB1/TOM1L1/DYNC1LI1/FBXO7/SETMAR/ORC1/CEP63/TRIP13/TPR/ATRIP | 48 |
| GO:0007098 | centrosome cycle | 24/1673 | 145/20772 | 0,000556811 | 0,015156042 | 0,014042869 | SASS6/TUBE1/HAUS1/CETN3/CEP44/CHEK1/CEP295/KIF11/BBS4/CNTLN/SSX2IP/SGO1/NPM1/PARD6B/GEN1/CCP110/CHORDC1/XRCC2/PLK4/KIF3A/PCM1/CHORDC1/MAP9/CEP63 | 24 |
| GO:0031125 | rRNA 3'-end processing | 5/1673 | 10/20772 | 0,000601587 | 0,016209422 | 0,015018881 | EXOSC8/ERI1/ERI2/RPS21/EXOSC9 | 5 |
| GO:2000234 | positive regulation of rRNA processing | 5/1673 | 10/20772 | 0,000601587 | 0,016209422 | 0,015018881 | UTP15/WDR75/HEATR1/WDR43/RIOK1 | 5 |
| GO:0007051 | spindle organization | 34/1673 | 235/20772 | 0,00061642 | 0,016444278 | 0,015236487 | TTK/SASS6/BORA/HAUS1/TRIM36/ASPM/SPATA17/NUF2/CENPH/DLGAP5/CLASP2/EFHC1/CENPE/KIF11/ABRAXAS1/LSM14A/SPAST/CEP126/KIF23/SMC3/BCCIP/INTS13/MAPRE1/ZNF207/RAN/CDCA8/STAG1/CCNB1/MAP9/TUBB/SENP6/CEP63/MAPRE3/TPR | 34 |
| GO:0021987 | cerebral cortex development | 24/1673 | 146/20772 | 0,000616468 | 0,016444278 | 0,015236487 | TBR1/EOMES/RELN/EMX2/SLIT2/CDON/FGF13/ASPM/PAX5/KIF14/EFHC1/BBS4/TRA2B/ARHGAP11B/PEX13/FAT4/FBXO45/YWHAE/GART/CDH2/CDK5R1/GLI3/NF1/KDM1A | 24 |

| | | | | | | | | |
|---|---|---|---|---|---|---|---|---|
| GO:0030705 | cytoskeleton-dependent intracellular transport | 33/1673 | 226/20772 | 0,000624445 | 0,016574188 | 0,015356856 | HOOK1/RASGRP1/HSBP1/BBS12/IFT70B/DYNC2H1/IFT70A/AGTPBP1/DYNLL1/WASF1/DYNLT2B/BLOC1S5/IFT88/LCA5/MAP6/SNAPIN/MAP2/LRPPRC/IFT56/SSX2IP/SPAST/AGBL4/RPGR/WDR35/IFT52/TTC21B/MYO6/KIF3A/PCM1/TUBB/HSPA8/INTU/IFT22 | 33 |
| GO:0000460 | maturation of 5.8S rRNA | 10/1673 | 38/20772 | 0,000650746 | 0,017186794 | 0,015924468 | FCF1/EXOSC8/RPF1/ERI1/ERI2/RPS21/UTP20/EXOSC9/MPHOSPH6/ABT1 | 10 |
| GO:0006367 | transcription initiation at RNA polymerase II promoter | 22/1673 | 130/20772 | 0,000682762 | 0,017943513 | 0,016625608 | DHX36/ELOC/CAND1/MED23/GTF2F2/TAF5/MED6/MED7/PSMC6/CCNH/TAF12/TAF13/POLR2I/E2F3/MED14/GTF2A1/GTF2H2/TBP/POLR2G/ZNF451/PAXIP1/POLR2D | 22 |
| GO:0035825 | homologous recombination | 15/1673 | 74/20772 | 0,000710814 | 0,018549633 | 0,01718721 | MND1/RMI1/RAD51C/RBBP8/RAD51AP1/RAD50/ATM/CENPS/TOP2B/MRE11/FANCM/CCNB1IP1/TOP2A/BRIP1/TRIP13 | 15 |
| GO:0006167 | AMP biosynthetic process | 6/1673 | 15/20772 | 0,000716256 | 0,018549633 | 0,01718721 | HPRT1/PPAT/ADSS2/GART/ADSL/PAICS | 6 |
| GO:0006177 | GMP biosynthetic process | 6/1673 | 15/20772 | 0,000716256 | 0,018549633 | 0,01718721 | HPRT1/GMPS/PPAT/GART/ADSL/PAICS | 6 |
| GO:0009060 | aerobic respiration | 32/1673 | 219/20772 | 0,00073946 | 0,019058053 | 0,017658288 | COX7C/TMEM135/DLD/MDH1B/NDUFA12/NDUFAB1/MSH2/SLC25A14/NDUFA4/DNAJC15/ATP5PB/DLAT/NDUFB3/NDUFC1/FH/NDUFB5/GHITM/ATP5PO/ATPSCKMT/CYCS/FXN/ADSL/UQCRH/CCNB1/NDUFS1/COX7A2L/UQCRC2/DGUOK/SLC25A33/AFG1L/UQCRFS1/ATP5F1B | 32 |
| GO:0006354 | DNA-templated transcription elongation | 24/1673 | 148/20772 | 0,00075268 | 0,019305515 | 0,017887574 | CCNT2/THOC1/MED23/GTF2F2/ENY2/MED6/MED7/ELP2/INTS2/ELP4/INTS7/EZH2/EAPP/INTS8/CCNT1/INTS13/POLR2I/MED14/CDK13/NCBP1/WDR43/LARP7/NCBP2/EAF1 | 24 |
| GO:0045132 | meiotic chromosome segregation | 22/1673 | 131/20772 | 0,000759377 | 0,019314833 | 0,017896208 | MND1/TTK/NDC1/RMI1/RAD51C/ASPM/NUF2/ATM/SGO1/CENPS/TOP2B/MRE11/SMC3/BUB1/FANCM/SGO2/TERF1/CCNB1IP1/TOP2A/BRIP1/CCNE1/TRIP13 | 22 |

| | | | | | | | | |
|---|---|---|---|---|---|---|---|---|
| GO:1900182 | positive regulation of protein localization to nucleus | 17/1673 | 90/20772 | 0,000760284 | 0,019314833 | 0,017896208 | EFCAB7/CCT8/CD2AP/RBM22/PIK3R1/ZIC1/EIF2AK3/CCT2/CCT6A/IPO5/CCT5/GLI3/RAN/CCT4/LARP7/TPR/CCT3 | 17 |
| GO:0007131 | reciprocal meiotic recombination | 14/1673 | 67/20772 | 0,000770877 | 0,019399191 | 0,01797437 | MND1/RMI1/RAD51C/RAD51AP1/RAD50/ATM/CENPS/TOP2B/MRE11/FANCM/CCNB1IP1/TOP2A/BRIP1/TRIP13 | 14 |
| GO:0140527 | reciprocal homologous recombination | 14/1673 | 67/20772 | 0,000770877 | 0,019399191 | 0,01797437 | MND1/RMI1/RAD51C/RAD51AP1/RAD50/ATM/CENPS/TOP2B/MRE11/FANCM/CCNB1IP1/TOP2A/BRIP1/TRIP13 | 14 |
| GO:0000959 | mitochondrial RNA metabolic process | 12/1673 | 53/20772 | 0,000852286 | 0,021297016 | 0,019732805 | FASTKD1/MTERF3/FASTKD2/YARS2/SLIRP/LRPPRC/TFAM/TRNT1/DARS2/CDK5RAP1/TFB2M/SLC25A33 | 12 |
| GO:0009126 | purine nucleoside monophosphate metabolic process | 11/1673 | 46/20772 | 0,000854276 | 0,021297016 | 0,019732805 | NT5C2/HPRT1/GMPS/PPAT/ADSS2/GART/ADSL/PAICS/GDA/DGUOK/MAGI3 | 11 |
| GO:0006520 | amino acid metabolic process | 43/1673 | 325/20772 | 0,000877623 | 0,021777306 | 0,020177818 | RIDA/BHMT/DDC/KYAT3/MTHFD2L/CDO1/DLD/PTS/SLC38A1/YARS2/AASS/AASDH/GMPS/DHFR/FARSB/CTPS2/DBT/PPAT/NARS2/ASNS/ARHGAP11B/ODC1/RARS1/SLC25A13/ASNSD1/GLUD2/GLUD2/QRSL1/IARS2/ADSS2/SMS/HIBCH/EPRS1/GART/MCCC2/MTR/ALDH5A1/SLC25A12/AASDHPPT/DARS2/PSAT1/IARS1/LARS1 | 43 |
| GO:0006352 | DNA-templated transcription initiation | 27/1673 | 176/20772 | 0,00088194 | 0,021783109 | 0,020183196 | DHX36/TBPL1/TAF9B/ELOC/SMARCA5/CAND1/MED23/GTF2F2/TAF5/MED6/MED7/PSMC6/TFAM/CCNH/TAF12/TAF13/POLR2I/E2F3/MED14/GTF2A1/GTF2H2/TBP/POLR2G/ZNF451/TFB2M/PAXIP1/POLR2D | 27 |
| GO:1903313 | positive regulation of mRNA metabolic process | 28/1673 | 185/20772 | 0,000887233 | 0,021812843 | 0,020210745 | DHX36/RIDA/CNOT7/EXOSC8/TUT4/TENT4B/TRA2B/RBMX/TRIM71/CLNS1A/EXOSC9/TRA2A/NCBP1/DCP2/NANOS1/NT5C3B/CNOT1/NCBP2/POLR2G/PARN/HSPA8/RC3H1/TNRC6C/NUDT21/CNOT9/DHX9/POLR2D/SF3B4 | 28 |

| | | | | | | | | |
|---|---|---|---|---|---|---|---|---|
| GO:0009168 | purine ribonucleoside monophosphate biosynthetic process | 7/1673 | 21/20772 | 0,000919791 | 0,022406769 | 0,020761049 | HPRT1/GMPS/PPAT/ADSS2/GART/ADSL/PAICS | 7 |
| GO:0016024 | CDP-diacylglycerol biosynthetic process | 7/1673 | 21/20772 | 0,000919791 | 0,022406769 | 0,020761049 | TAMM41/GPAM/AGPAT5/AGPAT5/AGPAT4/LCLAT1/CDS2 | 7 |
| GO:0140013 | meiotic nuclear division | 32/1673 | 222/20772 | 0,000934244 | 0,022655412 | 0,02099143 | TESMIN/MND1/TTK/CCNA1/NDC1/RMI1/NPM2/FIGNL1/KIF18A/RAD51C/ASPM/NUF2/RAD51AP1/ORC4/RAD50/ATM/SGO1/CENPS/TOP2B/MRE11/TDRKH/SMC3/BUB1/FANCM/SGO2/TERF1/CCNB1IP1/TOP2A/BRIP1/PDE3A/CCNE1/TRIP13 | 32 |
| GO:0000466 | maturation of 5.8S rRNA from tricistronic rRNA transcript (SSU-rRNA, 5.8S rRNA, LSU-rRNA) | 8/1673 | 27/20772 | 0,000962218 | 0,023228208 | 0,021522156 | FCF1/EXOSC8/ERI1/ERI2/RPS21/UTP20/EXOSC9/ABT1 | 8 |
| GO:0032206 | positive regulation of telomere maintenance | 17/1673 | 92/20772 | 0,00098407 | 0,023648699 | 0,021911762 | DHX36/ATR/CCT8/RAD50/ATM/MRE11/CCT2/CCT6A/TERF1/CCT5/CCT4/GNL3/PARN/RUVBL1/NAF1/CCT3/XRCC5 | 17 |
| GO:0006338 | chromatin remodeling | 55/1673 | 446/20772 | 0,001028336 | 0,024527931 | 0,022726418 | TBR1/EOMES/H1-6/ACTL6B/NPM2/H3-3A/H1-1/H4C3/SHPRH/HAT1/HDAC2/TASOR/SMARCAD1/RESF1/H4C11/ERCC6L/SMARCA5/H4C4/KDM4D/CHEK1/CHD1/RNF2/HMGA2/ASF1A/BCL7A/OIP5/ERCC6L2/RIF1/SS18/H4C8/NPM1/EZH2/CENPW/SMYD3/SETDB2/NAP1L5/ZRANB3/PHF10/MIS18A/KDM6A/GATAD1/CBX3/HJURP/ITGB3BP/TSPYL4/TTI1/H4C9/H4C5/H4C2/DDX21/SMARCC1/RUVBL1/HMGB2/TPR/KDM1A | 55 |

| | | | | | | | | |
|---|---|---|---|---|---|---|---|---|
| GO:0001731 | formation of translation preinitiation complex | 5/1673 | 11/20772 | 0,001029851 | 0,024527931 | 0,022726418 | DHX29/EIF2S3/MCTS1/DENR/EIF2S2 | 5 |
| GO:2000232 | regulation of rRNA processing | 6/1673 | 16/20772 | 0,00106828 | 0,025330099 | 0,023469668 | UTP15/METTL18/WDR75/HEATR1/WDR43/RIOK1 | 6 |
| GO:0045739 | positive regulation of DNA repair | 23/1673 | 144/20772 | 0,001184301 | 0,027956838 | 0,02590348 | ACTL6B/EYA4/UBE2V2/MBTD1/USP1/RBBP8/UBE2N/KDM4D/WDR48/RAD51AP1/BCL7A/ABRAXAS1/RIF1/CEBPG/PHF10/MARCHF6-DT/EPC1/MEAF6/SETMAR/EYA3/SMARCC1/RUVBL1/DHX9 | 23 |
| GO:0071168 | protein localization to chromatin | 12/1673 | 55/20772 | 0,001206539 | 0,028356315 | 0,026273616 | H4C3/TASOR/MSH2/H4C11/H4C4/H4C8/EZH2/WAPL/H4C9/MCM8/H4C5/H4C2 | 12 |
| GO:0000184 | nuclear-transcribed mRNA catabolic process, nonsense-mediated decay | 11/1673 | 48/20772 | 0,00124669 | 0,029007523 | 0,026876995 | MAGOHB/MAGOH/ETF1/SMG7/NCBP1/NBAS/SECISBP2/DCP2/UPF2/NCBP2/PARN | 11 |
| GO:1900151 | regulation of nuclear-transcribed mRNA catabolic process, deadenylation-dependent decay | 8/1673 | 28/20772 | 0,001252969 | 0,029007523 | 0,026876995 | DHX36/CNOT7/TENT4B/NANOS1/CNOT1/RC3H1/TNRC6C/DHX9 | 8 |
| GO:0009067 | aspartate family amino acid biosynthetic process | 7/1673 | 22/20772 | 0,001255996 | 0,029007523 | 0,026876995 | BHMT/MTHFD2L/AASS/ASNS/ASNSD1/MTR/AASDHPPT | 7 |
| GO:0046341 | CDP-diacylglycerol metabolic process | 7/1673 | 22/20772 | 0,001255996 | 0,029007523 | 0,026876995 | TAMM41/GPAM/AGPAT5/AGPAT5/AGPAT4/LCLAT1/CDS2 | 7 |

| | | | | | | | | |
|---|---|---|---|---|---|---|---|---|
| GO:0021537 | telencephalon development | 40/1673 | 302/20772 | 0,00127482 | 0,029315359 | 0,027162221 | TBR1/EOMES/RELN/EMX2/NEUROD1/SLIT2/CDON/FGF13/AGTPBP1/WDR89/ASPM/ZIC3/PAX5/HPRT1/ANXA3/KIF14/EFHC1/CASP3/ERBB4/BBS4/TENT2/TRA2B/EZH2/ARHGAP11B/PEX13/FAT4/ZIC1/FBXO45/BMP4/YWHAE/GART/CDH2/CDK5R1/GLI3/RAN/SECISBP2/KCNC1/NF1/NRG3/KDM1A | 40 |
| GO:0006414 | translational elongation | 13/1673 | 63/20772 | 0,001326617 | 0,030375533 | 0,028144528 | DPH3/SRP9/METTL18/EEF1B2/EEF1B2/DPH5/SEPSECS/NEMF/C6orf52/C6orf52/SECISBP2/EEF1A1/MRPL44 | 13 |
| GO:1902850 | microtubule cytoskeleton organization involved in mitosis | 27/1673 | 181/20772 | 0,001352513 | 0,030721121 | 0,028464733 | TTK/SASS6/BORA/NUF2/CENPH/DLGAP5/CLASP2/EFHC1/CENPE/KIF11/ABRAXAS1/LSM14A/SPDL1/SPAST/CEP126/KIF23/SMC3/BCCIP/INTS13/MAPRE1/ZNF207/RAN/CDCA8/STAG1/CCNB1/MAP9/TPR | 27 |
| GO:0006606 | protein import into nucleus | 26/1673 | 172/20772 | 0,001353227 | 0,030721121 | 0,028464733 | NXT2/RGPD6/EFCAB7/KPNA3/CSE1L/RPL23/NUP155/ATF2/NUP107/KPNA2/NUP153/RANBP2/RBM22/KPNA4/PIK3R1/ZIC1/BMP4/E2F3/IPO5/STK4/GLI3/TNPO1/RAN/IPO11/NF1/TPR | 26 |
| GO:0032204 | regulation of telomere maintenance | 21/1673 | 129/20772 | 0,00148544 | 0,033561695 | 0,031096674 | DHX36/ATR/CCT8/RAD50/ATM/TENT4B/NSMCE4A/MRE11/NSMCE2/CCT2/CCT6A/TERF1/CCT5/DCP2/CCT4/GNL3/PARN/RUVBL1/NAF1/CCT3/XRCC5 | 21 |
| GO:0048024 | regulation of mRNA splicing, via spliceosome | 19/1673 | 112/20772 | 0,001490932 | 0,033561695 | 0,031096674 | MAGOH/CELF6/RBM7/KHDRBS2/SRRM4/CWC22/RBFOX2/SRSF12/TRA2B/RBMX/WTAP/SRPK1/CLNS1A/TRA2A/NCBP1/SRSF7/LARP7/HSPA8/SF3B4 | 19 |
| GO:0043508 | negative regulation of JUN kinase activity | 6/1673 | 17/20772 | 0,001539191 | 0,034502453 | 0,031968336 | DUSP19/ZNF675/DNAJA1/AIDA/DUSP16/DUSP16 | 6 |
| GO:0019827 | stem cell population maintenance | 29/1673 | 201/20772 | 0,001556252 | 0,03473894 | 0,032187454 | EOMES/KIT/ACTL6B/PAX8/HDAC2/ASPM/LIG4/ZIC3/HMGA2/MED6/MED7/TUT4/BCL7A/RIF1/SS18/SINHCAF/MTF2/PHF10/SMC3/BRAF/SAV1/MED14/CDH2/WDR43/GNL3/PCM1/CNOT1/SMARCC1/FANCC | 29 |
| GO:0006450 | regulation of translational fidelity | 8/1673 | 29/20772 | 0,00160952 | 0,035778285 | 0,033150462 | IARS2/YRDC/IARS1/CDK5RAP1/DNAJC2/DTD1/LARS1/RPS5 | 8 |

| | | | | | | | | |
|---|---|---|---|---|---|---|---|---|
| GO:0140694 | non-membrane-bounded organelle assembly | 54/1673 | 445/20772 | 0,00163253 | 0,03604992 | 0,033402146 | TNNT1/PRKAA2/SASS6/SMAD4/HAUS1/CNOT7/ASPM/LSM3/CENPH/MTERF3/FASTKD2/CEP44/DLGAP5/MIS12/CLASP2/CEP295/CENPE/KIF11/MRPL22/DHX29/ABRAXAS1/LSM14A/NPM1/CENPW/KIF9/KNTC1/KIF23/BRIX1/SMC3/RPS23/RPL5/BCCIP/CCP110/SQLE/MAPRE1/ZNF207/CDCA8/NBDY/STAG1/PLK4/MAP9/CNOT1/TUBB/MRPS7/SENP6/CEP63/RPF2/RC3H1/MAPRE3/TPR/CDS2/ABT1/XRCC5/RPS5 | 54 |
| GO:0006188 | IMP biosynthetic process | 5/1673 | 12/20772 | 0,001648769 | 0,03604992 | 0,033402146 | HPRT1/PPAT/GART/ADSL/PAICS | 5 |
| GO:0009113 | purine nucleobase biosynthetic process | 5/1673 | 12/20772 | 0,001648769 | 0,03604992 | 0,033402146 | HPRT1/GMPS/PPAT/GART/PAICS | 5 |
| GO:0032790 | ribosome disassembly | 5/1673 | 12/20772 | 0,001648769 | 0,03604992 | 0,033402146 | GFM2/MRRF/MCTS1/MTIF2/DENR | 5 |
| GO:0006144 | purine nucleobase metabolic process | 7/1673 | 23/20772 | 0,001681162 | 0,036606186 | 0,033917555 | HPRT1/GMPS/PPAT/GART/PAICS/GDA/KDM1A | 7 |
| GO:0022904 | respiratory electron transport chain | 22/1673 | 139/20772 | 0,001687933 | 0,036606186 | 0,033917555 | COX7C/DLD/NDUFA12/NDUFAB1/NDUFA4/DNAJC15/NDUFB3/NDUFC1/NDUFB5/SLC25A13/GHITM/CYCS/GPD2/UQCRH/CCNB1/SLC25A12/NDUFS1/COX7A2L/UQCRC2/DGUOK/AFG1L/UQCRFS1 | 22 |
| GO:1903320 | regulation of protein modification by small protein conjugation or removal | 37/1673 | 278/20772 | 0,00173954 | 0,037521918 | 0,034766029 | UBE2V2/GLMN/SVBP/UBE2N/RWDD3/SKP2/RPL23/MTBP/WDR48/DNAJA1/RBX1/IVNS1ABP/DCUN1D4/CENPS/TBC1D7/UBA2/COPS4/RPS20/RPL5/MARCHF6-DT/FANCM/BTRC/COPS3/PPIA/SPOPL/PELI1/PLAA/UBQLN1/N4BP1/GNL3/COPS5/AMER1/OTUD4/PRMT3/PAXIP1/FANCI/KDM1A | 37 |
| GO:0034968 | histone lysine methylation | 20/1673 | 122/20772 | 0,001744224 | 0,037521918 | 0,034766029 | DYDC2/SMAD4/RLF/WDR5B/PAX5/N6AMT1/RIF1/EZH2/SMYD3/SETDB2/MTF2/PRDM5/KDM6A/NFYC/NFYA/SETMAR/BOD1/PAXIP1/KANSL2/KDM1A | 20 |

| | | | | | | | | |
|---|---|---|---|---|---|---|---|---|
| GO:0034243 | regulation of transcription elongation by RNA polymerase II | 17/1673 | 97/20772 | 0,001800275 | 0,038572155 | 0,035739129 | MED23/MED6/MED7/INTS2/INTS7/EZH2/EAPP/INTS8/CCNT1/INTS13/MED14/CDK13/NCBP1/WDR43/LARP7/NCBP2/EAF1 | 17 |
| GO:0045333 | cellular respiration | 36/1673 | 269/20772 | 0,001808806 | 0,038599927 | 0,035764861 | COX7C/TMEM135/DLD/MDH1B/NDUFA12/NDUFAB1/MSH2/SLC25A14/NDUFA4/DNAJC15/ATP5PB/DLAT/NDUFB3/NDUFC1/FH/NDUFB5/SLC25A13/GHITM/ATP5PO/ATPSCKMT/CYCS/GPD2/IFNAR1/FXN/ADSL/UQCRH/CCNB1/SLC25A12/NDUFS1/COX7A2L/UQCRC2/DGUOK/SLC25A33/AFG1L/UQCRFS1/ATP5F1B | 36 |
| GO:0002562 | somatic diversification of immune receptors via germline recombination within a single locus | 14/1673 | 73/20772 | 0,001850104 | 0,039167868 | 0,036291088 | THOC1/MSH2/LIG4/ATAD5/ATM/RIF1/XRCC4/EXO1/NHEJ1/POLB/MSH3/HMGB2/PAXIP1/DCAF1 | 14 |
| GO:0016444 | somatic cell DNA recombination | 14/1673 | 73/20772 | 0,001850104 | 0,039167868 | 0,036291088 | THOC1/MSH2/LIG4/ATAD5/ATM/RIF1/XRCC4/EXO1/NHEJ1/POLB/MSH3/HMGB2/PAXIP1/DCAF1 | 14 |
| GO:0016579 | protein deubiquitination | 25/1673 | 167/20772 | 0,001899136 | 0,040046994 | 0,037105645 | YOD1/USP12/TADA1/USP25/USP1/USP51/UCHL3/TAF5/ENY2/USP14/USP9X/ABRAXAS1/SUPT3H/PSMD14/OTUD3/USP37/STAMBP/TAF12/DESI2/OTUD6B/USP54/COPS5/OTUD4/USP8/USP38 | 25 |
| GO:0070979 | protein K11-linked ubiquitination | 8/1673 | 30/20772 | 0,002041793 | 0,0428857 | 0,039735855 | UBE2T/ANAPC13/ANAPC4/CDC16/CDC26/UBE2E3/ANAPC7/UBE2D3 | 8 |
| GO:0051170 | import into nucleus | 26/1673 | 177/20772 | 0,002049893 | 0,04288698 | 0,039737041 | NXT2/RGPD6/EFCAB7/KPNA3/CSE1L/RPL23/NUP155/ATF2/NUP107/KPNA2/NUP153/RANBP2/RBM22/KPNA4/PIK3R1/ZIC1/BMP4/E2F3/IPO5/STK4/GLI3/TNPO1/RAN/IPO11/NF1/TPR | 26 |
| GO:2000781 | positive regulation of double-strand break repair | 16/1673 | 90/20772 | 0,002080211 | 0,043351282 | 0,040167241 | ACTL6B/UBE2V2/MBTD1/RBBP8/UBE2N/KDM4D/WDR48/RAD51AP1/BCL7A/RIF1/PHF10/EPC1/MEAF6/SETMAR/SMARCC1/RUVBL1 | 16 |

| | | | | | | | | |
|---|---|---|---|---|---|---|---|---|
| GO:0010569 | regulation of double-strand break repair via homologous recombination | 14/1673 | 74/20772 | 0,002115938 | 0,043924236 | 0,040698114 | FIGNL1/MBTD1/USP51/PARPBP/RBBP8/CHEK1/WDR48/RAD51AP1/RPA2/RIF1/EPC1/MEAF6/RUVBL1/KDM1A | 14 |
| GO:0009167 | purine ribonucleoside monophosphate metabolic process | 10/1673 | 44/20772 | 0,002192926 | 0,044997147 | 0,041692221 | NT5C2/HPRT1/GMPS/PPAT/ADSS2/GART/ADSL/PAICS/GDA/MAGI3 | 10 |
| GO:0050685 | positive regulation of mRNA processing | 10/1673 | 44/20772 | 0,002192926 | 0,044997147 | 0,041692221 | DHX36/TRA2B/RBMX/CLNS1A/TRA2A/NCBP1/NCBP2/HSPA8/NUDT21/SF3B4 | 10 |
| GO:0090503 | RNA phosphodiester bond hydrolysis, exonucleolytic | 10/1673 | 44/20772 | 0,002192926 | 0,044997147 | 0,041692221 | CNOT7/EXOSC8/ANGEL2/ERI1/ERI2/EXOSC9/DCP2/CPSF3/CNOT1/PARN | 10 |
| GO:1903241 | U2-type prespliceosome assembly | 7/1673 | 24/20772 | 0,0022102 | 0,045177844 | 0,041859647 | SNRPG/SNRPF/SNRPE/SNRPB2/SNRPD2/SNRPA1/SF3B4 | 7 |
| GO:0042773 | ATP synthesis coupled electron transport | 19/1673 | 116/20772 | 0,002266601 | 0,045978381 | 0,042601386 | COX7C/DLD/NDUFA12/NDUFAB1/NDUFA4/DNAJC15/NDUFB3/NDUFC1/NDUFB5/GHITM/CYCS/UQCRH/CCNB1/NDUFS1/COX7A2L/UQCRC2/DGUOK/AFG1L/UQCRFS1 | 19 |
| GO:0042775 | mitochondrial ATP synthesis coupled electron transport | 19/1673 | 116/20772 | 0,002266601 | 0,045978381 | 0,042601386 | COX7C/DLD/NDUFA12/NDUFAB1/NDUFA4/DNAJC15/NDUFB3/NDUFC1/NDUFB5/GHITM/CYCS/UQCRH/CCNB1/NDUFS1/COX7A2L/UQCRC2/DGUOK/AFG1L/UQCRFS1 | 19 |

**decreasing genes specific for KO; MF**

| ID | Description | GeneRatio | BgRatio | pvalue | p.adjust | qvalue | geneID | Count |
|---|---|---|---|---|---|---|---|---|
| GO:0140097 | catalytic activity, acting on DNA | 53/1744 | 249/20696 | 2,3754E-10 | 2,4514E-07 | 2,2029E-07 | DHX36/DCLRE1A/TATDN1/FAN1/SHPRH/RFC3/SMARCAD1/RAD51C/N4BP2/RBBP8/MSH2/WRN/ERCC6L/LIG4/SMARCA5/DSCC1/DICER1/CHD1/ATAD5/RFC4/N6AMT1/HMGA2/NEIL3/PMS1/BLM/RAD50/TENT4B/ERCC6L2/TOP2B/MRE11/ZRANB3/GEN1/SMC3/EXO1/FANCM/WAPL/POLD3/RFC5/TERF1/XRCC2/TOP2A/MCM8/PTGES3/BRIP1/POLB/SETMAR/MSH3/RUVBL1/DHX9/MCM6/APEX1/XRCC5/XRCC6 | 53 |
| GO:0008094 | ATP-dependent activity, acting on DNA | 34/1744 | 132/20696 | 2,4394E-09 | 1,2587E-06 | 1,1311E-06 | DHX36/SHPRH/RFC3/SMARCAD1/RAD51C/MSH2/WRN/ERCC6L/SMARCA5/DSCC1/CHD1/ATAD5/RFC4/PMS1/BLM/RAD50/ERCC6L2/TOP2B/MRE11/ZRANB3/SMC3/FANCM/WAPL/RFC5/XRCC2/TOP2A/MCM8/BRIP1/MSH3/RUVBL1/DHX9/MCM6/XRCC5/XRCC6 | 34 |
| GO:0003735 | structural constituent of ribosome | 40/1744 | 194/20696 | 9,2445E-08 | 3,1801E-05 | 2,8577E-05 | RPS18/RSL24D1/RPL22L1/RPS29/MRPL42/MRPS6/RPS13/RPL35A/MRPS22/RPL30/RPL23/MRPL13/RPL9/MRPL47/MRPS31/MRPL22/RPS12/MRPS9/RPL27/MRPL1/RPS3A/RPL24/MRPS30/RPS20/RPS23/MRPS21/RPL5/RPS21/MRPL3/MRPL51/RPL36A/MRPL10/RPL21/MRPS15/RPL31/MRPS7/RPL10A/RPL41/MRPL44/RPS5 | 40 |
| GO:0000049 | tRNA binding | 22/1744 | 81/20696 | 5,805E-07 | 0,00014977 | 0,00013458 | RPL35A/YARS2/TRMT11/TYW5/XPOT/SSB/NSUN6/EIF2S3/RARS1/SEPSECS/IARS2/YRDC/NEMF/C6orf52/C6orf52/THUMPD3/ELP1/TRNT1/DARS2/IARS1/DTD1/EEF1A1 | 22 |

| | | | | | | | | |
|---|---|---|---|---|---|---|---|---|
| GO:0015631 | tubulin binding | 61/1744 | 390/20696 | 1,8136E-06 | 0,00037433 | 0,00033638 | HOOK1/STMN2/PIFO/DCX/EML5/CCDC181/FGF13/VBP1/AGTPBP1/KIF18A/STRBP/CEP70/TRIM36/PDCD5/KIF21A/MTUS2/SVBP/CETN3/CEP44/DLGAP5/MAP6/KIF14/CLASP2/EFHC1/CEP295/CENPE/KIF11/MAP2/NCALD/ABRAXAS1/DCDC1/BBS4/FNTA/LRPPRC/REEP1/CCDC66/CEP57/SPAST/AGBL4/JMY/KIF9/CACYBP/KIF23/SMC3/HSPH1/MAPRE1/CEP57L1/TBCEL/TERF1/ZNF207/CCT5/CDK5R1/KIF3A/KIF24/MAP9/POLB/KATNAL1/OFD1/MAPRE3/DNAL1/TPR | 61 |
| GO:0003678 | DNA helicase activity | 17/1744 | 58/20696 | 3,5115E-06 | 0,00060398 | 0,00054275 | DHX36/RFC3/WRN/DSCC1/RFC4/BLM/RAD50/MRE11/FANCM/RFC5/MCM8/BRIP1/RUVBL1/DHX9/MCM6/XRCC5/XRCC6 | 17 |
| GO:0000400 | four-way junction DNA binding | 8/1744 | 17/20696 | 3,0404E-05 | 0,00401024 | 0,00360366 | RAD51C/MSH2/WRN/BLM/GEN1/FANCM/XRCC2/HMGB2 | 8 |
| GO:0004520 | DNA endonuclease activity | 12/1744 | 37/20696 | 3,1087E-05 | 0,00401024 | 0,00360366 | FAN1/RAD51C/N4BP2/RBBP8/DICER1/RAD50/MRE11/ZRANB3/GEN1/EXO1/SETMAR/APEX1 | 12 |
| GO:0008168 | methyltransferase activity | 37/1744 | 219/20696 | 3,5173E-05 | 0,00403318 | 0,00362427 | BHMT/METTL5/FAM98B/EEF1AKMT1/ARMT1/METTL18/TRMT11/N6AMT1/EED/PRMT9/TRMT13/PCMTD2/PCMT1/CAMKMT/NSUN6/EZH2/SMYD3/SETDB2/DPH5/METTL15/PRDM5/TARBP1/HENMT1/ATPSCKMT/CMTR2/NDUFAF5/MTR/BMT2/THUMPD3/METTL25/METTL2A/SETMAR/PRMT3/TFB2M/METTL17/PRMT1/NSD1 | 37 |
| GO:0019783 | ubiquitin-like protein peptidase activity | 26/1744 | 134/20696 | 4,7267E-05 | 0,00487795 | 0,00438339 | YOD1/USP12/ZUP1/ATG4C/USP25/USP1/USP51/UCHL3/SENP1/USP14/USP9X/PSMD14/OTUD3/USP37/COPS4/STAMBP/DESI2/OTUD6B/USP54/COPS5/SENP6/OTUD4/USP8/USP38/ALG13/SENP5 | 26 |

| | | | | | | | | |
|---|---|---|---|---|---|---|---|---|
| GO:0140098 | catalytic activity, acting on RNA | 64/1744 | 462/20696 | 5,3913E-05 | 0,00505797 | 0,00454516 | DHX36/RIDA/NT5C3A/LACTB2/POLR2K/METTL5/DTWD1/CNOT7/AQR/YARS2/DICER1/TRMT11/RPP30/TYW5/POLR3B/ZC3H12B/PRIM2/TUT4/FARSB/DHX29/DHX15/CDKAL1/TRMT13/ANGEL2/NARS2/TENT2/ERI1/NSUN6/TSEN34/RARS1/QRSL1/ERI2/METTL15/SEPSECS/IARS2/EXO1/TARBP1/HENMT1/FANCM/DDX20/CMTR2/POLR2I/EPRS1/EXOSC9/N4BP1/DCP2/BRIP1/THUMPD3/CPSF3/TRNT1/RTCA/DDX52/METTL2A/CNOT1/DDX21/DARS2/IARS1/PARN/TFB2M/DHX9/APEX1/DTD1/LARS1/POP5 | 64 |
| GO:0061578 | K63-linked deubiquitinase activity | 7/1744 | 14/20696 | 6,0161E-05 | 0,00517381 | 0,00464925 | YOD1/USP14/USP9X/PSMD14/STAMBP/DESI2/OTUD4 | 7 |
| GO:0016741 | transferase activity, transferring one-carbon groups | 38/1744 | 235/20696 | 7,3919E-05 | 0,00586805 | 0,0052731 | BHMT/METTL5/FAM98B/EEF1AKMT1/ARMT1/METTL18/TRMT11/N6AMT1/EED/PRMT9/TRMT13/PCMTD2/PCMT1/CAMKMT/NSUN6/EZH2/SMYD3/SETDB2/DPH5/METTL15/PRDM5/TARBP1/HENMT1/ATPSCKMT/CMTR2/NDUFAF5/GART/MTR/BMT2/THUMPD3/METTL25/METTL2A/SETMAR/PRMT3/TFB2M/METTL17/PRMT1/NSD1 | 38 |
| GO:0003697 | single-stranded DNA binding | 24/1744 | 127/20696 | 0,00013968 | 0,00977721 | 0,00878593 | DHX36/MSH2/MMS22L/NEIL3/WDR48/RAD51AP1/RPA2/RBMS1/BLM/RAD50/LRPPRC/CNBP/ZRANB3/PURB/RAD23B/TERF1/SSBP1/MCM8/SETMAR/MSH3/POLR2G/HMGB2/DHX9/MCM6 | 24 |
| GO:0004386 | helicase activity | 30/1744 | 175/20696 | 0,00014211 | 0,00977721 | 0,00878593 | DHX36/SHPRH/RFC3/SMARCAD1/AQR/WRN/ERCC6L/SMARCA5/DSCC1/DICER1/CHD1/RFC4/BLM/DHX29/DHX15/RAD50/ERCC6L2/MRE11/FANCM/DDX20/RFC5/MCM8/BRIP1/DDX52/DDX21/RUVBL1/DHX9/MCM6/XRCC5/XRCC6 | 30 |
| GO:0000217 | DNA secondary structure binding | 11/1744 | 37/20696 | 0,00016081 | 0,01037225 | 0,00932063 | RAD51C/MSH2/WRN/HMGA2/NEIL3/RAD51AP1/BLM/GEN1/FANCM/XRCC2/HMGB2 | 11 |
| GO:0140142 | nucleocytoplasmic carrier activity | 10/1744 | 32/20696 | 0,00020157 | 0,01199566 | 0,01077946 | KPNA3/CSE1L/XPO7/KPNA2/XPO4/KPNA4/IPO5/TNPO1/RAN/IPO11 | 10 |
| GO:0042162 | telomeric DNA binding | 11/1744 | 38/20696 | 0,00020923 | 0,01199566 | 0,01077946 | WRN/ZBTB10/RPA2/BLM/RAD50/TERF1/SMG7/UPF2/APEX1/XRCC5/XRCC6 | 11 |

| | | | | | | | | |
|---|---|---|---|---|---|---|---|---|
| GO:0004536 | deoxyribonuclease activity | 14/1744 | 59/20696 | 0,00030751 | 0,01657759 | 0,01489683 | DCLRE1A/TATDN1/FAN1/RAD51C/N4BP2/RBBP8/DICER1/RAD50/MRE11/ZRANB3/GEN1/EXO1/SETMAR/APEX1 | 14 |
| GO:0048027 | mRNA 5'-UTR binding | 9/1744 | 28/20696 | 0,00033074 | 0,01657759 | 0,01489683 | DHX36/RPS13/RPS3A/UTP23/RPL5/CCT5/GNL3/RPL41/IGF2BP3 | 9 |
| GO:0004527 | exonuclease activity | 18/1744 | 88/20696 | 0,00033733 | 0,01657759 | 0,01489683 | DCLRE1A/TATDN1/FAN1/CNOT7/ENPP2/WRN/RAD50/ANGEL2/ERI1/MRE11/ERI2/EXO1/EXOSC9/DCP2/CPSF3/CNOT1/PARN/APEX1 | 18 |
| GO:0016796 | exonuclease activity, active with either ribo- or deoxyribonucleic acids and producing 5'-phosphomonoesters | 14/1744 | 60/20696 | 0,00037039 | 0,01663177 | 0,01494552 | DCLRE1A/TATDN1/CNOT7/ANGEL2/ERI1/MRE11/ERI2/EXO1/EXOSC9/DCP2/CPSF3/CNOT1/PARN/APEX1 | 14 |
| GO:0008017 | microtubule binding | 41/1744 | 281/20696 | 0,00038559 | 0,01663177 | 0,01494552 | HOOK1/DCX/EML5/CCDC181/FGF13/KIF18A/STRBP/KIF21A/MTUS2/SVBP/CETN3/CEP44/DLGAP5/MAP6/KIF14/CLASP2/CEP295/CENPE/KIF11/MAP2/ABRAXAS1/DCDC1/FNTA/LRPPRC/REEP1/CCDC66/CEP57/SPAST/JMY/KIF9/KIF23/MAPRE1/CEP57L1/TERF1/ZNF207/KIF3A/KIF24/MAP9/POLB/KATNAL1/MAPRE3 | 41 |
| GO:0004518 | nuclease activity | 35/1744 | 228/20696 | 0,00038679 | 0,01663177 | 0,01494552 | RIDA/DCLRE1A/LACTB2/TATDN1/FAN1/RAD51C/CNOT7/ENPP2/N4BP2/RBBP8/WRN/DICER1/RPP30/ZC3H12B/RAD50/ANGEL2/ASTE1/ERI1/TSEN34/MRE11/ZRANB3/ERI2/GEN1/EXO1/FANCM/EXOSC9/HARBI1/N4BP1/DCP2/CPSF3/CNOT1/SETMAR/PARN/APEX1/POP5 | 35 |
| GO:0019843 | rRNA binding | 17/1744 | 82/20696 | 0,00041368 | 0,01707662 | 0,01534527 | RPS18/MRPS6/RPS13/FASTKD2/RPL23/RPF1/RPL9/ERI1/NPM1/UTP23/BRIX1/PTCD3/RPL5/DDX21/MRPS7/RPF2/RPS5 | 17 |
| GO:0140662 | ATP-dependent protein folding chaperone | 12/1744 | 49/20696 | 0,00059955 | 0,02233182 | 0,02006766 | HSPA4L/HSPE1/HSPA13/CCT8/HSPH1/CCT2/CCT6A/CCT5/CCT4/HSPA9/HSPA8/CCT3 | 12 |

| | | | | | | | | |
|---|---|---|---|---|---|---|---|---|
| GO:0000979 | RNA polymerase II core promoter sequence-specific DNA binding | 7/1744 | 19/20696 | 0,0006059 | 0,02233182 | 0,02006766 | TBPL1/H3-3A/POU2F1/EZH2/H2AZ1/GTF2A1/TBP | 7 |
| GO:0046527 | glucosyltransferase activity | 7/1744 | 19/20696 | 0,0006059 | 0,02233182 | 0,02006766 | UGCG/ALG6/ALG10/ALG10B/POGLUT2/ALG8/POGLUT1 | 7 |
| GO:0043014 | alpha-tubulin binding | 11/1744 | 43/20696 | 0,00067589 | 0,0240525 | 0,02161388 | TRIM36/EFHC1/NCALD/BBS4/FNTA/SPAST/HSPH1/TBCEL/CDK5R1/OFD1/DNAL1 | 11 |
| GO:0140101 | catalytic activity, acting on a tRNA | 26/1744 | 158/20696 | 0,00072715 | 0,02501383 | 0,02247774 | NT5C3A/DTWD1/YARS2/TRMT11/RPP30/TYW5/FARSB/CDKAL1/TRMT13/NARS2/NSUN6/TSEN34/RARS1/QRSL1/SEPSECS/IARS2/TARBP1/EPRS1/THUMPD3/TRNT1/METTL2A/DARS2/IARS1/DTD1/LARS1/POP5 | 26 |
| GO:0101005 | deubiquitinase activity | 21/1744 | 119/20696 | 0,00091049 | 0,0303104 | 0,02723731 | YOD1/USP12/ZUP1/USP25/USP1/USP51/UCHL3/USP14/USP9X/PSMD14/OTUD3/USP37/STAMBP/DESI2/OTUD6B/USP54/COPS5/OTUD4/USP8/USP38/ALG13 | 21 |
| GO:0016888 | endodeoxyribonuclease activity, producing 5'-phosphomonoesters | 5/1744 | 11/20696 | 0,00126621 | 0,03959789 | 0,03558317 | FAN1/DICER1/GEN1/EXO1/APEX1 | 5 |
| GO:0051880 | G-quadruplex DNA binding | 5/1744 | 11/20696 | 0,00126621 | 0,03959789 | 0,03558317 | DHX36/WRN/BLM/RAD50/CNBP | 5 |
